# The phytopathogenic nature of *Dickeya aquatica* 174/2 and the dynamic early evolution of *Dickeya* pathogenicity

**DOI:** 10.1101/568105

**Authors:** Alexandre Duprey, Najwa Taib, Simon Leonard, Tiffany Garin, Jean-Pierre Flandrois, William Nasser, Céline Brochier-Armanet, Sylvie Reverchon

## Abstract

**Originality-Significance statement:** Although the reach of large-scale comparative studies has spread exponentially over the years, the phytopathogenic *Dickeya* group remains overlooked. In this work, we sequence the complete genome of *Dickeya aquatica* type strain, a species isolated from water that was first assumed to be non-phytopathogenic. We show that the proteome of *D. aquatica* contains a wide number of proteins involved in *Dickeya* virulence, including plant cell wall degrading enzymes, suggesting that this species could be in fact pathogenic. Using experimental approaches, we confirm this prediction and uncover the particular affinity of *D. aquatica* for acidic fruits. In-depth phylogenomic analyses reveal that *Dickeya* species display a great degree of genetic plasticity in the pathogenicity determinants, explaining how this bacterial group was able to colonize a wide variety of plants growing in different climates. These observations greatly advance our understanding of how bacteria adapt to new ecological niches.

**Summary:** *Dickeya* is a genus of phytopathogenic enterobacterales causing soft rot in a variety of plants (e.g. potato, chicory, maize). Among the species affiliated to this genus, *Dickeya aquatica*, described in 2014, remained particularly mysterious because it had no known host. Furthermore, while *D. aquatica* was proposed to represent a deep-branching species among *Dickeya* genus, its precise phylogenetic position remained elusive.

Here, we report the complete genome sequence of the *D. aquatica* type strain 174/2. We demonstrate the affinity of *D. aquatica*^T^ for acidic fruits such as tomato and cucumber, and show that exposure of this bacterium to acidic pH induces twitching motility. An in-depth phylogenomic analysis of all available *Dickeya* proteomes pinpoints *D. aquatica* as the second deepest branching lineage within this genus and reclassifies two lineages that likely correspond to new genomospecies (gs.): *Dickeya* gs. poaceaephila (*Dickeya* sp NCPPB 569) and *Dickeya* gs. undicola (*Dickeya* sp 2B12), together with a new putative genus, tentatively named *Prodigiosinella*. Finally, from comparative analyses of *Dickeya* proteomes we infer the complex evolutionary history of this genus, paving the way to study the adaptive patterns and processes of *Dickeya* to different environmental niches and hosts. In particular, we hypothetize that the lack of xylanases and xylose degradation pathways in *D. aquatica* could reflects adaptation to aquatic charophyte hosts which, in contrast to land plants, do not contain xyloglucans.

## Introduction

*Enterobacterales* represent one of the most studied orders of *Gammaproteobacteria*. According to current systematics, *Enterobacterales* are divided into eight families: *Enterobacteriaceae, Yersiniaceae, Thorselliaceae, Hafniaceae, Morganellaceae*, *Budviciaceae, Erwiniaceae*, and *Pectobacteriaceae. Enterobacterales* are widespread, being found in very different environments such as soils, fresh water, ocean, sediments, and many of them are associated with plants and animals, including insects and humans (Brenner and Farmer III, 2005). *Enterobacterales* include also important model organisms such as *Escherichia coli*, human pathogens such as *Salmonella, Shigella,* and *Yersinia* (Dekker and Frank, 2015), and plant pathogens such as *Erwiniaceae (e.g. Erwinia*, *Pantoea, Phaseolibacter)* and *Pectobacteriaceae* (*e.g. Pectobacterium, Dickeya, Brenneria, Lonsdalea)* (Hauben et al., 1998; Samson et al., 2005). These phytopathogens share virulence genes with zoopathogens such as type III secretion systems (T3SS) that inject effector proteins in eukaryotic cells to suppress the host innate immune defence (Buttner, 2016). They also produce specialized plant virulence factors such as pectinases found in pectinolytic bacteria (Hugouvieux-Cotte-Pattat et al., 2014).

Among pectinolytic enterobacterales, *Dickeya* is the causative agent of soft rot in a wide variety of plants including economically important crops (e.g. potato, chicory, maize, rice, tomato, sugar beet, pineapple, banana) and many ornamental plants (Ma et al., 2007). This causes substantial production losses amounting, for instance for potato, to tens of millions of Euros/year in Europe (Toth et al., 2011). The *Dickeya* genus was first described by Samson et al. (2005), who initially distinguished six species: *Dickeya dadantii, Dickeya dieffenbachiae, Dickeya chrysanthemi, Dickeya paradisiaca, Dickeya zeae*, and *Dickeya dianthicola*. Subsequently*, Dickeya dieffenbachiae* has been reclassified as *D. dadantii* subsp. *dieffenbachiae* (Brady et al., 2012).

More recently, three additional *Dickeya* species have been described: *D. solani*, a highly virulent species isolated from potatoes (Van der Wolf et al., 2014), *Dickeya aquatica* from freshwater rivers (Parkinson et al., 2014), and *Dickeya fangzhongdai* from pear trees displaying symptoms of bleeding canker in China (Tian et al., 2016). Thus, the *Dickeya* genus now comprises eight species with distinctive phenotypic features (Supplementary Table S1). Virulence mechanisms have been extensively studied in *Dickeya dadantii*. During infection, the bacterium enters the host using natural openings and multiplies in intercellular spaces without causing major damage. Then, it suddenly induces the production of aggression factors, such as pectinases, that break down the plant cell wall pectin, causing the macroscopic symptom called soft rot (Reverchon and Nasser, 2013). *Dickeya* populations were initially considered to be restricted to tropical and subtropical plant hosts and areas (Perombelon, 1990). This assumption was called into question following the identification of *D. dianthicola* strains from potato plants in Western Europe (Janse and Ruissen, 1988) and the first isolation of *D. solani* strains in France, Finland, Poland, the Netherlands, and Israel (Czajkowski et al., 2011; van der Wolf et al., 2014). It was postulated that the pathogen was introduced in Europe via the international trade of potato seeds (van der Wolf et al., 2014). Strains of *D. solani* have also been found in hyacinth, which was interpreted as a possible recent transfer from hyacinth to potato, possibly via contaminated irrigation waters (Sławiak *et al.*, 2009). This highlights the high capacity of adaptation and dissemination of *Dickeya* to new geographic areas and to new hosts.

According to a large phylogenomic analysis of 895 single copy gene families, *D. paradisiaca* was pinpointed as the first diverging species within *Dickeya*, while other species formed two groups, referred hereafter as to clusters I and II (Zhang et al., 2016). Cluster I encompassed *D. zeae* and *D. chrysanthemi*, while in cluster II, *D. solani* and *D. dadantii* formed the sister-lineage of *D. dianthicola* (Zhang et al., 2016). Species corresponding to these two clusters display different behaviours. For example, in temperate climate, *D. chrysanthemi* and *D. zeae* were frequently isolated from water and were rarely associated to potato infections, contrarily to *D. dianthicola* and *D. solani* (Potrykus et al. 2016). It is noteworthy, that in this study, strains M074 and M005, annotated as *D. chrysanthemi* and *D. solani*, respectively, branch in-between *D. dianthicola* and the *D. solani* / *D. dadantii* group within Cluster II, thus far away from the type strains of their species (Zhang et al., 2016). *D. fangzhongdai* branched at the base of the Cluster II in a tree based on seven housekeeping genes (Tian et al., 2016). Finally, according to a tree based on six housekeeping genes, *D. aquatica* emerge just after the divergence of *D. paradisiaca* but before the divergence of Cluster I and Cluster II (Parkinson et al., 2014). Yet, the associated bootstrap values were weak (Parkinson et al., 2014), meaning that the relative order of emergence of *D. paradisiaca* and *D. aquatica* remained to determine. Among *Dickeya*, *D. aquatica* is remarkable because all the three known strains 174/2^T^, 181/2, and Dw0440 have been isolated from waterways and, in contrast to other species, have no known vegetal host (Supplementary Table S1) (Parkinson et al., 2014; Ma et al., 2007). These features and its early-branching position within *Dickeya* make D. aquatica very interesting to study the origin and the early steps of the diversification of *Dickeya*, and in particular, the emergence of their virulence factors and capacity to infect plants.

These important questions were the focus of this study. For this purpose, we sequence the complete genome of the *D. aquatica* 174/2 type strain. Based on the phylogenetic analysis of 1,341 single copy core protein families, we show that *D. aquatica* represents the second deepest branching lineage within *Dickeya*, reclassify two lineages that likely correspond to new genomospecies (gs.): *Dickeya* gs. poaceaephila and *Dickeya* gs. undicola, and identify a new lineage, tentatively called *Prodigiosinella*, that likely represents the closest relative of *Dickeya*. We also highlight the presence of different virulence factors, suggesting that *D. aquatica* 174/2 could be pathogenic. Stimulated by these findings, we explore the pathogenic potential of *D. aquatica* 174/2. Surprisingly, we identify the acidic fruits tomato (pH 4.8) and cucumber (pH 5.1) as potential hosts for *D. aquatica*, reflecting the potential of this aquatic species to flexibly adapt to a new ecological niche provided by acidic fruits with a high water content. Accordingly, we show that this strain displays a specific induction of twitching motility under acidic conditions. Finally, using phylogenomic approaches, we trace back the origin and evolution of key factors associated with virulence and host specificity in *Dickeya*, including *D. aquatica*. From this study we infer the evolution of gene repertoires along the diversification of *Dickeya* and highlight a remarkable general tendency toward proteome reduction in all *Dickeya* species.

## Results and discussion

### General genomic features of D. aquatica 174/2 type strain

The *D. aquatica* 174/2^T^ genome consists in a single circular chromosome of 4,501,560 base pairs in size, with a GC content of 54.6%. The sequence was submitted to European Nucleotide Archive (accession GCA_900095885). This genome contains a total of 4,202 genes including 4,080 protein-coding DNA sequences (CDS), 22 ribosomal RNA-coding genes organized into seven operons, 76 tRNA-coding genes, and 23 non-coding RNA genes identified by sequence similarity with known functional RNAs entries in the RFAM database (Daub et al., 2015) (Figure 1). These features are typical of *Dickeya* species (Supplementary Table S2). The origin of chromosome replication (*oriC* position 1-8 bp) was predicted between *mnmG/gidA* and *mioC* as observed in *Escherichia coli* (Wolanski et al., 2014) and other enterobacterales, including *Dickeya* (Glasner et al., 2011; Zhou et al., 2015; Khayi et al., 2016). The terminus of replication is predicted between positions 2,220,802 and 2,220,829 bp with the *D. aquatica* 174/2^T^ Dif site (GGT<u>**T**</u>CGCATAATGTATATTATGTTAAAT) differing from the *E. coli* K-12 Dif site by only one nucleotide substitution (GGT<u>**G**</u>CGCATAATGTATATTATGTTAAAT). The distance from *oriC* to the predicted terminus was nearly equal for the two halves of the chromosome. It corresponds to a region near an inflection point in a slight strand-specific nucleotide compositional bias (Figure 1). The protein coding density is 85% of the genome with a slight preference for the leading strand utilization (57%). This density of predicted ORFs (slightly less than one per kilobase) is typical for *Enterobacterales*, including *Dickeya* species (Supplementary Table S2) (Glasner et al., 2011; Zhou et al., 2015; Khayi et al., 2016). Thus, despite its different ecological niche, *D. aquatica* shows genomic organisation very similar to other species of the *Dickeya* genus.

**Figure 1.**
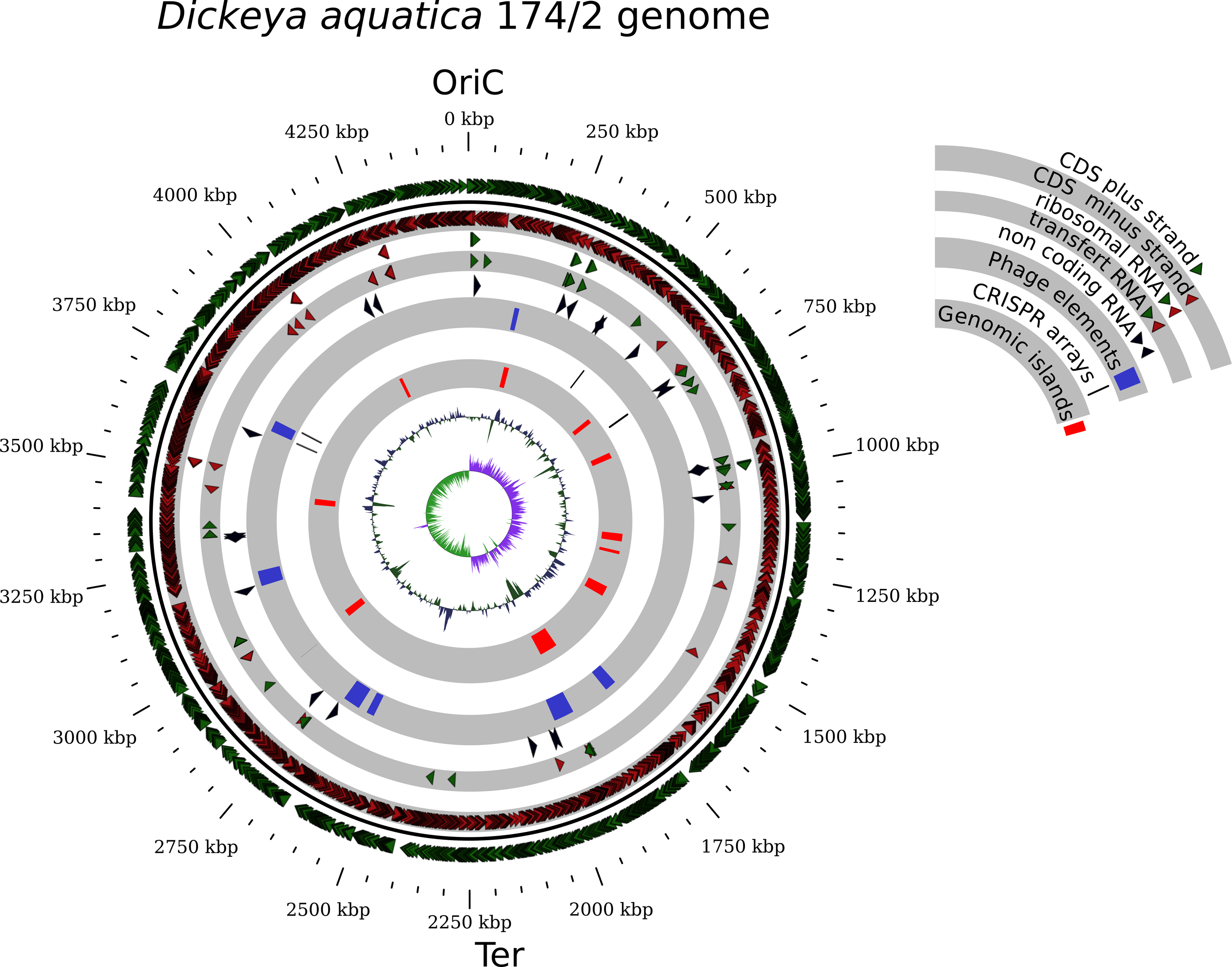
Genomic organisation of *Dickeya aquatica* 174/2^T^ chromosome. The chromosome is represented as a wheel and the origin of replication (OriC) and terminus (Ter) are indicated. The circles from outside to inside represent protein-coding sequences (CDS) on the forward strand, CDS on the reverse strand, distribution of ribosomal RNA operons (four on the right replichore and three on the left replichore), distribution of tRNA genes, and then distribution of ncRNA. The blue areas correspond to phage elements detected using PHAST. The thin black lines correspond to four CRISPR arrays and the red areas represent the ten genomic islands predicted using IslandViewer. The next circle (black) indicates the GC content and the central circle (green/purple) shows the GC-skew. The window size of the GC content and GC-skew is 100 nucleotides. Figure 1 was drawn using Gview https://server.gview.ca/.

#### Deciphering the early evolution of Dickeya

The massive release of *Dickeya* genomic sequences (7 complete genomes and 41 draft genomes) in public databases (Supplementary Table S3), in addition to the complete genome of *D. aquatica* 174/2^T^ reported in this study, provided an interesting resource to explore the evolutionary history of this genus. We investigated the phylogeny of *Dickeya* using 51 ribosomal proteins (rprots), as these were shown to be well suited to study the systematics of prokaryotes, especially *Proteobacteria* (Yutin et al., 2012; Ramulu et al., 2014), on the one hand, and 1,341 core protein families (core-pf), on the other hand. The maximum likelihood (ML) rprots tree, rooted with *Pectobacterium* (*Pectobacteriaceae*) and *Serratia* (*Yersiniaceae*), recovered the monophyly of *Dickeya* (Bootstrap Value (BV) = 100%, Supplementary Figure S1). Surprisingly, *Serratia* strains, used as outgroup to root the phylogeny of *Dickeya* together with *Pectobacterium*, did not form a monophyletic group, due to the robust clustering of *Serratia* sp. ATCC 39006 with *Dickeya* (BV = 99%). A similar grouping was also observed in trees based on the RNA components of the small and large subunits of the ribosome (SSU and LSU rRNA, respectively) (Supplementary Figure S2), suggesting that strain ATCC 39006 represents the sister-lineage of *Dickeya*, and was wrongly affiliated to the *Serratia*, based on its capacity to synthesize prodigiosin, a red pigment secondary metabolite with antimicrobial, anticancer, and immunosuppressant properties, characteristic of *Serratia* species (Thomson et al., 2000). The sister-relationship of ATCC 39006 and *Dickeya* is consistent with the close evolutionary link between the pectinolytic gene clusters (*pel*, *paeY*, *pemA*) shared by these taxa (Duprey et al., 2016b), and with the presence, in *Serratia* sp. ATCC 39006, of an homologue of the Vfm quorum sensing system previously reported as specific to *Dickeya* species (Supplementary Table S4 and Figure S3) (Nasser et al., 2013). The phylogenetic analysis of the genes involved in the biosynthesis of the prodigiosin showed that strain ATCC 39006 genes emerge in a cluster gathering sequences from unrelated taxa: *Serratia* (*Yersiniaceae*), *Hahella* (*Hahellaceae*), and *Streptomyces* (*Actinobacteria*), suggesting that the prodigiosin gene cluster spread among these lineages (including ATCC 39006 strain) through horizontal gene transfer (HGT) (Supplementary Figure S3). Interestingly, strain ATCC 39006, like the early-diverging *D. paradisiaca* species, was deprived of the *indABC* gene cluster responsible for the characteristic production of blue-pigmented indigoidine in *Dickeya* spp. (Supplementary Table S4) (Reverchon et al., 2002; Lee and Yu, 2006). Phylogenetic analyses showed that *Dickeya* IndA, IndB, and IndC proteins are closed to sequences from unrelated bacteria (Supplementary Figure S3), suggesting that indigoidine genes underwent HGT and have been secondarily acquired in *Dickeya* after the divergence with *D. paradisiaca*. The evolutionary link between strain ATCC 39006 and *Dickeya* is also consistent with comparison of their proteomes. In fact, strain ATCC 39006 share in average more proteins families with *Dickeya* (2,554/4,221 – 60.5%), compared to *Pectobacterium* (2,387/4,221 - 56.5%) or other *Serratia* species (2,162/4,221 - 51.2%) (Supplementary Table S5). Yet, the large evolutionary distance between ATCC 39006 strain and *Dickeya* (Supplementary Figure S1), its lower GC content (49.2% vs 52.6 - 56.9% in *Dickeya*) (Supplementary Table S3), and the lower number of protein families shared between ATCC 39006 and *Dickeya* (60.5%) compared to between *Dickeya* species (64.7% - 89.6%, Supplementary Table S5), suggested that strain ATCC 39006 does not belong to *Dickeya* and represents rather a close but distinct lineage, possibly a new genus, that we propose to call *Prodigiosinella*, with strain ATCC 39006 being reclassified as *Prodigiosinella confusarubida*. By using leBIBI^QBPP^ phylogenetic SSU rRNA-based positioning tool (Flandrois et al., 2015), we detected two *Erwinia* sp. Strains: MK01 (SSU accession number at the NCBI: AY690711.1) and MK09 (SSU accession number at the NCBI: AY690717.1), isolated from the rhizosphere of *Phragmites communis*, that were very closely related to ATCC 39006 and could belong to *P. confusarubida*.

According to these results and to decrease computation time, the core-pf phylogeny was inferred using *P. confusarubida* ATCC 39006 as outgroup. The ML core-pf and rprots trees displayed similar topologies (Figure 2 and Supplementary Figure S1), excepted regarding *D. chrysanthemi*, *D. sp.* NCPPB 569, and *D. solani* (see below). Worth to note, the rprots tree was overall less resolved than the core-pf tree (i.e. the former displayed lower BV than the latter). Both trees supported the monophyly of *D. paradisiaca*, *D. aquatica*, *D. zeae, D. dianthicola, D. dadantii*, and *D. solani* (BV ≥ 84%). The monophyly of *D. chrysanthemi* was recovered with core-pf (BV = 100%) but not with rprots. In fact, while the relationships among *D. chrysanthemi* strains were strongly supported in the core-pf tree, they were mostly unresolved (i.e. associated to weak BV) with rprots, meaning that these markers did not contain enough phylogenetic signal to resolved the *D. chrysanthemi* phylogeny. A composite group composed of strains annotated as *D. sp.*, *D. solani*, and *D. chrysanthemi*: M074, M005, B16, MK7, S1, NCPPB 3274, and ND14b (formerly misidentified as *Cedecea neteri* and transferred recently to *D. solani*) was present in both trees (BV ≥ 95%). By using leBIBI^QBPP^ tool (Flandrois et al., 2015), we identified strains B16, MK7, and S1 as *D. fangzhongdai*, suggesting that the whole clade could correspond to *D. fangzhongdai* (Alič et al., 2018).

**Figure 2.**
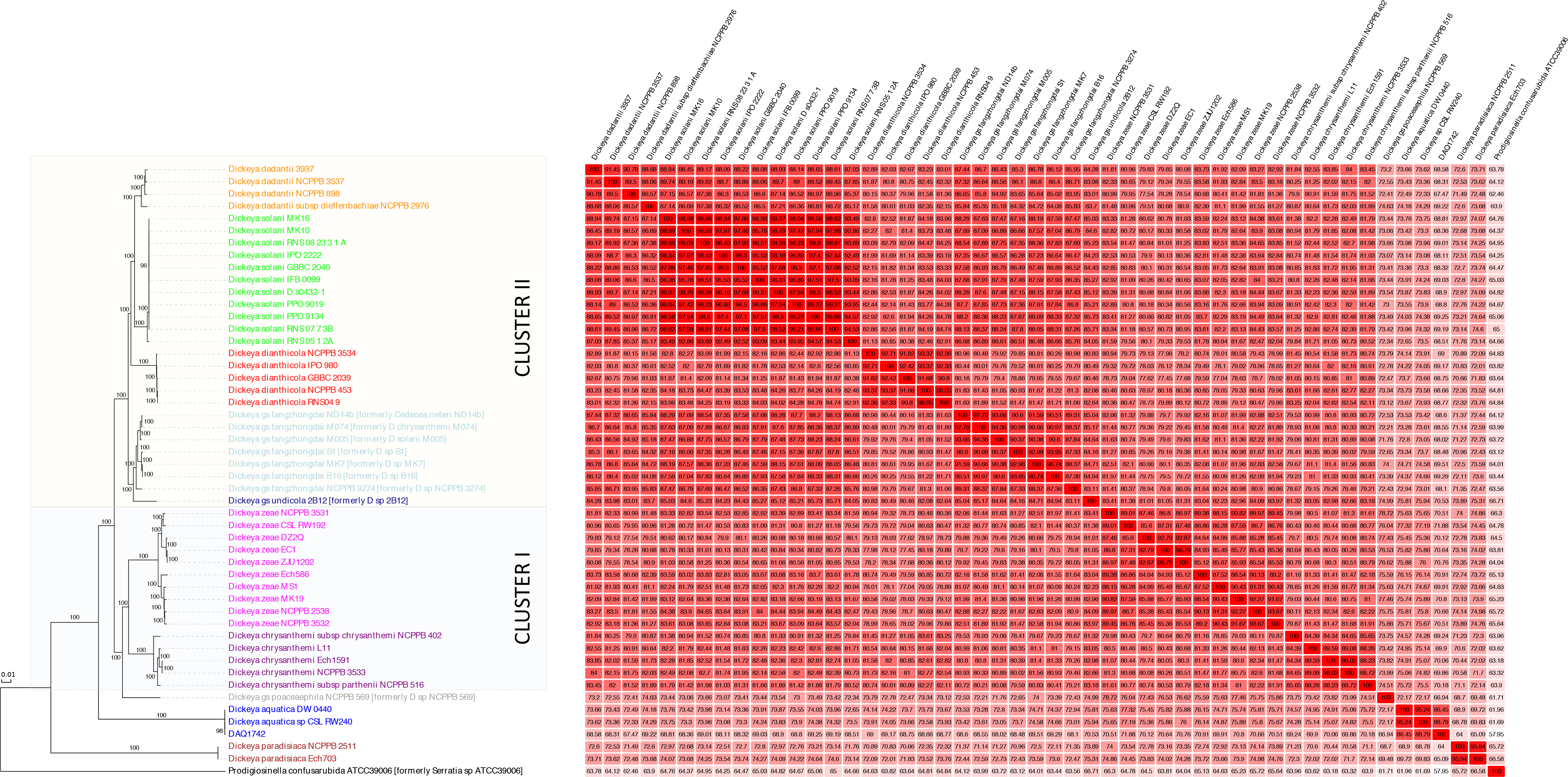
Phylogeny and proteome comparison of *Dickeya*. Maximal likelihood phylogeny (left) of 1,341 *Dickeya* single copy protein families (50 sequences, 414,696 amino acid positions). The tree was computed with IQ-TREE with the LG+C60 model and rooted using *Serratia* ATCC 39006. Numbers associated to branches correspond to ultrafast bootstrap values. The scale bar corresponds to evolutionary distance (i.e. the average number of the substitutions inferred per site). The table (right) corresponds to the *S*_AB_ association coefficient computed for each pair of strains as *S*_AB_=(100×2*N*_AB_)/(*N*_A_+*N*_B_), in which *N*_A_ is the number of protein families present in strain A, *N*_B_ is the number of protein families present in strain B and *N*_AB_ is the number of protein families shared by strain A and strain B. This coefficient ranged from 0 when both strains do not share any gene family to 100 when all the families present in strain A were also present in strain B. The figure was generated using Evolview (He et al., 2016).

Cluster I (*D. zeae* and *D. chrysanthemi*) and cluster II (*D. dianthicola*, *D. solani*, *D. dadantii*, *D. fangzhongdai* and *D. undicola*) were monophyletic (core-pf: both BV ≥ 100% and rprots: both BV ≥ 80%, respectively). Within cluster II, *D. fangzhongdai* diverged first (BV = 100%), while *D. solani* represented either the sister-group of *D. dadantii* (core-pf: BV = 100%) or *D. dianthicola* (rprots: BV = 98%). The conflicting position of *D. solani* was supported by high BV in both trees, indicating a real inconsistence between the two trees. To go further, AU tests performed on individual core-pf and rprots by using the core-pf and the rprots ML topologies (Supplementary Table S6). These tests showed that 1,063 out of 1,341 (79.3%) core-pf protein families reject the rprots topology, while 853 out of 1,341 (63.6%) core-pf protein families do not reject the core-pf topology. In contrast, 31.4% of the rprots do not reject the core-pf topology and 29.4% do not reject the rprots topology. Thus, a large majority of core-pf protein families favour the core-pf topology, while an equal number of rprots supports either one or the other topology. Accordingly, the rprots topology and in particular the grouping of *D. solani* with *D. dianthicola* could be questioned.

Regarding the first divergences within *Dickeya*, both trees pinpointed *D. paradisiaca* and *D. aquatica*, as early diverging lineages (core-pf: BV = 100%, rprots: BV = 86%), with *D. paradisiaca* emerging first (core-pf: BV = 100%, rprots: BV = 65%). Finally, the core-pf placed robustly *D. sp.* NCPPB 569, a strain isolated from sugarcane in Australia (Supplementary Table S3), at the base of Cluster I (BV = 100%), while its position was unresolved in the rprots tree (i.e. unsupported grouping with *D. aquatica*, BV = 46%). Irrespective of its position, the evolutionary distances between strain NCPPB 569 and the eight *Dickeya* species were roughly similar, suggesting NCPPB 569 could correspond to an independent lineage within *Dickeya*. This observation was in agreement with a previous work based on the phylogenetic analysis of *recA* (Parkinson et *al*. 2009), and led us to propose that strain NCPPB 569 could represent a distinct genomospecies, we proposed to name *Dickeya* gs. poaceaephila. This hypothesis was strengthened by the fact that strain NCPPB 569 could be distinguished from other species according to shared protein families (Figure 2 and Supplementary Figure S4). Among cluster II, *Dickeya* sp. 2B12 deserved attention.

In fact, this strain branched as a distant sister-lineage of *D. fangzhongdai* (core-pf: BV = 100% and rprots: BV = 95%). This suggested that this strain could also represent a new genomospecies that we have tentatively called *Dickeya* gs. undicola.

Altogether, the analysis of more than one thousand core protein families and ribosomal proteins allows to resolve most of the speciation events within *Dickeya* and provides a solid framework to investigate the origin and the evolution of important *Dickeya* biological features. As expected, both trees were largely consistent, even if the rprots tree was globally less resolved, in particular regarding *D. chrysanthemi* and *Dickeya* gs. poaceaephila. The main inconsistency concerned the relative order of divergence of *D. dadantii*, *D. dianthicola*, and *D. solani* within cluster II. AU tests suggested that rprots could contain a mix phylogenetic signal, and thus that the rprots tree could be less reliable than the core-pf tree. Determining the origin of the conflicting signal in rprots would require more investigations that are beyond the scope of the study. Because *D. solani* is a relatively late diverging lineage within *Dickeya*, we anticipated that this will not impact significantly inferences on the ancient *Dickeya* evolution. Nevertheless, in all analyses (e.g. evolution of virulence related systems, inference of ancestral gene repertoires, see below), we used the species tree based on core-pf, but by considering both possible placements for *D. solani*: either as the sister-group of *D. dadantii* or *D. dianthicola*. Regarding *D. aquatica*, the core-pf tree suggested that this species represent the second diverging lineage with *Dickeya*, while its position was unresolved according to rprots. Our analyses also reclassified strain ATCC 39006 as *Prodigiosinella confusarubida,* which could represent the closest relative of *Dickeya* genus. Better characterization of the biology and diversity of *Prodigiosinella* will provide important clues about their evolutionary links with *Dickeya* and the emergence of *Dickeya* as a genus.

#### Virulence-associated phenotypes of D. aquatica

While most *Dickeya* species have known vegetal hosts, all the identified strains of *D. aquatica* (174/2^T^, 181/2 and Dw0440) have been isolated from waterways (Supplementary Table S1) (Parkinson et al., 2014). To our knowledge, their pathogenicity has never been demonstrated. The *in silico* survey of the *D. aquatica* 174/2^T^ genome for transcriptional regulator binding sites revealed the presence of conserved key regulators of virulence (Supplementary Table S7). More precisely, all the regulators (KdgR, PecS, PecT, CRP, Fis, H-NS, GacA-GacS, RsmA-RsmB, MfbR) controlling virulence gene expression in *D. dadantii* (Reverchon et al., 2016) are present in *D. aquatica* 174/2^T^. The global regulators FNR and CRP displayed the highest number (approximately 80) of targets, followed by Fur, CpxR, ArcA, Lrp and KdgR each with more than 30 targets. The activator CRP and the repressor KdgR have a key role in *Dickeya* virulence (Duprey et al., 2016a), as they tighly control the pectin degradation pathway and are involved in coupling central metabolism to pectinase gene expression (Nasser et al., 1997), while Fur acts as a repressor of pectinase genes (Franza et al., 2002). In addition, Fur represses genes involved in the metabolism of iron, a metal that plays an important role as a virulence regulatory signal in *Dickeya* (Franza et al., 2002).

The presence of regulators controlling the virulence in *D. aquatica* 174/2^T^ was puzzling and prompted us to investigate its pathogenicity. For this, we compared the pathogenic potential of *D. aquatica* 174/2^T^ and *D. dadantii* 3937 on various infection models such as chicory leaves, potato tubers, cucumbers and tomato fruits (Figure 3). *D. aquatica* showed little efficiency in rotting potato and chicory. Indeed, only a small rotten area was observed near the entry point, opposed to *D. dadantii,* which spread inside the tuber or the leaf (Figure 3). Surprisingly, *D. aquatica* appeared particularly efficient on tomatoes and cucumbers, showing a generalised infection and fruit opening after 42 hours. By contrast, *D. dadantii* was much less efficient in infecting tomatoes and cucumbers. Importantly, tomatoes and cucumbers are acidic fruits (pH 4.8 and pH 5.1, respectively), whereas potato tubers and chicory leaves display higher pH (around 6.0 and 6.5, respectively), suggesting that *D. aquatica* could be more efficient on acidic fruits. Yet, neither *D. aquatica* nor *D. dadantii* are able to induce pathogenic symptoms on very acidic fruits such as pineapple (pH 4) or kiwi (pH 3) (Figure 3B). Previous studies have shown that some unrelated *Dickeya* species (i.e. *D. chrysanthemi*, *D. dianthicola*) are also capable of infecting tomatoes (Supplementary Table S1), supporting the hypothesis that recurrent adaptations to similar hosts occurred independently during the evolution of *Dickeya* genus.

**Figure 3.**
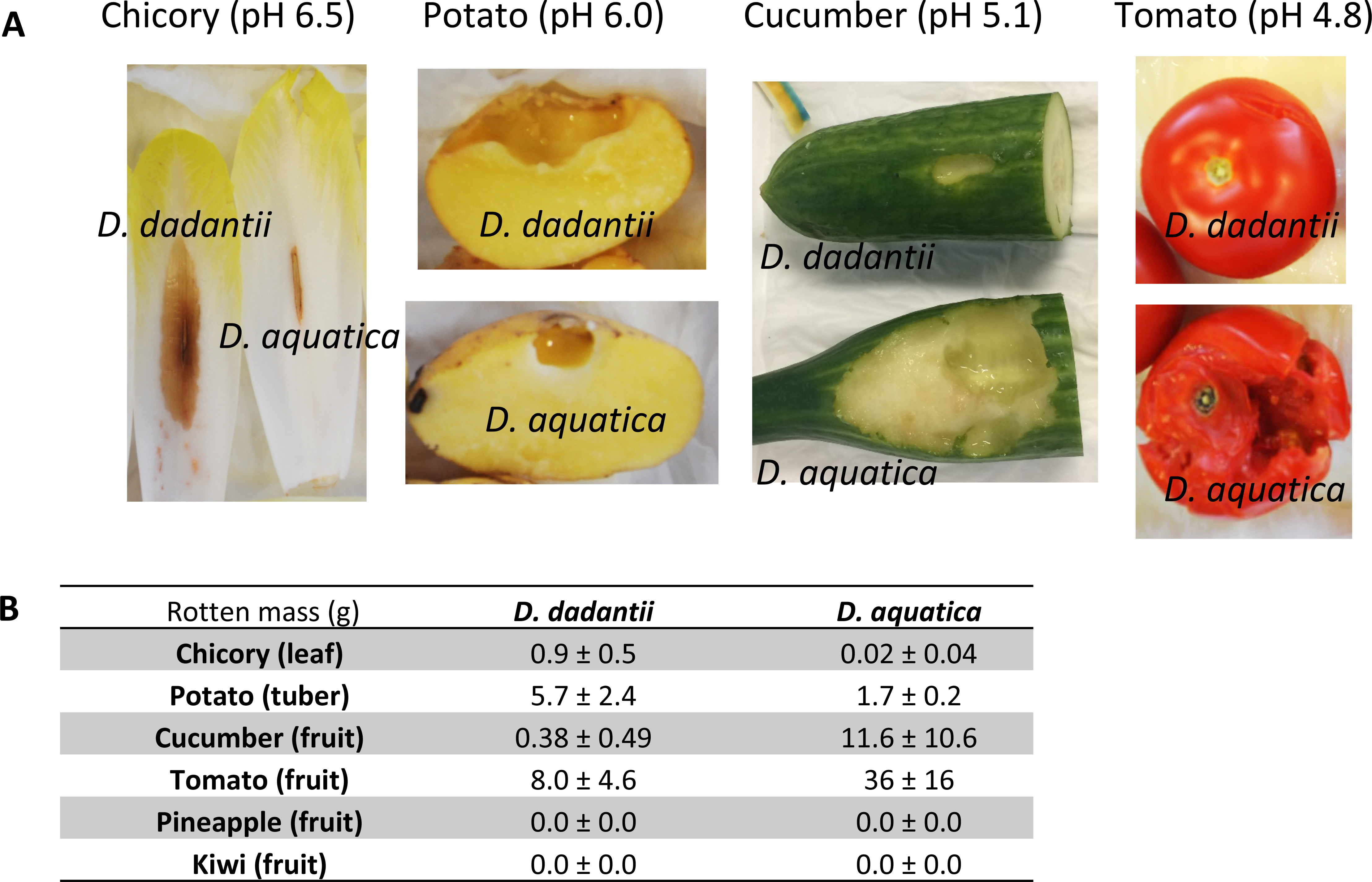
Virulence of *D. aquatica* on potato, chicory, cucumber and tomato. Bacterial cultures were grown in M63G minimal medium (M63 + 0.2% w/v glucose) and diluted to a given OD_600_ depending on the host: 0.2 (chicory) or 1 (potato, cucumber and tomato). For chicory, 5 μL of bacterial suspension were injected into a 2 cm incision at the center of the leaf. For potato, cucumber and tomato, 5, 100 or 200 μL of bacterial suspension were injected into the vegetable, respectively. Plants were incubated at 30°C with 100% humidity for 18 h (chicory) or 42 h (potato, cucumber and tomato). **A)** Picture of representative specimens of infected plants after incubation. Note that the rotten area was removed for potato. **B)** Quantification of the soft rot mass. Data is represented as mean +/− SD of 6 replicates. No soft rot symptoms were detected after 78 h for pineapple and kiwi fruits infected with 200 μL of bacterial suspension at OD_600_ 1.

#### Stress resistance phenotypes of D. aquatica

##### Osmotic stress

Depending on the plant host, *Dickeya* species encounter various stresses during the infectious process (Reverchon et al., 2016). We therefore assessed stress resistance of *D. aquatica* 174/2^T^ (Figure 4). This strain proved to be very sensitive to osmotic stress, as it displayed a 50% growth rate reduction on 0.3 M NaCl while *D. dadantii* was only slightly affected (20% growth rate reduction). This effect was even more pronounced on 0.5 M NaCl with a growth rate reduction of 90 % for *D. aquatica* and 43% for *D. dadantii* (Figure 4A). This sensitivity to osmotic stress could be linked to the lack of two osmoprotectant biosynthetic pathways in *D. aquatica*: glycine betaine biosynthesis (*betA-betB* gene cluster) and phosphoglycosyl glycerate biosynthesis (*pggS-pggP* gene cluster) (Jiang et al., 2016) (Supplementary Table S4). The *pggSP* gene cluster was present in *P. confusarubida* and all *Dickeya* species except *D. aquatica*. The phylogeny of the two genes was consistent with the phylogeny of species (Supplementary Figure S3), indicating that their absence in *D. aquatica* results from a specific and secondary loss in this taxon. In contrast, the *betABI* gene cluster was absent in *D. paradisiaca, D. aquatica*, *D. dianthicola* and Cluster I species (Supplementary Table S4). Their phylogenies suggested that the *betABI* gene cluster could have been present in the ancestor of *P. confusarubida* and *Dickeya*, and secondarily and independently lost in the *Dickeya* lineages mentioned above (Supplementary Figure S3). Finally, the *ousA* gene encoding the major osmoprotectant uptake system in *D. dadantii* is missing in *D. aquatica*, as well as in most other *Dickeya* species, suggesting either a recent acquisition by a few species or multiple losses of an ancestral system (Supplementary Table S4). Interestingly, *D. solani* and *D. dianthicola*, both lacking the *ousA* gene, are also more sensitive to osmotic stress than *D. dadantii* (Figure 4B).

**Figure 4.**
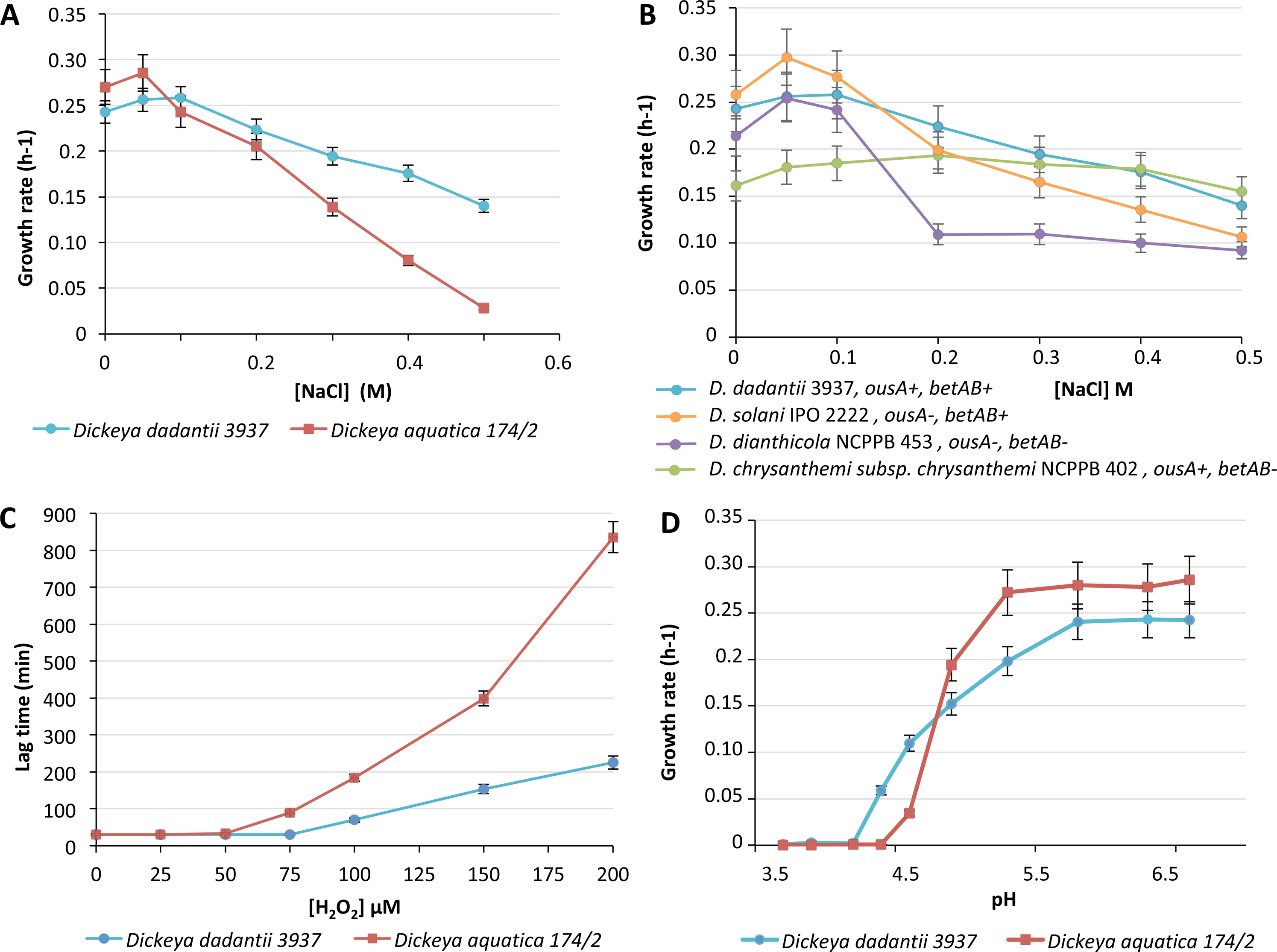
*D. aquatica* stress resistance. Bacteria were cultured at 30°C in 96 well plates using M63G (M63 + 0.2% w/v glucose) pH 7.0 as minimal medium. Bacterial growth (OD_600nm_) was monitored for 48 h using an Infinite^®^ 200 PRO - Tecan instrument. A and B) Resistance to osmotic stress was analysed using M63G enriched in 0.05 to 0.5 M NaCl (abscissa) and growth rates (ordinate) were determined. C) Resistance to oxidative stress was analysed in the same medium by adding H_2_O_2_ concentrations ranging from 25 to 200 μM (abscissa). The lag time (ordinate) is represented instead of the growth rate because after the degradation of H_2_O_2_ by bacterial catalases, the growth rates are similar. D) The pH effect on growth rate (ordinate) was analyzed using the same M63G medium buffered with malic acid at different pH ranging from 3.7 to 7.0 (abscissa).

##### Oxidative stress

*D. aquatica* 174/2^T^ was also very sensitive to oxidative stress, displaying longer lag time than *D. dadantii* in presence of 75 μM H_2_O_2_ (Figure 4C). This is consistent with the lack of the periplasmic superoxide dismutase SodC, whose gene was likely acquired in *Dickeya* after the divergence of *D. paradisiaca* and *D. aquatica* (Supplementary Figure S3), and the lack of the Fe-S cluster assembly SUF system (Supplementary Table S4). Iron-sulfur (Fe-S) clusters are fundamental to numerous biological processes in most organisms, but these protein cofactors can be prone to damage by various oxidants (e.g., O_2_, reactive oxygen species, and reactive nitrogen species) (Roche et al., 2013). In addition, the release of free iron in the bacterial cytoplasm amplifies the oxidative stress through the Fenton reaction, producing the highly toxic and reactive hydroxyl radical OH^.^. Most gammaproteobacteria have two Fe-S cluster biogenesis systems SUF and ISC (Roche et al., 2013). For example, *Escherichia coli* cells switch from the ISC to the SUF system under oxidative stress as OxyR, a sensor of oxidative stress, acts as an activator of *suf* operon expression (Lee et al., 2004). Both the SUF and ISC systems were shown to be required for *D. dadantii* growth in the plant host environment that is continually changing in terms of iron availability and redox conditions (Nachin et al., 2001; Expert et al., 2008; Rincon-Enriquez et al., 2008). While both the ISC and SUF systems were likely present in the ancestor of *Dickeya* and *P. confusarubida*, phylogeny and taxonomic distribution suggested that SUF genes were secondarily lost in *D. paradisiaca*, *D. aquatica*, *D*. gs. poaceaephila, and *D. zeae* (Supplementary Figure S3). The absence of both the SUF system and SodC in *D. aquatica* could be an important factor limiting its host range (Supplementary Figure S3). Finally, the presence of the DNA-binding protein Dps, which has a protective function against a wide range of stresses, in *P. confusarubida* and all *Dickeya* excepted *D. paradisiaca* suggested a specific loss in this latter lineage (Supplementary Figure S3).

Regarding other oxidative stress sources, the NorWVR system responsible for nitrite oxide detoxification was absent in the first diverging *Dickeya* species (Supplementary Table S4). The phylogenies of the corresponding proteins suggested secondary losses in *P. confusarubida*, *D. paradisiaca*, and *D. aquatica*, as these proteins were present in *Pectobacterium*, a closely related genus of *Dickeya* (Supplementary Figure S3). Finally, the ascorbate degradation pathway encoded by the *ula* genes was present in the *D. paradisiaca, D. aquatica, D. zeae*, and *D. chrysanthemi* species, suggesting that the pathway was also present in the ancestor of *Dickeya* and secondarily lost in *D.* gs. poaceaephila and in the ancestor of Cluster II (Supplementary Figure S3). Ascorbic acid, the major and probably the only antioxidant buffer in the plant apoplast, becomes oxidized during pathogen attack (Pignocchi and Foyer, 2003). Modification of the apoplastic redox state modulates receptor activity and signal transduction to regulate plant defence and growth (Pignocchi and Foyer, 2003). The capacity of some *Dickeya* species to catabolize ascorbate could be a strategy to weaken the plant defence.

##### Acidic stress

Regarding pH sensitivity, both *D. aquatica* 174/2^T^ and *D. dadantii* showed optimum growth at pH 7.0. *D. aquatica* 174/2^T^ displayed no significant growth rate reduction down to pH 5.3 while *D. dadantii* was slightly affected (20% growth rate reduction at pH 5.3 compared to optimum pH 7.0) (Figure 4D). Below pH 4.9, the growth of *D. aquatica* abruptly diminished compared to that of *D. dadantii* indicating that *D. aquatica* growth was much more impacted by low pH (Figure 4D). Bacterial response to acid stress involves evasion of cell damage and adaptation of the enzymatic profile by reducing reactions producing protons and promoting those consuming protons (Bearson et al., 1997). Concerning the protection of proteins against pH damage, *Dickeya* strains, including *D. aquatica* strain 174/2^T^, encode the lysine-rich protein Asr acid-inducible periplasmic chaperone, known to protect the proteins by sequestering protons due to its high basic amino acids composition (Seputiene et al., 2003). In *Dickeya*, the corresponding gene is located downstream the *rstAB* gene cluster, known to control the *asr* gene expression (Ogasawara et al., 2007). Yet, due to their atypical amino acid composition these proteins were often wrongly annotated as histone proteins (e.g. in *D. dadantii* NCPPB 898, *D. dianthicola* NCPPB 3534, *D. dianthicola* NCPPB 453, *D. dianthicola* GBBC 2039, and *D. chrysanthemi* ATCC 11663). In terms of metabolic adaptation, a major strategy for bacterial pH homeostasis consists in using decarboxylases to remove cytoplasmic protons (Krulwich et al., 2011). The glutamate-, arginine-, and lysine-inducible decarboxylases classically found in enteric bacteria (Foster, 2004) are absent in *Dickeya* species. However they contain some organic acid decarboxylases such as oxalate and malonate decarboxylases (Supplementary Table S4). Some *Dickeya* species, including *D. dadantii,* have two oxalate decarboxylation pathways: the frc-oxc pathway, being involved in the acid tolerance response in *E. coli* (Fontenot et al., 2013) and the OxdD decarboxylase pathway. Most *Dickeya* species harbour at least one of these oxalate decarboxylation pathways, except *D. aquatica*, *D.* gs. poaceaephila, *D. chrysanthemi*, and a few *D. zeae* strains, which were deprived of both pathways (Supplementary Table S4 and Supplementary Figure S3). The lack of the two oxalate-decarboxylation pathways in *D. aquatica* could contribute to its sensitivity to acidic pH. Finally, in contrast with *P. confusarubida* and most *Dickeya* species, *D. aquatica* and a few *D. zeae* strains were also devoid of malonate decarboxylation pathway (mdcABCDEFGHR) (Maderbocus *et al.*, 2017), which consumes protons (Supplementary Table S4), suggesting secondary gene losses in these strains, a hypothesis that was confirmed by phylogenetic analyses (Supplementary Figure S3).

##### Twitching motility

Interestingly, colony shape changes were observed when *D. aquatica* was grown at low pH in presence of malic acid (Figure 5). More precisely, colonies became wider (2-3 folds increase in diameter compared to unstressed *D. aquatica*) with a morphology characteristic of twitching motility (Henrichsen, 1972). Correspondingly, we detected genes encoding a complete type IV pilus assembly responsible for twitching motility in *D. aquatica* genome and named them *pil* genes according the *Pseudomonas aeruginosa* nomenclature (Maier and Wong, 2015). Among the collection of twenty-six *Dickeya* strains representing different species, only eight showed twitching at acidic pH (Figure 5). Interestingly, this phenotype appeared strain-dependent rather than species-dependent, as *D. solani*, strain RNS07.7.3B showed twitching motility at acidic pH, while *D. solani* strain PP09019 did not (Figure 5). As the *pil* genes, inherited by *Dickeya* from the common ancestor shared with *P. confusarubida,* were conserved in all species (Supplementary Table S4 and Figure S3), twitching was probably linked to strain-specific induction of these genes under acidic pH. It is noteworthy that twitching motility is strongly influenced by changes in the environment (Henrichsen, 1975). In plant pathogens, the contributions of type IV pilus to virulence have been investigated mainly in vascular pathogens, such as *Ralstonia* and *Xylella*, where they were proposed to contribute to bacterial colonization and spread in the xylem through cell attachment, biofilm formation, and twitching motility (Burdman et al., 2011). However, type IV piliation was also shown to be important for initial adhesion and colonization of leaves in a few non-vascular bacteria such as *Xanthomonas oryzae pv. oryzicola, Pseudomonas syringae pv. Tabaci* and *Pseudomonas syringae pv. syringae* (Burdman et al., 2011). The importance of type IV pilus in *Dickeya* pathogenicity is therefore an interesting question for the future studies.

**Figure 5.**
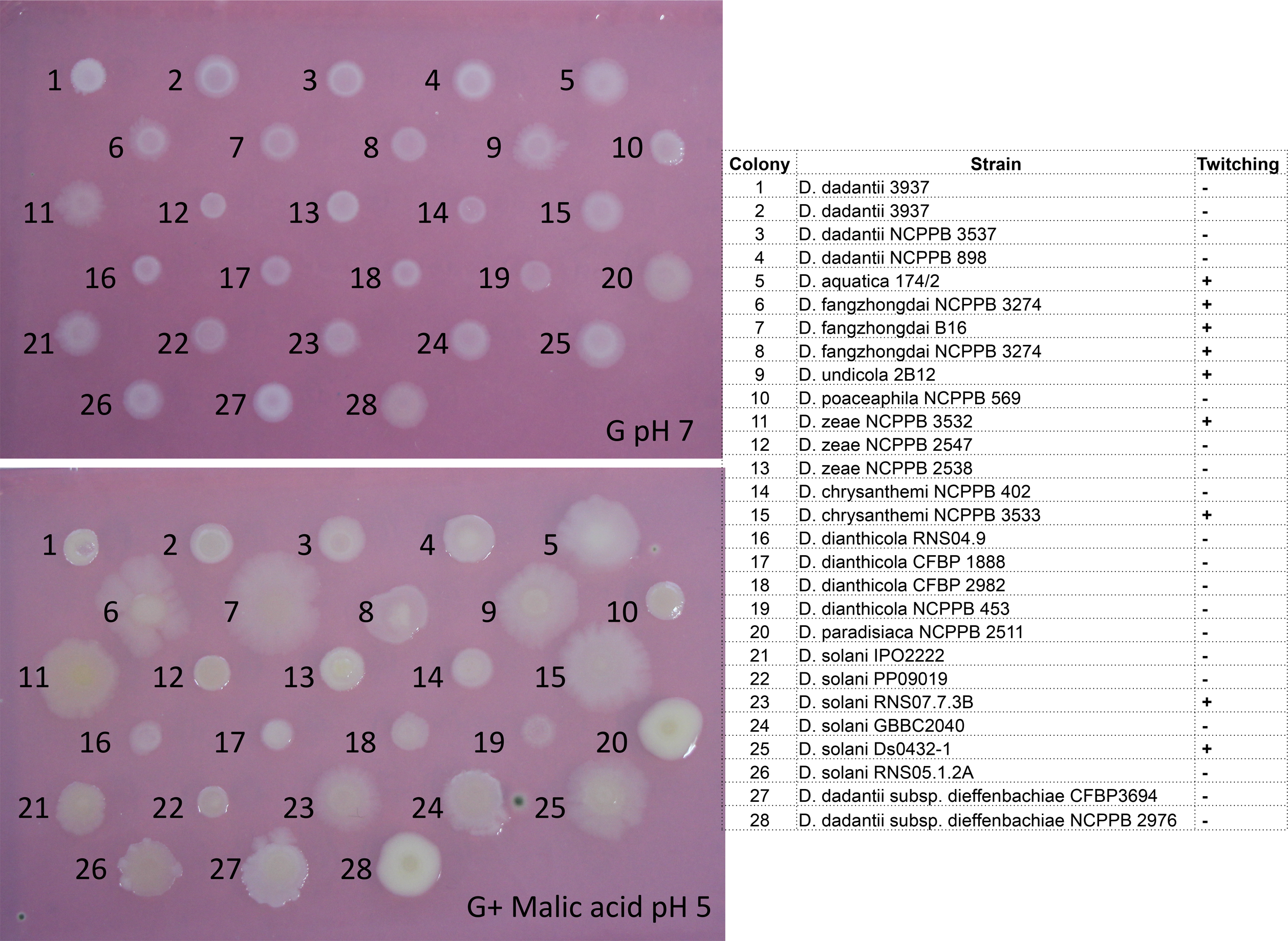
Twitching motility induced under acidic condition in *Dickeya* species. Colony morphologies of various *Dickeya* strains grown on M63G pH 7.0 and M63G buffered with malic acid at pH 5 in agar plates. The strains corresponding to colony numbers and their twitching phenotypes are indicated on the right.

##### Antimicrobial peptides

In response to infection, plants produce antimicrobial peptides (AMPs) to limit pathogen propagation. To overcome AMPs, bacteria remodel their envelope, and more precisely, they modify their LPS to decrease interaction with positively charged AMPs. Among the genes involved in LPS modification, the operons *arnABCDEFT* and *dltXABCD* turned out to be ancestral and conserved in all *Dickeya* species (Supplementary Figure S3), while other genes (i.e. *eptAB, pagP, lpxO* and *lpxT*) displayed different taxonomic distributions and evolutionary histories (Supplementary Table S4 and Figure S3). For example, *D. aquatica* 174/2^T^ is lacking *pagP*, *lpxO* and *lpxT* genes possibly contributing to a greater sensitivity to AMPs.

To conclude, our data indicate that, *D. aquatica* 174/2^T^ has an unanticipated phytopathogenic capacity similar to that of other *Dickeya* species. Particular stress resistance profiles and induction of twitching at acidic pH may contribute to its restricted host range. Based on this observation, we decided to compare the *D. aquatica* 174/2^T^ and other *Dickeya* species proteomes with a special focus on the virulence determinants including plant cell wall degrading enzymes, secretion systems, iron metabolism, plant adhesion elements and secondary metabolism.

#### Distribution of plant cell wall degrading enzymes in Dickeya

The plant cell wall is a complex and dynamic meshwork of polymers (cellulose, hemicellulose, pectin, structural glycoproteins) (Pauly and Keegstra, 2016). Among these polymers, pectin is the most complex and includes both linear regions composed of polygalacturonan and ramified regions (RGI and RGII, respectively). RGI contains a rhamnogalacturonan backbone and various lateral chains such as galactan, arabinan and galacturonan (Caffall and Mohnen, 2009). RGII contains a short galacturonan backbone, carrying four side chains, with a diversity of rare monosaccharides (O’Neill et al., 2004). The carboxylic groups of D-galacturonate residues are methyl-esterified to various degrees (up to 80%) and these residues are, to a lesser extent, acetylated at the C2 and/or C3 positions. Feruloyl esters are a type of modification commonly found in arabinan and galactan chains of ramified regions (Ishii, 1997). The virulence of *Dickeya* is correlated with their ability to synthesize and secrete plant cell wall degrading enzymes, including a full set of pectinases (Hugouvieux-Cotte-Pattat et al., 2014), xylanases and xylosidases (Keen et al., 1996) proteases PrtA, PrtB, PrtC, PrtG (Wandersman et al., 1987), and the cellulase Cel5Z (Py et al., 1991). The presence of these virulence factors varies depending on the species (Matsumoto et al., 2003; Duprey et al., 2016b). To obtain a comprehensive view of this phenomenon, we explored the “Plant cell wall degradosome” of *Dickeya* species including *D. aquatica* (**Figure** 6).

##### The pectinasome

The *Dickeya* pectinasome includes multiple pectate lyases (PelA, PelB, PelC, PelD, PelE, PelI, PelL, PelN, PelW, PelX, PelZ, Pel10), pectin lyases (PnlG, PnlH), polygalacturonases (PehK, PehN, PehV, PehW, PehX), pectin methyl esterases (PemA, PemB), pectin acetyl esterases (PaeX, PaeY), feruloyl esterases (FaeD, FaeT), rhamnogalacturonate lyases (RhiE, RhiF), and one periplasmic endogalactanase (GanA) (Figure 6) (Hugouvieux-Cotte-Pattat et al., 2014). *D.* gs. poaceaephila NCPPB 569 was the genospecies with the poorest pectinase content. According to their taxonomic distribution and phylogeny, most of these proteins could be inferred in the ancestor of *Dickeya* and in its sister lineage *P. confusarubida*. Regarding the *pelAED* cluster, while *P. confusarubida* has a single gene, the ancestor of *Dickeya* had two copies, *pelA* and an undifferentiated *pelDE* pectate lyase-coding gene (Duprey et al., 2016b), suggesting that a duplication event occurred in the stem of *Dickeya*. This undifferentiated *pelDE* has undergone a duplication event after the emergence of *D. paradisiaca*, giving rise to *pelD* and *pelE* (Supplementary Figure S5) (Duprey et al., 2016b). *D. aquatica* has then lost *pelE*, while *D. poaceaephila* has lost both *pelA* and *pelE*. The *pelA* gene was also lost in *D. dianthicola*, while *pelE* was lost in some *D. chrysanthemi* strains (Figure 6, Supplementary Figure S5). The *pelBC* cluster was also likely present in the *Dickeya* ancestor, and conserved in most *Dickeya* species. However, *pelB* was lost in *D.* gs. poaceaephila, and *D. dadantii subsp. dieffenbachiae* NCPPB 2976 (Supplementary Figure S3). The pectate lyase PelI was present in *Pectobacterium* and in all *Dickeya*, except *D. paradisiaca*, and in *P. confusarubida*, suggesting that it could be ancestral in *Dickeya*. Similarly, the phylogeny of pectin methyl esterase PemB indicated that the protein was present in *P. confusarubida* and *Pectobacterium*, suggesting an ancestral presence in *Dickeya*. Accordingly, its absence in *D. paradisiaca*, *D. aquatica*, *D.* gs. poaceaephila, and *D. zeae* strains likely reflected secondary losses. The phylogeny of this protein also suggested that the *pemB* gene found in *D. chrysanthemi* was acquired by HGT from Cluster II species (Supplementary Figure S3). Regarding the polygalacturonase PehN, a similar scenario could be inferred except that the corresponding gene was also lost in *P. confusarubida*, and that a few *D. zeae* strains seemed to have reacquired a PehN from Cluster II species by HGT (Supplementary Figure S3). Regarding the *pehVWX* cluster, the three genes were derived from a single *pehX* gene that was present in the ancestor of *P. confusarubida* and *Dickeya*. A gene duplication event occurred in the ancestor of *D. dianthicola*, *D. solani* and *D. dadantii* leading to PehV. A second event led to the divergence of PehW in the ancestor of *D. solani* and *D. dadantii* (Supplementary Figure S5). The polygalacturonase PehK was likely present in the ancestor of all *Dickeya* and secondarily lost in *D.* gs. poaceaephila*, D. solani*, and *D. dianthicola* (Supplementary Figure S3). The pectin lyase PnlG was likely present in the ancestor shared by *Dickeya* and *P. confusarubida*, and secondarily lost in *D. aquatica*, *D.* gs. poaceaephila, *D. chrysanthemi*, *D. dianthicola* and in some *D. zeae* strains (Supplementary Figure S3). Finally, the rare pectate lyase Pel10 and pectin lyase PnlH displayed patchy taxonomic distributions (Figure 6). Their phylogenies suggested that they spread through HGT in *Dickeya* (Supplementary Figure S3).

**Figure 6.**
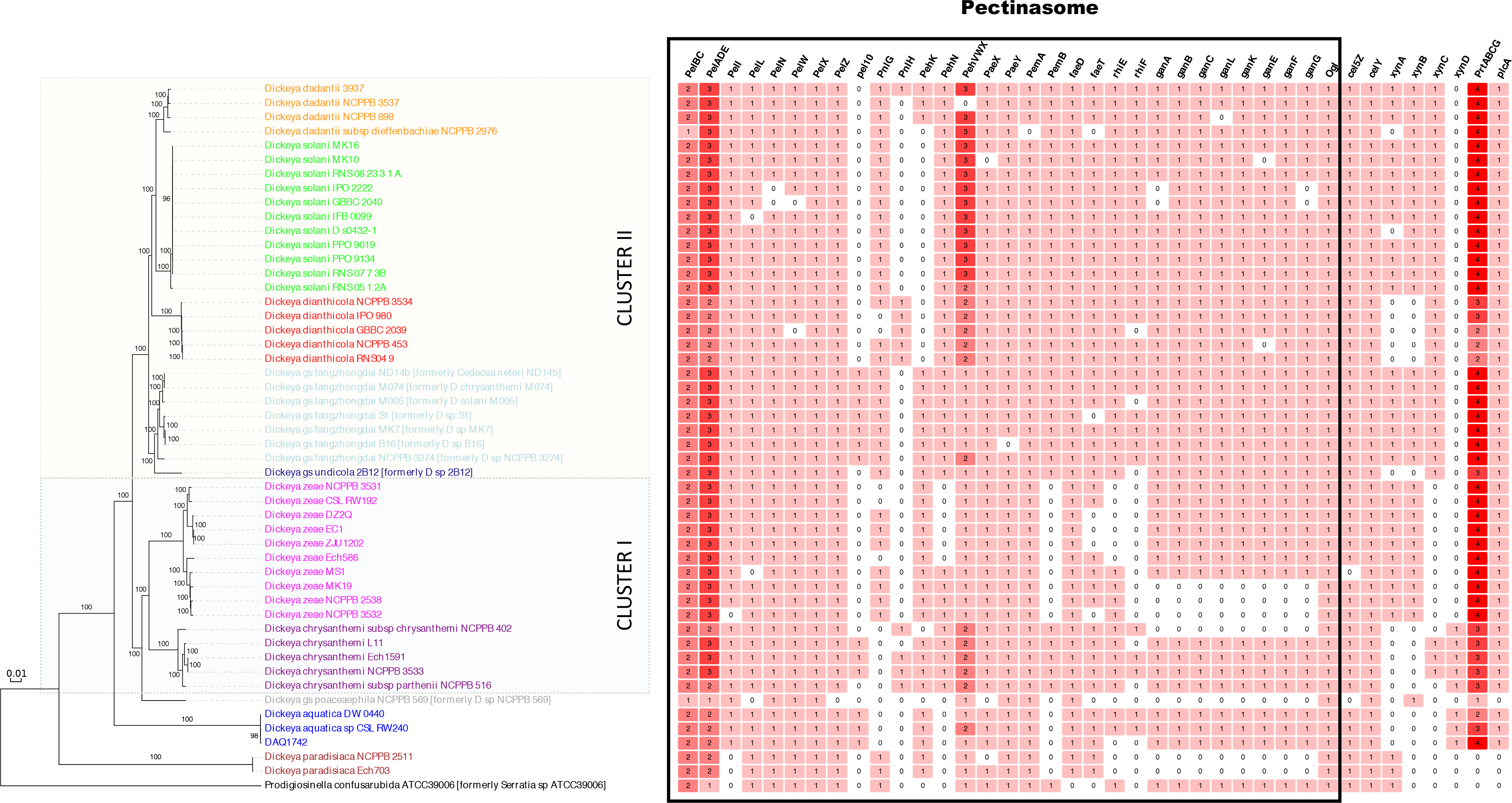
Distribution of plant cell wall degrading enzymes in *Dickeya*. The type of cell wall degrading enzyme and the number of homologues of each enzyme are indicated for each of the strains. The phylogeny on the left corresponds the core-pf ML tree. The figure was generated using Evolview (He et al., 2016).

The saturated and unsaturated digalacturonates resulting from pectin degradation by pectinases are converted into monogalacturonate and 5-keto-4-deoxyuronate by the oligogalacturonate lyase Ogl, which is present in all *Dickeya* species (Figure 6). The phylogenies of the *gan* gene cluster responsible for degradation of galactan chains in pectin-ramified regions and the rhamnogalacturonate lyase RhiE involved in degradation of RGI pectin-ramified regions indicated they were likely present in the ancestor of *P. confusarubida* and *Dickeya*, and secondarily lost in the basal *D. paradisiaca* species, in *D.* gs. poaceaephila, and in a few other strains (Figure 6). The ferulate esterases FaeT and FaeD were absent in *P. confusarubida* and were acquired in the basal *D. paradisiaca* and *D. aquatica* species. The FaeT enzyme was then secondarily lost in *D.* gs. poaceaephila, and *D. dadantii subsp dieffenbachiae* (Supplementary Figure S3). Altogether, the variability observed in the pectinasome among *Dickeya* species and even among different strains likely reflects the dynamic evolution of the corresponding gene families involving gene acquisitions and losses. More compellingly, this variability indicates that the pectinasome cannot be used to distinguish among various species.

##### Cellulases and Xylanases

Two cellulases Cel5Z and CelY were likely ancestral in *Dickeya* species except in *D.* gs. poaceaephila, which has lost Cel5Z. While Cel5Z was involved in cellulose degradation, CelY belonged to the *bcs* gene cluster responsible for cellulose fiber formation (Prigent-Combaret et al., 2012). Xylanases (XynA, XynB) and xylosidases (XynC, XynD) cleave xylan and xyloglucan, which belong to the hemicelluloses. The ß 1,4 xylan is mainly present in plant cell wall of monocots (Pena et al., 2016) and is further decorated, often by acetyl, arabinosyl, and glucuronosyl side-chain substitutions. Notably, the xylan substitution patterns depend on the plant species and are distinct in gymnosperms and angiosperms (Busse-Wicher et al., 2016). Xyloglucan is a β-1,4 glucan that can be substituted with a diverse array of glycosyl and nonglycosyl residues. The type and order of xyloglycan substituents depend on the plant species (Pauly and Keegstra, 2016). Xyloglucan polymers fall into one of two general types. In one type, three out of four backbone glucosyl residues are xylosylated, leading to an XXXG-type xyloglycan, which is predominant in most dicots. Another type of xyloglycan exhibits reduced xylosylation in that only two out of the four or more backbone glucosyl residues are xylosylated, resulting in the XXGGn-type xyloglucan, which is present in early land plants such as liverworts, mosses, lycophytes, and ferns of the order Polypodiales. This type of xyloglucan seems to be absent from the gymnosperms and angiosperms, with the exception of the grasses (Poales) and plants from the order Solanales, such as potato, tobacco and tomato (Pauly and Keegstra, 2016). Thus, it is tempting to hypothesize that the xylanase and xylosidase content of *Dickeya* species could be correlated with their plant host. Interestingly, all *Dickeya* species as well as *P. confusarubida* contained at least one XynA, XynB, XynC, or XynD coding gene (Figure 6). The phylogeny of XynA indicated that the corresponding gene was likely present in the ancestor of *P. confusarubida* and *Dickeya*, and secondarily lost in *D. dianthicola*, *D*. gs. undicola, *D. chrysanthemi*, *D*. gs. poaceaephila, and *D. aquatica* (Supplementary Figure S3). The XynB phylogeny suggested a secondary acquisition in *Dickeya* after the divergence of *P. confusarubida*, *D. paradisiaca*, and *D. aquatica*, followed by losses in *D. dianthicola*, *D. chrysanthemi*, and *D*. gs. undicola (Supplementary Figure S3). While the mode of action of the xylanase XynB was not studied, XynA is a glucuronoxylanase (CAZy family GH30) hydrolyzing the xylan backbone adjacent to each glucuronosyl side-chain (Urbanikova et al., 2011).

*D. aquatica, D. chrysanthemi, D*. gs. undicola, and *D. dianthicola* were deprived of both XynA and XynB (Figure 6) suggesting that these species would preferentially infect dicots as xylan is mainly present in plant cell wall of monocots. However, this is probably a tendency since *D. chrysanthemi* strain Ech1591 was isolated from maize, which is a monocot. The phylogeny of Xylosidase XynD suggested that the corresponding gene has been acquired by HGT in *D. aquatica* and *D. chrysanthemi*, (Supplementary Figure S3). A secondary acquisition via HGT could also be hypothesized for XynC (Supplementary Figure S3), as it is absent in *P. confusarubida* and the basal *Dickeya* species (i.e. *D. paradisiaca*, and *D. aquatica*), as well as in *D.* gs. poaceaephila, *D. zeae* and some *D. chrysanthemi* strains. Strikingly, the xylose degradation pathway (XylABFGHR), present in *P. confusarubida* and in all *Dickeya*, was absent in *D. aquatica*, indicating clearly a specific loss in this species (Supplementary Figure S3). The absence of the xylanases XynA and XynB, and xylosidase XynC as well as the xylose degradation pathway in *D. aquatica* could contribute to its restricted host range.

##### Proteases

Finally, the proteases PrtA, PrtB, PrtC, and PrtG, resulting from specific duplications that occurred during the diversification of *Dickeya*, and the associated type I protease secretion system PrtDEF were absent in *P. confusarubida* and the basal branching *D. paradisiaca*. Yet, *Pectobacterium* harbour closely related homologues of PrtDEF and a single protease-coding gene closely related to *Dickeya* PrtA, PrtB, PrtC and PrtG. This suggested that the whole system could have been present in the common ancestor they shared with *Dickeya*, and then secondarily lost in *P. confusarubida* and *D. paradisiaca* (Supplementary Figure S3). PrtG was specifically lost in *D. chrysanthemi, D. dianthicola D*. gs. undicola and *D.* gs. poaceaephila. This later genomospecies conserved only one protease PrtC whereas *D. dianthicola* strains RNS04-9 and NCPPB453 contained a *prtA* pseudogene and retained two proteases PrtC, PrtB (Figure 6).

##### Other factors

In addition to plant cell wall degrading enzymes, *Dickeya* use several other factors to colonize plant tissue and enhance the progression of disease. Such factors include the extracellular necrosis inducing protein NipE and the two paralogous proteins AvrL and AvrM. NipE and AvrL are conserved in *Pectobacterium* and in most *Dickeya* species, except in *D. paradisiaca*, *D.* gs. poaceaephila, and *P. confusarubida*, suggesting secondary losses in these lineages (Supplementary Figure S3). AvrM was also absent in *D. zeae*, some *D. chrysanthemi*, *D*. gs. undicola, and *D. dianthicola*. Interestingly *D. zeae* strains isolated from rice were the only *Dickeya* to be devoid of Avr proteins (Supplementary Table S4). Altogether, our data indicate that the host range specificity of the various *Dickeya* species is probably linked to the particular combination of plant cell wall degrading enzymes and accessory toxins they produce.

#### Distribution of secretion systems in Dickeya species

For all *Dickeya* species, possession of secretion systems allowing them to actively secrete virulence factors is of crucial importance. Unsurprisingly, different protein secretion systems (T1SS to T6SS) are present in *Dickeya* species (Supplementary Table S4). As previously mentioned, the type I protease secretion system PrtDEF was present in the ancestor of all *Dickeya* species and lost in the basal *D. paradisiaca* species and in *P. confusarubida*. All *Dickeya* species are equipped with the Out specific T2SS responsible for the secretion of most pectinases and the cellulase Cel5Z. The phylogeny of the components of the Out system indicated it was likely present in the ancestor shared by *Pectobacterium*, *P. confusarubida*, and *Dickeya*, and was conserved during the diversification of *Dickeya* (Supplementary Figure S3). A second Stt specific T2SS, allows secretion of the pectin lyase PnlH (Ferrandez and Condemine, 2008). Accordingly, the taxonomic distributions of the pectin lyase PnlH and the Stt T2SS components were similar, being present in *D. dianthicola, D. chrysanthemi,* and some *D. dadantii* strains, (Supplementary Table S4 and Figure 6). In *D.* gs. poaceaephila, the Stt T2SS was also present but not associated with PnlH, thus its function in this strain remains to be determined (Supplementary Table S4 and Figure 6). Interestingly, a T3SS is present in *Pectobacterium*, *P. confusarubida* and in all *Dickeya* species, except *D. paradisiaca* and *D.* gs. poaceaephila deprived of the T3SS and the associated DspE effector, and thus suggesting secondary losses (Supplementary Table S4). Therefore, these latter species are probably unable to suppress the plant immune response (see below). Interestingly, phylogenies of these proteins disclosed a close relationship with two bacterial phytopathogens, *Erwinia* and *Pseudomonas syringae*, suggesting that HGT occurred among these lineages (Supplementary Figure S3). In fact, the effector DspE belongs to the AvrE superfamily of Type III effectors (T3Es) (Degrave et al., 2015). The AvrE family is the only family of T3Es present in all type III-dependent, agriculturally important phytobacterial lineages that belong to the unrelated *Enterobacteriales*, *Xanthomonadales*, *Pseudomonadales* and *Ralstonia* taxa. This indicates that HGT of these effectors occurred in the ancestors of these important plant pathogen lineages (Jacobs et al., 2013). Recent studies indicated that AvrE-type effectors alter the sphingolipid pathway *in planta* by inhibiting the serine palmitoyl transferase (Siamer *et al.*, 2014). The sphingolipid biosynthetic pathway is induced during the plant hypersensitive response that blocks pathogen attack at the site of infection (Berkey *et al.*, 2012). Therefore, inhibition of this pathway delays hypersensitive response-dependent cell death and allows bacterial development *in planta* (Degrave et al., 2015). In *Dickeya* species, the T3SS genes are in synteny with the *plcA* gene encoding a phospholipase. These genes were probably acquired during the same event since *D. paradisiaca* and *D.* gs. poaceaephila that were deprived of the T3SS are also deprived of PlcA (Figure 6).

All *Dickeya* species and *P. confusarubida* were found to possess a two-partner secretion system (T5SS) CdiB-CdiA mediating bacterial intercellular competition. Their phylogenies clearly indicate an ancestral presence in both lineages (Supplementary Figure S3). CdiB is a transport protein that exports and presents CdiA proteins on the cell surface (Willett et al., 2015). The Cdi system is involved in contact-dependent growth inhibition (CDI) by delivering the C-terminal toxin domain of CdiA (CdiA-CT) to target bacteria (Aoki et al., 2010). Some *Dickeya* strains are equipped with two CdiA proteins, for example, *D. dadantii* 3937, which produces two different CdiA-CT toxins: the first one being a tRNase and the second one harbouring DNase activity (Aoki et al., 2010; Ruhe et al., 2013). Each Cdi system also encodes a specific CdiI antitoxin that interacts with the cognate CdiA-CT toxin and prevents auto-inhibition (Willett *et al.*, 2015).

The most striking feature in *D. aquatica* was the absence of both type IV (T4SS) and type VI (T6SS) secretion systems, a trait that was shared with *D. paradisiaca, D.* gs. poaceaephila and *P. confusarubida*. Yet, the presence of closely related T4SS and T6SS in *Pectobacterium* and other *Dickeya* species suggests that both systems were present in the ancestor of *Dickeya* and *Prodigiosinella*, and secondarily lost in the three mentioned species (Supplementary Figure S3). T4SS systems were used to transport a variety of biomolecules (DNA or proteins) across the bacterial envelope (Chandran Darbari and Waksman, 2015). Most T4SS detected in *Dickeya* species are associated with conjugal transfer proteins and thus, correspond likely to conjugative T4SS that transferred DNA (de la Cruz et al., 2010; Ilangovan et al., 2015). This process is instrumental in bacterial adaptation to environmental changes (Thomas and Nielsen, 2005). T6SS is used for interaction with the host and for inter-bacterial competition (Poole et al., 2011). Unfortunately, while effectors associated to the *Dickeya* T6SS carry C-terminal nuclease domains that degrade target cell DNA, little is known concerning their function in virulence (Ryu, 2015), limiting the interpretation of its absence in *D. aquatica*.

#### Distribution of plant adherence elements in Dickeya species

A chaperone-usher pilus assembly pathway, associated with type I fimbriae (*fimEAICDFGHB*), was present in *D. aquatica* but not in any other *Dickeya* species (Supplementary Table S4). Phylogenetic analyses suggested an acquisition from *Morganellaceae* or *Enterobacteriaceae*, through HGT (Supplementary Figure S3). The adhesin FimH, a two-domain protein at the tip of type I fimbriae is known to recognize mannoside structures and to be responsible for adhesion to both animal epithelial cells and plant surface (Haahtela et al., 1985; Sauer et al., 2000). Therefore, we can hypothesize that type I fimbriae could be involved in adherence of *D. aquatica* to plant surface. Interestingly, in *Xylella fastidiosa,* the Fim system is known to be an antagonist of twitching motility caused by type IV pili (De La Fuente et al., 2007). In *D. aquatica*, the mutually exclusive production of type I fimbriae and type IV pilus could be linked to the specific pH regulation of type IV pilus. When *D. aquatica* penetrates into the intercellular apoplast, which is an acidic compartment, twitching motility would be induced to favour bacteria dissemination in plant tissues, while the type I fimbriae would be no longer required during the colonization. This regulation of twitching would thus contribute to the efficiency of *D. aquatica* for infecting tomatoes, cucumbers, and probably other acidic fruits.

While the presence of type I fimbriae is a specific feature of *D. aquatica* among *Dickeya*, the Flp/Tad pilus, involved in plant surface adherence (Nykryri et al., 2013), is restricted to *D. chrysanthemi* likely as the consequence of a HGT from another proteobacterium (Supplementary Table S4 and Supplementary Figure S3). An operon encoding a multi-repeat adhesin (Dda3937_01477) associated to a T1SS secretion pathway was found in the genome of *D. dadantii* (Supplementary Table S4). This protein contains multiple cadherin-homologous domains and is likely involved in plant adhesion. Indeed, in *Pectobacterium atrosepticum*, such a multi-repeat adhesin secreted by a type I pathway was shown to be required for binding to the host plant (Perez-Mendoza et al., 2011). This adhesion and its secretion system were absent in the basal *Dickeya* species as well as in *D. zeae* and *D. chrysanthemi.* Phylogenetic analyses indicated an acquisition in the ancestor of Cluster II, followed by a secondary loss in *D. dianthicola* (Supplementary Figure S3).

From this analysis, it appears that the different *Dickeya* species retained distinct strategies to adhere to plant surfaces and that HGT played an essential role in the acquisition of the involved genes.

#### Distribution of iron assimilation systems in Dickeya species

Iron acquisition by *Dickeya* is required for the systemic progression of maceration symptoms in the plant hosts (Enard et al., 1988, Dellagi et al., 2005, Franza et al., 2005). To chelate iron from the surroundings, most *Dickeya* species synthesize and excrete two siderophores: the hydroxycarboxylate achromobactin encoded by the *acsABCDEF* cluster (Munzinger et al., 2000), and the catecholate chrysobactin encoded by the *cbsABCEFHP* genes (Persmark et al., 1989). The *Dickeya* strain EC16 produces dichrysobactin and linear/cyclic trichrysobactin in addition to the monomeric siderophore chrysobactin (Sandy and Butler, 2011). These siderophores form a complex with Fe(III) designated as ferric-siderophore (Franza and Expert, 2013). The ferric-siderophores are specifically recognized by outer membrane transporters (Acr for ferric-achromobactin; Fct for ferric-chrysobactin). These transporters are gated-channels energized by the cytoplasmic membrane-generated proton motive force transduced by the TonB protein and its auxiliary proteins ExbB and ExbD (Franza and Expert, 2013). Two pairs of ExbB and ExbD proteins are present in most *Dickeya* species. Transport of a ferric-siderophore across the inner membrane involves a specific ABC permease (CbrABCD for ferric achromobactin; CbuBCDG for ferric chrysobactin). Interestingly, the achromobactin genes (*acs* gene cluster) and related transport system (*cbr* gene cluster) were absent in *P. confusarubida* and *D. paradisiaca*, suggesting they were acquired by HGT in *Dickeya* after the divergence of these two lineages. *Dickeya* genes are closely related to *Pseudomonas fulva* and *P. syringae* sequences, suggesting an HGT between these plant-associated bacteria (Supplementary Figure S3). By contrast the chrysobactin genes (*cbs* gene cluster) and related transport system (*cbu* gene cluster) were likely present in the ancestor of *Dickeya* and *P. confusarubida* (Supplementary Figure S3), and then specifically lost in *D. dadantii* subspecies *dieffenbachiae.* In addition to ferric-siderophores, various other iron uptake systems are present in *Dickeya*. The ferrous iron transport systems FeoAB and EfeUOB can be inferred as ancestral in all *Dickeya* species and *P. confusarubida* (Supplementary Figure S3). In contrast, the taxonomic distribution and the phylogeny of the YfeABCD permease that can import both iron and manganese, suggested an acquisition through HGT by *D. chrysanthemi* and Cluster II species, except *D*. gs. undicola (Supplementary Table S4 and Supplementary Figure S3). The haem transport Hmu system was present in most *Dickeya*. Its phylogeny suggested an ancestral presence in *Dickeya*, followed by secondary losses in *D. dianthicola*, *D.* gs. undicola, *D. zeae*, and *D. aquatica* (Supplementary Figure S3, Supplementary Table S4). Although variable combinations of iron assimilation systems exist in *Dickeya* species, at least four systems were present in each species. This multiplicity underscores the fact that competition for this essential metal is critical for the outcome of the plant-*Dickeya* interaction.

#### Biosynthesis of secondary metabolites in Dickeya species

In addition to siderophores, some *Dickeya* species produce secondary metabolites such as the phytotoxin zeamine and the antifungal compound oocydin via non-ribosomal peptide synthases (NRPS) and polyketide synthases (PKS) (Zhou et al., 2011a; Matilla et al., 2012). To evaluate the diversity of secondary metabolites produced by the *Dickeya* genus, we screened the eight complete genomes (*D. paradisiaca* Ech703, *D. aquatica* 174/2, *D. zeae* EC1, *D. zeae* Ech586, *D. chrysanthemi* Ech1591, *D. solani* IPO2222, *D. fangzhongdai* N14b, *D. dadantii* 3937) and the three partial genomes *(D. dianthicola* RNS04.9, *D.* gs. poaceaephila Ech569, *D.* gs. undicola 2B12) for gene clusters encoding NRPS or/and PKS. Then, we analysed the evolutionary history of these genes clusters among the 49 *Dickeya* genomes. The *oocBCDEFGJKLMNOPQRSTUVW* gene cluster coding for oocydin biosynthesis proteins was present in *D. paradisiaca, D. zeae* strains isolated from rice, *D. chrysanthemi* subspecies *chrysanthemi, D. solani, D. dianthicola*, and *D. fangzhongdai* strain NCPPB 3274 (Supplementary Table S4). This could suggest an ancestral presence in *Dickeya* accompanied by losses in *D. aquatica*, *D*. gs. poaceaephila, *D. dadantii*, *D*. gs. undicola, most *D. fangzhongdai* strains, and some Cluster I strains (Supplementary Figure S3). However, the hypothesis of acquisition and spreading through HGT within *Dickeya* could not be excluded. The *zmsABCDEFGIJKLMNPQRS* gene cluster directing zeamine biosynthesis was restricted to *D. zeae* strains isolated from rice, *D. fangzhongdai* and *D. solani* (Supplementary Table S4), suggesting secondary acquisition by HGT (Supplementary Figure S3). In addition, genes involved in coronafic acid biosynthesis, a phytotoxin classically produced by *Pseudomonas syringae* (Bender et al., 1999), were present only in *D*. gs. poaceaephila and *D. dadantii* subspecies *dieffenbachiae* (Supplementary Table S4), suggesting specific acquisition via HGT. We detected four additional gene clusters encoding NRPS and PKS, (i) cluster 1 was specific to *D. paradisiaca* and *P. confusarubida*, (ii) cluster 2 was specific to *D. paradisiaca*, (iii) cluster 3 was specific to *D. aquatica*, suggesting recent acquisitions by these species, while (iv) cluster 4 was more widely distributed, being detected in *D. aquatica, D.* gs. poaceaephila*, D. fangzhongdai, D. solani*, some *D. dadantii* strains, and *D. zeae* except the strains isolated from rice (Supplementary Table S4). To conclude, each *Dickeya* species was characterized by a specific combination of large gene clusters possibly involved in the production and secretion of toxic secondary metabolites. These clusters were likely acquired from unrelated bacteria through HGT and could have been selected based on the constraints imposed by host or environmental factors. For example, the *D. zeae* strains can be subdivided in two groups, the strains isolated from rice, which produce both zeamine and oocydin, while the other strains infecting other crops produce the fourth-type metabolite encoded by cluster 4. This difference between *D. zeae* strains was used by Zhou et *al*. (2015) to define the distinct pathovar linked to rice as *D. zeae* subsp. *oryzae.*

#### Evolution of Dickeya gene repertoires

The 49 *Dickeya* proteomes used in this study contained in average 4,022 proteins, ranging in size from 3,533 (*D*. gs. poaceaephila) up-to 4,352 (*D. fangzhongdai* NCPPB 3274) proteins, representing a difference of 819 proteins (Supplementary Table S3). Comparison of the 197,073 proteins contained in the 49 *Dickeya* proteomes led to the delineation of 11,566 protein families (Figure 7). These protein families correspond to the pan-proteome of *Dickeya*. Among these protein families, 13.9% (1,604) were present at least in one copy in all *Dickeya* proteomes (Figure 7) and defined the core-proteome of this genus. Yet, the size of both pan- and core-proteomes could be slightly underestimated because the proteomes of some strains were deduced from draft genomes. Nevertheless, this meant that in average, ~39.9% of the proteins of any *Dickeya* proteome belonged to the core proteome, while a given *Dickeya* proteome encompassed only ~34.8% of the pan-proteome of this genus. Random taxonomic sampling-based rarefaction curves indicated that sequencing more *Dickeya* genomes will probably not change significantly the estimated size of the core-proteome, while it appears that the pan-proteome is far from being fully disclosed (Figure 7B). This highlights the high diversity and plasticity of the gene repertoires in *Dickeya*. This observation coupled to the great diversity of the virulence, stress resistance, metabolism, and secretion systems imply that none of the *Dickeya* strains could be regarded as a representative model for this genus. Core protein families could be punctually lost in a given strain, while some protein families can be present transiently in a few strains. Thus, it is also relevant to consider the persistent (i.e. protein families present in more than 90% of the strains) and volatile proteomes (i.e. protein families present in less than 10% of the strains) (Touchon et al., 2009). In *Dickeya*, most protein families could be classified either as persistent (2,714 protein families, 23.5%) or volatile (6,426 protein families, 55.6%) (Figure 7A). Unsurprisingly, the persistent proteome was enriched in proteins with known functions, while volatile proteome encompassed mostly hypothetical proteins, prophage elements, and transposases. It is tempting to consider the genes encoding for the volatile proteome as a reservoir of functional innovations, yet the adaptive potential of these genes remains a matter of debate (Touchon et al., 2009).

**Figure 7:**
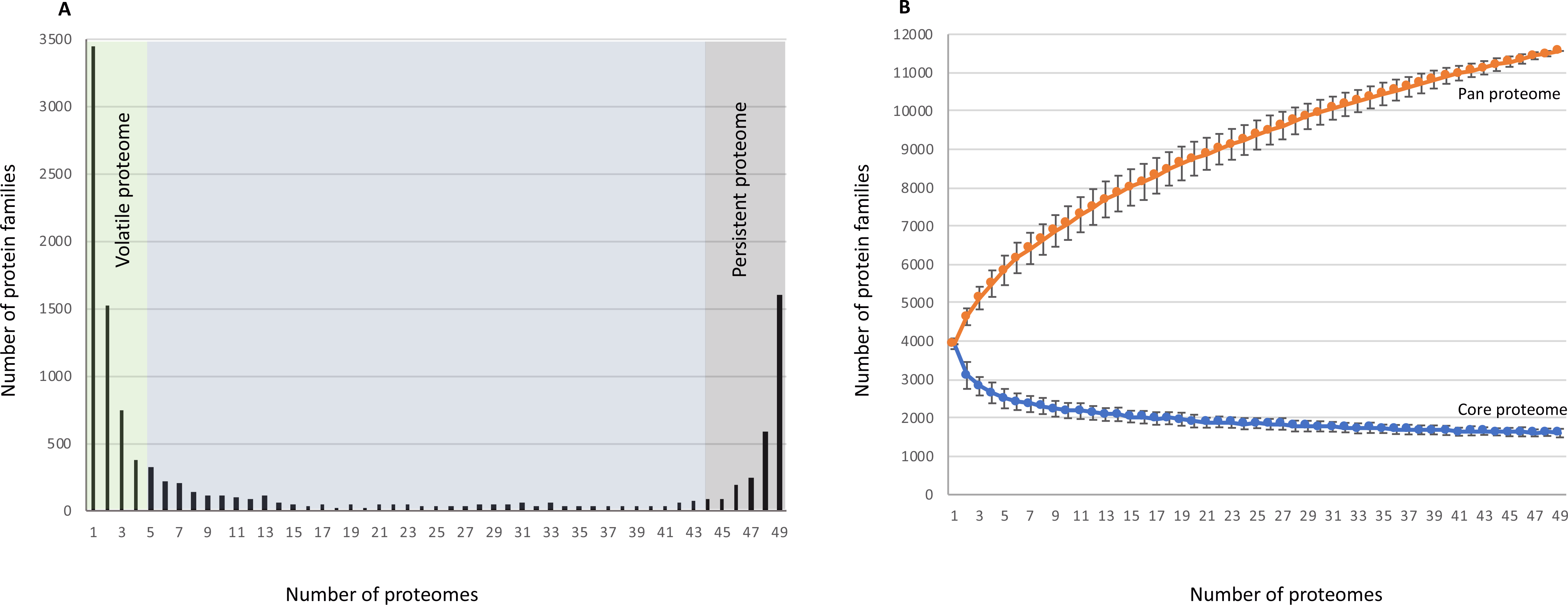
The core-, pan-, versatile-, and persistent-proteomes of *Dickeya* genus. The delineation of core-, pan-, versatile-, and persistant-proteomes was based on the taxonomic distribution of the 11,566 proteins families identified by analyzing the 49 proteomes of *Dickeya*. The core-proteome is defined by the protein families present at least in one copy in all 49 *Dickeya* proteomes, the pan-proteome is defined by the 11,566 protein families, while the versatile- and the persistent-proteomes are defined as the protein families present in less than 10% and in more than 90% of the 49 *Dickeya* proteomes, respectively. **(A) Distribution 11,566 protein families across the 49 *Dickeya* proteomes.** The 3,452 protein families present in a single *Dickeya* proteome (i.e. strain specific families) are on the left of the x-axis, while the 1,604 protein families defining the core-proteome are on the right of the x-axis. **(B) Estimation of the *Dickeya* core and pan-genomes** The graph shows the estimated sizes of the core- and pan-proteomes of *Dickeya* according to the number of considered strains. The curves were computed by calculating the core- and the pan-proteomes for an increasing number of strains randomly selected among the 49 *Dickeya* strains (100 replicates at each point). When all the 49 Dickeya strains were considered, the core- and the pan-proteomes encompasse 1,604 and 11,566 protein families.

Among the 11,566 protein families inferred in *Dickeya*, 3,452 corresponded to strain specific families (i.e. being present in a single proteome) (Figure 7A and Supplementary Table S8). The number of strain specific protein families ranged from 336 in *D. aquatica* 174/2^T^ and 304 in *D*. gs. Poaceaephila, down to zero in *Dickeya solani* strains MK16, PPO9134 and RNS0773B (Supplementary Table S8). To determine the origin of these strain specific protein families, we used them as seeds to query with BLASTP (e-value cut-off 10^−4^) a local database gathering 3,104 complete prokaryotic proteomes, including the 49 *Dickeya* and *P. confusarubida* ATCC 39006 proteomes. Results indicated that 1,051 (30.4%) *Dickeya* strain specific protein families displayed best hit in one of the other 48 *Dickeya* strains, meaning that those sequences were wrongly considered as strain specific because they did not satisfy the coverage and identity parameters used to delineate the protein families. This was not surprising because some protein families could be fast-evolving, meaning that applying uniform parameters can fail to delineate correctly these protein families and lead to an overestimation of the strain specific protein families. In contrast, 1,625 (47.1%) *Dickeya* strain specific protein families displayed best hits in non-*Dickeya* proteomes, meaning that the corresponding genes were likely acquired by HGT from non-*Dickeya* donors. The taxonomic distribution of the corresponding sequences pinpointed *Proteobacteria* (especially *Enterobacteriales*), and to a less extent *Firmicutes* as major donors (Supplementary Table S8 B-C). Yet, the contribution of these two phyla is likely overestimated due to their overrepresentation in sequence databases compared to other lineages. Accordingly, these results should be interpreted as general trends but additional data would be required to precisely estimate the real contribution of *Firmicutes* and *Proteobacteria*. Finally, 776 (22.5%) *Dickeya* strain specific protein families displayed no significant hits or no hits at all, indicating that the corresponding genes were truly strain specific or corresponded to annotation errors (false positives).

At the species level proteome size variation ranged from 435 (*D. aquatica*) to 120 (*D. paradisiaca*) proteins (Supplementary Table S8). The taxonomic distribution of the species-specific protein families displayed overall similar pictures, with most of them being present in all strains of the species (Supplementary Figure S6). This revealed a relative homogeneity of proteomes within species. Interestingly, a few strains diverged from this general trend, such as strain NCPBB 3274 within *D. fangzhongdai, D. aquatica* 174/2^T^, *D. chrysanthemi* NCPPB 402, *D. dadantii* NCPPB 2976, and *D. solani* RNS 0512A. This is consistent with some previous studies. For instance, *D. dadantii* NCPPB 2976 is part of the *dieffenbachiae* subspecies and has been shown to be clearly different from the other *D. dadantii* subsp *dadantii* strains based on ANI values (Zhang et al., 2016). Among, *D. solani*, strain RNS 0512A was proposed to define a novel *D. solani* sub-group based on the high number and wide distribution of nucleotide variations compared to other *D. solani* strains (Khayi et al., 2015).

Using COUNT (Csuros, 2010) on the 12,660 protein families built with SILIX and the topology of the core-pf tree, we inferred the ancestral protein repertoires at each node of the *Dickeya* core-pf phylogeny (Figure 8). COUNT provided similar results for different *Dickeya* species, irrespectively of the position of *D. solani*, as either the sister-group of *D. dadantii* or *D. dianthicola* (Supplementary Table S9), as suggested by core-pf and rprots phylogenetic analyses (see above). The main difference lies in the numbers of gains and losses at the base of *Dickeya*.

**Figure 8.**
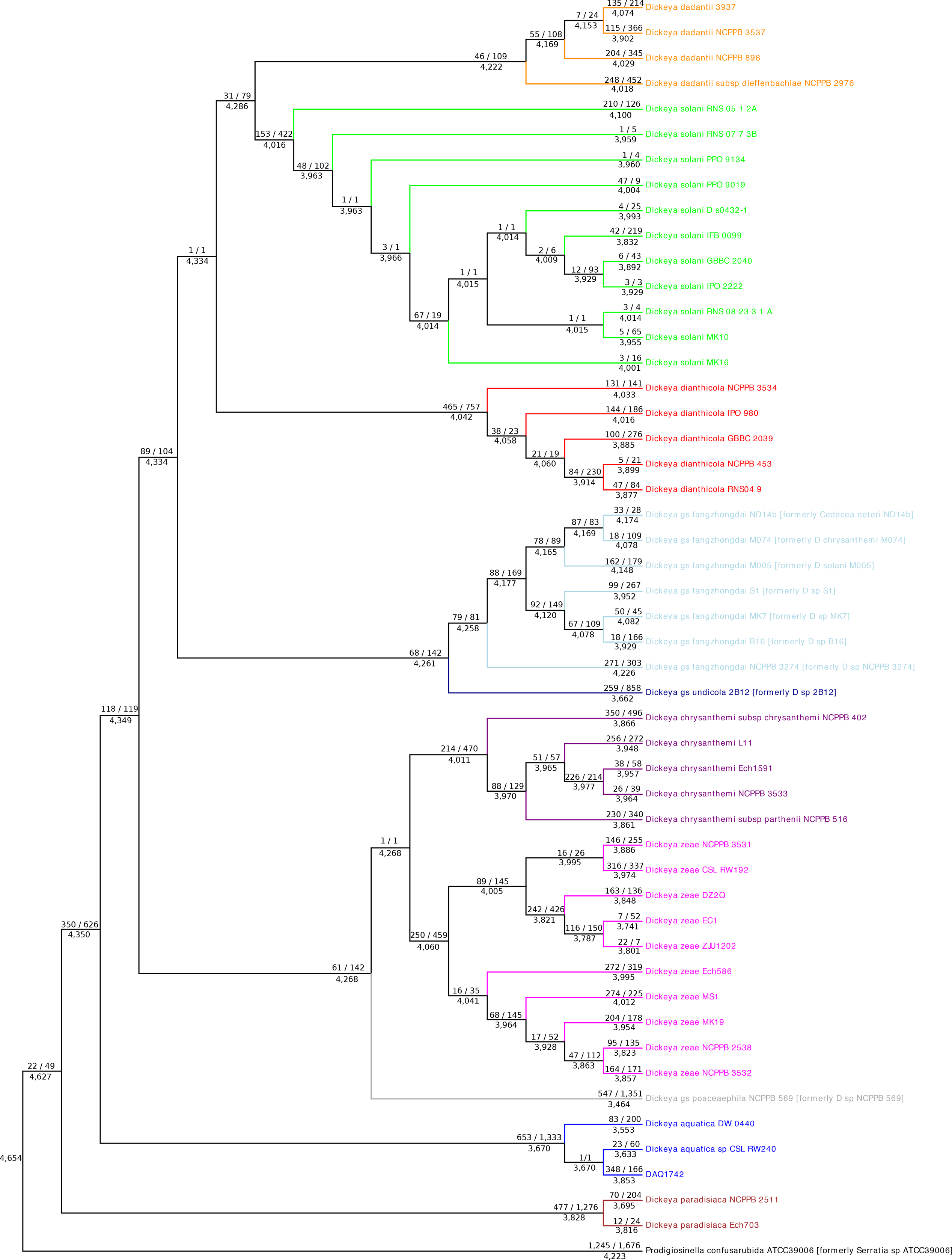
Evolution of protein family repertoires along the *Dickeya* phylogeny. The number of protein family repertoires, gains, and losses inferred by COUNT are mapped on the topology of the core-pf reference phylogeny of *Dickeya*. Values mapped above the branches correspond to gains / losses, while values below branches correspond to the number of protein families inferred. Considering the alternative placement of *D. solani* as sister-group of *D. dianthicola* provided very similar results (see Supplementary Table S9).

We inferred 4,627 protein families in the ancestor of *Dickeya*, while in average 3,921 protein families are contained in present-day *Dickeya* proteomes. This corresponds to a global loss of 18% of the protein families. Interestingly, loss of protein families dominated over gains and affected all *Dickeya* species (Figure 8). Highest protein losses were observed on the stems leading to *D*. gs. poaceaephila and *D. aquatica* and to a lesser extent in *D. paradisiaca*, *D*. gs. undicola, and *D. dianthicola*. These losses were only partially compensated by protein family gains. Surprisingly, more gains were observed in *D. aquatica* 174/2^T^, compared to the two other *D. aquatica* strains (Figure 8). We assume that this was not due to biases in the annotation process by RAST, because very similar results were obtained when using PROKKA (Seeman, 2014). In fact, this may reflect the fact that the genomes of DW 0440 and CSL RW240 strains were not completed, being reported as draft genomes. The general trends observed were robust irrespectly of the postion of *D. solani* relatively to *D. dadantii* and *D. dianthicola* (Supplementary Table S9).

### Characterization of the D. aquatica mobilome

Mobile Genetic Elements (MGEs) are the main actors of the HGT and include plasmids, viruses (phages and prophages) and transposons (Jackson et al., 2011). They are often localised within genomic islands on chromosomes. The mobilome of a strain is the repertoire of all the genes associated with MGEs. Using both PHAST and IslandViewer, we detected seven phage elements and ten genomic islands in *D. aquatica* 174/2^T^ (Figure 1, Supplementary Table S10). Most of the genomic islands contained genes of transposases, integrases or mobile elements that were likely remnants of HGT. They also contain 105 of the 336 ORFAN genes detected in *D. aquatica* 174/2^T^ strain. Among the seven detected prophages, P2, P6 and P7 were related to transposable Mu-phage. These were also present in some *D. zeae* strains isolated from river as well as *D. dianthicola* strains (Supplementary Table S10). P3 and P5 were specific to some *D. aquatica* strains, while P1, a defective prophage with only few conserved genes, and P4 were widespread in all *Dickeya* species (Supplementary Table S10). Among the ten genomic islands detected in *D. aquatica* 174/2^T^, seven (GI1, GI2, GI4, GI5, GI6, GI7, GI9) were mainly composed of mobile elements and small hypothetical proteins, GI2 also contained a type III restriction-modification system, whereas GI4 included a type I restriction-modification system and a toxin-antitoxin system (Supplementary Table S10). Similarly, GI6 contained a toxin-antitoxin system as well as an isolated non-ribosomal peptide synthase, which was also found in *D.* gs. poaceaephila and *D. zeae* (Supplementary Table S10). GI7 contained some metabolic proteins, including the previously mentioned cluster 3 encoding NRPS and PKS, as well as transporters, notably a cobalt/nickel ABC transporter (Supplementary Table S10). Excluding the mobile elements, GI1, GI2, GI4, GI5, GI6 GI7, and GI9 were specific to *D. aquatica* strains, even if a few genes composing these GIs can be punctually detected in some others strains (Supplementary Table S10). The three other genomic islands (GI3, GI8, GI10) were metabolic islands (Supplementary Table S10). GI3 has been laterally transferred between *Erwinia pyrofolia* and *D. aquatica* (Supplementary Figure S3). GI8 that included proteins related to fatty acid metabolism, was conserved in *D. zeae* strains isolated from rice or originated from China (Supplementary Table S10). GI10 contained proteins related to the complete carbapenem biosynthetic pathway CarABCDE and the associated resistance proteins CarFG (Supplementary Table S10). *P. confusarubida* contained genes coding for CarABCDE as well as CarFG, in agreement with its capacity to synthesize the carbapenem antibiotic (carbapen-2-em-3-carboxylic acid) (Thomson et al., 2000; Coulthurst et al., 2005). This cluster was conserved in *D. undicola, D. dadantii* subspecies *dieffenbachiae*, some *D. chrysanthemi* strains and in the *D. zeae* CSL-RW192 strain isolated from water (Supplementary Table S10). *D. paradisiaca* only contained the *carFG* resistance genes but was deprived of the biosynthetic genes (Supplementary Table S9). The phylogeny of CarABCDE and CarFG shows that relationships among the *Dickeya* sequences are inconsistent with the species phylogeny, suggesting a secondary acquisition through HGT in this genus (Supplementary Figure S3).

Overall, phages and genomic islands from *D. aquatica* are mostly species specific suggesting a wide genomic plasticity in the *Dickeya* genus.

### Concluding remarks

In this work, we sequenced the complete genome of *D. aquatica* 174/2 and showed that unlike initially supposed, this bacterium, similarly to other *Dickeya* species, is a phytopathogen. Who is the natural host of *D. aquatica* and why is this bacterium found in aquatic environments rather than associated with plants as are its close relatives? Althought the data at hand are not sufficient to answer these questions, we speculate that *D. aquatica* can be pathogenic for aquatic plants such as charophytes, which are a family of complex-structured algae living in a variety of wetland and freshwater habitats including those, from which the *D. aquatica* strains have been isolated. Charophytes are thought to be the closest ancestor of land plants. The cells of these algae are surrounded by polysaccharide-based cell walls. However, their cell walls are thin and cannot be distinguished as primary or secondary cell walls. Furthermore, they lack xyloglucans, which are common in most land plants (Sarkar et al., 2009). Intriguingly, *D. aquatica* lacks the xylanases and xylose degradation pathways and we hypothetize that this could reflect an evolutionary adaptation to charophyte hosts.

The comparison of *D. aquatica* 174/2 and *Dickeya* strain proteomes available in public databases showed that the *Dickeya* genus displays a remarkable diversity featuring many unique protein families and emphasizing that our knowledge of this genus is still limited. The real size of the pan-proteome is an open question encouraging further exploratory studies on these bacteria, the success of which will depend on the collection of new strains isolated from various environments and the sequencing of the genomes of newly identified representatives of the genus. Remarkably, we observed an enormous degree of genetic plasticity in the pathogenicity determinants that enable various *Dickeya* species to colonize a wide range of plant hosts. Furthermore, in this work, we have postulated the existence of a sister genus of *Dickeya*, *Prodigiosinella*, and characterized two new *Dickeya* genomospecies. The reconstruction of ancestral genomes allowed us to gain new insights into the evolutionary history of this genus and highlighted an evolutionary trajectory dominated by the loss of protein families.

## Experimental procedures

### Genome sequencing, assembly and annotation

DNA for sequencing of the *D. aquatica* type strain (174/2) genome was extracted from overnight broth culture using Promega bacterial genomic DNA kit. PacBio sequencing to > 350X coverage was performed by Eurofins Genomics (https://www.eurofinsgenomics.eu/). Reads were assembled using CANU (Koren et al., 2017). The annotation was performed automatically with RAST (Aziz et al., 2008), then expertly reviewed using MAGE (Vallenet et al., 2013) and literature data. The expert review allowed to assign 2,457 gene names, correct 547 annotations and add 5 CDS missed by RAST. The replication origin (oriC) was predicted by OriFinder (http://tubic.tju.edu.cn/Ori-Finder) (Gao and Zhang, 2008). The presence of mobile genetic elements in the *D. aquatica* 174/2 genome was investigated by the following online tools: IslandViewer (http://pathogenomics.sfu.ca/islandviewer) (Dhillon et al., 2015) for the GI regions, CRISPRDetect (http://brownlabtools.otago.ac.nz/CRISPRDetect/predict_crispr_array.html) (Biswas et al., 2016) for CRISPR arrays, whereas putative prophage sequences were identified by PHAST and PHASTER analysis (http://phast.wishartlab.com/) (Zhou et al., 2011b; Arndt et al., 2016). The genome sequence has been submitted to EMBL database under accession number GCA_900095885 (http://www.ebi.ac.uk/).

### Virulence assays on various hosts

Bacterial cultures were grown in M63G minimal medium (M63 + 0.2% w/v glucose) (Miller 1972) and diluted to a given OD_600_ depending on the host: 0.2 (chicory) or 1 (potato, cucumber, tomato, pineapple and kiwi). For chicory, 5 μL of bacterial suspension were injected into a 2 cm incision at the center of the leaf. For potato, cucumber and tomato, 5, 100 and 200 μL of bacterial suspension were injected into the vegetable, respectively. 200 μL of bacterial suspension were also injected into pineapple and kiwi fruits. Plants were incubated at 30°C with 100% humidity for 18 h (chicory) or 42 h (potato, cucumber and tomato) or 78 h (pineapple and kiwi). The soft rot mass is used to quantify virulence.

### Stress resistance assays

Bacteria were cultured at 30°C in 96 well plates using M63G (M63 + 0.2% w/v glucose) pH 7.0 as minimal medium. Bacterial growth (OD_600nm_) was monitored for 48 h using an Infinite^®^ 200 PRO - Tecan instrument. Resistance to osmotic stress was analysed using M63G enriched in 0.05 to 0.5 M NaCl. Resistance to oxidative stress was analysed in the same medium by adding H_2_O_2_ concentrations ranging from 25 to 200 μM. The pH effect was analyzed using the same M63G medium buffered with malic acid at different pH ranging from 3.7 to 7.0.

### Proteome database construction

We built a local database (DickeyaDB) gathering the proteomes of *Serratia* sp. ATCC 39006, 48 *Dickeya* available at the NCBI (by january 2017), and *D. aquatica* type strain 174/2 (Supplementary Table S3). A second database (prokaDB) containing 3,104 proteomes of prokaryotes, including the 50 proteomes of DickeyaDB, was also built.

### Assembly of Dickeya and Serratia ATCC 39006 protein families

*Dickeya* and *Serratia* ATCC 39006 protein families were assembled with SILIX version 1.2.9 (Miele et al., 2011). More precisely, pairwise comparisons of protein sequences contained in DickeyaDB were performed using the BLASTP program version 2.2.26 with default parameters (Altschul et al., 1997). Proteins in a pair providing HSP (High-scoring Segment Pairs) with identity over 60% and covering at least 80% of the protein lengths were gathered in the same family. This led to the assembly of 12,660 protein families, among which 1,493 were present in at least one copy in all DickeyaDB proteomes. In contrast, considering the 49 *Dickeya* proteomes without taking into account *Serratia* ATCC 39006 led to the assembly of 11,566 protein families, among which 1,604 were present at least in one copy in all *Dickeya* and *Serratia* ATCC 39006 proteomes, and 1,420 in exactly one copy.

### Inferrence of reference phylogenies of Dickeya

Reference phylogenies of *Dickeya* were inferred using ribosomal proteins (rprots) on the one hand and core protein families (core-pf) on the other hand. The rprots phylogenetic tree was rooted with sequences from *Serratia* ATCC 39006, together with three *Pectobacterium* species (*Pectobacterium carotovorum* PC1, *Pectobacterium atrosepticum* SCRI1043, and *Pectobacterium wasabia* WPP163) and five additional *Serratia* species (*Serratia marcescens* FGI94, *Serratia fonticola* DSMZ4576, *Serratia liquefaciens* ATCC27592, *Serratia proteamaculans* 568, and *Serratia plymuthica* AS9), while the core-pf phylogenetic tree was rooted with *Serratia* ATCC 39006 (see results).

Rprots sequences were extracted from the DickeyaDB using the engine of the riboDB database (Jauffrit et al., 2016). Briefly, the riboDB engine allows retrieving rprots sequences through a double approach combining reciprocal best-blast-hits and hidden Markov model (HMM) profiles searches.

Starting from the 1,420 core-pf protein families containing exactly one copy in each proteome of the DickeyaDB and 51 rprots, we applied several quality controls. First, within protein families extremely short sequences (<30% of the median length of the family) were discarded. Then, we used FastTreeMP (Price et al., 2010), and PhyloMCOA (de Vienne et al., 2012) to detect and discard outlier sequences, on the basis of nodal and patristic distances. At the end of the quality control process, 1,341 protein families present in more than 35 (70%) out of the 50 considered proteomes and 51 rprots were kept. For each of these protein families, multiple alignments were built using the CLUSTAL-Omega-1.1.0 program (Sievers et al., 2011) and trimmed using GBLOCKS (Castresana, 2000) with parameters set to a minimal trimming. The trimmed multiple alignments corresponding to 1,341 core-pf on the one hand and the 51 rprots on the other hand have been combined using Seaview-4.5.4 (Gouy et al., 2010) to build the core-pf and the rprots supermatrices containing 414,696 and 6,295 amino acid positions, respectively.

Maximum likelihood (ML) phylogenies of these supermatrices have been inferred with IQ-TREE-1.5.3 (Nguyen *et al.*, 2015) with a C60 profile mixture of the Le and Gascuel evolutionary model (Le and Gascuel 2008) and a gamma distribution with four site categories (Γ4) to model the heterogeneity of evolutionary rates across sites, as proposed by the model testing tool (Kalyaanamoorthy *et al.*, 2017) available in IQ-TREE. The robustness of the inferred ML trees was estimated with the non-parametric bootstrap procedure implemented in IQ-TREE-1.5.3 for the rprots supermatrix (100 replicates of the original alignments) and the ultrafast bootstrap approach for the core-pf supermatrix (1,000 replicates).

### Phylogenetic analysis of single markers

947 SSU rRNA and 125 LSU rRNA complete sequences from *Dickeya*, *Serratia* (including strain ATCC 39006), and *Pectobacterium* available in public databases were retrieved and aligned with MAFFT v7.222. The resulting multiple alignments were trimmed using BMGE-1.1 (default parameters). The phylogeny of the 947 SSU rRNA sequences was inferred using FastTree-2.1.9 (Price et al., 2010), with the GTR + gamma + cat 4 model, while the LSU rRNA tree was inferred with IQ-TREE with the TIM3+F+I+Γ4 model according to the BIC criterion, as suggested by the propose model tool available in IQ-TREE.

Homologues of proteins of interest were identified in the prokaDB with BLASTP. The first 75 HSP hits with evalue smaller than 10^−4^ were kept. The retrieved sequences, together with the seed were aligned using MAFFT v7.222, alignments were trimmed using BMGE-1.1 (default parameters). ML phylogenies were built using IQTREE-1.5.3 (Nguyen *et al.*, 2015) with the LG+Γ4+I+F model. In order to identify the origin and evolution of these proteins in *Dickeya*, their phylogenies were compared and reconciliated with the *Dickeya* species phylogenies, by considering the two alternative positions of *D. solani*: as either sister-group of *D. dadantii* or *D. dianthicola*.

### Identification of the D. fangzhongdai phylogenetic cluster

To determine whether some sequenced strains are related to *D. fangzhongdai,* we used all the *D. fangzhongdai* genes available in public databases SSU-rDNA (KT992690.1), *dnaX* (KT992713.1)*, fusA* (KT992697.1)*, purA* (KT992705.1)*, recA* (KT992693.1), *gapA* (KT992701.1) and *rplB* (KT992709.1) genes, we extracted the corresponding genes from the DickeyaDB database and then used these sequences as input for phylogenetic analyses using leBIBI^QBPP^ phylogenetic positioning tool (Flandrois et al., 2015). We found that the strains B16, MK7 and S1 were systematically affiliated with *D. fangzhongdai*.

### Ancestral gene content

The program COUNT (Csuros, 2010) was used for gene families evolutionary reconstruction in *Dickeya* species using the topology of the core-pf tree as reference. All the generated 12,660 families were submitted to COUNT, which can perform ancestral genome reconstruction by posterior probabilities in a phylogenetic birth-and-death model. Rates were optimized using a gain-loss-duplication model and three discrete gamma categories capturing rate variation across families, with other parameters set at default and allowing different gain-loss and duplication-loss rates for different branches. One hundred rounds of optimization were computed. COUNT was run twice: first with *D. solani* as sister-group of *D. dadantii* and then as sister-group of *D. dianthicola*.

## Acknowledgements

This work was supported by the Investissement d'Avenir grant (ANR-10-BINF-01-01), by the ANR Combicontrol grant (ANR-15-CE21-0003-01), by a grant from the FR BioEnviS and using the computing facilities of the Computing Cluster of the LBBE/PRABI. We would like to thank Georgi Muskhelishvili for critical reading the manuscript and Frederic Jauffrit as well as Mailys Dumet for providing the extraction of ribosomal proteins.

## Supplementary materials

**Supplementary Figure S1: Maximum Likelihood phylogenetic tree of *Dickeya* strains based on the Fr-protein supermatrix gathering 51 ribosomal proteins** (LG+C60, 58 sequences, 6,295 amino acid positions).

Numbers at branch correspond to bootstrap values (100 replicates of the original data set). The scale bar indicates the average number of substitutions per site. The figure was generated using Evolview (He et al., 2016).

**Supplementary Figure S2: Phylogenetic position of “ *Serratia* sp. ATCC 39006” based on LSU and SSU rDNA trees**

ML trees of the *Dickeya* genus inferred with a collection of **(A)**947 SSU rDNA and **(B)** 125 LSU rDNA sequences retrieved from public databases. The trees were rooted using *Pectobacterium* and *Serratia* sequences. The scale bars represent the estimated average number of substitution per site. Numbers at nodes represent ultrafast bootstrap values **(A)** and bootstrap values **(B)**. *Pectobacterium* sequences are in pink, *Serratia* sequences in purple, and *Dickeya* sequences in black. Worth to note, LSU and SSU rDNA sequences from *Serratia* sp. 39006 robustly branch with *Dickeya* sequences and not within the *Serratia* genus. The trees were drawn using the iToL webserver (Letunic and Bork, 2016).

**Supplementary Figure S3 - Single gene phylogenies**

Maximum Likelihood phylogenetic trees of 912 proteins of interest. Numbers associated with each branch correspond to ultrafast bootstrap values (1000 replicates of the original data set). The scale bars indicate the average number of substitutions per site. For each tree, the name and the annotation of the seed is provided in red.

**Supplementary Figure S4 - Non-metric multi-dimensional scaling (NMDS) plot of *Dickeya* genomes according to their gene content.** A Bray distance similarity matrix was calculated based on the presence/absence profiles of 8,115 gene families (present in at least 2 genomes) for the 49 analyzed genomes and used to generate NMDS coordinates for each strain. The shorter distance linking two genomes indicates higher similarities between these genomes. Genomes from different species are indicated in different colors.

**Supplementary Figure S5: Phylogenies and genetic organisation of *pelAED* and *pehVWX* clusters.**

Maximum likelihood trees of *pelAED* and *pehVWX* clusters together with genetic organisation of these clusters are shown. Numbers associated with each branch correspond to ultrafast bootstrap values (1000 replicates of the original data set). The scale bar indicates the average number of substitutions per site. For each tree, the name and the annotation of the seed is provided in red.

**Supplementary Figure S6: The number of shared protein families between and within *Dickeya* species**. Figures were generated using the package UpSetR (Conway et al. 2017).

**Supplementary Table S1: Phenotypic differentiation of species within the genus *Dickeya***. Table showing the phenotypic features of *Dickeya* species based on Samson et al. (2005), Parkinson et al. (2014), van der Wolf et al. (2014) and Tian et al. (2016).

**Supplementary Table S2: Genomic features of the *Dickeya* strains with completely sequenced genomes**

**Supplementary Table S3: List of the 49 *Dickeya* genomes used in this study indicating the plant host or habitat and the geographical origin of each strain**

**Supplementary Table S4: Distribution of genes with a role in pathogenicity or in adaptation to plant niche in *Dickeya* species**

Genes of *Dickeya* species were considered as present if identity of the encoded protein was higher than 60% of full-length amino acids sequence. If local alignments were too short with regard to the length of similar sequences, we performed a nucleotide BLAST on full-length DNA sequences with similar thresholds (10^−5^ e-value, 80% identity of full length sequence). This allowed us to eliminate false genes. Note that because some genomes represent only drafts, false negatives may occur.

**Supplementary Table S5: Protein family distribution within proteomes from *Dickeya, Pectobacterium* and *Serratia***.

Considered strains were *Dickeya aquatica* 174/2, *Dickeya dadantii* 3937, *Dickeya paradisiaca* NCPPB 2511, *Dickeya solani* IPO 2222, *Dickeya zeae* EC1, *Pectobacterium atrosepticum* SCRI1043, *Pectobacterium carotovorum* subsp carotovorum PC1, *Pectobacterium wasabiae* WPP163, *Serratia fonticola*, *Serratia liquefaciens* ATCC27592, *Serratia marcescens* FGI94, *Serratia plymuthica* AS9, *Serratia proteamaculans* 568 and *Serratia* sp ATCC39006. Families were built using SILIX version 1.2.9 with 60% of sequence identity and 80% of sequence coverage.

**Supplementary Table S6: Results of the AU tests performed on core-pf and rprots topologies using the 1,341 core-pf and the 51 rprots.**

**Supplementary Table S7: Distribution of Transcription factor-binding sites in *D. aquatica* 174/2 genome.** Binding sites for 56 transcriptional regulators were predicted in *D. aquatica* using nhmmer (Wheeler and Eddy, 2013).

**Supplementary Table S8: Protein family distribution within the proteomes of the 49 *Dickeya* strains.**

Families were computed with SILIX version 1.2.9 using 60% of sequence identity and 80% of sequence coverage.

**Supplementary Table S9: Comparison of inferences of ancestral gene repertoires in *Dickeya* considering the two possible alternative positions for *D. solani***

**Supplementary Table S10: Characterization of the mobilome of *D. aquatica* 174/2 and its conservation in other *Dickeya* species**

Genes of *Dickeya* species were considered as present if identity of the encoded protein was higher than 60% of full-length amino acids sequence. If local alignments were too short with regard to the length of similar sequences, we performed a nucleotide BLAST on full-length DNA sequences with similar thresholds (10^−5^ e-value, 80% identity of full length sequence). This allowed us to eliminate false genes. Note that because some genomes represent drafts, false negatives may occur.

