## Supplementary figures and images for "The phytopathogenic nature of *Dickeya aquatica* 174/2 and the dynamic early evolution of *Dickeya* pathogenicity"

### Supplementary Figure S1

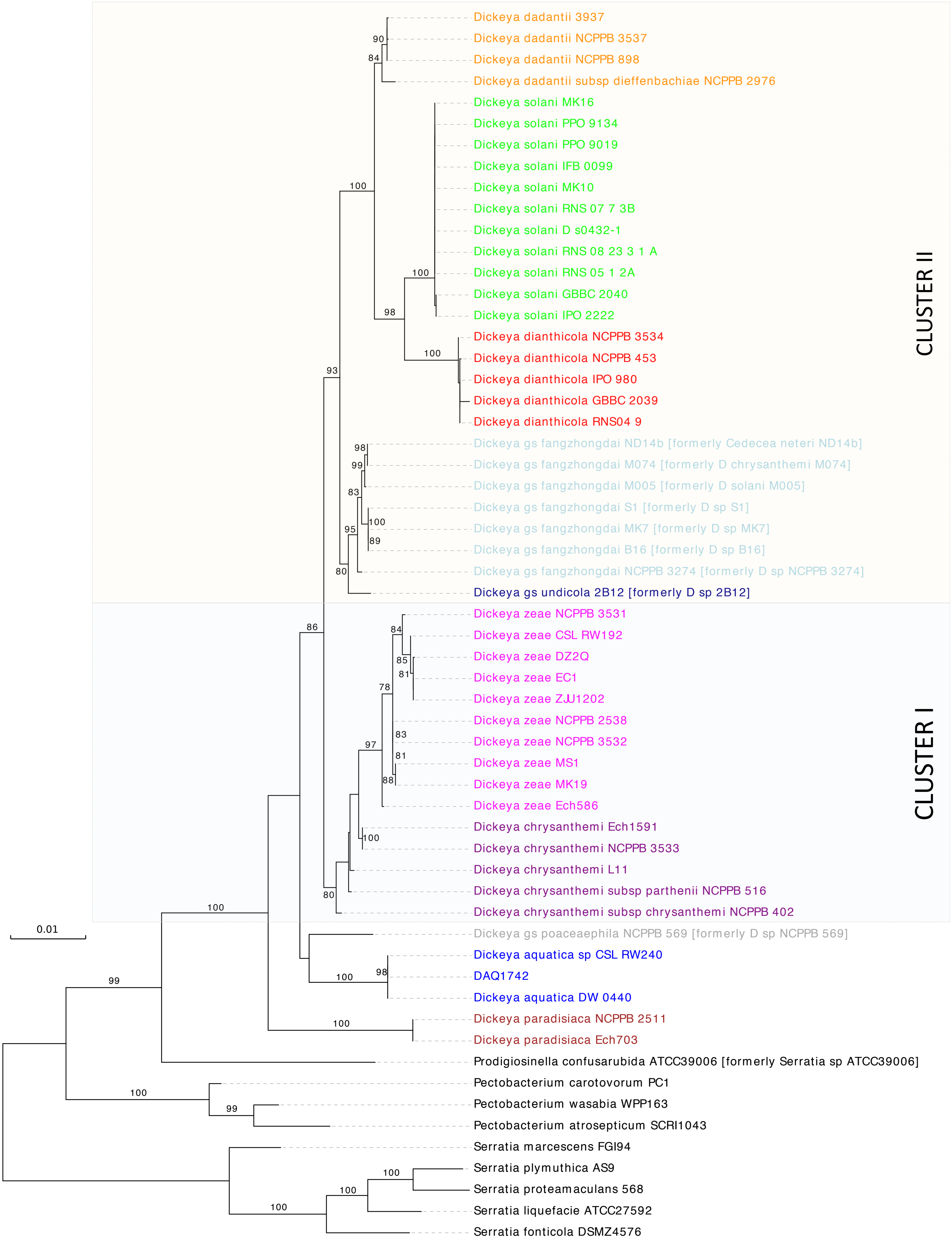

### Supplementary Figure S2

**SSU rDNA tree**

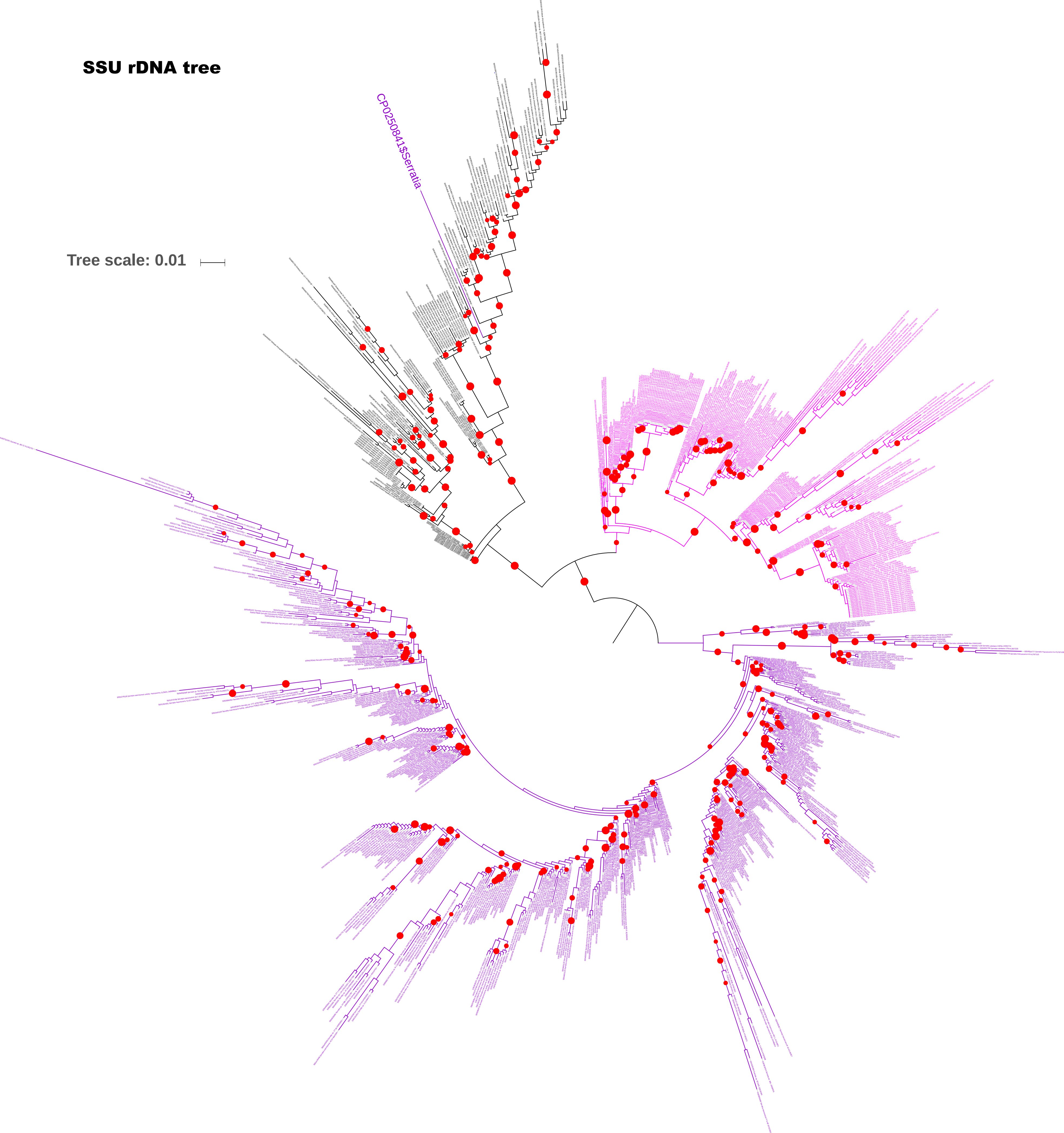

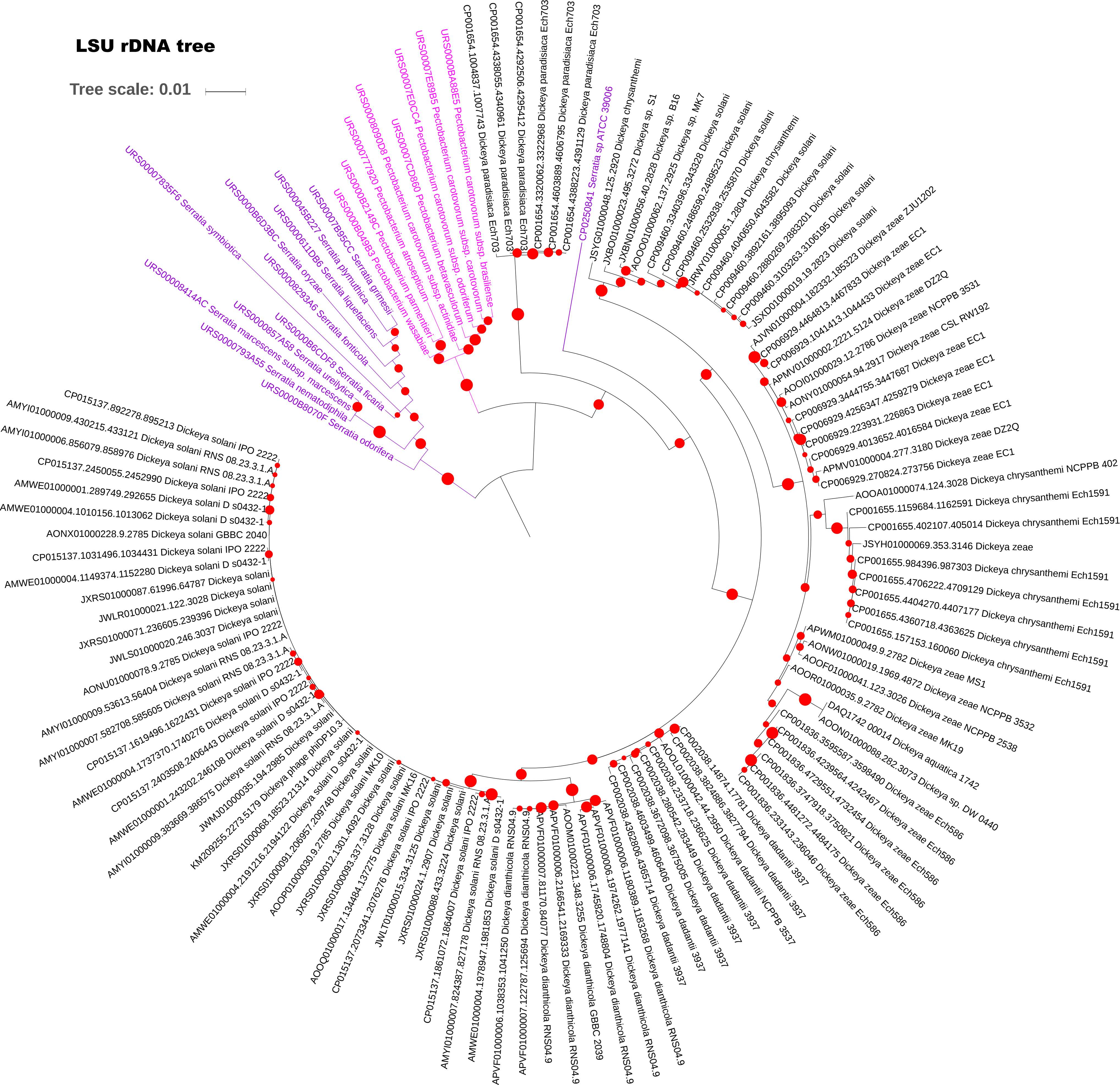

### Supplementary Figure S4

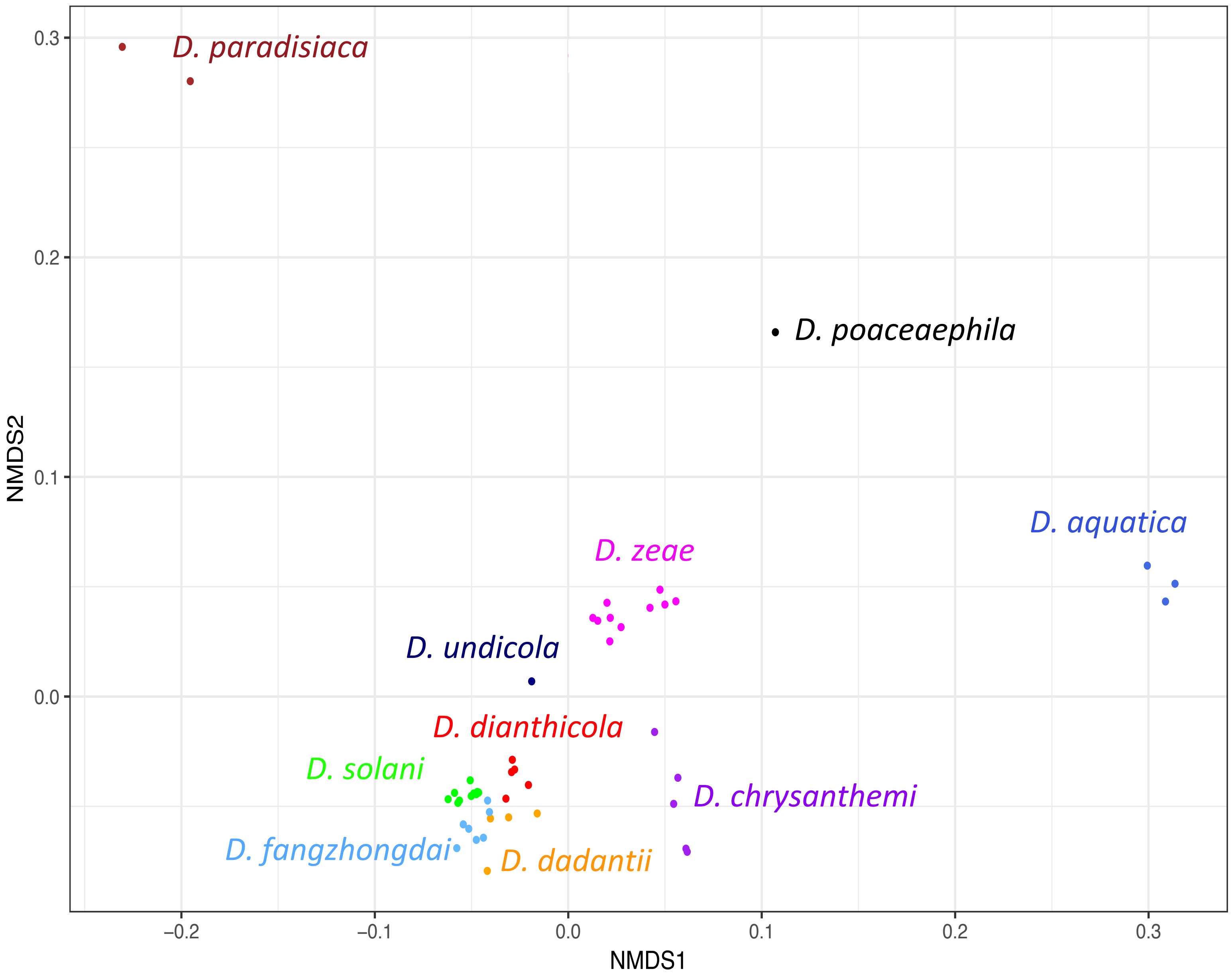

### Supplementary Figure S5

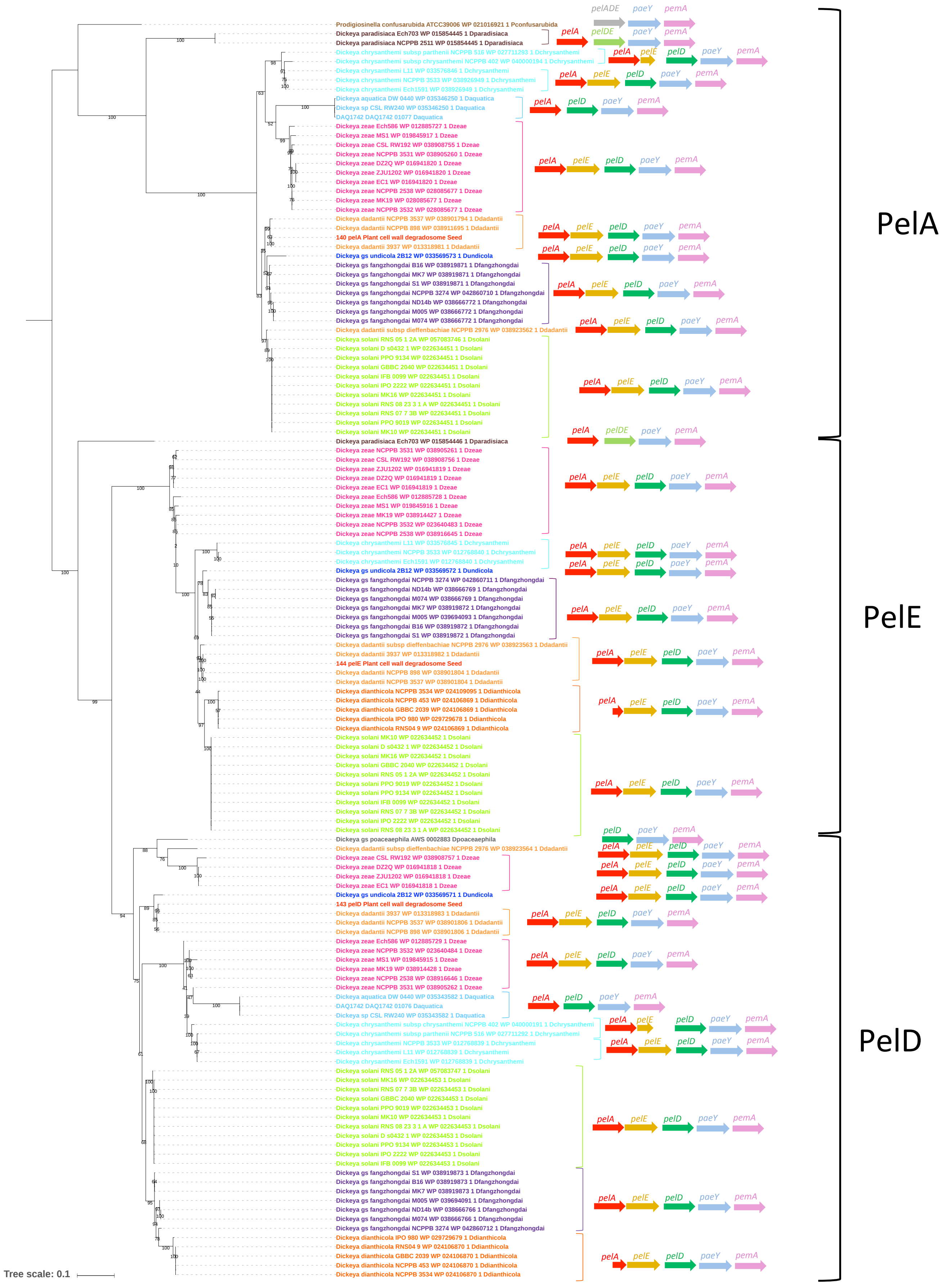

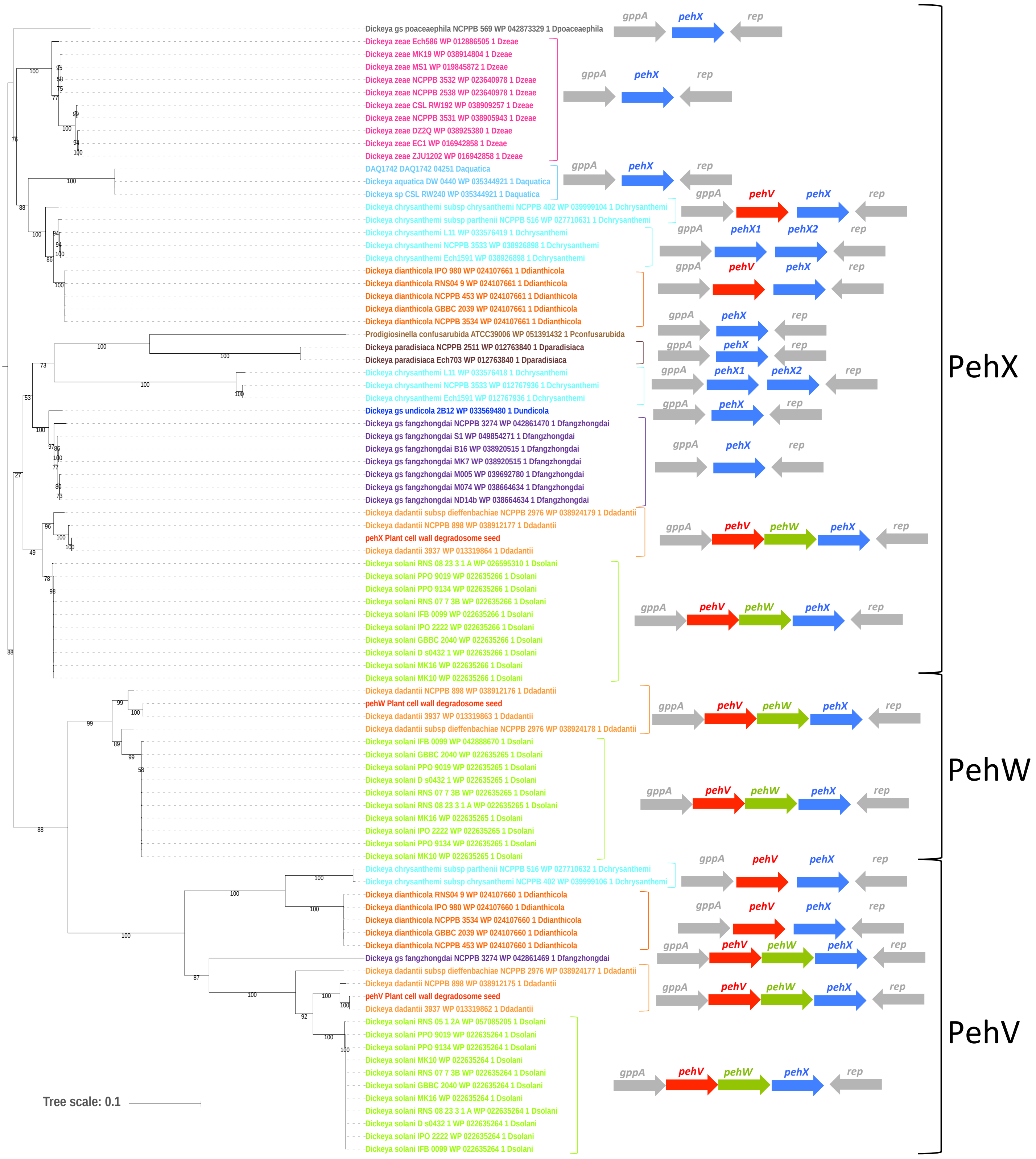

### Supplementary Figure S6

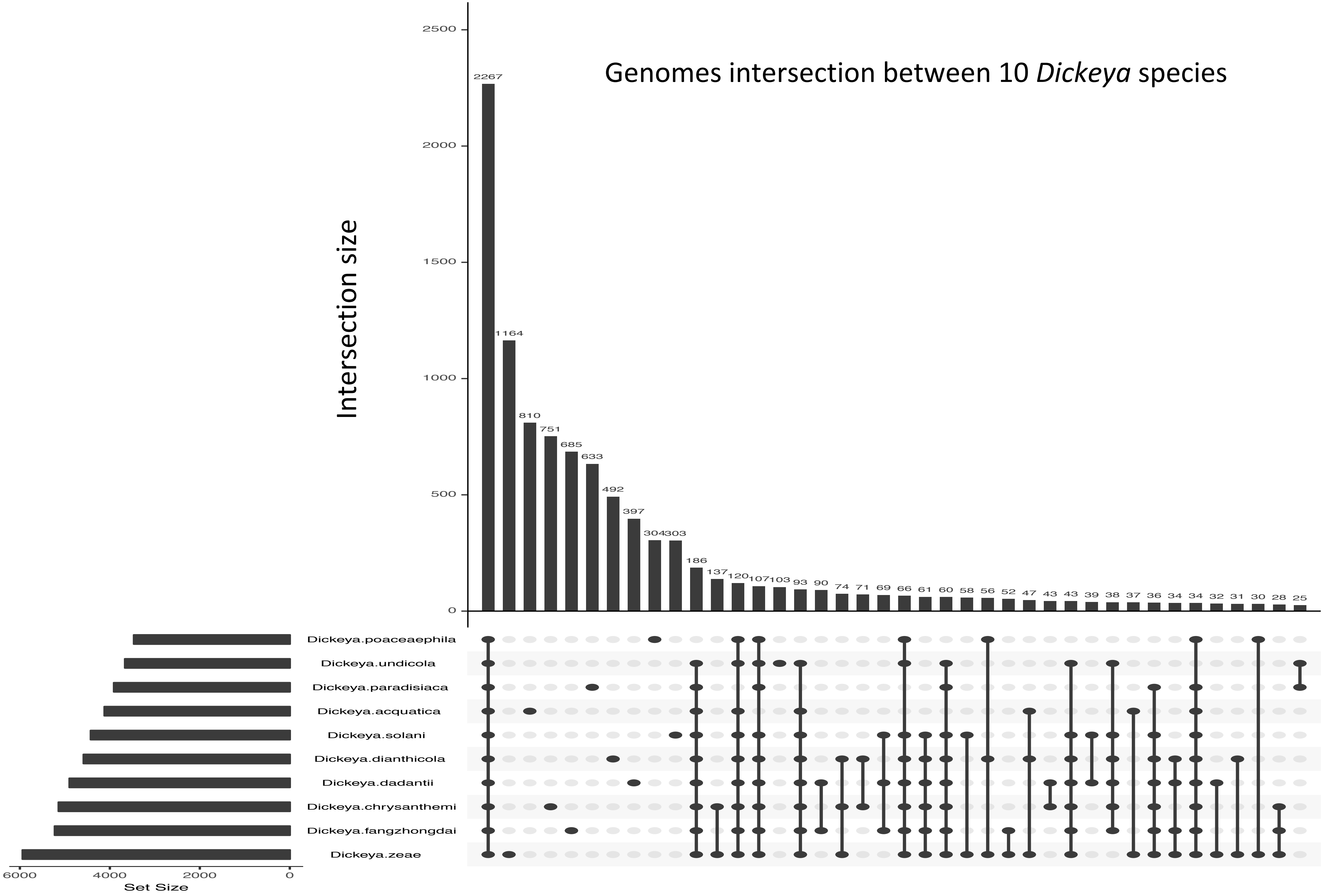

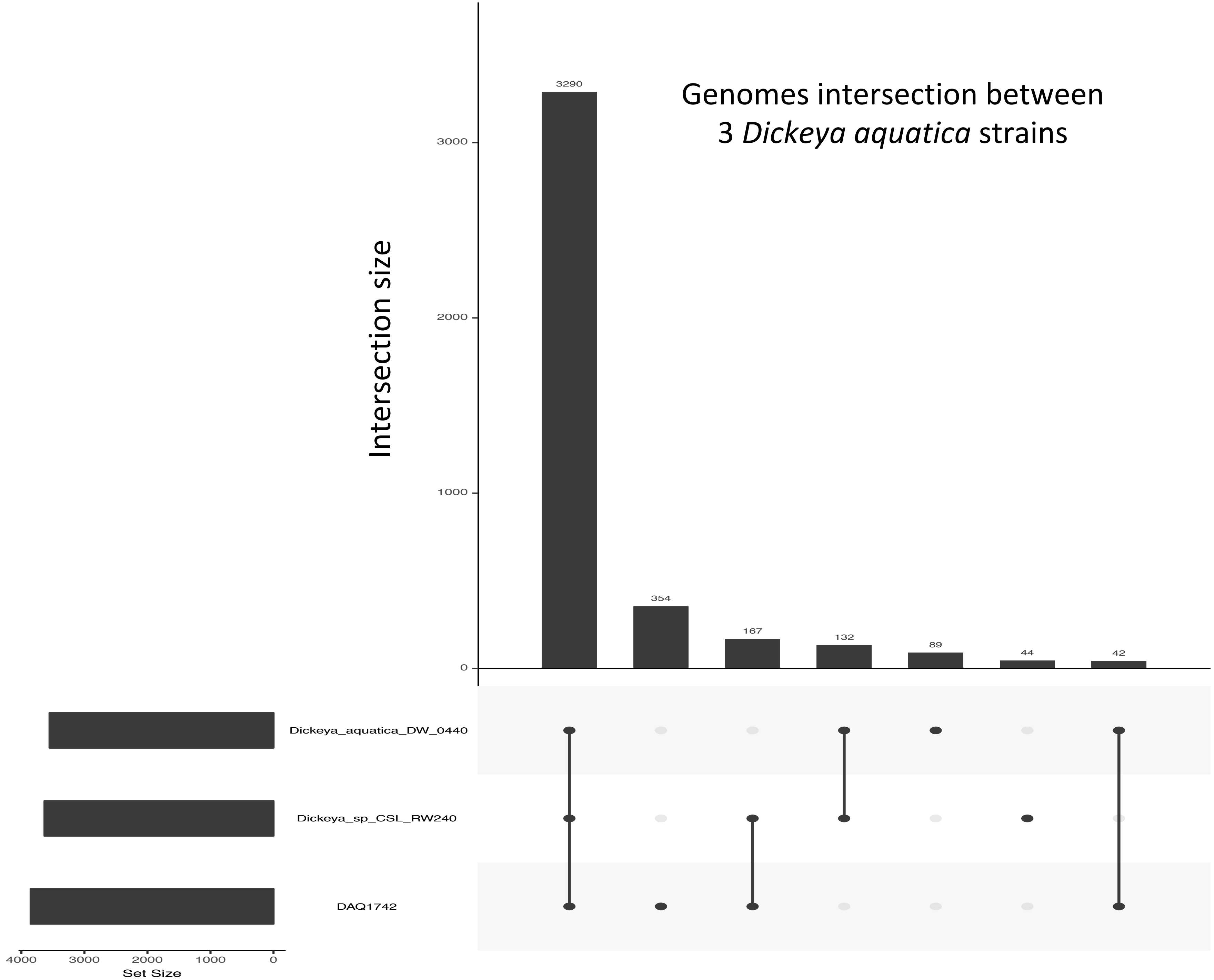

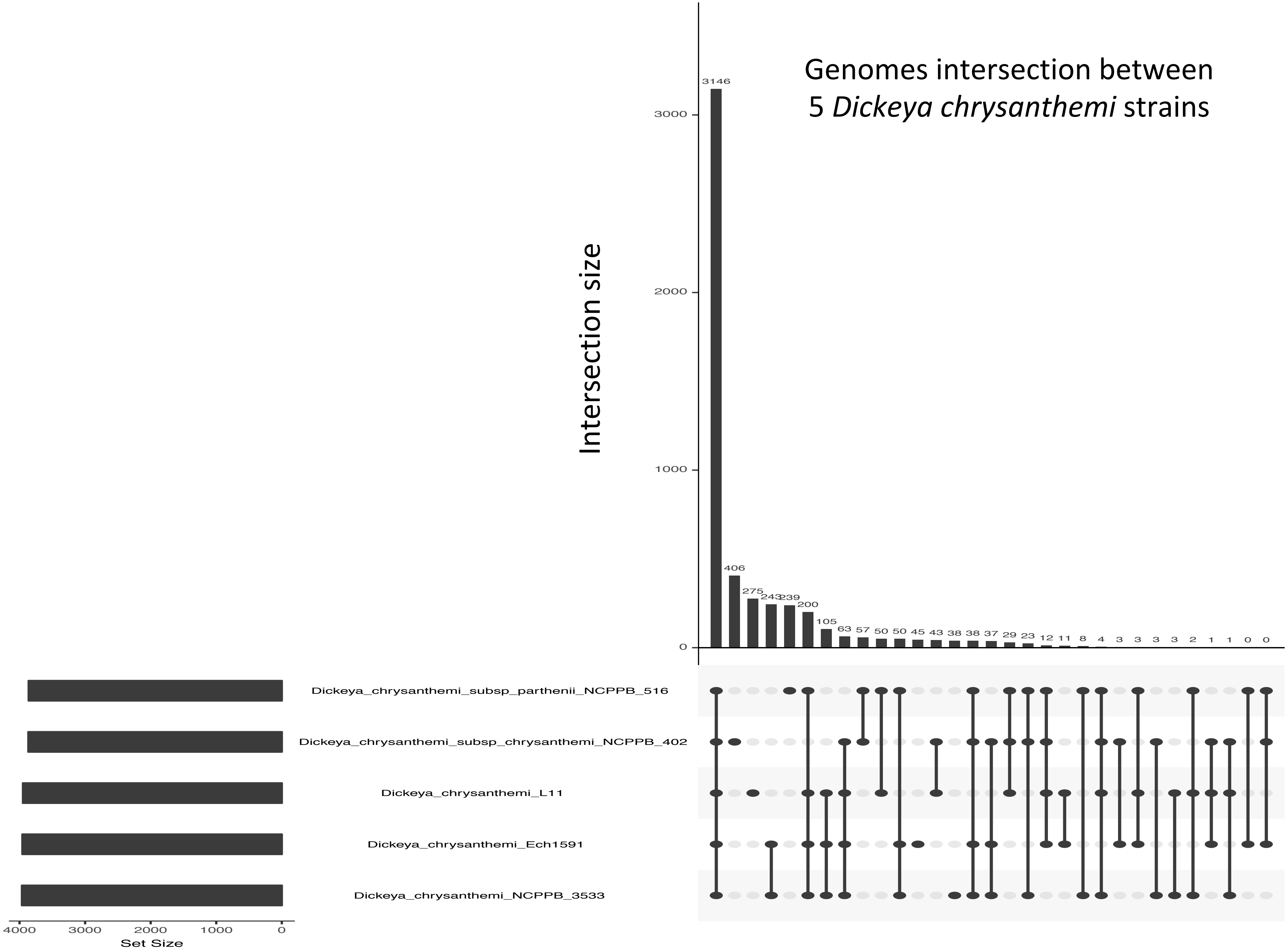

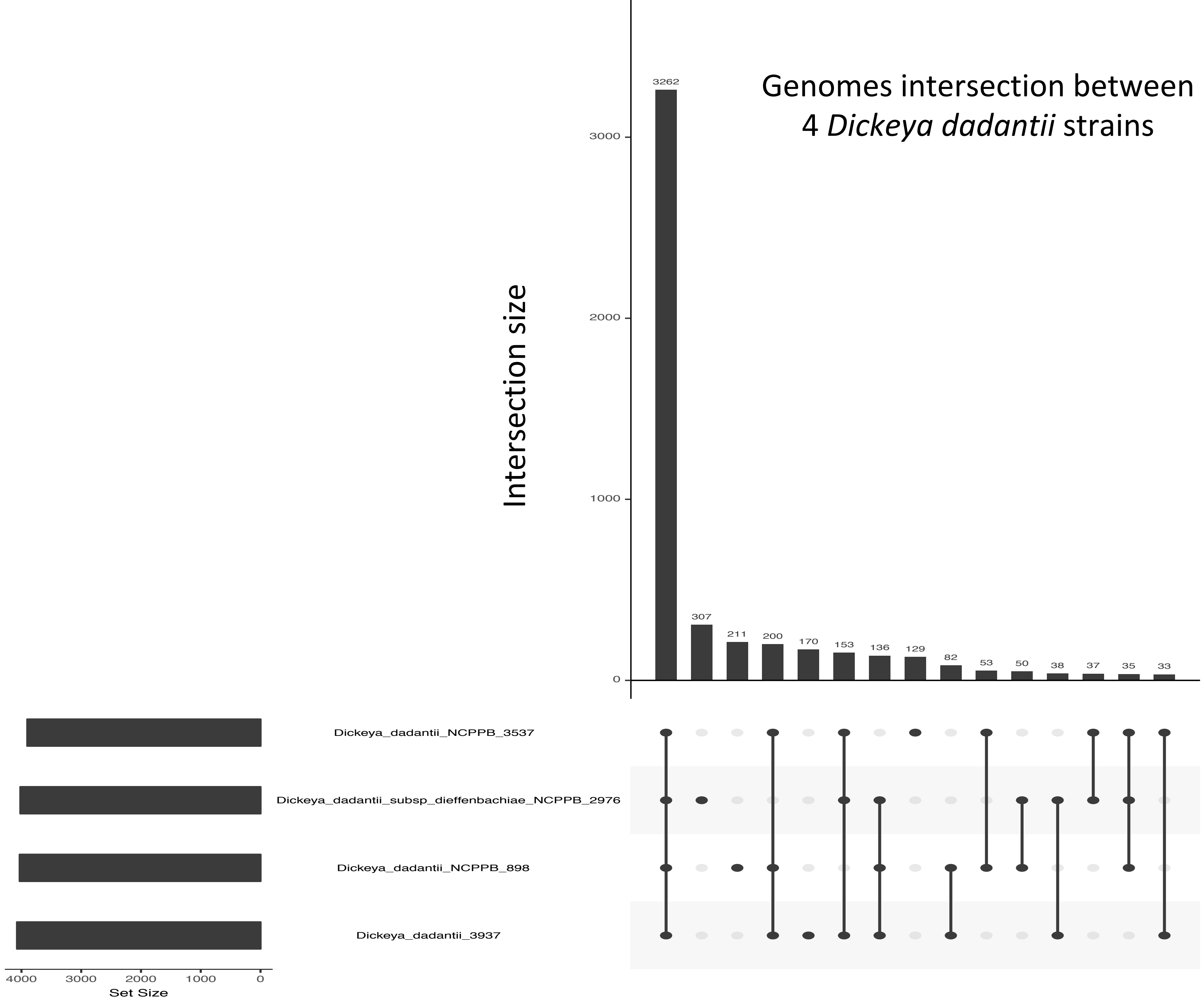

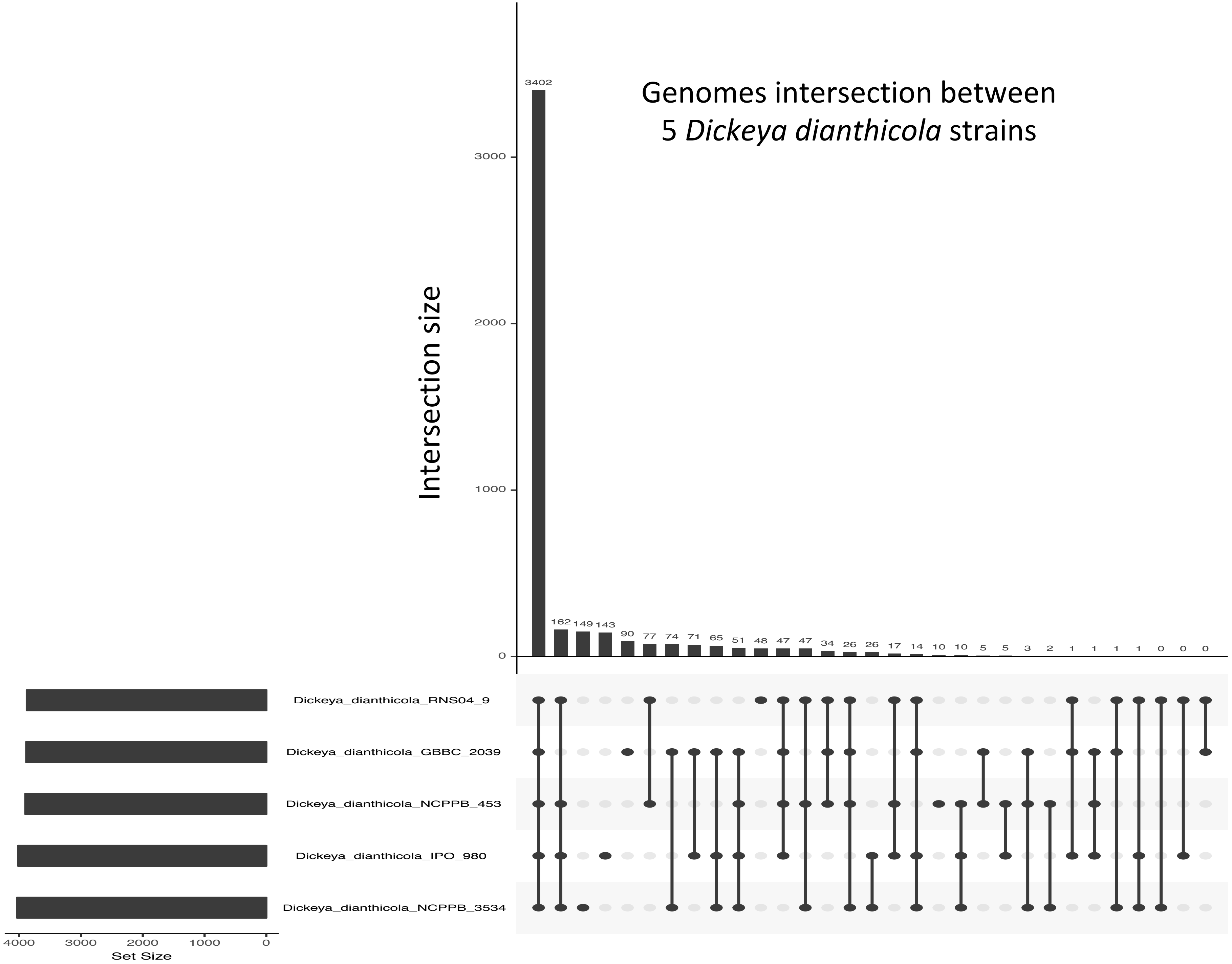

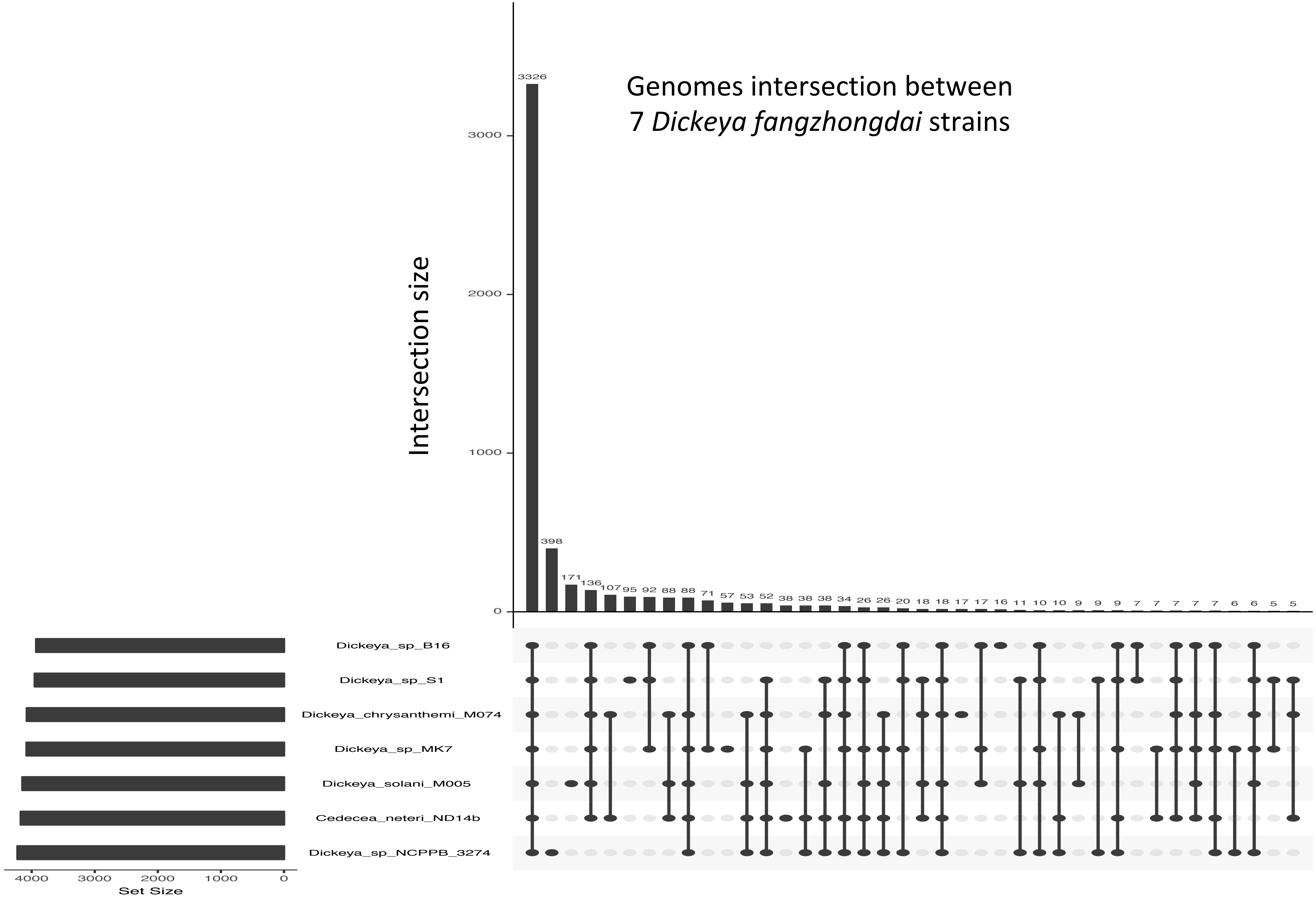

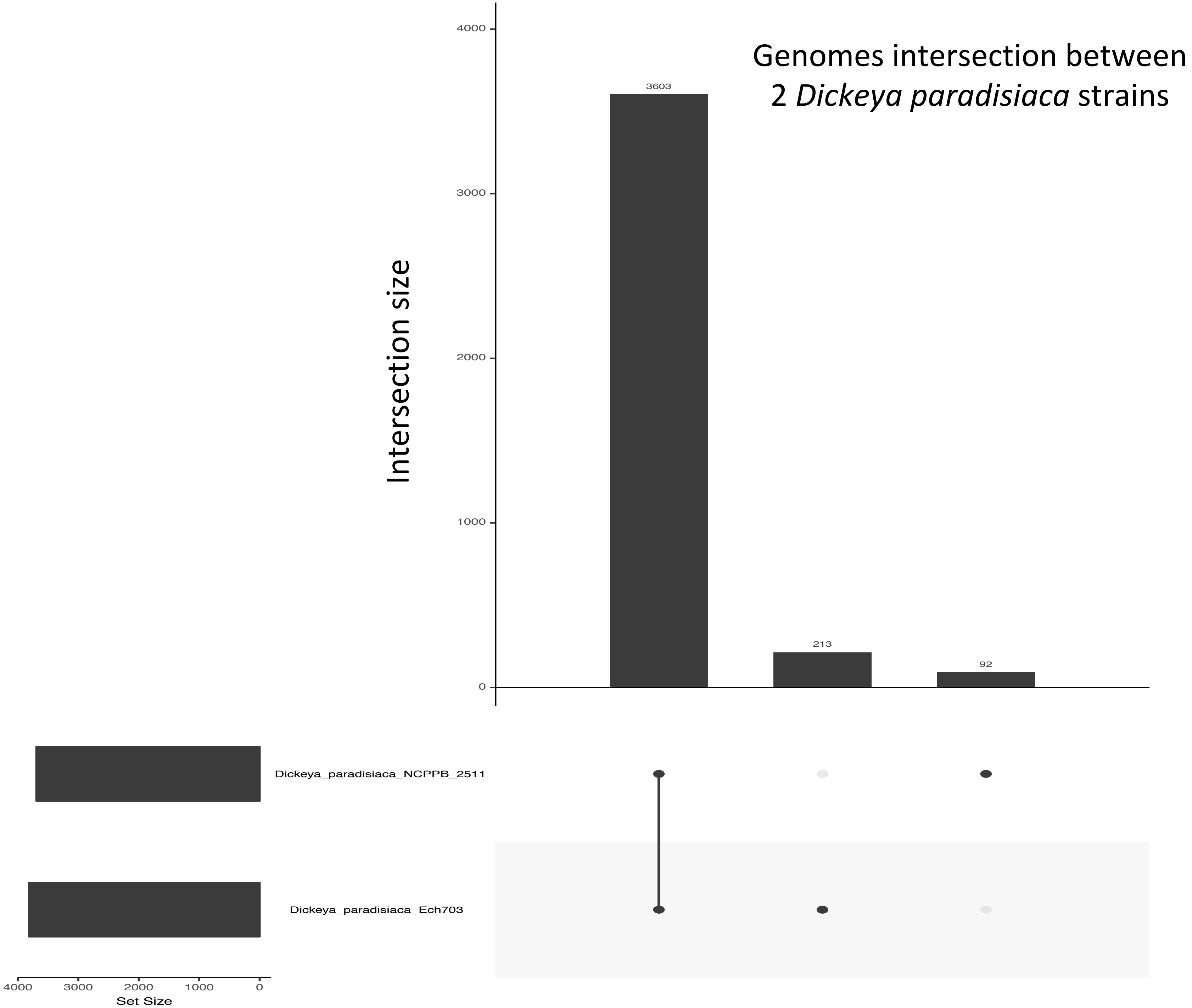

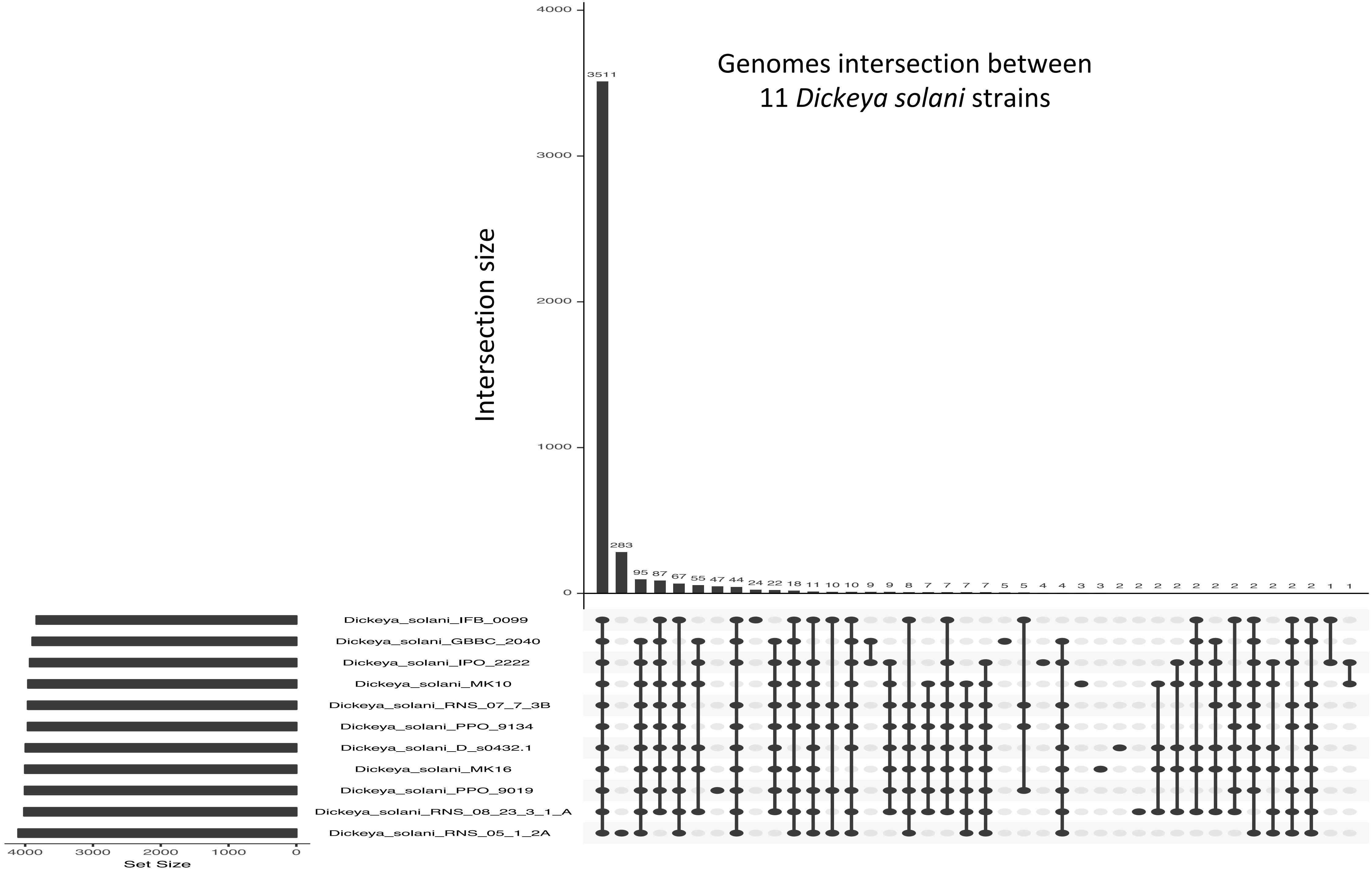

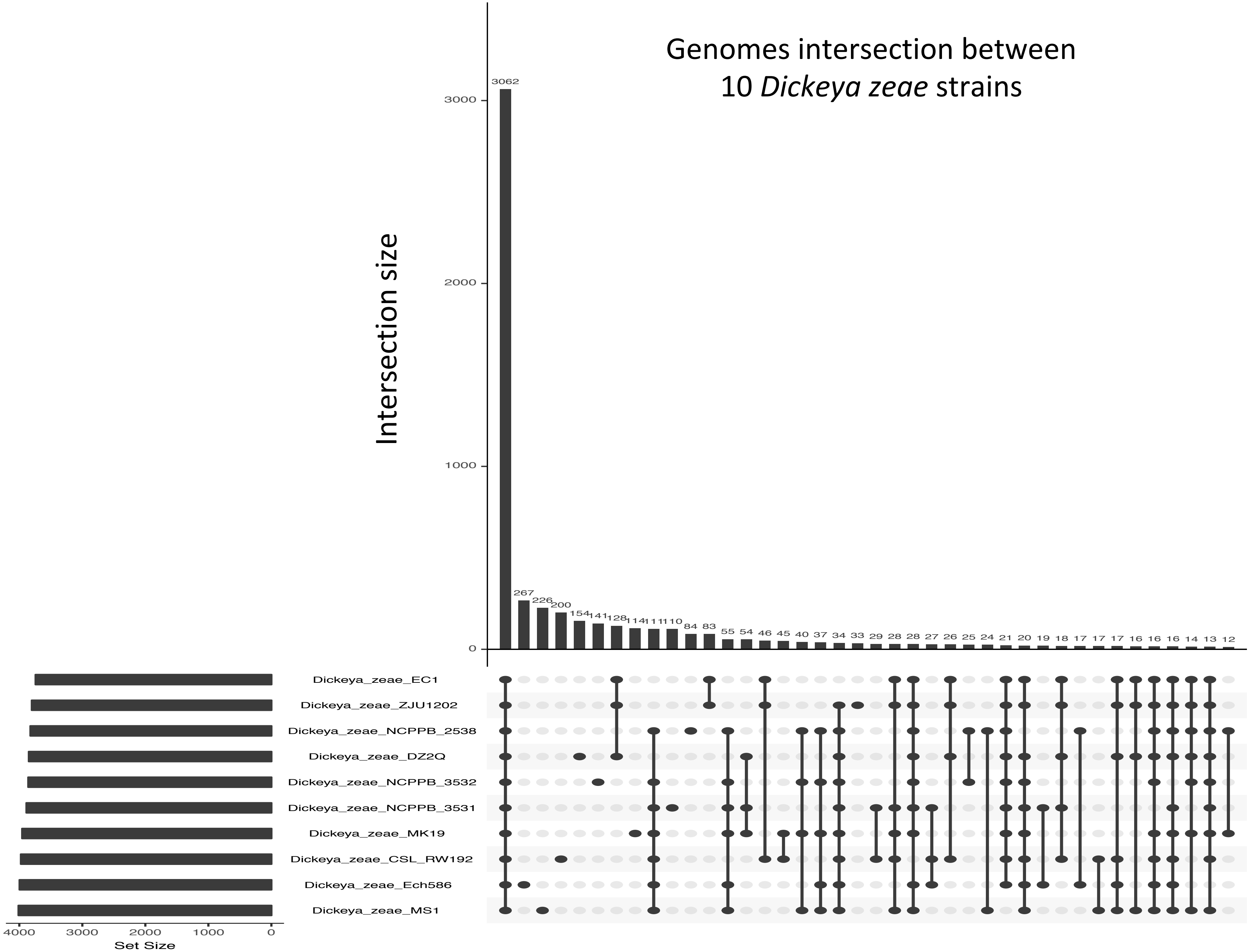
