## Supplementary Figure S3 for "The phytopathogenic nature of *Dickeya aquatica* 174/2 and the dynamic early evolution of *Dickeya* pathogenicity"

### Table of contents Figure S3


- **Plant adherence elements**
  - Flp pilus assembly (13 genes) 3-15
  - Fim pilus assembly (8 genes) 16-23
  - Type IV pilus assembly (10 genes) 24-33
  - Adhesin and associated secretion system (7 genes) 34-40
- **Secretion systems**
  - Type I protease secretion system Prt (3 genes) 41-43
  - Type II secretion system Out (14 genes) 44-57
  - Type II secretion system Stt (11 genes) 58-68
  - Type III secretion system Hrp and associated effectors or regulators (26 genes) 69-94
  - Type IV secretion system (10 genes) 95-104
  - Type V secretion system CdiA-B (3 genes) 105-107
  - TypeVI secretion system (24 genes) 108-131
- **Plant cell wall degradosome**
  - Pectinasome (35 genes) 132-166
  - Cellulases (2 genes) 167-168
  - Xyloglucan degradation (11 genes including xylose degradation) 169-179
  - Proteases (4 genes) 180-183
  - Phospholipase (1 gene) 184
- **Secondary metabolite biosynthesis**
  - Zeamine biosynthesis (17 genes) 185-201
  - Oocydin A biosynthesis (20 genes) 202-221
  - Coronafic acid biosynthesis (10 genes) 222-231
  - Cluster 1 unknown metabolite (9 genes) 232-240
  - Cluster 2 unknown metabolite (37 genes) 241-277
  - Cluster 3 unknown metabolite (16 genes) 278-293
  - Cluster 4 unknown metabolite (13 genes) 294-306
  - Pigment indigoidine synthesis (3 genes) 307-309
  - Pigment prodigiosin synthesis (15 genes) 310-324
- **Others virulence factors**
  - Insecticide toxins (4 genes) 325-328
  - Others toxins (3 genes) 329-331
- **Quorum sensing systems**
  - AHL-based ExpI-ExpR system (2 genes) 332-333
  - Vfm system (25 genes) 334-358
- **Metabolic pathways relevant for plant interaction**
  - Phosphonate utilisation (14 genes) 359-372
  - Opine degradation (7 genes) 373-379
  - Nitrogen fixation (20 genes) 380-399
  - Ascorbate metabolism (7 genes) 400-406
- **Response to stresses**
  - Response to antimicrobial peptides (21 genes) 407-427
  - Response to acidic stress (15 genes) 428-442
  - Response to osmotic stress (6 genes) 443-448
  - Response to oxidative stress (19 genes) 449-467
- **Iron metabolism**
  - Iron-binding proteins and iron storage (4 genes) 468-471
  - Achromobactin synthesis and transport (12 genes) 472-483
  - Chrysobactin synthesis and transport (14 genes) 484-497
  - Ferrous iron transport (5 genes) 498-502
  - Iron and Manganese transport (4 genes) 503-506
  - Hemin transport (4 genes) 507-510

#### - Phages and Genomics Islands


- Phage P1 (11 genes) 511-521
- Phages P2, P6, P7 related to Mu-like bacteriophage (49 genes) 522-570
- Phage P3 (67 genes) 571-637
- Phage P4 (27 genes) 638-664
- Phage P5 (55 genes) 665-719
- GI1 (10 genes) 720-729
- GI2 (24 genes) 730-753
- GI3 (19 genes) 754-772
- GI4 (22 genes) 773-794
- GI5 (14 genes) 795-808
- GI6 (10 genes) 809-818
- GI7 (61 genes) 819-879
- GI8 (9 genes) 880-888
- GI9 (18 genes) 889-906
- GI10 (8 genes) 907-914

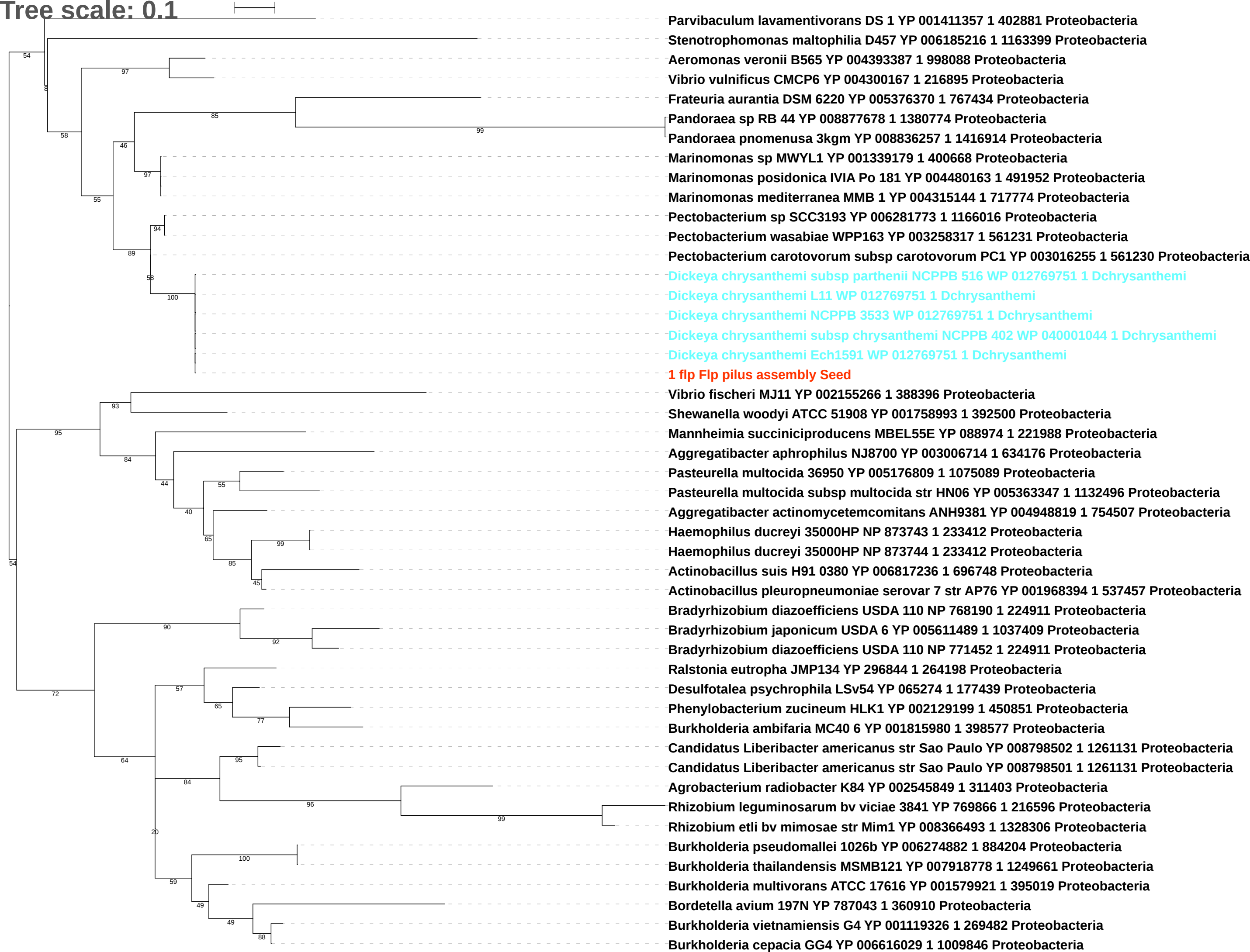
Tree scale: 0.1
Parvibaculum lavamentivorans DS 1 YP 001411357 1 402881 Proteobacteria
Stenotrophomonas maltophilia D457 YP 006185216 1 1163399 Proteobacteria
Aeromonas veronii B565 YP 004393387 1 998088 Proteobacteria
Vibrio vulnificus CMCP6 YP 004300167 1 216895 Proteobacteria
Frateuria aurantia DSM 6220 YP 005376370 1 767434 Proteobacteria
Pandoraea sp RB 44 YP 008877678 1 1380774 Proteobacteria
Pandoraea pnomenusa 3kgm YP 008836257 1 1416914 Proteobacteria
Marinomonas sp MWYL1 YP 001339179 1 400668 Proteobacteria
Marinomonas posidonica IVIA Po 181 YP 004480163 1 491952 Proteobacteria
Marinomonas mediterranea MMB 1 YP 004315144 1 717774 Proteobacteria
Pectobacterium sp SCC3193 YP 006281773 1 1166016 Proteobacteria
Pectobacterium wasabiae WPP163 YP 003258317 1 561231 Proteobacteria
Pectobacterium carotovorum subsp carotovorum PC1 YP 003016255 1 561230 Proteobacteria
Dickeya chrysanthemi subsp parthenii NCPPB 516 WP 012769751 1 Dchrysanthemi
Dickeya chrysanthemi L11 WP 012769751 1 Dchrysanthemi
Dickeya chrysanthemi NCPPB 3533 WP 012769751 1 Dchrysanthemi
Dickeya chrysanthemi subsp chrysanthemi NCPPB 402 WP 040001044 1 Dchrysanthemi
Dickeya chrysanthemi Ech1591 WP 012769751 1 Dchrysanthemi
1 flp Flp pilus assembly Seed
Vibrio fischeri MJ11 YP 002155266 1 388396 Proteobacteria
Shewanella woodyi ATCC 51908 YP 001758993 1 392500 Proteobacteria
Mannheimia succiniciproducens MBEL55E YP 088974 1 221988 Proteobacteria
Aggregatibacter aphrophilus NJ8700 YP 003006714 1 634176 Proteobacteria
Pasteurella multocida 36950 YP 005176809 1 1075089 Proteobacteria
Pasteurella multocida subsp multocida str HN06 YP 005363347 1 1132496 Proteobacteria
Aggregatibacter actinomycetemcomitans ANH9381 YP 004948819 1 754507 Proteobacteria
Haemophilus ducreyi 35000HP NP 873743 1 233412 Proteobacteria
Haemophilus ducreyi 35000HP NP 873744 1 233412 Proteobacteria
Actinobacillus suis H91 0380 YP 006817236 1 696748 Proteobacteria
Actinobacillus pleuropneumoniae serovar 7 str AP76 YP 001968394 1 537457 Proteobacteria
Bradyrhizobium diazoefficiens USDA 110 NP 768190 1 224911 Proteobacteria
Bradyrhizobium japonicum USDA 6 YP 005611489 1 1037409 Proteobacteria
Bradyrhizobium diazoefficiens USDA 110 NP 771452 1 224911 Proteobacteria
Ralstonia eutropha JMP134 YP 296844 1 264198 Proteobacteria
Desulfotalea psychrophila LSv54 YP 065274 1 177439 Proteobacteria
Phenylobacterium zucineum HLK1 YP 002129199 1 450851 Proteobacteria
Burkholderia ambifaria MC40 6 YP 001815980 1 398577 Proteobacteria
Candidatus Liberibacter americanus str Sao Paulo YP 008798502 1 1261131 Proteobacteria
Candidatus Liberibacter americanus str Sao Paulo YP 008798501 1 1261131 Proteobacteria
Agrobacterium radiobacter K84 YP 002545849 1 311403 Proteobacteria
Rhizobium leguminosarum bv viciae 3841 YP 769866 1 216596 Proteobacteria
Rhizobium etli bv mimosae str Mim1 YP 008366493 1 1328306 Proteobacteria
Burkholderia pseudomallei 1026b YP 006274882 1 884204 Proteobacteria
Burkholderia thailandensis MSMB121 YP 007918778 1 1249661 Proteobacteria
Burkholderia multivorans ATCC 17616 YP 001579921 1 395019 Proteobacteria
Bordetella avium 197N YP 787043 1 360910 Proteobacteria
Burkholderia vietnamiensis G4 YP 001119326 1 269482 Proteobacteria
Burkholderia cepacia GG4 YP 006616029 1 1009846 Proteobacteria

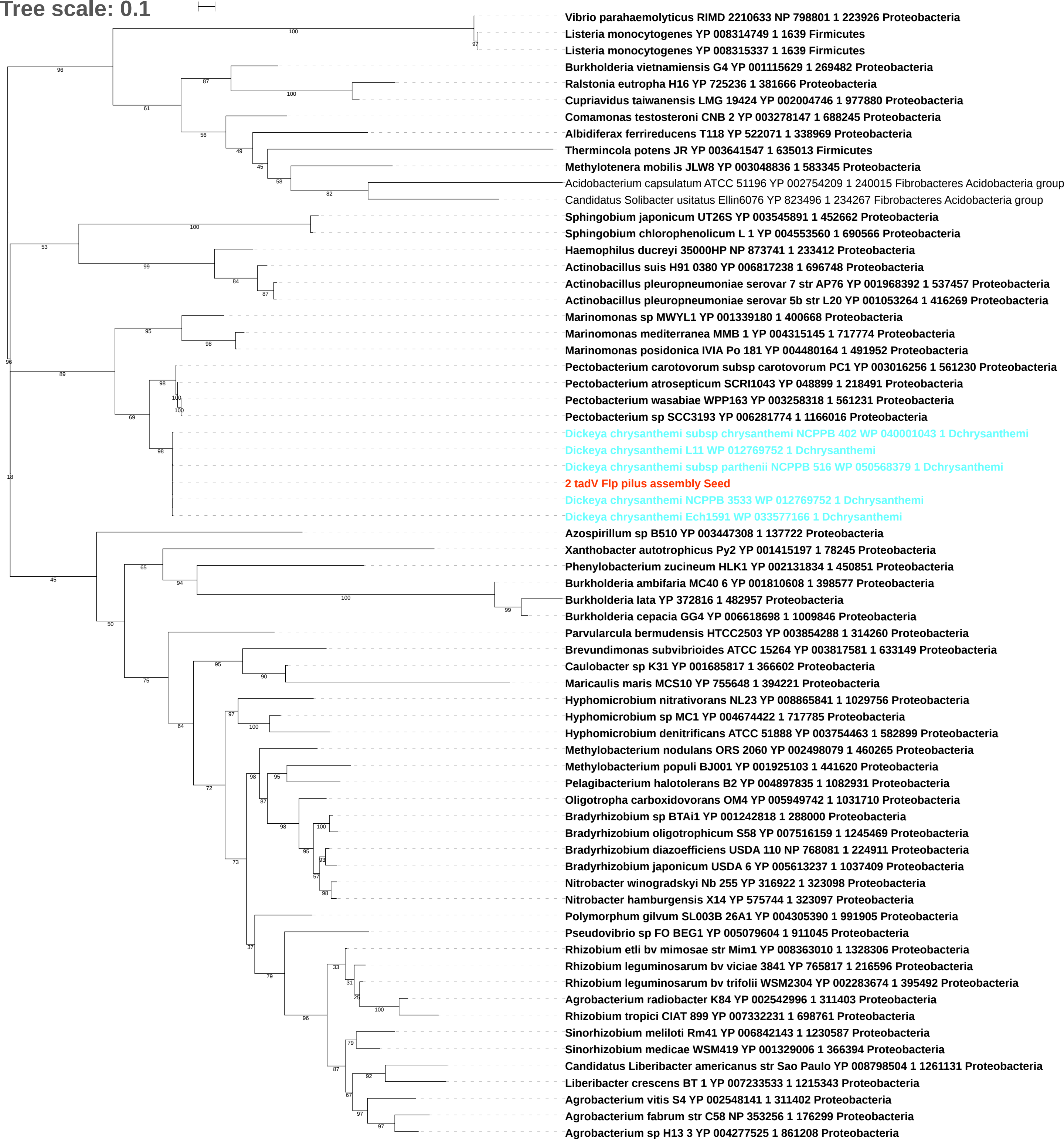
Tree scale: 0.1
Vibrio parahaemolyticus RIMD 2210633 NP 798801 1 223926 Proteobacteria
Listeria monocytogenes YP 008314749 1 1639 Firmicutes
Listeria monocytogenes YP 008315337 1 1639 Firmicutes
Burkholderia vietnamiensis G4 YP 001115629 1 269482 Proteobacteria
Ralstonia eutropha H16 YP 725236 1 381666 Proteobacteria
Cupriavidus taiwanensis LMG 19424 YP 002004746 1 977880 Proteobacteria
Comamonas testosteroni CNB 2 YP 003278147 1 688245 Proteobacteria
Albidiferax ferrireducens T118 YP 522071 1 338969 Proteobacteria
Thermincola potens JR YP 003641547 1 635013 Firmicutes
Methylotenera mobilis JLW8 YP 003048836 1 583345 Proteobacteria
Acidobacterium capsulatum ATCC 51196 YP 002754209 1 240015 Fibrobacteres Acidobacteria group
Candidatus Solibacter usitatus Ellin6076 YP 823496 1 234267 Fibrobacteres Acidobacteria group
Sphingobium japonicum UT26S YP 003545891 1 452662 Proteobacteria
Sphingobium chlorophenolicum L 1 YP 004553560 1 690566 Proteobacteria
Haemophilus ducreyi 35000HP NP 873741 1 233412 Proteobacteria
Actinobacillus suis H91 0380 YP 006817238 1 696748 Proteobacteria
Actinobacillus pleuropneumoniae serovar 7 str AP76 YP 001968392 1 537457 Proteobacteria
Actinobacillus pleuropneumoniae serovar 5b str L20 YP 001053264 1 416269 Proteobacteria
Marinomonas sp MWYL1 YP 001339180 1 400668 Proteobacteria
Marinomonas mediterranea MMB 1 YP 004315145 1 717774 Proteobacteria
Marinomonas posidonica IVIA Po 181 YP 004480164 1 491952 Proteobacteria
Pectobacterium carotovorum subsp carotovorum PC1 YP 003016256 1 561230 Proteobacteria
Pectobacterium atrosepticum SCRI1043 YP 048899 1 218491 Proteobacteria
Pectobacterium wasabiae WPP163 YP 003258318 1 561231 Proteobacteria
Pectobacterium sp SCC3193 YP 006281774 1 1166016 Proteobacteria
Dickeya chrysanthemi subsp chrysanthemi NCPPB 402 WP 040001043 1 Dchrysanthemi
Dickeya chrysanthemi L11 WP 012769752 1 Dchrysanthemi
Dickeya chrysanthemi subsp parthenii NCPPB 516 WP 050568379 1 Dchrysanthemi
2 tadV Flp pilus assembly Seed
Dickeya chrysanthemi NCPPB 3533 WP 012769752 1 Dchrysanthemi
Dickeya chrysanthemi Ech1591 WP 033577166 1 Dchrysanthemi
Azospirillum sp B510 YP 003447308 1 137722 Proteobacteria
Xanthobacter autotrophicus Py2 YP 001415197 1 78245 Proteobacteria
Phenylobacterium zucineum HLK1 YP 002131834 1 450851 Proteobacteria
Burkholderia ambifaria MC40 6 YP 001810608 1 398577 Proteobacteria
Burkholderia lata YP 372816 1 482957 Proteobacteria
Burkholderia cepacia GG4 YP 006618698 1 1009846 Proteobacteria
Parvularcula bermudensis HTCC2503 YP 003854288 1 314260 Proteobacteria
Brevundimonas subvibrioides ATCC 15264 YP 003817581 1 633149 Proteobacteria
Caulobacter sp K31 YP 001685817 1 366602 Proteobacteria
Maricaulis maris MCS10 YP 755648 1 394221 Proteobacteria
Hyphomicrobium nitrativorans NL23 YP 008865841 1 1029756 Proteobacteria
Hyphomicrobium sp MC1 YP 004674422 1 717785 Proteobacteria
Hyphomicrobium denitrificans ATCC 51888 YP 003754463 1 582899 Proteobacteria
Methylobacterium nodulans ORS 2060 YP 002498079 1 460265 Proteobacteria
Methylobacterium populi BJ001 YP 001925103 1 441620 Proteobacteria
Pelagibacterium halotolerans B2 YP 004897835 1 1082931 Proteobacteria
Oligotropha carboxidovorans OM4 YP 005949742 1 1031710 Proteobacteria
Bradyrhizobium sp BTAi1 YP 001242818 1 288000 Proteobacteria
Bradyrhizobium oligotrophicum S58 YP 007516159 1 1245469 Proteobacteria
Bradyrhizobium diazoefficiens USDA 110 NP 768081 1 224911 Proteobacteria
Bradyrhizobium japonicum USDA 6 YP 005613237 1 1037409 Proteobacteria
Nitrobacter winogradskyi Nb 255 YP 316922 1 323098 Proteobacteria
Nitrobacter hamburgensis X14 YP 575744 1 323097 Proteobacteria
Polymorphum gilvum SL003B 26A1 YP 004305390 1 991905 Proteobacteria
Pseudovibrio sp FO BEG1 YP 005079604 1 911045 Proteobacteria
Rhizobium etli bv mimosae str Mim1 YP 008363010 1 1328306 Proteobacteria
Rhizobium leguminosarum bv viciae 3841 YP 765817 1 216596 Proteobacteria
Rhizobium leguminosarum bv trifolii WSM2304 YP 002283674 1 395492 Proteobacteria
Agrobacterium radiobacter K84 YP 002542996 1 311403 Proteobacteria
Rhizobium tropici CIAT 899 YP 007332231 1 698761 Proteobacteria
Sinorhizobium meliloti Rm41 YP 006842143 1 1230587 Proteobacteria
Sinorhizobium medicae WSM419 YP 001329006 1 366394 Proteobacteria
Candidatus Liberibacter americanus str Sao Paulo YP 008798504 1 1261131 Proteobacteria
Liberibacter crescens BT 1 YP 007233533 1 1215343 Proteobacteria
Agrobacterium vitis S4 YP 002548141 1 311402 Proteobacteria
Agrobacterium fabrum str C58 NP 353256 1 176299 Proteobacteria
Agrobacterium sp H13 3 YP 004277525 1 861208 Proteobacteria

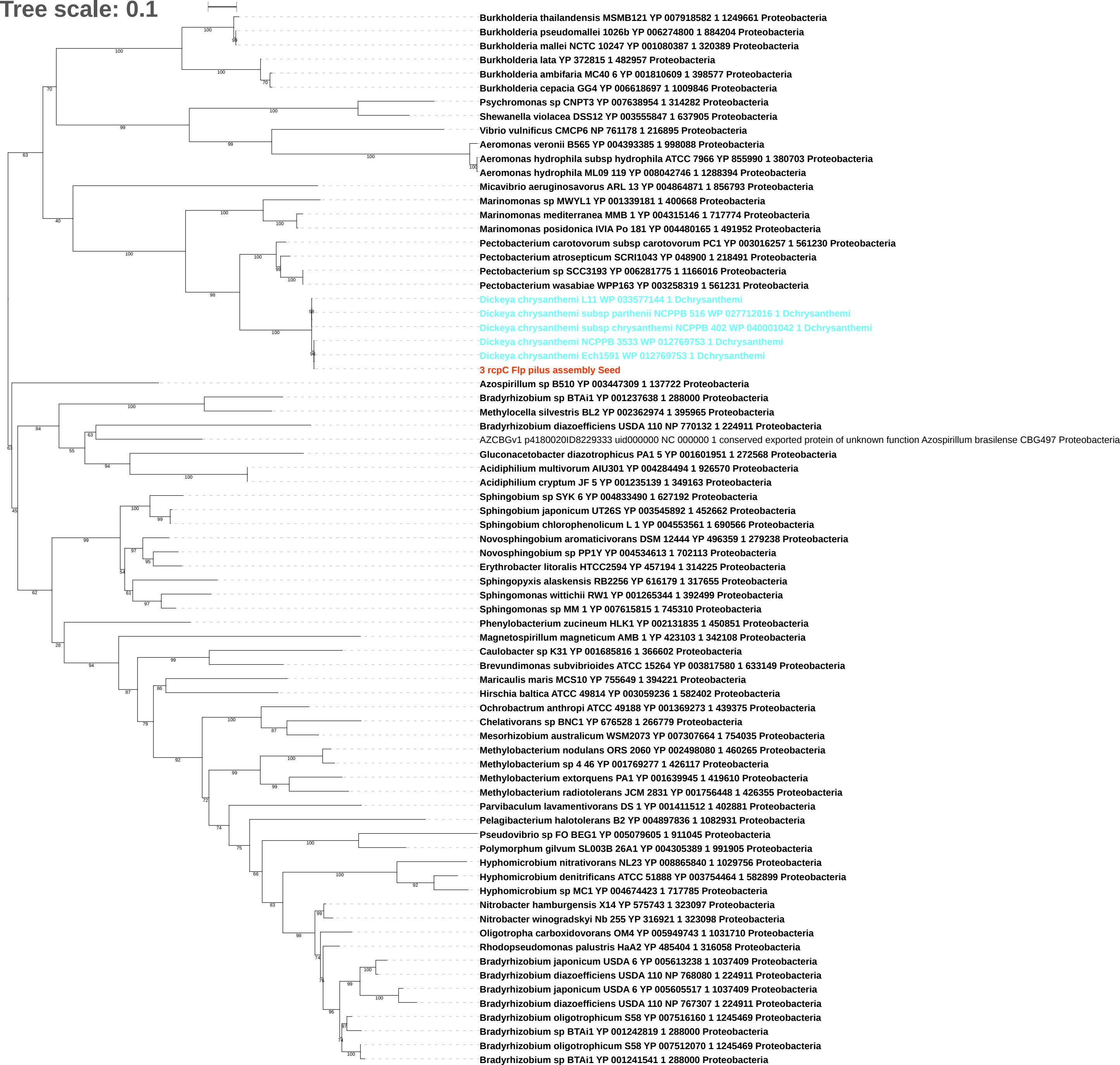
Tree scale: 0.1
Burkholderia thailandensis MSMB121 YP 007918582 1 1249661 Proteobacteria
Burkholderia pseudomallei 1026b YP 006274800 1 884204 Proteobacteria
Burkholderia mallei NCTC 10247 YP 001080387 1 320389 Proteobacteria
Burkholderia lata YP 372815 1 482957 Proteobacteria
Burkholderia ambifaria MC40 6 YP 001810609 1 398577 Proteobacteria
Burkholderia cepacia GG4 YP 006618697 1 1009846 Proteobacteria
Psychromonas sp CNPT3 YP 007638954 1 314282 Proteobacteria
Shewanella violacea DSS12 YP 003555847 1 637905 Proteobacteria
Vibrio vulnificus CMCP6 NP 761178 1 216895 Proteobacteria
Aeromonas veronii B565 YP 004393385 1 998088 Proteobacteria
Aeromonas hydrophila subsp hydrophila ATCC 7966 YP 855990 1 380703 Proteobacteria
Aeromonas hydrophila ML09 119 YP 008042746 1 1288394 Proteobacteria
Micavibrio aeruginosavorus ARL 13 YP 004864871 1 856793 Proteobacteria
Marinomonas sp MWYL1 YP 001339181 1 400668 Proteobacteria
Marinomonas mediterranea MMB 1 YP 004315146 1 717774 Proteobacteria
Marinomonas posidonica IVIA Po 181 YP 004480165 1 491952 Proteobacteria
Pectobacterium carotovorum subsp carotovorum PC1 YP 003016257 1 561230 Proteobacteria
Pectobacterium atrosepticum SCRI1043 YP 048900 1 218491 Proteobacteria
Pectobacterium sp SCC3193 YP 006281775 1 1166016 Proteobacteria
Pectobacterium wasabiae WPP163 YP 003258319 1 561231 Proteobacteria
Dickeya chrysanthemi L11 WP 033577144 1 Dchrysanthemi
Dickeya chrysanthemi subsp parthenii NCPPB 516 WP 027712016 1 Dchrysanthemi
Dickeya chrysanthemi subsp chrysanthemi NCPPB 402 WP 040001042 1 Dchrysanthemi
Dickeya chrysanthemi NCPPB 3533 WP 012769753 1 Dchrysanthemi
Dickeya chrysanthemi Ech1591 WP 012769753 1 Dchrysanthemi
3 rcpC Flp pilus assembly Seed
Azospirillum sp B510 YP 003447309 1 137722 Proteobacteria
Bradyrhizobium sp BTAi1 YP 001237638 1 288000 Proteobacteria
Methylocella silvestris BL2 YP 002362974 1 395965 Proteobacteria
Bradyrhizobium diazoefficiens USDA 110 NP 770132 1 224911 Proteobacteria
AZCBGv1 p4180020ID8229333 uid000000 NC 000000 1 conserved exported protein of unknown function Azospirillum brasilense CBG497 Proteobacteria
Gluconacetobacter diazotrophicus PA1 5 YP 001601951 1 272568 Proteobacteria
Acidiphilium multivorum AIU301 YP 004284494 1 926570 Proteobacteria
Acidiphilium cryptum JF 5 YP 001235139 1 349163 Proteobacteria
Sphingobium sp SYK 6 YP 004833490 1 627192 Proteobacteria
Sphingobium japonicum UT26S YP 003545892 1 452662 Proteobacteria
Sphingobium chlorophenolicum L 1 YP 004553561 1 690566 Proteobacteria
Novosphingobium aromaticivorans DSM 12444 YP 496359 1 279238 Proteobacteria
Novosphingobium sp PP1Y YP 004534613 1 702113 Proteobacteria
Erythrobacter litoralis HTCC2594 YP 457194 1 314225 Proteobacteria
Sphingopyxis alaskensis RB2256 YP 616179 1 317655 Proteobacteria
Sphingomonas wittichii RW1 YP 001265344 1 392499 Proteobacteria
Sphingomonas sp MM 1 YP 007615815 1 745310 Proteobacteria
Phenylobacterium zucineum HLK1 YP 002131835 1 450851 Proteobacteria
Magnetospirillum magneticum AMB 1 YP 423103 1 342108 Proteobacteria
Caulobacter sp K31 YP 001685816 1 366602 Proteobacteria
Brevundimonas subvibrioides ATCC 15264 YP 003817580 1 633149 Proteobacteria
Maricaulis maris MCS10 YP 755649 1 394221 Proteobacteria
Hirschia baltica ATCC 49814 YP 003059236 1 582402 Proteobacteria
Ochrobactrum anthropi ATCC 49188 YP 001369273 1 439375 Proteobacteria
Chelativorans sp BNC1 YP 676528 1 266779 Proteobacteria
Mesorhizobium australicum WSM2073 YP 007307664 1 754035 Proteobacteria
Methylobacterium nodulans ORS 2060 YP 002498080 1 460265 Proteobacteria
Methylobacterium sp 4 46 YP 001769277 1 426117 Proteobacteria
Methylobacterium extorquens PA1 YP 001639945 1 419610 Proteobacteria
Methylobacterium radiotolerans JCM 2831 YP 001756448 1 426355 Proteobacteria
Parvibaculum lavamentivorans DS 1 YP 001411512 1 402881 Proteobacteria
Pelagibacterium halotolerans B2 YP 004897836 1 1082931 Proteobacteria
Pseudovibrio sp FO BEG1 YP 005079605 1 911045 Proteobacteria
Polymorphum gilvum SL003B 26A1 YP 004305389 1 991905 Proteobacteria
Hyphomicrobium nitrativorans NL23 YP 008865840 1 1029756 Proteobacteria
Hyphomicrobium denitrificans ATCC 51888 YP 003754469 1 582899 Proteobacteria
Hyphomicrobium sp MC1 YP 004674423 1 717785 Proteobacteria
Nitrobacter hamburgensis X14 YP 575743 1 323097 Proteobacteria
Nitrobacter winogradskyi Nb 255 YP 316921 1 323098 Proteobacteria
Oligotropha carboxidovorans OM4 YP 005949743 1 1031710 Proteobacteria
Rhodopseudomonas palustris HaA2 YP 485404 1 316058 Proteobacteria
Bradyrhizobium japonicum USDA 6 YP 005613238 1 1037409 Proteobacteria
Bradyrhizobium diazoefficiens USDA 110 NP 768080 1 224911 Proteobacteria
Bradyrhizobium japonicum USDA 6 YP 005605517 1 1037409 Proteobacteria
Bradyrhizobium diazoefficiens USDA 110 NP 767307 1 224911 Proteobacteria
Bradyrhizobium oligotrophicum S58 YP 007516160 1 1245469 Proteobacteria
Bradyrhizobium sp BTAi1 YP 001242819 1 288000 Proteobacteria
Bradyrhizobium oligotrophicum S58 YP 007512070 1 1245469 Proteobacteria
Bradyrhizobium sp BTAi1 YP 001241541 1 288000 Proteobacteria

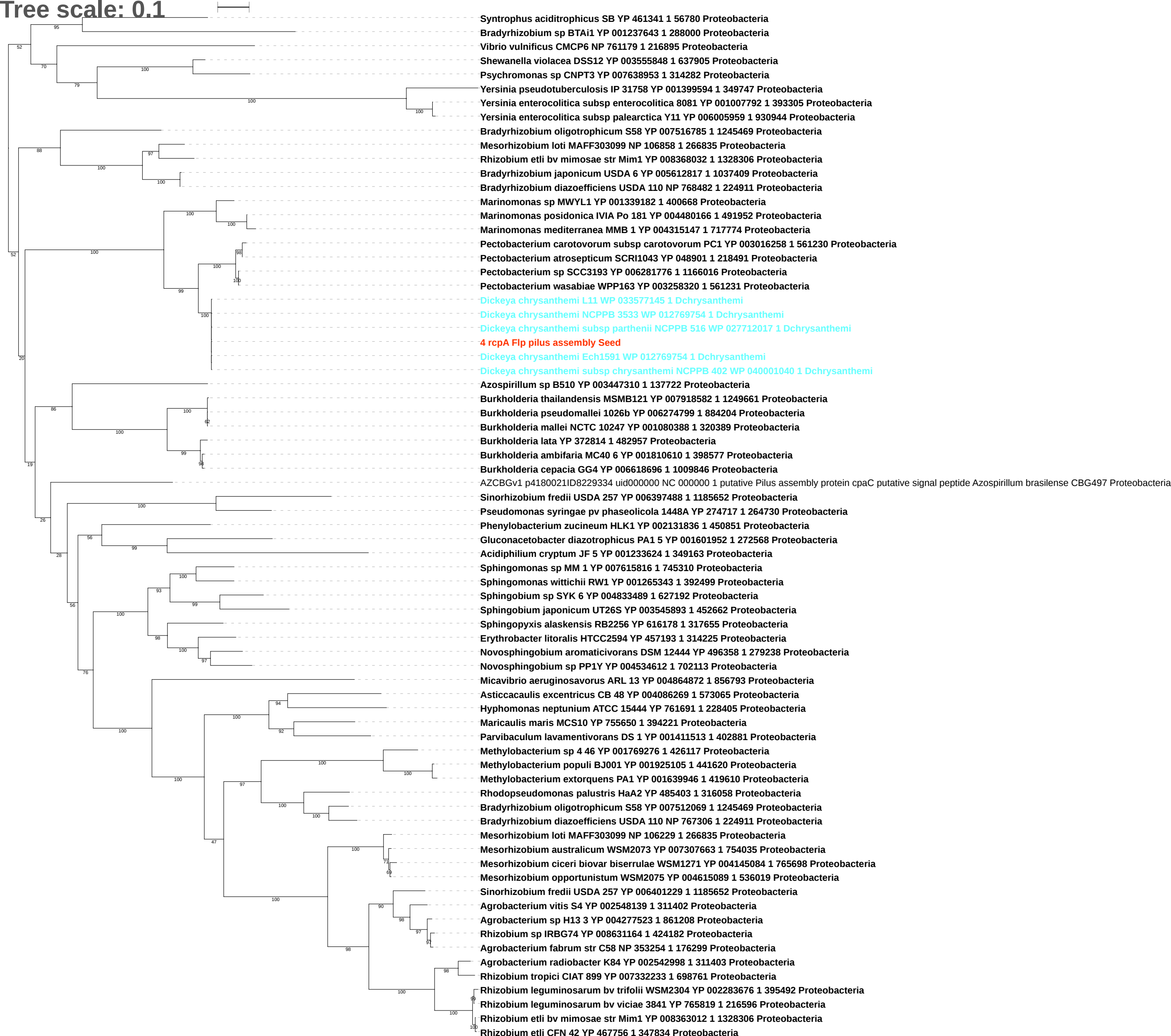

Tree scale: 0.1
Syntrophus aciditrophicus SB YP 461341 1 56780 Proteobacteria
Bradyrhizobium sp BTAi1 YP 001237643 1 288000 Proteobacteria
Vibrio vulnificus CMCP6 NP 761179 1 216895 Proteobacteria
Shewanella violacea DSS12 YP 003555848 1 637905 Proteobacteria
Psychromonas sp CNPT3 YP 007638953 1 314282 Proteobacteria
Yersinia pseudotuberculosis IP 31758 YP 001399594 1 349747 Proteobacteria
Yersinia enterocolitica subsp enterocolitica 8081 YP 001007792 1 393305 Proteobacteria
Yersinia enterocolitica subsp palearctica Y11 YP 006005959 1 930944 Proteobacteria
Bradyrhizobium oligotrophicum S58 YP 007516785 1 1245469 Proteobacteria
Mesorhizobium loti MAFF303099 NP 106858 1 266835 Proteobacteria
Rhizobium etli bv mimosae str Mim1 YP 008368032 1 1328306 Proteobacteria
Bradyrhizobium japonicum USDA 6 YP 005612817 1 1037409 Proteobacteria
Bradyrhizobium diazoefficiens USDA 110 NP 768482 1 224911 Proteobacteria
Marinomonas sp MWYL1 YP 001339182 1 400668 Proteobacteria
Marinomonas posidonica IVIA Po 181 YP 004480166 1 491952 Proteobacteria
Marinomonas mediterranea MMB 1 YP 004315147 1 717774 Proteobacteria
Pectobacterium carotovorum subsp carotovorum PC1 YP 003016258 1 561230 Proteobacteria
Pectobacterium atrosepticum SCRI1043 YP 048901 1 218491 Proteobacteria
Pectobacterium sp SCC3193 YP 006281776 1 1166016 Proteobacteria
Pectobacterium wasabiae WPP163 YP 003258320 1 561231 Proteobacteria
Dickeya chrysanthemi L11 WP 033577145 1 Dchrysanthemi
Dickeya chrysanthemi NCPPB 3533 WP 012769754 1 Dchrysanthemi
Dickeya chrysanthemi subsp parthenii NCPPB 516 WP 027712017 1 Dchrysanthemi
4 rcpA Flp pilus assembly Seed
Dickeya chrysanthemi Ech1591 WP 012769754 1 Dchrysanthemi
Dickeya chrysanthemi subsp chrysanthemi NCPPB 402 WP 040001040 1 Dchrysanthemi
Azospirillum sp B510 YP 003447310 1 137722 Proteobacteria
Burkholderia thailandensis MSMB121 YP 007918582 1 1249661 Proteobacteria
Burkholderia pseudomallei 1026b YP 006274799 1 884204 Proteobacteria
Burkholderia mallei NCTC 10247 YP 001080388 1 320389 Proteobacteria
Burkholderia lata YP 372814 1 482957 Proteobacteria
Burkholderia ambifaria MC40 6 YP 001810610 1 398577 Proteobacteria
Burkholderia cepacia GG4 YP 006618696 1 1009846 Proteobacteria
AZCBGv1 p4180021ID8229334 uid000000 NC 000000 1 putative Pilus assembly protein cpaC putative signal peptide Azospirillum brasilense CBG497 Proteobacteria
Sinorhizobium fredii USDA 257 YP 006397488 1 1185652 Proteobacteria
Pseudomonas syringae pv phaseolicola 1448A YP 274717 1 264730 Proteobacteria
Phenylobacterium zucineum HLK1 YP 002131836 1 450851 Proteobacteria
Gluconacetobacter diazotrophicus PA1 5 YP 001601952 1 272568 Proteobacteria
Acidiphilium cryptum JF 5 YP 001233624 1 349163 Proteobacteria
Sphingomonas sp MM 1 YP 007615816 1 745310 Proteobacteria
Sphingomonas wittichii RW1 YP 001265343 1 392499 Proteobacteria
Sphingobium sp SYK 6 YP 004833489 1 627192 Proteobacteria
Sphingobium japonicum UT26S YP 003545893 1 452662 Proteobacteria
Sphingopyxis alaskensis RB2256 YP 616178 1 317655 Proteobacteria
Erythrobacter litoralis HTCC2594 YP 457193 1 314225 Proteobacteria
Novosphingobium aromaticivorans DSM 12444 YP 496358 1 279238 Proteobacteria
Novosphingobium sp PP1Y YP 004534612 1 702113 Proteobacteria
Micavibrio aeruginosavorus ARL 13 YP 004864872 1 856793 Proteobacteria
Asticcacaulis excentricus CB 48 YP 004086269 1 573065 Proteobacteria
Hyphomonas neptunium ATCC 15444 YP 761691 1 228405 Proteobacteria
Maricaulis maris MCS10 YP 755650 1 394221 Proteobacteria
Parvibaculum lavamentivorans DS 1 YP 001411513 1 402881 Proteobacteria
Methylobacterium sp 4 46 YP 001769276 1 426117 Proteobacteria
Methylobacterium populi BJ001 YP 001925105 1 441620 Proteobacteria
Methylobacterium extorquens PA1 YP 001639946 1 419610 Proteobacteria
Rhodopseudomonas palustris HaA2 YP 485403 1 316058 Proteobacteria
Bradyrhizobium oligotrophicum S58 YP 007512069 1 1245469 Proteobacteria
Bradyrhizobium diazoefficiens USDA 110 NP 767306 1 224911 Proteobacteria
Mesorhizobium loti MAFF303099 NP 106229 1 266835 Proteobacteria
Mesorhizobium australicum WSM2073 YP 007307663 1 754035 Proteobacteria
Mesorhizobium ciceri biovar biserrulae WSM1271 YP 004145084 1 765698 Proteobacteria
Mesorhizobium opportunistum WSM2075 YP 004615089 1 536019 Proteobacteria
Sinorhizobium fredii USDA 257 YP 006401229 1 1185652 Proteobacteria
Agrobacterium vitis S4 YP 002548139 1 311402 Proteobacteria
Agrobacterium sp H13 3 YP 004277523 1 861208 Proteobacteria
Rhizobium sp IRBG74 YP 008631164 1 424182 Proteobacteria
Agrobacterium fabrum str C58 NP 353254 1 176299 Proteobacteria
Agrobacterium radiobacter K84 YP 002542998 1 311403 Proteobacteria
Rhizobium tropici CIAT 899 YP 007332233 1 698761 Proteobacteria
Rhizobium leguminosarum bv trifolii WSM2304 YP 002283676 1 395492 Proteobacteria
Rhizobium leguminosarum bv viciae 3841 YP 765819 1 216596 Proteobacteria
Rhizobium etli bv mimosae str Mim1 YP 008363012 1 1328306 Proteobacteria

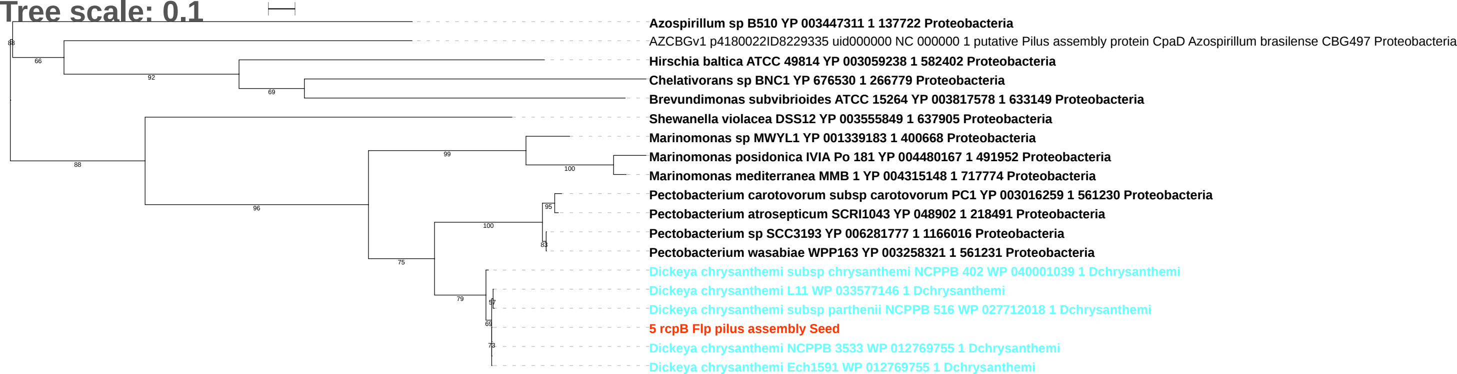

Tree scale: 0.1
Azospirillum sp B510 YP 003447311 1 137722 Proteobacteria
AZCBGv1 p4180022ID8229335 uid000000 NC 000000 1 putative Pilus assembly protein CpaD Azospirillum brasilense CBG497 Proteobacteria
Hirschia baltica ATCC 49814 YP 003059238 1 582402 Proteobacteria
Chelativorans sp BNC1 YP 676530 1 266779 Proteobacteria
Brevundimonas subvibrioides ATCC 15264 YP 003817578 1 633149 Proteobacteria
Shewanella violacea DSS12 YP 003555849 1 637905 Proteobacteria
Marinomonas sp MWYL1 YP 001339183 1 400668 Proteobacteria
Marinomonas posidonica IVIA Po 181 YP 004480167 1 491952 Proteobacteria
Marinomonas mediterranea MMB 1 YP 004315148 1 717774 Proteobacteria
Pectobacterium carotovorum subsp carotovorum PC1 YP 003016259 1 561230 Proteobacteria
Pectobacterium atrosepticum SCRI1043 YP 048902 1 218491 Proteobacteria
Pectobacterium sp SCC3193 YP 006281777 1 1166016 Proteobacteria
Pectobacterium wasabiae WPP163 YP 003258321 1 561231 Proteobacteria
Dickeya chrysanthemi subsp chrysanthemi NCPPB 402 WP 040001039 1 Dchrysanthemi
Dickeya chrysanthemi L11 WP 033577146 1 Dchrysanthemi
Dickeya chrysanthemi subsp parthenii NCPPB 516 WP 027712018 1 Dchrysanthemi
5 rcpB Flp pilus assembly Seed
Dickeya chrysanthemi NCPPB 3533 WP 012769755 1 Dchrysanthemi
Dickeya chrysanthemi Ech1591 WP 012769755 1 Dchrysanthemi

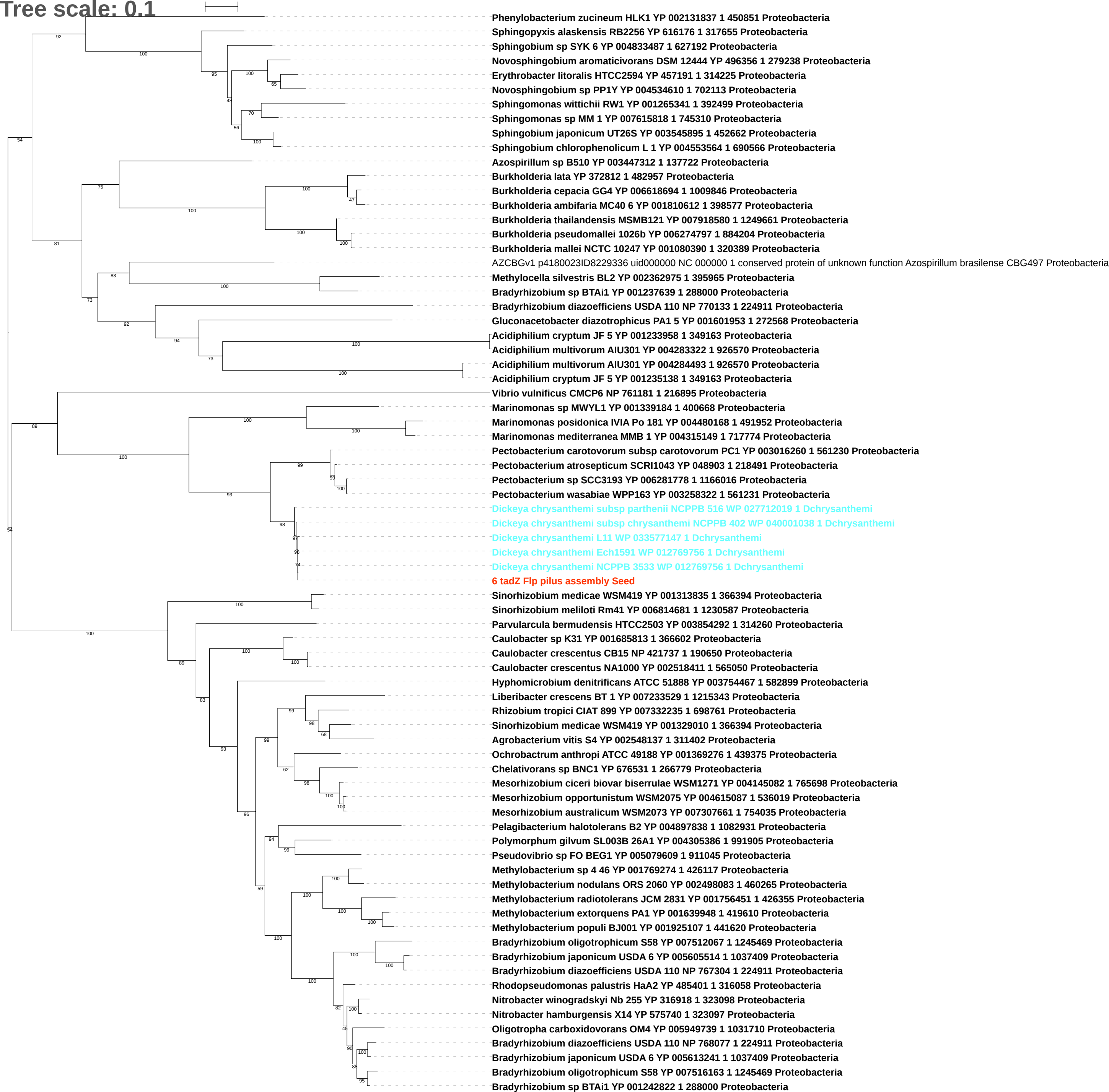

Tree scale: 0.1
Phenylobacterium zucineum HLK1 YP 002131837 1 450851 Proteobacteria
Sphingopyxis alaskensis RB2256 YP 616176 1 317655 Proteobacteria
Sphingobium sp SYK 6 YP 004833487 1 627192 Proteobacteria
Novosphingobium aromaticivorans DSM 12444 YP 496356 1 279238 Proteobacteria
Erythrobacter litoralis HTCC2594 YP 457191 1 314225 Proteobacteria
Novosphingobium sp PP1Y YP 004534610 1 702113 Proteobacteria
Sphingomonas wittichii RW1 YP 001265341 1 392499 Proteobacteria
Sphingomonas sp MM 1 YP 007615818 1 745310 Proteobacteria
Sphingobium japonicum UT26S YP 003545895 1 452662 Proteobacteria
Sphingobium chlorophenolicum L 1 YP 004553564 1 690566 Proteobacteria
Azospirillum sp B510 YP 003447312 1 137722 Proteobacteria
Burkholderia lata YP 372812 1 482957 Proteobacteria
Burkholderia cepacia GG4 YP 006618694 1 1009846 Proteobacteria
Burkholderia ambifaria MC40 6 YP 001810612 1 398577 Proteobacteria
Burkholderia thailandensis MSMB121 YP 007918580 1 1249661 Proteobacteria
Burkholderia pseudomallei 1026b YP 006274797 1 884204 Proteobacteria
Burkholderia mallei NCTC 10247 YP 001080390 1 320389 Proteobacteria
AZCBGv1 p4180023ID8229336 uid000000 NC 000000 1 conserved protein of unknown function Azospirillum brasilense CBG497 Proteobacteria
Methylocella silvestris BL2 YP 002362975 1 395965 Proteobacteria
Bradyrhizobium sp BTAi1 YP 001237639 1 288000 Proteobacteria
Bradyrhizobium diazoefficiens USDA 110 NP 770133 1 224911 Proteobacteria
Gluconacetobacter diazotrophicus PA1 5 YP 001601953 1 272568 Proteobacteria
Acidiphilium cryptum JF 5 YP 001233958 1 349163 Proteobacteria
Acidiphilium multivorum AIU301 YP 004283322 1 926570 Proteobacteria
Acidiphilium multivorum AIU301 YP 004284493 1 926570 Proteobacteria
Acidiphilium cryptum JF 5 YP 001235138 1 349163 Proteobacteria
Vibrio vulnificus CMCP6 NP 761181 1 216895 Proteobacteria
Marinomonas sp MWYL1 YP 001339184 1 400668 Proteobacteria
Marinomonas posidonica IVIA Po 181 YP 004480168 1 491952 Proteobacteria
Marinomonas mediterranea MMB 1 YP 004315149 1 717774 Proteobacteria
Pectobacterium carotovorum subsp carotovorum PC1 YP 003016260 1 561230 Proteobacteria
Pectobacterium atrosepticum SCRI1043 YP 048903 1 218491 Proteobacteria
Pectobacterium sp SCC3193 YP 006281778 1 1166016 Proteobacteria
Pectobacterium wasabiae WPP163 YP 003258322 1 561231 Proteobacteria
Dickeya chrysanthemi subsp parthenii NCPPB 516 WP 027712019 1 Dchrysanthemi
Dickeya chrysanthemi subsp chrysanthemi NCPPB 402 WP 040001038 1 Dchrysanthemi
Dickeya chrysanthemi L11 WP 033577147 1 Dchrysanthemi
Dickeya chrysanthemi Ech1591 WP 012769756 1 Dchrysanthemi
Dickeya chrysanthemi NCPPB 3533 WP 012769756 1 Dchrysanthemi
6 tadZ Flp pilus assembly Seed
Sinorhizobium medicae WSM419 YP 001313835 1 366394 Proteobacteria
Sinorhizobium meliloti Rm41 YP 006814681 1 1230587 Proteobacteria
Parvularcula bermudensis HTCC2503 YP 003854292 1 314260 Proteobacteria
Caulobacter sp K31 YP 001685813 1 366602 Proteobacteria
Caulobacter crescentus CB15 NP 421737 1 190650 Proteobacteria
Caulobacter crescentus NA1000 YP 002518411 1 565050 Proteobacteria
Hyphomicrobium denitrificans ATCC 51888 YP 003754467 1 582899 Proteobacteria
Liberibacter crescens BT 1 YP 007233529 1 1215343 Proteobacteria
Rhizobium tropici CIAT 899 YP 007332235 1 698761 Proteobacteria
Sinorhizobium medicae WSM419 YP 001329010 1 366394 Proteobacteria
Agrobacterium vitis S4 YP 002548137 1 311402 Proteobacteria
Ochrobactrum anthropi ATCC 49188 YP 001369276 1 439375 Proteobacteria
Chelativorans sp BNC1 YP 676531 1 266779 Proteobacteria
Mesorhizobium ciceri biovar biserrulae WSM1271 YP 004145082 1 765698 Proteobacteria
Mesorhizobium opportunistum WSM2075 YP 004615087 1 536019 Proteobacteria
Mesorhizobium australicum WSM2073 YP 007307661 1 754035 Proteobacteria
Pelagibacterium halotolerans B2 YP 004897838 1 1082931 Proteobacteria
Polymorphum gilvum SL003B 26A1 YP 004305386 1 991905 Proteobacteria
Pseudovibrio sp FO BEG1 YP 005079609 1 911045 Proteobacteria
Methylobacterium sp 4 46 YP 001769274 1 426117 Proteobacteria
Methylobacterium nodulans ORS 2060 YP 002498083 1 460265 Proteobacteria
Methylobacterium radiotolerans JCM 2831 YP 001756451 1 426355 Proteobacteria
Methylobacterium extorquens PA1 YP 001639948 1 419610 Proteobacteria
Methylobacterium populi BJ001 YP 001925107 1 441620 Proteobacteria
Bradyrhizobium oligotrophicum S58 YP 007512067 1 1245469 Proteobacteria
Bradyrhizobium japonicum USDA 6 YP 005605514 1 1037409 Proteobacteria
Bradyrhizobium diazoefficiens USDA 110 NP 767304 1 224911 Proteobacteria
Rhodopseudomonas palustris HaA2 YP 485401 1 316058 Proteobacteria
Nitrobacter winogradskyi Nb 255 YP 316918 1 323098 Proteobacteria
Nitrobacter hamburgensis X14 YP 575740 1 323097 Proteobacteria
Oligotropha carboxidovorans OM4 YP 005949739 1 1031710 Proteobacteria
Bradyrhizobium diazoefficiens USDA 110 NP 768077 1 224911 Proteobacteria
Bradyrhizobium japonicum USDA 6 YP 005613241 1 1037409 Proteobacteria
Bradyrhizobium oligotrophicum S58 YP 007516163 1 1245469 Proteobacteria
Bradyrhizobium sp BTAi1 YP 001242822 1 288000 Proteobacteria

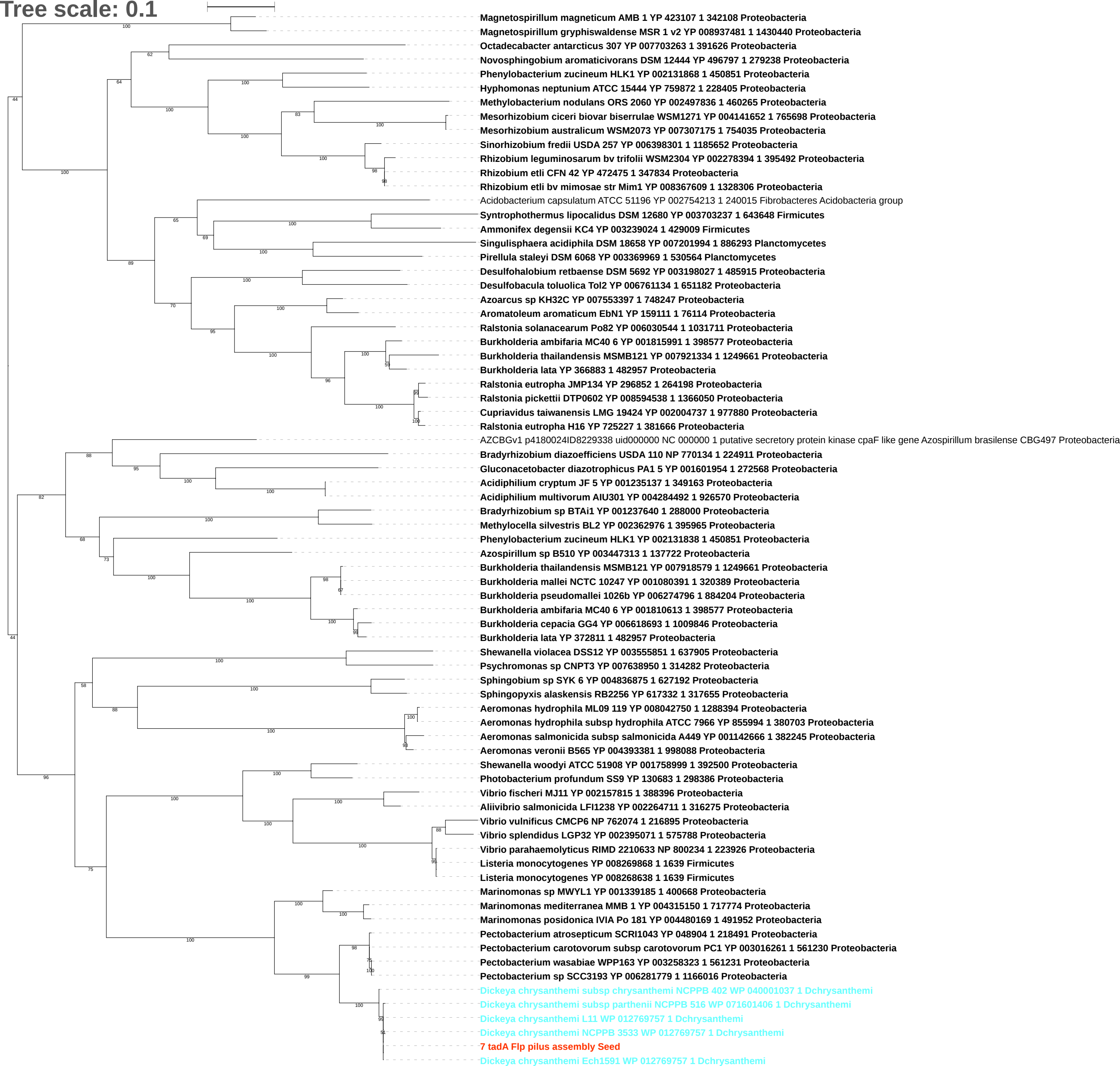
Tree scale: 0.1
Magnetospirillum magneticum AMB 1 YP 423107 1 342108 Proteobacteria
Magnetospirillum gryphiswaldense MSR 1 v2 YP 008937481 1 1430440 Proteobacteria
Octadecabacter antarcticus 307 YP 007703263 1 391626 Proteobacteria
Novosphingobium aromaticivorans DSM 12444 YP 496797 1 279238 Proteobacteria
Phenylobacterium zucineum HLK1 YP 002131868 1 450851 Proteobacteria
Hyphomonas neptunium ATCC 15444 YP 759872 1 228405 Proteobacteria
Methylobacterium nodulans ORS 2060 YP 002497836 1 460265 Proteobacteria
Mesorhizobium ciceri biovar biserrulae WSM1271 YP 004141652 1 765698 Proteobacteria
Mesorhizobium australicum WSM2073 YP 007307175 1 754035 Proteobacteria
Sinorhizobium fredii USDA 257 YP 006398301 1 1185652 Proteobacteria
Rhizobium leguminosarum bv trifolii WSM2304 YP 002278394 1 395492 Proteobacteria
Rhizobium etli CFN 42 YP 472475 1 347834 Proteobacteria
Rhizobium etli bv mimosae str Mim1 YP 008367609 1 1328306 Proteobacteria
Acidobacterium capsulatum ATCC 51196 YP 002754213 1 240015 Fibrobacteres Acidobacteria group
Syntrophothermus lipocalidus DSM 12680 YP 003703237 1 643648 Firmicutes
Ammonifex degensii KC4 YP 003239024 1 429009 Firmicutes
Singulisphaera acidiphila DSM 18658 YP 007201994 1 886293 Planctomycetes
Pirellula staleyi DSM 6068 YP 003369969 1 530564 Planctomycetes
Desulfohalobium retbaense DSM 5692 YP 003198027 1 485915 Proteobacteria
Desulfobacula toluolica Tol2 YP 006761134 1 651182 Proteobacteria
Azoarcus sp KH32C YP 007553397 1 748247 Proteobacteria
Aromatoleum aromaticum EbN1 YP 159111 1 76114 Proteobacteria
Ralstonia solanacearum Po82 YP 006030544 1 1031711 Proteobacteria
Burkholderia ambifaria MC40 6 YP 001815991 1 398577 Proteobacteria
Burkholderia thailandensis MSMB121 YP 007921334 1 1249661 Proteobacteria
Burkholderia lata YP 366883 1 482957 Proteobacteria
Ralstonia eutropha JMP134 YP 296852 1 264198 Proteobacteria
Ralstonia pickettii DTP0602 YP 008594538 1 1366050 Proteobacteria
Cupriavidus taiwanensis LMG 19424 YP 002004737 1 977880 Proteobacteria
Ralstonia eutropha H16 YP 725227 1 381666 Proteobacteria
AZCBGv1 p4180024ID8229338 uid000000 NC 000000 1 putative secretory protein kinase cpaF like gene Azospirillum brasilense CBG497 Proteobacteria
Bradyrhizobium diazoefficiens USDA 110 NP 770134 1 224911 Proteobacteria
Gluconacetobacter diazotrophicus PA1 5 YP 001601954 1 272568 Proteobacteria
Acidiphilium cryptum JF 5 YP 001235137 1 349163 Proteobacteria
Acidiphilium multivorum AIU301 YP 004284492 1 926570 Proteobacteria
Bradyrhizobium sp BTAi1 YP 001237640 1 288000 Proteobacteria
Methylocella silvestris BL2 YP 002362976 1 395965 Proteobacteria
Phenylobacterium zucineum HLK1 YP 002131838 1 450851 Proteobacteria
Azospirillum sp B510 YP 003447313 1 137722 Proteobacteria
Burkholderia thailandensis MSMB121 YP 007918579 1 1249661 Proteobacteria
Burkholderia mallei NCTC 10247 YP 001080391 1 320389 Proteobacteria
Burkholderia pseudomallei 1026b YP 006274796 1 884204 Proteobacteria
Burkholderia ambifaria MC40 6 YP 001810613 1 398577 Proteobacteria
Burkholderia cepacia GG4 YP 006618693 1 1009846 Proteobacteria
Burkholderia lata YP 372811 1 482957 Proteobacteria
Shewanella violacea DSS12 YP 003555851 1 637905 Proteobacteria
Psychromonas sp CNPT3 YP 007638950 1 314282 Proteobacteria
Sphingobium sp SYK 6 YP 004836875 1 627192 Proteobacteria
Sphingopyxis alaskensis RB2256 YP 617332 1 317655 Proteobacteria
Aeromonas hydrophila ML09 119 YP 008042750 1 1288394 Proteobacteria
Aeromonas hydrophila subsp hydrophila ATCC 7966 YP 855994 1 380703 Proteobacteria
Aeromonas salmonicida subsp salmonicida A449 YP 001142666 1 382245 Proteobacteria
Aeromonas veronii B565 YP 004393381 1 998088 Proteobacteria
Shewanella woodyi ATCC 51908 YP 001758999 1 392500 Proteobacteria
Photobacterium profundum SS9 YP 130683 1 298386 Proteobacteria
Vibrio fischeri MJ11 YP 002157815 1 388396 Proteobacteria
Aliivibrio salmonicida LFI1238 YP 002264711 1 316275 Proteobacteria
Vibrio vulnificus CMCP6 NP 762074 1 216895 Proteobacteria
Vibrio splendidus LGP32 YP 002395071 1 575788 Proteobacteria
Vibrio parahaemolyticus RIMD 2210633 NP 800234 1 223926 Proteobacteria
Listeria monocytogenes YP 008269868 1 1639 Firmicutes
Listeria monocytogenes YP 008268638 1 1639 Firmicutes
Marinomonas sp MWYL1 YP 001339185 1 400668 Proteobacteria
Marinomonas mediterranea MMB 1 YP 004315150 1 717774 Proteobacteria
Marinomonas posidonica IVIA Po 181 YP 004480169 1 491952 Proteobacteria
Pectobacterium atrosepticum SCRI1043 YP 048904 1 218491 Proteobacteria
Pectobacterium carotovorum subsp carotovorum PC1 YP 003016261 1 561230 Proteobacteria
Pectobacterium wasabiae WPP163 YP 003258323 1 561231 Proteobacteria
Pectobacterium sp SCC3193 YP 006281779 1 1166016 Proteobacteria
Dickeya chrysanthemi subsp chrysanthemi NCPPB 402 WP 040001037 1 Dchrysanthemi
Dickeya chrysanthemi subsp parthenii NCPPB 516 WP 071601406 1 Dchrysanthemi
Dickeya chrysanthemi L11 WP 012769757 1 Dchrysanthemi
Dickeya chrysanthemi NCPPB 3533 WP 012769757 1 Dchrysanthemi
7 tadA Flp pilus assembly Seed
Dickeya chrysanthemi Ech1591 WP 012769757 1 Dchrysanthemi

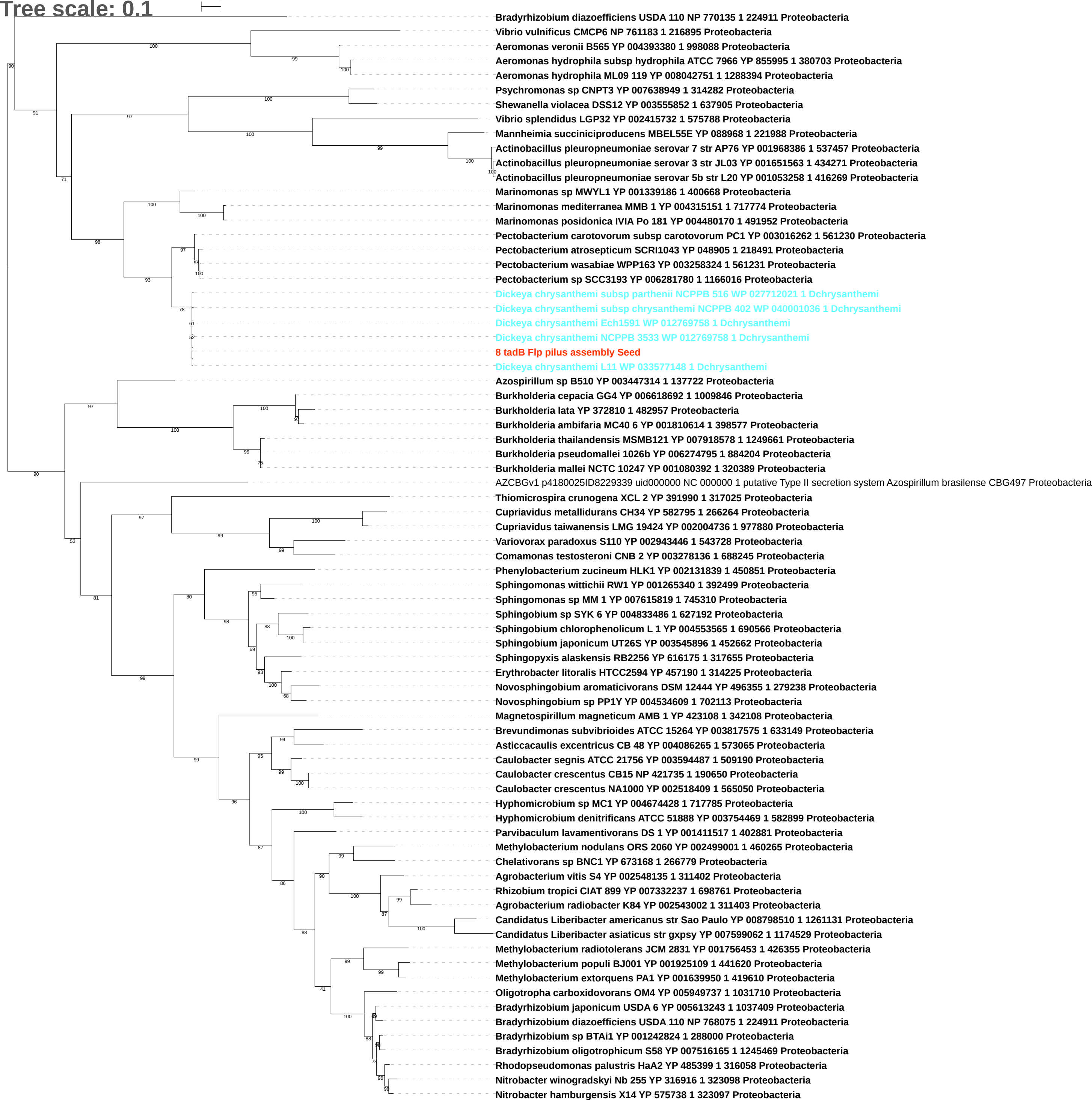
Tree scale: 0.1
Bradyrhizobium diazoefficiens USDA 110 NP 770135 1 224911 Proteobacteria
Vibrio vulnificus CMCP6 NP 761183 1 216895 Proteobacteria
Aeromonas veronii B565 YP 004393380 1 998088 Proteobacteria
Aeromonas hydrophila subsp hydrophila ATCC 7966 YP 855995 1 380703 Proteobacteria
Aeromonas hydrophila ML09 119 YP 008042751 1 1288394 Proteobacteria
Psychromonas sp CNPT3 YP 007638949 1 314282 Proteobacteria
Shewanella violacea DSS12 YP 003555852 1 637905 Proteobacteria
Vibrio splendidus LGP32 YP 002415732 1 575788 Proteobacteria
Mannheimia succiniciproducens MBEL55E YP 088968 1 221988 Proteobacteria
Actinobacillus pleuropneumoniae serovar 7 str AP76 YP 001968386 1 537457 Proteobacteria
Actinobacillus pleuropneumoniae serovar 3 str JL03 YP 001651563 1 434271 Proteobacteria
Actinobacillus pleuropneumoniae serovar 5b str L20 YP 001053258 1 416269 Proteobacteria
Marinomonas sp MWYL1 YP 001339186 1 400668 Proteobacteria
Marinomonas mediterranea MMB 1 YP 004315151 1 717774 Proteobacteria
Marinomonas posidonica IVIA Po 181 YP 004480170 1 491952 Proteobacteria
Pectobacterium carotovorum subsp carotovorum PC1 YP 003016262 1 561230 Proteobacteria
Pectobacterium atrosepticum SCRI1043 YP 048905 1 218491 Proteobacteria
Pectobacterium wasabiae WPP163 YP 003258324 1 561231 Proteobacteria
Pectobacterium sp SCC3193 YP 006281780 1 1166016 Proteobacteria
Dickeya chrysanthemi subsp parthenii NCPPB 516 WP 027712021 1 Dchrysanthemi
Dickeya chrysanthemi subsp chrysanthemi NCPPB 402 WP 040001036 1 Dchrysanthemi
Dickeya chrysanthemi Ech1591 WP 012769758 1 Dchrysanthemi
Dickeya chrysanthemi NCPPB 3533 WP 012769758 1 Dchrysanthemi
8 tadB Flp pilus assembly Seed
Dickeya chrysanthemi L11 WP 033577148 1 Dchrysanthemi
Azospirillum sp B510 YP 003447314 1 137722 Proteobacteria
Burkholderia cepacia GG4 YP 006618692 1 1009846 Proteobacteria
Burkholderia lata YP 372810 1 482957 Proteobacteria
Burkholderia ambifaria MC40 6 YP 001810614 1 398577 Proteobacteria
Burkholderia thailandensis MSMB121 YP 007918578 1 1249661 Proteobacteria
Burkholderia pseudomallei 1026b YP 006274795 1 884204 Proteobacteria
Burkholderia mallei NCTC 10247 YP 001080392 1 320389 Proteobacteria
AZCBGv1 p4180025ID8229339 uid000000 NC 000000 1 putative Type II secretion system Azospirillum brasilense CBG497 Proteobacteria
Thiomicrospira crunogena XCL 2 YP 391990 1 317025 Proteobacteria
Cupriavidus metallidurans CH34 YP 582795 1 266264 Proteobacteria
Cupriavidus taiwanensis LMG 19424 YP 002004736 1 977880 Proteobacteria
Variovorax paradoxus S110 YP 002943446 1 543728 Proteobacteria
Comamonas testosteroni CNB 2 YP 003278136 1 688245 Proteobacteria
Phenylobacterium zucineum HLK1 YP 002131839 1 450851 Proteobacteria
Sphingomonas wittichii RW1 YP 001265340 1 392499 Proteobacteria
Sphingomonas sp MM 1 YP 007615819 1 745310 Proteobacteria
Sphingobium sp SYK 6 YP 004833486 1 627192 Proteobacteria
Sphingobium chlorophenolicum L 1 YP 004553565 1 690566 Proteobacteria
Sphingobium japonicum UT26S YP 003545896 1 452662 Proteobacteria
Sphingopyxis alaskensis RB2256 YP 616175 1 317655 Proteobacteria
Erythrobacter litoralis HTCC2594 YP 457190 1 314225 Proteobacteria
Novosphingobium aromaticivorans DSM 12444 YP 496355 1 279238 Proteobacteria
Novosphingobium sp PP1Y YP 004534609 1 702113 Proteobacteria
Magnetospirillum magneticum AMB 1 YP 423108 1 342108 Proteobacteria
Brevundimonas subvibrioides ATCC 15264 YP 003817575 1 633149 Proteobacteria
Asticcacaulis excentricus CB 48 YP 004086265 1 573065 Proteobacteria
Caulobacter segnis ATCC 21756 YP 003594487 1 509190 Proteobacteria
Caulobacter crescentus CB15 NP 421735 1 190650 Proteobacteria
Caulobacter crescentus NA1000 YP 002518409 1 565050 Proteobacteria
Hyphomicrobium sp MC1 YP 004674428 1 717785 Proteobacteria
Hyphomicrobium denitrificans ATCC 51888 YP 003754469 1 582899 Proteobacteria
Parvibaculum lavamentivorans DS 1 YP 001411517 1 402881 Proteobacteria
Methylobacterium nodulans ORS 2060 YP 002499001 1 460265 Proteobacteria
Chelativorans sp BNC1 YP 673168 1 266779 Proteobacteria
Agrobacterium vitis S4 YP 002548135 1 311402 Proteobacteria
Rhizobium tropici CIAT 899 YP 007332237 1 698761 Proteobacteria
Agrobacterium radiobacter K84 YP 002543002 1 311403 Proteobacteria
Candidatus Liberibacter americanus str Sao Paulo YP 008798510 1 1261131 Proteobacteria
Candidatus Liberibacter asiaticus str gxpsy YP 007599062 1 1174529 Proteobacteria
Methylobacterium radiotolerans JCM 2831 YP 001756453 1 426355 Proteobacteria
Methylobacterium populi BJ001 YP 001925109 1 441620 Proteobacteria
Methylobacterium extorquens PA1 YP 001639950 1 419610 Proteobacteria
Oligotropha carboxidovorans OM4 YP 005949737 1 1031710 Proteobacteria
Bradyrhizobium japonicum USDA 6 YP 005613243 1 1037409 Proteobacteria
Bradyrhizobium diazoefficiens USDA 110 NP 768075 1 224911 Proteobacteria
Bradyrhizobium sp BTAi1 YP 001242824 1 288000 Proteobacteria
Bradyrhizobium oligotrophicum S58 YP 007516165 1 1245469 Proteobacteria
Rhodopseudomonas palustris HaA2 YP 485399 1 316058 Proteobacteria
Nitrobacter winogradskyi Nb 255 YP 316916 1 323098 Proteobacteria
Nitrobacter hamburgensis X14 YP 575738 1 323097 Proteobacteria

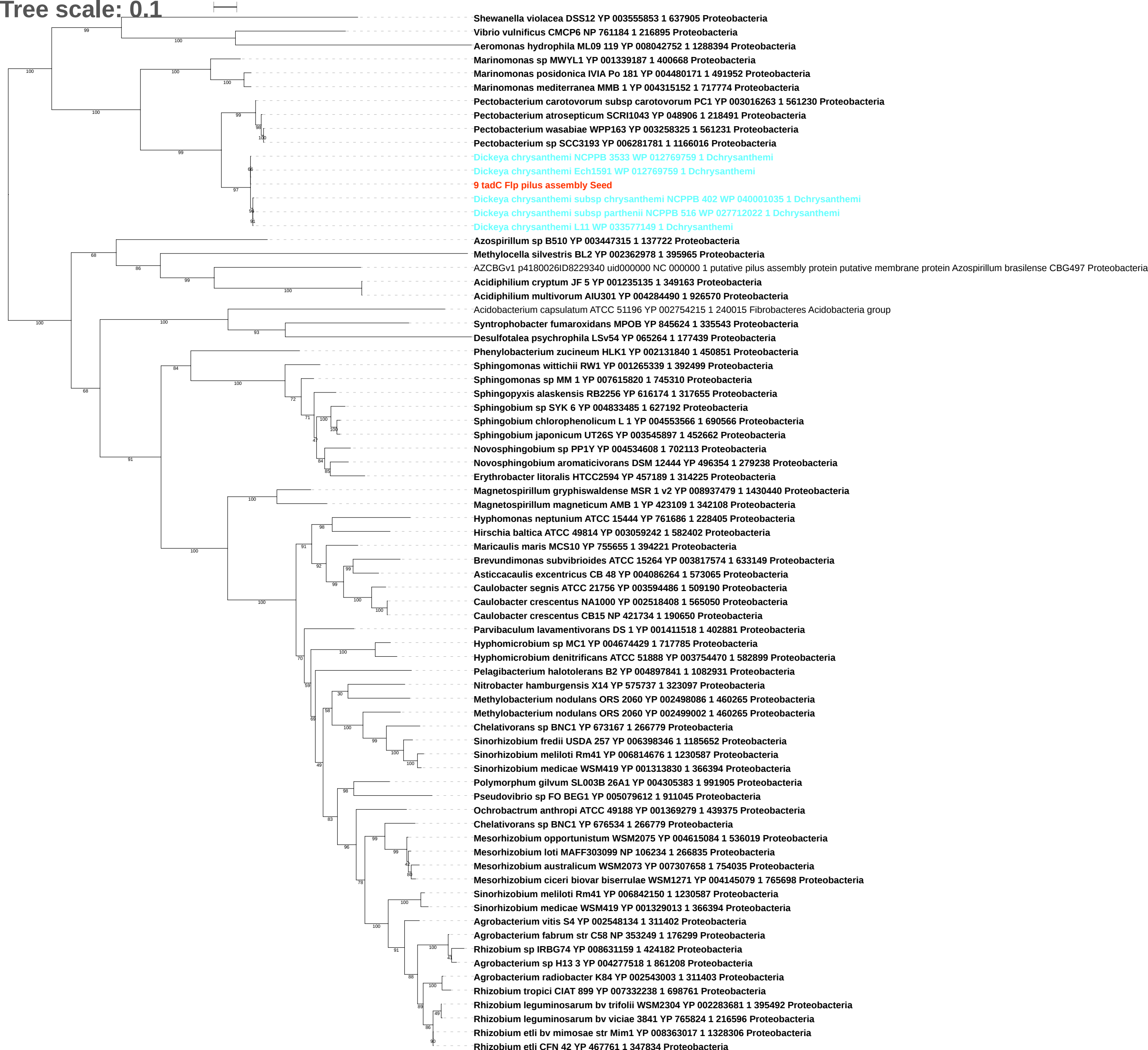
Tree scale: 0.1
Shewanella violacea DSS12 YP 003555853 1 637905 Proteobacteria
Vibrio vulnificus CMCP6 NP 761184 1 216895 Proteobacteria
Aeromonas hydrophila ML09 119 YP 008042752 1 1288394 Proteobacteria
Marinomonas sp MWYL1 YP 001339187 1 400668 Proteobacteria
Marinomonas posidonica IVIA Po 181 YP 004480171 1 491952 Proteobacteria
Marinomonas mediterranea MMB 1 YP 004315152 1 717774 Proteobacteria
Pectobacterium carotovorum subsp carotovorum PC1 YP 003016263 1 561230 Proteobacteria
Pectobacterium atrosepticum SCRI1043 YP 048906 1 218491 Proteobacteria
Pectobacterium wasabiae WPP163 YP 003258325 1 561231 Proteobacteria
Pectobacterium sp SCC3193 YP 006281781 1 1166016 Proteobacteria
Dickeya chrysanthemi NCPPB 3533 WP 012769759 1 Dchrysanthemi
Dickeya chrysanthemi Ech1591 WP 012769759 1 Dchrysanthemi
9 tadC Flp pilus assembly Seed
Dickeya chrysanthemi subsp chrysanthemi NCPPB 402 WP 040001035 1 Dchrysanthemi
Dickeya chrysanthemi subsp parthenii NCPPB 516 WP 027712022 1 Dchrysanthemi
Dickeya chrysanthemi L11 WP 033577149 1 Dchrysanthemi
Azospirillum sp B510 YP 003447315 1 137722 Proteobacteria
Methylocella silvestris BL2 YP 002362978 1 395965 Proteobacteria
AZCBGv1 p4180026ID8229340 uid000000 NC 000000 1 putative pilus assembly protein putative membrane protein Azospirillum brasilense CBG497 Proteobacteria
Acidiphilium cryptum JF 5 YP 001235135 1 349163 Proteobacteria
Acidiphilium multivorum AIU301 YP 004284490 1 926570 Proteobacteria
Acidobacterium capsulatum ATCC 51196 YP 002754215 1 240015 Fibrobacteres Acidobacteria group
Syntrophobacter fumaroxidans MPOB YP 845624 1 335543 Proteobacteria
Desulfotalea psychrophila LSv54 YP 065264 1 177439 Proteobacteria
Phenylobacterium zucineum HLK1 YP 002131840 1 450851 Proteobacteria
Sphingomonas wittichii RW1 YP 001265339 1 392499 Proteobacteria
Sphingomonas sp MM 1 YP 007615820 1 745310 Proteobacteria
Sphingopyxis alaskensis RB2256 YP 616174 1 317655 Proteobacteria
Sphingobium sp SYK 6 YP 004833485 1 627192 Proteobacteria
Sphingobium chlorophenolicum L 1 YP 004553566 1 690566 Proteobacteria
Sphingobium japonicum UT26S YP 003545897 1 452662 Proteobacteria
Novosphingobium sp PP1Y YP 004534608 1 702113 Proteobacteria
Novosphingobium aromaticivorans DSM 12444 YP 496354 1 279238 Proteobacteria
Erythrobacter litoralis HTCC2594 YP 457189 1 314225 Proteobacteria
Magnetospirillum gryphiswaldense MSR 1 v2 YP 008937479 1 1430440 Proteobacteria
Magnetospirillum magneticum AMB 1 YP 423109 1 342108 Proteobacteria
Hyphomonas neptunium ATCC 15444 YP 761686 1 228405 Proteobacteria
Hirschia baltica ATCC 49814 YP 003059242 1 582402 Proteobacteria
Maricaulis maris MCS10 YP 755655 1 394221 Proteobacteria
Brevundimonas subvibrioides ATCC 15264 YP 003817574 1 633149 Proteobacteria
Asticcacaulis excentricus CB 48 YP 004086264 1 573065 Proteobacteria
Caulobacter segnis ATCC 21756 YP 003594486 1 509190 Proteobacteria
Caulobacter crescentus NA1000 YP 002518408 1 565050 Proteobacteria
Caulobacter crescentus CB15 NP 421734 1 190650 Proteobacteria
Parvibaculum lavamentivorans DS 1 YP 001411518 1 402881 Proteobacteria
Hyphomicrobium sp MC1 YP 004674429 1 717785 Proteobacteria
Hyphomicrobium denitrificans ATCC 51888 YP 003754470 1 582899 Proteobacteria
Pelagibacterium halotolerans B2 YP 004897841 1 1082931 Proteobacteria
Nitrobacter hamburgensis X14 YP 575737 1 323097 Proteobacteria
Methylobacterium nodulans ORS 2060 YP 002498086 1 460265 Proteobacteria
Methylobacterium nodulans ORS 2060 YP 002499002 1 460265 Proteobacteria
Chelativorans sp BNC1 YP 673167 1 266779 Proteobacteria
Sinorhizobium fredii USDA 257 YP 006398346 1 1185652 Proteobacteria
Sinorhizobium meliloti Rm41 YP 006814676 1 1230587 Proteobacteria
Sinorhizobium medicae WSM419 YP 001313830 1 366394 Proteobacteria
Polymorphum gilvum SL003B 26A1 YP 004305383 1 991905 Proteobacteria
Pseudovibrio sp FO BEG1 YP 005079612 1 911045 Proteobacteria
Ochrobactrum anthropi ATCC 49188 YP 001369279 1 439375 Proteobacteria
Chelativorans sp BNC1 YP 676534 1 266779 Proteobacteria
Mesorhizobium opportunistum WSM2075 YP 004615084 1 536019 Proteobacteria
Mesorhizobium loti MAFF303099 NP 106234 1 266835 Proteobacteria
Mesorhizobium australicum WSM2073 YP 007307658 1 754035 Proteobacteria
Mesorhizobium ciceri biovar biserrulae WSM1271 YP 004145079 1 765698 Proteobacteria
Sinorhizobium meliloti Rm41 YP 006842150 1 1230587 Proteobacteria
Sinorhizobium medicae WSM419 YP 001329013 1 366394 Proteobacteria
Agrobacterium vitis S4 YP 002548134 1 311402 Proteobacteria
Agrobacterium fabrum str C58 NP 353249 1 176299 Proteobacteria
Rhizobium sp IRBG74 YP 008631159 1 424182 Proteobacteria
Agrobacterium sp H13 3 YP 004277518 1 861208 Proteobacteria
Agrobacterium radiobacter K84 YP 002543003 1 311403 Proteobacteria
Rhizobium tropici CIAT 899 YP 007332238 1 698761 Proteobacteria
Rhizobium leguminosarum bv trifolii WSM2304 YP 002283681 1 395492 Proteobacteria
Rhizobium leguminosarum bv viciae 3841 YP 765824 1 216596 Proteobacteria
Rhizobium etli bv mimosae str Mim1 YP 008363017 1 1328306 Proteobacteria
Rhizobium etli CFN 42 YP 467761 1 347834 Proteobacteria

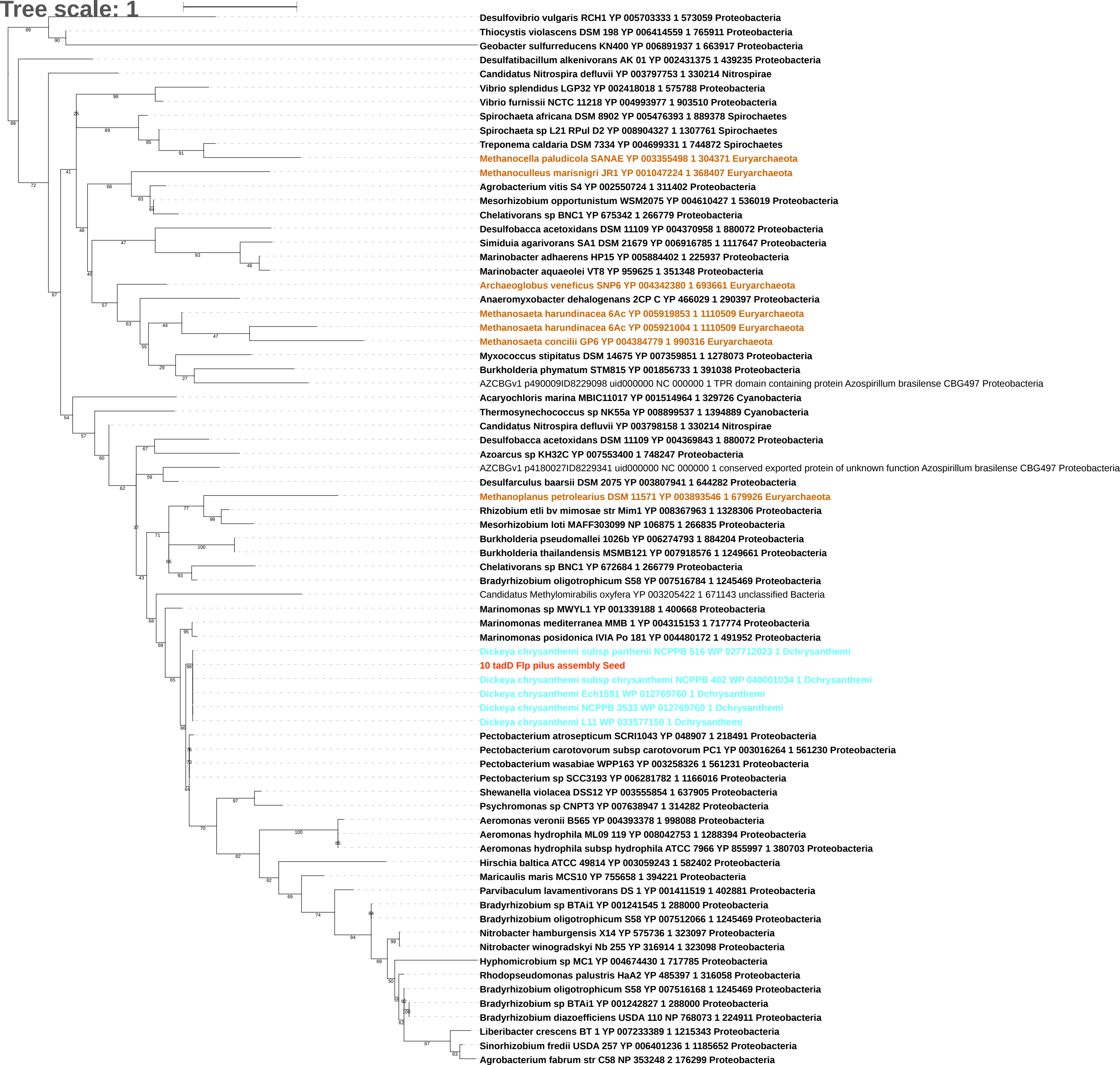
Tree scale: 1
Desulfovibrio vulgaris RCH1 YP 005703333 1 573059 Proteobacteria
Thiocystis violascens DSM 198 YP 006414559 1 765911 Proteobacteria
Geobacter sulfurreducens KN400 YP 006891937 1 663917 Proteobacteria
Desulfatibacillum alkenivorans AK 01 YP 002431375 1 439235 Proteobacteria
Candidatus Nitrospira defluvii YP 003797753 1 330214 Nitrospirae
Vibrio splendidus LGP32 YP 002418018 1 575788 Proteobacteria
Vibrio furnissii NCTC 11218 YP 004993977 1 903510 Proteobacteria
Spirochaeta africana DSM 8902 YP 005476393 1 889378 Spirochaetes
Spirochaeta sp L21 RPul D2 YP 008904327 1 1307761 Spirochaetes
Treponema caldaria DSM 7334 YP 004699331 1 744872 Spirochaetes
Methanocella paludicola SANAE YP 003355498 1 304371 Euryarchaeota
Methanoculleus marisnigri JR1 YP 001047224 1 368407 Euryarchaeota
Agrobacterium vitis S4 YP 002550724 1 311402 Proteobacteria
Mesorhizobium opportunistum WSM2075 YP 004610427 1 536019 Proteobacteria
Chelativorans sp BNC1 YP 675342 1 266779 Proteobacteria
Desulfobacca acetoxidans DSM 11109 YP 004370958 1 880072 Proteobacteria
Simiduia agarivorans SA1 DSM 21679 YP 006916785 1 1117647 Proteobacteria
Marinobacter adhaerens HP15 YP 005884402 1 225937 Proteobacteria
Marinobacter aquaeolei VT8 YP 959625 1 351348 Proteobacteria
Archaeoglobus veneficus SNP6 YP 004342380 1 693661 Euryarchaeota
Anaeromyxobacter dehalogenans 2CP C YP 466029 1 290397 Proteobacteria
Methanosaeta harundinacea 6Ac YP 005919853 1 1110509 Euryarchaeota
Methanosaeta harundinacea 6Ac YP 005921004 1 1110509 Euryarchaeota
Methanosaeta concilii GP6 YP 004384779 1 990316 Euryarchaeota
Myxococcus stipitatus DSM 14675 YP 007359851 1 1278073 Proteobacteria
Burkholderia phymatum STM815 YP 001856733 1 391038 Proteobacteria
AZCBGv1 p490009ID8229098 uid000000 NC 000000 1 TPR domain containing protein Azospirillum brasilense CBG497 Proteobacteria
Acaryochloris marina MBIC11017 YP 001514964 1 329726 Cyanobacteria
Thermosynechococcus sp NK55a YP 008899537 1 1394889 Cyanobacteria
Candidatus Nitrospira defluvii YP 003798158 1 330214 Nitrospirae
Desulfobacca acetoxidans DSM 11109 YP 004369843 1 880072 Proteobacteria
Azoarcus sp KH32C YP 007553400 1 748247 Proteobacteria
AZCBGv1 p4180027ID8229341 uid000000 NC 000000 1 conserved exported protein of unknown function Azospirillum brasilense CBG497 Proteobacteria
Desulfarculus baarsii DSM 2075 YP 003807941 1 644282 Proteobacteria
Methanoplanus petrolearius DSM 11571 YP 003893546 1 679926 Euryarchaeota
Rhizobium etli bv mimosae str Mim1 YP 008367963 1 1328306 Proteobacteria
Mesorhizobium loti MAFF303099 NP 106875 1 266835 Proteobacteria
Burkholderia pseudomallei 1026b YP 006274793 1 884204 Proteobacteria
Burkholderia thailandensis MSMB121 YP 007918576 1 1249661 Proteobacteria
Chelativorans sp BNC1 YP 672684 1 266779 Proteobacteria
Bradyrhizobium oligotrophicum S58 YP 007516784 1 1245469 Proteobacteria
Candidatus Methylomirabilis oxyfera YP 003205422 1 671143 unclassified Bacteria
Marinomonas sp MWYL1 YP 001339188 1 400668 Proteobacteria
Marinomonas mediterranea MMB 1 YP 004315153 1 717774 Proteobacteria
Marinomonas posidonica IVIA Po 181 YP 004480172 1 491952 Proteobacteria
Dickeya chrysanthemi subsp parthenii NCPPB 516 WP 027712023 1 Dchrysanthemi
10 tadD Flp pilus assembly Seed
Dickeya chrysanthemi subsp chrysanthemi NCPPB 402 WP 040001034 1 Dchrysanthemi
Dickeya chrysanthemi Ech1591 WP 012769760 1 Dchrysanthemi
Dickeya chrysanthemi NCPPB 3533 WP 012769760 1 Dchrysanthemi
Dickeya chrysanthemi L11 WP 033577150 1 Dchrysanthemi
Pectobacterium atrosepticum SCRI1043 YP 048907 1 218491 Proteobacteria
Pectobacterium carotovorum subsp carotovorum PC1 YP 003016264 1 561230 Proteobacteria
Pectobacterium wasabiae WPP163 YP 003258326 1 561231 Proteobacteria
Pectobacterium sp SCC3193 YP 006281782 1 1166016 Proteobacteria
Shewanella violacea DSS12 YP 003555854 1 637905 Proteobacteria
Psychromonas sp CNPT3 YP 007638947 1 314282 Proteobacteria
Aeromonas veronii B565 YP 004393378 1 998088 Proteobacteria
Aeromonas hydrophila ML09 119 YP 008042753 1 1288394 Proteobacteria
Aeromonas hydrophila subsp hydrophila ATCC 7966 YP 855997 1 380703 Proteobacteria
Hirschia baltica ATCC 49814 YP 003059243 1 582402 Proteobacteria
Maricaulis maris MCS10 YP 755658 1 394221 Proteobacteria
Parvibaculum lavamentivorans DS 1 YP 001411519 1 402881 Proteobacteria
Bradyrhizobium sp BTAi1 YP 001241545 1 288000 Proteobacteria
Bradyrhizobium oligotrophicum S58 YP 007512066 1 1245469 Proteobacteria
Nitrobacter hamburgensis X14 YP 575736 1 323097 Proteobacteria
Nitrobacter winogradskyi Nb 255 YP 316914 1 323098 Proteobacteria
Hyphomicrobium sp MC1 YP 004674430 1 717785 Proteobacteria
Rhodopseudomonas palustris HaA2 YP 485397 1 316058 Proteobacteria
Bradyrhizobium oligotrophicum S58 YP 007516168 1 1245469 Proteobacteria
Bradyrhizobium sp BTAi1 YP 001242827 1 288000 Proteobacteria
Bradyrhizobium diazoefficiens USDA 110 NP 768073 1 224911 Proteobacteria
Liberibacter crescens BT 1 YP 007233389 1 1215343 Proteobacteria
Sinorhizobium fredii USDA 257 YP 006401236 1 1185652 Proteobacteria
Agrobacterium fabrum str C58 NP 353248 2 176299 Proteobacteria

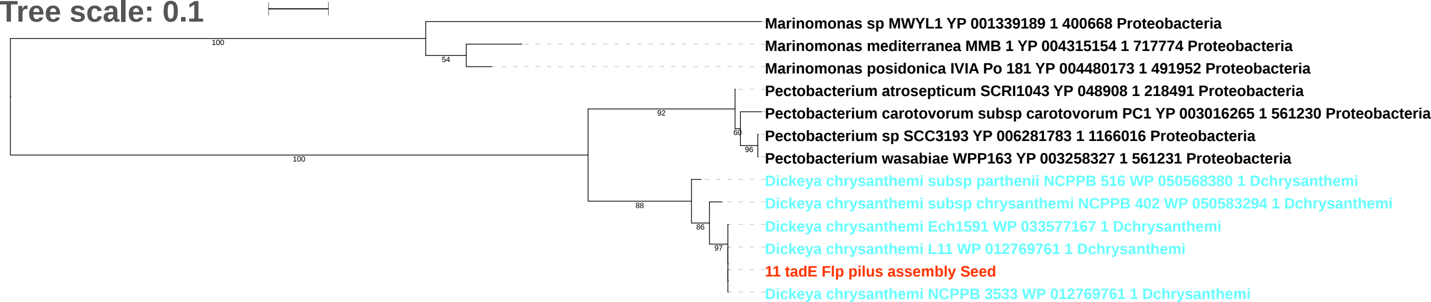

Tree scale: 0.1
Marinomonas sp MWYL1 YP 001339189 1 400668 Proteobacteria
Marinomonas mediterranea MMB 1 YP 004315154 1 717774 Proteobacteria
Marinomonas posidonica IVIA Po 181 YP 004480173 1 491952 Proteobacteria
Pectobacterium atrosepticum SCRI1043 YP 048908 1 218491 Proteobacteria
Pectobacterium carotovorum subsp carotovorum PC1 YP 003016265 1 561230 Proteobacteria
Pectobacterium sp SCC3193 YP 006281783 1 1166016 Proteobacteria
Pectobacterium wasabiae WPP163 YP 003258327 1 561231 Proteobacteria
Dickeya chrysanthemi subsp parthenii NCPPB 516 WP 050568380 1 Dchrysanthemi
Dickeya chrysanthemi subsp chrysanthemi NCPPB 402 WP 050583294 1 Dchrysanthemi
Dickeya chrysanthemi Ech1591 WP 033577167 1 Dchrysanthemi
Dickeya chrysanthemi L11 WP 012769761 1 Dchrysanthemi
11 tadE Flp pilus assembly Seed
Dickeya chrysanthemi NCPPB 3533 WP 012769761 1 Dchrysanthemi
100
54
100
92
60
96
88
86
97

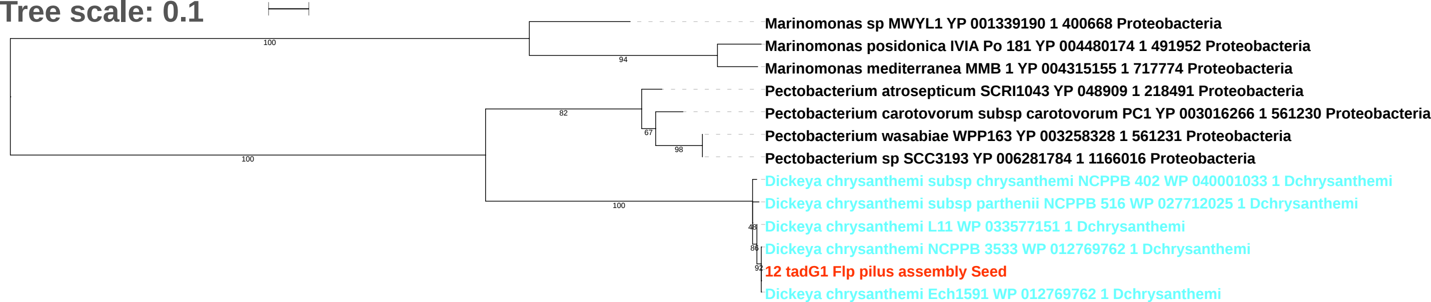
Tree scale: 0.1
Marinomonas sp MWYL1 YP 001339190 1 400668 Proteobacteria
Marinomonas posidonica IVIA Po 181 YP 004480174 1 491952 Proteobacteria
Marinomonas mediterranea MMB 1 YP 004315155 1 717774 Proteobacteria
Pectobacterium atrosepticum SCRI1043 YP 048909 1 218491 Proteobacteria
Pectobacterium carotovorum subsp carotovorum PC1 YP 003016266 1 561230 Proteobacteria
Pectobacterium wasabiae WPP163 YP 003258328 1 561231 Proteobacteria
Pectobacterium sp SCC3193 YP 006281784 1 1166016 Proteobacteria
Dickeya chrysanthemi subsp chrysanthemi NCPPB 402 WP 040001033 1 Dchrysanthemi
Dickeya chrysanthemi subsp parthenii NCPPB 516 WP 027712025 1 Dchrysanthemi
Dickeya chrysanthemi L11 WP 033577151 1 Dchrysanthemi
Dickeya chrysanthemi NCPPB 3533 WP 012769762 1 Dchrysanthemi
12 tadG1 Flp pilus assembly Seed
Dickeya chrysanthemi Ech1591 WP 012769762 1 Dchrysanthemi
100
94
82
67
98
100
100
48
86
92

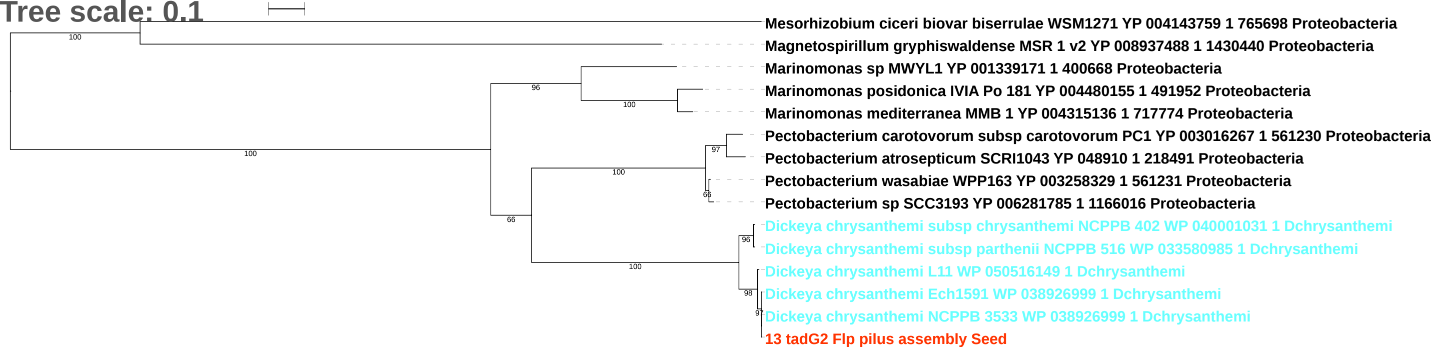

Tree scale: 0.1
Mesorhizobium ciceri biovar biserrulae WSM1271 YP 004143759 1 765698 Proteobacteria
Magnetospirillum gryphiswaldense MSR 1 v2 YP 008937488 1 1430440 Proteobacteria
Marinomonas sp MWYL1 YP 001339171 1 400668 Proteobacteria
Marinomonas posidonica IVIA Po 181 YP 004480155 1 491952 Proteobacteria
Marinomonas mediterranea MMB 1 YP 004315136 1 717774 Proteobacteria
Pectobacterium carotovorum subsp carotovorum PC1 YP 003016267 1 561230 Proteobacteria
Pectobacterium atrosepticum SCRI1043 YP 048910 1 218491 Proteobacteria
Pectobacterium wasabiae WPP163 YP 003258329 1 561231 Proteobacteria
Pectobacterium sp SCC3193 YP 006281785 1 1166016 Proteobacteria
Dickeya chrysanthemi subsp chrysanthemi NCPPB 402 WP 040001031 1 Dchrysanthemi
Dickeya chrysanthemi subsp parthenii NCPPB 516 WP 033580985 1 Dchrysanthemi
Dickeya chrysanthemi L11 WP 050516149 1 Dchrysanthemi
Dickeya chrysanthemi Ech1591 WP 038926999 1 Dchrysanthemi
Dickeya chrysanthemi NCPPB 3533 WP 038926999 1 Dchrysanthemi
13 tadG2 Flp pilus assembly Seed
100
100
96
100
97
100
66
66
96
100
98
97

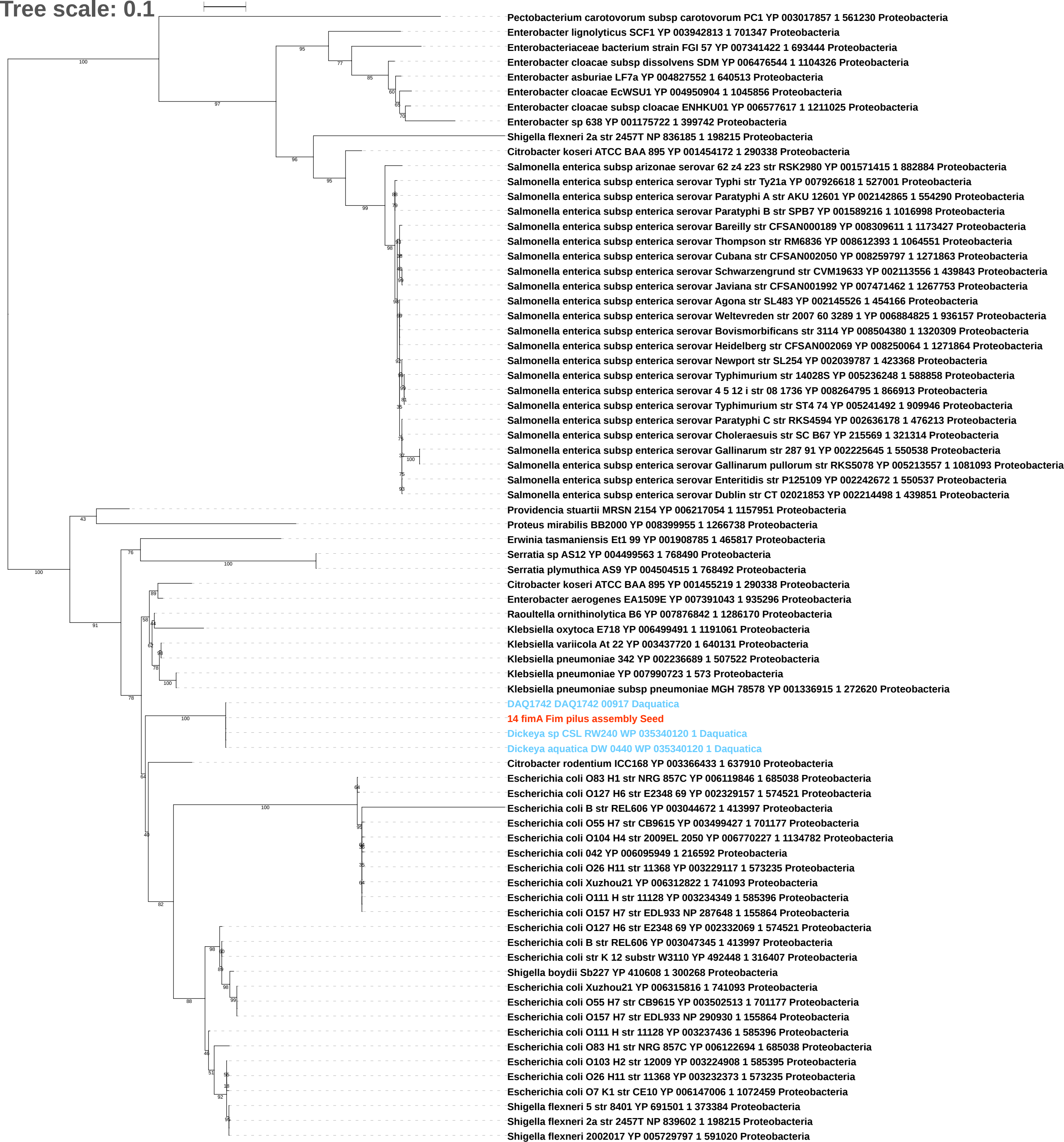
Tree scale: 0.1
Pectobacterium carotovorum subsp carotovorum PC1 YP 003017857 1 561230 Proteobacteria
Enterobacter lignolyticus SCF1 YP 003942813 1 701347 Proteobacteria
Enterobacteriaceae bacterium strain FGI 57 YP 007341422 1 693444 Proteobacteria
Enterobacter cloacae subsp dissolvens SDM YP 006476544 1 1104326 Proteobacteria
Enterobacter asburiae LF7a YP 004827552 1 640513 Proteobacteria
Enterobacter cloacae EcWSU1 YP 004950904 1 1045856 Proteobacteria
Enterobacter cloacae subsp cloacae ENHKU01 YP 006577617 1 1211025 Proteobacteria
Enterobacter sp 638 YP 001175722 1 399742 Proteobacteria
Shigella flexneri 2a str 2457T NP 836185 1 198215 Proteobacteria
Citrobacter koseri ATCC BAA 895 YP 001454172 1 290338 Proteobacteria
Salmonella enterica subsp arizonae serovar 62 z4 z23 str RSK2980 YP 001571415 1 882884 Proteobacteria
Salmonella enterica subsp enterica serovar Typhi str Ty21a YP 007926618 1 527001 Proteobacteria
Salmonella enterica subsp enterica serovar Paratyphi A str AKU 12601 YP 002142865 1 554290 Proteobacteria
Salmonella enterica subsp enterica serovar Paratyphi B str SPB7 YP 001589216 1 1016998 Proteobacteria
Salmonella enterica subsp enterica serovar Bareilly str CFSAN000189 YP 008309611 1 1173427 Proteobacteria
Salmonella enterica subsp enterica serovar Thompson str RM6836 YP 008612393 1 1064551 Proteobacteria
Salmonella enterica subsp enterica serovar Cubana str CFSAN002050 YP 008259797 1 1271863 Proteobacteria
Salmonella enterica subsp enterica serovar Schwarzengrund str CVM19633 YP 002113556 1 439843 Proteobacteria
Salmonella enterica subsp enterica serovar Javiana str CFSAN001992 YP 007471462 1 1267753 Proteobacteria
Salmonella enterica subsp enterica serovar Agona str SL483 YP 002145526 1 454166 Proteobacteria
Salmonella enterica subsp enterica serovar Weltevreden str 2007 60 3289 1 YP 006884825 1 936157 Proteobacteria
Salmonella enterica subsp enterica serovar Bovismorbificans str 3114 YP 008504380 1 1320309 Proteobacteria
Salmonella enterica subsp enterica serovar Heidelberg str CFSAN002069 YP 008250064 1 1271864 Proteobacteria
Salmonella enterica subsp enterica serovar Newport str SL254 YP 002039787 1 423368 Proteobacteria
Salmonella enterica subsp enterica serovar Typhimurium str 14028S YP 005236248 1 588858 Proteobacteria
Salmonella enterica subsp enterica serovar 4 5 12 i str 08 1736 YP 008264795 1 866913 Proteobacteria
Salmonella enterica subsp enterica serovar Typhimurium str ST4 74 YP 005241492 1 909946 Proteobacteria
Salmonella enterica subsp enterica serovar Paratyphi C str RKS4594 YP 002636178 1 476213 Proteobacteria
Salmonella enterica subsp enterica serovar Choleraesuis str SC B67 YP 215569 1 321314 Proteobacteria
Salmonella enterica subsp enterica serovar Gallinarum str 287 91 YP 002225645 1 550538 Proteobacteria
Salmonella enterica subsp enterica serovar Gallinarum pullorum str RKS5078 YP 005213557 1 1081093 Proteobacteria
Salmonella enterica subsp enterica serovar Enteritidis str P125109 YP 002242672 1 550537 Proteobacteria
Salmonella enterica subsp enterica serovar Dublin str CT 02021853 YP 002214498 1 439851 Proteobacteria
Providencia stuartii MRSN 2154 YP 006217054 1 1157951 Proteobacteria
Proteus mirabilis BB2000 YP 008399955 1 1266738 Proteobacteria
Erwinia tasmaniensis Et1 99 YP 001908785 1 465817 Proteobacteria
Serratia sp AS12 YP 004499563 1 768490 Proteobacteria
Serratia plymuthica AS9 YP 004504515 1 768492 Proteobacteria
Citrobacter koseri ATCC BAA 895 YP 001455219 1 290338 Proteobacteria
Enterobacter aerogenes EA1509E YP 007391043 1 935296 Proteobacteria
Raoultella ornithinolytica B6 YP 007876842 1 1286170 Proteobacteria
Klebsiella oxytoca E718 YP 006499491 1 1191061 Proteobacteria
Klebsiella variicola At 22 YP 003437720 1 640131 Proteobacteria
Klebsiella pneumoniae 342 YP 002236689 1 507522 Proteobacteria
Klebsiella pneumoniae YP 007990723 1 573 Proteobacteria
Klebsiella pneumoniae subsp pneumoniae MGH 78578 YP 001336915 1 272620 Proteobacteria
DAQ1742 DAQ1742 00917 Daquatica
14 fimA Fim pilus assembly Seed
Dickeya sp CSL RW240 WP 035340120 1 Daquatica
Dickeya aquatica DW 0440 WP 035340120 1 Daquatica
Citrobacter rodentium ICC168 YP 003366433 1 637910 Proteobacteria
Escherichia coli O83 H1 str NRG 857C YP 006119846 1 685038 Proteobacteria
Escherichia coli O127 H6 str E2348 69 YP 002329157 1 574521 Proteobacteria
Escherichia coli B str REL606 YP 003044672 1 413997 Proteobacteria
Escherichia coli O55 H7 str CB9615 YP 003499427 1 701177 Proteobacteria
Escherichia coli O104 H4 str 2009EL 2050 YP 006770227 1 1134782 Proteobacteria
Escherichia coli 042 YP 006095949 1 216592 Proteobacteria
Escherichia coli O26 H11 str 11368 YP 003229117 1 573235 Proteobacteria
Escherichia coli Xuzhou21 YP 006312822 1 741093 Proteobacteria
Escherichia coli O111 H str 11128 YP 003234349 1 585396 Proteobacteria
Escherichia coli O157 H7 str EDL933 NP 287648 1 155864 Proteobacteria
Escherichia coli O127 H6 str E2348 69 YP 002332069 1 574521 Proteobacteria
Escherichia coli B str REL606 YP 003047345 1 413997 Proteobacteria
Escherichia coli str K 12 substr W3110 YP 492448 1 316407 Proteobacteria
Shigella boydii Sb227 YP 410608 1 300268 Proteobacteria
Escherichia coli Xuzhou21 YP 006315816 1 741093 Proteobacteria
Escherichia coli O55 H7 str CB9615 YP 003502513 1 701177 Proteobacteria
Escherichia coli O157 H7 str EDL933 NP 290930 1 155864 Proteobacteria
Escherichia coli O111 H str 11128 YP 003237436 1 585396 Proteobacteria
Escherichia coli O83 H1 str NRG 857C YP 006122694 1 685038 Proteobacteria
Escherichia coli O103 H2 str 12009 YP 003224908 1 585395 Proteobacteria
Escherichia coli O26 H11 str 11368 YP 003232373 1 573235 Proteobacteria
Escherichia coli O7 K1 str CE10 YP 006147006 1 1072459 Proteobacteria
Shigella flexneri 5 str 8401 YP 691501 1 373384 Proteobacteria
Shigella flexneri 2a str 2457T NP 839602 1 198215 Proteobacteria
Shigella flexneri 2002017 YP 005729797 1 591020 Proteobacteria

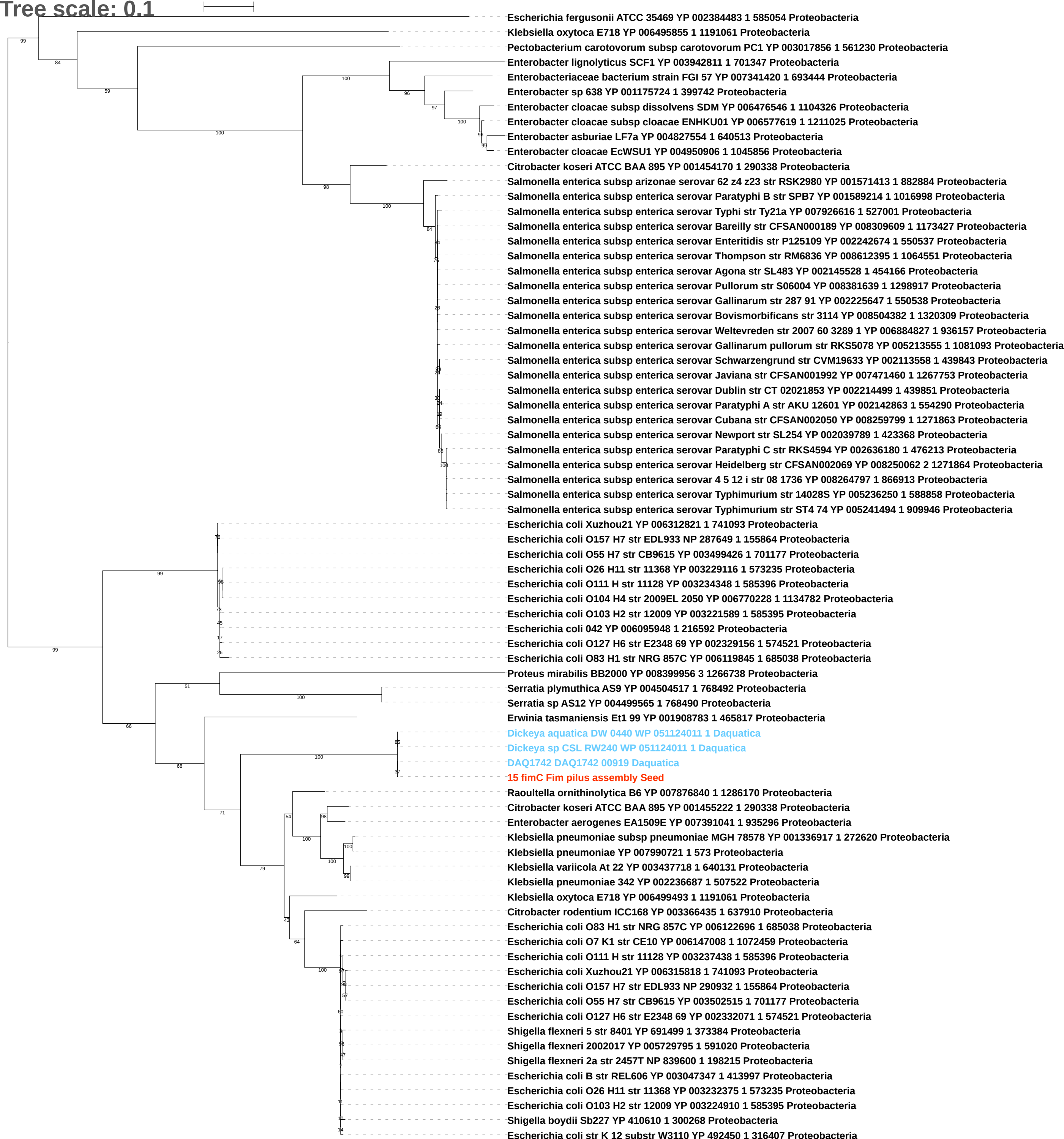

Tree scale: 0.1
Escherichia fergusonii ATCC 35469 YP 002384483 1 585054 Proteobacteria
Klebsiella oxytoca E718 YP 006495855 1 1191061 Proteobacteria
Pectobacterium carotovorum subsp carotovorum PC1 YP 003017856 1 561230 Proteobacteria
Enterobacter lignolyticus SCF1 YP 003942811 1 701347 Proteobacteria
Enterobacteriaceae bacterium strain FGI 57 YP 007341420 1 693444 Proteobacteria
Enterobacter sp 638 YP 001175724 1 399742 Proteobacteria
Enterobacter cloacae subsp dissolvens SDM YP 006476546 1 1104326 Proteobacteria
Enterobacter cloacae subsp cloacae ENHKU01 YP 006577619 1 1211025 Proteobacteria
Enterobacter asburiae LF7a YP 004827554 1 640513 Proteobacteria
Enterobacter cloacae EcWSU1 YP 004950906 1 1045856 Proteobacteria
Citrobacter koseri ATCC BAA 895 YP 001454170 1 290338 Proteobacteria
Salmonella enterica subsp arizonae serovar 62 z4 z23 str RSK2980 YP 001571413 1 882884 Proteobacteria
Salmonella enterica subsp enterica serovar Paratyphi B str SPB7 YP 001589214 1 1016998 Proteobacteria
Salmonella enterica subsp enterica serovar Typhi str Ty21a YP 007926616 1 527001 Proteobacteria
Salmonella enterica subsp enterica serovar Bareilly str CFSAN000189 YP 008309609 1 1173427 Proteobacteria
Salmonella enterica subsp enterica serovar Enteritidis str P125109 YP 002242674 1 550537 Proteobacteria
Salmonella enterica subsp enterica serovar Thompson str RM6836 YP 008612395 1 1064551 Proteobacteria
Salmonella enterica subsp enterica serovar Agona str SL483 YP 002145528 1 454166 Proteobacteria
Salmonella enterica subsp enterica serovar Pullorum str S06004 YP 008381639 1 1298917 Proteobacteria
Salmonella enterica subsp enterica serovar Gallinarum str 287 91 YP 002225647 1 550538 Proteobacteria
Salmonella enterica subsp enterica serovar Bovismorbificans str 3114 YP 008504382 1 1320309 Proteobacteria
Salmonella enterica subsp enterica serovar Weltevreden str 2007 60 3289 1 YP 006884827 1 936157 Proteobacteria
Salmonella enterica subsp enterica serovar Gallinarum pullorum str RKS5078 YP 005213555 1 1081093 Proteobacteria
Salmonella enterica subsp enterica serovar Schwarzengrund str CVM19633 YP 002113558 1 439843 Proteobacteria
Salmonella enterica subsp enterica serovar Javiana str CFSAN001992 YP 007471460 1 1267753 Proteobacteria
Salmonella enterica subsp enterica serovar Dublin str CT 02021853 YP 002214499 1 439851 Proteobacteria
Salmonella enterica subsp enterica serovar Paratyphi A str AKU 12601 YP 002142863 1 554290 Proteobacteria
Salmonella enterica subsp enterica serovar Cubana str CFSAN002050 YP 008259799 1 1271863 Proteobacteria
Salmonella enterica subsp enterica serovar Newport str SL254 YP 002039789 1 423368 Proteobacteria
Salmonella enterica subsp enterica serovar Paratyphi C str RKS4594 YP 002636180 1 476213 Proteobacteria
Salmonella enterica subsp enterica serovar Heidelberg str CFSAN002069 YP 008250062 2 1271864 Proteobacteria
Salmonella enterica subsp enterica serovar 4 5 12 i str 08 1736 YP 008264797 1 866913 Proteobacteria
Salmonella enterica subsp enterica serovar Typhimurium str 14028S YP 005236250 1 588858 Proteobacteria
Salmonella enterica subsp enterica serovar Typhimurium str ST4 74 YP 005241494 1 909946 Proteobacteria
Escherichia coli Xuzhou21 YP 006312821 1 741093 Proteobacteria
Escherichia coli O157 H7 str EDL933 NP 287649 1 155864 Proteobacteria
Escherichia coli O55 H7 str CB9615 YP 003499426 1 701177 Proteobacteria
Escherichia coli O26 H11 str 11368 YP 003229116 1 573235 Proteobacteria
Escherichia coli O111 H str 11128 YP 003234348 1 585396 Proteobacteria
Escherichia coli O104 H4 str 2009EL 2050 YP 006770228 1 1134782 Proteobacteria
Escherichia coli O103 H2 str 12009 YP 003221589 1 585395 Proteobacteria
Escherichia coli 042 YP 006095948 1 216592 Proteobacteria
Escherichia coli O127 H6 str E2348 69 YP 002329156 1 574521 Proteobacteria
Escherichia coli O83 H1 str NRG 857C YP 006119845 1 685038 Proteobacteria
Proteus mirabilis BB2000 YP 008399956 3 1266738 Proteobacteria
Serratia plymuthica AS9 YP 004504517 1 768492 Proteobacteria
Serratia sp AS12 YP 004499565 1 768490 Proteobacteria
Erwinia tasmaniensis Et1 99 YP 001908783 1 465817 Proteobacteria
Dickeya aquatica DW 0440 WP 051124011 1 Daquatica
Dickeya sp CSL RW240 WP 051124011 1 Daquatica
DAQ1742 DAQ1742 00919 Daquatica
15 fimC Fim pilus assembly Seed
Raoultella ornithinolytica B6 YP 007876840 1 1286170 Proteobacteria
Citrobacter koseri ATCC BAA 895 YP 001455222 1 290338 Proteobacteria
Enterobacter aerogenes EA1509E YP 007391041 1 935296 Proteobacteria
Klebsiella pneumoniae subsp pneumoniae MGH 78578 YP 001336917 1 272620 Proteobacteria
Klebsiella pneumoniae YP 007990721 1 573 Proteobacteria
Klebsiella variicola At 22 YP 003437718 1 640131 Proteobacteria
Klebsiella pneumoniae 342 YP 002236687 1 507522 Proteobacteria
Klebsiella oxytoca E718 YP 006499493 1 1191061 Proteobacteria
Citrobacter rodentium ICC168 YP 003366435 1 637910 Proteobacteria
Escherichia coli O83 H1 str NRG 857C YP 006122696 1 685038 Proteobacteria
Escherichia coli O7 K1 str CE10 YP 006147008 1 1072459 Proteobacteria
Escherichia coli O111 H str 11128 YP 003237438 1 585396 Proteobacteria
Escherichia coli Xuzhou21 YP 006315818 1 741093 Proteobacteria
Escherichia coli O157 H7 str EDL933 NP 290932 1 155864 Proteobacteria
Escherichia coli O55 H7 str CB9615 YP 003502515 1 701177 Proteobacteria
Escherichia coli O127 H6 str E2348 69 YP 002332071 1 574521 Proteobacteria
Shigella flexneri 5 str 8401 YP 691499 1 373384 Proteobacteria
Shigella flexneri 2002017 YP 005729795 1 591020 Proteobacteria
Shigella flexneri 2a str 2457T NP 839600 1 198215 Proteobacteria
Escherichia coli B str REL606 YP 003047347 1 413997 Proteobacteria
Escherichia coli O26 H11 str 11368 YP 003232375 1 573235 Proteobacteria
Escherichia coli O103 H2 str 12009 YP 003224910 1 585395 Proteobacteria
Shigella boydii Sb227 YP 410610 1 300268 Proteobacteria
Escherichia coli str K 12 substr W3110 YP 492450 1 316407 Proteobacteria

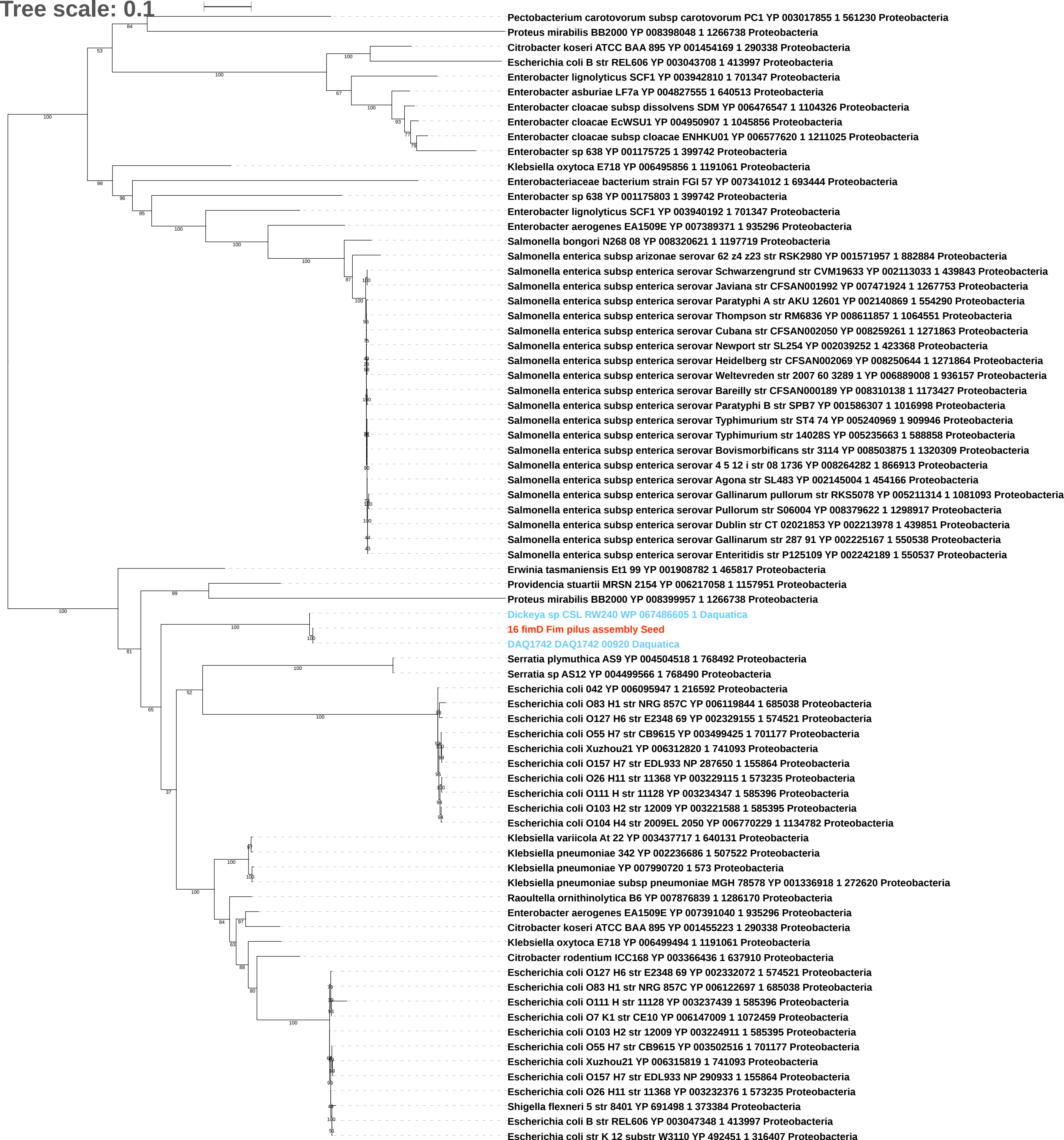
Tree scale: 0.1
Pectobacterium carotovorum subsp carotovorum PC1 YP 003017855 1 561230 Proteobacteria
Proteus mirabilis BB2000 YP 008398048 1 1266738 Proteobacteria
Citrobacter koseri ATCC BAA 895 YP 001454169 1 290338 Proteobacteria
Escherichia coli B str REL606 YP 003043708 1 413997 Proteobacteria
Enterobacter lignolyticus SCF1 YP 003942810 1 701347 Proteobacteria
Enterobacter asburiae LF7a YP 004827555 1 640513 Proteobacteria
Enterobacter cloacae subsp dissolvens SDM YP 006476547 1 1104326 Proteobacteria
Enterobacter cloacae EcWSU1 YP 004950907 1 1045856 Proteobacteria
Enterobacter cloacae subsp cloacae ENHKU01 YP 006577620 1 1211025 Proteobacteria
Enterobacter sp 638 YP 001175725 1 399742 Proteobacteria
Klebsiella oxytoca E718 YP 006495856 1 1191061 Proteobacteria
Enterobacteriaceae bacterium strain FGI 57 YP 007341012 1 693444 Proteobacteria
Enterobacter sp 638 YP 001175803 1 399742 Proteobacteria
Enterobacter lignolyticus SCF1 YP 003940192 1 701347 Proteobacteria
Enterobacter aerogenes EA1509E YP 007389371 1 935296 Proteobacteria
Salmonella bongori N268 08 YP 008320621 1 1197719 Proteobacteria
Salmonella enterica subsp arizonae serovar 62 z4 z23 str RSK2980 YP 001571957 1 882884 Proteobacteria
Salmonella enterica subsp enterica serovar Schwarzengrund str CVM19633 YP 002113033 1 439843 Proteobacteria
Salmonella enterica subsp enterica serovar Javiana str CFSAN001992 YP 007471924 1 1267753 Proteobacteria
Salmonella enterica subsp enterica serovar Paratyphi A str AKU 12601 YP 002140869 1 554290 Proteobacteria
Salmonella enterica subsp enterica serovar Thompson str RM6836 YP 008611857 1 1064551 Proteobacteria
Salmonella enterica subsp enterica serovar Cubana str CFSAN002050 YP 008259261 1 1271863 Proteobacteria
Salmonella enterica subsp enterica serovar Newport str SL254 YP 002039252 1 423368 Proteobacteria
Salmonella enterica subsp enterica serovar Heidelberg str CFSAN002069 YP 008250644 1 1271864 Proteobacteria
Salmonella enterica subsp enterica serovar Weltevreden str 2007 60 3289 1 YP 006889008 1 936157 Proteobacteria
Salmonella enterica subsp enterica serovar Bareilly str CFSAN000189 YP 008310138 1 1173427 Proteobacteria
Salmonella enterica subsp enterica serovar Paratyphi B str SPB7 YP 001586307 1 1016998 Proteobacteria
Salmonella enterica subsp enterica serovar Typhimurium str ST4 74 YP 005240969 1 909946 Proteobacteria
Salmonella enterica subsp enterica serovar Typhimurium str 14028S YP 005235663 1 588858 Proteobacteria
Salmonella enterica subsp enterica serovar Bovismorbificans str 3114 YP 008503875 1 1320309 Proteobacteria
Salmonella enterica subsp enterica serovar 4 5 12 i str 08 1736 YP 008264282 1 866913 Proteobacteria
Salmonella enterica subsp enterica serovar Agona str SL483 YP 002145004 1 454166 Proteobacteria
Salmonella enterica subsp enterica serovar Gallinarum pullorum str RKS5078 YP 005211314 1 1081093 Proteobacteria
Salmonella enterica subsp enterica serovar Pullorum str S06004 YP 008379622 1 1298917 Proteobacteria
Salmonella enterica subsp enterica serovar Dublin str CT 02021853 YP 002213978 1 439851 Proteobacteria
Salmonella enterica subsp enterica serovar Gallinarum str 287 91 YP 002225167 1 550538 Proteobacteria
Salmonella enterica subsp enterica serovar Enteritidis str P125109 YP 002242189 1 550537 Proteobacteria
Erwinia tasmaniensis Et1 99 YP 001908782 1 465817 Proteobacteria
Providencia stuartii MRSN 2154 YP 006217058 1 1157951 Proteobacteria
Proteus mirabilis BB2000 YP 008399957 1 1266738 Proteobacteria
Dickeya sp CSL RW240 WP 067486605 1 Daquatica
16 fimD Fim pilus assembly Seed
DAQ1742 DAQ1742 00920 Daquatica
Serratia plymuthica AS9 YP 004504518 1 768492 Proteobacteria
Serratia sp AS12 YP 004499566 1 768490 Proteobacteria
Escherichia coli 042 YP 006095947 1 216592 Proteobacteria
Escherichia coli O83 H1 str NRG 857C YP 006119844 1 685038 Proteobacteria
Escherichia coli O127 H6 str E2348 69 YP 002329155 1 574521 Proteobacteria
Escherichia coli O55 H7 str CB9615 YP 003499425 1 701177 Proteobacteria
Escherichia coli Xuzhou21 YP 006312820 1 741093 Proteobacteria
Escherichia coli O157 H7 str EDL933 NP 287650 1 155864 Proteobacteria
Escherichia coli O26 H11 str 11368 YP 003229115 1 573235 Proteobacteria
Escherichia coli O111 H str 11128 YP 003234347 1 585396 Proteobacteria
Escherichia coli O103 H2 str 12009 YP 003221588 1 585395 Proteobacteria
Escherichia coli O104 H4 str 2009EL 2050 YP 006770229 1 1134782 Proteobacteria
Klebsiella variicola At 22 YP 003437717 1 640131 Proteobacteria
Klebsiella pneumoniae 342 YP 002236686 1 507522 Proteobacteria
Klebsiella pneumoniae YP 007990720 1 573 Proteobacteria
Klebsiella pneumoniae subsp pneumoniae MGH 78578 YP 001336918 1 272620 Proteobacteria
Raoultella ornithinolytica B6 YP 007876839 1 1286170 Proteobacteria
Enterobacter aerogenes EA1509E YP 007391040 1 935296 Proteobacteria
Citrobacter koseri ATCC BAA 895 YP 001455223 1 290338 Proteobacteria
Klebsiella oxytoca E718 YP 006499494 1 1191061 Proteobacteria
Citrobacter rodentium ICC168 YP 003366436 1 637910 Proteobacteria
Escherichia coli O127 H6 str E2348 69 YP 002332072 1 574521 Proteobacteria
Escherichia coli O83 H1 str NRG 857C YP 006122697 1 685038 Proteobacteria
Escherichia coli O111 H str 11128 YP 003237439 1 585396 Proteobacteria
Escherichia coli O7 K1 str CE10 YP 006147009 1 1072459 Proteobacteria
Escherichia coli O103 H2 str 12009 YP 003224911 1 585395 Proteobacteria
Escherichia coli O55 H7 str CB9615 YP 003502516 1 701177 Proteobacteria
Escherichia coli Xuzhou21 YP 006315819 1 741093 Proteobacteria
Escherichia coli O157 H7 str EDL933 NP 290933 1 155864 Proteobacteria
Escherichia coli O26 H11 str 11368 YP 003232376 1 573235 Proteobacteria
Shigella flexneri 5 str 8401 YP 691498 1 373384 Proteobacteria
Escherichia coli B str REL606 YP 003047348 1 413997 Proteobacteria
Escherichia coli str K 12 substr W3110 YP 492451 1 316407 Proteobacteria

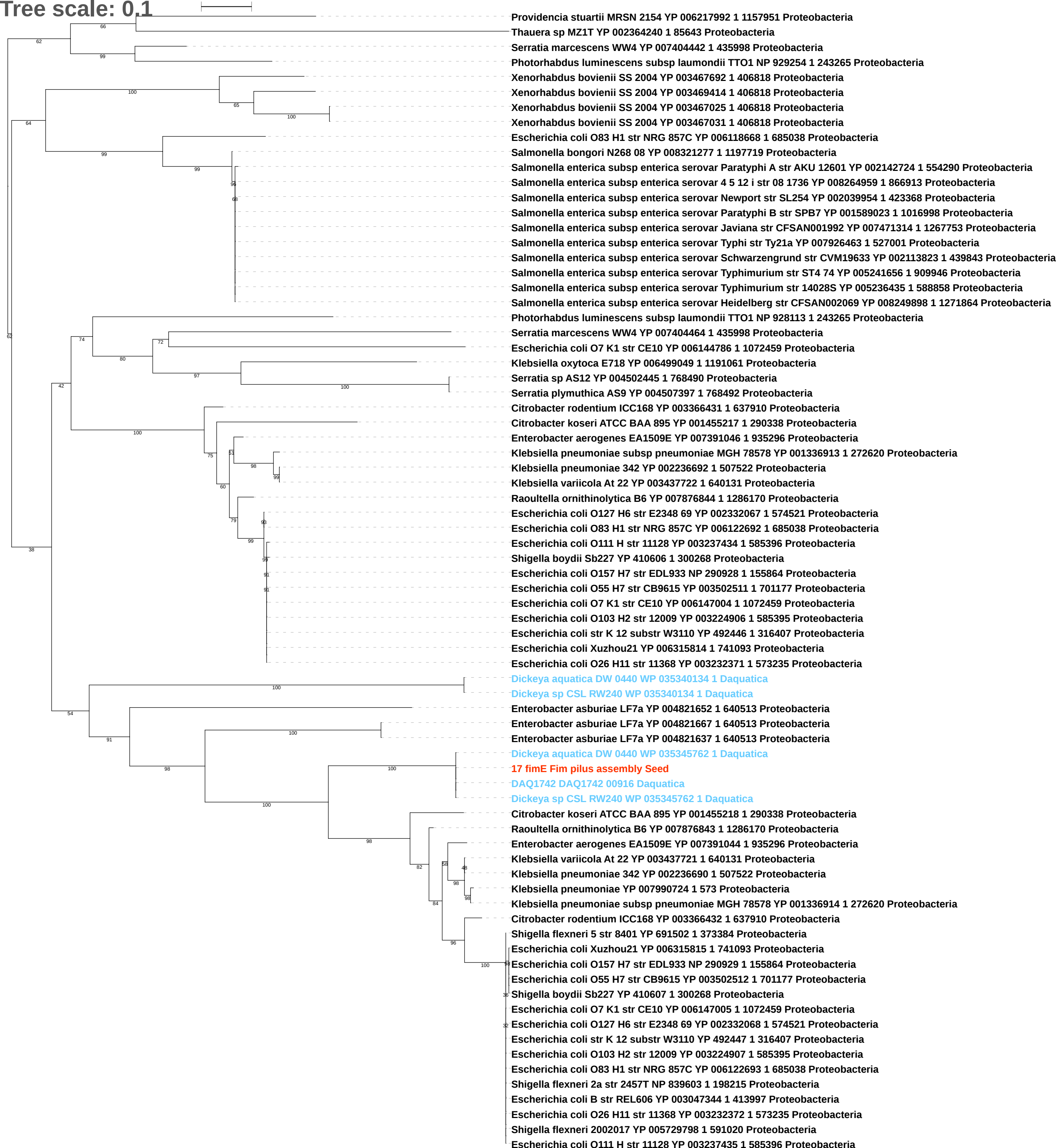
Tree scale: 0.1
Providencia stuartii MRSN 2154 YP 006217992 1 1157951 Proteobacteria
Thauera sp MZ1T YP 002364240 1 85643 Proteobacteria
Serratia marcescens WW4 YP 007404442 1 435998 Proteobacteria
Photorhabdus luminescens subsp laumondii TTO1 NP 929254 1 243265 Proteobacteria
Xenorhabdus bovienii SS 2004 YP 003467692 1 406818 Proteobacteria
Xenorhabdus bovienii SS 2004 YP 003469414 1 406818 Proteobacteria
Xenorhabdus bovienii SS 2004 YP 003467025 1 406818 Proteobacteria
Xenorhabdus bovienii SS 2004 YP 003467031 1 406818 Proteobacteria
Escherichia coli O83 H1 str NRG 857C YP 006118668 1 685038 Proteobacteria
Salmonella bongori N268 08 YP 008321277 1 1197719 Proteobacteria
Salmonella enterica subsp enterica serovar Paratyphi A str AKU 12601 YP 002142724 1 554290 Proteobacteria
Salmonella enterica subsp enterica serovar 4 5 12 i str 08 1736 YP 008264959 1 866913 Proteobacteria
Salmonella enterica subsp enterica serovar Newport str SL254 YP 002039954 1 423368 Proteobacteria
Salmonella enterica subsp enterica serovar Paratyphi B str SPB7 YP 001589023 1 1016998 Proteobacteria
Salmonella enterica subsp enterica serovar Javiana str CFSAN001992 YP 007471314 1 1267753 Proteobacteria
Salmonella enterica subsp enterica serovar Typhi str Ty21a YP 007926463 1 527001 Proteobacteria
Salmonella enterica subsp enterica serovar Schwarzengrund str CVM19633 YP 002113823 1 439843 Proteobacteria
Salmonella enterica subsp enterica serovar Typhimurium str ST4 74 YP 005241656 1 909946 Proteobacteria
Salmonella enterica subsp enterica serovar Typhimurium str 14028S YP 005236435 1 588858 Proteobacteria
Salmonella enterica subsp enterica serovar Heidelberg str CFSAN002069 YP 008249898 1 1271864 Proteobacteria
Photorhabdus luminescens subsp laumondii TTO1 NP 928113 1 243265 Proteobacteria
Serratia marcescens WW4 YP 007404464 1 435998 Proteobacteria
Escherichia coli O7 K1 str CE10 YP 006144786 1 1072459 Proteobacteria
Klebsiella oxytoca E718 YP 006499049 1 1191061 Proteobacteria
Serratia sp AS12 YP 004502445 1 768490 Proteobacteria
Serratia plymuthica AS9 YP 004507397 1 768492 Proteobacteria
Citrobacter rodentium ICC168 YP 003366431 1 637910 Proteobacteria
Citrobacter koseri ATCC BAA 895 YP 001455217 1 290338 Proteobacteria
Enterobacter aerogenes EA1509E YP 007391046 1 935296 Proteobacteria
Klebsiella pneumoniae subsp pneumoniae MGH 78578 YP 001336913 1 272620 Proteobacteria
Klebsiella pneumoniae 342 YP 002236692 1 507522 Proteobacteria
Klebsiella variicola At 22 YP 003437722 1 640131 Proteobacteria
Raoultella ornithinolytica B6 YP 007876844 1 1286170 Proteobacteria
Escherichia coli O127 H6 str E2348 69 YP 002332067 1 574521 Proteobacteria
Escherichia coli O83 H1 str NRG 857C YP 006122692 1 685038 Proteobacteria
Escherichia coli O111 H str 11128 YP 003237434 1 585396 Proteobacteria
Shigella boydii Sb227 YP 410606 1 300268 Proteobacteria
Escherichia coli O157 H7 str EDL933 NP 290928 1 155864 Proteobacteria
Escherichia coli O55 H7 str CB9615 YP 003502511 1 701177 Proteobacteria
Escherichia coli O7 K1 str CE10 YP 006147004 1 1072459 Proteobacteria
Escherichia coli O103 H2 str 12009 YP 003224906 1 585395 Proteobacteria
Escherichia coli str K 12 substr W3110 YP 492446 1 316407 Proteobacteria
Escherichia coli Xuzhou21 YP 006315814 1 741093 Proteobacteria
Escherichia coli O26 H11 str 11368 YP 003232371 1 573235 Proteobacteria
Dickeya aquatica DW 0440 WP 035340134 1 Daquatica
Dickeya sp CSL RW240 WP 035340134 1 Daquatica
Enterobacter asburiae LF7a YP 004821652 1 640513 Proteobacteria
Enterobacter asburiae LF7a YP 004821667 1 640513 Proteobacteria
Enterobacter asburiae LF7a YP 004821637 1 640513 Proteobacteria
Dickeya aquatica DW 0440 WP 035345762 1 Daquatica
17 fimE Fim pilus assembly Seed
DAQ1742 DAQ1742 00916 Daquatica
Dickeya sp CSL RW240 WP 035345762 1 Daquatica
Citrobacter koseri ATCC BAA 895 YP 001455218 1 290338 Proteobacteria
Raoultella ornithinolytica B6 YP 007876843 1 1286170 Proteobacteria
Enterobacter aerogenes EA1509E YP 007391044 1 935296 Proteobacteria
Klebsiella variicola At 22 YP 003437721 1 640131 Proteobacteria
Klebsiella pneumoniae 342 YP 002236690 1 507522 Proteobacteria
Klebsiella pneumoniae YP 007990724 1 573 Proteobacteria
Klebsiella pneumoniae subsp pneumoniae MGH 78578 YP 001336914 1 272620 Proteobacteria
Citrobacter rodentium ICC168 YP 003366432 1 637910 Proteobacteria
Shigella flexneri 5 str 8401 YP 691502 1 373384 Proteobacteria
Escherichia coli Xuzhou21 YP 006315815 1 741093 Proteobacteria
Escherichia coli O157 H7 str EDL933 NP 290929 1 155864 Proteobacteria
Escherichia coli O55 H7 str CB9615 YP 003502512 1 701177 Proteobacteria
Shigella boydii Sb227 YP 410607 1 300268 Proteobacteria
Escherichia coli O7 K1 str CE10 YP 006147005 1 1072459 Proteobacteria
Escherichia coli O127 H6 str E2348 69 YP 002332068 1 574521 Proteobacteria
Escherichia coli str K 12 substr W3110 YP 492447 1 316407 Proteobacteria
Escherichia coli O103 H2 str 12009 YP 003224907 1 585395 Proteobacteria
Escherichia coli O83 H1 str NRG 857C YP 006122693 1 685038 Proteobacteria
Shigella flexneri 2a str 2457T NP 839603 1 198215 Proteobacteria
Escherichia coli B str REL606 YP 003047344 1 413997 Proteobacteria
Escherichia coli O26 H11 str 11368 YP 003232372 1 573235 Proteobacteria
Shigella flexneri 2002017 YP 005729798 1 591020 Proteobacteria
Escherichia coli O111 H str 11128 YP 003237435 1 585396 Proteobacteria

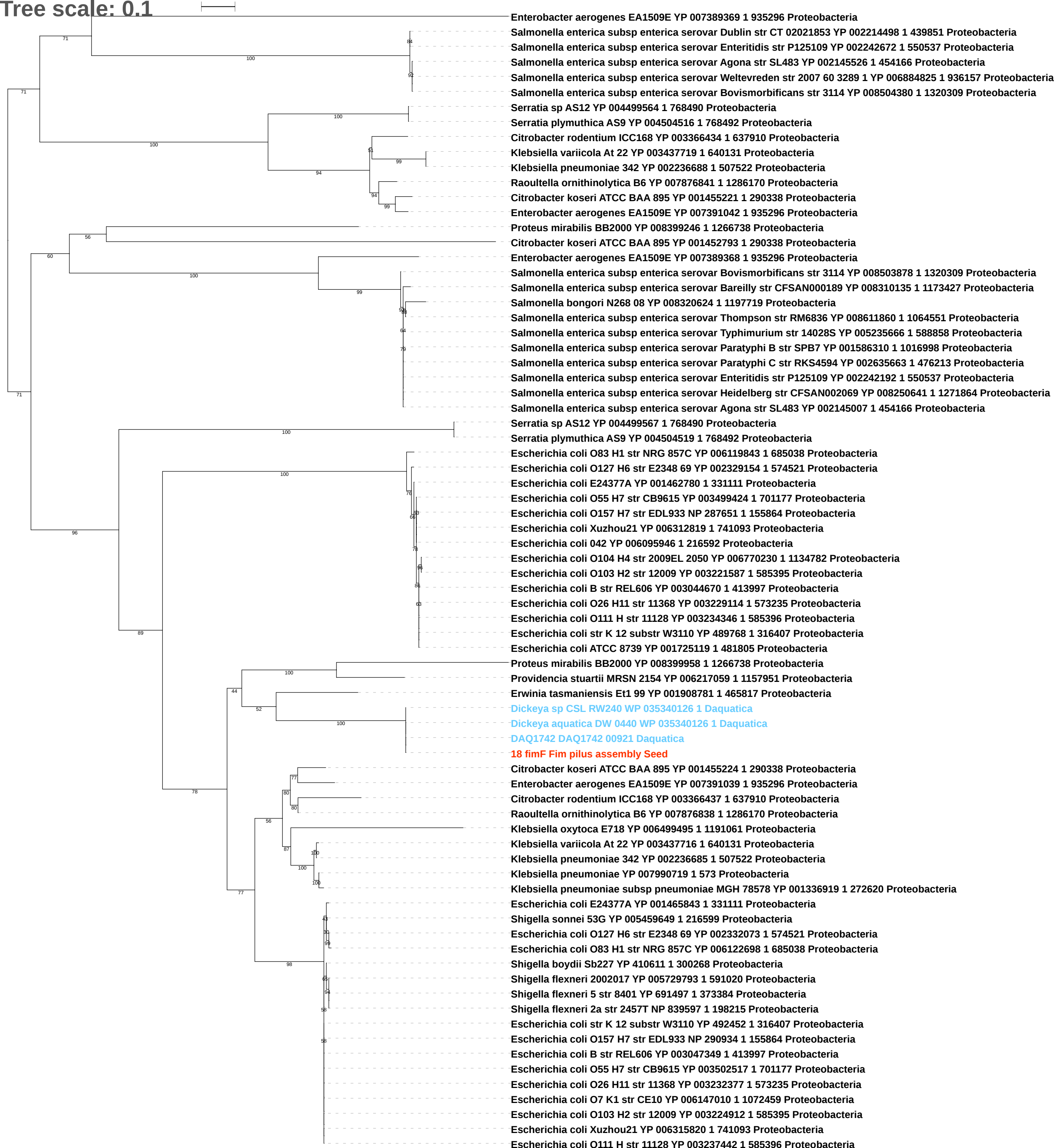
Tree scale: 0.1
Enterobacter aerogenes EA1509E YP 007389369 1 935296 Proteobacteria
Salmonella enterica subsp enterica serovar Dublin str CT 02021853 YP 002214498 1 439851 Proteobacteria
Salmonella enterica subsp enterica serovar Enteritidis str P125109 YP 002242672 1 550537 Proteobacteria
Salmonella enterica subsp enterica serovar Agona str SL483 YP 002145526 1 454166 Proteobacteria
Salmonella enterica subsp enterica serovar Weltevreden str 2007 60 3289 1 YP 006884825 1 936157 Proteobacteria
Salmonella enterica subsp enterica serovar Bovismorbificans str 3114 YP 008504380 1 1320309 Proteobacteria
Serratia sp AS12 YP 004499564 1 768490 Proteobacteria
Serratia plymuthica AS9 YP 004504516 1 768492 Proteobacteria
Citrobacter rodentium ICC168 YP 003366434 1 637910 Proteobacteria
Klebsiella variicola At 22 YP 003437719 1 640131 Proteobacteria
Klebsiella pneumoniae 342 YP 002236688 1 507522 Proteobacteria
Raoultella ornithinolytica B6 YP 007876841 1 1286170 Proteobacteria
Citrobacter koseri ATCC BAA 895 YP 001455221 1 290338 Proteobacteria
Enterobacter aerogenes EA1509E YP 007391042 1 935296 Proteobacteria
Proteus mirabilis BB2000 YP 008399246 1 1266738 Proteobacteria
Citrobacter koseri ATCC BAA 895 YP 001452793 1 290338 Proteobacteria
Enterobacter aerogenes EA1509E YP 007389368 1 935296 Proteobacteria
Salmonella enterica subsp enterica serovar Bovismorbificans str 3114 YP 008503878 1 1320309 Proteobacteria
Salmonella enterica subsp enterica serovar Bareilly str CFSAN000189 YP 008310135 1 1173427 Proteobacteria
Salmonella bongori N268 08 YP 008320624 1 1197719 Proteobacteria
Salmonella enterica subsp enterica serovar Thompson str RM6836 YP 008611860 1 1064551 Proteobacteria
Salmonella enterica subsp enterica serovar Typhimurium str 14028S YP 005235666 1 588858 Proteobacteria
Salmonella enterica subsp enterica serovar Paratyphi B str SPB7 YP 001586310 1 1016998 Proteobacteria
Salmonella enterica subsp enterica serovar Paratyphi C str RKS4594 YP 002635663 1 476213 Proteobacteria
Salmonella enterica subsp enterica serovar Enteritidis str P125109 YP 002242192 1 550537 Proteobacteria
Salmonella enterica subsp enterica serovar Heidelberg str CFSAN002069 YP 008250641 1 1271864 Proteobacteria
Salmonella enterica subsp enterica serovar Agona str SL483 YP 002145007 1 454166 Proteobacteria
Serratia sp AS12 YP 004499567 1 768490 Proteobacteria
Serratia plymuthica AS9 YP 004504519 1 768492 Proteobacteria
Escherichia coli O83 H1 str NRG 857C YP 006119843 1 685038 Proteobacteria
Escherichia coli O127 H6 str E2348 69 YP 002329154 1 574521 Proteobacteria
Escherichia coli E24377A YP 001462780 1 331111 Proteobacteria
Escherichia coli O55 H7 str CB9615 YP 003499424 1 701177 Proteobacteria
Escherichia coli O157 H7 str EDL933 NP 287651 1 155864 Proteobacteria
Escherichia coli Xuzhou21 YP 006312819 1 741093 Proteobacteria
Escherichia coli 042 YP 006095946 1 216592 Proteobacteria
Escherichia coli O104 H4 str 2009EL 2050 YP 006770230 1 1134782 Proteobacteria
Escherichia coli O103 H2 str 12009 YP 003221587 1 585395 Proteobacteria
Escherichia coli B str REL606 YP 003044670 1 413997 Proteobacteria
Escherichia coli O26 H11 str 11368 YP 003229114 1 573235 Proteobacteria
Escherichia coli O111 H str 11128 YP 003234346 1 585396 Proteobacteria
Escherichia coli str K 12 substr W3110 YP 489768 1 316407 Proteobacteria
Escherichia coli ATCC 8739 YP 001725119 1 481805 Proteobacteria
Proteus mirabilis BB2000 YP 008399958 1 1266738 Proteobacteria
Providencia stuartii MRSN 2154 YP 006217059 1 1157951 Proteobacteria
Erwinia tasmaniensis Et1 99 YP 001908781 1 465817 Proteobacteria
Dickeya sp CSL RW240 WP 035340126 1 Daquatica
Dickeya aquatica DW 0440 WP 035340126 1 Daquatica
DAQ1742 DAQ1742 00921 Daquatica
18 fimF Fim pilus assembly Seed
Citrobacter koseri ATCC BAA 895 YP 001455224 1 290338 Proteobacteria
Enterobacter aerogenes EA1509E YP 007391039 1 935296 Proteobacteria
Citrobacter rodentium ICC168 YP 003366437 1 637910 Proteobacteria
Raoultella ornithinolytica B6 YP 007876838 1 1286170 Proteobacteria
Klebsiella oxytoca E718 YP 006499495 1 1191061 Proteobacteria
Klebsiella variicola At 22 YP 003437716 1 640131 Proteobacteria
Klebsiella pneumoniae 342 YP 002236685 1 507522 Proteobacteria
Klebsiella pneumoniae YP 007990719 1 573 Proteobacteria
Klebsiella pneumoniae subsp pneumoniae MGH 78578 YP 001336919 1 272620 Proteobacteria
Escherichia coli E24377A YP 001465843 1 331111 Proteobacteria
Shigella sonnei 53G YP 005459649 1 216599 Proteobacteria
Escherichia coli O127 H6 str E2348 69 YP 002332073 1 574521 Proteobacteria
Escherichia coli O83 H1 str NRG 857C YP 006122698 1 685038 Proteobacteria
Shigella boydii Sb227 YP 410611 1 300268 Proteobacteria
Shigella flexneri 2002017 YP 005729793 1 591020 Proteobacteria
Shigella flexneri 5 str 8401 YP 691497 1 373384 Proteobacteria
Shigella flexneri 2a str 2457T NP 839597 1 198215 Proteobacteria
Escherichia coli str K 12 substr W3110 YP 492452 1 316407 Proteobacteria
Escherichia coli O157 H7 str EDL933 NP 290934 1 155864 Proteobacteria
Escherichia coli B str REL606 YP 003047349 1 413997 Proteobacteria
Escherichia coli O55 H7 str CB9615 YP 003502517 1 701177 Proteobacteria
Escherichia coli O26 H11 str 11368 YP 003232377 1 573235 Proteobacteria
Escherichia coli O7 K1 str CE10 YP 006147010 1 1072459 Proteobacteria
Escherichia coli O103 H2 str 12009 YP 003224912 1 585395 Proteobacteria
Escherichia coli Xuzhou21 YP 006315820 1 741093 Proteobacteria
Escherichia coli O111 H str 11128 YP 003237442 1 585396 Proteobacteria

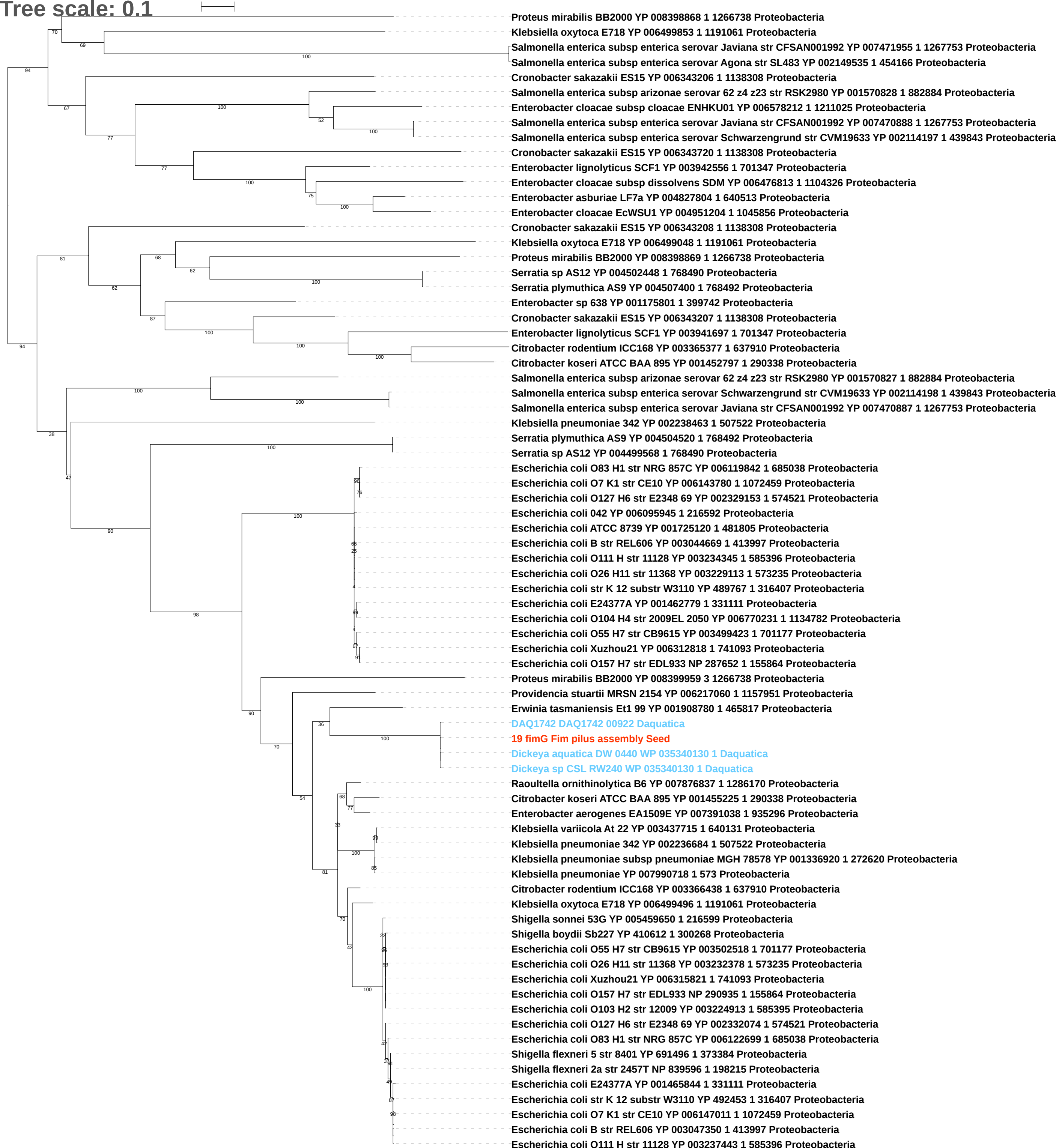

Tree scale: 0.1
Proteus mirabilis BB2000 YP 008398868 1 1266738 Proteobacteria
Klebsiella oxytoca E718 YP 006499853 1 1191061 Proteobacteria
Salmonella enterica subsp enterica serovar Javiana str CFSAN001992 YP 007471955 1 1267753 Proteobacteria
Salmonella enterica subsp enterica serovar Agona str SL483 YP 002149535 1 454166 Proteobacteria
Cronobacter sakazakii ES15 YP 006343206 1 1138308 Proteobacteria
Salmonella enterica subsp arizonae serovar 62 z4 z23 str RSK2980 YP 001570828 1 882884 Proteobacteria
Enterobacter cloacae subsp cloacae ENHKU01 YP 006578212 1 1211025 Proteobacteria
Salmonella enterica subsp enterica serovar Javiana str CFSAN001992 YP 007470888 1 1267753 Proteobacteria
Salmonella enterica subsp enterica serovar Schwarzengrund str CVM19633 YP 002114197 1 439843 Proteobacteria
Cronobacter sakazakii ES15 YP 006343720 1 1138308 Proteobacteria
Enterobacter lignolyticus SCF1 YP 003942556 1 701347 Proteobacteria
Enterobacter cloacae subsp dissolvens SDM YP 006476813 1 1104326 Proteobacteria
Enterobacter asburiae LF7a YP 004827804 1 640513 Proteobacteria
Enterobacter cloacae EcWSU1 YP 004951204 1 1045856 Proteobacteria
Cronobacter sakazakii ES15 YP 006343208 1 1138308 Proteobacteria
Klebsiella oxytoca E718 YP 006499048 1 1191061 Proteobacteria
Proteus mirabilis BB2000 YP 008398869 1 1266738 Proteobacteria
Serratia sp AS12 YP 004502448 1 768490 Proteobacteria
Serratia plymuthica AS9 YP 004507400 1 768492 Proteobacteria
Enterobacter sp 638 YP 001175801 1 399742 Proteobacteria
Cronobacter sakazakii ES15 YP 006343207 1 1138308 Proteobacteria
Enterobacter lignolyticus SCF1 YP 003941697 1 701347 Proteobacteria
Citrobacter rodentium ICC168 YP 003365377 1 637910 Proteobacteria
Citrobacter koseri ATCC BAA 895 YP 001452797 1 290338 Proteobacteria
Salmonella enterica subsp arizonae serovar 62 z4 z23 str RSK2980 YP 001570827 1 882884 Proteobacteria
Salmonella enterica subsp enterica serovar Schwarzengrund str CVM19633 YP 002114198 1 439843 Proteobacteria
Salmonella enterica subsp enterica serovar Javiana str CFSAN001992 YP 007470887 1 1267753 Proteobacteria
Klebsiella pneumoniae 342 YP 002238463 1 507522 Proteobacteria
Serratia plymuthica AS9 YP 004504520 1 768492 Proteobacteria
Serratia sp AS12 YP 004499568 1 768490 Proteobacteria
Escherichia coli O83 H1 str NRG 857C YP 006119842 1 685038 Proteobacteria
Escherichia coli O7 K1 str CE10 YP 006143780 1 1072459 Proteobacteria
Escherichia coli O127 H6 str E2348 69 YP 002329153 1 574521 Proteobacteria
Escherichia coli 042 YP 006095945 1 216592 Proteobacteria
Escherichia coli ATCC 8739 YP 001725120 1 481805 Proteobacteria
Escherichia coli B str REL606 YP 003044669 1 413997 Proteobacteria
Escherichia coli O111 H str 11128 YP 003234345 1 585396 Proteobacteria
Escherichia coli O26 H11 str 11368 YP 003229113 1 573235 Proteobacteria
Escherichia coli str K 12 substr W3110 YP 489767 1 316407 Proteobacteria
Escherichia coli E24377A YP 001462779 1 331111 Proteobacteria
Escherichia coli O104 H4 str 2009EL 2050 YP 006770231 1 1134782 Proteobacteria
Escherichia coli O55 H7 str CB9615 YP 003499423 1 701177 Proteobacteria
Escherichia coli Xuzhou21 YP 006312818 1 741093 Proteobacteria
Escherichia coli O157 H7 str EDL933 NP 287652 1 155864 Proteobacteria
Proteus mirabilis BB2000 YP 008399959 3 1266738 Proteobacteria
Providencia stuartii MRSN 2154 YP 006217060 1 1157951 Proteobacteria
Erwinia tasmaniensis Et1 99 YP 001908780 1 465817 Proteobacteria
DAQ1742 DAQ1742 00922 Daquatica
19 fimG Fim pilus assembly Seed
Dickeya aquatica DW 0440 WP 035340130 1 Daquatica
Dickeya sp CSL RW240 WP 035340130 1 Daquatica
Raoultella ornithinolytica B6 YP 007876837 1 1286170 Proteobacteria
Citrobacter koseri ATCC BAA 895 YP 001455225 1 290338 Proteobacteria
Enterobacter aerogenes EA1509E YP 007391038 1 935296 Proteobacteria
Klebsiella variicola At 22 YP 003437715 1 640131 Proteobacteria
Klebsiella pneumoniae 342 YP 002236684 1 507522 Proteobacteria
Klebsiella pneumoniae subsp pneumoniae MGH 78578 YP 001336920 1 272620 Proteobacteria
Klebsiella pneumoniae YP 007990718 1 573 Proteobacteria
Citrobacter rodentium ICC168 YP 003366438 1 637910 Proteobacteria
Klebsiella oxytoca E718 YP 006499496 1 1191061 Proteobacteria
Shigella sonnei 53G YP 005459650 1 216599 Proteobacteria
Shigella boydii Sb227 YP 410612 1 300268 Proteobacteria
Escherichia coli O55 H7 str CB9615 YP 003502518 1 701177 Proteobacteria
Escherichia coli O26 H11 str 11368 YP 003232378 1 573235 Proteobacteria
Escherichia coli Xuzhou21 YP 006315821 1 741093 Proteobacteria
Escherichia coli O157 H7 str EDL933 NP 290935 1 155864 Proteobacteria
Escherichia coli O103 H2 str 12009 YP 003224913 1 585395 Proteobacteria
Escherichia coli O127 H6 str E2348 69 YP 002332074 1 574521 Proteobacteria
Escherichia coli O83 H1 str NRG 857C YP 006122699 1 685038 Proteobacteria
Shigella flexneri 5 str 8401 YP 691496 1 373384 Proteobacteria
Shigella flexneri 2a str 2457T NP 839596 1 198215 Proteobacteria
Escherichia coli E24377A YP 001465844 1 331111 Proteobacteria
Escherichia coli str K 12 substr W3110 YP 492453 1 316407 Proteobacteria
Escherichia coli O7 K1 str CE10 YP 006147011 1 1072459 Proteobacteria
Escherichia coli B str REL606 YP 003047350 1 413997 Proteobacteria
Escherichia coli O111 H str 11128 YP 003237443 1 585396 Proteobacteria

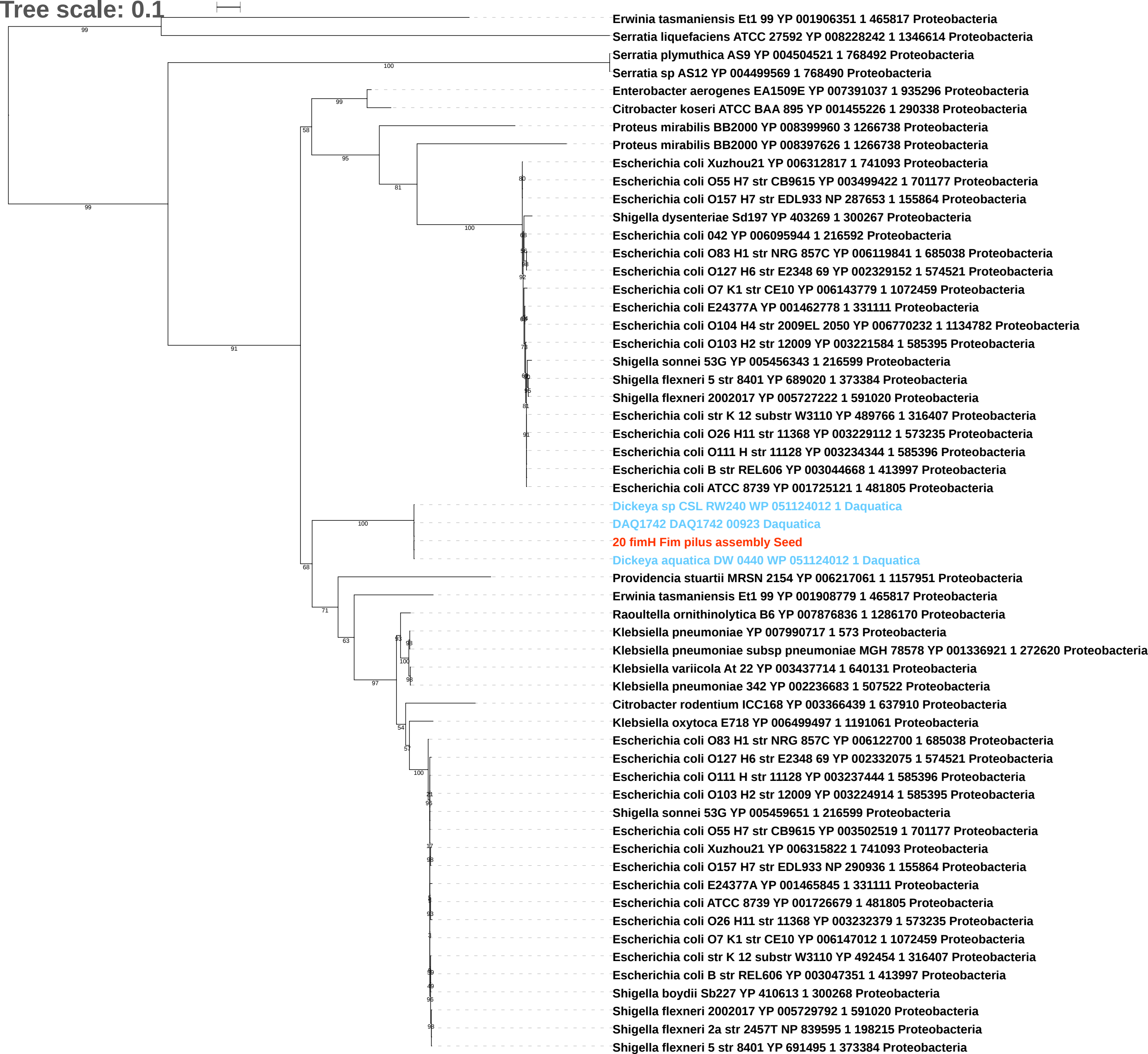
Tree scale: 0.1
Erwinia tasmaniensis Et1 99 YP 001906351 1 465817 Proteobacteria
Serratia liquefaciens ATCC 27592 YP 008228242 1 1346614 Proteobacteria
Serratia plymuthica AS9 YP 004504521 1 768492 Proteobacteria
Serratia sp AS12 YP 004499569 1 768490 Proteobacteria
Enterobacter aerogenes EA1509E YP 007391037 1 935296 Proteobacteria
Citrobacter koseri ATCC BAA 895 YP 001455226 1 290338 Proteobacteria
Proteus mirabilis BB2000 YP 008399960 3 1266738 Proteobacteria
Proteus mirabilis BB2000 YP 008397626 1 1266738 Proteobacteria
Escherichia coli Xuzhou21 YP 006312817 1 741093 Proteobacteria
Escherichia coli O55 H7 str CB9615 YP 003499422 1 701177 Proteobacteria
Escherichia coli O157 H7 str EDL933 NP 287653 1 155864 Proteobacteria
Shigella dysenteriae Sd197 YP 403269 1 300267 Proteobacteria
Escherichia coli 042 YP 006095944 1 216592 Proteobacteria
Escherichia coli O83 H1 str NRG 857C YP 006119841 1 685038 Proteobacteria
Escherichia coli O127 H6 str E2348 69 YP 002329152 1 574521 Proteobacteria
Escherichia coli O7 K1 str CE10 YP 006143779 1 1072459 Proteobacteria
Escherichia coli E24377A YP 001462778 1 331111 Proteobacteria
Escherichia coli O104 H4 str 2009EL 2050 YP 006770232 1 1134782 Proteobacteria
Escherichia coli O103 H2 str 12009 YP 003221584 1 585395 Proteobacteria
Shigella sonnei 53G YP 005456343 1 216599 Proteobacteria
Shigella flexneri 5 str 8401 YP 689020 1 373384 Proteobacteria
Shigella flexneri 2002017 YP 005727222 1 591020 Proteobacteria
Escherichia coli str K 12 substr W3110 YP 489766 1 316407 Proteobacteria
Escherichia coli O26 H11 str 11368 YP 003229112 1 573235 Proteobacteria
Escherichia coli O111 H str 11128 YP 003234344 1 585396 Proteobacteria
Escherichia coli B str REL606 YP 003044668 1 413997 Proteobacteria
Escherichia coli ATCC 8739 YP 001725121 1 481805 Proteobacteria
Dickeya sp CSL RW240 WP 051124012 1 Daquatica
DAQ1742 DAQ1742 00923 Daquatica
20 fimH Fim pilus assembly Seed
Dickeya aquatica DW 0440 WP 051124012 1 Daquatica
Providencia stuartii MRSN 2154 YP 006217061 1 1157951 Proteobacteria
Erwinia tasmaniensis Et1 99 YP 001908779 1 465817 Proteobacteria
Raoultella ornithinolytica B6 YP 007876836 1 1286170 Proteobacteria
Klebsiella pneumoniae YP 007990717 1 573 Proteobacteria
Klebsiella pneumoniae subsp pneumoniae MGH 78578 YP 001336921 1 272620 Proteobacteria
Klebsiella variicola At 22 YP 003437714 1 640131 Proteobacteria
Klebsiella pneumoniae 342 YP 002236683 1 507522 Proteobacteria
Citrobacter rodentium ICC168 YP 003366439 1 637910 Proteobacteria
Klebsiella oxytoca E718 YP 006499497 1 1191061 Proteobacteria
Escherichia coli O83 H1 str NRG 857C YP 006122700 1 685038 Proteobacteria
Escherichia coli O127 H6 str E2348 69 YP 002332075 1 574521 Proteobacteria
Escherichia coli O111 H str 11128 YP 003237444 1 585396 Proteobacteria
Escherichia coli O103 H2 str 12009 YP 003224914 1 585395 Proteobacteria
Shigella sonnei 53G YP 005459651 1 216599 Proteobacteria
Escherichia coli O55 H7 str CB9615 YP 003502519 1 701177 Proteobacteria
Escherichia coli Xuzhou21 YP 006315822 1 741093 Proteobacteria
Escherichia coli O157 H7 str EDL933 NP 290936 1 155864 Proteobacteria
Escherichia coli E24377A YP 001465845 1 331111 Proteobacteria
Escherichia coli ATCC 8739 YP 001726679 1 481805 Proteobacteria
Escherichia coli O26 H11 str 11368 YP 003232379 1 573235 Proteobacteria
Escherichia coli O7 K1 str CE10 YP 006147012 1 1072459 Proteobacteria
Escherichia coli str K 12 substr W3110 YP 492454 1 316407 Proteobacteria
Escherichia coli B str REL606 YP 003047351 1 413997 Proteobacteria
Shigella boydii Sb227 YP 410613 1 300268 Proteobacteria
Shigella flexneri 2002017 YP 005729792 1 591020 Proteobacteria
Shigella flexneri 2a str 2457T NP 839595 1 198215 Proteobacteria
Shigella flexneri 5 str 8401 YP 691495 1 373384 Proteobacteria

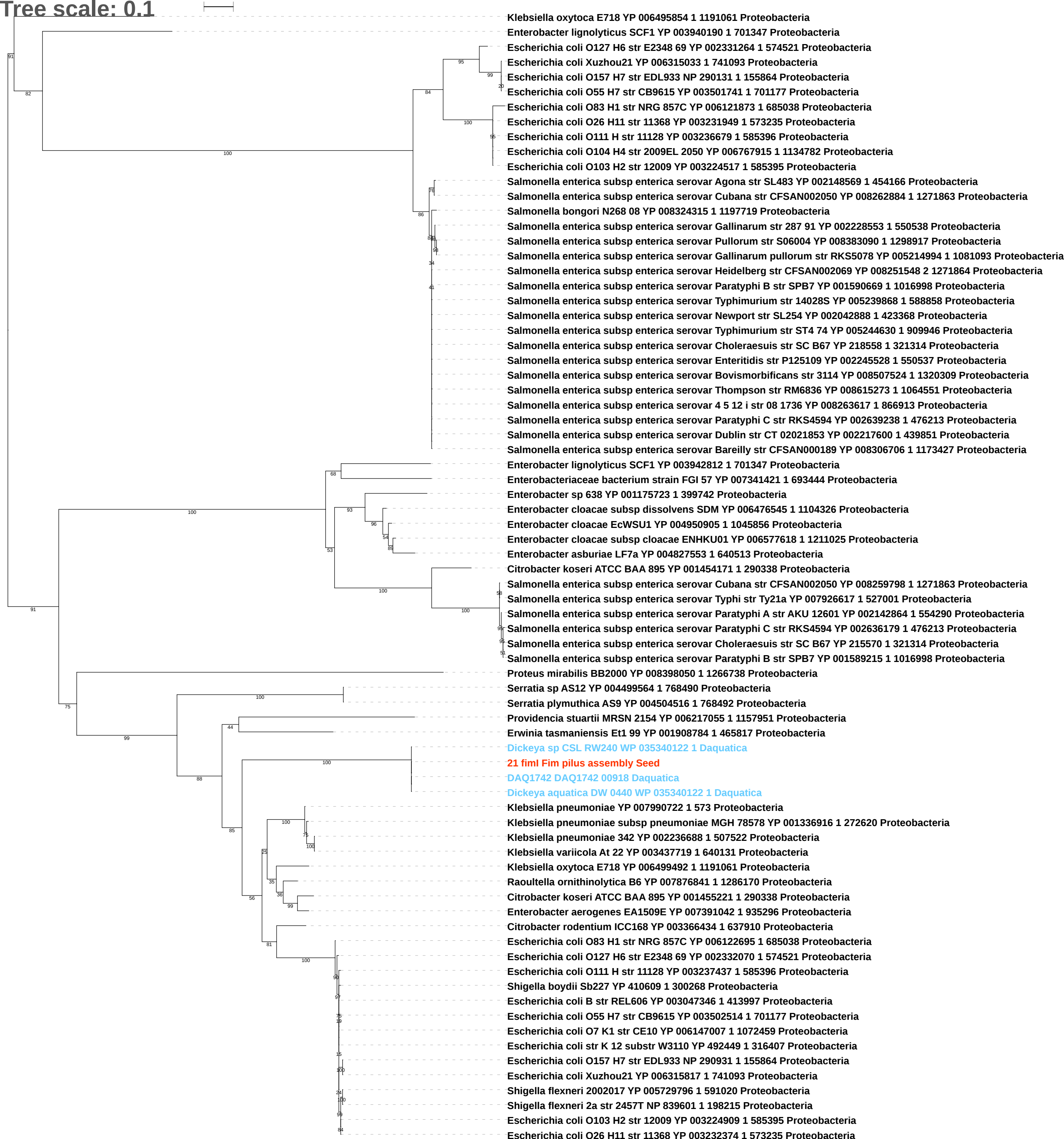
Tree scale: 0.1
Klebsiella oxytoca E718 YP 006495854 1 1191061 Proteobacteria
Enterobacter lignolyticus SCF1 YP 003940190 1 701347 Proteobacteria
Escherichia coli O127 H6 str E2348 69 YP 002331264 1 574521 Proteobacteria
Escherichia coli Xuzhou21 YP 006315033 1 741093 Proteobacteria
Escherichia coli O157 H7 str EDL933 NP 290131 1 155864 Proteobacteria
Escherichia coli O55 H7 str CB9615 YP 003501741 1 701177 Proteobacteria
Escherichia coli O83 H1 str NRG 857C YP 006121873 1 685038 Proteobacteria
Escherichia coli O26 H11 str 11368 YP 003231949 1 573235 Proteobacteria
Escherichia coli O111 H str 11128 YP 003236679 1 585396 Proteobacteria
Escherichia coli O104 H4 str 2009EL 2050 YP 006767915 1 1134782 Proteobacteria
Escherichia coli O103 H2 str 12009 YP 003224517 1 585395 Proteobacteria
Salmonella enterica subsp enterica serovar Agona str SL483 YP 002148569 1 454166 Proteobacteria
Salmonella enterica subsp enterica serovar Cubana str CFSAN002050 YP 008262884 1 1271863 Proteobacteria
Salmonella bongori N268 08 YP 008324315 1 1197719 Proteobacteria
Salmonella enterica subsp enterica serovar Gallinarum str 287 91 YP 002228553 1 550538 Proteobacteria
Salmonella enterica subsp enterica serovar Pullorum str S06004 YP 008383090 1 1298917 Proteobacteria
Salmonella enterica subsp enterica serovar Gallinarum pullorum str RKS5078 YP 005214994 1 1081093 Proteobacteria
Salmonella enterica subsp enterica serovar Heidelberg str CFSAN002069 YP 008251548 2 1271864 Proteobacteria
Salmonella enterica subsp enterica serovar Paratyphi B str SPB7 YP 001590669 1 1016998 Proteobacteria
Salmonella enterica subsp enterica serovar Typhimurium str 14028S YP 005239868 1 588858 Proteobacteria
Salmonella enterica subsp enterica serovar Newport str SL254 YP 002042888 1 423368 Proteobacteria
Salmonella enterica subsp enterica serovar Typhimurium str ST4 74 YP 005244630 1 909946 Proteobacteria
Salmonella enterica subsp enterica serovar Choleraesuis str SC B67 YP 218558 1 321314 Proteobacteria
Salmonella enterica subsp enterica serovar Enteritidis str P125109 YP 002245528 1 550537 Proteobacteria
Salmonella enterica subsp enterica serovar Bovismorbificans str 3114 YP 008507524 1 1320309 Proteobacteria
Salmonella enterica subsp enterica serovar Thompson str RM6836 YP 008615273 1 1064551 Proteobacteria
Salmonella enterica subsp enterica serovar 4 5 12 i str 08 1736 YP 008263617 1 866913 Proteobacteria
Salmonella enterica subsp enterica serovar Paratyphi C str RKS4594 YP 002639238 1 476213 Proteobacteria
Salmonella enterica subsp enterica serovar Dublin str CT 02021853 YP 002217600 1 439851 Proteobacteria
Salmonella enterica subsp enterica serovar Bareilly str CFSAN000189 YP 008306706 1 1173427 Proteobacteria
Enterobacter lignolyticus SCF1 YP 003942812 1 701347 Proteobacteria
Enterobacteriaceae bacterium strain FGI 57 YP 007341421 1 693444 Proteobacteria
Enterobacter sp 638 YP 001175723 1 399742 Proteobacteria
Enterobacter cloacae subsp dissolvens SDM YP 006476545 1 1104326 Proteobacteria
Enterobacter cloacae EcWSU1 YP 004950905 1 1045856 Proteobacteria
Enterobacter cloacae subsp cloacae ENHKU01 YP 006577618 1 1211025 Proteobacteria
Enterobacter asburiae LF7a YP 004827553 1 640513 Proteobacteria
Citrobacter koseri ATCC BAA 895 YP 001454171 1 290338 Proteobacteria
Salmonella enterica subsp enterica serovar Cubana str CFSAN002050 YP 008259798 1 1271863 Proteobacteria
Salmonella enterica subsp enterica serovar Typhi str Ty21a YP 007926617 1 527001 Proteobacteria
Salmonella enterica subsp enterica serovar Paratyphi A str AKU 12601 YP 002142864 1 554290 Proteobacteria
Salmonella enterica subsp enterica serovar Paratyphi C str RKS4594 YP 002636179 1 476213 Proteobacteria
Salmonella enterica subsp enterica serovar Choleraesuis str SC B67 YP 215570 1 321314 Proteobacteria
Salmonella enterica subsp enterica serovar Paratyphi B str SPB7 YP 001589215 1 1016998 Proteobacteria
Proteus mirabilis BB2000 YP 008398050 1 1266738 Proteobacteria
Serratia sp AS12 YP 004499564 1 768490 Proteobacteria
Serratia plymuthica AS9 YP 004504516 1 768492 Proteobacteria
Providencia stuartii MRSN 2154 YP 006217055 1 1157951 Proteobacteria
Erwinia tasmaniensis Et1 99 YP 001908784 1 465817 Proteobacteria
Dickeya sp CSL RW240 WP 035340122 1 Daquatica
21 fimI Fim pilus assembly Seed
DAQ1742 DAQ1742 00918 Daquatica
Dickeya aquatica DW 0440 WP 035340122 1 Daquatica
Klebsiella pneumoniae YP 007990722 1 573 Proteobacteria
Klebsiella pneumoniae subsp pneumoniae MGH 78578 YP 001336916 1 272620 Proteobacteria
Klebsiella pneumoniae 342 YP 002236688 1 507522 Proteobacteria
Klebsiella variicola At 22 YP 003437719 1 640131 Proteobacteria
Klebsiella oxytoca E718 YP 006499492 1 1191061 Proteobacteria
Raoultella ornithinolytica B6 YP 007876841 1 1286170 Proteobacteria
Citrobacter koseri ATCC BAA 895 YP 001455221 1 290338 Proteobacteria
Enterobacter aerogenes EA1509E YP 007391042 1 935296 Proteobacteria
Citrobacter rodentium ICC168 YP 003366434 1 637910 Proteobacteria
Escherichia coli O83 H1 str NRG 857C YP 006122695 1 685038 Proteobacteria
Escherichia coli O127 H6 str E2348 69 YP 002332070 1 574521 Proteobacteria
Escherichia coli O111 H str 11128 YP 003237437 1 585396 Proteobacteria
Shigella boydii Sb227 YP 410609 1 300268 Proteobacteria
Escherichia coli B str REL606 YP 003047346 1 413997 Proteobacteria
Escherichia coli O55 H7 str CB9615 YP 003502514 1 701177 Proteobacteria
Escherichia coli O7 K1 str CE10 YP 006147007 1 1072459 Proteobacteria
Escherichia coli str K 12 substr W3110 YP 492449 1 316407 Proteobacteria
Escherichia coli O157 H7 str EDL933 NP 290931 1 155864 Proteobacteria
Escherichia coli Xuzhou21 YP 006315817 1 741093 Proteobacteria
Shigella flexneri 2002017 YP 005729796 1 591020 Proteobacteria
Shigella flexneri 2a str 2457T NP 839601 1 198215 Proteobacteria
Escherichia coli O103 H2 str 12009 YP 003224909 1 585395 Proteobacteria
Escherichia coli O26 H11 str 11368 YP 003232374 1 573235 Proteobacteria

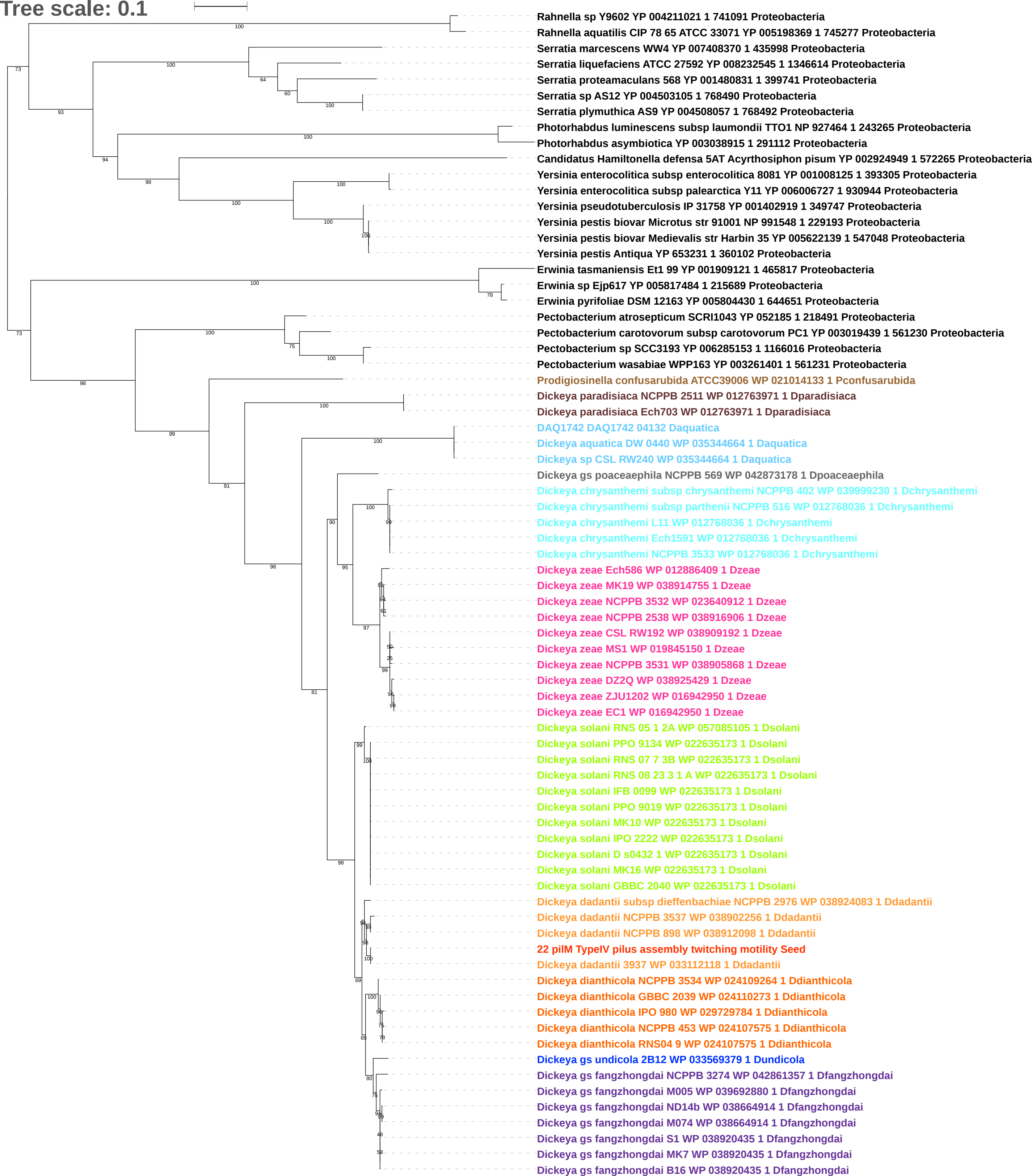

Tree scale: 0.1
Rahnella sp Y9602 YP 004211021 1 741091 Proteobacteria
Rahnella aquatilis CIP 78 65 ATCC 33071 YP 005198369 1 745277 Proteobacteria
Serratia marcescens WW4 YP 007408370 1 435998 Proteobacteria
Serratia liquefaciens ATCC 27592 YP 008232545 1 1346614 Proteobacteria
Serratia proteamaculans 568 YP 001480831 1 399741 Proteobacteria
Serratia sp AS12 YP 004503105 1 768490 Proteobacteria
Serratia plymuthica AS9 YP 004508057 1 768492 Proteobacteria
Photorhabdus luminescens subsp laumondii TTO1 NP 927464 1 243265 Proteobacteria
Photorhabdus asymbiotica YP 003038915 1 291112 Proteobacteria
Candidatus Hamiltonella defensa 5AT Acyrthosiphon pisum YP 002924949 1 572265 Proteobacteria
Yersinia enterocolitica subsp enterocolitica 8081 YP 001008125 1 393305 Proteobacteria
Yersinia enterocolitica subsp palearctica Y11 YP 006006727 1 930944 Proteobacteria
Yersinia pseudotuberculosis IP 31758 YP 001402919 1 349747 Proteobacteria
Yersinia pestis biovar Microtus str 91001 NP 991548 1 229193 Proteobacteria
Yersinia pestis biovar Medievalis str Harbin 35 YP 005622139 1 547048 Proteobacteria
Yersinia pestis Antiqua YP 653231 1 360102 Proteobacteria
Erwinia tasmaniensis Et1 99 YP 001909121 1 465817 Proteobacteria
Erwinia sp Ejp617 YP 005817484 1 215689 Proteobacteria
Erwinia pyrifoliae DSM 12163 YP 005804430 1 644651 Proteobacteria
Pectobacterium atrosepticum SCRI1043 YP 052185 1 218491 Proteobacteria
Pectobacterium carotovorum subsp carotovorum PC1 YP 003019439 1 561230 Proteobacteria
Pectobacterium sp SCC3193 YP 006285153 1 1166016 Proteobacteria
Pectobacterium wasabiae WPP163 YP 003261401 1 561231 Proteobacteria
Prodigiosinella confusarubida ATCC39006 WP 021014133 1 Pconfusarubida
Dickeya paradisiaca NCPPB 2511 WP 012763971 1 Dparadisiaca
Dickeya paradisiaca Ech703 WP 012763971 1 Dparadisiaca
DAQ1742 DAQ1742 04132 Daquatica
Dickeya aquatica DW 0440 WP 035344664 1 Daquatica
Dickeya sp CSL RW240 WP 035344664 1 Daquatica
Dickeya gs poaceaephila NCPPB 569 WP 042873178 1 Dpoaceaephila
Dickeya chrysanthemi subsp chrysanthemi NCPPB 402 WP 039999230 1 Dchrysanthemi
Dickeya chrysanthemi subsp parthenii NCPPB 516 WP 012768036 1 Dchrysanthemi
Dickeya chrysanthemi L11 WP 012768036 1 Dchrysanthemi
Dickeya chrysanthemi Ech1591 WP 012768036 1 Dchrysanthemi
Dickeya chrysanthemi NCPPB 3533 WP 012768036 1 Dchrysanthemi
Dickeya zeae Ech586 WP 012886409 1 Dzeae
Dickeya zeae MK19 WP 038914755 1 Dzeae
Dickeya zeae NCPPB 3532 WP 023640912 1 Dzeae
Dickeya zeae NCPPB 2538 WP 038916906 1 Dzeae
Dickeya zeae CSL RW192 WP 038909192 1 Dzeae
Dickeya zeae MS1 WP 019845150 1 Dzeae
Dickeya zeae NCPPB 3531 WP 038905868 1 Dzeae
Dickeya zeae DZ2Q WP 038925429 1 Dzeae
Dickeya zeae ZJU1202 WP 016942950 1 Dzeae
Dickeya zeae EC1 WP 016942950 1 Dzeae
Dickeya solani RNS 05 1 2A WP 057085105 1 Dsolani
Dickeya solani PPO 9134 WP 022635173 1 Dsolani
Dickeya solani RNS 07 7 3B WP 022635173 1 Dsolani
Dickeya solani RNS 08 23 3 1 A WP 022635173 1 Dsolani
Dickeya solani IFB 0099 WP 022635173 1 Dsolani
Dickeya solani PPO 9019 WP 022635173 1 Dsolani
Dickeya solani MK10 WP 022635173 1 Dsolani
Dickeya solani IPO 2222 WP 022635173 1 Dsolani
Dickeya solani D s0432 1 WP 022635173 1 Dsolani
Dickeya solani MK16 WP 022635173 1 Dsolani
Dickeya solani GBBC 2040 WP 022635173 1 Dsolani
Dickeya dadantii subsp dieffenbachiae NCPPB 2976 WP 038924083 1 Ddadantii
Dickeya dadantii NCPPB 3537 WP 038902256 1 Ddadantii
Dickeya dadantii NCPPB 898 WP 038912098 1 Ddadantii
22 pilM TypeIV pilus assembly twitching motility Seed
Dickeya dadantii 3937 WP 033112118 1 Ddadantii
Dickeya dianthicola NCPPB 3534 WP 024109264 1 Ddianthicola
Dickeya dianthicola GBBC 2039 WP 024110273 1 Ddianthicola
Dickeya dianthicola IPO 980 WP 029729784 1 Ddianthicola
Dickeya dianthicola NCPPB 453 WP 024107575 1 Ddianthicola
Dickeya dianthicola RNS04 9 WP 024107575 1 Ddianthicola
Dickeya gs undicola 2B12 WP 033569379 1 Dundicola
Dickeya gs fangzhongdai NCPPB 3274 WP 042861357 1 Dfangzhongdai
Dickeya gs fangzhongdai M005 WP 039692880 1 Dfangzhongdai
Dickeya gs fangzhongdai ND14b WP 038664914 1 Dfangzhongdai
Dickeya gs fangzhongdai M074 WP 038664914 1 Dfangzhongdai
Dickeya gs fangzhongdai S1 WP 038920435 1 Dfangzhongdai
Dickeya gs fangzhongdai MK7 WP 038920435 1 Dfangzhongdai
Dickeya gs fangzhongdai B16 WP 038920435 1 Dfangzhongdai

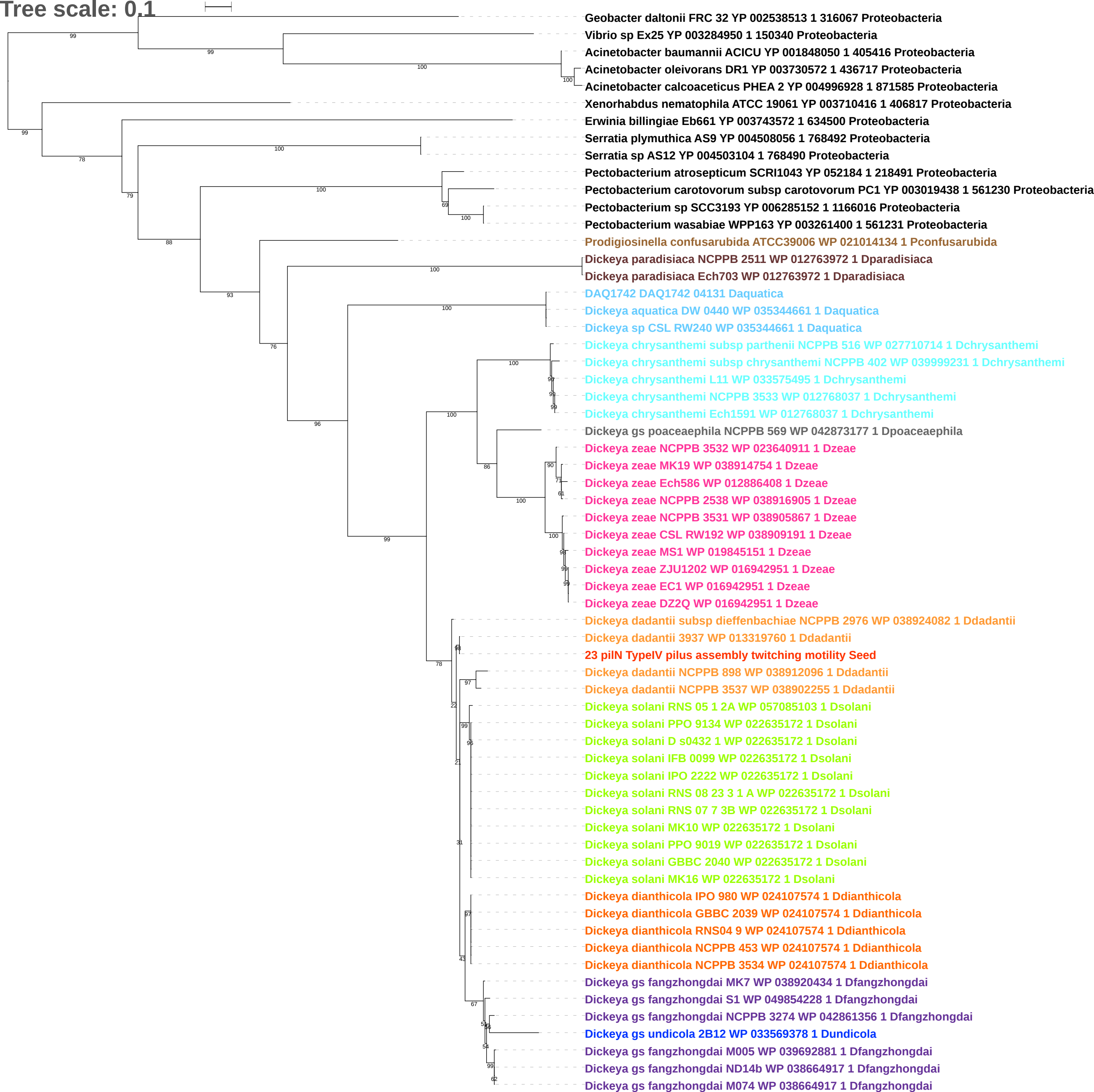
Tree scale: 0.1
Geobacter daltonii FRC 32 YP 002538513 1 316067 Proteobacteria
Vibrio sp Ex25 YP 003284950 1 150340 Proteobacteria
Acinetobacter baumannii ACICU YP 001848050 1 405416 Proteobacteria
Acinetobacter oleivorans DR1 YP 003730572 1 436717 Proteobacteria
Acinetobacter calcoaceticus PHEA 2 YP 004996928 1 871585 Proteobacteria
Xenorhabdus nematophila ATCC 19061 YP 003710416 1 406817 Proteobacteria
Erwinia billingiae Eb661 YP 003743572 1 634500 Proteobacteria
Serratia plymuthica AS9 YP 004508056 1 768492 Proteobacteria
Serratia sp AS12 YP 004503104 1 768490 Proteobacteria
Pectobacterium atrosepticum SCRI1043 YP 052184 1 218491 Proteobacteria
Pectobacterium carotovorum subsp carotovorum PC1 YP 003019438 1 561230 Proteobacteria
Pectobacterium sp SCC3193 YP 006285152 1 1166016 Proteobacteria
Pectobacterium wasabiae WPP163 YP 003261400 1 561231 Proteobacteria
Prodigiosinella confusarubida ATCC39006 WP 021014134 1 Pconfusarubida
Dickeya paradisiaca NCPPB 2511 WP 012763972 1 Dparadisiaca
Dickeya paradisiaca Ech703 WP 012763972 1 Dparadisiaca
DAQ1742 DAQ1742 04131 Daquatica
Dickeya aquatica DW 0440 WP 035344661 1 Daquatica
Dickeya sp CSL RW240 WP 035344661 1 Daquatica
Dickeya chrysanthemi subsp parthenii NCPPB 516 WP 027710714 1 Dchrysanthemi
Dickeya chrysanthemi subsp chrysanthemi NCPPB 402 WP 039999231 1 Dchrysanthemi
Dickeya chrysanthemi L11 WP 033575495 1 Dchrysanthemi
Dickeya chrysanthemi NCPPB 3533 WP 012768037 1 Dchrysanthemi
Dickeya chrysanthemi Ech1591 WP 012768037 1 Dchrysanthemi
Dickeya gs poaceaephila NCPPB 569 WP 042873177 1 Dpoaceaephila
Dickeya zeae NCPPB 3532 WP 023640911 1 Dzeae
Dickeya zeae MK19 WP 038914754 1 Dzeae
Dickeya zeae Ech586 WP 012886408 1 Dzeae
Dickeya zeae NCPPB 2538 WP 038916905 1 Dzeae
Dickeya zeae NCPPB 3531 WP 038905867 1 Dzeae
Dickeya zeae CSL RW192 WP 038909191 1 Dzeae
Dickeya zeae MS1 WP 019845151 1 Dzeae
Dickeya zeae ZJU1202 WP 016942951 1 Dzeae
Dickeya zeae EC1 WP 016942951 1 Dzeae
Dickeya zeae DZ2Q WP 016942951 1 Dzeae
Dickeya dadantii subsp dieffenbachiae NCPPB 2976 WP 038924082 1 Ddadantii
Dickeya dadantii 3937 WP 013319760 1 Ddadantii
23 pilN TypeIV pilus assembly twitching motility Seed
Dickeya dadantii NCPPB 898 WP 038912096 1 Ddadantii
Dickeya dadantii NCPPB 3537 WP 038902255 1 Ddadantii
Dickeya solani RNS 05 1 2A WP 057085103 1 Dsolani
Dickeya solani PPO 9134 WP 022635172 1 Dsolani
Dickeya solani D s0432 1 WP 022635172 1 Dsolani
Dickeya solani IFB 0099 WP 022635172 1 Dsolani
Dickeya solani IPO 2222 WP 022635172 1 Dsolani
Dickeya solani RNS 08 23 3 1 A WP 022635172 1 Dsolani
Dickeya solani RNS 07 7 3B WP 022635172 1 Dsolani
Dickeya solani MK10 WP 022635172 1 Dsolani
Dickeya solani PPO 9019 WP 022635172 1 Dsolani
Dickeya solani GBBC 2040 WP 022635172 1 Dsolani
Dickeya solani MK16 WP 022635172 1 Dsolani
Dickeya dianthicola IPO 980 WP 024107574 1 Ddianthicola
Dickeya dianthicola GBBC 2039 WP 024107574 1 Ddianthicola
Dickeya dianthicola RNS04 9 WP 024107574 1 Ddianthicola
Dickeya dianthicola NCPPB 453 WP 024107574 1 Ddianthicola
Dickeya dianthicola NCPPB 3534 WP 024107574 1 Ddianthicola
Dickeya gs fangzhongdai MK7 WP 038920434 1 Dfangzhongdai
Dickeya gs fangzhongdai S1 WP 049854228 1 Dfangzhongdai
Dickeya gs fangzhongdai NCPPB 3274 WP 042861356 1 Dfangzhongdai
Dickeya gs undicola 2B12 WP 033569378 1 Dundicola
Dickeya gs fangzhongdai M005 WP 039692881 1 Dfangzhongdai
Dickeya gs fangzhongdai ND14b WP 038664917 1 Dfangzhongdai
Dickeya gs fangzhongdai M074 WP 038664917 1 Dfangzhongdai

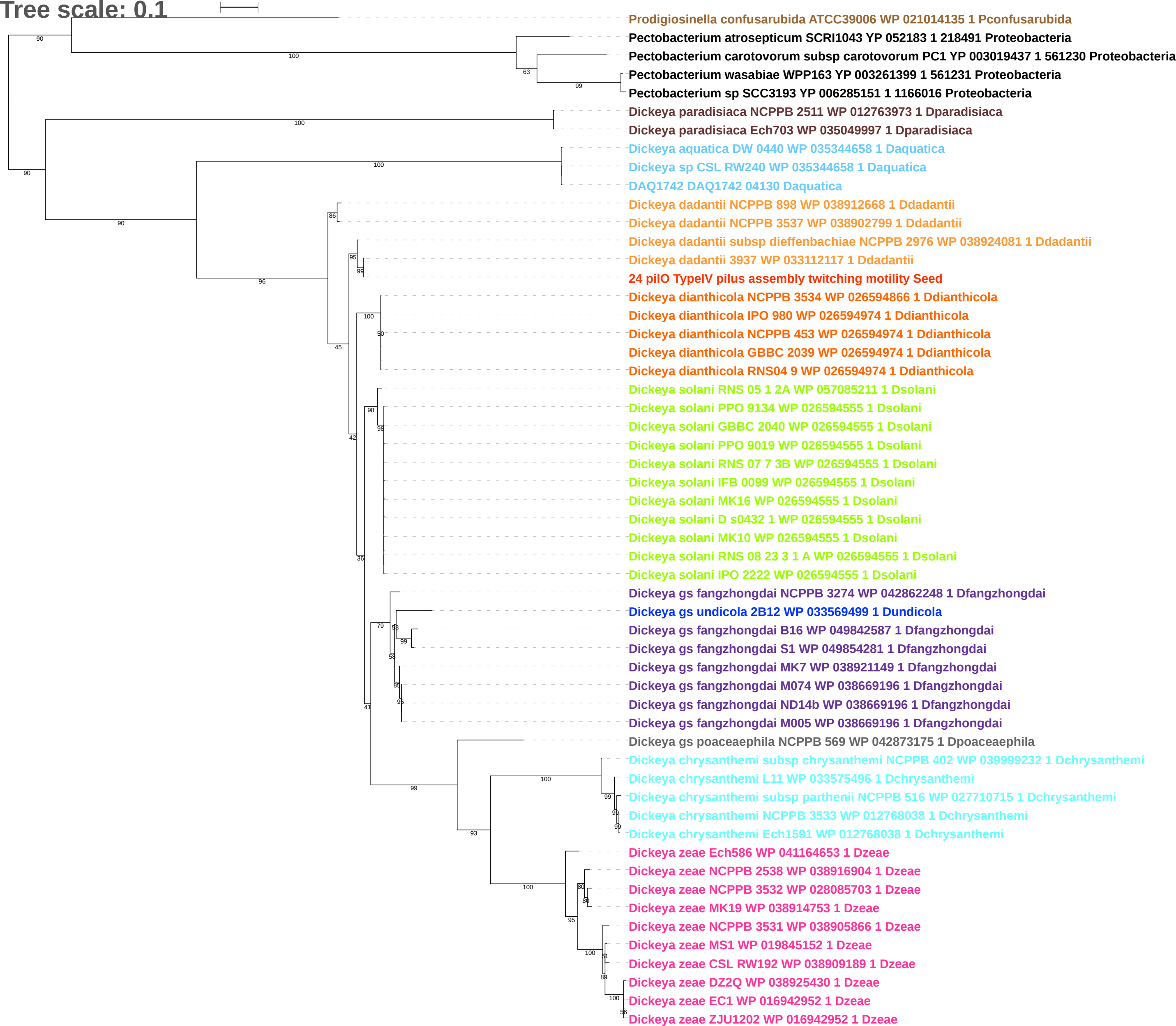
Tree scale: 0.1
Prodigiosinella confusarubida ATCC39006 WP 021014135 1 Pconfusarubida
Pectobacterium atrosepticum SCRI1043 YP 052183 1 218491 Proteobacteria
Pectobacterium carotovorum subsp carotovorum PC1 YP 003019437 1 561230 Proteobacteria
Pectobacterium wasabiae WPP163 YP 003261399 1 561231 Proteobacteria
Pectobacterium sp SCC3193 YP 006285151 1 1166016 Proteobacteria
Dickeya paradisiaca NCPPB 2511 WP 012763973 1 Dparadisiaca
Dickeya paradisiaca Ech703 WP 035049997 1 Dparadisiaca
Dickeya aquatica DW 0440 WP 035344658 1 Daquatica
Dickeya sp CSL RW240 WP 035344658 1 Daquatica
DAQ1742 DAQ1742 04130 Daquatica
Dickeya dadantii NCPPB 898 WP 038912668 1 Ddadantii
Dickeya dadantii NCPPB 3537 WP 038902799 1 Ddadantii
Dickeya dadantii subsp dieffenbachiae NCPPB 2976 WP 038924081 1 Ddadantii
Dickeya dadantii 3937 WP 033112117 1 Ddadantii
24 pilO TypeIV pilus assembly twitching motility Seed
Dickeya dianthicola NCPPB 3534 WP 026594866 1 Ddianthicola
Dickeya dianthicola IPO 980 WP 026594974 1 Ddianthicola
Dickeya dianthicola NCPPB 453 WP 026594974 1 Ddianthicola
Dickeya dianthicola GBBC 2039 WP 026594974 1 Ddianthicola
Dickeya dianthicola RNS04 9 WP 026594974 1 Ddianthicola
Dickeya solani RNS 05 1 2A WP 057085211 1 Dsolani
Dickeya solani PPO 9134 WP 026594555 1 Dsolani
Dickeya solani GBBC 2040 WP 026594555 1 Dsolani
Dickeya solani PPO 9019 WP 026594555 1 Dsolani
Dickeya solani RNS 07 7 3B WP 026594555 1 Dsolani
Dickeya solani IFB 0099 WP 026594555 1 Dsolani
Dickeya solani MK16 WP 026594555 1 Dsolani
Dickeya solani D s0432 1 WP 026594555 1 Dsolani
Dickeya solani MK10 WP 026594555 1 Dsolani
Dickeya solani RNS 08 23 3 1 A WP 026594555 1 Dsolani
Dickeya solani IPO 2222 WP 026594555 1 Dsolani
Dickeya gs fangzhongdai NCPPB 3274 WP 042862248 1 Dfangzhongdai
Dickeya gs undicola 2B12 WP 033569499 1 Dundicola
Dickeya gs fangzhongdai B16 WP 049842587 1 Dfangzhongdai
Dickeya gs fangzhongdai S1 WP 049854281 1 Dfangzhongdai
Dickeya gs fangzhongdai MK7 WP 038921149 1 Dfangzhongdai
Dickeya gs fangzhongdai M074 WP 038669196 1 Dfangzhongdai
Dickeya gs fangzhongdai ND14b WP 038669196 1 Dfangzhongdai
Dickeya gs fangzhongdai M005 WP 038669196 1 Dfangzhongdai
Dickeya gs poaceaephila NCPPB 569 WP 042873175 1 Dpoaceaephila
Dickeya chrysanthemi subsp chrysanthemi NCPPB 402 WP 039999232 1 Dchrysanthemi
Dickeya chrysanthemi L11 WP 033575496 1 Dchrysanthemi
Dickeya chrysanthemi subsp parthenii NCPPB 516 WP 027710715 1 Dchrysanthemi
Dickeya chrysanthemi NCPPB 3533 WP 012768038 1 Dchrysanthemi
Dickeya chrysanthemi Ech1591 WP 012768038 1 Dchrysanthemi
Dickeya zeae Ech586 WP 041164653 1 Dzeae
Dickeya zeae NCPPB 2538 WP 038916904 1 Dzeae
Dickeya zeae NCPPB 3532 WP 028085703 1 Dzeae
Dickeya zeae MK19 WP 038914753 1 Dzeae
Dickeya zeae NCPPB 3531 WP 038905866 1 Dzeae
Dickeya zeae MS1 WP 019845152 1 Dzeae
Dickeya zeae CSL RW192 WP 038909189 1 Dzeae
Dickeya zeae DZ2Q WP 038925430 1 Dzeae
Dickeya zeae EC1 WP 016942952 1 Dzeae
Dickeya zeae ZJU1202 WP 016942952 1 Dzeae

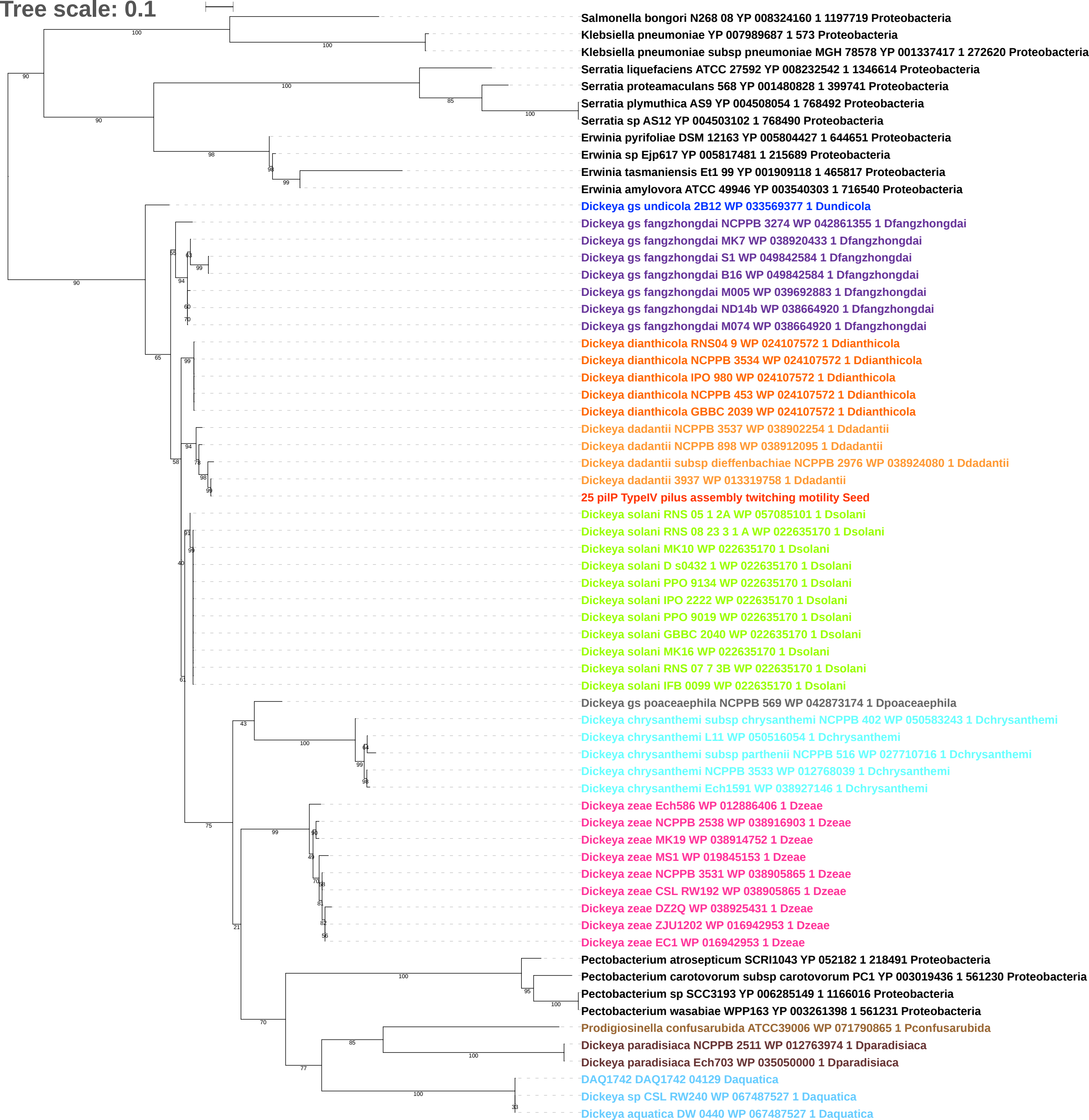
Tree scale: 0.1
Salmonella bongori N268 08 YP 008324160 1 1197719 Proteobacteria
Klebsiella pneumoniae YP 007989687 1 573 Proteobacteria
Klebsiella pneumoniae subsp pneumoniae MGH 78578 YP 001337417 1 272620 Proteobacteria
Serratia liquefaciens ATCC 27592 YP 008232542 1 1346614 Proteobacteria
Serratia proteamaculans 568 YP 001480828 1 399741 Proteobacteria
Serratia plymuthica AS9 YP 004508054 1 768492 Proteobacteria
Serratia sp AS12 YP 004503102 1 768490 Proteobacteria
Erwinia pyrifoliae DSM 12163 YP 005804427 1 644651 Proteobacteria
Erwinia sp Ejp617 YP 005817481 1 215689 Proteobacteria
Erwinia tasmaniensis Et1 99 YP 001909118 1 465817 Proteobacteria
Erwinia amylovora ATCC 49946 YP 003540303 1 716540 Proteobacteria
Dickeya gs undicola 2B12 WP 033569377 1 Dundicola
Dickeya gs fangzhongdai NCPPB 3274 WP 042861355 1 Dfangzhongdai
Dickeya gs fangzhongdai MK7 WP 038920433 1 Dfangzhongdai
Dickeya gs fangzhongdai S1 WP 049842584 1 Dfangzhongdai
Dickeya gs fangzhongdai B16 WP 049842584 1 Dfangzhongdai
Dickeya gs fangzhongdai M005 WP 039692883 1 Dfangzhongdai
Dickeya gs fangzhongdai ND14b WP 038664920 1 Dfangzhongdai
Dickeya gs fangzhongdai M074 WP 038664920 1 Dfangzhongdai
Dickeya dianthicola RNS04 9 WP 024107572 1 Ddianthicola
Dickeya dianthicola NCPPB 3534 WP 024107572 1 Ddianthicola
Dickeya dianthicola IPO 980 WP 024107572 1 Ddianthicola
Dickeya dianthicola NCPPB 453 WP 024107572 1 Ddianthicola
Dickeya dianthicola GBBC 2039 WP 024107572 1 Ddianthicola
Dickeya dadantii NCPPB 3537 WP 038902254 1 Ddadantii
Dickeya dadantii NCPPB 898 WP 038912095 1 Ddadantii
Dickeya dadantii subsp dieffenbachiae NCPPB 2976 WP 038924080 1 Ddadantii
Dickeya dadantii 3937 WP 013319758 1 Ddadantii
25 pilP TypeIV pilus assembly twitching motility Seed
Dickeya solani RNS 05 1 2A WP 057085101 1 Dsolani
Dickeya solani RNS 08 23 3 1 A WP 022635170 1 Dsolani
Dickeya solani MK10 WP 022635170 1 Dsolani
Dickeya solani D s0432 1 WP 022635170 1 Dsolani
Dickeya solani PPO 9134 WP 022635170 1 Dsolani
Dickeya solani IPO 2222 WP 022635170 1 Dsolani
Dickeya solani PPO 9019 WP 022635170 1 Dsolani
Dickeya solani GBBC 2040 WP 022635170 1 Dsolani
Dickeya solani MK16 WP 022635170 1 Dsolani
Dickeya solani RNS 07 7 3B WP 022635170 1 Dsolani
Dickeya solani IFB 0099 WP 022635170 1 Dsolani
Dickeya gs poaceaephila NCPPB 569 WP 042873174 1 Dpoaceaephila
Dickeya chrysanthemi subsp chrysanthemi NCPPB 402 WP 050583243 1 Dchrysanthemi
Dickeya chrysanthemi L11 WP 050516054 1 Dchrysanthemi
Dickeya chrysanthemi subsp parthenii NCPPB 516 WP 027710716 1 Dchrysanthemi
Dickeya chrysanthemi NCPPB 3533 WP 012768039 1 Dchrysanthemi
Dickeya chrysanthemi Ech1591 WP 038927146 1 Dchrysanthemi
Dickeya zeae Ech586 WP 012886406 1 Dzeae
Dickeya zeae NCPPB 2538 WP 038916903 1 Dzeae
Dickeya zeae MK19 WP 038914752 1 Dzeae
Dickeya zeae MS1 WP 019845153 1 Dzeae
Dickeya zeae NCPPB 3531 WP 038905865 1 Dzeae
Dickeya zeae CSL RW192 WP 038905865 1 Dzeae
Dickeya zeae DZ2Q WP 038925431 1 Dzeae
Dickeya zeae ZJU1202 WP 016942953 1 Dzeae
Dickeya zeae EC1 WP 016942953 1 Dzeae
Pectobacterium atrosepticum SCRI1043 YP 052182 1 218491 Proteobacteria
Pectobacterium carotovorum subsp carotovorum PC1 YP 003019436 1 561230 Proteobacteria
Pectobacterium sp SCC3193 YP 006285149 1 1166016 Proteobacteria
Pectobacterium wasabiae WPP163 YP 003261398 1 561231 Proteobacteria
Prodigiosinella confusarubida ATCC39006 WP 071790865 1 Pconfusarubida
Dickeya paradisiaca NCPPB 2511 WP 012763974 1 Dparadisiaca
Dickeya paradisiaca Ech703 WP 035050000 1 Dparadisiaca
DAQ1742 DAQ1742 04129 Daquatica
Dickeya sp CSL RW240 WP 067487527 1 Daquatica
Dickeya aquatica DW 0440 WP 067487527 1 Daquatica

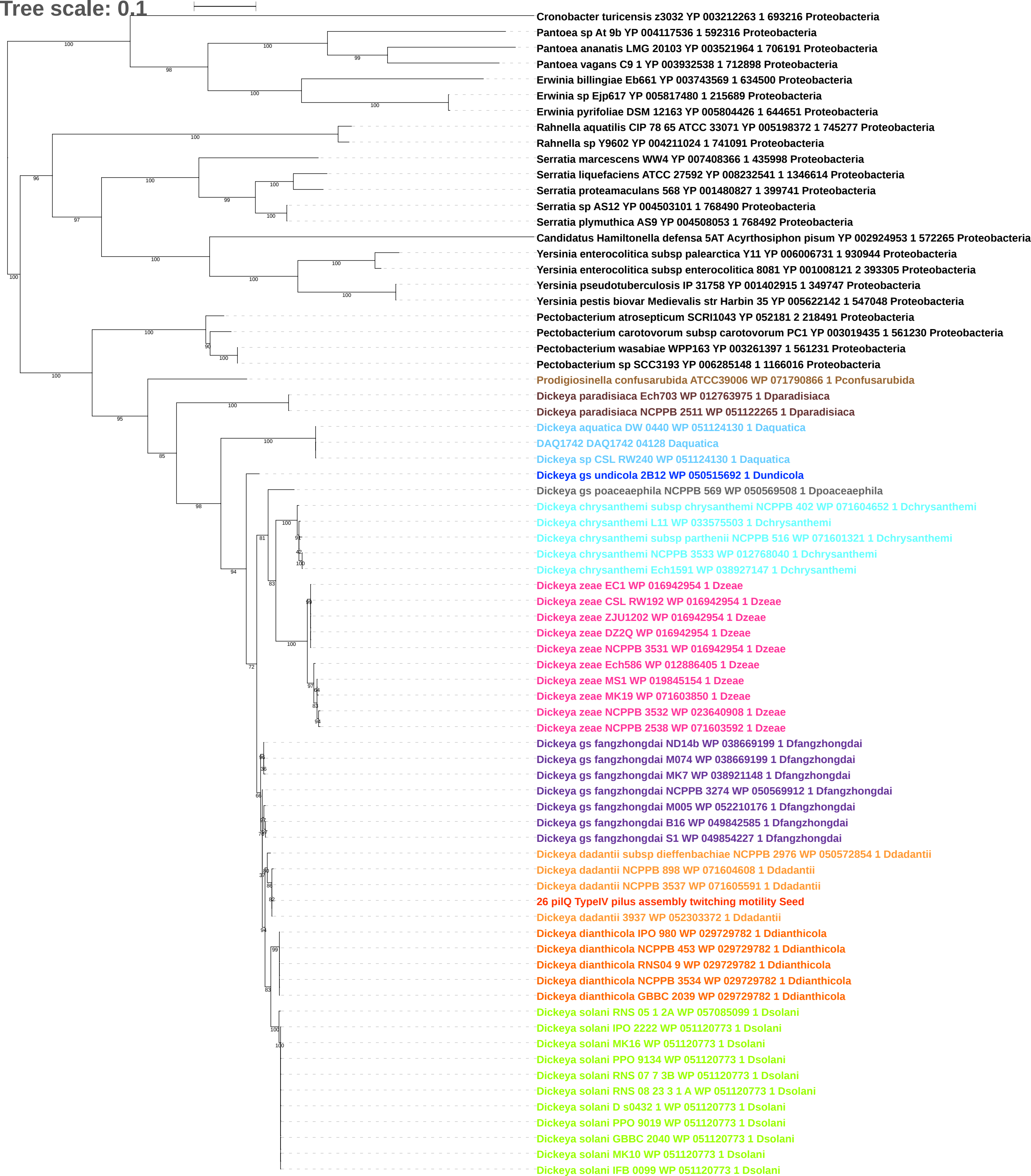
Tree scale: 0.1
Cronobacter turicensis z3032 YP 003212263 1 693216 Proteobacteria
Pantoea sp At 9b YP 004117536 1 592316 Proteobacteria
Pantoea ananatis LMG 20103 YP 003521964 1 706191 Proteobacteria
Pantoea vagans C9 1 YP 003932538 1 712898 Proteobacteria
Erwinia billingiae Eb661 YP 003743569 1 634500 Proteobacteria
Erwinia sp Ejp617 YP 005817480 1 215689 Proteobacteria
Erwinia pyrifoliae DSM 12163 YP 005804426 1 644651 Proteobacteria
Rahnella aquatilis CIP 78 65 ATCC 33071 YP 005198372 1 745277 Proteobacteria
Rahnella sp Y9602 YP 004211024 1 741091 Proteobacteria
Serratia marcescens WW4 YP 007408366 1 435998 Proteobacteria
Serratia liquefaciens ATCC 27592 YP 008232541 1 1346614 Proteobacteria
Serratia proteamaculans 568 YP 001480827 1 399741 Proteobacteria
Serratia sp AS12 YP 004503101 1 768490 Proteobacteria
Serratia plymuthica AS9 YP 004508053 1 768492 Proteobacteria
Candidatus Hamiltonella defensa 5AT Acyrthosiphon pisum YP 002924953 1 572265 Proteobacteria
Yersinia enterocolitica subsp palearctica Y11 YP 006006731 1 930944 Proteobacteria
Yersinia enterocolitica subsp enterocolitica 8081 YP 001008121 2 393305 Proteobacteria
Yersinia pseudotuberculosis IP 31758 YP 001402915 1 349747 Proteobacteria
Yersinia pestis biovar Medievalis str Harbin 35 YP 005622142 1 547048 Proteobacteria
Pectobacterium atrosepticum SCRI1043 YP 052181 2 218491 Proteobacteria
Pectobacterium carotovorum subsp carotovorum PC1 YP 003019435 1 561230 Proteobacteria
Pectobacterium wasabiae WPP163 YP 003261397 1 561231 Proteobacteria
Pectobacterium sp SCC3193 YP 006285148 1 1166016 Proteobacteria
Prodigiosinella confusarubida ATCC39006 WP 071790866 1 Pconfusarubida
Dickeya paradisiaca Ech703 WP 012763975 1 Dparadisiaca
Dickeya paradisiaca NCPPB 2511 WP 051122265 1 Dparadisiaca
Dickeya aquatica DW 0440 WP 051124130 1 Daquatica
DAQ1742 DAQ1742 04128 Daquatica
Dickeya sp CSL RW240 WP 051124130 1 Daquatica
Dickeya gs undicola 2B12 WP 050515692 1 Dundicola
Dickeya gs poaceaephila NCPPB 569 WP 050569508 1 Dpoaceaephila
Dickeya chrysanthemi subsp chrysanthemi NCPPB 402 WP 071604652 1 Dchrysanthemi
Dickeya chrysanthemi L11 WP 033575503 1 Dchrysanthemi
Dickeya chrysanthemi subsp parthenii NCPPB 516 WP 071601321 1 Dchrysanthemi
Dickeya chrysanthemi NCPPB 3533 WP 012768040 1 Dchrysanthemi
Dickeya chrysanthemi Ech1591 WP 038927147 1 Dchrysanthemi
Dickeya zeae EC1 WP 016942954 1 Dzeae
Dickeya zeae CSL RW192 WP 016942954 1 Dzeae
Dickeya zeae ZJU1202 WP 016942954 1 Dzeae
Dickeya zeae DZ2Q WP 016942954 1 Dzeae
Dickeya zeae NCPPB 3531 WP 016942954 1 Dzeae
Dickeya zeae Ech586 WP 012886405 1 Dzeae
Dickeya zeae MS1 WP 019845154 1 Dzeae
Dickeya zeae MK19 WP 071603850 1 Dzeae
Dickeya zeae NCPPB 3532 WP 023640908 1 Dzeae
Dickeya zeae NCPPB 2538 WP 071603592 1 Dzeae
Dickeya gs fangzhongdai ND14b WP 038669199 1 Dfangzhongdai
Dickeya gs fangzhongdai M074 WP 038669199 1 Dfangzhongdai
Dickeya gs fangzhongdai MK7 WP 038921148 1 Dfangzhongdai
Dickeya gs fangzhongdai NCPPB 3274 WP 050569912 1 Dfangzhongdai
Dickeya gs fangzhongdai M005 WP 052210176 1 Dfangzhongdai
Dickeya gs fangzhongdai B16 WP 049842585 1 Dfangzhongdai
Dickeya gs fangzhongdai S1 WP 049854227 1 Dfangzhongdai
Dickeya dadantii subsp dieffenbachiae NCPPB 2976 WP 050572854 1 Ddadantii
Dickeya dadantii NCPPB 898 WP 071604608 1 Ddadantii
Dickeya dadantii NCPPB 3537 WP 071605591 1 Ddadantii
26 pilQ TypeIV pilus assembly twitching motility Seed
Dickeya dadantii 3937 WP 052303372 1 Ddadantii
Dickeya dianthicola IPO 980 WP 029729782 1 Ddianthicola
Dickeya dianthicola NCPPB 453 WP 029729782 1 Ddianthicola
Dickeya dianthicola RNS04 9 WP 029729782 1 Ddianthicola
Dickeya dianthicola NCPPB 3534 WP 029729782 1 Ddianthicola
Dickeya dianthicola GBBC 2039 WP 029729782 1 Ddianthicola
Dickeya solani RNS 05 1 2A WP 057085099 1 Dsolani
Dickeya solani IPO 2222 WP 051120773 1 Dsolani
Dickeya solani MK16 WP 051120773 1 Dsolani
Dickeya solani PPO 9134 WP 051120773 1 Dsolani
Dickeya solani RNS 07 7 3B WP 051120773 1 Dsolani
Dickeya solani RNS 08 23 3 1 A WP 051120773 1 Dsolani
Dickeya solani D s0432 1 WP 051120773 1 Dsolani
Dickeya solani PPO 9019 WP 051120773 1 Dsolani
Dickeya solani GBBC 2040 WP 051120773 1 Dsolani
Dickeya solani MK10 WP 051120773 1 Dsolani
Dickeya solani IFB 0099 WP 051120773 1 Dsolani

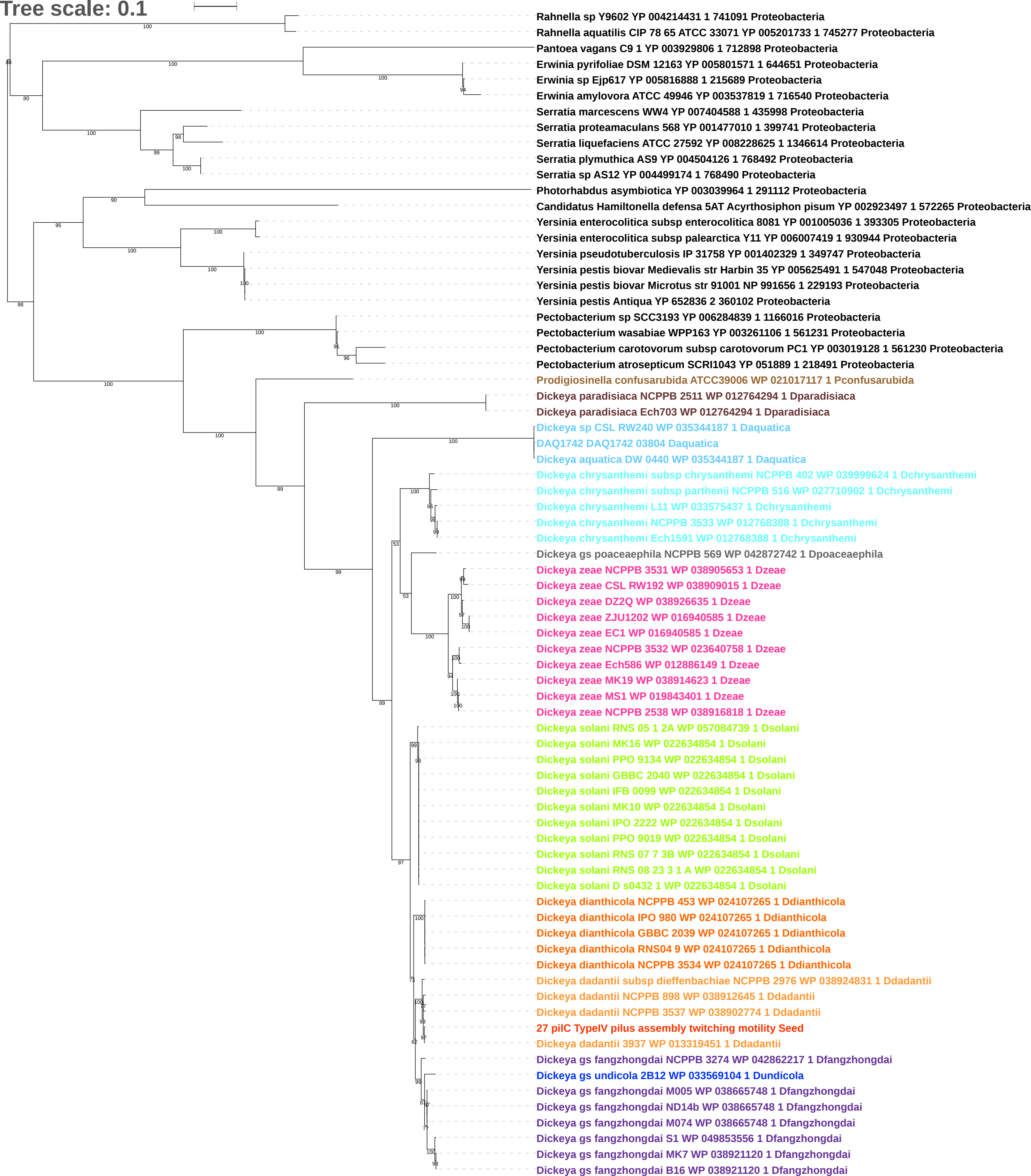
Tree scale: 0.1
Rahnella sp Y9602 YP 004214431 1 741091 Proteobacteria
Rahnella aquatilis CIP 78 65 ATCC 33071 YP 005201733 1 745277 Proteobacteria
Pantoea vagans C9 1 YP 003929806 1 712898 Proteobacteria
Erwinia pyrifoliae DSM 12163 YP 005801571 1 644651 Proteobacteria
Erwinia sp Ejp617 YP 005816888 1 215689 Proteobacteria
Erwinia amylovora ATCC 49946 YP 003537819 1 716540 Proteobacteria
Serratia marcescens WW4 YP 007404588 1 435998 Proteobacteria
Serratia proteamaculans 568 YP 001477010 1 399741 Proteobacteria
Serratia liquefaciens ATCC 27592 YP 008228625 1 1346614 Proteobacteria
Serratia plymuthica AS9 YP 004504126 1 768492 Proteobacteria
Serratia sp AS12 YP 004499174 1 768490 Proteobacteria
Photorhabdus asymbiotica YP 003039964 1 291112 Proteobacteria
Candidatus Hamiltonella defensa 5AT Acyrthosiphon pisum YP 002923497 1 572265 Proteobacteria
Yersinia enterocolitica subsp enterocolitica 8081 YP 001005036 1 393305 Proteobacteria
Yersinia enterocolitica subsp palearctica Y11 YP 006007419 1 930944 Proteobacteria
Yersinia pseudotuberculosis IP 31758 YP 001402329 1 349747 Proteobacteria
Yersinia pestis biovar Medievalis str Harbin 35 YP 005625491 1 547048 Proteobacteria
Yersinia pestis biovar Microtus str 91001 NP 991656 1 229193 Proteobacteria
Yersinia pestis Antiqua YP 652836 2 360102 Proteobacteria
Pectobacterium sp SCC3193 YP 006284839 1 1166016 Proteobacteria
Pectobacterium wasabiae WPP163 YP 003261106 1 561231 Proteobacteria
Pectobacterium carotovorum subsp carotovorum PC1 YP 003019128 1 561230 Proteobacteria
Pectobacterium atrosepticum SCRI1043 YP 051889 1 218491 Proteobacteria
Prodigiosinella confusarubida ATCC39006 WP 021017117 1 Pconfusarubida
Dickeya paradisiaca NCPPB 2511 WP 012764294 1 Dparadisiaca
Dickeya paradisiaca Ech703 WP 012764294 1 Dparadisiaca
Dickeya sp CSL RW240 WP 035344187 1 Daquatica
DAQ1742 DAQ1742 03804 Daquatica
Dickeya aquatica DW 0440 WP 035344187 1 Daquatica
Dickeya chrysanthemi subsp chrysanthemi NCPPB 402 WP 039999624 1 Dchrysanthemi
Dickeya chrysanthemi subsp parthenii NCPPB 516 WP 027710902 1 Dchrysanthemi
Dickeya chrysanthemi L11 WP 033575437 1 Dchrysanthemi
Dickeya chrysanthemi NCPPB 3533 WP 012768388 1 Dchrysanthemi
Dickeya chrysanthemi Ech1591 WP 012768388 1 Dchrysanthemi
Dickeya gs poaceaephila NCPPB 569 WP 042872742 1 Dpoaceaephila
Dickeya zeae NCPPB 3531 WP 038905653 1 Dzeae
Dickeya zeae CSL RW192 WP 038909015 1 Dzeae
Dickeya zeae DZ2Q WP 038926635 1 Dzeae
Dickeya zeae ZJU1202 WP 016940585 1 Dzeae
Dickeya zeae EC1 WP 016940585 1 Dzeae
Dickeya zeae NCPPB 3532 WP 023640758 1 Dzeae
Dickeya zeae Ech586 WP 012886149 1 Dzeae
Dickeya zeae MK19 WP 038914623 1 Dzeae
Dickeya zeae MS1 WP 019843401 1 Dzeae
Dickeya zeae NCPPB 2538 WP 038916818 1 Dzeae
Dickeya solani RNS 05 1 2A WP 057084739 1 Dsolani
Dickeya solani MK16 WP 022634854 1 Dsolani
Dickeya solani PPO 9134 WP 022634854 1 Dsolani
Dickeya solani GBBC 2040 WP 022634854 1 Dsolani
Dickeya solani IFB 0099 WP 022634854 1 Dsolani
Dickeya solani MK10 WP 022634854 1 Dsolani
Dickeya solani IPO 2222 WP 022634854 1 Dsolani
Dickeya solani PPO 9019 WP 022634854 1 Dsolani
Dickeya solani RNS 07 7 3B WP 022634854 1 Dsolani
Dickeya solani RNS 08 23 3 1 A WP 022634854 1 Dsolani
Dickeya solani D s0432 1 WP 022634854 1 Dsolani
Dickeya dianthicola NCPPB 453 WP 024107265 1 Ddianthicola
Dickeya dianthicola IPO 980 WP 024107265 1 Ddianthicola
Dickeya dianthicola GBBC 2039 WP 024107265 1 Ddianthicola
Dickeya dianthicola RNS04 9 WP 024107265 1 Ddianthicola
Dickeya dianthicola NCPPB 3534 WP 024107265 1 Ddianthicola
Dickeya dadantii subsp dieffenbachiae NCPPB 2976 WP 038924831 1 Ddadantii
Dickeya dadantii NCPPB 898 WP 038912645 1 Ddadantii
Dickeya dadantii NCPPB 3537 WP 038902774 1 Ddadantii
27 pilC TypeIV pilus assembly twitching motility Seed
Dickeya dadantii 3937 WP 013319451 1 Ddadantii
Dickeya gs fangzhongdai NCPPB 3274 WP 042862217 1 Dfangzhongdai
Dickeya gs undicola 2B12 WP 033569104 1 Dundicola
Dickeya gs fangzhongdai M005 WP 038665748 1 Dfangzhongdai
Dickeya gs fangzhongdai ND14b WP 038665748 1 Dfangzhongdai
Dickeya gs fangzhongdai M074 WP 038665748 1 Dfangzhongdai
Dickeya gs fangzhongdai S1 WP 049853556 1 Dfangzhongdai
Dickeya gs fangzhongdai MK7 WP 038921120 1 Dfangzhongdai
Dickeya gs fangzhongdai B16 WP 038921120 1 Dfangzhongdai

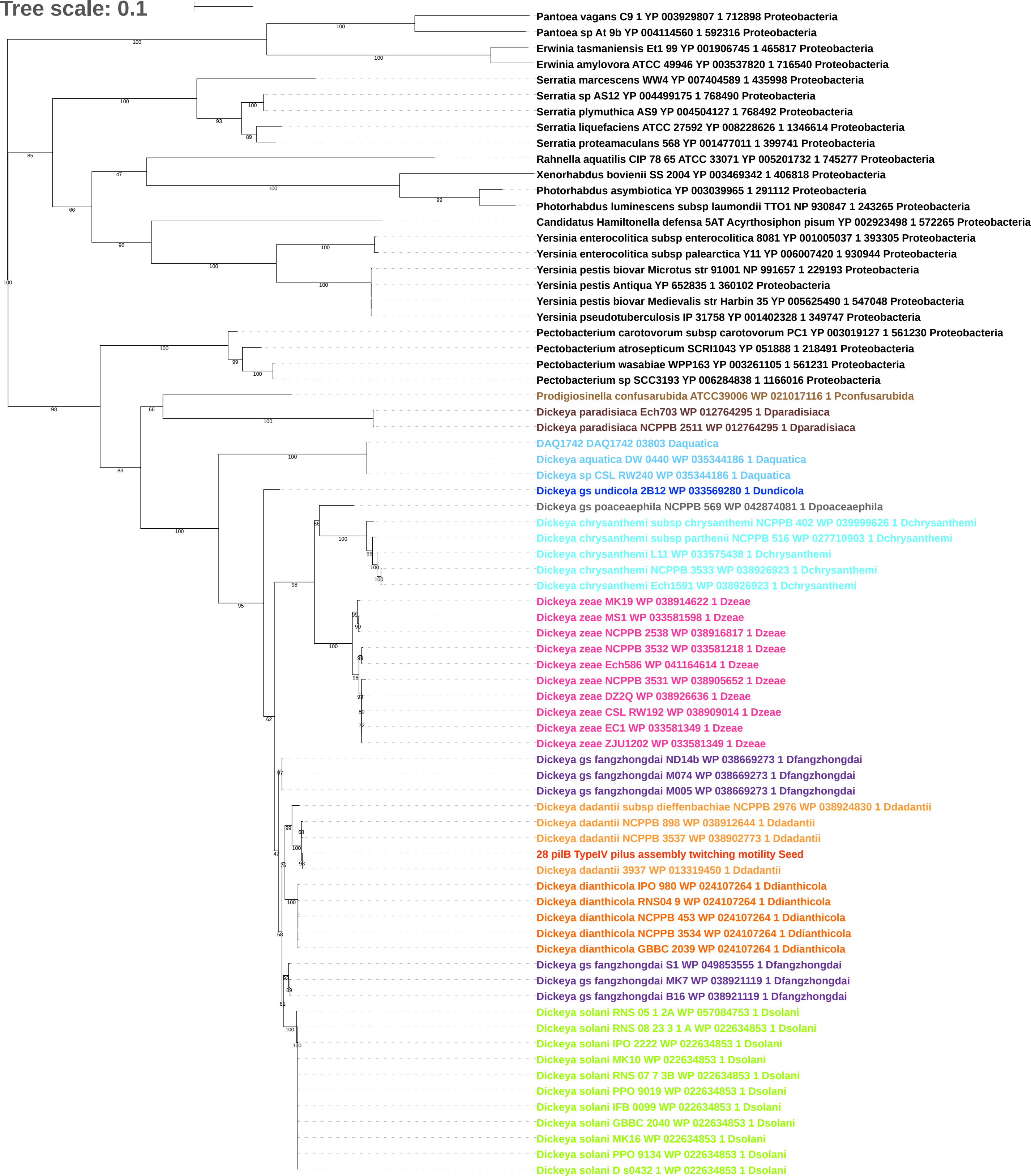
Tree scale: 0.1
Pantoea vagans C9 1 YP 003929807 1 712898 Proteobacteria
Pantoea sp At 9b YP 004114560 1 592316 Proteobacteria
Erwinia tasmaniensis Et1 99 YP 001906745 1 465817 Proteobacteria
Erwinia amylovora ATCC 49946 YP 003537820 1 716540 Proteobacteria
Serratia marcescens WW4 YP 007404589 1 435998 Proteobacteria
Serratia sp AS12 YP 004499175 1 768490 Proteobacteria
Serratia plymuthica AS9 YP 004504127 1 768492 Proteobacteria
Serratia liquefaciens ATCC 27592 YP 008228626 1 1346614 Proteobacteria
Serratia proteamaculans 568 YP 001477011 1 399741 Proteobacteria
Rahnella aquatilis CIP 78 65 ATCC 33071 YP 005201732 1 745277 Proteobacteria
Xenorhabdus bovienii SS 2004 YP 003469342 1 406818 Proteobacteria
Photorhabdus asymbiotica YP 003039965 1 291112 Proteobacteria
Photorhabdus luminescens subsp laumondii TTO1 NP 930847 1 243265 Proteobacteria
Candidatus Hamiltonella defensa 5AT Acyrthosiphon pisum YP 002923498 1 572265 Proteobacteria
Yersinia enterocolitica subsp enterocolitica 8081 YP 001005037 1 393305 Proteobacteria
Yersinia enterocolitica subsp palearctica Y11 YP 006007420 1 930944 Proteobacteria
Yersinia pestis biovar Microtus str 91001 NP 991657 1 229193 Proteobacteria
Yersinia pestis Antiqua YP 652835 1 360102 Proteobacteria
Yersinia pestis biovar Medievalis str Harbin 35 YP 005625490 1 547048 Proteobacteria
Yersinia pseudotuberculosis IP 31758 YP 001402328 1 349747 Proteobacteria
Pectobacterium carotovorum subsp carotovorum PC1 YP 003019127 1 561230 Proteobacteria
Pectobacterium atrosepticum SCRI1043 YP 051888 1 218491 Proteobacteria
Pectobacterium wasabiae WPP163 YP 003261105 1 561231 Proteobacteria
Pectobacterium sp SCC3193 YP 006284838 1 1166016 Proteobacteria
Prodigiosinella confusarubida ATCC39006 WP 021017116 1 Pconfusarubida
Dickeya paradisiaca Ech703 WP 012764295 1 Dparadisiaca
Dickeya paradisiaca NCPPB 2511 WP 012764295 1 Dparadisiaca
DAQ1742 DAQ1742 03803 Daquatica
Dickeya aquatica DW 0440 WP 035344186 1 Daquatica
Dickeya sp CSL RW240 WP 035344186 1 Daquatica
Dickeya gs undicola 2B12 WP 033569280 1 Dundicola
Dickeya gs poaceaephila NCPPB 569 WP 042874081 1 Dpoaceaephila
Dickeya chrysanthemi subsp chrysanthemi NCPPB 402 WP 039999626 1 Dchrysanthemi
Dickeya chrysanthemi subsp parthenii NCPPB 516 WP 027710903 1 Dchrysanthemi
Dickeya chrysanthemi L11 WP 033575438 1 Dchrysanthemi
Dickeya chrysanthemi NCPPB 3533 WP 038926923 1 Dchrysanthemi
Dickeya chrysanthemi Ech1591 WP 038926923 1 Dchrysanthemi
Dickeya zeae MK19 WP 038914622 1 Dzeae
Dickeya zeae MS1 WP 033581598 1 Dzeae
Dickeya zeae NCPPB 2538 WP 038916817 1 Dzeae
Dickeya zeae NCPPB 3532 WP 033581218 1 Dzeae
Dickeya zeae Ech586 WP 041164614 1 Dzeae
Dickeya zeae NCPPB 3531 WP 038905652 1 Dzeae
Dickeya zeae DZ2Q WP 038926636 1 Dzeae
Dickeya zeae CSL RW192 WP 038909014 1 Dzeae
Dickeya zeae EC1 WP 033581349 1 Dzeae
Dickeya zeae ZJU1202 WP 033581349 1 Dzeae
Dickeya gs fangzhongdai ND14b WP 038669273 1 Dfangzhongdai
Dickeya gs fangzhongdai M074 WP 038669273 1 Dfangzhongdai
Dickeya gs fangzhongdai M005 WP 038669273 1 Dfangzhongdai
Dickeya dadantii subsp dieffenbachiae NCPPB 2976 WP 038924830 1 Ddadantii
Dickeya dadantii NCPPB 898 WP 038912644 1 Ddadantii
Dickeya dadantii NCPPB 3537 WP 038902773 1 Ddadantii
28 pilB TypeIV pilus assembly twitching motility Seed
Dickeya dadantii 3937 WP 013319450 1 Ddadantii
Dickeya dianthicola IPO 980 WP 024107264 1 Ddianthicola
Dickeya dianthicola RNS04 9 WP 024107264 1 Ddianthicola
Dickeya dianthicola NCPPB 453 WP 024107264 1 Ddianthicola
Dickeya dianthicola NCPPB 3534 WP 024107264 1 Ddianthicola
Dickeya dianthicola GBBC 2039 WP 024107264 1 Ddianthicola
Dickeya gs fangzhongdai S1 WP 049853555 1 Dfangzhongdai
Dickeya gs fangzhongdai MK7 WP 038921119 1 Dfangzhongdai
Dickeya gs fangzhongdai B16 WP 038921119 1 Dfangzhongdai
Dickeya solani RNS 05 1 2A WP 057084753 1 Dsolani
Dickeya solani RNS 08 23 3 1 A WP 022634853 1 Dsolani
Dickeya solani IPO 2222 WP 022634853 1 Dsolani
Dickeya solani MK10 WP 022634853 1 Dsolani
Dickeya solani RNS 07 7 3B WP 022634853 1 Dsolani
Dickeya solani PPO 9019 WP 022634853 1 Dsolani
Dickeya solani IFB 0099 WP 022634853 1 Dsolani
Dickeya solani GBBC 2040 WP 022634853 1 Dsolani
Dickeya solani MK16 WP 022634853 1 Dsolani
Dickeya solani PPO 9134 WP 022634853 1 Dsolani
Dickeya solani D s0432 1 WP 022634853 1 Dsolani

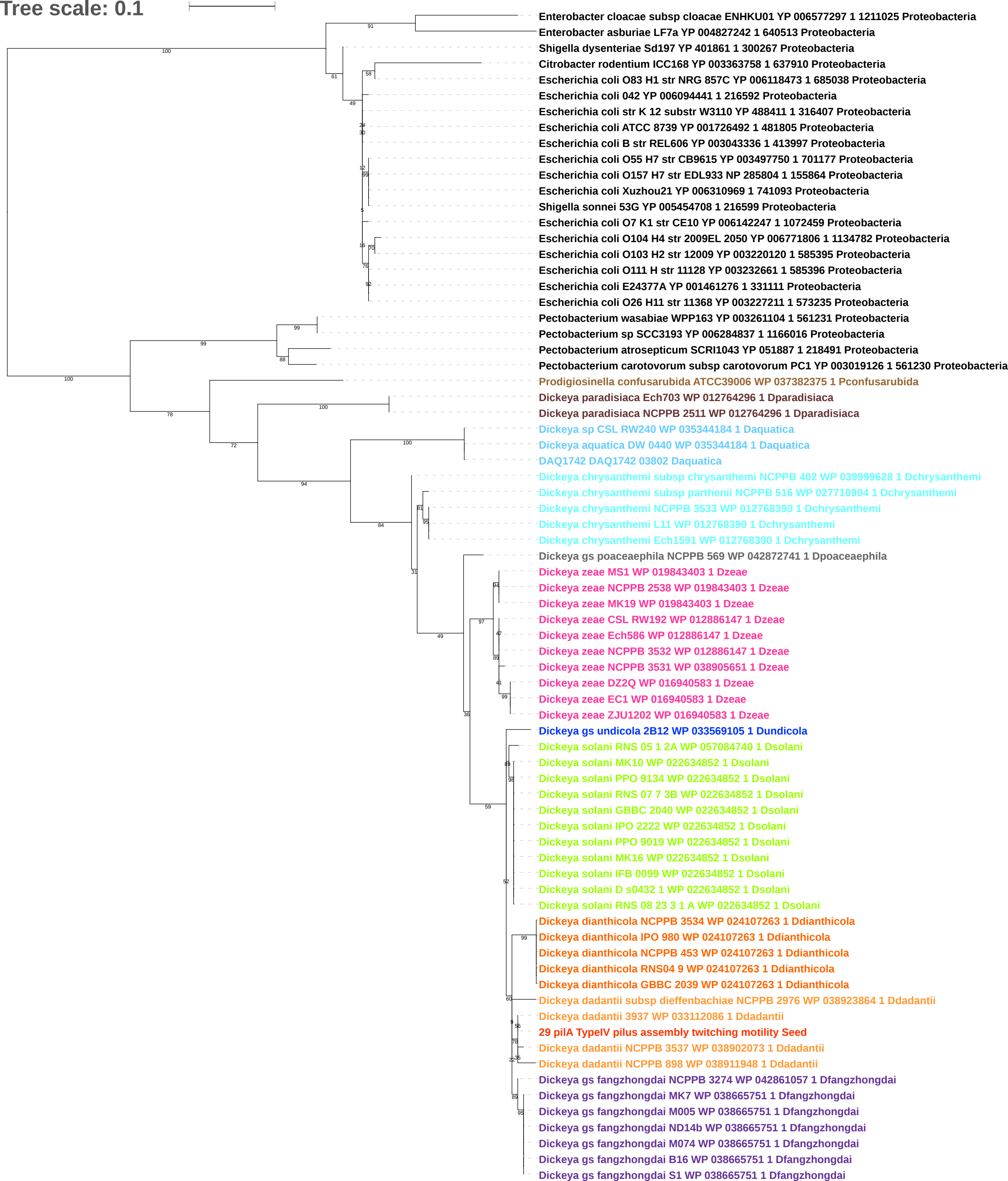
Tree scale: 0.1
Enterobacter cloacae subsp cloacae ENHKU01 YP 006577297 1 1211025 Proteobacteria
Enterobacter asburiae LF7a YP 004827242 1 640513 Proteobacteria
Shigella dysenteriae Sd197 YP 401861 1 300267 Proteobacteria
Citrobacter rodentium ICC168 YP 003363758 1 637910 Proteobacteria
Escherichia coli O83 H1 str NRG 857C YP 006118473 1 685038 Proteobacteria
Escherichia coli 042 YP 006094441 1 216592 Proteobacteria
Escherichia coli str K 12 substr W3110 YP 488411 1 316407 Proteobacteria
Escherichia coli ATCC 8739 YP 001726492 1 481805 Proteobacteria
Escherichia coli B str REL606 YP 003043336 1 413997 Proteobacteria
Escherichia coli O55 H7 str CB9615 YP 003497750 1 701177 Proteobacteria
Escherichia coli O157 H7 str EDL933 NP 285804 1 155864 Proteobacteria
Escherichia coli Xuzhou21 YP 006310969 1 741093 Proteobacteria
Shigella sonnei 53G YP 005454708 1 216599 Proteobacteria
Escherichia coli O7 K1 str CE10 YP 006142247 1 1072459 Proteobacteria
Escherichia coli O104 H4 str 2009EL 2050 YP 006771806 1 1134782 Proteobacteria
Escherichia coli O103 H2 str 12009 YP 003220120 1 585395 Proteobacteria
Escherichia coli O111 H str 11128 YP 003232661 1 585396 Proteobacteria
Escherichia coli E24377A YP 001461276 1 331111 Proteobacteria
Escherichia coli O26 H11 str 11368 YP 003227211 1 573235 Proteobacteria
Pectobacterium wasabiae WPP163 YP 003261104 1 561231 Proteobacteria
Pectobacterium sp SCC3193 YP 006284837 1 1166016 Proteobacteria
Pectobacterium atrosepticum SCRI1043 YP 051887 1 218491 Proteobacteria
Pectobacterium carotovorum subsp carotovorum PC1 YP 003019126 1 561230 Proteobacteria
Prodigiosinella confusarubida ATCC39006 WP 037382375 1 Pconfusarubida
Dickeya paradisiaca Ech703 WP 012764296 1 Dparadisiaca
Dickeya paradisiaca NCPPB 2511 WP 012764296 1 Dparadisiaca
Dickeya sp CSL RW240 WP 035344184 1 Daquatica
Dickeya aquatica DW 0440 WP 035344184 1 Daquatica
DAQ1742 DAQ1742 03802 Daquatica
Dickeya chrysanthemi subsp chrysanthemi NCPPB 402 WP 039999628 1 Dchrysanthemi
Dickeya chrysanthemi subsp parthenii NCPPB 516 WP 027710904 1 Dchrysanthemi
Dickeya chrysanthemi NCPPB 3533 WP 012768390 1 Dchrysanthemi
Dickeya chrysanthemi L11 WP 012768390 1 Dchrysanthemi
Dickeya chrysanthemi Ech1591 WP 012768390 1 Dchrysanthemi
Dickeya gs poaceaephila NCPPB 569 WP 042872741 1 Dpoaceaephila
Dickeya zeae MS1 WP 019843403 1 Dzeae
Dickeya zeae NCPPB 2538 WP 019843403 1 Dzeae
Dickeya zeae MK19 WP 019843403 1 Dzeae
Dickeya zeae CSL RW192 WP 012886147 1 Dzeae
Dickeya zeae Ech586 WP 012886147 1 Dzeae
Dickeya zeae NCPPB 3532 WP 012886147 1 Dzeae
Dickeya zeae NCPPB 3531 WP 038905651 1 Dzeae
Dickeya zeae DZ2Q WP 016940583 1 Dzeae
Dickeya zeae EC1 WP 016940583 1 Dzeae
Dickeya zeae ZJU1202 WP 016940583 1 Dzeae
Dickeya gs undicola 2B12 WP 033569105 1 Dundicola
Dickeya solani RNS 05 1 2A WP 057084740 1 Dsolani
Dickeya solani MK10 WP 022634852 1 Dsolani
Dickeya solani PPO 9134 WP 022634852 1 Dsolani
Dickeya solani RNS 07 7 3B WP 022634852 1 Dsolani
Dickeya solani GBBC 2040 WP 022634852 1 Dsolani
Dickeya solani IPO 2222 WP 022634852 1 Dsolani
Dickeya solani PPO 9019 WP 022634852 1 Dsolani
Dickeya solani MK16 WP 022634852 1 Dsolani
Dickeya solani IFB 0099 WP 022634852 1 Dsolani
Dickeya solani D s0432 1 WP 022634852 1 Dsolani
Dickeya solani RNS 08 23 3 1 A WP 022634852 1 Dsolani
Dickeya dianthicola NCPPB 3534 WP 024107263 1 Ddianthicola
Dickeya dianthicola IPO 980 WP 024107263 1 Ddianthicola
Dickeya dianthicola NCPPB 453 WP 024107263 1 Ddianthicola
Dickeya dianthicola RNS04 9 WP 024107263 1 Ddianthicola
Dickeya dianthicola GBBC 2039 WP 024107263 1 Ddianthicola
Dickeya dadantii subsp dieffenbachiae NCPPB 2976 WP 038923864 1 Ddadantii
Dickeya dadantii 3937 WP 033112086 1 Ddadantii
29 pilA TypeIV pilus assembly twitching motility Seed
Dickeya dadantii NCPPB 3537 WP 038902073 1 Ddadantii
Dickeya dadantii NCPPB 898 WP 038911948 1 Ddadantii
Dickeya gs fangzhongdai NCPPB 3274 WP 042861057 1 Dfangzhongdai
Dickeya gs fangzhongdai MK7 WP 038665751 1 Dfangzhongdai
Dickeya gs fangzhongdai M005 WP 038665751 1 Dfangzhongdai
Dickeya gs fangzhongdai ND14b WP 038665751 1 Dfangzhongdai
Dickeya gs fangzhongdai M074 WP 038665751 1 Dfangzhongdai
Dickeya gs fangzhongdai B16 WP 038665751 1 Dfangzhongdai
Dickeya gs fangzhongdai S1 WP 038665751 1 Dfangzhongdai

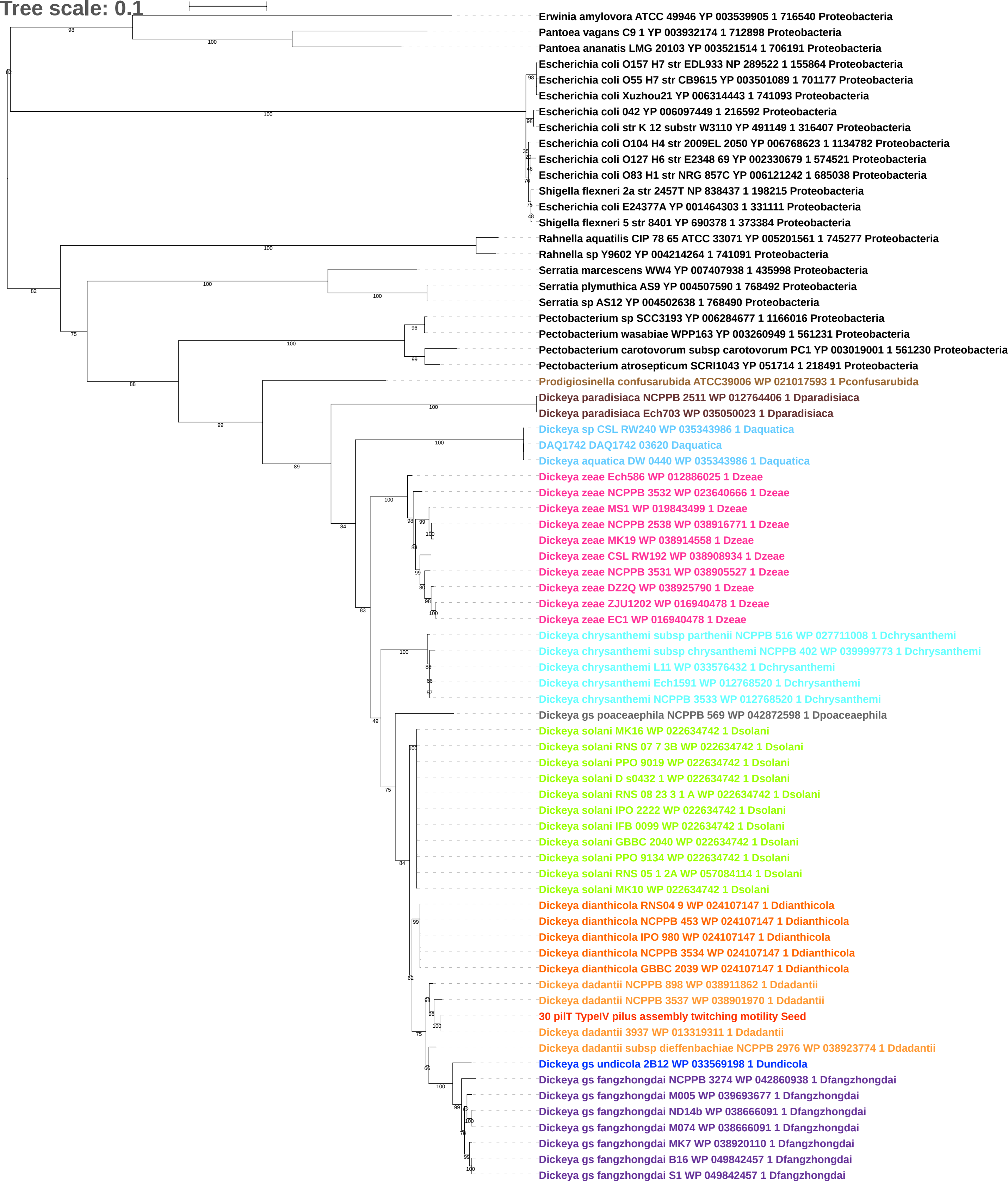
Tree scale: 0.1
Erwinia amylovora ATCC 49946 YP 003539905 1 716540 Proteobacteria
Pantoea vagans C9 1 YP 003932174 1 712898 Proteobacteria
Pantoea ananatis LMG 20103 YP 003521514 1 706191 Proteobacteria
Escherichia coli O157 H7 str EDL933 NP 289522 1 155864 Proteobacteria
Escherichia coli O55 H7 str CB9615 YP 003501089 1 701177 Proteobacteria
Escherichia coli Xuzhou21 YP 006314443 1 741093 Proteobacteria
Escherichia coli 042 YP 006097449 1 216592 Proteobacteria
Escherichia coli str K 12 substr W3110 YP 491149 1 316407 Proteobacteria
Escherichia coli O104 H4 str 2009EL 2050 YP 006768623 1 1134782 Proteobacteria
Escherichia coli O127 H6 str E2348 69 YP 002330679 1 574521 Proteobacteria
Escherichia coli O83 H1 str NRG 857C YP 006121242 1 685038 Proteobacteria
Shigella flexneri 2a str 2457T NP 838437 1 198215 Proteobacteria
Escherichia coli E24377A YP 001464303 1 331111 Proteobacteria
Shigella flexneri 5 str 8401 YP 690378 1 373384 Proteobacteria
Rahnella aquatilis CIP 78 65 ATCC 33071 YP 005201561 1 745277 Proteobacteria
Rahnella sp Y9602 YP 004214264 1 741091 Proteobacteria
Serratia marcescens WW4 YP 007407938 1 435998 Proteobacteria
Serratia plymuthica AS9 YP 004507590 1 768492 Proteobacteria
Serratia sp AS12 YP 004502638 1 768490 Proteobacteria
Pectobacterium sp SCC3193 YP 006284677 1 1166016 Proteobacteria
Pectobacterium wasabiae WPP163 YP 003260949 1 561231 Proteobacteria
Pectobacterium carotovorum subsp carotovorum PC1 YP 003019001 1 561230 Proteobacteria
Pectobacterium atrosepticum SCRI1043 YP 051714 1 218491 Proteobacteria
Prodigiosinella confusarubida ATCC39006 WP 021017593 1 Pconfusarubida
Dickeya paradisiaca NCPPB 2511 WP 012764406 1 Dparadisiaca
Dickeya paradisiaca Ech703 WP 035050023 1 Dparadisiaca
Dickeya sp CSL RW240 WP 035343986 1 Daquatica
DAQ1742 DAQ1742 03620 Daquatica
Dickeya aquatica DW 0440 WP 035343986 1 Daquatica
Dickeya zeae Ech586 WP 012886025 1 Dzeae
Dickeya zeae NCPPB 3532 WP 023640666 1 Dzeae
Dickeya zeae MS1 WP 019843499 1 Dzeae
Dickeya zeae NCPPB 2538 WP 038916771 1 Dzeae
Dickeya zeae MK19 WP 038914558 1 Dzeae
Dickeya zeae CSL RW192 WP 038908934 1 Dzeae
Dickeya zeae NCPPB 3531 WP 038905527 1 Dzeae
Dickeya zeae DZ2Q WP 038925790 1 Dzeae
Dickeya zeae ZJU1202 WP 016940478 1 Dzeae
Dickeya zeae EC1 WP 016940478 1 Dzeae
Dickeya chrysanthemi subsp parthenii NCPPB 516 WP 027711008 1 Dchrysanthemi
Dickeya chrysanthemi subsp chrysanthemi NCPPB 402 WP 039999773 1 Dchrysanthemi
Dickeya chrysanthemi L11 WP 033576432 1 Dchrysanthemi
Dickeya chrysanthemi Ech1591 WP 012768520 1 Dchrysanthemi
Dickeya chrysanthemi NCPPB 3533 WP 012768520 1 Dchrysanthemi
Dickeya gs poaceaephila NCPPB 569 WP 042872598 1 Dpoaceaephila
Dickeya solani MK16 WP 022634742 1 Dsolani
Dickeya solani RNS 07 7 3B WP 022634742 1 Dsolani
Dickeya solani PPO 9019 WP 022634742 1 Dsolani
Dickeya solani D s0432 1 WP 022634742 1 Dsolani
Dickeya solani RNS 08 23 3 1 A WP 022634742 1 Dsolani
Dickeya solani IPO 2222 WP 022634742 1 Dsolani
Dickeya solani IFB 0099 WP 022634742 1 Dsolani
Dickeya solani GBBC 2040 WP 022634742 1 Dsolani
Dickeya solani PPO 9134 WP 022634742 1 Dsolani
Dickeya solani RNS 05 1 2A WP 057084114 1 Dsolani
Dickeya solani MK10 WP 022634742 1 Dsolani
Dickeya dianthicola RNS04 9 WP 024107147 1 Ddianthicola
Dickeya dianthicola NCPPB 453 WP 024107147 1 Ddianthicola
Dickeya dianthicola IPO 980 WP 024107147 1 Ddianthicola
Dickeya dianthicola NCPPB 3534 WP 024107147 1 Ddianthicola
Dickeya dianthicola GBBC 2039 WP 024107147 1 Ddianthicola
Dickeya dadantii NCPPB 898 WP 038911862 1 Ddadantii
Dickeya dadantii NCPPB 3537 WP 038901970 1 Ddadantii
30 pilT TypeIV pilus assembly twitching motility Seed
Dickeya dadantii 3937 WP 013319311 1 Ddadantii
Dickeya dadantii subsp dieffenbachiae NCPPB 2976 WP 038923774 1 Ddadantii
Dickeya gs undicola 2B12 WP 033569198 1 Dundicola
Dickeya gs fangzhongdai NCPPB 3274 WP 042860938 1 Dfangzhongdai
Dickeya gs fangzhongdai M005 WP 039693677 1 Dfangzhongdai
Dickeya gs fangzhongdai ND14b WP 038666091 1 Dfangzhongdai
Dickeya gs fangzhongdai M074 WP 038666091 1 Dfangzhongdai
Dickeya gs fangzhongdai MK7 WP 038920110 1 Dfangzhongdai
Dickeya gs fangzhongdai B16 WP 049842457 1 Dfangzhongdai
Dickeya gs fangzhongdai S1 WP 049842457 1 Dfangzhongdai

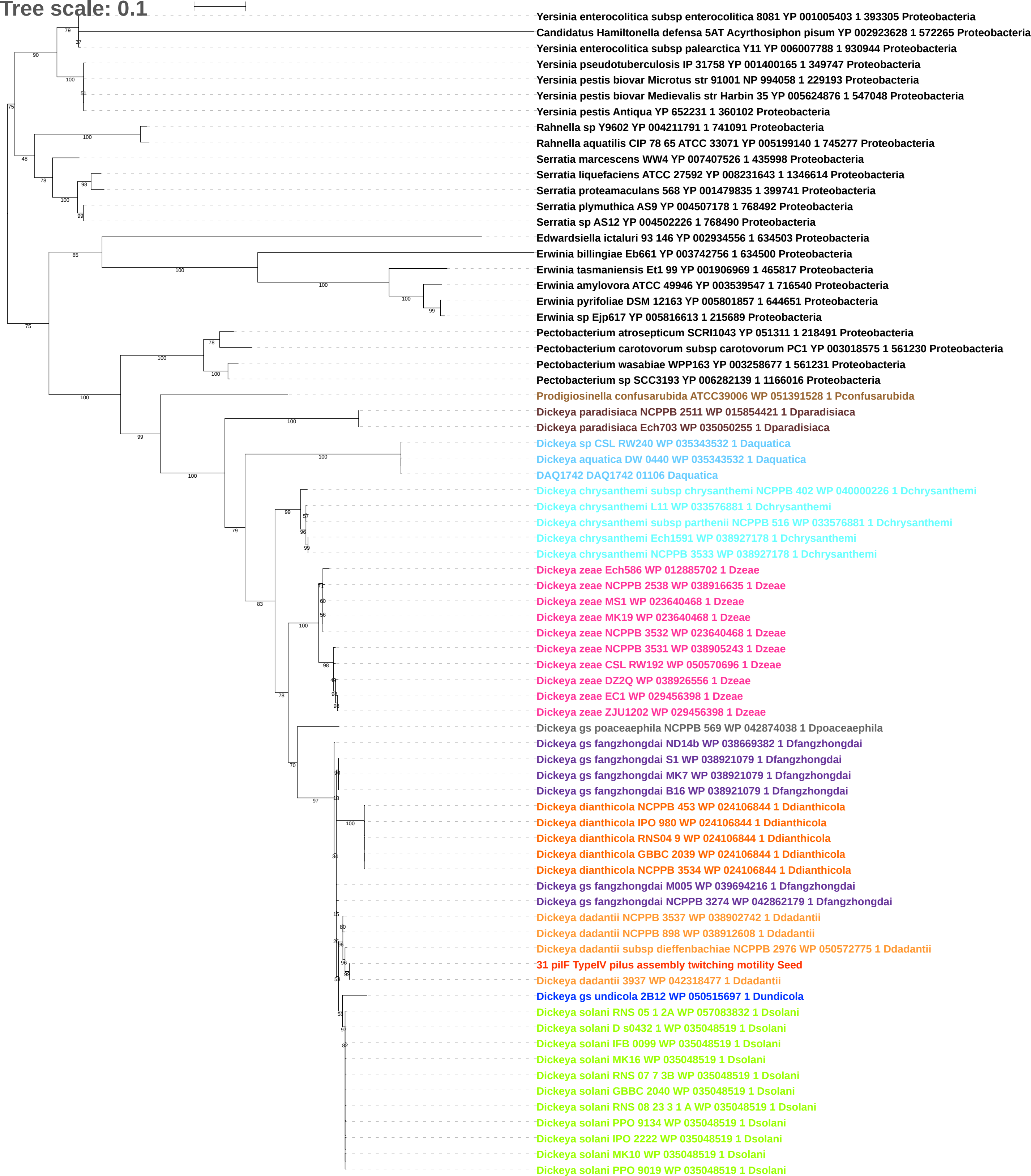
Tree scale: 0.1
Yersinia enterocolitica subsp enterocolitica 8081 YP 001005403 1 393305 Proteobacteria
Candidatus Hamiltonella defensa 5AT Acyrthosiphon pisum YP 002923628 1 572265 Proteobacteria
Yersinia enterocolitica subsp palearctica Y11 YP 006007788 1 930944 Proteobacteria
Yersinia pseudotuberculosis IP 31758 YP 001400165 1 349747 Proteobacteria
Yersinia pestis biovar Microtus str 91001 NP 994058 1 229193 Proteobacteria
Yersinia pestis biovar Medievalis str Harbin 35 YP 005624876 1 547048 Proteobacteria
Yersinia pestis Antiqua YP 652231 1 360102 Proteobacteria
Rahnella sp Y9602 YP 004211791 1 741091 Proteobacteria
Rahnella aquatilis CIP 78 65 ATCC 33071 YP 005199140 1 745277 Proteobacteria
Serratia marcescens WW4 YP 007407526 1 435998 Proteobacteria
Serratia liquefaciens ATCC 27592 YP 008231643 1 1346614 Proteobacteria
Serratia proteamaculans 568 YP 001479835 1 399741 Proteobacteria
Serratia plymuthica AS9 YP 004507178 1 768492 Proteobacteria
Serratia sp AS12 YP 004502226 1 768490 Proteobacteria
Edwardsiella ictaluri 93 146 YP 002934556 1 634503 Proteobacteria
Erwinia billingiae Eb661 YP 003742756 1 634500 Proteobacteria
Erwinia tasmaniensis Et1 99 YP 001906969 1 465817 Proteobacteria
Erwinia amylovora ATCC 49946 YP 003539547 1 716540 Proteobacteria
Erwinia pyrifoliae DSM 12163 YP 005801857 1 644651 Proteobacteria
Erwinia sp Ejp617 YP 005816613 1 215689 Proteobacteria
Pectobacterium atrosepticum SCRI1043 YP 051311 1 218491 Proteobacteria
Pectobacterium carotovorum subsp carotovorum PC1 YP 003018575 1 561230 Proteobacteria
Pectobacterium wasabiae WPP163 YP 003258677 1 561231 Proteobacteria
Pectobacterium sp SCC3193 YP 006282139 1 1166016 Proteobacteria
Prodigiosinella confusarubida ATCC39006 WP 051391528 1 Pconfusarubida
Dickeya paradisiaca NCPPB 2511 WP 015854421 1 Dparadisiaca
Dickeya paradisiaca Ech703 WP 035050255 1 Dparadisiaca
Dickeya sp CSL RW240 WP 035343532 1 Daquatica
Dickeya aquatica DW 0440 WP 035343532 1 Daquatica
DAQ1742 DAQ1742 01106 Daquatica
Dickeya chrysanthemi subsp chrysanthemi NCPPB 402 WP 040000226 1 Dchrysanthemi
Dickeya chrysanthemi L11 WP 033576881 1 Dchrysanthemi
Dickeya chrysanthemi subsp parthenii NCPPB 516 WP 033576881 1 Dchrysanthemi
Dickeya chrysanthemi Ech1591 WP 038927178 1 Dchrysanthemi
Dickeya chrysanthemi NCPPB 3533 WP 038927178 1 Dchrysanthemi
Dickeya zeae Ech586 WP 012885702 1 Dzeae
Dickeya zeae NCPPB 2538 WP 038916635 1 Dzeae
Dickeya zeae MS1 WP 023640468 1 Dzeae
Dickeya zeae MK19 WP 023640468 1 Dzeae
Dickeya zeae NCPPB 3532 WP 023640468 1 Dzeae
Dickeya zeae NCPPB 3531 WP 038905243 1 Dzeae
Dickeya zeae CSL RW192 WP 050570696 1 Dzeae
Dickeya zeae DZ2Q WP 038926556 1 Dzeae
Dickeya zeae EC1 WP 029456398 1 Dzeae
Dickeya zeae ZJU1202 WP 029456398 1 Dzeae
Dickeya gs poaceaephila NCPPB 569 WP 042874038 1 Dpoaceaephila
Dickeya gs fangzhongdai ND14b WP 038669382 1 Dfangzhongdai
Dickeya gs fangzhongdai S1 WP 038921079 1 Dfangzhongdai
Dickeya gs fangzhongdai MK7 WP 038921079 1 Dfangzhongdai
Dickeya gs fangzhongdai B16 WP 038921079 1 Dfangzhongdai
Dickeya dianthicola NCPPB 453 WP 024106844 1 Ddianthicola
Dickeya dianthicola IPO 980 WP 024106844 1 Ddianthicola
Dickeya dianthicola RNS04 9 WP 024106844 1 Ddianthicola
Dickeya dianthicola GBBC 2039 WP 024106844 1 Ddianthicola
Dickeya dianthicola NCPPB 3534 WP 024106844 1 Ddianthicola
Dickeya gs fangzhongdai M005 WP 039694216 1 Dfangzhongdai
Dickeya gs fangzhongdai NCPPB 3274 WP 042862179 1 Dfangzhongdai
Dickeya dadantii NCPPB 3537 WP 038902742 1 Ddadantii
Dickeya dadantii NCPPB 898 WP 038912608 1 Ddadantii
Dickeya dadantii subsp dieffenbachiae NCPPB 2976 WP 050572775 1 Ddadantii
31 pilF TypeIV pilus assembly twitching motility Seed
Dickeya dadantii 3937 WP 042318477 1 Ddadantii
Dickeya gs undicola 2B12 WP 050515697 1 Dundicola
Dickeya solani RNS 05 1 2A WP 057083832 1 Dsolani
Dickeya solani D s0432 1 WP 035048519 1 Dsolani
Dickeya solani IFB 0099 WP 035048519 1 Dsolani
Dickeya solani MK16 WP 035048519 1 Dsolani
Dickeya solani RNS 07 7 3B WP 035048519 1 Dsolani
Dickeya solani GBBC 2040 WP 035048519 1 Dsolani
Dickeya solani RNS 08 23 3 1 A WP 035048519 1 Dsolani
Dickeya solani PPO 9134 WP 035048519 1 Dsolani
Dickeya solani IPO 2222 WP 035048519 1 Dsolani
Dickeya solani MK10 WP 035048519 1 Dsolani
Dickeya solani PPO 9019 WP 035048519 1 Dsolani

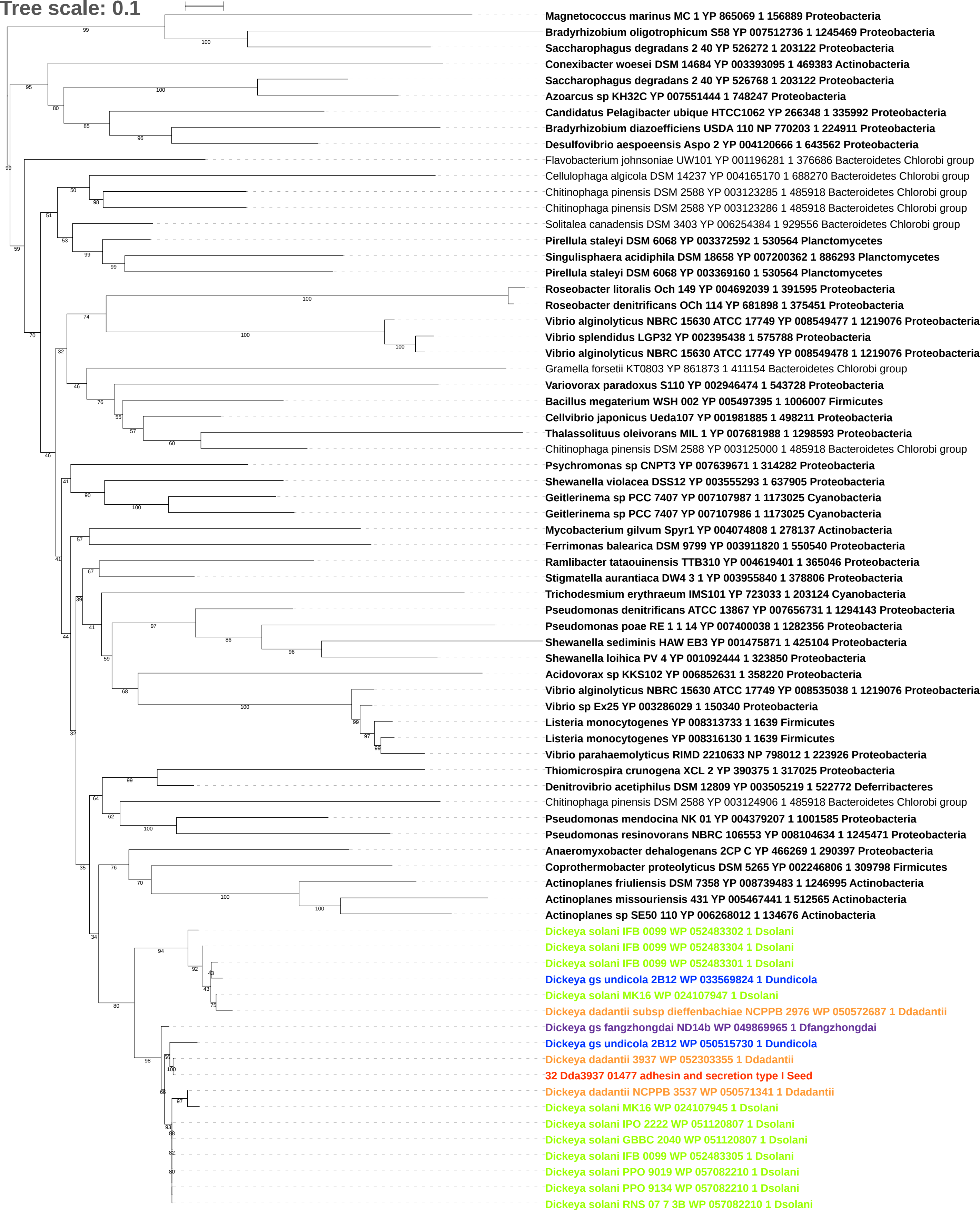
Tree scale: 0.1
Magnetococcus marinus MC 1 YP 865069 1 156889 Proteobacteria
Bradyrhizobium oligotrophicum S58 YP 007512736 1 1245469 Proteobacteria
Saccharophagus degradans 2 40 YP 526272 1 203122 Proteobacteria
Conexibacter woesei DSM 14684 YP 003393095 1 469383 Actinobacteria
Saccharophagus degradans 2 40 YP 526768 1 203122 Proteobacteria
Azoarcus sp KH32C YP 007551444 1 748247 Proteobacteria
Candidatus Pelagibacter ubique HTCC1062 YP 266348 1 335992 Proteobacteria
Bradyrhizobium diazoefficiens USDA 110 NP 770203 1 224911 Proteobacteria
Desulfovibrio aespoeensis Aspo 2 YP 004120666 1 643562 Proteobacteria
Flavobacterium johnsoniae UW101 YP 001196281 1 376686 Bacteroidetes Chlorobi group
Cellulophaga algicola DSM 14237 YP 004165170 1 688270 Bacteroidetes Chlorobi group
Chitinophaga pinensis DSM 2588 YP 003123285 1 485918 Bacteroidetes Chlorobi group
Chitinophaga pinensis DSM 2588 YP 003123286 1 485918 Bacteroidetes Chlorobi group
Solitalea canadensis DSM 3403 YP 006254384 1 929556 Bacteroidetes Chlorobi group
Pirellula staleyi DSM 6068 YP 003372592 1 530564 Planctomycetes
Singulisphaera acidiphila DSM 18658 YP 007200362 1 886293 Planctomycetes
Pirellula staleyi DSM 6068 YP 003369160 1 530564 Planctomycetes
Roseobacter litoralis Och 149 YP 004692039 1 391595 Proteobacteria
Roseobacter denitrificans OCh 114 YP 681898 1 375451 Proteobacteria
Vibrio alginolyticus NBRC 15630 ATCC 17749 YP 008549477 1 1219076 Proteobacteria
Vibrio splendidus LGP32 YP 002395438 1 575788 Proteobacteria
Vibrio alginolyticus NBRC 15630 ATCC 17749 YP 008549478 1 1219076 Proteobacteria
Gramella forsetii KT0803 YP 861873 1 411154 Bacteroidetes Chlorobi group
Variovorax paradoxus S110 YP 002946474 1 543728 Proteobacteria
Bacillus megaterium WSH 002 YP 005497395 1 1006007 Firmicutes
Cellvibrio japonicus Ueda107 YP 001981885 1 498211 Proteobacteria
Thalassolituus oleivorans MIL 1 YP 007681988 1 1298593 Proteobacteria
Chitinophaga pinensis DSM 2588 YP 003125000 1 485918 Bacteroidetes Chlorobi group
Psychromonas sp CNPT3 YP 007639671 1 314282 Proteobacteria
Shewanella violacea DSS12 YP 003555293 1 637905 Proteobacteria
Geitlerinema sp PCC 7407 YP 007107987 1 1173025 Cyanobacteria
Geitlerinema sp PCC 7407 YP 007107986 1 1173025 Cyanobacteria
Mycobacterium gilvum Spyr1 YP 004074808 1 278137 Actinobacteria
Ferrimonas balearica DSM 9799 YP 003911820 1 550540 Proteobacteria
Ramlibacter tataouinensis TTB310 YP 004619401 1 365046 Proteobacteria
Stigmatella aurantiaca DW4 3 1 YP 003955840 1 378806 Proteobacteria
Trichodesmium erythraeum IMS101 YP 723033 1 203124 Cyanobacteria
Pseudomonas denitrificans ATCC 13867 YP 007656731 1 1294143 Proteobacteria
Pseudomonas poae RE 1 1 14 YP 007400038 1 1282356 Proteobacteria
Shewanella sediminis HAW EB3 YP 001475871 1 425104 Proteobacteria
Shewanella loihica PV 4 YP 001092444 1 323850 Proteobacteria
Acidovorax sp KKS102 YP 006852631 1 358220 Proteobacteria
Vibrio alginolyticus NBRC 15630 ATCC 17749 YP 008535038 1 1219076 Proteobacteria
Vibrio sp Ex25 YP 003286029 1 150340 Proteobacteria
Listeria monocytogenes YP 008313733 1 1639 Firmicutes
Listeria monocytogenes YP 008316130 1 1639 Firmicutes
Vibrio parahaemolyticus RIMD 2210633 NP 798012 1 223926 Proteobacteria
Thiomicrospira crunogena XCL 2 YP 390375 1 317025 Proteobacteria
Denitrovibrio acetiphilus DSM 12809 YP 003505219 1 522772 Deferribacteres
Chitinophaga pinensis DSM 2588 YP 003124906 1 485918 Bacteroidetes Chlorobi group
Pseudomonas mendocina NK 01 YP 004379207 1 1001585 Proteobacteria
Pseudomonas resinovorans NBRC 106553 YP 008104634 1 1245471 Proteobacteria
Anaeromyxobacter dehalogenans 2CP C YP 466269 1 290397 Proteobacteria
Coprothermobacter proteolyticus DSM 5265 YP 002246806 1 309798 Firmicutes
Actinoplanes friuliensis DSM 7358 YP 008739483 1 1246995 Actinobacteria
Actinoplanes missouriensis 431 YP 005467441 1 512565 Actinobacteria
Actinoplanes sp SE50 110 YP 006268012 1 134676 Actinobacteria
Dickeya solani IFB 0099 WP 052483302 1 Dsolani
Dickeya solani IFB 0099 WP 052483304 1 Dsolani
Dickeya solani IFB 0099 WP 052483301 1 Dsolani
Dickeya gs undicola 2B12 WP 033569824 1 Dundicola
Dickeya solani MK16 WP 024107947 1 Dsolani
Dickeya dadantii subsp dieffenbachiae NCPPB 2976 WP 050572687 1 Ddadantii
Dickeya gs fangzhongdai ND14b WP 049869965 1 Dfangzhongdai
Dickeya gs undicola 2B12 WP 050515730 1 Dundicola
Dickeya dadantii 3937 WP 052303355 1 Ddadantii
32 Dda3937 01477 adhesin and secretion type I Seed
Dickeya dadantii NCPPB 3537 WP 050571341 1 Ddadantii
Dickeya solani MK16 WP 024107945 1 Dsolani
Dickeya solani IPO 2222 WP 051120807 1 Dsolani
Dickeya solani GBBC 2040 WP 051120807 1 Dsolani
Dickeya solani IFB 0099 WP 052483305 1 Dsolani
Dickeya solani PPO 9019 WP 057082210 1 Dsolani
Dickeya solani PPO 9134 WP 057082210 1 Dsolani
Dickeya solani RNS 07 7 3B WP 057082210 1 Dsolani

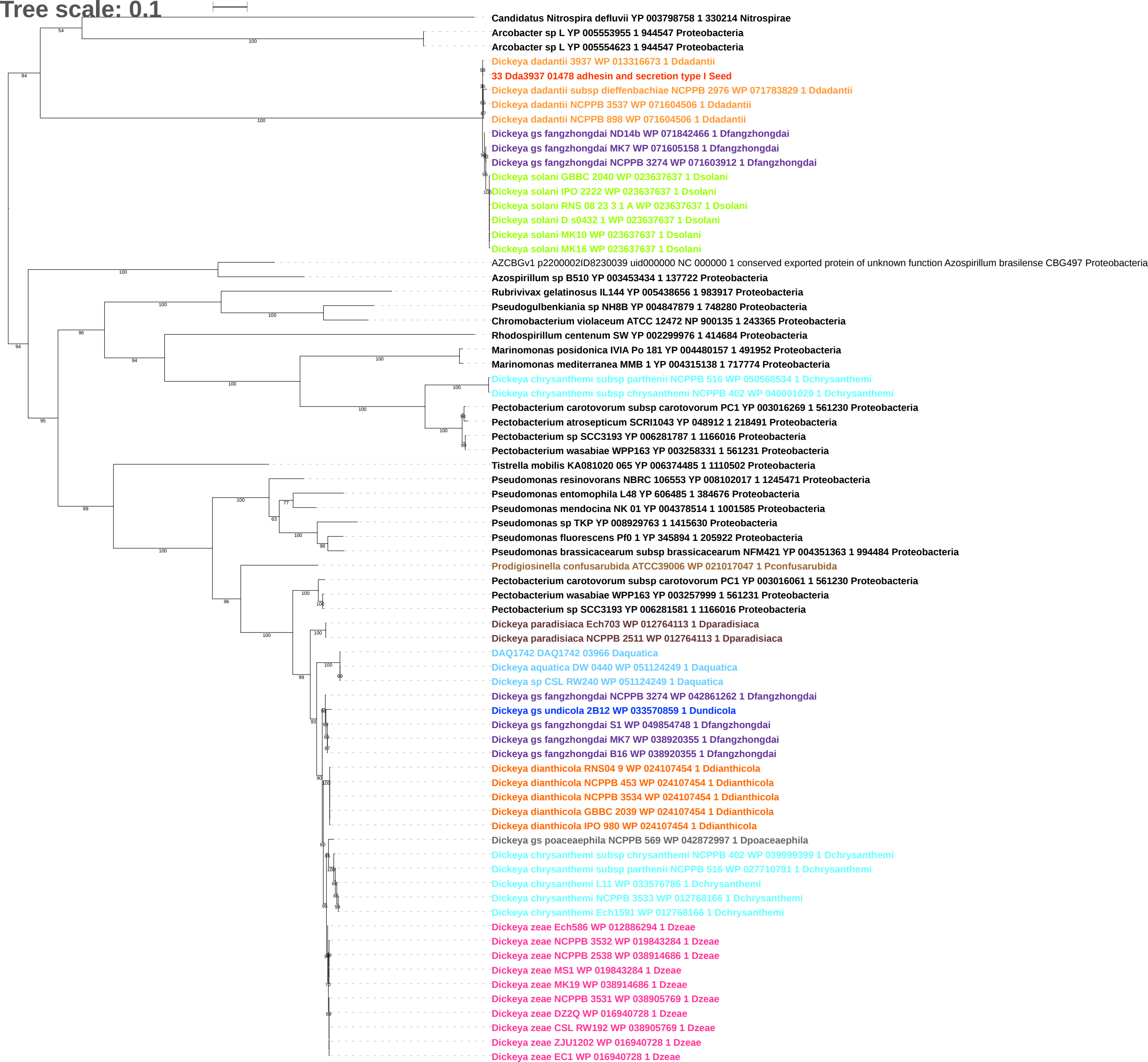
Tree scale: 0.1
Candidatus Nitrospira defluvii YP 003798758 1 330214 Nitrospirae
Arcobacter sp L YP 005553955 1 944547 Proteobacteria
Arcobacter sp L YP 005554623 1 944547 Proteobacteria
Dickeya dadantii 3937 WP 013316673 1 Ddadantii
33 Dda3937 01478 adhesin and secretion type I Seed
Dickeya dadantii subsp dieffenbachiae NCPPB 2976 WP 071783829 1 Ddadantii
Dickeya dadantii NCPPB 3537 WP 071604506 1 Ddadantii
Dickeya dadantii NCPPB 898 WP 071604506 1 Ddadantii
Dickeya gs fangzhongdai ND14b WP 071842466 1 Dfangzhongdai
Dickeya gs fangzhongdai MK7 WP 071605158 1 Dfangzhongdai
Dickeya gs fangzhongdai NCPPB 3274 WP 071603912 1 Dfangzhongdai
Dickeya solani GBBC 2040 WP 023637637 1 Dsolani
Dickeya solani IPO 2222 WP 023637637 1 Dsolani
Dickeya solani RNS 08 23 3 1 A WP 023637637 1 Dsolani
Dickeya solani D s0432 1 WP 023637637 1 Dsolani
Dickeya solani MK10 WP 023637637 1 Dsolani
Dickeya solani MK16 WP 023637637 1 Dsolani
AZCBGv1 p2200002ID8230039 uid000000 NC 000000 1 conserved exported protein of unknown function Azospirillum brasilense CBG497 Proteobacteria
Azospirillum sp B510 YP 003453434 1 137722 Proteobacteria
Rubrivivax gelatinosus IL144 YP 005438656 1 983917 Proteobacteria
Pseudogulbenkiania sp NH8B YP 004847879 1 748280 Proteobacteria
Chromobacterium violaceum ATCC 12472 NP 900135 1 243365 Proteobacteria
Rhodospirillum centenum SW YP 002299976 1 414684 Proteobacteria
Marinomonas posidonica IVIA Po 181 YP 004480157 1 491952 Proteobacteria
Marinomonas mediterranea MMB 1 YP 004315138 1 717774 Proteobacteria
Dickeya chrysanthemi subsp parthenii NCPPB 516 WP 050568534 1 Dchrysanthemi
Dickeya chrysanthemi subsp chrysanthemi NCPPB 402 WP 040001029 1 Dchrysanthemi
Pectobacterium carotovorum subsp carotovorum PC1 YP 003016269 1 561230 Proteobacteria
Pectobacterium atrosepticum SCRI1043 YP 048912 1 218491 Proteobacteria
Pectobacterium sp SCC3193 YP 006281787 1 1166016 Proteobacteria
Pectobacterium wasabiae WPP163 YP 003258331 1 561231 Proteobacteria
Tistrella mobilis KA081020 065 YP 006374485 1 1110502 Proteobacteria
Pseudomonas resinovorans NBRC 106553 YP 008102017 1 1245471 Proteobacteria
Pseudomonas entomophila L48 YP 606485 1 384676 Proteobacteria
Pseudomonas mendocina NK 01 YP 004378514 1 1001585 Proteobacteria
Pseudomonas sp TKP YP 008929763 1 1415630 Proteobacteria
Pseudomonas fluorescens Pf0 1 YP 345894 1 205922 Proteobacteria
Pseudomonas brassicacearum subsp brassicacearum NFM421 YP 004351363 1 994484 Proteobacteria
Prodigiosinella confusarubida ATCC39006 WP 021017047 1 Pconfusarubida
Pectobacterium carotovorum subsp carotovorum PC1 YP 003016061 1 561230 Proteobacteria
Pectobacterium wasabiae WPP163 YP 003257999 1 561231 Proteobacteria
Pectobacterium sp SCC3193 YP 006281581 1 1166016 Proteobacteria
Dickeya paradisiaca Ech703 WP 012764113 1 Dparadisiaca
Dickeya paradisiaca NCPPB 2511 WP 012764113 1 Dparadisiaca
DAQ1742 DAQ1742 03966 Daquatica
Dickeya aquatica DW 0440 WP 051124249 1 Daquatica
Dickeya sp CSL RW240 WP 051124249 1 Daquatica
Dickeya gs fangzhongdai NCPPB 3274 WP 042861262 1 Dfangzhongdai
Dickeya gs undicola 2B12 WP 033570859 1 Dundicola
Dickeya gs fangzhongdai S1 WP 049854748 1 Dfangzhongdai
Dickeya gs fangzhongdai MK7 WP 038920355 1 Dfangzhongdai
Dickeya gs fangzhongdai B16 WP 038920355 1 Dfangzhongdai
Dickeya dianthicola RNS04 9 WP 024107454 1 Ddianthicola
Dickeya dianthicola NCPPB 453 WP 024107454 1 Ddianthicola
Dickeya dianthicola NCPPB 3534 WP 024107454 1 Ddianthicola
Dickeya dianthicola GBBC 2039 WP 024107454 1 Ddianthicola
Dickeya dianthicola IPO 980 WP 024107454 1 Ddianthicola
Dickeya gs poaceaephila NCPPB 569 WP 042872997 1 Dpoaceaephila
Dickeya chrysanthemi subsp chrysanthemi NCPPB 402 WP 039999399 1 Dchrysanthemi
Dickeya chrysanthemi subsp parthenii NCPPB 516 WP 027710791 1 Dchrysanthemi
Dickeya chrysanthemi L11 WP 033576786 1 Dchrysanthemi
Dickeya chrysanthemi NCPPB 3533 WP 012768166 1 Dchrysanthemi
Dickeya chrysanthemi Ech1591 WP 012768166 1 Dchrysanthemi
Dickeya zeae Ech586 WP 012886294 1 Dzeae
Dickeya zeae NCPPB 3532 WP 019843284 1 Dzeae
Dickeya zeae NCPPB 2538 WP 038914686 1 Dzeae
Dickeya zeae MS1 WP 019843284 1 Dzeae
Dickeya zeae MK19 WP 038914686 1 Dzeae
Dickeya zeae NCPPB 3531 WP 038905769 1 Dzeae
Dickeya zeae DZ2Q WP 016940728 1 Dzeae
Dickeya zeae CSL RW192 WP 038905769 1 Dzeae
Dickeya zeae ZJU1202 WP 016940728 1 Dzeae
Dickeya zeae EC1 WP 016940728 1 Dzeae

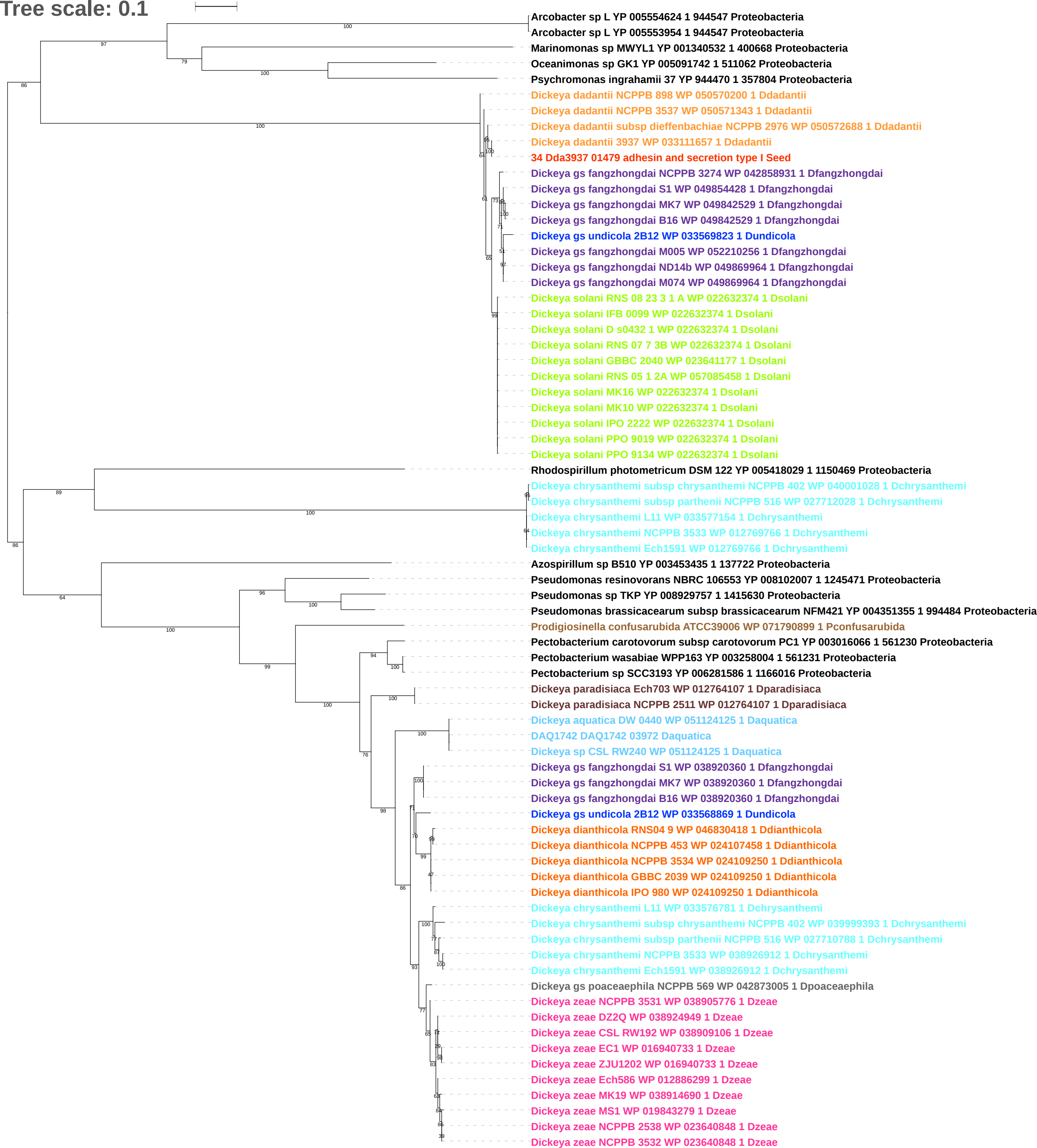
Tree scale: 0.1
Arcobacter sp L YP 005554624 1 944547 Proteobacteria
Arcobacter sp L YP 005553954 1 944547 Proteobacteria
Marinomonas sp MWYL1 YP 001340532 1 400668 Proteobacteria
Oceanimonas sp GK1 YP 005091742 1 511062 Proteobacteria
Psychromonas ingrahamii 37 YP 944470 1 357804 Proteobacteria
Dickeya dadantii NCPPB 898 WP 050570200 1 Ddadantii
Dickeya dadantii NCPPB 3537 WP 050571343 1 Ddadantii
Dickeya dadantii subsp dieffenbachiae NCPPB 2976 WP 050572688 1 Ddadantii
Dickeya dadantii 3937 WP 033111657 1 Ddadantii
34 Dda3937 01479 adhesin and secretion type I Seed
Dickeya gs fangzhongdai NCPPB 3274 WP 042858931 1 Dfangzhongdai
Dickeya gs fangzhongdai S1 WP 049854428 1 Dfangzhongdai
Dickeya gs fangzhongdai MK7 WP 049842529 1 Dfangzhongdai
Dickeya gs fangzhongdai B16 WP 049842529 1 Dfangzhongdai
Dickeya gs undicola 2B12 WP 033569823 1 Dundicola
Dickeya gs fangzhongdai M005 WP 052210256 1 Dfangzhongdai
Dickeya gs fangzhongdai ND14b WP 049869964 1 Dfangzhongdai
Dickeya gs fangzhongdai M074 WP 049869964 1 Dfangzhongdai
Dickeya solani RNS 08 23 3 1 A WP 022632374 1 Dsolani
Dickeya solani IFB 0099 WP 022632374 1 Dsolani
Dickeya solani D s0432 1 WP 022632374 1 Dsolani
Dickeya solani RNS 07 7 3B WP 022632374 1 Dsolani
Dickeya solani GBBC 2040 WP 023641177 1 Dsolani
Dickeya solani RNS 05 1 2A WP 057085458 1 Dsolani
Dickeya solani MK16 WP 022632374 1 Dsolani
Dickeya solani MK10 WP 022632374 1 Dsolani
Dickeya solani IPO 2222 WP 022632374 1 Dsolani
Dickeya solani PPO 9019 WP 022632374 1 Dsolani
Dickeya solani PPO 9134 WP 022632374 1 Dsolani
Rhodospirillum photometricum DSM 122 YP 005418029 1 1150469 Proteobacteria
Dickeya chrysanthemi subsp chrysanthemi NCPPB 402 WP 040001028 1 Dchrysanthemi
Dickeya chrysanthemi subsp parthenii NCPPB 516 WP 027712028 1 Dchrysanthemi
Dickeya chrysanthemi L11 WP 033577154 1 Dchrysanthemi
Dickeya chrysanthemi NCPPB 3533 WP 012769766 1 Dchrysanthemi
Dickeya chrysanthemi Ech1591 WP 012769766 1 Dchrysanthemi
Azospirillum sp B510 YP 003453435 1 137722 Proteobacteria
Pseudomonas resinovorans NBRC 106553 YP 008102007 1 1245471 Proteobacteria
Pseudomonas sp TKP YP 008929757 1 1415630 Proteobacteria
Pseudomonas brassicacearum subsp brassicacearum NFM421 YP 004351355 1 994484 Proteobacteria
Prodigiosinella confusarubida ATCC39006 WP 071790899 1 Pconfusarubida
Pectobacterium carotovorum subsp carotovorum PC1 YP 003016066 1 561230 Proteobacteria
Pectobacterium wasabiae WPP163 YP 003258004 1 561231 Proteobacteria
Pectobacterium sp SCC3193 YP 006281586 1 1166016 Proteobacteria
Dickeya paradisiaca Ech703 WP 012764107 1 Dparadisiaca
Dickeya paradisiaca NCPPB 2511 WP 012764107 1 Dparadisiaca
Dickeya aquatica DW 0440 WP 051124125 1 Daquatica
DAQ1742 DAQ1742 03972 Daquatica
Dickeya sp CSL RW240 WP 051124125 1 Daquatica
Dickeya gs fangzhongdai S1 WP 038920360 1 Dfangzhongdai
Dickeya gs fangzhongdai MK7 WP 038920360 1 Dfangzhongdai
Dickeya gs fangzhongdai B16 WP 038920360 1 Dfangzhongdai
Dickeya gs undicola 2B12 WP 033568869 1 Dundicola
Dickeya dianthicola RNS04 9 WP 046830418 1 Ddianthicola
Dickeya dianthicola NCPPB 453 WP 024107458 1 Ddianthicola
Dickeya dianthicola NCPPB 3534 WP 024109250 1 Ddianthicola
Dickeya dianthicola GBBC 2039 WP 024109250 1 Ddianthicola
Dickeya dianthicola IPO 980 WP 024109250 1 Ddianthicola
Dickeya chrysanthemi L11 WP 033576781 1 Dchrysanthemi
Dickeya chrysanthemi subsp chrysanthemi NCPPB 402 WP 039999393 1 Dchrysanthemi
Dickeya chrysanthemi subsp parthenii NCPPB 516 WP 027710788 1 Dchrysanthemi
Dickeya chrysanthemi NCPPB 3533 WP 038926912 1 Dchrysanthemi
Dickeya chrysanthemi Ech1591 WP 038926912 1 Dchrysanthemi
Dickeya gs poaceaephila NCPPB 569 WP 042873005 1 Dpoaceaephila
Dickeya zeae NCPPB 3531 WP 038905776 1 Dzeae
Dickeya zeae DZ2Q WP 038924949 1 Dzeae
Dickeya zeae CSL RW192 WP 038909106 1 Dzeae
Dickeya zeae EC1 WP 016940733 1 Dzeae
Dickeya zeae ZJU1202 WP 016940733 1 Dzeae
Dickeya zeae Ech586 WP 012886299 1 Dzeae
Dickeya zeae MK19 WP 038914690 1 Dzeae
Dickeya zeae MS1 WP 019843279 1 Dzeae
Dickeya zeae NCPPB 2538 WP 023640848 1 Dzeae
Dickeya zeae NCPPB 3532 WP 023640848 1 Dzeae

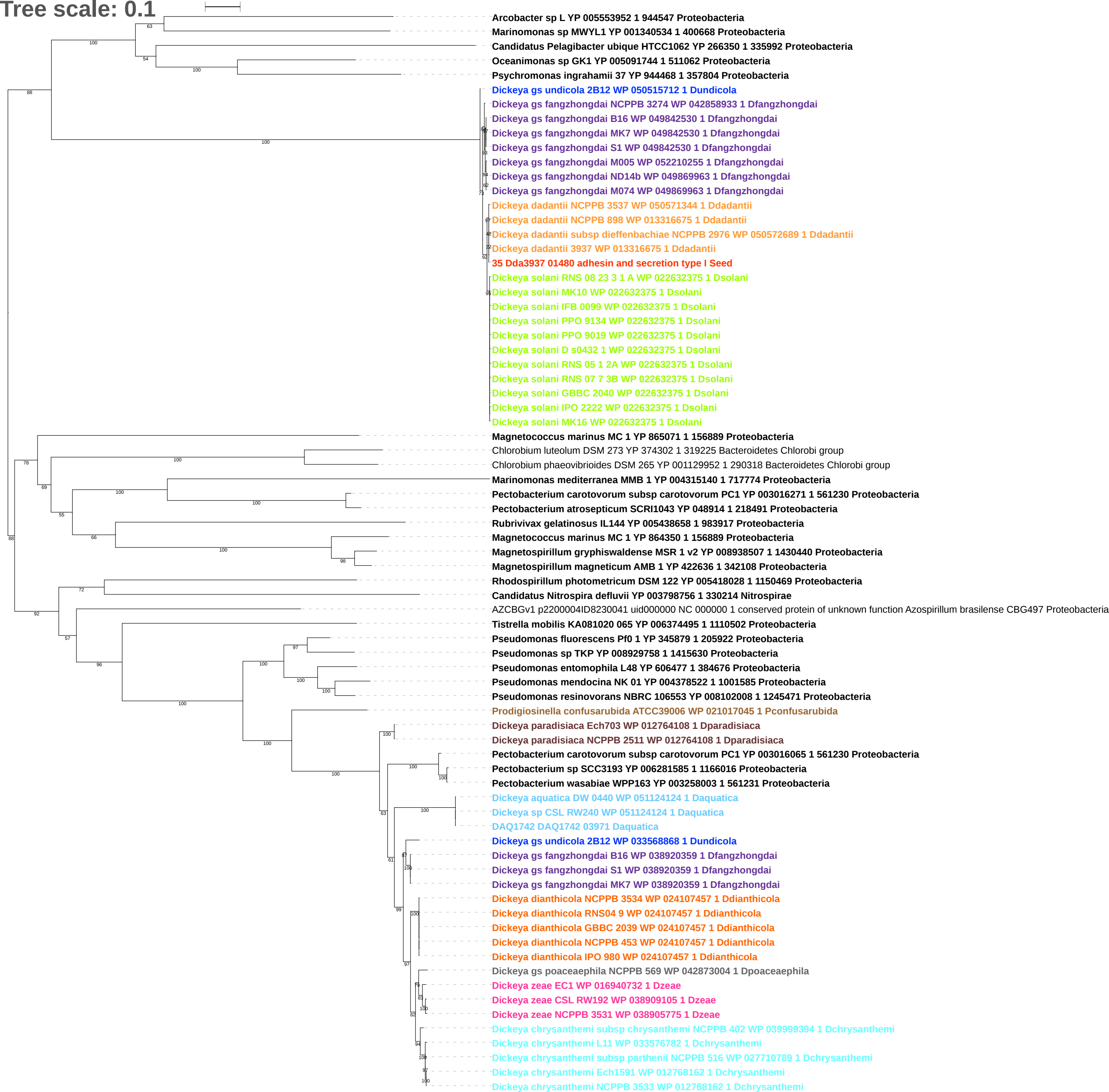
Tree scale: 0.1
Arcobacter sp L YP 005553952 1 944547 Proteobacteria
Marinomonas sp MWYL1 YP 001340534 1 400668 Proteobacteria
Candidatus Pelagibacter ubique HTCC1062 YP 266350 1 335992 Proteobacteria
Oceanimonas sp GK1 YP 005091744 1 511062 Proteobacteria
Psychromonas ingrahamii 37 YP 944468 1 357804 Proteobacteria
Dickeya gs undicola 2B12 WP 050515712 1 Dundicola
Dickeya gs fangzhongdai NCPPB 3274 WP 042858933 1 Dfangzhongdai
Dickeya gs fangzhongdai B16 WP 049842530 1 Dfangzhongdai
Dickeya gs fangzhongdai MK7 WP 049842530 1 Dfangzhongdai
Dickeya gs fangzhongdai S1 WP 049842530 1 Dfangzhongdai
Dickeya gs fangzhongdai M005 WP 052210255 1 Dfangzhongdai
Dickeya gs fangzhongdai ND14b WP 049869963 1 Dfangzhongdai
Dickeya gs fangzhongdai M074 WP 049869963 1 Dfangzhongdai
Dickeya dadantii NCPPB 3537 WP 050571344 1 Ddadantii
Dickeya dadantii NCPPB 898 WP 013316675 1 Ddadantii
Dickeya dadantii subsp dieffenbachiae NCPPB 2976 WP 050572689 1 Ddadantii
Dickeya dadantii 3937 WP 013316675 1 Ddadantii
35 Dda3937 01480 adhesin and secretion type I Seed
Dickeya solani RNS 08 23 3 1 A WP 022632375 1 Dsolani
Dickeya solani MK10 WP 022632375 1 Dsolani
Dickeya solani IFB 0099 WP 022632375 1 Dsolani
Dickeya solani PPO 9134 WP 022632375 1 Dsolani
Dickeya solani PPO 9019 WP 022632375 1 Dsolani
Dickeya solani D s0432 1 WP 022632375 1 Dsolani
Dickeya solani RNS 05 1 2A WP 022632375 1 Dsolani
Dickeya solani RNS 07 7 3B WP 022632375 1 Dsolani
Dickeya solani GBBC 2040 WP 022632375 1 Dsolani
Dickeya solani IPO 2222 WP 022632375 1 Dsolani
Dickeya solani MK16 WP 022632375 1 Dsolani
Magnetococcus marinus MC 1 YP 865071 1 156889 Proteobacteria
Chlorobium luteolum DSM 273 YP 374302 1 319225 Bacteroidetes Chlorobi group
Chlorobium phaeovibrioides DSM 265 YP 001129952 1 290318 Bacteroidetes Chlorobi group
Marinomonas mediterranea MMB 1 YP 004315140 1 717774 Proteobacteria
Pectobacterium carotovorum subsp carotovorum PC1 YP 003016271 1 561230 Proteobacteria
Pectobacterium atrosepticum SCRI1043 YP 048914 1 218491 Proteobacteria
Rubrivivax gelatinosus IL144 YP 005438658 1 983917 Proteobacteria
Magnetococcus marinus MC 1 YP 864350 1 156889 Proteobacteria
Magnetospirillum gryphiswaldense MSR 1 v2 YP 008938507 1 1430440 Proteobacteria
Magnetospirillum magneticum AMB 1 YP 422636 1 342108 Proteobacteria
Rhodospirillum photometricum DSM 122 YP 005418028 1 1150469 Proteobacteria
Candidatus Nitrospira defluvii YP 003798756 1 330214 Nitrospirae
AZCBGv1 p2200004ID8230041 uid000000 NC 000000 1 conserved protein of unknown function Azospirillum brasilense CBG497 Proteobacteria
Tistrella mobilis KA081020 065 YP 006374495 1 1110502 Proteobacteria
Pseudomonas fluorescens Pf0 1 YP 345879 1 205922 Proteobacteria
Pseudomonas sp TKP YP 008929758 1 1415630 Proteobacteria
Pseudomonas entomophila L48 YP 606477 1 384676 Proteobacteria
Pseudomonas mendocina NK 01 YP 004378522 1 1001585 Proteobacteria
Pseudomonas resinovorans NBRC 106553 YP 008102008 1 1245471 Proteobacteria
Prodigiosinella confusarubida ATCC39006 WP 021017045 1 Pconfusarubida
Dickeya paradisiaca Ech703 WP 012764108 1 Dparadisiaca
Dickeya paradisiaca NCPPB 2511 WP 012764108 1 Dparadisiaca
Pectobacterium carotovorum subsp carotovorum PC1 YP 003016065 1 561230 Proteobacteria
Pectobacterium sp SCC3193 YP 006281585 1 1166016 Proteobacteria
Pectobacterium wasabiae WPP163 YP 003258003 1 561231 Proteobacteria
Dickeya aquatica DW 0440 WP 051124124 1 Daquatica
Dickeya sp CSL RW240 WP 051124124 1 Daquatica
DAQ1742 DAQ1742 03971 Daquatica
Dickeya gs undicola 2B12 WP 033568868 1 Dundicola
Dickeya gs fangzhongdai B16 WP 038920359 1 Dfangzhongdai
Dickeya gs fangzhongdai S1 WP 038920359 1 Dfangzhongdai
Dickeya gs fangzhongdai MK7 WP 038920359 1 Dfangzhongdai
Dickeya dianthicola NCPPB 3534 WP 024107457 1 Ddianthicola
Dickeya dianthicola RNS04 9 WP 024107457 1 Ddianthicola
Dickeya dianthicola GBBC 2039 WP 024107457 1 Ddianthicola
Dickeya dianthicola NCPPB 453 WP 024107457 1 Ddianthicola
Dickeya dianthicola IPO 980 WP 024107457 1 Ddianthicola
Dickeya gs poaceaephila NCPPB 569 WP 042873004 1 Dpoaceaephila
Dickeya zeae EC1 WP 016940732 1 Dzeae
Dickeya zeae CSL RW192 WP 038909105 1 Dzeae
Dickeya zeae NCPPB 3531 WP 038905775 1 Dzeae
Dickeya chrysanthemi subsp chrysanthemi NCPPB 402 WP 039999394 1 Dchrysanthemi
Dickeya chrysanthemi L11 WP 033576782 1 Dchrysanthemi
Dickeya chrysanthemi subsp parthenii NCPPB 516 WP 027710789 1 Dchrysanthemi
Dickeya chrysanthemi Ech1591 WP 012768162 1 Dchrysanthemi
Dickeya chrysanthemi NCPPB 3533 WP 012768162 1 Dchrysanthemi

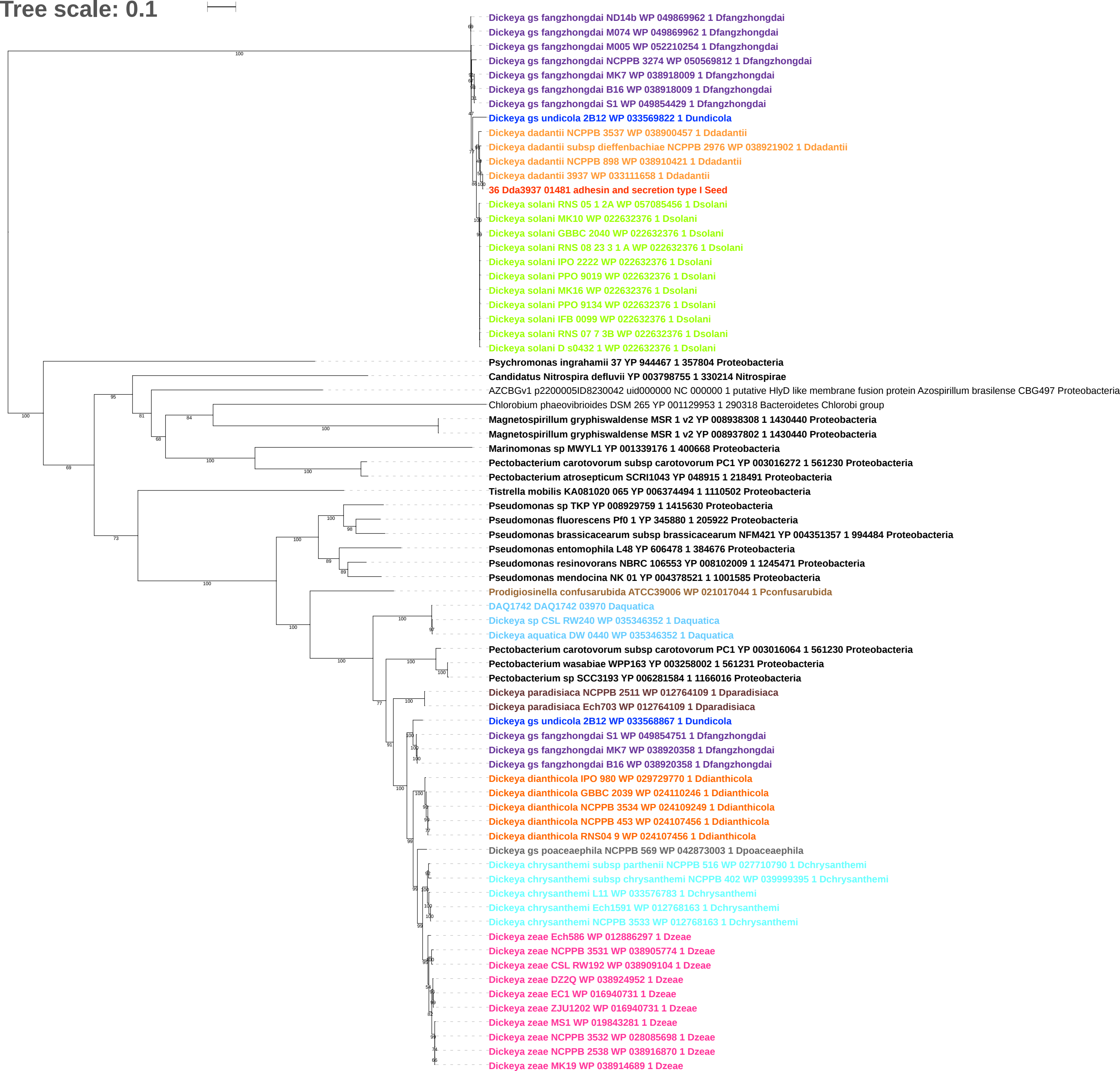
Tree scale: 0.1
Dickeya gs fangzhongdai ND14b WP 049869962 1 Dfangzhongdai
Dickeya gs fangzhongdai M074 WP 049869962 1 Dfangzhongdai
Dickeya gs fangzhongdai M005 WP 052210254 1 Dfangzhongdai
Dickeya gs fangzhongdai NCPPB 3274 WP 050569812 1 Dfangzhongdai
Dickeya gs fangzhongdai MK7 WP 038918009 1 Dfangzhongdai
Dickeya gs fangzhongdai B16 WP 038918009 1 Dfangzhongdai
Dickeya gs fangzhongdai S1 WP 049854429 1 Dfangzhongdai
Dickeya gs undicola 2B12 WP 033569822 1 Dundicola
Dickeya dadantii NCPPB 3537 WP 038900457 1 Ddadantii
Dickeya dadantii subsp dieffenbachiae NCPPB 2976 WP 038921902 1 Ddadantii
Dickeya dadantii NCPPB 898 WP 038910421 1 Ddadantii
Dickeya dadantii 3937 WP 033111658 1 Ddadantii
36 Dda3937 01481 adhesin and secretion type I Seed
Dickeya solani RNS 05 1 2A WP 057085456 1 Dsolani
Dickeya solani MK10 WP 022632376 1 Dsolani
Dickeya solani GBBC 2040 WP 022632376 1 Dsolani
Dickeya solani RNS 08 23 3 1 A WP 022632376 1 Dsolani
Dickeya solani IPO 2222 WP 022632376 1 Dsolani
Dickeya solani PPO 9019 WP 022632376 1 Dsolani
Dickeya solani MK16 WP 022632376 1 Dsolani
Dickeya solani PPO 9134 WP 022632376 1 Dsolani
Dickeya solani IFB 0099 WP 022632376 1 Dsolani
Dickeya solani RNS 07 7 3B WP 022632376 1 Dsolani
Dickeya solani D s0432 1 WP 022632376 1 Dsolani
Psychromonas ingrahamii 37 YP 944467 1 357804 Proteobacteria
Candidatus Nitrospira defluvii YP 003798755 1 330214 Nitrospirae
AZCBGv1 p2200005ID8230042 uid000000 NC 000000 1 putative HlyD like membrane fusion protein Azospirillum brasilense CBG497 Proteobacteria
Chlorobium phaeovibrioides DSM 265 YP 001129953 1 290318 Bacteroidetes Chlorobi group
Magnetospirillum gryphiswaldense MSR 1 v2 YP 008938308 1 1430440 Proteobacteria
Magnetospirillum gryphiswaldense MSR 1 v2 YP 008937802 1 1430440 Proteobacteria
Marinomonas sp MWYL1 YP 001339176 1 400668 Proteobacteria
Pectobacterium carotovorum subsp carotovorum PC1 YP 003016272 1 561230 Proteobacteria
Pectobacterium atrosepticum SCRI1043 YP 048915 1 218491 Proteobacteria
Tistrella mobilis KA081020 065 YP 006374494 1 1110502 Proteobacteria
Pseudomonas sp TKP YP 008929759 1 1415630 Proteobacteria
Pseudomonas fluorescens Pf0 1 YP 345880 1 205922 Proteobacteria
Pseudomonas brassicacearum subsp brassicacearum NFM421 YP 004351357 1 994484 Proteobacteria
Pseudomonas entomophila L48 YP 606478 1 384676 Proteobacteria
Pseudomonas resinovorans NBRC 106553 YP 008102009 1 1245471 Proteobacteria
Pseudomonas mendocina NK 01 YP 004378521 1 1001585 Proteobacteria
Prodigiosinella confusarubida ATCC39006 WP 021017044 1 Pconfusarubida
DAQ1742 DAQ1742 03970 Daquatica
Dickeya sp CSL RW240 WP 035346352 1 Daquatica
Dickeya aquatica DW 0440 WP 035346352 1 Daquatica
Pectobacterium carotovorum subsp carotovorum PC1 YP 003016064 1 561230 Proteobacteria
Pectobacterium wasabiae WPP163 YP 003258002 1 561231 Proteobacteria
Pectobacterium sp SCC3193 YP 006281584 1 1166016 Proteobacteria
Dickeya paradisiaca NCPPB 2511 WP 012764109 1 Dparadisiaca
Dickeya paradisiaca Ech703 WP 012764109 1 Dparadisiaca
Dickeya gs undicola 2B12 WP 033568867 1 Dundicola
Dickeya gs fangzhongdai S1 WP 049854751 1 Dfangzhongdai
Dickeya gs fangzhongdai MK7 WP 038920358 1 Dfangzhongdai
Dickeya gs fangzhongdai B16 WP 038920358 1 Dfangzhongdai
Dickeya dianthicola IPO 980 WP 029729770 1 Ddianthicola
Dickeya dianthicola GBBC 2039 WP 024110246 1 Ddianthicola
Dickeya dianthicola NCPPB 3534 WP 024109249 1 Ddianthicola
Dickeya dianthicola NCPPB 453 WP 024107456 1 Ddianthicola
Dickeya dianthicola RNS04 9 WP 024107456 1 Ddianthicola
Dickeya gs poaceaephila NCPPB 569 WP 042873003 1 Dpoaceaephila
Dickeya chrysanthemi subsp parthenii NCPPB 516 WP 027710790 1 Dchrysanthemi
Dickeya chrysanthemi subsp chrysanthemi NCPPB 402 WP 039999395 1 Dchrysanthemi
Dickeya chrysanthemi L11 WP 033576783 1 Dchrysanthemi
Dickeya chrysanthemi Ech1591 WP 012768163 1 Dchrysanthemi
Dickeya chrysanthemi NCPPB 3533 WP 012768163 1 Dchrysanthemi
Dickeya zeae Ech586 WP 012886297 1 Dzeae
Dickeya zeae NCPPB 3531 WP 038905774 1 Dzeae
Dickeya zeae CSL RW192 WP 038909104 1 Dzeae
Dickeya zeae DZ2Q WP 038924952 1 Dzeae
Dickeya zeae EC1 WP 016940731 1 Dzeae
Dickeya zeae ZJU1202 WP 016940731 1 Dzeae
Dickeya zeae MS1 WP 019843281 1 Dzeae
Dickeya zeae NCPPB 3532 WP 028085698 1 Dzeae
Dickeya zeae NCPPB 2538 WP 038916870 1 Dzeae
Dickeya zeae MK19 WP 038914689 1 Dzeae

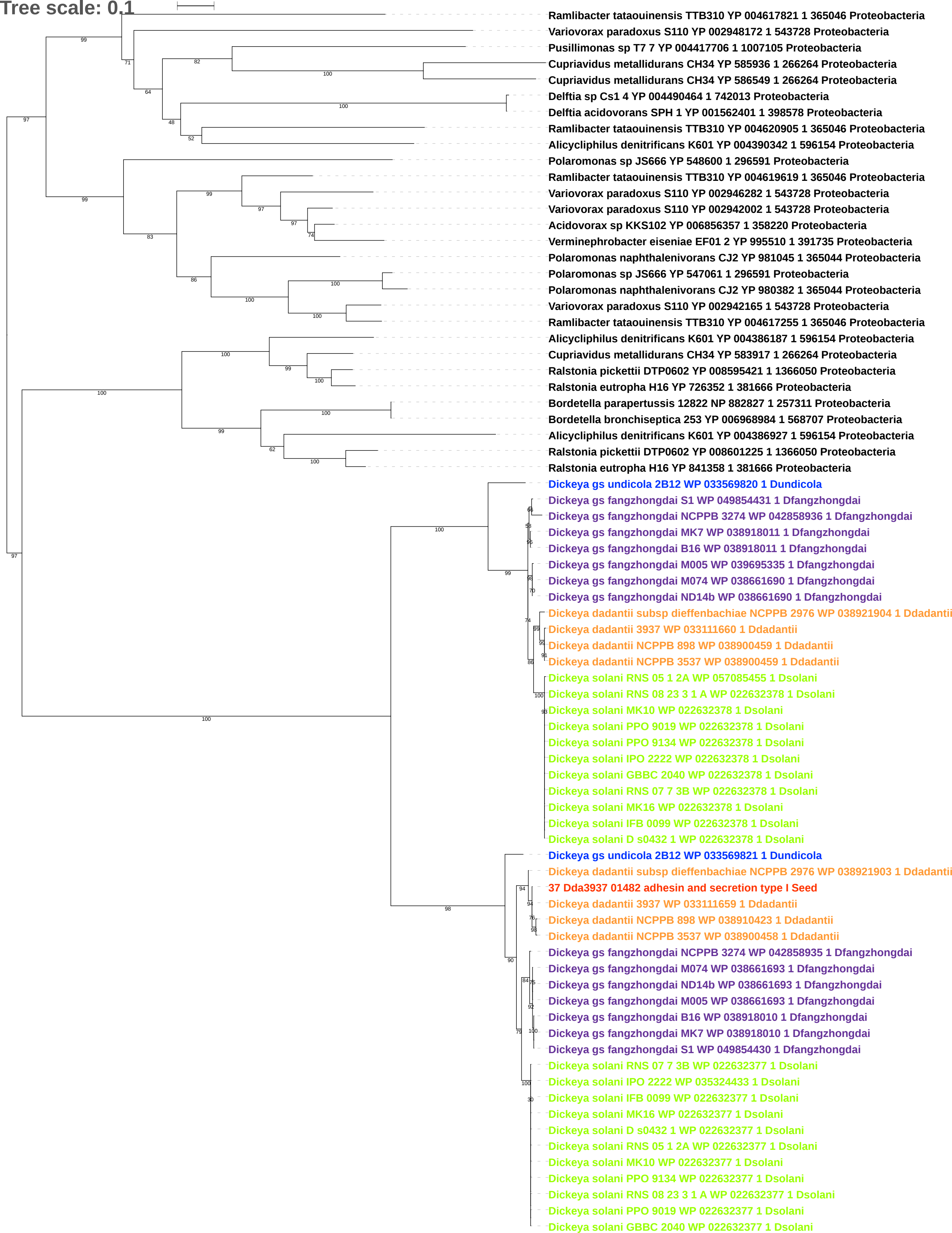
Tree scale: 0.1
Ramlibacter tataouinensis TTB310 YP 004617821 1 365046 Proteobacteria
Variovorax paradoxus S110 YP 002948172 1 543728 Proteobacteria
Pusillimonas sp T7 7 YP 004417706 1 1007105 Proteobacteria
Cupriavidus metallidurans CH34 YP 585936 1 266264 Proteobacteria
Cupriavidus metallidurans CH34 YP 586549 1 266264 Proteobacteria
Delftia sp Cs1 4 YP 004490464 1 742013 Proteobacteria
Delftia acidovorans SPH 1 YP 001562401 1 398578 Proteobacteria
Ramlibacter tataouinensis TTB310 YP 004620905 1 365046 Proteobacteria
Alicycliphilus denitrificans K601 YP 004390342 1 596154 Proteobacteria
Polaromonas sp JS666 YP 548600 1 296591 Proteobacteria
Ramlibacter tataouinensis TTB310 YP 004619619 1 365046 Proteobacteria
Variovorax paradoxus S110 YP 002946282 1 543728 Proteobacteria
Variovorax paradoxus S110 YP 002942002 1 543728 Proteobacteria
Acidovorax sp KKS102 YP 006856357 1 358220 Proteobacteria
Verminephrobacter eiseniae EF01 2 YP 995510 1 391735 Proteobacteria
Polaromonas naphthalenivorans CJ2 YP 981045 1 365044 Proteobacteria
Polaromonas sp JS666 YP 547061 1 296591 Proteobacteria
Polaromonas naphthalenivorans CJ2 YP 980382 1 365044 Proteobacteria
Variovorax paradoxus S110 YP 002942165 1 543728 Proteobacteria
Ramlibacter tataouinensis TTB310 YP 004617255 1 365046 Proteobacteria
Alicycliphilus denitrificans K601 YP 004386187 1 596154 Proteobacteria
Cupriavidus metallidurans CH34 YP 583917 1 266264 Proteobacteria
Ralstonia pickettii DTP0602 YP 008595421 1 1366050 Proteobacteria
Ralstonia eutropha H16 YP 726352 1 381666 Proteobacteria
Bordetella parapertussis 12822 NP 882827 1 257311 Proteobacteria
Bordetella bronchiseptica 253 YP 006968984 1 568707 Proteobacteria
Alicycliphilus denitrificans K601 YP 004386927 1 596154 Proteobacteria
Ralstonia pickettii DTP0602 YP 008601225 1 1366050 Proteobacteria
Ralstonia eutropha H16 YP 841358 1 381666 Proteobacteria
Dickeya gs undicola 2B12 WP 033569820 1 Dundicola
Dickeya gs fangzhongdai S1 WP 049854431 1 Dfangzhongdai
Dickeya gs fangzhongdai NCPPB 3274 WP 042858936 1 Dfangzhongdai
Dickeya gs fangzhongdai MK7 WP 038918011 1 Dfangzhongdai
Dickeya gs fangzhongdai B16 WP 038918011 1 Dfangzhongdai
Dickeya gs fangzhongdai M005 WP 039695335 1 Dfangzhongdai
Dickeya gs fangzhongdai M074 WP 038661690 1 Dfangzhongdai
Dickeya gs fangzhongdai ND14b WP 038661690 1 Dfangzhongdai
Dickeya dadantii subsp dieffenbachiae NCPPB 2976 WP 038921904 1 Ddadantii
Dickeya dadantii 3937 WP 033111660 1 Ddadantii
Dickeya dadantii NCPPB 898 WP 038900459 1 Ddadantii
Dickeya dadantii NCPPB 3537 WP 038900459 1 Ddadantii
Dickeya solani RNS 05 1 2A WP 057085455 1 Dsolani
Dickeya solani RNS 08 23 3 1 A WP 022632378 1 Dsolani
Dickeya solani MK10 WP 022632378 1 Dsolani
Dickeya solani PPO 9019 WP 022632378 1 Dsolani
Dickeya solani PPO 9134 WP 022632378 1 Dsolani
Dickeya solani IPO 2222 WP 022632378 1 Dsolani
Dickeya solani GBBC 2040 WP 022632378 1 Dsolani
Dickeya solani RNS 07 7 3B WP 022632378 1 Dsolani
Dickeya solani MK16 WP 022632378 1 Dsolani
Dickeya solani IFB 0099 WP 022632378 1 Dsolani
Dickeya solani D s0432 1 WP 022632378 1 Dsolani
Dickeya gs undicola 2B12 WP 033569821 1 Dundicola
Dickeya dadantii subsp dieffenbachiae NCPPB 2976 WP 038921903 1 Ddadantii
37 Dda3937 01482 adhesin and secretion type I Seed
Dickeya dadantii 3937 WP 033111659 1 Ddadantii
Dickeya dadantii NCPPB 898 WP 038910423 1 Ddadantii
Dickeya dadantii NCPPB 3537 WP 038900458 1 Ddadantii
Dickeya gs fangzhongdai NCPPB 3274 WP 042858935 1 Dfangzhongdai
Dickeya gs fangzhongdai M074 WP 038661693 1 Dfangzhongdai
Dickeya gs fangzhongdai ND14b WP 038661693 1 Dfangzhongdai
Dickeya gs fangzhongdai M005 WP 038661693 1 Dfangzhongdai
Dickeya gs fangzhongdai B16 WP 038918010 1 Dfangzhongdai
Dickeya gs fangzhongdai MK7 WP 038918010 1 Dfangzhongdai
Dickeya gs fangzhongdai S1 WP 049854430 1 Dfangzhongdai
Dickeya solani RNS 07 7 3B WP 022632377 1 Dsolani
Dickeya solani IPO 2222 WP 035324433 1 Dsolani
Dickeya solani IFB 0099 WP 022632377 1 Dsolani
Dickeya solani MK16 WP 022632377 1 Dsolani
Dickeya solani D s0432 1 WP 022632377 1 Dsolani
Dickeya solani RNS 05 1 2A WP 022632377 1 Dsolani
Dickeya solani MK10 WP 022632377 1 Dsolani
Dickeya solani PPO 9134 WP 022632377 1 Dsolani
Dickeya solani RNS 08 23 3 1 A WP 022632377 1 Dsolani
Dickeya solani PPO 9019 WP 022632377 1 Dsolani
Dickeya solani GBBC 2040 WP 022632377 1 Dsolani

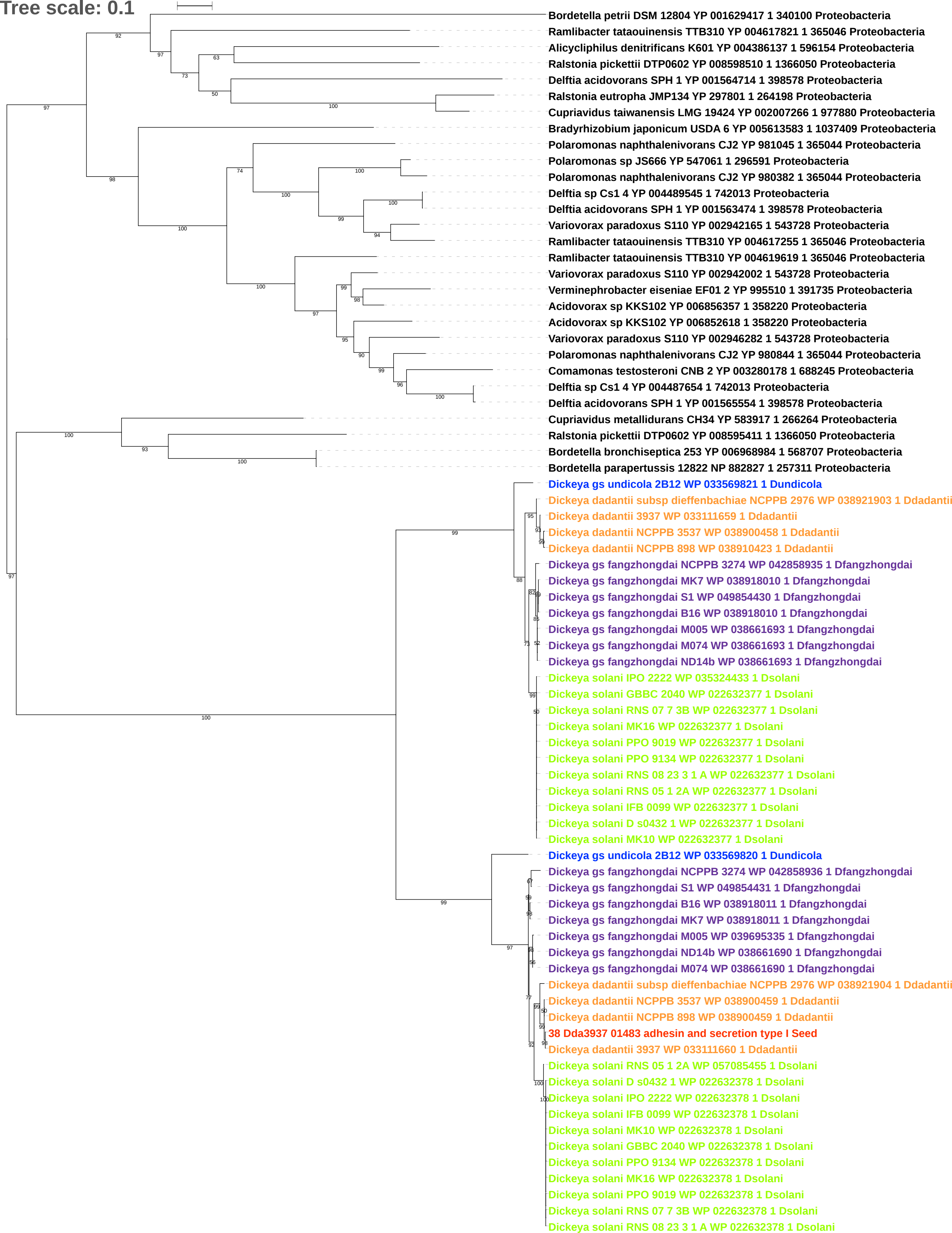
Tree scale: 0.1
Bordetella petrii DSM 12804 YP 001629417 1 340100 Proteobacteria
Ramlibacter tataouinensis TTB310 YP 004617821 1 365046 Proteobacteria
Alicycliphilus denitrificans K601 YP 004386137 1 596154 Proteobacteria
Ralstonia pickettii DTP0602 YP 008598510 1 1366050 Proteobacteria
Delftia acidovorans SPH 1 YP 001564714 1 398578 Proteobacteria
Ralstonia eutropha JMP134 YP 297801 1 264198 Proteobacteria
Cupriavidus taiwanensis LMG 19424 YP 002007266 1 977880 Proteobacteria
Bradyrhizobium japonicum USDA 6 YP 005613583 1 1037409 Proteobacteria
Polaromonas naphthalenivorans CJ2 YP 981045 1 365044 Proteobacteria
Polaromonas sp JS666 YP 547061 1 296591 Proteobacteria
Polaromonas naphthalenivorans CJ2 YP 980382 1 365044 Proteobacteria
Delftia sp Cs1 4 YP 004489545 1 742013 Proteobacteria
Delftia acidovorans SPH 1 YP 001563474 1 398578 Proteobacteria
Variovorax paradoxus S110 YP 002942165 1 543728 Proteobacteria
Ramlibacter tataouinensis TTB310 YP 004617255 1 365046 Proteobacteria
Ramlibacter tataouinensis TTB310 YP 004619619 1 365046 Proteobacteria
Variovorax paradoxus S110 YP 002942002 1 543728 Proteobacteria
Verminephrobacter eiseniae EF01 2 YP 995510 1 391735 Proteobacteria
Acidovorax sp KKS102 YP 006856357 1 358220 Proteobacteria
Acidovorax sp KKS102 YP 006852618 1 358220 Proteobacteria
Variovorax paradoxus S110 YP 002946282 1 543728 Proteobacteria
Polaromonas naphthalenivorans CJ2 YP 980844 1 365044 Proteobacteria
Comamonas testosteroni CNB 2 YP 003280178 1 688245 Proteobacteria
Delftia sp Cs1 4 YP 004487654 1 742013 Proteobacteria
Delftia acidovorans SPH 1 YP 001565554 1 398578 Proteobacteria
Cupriavidus metallidurans CH34 YP 583917 1 266264 Proteobacteria
Ralstonia pickettii DTP0602 YP 008595411 1 1366050 Proteobacteria
Bordetella bronchiseptica 253 YP 006968984 1 568707 Proteobacteria
Bordetella parapertussis 12822 NP 882827 1 257311 Proteobacteria
Dickeya gs undicola 2B12 WP 033569821 1 Dundicola
Dickeya dadantii subsp dieffenbachiae NCPPB 2976 WP 038921903 1 Ddadantii
Dickeya dadantii 3937 WP 033111659 1 Ddadantii
Dickeya dadantii NCPPB 3537 WP 038900458 1 Ddadantii
Dickeya dadantii NCPPB 898 WP 038910423 1 Ddadantii
Dickeya gs fangzhongdai NCPPB 3274 WP 042858935 1 Dfangzhongdai
Dickeya gs fangzhongdai MK7 WP 038918010 1 Dfangzhongdai
Dickeya gs fangzhongdai S1 WP 049854430 1 Dfangzhongdai
Dickeya gs fangzhongdai B16 WP 038918010 1 Dfangzhongdai
Dickeya gs fangzhongdai M005 WP 038661693 1 Dfangzhongdai
Dickeya gs fangzhongdai M074 WP 038661693 1 Dfangzhongdai
Dickeya gs fangzhongdai ND14b WP 038661693 1 Dfangzhongdai
Dickeya solani IPO 2222 WP 035324433 1 Dsolani
Dickeya solani GBBC 2040 WP 022632377 1 Dsolani
Dickeya solani RNS 07 7 3B WP 022632377 1 Dsolani
Dickeya solani MK16 WP 022632377 1 Dsolani
Dickeya solani PPO 9019 WP 022632377 1 Dsolani
Dickeya solani PPO 9134 WP 022632377 1 Dsolani
Dickeya solani RNS 08 23 3 1 A WP 022632377 1 Dsolani
Dickeya solani RNS 05 1 2A WP 022632377 1 Dsolani
Dickeya solani IFB 0099 WP 022632377 1 Dsolani
Dickeya solani D s0432 1 WP 022632377 1 Dsolani
Dickeya solani MK10 WP 022632377 1 Dsolani
Dickeya gs undicola 2B12 WP 033569820 1 Dundicola
Dickeya gs fangzhongdai NCPPB 3274 WP 042858936 1 Dfangzhongdai
Dickeya gs fangzhongdai S1 WP 049854431 1 Dfangzhongdai
Dickeya gs fangzhongdai B16 WP 038918011 1 Dfangzhongdai
Dickeya gs fangzhongdai MK7 WP 038918011 1 Dfangzhongdai
Dickeya gs fangzhongdai M005 WP 039695335 1 Dfangzhongdai
Dickeya gs fangzhongdai ND14b WP 038661690 1 Dfangzhongdai
Dickeya gs fangzhongdai M074 WP 038661690 1 Dfangzhongdai
Dickeya dadantii subsp dieffenbachiae NCPPB 2976 WP 038921904 1 Ddadantii
Dickeya dadantii NCPPB 3537 WP 038900459 1 Ddadantii
Dickeya dadantii NCPPB 898 WP 038900459 1 Ddadantii
38 Dda3937 01483 adhesin and secretion type I Seed
Dickeya dadantii 3937 WP 033111660 1 Ddadantii
Dickeya solani RNS 05 1 2A WP 057085455 1 Dsolani
Dickeya solani D s0432 1 WP 022632378 1 Dsolani
Dickeya solani IPO 2222 WP 022632378 1 Dsolani
Dickeya solani IFB 0099 WP 022632378 1 Dsolani
Dickeya solani MK10 WP 022632378 1 Dsolani
Dickeya solani GBBC 2040 WP 022632378 1 Dsolani
Dickeya solani PPO 9134 WP 022632378 1 Dsolani
Dickeya solani MK16 WP 022632378 1 Dsolani
Dickeya solani PPO 9019 WP 022632378 1 Dsolani
Dickeya solani RNS 07 7 3B WP 022632378 1 Dsolani
Dickeya solani RNS 08 23 3 1 A WP 022632378 1 Dsolani

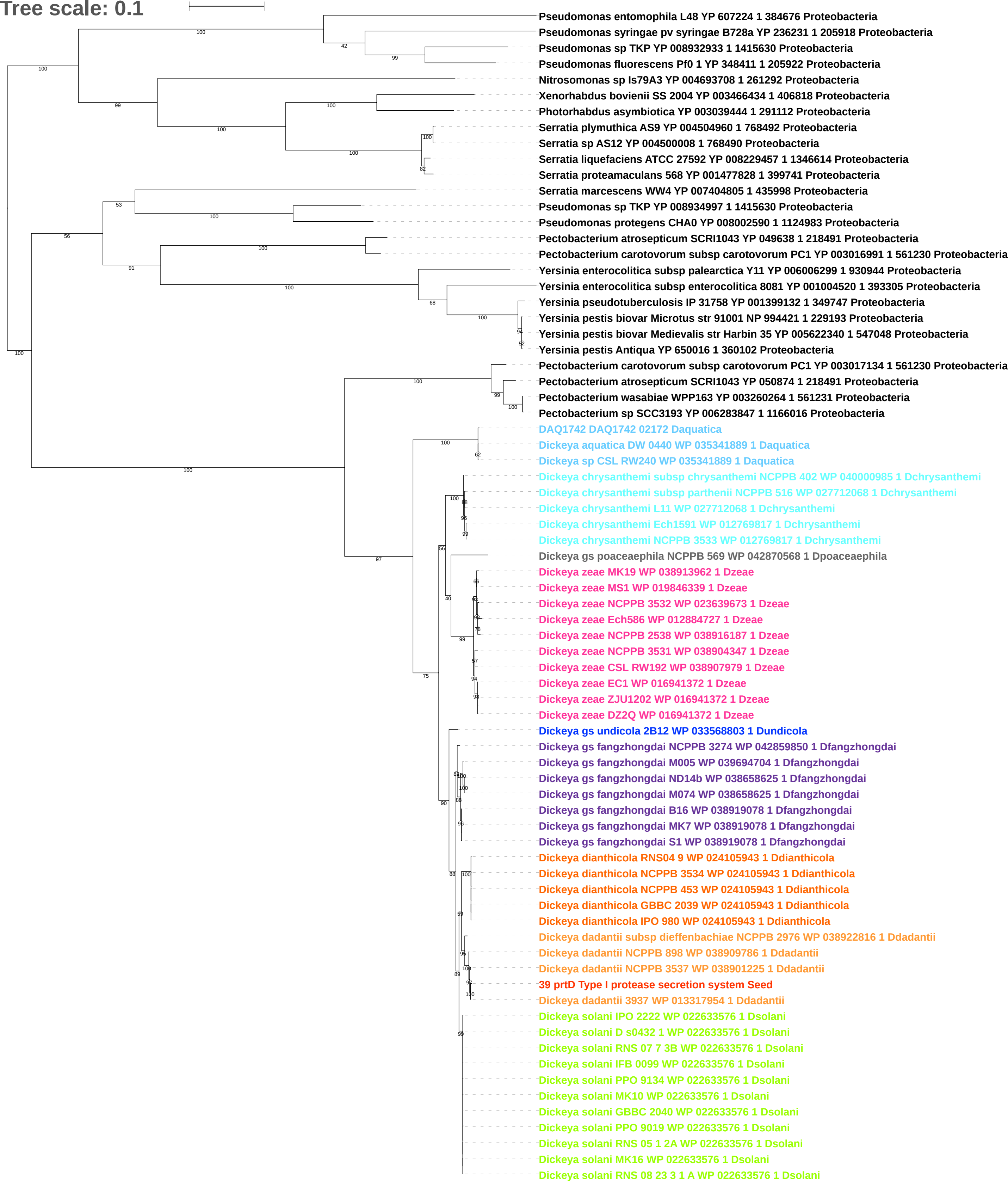
Tree scale: 0.1
Pseudomonas entomophila L48 YP 607224 1 384676 Proteobacteria
Pseudomonas syringae pv syringae B728a YP 236231 1 205918 Proteobacteria
Pseudomonas sp TKP YP 008932933 1 1415630 Proteobacteria
Pseudomonas fluorescens Pf0 1 YP 348411 1 205922 Proteobacteria
Nitrosomonas sp Is79A3 YP 004693708 1 261292 Proteobacteria
Xenorhabdus bovienii SS 2004 YP 003466434 1 406818 Proteobacteria
Photorhabdus asymbiotica YP 003039444 1 291112 Proteobacteria
Serratia plymuthica AS9 YP 004504960 1 768492 Proteobacteria
Serratia sp AS12 YP 004500008 1 768490 Proteobacteria
Serratia liquefaciens ATCC 27592 YP 008229457 1 1346614 Proteobacteria
Serratia proteamaculans 568 YP 001477828 1 399741 Proteobacteria
Serratia marcescens WW4 YP 007404805 1 435998 Proteobacteria
Pseudomonas sp TKP YP 008934997 1 1415630 Proteobacteria
Pseudomonas protegens CHA0 YP 008002590 1 1124983 Proteobacteria
Pectobacterium atrosepticum SCRI1043 YP 049638 1 218491 Proteobacteria
Pectobacterium carotovorum subsp carotovorum PC1 YP 003016991 1 561230 Proteobacteria
Yersinia enterocolitica subsp palearctica Y11 YP 006006299 1 930944 Proteobacteria
Yersinia enterocolitica subsp enterocolitica 8081 YP 001004520 1 393305 Proteobacteria
Yersinia pseudotuberculosis IP 31758 YP 001399132 1 349747 Proteobacteria
Yersinia pestis biovar Microtus str 91001 NP 994421 1 229193 Proteobacteria
Yersinia pestis biovar Medievalis str Harbin 35 YP 005622340 1 547048 Proteobacteria
Yersinia pestis Antiqua YP 650016 1 360102 Proteobacteria
Pectobacterium carotovorum subsp carotovorum PC1 YP 003017134 1 561230 Proteobacteria
Pectobacterium atrosepticum SCRI1043 YP 050874 1 218491 Proteobacteria
Pectobacterium wasabiae WPP163 YP 003260264 1 561231 Proteobacteria
Pectobacterium sp SCC3193 YP 006283847 1 1166016 Proteobacteria
DAQ1742 DAQ1742 02172 Daquatica
Dickeya aquatica DW 0440 WP 035341889 1 Daquatica
Dickeya sp CSL RW240 WP 035341889 1 Daquatica
Dickeya chrysanthemi subsp chrysanthemi NCPPB 402 WP 040000985 1 Dchrysanthemi
Dickeya chrysanthemi subsp parthenii NCPPB 516 WP 027712068 1 Dchrysanthemi
Dickeya chrysanthemi L11 WP 027712068 1 Dchrysanthemi
Dickeya chrysanthemi Ech1591 WP 012769817 1 Dchrysanthemi
Dickeya chrysanthemi NCPPB 3533 WP 012769817 1 Dchrysanthemi
Dickeya gs poaceaephila NCPPB 569 WP 042870568 1 Dpoaceaephila
Dickeya zeae MK19 WP 038913962 1 Dzeae
Dickeya zeae MS1 WP 019846339 1 Dzeae
Dickeya zeae NCPPB 3532 WP 023639673 1 Dzeae
Dickeya zeae Ech586 WP 012884727 1 Dzeae
Dickeya zeae NCPPB 2538 WP 038916187 1 Dzeae
Dickeya zeae NCPPB 3531 WP 038904347 1 Dzeae
Dickeya zeae CSL RW192 WP 038907979 1 Dzeae
Dickeya zeae EC1 WP 016941372 1 Dzeae
Dickeya zeae ZJU1202 WP 016941372 1 Dzeae
Dickeya zeae DZ2Q WP 016941372 1 Dzeae
Dickeya gs undicola 2B12 WP 033568803 1 Dundicola
Dickeya gs fangzhongdai NCPPB 3274 WP 042859850 1 Dfangzhongdai
Dickeya gs fangzhongdai M005 WP 039694704 1 Dfangzhongdai
Dickeya gs fangzhongdai ND14b WP 038658625 1 Dfangzhongdai
Dickeya gs fangzhongdai M074 WP 038658625 1 Dfangzhongdai
Dickeya gs fangzhongdai B16 WP 038919078 1 Dfangzhongdai
Dickeya gs fangzhongdai MK7 WP 038919078 1 Dfangzhongdai
Dickeya gs fangzhongdai S1 WP 038919078 1 Dfangzhongdai
Dickeya dianthicola RNS04 9 WP 024105943 1 Ddianthicola
Dickeya dianthicola NCPPB 3534 WP 024105943 1 Ddianthicola
Dickeya dianthicola NCPPB 453 WP 024105943 1 Ddianthicola
Dickeya dianthicola GBBC 2039 WP 024105943 1 Ddianthicola
Dickeya dianthicola IPO 980 WP 024105943 1 Ddianthicola
Dickeya dadantii subsp dieffenbachiae NCPPB 2976 WP 038922816 1 Ddadantii
Dickeya dadantii NCPPB 898 WP 038909786 1 Ddadantii
Dickeya dadantii NCPPB 3537 WP 038901225 1 Ddadantii
39 prtD Type I protease secretion system Seed
Dickeya dadantii 3937 WP 013317954 1 Ddadantii
Dickeya solani IPO 2222 WP 022633576 1 Dsolani
Dickeya solani D s0432 1 WP 022633576 1 Dsolani
Dickeya solani RNS 07 7 3B WP 022633576 1 Dsolani
Dickeya solani IFB 0099 WP 022633576 1 Dsolani
Dickeya solani PPO 9134 WP 022633576 1 Dsolani
Dickeya solani MK10 WP 022633576 1 Dsolani
Dickeya solani GBBC 2040 WP 022633576 1 Dsolani
Dickeya solani PPO 9019 WP 022633576 1 Dsolani
Dickeya solani RNS 05 1 2A WP 022633576 1 Dsolani
Dickeya solani MK16 WP 022633576 1 Dsolani
Dickeya solani RNS 08 23 3 1 A WP 022633576 1 Dsolani

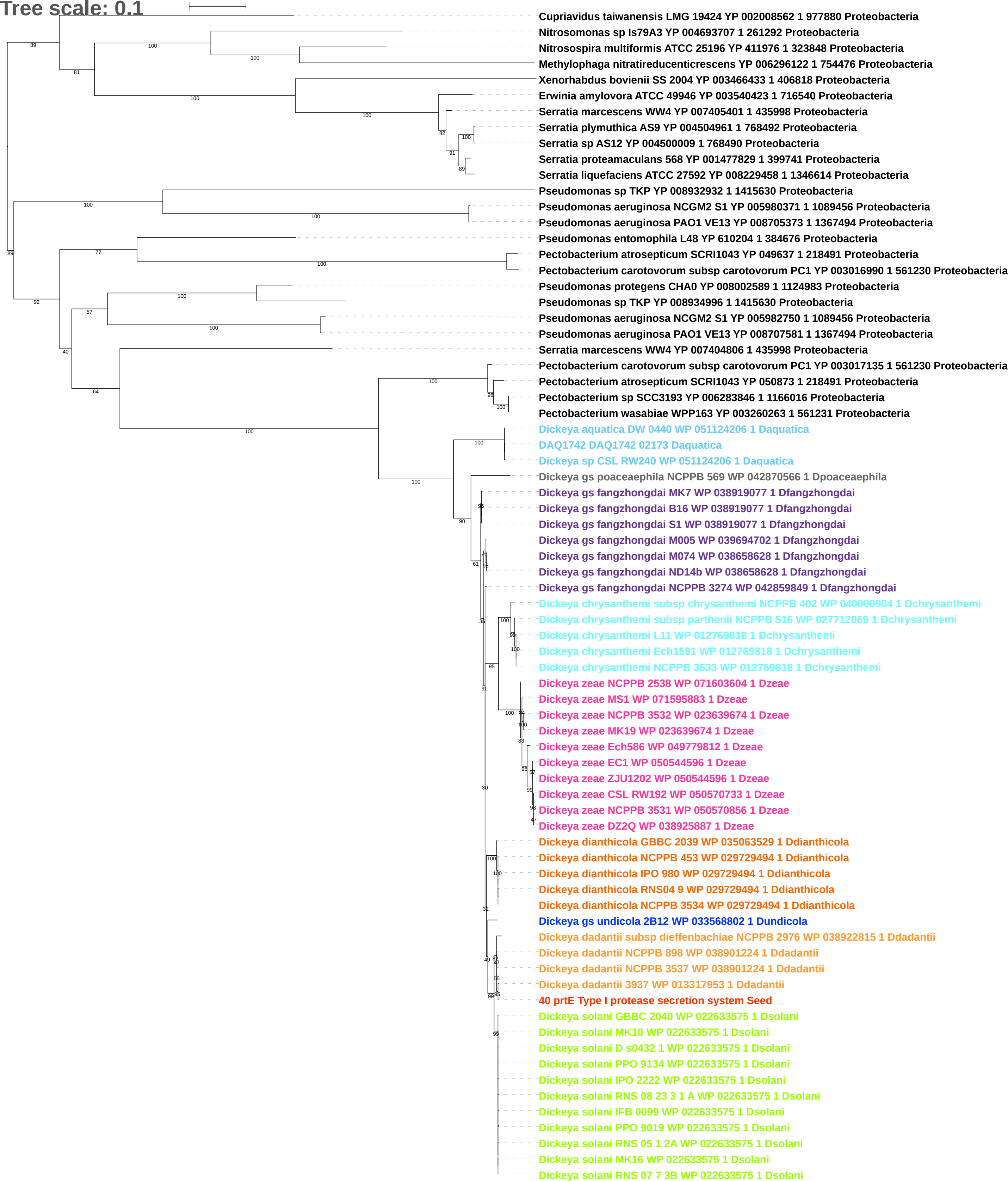
Tree scale: 0.1
Cupriavidus taiwanensis LMG 19424 YP 002008562 1 977880 Proteobacteria
Nitrosomonas sp Is79A3 YP 004693707 1 261292 Proteobacteria
Nitrosospira multiformis ATCC 25196 YP 411976 1 323848 Proteobacteria
Methylophaga nitratireducenticrescens YP 006296122 1 754476 Proteobacteria
Xenorhabdus bovienii SS 2004 YP 003466433 1 406818 Proteobacteria
Erwinia amylovora ATCC 49946 YP 003540423 1 716540 Proteobacteria
Serratia marcescens WW4 YP 007405401 1 435998 Proteobacteria
Serratia plymuthica AS9 YP 004504961 1 768492 Proteobacteria
Serratia sp AS12 YP 004500009 1 768490 Proteobacteria
Serratia proteamaculans 568 YP 001477829 1 399741 Proteobacteria
Serratia liquefaciens ATCC 27592 YP 008229458 1 1346614 Proteobacteria
Pseudomonas sp TKP YP 008932932 1 1415630 Proteobacteria
Pseudomonas aeruginosa NCGM2 S1 YP 005980371 1 1089456 Proteobacteria
Pseudomonas aeruginosa PAO1 VE13 YP 008705373 1 1367494 Proteobacteria
Pseudomonas entomophila L48 YP 610204 1 384676 Proteobacteria
Pectobacterium atrosepticum SCRI1043 YP 049637 1 218491 Proteobacteria
Pectobacterium carotovorum subsp carotovorum PC1 YP 003016990 1 561230 Proteobacteria
Pseudomonas protegens CHA0 YP 008002589 1 1124983 Proteobacteria
Pseudomonas sp TKP YP 008934996 1 1415630 Proteobacteria
Pseudomonas aeruginosa NCGM2 S1 YP 005982750 1 1089456 Proteobacteria
Pseudomonas aeruginosa PAO1 VE13 YP 008707581 1 1367494 Proteobacteria
Serratia marcescens WW4 YP 007404806 1 435998 Proteobacteria
Pectobacterium carotovorum subsp carotovorum PC1 YP 003017135 1 561230 Proteobacteria
Pectobacterium atrosepticum SCRI1043 YP 050873 1 218491 Proteobacteria
Pectobacterium sp SCC3193 YP 006283846 1 1166016 Proteobacteria
Pectobacterium wasabiae WPP163 YP 003260263 1 561231 Proteobacteria
Dickeya aquatica DW 0440 WP 051124206 1 Daquatica
DAQ1742 DAQ1742 02173 Daquatica
Dickeya sp CSL RW240 WP 051124206 1 Daquatica
Dickeya gs poaceaephila NCPPB 569 WP 042870566 1 Dpoaceaephila
Dickeya gs fangzhongdai MK7 WP 038919077 1 Dfangzhongdai
Dickeya gs fangzhongdai B16 WP 038919077 1 Dfangzhongdai
Dickeya gs fangzhongdai S1 WP 038919077 1 Dfangzhongdai
Dickeya gs fangzhongdai M005 WP 039694702 1 Dfangzhongdai
Dickeya gs fangzhongdai M074 WP 038658628 1 Dfangzhongdai
Dickeya gs fangzhongdai ND14b WP 038658628 1 Dfangzhongdai
Dickeya gs fangzhongdai NCPPB 3274 WP 042859849 1 Dfangzhongdai
Dickeya chrysanthemi subsp chrysanthemi NCPPB 402 WP 040000984 1 Dchrysanthemi
Dickeya chrysanthemi subsp parthenii NCPPB 516 WP 027712069 1 Dchrysanthemi
Dickeya chrysanthemi L11 WP 012769818 1 Dchrysanthemi
Dickeya chrysanthemi Ech1591 WP 012769818 1 Dchrysanthemi
Dickeya chrysanthemi NCPPB 3533 WP 012769818 1 Dchrysanthemi
Dickeya zeae NCPPB 2538 WP 071603604 1 Dzeae
Dickeya zeae MS1 WP 071595883 1 Dzeae
Dickeya zeae NCPPB 3532 WP 023639674 1 Dzeae
Dickeya zeae MK19 WP 023639674 1 Dzeae
Dickeya zeae Ech586 WP 049779812 1 Dzeae
Dickeya zeae EC1 WP 050544596 1 Dzeae
Dickeya zeae ZJU1202 WP 050544596 1 Dzeae
Dickeya zeae CSL RW192 WP 050570733 1 Dzeae
Dickeya zeae NCPPB 3531 WP 050570856 1 Dzeae
Dickeya zeae DZ2Q WP 038925887 1 Dzeae
Dickeya dianthicola GBBC 2039 WP 035063529 1 Ddianthicola
Dickeya dianthicola NCPPB 453 WP 029729494 1 Ddianthicola
Dickeya dianthicola IPO 980 WP 029729494 1 Ddianthicola
Dickeya dianthicola RNS04 9 WP 029729494 1 Ddianthicola
Dickeya dianthicola NCPPB 3534 WP 029729494 1 Ddianthicola
Dickeya gs undicola 2B12 WP 033568802 1 Dundicola
Dickeya dadantii subsp dieffenbachiae NCPPB 2976 WP 038922815 1 Ddadantii
Dickeya dadantii NCPPB 898 WP 038901224 1 Ddadantii
Dickeya dadantii NCPPB 3537 WP 038901224 1 Ddadantii
Dickeya dadantii 3937 WP 013317953 1 Ddadantii
40 prtE Type I protease secretion system Seed
Dickeya solani GBBC 2040 WP 022633575 1 Dsolani
Dickeya solani MK10 WP 022633575 1 Dsolani
Dickeya solani D s0432 1 WP 022633575 1 Dsolani
Dickeya solani PPO 9134 WP 022633575 1 Dsolani
Dickeya solani IPO 2222 WP 022633575 1 Dsolani
Dickeya solani RNS 08 23 3 1 A WP 022633575 1 Dsolani
Dickeya solani IFB 0099 WP 022633575 1 Dsolani
Dickeya solani PPO 9019 WP 022633575 1 Dsolani
Dickeya solani RNS 05 1 2A WP 022633575 1 Dsolani
Dickeya solani MK16 WP 022633575 1 Dsolani
Dickeya solani RNS 07 7 3B WP 022633575 1 Dsolani

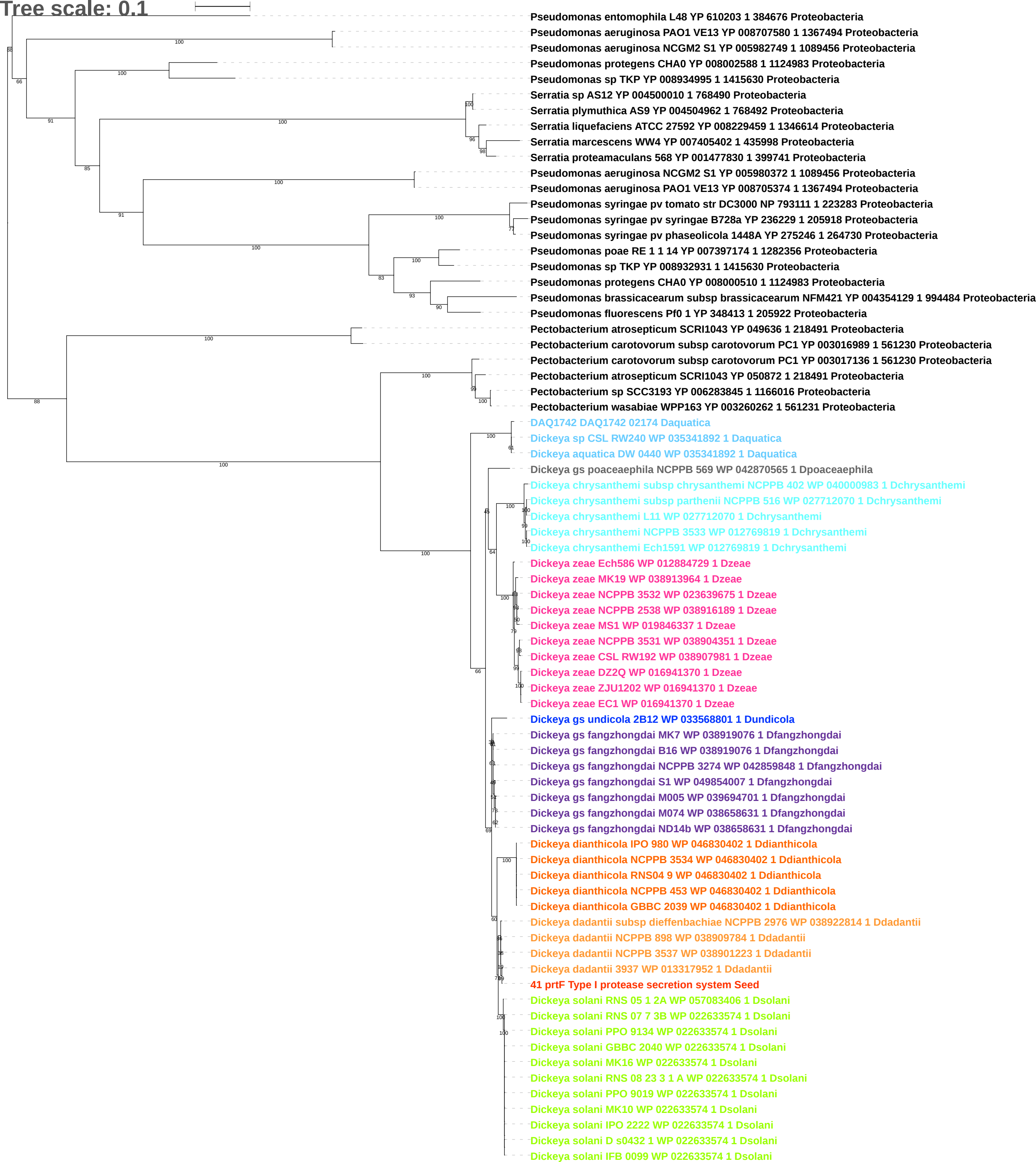
Tree scale: 0.1
Pseudomonas entomophila L48 YP 610203 1 384676 Proteobacteria
Pseudomonas aeruginosa PAO1 VE13 YP 008707580 1 1367494 Proteobacteria
Pseudomonas aeruginosa NCGM2 S1 YP 005982749 1 1089456 Proteobacteria
Pseudomonas protegens CHA0 YP 008002588 1 1124983 Proteobacteria
Pseudomonas sp TKP YP 008934995 1 1415630 Proteobacteria
Serratia sp AS12 YP 004500010 1 768490 Proteobacteria
Serratia plymuthica AS9 YP 004504962 1 768492 Proteobacteria
Serratia liquefaciens ATCC 27592 YP 008229459 1 1346614 Proteobacteria
Serratia marcescens WW4 YP 007405402 1 435998 Proteobacteria
Serratia proteamaculans 568 YP 001477830 1 399741 Proteobacteria
Pseudomonas aeruginosa NCGM2 S1 YP 005980372 1 1089456 Proteobacteria
Pseudomonas aeruginosa PAO1 VE13 YP 008705374 1 1367494 Proteobacteria
Pseudomonas syringae pv tomato str DC3000 NP 793111 1 223283 Proteobacteria
Pseudomonas syringae pv syringae B728a YP 236229 1 205918 Proteobacteria
Pseudomonas syringae pv phaseolicola 1448A YP 275246 1 264730 Proteobacteria
Pseudomonas poae RE 1 1 14 YP 007397174 1 1282356 Proteobacteria
Pseudomonas sp TKP YP 008932931 1 1415630 Proteobacteria
Pseudomonas protegens CHA0 YP 008000510 1 1124983 Proteobacteria
Pseudomonas brassicacearum subsp brassicacearum NFM421 YP 004354129 1 994484 Proteobacteria
Pseudomonas fluorescens Pf0 1 YP 348413 1 205922 Proteobacteria
Pectobacterium atrosepticum SCRI1043 YP 049636 1 218491 Proteobacteria
Pectobacterium carotovorum subsp carotovorum PC1 YP 003016989 1 561230 Proteobacteria
Pectobacterium carotovorum subsp carotovorum PC1 YP 003017136 1 561230 Proteobacteria
Pectobacterium atrosepticum SCRI1043 YP 050872 1 218491 Proteobacteria
Pectobacterium sp SCC3193 YP 006283845 1 1166016 Proteobacteria
Pectobacterium wasabiae WPP163 YP 003260262 1 561231 Proteobacteria
DAQ1742 DAQ1742 02174 Daquatica
Dickeya sp CSL RW240 WP 035341892 1 Daquatica
Dickeya aquatica DW 0440 WP 035341892 1 Daquatica
Dickeya gs poaceaephila NCPPB 569 WP 042870565 1 Dpoaceaephila
Dickeya chrysanthemi subsp chrysanthemi NCPPB 402 WP 040000983 1 Dchrysanthemi
Dickeya chrysanthemi subsp parthenii NCPPB 516 WP 027712070 1 Dchrysanthemi
Dickeya chrysanthemi L11 WP 027712070 1 Dchrysanthemi
Dickeya chrysanthemi NCPPB 3533 WP 012769819 1 Dchrysanthemi
Dickeya chrysanthemi Ech1591 WP 012769819 1 Dchrysanthemi
Dickeya zeae Ech586 WP 012884729 1 Dzeae
Dickeya zeae MK19 WP 038913964 1 Dzeae
Dickeya zeae NCPPB 3532 WP 023639675 1 Dzeae
Dickeya zeae NCPPB 2538 WP 038916189 1 Dzeae
Dickeya zeae MS1 WP 019846337 1 Dzeae
Dickeya zeae NCPPB 3531 WP 038904351 1 Dzeae
Dickeya zeae CSL RW192 WP 038907981 1 Dzeae
Dickeya zeae DZ2Q WP 016941370 1 Dzeae
Dickeya zeae ZJU1202 WP 016941370 1 Dzeae
Dickeya zeae EC1 WP 016941370 1 Dzeae
Dickeya gs undicola 2B12 WP 033568801 1 Dundicola
Dickeya gs fangzhongdai MK7 WP 038919076 1 Dfangzhongdai
Dickeya gs fangzhongdai B16 WP 038919076 1 Dfangzhongdai
Dickeya gs fangzhongdai NCPPB 3274 WP 042859848 1 Dfangzhongdai
Dickeya gs fangzhongdai S1 WP 049854007 1 Dfangzhongdai
Dickeya gs fangzhongdai M005 WP 039694701 1 Dfangzhongdai
Dickeya gs fangzhongdai M074 WP 038658631 1 Dfangzhongdai
Dickeya gs fangzhongdai ND14b WP 038658631 1 Dfangzhongdai
Dickeya dianthicola IPO 980 WP 046830402 1 Ddianthicola
Dickeya dianthicola NCPPB 3534 WP 046830402 1 Ddianthicola
Dickeya dianthicola RNS04 9 WP 046830402 1 Ddianthicola
Dickeya dianthicola NCPPB 453 WP 046830402 1 Ddianthicola
Dickeya dianthicola GBBC 2039 WP 046830402 1 Ddianthicola
Dickeya dadantii subsp dieffenbachiae NCPPB 2976 WP 038922814 1 Ddadantii
Dickeya dadantii NCPPB 898 WP 038909784 1 Ddadantii
Dickeya dadantii NCPPB 3537 WP 038901223 1 Ddadantii
Dickeya dadantii 3937 WP 013317952 1 Ddadantii
41 prtF Type I protease secretion system Seed
Dickeya solani RNS 05 1 2A WP 057083406 1 Dsolani
Dickeya solani RNS 07 7 3B WP 022633574 1 Dsolani
Dickeya solani PPO 9134 WP 022633574 1 Dsolani
Dickeya solani GBBC 2040 WP 022633574 1 Dsolani
Dickeya solani MK16 WP 022633574 1 Dsolani
Dickeya solani RNS 08 23 3 1 A WP 022633574 1 Dsolani
Dickeya solani PPO 9019 WP 022633574 1 Dsolani
Dickeya solani MK10 WP 022633574 1 Dsolani
Dickeya solani IPO 2222 WP 022633574 1 Dsolani
Dickeya solani D s0432 1 WP 022633574 1 Dsolani
Dickeya solani IFB 0099 WP 022633574 1 Dsolani

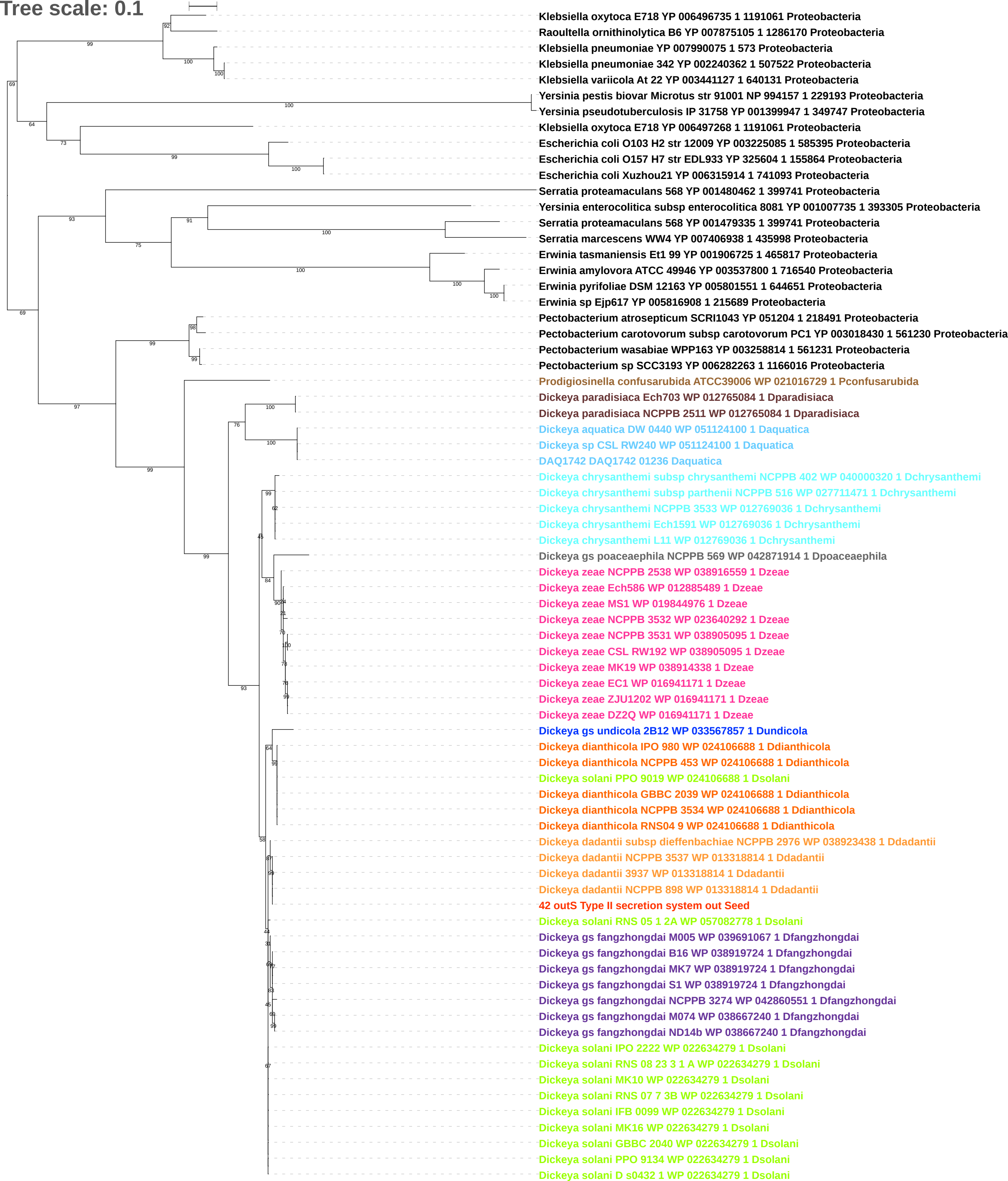

Tree scale: 0.1
Klebsiella oxytoca E718 YP 006496735 1 1191061 Proteobacteria
Raoultella ornithinolytica B6 YP 007875105 1 1286170 Proteobacteria
Klebsiella pneumoniae YP 007990075 1 573 Proteobacteria
Klebsiella pneumoniae 342 YP 002240362 1 507522 Proteobacteria
Klebsiella variicola At 22 YP 003441127 1 640131 Proteobacteria
Yersinia pestis biovar Microtus str 91001 NP 994157 1 229193 Proteobacteria
Yersinia pseudotuberculosis IP 31758 YP 001399947 1 349747 Proteobacteria
Klebsiella oxytoca E718 YP 006497268 1 1191061 Proteobacteria
Escherichia coli O103 H2 str 12009 YP 003225085 1 585395 Proteobacteria
Escherichia coli O157 H7 str EDL933 YP 325604 1 155864 Proteobacteria
Escherichia coli Xuzhou21 YP 006315914 1 741093 Proteobacteria
Serratia proteamaculans 568 YP 001480462 1 399741 Proteobacteria
Yersinia enterocolitica subsp enterocolitica 8081 YP 001007735 1 393305 Proteobacteria
Serratia proteamaculans 568 YP 001479335 1 399741 Proteobacteria
Serratia marcescens WW4 YP 007406938 1 435998 Proteobacteria
Erwinia tasmaniensis Et1 99 YP 001906725 1 465817 Proteobacteria
Erwinia amylovora ATCC 49946 YP 003537800 1 716540 Proteobacteria
Erwinia pyrifoliae DSM 12163 YP 005801551 1 644651 Proteobacteria
Erwinia sp Ejp617 YP 005816908 1 215689 Proteobacteria
Pectobacterium atrosepticum SCRI1043 YP 051204 1 218491 Proteobacteria
Pectobacterium carotovorum subsp carotovorum PC1 YP 003018430 1 561230 Proteobacteria
Pectobacterium wasabiae WPP163 YP 003258814 1 561231 Proteobacteria
Pectobacterium sp SCC3193 YP 006282263 1 1166016 Proteobacteria
Prodigiosinella confusarubida ATCC39006 WP 021016729 1 Pconfusarubida
Dickeya paradisiaca Ech703 WP 012765084 1 Dparadisiaca
Dickeya paradisiaca NCPPB 2511 WP 012765084 1 Dparadisiaca
Dickeya aquatica DW 0440 WP 051124100 1 Daquatica
Dickeya sp CSL RW240 WP 051124100 1 Daquatica
DAQ1742 DAQ1742 01236 Daquatica
Dickeya chrysanthemi subsp chrysanthemi NCPPB 402 WP 040000320 1 Dchrysanthemi
Dickeya chrysanthemi subsp parthenii NCPPB 516 WP 027711471 1 Dchrysanthemi
Dickeya chrysanthemi NCPPB 3533 WP 012769036 1 Dchrysanthemi
Dickeya chrysanthemi Ech1591 WP 012769036 1 Dchrysanthemi
Dickeya chrysanthemi L11 WP 012769036 1 Dchrysanthemi
Dickeya gs poaceaephila NCPPB 569 WP 042871914 1 Dpoaceaephila
Dickeya zeae NCPPB 2538 WP 038916559 1 Dzeae
Dickeya zeae Ech586 WP 012885489 1 Dzeae
Dickeya zeae MS1 WP 019844976 1 Dzeae
Dickeya zeae NCPPB 3532 WP 023640292 1 Dzeae
Dickeya zeae NCPPB 3531 WP 038905095 1 Dzeae
Dickeya zeae CSL RW192 WP 038905095 1 Dzeae
Dickeya zeae MK19 WP 038914338 1 Dzeae
Dickeya zeae EC1 WP 016941171 1 Dzeae
Dickeya zeae ZJU1202 WP 016941171 1 Dzeae
Dickeya zeae DZ2Q WP 016941171 1 Dzeae
Dickeya gs undicola 2B12 WP 033567857 1 Dundicola
Dickeya dianthicola IPO 980 WP 024106688 1 Ddianthicola
Dickeya dianthicola NCPPB 453 WP 024106688 1 Ddianthicola
Dickeya solani PPO 9019 WP 024106688 1 Dsolani
Dickeya dianthicola GBBC 2039 WP 024106688 1 Ddianthicola
Dickeya dianthicola NCPPB 3534 WP 024106688 1 Ddianthicola
Dickeya dianthicola RNS04 9 WP 024106688 1 Ddianthicola
Dickeya dadantii subsp dieffenbachiae NCPPB 2976 WP 038923438 1 Ddadantii
Dickeya dadantii NCPPB 3537 WP 013318814 1 Ddadantii
Dickeya dadantii 3937 WP 013318814 1 Ddadantii
Dickeya dadantii NCPPB 898 WP 013318814 1 Ddadantii
42 outS Type II secretion system out Seed
Dickeya solani RNS 05 1 2A WP 057082778 1 Dsolani
Dickeya gs fangzhongdai M005 WP 039691067 1 Dfangzhongdai
Dickeya gs fangzhongdai B16 WP 038919724 1 Dfangzhongdai
Dickeya gs fangzhongdai MK7 WP 038919724 1 Dfangzhongdai
Dickeya gs fangzhongdai S1 WP 038919724 1 Dfangzhongdai
Dickeya gs fangzhongdai NCPPB 3274 WP 042860551 1 Dfangzhongdai
Dickeya gs fangzhongdai M074 WP 038667240 1 Dfangzhongdai
Dickeya gs fangzhongdai ND14b WP 038667240 1 Dfangzhongdai
Dickeya solani IPO 2222 WP 022634279 1 Dsolani
Dickeya solani RNS 08 23 3 1 A WP 022634279 1 Dsolani
Dickeya solani MK10 WP 022634279 1 Dsolani
Dickeya solani RNS 07 7 3B WP 022634279 1 Dsolani
Dickeya solani IFB 0099 WP 022634279 1 Dsolani
Dickeya solani MK16 WP 022634279 1 Dsolani
Dickeya solani GBBC 2040 WP 022634279 1 Dsolani
Dickeya solani PPO 9134 WP 022634279 1 Dsolani
Dickeya solani D s0432 1 WP 022634279 1 Dsolani

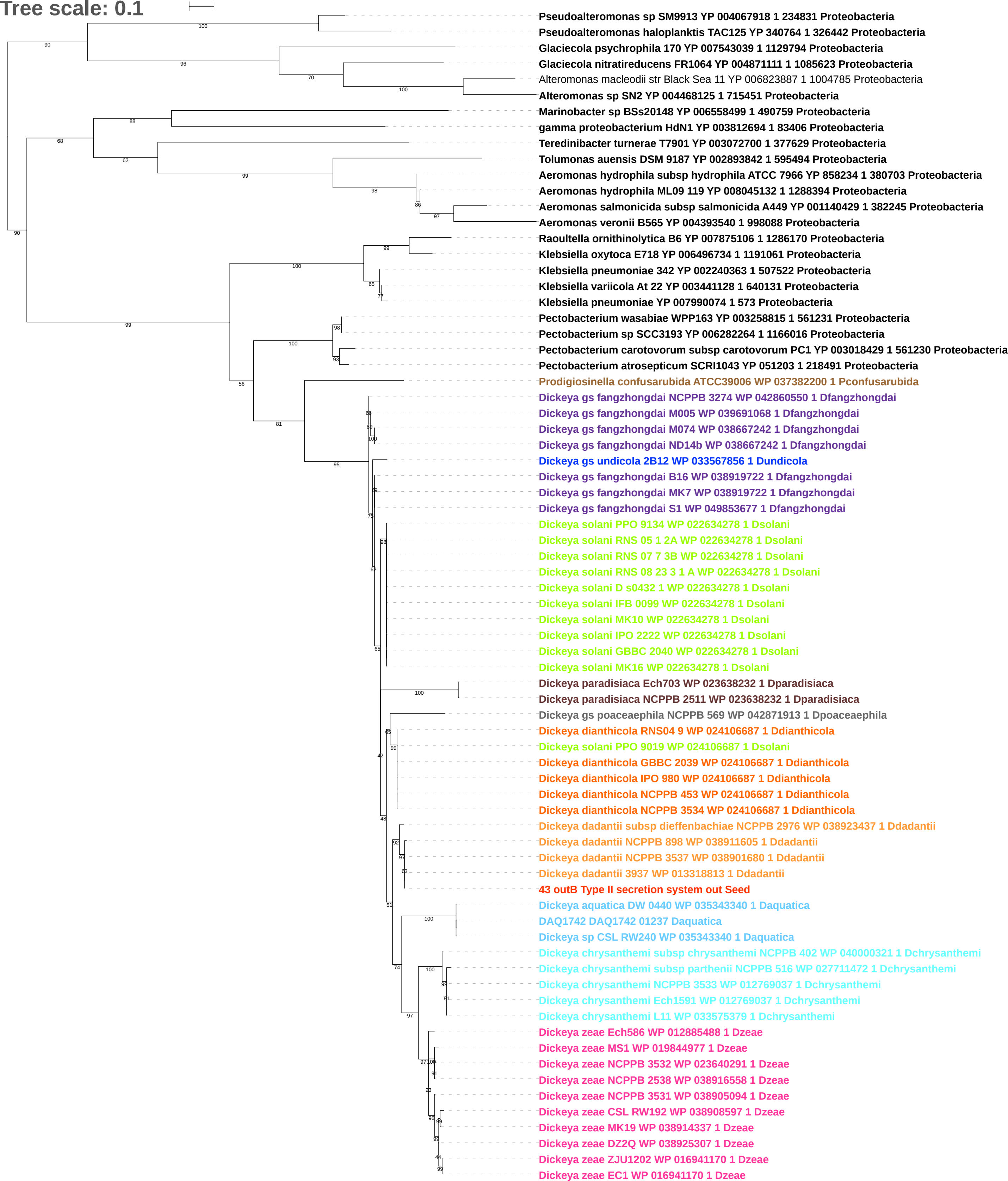
Tree scale: 0.1
Pseudoalteromonas sp SM9913 YP 004067918 1 234831 Proteobacteria
Pseudoalteromonas haloplanktis TAC125 YP 340764 1 326442 Proteobacteria
Glaciecola psychrophila 170 YP 007543039 1 1129794 Proteobacteria
Glaciecola nitratireducens FR1064 YP 004871111 1 1085623 Proteobacteria
Alteromonas macleodii str Black Sea 11 YP 006823887 1 1004785 Proteobacteria
Alteromonas sp SN2 YP 004468125 1 715451 Proteobacteria
Marinobacter sp BSs20148 YP 006558499 1 490759 Proteobacteria
gamma proteobacterium HdN1 YP 003812694 1 83406 Proteobacteria
Teredinibacter turnerae T7901 YP 003072700 1 377629 Proteobacteria
Tolumonas auensis DSM 9187 YP 002893842 1 595494 Proteobacteria
Aeromonas hydrophila subsp hydrophila ATCC 7966 YP 858234 1 380703 Proteobacteria
Aeromonas hydrophila ML09 119 YP 008045132 1 1288394 Proteobacteria
Aeromonas salmonicida subsp salmonicida A449 YP 001140429 1 382245 Proteobacteria
Aeromonas veronii B565 YP 004393540 1 998088 Proteobacteria
Raoultella ornithinolytica B6 YP 007875106 1 1286170 Proteobacteria
Klebsiella oxytoca E718 YP 006496734 1 1191061 Proteobacteria
Klebsiella pneumoniae 342 YP 002240363 1 507522 Proteobacteria
Klebsiella variicola At 22 YP 003441128 1 640131 Proteobacteria
Klebsiella pneumoniae YP 007990074 1 573 Proteobacteria
Pectobacterium wasabiae WPP163 YP 003258815 1 561231 Proteobacteria
Pectobacterium sp SCC3193 YP 006282264 1 1166016 Proteobacteria
Pectobacterium carotovorum subsp carotovorum PC1 YP 003018429 1 561230 Proteobacteria
Pectobacterium atrosepticum SCRI1043 YP 051203 1 218491 Proteobacteria
Prodigiosinella confusarubida ATCC39006 WP 037382200 1 Pconfusarubida
Dickeya gs fangzhongdai NCPPB 3274 WP 042860550 1 Dfangzhongdai
Dickeya gs fangzhongdai M005 WP 039691068 1 Dfangzhongdai
Dickeya gs fangzhongdai M074 WP 038667242 1 Dfangzhongdai
Dickeya gs fangzhongdai ND14b WP 038667242 1 Dfangzhongdai
Dickeya gs undicola 2B12 WP 033567856 1 Dundicola
Dickeya gs fangzhongdai B16 WP 038919722 1 Dfangzhongdai
Dickeya gs fangzhongdai MK7 WP 038919722 1 Dfangzhongdai
Dickeya gs fangzhongdai S1 WP 049853677 1 Dfangzhongdai
Dickeya solani PPO 9134 WP 022634278 1 Dsolani
Dickeya solani RNS 05 1 2A WP 022634278 1 Dsolani
Dickeya solani RNS 07 7 3B WP 022634278 1 Dsolani
Dickeya solani RNS 08 23 3 1 A WP 022634278 1 Dsolani
Dickeya solani D s0432 1 WP 022634278 1 Dsolani
Dickeya solani IFB 0099 WP 022634278 1 Dsolani
Dickeya solani MK10 WP 022634278 1 Dsolani
Dickeya solani IPO 2222 WP 022634278 1 Dsolani
Dickeya solani GBBC 2040 WP 022634278 1 Dsolani
Dickeya solani MK16 WP 022634278 1 Dsolani
Dickeya paradisiaca Ech703 WP 023638232 1 Dparadisiaca
Dickeya paradisiaca NCPPB 2511 WP 023638232 1 Dparadisiaca
Dickeya gs poaceaephila NCPPB 569 WP 042871913 1 Dpoaceaephila
Dickeya dianthicola RNS04 9 WP 024106687 1 Ddianthicola
Dickeya solani PPO 9019 WP 024106687 1 Dsolani
Dickeya dianthicola GBBC 2039 WP 024106687 1 Ddianthicola
Dickeya dianthicola IPO 980 WP 024106687 1 Ddianthicola
Dickeya dianthicola NCPPB 453 WP 024106687 1 Ddianthicola
Dickeya dianthicola NCPPB 3534 WP 024106687 1 Ddianthicola
Dickeya dadantii subsp dieffenbachiae NCPPB 2976 WP 038923437 1 Ddadantii
Dickeya dadantii NCPPB 898 WP 038911605 1 Ddadantii
Dickeya dadantii NCPPB 3537 WP 038901680 1 Ddadantii
Dickeya dadantii 3937 WP 013318813 1 Ddadantii
43 outB Type II secretion system out Seed
Dickeya aquatica DW 0440 WP 035343340 1 Daquatica
DAQ1742 DAQ1742 01237 Daquatica
Dickeya sp CSL RW240 WP 035343340 1 Daquatica
Dickeya chrysanthemi subsp chrysanthemi NCPPB 402 WP 040000321 1 Dchrysanthemi
Dickeya chrysanthemi subsp parthenii NCPPB 516 WP 027711472 1 Dchrysanthemi
Dickeya chrysanthemi NCPPB 3533 WP 012769037 1 Dchrysanthemi
Dickeya chrysanthemi Ech1591 WP 012769037 1 Dchrysanthemi
Dickeya chrysanthemi L11 WP 033575379 1 Dchrysanthemi
Dickeya zeae Ech586 WP 012885488 1 Dzeae
Dickeya zeae MS1 WP 019844977 1 Dzeae
Dickeya zeae NCPPB 3532 WP 023640291 1 Dzeae
Dickeya zeae NCPPB 2538 WP 038916558 1 Dzeae
Dickeya zeae NCPPB 3531 WP 038905094 1 Dzeae
Dickeya zeae CSL RW192 WP 038908597 1 Dzeae
Dickeya zeae MK19 WP 038914337 1 Dzeae
Dickeya zeae DZ2Q WP 038925307 1 Dzeae
Dickeya zeae ZJU1202 WP 016941170 1 Dzeae
Dickeya zeae EC1 WP 016941170 1 Dzeae

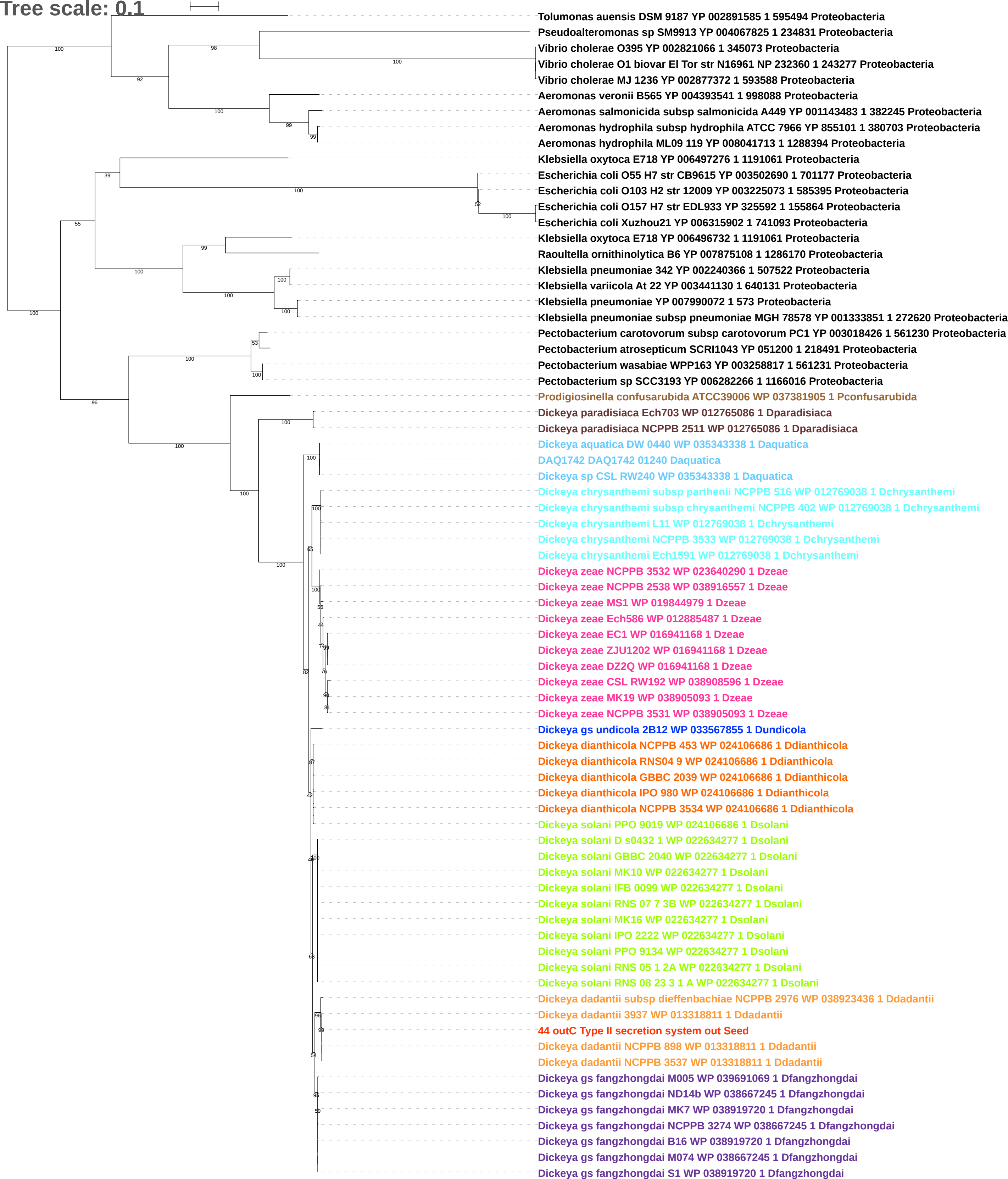
Tree scale: 0.1
Tolumonas auensis DSM 9187 YP 002891585 1 595494 Proteobacteria
Pseudoalteromonas sp SM9913 YP 004067825 1 234831 Proteobacteria
Vibrio cholerae O395 YP 002821066 1 345073 Proteobacteria
Vibrio cholerae O1 biovar El Tor str N16961 NP 232360 1 243277 Proteobacteria
Vibrio cholerae MJ 1236 YP 002877372 1 593588 Proteobacteria
Aeromonas veronii B565 YP 004393541 1 998088 Proteobacteria
Aeromonas salmonicida subsp salmonicida A449 YP 001143483 1 382245 Proteobacteria
Aeromonas hydrophila subsp hydrophila ATCC 7966 YP 855101 1 380703 Proteobacteria
Aeromonas hydrophila ML09 119 YP 008041713 1 1288394 Proteobacteria
Klebsiella oxytoca E718 YP 006497276 1 1191061 Proteobacteria
Escherichia coli O55 H7 str CB9615 YP 003502690 1 701177 Proteobacteria
Escherichia coli O103 H2 str 12009 YP 003225073 1 585395 Proteobacteria
Escherichia coli O157 H7 str EDL933 YP 325592 1 155864 Proteobacteria
Escherichia coli Xuzhou21 YP 006315902 1 741093 Proteobacteria
Klebsiella oxytoca E718 YP 006496732 1 1191061 Proteobacteria
Raoultella ornithinolytica B6 YP 007875108 1 1286170 Proteobacteria
Klebsiella pneumoniae 342 YP 002240366 1 507522 Proteobacteria
Klebsiella variicola At 22 YP 003441130 1 640131 Proteobacteria
Klebsiella pneumoniae YP 007990072 1 573 Proteobacteria
Klebsiella pneumoniae subsp pneumoniae MGH 78578 YP 001333851 1 272620 Proteobacteria
Pectobacterium carotovorum subsp carotovorum PC1 YP 003018426 1 561230 Proteobacteria
Pectobacterium atrosepticum SCRI1043 YP 051200 1 218491 Proteobacteria
Pectobacterium wasabiae WPP163 YP 003258817 1 561231 Proteobacteria
Pectobacterium sp SCC3193 YP 006282266 1 1166016 Proteobacteria
Prodigiosinella confusarubida ATCC39006 WP 037381905 1 Pconfusarubida
Dickeya paradisiaca Ech703 WP 012765086 1 Dparadisiaca
Dickeya paradisiaca NCPPB 2511 WP 012765086 1 Dparadisiaca
Dickeya aquatica DW 0440 WP 035343338 1 Daquatica
DAQ1742 DAQ1742 01240 Daquatica
Dickeya sp CSL RW240 WP 035343338 1 Daquatica
Dickeya chrysanthemi subsp parthenii NCPPB 516 WP 012769038 1 Dchrysanthemi
Dickeya chrysanthemi subsp chrysanthemi NCPPB 402 WP 012769038 1 Dchrysanthemi
Dickeya chrysanthemi L11 WP 012769038 1 Dchrysanthemi
Dickeya chrysanthemi NCPPB 3533 WP 012769038 1 Dchrysanthemi
Dickeya chrysanthemi Ech1591 WP 012769038 1 Dchrysanthemi
Dickeya zeae NCPPB 3532 WP 023640290 1 Dzeae
Dickeya zeae NCPPB 2538 WP 038916557 1 Dzeae
Dickeya zeae MS1 WP 019844979 1 Dzeae
Dickeya zeae Ech586 WP 012885487 1 Dzeae
Dickeya zeae EC1 WP 016941168 1 Dzeae
Dickeya zeae ZJU1202 WP 016941168 1 Dzeae
Dickeya zeae DZ2Q WP 016941168 1 Dzeae
Dickeya zeae CSL RW192 WP 038908596 1 Dzeae
Dickeya zeae MK19 WP 038905093 1 Dzeae
Dickeya zeae NCPPB 3531 WP 038905093 1 Dzeae
Dickeya gs undicola 2B12 WP 033567855 1 Dundicola
Dickeya dianthicola NCPPB 453 WP 024106686 1 Ddianthicola
Dickeya dianthicola RNS04 9 WP 024106686 1 Ddianthicola
Dickeya dianthicola GBBC 2039 WP 024106686 1 Ddianthicola
Dickeya dianthicola IPO 980 WP 024106686 1 Ddianthicola
Dickeya dianthicola NCPPB 3534 WP 024106686 1 Ddianthicola
Dickeya solani PPO 9019 WP 024106686 1 Dsolani
Dickeya solani D s0432 1 WP 022634277 1 Dsolani
Dickeya solani GBBC 2040 WP 022634277 1 Dsolani
Dickeya solani MK10 WP 022634277 1 Dsolani
Dickeya solani IFB 0099 WP 022634277 1 Dsolani
Dickeya solani RNS 07 7 3B WP 022634277 1 Dsolani
Dickeya solani MK16 WP 022634277 1 Dsolani
Dickeya solani IPO 2222 WP 022634277 1 Dsolani
Dickeya solani PPO 9134 WP 022634277 1 Dsolani
Dickeya solani RNS 05 1 2A WP 022634277 1 Dsolani
Dickeya solani RNS 08 23 3 1 A WP 022634277 1 Dsolani
Dickeya dadantii subsp dieffenbachiae NCPPB 2976 WP 038923436 1 Ddadantii
Dickeya dadantii 3937 WP 013318811 1 Ddadantii
44 outC Type II secretion system out Seed
Dickeya dadantii NCPPB 898 WP 013318811 1 Ddadantii
Dickeya dadantii NCPPB 3537 WP 013318811 1 Ddadantii
Dickeya gs fangzhongdai M005 WP 039691069 1 Dfangzhongdai
Dickeya gs fangzhongdai ND14b WP 038667245 1 Dfangzhongdai
Dickeya gs fangzhongdai MK7 WP 038919720 1 Dfangzhongdai
Dickeya gs fangzhongdai NCPPB 3274 WP 038667245 1 Dfangzhongdai
Dickeya gs fangzhongdai B16 WP 038919720 1 Dfangzhongdai
Dickeya gs fangzhongdai M074 WP 038667245 1 Dfangzhongdai
Dickeya gs fangzhongdai S1 WP 038919720 1 Dfangzhongdai

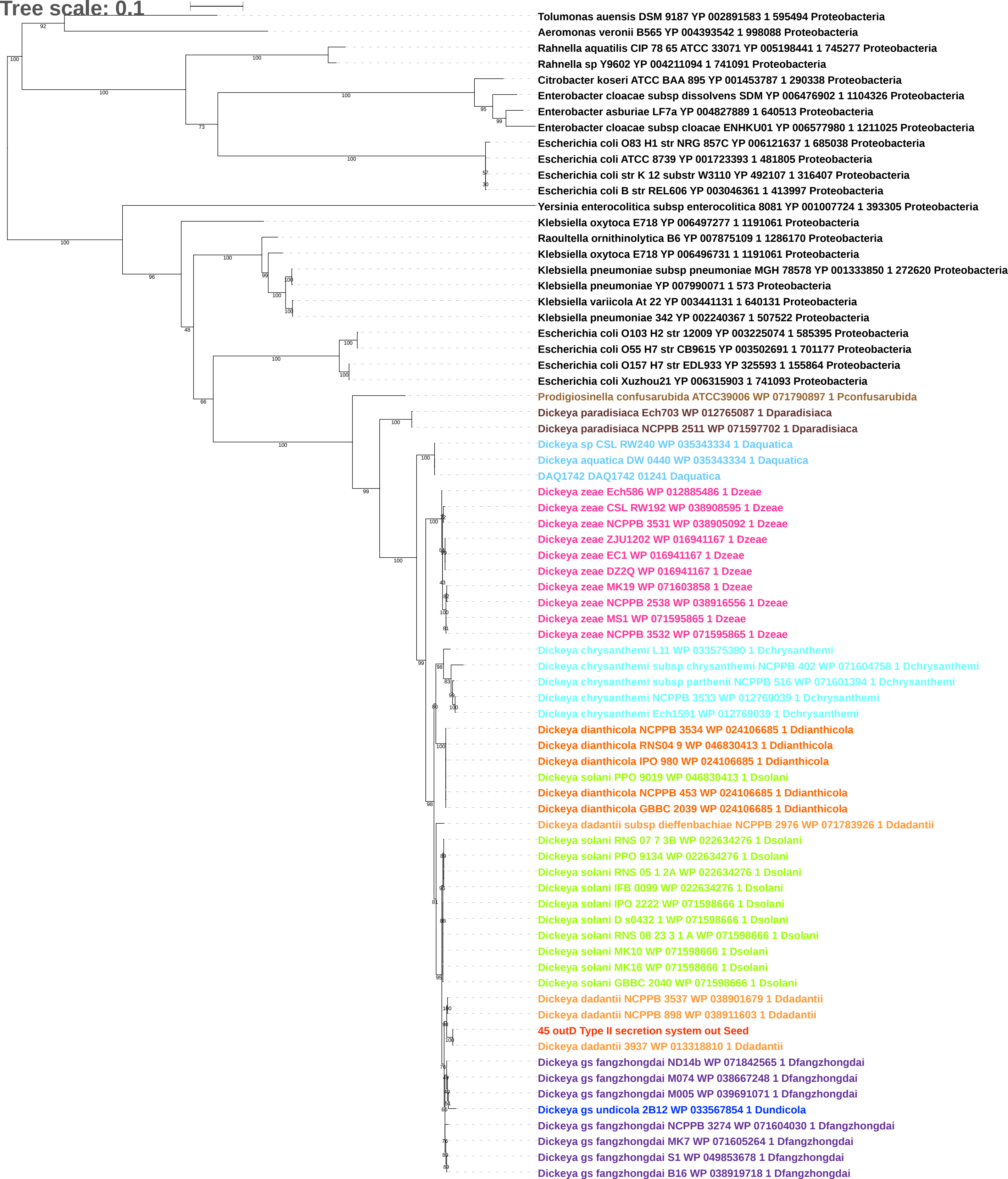
Tree scale: 0.1
Tolumonas auensis DSM 9187 YP 002891583 1 595494 Proteobacteria
Aeromonas veronii B565 YP 004393542 1 998088 Proteobacteria
Rahnella aquatilis CIP 78 65 ATCC 33071 YP 005198441 1 745277 Proteobacteria
Rahnella sp Y9602 YP 004211094 1 741091 Proteobacteria
Citrobacter koseri ATCC BAA 895 YP 001453787 1 290338 Proteobacteria
Enterobacter cloacae subsp dissolvens SDM YP 006476902 1 1104326 Proteobacteria
Enterobacter asburiae LF7a YP 004827889 1 640513 Proteobacteria
Enterobacter cloacae subsp cloacae ENHKU01 YP 006577980 1 1211025 Proteobacteria
Escherichia coli O83 H1 str NRG 857C YP 006121637 1 685038 Proteobacteria
Escherichia coli ATCC 8739 YP 001723393 1 481805 Proteobacteria
Escherichia coli str K 12 substr W3110 YP 492107 1 316407 Proteobacteria
Escherichia coli B str REL606 YP 003046361 1 413997 Proteobacteria
Yersinia enterocolitica subsp enterocolitica 8081 YP 001007724 1 393305 Proteobacteria
Klebsiella oxytoca E718 YP 006497277 1 1191061 Proteobacteria
Raoultella ornithinolytica B6 YP 007875109 1 1286170 Proteobacteria
Klebsiella oxytoca E718 YP 006496731 1 1191061 Proteobacteria
Klebsiella pneumoniae subsp pneumoniae MGH 78578 YP 001333850 1 272620 Proteobacteria
Klebsiella pneumoniae YP 007990071 1 573 Proteobacteria
Klebsiella variicola At 22 YP 003441131 1 640131 Proteobacteria
Klebsiella pneumoniae 342 YP 002240367 1 507522 Proteobacteria
Escherichia coli O103 H2 str 12009 YP 003225074 1 585395 Proteobacteria
Escherichia coli O55 H7 str CB9615 YP 003502691 1 701177 Proteobacteria
Escherichia coli O157 H7 str EDL933 YP 325593 1 155864 Proteobacteria
Escherichia coli Xuzhou21 YP 006315903 1 741093 Proteobacteria
Prodigiosinella confusarubida ATCC39006 WP 071790897 1 Pconfusarubida
Dickeya paradisiaca Ech703 WP 012765087 1 Dparadisiaca
Dickeya paradisiaca NCPPB 2511 WP 071597702 1 Dparadisiaca
Dickeya sp CSL RW240 WP 035343334 1 Daquatica
Dickeya aquatica DW 0440 WP 035343334 1 Daquatica
DAQ1742 DAQ1742 01241 Daquatica
Dickeya zeae Ech586 WP 012885486 1 Dzeae
Dickeya zeae CSL RW192 WP 038908595 1 Dzeae
Dickeya zeae NCPPB 3531 WP 038905092 1 Dzeae
Dickeya zeae ZJU1202 WP 016941167 1 Dzeae
Dickeya zeae EC1 WP 016941167 1 Dzeae
Dickeya zeae DZ2Q WP 016941167 1 Dzeae
Dickeya zeae MK19 WP 071603858 1 Dzeae
Dickeya zeae NCPPB 2538 WP 038916556 1 Dzeae
Dickeya zeae MS1 WP 071595865 1 Dzeae
Dickeya zeae NCPPB 3532 WP 071595865 1 Dzeae
Dickeya chrysanthemi L11 WP 033575380 1 Dchrysanthemi
Dickeya chrysanthemi subsp chrysanthemi NCPPB 402 WP 071604758 1 Dchrysanthemi
Dickeya chrysanthemi subsp parthenii NCPPB 516 WP 071601394 1 Dchrysanthemi
Dickeya chrysanthemi NCPPB 3533 WP 012769039 1 Dchrysanthemi
Dickeya chrysanthemi Ech1591 WP 012769039 1 Dchrysanthemi
Dickeya dianthicola NCPPB 3534 WP 024106685 1 Ddianthicola
Dickeya dianthicola RNS04 9 WP 046830413 1 Ddianthicola
Dickeya dianthicola IPO 980 WP 024106685 1 Ddianthicola
Dickeya solani PPO 9019 WP 046830413 1 Dsolani
Dickeya dianthicola NCPPB 453 WP 024106685 1 Ddianthicola
Dickeya dianthicola GBBC 2039 WP 024106685 1 Ddianthicola
Dickeya dadantii subsp dieffenbachiae NCPPB 2976 WP 071783926 1 Ddadantii
Dickeya solani RNS 07 7 3B WP 022634276 1 Dsolani
Dickeya solani PPO 9134 WP 022634276 1 Dsolani
Dickeya solani RNS 05 1 2A WP 022634276 1 Dsolani
Dickeya solani IFB 0099 WP 022634276 1 Dsolani
Dickeya solani IPO 2222 WP 071598666 1 Dsolani
Dickeya solani D s0432 1 WP 071598666 1 Dsolani
Dickeya solani RNS 08 23 3 1 A WP 071598666 1 Dsolani
Dickeya solani MK10 WP 071598666 1 Dsolani
Dickeya solani MK16 WP 071598666 1 Dsolani
Dickeya solani GBBC 2040 WP 071598666 1 Dsolani
Dickeya dadantii NCPPB 3537 WP 038901679 1 Ddadantii
Dickeya dadantii NCPPB 898 WP 038911603 1 Ddadantii
45 outD Type II secretion system out Seed
Dickeya dadantii 3937 WP 013318810 1 Ddadantii
Dickeya gs fangzhongdai ND14b WP 071842565 1 Dfangzhongdai
Dickeya gs fangzhongdai M074 WP 038667248 1 Dfangzhongdai
Dickeya gs fangzhongdai M005 WP 039691071 1 Dfangzhongdai
Dickeya gs undicola 2B12 WP 033567854 1 Dundicola
Dickeya gs fangzhongdai NCPPB 3274 WP 071604030 1 Dfangzhongdai
Dickeya gs fangzhongdai MK7 WP 071605264 1 Dfangzhongdai
Dickeya gs fangzhongdai S1 WP 049853678 1 Dfangzhongdai
Dickeya gs fangzhongdai B16 WP 038919718 1 Dfangzhongdai

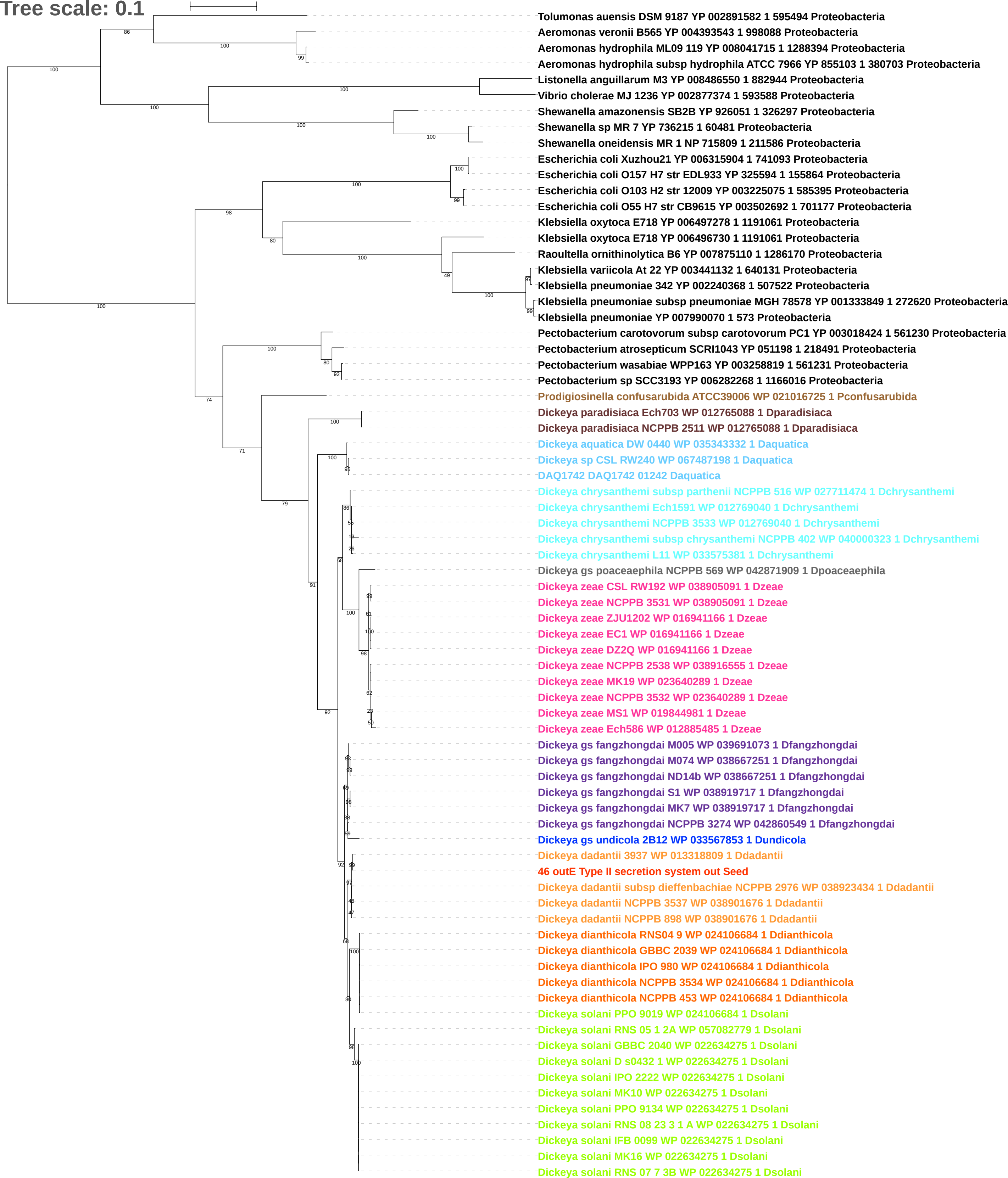
Tree scale: 0.1
Tolumonas auensis DSM 9187 YP 002891582 1 595494 Proteobacteria
Aeromonas veronii B565 YP 004393543 1 998088 Proteobacteria
Aeromonas hydrophila ML09 119 YP 008041715 1 1288394 Proteobacteria
Aeromonas hydrophila subsp hydrophila ATCC 7966 YP 855103 1 380703 Proteobacteria
Listonella anguillarum M3 YP 008486550 1 882944 Proteobacteria
Vibrio cholerae MJ 1236 YP 002877374 1 593588 Proteobacteria
Shewanella amazonensis SB2B YP 926051 1 326297 Proteobacteria
Shewanella sp MR 7 YP 736215 1 60481 Proteobacteria
Shewanella oneidensis MR 1 NP 715809 1 211586 Proteobacteria
Escherichia coli Xuzhou21 YP 006315904 1 741093 Proteobacteria
Escherichia coli O157 H7 str EDL933 YP 325594 1 155864 Proteobacteria
Escherichia coli O103 H2 str 12009 YP 003225075 1 585395 Proteobacteria
Escherichia coli O55 H7 str CB9615 YP 003502692 1 701177 Proteobacteria
Klebsiella oxytoca E718 YP 006497278 1 1191061 Proteobacteria
Klebsiella oxytoca E718 YP 006496730 1 1191061 Proteobacteria
Raoultella ornithinolytica B6 YP 007875110 1 1286170 Proteobacteria
Klebsiella variicola At 22 YP 003441132 1 640131 Proteobacteria
Klebsiella pneumoniae 342 YP 002240368 1 507522 Proteobacteria
Klebsiella pneumoniae subsp pneumoniae MGH 78578 YP 001333849 1 272620 Proteobacteria
Klebsiella pneumoniae YP 007990070 1 573 Proteobacteria
Pectobacterium carotovorum subsp carotovorum PC1 YP 003018424 1 561230 Proteobacteria
Pectobacterium atrosepticum SCRI1043 YP 051198 1 218491 Proteobacteria
Pectobacterium wasabiae WPP163 YP 003258819 1 561231 Proteobacteria
Pectobacterium sp SCC3193 YP 006282268 1 1166016 Proteobacteria
Prodigiosinella confusarubida ATCC39006 WP 021016725 1 Pconfusarubida
Dickeya paradisiaca Ech703 WP 012765088 1 Dparadisiaca
Dickeya paradisiaca NCPPB 2511 WP 012765088 1 Dparadisiaca
Dickeya aquatica DW 0440 WP 035343332 1 Daquatica
Dickeya sp CSL RW240 WP 067487198 1 Daquatica
DAQ1742 DAQ1742 01242 Daquatica
Dickeya chrysanthemi subsp parthenii NCPPB 516 WP 027711474 1 Dchrysanthemi
Dickeya chrysanthemi Ech1591 WP 012769040 1 Dchrysanthemi
Dickeya chrysanthemi NCPPB 3533 WP 012769040 1 Dchrysanthemi
Dickeya chrysanthemi subsp chrysanthemi NCPPB 402 WP 040000323 1 Dchrysanthemi
Dickeya chrysanthemi L11 WP 033575381 1 Dchrysanthemi
Dickeya gs poaceaephila NCPPB 569 WP 042871909 1 Dpoaceaephila
Dickeya zeae CSL RW192 WP 038905091 1 Dzeae
Dickeya zeae NCPPB 3531 WP 038905091 1 Dzeae
Dickeya zeae ZJU1202 WP 016941166 1 Dzeae
Dickeya zeae EC1 WP 016941166 1 Dzeae
Dickeya zeae DZ2Q WP 016941166 1 Dzeae
Dickeya zeae NCPPB 2538 WP 038916555 1 Dzeae
Dickeya zeae MK19 WP 023640289 1 Dzeae
Dickeya zeae NCPPB 3532 WP 023640289 1 Dzeae
Dickeya zeae MS1 WP 019844981 1 Dzeae
Dickeya zeae Ech586 WP 012885485 1 Dzeae
Dickeya gs fangzhongdai M005 WP 039691073 1 Dfangzhongdai
Dickeya gs fangzhongdai M074 WP 038667251 1 Dfangzhongdai
Dickeya gs fangzhongdai ND14b WP 038667251 1 Dfangzhongdai
Dickeya gs fangzhongdai S1 WP 038919717 1 Dfangzhongdai
Dickeya gs fangzhongdai MK7 WP 038919717 1 Dfangzhongdai
Dickeya gs fangzhongdai NCPPB 3274 WP 042860549 1 Dfangzhongdai
Dickeya gs undicola 2B12 WP 033567853 1 Dundicola
Dickeya dadantii 3937 WP 013318809 1 Ddadantii
46 outE Type II secretion system out Seed
Dickeya dadantii subsp dieffenbachiae NCPPB 2976 WP 038923434 1 Ddadantii
Dickeya dadantii NCPPB 3537 WP 038901676 1 Ddadantii
Dickeya dadantii NCPPB 898 WP 038901676 1 Ddadantii
Dickeya dianthicola RNS04 9 WP 024106684 1 Ddianthicola
Dickeya dianthicola GBBC 2039 WP 024106684 1 Ddianthicola
Dickeya dianthicola IPO 980 WP 024106684 1 Ddianthicola
Dickeya dianthicola NCPPB 3534 WP 024106684 1 Ddianthicola
Dickeya dianthicola NCPPB 453 WP 024106684 1 Ddianthicola
Dickeya solani PPO 9019 WP 024106684 1 Dsolani
Dickeya solani RNS 05 1 2A WP 057082779 1 Dsolani
Dickeya solani GBBC 2040 WP 022634275 1 Dsolani
Dickeya solani D s0432 1 WP 022634275 1 Dsolani
Dickeya solani IPO 2222 WP 022634275 1 Dsolani
Dickeya solani MK10 WP 022634275 1 Dsolani
Dickeya solani PPO 9134 WP 022634275 1 Dsolani
Dickeya solani RNS 08 23 3 1 A WP 022634275 1 Dsolani
Dickeya solani IFB 0099 WP 022634275 1 Dsolani
Dickeya solani MK16 WP 022634275 1 Dsolani
Dickeya solani RNS 07 7 3B WP 022634275 1 Dsolani

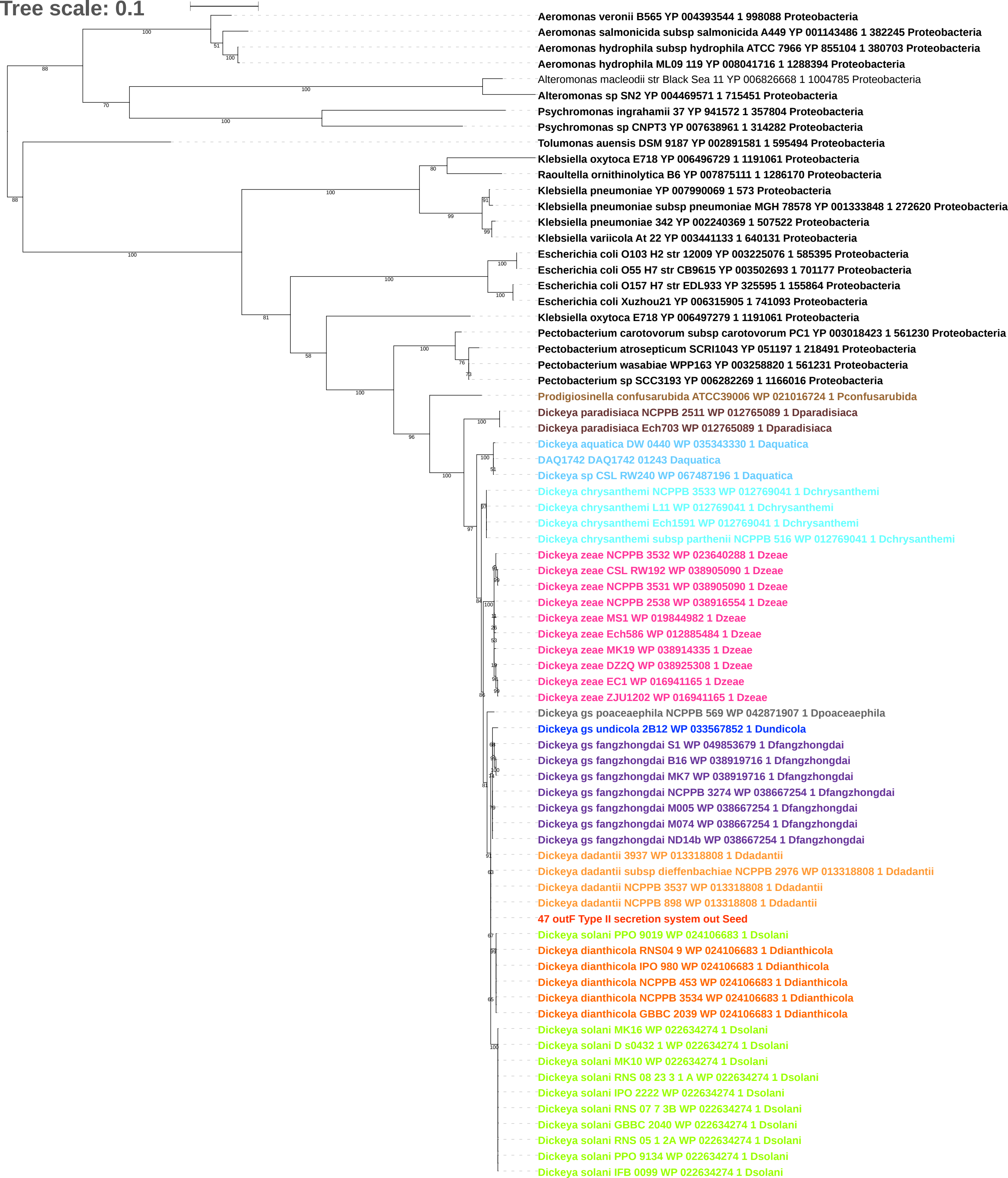
Tree scale: 0.1
Aeromonas veronii B565 YP 004393544 1 998088 Proteobacteria
Aeromonas salmonicida subsp salmonicida A449 YP 001143486 1 382245 Proteobacteria
Aeromonas hydrophila subsp hydrophila ATCC 7966 YP 855104 1 380703 Proteobacteria
Aeromonas hydrophila ML09 119 YP 008041716 1 1288394 Proteobacteria
Alteromonas macleodii str Black Sea 11 YP 006826668 1 1004785 Proteobacteria
Alteromonas sp SN2 YP 004469571 1 715451 Proteobacteria
Psychromonas ingrahamii 37 YP 941572 1 357804 Proteobacteria
Psychromonas sp CNPT3 YP 007638961 1 314282 Proteobacteria
Tolumonas auensis DSM 9187 YP 002891581 1 595494 Proteobacteria
Klebsiella oxytoca E718 YP 006496729 1 1191061 Proteobacteria
Raoultella ornithinolytica B6 YP 007875111 1 1286170 Proteobacteria
Klebsiella pneumoniae YP 007990069 1 573 Proteobacteria
Klebsiella pneumoniae subsp pneumoniae MGH 78578 YP 001333848 1 272620 Proteobacteria
Klebsiella pneumoniae 342 YP 002240369 1 507522 Proteobacteria
Klebsiella variicola At 22 YP 003441133 1 640131 Proteobacteria
Escherichia coli O103 H2 str 12009 YP 003225076 1 585395 Proteobacteria
Escherichia coli O55 H7 str CB9615 YP 003502693 1 701177 Proteobacteria
Escherichia coli O157 H7 str EDL933 YP 325595 1 155864 Proteobacteria
Escherichia coli Xuzhou21 YP 006315905 1 741093 Proteobacteria
Klebsiella oxytoca E718 YP 006497279 1 1191061 Proteobacteria
Pectobacterium carotovorum subsp carotovorum PC1 YP 003018423 1 561230 Proteobacteria
Pectobacterium atrosepticum SCRI1043 YP 051197 1 218491 Proteobacteria
Pectobacterium wasabiae WPP163 YP 003258820 1 561231 Proteobacteria
Pectobacterium sp SCC3193 YP 006282269 1 1166016 Proteobacteria
Prodigiosinella confusarubida ATCC39006 WP 021016724 1 Pconfusarubida
Dickeya paradisiaca NCPPB 2511 WP 012765089 1 Dparadisiaca
Dickeya paradisiaca Ech703 WP 012765089 1 Dparadisiaca
Dickeya aquatica DW 0440 WP 035343330 1 Daquatica
DAQ1742 DAQ1742 01243 Daquatica
Dickeya sp CSL RW240 WP 067487196 1 Daquatica
Dickeya chrysanthemi NCPPB 3533 WP 012769041 1 Dchrysanthemi
Dickeya chrysanthemi L11 WP 012769041 1 Dchrysanthemi
Dickeya chrysanthemi Ech1591 WP 012769041 1 Dchrysanthemi
Dickeya chrysanthemi subsp parthenii NCPPB 516 WP 012769041 1 Dchrysanthemi
Dickeya zeae NCPPB 3532 WP 023640288 1 Dzeae
Dickeya zeae CSL RW192 WP 038905090 1 Dzeae
Dickeya zeae NCPPB 3531 WP 038905090 1 Dzeae
Dickeya zeae NCPPB 2538 WP 038916554 1 Dzeae
Dickeya zeae MS1 WP 019844982 1 Dzeae
Dickeya zeae Ech586 WP 012885484 1 Dzeae
Dickeya zeae MK19 WP 038914335 1 Dzeae
Dickeya zeae DZ2Q WP 038925308 1 Dzeae
Dickeya zeae EC1 WP 016941165 1 Dzeae
Dickeya zeae ZJU1202 WP 016941165 1 Dzeae
Dickeya gs poaceaephila NCPPB 569 WP 042871907 1 Dpoaceaephila
Dickeya gs undicola 2B12 WP 033567852 1 Dundicola
Dickeya gs fangzhongdai S1 WP 049853679 1 Dfangzhongdai
Dickeya gs fangzhongdai B16 WP 038919716 1 Dfangzhongdai
Dickeya gs fangzhongdai MK7 WP 038919716 1 Dfangzhongdai
Dickeya gs fangzhongdai NCPPB 3274 WP 038667254 1 Dfangzhongdai
Dickeya gs fangzhongdai M005 WP 038667254 1 Dfangzhongdai
Dickeya gs fangzhongdai M074 WP 038667254 1 Dfangzhongdai
Dickeya gs fangzhongdai ND14b WP 038667254 1 Dfangzhongdai
Dickeya dadantii 3937 WP 013318808 1 Ddadantii
Dickeya dadantii subsp dieffenbachiae NCPPB 2976 WP 013318808 1 Ddadantii
Dickeya dadantii NCPPB 3537 WP 013318808 1 Ddadantii
Dickeya dadantii NCPPB 898 WP 013318808 1 Ddadantii
47 outF Type II secretion system out Seed
Dickeya solani PPO 9019 WP 024106683 1 Dsolani
Dickeya dianthicola RNS04 9 WP 024106683 1 Ddianthicola
Dickeya dianthicola IPO 980 WP 024106683 1 Ddianthicola
Dickeya dianthicola NCPPB 453 WP 024106683 1 Ddianthicola
Dickeya dianthicola NCPPB 3534 WP 024106683 1 Ddianthicola
Dickeya dianthicola GBBC 2039 WP 024106683 1 Ddianthicola
Dickeya solani MK16 WP 022634274 1 Dsolani
Dickeya solani D s0432 1 WP 022634274 1 Dsolani
Dickeya solani MK10 WP 022634274 1 Dsolani
Dickeya solani RNS 08 23 3 1 A WP 022634274 1 Dsolani
Dickeya solani IPO 2222 WP 022634274 1 Dsolani
Dickeya solani RNS 07 7 3B WP 022634274 1 Dsolani
Dickeya solani GBBC 2040 WP 022634274 1 Dsolani
Dickeya solani RNS 05 1 2A WP 022634274 1 Dsolani
Dickeya solani PPO 9134 WP 022634274 1 Dsolani
Dickeya solani IFB 0099 WP 022634274 1 Dsolani

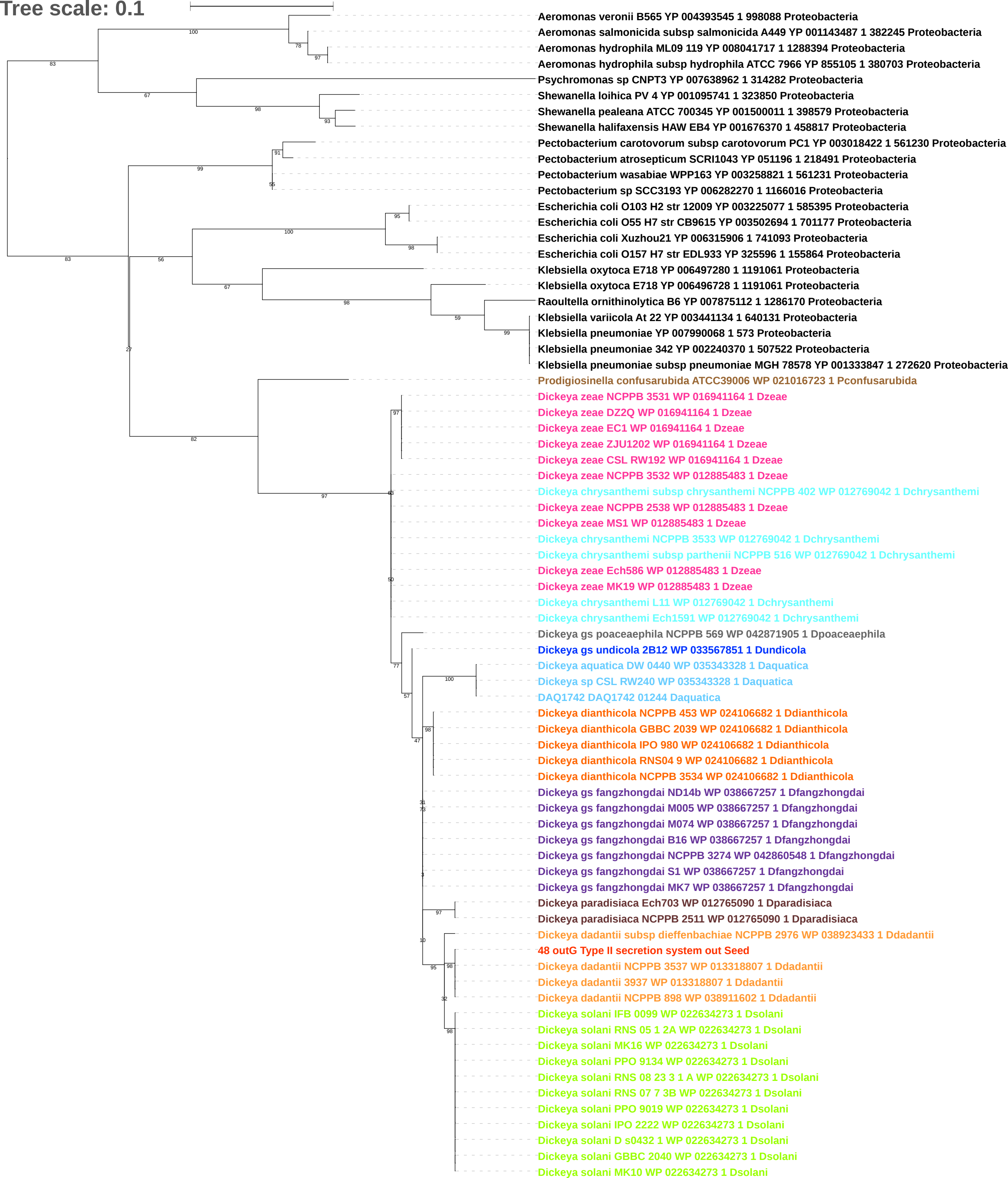
Tree scale: 0.1
Aeromonas veronii B565 YP 004393545 1 998088 Proteobacteria
Aeromonas salmonicida subsp salmonicida A449 YP 001143487 1 382245 Proteobacteria
Aeromonas hydrophila ML09 119 YP 008041717 1 1288394 Proteobacteria
Aeromonas hydrophila subsp hydrophila ATCC 7966 YP 855105 1 380703 Proteobacteria
Psychromonas sp CNPT3 YP 007638962 1 314282 Proteobacteria
Shewanella loihica PV 4 YP 001095741 1 323850 Proteobacteria
Shewanella pealeana ATCC 700345 YP 001500011 1 398579 Proteobacteria
Shewanella halifaxensis HAW EB4 YP 001676370 1 458817 Proteobacteria
Pectobacterium carotovorum subsp carotovorum PC1 YP 003018422 1 561230 Proteobacteria
Pectobacterium atrosepticum SCRI1043 YP 051196 1 218491 Proteobacteria
Pectobacterium wasabiae WPP163 YP 003258821 1 561231 Proteobacteria
Pectobacterium sp SCC3193 YP 006282270 1 1166016 Proteobacteria
Escherichia coli O103 H2 str 12009 YP 003225077 1 585395 Proteobacteria
Escherichia coli O55 H7 str CB9615 YP 003502694 1 701177 Proteobacteria
Escherichia coli Xuzhou21 YP 006315906 1 741093 Proteobacteria
Escherichia coli O157 H7 str EDL933 YP 325596 1 155864 Proteobacteria
Klebsiella oxytoca E718 YP 006497280 1 1191061 Proteobacteria
Klebsiella oxytoca E718 YP 006496728 1 1191061 Proteobacteria
Raoultella ornithinolytica B6 YP 007875112 1 1286170 Proteobacteria
Klebsiella variicola At 22 YP 003441134 1 640131 Proteobacteria
Klebsiella pneumoniae YP 007990068 1 573 Proteobacteria
Klebsiella pneumoniae 342 YP 002240370 1 507522 Proteobacteria
Klebsiella pneumoniae subsp pneumoniae MGH 78578 YP 001333847 1 272620 Proteobacteria
Prodigiosinella confusarubida ATCC39006 WP 021016723 1 Pconfusarubida
Dickeya zeae NCPPB 3531 WP 016941164 1 Dzeae
Dickeya zeae DZ2Q WP 016941164 1 Dzeae
Dickeya zeae EC1 WP 016941164 1 Dzeae
Dickeya zeae ZJU1202 WP 016941164 1 Dzeae
Dickeya zeae CSL RW192 WP 016941164 1 Dzeae
Dickeya zeae NCPPB 3532 WP 012885483 1 Dzeae
Dickeya chrysanthemi subsp chrysanthemi NCPPB 402 WP 012769042 1 Dchrysanthemi
Dickeya zeae NCPPB 2538 WP 012885483 1 Dzeae
Dickeya zeae MS1 WP 012885483 1 Dzeae
Dickeya chrysanthemi NCPPB 3533 WP 012769042 1 Dchrysanthemi
Dickeya chrysanthemi subsp parthenii NCPPB 516 WP 012769042 1 Dchrysanthemi
Dickeya zeae Ech586 WP 012885483 1 Dzeae
Dickeya zeae MK19 WP 012885483 1 Dzeae
Dickeya chrysanthemi L11 WP 012769042 1 Dchrysanthemi
Dickeya chrysanthemi Ech1591 WP 012769042 1 Dchrysanthemi
Dickeya gs poaceaephila NCPPB 569 WP 042871905 1 Dpoaceaephila
Dickeya gs undicola 2B12 WP 033567851 1 Dundicola
Dickeya aquatica DW 0440 WP 035343328 1 Daquatica
Dickeya sp CSL RW240 WP 035343328 1 Daquatica
DAQ1742 DAQ1742 01244 Daquatica
Dickeya dianthicola NCPPB 453 WP 024106682 1 Ddianthicola
Dickeya dianthicola GBBC 2039 WP 024106682 1 Ddianthicola
Dickeya dianthicola IPO 980 WP 024106682 1 Ddianthicola
Dickeya dianthicola RNS04 9 WP 024106682 1 Ddianthicola
Dickeya dianthicola NCPPB 3534 WP 024106682 1 Ddianthicola
Dickeya gs fangzhongdai ND14b WP 038667257 1 Dfangzhongdai
Dickeya gs fangzhongdai M005 WP 038667257 1 Dfangzhongdai
Dickeya gs fangzhongdai M074 WP 038667257 1 Dfangzhongdai
Dickeya gs fangzhongdai B16 WP 038667257 1 Dfangzhongdai
Dickeya gs fangzhongdai NCPPB 3274 WP 042860548 1 Dfangzhongdai
Dickeya gs fangzhongdai S1 WP 038667257 1 Dfangzhongdai
Dickeya gs fangzhongdai MK7 WP 038667257 1 Dfangzhongdai
Dickeya paradisiaca Ech703 WP 012765090 1 Dparadisiaca
Dickeya paradisiaca NCPPB 2511 WP 012765090 1 Dparadisiaca
Dickeya dadantii subsp dieffenbachiae NCPPB 2976 WP 038923433 1 Ddadantii
48 outG Type II secretion system out Seed
Dickeya dadantii NCPPB 3537 WP 013318807 1 Ddadantii
Dickeya dadantii 3937 WP 013318807 1 Ddadantii
Dickeya dadantii NCPPB 898 WP 038911602 1 Ddadantii
Dickeya solani IFB 0099 WP 022634273 1 Dsolani
Dickeya solani RNS 05 1 2A WP 022634273 1 Dsolani
Dickeya solani MK16 WP 022634273 1 Dsolani
Dickeya solani PPO 9134 WP 022634273 1 Dsolani
Dickeya solani RNS 08 23 3 1 A WP 022634273 1 Dsolani
Dickeya solani RNS 07 7 3B WP 022634273 1 Dsolani
Dickeya solani PPO 9019 WP 022634273 1 Dsolani
Dickeya solani IPO 2222 WP 022634273 1 Dsolani
Dickeya solani D s0432 1 WP 022634273 1 Dsolani
Dickeya solani GBBC 2040 WP 022634273 1 Dsolani
Dickeya solani MK10 WP 022634273 1 Dsolani

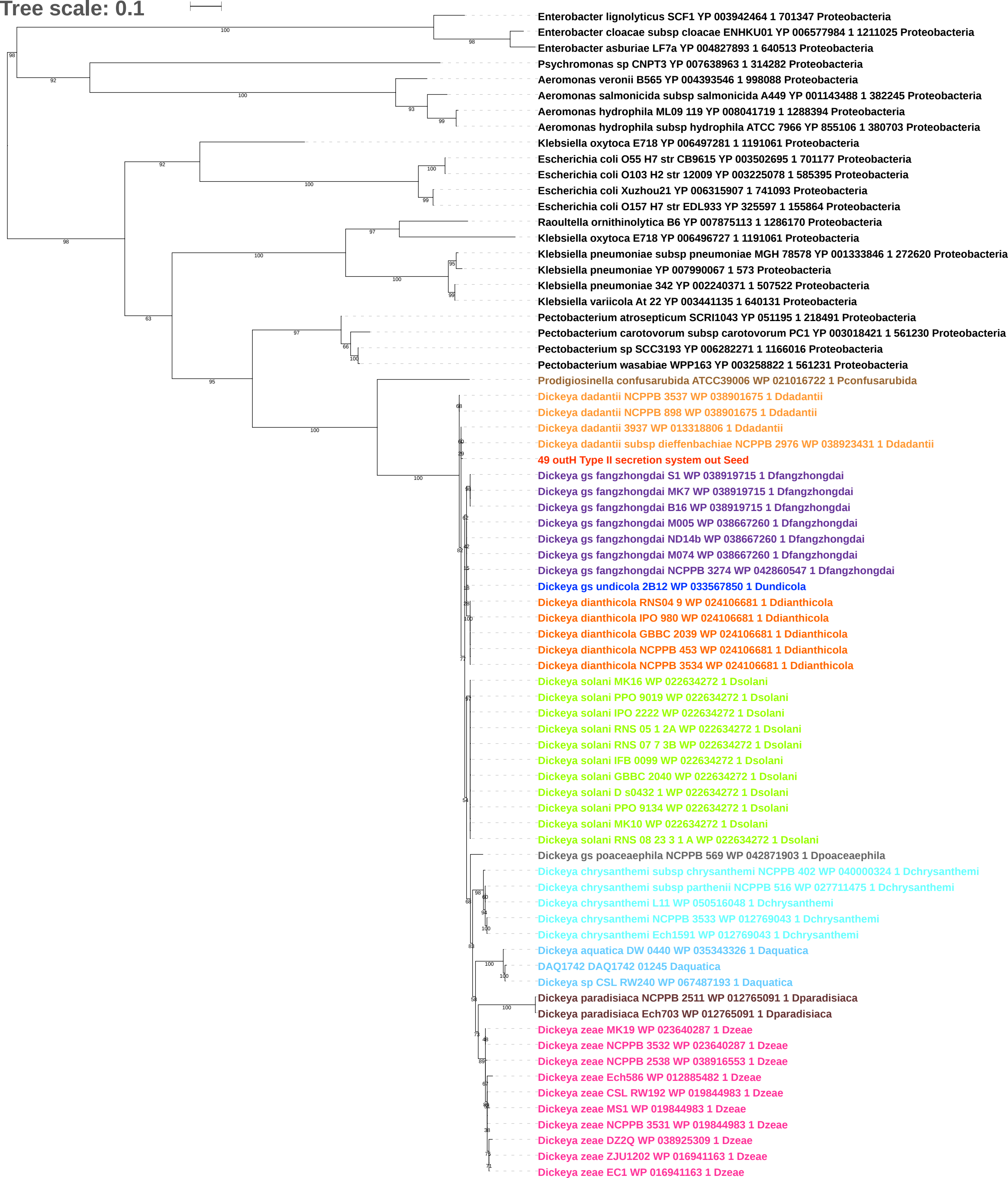
Tree scale: 0.1
Enterobacter lignolyticus SCF1 YP 003942464 1 701347 Proteobacteria
Enterobacter cloacae subsp cloacae ENHKU01 YP 006577984 1 1211025 Proteobacteria
Enterobacter asburiae LF7a YP 004827893 1 640513 Proteobacteria
Psychromonas sp CNPT3 YP 007638963 1 314282 Proteobacteria
Aeromonas veronii B565 YP 004393546 1 998088 Proteobacteria
Aeromonas salmonicida subsp salmonicida A449 YP 001143488 1 382245 Proteobacteria
Aeromonas hydrophila ML09 119 YP 008041719 1 1288394 Proteobacteria
Aeromonas hydrophila subsp hydrophila ATCC 7966 YP 855106 1 380703 Proteobacteria
Klebsiella oxytoca E718 YP 006497281 1 1191061 Proteobacteria
Escherichia coli O55 H7 str CB9615 YP 003502695 1 701177 Proteobacteria
Escherichia coli O103 H2 str 12009 YP 003225078 1 585395 Proteobacteria
Escherichia coli Xuzhou21 YP 006315907 1 741093 Proteobacteria
Escherichia coli O157 H7 str EDL933 YP 325597 1 155864 Proteobacteria
Raoultella ornithinolytica B6 YP 007875113 1 1286170 Proteobacteria
Klebsiella oxytoca E718 YP 006496727 1 1191061 Proteobacteria
Klebsiella pneumoniae subsp pneumoniae MGH 78578 YP 001333846 1 272620 Proteobacteria
Klebsiella pneumoniae YP 007990067 1 573 Proteobacteria
Klebsiella pneumoniae 342 YP 002240371 1 507522 Proteobacteria
Klebsiella variicola At 22 YP 003441135 1 640131 Proteobacteria
Pectobacterium atrosepticum SCRI1043 YP 051195 1 218491 Proteobacteria
Pectobacterium carotovorum subsp carotovorum PC1 YP 003018421 1 561230 Proteobacteria
Pectobacterium sp SCC3193 YP 006282271 1 1166016 Proteobacteria
Pectobacterium wasabiae WPP163 YP 003258822 1 561231 Proteobacteria
Prodigiosinella confusarubida ATCC39006 WP 021016722 1 Pconfusarubida
Dickeya dadantii NCPPB 3537 WP 038901675 1 Ddadantii
Dickeya dadantii NCPPB 898 WP 038901675 1 Ddadantii
Dickeya dadantii 3937 WP 013318806 1 Ddadantii
Dickeya dadantii subsp dieffenbachiae NCPPB 2976 WP 038923431 1 Ddadantii
49 outH Type II secretion system out Seed
Dickeya gs fangzhongdai S1 WP 038919715 1 Dfangzhongdai
Dickeya gs fangzhongdai MK7 WP 038919715 1 Dfangzhongdai
Dickeya gs fangzhongdai B16 WP 038919715 1 Dfangzhongdai
Dickeya gs fangzhongdai M005 WP 038667260 1 Dfangzhongdai
Dickeya gs fangzhongdai ND14b WP 038667260 1 Dfangzhongdai
Dickeya gs fangzhongdai M074 WP 038667260 1 Dfangzhongdai
Dickeya gs fangzhongdai NCPPB 3274 WP 042860547 1 Dfangzhongdai
Dickeya gs undicola 2B12 WP 033567850 1 Dundicola
Dickeya dianthicola RNS04 9 WP 024106681 1 Ddianthicola
Dickeya dianthicola IPO 980 WP 024106681 1 Ddianthicola
Dickeya dianthicola GBBC 2039 WP 024106681 1 Ddianthicola
Dickeya dianthicola NCPPB 453 WP 024106681 1 Ddianthicola
Dickeya dianthicola NCPPB 3534 WP 024106681 1 Ddianthicola
Dickeya solani MK16 WP 022634272 1 Dsolani
Dickeya solani PPO 9019 WP 022634272 1 Dsolani
Dickeya solani IPO 2222 WP 022634272 1 Dsolani
Dickeya solani RNS 05 1 2A WP 022634272 1 Dsolani
Dickeya solani RNS 07 7 3B WP 022634272 1 Dsolani
Dickeya solani IFB 0099 WP 022634272 1 Dsolani
Dickeya solani GBBC 2040 WP 022634272 1 Dsolani
Dickeya solani D s0432 1 WP 022634272 1 Dsolani
Dickeya solani PPO 9134 WP 022634272 1 Dsolani
Dickeya solani MK10 WP 022634272 1 Dsolani
Dickeya solani RNS 08 23 3 1 A WP 022634272 1 Dsolani
Dickeya gs poaceaephila NCPPB 569 WP 042871903 1 Dpoaceaephila
Dickeya chrysanthemi subsp chrysanthemi NCPPB 402 WP 040000324 1 Dchrysanthemi
Dickeya chrysanthemi subsp parthenii NCPPB 516 WP 027711475 1 Dchrysanthemi
Dickeya chrysanthemi L11 WP 050516048 1 Dchrysanthemi
Dickeya chrysanthemi NCPPB 3533 WP 012769043 1 Dchrysanthemi
Dickeya chrysanthemi Ech1591 WP 012769043 1 Dchrysanthemi
Dickeya aquatica DW 0440 WP 035343326 1 Daquatica
DAQ1742 DAQ1742 01245 Daquatica
Dickeya sp CSL RW240 WP 067487193 1 Daquatica
Dickeya paradisiaca NCPPB 2511 WP 012765091 1 Dparadisiaca
Dickeya paradisiaca Ech703 WP 012765091 1 Dparadisiaca
Dickeya zeae MK19 WP 023640287 1 Dzeae
Dickeya zeae NCPPB 3532 WP 023640287 1 Dzeae
Dickeya zeae NCPPB 2538 WP 038916553 1 Dzeae
Dickeya zeae Ech586 WP 012885482 1 Dzeae
Dickeya zeae CSL RW192 WP 019844983 1 Dzeae
Dickeya zeae MS1 WP 019844983 1 Dzeae
Dickeya zeae NCPPB 3531 WP 019844983 1 Dzeae
Dickeya zeae DZ2Q WP 038925309 1 Dzeae
Dickeya zeae ZJU1202 WP 016941163 1 Dzeae
Dickeya zeae EC1 WP 016941163 1 Dzeae

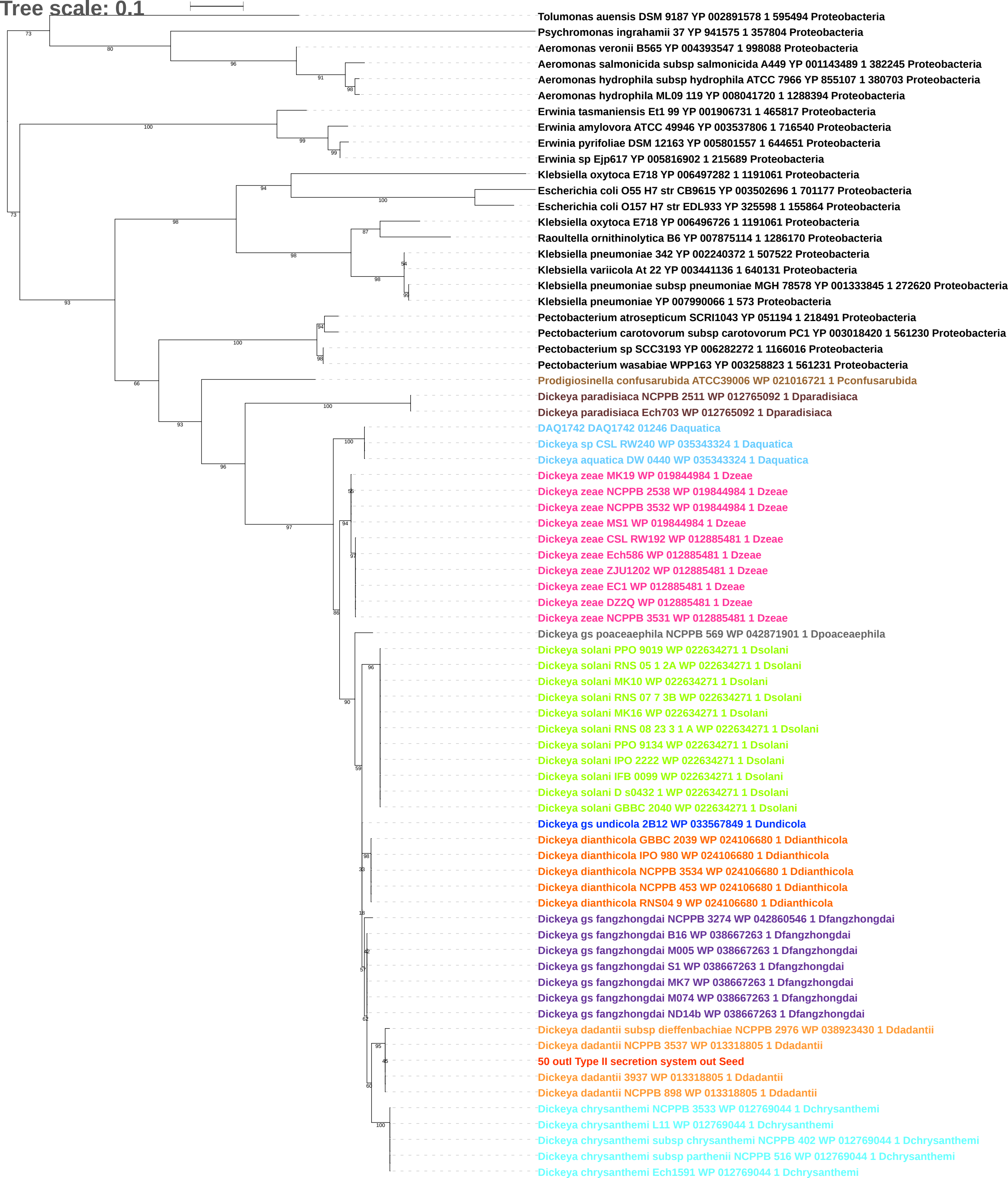
Tree scale: 0.1
Tolumonas auensis DSM 9187 YP 002891578 1 595494 Proteobacteria
Psychromonas ingrahamii 37 YP 941575 1 357804 Proteobacteria
Aeromonas veronii B565 YP 004393547 1 998088 Proteobacteria
Aeromonas salmonicida subsp salmonicida A449 YP 001143489 1 382245 Proteobacteria
Aeromonas hydrophila subsp hydrophila ATCC 7966 YP 855107 1 380703 Proteobacteria
Aeromonas hydrophila ML09 119 YP 008041720 1 1288394 Proteobacteria
Erwinia tasmaniensis Et1 99 YP 001906731 1 465817 Proteobacteria
Erwinia amylovora ATCC 49946 YP 003537806 1 716540 Proteobacteria
Erwinia pyrifoliae DSM 12163 YP 005801557 1 644651 Proteobacteria
Erwinia sp Ejp617 YP 005816902 1 215689 Proteobacteria
Klebsiella oxytoca E718 YP 006497282 1 1191061 Proteobacteria
Escherichia coli O55 H7 str CB9615 YP 003502696 1 701177 Proteobacteria
Escherichia coli O157 H7 str EDL933 YP 325598 1 155864 Proteobacteria
Klebsiella oxytoca E718 YP 006496726 1 1191061 Proteobacteria
Raoultella ornithinolytica B6 YP 007875114 1 1286170 Proteobacteria
Klebsiella pneumoniae 342 YP 002240372 1 507522 Proteobacteria
Klebsiella variicola At 22 YP 003441136 1 640131 Proteobacteria
Klebsiella pneumoniae subsp pneumoniae MGH 78578 YP 001333845 1 272620 Proteobacteria
Klebsiella pneumoniae YP 007990066 1 573 Proteobacteria
Pectobacterium atrosepticum SCRI1043 YP 051194 1 218491 Proteobacteria
Pectobacterium carotovorum subsp carotovorum PC1 YP 003018420 1 561230 Proteobacteria
Pectobacterium sp SCC3193 YP 006282272 1 1166016 Proteobacteria
Pectobacterium wasabiae WPP163 YP 003258823 1 561231 Proteobacteria
Prodigiosinella confusarubida ATCC39006 WP 021016721 1 Pconfusarubida
Dickeya paradisiaca NCPPB 2511 WP 012765092 1 Dparadisiaca
Dickeya paradisiaca Ech703 WP 012765092 1 Dparadisiaca
DAQ1742 DAQ1742 01246 Daquatica
Dickeya sp CSL RW240 WP 035343324 1 Daquatica
Dickeya aquatica DW 0440 WP 035343324 1 Daquatica
Dickeya zeae MK19 WP 019844984 1 Dzeae
Dickeya zeae NCPPB 2538 WP 019844984 1 Dzeae
Dickeya zeae NCPPB 3532 WP 019844984 1 Dzeae
Dickeya zeae MS1 WP 019844984 1 Dzeae
Dickeya zeae CSL RW192 WP 012885481 1 Dzeae
Dickeya zeae Ech586 WP 012885481 1 Dzeae
Dickeya zeae ZJU1202 WP 012885481 1 Dzeae
Dickeya zeae EC1 WP 012885481 1 Dzeae
Dickeya zeae DZ2Q WP 012885481 1 Dzeae
Dickeya zeae NCPPB 3531 WP 012885481 1 Dzeae
Dickeya gs poaceaephila NCPPB 569 WP 042871901 1 Dpoaceaephila
Dickeya solani PPO 9019 WP 022634271 1 Dsolani
Dickeya solani RNS 05 1 2A WP 022634271 1 Dsolani
Dickeya solani MK10 WP 022634271 1 Dsolani
Dickeya solani RNS 07 7 3B WP 022634271 1 Dsolani
Dickeya solani MK16 WP 022634271 1 Dsolani
Dickeya solani RNS 08 23 3 1 A WP 022634271 1 Dsolani
Dickeya solani PPO 9134 WP 022634271 1 Dsolani
Dickeya solani IPO 2222 WP 022634271 1 Dsolani
Dickeya solani IFB 0099 WP 022634271 1 Dsolani
Dickeya solani D s0432 1 WP 022634271 1 Dsolani
Dickeya solani GBBC 2040 WP 022634271 1 Dsolani
Dickeya gs undicola 2B12 WP 033567849 1 Dundicola
Dickeya dianthicola GBBC 2039 WP 024106680 1 Ddianthicola
Dickeya dianthicola IPO 980 WP 024106680 1 Ddianthicola
Dickeya dianthicola NCPPB 3534 WP 024106680 1 Ddianthicola
Dickeya dianthicola NCPPB 453 WP 024106680 1 Ddianthicola
Dickeya dianthicola RNS04 9 WP 024106680 1 Ddianthicola
Dickeya gs fangzhongdai NCPPB 3274 WP 042860546 1 Dfangzhongdai
Dickeya gs fangzhongdai B16 WP 038667263 1 Dfangzhongdai
Dickeya gs fangzhongdai M005 WP 038667263 1 Dfangzhongdai
Dickeya gs fangzhongdai S1 WP 038667263 1 Dfangzhongdai
Dickeya gs fangzhongdai MK7 WP 038667263 1 Dfangzhongdai
Dickeya gs fangzhongdai M074 WP 038667263 1 Dfangzhongdai
Dickeya gs fangzhongdai ND14b WP 038667263 1 Dfangzhongdai
Dickeya dadantii subsp dieffenbachiae NCPPB 2976 WP 038923430 1 Ddadantii
Dickeya dadantii NCPPB 3537 WP 013318805 1 Ddadantii
50 outI Type II secretion system out Seed
Dickeya dadantii 3937 WP 013318805 1 Ddadantii
Dickeya dadantii NCPPB 898 WP 013318805 1 Ddadantii
Dickeya chrysanthemi NCPPB 3533 WP 012769044 1 Dchrysanthemi
Dickeya chrysanthemi L11 WP 012769044 1 Dchrysanthemi
Dickeya chrysanthemi subsp chrysanthemi NCPPB 402 WP 012769044 1 Dchrysanthemi
Dickeya chrysanthemi subsp parthenii NCPPB 516 WP 012769044 1 Dchrysanthemi
Dickeya chrysanthemi Ech1591 WP 012769044 1 Dchrysanthemi

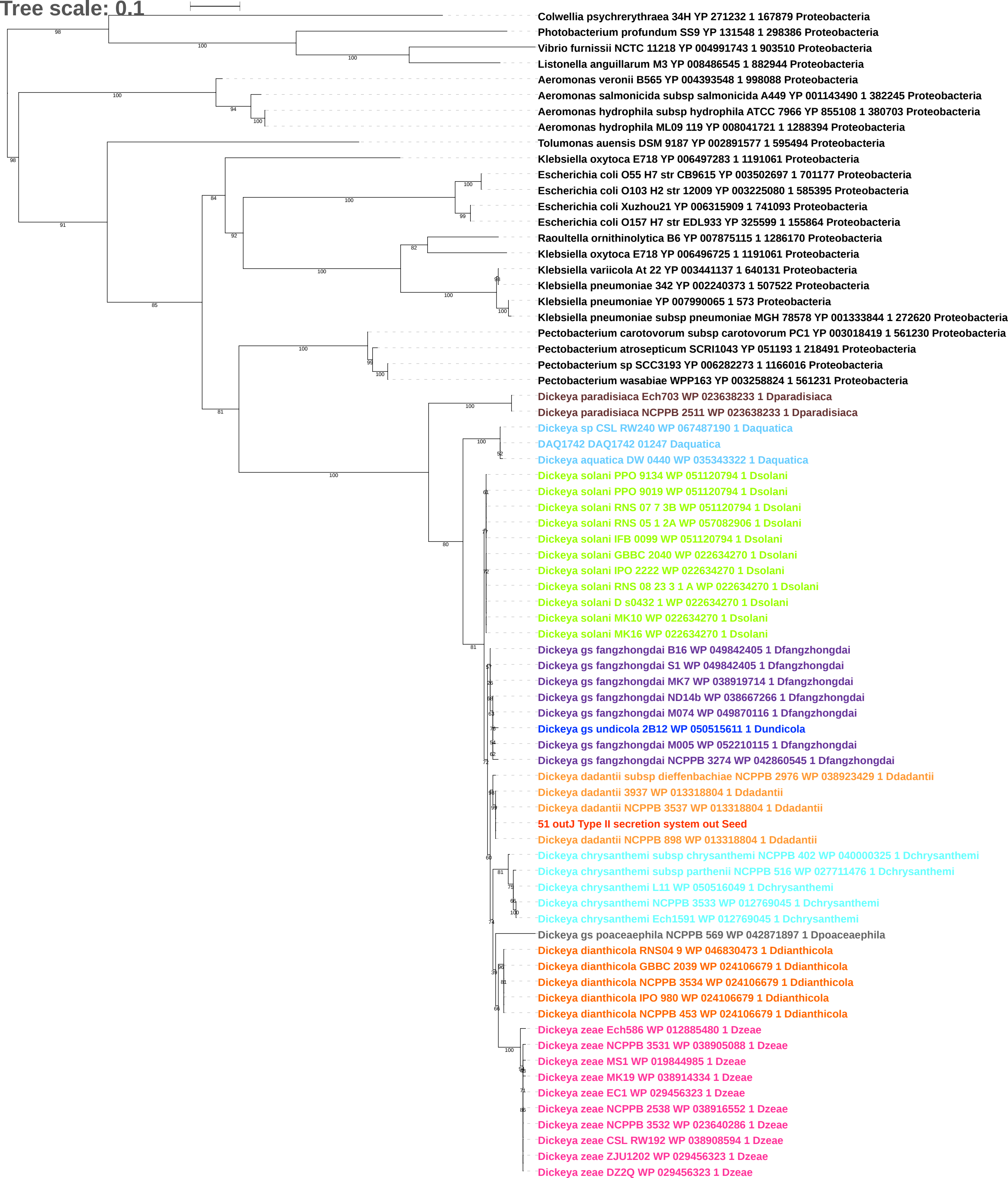
Tree scale: 0.1
Colwellia psychrerythraea 34H YP 271232 1 167879 Proteobacteria
Photobacterium profundum SS9 YP 131548 1 298386 Proteobacteria
Vibrio furnissii NCTC 11218 YP 004991743 1 903510 Proteobacteria
Listonella anguillarum M3 YP 008486545 1 882944 Proteobacteria
Aeromonas veronii B565 YP 004393548 1 998088 Proteobacteria
Aeromonas salmonicida subsp salmonicida A449 YP 001143490 1 382245 Proteobacteria
Aeromonas hydrophila subsp hydrophila ATCC 7966 YP 855108 1 380703 Proteobacteria
Aeromonas hydrophila ML09 119 YP 008041721 1 1288394 Proteobacteria
Tolumonas auensis DSM 9187 YP 002891577 1 595494 Proteobacteria
Klebsiella oxytoca E718 YP 006497283 1 1191061 Proteobacteria
Escherichia coli O55 H7 str CB9615 YP 003502697 1 701177 Proteobacteria
Escherichia coli O103 H2 str 12009 YP 003225080 1 585395 Proteobacteria
Escherichia coli Xuzhou21 YP 006315909 1 741093 Proteobacteria
Escherichia coli O157 H7 str EDL933 YP 325599 1 155864 Proteobacteria
Raoultella ornithinolytica B6 YP 007875115 1 1286170 Proteobacteria
Klebsiella oxytoca E718 YP 006496725 1 1191061 Proteobacteria
Klebsiella variicola At 22 YP 003441137 1 640131 Proteobacteria
Klebsiella pneumoniae 342 YP 002240373 1 507522 Proteobacteria
Klebsiella pneumoniae YP 007990065 1 573 Proteobacteria
Klebsiella pneumoniae subsp pneumoniae MGH 78578 YP 001333844 1 272620 Proteobacteria
Pectobacterium carotovorum subsp carotovorum PC1 YP 003018419 1 561230 Proteobacteria
Pectobacterium atrosepticum SCRI1043 YP 051193 1 218491 Proteobacteria
Pectobacterium sp SCC3193 YP 006282273 1 1166016 Proteobacteria
Pectobacterium wasabiae WPP163 YP 003258824 1 561231 Proteobacteria
Dickeya paradisiaca Ech703 WP 023638233 1 Dparadisiaca
Dickeya paradisiaca NCPPB 2511 WP 023638233 1 Dparadisiaca
Dickeya sp CSL RW240 WP 067487190 1 Daquatica
DAQ1742 DAQ1742 01247 Daquatica
Dickeya aquatica DW 0440 WP 035343322 1 Daquatica
Dickeya solani PPO 9134 WP 051120794 1 Dsolani
Dickeya solani PPO 9019 WP 051120794 1 Dsolani
Dickeya solani RNS 07 7 3B WP 051120794 1 Dsolani
Dickeya solani RNS 05 1 2A WP 057082906 1 Dsolani
Dickeya solani IFB 0099 WP 051120794 1 Dsolani
Dickeya solani GBBC 2040 WP 022634270 1 Dsolani
Dickeya solani IPO 2222 WP 022634270 1 Dsolani
Dickeya solani RNS 08 23 3 1 A WP 022634270 1 Dsolani
Dickeya solani D s0432 1 WP 022634270 1 Dsolani
Dickeya solani MK10 WP 022634270 1 Dsolani
Dickeya solani MK16 WP 022634270 1 Dsolani
Dickeya gs fangzhongdai B16 WP 049842405 1 Dfangzhongdai
Dickeya gs fangzhongdai S1 WP 049842405 1 Dfangzhongdai
Dickeya gs fangzhongdai MK7 WP 038919714 1 Dfangzhongdai
Dickeya gs fangzhongdai ND14b WP 038667266 1 Dfangzhongdai
Dickeya gs fangzhongdai M074 WP 049870116 1 Dfangzhongdai
Dickeya gs undicola 2B12 WP 050515611 1 Dundicola
Dickeya gs fangzhongdai M005 WP 052210115 1 Dfangzhongdai
Dickeya gs fangzhongdai NCPPB 3274 WP 042860545 1 Dfangzhongdai
Dickeya dadantii subsp dieffenbachiae NCPPB 2976 WP 038923429 1 Ddadantii
Dickeya dadantii 3937 WP 013318804 1 Ddadantii
Dickeya dadantii NCPPB 3537 WP 013318804 1 Ddadantii
51 outJ Type II secretion system out Seed
Dickeya dadantii NCPPB 898 WP 013318804 1 Ddadantii
Dickeya chrysanthemi subsp chrysanthemi NCPPB 402 WP 040000325 1 Dchrysanthemi
Dickeya chrysanthemi subsp parthenii NCPPB 516 WP 027711476 1 Dchrysanthemi
Dickeya chrysanthemi L11 WP 050516049 1 Dchrysanthemi
Dickeya chrysanthemi NCPPB 3533 WP 012769045 1 Dchrysanthemi
Dickeya chrysanthemi Ech1591 WP 012769045 1 Dchrysanthemi
Dickeya gs poaceaephila NCPPB 569 WP 042871897 1 Dpoaceaephila
Dickeya dianthicola RNS04 9 WP 046830473 1 Ddianthicola
Dickeya dianthicola GBBC 2039 WP 024106679 1 Ddianthicola
Dickeya dianthicola NCPPB 3534 WP 024106679 1 Ddianthicola
Dickeya dianthicola IPO 980 WP 024106679 1 Ddianthicola
Dickeya dianthicola NCPPB 453 WP 024106679 1 Ddianthicola
Dickeya zeae Ech586 WP 012885480 1 Dzeae
Dickeya zeae NCPPB 3531 WP 038905088 1 Dzeae
Dickeya zeae MS1 WP 019844985 1 Dzeae
Dickeya zeae MK19 WP 038914334 1 Dzeae
Dickeya zeae EC1 WP 029456323 1 Dzeae
Dickeya zeae NCPPB 2538 WP 038916552 1 Dzeae
Dickeya zeae NCPPB 3532 WP 023640286 1 Dzeae
Dickeya zeae CSL RW192 WP 038908594 1 Dzeae
Dickeya zeae ZJU1202 WP 029456323 1 Dzeae
Dickeya zeae DZ2Q WP 029456323 1 Dzeae

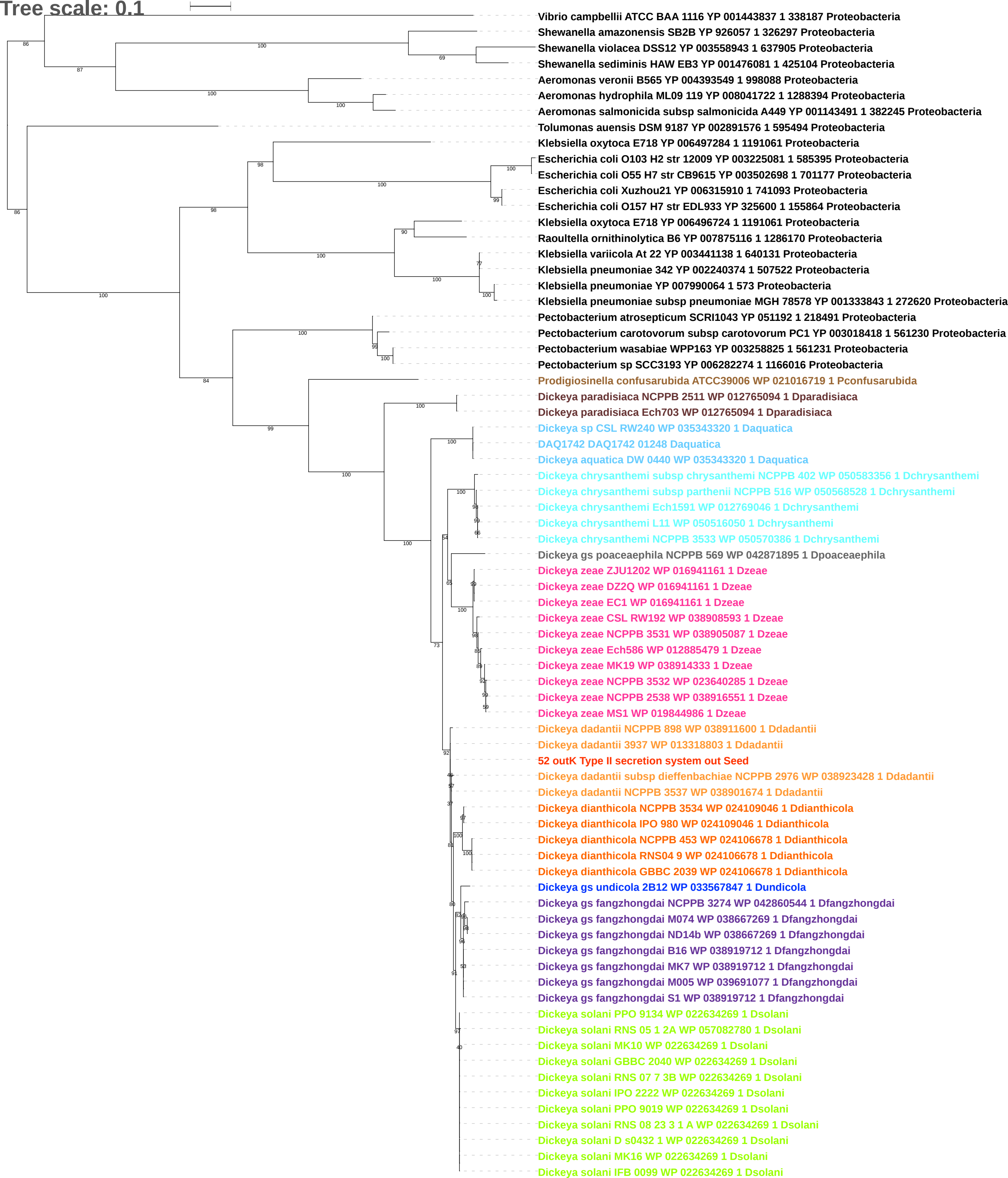
Tree scale: 0.1
Vibrio campbellii ATCC BAA 1116 YP 001443837 1 338187 Proteobacteria
Shewanella amazonensis SB2B YP 926057 1 326297 Proteobacteria
Shewanella violacea DSS12 YP 003558943 1 637905 Proteobacteria
Shewanella sediminis HAW EB3 YP 001476081 1 425104 Proteobacteria
Aeromonas veronii B565 YP 004393549 1 998088 Proteobacteria
Aeromonas hydrophila ML09 119 YP 008041722 1 1288394 Proteobacteria
Aeromonas salmonicida subsp salmonicida A449 YP 001143491 1 382245 Proteobacteria
Tolumonas auensis DSM 9187 YP 002891576 1 595494 Proteobacteria
Klebsiella oxytoca E718 YP 006497284 1 1191061 Proteobacteria
Escherichia coli O103 H2 str 12009 YP 003225081 1 585395 Proteobacteria
Escherichia coli O55 H7 str CB9615 YP 003502698 1 701177 Proteobacteria
Escherichia coli Xuzhou21 YP 006315910 1 741093 Proteobacteria
Escherichia coli O157 H7 str EDL933 YP 325600 1 155864 Proteobacteria
Klebsiella oxytoca E718 YP 006496724 1 1191061 Proteobacteria
Raoultella ornithinolytica B6 YP 007875116 1 1286170 Proteobacteria
Klebsiella variicola At 22 YP 003441138 1 640131 Proteobacteria
Klebsiella pneumoniae 342 YP 002240374 1 507522 Proteobacteria
Klebsiella pneumoniae YP 007990064 1 573 Proteobacteria
Klebsiella pneumoniae subsp pneumoniae MGH 78578 YP 001333843 1 272620 Proteobacteria
Pectobacterium atrosepticum SCRI1043 YP 051192 1 218491 Proteobacteria
Pectobacterium carotovorum subsp carotovorum PC1 YP 003018418 1 561230 Proteobacteria
Pectobacterium wasabiae WPP163 YP 003258825 1 561231 Proteobacteria
Pectobacterium sp SCC3193 YP 006282274 1 1166016 Proteobacteria
Prodigiosinella confusarubida ATCC39006 WP 021016719 1 Pconfusarubida
Dickeya paradisiaca NCPPB 2511 WP 012765094 1 Dparadisiaca
Dickeya paradisiaca Ech703 WP 012765094 1 Dparadisiaca
Dickeya sp CSL RW240 WP 035343320 1 Daquatica
DAQ1742 DAQ1742 01248 Daquatica
Dickeya aquatica DW 0440 WP 035343320 1 Daquatica
Dickeya chrysanthemi subsp chrysanthemi NCPPB 402 WP 050583356 1 Dchrysanthemi
Dickeya chrysanthemi subsp parthenii NCPPB 516 WP 050568528 1 Dchrysanthemi
Dickeya chrysanthemi Ech1591 WP 012769046 1 Dchrysanthemi
Dickeya chrysanthemi L11 WP 050516050 1 Dchrysanthemi
Dickeya chrysanthemi NCPPB 3533 WP 050570386 1 Dchrysanthemi
Dickeya gs poaceaephila NCPPB 569 WP 042871895 1 Dpoaceaephila
Dickeya zeae ZJU1202 WP 016941161 1 Dzeae
Dickeya zeae DZ2Q WP 016941161 1 Dzeae
Dickeya zeae EC1 WP 016941161 1 Dzeae
Dickeya zeae CSL RW192 WP 038908593 1 Dzeae
Dickeya zeae NCPPB 3531 WP 038905087 1 Dzeae
Dickeya zeae Ech586 WP 012885479 1 Dzeae
Dickeya zeae MK19 WP 038914333 1 Dzeae
Dickeya zeae NCPPB 3532 WP 023640285 1 Dzeae
Dickeya zeae NCPPB 2538 WP 038916551 1 Dzeae
Dickeya zeae MS1 WP 019844986 1 Dzeae
Dickeya dadantii NCPPB 898 WP 038911600 1 Ddadantii
Dickeya dadantii 3937 WP 013318803 1 Ddadantii
52 outK Type II secretion system out Seed
Dickeya dadantii subsp dieffenbachiae NCPPB 2976 WP 038923428 1 Ddadantii
Dickeya dadantii NCPPB 3537 WP 038901674 1 Ddadantii
Dickeya dianthicola NCPPB 3534 WP 024109046 1 Ddianthicola
Dickeya dianthicola IPO 980 WP 024109046 1 Ddianthicola
Dickeya dianthicola NCPPB 453 WP 024106678 1 Ddianthicola
Dickeya dianthicola RNS04 9 WP 024106678 1 Ddianthicola
Dickeya dianthicola GBBC 2039 WP 024106678 1 Ddianthicola
Dickeya gs undicola 2B12 WP 033567847 1 Dundicola
Dickeya gs fangzhongdai NCPPB 3274 WP 042860544 1 Dfangzhongdai
Dickeya gs fangzhongdai M074 WP 038667269 1 Dfangzhongdai
Dickeya gs fangzhongdai ND14b WP 038667269 1 Dfangzhongdai
Dickeya gs fangzhongdai B16 WP 038919712 1 Dfangzhongdai
Dickeya gs fangzhongdai MK7 WP 038919712 1 Dfangzhongdai
Dickeya gs fangzhongdai M005 WP 039691077 1 Dfangzhongdai
Dickeya gs fangzhongdai S1 WP 038919712 1 Dfangzhongdai
Dickeya solani PPO 9134 WP 022634269 1 Dsolani
Dickeya solani RNS 05 1 2A WP 057082780 1 Dsolani
Dickeya solani MK10 WP 022634269 1 Dsolani
Dickeya solani GBBC 2040 WP 022634269 1 Dsolani
Dickeya solani RNS 07 7 3B WP 022634269 1 Dsolani
Dickeya solani IPO 2222 WP 022634269 1 Dsolani
Dickeya solani PPO 9019 WP 022634269 1 Dsolani
Dickeya solani RNS 08 23 3 1 A WP 022634269 1 Dsolani
Dickeya solani D s0432 1 WP 022634269 1 Dsolani
Dickeya solani MK16 WP 022634269 1 Dsolani
Dickeya solani IFB 0099 WP 022634269 1 Dsolani

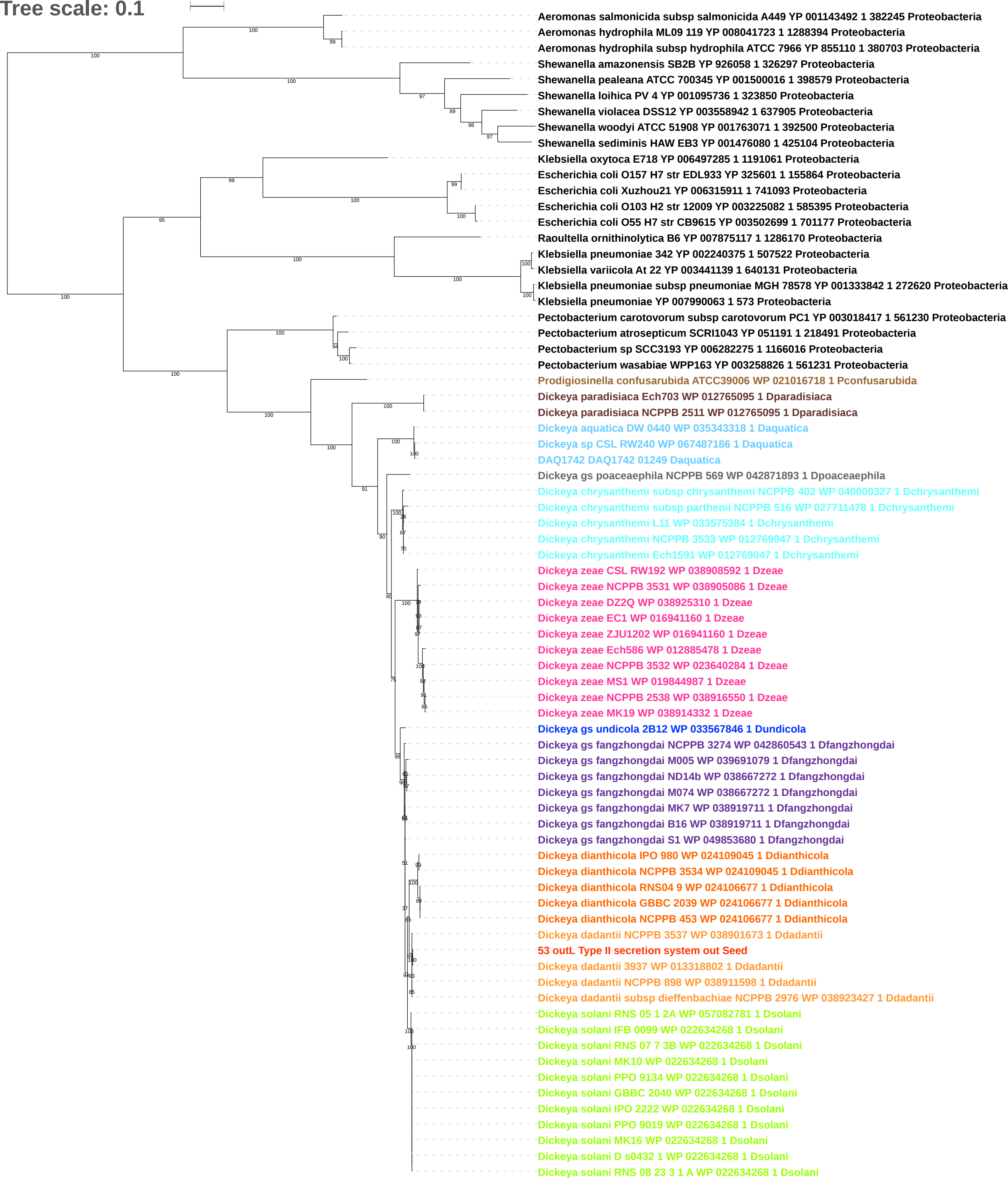

Tree scale: 0.1
Aeromonas salmonicida subsp salmonicida A449 YP 001143492 1 382245 Proteobacteria
Aeromonas hydrophila ML09 119 YP 008041723 1 1288394 Proteobacteria
Aeromonas hydrophila subsp hydrophila ATCC 7966 YP 855110 1 380703 Proteobacteria
Shewanella amazonensis SB2B YP 926058 1 326297 Proteobacteria
Shewanella pealeana ATCC 700345 YP 001500016 1 398579 Proteobacteria
Shewanella loihica PV 4 YP 001095736 1 323850 Proteobacteria
Shewanella violacea DSS12 YP 003558942 1 637905 Proteobacteria
Shewanella woodyi ATCC 51908 YP 001763071 1 392500 Proteobacteria
Shewanella sediminis HAW EB3 YP 001476080 1 425104 Proteobacteria
Klebsiella oxytoca E718 YP 006497285 1 1191061 Proteobacteria
Escherichia coli O157 H7 str EDL933 YP 325601 1 155864 Proteobacteria
Escherichia coli Xuzhou21 YP 006315911 1 741093 Proteobacteria
Escherichia coli O103 H2 str 12009 YP 003225082 1 585395 Proteobacteria
Escherichia coli O55 H7 str CB9615 YP 003502699 1 701177 Proteobacteria
Raoultella ornithinolytica B6 YP 007875117 1 1286170 Proteobacteria
Klebsiella pneumoniae 342 YP 002240375 1 507522 Proteobacteria
Klebsiella variicola At 22 YP 003441139 1 640131 Proteobacteria
Klebsiella pneumoniae subsp pneumoniae MGH 78578 YP 001333842 1 272620 Proteobacteria
Klebsiella pneumoniae YP 007990063 1 573 Proteobacteria
Pectobacterium carotovorum subsp carotovorum PC1 YP 003018417 1 561230 Proteobacteria
Pectobacterium atrosepticum SCRI1043 YP 051191 1 218491 Proteobacteria
Pectobacterium sp SCC3193 YP 006282275 1 1166016 Proteobacteria
Pectobacterium wasabiae WPP163 YP 003258826 1 561231 Proteobacteria
Prodigiosinella confusarubida ATCC39006 WP 021016718 1 Pconfusarubida
Dickeya paradisiaca Ech703 WP 012765095 1 Dparadisiaca
Dickeya paradisiaca NCPPB 2511 WP 012765095 1 Dparadisiaca
Dickeya aquatica DW 0440 WP 035343318 1 Daquatica
Dickeya sp CSL RW240 WP 067487186 1 Daquatica
DAQ1742 DAQ1742 01249 Daquatica
Dickeya gs poaceaephila NCPPB 569 WP 042871893 1 Dpoaceaephila
Dickeya chrysanthemi subsp chrysanthemi NCPPB 402 WP 040000327 1 Dchrysanthemi
Dickeya chrysanthemi subsp parthenii NCPPB 516 WP 027711478 1 Dchrysanthemi
Dickeya chrysanthemi L11 WP 033575384 1 Dchrysanthemi
Dickeya chrysanthemi NCPPB 3533 WP 012769047 1 Dchrysanthemi
Dickeya chrysanthemi Ech1591 WP 012769047 1 Dchrysanthemi
Dickeya zeae CSL RW192 WP 038908592 1 Dzeae
Dickeya zeae NCPPB 3531 WP 038905086 1 Dzeae
Dickeya zeae DZ2Q WP 038925310 1 Dzeae
Dickeya zeae EC1 WP 016941160 1 Dzeae
Dickeya zeae ZJU1202 WP 016941160 1 Dzeae
Dickeya zeae Ech586 WP 012885478 1 Dzeae
Dickeya zeae NCPPB 3532 WP 023640284 1 Dzeae
Dickeya zeae MS1 WP 019844987 1 Dzeae
Dickeya zeae NCPPB 2538 WP 038916550 1 Dzeae
Dickeya zeae MK19 WP 038914332 1 Dzeae
Dickeya gs undicola 2B12 WP 033567846 1 Dundicola
Dickeya gs fangzhongdai NCPPB 3274 WP 042860543 1 Dfangzhongdai
Dickeya gs fangzhongdai M005 WP 039691079 1 Dfangzhongdai
Dickeya gs fangzhongdai ND14b WP 038667272 1 Dfangzhongdai
Dickeya gs fangzhongdai M074 WP 038667272 1 Dfangzhongdai
Dickeya gs fangzhongdai MK7 WP 038919711 1 Dfangzhongdai
Dickeya gs fangzhongdai B16 WP 038919711 1 Dfangzhongdai
Dickeya gs fangzhongdai S1 WP 049853680 1 Dfangzhongdai
Dickeya dianthicola IPO 980 WP 024109045 1 Ddianthicola
Dickeya dianthicola NCPPB 3534 WP 024109045 1 Ddianthicola
Dickeya dianthicola RNS04 9 WP 024106677 1 Ddianthicola
Dickeya dianthicola GBBC 2039 WP 024106677 1 Ddianthicola
Dickeya dianthicola NCPPB 453 WP 024106677 1 Ddianthicola
Dickeya dadantii NCPPB 3537 WP 038901673 1 Ddadantii
53 outL Type II secretion system out Seed
Dickeya dadantii 3937 WP 013318802 1 Ddadantii
Dickeya dadantii NCPPB 898 WP 038911598 1 Ddadantii
Dickeya dadantii subsp dieffenbachiae NCPPB 2976 WP 038923427 1 Ddadantii
Dickeya solani RNS 05 1 2A WP 057082781 1 Dsolani
Dickeya solani IFB 0099 WP 022634268 1 Dsolani
Dickeya solani RNS 07 7 3B WP 022634268 1 Dsolani
Dickeya solani MK10 WP 022634268 1 Dsolani
Dickeya solani PPO 9134 WP 022634268 1 Dsolani
Dickeya solani GBBC 2040 WP 022634268 1 Dsolani
Dickeya solani IPO 2222 WP 022634268 1 Dsolani
Dickeya solani PPO 9019 WP 022634268 1 Dsolani
Dickeya solani MK16 WP 022634268 1 Dsolani
Dickeya solani D s0432 1 WP 022634268 1 Dsolani
Dickeya solani RNS 08 23 3 1 A WP 022634268 1 Dsolani

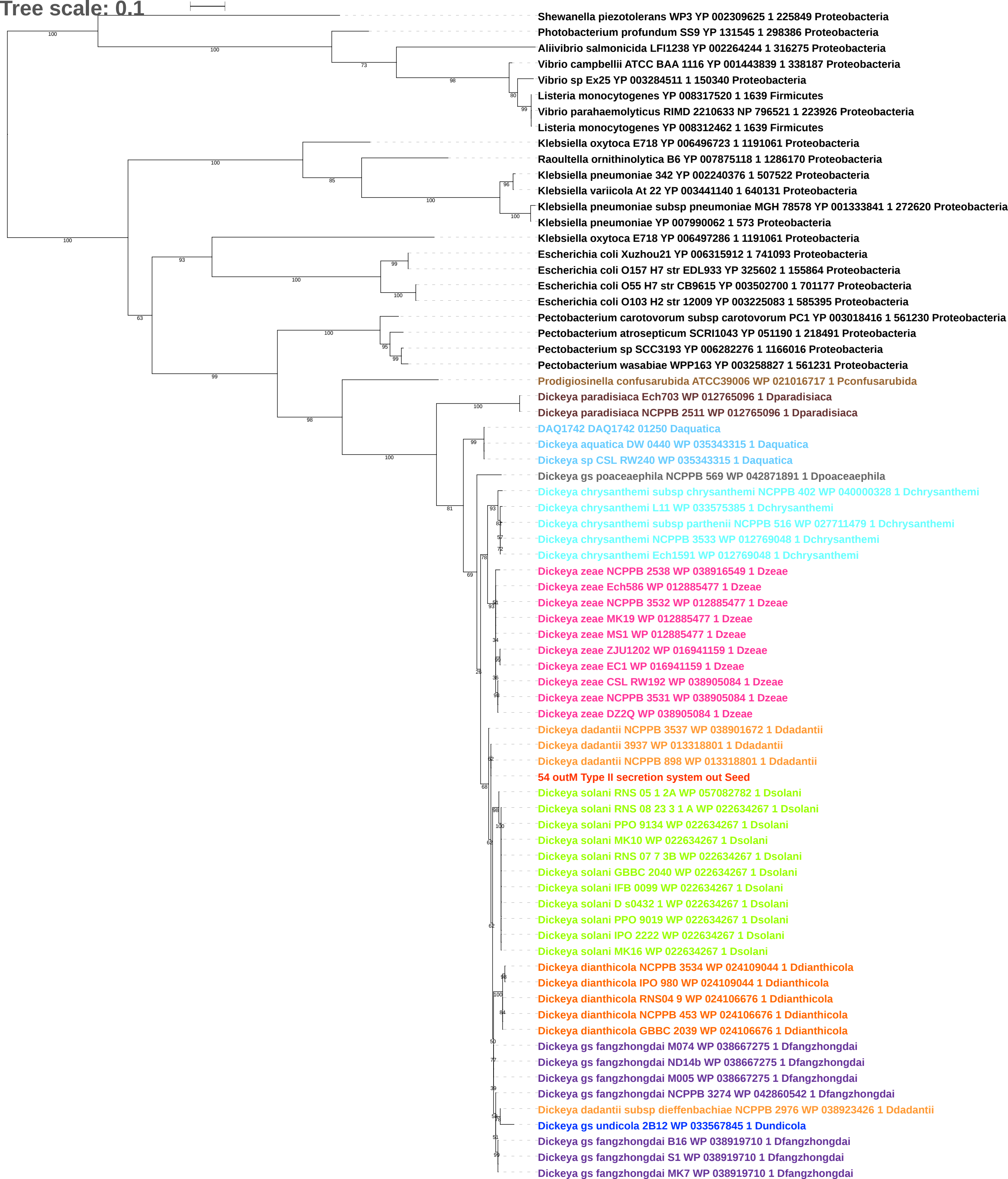
Tree scale: 0.1
Shewanella piezotolerans WP3 YP 002309625 1 225849 Proteobacteria
Photobacterium profundum SS9 YP 131545 1 298386 Proteobacteria
Aliivibrio salmonicida LFI1238 YP 002264244 1 316275 Proteobacteria
Vibrio campbellii ATCC BAA 1116 YP 001443839 1 338187 Proteobacteria
Vibrio sp Ex25 YP 003284511 1 150340 Proteobacteria
Listeria monocytogenes YP 008317520 1 1639 Firmicutes
Vibrio parahaemolyticus RIMD 2210633 NP 796521 1 223926 Proteobacteria
Listeria monocytogenes YP 008312462 1 1639 Firmicutes
Klebsiella oxytoca E718 YP 006496723 1 1191061 Proteobacteria
Raoultella ornithinolytica B6 YP 007875118 1 1286170 Proteobacteria
Klebsiella pneumoniae 342 YP 002240376 1 507522 Proteobacteria
Klebsiella variicola At 22 YP 003441140 1 640131 Proteobacteria
Klebsiella pneumoniae subsp pneumoniae MGH 78578 YP 001333841 1 272620 Proteobacteria
Klebsiella pneumoniae YP 007990062 1 573 Proteobacteria
Klebsiella oxytoca E718 YP 006497286 1 1191061 Proteobacteria
Escherichia coli Xuzhou21 YP 006315912 1 741093 Proteobacteria
Escherichia coli O157 H7 str EDL933 YP 325602 1 155864 Proteobacteria
Escherichia coli O55 H7 str CB9615 YP 003502700 1 701177 Proteobacteria
Escherichia coli O103 H2 str 12009 YP 003225083 1 585395 Proteobacteria
Pectobacterium carotovorum subsp carotovorum PC1 YP 003018416 1 561230 Proteobacteria
Pectobacterium atrosepticum SCRI1043 YP 051190 1 218491 Proteobacteria
Pectobacterium sp SCC3193 YP 006282276 1 1166016 Proteobacteria
Pectobacterium wasabiae WPP163 YP 003258827 1 561231 Proteobacteria
Prodigiosinella confusarubida ATCC39006 WP 021016717 1 Pconfusarubida
Dickeya paradisiaca Ech703 WP 012765096 1 Dparadisiaca
Dickeya paradisiaca NCPPB 2511 WP 012765096 1 Dparadisiaca
DAQ1742 DAQ1742 01250 Daquatica
Dickeya aquatica DW 0440 WP 035343315 1 Daquatica
Dickeya sp CSL RW240 WP 035343315 1 Daquatica
Dickeya gs poaceaephila NCPPB 569 WP 042871891 1 Dpoaceaephila
Dickeya chrysanthemi subsp chrysanthemi NCPPB 402 WP 040000328 1 Dchrysanthemi
Dickeya chrysanthemi L11 WP 033575385 1 Dchrysanthemi
Dickeya chrysanthemi subsp parthenii NCPPB 516 WP 027711479 1 Dchrysanthemi
Dickeya chrysanthemi NCPPB 3533 WP 012769048 1 Dchrysanthemi
Dickeya chrysanthemi Ech1591 WP 012769048 1 Dchrysanthemi
Dickeya zeae NCPPB 2538 WP 038916549 1 Dzeae
Dickeya zeae Ech586 WP 012885477 1 Dzeae
Dickeya zeae NCPPB 3532 WP 012885477 1 Dzeae
Dickeya zeae MK19 WP 012885477 1 Dzeae
Dickeya zeae MS1 WP 012885477 1 Dzeae
Dickeya zeae ZJU1202 WP 016941159 1 Dzeae
Dickeya zeae EC1 WP 016941159 1 Dzeae
Dickeya zeae CSL RW192 WP 038905084 1 Dzeae
Dickeya zeae NCPPB 3531 WP 038905084 1 Dzeae
Dickeya zeae DZ2Q WP 038905084 1 Dzeae
Dickeya dadantii NCPPB 3537 WP 038901672 1 Ddadantii
Dickeya dadantii 3937 WP 013318801 1 Ddadantii
Dickeya dadantii NCPPB 898 WP 013318801 1 Ddadantii
54 outM Type II secretion system out Seed
Dickeya solani RNS 05 1 2A WP 057082782 1 Dsolani
Dickeya solani RNS 08 23 3 1 A WP 022634267 1 Dsolani
Dickeya solani PPO 9134 WP 022634267 1 Dsolani
Dickeya solani MK10 WP 022634267 1 Dsolani
Dickeya solani RNS 07 7 3B WP 022634267 1 Dsolani
Dickeya solani GBBC 2040 WP 022634267 1 Dsolani
Dickeya solani IFB 0099 WP 022634267 1 Dsolani
Dickeya solani D s0432 1 WP 022634267 1 Dsolani
Dickeya solani PPO 9019 WP 022634267 1 Dsolani
Dickeya solani IPO 2222 WP 022634267 1 Dsolani
Dickeya solani MK16 WP 022634267 1 Dsolani
Dickeya dianthicola NCPPB 3534 WP 024109044 1 Ddianthicola
Dickeya dianthicola IPO 980 WP 024109044 1 Ddianthicola
Dickeya dianthicola RNS04 9 WP 024106676 1 Ddianthicola
Dickeya dianthicola NCPPB 453 WP 024106676 1 Ddianthicola
Dickeya dianthicola GBBC 2039 WP 024106676 1 Ddianthicola
Dickeya gs fangzhongdai M074 WP 038667275 1 Dfangzhongdai
Dickeya gs fangzhongdai ND14b WP 038667275 1 Dfangzhongdai
Dickeya gs fangzhongdai M005 WP 038667275 1 Dfangzhongdai
Dickeya gs fangzhongdai NCPPB 3274 WP 042860542 1 Dfangzhongdai
Dickeya dadantii subsp dieffenbachiae NCPPB 2976 WP 038923426 1 Ddadantii
Dickeya gs undicola 2B12 WP 033567845 1 Dundicola
Dickeya gs fangzhongdai B16 WP 038919710 1 Dfangzhongdai
Dickeya gs fangzhongdai S1 WP 038919710 1 Dfangzhongdai
Dickeya gs fangzhongdai MK7 WP 038919710 1 Dfangzhongdai

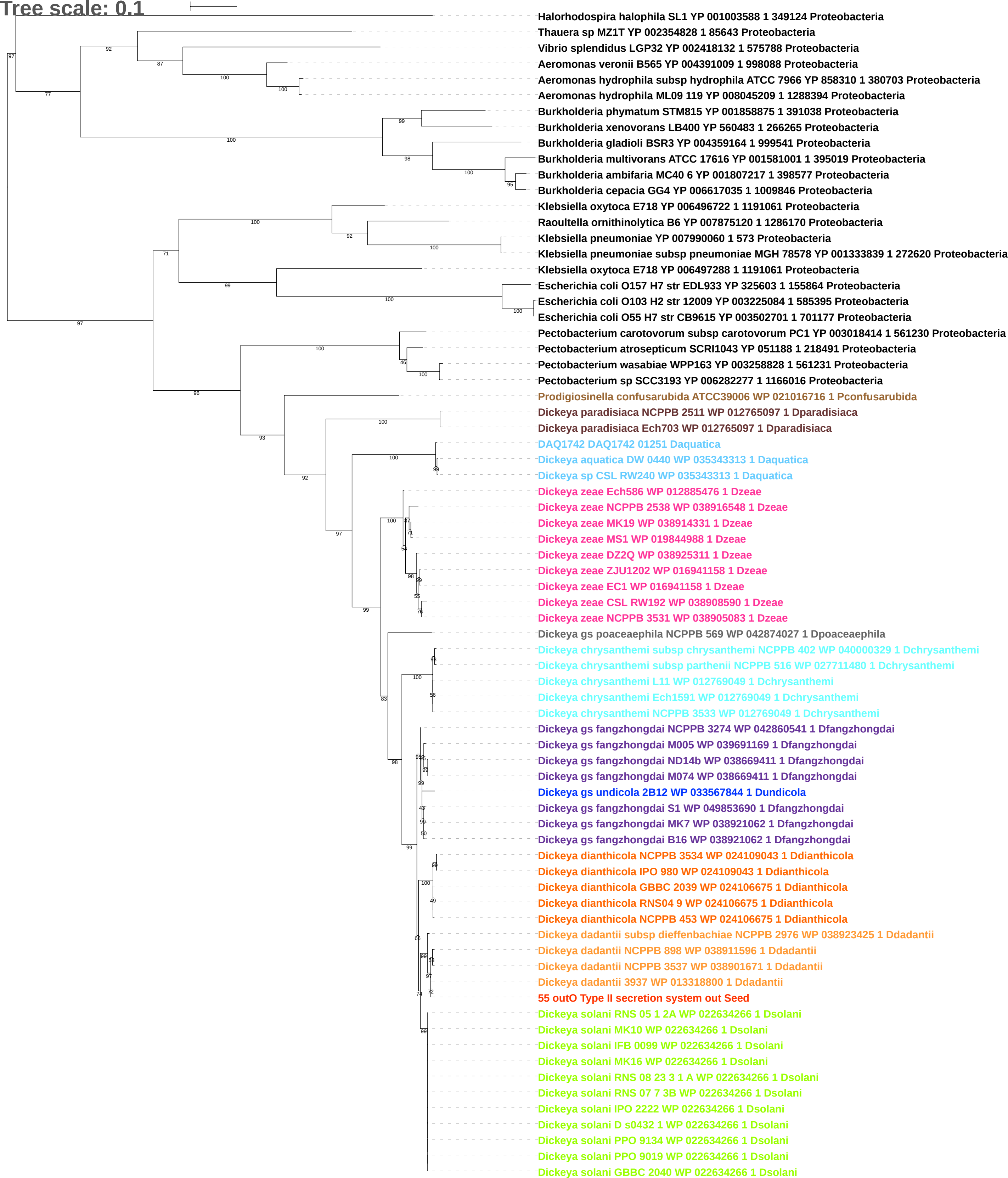
Tree scale: 0.1
Halorhodospira halophila SL1 YP 001003588 1 349124 Proteobacteria
Thauera sp MZ1T YP 002354828 1 85643 Proteobacteria
Vibrio splendidus LGP32 YP 002418132 1 575788 Proteobacteria
Aeromonas veronii B565 YP 004391009 1 998088 Proteobacteria
Aeromonas hydrophila subsp hydrophila ATCC 7966 YP 858310 1 380703 Proteobacteria
Aeromonas hydrophila ML09 119 YP 008045209 1 1288394 Proteobacteria
Burkholderia phymatum STM815 YP 001858875 1 391038 Proteobacteria
Burkholderia xenovorans LB400 YP 560483 1 266265 Proteobacteria
Burkholderia gladioli BSR3 YP 004359164 1 999541 Proteobacteria
Burkholderia multivorans ATCC 17616 YP 001581001 1 395019 Proteobacteria
Burkholderia ambifaria MC40 6 YP 001807217 1 398577 Proteobacteria
Burkholderia cepacia GG4 YP 006617035 1 1009846 Proteobacteria
Klebsiella oxytoca E718 YP 006496722 1 1191061 Proteobacteria
Raoultella ornithinolytica B6 YP 007875120 1 1286170 Proteobacteria
Klebsiella pneumoniae YP 007990060 1 573 Proteobacteria
Klebsiella pneumoniae subsp pneumoniae MGH 78578 YP 001333839 1 272620 Proteobacteria
Klebsiella oxytoca E718 YP 006497288 1 1191061 Proteobacteria
Escherichia coli O157 H7 str EDL933 YP 325603 1 155864 Proteobacteria
Escherichia coli O103 H2 str 12009 YP 003225084 1 585395 Proteobacteria
Escherichia coli O55 H7 str CB9615 YP 003502701 1 701177 Proteobacteria
Pectobacterium carotovorum subsp carotovorum PC1 YP 003018414 1 561230 Proteobacteria
Pectobacterium atrosepticum SCRI1043 YP 051188 1 218491 Proteobacteria
Pectobacterium wasabiae WPP163 YP 003258828 1 561231 Proteobacteria
Pectobacterium sp SCC3193 YP 006282277 1 1166016 Proteobacteria
Prodigiosinella confusarubida ATCC39006 WP 021016716 1 Pconfusarubida
Dickeya paradisiaca NCPPB 2511 WP 012765097 1 Dparadisiaca
Dickeya paradisiaca Ech703 WP 012765097 1 Dparadisiaca
DAQ1742 DAQ1742 01251 Daquatica
Dickeya aquatica DW 0440 WP 035343313 1 Daquatica
Dickeya sp CSL RW240 WP 035343313 1 Daquatica
Dickeya zeae Ech586 WP 012885476 1 Dzeae
Dickeya zeae NCPPB 2538 WP 038916548 1 Dzeae
Dickeya zeae MK19 WP 038914331 1 Dzeae
Dickeya zeae MS1 WP 019844988 1 Dzeae
Dickeya zeae DZ2Q WP 038925311 1 Dzeae
Dickeya zeae ZJU1202 WP 016941158 1 Dzeae
Dickeya zeae EC1 WP 016941158 1 Dzeae
Dickeya zeae CSL RW192 WP 038908590 1 Dzeae
Dickeya zeae NCPPB 3531 WP 038905083 1 Dzeae
Dickeya gs poaceaephila NCPPB 569 WP 042874027 1 Dpoaceaephila
Dickeya chrysanthemi subsp chrysanthemi NCPPB 402 WP 040000329 1 Dchrysanthemi
Dickeya chrysanthemi subsp parthenii NCPPB 516 WP 027711480 1 Dchrysanthemi
Dickeya chrysanthemi L11 WP 012769049 1 Dchrysanthemi
Dickeya chrysanthemi Ech1591 WP 012769049 1 Dchrysanthemi
Dickeya chrysanthemi NCPPB 3533 WP 012769049 1 Dchrysanthemi
Dickeya gs fangzhongdai NCPPB 3274 WP 042860541 1 Dfangzhongdai
Dickeya gs fangzhongdai M005 WP 039691169 1 Dfangzhongdai
Dickeya gs fangzhongdai ND14b WP 038669411 1 Dfangzhongdai
Dickeya gs fangzhongdai M074 WP 038669411 1 Dfangzhongdai
Dickeya gs undicola 2B12 WP 033567844 1 Dundicola
Dickeya gs fangzhongdai S1 WP 049853690 1 Dfangzhongdai
Dickeya gs fangzhongdai MK7 WP 038921062 1 Dfangzhongdai
Dickeya gs fangzhongdai B16 WP 038921062 1 Dfangzhongdai
Dickeya dianthicola NCPPB 3534 WP 024109043 1 Ddianthicola
Dickeya dianthicola IPO 980 WP 024109043 1 Ddianthicola
Dickeya dianthicola GBBC 2039 WP 024106675 1 Ddianthicola
Dickeya dianthicola RNS04 9 WP 024106675 1 Ddianthicola
Dickeya dianthicola NCPPB 453 WP 024106675 1 Ddianthicola
Dickeya dadantii subsp dieffenbachiae NCPPB 2976 WP 038923425 1 Ddadantii
Dickeya dadantii NCPPB 898 WP 038911596 1 Ddadantii
Dickeya dadantii NCPPB 3537 WP 038901671 1 Ddadantii
Dickeya dadantii 3937 WP 013318800 1 Ddadantii
55 outO Type II secretion system out Seed
Dickeya solani RNS 05 1 2A WP 022634266 1 Dsolani
Dickeya solani MK10 WP 022634266 1 Dsolani
Dickeya solani IFB 0099 WP 022634266 1 Dsolani
Dickeya solani MK16 WP 022634266 1 Dsolani
Dickeya solani RNS 08 23 3 1 A WP 022634266 1 Dsolani
Dickeya solani RNS 07 7 3B WP 022634266 1 Dsolani
Dickeya solani IPO 2222 WP 022634266 1 Dsolani
Dickeya solani D s0432 1 WP 022634266 1 Dsolani
Dickeya solani PPO 9134 WP 022634266 1 Dsolani
Dickeya solani PPO 9019 WP 022634266 1 Dsolani
Dickeya solani GBBC 2040 WP 022634266 1 Dsolani

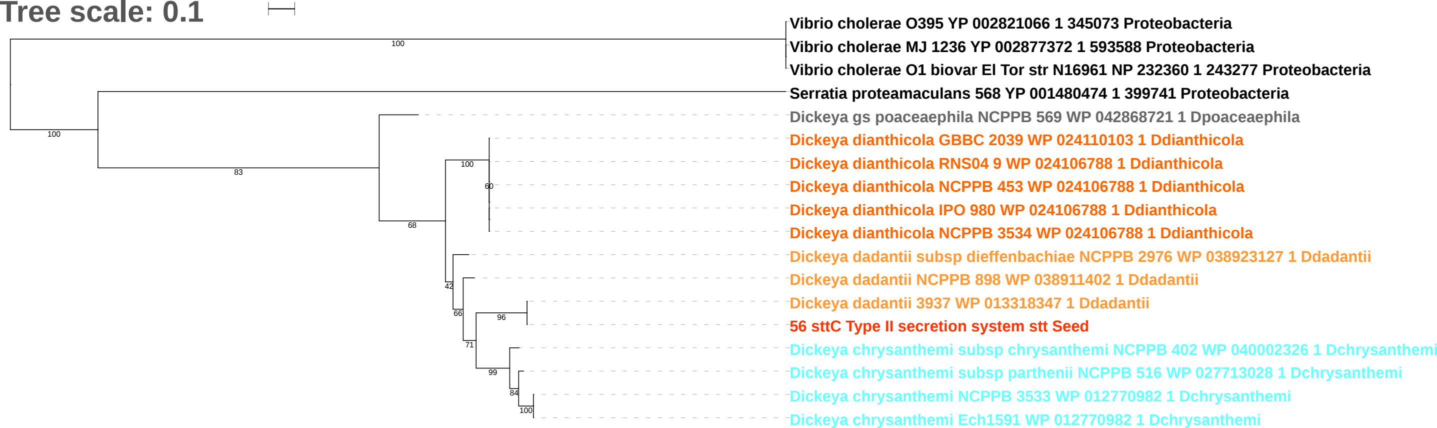

Tree scale: 0.1
Vibrio cholerae O395 YP 002821066 1 345073 Proteobacteria
Vibrio cholerae MJ 1236 YP 002877372 1 593588 Proteobacteria
Vibrio cholerae O1 biovar El Tor str N16961 NP 232360 1 243277 Proteobacteria
Serratia proteamaculans 568 YP 001480474 1 399741 Proteobacteria
Dickeya gs poaceaephila NCPPB 569 WP 042868721 1 Dpoaceaephila
Dickeya dianthicola GBBC 2039 WP 024110103 1 Ddianthicola
Dickeya dianthicola RNS04 9 WP 024106788 1 Ddianthicola
Dickeya dianthicola NCPPB 453 WP 024106788 1 Ddianthicola
Dickeya dianthicola IPO 980 WP 024106788 1 Ddianthicola
Dickeya dianthicola NCPPB 3534 WP 024106788 1 Ddianthicola
Dickeya dadantii subsp dieffenbachiae NCPPB 2976 WP 038923127 1 Ddadantii
Dickeya dadantii NCPPB 898 WP 038911402 1 Ddadantii
Dickeya dadantii 3937 WP 013318347 1 Ddadantii
56 sttC Type II secretion system stt Seed
Dickeya chrysanthemi subsp chrysanthemi NCPPB 402 WP 040002326 1 Dchrysanthemi
Dickeya chrysanthemi subsp parthenii NCPPB 516 WP 027713028 1 Dchrysanthemi
Dickeya chrysanthemi NCPPB 3533 WP 012770982 1 Dchrysanthemi
Dickeya chrysanthemi Ech1591 WP 012770982 1 Dchrysanthemi
100
100
83
68
100
60
42
66
96
71
99
84
100

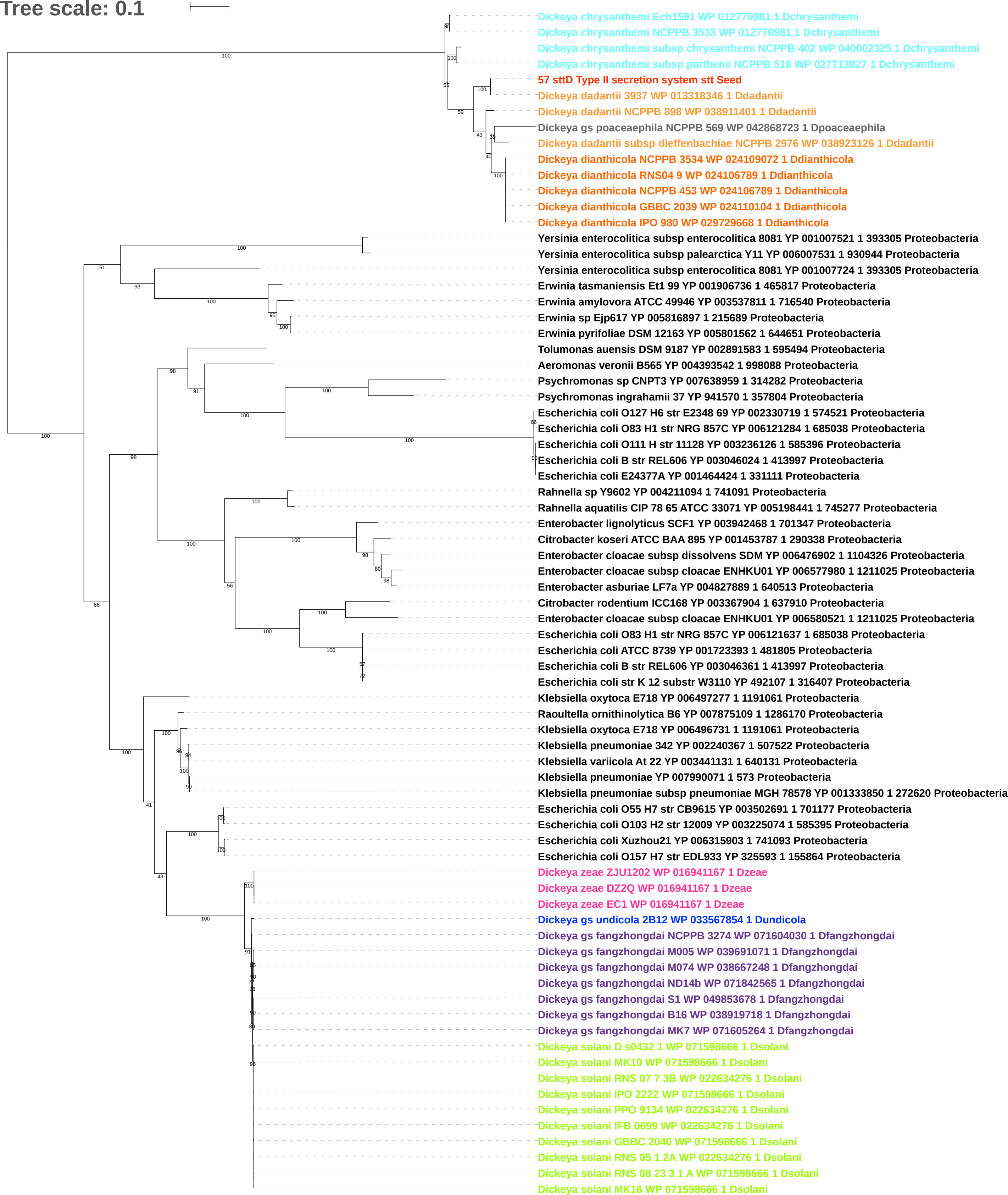
Tree scale: 0.1
Dickeya chrysanthemi Ech1591 WP 012770981 1 Dchrysanthemi
Dickeya chrysanthemi NCPPB 3533 WP 012770981 1 Dchrysanthemi
Dickeya chrysanthemi subsp chrysanthemi NCPPB 402 WP 040002325 1 Dchrysanthemi
Dickeya chrysanthemi subsp parthenii NCPPB 516 WP 027713027 1 Dchrysanthemi
57 sttD Type II secretion system stt Seed
Dickeya dadantii 3937 WP 013318346 1 Ddadantii
Dickeya dadantii NCPPB 898 WP 038911401 1 Ddadantii
Dickeya gs poaceaephila NCPPB 569 WP 042868723 1 Dpoaceaephila
Dickeya dadantii subsp dieffenbachiae NCPPB 2976 WP 038923126 1 Ddadantii
Dickeya dianthicola NCPPB 3534 WP 024109072 1 Ddianthicola
Dickeya dianthicola RNS04 9 WP 024106789 1 Ddianthicola
Dickeya dianthicola NCPPB 453 WP 024106789 1 Ddianthicola
Dickeya dianthicola GBBC 2039 WP 024110104 1 Ddianthicola
Dickeya dianthicola IPO 980 WP 029729668 1 Ddianthicola
Yersinia enterocolitica subsp enterocolitica 8081 YP 001007521 1 393305 Proteobacteria
Yersinia enterocolitica subsp palearctica Y11 YP 006007531 1 930944 Proteobacteria
Yersinia enterocolitica subsp enterocolitica 8081 YP 001007724 1 393305 Proteobacteria
Erwinia tasmaniensis Et1 99 YP 001906736 1 465817 Proteobacteria
Erwinia amylovora ATCC 49946 YP 003537811 1 716540 Proteobacteria
Erwinia sp Ejp617 YP 005816897 1 215689 Proteobacteria
Erwinia pyrifoliae DSM 12163 YP 005801562 1 644651 Proteobacteria
Tolumonas auensis DSM 9187 YP 002891583 1 595494 Proteobacteria
Aeromonas veronii B565 YP 004393542 1 998088 Proteobacteria
Psychromonas sp CNPT3 YP 007638959 1 314282 Proteobacteria
Psychromonas ingrahamii 37 YP 941570 1 357804 Proteobacteria
Escherichia coli O127 H6 str E2348 69 YP 002330719 1 574521 Proteobacteria
Escherichia coli O83 H1 str NRG 857C YP 006121284 1 685038 Proteobacteria
Escherichia coli O111 H str 11128 YP 003236126 1 585396 Proteobacteria
Escherichia coli B str REL606 YP 003046024 1 413997 Proteobacteria
Escherichia coli E24377A YP 001464424 1 331111 Proteobacteria
Rahnella sp Y9602 YP 004211094 1 741091 Proteobacteria
Rahnella aquatilis CIP 78 65 ATCC 33071 YP 005198441 1 745277 Proteobacteria
Enterobacter lignolyticus SCF1 YP 003942468 1 701347 Proteobacteria
Citrobacter koseri ATCC BAA 895 YP 001453787 1 290338 Proteobacteria
Enterobacter cloacae subsp dissolvens SDM YP 006476902 1 1104326 Proteobacteria
Enterobacter cloacae subsp cloacae ENHKU01 YP 006577980 1 1211025 Proteobacteria
Enterobacter asburiae LF7a YP 004827889 1 640513 Proteobacteria
Citrobacter rodentium ICC168 YP 003367904 1 637910 Proteobacteria
Enterobacter cloacae subsp cloacae ENHKU01 YP 006580521 1 1211025 Proteobacteria
Escherichia coli O83 H1 str NRG 857C YP 006121637 1 685038 Proteobacteria
Escherichia coli ATCC 8739 YP 001723393 1 481805 Proteobacteria
Escherichia coli B str REL606 YP 003046361 1 413997 Proteobacteria
Escherichia coli str K 12 substr W3110 YP 492107 1 316407 Proteobacteria
Klebsiella oxytoca E718 YP 006497277 1 1191061 Proteobacteria
Raoultella ornithinolytica B6 YP 007875109 1 1286170 Proteobacteria
Klebsiella oxytoca E718 YP 006496731 1 1191061 Proteobacteria
Klebsiella pneumoniae 342 YP 002240367 1 507522 Proteobacteria
Klebsiella variicola At 22 YP 003441131 1 640131 Proteobacteria
Klebsiella pneumoniae YP 007990071 1 573 Proteobacteria
Klebsiella pneumoniae subsp pneumoniae MGH 78578 YP 001333850 1 272620 Proteobacteria
Escherichia coli O55 H7 str CB9615 YP 003502691 1 701177 Proteobacteria
Escherichia coli O103 H2 str 12009 YP 003225074 1 585395 Proteobacteria
Escherichia coli Xuzhou21 YP 006315903 1 741093 Proteobacteria
Escherichia coli O157 H7 str EDL933 YP 325593 1 155864 Proteobacteria
Dickeya zeae ZJU1202 WP 016941167 1 Dzeae
Dickeya zeae DZ2Q WP 016941167 1 Dzeae
Dickeya zeae EC1 WP 016941167 1 Dzeae
Dickeya gs undicola 2B12 WP 033567854 1 Dundicola
Dickeya gs fangzhongdai NCPPB 3274 WP 071604030 1 Dfangzhongdai
Dickeya gs fangzhongdai M005 WP 039691071 1 Dfangzhongdai
Dickeya gs fangzhongdai M074 WP 038667248 1 Dfangzhongdai
Dickeya gs fangzhongdai ND14b WP 071842565 1 Dfangzhongdai
Dickeya gs fangzhongdai S1 WP 049853678 1 Dfangzhongdai
Dickeya gs fangzhongdai B16 WP 038919718 1 Dfangzhongdai
Dickeya gs fangzhongdai MK7 WP 071605264 1 Dfangzhongdai
Dickeya solani D s0432 1 WP 071598666 1 Dsolani
Dickeya solani MK10 WP 071598666 1 Dsolani
Dickeya solani RNS 07 7 3B WP 022634276 1 Dsolani
Dickeya solani IPO 2222 WP 071598666 1 Dsolani
Dickeya solani PPO 9134 WP 022634276 1 Dsolani
Dickeya solani IFB 0099 WP 022634276 1 Dsolani
Dickeya solani GBBC 2040 WP 071598666 1 Dsolani
Dickeya solani RNS 05 1 2A WP 022634276 1 Dsolani
Dickeya solani RNS 08 23 3 1 A WP 071598666 1 Dsolani
Dickeya solani MK16 WP 071598666 1 Dsolani

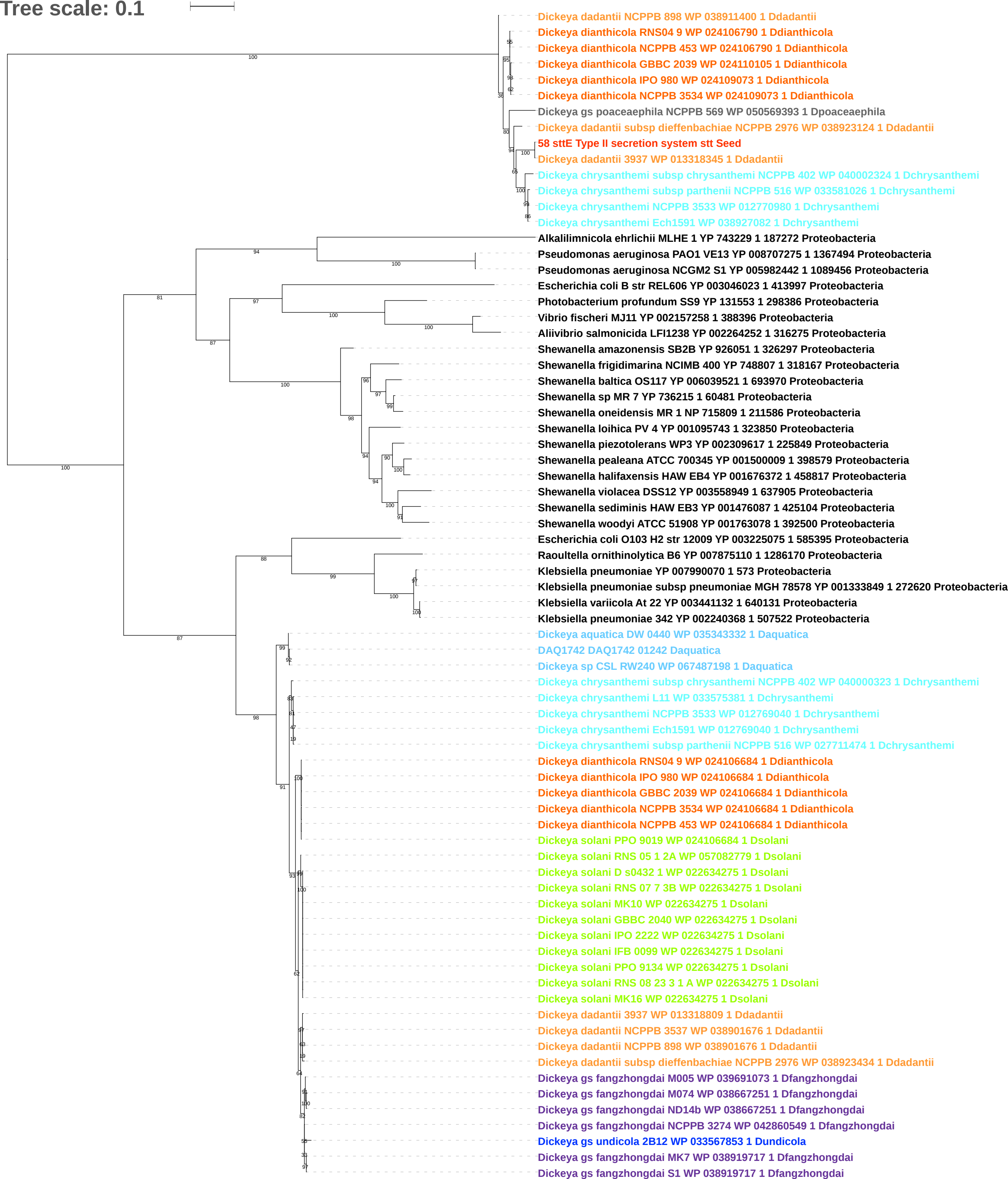
Tree scale: 0.1
Dickeya dadantii NCPPB 898 WP 038911400 1 Ddadantii
Dickeya dianthicola RNS04 9 WP 024106790 1 Ddianthicola
Dickeya dianthicola NCPPB 453 WP 024106790 1 Ddianthicola
Dickeya dianthicola GBBC 2039 WP 024110105 1 Ddianthicola
Dickeya dianthicola IPO 980 WP 024109073 1 Ddianthicola
Dickeya dianthicola NCPPB 3534 WP 024109073 1 Ddianthicola
Dickeya gs poaceaephila NCPPB 569 WP 050569393 1 Dpoaceaephila
Dickeya dadantii subsp dieffenbachiae NCPPB 2976 WP 038923124 1 Ddadantii
58 sttE Type II secretion system stt Seed
Dickeya dadantii 3937 WP 013318345 1 Ddadantii
Dickeya chrysanthemi subsp chrysanthemi NCPPB 402 WP 040002324 1 Dchrysanthemi
Dickeya chrysanthemi subsp parthenii NCPPB 516 WP 033581026 1 Dchrysanthemi
Dickeya chrysanthemi NCPPB 3533 WP 012770980 1 Dchrysanthemi
Dickeya chrysanthemi Ech1591 WP 038927082 1 Dchrysanthemi
Alkalilimnicola ehrlichii MLHE 1 YP 743229 1 187272 Proteobacteria
Pseudomonas aeruginosa PAO1 VE13 YP 008707275 1 1367494 Proteobacteria
Pseudomonas aeruginosa NCGM2 S1 YP 005982442 1 1089456 Proteobacteria
Escherichia coli B str REL606 YP 003046023 1 413997 Proteobacteria
Photobacterium profundum SS9 YP 131553 1 298386 Proteobacteria
Vibrio fischeri MJ11 YP 002157258 1 388396 Proteobacteria
Aliivibrio salmonicida LFI1238 YP 002264252 1 316275 Proteobacteria
Shewanella amazonensis SB2B YP 926051 1 326297 Proteobacteria
Shewanella frigidimarina NCIMB 400 YP 748807 1 318167 Proteobacteria
Shewanella baltica OS117 YP 006039521 1 693970 Proteobacteria
Shewanella sp MR 7 YP 736215 1 60481 Proteobacteria
Shewanella oneidensis MR 1 NP 715809 1 211586 Proteobacteria
Shewanella loihica PV 4 YP 001095743 1 323850 Proteobacteria
Shewanella piezotolerans WP3 YP 002309617 1 225849 Proteobacteria
Shewanella pealeana ATCC 700345 YP 001500009 1 398579 Proteobacteria
Shewanella halifaxensis HAW EB4 YP 001676372 1 458817 Proteobacteria
Shewanella violacea DSS12 YP 003558949 1 637905 Proteobacteria
Shewanella sediminis HAW EB3 YP 001476087 1 425104 Proteobacteria
Shewanella woodyi ATCC 51908 YP 001763078 1 392500 Proteobacteria
Escherichia coli O103 H2 str 12009 YP 003225075 1 585395 Proteobacteria
Raoultella ornithinolytica B6 YP 007875110 1 1286170 Proteobacteria
Klebsiella pneumoniae YP 007990070 1 573 Proteobacteria
Klebsiella pneumoniae subsp pneumoniae MGH 78578 YP 001333849 1 272620 Proteobacteria
Klebsiella variicola At 22 YP 003441132 1 640131 Proteobacteria
Klebsiella pneumoniae 342 YP 002240368 1 507522 Proteobacteria
Dickeya aquatica DW 0440 WP 035343332 1 Daquatica
DAQ1742 DAQ1742 01242 Daquatica
Dickeya sp CSL RW240 WP 067487198 1 Daquatica
Dickeya chrysanthemi subsp chrysanthemi NCPPB 402 WP 040000323 1 Dchrysanthemi
Dickeya chrysanthemi L11 WP 033575381 1 Dchrysanthemi
Dickeya chrysanthemi NCPPB 3533 WP 012769040 1 Dchrysanthemi
Dickeya chrysanthemi Ech1591 WP 012769040 1 Dchrysanthemi
Dickeya chrysanthemi subsp parthenii NCPPB 516 WP 027711474 1 Dchrysanthemi
Dickeya dianthicola RNS04 9 WP 024106684 1 Ddianthicola
Dickeya dianthicola IPO 980 WP 024106684 1 Ddianthicola
Dickeya dianthicola GBBC 2039 WP 024106684 1 Ddianthicola
Dickeya dianthicola NCPPB 3534 WP 024106684 1 Ddianthicola
Dickeya dianthicola NCPPB 453 WP 024106684 1 Ddianthicola
Dickeya solani PPO 9019 WP 024106684 1 Dsolani
Dickeya solani RNS 05 1 2A WP 057082779 1 Dsolani
Dickeya solani D s0432 1 WP 022634275 1 Dsolani
Dickeya solani RNS 07 7 3B WP 022634275 1 Dsolani
Dickeya solani MK10 WP 022634275 1 Dsolani
Dickeya solani GBBC 2040 WP 022634275 1 Dsolani
Dickeya solani IPO 2222 WP 022634275 1 Dsolani
Dickeya solani IFB 0099 WP 022634275 1 Dsolani
Dickeya solani PPO 9134 WP 022634275 1 Dsolani
Dickeya solani RNS 08 23 3 1 A WP 022634275 1 Dsolani
Dickeya solani MK16 WP 022634275 1 Dsolani
Dickeya dadantii 3937 WP 013318809 1 Ddadantii
Dickeya dadantii NCPPB 3537 WP 038901676 1 Ddadantii
Dickeya dadantii NCPPB 898 WP 038901676 1 Ddadantii
Dickeya dadantii subsp dieffenbachiae NCPPB 2976 WP 038923434 1 Ddadantii
Dickeya gs fangzhongdai M005 WP 039691073 1 Dfangzhongdai
Dickeya gs fangzhongdai M074 WP 038667251 1 Dfangzhongdai
Dickeya gs fangzhongdai ND14b WP 038667251 1 Dfangzhongdai
Dickeya gs fangzhongdai NCPPB 3274 WP 042860549 1 Dfangzhongdai
Dickeya gs undicola 2B12 WP 033567853 1 Dundicola
Dickeya gs fangzhongdai MK7 WP 038919717 1 Dfangzhongdai
Dickeya gs fangzhongdai S1 WP 038919717 1 Dfangzhongdai

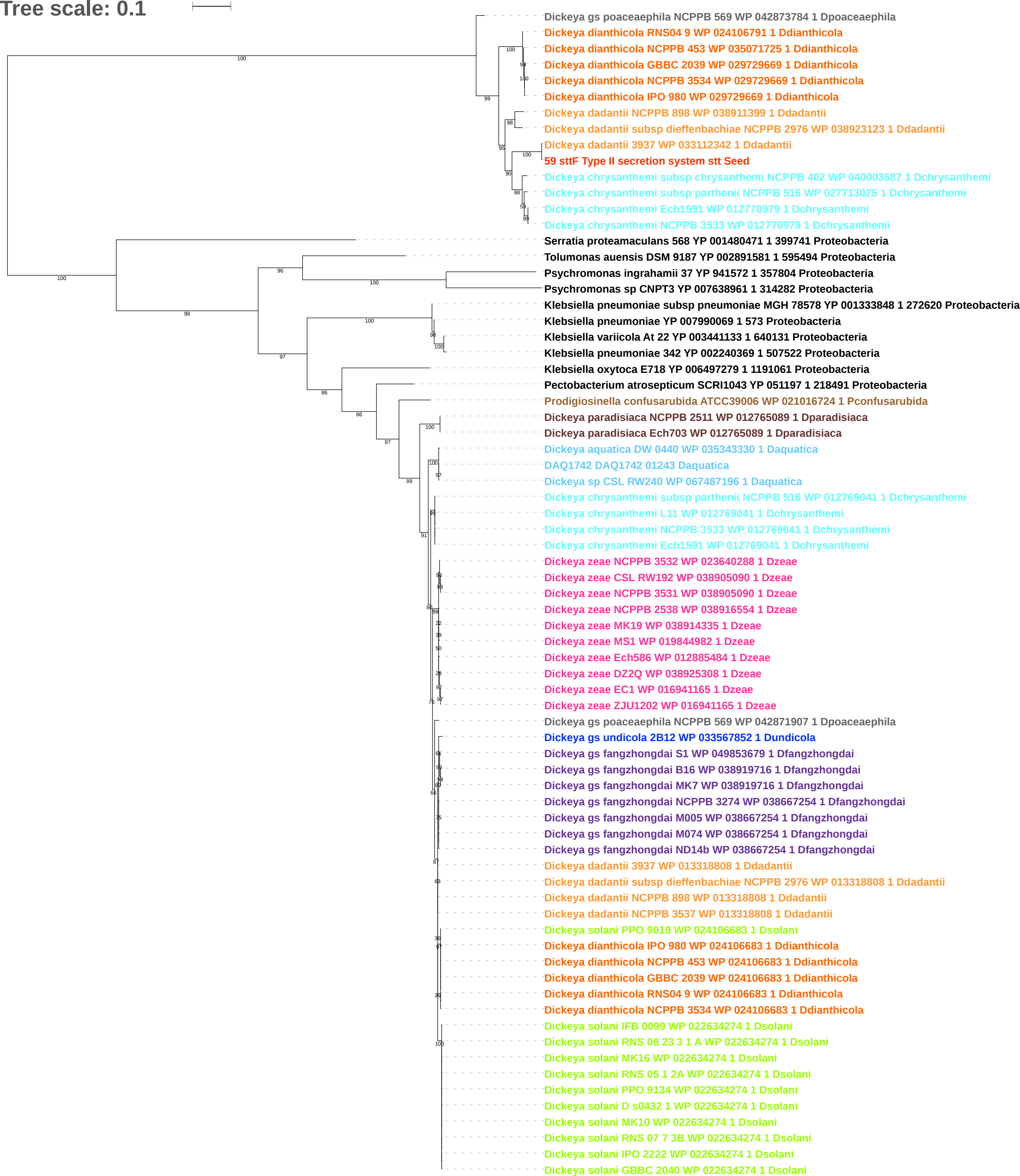
Tree scale: 0.1
Dickeya gs poaceaephila NCPPB 569 WP 042873784 1 Dpoaceaephila
Dickeya dianthicola RNS04 9 WP 024106791 1 Ddianthicola
Dickeya dianthicola NCPPB 453 WP 035071725 1 Ddianthicola
Dickeya dianthicola GBBC 2039 WP 029729669 1 Ddianthicola
Dickeya dianthicola NCPPB 3534 WP 029729669 1 Ddianthicola
Dickeya dianthicola IPO 980 WP 029729669 1 Ddianthicola
Dickeya dadantii NCPPB 898 WP 038911399 1 Ddadantii
Dickeya dadantii subsp dieffenbachiae NCPPB 2976 WP 038923123 1 Ddadantii
Dickeya dadantii 3937 WP 033112342 1 Ddadantii
59 sttF Type II secretion system stt Seed
Dickeya chrysanthemi subsp chrysanthemi NCPPB 402 WP 040003687 1 Dchrysanthemi
Dickeya chrysanthemi subsp parthenii NCPPB 516 WP 027713025 1 Dchrysanthemi
Dickeya chrysanthemi Ech1591 WP 012770979 1 Dchrysanthemi
Dickeya chrysanthemi NCPPB 3533 WP 012770979 1 Dchrysanthemi
Serratia proteamaculans 568 YP 001480471 1 399741 Proteobacteria
Tolumonas auensis DSM 9187 YP 002891581 1 595494 Proteobacteria
Psychromonas ingrahamii 37 YP 941572 1 357804 Proteobacteria
Psychromonas sp CNPT3 YP 007638961 1 314282 Proteobacteria
Klebsiella pneumoniae subsp pneumoniae MGH 78578 YP 001333848 1 272620 Proteobacteria
Klebsiella pneumoniae YP 007990069 1 573 Proteobacteria
Klebsiella variicola At 22 YP 003441133 1 640131 Proteobacteria
Klebsiella pneumoniae 342 YP 002240369 1 507522 Proteobacteria
Klebsiella oxytoca E718 YP 006497279 1 1191061 Proteobacteria
Pectobacterium atrosepticum SCRI1043 YP 051197 1 218491 Proteobacteria
Prodigiosinella confusarubida ATCC39006 WP 021016724 1 Pconfusarubida
Dickeya paradisiaca NCPPB 2511 WP 012765089 1 Dparadisiaca
Dickeya paradisiaca Ech703 WP 012765089 1 Dparadisiaca
Dickeya aquatica DW 0440 WP 035343330 1 Daquatica
DAQ1742 DAQ1742 01243 Daquatica
Dickeya sp CSL RW240 WP 067487196 1 Daquatica
Dickeya chrysanthemi subsp parthenii NCPPB 516 WP 012769041 1 Dchrysanthemi
Dickeya chrysanthemi L11 WP 012769041 1 Dchrysanthemi
Dickeya chrysanthemi NCPPB 3533 WP 012769041 1 Dchrysanthemi
Dickeya chrysanthemi Ech1591 WP 012769041 1 Dchrysanthemi
Dickeya zeae NCPPB 3532 WP 023640288 1 Dzeae
Dickeya zeae CSL RW192 WP 038905090 1 Dzeae
Dickeya zeae NCPPB 3531 WP 038905090 1 Dzeae
Dickeya zeae NCPPB 2538 WP 038916554 1 Dzeae
Dickeya zeae MK19 WP 038914335 1 Dzeae
Dickeya zeae MS1 WP 019844982 1 Dzeae
Dickeya zeae Ech586 WP 012885484 1 Dzeae
Dickeya zeae DZ2Q WP 038925308 1 Dzeae
Dickeya zeae EC1 WP 016941165 1 Dzeae
Dickeya zeae ZJU1202 WP 016941165 1 Dzeae
Dickeya gs poaceaephila NCPPB 569 WP 042871907 1 Dpoaceaephila
Dickeya gs undicola 2B12 WP 033567852 1 Dundicola
Dickeya gs fangzhongdai S1 WP 049853679 1 Dfangzhongdai
Dickeya gs fangzhongdai B16 WP 038919716 1 Dfangzhongdai
Dickeya gs fangzhongdai MK7 WP 038919716 1 Dfangzhongdai
Dickeya gs fangzhongdai NCPPB 3274 WP 038667254 1 Dfangzhongdai
Dickeya gs fangzhongdai M005 WP 038667254 1 Dfangzhongdai
Dickeya gs fangzhongdai M074 WP 038667254 1 Dfangzhongdai
Dickeya gs fangzhongdai ND14b WP 038667254 1 Dfangzhongdai
Dickeya dadantii 3937 WP 013318808 1 Ddadantii
Dickeya dadantii subsp dieffenbachiae NCPPB 2976 WP 013318808 1 Ddadantii
Dickeya dadantii NCPPB 898 WP 013318808 1 Ddadantii
Dickeya dadantii NCPPB 3537 WP 013318808 1 Ddadantii
Dickeya solani PPO 9019 WP 024106683 1 Dsolani
Dickeya dianthicola IPO 980 WP 024106683 1 Ddianthicola
Dickeya dianthicola NCPPB 453 WP 024106683 1 Ddianthicola
Dickeya dianthicola GBBC 2039 WP 024106683 1 Ddianthicola
Dickeya dianthicola RNS04 9 WP 024106683 1 Ddianthicola
Dickeya dianthicola NCPPB 3534 WP 024106683 1 Ddianthicola
Dickeya solani IFB 0099 WP 022634274 1 Dsolani
Dickeya solani RNS 08 23 3 1 A WP 022634274 1 Dsolani
Dickeya solani MK16 WP 022634274 1 Dsolani
Dickeya solani RNS 05 1 2A WP 022634274 1 Dsolani
Dickeya solani PPO 9134 WP 022634274 1 Dsolani
Dickeya solani D s0432 1 WP 022634274 1 Dsolani
Dickeya solani MK10 WP 022634274 1 Dsolani
Dickeya solani RNS 07 7 3B WP 022634274 1 Dsolani
Dickeya solani IPO 2222 WP 022634274 1 Dsolani
Dickeya solani GBBC 2040 WP 022634274 1 Dsolani

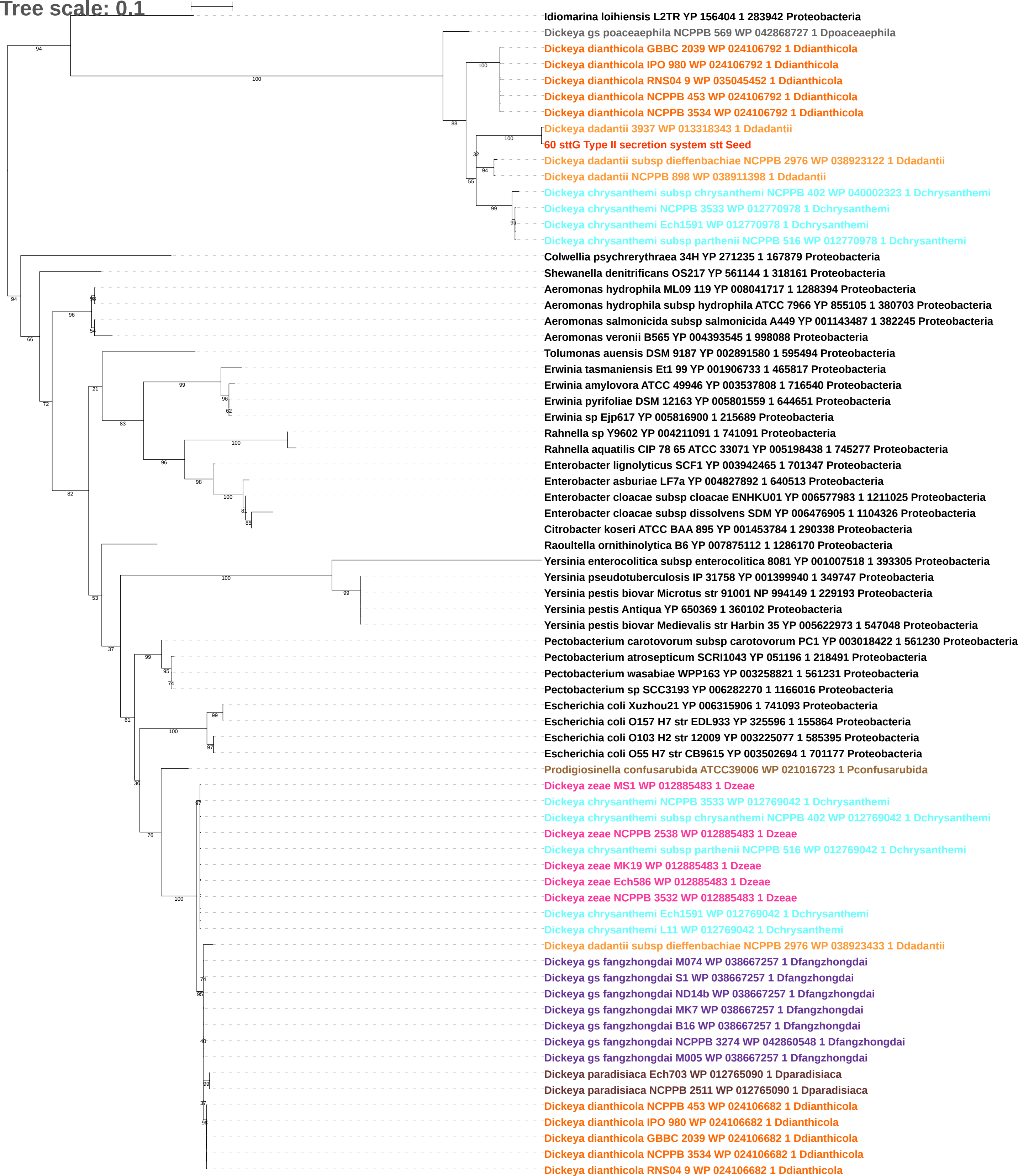
Tree scale: 0.1
Idiomarina loihiensis L2TR YP 156404 1 283942 Proteobacteria
Dickeya gs poaceaephila NCPPB 569 WP 042868727 1 Dpoaceaephila
Dickeya dianthicola GBBC 2039 WP 024106792 1 Ddianthicola
Dickeya dianthicola IPO 980 WP 024106792 1 Ddianthicola
Dickeya dianthicola RNS04 9 WP 035045452 1 Ddianthicola
Dickeya dianthicola NCPPB 453 WP 024106792 1 Ddianthicola
Dickeya dianthicola NCPPB 3534 WP 024106792 1 Ddianthicola
Dickeya dadantii 3937 WP 013318343 1 Ddadantii
60 sttG Type II secretion system stt Seed
Dickeya dadantii subsp dieffenbachiae NCPPB 2976 WP 038923122 1 Ddadantii
Dickeya dadantii NCPPB 898 WP 038911398 1 Ddadantii
Dickeya chrysanthemi subsp chrysanthemi NCPPB 402 WP 040002323 1 Dchrysanthemi
Dickeya chrysanthemi NCPPB 3533 WP 012770978 1 Dchrysanthemi
Dickeya chrysanthemi Ech1591 WP 012770978 1 Dchrysanthemi
Dickeya chrysanthemi subsp parthenii NCPPB 516 WP 012770978 1 Dchrysanthemi
Colwellia psychrerythraea 34H YP 271235 1 167879 Proteobacteria
Shewanella denitrificans OS217 YP 561144 1 318161 Proteobacteria
Aeromonas hydrophila ML09 119 YP 008041717 1 1288394 Proteobacteria
Aeromonas hydrophila subsp hydrophila ATCC 7966 YP 855105 1 380703 Proteobacteria
Aeromonas salmonicida subsp salmonicida A449 YP 001143487 1 382245 Proteobacteria
Aeromonas veronii B565 YP 004393545 1 998088 Proteobacteria
Tolumonas auensis DSM 9187 YP 002891580 1 595494 Proteobacteria
Erwinia tasmaniensis Et1 99 YP 001906733 1 465817 Proteobacteria
Erwinia amylovora ATCC 49946 YP 003537808 1 716540 Proteobacteria
Erwinia pyrifoliae DSM 12163 YP 005801559 1 644651 Proteobacteria
Erwinia sp Ejp617 YP 005816900 1 215689 Proteobacteria
Rahnella sp Y9602 YP 004211091 1 741091 Proteobacteria
Rahnella aquatilis CIP 78 65 ATCC 33071 YP 005198438 1 745277 Proteobacteria
Enterobacter lignolyticus SCF1 YP 003942465 1 701347 Proteobacteria
Enterobacter asburiae LF7a YP 004827892 1 640513 Proteobacteria
Enterobacter cloacae subsp cloacae ENHKU01 YP 006577983 1 1211025 Proteobacteria
Enterobacter cloacae subsp dissolvens SDM YP 006476905 1 1104326 Proteobacteria
Citrobacter koseri ATCC BAA 895 YP 001453784 1 290338 Proteobacteria
Raoultella ornithinolytica B6 YP 007875112 1 1286170 Proteobacteria
Yersinia enterocolitica subsp enterocolitica 8081 YP 001007518 1 393305 Proteobacteria
Yersinia pseudotuberculosis IP 31758 YP 001399940 1 349747 Proteobacteria
Yersinia pestis biovar Microtus str 91001 NP 994149 1 229193 Proteobacteria
Yersinia pestis Antiqua YP 650369 1 360102 Proteobacteria
Yersinia pestis biovar Medievalis str Harbin 35 YP 005622973 1 547048 Proteobacteria
Pectobacterium carotovorum subsp carotovorum PC1 YP 003018422 1 561230 Proteobacteria
Pectobacterium atrosepticum SCRI1043 YP 051196 1 218491 Proteobacteria
Pectobacterium wasabiae WPP163 YP 003258821 1 561231 Proteobacteria
Pectobacterium sp SCC3193 YP 006282270 1 1166016 Proteobacteria
Escherichia coli Xuzhou21 YP 006315906 1 741093 Proteobacteria
Escherichia coli O157 H7 str EDL933 YP 325596 1 155864 Proteobacteria
Escherichia coli O103 H2 str 12009 YP 003225077 1 585395 Proteobacteria
Escherichia coli O55 H7 str CB9615 YP 003502694 1 701177 Proteobacteria
Prodigiosinella confusarubida ATCC39006 WP 021016723 1 Pconfusarubida
Dickeya zeae MS1 WP 012885483 1 Dzeae
Dickeya chrysanthemi NCPPB 3533 WP 012769042 1 Dchrysanthemi
Dickeya chrysanthemi subsp chrysanthemi NCPPB 402 WP 012769042 1 Dchrysanthemi
Dickeya zeae NCPPB 2538 WP 012885483 1 Dzeae
Dickeya chrysanthemi subsp parthenii NCPPB 516 WP 012769042 1 Dchrysanthemi
Dickeya zeae MK19 WP 012885483 1 Dzeae
Dickeya zeae Ech586 WP 012885483 1 Dzeae
Dickeya zeae NCPPB 3532 WP 012885483 1 Dzeae
Dickeya chrysanthemi Ech1591 WP 012769042 1 Dchrysanthemi
Dickeya chrysanthemi L11 WP 012769042 1 Dchrysanthemi
Dickeya dadantii subsp dieffenbachiae NCPPB 2976 WP 038923433 1 Ddadantii
Dickeya gs fangzhongdai M074 WP 038667257 1 Dfangzhongdai
Dickeya gs fangzhongdai S1 WP 038667257 1 Dfangzhongdai
Dickeya gs fangzhongdai ND14b WP 038667257 1 Dfangzhongdai
Dickeya gs fangzhongdai MK7 WP 038667257 1 Dfangzhongdai
Dickeya gs fangzhongdai B16 WP 038667257 1 Dfangzhongdai
Dickeya gs fangzhongdai NCPPB 3274 WP 042860548 1 Dfangzhongdai
Dickeya gs fangzhongdai M005 WP 038667257 1 Dfangzhongdai
Dickeya paradisiaca Ech703 WP 012765090 1 Dparadisiaca
Dickeya paradisiaca NCPPB 2511 WP 012765090 1 Dparadisiaca
Dickeya dianthicola NCPPB 453 WP 024106682 1 Ddianthicola
Dickeya dianthicola IPO 980 WP 024106682 1 Ddianthicola
Dickeya dianthicola GBBC 2039 WP 024106682 1 Ddianthicola
Dickeya dianthicola NCPPB 3534 WP 024106682 1 Ddianthicola
Dickeya dianthicola RNS04 9 WP 024106682 1 Ddianthicola

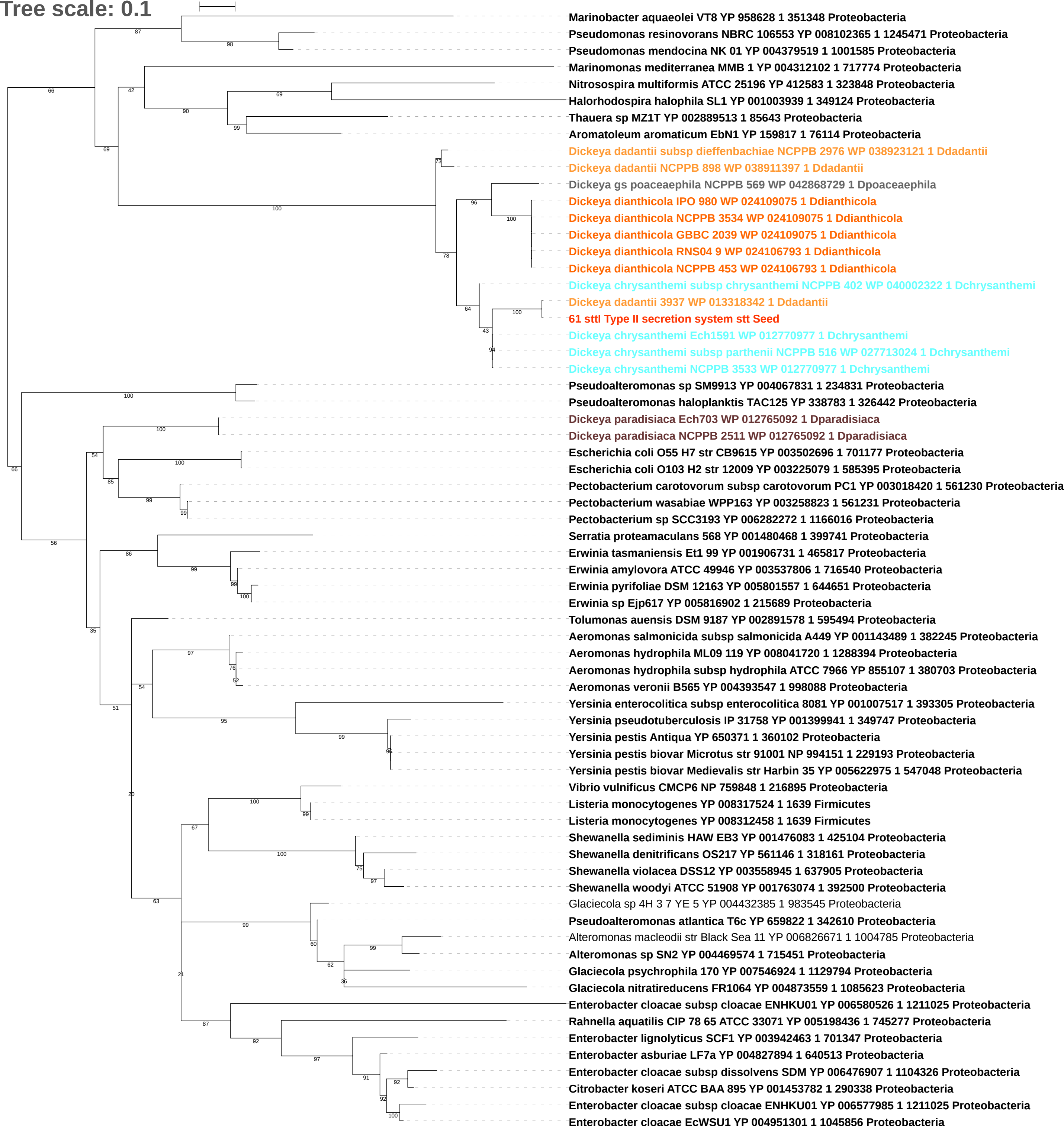
Tree scale: 0.1
Marinobacter aquaeolei VT8 YP 958628 1 351348 Proteobacteria
Pseudomonas resinovorans NBRC 106553 YP 008102365 1 1245471 Proteobacteria
Pseudomonas mendocina NK 01 YP 004379519 1 1001585 Proteobacteria
Marinomonas mediterranea MMB 1 YP 004312102 1 717774 Proteobacteria
Nitrosospira multiformis ATCC 25196 YP 412583 1 323848 Proteobacteria
Halorhodospira halophila SL1 YP 001003939 1 349124 Proteobacteria
Thauera sp MZ1T YP 002889513 1 85643 Proteobacteria
Aromatoleum aromaticum EbN1 YP 159817 1 76114 Proteobacteria
Dickeya dadantii subsp dieffenbachiae NCPPB 2976 WP 038923121 1 Ddadantii
Dickeya dadantii NCPPB 898 WP 038911397 1 Ddadantii
Dickeya gs poaceaephila NCPPB 569 WP 042868729 1 Dpoaceaephila
Dickeya dianthicola IPO 980 WP 024109075 1 Ddianthicola
Dickeya dianthicola NCPPB 3534 WP 024109075 1 Ddianthicola
Dickeya dianthicola GBBC 2039 WP 024109075 1 Ddianthicola
Dickeya dianthicola RNS04 9 WP 024106793 1 Ddianthicola
Dickeya dianthicola NCPPB 453 WP 024106793 1 Ddianthicola
Dickeya chrysanthemi subsp chrysanthemi NCPPB 402 WP 040002322 1 Dchrysanthemi
Dickeya dadantii 3937 WP 013318342 1 Ddadantii
61 sttI Type II secretion system stt Seed
Dickeya chrysanthemi Ech1591 WP 012770977 1 Dchrysanthemi
Dickeya chrysanthemi subsp parthenii NCPPB 516 WP 027713024 1 Dchrysanthemi
Dickeya chrysanthemi NCPPB 3533 WP 012770977 1 Dchrysanthemi
Pseudoalteromonas sp SM9913 YP 004067831 1 234831 Proteobacteria
Pseudoalteromonas haloplanktis TAC125 YP 338783 1 326442 Proteobacteria
Dickeya paradisiaca Ech703 WP 012765092 1 Dparadisiaca
Dickeya paradisiaca NCPPB 2511 WP 012765092 1 Dparadisiaca
Escherichia coli O55 H7 str CB9615 YP 003502696 1 701177 Proteobacteria
Escherichia coli O103 H2 str 12009 YP 003225079 1 585395 Proteobacteria
Pectobacterium carotovorum subsp carotovorum PC1 YP 003018420 1 561230 Proteobacteria
Pectobacterium wasabiae WPP163 YP 003258823 1 561231 Proteobacteria
Pectobacterium sp SCC3193 YP 006282272 1 1166016 Proteobacteria
Serratia proteamaculans 568 YP 001480468 1 399741 Proteobacteria
Erwinia tasmaniensis Et1 99 YP 001906731 1 465817 Proteobacteria
Erwinia amylovora ATCC 49946 YP 003537806 1 716540 Proteobacteria
Erwinia pyrifoliae DSM 12163 YP 005801557 1 644651 Proteobacteria
Erwinia sp Ejp617 YP 005816902 1 215689 Proteobacteria
Tolumonas auensis DSM 9187 YP 002891578 1 595494 Proteobacteria
Aeromonas salmonicida subsp salmonicida A449 YP 001143489 1 382245 Proteobacteria
Aeromonas hydrophila ML09 119 YP 008041720 1 1288394 Proteobacteria
Aeromonas hydrophila subsp hydrophila ATCC 7966 YP 855107 1 380703 Proteobacteria
Aeromonas veronii B565 YP 004393547 1 998088 Proteobacteria
Yersinia enterocolitica subsp enterocolitica 8081 YP 001007517 1 393305 Proteobacteria
Yersinia pseudotuberculosis IP 31758 YP 001399941 1 349747 Proteobacteria
Yersinia pestis Antiqua YP 650371 1 360102 Proteobacteria
Yersinia pestis biovar Microtus str 91001 NP 994151 1 229193 Proteobacteria
Yersinia pestis biovar Medievalis str Harbin 35 YP 005622975 1 547048 Proteobacteria
Vibrio vulnificus CMCP6 NP 759848 1 216895 Proteobacteria
Listeria monocytogenes YP 008317524 1 1639 Firmicutes
Listeria monocytogenes YP 008312458 1 1639 Firmicutes
Shewanella sediminis HAW EB3 YP 001476083 1 425104 Proteobacteria
Shewanella denitrificans OS217 YP 561146 1 318161 Proteobacteria
Shewanella violacea DSS12 YP 003558945 1 637905 Proteobacteria
Shewanella woodyi ATCC 51908 YP 001763074 1 392500 Proteobacteria
Glaciecola sp 4H 3 7 YE 5 YP 004432385 1 983545 Proteobacteria
Pseudoalteromonas atlantica T6c YP 659822 1 342610 Proteobacteria
Alteromonas macleodii str Black Sea 11 YP 006826671 1 1004785 Proteobacteria
Alteromonas sp SN2 YP 004469574 1 715451 Proteobacteria
Glaciecola psychrophila 170 YP 007546924 1 1129794 Proteobacteria
Glaciecola nitratireducens FR1064 YP 004873559 1 1085623 Proteobacteria
Enterobacter cloacae subsp cloacae ENHKU01 YP 006580526 1 1211025 Proteobacteria
Rahnella aquatilis CIP 78 65 ATCC 33071 YP 005198436 1 745277 Proteobacteria
Enterobacter lignolyticus SCF1 YP 003942463 1 701347 Proteobacteria
Enterobacter asburiae LF7a YP 004827894 1 640513 Proteobacteria
Enterobacter cloacae subsp dissolvens SDM YP 006476907 1 1104326 Proteobacteria
Citrobacter koseri ATCC BAA 895 YP 001453782 1 290338 Proteobacteria
Enterobacter cloacae subsp cloacae ENHKU01 YP 006577985 1 1211025 Proteobacteria
Enterobacter cloacae EcWSU1 YP 004951301 1 1045856 Proteobacteria

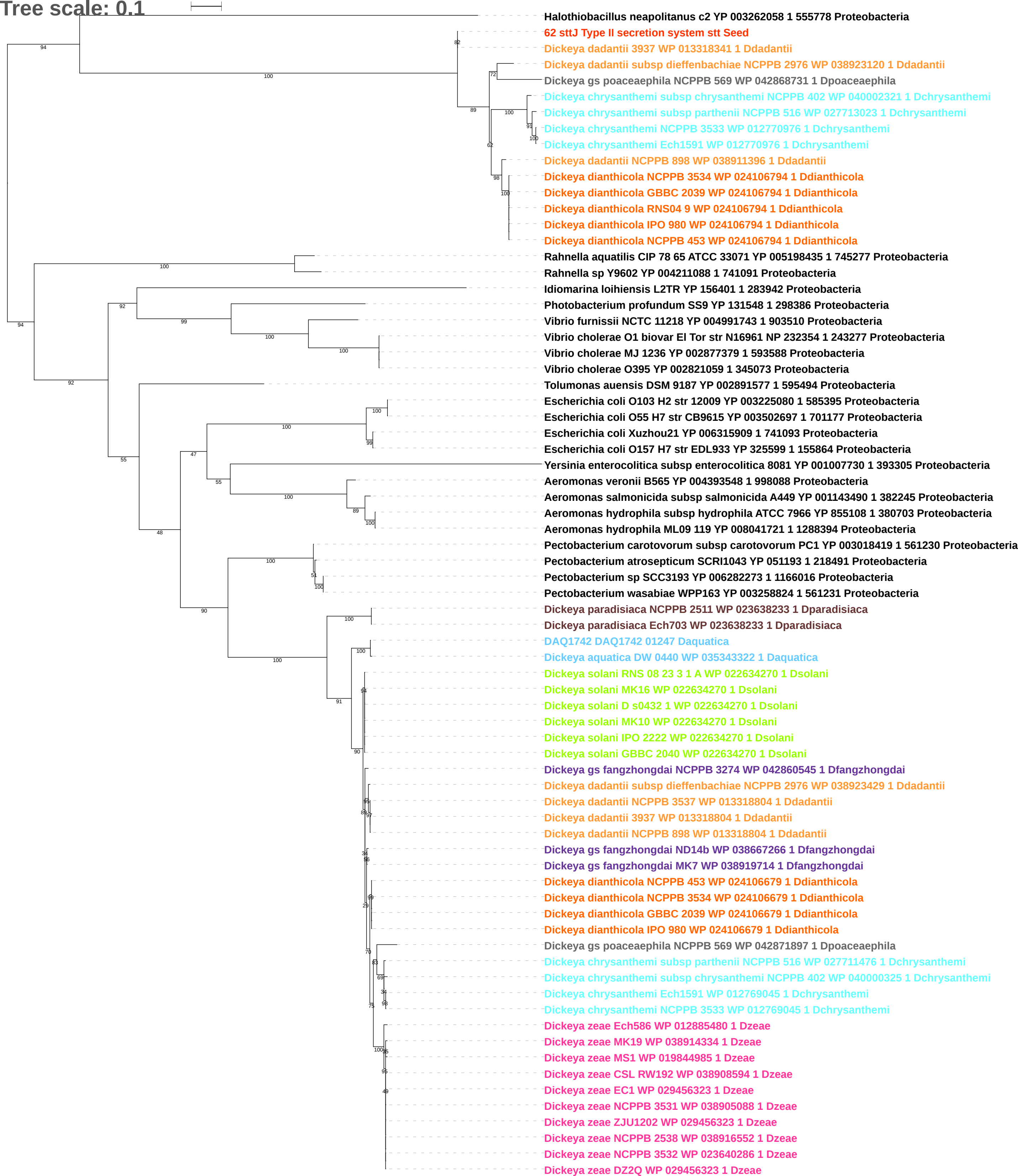
Tree scale: 0.1
Halothiobacillus neapolitanus c2 YP 003262058 1 555778 Proteobacteria
62 sttJ Type II secretion system stt Seed
Dickeya dadantii 3937 WP 013318341 1 Ddadantii
Dickeya dadantii subsp dieffenbachiae NCPPB 2976 WP 038923120 1 Ddadantii
Dickeya gs poaceaephila NCPPB 569 WP 042868731 1 Dpoaceaephila
Dickeya chrysanthemi subsp chrysanthemi NCPPB 402 WP 040002321 1 Dchrysanthemi
Dickeya chrysanthemi subsp parthenii NCPPB 516 WP 027713023 1 Dchrysanthemi
Dickeya chrysanthemi NCPPB 3533 WP 012770976 1 Dchrysanthemi
Dickeya chrysanthemi Ech1591 WP 012770976 1 Dchrysanthemi
Dickeya dadantii NCPPB 898 WP 038911396 1 Ddadantii
Dickeya dianthicola NCPPB 3534 WP 024106794 1 Ddianthicola
Dickeya dianthicola GBBC 2039 WP 024106794 1 Ddianthicola
Dickeya dianthicola RNS04 9 WP 024106794 1 Ddianthicola
Dickeya dianthicola IPO 980 WP 024106794 1 Ddianthicola
Dickeya dianthicola NCPPB 453 WP 024106794 1 Ddianthicola
Rahnella aquatilis CIP 78 65 ATCC 33071 YP 005198435 1 745277 Proteobacteria
Rahnella sp Y9602 YP 004211088 1 741091 Proteobacteria
Idiomarina loihiensis L2TR YP 156401 1 283942 Proteobacteria
Photobacterium profundum SS9 YP 131548 1 298386 Proteobacteria
Vibrio furnissii NCTC 11218 YP 004991743 1 903510 Proteobacteria
Vibrio cholerae O1 biovar El Tor str N16961 NP 232354 1 243277 Proteobacteria
Vibrio cholerae MJ 1236 YP 002877379 1 593588 Proteobacteria
Vibrio cholerae O395 YP 002821059 1 345073 Proteobacteria
Tolumonas auensis DSM 9187 YP 002891577 1 595494 Proteobacteria
Escherichia coli O103 H2 str 12009 YP 003225080 1 585395 Proteobacteria
Escherichia coli O55 H7 str CB9615 YP 003502697 1 701177 Proteobacteria
Escherichia coli Xuzhou21 YP 006315909 1 741093 Proteobacteria
Escherichia coli O157 H7 str EDL933 YP 325599 1 155864 Proteobacteria
Yersinia enterocolitica subsp enterocolitica 8081 YP 001007730 1 393305 Proteobacteria
Aeromonas veronii B565 YP 004393548 1 998088 Proteobacteria
Aeromonas salmonicida subsp salmonicida A449 YP 001143490 1 382245 Proteobacteria
Aeromonas hydrophila subsp hydrophila ATCC 7966 YP 855108 1 380703 Proteobacteria
Aeromonas hydrophila ML09 119 YP 008041721 1 1288394 Proteobacteria
Pectobacterium carotovorum subsp carotovorum PC1 YP 003018419 1 561230 Proteobacteria
Pectobacterium atrosepticum SCRI1043 YP 051193 1 218491 Proteobacteria
Pectobacterium sp SCC3193 YP 006282273 1 1166016 Proteobacteria
Pectobacterium wasabiae WPP163 YP 003258824 1 561231 Proteobacteria
Dickeya paradisiaca NCPPB 2511 WP 023638233 1 Dparadisiaca
Dickeya paradisiaca Ech703 WP 023638233 1 Dparadisiaca
DAQ1742 DAQ1742 01247 Daquatica
Dickeya aquatica DW 0440 WP 035343322 1 Daquatica
Dickeya solani RNS 08 23 3 1 A WP 022634270 1 Dsolani
Dickeya solani MK16 WP 022634270 1 Dsolani
Dickeya solani D s0432 1 WP 022634270 1 Dsolani
Dickeya solani MK10 WP 022634270 1 Dsolani
Dickeya solani IPO 2222 WP 022634270 1 Dsolani
Dickeya solani GBBC 2040 WP 022634270 1 Dsolani
Dickeya gs fangzhongdai NCPPB 3274 WP 042860545 1 Dfangzhongdai
Dickeya dadantii subsp dieffenbachiae NCPPB 2976 WP 038923429 1 Ddadantii
Dickeya dadantii NCPPB 3537 WP 013318804 1 Ddadantii
Dickeya dadantii 3937 WP 013318804 1 Ddadantii
Dickeya dadantii NCPPB 898 WP 013318804 1 Ddadantii
Dickeya gs fangzhongdai ND14b WP 038667266 1 Dfangzhongdai
Dickeya gs fangzhongdai MK7 WP 038919714 1 Dfangzhongdai
Dickeya dianthicola NCPPB 453 WP 024106679 1 Ddianthicola
Dickeya dianthicola NCPPB 3534 WP 024106679 1 Ddianthicola
Dickeya dianthicola GBBC 2039 WP 024106679 1 Ddianthicola
Dickeya dianthicola IPO 980 WP 024106679 1 Ddianthicola
Dickeya gs poaceaephila NCPPB 569 WP 042871897 1 Dpoaceaephila
Dickeya chrysanthemi subsp parthenii NCPPB 516 WP 027711476 1 Dchrysanthemi
Dickeya chrysanthemi subsp chrysanthemi NCPPB 402 WP 040000325 1 Dchrysanthemi
Dickeya chrysanthemi Ech1591 WP 012769045 1 Dchrysanthemi
Dickeya chrysanthemi NCPPB 3533 WP 012769045 1 Dchrysanthemi
Dickeya zeae Ech586 WP 012885480 1 Dzeae
Dickeya zeae MK19 WP 038914334 1 Dzeae
Dickeya zeae MS1 WP 019844985 1 Dzeae
Dickeya zeae CSL RW192 WP 038908594 1 Dzeae
Dickeya zeae EC1 WP 029456323 1 Dzeae
Dickeya zeae NCPPB 3531 WP 038905088 1 Dzeae
Dickeya zeae ZJU1202 WP 029456323 1 Dzeae
Dickeya zeae NCPPB 2538 WP 038916552 1 Dzeae
Dickeya zeae NCPPB 3532 WP 023640286 1 Dzeae
Dickeya zeae DZ2Q WP 029456323 1 Dzeae

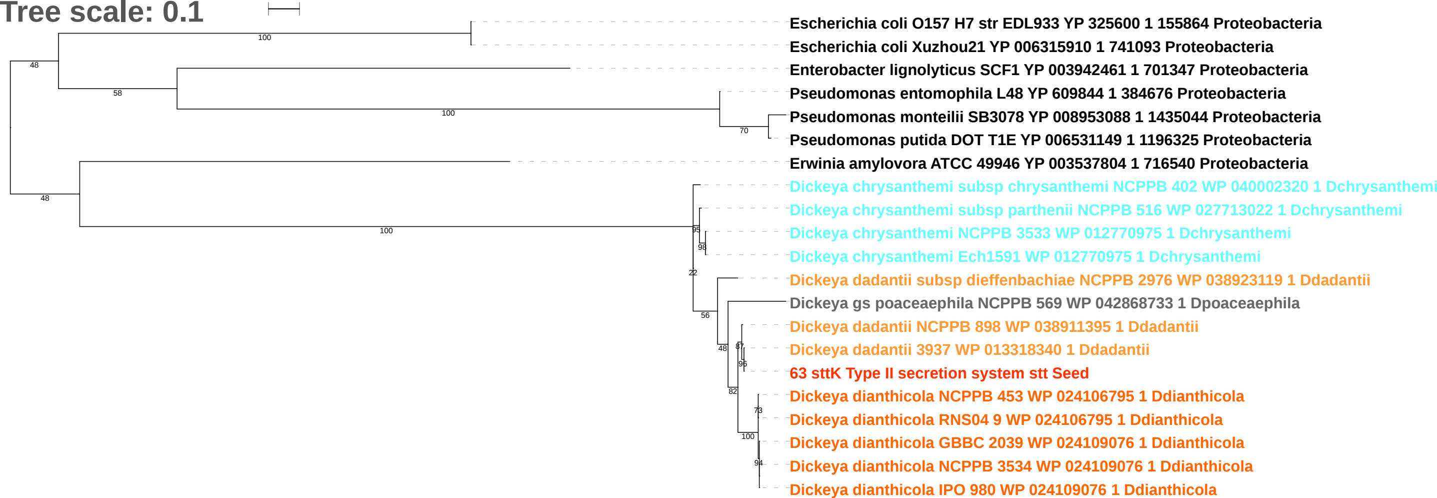
Tree scale: 0.1
Escherichia coli O157 H7 str EDL933 YP 325600 1 155864 Proteobacteria
Escherichia coli Xuzhou21 YP 006315910 1 741093 Proteobacteria
Enterobacter lignolyticus SCF1 YP 003942461 1 701347 Proteobacteria
Pseudomonas entomophila L48 YP 609844 1 384676 Proteobacteria
Pseudomonas monteilii SB3078 YP 008953088 1 1435044 Proteobacteria
Pseudomonas putida DOT T1E YP 006531149 1 1196325 Proteobacteria
Erwinia amylovora ATCC 49946 YP 003537804 1 716540 Proteobacteria
Dickeya chrysanthemi subsp chrysanthemi NCPPB 402 WP 040002320 1 Dchrysanthemi
Dickeya chrysanthemi subsp parthenii NCPPB 516 WP 027713022 1 Dchrysanthemi
Dickeya chrysanthemi NCPPB 3533 WP 012770975 1 Dchrysanthemi
Dickeya chrysanthemi Ech1591 WP 012770975 1 Dchrysanthemi
Dickeya dadantii subsp dieffenbachiae NCPPB 2976 WP 038923119 1 Ddadantii
Dickeya gs poaceaephila NCPPB 569 WP 042868733 1 Dpoaceaephila
Dickeya dadantii NCPPB 898 WP 038911395 1 Ddadantii
Dickeya dadantii 3937 WP 013318340 1 Ddadantii
63 sttK Type II secretion system stt Seed
Dickeya dianthicola NCPPB 453 WP 024106795 1 Ddianthicola
Dickeya dianthicola RNS04 9 WP 024106795 1 Ddianthicola
Dickeya dianthicola GBBC 2039 WP 024109076 1 Ddianthicola
Dickeya dianthicola NCPPB 3534 WP 024109076 1 Ddianthicola
Dickeya dianthicola IPO 980 WP 024109076 1 Ddianthicola

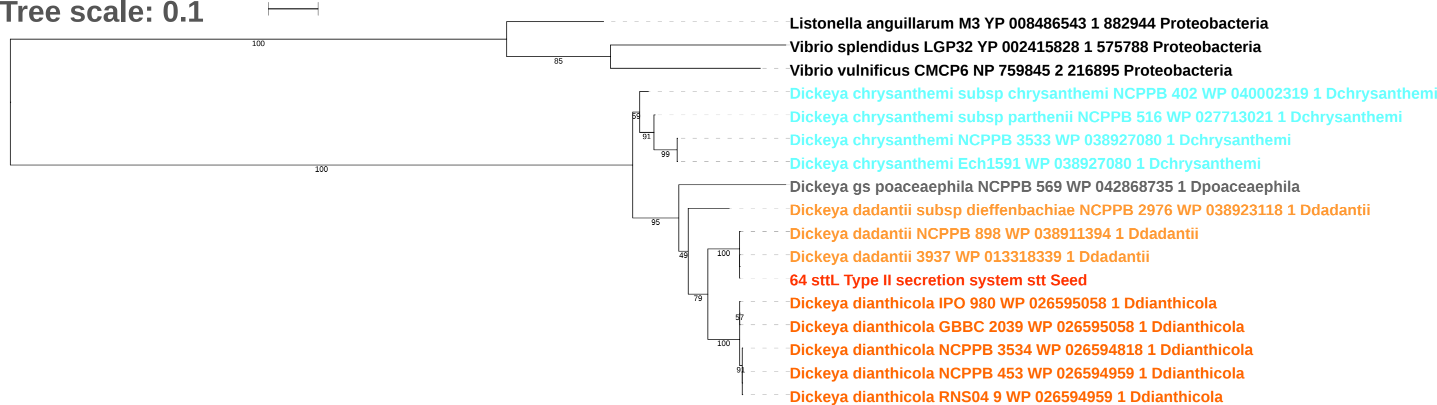

Tree scale: 0.1
Listonella anguillarum M3 YP 008486543 1 882944 Proteobacteria
Vibrio splendidus LGP32 YP 002415828 1 575788 Proteobacteria
Vibrio vulnificus CMCP6 NP 759845 2 216895 Proteobacteria
Dickeya chrysanthemi subsp chrysanthemi NCPPB 402 WP 040002319 1 Dchrysanthemi
Dickeya chrysanthemi subsp parthenii NCPPB 516 WP 027713021 1 Dchrysanthemi
Dickeya chrysanthemi NCPPB 3533 WP 038927080 1 Dchrysanthemi
Dickeya chrysanthemi Ech1591 WP 038927080 1 Dchrysanthemi
Dickeya gs poaceaephila NCPPB 569 WP 042868735 1 Dpoaceaephila
Dickeya dadantii subsp dieffenbachiae NCPPB 2976 WP 038923118 1 Ddadantii
Dickeya dadantii NCPPB 898 WP 038911394 1 Ddadantii
Dickeya dadantii 3937 WP 013318339 1 Ddadantii
64 sttL Type II secretion system stt Seed
Dickeya dianthicola IPO 980 WP 026595058 1 Ddianthicola
Dickeya dianthicola GBBC 2039 WP 026595058 1 Ddianthicola
Dickeya dianthicola NCPPB 3534 WP 026594818 1 Ddianthicola
Dickeya dianthicola NCPPB 453 WP 026594959 1 Ddianthicola
Dickeya dianthicola RNS04 9 WP 026594959 1 Ddianthicola
100
85
100
59
91
99
95
49
100
79
57
100
91

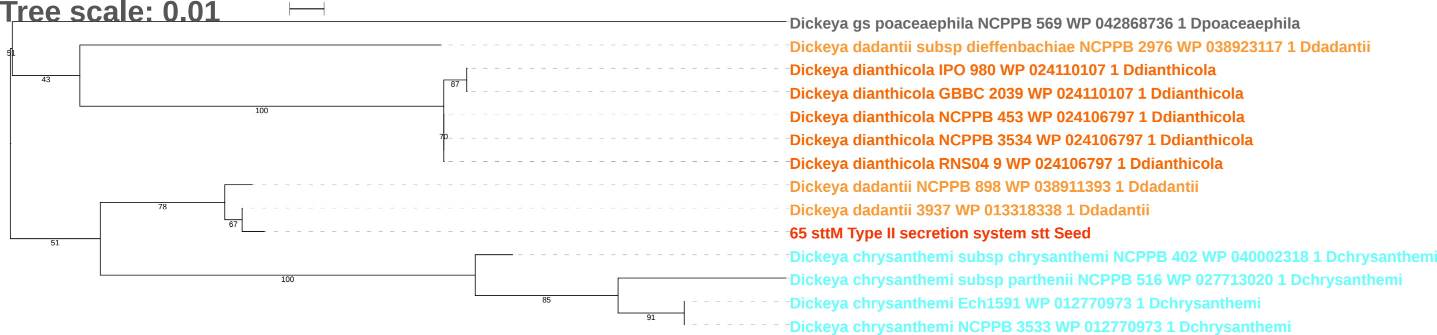

Tree scale: 0.01
Dickeya gs poaceaephila NCPPB 569 WP 042868736 1 Dpoaceaephila
Dickeya dadantii subsp dieffenbachiae NCPPB 2976 WP 038923117 1 Ddadantii
Dickeya dianthicola IPO 980 WP 024110107 1 Ddianthicola
Dickeya dianthicola GBBC 2039 WP 024110107 1 Ddianthicola
Dickeya dianthicola NCPPB 453 WP 024106797 1 Ddianthicola
Dickeya dianthicola NCPPB 3534 WP 024106797 1 Ddianthicola
Dickeya dianthicola RNS04 9 WP 024106797 1 Ddianthicola
Dickeya dadantii NCPPB 898 WP 038911393 1 Ddadantii
Dickeya dadantii 3937 WP 013318338 1 Ddadantii
65 sttM Type II secretion system stt Seed
Dickeya chrysanthemi subsp chrysanthemi NCPPB 402 WP 040002318 1 Dchrysanthemi
Dickeya chrysanthemi subsp parthenii NCPPB 516 WP 027713020 1 Dchrysanthemi
Dickeya chrysanthemi Ech1591 WP 012770973 1 Dchrysanthemi
Dickeya chrysanthemi NCPPB 3533 WP 012770973 1 Dchrysanthemi
51
43
100
87
70
78
67
51
100
85
91

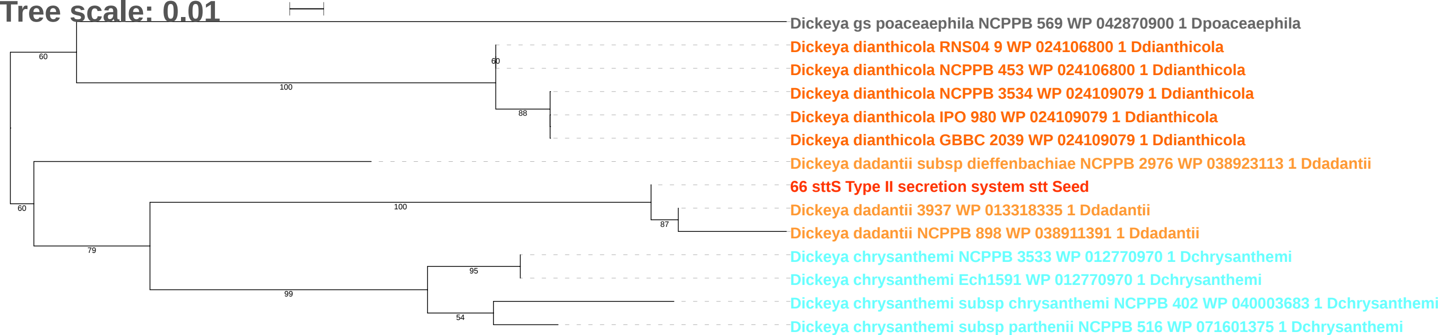

Tree scale: 0.01
Dickeya gs poaceaephila NCPPB 569 WP 042870900 1 Dpoaceaephila
Dickeya dianthicola RNS04 9 WP 024106800 1 Ddianthicola
Dickeya dianthicola NCPPB 453 WP 024106800 1 Ddianthicola
Dickeya dianthicola NCPPB 3534 WP 024109079 1 Ddianthicola
Dickeya dianthicola IPO 980 WP 024109079 1 Ddianthicola
Dickeya dianthicola GBBC 2039 WP 024109079 1 Ddianthicola
Dickeya dadantii subsp dieffenbachiae NCPPB 2976 WP 038923113 1 Ddadantii
66 sttS Type II secretion system stt Seed
Dickeya dadantii 3937 WP 013318335 1 Ddadantii
Dickeya dadantii NCPPB 898 WP 038911391 1 Ddadantii
Dickeya chrysanthemi NCPPB 3533 WP 012770970 1 Dchrysanthemi
Dickeya chrysanthemi Ech1591 WP 012770970 1 Dchrysanthemi
Dickeya chrysanthemi subsp chrysanthemi NCPPB 402 WP 040003683 1 Dchrysanthemi
Dickeya chrysanthemi subsp parthenii NCPPB 516 WP 071601375 1 Dchrysanthemi
60
100
60
88
60
100
87
79
95
99
54

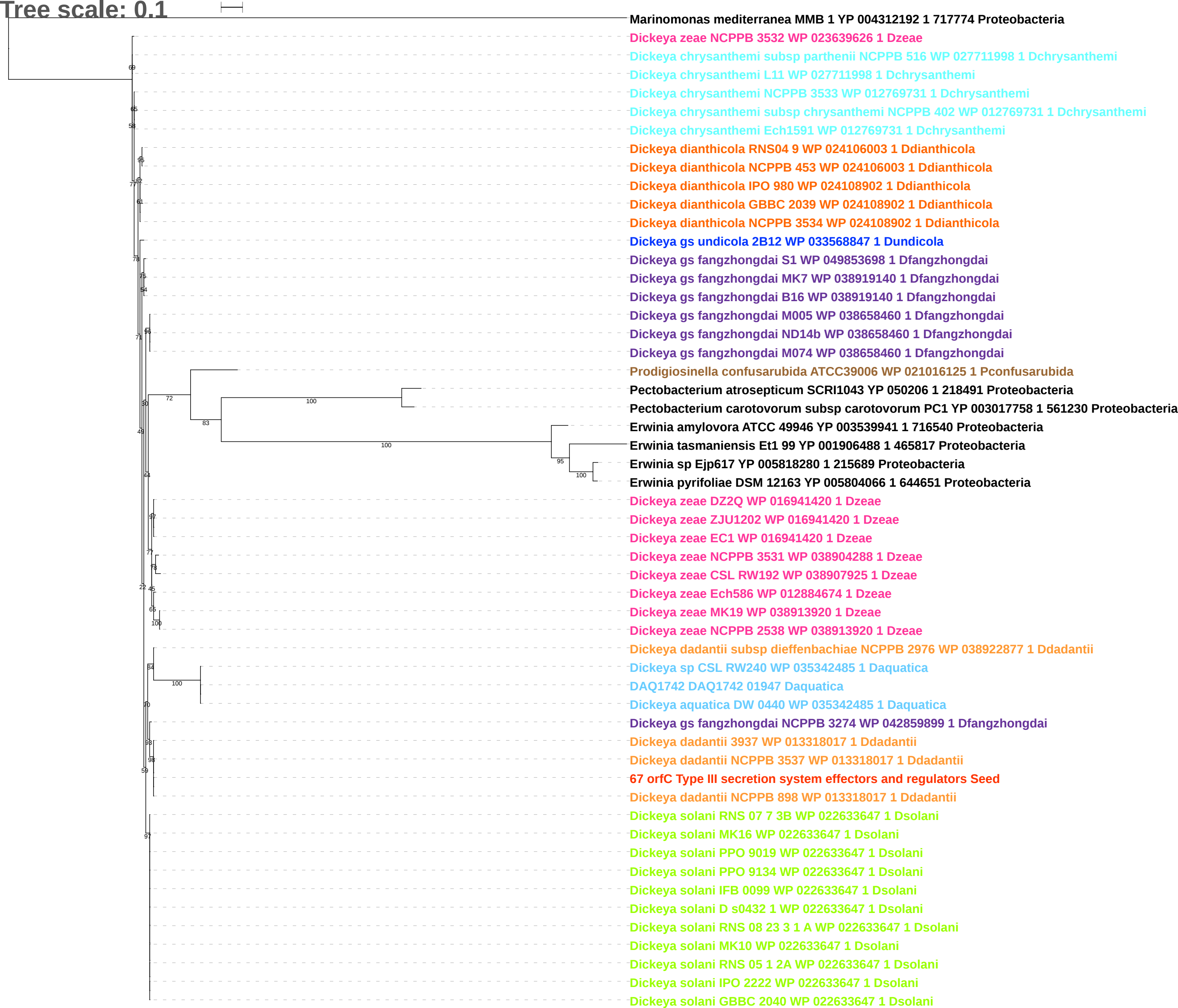
Tree scale: 0.1
Marinomonas mediterranea MMB 1 YP 004312192 1 717774 Proteobacteria
Dickeya zeae NCPPB 3532 WP 023639626 1 Dzeae
Dickeya chrysanthemi subsp parthenii NCPPB 516 WP 027711998 1 Dchrysanthemi
Dickeya chrysanthemi L11 WP 027711998 1 Dchrysanthemi
Dickeya chrysanthemi NCPPB 3533 WP 012769731 1 Dchrysanthemi
Dickeya chrysanthemi subsp chrysanthemi NCPPB 402 WP 012769731 1 Dchrysanthemi
Dickeya chrysanthemi Ech1591 WP 012769731 1 Dchrysanthemi
Dickeya dianthicola RNS04 9 WP 024106003 1 Ddianthicola
Dickeya dianthicola NCPPB 453 WP 024106003 1 Ddianthicola
Dickeya dianthicola IPO 980 WP 024108902 1 Ddianthicola
Dickeya dianthicola GBBC 2039 WP 024108902 1 Ddianthicola
Dickeya dianthicola NCPPB 3534 WP 024108902 1 Ddianthicola
Dickeya gs undicola 2B12 WP 033568847 1 Dundicola
Dickeya gs fangzhongdai S1 WP 049853698 1 Dfangzhongdai
Dickeya gs fangzhongdai MK7 WP 038919140 1 Dfangzhongdai
Dickeya gs fangzhongdai B16 WP 038919140 1 Dfangzhongdai
Dickeya gs fangzhongdai M005 WP 038658460 1 Dfangzhongdai
Dickeya gs fangzhongdai ND14b WP 038658460 1 Dfangzhongdai
Dickeya gs fangzhongdai M074 WP 038658460 1 Dfangzhongdai
Prodigiosinella confusarubida ATCC39006 WP 021016125 1 Pconfusarubida
Pectobacterium atrosepticum SCRI1043 YP 050206 1 218491 Proteobacteria
Pectobacterium carotovorum subsp carotovorum PC1 YP 003017758 1 561230 Proteobacteria
Erwinia amylovora ATCC 49946 YP 003539941 1 716540 Proteobacteria
Erwinia tasmaniensis Et1 99 YP 001906488 1 465817 Proteobacteria
Erwinia sp Ejp617 YP 005818280 1 215689 Proteobacteria
Erwinia pyrifoliae DSM 12163 YP 005804066 1 644651 Proteobacteria
Dickeya zeae DZ2Q WP 016941420 1 Dzeae
Dickeya zeae ZJU1202 WP 016941420 1 Dzeae
Dickeya zeae EC1 WP 016941420 1 Dzeae
Dickeya zeae NCPPB 3531 WP 038904288 1 Dzeae
Dickeya zeae CSL RW192 WP 038907925 1 Dzeae
Dickeya zeae Ech586 WP 012884674 1 Dzeae
Dickeya zeae MK19 WP 038913920 1 Dzeae
Dickeya zeae NCPPB 2538 WP 038913920 1 Dzeae
Dickeya dadantii subsp dieffenbachiae NCPPB 2976 WP 038922877 1 Ddadantii
Dickeya sp CSL RW240 WP 035342485 1 Daquatica
DAQ1742 DAQ1742 01947 Daquatica
Dickeya aquatica DW 0440 WP 035342485 1 Daquatica
Dickeya gs fangzhongdai NCPPB 3274 WP 042859899 1 Dfangzhongdai
Dickeya dadantii 3937 WP 013318017 1 Ddadantii
Dickeya dadantii NCPPB 3537 WP 013318017 1 Ddadantii
67 orfC Type III secretion system effectors and regulators Seed
Dickeya dadantii NCPPB 898 WP 013318017 1 Ddadantii
Dickeya solani RNS 07 7 3B WP 022633647 1 Dsolani
Dickeya solani MK16 WP 022633647 1 Dsolani
Dickeya solani PPO 9019 WP 022633647 1 Dsolani
Dickeya solani PPO 9134 WP 022633647 1 Dsolani
Dickeya solani IFB 0099 WP 022633647 1 Dsolani
Dickeya solani D s0432 1 WP 022633647 1 Dsolani
Dickeya solani RNS 08 23 3 1 A WP 022633647 1 Dsolani
Dickeya solani MK10 WP 022633647 1 Dsolani
Dickeya solani RNS 05 1 2A WP 022633647 1 Dsolani
Dickeya solani IPO 2222 WP 022633647 1 Dsolani
Dickeya solani GBBC 2040 WP 022633647 1 Dsolani

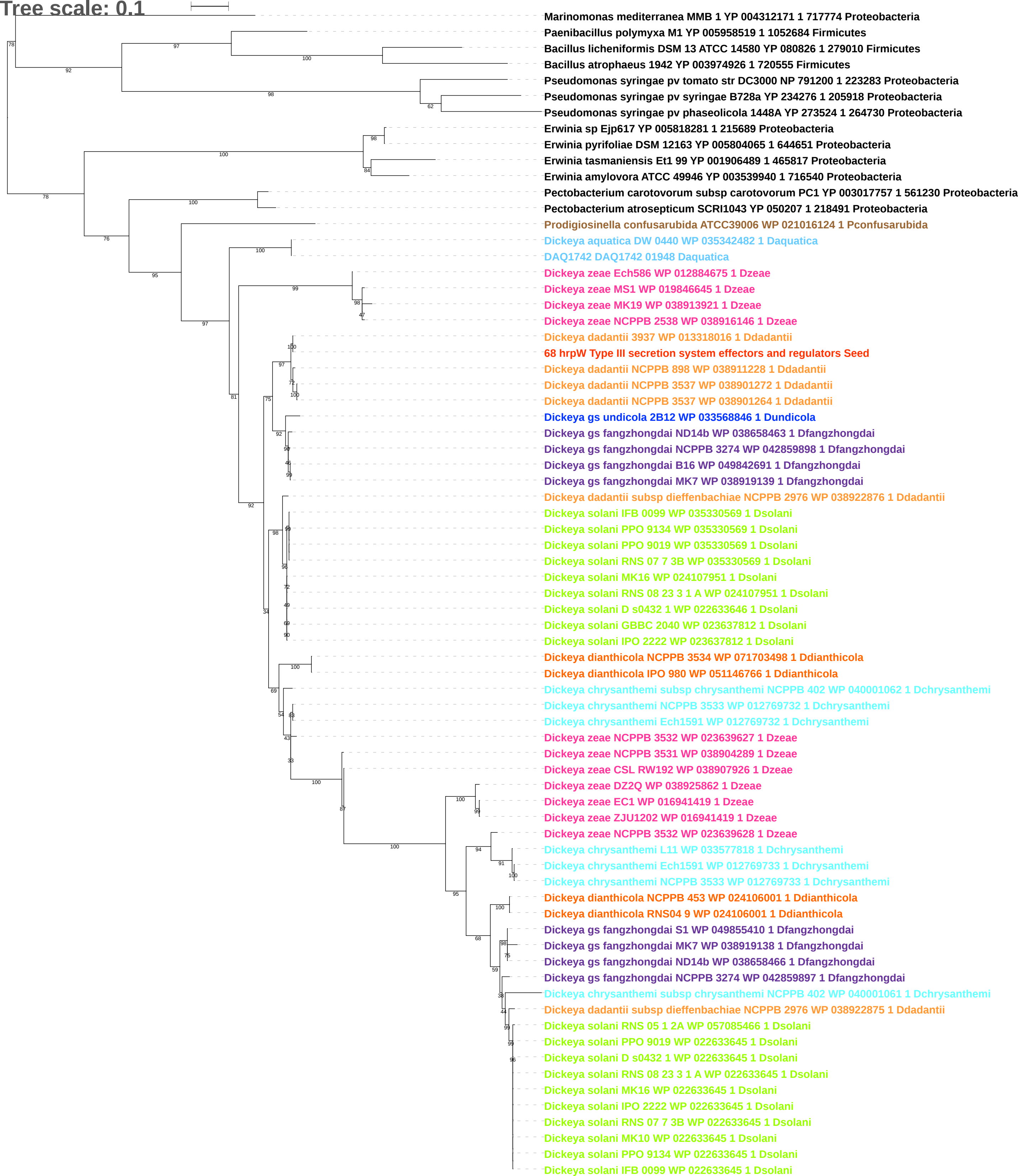
Tree scale: 0.1
Marinomonas mediterranea MMB 1 YP 004312171 1 717774 Proteobacteria
Paenibacillus polymyxa M1 YP 005958519 1 1052684 Firmicutes
Bacillus licheniformis DSM 13 ATCC 14580 YP 080826 1 279010 Firmicutes
Bacillus atrophaeus 1942 YP 003974926 1 720555 Firmicutes
Pseudomonas syringae pv tomato str DC3000 NP 791200 1 223283 Proteobacteria
Pseudomonas syringae pv syringae B728a YP 234276 1 205918 Proteobacteria
Pseudomonas syringae pv phaseolicola 1448A YP 273524 1 264730 Proteobacteria
Erwinia sp Ejp617 YP 005818281 1 215689 Proteobacteria
Erwinia pyrifoliae DSM 12163 YP 005804065 1 644651 Proteobacteria
Erwinia tasmaniensis Et1 99 YP 001906489 1 465817 Proteobacteria
Erwinia amylovora ATCC 49946 YP 003539940 1 716540 Proteobacteria
Pectobacterium carotovorum subsp carotovorum PC1 YP 003017757 1 561230 Proteobacteria
Pectobacterium atrosepticum SCRI1043 YP 050207 1 218491 Proteobacteria
Prodigiosinella confusarubida ATCC39006 WP 021016124 1 Pconfusarubida
Dickeya aquatica DW 0440 WP 035342482 1 Daquatica
DAQ1742 DAQ1742 01948 Daquatica
Dickeya zeae Ech586 WP 012884675 1 Dzeae
Dickeya zeae MS1 WP 019846645 1 Dzeae
Dickeya zeae MK19 WP 038913921 1 Dzeae
Dickeya zeae NCPPB 2538 WP 038916146 1 Dzeae
Dickeya dadantii 3937 WP 013318016 1 Ddadantii
68 hrpW Type III secretion system effectors and regulators Seed
Dickeya dadantii NCPPB 898 WP 038911228 1 Ddadantii
Dickeya dadantii NCPPB 3537 WP 038901272 1 Ddadantii
Dickeya dadantii NCPPB 3537 WP 038901264 1 Ddadantii
Dickeya gs undicola 2B12 WP 033568846 1 Dundicola
Dickeya gs fangzhongdai ND14b WP 038658463 1 Dfangzhongdai
Dickeya gs fangzhongdai NCPPB 3274 WP 042859898 1 Dfangzhongdai
Dickeya gs fangzhongdai B16 WP 049842691 1 Dfangzhongdai
Dickeya gs fangzhongdai MK7 WP 038919139 1 Dfangzhongdai
Dickeya dadantii subsp dieffenbachiae NCPPB 2976 WP 038922876 1 Ddadantii
Dickeya solani IFB 0099 WP 035330569 1 Dsolani
Dickeya solani PPO 9134 WP 035330569 1 Dsolani
Dickeya solani PPO 9019 WP 035330569 1 Dsolani
Dickeya solani RNS 07 7 3B WP 035330569 1 Dsolani
Dickeya solani MK16 WP 024107951 1 Dsolani
Dickeya solani RNS 08 23 3 1 A WP 024107951 1 Dsolani
Dickeya solani D s0432 1 WP 022633646 1 Dsolani
Dickeya solani GBBC 2040 WP 023637812 1 Dsolani
Dickeya solani IPO 2222 WP 023637812 1 Dsolani
Dickeya dianthicola NCPPB 3534 WP 071703498 1 Ddianthicola
Dickeya dianthicola IPO 980 WP 051146766 1 Ddianthicola
Dickeya chrysanthemi subsp chrysanthemi NCPPB 402 WP 040001062 1 Dchrysanthemi
Dickeya chrysanthemi NCPPB 3533 WP 012769732 1 Dchrysanthemi
Dickeya chrysanthemi Ech1591 WP 012769732 1 Dchrysanthemi
Dickeya zeae NCPPB 3532 WP 023639627 1 Dzeae
Dickeya zeae NCPPB 3531 WP 038904289 1 Dzeae
Dickeya zeae CSL RW192 WP 038907926 1 Dzeae
Dickeya zeae DZ2Q WP 038925862 1 Dzeae
Dickeya zeae EC1 WP 016941419 1 Dzeae
Dickeya zeae ZJU1202 WP 016941419 1 Dzeae
Dickeya zeae NCPPB 3532 WP 023639628 1 Dzeae
Dickeya chrysanthemi L11 WP 033577818 1 Dchrysanthemi
Dickeya chrysanthemi Ech1591 WP 012769733 1 Dchrysanthemi
Dickeya chrysanthemi NCPPB 3533 WP 012769733 1 Dchrysanthemi
Dickeya dianthicola NCPPB 453 WP 024106001 1 Ddianthicola
Dickeya dianthicola RNS04 9 WP 024106001 1 Ddianthicola
Dickeya gs fangzhongdai S1 WP 049855410 1 Dfangzhongdai
Dickeya gs fangzhongdai MK7 WP 038919138 1 Dfangzhongdai
Dickeya gs fangzhongdai ND14b WP 038658466 1 Dfangzhongdai
Dickeya gs fangzhongdai NCPPB 3274 WP 042859897 1 Dfangzhongdai
Dickeya chrysanthemi subsp chrysanthemi NCPPB 402 WP 040001061 1 Dchrysanthemi
Dickeya dadantii subsp dieffenbachiae NCPPB 2976 WP 038922875 1 Ddadantii
Dickeya solani RNS 05 1 2A WP 057085466 1 Dsolani
Dickeya solani PPO 9019 WP 022633645 1 Dsolani
Dickeya solani D s0432 1 WP 022633645 1 Dsolani
Dickeya solani RNS 08 23 3 1 A WP 022633645 1 Dsolani
Dickeya solani MK16 WP 022633645 1 Dsolani
Dickeya solani IPO 2222 WP 022633645 1 Dsolani
Dickeya solani RNS 07 7 3B WP 022633645 1 Dsolani
Dickeya solani MK10 WP 022633645 1 Dsolani
Dickeya solani PPO 9134 WP 022633645 1 Dsolani
Dickeya solani IFB 0099 WP 022633645 1 Dsolani

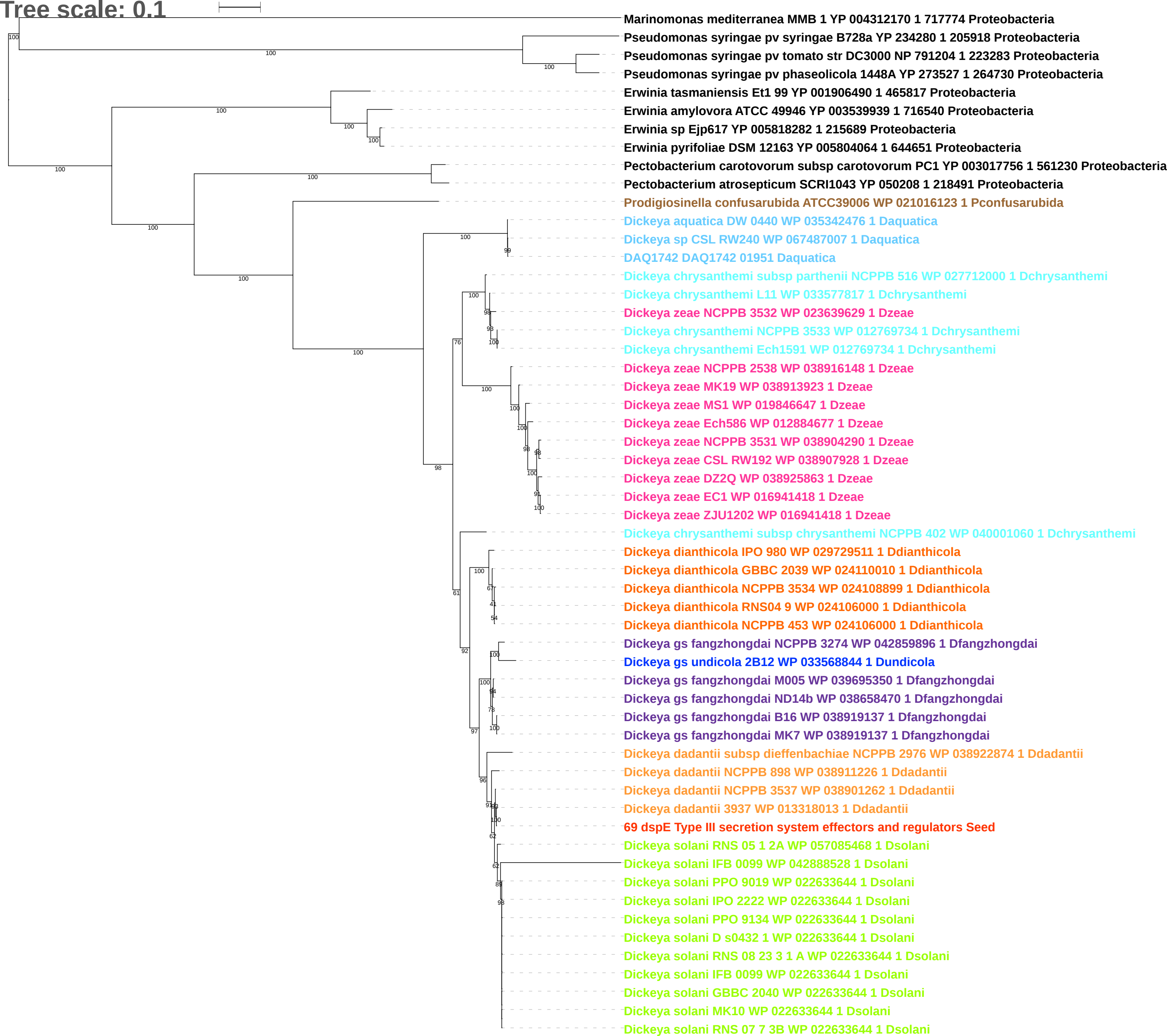
Tree scale: 0.1
Marinomonas mediterranea MMB 1 YP 004312170 1 717774 Proteobacteria
Pseudomonas syringae pv syringae B728a YP 234280 1 205918 Proteobacteria
Pseudomonas syringae pv tomato str DC3000 NP 791204 1 223283 Proteobacteria
Pseudomonas syringae pv phaseolicola 1448A YP 273527 1 264730 Proteobacteria
Erwinia tasmaniensis Et1 99 YP 001906490 1 465817 Proteobacteria
Erwinia amylovora ATCC 49946 YP 003539939 1 716540 Proteobacteria
Erwinia sp Ejp617 YP 005818282 1 215689 Proteobacteria
Erwinia pyrifoliae DSM 12163 YP 005804064 1 644651 Proteobacteria
Pectobacterium carotovorum subsp carotovorum PC1 YP 003017756 1 561230 Proteobacteria
Pectobacterium atrosepticum SCRI1043 YP 050208 1 218491 Proteobacteria
Prodigiosinella confusarubida ATCC39006 WP 021016123 1 Pconfusarubida
Dickeya aquatica DW 0440 WP 035342476 1 Daquatica
Dickeya sp CSL RW240 WP 067487007 1 Daquatica
DAQ1742 DAQ1742 01951 Daquatica
Dickeya chrysanthemi subsp parthenii NCPPB 516 WP 027712000 1 Dchrysanthemi
Dickeya chrysanthemi L11 WP 033577817 1 Dchrysanthemi
Dickeya zeae NCPPB 3532 WP 023639629 1 Dzeae
Dickeya chrysanthemi NCPPB 3533 WP 012769734 1 Dchrysanthemi
Dickeya chrysanthemi Ech1591 WP 012769734 1 Dchrysanthemi
Dickeya zeae NCPPB 2538 WP 038916148 1 Dzeae
Dickeya zeae MK19 WP 038913923 1 Dzeae
Dickeya zeae MS1 WP 019846647 1 Dzeae
Dickeya zeae Ech586 WP 012884677 1 Dzeae
Dickeya zeae NCPPB 3531 WP 038904290 1 Dzeae
Dickeya zeae CSL RW192 WP 038907928 1 Dzeae
Dickeya zeae DZ2Q WP 038925863 1 Dzeae
Dickeya zeae EC1 WP 016941418 1 Dzeae
Dickeya zeae ZJU1202 WP 016941418 1 Dzeae
Dickeya chrysanthemi subsp chrysanthemi NCPPB 402 WP 040001060 1 Dchrysanthemi
Dickeya dianthicola IPO 980 WP 029729511 1 Ddianthicola
Dickeya dianthicola GBBC 2039 WP 024110010 1 Ddianthicola
Dickeya dianthicola NCPPB 3534 WP 024108899 1 Ddianthicola
Dickeya dianthicola RNS04 9 WP 024106000 1 Ddianthicola
Dickeya dianthicola NCPPB 453 WP 024106000 1 Ddianthicola
Dickeya gs fangzhongdai NCPPB 3274 WP 042859896 1 Dfangzhongdai
Dickeya gs undicola 2B12 WP 033568844 1 Dundicola
Dickeya gs fangzhongdai M005 WP 039695350 1 Dfangzhongdai
Dickeya gs fangzhongdai ND14b WP 038658470 1 Dfangzhongdai
Dickeya gs fangzhongdai B16 WP 038919137 1 Dfangzhongdai
Dickeya gs fangzhongdai MK7 WP 038919137 1 Dfangzhongdai
Dickeya dadantii subsp dieffenbachiae NCPPB 2976 WP 038922874 1 Ddadantii
Dickeya dadantii NCPPB 898 WP 038911226 1 Ddadantii
Dickeya dadantii NCPPB 3537 WP 038901262 1 Ddadantii
Dickeya dadantii 3937 WP 013318013 1 Ddadantii
69 dspE Type III secretion system effectors and regulators Seed
Dickeya solani RNS 05 1 2A WP 057085468 1 Dsolani
Dickeya solani IFB 0099 WP 042888528 1 Dsolani
Dickeya solani PPO 9019 WP 022633644 1 Dsolani
Dickeya solani IPO 2222 WP 022633644 1 Dsolani
Dickeya solani PPO 9134 WP 022633644 1 Dsolani
Dickeya solani D s0432 1 WP 022633644 1 Dsolani
Dickeya solani RNS 08 23 3 1 A WP 022633644 1 Dsolani
Dickeya solani IFB 0099 WP 022633644 1 Dsolani
Dickeya solani GBBC 2040 WP 022633644 1 Dsolani
Dickeya solani MK10 WP 022633644 1 Dsolani
Dickeya solani RNS 07 7 3B WP 022633644 1 Dsolani

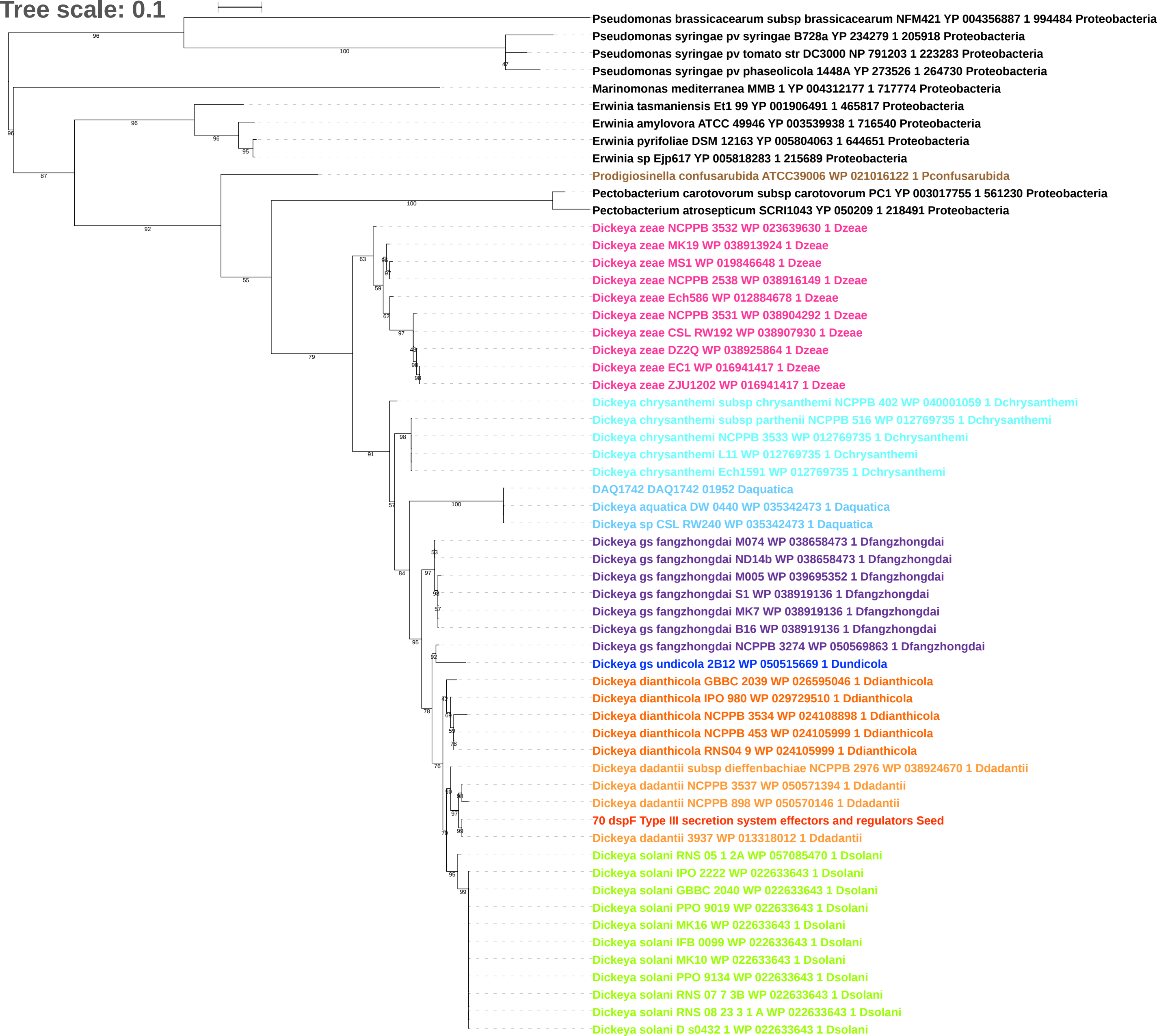

Tree scale: 0.1
Pseudomonas brassicacearum subsp brassicacearum NFM421 YP 004356887 1 994484 Proteobacteria
Pseudomonas syringae pv syringae B728a YP 234279 1 205918 Proteobacteria
Pseudomonas syringae pv tomato str DC3000 NP 791203 1 223283 Proteobacteria
Pseudomonas syringae pv phaseolicola 1448A YP 273526 1 264730 Proteobacteria
Marinomonas mediterranea MMB 1 YP 004312177 1 717774 Proteobacteria
Erwinia tasmaniensis Et1 99 YP 001906491 1 465817 Proteobacteria
Erwinia amylovora ATCC 49946 YP 003539938 1 716540 Proteobacteria
Erwinia pyrifoliae DSM 12163 YP 005804063 1 644651 Proteobacteria
Erwinia sp Ejp617 YP 005818283 1 215689 Proteobacteria
Prodigiosinella confusarubida ATCC39006 WP 021016122 1 Pconfusarubida
Pectobacterium carotovorum subsp carotovorum PC1 YP 003017755 1 561230 Proteobacteria
Pectobacterium atrosepticum SCRI1043 YP 050209 1 218491 Proteobacteria
Dickeya zeae NCPPB 3532 WP 023639630 1 Dzeae
Dickeya zeae MK19 WP 038913924 1 Dzeae
Dickeya zeae MS1 WP 019846648 1 Dzeae
Dickeya zeae NCPPB 2538 WP 038916149 1 Dzeae
Dickeya zeae Ech586 WP 012884678 1 Dzeae
Dickeya zeae NCPPB 3531 WP 038904292 1 Dzeae
Dickeya zeae CSL RW192 WP 038907930 1 Dzeae
Dickeya zeae DZ2Q WP 038925864 1 Dzeae
Dickeya zeae EC1 WP 016941417 1 Dzeae
Dickeya zeae ZJU1202 WP 016941417 1 Dzeae
Dickeya chrysanthemi subsp chrysanthemi NCPPB 402 WP 040001059 1 Dchrysanthemi
Dickeya chrysanthemi subsp parthenii NCPPB 516 WP 012769735 1 Dchrysanthemi
Dickeya chrysanthemi NCPPB 3533 WP 012769735 1 Dchrysanthemi
Dickeya chrysanthemi L11 WP 012769735 1 Dchrysanthemi
Dickeya chrysanthemi Ech1591 WP 012769735 1 Dchrysanthemi
DAQ1742 DAQ1742 01952 Daquatica
Dickeya aquatica DW 0440 WP 035342473 1 Daquatica
Dickeya sp CSL RW240 WP 035342473 1 Daquatica
Dickeya gs fangzhongdai M074 WP 038658473 1 Dfangzhongdai
Dickeya gs fangzhongdai ND14b WP 038658473 1 Dfangzhongdai
Dickeya gs fangzhongdai M005 WP 039695352 1 Dfangzhongdai
Dickeya gs fangzhongdai S1 WP 038919136 1 Dfangzhongdai
Dickeya gs fangzhongdai MK7 WP 038919136 1 Dfangzhongdai
Dickeya gs fangzhongdai B16 WP 038919136 1 Dfangzhongdai
Dickeya gs fangzhongdai NCPPB 3274 WP 050569863 1 Dfangzhongdai
Dickeya gs undicola 2B12 WP 050515669 1 Dundicola
Dickeya dianthicola GBBC 2039 WP 026595046 1 Ddianthicola
Dickeya dianthicola IPO 980 WP 029729510 1 Ddianthicola
Dickeya dianthicola NCPPB 3534 WP 024108898 1 Ddianthicola
Dickeya dianthicola NCPPB 453 WP 024105999 1 Ddianthicola
Dickeya dianthicola RNS04 9 WP 024105999 1 Ddianthicola
Dickeya dadantii subsp dieffenbachiae NCPPB 2976 WP 038924670 1 Ddadantii
Dickeya dadantii NCPPB 3537 WP 050571394 1 Ddadantii
Dickeya dadantii NCPPB 898 WP 050570146 1 Ddadantii
70 dspF Type III secretion system effectors and regulators Seed
Dickeya dadantii 3937 WP 013318012 1 Ddadantii
Dickeya solani RNS 05 1 2A WP 057085470 1 Dsolani
Dickeya solani IPO 2222 WP 022633643 1 Dsolani
Dickeya solani GBBC 2040 WP 022633643 1 Dsolani
Dickeya solani PPO 9019 WP 022633643 1 Dsolani
Dickeya solani MK16 WP 022633643 1 Dsolani
Dickeya solani IFB 0099 WP 022633643 1 Dsolani
Dickeya solani MK10 WP 022633643 1 Dsolani
Dickeya solani PPO 9134 WP 022633643 1 Dsolani
Dickeya solani RNS 07 7 3B WP 022633643 1 Dsolani
Dickeya solani RNS 08 23 3 1 A WP 022633643 1 Dsolani
Dickeya solani D s0432 1 WP 022633643 1 Dsolani

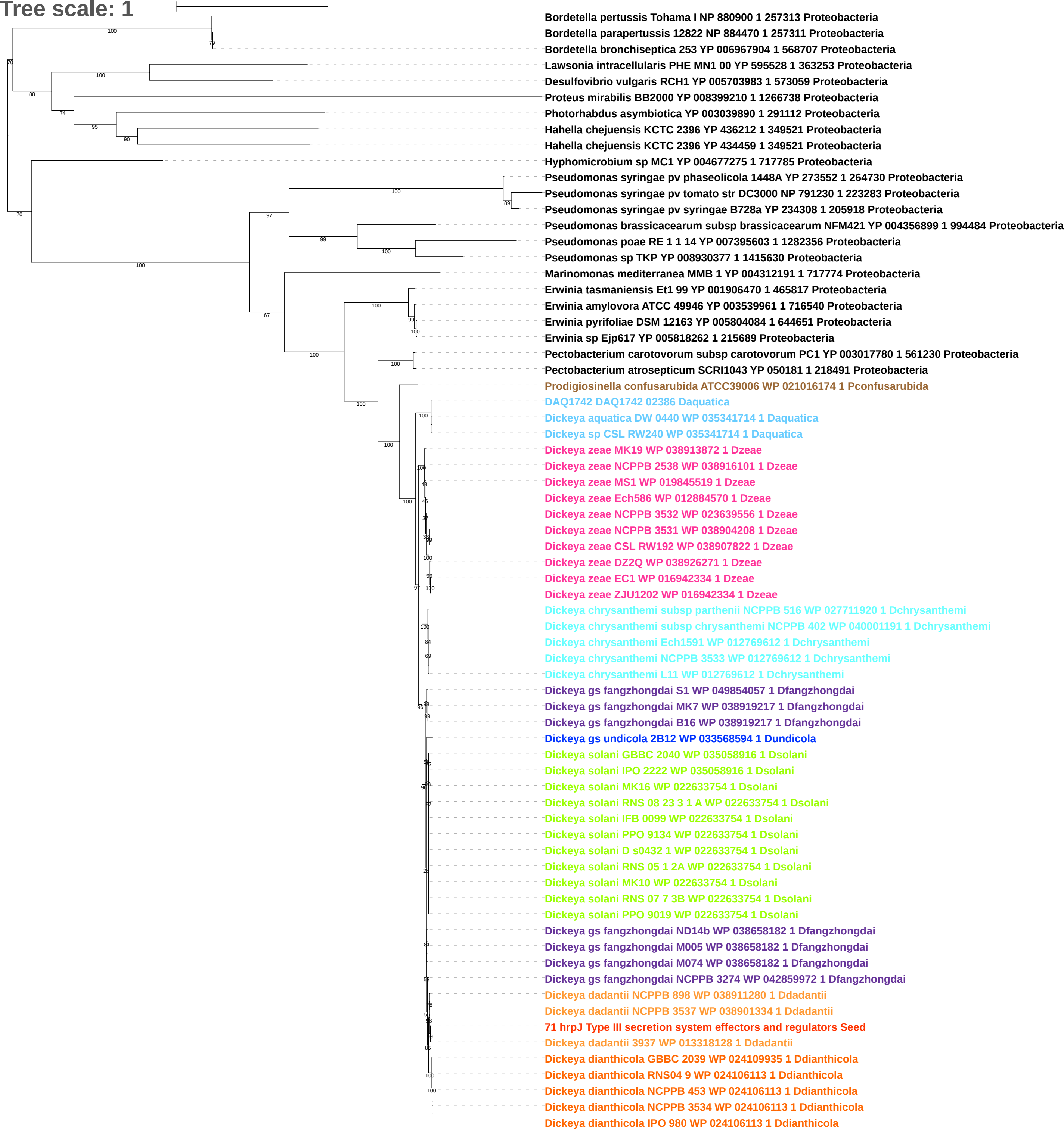

Tree scale: 1
Bordetella pertussis Tohama I NP 880900 1 257313 Proteobacteria
Bordetella parapertussis 12822 NP 884470 1 257311 Proteobacteria
Bordetella bronchiseptica 253 YP 006967904 1 568707 Proteobacteria
Lawsonia intracellularis PHE MN1 00 YP 595528 1 363253 Proteobacteria
Desulfovibrio vulgaris RCH1 YP 005703983 1 573059 Proteobacteria
Proteus mirabilis BB2000 YP 008399210 1 1266738 Proteobacteria
Photorhabdus asymbiotica YP 003039890 1 291112 Proteobacteria
Hahella chejuensis KCTC 2396 YP 436212 1 349521 Proteobacteria
Hahella chejuensis KCTC 2396 YP 434459 1 349521 Proteobacteria
Hyphomicrobium sp MC1 YP 004677275 1 717785 Proteobacteria
Pseudomonas syringae pv phaseolicola 1448A YP 273552 1 264730 Proteobacteria
Pseudomonas syringae pv tomato str DC3000 NP 791230 1 223283 Proteobacteria
Pseudomonas syringae pv syringae B728a YP 234308 1 205918 Proteobacteria
Pseudomonas brassicacearum subsp brassicacearum NFM421 YP 004356899 1 994484 Proteobacteria
Pseudomonas poae RE 1 1 14 YP 007395603 1 1282356 Proteobacteria
Pseudomonas sp TKP YP 008930377 1 1415630 Proteobacteria
Marinomonas mediterranea MMB 1 YP 004312191 1 717774 Proteobacteria
Erwinia tasmaniensis Et1 99 YP 001906470 1 465817 Proteobacteria
Erwinia amylovora ATCC 49946 YP 003539961 1 716540 Proteobacteria
Erwinia pyrifoliae DSM 12163 YP 005804084 1 644651 Proteobacteria
Erwinia sp Ejp617 YP 005818262 1 215689 Proteobacteria
Pectobacterium carotovorum subsp carotovorum PC1 YP 003017780 1 561230 Proteobacteria
Pectobacterium atrosepticum SCRI1043 YP 050181 1 218491 Proteobacteria
Prodigiosinella confusarubida ATCC39006 WP 021016174 1 Pconfusarubida
DAQ1742 DAQ1742 02386 Daquatica
Dickeya aquatica DW 0440 WP 035341714 1 Daquatica
Dickeya sp CSL RW240 WP 035341714 1 Daquatica
Dickeya zeae MK19 WP 038913872 1 Dzeae
Dickeya zeae NCPPB 2538 WP 038916101 1 Dzeae
Dickeya zeae MS1 WP 019845519 1 Dzeae
Dickeya zeae Ech586 WP 012884570 1 Dzeae
Dickeya zeae NCPPB 3532 WP 023639556 1 Dzeae
Dickeya zeae NCPPB 3531 WP 038904208 1 Dzeae
Dickeya zeae CSL RW192 WP 038907822 1 Dzeae
Dickeya zeae DZ2Q WP 038926271 1 Dzeae
Dickeya zeae EC1 WP 016942334 1 Dzeae
Dickeya zeae ZJU1202 WP 016942334 1 Dzeae
Dickeya chrysanthemi subsp parthenii NCPPB 516 WP 027711920 1 Dchrysanthemi
Dickeya chrysanthemi subsp chrysanthemi NCPPB 402 WP 040001191 1 Dchrysanthemi
Dickeya chrysanthemi Ech1591 WP 012769612 1 Dchrysanthemi
Dickeya chrysanthemi NCPPB 3533 WP 012769612 1 Dchrysanthemi
Dickeya chrysanthemi L11 WP 012769612 1 Dchrysanthemi
Dickeya gs fangzhongdai S1 WP 049854057 1 Dfangzhongdai
Dickeya gs fangzhongdai MK7 WP 038919217 1 Dfangzhongdai
Dickeya gs fangzhongdai B16 WP 038919217 1 Dfangzhongdai
Dickeya gs undicola 2B12 WP 033568594 1 Dundicola
Dickeya solani GBBC 2040 WP 035058916 1 Dsolani
Dickeya solani IPO 2222 WP 035058916 1 Dsolani
Dickeya solani MK16 WP 022633754 1 Dsolani
Dickeya solani RNS 08 23 3 1 A WP 022633754 1 Dsolani
Dickeya solani IFB 0099 WP 022633754 1 Dsolani
Dickeya solani PPO 9134 WP 022633754 1 Dsolani
Dickeya solani D s0432 1 WP 022633754 1 Dsolani
Dickeya solani RNS 05 1 2A WP 022633754 1 Dsolani
Dickeya solani MK10 WP 022633754 1 Dsolani
Dickeya solani RNS 07 7 3B WP 022633754 1 Dsolani
Dickeya solani PPO 9019 WP 022633754 1 Dsolani
Dickeya gs fangzhongdai ND14b WP 038658182 1 Dfangzhongdai
Dickeya gs fangzhongdai M005 WP 038658182 1 Dfangzhongdai
Dickeya gs fangzhongdai M074 WP 038658182 1 Dfangzhongdai
Dickeya gs fangzhongdai NCPPB 3274 WP 042859972 1 Dfangzhongdai
Dickeya dadantii NCPPB 898 WP 038911280 1 Ddadantii
Dickeya dadantii NCPPB 3537 WP 038901334 1 Ddadantii
71 hrpJ Type III secretion system effectors and regulators Seed
Dickeya dadantii 3937 WP 013318128 1 Ddadantii
Dickeya dianthicola GBBC 2039 WP 024109935 1 Ddianthicola
Dickeya dianthicola RNS04 9 WP 024106113 1 Ddianthicola
Dickeya dianthicola NCPPB 453 WP 024106113 1 Ddianthicola
Dickeya dianthicola NCPPB 3534 WP 024106113 1 Ddianthicola
Dickeya dianthicola IPO 980 WP 024106113 1 Ddianthicola

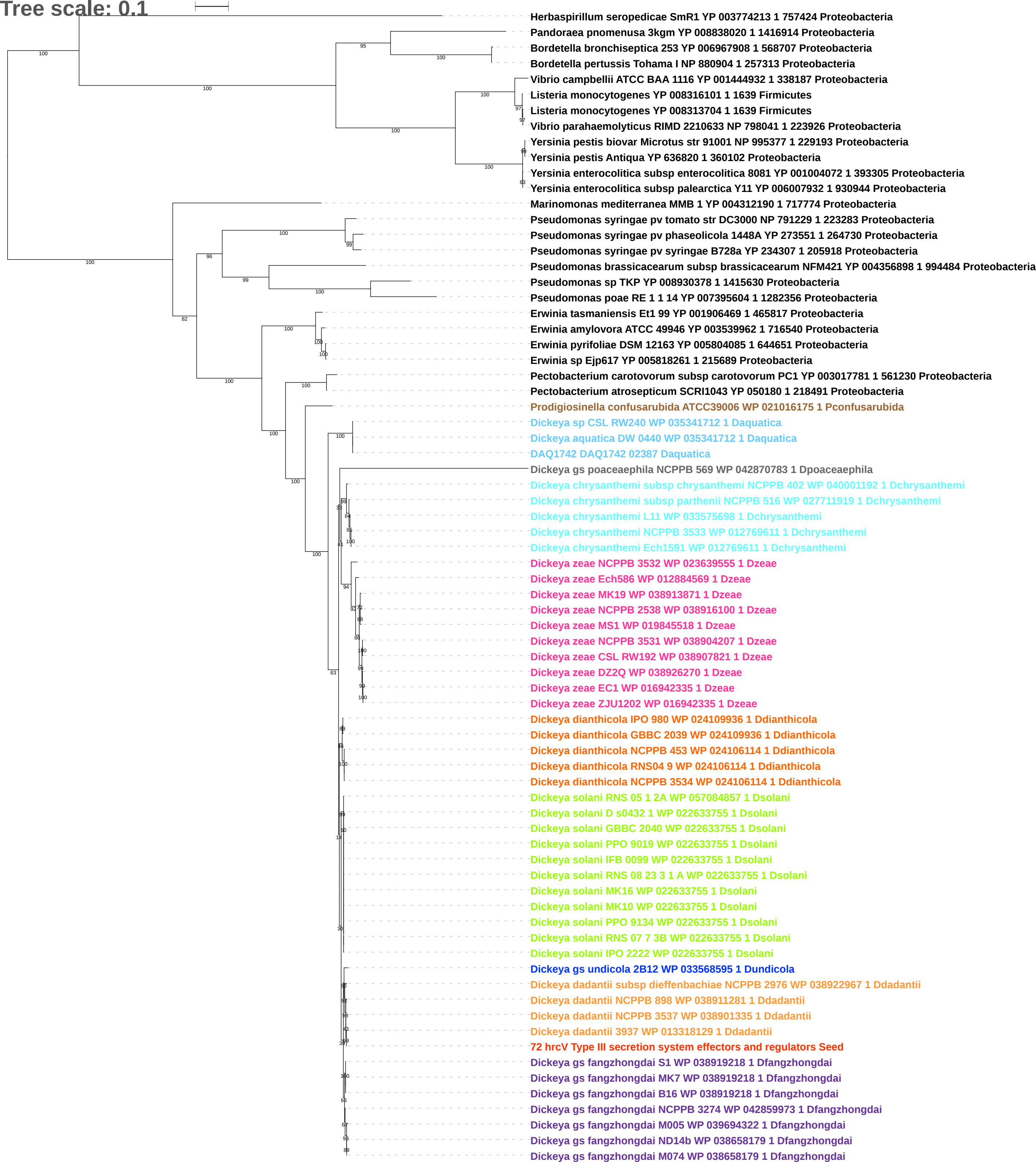

Tree scale: 0.1
Herbaspirillum seropedicae SmR1 YP 003774213 1 757424 Proteobacteria
Pandoraea pnomenusa 3kgm YP 008838020 1 1416914 Proteobacteria
Bordetella bronchiseptica 253 YP 006967908 1 568707 Proteobacteria
Bordetella pertussis Tohama I NP 880904 1 257313 Proteobacteria
Vibrio campbellii ATCC BAA 1116 YP 001444932 1 338187 Proteobacteria
Listeria monocytogenes YP 008316101 1 1639 Firmicutes
Listeria monocytogenes YP 008313704 1 1639 Firmicutes
Vibrio parahaemolyticus RIMD 2210633 NP 798041 1 223926 Proteobacteria
Yersinia pestis biovar Microtus str 91001 NP 995377 1 229193 Proteobacteria
Yersinia pestis Antiqua YP 636820 1 360102 Proteobacteria
Yersinia enterocolitica subsp enterocolitica 8081 YP 001004072 1 393305 Proteobacteria
Yersinia enterocolitica subsp palearctica Y11 YP 006007932 1 930944 Proteobacteria
Marinomonas mediterranea MMB 1 YP 004312190 1 717774 Proteobacteria
Pseudomonas syringae pv tomato str DC3000 NP 791229 1 223283 Proteobacteria
Pseudomonas syringae pv phaseolicola 1448A YP 273551 1 264730 Proteobacteria
Pseudomonas syringae pv syringae B728a YP 234307 1 205918 Proteobacteria
Pseudomonas brassicacearum subsp brassicacearum NFM421 YP 004356898 1 994484 Proteobacteria
Pseudomonas sp TKP YP 008930378 1 1415630 Proteobacteria
Pseudomonas poae RE 1 1 14 YP 007395604 1 1282356 Proteobacteria
Erwinia tasmaniensis Et1 99 YP 001906469 1 465817 Proteobacteria
Erwinia amylovora ATCC 49946 YP 003539962 1 716540 Proteobacteria
Erwinia pyrifoliae DSM 12163 YP 005804085 1 644651 Proteobacteria
Erwinia sp Ejp617 YP 005818261 1 215689 Proteobacteria
Pectobacterium carotovorum subsp carotovorum PC1 YP 003017781 1 561230 Proteobacteria
Pectobacterium atrosepticum SCRI1043 YP 050180 1 218491 Proteobacteria
Prodigiosinella confusarubida ATCC39006 WP 021016175 1 Pconfusarubida
Dickeya sp CSL RW240 WP 035341712 1 Daquatica
Dickeya aquatica DW 0440 WP 035341712 1 Daquatica
DAQ1742 DAQ1742 02387 Daquatica
Dickeya gs poaceaephila NCPPB 569 WP 042870783 1 Dpoaceaephila
Dickeya chrysanthemi subsp chrysanthemi NCPPB 402 WP 040001192 1 Dchrysanthemi
Dickeya chrysanthemi subsp parthenii NCPPB 516 WP 027711919 1 Dchrysanthemi
Dickeya chrysanthemi L11 WP 033575698 1 Dchrysanthemi
Dickeya chrysanthemi NCPPB 3533 WP 012769611 1 Dchrysanthemi
Dickeya chrysanthemi Ech1591 WP 012769611 1 Dchrysanthemi
Dickeya zeae NCPPB 3532 WP 023639555 1 Dzeae
Dickeya zeae Ech586 WP 012884569 1 Dzeae
Dickeya zeae MK19 WP 038913871 1 Dzeae
Dickeya zeae NCPPB 2538 WP 038916100 1 Dzeae
Dickeya zeae MS1 WP 019845518 1 Dzeae
Dickeya zeae NCPPB 3531 WP 038904207 1 Dzeae
Dickeya zeae CSL RW192 WP 038907821 1 Dzeae
Dickeya zeae DZ2Q WP 038926270 1 Dzeae
Dickeya zeae EC1 WP 016942335 1 Dzeae
Dickeya zeae ZJU1202 WP 016942335 1 Dzeae
Dickeya dianthicola IPO 980 WP 024109936 1 Ddianthicola
Dickeya dianthicola GBBC 2039 WP 024109936 1 Ddianthicola
Dickeya dianthicola NCPPB 453 WP 024106114 1 Ddianthicola
Dickeya dianthicola RNS04 9 WP 024106114 1 Ddianthicola
Dickeya dianthicola NCPPB 3534 WP 024106114 1 Ddianthicola
Dickeya solani RNS 05 1 2A WP 057084857 1 Dsolani
Dickeya solani D s0432 1 WP 022633755 1 Dsolani
Dickeya solani GBBC 2040 WP 022633755 1 Dsolani
Dickeya solani PPO 9019 WP 022633755 1 Dsolani
Dickeya solani IFB 0099 WP 022633755 1 Dsolani
Dickeya solani RNS 08 23 3 1 A WP 022633755 1 Dsolani
Dickeya solani MK16 WP 022633755 1 Dsolani
Dickeya solani MK10 WP 022633755 1 Dsolani
Dickeya solani PPO 9134 WP 022633755 1 Dsolani
Dickeya solani RNS 07 7 3B WP 022633755 1 Dsolani
Dickeya solani IPO 2222 WP 022633755 1 Dsolani
Dickeya gs undicola 2B12 WP 033568595 1 Dundicola
Dickeya dadantii subsp dieffenbachiae NCPPB 2976 WP 038922967 1 Ddadantii
Dickeya dadantii NCPPB 898 WP 038911281 1 Ddadantii
Dickeya dadantii NCPPB 3537 WP 038901335 1 Ddadantii
Dickeya dadantii 3937 WP 013318129 1 Ddadantii
72 hrcV Type III secretion system effectors and regulators Seed
Dickeya gs fangzhongdai S1 WP 038919218 1 Dfangzhongdai
Dickeya gs fangzhongdai MK7 WP 038919218 1 Dfangzhongdai
Dickeya gs fangzhongdai B16 WP 038919218 1 Dfangzhongdai
Dickeya gs fangzhongdai NCPPB 3274 WP 042859973 1 Dfangzhongdai
Dickeya gs fangzhongdai M005 WP 039694322 1 Dfangzhongdai
Dickeya gs fangzhongdai ND14b WP 038658179 1 Dfangzhongdai
Dickeya gs fangzhongdai M074 WP 038658179 1 Dfangzhongdai

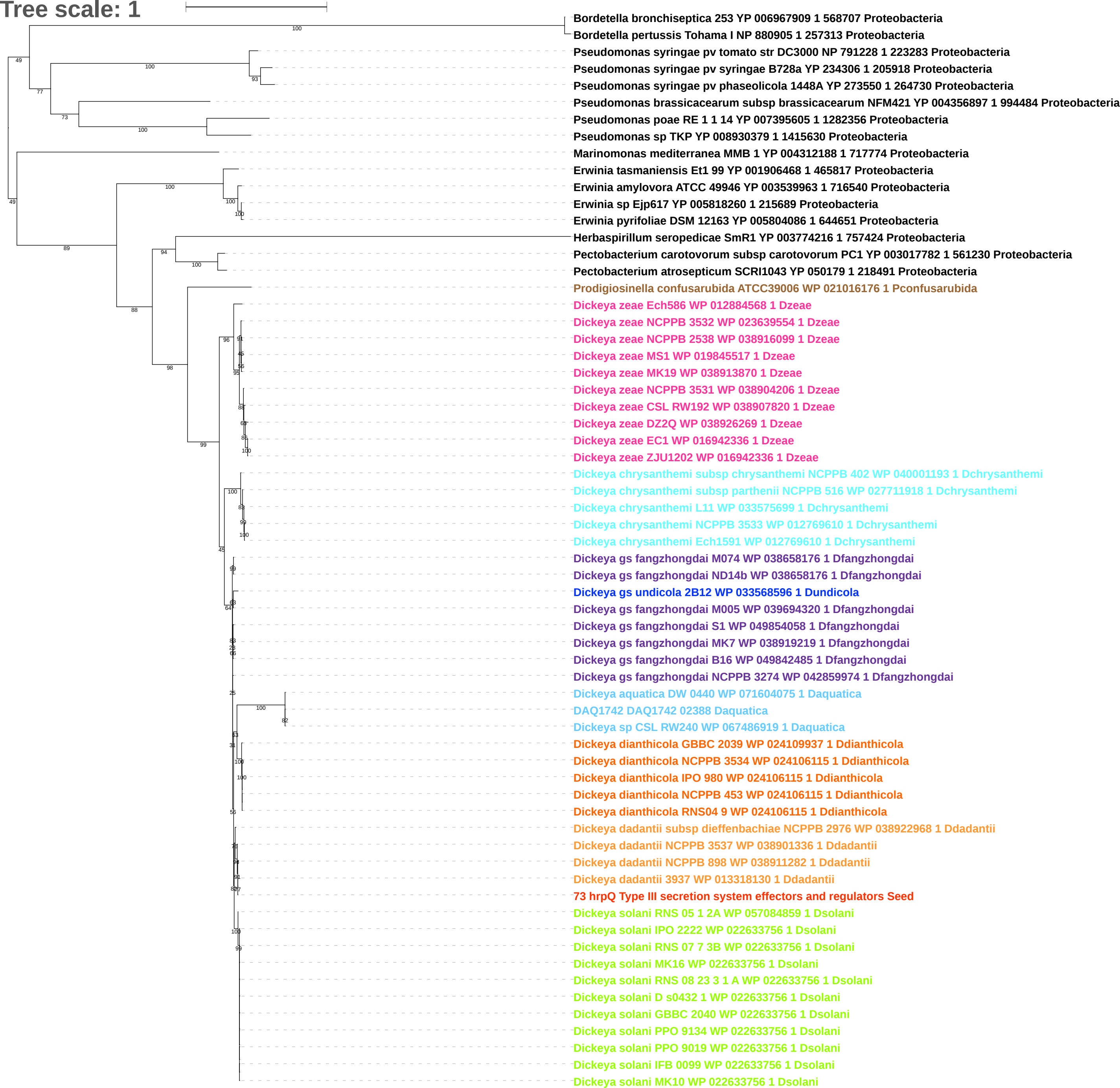

Tree scale: 1
Bordetella bronchiseptica 253 YP 006967909 1 568707 Proteobacteria
Bordetella pertussis Tohama I NP 880905 1 257313 Proteobacteria
Pseudomonas syringae pv tomato str DC3000 NP 791228 1 223283 Proteobacteria
Pseudomonas syringae pv syringae B728a YP 234306 1 205918 Proteobacteria
Pseudomonas syringae pv phaseolicola 1448A YP 273550 1 264730 Proteobacteria
Pseudomonas brassicacearum subsp brassicacearum NFM421 YP 004356897 1 994484 Proteobacteria
Pseudomonas poae RE 1 1 14 YP 007395605 1 1282356 Proteobacteria
Pseudomonas sp TKP YP 008930379 1 1415630 Proteobacteria
Marinomonas mediterranea MMB 1 YP 004312188 1 717774 Proteobacteria
Erwinia tasmaniensis Et1 99 YP 001906468 1 465817 Proteobacteria
Erwinia amylovora ATCC 49946 YP 003539963 1 716540 Proteobacteria
Erwinia sp Ejp617 YP 005818260 1 215689 Proteobacteria
Erwinia pyrifoliae DSM 12163 YP 005804086 1 644651 Proteobacteria
Herbaspirillum seropedicae SmR1 YP 003774216 1 757424 Proteobacteria
Pectobacterium carotovorum subsp carotovorum PC1 YP 003017782 1 561230 Proteobacteria
Pectobacterium atrosepticum SCRI1043 YP 050179 1 218491 Proteobacteria
Prodigiosinella confusarubida ATCC39006 WP 021016176 1 Pconfusarubida
Dickeya zeae Ech586 WP 012884568 1 Dzeae
Dickeya zeae NCPPB 3532 WP 023639554 1 Dzeae
Dickeya zeae NCPPB 2538 WP 038916099 1 Dzeae
Dickeya zeae MS1 WP 019845517 1 Dzeae
Dickeya zeae MK19 WP 038913870 1 Dzeae
Dickeya zeae NCPPB 3531 WP 038904206 1 Dzeae
Dickeya zeae CSL RW192 WP 038907820 1 Dzeae
Dickeya zeae DZ2Q WP 038926269 1 Dzeae
Dickeya zeae EC1 WP 016942336 1 Dzeae
Dickeya zeae ZJU1202 WP 016942336 1 Dzeae
Dickeya chrysanthemi subsp chrysanthemi NCPPB 402 WP 040001193 1 Dchrysanthemi
Dickeya chrysanthemi subsp parthenii NCPPB 516 WP 027711918 1 Dchrysanthemi
Dickeya chrysanthemi L11 WP 033575699 1 Dchrysanthemi
Dickeya chrysanthemi NCPPB 3533 WP 012769610 1 Dchrysanthemi
Dickeya chrysanthemi Ech1591 WP 012769610 1 Dchrysanthemi
Dickeya gs fangzhongdai M074 WP 038658176 1 Dfangzhongdai
Dickeya gs fangzhongdai ND14b WP 038658176 1 Dfangzhongdai
Dickeya gs undicola 2B12 WP 033568596 1 Dundicola
Dickeya gs fangzhongdai M005 WP 039694320 1 Dfangzhongdai
Dickeya gs fangzhongdai S1 WP 049854058 1 Dfangzhongdai
Dickeya gs fangzhongdai MK7 WP 038919219 1 Dfangzhongdai
Dickeya gs fangzhongdai B16 WP 049842485 1 Dfangzhongdai
Dickeya gs fangzhongdai NCPPB 3274 WP 042859974 1 Dfangzhongdai
Dickeya aquatica DW 0440 WP 071604075 1 Daquatica
DAQ1742 DAQ1742 02388 Daquatica
Dickeya sp CSL RW240 WP 067486919 1 Daquatica
Dickeya dianthicola GBBC 2039 WP 024109937 1 Ddianthicola
Dickeya dianthicola NCPPB 3534 WP 024106115 1 Ddianthicola
Dickeya dianthicola IPO 980 WP 024106115 1 Ddianthicola
Dickeya dianthicola NCPPB 453 WP 024106115 1 Ddianthicola
Dickeya dianthicola RNS04 9 WP 024106115 1 Ddianthicola
Dickeya dadantii subsp dieffenbachiae NCPPB 2976 WP 038922968 1 Ddadantii
Dickeya dadantii NCPPB 3537 WP 038901336 1 Ddadantii
Dickeya dadantii NCPPB 898 WP 038911282 1 Ddadantii
Dickeya dadantii 3937 WP 013318130 1 Ddadantii
73 hrpQ Type III secretion system effectors and regulators Seed
Dickeya solani RNS 05 1 2A WP 057084859 1 Dsolani
Dickeya solani IPO 2222 WP 022633756 1 Dsolani
Dickeya solani RNS 07 7 3B WP 022633756 1 Dsolani
Dickeya solani MK16 WP 022633756 1 Dsolani
Dickeya solani RNS 08 23 3 1 A WP 022633756 1 Dsolani
Dickeya solani D s0432 1 WP 022633756 1 Dsolani
Dickeya solani GBBC 2040 WP 022633756 1 Dsolani
Dickeya solani PPO 9134 WP 022633756 1 Dsolani
Dickeya solani PPO 9019 WP 022633756 1 Dsolani
Dickeya solani IFB 0099 WP 022633756 1 Dsolani
Dickeya solani MK10 WP 022633756 1 Dsolani

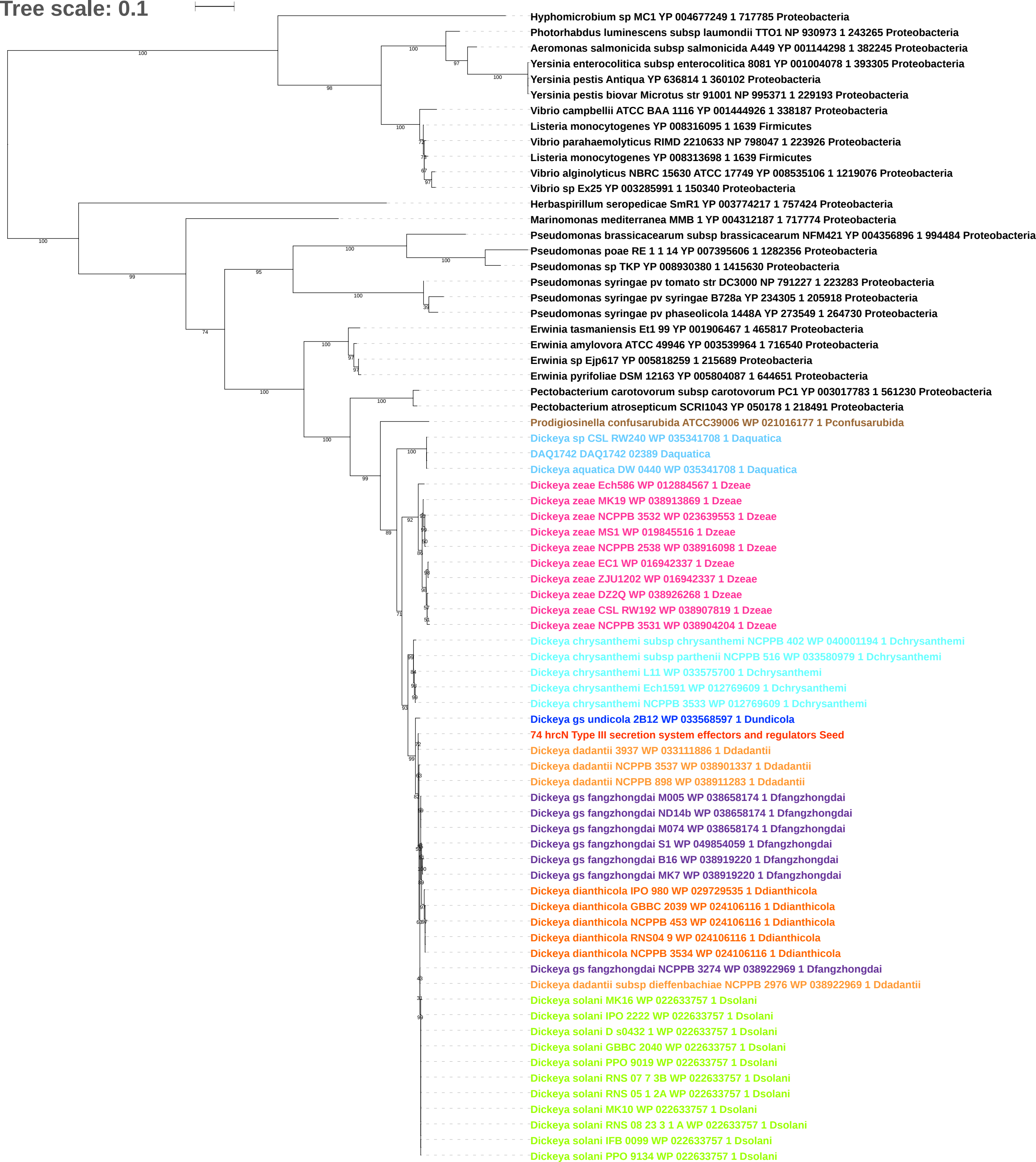
Tree scale: 0.1
Hyphomicrobium sp MC1 YP 004677249 1 717785 Proteobacteria
Photorhabdus luminescens subsp laumondii TTO1 NP 930973 1 243265 Proteobacteria
Aeromonas salmonicida subsp salmonicida A449 YP 001144298 1 382245 Proteobacteria
Yersinia enterocolitica subsp enterocolitica 8081 YP 001004078 1 393305 Proteobacteria
Yersinia pestis Antiqua YP 636814 1 360102 Proteobacteria
Yersinia pestis biovar Microtus str 91001 NP 995371 1 229193 Proteobacteria
Vibrio campbellii ATCC BAA 1116 YP 001444926 1 338187 Proteobacteria
Listeria monocytogenes YP 008316095 1 1639 Firmicutes
Vibrio parahaemolyticus RIMD 2210633 NP 798047 1 223926 Proteobacteria
Listeria monocytogenes YP 008313698 1 1639 Firmicutes
Vibrio alginolyticus NBRC 15630 ATCC 17749 YP 008535106 1 1219076 Proteobacteria
Vibrio sp Ex25 YP 003285991 1 150340 Proteobacteria
Herbaspirillum seropedicae SmR1 YP 003774217 1 757424 Proteobacteria
Marinomonas mediterranea MMB 1 YP 004312187 1 717774 Proteobacteria
Pseudomonas brassicacearum subsp brassicacearum NFM421 YP 004356896 1 994484 Proteobacteria
Pseudomonas poae RE 1 1 14 YP 007395606 1 1282356 Proteobacteria
Pseudomonas sp TKP YP 008930380 1 1415630 Proteobacteria
Pseudomonas syringae pv tomato str DC3000 NP 791227 1 223283 Proteobacteria
Pseudomonas syringae pv syringae B728a YP 234305 1 205918 Proteobacteria
Pseudomonas syringae pv phaseolicola 1448A YP 273549 1 264730 Proteobacteria
Erwinia tasmaniensis Et1 99 YP 001906467 1 465817 Proteobacteria
Erwinia amylovora ATCC 49946 YP 003539964 1 716540 Proteobacteria
Erwinia sp Ejp617 YP 005818259 1 215689 Proteobacteria
Erwinia pyrifoliae DSM 12163 YP 005804087 1 644651 Proteobacteria
Pectobacterium carotovorum subsp carotovorum PC1 YP 003017783 1 561230 Proteobacteria
Pectobacterium atrosepticum SCRI1043 YP 050178 1 218491 Proteobacteria
Prodigiosinella confusarubida ATCC39006 WP 021016177 1 Pconfusarubida
Dickeya sp CSL RW240 WP 035341708 1 Daquatica
DAQ1742 DAQ1742 02389 Daquatica
Dickeya aquatica DW 0440 WP 035341708 1 Daquatica
Dickeya zeae Ech586 WP 012884567 1 Dzeae
Dickeya zeae MK19 WP 038913869 1 Dzeae
Dickeya zeae NCPPB 3532 WP 023639553 1 Dzeae
Dickeya zeae MS1 WP 019845516 1 Dzeae
Dickeya zeae NCPPB 2538 WP 038916098 1 Dzeae
Dickeya zeae EC1 WP 016942337 1 Dzeae
Dickeya zeae ZJU1202 WP 016942337 1 Dzeae
Dickeya zeae DZ2Q WP 038926268 1 Dzeae
Dickeya zeae CSL RW192 WP 038907819 1 Dzeae
Dickeya zeae NCPPB 3531 WP 038904204 1 Dzeae
Dickeya chrysanthemi subsp chrysanthemi NCPPB 402 WP 040001194 1 Dchrysanthemi
Dickeya chrysanthemi subsp parthenii NCPPB 516 WP 033580979 1 Dchrysanthemi
Dickeya chrysanthemi L11 WP 033575700 1 Dchrysanthemi
Dickeya chrysanthemi Ech1591 WP 012769609 1 Dchrysanthemi
Dickeya chrysanthemi NCPPB 3533 WP 012769609 1 Dchrysanthemi
Dickeya gs undicola 2B12 WP 033568597 1 Dundicola
74 hrcN Type III secretion system effectors and regulators Seed
Dickeya dadantii 3937 WP 033111886 1 Ddadantii
Dickeya dadantii NCPPB 3537 WP 038901337 1 Ddadantii
Dickeya dadantii NCPPB 898 WP 038911283 1 Ddadantii
Dickeya gs fangzhongdai M005 WP 038658174 1 Dfangzhongdai
Dickeya gs fangzhongdai ND14b WP 038658174 1 Dfangzhongdai
Dickeya gs fangzhongdai M074 WP 038658174 1 Dfangzhongdai
Dickeya gs fangzhongdai S1 WP 049854059 1 Dfangzhongdai
Dickeya gs fangzhongdai B16 WP 038919220 1 Dfangzhongdai
Dickeya gs fangzhongdai MK7 WP 038919220 1 Dfangzhongdai
Dickeya dianthicola IPO 980 WP 029729535 1 Ddianthicola
Dickeya dianthicola GBBC 2039 WP 024106116 1 Ddianthicola
Dickeya dianthicola NCPPB 453 WP 024106116 1 Ddianthicola
Dickeya dianthicola RNS04 9 WP 024106116 1 Ddianthicola
Dickeya dianthicola NCPPB 3534 WP 024106116 1 Ddianthicola
Dickeya gs fangzhongdai NCPPB 3274 WP 038922969 1 Dfangzhongdai
Dickeya dadantii subsp dieffenbachiae NCPPB 2976 WP 038922969 1 Ddadantii
Dickeya solani MK16 WP 022633757 1 Dsolani
Dickeya solani IPO 2222 WP 022633757 1 Dsolani
Dickeya solani D s0432 1 WP 022633757 1 Dsolani
Dickeya solani GBBC 2040 WP 022633757 1 Dsolani
Dickeya solani PPO 9019 WP 022633757 1 Dsolani
Dickeya solani RNS 07 7 3B WP 022633757 1 Dsolani
Dickeya solani RNS 05 1 2A WP 022633757 1 Dsolani
Dickeya solani MK10 WP 022633757 1 Dsolani
Dickeya solani RNS 08 23 3 1 A WP 022633757 1 Dsolani
Dickeya solani IFB 0099 WP 022633757 1 Dsolani
Dickeya solani PPO 9134 WP 022633757 1 Dsolani

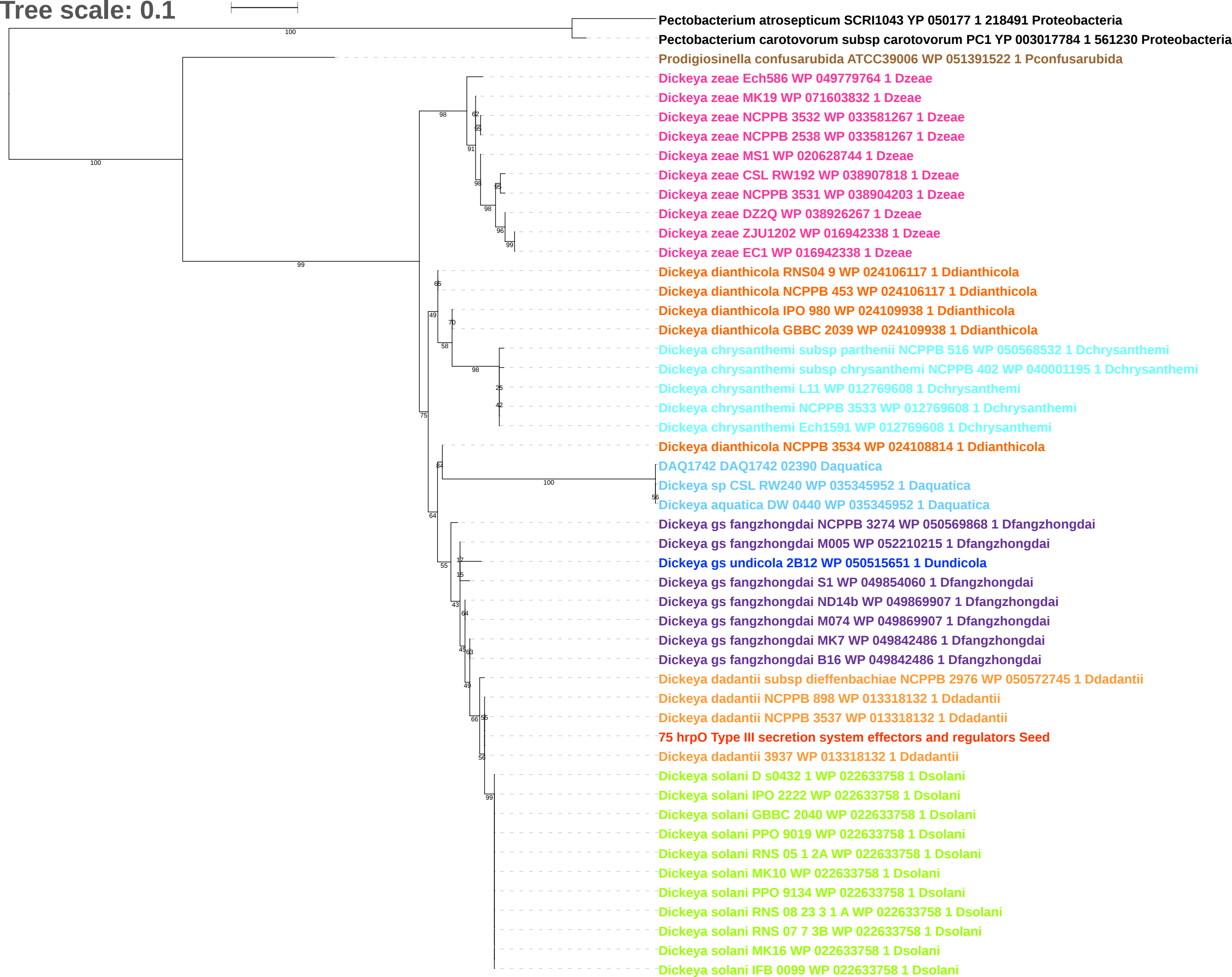
Tree scale: 0.1
Pectobacterium atrosepticum SCRI1043 YP 050177 1 218491 Proteobacteria
Pectobacterium carotovorum subsp carotovorum PC1 YP 003017784 1 561230 Proteobacteria
Prodigiosinella confusarubida ATCC39006 WP 051391522 1 Pconfusarubida
Dickeya zeae Ech586 WP 049779764 1 Dzeae
Dickeya zeae MK19 WP 071603832 1 Dzeae
Dickeya zeae NCPPB 3532 WP 033581267 1 Dzeae
Dickeya zeae NCPPB 2538 WP 033581267 1 Dzeae
Dickeya zeae MS1 WP 020628744 1 Dzeae
Dickeya zeae CSL RW192 WP 038907818 1 Dzeae
Dickeya zeae NCPPB 3531 WP 038904203 1 Dzeae
Dickeya zeae DZ2Q WP 038926267 1 Dzeae
Dickeya zeae ZJU1202 WP 016942338 1 Dzeae
Dickeya zeae EC1 WP 016942338 1 Dzeae
Dickeya dianthicola RNS04 9 WP 024106117 1 Ddianthicola
Dickeya dianthicola NCPPB 453 WP 024106117 1 Ddianthicola
Dickeya dianthicola IPO 980 WP 024109938 1 Ddianthicola
Dickeya dianthicola GBBC 2039 WP 024109938 1 Ddianthicola
Dickeya chrysanthemi subsp parthenii NCPPB 516 WP 050568532 1 Dchrysanthemi
Dickeya chrysanthemi subsp chrysanthemi NCPPB 402 WP 040001195 1 Dchrysanthemi
Dickeya chrysanthemi L11 WP 012769608 1 Dchrysanthemi
Dickeya chrysanthemi NCPPB 3533 WP 012769608 1 Dchrysanthemi
Dickeya chrysanthemi Ech1591 WP 012769608 1 Dchrysanthemi
Dickeya dianthicola NCPPB 3534 WP 024108814 1 Ddianthicola
DAQ1742 DAQ1742 02390 Daquatica
Dickeya sp CSL RW240 WP 035345952 1 Daquatica
Dickeya aquatica DW 0440 WP 035345952 1 Daquatica
Dickeya gs fangzhongdai NCPPB 3274 WP 050569868 1 Dfangzhongdai
Dickeya gs fangzhongdai M005 WP 052210215 1 Dfangzhongdai
Dickeya gs undicola 2B12 WP 050515651 1 Dundicola
Dickeya gs fangzhongdai S1 WP 049854060 1 Dfangzhongdai
Dickeya gs fangzhongdai ND14b WP 049869907 1 Dfangzhongdai
Dickeya gs fangzhongdai M074 WP 049869907 1 Dfangzhongdai
Dickeya gs fangzhongdai MK7 WP 049842486 1 Dfangzhongdai
Dickeya gs fangzhongdai B16 WP 049842486 1 Dfangzhongdai
Dickeya dadantii subsp dieffenbachiae NCPPB 2976 WP 050572745 1 Ddadantii
Dickeya dadantii NCPPB 898 WP 013318132 1 Ddadantii
Dickeya dadantii NCPPB 3537 WP 013318132 1 Ddadantii
75 hrpO Type III secretion system effectors and regulators Seed
Dickeya dadantii 3937 WP 013318132 1 Ddadantii
Dickeya solani D s0432 1 WP 022633758 1 Dsolani
Dickeya solani IPO 2222 WP 022633758 1 Dsolani
Dickeya solani GBBC 2040 WP 022633758 1 Dsolani
Dickeya solani PPO 9019 WP 022633758 1 Dsolani
Dickeya solani RNS 05 1 2A WP 022633758 1 Dsolani
Dickeya solani MK10 WP 022633758 1 Dsolani
Dickeya solani PPO 9134 WP 022633758 1 Dsolani
Dickeya solani RNS 08 23 3 1 A WP 022633758 1 Dsolani
Dickeya solani RNS 07 7 3B WP 022633758 1 Dsolani
Dickeya solani MK16 WP 022633758 1 Dsolani
Dickeya solani IFB 0099 WP 022633758 1 Dsolani

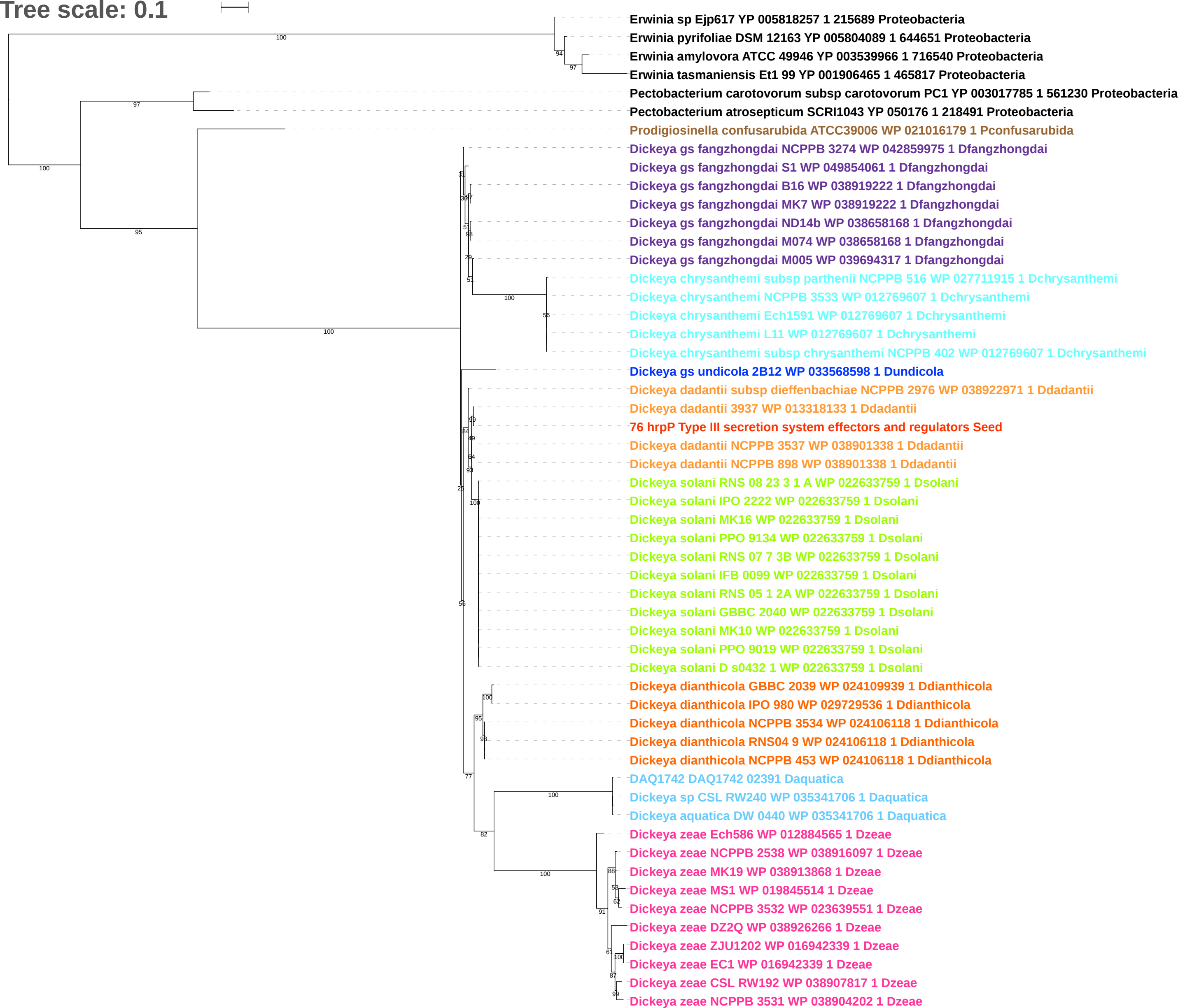
Tree scale: 0.1
Erwinia sp Ejp617 YP 005818257 1 215689 Proteobacteria
Erwinia pyrifoliae DSM 12163 YP 005804089 1 644651 Proteobacteria
Erwinia amylovora ATCC 49946 YP 003539966 1 716540 Proteobacteria
Erwinia tasmaniensis Et1 99 YP 001906465 1 465817 Proteobacteria
Pectobacterium carotovorum subsp carotovorum PC1 YP 003017785 1 561230 Proteobacteria
Pectobacterium atrosepticum SCRI1043 YP 050176 1 218491 Proteobacteria
Prodigiosinella confusarubida ATCC39006 WP 021016179 1 Pconfusarubida
Dickeya gs fangzhongdai NCPPB 3274 WP 042859975 1 Dfangzhongdai
Dickeya gs fangzhongdai S1 WP 049854061 1 Dfangzhongdai
Dickeya gs fangzhongdai B16 WP 038919222 1 Dfangzhongdai
Dickeya gs fangzhongdai MK7 WP 038919222 1 Dfangzhongdai
Dickeya gs fangzhongdai ND14b WP 038658168 1 Dfangzhongdai
Dickeya gs fangzhongdai M074 WP 038658168 1 Dfangzhongdai
Dickeya gs fangzhongdai M005 WP 039694317 1 Dfangzhongdai
Dickeya chrysanthemi subsp parthenii NCPPB 516 WP 027711915 1 Dchrysanthemi
Dickeya chrysanthemi NCPPB 3533 WP 012769607 1 Dchrysanthemi
Dickeya chrysanthemi Ech1591 WP 012769607 1 Dchrysanthemi
Dickeya chrysanthemi L11 WP 012769607 1 Dchrysanthemi
Dickeya chrysanthemi subsp chrysanthemi NCPPB 402 WP 012769607 1 Dchrysanthemi
Dickeya gs undicola 2B12 WP 033568598 1 Dundicola
Dickeya dadantii subsp dieffenbachiae NCPPB 2976 WP 038922971 1 Ddadantii
Dickeya dadantii 3937 WP 013318133 1 Ddadantii
76 hrpP Type III secretion system effectors and regulators Seed
Dickeya dadantii NCPPB 3537 WP 038901338 1 Ddadantii
Dickeya dadantii NCPPB 898 WP 038901338 1 Ddadantii
Dickeya solani RNS 08 23 3 1 A WP 022633759 1 Dsolani
Dickeya solani IPO 2222 WP 022633759 1 Dsolani
Dickeya solani MK16 WP 022633759 1 Dsolani
Dickeya solani PPO 9134 WP 022633759 1 Dsolani
Dickeya solani RNS 07 7 3B WP 022633759 1 Dsolani
Dickeya solani IFB 0099 WP 022633759 1 Dsolani
Dickeya solani RNS 05 1 2A WP 022633759 1 Dsolani
Dickeya solani GBBC 2040 WP 022633759 1 Dsolani
Dickeya solani MK10 WP 022633759 1 Dsolani
Dickeya solani PPO 9019 WP 022633759 1 Dsolani
Dickeya solani D s0432 1 WP 022633759 1 Dsolani
Dickeya dianthicola GBBC 2039 WP 024109939 1 Ddianthicola
Dickeya dianthicola IPO 980 WP 029729536 1 Ddianthicola
Dickeya dianthicola NCPPB 3534 WP 024106118 1 Ddianthicola
Dickeya dianthicola RNS04 9 WP 024106118 1 Ddianthicola
Dickeya dianthicola NCPPB 453 WP 024106118 1 Ddianthicola
DAQ1742 DAQ1742 02391 Daquatica
Dickeya sp CSL RW240 WP 035341706 1 Daquatica
Dickeya aquatica DW 0440 WP 035341706 1 Daquatica
Dickeya zeae Ech586 WP 012884565 1 Dzeae
Dickeya zeae NCPPB 2538 WP 038916097 1 Dzeae
Dickeya zeae MK19 WP 038913868 1 Dzeae
Dickeya zeae MS1 WP 019845514 1 Dzeae
Dickeya zeae NCPPB 3532 WP 023639551 1 Dzeae
Dickeya zeae DZ2Q WP 038926266 1 Dzeae
Dickeya zeae ZJU1202 WP 016942339 1 Dzeae
Dickeya zeae EC1 WP 016942339 1 Dzeae
Dickeya zeae CSL RW192 WP 038907817 1 Dzeae
Dickeya zeae NCPPB 3531 WP 038904202 1 Dzeae

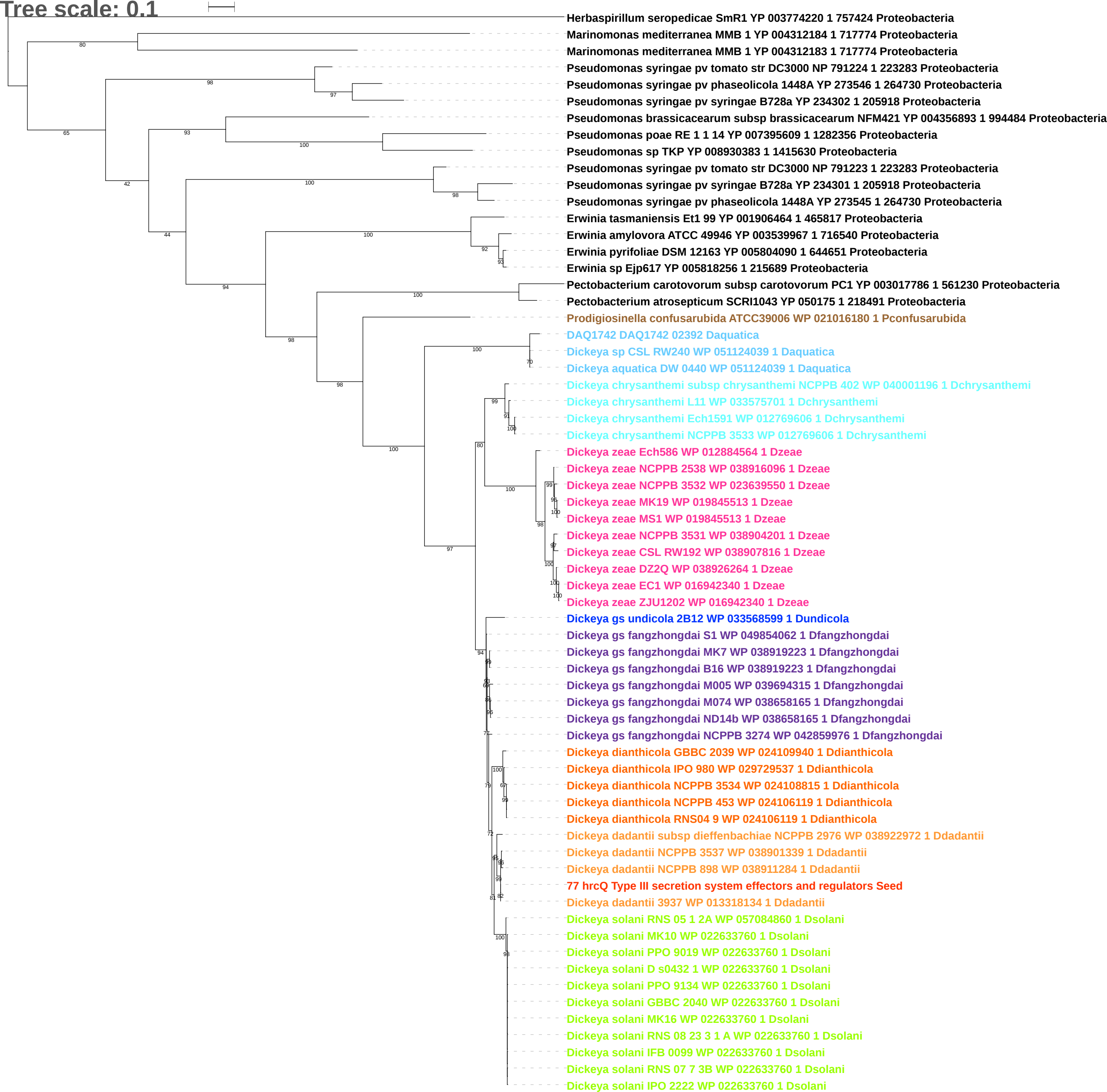
Tree scale: 0.1
Herbaspirillum seropedicae SmR1 YP 003774220 1 757424 Proteobacteria
Marinomonas mediterranea MMB 1 YP 004312184 1 717774 Proteobacteria
Marinomonas mediterranea MMB 1 YP 004312183 1 717774 Proteobacteria
Pseudomonas syringae pv tomato str DC3000 NP 791224 1 223283 Proteobacteria
Pseudomonas syringae pv phaseolicola 1448A YP 273546 1 264730 Proteobacteria
Pseudomonas syringae pv syringae B728a YP 234302 1 205918 Proteobacteria
Pseudomonas brassicacearum subsp brassicacearum NFM421 YP 004356893 1 994484 Proteobacteria
Pseudomonas poae RE 1 1 14 YP 007395609 1 1282356 Proteobacteria
Pseudomonas sp TKP YP 008930383 1 1415630 Proteobacteria
Pseudomonas syringae pv tomato str DC3000 NP 791223 1 223283 Proteobacteria
Pseudomonas syringae pv syringae B728a YP 234301 1 205918 Proteobacteria
Pseudomonas syringae pv phaseolicola 1448A YP 273545 1 264730 Proteobacteria
Erwinia tasmaniensis Et1 99 YP 001906464 1 465817 Proteobacteria
Erwinia amylovora ATCC 49946 YP 003539967 1 716540 Proteobacteria
Erwinia pyrifoliae DSM 12163 YP 005804090 1 644651 Proteobacteria
Erwinia sp Ejp617 YP 005818256 1 215689 Proteobacteria
Pectobacterium carotovorum subsp carotovorum PC1 YP 003017786 1 561230 Proteobacteria
Pectobacterium atrosepticum SCRI1043 YP 050175 1 218491 Proteobacteria
Prodigiosinella confusarubida ATCC39006 WP 021016180 1 Pconfusarubida
DAQ1742 DAQ1742 02392 Daquatica
Dickeya sp CSL RW240 WP 051124039 1 Daquatica
Dickeya aquatica DW 0440 WP 051124039 1 Daquatica
Dickeya chrysanthemi subsp chrysanthemi NCPPB 402 WP 040001196 1 Dchrysanthemi
Dickeya chrysanthemi L11 WP 033575701 1 Dchrysanthemi
Dickeya chrysanthemi Ech1591 WP 012769606 1 Dchrysanthemi
Dickeya chrysanthemi NCPPB 3533 WP 012769606 1 Dchrysanthemi
Dickeya zeae Ech586 WP 012884564 1 Dzeae
Dickeya zeae NCPPB 2538 WP 038916096 1 Dzeae
Dickeya zeae NCPPB 3532 WP 023639550 1 Dzeae
Dickeya zeae MK19 WP 019845513 1 Dzeae
Dickeya zeae MS1 WP 019845513 1 Dzeae
Dickeya zeae NCPPB 3531 WP 038904201 1 Dzeae
Dickeya zeae CSL RW192 WP 038907816 1 Dzeae
Dickeya zeae DZ2Q WP 038926264 1 Dzeae
Dickeya zeae EC1 WP 016942340 1 Dzeae
Dickeya zeae ZJU1202 WP 016942340 1 Dzeae
Dickeya gs undicola 2B12 WP 033568599 1 Dundicola
Dickeya gs fangzhongdai S1 WP 049854062 1 Dfangzhongdai
Dickeya gs fangzhongdai MK7 WP 038919223 1 Dfangzhongdai
Dickeya gs fangzhongdai B16 WP 038919223 1 Dfangzhongdai
Dickeya gs fangzhongdai M005 WP 039694315 1 Dfangzhongdai
Dickeya gs fangzhongdai M074 WP 038658165 1 Dfangzhongdai
Dickeya gs fangzhongdai ND14b WP 038658165 1 Dfangzhongdai
Dickeya gs fangzhongdai NCPPB 3274 WP 042859976 1 Dfangzhongdai
Dickeya dianthicola GBBC 2039 WP 024109940 1 Ddianthicola
Dickeya dianthicola IPO 980 WP 029729537 1 Ddianthicola
Dickeya dianthicola NCPPB 3534 WP 024108815 1 Ddianthicola
Dickeya dianthicola NCPPB 453 WP 024106119 1 Ddianthicola
Dickeya dianthicola RNS04 9 WP 024106119 1 Ddianthicola
Dickeya dadantii subsp dieffenbachiae NCPPB 2976 WP 038922972 1 Ddadantii
Dickeya dadantii NCPPB 3537 WP 038901339 1 Ddadantii
Dickeya dadantii NCPPB 898 WP 038911284 1 Ddadantii
77 hrcQ Type III secretion system effectors and regulators Seed
Dickeya dadantii 3937 WP 013318134 1 Ddadantii
Dickeya solani RNS 05 1 2A WP 057084860 1 Dsolani
Dickeya solani MK10 WP 022633760 1 Dsolani
Dickeya solani PPO 9019 WP 022633760 1 Dsolani
Dickeya solani D s0432 1 WP 022633760 1 Dsolani
Dickeya solani PPO 9134 WP 022633760 1 Dsolani
Dickeya solani GBBC 2040 WP 022633760 1 Dsolani
Dickeya solani MK16 WP 022633760 1 Dsolani
Dickeya solani RNS 08 23 3 1 A WP 022633760 1 Dsolani
Dickeya solani IFB 0099 WP 022633760 1 Dsolani
Dickeya solani RNS 07 7 3B WP 022633760 1 Dsolani
Dickeya solani IPO 2222 WP 022633760 1 Dsolani

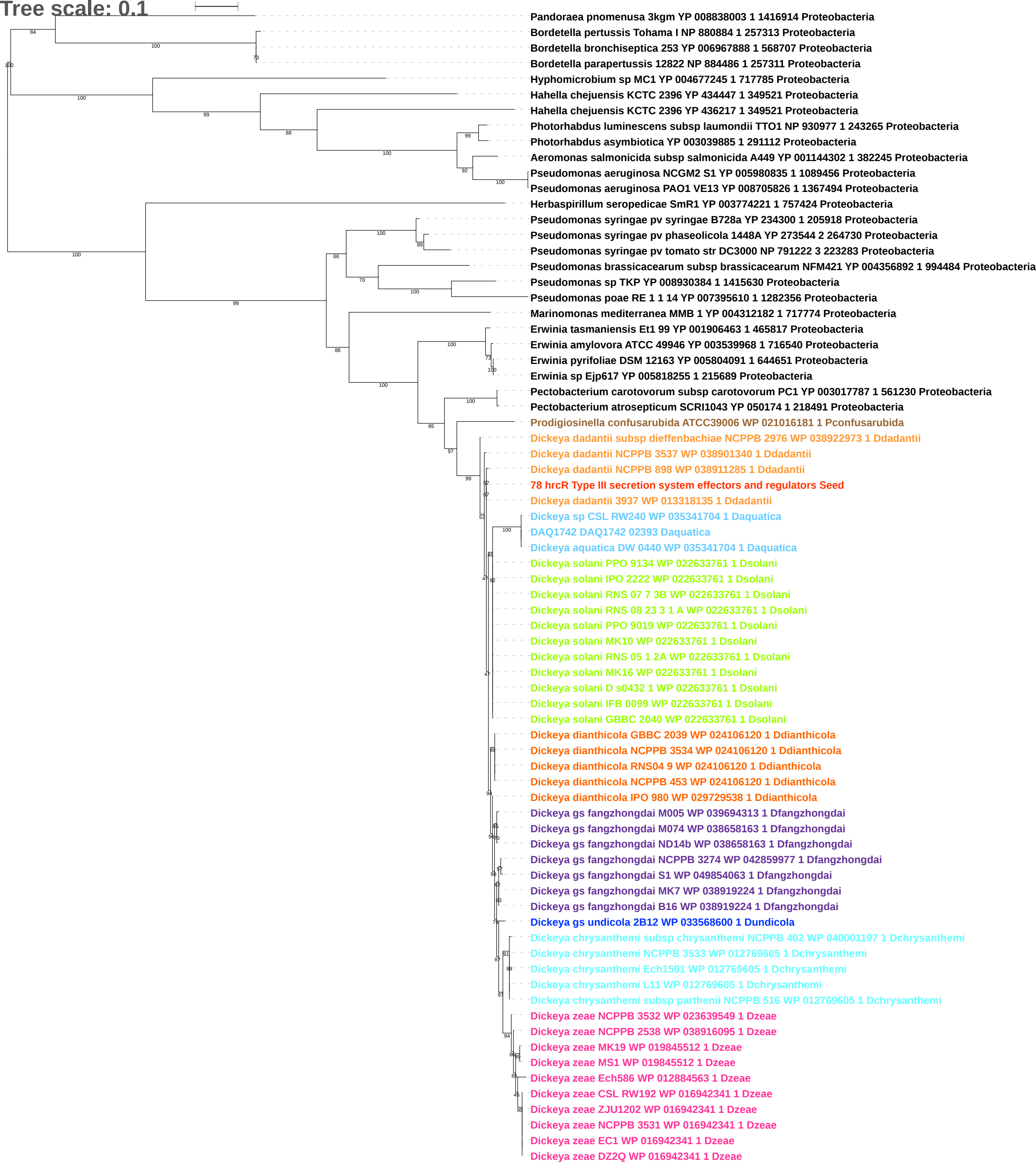
Tree scale: 0.1
Pandoraea pnomenusa 3kgm YP 008838003 1 1416914 Proteobacteria
Bordetella pertussis Tohama I NP 880884 1 257313 Proteobacteria
Bordetella bronchiseptica 253 YP 006967888 1 568707 Proteobacteria
Bordetella parapertussis 12822 NP 884486 1 257311 Proteobacteria
Hyphomicrobium sp MC1 YP 004677245 1 717785 Proteobacteria
Hahella chejuensis KCTC 2396 YP 434447 1 349521 Proteobacteria
Hahella chejuensis KCTC 2396 YP 436217 1 349521 Proteobacteria
Photorhabdus luminescens subsp laumondii TTO1 NP 930977 1 243265 Proteobacteria
Photorhabdus asymbiotica YP 003039885 1 291112 Proteobacteria
Aeromonas salmonicida subsp salmonicida A449 YP 001144302 1 382245 Proteobacteria
Pseudomonas aeruginosa NCGM2 S1 YP 005980835 1 1089456 Proteobacteria
Pseudomonas aeruginosa PAO1 VE13 YP 008705826 1 1367494 Proteobacteria
Herbaspirillum seropedicae SmR1 YP 003774221 1 757424 Proteobacteria
Pseudomonas syringae pv syringae B728a YP 234300 1 205918 Proteobacteria
Pseudomonas syringae pv phaseolicola 1448A YP 273544 2 264730 Proteobacteria
Pseudomonas syringae pv tomato str DC3000 NP 791222 3 223283 Proteobacteria
Pseudomonas brassicacearum subsp brassicacearum NFM421 YP 004356892 1 994484 Proteobacteria
Pseudomonas sp TKP YP 008930384 1 1415630 Proteobacteria
Pseudomonas poae RE 1 1 14 YP 007395610 1 1282356 Proteobacteria
Marinomonas mediterranea MMB 1 YP 004312182 1 717774 Proteobacteria
Erwinia tasmaniensis Et1 99 YP 001906463 1 465817 Proteobacteria
Erwinia amylovora ATCC 49946 YP 003539968 1 716540 Proteobacteria
Erwinia pyrifoliae DSM 12163 YP 005804091 1 644651 Proteobacteria
Erwinia sp Ejp617 YP 005818255 1 215689 Proteobacteria
Pectobacterium carotovorum subsp carotovorum PC1 YP 003017787 1 561230 Proteobacteria
Pectobacterium atrosepticum SCRI1043 YP 050174 1 218491 Proteobacteria
Prodigiosinella confusarubida ATCC39006 WP 021016181 1 Pconfusarubida
Dickeya dadantii subsp dieffenbachiae NCPPB 2976 WP 038922973 1 Ddadantii
Dickeya dadantii NCPPB 3537 WP 038901340 1 Ddadantii
Dickeya dadantii NCPPB 898 WP 038911285 1 Ddadantii
78 hrcR Type III secretion system effectors and regulators Seed
Dickeya dadantii 3937 WP 013318135 1 Ddadantii
Dickeya sp CSL RW240 WP 035341704 1 Daquatica
DAQ1742 DAQ1742 02393 Daquatica
Dickeya aquatica DW 0440 WP 035341704 1 Daquatica
Dickeya solani PPO 9134 WP 022633761 1 Dsolani
Dickeya solani IPO 2222 WP 022633761 1 Dsolani
Dickeya solani RNS 07 7 3B WP 022633761 1 Dsolani
Dickeya solani RNS 08 23 3 1 A WP 022633761 1 Dsolani
Dickeya solani PPO 9019 WP 022633761 1 Dsolani
Dickeya solani MK10 WP 022633761 1 Dsolani
Dickeya solani RNS 05 1 2A WP 022633761 1 Dsolani
Dickeya solani MK16 WP 022633761 1 Dsolani
Dickeya solani D s0432 1 WP 022633761 1 Dsolani
Dickeya solani IFB 0099 WP 022633761 1 Dsolani
Dickeya solani GBBC 2040 WP 022633761 1 Dsolani
Dickeya dianthicola GBBC 2039 WP 024106120 1 Ddianthicola
Dickeya dianthicola NCPPB 3534 WP 024106120 1 Ddianthicola
Dickeya dianthicola RNS04 9 WP 024106120 1 Ddianthicola
Dickeya dianthicola NCPPB 453 WP 024106120 1 Ddianthicola
Dickeya dianthicola IPO 980 WP 029729538 1 Ddianthicola
Dickeya gs fangzhongdai M005 WP 039694313 1 Dfangzhongdai
Dickeya gs fangzhongdai M074 WP 038658163 1 Dfangzhongdai
Dickeya gs fangzhongdai ND14b WP 038658163 1 Dfangzhongdai
Dickeya gs fangzhongdai NCPPB 3274 WP 042859977 1 Dfangzhongdai
Dickeya gs fangzhongdai S1 WP 049854063 1 Dfangzhongdai
Dickeya gs fangzhongdai MK7 WP 038919224 1 Dfangzhongdai
Dickeya gs fangzhongdai B16 WP 038919224 1 Dfangzhongdai
Dickeya gs undicola 2B12 WP 033568600 1 Dundicola
Dickeya chrysanthemi subsp chrysanthemi NCPPB 402 WP 040001197 1 Dchrysanthemi
Dickeya chrysanthemi NCPPB 3533 WP 012769605 1 Dchrysanthemi
Dickeya chrysanthemi Ech1591 WP 012769605 1 Dchrysanthemi
Dickeya chrysanthemi L11 WP 012769605 1 Dchrysanthemi
Dickeya chrysanthemi subsp parthenii NCPPB 516 WP 012769605 1 Dchrysanthemi
Dickeya zeae NCPPB 3532 WP 023639549 1 Dzeae
Dickeya zeae NCPPB 2538 WP 038916095 1 Dzeae
Dickeya zeae MK19 WP 019845512 1 Dzeae
Dickeya zeae MS1 WP 019845512 1 Dzeae
Dickeya zeae Ech586 WP 012884563 1 Dzeae
Dickeya zeae CSL RW192 WP 016942341 1 Dzeae
Dickeya zeae ZJU1202 WP 016942341 1 Dzeae
Dickeya zeae NCPPB 3531 WP 016942341 1 Dzeae
Dickeya zeae EC1 WP 016942341 1 Dzeae
Dickeya zeae DZ2Q WP 016942341 1 Dzeae

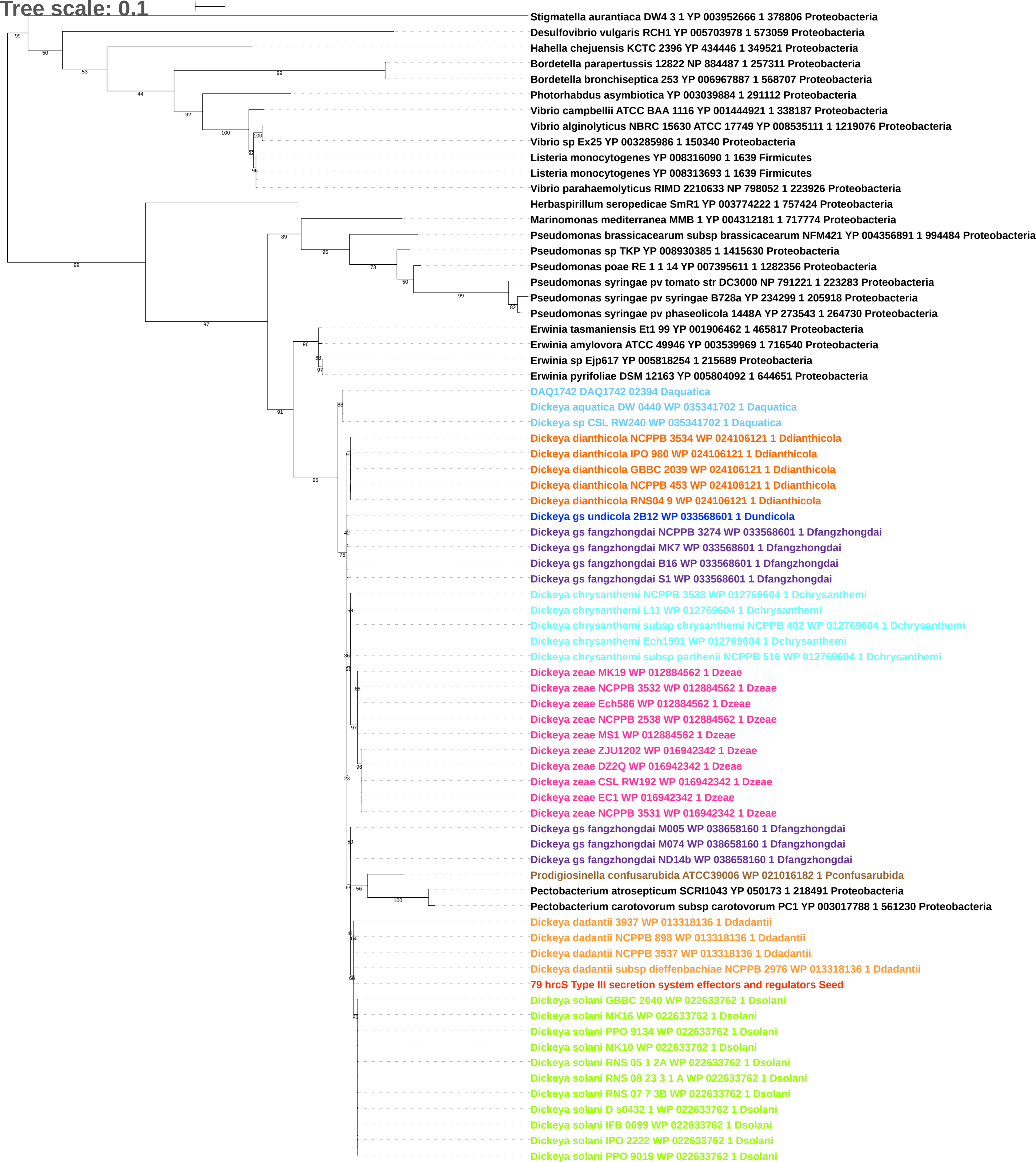
Tree scale: 0.1
Stigmatella aurantiaca DW4 3 1 YP 003952666 1 378806 Proteobacteria
Desulfovibrio vulgaris RCH1 YP 005703978 1 573059 Proteobacteria
Hahella chejuensis KCTC 2396 YP 434446 1 349521 Proteobacteria
Bordetella parapertussis 12822 NP 884487 1 257311 Proteobacteria
Bordetella bronchiseptica 253 YP 006967887 1 568707 Proteobacteria
Photorhabdus asymbiotica YP 003039884 1 291112 Proteobacteria
Vibrio campbellii ATCC BAA 1116 YP 001444921 1 338187 Proteobacteria
Vibrio alginolyticus NBRC 15630 ATCC 17749 YP 008535111 1 1219076 Proteobacteria
Vibrio sp Ex25 YP 003285986 1 150340 Proteobacteria
Listeria monocytogenes YP 008316090 1 1639 Firmicutes
Listeria monocytogenes YP 008313693 1 1639 Firmicutes
Vibrio parahaemolyticus RIMD 2210633 NP 798052 1 223926 Proteobacteria
Herbaspirillum seropedicae SmR1 YP 003774222 1 757424 Proteobacteria
Marinomonas mediterranea MMB 1 YP 004312181 1 717774 Proteobacteria
Pseudomonas brassicacearum subsp brassicacearum NFM421 YP 004356891 1 994484 Proteobacteria
Pseudomonas sp TKP YP 008930385 1 1415630 Proteobacteria
Pseudomonas poae RE 1 1 14 YP 007395611 1 1282356 Proteobacteria
Pseudomonas syringae pv tomato str DC3000 NP 791221 1 223283 Proteobacteria
Pseudomonas syringae pv syringae B728a YP 234299 1 205918 Proteobacteria
Pseudomonas syringae pv phaseolicola 1448A YP 273543 1 264730 Proteobacteria
Erwinia tasmaniensis Et1 99 YP 001906462 1 465817 Proteobacteria
Erwinia amylovora ATCC 49946 YP 003539969 1 716540 Proteobacteria
Erwinia sp Ejp617 YP 005818254 1 215689 Proteobacteria
Erwinia pyrifoliae DSM 12163 YP 005804092 1 644651 Proteobacteria
DAQ1742 DAQ1742 02394 Daquatica
Dickeya aquatica DW 0440 WP 035341702 1 Daquatica
Dickeya sp CSL RW240 WP 035341702 1 Daquatica
Dickeya dianthicola NCPPB 3534 WP 024106121 1 Ddianthicola
Dickeya dianthicola IPO 980 WP 024106121 1 Ddianthicola
Dickeya dianthicola GBBC 2039 WP 024106121 1 Ddianthicola
Dickeya dianthicola NCPPB 453 WP 024106121 1 Ddianthicola
Dickeya dianthicola RNS04 9 WP 024106121 1 Ddianthicola
Dickeya gs undicola 2B12 WP 033568601 1 Dundicola
Dickeya gs fangzhongdai NCPPB 3274 WP 033568601 1 Dfangzhongdai
Dickeya gs fangzhongdai MK7 WP 033568601 1 Dfangzhongdai
Dickeya gs fangzhongdai B16 WP 033568601 1 Dfangzhongdai
Dickeya gs fangzhongdai S1 WP 033568601 1 Dfangzhongdai
Dickeya chrysanthemi NCPPB 3533 WP 012769604 1 Dchrysanthemi
Dickeya chrysanthemi L11 WP 012769604 1 Dchrysanthemi
Dickeya chrysanthemi subsp chrysanthemi NCPPB 402 WP 012769604 1 Dchrysanthemi
Dickeya chrysanthemi Ech1591 WP 012769604 1 Dchrysanthemi
Dickeya chrysanthemi subsp parthenii NCPPB 516 WP 012769604 1 Dchrysanthemi
Dickeya zeae MK19 WP 012884562 1 Dzeae
Dickeya zeae NCPPB 3532 WP 012884562 1 Dzeae
Dickeya zeae Ech586 WP 012884562 1 Dzeae
Dickeya zeae NCPPB 2538 WP 012884562 1 Dzeae
Dickeya zeae MS1 WP 012884562 1 Dzeae
Dickeya zeae ZJU1202 WP 016942342 1 Dzeae
Dickeya zeae DZ2Q WP 016942342 1 Dzeae
Dickeya zeae CSL RW192 WP 016942342 1 Dzeae
Dickeya zeae EC1 WP 016942342 1 Dzeae
Dickeya zeae NCPPB 3531 WP 016942342 1 Dzeae
Dickeya gs fangzhongdai M005 WP 038658160 1 Dfangzhongdai
Dickeya gs fangzhongdai M074 WP 038658160 1 Dfangzhongdai
Dickeya gs fangzhongdai ND14b WP 038658160 1 Dfangzhongdai
Prodigiosinella confusarubida ATCC39006 WP 021016182 1 Pconfusarubida
Pectobacterium atrosepticum SCRI1043 YP 050173 1 218491 Proteobacteria
Pectobacterium carotovorum subsp carotovorum PC1 YP 003017788 1 561230 Proteobacteria
Dickeya dadantii 3937 WP 013318136 1 Ddadantii
Dickeya dadantii NCPPB 898 WP 013318136 1 Ddadantii
Dickeya dadantii NCPPB 3537 WP 013318136 1 Ddadantii
Dickeya dadantii subsp dieffenbachiae NCPPB 2976 WP 013318136 1 Ddadantii
79 hrcS Type III secretion system effectors and regulators Seed
Dickeya solani GBBC 2040 WP 022633762 1 Dsolani
Dickeya solani MK16 WP 022633762 1 Dsolani
Dickeya solani PPO 9134 WP 022633762 1 Dsolani
Dickeya solani MK10 WP 022633762 1 Dsolani
Dickeya solani RNS 05 1 2A WP 022633762 1 Dsolani
Dickeya solani RNS 08 23 3 1 A WP 022633762 1 Dsolani
Dickeya solani RNS 07 7 3B WP 022633762 1 Dsolani
Dickeya solani D s0432 1 WP 022633762 1 Dsolani
Dickeya solani IFB 0099 WP 022633762 1 Dsolani
Dickeya solani IPO 2222 WP 022633762 1 Dsolani
Dickeya solani PPO 9019 WP 022633762 1 Dsolani

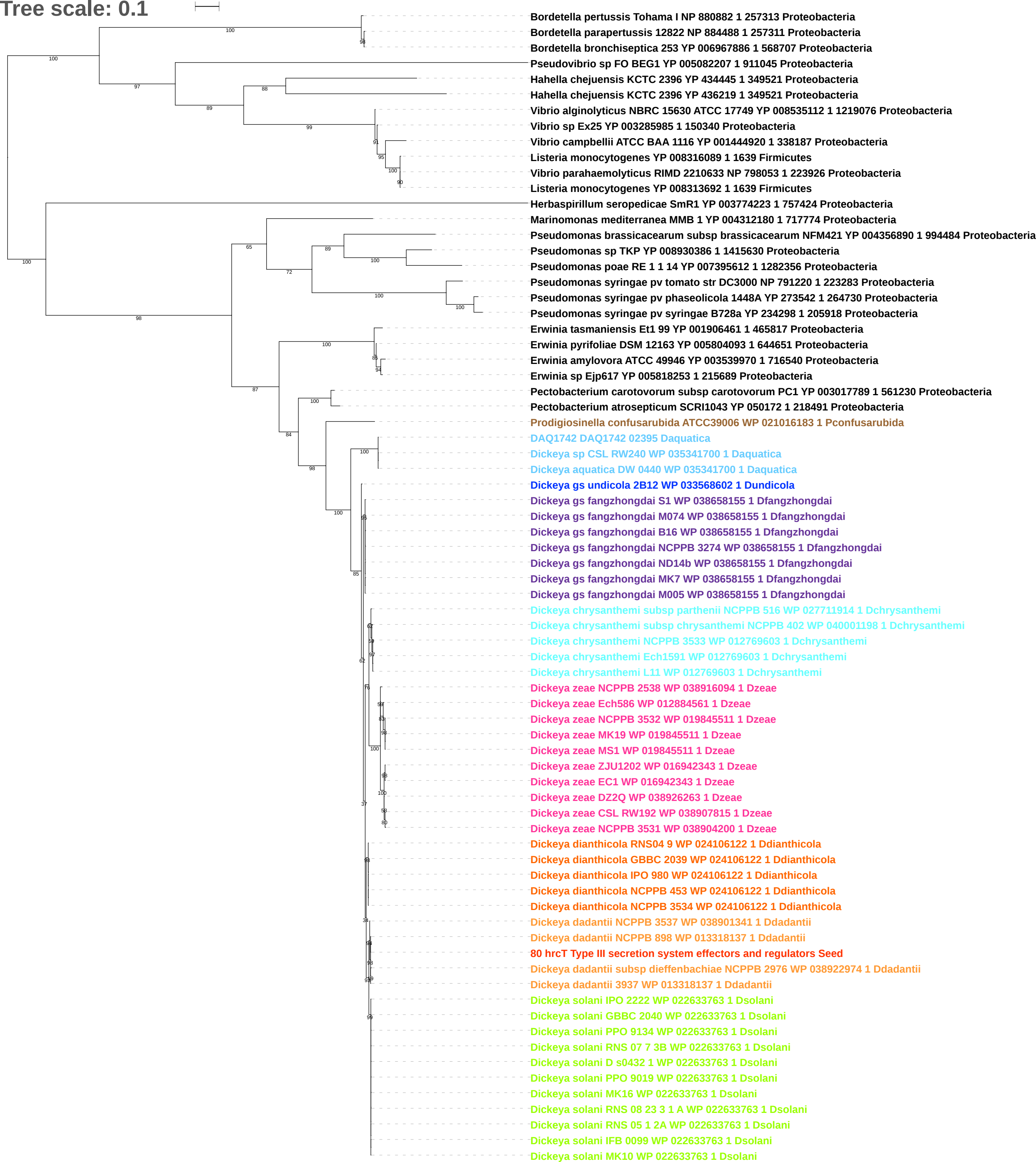
Tree scale: 0.1
Bordetella pertussis Tohama I NP 880882 1 257313 Proteobacteria
Bordetella parapertussis 12822 NP 884488 1 257311 Proteobacteria
Bordetella bronchiseptica 253 YP 006967886 1 568707 Proteobacteria
Pseudovibrio sp FO BEG1 YP 005082207 1 911045 Proteobacteria
Hahella chejuensis KCTC 2396 YP 434445 1 349521 Proteobacteria
Hahella chejuensis KCTC 2396 YP 436219 1 349521 Proteobacteria
Vibrio alginolyticus NBRC 15630 ATCC 17749 YP 008535112 1 1219076 Proteobacteria
Vibrio sp Ex25 YP 003285985 1 150340 Proteobacteria
Vibrio campbellii ATCC BAA 1116 YP 001444920 1 338187 Proteobacteria
Listeria monocytogenes YP 008316089 1 1639 Firmicutes
Vibrio parahaemolyticus RIMD 2210633 NP 798053 1 223926 Proteobacteria
Listeria monocytogenes YP 008313692 1 1639 Firmicutes
Herbaspirillum seropedicae SmR1 YP 003774223 1 757424 Proteobacteria
Marinomonas mediterranea MMB 1 YP 004312180 1 717774 Proteobacteria
Pseudomonas brassicacearum subsp brassicacearum NFM421 YP 004356890 1 994484 Proteobacteria
Pseudomonas sp TKP YP 008930386 1 1415630 Proteobacteria
Pseudomonas poae RE 1 1 14 YP 007395612 1 1282356 Proteobacteria
Pseudomonas syringae pv tomato str DC3000 NP 791220 1 223283 Proteobacteria
Pseudomonas syringae pv phaseolicola 1448A YP 273542 1 264730 Proteobacteria
Pseudomonas syringae pv syringae B728a YP 234298 1 205918 Proteobacteria
Erwinia tasmaniensis Et1 99 YP 001906461 1 465817 Proteobacteria
Erwinia pyrifoliae DSM 12163 YP 005804093 1 644651 Proteobacteria
Erwinia amylovora ATCC 49946 YP 003539970 1 716540 Proteobacteria
Erwinia sp Ejp617 YP 005818253 1 215689 Proteobacteria
Pectobacterium carotovorum subsp carotovorum PC1 YP 003017789 1 561230 Proteobacteria
Pectobacterium atrosepticum SCRI1043 YP 050172 1 218491 Proteobacteria
Prodigiosinella confusarubida ATCC39006 WP 021016183 1 Pconfusarubida
DAQ1742 DAQ1742 02395 Daquatica
Dickeya sp CSL RW240 WP 035341700 1 Daquatica
Dickeya aquatica DW 0440 WP 035341700 1 Daquatica
Dickeya gs undicola 2B12 WP 033568602 1 Dundicola
Dickeya gs fangzhongdai S1 WP 038658155 1 Dfangzhongdai
Dickeya gs fangzhongdai M074 WP 038658155 1 Dfangzhongdai
Dickeya gs fangzhongdai B16 WP 038658155 1 Dfangzhongdai
Dickeya gs fangzhongdai NCPPB 3274 WP 038658155 1 Dfangzhongdai
Dickeya gs fangzhongdai ND14b WP 038658155 1 Dfangzhongdai
Dickeya gs fangzhongdai MK7 WP 038658155 1 Dfangzhongdai
Dickeya gs fangzhongdai M005 WP 038658155 1 Dfangzhongdai
Dickeya chrysanthemi subsp parthenii NCPPB 516 WP 027711914 1 Dchrysanthemi
Dickeya chrysanthemi subsp chrysanthemi NCPPB 402 WP 040001198 1 Dchrysanthemi
Dickeya chrysanthemi NCPPB 3533 WP 012769603 1 Dchrysanthemi
Dickeya chrysanthemi Ech1591 WP 012769603 1 Dchrysanthemi
Dickeya chrysanthemi L11 WP 012769603 1 Dchrysanthemi
Dickeya zeae NCPPB 2538 WP 038916094 1 Dzeae
Dickeya zeae Ech586 WP 012884561 1 Dzeae
Dickeya zeae NCPPB 3532 WP 019845511 1 Dzeae
Dickeya zeae MK19 WP 019845511 1 Dzeae
Dickeya zeae MS1 WP 019845511 1 Dzeae
Dickeya zeae ZJU1202 WP 016942343 1 Dzeae
Dickeya zeae EC1 WP 016942343 1 Dzeae
Dickeya zeae DZ2Q WP 038926263 1 Dzeae
Dickeya zeae CSL RW192 WP 038907815 1 Dzeae
Dickeya zeae NCPPB 3531 WP 038904200 1 Dzeae
Dickeya dianthicola RNS04 9 WP 024106122 1 Ddianthicola
Dickeya dianthicola GBBC 2039 WP 024106122 1 Ddianthicola
Dickeya dianthicola IPO 980 WP 024106122 1 Ddianthicola
Dickeya dianthicola NCPPB 453 WP 024106122 1 Ddianthicola
Dickeya dianthicola NCPPB 3534 WP 024106122 1 Ddianthicola
Dickeya dadantii NCPPB 3537 WP 038901341 1 Ddadantii
Dickeya dadantii NCPPB 898 WP 013318137 1 Ddadantii
80 hrcT Type III secretion system effectors and regulators Seed
Dickeya dadantii subsp dieffenbachiae NCPPB 2976 WP 038922974 1 Ddadantii
Dickeya dadantii 3937 WP 013318137 1 Ddadantii
Dickeya solani IPO 2222 WP 022633763 1 Dsolani
Dickeya solani GBBC 2040 WP 022633763 1 Dsolani
Dickeya solani PPO 9134 WP 022633763 1 Dsolani
Dickeya solani RNS 07 7 3B WP 022633763 1 Dsolani
Dickeya solani D s0432 1 WP 022633763 1 Dsolani
Dickeya solani PPO 9019 WP 022633763 1 Dsolani
Dickeya solani MK16 WP 022633763 1 Dsolani
Dickeya solani RNS 08 23 3 1 A WP 022633763 1 Dsolani
Dickeya solani RNS 05 1 2A WP 022633763 1 Dsolani
Dickeya solani IFB 0099 WP 022633763 1 Dsolani
Dickeya solani MK10 WP 022633763 1 Dsolani

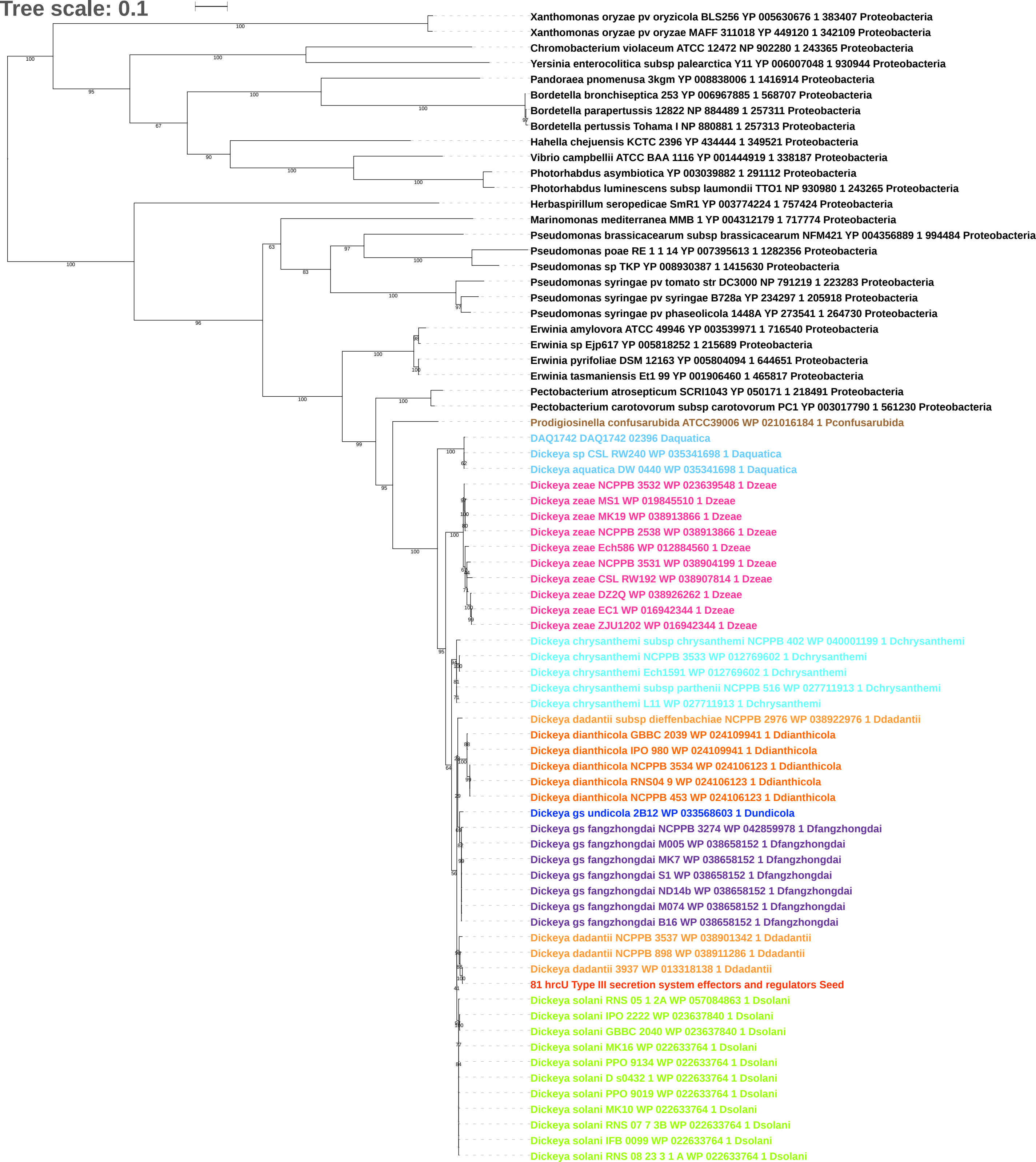

Tree scale: 0.1
Xanthomonas oryzae pv oryzicola BLS256 YP 005630676 1 383407 Proteobacteria
Xanthomonas oryzae pv oryzae MAFF 311018 YP 449120 1 342109 Proteobacteria
Chromobacterium violaceum ATCC 12472 NP 902280 1 243365 Proteobacteria
Yersinia enterocolitica subsp palearctica Y11 YP 006007048 1 930944 Proteobacteria
Pandoraea pnomenusa 3kgm YP 008838006 1 1416914 Proteobacteria
Bordetella bronchiseptica 253 YP 006967885 1 568707 Proteobacteria
Bordetella parapertussis 12822 NP 884489 1 257311 Proteobacteria
Bordetella pertussis Tohama I NP 880881 1 257313 Proteobacteria
Hahella chejuensis KCTC 2396 YP 434444 1 349521 Proteobacteria
Vibrio campbellii ATCC BAA 1116 YP 001444919 1 338187 Proteobacteria
Photorhabdus asymbiotica YP 003039882 1 291112 Proteobacteria
Photorhabdus luminescens subsp laumondii TTO1 NP 930980 1 243265 Proteobacteria
Herbaspirillum seropedicae SmR1 YP 003774224 1 757424 Proteobacteria
Marinomonas mediterranea MMB 1 YP 004312179 1 717774 Proteobacteria
Pseudomonas brassicacearum subsp brassicacearum NFM421 YP 004356889 1 994484 Proteobacteria
Pseudomonas poae RE 1 1 14 YP 007395613 1 1282356 Proteobacteria
Pseudomonas sp TKP YP 008930387 1 1415630 Proteobacteria
Pseudomonas syringae pv tomato str DC3000 NP 791219 1 223283 Proteobacteria
Pseudomonas syringae pv syringae B728a YP 234297 1 205918 Proteobacteria
Pseudomonas syringae pv phaseolicola 1448A YP 273541 1 264730 Proteobacteria
Erwinia amylovora ATCC 49946 YP 003539971 1 716540 Proteobacteria
Erwinia sp Ejp617 YP 005818252 1 215689 Proteobacteria
Erwinia pyrifoliae DSM 12163 YP 005804094 1 644651 Proteobacteria
Erwinia tasmaniensis Et1 99 YP 001906460 1 465817 Proteobacteria
Pectobacterium atrosepticum SCRI1043 YP 050171 1 218491 Proteobacteria
Pectobacterium carotovorum subsp carotovorum PC1 YP 003017790 1 561230 Proteobacteria
Prodigiosinella confusarubida ATCC39006 WP 021016184 1 Pconfusarubida
DAQ1742 DAQ1742 02396 Daquatica
Dickeya sp CSL RW240 WP 035341698 1 Daquatica
Dickeya aquatica DW 0440 WP 035341698 1 Daquatica
Dickeya zeae NCPPB 3532 WP 023639548 1 Dzeae
Dickeya zeae MS1 WP 019845510 1 Dzeae
Dickeya zeae MK19 WP 038913866 1 Dzeae
Dickeya zeae NCPPB 2538 WP 038913866 1 Dzeae
Dickeya zeae Ech586 WP 012884560 1 Dzeae
Dickeya zeae NCPPB 3531 WP 038904199 1 Dzeae
Dickeya zeae CSL RW192 WP 038907814 1 Dzeae
Dickeya zeae DZ2Q WP 038926262 1 Dzeae
Dickeya zeae EC1 WP 016942344 1 Dzeae
Dickeya zeae ZJU1202 WP 016942344 1 Dzeae
Dickeya chrysanthemi subsp chrysanthemi NCPPB 402 WP 040001199 1 Dchrysanthemi
Dickeya chrysanthemi NCPPB 3533 WP 012769602 1 Dchrysanthemi
Dickeya chrysanthemi Ech1591 WP 012769602 1 Dchrysanthemi
Dickeya chrysanthemi subsp parthenii NCPPB 516 WP 027711913 1 Dchrysanthemi
Dickeya chrysanthemi L11 WP 027711913 1 Dchrysanthemi
Dickeya dadantii subsp dieffenbachiae NCPPB 2976 WP 038922976 1 Ddadantii
Dickeya dianthicola GBBC 2039 WP 024109941 1 Ddianthicola
Dickeya dianthicola IPO 980 WP 024109941 1 Ddianthicola
Dickeya dianthicola NCPPB 3534 WP 024106123 1 Ddianthicola
Dickeya dianthicola RNS04 9 WP 024106123 1 Ddianthicola
Dickeya dianthicola NCPPB 453 WP 024106123 1 Ddianthicola
Dickeya gs undicola 2B12 WP 033568603 1 Dundicola
Dickeya gs fangzhongdai NCPPB 3274 WP 042859978 1 Dfangzhongdai
Dickeya gs fangzhongdai M005 WP 038658152 1 Dfangzhongdai
Dickeya gs fangzhongdai MK7 WP 038658152 1 Dfangzhongdai
Dickeya gs fangzhongdai S1 WP 038658152 1 Dfangzhongdai
Dickeya gs fangzhongdai ND14b WP 038658152 1 Dfangzhongdai
Dickeya gs fangzhongdai M074 WP 038658152 1 Dfangzhongdai
Dickeya gs fangzhongdai B16 WP 038658152 1 Dfangzhongdai
Dickeya dadantii NCPPB 3537 WP 038901342 1 Ddadantii
Dickeya dadantii NCPPB 898 WP 038911286 1 Ddadantii
Dickeya dadantii 3937 WP 013318138 1 Ddadantii
81 hrcU Type III secretion system effectors and regulators Seed
Dickeya solani RNS 05 1 2A WP 057084863 1 Dsolani
Dickeya solani IPO 2222 WP 023637840 1 Dsolani
Dickeya solani GBBC 2040 WP 023637840 1 Dsolani
Dickeya solani MK16 WP 022633764 1 Dsolani
Dickeya solani PPO 9134 WP 022633764 1 Dsolani
Dickeya solani D s0432 1 WP 022633764 1 Dsolani
Dickeya solani PPO 9019 WP 022633764 1 Dsolani
Dickeya solani MK10 WP 022633764 1 Dsolani
Dickeya solani RNS 07 7 3B WP 022633764 1 Dsolani
Dickeya solani IFB 0099 WP 022633764 1 Dsolani
Dickeya solani RNS 08 23 3 1 A WP 022633764 1 Dsolani

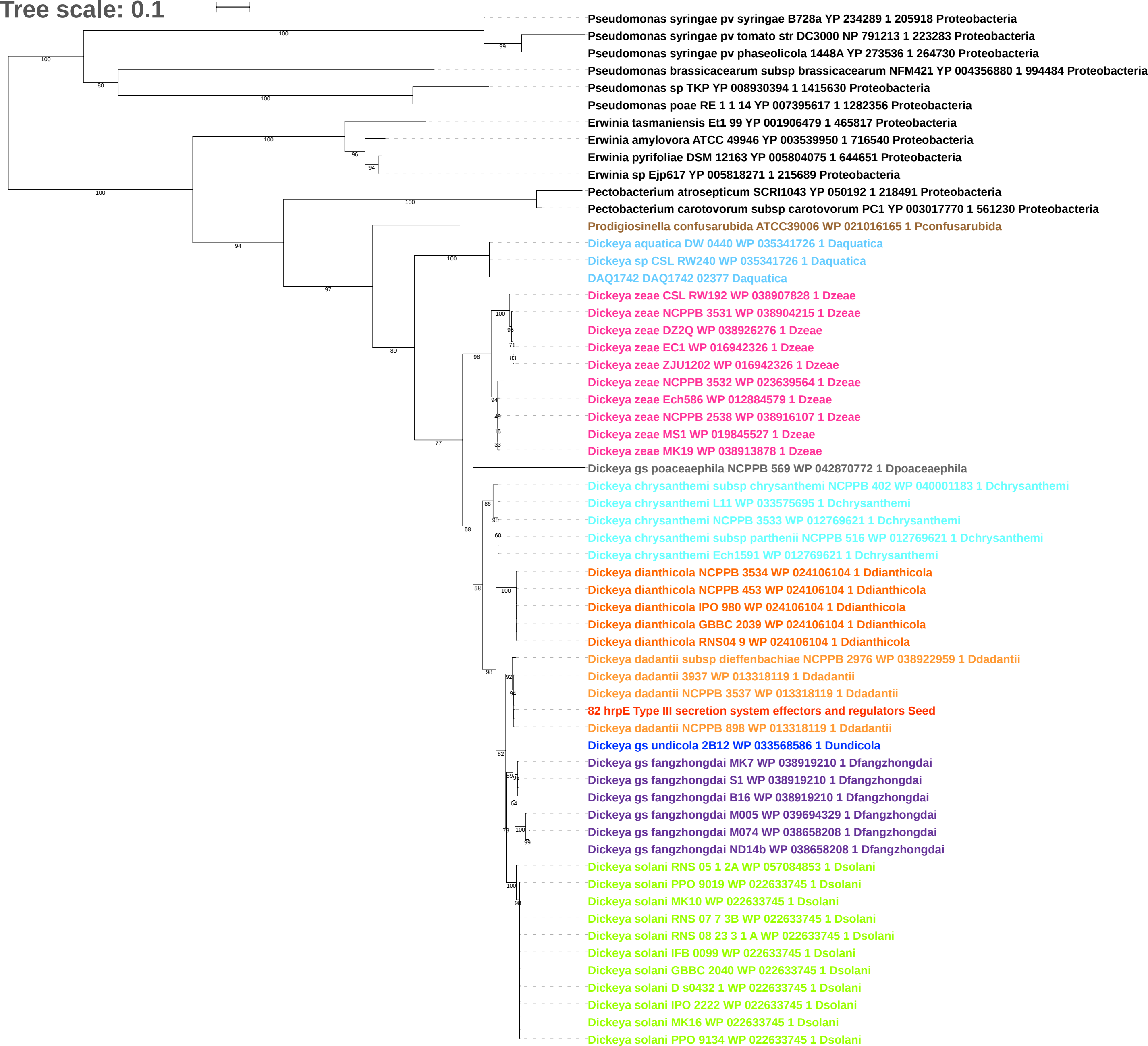
Tree scale: 0.1
Pseudomonas syringae pv syringae B728a YP 234289 1 205918 Proteobacteria
Pseudomonas syringae pv tomato str DC3000 NP 791213 1 223283 Proteobacteria
Pseudomonas syringae pv phaseolicola 1448A YP 273536 1 264730 Proteobacteria
Pseudomonas brassicacearum subsp brassicacearum NFM421 YP 004356880 1 994484 Proteobacteria
Pseudomonas sp TKP YP 008930394 1 1415630 Proteobacteria
Pseudomonas poae RE 1 1 14 YP 007395617 1 1282356 Proteobacteria
Erwinia tasmaniensis Et1 99 YP 001906479 1 465817 Proteobacteria
Erwinia amylovora ATCC 49946 YP 003539950 1 716540 Proteobacteria
Erwinia pyrifoliae DSM 12163 YP 005804075 1 644651 Proteobacteria
Erwinia sp Ejp617 YP 005818271 1 215689 Proteobacteria
Pectobacterium atrosepticum SCRI1043 YP 050192 1 218491 Proteobacteria
Pectobacterium carotovorum subsp carotovorum PC1 YP 003017770 1 561230 Proteobacteria
Prodigiosinella confusarubida ATCC39006 WP 021016165 1 Pconfusarubida
Dickeya aquatica DW 0440 WP 035341726 1 Daquatica
Dickeya sp CSL RW240 WP 035341726 1 Daquatica
DAQ1742 DAQ1742 02377 Daquatica
Dickeya zeae CSL RW192 WP 038907828 1 Dzeae
Dickeya zeae NCPPB 3531 WP 038904215 1 Dzeae
Dickeya zeae DZ2Q WP 038926276 1 Dzeae
Dickeya zeae EC1 WP 016942326 1 Dzeae
Dickeya zeae ZJU1202 WP 016942326 1 Dzeae
Dickeya zeae NCPPB 3532 WP 023639564 1 Dzeae
Dickeya zeae Ech586 WP 012884579 1 Dzeae
Dickeya zeae NCPPB 2538 WP 038916107 1 Dzeae
Dickeya zeae MS1 WP 019845527 1 Dzeae
Dickeya zeae MK19 WP 038913878 1 Dzeae
Dickeya gs poaceaephila NCPPB 569 WP 042870772 1 Dpoaceaephila
Dickeya chrysanthemi subsp chrysanthemi NCPPB 402 WP 040001183 1 Dchrysanthemi
Dickeya chrysanthemi L11 WP 033575695 1 Dchrysanthemi
Dickeya chrysanthemi NCPPB 3533 WP 012769621 1 Dchrysanthemi
Dickeya chrysanthemi subsp parthenii NCPPB 516 WP 012769621 1 Dchrysanthemi
Dickeya chrysanthemi Ech1591 WP 012769621 1 Dchrysanthemi
Dickeya dianthicola NCPPB 3534 WP 024106104 1 Ddianthicola
Dickeya dianthicola NCPPB 453 WP 024106104 1 Ddianthicola
Dickeya dianthicola IPO 980 WP 024106104 1 Ddianthicola
Dickeya dianthicola GBBC 2039 WP 024106104 1 Ddianthicola
Dickeya dianthicola RNS04 9 WP 024106104 1 Ddianthicola
Dickeya dadantii subsp dieffenbachiae NCPPB 2976 WP 038922959 1 Ddadantii
Dickeya dadantii 3937 WP 013318119 1 Ddadantii
Dickeya dadantii NCPPB 3537 WP 013318119 1 Ddadantii
82 hrpE Type III secretion system effectors and regulators Seed
Dickeya dadantii NCPPB 898 WP 013318119 1 Ddadantii
Dickeya gs undicola 2B12 WP 033568586 1 Dundicola
Dickeya gs fangzhongdai MK7 WP 038919210 1 Dfangzhongdai
Dickeya gs fangzhongdai S1 WP 038919210 1 Dfangzhongdai
Dickeya gs fangzhongdai B16 WP 038919210 1 Dfangzhongdai
Dickeya gs fangzhongdai M005 WP 039694329 1 Dfangzhongdai
Dickeya gs fangzhongdai M074 WP 038658208 1 Dfangzhongdai
Dickeya gs fangzhongdai ND14b WP 038658208 1 Dfangzhongdai
Dickeya solani RNS 05 1 2A WP 057084853 1 Dsolani
Dickeya solani PPO 9019 WP 022633745 1 Dsolani
Dickeya solani MK10 WP 022633745 1 Dsolani
Dickeya solani RNS 07 7 3B WP 022633745 1 Dsolani
Dickeya solani RNS 08 23 3 1 A WP 022633745 1 Dsolani
Dickeya solani IFB 0099 WP 022633745 1 Dsolani
Dickeya solani GBBC 2040 WP 022633745 1 Dsolani
Dickeya solani D s0432 1 WP 022633745 1 Dsolani
Dickeya solani IPO 2222 WP 022633745 1 Dsolani
Dickeya solani MK16 WP 022633745 1 Dsolani
Dickeya solani PPO 9134 WP 022633745 1 Dsolani

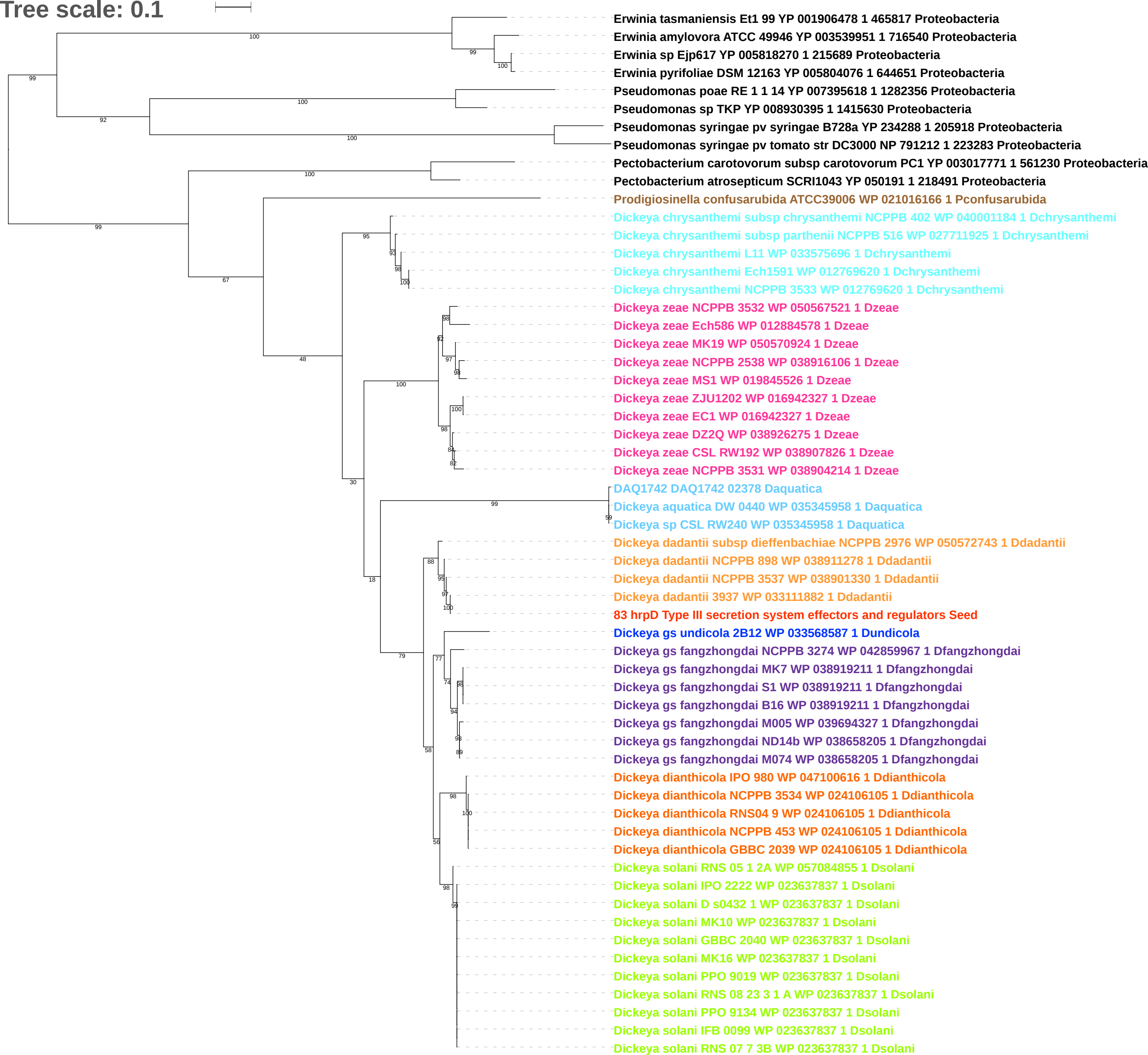
Tree scale: 0.1
Erwinia tasmaniensis Et1 99 YP 001906478 1 465817 Proteobacteria
Erwinia amylovora ATCC 49946 YP 003539951 1 716540 Proteobacteria
Erwinia sp Ejp617 YP 005818270 1 215689 Proteobacteria
Erwinia pyrifoliae DSM 12163 YP 005804076 1 644651 Proteobacteria
Pseudomonas poae RE 1 1 14 YP 007395618 1 1282356 Proteobacteria
Pseudomonas sp TKP YP 008930395 1 1415630 Proteobacteria
Pseudomonas syringae pv syringae B728a YP 234288 1 205918 Proteobacteria
Pseudomonas syringae pv tomato str DC3000 NP 791212 1 223283 Proteobacteria
Pectobacterium carotovorum subsp carotovorum PC1 YP 003017771 1 561230 Proteobacteria
Pectobacterium atrosepticum SCRI1043 YP 050191 1 218491 Proteobacteria
Prodigiosinella confusarubida ATCC39006 WP 021016166 1 Pconfusarubida
Dickeya chrysanthemi subsp chrysanthemi NCPPB 402 WP 040001184 1 Dchrysanthemi
Dickeya chrysanthemi subsp parthenii NCPPB 516 WP 027711925 1 Dchrysanthemi
Dickeya chrysanthemi L11 WP 033575696 1 Dchrysanthemi
Dickeya chrysanthemi Ech1591 WP 012769620 1 Dchrysanthemi
Dickeya chrysanthemi NCPPB 3533 WP 012769620 1 Dchrysanthemi
Dickeya zeae NCPPB 3532 WP 050567521 1 Dzeae
Dickeya zeae Ech586 WP 012884578 1 Dzeae
Dickeya zeae MK19 WP 050570924 1 Dzeae
Dickeya zeae NCPPB 2538 WP 038916106 1 Dzeae
Dickeya zeae MS1 WP 019845526 1 Dzeae
Dickeya zeae ZJU1202 WP 016942327 1 Dzeae
Dickeya zeae EC1 WP 016942327 1 Dzeae
Dickeya zeae DZ2Q WP 038926275 1 Dzeae
Dickeya zeae CSL RW192 WP 038907826 1 Dzeae
Dickeya zeae NCPPB 3531 WP 038904214 1 Dzeae
DAQ1742 DAQ1742 02378 Daquatica
Dickeya aquatica DW 0440 WP 035345958 1 Daquatica
Dickeya sp CSL RW240 WP 035345958 1 Daquatica
Dickeya dadantii subsp dieffenbachiae NCPPB 2976 WP 050572743 1 Ddadantii
Dickeya dadantii NCPPB 898 WP 038911278 1 Ddadantii
Dickeya dadantii NCPPB 3537 WP 038901330 1 Ddadantii
Dickeya dadantii 3937 WP 033111882 1 Ddadantii
83 hrpD Type III secretion system effectors and regulators Seed
Dickeya gs undicola 2B12 WP 033568587 1 Dundicola
Dickeya gs fangzhongdai NCPPB 3274 WP 042859967 1 Dfangzhongdai
Dickeya gs fangzhongdai MK7 WP 038919211 1 Dfangzhongdai
Dickeya gs fangzhongdai S1 WP 038919211 1 Dfangzhongdai
Dickeya gs fangzhongdai B16 WP 038919211 1 Dfangzhongdai
Dickeya gs fangzhongdai M005 WP 039694327 1 Dfangzhongdai
Dickeya gs fangzhongdai ND14b WP 038658205 1 Dfangzhongdai
Dickeya gs fangzhongdai M074 WP 038658205 1 Dfangzhongdai
Dickeya dianthicola IPO 980 WP 047100616 1 Ddianthicola
Dickeya dianthicola NCPPB 3534 WP 024106105 1 Ddianthicola
Dickeya dianthicola RNS04 9 WP 024106105 1 Ddianthicola
Dickeya dianthicola NCPPB 453 WP 024106105 1 Ddianthicola
Dickeya dianthicola GBBC 2039 WP 024106105 1 Ddianthicola
Dickeya solani RNS 05 1 2A WP 057084855 1 Dsolani
Dickeya solani IPO 2222 WP 023637837 1 Dsolani
Dickeya solani D s0432 1 WP 023637837 1 Dsolani
Dickeya solani MK10 WP 023637837 1 Dsolani
Dickeya solani GBBC 2040 WP 023637837 1 Dsolani
Dickeya solani MK16 WP 023637837 1 Dsolani
Dickeya solani PPO 9019 WP 023637837 1 Dsolani
Dickeya solani RNS 08 23 3 1 A WP 023637837 1 Dsolani
Dickeya solani PPO 9134 WP 023637837 1 Dsolani
Dickeya solani IFB 0099 WP 023637837 1 Dsolani
Dickeya solani RNS 07 7 3B WP 023637837 1 Dsolani

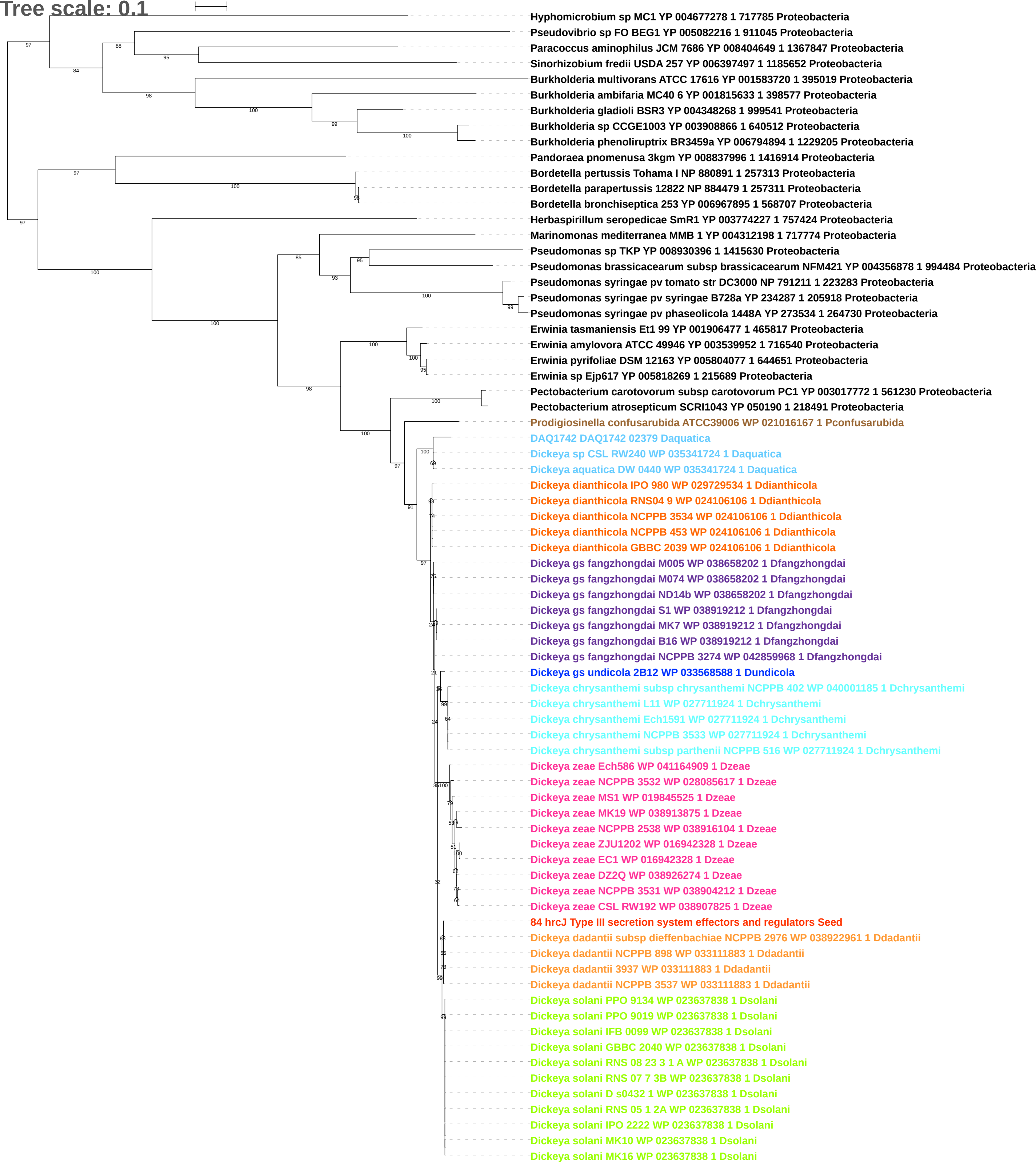
Tree scale: 0.1
Hyphomicrobium sp MC1 YP 004677278 1 717785 Proteobacteria
Pseudovibrio sp FO BEG1 YP 005082216 1 911045 Proteobacteria
Paracoccus aminophilus JCM 7686 YP 008404649 1 1367847 Proteobacteria
Sinorhizobium fredii USDA 257 YP 006397497 1 1185652 Proteobacteria
Burkholderia multivorans ATCC 17616 YP 001583720 1 395019 Proteobacteria
Burkholderia ambifaria MC40 6 YP 001815633 1 398577 Proteobacteria
Burkholderia gladioli BSR3 YP 004348268 1 999541 Proteobacteria
Burkholderia sp CCGE1003 YP 003908866 1 640512 Proteobacteria
Burkholderia phenoliruptrix BR3459a YP 006794894 1 1229205 Proteobacteria
Pandoraea pnomenusa 3kgm YP 008837996 1 1416914 Proteobacteria
Bordetella pertussis Tohama I NP 880891 1 257313 Proteobacteria
Bordetella parapertussis 12822 NP 884479 1 257311 Proteobacteria
Bordetella bronchiseptica 253 YP 006967895 1 568707 Proteobacteria
Herbaspirillum seropedicae SmR1 YP 003774227 1 757424 Proteobacteria
Marinomonas mediterranea MMB 1 YP 004312198 1 717774 Proteobacteria
Pseudomonas sp TKP YP 008930396 1 1415630 Proteobacteria
Pseudomonas brassicacearum subsp brassicacearum NFM421 YP 004356878 1 994484 Proteobacteria
Pseudomonas syringae pv tomato str DC3000 NP 791211 1 223283 Proteobacteria
Pseudomonas syringae pv syringae B728a YP 234287 1 205918 Proteobacteria
Pseudomonas syringae pv phaseolicola 1448A YP 273534 1 264730 Proteobacteria
Erwinia tasmaniensis Et1 99 YP 001906477 1 465817 Proteobacteria
Erwinia amylovora ATCC 49946 YP 003539952 1 716540 Proteobacteria
Erwinia pyrifoliae DSM 12163 YP 005804077 1 644651 Proteobacteria
Erwinia sp Ejp617 YP 005818269 1 215689 Proteobacteria
Pectobacterium carotovorum subsp carotovorum PC1 YP 003017772 1 561230 Proteobacteria
Pectobacterium atrosepticum SCRI1043 YP 050190 1 218491 Proteobacteria
Prodigiosinella confusarubida ATCC39006 WP 021016167 1 Pconfusarubida
DAQ1742 DAQ1742 02379 Daquatica
Dickeya sp CSL RW240 WP 035341724 1 Daquatica
Dickeya aquatica DW 0440 WP 035341724 1 Daquatica
Dickeya dianthicola IPO 980 WP 029729534 1 Ddianthicola
Dickeya dianthicola RNS04 9 WP 024106106 1 Ddianthicola
Dickeya dianthicola NCPPB 3534 WP 024106106 1 Ddianthicola
Dickeya dianthicola NCPPB 453 WP 024106106 1 Ddianthicola
Dickeya dianthicola GBBC 2039 WP 024106106 1 Ddianthicola
Dickeya gs fangzhongdai M005 WP 038658202 1 Dfangzhongdai
Dickeya gs fangzhongdai M074 WP 038658202 1 Dfangzhongdai
Dickeya gs fangzhongdai ND14b WP 038658202 1 Dfangzhongdai
Dickeya gs fangzhongdai S1 WP 038919212 1 Dfangzhongdai
Dickeya gs fangzhongdai MK7 WP 038919212 1 Dfangzhongdai
Dickeya gs fangzhongdai B16 WP 038919212 1 Dfangzhongdai
Dickeya gs fangzhongdai NCPPB 3274 WP 042859968 1 Dfangzhongdai
Dickeya gs undicola 2B12 WP 033568588 1 Dundicola
Dickeya chrysanthemi subsp chrysanthemi NCPPB 402 WP 040001185 1 Dchrysanthemi
Dickeya chrysanthemi L11 WP 027711924 1 Dchrysanthemi
Dickeya chrysanthemi Ech1591 WP 027711924 1 Dchrysanthemi
Dickeya chrysanthemi NCPPB 3533 WP 027711924 1 Dchrysanthemi
Dickeya chrysanthemi subsp parthenii NCPPB 516 WP 027711924 1 Dchrysanthemi
Dickeya zeae Ech586 WP 041164909 1 Dzeae
Dickeya zeae NCPPB 3532 WP 028085617 1 Dzeae
Dickeya zeae MS1 WP 019845525 1 Dzeae
Dickeya zeae MK19 WP 038913875 1 Dzeae
Dickeya zeae NCPPB 2538 WP 038916104 1 Dzeae
Dickeya zeae ZJU1202 WP 016942328 1 Dzeae
Dickeya zeae EC1 WP 016942328 1 Dzeae
Dickeya zeae DZ2Q WP 038926274 1 Dzeae
Dickeya zeae NCPPB 3531 WP 038904212 1 Dzeae
Dickeya zeae CSL RW192 WP 038907825 1 Dzeae
84 hrcJ Type III secretion system effectors and regulators Seed
Dickeya dadantii subsp dieffenbachiae NCPPB 2976 WP 038922961 1 Ddadantii
Dickeya dadantii NCPPB 898 WP 033111883 1 Ddadantii
Dickeya dadantii 3937 WP 033111883 1 Ddadantii
Dickeya dadantii NCPPB 3537 WP 033111883 1 Ddadantii
Dickeya solani PPO 9134 WP 023637838 1 Dsolani
Dickeya solani PPO 9019 WP 023637838 1 Dsolani
Dickeya solani IFB 0099 WP 023637838 1 Dsolani
Dickeya solani GBBC 2040 WP 023637838 1 Dsolani
Dickeya solani RNS 08 23 3 1 A WP 023637838 1 Dsolani
Dickeya solani RNS 07 7 3B WP 023637838 1 Dsolani
Dickeya solani D s0432 1 WP 023637838 1 Dsolani
Dickeya solani RNS 05 1 2A WP 023637838 1 Dsolani
Dickeya solani IPO 2222 WP 023637838 1 Dsolani
Dickeya solani MK10 WP 023637838 1 Dsolani
Dickeya solani MK16 WP 023637838 1 Dsolani

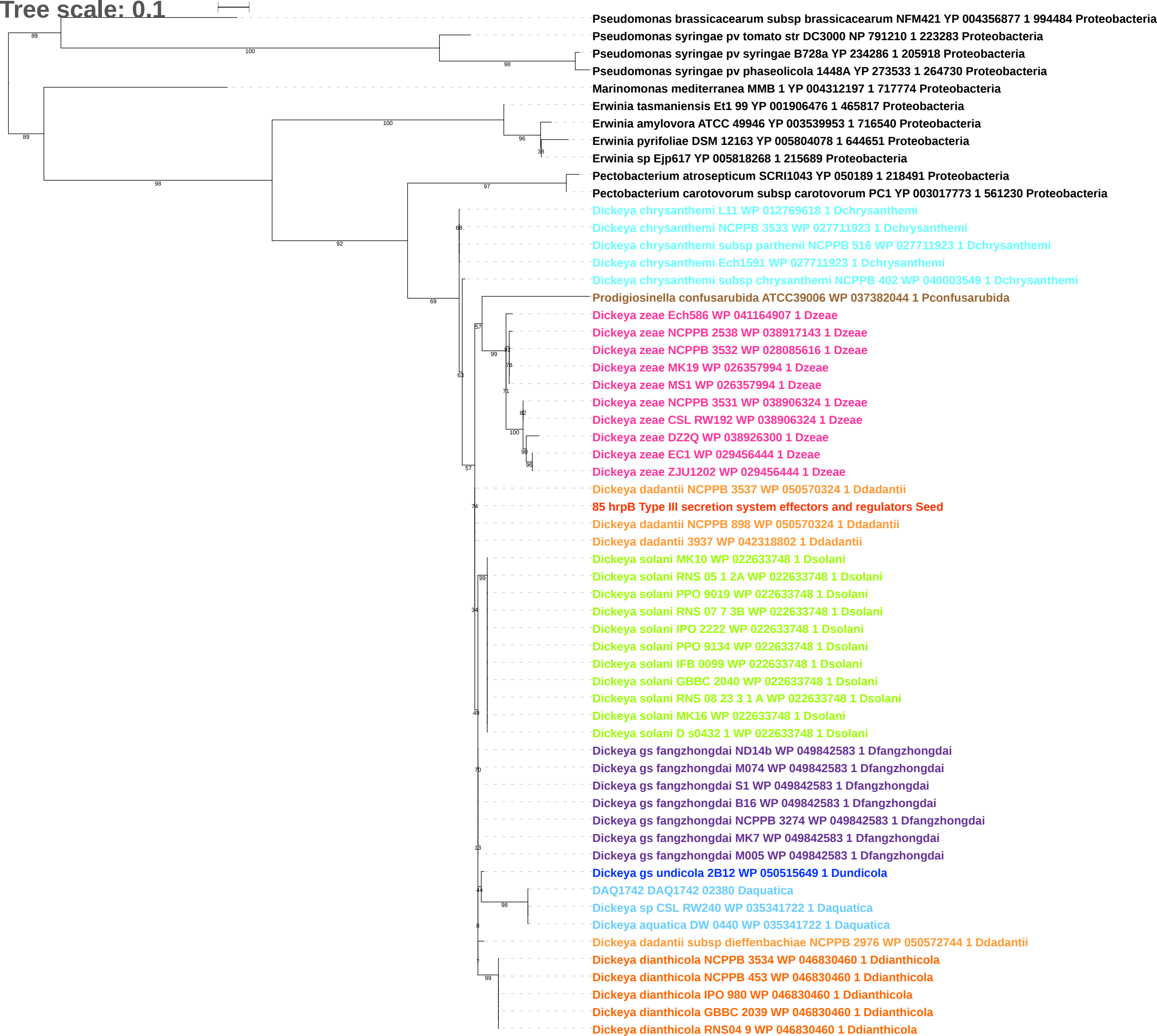
Tree scale: 0.1
Pseudomonas brassicacearum subsp brassicacearum NFM421 YP 004356877 1 994484 Proteobacteria
Pseudomonas syringae pv tomato str DC3000 NP 791210 1 223283 Proteobacteria
Pseudomonas syringae pv syringae B728a YP 234286 1 205918 Proteobacteria
Pseudomonas syringae pv phaseolicola 1448A YP 273533 1 264730 Proteobacteria
Marinomonas mediterranea MMB 1 YP 004312197 1 717774 Proteobacteria
Erwinia tasmaniensis Et1 99 YP 001906476 1 465817 Proteobacteria
Erwinia amylovora ATCC 49946 YP 003539953 1 716540 Proteobacteria
Erwinia pyrifoliae DSM 12163 YP 005804078 1 644651 Proteobacteria
Erwinia sp Ejp617 YP 005818268 1 215689 Proteobacteria
Pectobacterium atrosepticum SCRI1043 YP 050189 1 218491 Proteobacteria
Pectobacterium carotovorum subsp carotovorum PC1 YP 003017773 1 561230 Proteobacteria
Dickeya chrysanthemi L11 WP 012769618 1 Dchrysanthemi
Dickeya chrysanthemi NCPPB 3533 WP 027711923 1 Dchrysanthemi
Dickeya chrysanthemi subsp parthenii NCPPB 516 WP 027711923 1 Dchrysanthemi
Dickeya chrysanthemi Ech1591 WP 027711923 1 Dchrysanthemi
Dickeya chrysanthemi subsp chrysanthemi NCPPB 402 WP 040003549 1 Dchrysanthemi
Prodigiosinella confusarubida ATCC39006 WP 037382044 1 Pconfusarubida
Dickeya zeae Ech586 WP 041164907 1 Dzeae
Dickeya zeae NCPPB 2538 WP 038917143 1 Dzeae
Dickeya zeae NCPPB 3532 WP 028085616 1 Dzeae
Dickeya zeae MK19 WP 026357994 1 Dzeae
Dickeya zeae MS1 WP 026357994 1 Dzeae
Dickeya zeae NCPPB 3531 WP 038906324 1 Dzeae
Dickeya zeae CSL RW192 WP 038906324 1 Dzeae
Dickeya zeae DZ2Q WP 038926300 1 Dzeae
Dickeya zeae EC1 WP 029456444 1 Dzeae
Dickeya zeae ZJU1202 WP 029456444 1 Dzeae
Dickeya dadantii NCPPB 3537 WP 050570324 1 Ddadantii
85 hrpB Type III secretion system effectors and regulators Seed
Dickeya dadantii NCPPB 898 WP 050570324 1 Ddadantii
Dickeya dadantii 3937 WP 042318802 1 Ddadantii
Dickeya solani MK10 WP 022633748 1 Dsolani
Dickeya solani RNS 05 1 2A WP 022633748 1 Dsolani
Dickeya solani PPO 9019 WP 022633748 1 Dsolani
Dickeya solani RNS 07 7 3B WP 022633748 1 Dsolani
Dickeya solani IPO 2222 WP 022633748 1 Dsolani
Dickeya solani PPO 9134 WP 022633748 1 Dsolani
Dickeya solani IFB 0099 WP 022633748 1 Dsolani
Dickeya solani GBBC 2040 WP 022633748 1 Dsolani
Dickeya solani RNS 08 23 3 1 A WP 022633748 1 Dsolani
Dickeya solani MK16 WP 022633748 1 Dsolani
Dickeya solani D s0432 1 WP 022633748 1 Dsolani
Dickeya gs fangzhongdai ND14b WP 049842583 1 Dfangzhongdai
Dickeya gs fangzhongdai M074 WP 049842583 1 Dfangzhongdai
Dickeya gs fangzhongdai S1 WP 049842583 1 Dfangzhongdai
Dickeya gs fangzhongdai B16 WP 049842583 1 Dfangzhongdai
Dickeya gs fangzhongdai NCPPB 3274 WP 049842583 1 Dfangzhongdai
Dickeya gs fangzhongdai MK7 WP 049842583 1 Dfangzhongdai
Dickeya gs fangzhongdai M005 WP 049842583 1 Dfangzhongdai
Dickeya gs undicola 2B12 WP 050515649 1 Dundicola
DAQ1742 DAQ1742 02380 Daquatica
Dickeya sp CSL RW240 WP 035341722 1 Daquatica
Dickeya aquatica DW 0440 WP 035341722 1 Daquatica
Dickeya dadantii subsp dieffenbachiae NCPPB 2976 WP 050572744 1 Ddadantii
Dickeya dianthicola NCPPB 3534 WP 046830460 1 Ddianthicola
Dickeya dianthicola NCPPB 453 WP 046830460 1 Ddianthicola
Dickeya dianthicola IPO 980 WP 046830460 1 Ddianthicola
Dickeya dianthicola GBBC 2039 WP 046830460 1 Ddianthicola
Dickeya dianthicola RNS04 9 WP 046830460 1 Ddianthicola

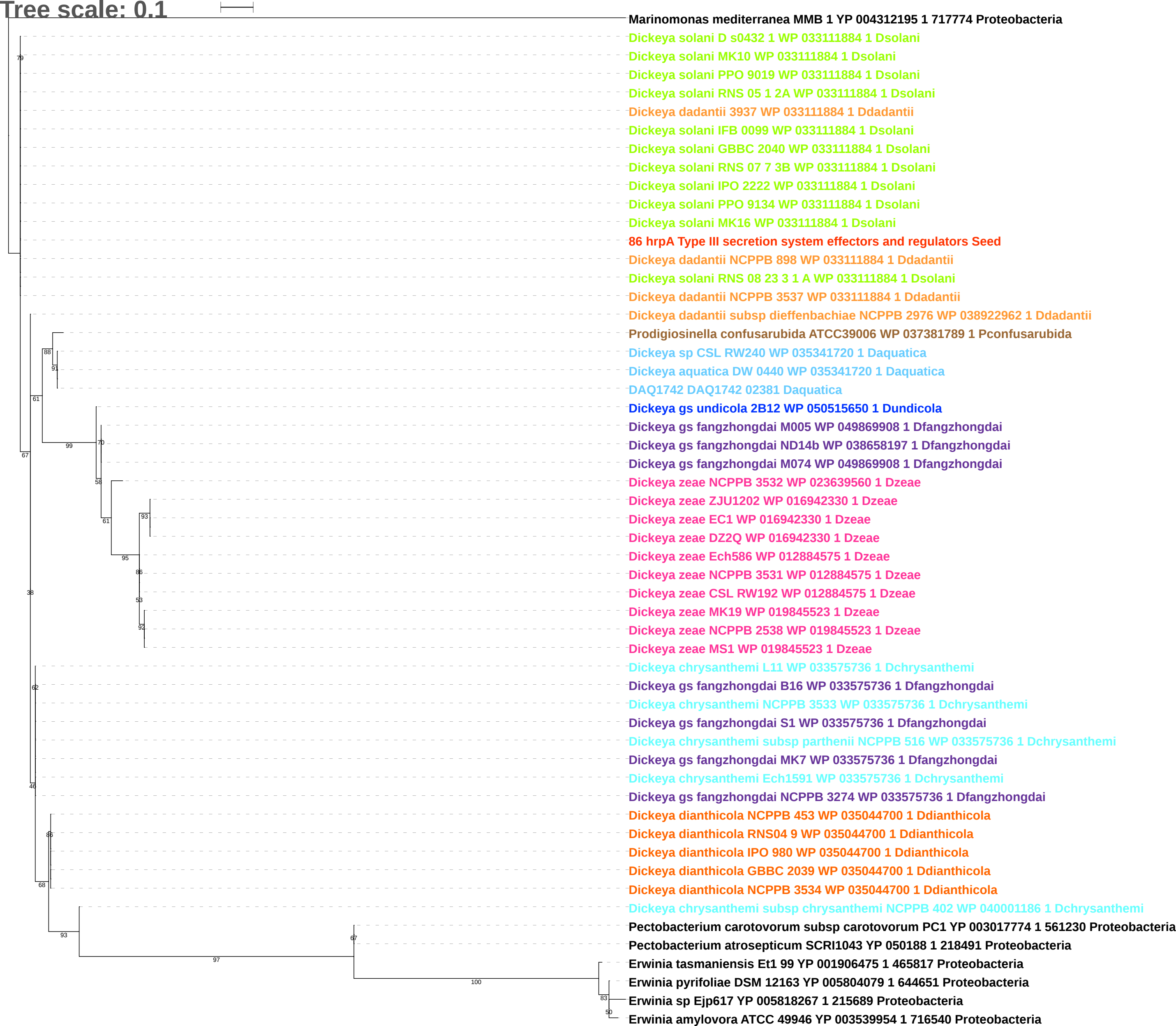
Tree scale: 0.1
Marinomonas mediterranea MMB 1 YP 004312195 1 717774 Proteobacteria
Dickeya solani D s0432 1 WP 033111884 1 Dsolani
Dickeya solani MK10 WP 033111884 1 Dsolani
Dickeya solani PPO 9019 WP 033111884 1 Dsolani
Dickeya solani RNS 05 1 2A WP 033111884 1 Dsolani
Dickeya dadantii 3937 WP 033111884 1 Ddadantii
Dickeya solani IFB 0099 WP 033111884 1 Dsolani
Dickeya solani GBBC 2040 WP 033111884 1 Dsolani
Dickeya solani RNS 07 7 3B WP 033111884 1 Dsolani
Dickeya solani IPO 2222 WP 033111884 1 Dsolani
Dickeya solani PPO 9134 WP 033111884 1 Dsolani
Dickeya solani MK16 WP 033111884 1 Dsolani
86 hrpA Type III secretion system effectors and regulators Seed
Dickeya dadantii NCPPB 898 WP 033111884 1 Ddadantii
Dickeya solani RNS 08 23 3 1 A WP 033111884 1 Dsolani
Dickeya dadantii NCPPB 3537 WP 033111884 1 Ddadantii
Dickeya dadantii subsp dieffenbachiae NCPPB 2976 WP 038922962 1 Ddadantii
Prodigiosinella confusarubida ATCC39006 WP 037381789 1 Pconfusarubida
Dickeya sp CSL RW240 WP 035341720 1 Daquatica
Dickeya aquatica DW 0440 WP 035341720 1 Daquatica
DAQ1742 DAQ1742 02381 Daquatica
Dickeya gs undicola 2B12 WP 050515650 1 Dundicola
Dickeya gs fangzhongdai M005 WP 049869908 1 Dfangzhongdai
Dickeya gs fangzhongdai ND14b WP 038658197 1 Dfangzhongdai
Dickeya gs fangzhongdai M074 WP 049869908 1 Dfangzhongdai
Dickeya zeae NCPPB 3532 WP 023639560 1 Dzeae
Dickeya zeae ZJU1202 WP 016942330 1 Dzeae
Dickeya zeae EC1 WP 016942330 1 Dzeae
Dickeya zeae DZ2Q WP 016942330 1 Dzeae
Dickeya zeae Ech586 WP 012884575 1 Dzeae
Dickeya zeae NCPPB 3531 WP 012884575 1 Dzeae
Dickeya zeae CSL RW192 WP 012884575 1 Dzeae
Dickeya zeae MK19 WP 019845523 1 Dzeae
Dickeya zeae NCPPB 2538 WP 019845523 1 Dzeae
Dickeya zeae MS1 WP 019845523 1 Dzeae
Dickeya chrysanthemi L11 WP 033575736 1 Dchrysanthemi
Dickeya gs fangzhongdai B16 WP 033575736 1 Dfangzhongdai
Dickeya chrysanthemi NCPPB 3533 WP 033575736 1 Dchrysanthemi
Dickeya gs fangzhongdai S1 WP 033575736 1 Dfangzhongdai
Dickeya chrysanthemi subsp parthenii NCPPB 516 WP 033575736 1 Dchrysanthemi
Dickeya gs fangzhongdai MK7 WP 033575736 1 Dfangzhongdai
Dickeya chrysanthemi Ech1591 WP 033575736 1 Dchrysanthemi
Dickeya gs fangzhongdai NCPPB 3274 WP 033575736 1 Dfangzhongdai
Dickeya dianthicola NCPPB 453 WP 035044700 1 Ddianthicola
Dickeya dianthicola RNS04 9 WP 035044700 1 Ddianthicola
Dickeya dianthicola IPO 980 WP 035044700 1 Ddianthicola
Dickeya dianthicola GBBC 2039 WP 035044700 1 Ddianthicola
Dickeya dianthicola NCPPB 3534 WP 035044700 1 Ddianthicola
Dickeya chrysanthemi subsp chrysanthemi NCPPB 402 WP 040001186 1 Dchrysanthemi
Pectobacterium carotovorum subsp carotovorum PC1 YP 003017774 1 561230 Proteobacteria
Pectobacterium atrosepticum SCRI1043 YP 050188 1 218491 Proteobacteria
Erwinia tasmaniensis Et1 99 YP 001906475 1 465817 Proteobacteria
Erwinia pyrifoliae DSM 12163 YP 005804079 1 644651 Proteobacteria
Erwinia sp Ejp617 YP 005818267 1 215689 Proteobacteria
Erwinia amylovora ATCC 49946 YP 003539954 1 716540 Proteobacteria

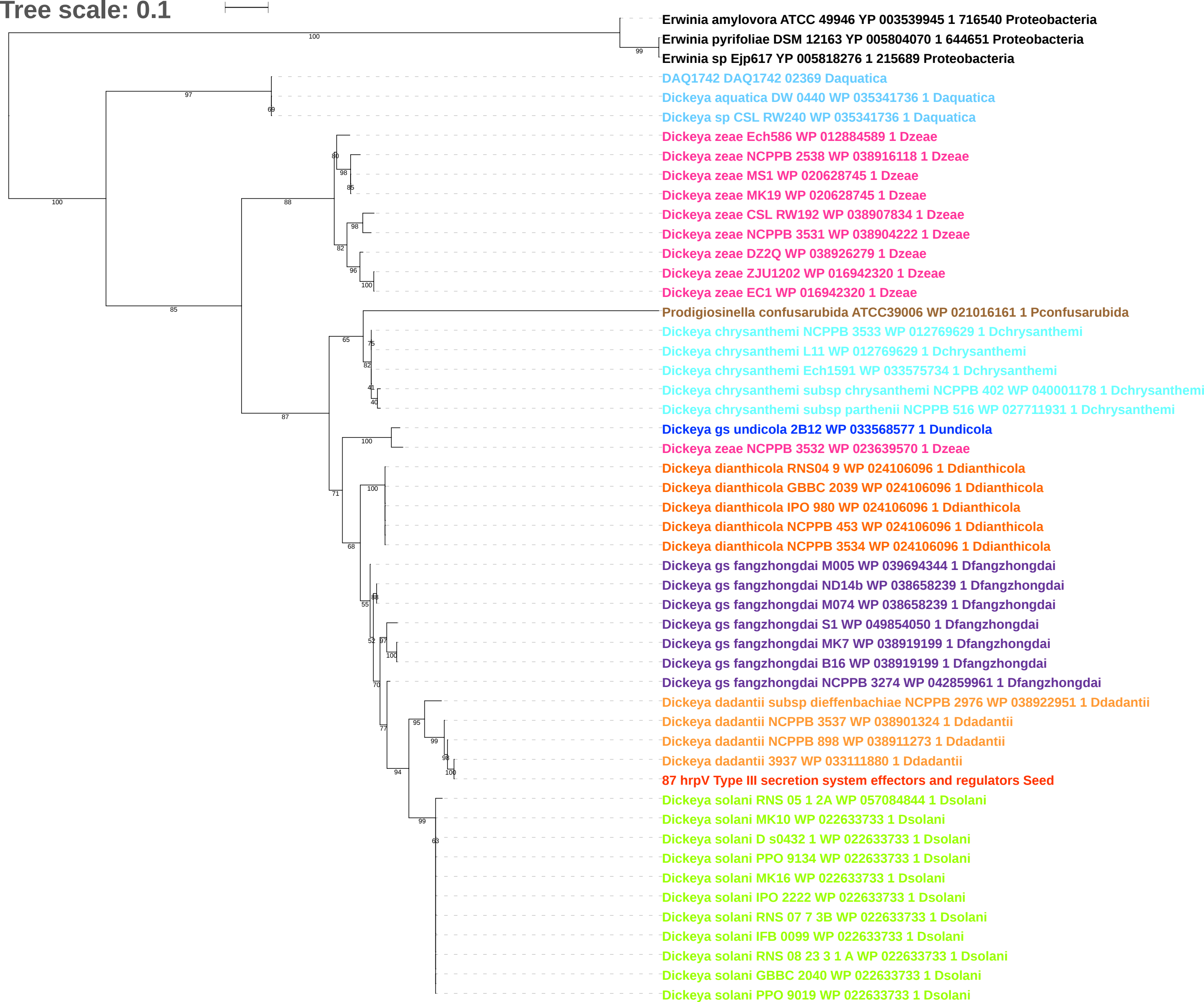
Tree scale: 0.1
Erwinia amylovora ATCC 49946 YP 003539945 1 716540 Proteobacteria
Erwinia pyrifoliae DSM 12163 YP 005804070 1 644651 Proteobacteria
Erwinia sp Ejp617 YP 005818276 1 215689 Proteobacteria
DAQ1742 DAQ1742 02369 Daquatica
Dickeya aquatica DW 0440 WP 035341736 1 Daquatica
Dickeya sp CSL RW240 WP 035341736 1 Daquatica
Dickeya zeae Ech586 WP 012884589 1 Dzeae
Dickeya zeae NCPPB 2538 WP 038916118 1 Dzeae
Dickeya zeae MS1 WP 020628745 1 Dzeae
Dickeya zeae MK19 WP 020628745 1 Dzeae
Dickeya zeae CSL RW192 WP 038907834 1 Dzeae
Dickeya zeae NCPPB 3531 WP 038904222 1 Dzeae
Dickeya zeae DZ2Q WP 038926279 1 Dzeae
Dickeya zeae ZJU1202 WP 016942320 1 Dzeae
Dickeya zeae EC1 WP 016942320 1 Dzeae
Prodigiosinella confusarubida ATCC39006 WP 021016161 1 Pconfusarubida
Dickeya chrysanthemi NCPPB 3533 WP 012769629 1 Dchrysanthemi
Dickeya chrysanthemi L11 WP 012769629 1 Dchrysanthemi
Dickeya chrysanthemi Ech1591 WP 033575734 1 Dchrysanthemi
Dickeya chrysanthemi subsp chrysanthemi NCPPB 402 WP 040001178 1 Dchrysanthemi
Dickeya chrysanthemi subsp parthenii NCPPB 516 WP 027711931 1 Dchrysanthemi
Dickeya gs undicola 2B12 WP 033568577 1 Dundicola
Dickeya zeae NCPPB 3532 WP 023639570 1 Dzeae
Dickeya dianthicola RNS04 9 WP 024106096 1 Ddianthicola
Dickeya dianthicola GBBC 2039 WP 024106096 1 Ddianthicola
Dickeya dianthicola IPO 980 WP 024106096 1 Ddianthicola
Dickeya dianthicola NCPPB 453 WP 024106096 1 Ddianthicola
Dickeya dianthicola NCPPB 3534 WP 024106096 1 Ddianthicola
Dickeya gs fangzhongdai M005 WP 039694344 1 Dfangzhongdai
Dickeya gs fangzhongdai ND14b WP 038658239 1 Dfangzhongdai
Dickeya gs fangzhongdai M074 WP 038658239 1 Dfangzhongdai
Dickeya gs fangzhongdai S1 WP 049854050 1 Dfangzhongdai
Dickeya gs fangzhongdai MK7 WP 038919199 1 Dfangzhongdai
Dickeya gs fangzhongdai B16 WP 038919199 1 Dfangzhongdai
Dickeya gs fangzhongdai NCPPB 3274 WP 042859961 1 Dfangzhongdai
Dickeya dadantii subsp dieffenbachiae NCPPB 2976 WP 038922951 1 Ddadantii
Dickeya dadantii NCPPB 3537 WP 038901324 1 Ddadantii
Dickeya dadantii NCPPB 898 WP 038911273 1 Ddadantii
Dickeya dadantii 3937 WP 033111880 1 Ddadantii
87 hrpV Type III secretion system effectors and regulators Seed
Dickeya solani RNS 05 1 2A WP 057084844 1 Dsolani
Dickeya solani MK10 WP 022633733 1 Dsolani
Dickeya solani D s0432 1 WP 022633733 1 Dsolani
Dickeya solani PPO 9134 WP 022633733 1 Dsolani
Dickeya solani MK16 WP 022633733 1 Dsolani
Dickeya solani IPO 2222 WP 022633733 1 Dsolani
Dickeya solani RNS 07 7 3B WP 022633733 1 Dsolani
Dickeya solani IFB 0099 WP 022633733 1 Dsolani
Dickeya solani RNS 08 23 3 1 A WP 022633733 1 Dsolani
Dickeya solani GBBC 2040 WP 022633733 1 Dsolani
Dickeya solani PPO 9019 WP 022633733 1 Dsolani

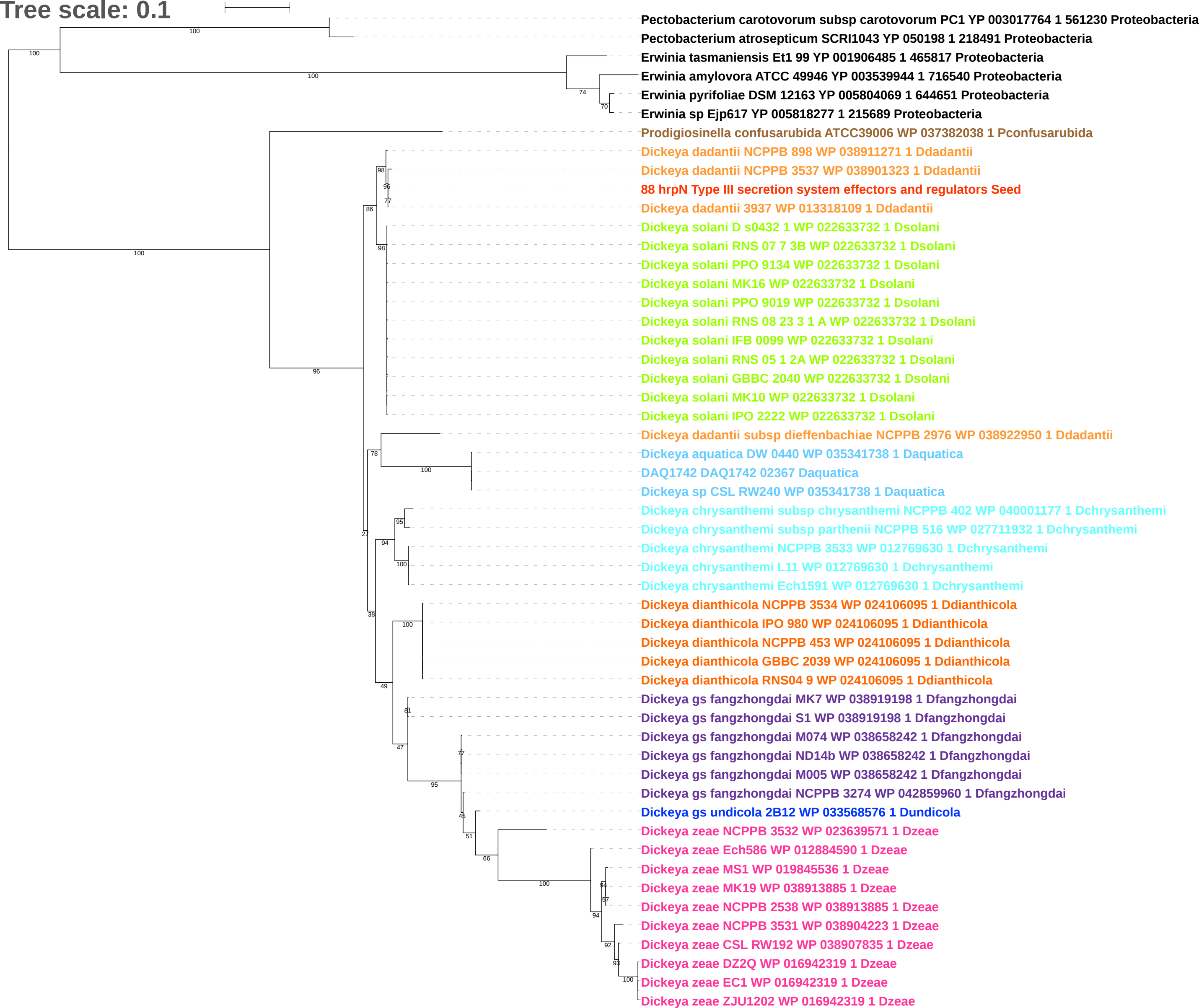
Tree scale: 0.1
Pectobacterium carotovorum subsp carotovorum PC1 YP 003017764 1 561230 Proteobacteria
Pectobacterium atrosepticum SCRI1043 YP 050198 1 218491 Proteobacteria
Erwinia tasmaniensis Et1 99 YP 001906485 1 465817 Proteobacteria
Erwinia amylovora ATCC 49946 YP 003539944 1 716540 Proteobacteria
Erwinia pyrifoliae DSM 12163 YP 005804069 1 644651 Proteobacteria
Erwinia sp Ejp617 YP 005818277 1 215689 Proteobacteria
Prodigiosinella confusarubida ATCC39006 WP 037382038 1 Pconfusarubida
Dickeya dadantii NCPPB 898 WP 038911271 1 Ddadantii
Dickeya dadantii NCPPB 3537 WP 038901323 1 Ddadantii
88 hrpN Type III secretion system effectors and regulators Seed
Dickeya dadantii 3937 WP 013318109 1 Ddadantii
Dickeya solani D s0432 1 WP 022633732 1 Dsolani
Dickeya solani RNS 07 7 3B WP 022633732 1 Dsolani
Dickeya solani PPO 9134 WP 022633732 1 Dsolani
Dickeya solani MK16 WP 022633732 1 Dsolani
Dickeya solani PPO 9019 WP 022633732 1 Dsolani
Dickeya solani RNS 08 23 3 1 A WP 022633732 1 Dsolani
Dickeya solani IFB 0099 WP 022633732 1 Dsolani
Dickeya solani RNS 05 1 2A WP 022633732 1 Dsolani
Dickeya solani GBBC 2040 WP 022633732 1 Dsolani
Dickeya solani MK10 WP 022633732 1 Dsolani
Dickeya solani IPO 2222 WP 022633732 1 Dsolani
Dickeya dadantii subsp dieffenbachiae NCPPB 2976 WP 038922950 1 Ddadantii
Dickeya aquatica DW 0440 WP 035341738 1 Daquatica
DAQ1742 DAQ1742 02367 Daquatica
Dickeya sp CSL RW240 WP 035341738 1 Daquatica
Dickeya chrysanthemi subsp chrysanthemi NCPPB 402 WP 040001177 1 Dchrysanthemi
Dickeya chrysanthemi subsp parthenii NCPPB 516 WP 027711932 1 Dchrysanthemi
Dickeya chrysanthemi NCPPB 3533 WP 012769630 1 Dchrysanthemi
Dickeya chrysanthemi L11 WP 012769630 1 Dchrysanthemi
Dickeya chrysanthemi Ech1591 WP 012769630 1 Dchrysanthemi
Dickeya dianthicola NCPPB 3534 WP 024106095 1 Ddianthicola
Dickeya dianthicola IPO 980 WP 024106095 1 Ddianthicola
Dickeya dianthicola NCPPB 453 WP 024106095 1 Ddianthicola
Dickeya dianthicola GBBC 2039 WP 024106095 1 Ddianthicola
Dickeya dianthicola RNS04 9 WP 024106095 1 Ddianthicola
Dickeya gs fangzhongdai MK7 WP 038919198 1 Dfangzhongdai
Dickeya gs fangzhongdai S1 WP 038919198 1 Dfangzhongdai
Dickeya gs fangzhongdai M074 WP 038658242 1 Dfangzhongdai
Dickeya gs fangzhongdai ND14b WP 038658242 1 Dfangzhongdai
Dickeya gs fangzhongdai M005 WP 038658242 1 Dfangzhongdai
Dickeya gs fangzhongdai NCPPB 3274 WP 042859960 1 Dfangzhongdai
Dickeya gs undicola 2B12 WP 033568576 1 Dundicola
Dickeya zeae NCPPB 3532 WP 023639571 1 Dzeae
Dickeya zeae Ech586 WP 012884590 1 Dzeae
Dickeya zeae MS1 WP 019845536 1 Dzeae
Dickeya zeae MK19 WP 038913885 1 Dzeae
Dickeya zeae NCPPB 2538 WP 038913885 1 Dzeae
Dickeya zeae NCPPB 3531 WP 038904223 1 Dzeae
Dickeya zeae CSL RW192 WP 038907835 1 Dzeae
Dickeya zeae DZ2Q WP 016942319 1 Dzeae
Dickeya zeae EC1 WP 016942319 1 Dzeae
Dickeya zeae ZJU1202 WP 016942319 1 Dzeae

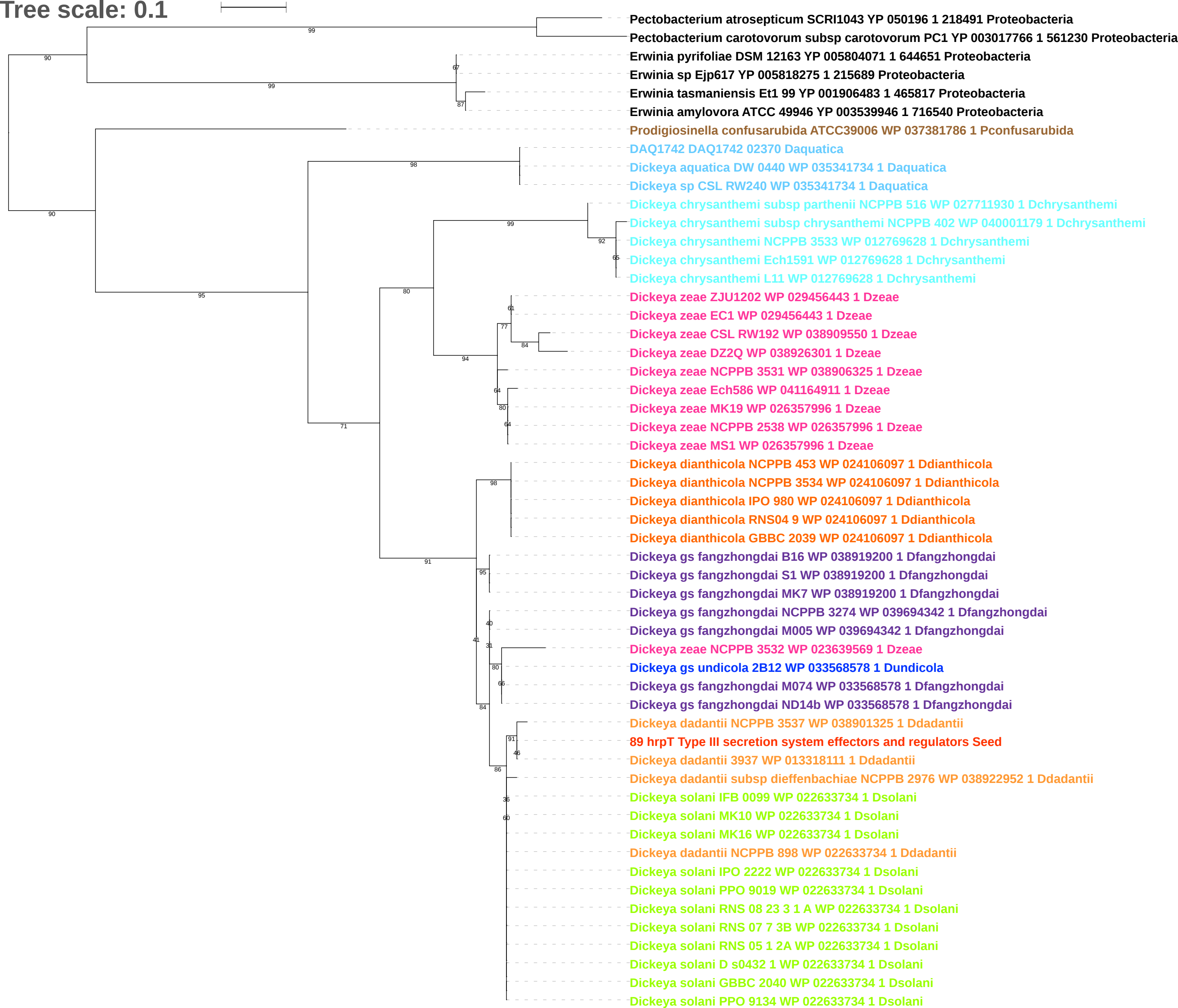
Tree scale: 0.1
Pectobacterium atrosepticum SCRI1043 YP 050196 1 218491 Proteobacteria
Pectobacterium carotovorum subsp carotovorum PC1 YP 003017766 1 561230 Proteobacteria
Erwinia pyrifoliae DSM 12163 YP 005804071 1 644651 Proteobacteria
Erwinia sp Ejp617 YP 005818275 1 215689 Proteobacteria
Erwinia tasmaniensis Et1 99 YP 001906483 1 465817 Proteobacteria
Erwinia amylovora ATCC 49946 YP 003539946 1 716540 Proteobacteria
Prodigiosinella confusarubida ATCC39006 WP 037381786 1 Pconfusarubida
DAQ1742 DAQ1742 02370 Daquatica
Dickeya aquatica DW 0440 WP 035341734 1 Daquatica
Dickeya sp CSL RW240 WP 035341734 1 Daquatica
Dickeya chrysanthemi subsp parthenii NCPPB 516 WP 027711930 1 Dchrysanthemi
Dickeya chrysanthemi subsp chrysanthemi NCPPB 402 WP 040001179 1 Dchrysanthemi
Dickeya chrysanthemi NCPPB 3533 WP 012769628 1 Dchrysanthemi
Dickeya chrysanthemi Ech1591 WP 012769628 1 Dchrysanthemi
Dickeya chrysanthemi L11 WP 012769628 1 Dchrysanthemi
Dickeya zeae ZJU1202 WP 029456443 1 Dzeae
Dickeya zeae EC1 WP 029456443 1 Dzeae
Dickeya zeae CSL RW192 WP 038909550 1 Dzeae
Dickeya zeae DZ2Q WP 038926301 1 Dzeae
Dickeya zeae NCPPB 3531 WP 038906325 1 Dzeae
Dickeya zeae Ech586 WP 041164911 1 Dzeae
Dickeya zeae MK19 WP 026357996 1 Dzeae
Dickeya zeae NCPPB 2538 WP 026357996 1 Dzeae
Dickeya zeae MS1 WP 026357996 1 Dzeae
Dickeya dianthicola NCPPB 453 WP 024106097 1 Ddianthicola
Dickeya dianthicola NCPPB 3534 WP 024106097 1 Ddianthicola
Dickeya dianthicola IPO 980 WP 024106097 1 Ddianthicola
Dickeya dianthicola RNS04 9 WP 024106097 1 Ddianthicola
Dickeya dianthicola GBBC 2039 WP 024106097 1 Ddianthicola
Dickeya gs fangzhongdai B16 WP 038919200 1 Dfangzhongdai
Dickeya gs fangzhongdai S1 WP 038919200 1 Dfangzhongdai
Dickeya gs fangzhongdai MK7 WP 038919200 1 Dfangzhongdai
Dickeya gs fangzhongdai NCPPB 3274 WP 039694342 1 Dfangzhongdai
Dickeya gs fangzhongdai M005 WP 039694342 1 Dfangzhongdai
Dickeya zeae NCPPB 3532 WP 023639569 1 Dzeae
Dickeya gs undicola 2B12 WP 033568578 1 Dundicola
Dickeya gs fangzhongdai M074 WP 033568578 1 Dfangzhongdai
Dickeya gs fangzhongdai ND14b WP 033568578 1 Dfangzhongdai
Dickeya dadantii NCPPB 3537 WP 038901325 1 Ddadantii
89 hrpT Type III secretion system effectors and regulators Seed
Dickeya dadantii 3937 WP 013318111 1 Ddadantii
Dickeya dadantii subsp dieffenbachiae NCPPB 2976 WP 038922952 1 Ddadantii
Dickeya solani IFB 0099 WP 022633734 1 Dsolani
Dickeya solani MK10 WP 022633734 1 Dsolani
Dickeya solani MK16 WP 022633734 1 Dsolani
Dickeya dadantii NCPPB 898 WP 022633734 1 Ddadantii
Dickeya solani IPO 2222 WP 022633734 1 Dsolani
Dickeya solani PPO 9019 WP 022633734 1 Dsolani
Dickeya solani RNS 08 23 3 1 A WP 022633734 1 Dsolani
Dickeya solani RNS 07 7 3B WP 022633734 1 Dsolani
Dickeya solani RNS 05 1 2A WP 022633734 1 Dsolani
Dickeya solani D s0432 1 WP 022633734 1 Dsolani
Dickeya solani GBBC 2040 WP 022633734 1 Dsolani
Dickeya solani PPO 9134 WP 022633734 1 Dsolani

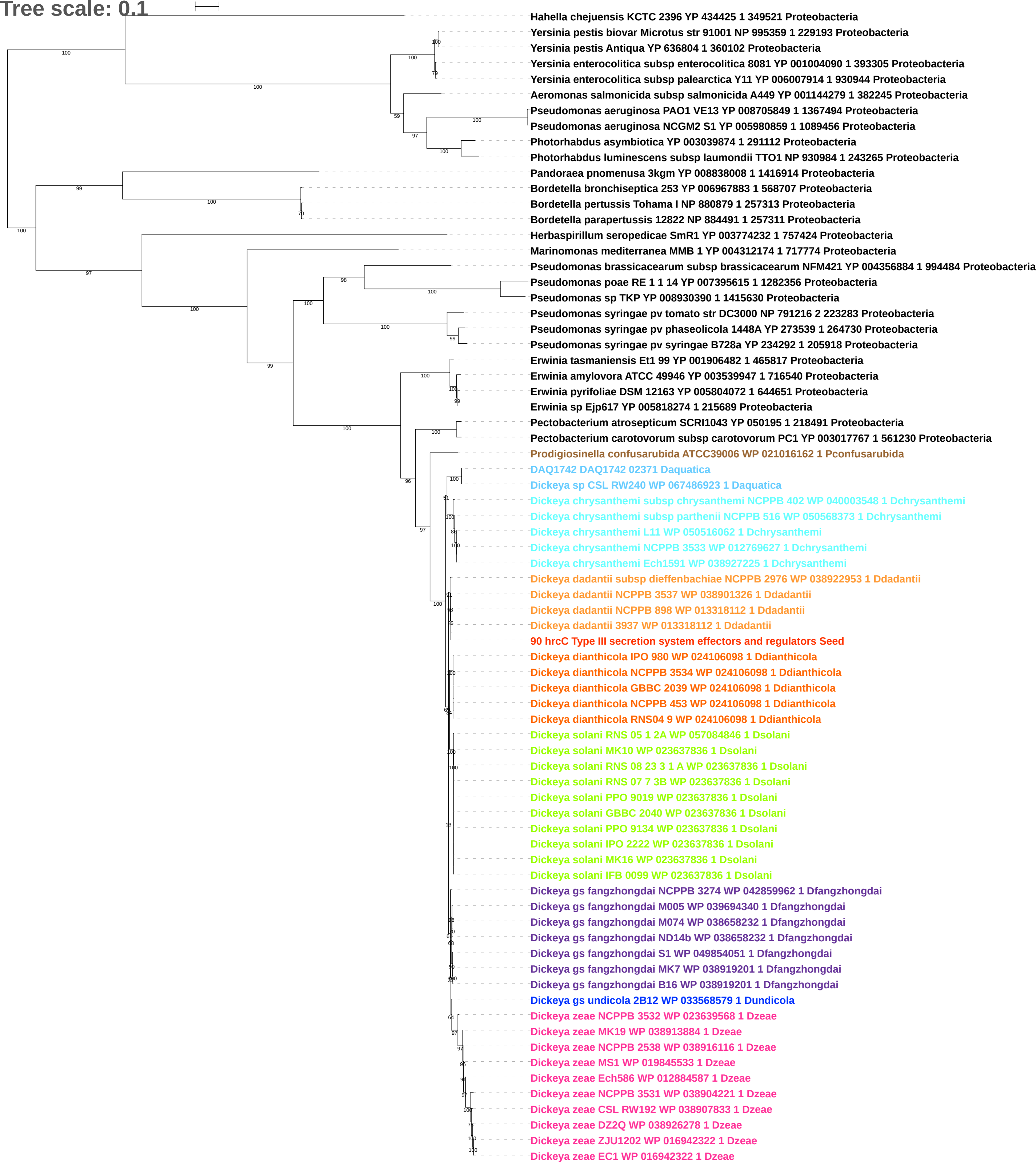
Tree scale: 0.1
Hahella chejuensis KCTC 2396 YP 434425 1 349521 Proteobacteria
Yersinia pestis biovar Microtus str 91001 NP 995359 1 229193 Proteobacteria
Yersinia pestis Antiqua YP 636804 1 360102 Proteobacteria
Yersinia enterocolitica subsp enterocolitica 8081 YP 001004090 1 393305 Proteobacteria
Yersinia enterocolitica subsp palearctica Y11 YP 006007914 1 930944 Proteobacteria
Aeromonas salmonicida subsp salmonicida A449 YP 001144279 1 382245 Proteobacteria
Pseudomonas aeruginosa PAO1 VE13 YP 008705849 1 1367494 Proteobacteria
Pseudomonas aeruginosa NCGM2 S1 YP 005980859 1 1089456 Proteobacteria
Photorhabdus asymbiotica YP 003039874 1 291112 Proteobacteria
Photorhabdus luminescens subsp laumondii TTO1 NP 930984 1 243265 Proteobacteria
Pandoraea pnomenusa 3kgm YP 008838008 1 1416914 Proteobacteria
Bordetella bronchiseptica 253 YP 006967883 1 568707 Proteobacteria
Bordetella pertussis Tohama I NP 880879 1 257313 Proteobacteria
Bordetella parapertussis 12822 NP 884491 1 257311 Proteobacteria
Herbaspirillum seropedicae SmR1 YP 003774232 1 757424 Proteobacteria
Marinomonas mediterranea MMB 1 YP 004312174 1 717774 Proteobacteria
Pseudomonas brassicacearum subsp brassicacearum NFM421 YP 004356884 1 994484 Proteobacteria
Pseudomonas poae RE 1 1 14 YP 007395615 1 1282356 Proteobacteria
Pseudomonas sp TKP YP 008930390 1 1415630 Proteobacteria
Pseudomonas syringae pv tomato str DC3000 NP 791216 2 223283 Proteobacteria
Pseudomonas syringae pv phaseolicola 1448A YP 273539 1 264730 Proteobacteria
Pseudomonas syringae pv syringae B728a YP 234292 1 205918 Proteobacteria
Erwinia tasmaniensis Et1 99 YP 001906482 1 465817 Proteobacteria
Erwinia amylovora ATCC 49946 YP 003539947 1 716540 Proteobacteria
Erwinia pyrifoliae DSM 12163 YP 005804072 1 644651 Proteobacteria
Erwinia sp Ejp617 YP 005818274 1 215689 Proteobacteria
Pectobacterium atrosepticum SCRI1043 YP 050195 1 218491 Proteobacteria
Pectobacterium carotovorum subsp carotovorum PC1 YP 003017767 1 561230 Proteobacteria
Prodigiosinella confusarubida ATCC39006 WP 021016162 1 Pconfusarubida
DAQ1742 DAQ1742 02371 Daquatica
Dickeya sp CSL RW240 WP 067486923 1 Daquatica
Dickeya chrysanthemi subsp chrysanthemi NCPPB 402 WP 040003548 1 Dchrysanthemi
Dickeya chrysanthemi subsp parthenii NCPPB 516 WP 050568373 1 Dchrysanthemi
Dickeya chrysanthemi L11 WP 050516062 1 Dchrysanthemi
Dickeya chrysanthemi NCPPB 3533 WP 012769627 1 Dchrysanthemi
Dickeya chrysanthemi Ech1591 WP 038927225 1 Dchrysanthemi
Dickeya dadantii subsp dieffenbachiae NCPPB 2976 WP 038922953 1 Ddadantii
Dickeya dadantii NCPPB 3537 WP 038901326 1 Ddadantii
Dickeya dadantii NCPPB 898 WP 013318112 1 Ddadantii
Dickeya dadantii 3937 WP 013318112 1 Ddadantii
90 hrcC Type III secretion system effectors and regulators Seed
Dickeya dianthicola IPO 980 WP 024106098 1 Ddianthicola
Dickeya dianthicola NCPPB 3534 WP 024106098 1 Ddianthicola
Dickeya dianthicola GBBC 2039 WP 024106098 1 Ddianthicola
Dickeya dianthicola NCPPB 453 WP 024106098 1 Ddianthicola
Dickeya dianthicola RNS04 9 WP 024106098 1 Ddianthicola
Dickeya solani RNS 05 1 2A WP 057084846 1 Dsolani
Dickeya solani MK10 WP 023637836 1 Dsolani
Dickeya solani RNS 08 23 3 1 A WP 023637836 1 Dsolani
Dickeya solani RNS 07 7 3B WP 023637836 1 Dsolani
Dickeya solani PPO 9019 WP 023637836 1 Dsolani
Dickeya solani GBBC 2040 WP 023637836 1 Dsolani
Dickeya solani PPO 9134 WP 023637836 1 Dsolani
Dickeya solani IPO 2222 WP 023637836 1 Dsolani
Dickeya solani MK16 WP 023637836 1 Dsolani
Dickeya solani IFB 0099 WP 023637836 1 Dsolani
Dickeya gs fangzhongdai NCPPB 3274 WP 042859962 1 Dfangzhongdai
Dickeya gs fangzhongdai M005 WP 039694340 1 Dfangzhongdai
Dickeya gs fangzhongdai M074 WP 038658232 1 Dfangzhongdai
Dickeya gs fangzhongdai ND14b WP 038658232 1 Dfangzhongdai
Dickeya gs fangzhongdai S1 WP 049854051 1 Dfangzhongdai
Dickeya gs fangzhongdai MK7 WP 038919201 1 Dfangzhongdai
Dickeya gs fangzhongdai B16 WP 038919201 1 Dfangzhongdai
Dickeya gs undicola 2B12 WP 033568579 1 Dundicola
Dickeya zeae NCPPB 3532 WP 023639568 1 Dzeae
Dickeya zeae MK19 WP 038913884 1 Dzeae
Dickeya zeae NCPPB 2538 WP 038916116 1 Dzeae
Dickeya zeae MS1 WP 019845533 1 Dzeae
Dickeya zeae Ech586 WP 012884587 1 Dzeae
Dickeya zeae NCPPB 3531 WP 038904221 1 Dzeae
Dickeya zeae CSL RW192 WP 038907833 1 Dzeae
Dickeya zeae DZ2Q WP 038926278 1 Dzeae
Dickeya zeae ZJU1202 WP 016942322 1 Dzeae
Dickeya zeae EC1 WP 016942322 1 Dzeae

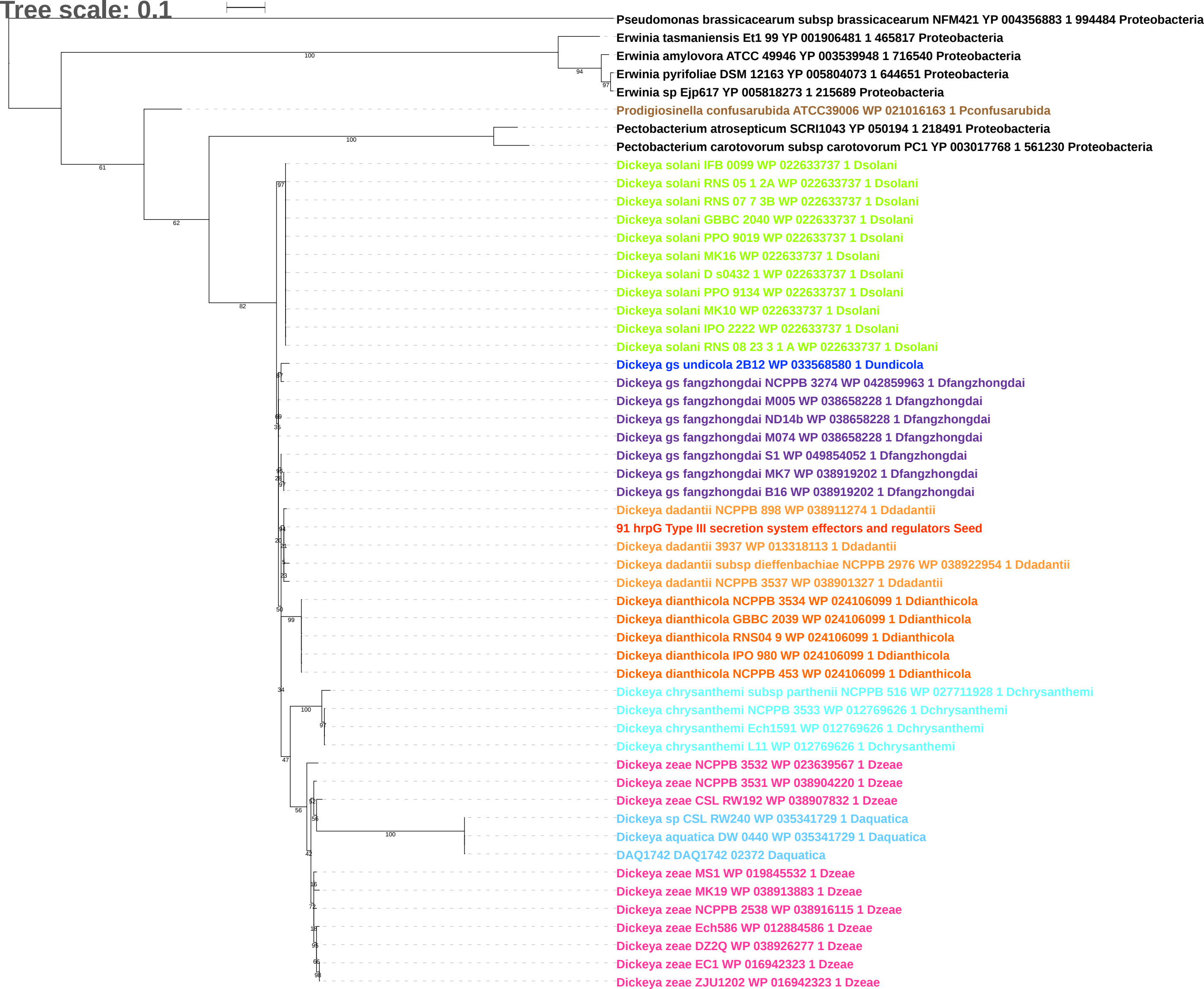

Tree scale: 0.1
Pseudomonas brassicacearum subsp brassicacearum NFM421 YP 004356883 1 994484 Proteobacteria
Erwinia tasmaniensis Et1 99 YP 001906481 1 465817 Proteobacteria
Erwinia amylovora ATCC 49946 YP 003539948 1 716540 Proteobacteria
Erwinia pyrifoliae DSM 12163 YP 005804073 1 644651 Proteobacteria
Erwinia sp Ejp617 YP 005818273 1 215689 Proteobacteria
Prodigiosinella confusarubida ATCC39006 WP 021016163 1 Pconfusarubida
Pectobacterium atrosepticum SCRI1043 YP 050194 1 218491 Proteobacteria
Pectobacterium carotovorum subsp carotovorum PC1 YP 003017768 1 561230 Proteobacteria
Dickeya solani IFB 0099 WP 022633737 1 Dsolani
Dickeya solani RNS 05 1 2A WP 022633737 1 Dsolani
Dickeya solani RNS 07 7 3B WP 022633737 1 Dsolani
Dickeya solani GBBC 2040 WP 022633737 1 Dsolani
Dickeya solani PPO 9019 WP 022633737 1 Dsolani
Dickeya solani MK16 WP 022633737 1 Dsolani
Dickeya solani D s0432 1 WP 022633737 1 Dsolani
Dickeya solani PPO 9134 WP 022633737 1 Dsolani
Dickeya solani MK10 WP 022633737 1 Dsolani
Dickeya solani IPO 2222 WP 022633737 1 Dsolani
Dickeya solani RNS 08 23 3 1 A WP 022633737 1 Dsolani
Dickeya gs undicola 2B12 WP 033568580 1 Dundicola
Dickeya gs fangzhongdai NCPPB 3274 WP 042859963 1 Dfangzhongdai
Dickeya gs fangzhongdai M005 WP 038658228 1 Dfangzhongdai
Dickeya gs fangzhongdai ND14b WP 038658228 1 Dfangzhongdai
Dickeya gs fangzhongdai M074 WP 038658228 1 Dfangzhongdai
Dickeya gs fangzhongdai S1 WP 049854052 1 Dfangzhongdai
Dickeya gs fangzhongdai MK7 WP 038919202 1 Dfangzhongdai
Dickeya gs fangzhongdai B16 WP 038919202 1 Dfangzhongdai
Dickeya dadantii NCPPB 898 WP 038911274 1 Ddadantii
91 hrpG Type III secretion system effectors and regulators Seed
Dickeya dadantii 3937 WP 013318113 1 Ddadantii
Dickeya dadantii subsp dieffenbachiae NCPPB 2976 WP 038922954 1 Ddadantii
Dickeya dadantii NCPPB 3537 WP 038901327 1 Ddadantii
Dickeya dianthicola NCPPB 3534 WP 024106099 1 Ddianthicola
Dickeya dianthicola GBBC 2039 WP 024106099 1 Ddianthicola
Dickeya dianthicola RNS04 9 WP 024106099 1 Ddianthicola
Dickeya dianthicola IPO 980 WP 024106099 1 Ddianthicola
Dickeya dianthicola NCPPB 453 WP 024106099 1 Ddianthicola
Dickeya chrysanthemi subsp parthenii NCPPB 516 WP 027711928 1 Dchrysanthemi
Dickeya chrysanthemi NCPPB 3533 WP 012769626 1 Dchrysanthemi
Dickeya chrysanthemi Ech1591 WP 012769626 1 Dchrysanthemi
Dickeya chrysanthemi L11 WP 012769626 1 Dchrysanthemi
Dickeya zeae NCPPB 3532 WP 023639567 1 Dzeae
Dickeya zeae NCPPB 3531 WP 038904220 1 Dzeae
Dickeya zeae CSL RW192 WP 038907832 1 Dzeae
Dickeya sp CSL RW240 WP 035341729 1 Daquatica
Dickeya aquatica DW 0440 WP 035341729 1 Daquatica
DAQ1742 DAQ1742 02372 Daquatica
Dickeya zeae MS1 WP 019845532 1 Dzeae
Dickeya zeae MK19 WP 038913883 1 Dzeae
Dickeya zeae NCPPB 2538 WP 038916115 1 Dzeae
Dickeya zeae Ech586 WP 012884586 1 Dzeae
Dickeya zeae DZ2Q WP 038926277 1 Dzeae
Dickeya zeae EC1 WP 016942323 1 Dzeae
Dickeya zeae ZJU1202 WP 016942323 1 Dzeae

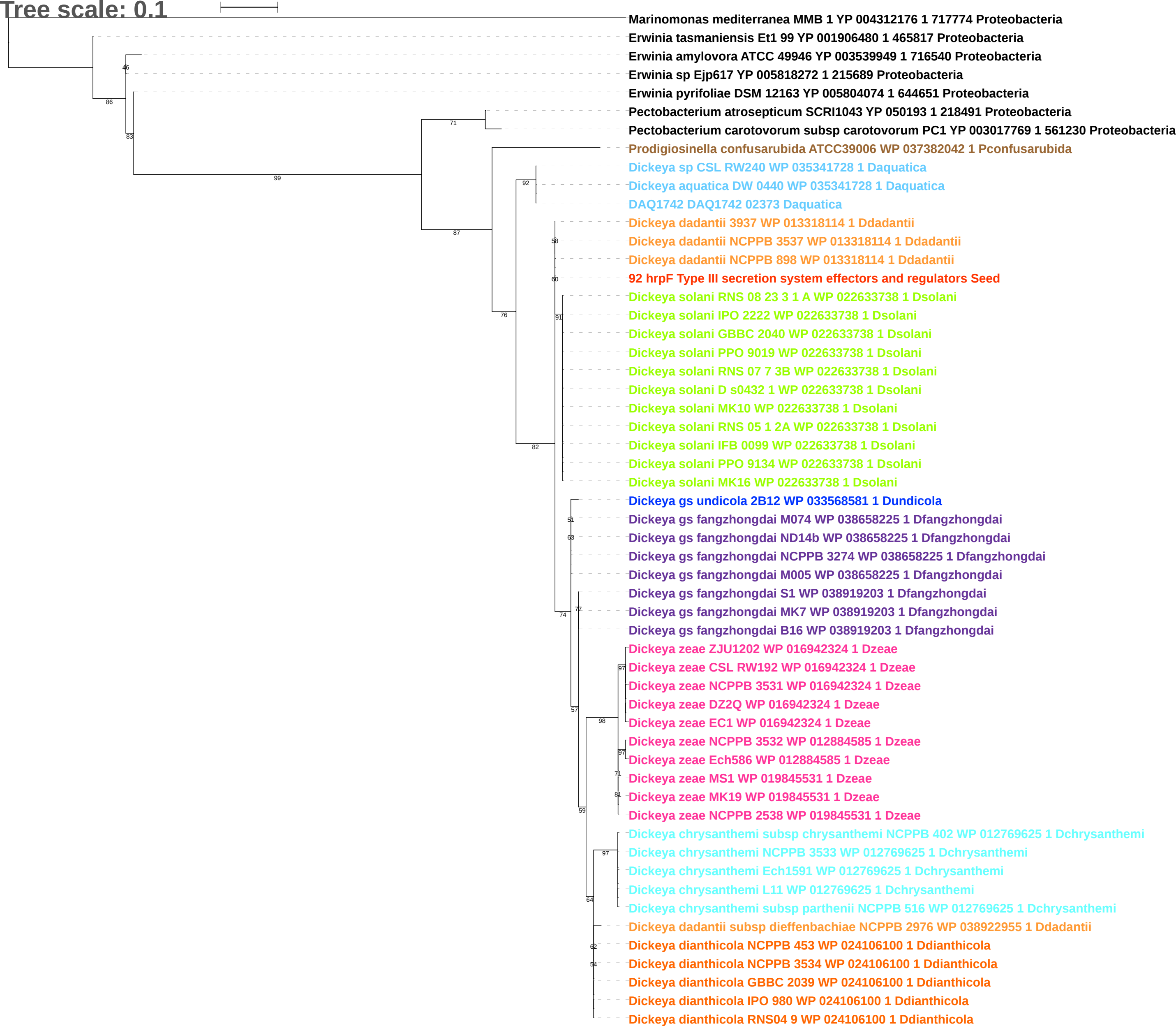
Tree scale: 0.1
Marinomonas mediterranea MMB 1 YP 004312176 1 717774 Proteobacteria
Erwinia tasmaniensis Et1 99 YP 001906480 1 465817 Proteobacteria
Erwinia amylovora ATCC 49946 YP 003539949 1 716540 Proteobacteria
Erwinia sp Ejp617 YP 005818272 1 215689 Proteobacteria
Erwinia pyrifoliae DSM 12163 YP 005804074 1 644651 Proteobacteria
Pectobacterium atrosepticum SCRI1043 YP 050193 1 218491 Proteobacteria
Pectobacterium carotovorum subsp carotovorum PC1 YP 003017769 1 561230 Proteobacteria
Prodigiosinella confusarubida ATCC39006 WP 037382042 1 Pconfusarubida
Dickeya sp CSL RW240 WP 035341728 1 Daquatica
Dickeya aquatica DW 0440 WP 035341728 1 Daquatica
DAQ1742 DAQ1742 02373 Daquatica
Dickeya dadantii 3937 WP 013318114 1 Ddadantii
Dickeya dadantii NCPPB 3537 WP 013318114 1 Ddadantii
Dickeya dadantii NCPPB 898 WP 013318114 1 Ddadantii
92 hrpF Type III secretion system effectors and regulators Seed
Dickeya solani RNS 08 23 3 1 A WP 022633738 1 Dsolani
Dickeya solani IPO 2222 WP 022633738 1 Dsolani
Dickeya solani GBBC 2040 WP 022633738 1 Dsolani
Dickeya solani PPO 9019 WP 022633738 1 Dsolani
Dickeya solani RNS 07 7 3B WP 022633738 1 Dsolani
Dickeya solani D s0432 1 WP 022633738 1 Dsolani
Dickeya solani MK10 WP 022633738 1 Dsolani
Dickeya solani RNS 05 1 2A WP 022633738 1 Dsolani
Dickeya solani IFB 0099 WP 022633738 1 Dsolani
Dickeya solani PPO 9134 WP 022633738 1 Dsolani
Dickeya solani MK16 WP 022633738 1 Dsolani
Dickeya gs undicola 2B12 WP 033568581 1 Dundicola
Dickeya gs fangzhongdai M074 WP 038658225 1 Dfangzhongdai
Dickeya gs fangzhongdai ND14b WP 038658225 1 Dfangzhongdai
Dickeya gs fangzhongdai NCPPB 3274 WP 038658225 1 Dfangzhongdai
Dickeya gs fangzhongdai M005 WP 038658225 1 Dfangzhongdai
Dickeya gs fangzhongdai S1 WP 038919203 1 Dfangzhongdai
Dickeya gs fangzhongdai MK7 WP 038919203 1 Dfangzhongdai
Dickeya gs fangzhongdai B16 WP 038919203 1 Dfangzhongdai
Dickeya zeae ZJU1202 WP 016942324 1 Dzeae
Dickeya zeae CSL RW192 WP 016942324 1 Dzeae
Dickeya zeae NCPPB 3531 WP 016942324 1 Dzeae
Dickeya zeae DZ2Q WP 016942324 1 Dzeae
Dickeya zeae EC1 WP 016942324 1 Dzeae
Dickeya zeae NCPPB 3532 WP 012884585 1 Dzeae
Dickeya zeae Ech586 WP 012884585 1 Dzeae
Dickeya zeae MS1 WP 019845531 1 Dzeae
Dickeya zeae MK19 WP 019845531 1 Dzeae
Dickeya zeae NCPPB 2538 WP 019845531 1 Dzeae
Dickeya chrysanthemi subsp chrysanthemi NCPPB 402 WP 012769625 1 Dchrysanthemi
Dickeya chrysanthemi NCPPB 3533 WP 012769625 1 Dchrysanthemi
Dickeya chrysanthemi Ech1591 WP 012769625 1 Dchrysanthemi
Dickeya chrysanthemi L11 WP 012769625 1 Dchrysanthemi
Dickeya chrysanthemi subsp parthenii NCPPB 516 WP 012769625 1 Dchrysanthemi
Dickeya dadantii subsp dieffenbachiae NCPPB 2976 WP 038922955 1 Ddadantii
Dickeya dianthicola NCPPB 453 WP 024106100 1 Ddianthicola
Dickeya dianthicola NCPPB 3534 WP 024106100 1 Ddianthicola
Dickeya dianthicola GBBC 2039 WP 024106100 1 Ddianthicola
Dickeya dianthicola IPO 980 WP 024106100 1 Ddianthicola
Dickeya dianthicola RNS04 9 WP 024106100 1 Ddianthicola

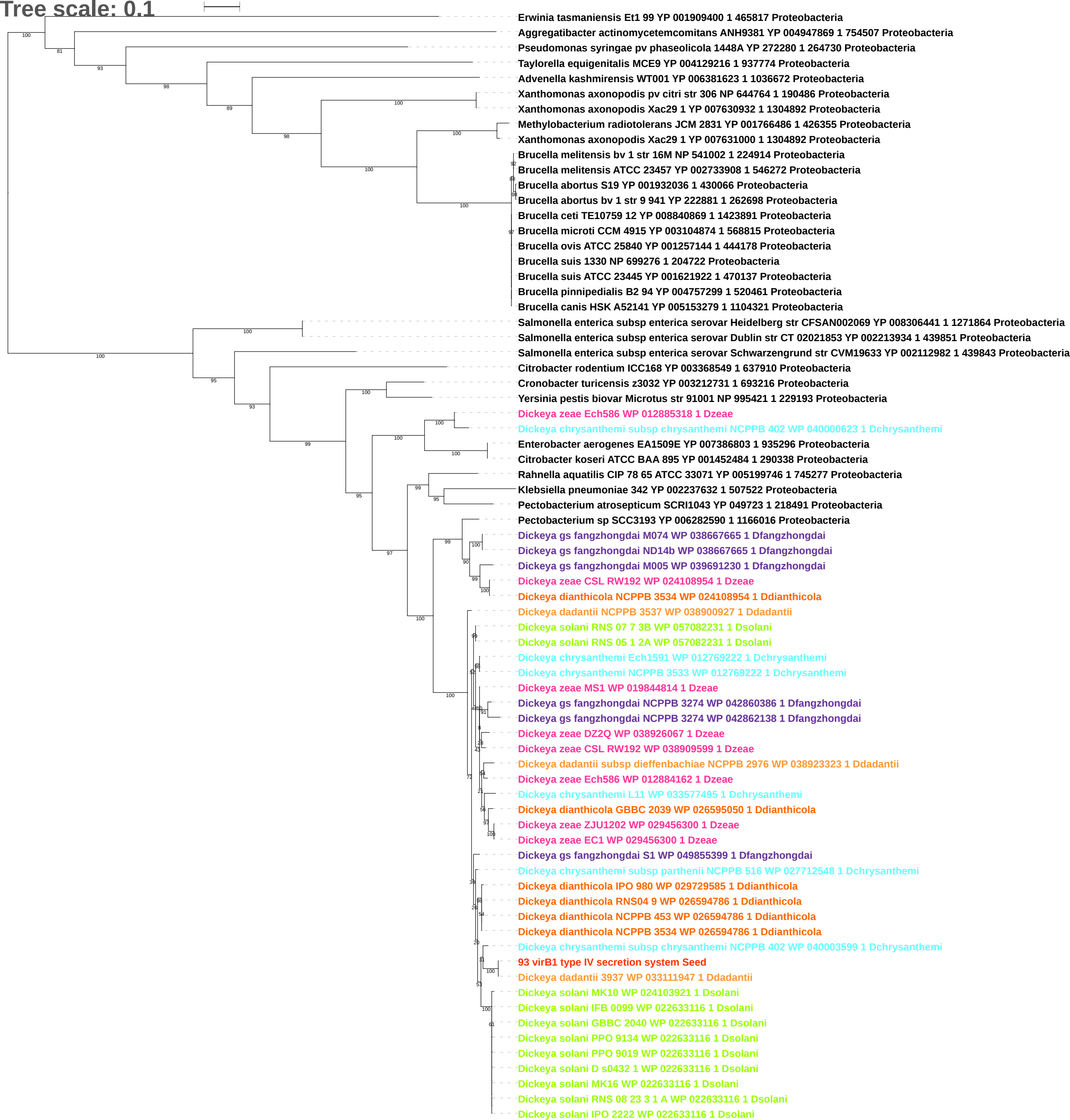
Tree scale: 0.1
Erwinia tasmaniensis Et1 99 YP 001909400 1 465817 Proteobacteria
Aggregatibacter actinomycetemcomitans ANH9381 YP 004947869 1 754507 Proteobacteria
Pseudomonas syringae pv phaseolicola 1448A YP 272280 1 264730 Proteobacteria
Taylorella equigenitalis MCE9 YP 004129216 1 937774 Proteobacteria
Advenella kashmirensis WT001 YP 006381623 1 1036672 Proteobacteria
Xanthomonas axonopodis pv citri str 306 NP 644764 1 190486 Proteobacteria
Xanthomonas axonopodis Xac29 1 YP 007630932 1 1304892 Proteobacteria
Methylobacterium radiotolerans JCM 2831 YP 001766486 1 426355 Proteobacteria
Xanthomonas axonopodis Xac29 1 YP 007631000 1 1304892 Proteobacteria
Brucella melitensis bv 1 str 16M NP 541002 1 224914 Proteobacteria
Brucella melitensis ATCC 23457 YP 002733908 1 546272 Proteobacteria
Brucella abortus S19 YP 001932036 1 430066 Proteobacteria
Brucella abortus bv 1 str 9 941 YP 222881 1 262698 Proteobacteria
Brucella ceti TE10759 12 YP 008840869 1 1423891 Proteobacteria
Brucella microti CCM 4915 YP 003104874 1 568815 Proteobacteria
Brucella ovis ATCC 25840 YP 001257144 1 444178 Proteobacteria
Brucella suis 1330 NP 699276 1 204722 Proteobacteria
Brucella suis ATCC 23445 YP 001621922 1 470137 Proteobacteria
Brucella pinnipedialis B2 94 YP 004757299 1 520461 Proteobacteria
Brucella canis HSK A52141 YP 005153279 1 1104321 Proteobacteria
Salmonella enterica subsp enterica serovar Heidelberg str CFSAN002069 YP 008306441 1 1271864 Proteobacteria
Salmonella enterica subsp enterica serovar Dublin str CT 02021853 YP 002213934 1 439851 Proteobacteria
Salmonella enterica subsp enterica serovar Schwarzengrund str CVM19633 YP 002112982 1 439843 Proteobacteria
Citrobacter rodentium ICC168 YP 003368549 1 637910 Proteobacteria
Cronobacter turicensis z3032 YP 003212731 1 693216 Proteobacteria
Yersinia pestis biovar Microtus str 91001 NP 995421 1 229193 Proteobacteria
Dickeya zeae Ech586 WP 012885318 1 Dzeae
Dickeya chrysanthemi subsp chrysanthemi NCPPB 402 WP 040000623 1 Dchrysanthemi
Enterobacter aerogenes EA1509E YP 007386803 1 935296 Proteobacteria
Citrobacter koseri ATCC BAA 895 YP 001452484 1 290338 Proteobacteria
Rahnella aquatilis CIP 78 65 ATCC 33071 YP 005199746 1 745277 Proteobacteria
Klebsiella pneumoniae 342 YP 002237632 1 507522 Proteobacteria
Pectobacterium atrosepticum SCRI1043 YP 049723 1 218491 Proteobacteria
Pectobacterium sp SCC3193 YP 006282590 1 1166016 Proteobacteria
Dickeya gs fangzhongdai M074 WP 038667665 1 Dfangzhongdai
Dickeya gs fangzhongdai ND14b WP 038667665 1 Dfangzhongdai
Dickeya gs fangzhongdai M005 WP 039691230 1 Dfangzhongdai
Dickeya zeae CSL RW192 WP 024108954 1 Dzeae
Dickeya dianthicola NCPPB 3534 WP 024108954 1 Ddianthicola
Dickeya dadantii NCPPB 3537 WP 038900927 1 Ddadantii
Dickeya solani RNS 07 7 3B WP 057082231 1 Dsolani
Dickeya solani RNS 05 1 2A WP 057082231 1 Dsolani
Dickeya chrysanthemi Ech1591 WP 012769222 1 Dchrysanthemi
Dickeya chrysanthemi NCPPB 3533 WP 012769222 1 Dchrysanthemi
Dickeya zeae MS1 WP 019844814 1 Dzeae
Dickeya gs fangzhongdai NCPPB 3274 WP 042860386 1 Dfangzhongdai
Dickeya gs fangzhongdai NCPPB 3274 WP 042862138 1 Dfangzhongdai
Dickeya zeae DZ2Q WP 038926067 1 Dzeae
Dickeya zeae CSL RW192 WP 038909599 1 Dzeae
Dickeya dadantii subsp dieffenbachiae NCPPB 2976 WP 038923323 1 Ddadantii
Dickeya zeae Ech586 WP 012884162 1 Dzeae
Dickeya chrysanthemi L11 WP 033577495 1 Dchrysanthemi
Dickeya dianthicola GBBC 2039 WP 026595050 1 Ddianthicola
Dickeya zeae ZJU1202 WP 029456300 1 Dzeae
Dickeya zeae EC1 WP 029456300 1 Dzeae
Dickeya gs fangzhongdai S1 WP 049855399 1 Dfangzhongdai
Dickeya chrysanthemi subsp parthenii NCPPB 516 WP 027712548 1 Dchrysanthemi
Dickeya dianthicola IPO 980 WP 029729585 1 Ddianthicola
Dickeya dianthicola RNS04 9 WP 026594786 1 Ddianthicola
Dickeya dianthicola NCPPB 453 WP 026594786 1 Ddianthicola
Dickeya dianthicola NCPPB 3534 WP 026594786 1 Ddianthicola
Dickeya chrysanthemi subsp chrysanthemi NCPPB 402 WP 040003599 1 Dchrysanthemi
93 virB1 type IV secretion system Seed
Dickeya dadantii 3937 WP 033111947 1 Ddadantii
Dickeya solani MK10 WP 024103921 1 Dsolani
Dickeya solani IFB 0099 WP 022633116 1 Dsolani
Dickeya solani GBBC 2040 WP 022633116 1 Dsolani
Dickeya solani PPO 9134 WP 022633116 1 Dsolani
Dickeya solani PPO 9019 WP 022633116 1 Dsolani
Dickeya solani D s0432 1 WP 022633116 1 Dsolani
Dickeya solani MK16 WP 022633116 1 Dsolani
Dickeya solani RNS 08 23 3 1 A WP 022633116 1 Dsolani
Dickeya solani IPO 2222 WP 022633116 1 Dsolani

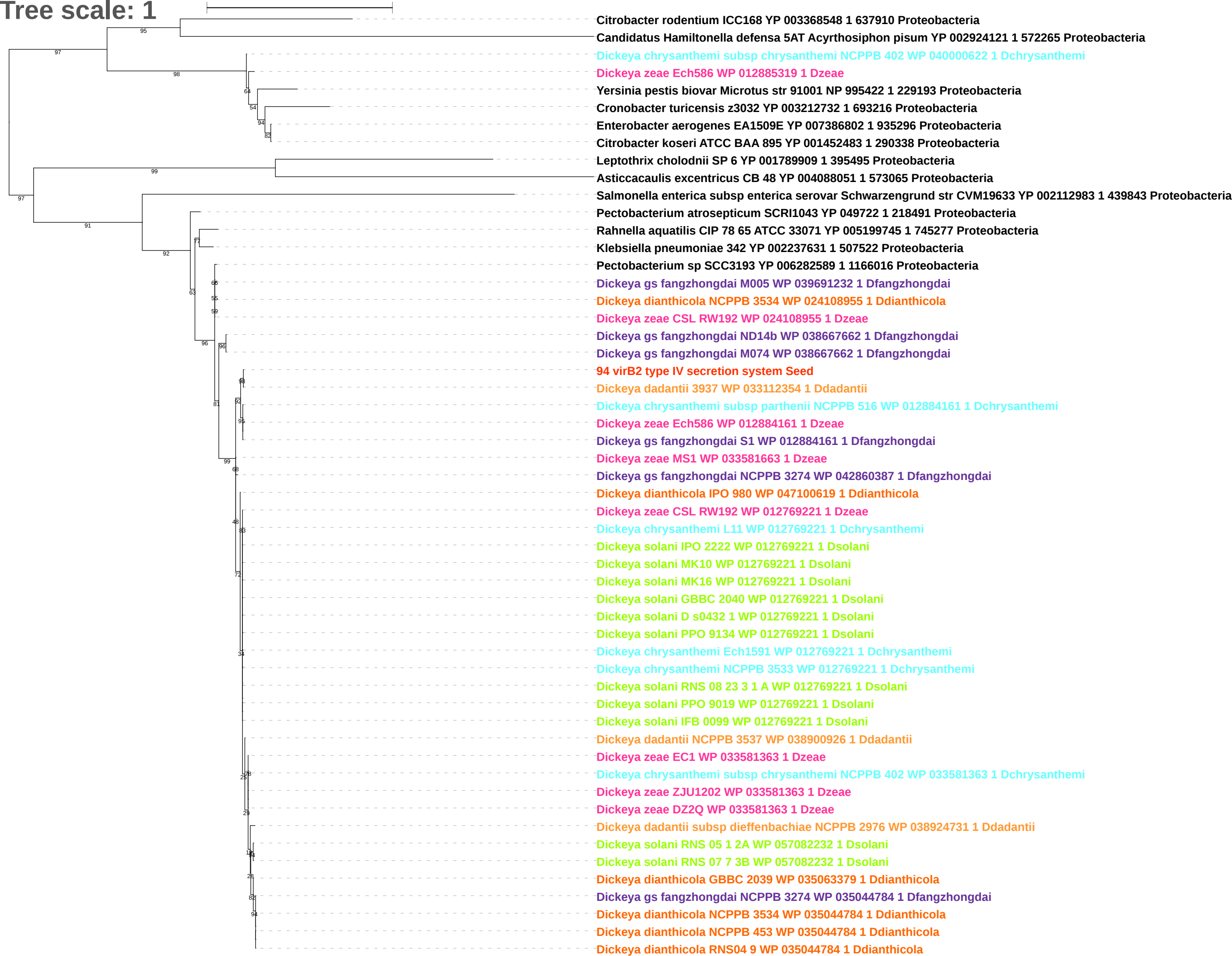
Tree scale: 1
Citrobacter rodentium ICC168 YP 003368548 1 637910 Proteobacteria
Candidatus Hamiltonella defensa 5AT Acyrthosiphon pisum YP 002924121 1 572265 Proteobacteria
Dickeya chrysanthemi subsp chrysanthemi NCPPB 402 WP 040000622 1 Dchrysanthemi
Dickeya zeae Ech586 WP 012885319 1 Dzeae
Yersinia pestis biovar Microtus str 91001 NP 995422 1 229193 Proteobacteria
Cronobacter turicensis z3032 YP 003212732 1 693216 Proteobacteria
Enterobacter aerogenes EA1509E YP 007386802 1 935296 Proteobacteria
Citrobacter koseri ATCC BAA 895 YP 001452483 1 290338 Proteobacteria
Leptothrix cholodnii SP 6 YP 001789909 1 395495 Proteobacteria
Asticcacaulis excentricus CB 48 YP 004088051 1 573065 Proteobacteria
Salmonella enterica subsp enterica serovar Schwarzengrund str CVM19633 YP 002112983 1 439843 Proteobacteria
Pectobacterium atrosepticum SCRI1043 YP 049722 1 218491 Proteobacteria
Rahnella aquatilis CIP 78 65 ATCC 33071 YP 005199745 1 745277 Proteobacteria
Klebsiella pneumoniae 342 YP 002237631 1 507522 Proteobacteria
Pectobacterium sp SCC3193 YP 006282589 1 1166016 Proteobacteria
Dickeya gs fangzhongdai M005 WP 039691232 1 Dfangzhongdai
Dickeya dianthicola NCPPB 3534 WP 024108955 1 Ddianthicola
Dickeya zeae CSL RW192 WP 024108955 1 Dzeae
Dickeya gs fangzhongdai ND14b WP 038667662 1 Dfangzhongdai
Dickeya gs fangzhongdai M074 WP 038667662 1 Dfangzhongdai
94 virB2 type IV secretion system Seed
Dickeya dadantii 3937 WP 033112354 1 Ddadantii
Dickeya chrysanthemi subsp parthenii NCPPB 516 WP 012884161 1 Dchrysanthemi
Dickeya zeae Ech586 WP 012884161 1 Dzeae
Dickeya gs fangzhongdai S1 WP 012884161 1 Dfangzhongdai
Dickeya zeae MS1 WP 033581663 1 Dzeae
Dickeya gs fangzhongdai NCPPB 3274 WP 042860387 1 Dfangzhongdai
Dickeya dianthicola IPO 980 WP 047100619 1 Ddianthicola
Dickeya zeae CSL RW192 WP 012769221 1 Dzeae
Dickeya chrysanthemi L11 WP 012769221 1 Dchrysanthemi
Dickeya solani IPO 2222 WP 012769221 1 Dsolani
Dickeya solani MK10 WP 012769221 1 Dsolani
Dickeya solani MK16 WP 012769221 1 Dsolani
Dickeya solani GBBC 2040 WP 012769221 1 Dsolani
Dickeya solani D s0432 1 WP 012769221 1 Dsolani
Dickeya solani PPO 9134 WP 012769221 1 Dsolani
Dickeya chrysanthemi Ech1591 WP 012769221 1 Dchrysanthemi
Dickeya chrysanthemi NCPPB 3533 WP 012769221 1 Dchrysanthemi
Dickeya solani RNS 08 23 3 1 A WP 012769221 1 Dsolani
Dickeya solani PPO 9019 WP 012769221 1 Dsolani
Dickeya solani IFB 0099 WP 012769221 1 Dsolani
Dickeya dadantii NCPPB 3537 WP 038900926 1 Ddadantii
Dickeya zeae EC1 WP 033581363 1 Dzeae
Dickeya chrysanthemi subsp chrysanthemi NCPPB 402 WP 033581363 1 Dchrysanthemi
Dickeya zeae ZJU1202 WP 033581363 1 Dzeae
Dickeya zeae DZ2Q WP 033581363 1 Dzeae
Dickeya dadantii subsp dieffenbachiae NCPPB 2976 WP 038924731 1 Ddadantii
Dickeya solani RNS 05 1 2A WP 057082232 1 Dsolani
Dickeya solani RNS 07 7 3B WP 057082232 1 Dsolani
Dickeya dianthicola GBBC 2039 WP 035063379 1 Ddianthicola
Dickeya gs fangzhongdai NCPPB 3274 WP 035044784 1 Dfangzhongdai
Dickeya dianthicola NCPPB 3534 WP 035044784 1 Ddianthicola
Dickeya dianthicola NCPPB 453 WP 035044784 1 Ddianthicola
Dickeya dianthicola RNS04 9 WP 035044784 1 Ddianthicola

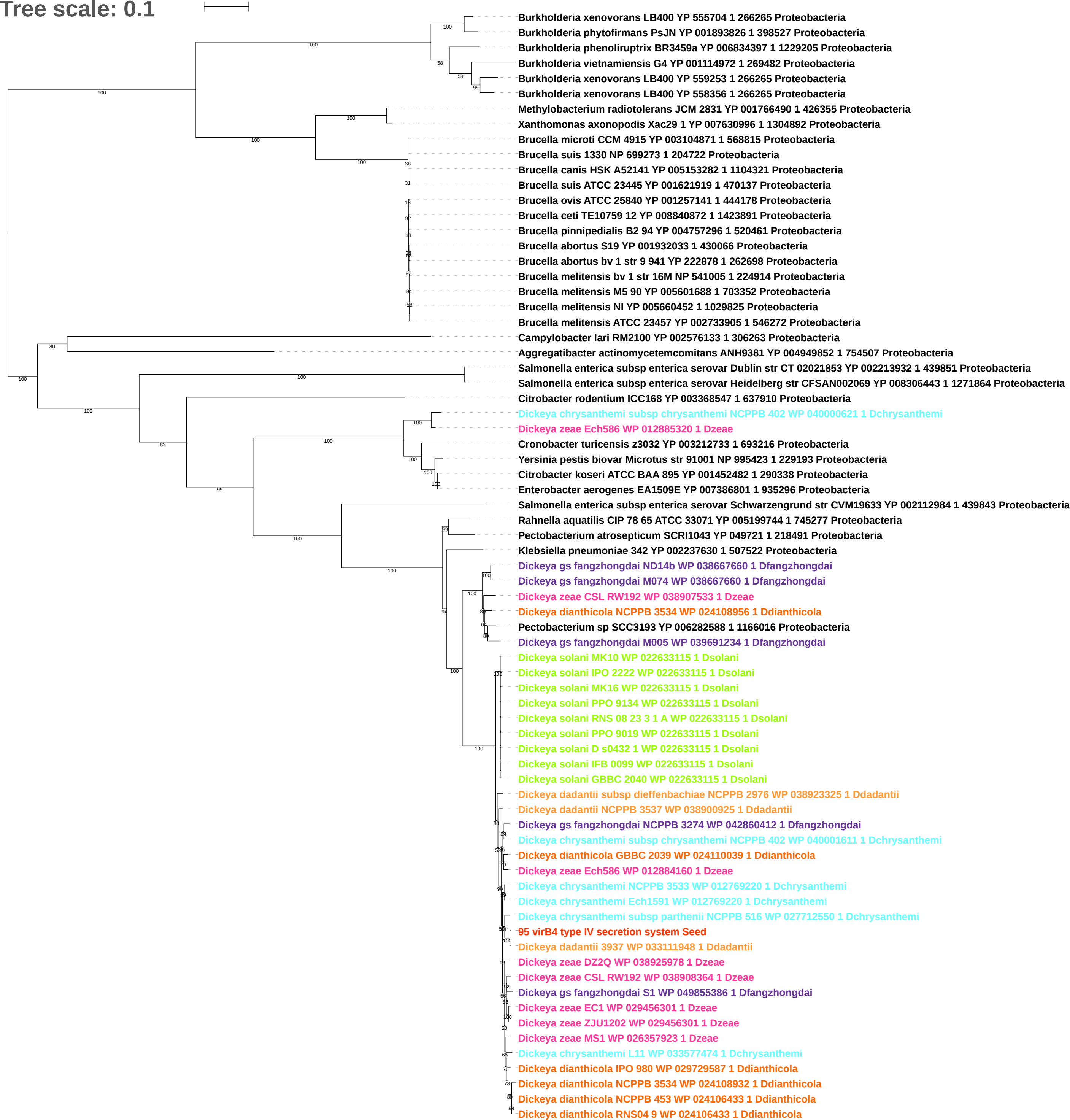
Tree scale: 0.1
Burkholderia xenovorans LB400 YP 555704 1 266265 Proteobacteria
Burkholderia phytofirmans PsJN YP 001893826 1 398527 Proteobacteria
Burkholderia phenoliruptrix BR3459a YP 006834397 1 1229205 Proteobacteria
Burkholderia vietnamiensis G4 YP 001114972 1 269482 Proteobacteria
Burkholderia xenovorans LB400 YP 559253 1 266265 Proteobacteria
Burkholderia xenovorans LB400 YP 558356 1 266265 Proteobacteria
Methylobacterium radiotolerans JCM 2831 YP 001766490 1 426355 Proteobacteria
Xanthomonas axonopodis Xac29 1 YP 007630996 1 1304892 Proteobacteria
Brucella microti CCM 4915 YP 003104871 1 568815 Proteobacteria
Brucella suis 1330 NP 699273 1 204722 Proteobacteria
Brucella canis HSK A52141 YP 005153282 1 1104321 Proteobacteria
Brucella suis ATCC 23445 YP 001621919 1 470137 Proteobacteria
Brucella ovis ATCC 25840 YP 001257141 1 444178 Proteobacteria
Brucella ceti TE10759 12 YP 008840872 1 1423891 Proteobacteria
Brucella pinnipedialis B2 94 YP 004757296 1 520461 Proteobacteria
Brucella abortus S19 YP 001932033 1 430066 Proteobacteria
Brucella abortus bv 1 str 9 941 YP 222878 1 262698 Proteobacteria
Brucella melitensis bv 1 str 16M NP 541005 1 224914 Proteobacteria
Brucella melitensis M5 90 YP 005601688 1 703352 Proteobacteria
Brucella melitensis NI YP 005660452 1 1029825 Proteobacteria
Brucella melitensis ATCC 23457 YP 002733905 1 546272 Proteobacteria
Campylobacter lari RM2100 YP 002576133 1 306263 Proteobacteria
Aggregatibacter actinomycetemcomitans ANH9381 YP 004949852 1 754507 Proteobacteria
Salmonella enterica subsp enterica serovar Dublin str CT 02021853 YP 002213932 1 439851 Proteobacteria
Salmonella enterica subsp enterica serovar Heidelberg str CFSAN002069 YP 008306443 1 1271864 Proteobacteria
Citrobacter rodentium ICC168 YP 003368547 1 637910 Proteobacteria
Dickeya chrysanthemi subsp chrysanthemi NCPPB 402 WP 040000621 1 Dchrysanthemi
Dickeya zeae Ech586 WP 012885320 1 Dzeae
Cronobacter turicensis z3032 YP 003212733 1 693216 Proteobacteria
Yersinia pestis biovar Microtus str 91001 NP 995423 1 229193 Proteobacteria
Citrobacter koseri ATCC BAA 895 YP 001452482 1 290338 Proteobacteria
Enterobacter aerogenes EA1509E YP 007386801 1 935296 Proteobacteria
Salmonella enterica subsp enterica serovar Schwarzengrund str CVM19633 YP 002112984 1 439843 Proteobacteria
Rahnella aquatilis CIP 78 65 ATCC 33071 YP 005199744 1 745277 Proteobacteria
Pectobacterium atrosepticum SCRI1043 YP 049721 1 218491 Proteobacteria
Klebsiella pneumoniae 342 YP 002237630 1 507522 Proteobacteria
Dickeya gs fangzhongdai ND14b WP 038667660 1 Dfangzhongdai
Dickeya gs fangzhongdai M074 WP 038667660 1 Dfangzhongdai
Dickeya zeae CSL RW192 WP 038907533 1 Dzeae
Dickeya dianthicola NCPPB 3534 WP 024108956 1 Ddianthicola
Pectobacterium sp SCC3193 YP 006282588 1 1166016 Proteobacteria
Dickeya gs fangzhongdai M005 WP 039691234 1 Dfangzhongdai
Dickeya solani MK10 WP 022633115 1 Dsolani
Dickeya solani IPO 2222 WP 022633115 1 Dsolani
Dickeya solani MK16 WP 022633115 1 Dsolani
Dickeya solani PPO 9134 WP 022633115 1 Dsolani
Dickeya solani RNS 08 23 3 1 A WP 022633115 1 Dsolani
Dickeya solani PPO 9019 WP 022633115 1 Dsolani
Dickeya solani D s0432 1 WP 022633115 1 Dsolani
Dickeya solani IFB 0099 WP 022633115 1 Dsolani
Dickeya solani GBBC 2040 WP 022633115 1 Dsolani
Dickeya dadantii subsp dieffenbachiae NCPPB 2976 WP 038923325 1 Ddadantii
Dickeya dadantii NCPPB 3537 WP 038900925 1 Ddadantii
Dickeya gs fangzhongdai NCPPB 3274 WP 042860412 1 Dfangzhongdai
Dickeya chrysanthemi subsp chrysanthemi NCPPB 402 WP 040001611 1 Dchrysanthemi
Dickeya dianthicola GBBC 2039 WP 024110039 1 Ddianthicola
Dickeya zeae Ech586 WP 012884160 1 Dzeae
Dickeya chrysanthemi NCPPB 3533 WP 012769220 1 Dchrysanthemi
Dickeya chrysanthemi Ech1591 WP 012769220 1 Dchrysanthemi
Dickeya chrysanthemi subsp parthenii NCPPB 516 WP 027712550 1 Dchrysanthemi
95 virB4 type IV secretion system Seed
Dickeya dadantii 3937 WP 033111948 1 Ddadantii
Dickeya zeae DZ2Q WP 038925978 1 Dzeae
Dickeya zeae CSL RW192 WP 038908364 1 Dzeae
Dickeya gs fangzhongdai S1 WP 049855386 1 Dfangzhongdai
Dickeya zeae EC1 WP 029456301 1 Dzeae
Dickeya zeae ZJU1202 WP 029456301 1 Dzeae
Dickeya zeae MS1 WP 026357923 1 Dzeae
Dickeya chrysanthemi L11 WP 033577474 1 Dchrysanthemi
Dickeya dianthicola IPO 980 WP 029729587 1 Ddianthicola
Dickeya dianthicola NCPPB 3534 WP 024108932 1 Ddianthicola
Dickeya dianthicola NCPPB 453 WP 024106433 1 Ddianthicola
Dickeya dianthicola RNS04 9 WP 024106433 1 Ddianthicola

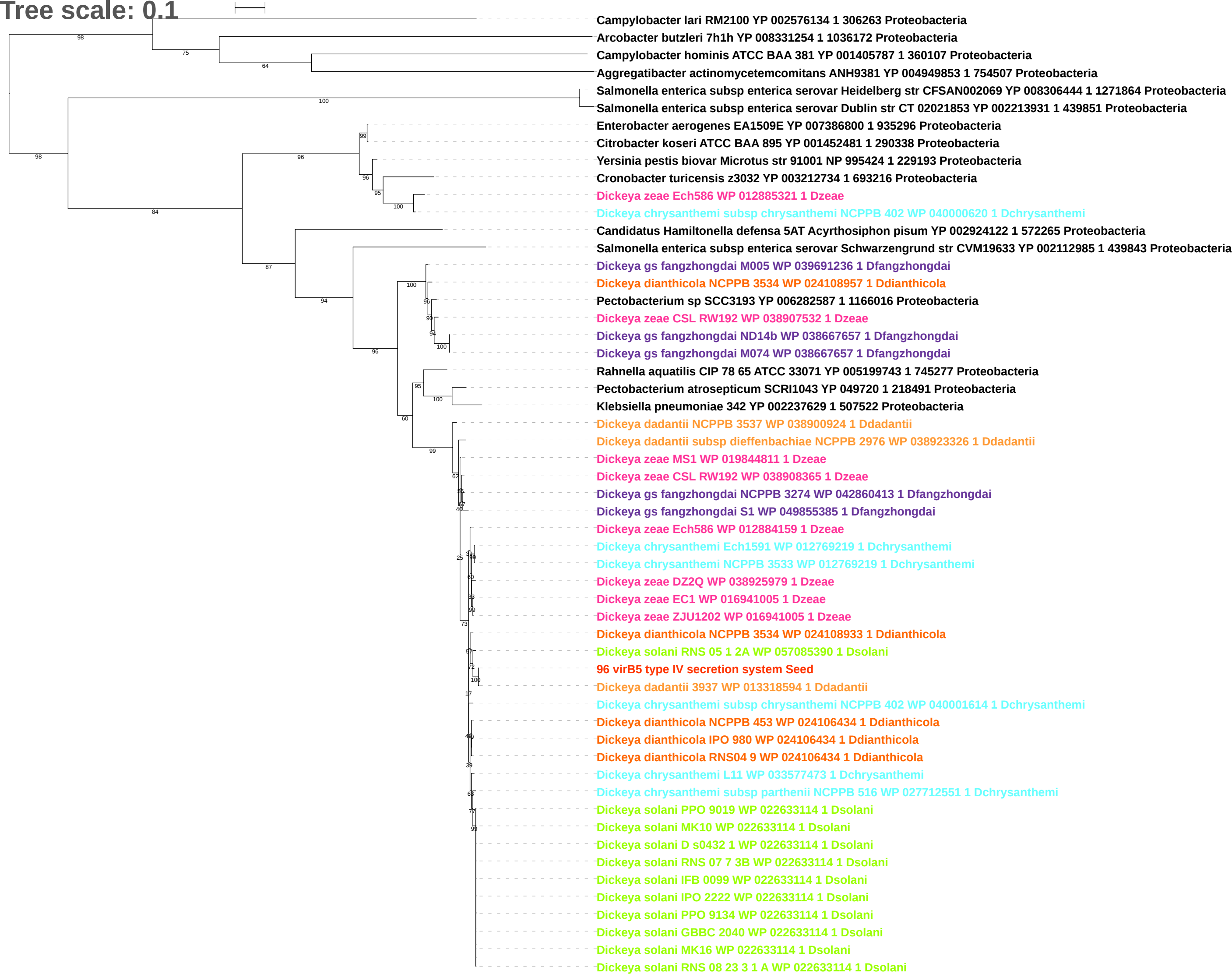

Tree scale: 0.1
Campylobacter lari RM2100 YP 002576134 1 306263 Proteobacteria
Arcobacter butzleri 7h1h YP 008331254 1 1036172 Proteobacteria
Campylobacter hominis ATCC BAA 381 YP 001405787 1 360107 Proteobacteria
Aggregatibacter actinomycetemcomitans ANH9381 YP 004949853 1 754507 Proteobacteria
Salmonella enterica subsp enterica serovar Heidelberg str CFSAN002069 YP 008306444 1 1271864 Proteobacteria
Salmonella enterica subsp enterica serovar Dublin str CT 02021853 YP 002213931 1 439851 Proteobacteria
Enterobacter aerogenes EA1509E YP 007386800 1 935296 Proteobacteria
Citrobacter koseri ATCC BAA 895 YP 001452481 1 290338 Proteobacteria
Yersinia pestis biovar Microtus str 91001 NP 995424 1 229193 Proteobacteria
Cronobacter turicensis z3032 YP 003212734 1 693216 Proteobacteria
Dickeya zeae Ech586 WP 012885321 1 Dzeae
Dickeya chrysanthemi subsp chrysanthemi NCPPB 402 WP 040000620 1 Dchrysanthemi
Candidatus Hamiltonella defensa 5AT Acyrthosiphon pisum YP 002924122 1 572265 Proteobacteria
Salmonella enterica subsp enterica serovar Schwarzengrund str CVM19633 YP 002112985 1 439843 Proteobacteria
Dickeya gs fangzhongdai M005 WP 039691236 1 Dfangzhongdai
Dickeya dianthicola NCPPB 3534 WP 024108957 1 Ddianthicola
Pectobacterium sp SCC3193 YP 006282587 1 1166016 Proteobacteria
Dickeya zeae CSL RW192 WP 038907532 1 Dzeae
Dickeya gs fangzhongdai ND14b WP 038667657 1 Dfangzhongdai
Dickeya gs fangzhongdai M074 WP 038667657 1 Dfangzhongdai
Rahnella aquatilis CIP 78 65 ATCC 33071 YP 005199743 1 745277 Proteobacteria
Pectobacterium atrosepticum SCRI1043 YP 049720 1 218491 Proteobacteria
Klebsiella pneumoniae 342 YP 002237629 1 507522 Proteobacteria
Dickeya dadantii NCPPB 3537 WP 038900924 1 Ddadantii
Dickeya dadantii subsp dieffenbachiae NCPPB 2976 WP 038923326 1 Ddadantii
Dickeya zeae MS1 WP 019844811 1 Dzeae
Dickeya zeae CSL RW192 WP 038908365 1 Dzeae
Dickeya gs fangzhongdai NCPPB 3274 WP 042860413 1 Dfangzhongdai
Dickeya gs fangzhongdai S1 WP 049855385 1 Dfangzhongdai
Dickeya zeae Ech586 WP 012884159 1 Dzeae
Dickeya chrysanthemi Ech1591 WP 012769219 1 Dchrysanthemi
Dickeya chrysanthemi NCPPB 3533 WP 012769219 1 Dchrysanthemi
Dickeya zeae DZ2Q WP 038925979 1 Dzeae
Dickeya zeae EC1 WP 016941005 1 Dzeae
Dickeya zeae ZJU1202 WP 016941005 1 Dzeae
Dickeya dianthicola NCPPB 3534 WP 024108933 1 Ddianthicola
Dickeya solani RNS 05 1 2A WP 057085390 1 Dsolani
96 virB5 type IV secretion system Seed
Dickeya dadantii 3937 WP 013318594 1 Ddadantii
Dickeya chrysanthemi subsp chrysanthemi NCPPB 402 WP 040001614 1 Dchrysanthemi
Dickeya dianthicola NCPPB 453 WP 024106434 1 Ddianthicola
Dickeya dianthicola IPO 980 WP 024106434 1 Ddianthicola
Dickeya dianthicola RNS04 9 WP 024106434 1 Ddianthicola
Dickeya chrysanthemi L11 WP 033577473 1 Dchrysanthemi
Dickeya chrysanthemi subsp parthenii NCPPB 516 WP 027712551 1 Dchrysanthemi
Dickeya solani PPO 9019 WP 022633114 1 Dsolani
Dickeya solani MK10 WP 022633114 1 Dsolani
Dickeya solani D s0432 1 WP 022633114 1 Dsolani
Dickeya solani RNS 07 7 3B WP 022633114 1 Dsolani
Dickeya solani IFB 0099 WP 022633114 1 Dsolani
Dickeya solani IPO 2222 WP 022633114 1 Dsolani
Dickeya solani PPO 9134 WP 022633114 1 Dsolani
Dickeya solani GBBC 2040 WP 022633114 1 Dsolani
Dickeya solani MK16 WP 022633114 1 Dsolani
Dickeya solani RNS 08 23 3 1 A WP 022633114 1 Dsolani

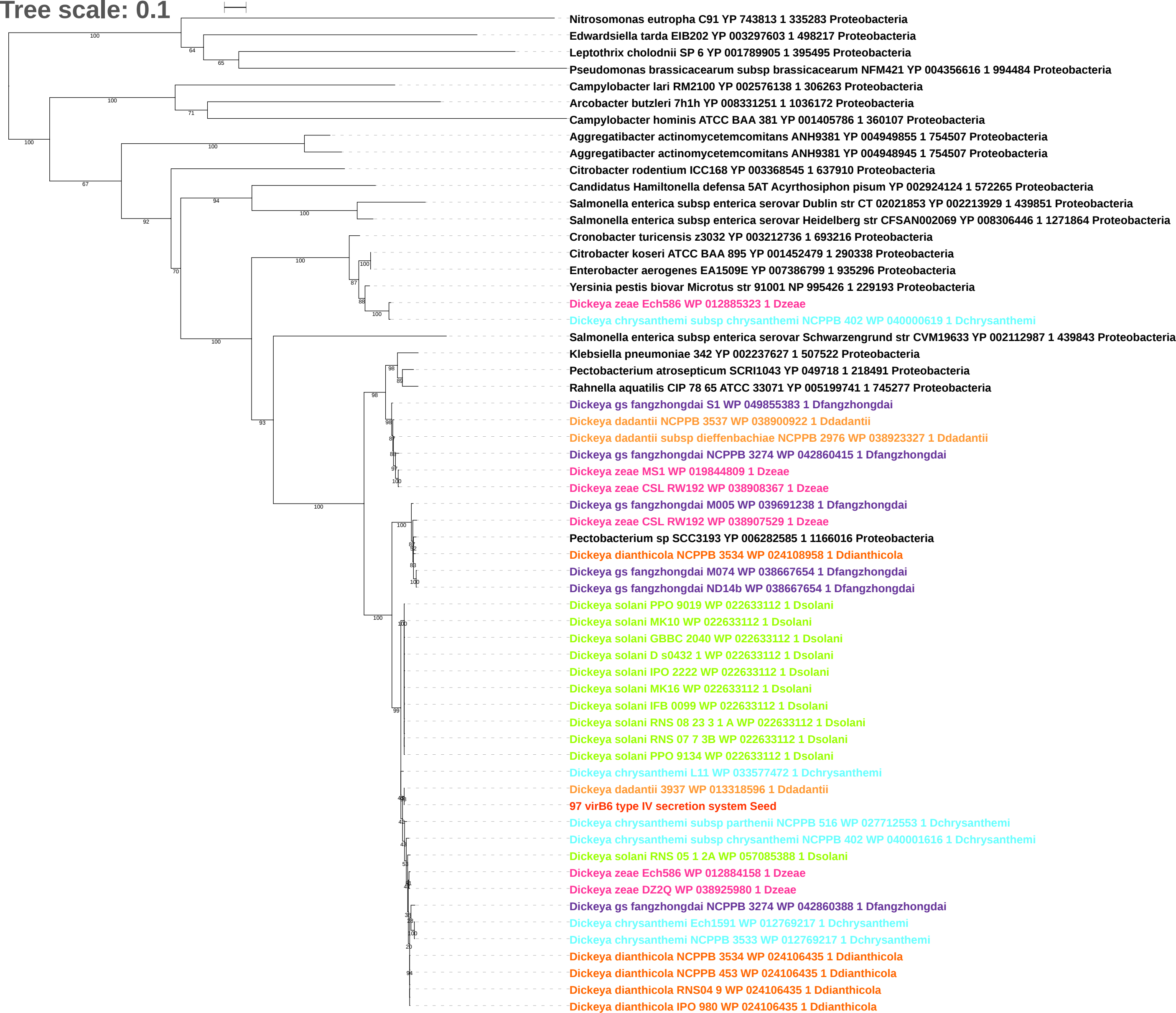
Tree scale: 0.1
Nitrosomonas eutropha C91 YP 743813 1 335283 Proteobacteria
Edwardsiella tarda EIB202 YP 003297603 1 498217 Proteobacteria
Leptothrix cholodnii SP 6 YP 001789905 1 395495 Proteobacteria
Pseudomonas brassicacearum subsp brassicacearum NFM421 YP 004356616 1 994484 Proteobacteria
Campylobacter lari RM2100 YP 002576138 1 306263 Proteobacteria
Arcobacter butzleri 7h1h YP 008331251 1 1036172 Proteobacteria
Campylobacter hominis ATCC BAA 381 YP 001405786 1 360107 Proteobacteria
Aggregatibacter actinomycetemcomitans ANH9381 YP 004949855 1 754507 Proteobacteria
Aggregatibacter actinomycetemcomitans ANH9381 YP 004948945 1 754507 Proteobacteria
Citrobacter rodentium ICC168 YP 003368545 1 637910 Proteobacteria
Candidatus Hamiltonella defensa 5AT Acyrthosiphon pisum YP 002924124 1 572265 Proteobacteria
Salmonella enterica subsp enterica serovar Dublin str CT 02021853 YP 002213929 1 439851 Proteobacteria
Salmonella enterica subsp enterica serovar Heidelberg str CFSAN002069 YP 008306446 1 1271864 Proteobacteria
Cronobacter turicensis z3032 YP 003212736 1 693216 Proteobacteria
Citrobacter koseri ATCC BAA 895 YP 001452479 1 290338 Proteobacteria
Enterobacter aerogenes EA1509E YP 007386799 1 935296 Proteobacteria
Yersinia pestis biovar Microtus str 91001 NP 995426 1 229193 Proteobacteria
Dickeya zeae Ech586 WP 012885323 1 Dzeae
Dickeya chrysanthemi subsp chrysanthemi NCPPB 402 WP 040000619 1 Dchrysanthemi
Salmonella enterica subsp enterica serovar Schwarzengrund str CVM19633 YP 002112987 1 439843 Proteobacteria
Klebsiella pneumoniae 342 YP 002237627 1 507522 Proteobacteria
Pectobacterium atrosepticum SCRI1043 YP 049718 1 218491 Proteobacteria
Rahnella aquatilis CIP 78 65 ATCC 33071 YP 005199741 1 745277 Proteobacteria
Dickeya gs fangzhongdai S1 WP 049855383 1 Dfangzhongdai
Dickeya dadantii NCPPB 3537 WP 038900922 1 Ddadantii
Dickeya dadantii subsp dieffenbachiae NCPPB 2976 WP 038923327 1 Ddadantii
Dickeya gs fangzhongdai NCPPB 3274 WP 042860415 1 Dfangzhongdai
Dickeya zeae MS1 WP 019844809 1 Dzeae
Dickeya zeae CSL RW192 WP 038908367 1 Dzeae
Dickeya gs fangzhongdai M005 WP 039691238 1 Dfangzhongdai
Dickeya zeae CSL RW192 WP 038907529 1 Dzeae
Pectobacterium sp SCC3193 YP 006282585 1 1166016 Proteobacteria
Dickeya dianthicola NCPPB 3534 WP 024108958 1 Ddianthicola
Dickeya gs fangzhongdai M074 WP 038667654 1 Dfangzhongdai
Dickeya gs fangzhongdai ND14b WP 038667654 1 Dfangzhongdai
Dickeya solani PPO 9019 WP 022633112 1 Dsolani
Dickeya solani MK10 WP 022633112 1 Dsolani
Dickeya solani GBBC 2040 WP 022633112 1 Dsolani
Dickeya solani D s0432 1 WP 022633112 1 Dsolani
Dickeya solani IPO 2222 WP 022633112 1 Dsolani
Dickeya solani MK16 WP 022633112 1 Dsolani
Dickeya solani IFB 0099 WP 022633112 1 Dsolani
Dickeya solani RNS 08 23 3 1 A WP 022633112 1 Dsolani
Dickeya solani RNS 07 7 3B WP 022633112 1 Dsolani
Dickeya solani PPO 9134 WP 022633112 1 Dsolani
Dickeya chrysanthemi L11 WP 033577472 1 Dchrysanthemi
Dickeya dadantii 3937 WP 013318596 1 Ddadantii
97 virB6 type IV secretion system Seed
Dickeya chrysanthemi subsp parthenii NCPPB 516 WP 027712553 1 Dchrysanthemi
Dickeya chrysanthemi subsp chrysanthemi NCPPB 402 WP 040001616 1 Dchrysanthemi
Dickeya solani RNS 05 1 2A WP 057085388 1 Dsolani
Dickeya zeae Ech586 WP 012884158 1 Dzeae
Dickeya zeae DZ2Q WP 038925980 1 Dzeae
Dickeya gs fangzhongdai NCPPB 3274 WP 042860388 1 Dfangzhongdai
Dickeya chrysanthemi Ech1591 WP 012769217 1 Dchrysanthemi
Dickeya chrysanthemi NCPPB 3533 WP 012769217 1 Dchrysanthemi
Dickeya dianthicola NCPPB 3534 WP 024106435 1 Ddianthicola
Dickeya dianthicola NCPPB 453 WP 024106435 1 Ddianthicola
Dickeya dianthicola RNS04 9 WP 024106435 1 Ddianthicola
Dickeya dianthicola IPO 980 WP 024106435 1 Ddianthicola

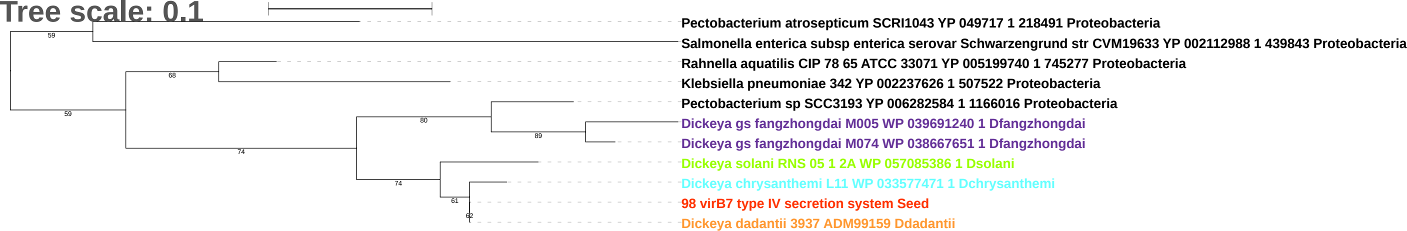

Tree scale: 0.1
Pectobacterium atrosepticum SCRI1043 YP 049717 1 218491 Proteobacteria
Salmonella enterica subsp enterica serovar Schwarzengrund str CVM19633 YP 002112988 1 439843 Proteobacteria
Rahnella aquatilis CIP 78 65 ATCC 33071 YP 005199740 1 745277 Proteobacteria
Klebsiella pneumoniae 342 YP 002237626 1 507522 Proteobacteria
Pectobacterium sp SCC3193 YP 006282584 1 1166016 Proteobacteria
Dickeya gs fangzhongdai M005 WP 039691240 1 Dfangzhongdai
Dickeya gs fangzhongdai M074 WP 038667651 1 Dfangzhongdai
Dickeya solani RNS 05 1 2A WP 057085386 1 Dsolani
Dickeya chrysanthemi L11 WP 033577471 1 Dchrysanthemi
98 virB7 type IV secretion system Seed
Dickeya dadantii 3937 ADM99159 Ddadantii
59
68
59
80
89
74
74
61
62

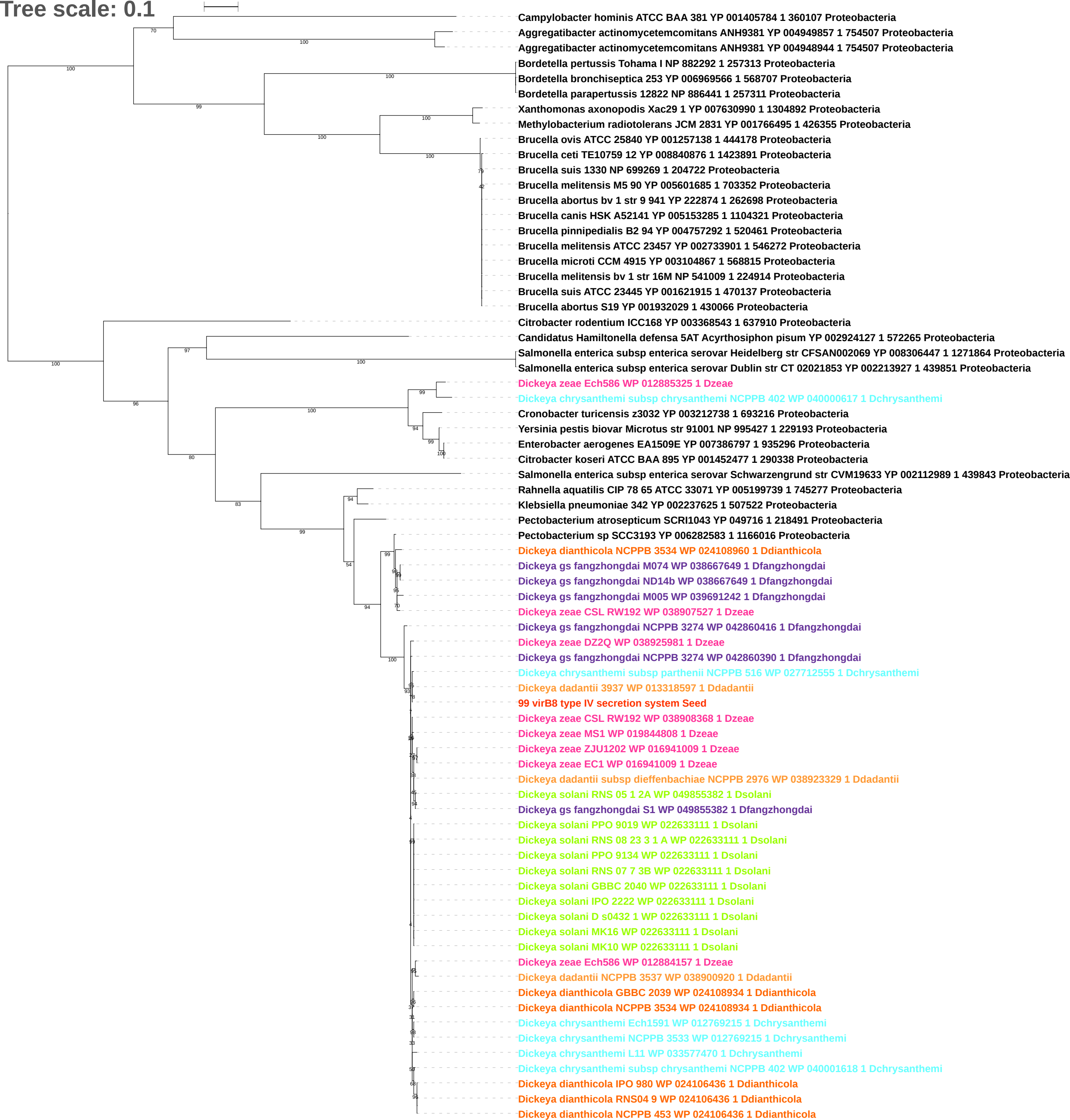
Tree scale: 0.1
Campylobacter hominis ATCC BAA 381 YP 001405784 1 360107 Proteobacteria
Aggregatibacter actinomycetemcomitans ANH9381 YP 004949857 1 754507 Proteobacteria
Aggregatibacter actinomycetemcomitans ANH9381 YP 004948944 1 754507 Proteobacteria
Bordetella pertussis Tohama I NP 882292 1 257313 Proteobacteria
Bordetella bronchiseptica 253 YP 006969566 1 568707 Proteobacteria
Bordetella parapertussis 12822 NP 886441 1 257311 Proteobacteria
Xanthomonas axonopodis Xac29 1 YP 007630990 1 1304892 Proteobacteria
Methylobacterium radiotolerans JCM 2831 YP 001766495 1 426355 Proteobacteria
Brucella ovis ATCC 25840 YP 001257138 1 444178 Proteobacteria
Brucella ceti TE10759 12 YP 008840876 1 1423891 Proteobacteria
Brucella suis 1330 NP 699269 1 204722 Proteobacteria
Brucella melitensis M5 90 YP 005601685 1 703352 Proteobacteria
Brucella abortus bv 1 str 9 941 YP 222874 1 262698 Proteobacteria
Brucella canis HSK A52141 YP 005153285 1 1104321 Proteobacteria
Brucella pinnipedialis B2 94 YP 004757292 1 520461 Proteobacteria
Brucella melitensis ATCC 23457 YP 002733901 1 546272 Proteobacteria
Brucella microti CCM 4915 YP 003104867 1 568815 Proteobacteria
Brucella melitensis bv 1 str 16M NP 541009 1 224914 Proteobacteria
Brucella suis ATCC 23445 YP 001621915 1 470137 Proteobacteria
Brucella abortus S19 YP 001932029 1 430066 Proteobacteria
Citrobacter rodentium ICC168 YP 003368543 1 637910 Proteobacteria
Candidatus Hamiltonella defensa 5AT Acyrthosiphon pisum YP 002924127 1 572265 Proteobacteria
Salmonella enterica subsp enterica serovar Heidelberg str CFSAN002069 YP 008306447 1 1271864 Proteobacteria
Salmonella enterica subsp enterica serovar Dublin str CT 02021853 YP 002213927 1 439851 Proteobacteria
Dickeya zeae Ech586 WP 012885325 1 Dzeae
Dickeya chrysanthemi subsp chrysanthemi NCPPB 402 WP 040000617 1 Dchrysanthemi
Cronobacter turicensis z3032 YP 003212738 1 693216 Proteobacteria
Yersinia pestis biovar Microtus str 91001 NP 995427 1 229193 Proteobacteria
Enterobacter aerogenes EA1509E YP 007386797 1 935296 Proteobacteria
Citrobacter koseri ATCC BAA 895 YP 001452477 1 290338 Proteobacteria
Salmonella enterica subsp enterica serovar Schwarzengrund str CVM19633 YP 002112989 1 439843 Proteobacteria
Rahnella aquatilis CIP 78 65 ATCC 33071 YP 005199739 1 745277 Proteobacteria
Klebsiella pneumoniae 342 YP 002237625 1 507522 Proteobacteria
Pectobacterium atrosepticum SCRI1043 YP 049716 1 218491 Proteobacteria
Pectobacterium sp SCC3193 YP 006282583 1 1166016 Proteobacteria
Dickeya dianthicola NCPPB 3534 WP 024108960 1 Ddianthicola
Dickeya gs fangzhongdai M074 WP 038667649 1 Dfangzhongdai
Dickeya gs fangzhongdai ND14b WP 038667649 1 Dfangzhongdai
Dickeya gs fangzhongdai M005 WP 039691242 1 Dfangzhongdai
Dickeya zeae CSL RW192 WP 038907527 1 Dzeae
Dickeya gs fangzhongdai NCPPB 3274 WP 042860416 1 Dfangzhongdai
Dickeya zeae DZ2Q WP 038925981 1 Dzeae
Dickeya gs fangzhongdai NCPPB 3274 WP 042860390 1 Dfangzhongdai
Dickeya chrysanthemi subsp parthenii NCPPB 516 WP 027712555 1 Dchrysanthemi
Dickeya dadantii 3937 WP 013318597 1 Ddadantii
99 virB8 type IV secretion system Seed
Dickeya zeae CSL RW192 WP 038908368 1 Dzeae
Dickeya zeae MS1 WP 019844808 1 Dzeae
Dickeya zeae ZJU1202 WP 016941009 1 Dzeae
Dickeya zeae EC1 WP 016941009 1 Dzeae
Dickeya dadantii subsp dieffenbachiae NCPPB 2976 WP 038923329 1 Ddadantii
Dickeya solani RNS 05 1 2A WP 049855382 1 Dsolani
Dickeya gs fangzhongdai S1 WP 049855382 1 Dfangzhongdai
Dickeya solani PPO 9019 WP 022633111 1 Dsolani
Dickeya solani RNS 08 23 3 1 A WP 022633111 1 Dsolani
Dickeya solani PPO 9134 WP 022633111 1 Dsolani
Dickeya solani RNS 07 7 3B WP 022633111 1 Dsolani
Dickeya solani GBBC 2040 WP 022633111 1 Dsolani
Dickeya solani IPO 2222 WP 022633111 1 Dsolani
Dickeya solani D s0432 1 WP 022633111 1 Dsolani
Dickeya solani MK16 WP 022633111 1 Dsolani
Dickeya solani MK10 WP 022633111 1 Dsolani
Dickeya zeae Ech586 WP 012884157 1 Dzeae
Dickeya dadantii NCPPB 3537 WP 038900920 1 Ddadantii
Dickeya dianthicola GBBC 2039 WP 024108934 1 Ddianthicola
Dickeya dianthicola NCPPB 3534 WP 024108934 1 Ddianthicola
Dickeya chrysanthemi Ech1591 WP 012769215 1 Dchrysanthemi
Dickeya chrysanthemi NCPPB 3533 WP 012769215 1 Dchrysanthemi
Dickeya chrysanthemi L11 WP 033577470 1 Dchrysanthemi
Dickeya chrysanthemi subsp chrysanthemi NCPPB 402 WP 040001618 1 Dchrysanthemi
Dickeya dianthicola IPO 980 WP 024106436 1 Ddianthicola
Dickeya dianthicola RNS04 9 WP 024106436 1 Ddianthicola
Dickeya dianthicola NCPPB 453 WP 024106436 1 Ddianthicola

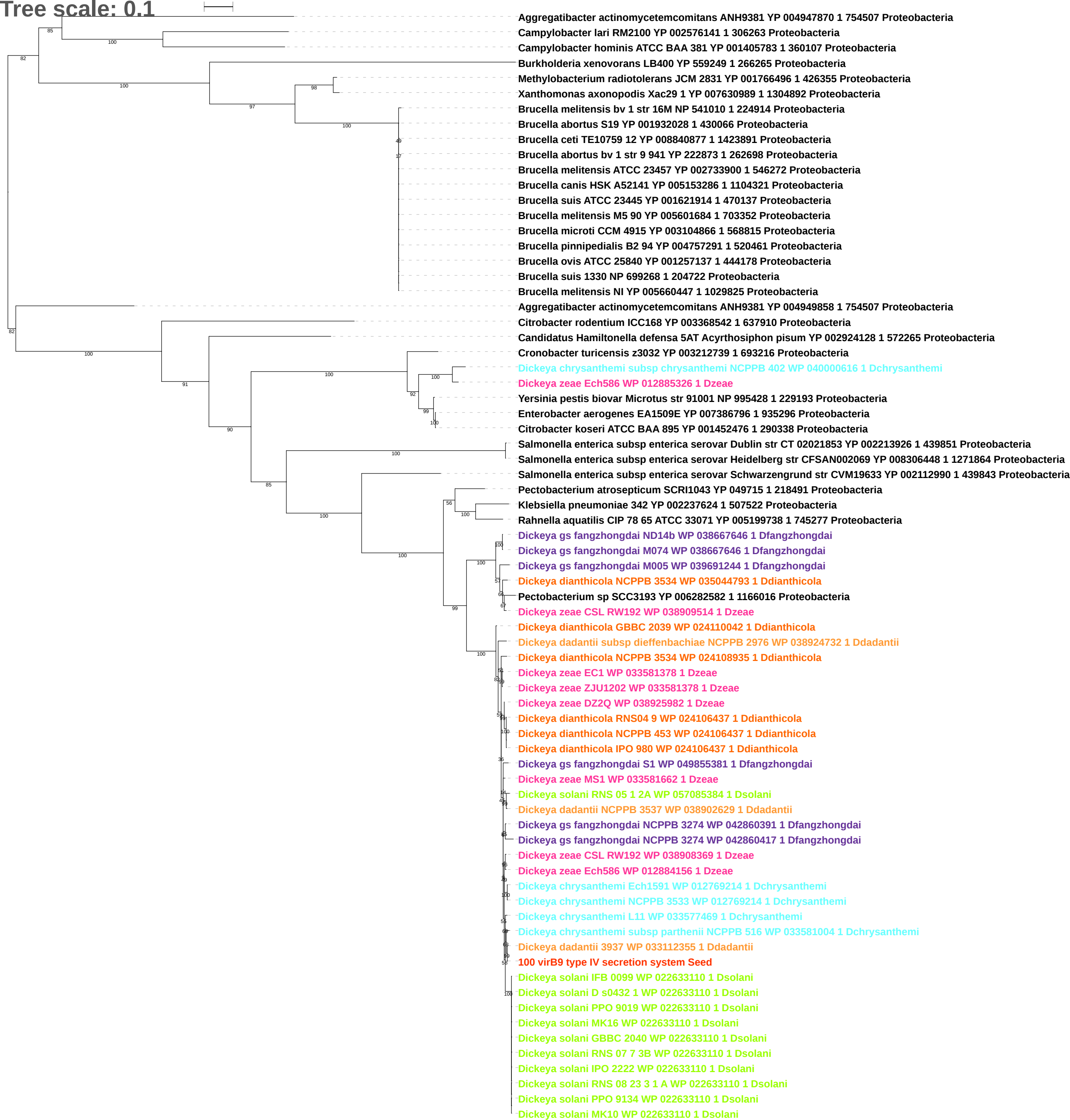
Tree scale: 0.1
Aggregatibacter actinomycetemcomitans ANH9381 YP 004947870 1 754507 Proteobacteria
Campylobacter lari RM2100 YP 002576141 1 306263 Proteobacteria
Campylobacter hominis ATCC BAA 381 YP 001405783 1 360107 Proteobacteria
Burkholderia xenovorans LB400 YP 559249 1 266265 Proteobacteria
Methylobacterium radiotolerans JCM 2831 YP 001766496 1 426355 Proteobacteria
Xanthomonas axonopodis Xac29 1 YP 007630989 1 1304892 Proteobacteria
Brucella melitensis bv 1 str 16M NP 541010 1 224914 Proteobacteria
Brucella abortus S19 YP 001932028 1 430066 Proteobacteria
Brucella ceti TE10759 12 YP 008840877 1 1423891 Proteobacteria
Brucella abortus bv 1 str 9 941 YP 222873 1 262698 Proteobacteria
Brucella melitensis ATCC 23457 YP 002733900 1 546272 Proteobacteria
Brucella canis HSK A52141 YP 005153286 1 1104321 Proteobacteria
Brucella suis ATCC 23445 YP 001621914 1 470137 Proteobacteria
Brucella melitensis M5 90 YP 005601684 1 703352 Proteobacteria
Brucella microti CCM 4915 YP 003104866 1 568815 Proteobacteria
Brucella pinnipedialis B2 94 YP 004757291 1 520461 Proteobacteria
Brucella ovis ATCC 25840 YP 001257137 1 444178 Proteobacteria
Brucella suis 1330 NP 699268 1 204722 Proteobacteria
Brucella melitensis NI YP 005660447 1 1029825 Proteobacteria
Aggregatibacter actinomycetemcomitans ANH9381 YP 004949858 1 754507 Proteobacteria
Citrobacter rodentium ICC168 YP 003368542 1 637910 Proteobacteria
Candidatus Hamiltonella defensa 5AT Acyrthosiphon pisum YP 002924128 1 572265 Proteobacteria
Cronobacter turicensis z3032 YP 003212739 1 693216 Proteobacteria
Dickeya chrysanthemi subsp chrysanthemi NCPPB 402 WP 040000616 1 Dchrysanthemi
Dickeya zeae Ech586 WP 012885326 1 Dzeae
Yersinia pestis biovar Microtus str 91001 NP 995428 1 229193 Proteobacteria
Enterobacter aerogenes EA1509E YP 007386796 1 935296 Proteobacteria
Citrobacter koseri ATCC BAA 895 YP 001452476 1 290338 Proteobacteria
Salmonella enterica subsp enterica serovar Dublin str CT 02021853 YP 002213926 1 439851 Proteobacteria
Salmonella enterica subsp enterica serovar Heidelberg str CFSAN002069 YP 008306448 1 1271864 Proteobacteria
Salmonella enterica subsp enterica serovar Schwarzengrund str CVM19633 YP 002112990 1 439843 Proteobacteria
Pectobacterium atrosepticum SCRI1043 YP 049715 1 218491 Proteobacteria
Klebsiella pneumoniae 342 YP 002237624 1 507522 Proteobacteria
Rahnella aquatilis CIP 78 65 ATCC 33071 YP 005199738 1 745277 Proteobacteria
Dickeya gs fangzhongdai ND14b WP 038667646 1 Dfangzhongdai
Dickeya gs fangzhongdai M074 WP 038667646 1 Dfangzhongdai
Dickeya gs fangzhongdai M005 WP 039691244 1 Dfangzhongdai
Dickeya dianthicola NCPPB 3534 WP 035044793 1 Ddianthicola
Pectobacterium sp SCC3193 YP 006282582 1 1166016 Proteobacteria
Dickeya zeae CSL RW192 WP 038909514 1 Dzeae
Dickeya dianthicola GBBC 2039 WP 024110042 1 Ddianthicola
Dickeya dadantii subsp dieffenbachiae NCPPB 2976 WP 038924732 1 Ddadantii
Dickeya dianthicola NCPPB 3534 WP 024108935 1 Ddianthicola
Dickeya zeae EC1 WP 033581378 1 Dzeae
Dickeya zeae ZJU1202 WP 033581378 1 Dzeae
Dickeya zeae DZ2Q WP 038925982 1 Dzeae
Dickeya dianthicola RNS04 9 WP 024106437 1 Ddianthicola
Dickeya dianthicola NCPPB 453 WP 024106437 1 Ddianthicola
Dickeya dianthicola IPO 980 WP 024106437 1 Ddianthicola
Dickeya gs fangzhongdai S1 WP 049855381 1 Dfangzhongdai
Dickeya zeae MS1 WP 033581662 1 Dzeae
Dickeya solani RNS 05 1 2A WP 057085384 1 Dsolani
Dickeya dadantii NCPPB 3537 WP 038902629 1 Ddadantii
Dickeya gs fangzhongdai NCPPB 3274 WP 042860391 1 Dfangzhongdai
Dickeya gs fangzhongdai NCPPB 3274 WP 042860417 1 Dfangzhongdai
Dickeya zeae CSL RW192 WP 038908369 1 Dzeae
Dickeya zeae Ech586 WP 012884156 1 Dzeae
Dickeya chrysanthemi Ech1591 WP 012769214 1 Dchrysanthemi
Dickeya chrysanthemi NCPPB 3533 WP 012769214 1 Dchrysanthemi
Dickeya chrysanthemi L11 WP 033577469 1 Dchrysanthemi
Dickeya chrysanthemi subsp parthenii NCPPB 516 WP 033581004 1 Dchrysanthemi
Dickeya dadantii 3937 WP 033112355 1 Ddadantii
100 virB9 type IV secretion system Seed
Dickeya solani IFB 0099 WP 022633110 1 Dsolani
Dickeya solani D s0432 1 WP 022633110 1 Dsolani
Dickeya solani PPO 9019 WP 022633110 1 Dsolani
Dickeya solani MK16 WP 022633110 1 Dsolani
Dickeya solani GBBC 2040 WP 022633110 1 Dsolani
Dickeya solani RNS 07 7 3B WP 022633110 1 Dsolani
Dickeya solani IPO 2222 WP 022633110 1 Dsolani
Dickeya solani RNS 08 23 3 1 A WP 022633110 1 Dsolani
Dickeya solani PPO 9134 WP 022633110 1 Dsolani
Dickeya solani MK10 WP 022633110 1 Dsolani

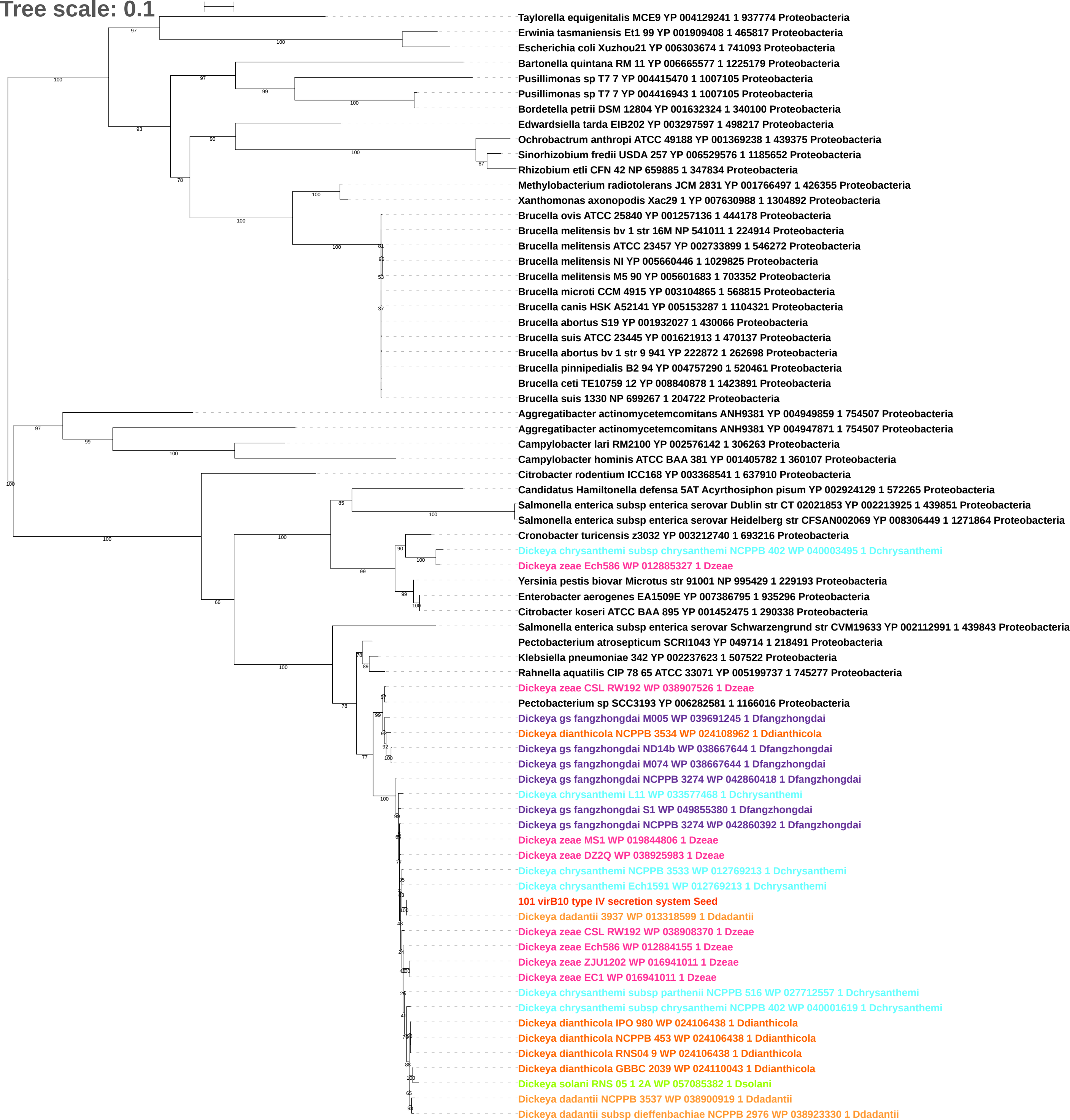

Tree scale: 0.1
Taylorella equigenitalis MCE9 YP 004129241 1 937774 Proteobacteria
Erwinia tasmaniensis Et1 99 YP 001909408 1 465817 Proteobacteria
Escherichia coli Xuzhou21 YP 006303674 1 741093 Proteobacteria
Bartonella quintana RM 11 YP 006665577 1 1225179 Proteobacteria
Pusillimonas sp T7 7 YP 004415470 1 1007105 Proteobacteria
Pusillimonas sp T7 7 YP 004416943 1 1007105 Proteobacteria
Bordetella petrii DSM 12804 YP 001632324 1 340100 Proteobacteria
Edwardsiella tarda EIB202 YP 003297597 1 498217 Proteobacteria
Ochrobactrum anthropi ATCC 49188 YP 001369238 1 439375 Proteobacteria
Sinorhizobium fredii USDA 257 YP 006529576 1 1185652 Proteobacteria
Rhizobium etli CFN 42 NP 659885 1 347834 Proteobacteria
Methylobacterium radiotolerans JCM 2831 YP 001766497 1 426355 Proteobacteria
Xanthomonas axonopodis Xac29 1 YP 007630988 1 1304892 Proteobacteria
Brucella ovis ATCC 25840 YP 001257136 1 444178 Proteobacteria
Brucella melitensis bv 1 str 16M NP 541011 1 224914 Proteobacteria
Brucella melitensis ATCC 23457 YP 002733899 1 546272 Proteobacteria
Brucella melitensis NI YP 005660446 1 1029825 Proteobacteria
Brucella melitensis M5 90 YP 005601683 1 703352 Proteobacteria
Brucella microti CCM 4915 YP 003104865 1 568815 Proteobacteria
Brucella canis HSK A52141 YP 005153287 1 1104321 Proteobacteria
Brucella abortus S19 YP 001932027 1 430066 Proteobacteria
Brucella suis ATCC 23445 YP 001621913 1 470137 Proteobacteria
Brucella abortus bv 1 str 9 941 YP 222872 1 262698 Proteobacteria
Brucella pinnipedialis B2 94 YP 004757290 1 520461 Proteobacteria
Brucella ceti TE10759 12 YP 008840878 1 1423891 Proteobacteria
Brucella suis 1330 NP 699267 1 204722 Proteobacteria
Aggregatibacter actinomycetemcomitans ANH9381 YP 004949859 1 754507 Proteobacteria
Aggregatibacter actinomycetemcomitans ANH9381 YP 004947871 1 754507 Proteobacteria
Campylobacter lari RM2100 YP 002576142 1 306263 Proteobacteria
Campylobacter hominis ATCC BAA 381 YP 001405782 1 360107 Proteobacteria
Citrobacter rodentium ICC168 YP 003368541 1 637910 Proteobacteria
Candidatus Hamiltonella defensa 5AT Acyrthosiphon pisum YP 002924129 1 572265 Proteobacteria
Salmonella enterica subsp enterica serovar Dublin str CT 02021853 YP 002213925 1 439851 Proteobacteria
Salmonella enterica subsp enterica serovar Heidelberg str CFSAN002069 YP 008306449 1 1271864 Proteobacteria
Cronobacter turicensis z3032 YP 003212740 1 693216 Proteobacteria
Dickeya chrysanthemi subsp chrysanthemi NCPPB 402 WP 040003495 1 Dchrysanthemi
Dickeya zeae Ech586 WP 012885327 1 Dzeae
Yersinia pestis biovar Microtus str 91001 NP 995429 1 229193 Proteobacteria
Enterobacter aerogenes EA1509E YP 007386795 1 935296 Proteobacteria
Citrobacter koseri ATCC BAA 895 YP 001452475 1 290338 Proteobacteria
Salmonella enterica subsp enterica serovar Schwarzengrund str CVM19633 YP 002112991 1 439843 Proteobacteria
Pectobacterium atrosepticum SCRI1043 YP 049714 1 218491 Proteobacteria
Klebsiella pneumoniae 342 YP 002237623 1 507522 Proteobacteria
Rahnella aquatilis CIP 78 65 ATCC 33071 YP 005199737 1 745277 Proteobacteria
Dickeya zeae CSL RW192 WP 038907526 1 Dzeae
Pectobacterium sp SCC3193 YP 006282581 1 1166016 Proteobacteria
Dickeya gs fangzhongdai M005 WP 039691245 1 Dfangzhongdai
Dickeya dianthicola NCPPB 3534 WP 024108962 1 Ddianthicola
Dickeya gs fangzhongdai ND14b WP 038667644 1 Dfangzhongdai
Dickeya gs fangzhongdai M074 WP 038667644 1 Dfangzhongdai
Dickeya gs fangzhongdai NCPPB 3274 WP 042860418 1 Dfangzhongdai
Dickeya chrysanthemi L11 WP 033577468 1 Dchrysanthemi
Dickeya gs fangzhongdai S1 WP 049855380 1 Dfangzhongdai
Dickeya gs fangzhongdai NCPPB 3274 WP 042860392 1 Dfangzhongdai
Dickeya zeae MS1 WP 019844806 1 Dzeae
Dickeya zeae DZ2Q WP 038925983 1 Dzeae
Dickeya chrysanthemi NCPPB 3533 WP 012769213 1 Dchrysanthemi
Dickeya chrysanthemi Ech1591 WP 012769213 1 Dchrysanthemi
101 virB10 type IV secretion system Seed
Dickeya dadantii 3937 WP 013318599 1 Ddadantii
Dickeya zeae CSL RW192 WP 038908370 1 Dzeae
Dickeya zeae Ech586 WP 012884155 1 Dzeae
Dickeya zeae ZJU1202 WP 016941011 1 Dzeae
Dickeya zeae EC1 WP 016941011 1 Dzeae
Dickeya chrysanthemi subsp parthenii NCPPB 516 WP 027712557 1 Dchrysanthemi
Dickeya chrysanthemi subsp chrysanthemi NCPPB 402 WP 040001619 1 Dchrysanthemi
Dickeya dianthicola IPO 980 WP 024106438 1 Ddianthicola
Dickeya dianthicola NCPPB 453 WP 024106438 1 Ddianthicola
Dickeya dianthicola RNS04 9 WP 024106438 1 Ddianthicola
Dickeya dianthicola GBBC 2039 WP 024110043 1 Ddianthicola
Dickeya solani RNS 05 1 2A WP 057085382 1 Dsolani
Dickeya dadantii NCPPB 3537 WP 038900919 1 Ddadantii
Dickeya dadantii subsp dieffenbachiae NCPPB 2976 WP 038923330 1 Ddadantii

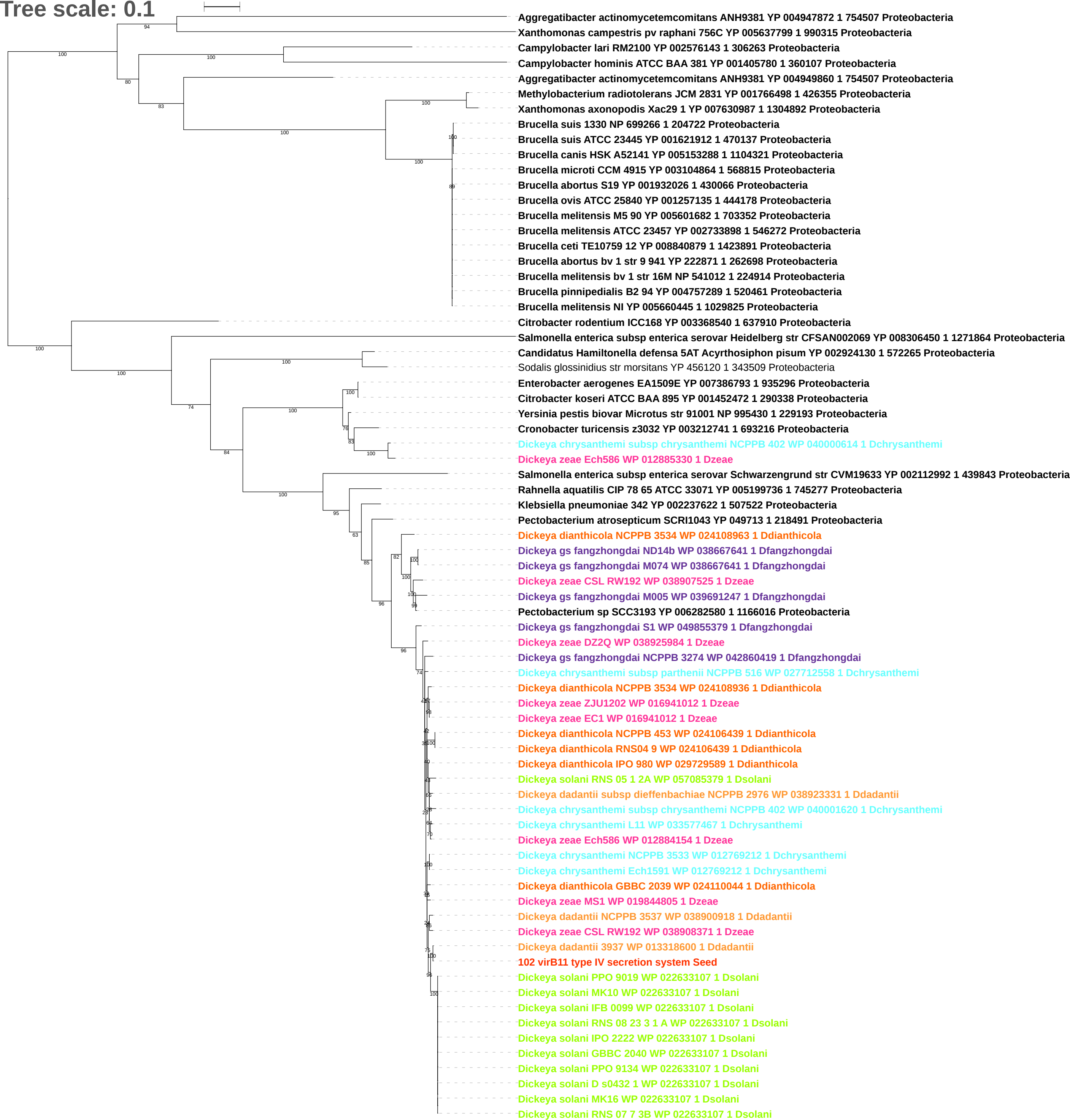
Tree scale: 0.1
Aggregatibacter actinomycetemcomitans ANH9381 YP 004947872 1 754507 Proteobacteria
Xanthomonas campestris pv raphani 756C YP 005637799 1 990315 Proteobacteria
Campylobacter lari RM2100 YP 002576143 1 306263 Proteobacteria
Campylobacter hominis ATCC BAA 381 YP 001405780 1 360107 Proteobacteria
Aggregatibacter actinomycetemcomitans ANH9381 YP 004949860 1 754507 Proteobacteria
Methylobacterium radiotolerans JCM 2831 YP 001766498 1 426355 Proteobacteria
Xanthomonas axonopodis Xac29 1 YP 007630987 1 1304892 Proteobacteria
Brucella suis 1330 NP 699266 1 204722 Proteobacteria
Brucella suis ATCC 23445 YP 001621912 1 470137 Proteobacteria
Brucella canis HSK A52141 YP 005153288 1 1104321 Proteobacteria
Brucella microti CCM 4915 YP 003104864 1 568815 Proteobacteria
Brucella abortus S19 YP 001932026 1 430066 Proteobacteria
Brucella ovis ATCC 25840 YP 001257135 1 444178 Proteobacteria
Brucella melitensis M5 90 YP 005601682 1 703352 Proteobacteria
Brucella melitensis ATCC 23457 YP 002733898 1 546272 Proteobacteria
Brucella ceti TE10759 12 YP 008840879 1 1423891 Proteobacteria
Brucella abortus bv 1 str 9 941 YP 222871 1 262698 Proteobacteria
Brucella melitensis bv 1 str 16M NP 541012 1 224914 Proteobacteria
Brucella pinnipedialis B2 94 YP 004757289 1 520461 Proteobacteria
Brucella melitensis NI YP 005660445 1 1029825 Proteobacteria
Citrobacter rodentium ICC168 YP 003368540 1 637910 Proteobacteria
Salmonella enterica subsp enterica serovar Heidelberg str CFSAN002069 YP 008306450 1 1271864 Proteobacteria
Candidatus Hamiltonella defensa 5AT Acyrthosiphon pisum YP 002924130 1 572265 Proteobacteria
Sodalis glossinidius str morsitans YP 456120 1 343509 Proteobacteria
Enterobacter aerogenes EA1509E YP 007386793 1 935296 Proteobacteria
Citrobacter koseri ATCC BAA 895 YP 001452472 1 290338 Proteobacteria
Yersinia pestis biovar Microtus str 91001 NP 995430 1 229193 Proteobacteria
Cronobacter turicensis z3032 YP 003212741 1 693216 Proteobacteria
Dickeya chrysanthemi subsp chrysanthemi NCPPB 402 WP 040000614 1 Dchrysanthemi
Dickeya zeae Ech586 WP 012885330 1 Dzeae
Salmonella enterica subsp enterica serovar Schwarzengrund str CVM19633 YP 002112992 1 439843 Proteobacteria
Rahnella aquatilis CIP 78 65 ATCC 33071 YP 005199736 1 745277 Proteobacteria
Klebsiella pneumoniae 342 YP 002237622 1 507522 Proteobacteria
Pectobacterium atrosepticum SCRI1043 YP 049713 1 218491 Proteobacteria
Dickeya dianthicola NCPPB 3534 WP 024108963 1 Ddianthicola
Dickeya gs fangzhongdai ND14b WP 038667641 1 Dfangzhongdai
Dickeya gs fangzhongdai M074 WP 038667641 1 Dfangzhongdai
Dickeya zeae CSL RW192 WP 038907525 1 Dzeae
Dickeya gs fangzhongdai M005 WP 039691247 1 Dfangzhongdai
Pectobacterium sp SCC3193 YP 006282580 1 1166016 Proteobacteria
Dickeya gs fangzhongdai S1 WP 049855379 1 Dfangzhongdai
Dickeya zeae DZ2Q WP 038925984 1 Dzeae
Dickeya gs fangzhongdai NCPPB 3274 WP 042860419 1 Dfangzhongdai
Dickeya chrysanthemi subsp parthenii NCPPB 516 WP 027712558 1 Dchrysanthemi
Dickeya dianthicola NCPPB 3534 WP 024108936 1 Ddianthicola
Dickeya zeae ZJU1202 WP 016941012 1 Dzeae
Dickeya zeae EC1 WP 016941012 1 Dzeae
Dickeya dianthicola NCPPB 453 WP 024106439 1 Ddianthicola
Dickeya dianthicola RNS04 9 WP 024106439 1 Ddianthicola
Dickeya dianthicola IPO 980 WP 029729589 1 Ddianthicola
Dickeya solani RNS 05 1 2A WP 057085379 1 Dsolani
Dickeya dadantii subsp dieffenbachiae NCPPB 2976 WP 038923331 1 Ddadantii
Dickeya chrysanthemi subsp chrysanthemi NCPPB 402 WP 040001620 1 Dchrysanthemi
Dickeya chrysanthemi L11 WP 033577467 1 Dchrysanthemi
Dickeya zeae Ech586 WP 012884154 1 Dzeae
Dickeya chrysanthemi NCPPB 3533 WP 012769212 1 Dchrysanthemi
Dickeya chrysanthemi Ech1591 WP 012769212 1 Dchrysanthemi
Dickeya dianthicola GBBC 2039 WP 024110044 1 Ddianthicola
Dickeya zeae MS1 WP 019844805 1 Dzeae
Dickeya dadantii NCPPB 3537 WP 038900918 1 Ddadantii
Dickeya zeae CSL RW192 WP 038908371 1 Dzeae
Dickeya dadantii 3937 WP 013318600 1 Ddadantii
102 virB11 type IV secretion system Seed
Dickeya solani PPO 9019 WP 022633107 1 Dsolani
Dickeya solani MK10 WP 022633107 1 Dsolani
Dickeya solani IFB 0099 WP 022633107 1 Dsolani
Dickeya solani RNS 08 23 3 1 A WP 022633107 1 Dsolani
Dickeya solani IPO 2222 WP 022633107 1 Dsolani
Dickeya solani GBBC 2040 WP 022633107 1 Dsolani
Dickeya solani PPO 9134 WP 022633107 1 Dsolani
Dickeya solani D s0432 1 WP 022633107 1 Dsolani
Dickeya solani MK16 WP 022633107 1 Dsolani
Dickeya solani RNS 07 7 3B WP 022633107 1 Dsolani

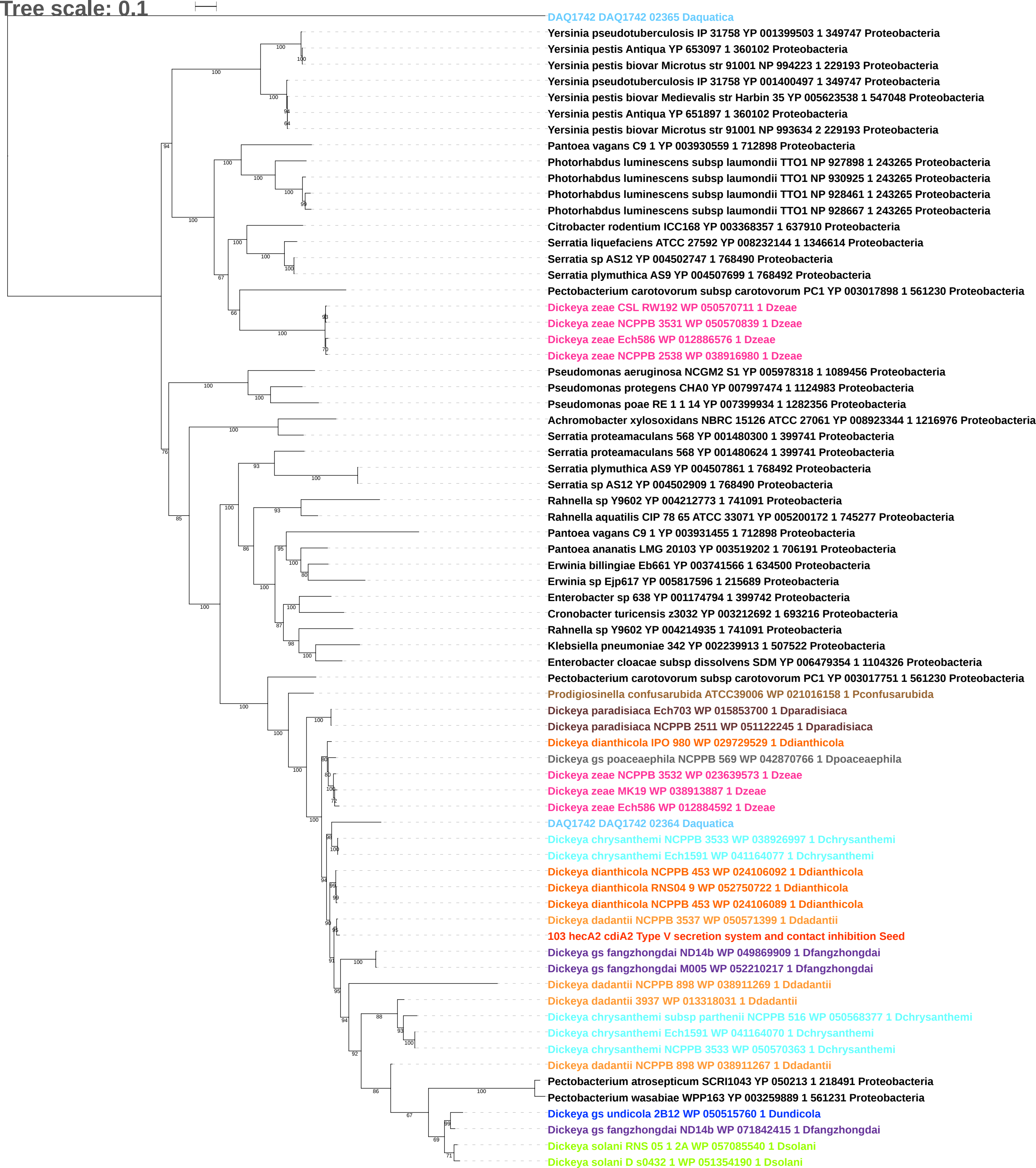
Tree scale: 0.1
DAQ1742 DAQ1742 02365 Daquatica
Yersinia pseudotuberculosis IP 31758 YP 001399503 1 349747 Proteobacteria
Yersinia pestis Antiqua YP 653097 1 360102 Proteobacteria
Yersinia pestis biovar Microtus str 91001 NP 994223 1 229193 Proteobacteria
Yersinia pseudotuberculosis IP 31758 YP 001400497 1 349747 Proteobacteria
Yersinia pestis biovar Medievalis str Harbin 35 YP 005623538 1 547048 Proteobacteria
Yersinia pestis Antiqua YP 651897 1 360102 Proteobacteria
Yersinia pestis biovar Microtus str 91001 NP 993634 2 229193 Proteobacteria
Pantoea vagans C9 1 YP 003930559 1 712898 Proteobacteria
Photorhabdus luminescens subsp laumondii TTO1 NP 927898 1 243265 Proteobacteria
Photorhabdus luminescens subsp laumondii TTO1 NP 930925 1 243265 Proteobacteria
Photorhabdus luminescens subsp laumondii TTO1 NP 928461 1 243265 Proteobacteria
Photorhabdus luminescens subsp laumondii TTO1 NP 928667 1 243265 Proteobacteria
Citrobacter rodentium ICC168 YP 003368357 1 637910 Proteobacteria
Serratia liquefaciens ATCC 27592 YP 008232144 1 1346614 Proteobacteria
Serratia sp AS12 YP 004502747 1 768490 Proteobacteria
Serratia plymuthica AS9 YP 004507699 1 768492 Proteobacteria
Pectobacterium carotovorum subsp carotovorum PC1 YP 003017898 1 561230 Proteobacteria
Dickeya zeae CSL RW192 WP 050570711 1 Dzeae
Dickeya zeae NCPPB 3531 WP 050570839 1 Dzeae
Dickeya zeae Ech586 WP 012886576 1 Dzeae
Dickeya zeae NCPPB 2538 WP 038916980 1 Dzeae
Pseudomonas aeruginosa NCGM2 S1 YP 005978318 1 1089456 Proteobacteria
Pseudomonas protegens CHA0 YP 007997474 1 1124983 Proteobacteria
Pseudomonas poae RE 1 1 14 YP 007399934 1 1282356 Proteobacteria
Achromobacter xylosoxidans NBRC 15126 ATCC 27061 YP 008923344 1 1216976 Proteobacteria
Serratia proteamaculans 568 YP 001480300 1 399741 Proteobacteria
Serratia proteamaculans 568 YP 001480624 1 399741 Proteobacteria
Serratia plymuthica AS9 YP 004507861 1 768492 Proteobacteria
Serratia sp AS12 YP 004502909 1 768490 Proteobacteria
Rahnella sp Y9602 YP 004212773 1 741091 Proteobacteria
Rahnella aquatilis CIP 78 65 ATCC 33071 YP 005200172 1 745277 Proteobacteria
Pantoea vagans C9 1 YP 003931455 1 712898 Proteobacteria
Pantoea ananatis LMG 20103 YP 003519202 1 706191 Proteobacteria
Erwinia billingiae Eb661 YP 003741566 1 634500 Proteobacteria
Erwinia sp Ejp617 YP 005817596 1 215689 Proteobacteria
Enterobacter sp 638 YP 001174794 1 399742 Proteobacteria
Cronobacter turicensis z3032 YP 003212692 1 693216 Proteobacteria
Rahnella sp Y9602 YP 004214935 1 741091 Proteobacteria
Klebsiella pneumoniae 342 YP 002239913 1 507522 Proteobacteria
Enterobacter cloacae subsp dissolvens SDM YP 006479354 1 1104326 Proteobacteria
Pectobacterium carotovorum subsp carotovorum PC1 YP 003017751 1 561230 Proteobacteria
Prodigiosinella confusarubida ATCC39006 WP 021016158 1 Pconfusarubida
Dickeya paradisiaca Ech703 WP 015853700 1 Dparadisiaca
Dickeya paradisiaca NCPPB 2511 WP 051122245 1 Dparadisiaca
Dickeya dianthicola IPO 980 WP 029729529 1 Ddianthicola
Dickeya gs poaceaephila NCPPB 569 WP 042870766 1 Dpoaceaephila
Dickeya zeae NCPPB 3532 WP 023639573 1 Dzeae
Dickeya zeae MK19 WP 038913887 1 Dzeae
Dickeya zeae Ech586 WP 012884592 1 Dzeae
DAQ1742 DAQ1742 02364 Daquatica
Dickeya chrysanthemi NCPPB 3533 WP 038926997 1 Dchrysanthemi
Dickeya chrysanthemi Ech1591 WP 041164077 1 Dchrysanthemi
Dickeya dianthicola NCPPB 453 WP 024106092 1 Ddianthicola
Dickeya dianthicola RNS04 9 WP 052750722 1 Ddianthicola
Dickeya dianthicola NCPPB 453 WP 024106089 1 Ddianthicola
Dickeya dadantii NCPPB 3537 WP 050571399 1 Ddadantii
103 hecA2 cdiA2 Type V secretion system and contact inhibition Seed
Dickeya gs fangzhongdai ND14b WP 049869909 1 Dfangzhongdai
Dickeya gs fangzhongdai M005 WP 052210217 1 Dfangzhongdai
Dickeya dadantii NCPPB 898 WP 038911269 1 Ddadantii
Dickeya dadantii 3937 WP 013318031 1 Ddadantii
Dickeya chrysanthemi subsp parthenii NCPPB 516 WP 050568377 1 Dchrysanthemi
Dickeya chrysanthemi Ech1591 WP 041164070 1 Dchrysanthemi
Dickeya chrysanthemi NCPPB 3533 WP 050570363 1 Dchrysanthemi
Dickeya dadantii NCPPB 898 WP 038911267 1 Ddadantii
Pectobacterium atrosepticum SCRI1043 YP 050213 1 218491 Proteobacteria
Pectobacterium wasabiae WPP163 YP 003259889 1 561231 Proteobacteria
Dickeya gs undicola 2B12 WP 050515760 1 Dundicola
Dickeya gs fangzhongdai ND14b WP 071842415 1 Dfangzhongdai
Dickeya solani RNS 05 1 2A WP 057085540 1 Dsolani
Dickeya solani D s0432 1 WP 051354190 1 Dsolani

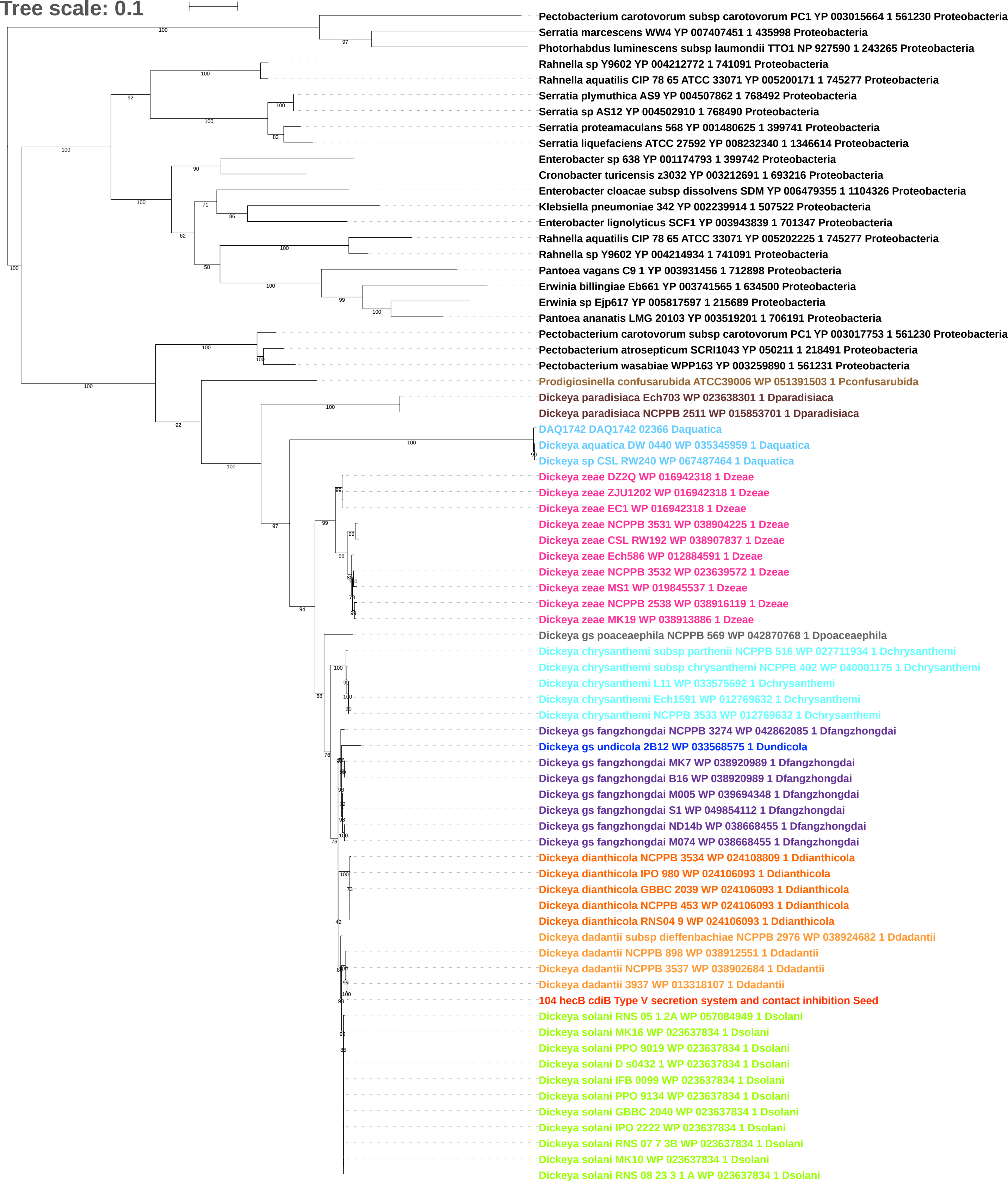
Tree scale: 0.1
Pectobacterium carotovorum subsp carotovorum PC1 YP 003015664 1 561230 Proteobacteria
Serratia marcescens WW4 YP 007407451 1 435998 Proteobacteria
Photorhabdus luminescens subsp laumondii TTO1 NP 927590 1 243265 Proteobacteria
Rahnella sp Y9602 YP 004212772 1 741091 Proteobacteria
Rahnella aquatilis CIP 78 65 ATCC 33071 YP 005200171 1 745277 Proteobacteria
Serratia plymuthica AS9 YP 004507862 1 768492 Proteobacteria
Serratia sp AS12 YP 004502910 1 768490 Proteobacteria
Serratia proteamaculans 568 YP 001480625 1 399741 Proteobacteria
Serratia liquefaciens ATCC 27592 YP 008232340 1 1346614 Proteobacteria
Enterobacter sp 638 YP 001174793 1 399742 Proteobacteria
Cronobacter turicensis z3032 YP 003212691 1 693216 Proteobacteria
Enterobacter cloacae subsp dissolvens SDM YP 006479355 1 1104326 Proteobacteria
Klebsiella pneumoniae 342 YP 002239914 1 507522 Proteobacteria
Enterobacter lignolyticus SCF1 YP 003943839 1 701347 Proteobacteria
Rahnella aquatilis CIP 78 65 ATCC 33071 YP 005202225 1 745277 Proteobacteria
Rahnella sp Y9602 YP 004214934 1 741091 Proteobacteria
Pantoea vagans C9 1 YP 003931456 1 712898 Proteobacteria
Erwinia billingiae Eb661 YP 003741565 1 634500 Proteobacteria
Erwinia sp Ejp617 YP 005817597 1 215689 Proteobacteria
Pantoea ananatis LMG 20103 YP 003519201 1 706191 Proteobacteria
Pectobacterium carotovorum subsp carotovorum PC1 YP 003017753 1 561230 Proteobacteria
Pectobacterium atrosepticum SCRI1043 YP 050211 1 218491 Proteobacteria
Pectobacterium wasabiae WPP163 YP 003259890 1 561231 Proteobacteria
Prodigiosinella confusarubida ATCC39006 WP 051391503 1 Pconfusarubida
Dickeya paradisiaca Ech703 WP 023638301 1 Dparadisiaca
Dickeya paradisiaca NCPPB 2511 WP 015853701 1 Dparadisiaca
DAQ1742 DAQ1742 02366 Daquatica
Dickeya aquatica DW 0440 WP 035345959 1 Daquatica
Dickeya sp CSL RW240 WP 067487464 1 Daquatica
Dickeya zeae DZ2Q WP 016942318 1 Dzeae
Dickeya zeae ZJU1202 WP 016942318 1 Dzeae
Dickeya zeae EC1 WP 016942318 1 Dzeae
Dickeya zeae NCPPB 3531 WP 038904225 1 Dzeae
Dickeya zeae CSL RW192 WP 038907837 1 Dzeae
Dickeya zeae Ech586 WP 012884591 1 Dzeae
Dickeya zeae NCPPB 3532 WP 023639572 1 Dzeae
Dickeya zeae MS1 WP 019845537 1 Dzeae
Dickeya zeae NCPPB 2538 WP 038916119 1 Dzeae
Dickeya zeae MK19 WP 038913886 1 Dzeae
Dickeya gs poaceaephila NCPPB 569 WP 042870768 1 Dpoaceaephila
Dickeya chrysanthemi subsp parthenii NCPPB 516 WP 027711934 1 Dchrysanthemi
Dickeya chrysanthemi subsp chrysanthemi NCPPB 402 WP 040001175 1 Dchrysanthemi
Dickeya chrysanthemi L11 WP 033575692 1 Dchrysanthemi
Dickeya chrysanthemi Ech1591 WP 012769632 1 Dchrysanthemi
Dickeya chrysanthemi NCPPB 3533 WP 012769632 1 Dchrysanthemi
Dickeya gs fangzhongdai NCPPB 3274 WP 042862085 1 Dfangzhongdai
Dickeya gs undicola 2B12 WP 033568575 1 Dundicola
Dickeya gs fangzhongdai MK7 WP 038920989 1 Dfangzhongdai
Dickeya gs fangzhongdai B16 WP 038920989 1 Dfangzhongdai
Dickeya gs fangzhongdai M005 WP 039694348 1 Dfangzhongdai
Dickeya gs fangzhongdai S1 WP 049854112 1 Dfangzhongdai
Dickeya gs fangzhongdai ND14b WP 038668455 1 Dfangzhongdai
Dickeya gs fangzhongdai M074 WP 038668455 1 Dfangzhongdai
Dickeya dianthicola NCPPB 3534 WP 024108809 1 Ddianthicola
Dickeya dianthicola IPO 980 WP 024106093 1 Ddianthicola
Dickeya dianthicola GBBC 2039 WP 024106093 1 Ddianthicola
Dickeya dianthicola NCPPB 453 WP 024106093 1 Ddianthicola
Dickeya dianthicola RNS04 9 WP 024106093 1 Ddianthicola
Dickeya dadantii subsp dieffenbachiae NCPPB 2976 WP 038924682 1 Ddadantii
Dickeya dadantii NCPPB 898 WP 038912551 1 Ddadantii
Dickeya dadantii NCPPB 3537 WP 038902684 1 Ddadantii
Dickeya dadantii 3937 WP 013318107 1 Ddadantii
104 hecB cdiB Type V secretion system and contact inhibition Seed
Dickeya solani RNS 05 1 2A WP 057084949 1 Dsolani
Dickeya solani MK16 WP 023637834 1 Dsolani
Dickeya solani PPO 9019 WP 023637834 1 Dsolani
Dickeya solani D s0432 1 WP 023637834 1 Dsolani
Dickeya solani IFB 0099 WP 023637834 1 Dsolani
Dickeya solani PPO 9134 WP 023637834 1 Dsolani
Dickeya solani GBBC 2040 WP 023637834 1 Dsolani
Dickeya solani IPO 2222 WP 023637834 1 Dsolani
Dickeya solani RNS 07 7 3B WP 023637834 1 Dsolani
Dickeya solani MK10 WP 023637834 1 Dsolani
Dickeya solani RNS 08 23 3 1 A WP 023637834 1 Dsolani

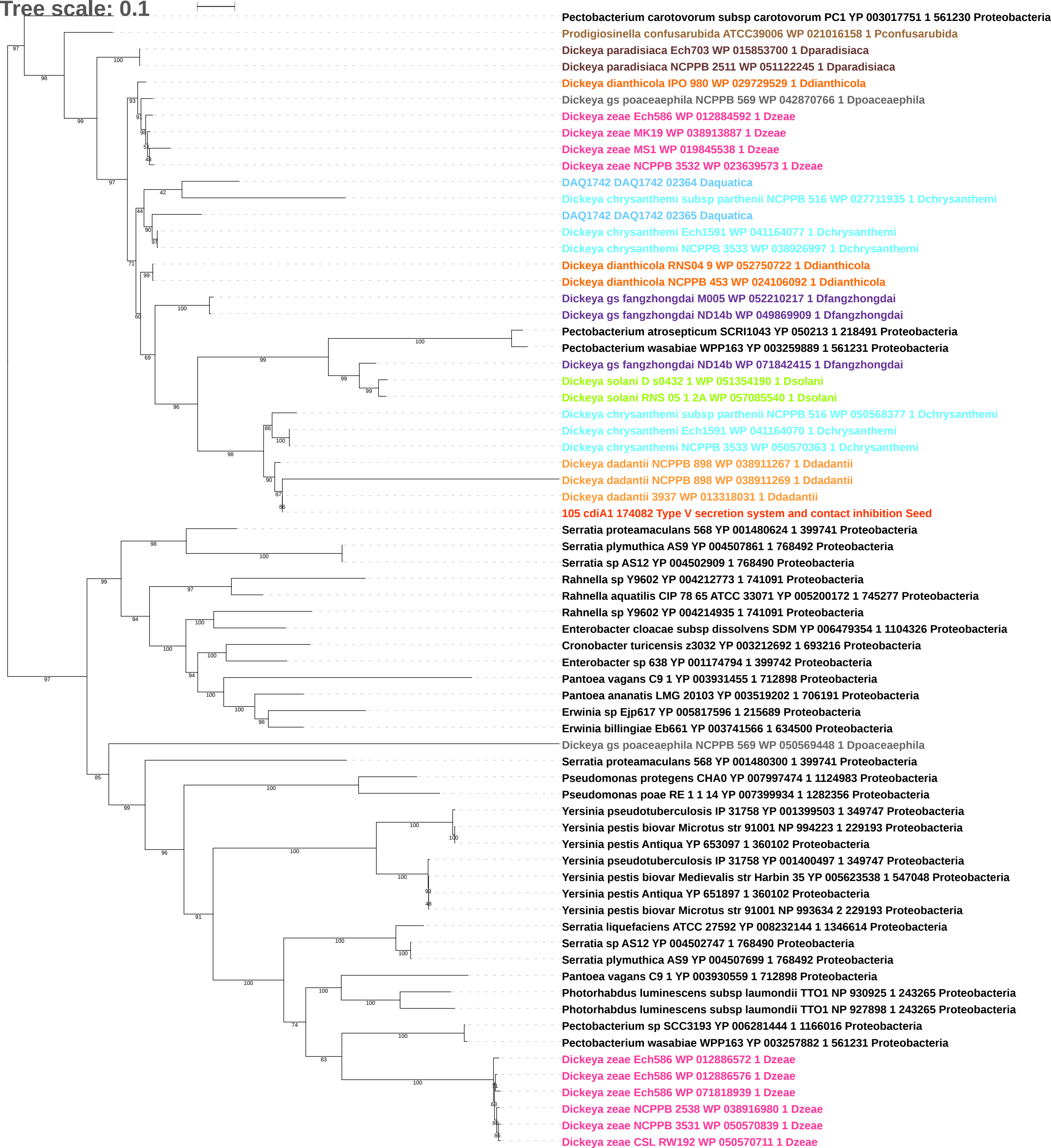
Tree scale: 0.1
Pectobacterium carotovorum subsp carotovorum PC1 YP 003017751 1 561230 Proteobacteria
Prodigiosinella confusarubida ATCC39006 WP 021016158 1 Pconfusarubida
Dickeya paradisiaca Ech703 WP 015853700 1 Dparadisiaca
Dickeya paradisiaca NCPPB 2511 WP 051122245 1 Dparadisiaca
Dickeya dianthicola IPO 980 WP 029729529 1 Ddianthicola
Dickeya gs poaceaephila NCPPB 569 WP 042870766 1 Dpoaceaephila
Dickeya zeae Ech586 WP 012884592 1 Dzeae
Dickeya zeae MK19 WP 038913887 1 Dzeae
Dickeya zeae MS1 WP 019845538 1 Dzeae
Dickeya zeae NCPPB 3532 WP 023639573 1 Dzeae
DAQ1742 DAQ1742 02364 Daquatica
Dickeya chrysanthemi subsp parthenii NCPPB 516 WP 027711935 1 Dchrysanthemi
DAQ1742 DAQ1742 02365 Daquatica
Dickeya chrysanthemi Ech1591 WP 041164077 1 Dchrysanthemi
Dickeya chrysanthemi NCPPB 3533 WP 038926997 1 Dchrysanthemi
Dickeya dianthicola RNS04 9 WP 052750722 1 Ddianthicola
Dickeya dianthicola NCPPB 453 WP 024106092 1 Ddianthicola
Dickeya gs fangzhongdai M005 WP 052210217 1 Dfangzhongdai
Dickeya gs fangzhongdai ND14b WP 049869909 1 Dfangzhongdai
Pectobacterium atrosepticum SCRI1043 YP 050213 1 218491 Proteobacteria
Pectobacterium wasabiae WPP163 YP 003259889 1 561231 Proteobacteria
Dickeya gs fangzhongdai ND14b WP 071842415 1 Dfangzhongdai
Dickeya solani D s0432 1 WP 051354190 1 Dsolani
Dickeya solani RNS 05 1 2A WP 057085540 1 Dsolani
Dickeya chrysanthemi subsp parthenii NCPPB 516 WP 050568377 1 Dchrysanthemi
Dickeya chrysanthemi Ech1591 WP 041164070 1 Dchrysanthemi
Dickeya chrysanthemi NCPPB 3533 WP 050570363 1 Dchrysanthemi
Dickeya dadantii NCPPB 898 WP 038911267 1 Ddadantii
Dickeya dadantii NCPPB 898 WP 038911269 1 Ddadantii
Dickeya dadantii 3937 WP 013318031 1 Ddadantii
105 cdiA1 174082 Type V secretion system and contact inhibition Seed
Serratia proteamaculans 568 YP 001480624 1 399741 Proteobacteria
Serratia plymuthica AS9 YP 004507861 1 768492 Proteobacteria
Serratia sp AS12 YP 004502909 1 768490 Proteobacteria
Rahnella sp Y9602 YP 004212773 1 741091 Proteobacteria
Rahnella aquatilis CIP 78 65 ATCC 33071 YP 005200172 1 745277 Proteobacteria
Rahnella sp Y9602 YP 004214935 1 741091 Proteobacteria
Enterobacter cloacae subsp dissolvens SDM YP 006479354 1 1104326 Proteobacteria
Cronobacter turicensis z3032 YP 003212692 1 693216 Proteobacteria
Enterobacter sp 638 YP 001174794 1 399742 Proteobacteria
Pantoea vagans C9 1 YP 003931455 1 712898 Proteobacteria
Pantoea ananatis LMG 20103 YP 003519202 1 706191 Proteobacteria
Erwinia sp Ejp617 YP 005817596 1 215689 Proteobacteria
Erwinia billingiae Eb661 YP 003741566 1 634500 Proteobacteria
Dickeya gs poaceaephila NCPPB 569 WP 050569448 1 Dpoaceaephila
Serratia proteamaculans 568 YP 001480300 1 399741 Proteobacteria
Pseudomonas protegens CHA0 YP 007997474 1 1124983 Proteobacteria
Pseudomonas poae RE 1 1 14 YP 007399934 1 1282356 Proteobacteria
Yersinia pseudotuberculosis IP 31758 YP 001399503 1 349747 Proteobacteria
Yersinia pestis biovar Microtus str 91001 NP 994223 1 229193 Proteobacteria
Yersinia pestis Antiqua YP 653097 1 360102 Proteobacteria
Yersinia pseudotuberculosis IP 31758 YP 001400497 1 349747 Proteobacteria
Yersinia pestis biovar Medievalis str Harbin 35 YP 005623538 1 547048 Proteobacteria
Yersinia pestis Antiqua YP 651897 1 360102 Proteobacteria
Yersinia pestis biovar Microtus str 91001 NP 993634 2 229193 Proteobacteria
Serratia liquefaciens ATCC 27592 YP 008232144 1 1346614 Proteobacteria
Serratia sp AS12 YP 004502747 1 768490 Proteobacteria
Serratia plymuthica AS9 YP 004507699 1 768492 Proteobacteria
Pantoea vagans C9 1 YP 003930559 1 712898 Proteobacteria
Photorhabdus luminescens subsp laumondii TTO1 NP 930925 1 243265 Proteobacteria
Photorhabdus luminescens subsp laumondii TTO1 NP 927898 1 243265 Proteobacteria
Pectobacterium sp SCC3193 YP 006281444 1 1166016 Proteobacteria
Pectobacterium wasabiae WPP163 YP 003257882 1 561231 Proteobacteria
Dickeya zeae Ech586 WP 012886572 1 Dzeae
Dickeya zeae Ech586 WP 012886576 1 Dzeae
Dickeya zeae Ech586 WP 071818939 1 Dzeae
Dickeya zeae NCPPB 2538 WP 038916980 1 Dzeae
Dickeya zeae NCPPB 3531 WP 050570839 1 Dzeae
Dickeya zeae CSL RW192 WP 050570711 1 Dzeae

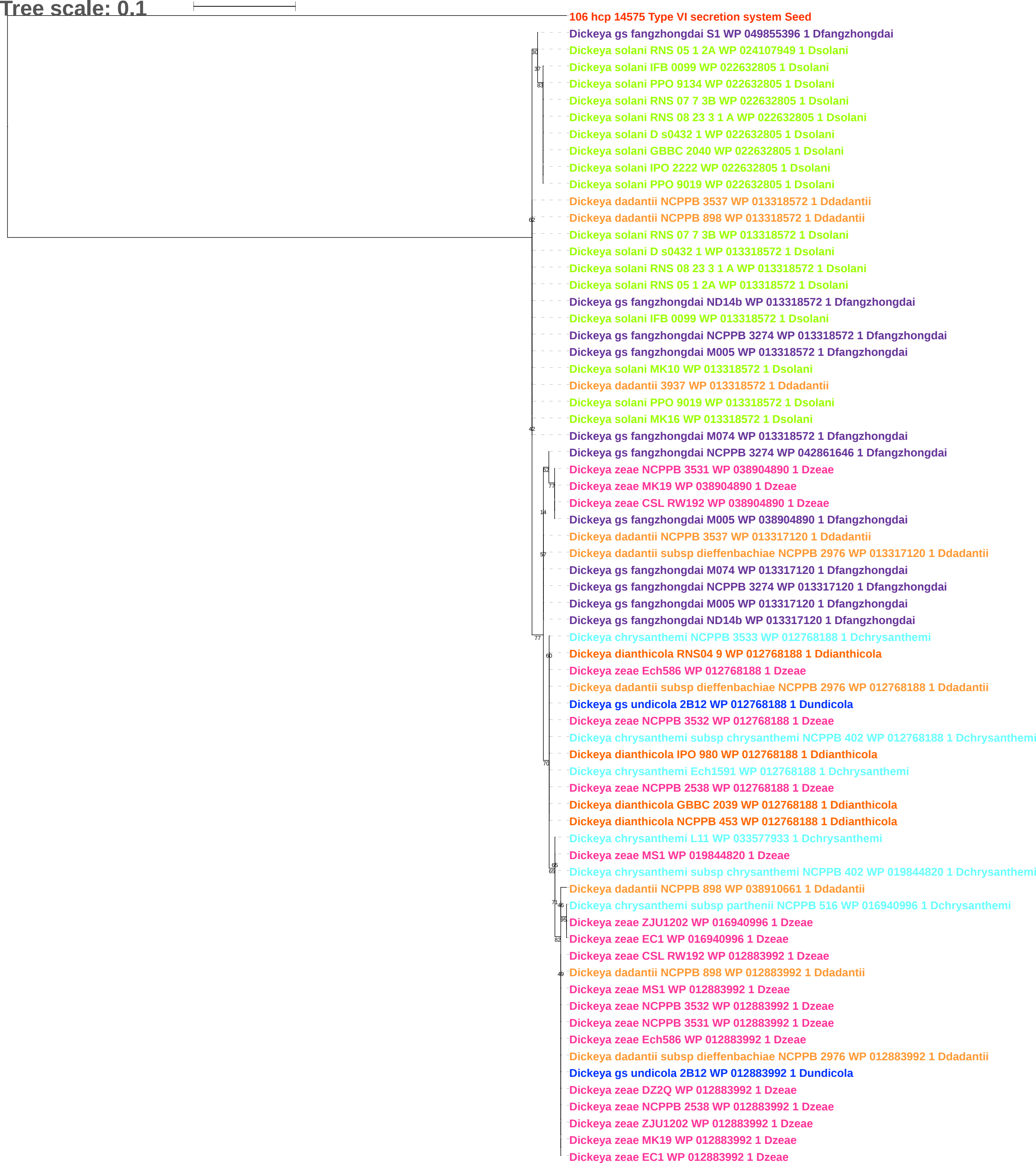
Tree scale: 0.1
106 hcp 14575 Type VI secretion system Seed
Dickeya gs fangzhongdai S1 WP 049855396 1 Dfangzhongdai
Dickeya solani RNS 05 1 2A WP 024107949 1 Dsolani
Dickeya solani IFB 0099 WP 022632805 1 Dsolani
Dickeya solani PPO 9134 WP 022632805 1 Dsolani
Dickeya solani RNS 07 7 3B WP 022632805 1 Dsolani
Dickeya solani RNS 08 23 3 1 A WP 022632805 1 Dsolani
Dickeya solani D s0432 1 WP 022632805 1 Dsolani
Dickeya solani GBBC 2040 WP 022632805 1 Dsolani
Dickeya solani IPO 2222 WP 022632805 1 Dsolani
Dickeya solani PPO 9019 WP 022632805 1 Dsolani
Dickeya dadantii NCPPB 3537 WP 013318572 1 Ddadantii
Dickeya dadantii NCPPB 898 WP 013318572 1 Ddadantii
Dickeya solani RNS 07 7 3B WP 013318572 1 Dsolani
Dickeya solani D s0432 1 WP 013318572 1 Dsolani
Dickeya solani RNS 08 23 3 1 A WP 013318572 1 Dsolani
Dickeya solani RNS 05 1 2A WP 013318572 1 Dsolani
Dickeya gs fangzhongdai ND14b WP 013318572 1 Dfangzhongdai
Dickeya solani IFB 0099 WP 013318572 1 Dsolani
Dickeya gs fangzhongdai NCPPB 3274 WP 013318572 1 Dfangzhongdai
Dickeya gs fangzhongdai M005 WP 013318572 1 Dfangzhongdai
Dickeya solani MK10 WP 013318572 1 Dsolani
Dickeya dadantii 3937 WP 013318572 1 Ddadantii
Dickeya solani PPO 9019 WP 013318572 1 Dsolani
Dickeya solani MK16 WP 013318572 1 Dsolani
Dickeya gs fangzhongdai M074 WP 013318572 1 Dfangzhongdai
Dickeya gs fangzhongdai NCPPB 3274 WP 042861646 1 Dfangzhongdai
Dickeya zeae NCPPB 3531 WP 038904890 1 Dzeae
Dickeya zeae MK19 WP 038904890 1 Dzeae
Dickeya zeae CSL RW192 WP 038904890 1 Dzeae
Dickeya gs fangzhongdai M005 WP 038904890 1 Dfangzhongdai
Dickeya dadantii NCPPB 3537 WP 013317120 1 Ddadantii
Dickeya dadantii subsp dieffenbachiae NCPPB 2976 WP 013317120 1 Ddadantii
Dickeya gs fangzhongdai M074 WP 013317120 1 Dfangzhongdai
Dickeya gs fangzhongdai NCPPB 3274 WP 013317120 1 Dfangzhongdai
Dickeya gs fangzhongdai M005 WP 013317120 1 Dfangzhongdai
Dickeya gs fangzhongdai ND14b WP 013317120 1 Dfangzhongdai
Dickeya chrysanthemi NCPPB 3533 WP 012768188 1 Dchrysanthemi
Dickeya dianthicola RNS04 9 WP 012768188 1 Ddianthicola
Dickeya zeae Ech586 WP 012768188 1 Dzeae
Dickeya dadantii subsp dieffenbachiae NCPPB 2976 WP 012768188 1 Ddadantii
Dickeya gs undicola 2B12 WP 012768188 1 Dundicola
Dickeya zeae NCPPB 3532 WP 012768188 1 Dzeae
Dickeya chrysanthemi subsp chrysanthemi NCPPB 402 WP 012768188 1 Dchrysanthemi
Dickeya dianthicola IPO 980 WP 012768188 1 Ddianthicola
Dickeya chrysanthemi Ech1591 WP 012768188 1 Dchrysanthemi
Dickeya zeae NCPPB 2538 WP 012768188 1 Dzeae
Dickeya dianthicola GBBC 2039 WP 012768188 1 Ddianthicola
Dickeya dianthicola NCPPB 453 WP 012768188 1 Ddianthicola
Dickeya chrysanthemi L11 WP 033577933 1 Dchrysanthemi
Dickeya zeae MS1 WP 019844820 1 Dzeae
Dickeya chrysanthemi subsp chrysanthemi NCPPB 402 WP 019844820 1 Dchrysanthemi
Dickeya dadantii NCPPB 898 WP 038910661 1 Ddadantii
Dickeya chrysanthemi subsp parthenii NCPPB 516 WP 016940996 1 Dchrysanthemi
Dickeya zeae ZJU1202 WP 016940996 1 Dzeae
Dickeya zeae EC1 WP 016940996 1 Dzeae
Dickeya zeae CSL RW192 WP 012883992 1 Dzeae
Dickeya dadantii NCPPB 898 WP 012883992 1 Ddadantii
Dickeya zeae MS1 WP 012883992 1 Dzeae
Dickeya zeae NCPPB 3532 WP 012883992 1 Dzeae
Dickeya zeae NCPPB 3531 WP 012883992 1 Dzeae
Dickeya zeae Ech586 WP 012883992 1 Dzeae
Dickeya dadantii subsp dieffenbachiae NCPPB 2976 WP 012883992 1 Ddadantii
Dickeya gs undicola 2B12 WP 012883992 1 Dundicola
Dickeya zeae DZ2Q WP 012883992 1 Dzeae
Dickeya zeae NCPPB 2538 WP 012883992 1 Dzeae
Dickeya zeae ZJU1202 WP 012883992 1 Dzeae
Dickeya zeae MK19 WP 012883992 1 Dzeae
Dickeya zeae EC1 WP 012883992 1 Dzeae

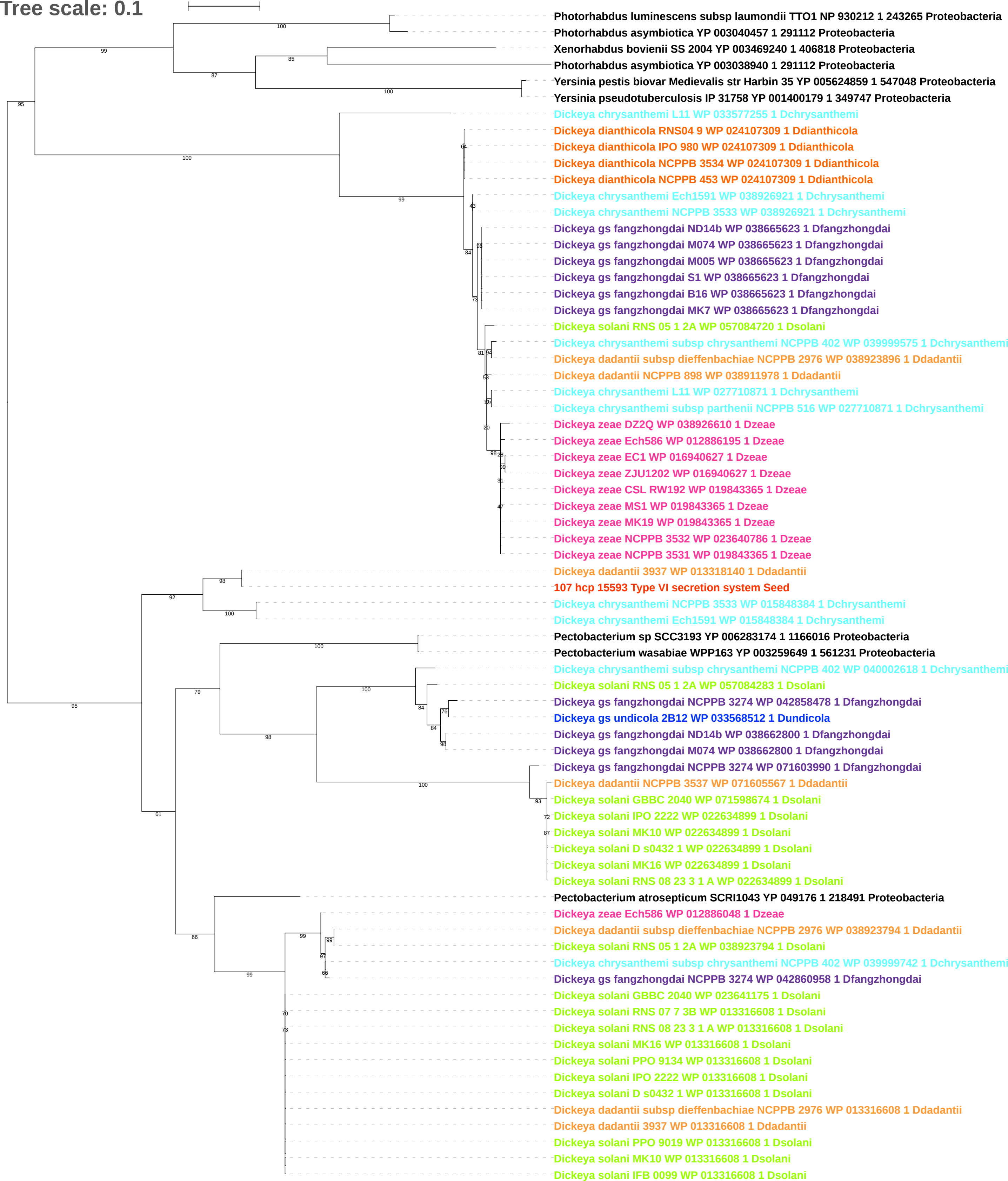

Tree scale: 0.1
Photorhabdus luminescens subsp laumondii TTO1 NP 930212 1 243265 Proteobacteria
Photorhabdus asymbiotica YP 003040457 1 291112 Proteobacteria
Xenorhabdus bovienii SS 2004 YP 003469240 1 406818 Proteobacteria
Photorhabdus asymbiotica YP 003038940 1 291112 Proteobacteria
Yersinia pestis biovar Medievalis str Harbin 35 YP 005624859 1 547048 Proteobacteria
Yersinia pseudotuberculosis IP 31758 YP 001400179 1 349747 Proteobacteria
Dickeya chrysanthemi L11 WP 033577255 1 Dchrysanthemi
Dickeya dianthicola RNS04 9 WP 024107309 1 Ddianthicola
Dickeya dianthicola IPO 980 WP 024107309 1 Ddianthicola
Dickeya dianthicola NCPPB 3534 WP 024107309 1 Ddianthicola
Dickeya dianthicola NCPPB 453 WP 024107309 1 Ddianthicola
Dickeya chrysanthemi Ech1591 WP 038926921 1 Dchrysanthemi
Dickeya chrysanthemi NCPPB 3533 WP 038926921 1 Dchrysanthemi
Dickeya gs fangzhongdai ND14b WP 038665623 1 Dfangzhongdai
Dickeya gs fangzhongdai M074 WP 038665623 1 Dfangzhongdai
Dickeya gs fangzhongdai M005 WP 038665623 1 Dfangzhongdai
Dickeya gs fangzhongdai S1 WP 038665623 1 Dfangzhongdai
Dickeya gs fangzhongdai B16 WP 038665623 1 Dfangzhongdai
Dickeya gs fangzhongdai MK7 WP 038665623 1 Dfangzhongdai
Dickeya solani RNS 05 1 2A WP 057084720 1 Dsolani
Dickeya chrysanthemi subsp chrysanthemi NCPPB 402 WP 039999575 1 Dchrysanthemi
Dickeya dadantii subsp dieffenbachiae NCPPB 2976 WP 038923896 1 Ddadantii
Dickeya dadantii NCPPB 898 WP 038911978 1 Ddadantii
Dickeya chrysanthemi L11 WP 027710871 1 Dchrysanthemi
Dickeya chrysanthemi subsp parthenii NCPPB 516 WP 027710871 1 Dchrysanthemi
Dickeya zeae DZ2Q WP 038926610 1 Dzeae
Dickeya zeae Ech586 WP 012886195 1 Dzeae
Dickeya zeae EC1 WP 016940627 1 Dzeae
Dickeya zeae ZJU1202 WP 016940627 1 Dzeae
Dickeya zeae CSL RW192 WP 019843365 1 Dzeae
Dickeya zeae MS1 WP 019843365 1 Dzeae
Dickeya zeae MK19 WP 019843365 1 Dzeae
Dickeya zeae NCPPB 3532 WP 023640786 1 Dzeae
Dickeya zeae NCPPB 3531 WP 019843365 1 Dzeae
Dickeya dadantii 3937 WP 013318140 1 Ddadantii
107 hcp 15593 Type VI secretion system Seed
Dickeya chrysanthemi NCPPB 3533 WP 015848384 1 Dchrysanthemi
Dickeya chrysanthemi Ech1591 WP 015848384 1 Dchrysanthemi
Pectobacterium sp SCC3193 YP 006283174 1 1166016 Proteobacteria
Pectobacterium wasabiae WPP163 YP 003259649 1 561231 Proteobacteria
Dickeya chrysanthemi subsp chrysanthemi NCPPB 402 WP 040002618 1 Dchrysanthemi
Dickeya solani RNS 05 1 2A WP 057084283 1 Dsolani
Dickeya gs fangzhongdai NCPPB 3274 WP 042858478 1 Dfangzhongdai
Dickeya gs undicola 2B12 WP 033568512 1 Dundicola
Dickeya gs fangzhongdai ND14b WP 038662800 1 Dfangzhongdai
Dickeya gs fangzhongdai M074 WP 038662800 1 Dfangzhongdai
Dickeya gs fangzhongdai NCPPB 3274 WP 071603990 1 Dfangzhongdai
Dickeya dadantii NCPPB 3537 WP 071605567 1 Ddadantii
Dickeya solani GBBC 2040 WP 071598674 1 Dsolani
Dickeya solani IPO 2222 WP 022634899 1 Dsolani
Dickeya solani MK10 WP 022634899 1 Dsolani
Dickeya solani D s0432 1 WP 022634899 1 Dsolani
Dickeya solani MK16 WP 022634899 1 Dsolani
Dickeya solani RNS 08 23 3 1 A WP 022634899 1 Dsolani
Pectobacterium atrosepticum SCRI1043 YP 049176 1 218491 Proteobacteria
Dickeya zeae Ech586 WP 012886048 1 Dzeae
Dickeya dadantii subsp dieffenbachiae NCPPB 2976 WP 038923794 1 Ddadantii
Dickeya solani RNS 05 1 2A WP 038923794 1 Dsolani
Dickeya chrysanthemi subsp chrysanthemi NCPPB 402 WP 039999742 1 Dchrysanthemi
Dickeya gs fangzhongdai NCPPB 3274 WP 042860958 1 Dfangzhongdai
Dickeya solani GBBC 2040 WP 023641175 1 Dsolani
Dickeya solani RNS 07 7 3B WP 013316608 1 Dsolani
Dickeya solani RNS 08 23 3 1 A WP 013316608 1 Dsolani
Dickeya solani MK16 WP 013316608 1 Dsolani
Dickeya solani PPO 9134 WP 013316608 1 Dsolani
Dickeya solani IPO 2222 WP 013316608 1 Dsolani
Dickeya solani D s0432 1 WP 013316608 1 Dsolani
Dickeya dadantii subsp dieffenbachiae NCPPB 2976 WP 013316608 1 Ddadantii
Dickeya dadantii 3937 WP 013316608 1 Ddadantii
Dickeya solani PPO 9019 WP 013316608 1 Dsolani
Dickeya solani MK10 WP 013316608 1 Dsolani
Dickeya solani IFB 0099 WP 013316608 1 Dsolani

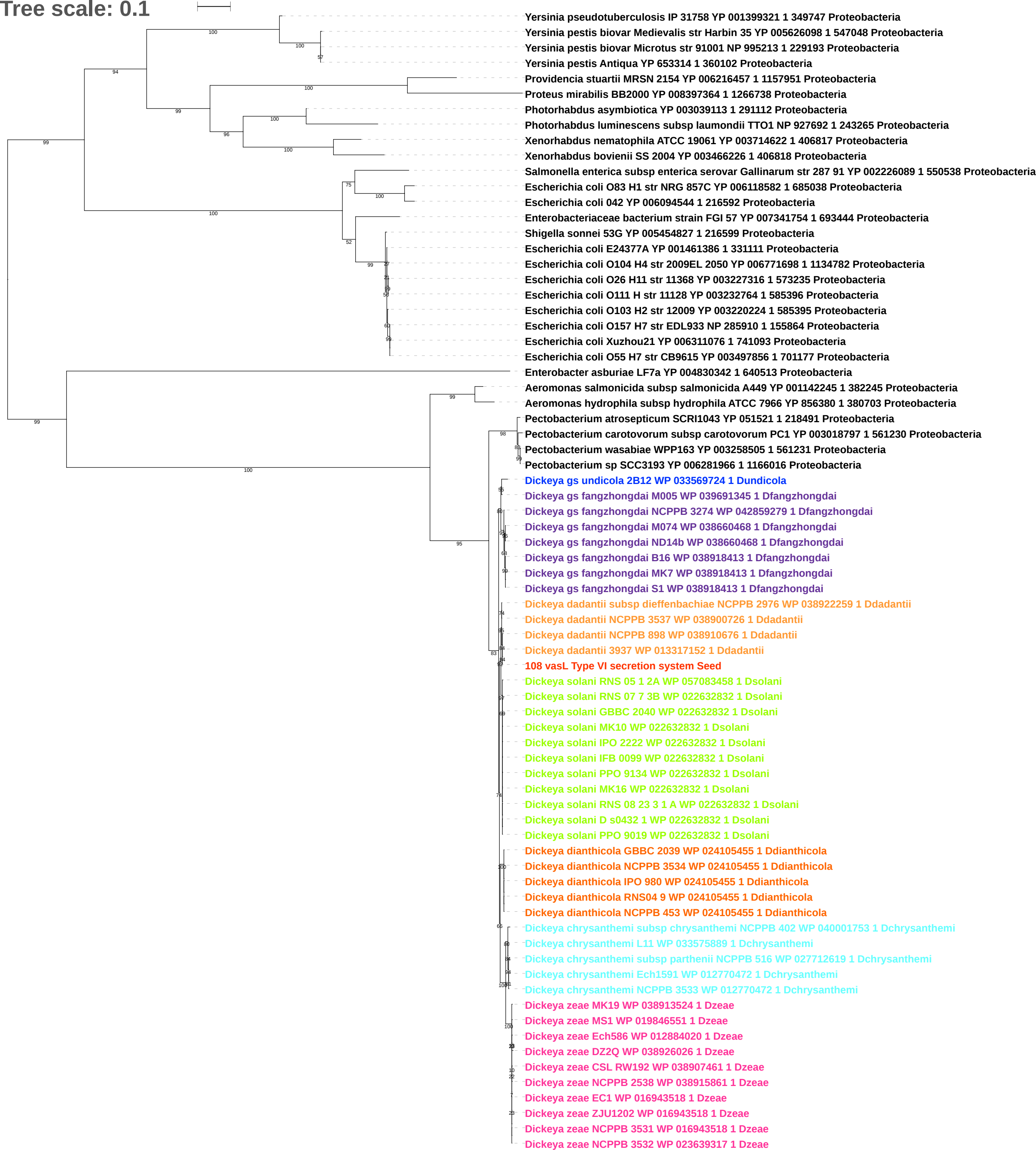
Tree scale: 0.1
Yersinia pseudotuberculosis IP 31758 YP 001399321 1 349747 Proteobacteria
Yersinia pestis biovar Medievalis str Harbin 35 YP 005626098 1 547048 Proteobacteria
Yersinia pestis biovar Microtus str 91001 NP 995213 1 229193 Proteobacteria
Yersinia pestis Antiqua YP 653314 1 360102 Proteobacteria
Providencia stuartii MRSN 2154 YP 006216457 1 1157951 Proteobacteria
Proteus mirabilis BB2000 YP 008397364 1 1266738 Proteobacteria
Photorhabdus asymbiotica YP 003039113 1 291112 Proteobacteria
Photorhabdus luminescens subsp laumondii TTO1 NP 927692 1 243265 Proteobacteria
Xenorhabdus nematophila ATCC 19061 YP 003714622 1 406817 Proteobacteria
Xenorhabdus bovienii SS 2004 YP 003466226 1 406818 Proteobacteria
Salmonella enterica subsp enterica serovar Gallinarum str 287 91 YP 002226089 1 550538 Proteobacteria
Escherichia coli O83 H1 str NRG 857C YP 006118582 1 685038 Proteobacteria
Escherichia coli 042 YP 006094544 1 216592 Proteobacteria
Enterobacteriaceae bacterium strain FGI 57 YP 007341754 1 693444 Proteobacteria
Shigella sonnei 53G YP 005454827 1 216599 Proteobacteria
Escherichia coli E24377A YP 001461386 1 331111 Proteobacteria
Escherichia coli O104 H4 str 2009EL 2050 YP 006771698 1 1134782 Proteobacteria
Escherichia coli O26 H11 str 11368 YP 003227316 1 573235 Proteobacteria
Escherichia coli O111 H str 11128 YP 003232764 1 585396 Proteobacteria
Escherichia coli O103 H2 str 12009 YP 003220224 1 585395 Proteobacteria
Escherichia coli O157 H7 str EDL933 NP 285910 1 155864 Proteobacteria
Escherichia coli Xuzhou21 YP 006311076 1 741093 Proteobacteria
Escherichia coli O55 H7 str CB9615 YP 003497856 1 701177 Proteobacteria
Enterobacter asburiae LF7a YP 004830342 1 640513 Proteobacteria
Aeromonas salmonicida subsp salmonicida A449 YP 001142245 1 382245 Proteobacteria
Aeromonas hydrophila subsp hydrophila ATCC 7966 YP 856380 1 380703 Proteobacteria
Pectobacterium atrosepticum SCRI1043 YP 051521 1 218491 Proteobacteria
Pectobacterium carotovorum subsp carotovorum PC1 YP 003018797 1 561230 Proteobacteria
Pectobacterium wasabiae WPP163 YP 003258505 1 561231 Proteobacteria
Pectobacterium sp SCC3193 YP 006281966 1 1166016 Proteobacteria
Dickeya gs undicola 2B12 WP 033569724 1 Dundicola
Dickeya gs fangzhongdai M005 WP 039691345 1 Dfangzhongdai
Dickeya gs fangzhongdai NCPPB 3274 WP 042859279 1 Dfangzhongdai
Dickeya gs fangzhongdai M074 WP 038660468 1 Dfangzhongdai
Dickeya gs fangzhongdai ND14b WP 038660468 1 Dfangzhongdai
Dickeya gs fangzhongdai B16 WP 038918413 1 Dfangzhongdai
Dickeya gs fangzhongdai MK7 WP 038918413 1 Dfangzhongdai
Dickeya gs fangzhongdai S1 WP 038918413 1 Dfangzhongdai
Dickeya dadantii subsp dieffenbachiae NCPPB 2976 WP 038922259 1 Ddadantii
Dickeya dadantii NCPPB 3537 WP 038900726 1 Ddadantii
Dickeya dadantii NCPPB 898 WP 038910676 1 Ddadantii
Dickeya dadantii 3937 WP 013317152 1 Ddadantii
108 vasL Type VI secretion system Seed
Dickeya solani RNS 05 1 2A WP 057083458 1 Dsolani
Dickeya solani RNS 07 7 3B WP 022632832 1 Dsolani
Dickeya solani GBBC 2040 WP 022632832 1 Dsolani
Dickeya solani MK10 WP 022632832 1 Dsolani
Dickeya solani IPO 2222 WP 022632832 1 Dsolani
Dickeya solani IFB 0099 WP 022632832 1 Dsolani
Dickeya solani PPO 9134 WP 022632832 1 Dsolani
Dickeya solani MK16 WP 022632832 1 Dsolani
Dickeya solani RNS 08 23 3 1 A WP 022632832 1 Dsolani
Dickeya solani D s0432 1 WP 022632832 1 Dsolani
Dickeya solani PPO 9019 WP 022632832 1 Dsolani
Dickeya dianthicola GBBC 2039 WP 024105455 1 Ddianthicola
Dickeya dianthicola NCPPB 3534 WP 024105455 1 Ddianthicola
Dickeya dianthicola IPO 980 WP 024105455 1 Ddianthicola
Dickeya dianthicola RNS04 9 WP 024105455 1 Ddianthicola
Dickeya dianthicola NCPPB 453 WP 024105455 1 Ddianthicola
Dickeya chrysanthemi subsp chrysanthemi NCPPB 402 WP 040001753 1 Dchrysanthemi
Dickeya chrysanthemi L11 WP 033575889 1 Dchrysanthemi
Dickeya chrysanthemi subsp parthenii NCPPB 516 WP 027712619 1 Dchrysanthemi
Dickeya chrysanthemi Ech1591 WP 012770472 1 Dchrysanthemi
Dickeya chrysanthemi NCPPB 3533 WP 012770472 1 Dchrysanthemi
Dickeya zeae MK19 WP 038913524 1 Dzeae
Dickeya zeae MS1 WP 019846551 1 Dzeae
Dickeya zeae Ech586 WP 012884020 1 Dzeae
Dickeya zeae DZ2Q WP 038926026 1 Dzeae
Dickeya zeae CSL RW192 WP 038907461 1 Dzeae
Dickeya zeae NCPPB 2538 WP 038915861 1 Dzeae
Dickeya zeae EC1 WP 016943518 1 Dzeae
Dickeya zeae ZJU1202 WP 016943518 1 Dzeae
Dickeya zeae NCPPB 3531 WP 016943518 1 Dzeae
Dickeya zeae NCPPB 3532 WP 023639317 1 Dzeae

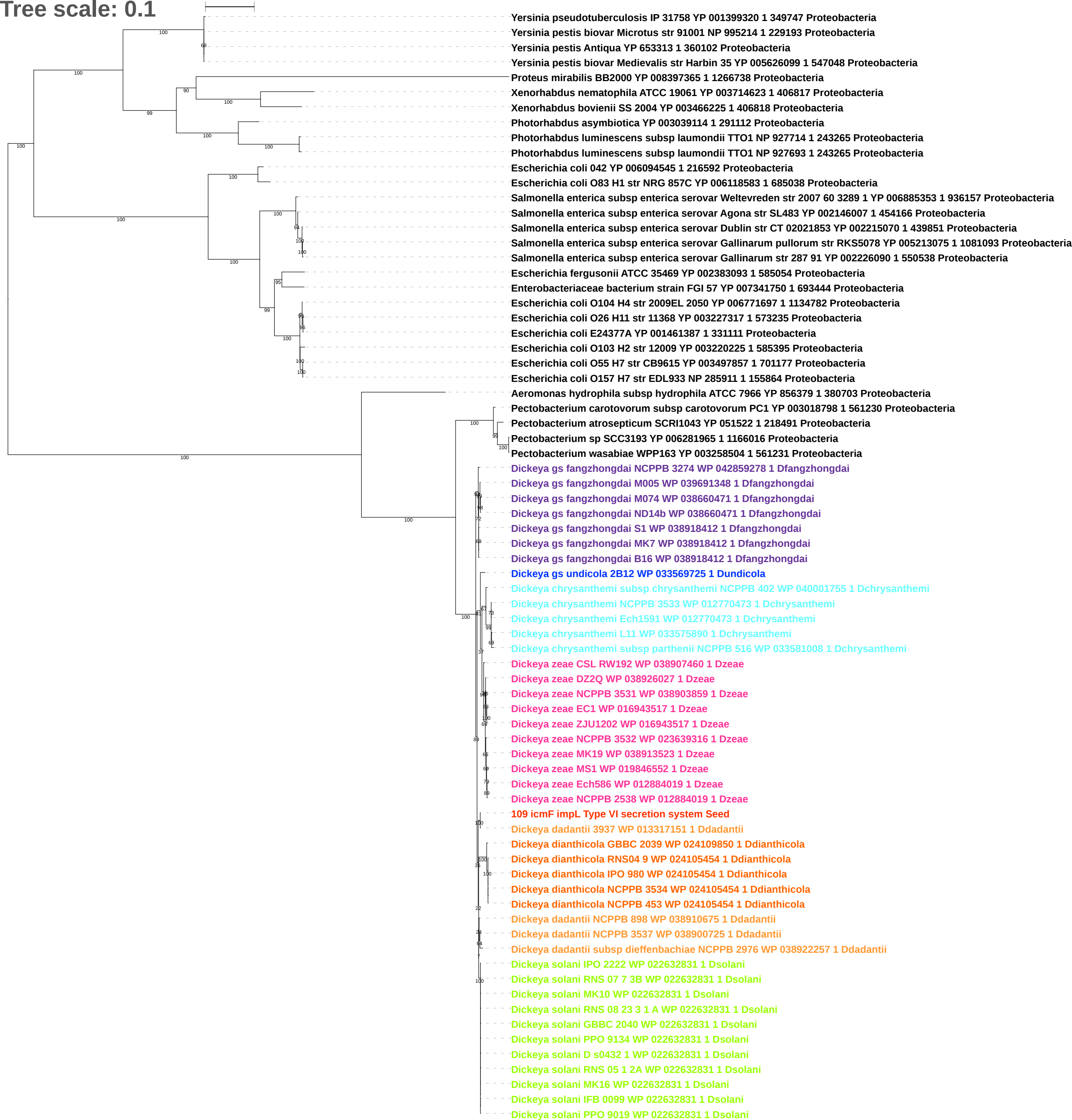
Tree scale: 0.1
Yersinia pseudotuberculosis IP 31758 YP 001399320 1 349747 Proteobacteria
Yersinia pestis biovar Microtus str 91001 NP 995214 1 229193 Proteobacteria
Yersinia pestis Antiqua YP 653313 1 360102 Proteobacteria
Yersinia pestis biovar Medievalis str Harbin 35 YP 005626099 1 547048 Proteobacteria
Proteus mirabilis BB2000 YP 008397365 1 1266738 Proteobacteria
Xenorhabdus nematophila ATCC 19061 YP 003714623 1 406817 Proteobacteria
Xenorhabdus bovienii SS 2004 YP 003466225 1 406818 Proteobacteria
Photorhabdus asymbiotica YP 003039114 1 291112 Proteobacteria
Photorhabdus luminescens subsp laumondii TTO1 NP 927714 1 243265 Proteobacteria
Photorhabdus luminescens subsp laumondii TTO1 NP 927693 1 243265 Proteobacteria
Escherichia coli 042 YP 006094545 1 216592 Proteobacteria
Escherichia coli O83 H1 str NRG 857C YP 006118583 1 685038 Proteobacteria
Salmonella enterica subsp enterica serovar Weltevreden str 2007 60 3289 1 YP 006885353 1 936157 Proteobacteria
Salmonella enterica subsp enterica serovar Agona str SL483 YP 002146007 1 454166 Proteobacteria
Salmonella enterica subsp enterica serovar Dublin str CT 02021853 YP 002215070 1 439851 Proteobacteria
Salmonella enterica subsp enterica serovar Gallinarum pullorum str RKS5078 YP 005213075 1 1081093 Proteobacteria
Salmonella enterica subsp enterica serovar Gallinarum str 287 91 YP 002226090 1 550538 Proteobacteria
Escherichia fergusonii ATCC 35469 YP 002383093 1 585054 Proteobacteria
Enterobacteriaceae bacterium strain FGI 57 YP 007341750 1 693444 Proteobacteria
Escherichia coli O104 H4 str 2009EL 2050 YP 006771697 1 1134782 Proteobacteria
Escherichia coli O26 H11 str 11368 YP 003227317 1 573235 Proteobacteria
Escherichia coli E24377A YP 001461387 1 331111 Proteobacteria
Escherichia coli O103 H2 str 12009 YP 003220225 1 585395 Proteobacteria
Escherichia coli O55 H7 str CB9615 YP 003497857 1 701177 Proteobacteria
Escherichia coli O157 H7 str EDL933 NP 285911 1 155864 Proteobacteria
Aeromonas hydrophila subsp hydrophila ATCC 7966 YP 856379 1 380703 Proteobacteria
Pectobacterium carotovorum subsp carotovorum PC1 YP 003018798 1 561230 Proteobacteria
Pectobacterium atrosepticum SCRI1043 YP 051522 1 218491 Proteobacteria
Pectobacterium sp SCC3193 YP 006281965 1 1166016 Proteobacteria
Pectobacterium wasabiae WPP163 YP 003258504 1 561231 Proteobacteria
Dickeya gs fangzhongdai NCPPB 3274 WP 042859278 1 Dfangzhongdai
Dickeya gs fangzhongdai M005 WP 039691348 1 Dfangzhongdai
Dickeya gs fangzhongdai M074 WP 038660471 1 Dfangzhongdai
Dickeya gs fangzhongdai ND14b WP 038660471 1 Dfangzhongdai
Dickeya gs fangzhongdai S1 WP 038918412 1 Dfangzhongdai
Dickeya gs fangzhongdai MK7 WP 038918412 1 Dfangzhongdai
Dickeya gs fangzhongdai B16 WP 038918412 1 Dfangzhongdai
Dickeya gs undicola 2B12 WP 033569725 1 Dundicola
Dickeya chrysanthemi subsp chrysanthemi NCPPB 402 WP 040001755 1 Dchrysanthemi
Dickeya chrysanthemi NCPPB 3533 WP 012770473 1 Dchrysanthemi
Dickeya chrysanthemi Ech1591 WP 012770473 1 Dchrysanthemi
Dickeya chrysanthemi L11 WP 033575890 1 Dchrysanthemi
Dickeya chrysanthemi subsp parthenii NCPPB 516 WP 033581008 1 Dchrysanthemi
Dickeya zeae CSL RW192 WP 038907460 1 Dzeae
Dickeya zeae DZ2Q WP 038926027 1 Dzeae
Dickeya zeae NCPPB 3531 WP 038903859 1 Dzeae
Dickeya zeae EC1 WP 016943517 1 Dzeae
Dickeya zeae ZJU1202 WP 016943517 1 Dzeae
Dickeya zeae NCPPB 3532 WP 023639316 1 Dzeae
Dickeya zeae MK19 WP 038913523 1 Dzeae
Dickeya zeae MS1 WP 019846552 1 Dzeae
Dickeya zeae Ech586 WP 012884019 1 Dzeae
Dickeya zeae NCPPB 2538 WP 012884019 1 Dzeae
109 icmF impL Type VI secretion system Seed
Dickeya dadantii 3937 WP 013317151 1 Ddadantii
Dickeya dianthicola GBBC 2039 WP 024109850 1 Ddianthicola
Dickeya dianthicola RNS04 9 WP 024105454 1 Ddianthicola
Dickeya dianthicola IPO 980 WP 024105454 1 Ddianthicola
Dickeya dianthicola NCPPB 3534 WP 024105454 1 Ddianthicola
Dickeya dianthicola NCPPB 453 WP 024105454 1 Ddianthicola
Dickeya dadantii NCPPB 898 WP 038910675 1 Ddadantii
Dickeya dadantii NCPPB 3537 WP 038900725 1 Ddadantii
Dickeya dadantii subsp dieffenbachiae NCPPB 2976 WP 038922257 1 Ddadantii
Dickeya solani IPO 2222 WP 022632831 1 Dsolani
Dickeya solani RNS 07 7 3B WP 022632831 1 Dsolani
Dickeya solani MK10 WP 022632831 1 Dsolani
Dickeya solani RNS 08 23 3 1 A WP 022632831 1 Dsolani
Dickeya solani GBBC 2040 WP 022632831 1 Dsolani
Dickeya solani PPO 9134 WP 022632831 1 Dsolani
Dickeya solani D s0432 1 WP 022632831 1 Dsolani
Dickeya solani RNS 05 1 2A WP 022632831 1 Dsolani
Dickeya solani MK16 WP 022632831 1 Dsolani
Dickeya solani IFB 0099 WP 022632831 1 Dsolani
Dickeya solani PPO 9019 WP 022632831 1 Dsolani

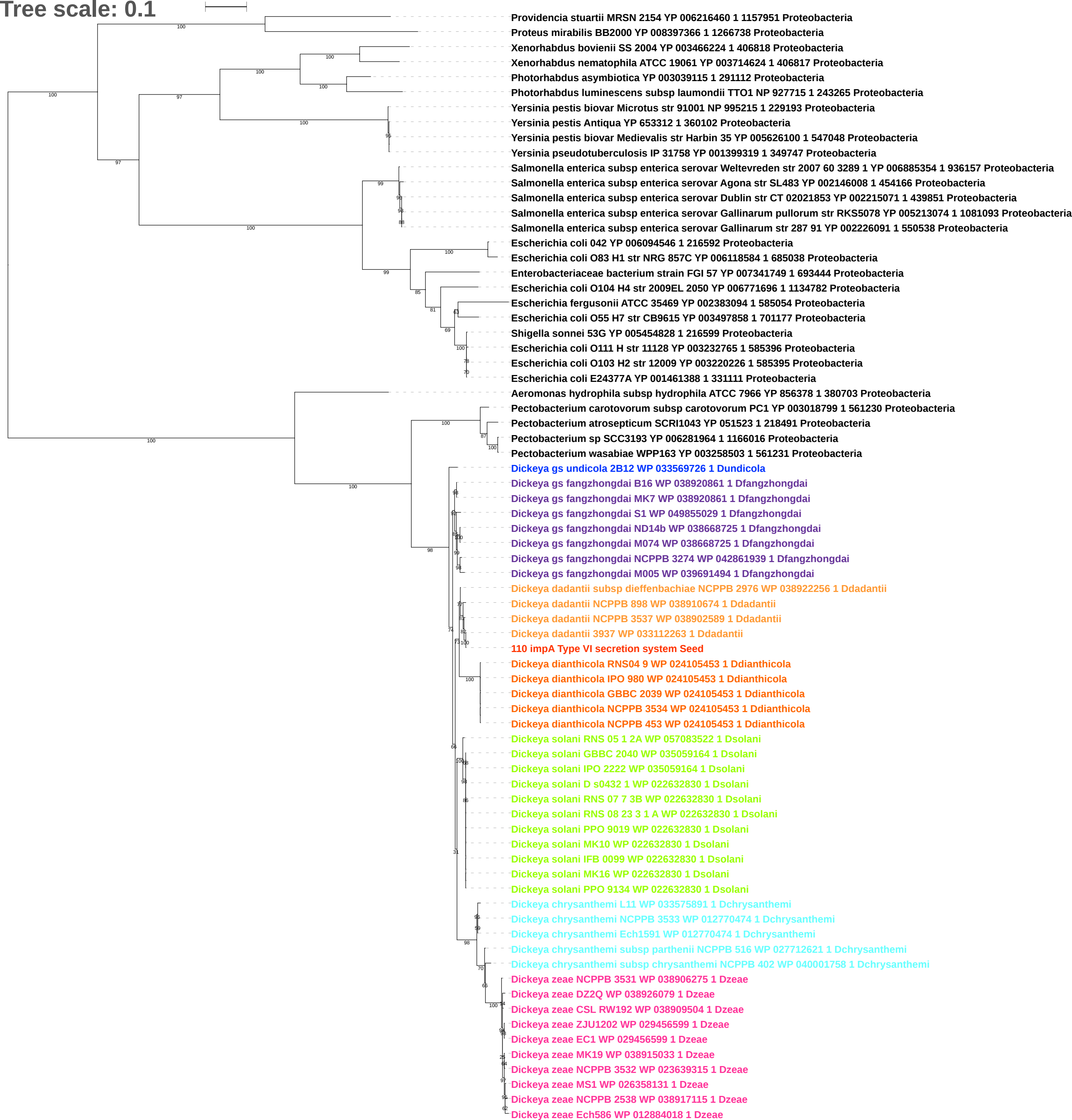
Tree scale: 0.1
Providencia stuartii MRSN 2154 YP 006216460 1 1157951 Proteobacteria
Proteus mirabilis BB2000 YP 008397366 1 1266738 Proteobacteria
Xenorhabdus bovienii SS 2004 YP 003466224 1 406818 Proteobacteria
Xenorhabdus nematophila ATCC 19061 YP 003714624 1 406817 Proteobacteria
Photorhabdus asymbiotica YP 003039115 1 291112 Proteobacteria
Photorhabdus luminescens subsp laumondii TTO1 NP 927715 1 243265 Proteobacteria
Yersinia pestis biovar Microtus str 91001 NP 995215 1 229193 Proteobacteria
Yersinia pestis Antiqua YP 653312 1 360102 Proteobacteria
Yersinia pestis biovar Medievalis str Harbin 35 YP 005626100 1 547048 Proteobacteria
Yersinia pseudotuberculosis IP 31758 YP 001399319 1 349747 Proteobacteria
Salmonella enterica subsp enterica serovar Weltevreden str 2007 60 3289 1 YP 006885354 1 936157 Proteobacteria
Salmonella enterica subsp enterica serovar Agona str SL483 YP 002146008 1 454166 Proteobacteria
Salmonella enterica subsp enterica serovar Dublin str CT 02021853 YP 002215071 1 439851 Proteobacteria
Salmonella enterica subsp enterica serovar Gallinarum pullorum str RKS5078 YP 005213074 1 1081093 Proteobacteria
Salmonella enterica subsp enterica serovar Gallinarum str 287 91 YP 002226091 1 550538 Proteobacteria
Escherichia coli 042 YP 006094546 1 216592 Proteobacteria
Escherichia coli O83 H1 str NRG 857C YP 006118584 1 685038 Proteobacteria
Enterobacteriaceae bacterium strain FGI 57 YP 007341749 1 693444 Proteobacteria
Escherichia coli O104 H4 str 2009EL 2050 YP 006771696 1 1134782 Proteobacteria
Escherichia fergusonii ATCC 35469 YP 002383094 1 585054 Proteobacteria
Escherichia coli O55 H7 str CB9615 YP 003497858 1 701177 Proteobacteria
Shigella sonnei 53G YP 005454828 1 216599 Proteobacteria
Escherichia coli O111 H str 11128 YP 003232765 1 585396 Proteobacteria
Escherichia coli O103 H2 str 12009 YP 003220226 1 585395 Proteobacteria
Escherichia coli E24377A YP 001461388 1 331111 Proteobacteria
Aeromonas hydrophila subsp hydrophila ATCC 7966 YP 856378 1 380703 Proteobacteria
Pectobacterium carotovorum subsp carotovorum PC1 YP 003018799 1 561230 Proteobacteria
Pectobacterium atrosepticum SCRI1043 YP 051523 1 218491 Proteobacteria
Pectobacterium sp SCC3193 YP 006281964 1 1166016 Proteobacteria
Pectobacterium wasabiae WPP163 YP 003258503 1 561231 Proteobacteria
Dickeya gs undicola 2B12 WP 033569726 1 Dundicola
Dickeya gs fangzhongdai B16 WP 038920861 1 Dfangzhongdai
Dickeya gs fangzhongdai MK7 WP 038920861 1 Dfangzhongdai
Dickeya gs fangzhongdai S1 WP 049855029 1 Dfangzhongdai
Dickeya gs fangzhongdai ND14b WP 038668725 1 Dfangzhongdai
Dickeya gs fangzhongdai M074 WP 038668725 1 Dfangzhongdai
Dickeya gs fangzhongdai NCPPB 3274 WP 042861939 1 Dfangzhongdai
Dickeya gs fangzhongdai M005 WP 039691494 1 Dfangzhongdai
Dickeya dadantii subsp dieffenbachiae NCPPB 2976 WP 038922256 1 Ddadantii
Dickeya dadantii NCPPB 898 WP 038910674 1 Ddadantii
Dickeya dadantii NCPPB 3537 WP 038902589 1 Ddadantii
Dickeya dadantii 3937 WP 033112263 1 Ddadantii
110 impA Type VI secretion system Seed
Dickeya dianthicola RNS04 9 WP 024105453 1 Ddianthicola
Dickeya dianthicola IPO 980 WP 024105453 1 Ddianthicola
Dickeya dianthicola GBBC 2039 WP 024105453 1 Ddianthicola
Dickeya dianthicola NCPPB 3534 WP 024105453 1 Ddianthicola
Dickeya dianthicola NCPPB 453 WP 024105453 1 Ddianthicola
Dickeya solani RNS 05 1 2A WP 057083522 1 Dsolani
Dickeya solani GBBC 2040 WP 035059164 1 Dsolani
Dickeya solani IPO 2222 WP 035059164 1 Dsolani
Dickeya solani D s0432 1 WP 022632830 1 Dsolani
Dickeya solani RNS 07 7 3B WP 022632830 1 Dsolani
Dickeya solani RNS 08 23 3 1 A WP 022632830 1 Dsolani
Dickeya solani PPO 9019 WP 022632830 1 Dsolani
Dickeya solani MK10 WP 022632830 1 Dsolani
Dickeya solani IFB 0099 WP 022632830 1 Dsolani
Dickeya solani MK16 WP 022632830 1 Dsolani
Dickeya solani PPO 9134 WP 022632830 1 Dsolani
Dickeya chrysanthemi L11 WP 033575891 1 Dchrysanthemi
Dickeya chrysanthemi NCPPB 3533 WP 012770474 1 Dchrysanthemi
Dickeya chrysanthemi Ech1591 WP 012770474 1 Dchrysanthemi
Dickeya chrysanthemi subsp parthenii NCPPB 516 WP 027712621 1 Dchrysanthemi
Dickeya chrysanthemi subsp chrysanthemi NCPPB 402 WP 040001758 1 Dchrysanthemi
Dickeya zeae NCPPB 3531 WP 038906275 1 Dzeae
Dickeya zeae DZ2Q WP 038926079 1 Dzeae
Dickeya zeae CSL RW192 WP 038909504 1 Dzeae
Dickeya zeae ZJU1202 WP 029456599 1 Dzeae
Dickeya zeae EC1 WP 029456599 1 Dzeae
Dickeya zeae MK19 WP 038915033 1 Dzeae
Dickeya zeae NCPPB 3532 WP 023639315 1 Dzeae
Dickeya zeae MS1 WP 026358131 1 Dzeae
Dickeya zeae NCPPB 2538 WP 038917115 1 Dzeae
Dickeya zeae Ech586 WP 012884018 1 Dzeae

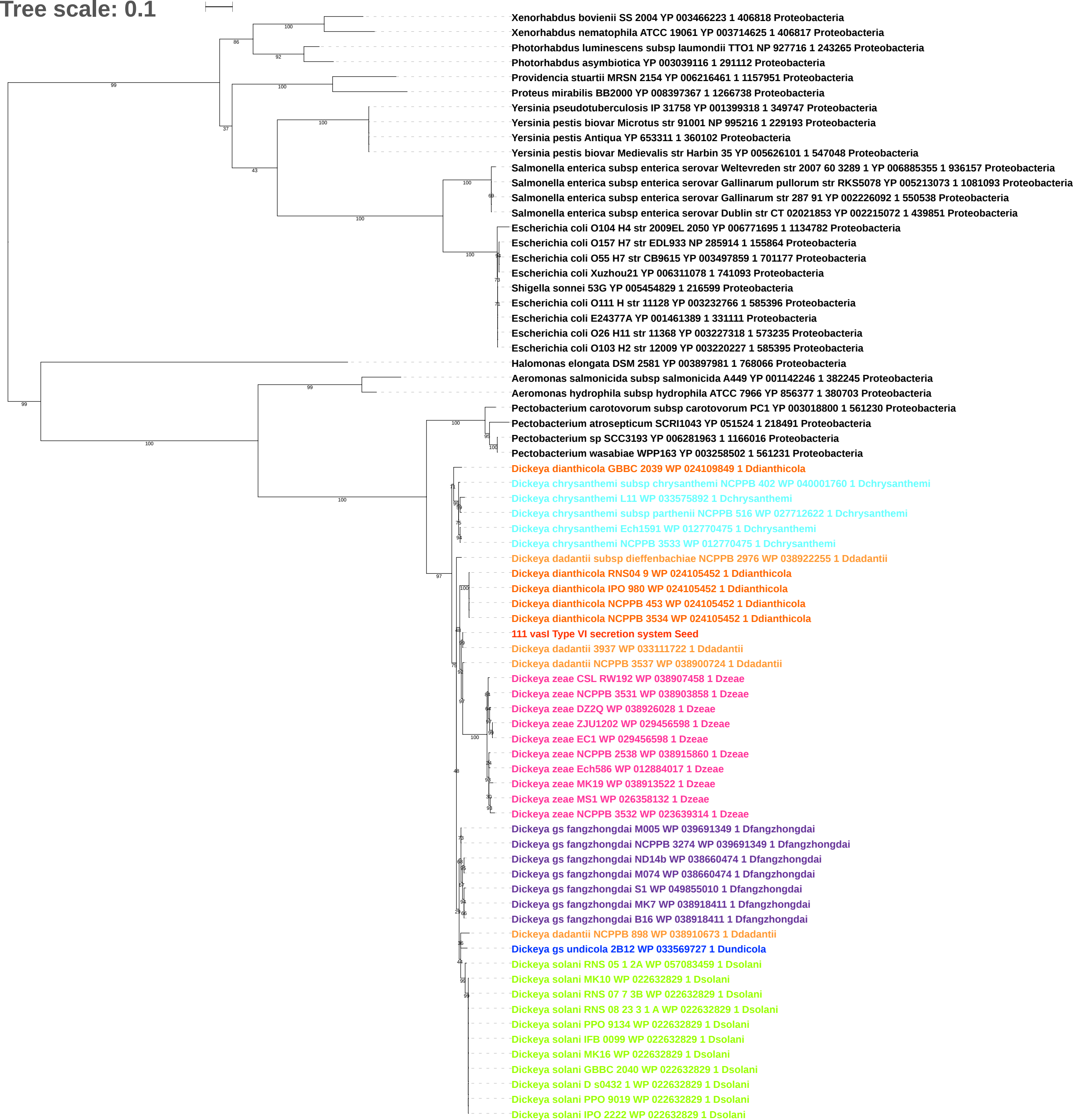
Tree scale: 0.1
Xenorhabdus bovienii SS 2004 YP 003466223 1 406818 Proteobacteria
Xenorhabdus nematophila ATCC 19061 YP 003714625 1 406817 Proteobacteria
Photorhabdus luminescens subsp laumondii TTO1 NP 927716 1 243265 Proteobacteria
Photorhabdus asymbiotica YP 003039116 1 291112 Proteobacteria
Providencia stuartii MRSN 2154 YP 006216461 1 1157951 Proteobacteria
Proteus mirabilis BB2000 YP 008397367 1 1266738 Proteobacteria
Yersinia pseudotuberculosis IP 31758 YP 001399318 1 349747 Proteobacteria
Yersinia pestis biovar Microtus str 91001 NP 995216 1 229193 Proteobacteria
Yersinia pestis Antiqua YP 653311 1 360102 Proteobacteria
Yersinia pestis biovar Medievalis str Harbin 35 YP 005626101 1 547048 Proteobacteria
Salmonella enterica subsp enterica serovar Weltevreden str 2007 60 3289 1 YP 006885355 1 936157 Proteobacteria
Salmonella enterica subsp enterica serovar Gallinarum pullorum str RKS5078 YP 005213073 1 1081093 Proteobacteria
Salmonella enterica subsp enterica serovar Gallinarum str 287 91 YP 002226092 1 550538 Proteobacteria
Salmonella enterica subsp enterica serovar Dublin str CT 02021853 YP 002215072 1 439851 Proteobacteria
Escherichia coli O104 H4 str 2009EL 2050 YP 006771695 1 1134782 Proteobacteria
Escherichia coli O157 H7 str EDL933 NP 285914 1 155864 Proteobacteria
Escherichia coli O55 H7 str CB9615 YP 003497859 1 701177 Proteobacteria
Escherichia coli Xuzhou21 YP 006311078 1 741093 Proteobacteria
Shigella sonnei 53G YP 005454829 1 216599 Proteobacteria
Escherichia coli O111 H str 11128 YP 003232766 1 585396 Proteobacteria
Escherichia coli E24377A YP 001461389 1 331111 Proteobacteria
Escherichia coli O26 H11 str 11368 YP 003227318 1 573235 Proteobacteria
Escherichia coli O103 H2 str 12009 YP 003220227 1 585395 Proteobacteria
Halomonas elongata DSM 2581 YP 003897981 1 768066 Proteobacteria
Aeromonas salmonicida subsp salmonicida A449 YP 001142246 1 382245 Proteobacteria
Aeromonas hydrophila subsp hydrophila ATCC 7966 YP 856377 1 380703 Proteobacteria
Pectobacterium carotovorum subsp carotovorum PC1 YP 003018800 1 561230 Proteobacteria
Pectobacterium atrosepticum SCRI1043 YP 051524 1 218491 Proteobacteria
Pectobacterium sp SCC3193 YP 006281963 1 1166016 Proteobacteria
Pectobacterium wasabiae WPP163 YP 003258502 1 561231 Proteobacteria
Dickeya dianthicola GBBC 2039 WP 024109849 1 Ddianthicola
Dickeya chrysanthemi subsp chrysanthemi NCPPB 402 WP 040001760 1 Dchrysanthemi
Dickeya chrysanthemi L11 WP 033575892 1 Dchrysanthemi
Dickeya chrysanthemi subsp parthenii NCPPB 516 WP 027712622 1 Dchrysanthemi
Dickeya chrysanthemi Ech1591 WP 012770475 1 Dchrysanthemi
Dickeya chrysanthemi NCPPB 3533 WP 012770475 1 Dchrysanthemi
Dickeya dadantii subsp dieffenbachiae NCPPB 2976 WP 038922255 1 Ddadantii
Dickeya dianthicola RNS04 9 WP 024105452 1 Ddianthicola
Dickeya dianthicola IPO 980 WP 024105452 1 Ddianthicola
Dickeya dianthicola NCPPB 453 WP 024105452 1 Ddianthicola
Dickeya dianthicola NCPPB 3534 WP 024105452 1 Ddianthicola
111 vasI Type VI secretion system Seed
Dickeya dadantii 3937 WP 033111722 1 Ddadantii
Dickeya dadantii NCPPB 3537 WP 038900724 1 Ddadantii
Dickeya zeae CSL RW192 WP 038907458 1 Dzeae
Dickeya zeae NCPPB 3531 WP 038903858 1 Dzeae
Dickeya zeae DZ2Q WP 038926028 1 Dzeae
Dickeya zeae ZJU1202 WP 029456598 1 Dzeae
Dickeya zeae EC1 WP 029456598 1 Dzeae
Dickeya zeae NCPPB 2538 WP 038915860 1 Dzeae
Dickeya zeae Ech586 WP 012884017 1 Dzeae
Dickeya zeae MK19 WP 038913522 1 Dzeae
Dickeya zeae MS1 WP 026358132 1 Dzeae
Dickeya zeae NCPPB 3532 WP 023639314 1 Dzeae
Dickeya gs fangzhongdai M005 WP 039691349 1 Dfangzhongdai
Dickeya gs fangzhongdai NCPPB 3274 WP 039691349 1 Dfangzhongdai
Dickeya gs fangzhongdai ND14b WP 038660474 1 Dfangzhongdai
Dickeya gs fangzhongdai M074 WP 038660474 1 Dfangzhongdai
Dickeya gs fangzhongdai S1 WP 049855010 1 Dfangzhongdai
Dickeya gs fangzhongdai MK7 WP 038918411 1 Dfangzhongdai
Dickeya gs fangzhongdai B16 WP 038918411 1 Dfangzhongdai
Dickeya dadantii NCPPB 898 WP 038910673 1 Ddadantii
Dickeya gs undicola 2B12 WP 033569727 1 Dundicola
Dickeya solani RNS 05 1 2A WP 057083459 1 Dsolani
Dickeya solani MK10 WP 022632829 1 Dsolani
Dickeya solani RNS 07 7 3B WP 022632829 1 Dsolani
Dickeya solani RNS 08 23 3 1 A WP 022632829 1 Dsolani
Dickeya solani PPO 9134 WP 022632829 1 Dsolani
Dickeya solani IFB 0099 WP 022632829 1 Dsolani
Dickeya solani MK16 WP 022632829 1 Dsolani
Dickeya solani GBBC 2040 WP 022632829 1 Dsolani
Dickeya solani D s0432 1 WP 022632829 1 Dsolani
Dickeya solani PPO 9019 WP 022632829 1 Dsolani
Dickeya solani IPO 2222 WP 022632829 1 Dsolani

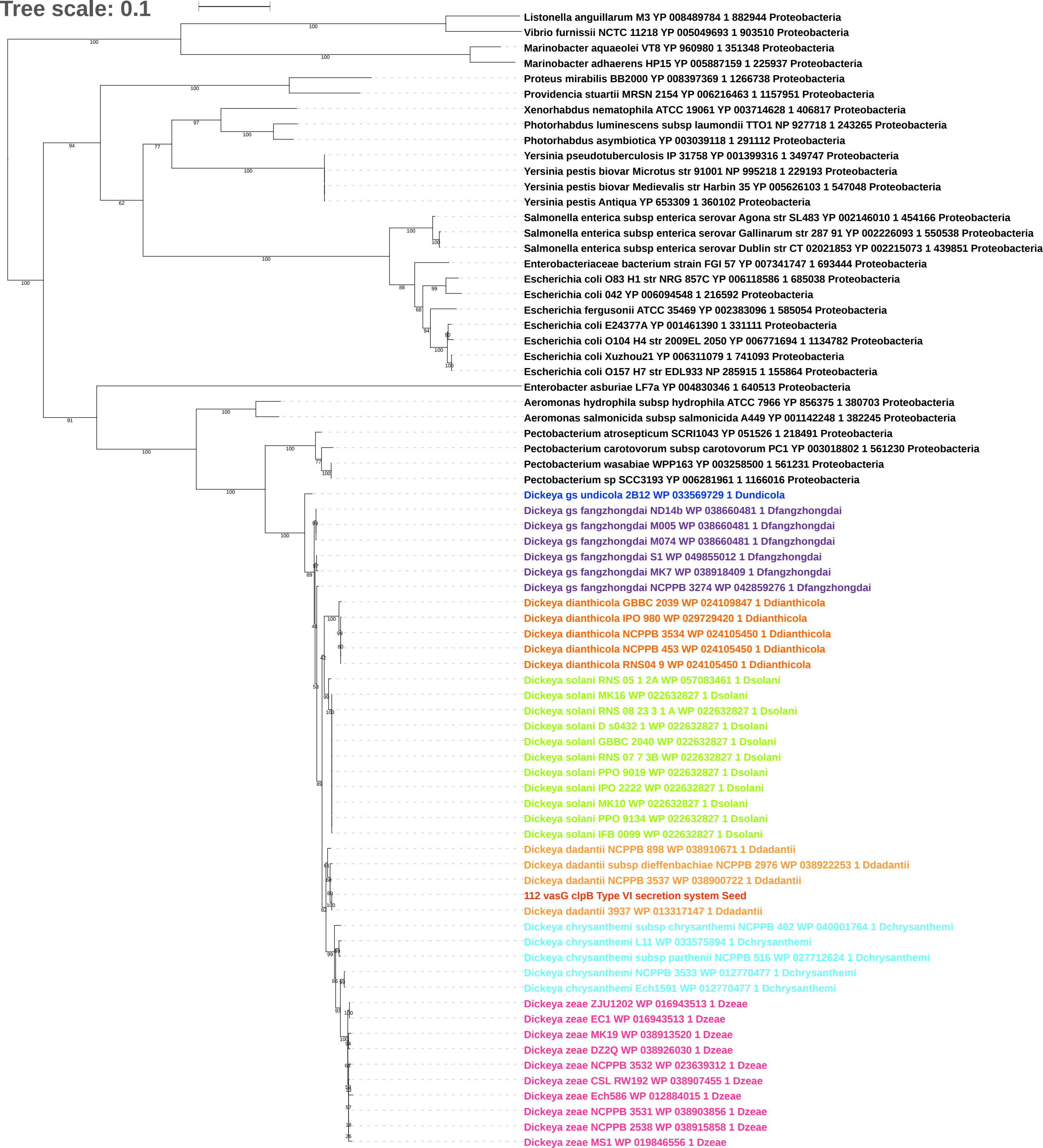
Tree scale: 0.1
Listonella anguillarum M3 YP 008489784 1 882944 Proteobacteria
Vibrio furnissii NCTC 11218 YP 005049693 1 903510 Proteobacteria
Marinobacter aquaeolei VT8 YP 960980 1 351348 Proteobacteria
Marinobacter adhaerens HP15 YP 005887159 1 225937 Proteobacteria
Proteus mirabilis BB2000 YP 008397369 1 1266738 Proteobacteria
Providencia stuartii MRSN 2154 YP 006216463 1 1157951 Proteobacteria
Xenorhabdus nematophila ATCC 19061 YP 003714628 1 406817 Proteobacteria
Photorhabdus luminescens subsp laumondii TTO1 NP 927718 1 243265 Proteobacteria
Photorhabdus asymbiotica YP 003039118 1 291112 Proteobacteria
Yersinia pseudotuberculosis IP 31758 YP 001399316 1 349747 Proteobacteria
Yersinia pestis biovar Microtus str 91001 NP 995218 1 229193 Proteobacteria
Yersinia pestis biovar Medievalis str Harbin 35 YP 005626103 1 547048 Proteobacteria
Yersinia pestis Antiqua YP 653309 1 360102 Proteobacteria
Salmonella enterica subsp enterica serovar Agona str SL483 YP 002146010 1 454166 Proteobacteria
Salmonella enterica subsp enterica serovar Gallinarum str 287 91 YP 002226093 1 550538 Proteobacteria
Salmonella enterica subsp enterica serovar Dublin str CT 02021853 YP 002215073 1 439851 Proteobacteria
Enterobacteriaceae bacterium strain FGI 57 YP 007341747 1 693444 Proteobacteria
Escherichia coli O83 H1 str NRG 857C YP 006118586 1 685038 Proteobacteria
Escherichia coli 042 YP 006094548 1 216592 Proteobacteria
Escherichia fergusonii ATCC 35469 YP 002383096 1 585054 Proteobacteria
Escherichia coli E24377A YP 001461390 1 331111 Proteobacteria
Escherichia coli O104 H4 str 2009EL 2050 YP 006771694 1 1134782 Proteobacteria
Escherichia coli Xuzhou21 YP 006311079 1 741093 Proteobacteria
Escherichia coli O157 H7 str EDL933 NP 285915 1 155864 Proteobacteria
Enterobacter asburiae LF7a YP 004830346 1 640513 Proteobacteria
Aeromonas hydrophila subsp hydrophila ATCC 7966 YP 856375 1 380703 Proteobacteria
Aeromonas salmonicida subsp salmonicida A449 YP 001142248 1 382245 Proteobacteria
Pectobacterium atrosepticum SCRI1043 YP 051526 1 218491 Proteobacteria
Pectobacterium carotovorum subsp carotovorum PC1 YP 003018802 1 561230 Proteobacteria
Pectobacterium wasabiae WPP163 YP 003258500 1 561231 Proteobacteria
Pectobacterium sp SCC3193 YP 006281961 1 1166016 Proteobacteria
Dickeya gs undicola 2B12 WP 033569729 1 Dundicola
Dickeya gs fangzhongdai ND14b WP 038660481 1 Dfangzhongdai
Dickeya gs fangzhongdai M005 WP 038660481 1 Dfangzhongdai
Dickeya gs fangzhongdai M074 WP 038660481 1 Dfangzhongdai
Dickeya gs fangzhongdai S1 WP 049855012 1 Dfangzhongdai
Dickeya gs fangzhongdai MK7 WP 038918409 1 Dfangzhongdai
Dickeya gs fangzhongdai NCPPB 3274 WP 042859276 1 Dfangzhongdai
Dickeya dianthicola GBBC 2039 WP 024109847 1 Ddianthicola
Dickeya dianthicola IPO 980 WP 029729420 1 Ddianthicola
Dickeya dianthicola NCPPB 3534 WP 024105450 1 Ddianthicola
Dickeya dianthicola NCPPB 453 WP 024105450 1 Ddianthicola
Dickeya dianthicola RNS04 9 WP 024105450 1 Ddianthicola
Dickeya solani RNS 05 1 2A WP 057083461 1 Dsolani
Dickeya solani MK16 WP 022632827 1 Dsolani
Dickeya solani RNS 08 23 3 1 A WP 022632827 1 Dsolani
Dickeya solani D s0432 1 WP 022632827 1 Dsolani
Dickeya solani GBBC 2040 WP 022632827 1 Dsolani
Dickeya solani RNS 07 7 3B WP 022632827 1 Dsolani
Dickeya solani PPO 9019 WP 022632827 1 Dsolani
Dickeya solani IPO 2222 WP 022632827 1 Dsolani
Dickeya solani MK10 WP 022632827 1 Dsolani
Dickeya solani PPO 9134 WP 022632827 1 Dsolani
Dickeya solani IFB 0099 WP 022632827 1 Dsolani
Dickeya dadantii NCPPB 898 WP 038910671 1 Ddadantii
Dickeya dadantii subsp dieffenbachiae NCPPB 2976 WP 038922253 1 Ddadantii
Dickeya dadantii NCPPB 3537 WP 038900722 1 Ddadantii
112 vasG clpB Type VI secretion system Seed
Dickeya dadantii 3937 WP 013317147 1 Ddadantii
Dickeya chrysanthemi subsp chrysanthemi NCPPB 402 WP 040001764 1 Dchrysanthemi
Dickeya chrysanthemi L11 WP 033575894 1 Dchrysanthemi
Dickeya chrysanthemi subsp parthenii NCPPB 516 WP 027712624 1 Dchrysanthemi
Dickeya chrysanthemi NCPPB 3533 WP 012770477 1 Dchrysanthemi
Dickeya chrysanthemi Ech1591 WP 012770477 1 Dchrysanthemi
Dickeya zeae ZJU1202 WP 016943513 1 Dzeae
Dickeya zeae EC1 WP 016943513 1 Dzeae
Dickeya zeae MK19 WP 038913520 1 Dzeae
Dickeya zeae DZ2Q WP 038926030 1 Dzeae
Dickeya zeae NCPPB 3532 WP 023639312 1 Dzeae
Dickeya zeae CSL RW192 WP 038907455 1 Dzeae
Dickeya zeae Ech586 WP 012884015 1 Dzeae
Dickeya zeae NCPPB 3531 WP 038903856 1 Dzeae
Dickeya zeae NCPPB 2538 WP 038915858 1 Dzeae
Dickeya zeae MS1 WP 019846556 1 Dzeae

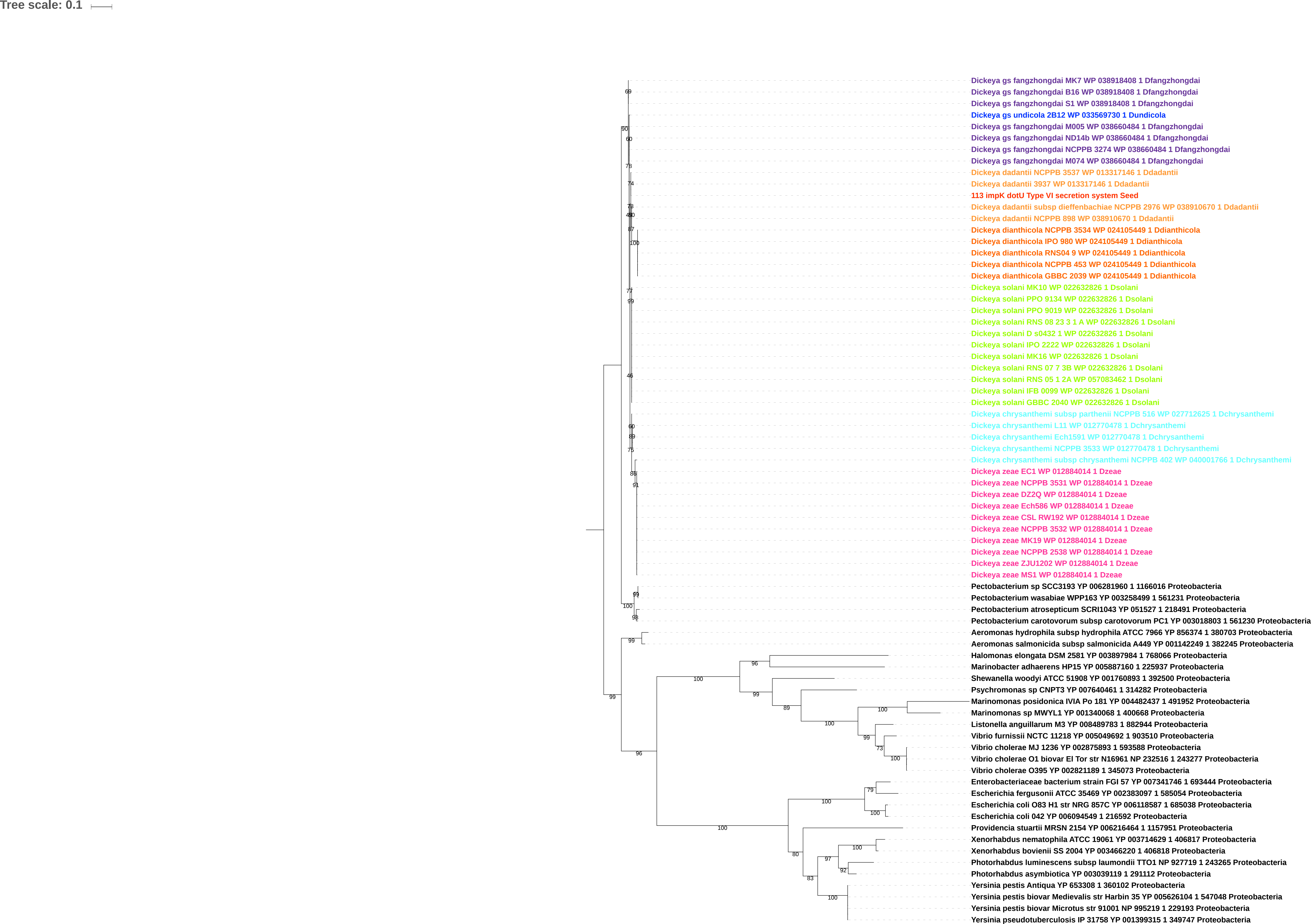
Tree scale: 0.1
Dickeya gs fangzhongdai MK7 WP 038918408 1 Dfangzhongdai
Dickeya gs fangzhongdai B16 WP 038918408 1 Dfangzhongdai
Dickeya gs fangzhongdai S1 WP 038918408 1 Dfangzhongdai
Dickeya gs undicola 2B12 WP 033569730 1 Dundicola
Dickeya gs fangzhongdai M005 WP 038660484 1 Dfangzhongdai
Dickeya gs fangzhongdai ND14b WP 038660484 1 Dfangzhongdai
Dickeya gs fangzhongdai NCPPB 3274 WP 038660484 1 Dfangzhongdai
Dickeya gs fangzhongdai M074 WP 038660484 1 Dfangzhongdai
Dickeya dadantii NCPPB 3537 WP 013317146 1 Ddadantii
Dickeya dadantii 3937 WP 013317146 1 Ddadantii
113 impK dotU Type VI secretion system Seed
Dickeya dadantii subsp dieffenbachiae NCPPB 2976 WP 038910670 1 Ddadantii
Dickeya dadantii NCPPB 898 WP 038910670 1 Ddadantii
Dickeya dianthicola NCPPB 3534 WP 024105449 1 Ddianthicola
Dickeya dianthicola IPO 980 WP 024105449 1 Ddianthicola
Dickeya dianthicola RNS04 9 WP 024105449 1 Ddianthicola
Dickeya dianthicola NCPPB 453 WP 024105449 1 Ddianthicola
Dickeya dianthicola GBBC 2039 WP 024105449 1 Ddianthicola
Dickeya solani MK10 WP 022632826 1 Dsolani
Dickeya solani PPO 9134 WP 022632826 1 Dsolani
Dickeya solani PPO 9019 WP 022632826 1 Dsolani
Dickeya solani RNS 08 23 3 1 A WP 022632826 1 Dsolani
Dickeya solani D s0432 1 WP 022632826 1 Dsolani
Dickeya solani IPO 2222 WP 022632826 1 Dsolani
Dickeya solani MK16 WP 022632826 1 Dsolani
Dickeya solani RNS 07 7 3B WP 022632826 1 Dsolani
Dickeya solani RNS 05 1 2A WP 057083462 1 Dsolani
Dickeya solani IFB 0099 WP 022632826 1 Dsolani
Dickeya solani GBBC 2040 WP 022632826 1 Dsolani
Dickeya chrysanthemi subsp parthenii NCPPB 516 WP 027712625 1 Dchrysanthemi
Dickeya chrysanthemi L11 WP 012770478 1 Dchrysanthemi
Dickeya chrysanthemi Ech1591 WP 012770478 1 Dchrysanthemi
Dickeya chrysanthemi NCPPB 3533 WP 012770478 1 Dchrysanthemi
Dickeya chrysanthemi subsp chrysanthemi NCPPB 402 WP 040001766 1 Dchrysanthemi
Dickeya zeae EC1 WP 012884014 1 Dzeae
Dickeya zeae NCPPB 3531 WP 012884014 1 Dzeae
Dickeya zeae DZ2Q WP 012884014 1 Dzeae
Dickeya zeae Ech586 WP 012884014 1 Dzeae
Dickeya zeae CSL RW192 WP 012884014 1 Dzeae
Dickeya zeae NCPPB 3532 WP 012884014 1 Dzeae
Dickeya zeae MK19 WP 012884014 1 Dzeae
Dickeya zeae NCPPB 2538 WP 012884014 1 Dzeae
Dickeya zeae ZJU1202 WP 012884014 1 Dzeae
Dickeya zeae MS1 WP 012884014 1 Dzeae
Pectobacterium sp SCC3193 YP 006281960 1 1166016 Proteobacteria
Pectobacterium wasabiae WPP163 YP 003258499 1 561231 Proteobacteria
Pectobacterium atrosepticum SCRI1043 YP 051527 1 218491 Proteobacteria
Pectobacterium carotovorum subsp carotovorum PC1 YP 003018803 1 561230 Proteobacteria
Aeromonas hydrophila subsp hydrophila ATCC 7966 YP 856374 1 380703 Proteobacteria
Aeromonas salmonicida subsp salmonicida A449 YP 001142249 1 382245 Proteobacteria
Halomonas elongata DSM 2581 YP 003897984 1 768066 Proteobacteria
Marinobacter adhaerens HP15 YP 005887160 1 225937 Proteobacteria
Shewanella woodyi ATCC 51908 YP 001760893 1 392500 Proteobacteria
Psychromonas sp CNPT3 YP 007640461 1 314282 Proteobacteria
Marinomonas posidonica IVIA Po 181 YP 004482437 1 491952 Proteobacteria
Marinomonas sp MWYL1 YP 001340068 1 400668 Proteobacteria
Listonella anguillarum M3 YP 008489783 1 882944 Proteobacteria
Vibrio furnissii NCTC 11218 YP 005049692 1 903510 Proteobacteria
Vibrio cholerae MJ 1236 YP 002875893 1 593588 Proteobacteria
Vibrio cholerae O1 biovar El Tor str N16961 NP 232516 1 243277 Proteobacteria
Vibrio cholerae O395 YP 002821189 1 345073 Proteobacteria
Enterobacteriaceae bacterium strain FGI 57 YP 007341746 1 693444 Proteobacteria
Escherichia fergusonii ATCC 35469 YP 002383097 1 585054 Proteobacteria
Escherichia coli O83 H1 str NRG 857C YP 006118587 1 685038 Proteobacteria
Escherichia coli 042 YP 006094549 1 216592 Proteobacteria
Providencia stuartii MRSN 2154 YP 006216464 1 1157951 Proteobacteria
Xenorhabdus nematophila ATCC 19061 YP 003714629 1 406817 Proteobacteria
Xenorhabdus bovienii SS 2004 YP 003466220 1 406818 Proteobacteria
Photorhabdus luminescens subsp laumondii TTO1 NP 927719 1 243265 Proteobacteria
Photorhabdus asymbiotica YP 003039119 1 291112 Proteobacteria
Yersinia pestis Antiqua YP 653308 1 360102 Proteobacteria
Yersinia pestis biovar Medievalis str Harbin 35 YP 005626104 1 547048 Proteobacteria
Yersinia pestis biovar Microtus str 91001 NP 995219 1 229193 Proteobacteria
Yersinia pseudotuberculosis IP 31758 YP 001399315 1 349747 Proteobacteria

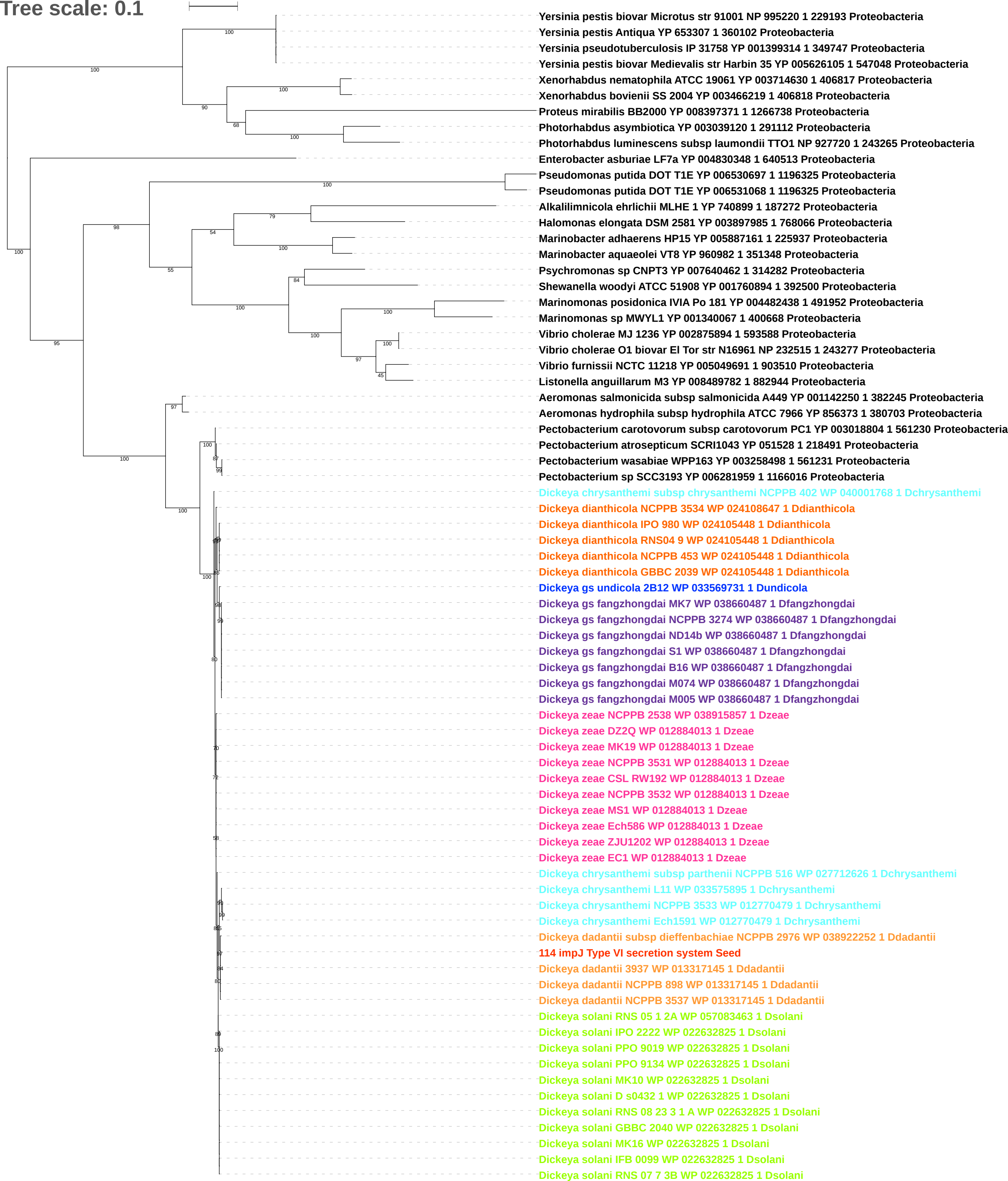
Tree scale: 0.1
Yersinia pestis biovar Microtus str 91001 NP 995220 1 229193 Proteobacteria
Yersinia pestis Antiqua YP 653307 1 360102 Proteobacteria
Yersinia pseudotuberculosis IP 31758 YP 001399314 1 349747 Proteobacteria
Yersinia pestis biovar Medievalis str Harbin 35 YP 005626105 1 547048 Proteobacteria
Xenorhabdus nematophila ATCC 19061 YP 003714630 1 406817 Proteobacteria
Xenorhabdus bovienii SS 2004 YP 003466219 1 406818 Proteobacteria
Proteus mirabilis BB2000 YP 008397371 1 1266738 Proteobacteria
Photorhabdus asymbiotica YP 003039120 1 291112 Proteobacteria
Photorhabdus luminescens subsp laumondii TTO1 NP 927720 1 243265 Proteobacteria
Enterobacter asburiae LF7a YP 004830348 1 640513 Proteobacteria
Pseudomonas putida DOT T1E YP 006530697 1 1196325 Proteobacteria
Pseudomonas putida DOT T1E YP 006531068 1 1196325 Proteobacteria
Alkalilimnicola ehrlichii MLHE 1 YP 740899 1 187272 Proteobacteria
Halomonas elongata DSM 2581 YP 003897985 1 768066 Proteobacteria
Marinobacter adhaerens HP15 YP 005887161 1 225937 Proteobacteria
Marinobacter aquaeolei VT8 YP 960982 1 351348 Proteobacteria
Psychromonas sp CNPT3 YP 007640462 1 314282 Proteobacteria
Shewanella woodyi ATCC 51908 YP 001760894 1 392500 Proteobacteria
Marinomonas posidonica IVIA Po 181 YP 004482438 1 491952 Proteobacteria
Marinomonas sp MWYL1 YP 001340067 1 400668 Proteobacteria
Vibrio cholerae MJ 1236 YP 002875894 1 593588 Proteobacteria
Vibrio cholerae O1 biovar El Tor str N16961 NP 232515 1 243277 Proteobacteria
Vibrio furnissii NCTC 11218 YP 005049691 1 903510 Proteobacteria
Listonella anguillarum M3 YP 008489782 1 882944 Proteobacteria
Aeromonas salmonicida subsp salmonicida A449 YP 001142250 1 382245 Proteobacteria
Aeromonas hydrophila subsp hydrophila ATCC 7966 YP 856373 1 380703 Proteobacteria
Pectobacterium carotovorum subsp carotovorum PC1 YP 003018804 1 561230 Proteobacteria
Pectobacterium atrosepticum SCRI1043 YP 051528 1 218491 Proteobacteria
Pectobacterium wasabiae WPP163 YP 003258498 1 561231 Proteobacteria
Pectobacterium sp SCC3193 YP 006281959 1 1166016 Proteobacteria
Dickeya chrysanthemi subsp chrysanthemi NCPPB 402 WP 040001768 1 Dchrysanthemi
Dickeya dianthicola NCPPB 3534 WP 024108647 1 Ddianthicola
Dickeya dianthicola IPO 980 WP 024105448 1 Ddianthicola
Dickeya dianthicola RNS04 9 WP 024105448 1 Ddianthicola
Dickeya dianthicola NCPPB 453 WP 024105448 1 Ddianthicola
Dickeya dianthicola GBBC 2039 WP 024105448 1 Ddianthicola
Dickeya gs undicola 2B12 WP 033569731 1 Dundicola
Dickeya gs fangzhongdai MK7 WP 038660487 1 Dfangzhongdai
Dickeya gs fangzhongdai NCPPB 3274 WP 038660487 1 Dfangzhongdai
Dickeya gs fangzhongdai ND14b WP 038660487 1 Dfangzhongdai
Dickeya gs fangzhongdai S1 WP 038660487 1 Dfangzhongdai
Dickeya gs fangzhongdai B16 WP 038660487 1 Dfangzhongdai
Dickeya gs fangzhongdai M074 WP 038660487 1 Dfangzhongdai
Dickeya gs fangzhongdai M005 WP 038660487 1 Dfangzhongdai
Dickeya zeae NCPPB 2538 WP 038915857 1 Dzeae
Dickeya zeae DZ2Q WP 012884013 1 Dzeae
Dickeya zeae MK19 WP 012884013 1 Dzeae
Dickeya zeae NCPPB 3531 WP 012884013 1 Dzeae
Dickeya zeae CSL RW192 WP 012884013 1 Dzeae
Dickeya zeae NCPPB 3532 WP 012884013 1 Dzeae
Dickeya zeae MS1 WP 012884013 1 Dzeae
Dickeya zeae Ech586 WP 012884013 1 Dzeae
Dickeya zeae ZJU1202 WP 012884013 1 Dzeae
Dickeya zeae EC1 WP 012884013 1 Dzeae
Dickeya chrysanthemi subsp parthenii NCPPB 516 WP 027712626 1 Dchrysanthemi
Dickeya chrysanthemi L11 WP 033575895 1 Dchrysanthemi
Dickeya chrysanthemi NCPPB 3533 WP 012770479 1 Dchrysanthemi
Dickeya chrysanthemi Ech1591 WP 012770479 1 Dchrysanthemi
Dickeya dadantii subsp dieffenbachiae NCPPB 2976 WP 038922252 1 Ddadantii
114 impJ Type VI secretion system Seed
Dickeya dadantii 3937 WP 013317145 1 Ddadantii
Dickeya dadantii NCPPB 898 WP 013317145 1 Ddadantii
Dickeya dadantii NCPPB 3537 WP 013317145 1 Ddadantii
Dickeya solani RNS 05 1 2A WP 057083463 1 Dsolani
Dickeya solani IPO 2222 WP 022632825 1 Dsolani
Dickeya solani PPO 9019 WP 022632825 1 Dsolani
Dickeya solani PPO 9134 WP 022632825 1 Dsolani
Dickeya solani MK10 WP 022632825 1 Dsolani
Dickeya solani D s0432 1 WP 022632825 1 Dsolani
Dickeya solani RNS 08 23 3 1 A WP 022632825 1 Dsolani
Dickeya solani GBBC 2040 WP 022632825 1 Dsolani
Dickeya solani MK16 WP 022632825 1 Dsolani
Dickeya solani IFB 0099 WP 022632825 1 Dsolani
Dickeya solani RNS 07 7 3B WP 022632825 1 Dsolani

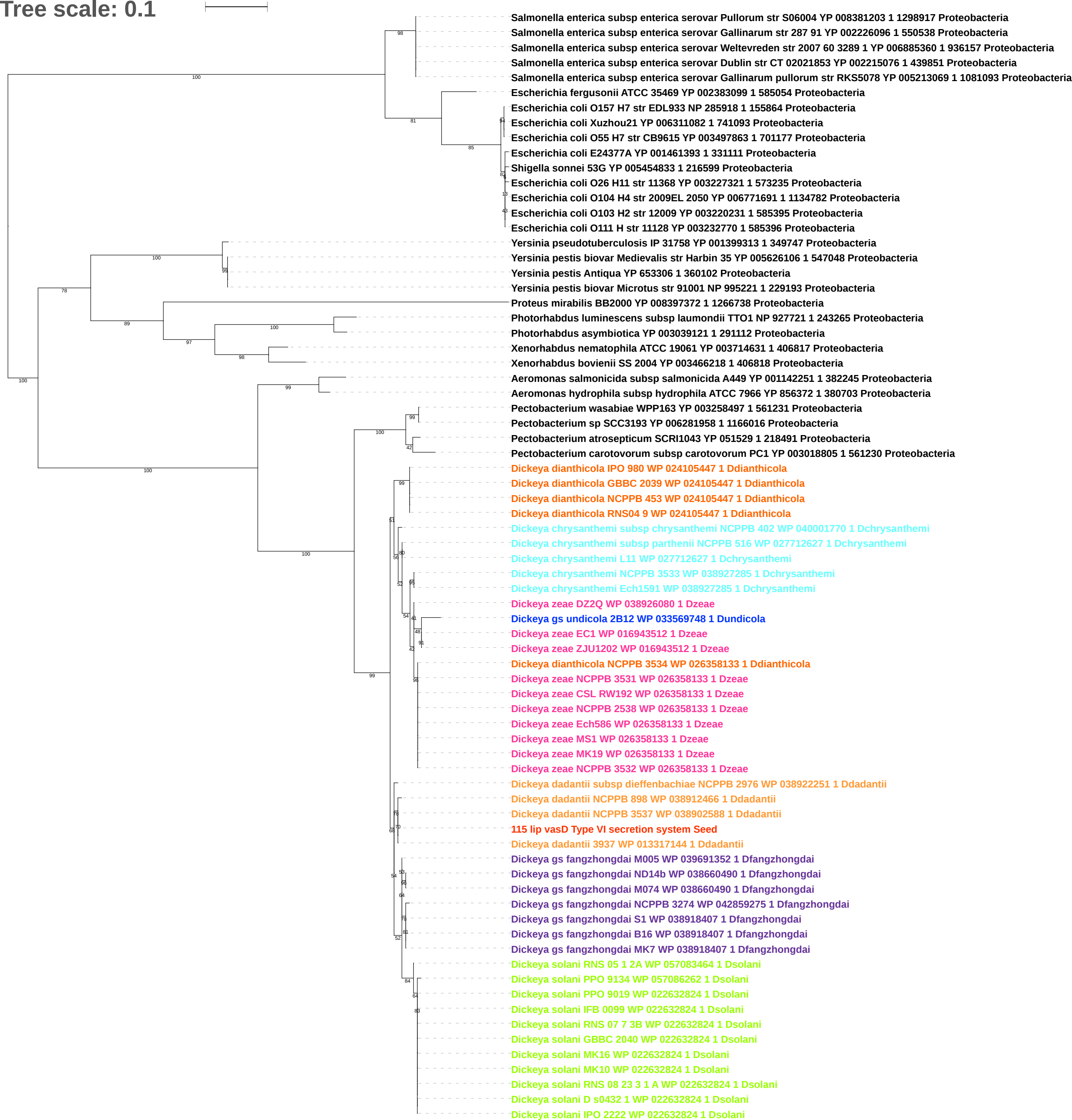
Tree scale: 0.1
Salmonella enterica subsp enterica serovar Pullorum str S06004 YP 008381203 1 1298917 Proteobacteria
Salmonella enterica subsp enterica serovar Gallinarum str 287 91 YP 002226096 1 550538 Proteobacteria
Salmonella enterica subsp enterica serovar Weltevreden str 2007 60 3289 1 YP 006885360 1 936157 Proteobacteria
Salmonella enterica subsp enterica serovar Dublin str CT 02021853 YP 002215076 1 439851 Proteobacteria
Salmonella enterica subsp enterica serovar Gallinarum pullorum str RKS5078 YP 005213069 1 1081093 Proteobacteria
Escherichia fergusonii ATCC 35469 YP 002383099 1 585054 Proteobacteria
Escherichia coli O157 H7 str EDL933 NP 285918 1 155864 Proteobacteria
Escherichia coli Xuzhou21 YP 006311082 1 741093 Proteobacteria
Escherichia coli O55 H7 str CB9615 YP 003497863 1 701177 Proteobacteria
Escherichia coli E24377A YP 001461393 1 331111 Proteobacteria
Shigella sonnei 53G YP 005454833 1 216599 Proteobacteria
Escherichia coli O26 H11 str 11368 YP 003227321 1 573235 Proteobacteria
Escherichia coli O104 H4 str 2009EL 2050 YP 006771691 1 1134782 Proteobacteria
Escherichia coli O103 H2 str 12009 YP 003220231 1 585395 Proteobacteria
Escherichia coli O111 H str 11128 YP 003232770 1 585396 Proteobacteria
Yersinia pseudotuberculosis IP 31758 YP 001399313 1 349747 Proteobacteria
Yersinia pestis biovar Medievalis str Harbin 35 YP 005626106 1 547048 Proteobacteria
Yersinia pestis Antiqua YP 653306 1 360102 Proteobacteria
Yersinia pestis biovar Microtus str 91001 NP 995221 1 229193 Proteobacteria
Proteus mirabilis BB2000 YP 008397372 1 1266738 Proteobacteria
Photorhabdus luminescens subsp laumondii TTO1 NP 927721 1 243265 Proteobacteria
Photorhabdus asymbiotica YP 003039121 1 291112 Proteobacteria
Xenorhabdus nematophila ATCC 19061 YP 003714631 1 406817 Proteobacteria
Xenorhabdus bovienii SS 2004 YP 003466218 1 406818 Proteobacteria
Aeromonas salmonicida subsp salmonicida A449 YP 001142251 1 382245 Proteobacteria
Aeromonas hydrophila subsp hydrophila ATCC 7966 YP 856372 1 380703 Proteobacteria
Pectobacterium wasabiae WPP163 YP 003258497 1 561231 Proteobacteria
Pectobacterium sp SCC3193 YP 006281958 1 1166016 Proteobacteria
Pectobacterium atrosepticum SCRI1043 YP 051529 1 218491 Proteobacteria
Pectobacterium carotovorum subsp carotovorum PC1 YP 003018805 1 561230 Proteobacteria
Dickeya dianthicola IPO 980 WP 024105447 1 Ddianthicola
Dickeya dianthicola GBBC 2039 WP 024105447 1 Ddianthicola
Dickeya dianthicola NCPPB 453 WP 024105447 1 Ddianthicola
Dickeya dianthicola RNS04 9 WP 024105447 1 Ddianthicola
Dickeya chrysanthemi subsp chrysanthemi NCPPB 402 WP 040001770 1 Dchrysanthemi
Dickeya chrysanthemi subsp parthenii NCPPB 516 WP 027712627 1 Dchrysanthemi
Dickeya chrysanthemi L11 WP 027712627 1 Dchrysanthemi
Dickeya chrysanthemi NCPPB 3533 WP 038927285 1 Dchrysanthemi
Dickeya chrysanthemi Ech1591 WP 038927285 1 Dchrysanthemi
Dickeya zeae DZ2Q WP 038926080 1 Dzeae
Dickeya gs undicola 2B12 WP 033569748 1 Dundicola
Dickeya zeae EC1 WP 016943512 1 Dzeae
Dickeya zeae ZJU1202 WP 016943512 1 Dzeae
Dickeya dianthicola NCPPB 3534 WP 026358133 1 Ddianthicola
Dickeya zeae NCPPB 3531 WP 026358133 1 Dzeae
Dickeya zeae CSL RW192 WP 026358133 1 Dzeae
Dickeya zeae NCPPB 2538 WP 026358133 1 Dzeae
Dickeya zeae Ech586 WP 026358133 1 Dzeae
Dickeya zeae MS1 WP 026358133 1 Dzeae
Dickeya zeae MK19 WP 026358133 1 Dzeae
Dickeya zeae NCPPB 3532 WP 026358133 1 Dzeae
Dickeya dadantii subsp dieffenbachiae NCPPB 2976 WP 038922251 1 Ddadantii
Dickeya dadantii NCPPB 898 WP 038912466 1 Ddadantii
Dickeya dadantii NCPPB 3537 WP 038902588 1 Ddadantii
115 lip vasD Type VI secretion system Seed
Dickeya dadantii 3937 WP 013317144 1 Ddadantii
Dickeya gs fangzhongdai M005 WP 039691352 1 Dfangzhongdai
Dickeya gs fangzhongdai ND14b WP 038660490 1 Dfangzhongdai
Dickeya gs fangzhongdai M074 WP 038660490 1 Dfangzhongdai
Dickeya gs fangzhongdai NCPPB 3274 WP 042859275 1 Dfangzhongdai
Dickeya gs fangzhongdai S1 WP 038918407 1 Dfangzhongdai
Dickeya gs fangzhongdai B16 WP 038918407 1 Dfangzhongdai
Dickeya gs fangzhongdai MK7 WP 038918407 1 Dfangzhongdai
Dickeya solani RNS 05 1 2A WP 057083464 1 Dsolani
Dickeya solani PPO 9134 WP 057086262 1 Dsolani
Dickeya solani PPO 9019 WP 022632824 1 Dsolani
Dickeya solani IFB 0099 WP 022632824 1 Dsolani
Dickeya solani RNS 07 7 3B WP 022632824 1 Dsolani
Dickeya solani GBBC 2040 WP 022632824 1 Dsolani
Dickeya solani MK16 WP 022632824 1 Dsolani
Dickeya solani MK10 WP 022632824 1 Dsolani
Dickeya solani RNS 08 23 3 1 A WP 022632824 1 Dsolani
Dickeya solani D s0432 1 WP 022632824 1 Dsolani
Dickeya solani IPO 2222 WP 022632824 1 Dsolani

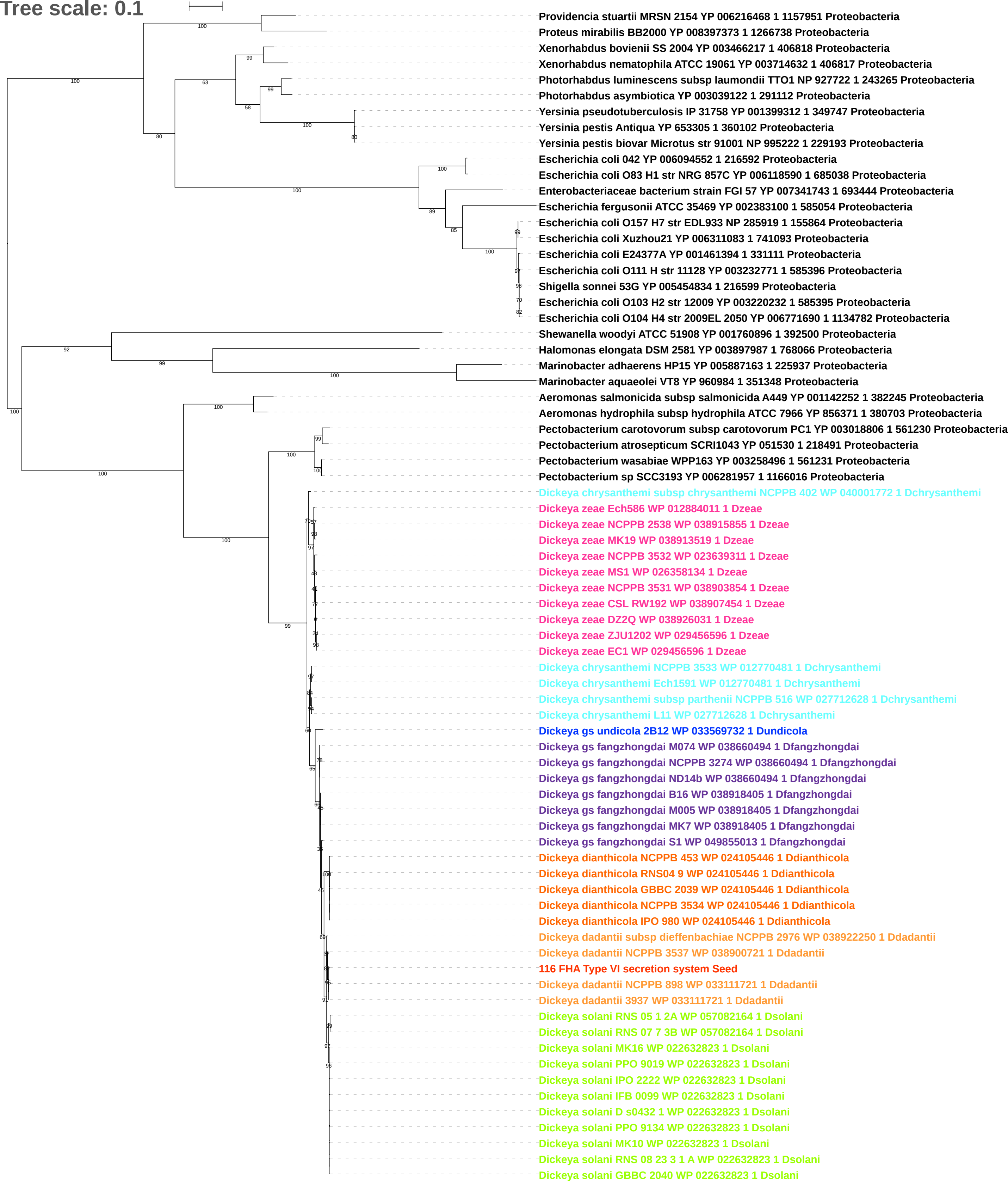
Tree scale: 0.1
Providencia stuartii MRSN 2154 YP 006216468 1 1157951 Proteobacteria
Proteus mirabilis BB2000 YP 008397373 1 1266738 Proteobacteria
Xenorhabdus bovienii SS 2004 YP 003466217 1 406818 Proteobacteria
Xenorhabdus nematophila ATCC 19061 YP 003714632 1 406817 Proteobacteria
Photorhabdus luminescens subsp laumondii TTO1 NP 927722 1 243265 Proteobacteria
Photorhabdus asymbiotica YP 003039122 1 291112 Proteobacteria
Yersinia pseudotuberculosis IP 31758 YP 001399312 1 349747 Proteobacteria
Yersinia pestis Antiqua YP 653305 1 360102 Proteobacteria
Yersinia pestis biovar Microtus str 91001 NP 995222 1 229193 Proteobacteria
Escherichia coli 042 YP 006094552 1 216592 Proteobacteria
Escherichia coli O83 H1 str NRG 857C YP 006118590 1 685038 Proteobacteria
Enterobacteriaceae bacterium strain FGI 57 YP 007341743 1 693444 Proteobacteria
Escherichia fergusonii ATCC 35469 YP 002383100 1 585054 Proteobacteria
Escherichia coli O157 H7 str EDL933 NP 285919 1 155864 Proteobacteria
Escherichia coli Xuzhou21 YP 006311083 1 741093 Proteobacteria
Escherichia coli E24377A YP 001461394 1 331111 Proteobacteria
Escherichia coli O111 H str 11128 YP 003232771 1 585396 Proteobacteria
Shigella sonnei 53G YP 005454834 1 216599 Proteobacteria
Escherichia coli O103 H2 str 12009 YP 003220232 1 585395 Proteobacteria
Escherichia coli O104 H4 str 2009EL 2050 YP 006771690 1 1134782 Proteobacteria
Shewanella woodyi ATCC 51908 YP 001760896 1 392500 Proteobacteria
Halomonas elongata DSM 2581 YP 003897987 1 768066 Proteobacteria
Marinobacter adhaerens HP15 YP 005887163 1 225937 Proteobacteria
Marinobacter aquaeolei VT8 YP 960984 1 351348 Proteobacteria
Aeromonas salmonicida subsp salmonicida A449 YP 001142252 1 382245 Proteobacteria
Aeromonas hydrophila subsp hydrophila ATCC 7966 YP 856371 1 380703 Proteobacteria
Pectobacterium carotovorum subsp carotovorum PC1 YP 003018806 1 561230 Proteobacteria
Pectobacterium atrosepticum SCRI1043 YP 051530 1 218491 Proteobacteria
Pectobacterium wasabiae WPP163 YP 003258496 1 561231 Proteobacteria
Pectobacterium sp SCC3193 YP 006281957 1 1166016 Proteobacteria
Dickeya chrysanthemi subsp chrysanthemi NCPPB 402 WP 040001772 1 Dchrysanthemi
Dickeya zeae Ech586 WP 012884011 1 Dzeae
Dickeya zeae NCPPB 2538 WP 038915855 1 Dzeae
Dickeya zeae MK19 WP 038913519 1 Dzeae
Dickeya zeae NCPPB 3532 WP 023639311 1 Dzeae
Dickeya zeae MS1 WP 026358134 1 Dzeae
Dickeya zeae NCPPB 3531 WP 038903854 1 Dzeae
Dickeya zeae CSL RW192 WP 038907454 1 Dzeae
Dickeya zeae DZ2Q WP 038926031 1 Dzeae
Dickeya zeae ZJU1202 WP 029456596 1 Dzeae
Dickeya zeae EC1 WP 029456596 1 Dzeae
Dickeya chrysanthemi NCPPB 3533 WP 012770481 1 Dchrysanthemi
Dickeya chrysanthemi Ech1591 WP 012770481 1 Dchrysanthemi
Dickeya chrysanthemi subsp parthenii NCPPB 516 WP 027712628 1 Dchrysanthemi
Dickeya chrysanthemi L11 WP 027712628 1 Dchrysanthemi
Dickeya gs undicola 2B12 WP 033569732 1 Dundicola
Dickeya gs fangzhongdai M074 WP 038660494 1 Dfangzhongdai
Dickeya gs fangzhongdai NCPPB 3274 WP 038660494 1 Dfangzhongdai
Dickeya gs fangzhongdai ND14b WP 038660494 1 Dfangzhongdai
Dickeya gs fangzhongdai B16 WP 038918405 1 Dfangzhongdai
Dickeya gs fangzhongdai M005 WP 038918405 1 Dfangzhongdai
Dickeya gs fangzhongdai MK7 WP 038918405 1 Dfangzhongdai
Dickeya gs fangzhongdai S1 WP 049855013 1 Dfangzhongdai
Dickeya dianthicola NCPPB 453 WP 024105446 1 Ddianthicola
Dickeya dianthicola RNS04 9 WP 024105446 1 Ddianthicola
Dickeya dianthicola GBBC 2039 WP 024105446 1 Ddianthicola
Dickeya dianthicola NCPPB 3534 WP 024105446 1 Ddianthicola
Dickeya dianthicola IPO 980 WP 024105446 1 Ddianthicola
Dickeya dadantii subsp dieffenbachiae NCPPB 2976 WP 038922250 1 Ddadantii
Dickeya dadantii NCPPB 3537 WP 038900721 1 Ddadantii
116 FHA Type VI secretion system Seed
Dickeya dadantii NCPPB 898 WP 033111721 1 Ddadantii
Dickeya dadantii 3937 WP 033111721 1 Ddadantii
Dickeya solani RNS 05 1 2A WP 057082164 1 Dsolani
Dickeya solani RNS 07 7 3B WP 057082164 1 Dsolani
Dickeya solani MK16 WP 022632823 1 Dsolani
Dickeya solani PPO 9019 WP 022632823 1 Dsolani
Dickeya solani IPO 2222 WP 022632823 1 Dsolani
Dickeya solani IFB 0099 WP 022632823 1 Dsolani
Dickeya solani D s0432 1 WP 022632823 1 Dsolani
Dickeya solani PPO 9134 WP 022632823 1 Dsolani
Dickeya solani MK10 WP 022632823 1 Dsolani
Dickeya solani RNS 08 23 3 1 A WP 022632823 1 Dsolani
Dickeya solani GBBC 2040 WP 022632823 1 Dsolani

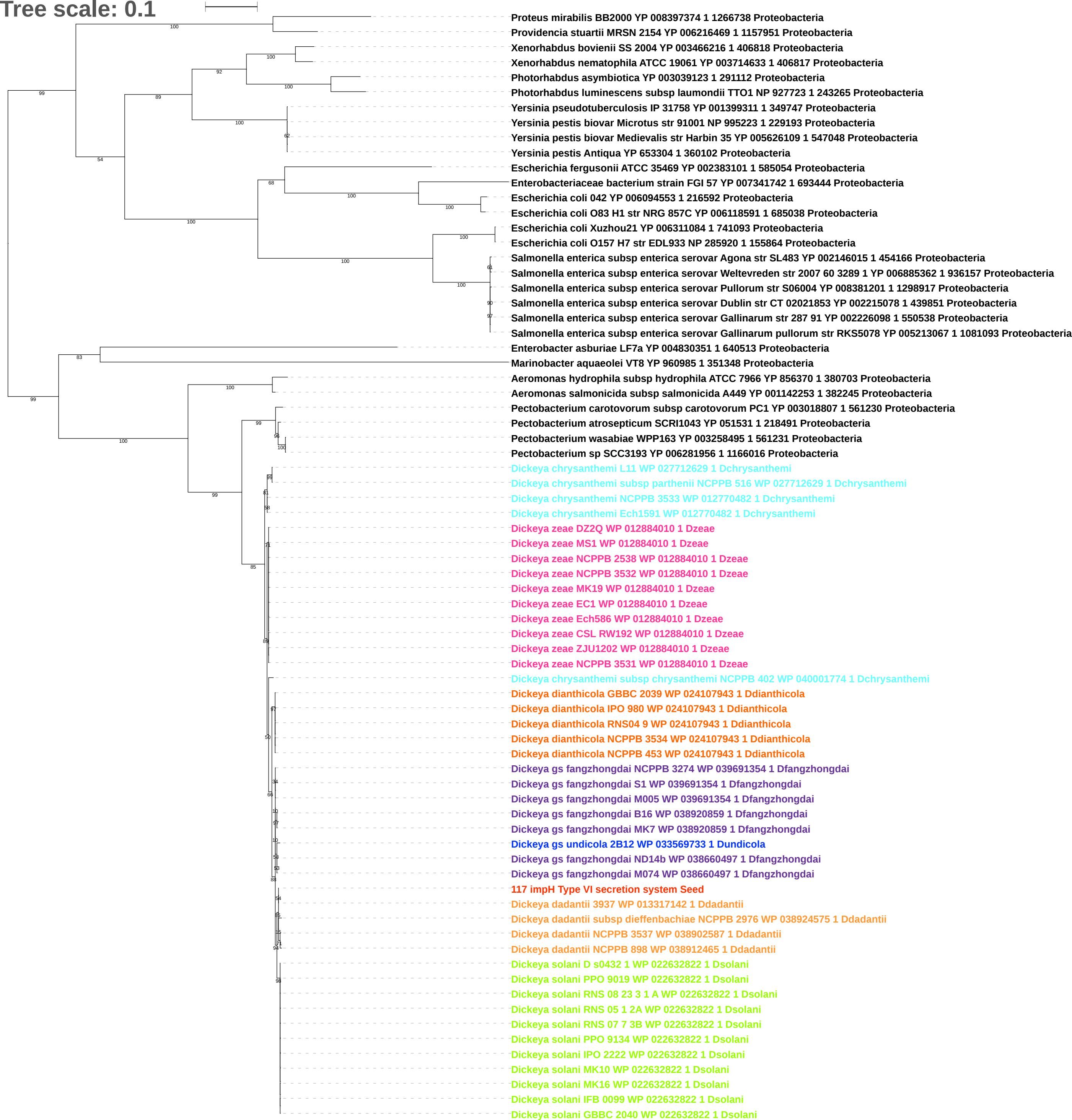
Tree scale: 0.1
Proteus mirabilis BB2000 YP 008397374 1 1266738 Proteobacteria
Providencia stuartii MRSN 2154 YP 006216469 1 1157951 Proteobacteria
Xenorhabdus bovienii SS 2004 YP 003466216 1 406818 Proteobacteria
Xenorhabdus nematophila ATCC 19061 YP 003714633 1 406817 Proteobacteria
Photorhabdus asymbiotica YP 003039123 1 291112 Proteobacteria
Photorhabdus luminescens subsp laumondii TTO1 NP 927723 1 243265 Proteobacteria
Yersinia pseudotuberculosis IP 31758 YP 001399311 1 349747 Proteobacteria
Yersinia pestis biovar Microtus str 91001 NP 995223 1 229193 Proteobacteria
Yersinia pestis biovar Medievalis str Harbin 35 YP 005626109 1 547048 Proteobacteria
Yersinia pestis Antiqua YP 653304 1 360102 Proteobacteria
Escherichia fergusonii ATCC 35469 YP 002383101 1 585054 Proteobacteria
Enterobacteriaceae bacterium strain FGI 57 YP 007341742 1 693444 Proteobacteria
Escherichia coli 042 YP 006094553 1 216592 Proteobacteria
Escherichia coli O83 H1 str NRG 857C YP 006118591 1 685038 Proteobacteria
Escherichia coli Xuzhou21 YP 006311084 1 741093 Proteobacteria
Escherichia coli O157 H7 str EDL933 NP 285920 1 155864 Proteobacteria
Salmonella enterica subsp enterica serovar Agona str SL483 YP 002146015 1 454166 Proteobacteria
Salmonella enterica subsp enterica serovar Weltevreden str 2007 60 3289 1 YP 006885362 1 936157 Proteobacteria
Salmonella enterica subsp enterica serovar Pullorum str S06004 YP 008381201 1 1298917 Proteobacteria
Salmonella enterica subsp enterica serovar Dublin str CT 02021853 YP 002215078 1 439851 Proteobacteria
Salmonella enterica subsp enterica serovar Gallinarum str 287 91 YP 002226098 1 550538 Proteobacteria
Salmonella enterica subsp enterica serovar Gallinarum pullorum str RKS5078 YP 005213067 1 1081093 Proteobacteria
Enterobacter asburiae LF7a YP 004830351 1 640513 Proteobacteria
Marinobacter aquaeolei VT8 YP 960985 1 351348 Proteobacteria
Aeromonas hydrophila subsp hydrophila ATCC 7966 YP 856370 1 380703 Proteobacteria
Aeromonas salmonicida subsp salmonicida A449 YP 001142253 1 382245 Proteobacteria
Pectobacterium carotovorum subsp carotovorum PC1 YP 003018807 1 561230 Proteobacteria
Pectobacterium atrosepticum SCRI1043 YP 051531 1 218491 Proteobacteria
Pectobacterium wasabiae WPP163 YP 003258495 1 561231 Proteobacteria
Pectobacterium sp SCC3193 YP 006281956 1 1166016 Proteobacteria
Dickeya chrysanthemi L11 WP 027712629 1 Dchrysanthemi
Dickeya chrysanthemi subsp parthenii NCPPB 516 WP 027712629 1 Dchrysanthemi
Dickeya chrysanthemi NCPPB 3533 WP 012770482 1 Dchrysanthemi
Dickeya chrysanthemi Ech1591 WP 012770482 1 Dchrysanthemi
Dickeya zeae DZ2Q WP 012884010 1 Dzeae
Dickeya zeae MS1 WP 012884010 1 Dzeae
Dickeya zeae NCPPB 2538 WP 012884010 1 Dzeae
Dickeya zeae NCPPB 3532 WP 012884010 1 Dzeae
Dickeya zeae MK19 WP 012884010 1 Dzeae
Dickeya zeae EC1 WP 012884010 1 Dzeae
Dickeya zeae Ech586 WP 012884010 1 Dzeae
Dickeya zeae CSL RW192 WP 012884010 1 Dzeae
Dickeya zeae ZJU1202 WP 012884010 1 Dzeae
Dickeya zeae NCPPB 3531 WP 012884010 1 Dzeae
Dickeya chrysanthemi subsp chrysanthemi NCPPB 402 WP 040001774 1 Dchrysanthemi
Dickeya dianthicola GBBC 2039 WP 024107943 1 Ddianthicola
Dickeya dianthicola IPO 980 WP 024107943 1 Ddianthicola
Dickeya dianthicola RNS04 9 WP 024107943 1 Ddianthicola
Dickeya dianthicola NCPPB 3534 WP 024107943 1 Ddianthicola
Dickeya dianthicola NCPPB 453 WP 024107943 1 Ddianthicola
Dickeya gs fangzhongdai NCPPB 3274 WP 039691354 1 Dfangzhongdai
Dickeya gs fangzhongdai S1 WP 039691354 1 Dfangzhongdai
Dickeya gs fangzhongdai M005 WP 039691354 1 Dfangzhongdai
Dickeya gs fangzhongdai B16 WP 038920859 1 Dfangzhongdai
Dickeya gs fangzhongdai MK7 WP 038920859 1 Dfangzhongdai
Dickeya gs undicola 2B12 WP 033569733 1 Dundicola
Dickeya gs fangzhongdai ND14b WP 038660497 1 Dfangzhongdai
Dickeya gs fangzhongdai M074 WP 038660497 1 Dfangzhongdai
117 impH Type VI secretion system Seed
Dickeya dadantii 3937 WP 013317142 1 Ddadantii
Dickeya dadantii subsp dieffenbachiae NCPPB 2976 WP 038924575 1 Ddadantii
Dickeya dadantii NCPPB 3537 WP 038902587 1 Ddadantii
Dickeya dadantii NCPPB 898 WP 038912465 1 Ddadantii
Dickeya solani D s0432 1 WP 022632822 1 Dsolani
Dickeya solani PPO 9019 WP 022632822 1 Dsolani
Dickeya solani RNS 08 23 3 1 A WP 022632822 1 Dsolani
Dickeya solani RNS 05 1 2A WP 022632822 1 Dsolani
Dickeya solani RNS 07 7 3B WP 022632822 1 Dsolani
Dickeya solani PPO 9134 WP 022632822 1 Dsolani
Dickeya solani IPO 2222 WP 022632822 1 Dsolani
Dickeya solani MK10 WP 022632822 1 Dsolani
Dickeya solani MK16 WP 022632822 1 Dsolani
Dickeya solani IFB 0099 WP 022632822 1 Dsolani
Dickeya solani GBBC 2040 WP 022632822 1 Dsolani

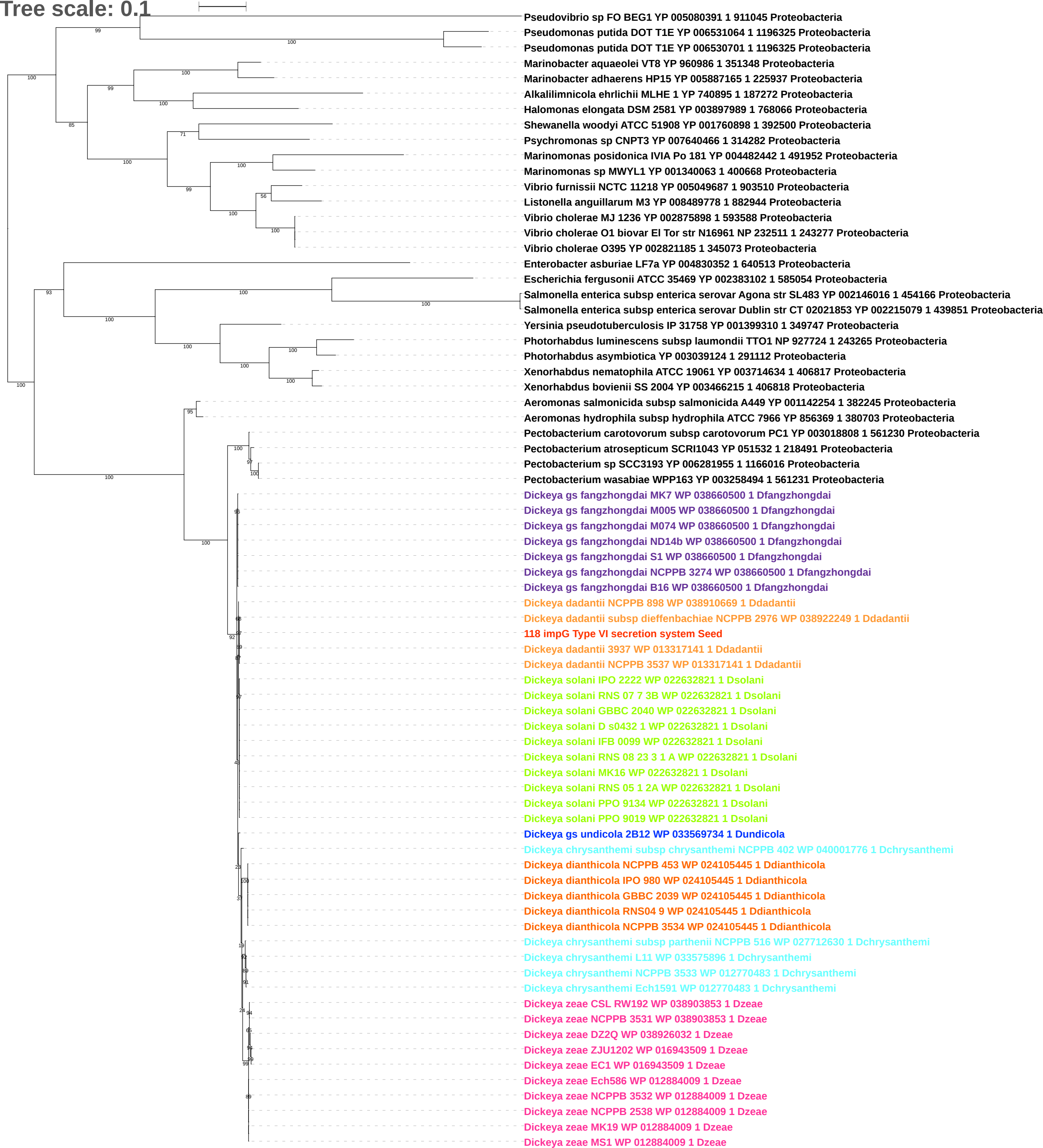
Tree scale: 0.1
Pseudovibrio sp FO BEG1 YP 005080391 1 911045 Proteobacteria
Pseudomonas putida DOT T1E YP 006531064 1 1196325 Proteobacteria
Pseudomonas putida DOT T1E YP 006530701 1 1196325 Proteobacteria
Marinobacter aquaeolei VT8 YP 960986 1 351348 Proteobacteria
Marinobacter adhaerens HP15 YP 005887165 1 225937 Proteobacteria
Alkalilimnicola ehrlichii MLHE 1 YP 740895 1 187272 Proteobacteria
Halomonas elongata DSM 2581 YP 003897989 1 768066 Proteobacteria
Shewanella woodyi ATCC 51908 YP 001760898 1 392500 Proteobacteria
Psychromonas sp CNPT3 YP 007640466 1 314282 Proteobacteria
Marinomonas posidonica IVIA Po 181 YP 004482442 1 491952 Proteobacteria
Marinomonas sp MWYL1 YP 001340063 1 400668 Proteobacteria
Vibrio furnissii NCTC 11218 YP 005049687 1 903510 Proteobacteria
Listonella anguillarum M3 YP 008489778 1 882944 Proteobacteria
Vibrio cholerae MJ 1236 YP 002875898 1 593588 Proteobacteria
Vibrio cholerae O1 biovar El Tor str N16961 NP 232511 1 243277 Proteobacteria
Vibrio cholerae O395 YP 002821185 1 345073 Proteobacteria
Enterobacter asburiae LF7a YP 004830352 1 640513 Proteobacteria
Escherichia fergusonii ATCC 35469 YP 002383102 1 585054 Proteobacteria
Salmonella enterica subsp enterica serovar Agona str SL483 YP 002146016 1 454166 Proteobacteria
Salmonella enterica subsp enterica serovar Dublin str CT 02021853 YP 002215079 1 439851 Proteobacteria
Yersinia pseudotuberculosis IP 31758 YP 001399310 1 349747 Proteobacteria
Photorhabdus luminescens subsp laumondii TTO1 NP 927724 1 243265 Proteobacteria
Photorhabdus asymbiotica YP 003039124 1 291112 Proteobacteria
Xenorhabdus nematophila ATCC 19061 YP 003714634 1 406817 Proteobacteria
Xenorhabdus bovienii SS 2004 YP 003466215 1 406818 Proteobacteria
Aeromonas salmonicida subsp salmonicida A449 YP 001142254 1 382245 Proteobacteria
Aeromonas hydrophila subsp hydrophila ATCC 7966 YP 856369 1 380703 Proteobacteria
Pectobacterium carotovorum subsp carotovorum PC1 YP 003018808 1 561230 Proteobacteria
Pectobacterium atrosepticum SCRI1043 YP 051532 1 218491 Proteobacteria
Pectobacterium sp SCC3193 YP 006281955 1 1166016 Proteobacteria
Pectobacterium wasabiae WPP163 YP 003258494 1 561231 Proteobacteria
Dickeya gs fangzhongdai MK7 WP 038660500 1 Dfangzhongdai
Dickeya gs fangzhongdai M005 WP 038660500 1 Dfangzhongdai
Dickeya gs fangzhongdai M074 WP 038660500 1 Dfangzhongdai
Dickeya gs fangzhongdai ND14b WP 038660500 1 Dfangzhongdai
Dickeya gs fangzhongdai S1 WP 038660500 1 Dfangzhongdai
Dickeya gs fangzhongdai NCPPB 3274 WP 038660500 1 Dfangzhongdai
Dickeya gs fangzhongdai B16 WP 038660500 1 Dfangzhongdai
Dickeya dadantii NCPPB 898 WP 038910669 1 Ddadantii
Dickeya dadantii subsp dieffenbachiae NCPPB 2976 WP 038922249 1 Ddadantii
118 impG Type VI secretion system Seed
Dickeya dadantii 3937 WP 013317141 1 Ddadantii
Dickeya dadantii NCPPB 3537 WP 013317141 1 Ddadantii
Dickeya solani IPO 2222 WP 022632821 1 Dsolani
Dickeya solani RNS 07 7 3B WP 022632821 1 Dsolani
Dickeya solani GBBC 2040 WP 022632821 1 Dsolani
Dickeya solani D s0432 1 WP 022632821 1 Dsolani
Dickeya solani IFB 0099 WP 022632821 1 Dsolani
Dickeya solani RNS 08 23 3 1 A WP 022632821 1 Dsolani
Dickeya solani MK16 WP 022632821 1 Dsolani
Dickeya solani RNS 05 1 2A WP 022632821 1 Dsolani
Dickeya solani PPO 9134 WP 022632821 1 Dsolani
Dickeya solani PPO 9019 WP 022632821 1 Dsolani
Dickeya gs undicola 2B12 WP 033569734 1 Dundicola
Dickeya chrysanthemi subsp chrysanthemi NCPPB 402 WP 040001776 1 Dchrysanthemi
Dickeya dianthicola NCPPB 453 WP 024105445 1 Ddianthicola
Dickeya dianthicola IPO 980 WP 024105445 1 Ddianthicola
Dickeya dianthicola GBBC 2039 WP 024105445 1 Ddianthicola
Dickeya dianthicola RNS04 9 WP 024105445 1 Ddianthicola
Dickeya dianthicola NCPPB 3534 WP 024105445 1 Ddianthicola
Dickeya chrysanthemi subsp parthenii NCPPB 516 WP 027712630 1 Dchrysanthemi
Dickeya chrysanthemi L11 WP 033575896 1 Dchrysanthemi
Dickeya chrysanthemi NCPPB 3533 WP 012770483 1 Dchrysanthemi
Dickeya chrysanthemi Ech1591 WP 012770483 1 Dchrysanthemi
Dickeya zeae CSL RW192 WP 038903853 1 Dzeae
Dickeya zeae NCPPB 3531 WP 038903853 1 Dzeae
Dickeya zeae DZ2Q WP 038926032 1 Dzeae
Dickeya zeae ZJU1202 WP 016943509 1 Dzeae
Dickeya zeae EC1 WP 016943509 1 Dzeae
Dickeya zeae Ech586 WP 012884009 1 Dzeae
Dickeya zeae NCPPB 3532 WP 012884009 1 Dzeae
Dickeya zeae NCPPB 2538 WP 012884009 1 Dzeae
Dickeya zeae MK19 WP 012884009 1 Dzeae
Dickeya zeae MS1 WP 012884009 1 Dzeae

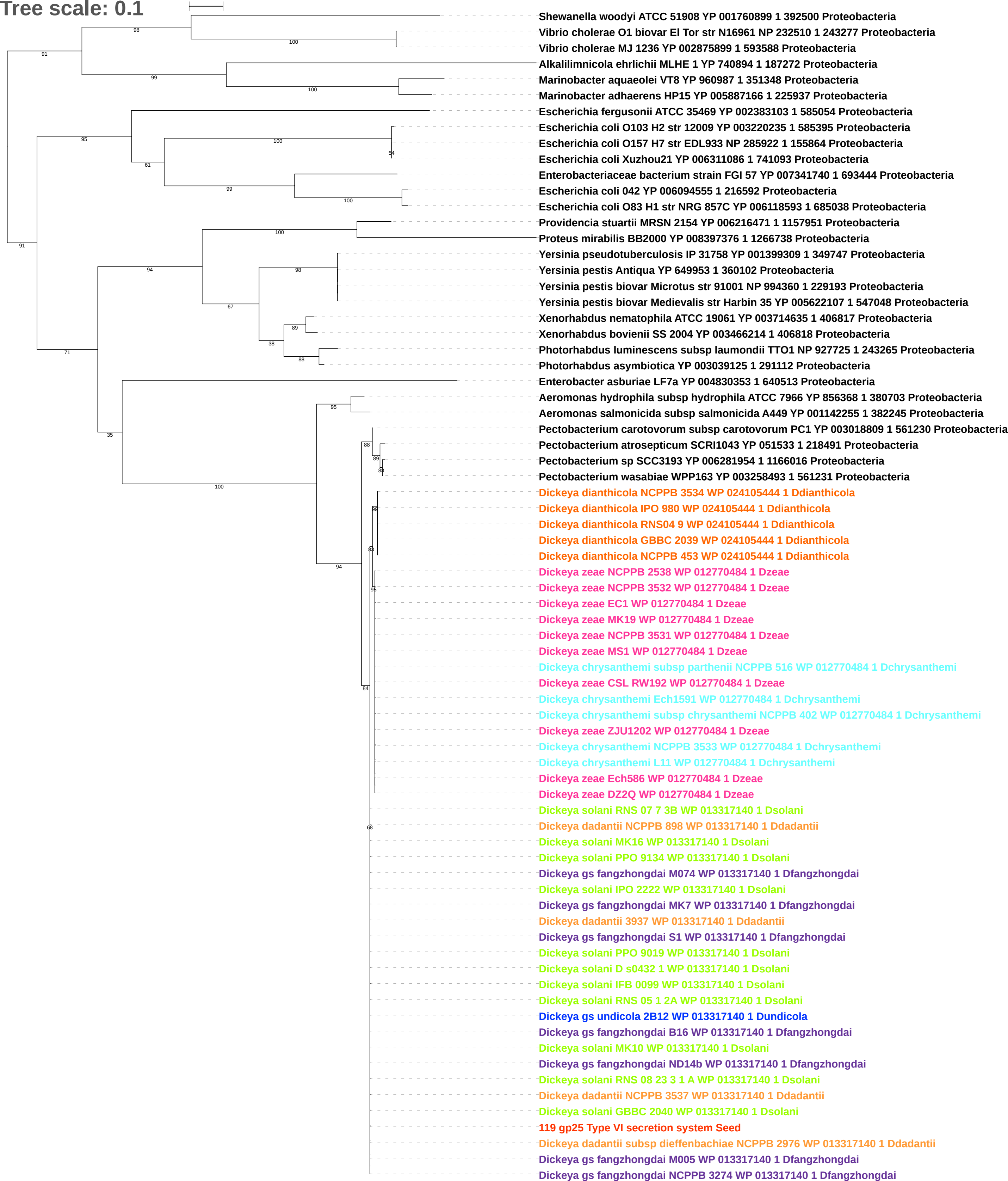
Tree scale: 0.1
Shewanella woodyi ATCC 51908 YP 001760899 1 392500 Proteobacteria
Vibrio cholerae O1 biovar El Tor str N16961 NP 232510 1 243277 Proteobacteria
Vibrio cholerae MJ 1236 YP 002875899 1 593588 Proteobacteria
Alkalilimnicola ehrlichii MLHE 1 YP 740894 1 187272 Proteobacteria
Marinobacter aquaeolei VT8 YP 960987 1 351348 Proteobacteria
Marinobacter adhaerens HP15 YP 005887166 1 225937 Proteobacteria
Escherichia fergusonii ATCC 35469 YP 002383103 1 585054 Proteobacteria
Escherichia coli O103 H2 str 12009 YP 003220235 1 585395 Proteobacteria
Escherichia coli O157 H7 str EDL933 NP 285922 1 155864 Proteobacteria
Escherichia coli Xuzhou21 YP 006311086 1 741093 Proteobacteria
Enterobacteriaceae bacterium strain FGI 57 YP 007341740 1 693444 Proteobacteria
Escherichia coli 042 YP 006094555 1 216592 Proteobacteria
Escherichia coli O83 H1 str NRG 857C YP 006118593 1 685038 Proteobacteria
Providencia stuartii MRSN 2154 YP 006216471 1 1157951 Proteobacteria
Proteus mirabilis BB2000 YP 008397376 1 1266738 Proteobacteria
Yersinia pseudotuberculosis IP 31758 YP 001399309 1 349747 Proteobacteria
Yersinia pestis Antiqua YP 649953 1 360102 Proteobacteria
Yersinia pestis biovar Microtus str 91001 NP 994360 1 229193 Proteobacteria
Yersinia pestis biovar Medievalis str Harbin 35 YP 005622107 1 547048 Proteobacteria
Xenorhabdus nematophila ATCC 19061 YP 003714635 1 406817 Proteobacteria
Xenorhabdus bovienii SS 2004 YP 003466214 1 406818 Proteobacteria
Photorhabdus luminescens subsp laumondii TTO1 NP 927725 1 243265 Proteobacteria
Photorhabdus asymbiotica YP 003039125 1 291112 Proteobacteria
Enterobacter asburiae LF7a YP 004830353 1 640513 Proteobacteria
Aeromonas hydrophila subsp hydrophila ATCC 7966 YP 856368 1 380703 Proteobacteria
Aeromonas salmonicida subsp salmonicida A449 YP 001142255 1 382245 Proteobacteria
Pectobacterium carotovorum subsp carotovorum PC1 YP 003018809 1 561230 Proteobacteria
Pectobacterium atrosepticum SCRI1043 YP 051533 1 218491 Proteobacteria
Pectobacterium sp SCC3193 YP 006281954 1 1166016 Proteobacteria
Pectobacterium wasabiae WPP163 YP 003258493 1 561231 Proteobacteria
Dickeya dianthicola NCPPB 3534 WP 024105444 1 Ddianthicola
Dickeya dianthicola IPO 980 WP 024105444 1 Ddianthicola
Dickeya dianthicola RNS04 9 WP 024105444 1 Ddianthicola
Dickeya dianthicola GBBC 2039 WP 024105444 1 Ddianthicola
Dickeya dianthicola NCPPB 453 WP 024105444 1 Ddianthicola
Dickeya zeae NCPPB 2538 WP 012770484 1 Dzeae
Dickeya zeae NCPPB 3532 WP 012770484 1 Dzeae
Dickeya zeae EC1 WP 012770484 1 Dzeae
Dickeya zeae MK19 WP 012770484 1 Dzeae
Dickeya zeae NCPPB 3531 WP 012770484 1 Dzeae
Dickeya zeae MS1 WP 012770484 1 Dzeae
Dickeya chrysanthemi subsp parthenii NCPPB 516 WP 012770484 1 Dchrysanthemi
Dickeya zeae CSL RW192 WP 012770484 1 Dzeae
Dickeya chrysanthemi Ech1591 WP 012770484 1 Dchrysanthemi
Dickeya chrysanthemi subsp chrysanthemi NCPPB 402 WP 012770484 1 Dchrysanthemi
Dickeya zeae ZJU1202 WP 012770484 1 Dzeae
Dickeya chrysanthemi NCPPB 3533 WP 012770484 1 Dchrysanthemi
Dickeya chrysanthemi L11 WP 012770484 1 Dchrysanthemi
Dickeya zeae Ech586 WP 012770484 1 Dzeae
Dickeya zeae DZ2Q WP 012770484 1 Dzeae
Dickeya solani RNS 07 7 3B WP 013317140 1 Dsolani
Dickeya dadantii NCPPB 898 WP 013317140 1 Ddadantii
Dickeya solani MK16 WP 013317140 1 Dsolani
Dickeya solani PPO 9134 WP 013317140 1 Dsolani
Dickeya gs fangzhongdai M074 WP 013317140 1 Dfangzhongdai
Dickeya solani IPO 2222 WP 013317140 1 Dsolani
Dickeya gs fangzhongdai MK7 WP 013317140 1 Dfangzhongdai
Dickeya dadantii 3937 WP 013317140 1 Ddadantii
Dickeya gs fangzhongdai S1 WP 013317140 1 Dfangzhongdai
Dickeya solani PPO 9019 WP 013317140 1 Dsolani
Dickeya solani D s0432 1 WP 013317140 1 Dsolani
Dickeya solani IFB 0099 WP 013317140 1 Dsolani
Dickeya solani RNS 05 1 2A WP 013317140 1 Dsolani
Dickeya gs undicola 2B12 WP 013317140 1 Dundicola
Dickeya gs fangzhongdai B16 WP 013317140 1 Dfangzhongdai
Dickeya solani MK10 WP 013317140 1 Dsolani
Dickeya gs fangzhongdai ND14b WP 013317140 1 Dfangzhongdai
Dickeya solani RNS 08 23 3 1 A WP 013317140 1 Dsolani
Dickeya dadantii NCPPB 3537 WP 013317140 1 Ddadantii
Dickeya solani GBBC 2040 WP 013317140 1 Dsolani
119 gp25 Type VI secretion system Seed
Dickeya dadantii subsp dieffenbachiae NCPPB 2976 WP 013317140 1 Ddadantii
Dickeya gs fangzhongdai M005 WP 013317140 1 Dfangzhongdai
Dickeya gs fangzhongdai NCPPB 3274 WP 013317140 1 Dfangzhongdai

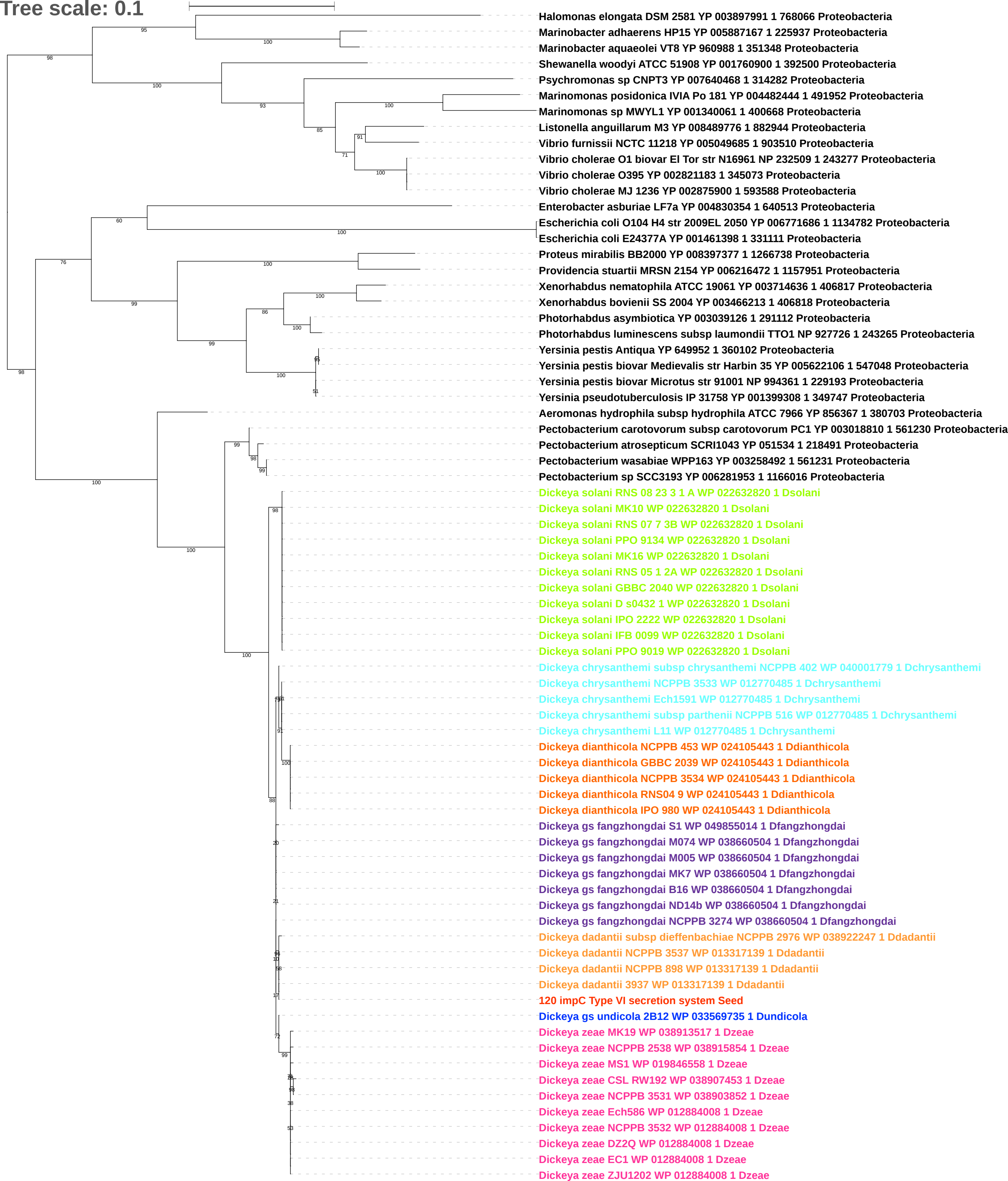
Tree scale: 0.1
Halomonas elongata DSM 2581 YP 003897991 1 768066 Proteobacteria
Marinobacter adhaerens HP15 YP 005887167 1 225937 Proteobacteria
Marinobacter aquaeolei VT8 YP 960988 1 351348 Proteobacteria
Shewanella woodyi ATCC 51908 YP 001760900 1 392500 Proteobacteria
Psychromonas sp CNPT3 YP 007640468 1 314282 Proteobacteria
Marinomonas posidonica IVIA Po 181 YP 004482444 1 491952 Proteobacteria
Marinomonas sp MWYL1 YP 001340061 1 400668 Proteobacteria
Listonella anguillarum M3 YP 008489776 1 882944 Proteobacteria
Vibrio furnissii NCTC 11218 YP 005049685 1 903510 Proteobacteria
Vibrio cholerae O1 biovar El Tor str N16961 NP 232509 1 243277 Proteobacteria
Vibrio cholerae O395 YP 002821183 1 345073 Proteobacteria
Vibrio cholerae MJ 1236 YP 002875900 1 593588 Proteobacteria
Enterobacter asburiae LF7a YP 004830354 1 640513 Proteobacteria
Escherichia coli O104 H4 str 2009EL 2050 YP 006771686 1 1134782 Proteobacteria
Escherichia coli E24377A YP 001461398 1 331111 Proteobacteria
Proteus mirabilis BB2000 YP 008397377 1 1266738 Proteobacteria
Providencia stuartii MRSN 2154 YP 006216472 1 1157951 Proteobacteria
Xenorhabdus nematophila ATCC 19061 YP 003714636 1 406817 Proteobacteria
Xenorhabdus bovienii SS 2004 YP 003466213 1 406818 Proteobacteria
Photorhabdus asymbiotica YP 003039126 1 291112 Proteobacteria
Photorhabdus luminescens subsp laumondii TTO1 NP 927726 1 243265 Proteobacteria
Yersinia pestis Antiqua YP 649952 1 360102 Proteobacteria
Yersinia pestis biovar Medievalis str Harbin 35 YP 005622106 1 547048 Proteobacteria
Yersinia pestis biovar Microtus str 91001 NP 994361 1 229193 Proteobacteria
Yersinia pseudotuberculosis IP 31758 YP 001399308 1 349747 Proteobacteria
Aeromonas hydrophila subsp hydrophila ATCC 7966 YP 856367 1 380703 Proteobacteria
Pectobacterium carotovorum subsp carotovorum PC1 YP 003018810 1 561230 Proteobacteria
Pectobacterium atrosepticum SCRI1043 YP 051534 1 218491 Proteobacteria
Pectobacterium wasabiae WPP163 YP 003258492 1 561231 Proteobacteria
Pectobacterium sp SCC3193 YP 006281953 1 1166016 Proteobacteria
Dickeya solani RNS 08 23 3 1 A WP 022632820 1 Dsolani
Dickeya solani MK10 WP 022632820 1 Dsolani
Dickeya solani RNS 07 7 3B WP 022632820 1 Dsolani
Dickeya solani PPO 9134 WP 022632820 1 Dsolani
Dickeya solani MK16 WP 022632820 1 Dsolani
Dickeya solani RNS 05 1 2A WP 022632820 1 Dsolani
Dickeya solani GBBC 2040 WP 022632820 1 Dsolani
Dickeya solani D s0432 1 WP 022632820 1 Dsolani
Dickeya solani IPO 2222 WP 022632820 1 Dsolani
Dickeya solani IFB 0099 WP 022632820 1 Dsolani
Dickeya solani PPO 9019 WP 022632820 1 Dsolani
Dickeya chrysanthemi subsp chrysanthemi NCPPB 402 WP 040001779 1 Dchrysanthemi
Dickeya chrysanthemi NCPPB 3533 WP 012770485 1 Dchrysanthemi
Dickeya chrysanthemi Ech1591 WP 012770485 1 Dchrysanthemi
Dickeya chrysanthemi subsp parthenii NCPPB 516 WP 012770485 1 Dchrysanthemi
Dickeya chrysanthemi L11 WP 012770485 1 Dchrysanthemi
Dickeya dianthicola NCPPB 453 WP 024105443 1 Ddianthicola
Dickeya dianthicola GBBC 2039 WP 024105443 1 Ddianthicola
Dickeya dianthicola NCPPB 3534 WP 024105443 1 Ddianthicola
Dickeya dianthicola RNS04 9 WP 024105443 1 Ddianthicola
Dickeya dianthicola IPO 980 WP 024105443 1 Ddianthicola
Dickeya gs fangzhongdai S1 WP 049855014 1 Dfangzhongdai
Dickeya gs fangzhongdai M074 WP 038660504 1 Dfangzhongdai
Dickeya gs fangzhongdai M005 WP 038660504 1 Dfangzhongdai
Dickeya gs fangzhongdai MK7 WP 038660504 1 Dfangzhongdai
Dickeya gs fangzhongdai B16 WP 038660504 1 Dfangzhongdai
Dickeya gs fangzhongdai ND14b WP 038660504 1 Dfangzhongdai
Dickeya gs fangzhongdai NCPPB 3274 WP 038660504 1 Dfangzhongdai
Dickeya dadantii subsp dieffenbachiae NCPPB 2976 WP 038922247 1 Ddadantii
Dickeya dadantii NCPPB 3537 WP 013317139 1 Ddadantii
Dickeya dadantii NCPPB 898 WP 013317139 1 Ddadantii
Dickeya dadantii 3937 WP 013317139 1 Ddadantii
120 impC Type VI secretion system Seed
Dickeya gs undicola 2B12 WP 033569735 1 Dundicola
Dickeya zeae MK19 WP 038913517 1 Dzeae
Dickeya zeae NCPPB 2538 WP 038915854 1 Dzeae
Dickeya zeae MS1 WP 019846558 1 Dzeae
Dickeya zeae CSL RW192 WP 038907453 1 Dzeae
Dickeya zeae NCPPB 3531 WP 038903852 1 Dzeae
Dickeya zeae Ech586 WP 012884008 1 Dzeae
Dickeya zeae NCPPB 3532 WP 012884008 1 Dzeae
Dickeya zeae DZ2Q WP 012884008 1 Dzeae
Dickeya zeae EC1 WP 012884008 1 Dzeae
Dickeya zeae ZJU1202 WP 012884008 1 Dzeae

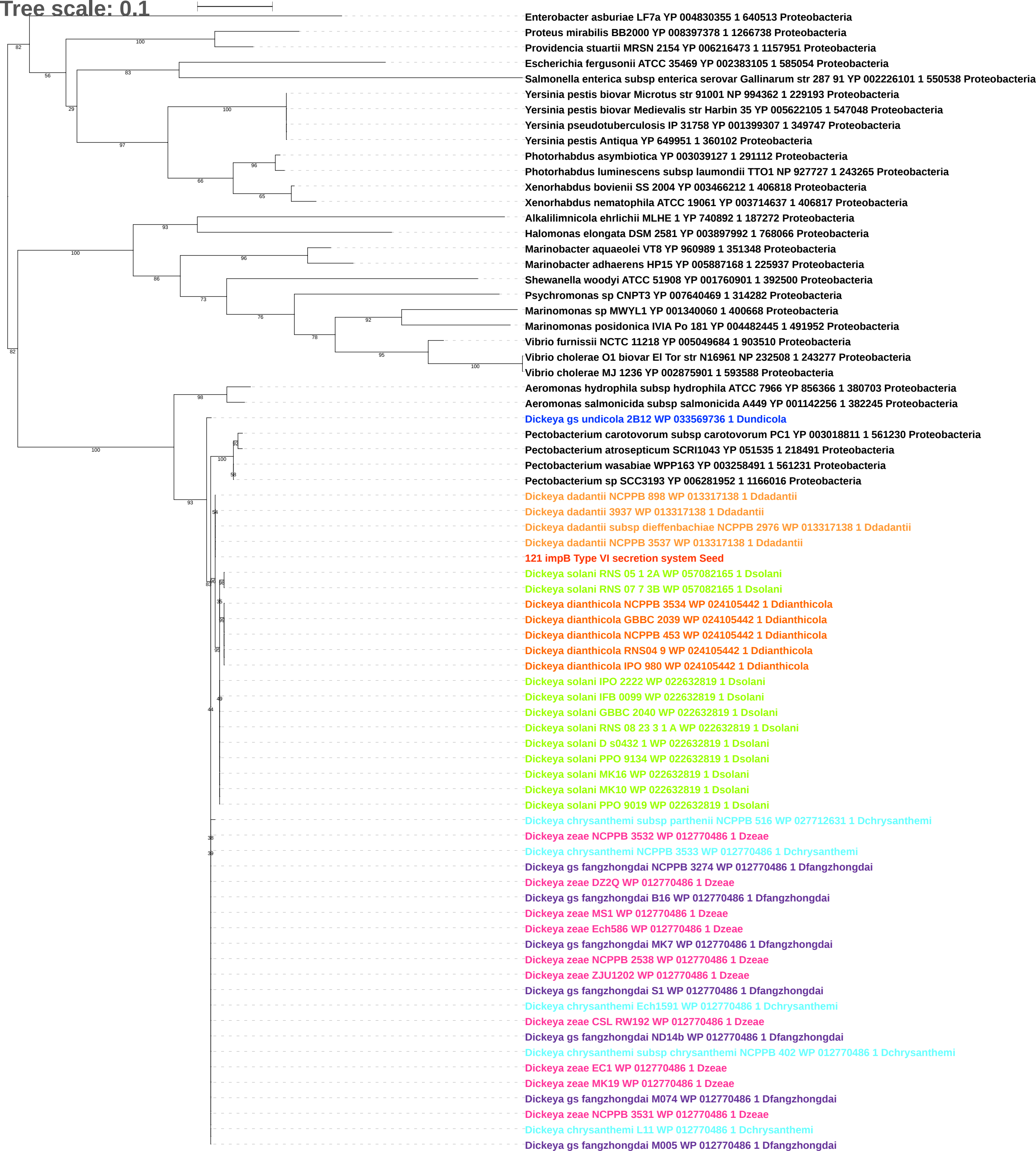
Tree scale: 0.1
Enterobacter asburiae LF7a YP 004830355 1 640513 Proteobacteria
Proteus mirabilis BB2000 YP 008397378 1 1266738 Proteobacteria
Providencia stuartii MRSN 2154 YP 006216473 1 1157951 Proteobacteria
Escherichia fergusonii ATCC 35469 YP 002383105 1 585054 Proteobacteria
Salmonella enterica subsp enterica serovar Gallinarum str 287 91 YP 002226101 1 550538 Proteobacteria
Yersinia pestis biovar Microtus str 91001 NP 994362 1 229193 Proteobacteria
Yersinia pestis biovar Medievalis str Harbin 35 YP 005622105 1 547048 Proteobacteria
Yersinia pseudotuberculosis IP 31758 YP 001399307 1 349747 Proteobacteria
Yersinia pestis Antiqua YP 649951 1 360102 Proteobacteria
Photorhabdus asymbiotica YP 003039127 1 291112 Proteobacteria
Photorhabdus luminescens subsp laumondii TTO1 NP 927727 1 243265 Proteobacteria
Xenorhabdus bovienii SS 2004 YP 003466212 1 406818 Proteobacteria
Xenorhabdus nematophila ATCC 19061 YP 003714637 1 406817 Proteobacteria
Alkalilimnicola ehrlichii MLHE 1 YP 740892 1 187272 Proteobacteria
Halomonas elongata DSM 2581 YP 003897992 1 768066 Proteobacteria
Marinobacter aquaeolei VT8 YP 960989 1 351348 Proteobacteria
Marinobacter adhaerens HP15 YP 005887168 1 225937 Proteobacteria
Shewanella woodyi ATCC 51908 YP 001760901 1 392500 Proteobacteria
Psychromonas sp CNPT3 YP 007640469 1 314282 Proteobacteria
Marinomonas sp MWYL1 YP 001340060 1 400668 Proteobacteria
Marinomonas posidonica IVIA Po 181 YP 004482445 1 491952 Proteobacteria
Vibrio furnissii NCTC 11218 YP 005049684 1 903510 Proteobacteria
Vibrio cholerae O1 biovar El Tor str N16961 NP 232508 1 243277 Proteobacteria
Vibrio cholerae MJ 1236 YP 002875901 1 593588 Proteobacteria
Aeromonas hydrophila subsp hydrophila ATCC 7966 YP 856366 1 380703 Proteobacteria
Aeromonas salmonicida subsp salmonicida A449 YP 001142256 1 382245 Proteobacteria
Dickeya gs undicola 2B12 WP 033569736 1 Dundicola
Pectobacterium carotovorum subsp carotovorum PC1 YP 003018811 1 561230 Proteobacteria
Pectobacterium atrosepticum SCRI1043 YP 051535 1 218491 Proteobacteria
Pectobacterium wasabiae WPP163 YP 003258491 1 561231 Proteobacteria
Pectobacterium sp SCC3193 YP 006281952 1 1166016 Proteobacteria
Dickeya dadantii NCPPB 898 WP 013317138 1 Ddadantii
Dickeya dadantii 3937 WP 013317138 1 Ddadantii
Dickeya dadantii subsp dieffenbachiae NCPPB 2976 WP 013317138 1 Ddadantii
Dickeya dadantii NCPPB 3537 WP 013317138 1 Ddadantii
121 impB Type VI secretion system Seed
Dickeya solani RNS 05 1 2A WP 057082165 1 Dsolani
Dickeya solani RNS 07 7 3B WP 057082165 1 Dsolani
Dickeya dianthicola NCPPB 3534 WP 024105442 1 Ddianthicola
Dickeya dianthicola GBBC 2039 WP 024105442 1 Ddianthicola
Dickeya dianthicola NCPPB 453 WP 024105442 1 Ddianthicola
Dickeya dianthicola RNS04 9 WP 024105442 1 Ddianthicola
Dickeya dianthicola IPO 980 WP 024105442 1 Ddianthicola
Dickeya solani IPO 2222 WP 022632819 1 Dsolani
Dickeya solani IFB 0099 WP 022632819 1 Dsolani
Dickeya solani GBBC 2040 WP 022632819 1 Dsolani
Dickeya solani RNS 08 23 3 1 A WP 022632819 1 Dsolani
Dickeya solani D s0432 1 WP 022632819 1 Dsolani
Dickeya solani PPO 9134 WP 022632819 1 Dsolani
Dickeya solani MK16 WP 022632819 1 Dsolani
Dickeya solani MK10 WP 022632819 1 Dsolani
Dickeya solani PPO 9019 WP 022632819 1 Dsolani
Dickeya chrysanthemi subsp parthenii NCPPB 516 WP 027712631 1 Dchrysanthemi
Dickeya zeae NCPPB 3532 WP 012770486 1 Dzeae
Dickeya chrysanthemi NCPPB 3533 WP 012770486 1 Dchrysanthemi
Dickeya gs fangzhongdai NCPPB 3274 WP 012770486 1 Dfangzhongdai
Dickeya zeae DZ2Q WP 012770486 1 Dzeae
Dickeya gs fangzhongdai B16 WP 012770486 1 Dfangzhongdai
Dickeya zeae MS1 WP 012770486 1 Dzeae
Dickeya zeae Ech586 WP 012770486 1 Dzeae
Dickeya gs fangzhongdai MK7 WP 012770486 1 Dfangzhongdai
Dickeya zeae NCPPB 2538 WP 012770486 1 Dzeae
Dickeya zeae ZJU1202 WP 012770486 1 Dzeae
Dickeya gs fangzhongdai S1 WP 012770486 1 Dfangzhongdai
Dickeya chrysanthemi Ech1591 WP 012770486 1 Dchrysanthemi
Dickeya zeae CSL RW192 WP 012770486 1 Dzeae
Dickeya gs fangzhongdai ND14b WP 012770486 1 Dfangzhongdai
Dickeya chrysanthemi subsp chrysanthemi NCPPB 402 WP 012770486 1 Dchrysanthemi
Dickeya zeae EC1 WP 012770486 1 Dzeae
Dickeya zeae MK19 WP 012770486 1 Dzeae
Dickeya gs fangzhongdai M074 WP 012770486 1 Dfangzhongdai
Dickeya zeae NCPPB 3531 WP 012770486 1 Dzeae
Dickeya chrysanthemi L11 WP 012770486 1 Dchrysanthemi
Dickeya gs fangzhongdai M005 WP 012770486 1 Dfangzhongdai

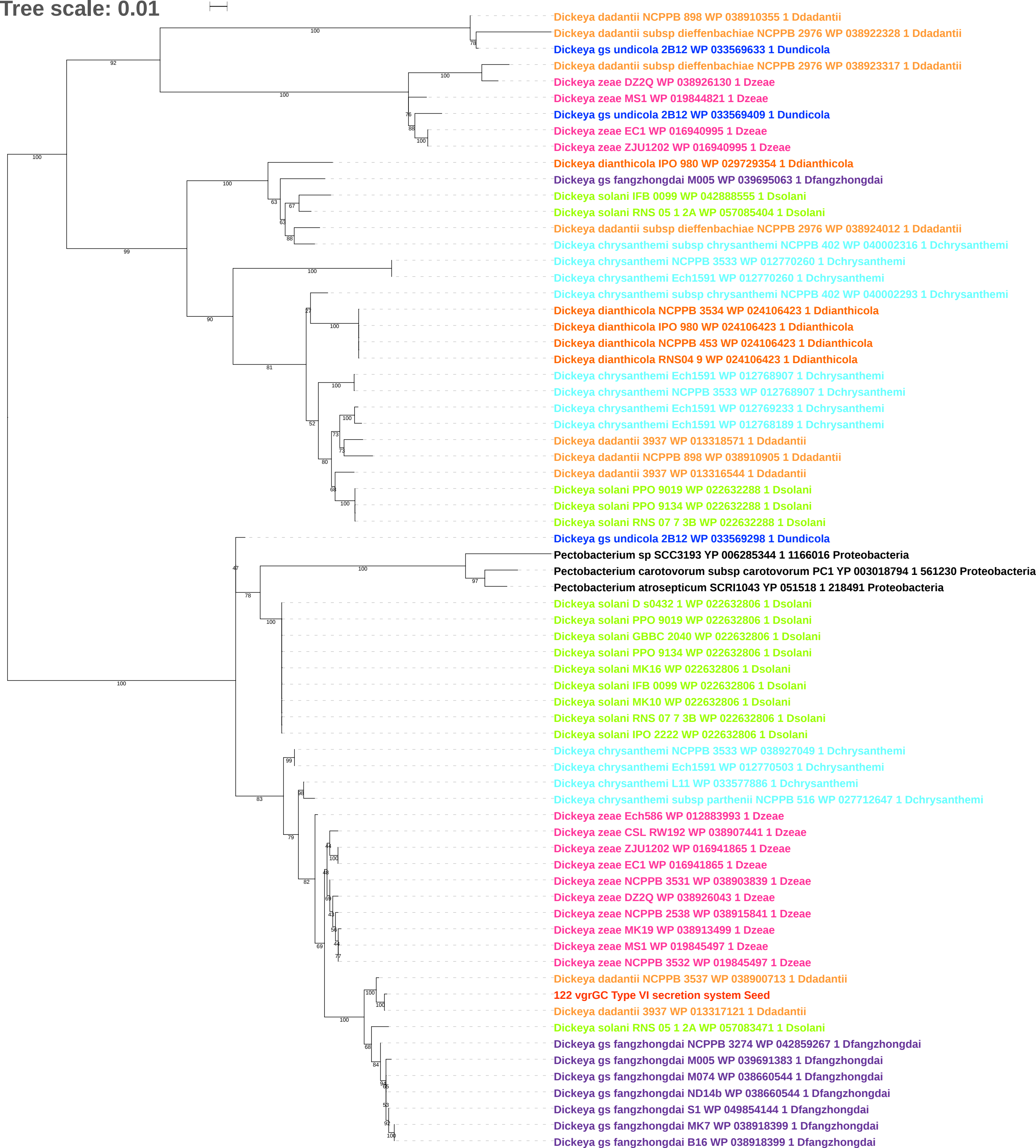
Tree scale: 0.01
Dickeya dadantii NCPPB 898 WP 038910355 1 Ddadantii
Dickeya dadantii subsp dieffenbachiae NCPPB 2976 WP 038922328 1 Ddadantii
Dickeya gs undicola 2B12 WP 033569633 1 Dundicola
Dickeya dadantii subsp dieffenbachiae NCPPB 2976 WP 038923317 1 Ddadantii
Dickeya zeae DZ2Q WP 038926130 1 Dzeae
Dickeya zeae MS1 WP 019844821 1 Dzeae
Dickeya gs undicola 2B12 WP 033569409 1 Dundicola
Dickeya zeae EC1 WP 016940995 1 Dzeae
Dickeya zeae ZJU1202 WP 016940995 1 Dzeae
Dickeya dianthicola IPO 980 WP 029729354 1 Ddianthicola
Dickeya gs fangzhongdai M005 WP 039695063 1 Dfangzhongdai
Dickeya solani IFB 0099 WP 042888555 1 Dsolani
Dickeya solani RNS 05 1 2A WP 057085404 1 Dsolani
Dickeya dadantii subsp dieffenbachiae NCPPB 2976 WP 038924012 1 Ddadantii
Dickeya chrysanthemi subsp chrysanthemi NCPPB 402 WP 040002316 1 Dchrysanthemi
Dickeya chrysanthemi NCPPB 3533 WP 012770260 1 Dchrysanthemi
Dickeya chrysanthemi Ech1591 WP 012770260 1 Dchrysanthemi
Dickeya chrysanthemi subsp chrysanthemi NCPPB 402 WP 040002293 1 Dchrysanthemi
Dickeya dianthicola NCPPB 3534 WP 024106423 1 Ddianthicola
Dickeya dianthicola IPO 980 WP 024106423 1 Ddianthicola
Dickeya dianthicola NCPPB 453 WP 024106423 1 Ddianthicola
Dickeya dianthicola RNS04 9 WP 024106423 1 Ddianthicola
Dickeya chrysanthemi Ech1591 WP 012768907 1 Dchrysanthemi
Dickeya chrysanthemi NCPPB 3533 WP 012768907 1 Dchrysanthemi
Dickeya chrysanthemi Ech1591 WP 012769233 1 Dchrysanthemi
Dickeya chrysanthemi Ech1591 WP 012768189 1 Dchrysanthemi
Dickeya dadantii 3937 WP 013318571 1 Ddadantii
Dickeya dadantii NCPPB 898 WP 038910905 1 Ddadantii
Dickeya dadantii 3937 WP 013316544 1 Ddadantii
Dickeya solani PPO 9019 WP 022632288 1 Dsolani
Dickeya solani PPO 9134 WP 022632288 1 Dsolani
Dickeya solani RNS 07 7 3B WP 022632288 1 Dsolani
Dickeya gs undicola 2B12 WP 033569298 1 Dundicola
Pectobacterium sp SCC3193 YP 006285344 1 1166016 Proteobacteria
Pectobacterium carotovorum subsp carotovorum PC1 YP 003018794 1 561230 Proteobacteria
Pectobacterium atrosepticum SCRI1043 YP 051518 1 218491 Proteobacteria
Dickeya solani D s0432 1 WP 022632806 1 Dsolani
Dickeya solani PPO 9019 WP 022632806 1 Dsolani
Dickeya solani GBBC 2040 WP 022632806 1 Dsolani
Dickeya solani PPO 9134 WP 022632806 1 Dsolani
Dickeya solani MK16 WP 022632806 1 Dsolani
Dickeya solani IFB 0099 WP 022632806 1 Dsolani
Dickeya solani MK10 WP 022632806 1 Dsolani
Dickeya solani RNS 07 7 3B WP 022632806 1 Dsolani
Dickeya solani IPO 2222 WP 022632806 1 Dsolani
Dickeya chrysanthemi NCPPB 3533 WP 038927049 1 Dchrysanthemi
Dickeya chrysanthemi Ech1591 WP 012770503 1 Dchrysanthemi
Dickeya chrysanthemi L11 WP 033577886 1 Dchrysanthemi
Dickeya chrysanthemi subsp parthenii NCPPB 516 WP 027712647 1 Dchrysanthemi
Dickeya zeae Ech586 WP 012883993 1 Dzeae
Dickeya zeae CSL RW192 WP 038907441 1 Dzeae
Dickeya zeae ZJU1202 WP 016941865 1 Dzeae
Dickeya zeae EC1 WP 016941865 1 Dzeae
Dickeya zeae NCPPB 3531 WP 038903839 1 Dzeae
Dickeya zeae DZ2Q WP 038926043 1 Dzeae
Dickeya zeae NCPPB 2538 WP 038915841 1 Dzeae
Dickeya zeae MK19 WP 038913499 1 Dzeae
Dickeya zeae MS1 WP 019845497 1 Dzeae
Dickeya zeae NCPPB 3532 WP 019845497 1 Dzeae
Dickeya dadantii NCPPB 3537 WP 038900713 1 Ddadantii
122 vgrGC Type VI secretion system Seed
Dickeya dadantii 3937 WP 013317121 1 Ddadantii
Dickeya solani RNS 05 1 2A WP 057083471 1 Dsolani
Dickeya gs fangzhongdai NCPPB 3274 WP 042859267 1 Dfangzhongdai
Dickeya gs fangzhongdai M005 WP 039691383 1 Dfangzhongdai
Dickeya gs fangzhongdai M074 WP 038660544 1 Dfangzhongdai
Dickeya gs fangzhongdai ND14b WP 038660544 1 Dfangzhongdai
Dickeya gs fangzhongdai S1 WP 049854144 1 Dfangzhongdai
Dickeya gs fangzhongdai MK7 WP 038918399 1 Dfangzhongdai
Dickeya gs fangzhongdai B16 WP 038918399 1 Dfangzhongdai

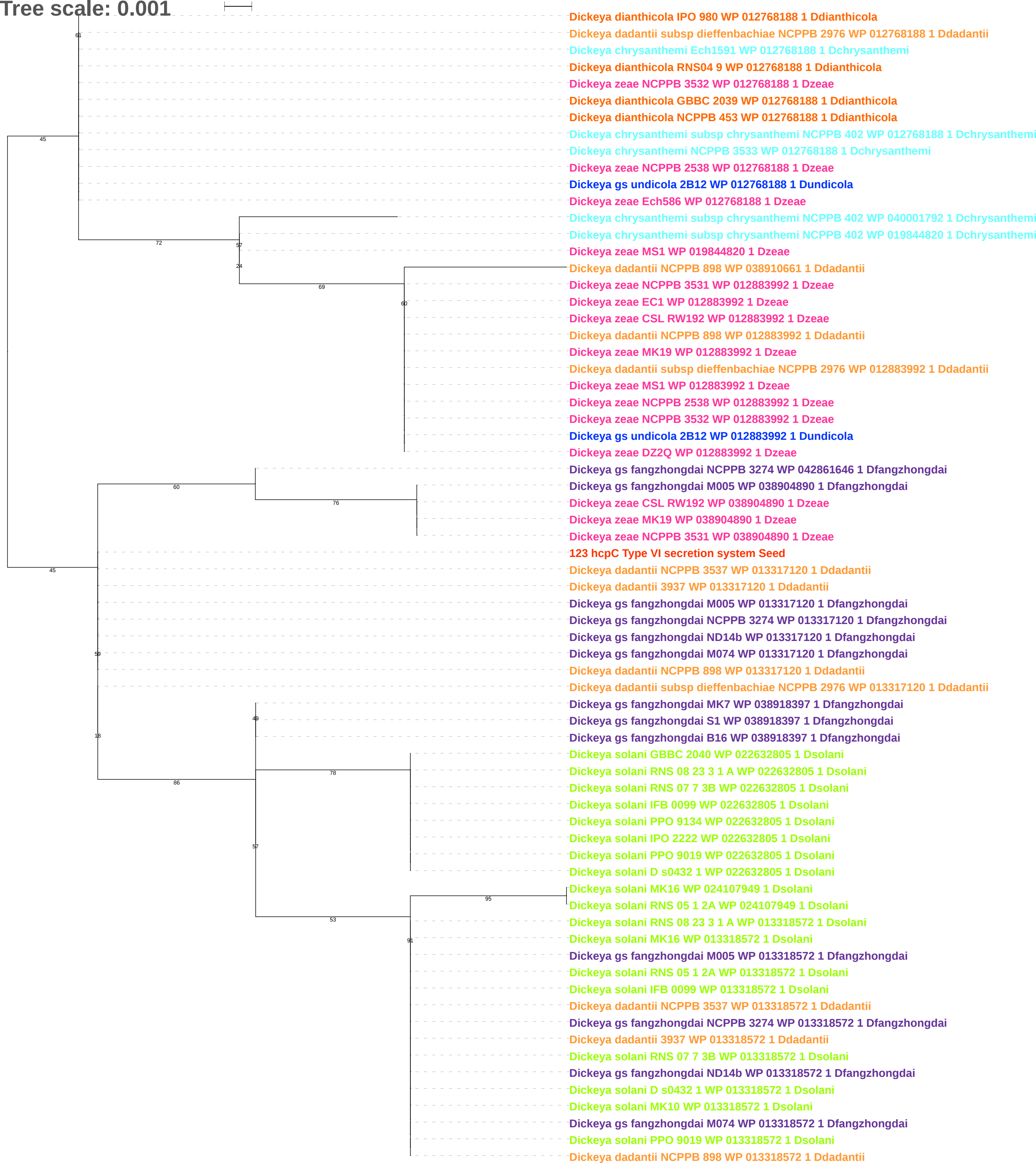
Tree scale: 0.001
Dickeya dianthicola IPO 980 WP 012768188 1 Ddianthicola
Dickeya dadantii subsp dieffenbachiae NCPPB 2976 WP 012768188 1 Ddadantii
Dickeya chrysanthemi Ech1591 WP 012768188 1 Dchrysanthemi
Dickeya dianthicola RNS04 9 WP 012768188 1 Ddianthicola
Dickeya zeae NCPPB 3532 WP 012768188 1 Dzeae
Dickeya dianthicola GBBC 2039 WP 012768188 1 Ddianthicola
Dickeya dianthicola NCPPB 453 WP 012768188 1 Ddianthicola
Dickeya chrysanthemi subsp chrysanthemi NCPPB 402 WP 012768188 1 Dchrysanthemi
Dickeya chrysanthemi NCPPB 3533 WP 012768188 1 Dchrysanthemi
Dickeya zeae NCPPB 2538 WP 012768188 1 Dzeae
Dickeya gs undicola 2B12 WP 012768188 1 Dundicola
Dickeya zeae Ech586 WP 012768188 1 Dzeae
Dickeya chrysanthemi subsp chrysanthemi NCPPB 402 WP 040001792 1 Dchrysanthemi
Dickeya chrysanthemi subsp chrysanthemi NCPPB 402 WP 019844820 1 Dchrysanthemi
Dickeya zeae MS1 WP 019844820 1 Dzeae
Dickeya dadantii NCPPB 898 WP 038910661 1 Ddadantii
Dickeya zeae NCPPB 3531 WP 012883992 1 Dzeae
Dickeya zeae EC1 WP 012883992 1 Dzeae
Dickeya zeae CSL RW192 WP 012883992 1 Dzeae
Dickeya dadantii NCPPB 898 WP 012883992 1 Ddadantii
Dickeya zeae MK19 WP 012883992 1 Dzeae
Dickeya dadantii subsp dieffenbachiae NCPPB 2976 WP 012883992 1 Ddadantii
Dickeya zeae MS1 WP 012883992 1 Dzeae
Dickeya zeae NCPPB 2538 WP 012883992 1 Dzeae
Dickeya zeae NCPPB 3532 WP 012883992 1 Dzeae
Dickeya gs undicola 2B12 WP 012883992 1 Dundicola
Dickeya zeae DZ2Q WP 012883992 1 Dzeae
Dickeya gs fangzhongdai NCPPB 3274 WP 042861646 1 Dfangzhongdai
Dickeya gs fangzhongdai M005 WP 038904890 1 Dfangzhongdai
Dickeya zeae CSL RW192 WP 038904890 1 Dzeae
Dickeya zeae MK19 WP 038904890 1 Dzeae
Dickeya zeae NCPPB 3531 WP 038904890 1 Dzeae
123 hcpC Type VI secretion system Seed
Dickeya dadantii NCPPB 3537 WP 013317120 1 Ddadantii
Dickeya dadantii 3937 WP 013317120 1 Ddadantii
Dickeya gs fangzhongdai M005 WP 013317120 1 Dfangzhongdai
Dickeya gs fangzhongdai NCPPB 3274 WP 013317120 1 Dfangzhongdai
Dickeya gs fangzhongdai ND14b WP 013317120 1 Dfangzhongdai
Dickeya gs fangzhongdai M074 WP 013317120 1 Dfangzhongdai
Dickeya dadantii NCPPB 898 WP 013317120 1 Ddadantii
Dickeya dadantii subsp dieffenbachiae NCPPB 2976 WP 013317120 1 Ddadantii
Dickeya gs fangzhongdai MK7 WP 038918397 1 Dfangzhongdai
Dickeya gs fangzhongdai S1 WP 038918397 1 Dfangzhongdai
Dickeya gs fangzhongdai B16 WP 038918397 1 Dfangzhongdai
Dickeya solani GBBC 2040 WP 022632805 1 Dsolani
Dickeya solani RNS 08 23 3 1 A WP 022632805 1 Dsolani
Dickeya solani RNS 07 7 3B WP 022632805 1 Dsolani
Dickeya solani IFB 0099 WP 022632805 1 Dsolani
Dickeya solani PPO 9134 WP 022632805 1 Dsolani
Dickeya solani IPO 2222 WP 022632805 1 Dsolani
Dickeya solani PPO 9019 WP 022632805 1 Dsolani
Dickeya solani D s0432 1 WP 022632805 1 Dsolani
Dickeya solani MK16 WP 024107949 1 Dsolani
Dickeya solani RNS 05 1 2A WP 024107949 1 Dsolani
Dickeya solani RNS 08 23 3 1 A WP 013318572 1 Dsolani
Dickeya solani MK16 WP 013318572 1 Dsolani
Dickeya gs fangzhongdai M005 WP 013318572 1 Dfangzhongdai
Dickeya solani RNS 05 1 2A WP 013318572 1 Dsolani
Dickeya solani IFB 0099 WP 013318572 1 Dsolani
Dickeya dadantii NCPPB 3537 WP 013318572 1 Ddadantii
Dickeya gs fangzhongdai NCPPB 3274 WP 013318572 1 Dfangzhongdai
Dickeya dadantii 3937 WP 013318572 1 Ddadantii
Dickeya solani RNS 07 7 3B WP 013318572 1 Dsolani
Dickeya gs fangzhongdai ND14b WP 013318572 1 Dfangzhongdai
Dickeya solani D s0432 1 WP 013318572 1 Dsolani
Dickeya solani MK10 WP 013318572 1 Dsolani
Dickeya gs fangzhongdai M074 WP 013318572 1 Dfangzhongdai
Dickeya solani PPO 9019 WP 013318572 1 Dsolani
Dickeya dadantii NCPPB 898 WP 013318572 1 Ddadantii

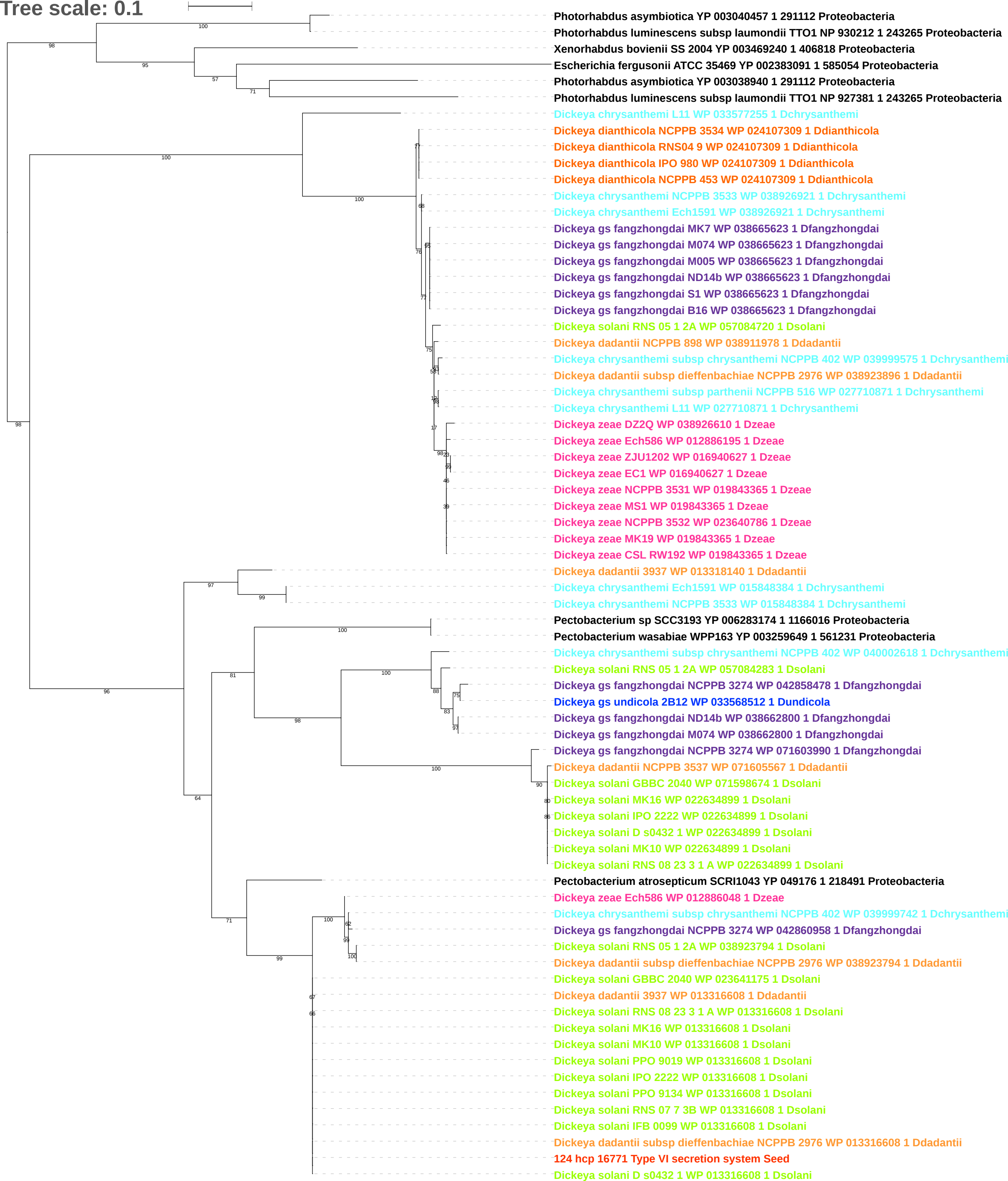
Tree scale: 0.1
Photorhabdus asymbiotica YP 003040457 1 291112 Proteobacteria
Photorhabdus luminescens subsp laumondii TTO1 NP 930212 1 243265 Proteobacteria
Xenorhabdus bovienii SS 2004 YP 003469240 1 406818 Proteobacteria
Escherichia fergusonii ATCC 35469 YP 002383091 1 585054 Proteobacteria
Photorhabdus asymbiotica YP 003038940 1 291112 Proteobacteria
Photorhabdus luminescens subsp laumondii TTO1 NP 927381 1 243265 Proteobacteria
Dickeya chrysanthemi L11 WP 033577255 1 Dchrysanthemi
Dickeya dianthicola NCPPB 3534 WP 024107309 1 Ddianthicola
Dickeya dianthicola RNS04 9 WP 024107309 1 Ddianthicola
Dickeya dianthicola IPO 980 WP 024107309 1 Ddianthicola
Dickeya dianthicola NCPPB 453 WP 024107309 1 Ddianthicola
Dickeya chrysanthemi NCPPB 3533 WP 038926921 1 Dchrysanthemi
Dickeya chrysanthemi Ech1591 WP 038926921 1 Dchrysanthemi
Dickeya gs fangzhongdai MK7 WP 038665623 1 Dfangzhongdai
Dickeya gs fangzhongdai M074 WP 038665623 1 Dfangzhongdai
Dickeya gs fangzhongdai M005 WP 038665623 1 Dfangzhongdai
Dickeya gs fangzhongdai ND14b WP 038665623 1 Dfangzhongdai
Dickeya gs fangzhongdai S1 WP 038665623 1 Dfangzhongdai
Dickeya gs fangzhongdai B16 WP 038665623 1 Dfangzhongdai
Dickeya solani RNS 05 1 2A WP 057084720 1 Dsolani
Dickeya dadantii NCPPB 898 WP 038911978 1 Ddadantii
Dickeya chrysanthemi subsp chrysanthemi NCPPB 402 WP 039999575 1 Dchrysanthemi
Dickeya dadantii subsp dieffenbachiae NCPPB 2976 WP 038923896 1 Ddadantii
Dickeya chrysanthemi subsp parthenii NCPPB 516 WP 027710871 1 Dchrysanthemi
Dickeya chrysanthemi L11 WP 027710871 1 Dchrysanthemi
Dickeya zeae DZ2Q WP 038926610 1 Dzeae
Dickeya zeae Ech586 WP 012886195 1 Dzeae
Dickeya zeae ZJU1202 WP 016940627 1 Dzeae
Dickeya zeae EC1 WP 016940627 1 Dzeae
Dickeya zeae NCPPB 3531 WP 019843365 1 Dzeae
Dickeya zeae MS1 WP 019843365 1 Dzeae
Dickeya zeae NCPPB 3532 WP 023640786 1 Dzeae
Dickeya zeae MK19 WP 019843365 1 Dzeae
Dickeya zeae CSL RW192 WP 019843365 1 Dzeae
Dickeya dadantii 3937 WP 013318140 1 Ddadantii
Dickeya chrysanthemi Ech1591 WP 015848384 1 Dchrysanthemi
Dickeya chrysanthemi NCPPB 3533 WP 015848384 1 Dchrysanthemi
Pectobacterium sp SCC3193 YP 006283174 1 1166016 Proteobacteria
Pectobacterium wasabiae WPP163 YP 003259649 1 561231 Proteobacteria
Dickeya chrysanthemi subsp chrysanthemi NCPPB 402 WP 040002618 1 Dchrysanthemi
Dickeya solani RNS 05 1 2A WP 057084283 1 Dsolani
Dickeya gs fangzhongdai NCPPB 3274 WP 042858478 1 Dfangzhongdai
Dickeya gs undicola 2B12 WP 033568512 1 Dundicola
Dickeya gs fangzhongdai ND14b WP 038662800 1 Dfangzhongdai
Dickeya gs fangzhongdai M074 WP 038662800 1 Dfangzhongdai
Dickeya gs fangzhongdai NCPPB 3274 WP 071603990 1 Dfangzhongdai
Dickeya dadantii NCPPB 3537 WP 071605567 1 Ddadantii
Dickeya solani GBBC 2040 WP 071598674 1 Dsolani
Dickeya solani MK16 WP 022634899 1 Dsolani
Dickeya solani IPO 2222 WP 022634899 1 Dsolani
Dickeya solani D s0432 1 WP 022634899 1 Dsolani
Dickeya solani MK10 WP 022634899 1 Dsolani
Dickeya solani RNS 08 23 3 1 A WP 022634899 1 Dsolani
Pectobacterium atrosepticum SCRI1043 YP 049176 1 218491 Proteobacteria
Dickeya zeae Ech586 WP 012886048 1 Dzeae
Dickeya chrysanthemi subsp chrysanthemi NCPPB 402 WP 039999742 1 Dchrysanthemi
Dickeya gs fangzhongdai NCPPB 3274 WP 042860958 1 Dfangzhongdai
Dickeya solani RNS 05 1 2A WP 038923794 1 Dsolani
Dickeya dadantii subsp dieffenbachiae NCPPB 2976 WP 038923794 1 Ddadantii
Dickeya solani GBBC 2040 WP 023641175 1 Dsolani
Dickeya dadantii 3937 WP 013316608 1 Ddadantii
Dickeya solani RNS 08 23 3 1 A WP 013316608 1 Dsolani
Dickeya solani MK16 WP 013316608 1 Dsolani
Dickeya solani MK10 WP 013316608 1 Dsolani
Dickeya solani PPO 9019 WP 013316608 1 Dsolani
Dickeya solani IPO 2222 WP 013316608 1 Dsolani
Dickeya solani PPO 9134 WP 013316608 1 Dsolani
Dickeya solani RNS 07 7 3B WP 013316608 1 Dsolani
Dickeya solani IFB 0099 WP 013316608 1 Dsolani
Dickeya dadantii subsp dieffenbachiae NCPPB 2976 WP 013316608 1 Ddadantii
124 hcp 16771 Type VI secretion system Seed
Dickeya solani D s0432 1 WP 013316608 1 Dsolani

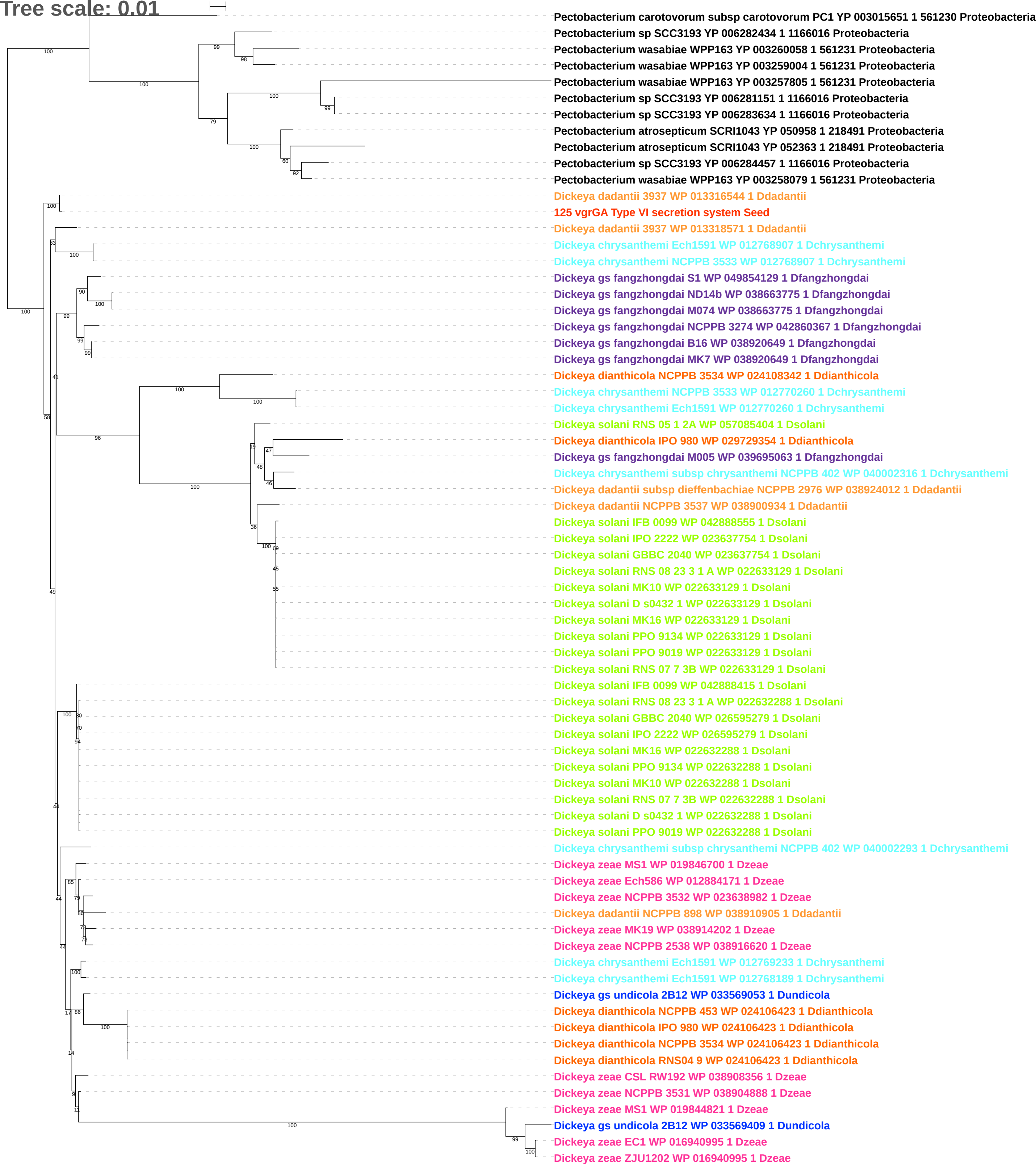
Tree scale: 0.01
Pectobacterium carotovorum subsp carotovorum PC1 YP 003015651 1 561230 Proteobacteria
Pectobacterium sp SCC3193 YP 006282434 1 1166016 Proteobacteria
Pectobacterium wasabiae WPP163 YP 003260058 1 561231 Proteobacteria
Pectobacterium wasabiae WPP163 YP 003259004 1 561231 Proteobacteria
Pectobacterium wasabiae WPP163 YP 003257805 1 561231 Proteobacteria
Pectobacterium sp SCC3193 YP 006281151 1 1166016 Proteobacteria
Pectobacterium sp SCC3193 YP 006283634 1 1166016 Proteobacteria
Pectobacterium atrosepticum SCRI1043 YP 050958 1 218491 Proteobacteria
Pectobacterium atrosepticum SCRI1043 YP 052363 1 218491 Proteobacteria
Pectobacterium sp SCC3193 YP 006284457 1 1166016 Proteobacteria
Pectobacterium wasabiae WPP163 YP 003258079 1 561231 Proteobacteria
Dickeya dadantii 3937 WP 013316544 1 Ddadantii
125 vgrGA Type VI secretion system Seed
Dickeya dadantii 3937 WP 013318571 1 Ddadantii
Dickeya chrysanthemi Ech1591 WP 012768907 1 Dchrysanthemi
Dickeya chrysanthemi NCPPB 3533 WP 012768907 1 Dchrysanthemi
Dickeya gs fangzhongdai S1 WP 049854129 1 Dfangzhongdai
Dickeya gs fangzhongdai ND14b WP 038663775 1 Dfangzhongdai
Dickeya gs fangzhongdai M074 WP 038663775 1 Dfangzhongdai
Dickeya gs fangzhongdai NCPPB 3274 WP 042860367 1 Dfangzhongdai
Dickeya gs fangzhongdai B16 WP 038920649 1 Dfangzhongdai
Dickeya gs fangzhongdai MK7 WP 038920649 1 Dfangzhongdai
Dickeya dianthicola NCPPB 3534 WP 024108342 1 Ddianthicola
Dickeya chrysanthemi NCPPB 3533 WP 012770260 1 Dchrysanthemi
Dickeya chrysanthemi Ech1591 WP 012770260 1 Dchrysanthemi
Dickeya solani RNS 05 1 2A WP 057085404 1 Dsolani
Dickeya dianthicola IPO 980 WP 029729354 1 Ddianthicola
Dickeya gs fangzhongdai M005 WP 039695063 1 Dfangzhongdai
Dickeya chrysanthemi subsp chrysanthemi NCPPB 402 WP 040002316 1 Dchrysanthemi
Dickeya dadantii subsp dieffenbachiae NCPPB 2976 WP 038924012 1 Ddadantii
Dickeya dadantii NCPPB 3537 WP 038900934 1 Ddadantii
Dickeya solani IFB 0099 WP 042888555 1 Dsolani
Dickeya solani IPO 2222 WP 023637754 1 Dsolani
Dickeya solani GBBC 2040 WP 023637754 1 Dsolani
Dickeya solani RNS 08 23 3 1 A WP 022633129 1 Dsolani
Dickeya solani MK10 WP 022633129 1 Dsolani
Dickeya solani D s0432 1 WP 022633129 1 Dsolani
Dickeya solani MK16 WP 022633129 1 Dsolani
Dickeya solani PPO 9134 WP 022633129 1 Dsolani
Dickeya solani PPO 9019 WP 022633129 1 Dsolani
Dickeya solani RNS 07 7 3B WP 022633129 1 Dsolani
Dickeya solani IFB 0099 WP 042888415 1 Dsolani
Dickeya solani RNS 08 23 3 1 A WP 022632288 1 Dsolani
Dickeya solani GBBC 2040 WP 026595279 1 Dsolani
Dickeya solani IPO 2222 WP 026595279 1 Dsolani
Dickeya solani MK16 WP 022632288 1 Dsolani
Dickeya solani PPO 9134 WP 022632288 1 Dsolani
Dickeya solani MK10 WP 022632288 1 Dsolani
Dickeya solani RNS 07 7 3B WP 022632288 1 Dsolani
Dickeya solani D s0432 1 WP 022632288 1 Dsolani
Dickeya solani PPO 9019 WP 022632288 1 Dsolani
Dickeya chrysanthemi subsp chrysanthemi NCPPB 402 WP 040002293 1 Dchrysanthemi
Dickeya zeae MS1 WP 019846700 1 Dzeae
Dickeya zeae Ech586 WP 012884171 1 Dzeae
Dickeya zeae NCPPB 3532 WP 023638982 1 Dzeae
Dickeya dadantii NCPPB 898 WP 038910905 1 Ddadantii
Dickeya zeae MK19 WP 038914202 1 Dzeae
Dickeya zeae NCPPB 2538 WP 038916620 1 Dzeae
Dickeya chrysanthemi Ech1591 WP 012769233 1 Dchrysanthemi
Dickeya chrysanthemi Ech1591 WP 012768189 1 Dchrysanthemi
Dickeya gs undicola 2B12 WP 033569053 1 Dundicola
Dickeya dianthicola NCPPB 453 WP 024106423 1 Ddianthicola
Dickeya dianthicola IPO 980 WP 024106423 1 Ddianthicola
Dickeya dianthicola NCPPB 3534 WP 024106423 1 Ddianthicola
Dickeya dianthicola RNS04 9 WP 024106423 1 Ddianthicola
Dickeya zeae CSL RW192 WP 038908356 1 Dzeae
Dickeya zeae NCPPB 3531 WP 038904888 1 Dzeae
Dickeya zeae MS1 WP 019844821 1 Dzeae
Dickeya gs undicola 2B12 WP 033569409 1 Dundicola
Dickeya zeae EC1 WP 016940995 1 Dzeae
Dickeya zeae ZJU1202 WP 016940995 1 Dzeae

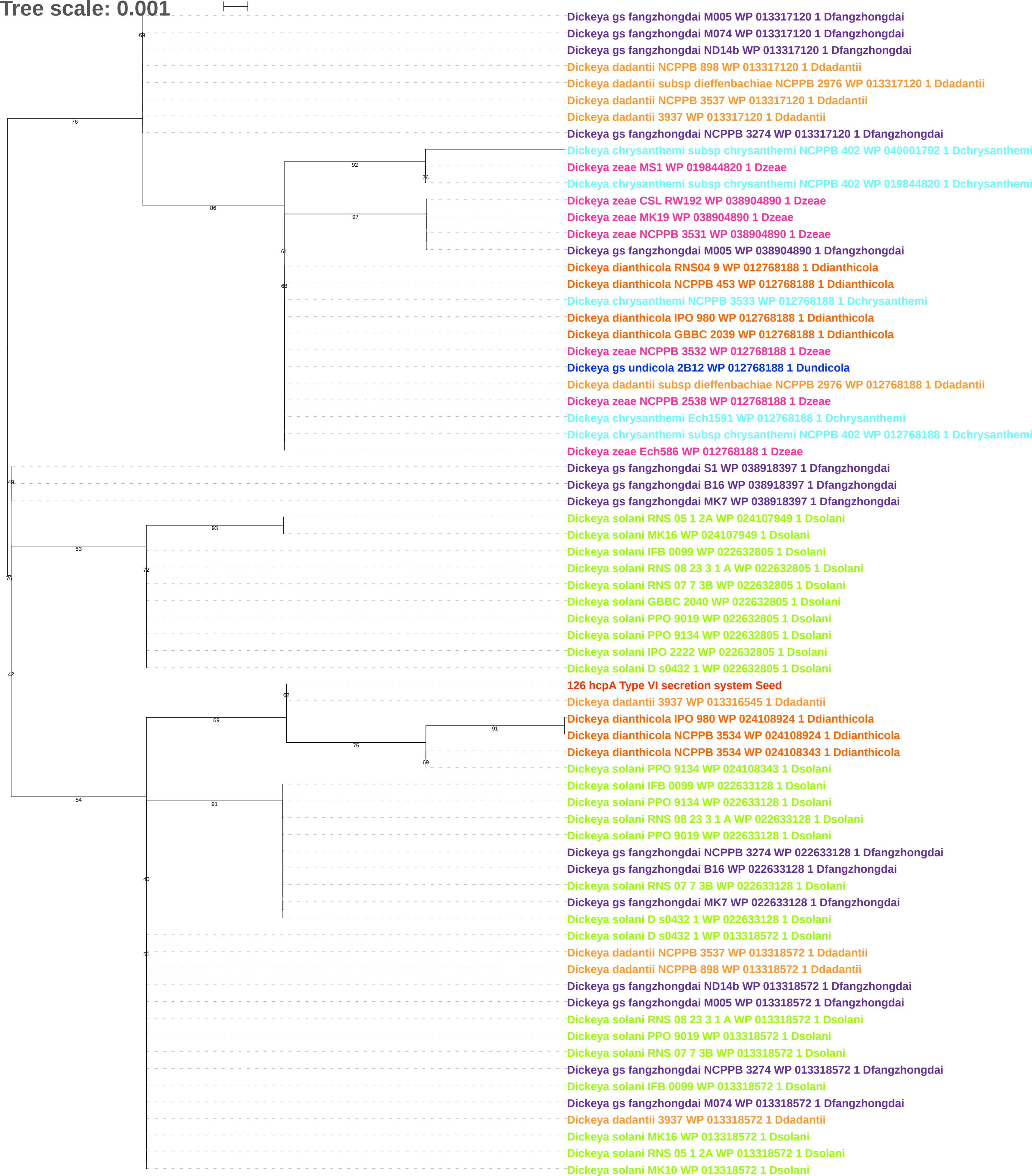
Tree scale: 0.001
Dickeya gs fangzhongdai M005 WP 013317120 1 Dfangzhongdai
Dickeya gs fangzhongdai M074 WP 013317120 1 Dfangzhongdai
Dickeya gs fangzhongdai ND14b WP 013317120 1 Dfangzhongdai
Dickeya dadantii NCPPB 898 WP 013317120 1 Ddadantii
Dickeya dadantii subsp dieffenbachiae NCPPB 2976 WP 013317120 1 Ddadantii
Dickeya dadantii NCPPB 3537 WP 013317120 1 Ddadantii
Dickeya dadantii 3937 WP 013317120 1 Ddadantii
Dickeya gs fangzhongdai NCPPB 3274 WP 013317120 1 Dfangzhongdai
Dickeya chrysanthemi subsp chrysanthemi NCPPB 402 WP 040001792 1 Dchrysanthemi
Dickeya zeae MS1 WP 019844820 1 Dzeae
Dickeya chrysanthemi subsp chrysanthemi NCPPB 402 WP 019844820 1 Dchrysanthemi
Dickeya zeae CSL RW192 WP 038904890 1 Dzeae
Dickeya zeae MK19 WP 038904890 1 Dzeae
Dickeya zeae NCPPB 3531 WP 038904890 1 Dzeae
Dickeya gs fangzhongdai M005 WP 038904890 1 Dfangzhongdai
Dickeya dianthicola RNS04 9 WP 012768188 1 Ddianthicola
Dickeya dianthicola NCPPB 453 WP 012768188 1 Ddianthicola
Dickeya chrysanthemi NCPPB 3533 WP 012768188 1 Dchrysanthemi
Dickeya dianthicola IPO 980 WP 012768188 1 Ddianthicola
Dickeya dianthicola GBBC 2039 WP 012768188 1 Ddianthicola
Dickeya zeae NCPPB 3532 WP 012768188 1 Dzeae
Dickeya gs undicola 2B12 WP 012768188 1 Dundicola
Dickeya dadantii subsp dieffenbachiae NCPPB 2976 WP 012768188 1 Ddadantii
Dickeya zeae NCPPB 2538 WP 012768188 1 Dzeae
Dickeya chrysanthemi Ech1591 WP 012768188 1 Dchrysanthemi
Dickeya chrysanthemi subsp chrysanthemi NCPPB 402 WP 012768188 1 Dchrysanthemi
Dickeya zeae Ech586 WP 012768188 1 Dzeae
Dickeya gs fangzhongdai S1 WP 038918397 1 Dfangzhongdai
Dickeya gs fangzhongdai B16 WP 038918397 1 Dfangzhongdai
Dickeya gs fangzhongdai MK7 WP 038918397 1 Dfangzhongdai
Dickeya solani RNS 05 1 2A WP 024107949 1 Dsolani
Dickeya solani MK16 WP 024107949 1 Dsolani
Dickeya solani IFB 0099 WP 022632805 1 Dsolani
Dickeya solani RNS 08 23 3 1 A WP 022632805 1 Dsolani
Dickeya solani RNS 07 7 3B WP 022632805 1 Dsolani
Dickeya solani GBBC 2040 WP 022632805 1 Dsolani
Dickeya solani PPO 9019 WP 022632805 1 Dsolani
Dickeya solani PPO 9134 WP 022632805 1 Dsolani
Dickeya solani IPO 2222 WP 022632805 1 Dsolani
Dickeya solani D s0432 1 WP 022632805 1 Dsolani
126 hcpA Type VI secretion system Seed
Dickeya dadantii 3937 WP 013316545 1 Ddadantii
Dickeya dianthicola IPO 980 WP 024108924 1 Ddianthicola
Dickeya dianthicola NCPPB 3534 WP 024108924 1 Ddianthicola
Dickeya dianthicola NCPPB 3534 WP 024108343 1 Ddianthicola
Dickeya solani PPO 9134 WP 024108343 1 Dsolani
Dickeya solani IFB 0099 WP 022633128 1 Dsolani
Dickeya solani PPO 9134 WP 022633128 1 Dsolani
Dickeya solani RNS 08 23 3 1 A WP 022633128 1 Dsolani
Dickeya solani PPO 9019 WP 022633128 1 Dsolani
Dickeya gs fangzhongdai NCPPB 3274 WP 022633128 1 Dfangzhongdai
Dickeya gs fangzhongdai B16 WP 022633128 1 Dfangzhongdai
Dickeya solani RNS 07 7 3B WP 022633128 1 Dsolani
Dickeya gs fangzhongdai MK7 WP 022633128 1 Dfangzhongdai
Dickeya solani D s0432 1 WP 022633128 1 Dsolani
Dickeya solani D s0432 1 WP 013318572 1 Dsolani
Dickeya dadantii NCPPB 3537 WP 013318572 1 Ddadantii
Dickeya dadantii NCPPB 898 WP 013318572 1 Ddadantii
Dickeya gs fangzhongdai ND14b WP 013318572 1 Dfangzhongdai
Dickeya gs fangzhongdai M005 WP 013318572 1 Dfangzhongdai
Dickeya solani RNS 08 23 3 1 A WP 013318572 1 Dsolani
Dickeya solani PPO 9019 WP 013318572 1 Dsolani
Dickeya solani RNS 07 7 3B WP 013318572 1 Dsolani
Dickeya gs fangzhongdai NCPPB 3274 WP 013318572 1 Dfangzhongdai
Dickeya solani IFB 0099 WP 013318572 1 Dsolani
Dickeya gs fangzhongdai M074 WP 013318572 1 Dfangzhongdai
Dickeya dadantii 3937 WP 013318572 1 Ddadantii
Dickeya solani MK16 WP 013318572 1 Dsolani
Dickeya solani RNS 05 1 2A WP 013318572 1 Dsolani
Dickeya solani MK10 WP 013318572 1 Dsolani

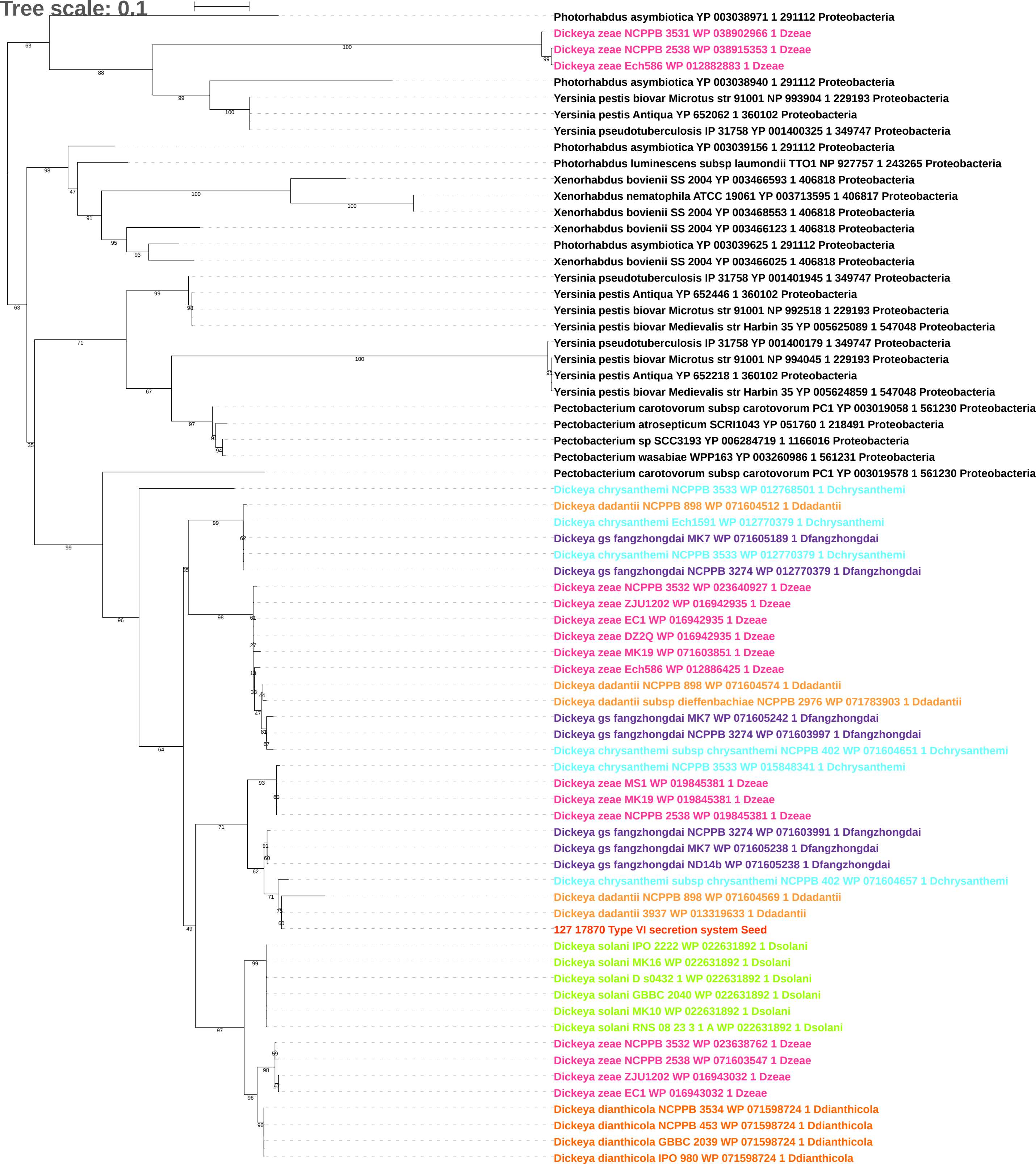
Tree scale: 0.1
Photorhabdus asymbiotica YP 003038971 1 291112 Proteobacteria
Dickeya zeae NCPPB 3531 WP 038902966 1 Dzeae
Dickeya zeae NCPPB 2538 WP 038915353 1 Dzeae
Dickeya zeae Ech586 WP 012882883 1 Dzeae
Photorhabdus asymbiotica YP 003038940 1 291112 Proteobacteria
Yersinia pestis biovar Microtus str 91001 NP 993904 1 229193 Proteobacteria
Yersinia pestis Antiqua YP 652062 1 360102 Proteobacteria
Yersinia pseudotuberculosis IP 31758 YP 001400325 1 349747 Proteobacteria
Photorhabdus asymbiotica YP 003039156 1 291112 Proteobacteria
Photorhabdus luminescens subsp laumondii TTO1 NP 927757 1 243265 Proteobacteria
Xenorhabdus bovienii SS 2004 YP 003466593 1 406818 Proteobacteria
Xenorhabdus nematophila ATCC 19061 YP 003713595 1 406817 Proteobacteria
Xenorhabdus bovienii SS 2004 YP 003468553 1 406818 Proteobacteria
Xenorhabdus bovienii SS 2004 YP 003466123 1 406818 Proteobacteria
Photorhabdus asymbiotica YP 003039625 1 291112 Proteobacteria
Xenorhabdus bovienii SS 2004 YP 003466025 1 406818 Proteobacteria
Yersinia pseudotuberculosis IP 31758 YP 001401945 1 349747 Proteobacteria
Yersinia pestis Antiqua YP 652446 1 360102 Proteobacteria
Yersinia pestis biovar Microtus str 91001 NP 992518 1 229193 Proteobacteria
Yersinia pestis biovar Medievalis str Harbin 35 YP 005625089 1 547048 Proteobacteria
Yersinia pseudotuberculosis IP 31758 YP 001400179 1 349747 Proteobacteria
Yersinia pestis biovar Microtus str 91001 NP 994045 1 229193 Proteobacteria
Yersinia pestis Antiqua YP 652218 1 360102 Proteobacteria
Yersinia pestis biovar Medievalis str Harbin 35 YP 005624859 1 547048 Proteobacteria
Pectobacterium carotovorum subsp carotovorum PC1 YP 003019058 1 561230 Proteobacteria
Pectobacterium atrosepticum SCRI1043 YP 051760 1 218491 Proteobacteria
Pectobacterium sp SCC3193 YP 006284719 1 1166016 Proteobacteria
Pectobacterium wasabiae WPP163 YP 003260986 1 561231 Proteobacteria
Pectobacterium carotovorum subsp carotovorum PC1 YP 003019578 1 561230 Proteobacteria
Dickeya chrysanthemi NCPPB 3533 WP 012768501 1 Dchrysanthemi
Dickeya dadantii NCPPB 898 WP 071604512 1 Ddadantii
Dickeya chrysanthemi Ech1591 WP 012770379 1 Dchrysanthemi
Dickeya gs fangzhongdai MK7 WP 071605189 1 Dfangzhongdai
Dickeya chrysanthemi NCPPB 3533 WP 012770379 1 Dchrysanthemi
Dickeya gs fangzhongdai NCPPB 3274 WP 012770379 1 Dfangzhongdai
Dickeya zeae NCPPB 3532 WP 023640927 1 Dzeae
Dickeya zeae ZJU1202 WP 016942935 1 Dzeae
Dickeya zeae EC1 WP 016942935 1 Dzeae
Dickeya zeae DZ2Q WP 016942935 1 Dzeae
Dickeya zeae MK19 WP 071603851 1 Dzeae
Dickeya zeae Ech586 WP 012886425 1 Dzeae
Dickeya dadantii NCPPB 898 WP 071604574 1 Ddadantii
Dickeya dadantii subsp dieffenbachiae NCPPB 2976 WP 071783903 1 Ddadantii
Dickeya gs fangzhongdai MK7 WP 071605242 1 Dfangzhongdai
Dickeya gs fangzhongdai NCPPB 3274 WP 071603997 1 Dfangzhongdai
Dickeya chrysanthemi subsp chrysanthemi NCPPB 402 WP 071604651 1 Dchrysanthemi
Dickeya chrysanthemi NCPPB 3533 WP 015848341 1 Dchrysanthemi
Dickeya zeae MS1 WP 019845381 1 Dzeae
Dickeya zeae MK19 WP 019845381 1 Dzeae
Dickeya zeae NCPPB 2538 WP 019845381 1 Dzeae
Dickeya gs fangzhongdai NCPPB 3274 WP 071603991 1 Dfangzhongdai
Dickeya gs fangzhongdai MK7 WP 071605238 1 Dfangzhongdai
Dickeya gs fangzhongdai ND14b WP 071605238 1 Dfangzhongdai
Dickeya chrysanthemi subsp chrysanthemi NCPPB 402 WP 071604657 1 Dchrysanthemi
Dickeya dadantii NCPPB 898 WP 071604569 1 Ddadantii
Dickeya dadantii 3937 WP 013319633 1 Ddadantii
127 17870 Type VI secretion system Seed
Dickeya solani IPO 2222 WP 022631892 1 Dsolani
Dickeya solani MK16 WP 022631892 1 Dsolani
Dickeya solani D s0432 1 WP 022631892 1 Dsolani
Dickeya solani GBBC 2040 WP 022631892 1 Dsolani
Dickeya solani MK10 WP 022631892 1 Dsolani
Dickeya solani RNS 08 23 3 1 A WP 022631892 1 Dsolani
Dickeya zeae NCPPB 3532 WP 023638762 1 Dzeae
Dickeya zeae NCPPB 2538 WP 071603547 1 Dzeae
Dickeya zeae ZJU1202 WP 016943032 1 Dzeae
Dickeya zeae EC1 WP 016943032 1 Dzeae
Dickeya dianthicola NCPPB 3534 WP 071598724 1 Ddianthicola
Dickeya dianthicola NCPPB 453 WP 071598724 1 Ddianthicola
Dickeya dianthicola GBBC 2039 WP 071598724 1 Ddianthicola
Dickeya dianthicola IPO 980 WP 071598724 1 Ddianthicola

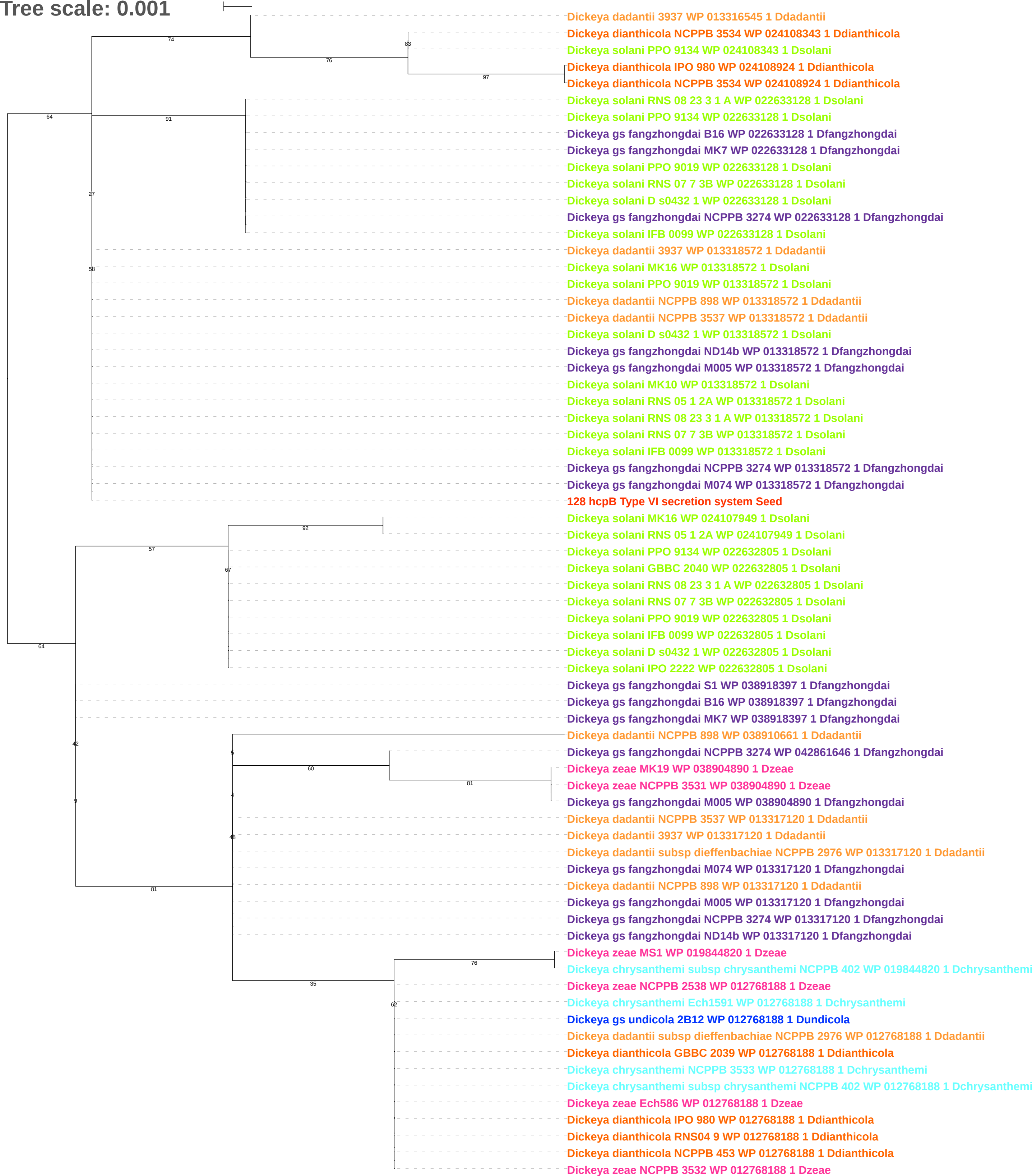
Tree scale: 0.001
Dickeya dadantii 3937 WP 013316545 1 Ddadantii
Dickeya dianthicola NCPPB 3534 WP 024108343 1 Ddianthicola
Dickeya solani PPO 9134 WP 024108343 1 Dsolani
Dickeya dianthicola IPO 980 WP 024108924 1 Ddianthicola
Dickeya dianthicola NCPPB 3534 WP 024108924 1 Ddianthicola
Dickeya solani RNS 08 23 3 1 A WP 022633128 1 Dsolani
Dickeya solani PPO 9134 WP 022633128 1 Dsolani
Dickeya gs fangzhongdai B16 WP 022633128 1 Dfangzhongdai
Dickeya gs fangzhongdai MK7 WP 022633128 1 Dfangzhongdai
Dickeya solani PPO 9019 WP 022633128 1 Dsolani
Dickeya solani RNS 07 7 3B WP 022633128 1 Dsolani
Dickeya solani D s0432 1 WP 022633128 1 Dsolani
Dickeya gs fangzhongdai NCPPB 3274 WP 022633128 1 Dfangzhongdai
Dickeya solani IFB 0099 WP 022633128 1 Dsolani
Dickeya dadantii 3937 WP 013318572 1 Ddadantii
Dickeya solani MK16 WP 013318572 1 Dsolani
Dickeya solani PPO 9019 WP 013318572 1 Dsolani
Dickeya dadantii NCPPB 898 WP 013318572 1 Ddadantii
Dickeya dadantii NCPPB 3537 WP 013318572 1 Ddadantii
Dickeya solani D s0432 1 WP 013318572 1 Dsolani
Dickeya gs fangzhongdai ND14b WP 013318572 1 Dfangzhongdai
Dickeya gs fangzhongdai M005 WP 013318572 1 Dfangzhongdai
Dickeya solani MK10 WP 013318572 1 Dsolani
Dickeya solani RNS 05 1 2A WP 013318572 1 Dsolani
Dickeya solani RNS 08 23 3 1 A WP 013318572 1 Dsolani
Dickeya solani RNS 07 7 3B WP 013318572 1 Dsolani
Dickeya solani IFB 0099 WP 013318572 1 Dsolani
Dickeya gs fangzhongdai NCPPB 3274 WP 013318572 1 Dfangzhongdai
Dickeya gs fangzhongdai M074 WP 013318572 1 Dfangzhongdai
128 hcpB Type VI secretion system Seed
Dickeya solani MK16 WP 024107949 1 Dsolani
Dickeya solani RNS 05 1 2A WP 024107949 1 Dsolani
Dickeya solani PPO 9134 WP 022632805 1 Dsolani
Dickeya solani GBBC 2040 WP 022632805 1 Dsolani
Dickeya solani RNS 08 23 3 1 A WP 022632805 1 Dsolani
Dickeya solani RNS 07 7 3B WP 022632805 1 Dsolani
Dickeya solani PPO 9019 WP 022632805 1 Dsolani
Dickeya solani IFB 0099 WP 022632805 1 Dsolani
Dickeya solani D s0432 1 WP 022632805 1 Dsolani
Dickeya solani IPO 2222 WP 022632805 1 Dsolani
Dickeya gs fangzhongdai S1 WP 038918397 1 Dfangzhongdai
Dickeya gs fangzhongdai B16 WP 038918397 1 Dfangzhongdai
Dickeya gs fangzhongdai MK7 WP 038918397 1 Dfangzhongdai
Dickeya dadantii NCPPB 898 WP 038910661 1 Ddadantii
Dickeya gs fangzhongdai NCPPB 3274 WP 042861646 1 Dfangzhongdai
Dickeya zeae MK19 WP 038904890 1 Dzeae
Dickeya zeae NCPPB 3531 WP 038904890 1 Dzeae
Dickeya gs fangzhongdai M005 WP 038904890 1 Dfangzhongdai
Dickeya dadantii NCPPB 3537 WP 013317120 1 Ddadantii
Dickeya dadantii 3937 WP 013317120 1 Ddadantii
Dickeya dadantii subsp dieffenbachiae NCPPB 2976 WP 013317120 1 Ddadantii
Dickeya gs fangzhongdai M074 WP 013317120 1 Dfangzhongdai
Dickeya dadantii NCPPB 898 WP 013317120 1 Ddadantii
Dickeya gs fangzhongdai M005 WP 013317120 1 Dfangzhongdai
Dickeya gs fangzhongdai NCPPB 3274 WP 013317120 1 Dfangzhongdai
Dickeya gs fangzhongdai ND14b WP 013317120 1 Dfangzhongdai
Dickeya zeae MS1 WP 019844820 1 Dzeae
Dickeya chrysanthemi subsp chrysanthemi NCPPB 402 WP 019844820 1 Dchrysanthemi
Dickeya zeae NCPPB 2538 WP 012768188 1 Dzeae
Dickeya chrysanthemi Ech1591 WP 012768188 1 Dchrysanthemi
Dickeya gs undicola 2B12 WP 012768188 1 Dundicola
Dickeya dadantii subsp dieffenbachiae NCPPB 2976 WP 012768188 1 Ddadantii
Dickeya dianthicola GBBC 2039 WP 012768188 1 Ddianthicola
Dickeya chrysanthemi NCPPB 3533 WP 012768188 1 Dchrysanthemi
Dickeya chrysanthemi subsp chrysanthemi NCPPB 402 WP 012768188 1 Dchrysanthemi
Dickeya zeae Ech586 WP 012768188 1 Dzeae
Dickeya dianthicola IPO 980 WP 012768188 1 Ddianthicola
Dickeya dianthicola RNS04 9 WP 012768188 1 Ddianthicola
Dickeya dianthicola NCPPB 453 WP 012768188 1 Ddianthicola
Dickeya zeae NCPPB 3532 WP 012768188 1 Dzeae

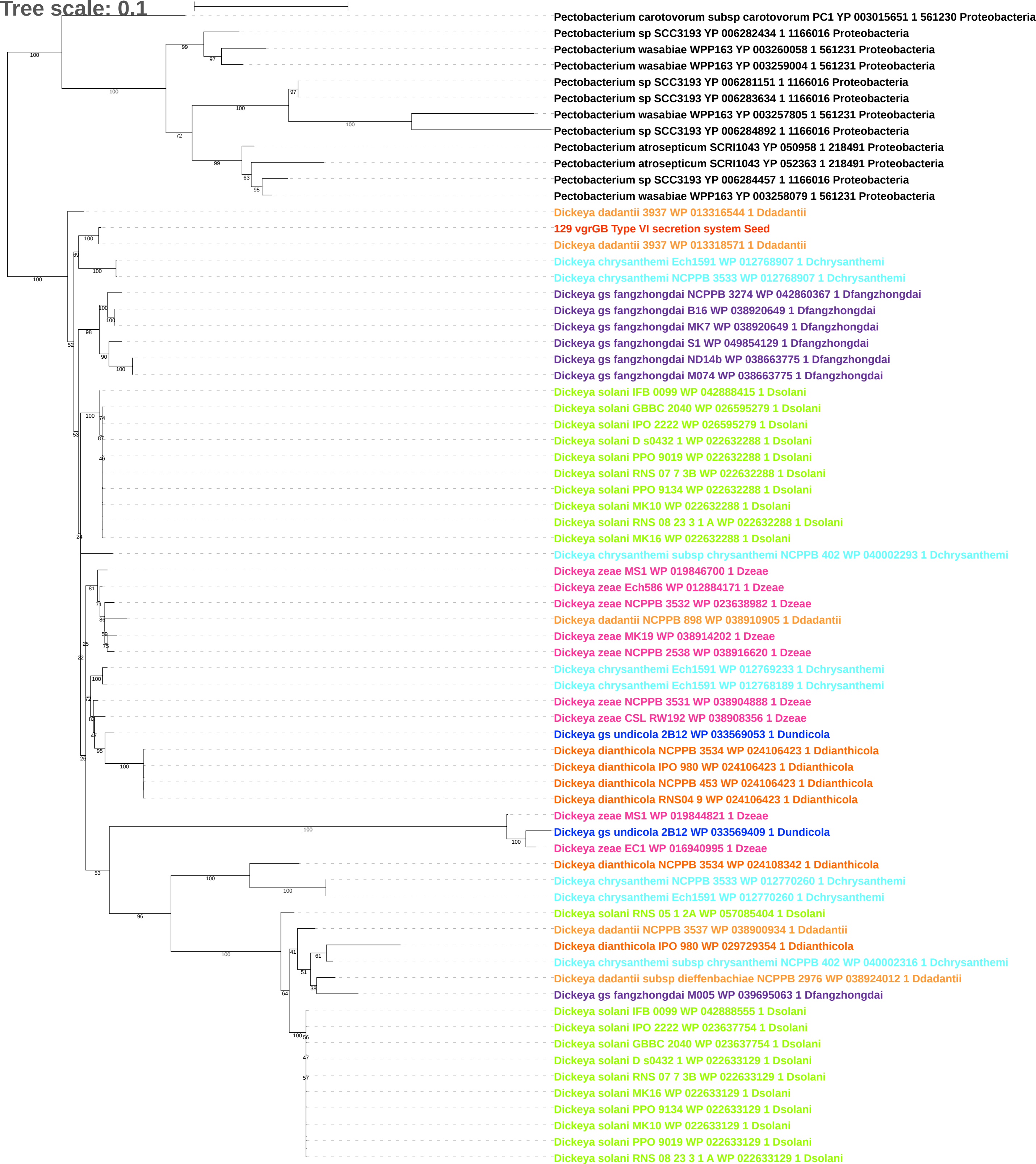
Tree scale: 0.1
Pectobacterium carotovorum subsp carotovorum PC1 YP 003015651 1 561230 Proteobacteria
Pectobacterium sp SCC3193 YP 006282434 1 1166016 Proteobacteria
Pectobacterium wasabiae WPP163 YP 003260058 1 561231 Proteobacteria
Pectobacterium wasabiae WPP163 YP 003259004 1 561231 Proteobacteria
Pectobacterium sp SCC3193 YP 006281151 1 1166016 Proteobacteria
Pectobacterium sp SCC3193 YP 006283634 1 1166016 Proteobacteria
Pectobacterium wasabiae WPP163 YP 003257805 1 561231 Proteobacteria
Pectobacterium sp SCC3193 YP 006284892 1 1166016 Proteobacteria
Pectobacterium atrosepticum SCRI1043 YP 050958 1 218491 Proteobacteria
Pectobacterium atrosepticum SCRI1043 YP 052363 1 218491 Proteobacteria
Pectobacterium sp SCC3193 YP 006284457 1 1166016 Proteobacteria
Pectobacterium wasabiae WPP163 YP 003258079 1 561231 Proteobacteria
Dickeya dadantii 3937 WP 013316544 1 Ddadantii
129 vgrGB Type VI secretion system Seed
Dickeya dadantii 3937 WP 013318571 1 Ddadantii
Dickeya chrysanthemi Ech1591 WP 012768907 1 Dchrysanthemi
Dickeya chrysanthemi NCPPB 3533 WP 012768907 1 Dchrysanthemi
Dickeya gs fangzhongdai NCPPB 3274 WP 042860367 1 Dfangzhongdai
Dickeya gs fangzhongdai B16 WP 038920649 1 Dfangzhongdai
Dickeya gs fangzhongdai MK7 WP 038920649 1 Dfangzhongdai
Dickeya gs fangzhongdai S1 WP 049854129 1 Dfangzhongdai
Dickeya gs fangzhongdai ND14b WP 038663775 1 Dfangzhongdai
Dickeya gs fangzhongdai M074 WP 038663775 1 Dfangzhongdai
Dickeya solani IFB 0099 WP 042888415 1 Dsolani
Dickeya solani GBBC 2040 WP 026595279 1 Dsolani
Dickeya solani IPO 2222 WP 026595279 1 Dsolani
Dickeya solani D s0432 1 WP 022632288 1 Dsolani
Dickeya solani PPO 9019 WP 022632288 1 Dsolani
Dickeya solani RNS 07 7 3B WP 022632288 1 Dsolani
Dickeya solani PPO 9134 WP 022632288 1 Dsolani
Dickeya solani MK10 WP 022632288 1 Dsolani
Dickeya solani RNS 08 23 3 1 A WP 022632288 1 Dsolani
Dickeya solani MK16 WP 022632288 1 Dsolani
Dickeya chrysanthemi subsp chrysanthemi NCPPB 402 WP 040002293 1 Dchrysanthemi
Dickeya zeae MS1 WP 019846700 1 Dzeae
Dickeya zeae Ech586 WP 012884171 1 Dzeae
Dickeya zeae NCPPB 3532 WP 023638982 1 Dzeae
Dickeya dadantii NCPPB 898 WP 038910905 1 Ddadantii
Dickeya zeae MK19 WP 038914202 1 Dzeae
Dickeya zeae NCPPB 2538 WP 038916620 1 Dzeae
Dickeya chrysanthemi Ech1591 WP 012769233 1 Dchrysanthemi
Dickeya chrysanthemi Ech1591 WP 012768189 1 Dchrysanthemi
Dickeya zeae NCPPB 3531 WP 038904888 1 Dzeae
Dickeya zeae CSL RW192 WP 038908356 1 Dzeae
Dickeya gs undicola 2B12 WP 033569053 1 Dundicola
Dickeya dianthicola NCPPB 3534 WP 024106423 1 Ddianthicola
Dickeya dianthicola IPO 980 WP 024106423 1 Ddianthicola
Dickeya dianthicola NCPPB 453 WP 024106423 1 Ddianthicola
Dickeya dianthicola RNS04 9 WP 024106423 1 Ddianthicola
Dickeya zeae MS1 WP 019844821 1 Dzeae
Dickeya gs undicola 2B12 WP 033569409 1 Dundicola
Dickeya zeae EC1 WP 016940995 1 Dzeae
Dickeya dianthicola NCPPB 3534 WP 024108342 1 Ddianthicola
Dickeya chrysanthemi NCPPB 3533 WP 012770260 1 Dchrysanthemi
Dickeya chrysanthemi Ech1591 WP 012770260 1 Dchrysanthemi
Dickeya solani RNS 05 1 2A WP 057085404 1 Dsolani
Dickeya dadantii NCPPB 3537 WP 038900934 1 Ddadantii
Dickeya dianthicola IPO 980 WP 029729354 1 Ddianthicola
Dickeya chrysanthemi subsp chrysanthemi NCPPB 402 WP 040002316 1 Dchrysanthemi
Dickeya dadantii subsp dieffenbachiae NCPPB 2976 WP 038924012 1 Ddadantii
Dickeya gs fangzhongdai M005 WP 039695063 1 Dfangzhongdai
Dickeya solani IFB 0099 WP 042888555 1 Dsolani
Dickeya solani IPO 2222 WP 023637754 1 Dsolani
Dickeya solani GBBC 2040 WP 023637754 1 Dsolani
Dickeya solani D s0432 1 WP 022633129 1 Dsolani
Dickeya solani RNS 07 7 3B WP 022633129 1 Dsolani
Dickeya solani MK16 WP 022633129 1 Dsolani
Dickeya solani PPO 9134 WP 022633129 1 Dsolani
Dickeya solani MK10 WP 022633129 1 Dsolani
Dickeya solani PPO 9019 WP 022633129 1 Dsolani
Dickeya solani RNS 08 23 3 1 A WP 022633129 1 Dsolani

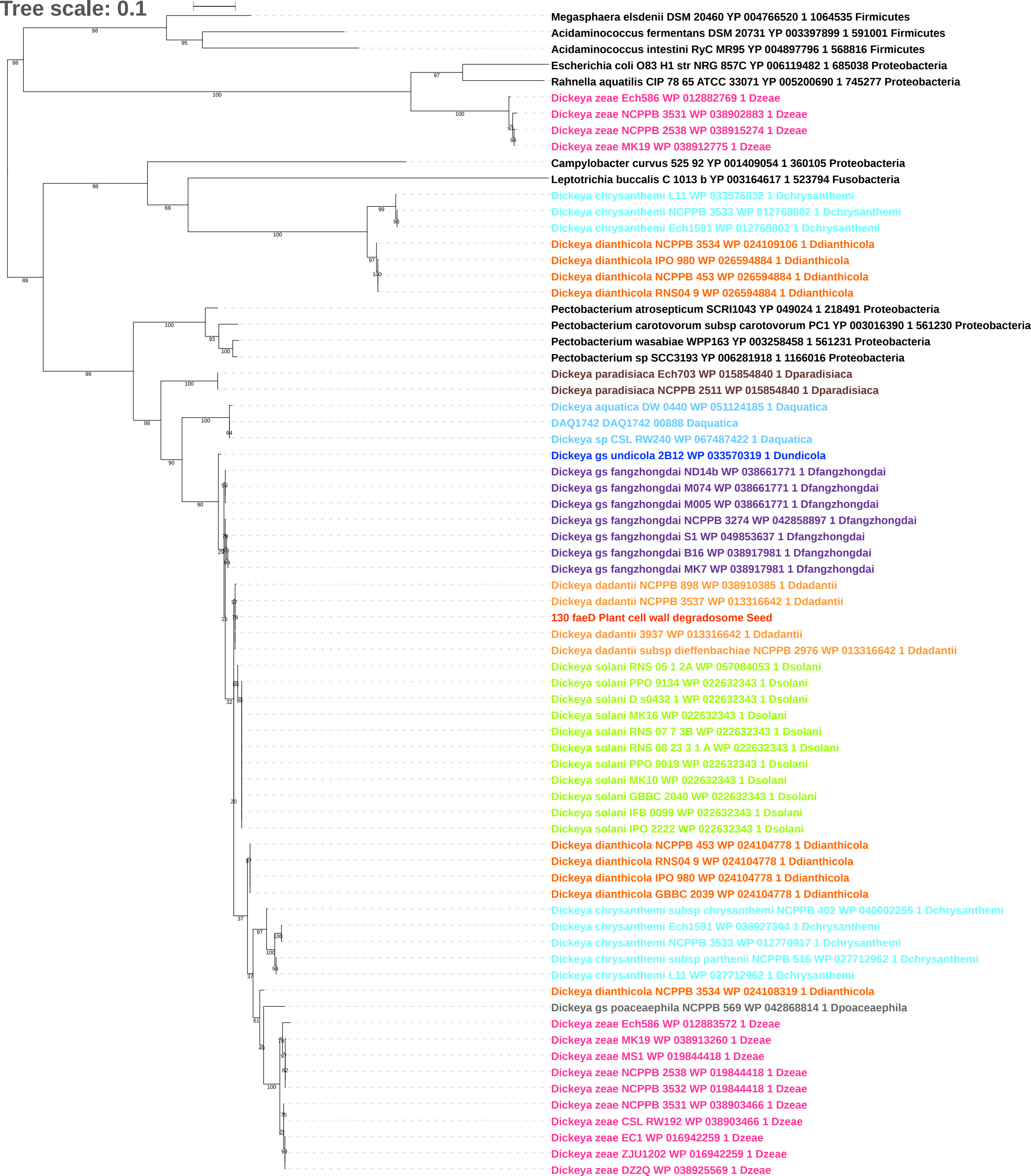
Tree scale: 0.1
Megasphaera elsdenii DSM 20460 YP 004766520 1 1064535 Firmicutes
Acidaminococcus fermentans DSM 20731 YP 003397899 1 591001 Firmicutes
Acidaminococcus intestini RyC MR95 YP 004897796 1 568816 Firmicutes
Escherichia coli O83 H1 str NRG 857C YP 006119482 1 685038 Proteobacteria
Rahnella aquatilis CIP 78 65 ATCC 33071 YP 005200690 1 745277 Proteobacteria
Dickeya zeae Ech586 WP 012882769 1 Dzeae
Dickeya zeae NCPPB 3531 WP 038902883 1 Dzeae
Dickeya zeae NCPPB 2538 WP 038915274 1 Dzeae
Dickeya zeae MK19 WP 038912775 1 Dzeae
Campylobacter curvus 525 92 YP 001409054 1 360105 Proteobacteria
Leptotrichia buccalis C 1013 b YP 003164617 1 523794 Fusobacteria
Dickeya chrysanthemi L11 WP 033576832 1 Dchrysanthemi
Dickeya chrysanthemi NCPPB 3533 WP 012768802 1 Dchrysanthemi
Dickeya chrysanthemi Ech1591 WP 012768802 1 Dchrysanthemi
Dickeya dianthicola NCPPB 3534 WP 024109106 1 Ddianthicola
Dickeya dianthicola IPO 980 WP 026594884 1 Ddianthicola
Dickeya dianthicola NCPPB 453 WP 026594884 1 Ddianthicola
Dickeya dianthicola RNS04 9 WP 026594884 1 Ddianthicola
Pectobacterium atrosepticum SCRI1043 YP 049024 1 218491 Proteobacteria
Pectobacterium carotovorum subsp carotovorum PC1 YP 003016390 1 561230 Proteobacteria
Pectobacterium wasabiae WPP163 YP 003258458 1 561231 Proteobacteria
Pectobacterium sp SCC3193 YP 006281918 1 1166016 Proteobacteria
Dickeya paradisiaca Ech703 WP 015854840 1 Dparadisiaca
Dickeya paradisiaca NCPPB 2511 WP 015854840 1 Dparadisiaca
Dickeya aquatica DW 0440 WP 051124185 1 Daquatica
DAQ1742 DAQ1742 00888 Daquatica
Dickeya sp CSL RW240 WP 067487422 1 Daquatica
Dickeya gs undicola 2B12 WP 033570319 1 Dundicola
Dickeya gs fangzhongdai ND14b WP 038661771 1 Dfangzhongdai
Dickeya gs fangzhongdai M074 WP 038661771 1 Dfangzhongdai
Dickeya gs fangzhongdai M005 WP 038661771 1 Dfangzhongdai
Dickeya gs fangzhongdai NCPPB 3274 WP 042858897 1 Dfangzhongdai
Dickeya gs fangzhongdai S1 WP 049853637 1 Dfangzhongdai
Dickeya gs fangzhongdai B16 WP 038917981 1 Dfangzhongdai
Dickeya gs fangzhongdai MK7 WP 038917981 1 Dfangzhongdai
Dickeya dadantii NCPPB 898 WP 038910385 1 Ddadantii
Dickeya dadantii NCPPB 3537 WP 013316642 1 Ddadantii
130 faeD Plant cell wall degradosome Seed
Dickeya dadantii 3937 WP 013316642 1 Ddadantii
Dickeya dadantii subsp dieffenbachiae NCPPB 2976 WP 013316642 1 Ddadantii
Dickeya solani RNS 05 1 2A WP 057084053 1 Dsolani
Dickeya solani PPO 9134 WP 022632343 1 Dsolani
Dickeya solani D s0432 1 WP 022632343 1 Dsolani
Dickeya solani MK16 WP 022632343 1 Dsolani
Dickeya solani RNS 07 7 3B WP 022632343 1 Dsolani
Dickeya solani RNS 08 23 3 1 A WP 022632343 1 Dsolani
Dickeya solani PPO 9019 WP 022632343 1 Dsolani
Dickeya solani MK10 WP 022632343 1 Dsolani
Dickeya solani GBBC 2040 WP 022632343 1 Dsolani
Dickeya solani IFB 0099 WP 022632343 1 Dsolani
Dickeya solani IPO 2222 WP 022632343 1 Dsolani
Dickeya dianthicola NCPPB 453 WP 024104778 1 Ddianthicola
Dickeya dianthicola RNS04 9 WP 024104778 1 Ddianthicola
Dickeya dianthicola IPO 980 WP 024104778 1 Ddianthicola
Dickeya dianthicola GBBC 2039 WP 024104778 1 Ddianthicola
Dickeya chrysanthemi subsp chrysanthemi NCPPB 402 WP 040002255 1 Dchrysanthemi
Dickeya chrysanthemi Ech1591 WP 038927304 1 Dchrysanthemi
Dickeya chrysanthemi NCPPB 3533 WP 012770917 1 Dchrysanthemi
Dickeya chrysanthemi subsp parthenii NCPPB 516 WP 027712962 1 Dchrysanthemi
Dickeya chrysanthemi L11 WP 027712962 1 Dchrysanthemi
Dickeya dianthicola NCPPB 3534 WP 024108319 1 Ddianthicola
Dickeya gs poaceaephila NCPPB 569 WP 042868814 1 Dpoaceaephila
Dickeya zeae Ech586 WP 012883572 1 Dzeae
Dickeya zeae MK19 WP 038913260 1 Dzeae
Dickeya zeae MS1 WP 019844418 1 Dzeae
Dickeya zeae NCPPB 2538 WP 019844418 1 Dzeae
Dickeya zeae NCPPB 3532 WP 019844418 1 Dzeae
Dickeya zeae NCPPB 3531 WP 038903466 1 Dzeae
Dickeya zeae CSL RW192 WP 038903466 1 Dzeae
Dickeya zeae EC1 WP 016942259 1 Dzeae
Dickeya zeae ZJU1202 WP 016942259 1 Dzeae
Dickeya zeae DZ2Q WP 038925569 1 Dzeae

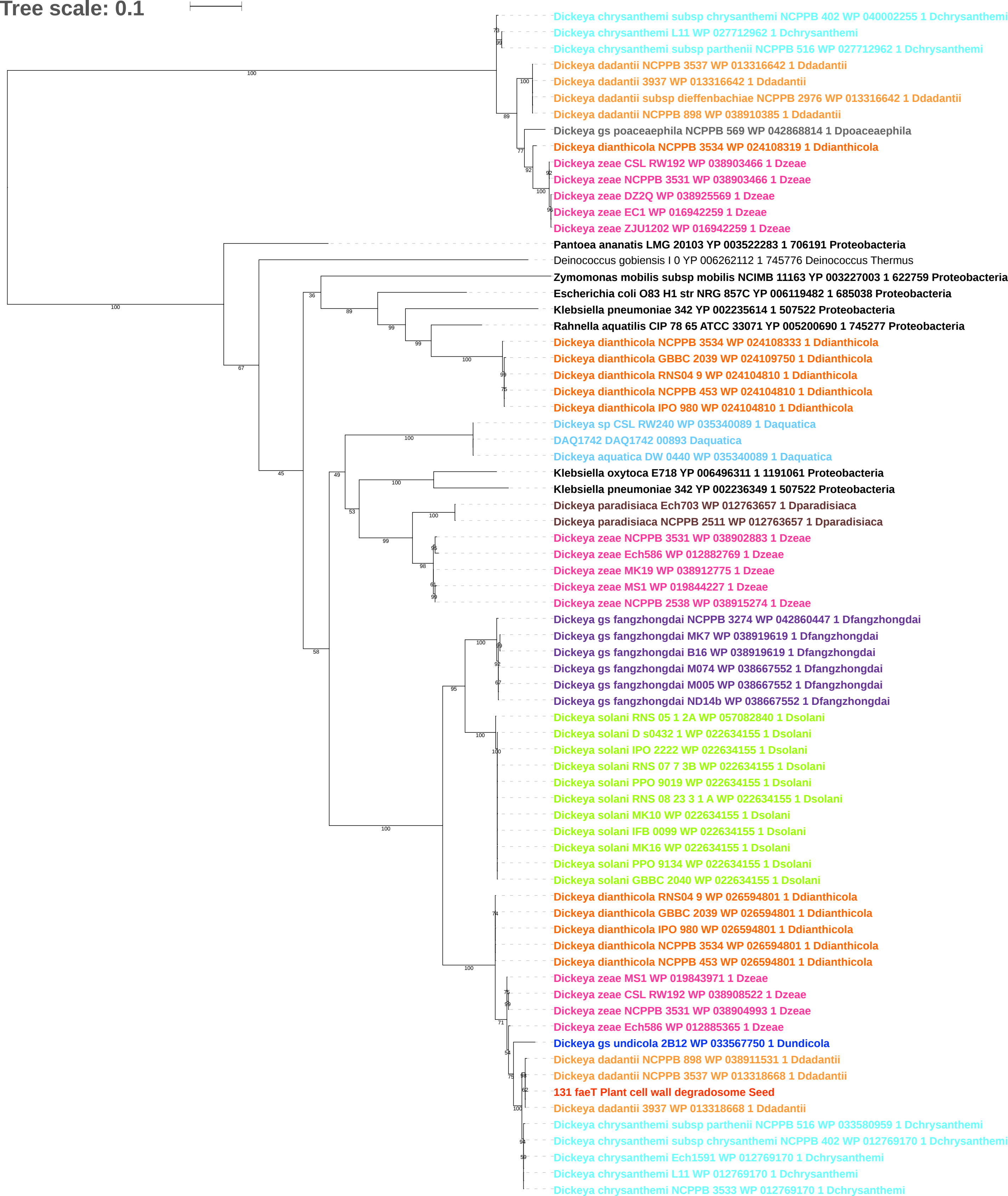
Tree scale: 0.1
Dickeya chrysanthemi subsp chrysanthemi NCPPB 402 WP 040002255 1 Dchrysanthemi
Dickeya chrysanthemi L11 WP 027712962 1 Dchrysanthemi
Dickeya chrysanthemi subsp parthenii NCPPB 516 WP 027712962 1 Dchrysanthemi
Dickeya dadantii NCPPB 3537 WP 013316642 1 Ddadantii
Dickeya dadantii 3937 WP 013316642 1 Ddadantii
Dickeya dadantii subsp dieffenbachiae NCPPB 2976 WP 013316642 1 Ddadantii
Dickeya dadantii NCPPB 898 WP 038910385 1 Ddadantii
Dickeya gs poaceaephila NCPPB 569 WP 042868814 1 Dpoaceaephila
Dickeya dianthicola NCPPB 3534 WP 024108319 1 Ddianthicola
Dickeya zeae CSL RW192 WP 038903466 1 Dzeae
Dickeya zeae NCPPB 3531 WP 038903466 1 Dzeae
Dickeya zeae DZ2Q WP 038925569 1 Dzeae
Dickeya zeae EC1 WP 016942259 1 Dzeae
Dickeya zeae ZJU1202 WP 016942259 1 Dzeae
Pantoea ananatis LMG 20103 YP 003522283 1 706191 Proteobacteria
Deinococcus gobiensis I 0 YP 006262112 1 745776 Deinococcus Thermus
Zymomonas mobilis subsp mobilis NCIMB 11163 YP 003227003 1 622759 Proteobacteria
Escherichia coli O83 H1 str NRG 857C YP 006119482 1 685038 Proteobacteria
Klebsiella pneumoniae 342 YP 002235614 1 507522 Proteobacteria
Rahnella aquatilis CIP 78 65 ATCC 33071 YP 005200690 1 745277 Proteobacteria
Dickeya dianthicola NCPPB 3534 WP 024108333 1 Ddianthicola
Dickeya dianthicola GBBC 2039 WP 024109750 1 Ddianthicola
Dickeya dianthicola RNS04 9 WP 024104810 1 Ddianthicola
Dickeya dianthicola NCPPB 453 WP 024104810 1 Ddianthicola
Dickeya dianthicola IPO 980 WP 024104810 1 Ddianthicola
Dickeya sp CSL RW240 WP 035340089 1 Daquatica
DAQ1742 DAQ1742 00893 Daquatica
Dickeya aquatica DW 0440 WP 035340089 1 Daquatica
Klebsiella oxytoca E718 YP 006496311 1 1191061 Proteobacteria
Klebsiella pneumoniae 342 YP 002236349 1 507522 Proteobacteria
Dickeya paradisiaca Ech703 WP 012763657 1 Dparadisiaca
Dickeya paradisiaca NCPPB 2511 WP 012763657 1 Dparadisiaca
Dickeya zeae NCPPB 3531 WP 038902883 1 Dzeae
Dickeya zeae Ech586 WP 012882769 1 Dzeae
Dickeya zeae MK19 WP 038912775 1 Dzeae
Dickeya zeae MS1 WP 019844227 1 Dzeae
Dickeya zeae NCPPB 2538 WP 038915274 1 Dzeae
Dickeya gs fangzhongdai NCPPB 3274 WP 042860447 1 Dfangzhongdai
Dickeya gs fangzhongdai MK7 WP 038919619 1 Dfangzhongdai
Dickeya gs fangzhongdai B16 WP 038919619 1 Dfangzhongdai
Dickeya gs fangzhongdai M074 WP 038667552 1 Dfangzhongdai
Dickeya gs fangzhongdai M005 WP 038667552 1 Dfangzhongdai
Dickeya gs fangzhongdai ND14b WP 038667552 1 Dfangzhongdai
Dickeya solani RNS 05 1 2A WP 057082840 1 Dsolani
Dickeya solani D s0432 1 WP 022634155 1 Dsolani
Dickeya solani IPO 2222 WP 022634155 1 Dsolani
Dickeya solani RNS 07 7 3B WP 022634155 1 Dsolani
Dickeya solani PPO 9019 WP 022634155 1 Dsolani
Dickeya solani RNS 08 23 3 1 A WP 022634155 1 Dsolani
Dickeya solani MK10 WP 022634155 1 Dsolani
Dickeya solani IFB 0099 WP 022634155 1 Dsolani
Dickeya solani MK16 WP 022634155 1 Dsolani
Dickeya solani PPO 9134 WP 022634155 1 Dsolani
Dickeya solani GBBC 2040 WP 022634155 1 Dsolani
Dickeya dianthicola RNS04 9 WP 026594801 1 Ddianthicola
Dickeya dianthicola GBBC 2039 WP 026594801 1 Ddianthicola
Dickeya dianthicola IPO 980 WP 026594801 1 Ddianthicola
Dickeya dianthicola NCPPB 3534 WP 026594801 1 Ddianthicola
Dickeya dianthicola NCPPB 453 WP 026594801 1 Ddianthicola
Dickeya zeae MS1 WP 019843971 1 Dzeae
Dickeya zeae CSL RW192 WP 038908522 1 Dzeae
Dickeya zeae NCPPB 3531 WP 038904993 1 Dzeae
Dickeya zeae Ech586 WP 012885365 1 Dzeae
Dickeya gs undicola 2B12 WP 033567750 1 Dundicola
Dickeya dadantii NCPPB 898 WP 038911531 1 Ddadantii
Dickeya dadantii NCPPB 3537 WP 013318668 1 Ddadantii
131 faeT Plant cell wall degradosome Seed
Dickeya dadantii 3937 WP 013318668 1 Ddadantii
Dickeya chrysanthemi subsp parthenii NCPPB 516 WP 033580959 1 Dchrysanthemi
Dickeya chrysanthemi subsp chrysanthemi NCPPB 402 WP 012769170 1 Dchrysanthemi
Dickeya chrysanthemi Ech1591 WP 012769170 1 Dchrysanthemi
Dickeya chrysanthemi L11 WP 012769170 1 Dchrysanthemi
Dickeya chrysanthemi NCPPB 3533 WP 012769170 1 Dchrysanthemi

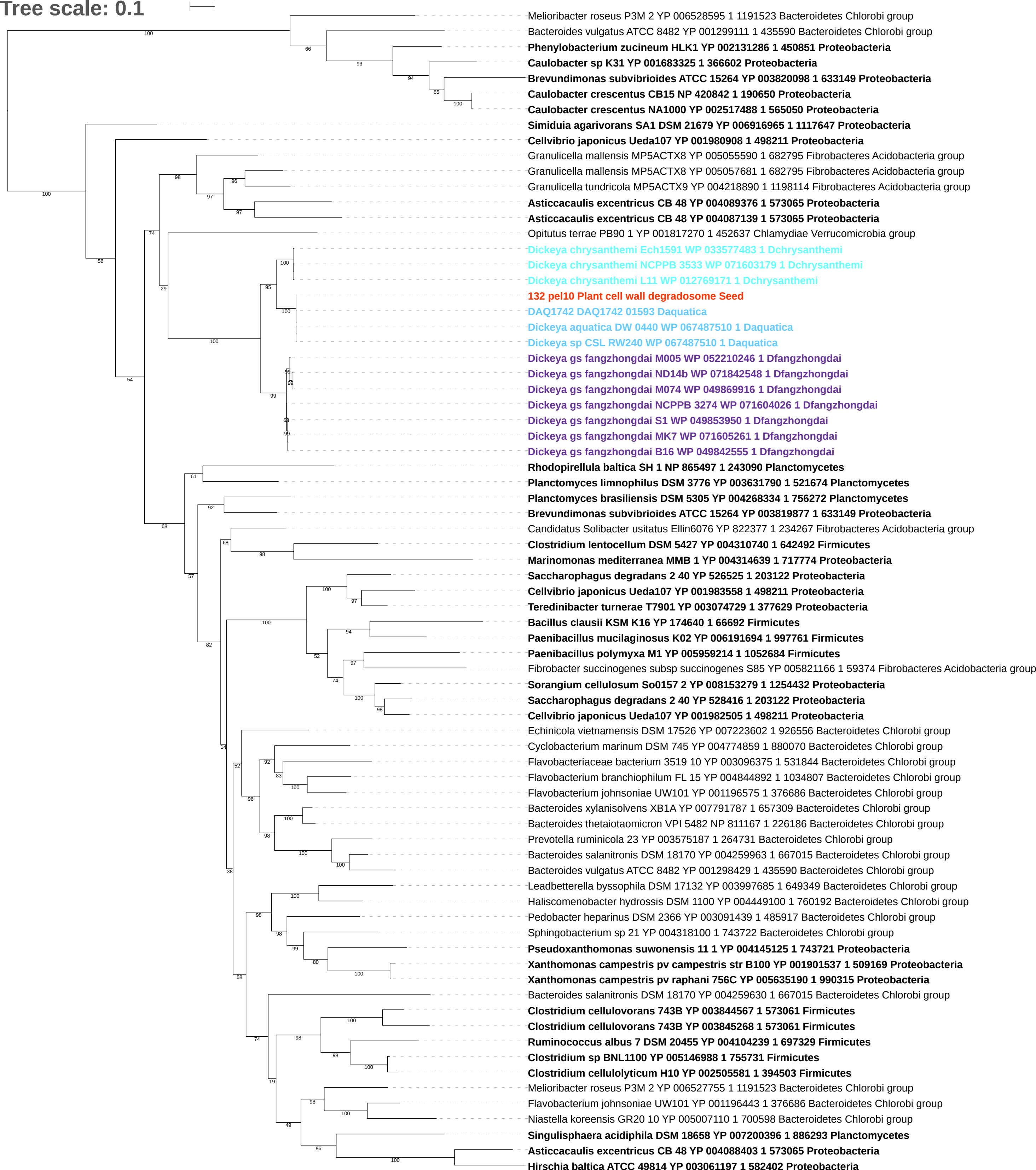

Tree scale: 0.1
Melioribacter roseus P3M 2 YP 006528595 1 1191523 Bacteroidetes Chlorobi group
Bacteroides vulgatus ATCC 8482 YP 001299111 1 435590 Bacteroidetes Chlorobi group
Phenylobacterium zucineum HLK1 YP 002131286 1 450851 Proteobacteria
Caulobacter sp K31 YP 001683325 1 366602 Proteobacteria
Brevundimonas subvibrioides ATCC 15264 YP 003820098 1 633149 Proteobacteria
Caulobacter crescentus CB15 NP 420842 1 190650 Proteobacteria
Caulobacter crescentus NA1000 YP 002517488 1 565050 Proteobacteria
Simiduia agarivorans SA1 DSM 21679 YP 006916965 1 1117647 Proteobacteria
Cellvibrio japonicus Ueda107 YP 001980908 1 498211 Proteobacteria
Granulicella mallensis MP5ACTX8 YP 005055590 1 682795 Fibrobacteres Acidobacteria group
Granulicella mallensis MP5ACTX8 YP 005057681 1 682795 Fibrobacteres Acidobacteria group
Granulicella tundricola MP5ACTX9 YP 004218890 1 1198114 Fibrobacteres Acidobacteria group
Asticcacaulis excentricus CB 48 YP 004089376 1 573065 Proteobacteria
Asticcacaulis excentricus CB 48 YP 004087139 1 573065 Proteobacteria
Opitutus terrae PB90 1 YP 001817270 1 452637 Chlamydiae Verrucomicrobia group
Dickeya chrysanthemi Ech1591 WP 033577483 1 Dchrysanthemi
Dickeya chrysanthemi NCPPB 3533 WP 071603179 1 Dchrysanthemi
Dickeya chrysanthemi L11 WP 012769171 1 Dchrysanthemi
132 pel10 Plant cell wall degradosome Seed
DAQ1742 DAQ1742 01593 Daquatica
Dickeya aquatica DW 0440 WP 067487510 1 Daquatica
Dickeya sp CSL RW240 WP 067487510 1 Daquatica
Dickeya gs fangzhongdai M005 WP 052210246 1 Dfangzhongdai
Dickeya gs fangzhongdai ND14b WP 071842548 1 Dfangzhongdai
Dickeya gs fangzhongdai M074 WP 049869916 1 Dfangzhongdai
Dickeya gs fangzhongdai NCPPB 3274 WP 071604026 1 Dfangzhongdai
Dickeya gs fangzhongdai S1 WP 049853950 1 Dfangzhongdai
Dickeya gs fangzhongdai MK7 WP 071605261 1 Dfangzhongdai
Dickeya gs fangzhongdai B16 WP 049842555 1 Dfangzhongdai
Rhodopirellula baltica SH 1 NP 865497 1 243090 Planctomycetes
Planctomyces limnophilus DSM 3776 YP 003631790 1 521674 Planctomycetes
Planctomyces brasiliensis DSM 5305 YP 004268334 1 756272 Planctomycetes
Brevundimonas subvibrioides ATCC 15264 YP 003819877 1 633149 Proteobacteria
Candidatus Solibacter usitatus Ellin6076 YP 822377 1 234267 Fibrobacteres Acidobacteria group
Clostridium lentocellum DSM 5427 YP 004310740 1 642492 Firmicutes
Marinomonas mediterranea MMB 1 YP 004314639 1 717774 Proteobacteria
Saccharophagus degradans 2 40 YP 526525 1 203122 Proteobacteria
Cellvibrio japonicus Ueda107 YP 001983558 1 498211 Proteobacteria
Teredinibacter turnerae T7901 YP 003074729 1 377629 Proteobacteria
Bacillus clausii KSM K16 YP 174640 1 66692 Firmicutes
Paenibacillus mucilaginosus K02 YP 006191694 1 997761 Firmicutes
Paenibacillus polymyxa M1 YP 005959214 1 1052684 Firmicutes
Fibrobacter succinogenes subsp succinogenes S85 YP 005821166 1 59374 Fibrobacteres Acidobacteria group
Sorangium cellulosum So0157 2 YP 008153279 1 1254432 Proteobacteria
Saccharophagus degradans 2 40 YP 528416 1 203122 Proteobacteria
Cellvibrio japonicus Ueda107 YP 001982505 1 498211 Proteobacteria
Echinicola vietnamensis DSM 17526 YP 007223602 1 926556 Bacteroidetes Chlorobi group
Cyclobacterium marinum DSM 745 YP 004774859 1 880070 Bacteroidetes Chlorobi group
Flavobacteriaceae bacterium 3519 10 YP 003096375 1 531844 Bacteroidetes Chlorobi group
Flavobacterium branchiophilum FL 15 YP 004844892 1 1034807 Bacteroidetes Chlorobi group
Flavobacterium johnsoniae UW101 YP 001196575 1 376686 Bacteroidetes Chlorobi group
Bacteroides xylanisolvens XB1A YP 007791787 1 657309 Bacteroidetes Chlorobi group
Bacteroides thetaiotaomicron VPI 5482 NP 811167 1 226186 Bacteroidetes Chlorobi group
Prevotella ruminicola 23 YP 003575187 1 264731 Bacteroidetes Chlorobi group
Bacteroides salanitronis DSM 18170 YP 004259963 1 667015 Bacteroidetes Chlorobi group
Bacteroides vulgatus ATCC 8482 YP 001298429 1 435590 Bacteroidetes Chlorobi group
Leadbetterella byssophila DSM 17132 YP 003997685 1 649349 Bacteroidetes Chlorobi group
Haliscomenobacter hydrossis DSM 1100 YP 004449100 1 760192 Bacteroidetes Chlorobi group
Pedobacter heparinus DSM 2366 YP 003091439 1 485917 Bacteroidetes Chlorobi group
Sphingobacterium sp 21 YP 004318100 1 743722 Bacteroidetes Chlorobi group
Pseudoxanthomonas suwonensis 11 1 YP 004145125 1 743721 Proteobacteria
Xanthomonas campestris pv campestris str B100 YP 001901537 1 509169 Proteobacteria
Xanthomonas campestris pv raphani 756C YP 005635190 1 990315 Proteobacteria
Bacteroides salanitronis DSM 18170 YP 004259630 1 667015 Bacteroidetes Chlorobi group
Clostridium cellulovorans 743B YP 003844567 1 573061 Firmicutes
Clostridium cellulovorans 743B YP 003845268 1 573061 Firmicutes
Ruminococcus albus 7 DSM 20455 YP 004104239 1 697329 Firmicutes
Clostridium sp BNL1100 YP 005146988 1 755731 Firmicutes
Clostridium cellulolyticum H10 YP 002505581 1 394503 Firmicutes
Melioribacter roseus P3M 2 YP 006527755 1 1191523 Bacteroidetes Chlorobi group
Flavobacterium johnsoniae UW101 YP 001196443 1 376686 Bacteroidetes Chlorobi group
Niastella koreensis GR20 10 YP 005007110 1 700598 Bacteroidetes Chlorobi group
Singulisphaera acidiphila DSM 18658 YP 007200396 1 886293 Planctomycetes
Asticcacaulis excentricus CB 48 YP 004088403 1 573065 Proteobacteria
Hirschia baltica ATCC 49814 YP 003061197 1 582402 Proteobacteria

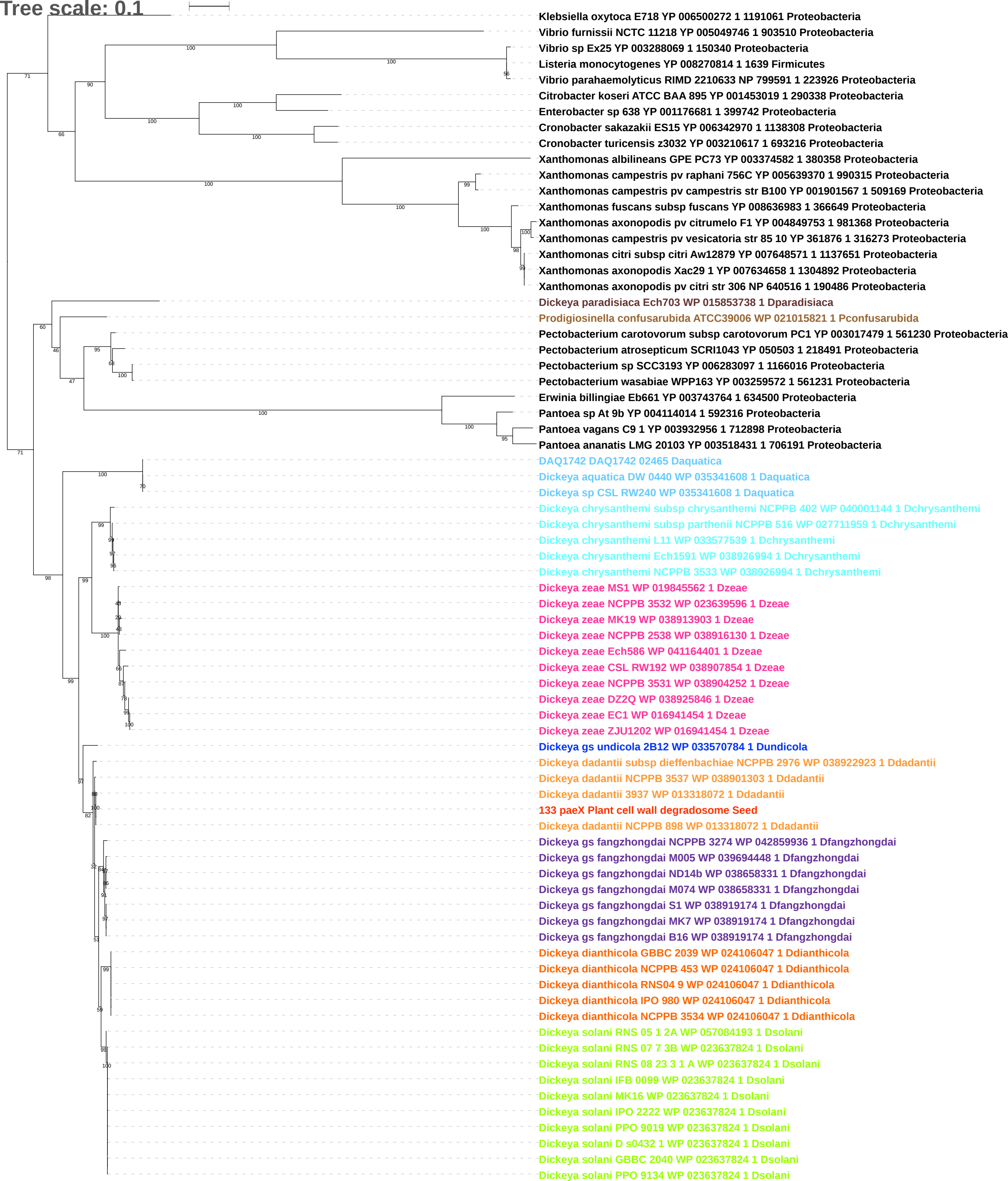
Tree scale: 0.1
Klebsiella oxytoca E718 YP 006500272 1 1191061 Proteobacteria
Vibrio furnissii NCTC 11218 YP 005049746 1 903510 Proteobacteria
Vibrio sp Ex25 YP 003288069 1 150340 Proteobacteria
Listeria monocytogenes YP 008270814 1 1639 Firmicutes
Vibrio parahaemolyticus RIMD 2210633 NP 799591 1 223926 Proteobacteria
Citrobacter koseri ATCC BAA 895 YP 001453019 1 290338 Proteobacteria
Enterobacter sp 638 YP 001176681 1 399742 Proteobacteria
Cronobacter sakazakii ES15 YP 006342970 1 1138308 Proteobacteria
Cronobacter turicensis z3032 YP 003210617 1 693216 Proteobacteria
Xanthomonas albilineans GPE PC73 YP 003374582 1 380358 Proteobacteria
Xanthomonas campestris pv raphani 756C YP 005639370 1 990315 Proteobacteria
Xanthomonas campestris pv campestris str B100 YP 001901567 1 509169 Proteobacteria
Xanthomonas fuscans subsp fuscans YP 008636983 1 366649 Proteobacteria
Xanthomonas axonopodis pv citrumelo F1 YP 004849753 1 981368 Proteobacteria
Xanthomonas campestris pv vesicatoria str 85 10 YP 361876 1 316273 Proteobacteria
Xanthomonas citri subsp citri Aw12879 YP 007648571 1 1137651 Proteobacteria
Xanthomonas axonopodis Xac29 1 YP 007634658 1 1304892 Proteobacteria
Xanthomonas axonopodis pv citri str 306 NP 640516 1 190486 Proteobacteria
Dickeya paradisiaca Ech703 WP 015853738 1 Dparadisiaca
Prodigiosinella confusarubida ATCC39006 WP 021015821 1 Pconfusarubida
Pectobacterium carotovorum subsp carotovorum PC1 YP 003017479 1 561230 Proteobacteria
Pectobacterium atrosepticum SCRI1043 YP 050503 1 218491 Proteobacteria
Pectobacterium sp SCC3193 YP 006283097 1 1166016 Proteobacteria
Pectobacterium wasabiae WPP163 YP 003259572 1 561231 Proteobacteria
Erwinia billingiae Eb661 YP 003743764 1 634500 Proteobacteria
Pantoea sp At 9b YP 004114014 1 592316 Proteobacteria
Pantoea vagans C9 1 YP 003932956 1 712898 Proteobacteria
Pantoea ananatis LMG 20103 YP 003518431 1 706191 Proteobacteria
DAQ1742 DAQ1742 02465 Daquatica
Dickeya aquatica DW 0440 WP 035341608 1 Daquatica
Dickeya sp CSL RW240 WP 035341608 1 Daquatica
Dickeya chrysanthemi subsp chrysanthemi NCPPB 402 WP 040001144 1 Dchrysanthemi
Dickeya chrysanthemi subsp parthenii NCPPB 516 WP 027711959 1 Dchrysanthemi
Dickeya chrysanthemi L11 WP 033577539 1 Dchrysanthemi
Dickeya chrysanthemi Ech1591 WP 038926994 1 Dchrysanthemi
Dickeya chrysanthemi NCPPB 3533 WP 038926994 1 Dchrysanthemi
Dickeya zeae MS1 WP 019845562 1 Dzeae
Dickeya zeae NCPPB 3532 WP 023639596 1 Dzeae
Dickeya zeae MK19 WP 038913903 1 Dzeae
Dickeya zeae NCPPB 2538 WP 038916130 1 Dzeae
Dickeya zeae Ech586 WP 041164401 1 Dzeae
Dickeya zeae CSL RW192 WP 038907854 1 Dzeae
Dickeya zeae NCPPB 3531 WP 038904252 1 Dzeae
Dickeya zeae DZ2Q WP 038925846 1 Dzeae
Dickeya zeae EC1 WP 016941454 1 Dzeae
Dickeya zeae ZJU1202 WP 016941454 1 Dzeae
Dickeya gs undicola 2B12 WP 033570784 1 Dundicola
Dickeya dadantii subsp dieffenbachiae NCPPB 2976 WP 038922923 1 Ddadantii
Dickeya dadantii NCPPB 3537 WP 038901303 1 Ddadantii
Dickeya dadantii 3937 WP 013318072 1 Ddadantii
133 paeX Plant cell wall degradosome Seed
Dickeya dadantii NCPPB 898 WP 013318072 1 Ddadantii
Dickeya gs fangzhongdai NCPPB 3274 WP 042859936 1 Dfangzhongdai
Dickeya gs fangzhongdai M005 WP 039694448 1 Dfangzhongdai
Dickeya gs fangzhongdai ND14b WP 038658331 1 Dfangzhongdai
Dickeya gs fangzhongdai M074 WP 038658331 1 Dfangzhongdai
Dickeya gs fangzhongdai S1 WP 038919174 1 Dfangzhongdai
Dickeya gs fangzhongdai MK7 WP 038919174 1 Dfangzhongdai
Dickeya gs fangzhongdai B16 WP 038919174 1 Dfangzhongdai
Dickeya dianthicola GBBC 2039 WP 024106047 1 Ddianthicola
Dickeya dianthicola NCPPB 453 WP 024106047 1 Ddianthicola
Dickeya dianthicola RNS04 9 WP 024106047 1 Ddianthicola
Dickeya dianthicola IPO 980 WP 024106047 1 Ddianthicola
Dickeya dianthicola NCPPB 3534 WP 024106047 1 Ddianthicola
Dickeya solani RNS 05 1 2A WP 057084193 1 Dsolani
Dickeya solani RNS 07 7 3B WP 023637824 1 Dsolani
Dickeya solani RNS 08 23 3 1 A WP 023637824 1 Dsolani
Dickeya solani IFB 0099 WP 023637824 1 Dsolani
Dickeya solani MK16 WP 023637824 1 Dsolani
Dickeya solani IPO 2222 WP 023637824 1 Dsolani
Dickeya solani PPO 9019 WP 023637824 1 Dsolani
Dickeya solani D s0432 1 WP 023637824 1 Dsolani
Dickeya solani GBBC 2040 WP 023637824 1 Dsolani
Dickeya solani PPO 9134 WP 023637824 1 Dsolani

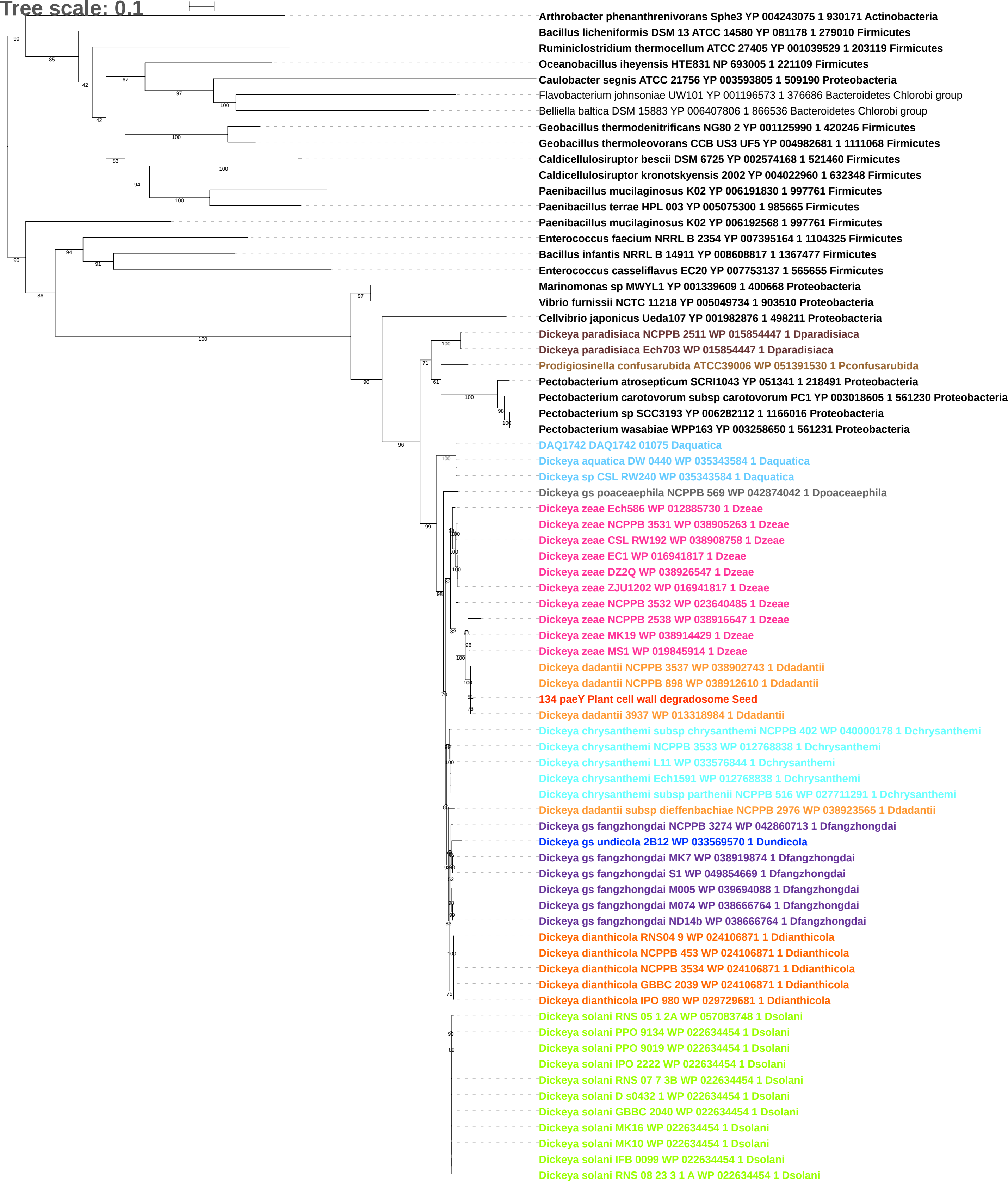

Tree scale: 0.1
Arthrobacter phenanthrenivorans Sphe3 YP 004243075 1 930171 Actinobacteria
Bacillus licheniformis DSM 13 ATCC 14580 YP 081178 1 279010 Firmicutes
Ruminiclostridium thermocellum ATCC 27405 YP 001039529 1 203119 Firmicutes
Oceanobacillus iheyensis HTE831 NP 693005 1 221109 Firmicutes
Caulobacter segnis ATCC 21756 YP 003593805 1 509190 Proteobacteria
Flavobacterium johnsoniae UW101 YP 001196573 1 376686 Bacteroidetes Chlorobi group
Belliella baltica DSM 15883 YP 006407806 1 866536 Bacteroidetes Chlorobi group
Geobacillus thermodenitrificans NG80 2 YP 001125990 1 420246 Firmicutes
Geobacillus thermoleovorans CCB US3 UF5 YP 004982681 1 1111068 Firmicutes
Caldicellulosiruptor bescii DSM 6725 YP 002574168 1 521460 Firmicutes
Caldicellulosiruptor kronotskyensis 2002 YP 004022960 1 632348 Firmicutes
Paenibacillus mucilaginosus K02 YP 006191830 1 997761 Firmicutes
Paenibacillus terrae HPL 003 YP 005075300 1 985665 Firmicutes
Paenibacillus mucilaginosus K02 YP 006192568 1 997761 Firmicutes
Enterococcus faecium NRRL B 2354 YP 007395164 1 1104325 Firmicutes
Bacillus infantis NRRL B 14911 YP 008608817 1 1367477 Firmicutes
Enterococcus casseliflavus EC20 YP 007753137 1 565655 Firmicutes
Marinomonas sp MWYL1 YP 001339609 1 400668 Proteobacteria
Vibrio furnissii NCTC 11218 YP 005049734 1 903510 Proteobacteria
Cellvibrio japonicus Ueda107 YP 001982876 1 498211 Proteobacteria
Dickeya paradisiaca NCPPB 2511 WP 015854447 1 Dparadisiaca
Dickeya paradisiaca Ech703 WP 015854447 1 Dparadisiaca
Prodigiosinella confusarubida ATCC39006 WP 051391530 1 Pconfusarubida
Pectobacterium atrosepticum SCRI1043 YP 051341 1 218491 Proteobacteria
Pectobacterium carotovorum subsp carotovorum PC1 YP 003018605 1 561230 Proteobacteria
Pectobacterium sp SCC3193 YP 006282112 1 1166016 Proteobacteria
Pectobacterium wasabiae WPP163 YP 003258650 1 561231 Proteobacteria
DAQ1742 DAQ1742 01075 Daquatica
Dickeya aquatica DW 0440 WP 035343584 1 Daquatica
Dickeya sp CSL RW240 WP 035343584 1 Daquatica
Dickeya gs poaceaephila NCPPB 569 WP 042874042 1 Dpoaceaephila
Dickeya zeae Ech586 WP 012885730 1 Dzeae
Dickeya zeae NCPPB 3531 WP 038905263 1 Dzeae
Dickeya zeae CSL RW192 WP 038908758 1 Dzeae
Dickeya zeae EC1 WP 016941817 1 Dzeae
Dickeya zeae DZ2Q WP 038926547 1 Dzeae
Dickeya zeae ZJU1202 WP 016941817 1 Dzeae
Dickeya zeae NCPPB 3532 WP 023640485 1 Dzeae
Dickeya zeae NCPPB 2538 WP 038916647 1 Dzeae
Dickeya zeae MK19 WP 038914429 1 Dzeae
Dickeya zeae MS1 WP 019845914 1 Dzeae
Dickeya dadantii NCPPB 3537 WP 038902743 1 Ddadantii
Dickeya dadantii NCPPB 898 WP 038912610 1 Ddadantii
134 paeY Plant cell wall degradosome Seed
Dickeya dadantii 3937 WP 013318984 1 Ddadantii
Dickeya chrysanthemi subsp chrysanthemi NCPPB 402 WP 040000178 1 Dchrysanthemi
Dickeya chrysanthemi NCPPB 3533 WP 012768838 1 Dchrysanthemi
Dickeya chrysanthemi L11 WP 033576844 1 Dchrysanthemi
Dickeya chrysanthemi Ech1591 WP 012768838 1 Dchrysanthemi
Dickeya chrysanthemi subsp parthenii NCPPB 516 WP 027711291 1 Dchrysanthemi
Dickeya dadantii subsp dieffenbachiae NCPPB 2976 WP 038923565 1 Ddadantii
Dickeya gs fangzhongdai NCPPB 3274 WP 042860713 1 Dfangzhongdai
Dickeya gs undicola 2B12 WP 033569570 1 Dundicola
Dickeya gs fangzhongdai MK7 WP 038919874 1 Dfangzhongdai
Dickeya gs fangzhongdai S1 WP 049854669 1 Dfangzhongdai
Dickeya gs fangzhongdai M005 WP 039694088 1 Dfangzhongdai
Dickeya gs fangzhongdai M074 WP 038666764 1 Dfangzhongdai
Dickeya gs fangzhongdai ND14b WP 038666764 1 Dfangzhongdai
Dickeya dianthicola RNS04 9 WP 024106871 1 Ddianthicola
Dickeya dianthicola NCPPB 453 WP 024106871 1 Ddianthicola
Dickeya dianthicola NCPPB 3534 WP 024106871 1 Ddianthicola
Dickeya dianthicola GBBC 2039 WP 024106871 1 Ddianthicola
Dickeya dianthicola IPO 980 WP 029729681 1 Ddianthicola
Dickeya solani RNS 05 1 2A WP 057083748 1 Dsolani
Dickeya solani PPO 9134 WP 022634454 1 Dsolani
Dickeya solani PPO 9019 WP 022634454 1 Dsolani
Dickeya solani IPO 2222 WP 022634454 1 Dsolani
Dickeya solani RNS 07 7 3B WP 022634454 1 Dsolani
Dickeya solani D s0432 1 WP 022634454 1 Dsolani
Dickeya solani GBBC 2040 WP 022634454 1 Dsolani
Dickeya solani MK16 WP 022634454 1 Dsolani
Dickeya solani MK10 WP 022634454 1 Dsolani
Dickeya solani IFB 0099 WP 022634454 1 Dsolani
Dickeya solani RNS 08 23 3 1 A WP 022634454 1 Dsolani

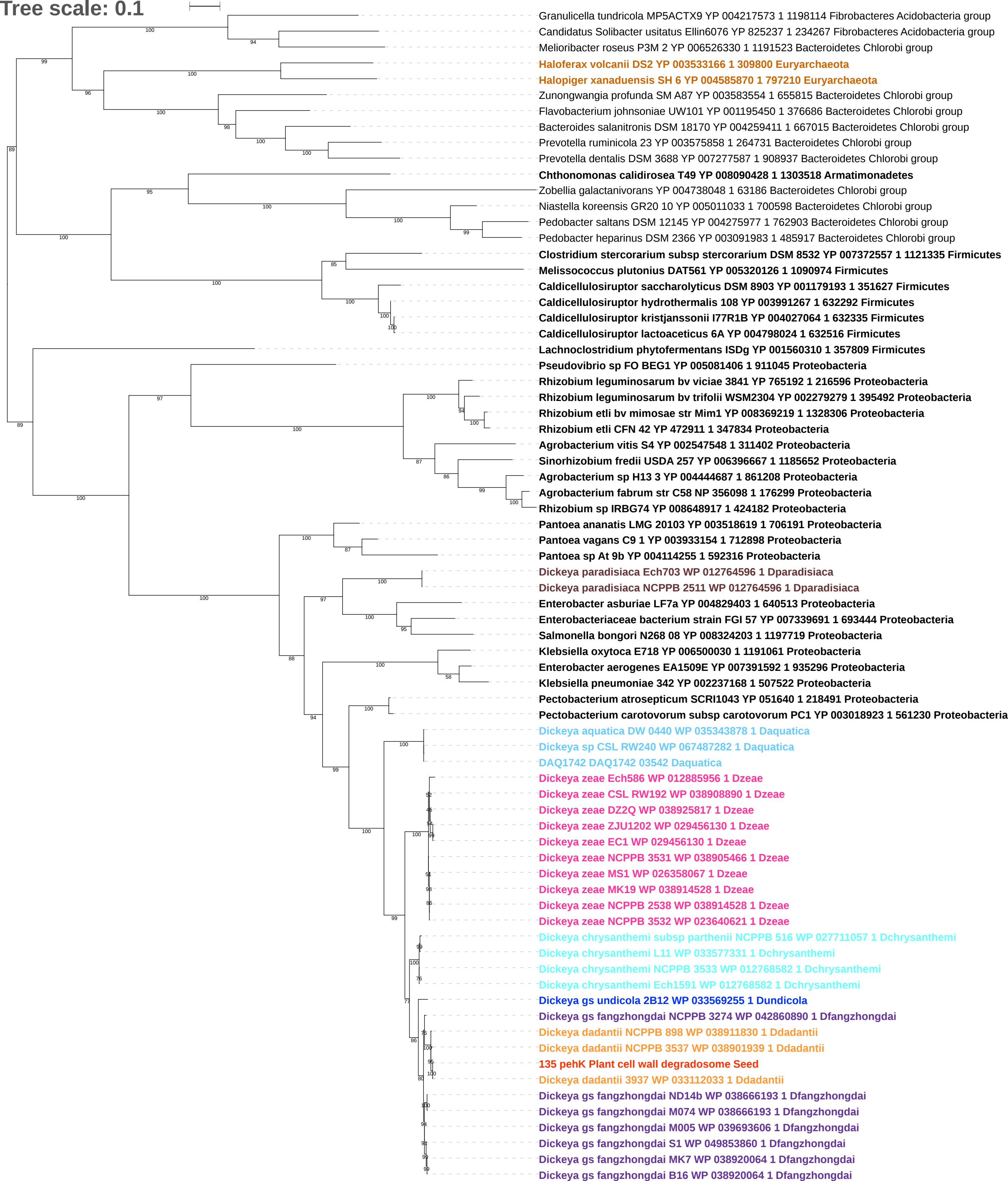
Tree scale: 0.1
Granulicella tundricola MP5ACTX9 YP 004217573 1 1198114 Fibrobacteres Acidobacteria group
Candidatus Solibacter usitatus Ellin6076 YP 825237 1 234267 Fibrobacteres Acidobacteria group
Melioribacter roseus P3M 2 YP 006526330 1 1191523 Bacteroidetes Chlorobi group
Haloferax volcanii DS2 YP 003533166 1 309800 Euryarchaeota
Halopiger xanaduensis SH 6 YP 004585870 1 797210 Euryarchaeota
Zunongwangia profunda SM A87 YP 003583554 1 655815 Bacteroidetes Chlorobi group
Flavobacterium johnsoniae UW101 YP 001195450 1 376686 Bacteroidetes Chlorobi group
Bacteroides salanitronis DSM 18170 YP 004259411 1 667015 Bacteroidetes Chlorobi group
Prevotella ruminicola 23 YP 003575858 1 264731 Bacteroidetes Chlorobi group
Prevotella dentalis DSM 3688 YP 007277587 1 908937 Bacteroidetes Chlorobi group
Chthonomonas calidirosea T49 YP 008090428 1 1303518 Armatimonadetes
Zobellia galactanivorans YP 004738048 1 63186 Bacteroidetes Chlorobi group
Niastella koreensis GR20 10 YP 005011033 1 700598 Bacteroidetes Chlorobi group
Pedobacter saltans DSM 12145 YP 004275977 1 762903 Bacteroidetes Chlorobi group
Pedobacter heparinus DSM 2366 YP 003091983 1 485917 Bacteroidetes Chlorobi group
Clostridium stercorarium subsp stercorarium DSM 8532 YP 007372557 1 1121335 Firmicutes
Melissococcus plutonius DAT561 YP 005320126 1 1090974 Firmicutes
Caldicellulosiruptor saccharolyticus DSM 8903 YP 001179193 1 351627 Firmicutes
Caldicellulosiruptor hydrothermalis 108 YP 003991267 1 632292 Firmicutes
Caldicellulosiruptor kristjanssonii I77R1B YP 004027064 1 632335 Firmicutes
Caldicellulosiruptor lactoaceticus 6A YP 004798024 1 632516 Firmicutes
Lachnoclostridium phytofermentans ISDg YP 001560310 1 357809 Firmicutes
Pseudovibrio sp FO BEG1 YP 005081406 1 911045 Proteobacteria
Rhizobium leguminosarum bv viciae 3841 YP 765192 1 216596 Proteobacteria
Rhizobium leguminosarum bv trifolii WSM2304 YP 002279279 1 395492 Proteobacteria
Rhizobium etli bv mimosae str Mim1 YP 008369219 1 1328306 Proteobacteria
Rhizobium etli CFN 42 YP 472911 1 347834 Proteobacteria
Agrobacterium vitis S4 YP 002547548 1 311402 Proteobacteria
Sinorhizobium fredii USDA 257 YP 006396667 1 1185652 Proteobacteria
Agrobacterium sp H13 3 YP 004444687 1 861208 Proteobacteria
Agrobacterium fabrum str C58 NP 356098 1 176299 Proteobacteria
Rhizobium sp IRBG74 YP 008648917 1 424182 Proteobacteria
Pantoea ananatis LMG 20103 YP 003518619 1 706191 Proteobacteria
Pantoea vagans C9 1 YP 003933154 1 712898 Proteobacteria
Pantoea sp At 9b YP 004114255 1 592316 Proteobacteria
Dickeya paradisiaca Ech703 WP 012764596 1 Dparadisiaca
Dickeya paradisiaca NCPPB 2511 WP 012764596 1 Dparadisiaca
Enterobacter asburiae LF7a YP 004829403 1 640513 Proteobacteria
Enterobacteriaceae bacterium strain FGI 57 YP 007339691 1 693444 Proteobacteria
Salmonella bongori N268 08 YP 008324203 1 1197719 Proteobacteria
Klebsiella oxytoca E718 YP 006500030 1 1191061 Proteobacteria
Enterobacter aerogenes EA1509E YP 007391592 1 935296 Proteobacteria
Klebsiella pneumoniae 342 YP 002237168 1 507522 Proteobacteria
Pectobacterium atrosepticum SCRI1043 YP 051640 1 218491 Proteobacteria
Pectobacterium carotovorum subsp carotovorum PC1 YP 003018923 1 561230 Proteobacteria
Dickeya aquatica DW 0440 WP 035343878 1 Daquatica
Dickeya sp CSL RW240 WP 067487282 1 Daquatica
DAQ1742 DAQ1742 03542 Daquatica
Dickeya zeae Ech586 WP 012885956 1 Dzeae
Dickeya zeae CSL RW192 WP 038908890 1 Dzeae
Dickeya zeae DZ2Q WP 038925817 1 Dzeae
Dickeya zeae ZJU1202 WP 029456130 1 Dzeae
Dickeya zeae EC1 WP 029456130 1 Dzeae
Dickeya zeae NCPPB 3531 WP 038905466 1 Dzeae
Dickeya zeae MS1 WP 026358067 1 Dzeae
Dickeya zeae MK19 WP 038914528 1 Dzeae
Dickeya zeae NCPPB 2538 WP 038914528 1 Dzeae
Dickeya zeae NCPPB 3532 WP 023640621 1 Dzeae
Dickeya chrysanthemi subsp parthenii NCPPB 516 WP 027711057 1 Dchrysanthemi
Dickeya chrysanthemi L11 WP 033577331 1 Dchrysanthemi
Dickeya chrysanthemi NCPPB 3533 WP 012768582 1 Dchrysanthemi
Dickeya chrysanthemi Ech1591 WP 012768582 1 Dchrysanthemi
Dickeya gs undicola 2B12 WP 033569255 1 Dundicola
Dickeya gs fangzhongdai NCPPB 3274 WP 042860890 1 Dfangzhongdai
Dickeya dadantii NCPPB 898 WP 038911830 1 Ddadantii
Dickeya dadantii NCPPB 3537 WP 038901939 1 Ddadantii
135 pehK Plant cell wall degradosome Seed
Dickeya dadantii 3937 WP 033112033 1 Ddadantii
Dickeya gs fangzhongdai ND14b WP 038666193 1 Dfangzhongdai
Dickeya gs fangzhongdai M074 WP 038666193 1 Dfangzhongdai
Dickeya gs fangzhongdai M005 WP 039693606 1 Dfangzhongdai
Dickeya gs fangzhongdai S1 WP 049853860 1 Dfangzhongdai
Dickeya gs fangzhongdai MK7 WP 038920064 1 Dfangzhongdai
Dickeya gs fangzhongdai B16 WP 038920064 1 Dfangzhongdai

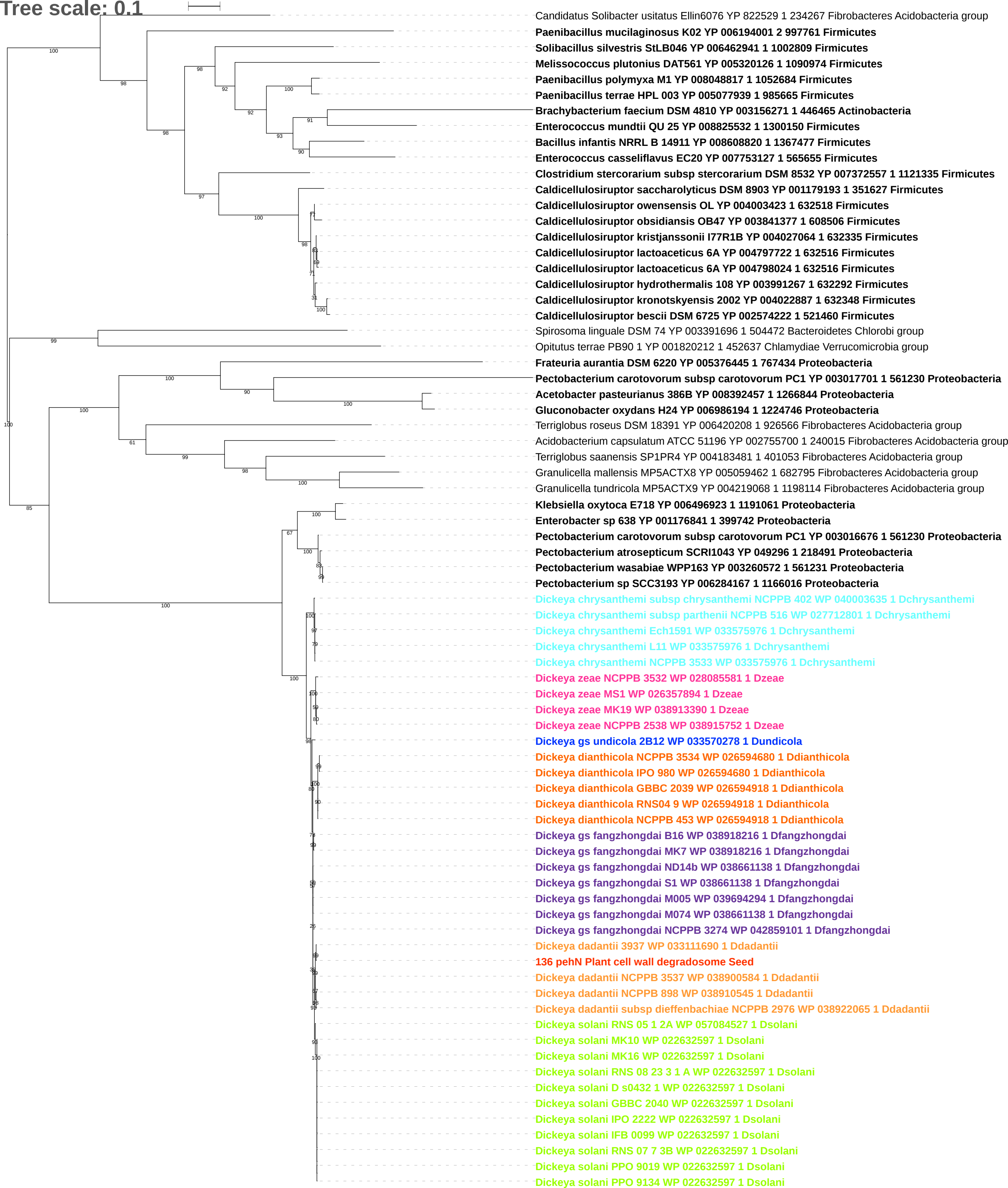
Tree scale: 0.1
Candidatus Solibacter usitatus Ellin6076 YP 822529 1 234267 Fibrobacteres Acidobacteria group
Paenibacillus mucilaginosus K02 YP 006194001 2 997761 Firmicutes
Solibacillus silvestris StLB046 YP 006462941 1 1002809 Firmicutes
Melissococcus plutonius DAT561 YP 005320126 1 1090974 Firmicutes
Paenibacillus polymyxa M1 YP 008048817 1 1052684 Firmicutes
Paenibacillus terrae HPL 003 YP 005077939 1 985665 Firmicutes
Brachybacterium faecium DSM 4810 YP 003156271 1 446465 Actinobacteria
Enterococcus mundtii QU 25 YP 008825532 1 1300150 Firmicutes
Bacillus infantis NRRL B 14911 YP 008608820 1 1367477 Firmicutes
Enterococcus casseliflavus EC20 YP 007753127 1 565655 Firmicutes
Clostridium stercorarium subsp stercorarium DSM 8532 YP 007372557 1 1121335 Firmicutes
Caldicellulosiruptor saccharolyticus DSM 8903 YP 001179193 1 351627 Firmicutes
Caldicellulosiruptor owensensis OL YP 004003423 1 632518 Firmicutes
Caldicellulosiruptor obsidiansis OB47 YP 003841377 1 608506 Firmicutes
Caldicellulosiruptor kristjanssonii I77R1B YP 004027064 1 632335 Firmicutes
Caldicellulosiruptor lactoaceticus 6A YP 004797722 1 632516 Firmicutes
Caldicellulosiruptor lactoaceticus 6A YP 004798024 1 632516 Firmicutes
Caldicellulosiruptor hydrothermalis 108 YP 003991267 1 632292 Firmicutes
Caldicellulosiruptor kronotskyensis 2002 YP 004022887 1 632348 Firmicutes
Caldicellulosiruptor bescii DSM 6725 YP 002574222 1 521460 Firmicutes
Spirosoma linguale DSM 74 YP 003391696 1 504472 Bacteroidetes Chlorobi group
Opitutus terrae PB90 1 YP 001820212 1 452637 Chlamydiae Verrucomicrobia group
Frateuria aurantia DSM 6220 YP 005376445 1 767434 Proteobacteria
Pectobacterium carotovorum subsp carotovorum PC1 YP 003017701 1 561230 Proteobacteria
Acetobacter pasteurianus 386B YP 008392457 1 1266844 Proteobacteria
Gluconobacter oxydans H24 YP 006986194 1 1224746 Proteobacteria
Terriglobus roseus DSM 18391 YP 006420208 1 926566 Fibrobacteres Acidobacteria group
Acidobacterium capsulatum ATCC 51196 YP 002755700 1 240015 Fibrobacteres Acidobacteria group
Terriglobus saanensis SP1PR4 YP 004183481 1 401053 Fibrobacteres Acidobacteria group
Granulicella mallensis MP5ACTX8 YP 005059462 1 682795 Fibrobacteres Acidobacteria group
Granulicella tundricola MP5ACTX9 YP 004219068 1 1198114 Fibrobacteres Acidobacteria group
Klebsiella oxytoca E718 YP 006496923 1 1191061 Proteobacteria
Enterobacter sp 638 YP 001176841 1 399742 Proteobacteria
Pectobacterium carotovorum subsp carotovorum PC1 YP 003016676 1 561230 Proteobacteria
Pectobacterium atrosepticum SCRI1043 YP 049296 1 218491 Proteobacteria
Pectobacterium wasabiae WPP163 YP 003260572 1 561231 Proteobacteria
Pectobacterium sp SCC3193 YP 006284167 1 1166016 Proteobacteria
Dickeya chrysanthemi subsp chrysanthemi NCPPB 402 WP 040003635 1 Dchrysanthemi
Dickeya chrysanthemi subsp parthenii NCPPB 516 WP 027712801 1 Dchrysanthemi
Dickeya chrysanthemi Ech1591 WP 033575976 1 Dchrysanthemi
Dickeya chrysanthemi L11 WP 033575976 1 Dchrysanthemi
Dickeya chrysanthemi NCPPB 3533 WP 033575976 1 Dchrysanthemi
Dickeya zeae NCPPB 3532 WP 028085581 1 Dzeae
Dickeya zeae MS1 WP 026357894 1 Dzeae
Dickeya zeae MK19 WP 038913390 1 Dzeae
Dickeya zeae NCPPB 2538 WP 038915752 1 Dzeae
Dickeya gs undicola 2B12 WP 033570278 1 Dundicola
Dickeya dianthicola NCPPB 3534 WP 026594680 1 Ddianthicola
Dickeya dianthicola IPO 980 WP 026594680 1 Ddianthicola
Dickeya dianthicola GBBC 2039 WP 026594918 1 Ddianthicola
Dickeya dianthicola RNS04 9 WP 026594918 1 Ddianthicola
Dickeya dianthicola NCPPB 453 WP 026594918 1 Ddianthicola
Dickeya gs fangzhongdai B16 WP 038918216 1 Dfangzhongdai
Dickeya gs fangzhongdai MK7 WP 038918216 1 Dfangzhongdai
Dickeya gs fangzhongdai ND14b WP 038661138 1 Dfangzhongdai
Dickeya gs fangzhongdai S1 WP 038661138 1 Dfangzhongdai
Dickeya gs fangzhongdai M005 WP 039694294 1 Dfangzhongdai
Dickeya gs fangzhongdai M074 WP 038661138 1 Dfangzhongdai
Dickeya gs fangzhongdai NCPPB 3274 WP 042859101 1 Dfangzhongdai
Dickeya dadantii 3937 WP 033111690 1 Ddadantii
136 pehN Plant cell wall degradosome Seed
Dickeya dadantii NCPPB 3537 WP 038900584 1 Ddadantii
Dickeya dadantii NCPPB 898 WP 038910545 1 Ddadantii
Dickeya dadantii subsp dieffenbachiae NCPPB 2976 WP 038922065 1 Ddadantii
Dickeya solani RNS 05 1 2A WP 057084527 1 Dsolani
Dickeya solani MK10 WP 022632597 1 Dsolani
Dickeya solani MK16 WP 022632597 1 Dsolani
Dickeya solani RNS 08 23 3 1 A WP 022632597 1 Dsolani
Dickeya solani D s0432 1 WP 022632597 1 Dsolani
Dickeya solani GBBC 2040 WP 022632597 1 Dsolani
Dickeya solani IPO 2222 WP 022632597 1 Dsolani
Dickeya solani IFB 0099 WP 022632597 1 Dsolani
Dickeya solani RNS 07 7 3B WP 022632597 1 Dsolani
Dickeya solani PPO 9019 WP 022632597 1 Dsolani
Dickeya solani PPO 9134 WP 022632597 1 Dsolani

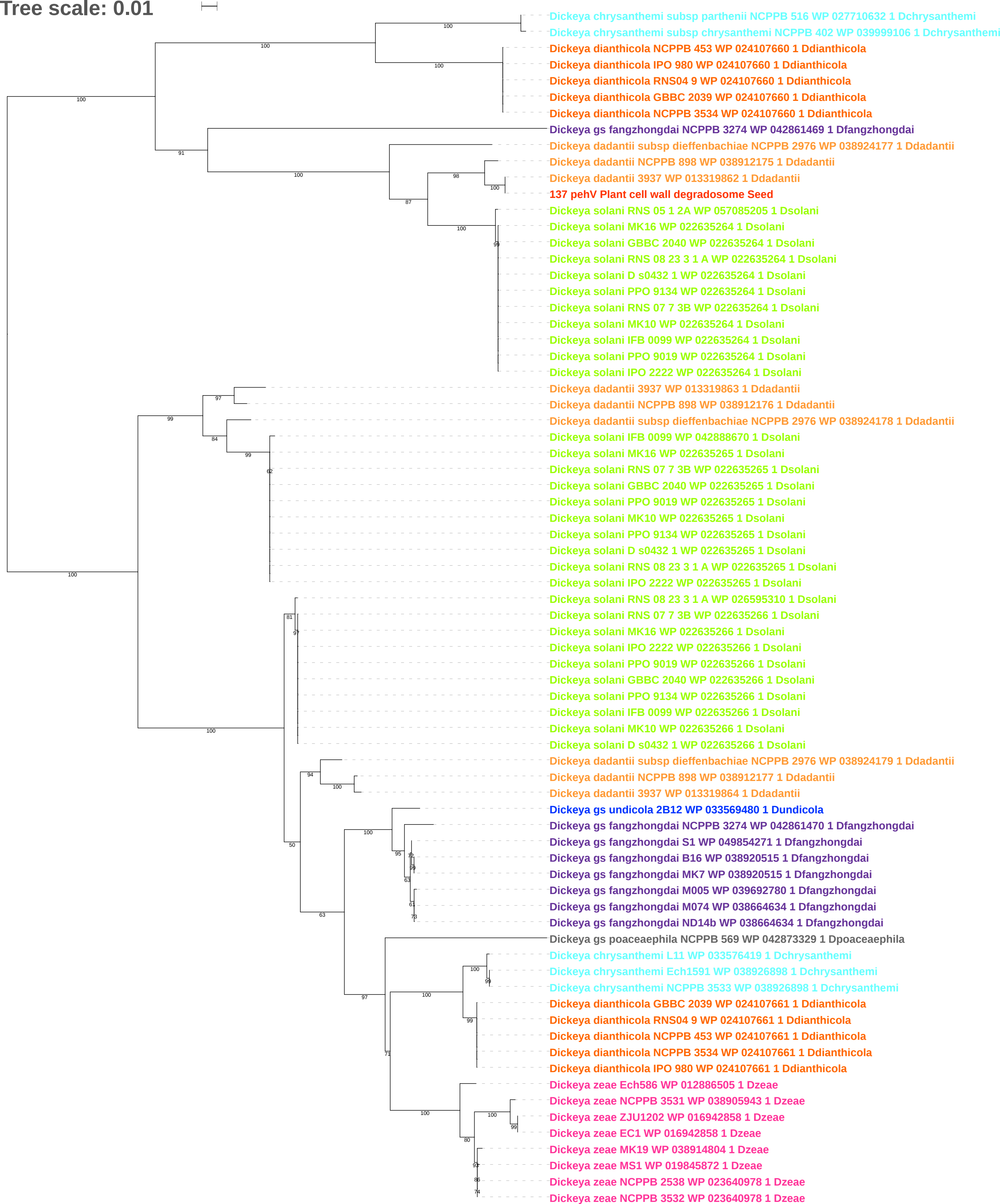
Tree scale: 0.01
Dickeya chrysanthemi subsp parthenii NCPPB 516 WP 027710632 1 Dchrysanthemi
Dickeya chrysanthemi subsp chrysanthemi NCPPB 402 WP 039999106 1 Dchrysanthemi
Dickeya dianthicola NCPPB 453 WP 024107660 1 Ddianthicola
Dickeya dianthicola IPO 980 WP 024107660 1 Ddianthicola
Dickeya dianthicola RNS04 9 WP 024107660 1 Ddianthicola
Dickeya dianthicola GBBC 2039 WP 024107660 1 Ddianthicola
Dickeya dianthicola NCPPB 3534 WP 024107660 1 Ddianthicola
Dickeya gs fangzhongdai NCPPB 3274 WP 042861469 1 Dfangzhongdai
Dickeya dadantii subsp dieffenbachiae NCPPB 2976 WP 038924177 1 Ddadantii
Dickeya dadantii NCPPB 898 WP 038912175 1 Ddadantii
Dickeya dadantii 3937 WP 013319862 1 Ddadantii
137 pehV Plant cell wall degradosome Seed
Dickeya solani RNS 05 1 2A WP 057085205 1 Dsolani
Dickeya solani MK16 WP 022635264 1 Dsolani
Dickeya solani GBBC 2040 WP 022635264 1 Dsolani
Dickeya solani RNS 08 23 3 1 A WP 022635264 1 Dsolani
Dickeya solani D s0432 1 WP 022635264 1 Dsolani
Dickeya solani PPO 9134 WP 022635264 1 Dsolani
Dickeya solani RNS 07 7 3B WP 022635264 1 Dsolani
Dickeya solani MK10 WP 022635264 1 Dsolani
Dickeya solani IFB 0099 WP 022635264 1 Dsolani
Dickeya solani PPO 9019 WP 022635264 1 Dsolani
Dickeya solani IPO 2222 WP 022635264 1 Dsolani
Dickeya dadantii 3937 WP 013319863 1 Ddadantii
Dickeya dadantii NCPPB 898 WP 038912176 1 Ddadantii
Dickeya dadantii subsp dieffenbachiae NCPPB 2976 WP 038924178 1 Ddadantii
Dickeya solani IFB 0099 WP 042888670 1 Dsolani
Dickeya solani MK16 WP 022635265 1 Dsolani
Dickeya solani RNS 07 7 3B WP 022635265 1 Dsolani
Dickeya solani GBBC 2040 WP 022635265 1 Dsolani
Dickeya solani PPO 9019 WP 022635265 1 Dsolani
Dickeya solani MK10 WP 022635265 1 Dsolani
Dickeya solani PPO 9134 WP 022635265 1 Dsolani
Dickeya solani D s0432 1 WP 022635265 1 Dsolani
Dickeya solani RNS 08 23 3 1 A WP 022635265 1 Dsolani
Dickeya solani IPO 2222 WP 022635265 1 Dsolani
Dickeya solani RNS 08 23 3 1 A WP 026595310 1 Dsolani
Dickeya solani RNS 07 7 3B WP 022635266 1 Dsolani
Dickeya solani MK16 WP 022635266 1 Dsolani
Dickeya solani IPO 2222 WP 022635266 1 Dsolani
Dickeya solani PPO 9019 WP 022635266 1 Dsolani
Dickeya solani GBBC 2040 WP 022635266 1 Dsolani
Dickeya solani PPO 9134 WP 022635266 1 Dsolani
Dickeya solani IFB 0099 WP 022635266 1 Dsolani
Dickeya solani MK10 WP 022635266 1 Dsolani
Dickeya solani D s0432 1 WP 022635266 1 Dsolani
Dickeya dadantii subsp dieffenbachiae NCPPB 2976 WP 038924179 1 Ddadantii
Dickeya dadantii NCPPB 898 WP 038912177 1 Ddadantii
Dickeya dadantii 3937 WP 013319864 1 Ddadantii
Dickeya gs undicola 2B12 WP 033569480 1 Dundicola
Dickeya gs fangzhongdai NCPPB 3274 WP 042861470 1 Dfangzhongdai
Dickeya gs fangzhongdai S1 WP 049854271 1 Dfangzhongdai
Dickeya gs fangzhongdai B16 WP 038920515 1 Dfangzhongdai
Dickeya gs fangzhongdai MK7 WP 038920515 1 Dfangzhongdai
Dickeya gs fangzhongdai M005 WP 039692780 1 Dfangzhongdai
Dickeya gs fangzhongdai M074 WP 038664634 1 Dfangzhongdai
Dickeya gs fangzhongdai ND14b WP 038664634 1 Dfangzhongdai
Dickeya gs poaceaephila NCPPB 569 WP 042873329 1 Dpoaceaephila
Dickeya chrysanthemi L11 WP 033576419 1 Dchrysanthemi
Dickeya chrysanthemi Ech1591 WP 038926898 1 Dchrysanthemi
Dickeya chrysanthemi NCPPB 3533 WP 038926898 1 Dchrysanthemi
Dickeya dianthicola GBBC 2039 WP 024107661 1 Ddianthicola
Dickeya dianthicola RNS04 9 WP 024107661 1 Ddianthicola
Dickeya dianthicola NCPPB 453 WP 024107661 1 Ddianthicola
Dickeya dianthicola NCPPB 3534 WP 024107661 1 Ddianthicola
Dickeya dianthicola IPO 980 WP 024107661 1 Ddianthicola
Dickeya zeae Ech586 WP 012886505 1 Dzeae
Dickeya zeae NCPPB 3531 WP 038905943 1 Dzeae
Dickeya zeae ZJU1202 WP 016942858 1 Dzeae
Dickeya zeae EC1 WP 016942858 1 Dzeae
Dickeya zeae MK19 WP 038914804 1 Dzeae
Dickeya zeae MS1 WP 019845872 1 Dzeae
Dickeya zeae NCPPB 2538 WP 023640978 1 Dzeae
Dickeya zeae NCPPB 3532 WP 023640978 1 Dzeae

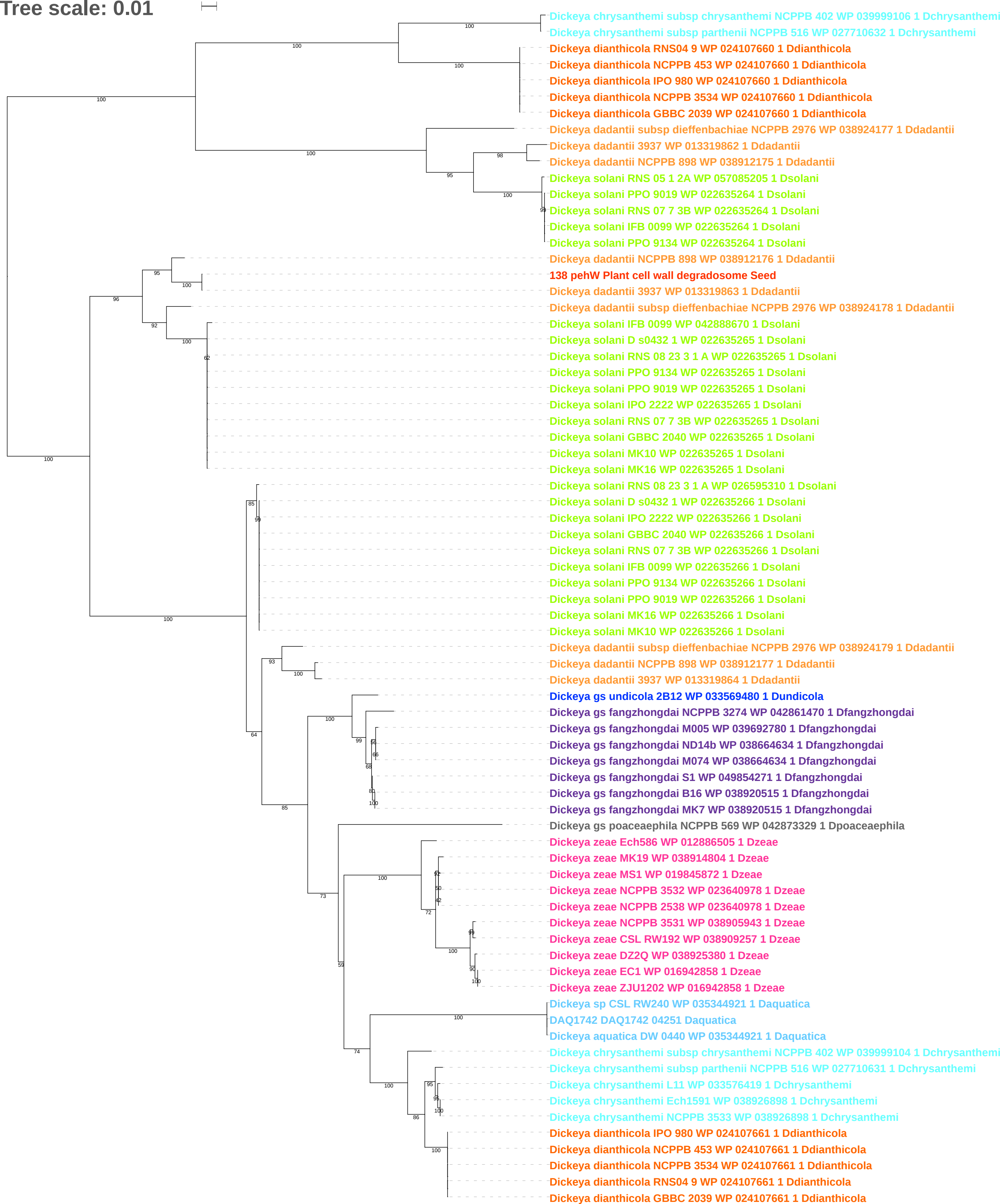
Tree scale: 0.01
Dickeya chrysanthemi subsp chrysanthemi NCPPB 402 WP 039999106 1 Dchrysanthemi
Dickeya chrysanthemi subsp parthenii NCPPB 516 WP 027710632 1 Dchrysanthemi
Dickeya dianthicola RNS04 9 WP 024107660 1 Ddianthicola
Dickeya dianthicola NCPPB 453 WP 024107660 1 Ddianthicola
Dickeya dianthicola IPO 980 WP 024107660 1 Ddianthicola
Dickeya dianthicola NCPPB 3534 WP 024107660 1 Ddianthicola
Dickeya dianthicola GBBC 2039 WP 024107660 1 Ddianthicola
Dickeya dadantii subsp dieffenbachiae NCPPB 2976 WP 038924177 1 Ddadantii
Dickeya dadantii 3937 WP 013319862 1 Ddadantii
Dickeya dadantii NCPPB 898 WP 038912175 1 Ddadantii
Dickeya solani RNS 05 1 2A WP 057085205 1 Dsolani
Dickeya solani PPO 9019 WP 022635264 1 Dsolani
Dickeya solani RNS 07 7 3B WP 022635264 1 Dsolani
Dickeya solani IFB 0099 WP 022635264 1 Dsolani
Dickeya solani PPO 9134 WP 022635264 1 Dsolani
Dickeya dadantii NCPPB 898 WP 038912176 1 Ddadantii
138 pehW Plant cell wall degradosome Seed
Dickeya dadantii 3937 WP 013319863 1 Ddadantii
Dickeya dadantii subsp dieffenbachiae NCPPB 2976 WP 038924178 1 Ddadantii
Dickeya solani IFB 0099 WP 042888670 1 Dsolani
Dickeya solani D s0432 1 WP 022635265 1 Dsolani
Dickeya solani RNS 08 23 3 1 A WP 022635265 1 Dsolani
Dickeya solani PPO 9134 WP 022635265 1 Dsolani
Dickeya solani PPO 9019 WP 022635265 1 Dsolani
Dickeya solani IPO 2222 WP 022635265 1 Dsolani
Dickeya solani RNS 07 7 3B WP 022635265 1 Dsolani
Dickeya solani GBBC 2040 WP 022635265 1 Dsolani
Dickeya solani MK10 WP 022635265 1 Dsolani
Dickeya solani MK16 WP 022635265 1 Dsolani
Dickeya solani RNS 08 23 3 1 A WP 026595310 1 Dsolani
Dickeya solani D s0432 1 WP 022635266 1 Dsolani
Dickeya solani IPO 2222 WP 022635266 1 Dsolani
Dickeya solani GBBC 2040 WP 022635266 1 Dsolani
Dickeya solani RNS 07 7 3B WP 022635266 1 Dsolani
Dickeya solani IFB 0099 WP 022635266 1 Dsolani
Dickeya solani PPO 9134 WP 022635266 1 Dsolani
Dickeya solani PPO 9019 WP 022635266 1 Dsolani
Dickeya solani MK16 WP 022635266 1 Dsolani
Dickeya solani MK10 WP 022635266 1 Dsolani
Dickeya dadantii subsp dieffenbachiae NCPPB 2976 WP 038924179 1 Ddadantii
Dickeya dadantii NCPPB 898 WP 038912177 1 Ddadantii
Dickeya dadantii 3937 WP 013319864 1 Ddadantii
Dickeya gs undicola 2B12 WP 033569480 1 Dundicola
Dickeya gs fangzhongdai NCPPB 3274 WP 042861470 1 Dfangzhongdai
Dickeya gs fangzhongdai M005 WP 039692780 1 Dfangzhongdai
Dickeya gs fangzhongdai ND14b WP 038664634 1 Dfangzhongdai
Dickeya gs fangzhongdai M074 WP 038664634 1 Dfangzhongdai
Dickeya gs fangzhongdai S1 WP 049854271 1 Dfangzhongdai
Dickeya gs fangzhongdai B16 WP 038920515 1 Dfangzhongdai
Dickeya gs fangzhongdai MK7 WP 038920515 1 Dfangzhongdai
Dickeya gs poaceaephila NCPPB 569 WP 042873329 1 Dpoaceaephila
Dickeya zeae Ech586 WP 012886505 1 Dzeae
Dickeya zeae MK19 WP 038914804 1 Dzeae
Dickeya zeae MS1 WP 019845872 1 Dzeae
Dickeya zeae NCPPB 3532 WP 023640978 1 Dzeae
Dickeya zeae NCPPB 2538 WP 023640978 1 Dzeae
Dickeya zeae NCPPB 3531 WP 038905943 1 Dzeae
Dickeya zeae CSL RW192 WP 038909257 1 Dzeae
Dickeya zeae DZ2Q WP 038925380 1 Dzeae
Dickeya zeae EC1 WP 016942858 1 Dzeae
Dickeya zeae ZJU1202 WP 016942858 1 Dzeae
Dickeya sp CSL RW240 WP 035344921 1 Daquatica
DAQ1742 DAQ1742 04251 Daquatica
Dickeya aquatica DW 0440 WP 035344921 1 Daquatica
Dickeya chrysanthemi subsp chrysanthemi NCPPB 402 WP 039999104 1 Dchrysanthemi
Dickeya chrysanthemi subsp parthenii NCPPB 516 WP 027710631 1 Dchrysanthemi
Dickeya chrysanthemi L11 WP 033576419 1 Dchrysanthemi
Dickeya chrysanthemi Ech1591 WP 038926898 1 Dchrysanthemi
Dickeya chrysanthemi NCPPB 3533 WP 038926898 1 Dchrysanthemi
Dickeya dianthicola IPO 980 WP 024107661 1 Ddianthicola
Dickeya dianthicola NCPPB 453 WP 024107661 1 Ddianthicola
Dickeya dianthicola NCPPB 3534 WP 024107661 1 Ddianthicola
Dickeya dianthicola RNS04 9 WP 024107661 1 Ddianthicola
Dickeya dianthicola GBBC 2039 WP 024107661 1 Ddianthicola

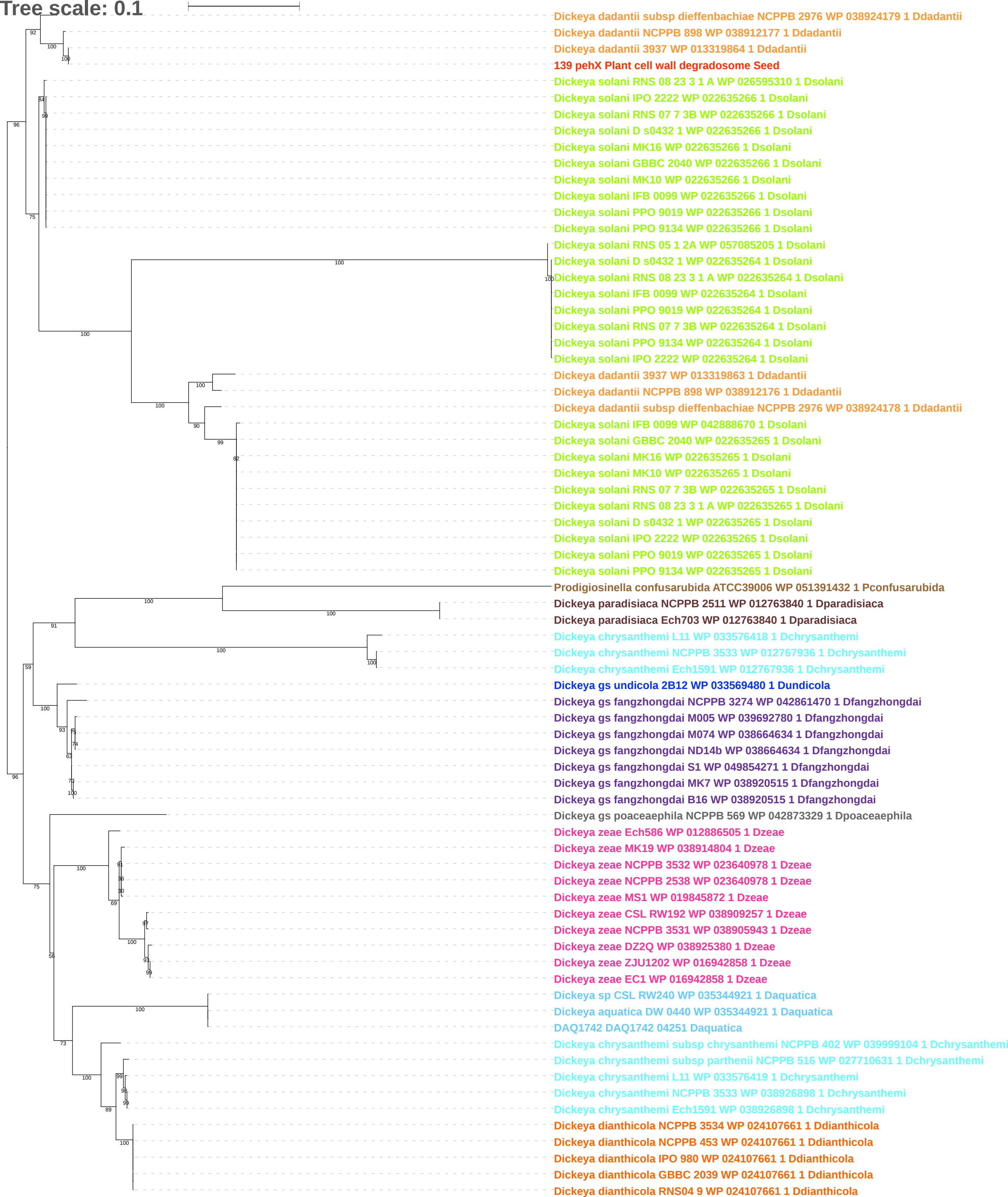
Tree scale: 0.1
Dickeya dadantii subsp dieffenbachiae NCPPB 2976 WP 038924179 1 Ddadantii
Dickeya dadantii NCPPB 898 WP 038912177 1 Ddadantii
Dickeya dadantii 3937 WP 013319864 1 Ddadantii
139 pehX Plant cell wall degradosome Seed
Dickeya solani RNS 08 23 3 1 A WP 026595310 1 Dsolani
Dickeya solani IPO 2222 WP 022635266 1 Dsolani
Dickeya solani RNS 07 7 3B WP 022635266 1 Dsolani
Dickeya solani D s0432 1 WP 022635266 1 Dsolani
Dickeya solani MK16 WP 022635266 1 Dsolani
Dickeya solani GBBC 2040 WP 022635266 1 Dsolani
Dickeya solani MK10 WP 022635266 1 Dsolani
Dickeya solani IFB 0099 WP 022635266 1 Dsolani
Dickeya solani PPO 9019 WP 022635266 1 Dsolani
Dickeya solani PPO 9134 WP 022635266 1 Dsolani
Dickeya solani RNS 05 1 2A WP 057085205 1 Dsolani
Dickeya solani D s0432 1 WP 022635264 1 Dsolani
Dickeya solani RNS 08 23 3 1 A WP 022635264 1 Dsolani
Dickeya solani IFB 0099 WP 022635264 1 Dsolani
Dickeya solani PPO 9019 WP 022635264 1 Dsolani
Dickeya solani RNS 07 7 3B WP 022635264 1 Dsolani
Dickeya solani PPO 9134 WP 022635264 1 Dsolani
Dickeya solani IPO 2222 WP 022635264 1 Dsolani
Dickeya dadantii 3937 WP 013319863 1 Ddadantii
Dickeya dadantii NCPPB 898 WP 038912176 1 Ddadantii
Dickeya dadantii subsp dieffenbachiae NCPPB 2976 WP 038924178 1 Ddadantii
Dickeya solani IFB 0099 WP 042888670 1 Dsolani
Dickeya solani GBBC 2040 WP 022635265 1 Dsolani
Dickeya solani MK16 WP 022635265 1 Dsolani
Dickeya solani MK10 WP 022635265 1 Dsolani
Dickeya solani RNS 07 7 3B WP 022635265 1 Dsolani
Dickeya solani RNS 08 23 3 1 A WP 022635265 1 Dsolani
Dickeya solani D s0432 1 WP 022635265 1 Dsolani
Dickeya solani IPO 2222 WP 022635265 1 Dsolani
Dickeya solani PPO 9019 WP 022635265 1 Dsolani
Dickeya solani PPO 9134 WP 022635265 1 Dsolani
Prodigiosinella confusarubida ATCC39006 WP 051391432 1 Pconfusarubida
Dickeya paradisiaca NCPPB 2511 WP 012763840 1 Dparadisiaca
Dickeya paradisiaca Ech703 WP 012763840 1 Dparadisiaca
Dickeya chrysanthemi L11 WP 033576418 1 Dchrysanthemi
Dickeya chrysanthemi NCPPB 3533 WP 012767936 1 Dchrysanthemi
Dickeya chrysanthemi Ech1591 WP 012767936 1 Dchrysanthemi
Dickeya gs undicola 2B12 WP 033569480 1 Dundicola
Dickeya gs fangzhongdai NCPPB 3274 WP 042861470 1 Dfangzhongdai
Dickeya gs fangzhongdai M005 WP 039692780 1 Dfangzhongdai
Dickeya gs fangzhongdai M074 WP 038664634 1 Dfangzhongdai
Dickeya gs fangzhongdai ND14b WP 038664634 1 Dfangzhongdai
Dickeya gs fangzhongdai S1 WP 049854271 1 Dfangzhongdai
Dickeya gs fangzhongdai MK7 WP 038920515 1 Dfangzhongdai
Dickeya gs fangzhongdai B16 WP 038920515 1 Dfangzhongdai
Dickeya gs poaceaephila NCPPB 569 WP 042873329 1 Dpoaceaephila
Dickeya zeae Ech586 WP 012886505 1 Dzeae
Dickeya zeae MK19 WP 038914804 1 Dzeae
Dickeya zeae NCPPB 3532 WP 023640978 1 Dzeae
Dickeya zeae NCPPB 2538 WP 023640978 1 Dzeae
Dickeya zeae MS1 WP 019845872 1 Dzeae
Dickeya zeae CSL RW192 WP 038909257 1 Dzeae
Dickeya zeae NCPPB 3531 WP 038905943 1 Dzeae
Dickeya zeae DZ2Q WP 038925380 1 Dzeae
Dickeya zeae ZJU1202 WP 016942858 1 Dzeae
Dickeya zeae EC1 WP 016942858 1 Dzeae
Dickeya sp CSL RW240 WP 035344921 1 Daquatica
Dickeya aquatica DW 0440 WP 035344921 1 Daquatica
DAQ1742 DAQ1742 04251 Daquatica
Dickeya chrysanthemi subsp chrysanthemi NCPPB 402 WP 039999104 1 Dchrysanthemi
Dickeya chrysanthemi subsp parthenii NCPPB 516 WP 027710631 1 Dchrysanthemi
Dickeya chrysanthemi L11 WP 033576419 1 Dchrysanthemi
Dickeya chrysanthemi NCPPB 3533 WP 038926898 1 Dchrysanthemi
Dickeya chrysanthemi Ech1591 WP 038926898 1 Dchrysanthemi
Dickeya dianthicola NCPPB 3534 WP 024107661 1 Ddianthicola
Dickeya dianthicola NCPPB 453 WP 024107661 1 Ddianthicola
Dickeya dianthicola IPO 980 WP 024107661 1 Ddianthicola
Dickeya dianthicola GBBC 2039 WP 024107661 1 Ddianthicola
Dickeya dianthicola RNS04 9 WP 024107661 1 Ddianthicola

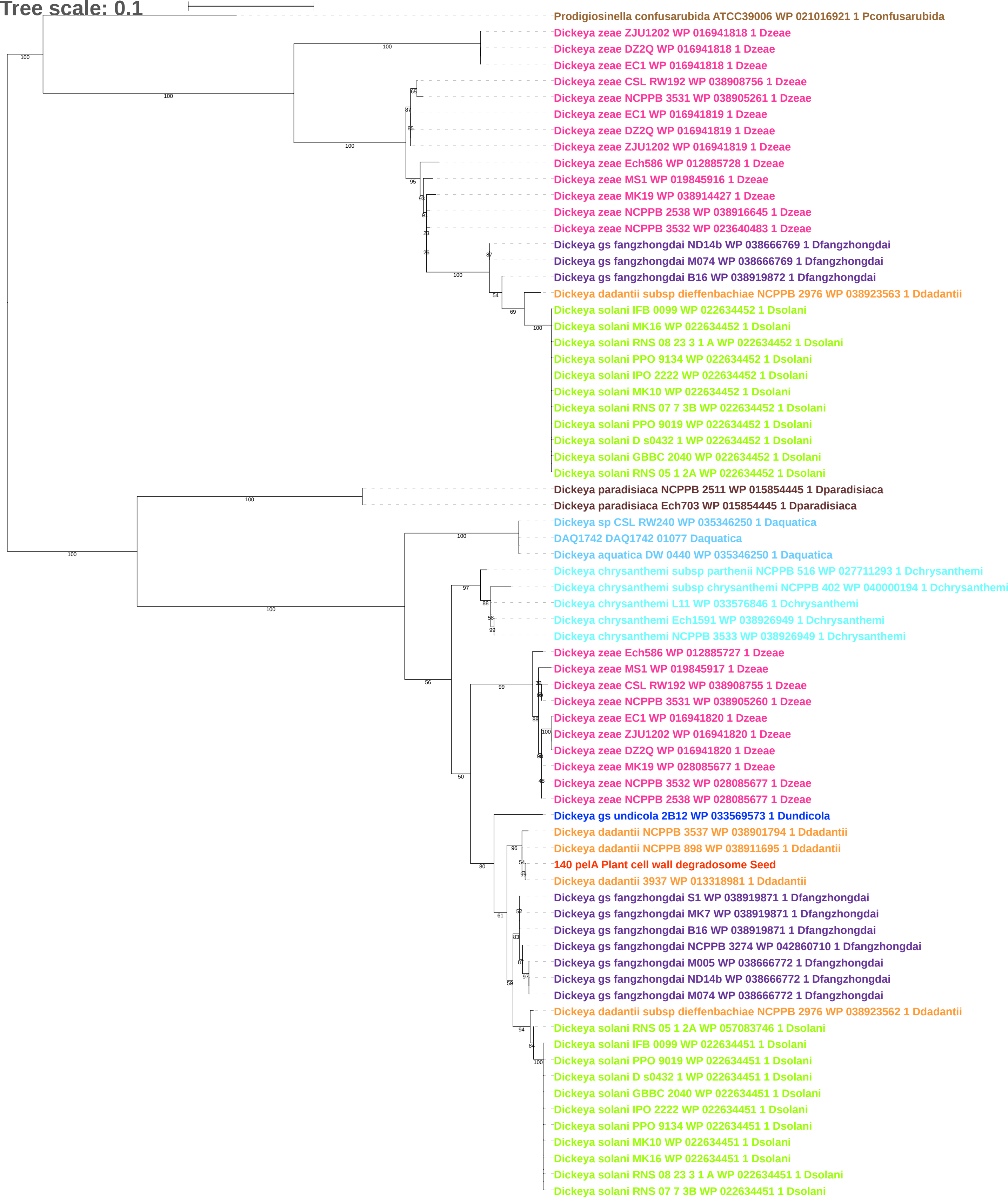
Tree scale: 0.1
Prodigiosinella confusarubida ATCC39006 WP 021016921 1 Pconfusarubida
Dickeya zeae ZJU1202 WP 016941818 1 Dzeae
Dickeya zeae DZ2Q WP 016941818 1 Dzeae
Dickeya zeae EC1 WP 016941818 1 Dzeae
Dickeya zeae CSL RW192 WP 038908756 1 Dzeae
Dickeya zeae NCPPB 3531 WP 038905261 1 Dzeae
Dickeya zeae EC1 WP 016941819 1 Dzeae
Dickeya zeae DZ2Q WP 016941819 1 Dzeae
Dickeya zeae ZJU1202 WP 016941819 1 Dzeae
Dickeya zeae Ech586 WP 012885728 1 Dzeae
Dickeya zeae MS1 WP 019845916 1 Dzeae
Dickeya zeae MK19 WP 038914427 1 Dzeae
Dickeya zeae NCPPB 2538 WP 038916645 1 Dzeae
Dickeya zeae NCPPB 3532 WP 023640483 1 Dzeae
Dickeya gs fangzhongdai ND14b WP 038666769 1 Dfangzhongdai
Dickeya gs fangzhongdai M074 WP 038666769 1 Dfangzhongdai
Dickeya gs fangzhongdai B16 WP 038919872 1 Dfangzhongdai
Dickeya dadantii subsp dieffenbachiae NCPPB 2976 WP 038923563 1 Ddadantii
Dickeya solani IFB 0099 WP 022634452 1 Dsolani
Dickeya solani MK16 WP 022634452 1 Dsolani
Dickeya solani RNS 08 23 3 1 A WP 022634452 1 Dsolani
Dickeya solani PPO 9134 WP 022634452 1 Dsolani
Dickeya solani IPO 2222 WP 022634452 1 Dsolani
Dickeya solani MK10 WP 022634452 1 Dsolani
Dickeya solani RNS 07 7 3B WP 022634452 1 Dsolani
Dickeya solani PPO 9019 WP 022634452 1 Dsolani
Dickeya solani D s0432 1 WP 022634452 1 Dsolani
Dickeya solani GBBC 2040 WP 022634452 1 Dsolani
Dickeya solani RNS 05 1 2A WP 022634452 1 Dsolani
Dickeya paradisiaca NCPPB 2511 WP 015854445 1 Dparadisiaca
Dickeya paradisiaca Ech703 WP 015854445 1 Dparadisiaca
Dickeya sp CSL RW240 WP 035346250 1 Daquatica
DAQ1742 DAQ1742 01077 Daquatica
Dickeya aquatica DW 0440 WP 035346250 1 Daquatica
Dickeya chrysanthemi subsp parthenii NCPPB 516 WP 027711293 1 Dchrysanthemi
Dickeya chrysanthemi subsp chrysanthemi NCPPB 402 WP 040000194 1 Dchrysanthemi
Dickeya chrysanthemi L11 WP 033576846 1 Dchrysanthemi
Dickeya chrysanthemi Ech1591 WP 038926949 1 Dchrysanthemi
Dickeya chrysanthemi NCPPB 3533 WP 038926949 1 Dchrysanthemi
Dickeya zeae Ech586 WP 012885727 1 Dzeae
Dickeya zeae MS1 WP 019845917 1 Dzeae
Dickeya zeae CSL RW192 WP 038908755 1 Dzeae
Dickeya zeae NCPPB 3531 WP 038905260 1 Dzeae
Dickeya zeae EC1 WP 016941820 1 Dzeae
Dickeya zeae ZJU1202 WP 016941820 1 Dzeae
Dickeya zeae DZ2Q WP 016941820 1 Dzeae
Dickeya zeae MK19 WP 028085677 1 Dzeae
Dickeya zeae NCPPB 3532 WP 028085677 1 Dzeae
Dickeya zeae NCPPB 2538 WP 028085677 1 Dzeae
Dickeya gs undicola 2B12 WP 033569573 1 Dundicola
Dickeya dadantii NCPPB 3537 WP 038901794 1 Ddadantii
Dickeya dadantii NCPPB 898 WP 038911695 1 Ddadantii
140 pelA Plant cell wall degradosome Seed
Dickeya dadantii 3937 WP 013318981 1 Ddadantii
Dickeya gs fangzhongdai S1 WP 038919871 1 Dfangzhongdai
Dickeya gs fangzhongdai MK7 WP 038919871 1 Dfangzhongdai
Dickeya gs fangzhongdai B16 WP 038919871 1 Dfangzhongdai
Dickeya gs fangzhongdai NCPPB 3274 WP 042860710 1 Dfangzhongdai
Dickeya gs fangzhongdai M005 WP 038666772 1 Dfangzhongdai
Dickeya gs fangzhongdai ND14b WP 038666772 1 Dfangzhongdai
Dickeya gs fangzhongdai M074 WP 038666772 1 Dfangzhongdai
Dickeya dadantii subsp dieffenbachiae NCPPB 2976 WP 038923562 1 Ddadantii
Dickeya solani RNS 05 1 2A WP 057083746 1 Dsolani
Dickeya solani IFB 0099 WP 022634451 1 Dsolani
Dickeya solani PPO 9019 WP 022634451 1 Dsolani
Dickeya solani D s0432 1 WP 022634451 1 Dsolani
Dickeya solani GBBC 2040 WP 022634451 1 Dsolani
Dickeya solani IPO 2222 WP 022634451 1 Dsolani
Dickeya solani PPO 9134 WP 022634451 1 Dsolani
Dickeya solani MK10 WP 022634451 1 Dsolani
Dickeya solani MK16 WP 022634451 1 Dsolani
Dickeya solani RNS 08 23 3 1 A WP 022634451 1 Dsolani
Dickeya solani RNS 07 7 3B WP 022634451 1 Dsolani

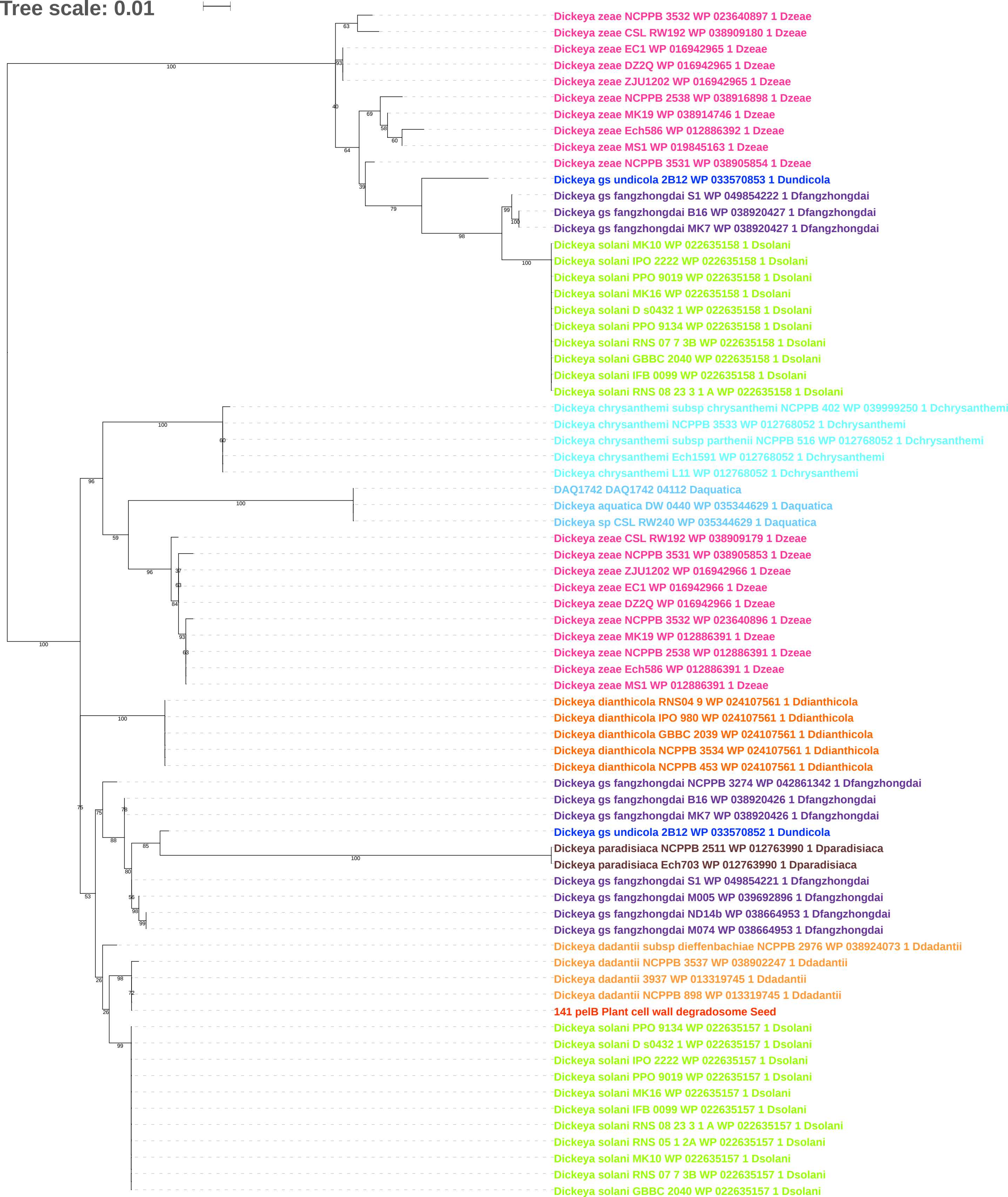
Tree scale: 0.01
Dickeya zeae NCPPB 3532 WP 023640897 1 Dzeae
Dickeya zeae CSL RW192 WP 038909180 1 Dzeae
Dickeya zeae EC1 WP 016942965 1 Dzeae
Dickeya zeae DZ2Q WP 016942965 1 Dzeae
Dickeya zeae ZJU1202 WP 016942965 1 Dzeae
Dickeya zeae NCPPB 2538 WP 038916898 1 Dzeae
Dickeya zeae MK19 WP 038914746 1 Dzeae
Dickeya zeae Ech586 WP 012886392 1 Dzeae
Dickeya zeae MS1 WP 019845163 1 Dzeae
Dickeya zeae NCPPB 3531 WP 038905854 1 Dzeae
Dickeya gs undicola 2B12 WP 033570853 1 Dundicola
Dickeya gs fangzhongdai S1 WP 049854222 1 Dfangzhongdai
Dickeya gs fangzhongdai B16 WP 038920427 1 Dfangzhongdai
Dickeya gs fangzhongdai MK7 WP 038920427 1 Dfangzhongdai
Dickeya solani MK10 WP 022635158 1 Dsolani
Dickeya solani IPO 2222 WP 022635158 1 Dsolani
Dickeya solani PPO 9019 WP 022635158 1 Dsolani
Dickeya solani MK16 WP 022635158 1 Dsolani
Dickeya solani D s0432 1 WP 022635158 1 Dsolani
Dickeya solani PPO 9134 WP 022635158 1 Dsolani
Dickeya solani RNS 07 7 3B WP 022635158 1 Dsolani
Dickeya solani GBBC 2040 WP 022635158 1 Dsolani
Dickeya solani IFB 0099 WP 022635158 1 Dsolani
Dickeya solani RNS 08 23 3 1 A WP 022635158 1 Dsolani
Dickeya chrysanthemi subsp chrysanthemi NCPPB 402 WP 039999250 1 Dchrysanthemi
Dickeya chrysanthemi NCPPB 3533 WP 012768052 1 Dchrysanthemi
Dickeya chrysanthemi subsp parthenii NCPPB 516 WP 012768052 1 Dchrysanthemi
Dickeya chrysanthemi Ech1591 WP 012768052 1 Dchrysanthemi
Dickeya chrysanthemi L11 WP 012768052 1 Dchrysanthemi
DAQ1742 DAQ1742 04112 Daquatica
Dickeya aquatica DW 0440 WP 035344629 1 Daquatica
Dickeya sp CSL RW240 WP 035344629 1 Daquatica
Dickeya zeae CSL RW192 WP 038909179 1 Dzeae
Dickeya zeae NCPPB 3531 WP 038905853 1 Dzeae
Dickeya zeae ZJU1202 WP 016942966 1 Dzeae
Dickeya zeae EC1 WP 016942966 1 Dzeae
Dickeya zeae DZ2Q WP 016942966 1 Dzeae
Dickeya zeae NCPPB 3532 WP 023640896 1 Dzeae
Dickeya zeae MK19 WP 012886391 1 Dzeae
Dickeya zeae NCPPB 2538 WP 012886391 1 Dzeae
Dickeya zeae Ech586 WP 012886391 1 Dzeae
Dickeya zeae MS1 WP 012886391 1 Dzeae
Dickeya dianthicola RNS04 9 WP 024107561 1 Ddianthicola
Dickeya dianthicola IPO 980 WP 024107561 1 Ddianthicola
Dickeya dianthicola GBBC 2039 WP 024107561 1 Ddianthicola
Dickeya dianthicola NCPPB 3534 WP 024107561 1 Ddianthicola
Dickeya dianthicola NCPPB 453 WP 024107561 1 Ddianthicola
Dickeya gs fangzhongdai NCPPB 3274 WP 042861342 1 Dfangzhongdai
Dickeya gs fangzhongdai B16 WP 038920426 1 Dfangzhongdai
Dickeya gs fangzhongdai MK7 WP 038920426 1 Dfangzhongdai
Dickeya gs undicola 2B12 WP 033570852 1 Dundicola
Dickeya paradisiaca NCPPB 2511 WP 012763990 1 Dparadisiaca
Dickeya paradisiaca Ech703 WP 012763990 1 Dparadisiaca
Dickeya gs fangzhongdai S1 WP 049854221 1 Dfangzhongdai
Dickeya gs fangzhongdai M005 WP 039692896 1 Dfangzhongdai
Dickeya gs fangzhongdai ND14b WP 038664953 1 Dfangzhongdai
Dickeya gs fangzhongdai M074 WP 038664953 1 Dfangzhongdai
Dickeya dadantii subsp dieffenbachiae NCPPB 2976 WP 038924073 1 Ddadantii
Dickeya dadantii NCPPB 3537 WP 038902247 1 Ddadantii
Dickeya dadantii 3937 WP 013319745 1 Ddadantii
Dickeya dadantii NCPPB 898 WP 013319745 1 Ddadantii
141 pelB Plant cell wall degradosome Seed
Dickeya solani PPO 9134 WP 022635157 1 Dsolani
Dickeya solani D s0432 1 WP 022635157 1 Dsolani
Dickeya solani IPO 2222 WP 022635157 1 Dsolani
Dickeya solani PPO 9019 WP 022635157 1 Dsolani
Dickeya solani MK16 WP 022635157 1 Dsolani
Dickeya solani IFB 0099 WP 022635157 1 Dsolani
Dickeya solani RNS 08 23 3 1 A WP 022635157 1 Dsolani
Dickeya solani RNS 05 1 2A WP 022635157 1 Dsolani
Dickeya solani MK10 WP 022635157 1 Dsolani
Dickeya solani RNS 07 7 3B WP 022635157 1 Dsolani
Dickeya solani GBBC 2040 WP 022635157 1 Dsolani

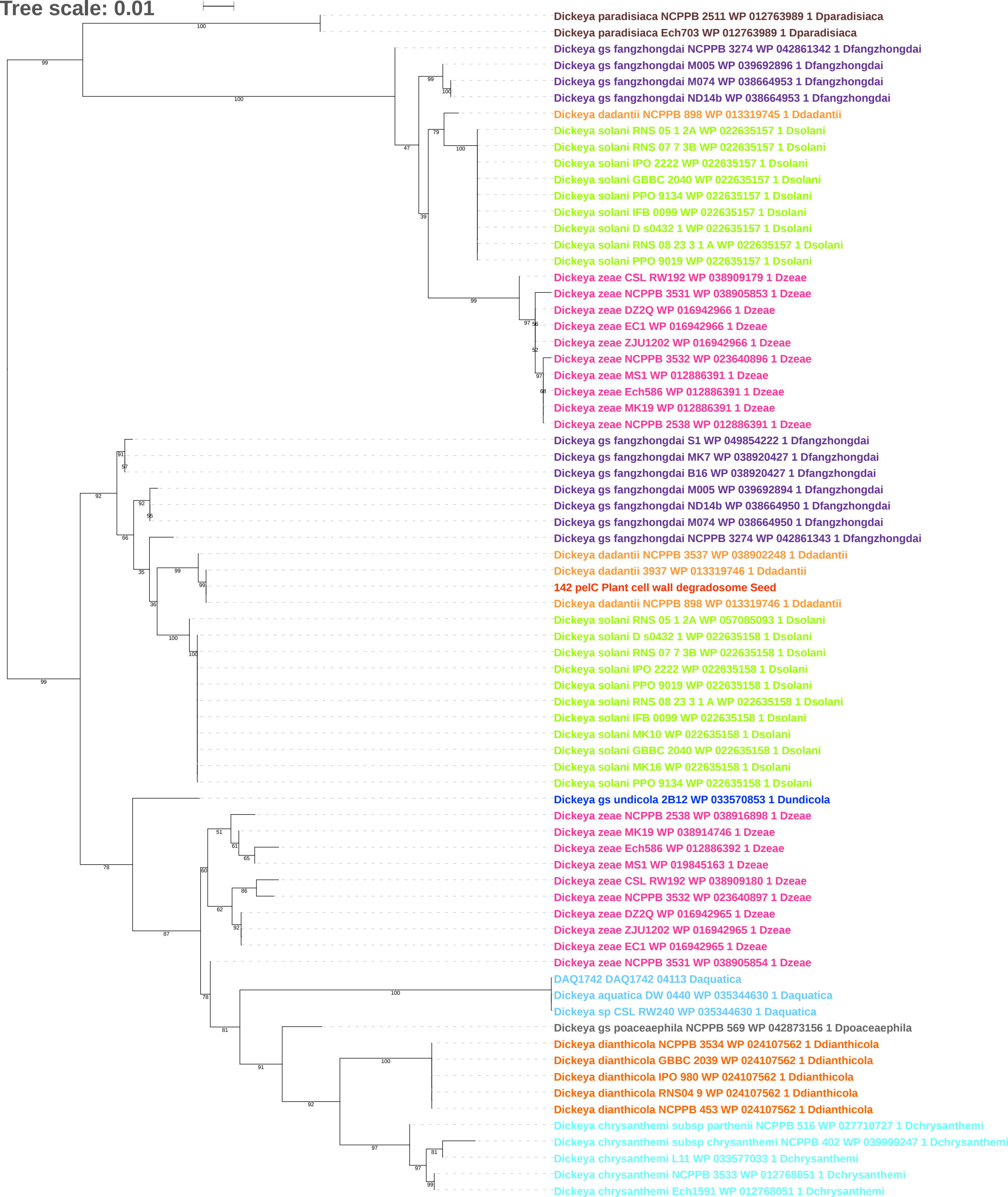
Tree scale: 0.01
Dickeya paradisiaca NCPPB 2511 WP 012763989 1 Dparadisiaca
Dickeya paradisiaca Ech703 WP 012763989 1 Dparadisiaca
Dickeya gs fangzhongdai NCPPB 3274 WP 042861342 1 Dfangzhongdai
Dickeya gs fangzhongdai M005 WP 039692896 1 Dfangzhongdai
Dickeya gs fangzhongdai M074 WP 038664953 1 Dfangzhongdai
Dickeya gs fangzhongdai ND14b WP 038664953 1 Dfangzhongdai
Dickeya dadantii NCPPB 898 WP 013319745 1 Ddadantii
Dickeya solani RNS 05 1 2A WP 022635157 1 Dsolani
Dickeya solani RNS 07 7 3B WP 022635157 1 Dsolani
Dickeya solani IPO 2222 WP 022635157 1 Dsolani
Dickeya solani GBBC 2040 WP 022635157 1 Dsolani
Dickeya solani PPO 9134 WP 022635157 1 Dsolani
Dickeya solani IFB 0099 WP 022635157 1 Dsolani
Dickeya solani D s0432 1 WP 022635157 1 Dsolani
Dickeya solani RNS 08 23 3 1 A WP 022635157 1 Dsolani
Dickeya solani PPO 9019 WP 022635157 1 Dsolani
Dickeya zeae CSL RW192 WP 038909179 1 Dzeae
Dickeya zeae NCPPB 3531 WP 038905853 1 Dzeae
Dickeya zeae DZ2Q WP 016942966 1 Dzeae
Dickeya zeae EC1 WP 016942966 1 Dzeae
Dickeya zeae ZJU1202 WP 016942966 1 Dzeae
Dickeya zeae NCPPB 3532 WP 023640896 1 Dzeae
Dickeya zeae MS1 WP 012886391 1 Dzeae
Dickeya zeae Ech586 WP 012886391 1 Dzeae
Dickeya zeae MK19 WP 012886391 1 Dzeae
Dickeya zeae NCPPB 2538 WP 012886391 1 Dzeae
Dickeya gs fangzhongdai S1 WP 049854222 1 Dfangzhongdai
Dickeya gs fangzhongdai MK7 WP 038920427 1 Dfangzhongdai
Dickeya gs fangzhongdai B16 WP 038920427 1 Dfangzhongdai
Dickeya gs fangzhongdai M005 WP 039692894 1 Dfangzhongdai
Dickeya gs fangzhongdai ND14b WP 038664950 1 Dfangzhongdai
Dickeya gs fangzhongdai M074 WP 038664950 1 Dfangzhongdai
Dickeya gs fangzhongdai NCPPB 3274 WP 042861343 1 Dfangzhongdai
Dickeya dadantii NCPPB 3537 WP 038902248 1 Ddadantii
Dickeya dadantii 3937 WP 013319746 1 Ddadantii
142 pelC Plant cell wall degradosome Seed
Dickeya dadantii NCPPB 898 WP 013319746 1 Ddadantii
Dickeya solani RNS 05 1 2A WP 057085093 1 Dsolani
Dickeya solani D s0432 1 WP 022635158 1 Dsolani
Dickeya solani RNS 07 7 3B WP 022635158 1 Dsolani
Dickeya solani IPO 2222 WP 022635158 1 Dsolani
Dickeya solani PPO 9019 WP 022635158 1 Dsolani
Dickeya solani RNS 08 23 3 1 A WP 022635158 1 Dsolani
Dickeya solani IFB 0099 WP 022635158 1 Dsolani
Dickeya solani MK10 WP 022635158 1 Dsolani
Dickeya solani GBBC 2040 WP 022635158 1 Dsolani
Dickeya solani MK16 WP 022635158 1 Dsolani
Dickeya solani PPO 9134 WP 022635158 1 Dsolani
Dickeya gs undicola 2B12 WP 033570853 1 Dundicola
Dickeya zeae NCPPB 2538 WP 038916898 1 Dzeae
Dickeya zeae MK19 WP 038914746 1 Dzeae
Dickeya zeae Ech586 WP 012886392 1 Dzeae
Dickeya zeae MS1 WP 019845163 1 Dzeae
Dickeya zeae CSL RW192 WP 038909180 1 Dzeae
Dickeya zeae NCPPB 3532 WP 023640897 1 Dzeae
Dickeya zeae DZ2Q WP 016942965 1 Dzeae
Dickeya zeae ZJU1202 WP 016942965 1 Dzeae
Dickeya zeae EC1 WP 016942965 1 Dzeae
Dickeya zeae NCPPB 3531 WP 038905854 1 Dzeae
DAQ1742 DAQ1742 04113 Daquatica
Dickeya aquatica DW 0440 WP 035344630 1 Daquatica
Dickeya sp CSL RW240 WP 035344630 1 Daquatica
Dickeya gs poaceaephila NCPPB 569 WP 042873156 1 Dpoaceaephila
Dickeya dianthicola NCPPB 3534 WP 024107562 1 Ddianthicola
Dickeya dianthicola GBBC 2039 WP 024107562 1 Ddianthicola
Dickeya dianthicola IPO 980 WP 024107562 1 Ddianthicola
Dickeya dianthicola RNS04 9 WP 024107562 1 Ddianthicola
Dickeya dianthicola NCPPB 453 WP 024107562 1 Ddianthicola
Dickeya chrysanthemi subsp parthenii NCPPB 516 WP 027710727 1 Dchrysanthemi
Dickeya chrysanthemi subsp chrysanthemi NCPPB 402 WP 039999247 1 Dchrysanthemi
Dickeya chrysanthemi L11 WP 033577033 1 Dchrysanthemi
Dickeya chrysanthemi NCPPB 3533 WP 012768051 1 Dchrysanthemi
Dickeya chrysanthemi Ech1591 WP 012768051 1 Dchrysanthemi

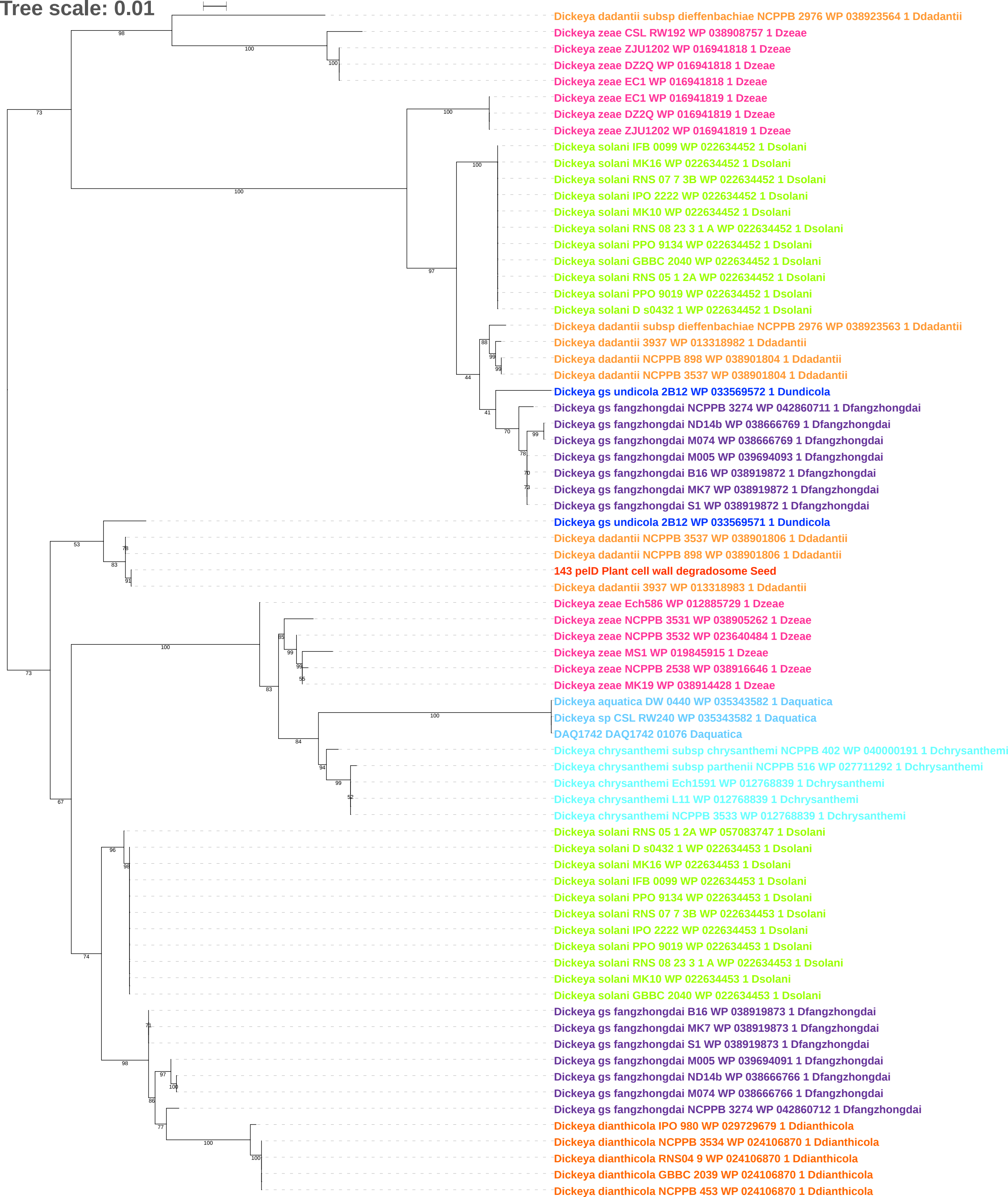
Tree scale: 0.01
Dickeya dadantii subsp dieffenbachiae NCPPB 2976 WP 038923564 1 Ddadantii
Dickeya zeae CSL RW192 WP 038908757 1 Dzeae
Dickeya zeae ZJU1202 WP 016941818 1 Dzeae
Dickeya zeae DZ2Q WP 016941818 1 Dzeae
Dickeya zeae EC1 WP 016941818 1 Dzeae
Dickeya zeae EC1 WP 016941819 1 Dzeae
Dickeya zeae DZ2Q WP 016941819 1 Dzeae
Dickeya zeae ZJU1202 WP 016941819 1 Dzeae
Dickeya solani IFB 0099 WP 022634452 1 Dsolani
Dickeya solani MK16 WP 022634452 1 Dsolani
Dickeya solani RNS 07 7 3B WP 022634452 1 Dsolani
Dickeya solani IPO 2222 WP 022634452 1 Dsolani
Dickeya solani MK10 WP 022634452 1 Dsolani
Dickeya solani RNS 08 23 3 1 A WP 022634452 1 Dsolani
Dickeya solani PPO 9134 WP 022634452 1 Dsolani
Dickeya solani GBBC 2040 WP 022634452 1 Dsolani
Dickeya solani RNS 05 1 2A WP 022634452 1 Dsolani
Dickeya solani PPO 9019 WP 022634452 1 Dsolani
Dickeya solani D s0432 1 WP 022634452 1 Dsolani
Dickeya dadantii subsp dieffenbachiae NCPPB 2976 WP 038923563 1 Ddadantii
Dickeya dadantii 3937 WP 013318982 1 Ddadantii
Dickeya dadantii NCPPB 898 WP 038901804 1 Ddadantii
Dickeya dadantii NCPPB 3537 WP 038901804 1 Ddadantii
Dickeya gs undicola 2B12 WP 033569572 1 Dundicola
Dickeya gs fangzhongdai NCPPB 3274 WP 042860711 1 Dfangzhongdai
Dickeya gs fangzhongdai ND14b WP 038666769 1 Dfangzhongdai
Dickeya gs fangzhongdai M074 WP 038666769 1 Dfangzhongdai
Dickeya gs fangzhongdai M005 WP 039694093 1 Dfangzhongdai
Dickeya gs fangzhongdai B16 WP 038919872 1 Dfangzhongdai
Dickeya gs fangzhongdai MK7 WP 038919872 1 Dfangzhongdai
Dickeya gs fangzhongdai S1 WP 038919872 1 Dfangzhongdai
Dickeya gs undicola 2B12 WP 033569571 1 Dundicola
Dickeya dadantii NCPPB 3537 WP 038901806 1 Ddadantii
Dickeya dadantii NCPPB 898 WP 038901806 1 Ddadantii
143 pelD Plant cell wall degradosome Seed
Dickeya dadantii 3937 WP 013318983 1 Ddadantii
Dickeya zeae Ech586 WP 012885729 1 Dzeae
Dickeya zeae NCPPB 3531 WP 038905262 1 Dzeae
Dickeya zeae NCPPB 3532 WP 023640484 1 Dzeae
Dickeya zeae MS1 WP 019845915 1 Dzeae
Dickeya zeae NCPPB 2538 WP 038916646 1 Dzeae
Dickeya zeae MK19 WP 038914428 1 Dzeae
Dickeya aquatica DW 0440 WP 035343582 1 Daquatica
Dickeya sp CSL RW240 WP 035343582 1 Daquatica
DAQ1742 DAQ1742 01076 Daquatica
Dickeya chrysanthemi subsp chrysanthemi NCPPB 402 WP 040000191 1 Dchrysanthemi
Dickeya chrysanthemi subsp parthenii NCPPB 516 WP 027711292 1 Dchrysanthemi
Dickeya chrysanthemi Ech1591 WP 012768839 1 Dchrysanthemi
Dickeya chrysanthemi L11 WP 012768839 1 Dchrysanthemi
Dickeya chrysanthemi NCPPB 3533 WP 012768839 1 Dchrysanthemi
Dickeya solani RNS 05 1 2A WP 057083747 1 Dsolani
Dickeya solani D s0432 1 WP 022634453 1 Dsolani
Dickeya solani MK16 WP 022634453 1 Dsolani
Dickeya solani IFB 0099 WP 022634453 1 Dsolani
Dickeya solani PPO 9134 WP 022634453 1 Dsolani
Dickeya solani RNS 07 7 3B WP 022634453 1 Dsolani
Dickeya solani IPO 2222 WP 022634453 1 Dsolani
Dickeya solani PPO 9019 WP 022634453 1 Dsolani
Dickeya solani RNS 08 23 3 1 A WP 022634453 1 Dsolani
Dickeya solani MK10 WP 022634453 1 Dsolani
Dickeya solani GBBC 2040 WP 022634453 1 Dsolani
Dickeya gs fangzhongdai B16 WP 038919873 1 Dfangzhongdai
Dickeya gs fangzhongdai MK7 WP 038919873 1 Dfangzhongdai
Dickeya gs fangzhongdai S1 WP 038919873 1 Dfangzhongdai
Dickeya gs fangzhongdai M005 WP 039694091 1 Dfangzhongdai
Dickeya gs fangzhongdai ND14b WP 038666766 1 Dfangzhongdai
Dickeya gs fangzhongdai M074 WP 038666766 1 Dfangzhongdai
Dickeya gs fangzhongdai NCPPB 3274 WP 042860712 1 Dfangzhongdai
Dickeya dianthicola IPO 980 WP 029729679 1 Ddianthicola
Dickeya dianthicola NCPPB 3534 WP 024106870 1 Ddianthicola
Dickeya dianthicola RNS04 9 WP 024106870 1 Ddianthicola
Dickeya dianthicola GBBC 2039 WP 024106870 1 Ddianthicola
Dickeya dianthicola NCPPB 453 WP 024106870 1 Ddianthicola

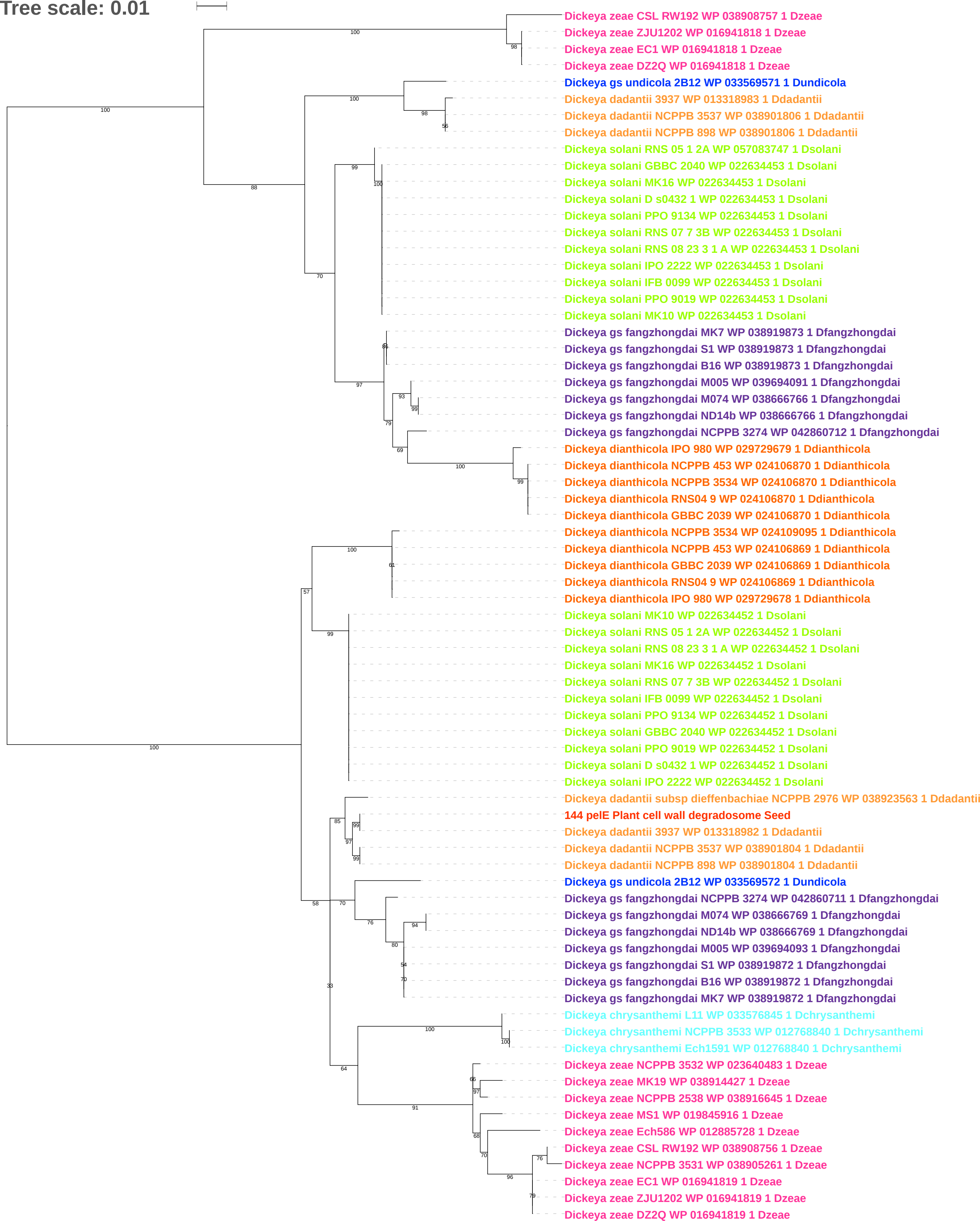
Tree scale: 0.01
Dickeya zeae CSL RW192 WP 038908757 1 Dzeae
Dickeya zeae ZJU1202 WP 016941818 1 Dzeae
Dickeya zeae EC1 WP 016941818 1 Dzeae
Dickeya zeae DZ2Q WP 016941818 1 Dzeae
Dickeya gs undicola 2B12 WP 033569571 1 Dundicola
Dickeya dadantii 3937 WP 013318983 1 Ddadantii
Dickeya dadantii NCPPB 3537 WP 038901806 1 Ddadantii
Dickeya dadantii NCPPB 898 WP 038901806 1 Ddadantii
Dickeya solani RNS 05 1 2A WP 057083747 1 Dsolani
Dickeya solani GBBC 2040 WP 022634453 1 Dsolani
Dickeya solani MK16 WP 022634453 1 Dsolani
Dickeya solani D s0432 1 WP 022634453 1 Dsolani
Dickeya solani PPO 9134 WP 022634453 1 Dsolani
Dickeya solani RNS 07 7 3B WP 022634453 1 Dsolani
Dickeya solani RNS 08 23 3 1 A WP 022634453 1 Dsolani
Dickeya solani IPO 2222 WP 022634453 1 Dsolani
Dickeya solani IFB 0099 WP 022634453 1 Dsolani
Dickeya solani PPO 9019 WP 022634453 1 Dsolani
Dickeya solani MK10 WP 022634453 1 Dsolani
Dickeya gs fangzhongdai MK7 WP 038919873 1 Dfangzhongdai
Dickeya gs fangzhongdai S1 WP 038919873 1 Dfangzhongdai
Dickeya gs fangzhongdai B16 WP 038919873 1 Dfangzhongdai
Dickeya gs fangzhongdai M005 WP 039694091 1 Dfangzhongdai
Dickeya gs fangzhongdai M074 WP 038666766 1 Dfangzhongdai
Dickeya gs fangzhongdai ND14b WP 038666766 1 Dfangzhongdai
Dickeya gs fangzhongdai NCPPB 3274 WP 042860712 1 Dfangzhongdai
Dickeya dianthicola IPO 980 WP 029729679 1 Ddianthicola
Dickeya dianthicola NCPPB 453 WP 024106870 1 Ddianthicola
Dickeya dianthicola NCPPB 3534 WP 024106870 1 Ddianthicola
Dickeya dianthicola RNS04 9 WP 024106870 1 Ddianthicola
Dickeya dianthicola GBBC 2039 WP 024106870 1 Ddianthicola
Dickeya dianthicola NCPPB 3534 WP 024109095 1 Ddianthicola
Dickeya dianthicola NCPPB 453 WP 024106869 1 Ddianthicola
Dickeya dianthicola GBBC 2039 WP 024106869 1 Ddianthicola
Dickeya dianthicola RNS04 9 WP 024106869 1 Ddianthicola
Dickeya dianthicola IPO 980 WP 029729678 1 Ddianthicola
Dickeya solani MK10 WP 022634452 1 Dsolani
Dickeya solani RNS 05 1 2A WP 022634452 1 Dsolani
Dickeya solani RNS 08 23 3 1 A WP 022634452 1 Dsolani
Dickeya solani MK16 WP 022634452 1 Dsolani
Dickeya solani RNS 07 7 3B WP 022634452 1 Dsolani
Dickeya solani IFB 0099 WP 022634452 1 Dsolani
Dickeya solani PPO 9134 WP 022634452 1 Dsolani
Dickeya solani GBBC 2040 WP 022634452 1 Dsolani
Dickeya solani PPO 9019 WP 022634452 1 Dsolani
Dickeya solani D s0432 1 WP 022634452 1 Dsolani
Dickeya solani IPO 2222 WP 022634452 1 Dsolani
Dickeya dadantii subsp dieffenbachiae NCPPB 2976 WP 038923563 1 Ddadantii
144 pelE Plant cell wall degradosome Seed
Dickeya dadantii 3937 WP 013318982 1 Ddadantii
Dickeya dadantii NCPPB 3537 WP 038901804 1 Ddadantii
Dickeya dadantii NCPPB 898 WP 038901804 1 Ddadantii
Dickeya gs undicola 2B12 WP 033569572 1 Dundicola
Dickeya gs fangzhongdai NCPPB 3274 WP 042860711 1 Dfangzhongdai
Dickeya gs fangzhongdai M074 WP 038666769 1 Dfangzhongdai
Dickeya gs fangzhongdai ND14b WP 038666769 1 Dfangzhongdai
Dickeya gs fangzhongdai M005 WP 039694093 1 Dfangzhongdai
Dickeya gs fangzhongdai S1 WP 038919872 1 Dfangzhongdai
Dickeya gs fangzhongdai B16 WP 038919872 1 Dfangzhongdai
Dickeya gs fangzhongdai MK7 WP 038919872 1 Dfangzhongdai
Dickeya chrysanthemi L11 WP 033576845 1 Dchrysanthemi
Dickeya chrysanthemi NCPPB 3533 WP 012768840 1 Dchrysanthemi
Dickeya chrysanthemi Ech1591 WP 012768840 1 Dchrysanthemi
Dickeya zeae NCPPB 3532 WP 023640483 1 Dzeae
Dickeya zeae MK19 WP 038914427 1 Dzeae
Dickeya zeae NCPPB 2538 WP 038916645 1 Dzeae
Dickeya zeae MS1 WP 019845916 1 Dzeae
Dickeya zeae Ech586 WP 012885728 1 Dzeae
Dickeya zeae CSL RW192 WP 038908756 1 Dzeae
Dickeya zeae NCPPB 3531 WP 038905261 1 Dzeae
Dickeya zeae EC1 WP 016941819 1 Dzeae
Dickeya zeae ZJU1202 WP 016941819 1 Dzeae
Dickeya zeae DZ2Q WP 016941819 1 Dzeae

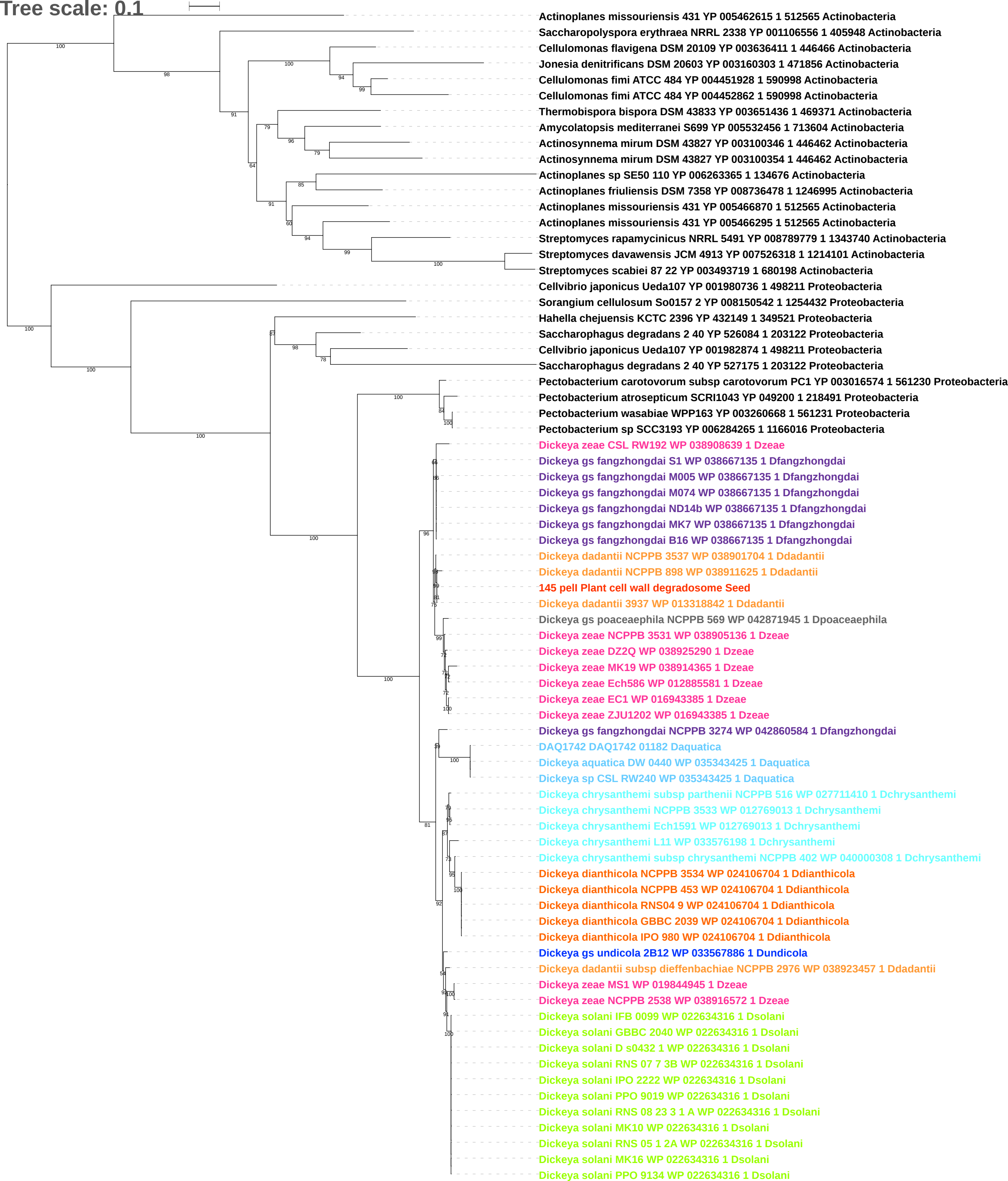
Tree scale: 0.1
Actinoplanes missouriensis 431 YP 005462615 1 512565 Actinobacteria
Saccharopolyspora erythraea NRRL 2338 YP 001106556 1 405948 Actinobacteria
Cellulomonas flavigena DSM 20109 YP 003636411 1 446466 Actinobacteria
Jonesia denitrificans DSM 20603 YP 003160303 1 471856 Actinobacteria
Cellulomonas fimi ATCC 484 YP 004451928 1 590998 Actinobacteria
Cellulomonas fimi ATCC 484 YP 004452862 1 590998 Actinobacteria
Thermobispora bispora DSM 43833 YP 003651436 1 469371 Actinobacteria
Amycolatopsis mediterranei S699 YP 005532456 1 713604 Actinobacteria
Actinosynnema mirum DSM 43827 YP 003100346 1 446462 Actinobacteria
Actinosynnema mirum DSM 43827 YP 003100354 1 446462 Actinobacteria
Actinoplanes sp SE50 110 YP 006263365 1 134676 Actinobacteria
Actinoplanes friuliensis DSM 7358 YP 008736478 1 1246995 Actinobacteria
Actinoplanes missouriensis 431 YP 005466870 1 512565 Actinobacteria
Actinoplanes missouriensis 431 YP 005466295 1 512565 Actinobacteria
Streptomyces rapamycinicus NRRL 5491 YP 008789779 1 1343740 Actinobacteria
Streptomyces davawensis JCM 4913 YP 007526318 1 1214101 Actinobacteria
Streptomyces scabiei 87 22 YP 003493719 1 680198 Actinobacteria
Cellvibrio japonicus Ueda107 YP 001980736 1 498211 Proteobacteria
Sorangium cellulosum So0157 2 YP 008150542 1 1254432 Proteobacteria
Hahella chejuensis KCTC 2396 YP 432149 1 349521 Proteobacteria
Saccharophagus degradans 2 40 YP 526084 1 203122 Proteobacteria
Cellvibrio japonicus Ueda107 YP 001982874 1 498211 Proteobacteria
Saccharophagus degradans 2 40 YP 527175 1 203122 Proteobacteria
Pectobacterium carotovorum subsp carotovorum PC1 YP 003016574 1 561230 Proteobacteria
Pectobacterium atrosepticum SCRI1043 YP 049200 1 218491 Proteobacteria
Pectobacterium wasabiae WPP163 YP 003260668 1 561231 Proteobacteria
Pectobacterium sp SCC3193 YP 006284265 1 1166016 Proteobacteria
Dickeya zeae CSL RW192 WP 038908639 1 Dzeae
Dickeya gs fangzhongdai S1 WP 038667135 1 Dfangzhongdai
Dickeya gs fangzhongdai M005 WP 038667135 1 Dfangzhongdai
Dickeya gs fangzhongdai M074 WP 038667135 1 Dfangzhongdai
Dickeya gs fangzhongdai ND14b WP 038667135 1 Dfangzhongdai
Dickeya gs fangzhongdai MK7 WP 038667135 1 Dfangzhongdai
Dickeya gs fangzhongdai B16 WP 038667135 1 Dfangzhongdai
Dickeya dadantii NCPPB 3537 WP 038901704 1 Ddadantii
Dickeya dadantii NCPPB 898 WP 038911625 1 Ddadantii
145 pelI Plant cell wall degradosome Seed
Dickeya dadantii 3937 WP 013318842 1 Ddadantii
Dickeya gs poaceaephila NCPPB 569 WP 042871945 1 Dpoaceaephila
Dickeya zeae NCPPB 3531 WP 038905136 1 Dzeae
Dickeya zeae DZ2Q WP 038925290 1 Dzeae
Dickeya zeae MK19 WP 038914365 1 Dzeae
Dickeya zeae Ech586 WP 012885581 1 Dzeae
Dickeya zeae EC1 WP 016943385 1 Dzeae
Dickeya zeae ZJU1202 WP 016943385 1 Dzeae
Dickeya gs fangzhongdai NCPPB 3274 WP 042860584 1 Dfangzhongdai
DAQ1742 DAQ1742 01182 Daquatica
Dickeya aquatica DW 0440 WP 035343425 1 Daquatica
Dickeya sp CSL RW240 WP 035343425 1 Daquatica
Dickeya chrysanthemi subsp parthenii NCPPB 516 WP 027711410 1 Dchrysanthemi
Dickeya chrysanthemi NCPPB 3533 WP 012769013 1 Dchrysanthemi
Dickeya chrysanthemi Ech1591 WP 012769013 1 Dchrysanthemi
Dickeya chrysanthemi L11 WP 033576198 1 Dchrysanthemi
Dickeya chrysanthemi subsp chrysanthemi NCPPB 402 WP 040000308 1 Dchrysanthemi
Dickeya dianthicola NCPPB 3534 WP 024106704 1 Ddianthicola
Dickeya dianthicola NCPPB 453 WP 024106704 1 Ddianthicola
Dickeya dianthicola RNS04 9 WP 024106704 1 Ddianthicola
Dickeya dianthicola GBBC 2039 WP 024106704 1 Ddianthicola
Dickeya dianthicola IPO 980 WP 024106704 1 Ddianthicola
Dickeya gs undicola 2B12 WP 033567886 1 Dundicola
Dickeya dadantii subsp dieffenbachiae NCPPB 2976 WP 038923457 1 Ddadantii
Dickeya zeae MS1 WP 019844945 1 Dzeae
Dickeya zeae NCPPB 2538 WP 038916572 1 Dzeae
Dickeya solani IFB 0099 WP 022634316 1 Dsolani
Dickeya solani GBBC 2040 WP 022634316 1 Dsolani
Dickeya solani D s0432 1 WP 022634316 1 Dsolani
Dickeya solani RNS 07 7 3B WP 022634316 1 Dsolani
Dickeya solani IPO 2222 WP 022634316 1 Dsolani
Dickeya solani PPO 9019 WP 022634316 1 Dsolani
Dickeya solani RNS 08 23 3 1 A WP 022634316 1 Dsolani
Dickeya solani MK10 WP 022634316 1 Dsolani
Dickeya solani RNS 05 1 2A WP 022634316 1 Dsolani
Dickeya solani MK16 WP 022634316 1 Dsolani
Dickeya solani PPO 9134 WP 022634316 1 Dsolani

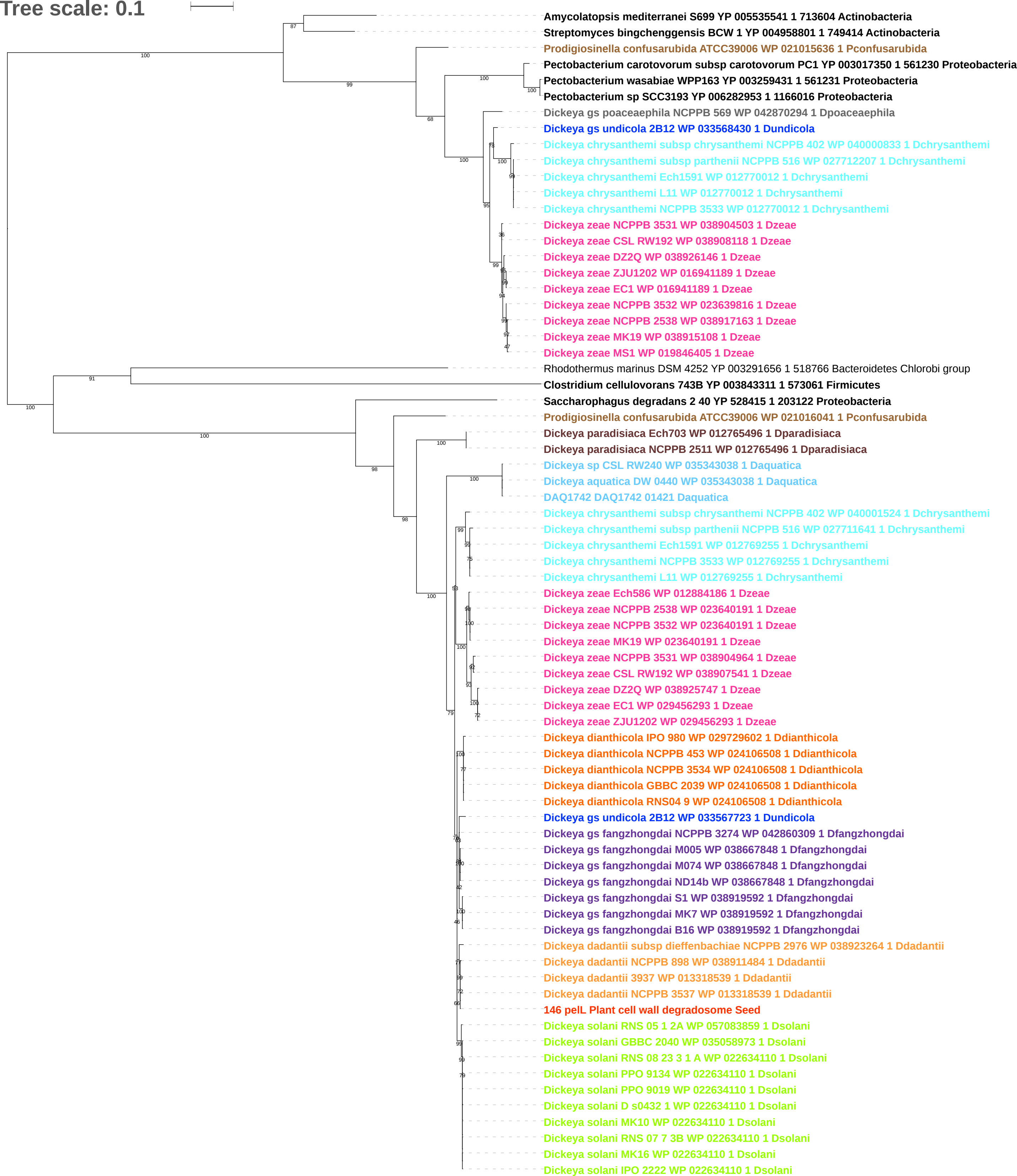
Tree scale: 0.1
Amycolatopsis mediterranei S699 YP 005535541 1 713604 Actinobacteria
Streptomyces bingchenggensis BCW 1 YP 004958801 1 749414 Actinobacteria
Prodigiosinella confusarubida ATCC39006 WP 021015636 1 Pconfusarubida
Pectobacterium carotovorum subsp carotovorum PC1 YP 003017350 1 561230 Proteobacteria
Pectobacterium wasabiae WPP163 YP 003259431 1 561231 Proteobacteria
Pectobacterium sp SCC3193 YP 006282953 1 1166016 Proteobacteria
Dickeya gs poaceaephila NCPPB 569 WP 042870294 1 Dpoaceaephila
Dickeya gs undicola 2B12 WP 033568430 1 Dundicola
Dickeya chrysanthemi subsp chrysanthemi NCPPB 402 WP 040000833 1 Dchrysanthemi
Dickeya chrysanthemi subsp parthenii NCPPB 516 WP 027712207 1 Dchrysanthemi
Dickeya chrysanthemi Ech1591 WP 012770012 1 Dchrysanthemi
Dickeya chrysanthemi L11 WP 012770012 1 Dchrysanthemi
Dickeya chrysanthemi NCPPB 3533 WP 012770012 1 Dchrysanthemi
Dickeya zeae NCPPB 3531 WP 038904503 1 Dzeae
Dickeya zeae CSL RW192 WP 038908118 1 Dzeae
Dickeya zeae DZ2Q WP 038926146 1 Dzeae
Dickeya zeae ZJU1202 WP 016941189 1 Dzeae
Dickeya zeae EC1 WP 016941189 1 Dzeae
Dickeya zeae NCPPB 3532 WP 023639816 1 Dzeae
Dickeya zeae NCPPB 2538 WP 038917163 1 Dzeae
Dickeya zeae MK19 WP 038915108 1 Dzeae
Dickeya zeae MS1 WP 019846405 1 Dzeae
Rhodothermus marinus DSM 4252 YP 003291656 1 518766 Bacteroidetes Chlorobi group
Clostridium cellulovorans 743B YP 003843311 1 573061 Firmicutes
Saccharophagus degradans 2 40 YP 528415 1 203122 Proteobacteria
Prodigiosinella confusarubida ATCC39006 WP 021016041 1 Pconfusarubida
Dickeya paradisiaca Ech703 WP 012765496 1 Dparadisiaca
Dickeya paradisiaca NCPPB 2511 WP 012765496 1 Dparadisiaca
Dickeya sp CSL RW240 WP 035343038 1 Daquatica
Dickeya aquatica DW 0440 WP 035343038 1 Daquatica
DAQ1742 DAQ1742 01421 Daquatica
Dickeya chrysanthemi subsp chrysanthemi NCPPB 402 WP 040001524 1 Dchrysanthemi
Dickeya chrysanthemi subsp parthenii NCPPB 516 WP 027711641 1 Dchrysanthemi
Dickeya chrysanthemi Ech1591 WP 012769255 1 Dchrysanthemi
Dickeya chrysanthemi NCPPB 3533 WP 012769255 1 Dchrysanthemi
Dickeya chrysanthemi L11 WP 012769255 1 Dchrysanthemi
Dickeya zeae Ech586 WP 012884186 1 Dzeae
Dickeya zeae NCPPB 2538 WP 023640191 1 Dzeae
Dickeya zeae NCPPB 3532 WP 023640191 1 Dzeae
Dickeya zeae MK19 WP 023640191 1 Dzeae
Dickeya zeae NCPPB 3531 WP 038904964 1 Dzeae
Dickeya zeae CSL RW192 WP 038907541 1 Dzeae
Dickeya zeae DZ2Q WP 038925747 1 Dzeae
Dickeya zeae EC1 WP 029456293 1 Dzeae
Dickeya zeae ZJU1202 WP 029456293 1 Dzeae
Dickeya dianthicola IPO 980 WP 029729602 1 Ddianthicola
Dickeya dianthicola NCPPB 453 WP 024106508 1 Ddianthicola
Dickeya dianthicola NCPPB 3534 WP 024106508 1 Ddianthicola
Dickeya dianthicola GBBC 2039 WP 024106508 1 Ddianthicola
Dickeya dianthicola RNS04 9 WP 024106508 1 Ddianthicola
Dickeya gs undicola 2B12 WP 033567723 1 Dundicola
Dickeya gs fangzhongdai NCPPB 3274 WP 042860309 1 Dfangzhongdai
Dickeya gs fangzhongdai M005 WP 038667848 1 Dfangzhongdai
Dickeya gs fangzhongdai M074 WP 038667848 1 Dfangzhongdai
Dickeya gs fangzhongdai ND14b WP 038667848 1 Dfangzhongdai
Dickeya gs fangzhongdai S1 WP 038919592 1 Dfangzhongdai
Dickeya gs fangzhongdai MK7 WP 038919592 1 Dfangzhongdai
Dickeya gs fangzhongdai B16 WP 038919592 1 Dfangzhongdai
Dickeya dadantii subsp dieffenbachiae NCPPB 2976 WP 038923264 1 Ddadantii
Dickeya dadantii NCPPB 898 WP 038911484 1 Ddadantii
Dickeya dadantii 3937 WP 013318539 1 Ddadantii
Dickeya dadantii NCPPB 3537 WP 013318539 1 Ddadantii
146 pelL Plant cell wall degradosome Seed
Dickeya solani RNS 05 1 2A WP 057083859 1 Dsolani
Dickeya solani GBBC 2040 WP 035058973 1 Dsolani
Dickeya solani RNS 08 23 3 1 A WP 022634110 1 Dsolani
Dickeya solani PPO 9134 WP 022634110 1 Dsolani
Dickeya solani PPO 9019 WP 022634110 1 Dsolani
Dickeya solani D s0432 1 WP 022634110 1 Dsolani
Dickeya solani MK10 WP 022634110 1 Dsolani
Dickeya solani RNS 07 7 3B WP 022634110 1 Dsolani
Dickeya solani MK16 WP 022634110 1 Dsolani
Dickeya solani IPO 2222 WP 022634110 1 Dsolani

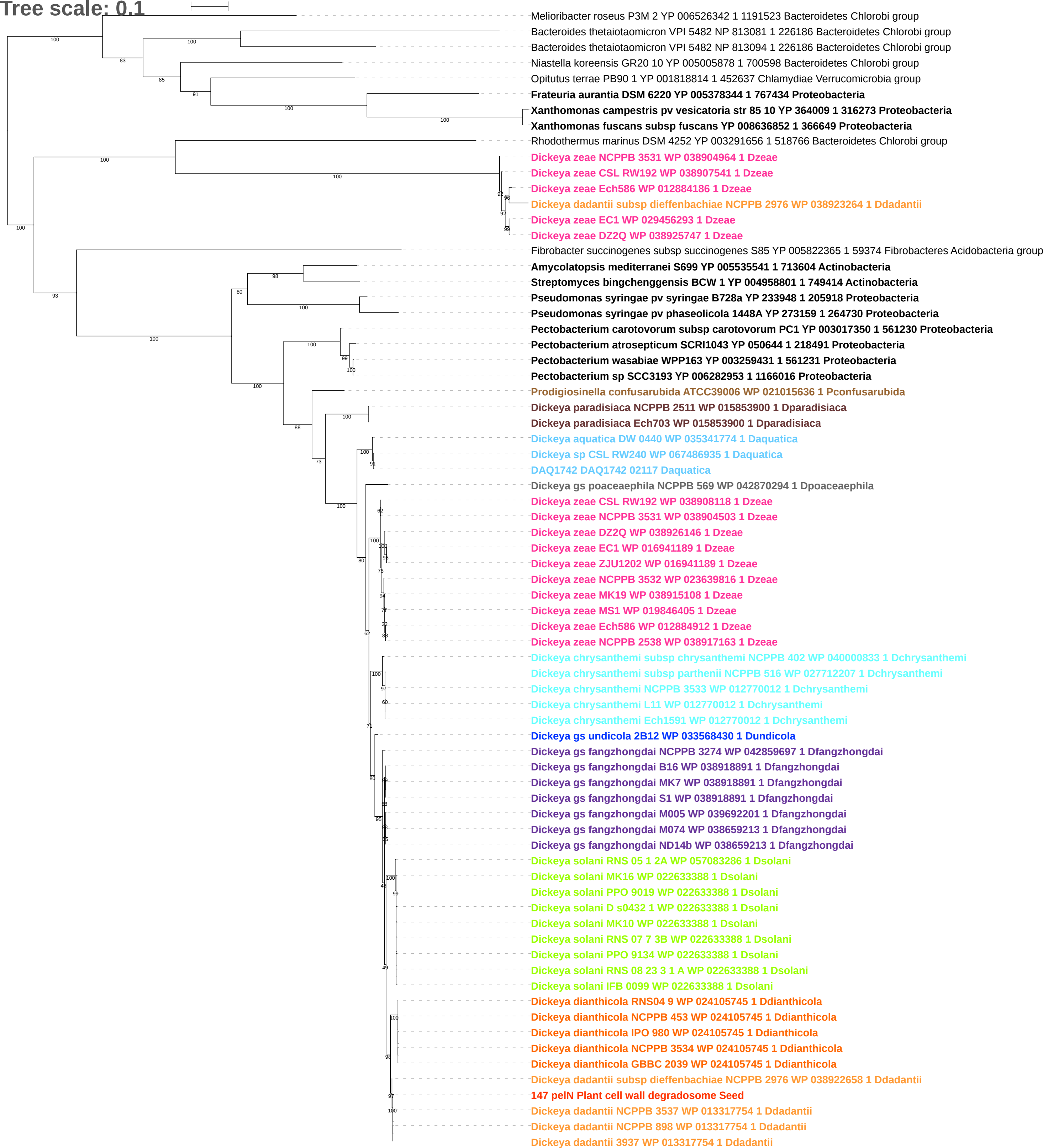
Tree scale: 0.1
Melioribacter roseus P3M 2 YP 006526342 1 1191523 Bacteroidetes Chlorobi group
Bacteroides thetaiotaomicron VPI 5482 NP 813081 1 226186 Bacteroidetes Chlorobi group
Bacteroides thetaiotaomicron VPI 5482 NP 813094 1 226186 Bacteroidetes Chlorobi group
Niastella koreensis GR20 10 YP 005005878 1 700598 Bacteroidetes Chlorobi group
Opitutus terrae PB90 1 YP 001818814 1 452637 Chlamydiae Verrucomicrobia group
Frateuria aurantia DSM 6220 YP 005378344 1 767434 Proteobacteria
Xanthomonas campestris pv vesicatoria str 85 10 YP 364009 1 316273 Proteobacteria
Xanthomonas fuscans subsp fuscans YP 008636852 1 366649 Proteobacteria
Rhodothermus marinus DSM 4252 YP 003291656 1 518766 Bacteroidetes Chlorobi group
Dickeya zeae NCPPB 3531 WP 038904964 1 Dzeae
Dickeya zeae CSL RW192 WP 038907541 1 Dzeae
Dickeya zeae Ech586 WP 012884186 1 Dzeae
Dickeya dadantii subsp dieffenbachiae NCPPB 2976 WP 038923264 1 Ddadantii
Dickeya zeae EC1 WP 029456293 1 Dzeae
Dickeya zeae DZ2Q WP 038925747 1 Dzeae
Fibrobacter succinogenes subsp succinogenes S85 YP 005822365 1 59374 Fibrobacteres Acidobacteria group
Amycolatopsis mediterranei S699 YP 005535541 1 713604 Actinobacteria
Streptomyces bingchenggensis BCW 1 YP 004958801 1 749414 Actinobacteria
Pseudomonas syringae pv syringae B728a YP 233948 1 205918 Proteobacteria
Pseudomonas syringae pv phaseolicola 1448A YP 273159 1 264730 Proteobacteria
Pectobacterium carotovorum subsp carotovorum PC1 YP 003017350 1 561230 Proteobacteria
Pectobacterium atrosepticum SCRI1043 YP 050644 1 218491 Proteobacteria
Pectobacterium wasabiae WPP163 YP 003259431 1 561231 Proteobacteria
Pectobacterium sp SCC3193 YP 006282953 1 1166016 Proteobacteria
Prodigiosinella confusarubida ATCC39006 WP 021015636 1 Pconfusarubida
Dickeya paradisiaca NCPPB 2511 WP 015853900 1 Dparadisiaca
Dickeya paradisiaca Ech703 WP 015853900 1 Dparadisiaca
Dickeya aquatica DW 0440 WP 035341774 1 Daquatica
Dickeya sp CSL RW240 WP 067486935 1 Daquatica
DAQ1742 DAQ1742 02117 Daquatica
Dickeya gs poaceaephila NCPPB 569 WP 042870294 1 Dpoaceaephila
Dickeya zeae CSL RW192 WP 038908118 1 Dzeae
Dickeya zeae NCPPB 3531 WP 038904503 1 Dzeae
Dickeya zeae DZ2Q WP 038926146 1 Dzeae
Dickeya zeae EC1 WP 016941189 1 Dzeae
Dickeya zeae ZJU1202 WP 016941189 1 Dzeae
Dickeya zeae NCPPB 3532 WP 023639816 1 Dzeae
Dickeya zeae MK19 WP 038915108 1 Dzeae
Dickeya zeae MS1 WP 019846405 1 Dzeae
Dickeya zeae Ech586 WP 012884912 1 Dzeae
Dickeya zeae NCPPB 2538 WP 038917163 1 Dzeae
Dickeya chrysanthemi subsp chrysanthemi NCPPB 402 WP 040000833 1 Dchrysanthemi
Dickeya chrysanthemi subsp parthenii NCPPB 516 WP 027712207 1 Dchrysanthemi
Dickeya chrysanthemi NCPPB 3533 WP 012770012 1 Dchrysanthemi
Dickeya chrysanthemi L11 WP 012770012 1 Dchrysanthemi
Dickeya chrysanthemi Ech1591 WP 012770012 1 Dchrysanthemi
Dickeya gs undicola 2B12 WP 033568430 1 Dundicola
Dickeya gs fangzhongdai NCPPB 3274 WP 042859697 1 Dfangzhongdai
Dickeya gs fangzhongdai B16 WP 038918891 1 Dfangzhongdai
Dickeya gs fangzhongdai MK7 WP 038918891 1 Dfangzhongdai
Dickeya gs fangzhongdai S1 WP 038918891 1 Dfangzhongdai
Dickeya gs fangzhongdai M005 WP 039692201 1 Dfangzhongdai
Dickeya gs fangzhongdai M074 WP 038659213 1 Dfangzhongdai
Dickeya gs fangzhongdai ND14b WP 038659213 1 Dfangzhongdai
Dickeya solani RNS 05 1 2A WP 057083286 1 Dsolani
Dickeya solani MK16 WP 022633388 1 Dsolani
Dickeya solani PPO 9019 WP 022633388 1 Dsolani
Dickeya solani D s0432 1 WP 022633388 1 Dsolani
Dickeya solani MK10 WP 022633388 1 Dsolani
Dickeya solani RNS 07 7 3B WP 022633388 1 Dsolani
Dickeya solani PPO 9134 WP 022633388 1 Dsolani
Dickeya solani RNS 08 23 3 1 A WP 022633388 1 Dsolani
Dickeya solani IFB 0099 WP 022633388 1 Dsolani
Dickeya dianthicola RNS04 9 WP 024105745 1 Ddianthicola
Dickeya dianthicola NCPPB 453 WP 024105745 1 Ddianthicola
Dickeya dianthicola IPO 980 WP 024105745 1 Ddianthicola
Dickeya dianthicola NCPPB 3534 WP 024105745 1 Ddianthicola
Dickeya dianthicola GBBC 2039 WP 024105745 1 Ddianthicola
Dickeya dadantii subsp dieffenbachiae NCPPB 2976 WP 038922658 1 Ddadantii
147 pelN Plant cell wall degradosome Seed
Dickeya dadantii NCPPB 3537 WP 013317754 1 Ddadantii
Dickeya dadantii NCPPB 898 WP 013317754 1 Ddadantii
Dickeya dadantii 3937 WP 013317754 1 Ddadantii

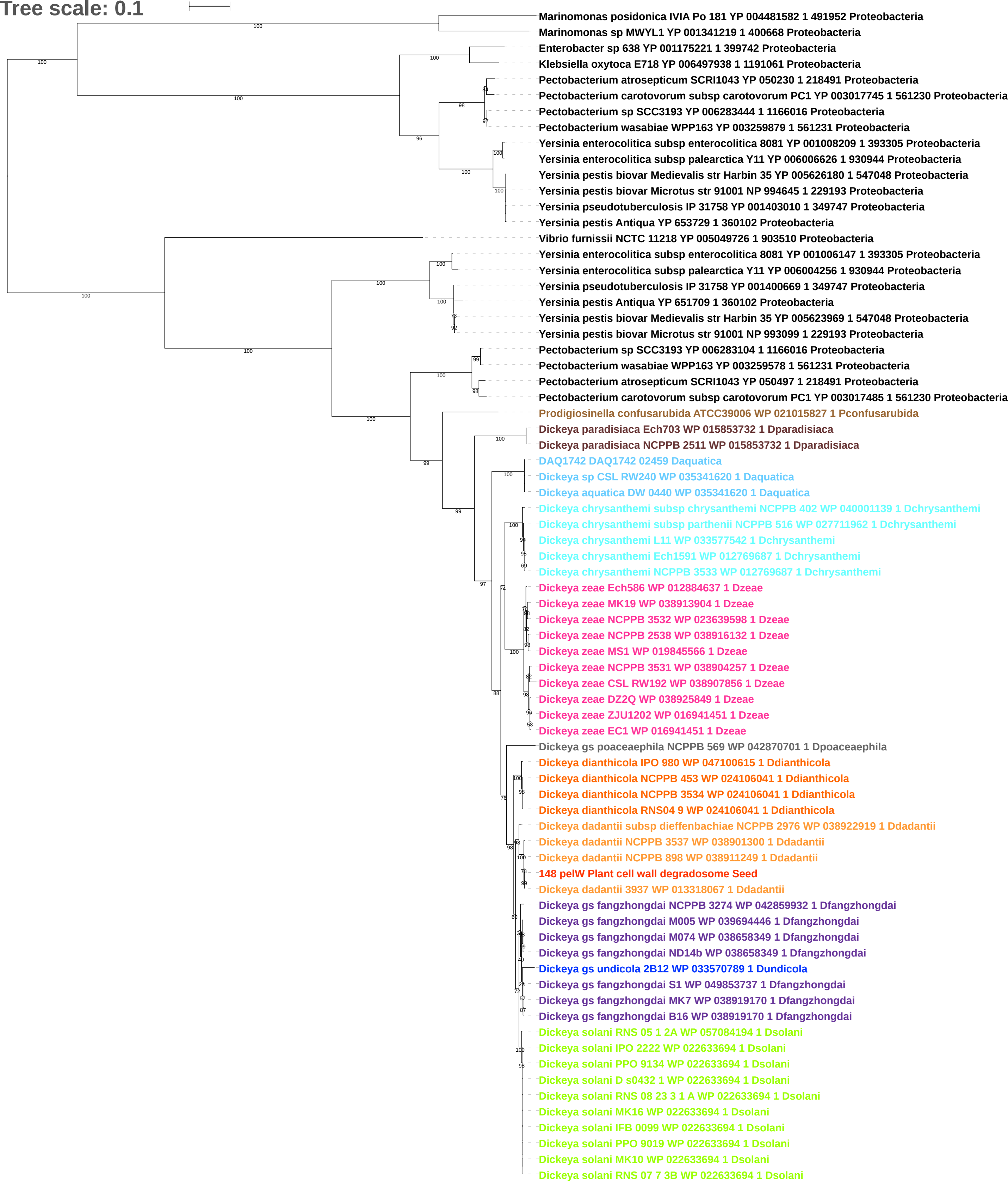
Tree scale: 0.1
Marinomonas posidonica IVIA Po 181 YP 004481582 1 491952 Proteobacteria
Marinomonas sp MWYL1 YP 001341219 1 400668 Proteobacteria
Enterobacter sp 638 YP 001175221 1 399742 Proteobacteria
Klebsiella oxytoca E718 YP 006497938 1 1191061 Proteobacteria
Pectobacterium atrosepticum SCRI1043 YP 050230 1 218491 Proteobacteria
Pectobacterium carotovorum subsp carotovorum PC1 YP 003017745 1 561230 Proteobacteria
Pectobacterium sp SCC3193 YP 006283444 1 1166016 Proteobacteria
Pectobacterium wasabiae WPP163 YP 003259879 1 561231 Proteobacteria
Yersinia enterocolitica subsp enterocolitica 8081 YP 001008209 1 393305 Proteobacteria
Yersinia enterocolitica subsp palearctica Y11 YP 006006626 1 930944 Proteobacteria
Yersinia pestis biovar Medievalis str Harbin 35 YP 005626180 1 547048 Proteobacteria
Yersinia pestis biovar Microtus str 91001 NP 994645 1 229193 Proteobacteria
Yersinia pseudotuberculosis IP 31758 YP 001403010 1 349747 Proteobacteria
Yersinia pestis Antiqua YP 653729 1 360102 Proteobacteria
Vibrio furnissii NCTC 11218 YP 005049726 1 903510 Proteobacteria
Yersinia enterocolitica subsp enterocolitica 8081 YP 001006147 1 393305 Proteobacteria
Yersinia enterocolitica subsp palearctica Y11 YP 006004256 1 930944 Proteobacteria
Yersinia pseudotuberculosis IP 31758 YP 001400669 1 349747 Proteobacteria
Yersinia pestis Antiqua YP 651709 1 360102 Proteobacteria
Yersinia pestis biovar Medievalis str Harbin 35 YP 005623969 1 547048 Proteobacteria
Yersinia pestis biovar Microtus str 91001 NP 993099 1 229193 Proteobacteria
Pectobacterium sp SCC3193 YP 006283104 1 1166016 Proteobacteria
Pectobacterium wasabiae WPP163 YP 003259578 1 561231 Proteobacteria
Pectobacterium atrosepticum SCRI1043 YP 050497 1 218491 Proteobacteria
Pectobacterium carotovorum subsp carotovorum PC1 YP 003017485 1 561230 Proteobacteria
Prodigiosinella confusarubida ATCC39006 WP 021015827 1 Pconfusarubida
Dickeya paradisiaca Ech703 WP 015853732 1 Dparadisiaca
Dickeya paradisiaca NCPPB 2511 WP 015853732 1 Dparadisiaca
DAQ1742 DAQ1742 02459 Daquatica
Dickeya sp CSL RW240 WP 035341620 1 Daquatica
Dickeya aquatica DW 0440 WP 035341620 1 Daquatica
Dickeya chrysanthemi subsp chrysanthemi NCPPB 402 WP 040001139 1 Dchrysanthemi
Dickeya chrysanthemi subsp parthenii NCPPB 516 WP 027711962 1 Dchrysanthemi
Dickeya chrysanthemi L11 WP 033577542 1 Dchrysanthemi
Dickeya chrysanthemi Ech1591 WP 012769687 1 Dchrysanthemi
Dickeya chrysanthemi NCPPB 3533 WP 012769687 1 Dchrysanthemi
Dickeya zeae Ech586 WP 012884637 1 Dzeae
Dickeya zeae MK19 WP 038913904 1 Dzeae
Dickeya zeae NCPPB 3532 WP 023639598 1 Dzeae
Dickeya zeae NCPPB 2538 WP 038916132 1 Dzeae
Dickeya zeae MS1 WP 019845566 1 Dzeae
Dickeya zeae NCPPB 3531 WP 038904257 1 Dzeae
Dickeya zeae CSL RW192 WP 038907856 1 Dzeae
Dickeya zeae DZ2Q WP 038925849 1 Dzeae
Dickeya zeae ZJU1202 WP 016941451 1 Dzeae
Dickeya zeae EC1 WP 016941451 1 Dzeae
Dickeya gs poaceaephila NCPPB 569 WP 042870701 1 Dpoaceaephila
Dickeya dianthicola IPO 980 WP 047100615 1 Ddianthicola
Dickeya dianthicola NCPPB 453 WP 024106041 1 Ddianthicola
Dickeya dianthicola NCPPB 3534 WP 024106041 1 Ddianthicola
Dickeya dianthicola RNS04 9 WP 024106041 1 Ddianthicola
Dickeya dadantii subsp dieffenbachiae NCPPB 2976 WP 038922919 1 Ddadantii
Dickeya dadantii NCPPB 3537 WP 038901300 1 Ddadantii
Dickeya dadantii NCPPB 898 WP 038911249 1 Ddadantii
148 pelW Plant cell wall degradosome Seed
Dickeya dadantii 3937 WP 013318067 1 Ddadantii
Dickeya gs fangzhongdai NCPPB 3274 WP 042859932 1 Dfangzhongdai
Dickeya gs fangzhongdai M005 WP 039694446 1 Dfangzhongdai
Dickeya gs fangzhongdai M074 WP 038658349 1 Dfangzhongdai
Dickeya gs fangzhongdai ND14b WP 038658349 1 Dfangzhongdai
Dickeya gs undicola 2B12 WP 033570789 1 Dundicola
Dickeya gs fangzhongdai S1 WP 049853737 1 Dfangzhongdai
Dickeya gs fangzhongdai MK7 WP 038919170 1 Dfangzhongdai
Dickeya gs fangzhongdai B16 WP 038919170 1 Dfangzhongdai
Dickeya solani RNS 05 1 2A WP 057084194 1 Dsolani
Dickeya solani IPO 2222 WP 022633694 1 Dsolani
Dickeya solani PPO 9134 WP 022633694 1 Dsolani
Dickeya solani D s0432 1 WP 022633694 1 Dsolani
Dickeya solani RNS 08 23 3 1 A WP 022633694 1 Dsolani
Dickeya solani MK16 WP 022633694 1 Dsolani
Dickeya solani IFB 0099 WP 022633694 1 Dsolani
Dickeya solani PPO 9019 WP 022633694 1 Dsolani
Dickeya solani MK10 WP 022633694 1 Dsolani
Dickeya solani RNS 07 7 3B WP 022633694 1 Dsolani

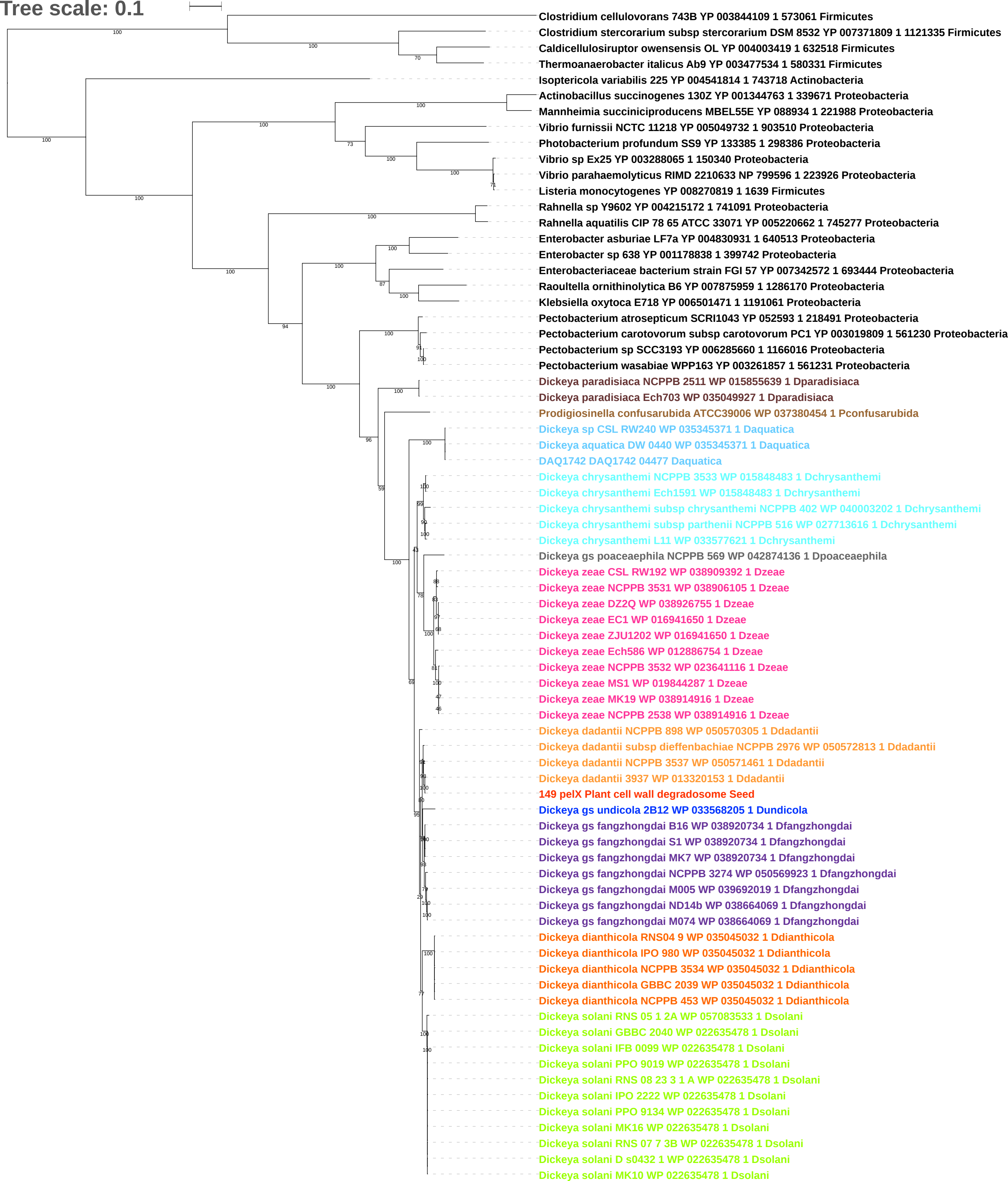
Tree scale: 0.1
Clostridium cellulovorans 743B YP 003844109 1 573061 Firmicutes
Clostridium stercorarium subsp stercorarium DSM 8532 YP 007371809 1 1121335 Firmicutes
Caldicellulosiruptor owensensis OL YP 004003419 1 632518 Firmicutes
Thermoanaerobacter italicus Ab9 YP 003477534 1 580331 Firmicutes
Isoptericola variabilis 225 YP 004541814 1 743718 Actinobacteria
Actinobacillus succinogenes 130Z YP 001344763 1 339671 Proteobacteria
Mannheimia succiniciproducens MBEL55E YP 088934 1 221988 Proteobacteria
Vibrio furnissii NCTC 11218 YP 005049732 1 903510 Proteobacteria
Photobacterium profundum SS9 YP 133385 1 298386 Proteobacteria
Vibrio sp Ex25 YP 003288065 1 150340 Proteobacteria
Vibrio parahaemolyticus RIMD 2210633 NP 799596 1 223926 Proteobacteria
Listeria monocytogenes YP 008270819 1 1639 Firmicutes
Rahnella sp Y9602 YP 004215172 1 741091 Proteobacteria
Rahnella aquatilis CIP 78 65 ATCC 33071 YP 005220662 1 745277 Proteobacteria
Enterobacter asburiae LF7a YP 004830931 1 640513 Proteobacteria
Enterobacter sp 638 YP 001178838 1 399742 Proteobacteria
Enterobacteriaceae bacterium strain FGI 57 YP 007342572 1 693444 Proteobacteria
Raoultella ornithinolytica B6 YP 007875959 1 1286170 Proteobacteria
Klebsiella oxytoca E718 YP 006501471 1 1191061 Proteobacteria
Pectobacterium atrosepticum SCRI1043 YP 052593 1 218491 Proteobacteria
Pectobacterium carotovorum subsp carotovorum PC1 YP 003019809 1 561230 Proteobacteria
Pectobacterium sp SCC3193 YP 006285660 1 1166016 Proteobacteria
Pectobacterium wasabiae WPP163 YP 003261857 1 561231 Proteobacteria
Dickeya paradisiaca NCPPB 2511 WP 015855639 1 Dparadisiaca
Dickeya paradisiaca Ech703 WP 035049927 1 Dparadisiaca
Prodigiosinella confusarubida ATCC39006 WP 037380454 1 Pconfusarubida
Dickeya sp CSL RW240 WP 035345371 1 Daquatica
Dickeya aquatica DW 0440 WP 035345371 1 Daquatica
DAQ1742 DAQ1742 04477 Daquatica
Dickeya chrysanthemi NCPPB 3533 WP 015848483 1 Dchrysanthemi
Dickeya chrysanthemi Ech1591 WP 015848483 1 Dchrysanthemi
Dickeya chrysanthemi subsp chrysanthemi NCPPB 402 WP 040003202 1 Dchrysanthemi
Dickeya chrysanthemi subsp parthenii NCPPB 516 WP 027713616 1 Dchrysanthemi
Dickeya chrysanthemi L11 WP 033577621 1 Dchrysanthemi
Dickeya gs poaceaephila NCPPB 569 WP 042874136 1 Dpoaceaephila
Dickeya zeae CSL RW192 WP 038909392 1 Dzeae
Dickeya zeae NCPPB 3531 WP 038906105 1 Dzeae
Dickeya zeae DZ2Q WP 038926755 1 Dzeae
Dickeya zeae EC1 WP 016941650 1 Dzeae
Dickeya zeae ZJU1202 WP 016941650 1 Dzeae
Dickeya zeae Ech586 WP 012886754 1 Dzeae
Dickeya zeae NCPPB 3532 WP 023641116 1 Dzeae
Dickeya zeae MS1 WP 019844287 1 Dzeae
Dickeya zeae MK19 WP 038914916 1 Dzeae
Dickeya zeae NCPPB 2538 WP 038914916 1 Dzeae
Dickeya dadantii NCPPB 898 WP 050570305 1 Ddadantii
Dickeya dadantii subsp dieffenbachiae NCPPB 2976 WP 050572813 1 Ddadantii
Dickeya dadantii NCPPB 3537 WP 050571461 1 Ddadantii
Dickeya dadantii 3937 WP 013320153 1 Ddadantii
149 pelX Plant cell wall degradosome Seed
Dickeya gs undicola 2B12 WP 033568205 1 Dundicola
Dickeya gs fangzhongdai B16 WP 038920734 1 Dfangzhongdai
Dickeya gs fangzhongdai S1 WP 038920734 1 Dfangzhongdai
Dickeya gs fangzhongdai MK7 WP 038920734 1 Dfangzhongdai
Dickeya gs fangzhongdai NCPPB 3274 WP 050569923 1 Dfangzhongdai
Dickeya gs fangzhongdai M005 WP 039692019 1 Dfangzhongdai
Dickeya gs fangzhongdai ND14b WP 038664069 1 Dfangzhongdai
Dickeya gs fangzhongdai M074 WP 038664069 1 Dfangzhongdai
Dickeya dianthicola RNS04 9 WP 035045032 1 Ddianthicola
Dickeya dianthicola IPO 980 WP 035045032 1 Ddianthicola
Dickeya dianthicola NCPPB 3534 WP 035045032 1 Ddianthicola
Dickeya dianthicola GBBC 2039 WP 035045032 1 Ddianthicola
Dickeya dianthicola NCPPB 453 WP 035045032 1 Ddianthicola
Dickeya solani RNS 05 1 2A WP 057083533 1 Dsolani
Dickeya solani GBBC 2040 WP 022635478 1 Dsolani
Dickeya solani IFB 0099 WP 022635478 1 Dsolani
Dickeya solani PPO 9019 WP 022635478 1 Dsolani
Dickeya solani RNS 08 23 3 1 A WP 022635478 1 Dsolani
Dickeya solani IPO 2222 WP 022635478 1 Dsolani
Dickeya solani PPO 9134 WP 022635478 1 Dsolani
Dickeya solani MK16 WP 022635478 1 Dsolani
Dickeya solani RNS 07 7 3B WP 022635478 1 Dsolani
Dickeya solani D s0432 1 WP 022635478 1 Dsolani
Dickeya solani MK10 WP 022635478 1 Dsolani

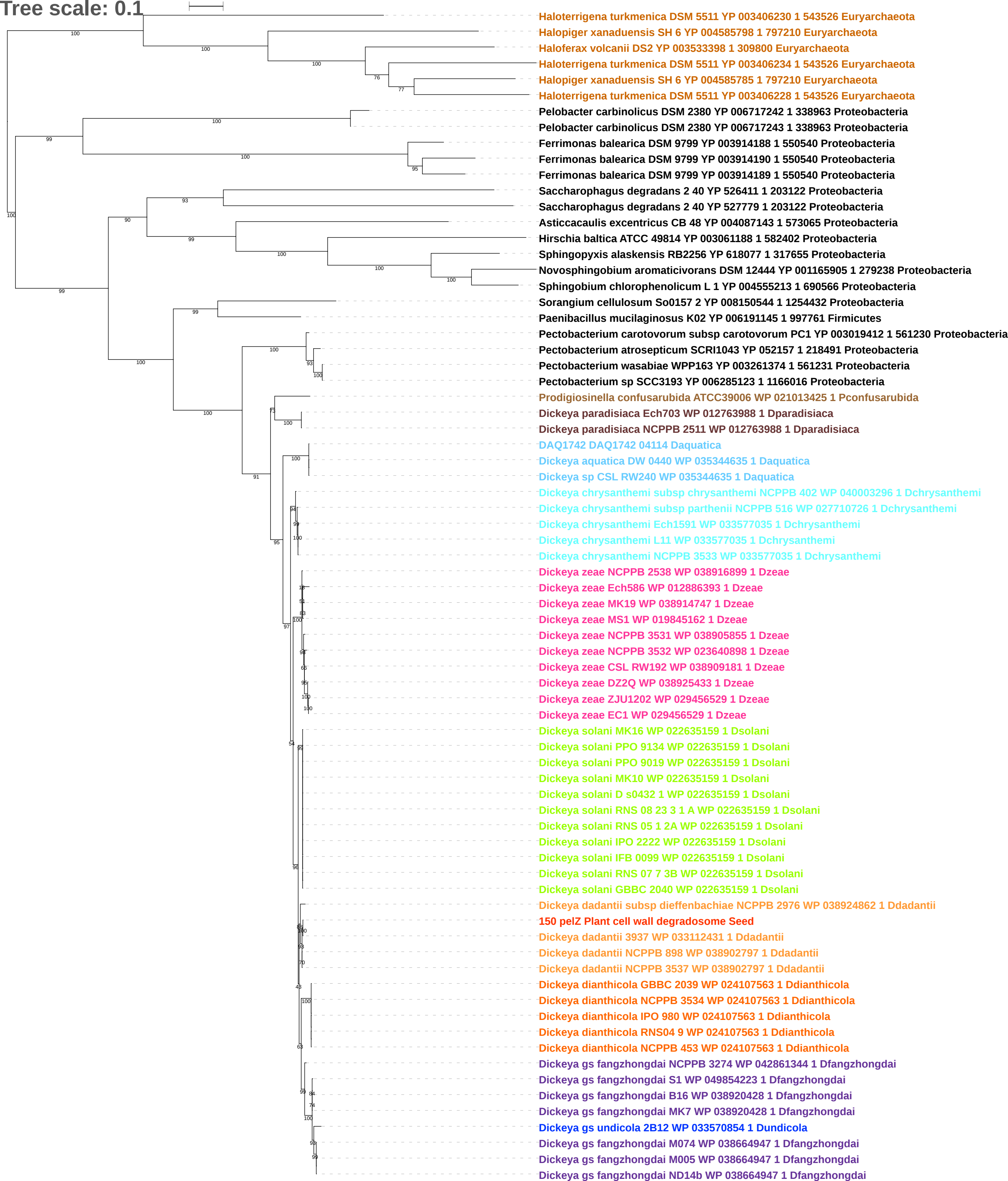
Tree scale: 0.1
Haloterrigena turkmenica DSM 5511 YP 003406230 1 543526 Euryarchaeota
Halopiger xanaduensis SH 6 YP 004585798 1 797210 Euryarchaeota
Haloferax volcanii DS2 YP 003533398 1 309800 Euryarchaeota
Haloterrigena turkmenica DSM 5511 YP 003406234 1 543526 Euryarchaeota
Halopiger xanaduensis SH 6 YP 004585785 1 797210 Euryarchaeota
Haloterrigena turkmenica DSM 5511 YP 003406228 1 543526 Euryarchaeota
Pelobacter carbinolicus DSM 2380 YP 006717242 1 338963 Proteobacteria
Pelobacter carbinolicus DSM 2380 YP 006717243 1 338963 Proteobacteria
Ferrimonas balearica DSM 9799 YP 003914188 1 550540 Proteobacteria
Ferrimonas balearica DSM 9799 YP 003914190 1 550540 Proteobacteria
Ferrimonas balearica DSM 9799 YP 003914189 1 550540 Proteobacteria
Saccharophagus degradans 2 40 YP 526411 1 203122 Proteobacteria
Saccharophagus degradans 2 40 YP 527779 1 203122 Proteobacteria
Asticcacaulis excentricus CB 48 YP 004087143 1 573065 Proteobacteria
Hirschia baltica ATCC 49814 YP 003061188 1 582402 Proteobacteria
Sphingopyxis alaskensis RB2256 YP 618077 1 317655 Proteobacteria
Novosphingobium aromaticivorans DSM 12444 YP 001165905 1 279238 Proteobacteria
Sphingobium chlorophenolicum L 1 YP 004555213 1 690566 Proteobacteria
Sorangium cellulosum So0157 2 YP 008150544 1 1254432 Proteobacteria
Paenibacillus mucilaginosus K02 YP 006191145 1 997761 Firmicutes
Pectobacterium carotovorum subsp carotovorum PC1 YP 003019412 1 561230 Proteobacteria
Pectobacterium atrosepticum SCRI1043 YP 052157 1 218491 Proteobacteria
Pectobacterium wasabiae WPP163 YP 003261374 1 561231 Proteobacteria
Pectobacterium sp SCC3193 YP 006285123 1 1166016 Proteobacteria
Prodigiosinella confusarubida ATCC39006 WP 021013425 1 Pconfusarubida
Dickeya paradisiaca Ech703 WP 012763988 1 Dparadisiaca
Dickeya paradisiaca NCPPB 2511 WP 012763988 1 Dparadisiaca
DAQ1742 DAQ1742 04114 Daquatica
Dickeya aquatica DW 0440 WP 035344635 1 Daquatica
Dickeya sp CSL RW240 WP 035344635 1 Daquatica
Dickeya chrysanthemi subsp chrysanthemi NCPPB 402 WP 040003296 1 Dchrysanthemi
Dickeya chrysanthemi subsp parthenii NCPPB 516 WP 027710726 1 Dchrysanthemi
Dickeya chrysanthemi Ech1591 WP 033577035 1 Dchrysanthemi
Dickeya chrysanthemi L11 WP 033577035 1 Dchrysanthemi
Dickeya chrysanthemi NCPPB 3533 WP 033577035 1 Dchrysanthemi
Dickeya zeae NCPPB 2538 WP 038916899 1 Dzeae
Dickeya zeae Ech586 WP 012886393 1 Dzeae
Dickeya zeae MK19 WP 038914747 1 Dzeae
Dickeya zeae MS1 WP 019845162 1 Dzeae
Dickeya zeae NCPPB 3531 WP 038905855 1 Dzeae
Dickeya zeae NCPPB 3532 WP 023640898 1 Dzeae
Dickeya zeae CSL RW192 WP 038909181 1 Dzeae
Dickeya zeae DZ2Q WP 038925433 1 Dzeae
Dickeya zeae ZJU1202 WP 029456529 1 Dzeae
Dickeya zeae EC1 WP 029456529 1 Dzeae
Dickeya solani MK16 WP 022635159 1 Dsolani
Dickeya solani PPO 9134 WP 022635159 1 Dsolani
Dickeya solani PPO 9019 WP 022635159 1 Dsolani
Dickeya solani MK10 WP 022635159 1 Dsolani
Dickeya solani D s0432 1 WP 022635159 1 Dsolani
Dickeya solani RNS 08 23 3 1 A WP 022635159 1 Dsolani
Dickeya solani RNS 05 1 2A WP 022635159 1 Dsolani
Dickeya solani IPO 2222 WP 022635159 1 Dsolani
Dickeya solani IFB 0099 WP 022635159 1 Dsolani
Dickeya solani RNS 07 7 3B WP 022635159 1 Dsolani
Dickeya solani GBBC 2040 WP 022635159 1 Dsolani
Dickeya dadantii subsp dieffenbachiae NCPPB 2976 WP 038924862 1 Ddadantii
150 pelZ Plant cell wall degradosome Seed
Dickeya dadantii 3937 WP 033112431 1 Ddadantii
Dickeya dadantii NCPPB 898 WP 038902797 1 Ddadantii
Dickeya dadantii NCPPB 3537 WP 038902797 1 Ddadantii
Dickeya dianthicola GBBC 2039 WP 024107563 1 Ddianthicola
Dickeya dianthicola NCPPB 3534 WP 024107563 1 Ddianthicola
Dickeya dianthicola IPO 980 WP 024107563 1 Ddianthicola
Dickeya dianthicola RNS04 9 WP 024107563 1 Ddianthicola
Dickeya dianthicola NCPPB 453 WP 024107563 1 Ddianthicola
Dickeya gs fangzhongdai NCPPB 3274 WP 042861344 1 Dfangzhongdai
Dickeya gs fangzhongdai S1 WP 049854223 1 Dfangzhongdai
Dickeya gs fangzhongdai B16 WP 038920428 1 Dfangzhongdai
Dickeya gs fangzhongdai MK7 WP 038920428 1 Dfangzhongdai
Dickeya gs undicola 2B12 WP 033570854 1 Dundicola
Dickeya gs fangzhongdai M074 WP 038664947 1 Dfangzhongdai
Dickeya gs fangzhongdai M005 WP 038664947 1 Dfangzhongdai
Dickeya gs fangzhongdai ND14b WP 038664947 1 Dfangzhongdai

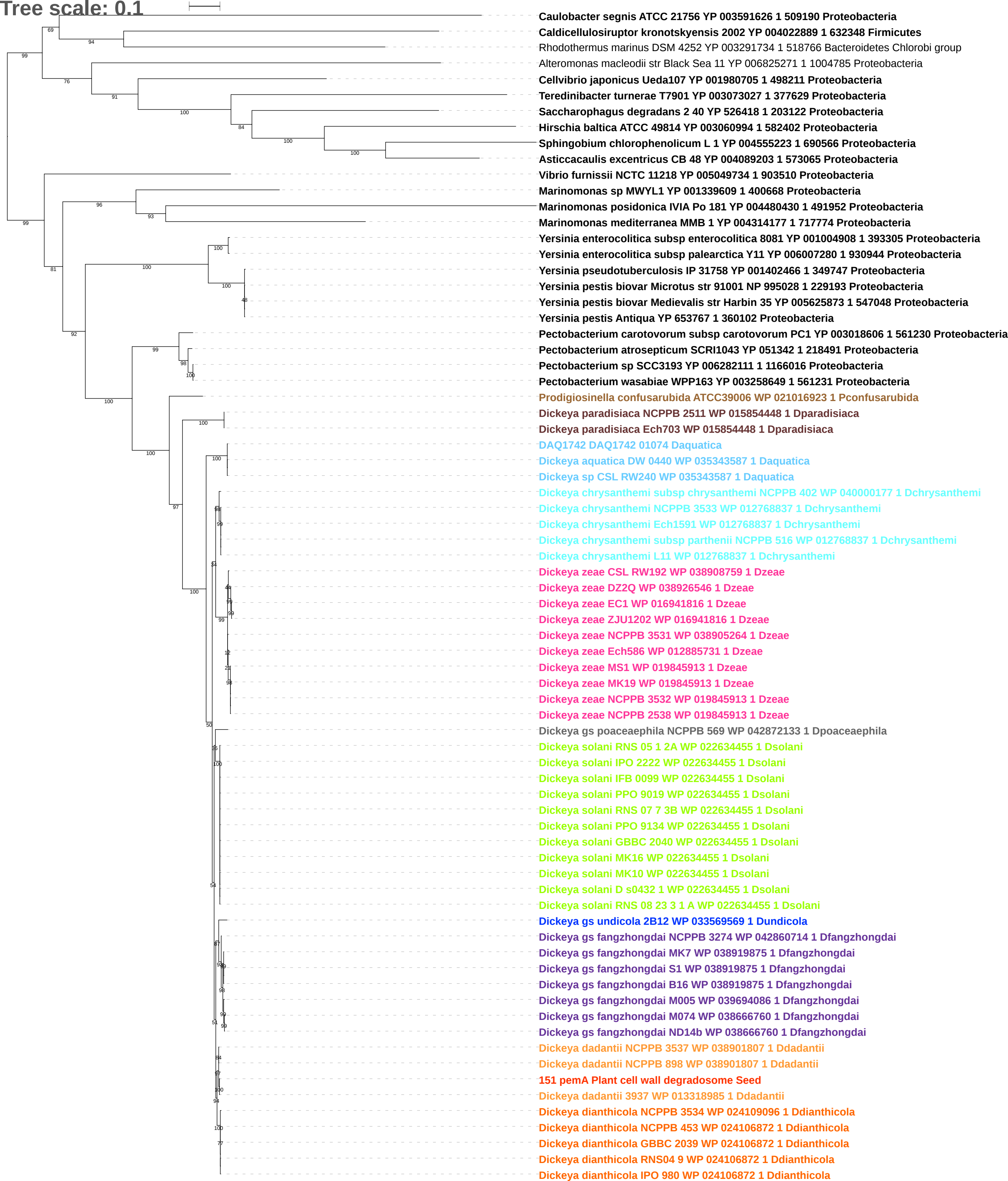
Tree scale: 0.1
Caulobacter segnis ATCC 21756 YP 003591626 1 509190 Proteobacteria
Caldicellulosiruptor kronotskyensis 2002 YP 004022889 1 632348 Firmicutes
Rhodothermus marinus DSM 4252 YP 003291734 1 518766 Bacteroidetes Chlorobi group
Alteromonas macleodii str Black Sea 11 YP 006825271 1 1004785 Proteobacteria
Cellvibrio japonicus Ueda107 YP 001980705 1 498211 Proteobacteria
Teredinibacter turnerae T7901 YP 003073027 1 377629 Proteobacteria
Saccharophagus degradans 2 40 YP 526418 1 203122 Proteobacteria
Hirschia baltica ATCC 49814 YP 003060994 1 582402 Proteobacteria
Sphingobium chlorophenolicum L 1 YP 004555223 1 690566 Proteobacteria
Asticcacaulis excentricus CB 48 YP 004089203 1 573065 Proteobacteria
Vibrio furnissii NCTC 11218 YP 005049734 1 903510 Proteobacteria
Marinomonas sp MWYL1 YP 001339609 1 400668 Proteobacteria
Marinomonas posidonica IVIA Po 181 YP 004480430 1 491952 Proteobacteria
Marinomonas mediterranea MMB 1 YP 004314177 1 717774 Proteobacteria
Yersinia enterocolitica subsp enterocolitica 8081 YP 001004908 1 393305 Proteobacteria
Yersinia enterocolitica subsp palearctica Y11 YP 006007280 1 930944 Proteobacteria
Yersinia pseudotuberculosis IP 31758 YP 001402466 1 349747 Proteobacteria
Yersinia pestis biovar Microtus str 91001 NP 995028 1 229193 Proteobacteria
Yersinia pestis biovar Medievalis str Harbin 35 YP 005625873 1 547048 Proteobacteria
Yersinia pestis Antiqua YP 653767 1 360102 Proteobacteria
Pectobacterium carotovorum subsp carotovorum PC1 YP 003018606 1 561230 Proteobacteria
Pectobacterium atrosepticum SCRI1043 YP 051342 1 218491 Proteobacteria
Pectobacterium sp SCC3193 YP 006282111 1 1166016 Proteobacteria
Pectobacterium wasabiae WPP163 YP 003258649 1 561231 Proteobacteria
Prodigiosinella confusarubida ATCC39006 WP 021016923 1 Pconfusarubida
Dickeya paradisiaca NCPPB 2511 WP 015854448 1 Dparadisiaca
Dickeya paradisiaca Ech703 WP 015854448 1 Dparadisiaca
DAQ1742 DAQ1742 01074 Daquatica
Dickeya aquatica DW 0440 WP 035343587 1 Daquatica
Dickeya sp CSL RW240 WP 035343587 1 Daquatica
Dickeya chrysanthemi subsp chrysanthemi NCPPB 402 WP 040000177 1 Dchrysanthemi
Dickeya chrysanthemi NCPPB 3533 WP 012768837 1 Dchrysanthemi
Dickeya chrysanthemi Ech1591 WP 012768837 1 Dchrysanthemi
Dickeya chrysanthemi subsp parthenii NCPPB 516 WP 012768837 1 Dchrysanthemi
Dickeya chrysanthemi L11 WP 012768837 1 Dchrysanthemi
Dickeya zeae CSL RW192 WP 038908759 1 Dzeae
Dickeya zeae DZ2Q WP 038926546 1 Dzeae
Dickeya zeae EC1 WP 016941816 1 Dzeae
Dickeya zeae ZJU1202 WP 016941816 1 Dzeae
Dickeya zeae NCPPB 3531 WP 038905264 1 Dzeae
Dickeya zeae Ech586 WP 012885731 1 Dzeae
Dickeya zeae MS1 WP 019845913 1 Dzeae
Dickeya zeae MK19 WP 019845913 1 Dzeae
Dickeya zeae NCPPB 3532 WP 019845913 1 Dzeae
Dickeya zeae NCPPB 2538 WP 019845913 1 Dzeae
Dickeya gs poaceaephila NCPPB 569 WP 042872133 1 Dpoaceaephila
Dickeya solani RNS 05 1 2A WP 022634455 1 Dsolani
Dickeya solani IPO 2222 WP 022634455 1 Dsolani
Dickeya solani IFB 0099 WP 022634455 1 Dsolani
Dickeya solani PPO 9019 WP 022634455 1 Dsolani
Dickeya solani RNS 07 7 3B WP 022634455 1 Dsolani
Dickeya solani PPO 9134 WP 022634455 1 Dsolani
Dickeya solani GBBC 2040 WP 022634455 1 Dsolani
Dickeya solani MK16 WP 022634455 1 Dsolani
Dickeya solani MK10 WP 022634455 1 Dsolani
Dickeya solani D s0432 1 WP 022634455 1 Dsolani
Dickeya solani RNS 08 23 3 1 A WP 022634455 1 Dsolani
Dickeya gs undicola 2B12 WP 033569569 1 Dundicola
Dickeya gs fangzhongdai NCPPB 3274 WP 042860714 1 Dfangzhongdai
Dickeya gs fangzhongdai MK7 WP 038919875 1 Dfangzhongdai
Dickeya gs fangzhongdai S1 WP 038919875 1 Dfangzhongdai
Dickeya gs fangzhongdai B16 WP 038919875 1 Dfangzhongdai
Dickeya gs fangzhongdai M005 WP 039694086 1 Dfangzhongdai
Dickeya gs fangzhongdai M074 WP 038666760 1 Dfangzhongdai
Dickeya gs fangzhongdai ND14b WP 038666760 1 Dfangzhongdai
Dickeya dadantii NCPPB 3537 WP 038901807 1 Ddadantii
Dickeya dadantii NCPPB 898 WP 038901807 1 Ddadantii
151 pemA Plant cell wall degradosome Seed
Dickeya dadantii 3937 WP 013318985 1 Ddadantii
Dickeya dianthicola NCPPB 3534 WP 024109096 1 Ddianthicola
Dickeya dianthicola NCPPB 453 WP 024106872 1 Ddianthicola
Dickeya dianthicola GBBC 2039 WP 024106872 1 Ddianthicola
Dickeya dianthicola RNS04 9 WP 024106872 1 Ddianthicola
Dickeya dianthicola IPO 980 WP 024106872 1 Ddianthicola

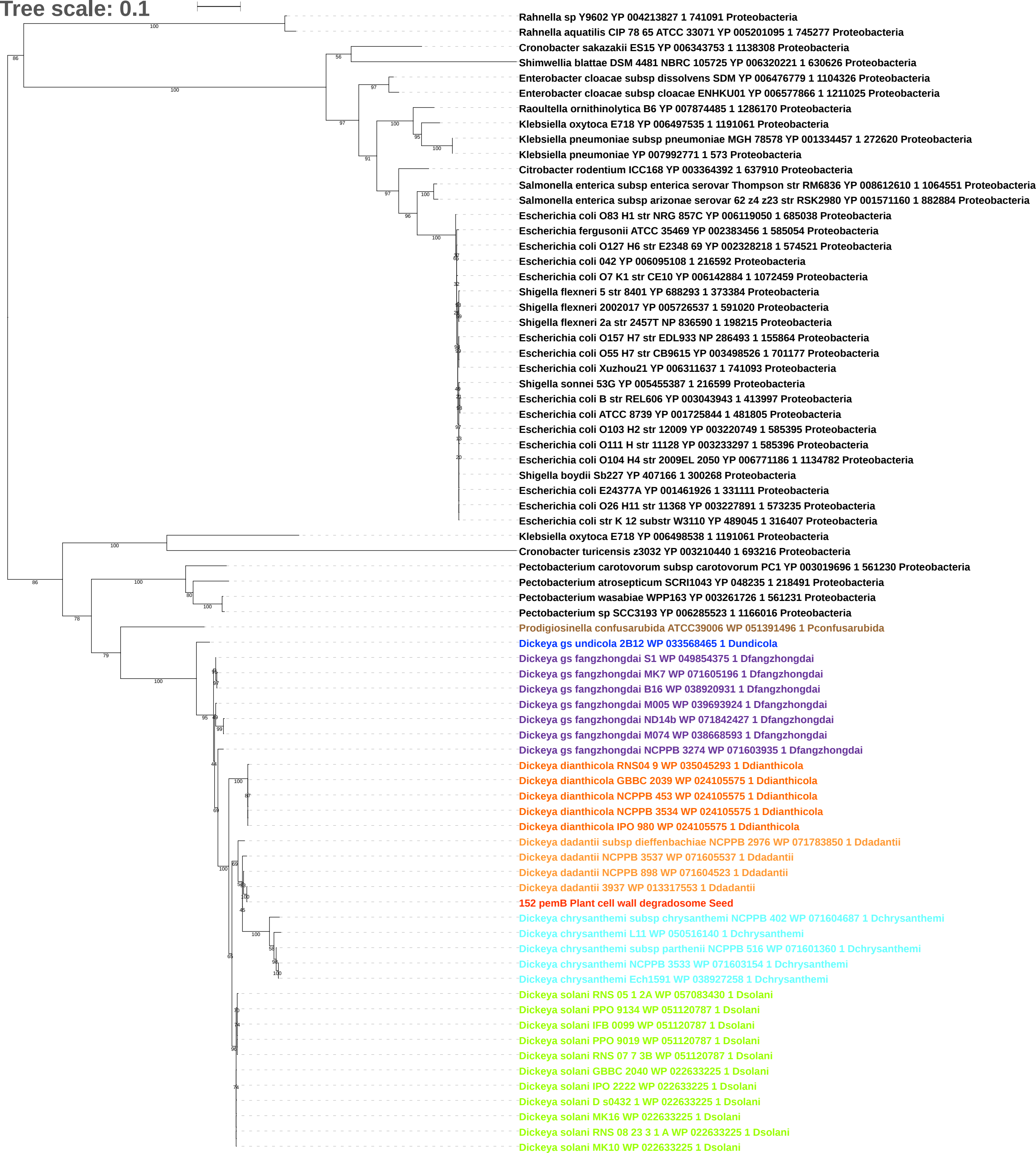
Tree scale: 0.1
Rahnella sp Y9602 YP 004213827 1 741091 Proteobacteria
Rahnella aquatilis CIP 78 65 ATCC 33071 YP 005201095 1 745277 Proteobacteria
Cronobacter sakazakii ES15 YP 006343753 1 1138308 Proteobacteria
Shimwellia blattae DSM 4481 NBRC 105725 YP 006320221 1 630626 Proteobacteria
Enterobacter cloacae subsp dissolvens SDM YP 006476779 1 1104326 Proteobacteria
Enterobacter cloacae subsp cloacae ENHKU01 YP 006577866 1 1211025 Proteobacteria
Raoultella ornithinolytica B6 YP 007874485 1 1286170 Proteobacteria
Klebsiella oxytoca E718 YP 006497535 1 1191061 Proteobacteria
Klebsiella pneumoniae subsp pneumoniae MGH 78578 YP 001334457 1 272620 Proteobacteria
Klebsiella pneumoniae YP 007992771 1 573 Proteobacteria
Citrobacter rodentium ICC168 YP 003364392 1 637910 Proteobacteria
Salmonella enterica subsp enterica serovar Thompson str RM6836 YP 008612610 1 1064551 Proteobacteria
Salmonella enterica subsp arizonae serovar 62 z4 z23 str RSK2980 YP 001571160 1 882884 Proteobacteria
Escherichia coli O83 H1 str NRG 857C YP 006119050 1 685038 Proteobacteria
Escherichia fergusonii ATCC 35469 YP 002383456 1 585054 Proteobacteria
Escherichia coli O127 H6 str E2348 69 YP 002328218 1 574521 Proteobacteria
Escherichia coli 042 YP 006095108 1 216592 Proteobacteria
Escherichia coli O7 K1 str CE10 YP 006142884 1 1072459 Proteobacteria
Shigella flexneri 5 str 8401 YP 688293 1 373384 Proteobacteria
Shigella flexneri 2002017 YP 005726537 1 591020 Proteobacteria
Shigella flexneri 2a str 2457T NP 836590 1 198215 Proteobacteria
Escherichia coli O157 H7 str EDL933 NP 286493 1 155864 Proteobacteria
Escherichia coli O55 H7 str CB9615 YP 003498526 1 701177 Proteobacteria
Escherichia coli Xuzhou21 YP 006311637 1 741093 Proteobacteria
Shigella sonnei 53G YP 005455387 1 216599 Proteobacteria
Escherichia coli B str REL606 YP 003043943 1 413997 Proteobacteria
Escherichia coli ATCC 8739 YP 001725844 1 481805 Proteobacteria
Escherichia coli O103 H2 str 12009 YP 003220749 1 585395 Proteobacteria
Escherichia coli O111 H str 11128 YP 003233297 1 585396 Proteobacteria
Escherichia coli O104 H4 str 2009EL 2050 YP 006771186 1 1134782 Proteobacteria
Shigella boydii Sb227 YP 407166 1 300268 Proteobacteria
Escherichia coli E24377A YP 001461926 1 331111 Proteobacteria
Escherichia coli O26 H11 str 11368 YP 003227891 1 573235 Proteobacteria
Escherichia coli str K 12 substr W3110 YP 489045 1 316407 Proteobacteria
Klebsiella oxytoca E718 YP 006498538 1 1191061 Proteobacteria
Cronobacter turicensis z3032 YP 003210440 1 693216 Proteobacteria
Pectobacterium carotovorum subsp carotovorum PC1 YP 003019696 1 561230 Proteobacteria
Pectobacterium atrosepticum SCRI1043 YP 048235 1 218491 Proteobacteria
Pectobacterium wasabiae WPP163 YP 003261726 1 561231 Proteobacteria
Pectobacterium sp SCC3193 YP 006285523 1 1166016 Proteobacteria
Prodigiosinella confusarubida ATCC39006 WP 051391496 1 Pconfusarubida
Dickeya gs undicola 2B12 WP 033568465 1 Dundicola
Dickeya gs fangzhongdai S1 WP 049854375 1 Dfangzhongdai
Dickeya gs fangzhongdai MK7 WP 071605196 1 Dfangzhongdai
Dickeya gs fangzhongdai B16 WP 038920931 1 Dfangzhongdai
Dickeya gs fangzhongdai M005 WP 039693924 1 Dfangzhongdai
Dickeya gs fangzhongdai ND14b WP 071842427 1 Dfangzhongdai
Dickeya gs fangzhongdai M074 WP 038668593 1 Dfangzhongdai
Dickeya gs fangzhongdai NCPPB 3274 WP 071603935 1 Dfangzhongdai
Dickeya dianthicola RNS04 9 WP 035045293 1 Ddianthicola
Dickeya dianthicola GBBC 2039 WP 024105575 1 Ddianthicola
Dickeya dianthicola NCPPB 453 WP 024105575 1 Ddianthicola
Dickeya dianthicola NCPPB 3534 WP 024105575 1 Ddianthicola
Dickeya dianthicola IPO 980 WP 024105575 1 Ddianthicola
Dickeya dadantii subsp dieffenbachiae NCPPB 2976 WP 071783850 1 Ddadantii
Dickeya dadantii NCPPB 3537 WP 071605537 1 Ddadantii
Dickeya dadantii NCPPB 898 WP 071604523 1 Ddadantii
Dickeya dadantii 3937 WP 013317553 1 Ddadantii
152 pemB Plant cell wall degradosome Seed
Dickeya chrysanthemi subsp chrysanthemi NCPPB 402 WP 071604687 1 Dchrysanthemi
Dickeya chrysanthemi L11 WP 050516140 1 Dchrysanthemi
Dickeya chrysanthemi subsp parthenii NCPPB 516 WP 071601360 1 Dchrysanthemi
Dickeya chrysanthemi NCPPB 3533 WP 071603154 1 Dchrysanthemi
Dickeya chrysanthemi Ech1591 WP 038927258 1 Dchrysanthemi
Dickeya solani RNS 05 1 2A WP 057083430 1 Dsolani
Dickeya solani PPO 9134 WP 051120787 1 Dsolani
Dickeya solani IFB 0099 WP 051120787 1 Dsolani
Dickeya solani PPO 9019 WP 051120787 1 Dsolani
Dickeya solani RNS 07 7 3B WP 051120787 1 Dsolani
Dickeya solani GBBC 2040 WP 022633225 1 Dsolani
Dickeya solani IPO 2222 WP 022633225 1 Dsolani
Dickeya solani D s0432 1 WP 022633225 1 Dsolani
Dickeya solani MK16 WP 022633225 1 Dsolani
Dickeya solani RNS 08 23 3 1 A WP 022633225 1 Dsolani
Dickeya solani MK10 WP 022633225 1 Dsolani

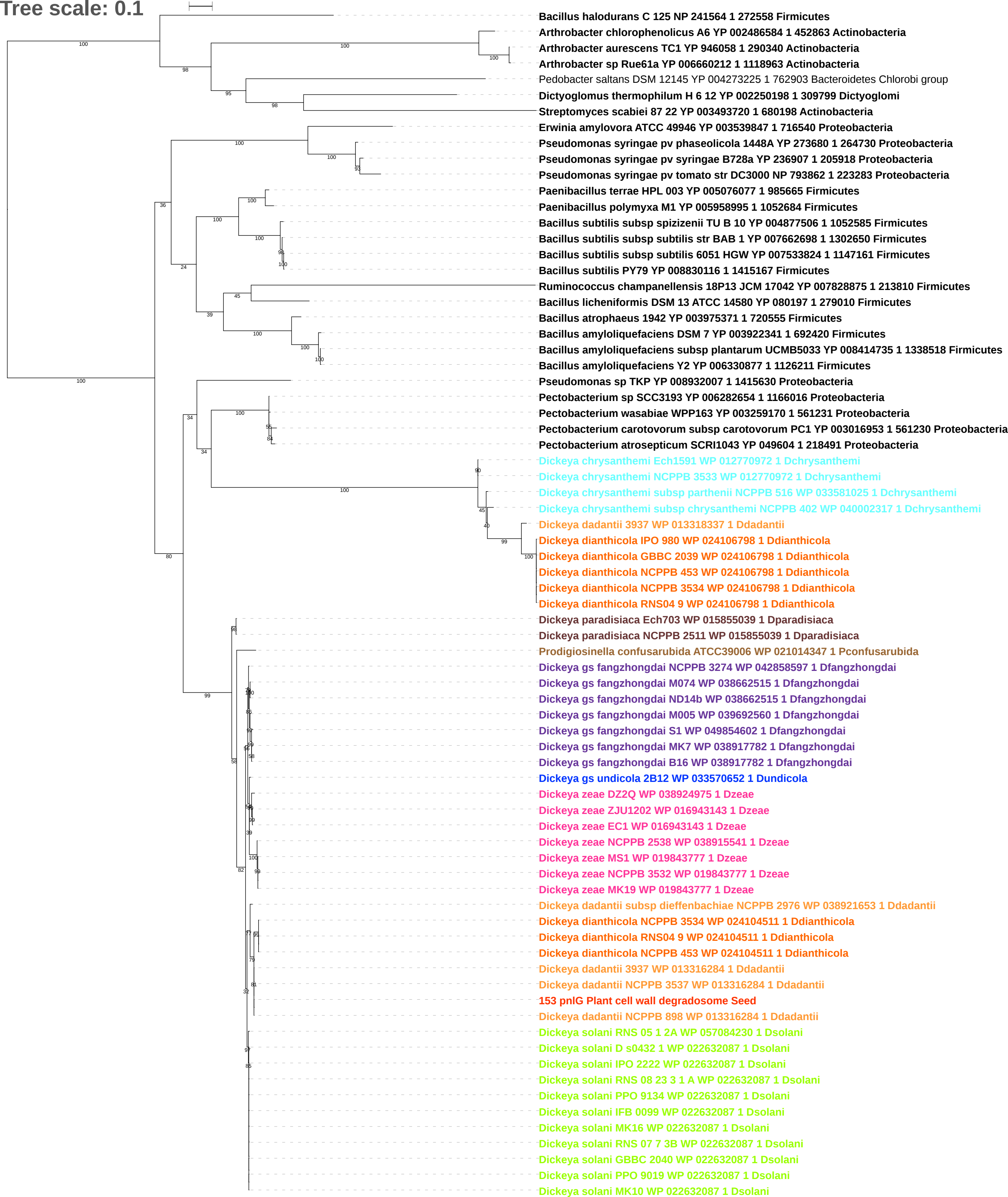
Tree scale: 0.1
Bacillus halodurans C 125 NP 241564 1 272558 Firmicutes
Arthrobacter chlorophenolicus A6 YP 002486584 1 452863 Actinobacteria
Arthrobacter aurescens TC1 YP 946058 1 290340 Actinobacteria
Arthrobacter sp Rue61a YP 006660212 1 1118963 Actinobacteria
Pedobacter saltans DSM 12145 YP 004273225 1 762903 Bacteroidetes Chlorobi group
Dictyoglomus thermophilum H 6 12 YP 002250198 1 309799 Dictyoglomi
Streptomyces scabiei 87 22 YP 003493720 1 680198 Actinobacteria
Erwinia amylovora ATCC 49946 YP 003539847 1 716540 Proteobacteria
Pseudomonas syringae pv phaseolicola 1448A YP 273680 1 264730 Proteobacteria
Pseudomonas syringae pv syringae B728a YP 236907 1 205918 Proteobacteria
Pseudomonas syringae pv tomato str DC3000 NP 793862 1 223283 Proteobacteria
Paenibacillus terrae HPL 003 YP 005076077 1 985665 Firmicutes
Paenibacillus polymyxa M1 YP 005958995 1 1052684 Firmicutes
Bacillus subtilis subsp spizizenii TU B 10 YP 004877506 1 1052585 Firmicutes
Bacillus subtilis subsp subtilis str BAB 1 YP 007662698 1 1302650 Firmicutes
Bacillus subtilis subsp subtilis 6051 HGW YP 007533824 1 1147161 Firmicutes
Bacillus subtilis PY79 YP 008830116 1 1415167 Firmicutes
Ruminococcus champanellensis 18P13 JCM 17042 YP 007828875 1 213810 Firmicutes
Bacillus licheniformis DSM 13 ATCC 14580 YP 080197 1 279010 Firmicutes
Bacillus atrophaeus 1942 YP 003975371 1 720555 Firmicutes
Bacillus amyloliquefaciens DSM 7 YP 003922341 1 692420 Firmicutes
Bacillus amyloliquefaciens subsp plantarum UCMB5033 YP 008414735 1 1338518 Firmicutes
Bacillus amyloliquefaciens Y2 YP 006330877 1 1126211 Firmicutes
Pseudomonas sp TKP YP 008932007 1 1415630 Proteobacteria
Pectobacterium sp SCC3193 YP 006282654 1 1166016 Proteobacteria
Pectobacterium wasabiae WPP163 YP 003259170 1 561231 Proteobacteria
Pectobacterium carotovorum subsp carotovorum PC1 YP 003016953 1 561230 Proteobacteria
Pectobacterium atrosepticum SCRI1043 YP 049604 1 218491 Proteobacteria
Dickeya chrysanthemi Ech1591 WP 012770972 1 Dchrysanthemi
Dickeya chrysanthemi NCPPB 3533 WP 012770972 1 Dchrysanthemi
Dickeya chrysanthemi subsp parthenii NCPPB 516 WP 033581025 1 Dchrysanthemi
Dickeya chrysanthemi subsp chrysanthemi NCPPB 402 WP 040002317 1 Dchrysanthemi
Dickeya dadantii 3937 WP 013318337 1 Ddadantii
Dickeya dianthicola IPO 980 WP 024106798 1 Ddianthicola
Dickeya dianthicola GBBC 2039 WP 024106798 1 Ddianthicola
Dickeya dianthicola NCPPB 453 WP 024106798 1 Ddianthicola
Dickeya dianthicola NCPPB 3534 WP 024106798 1 Ddianthicola
Dickeya dianthicola RNS04 9 WP 024106798 1 Ddianthicola
Dickeya paradisiaca Ech703 WP 015855039 1 Dparadisiaca
Dickeya paradisiaca NCPPB 2511 WP 015855039 1 Dparadisiaca
Prodigiosinella confusarubida ATCC39006 WP 021014347 1 Pconfusarubida
Dickeya gs fangzhongdai NCPPB 3274 WP 042858597 1 Dfangzhongdai
Dickeya gs fangzhongdai M074 WP 038662515 1 Dfangzhongdai
Dickeya gs fangzhongdai ND14b WP 038662515 1 Dfangzhongdai
Dickeya gs fangzhongdai M005 WP 039692560 1 Dfangzhongdai
Dickeya gs fangzhongdai S1 WP 049854602 1 Dfangzhongdai
Dickeya gs fangzhongdai MK7 WP 038917782 1 Dfangzhongdai
Dickeya gs fangzhongdai B16 WP 038917782 1 Dfangzhongdai
Dickeya gs undicola 2B12 WP 033570652 1 Dundicola
Dickeya zeae DZ2Q WP 038924975 1 Dzeae
Dickeya zeae ZJU1202 WP 016943143 1 Dzeae
Dickeya zeae EC1 WP 016943143 1 Dzeae
Dickeya zeae NCPPB 2538 WP 038915541 1 Dzeae
Dickeya zeae MS1 WP 019843777 1 Dzeae
Dickeya zeae NCPPB 3532 WP 019843777 1 Dzeae
Dickeya zeae MK19 WP 019843777 1 Dzeae
Dickeya dadantii subsp dieffenbachiae NCPPB 2976 WP 038921653 1 Ddadantii
Dickeya dianthicola NCPPB 3534 WP 024104511 1 Ddianthicola
Dickeya dianthicola RNS04 9 WP 024104511 1 Ddianthicola
Dickeya dianthicola NCPPB 453 WP 024104511 1 Ddianthicola
Dickeya dadantii 3937 WP 013316284 1 Ddadantii
Dickeya dadantii NCPPB 3537 WP 013316284 1 Ddadantii
153 pnlG Plant cell wall degradosome Seed
Dickeya dadantii NCPPB 898 WP 013316284 1 Ddadantii
Dickeya solani RNS 05 1 2A WP 057084230 1 Dsolani
Dickeya solani D s0432 1 WP 022632087 1 Dsolani
Dickeya solani IPO 2222 WP 022632087 1 Dsolani
Dickeya solani RNS 08 23 3 1 A WP 022632087 1 Dsolani
Dickeya solani PPO 9134 WP 022632087 1 Dsolani
Dickeya solani IFB 0099 WP 022632087 1 Dsolani
Dickeya solani MK16 WP 022632087 1 Dsolani
Dickeya solani RNS 07 7 3B WP 022632087 1 Dsolani
Dickeya solani GBBC 2040 WP 022632087 1 Dsolani
Dickeya solani PPO 9019 WP 022632087 1 Dsolani
Dickeya solani MK10 WP 022632087 1 Dsolani

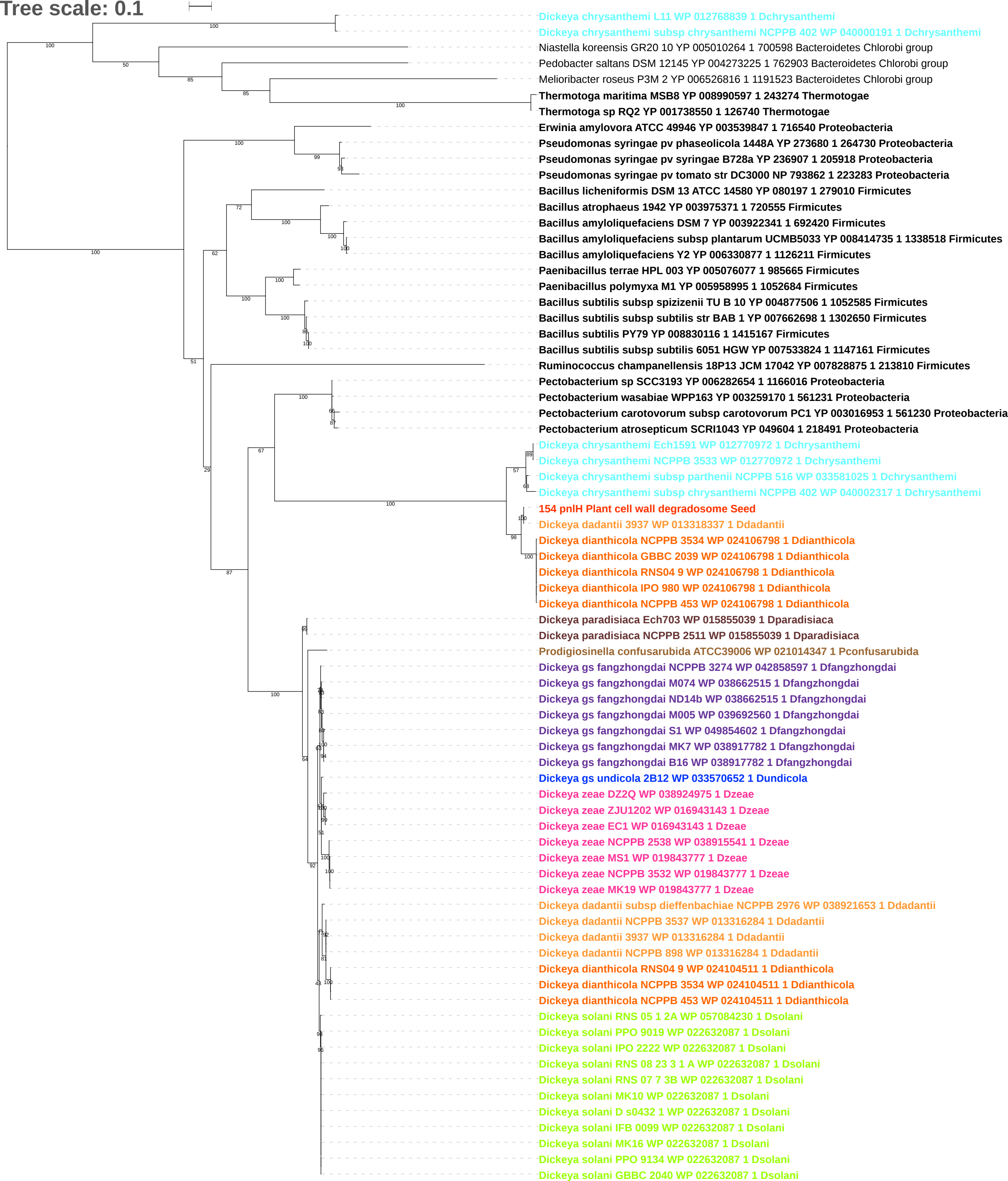
Tree scale: 0.1
Dickeya chrysanthemi L11 WP 012768839 1 Dchrysanthemi
Dickeya chrysanthemi subsp chrysanthemi NCPPB 402 WP 040000191 1 Dchrysanthemi
Niastella koreensis GR20 10 YP 005010264 1 700598 Bacteroidetes Chlorobi group
Pedobacter saltans DSM 12145 YP 004273225 1 762903 Bacteroidetes Chlorobi group
Melioribacter roseus P3M 2 YP 006526816 1 1191523 Bacteroidetes Chlorobi group
Thermotoga maritima MSB8 YP 008990597 1 243274 Thermotogae
Thermotoga sp RQ2 YP 001738550 1 126740 Thermotogae
Erwinia amylovora ATCC 49946 YP 003539847 1 716540 Proteobacteria
Pseudomonas syringae pv phaseolicola 1448A YP 273680 1 264730 Proteobacteria
Pseudomonas syringae pv syringae B728a YP 236907 1 205918 Proteobacteria
Pseudomonas syringae pv tomato str DC3000 NP 793862 1 223283 Proteobacteria
Bacillus licheniformis DSM 13 ATCC 14580 YP 080197 1 279010 Firmicutes
Bacillus atrophaeus 1942 YP 003975371 1 720555 Firmicutes
Bacillus amyloliquefaciens DSM 7 YP 003922341 1 692420 Firmicutes
Bacillus amyloliquefaciens subsp plantarum UCMB5033 YP 008414735 1 1338518 Firmicutes
Bacillus amyloliquefaciens Y2 YP 006330877 1 1126211 Firmicutes
Paenibacillus terrae HPL 003 YP 005076077 1 985665 Firmicutes
Paenibacillus polymyxa M1 YP 005958995 1 1052684 Firmicutes
Bacillus subtilis subsp spizizenii TU B 10 YP 004877506 1 1052585 Firmicutes
Bacillus subtilis subsp subtilis str BAB 1 YP 007662698 1 1302650 Firmicutes
Bacillus subtilis PY79 YP 008830116 1 1415167 Firmicutes
Bacillus subtilis subsp subtilis 6051 HGW YP 007533824 1 1147161 Firmicutes
Ruminococcus champanellensis 18P13 JCM 17042 YP 007828875 1 213810 Firmicutes
Pectobacterium sp SCC3193 YP 006282654 1 1166016 Proteobacteria
Pectobacterium wasabiae WPP163 YP 003259170 1 561231 Proteobacteria
Pectobacterium carotovorum subsp carotovorum PC1 YP 003016953 1 561230 Proteobacteria
Pectobacterium atrosepticum SCRI1043 YP 049604 1 218491 Proteobacteria
Dickeya chrysanthemi Ech1591 WP 012770972 1 Dchrysanthemi
Dickeya chrysanthemi NCPPB 3533 WP 012770972 1 Dchrysanthemi
Dickeya chrysanthemi subsp parthenii NCPPB 516 WP 033581025 1 Dchrysanthemi
Dickeya chrysanthemi subsp chrysanthemi NCPPB 402 WP 040002317 1 Dchrysanthemi
154 pnlH Plant cell wall degradosome Seed
Dickeya dadantii 3937 WP 013318337 1 Ddadantii
Dickeya dianthicola NCPPB 3534 WP 024106798 1 Ddianthicola
Dickeya dianthicola GBBC 2039 WP 024106798 1 Ddianthicola
Dickeya dianthicola RNS04 9 WP 024106798 1 Ddianthicola
Dickeya dianthicola IPO 980 WP 024106798 1 Ddianthicola
Dickeya dianthicola NCPPB 453 WP 024106798 1 Ddianthicola
Dickeya paradisiaca Ech703 WP 015855039 1 Dparadisiaca
Dickeya paradisiaca NCPPB 2511 WP 015855039 1 Dparadisiaca
Prodigiosinella confusarubida ATCC39006 WP 021014347 1 Pconfusarubida
Dickeya gs fangzhongdai NCPPB 3274 WP 042858597 1 Dfangzhongdai
Dickeya gs fangzhongdai M074 WP 038662515 1 Dfangzhongdai
Dickeya gs fangzhongdai ND14b WP 038662515 1 Dfangzhongdai
Dickeya gs fangzhongdai M005 WP 039692560 1 Dfangzhongdai
Dickeya gs fangzhongdai S1 WP 049854602 1 Dfangzhongdai
Dickeya gs fangzhongdai MK7 WP 038917782 1 Dfangzhongdai
Dickeya gs fangzhongdai B16 WP 038917782 1 Dfangzhongdai
Dickeya gs undicola 2B12 WP 033570652 1 Dundicola
Dickeya zeae DZ2Q WP 038924975 1 Dzeae
Dickeya zeae ZJU1202 WP 016943143 1 Dzeae
Dickeya zeae EC1 WP 016943143 1 Dzeae
Dickeya zeae NCPPB 2538 WP 038915541 1 Dzeae
Dickeya zeae MS1 WP 019843777 1 Dzeae
Dickeya zeae NCPPB 3532 WP 019843777 1 Dzeae
Dickeya zeae MK19 WP 019843777 1 Dzeae
Dickeya dadantii subsp dieffenbachiae NCPPB 2976 WP 038921653 1 Ddadantii
Dickeya dadantii NCPPB 3537 WP 013316284 1 Ddadantii
Dickeya dadantii 3937 WP 013316284 1 Ddadantii
Dickeya dadantii NCPPB 898 WP 013316284 1 Ddadantii
Dickeya dianthicola RNS04 9 WP 024104511 1 Ddianthicola
Dickeya dianthicola NCPPB 3534 WP 024104511 1 Ddianthicola
Dickeya dianthicola NCPPB 453 WP 024104511 1 Ddianthicola
Dickeya solani RNS 05 1 2A WP 057084230 1 Dsolani
Dickeya solani PPO 9019 WP 022632087 1 Dsolani
Dickeya solani IPO 2222 WP 022632087 1 Dsolani
Dickeya solani RNS 08 23 3 1 A WP 022632087 1 Dsolani
Dickeya solani RNS 07 7 3B WP 022632087 1 Dsolani
Dickeya solani MK10 WP 022632087 1 Dsolani
Dickeya solani D s0432 1 WP 022632087 1 Dsolani
Dickeya solani IFB 0099 WP 022632087 1 Dsolani
Dickeya solani MK16 WP 022632087 1 Dsolani
Dickeya solani PPO 9134 WP 022632087 1 Dsolani
Dickeya solani GBBC 2040 WP 022632087 1 Dsolani

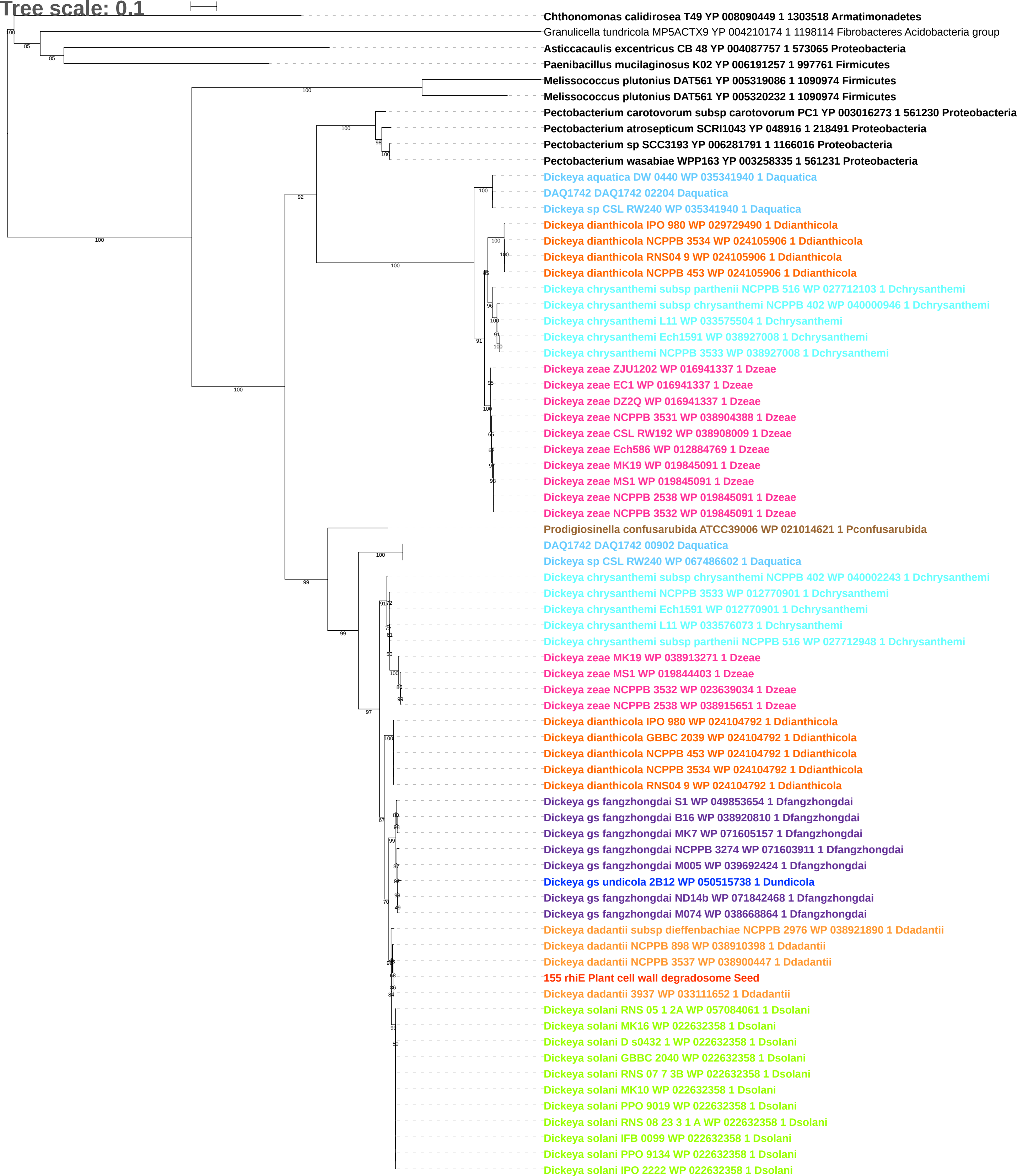
Tree scale: 0.1
Chthonomonas calidirosea T49 YP 008090449 1 1303518 Armatimonadetes
Granulicella tundricola MP5ACTX9 YP 004210174 1 1198114 Fibrobacteres Acidobacteria group
Asticcacaulis excentricus CB 48 YP 004087757 1 573065 Proteobacteria
Paenibacillus mucilaginosus K02 YP 006191257 1 997761 Firmicutes
Melissococcus plutonius DAT561 YP 005319086 1 1090974 Firmicutes
Melissococcus plutonius DAT561 YP 005320232 1 1090974 Firmicutes
Pectobacterium carotovorum subsp carotovorum PC1 YP 003016273 1 561230 Proteobacteria
Pectobacterium atrosepticum SCRI1043 YP 048916 1 218491 Proteobacteria
Pectobacterium sp SCC3193 YP 006281791 1 1166016 Proteobacteria
Pectobacterium wasabiae WPP163 YP 003258335 1 561231 Proteobacteria
Dickeya aquatica DW 0440 WP 035341940 1 Daquatica
DAQ1742 DAQ1742 02204 Daquatica
Dickeya sp CSL RW240 WP 035341940 1 Daquatica
Dickeya dianthicola IPO 980 WP 029729490 1 Ddianthicola
Dickeya dianthicola NCPPB 3534 WP 024105906 1 Ddianthicola
Dickeya dianthicola RNS04 9 WP 024105906 1 Ddianthicola
Dickeya dianthicola NCPPB 453 WP 024105906 1 Ddianthicola
Dickeya chrysanthemi subsp parthenii NCPPB 516 WP 027712103 1 Dchrysanthemi
Dickeya chrysanthemi subsp chrysanthemi NCPPB 402 WP 040000946 1 Dchrysanthemi
Dickeya chrysanthemi L11 WP 033575504 1 Dchrysanthemi
Dickeya chrysanthemi Ech1591 WP 038927008 1 Dchrysanthemi
Dickeya chrysanthemi NCPPB 3533 WP 038927008 1 Dchrysanthemi
Dickeya zeae ZJU1202 WP 016941337 1 Dzeae
Dickeya zeae EC1 WP 016941337 1 Dzeae
Dickeya zeae DZ2Q WP 016941337 1 Dzeae
Dickeya zeae NCPPB 3531 WP 038904388 1 Dzeae
Dickeya zeae CSL RW192 WP 038908009 1 Dzeae
Dickeya zeae Ech586 WP 012884769 1 Dzeae
Dickeya zeae MK19 WP 019845091 1 Dzeae
Dickeya zeae MS1 WP 019845091 1 Dzeae
Dickeya zeae NCPPB 2538 WP 019845091 1 Dzeae
Dickeya zeae NCPPB 3532 WP 019845091 1 Dzeae
Prodigiosinella confusarubida ATCC39006 WP 021014621 1 Pconfusarubida
DAQ1742 DAQ1742 00902 Daquatica
Dickeya sp CSL RW240 WP 067486602 1 Daquatica
Dickeya chrysanthemi subsp chrysanthemi NCPPB 402 WP 040002243 1 Dchrysanthemi
Dickeya chrysanthemi NCPPB 3533 WP 012770901 1 Dchrysanthemi
Dickeya chrysanthemi Ech1591 WP 012770901 1 Dchrysanthemi
Dickeya chrysanthemi L11 WP 033576073 1 Dchrysanthemi
Dickeya chrysanthemi subsp parthenii NCPPB 516 WP 027712948 1 Dchrysanthemi
Dickeya zeae MK19 WP 038913271 1 Dzeae
Dickeya zeae MS1 WP 019844403 1 Dzeae
Dickeya zeae NCPPB 3532 WP 023639034 1 Dzeae
Dickeya zeae NCPPB 2538 WP 038915651 1 Dzeae
Dickeya dianthicola IPO 980 WP 024104792 1 Ddianthicola
Dickeya dianthicola GBBC 2039 WP 024104792 1 Ddianthicola
Dickeya dianthicola NCPPB 453 WP 024104792 1 Ddianthicola
Dickeya dianthicola NCPPB 3534 WP 024104792 1 Ddianthicola
Dickeya dianthicola RNS04 9 WP 024104792 1 Ddianthicola
Dickeya gs fangzhongdai S1 WP 049853654 1 Dfangzhongdai
Dickeya gs fangzhongdai B16 WP 038920810 1 Dfangzhongdai
Dickeya gs fangzhongdai MK7 WP 071605157 1 Dfangzhongdai
Dickeya gs fangzhongdai NCPPB 3274 WP 071603911 1 Dfangzhongdai
Dickeya gs fangzhongdai M005 WP 039692424 1 Dfangzhongdai
Dickeya gs undicola 2B12 WP 050515738 1 Dundicola
Dickeya gs fangzhongdai ND14b WP 071842468 1 Dfangzhongdai
Dickeya gs fangzhongdai M074 WP 038668864 1 Dfangzhongdai
Dickeya dadantii subsp dieffenbachiae NCPPB 2976 WP 038921890 1 Ddadantii
Dickeya dadantii NCPPB 898 WP 038910398 1 Ddadantii
Dickeya dadantii NCPPB 3537 WP 038900447 1 Ddadantii
155 rhiE Plant cell wall degradosome Seed
Dickeya dadantii 3937 WP 033111652 1 Ddadantii
Dickeya solani RNS 05 1 2A WP 057084061 1 Dsolani
Dickeya solani MK16 WP 022632358 1 Dsolani
Dickeya solani D s0432 1 WP 022632358 1 Dsolani
Dickeya solani GBBC 2040 WP 022632358 1 Dsolani
Dickeya solani RNS 07 7 3B WP 022632358 1 Dsolani
Dickeya solani MK10 WP 022632358 1 Dsolani
Dickeya solani PPO 9019 WP 022632358 1 Dsolani
Dickeya solani RNS 08 23 3 1 A WP 022632358 1 Dsolani
Dickeya solani IFB 0099 WP 022632358 1 Dsolani
Dickeya solani PPO 9134 WP 022632358 1 Dsolani
Dickeya solani IPO 2222 WP 022632358 1 Dsolani

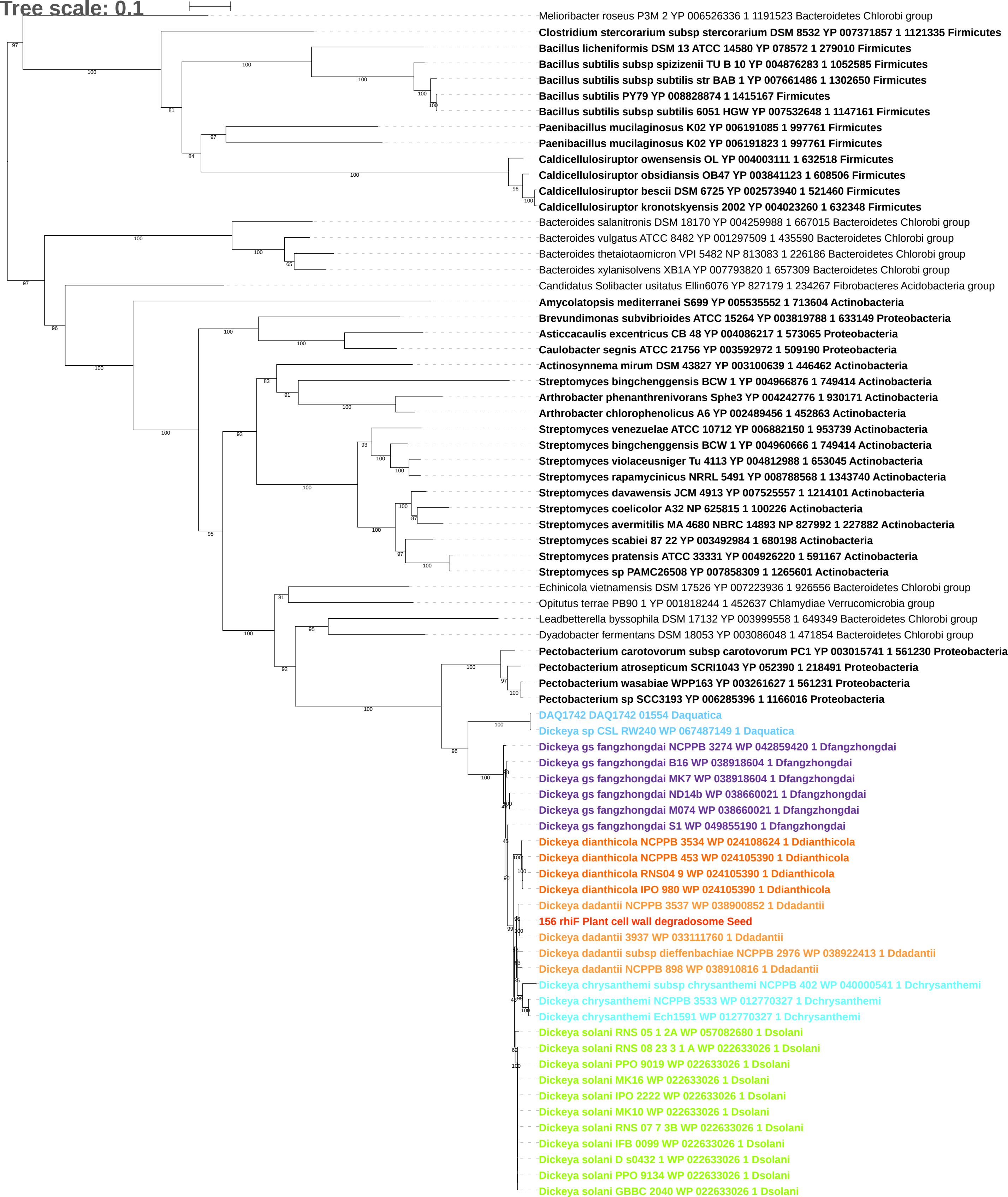
Tree scale: 0.1
Melioribacter roseus P3M 2 YP 006526336 1 1191523 Bacteroidetes Chlorobi group
Clostridium stercorarium subsp stercorarium DSM 8532 YP 007371857 1 1121335 Firmicutes
Bacillus licheniformis DSM 13 ATCC 14580 YP 078572 1 279010 Firmicutes
Bacillus subtilis subsp spizizenii TU B 10 YP 004876283 1 1052585 Firmicutes
Bacillus subtilis subsp subtilis str BAB 1 YP 007661486 1 1302650 Firmicutes
Bacillus subtilis PY79 YP 008828874 1 1415167 Firmicutes
Bacillus subtilis subsp subtilis 6051 HGW YP 007532648 1 1147161 Firmicutes
Paenibacillus mucilaginosus K02 YP 006191085 1 997761 Firmicutes
Paenibacillus mucilaginosus K02 YP 006191823 1 997761 Firmicutes
Caldicellulosiruptor owensensis OL YP 004003111 1 632518 Firmicutes
Caldicellulosiruptor obsidiansis OB47 YP 003841123 1 608506 Firmicutes
Caldicellulosiruptor bescii DSM 6725 YP 002573940 1 521460 Firmicutes
Caldicellulosiruptor kronotskyensis 2002 YP 004023260 1 632348 Firmicutes
Bacteroides salanitronis DSM 18170 YP 004259988 1 667015 Bacteroidetes Chlorobi group
Bacteroides vulgatus ATCC 8482 YP 001297509 1 435590 Bacteroidetes Chlorobi group
Bacteroides thetaiotaomicron VPI 5482 NP 813083 1 226186 Bacteroidetes Chlorobi group
Bacteroides xylanisolvens XB1A YP 007793820 1 657309 Bacteroidetes Chlorobi group
Candidatus Solibacter usitatus Ellin6076 YP 827179 1 234267 Fibrobacteres Acidobacteria group
Amycolatopsis mediterranei S699 YP 005535552 1 713604 Actinobacteria
Brevundimonas subvibrioides ATCC 15264 YP 003819788 1 633149 Proteobacteria
Asticcacaulis excentricus CB 48 YP 004086217 1 573065 Proteobacteria
Caulobacter segnis ATCC 21756 YP 003592972 1 509190 Proteobacteria
Actinosynnema mirum DSM 43827 YP 003100639 1 446462 Actinobacteria
Streptomyces bingchenggensis BCW 1 YP 004966876 1 749414 Actinobacteria
Arthrobacter phenanthrenivorans Sphe3 YP 004242776 1 930171 Actinobacteria
Arthrobacter chlorophenolicus A6 YP 002489456 1 452863 Actinobacteria
Streptomyces venezuelae ATCC 10712 YP 006882150 1 953739 Actinobacteria
Streptomyces bingchenggensis BCW 1 YP 004960666 1 749414 Actinobacteria
Streptomyces violaceusniger Tu 4113 YP 004812988 1 653045 Actinobacteria
Streptomyces rapamycinicus NRRL 5491 YP 008788568 1 1343740 Actinobacteria
Streptomyces davawensis JCM 4913 YP 007525557 1 1214101 Actinobacteria
Streptomyces coelicolor A32 NP 625815 1 100226 Actinobacteria
Streptomyces avermitilis MA 4680 NBRC 14893 NP 827992 1 227882 Actinobacteria
Streptomyces scabiei 87 22 YP 003492984 1 680198 Actinobacteria
Streptomyces pratensis ATCC 33331 YP 004926220 1 591167 Actinobacteria
Streptomyces sp PAMC26508 YP 007858309 1 1265601 Actinobacteria
Echinicola vietnamensis DSM 17526 YP 007223936 1 926556 Bacteroidetes Chlorobi group
Opitutus terrae PB90 1 YP 001818244 1 452637 Chlamydiae Verrucomicrobia group
Leadbetterella byssophila DSM 17132 YP 003999558 1 649349 Bacteroidetes Chlorobi group
Dyadobacter fermentans DSM 18053 YP 003086048 1 471854 Bacteroidetes Chlorobi group
Pectobacterium carotovorum subsp carotovorum PC1 YP 003015741 1 561230 Proteobacteria
Pectobacterium atrosepticum SCRI1043 YP 052390 1 218491 Proteobacteria
Pectobacterium wasabiae WPP163 YP 003261627 1 561231 Proteobacteria
Pectobacterium sp SCC3193 YP 006285396 1 1166016 Proteobacteria
DAQ1742 DAQ1742 01554 Daquatica
Dickeya sp CSL RW240 WP 067487149 1 Daquatica
Dickeya gs fangzhongdai NCPPB 3274 WP 042859420 1 Dfangzhongdai
Dickeya gs fangzhongdai B16 WP 038918604 1 Dfangzhongdai
Dickeya gs fangzhongdai MK7 WP 038918604 1 Dfangzhongdai
Dickeya gs fangzhongdai ND14b WP 038660021 1 Dfangzhongdai
Dickeya gs fangzhongdai M074 WP 038660021 1 Dfangzhongdai
Dickeya gs fangzhongdai S1 WP 049855190 1 Dfangzhongdai
Dickeya dianthicola NCPPB 3534 WP 024108624 1 Ddianthicola
Dickeya dianthicola NCPPB 453 WP 024105390 1 Ddianthicola
Dickeya dianthicola RNS04 9 WP 024105390 1 Ddianthicola
Dickeya dianthicola IPO 980 WP 024105390 1 Ddianthicola
Dickeya dadantii NCPPB 3537 WP 038900852 1 Ddadantii
156 rhiF Plant cell wall degradosome Seed
Dickeya dadantii 3937 WP 033111760 1 Ddadantii
Dickeya dadantii subsp dieffenbachiae NCPPB 2976 WP 038922413 1 Ddadantii
Dickeya dadantii NCPPB 898 WP 038910816 1 Ddadantii
Dickeya chrysanthemi subsp chrysanthemi NCPPB 402 WP 040000541 1 Dchrysanthemi
Dickeya chrysanthemi NCPPB 3533 WP 012770327 1 Dchrysanthemi
Dickeya chrysanthemi Ech1591 WP 012770327 1 Dchrysanthemi
Dickeya solani RNS 05 1 2A WP 057082680 1 Dsolani
Dickeya solani RNS 08 23 3 1 A WP 022633026 1 Dsolani
Dickeya solani PPO 9019 WP 022633026 1 Dsolani
Dickeya solani MK16 WP 022633026 1 Dsolani
Dickeya solani IPO 2222 WP 022633026 1 Dsolani
Dickeya solani MK10 WP 022633026 1 Dsolani
Dickeya solani RNS 07 7 3B WP 022633026 1 Dsolani
Dickeya solani IFB 0099 WP 022633026 1 Dsolani
Dickeya solani D s0432 1 WP 022633026 1 Dsolani
Dickeya solani PPO 9134 WP 022633026 1 Dsolani
Dickeya solani GBBC 2040 WP 022633026 1 Dsolani

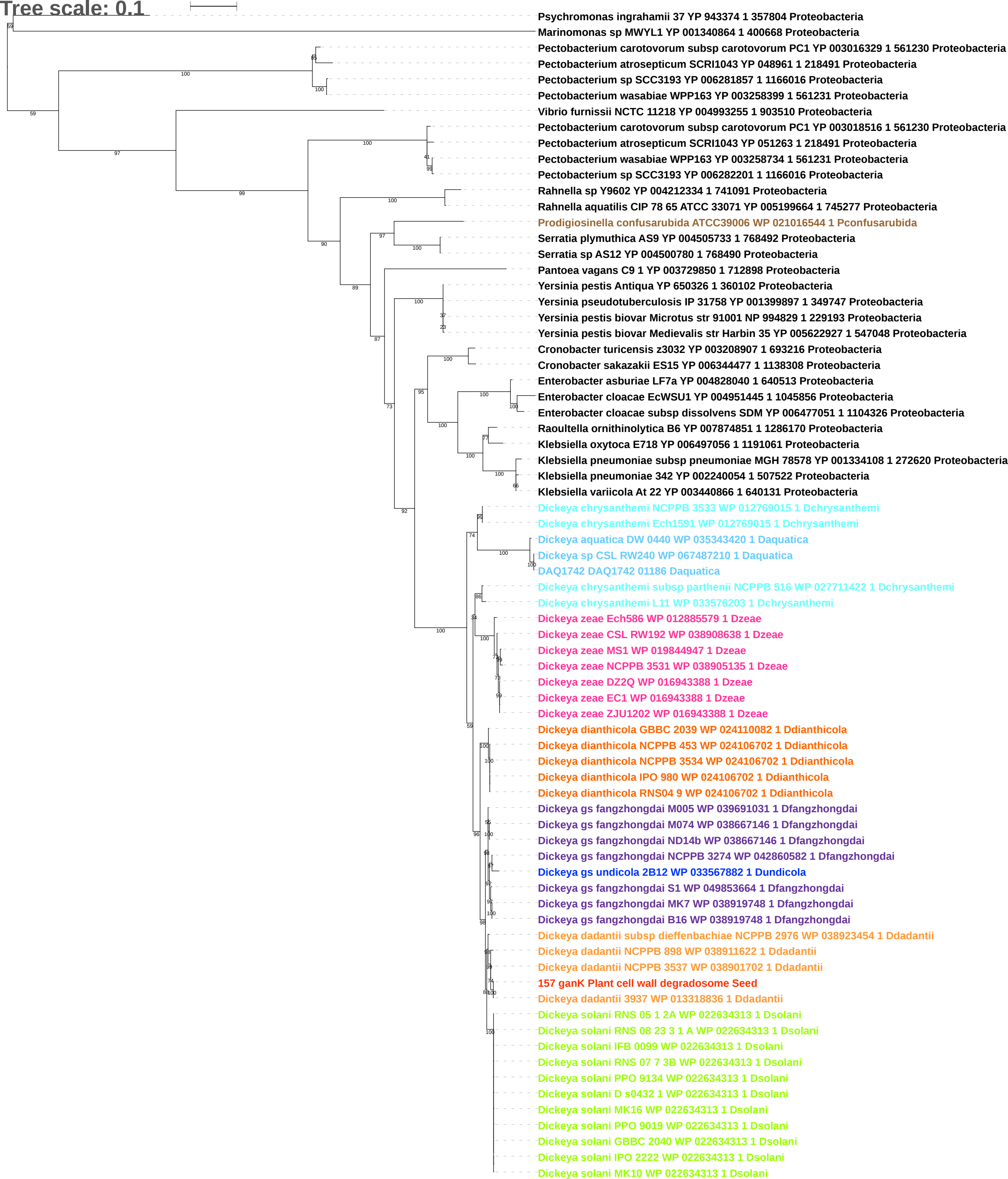
Tree scale: 0.1
Psychromonas ingrahamii 37 YP 943374 1 357804 Proteobacteria
Marinomonas sp MWYL1 YP 001340864 1 400668 Proteobacteria
Pectobacterium carotovorum subsp carotovorum PC1 YP 003016329 1 561230 Proteobacteria
Pectobacterium atrosepticum SCRI1043 YP 048961 1 218491 Proteobacteria
Pectobacterium sp SCC3193 YP 006281857 1 1166016 Proteobacteria
Pectobacterium wasabiae WPP163 YP 003258399 1 561231 Proteobacteria
Vibrio furnissii NCTC 11218 YP 004993255 1 903510 Proteobacteria
Pectobacterium carotovorum subsp carotovorum PC1 YP 003018516 1 561230 Proteobacteria
Pectobacterium atrosepticum SCRI1043 YP 051263 1 218491 Proteobacteria
Pectobacterium wasabiae WPP163 YP 003258734 1 561231 Proteobacteria
Pectobacterium sp SCC3193 YP 006282201 1 1166016 Proteobacteria
Rahnella sp Y9602 YP 004212334 1 741091 Proteobacteria
Rahnella aquatilis CIP 78 65 ATCC 33071 YP 005199664 1 745277 Proteobacteria
Prodigiosinella confusarubida ATCC39006 WP 021016544 1 Pconfusarubida
Serratia plymuthica AS9 YP 004505733 1 768492 Proteobacteria
Serratia sp AS12 YP 004500780 1 768490 Proteobacteria
Pantoea vagans C9 1 YP 003729850 1 712898 Proteobacteria
Yersinia pestis Antiqua YP 650326 1 360102 Proteobacteria
Yersinia pseudotuberculosis IP 31758 YP 001399897 1 349747 Proteobacteria
Yersinia pestis biovar Microtus str 91001 NP 994829 1 229193 Proteobacteria
Yersinia pestis biovar Medievalis str Harbin 35 YP 005622927 1 547048 Proteobacteria
Cronobacter turicensis z3032 YP 003208907 1 693216 Proteobacteria
Cronobacter sakazakii ES15 YP 006344477 1 1138308 Proteobacteria
Enterobacter asburiae LF7a YP 004828040 1 640513 Proteobacteria
Enterobacter cloacae EcWSU1 YP 004951445 1 1045856 Proteobacteria
Enterobacter cloacae subsp dissolvens SDM YP 006477051 1 1104326 Proteobacteria
Raoultella ornithinolytica B6 YP 007874851 1 1286170 Proteobacteria
Klebsiella oxytoca E718 YP 006497056 1 1191061 Proteobacteria
Klebsiella pneumoniae subsp pneumoniae MGH 78578 YP 001334108 1 272620 Proteobacteria
Klebsiella pneumoniae 342 YP 002240054 1 507522 Proteobacteria
Klebsiella variicola At 22 YP 003440866 1 640131 Proteobacteria
Dickeya chrysanthemi NCPPB 3533 WP 012769015 1 Dchrysanthemi
Dickeya chrysanthemi Ech1591 WP 012769015 1 Dchrysanthemi
Dickeya aquatica DW 0440 WP 035343420 1 Daquatica
Dickeya sp CSL RW240 WP 067487210 1 Daquatica
DAQ1742 DAQ1742 01186 Daquatica
Dickeya chrysanthemi subsp parthenii NCPPB 516 WP 027711422 1 Dchrysanthemi
Dickeya chrysanthemi L11 WP 033576203 1 Dchrysanthemi
Dickeya zeae Ech586 WP 012885579 1 Dzeae
Dickeya zeae CSL RW192 WP 038908638 1 Dzeae
Dickeya zeae MS1 WP 019844947 1 Dzeae
Dickeya zeae NCPPB 3531 WP 038905135 1 Dzeae
Dickeya zeae DZ2Q WP 016943388 1 Dzeae
Dickeya zeae EC1 WP 016943388 1 Dzeae
Dickeya zeae ZJU1202 WP 016943388 1 Dzeae
Dickeya dianthicola GBBC 2039 WP 024110082 1 Ddianthicola
Dickeya dianthicola NCPPB 453 WP 024106702 1 Ddianthicola
Dickeya dianthicola NCPPB 3534 WP 024106702 1 Ddianthicola
Dickeya dianthicola IPO 980 WP 024106702 1 Ddianthicola
Dickeya dianthicola RNS04 9 WP 024106702 1 Ddianthicola
Dickeya gs fangzhongdai M005 WP 039691031 1 Dfangzhongdai
Dickeya gs fangzhongdai M074 WP 038667146 1 Dfangzhongdai
Dickeya gs fangzhongdai ND14b WP 038667146 1 Dfangzhongdai
Dickeya gs fangzhongdai NCPPB 3274 WP 042860582 1 Dfangzhongdai
Dickeya gs undicola 2B12 WP 033567882 1 Dundicola
Dickeya gs fangzhongdai S1 WP 049853664 1 Dfangzhongdai
Dickeya gs fangzhongdai MK7 WP 038919748 1 Dfangzhongdai
Dickeya gs fangzhongdai B16 WP 038919748 1 Dfangzhongdai
Dickeya dadantii subsp dieffenbachiae NCPPB 2976 WP 038923454 1 Ddadantii
Dickeya dadantii NCPPB 898 WP 038911622 1 Ddadantii
Dickeya dadantii NCPPB 3537 WP 038901702 1 Ddadantii
157 ganK Plant cell wall degradosome Seed
Dickeya dadantii 3937 WP 013318836 1 Ddadantii
Dickeya solani RNS 05 1 2A WP 022634313 1 Dsolani
Dickeya solani RNS 08 23 3 1 A WP 022634313 1 Dsolani
Dickeya solani IFB 0099 WP 022634313 1 Dsolani
Dickeya solani RNS 07 7 3B WP 022634313 1 Dsolani
Dickeya solani PPO 9134 WP 022634313 1 Dsolani
Dickeya solani D s0432 1 WP 022634313 1 Dsolani
Dickeya solani MK16 WP 022634313 1 Dsolani
Dickeya solani PPO 9019 WP 022634313 1 Dsolani
Dickeya solani GBBC 2040 WP 022634313 1 Dsolani
Dickeya solani IPO 2222 WP 022634313 1 Dsolani
Dickeya solani MK10 WP 022634313 1 Dsolani

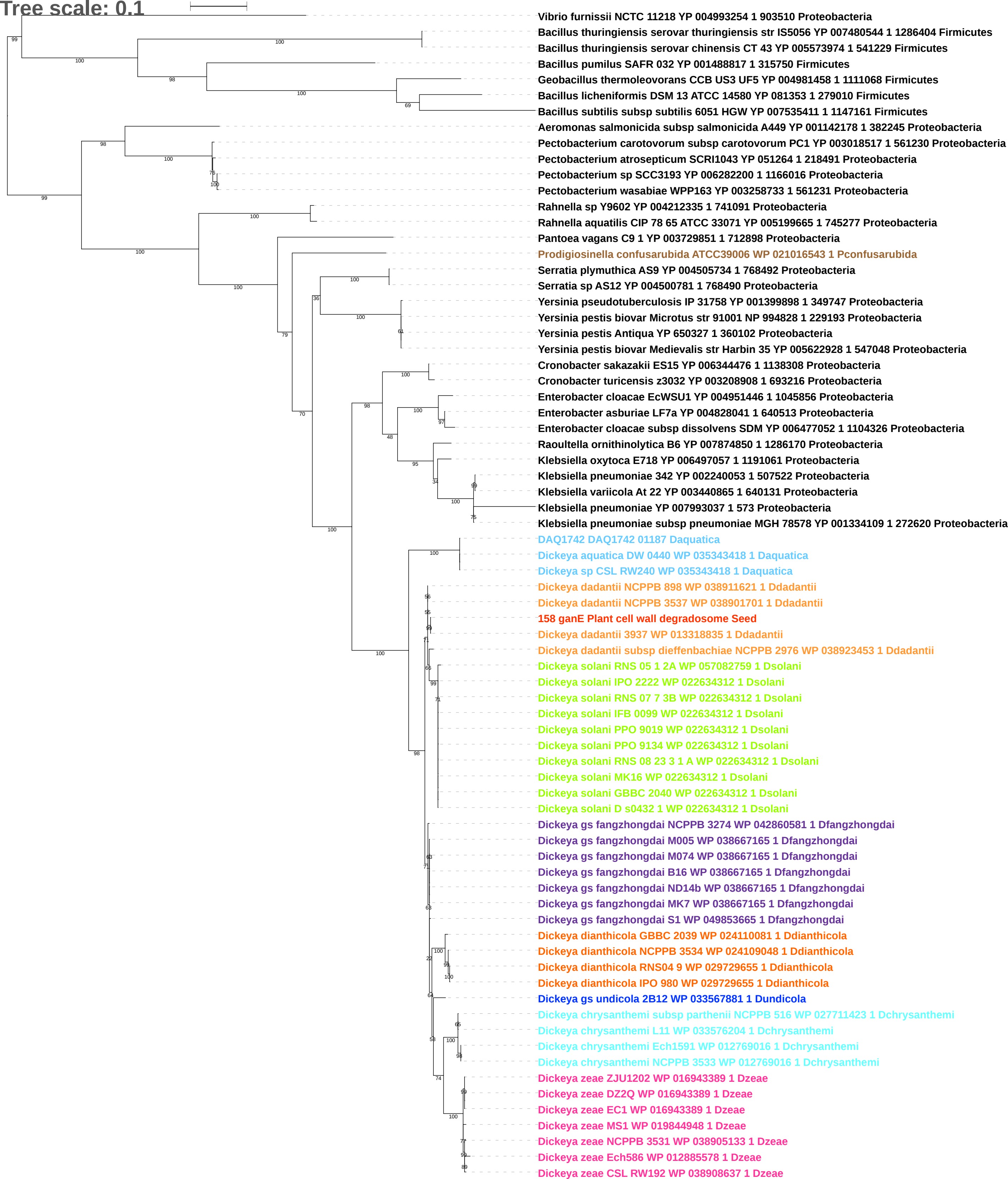
Tree scale: 0.1
Vibrio furnissii NCTC 11218 YP 004993254 1 903510 Proteobacteria
Bacillus thuringiensis serovar thuringiensis str IS5056 YP 007480544 1 1286404 Firmicutes
Bacillus thuringiensis serovar chinensis CT 43 YP 005573974 1 541229 Firmicutes
Bacillus pumilus SAFR 032 YP 001488817 1 315750 Firmicutes
Geobacillus thermoleovorans CCB US3 UF5 YP 004981458 1 1111068 Firmicutes
Bacillus licheniformis DSM 13 ATCC 14580 YP 081353 1 279010 Firmicutes
Bacillus subtilis subsp subtilis 6051 HGW YP 007535411 1 1147161 Firmicutes
Aeromonas salmonicida subsp salmonicida A449 YP 001142178 1 382245 Proteobacteria
Pectobacterium carotovorum subsp carotovorum PC1 YP 003018517 1 561230 Proteobacteria
Pectobacterium atrosepticum SCRI1043 YP 051264 1 218491 Proteobacteria
Pectobacterium sp SCC3193 YP 006282200 1 1166016 Proteobacteria
Pectobacterium wasabiae WPP163 YP 003258733 1 561231 Proteobacteria
Rahnella sp Y9602 YP 004212335 1 741091 Proteobacteria
Rahnella aquatilis CIP 78 65 ATCC 33071 YP 005199665 1 745277 Proteobacteria
Pantoea vagans C9 1 YP 003729851 1 712898 Proteobacteria
Prodigiosinella confusarubida ATCC39006 WP 021016543 1 Pconfusarubida
Serratia plymuthica AS9 YP 004505734 1 768492 Proteobacteria
Serratia sp AS12 YP 004500781 1 768490 Proteobacteria
Yersinia pseudotuberculosis IP 31758 YP 001399898 1 349747 Proteobacteria
Yersinia pestis biovar Microtus str 91001 NP 994828 1 229193 Proteobacteria
Yersinia pestis Antiqua YP 650327 1 360102 Proteobacteria
Yersinia pestis biovar Medievalis str Harbin 35 YP 005622928 1 547048 Proteobacteria
Cronobacter sakazakii ES15 YP 006344476 1 1138308 Proteobacteria
Cronobacter turicensis z3032 YP 003208908 1 693216 Proteobacteria
Enterobacter cloacae EcWSU1 YP 004951446 1 1045856 Proteobacteria
Enterobacter asburiae LF7a YP 004828041 1 640513 Proteobacteria
Enterobacter cloacae subsp dissolvens SDM YP 006477052 1 1104326 Proteobacteria
Raoultella ornithinolytica B6 YP 007874850 1 1286170 Proteobacteria
Klebsiella oxytoca E718 YP 006497057 1 1191061 Proteobacteria
Klebsiella pneumoniae 342 YP 002240053 1 507522 Proteobacteria
Klebsiella variicola At 22 YP 003440865 1 640131 Proteobacteria
Klebsiella pneumoniae YP 007993037 1 573 Proteobacteria
Klebsiella pneumoniae subsp pneumoniae MGH 78578 YP 001334109 1 272620 Proteobacteria
DAQ1742 DAQ1742 01187 Daquatica
Dickeya aquatica DW 0440 WP 035343418 1 Daquatica
Dickeya sp CSL RW240 WP 035343418 1 Daquatica
Dickeya dadantii NCPPB 898 WP 038911621 1 Ddadantii
Dickeya dadantii NCPPB 3537 WP 038901701 1 Ddadantii
158 ganE Plant cell wall degradosome Seed
Dickeya dadantii 3937 WP 013318835 1 Ddadantii
Dickeya dadantii subsp dieffenbachiae NCPPB 2976 WP 038923453 1 Ddadantii
Dickeya solani RNS 05 1 2A WP 057082759 1 Dsolani
Dickeya solani IPO 2222 WP 022634312 1 Dsolani
Dickeya solani RNS 07 7 3B WP 022634312 1 Dsolani
Dickeya solani IFB 0099 WP 022634312 1 Dsolani
Dickeya solani PPO 9019 WP 022634312 1 Dsolani
Dickeya solani PPO 9134 WP 022634312 1 Dsolani
Dickeya solani RNS 08 23 3 1 A WP 022634312 1 Dsolani
Dickeya solani MK16 WP 022634312 1 Dsolani
Dickeya solani GBBC 2040 WP 022634312 1 Dsolani
Dickeya solani D s0432 1 WP 022634312 1 Dsolani
Dickeya gs fangzhongdai NCPPB 3274 WP 042860581 1 Dfangzhongdai
Dickeya gs fangzhongdai M005 WP 038667165 1 Dfangzhongdai
Dickeya gs fangzhongdai M074 WP 038667165 1 Dfangzhongdai
Dickeya gs fangzhongdai B16 WP 038667165 1 Dfangzhongdai
Dickeya gs fangzhongdai ND14b WP 038667165 1 Dfangzhongdai
Dickeya gs fangzhongdai MK7 WP 038667165 1 Dfangzhongdai
Dickeya gs fangzhongdai S1 WP 049853665 1 Dfangzhongdai
Dickeya dianthicola GBBC 2039 WP 024110081 1 Ddianthicola
Dickeya dianthicola NCPPB 3534 WP 024109048 1 Ddianthicola
Dickeya dianthicola RNS04 9 WP 029729655 1 Ddianthicola
Dickeya dianthicola IPO 980 WP 029729655 1 Ddianthicola
Dickeya gs undicola 2B12 WP 033567881 1 Dundicola
Dickeya chrysanthemi subsp parthenii NCPPB 516 WP 027711423 1 Dchrysanthemi
Dickeya chrysanthemi L11 WP 033576204 1 Dchrysanthemi
Dickeya chrysanthemi Ech1591 WP 012769016 1 Dchrysanthemi
Dickeya chrysanthemi NCPPB 3533 WP 012769016 1 Dchrysanthemi
Dickeya zeae ZJU1202 WP 016943389 1 Dzeae
Dickeya zeae DZ2Q WP 016943389 1 Dzeae
Dickeya zeae EC1 WP 016943389 1 Dzeae
Dickeya zeae MS1 WP 019844948 1 Dzeae
Dickeya zeae NCPPB 3531 WP 038905133 1 Dzeae
Dickeya zeae Ech586 WP 012885578 1 Dzeae
Dickeya zeae CSL RW192 WP 038908637 1 Dzeae

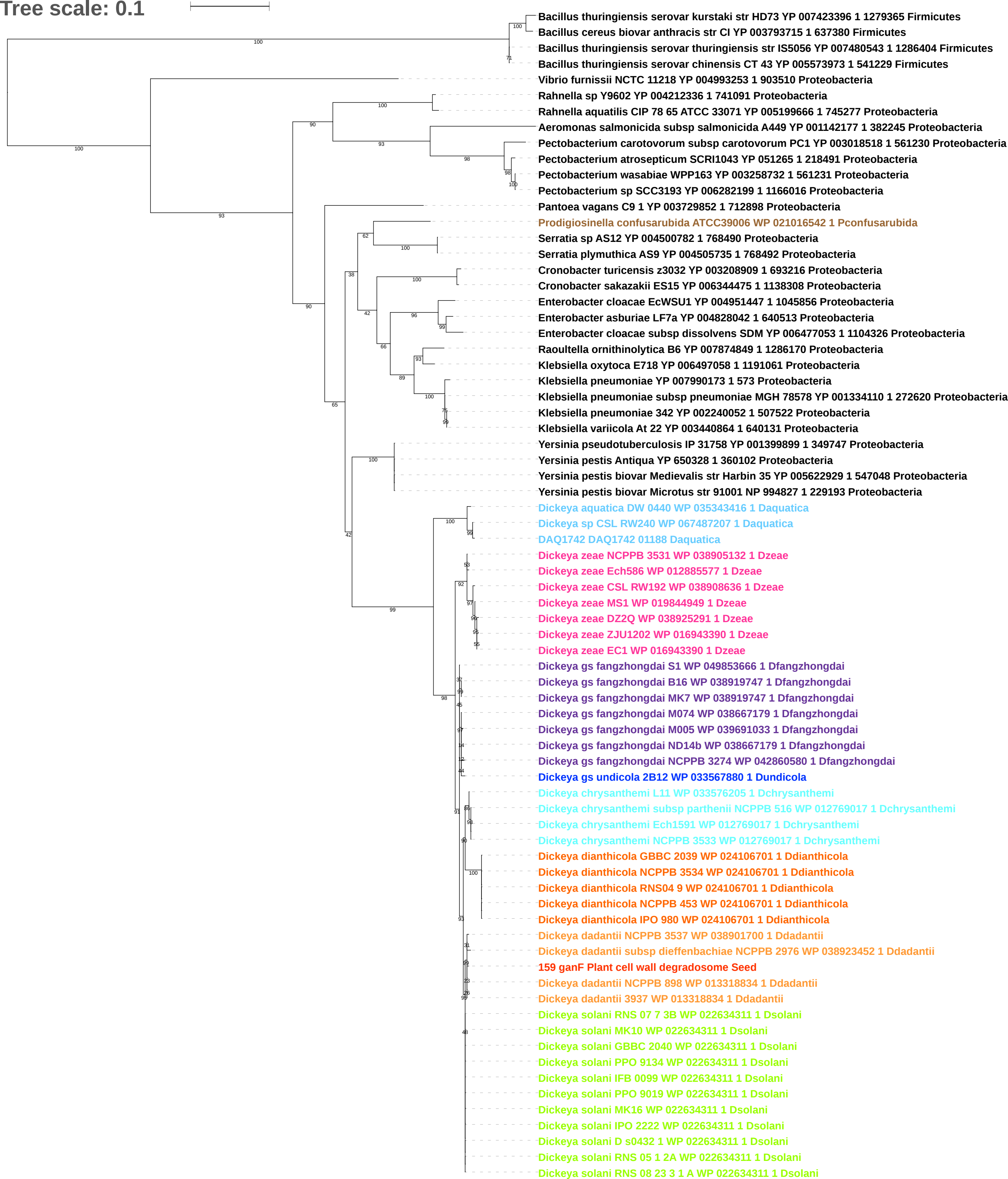
Tree scale: 0.1
Bacillus thuringiensis serovar kurstaki str HD73 YP 007423396 1 1279365 Firmicutes
Bacillus cereus biovar anthracis str CI YP 003793715 1 637380 Firmicutes
Bacillus thuringiensis serovar thuringiensis str IS5056 YP 007480543 1 1286404 Firmicutes
Bacillus thuringiensis serovar chinensis CT 43 YP 005573973 1 541229 Firmicutes
Vibrio furnissii NCTC 11218 YP 004993253 1 903510 Proteobacteria
Rahnella sp Y9602 YP 004212336 1 741091 Proteobacteria
Rahnella aquatilis CIP 78 65 ATCC 33071 YP 005199666 1 745277 Proteobacteria
Aeromonas salmonicida subsp salmonicida A449 YP 001142177 1 382245 Proteobacteria
Pectobacterium carotovorum subsp carotovorum PC1 YP 003018518 1 561230 Proteobacteria
Pectobacterium atrosepticum SCRI1043 YP 051265 1 218491 Proteobacteria
Pectobacterium wasabiae WPP163 YP 003258732 1 561231 Proteobacteria
Pectobacterium sp SCC3193 YP 006282199 1 1166016 Proteobacteria
Pantoea vagans C9 1 YP 003729852 1 712898 Proteobacteria
Prodigiosinella confusarubida ATCC39006 WP 021016542 1 Pconfusarubida
Serratia sp AS12 YP 004500782 1 768490 Proteobacteria
Serratia plymuthica AS9 YP 004505735 1 768492 Proteobacteria
Cronobacter turicensis z3032 YP 003208909 1 693216 Proteobacteria
Cronobacter sakazakii ES15 YP 006344475 1 1138308 Proteobacteria
Enterobacter cloacae EcWSU1 YP 004951447 1 1045856 Proteobacteria
Enterobacter asburiae LF7a YP 004828042 1 640513 Proteobacteria
Enterobacter cloacae subsp dissolvens SDM YP 006477053 1 1104326 Proteobacteria
Raoultella ornithinolytica B6 YP 007874849 1 1286170 Proteobacteria
Klebsiella oxytoca E718 YP 006497058 1 1191061 Proteobacteria
Klebsiella pneumoniae YP 007990173 1 573 Proteobacteria
Klebsiella pneumoniae subsp pneumoniae MGH 78578 YP 001334110 1 272620 Proteobacteria
Klebsiella pneumoniae 342 YP 002240052 1 507522 Proteobacteria
Klebsiella variicola At 22 YP 003440864 1 640131 Proteobacteria
Yersinia pseudotuberculosis IP 31758 YP 001399899 1 349747 Proteobacteria
Yersinia pestis Antiqua YP 650328 1 360102 Proteobacteria
Yersinia pestis biovar Medievalis str Harbin 35 YP 005622929 1 547048 Proteobacteria
Yersinia pestis biovar Microtus str 91001 NP 994827 1 229193 Proteobacteria
Dickeya aquatica DW 0440 WP 035343416 1 Daquatica
Dickeya sp CSL RW240 WP 067487207 1 Daquatica
DAQ1742 DAQ1742 01188 Daquatica
Dickeya zeae NCPPB 3531 WP 038905132 1 Dzeae
Dickeya zeae Ech586 WP 012885577 1 Dzeae
Dickeya zeae CSL RW192 WP 038908636 1 Dzeae
Dickeya zeae MS1 WP 019844949 1 Dzeae
Dickeya zeae DZ2Q WP 038925291 1 Dzeae
Dickeya zeae ZJU1202 WP 016943390 1 Dzeae
Dickeya zeae EC1 WP 016943390 1 Dzeae
Dickeya gs fangzhongdai S1 WP 049853666 1 Dfangzhongdai
Dickeya gs fangzhongdai B16 WP 038919747 1 Dfangzhongdai
Dickeya gs fangzhongdai MK7 WP 038919747 1 Dfangzhongdai
Dickeya gs fangzhongdai M074 WP 038667179 1 Dfangzhongdai
Dickeya gs fangzhongdai M005 WP 039691033 1 Dfangzhongdai
Dickeya gs fangzhongdai ND14b WP 038667179 1 Dfangzhongdai
Dickeya gs fangzhongdai NCPPB 3274 WP 042860580 1 Dfangzhongdai
Dickeya gs undicola 2B12 WP 033567880 1 Dundicola
Dickeya chrysanthemi L11 WP 033576205 1 Dchrysanthemi
Dickeya chrysanthemi subsp parthenii NCPPB 516 WP 012769017 1 Dchrysanthemi
Dickeya chrysanthemi Ech1591 WP 012769017 1 Dchrysanthemi
Dickeya chrysanthemi NCPPB 3533 WP 012769017 1 Dchrysanthemi
Dickeya dianthicola GBBC 2039 WP 024106701 1 Ddianthicola
Dickeya dianthicola NCPPB 3534 WP 024106701 1 Ddianthicola
Dickeya dianthicola RNS04 9 WP 024106701 1 Ddianthicola
Dickeya dianthicola NCPPB 453 WP 024106701 1 Ddianthicola
Dickeya dianthicola IPO 980 WP 024106701 1 Ddianthicola
Dickeya dadantii NCPPB 3537 WP 038901700 1 Ddadantii
Dickeya dadantii subsp dieffenbachiae NCPPB 2976 WP 038923452 1 Ddadantii
159 ganF Plant cell wall degradosome Seed
Dickeya dadantii NCPPB 898 WP 013318834 1 Ddadantii
Dickeya dadantii 3937 WP 013318834 1 Ddadantii
Dickeya solani RNS 07 7 3B WP 022634311 1 Dsolani
Dickeya solani MK10 WP 022634311 1 Dsolani
Dickeya solani GBBC 2040 WP 022634311 1 Dsolani
Dickeya solani PPO 9134 WP 022634311 1 Dsolani
Dickeya solani IFB 0099 WP 022634311 1 Dsolani
Dickeya solani PPO 9019 WP 022634311 1 Dsolani
Dickeya solani MK16 WP 022634311 1 Dsolani
Dickeya solani IPO 2222 WP 022634311 1 Dsolani
Dickeya solani D s0432 1 WP 022634311 1 Dsolani
Dickeya solani RNS 05 1 2A WP 022634311 1 Dsolani
Dickeya solani RNS 08 23 3 1 A WP 022634311 1 Dsolani

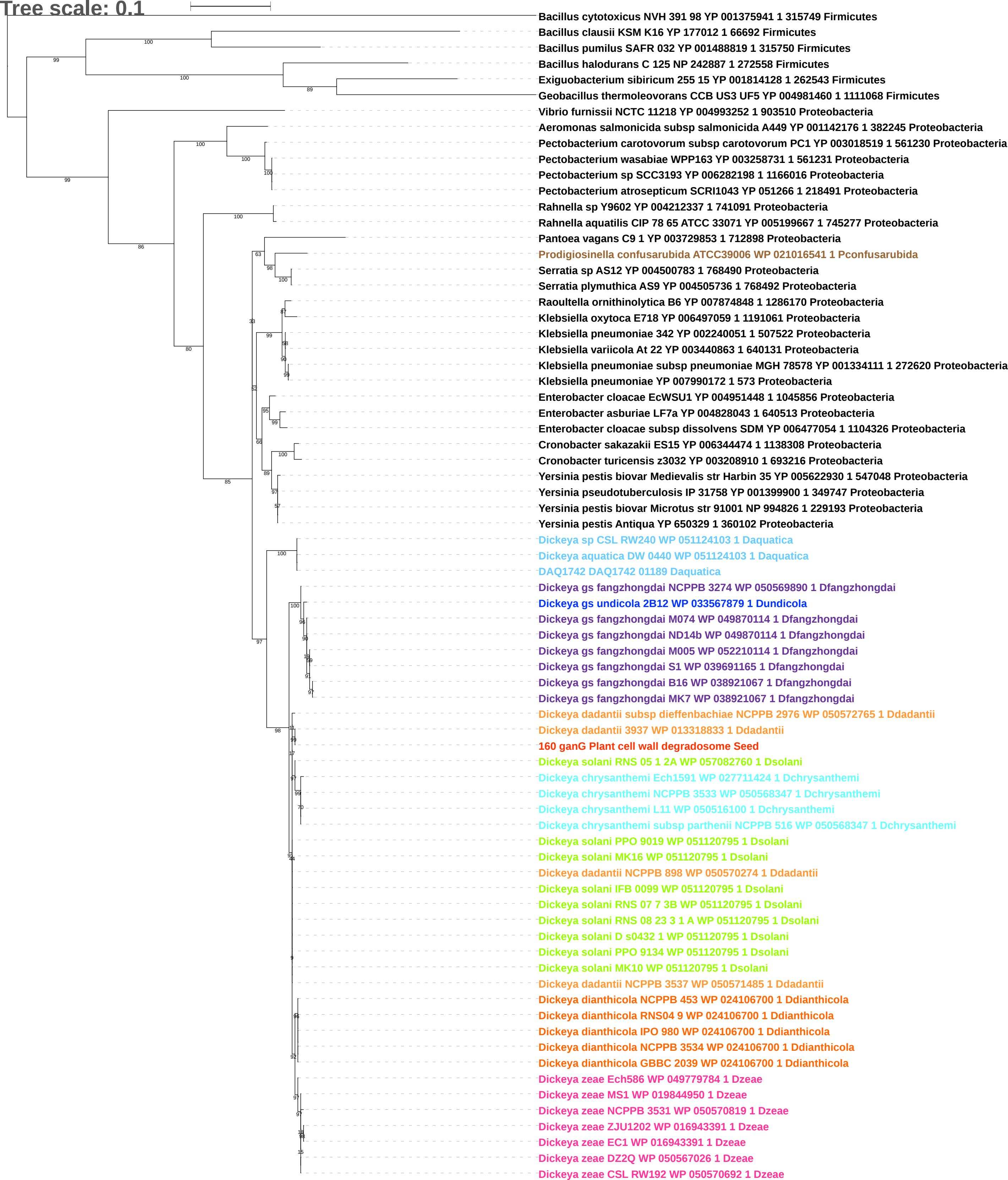
Tree scale: 0.1
Bacillus cytotoxicus NVH 391 98 YP 001375941 1 315749 Firmicutes
Bacillus clausii KSM K16 YP 177012 1 66692 Firmicutes
Bacillus pumilus SAFR 032 YP 001488819 1 315750 Firmicutes
Bacillus halodurans C 125 NP 242887 1 272558 Firmicutes
Exiguobacterium sibiricum 255 15 YP 001814128 1 262543 Firmicutes
Geobacillus thermoleovorans CCB US3 UF5 YP 004981460 1 1111068 Firmicutes
Vibrio furnissii NCTC 11218 YP 004993252 1 903510 Proteobacteria
Aeromonas salmonicida subsp salmonicida A449 YP 001142176 1 382245 Proteobacteria
Pectobacterium carotovorum subsp carotovorum PC1 YP 003018519 1 561230 Proteobacteria
Pectobacterium wasabiae WPP163 YP 003258731 1 561231 Proteobacteria
Pectobacterium sp SCC3193 YP 006282198 1 1166016 Proteobacteria
Pectobacterium atrosepticum SCRI1043 YP 051266 1 218491 Proteobacteria
Rahnella sp Y9602 YP 004212337 1 741091 Proteobacteria
Rahnella aquatilis CIP 78 65 ATCC 33071 YP 005199667 1 745277 Proteobacteria
Pantoea vagans C9 1 YP 003729853 1 712898 Proteobacteria
Prodigiosinella confusarubida ATCC39006 WP 021016541 1 Pconfusarubida
Serratia sp AS12 YP 004500783 1 768490 Proteobacteria
Serratia plymuthica AS9 YP 004505736 1 768492 Proteobacteria
Raoultella ornithinolytica B6 YP 007874848 1 1286170 Proteobacteria
Klebsiella oxytoca E718 YP 006497059 1 1191061 Proteobacteria
Klebsiella pneumoniae 342 YP 002240051 1 507522 Proteobacteria
Klebsiella variicola At 22 YP 003440863 1 640131 Proteobacteria
Klebsiella pneumoniae subsp pneumoniae MGH 78578 YP 001334111 1 272620 Proteobacteria
Klebsiella pneumoniae YP 007990172 1 573 Proteobacteria
Enterobacter cloacae EcWSU1 YP 004951448 1 1045856 Proteobacteria
Enterobacter asburiae LF7a YP 004828043 1 640513 Proteobacteria
Enterobacter cloacae subsp dissolvens SDM YP 006477054 1 1104326 Proteobacteria
Cronobacter sakazakii ES15 YP 006344474 1 1138308 Proteobacteria
Cronobacter turicensis z3032 YP 003208910 1 693216 Proteobacteria
Yersinia pestis biovar Medievalis str Harbin 35 YP 005622930 1 547048 Proteobacteria
Yersinia pseudotuberculosis IP 31758 YP 001399900 1 349747 Proteobacteria
Yersinia pestis biovar Microtus str 91001 NP 994826 1 229193 Proteobacteria
Yersinia pestis Antiqua YP 650329 1 360102 Proteobacteria
Dickeya sp CSL RW240 WP 051124103 1 Daquatica
Dickeya aquatica DW 0440 WP 051124103 1 Daquatica
DAQ1742 DAQ1742 01189 Daquatica
Dickeya gs fangzhongdai NCPPB 3274 WP 050569890 1 Dfangzhongdai
Dickeya gs undicola 2B12 WP 033567879 1 Dundicola
Dickeya gs fangzhongdai M074 WP 049870114 1 Dfangzhongdai
Dickeya gs fangzhongdai ND14b WP 049870114 1 Dfangzhongdai
Dickeya gs fangzhongdai M005 WP 052210114 1 Dfangzhongdai
Dickeya gs fangzhongdai S1 WP 039691165 1 Dfangzhongdai
Dickeya gs fangzhongdai B16 WP 038921067 1 Dfangzhongdai
Dickeya gs fangzhongdai MK7 WP 038921067 1 Dfangzhongdai
Dickeya dadantii subsp dieffenbachiae NCPPB 2976 WP 050572765 1 Ddadantii
Dickeya dadantii 3937 WP 013318833 1 Ddadantii
160 ganG Plant cell wall degradosome Seed
Dickeya solani RNS 05 1 2A WP 057082760 1 Dsolani
Dickeya chrysanthemi Ech1591 WP 027711424 1 Dchrysanthemi
Dickeya chrysanthemi NCPPB 3533 WP 050568347 1 Dchrysanthemi
Dickeya chrysanthemi L11 WP 050516100 1 Dchrysanthemi
Dickeya chrysanthemi subsp parthenii NCPPB 516 WP 050568347 1 Dchrysanthemi
Dickeya solani PPO 9019 WP 051120795 1 Dsolani
Dickeya solani MK16 WP 051120795 1 Dsolani
Dickeya dadantii NCPPB 898 WP 050570274 1 Ddadantii
Dickeya solani IFB 0099 WP 051120795 1 Dsolani
Dickeya solani RNS 07 7 3B WP 051120795 1 Dsolani
Dickeya solani RNS 08 23 3 1 A WP 051120795 1 Dsolani
Dickeya solani D s0432 1 WP 051120795 1 Dsolani
Dickeya solani PPO 9134 WP 051120795 1 Dsolani
Dickeya solani MK10 WP 051120795 1 Dsolani
Dickeya dadantii NCPPB 3537 WP 050571485 1 Ddadantii
Dickeya dianthicola NCPPB 453 WP 024106700 1 Ddianthicola
Dickeya dianthicola RNS04 9 WP 024106700 1 Ddianthicola
Dickeya dianthicola IPO 980 WP 024106700 1 Ddianthicola
Dickeya dianthicola NCPPB 3534 WP 024106700 1 Ddianthicola
Dickeya dianthicola GBBC 2039 WP 024106700 1 Ddianthicola
Dickeya zeae Ech586 WP 049779784 1 Dzeae
Dickeya zeae MS1 WP 019844950 1 Dzeae
Dickeya zeae NCPPB 3531 WP 050570819 1 Dzeae
Dickeya zeae ZJU1202 WP 016943391 1 Dzeae
Dickeya zeae EC1 WP 016943391 1 Dzeae
Dickeya zeae DZ2Q WP 050567026 1 Dzeae
Dickeya zeae CSL RW192 WP 050570692 1 Dzeae

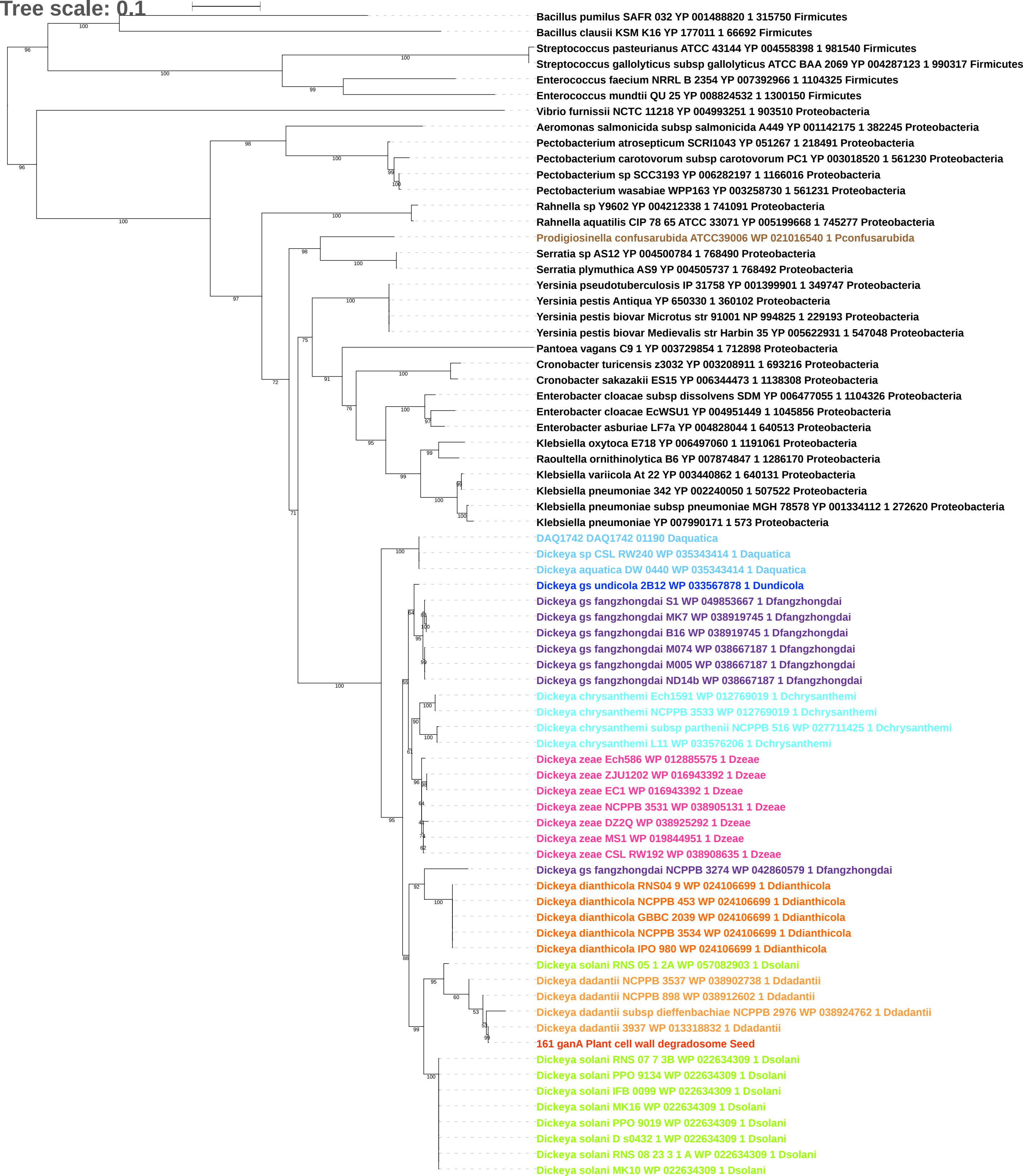
Tree scale: 0.1
Bacillus pumilus SAFR 032 YP 001488820 1 315750 Firmicutes
Bacillus clausii KSM K16 YP 177011 1 66692 Firmicutes
Streptococcus pasteurianus ATCC 43144 YP 004558398 1 981540 Firmicutes
Streptococcus gallolyticus subsp gallolyticus ATCC BAA 2069 YP 004287123 1 990317 Firmicutes
Enterococcus faecium NRRL B 2354 YP 007392966 1 1104325 Firmicutes
Enterococcus mundtii QU 25 YP 008824532 1 1300150 Firmicutes
Vibrio furnissii NCTC 11218 YP 004993251 1 903510 Proteobacteria
Aeromonas salmonicida subsp salmonicida A449 YP 001142175 1 382245 Proteobacteria
Pectobacterium atrosepticum SCRI1043 YP 051267 1 218491 Proteobacteria
Pectobacterium carotovorum subsp carotovorum PC1 YP 003018520 1 561230 Proteobacteria
Pectobacterium sp SCC3193 YP 006282197 1 1166016 Proteobacteria
Pectobacterium wasabiae WPP163 YP 003258730 1 561231 Proteobacteria
Rahnella sp Y9602 YP 004212338 1 741091 Proteobacteria
Rahnella aquatilis CIP 78 65 ATCC 33071 YP 005199668 1 745277 Proteobacteria
Prodigiosinella confusarubida ATCC39006 WP 021016540 1 Pconfusarubida
Serratia sp AS12 YP 004500784 1 768490 Proteobacteria
Serratia plymuthica AS9 YP 004505737 1 768492 Proteobacteria
Yersinia pseudotuberculosis IP 31758 YP 001399901 1 349747 Proteobacteria
Yersinia pestis Antiqua YP 650330 1 360102 Proteobacteria
Yersinia pestis biovar Microtus str 91001 NP 994825 1 229193 Proteobacteria
Yersinia pestis biovar Medievalis str Harbin 35 YP 005622931 1 547048 Proteobacteria
Pantoea vagans C9 1 YP 003729854 1 712898 Proteobacteria
Cronobacter turicensis z3032 YP 003208911 1 693216 Proteobacteria
Cronobacter sakazakii ES15 YP 006344473 1 1138308 Proteobacteria
Enterobacter cloacae subsp dissolvens SDM YP 006477055 1 1104326 Proteobacteria
Enterobacter cloacae EcWSU1 YP 004951449 1 1045856 Proteobacteria
Enterobacter asburiae LF7a YP 004828044 1 640513 Proteobacteria
Klebsiella oxytoca E718 YP 006497060 1 1191061 Proteobacteria
Raoultella ornithinolytica B6 YP 007874847 1 1286170 Proteobacteria
Klebsiella variicola At 22 YP 003440862 1 640131 Proteobacteria
Klebsiella pneumoniae 342 YP 002240050 1 507522 Proteobacteria
Klebsiella pneumoniae subsp pneumoniae MGH 78578 YP 001334112 1 272620 Proteobacteria
Klebsiella pneumoniae YP 007990171 1 573 Proteobacteria
DAQ1742 DAQ1742 01190 Daquatica
Dickeya sp CSL RW240 WP 035343414 1 Daquatica
Dickeya aquatica DW 0440 WP 035343414 1 Daquatica
Dickeya gs undicola 2B12 WP 033567878 1 Dundicola
Dickeya gs fangzhongdai S1 WP 049853667 1 Dfangzhongdai
Dickeya gs fangzhongdai MK7 WP 038919745 1 Dfangzhongdai
Dickeya gs fangzhongdai B16 WP 038919745 1 Dfangzhongdai
Dickeya gs fangzhongdai M074 WP 038667187 1 Dfangzhongdai
Dickeya gs fangzhongdai M005 WP 038667187 1 Dfangzhongdai
Dickeya gs fangzhongdai ND14b WP 038667187 1 Dfangzhongdai
Dickeya chrysanthemi Ech1591 WP 012769019 1 Dchrysanthemi
Dickeya chrysanthemi NCPPB 3533 WP 012769019 1 Dchrysanthemi
Dickeya chrysanthemi subsp parthenii NCPPB 516 WP 027711425 1 Dchrysanthemi
Dickeya chrysanthemi L11 WP 033576206 1 Dchrysanthemi
Dickeya zeae Ech586 WP 012885575 1 Dzeae
Dickeya zeae ZJU1202 WP 016943392 1 Dzeae
Dickeya zeae EC1 WP 016943392 1 Dzeae
Dickeya zeae NCPPB 3531 WP 038905131 1 Dzeae
Dickeya zeae DZ2Q WP 038925292 1 Dzeae
Dickeya zeae MS1 WP 019844951 1 Dzeae
Dickeya zeae CSL RW192 WP 038908635 1 Dzeae
Dickeya gs fangzhongdai NCPPB 3274 WP 042860579 1 Dfangzhongdai
Dickeya dianthicola RNS04 9 WP 024106699 1 Ddianthicola
Dickeya dianthicola NCPPB 453 WP 024106699 1 Ddianthicola
Dickeya dianthicola GBBC 2039 WP 024106699 1 Ddianthicola
Dickeya dianthicola NCPPB 3534 WP 024106699 1 Ddianthicola
Dickeya dianthicola IPO 980 WP 024106699 1 Ddianthicola
Dickeya solani RNS 05 1 2A WP 057082903 1 Dsolani
Dickeya dadantii NCPPB 3537 WP 038902738 1 Ddadantii
Dickeya dadantii NCPPB 898 WP 038912602 1 Ddadantii
Dickeya dadantii subsp dieffenbachiae NCPPB 2976 WP 038924762 1 Ddadantii
Dickeya dadantii 3937 WP 013318832 1 Ddadantii
161 ganA Plant cell wall degradosome Seed
Dickeya solani RNS 07 7 3B WP 022634309 1 Dsolani
Dickeya solani PPO 9134 WP 022634309 1 Dsolani
Dickeya solani IFB 0099 WP 022634309 1 Dsolani
Dickeya solani MK16 WP 022634309 1 Dsolani
Dickeya solani PPO 9019 WP 022634309 1 Dsolani
Dickeya solani D s0432 1 WP 022634309 1 Dsolani
Dickeya solani RNS 08 23 3 1 A WP 022634309 1 Dsolani
Dickeya solani MK10 WP 022634309 1 Dsolani

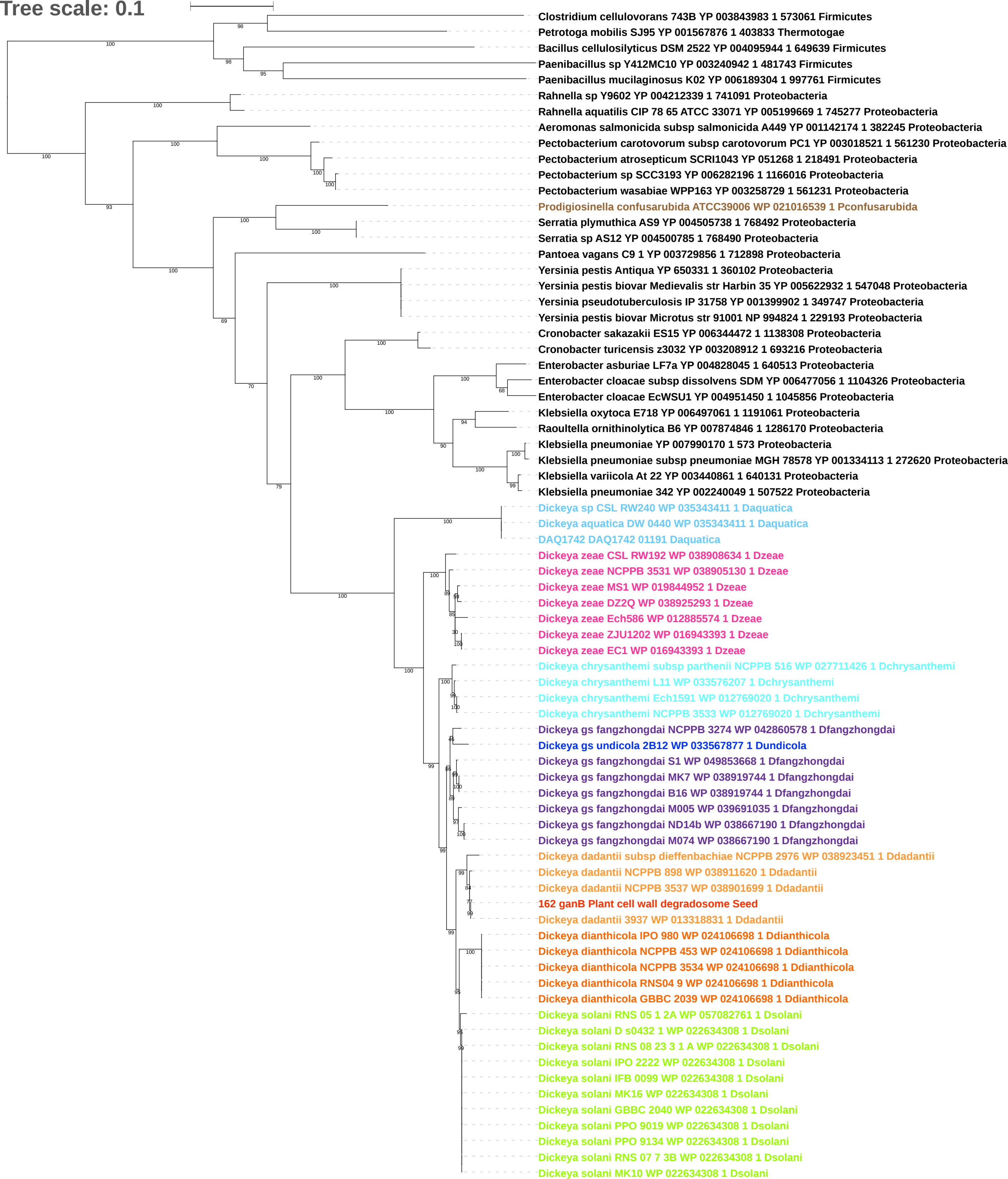
Tree scale: 0.1
Clostridium cellulovorans 743B YP 003843983 1 573061 Firmicutes
Petrotoga mobilis SJ95 YP 001567876 1 403833 Thermotogae
Bacillus cellulosilyticus DSM 2522 YP 004095944 1 649639 Firmicutes
Paenibacillus sp Y412MC10 YP 003240942 1 481743 Firmicutes
Paenibacillus mucilaginosus K02 YP 006189304 1 997761 Firmicutes
Rahnella sp Y9602 YP 004212339 1 741091 Proteobacteria
Rahnella aquatilis CIP 78 65 ATCC 33071 YP 005199669 1 745277 Proteobacteria
Aeromonas salmonicida subsp salmonicida A449 YP 001142174 1 382245 Proteobacteria
Pectobacterium carotovorum subsp carotovorum PC1 YP 003018521 1 561230 Proteobacteria
Pectobacterium atrosepticum SCRI1043 YP 051268 1 218491 Proteobacteria
Pectobacterium sp SCC3193 YP 006282196 1 1166016 Proteobacteria
Pectobacterium wasabiae WPP163 YP 003258729 1 561231 Proteobacteria
Prodigiosinella confusarubida ATCC39006 WP 021016539 1 Pconfusarubida
Serratia plymuthica AS9 YP 004505738 1 768492 Proteobacteria
Serratia sp AS12 YP 004500785 1 768490 Proteobacteria
Pantoea vagans C9 1 YP 003729856 1 712898 Proteobacteria
Yersinia pestis Antiqua YP 650331 1 360102 Proteobacteria
Yersinia pestis biovar Medievalis str Harbin 35 YP 005622932 1 547048 Proteobacteria
Yersinia pseudotuberculosis IP 31758 YP 001399902 1 349747 Proteobacteria
Yersinia pestis biovar Microtus str 91001 NP 994824 1 229193 Proteobacteria
Cronobacter sakazakii ES15 YP 006344472 1 1138308 Proteobacteria
Cronobacter turicensis z3032 YP 003208912 1 693216 Proteobacteria
Enterobacter asburiae LF7a YP 004828045 1 640513 Proteobacteria
Enterobacter cloacae subsp dissolvens SDM YP 006477056 1 1104326 Proteobacteria
Enterobacter cloacae EcWSU1 YP 004951450 1 1045856 Proteobacteria
Klebsiella oxytoca E718 YP 006497061 1 1191061 Proteobacteria
Raoultella ornithinolytica B6 YP 007874846 1 1286170 Proteobacteria
Klebsiella pneumoniae YP 007990170 1 573 Proteobacteria
Klebsiella pneumoniae subsp pneumoniae MGH 78578 YP 001334113 1 272620 Proteobacteria
Klebsiella variicola At 22 YP 003440861 1 640131 Proteobacteria
Klebsiella pneumoniae 342 YP 002240049 1 507522 Proteobacteria
Dickeya sp CSL RW240 WP 035343411 1 Daquatica
Dickeya aquatica DW 0440 WP 035343411 1 Daquatica
DAQ1742 DAQ1742 01191 Daquatica
Dickeya zeae CSL RW192 WP 038908634 1 Dzeae
Dickeya zeae NCPPB 3531 WP 038905130 1 Dzeae
Dickeya zeae MS1 WP 019844952 1 Dzeae
Dickeya zeae DZ2Q WP 038925293 1 Dzeae
Dickeya zeae Ech586 WP 012885574 1 Dzeae
Dickeya zeae ZJU1202 WP 016943393 1 Dzeae
Dickeya zeae EC1 WP 016943393 1 Dzeae
Dickeya chrysanthemi subsp parthenii NCPPB 516 WP 027711426 1 Dchrysanthemi
Dickeya chrysanthemi L11 WP 033576207 1 Dchrysanthemi
Dickeya chrysanthemi Ech1591 WP 012769020 1 Dchrysanthemi
Dickeya chrysanthemi NCPPB 3533 WP 012769020 1 Dchrysanthemi
Dickeya gs fangzhongdai NCPPB 3274 WP 042860578 1 Dfangzhongdai
Dickeya gs undicola 2B12 WP 033567877 1 Dundicola
Dickeya gs fangzhongdai S1 WP 049853668 1 Dfangzhongdai
Dickeya gs fangzhongdai MK7 WP 038919744 1 Dfangzhongdai
Dickeya gs fangzhongdai B16 WP 038919744 1 Dfangzhongdai
Dickeya gs fangzhongdai M005 WP 039691035 1 Dfangzhongdai
Dickeya gs fangzhongdai ND14b WP 038667190 1 Dfangzhongdai
Dickeya gs fangzhongdai M074 WP 038667190 1 Dfangzhongdai
Dickeya dadantii subsp dieffenbachiae NCPPB 2976 WP 038923451 1 Ddadantii
Dickeya dadantii NCPPB 898 WP 038911620 1 Ddadantii
Dickeya dadantii NCPPB 3537 WP 038901699 1 Ddadantii
162 ganB Plant cell wall degradosome Seed
Dickeya dadantii 3937 WP 013318831 1 Ddadantii
Dickeya dianthicola IPO 980 WP 024106698 1 Ddianthicola
Dickeya dianthicola NCPPB 453 WP 024106698 1 Ddianthicola
Dickeya dianthicola NCPPB 3534 WP 024106698 1 Ddianthicola
Dickeya dianthicola RNS04 9 WP 024106698 1 Ddianthicola
Dickeya dianthicola GBBC 2039 WP 024106698 1 Ddianthicola
Dickeya solani RNS 05 1 2A WP 057082761 1 Dsolani
Dickeya solani D s0432 1 WP 022634308 1 Dsolani
Dickeya solani RNS 08 23 3 1 A WP 022634308 1 Dsolani
Dickeya solani IPO 2222 WP 022634308 1 Dsolani
Dickeya solani IFB 0099 WP 022634308 1 Dsolani
Dickeya solani MK16 WP 022634308 1 Dsolani
Dickeya solani GBBC 2040 WP 022634308 1 Dsolani
Dickeya solani PPO 9019 WP 022634308 1 Dsolani
Dickeya solani PPO 9134 WP 022634308 1 Dsolani
Dickeya solani RNS 07 7 3B WP 022634308 1 Dsolani
Dickeya solani MK10 WP 022634308 1 Dsolani

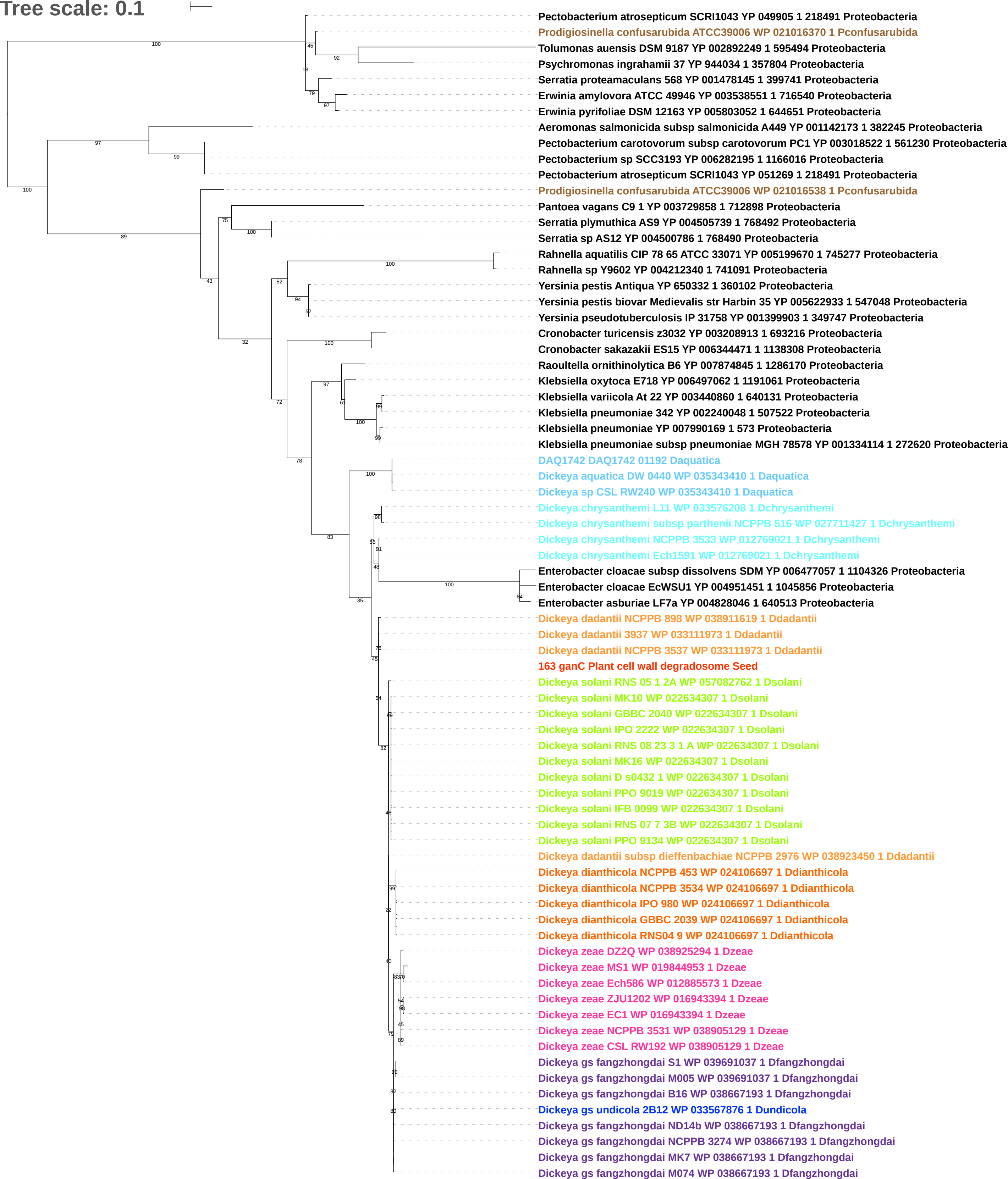
Tree scale: 0.1
Pectobacterium atrosepticum SCRI1043 YP 049905 1 218491 Proteobacteria
Prodigiosinella confusarubida ATCC39006 WP 021016370 1 Pconfusarubida
Tolumonas auensis DSM 9187 YP 002892249 1 595494 Proteobacteria
Psychromonas ingrahamii 37 YP 944034 1 357804 Proteobacteria
Serratia proteamaculans 568 YP 001478145 1 399741 Proteobacteria
Erwinia amylovora ATCC 49946 YP 003538551 1 716540 Proteobacteria
Erwinia pyrifoliae DSM 12163 YP 005803052 1 644651 Proteobacteria
Aeromonas salmonicida subsp salmonicida A449 YP 001142173 1 382245 Proteobacteria
Pectobacterium carotovorum subsp carotovorum PC1 YP 003018522 1 561230 Proteobacteria
Pectobacterium sp SCC3193 YP 006282195 1 1166016 Proteobacteria
Pectobacterium atrosepticum SCRI1043 YP 051269 1 218491 Proteobacteria
Prodigiosinella confusarubida ATCC39006 WP 021016538 1 Pconfusarubida
Pantoea vagans C9 1 YP 003729858 1 712898 Proteobacteria
Serratia plymuthica AS9 YP 004505739 1 768492 Proteobacteria
Serratia sp AS12 YP 004500786 1 768490 Proteobacteria
Rahnella aquatilis CIP 78 65 ATCC 33071 YP 005199670 1 745277 Proteobacteria
Rahnella sp Y9602 YP 004212340 1 741091 Proteobacteria
Yersinia pestis Antiqua YP 650332 1 360102 Proteobacteria
Yersinia pestis biovar Medievalis str Harbin 35 YP 005622933 1 547048 Proteobacteria
Yersinia pseudotuberculosis IP 31758 YP 001399903 1 349747 Proteobacteria
Cronobacter turicensis z3032 YP 003208913 1 693216 Proteobacteria
Cronobacter sakazakii ES15 YP 006344471 1 1138308 Proteobacteria
Raoultella ornithinolytica B6 YP 007874845 1 1286170 Proteobacteria
Klebsiella oxytoca E718 YP 006497062 1 1191061 Proteobacteria
Klebsiella variicola At 22 YP 003440860 1 640131 Proteobacteria
Klebsiella pneumoniae 342 YP 002240048 1 507522 Proteobacteria
Klebsiella pneumoniae YP 007990169 1 573 Proteobacteria
Klebsiella pneumoniae subsp pneumoniae MGH 78578 YP 001334114 1 272620 Proteobacteria
DAQ1742 DAQ1742 01192 Daquatica
Dickeya aquatica DW 0440 WP 035343410 1 Daquatica
Dickeya sp CSL RW240 WP 035343410 1 Daquatica
Dickeya chrysanthemi L11 WP 033576208 1 Dchrysanthemi
Dickeya chrysanthemi subsp parthenii NCPPB 516 WP 027711427 1 Dchrysanthemi
Dickeya chrysanthemi NCPPB 3533 WP 012769021 1 Dchrysanthemi
Dickeya chrysanthemi Ech1591 WP 012769021 1 Dchrysanthemi
Enterobacter cloacae subsp dissolvens SDM YP 006477057 1 1104326 Proteobacteria
Enterobacter cloacae EcWSU1 YP 004951451 1 1045856 Proteobacteria
Enterobacter asburiae LF7a YP 004828046 1 640513 Proteobacteria
Dickeya dadantii NCPPB 898 WP 038911619 1 Ddadantii
Dickeya dadantii 3937 WP 033111973 1 Ddadantii
Dickeya dadantii NCPPB 3537 WP 033111973 1 Ddadantii
163 ganC Plant cell wall degradosome Seed
Dickeya solani RNS 05 1 2A WP 057082762 1 Dsolani
Dickeya solani MK10 WP 022634307 1 Dsolani
Dickeya solani GBBC 2040 WP 022634307 1 Dsolani
Dickeya solani IPO 2222 WP 022634307 1 Dsolani
Dickeya solani RNS 08 23 3 1 A WP 022634307 1 Dsolani
Dickeya solani MK16 WP 022634307 1 Dsolani
Dickeya solani D s0432 1 WP 022634307 1 Dsolani
Dickeya solani PPO 9019 WP 022634307 1 Dsolani
Dickeya solani IFB 0099 WP 022634307 1 Dsolani
Dickeya solani RNS 07 7 3B WP 022634307 1 Dsolani
Dickeya solani PPO 9134 WP 022634307 1 Dsolani
Dickeya dadantii subsp dieffenbachiae NCPPB 2976 WP 038923450 1 Ddadantii
Dickeya dianthicola NCPPB 453 WP 024106697 1 Ddianthicola
Dickeya dianthicola NCPPB 3534 WP 024106697 1 Ddianthicola
Dickeya dianthicola IPO 980 WP 024106697 1 Ddianthicola
Dickeya dianthicola GBBC 2039 WP 024106697 1 Ddianthicola
Dickeya dianthicola RNS04 9 WP 024106697 1 Ddianthicola
Dickeya zeae DZ2Q WP 038925294 1 Dzeae
Dickeya zeae MS1 WP 019844953 1 Dzeae
Dickeya zeae Ech586 WP 012885573 1 Dzeae
Dickeya zeae ZJU1202 WP 016943394 1 Dzeae
Dickeya zeae EC1 WP 016943394 1 Dzeae
Dickeya zeae NCPPB 3531 WP 038905129 1 Dzeae
Dickeya zeae CSL RW192 WP 038905129 1 Dzeae
Dickeya gs fangzhongdai S1 WP 039691037 1 Dfangzhongdai
Dickeya gs fangzhongdai M005 WP 039691037 1 Dfangzhongdai
Dickeya gs fangzhongdai B16 WP 038667193 1 Dfangzhongdai
Dickeya gs undicola 2B12 WP 033567876 1 Dundicola
Dickeya gs fangzhongdai ND14b WP 038667193 1 Dfangzhongdai
Dickeya gs fangzhongdai NCPPB 3274 WP 038667193 1 Dfangzhongdai
Dickeya gs fangzhongdai MK7 WP 038667193 1 Dfangzhongdai
Dickeya gs fangzhongdai M074 WP 038667193 1 Dfangzhongdai

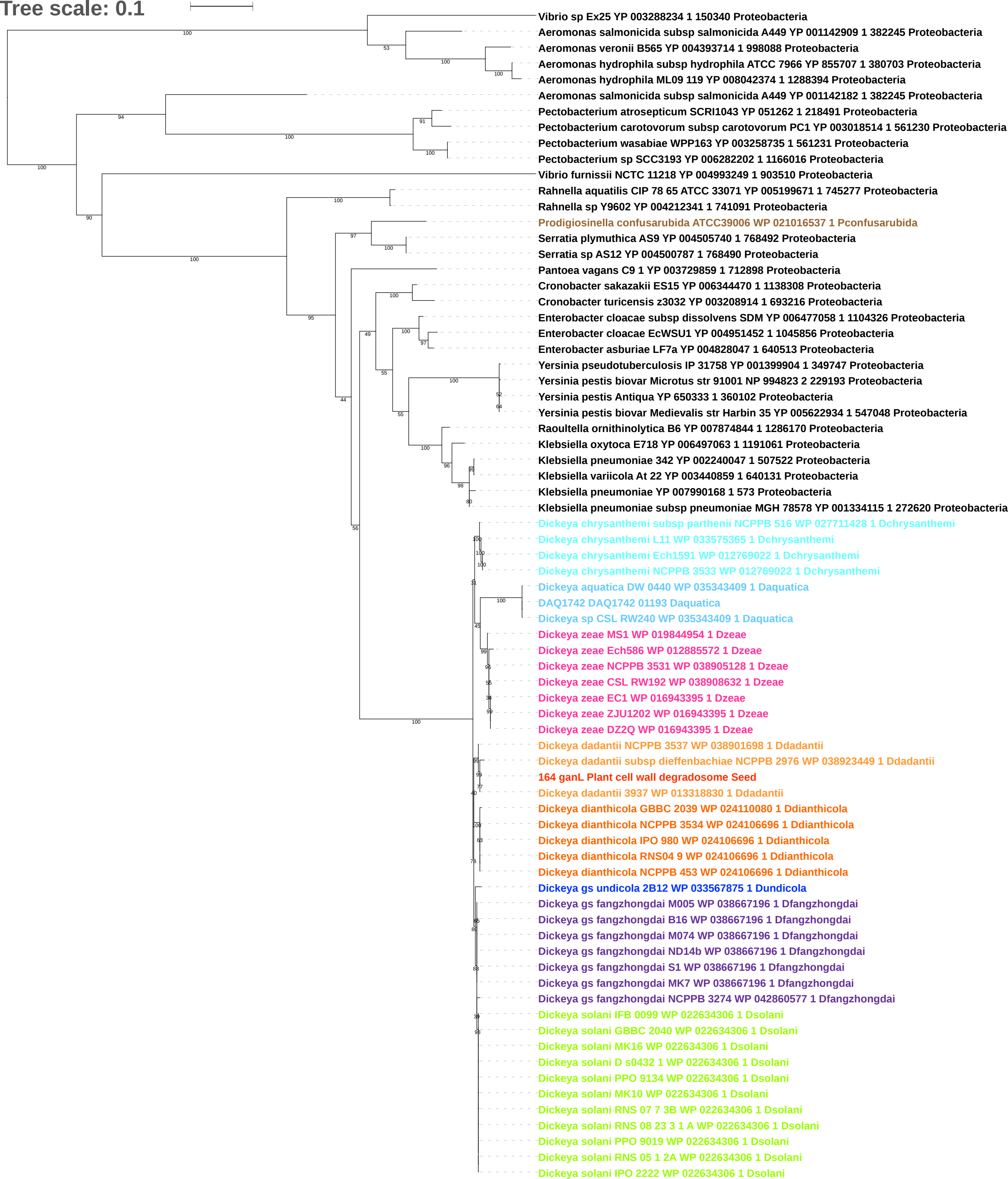
Tree scale: 0.1
Vibrio sp Ex25 YP 003288234 1 150340 Proteobacteria
Aeromonas salmonicida subsp salmonicida A449 YP 001142909 1 382245 Proteobacteria
Aeromonas veronii B565 YP 004393714 1 998088 Proteobacteria
Aeromonas hydrophila subsp hydrophila ATCC 7966 YP 855707 1 380703 Proteobacteria
Aeromonas hydrophila ML09 119 YP 008042374 1 1288394 Proteobacteria
Aeromonas salmonicida subsp salmonicida A449 YP 001142182 1 382245 Proteobacteria
Pectobacterium atrosepticum SCRI1043 YP 051262 1 218491 Proteobacteria
Pectobacterium carotovorum subsp carotovorum PC1 YP 003018514 1 561230 Proteobacteria
Pectobacterium wasabiae WPP163 YP 003258735 1 561231 Proteobacteria
Pectobacterium sp SCC3193 YP 006282202 1 1166016 Proteobacteria
Vibrio furnissii NCTC 11218 YP 004993249 1 903510 Proteobacteria
Rahnella aquatilis CIP 78 65 ATCC 33071 YP 005199671 1 745277 Proteobacteria
Rahnella sp Y9602 YP 004212341 1 741091 Proteobacteria
Prodigiosinella confusarubida ATCC39006 WP 021016537 1 Pconfusarubida
Serratia plymuthica AS9 YP 004505740 1 768492 Proteobacteria
Serratia sp AS12 YP 004500787 1 768490 Proteobacteria
Pantoea vagans C9 1 YP 003729859 1 712898 Proteobacteria
Cronobacter sakazakii ES15 YP 006344470 1 1138308 Proteobacteria
Cronobacter turicensis z3032 YP 003208914 1 693216 Proteobacteria
Enterobacter cloacae subsp dissolvens SDM YP 006477058 1 1104326 Proteobacteria
Enterobacter cloacae EcWSU1 YP 004951452 1 1045856 Proteobacteria
Enterobacter asburiae LF7a YP 004828047 1 640513 Proteobacteria
Yersinia pseudotuberculosis IP 31758 YP 001399904 1 349747 Proteobacteria
Yersinia pestis biovar Microtus str 91001 NP 994823 2 229193 Proteobacteria
Yersinia pestis Antiqua YP 650333 1 360102 Proteobacteria
Yersinia pestis biovar Medievalis str Harbin 35 YP 005622934 1 547048 Proteobacteria
Raoultella ornithinolytica B6 YP 007874844 1 1286170 Proteobacteria
Klebsiella oxytoca E718 YP 006497063 1 1191061 Proteobacteria
Klebsiella pneumoniae 342 YP 002240047 1 507522 Proteobacteria
Klebsiella variicola At 22 YP 003440859 1 640131 Proteobacteria
Klebsiella pneumoniae YP 007990168 1 573 Proteobacteria
Klebsiella pneumoniae subsp pneumoniae MGH 78578 YP 001334115 1 272620 Proteobacteria
Dickeya chrysanthemi subsp parthenii NCPPB 516 WP 027711428 1 Dchrysanthemi
Dickeya chrysanthemi L11 WP 033575365 1 Dchrysanthemi
Dickeya chrysanthemi Ech1591 WP 012769022 1 Dchrysanthemi
Dickeya chrysanthemi NCPPB 3533 WP 012769022 1 Dchrysanthemi
Dickeya aquatica DW 0440 WP 035343409 1 Daquatica
DAQ1742 DAQ1742 01193 Daquatica
Dickeya sp CSL RW240 WP 035343409 1 Daquatica
Dickeya zeae MS1 WP 019844954 1 Dzeae
Dickeya zeae Ech586 WP 012885572 1 Dzeae
Dickeya zeae NCPPB 3531 WP 038905128 1 Dzeae
Dickeya zeae CSL RW192 WP 038908632 1 Dzeae
Dickeya zeae EC1 WP 016943395 1 Dzeae
Dickeya zeae ZJU1202 WP 016943395 1 Dzeae
Dickeya zeae DZ2Q WP 016943395 1 Dzeae
Dickeya dadantii NCPPB 3537 WP 038901698 1 Ddadantii
Dickeya dadantii subsp dieffenbachiae NCPPB 2976 WP 038923449 1 Ddadantii
164 ganL Plant cell wall degradosome Seed
Dickeya dadantii 3937 WP 013318830 1 Ddadantii
Dickeya dianthicola GBBC 2039 WP 024110080 1 Ddianthicola
Dickeya dianthicola NCPPB 3534 WP 024106696 1 Ddianthicola
Dickeya dianthicola IPO 980 WP 024106696 1 Ddianthicola
Dickeya dianthicola RNS04 9 WP 024106696 1 Ddianthicola
Dickeya dianthicola NCPPB 453 WP 024106696 1 Ddianthicola
Dickeya gs undicola 2B12 WP 033567875 1 Dundicola
Dickeya gs fangzhongdai M005 WP 038667196 1 Dfangzhongdai
Dickeya gs fangzhongdai B16 WP 038667196 1 Dfangzhongdai
Dickeya gs fangzhongdai M074 WP 038667196 1 Dfangzhongdai
Dickeya gs fangzhongdai ND14b WP 038667196 1 Dfangzhongdai
Dickeya gs fangzhongdai S1 WP 038667196 1 Dfangzhongdai
Dickeya gs fangzhongdai MK7 WP 038667196 1 Dfangzhongdai
Dickeya gs fangzhongdai NCPPB 3274 WP 042860577 1 Dfangzhongdai
Dickeya solani IFB 0099 WP 022634306 1 Dsolani
Dickeya solani GBBC 2040 WP 022634306 1 Dsolani
Dickeya solani MK16 WP 022634306 1 Dsolani
Dickeya solani D s0432 1 WP 022634306 1 Dsolani
Dickeya solani PPO 9134 WP 022634306 1 Dsolani
Dickeya solani MK10 WP 022634306 1 Dsolani
Dickeya solani RNS 07 7 3B WP 022634306 1 Dsolani
Dickeya solani RNS 08 23 3 1 A WP 022634306 1 Dsolani
Dickeya solani PPO 9019 WP 022634306 1 Dsolani
Dickeya solani RNS 05 1 2A WP 022634306 1 Dsolani
Dickeya solani IPO 2222 WP 022634306 1 Dsolani

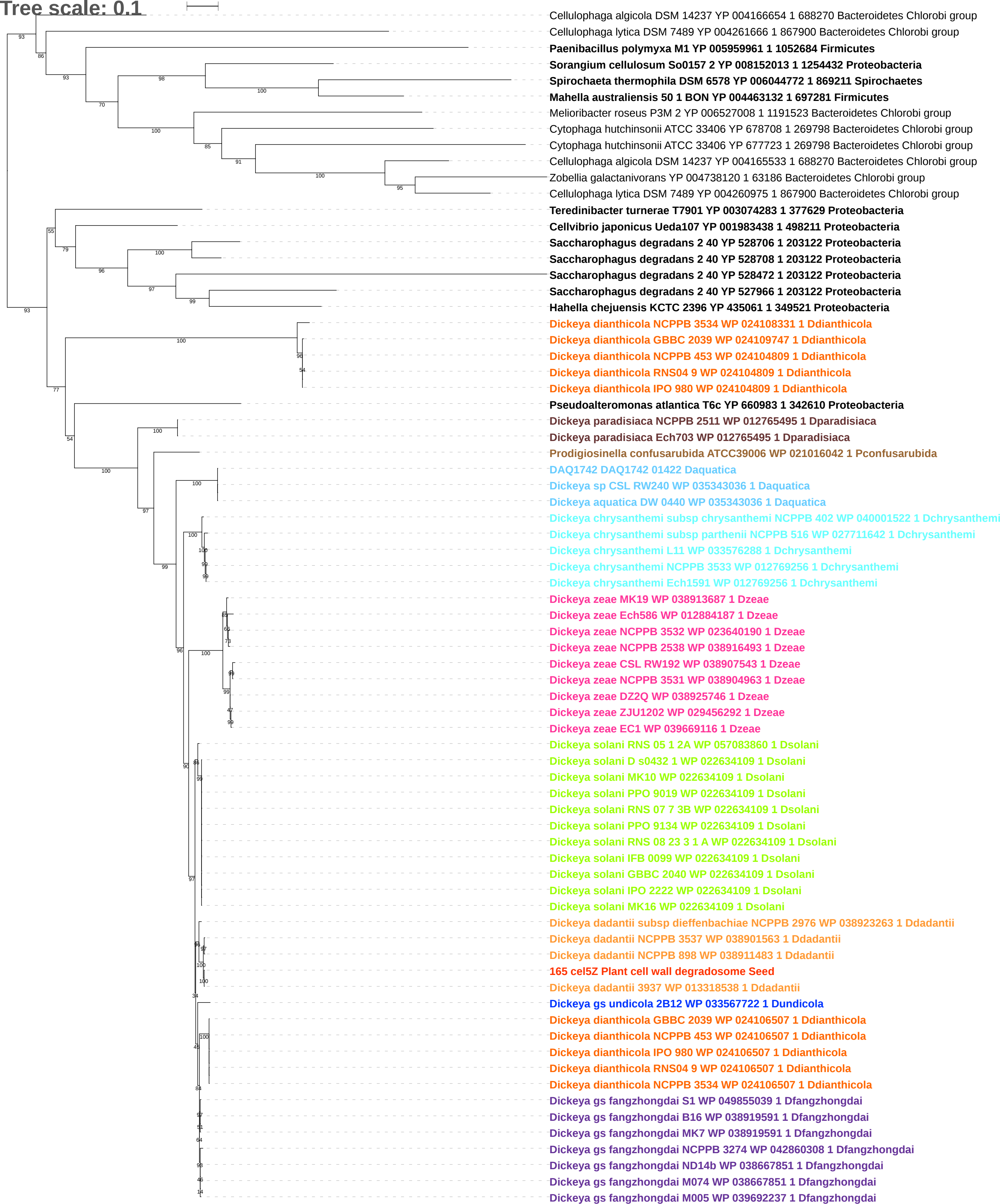
Tree scale: 0.1
Cellulophaga algicola DSM 14237 YP 004166654 1 688270 Bacteroidetes Chlorobi group
Cellulophaga lytica DSM 7489 YP 004261666 1 867900 Bacteroidetes Chlorobi group
Paenibacillus polymyxa M1 YP 005959961 1 1052684 Firmicutes
Sorangium cellulosum So0157 2 YP 008152013 1 1254432 Proteobacteria
Spirochaeta thermophila DSM 6578 YP 006044772 1 869211 Spirochaetes
Mahella australiensis 50 1 BON YP 004463132 1 697281 Firmicutes
Melioribacter roseus P3M 2 YP 006527008 1 1191523 Bacteroidetes Chlorobi group
Cytophaga hutchinsonii ATCC 33406 YP 678708 1 269798 Bacteroidetes Chlorobi group
Cytophaga hutchinsonii ATCC 33406 YP 677723 1 269798 Bacteroidetes Chlorobi group
Cellulophaga algicola DSM 14237 YP 004165533 1 688270 Bacteroidetes Chlorobi group
Zobellia galactanivorans YP 004738120 1 63186 Bacteroidetes Chlorobi group
Cellulophaga lytica DSM 7489 YP 004260975 1 867900 Bacteroidetes Chlorobi group
Teredinibacter turnerae T7901 YP 003074283 1 377629 Proteobacteria
Cellvibrio japonicus Ueda107 YP 001983438 1 498211 Proteobacteria
Saccharophagus degradans 2 40 YP 528706 1 203122 Proteobacteria
Saccharophagus degradans 2 40 YP 528708 1 203122 Proteobacteria
Saccharophagus degradans 2 40 YP 528472 1 203122 Proteobacteria
Saccharophagus degradans 2 40 YP 527966 1 203122 Proteobacteria
Hahella chejuensis KCTC 2396 YP 435061 1 349521 Proteobacteria
Dickeya dianthicola NCPPB 3534 WP 024108331 1 Ddianthicola
Dickeya dianthicola GBBC 2039 WP 024109747 1 Ddianthicola
Dickeya dianthicola NCPPB 453 WP 024104809 1 Ddianthicola
Dickeya dianthicola RNS04 9 WP 024104809 1 Ddianthicola
Dickeya dianthicola IPO 980 WP 024104809 1 Ddianthicola
Pseudoalteromonas atlantica T6c YP 660983 1 342610 Proteobacteria
Dickeya paradisiaca NCPPB 2511 WP 012765495 1 Dparadisiaca
Dickeya paradisiaca Ech703 WP 012765495 1 Dparadisiaca
Prodigiosinella confusarubida ATCC39006 WP 021016042 1 Pconfusarubida
DAQ1742 DAQ1742 01422 Daquatica
Dickeya sp CSL RW240 WP 035343036 1 Daquatica
Dickeya aquatica DW 0440 WP 035343036 1 Daquatica
Dickeya chrysanthemi subsp chrysanthemi NCPPB 402 WP 040001522 1 Dchrysanthemi
Dickeya chrysanthemi subsp parthenii NCPPB 516 WP 027711642 1 Dchrysanthemi
Dickeya chrysanthemi L11 WP 033576288 1 Dchrysanthemi
Dickeya chrysanthemi NCPPB 3533 WP 012769256 1 Dchrysanthemi
Dickeya chrysanthemi Ech1591 WP 012769256 1 Dchrysanthemi
Dickeya zeae MK19 WP 038913687 1 Dzeae
Dickeya zeae Ech586 WP 012884187 1 Dzeae
Dickeya zeae NCPPB 3532 WP 023640190 1 Dzeae
Dickeya zeae NCPPB 2538 WP 038916493 1 Dzeae
Dickeya zeae CSL RW192 WP 038907543 1 Dzeae
Dickeya zeae NCPPB 3531 WP 038904963 1 Dzeae
Dickeya zeae DZ2Q WP 038925746 1 Dzeae
Dickeya zeae ZJU1202 WP 029456292 1 Dzeae
Dickeya zeae EC1 WP 039669116 1 Dzeae
Dickeya solani RNS 05 1 2A WP 057083860 1 Dsolani
Dickeya solani D s0432 1 WP 022634109 1 Dsolani
Dickeya solani MK10 WP 022634109 1 Dsolani
Dickeya solani PPO 9019 WP 022634109 1 Dsolani
Dickeya solani RNS 07 7 3B WP 022634109 1 Dsolani
Dickeya solani PPO 9134 WP 022634109 1 Dsolani
Dickeya solani RNS 08 23 3 1 A WP 022634109 1 Dsolani
Dickeya solani IFB 0099 WP 022634109 1 Dsolani
Dickeya solani GBBC 2040 WP 022634109 1 Dsolani
Dickeya solani IPO 2222 WP 022634109 1 Dsolani
Dickeya solani MK16 WP 022634109 1 Dsolani
Dickeya dadantii subsp dieffenbachiae NCPPB 2976 WP 038923263 1 Ddadantii
Dickeya dadantii NCPPB 3537 WP 038901563 1 Ddadantii
Dickeya dadantii NCPPB 898 WP 038911483 1 Ddadantii
165 cel5Z Plant cell wall degradosome Seed
Dickeya dadantii 3937 WP 013318538 1 Ddadantii
Dickeya gs undicola 2B12 WP 033567722 1 Dundicola
Dickeya dianthicola GBBC 2039 WP 024106507 1 Ddianthicola
Dickeya dianthicola NCPPB 453 WP 024106507 1 Ddianthicola
Dickeya dianthicola IPO 980 WP 024106507 1 Ddianthicola
Dickeya dianthicola RNS04 9 WP 024106507 1 Ddianthicola
Dickeya dianthicola NCPPB 3534 WP 024106507 1 Ddianthicola
Dickeya gs fangzhongdai S1 WP 049855039 1 Dfangzhongdai
Dickeya gs fangzhongdai B16 WP 038919591 1 Dfangzhongdai
Dickeya gs fangzhongdai MK7 WP 038919591 1 Dfangzhongdai
Dickeya gs fangzhongdai NCPPB 3274 WP 042860308 1 Dfangzhongdai
Dickeya gs fangzhongdai ND14b WP 038667851 1 Dfangzhongdai
Dickeya gs fangzhongdai M074 WP 038667851 1 Dfangzhongdai
Dickeya gs fangzhongdai M005 WP 039692237 1 Dfangzhongdai

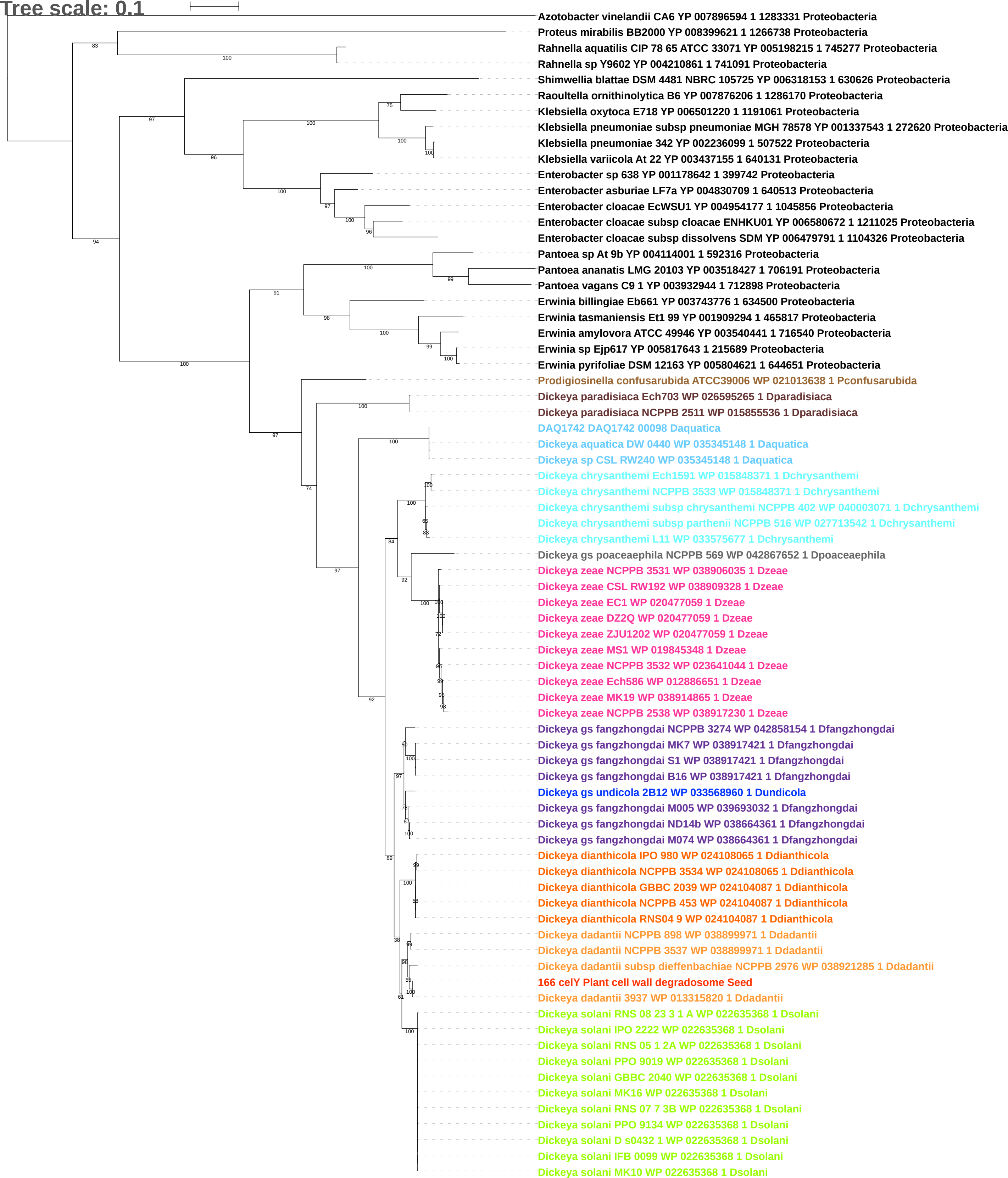
Tree scale: 0.1
Azotobacter vinelandii CA6 YP 007896594 1 1283331 Proteobacteria
Proteus mirabilis BB2000 YP 008399621 1 1266738 Proteobacteria
Rahnella aquatilis CIP 78 65 ATCC 33071 YP 005198215 1 745277 Proteobacteria
Rahnella sp Y9602 YP 004210861 1 741091 Proteobacteria
Shimwellia blattae DSM 4481 NBRC 105725 YP 006318153 1 630626 Proteobacteria
Raoultella ornithinolytica B6 YP 007876206 1 1286170 Proteobacteria
Klebsiella oxytoca E718 YP 006501220 1 1191061 Proteobacteria
Klebsiella pneumoniae subsp pneumoniae MGH 78578 YP 001337543 1 272620 Proteobacteria
Klebsiella pneumoniae 342 YP 002236099 1 507522 Proteobacteria
Klebsiella variicola At 22 YP 003437155 1 640131 Proteobacteria
Enterobacter sp 638 YP 001178642 1 399742 Proteobacteria
Enterobacter asburiae LF7a YP 004830709 1 640513 Proteobacteria
Enterobacter cloacae EcWSU1 YP 004954177 1 1045856 Proteobacteria
Enterobacter cloacae subsp cloacae ENHKU01 YP 006580672 1 1211025 Proteobacteria
Enterobacter cloacae subsp dissolvens SDM YP 006479791 1 1104326 Proteobacteria
Pantoea sp At 9b YP 004114001 1 592316 Proteobacteria
Pantoea ananatis LMG 20103 YP 003518427 1 706191 Proteobacteria
Pantoea vagans C9 1 YP 003932944 1 712898 Proteobacteria
Erwinia billingiae Eb661 YP 003743776 1 634500 Proteobacteria
Erwinia tasmaniensis Et1 99 YP 001909294 1 465817 Proteobacteria
Erwinia amylovora ATCC 49946 YP 003540441 1 716540 Proteobacteria
Erwinia sp Ejp617 YP 005817643 1 215689 Proteobacteria
Erwinia pyrifoliae DSM 12163 YP 005804621 1 644651 Proteobacteria
Prodigiosinella confusarubida ATCC39006 WP 021013638 1 Pconfusarubida
Dickeya paradisiaca Ech703 WP 026595265 1 Dparadisiaca
Dickeya paradisiaca NCPPB 2511 WP 015855536 1 Dparadisiaca
DAQ1742 DAQ1742 00098 Daquatica
Dickeya aquatica DW 0440 WP 035345148 1 Daquatica
Dickeya sp CSL RW240 WP 035345148 1 Daquatica
Dickeya chrysanthemi Ech1591 WP 015848371 1 Dchrysanthemi
Dickeya chrysanthemi NCPPB 3533 WP 015848371 1 Dchrysanthemi
Dickeya chrysanthemi subsp chrysanthemi NCPPB 402 WP 040003071 1 Dchrysanthemi
Dickeya chrysanthemi subsp parthenii NCPPB 516 WP 027713542 1 Dchrysanthemi
Dickeya chrysanthemi L11 WP 033575677 1 Dchrysanthemi
Dickeya gs poaceaephila NCPPB 569 WP 042867652 1 Dpoaceaephila
Dickeya zeae NCPPB 3531 WP 038906035 1 Dzeae
Dickeya zeae CSL RW192 WP 038909328 1 Dzeae
Dickeya zeae EC1 WP 020477059 1 Dzeae
Dickeya zeae DZ2Q WP 020477059 1 Dzeae
Dickeya zeae ZJU1202 WP 020477059 1 Dzeae
Dickeya zeae MS1 WP 019845348 1 Dzeae
Dickeya zeae NCPPB 3532 WP 023641044 1 Dzeae
Dickeya zeae Ech586 WP 012886651 1 Dzeae
Dickeya zeae MK19 WP 038914865 1 Dzeae
Dickeya zeae NCPPB 2538 WP 038917230 1 Dzeae
Dickeya gs fangzhongdai NCPPB 3274 WP 042858154 1 Dfangzhongdai
Dickeya gs fangzhongdai MK7 WP 038917421 1 Dfangzhongdai
Dickeya gs fangzhongdai S1 WP 038917421 1 Dfangzhongdai
Dickeya gs fangzhongdai B16 WP 038917421 1 Dfangzhongdai
Dickeya gs undicola 2B12 WP 033568960 1 Dundicola
Dickeya gs fangzhongdai M005 WP 039693032 1 Dfangzhongdai
Dickeya gs fangzhongdai ND14b WP 038664361 1 Dfangzhongdai
Dickeya gs fangzhongdai M074 WP 038664361 1 Dfangzhongdai
Dickeya dianthicola IPO 980 WP 024108065 1 Ddianthicola
Dickeya dianthicola NCPPB 3534 WP 024108065 1 Ddianthicola
Dickeya dianthicola GBBC 2039 WP 024104087 1 Ddianthicola
Dickeya dianthicola NCPPB 453 WP 024104087 1 Ddianthicola
Dickeya dianthicola RNS04 9 WP 024104087 1 Ddianthicola
Dickeya dadantii NCPPB 898 WP 038899971 1 Ddadantii
Dickeya dadantii NCPPB 3537 WP 038899971 1 Ddadantii
Dickeya dadantii subsp dieffenbachiae NCPPB 2976 WP 038921285 1 Ddadantii
166 celY Plant cell wall degradosome Seed
Dickeya dadantii 3937 WP 013315820 1 Ddadantii
Dickeya solani RNS 08 23 3 1 A WP 022635368 1 Dsolani
Dickeya solani IPO 2222 WP 022635368 1 Dsolani
Dickeya solani RNS 05 1 2A WP 022635368 1 Dsolani
Dickeya solani PPO 9019 WP 022635368 1 Dsolani
Dickeya solani GBBC 2040 WP 022635368 1 Dsolani
Dickeya solani MK16 WP 022635368 1 Dsolani
Dickeya solani RNS 07 7 3B WP 022635368 1 Dsolani
Dickeya solani PPO 9134 WP 022635368 1 Dsolani
Dickeya solani D s0432 1 WP 022635368 1 Dsolani
Dickeya solani IFB 0099 WP 022635368 1 Dsolani
Dickeya solani MK10 WP 022635368 1 Dsolani

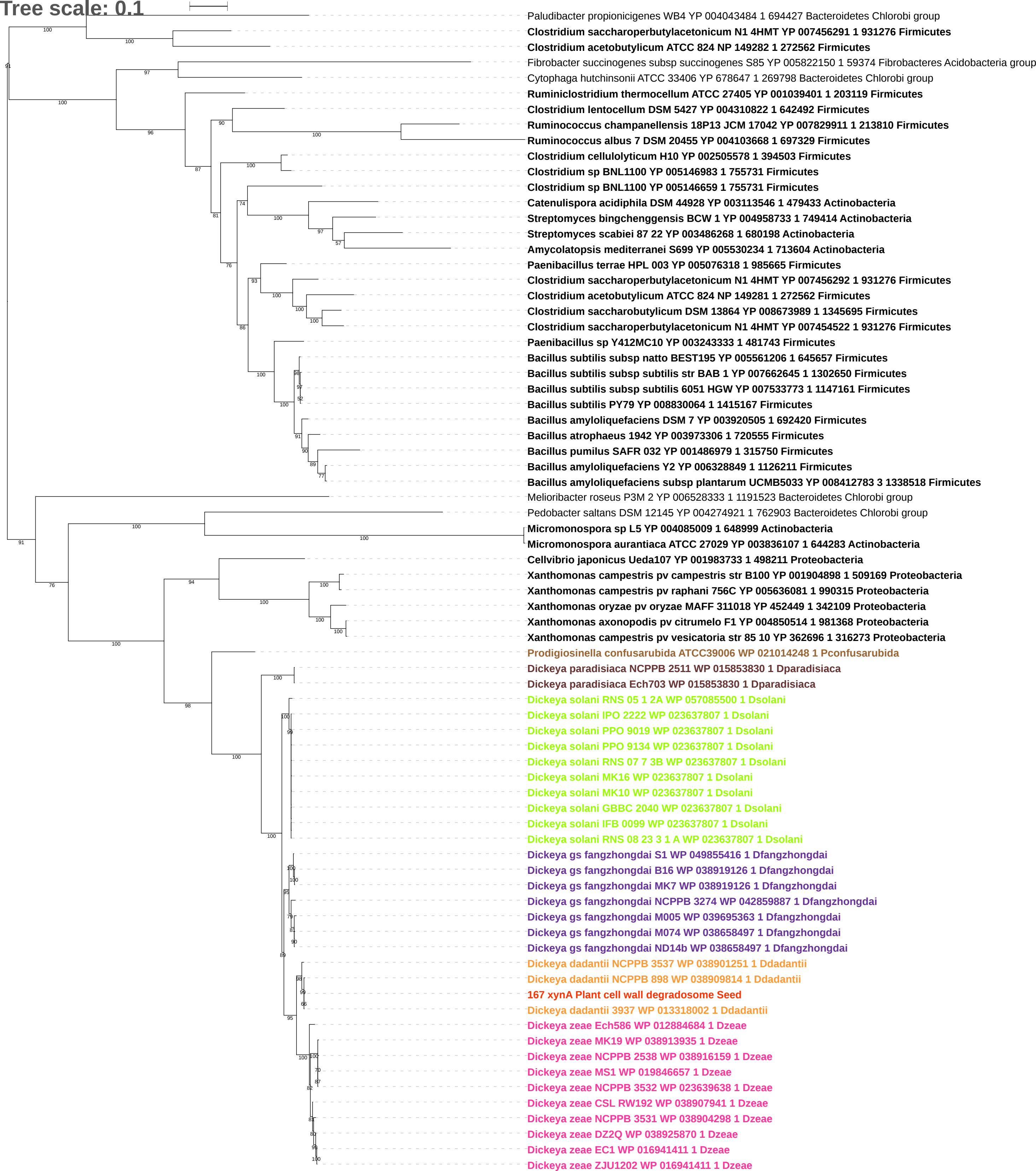
Tree scale: 0.1
Paludibacter propionicigenes WB4 YP 004043484 1 694427 Bacteroidetes Chlorobi group
Clostridium saccharoperbutylacetonicum N1 4HMT YP 007456291 1 931276 Firmicutes
Clostridium acetobutylicum ATCC 824 NP 149282 1 272562 Firmicutes
Fibrobacter succinogenes subsp succinogenes S85 YP 005822150 1 59374 Fibrobacteres Acidobacteria group
Cytophaga hutchinsonii ATCC 33406 YP 678647 1 269798 Bacteroidetes Chlorobi group
Ruminiclostridium thermocellum ATCC 27405 YP 001039401 1 203119 Firmicutes
Clostridium lentocellum DSM 5427 YP 004310822 1 642492 Firmicutes
Ruminococcus champanellensis 18P13 JCM 17042 YP 007829911 1 213810 Firmicutes
Ruminococcus albus 7 DSM 20455 YP 004103668 1 697329 Firmicutes
Clostridium cellulolyticum H10 YP 002505578 1 394503 Firmicutes
Clostridium sp BNL1100 YP 005146983 1 755731 Firmicutes
Clostridium sp BNL1100 YP 005146659 1 755731 Firmicutes
Catenulispora acidiphila DSM 44928 YP 003113546 1 479433 Actinobacteria
Streptomyces bingchenggensis BCW 1 YP 004958733 1 749414 Actinobacteria
Streptomyces scabiei 87 22 YP 003486268 1 680198 Actinobacteria
Amycolatopsis mediterranei S699 YP 005530234 1 713604 Actinobacteria
Paenibacillus terrae HPL 003 YP 005076318 1 985665 Firmicutes
Clostridium saccharoperbutylacetonicum N1 4HMT YP 007456292 1 931276 Firmicutes
Clostridium acetobutylicum ATCC 824 NP 149281 1 272562 Firmicutes
Clostridium saccharobutylicum DSM 13864 YP 008673989 1 1345695 Firmicutes
Clostridium saccharoperbutylacetonicum N1 4HMT YP 007454522 1 931276 Firmicutes
Paenibacillus sp Y412MC10 YP 003243333 1 481743 Firmicutes
Bacillus subtilis subsp natto BEST195 YP 005561206 1 645657 Firmicutes
Bacillus subtilis subsp subtilis str BAB 1 YP 007662645 1 1302650 Firmicutes
Bacillus subtilis subsp subtilis 6051 HGW YP 007533773 1 1147161 Firmicutes
Bacillus subtilis PY79 YP 008830064 1 1415167 Firmicutes
Bacillus amyloliquefaciens DSM 7 YP 003920505 1 692420 Firmicutes
Bacillus atrophaeus 1942 YP 003973306 1 720555 Firmicutes
Bacillus pumilus SAFR 032 YP 001486979 1 315750 Firmicutes
Bacillus amyloliquefaciens Y2 YP 006328849 1 1126211 Firmicutes
Bacillus amyloliquefaciens subsp plantarum UCMB5033 YP 008412783 3 1338518 Firmicutes
Melioribacter roseus P3M 2 YP 006528333 1 1191523 Bacteroidetes Chlorobi group
Pedobacter saltans DSM 12145 YP 004274921 1 762903 Bacteroidetes Chlorobi group
Micromonospora sp L5 YP 004085009 1 648999 Actinobacteria
Micromonospora aurantiaca ATCC 27029 YP 003836107 1 644283 Actinobacteria
Cellvibrio japonicus Ueda107 YP 001983733 1 498211 Proteobacteria
Xanthomonas campestris pv campestris str B100 YP 001904898 1 509169 Proteobacteria
Xanthomonas campestris pv raphani 756C YP 005636081 1 990315 Proteobacteria
Xanthomonas oryzae pv oryzae MAFF 311018 YP 452449 1 342109 Proteobacteria
Xanthomonas axonopodis pv citrumelo F1 YP 004850514 1 981368 Proteobacteria
Xanthomonas campestris pv vesicatoria str 85 10 YP 362696 1 316273 Proteobacteria
Prodigiosinella confusarubida ATCC39006 WP 021014248 1 Pconfusarubida
Dickeya paradisiaca NCPPB 2511 WP 015853830 1 Dparadisiaca
Dickeya paradisiaca Ech703 WP 015853830 1 Dparadisiaca
Dickeya solani RNS 05 1 2A WP 057085500 1 Dsolani
Dickeya solani IPO 2222 WP 023637807 1 Dsolani
Dickeya solani PPO 9019 WP 023637807 1 Dsolani
Dickeya solani PPO 9134 WP 023637807 1 Dsolani
Dickeya solani RNS 07 7 3B WP 023637807 1 Dsolani
Dickeya solani MK16 WP 023637807 1 Dsolani
Dickeya solani MK10 WP 023637807 1 Dsolani
Dickeya solani GBBC 2040 WP 023637807 1 Dsolani
Dickeya solani IFB 0099 WP 023637807 1 Dsolani
Dickeya solani RNS 08 23 3 1 A WP 023637807 1 Dsolani
Dickeya gs fangzhongdai S1 WP 049855416 1 Dfangzhongdai
Dickeya gs fangzhongdai B16 WP 038919126 1 Dfangzhongdai
Dickeya gs fangzhongdai MK7 WP 038919126 1 Dfangzhongdai
Dickeya gs fangzhongdai NCPPB 3274 WP 042859887 1 Dfangzhongdai
Dickeya gs fangzhongdai M005 WP 039695363 1 Dfangzhongdai
Dickeya gs fangzhongdai M074 WP 038658497 1 Dfangzhongdai
Dickeya gs fangzhongdai ND14b WP 038658497 1 Dfangzhongdai
Dickeya dadantii NCPPB 3537 WP 038901251 1 Ddadantii
Dickeya dadantii NCPPB 898 WP 038909814 1 Ddadantii
167 xynA Plant cell wall degradosome Seed
Dickeya dadantii 3937 WP 013318002 1 Ddadantii
Dickeya zeae Ech586 WP 012884684 1 Dzeae
Dickeya zeae MK19 WP 038913935 1 Dzeae
Dickeya zeae NCPPB 2538 WP 038916159 1 Dzeae
Dickeya zeae MS1 WP 019846657 1 Dzeae
Dickeya zeae NCPPB 3532 WP 023639638 1 Dzeae
Dickeya zeae CSL RW192 WP 038907941 1 Dzeae
Dickeya zeae NCPPB 3531 WP 038904298 1 Dzeae
Dickeya zeae DZ2Q WP 038925870 1 Dzeae
Dickeya zeae EC1 WP 016941411 1 Dzeae
Dickeya zeae ZJU1202 WP 016941411 1 Dzeae

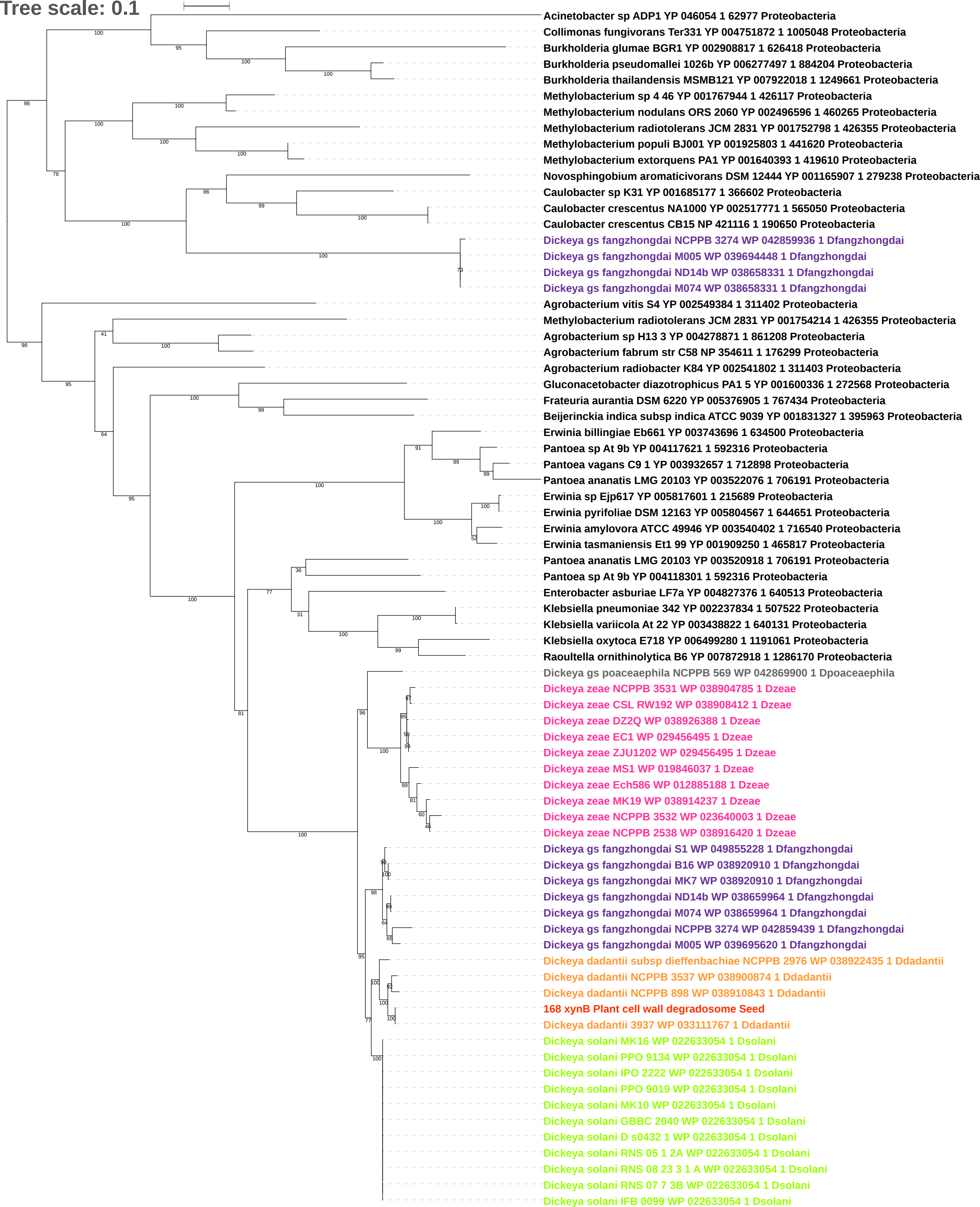
Tree scale: 0.1
Acinetobacter sp ADP1 YP 046054 1 62977 Proteobacteria
Collimonas fungivorans Ter331 YP 004751872 1 1005048 Proteobacteria
Burkholderia glumae BGR1 YP 002908817 1 626418 Proteobacteria
Burkholderia pseudomallei 1026b YP 006277497 1 884204 Proteobacteria
Burkholderia thailandensis MSMB121 YP 007922018 1 1249661 Proteobacteria
Methylobacterium sp 4 46 YP 001767944 1 426117 Proteobacteria
Methylobacterium nodulans ORS 2060 YP 002496596 1 460265 Proteobacteria
Methylobacterium radiotolerans JCM 2831 YP 001752798 1 426355 Proteobacteria
Methylobacterium populi BJ001 YP 001925803 1 441620 Proteobacteria
Methylobacterium extorquens PA1 YP 001640393 1 419610 Proteobacteria
Novosphingobium aromaticivorans DSM 12444 YP 001165907 1 279238 Proteobacteria
Caulobacter sp K31 YP 001685177 1 366602 Proteobacteria
Caulobacter crescentus NA1000 YP 002517771 1 565050 Proteobacteria
Caulobacter crescentus CB15 NP 421116 1 190650 Proteobacteria
Dickeya gs fangzhongdai NCPPB 3274 WP 042859936 1 Dfangzhongdai
Dickeya gs fangzhongdai M005 WP 039694448 1 Dfangzhongdai
Dickeya gs fangzhongdai ND14b WP 038658331 1 Dfangzhongdai
Dickeya gs fangzhongdai M074 WP 038658331 1 Dfangzhongdai
Agrobacterium vitis S4 YP 002549384 1 311402 Proteobacteria
Methylobacterium radiotolerans JCM 2831 YP 001754214 1 426355 Proteobacteria
Agrobacterium sp H13 3 YP 004278871 1 861208 Proteobacteria
Agrobacterium fabrum str C58 NP 354611 1 176299 Proteobacteria
Agrobacterium radiobacter K84 YP 002541802 1 311403 Proteobacteria
Gluconacetobacter diazotrophicus PA1 5 YP 001600336 1 272568 Proteobacteria
Frateuria aurantia DSM 6220 YP 005376905 1 767434 Proteobacteria
Beijerinckia indica subsp indica ATCC 9039 YP 001831327 1 395963 Proteobacteria
Erwinia billingiae Eb661 YP 003743696 1 634500 Proteobacteria
Pantoea sp At 9b YP 004117621 1 592316 Proteobacteria
Pantoea vagans C9 1 YP 003932657 1 712898 Proteobacteria
Pantoea ananatis LMG 20103 YP 003522076 1 706191 Proteobacteria
Erwinia sp Ejp617 YP 005817601 1 215689 Proteobacteria
Erwinia pyrifoliae DSM 12163 YP 005804567 1 644651 Proteobacteria
Erwinia amylovora ATCC 49946 YP 003540402 1 716540 Proteobacteria
Erwinia tasmaniensis Et1 99 YP 001909250 1 465817 Proteobacteria
Pantoea ananatis LMG 20103 YP 003520918 1 706191 Proteobacteria
Pantoea sp At 9b YP 004118301 1 592316 Proteobacteria
Enterobacter asburiae LF7a YP 004827376 1 640513 Proteobacteria
Klebsiella pneumoniae 342 YP 002237834 1 507522 Proteobacteria
Klebsiella variicola At 22 YP 003438822 1 640131 Proteobacteria
Klebsiella oxytoca E718 YP 006499280 1 1191061 Proteobacteria
Raoultella ornithinolytica B6 YP 007872918 1 1286170 Proteobacteria
Dickeya gs poaceaephila NCPPB 569 WP 042869900 1 Dpoaceaephila
Dickeya zeae NCPPB 3531 WP 038904785 1 Dzeae
Dickeya zeae CSL RW192 WP 038908412 1 Dzeae
Dickeya zeae DZ2Q WP 038926388 1 Dzeae
Dickeya zeae EC1 WP 029456495 1 Dzeae
Dickeya zeae ZJU1202 WP 029456495 1 Dzeae
Dickeya zeae MS1 WP 019846037 1 Dzeae
Dickeya zeae Ech586 WP 012885188 1 Dzeae
Dickeya zeae MK19 WP 038914237 1 Dzeae
Dickeya zeae NCPPB 3532 WP 023640003 1 Dzeae
Dickeya zeae NCPPB 2538 WP 038916420 1 Dzeae
Dickeya gs fangzhongdai S1 WP 049855228 1 Dfangzhongdai
Dickeya gs fangzhongdai B16 WP 038920910 1 Dfangzhongdai
Dickeya gs fangzhongdai MK7 WP 038920910 1 Dfangzhongdai
Dickeya gs fangzhongdai ND14b WP 038659964 1 Dfangzhongdai
Dickeya gs fangzhongdai M074 WP 038659964 1 Dfangzhongdai
Dickeya gs fangzhongdai NCPPB 3274 WP 042859439 1 Dfangzhongdai
Dickeya gs fangzhongdai M005 WP 039695620 1 Dfangzhongdai
Dickeya dadantii subsp dieffenbachiae NCPPB 2976 WP 038922435 1 Ddadantii
Dickeya dadantii NCPPB 3537 WP 038900874 1 Ddadantii
Dickeya dadantii NCPPB 898 WP 038910843 1 Ddadantii
168 xynB Plant cell wall degradosome Seed
Dickeya dadantii 3937 WP 033111767 1 Ddadantii
Dickeya solani MK16 WP 022633054 1 Dsolani
Dickeya solani PPO 9134 WP 022633054 1 Dsolani
Dickeya solani IPO 2222 WP 022633054 1 Dsolani
Dickeya solani PPO 9019 WP 022633054 1 Dsolani
Dickeya solani MK10 WP 022633054 1 Dsolani
Dickeya solani GBBC 2040 WP 022633054 1 Dsolani
Dickeya solani D s0432 1 WP 022633054 1 Dsolani
Dickeya solani RNS 05 1 2A WP 022633054 1 Dsolani
Dickeya solani RNS 08 23 3 1 A WP 022633054 1 Dsolani
Dickeya solani RNS 07 7 3B WP 022633054 1 Dsolani
Dickeya solani IFB 0099 WP 022633054 1 Dsolani

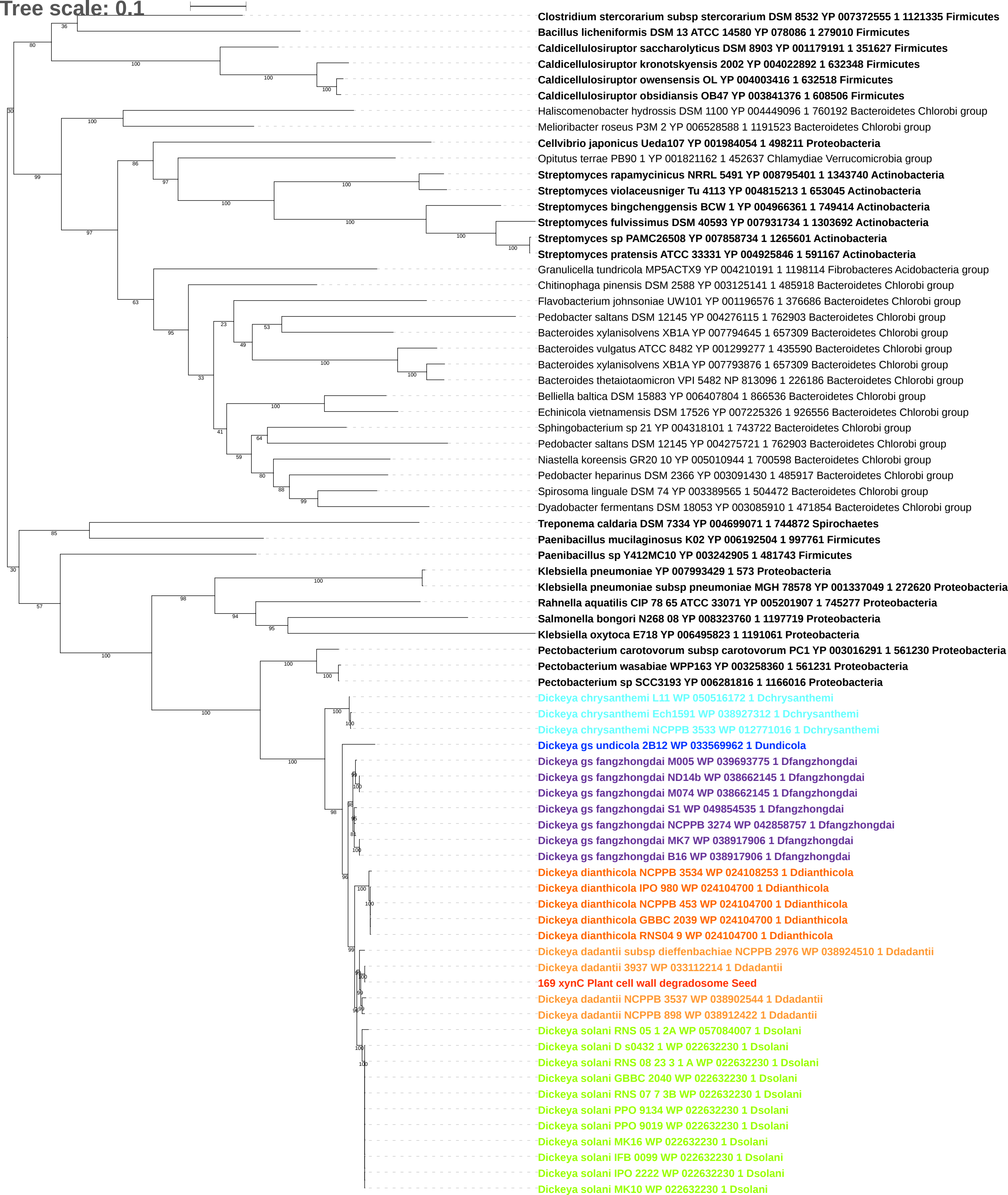
Tree scale: 0.1
Clostridium stercorarium subsp stercorarium DSM 8532 YP 007372555 1 1121335 Firmicutes
Bacillus licheniformis DSM 13 ATCC 14580 YP 078086 1 279010 Firmicutes
Caldicellulosiruptor saccharolyticus DSM 8903 YP 001179191 1 351627 Firmicutes
Caldicellulosiruptor kronotskyensis 2002 YP 004022892 1 632348 Firmicutes
Caldicellulosiruptor owensensis OL YP 004003416 1 632518 Firmicutes
Caldicellulosiruptor obsidiansis OB47 YP 003841376 1 608506 Firmicutes
Haliscomenobacter hydrossis DSM 1100 YP 004449096 1 760192 Bacteroidetes Chlorobi group
Melioribacter roseus P3M 2 YP 006528588 1 1191523 Bacteroidetes Chlorobi group
Cellvibrio japonicus Ueda107 YP 001984054 1 498211 Proteobacteria
Opitutus terrae PB90 1 YP 001821162 1 452637 Chlamydiae Verrucomicrobia group
Streptomyces rapamycinicus NRRL 5491 YP 008795401 1 1343740 Actinobacteria
Streptomyces violaceusniger Tu 4113 YP 004815213 1 653045 Actinobacteria
Streptomyces bingchenggensis BCW 1 YP 004966361 1 749414 Actinobacteria
Streptomyces fulvissimus DSM 40593 YP 007931734 1 1303692 Actinobacteria
Streptomyces sp PAMC26508 YP 007858734 1 1265601 Actinobacteria
Streptomyces pratensis ATCC 33331 YP 004925846 1 591167 Actinobacteria
Granulicella tundricola MP5ACTX9 YP 004210191 1 1198114 Fibrobacteres Acidobacteria group
Chitinophaga pinensis DSM 2588 YP 003125141 1 485918 Bacteroidetes Chlorobi group
Flavobacterium johnsoniae UW101 YP 001196576 1 376686 Bacteroidetes Chlorobi group
Pedobacter saltans DSM 12145 YP 004276115 1 762903 Bacteroidetes Chlorobi group
Bacteroides xylanisolvens XB1A YP 007794645 1 657309 Bacteroidetes Chlorobi group
Bacteroides vulgatus ATCC 8482 YP 001299277 1 435590 Bacteroidetes Chlorobi group
Bacteroides xylanisolvens XB1A YP 007793876 1 657309 Bacteroidetes Chlorobi group
Bacteroides thetaiotaomicron VPI 5482 NP 813096 1 226186 Bacteroidetes Chlorobi group
Belliella baltica DSM 15883 YP 006407804 1 866536 Bacteroidetes Chlorobi group
Echinicola vietnamensis DSM 17526 YP 007225326 1 926556 Bacteroidetes Chlorobi group
Sphingobacterium sp 21 YP 004318101 1 743722 Bacteroidetes Chlorobi group
Pedobacter saltans DSM 12145 YP 004275721 1 762903 Bacteroidetes Chlorobi group
Niastella koreensis GR20 10 YP 005010944 1 700598 Bacteroidetes Chlorobi group
Pedobacter heparinus DSM 2366 YP 003091430 1 485917 Bacteroidetes Chlorobi group
Spirosoma linguale DSM 74 YP 003389565 1 504472 Bacteroidetes Chlorobi group
Dyadobacter fermentans DSM 18053 YP 003085910 1 471854 Bacteroidetes Chlorobi group
Treponema caldaria DSM 7334 YP 004699071 1 744872 Spirochaetes
Paenibacillus mucilaginosus K02 YP 006192504 1 997761 Firmicutes
Paenibacillus sp Y412MC10 YP 003242905 1 481743 Firmicutes
Klebsiella pneumoniae YP 007993429 1 573 Proteobacteria
Klebsiella pneumoniae subsp pneumoniae MGH 78578 YP 001337049 1 272620 Proteobacteria
Rahnella aquatilis CIP 78 65 ATCC 33071 YP 005201907 1 745277 Proteobacteria
Salmonella bongori N268 08 YP 008323760 1 1197719 Proteobacteria
Klebsiella oxytoca E718 YP 006495823 1 1191061 Proteobacteria
Pectobacterium carotovorum subsp carotovorum PC1 YP 003016291 1 561230 Proteobacteria
Pectobacterium wasabiae WPP163 YP 003258360 1 561231 Proteobacteria
Pectobacterium sp SCC3193 YP 006281816 1 1166016 Proteobacteria
Dickeya chrysanthemi L11 WP 050516172 1 Dchrysanthemi
Dickeya chrysanthemi Ech1591 WP 038927312 1 Dchrysanthemi
Dickeya chrysanthemi NCPPB 3533 WP 012771016 1 Dchrysanthemi
Dickeya gs undicola 2B12 WP 033569962 1 Dundicola
Dickeya gs fangzhongdai M005 WP 039693775 1 Dfangzhongdai
Dickeya gs fangzhongdai ND14b WP 038662145 1 Dfangzhongdai
Dickeya gs fangzhongdai M074 WP 038662145 1 Dfangzhongdai
Dickeya gs fangzhongdai S1 WP 049854535 1 Dfangzhongdai
Dickeya gs fangzhongdai NCPPB 3274 WP 042858757 1 Dfangzhongdai
Dickeya gs fangzhongdai MK7 WP 038917906 1 Dfangzhongdai
Dickeya gs fangzhongdai B16 WP 038917906 1 Dfangzhongdai
Dickeya dianthicola NCPPB 3534 WP 024108253 1 Ddianthicola
Dickeya dianthicola IPO 980 WP 024104700 1 Ddianthicola
Dickeya dianthicola NCPPB 453 WP 024104700 1 Ddianthicola
Dickeya dianthicola GBBC 2039 WP 024104700 1 Ddianthicola
Dickeya dianthicola RNS04 9 WP 024104700 1 Ddianthicola
Dickeya dadantii subsp dieffenbachiae NCPPB 2976 WP 038924510 1 Ddadantii
Dickeya dadantii 3937 WP 033112214 1 Ddadantii
169 xynC Plant cell wall degradosome Seed
Dickeya dadantii NCPPB 3537 WP 038902544 1 Ddadantii
Dickeya dadantii NCPPB 898 WP 038912422 1 Ddadantii
Dickeya solani RNS 05 1 2A WP 057084007 1 Dsolani
Dickeya solani D s0432 1 WP 022632230 1 Dsolani
Dickeya solani RNS 08 23 3 1 A WP 022632230 1 Dsolani
Dickeya solani GBBC 2040 WP 022632230 1 Dsolani
Dickeya solani RNS 07 7 3B WP 022632230 1 Dsolani
Dickeya solani PPO 9134 WP 022632230 1 Dsolani
Dickeya solani PPO 9019 WP 022632230 1 Dsolani
Dickeya solani MK16 WP 022632230 1 Dsolani
Dickeya solani IFB 0099 WP 022632230 1 Dsolani
Dickeya solani IPO 2222 WP 022632230 1 Dsolani
Dickeya solani MK10 WP 022632230 1 Dsolani

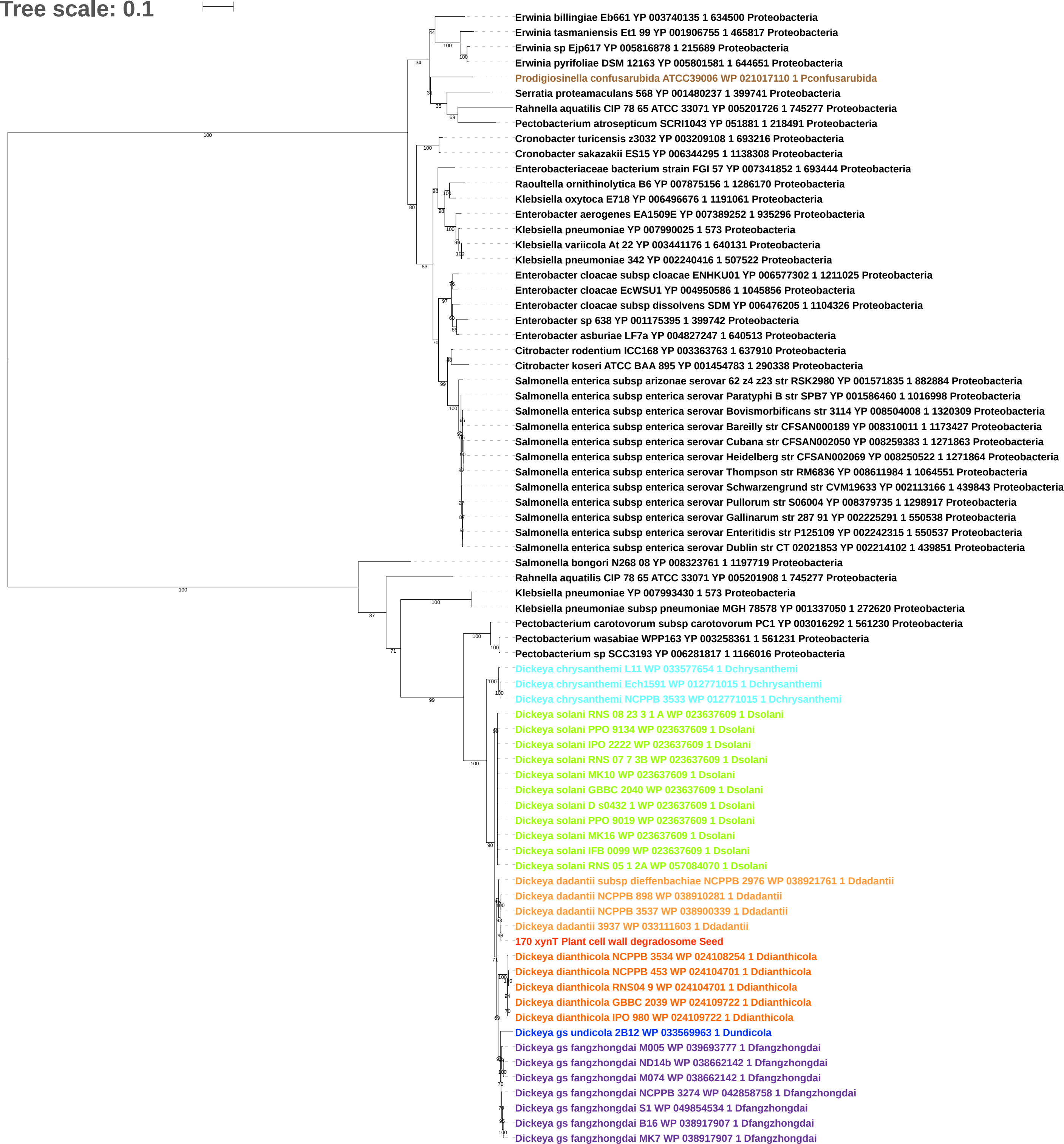
Tree scale: 0.1
Erwinia billingiae Eb661 YP 003740135 1 634500 Proteobacteria
Erwinia tasmaniensis Et1 99 YP 001906755 1 465817 Proteobacteria
Erwinia sp Ejp617 YP 005816878 1 215689 Proteobacteria
Erwinia pyrifoliae DSM 12163 YP 005801581 1 644651 Proteobacteria
Prodigiosinella confusarubida ATCC39006 WP 021017110 1 Pconfusarubida
Serratia proteamaculans 568 YP 001480237 1 399741 Proteobacteria
Rahnella aquatilis CIP 78 65 ATCC 33071 YP 005201726 1 745277 Proteobacteria
Pectobacterium atrosepticum SCRI1043 YP 051881 1 218491 Proteobacteria
Cronobacter turicensis z3032 YP 003209108 1 693216 Proteobacteria
Cronobacter sakazakii ES15 YP 006344295 1 1138308 Proteobacteria
Enterobacteriaceae bacterium strain FGI 57 YP 007341852 1 693444 Proteobacteria
Raoultella ornithinolytica B6 YP 007875156 1 1286170 Proteobacteria
Klebsiella oxytoca E718 YP 006496676 1 1191061 Proteobacteria
Enterobacter aerogenes EA1509E YP 007389252 1 935296 Proteobacteria
Klebsiella pneumoniae YP 007990025 1 573 Proteobacteria
Klebsiella variicola At 22 YP 003441176 1 640131 Proteobacteria
Klebsiella pneumoniae 342 YP 002240416 1 507522 Proteobacteria
Enterobacter cloacae subsp cloacae ENHKU01 YP 006577302 1 1211025 Proteobacteria
Enterobacter cloacae EcWSU1 YP 004950586 1 1045856 Proteobacteria
Enterobacter cloacae subsp dissolvens SDM YP 006476205 1 1104326 Proteobacteria
Enterobacter sp 638 YP 001175395 1 399742 Proteobacteria
Enterobacter asburiae LF7a YP 004827247 1 640513 Proteobacteria
Citrobacter rodentium ICC168 YP 003363763 1 637910 Proteobacteria
Citrobacter koseri ATCC BAA 895 YP 001454783 1 290338 Proteobacteria
Salmonella enterica subsp arizonae serovar 62 z4 z23 str RSK2980 YP 001571835 1 882884 Proteobacteria
Salmonella enterica subsp enterica serovar Paratyphi B str SPB7 YP 001586460 1 1016998 Proteobacteria
Salmonella enterica subsp enterica serovar Bovismorbificans str 3114 YP 008504008 1 1320309 Proteobacteria
Salmonella enterica subsp enterica serovar Bareilly str CFSAN000189 YP 008310011 1 1173427 Proteobacteria
Salmonella enterica subsp enterica serovar Cubana str CFSAN002050 YP 008259383 1 1271863 Proteobacteria
Salmonella enterica subsp enterica serovar Heidelberg str CFSAN002069 YP 008250522 1 1271864 Proteobacteria
Salmonella enterica subsp enterica serovar Thompson str RM6836 YP 008611984 1 1064551 Proteobacteria
Salmonella enterica subsp enterica serovar Schwarzengrund str CVM19633 YP 002113166 1 439843 Proteobacteria
Salmonella enterica subsp enterica serovar Pullorum str S06004 YP 008379735 1 1298917 Proteobacteria
Salmonella enterica subsp enterica serovar Gallinarum str 287 91 YP 002225291 1 550538 Proteobacteria
Salmonella enterica subsp enterica serovar Enteritidis str P125109 YP 002242315 1 550537 Proteobacteria
Salmonella enterica subsp enterica serovar Dublin str CT 02021853 YP 002214102 1 439851 Proteobacteria
Salmonella bongori N268 08 YP 008323761 1 1197719 Proteobacteria
Rahnella aquatilis CIP 78 65 ATCC 33071 YP 005201908 1 745277 Proteobacteria
Klebsiella pneumoniae YP 007993430 1 573 Proteobacteria
Klebsiella pneumoniae subsp pneumoniae MGH 78578 YP 001337050 1 272620 Proteobacteria
Pectobacterium carotovorum subsp carotovorum PC1 YP 003016292 1 561230 Proteobacteria
Pectobacterium wasabiae WPP163 YP 003258361 1 561231 Proteobacteria
Pectobacterium sp SCC3193 YP 006281817 1 1166016 Proteobacteria
Dickeya chrysanthemi L11 WP 033577654 1 Dchrysanthemi
Dickeya chrysanthemi Ech1591 WP 012771015 1 Dchrysanthemi
Dickeya chrysanthemi NCPPB 3533 WP 012771015 1 Dchrysanthemi
Dickeya solani RNS 08 23 3 1 A WP 023637609 1 Dsolani
Dickeya solani PPO 9134 WP 023637609 1 Dsolani
Dickeya solani IPO 2222 WP 023637609 1 Dsolani
Dickeya solani RNS 07 7 3B WP 023637609 1 Dsolani
Dickeya solani MK10 WP 023637609 1 Dsolani
Dickeya solani GBBC 2040 WP 023637609 1 Dsolani
Dickeya solani D s0432 1 WP 023637609 1 Dsolani
Dickeya solani PPO 9019 WP 023637609 1 Dsolani
Dickeya solani MK16 WP 023637609 1 Dsolani
Dickeya solani IFB 0099 WP 023637609 1 Dsolani
Dickeya solani RNS 05 1 2A WP 057084070 1 Dsolani
Dickeya dadantii subsp dieffenbachiae NCPPB 2976 WP 038921761 1 Ddadantii
Dickeya dadantii NCPPB 898 WP 038910281 1 Ddadantii
Dickeya dadantii NCPPB 3537 WP 038900339 1 Ddadantii
Dickeya dadantii 3937 WP 033111603 1 Ddadantii
170 xynT Plant cell wall degradosome Seed
Dickeya dianthicola NCPPB 3534 WP 024108254 1 Ddianthicola
Dickeya dianthicola NCPPB 453 WP 024104701 1 Ddianthicola
Dickeya dianthicola RNS04 9 WP 024104701 1 Ddianthicola
Dickeya dianthicola GBBC 2039 WP 024109722 1 Ddianthicola
Dickeya dianthicola IPO 980 WP 024109722 1 Ddianthicola
Dickeya gs undicola 2B12 WP 033569963 1 Dundicola
Dickeya gs fangzhongdai M005 WP 039693777 1 Dfangzhongdai
Dickeya gs fangzhongdai ND14b WP 038662142 1 Dfangzhongdai
Dickeya gs fangzhongdai M074 WP 038662142 1 Dfangzhongdai
Dickeya gs fangzhongdai NCPPB 3274 WP 042858758 1 Dfangzhongdai
Dickeya gs fangzhongdai S1 WP 049854534 1 Dfangzhongdai
Dickeya gs fangzhongdai B16 WP 038917907 1 Dfangzhongdai
Dickeya gs fangzhongdai MK7 WP 038917907 1 Dfangzhongdai

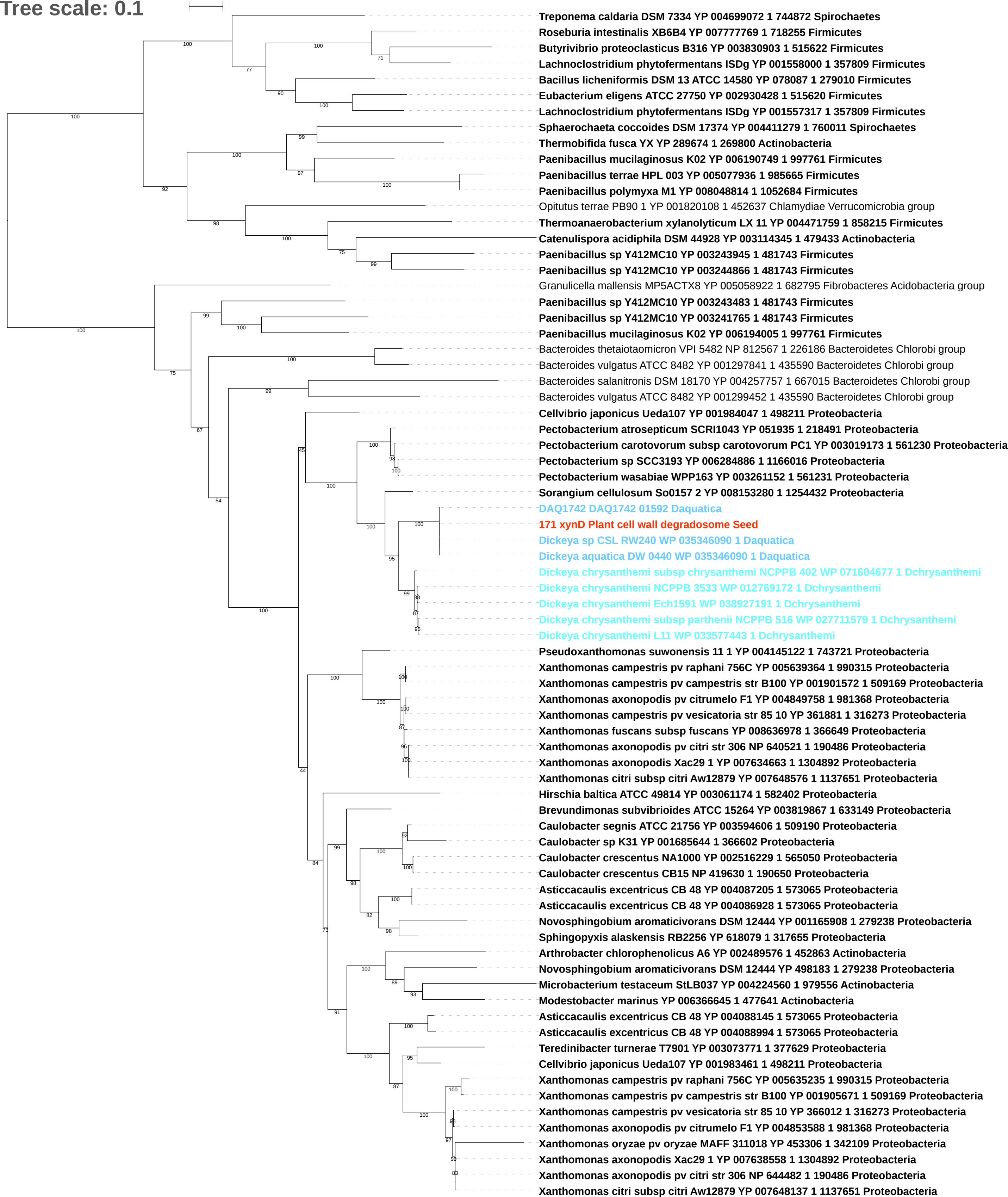
Tree scale: 0.1
Treponema caldaria DSM 7334 YP 004699072 1 744872 Spirochaetes
Roseburia intestinalis XB6B4 YP 007777769 1 718255 Firmicutes
Butyrivibrio proteoclasticus B316 YP 003830903 1 515622 Firmicutes
Lachnoclostridium phytofermentans ISDg YP 001558000 1 357809 Firmicutes
Bacillus licheniformis DSM 13 ATCC 14580 YP 078087 1 279010 Firmicutes
Eubacterium eligens ATCC 27750 YP 002930428 1 515620 Firmicutes
Lachnoclostridium phytofermentans ISDg YP 001557317 1 357809 Firmicutes
Sphaerochaeta coccoides DSM 17374 YP 004411279 1 760011 Spirochaetes
Thermobifida fusca YX YP 289674 1 269800 Actinobacteria
Paenibacillus mucilaginosus K02 YP 006190749 1 997761 Firmicutes
Paenibacillus terrae HPL 003 YP 005077936 1 985665 Firmicutes
Paenibacillus polymyxa M1 YP 008048814 1 1052684 Firmicutes
Opitutus terrae PB90 1 YP 001820108 1 452637 Chlamydiae Verrucomicrobia group
Thermoanaerobacterium xylanolyticum LX 11 YP 004471759 1 858215 Firmicutes
Catenulispora acidiphila DSM 44928 YP 003114345 1 479433 Actinobacteria
Paenibacillus sp Y412MC10 YP 003243945 1 481743 Firmicutes
Paenibacillus sp Y412MC10 YP 003244866 1 481743 Firmicutes
Granulicella mallensis MP5ACTX8 YP 005058922 1 682795 Fibrobacteres Acidobacteria group
Paenibacillus sp Y412MC10 YP 003243483 1 481743 Firmicutes
Paenibacillus sp Y412MC10 YP 003241765 1 481743 Firmicutes
Paenibacillus mucilaginosus K02 YP 006194005 1 997761 Firmicutes
Bacteroides thetaiotaomicron VPI 5482 NP 812567 1 226186 Bacteroidetes Chlorobi group
Bacteroides vulgatus ATCC 8482 YP 001297841 1 435590 Bacteroidetes Chlorobi group
Bacteroides salanitronis DSM 18170 YP 004257757 1 667015 Bacteroidetes Chlorobi group
Bacteroides vulgatus ATCC 8482 YP 001299452 1 435590 Bacteroidetes Chlorobi group
Cellvibrio japonicus Ueda107 YP 001984047 1 498211 Proteobacteria
Pectobacterium atrosepticum SCRI1043 YP 051935 1 218491 Proteobacteria
Pectobacterium carotovorum subsp carotovorum PC1 YP 003019173 1 561230 Proteobacteria
Pectobacterium sp SCC3193 YP 006284886 1 1166016 Proteobacteria
Pectobacterium wasabiae WPP163 YP 003261152 1 561231 Proteobacteria
Sorangium cellulosum So0157 2 YP 008153280 1 1254432 Proteobacteria
DAQ1742 DAQ1742 01592 Daquatica
171 xynD Plant cell wall degradosome Seed
Dickeya sp CSL RW240 WP 035346090 1 Daquatica
Dickeya aquatica DW 0440 WP 035346090 1 Daquatica
Dickeya chrysanthemi subsp chrysanthemi NCPPB 402 WP 071604677 1 Dchrysanthemi
Dickeya chrysanthemi NCPPB 3533 WP 012769172 1 Dchrysanthemi
Dickeya chrysanthemi Ech1591 WP 038927191 1 Dchrysanthemi
Dickeya chrysanthemi subsp parthenii NCPPB 516 WP 027711579 1 Dchrysanthemi
Dickeya chrysanthemi L11 WP 033577443 1 Dchrysanthemi
Pseudoxanthomonas suwonensis 11 1 YP 004145122 1 743721 Proteobacteria
Xanthomonas campestris pv raphani 756C YP 005639364 1 990315 Proteobacteria
Xanthomonas campestris pv campestris str B100 YP 001901572 1 509169 Proteobacteria
Xanthomonas axonopodis pv citrumelo F1 YP 004849758 1 981368 Proteobacteria
Xanthomonas campestris pv vesicatoria str 85 10 YP 361881 1 316273 Proteobacteria
Xanthomonas fuscans subsp fuscans YP 008636978 1 366649 Proteobacteria
Xanthomonas axonopodis pv citri str 306 NP 640521 1 190486 Proteobacteria
Xanthomonas axonopodis Xac29 1 YP 007634663 1 1304892 Proteobacteria
Xanthomonas citri subsp citri Aw12879 YP 007648576 1 1137651 Proteobacteria
Hirschia baltica ATCC 49814 YP 003061174 1 582402 Proteobacteria
Brevundimonas subvibrioides ATCC 15264 YP 003819867 1 633149 Proteobacteria
Caulobacter segnis ATCC 21756 YP 003594606 1 509190 Proteobacteria
Caulobacter sp K31 YP 001685644 1 366602 Proteobacteria
Caulobacter crescentus NA1000 YP 002516229 1 565050 Proteobacteria
Caulobacter crescentus CB15 NP 419630 1 190650 Proteobacteria
Asticcacaulis excentricus CB 48 YP 004087205 1 573065 Proteobacteria
Asticcacaulis excentricus CB 48 YP 004086928 1 573065 Proteobacteria
Novosphingobium aromaticivorans DSM 12444 YP 001165908 1 279238 Proteobacteria
Sphingopyxis alaskensis RB2256 YP 618079 1 317655 Proteobacteria
Arthrobacter chlorophenolicus A6 YP 002489576 1 452863 Actinobacteria
Novosphingobium aromaticivorans DSM 12444 YP 498183 1 279238 Proteobacteria
Microbacterium testaceum StLB037 YP 004224560 1 979556 Actinobacteria
Modestobacter marinus YP 006366645 1 477641 Actinobacteria
Asticcacaulis excentricus CB 48 YP 004088145 1 573065 Proteobacteria
Asticcacaulis excentricus CB 48 YP 004088994 1 573065 Proteobacteria
Teredinibacter turnerae T7901 YP 003073771 1 377629 Proteobacteria
Cellvibrio japonicus Ueda107 YP 001983461 1 498211 Proteobacteria
Xanthomonas campestris pv raphani 756C YP 005635235 1 990315 Proteobacteria
Xanthomonas campestris pv campestris str B100 YP 001905671 1 509169 Proteobacteria
Xanthomonas campestris pv vesicatoria str 85 10 YP 366012 1 316273 Proteobacteria
Xanthomonas axonopodis pv citrumelo F1 YP 004853588 1 981368 Proteobacteria
Xanthomonas oryzae pv oryzae MAFF 311018 YP 453306 1 342109 Proteobacteria
Xanthomonas axonopodis Xac29 1 YP 007638558 1 1304892 Proteobacteria
Xanthomonas axonopodis pv citri str 306 NP 644482 1 190486 Proteobacteria
Xanthomonas citri subsp citri Aw12879 YP 007648137 1 1137651 Proteobacteria

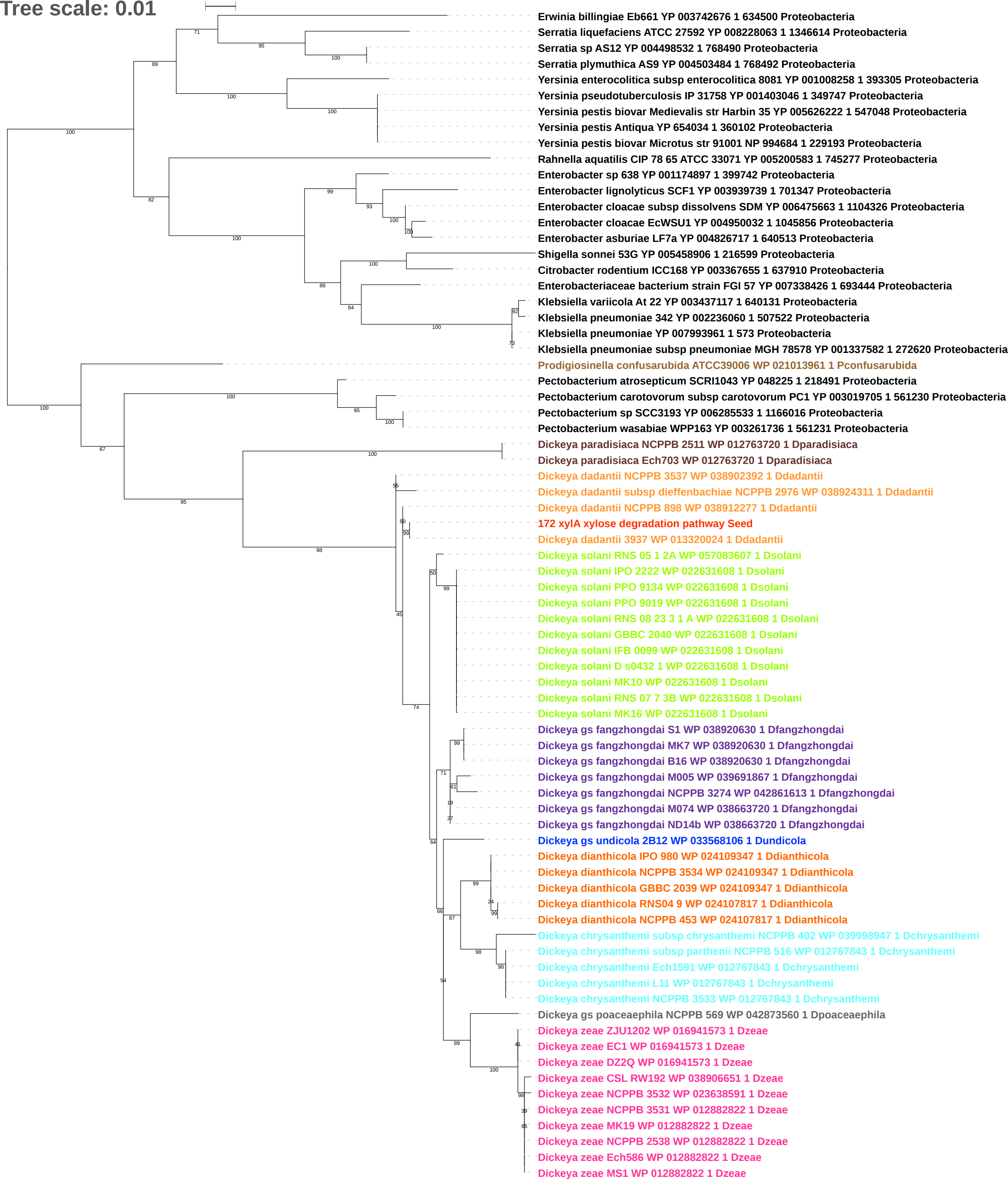
Tree scale: 0.01
Erwinia billingiae Eb661 YP 003742676 1 634500 Proteobacteria
Serratia liquefaciens ATCC 27592 YP 008228063 1 1346614 Proteobacteria
Serratia sp AS12 YP 004498532 1 768490 Proteobacteria
Serratia plymuthica AS9 YP 004503484 1 768492 Proteobacteria
Yersinia enterocolitica subsp enterocolitica 8081 YP 001008258 1 393305 Proteobacteria
Yersinia pseudotuberculosis IP 31758 YP 001403046 1 349747 Proteobacteria
Yersinia pestis biovar Medievalis str Harbin 35 YP 005626222 1 547048 Proteobacteria
Yersinia pestis Antiqua YP 654034 1 360102 Proteobacteria
Yersinia pestis biovar Microtus str 91001 NP 994684 1 229193 Proteobacteria
Rahnella aquatilis CIP 78 65 ATCC 33071 YP 005200583 1 745277 Proteobacteria
Enterobacter sp 638 YP 001174897 1 399742 Proteobacteria
Enterobacter lignolyticus SCF1 YP 003939739 1 701347 Proteobacteria
Enterobacter cloacae subsp dissolvens SDM YP 006475663 1 1104326 Proteobacteria
Enterobacter cloacae EcWSU1 YP 004950032 1 1045856 Proteobacteria
Enterobacter asburiae LF7a YP 004826717 1 640513 Proteobacteria
Shigella sonnei 53G YP 005458906 1 216599 Proteobacteria
Citrobacter rodentium ICC168 YP 003367655 1 637910 Proteobacteria
Enterobacteriaceae bacterium strain FGI 57 YP 007338426 1 693444 Proteobacteria
Klebsiella variicola At 22 YP 003437117 1 640131 Proteobacteria
Klebsiella pneumoniae 342 YP 002236060 1 507522 Proteobacteria
Klebsiella pneumoniae YP 007993961 1 573 Proteobacteria
Klebsiella pneumoniae subsp pneumoniae MGH 78578 YP 001337582 1 272620 Proteobacteria
Prodigiosinella confusarubida ATCC39006 WP 021013961 1 Pconfusarubida
Pectobacterium atrosepticum SCRI1043 YP 048225 1 218491 Proteobacteria
Pectobacterium carotovorum subsp carotovorum PC1 YP 003019705 1 561230 Proteobacteria
Pectobacterium sp SCC3193 YP 006285533 1 1166016 Proteobacteria
Pectobacterium wasabiae WPP163 YP 003261736 1 561231 Proteobacteria
Dickeya paradisiaca NCPPB 2511 WP 012763720 1 Dparadisiaca
Dickeya paradisiaca Ech703 WP 012763720 1 Dparadisiaca
Dickeya dadantii NCPPB 3537 WP 038902392 1 Ddadantii
Dickeya dadantii subsp dieffenbachiae NCPPB 2976 WP 038924311 1 Ddadantii
Dickeya dadantii NCPPB 898 WP 038912277 1 Ddadantii
172 xylA xylose degradation pathway Seed
Dickeya dadantii 3937 WP 013320024 1 Ddadantii
Dickeya solani RNS 05 1 2A WP 057083607 1 Dsolani
Dickeya solani IPO 2222 WP 022631608 1 Dsolani
Dickeya solani PPO 9134 WP 022631608 1 Dsolani
Dickeya solani PPO 9019 WP 022631608 1 Dsolani
Dickeya solani RNS 08 23 3 1 A WP 022631608 1 Dsolani
Dickeya solani GBBC 2040 WP 022631608 1 Dsolani
Dickeya solani IFB 0099 WP 022631608 1 Dsolani
Dickeya solani D s0432 1 WP 022631608 1 Dsolani
Dickeya solani MK10 WP 022631608 1 Dsolani
Dickeya solani RNS 07 7 3B WP 022631608 1 Dsolani
Dickeya solani MK16 WP 022631608 1 Dsolani
Dickeya gs fangzhongdai S1 WP 038920630 1 Dfangzhongdai
Dickeya gs fangzhongdai MK7 WP 038920630 1 Dfangzhongdai
Dickeya gs fangzhongdai B16 WP 038920630 1 Dfangzhongdai
Dickeya gs fangzhongdai M005 WP 039691867 1 Dfangzhongdai
Dickeya gs fangzhongdai NCPPB 3274 WP 042861613 1 Dfangzhongdai
Dickeya gs fangzhongdai M074 WP 038663720 1 Dfangzhongdai
Dickeya gs fangzhongdai ND14b WP 038663720 1 Dfangzhongdai
Dickeya gs undicola 2B12 WP 033568106 1 Dundicola
Dickeya dianthicola IPO 980 WP 024109347 1 Ddianthicola
Dickeya dianthicola NCPPB 3534 WP 024109347 1 Ddianthicola
Dickeya dianthicola GBBC 2039 WP 024109347 1 Ddianthicola
Dickeya dianthicola RNS04 9 WP 024107817 1 Ddianthicola
Dickeya dianthicola NCPPB 453 WP 024107817 1 Ddianthicola
Dickeya chrysanthemi subsp chrysanthemi NCPPB 402 WP 039998947 1 Dchrysanthemi
Dickeya chrysanthemi subsp parthenii NCPPB 516 WP 012767843 1 Dchrysanthemi
Dickeya chrysanthemi Ech1591 WP 012767843 1 Dchrysanthemi
Dickeya chrysanthemi L11 WP 012767843 1 Dchrysanthemi
Dickeya chrysanthemi NCPPB 3533 WP 012767843 1 Dchrysanthemi
Dickeya gs poaceaephila NCPPB 569 WP 042873560 1 Dpoaceaephila
Dickeya zeae ZJU1202 WP 016941573 1 Dzeae
Dickeya zeae EC1 WP 016941573 1 Dzeae
Dickeya zeae DZ2Q WP 016941573 1 Dzeae
Dickeya zeae CSL RW192 WP 038906651 1 Dzeae
Dickeya zeae NCPPB 3532 WP 023638591 1 Dzeae
Dickeya zeae NCPPB 3531 WP 012882822 1 Dzeae
Dickeya zeae MK19 WP 012882822 1 Dzeae
Dickeya zeae NCPPB 2538 WP 012882822 1 Dzeae
Dickeya zeae Ech586 WP 012882822 1 Dzeae
Dickeya zeae MS1 WP 012882822 1 Dzeae

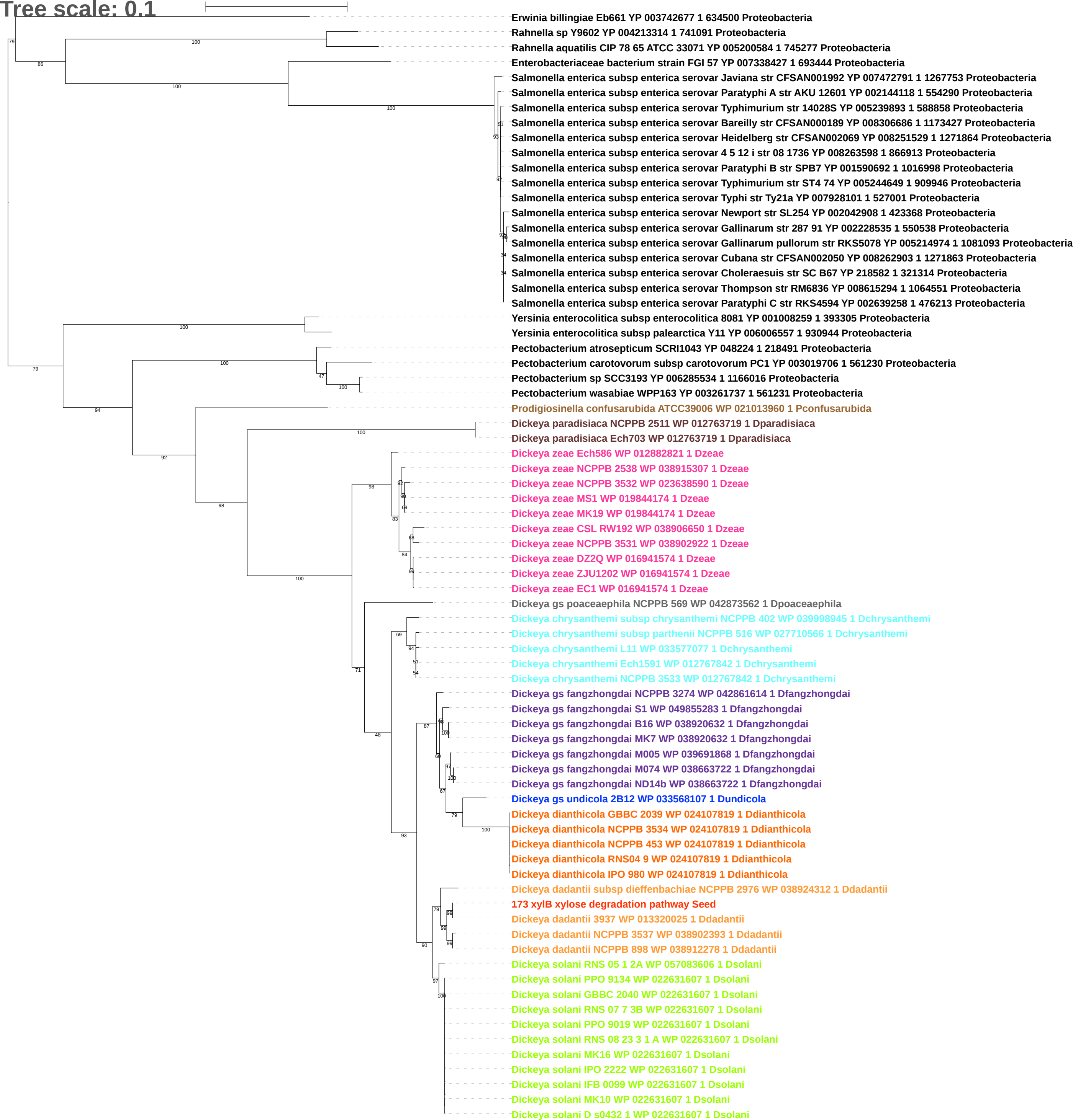
Tree scale: 0.1
Erwinia billingiae Eb661 YP 003742677 1 634500 Proteobacteria
Rahnella sp Y9602 YP 004213314 1 741091 Proteobacteria
Rahnella aquatilis CIP 78 65 ATCC 33071 YP 005200584 1 745277 Proteobacteria
Enterobacteriaceae bacterium strain FGI 57 YP 007338427 1 693444 Proteobacteria
Salmonella enterica subsp enterica serovar Javiana str CFSAN001992 YP 007472791 1 1267753 Proteobacteria
Salmonella enterica subsp enterica serovar Paratyphi A str AKU 12601 YP 002144118 1 554290 Proteobacteria
Salmonella enterica subsp enterica serovar Typhimurium str 14028S YP 005239893 1 588858 Proteobacteria
Salmonella enterica subsp enterica serovar Bareilly str CFSAN000189 YP 008306686 1 1173427 Proteobacteria
Salmonella enterica subsp enterica serovar Heidelberg str CFSAN002069 YP 008251529 1 1271864 Proteobacteria
Salmonella enterica subsp enterica serovar 4 5 12 i str 08 1736 YP 008263598 1 866913 Proteobacteria
Salmonella enterica subsp enterica serovar Paratyphi B str SPB7 YP 001590692 1 1016998 Proteobacteria
Salmonella enterica subsp enterica serovar Typhimurium str ST4 74 YP 005244649 1 909946 Proteobacteria
Salmonella enterica subsp enterica serovar Typhi str Ty21a YP 007928101 1 527001 Proteobacteria
Salmonella enterica subsp enterica serovar Newport str SL254 YP 002042908 1 423368 Proteobacteria
Salmonella enterica subsp enterica serovar Gallinarum str 287 91 YP 002228535 1 550538 Proteobacteria
Salmonella enterica subsp enterica serovar Gallinarum pullorum str RKS5078 YP 005214974 1 1081093 Proteobacteria
Salmonella enterica subsp enterica serovar Cubana str CFSAN002050 YP 008262903 1 1271863 Proteobacteria
Salmonella enterica subsp enterica serovar Choleraesuis str SC B67 YP 218582 1 321314 Proteobacteria
Salmonella enterica subsp enterica serovar Thompson str RM6836 YP 008615294 1 1064551 Proteobacteria
Salmonella enterica subsp enterica serovar Paratyphi C str RKS4594 YP 002639258 1 476213 Proteobacteria
Yersinia enterocolitica subsp enterocolitica 8081 YP 001008259 1 393305 Proteobacteria
Yersinia enterocolitica subsp palearctica Y11 YP 006006557 1 930944 Proteobacteria
Pectobacterium atrosepticum SCRI1043 YP 048224 1 218491 Proteobacteria
Pectobacterium carotovorum subsp carotovorum PC1 YP 003019706 1 561230 Proteobacteria
Pectobacterium sp SCC3193 YP 006285534 1 1166016 Proteobacteria
Pectobacterium wasabiae WPP163 YP 003261737 1 561231 Proteobacteria
Prodigiosinella confusarubida ATCC39006 WP 021013960 1 Pconfusarubida
Dickeya paradisiaca NCPPB 2511 WP 012763719 1 Dparadisiaca
Dickeya paradisiaca Ech703 WP 012763719 1 Dparadisiaca
Dickeya zeae Ech586 WP 012882821 1 Dzeae
Dickeya zeae NCPPB 2538 WP 038915307 1 Dzeae
Dickeya zeae NCPPB 3532 WP 023638590 1 Dzeae
Dickeya zeae MS1 WP 019844174 1 Dzeae
Dickeya zeae MK19 WP 019844174 1 Dzeae
Dickeya zeae CSL RW192 WP 038906650 1 Dzeae
Dickeya zeae NCPPB 3531 WP 038902922 1 Dzeae
Dickeya zeae DZ2Q WP 016941574 1 Dzeae
Dickeya zeae ZJU1202 WP 016941574 1 Dzeae
Dickeya zeae EC1 WP 016941574 1 Dzeae
Dickeya gs poaceaephila NCPPB 569 WP 042873562 1 Dpoaceaephila
Dickeya chrysanthemi subsp chrysanthemi NCPPB 402 WP 039998945 1 Dchrysanthemi
Dickeya chrysanthemi subsp parthenii NCPPB 516 WP 027710566 1 Dchrysanthemi
Dickeya chrysanthemi L11 WP 033577077 1 Dchrysanthemi
Dickeya chrysanthemi Ech1591 WP 012767842 1 Dchrysanthemi
Dickeya chrysanthemi NCPPB 3533 WP 012767842 1 Dchrysanthemi
Dickeya gs fangzhongdai NCPPB 3274 WP 042861614 1 Dfangzhongdai
Dickeya gs fangzhongdai S1 WP 049855283 1 Dfangzhongdai
Dickeya gs fangzhongdai B16 WP 038920632 1 Dfangzhongdai
Dickeya gs fangzhongdai MK7 WP 038920632 1 Dfangzhongdai
Dickeya gs fangzhongdai M005 WP 039691868 1 Dfangzhongdai
Dickeya gs fangzhongdai M074 WP 038663722 1 Dfangzhongdai
Dickeya gs fangzhongdai ND14b WP 038663722 1 Dfangzhongdai
Dickeya gs undicola 2B12 WP 033568107 1 Dundicola
Dickeya dianthicola GBBC 2039 WP 024107819 1 Ddianthicola
Dickeya dianthicola NCPPB 3534 WP 024107819 1 Ddianthicola
Dickeya dianthicola NCPPB 453 WP 024107819 1 Ddianthicola
Dickeya dianthicola RNS04 9 WP 024107819 1 Ddianthicola
Dickeya dianthicola IPO 980 WP 024107819 1 Ddianthicola
Dickeya dadantii subsp dieffenbachiae NCPPB 2976 WP 038924312 1 Ddadantii
173 xylB xylose degradation pathway Seed
Dickeya dadantii 3937 WP 013320025 1 Ddadantii
Dickeya dadantii NCPPB 3537 WP 038902393 1 Ddadantii
Dickeya dadantii NCPPB 898 WP 038912278 1 Ddadantii
Dickeya solani RNS 05 1 2A WP 057083606 1 Dsolani
Dickeya solani PPO 9134 WP 022631607 1 Dsolani
Dickeya solani GBBC 2040 WP 022631607 1 Dsolani
Dickeya solani RNS 07 7 3B WP 022631607 1 Dsolani
Dickeya solani PPO 9019 WP 022631607 1 Dsolani
Dickeya solani RNS 08 23 3 1 A WP 022631607 1 Dsolani
Dickeya solani MK16 WP 022631607 1 Dsolani
Dickeya solani IPO 2222 WP 022631607 1 Dsolani
Dickeya solani IFB 0099 WP 022631607 1 Dsolani
Dickeya solani MK10 WP 022631607 1 Dsolani
Dickeya solani D s0432 1 WP 022631607 1 Dsolani

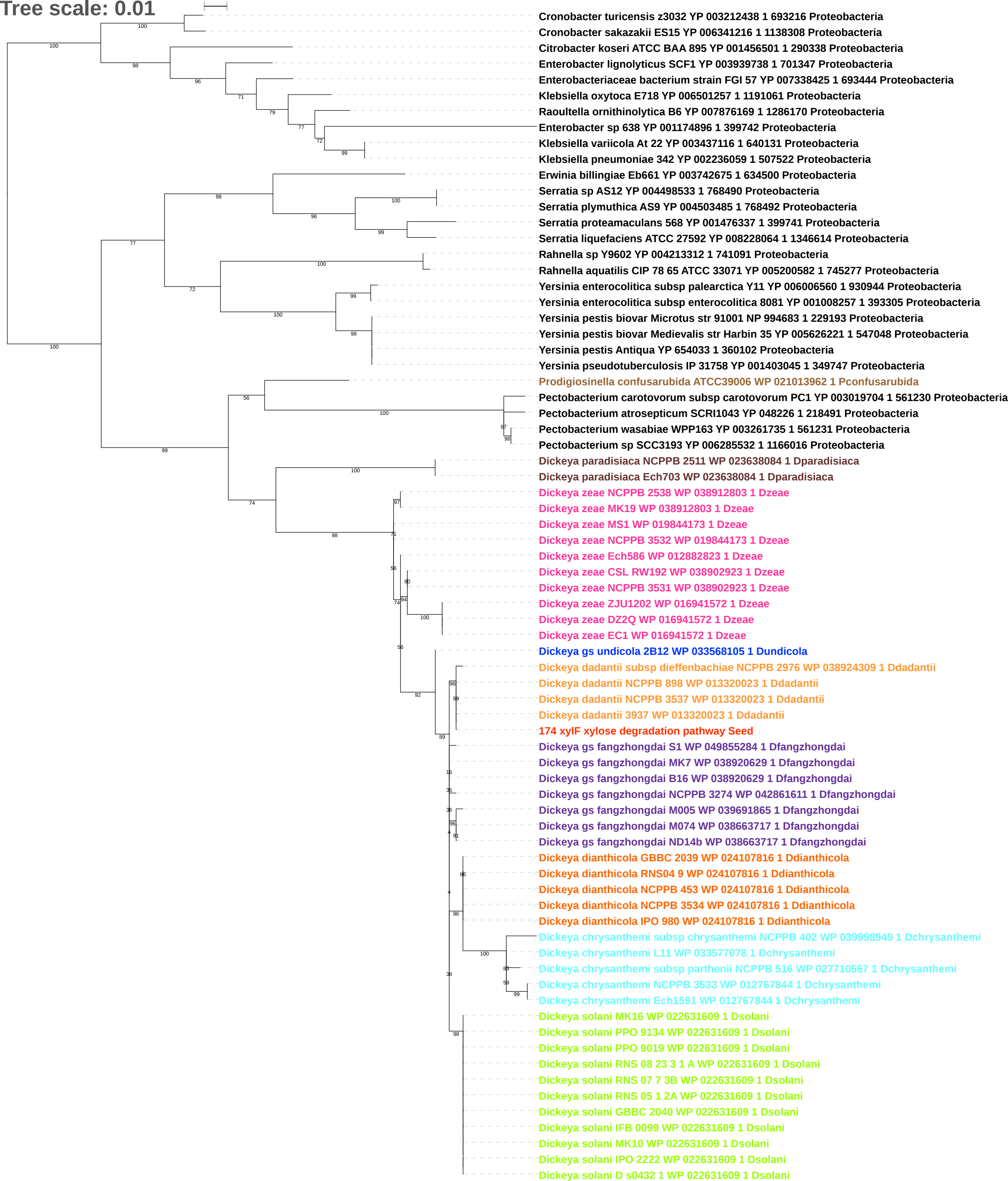
Tree scale: 0.01
Cronobacter turicensis z3032 YP 003212438 1 693216 Proteobacteria
Cronobacter sakazakii ES15 YP 006341216 1 1138308 Proteobacteria
Citrobacter koseri ATCC BAA 895 YP 001456501 1 290338 Proteobacteria
Enterobacter lignolyticus SCF1 YP 003939738 1 701347 Proteobacteria
Enterobacteriaceae bacterium strain FGI 57 YP 007338425 1 693444 Proteobacteria
Klebsiella oxytoca E718 YP 006501257 1 1191061 Proteobacteria
Raoultella ornithinolytica B6 YP 007876169 1 1286170 Proteobacteria
Enterobacter sp 638 YP 001174896 1 399742 Proteobacteria
Klebsiella variicola At 22 YP 003437116 1 640131 Proteobacteria
Klebsiella pneumoniae 342 YP 002236059 1 507522 Proteobacteria
Erwinia billingiae Eb661 YP 003742675 1 634500 Proteobacteria
Serratia sp AS12 YP 004498533 1 768490 Proteobacteria
Serratia plymuthica AS9 YP 004503485 1 768492 Proteobacteria
Serratia proteamaculans 568 YP 001476337 1 399741 Proteobacteria
Serratia liquefaciens ATCC 27592 YP 008228064 1 1346614 Proteobacteria
Rahnella sp Y9602 YP 004213312 1 741091 Proteobacteria
Rahnella aquatilis CIP 78 65 ATCC 33071 YP 005200582 1 745277 Proteobacteria
Yersinia enterocolitica subsp palearctica Y11 YP 006006560 1 930944 Proteobacteria
Yersinia enterocolitica subsp enterocolitica 8081 YP 001008257 1 393305 Proteobacteria
Yersinia pestis biovar Microtus str 91001 NP 994683 1 229193 Proteobacteria
Yersinia pestis biovar Medievalis str Harbin 35 YP 005626221 1 547048 Proteobacteria
Yersinia pestis Antiqua YP 654033 1 360102 Proteobacteria
Yersinia pseudotuberculosis IP 31758 YP 001403045 1 349747 Proteobacteria
Prodigiosinella confusarubida ATCC39006 WP 021013962 1 Pconfusarubida
Pectobacterium carotovorum subsp carotovorum PC1 YP 003019704 1 561230 Proteobacteria
Pectobacterium atrosepticum SCRI1043 YP 048226 1 218491 Proteobacteria
Pectobacterium wasabiae WPP163 YP 003261735 1 561231 Proteobacteria
Pectobacterium sp SCC3193 YP 006285532 1 1166016 Proteobacteria
Dickeya paradisiaca NCPPB 2511 WP 023638084 1 Dparadisiaca
Dickeya paradisiaca Ech703 WP 023638084 1 Dparadisiaca
Dickeya zeae NCPPB 2538 WP 038912803 1 Dzeae
Dickeya zeae MK19 WP 038912803 1 Dzeae
Dickeya zeae MS1 WP 019844173 1 Dzeae
Dickeya zeae NCPPB 3532 WP 019844173 1 Dzeae
Dickeya zeae Ech586 WP 012882823 1 Dzeae
Dickeya zeae CSL RW192 WP 038902923 1 Dzeae
Dickeya zeae NCPPB 3531 WP 038902923 1 Dzeae
Dickeya zeae ZJU1202 WP 016941572 1 Dzeae
Dickeya zeae DZ2Q WP 016941572 1 Dzeae
Dickeya zeae EC1 WP 016941572 1 Dzeae
Dickeya gs undicola 2B12 WP 033568105 1 Dundicola
Dickeya dadantii subsp dieffenbachiae NCPPB 2976 WP 038924309 1 Ddadantii
Dickeya dadantii NCPPB 898 WP 013320023 1 Ddadantii
Dickeya dadantii NCPPB 3537 WP 013320023 1 Ddadantii
Dickeya dadantii 3937 WP 013320023 1 Ddadantii
174 xylF xylose degradation pathway Seed
Dickeya gs fangzhongdai S1 WP 049855284 1 Dfangzhongdai
Dickeya gs fangzhongdai MK7 WP 038920629 1 Dfangzhongdai
Dickeya gs fangzhongdai B16 WP 038920629 1 Dfangzhongdai
Dickeya gs fangzhongdai NCPPB 3274 WP 042861611 1 Dfangzhongdai
Dickeya gs fangzhongdai M005 WP 039691865 1 Dfangzhongdai
Dickeya gs fangzhongdai M074 WP 038663717 1 Dfangzhongdai
Dickeya gs fangzhongdai ND14b WP 038663717 1 Dfangzhongdai
Dickeya dianthicola GBBC 2039 WP 024107816 1 Ddianthicola
Dickeya dianthicola RNS04 9 WP 024107816 1 Ddianthicola
Dickeya dianthicola NCPPB 453 WP 024107816 1 Ddianthicola
Dickeya dianthicola NCPPB 3534 WP 024107816 1 Ddianthicola
Dickeya dianthicola IPO 980 WP 024107816 1 Ddianthicola
Dickeya chrysanthemi subsp chrysanthemi NCPPB 402 WP 039998949 1 Dchrysanthemi
Dickeya chrysanthemi L11 WP 033577078 1 Dchrysanthemi
Dickeya chrysanthemi subsp parthenii NCPPB 516 WP 027710567 1 Dchrysanthemi
Dickeya chrysanthemi NCPPB 3533 WP 012767844 1 Dchrysanthemi
Dickeya chrysanthemi Ech1591 WP 012767844 1 Dchrysanthemi
Dickeya solani MK16 WP 022631609 1 Dsolani
Dickeya solani PPO 9134 WP 022631609 1 Dsolani
Dickeya solani PPO 9019 WP 022631609 1 Dsolani
Dickeya solani RNS 08 23 3 1 A WP 022631609 1 Dsolani
Dickeya solani RNS 07 7 3B WP 022631609 1 Dsolani
Dickeya solani RNS 05 1 2A WP 022631609 1 Dsolani
Dickeya solani GBBC 2040 WP 022631609 1 Dsolani
Dickeya solani IFB 0099 WP 022631609 1 Dsolani
Dickeya solani MK10 WP 022631609 1 Dsolani
Dickeya solani IPO 2222 WP 022631609 1 Dsolani
Dickeya solani D s0432 1 WP 022631609 1 Dsolani

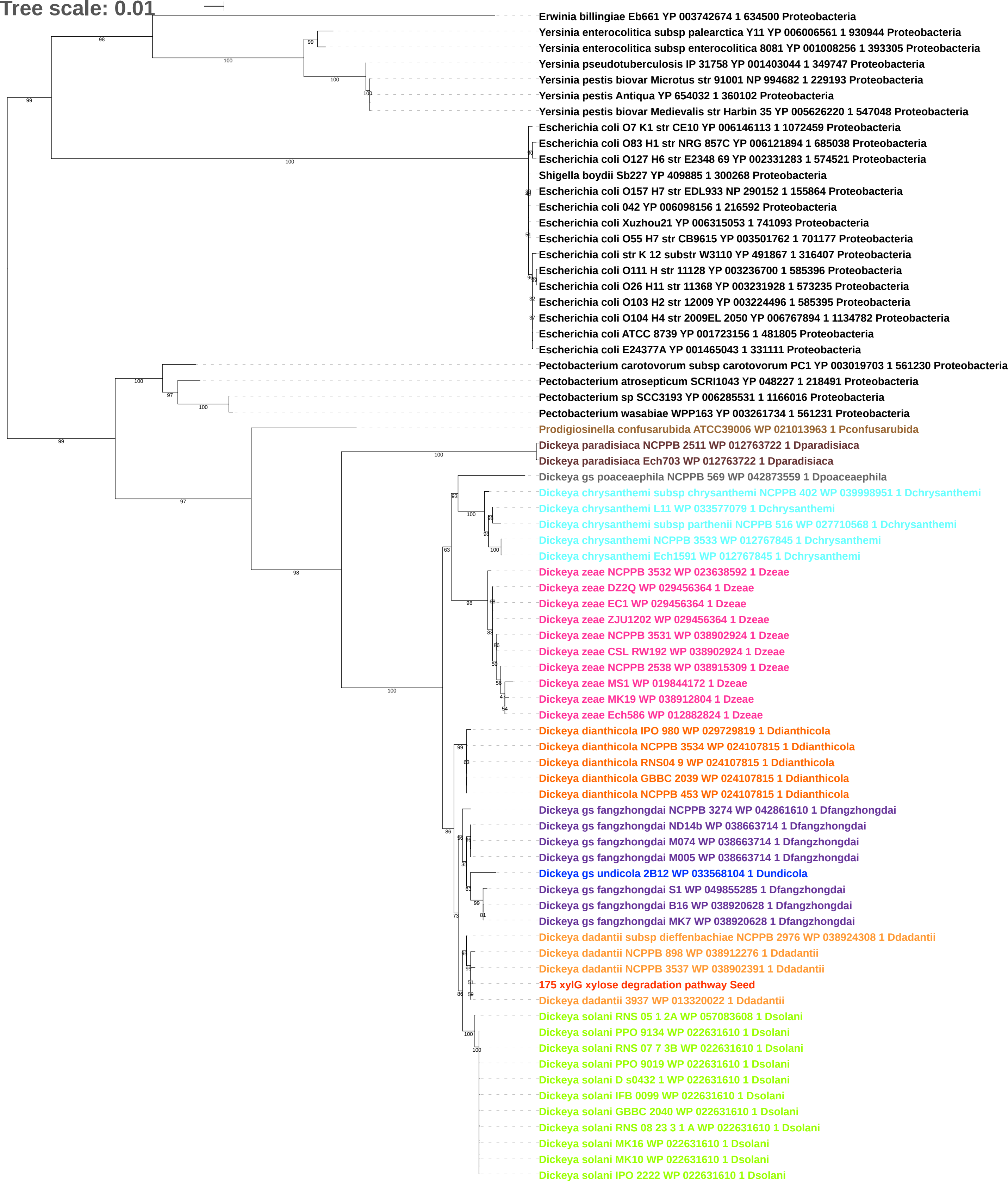

Tree scale: 0.01
Erwinia billingiae Eb661 YP 003742674 1 634500 Proteobacteria
Yersinia enterocolitica subsp palearctica Y11 YP 006006561 1 930944 Proteobacteria
Yersinia enterocolitica subsp enterocolitica 8081 YP 001008256 1 393305 Proteobacteria
Yersinia pseudotuberculosis IP 31758 YP 001403044 1 349747 Proteobacteria
Yersinia pestis biovar Microtus str 91001 NP 994682 1 229193 Proteobacteria
Yersinia pestis Antiqua YP 654032 1 360102 Proteobacteria
Yersinia pestis biovar Medievalis str Harbin 35 YP 005626220 1 547048 Proteobacteria
Escherichia coli O7 K1 str CE10 YP 006146113 1 1072459 Proteobacteria
Escherichia coli O83 H1 str NRG 857C YP 006121894 1 685038 Proteobacteria
Escherichia coli O127 H6 str E2348 69 YP 002331283 1 574521 Proteobacteria
Shigella boydii Sb227 YP 409885 1 300268 Proteobacteria
Escherichia coli O157 H7 str EDL933 NP 290152 1 155864 Proteobacteria
Escherichia coli 042 YP 006098156 1 216592 Proteobacteria
Escherichia coli Xuzhou21 YP 006315053 1 741093 Proteobacteria
Escherichia coli O55 H7 str CB9615 YP 003501762 1 701177 Proteobacteria
Escherichia coli str K 12 substr W3110 YP 491867 1 316407 Proteobacteria
Escherichia coli O111 H str 11128 YP 003236700 1 585396 Proteobacteria
Escherichia coli O26 H11 str 11368 YP 003231928 1 573235 Proteobacteria
Escherichia coli O103 H2 str 12009 YP 003224496 1 585395 Proteobacteria
Escherichia coli O104 H4 str 2009EL 2050 YP 006767894 1 1134782 Proteobacteria
Escherichia coli ATCC 8739 YP 001723156 1 481805 Proteobacteria
Escherichia coli E24377A YP 001465043 1 331111 Proteobacteria
Pectobacterium carotovorum subsp carotovorum PC1 YP 003019703 1 561230 Proteobacteria
Pectobacterium atrosepticum SCRI1043 YP 048227 1 218491 Proteobacteria
Pectobacterium sp SCC3193 YP 006285531 1 1166016 Proteobacteria
Pectobacterium wasabiae WPP163 YP 003261734 1 561231 Proteobacteria
Prodigiosinella confusarubida ATCC39006 WP 021013963 1 Pconfusarubida
Dickeya paradisiaca NCPPB 2511 WP 012763722 1 Dparadisiaca
Dickeya paradisiaca Ech703 WP 012763722 1 Dparadisiaca
Dickeya gs poaceaephila NCPPB 569 WP 042873559 1 Dpoaceaephila
Dickeya chrysanthemi subsp chrysanthemi NCPPB 402 WP 039998951 1 Dchrysanthemi
Dickeya chrysanthemi L11 WP 033577079 1 Dchrysanthemi
Dickeya chrysanthemi subsp parthenii NCPPB 516 WP 027710568 1 Dchrysanthemi
Dickeya chrysanthemi NCPPB 3533 WP 012767845 1 Dchrysanthemi
Dickeya chrysanthemi Ech1591 WP 012767845 1 Dchrysanthemi
Dickeya zeae NCPPB 3532 WP 023638592 1 Dzeae
Dickeya zeae DZ2Q WP 029456364 1 Dzeae
Dickeya zeae EC1 WP 029456364 1 Dzeae
Dickeya zeae ZJU1202 WP 029456364 1 Dzeae
Dickeya zeae NCPPB 3531 WP 038902924 1 Dzeae
Dickeya zeae CSL RW192 WP 038902924 1 Dzeae
Dickeya zeae NCPPB 2538 WP 038915309 1 Dzeae
Dickeya zeae MS1 WP 019844172 1 Dzeae
Dickeya zeae MK19 WP 038912804 1 Dzeae
Dickeya zeae Ech586 WP 012882824 1 Dzeae
Dickeya dianthicola IPO 980 WP 029729819 1 Ddianthicola
Dickeya dianthicola NCPPB 3534 WP 024107815 1 Ddianthicola
Dickeya dianthicola RNS04 9 WP 024107815 1 Ddianthicola
Dickeya dianthicola GBBC 2039 WP 024107815 1 Ddianthicola
Dickeya dianthicola NCPPB 453 WP 024107815 1 Ddianthicola
Dickeya gs fangzhongdai NCPPB 3274 WP 042861610 1 Dfangzhongdai
Dickeya gs fangzhongdai ND14b WP 038663714 1 Dfangzhongdai
Dickeya gs fangzhongdai M074 WP 038663714 1 Dfangzhongdai
Dickeya gs fangzhongdai M005 WP 038663714 1 Dfangzhongdai
Dickeya gs undicola 2B12 WP 033568104 1 Dundicola
Dickeya gs fangzhongdai S1 WP 049855285 1 Dfangzhongdai
Dickeya gs fangzhongdai B16 WP 038920628 1 Dfangzhongdai
Dickeya gs fangzhongdai MK7 WP 038920628 1 Dfangzhongdai
Dickeya dadantii subsp dieffenbachiae NCPPB 2976 WP 038924308 1 Ddadantii
Dickeya dadantii NCPPB 898 WP 038912276 1 Ddadantii
Dickeya dadantii NCPPB 3537 WP 038902391 1 Ddadantii
175 xylG xylose degradation pathway Seed
Dickeya dadantii 3937 WP 013320022 1 Ddadantii
Dickeya solani RNS 05 1 2A WP 057083608 1 Dsolani
Dickeya solani PPO 9134 WP 022631610 1 Dsolani
Dickeya solani RNS 07 7 3B WP 022631610 1 Dsolani
Dickeya solani PPO 9019 WP 022631610 1 Dsolani
Dickeya solani D s0432 1 WP 022631610 1 Dsolani
Dickeya solani IFB 0099 WP 022631610 1 Dsolani
Dickeya solani GBBC 2040 WP 022631610 1 Dsolani
Dickeya solani RNS 08 23 3 1 A WP 022631610 1 Dsolani
Dickeya solani MK16 WP 022631610 1 Dsolani
Dickeya solani MK10 WP 022631610 1 Dsolani
Dickeya solani IPO 2222 WP 022631610 1 Dsolani

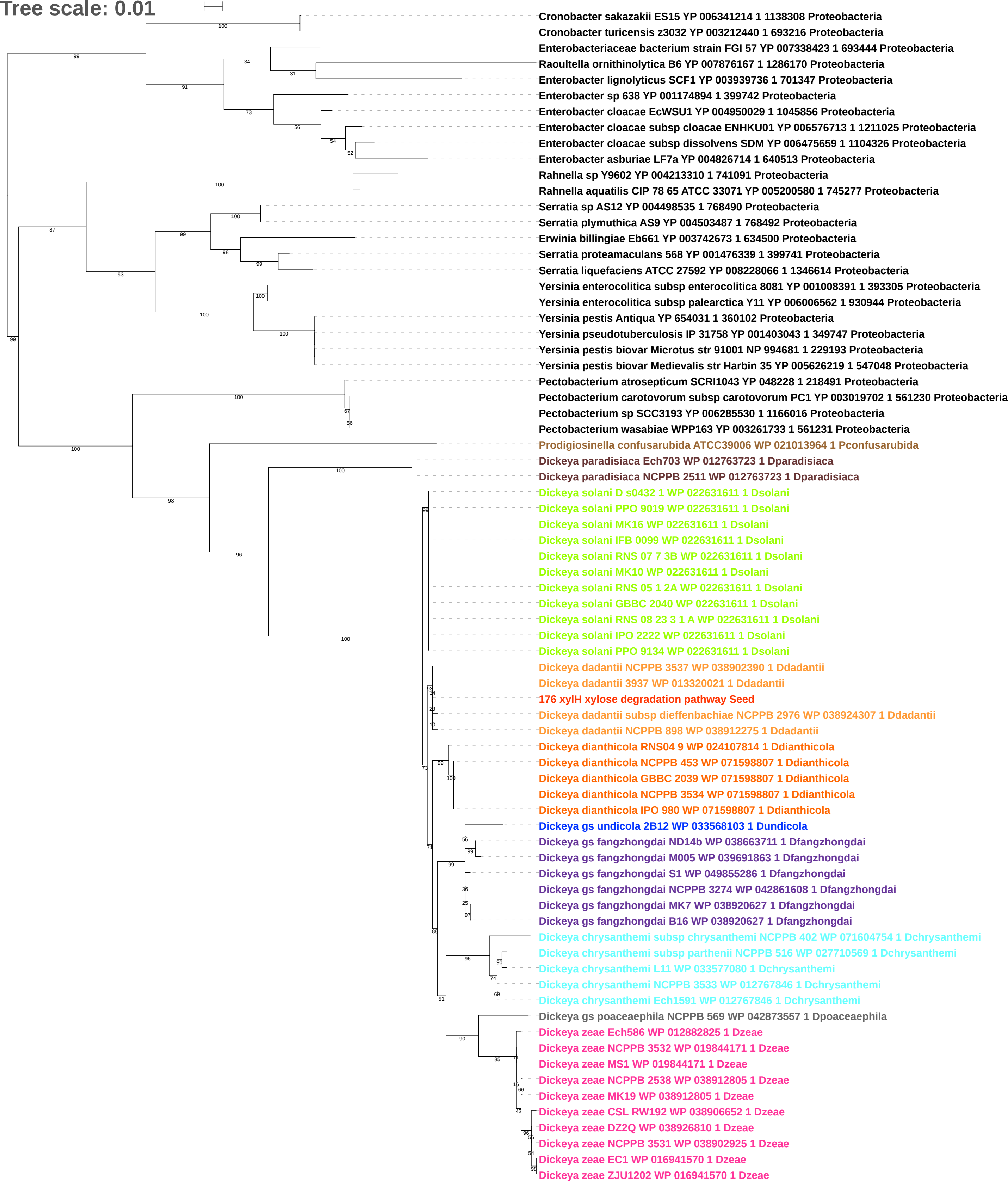
Tree scale: 0.01
Cronobacter sakazakii ES15 YP 006341214 1 1138308 Proteobacteria
Cronobacter turicensis z3032 YP 003212440 1 693216 Proteobacteria
Enterobacteriaceae bacterium strain FGI 57 YP 007338423 1 693444 Proteobacteria
Raoultella ornithinolytica B6 YP 007876167 1 1286170 Proteobacteria
Enterobacter lignolyticus SCF1 YP 003939736 1 701347 Proteobacteria
Enterobacter sp 638 YP 001174894 1 399742 Proteobacteria
Enterobacter cloacae EcWSU1 YP 004950029 1 1045856 Proteobacteria
Enterobacter cloacae subsp cloacae ENHKU01 YP 006576713 1 1211025 Proteobacteria
Enterobacter cloacae subsp dissolvens SDM YP 006475659 1 1104326 Proteobacteria
Enterobacter asburiae LF7a YP 004826714 1 640513 Proteobacteria
Rahnella sp Y9602 YP 004213310 1 741091 Proteobacteria
Rahnella aquatilis CIP 78 65 ATCC 33071 YP 005200580 1 745277 Proteobacteria
Serratia sp AS12 YP 004498535 1 768490 Proteobacteria
Serratia plymuthica AS9 YP 004503487 1 768492 Proteobacteria
Erwinia billingiae Eb661 YP 003742673 1 634500 Proteobacteria
Serratia proteamaculans 568 YP 001476339 1 399741 Proteobacteria
Serratia liquefaciens ATCC 27592 YP 008228066 1 1346614 Proteobacteria
Yersinia enterocolitica subsp enterocolitica 8081 YP 001008391 1 393305 Proteobacteria
Yersinia enterocolitica subsp palearctica Y11 YP 006006562 1 930944 Proteobacteria
Yersinia pestis Antiqua YP 654031 1 360102 Proteobacteria
Yersinia pseudotuberculosis IP 31758 YP 001403043 1 349747 Proteobacteria
Yersinia pestis biovar Microtus str 91001 NP 994681 1 229193 Proteobacteria
Yersinia pestis biovar Medievalis str Harbin 35 YP 005626219 1 547048 Proteobacteria
Pectobacterium atrosepticum SCRI1043 YP 048228 1 218491 Proteobacteria
Pectobacterium carotovorum subsp carotovorum PC1 YP 003019702 1 561230 Proteobacteria
Pectobacterium sp SCC3193 YP 006285530 1 1166016 Proteobacteria
Pectobacterium wasabiae WPP163 YP 003261733 1 561231 Proteobacteria
Prodigiosinella confusarubida ATCC39006 WP 021013964 1 Pconfusarubida
Dickeya paradisiaca Ech703 WP 012763723 1 Dparadisiaca
Dickeya paradisiaca NCPPB 2511 WP 012763723 1 Dparadisiaca
Dickeya solani D s0432 1 WP 022631611 1 Dsolani
Dickeya solani PPO 9019 WP 022631611 1 Dsolani
Dickeya solani MK16 WP 022631611 1 Dsolani
Dickeya solani IFB 0099 WP 022631611 1 Dsolani
Dickeya solani RNS 07 7 3B WP 022631611 1 Dsolani
Dickeya solani MK10 WP 022631611 1 Dsolani
Dickeya solani RNS 05 1 2A WP 022631611 1 Dsolani
Dickeya solani GBBC 2040 WP 022631611 1 Dsolani
Dickeya solani RNS 08 23 3 1 A WP 022631611 1 Dsolani
Dickeya solani IPO 2222 WP 022631611 1 Dsolani
Dickeya solani PPO 9134 WP 022631611 1 Dsolani
Dickeya dadantii NCPPB 3537 WP 038902390 1 Ddadantii
Dickeya dadantii 3937 WP 013320021 1 Ddadantii
176 xylH xylose degradation pathway Seed
Dickeya dadantii subsp dieffenbachiae NCPPB 2976 WP 038924307 1 Ddadantii
Dickeya dadantii NCPPB 898 WP 038912275 1 Ddadantii
Dickeya dianthicola RNS04 9 WP 024107814 1 Ddianthicola
Dickeya dianthicola NCPPB 453 WP 071598807 1 Ddianthicola
Dickeya dianthicola GBBC 2039 WP 071598807 1 Ddianthicola
Dickeya dianthicola NCPPB 3534 WP 071598807 1 Ddianthicola
Dickeya dianthicola IPO 980 WP 071598807 1 Ddianthicola
Dickeya gs undicola 2B12 WP 033568103 1 Dundicola
Dickeya gs fangzhongdai ND14b WP 038663711 1 Dfangzhongdai
Dickeya gs fangzhongdai M005 WP 039691863 1 Dfangzhongdai
Dickeya gs fangzhongdai S1 WP 049855286 1 Dfangzhongdai
Dickeya gs fangzhongdai NCPPB 3274 WP 042861608 1 Dfangzhongdai
Dickeya gs fangzhongdai MK7 WP 038920627 1 Dfangzhongdai
Dickeya gs fangzhongdai B16 WP 038920627 1 Dfangzhongdai
Dickeya chrysanthemi subsp chrysanthemi NCPPB 402 WP 071604754 1 Dchrysanthemi
Dickeya chrysanthemi subsp parthenii NCPPB 516 WP 027710569 1 Dchrysanthemi
Dickeya chrysanthemi L11 WP 033577080 1 Dchrysanthemi
Dickeya chrysanthemi NCPPB 3533 WP 012767846 1 Dchrysanthemi
Dickeya chrysanthemi Ech1591 WP 012767846 1 Dchrysanthemi
Dickeya gs poaceaephila NCPPB 569 WP 042873557 1 Dpoaceaephila
Dickeya zeae Ech586 WP 012882825 1 Dzeae
Dickeya zeae NCPPB 3532 WP 019844171 1 Dzeae
Dickeya zeae MS1 WP 019844171 1 Dzeae
Dickeya zeae NCPPB 2538 WP 038912805 1 Dzeae
Dickeya zeae MK19 WP 038912805 1 Dzeae
Dickeya zeae CSL RW192 WP 038906652 1 Dzeae
Dickeya zeae DZ2Q WP 038926810 1 Dzeae
Dickeya zeae NCPPB 3531 WP 038902925 1 Dzeae
Dickeya zeae EC1 WP 016941570 1 Dzeae
Dickeya zeae ZJU1202 WP 016941570 1 Dzeae

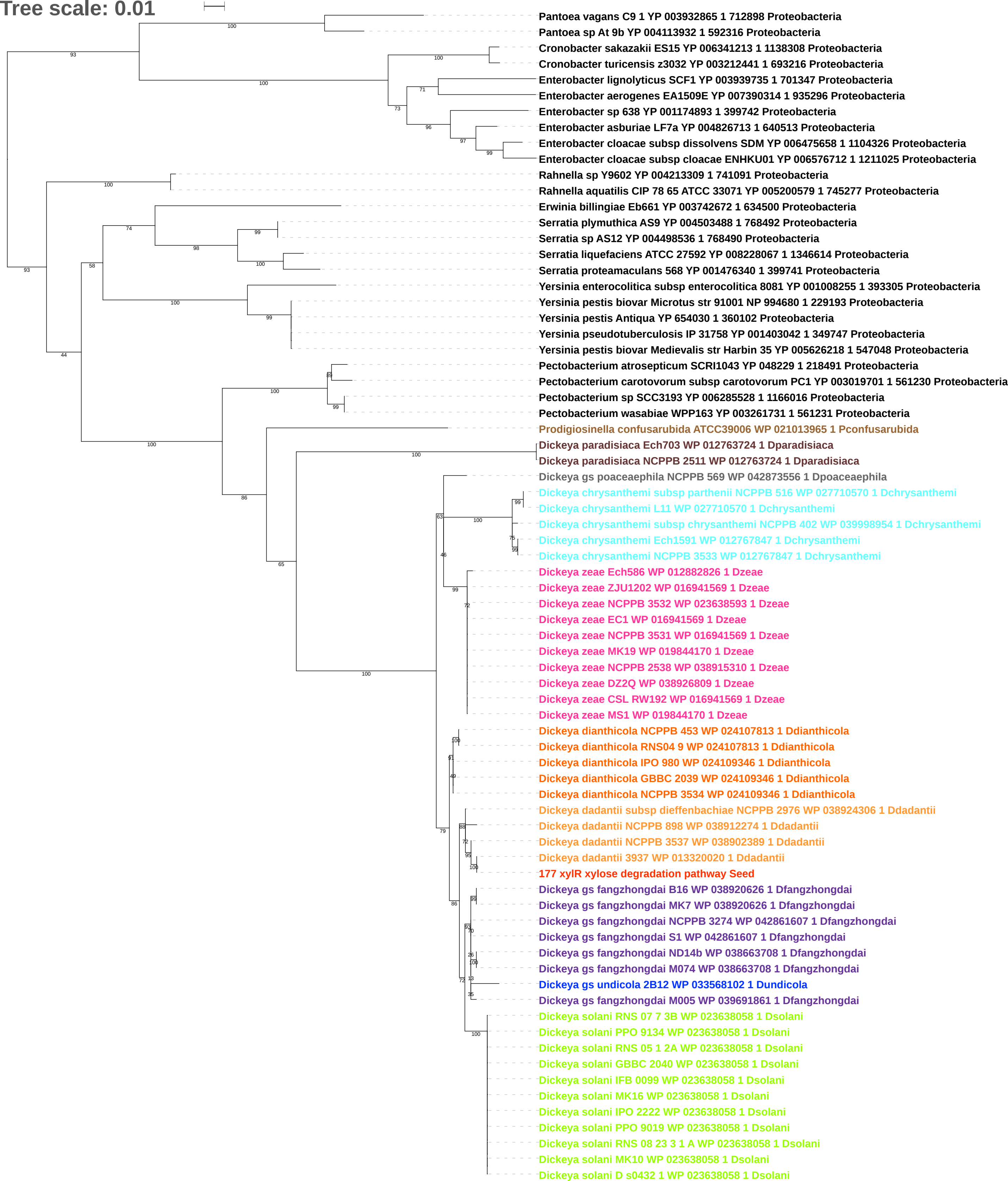
Tree scale: 0.01
Pantoea vagans C9 1 YP 003932865 1 712898 Proteobacteria
Pantoea sp At 9b YP 004113932 1 592316 Proteobacteria
Cronobacter sakazakii ES15 YP 006341213 1 1138308 Proteobacteria
Cronobacter turicensis z3032 YP 003212441 1 693216 Proteobacteria
Enterobacter lignolyticus SCF1 YP 003939735 1 701347 Proteobacteria
Enterobacter aerogenes EA1509E YP 007390314 1 935296 Proteobacteria
Enterobacter sp 638 YP 001174893 1 399742 Proteobacteria
Enterobacter asburiae LF7a YP 004826713 1 640513 Proteobacteria
Enterobacter cloacae subsp dissolvens SDM YP 006475658 1 1104326 Proteobacteria
Enterobacter cloacae subsp cloacae ENHKU01 YP 006576712 1 1211025 Proteobacteria
Rahnella sp Y9602 YP 004213309 1 741091 Proteobacteria
Rahnella aquatilis CIP 78 65 ATCC 33071 YP 005200579 1 745277 Proteobacteria
Erwinia billingiae Eb661 YP 003742672 1 634500 Proteobacteria
Serratia plymuthica AS9 YP 004503488 1 768492 Proteobacteria
Serratia sp AS12 YP 004498536 1 768490 Proteobacteria
Serratia liquefaciens ATCC 27592 YP 008228067 1 1346614 Proteobacteria
Serratia proteamaculans 568 YP 001476340 1 399741 Proteobacteria
Yersinia enterocolitica subsp enterocolitica 8081 YP 001008255 1 393305 Proteobacteria
Yersinia pestis biovar Microtus str 91001 NP 994680 1 229193 Proteobacteria
Yersinia pestis Antiqua YP 654030 1 360102 Proteobacteria
Yersinia pseudotuberculosis IP 31758 YP 001403042 1 349747 Proteobacteria
Yersinia pestis biovar Medievalis str Harbin 35 YP 005626218 1 547048 Proteobacteria
Pectobacterium atrosepticum SCRI1043 YP 048229 1 218491 Proteobacteria
Pectobacterium carotovorum subsp carotovorum PC1 YP 003019701 1 561230 Proteobacteria
Pectobacterium sp SCC3193 YP 006285528 1 1166016 Proteobacteria
Pectobacterium wasabiae WPP163 YP 003261731 1 561231 Proteobacteria
Prodigiosinella confusarubida ATCC39006 WP 021013965 1 Pconfusarubida
Dickeya paradisiaca Ech703 WP 012763724 1 Dparadisiaca
Dickeya paradisiaca NCPPB 2511 WP 012763724 1 Dparadisiaca
Dickeya gs poaceaephila NCPPB 569 WP 042873556 1 Dpoaceaephila
Dickeya chrysanthemi subsp parthenii NCPPB 516 WP 027710570 1 Dchrysanthemi
Dickeya chrysanthemi L11 WP 027710570 1 Dchrysanthemi
Dickeya chrysanthemi subsp chrysanthemi NCPPB 402 WP 039998954 1 Dchrysanthemi
Dickeya chrysanthemi Ech1591 WP 012767847 1 Dchrysanthemi
Dickeya chrysanthemi NCPPB 3533 WP 012767847 1 Dchrysanthemi
Dickeya zeae Ech586 WP 012882826 1 Dzeae
Dickeya zeae ZJU1202 WP 016941569 1 Dzeae
Dickeya zeae NCPPB 3532 WP 023638593 1 Dzeae
Dickeya zeae EC1 WP 016941569 1 Dzeae
Dickeya zeae NCPPB 3531 WP 016941569 1 Dzeae
Dickeya zeae MK19 WP 019844170 1 Dzeae
Dickeya zeae NCPPB 2538 WP 038915310 1 Dzeae
Dickeya zeae DZ2Q WP 038926809 1 Dzeae
Dickeya zeae CSL RW192 WP 016941569 1 Dzeae
Dickeya zeae MS1 WP 019844170 1 Dzeae
Dickeya dianthicola NCPPB 453 WP 024107813 1 Ddianthicola
Dickeya dianthicola RNS04 9 WP 024107813 1 Ddianthicola
Dickeya dianthicola IPO 980 WP 024109346 1 Ddianthicola
Dickeya dianthicola GBBC 2039 WP 024109346 1 Ddianthicola
Dickeya dianthicola NCPPB 3534 WP 024109346 1 Ddianthicola
Dickeya dadantii subsp dieffenbachiae NCPPB 2976 WP 038924306 1 Ddadantii
Dickeya dadantii NCPPB 898 WP 038912274 1 Ddadantii
Dickeya dadantii NCPPB 3537 WP 038902389 1 Ddadantii
Dickeya dadantii 3937 WP 013320020 1 Ddadantii
177 xylR xylose degradation pathway Seed
Dickeya gs fangzhongdai B16 WP 038920626 1 Dfangzhongdai
Dickeya gs fangzhongdai MK7 WP 038920626 1 Dfangzhongdai
Dickeya gs fangzhongdai NCPPB 3274 WP 042861607 1 Dfangzhongdai
Dickeya gs fangzhongdai S1 WP 042861607 1 Dfangzhongdai
Dickeya gs fangzhongdai ND14b WP 038663708 1 Dfangzhongdai
Dickeya gs fangzhongdai M074 WP 038663708 1 Dfangzhongdai
Dickeya gs undicola 2B12 WP 033568102 1 Dundicola
Dickeya gs fangzhongdai M005 WP 039691861 1 Dfangzhongdai
Dickeya solani RNS 07 7 3B WP 023638058 1 Dsolani
Dickeya solani PPO 9134 WP 023638058 1 Dsolani
Dickeya solani RNS 05 1 2A WP 023638058 1 Dsolani
Dickeya solani GBBC 2040 WP 023638058 1 Dsolani
Dickeya solani IFB 0099 WP 023638058 1 Dsolani
Dickeya solani MK16 WP 023638058 1 Dsolani
Dickeya solani IPO 2222 WP 023638058 1 Dsolani
Dickeya solani PPO 9019 WP 023638058 1 Dsolani
Dickeya solani RNS 08 23 3 1 A WP 023638058 1 Dsolani
Dickeya solani MK10 WP 023638058 1 Dsolani
Dickeya solani D s0432 1 WP 023638058 1 Dsolani

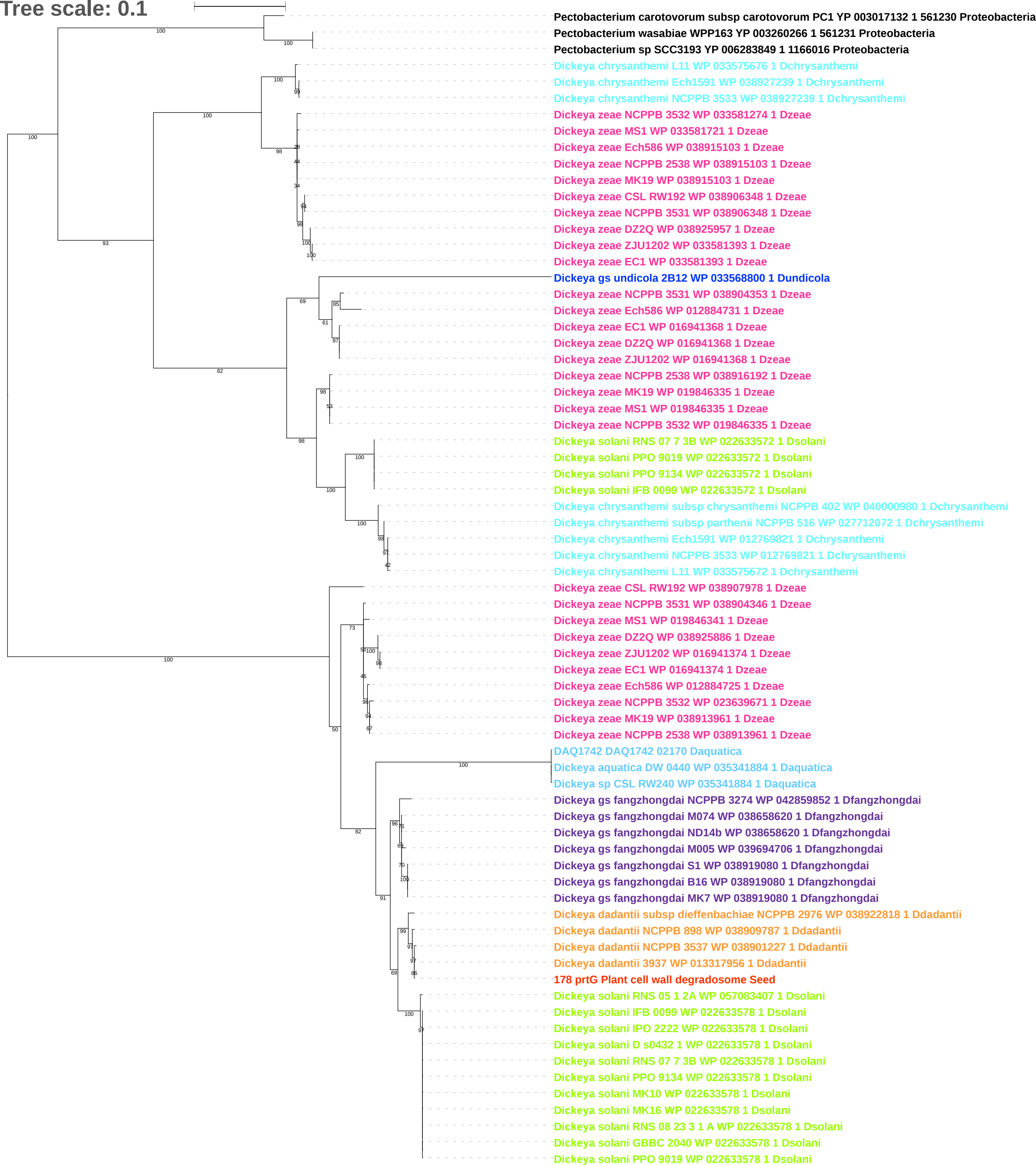
Tree scale: 0.1
Pectobacterium carotovorum subsp carotovorum PC1 YP 003017132 1 561230 Proteobacteria
Pectobacterium wasabiae WPP163 YP 003260266 1 561231 Proteobacteria
Pectobacterium sp SCC3193 YP 006283849 1 1166016 Proteobacteria
Dickeya chrysanthemi L11 WP 033575676 1 Dchrysanthemi
Dickeya chrysanthemi Ech1591 WP 038927239 1 Dchrysanthemi
Dickeya chrysanthemi NCPPB 3533 WP 038927239 1 Dchrysanthemi
Dickeya zeae NCPPB 3532 WP 033581274 1 Dzeae
Dickeya zeae MS1 WP 033581721 1 Dzeae
Dickeya zeae Ech586 WP 038915103 1 Dzeae
Dickeya zeae NCPPB 2538 WP 038915103 1 Dzeae
Dickeya zeae MK19 WP 038915103 1 Dzeae
Dickeya zeae CSL RW192 WP 038906348 1 Dzeae
Dickeya zeae NCPPB 3531 WP 038906348 1 Dzeae
Dickeya zeae DZ2Q WP 038925957 1 Dzeae
Dickeya zeae ZJU1202 WP 033581393 1 Dzeae
Dickeya zeae EC1 WP 033581393 1 Dzeae
Dickeya gs undicola 2B12 WP 033568800 1 Dundicola
Dickeya zeae NCPPB 3531 WP 038904353 1 Dzeae
Dickeya zeae Ech586 WP 012884731 1 Dzeae
Dickeya zeae EC1 WP 016941368 1 Dzeae
Dickeya zeae DZ2Q WP 016941368 1 Dzeae
Dickeya zeae ZJU1202 WP 016941368 1 Dzeae
Dickeya zeae NCPPB 2538 WP 038916192 1 Dzeae
Dickeya zeae MK19 WP 019846335 1 Dzeae
Dickeya zeae MS1 WP 019846335 1 Dzeae
Dickeya zeae NCPPB 3532 WP 019846335 1 Dzeae
Dickeya solani RNS 07 7 3B WP 022633572 1 Dsolani
Dickeya solani PPO 9019 WP 022633572 1 Dsolani
Dickeya solani PPO 9134 WP 022633572 1 Dsolani
Dickeya solani IFB 0099 WP 022633572 1 Dsolani
Dickeya chrysanthemi subsp chrysanthemi NCPPB 402 WP 040000980 1 Dchrysanthemi
Dickeya chrysanthemi subsp parthenii NCPPB 516 WP 027712072 1 Dchrysanthemi
Dickeya chrysanthemi Ech1591 WP 012769821 1 Dchrysanthemi
Dickeya chrysanthemi NCPPB 3533 WP 012769821 1 Dchrysanthemi
Dickeya chrysanthemi L11 WP 033575672 1 Dchrysanthemi
Dickeya zeae CSL RW192 WP 038907978 1 Dzeae
Dickeya zeae NCPPB 3531 WP 038904346 1 Dzeae
Dickeya zeae MS1 WP 019846341 1 Dzeae
Dickeya zeae DZ2Q WP 038925886 1 Dzeae
Dickeya zeae ZJU1202 WP 016941374 1 Dzeae
Dickeya zeae EC1 WP 016941374 1 Dzeae
Dickeya zeae Ech586 WP 012884725 1 Dzeae
Dickeya zeae NCPPB 3532 WP 023639671 1 Dzeae
Dickeya zeae MK19 WP 038913961 1 Dzeae
Dickeya zeae NCPPB 2538 WP 038913961 1 Dzeae
DAQ1742 DAQ1742 02170 Daquatica
Dickeya aquatica DW 0440 WP 035341884 1 Daquatica
Dickeya sp CSL RW240 WP 035341884 1 Daquatica
Dickeya gs fangzhongdai NCPPB 3274 WP 042859852 1 Dfangzhongdai
Dickeya gs fangzhongdai M074 WP 038658620 1 Dfangzhongdai
Dickeya gs fangzhongdai ND14b WP 038658620 1 Dfangzhongdai
Dickeya gs fangzhongdai M005 WP 039694706 1 Dfangzhongdai
Dickeya gs fangzhongdai S1 WP 038919080 1 Dfangzhongdai
Dickeya gs fangzhongdai B16 WP 038919080 1 Dfangzhongdai
Dickeya gs fangzhongdai MK7 WP 038919080 1 Dfangzhongdai
Dickeya dadantii subsp dieffenbachiae NCPPB 2976 WP 038922818 1 Ddadantii
Dickeya dadantii NCPPB 898 WP 038909787 1 Ddadantii
Dickeya dadantii NCPPB 3537 WP 038901227 1 Ddadantii
Dickeya dadantii 3937 WP 013317956 1 Ddadantii
178 prtG Plant cell wall degradosome Seed
Dickeya solani RNS 05 1 2A WP 057083407 1 Dsolani
Dickeya solani IFB 0099 WP 022633578 1 Dsolani
Dickeya solani IPO 2222 WP 022633578 1 Dsolani
Dickeya solani D s0432 1 WP 022633578 1 Dsolani
Dickeya solani RNS 07 7 3B WP 022633578 1 Dsolani
Dickeya solani PPO 9134 WP 022633578 1 Dsolani
Dickeya solani MK10 WP 022633578 1 Dsolani
Dickeya solani MK16 WP 022633578 1 Dsolani
Dickeya solani RNS 08 23 3 1 A WP 022633578 1 Dsolani
Dickeya solani GBBC 2040 WP 022633578 1 Dsolani
Dickeya solani PPO 9019 WP 022633578 1 Dsolani

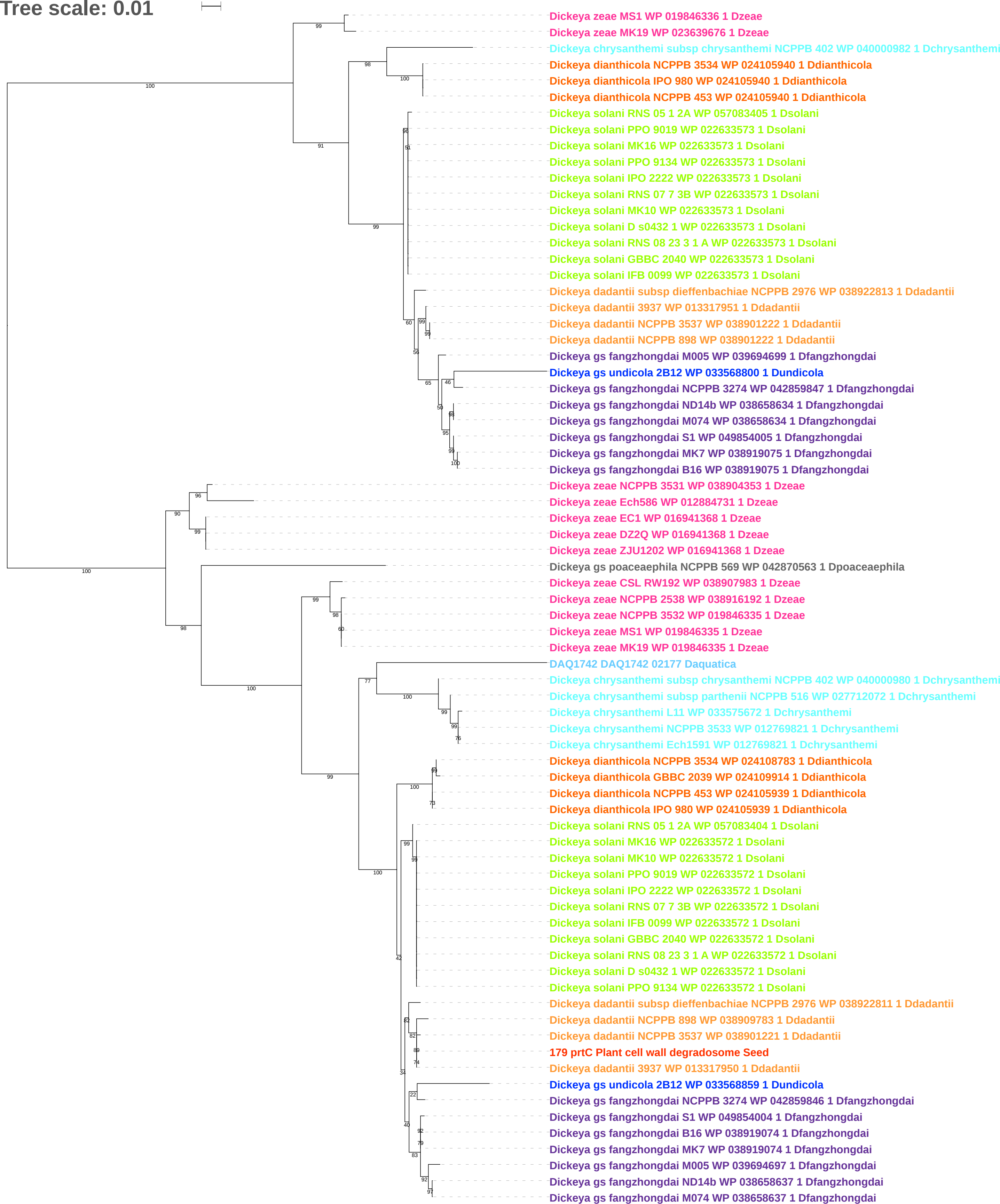
Tree scale: 0.01
Dickeya zeae MS1 WP 019846336 1 Dzeae
Dickeya zeae MK19 WP 023639676 1 Dzeae
Dickeya chrysanthemi subsp chrysanthemi NCPPB 402 WP 040000982 1 Dchrysanthemi
Dickeya dianthicola NCPPB 3534 WP 024105940 1 Ddianthicola
Dickeya dianthicola IPO 980 WP 024105940 1 Ddianthicola
Dickeya dianthicola NCPPB 453 WP 024105940 1 Ddianthicola
Dickeya solani RNS 05 1 2A WP 057083405 1 Dsolani
Dickeya solani PPO 9019 WP 022633573 1 Dsolani
Dickeya solani MK16 WP 022633573 1 Dsolani
Dickeya solani PPO 9134 WP 022633573 1 Dsolani
Dickeya solani IPO 2222 WP 022633573 1 Dsolani
Dickeya solani RNS 07 7 3B WP 022633573 1 Dsolani
Dickeya solani MK10 WP 022633573 1 Dsolani
Dickeya solani D s0432 1 WP 022633573 1 Dsolani
Dickeya solani RNS 08 23 3 1 A WP 022633573 1 Dsolani
Dickeya solani GBBC 2040 WP 022633573 1 Dsolani
Dickeya solani IFB 0099 WP 022633573 1 Dsolani
Dickeya dadantii subsp dieffenbachiae NCPPB 2976 WP 038922813 1 Ddadantii
Dickeya dadantii 3937 WP 013317951 1 Ddadantii
Dickeya dadantii NCPPB 3537 WP 038901222 1 Ddadantii
Dickeya dadantii NCPPB 898 WP 038901222 1 Ddadantii
Dickeya gs fangzhongdai M005 WP 039694699 1 Dfangzhongdai
Dickeya gs undicola 2B12 WP 033568800 1 Dundicola
Dickeya gs fangzhongdai NCPPB 3274 WP 042859847 1 Dfangzhongdai
Dickeya gs fangzhongdai ND14b WP 038658634 1 Dfangzhongdai
Dickeya gs fangzhongdai M074 WP 038658634 1 Dfangzhongdai
Dickeya gs fangzhongdai S1 WP 049854005 1 Dfangzhongdai
Dickeya gs fangzhongdai MK7 WP 038919075 1 Dfangzhongdai
Dickeya gs fangzhongdai B16 WP 038919075 1 Dfangzhongdai
Dickeya zeae NCPPB 3531 WP 038904353 1 Dzeae
Dickeya zeae Ech586 WP 012884731 1 Dzeae
Dickeya zeae EC1 WP 016941368 1 Dzeae
Dickeya zeae DZ2Q WP 016941368 1 Dzeae
Dickeya zeae ZJU1202 WP 016941368 1 Dzeae
Dickeya gs poaceaephila NCPPB 569 WP 042870563 1 Dpoaceaephila
Dickeya zeae CSL RW192 WP 038907983 1 Dzeae
Dickeya zeae NCPPB 2538 WP 038916192 1 Dzeae
Dickeya zeae NCPPB 3532 WP 019846335 1 Dzeae
Dickeya zeae MS1 WP 019846335 1 Dzeae
Dickeya zeae MK19 WP 019846335 1 Dzeae
DAQ1742 DAQ1742 02177 Daquatica
Dickeya chrysanthemi subsp chrysanthemi NCPPB 402 WP 040000980 1 Dchrysanthemi
Dickeya chrysanthemi subsp parthenii NCPPB 516 WP 027712072 1 Dchrysanthemi
Dickeya chrysanthemi L11 WP 033575672 1 Dchrysanthemi
Dickeya chrysanthemi NCPPB 3533 WP 012769821 1 Dchrysanthemi
Dickeya chrysanthemi Ech1591 WP 012769821 1 Dchrysanthemi
Dickeya dianthicola NCPPB 3534 WP 024108783 1 Ddianthicola
Dickeya dianthicola GBBC 2039 WP 024109914 1 Ddianthicola
Dickeya dianthicola NCPPB 453 WP 024105939 1 Ddianthicola
Dickeya dianthicola IPO 980 WP 024105939 1 Ddianthicola
Dickeya solani RNS 05 1 2A WP 057083404 1 Dsolani
Dickeya solani MK16 WP 022633572 1 Dsolani
Dickeya solani MK10 WP 022633572 1 Dsolani
Dickeya solani PPO 9019 WP 022633572 1 Dsolani
Dickeya solani IPO 2222 WP 022633572 1 Dsolani
Dickeya solani RNS 07 7 3B WP 022633572 1 Dsolani
Dickeya solani IFB 0099 WP 022633572 1 Dsolani
Dickeya solani GBBC 2040 WP 022633572 1 Dsolani
Dickeya solani RNS 08 23 3 1 A WP 022633572 1 Dsolani
Dickeya solani D s0432 1 WP 022633572 1 Dsolani
Dickeya solani PPO 9134 WP 022633572 1 Dsolani
Dickeya dadantii subsp dieffenbachiae NCPPB 2976 WP 038922811 1 Ddadantii
Dickeya dadantii NCPPB 898 WP 038909783 1 Ddadantii
Dickeya dadantii NCPPB 3537 WP 038901221 1 Ddadantii
179 prtC Plant cell wall degradosome Seed
Dickeya dadantii 3937 WP 013317950 1 Ddadantii
Dickeya gs undicola 2B12 WP 033568859 1 Dundicola
Dickeya gs fangzhongdai NCPPB 3274 WP 042859846 1 Dfangzhongdai
Dickeya gs fangzhongdai S1 WP 049854004 1 Dfangzhongdai
Dickeya gs fangzhongdai B16 WP 038919074 1 Dfangzhongdai
Dickeya gs fangzhongdai MK7 WP 038919074 1 Dfangzhongdai
Dickeya gs fangzhongdai M005 WP 039694697 1 Dfangzhongdai
Dickeya gs fangzhongdai ND14b WP 038658637 1 Dfangzhongdai
Dickeya gs fangzhongdai M074 WP 038658637 1 Dfangzhongdai

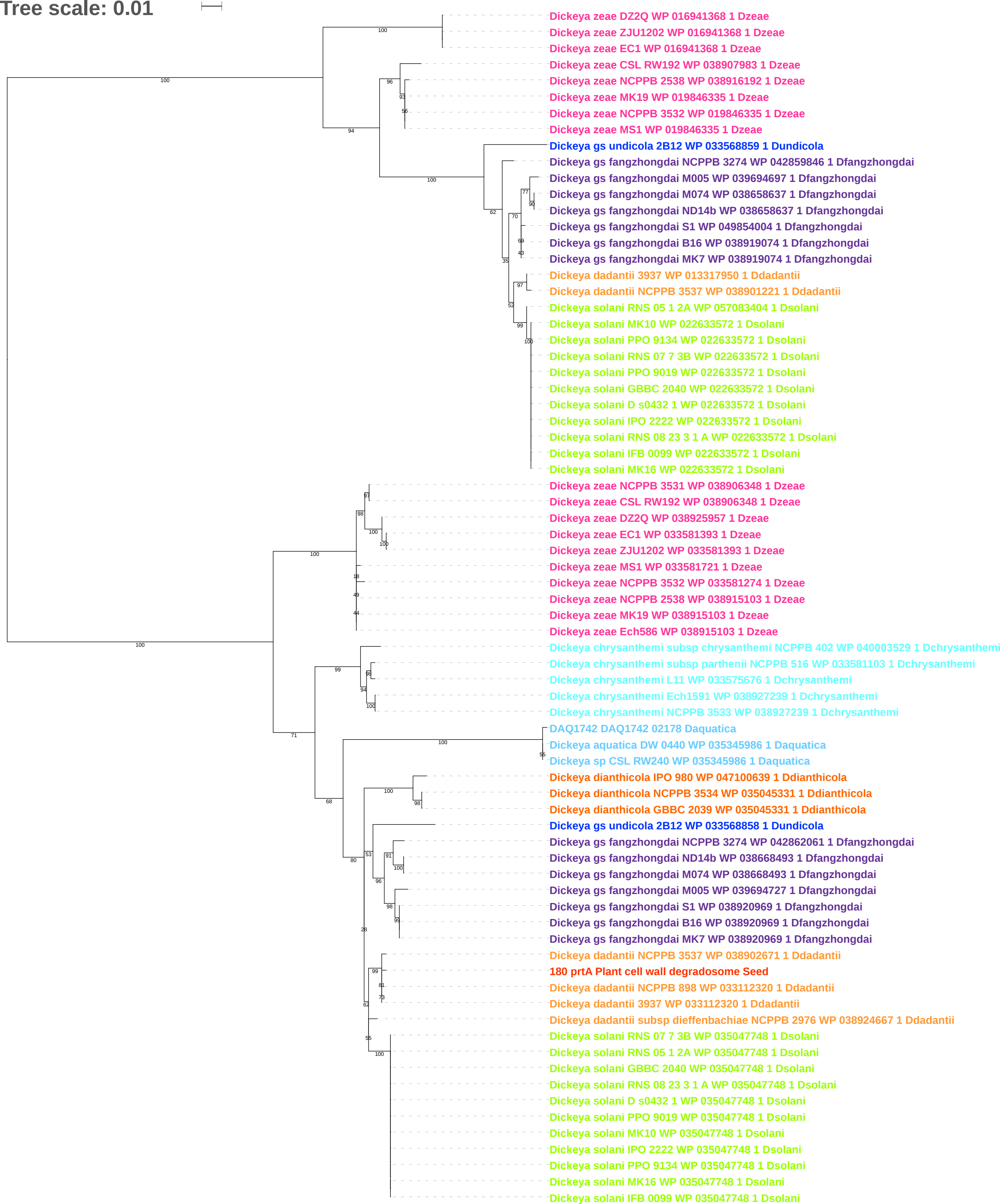
Tree scale: 0.01
Dickeya zeae DZ2Q WP 016941368 1 Dzeae
Dickeya zeae ZJU1202 WP 016941368 1 Dzeae
Dickeya zeae EC1 WP 016941368 1 Dzeae
Dickeya zeae CSL RW192 WP 038907983 1 Dzeae
Dickeya zeae NCPPB 2538 WP 038916192 1 Dzeae
Dickeya zeae MK19 WP 019846335 1 Dzeae
Dickeya zeae NCPPB 3532 WP 019846335 1 Dzeae
Dickeya zeae MS1 WP 019846335 1 Dzeae
Dickeya gs undicola 2B12 WP 033568859 1 Dundicola
Dickeya gs fangzhongdai NCPPB 3274 WP 042859846 1 Dfangzhongdai
Dickeya gs fangzhongdai M005 WP 039694697 1 Dfangzhongdai
Dickeya gs fangzhongdai M074 WP 038658637 1 Dfangzhongdai
Dickeya gs fangzhongdai ND14b WP 038658637 1 Dfangzhongdai
Dickeya gs fangzhongdai S1 WP 049854004 1 Dfangzhongdai
Dickeya gs fangzhongdai B16 WP 038919074 1 Dfangzhongdai
Dickeya gs fangzhongdai MK7 WP 038919074 1 Dfangzhongdai
Dickeya dadantii 3937 WP 013317950 1 Ddadantii
Dickeya dadantii NCPPB 3537 WP 038901221 1 Ddadantii
Dickeya solani RNS 05 1 2A WP 057083404 1 Dsolani
Dickeya solani MK10 WP 022633572 1 Dsolani
Dickeya solani PPO 9134 WP 022633572 1 Dsolani
Dickeya solani RNS 07 7 3B WP 022633572 1 Dsolani
Dickeya solani PPO 9019 WP 022633572 1 Dsolani
Dickeya solani GBBC 2040 WP 022633572 1 Dsolani
Dickeya solani D s0432 1 WP 022633572 1 Dsolani
Dickeya solani IPO 2222 WP 022633572 1 Dsolani
Dickeya solani RNS 08 23 3 1 A WP 022633572 1 Dsolani
Dickeya solani IFB 0099 WP 022633572 1 Dsolani
Dickeya solani MK16 WP 022633572 1 Dsolani
Dickeya zeae NCPPB 3531 WP 038906348 1 Dzeae
Dickeya zeae CSL RW192 WP 038906348 1 Dzeae
Dickeya zeae DZ2Q WP 038925957 1 Dzeae
Dickeya zeae EC1 WP 033581393 1 Dzeae
Dickeya zeae ZJU1202 WP 033581393 1 Dzeae
Dickeya zeae MS1 WP 033581721 1 Dzeae
Dickeya zeae NCPPB 3532 WP 033581274 1 Dzeae
Dickeya zeae NCPPB 2538 WP 038915103 1 Dzeae
Dickeya zeae MK19 WP 038915103 1 Dzeae
Dickeya zeae Ech586 WP 038915103 1 Dzeae
Dickeya chrysanthemi subsp chrysanthemi NCPPB 402 WP 040003529 1 Dchrysanthemi
Dickeya chrysanthemi subsp parthenii NCPPB 516 WP 033581103 1 Dchrysanthemi
Dickeya chrysanthemi L11 WP 033575676 1 Dchrysanthemi
Dickeya chrysanthemi Ech1591 WP 038927239 1 Dchrysanthemi
Dickeya chrysanthemi NCPPB 3533 WP 038927239 1 Dchrysanthemi
DAQ1742 DAQ1742 02178 Daquatica
Dickeya aquatica DW 0440 WP 035345986 1 Daquatica
Dickeya sp CSL RW240 WP 035345986 1 Daquatica
Dickeya dianthicola IPO 980 WP 047100639 1 Ddianthicola
Dickeya dianthicola NCPPB 3534 WP 035045331 1 Ddianthicola
Dickeya dianthicola GBBC 2039 WP 035045331 1 Ddianthicola
Dickeya gs undicola 2B12 WP 033568858 1 Dundicola
Dickeya gs fangzhongdai NCPPB 3274 WP 042862061 1 Dfangzhongdai
Dickeya gs fangzhongdai ND14b WP 038668493 1 Dfangzhongdai
Dickeya gs fangzhongdai M074 WP 038668493 1 Dfangzhongdai
Dickeya gs fangzhongdai M005 WP 039694727 1 Dfangzhongdai
Dickeya gs fangzhongdai S1 WP 038920969 1 Dfangzhongdai
Dickeya gs fangzhongdai B16 WP 038920969 1 Dfangzhongdai
Dickeya gs fangzhongdai MK7 WP 038920969 1 Dfangzhongdai
Dickeya dadantii NCPPB 3537 WP 038902671 1 Ddadantii
180 prtA Plant cell wall degradosome Seed
Dickeya dadantii NCPPB 898 WP 033112320 1 Ddadantii
Dickeya dadantii 3937 WP 033112320 1 Ddadantii
Dickeya dadantii subsp dieffenbachiae NCPPB 2976 WP 038924667 1 Ddadantii
Dickeya solani RNS 07 7 3B WP 035047748 1 Dsolani
Dickeya solani RNS 05 1 2A WP 035047748 1 Dsolani
Dickeya solani GBBC 2040 WP 035047748 1 Dsolani
Dickeya solani RNS 08 23 3 1 A WP 035047748 1 Dsolani
Dickeya solani D s0432 1 WP 035047748 1 Dsolani
Dickeya solani PPO 9019 WP 035047748 1 Dsolani
Dickeya solani MK10 WP 035047748 1 Dsolani
Dickeya solani IPO 2222 WP 035047748 1 Dsolani
Dickeya solani PPO 9134 WP 035047748 1 Dsolani
Dickeya solani MK16 WP 035047748 1 Dsolani
Dickeya solani IFB 0099 WP 035047748 1 Dsolani

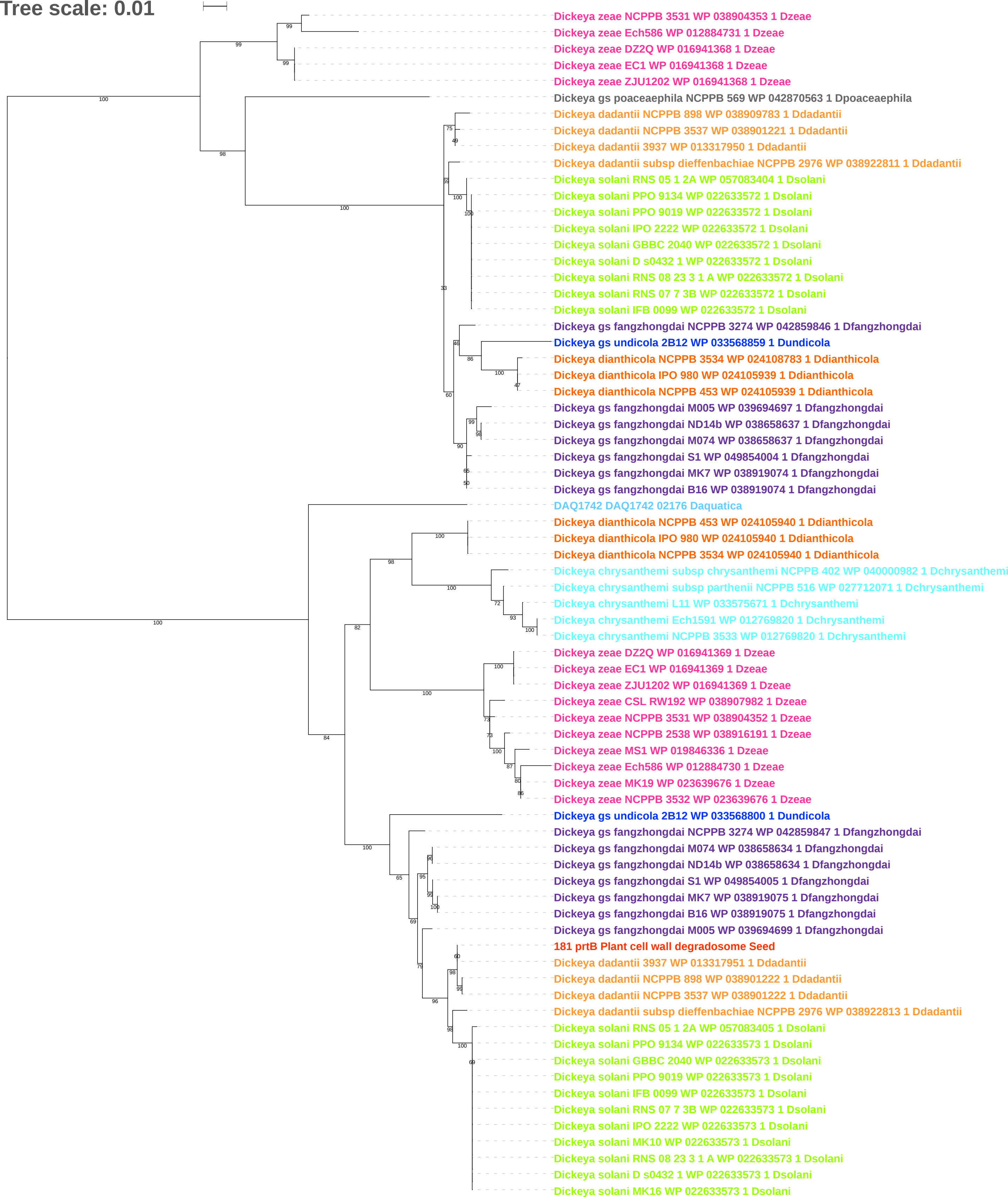
Tree scale: 0.01
Dickeya zeae NCPPB 3531 WP 038904353 1 Dzeae
Dickeya zeae Ech586 WP 012884731 1 Dzeae
Dickeya zeae DZ2Q WP 016941368 1 Dzeae
Dickeya zeae EC1 WP 016941368 1 Dzeae
Dickeya zeae ZJU1202 WP 016941368 1 Dzeae
Dickeya gs poaceaephila NCPPB 569 WP 042870563 1 Dpoaceaephila
Dickeya dadantii NCPPB 898 WP 038909783 1 Ddadantii
Dickeya dadantii NCPPB 3537 WP 038901221 1 Ddadantii
Dickeya dadantii 3937 WP 013317950 1 Ddadantii
Dickeya dadantii subsp dieffenbachiae NCPPB 2976 WP 038922811 1 Ddadantii
Dickeya solani RNS 05 1 2A WP 057083404 1 Dsolani
Dickeya solani PPO 9134 WP 022633572 1 Dsolani
Dickeya solani PPO 9019 WP 022633572 1 Dsolani
Dickeya solani IPO 2222 WP 022633572 1 Dsolani
Dickeya solani GBBC 2040 WP 022633572 1 Dsolani
Dickeya solani D s0432 1 WP 022633572 1 Dsolani
Dickeya solani RNS 08 23 3 1 A WP 022633572 1 Dsolani
Dickeya solani RNS 07 7 3B WP 022633572 1 Dsolani
Dickeya solani IFB 0099 WP 022633572 1 Dsolani
Dickeya gs fangzhongdai NCPPB 3274 WP 042859846 1 Dfangzhongdai
Dickeya gs undicola 2B12 WP 033568859 1 Dundicola
Dickeya dianthicola NCPPB 3534 WP 024108783 1 Ddianthicola
Dickeya dianthicola IPO 980 WP 024105939 1 Ddianthicola
Dickeya dianthicola NCPPB 453 WP 024105939 1 Ddianthicola
Dickeya gs fangzhongdai M005 WP 039694697 1 Dfangzhongdai
Dickeya gs fangzhongdai ND14b WP 038658637 1 Dfangzhongdai
Dickeya gs fangzhongdai M074 WP 038658637 1 Dfangzhongdai
Dickeya gs fangzhongdai S1 WP 049854004 1 Dfangzhongdai
Dickeya gs fangzhongdai MK7 WP 038919074 1 Dfangzhongdai
Dickeya gs fangzhongdai B16 WP 038919074 1 Dfangzhongdai
DAQ1742 DAQ1742 02176 Daquatica
Dickeya dianthicola NCPPB 453 WP 024105940 1 Ddianthicola
Dickeya dianthicola IPO 980 WP 024105940 1 Ddianthicola
Dickeya dianthicola NCPPB 3534 WP 024105940 1 Ddianthicola
Dickeya chrysanthemi subsp chrysanthemi NCPPB 402 WP 040000982 1 Dchrysanthemi
Dickeya chrysanthemi subsp parthenii NCPPB 516 WP 027712071 1 Dchrysanthemi
Dickeya chrysanthemi L11 WP 033575671 1 Dchrysanthemi
Dickeya chrysanthemi Ech1591 WP 012769820 1 Dchrysanthemi
Dickeya chrysanthemi NCPPB 3533 WP 012769820 1 Dchrysanthemi
Dickeya zeae DZ2Q WP 016941369 1 Dzeae
Dickeya zeae EC1 WP 016941369 1 Dzeae
Dickeya zeae ZJU1202 WP 016941369 1 Dzeae
Dickeya zeae CSL RW192 WP 038907982 1 Dzeae
Dickeya zeae NCPPB 3531 WP 038904352 1 Dzeae
Dickeya zeae NCPPB 2538 WP 038916191 1 Dzeae
Dickeya zeae MS1 WP 019846336 1 Dzeae
Dickeya zeae Ech586 WP 012884730 1 Dzeae
Dickeya zeae MK19 WP 023639676 1 Dzeae
Dickeya zeae NCPPB 3532 WP 023639676 1 Dzeae
Dickeya gs undicola 2B12 WP 033568800 1 Dundicola
Dickeya gs fangzhongdai NCPPB 3274 WP 042859847 1 Dfangzhongdai
Dickeya gs fangzhongdai M074 WP 038658634 1 Dfangzhongdai
Dickeya gs fangzhongdai ND14b WP 038658634 1 Dfangzhongdai
Dickeya gs fangzhongdai S1 WP 049854005 1 Dfangzhongdai
Dickeya gs fangzhongdai MK7 WP 038919075 1 Dfangzhongdai
Dickeya gs fangzhongdai B16 WP 038919075 1 Dfangzhongdai
Dickeya gs fangzhongdai M005 WP 039694699 1 Dfangzhongdai
181 prtB Plant cell wall degradosome Seed
Dickeya dadantii 3937 WP 013317951 1 Ddadantii
Dickeya dadantii NCPPB 898 WP 038901222 1 Ddadantii
Dickeya dadantii NCPPB 3537 WP 038901222 1 Ddadantii
Dickeya dadantii subsp dieffenbachiae NCPPB 2976 WP 038922813 1 Ddadantii
Dickeya solani RNS 05 1 2A WP 057083405 1 Dsolani
Dickeya solani PPO 9134 WP 022633573 1 Dsolani
Dickeya solani GBBC 2040 WP 022633573 1 Dsolani
Dickeya solani PPO 9019 WP 022633573 1 Dsolani
Dickeya solani IFB 0099 WP 022633573 1 Dsolani
Dickeya solani RNS 07 7 3B WP 022633573 1 Dsolani
Dickeya solani IPO 2222 WP 022633573 1 Dsolani
Dickeya solani MK10 WP 022633573 1 Dsolani
Dickeya solani RNS 08 23 3 1 A WP 022633573 1 Dsolani
Dickeya solani D s0432 1 WP 022633573 1 Dsolani
Dickeya solani MK16 WP 022633573 1 Dsolani

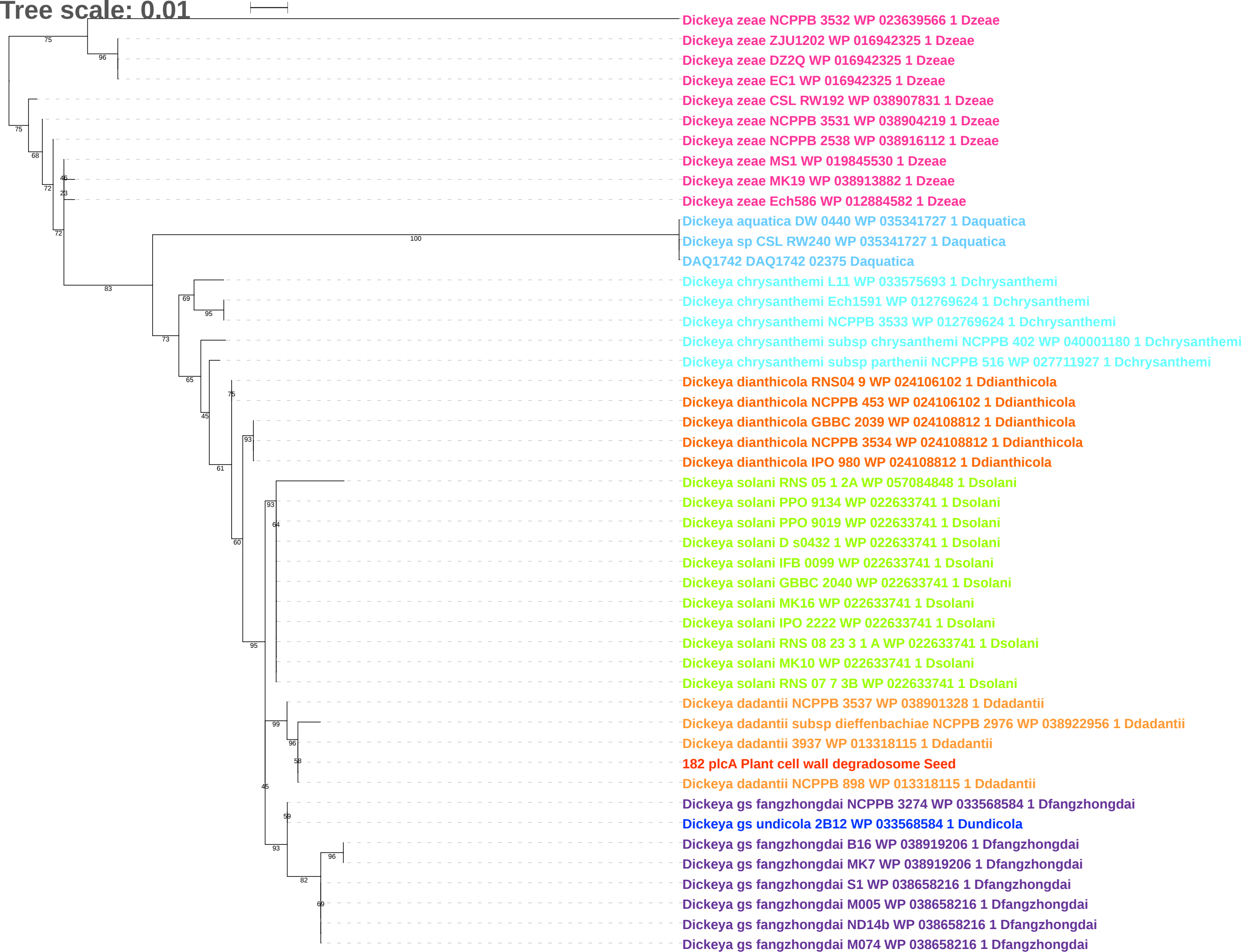
Tree scale: 0.01
Dickeya zeae NCPPB 3532 WP 023639566 1 Dzeae
Dickeya zeae ZJU1202 WP 016942325 1 Dzeae
Dickeya zeae DZ2Q WP 016942325 1 Dzeae
Dickeya zeae EC1 WP 016942325 1 Dzeae
Dickeya zeae CSL RW192 WP 038907831 1 Dzeae
Dickeya zeae NCPPB 3531 WP 038904219 1 Dzeae
Dickeya zeae NCPPB 2538 WP 038916112 1 Dzeae
Dickeya zeae MS1 WP 019845530 1 Dzeae
Dickeya zeae MK19 WP 038913882 1 Dzeae
Dickeya zeae Ech586 WP 012884582 1 Dzeae
Dickeya aquatica DW 0440 WP 035341727 1 Daquatica
Dickeya sp CSL RW240 WP 035341727 1 Daquatica
DAQ1742 DAQ1742 02375 Daquatica
Dickeya chrysanthemi L11 WP 033575693 1 Dchrysanthemi
Dickeya chrysanthemi Ech1591 WP 012769624 1 Dchrysanthemi
Dickeya chrysanthemi NCPPB 3533 WP 012769624 1 Dchrysanthemi
Dickeya chrysanthemi subsp chrysanthemi NCPPB 402 WP 040001180 1 Dchrysanthemi
Dickeya chrysanthemi subsp parthenii NCPPB 516 WP 027711927 1 Dchrysanthemi
Dickeya dianthicola RNS04 9 WP 024106102 1 Ddianthicola
Dickeya dianthicola NCPPB 453 WP 024106102 1 Ddianthicola
Dickeya dianthicola GBBC 2039 WP 024108812 1 Ddianthicola
Dickeya dianthicola NCPPB 3534 WP 024108812 1 Ddianthicola
Dickeya dianthicola IPO 980 WP 024108812 1 Ddianthicola
Dickeya solani RNS 05 1 2A WP 057084848 1 Dsolani
Dickeya solani PPO 9134 WP 022633741 1 Dsolani
Dickeya solani PPO 9019 WP 022633741 1 Dsolani
Dickeya solani D s0432 1 WP 022633741 1 Dsolani
Dickeya solani IFB 0099 WP 022633741 1 Dsolani
Dickeya solani GBBC 2040 WP 022633741 1 Dsolani
Dickeya solani MK16 WP 022633741 1 Dsolani
Dickeya solani IPO 2222 WP 022633741 1 Dsolani
Dickeya solani RNS 08 23 3 1 A WP 022633741 1 Dsolani
Dickeya solani MK10 WP 022633741 1 Dsolani
Dickeya solani RNS 07 7 3B WP 022633741 1 Dsolani
Dickeya dadantii NCPPB 3537 WP 038901328 1 Ddadantii
Dickeya dadantii subsp dieffenbachiae NCPPB 2976 WP 038922956 1 Ddadantii
Dickeya dadantii 3937 WP 013318115 1 Ddadantii
182 plcA Plant cell wall degradosome Seed
Dickeya dadantii NCPPB 898 WP 013318115 1 Ddadantii
Dickeya gs fangzhongdai NCPPB 3274 WP 033568584 1 Dfangzhongdai
Dickeya gs undicola 2B12 WP 033568584 1 Dundicola
Dickeya gs fangzhongdai B16 WP 038919206 1 Dfangzhongdai
Dickeya gs fangzhongdai MK7 WP 038919206 1 Dfangzhongdai
Dickeya gs fangzhongdai S1 WP 038658216 1 Dfangzhongdai
Dickeya gs fangzhongdai M005 WP 038658216 1 Dfangzhongdai
Dickeya gs fangzhongdai ND14b WP 038658216 1 Dfangzhongdai
Dickeya gs fangzhongdai M074 WP 038658216 1 Dfangzhongdai

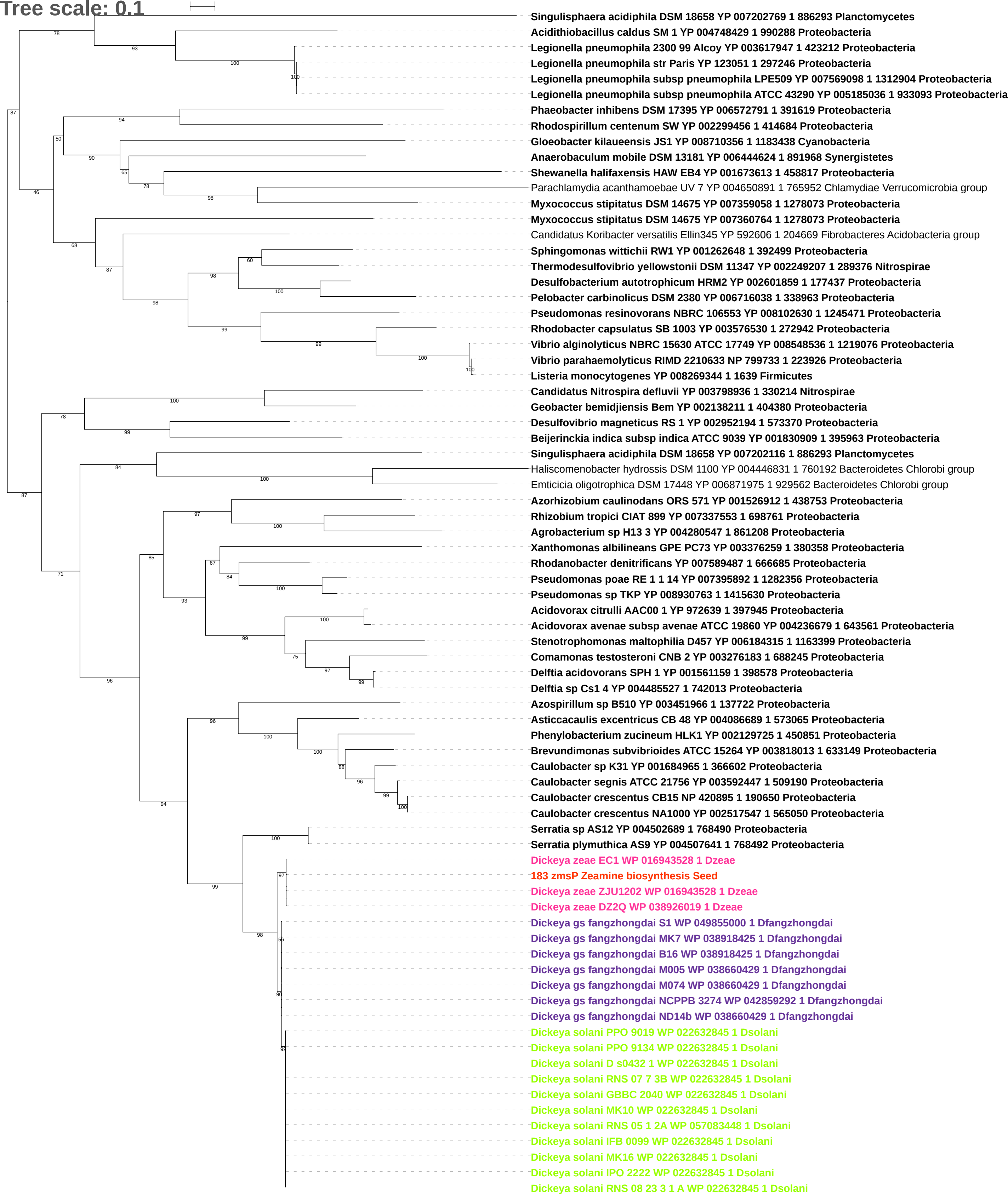
Tree scale: 0.1
Singulisphaera acidiphila DSM 18658 YP 007202769 1 886293 Planctomycetes
Acidithiobacillus caldus SM 1 YP 004748429 1 990288 Proteobacteria
Legionella pneumophila 2300 99 Alcoy YP 003617947 1 423212 Proteobacteria
Legionella pneumophila str Paris YP 123051 1 297246 Proteobacteria
Legionella pneumophila subsp pneumophila LPE509 YP 007569098 1 1312904 Proteobacteria
Legionella pneumophila subsp pneumophila ATCC 43290 YP 005185036 1 933093 Proteobacteria
Phaeobacter inhibens DSM 17395 YP 006572791 1 391619 Proteobacteria
Rhodospirillum centenum SW YP 002299456 1 414684 Proteobacteria
Gloeobacter kilaueensis JS1 YP 008710356 1 1183438 Cyanobacteria
Anaerobaculum mobile DSM 13181 YP 006444624 1 891968 Synergistetes
Shewanella halifaxensis HAW EB4 YP 001673613 1 458817 Proteobacteria
Parachlamydia acanthamoebae UV 7 YP 004650891 1 765952 Chlamydiae Verrucomicrobia group
Myxococcus stipitatus DSM 14675 YP 007359058 1 1278073 Proteobacteria
Myxococcus stipitatus DSM 14675 YP 007360764 1 1278073 Proteobacteria
Candidatus Koribacter versatilis Ellin345 YP 592606 1 204669 Fibrobacteres Acidobacteria group
Sphingomonas wittichii RW1 YP 001262648 1 392499 Proteobacteria
Thermodesulfovibrio yellowstonii DSM 11347 YP 002249207 1 289376 Nitrospirae
Desulfobacterium autotrophicum HRM2 YP 002601859 1 177437 Proteobacteria
Pelobacter carbinolicus DSM 2380 YP 006716038 1 338963 Proteobacteria
Pseudomonas resinovorans NBRC 106553 YP 008102630 1 1245471 Proteobacteria
Rhodobacter capsulatus SB 1003 YP 003576530 1 272942 Proteobacteria
Vibrio alginolyticus NBRC 15630 ATCC 17749 YP 008548536 1 1219076 Proteobacteria
Vibrio parahaemolyticus RIMD 2210633 NP 799733 1 223926 Proteobacteria
Listeria monocytogenes YP 008269344 1 1639 Firmicutes
Candidatus Nitrospira defluvii YP 003798936 1 330214 Nitrospirae
Geobacter bemidjiensis Bem YP 002138211 1 404380 Proteobacteria
Desulfovibrio magneticus RS 1 YP 002952194 1 573370 Proteobacteria
Beijerinckia indica subsp indica ATCC 9039 YP 001830909 1 395963 Proteobacteria
Singulisphaera acidiphila DSM 18658 YP 007202116 1 886293 Planctomycetes
Haliscomenobacter hydrossis DSM 1100 YP 004446831 1 760192 Bacteroidetes Chlorobi group
Emticicia oligotrophica DSM 17448 YP 006871975 1 929562 Bacteroidetes Chlorobi group
Azorhizobium caulinodans ORS 571 YP 001526912 1 438753 Proteobacteria
Rhizobium tropici CIAT 899 YP 007337553 1 698761 Proteobacteria
Agrobacterium sp H13 3 YP 004280547 1 861208 Proteobacteria
Xanthomonas albilineans GPE PC73 YP 003376259 1 380358 Proteobacteria
Rhodanobacter denitrificans YP 007589487 1 666685 Proteobacteria
Pseudomonas poae RE 1 1 14 YP 007395892 1 1282356 Proteobacteria
Pseudomonas sp TKP YP 008930763 1 1415630 Proteobacteria
Acidovorax citrulli AAC00 1 YP 972639 1 397945 Proteobacteria
Acidovorax avenae subsp avenae ATCC 19860 YP 004236679 1 643561 Proteobacteria
Stenotrophomonas maltophilia D457 YP 006184315 1 1163399 Proteobacteria
Comamonas testosteroni CNB 2 YP 003276183 1 688245 Proteobacteria
Delftia acidovorans SPH 1 YP 001561159 1 398578 Proteobacteria
Delftia sp Cs1 4 YP 004485527 1 742013 Proteobacteria
Azospirillum sp B510 YP 003451966 1 137722 Proteobacteria
Asticcacaulis excentricus CB 48 YP 004086689 1 573065 Proteobacteria
Phenylobacterium zucineum HLK1 YP 002129725 1 450851 Proteobacteria
Brevundimonas subvibrioides ATCC 15264 YP 003818013 1 633149 Proteobacteria
Caulobacter sp K31 YP 001684965 1 366602 Proteobacteria
Caulobacter segnis ATCC 21756 YP 003592447 1 509190 Proteobacteria
Caulobacter crescentus CB15 NP 420895 1 190650 Proteobacteria
Caulobacter crescentus NA1000 YP 002517547 1 565050 Proteobacteria
Serratia sp AS12 YP 004502689 1 768490 Proteobacteria
Serratia plymuthica AS9 YP 004507641 1 768492 Proteobacteria
Dickeya zeae EC1 WP 016943528 1 Dzeae
183 zmsP Zeamine biosynthesis Seed
Dickeya zeae ZJU1202 WP 016943528 1 Dzeae
Dickeya zeae DZ2Q WP 038926019 1 Dzeae
Dickeya gs fangzhongdai S1 WP 049855000 1 Dfangzhongdai
Dickeya gs fangzhongdai MK7 WP 038918425 1 Dfangzhongdai
Dickeya gs fangzhongdai B16 WP 038918425 1 Dfangzhongdai
Dickeya gs fangzhongdai M005 WP 038660429 1 Dfangzhongdai
Dickeya gs fangzhongdai M074 WP 038660429 1 Dfangzhongdai
Dickeya gs fangzhongdai NCPPB 3274 WP 042859292 1 Dfangzhongdai
Dickeya gs fangzhongdai ND14b WP 038660429 1 Dfangzhongdai
Dickeya solani PPO 9019 WP 022632845 1 Dsolani
Dickeya solani PPO 9134 WP 022632845 1 Dsolani
Dickeya solani D s0432 1 WP 022632845 1 Dsolani
Dickeya solani RNS 07 7 3B WP 022632845 1 Dsolani
Dickeya solani GBBC 2040 WP 022632845 1 Dsolani
Dickeya solani MK10 WP 022632845 1 Dsolani
Dickeya solani RNS 05 1 2A WP 057083448 1 Dsolani
Dickeya solani IFB 0099 WP 022632845 1 Dsolani
Dickeya solani MK16 WP 022632845 1 Dsolani
Dickeya solani IPO 2222 WP 022632845 1 Dsolani
Dickeya solani RNS 08 23 3 1 A WP 022632845 1 Dsolani

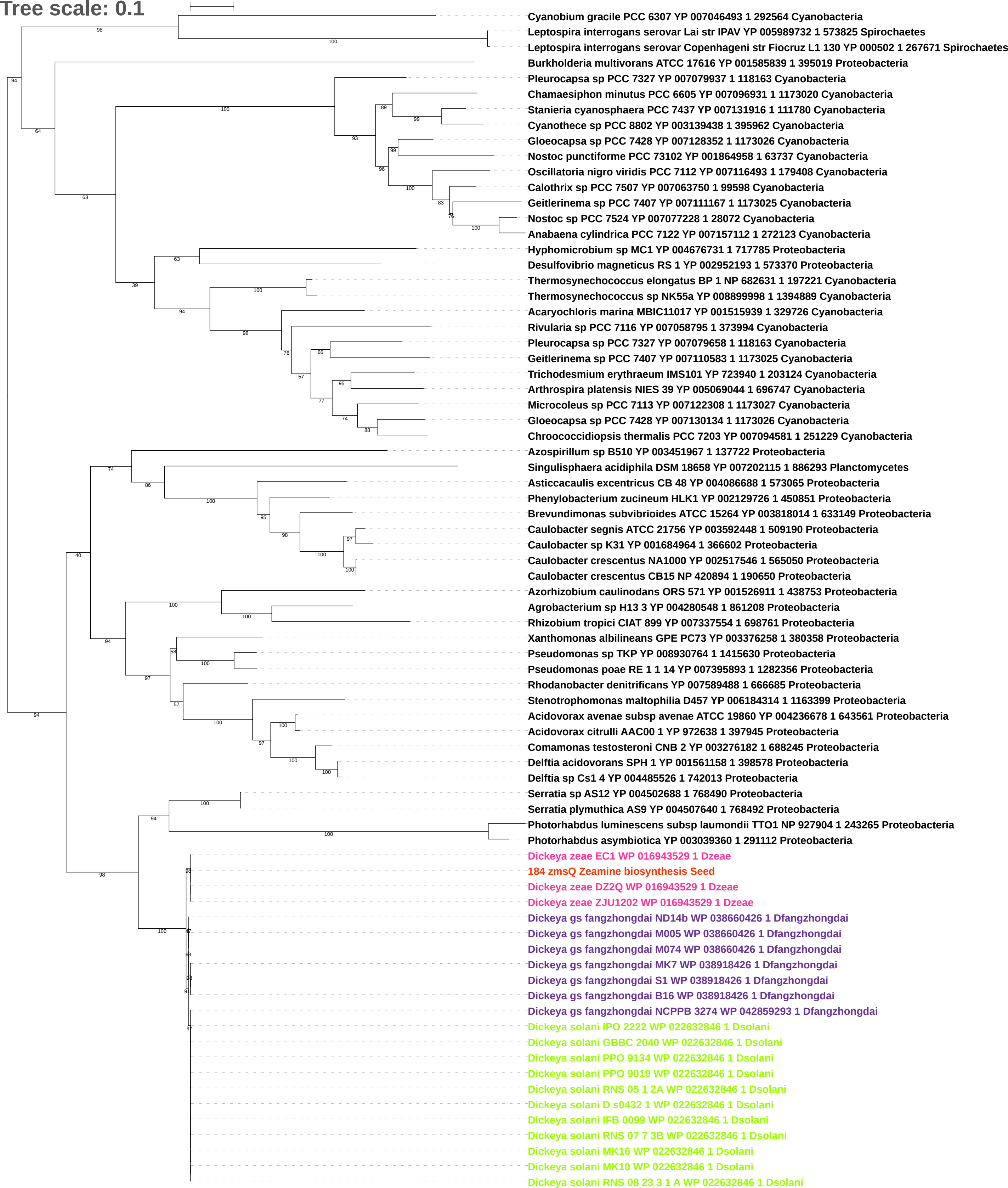
Tree scale: 0.1
Cyanobium gracile PCC 6307 YP 007046493 1 292564 Cyanobacteria
Leptospira interrogans serovar Lai str IPAV YP 005989732 1 573825 Spirochaetes
Leptospira interrogans serovar Copenhageni str Fiocruz L1 130 YP 000502 1 267671 Spirochaetes
Burkholderia multivorans ATCC 17616 YP 001585839 1 395019 Proteobacteria
Pleurocapsa sp PCC 7327 YP 007079937 1 118163 Cyanobacteria
Chamaesiphon minutus PCC 6605 YP 007096931 1 1173020 Cyanobacteria
Stanieria cyanosphaera PCC 7437 YP 007131916 1 111780 Cyanobacteria
Cyanothece sp PCC 8802 YP 003139438 1 395962 Cyanobacteria
Gloeocapsa sp PCC 7428 YP 007128352 1 1173026 Cyanobacteria
Nostoc punctiforme PCC 73102 YP 001864958 1 63737 Cyanobacteria
Oscillatoria nigro viridis PCC 7112 YP 007116493 1 179408 Cyanobacteria
Calothrix sp PCC 7507 YP 007063750 1 99598 Cyanobacteria
Geitlerinema sp PCC 7407 YP 007111167 1 1173025 Cyanobacteria
Nostoc sp PCC 7524 YP 007077228 1 28072 Cyanobacteria
Anabaena cylindrica PCC 7122 YP 007157112 1 272123 Cyanobacteria
Hyphomicrobium sp MC1 YP 004676731 1 717785 Proteobacteria
Desulfovibrio magneticus RS 1 YP 002952193 1 573370 Proteobacteria
Thermosynechococcus elongatus BP 1 NP 682631 1 197221 Cyanobacteria
Thermosynechococcus sp NK55a YP 008899998 1 1394889 Cyanobacteria
Acaryochloris marina MBIC11017 YP 001515939 1 329726 Cyanobacteria
Rivularia sp PCC 7116 YP 007058795 1 373994 Cyanobacteria
Pleurocapsa sp PCC 7327 YP 007079658 1 118163 Cyanobacteria
Geitlerinema sp PCC 7407 YP 007110583 1 1173025 Cyanobacteria
Trichodesmium erythraeum IMS101 YP 723940 1 203124 Cyanobacteria
Arthrospira platensis NIES 39 YP 005069044 1 696747 Cyanobacteria
Microcoleus sp PCC 7113 YP 007122308 1 1173027 Cyanobacteria
Gloeocapsa sp PCC 7428 YP 007130134 1 1173026 Cyanobacteria
Chroococcidiopsis thermalis PCC 7203 YP 007094581 1 251229 Cyanobacteria
Azospirillum sp B510 YP 003451967 1 137722 Proteobacteria
Singulisphaera acidiphila DSM 18658 YP 007202115 1 886293 Planctomycetes
Asticcacaulis excentricus CB 48 YP 004086688 1 573065 Proteobacteria
Phenylobacterium zucineum HLK1 YP 002129726 1 450851 Proteobacteria
Brevundimonas subvibrioides ATCC 15264 YP 003818014 1 633149 Proteobacteria
Caulobacter segnis ATCC 21756 YP 003592448 1 509190 Proteobacteria
Caulobacter sp K31 YP 001684964 1 366602 Proteobacteria
Caulobacter crescentus NA1000 YP 002517546 1 565050 Proteobacteria
Caulobacter crescentus CB15 NP 420894 1 190650 Proteobacteria
Azorhizobium caulinodans ORS 571 YP 001526911 1 438753 Proteobacteria
Agrobacterium sp H13 3 YP 004280548 1 861208 Proteobacteria
Rhizobium tropici CIAT 899 YP 007337554 1 698761 Proteobacteria
Xanthomonas albilineans GPE PC73 YP 003376258 1 380358 Proteobacteria
Pseudomonas sp TKP YP 008930764 1 1415630 Proteobacteria
Pseudomonas poae RE 1 1 14 YP 007395893 1 1282356 Proteobacteria
Rhodanobacter denitrificans YP 007589488 1 666685 Proteobacteria
Stenotrophomonas maltophilia D457 YP 006184314 1 1163399 Proteobacteria
Acidovorax avenae subsp avenae ATCC 19860 YP 004236678 1 643561 Proteobacteria
Acidovorax citrulli AAC00 1 YP 972638 1 397945 Proteobacteria
Comamonas testosteroni CNB 2 YP 003276182 1 688245 Proteobacteria
Delftia acidovorans SPH 1 YP 001561158 1 398578 Proteobacteria
Delftia sp Cs1 4 YP 004485526 1 742013 Proteobacteria
Serratia sp AS12 YP 004502688 1 768490 Proteobacteria
Serratia plymuthica AS9 YP 004507640 1 768492 Proteobacteria
Photorhabdus luminescens subsp laumondii TTO1 NP 927904 1 243265 Proteobacteria
Photorhabdus asymbiotica YP 003039360 1 291112 Proteobacteria
Dickeya zeae EC1 WP 016943529 1 Dzeae
184 zmsQ Zeamine biosynthesis Seed
Dickeya zeae DZ2Q WP 016943529 1 Dzeae
Dickeya zeae ZJU1202 WP 016943529 1 Dzeae
Dickeya gs fangzhongdai ND14b WP 038660426 1 Dfangzhongdai
Dickeya gs fangzhongdai M005 WP 038660426 1 Dfangzhongdai
Dickeya gs fangzhongdai M074 WP 038660426 1 Dfangzhongdai
Dickeya gs fangzhongdai MK7 WP 038918426 1 Dfangzhongdai
Dickeya gs fangzhongdai S1 WP 038918426 1 Dfangzhongdai
Dickeya gs fangzhongdai B16 WP 038918426 1 Dfangzhongdai
Dickeya gs fangzhongdai NCPPB 3274 WP 042859293 1 Dfangzhongdai
Dickeya solani IPO 2222 WP 022632846 1 Dsolani
Dickeya solani GBBC 2040 WP 022632846 1 Dsolani
Dickeya solani PPO 9134 WP 022632846 1 Dsolani
Dickeya solani PPO 9019 WP 022632846 1 Dsolani
Dickeya solani RNS 05 1 2A WP 022632846 1 Dsolani
Dickeya solani D s0432 1 WP 022632846 1 Dsolani
Dickeya solani IFB 0099 WP 022632846 1 Dsolani
Dickeya solani RNS 07 7 3B WP 022632846 1 Dsolani
Dickeya solani MK16 WP 022632846 1 Dsolani
Dickeya solani MK10 WP 022632846 1 Dsolani
Dickeya solani RNS 08 23 3 1 A WP 022632846 1 Dsolani

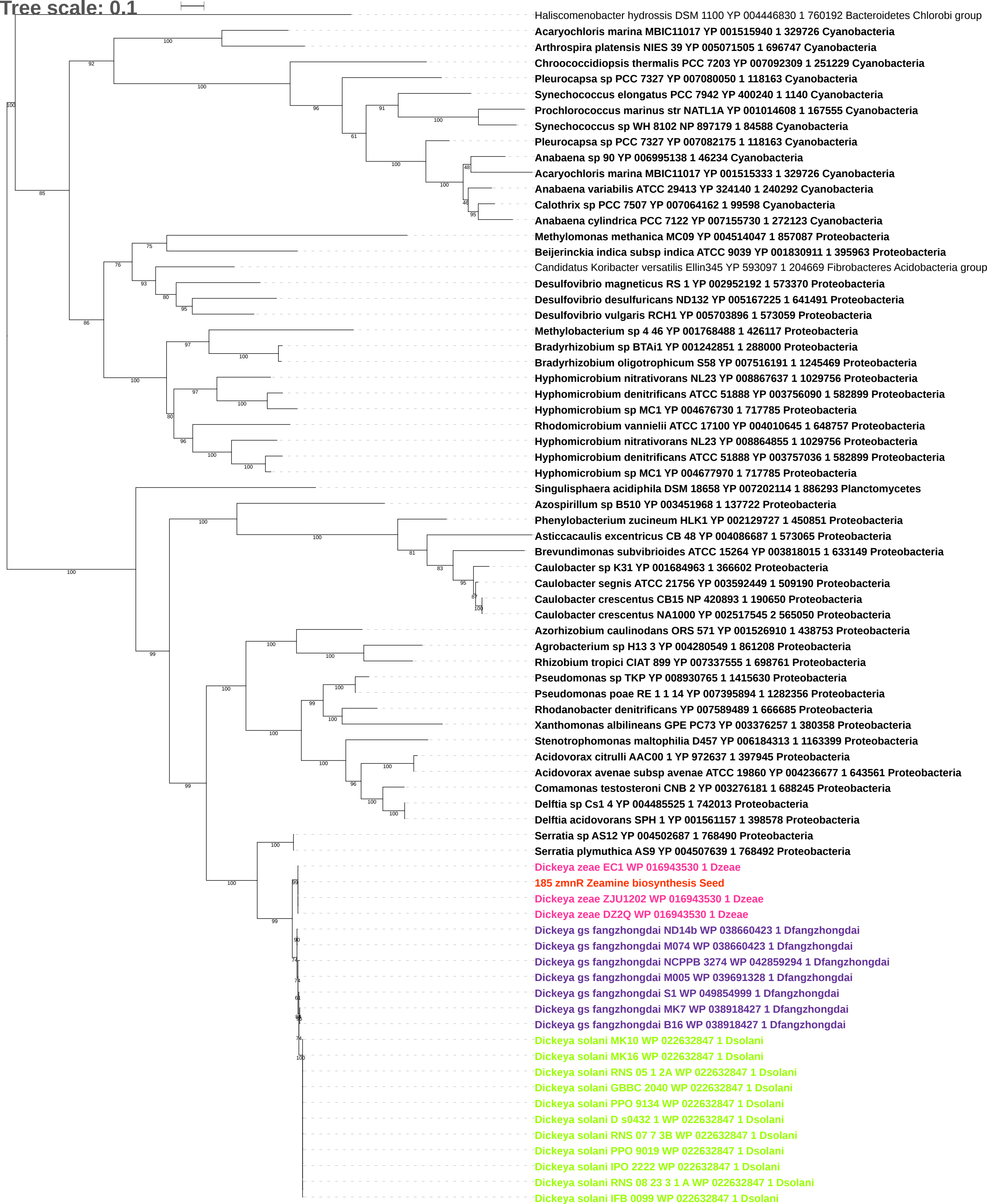
Tree scale: 0.1
Haliscomenobacter hydrossis DSM 1100 YP 004446830 1 760192 Bacteroidetes Chlorobi group
Acaryochloris marina MBIC11017 YP 001515940 1 329726 Cyanobacteria
Arthrospira platensis NIES 39 YP 005071505 1 696747 Cyanobacteria
Chroococcidiopsis thermalis PCC 7203 YP 007092309 1 251229 Cyanobacteria
Pleurocapsa sp PCC 7327 YP 007080050 1 118163 Cyanobacteria
Synechococcus elongatus PCC 7942 YP 400240 1 1140 Cyanobacteria
Prochlorococcus marinus str NATL1A YP 001014608 1 167555 Cyanobacteria
Synechococcus sp WH 8102 NP 897179 1 84588 Cyanobacteria
Pleurocapsa sp PCC 7327 YP 007082175 1 118163 Cyanobacteria
Anabaena sp 90 YP 006995138 1 46234 Cyanobacteria
Acaryochloris marina MBIC11017 YP 001515333 1 329726 Cyanobacteria
Anabaena variabilis ATCC 29413 YP 324140 1 240292 Cyanobacteria
Calothrix sp PCC 7507 YP 007064162 1 99598 Cyanobacteria
Anabaena cylindrica PCC 7122 YP 007155730 1 272123 Cyanobacteria
Methylomonas methanica MC09 YP 004514047 1 857087 Proteobacteria
Beijerinckia indica subsp indica ATCC 9039 YP 001830911 1 395963 Proteobacteria
Candidatus Koribacter versatilis Ellin345 YP 593097 1 204669 Fibrobacteres Acidobacteria group
Desulfovibrio magneticus RS 1 YP 002952192 1 573370 Proteobacteria
Desulfovibrio desulfuricans ND132 YP 005167225 1 641491 Proteobacteria
Desulfovibrio vulgaris RCH1 YP 005703896 1 573059 Proteobacteria
Methylobacterium sp 4 46 YP 001768488 1 426117 Proteobacteria
Bradyrhizobium sp BTAi1 YP 001242851 1 288000 Proteobacteria
Bradyrhizobium oligotrophicum S58 YP 007516191 1 1245469 Proteobacteria
Hyphomicrobium nitrativorans NL23 YP 008867637 1 1029756 Proteobacteria
Hyphomicrobium denitrificans ATCC 51888 YP 003756090 1 582899 Proteobacteria
Hyphomicrobium sp MC1 YP 004676730 1 717785 Proteobacteria
Rhodomicrobium vannielii ATCC 17100 YP 004010645 1 648757 Proteobacteria
Hyphomicrobium nitrativorans NL23 YP 008864855 1 1029756 Proteobacteria
Hyphomicrobium denitrificans ATCC 51888 YP 003757036 1 582899 Proteobacteria
Hyphomicrobium sp MC1 YP 004677970 1 717785 Proteobacteria
Singulisphaera acidiphila DSM 18658 YP 007202114 1 886293 Planctomycetes
Azospirillum sp B510 YP 003451968 1 137722 Proteobacteria
Phenylobacterium zucineum HLK1 YP 002129727 1 450851 Proteobacteria
Asticcacaulis excentricus CB 48 YP 004086687 1 573065 Proteobacteria
Brevundimonas subvibrioides ATCC 15264 YP 003818015 1 633149 Proteobacteria
Caulobacter sp K31 YP 001684963 1 366602 Proteobacteria
Caulobacter segnis ATCC 21756 YP 003592449 1 509190 Proteobacteria
Caulobacter crescentus CB15 NP 420893 1 190650 Proteobacteria
Caulobacter crescentus NA1000 YP 002517545 2 565050 Proteobacteria
Azorhizobium caulinodans ORS 571 YP 001526910 1 438753 Proteobacteria
Agrobacterium sp H13 3 YP 004280549 1 861208 Proteobacteria
Rhizobium tropici CIAT 899 YP 007337555 1 698761 Proteobacteria
Pseudomonas sp TKP YP 008930765 1 1415630 Proteobacteria
Pseudomonas poae RE 1 1 14 YP 007395894 1 1282356 Proteobacteria
Rhodanobacter denitrificans YP 007589489 1 666685 Proteobacteria
Xanthomonas albilineans GPE PC73 YP 003376257 1 380358 Proteobacteria
Stenotrophomonas maltophilia D457 YP 006184313 1 1163399 Proteobacteria
Acidovorax citrulli AAC00 1 YP 972637 1 397945 Proteobacteria
Acidovorax avenae subsp avenae ATCC 19860 YP 004236677 1 643561 Proteobacteria
Comamonas testosteroni CNB 2 YP 003276181 1 688245 Proteobacteria
Delftia sp Cs1 4 YP 004485525 1 742013 Proteobacteria
Delftia acidovorans SPH 1 YP 001561157 1 398578 Proteobacteria
Serratia sp AS12 YP 004502687 1 768490 Proteobacteria
Serratia plymuthica AS9 YP 004507639 1 768492 Proteobacteria
Dickeya zeae EC1 WP 016943530 1 Dzeae
185 zmnR Zeamine biosynthesis Seed
Dickeya zeae ZJU1202 WP 016943530 1 Dzeae
Dickeya zeae DZ2Q WP 016943530 1 Dzeae
Dickeya gs fangzhongdai ND14b WP 038660423 1 Dfangzhongdai
Dickeya gs fangzhongdai M074 WP 038660423 1 Dfangzhongdai
Dickeya gs fangzhongdai NCPPB 3274 WP 042859294 1 Dfangzhongdai
Dickeya gs fangzhongdai M005 WP 039691328 1 Dfangzhongdai
Dickeya gs fangzhongdai S1 WP 049854999 1 Dfangzhongdai
Dickeya gs fangzhongdai MK7 WP 038918427 1 Dfangzhongdai
Dickeya gs fangzhongdai B16 WP 038918427 1 Dfangzhongdai
Dickeya solani MK10 WP 022632847 1 Dsolani
Dickeya solani MK16 WP 022632847 1 Dsolani
Dickeya solani RNS 05 1 2A WP 022632847 1 Dsolani
Dickeya solani GBBC 2040 WP 022632847 1 Dsolani
Dickeya solani PPO 9134 WP 022632847 1 Dsolani
Dickeya solani D s0432 1 WP 022632847 1 Dsolani
Dickeya solani RNS 07 7 3B WP 022632847 1 Dsolani
Dickeya solani PPO 9019 WP 022632847 1 Dsolani
Dickeya solani IPO 2222 WP 022632847 1 Dsolani
Dickeya solani RNS 08 23 3 1 A WP 022632847 1 Dsolani
Dickeya solani IFB 0099 WP 022632847 1 Dsolani

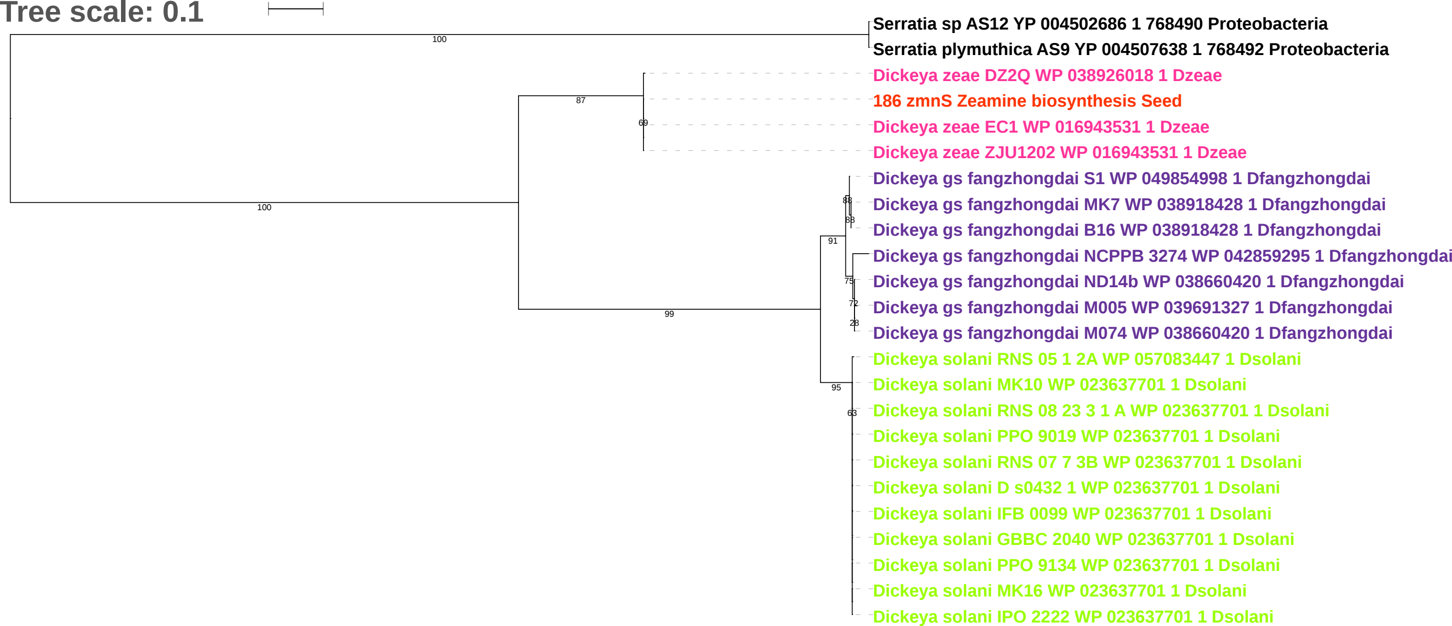
Tree scale: 0.1
100
100
87
69
99
91
95
88
88
75
72
28
63
Serratia sp AS12 YP 004502686 1 768490 Proteobacteria
Serratia plymuthica AS9 YP 004507638 1 768492 Proteobacteria
Dickeya zeae DZ2Q WP 038926018 1 Dzeae
186 zmnS Zeamine biosynthesis Seed
Dickeya zeae EC1 WP 016943531 1 Dzeae
Dickeya zeae ZJU1202 WP 016943531 1 Dzeae
Dickeya gs fangzhongdai S1 WP 049854998 1 Dfangzhongdai
Dickeya gs fangzhongdai MK7 WP 038918428 1 Dfangzhongdai
Dickeya gs fangzhongdai B16 WP 038918428 1 Dfangzhongdai
Dickeya gs fangzhongdai NCPPB 3274 WP 042859295 1 Dfangzhongdai
Dickeya gs fangzhongdai ND14b WP 038660420 1 Dfangzhongdai
Dickeya gs fangzhongdai M005 WP 039691327 1 Dfangzhongdai
Dickeya gs fangzhongdai M074 WP 038660420 1 Dfangzhongdai
Dickeya solani RNS 05 1 2A WP 057083447 1 Dsolani
Dickeya solani MK10 WP 023637701 1 Dsolani
Dickeya solani RNS 08 23 3 1 A WP 023637701 1 Dsolani
Dickeya solani PPO 9019 WP 023637701 1 Dsolani
Dickeya solani RNS 07 7 3B WP 023637701 1 Dsolani
Dickeya solani D s0432 1 WP 023637701 1 Dsolani
Dickeya solani IFB 0099 WP 023637701 1 Dsolani
Dickeya solani GBBC 2040 WP 023637701 1 Dsolani
Dickeya solani PPO 9134 WP 023637701 1 Dsolani
Dickeya solani MK16 WP 023637701 1 Dsolani
Dickeya solani IPO 2222 WP 023637701 1 Dsolani

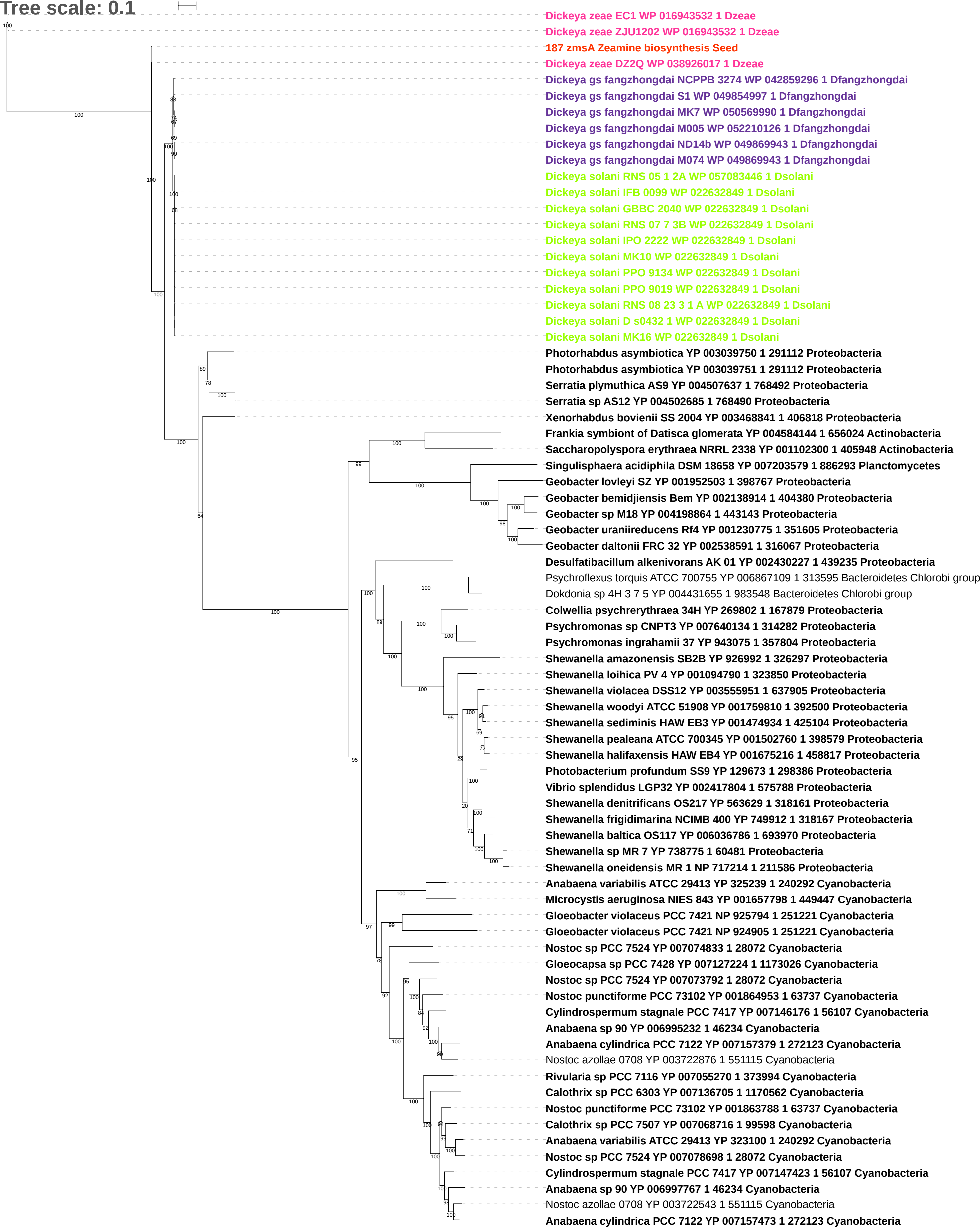
Tree scale: 0.1
Dickeya zeae EC1 WP 016943532 1 Dzeae
Dickeya zeae ZJU1202 WP 016943532 1 Dzeae
187 zmsA Zeamine biosynthesis Seed
Dickeya zeae DZ2Q WP 038926017 1 Dzeae
Dickeya gs fangzhongdai NCPPB 3274 WP 042859296 1 Dfangzhongdai
Dickeya gs fangzhongdai S1 WP 049854997 1 Dfangzhongdai
Dickeya gs fangzhongdai MK7 WP 050569990 1 Dfangzhongdai
Dickeya gs fangzhongdai M005 WP 052210126 1 Dfangzhongdai
Dickeya gs fangzhongdai ND14b WP 049869943 1 Dfangzhongdai
Dickeya gs fangzhongdai M074 WP 049869943 1 Dfangzhongdai
Dickeya solani RNS 05 1 2A WP 057083446 1 Dsolani
Dickeya solani IFB 0099 WP 022632849 1 Dsolani
Dickeya solani GBBC 2040 WP 022632849 1 Dsolani
Dickeya solani RNS 07 7 3B WP 022632849 1 Dsolani
Dickeya solani IPO 2222 WP 022632849 1 Dsolani
Dickeya solani MK10 WP 022632849 1 Dsolani
Dickeya solani PPO 9134 WP 022632849 1 Dsolani
Dickeya solani PPO 9019 WP 022632849 1 Dsolani
Dickeya solani RNS 08 23 3 1 A WP 022632849 1 Dsolani
Dickeya solani D s0432 1 WP 022632849 1 Dsolani
Dickeya solani MK16 WP 022632849 1 Dsolani
Photorhabdus asymbiotica YP 003039750 1 291112 Proteobacteria
Photorhabdus asymbiotica YP 003039751 1 291112 Proteobacteria
Serratia plymuthica AS9 YP 004507637 1 768492 Proteobacteria
Serratia sp AS12 YP 004502685 1 768490 Proteobacteria
Xenorhabdus bovienii SS 2004 YP 003468841 1 406818 Proteobacteria
Frankia symbiont of Datisca glomerata YP 004584144 1 656024 Actinobacteria
Saccharopolyspora erythraea NRRL 2338 YP 001102300 1 405948 Actinobacteria
Singulisphaera acidiphila DSM 18658 YP 007203579 1 886293 Planctomycetes
Geobacter lovleyi SZ YP 001952503 1 398767 Proteobacteria
Geobacter bemidjiensis Bem YP 002138914 1 404380 Proteobacteria
Geobacter sp M18 YP 004198864 1 443143 Proteobacteria
Geobacter uraniireducens Rf4 YP 001230775 1 351605 Proteobacteria
Geobacter daltonii FRC 32 YP 002538591 1 316067 Proteobacteria
Desulfatibacillum alkenivorans AK 01 YP 002430227 1 439235 Proteobacteria
Psychroflexus torquis ATCC 700755 YP 006867109 1 313595 Bacteroidetes Chlorobi group
Dokdonia sp 4H 3 7 5 YP 004431655 1 983548 Bacteroidetes Chlorobi group
Colwellia psychrerythraea 34H YP 269802 1 167879 Proteobacteria
Psychromonas sp CNPT3 YP 007640134 1 314282 Proteobacteria
Psychromonas ingrahamii 37 YP 943075 1 357804 Proteobacteria
Shewanella amazonensis SB2B YP 926992 1 326297 Proteobacteria
Shewanella loihica PV 4 YP 001094790 1 323850 Proteobacteria
Shewanella violacea DSS12 YP 003555951 1 637905 Proteobacteria
Shewanella woodyi ATCC 51908 YP 001759810 1 392500 Proteobacteria
Shewanella sediminis HAW EB3 YP 001474934 1 425104 Proteobacteria
Shewanella pealeana ATCC 700345 YP 001502760 1 398579 Proteobacteria
Shewanella halifaxensis HAW EB4 YP 001675216 1 458817 Proteobacteria
Photobacterium profundum SS9 YP 129673 1 298386 Proteobacteria
Vibrio splendidus LGP32 YP 002417804 1 575788 Proteobacteria
Shewanella denitrificans OS217 YP 563629 1 318161 Proteobacteria
Shewanella frigidimarina NCIMB 400 YP 749912 1 318167 Proteobacteria
Shewanella baltica OS117 YP 006036786 1 693970 Proteobacteria
Shewanella sp MR 7 YP 738775 1 60481 Proteobacteria
Shewanella oneidensis MR 1 NP 717214 1 211586 Proteobacteria
Anabaena variabilis ATCC 29413 YP 325239 1 240292 Cyanobacteria
Microcystis aeruginosa NIES 843 YP 001657798 1 449447 Cyanobacteria
Gloeobacter violaceus PCC 7421 NP 925794 1 251221 Cyanobacteria
Gloeobacter violaceus PCC 7421 NP 924905 1 251221 Cyanobacteria
Nostoc sp PCC 7524 YP 007074833 1 28072 Cyanobacteria
Gloeocapsa sp PCC 7428 YP 007127224 1 1173026 Cyanobacteria
Nostoc sp PCC 7524 YP 007073792 1 28072 Cyanobacteria
Nostoc punctiforme PCC 73102 YP 001864953 1 63737 Cyanobacteria
Cylindrospermum stagnale PCC 7417 YP 007146176 1 56107 Cyanobacteria
Anabaena sp 90 YP 006995232 1 46234 Cyanobacteria
Anabaena cylindrica PCC 7122 YP 007157379 1 272123 Cyanobacteria
Nostoc azollae 0708 YP 003722876 1 551115 Cyanobacteria
Rivularia sp PCC 7116 YP 007055270 1 373994 Cyanobacteria
Calothrix sp PCC 6303 YP 007136705 1 1170562 Cyanobacteria
Nostoc punctiforme PCC 73102 YP 001863788 1 63737 Cyanobacteria
Calothrix sp PCC 7507 YP 007068716 1 99598 Cyanobacteria
Anabaena variabilis ATCC 29413 YP 323100 1 240292 Cyanobacteria
Nostoc sp PCC 7524 YP 007078698 1 28072 Cyanobacteria
Cylindrospermum stagnale PCC 7417 YP 007147423 1 56107 Cyanobacteria
Anabaena sp 90 YP 006997767 1 46234 Cyanobacteria
Nostoc azollae 0708 YP 003722543 1 551115 Cyanobacteria
Anabaena cylindrica PCC 7122 YP 007157473 1 272123 Cyanobacteria

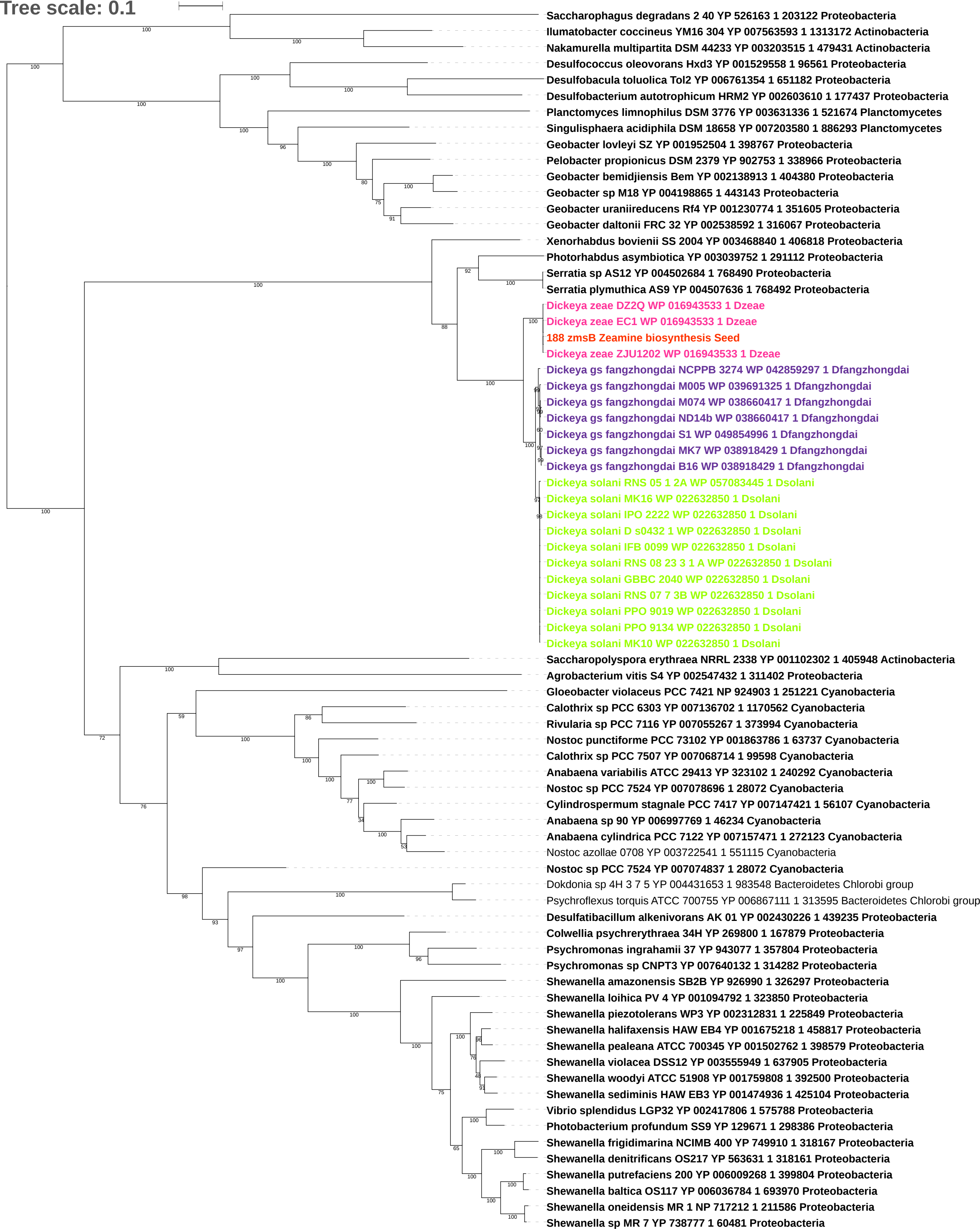
Tree scale: 0.1
Saccharophagus degradans 2 40 YP 526163 1 203122 Proteobacteria
Ilumatobacter coccineus YM16 304 YP 007563593 1 1313172 Actinobacteria
Nakamurella multipartita DSM 44233 YP 003203515 1 479431 Actinobacteria
Desulfococcus oleovorans Hxd3 YP 001529558 1 96561 Proteobacteria
Desulfobacula toluolica Tol2 YP 006761354 1 651182 Proteobacteria
Desulfobacterium autotrophicum HRM2 YP 002603610 1 177437 Proteobacteria
Planctomyces limnophilus DSM 3776 YP 003631336 1 521674 Planctomycetes
Singulisphaera acidiphila DSM 18658 YP 007203580 1 886293 Planctomycetes
Geobacter lovleyi SZ YP 001952504 1 398767 Proteobacteria
Pelobacter propionicus DSM 2379 YP 902753 1 338966 Proteobacteria
Geobacter bemidjiensis Bem YP 002138913 1 404380 Proteobacteria
Geobacter sp M18 YP 004198865 1 443143 Proteobacteria
Geobacter uraniireducens Rf4 YP 001230774 1 351605 Proteobacteria
Geobacter daltonii FRC 32 YP 002538592 1 316067 Proteobacteria
Xenorhabdus bovienii SS 2004 YP 003468840 1 406818 Proteobacteria
Photorhabdus asymbiotica YP 003039752 1 291112 Proteobacteria
Serratia sp AS12 YP 004502684 1 768490 Proteobacteria
Serratia plymuthica AS9 YP 004507636 1 768492 Proteobacteria
Dickeya zeae DZ2Q WP 016943533 1 Dzeae
Dickeya zeae EC1 WP 016943533 1 Dzeae
188 zmsB Zeamine biosynthesis Seed
Dickeya zeae ZJU1202 WP 016943533 1 Dzeae
Dickeya gs fangzhongdai NCPPB 3274 WP 042859297 1 Dfangzhongdai
Dickeya gs fangzhongdai M005 WP 039691325 1 Dfangzhongdai
Dickeya gs fangzhongdai M074 WP 038660417 1 Dfangzhongdai
Dickeya gs fangzhongdai ND14b WP 038660417 1 Dfangzhongdai
Dickeya gs fangzhongdai S1 WP 049854996 1 Dfangzhongdai
Dickeya gs fangzhongdai MK7 WP 038918429 1 Dfangzhongdai
Dickeya gs fangzhongdai B16 WP 038918429 1 Dfangzhongdai
Dickeya solani RNS 05 1 2A WP 057083445 1 Dsolani
Dickeya solani MK16 WP 022632850 1 Dsolani
Dickeya solani IPO 2222 WP 022632850 1 Dsolani
Dickeya solani D s0432 1 WP 022632850 1 Dsolani
Dickeya solani IFB 0099 WP 022632850 1 Dsolani
Dickeya solani RNS 08 23 3 1 A WP 022632850 1 Dsolani
Dickeya solani GBBC 2040 WP 022632850 1 Dsolani
Dickeya solani RNS 07 7 3B WP 022632850 1 Dsolani
Dickeya solani PPO 9019 WP 022632850 1 Dsolani
Dickeya solani PPO 9134 WP 022632850 1 Dsolani
Dickeya solani MK10 WP 022632850 1 Dsolani
Saccharopolyspora erythraea NRRL 2338 YP 001102302 1 405948 Actinobacteria
Agrobacterium vitis S4 YP 002547432 1 311402 Proteobacteria
Gloeobacter violaceus PCC 7421 NP 924903 1 251221 Cyanobacteria
Calothrix sp PCC 6303 YP 007136702 1 1170562 Cyanobacteria
Rivularia sp PCC 7116 YP 007055267 1 373994 Cyanobacteria
Nostoc punctiforme PCC 73102 YP 001863786 1 63737 Cyanobacteria
Calothrix sp PCC 7507 YP 007068714 1 99598 Cyanobacteria
Anabaena variabilis ATCC 29413 YP 323102 1 240292 Cyanobacteria
Nostoc sp PCC 7524 YP 007078696 1 28072 Cyanobacteria
Cylindrospermum stagnale PCC 7417 YP 007147421 1 56107 Cyanobacteria
Anabaena sp 90 YP 006997769 1 46234 Cyanobacteria
Anabaena cylindrica PCC 7122 YP 007157471 1 272123 Cyanobacteria
Nostoc azollae 0708 YP 003722541 1 551115 Cyanobacteria
Nostoc sp PCC 7524 YP 007074837 1 28072 Cyanobacteria
Dokdonia sp 4H 3 7 5 YP 004431653 1 983548 Bacteroidetes Chlorobi group
Psychroflexus torquis ATCC 700755 YP 006867111 1 313595 Bacteroidetes Chlorobi group
Desulfatibacillum alkenivorans AK 01 YP 002430226 1 439235 Proteobacteria
Colwellia psychrerythraea 34H YP 269800 1 167879 Proteobacteria
Psychromonas ingrahamii 37 YP 943077 1 357804 Proteobacteria
Psychromonas sp CNPT3 YP 007640132 1 314282 Proteobacteria
Shewanella amazonensis SB2B YP 926990 1 326297 Proteobacteria
Shewanella loihica PV 4 YP 001094792 1 323850 Proteobacteria
Shewanella piezotolerans WP3 YP 002312831 1 225849 Proteobacteria
Shewanella halifaxensis HAW EB4 YP 001675218 1 458817 Proteobacteria
Shewanella pealeana ATCC 700345 YP 001502762 1 398579 Proteobacteria
Shewanella violacea DSS12 YP 003555949 1 637905 Proteobacteria
Shewanella woodyi ATCC 51908 YP 001759808 1 392500 Proteobacteria
Shewanella sediminis HAW EB3 YP 001474936 1 425104 Proteobacteria
Vibrio splendidus LGP32 YP 002417806 1 575788 Proteobacteria
Photobacterium profundum SS9 YP 129671 1 298386 Proteobacteria
Shewanella frigidimarina NCIMB 400 YP 749910 1 318167 Proteobacteria
Shewanella denitrificans OS217 YP 563631 1 318161 Proteobacteria
Shewanella putrefaciens 200 YP 006009268 1 399804 Proteobacteria
Shewanella baltica OS117 YP 006036784 1 693970 Proteobacteria
Shewanella oneidensis MR 1 NP 717212 1 211586 Proteobacteria
Shewanella sp MR 7 YP 738777 1 60481 Proteobacteria

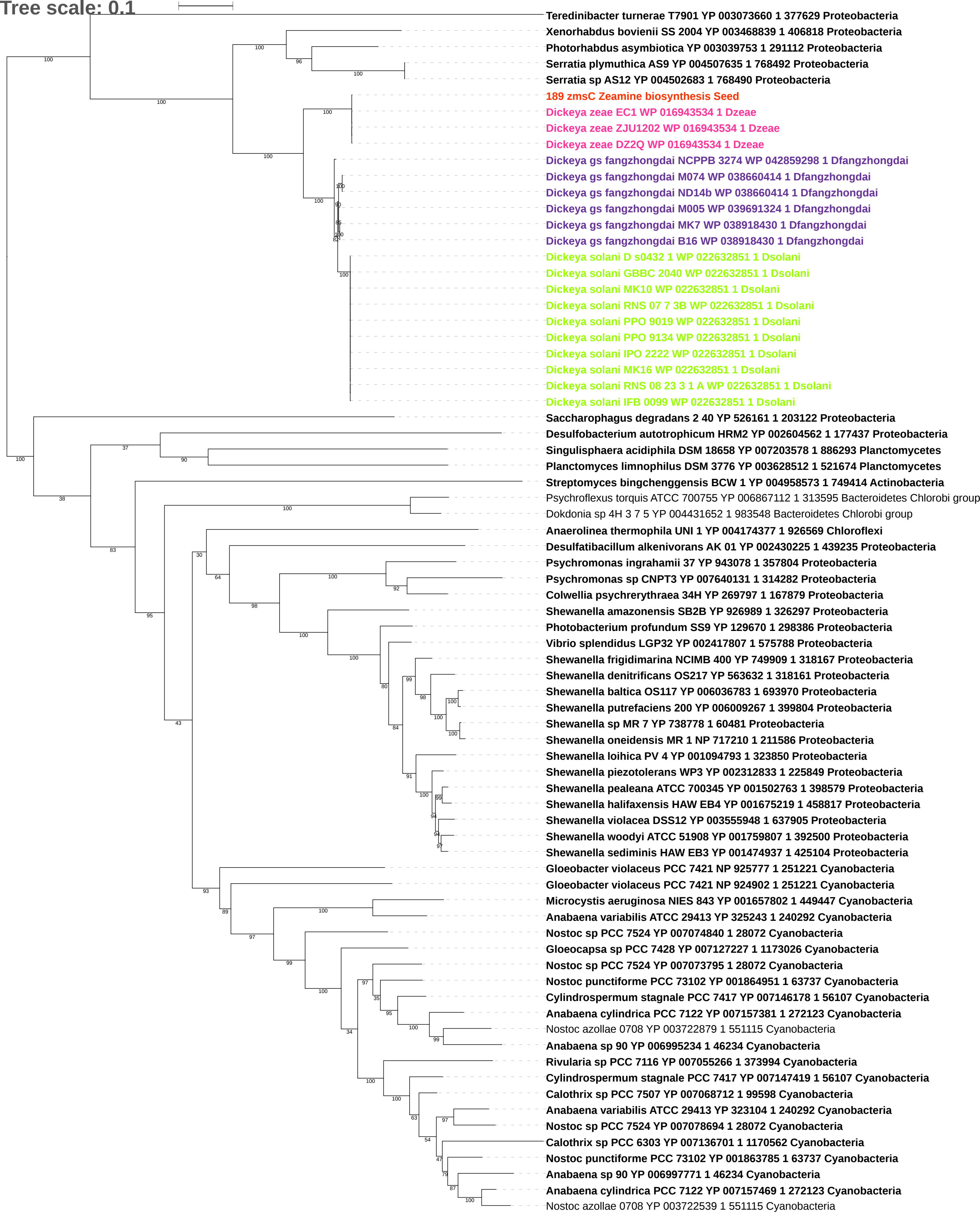
Tree scale: 0.1
Teredinibacter turnerae T7901 YP 003073660 1 377629 Proteobacteria
Xenorhabdus bovienii SS 2004 YP 003468839 1 406818 Proteobacteria
Photorhabdus asymbiotica YP 003039753 1 291112 Proteobacteria
Serratia plymuthica AS9 YP 004507635 1 768492 Proteobacteria
Serratia sp AS12 YP 004502683 1 768490 Proteobacteria
189 zmsC Zeamine biosynthesis Seed
Dickeya zeae EC1 WP 016943534 1 Dzeae
Dickeya zeae ZJU1202 WP 016943534 1 Dzeae
Dickeya zeae DZ2Q WP 016943534 1 Dzeae
Dickeya gs fangzhongdai NCPPB 3274 WP 042859298 1 Dfangzhongdai
Dickeya gs fangzhongdai M074 WP 038660414 1 Dfangzhongdai
Dickeya gs fangzhongdai ND14b WP 038660414 1 Dfangzhongdai
Dickeya gs fangzhongdai M005 WP 039691324 1 Dfangzhongdai
Dickeya gs fangzhongdai MK7 WP 038918430 1 Dfangzhongdai
Dickeya gs fangzhongdai B16 WP 038918430 1 Dfangzhongdai
Dickeya solani D s0432 1 WP 022632851 1 Dsolani
Dickeya solani GBBC 2040 WP 022632851 1 Dsolani
Dickeya solani MK10 WP 022632851 1 Dsolani
Dickeya solani RNS 07 7 3B WP 022632851 1 Dsolani
Dickeya solani PPO 9019 WP 022632851 1 Dsolani
Dickeya solani PPO 9134 WP 022632851 1 Dsolani
Dickeya solani IPO 2222 WP 022632851 1 Dsolani
Dickeya solani MK16 WP 022632851 1 Dsolani
Dickeya solani RNS 08 23 3 1 A WP 022632851 1 Dsolani
Dickeya solani IFB 0099 WP 022632851 1 Dsolani
Saccharophagus degradans 2 40 YP 526161 1 203122 Proteobacteria
Desulfobacterium autotrophicum HRM2 YP 002604562 1 177437 Proteobacteria
Singulisphaera acidiphila DSM 18658 YP 007203578 1 886293 Planctomycetes
Planctomyces limnophilus DSM 3776 YP 003628512 1 521674 Planctomycetes
Streptomyces bingchenggensis BCW 1 YP 004958573 1 749414 Actinobacteria
Psychroflexus torquis ATCC 700755 YP 006867112 1 313595 Bacteroidetes Chlorobi group
Dokdonia sp 4H 3 7 5 YP 004431652 1 983548 Bacteroidetes Chlorobi group
Anaerolinea thermophila UNI 1 YP 004174377 1 926569 Chloroflexi
Desulfatibacillum alkenivorans AK 01 YP 002430225 1 439235 Proteobacteria
Psychromonas ingrahamii 37 YP 943078 1 357804 Proteobacteria
Psychromonas sp CNPT3 YP 007640131 1 314282 Proteobacteria
Colwellia psychrerythraea 34H YP 269797 1 167879 Proteobacteria
Shewanella amazonensis SB2B YP 926989 1 326297 Proteobacteria
Photobacterium profundum SS9 YP 129670 1 298386 Proteobacteria
Vibrio splendidus LGP32 YP 002417807 1 575788 Proteobacteria
Shewanella frigidimarina NCIMB 400 YP 749909 1 318167 Proteobacteria
Shewanella denitrificans OS217 YP 563632 1 318161 Proteobacteria
Shewanella baltica OS117 YP 006036783 1 693970 Proteobacteria
Shewanella putrefaciens 200 YP 006009267 1 399804 Proteobacteria
Shewanella sp MR 7 YP 738778 1 60481 Proteobacteria
Shewanella oneidensis MR 1 NP 717210 1 211586 Proteobacteria
Shewanella loihica PV 4 YP 001094793 1 323850 Proteobacteria
Shewanella piezotolerans WP3 YP 002312833 1 225849 Proteobacteria
Shewanella pealeana ATCC 700345 YP 001502763 1 398579 Proteobacteria
Shewanella halifaxensis HAW EB4 YP 001675219 1 458817 Proteobacteria
Shewanella violacea DSS12 YP 003555948 1 637905 Proteobacteria
Shewanella woodyi ATCC 51908 YP 001759807 1 392500 Proteobacteria
Shewanella sediminis HAW EB3 YP 001474937 1 425104 Proteobacteria
Gloeobacter violaceus PCC 7421 NP 925777 1 251221 Cyanobacteria
Gloeobacter violaceus PCC 7421 NP 924902 1 251221 Cyanobacteria
Microcystis aeruginosa NIES 843 YP 001657802 1 449447 Cyanobacteria
Anabaena variabilis ATCC 29413 YP 325243 1 240292 Cyanobacteria
Nostoc sp PCC 7524 YP 007074840 1 28072 Cyanobacteria
Gloeocapsa sp PCC 7428 YP 007127227 1 1173026 Cyanobacteria
Nostoc sp PCC 7524 YP 007073795 1 28072 Cyanobacteria
Nostoc punctiforme PCC 73102 YP 001864951 1 63737 Cyanobacteria
Cylindrospermum stagnale PCC 7417 YP 007146178 1 56107 Cyanobacteria
Anabaena cylindrica PCC 7122 YP 007157381 1 272123 Cyanobacteria
Nostoc azollae 0708 YP 003722879 1 551115 Cyanobacteria
Anabaena sp 90 YP 006995234 1 46234 Cyanobacteria
Rivularia sp PCC 7116 YP 007055266 1 373994 Cyanobacteria
Cylindrospermum stagnale PCC 7417 YP 007147419 1 56107 Cyanobacteria
Calothrix sp PCC 7507 YP 007068712 1 99598 Cyanobacteria
Anabaena variabilis ATCC 29413 YP 323104 1 240292 Cyanobacteria
Nostoc sp PCC 7524 YP 007078694 1 28072 Cyanobacteria
Calothrix sp PCC 6303 YP 007136701 1 1170562 Cyanobacteria
Nostoc punctiforme PCC 73102 YP 001863785 1 63737 Cyanobacteria
Anabaena sp 90 YP 006997771 1 46234 Cyanobacteria
Anabaena cylindrica PCC 7122 YP 007157469 1 272123 Cyanobacteria
Nostoc azollae 0708 YP 003722539 1 551115 Cyanobacteria

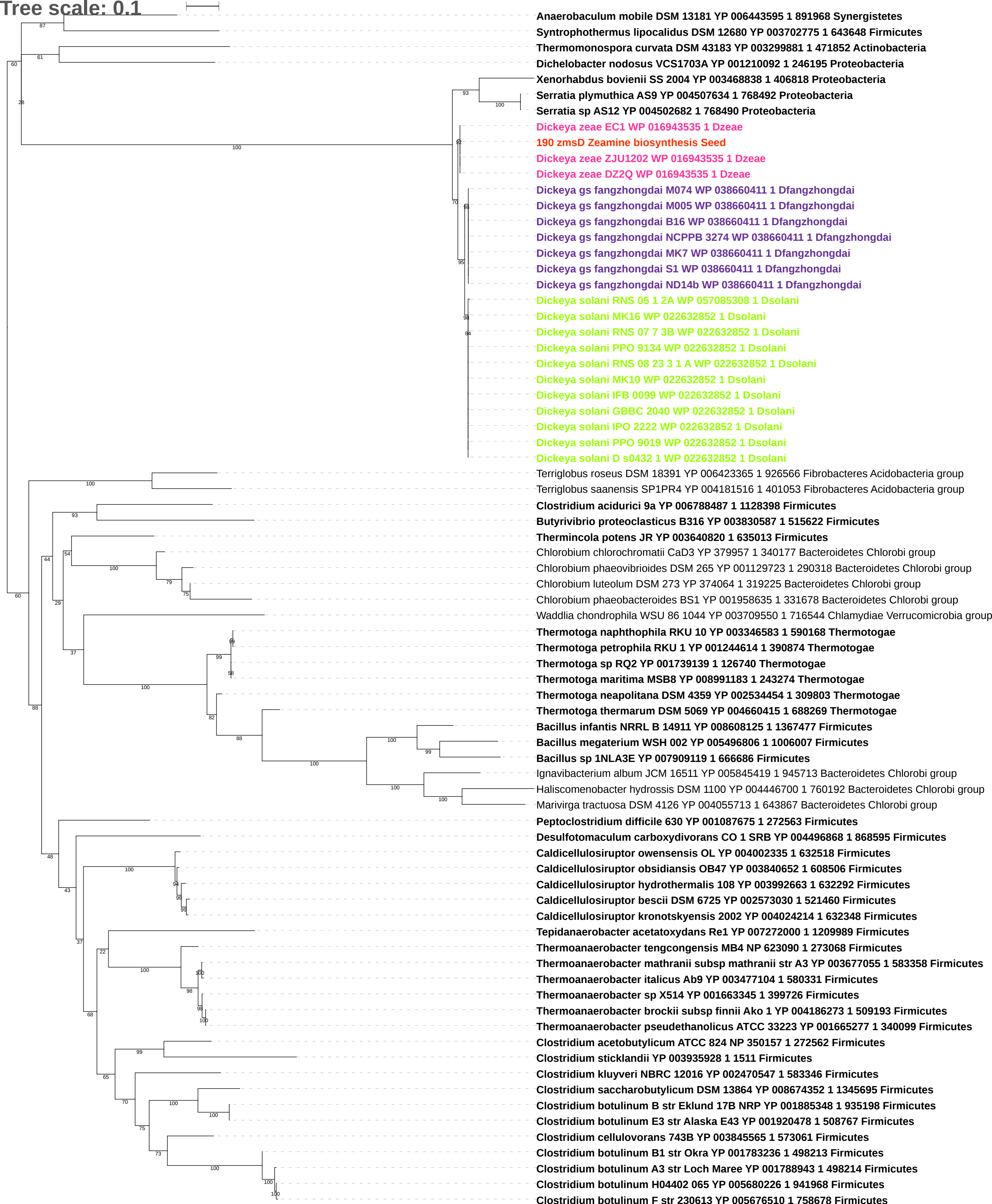
Tree scale: 0.1
Anaerobaculum mobile DSM 13181 YP 006443595 1 891968 Synergistetes
Syntrophothermus lipocalidus DSM 12680 YP 003702775 1 643648 Firmicutes
Thermomonospora curvata DSM 43183 YP 003299881 1 471852 Actinobacteria
Dichelobacter nodosus VCS1703A YP 001210092 1 246195 Proteobacteria
Xenorhabdus bovienii SS 2004 YP 003468838 1 406818 Proteobacteria
Serratia plymuthica AS9 YP 004507634 1 768492 Proteobacteria
Serratia sp AS12 YP 004502682 1 768490 Proteobacteria
Dickeya zeae EC1 WP 016943535 1 Dzeae
190 zmsD Zeamine biosynthesis Seed
Dickeya zeae ZJU1202 WP 016943535 1 Dzeae
Dickeya zeae DZ2Q WP 016943535 1 Dzeae
Dickeya gs fangzhongdai M074 WP 038660411 1 Dfangzhongdai
Dickeya gs fangzhongdai M005 WP 038660411 1 Dfangzhongdai
Dickeya gs fangzhongdai B16 WP 038660411 1 Dfangzhongdai
Dickeya gs fangzhongdai NCPPB 3274 WP 038660411 1 Dfangzhongdai
Dickeya gs fangzhongdai MK7 WP 038660411 1 Dfangzhongdai
Dickeya gs fangzhongdai S1 WP 038660411 1 Dfangzhongdai
Dickeya gs fangzhongdai ND14b WP 038660411 1 Dfangzhongdai
Dickeya solani RNS 05 1 2A WP 057085308 1 Dsolani
Dickeya solani MK16 WP 022632852 1 Dsolani
Dickeya solani RNS 07 7 3B WP 022632852 1 Dsolani
Dickeya solani PPO 9134 WP 022632852 1 Dsolani
Dickeya solani RNS 08 23 3 1 A WP 022632852 1 Dsolani
Dickeya solani MK10 WP 022632852 1 Dsolani
Dickeya solani IFB 0099 WP 022632852 1 Dsolani
Dickeya solani GBBC 2040 WP 022632852 1 Dsolani
Dickeya solani IPO 2222 WP 022632852 1 Dsolani
Dickeya solani PPO 9019 WP 022632852 1 Dsolani
Dickeya solani D s0432 1 WP 022632852 1 Dsolani
Terriglobus roseus DSM 18391 YP 006423365 1 926566 Fibrobacteres Acidobacteria group
Terriglobus saanensis SP1PR4 YP 004181516 1 401053 Fibrobacteres Acidobacteria group
Clostridium acidurici 9a YP 006788487 1 1128398 Firmicutes
Butyrivibrio proteoclasticus B316 YP 003830587 1 515622 Firmicutes
Thermincola potens JR YP 003640820 1 635013 Firmicutes
Chlorobium chlorochromatii CaD3 YP 379957 1 340177 Bacteroidetes Chlorobi group
Chlorobium phaeovibrioides DSM 265 YP 001129723 1 290318 Bacteroidetes Chlorobi group
Chlorobium luteolum DSM 273 YP 374064 1 319225 Bacteroidetes Chlorobi group
Chlorobium phaeobacteroides BS1 YP 001958635 1 331678 Bacteroidetes Chlorobi group
Waddlia chondrophila WSU 86 1044 YP 003709550 1 716544 Chlamydiae Verrucomicrobia group
Thermotoga naphthophila RKU 10 YP 003346583 1 590168 Thermotogae
Thermotoga petrophila RKU 1 YP 001244614 1 390874 Thermotogae
Thermotoga sp RQ2 YP 001739139 1 126740 Thermotogae
Thermotoga maritima MSB8 YP 008991183 1 243274 Thermotogae
Thermotoga neapolitana DSM 4359 YP 002534454 1 309803 Thermotogae
Thermotoga thermarum DSM 5069 YP 004660415 1 688269 Thermotogae
Bacillus infantis NRRL B 14911 YP 008608125 1 1367477 Firmicutes
Bacillus megaterium WSH 002 YP 005496806 1 1006007 Firmicutes
Bacillus sp 1NLA3E YP 007909119 1 666686 Firmicutes
Ignavibacterium album JCM 16511 YP 005845419 1 945713 Bacteroidetes Chlorobi group
Haliscomenobacter hydrossis DSM 1100 YP 004446700 1 760192 Bacteroidetes Chlorobi group
Marivirga tractuosa DSM 4126 YP 004055713 1 643867 Bacteroidetes Chlorobi group
Peptoclostridium difficile 630 YP 001087675 1 272563 Firmicutes
Desulfotomaculum carboxydivorans CO 1 SRB YP 004496868 1 868595 Firmicutes
Caldicellulosiruptor owensensis OL YP 004002335 1 632518 Firmicutes
Caldicellulosiruptor obsidiansis OB47 YP 003840652 1 608506 Firmicutes
Caldicellulosiruptor hydrothermalis 108 YP 003992663 1 632292 Firmicutes
Caldicellulosiruptor bescii DSM 6725 YP 002573030 1 521460 Firmicutes
Caldicellulosiruptor kronotskyensis 2002 YP 004024214 1 632348 Firmicutes
Tepidanaerobacter acetatoxydans Re1 YP 007272000 1 1209989 Firmicutes
Thermoanaerobacter tengcongensis MB4 NP 623090 1 273068 Firmicutes
Thermoanaerobacter mathranii subsp mathranii str A3 YP 003677055 1 583358 Firmicutes
Thermoanaerobacter italicus Ab9 YP 003477104 1 580331 Firmicutes
Thermoanaerobacter sp X514 YP 001663345 1 399726 Firmicutes
Thermoanaerobacter brockii subsp finnii Ako 1 YP 004186273 1 509193 Firmicutes
Thermoanaerobacter pseudethanolicus ATCC 33223 YP 001665277 1 340099 Firmicutes
Clostridium acetobutylicum ATCC 824 NP 350157 1 272562 Firmicutes
Clostridium sticklandii YP 003935928 1 1511 Firmicutes
Clostridium kluyveri NBRC 12016 YP 002470547 1 583346 Firmicutes
Clostridium saccharobutylicum DSM 13864 YP 008674352 1 1345695 Firmicutes
Clostridium botulinum B str Eklund 17B NRP YP 001885348 1 935198 Firmicutes
Clostridium botulinum E3 str Alaska E43 YP 001920478 1 508767 Firmicutes
Clostridium cellulovorans 743B YP 003845565 1 573061 Firmicutes
Clostridium botulinum B1 str Okra YP 001783236 1 498213 Firmicutes
Clostridium botulinum A3 str Loch Maree YP 001788943 1 498214 Firmicutes
Clostridium botulinum H04402 065 YP 005680226 1 941968 Firmicutes
Clostridium botulinum F str 230613 YP 005676510 1 758678 Firmicutes

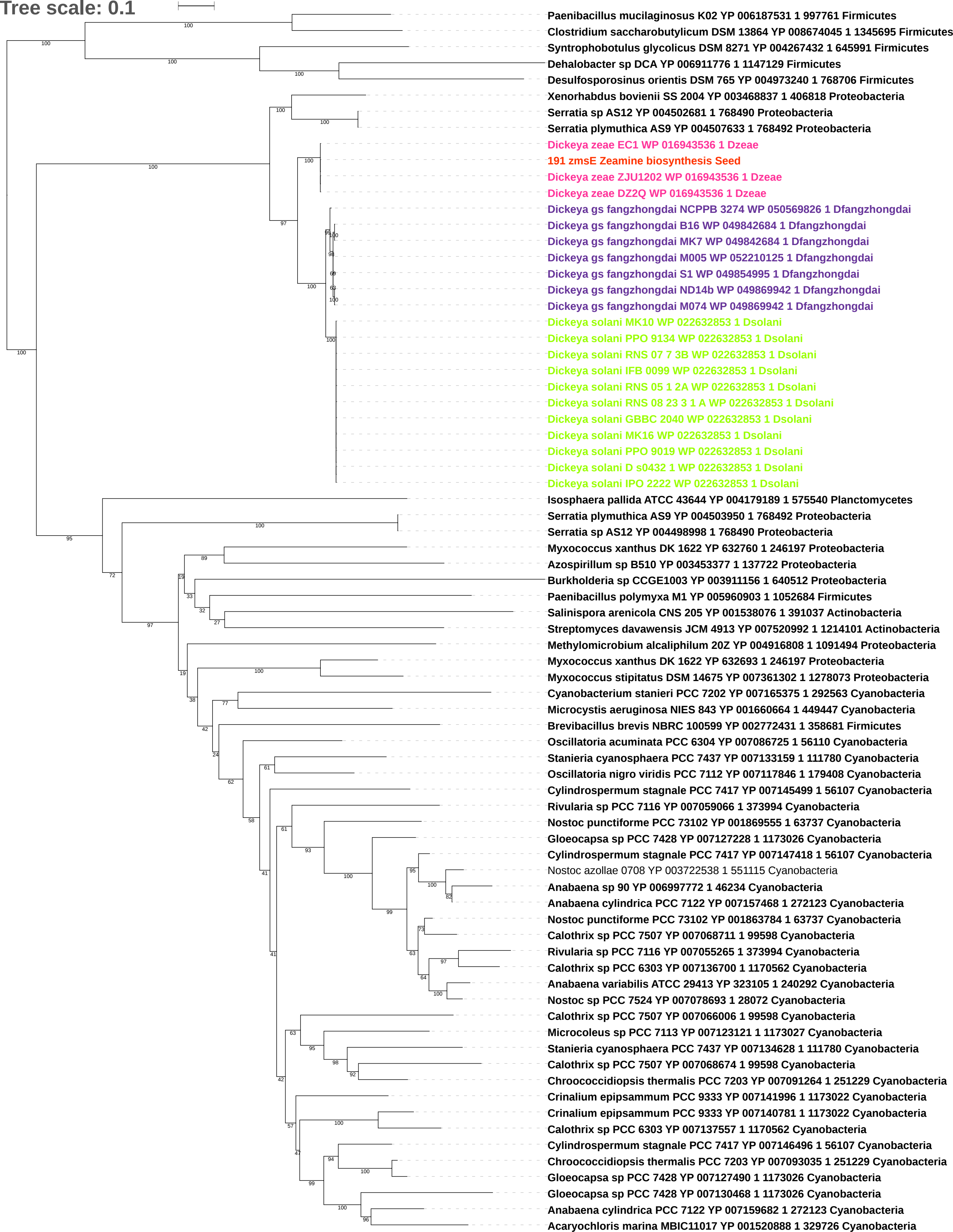
Tree scale: 0.1
Paenibacillus mucilaginosus K02 YP 006187531 1 997761 Firmicutes
Clostridium saccharobutylicum DSM 13864 YP 008674045 1 1345695 Firmicutes
Syntrophobotulus glycolicus DSM 8271 YP 004267432 1 645991 Firmicutes
Dehalobacter sp DCA YP 006911776 1 1147129 Firmicutes
Desulfosporosinus orientis DSM 765 YP 004973240 1 768706 Firmicutes
Xenorhabdus bovienii SS 2004 YP 003468837 1 406818 Proteobacteria
Serratia sp AS12 YP 004502681 1 768490 Proteobacteria
Serratia plymuthica AS9 YP 004507633 1 768492 Proteobacteria
Dickeya zeae EC1 WP 016943536 1 Dzeae
191 zmsE Zeamine biosynthesis Seed
Dickeya zeae ZJU1202 WP 016943536 1 Dzeae
Dickeya zeae DZ2Q WP 016943536 1 Dzeae
Dickeya gs fangzhongdai NCPPB 3274 WP 050569826 1 Dfangzhongdai
Dickeya gs fangzhongdai B16 WP 049842684 1 Dfangzhongdai
Dickeya gs fangzhongdai MK7 WP 049842684 1 Dfangzhongdai
Dickeya gs fangzhongdai M005 WP 052210125 1 Dfangzhongdai
Dickeya gs fangzhongdai S1 WP 049854995 1 Dfangzhongdai
Dickeya gs fangzhongdai ND14b WP 049869942 1 Dfangzhongdai
Dickeya gs fangzhongdai M074 WP 049869942 1 Dfangzhongdai
Dickeya solani MK10 WP 022632853 1 Dsolani
Dickeya solani PPO 9134 WP 022632853 1 Dsolani
Dickeya solani RNS 07 7 3B WP 022632853 1 Dsolani
Dickeya solani IFB 0099 WP 022632853 1 Dsolani
Dickeya solani RNS 05 1 2A WP 022632853 1 Dsolani
Dickeya solani RNS 08 23 3 1 A WP 022632853 1 Dsolani
Dickeya solani GBBC 2040 WP 022632853 1 Dsolani
Dickeya solani MK16 WP 022632853 1 Dsolani
Dickeya solani PPO 9019 WP 022632853 1 Dsolani
Dickeya solani D s0432 1 WP 022632853 1 Dsolani
Dickeya solani IPO 2222 WP 022632853 1 Dsolani
Isosphaera pallida ATCC 43644 YP 004179189 1 575540 Planctomycetes
Serratia plymuthica AS9 YP 004503950 1 768492 Proteobacteria
Serratia sp AS12 YP 004498998 1 768490 Proteobacteria
Myxococcus xanthus DK 1622 YP 632760 1 246197 Proteobacteria
Azospirillum sp B510 YP 003453377 1 137722 Proteobacteria
Burkholderia sp CCGE1003 YP 003911156 1 640512 Proteobacteria
Paenibacillus polymyxa M1 YP 005960903 1 1052684 Firmicutes
Salinispora arenicola CNS 205 YP 001538076 1 391037 Actinobacteria
Streptomyces davawensis JCM 4913 YP 007520992 1 1214101 Actinobacteria
Methylomicrobium alcaliphilum 20Z YP 004916808 1 1091494 Proteobacteria
Myxococcus xanthus DK 1622 YP 632693 1 246197 Proteobacteria
Myxococcus stipitatus DSM 14675 YP 007361302 1 1278073 Proteobacteria
Cyanobacterium stanieri PCC 7202 YP 007165375 1 292563 Cyanobacteria
Microcystis aeruginosa NIES 843 YP 001660664 1 449447 Cyanobacteria
Brevibacillus brevis NBRC 100599 YP 002772431 1 358681 Firmicutes
Oscillatoria acuminata PCC 6304 YP 007086725 1 56110 Cyanobacteria
Stanieria cyanosphaera PCC 7437 YP 007133159 1 111780 Cyanobacteria
Oscillatoria nigro viridis PCC 7112 YP 007117846 1 179408 Cyanobacteria
Cylindrospermum stagnale PCC 7417 YP 007145499 1 56107 Cyanobacteria
Rivularia sp PCC 7116 YP 007059066 1 373994 Cyanobacteria
Nostoc punctiforme PCC 73102 YP 001869555 1 63737 Cyanobacteria
Gloeocapsa sp PCC 7428 YP 007127228 1 1173026 Cyanobacteria
Cylindrospermum stagnale PCC 7417 YP 007147418 1 56107 Cyanobacteria
Nostoc azollae 0708 YP 003722538 1 551115 Cyanobacteria
Anabaena sp 90 YP 006997772 1 46234 Cyanobacteria
Anabaena cylindrica PCC 7122 YP 007157468 1 272123 Cyanobacteria
Nostoc punctiforme PCC 73102 YP 001863784 1 63737 Cyanobacteria
Calothrix sp PCC 7507 YP 007068711 1 99598 Cyanobacteria
Rivularia sp PCC 7116 YP 007055265 1 373994 Cyanobacteria
Calothrix sp PCC 6303 YP 007136700 1 1170562 Cyanobacteria
Anabaena variabilis ATCC 29413 YP 323105 1 240292 Cyanobacteria
Nostoc sp PCC 7524 YP 007078693 1 28072 Cyanobacteria
Calothrix sp PCC 7507 YP 007066006 1 99598 Cyanobacteria
Microcoleus sp PCC 7113 YP 007123121 1 1173027 Cyanobacteria
Stanieria cyanosphaera PCC 7437 YP 007134628 1 111780 Cyanobacteria
Calothrix sp PCC 7507 YP 007068674 1 99598 Cyanobacteria
Chroococcidiopsis thermalis PCC 7203 YP 007091264 1 251229 Cyanobacteria
Crinalium epipsammum PCC 9333 YP 007141996 1 1173022 Cyanobacteria
Crinalium epipsammum PCC 9333 YP 007140781 1 1173022 Cyanobacteria
Calothrix sp PCC 6303 YP 007137557 1 1170562 Cyanobacteria
Cylindrospermum stagnale PCC 7417 YP 007146496 1 56107 Cyanobacteria
Chroococcidiopsis thermalis PCC 7203 YP 007093035 1 251229 Cyanobacteria
Gloeocapsa sp PCC 7428 YP 007127490 1 1173026 Cyanobacteria
Gloeocapsa sp PCC 7428 YP 007130468 1 1173026 Cyanobacteria
Anabaena cylindrica PCC 7122 YP 007159682 1 272123 Cyanobacteria
Acaryochloris marina MBIC11017 YP 001520888 1 329726 Cyanobacteria

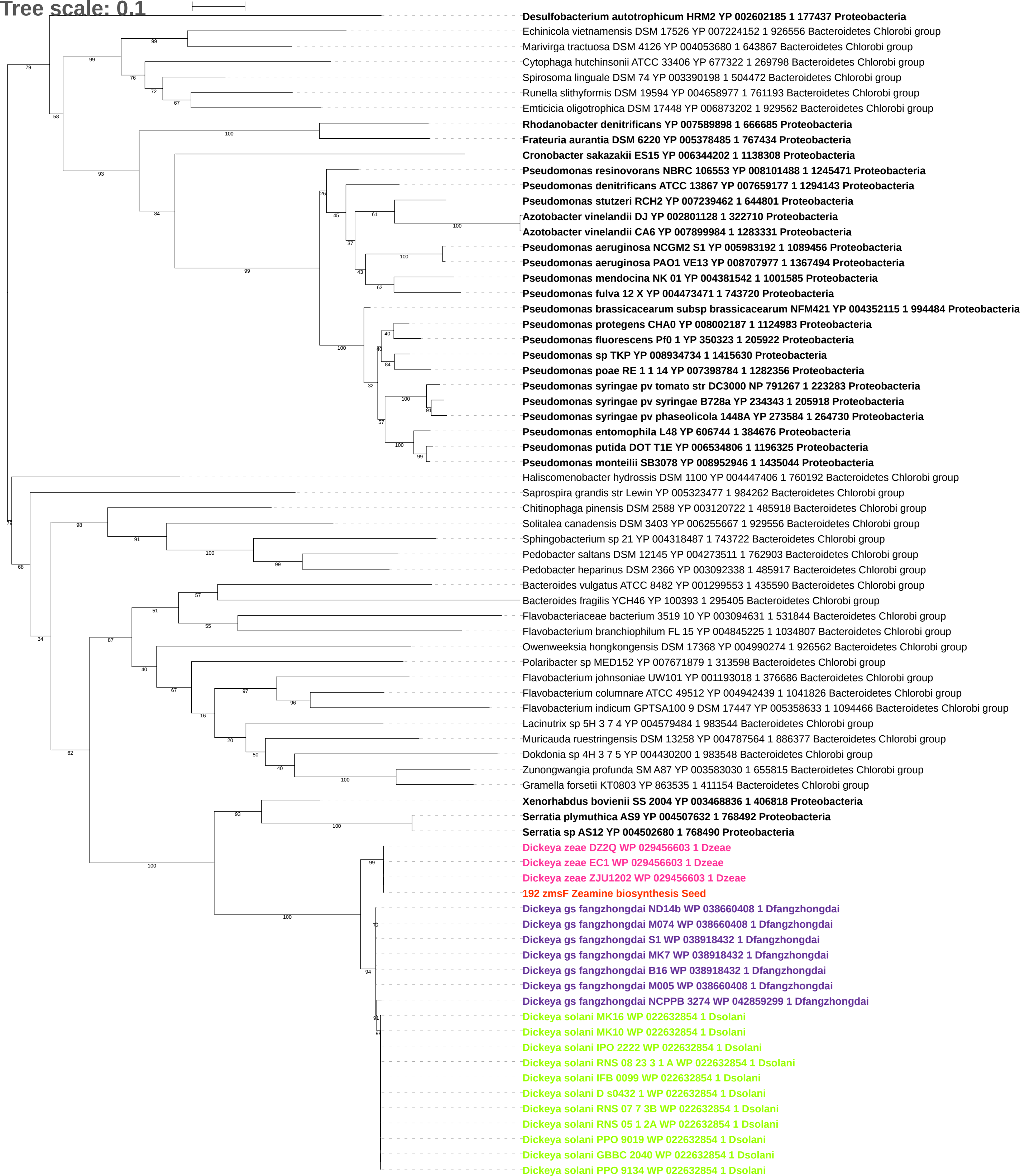
Tree scale: 0.1
Desulfobacterium autotrophicum HRM2 YP 002602185 1 177437 Proteobacteria
Echinicola vietnamensis DSM 17526 YP 007224152 1 926556 Bacteroidetes Chlorobi group
Marivirga tractuosa DSM 4126 YP 004053680 1 643867 Bacteroidetes Chlorobi group
Cytophaga hutchinsonii ATCC 33406 YP 677322 1 269798 Bacteroidetes Chlorobi group
Spirosoma linguale DSM 74 YP 003390198 1 504472 Bacteroidetes Chlorobi group
Runella slithyformis DSM 19594 YP 004658977 1 761193 Bacteroidetes Chlorobi group
Emticicia oligotrophica DSM 17448 YP 006873202 1 929562 Bacteroidetes Chlorobi group
Rhodanobacter denitrificans YP 007589898 1 666685 Proteobacteria
Frateuria aurantia DSM 6220 YP 005378485 1 767434 Proteobacteria
Cronobacter sakazakii ES15 YP 006344202 1 1138308 Proteobacteria
Pseudomonas resinovorans NBRC 106553 YP 008101488 1 1245471 Proteobacteria
Pseudomonas denitrificans ATCC 13867 YP 007659177 1 1294143 Proteobacteria
Pseudomonas stutzeri RCH2 YP 007239462 1 644801 Proteobacteria
Azotobacter vinelandii DJ YP 002801128 1 322710 Proteobacteria
Azotobacter vinelandii CA6 YP 007899984 1 1283331 Proteobacteria
Pseudomonas aeruginosa NCGM2 S1 YP 005983192 1 1089456 Proteobacteria
Pseudomonas aeruginosa PAO1 VE13 YP 008707977 1 1367494 Proteobacteria
Pseudomonas mendocina NK 01 YP 004381542 1 1001585 Proteobacteria
Pseudomonas fulva 12 X YP 004473471 1 743720 Proteobacteria
Pseudomonas brassicacearum subsp brassicacearum NFM421 YP 004352115 1 994484 Proteobacteria
Pseudomonas protegens CHA0 YP 008002187 1 1124983 Proteobacteria
Pseudomonas fluorescens Pf0 1 YP 350323 1 205922 Proteobacteria
Pseudomonas sp TKP YP 008934734 1 1415630 Proteobacteria
Pseudomonas poae RE 1 1 14 YP 007398784 1 1282356 Proteobacteria
Pseudomonas syringae pv tomato str DC3000 NP 791267 1 223283 Proteobacteria
Pseudomonas syringae pv syringae B728a YP 234343 1 205918 Proteobacteria
Pseudomonas syringae pv phaseolicola 1448A YP 273584 1 264730 Proteobacteria
Pseudomonas entomophila L48 YP 606744 1 384676 Proteobacteria
Pseudomonas putida DOT T1E YP 006534806 1 1196325 Proteobacteria
Pseudomonas monteilii SB3078 YP 008952946 1 1435044 Proteobacteria
Haliscomenobacter hydrossis DSM 1100 YP 004447406 1 760192 Bacteroidetes Chlorobi group
Saprospira grandis str Lewin YP 005323477 1 984262 Bacteroidetes Chlorobi group
Chitinophaga pinensis DSM 2588 YP 003120722 1 485918 Bacteroidetes Chlorobi group
Solitalea canadensis DSM 3403 YP 006255667 1 929556 Bacteroidetes Chlorobi group
Sphingobacterium sp 21 YP 004318487 1 743722 Bacteroidetes Chlorobi group
Pedobacter saltans DSM 12145 YP 004273511 1 762903 Bacteroidetes Chlorobi group
Pedobacter heparinus DSM 2366 YP 003092338 1 485917 Bacteroidetes Chlorobi group
Bacteroides vulgatus ATCC 8482 YP 001299553 1 435590 Bacteroidetes Chlorobi group
Bacteroides fragilis YCH46 YP 100393 1 295405 Bacteroidetes Chlorobi group
Flavobacteriaceae bacterium 3519 10 YP 003094631 1 531844 Bacteroidetes Chlorobi group
Flavobacterium branchiophilum FL 15 YP 004845225 1 1034807 Bacteroidetes Chlorobi group
Owenweeksia hongkongensis DSM 17368 YP 004990274 1 926562 Bacteroidetes Chlorobi group
Polaribacter sp MED152 YP 007671879 1 313598 Bacteroidetes Chlorobi group
Flavobacterium johnsoniae UW101 YP 001193018 1 376686 Bacteroidetes Chlorobi group
Flavobacterium columnare ATCC 49512 YP 004942439 1 1041826 Bacteroidetes Chlorobi group
Flavobacterium indicum GPTSA100 9 DSM 17447 YP 005358633 1 1094466 Bacteroidetes Chlorobi group
Lacinutrix sp 5H 3 7 4 YP 004579484 1 983544 Bacteroidetes Chlorobi group
Muricauda ruestringensis DSM 13258 YP 004787564 1 886377 Bacteroidetes Chlorobi group
Dokdonia sp 4H 3 7 5 YP 004430200 1 983548 Bacteroidetes Chlorobi group
Zunongwangia profunda SM A87 YP 003583030 1 655815 Bacteroidetes Chlorobi group
Gramella forsetii KT0803 YP 863535 1 411154 Bacteroidetes Chlorobi group
Xenorhabdus bovienii SS 2004 YP 003468836 1 406818 Proteobacteria
Serratia plymuthica AS9 YP 004507632 1 768492 Proteobacteria
Serratia sp AS12 YP 004502680 1 768490 Proteobacteria
Dickeya zeae DZ2Q WP 029456603 1 Dzeae
Dickeya zeae EC1 WP 029456603 1 Dzeae
Dickeya zeae ZJU1202 WP 029456603 1 Dzeae
192 zmsF Zeamine biosynthesis Seed
Dickeya gs fangzhongdai ND14b WP 038660408 1 Dfangzhongdai
Dickeya gs fangzhongdai M074 WP 038660408 1 Dfangzhongdai
Dickeya gs fangzhongdai S1 WP 038918432 1 Dfangzhongdai
Dickeya gs fangzhongdai MK7 WP 038918432 1 Dfangzhongdai
Dickeya gs fangzhongdai B16 WP 038918432 1 Dfangzhongdai
Dickeya gs fangzhongdai M005 WP 038660408 1 Dfangzhongdai
Dickeya gs fangzhongdai NCPPB 3274 WP 042859299 1 Dfangzhongdai
Dickeya solani MK16 WP 022632854 1 Dsolani
Dickeya solani MK10 WP 022632854 1 Dsolani
Dickeya solani IPO 2222 WP 022632854 1 Dsolani
Dickeya solani RNS 08 23 3 1 A WP 022632854 1 Dsolani
Dickeya solani IFB 0099 WP 022632854 1 Dsolani
Dickeya solani D s0432 1 WP 022632854 1 Dsolani
Dickeya solani RNS 07 7 3B WP 022632854 1 Dsolani
Dickeya solani RNS 05 1 2A WP 022632854 1 Dsolani
Dickeya solani PPO 9019 WP 022632854 1 Dsolani
Dickeya solani GBBC 2040 WP 022632854 1 Dsolani
Dickeya solani PPO 9134 WP 022632854 1 Dsolani

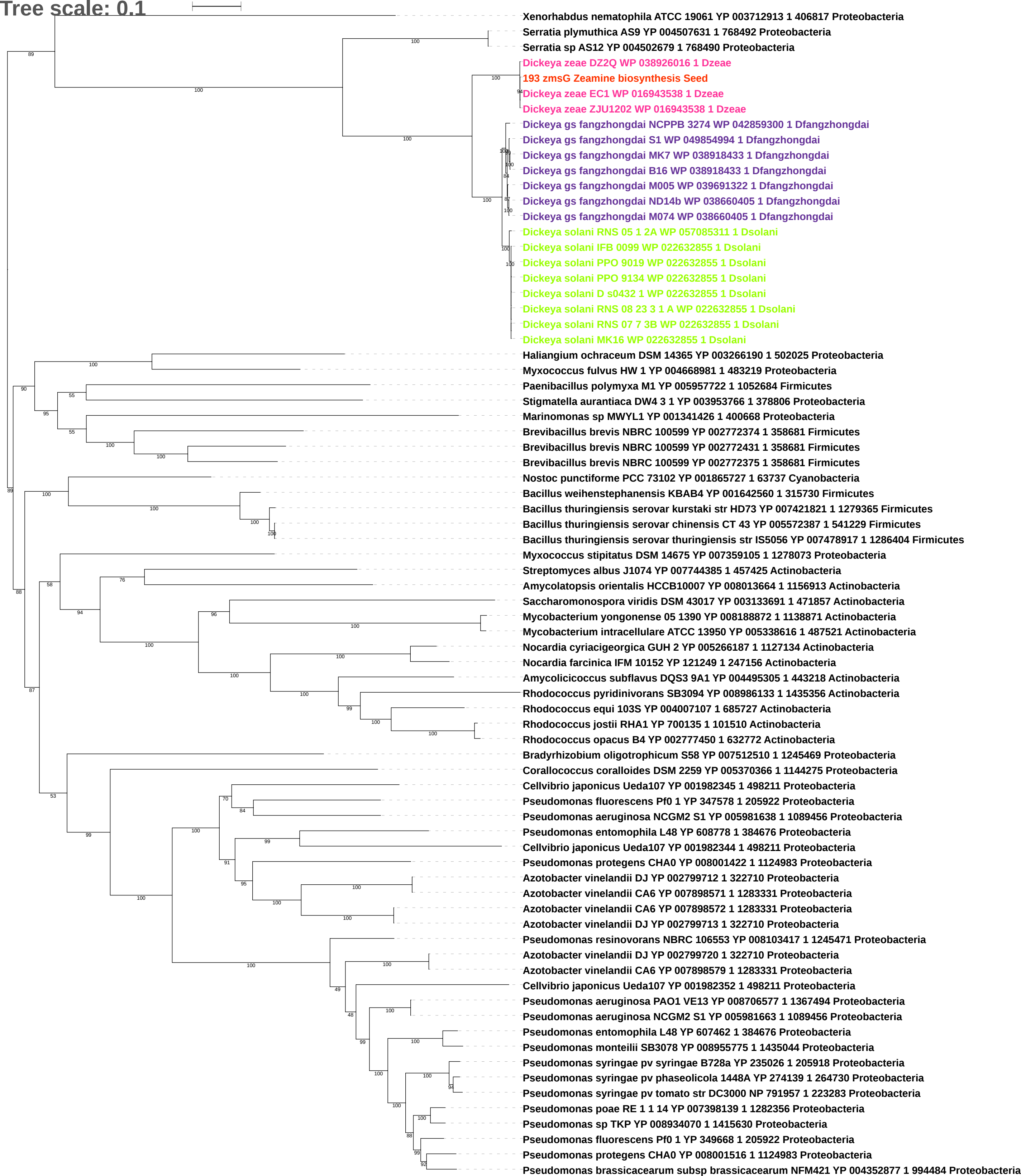
Tree scale: 0.1
Xenorhabdus nematophila ATCC 19061 YP 003712913 1 406817 Proteobacteria
Serratia plymuthica AS9 YP 004507631 1 768492 Proteobacteria
Serratia sp AS12 YP 004502679 1 768490 Proteobacteria
Dickeya zeae DZ2Q WP 038926016 1 Dzeae
193 zmsG Zeamine biosynthesis Seed
Dickeya zeae EC1 WP 016943538 1 Dzeae
Dickeya zeae ZJU1202 WP 016943538 1 Dzeae
Dickeya gs fangzhongdai NCPPB 3274 WP 042859300 1 Dfangzhongdai
Dickeya gs fangzhongdai S1 WP 049854994 1 Dfangzhongdai
Dickeya gs fangzhongdai MK7 WP 038918433 1 Dfangzhongdai
Dickeya gs fangzhongdai B16 WP 038918433 1 Dfangzhongdai
Dickeya gs fangzhongdai M005 WP 039691322 1 Dfangzhongdai
Dickeya gs fangzhongdai ND14b WP 038660405 1 Dfangzhongdai
Dickeya gs fangzhongdai M074 WP 038660405 1 Dfangzhongdai
Dickeya solani RNS 05 1 2A WP 057085311 1 Dsolani
Dickeya solani IFB 0099 WP 022632855 1 Dsolani
Dickeya solani PPO 9019 WP 022632855 1 Dsolani
Dickeya solani PPO 9134 WP 022632855 1 Dsolani
Dickeya solani D s0432 1 WP 022632855 1 Dsolani
Dickeya solani RNS 08 23 3 1 A WP 022632855 1 Dsolani
Dickeya solani RNS 07 7 3B WP 022632855 1 Dsolani
Dickeya solani MK16 WP 022632855 1 Dsolani
Haliangium ochraceum DSM 14365 YP 003266190 1 502025 Proteobacteria
Myxococcus fulvus HW 1 YP 004668981 1 483219 Proteobacteria
Paenibacillus polymyxa M1 YP 005957722 1 1052684 Firmicutes
Stigmatella aurantiaca DW4 3 1 YP 003953766 1 378806 Proteobacteria
Marinomonas sp MWYL1 YP 001341426 1 400668 Proteobacteria
Brevibacillus brevis NBRC 100599 YP 002772374 1 358681 Firmicutes
Brevibacillus brevis NBRC 100599 YP 002772431 1 358681 Firmicutes
Brevibacillus brevis NBRC 100599 YP 002772375 1 358681 Firmicutes
Nostoc punctiforme PCC 73102 YP 001865727 1 63737 Cyanobacteria
Bacillus weihenstephanensis KBAB4 YP 001642560 1 315730 Firmicutes
Bacillus thuringiensis serovar kurstaki str HD73 YP 007421821 1 1279365 Firmicutes
Bacillus thuringiensis serovar chinensis CT 43 YP 005572387 1 541229 Firmicutes
Bacillus thuringiensis serovar thuringiensis str IS5056 YP 007478917 1 1286404 Firmicutes
Myxococcus stipitatus DSM 14675 YP 007359105 1 1278073 Proteobacteria
Streptomyces albus J1074 YP 007744385 1 457425 Actinobacteria
Amycolatopsis orientalis HCCB10007 YP 008013664 1 1156913 Actinobacteria
Saccharomonospora viridis DSM 43017 YP 003133691 1 471857 Actinobacteria
Mycobacterium yongonense 05 1390 YP 008188872 1 1138871 Actinobacteria
Mycobacterium intracellulare ATCC 13950 YP 005338616 1 487521 Actinobacteria
Nocardia cyriacigeorgica GUH 2 YP 005266187 1 1127134 Actinobacteria
Nocardia farcinica IFM 10152 YP 121249 1 247156 Actinobacteria
Amycolicicoccus subflavus DQS3 9A1 YP 004495305 1 443218 Actinobacteria
Rhodococcus pyridinivorans SB3094 YP 008986133 1 1435356 Actinobacteria
Rhodococcus equi 103S YP 004007107 1 685727 Actinobacteria
Rhodococcus jostii RHA1 YP 700135 1 101510 Actinobacteria
Rhodococcus opacus B4 YP 002777450 1 632772 Actinobacteria
Bradyrhizobium oligotrophicum S58 YP 007512510 1 1245469 Proteobacteria
Corallococcus coralloides DSM 2259 YP 005370366 1 1144275 Proteobacteria
Cellvibrio japonicus Ueda107 YP 001982345 1 498211 Proteobacteria
Pseudomonas fluorescens Pf0 1 YP 347578 1 205922 Proteobacteria
Pseudomonas aeruginosa NCGM2 S1 YP 005981638 1 1089456 Proteobacteria
Pseudomonas entomophila L48 YP 608778 1 384676 Proteobacteria
Cellvibrio japonicus Ueda107 YP 001982344 1 498211 Proteobacteria
Pseudomonas protegens CHA0 YP 008001422 1 1124983 Proteobacteria
Azotobacter vinelandii DJ YP 002799712 1 322710 Proteobacteria
Azotobacter vinelandii CA6 YP 007898571 1 1283331 Proteobacteria
Azotobacter vinelandii CA6 YP 007898572 1 1283331 Proteobacteria
Azotobacter vinelandii DJ YP 002799713 1 322710 Proteobacteria
Pseudomonas resinovorans NBRC 106553 YP 008103417 1 1245471 Proteobacteria
Azotobacter vinelandii DJ YP 002799720 1 322710 Proteobacteria
Azotobacter vinelandii CA6 YP 007898579 1 1283331 Proteobacteria
Cellvibrio japonicus Ueda107 YP 001982352 1 498211 Proteobacteria
Pseudomonas aeruginosa PAO1 VE13 YP 008706577 1 1367494 Proteobacteria
Pseudomonas aeruginosa NCGM2 S1 YP 005981663 1 1089456 Proteobacteria
Pseudomonas entomophila L48 YP 607462 1 384676 Proteobacteria
Pseudomonas monteilii SB3078 YP 008955775 1 1435044 Proteobacteria
Pseudomonas syringae pv syringae B728a YP 235026 1 205918 Proteobacteria
Pseudomonas syringae pv phaseolicola 1448A YP 274139 1 264730 Proteobacteria
Pseudomonas syringae pv tomato str DC3000 NP 791957 1 223283 Proteobacteria
Pseudomonas poae RE 1 1 14 YP 007398139 1 1282356 Proteobacteria
Pseudomonas sp TKP YP 008934070 1 1415630 Proteobacteria
Pseudomonas fluorescens Pf0 1 YP 349668 1 205922 Proteobacteria
Pseudomonas protegens CHA0 YP 008001516 1 1124983 Proteobacteria
Pseudomonas brassicacearum subsp brassicacearum NFM421 YP 004352877 1 994484 Proteobacteria

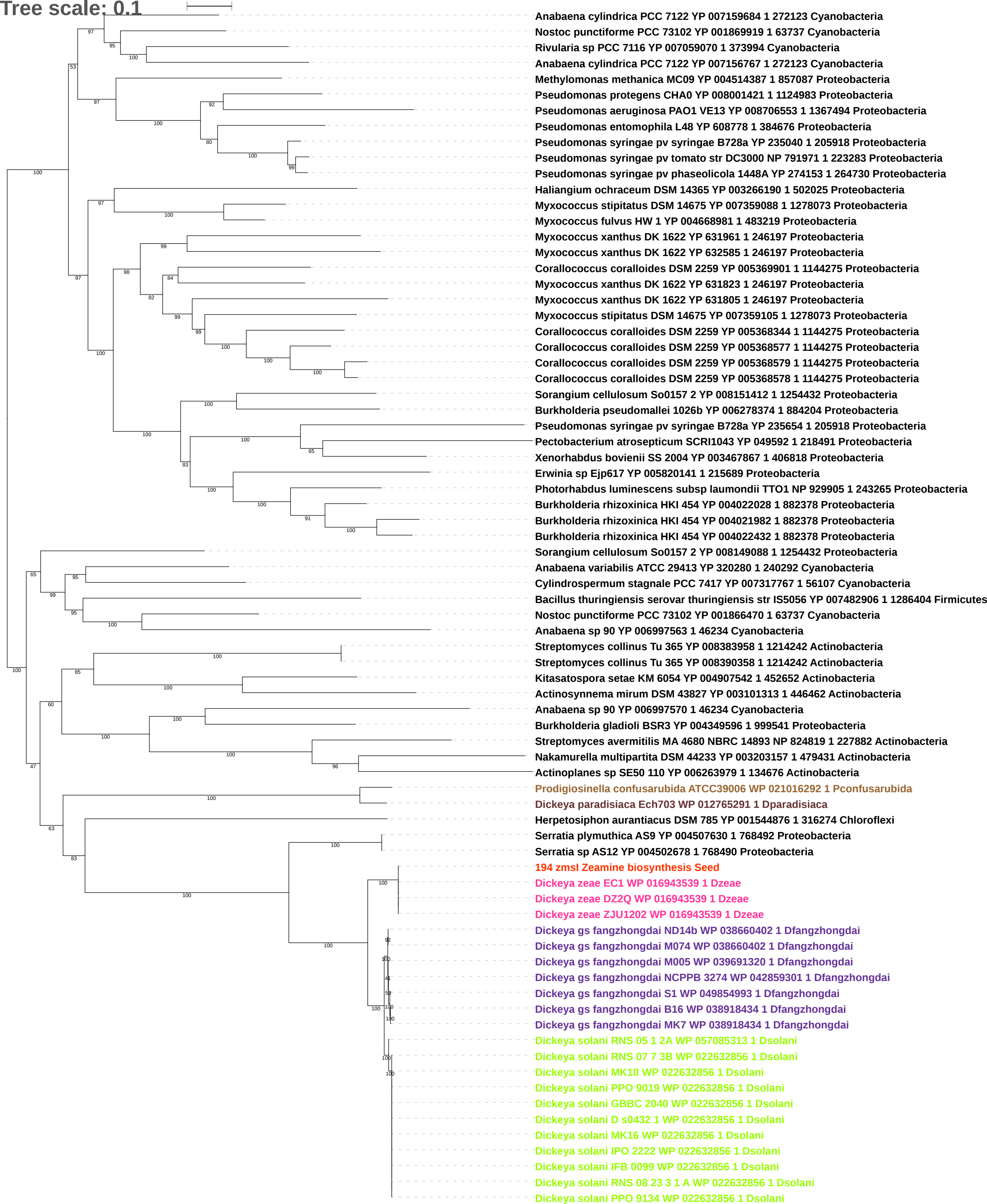
Tree scale: 0.1
Anabaena cylindrica PCC 7122 YP 007159684 1 272123 Cyanobacteria
Nostoc punctiforme PCC 73102 YP 001869919 1 63737 Cyanobacteria
Rivularia sp PCC 7116 YP 007059070 1 373994 Cyanobacteria
Anabaena cylindrica PCC 7122 YP 007156767 1 272123 Cyanobacteria
Methylomonas methanica MC09 YP 004514387 1 857087 Proteobacteria
Pseudomonas protegens CHA0 YP 008001421 1 1124983 Proteobacteria
Pseudomonas aeruginosa PAO1 VE13 YP 008706553 1 1367494 Proteobacteria
Pseudomonas entomophila L48 YP 608778 1 384676 Proteobacteria
Pseudomonas syringae pv syringae B728a YP 235040 1 205918 Proteobacteria
Pseudomonas syringae pv tomato str DC3000 NP 791971 1 223283 Proteobacteria
Pseudomonas syringae pv phaseolicola 1448A YP 274153 1 264730 Proteobacteria
Haliangium ochraceum DSM 14365 YP 003266190 1 502025 Proteobacteria
Myxococcus stipitatus DSM 14675 YP 007359088 1 1278073 Proteobacteria
Myxococcus fulvus HW 1 YP 004668981 1 483219 Proteobacteria
Myxococcus xanthus DK 1622 YP 631961 1 246197 Proteobacteria
Myxococcus xanthus DK 1622 YP 632585 1 246197 Proteobacteria
Corallococcus coralloides DSM 2259 YP 005369901 1 1144275 Proteobacteria
Myxococcus xanthus DK 1622 YP 631823 1 246197 Proteobacteria
Myxococcus xanthus DK 1622 YP 631805 1 246197 Proteobacteria
Myxococcus stipitatus DSM 14675 YP 007359105 1 1278073 Proteobacteria
Corallococcus coralloides DSM 2259 YP 005368344 1 1144275 Proteobacteria
Corallococcus coralloides DSM 2259 YP 005368577 1 1144275 Proteobacteria
Corallococcus coralloides DSM 2259 YP 005368579 1 1144275 Proteobacteria
Corallococcus coralloides DSM 2259 YP 005368578 1 1144275 Proteobacteria
Sorangium cellulosum So0157 2 YP 008151412 1 1254432 Proteobacteria
Burkholderia pseudomallei 1026b YP 006278374 1 884204 Proteobacteria
Pseudomonas syringae pv syringae B728a YP 235654 1 205918 Proteobacteria
Pectobacterium atrosepticum SCRI1043 YP 049592 1 218491 Proteobacteria
Xenorhabdus bovienii SS 2004 YP 003467867 1 406818 Proteobacteria
Erwinia sp Ejp617 YP 005820141 1 215689 Proteobacteria
Photorhabdus luminescens subsp laumondii TTO1 NP 929905 1 243265 Proteobacteria
Burkholderia rhizoxinica HKI 454 YP 004022028 1 882378 Proteobacteria
Burkholderia rhizoxinica HKI 454 YP 004021982 1 882378 Proteobacteria
Burkholderia rhizoxinica HKI 454 YP 004022432 1 882378 Proteobacteria
Sorangium cellulosum So0157 2 YP 008149088 1 1254432 Proteobacteria
Anabaena variabilis ATCC 29413 YP 320280 1 240292 Cyanobacteria
Cylindrospermum stagnale PCC 7417 YP 007317767 1 56107 Cyanobacteria
Bacillus thuringiensis serovar thuringiensis str IS5056 YP 007482906 1 1286404 Firmicutes
Nostoc punctiforme PCC 73102 YP 001866470 1 63737 Cyanobacteria
Anabaena sp 90 YP 006997563 1 46234 Cyanobacteria
Streptomyces collinus Tu 365 YP 008383958 1 1214242 Actinobacteria
Streptomyces collinus Tu 365 YP 008390358 1 1214242 Actinobacteria
Kitasatospora setae KM 6054 YP 004907542 1 452652 Actinobacteria
Actinosynnema mirum DSM 43827 YP 003101313 1 446462 Actinobacteria
Anabaena sp 90 YP 006997570 1 46234 Cyanobacteria
Burkholderia gladioli BSR3 YP 004349596 1 999541 Proteobacteria
Streptomyces avermitilis MA 4680 NBRC 14893 NP 824819 1 227882 Actinobacteria
Nakamurella multipartita DSM 44233 YP 003203157 1 479431 Actinobacteria
Actinoplanes sp SE50 110 YP 006263979 1 134676 Actinobacteria
Prodigiosinella confusarubida ATCC39006 WP 021016292 1 Pconfusarubida
Dickeya paradisiaca Ech703 WP 012765291 1 Dparadisiaca
Herpetosiphon aurantiacus DSM 785 YP 001544876 1 316274 Chloroflexi
Serratia plymuthica AS9 YP 004507630 1 768492 Proteobacteria
Serratia sp AS12 YP 004502678 1 768490 Proteobacteria
194 zmsI Zeamine biosynthesis Seed
Dickeya zeae EC1 WP 016943539 1 Dzeae
Dickeya zeae DZ2Q WP 016943539 1 Dzeae
Dickeya zeae ZJU1202 WP 016943539 1 Dzeae
Dickeya gs fangzhongdai ND14b WP 038660402 1 Dfangzhongdai
Dickeya gs fangzhongdai M074 WP 038660402 1 Dfangzhongdai
Dickeya gs fangzhongdai M005 WP 039691320 1 Dfangzhongdai
Dickeya gs fangzhongdai NCPPB 3274 WP 042859301 1 Dfangzhongdai
Dickeya gs fangzhongdai S1 WP 049854993 1 Dfangzhongdai
Dickeya gs fangzhongdai B16 WP 038918434 1 Dfangzhongdai
Dickeya gs fangzhongdai MK7 WP 038918434 1 Dfangzhongdai
Dickeya solani RNS 05 1 2A WP 057085313 1 Dsolani
Dickeya solani RNS 07 7 3B WP 022632856 1 Dsolani
Dickeya solani MK10 WP 022632856 1 Dsolani
Dickeya solani PPO 9019 WP 022632856 1 Dsolani
Dickeya solani GBBC 2040 WP 022632856 1 Dsolani
Dickeya solani D s0432 1 WP 022632856 1 Dsolani
Dickeya solani MK16 WP 022632856 1 Dsolani
Dickeya solani IPO 2222 WP 022632856 1 Dsolani
Dickeya solani IFB 0099 WP 022632856 1 Dsolani
Dickeya solani RNS 08 23 3 1 A WP 022632856 1 Dsolani
Dickeya solani PPO 9134 WP 022632856 1 Dsolani

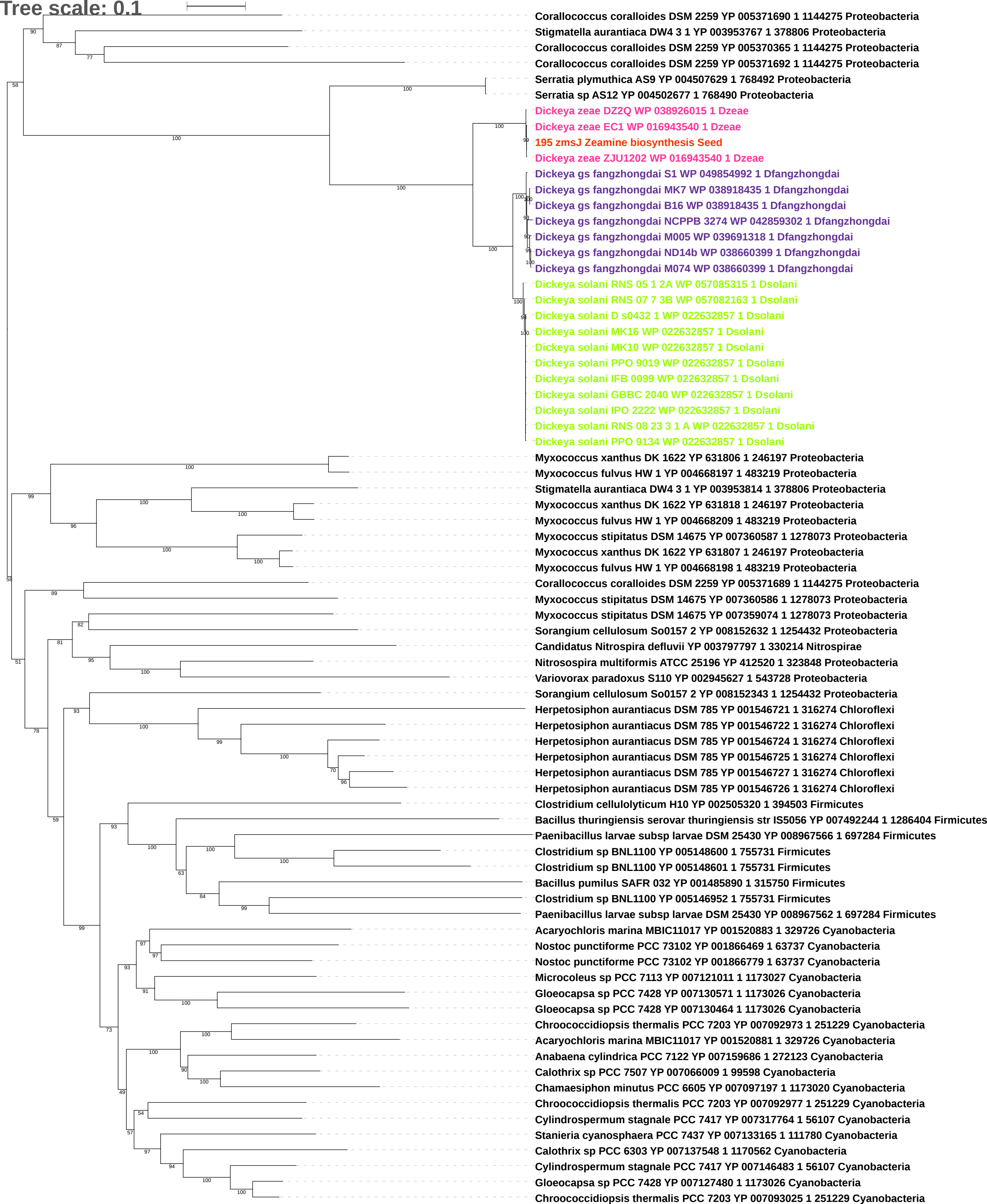

Tree scale: 0.1
Corallococcus coralloides DSM 2259 YP 005371690 1 1144275 Proteobacteria
Stigmatella aurantiaca DW4 3 1 YP 003953767 1 378806 Proteobacteria
Corallococcus coralloides DSM 2259 YP 005370365 1 1144275 Proteobacteria
Corallococcus coralloides DSM 2259 YP 005371692 1 1144275 Proteobacteria
Serratia plymuthica AS9 YP 004507629 1 768492 Proteobacteria
Serratia sp AS12 YP 004502677 1 768490 Proteobacteria
Dickeya zeae DZ2Q WP 038926015 1 Dzeae
Dickeya zeae EC1 WP 016943540 1 Dzeae
195 zmsJ Zeamine biosynthesis Seed
Dickeya zeae ZJU1202 WP 016943540 1 Dzeae
Dickeya gs fangzhongdai S1 WP 049854992 1 Dfangzhongdai
Dickeya gs fangzhongdai MK7 WP 038918435 1 Dfangzhongdai
Dickeya gs fangzhongdai B16 WP 038918435 1 Dfangzhongdai
Dickeya gs fangzhongdai NCPPB 3274 WP 042859302 1 Dfangzhongdai
Dickeya gs fangzhongdai M005 WP 039691318 1 Dfangzhongdai
Dickeya gs fangzhongdai ND14b WP 038660399 1 Dfangzhongdai
Dickeya gs fangzhongdai M074 WP 038660399 1 Dfangzhongdai
Dickeya solani RNS 05 1 2A WP 057085315 1 Dsolani
Dickeya solani RNS 07 7 3B WP 057082163 1 Dsolani
Dickeya solani D s0432 1 WP 022632857 1 Dsolani
Dickeya solani MK16 WP 022632857 1 Dsolani
Dickeya solani MK10 WP 022632857 1 Dsolani
Dickeya solani PPO 9019 WP 022632857 1 Dsolani
Dickeya solani IFB 0099 WP 022632857 1 Dsolani
Dickeya solani GBBC 2040 WP 022632857 1 Dsolani
Dickeya solani IPO 2222 WP 022632857 1 Dsolani
Dickeya solani RNS 08 23 3 1 A WP 022632857 1 Dsolani
Dickeya solani PPO 9134 WP 022632857 1 Dsolani
Myxococcus xanthus DK 1622 YP 631806 1 246197 Proteobacteria
Myxococcus fulvus HW 1 YP 004668197 1 483219 Proteobacteria
Stigmatella aurantiaca DW4 3 1 YP 003953814 1 378806 Proteobacteria
Myxococcus xanthus DK 1622 YP 631818 1 246197 Proteobacteria
Myxococcus fulvus HW 1 YP 004668209 1 483219 Proteobacteria
Myxococcus stipitatus DSM 14675 YP 007360587 1 1278073 Proteobacteria
Myxococcus xanthus DK 1622 YP 631807 1 246197 Proteobacteria
Myxococcus fulvus HW 1 YP 004668198 1 483219 Proteobacteria
Corallococcus coralloides DSM 2259 YP 005371689 1 1144275 Proteobacteria
Myxococcus stipitatus DSM 14675 YP 007360586 1 1278073 Proteobacteria
Myxococcus stipitatus DSM 14675 YP 007359074 1 1278073 Proteobacteria
Sorangium cellulosum So0157 2 YP 008152632 1 1254432 Proteobacteria
Candidatus Nitrospira defluvii YP 003797797 1 330214 Nitrospirae
Nitrosospira multiformis ATCC 25196 YP 412520 1 323848 Proteobacteria
Variovorax paradoxus S110 YP 002945627 1 543728 Proteobacteria
Sorangium cellulosum So0157 2 YP 008152343 1 1254432 Proteobacteria
Herpetosiphon aurantiacus DSM 785 YP 001546721 1 316274 Chloroflexi
Herpetosiphon aurantiacus DSM 785 YP 001546722 1 316274 Chloroflexi
Herpetosiphon aurantiacus DSM 785 YP 001546724 1 316274 Chloroflexi
Herpetosiphon aurantiacus DSM 785 YP 001546725 1 316274 Chloroflexi
Herpetosiphon aurantiacus DSM 785 YP 001546727 1 316274 Chloroflexi
Herpetosiphon aurantiacus DSM 785 YP 001546726 1 316274 Chloroflexi
Clostridium cellulolyticum H10 YP 002505320 1 394503 Firmicutes
Bacillus thuringiensis serovar thuringiensis str IS5056 YP 007492244 1 1286404 Firmicutes
Paenibacillus larvae subsp larvae DSM 25430 YP 008967566 1 697284 Firmicutes
Clostridium sp BNL1100 YP 005148600 1 755731 Firmicutes
Clostridium sp BNL1100 YP 005148601 1 755731 Firmicutes
Bacillus pumilus SAFR 032 YP 001485890 1 315750 Firmicutes
Clostridium sp BNL1100 YP 005146952 1 755731 Firmicutes
Paenibacillus larvae subsp larvae DSM 25430 YP 008967562 1 697284 Firmicutes
Acaryochloris marina MBIC11017 YP 001520883 1 329726 Cyanobacteria
Nostoc punctiforme PCC 73102 YP 001866469 1 63737 Cyanobacteria
Nostoc punctiforme PCC 73102 YP 001866779 1 63737 Cyanobacteria
Microcoleus sp PCC 7113 YP 007121011 1 1173027 Cyanobacteria
Gloeocapsa sp PCC 7428 YP 007130571 1 1173026 Cyanobacteria
Gloeocapsa sp PCC 7428 YP 007130464 1 1173026 Cyanobacteria
Chroococcidiopsis thermalis PCC 7203 YP 007092973 1 251229 Cyanobacteria
Acaryochloris marina MBIC11017 YP 001520881 1 329726 Cyanobacteria
Anabaena cylindrica PCC 7122 YP 007159686 1 272123 Cyanobacteria
Calothrix sp PCC 7507 YP 007066009 1 99598 Cyanobacteria
Chamaesiphon minutus PCC 6605 YP 007097197 1 1173020 Cyanobacteria
Chroococcidiopsis thermalis PCC 7203 YP 007092977 1 251229 Cyanobacteria
Cylindrospermum stagnale PCC 7417 YP 007317764 1 56107 Cyanobacteria
Stanieria cyanosphaera PCC 7437 YP 007133165 1 111780 Cyanobacteria
Calothrix sp PCC 6303 YP 007137548 1 1170562 Cyanobacteria
Cylindrospermum stagnale PCC 7417 YP 007146483 1 56107 Cyanobacteria
Gloeocapsa sp PCC 7428 YP 007127480 1 1173026 Cyanobacteria
Chroococcidiopsis thermalis PCC 7203 YP 007093025 1 251229 Cyanobacteria

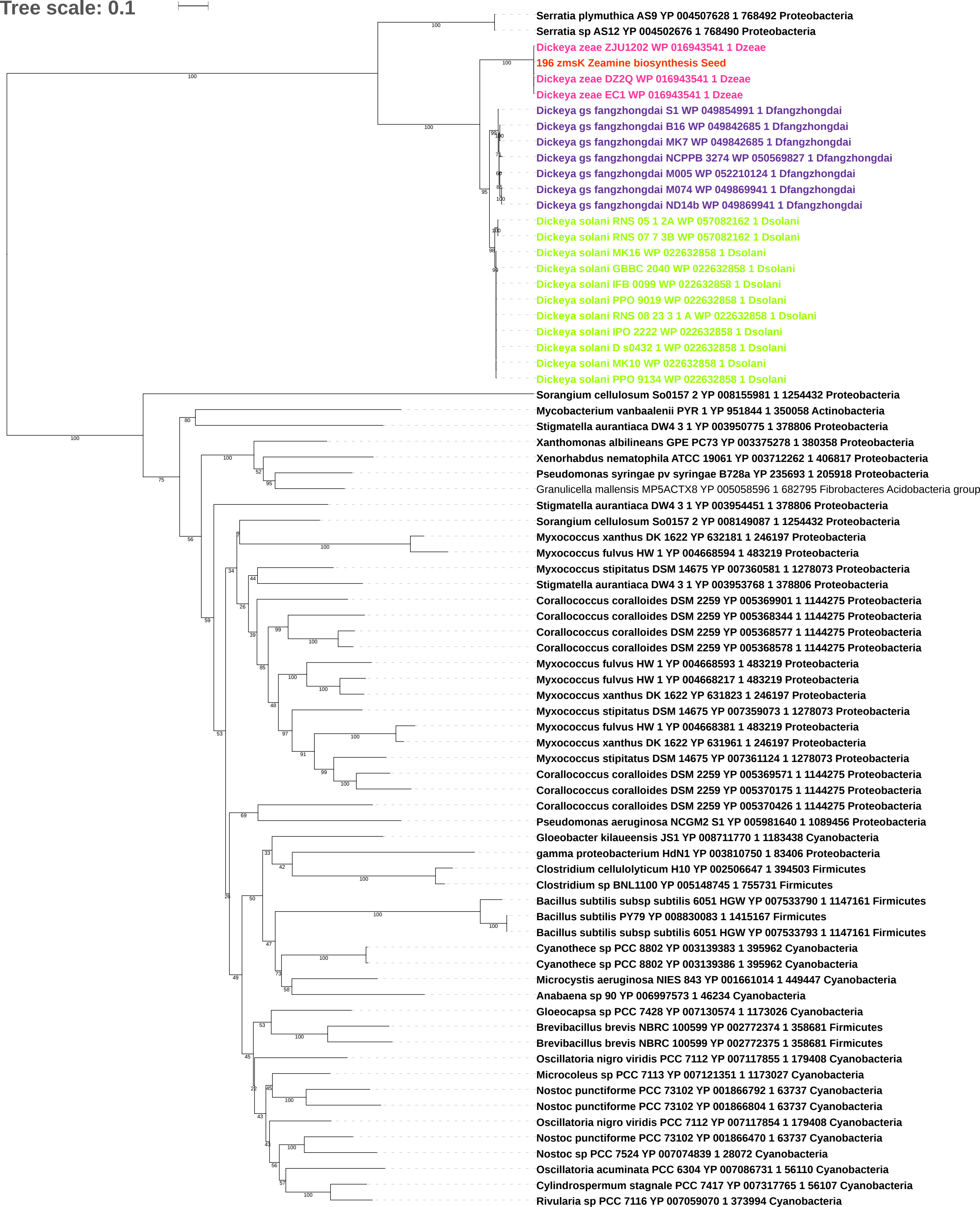

Tree scale: 0.1
Serratia plymuthica AS9 YP 004507628 1 768492 Proteobacteria
Serratia sp AS12 YP 004502676 1 768490 Proteobacteria
Dickeya zeae ZJU1202 WP 016943541 1 Dzeae
196 zmsK Zeamine biosynthesis Seed
Dickeya zeae DZ2Q WP 016943541 1 Dzeae
Dickeya zeae EC1 WP 016943541 1 Dzeae
Dickeya gs fangzhongdai S1 WP 049854991 1 Dfangzhongdai
Dickeya gs fangzhongdai B16 WP 049842685 1 Dfangzhongdai
Dickeya gs fangzhongdai MK7 WP 049842685 1 Dfangzhongdai
Dickeya gs fangzhongdai NCPPB 3274 WP 050569827 1 Dfangzhongdai
Dickeya gs fangzhongdai M005 WP 052210124 1 Dfangzhongdai
Dickeya gs fangzhongdai M074 WP 049869941 1 Dfangzhongdai
Dickeya gs fangzhongdai ND14b WP 049869941 1 Dfangzhongdai
Dickeya solani RNS 05 1 2A WP 057082162 1 Dsolani
Dickeya solani RNS 07 7 3B WP 057082162 1 Dsolani
Dickeya solani MK16 WP 022632858 1 Dsolani
Dickeya solani GBBC 2040 WP 022632858 1 Dsolani
Dickeya solani IFB 0099 WP 022632858 1 Dsolani
Dickeya solani PPO 9019 WP 022632858 1 Dsolani
Dickeya solani RNS 08 23 3 1 A WP 022632858 1 Dsolani
Dickeya solani IPO 2222 WP 022632858 1 Dsolani
Dickeya solani D s0432 1 WP 022632858 1 Dsolani
Dickeya solani MK10 WP 022632858 1 Dsolani
Dickeya solani PPO 9134 WP 022632858 1 Dsolani
Sorangium cellulosum So0157 2 YP 008155981 1 1254432 Proteobacteria
Mycobacterium vanbaalenii PYR 1 YP 951844 1 350058 Actinobacteria
Stigmatella aurantiaca DW4 3 1 YP 003950775 1 378806 Proteobacteria
Xanthomonas albilineans GPE PC73 YP 003375278 1 380358 Proteobacteria
Xenorhabdus nematophila ATCC 19061 YP 003712262 1 406817 Proteobacteria
Pseudomonas syringae pv syringae B728a YP 235693 1 205918 Proteobacteria
Granulicella mallensis MP5ACTX8 YP 005058596 1 682795 Fibrobacteres Acidobacteria group
Stigmatella aurantiaca DW4 3 1 YP 003954451 1 378806 Proteobacteria
Sorangium cellulosum So0157 2 YP 008149087 1 1254432 Proteobacteria
Myxococcus xanthus DK 1622 YP 632181 1 246197 Proteobacteria
Myxococcus fulvus HW 1 YP 004668594 1 483219 Proteobacteria
Myxococcus stipitatus DSM 14675 YP 007360581 1 1278073 Proteobacteria
Stigmatella aurantiaca DW4 3 1 YP 003953768 1 378806 Proteobacteria
Corallococcus coralloides DSM 2259 YP 005369901 1 1144275 Proteobacteria
Corallococcus coralloides DSM 2259 YP 005368344 1 1144275 Proteobacteria
Corallococcus coralloides DSM 2259 YP 005368577 1 1144275 Proteobacteria
Corallococcus coralloides DSM 2259 YP 005368578 1 1144275 Proteobacteria
Myxococcus fulvus HW 1 YP 004668593 1 483219 Proteobacteria
Myxococcus fulvus HW 1 YP 004668217 1 483219 Proteobacteria
Myxococcus xanthus DK 1622 YP 631823 1 246197 Proteobacteria
Myxococcus stipitatus DSM 14675 YP 007359073 1 1278073 Proteobacteria
Myxococcus fulvus HW 1 YP 004668381 1 483219 Proteobacteria
Myxococcus xanthus DK 1622 YP 631961 1 246197 Proteobacteria
Myxococcus stipitatus DSM 14675 YP 007361124 1 1278073 Proteobacteria
Corallococcus coralloides DSM 2259 YP 005369571 1 1144275 Proteobacteria
Corallococcus coralloides DSM 2259 YP 005370175 1 1144275 Proteobacteria
Corallococcus coralloides DSM 2259 YP 005370426 1 1144275 Proteobacteria
Pseudomonas aeruginosa NCGM2 S1 YP 005981640 1 1089456 Proteobacteria
Gloeobacter kilaueensis JS1 YP 008711770 1 1183438 Cyanobacteria
gamma proteobacterium HdN1 YP 003810750 1 83406 Proteobacteria
Clostridium cellulolyticum H10 YP 002506647 1 394503 Firmicutes
Clostridium sp BNL1100 YP 005148745 1 755731 Firmicutes
Bacillus subtilis subsp subtilis 6051 HGW YP 007533790 1 1147161 Firmicutes
Bacillus subtilis PY79 YP 008830083 1 1415167 Firmicutes
Bacillus subtilis subsp subtilis 6051 HGW YP 007533793 1 1147161 Firmicutes
Cyanothece sp PCC 8802 YP 003139383 1 395962 Cyanobacteria
Cyanothece sp PCC 8802 YP 003139386 1 395962 Cyanobacteria
Microcystis aeruginosa NIES 843 YP 001661014 1 449447 Cyanobacteria
Anabaena sp 90 YP 006997573 1 46234 Cyanobacteria
Gloeocapsa sp PCC 7428 YP 007130574 1 1173026 Cyanobacteria
Brevibacillus brevis NBRC 100599 YP 002772374 1 358681 Firmicutes
Brevibacillus brevis NBRC 100599 YP 002772375 1 358681 Firmicutes
Oscillatoria nigro viridis PCC 7112 YP 007117855 1 179408 Cyanobacteria
Microcoleus sp PCC 7113 YP 007121351 1 1173027 Cyanobacteria
Nostoc punctiforme PCC 73102 YP 001866792 1 63737 Cyanobacteria
Nostoc punctiforme PCC 73102 YP 001866804 1 63737 Cyanobacteria
Oscillatoria nigro viridis PCC 7112 YP 007117854 1 179408 Cyanobacteria
Nostoc punctiforme PCC 73102 YP 001866470 1 63737 Cyanobacteria
Nostoc sp PCC 7524 YP 007074839 1 28072 Cyanobacteria
Oscillatoria acuminata PCC 6304 YP 007086731 1 56110 Cyanobacteria
Cylindrospermum stagnale PCC 7417 YP 007317765 1 56107 Cyanobacteria
Rivularia sp PCC 7116 YP 007059070 1 373994 Cyanobacteria

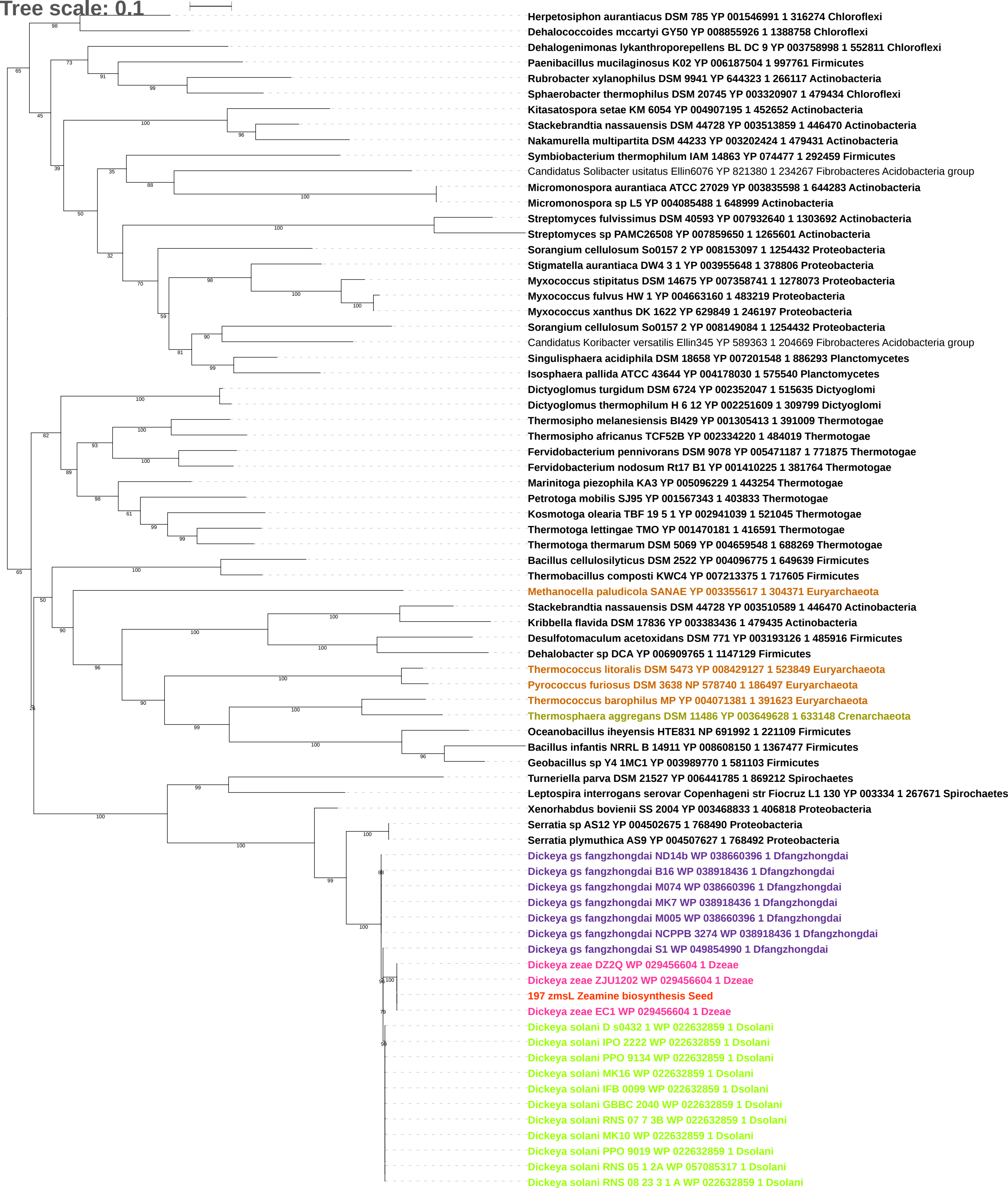
Tree scale: 0.1
Herpetosiphon aurantiacus DSM 785 YP 001546991 1 316274 Chloroflexi
Dehalococcoides mccartyi GY50 YP 008855926 1 1388758 Chloroflexi
Dehalogenimonas lykanthroporepellens BL DC 9 YP 003758998 1 552811 Chloroflexi
Paenibacillus mucilaginosus K02 YP 006187504 1 997761 Firmicutes
Rubrobacter xylanophilus DSM 9941 YP 644323 1 266117 Actinobacteria
Sphaerobacter thermophilus DSM 20745 YP 003320907 1 479434 Chloroflexi
Kitasatospora setae KM 6054 YP 004907195 1 452652 Actinobacteria
Stackebrandtia nassauensis DSM 44728 YP 003513859 1 446470 Actinobacteria
Nakamurella multipartita DSM 44233 YP 003202424 1 479431 Actinobacteria
Symbiobacterium thermophilum IAM 14863 YP 074477 1 292459 Firmicutes
Candidatus Solibacter usitatus Ellin6076 YP 821380 1 234267 Fibrobacteres Acidobacteria group
Micromonospora aurantiaca ATCC 27029 YP 003835598 1 644283 Actinobacteria
Micromonospora sp L5 YP 004085488 1 648999 Actinobacteria
Streptomyces fulvissimus DSM 40593 YP 007932640 1 1303692 Actinobacteria
Streptomyces sp PAMC26508 YP 007859650 1 1265601 Actinobacteria
Sorangium cellulosum So0157 2 YP 008153097 1 1254432 Proteobacteria
Stigmatella aurantiaca DW4 3 1 YP 003955648 1 378806 Proteobacteria
Myxococcus stipitatus DSM 14675 YP 007358741 1 1278073 Proteobacteria
Myxococcus fulvus HW 1 YP 004663160 1 483219 Proteobacteria
Myxococcus xanthus DK 1622 YP 629849 1 246197 Proteobacteria
Sorangium cellulosum So0157 2 YP 008149084 1 1254432 Proteobacteria
Candidatus Koribacter versatilis Ellin345 YP 589363 1 204669 Fibrobacteres Acidobacteria group
Singulisphaera acidiphila DSM 18658 YP 007201548 1 886293 Planctomycetes
Isosphaera pallida ATCC 43644 YP 004178030 1 575540 Planctomycetes
Dictyoglomus turgidum DSM 6724 YP 002352047 1 515635 Dictyoglomi
Dictyoglomus thermophilum H 6 12 YP 002251609 1 309799 Dictyoglomi
Thermosipho melanesiensis BI429 YP 001305413 1 391009 Thermotogae
Thermosipho africanus TCF52B YP 002334220 1 484019 Thermotogae
Fervidobacterium pennivorans DSM 9078 YP 005471187 1 771875 Thermotogae
Fervidobacterium nodosum Rt17 B1 YP 001410225 1 381764 Thermotogae
Marinitoga piezophila KA3 YP 005096229 1 443254 Thermotogae
Petrotoga mobilis SJ95 YP 001567343 1 403833 Thermotogae
Kosmotoga olearia TBF 19 5 1 YP 002941039 1 521045 Thermotogae
Thermotoga lettingae TMO YP 001470181 1 416591 Thermotogae
Thermotoga thermarum DSM 5069 YP 004659548 1 688269 Thermotogae
Bacillus cellulosilyticus DSM 2522 YP 004096775 1 649639 Firmicutes
Thermobacillus composti KWC4 YP 007213375 1 717605 Firmicutes
Methanocella paludicola SANAE YP 003355617 1 304371 Euryarchaeota
Stackebrandtia nassauensis DSM 44728 YP 003510589 1 446470 Actinobacteria
Kribbella flavida DSM 17836 YP 003383436 1 479435 Actinobacteria
Desulfotomaculum acetoxidans DSM 771 YP 003193126 1 485916 Firmicutes
Dehalobacter sp DCA YP 006909765 1 1147129 Firmicutes
Thermococcus litoralis DSM 5473 YP 008429127 1 523849 Euryarchaeota
Pyrococcus furiosus DSM 3638 NP 578740 1 186497 Euryarchaeota
Thermococcus barophilus MP YP 004071381 1 391623 Euryarchaeota
Thermosphaera aggregans DSM 11486 YP 003649628 1 633148 Crenarchaeota
Oceanobacillus iheyensis HTE831 NP 691992 1 221109 Firmicutes
Bacillus infantis NRRL B 14911 YP 008608150 1 1367477 Firmicutes
Geobacillus sp Y4 1MC1 YP 003989770 1 581103 Firmicutes
Turneriella parva DSM 21527 YP 006441785 1 869212 Spirochaetes
Leptospira interrogans serovar Copenhageni str Fiocruz L1 130 YP 003334 1 267671 Spirochaetes
Xenorhabdus bovienii SS 2004 YP 003468833 1 406818 Proteobacteria
Serratia sp AS12 YP 004502675 1 768490 Proteobacteria
Serratia plymuthica AS9 YP 004507627 1 768492 Proteobacteria
Dickeya gs fangzhongdai ND14b WP 038660396 1 Dfangzhongdai
Dickeya gs fangzhongdai B16 WP 038918436 1 Dfangzhongdai
Dickeya gs fangzhongdai M074 WP 038660396 1 Dfangzhongdai
Dickeya gs fangzhongdai MK7 WP 038918436 1 Dfangzhongdai
Dickeya gs fangzhongdai M005 WP 038660396 1 Dfangzhongdai
Dickeya gs fangzhongdai NCPPB 3274 WP 038918436 1 Dfangzhongdai
Dickeya gs fangzhongdai S1 WP 049854990 1 Dfangzhongdai
Dickeya zeae DZ2Q WP 029456604 1 Dzeae
Dickeya zeae ZJU1202 WP 029456604 1 Dzeae
197 zmsL Zeamine biosynthesis Seed
Dickeya zeae EC1 WP 029456604 1 Dzeae
Dickeya solani D s0432 1 WP 022632859 1 Dsolani
Dickeya solani IPO 2222 WP 022632859 1 Dsolani
Dickeya solani PPO 9134 WP 022632859 1 Dsolani
Dickeya solani MK16 WP 022632859 1 Dsolani
Dickeya solani IFB 0099 WP 022632859 1 Dsolani
Dickeya solani GBBC 2040 WP 022632859 1 Dsolani
Dickeya solani RNS 07 7 3B WP 022632859 1 Dsolani
Dickeya solani MK10 WP 022632859 1 Dsolani
Dickeya solani PPO 9019 WP 022632859 1 Dsolani
Dickeya solani RNS 05 1 2A WP 057085317 1 Dsolani
Dickeya solani RNS 08 23 3 1 A WP 022632859 1 Dsolani

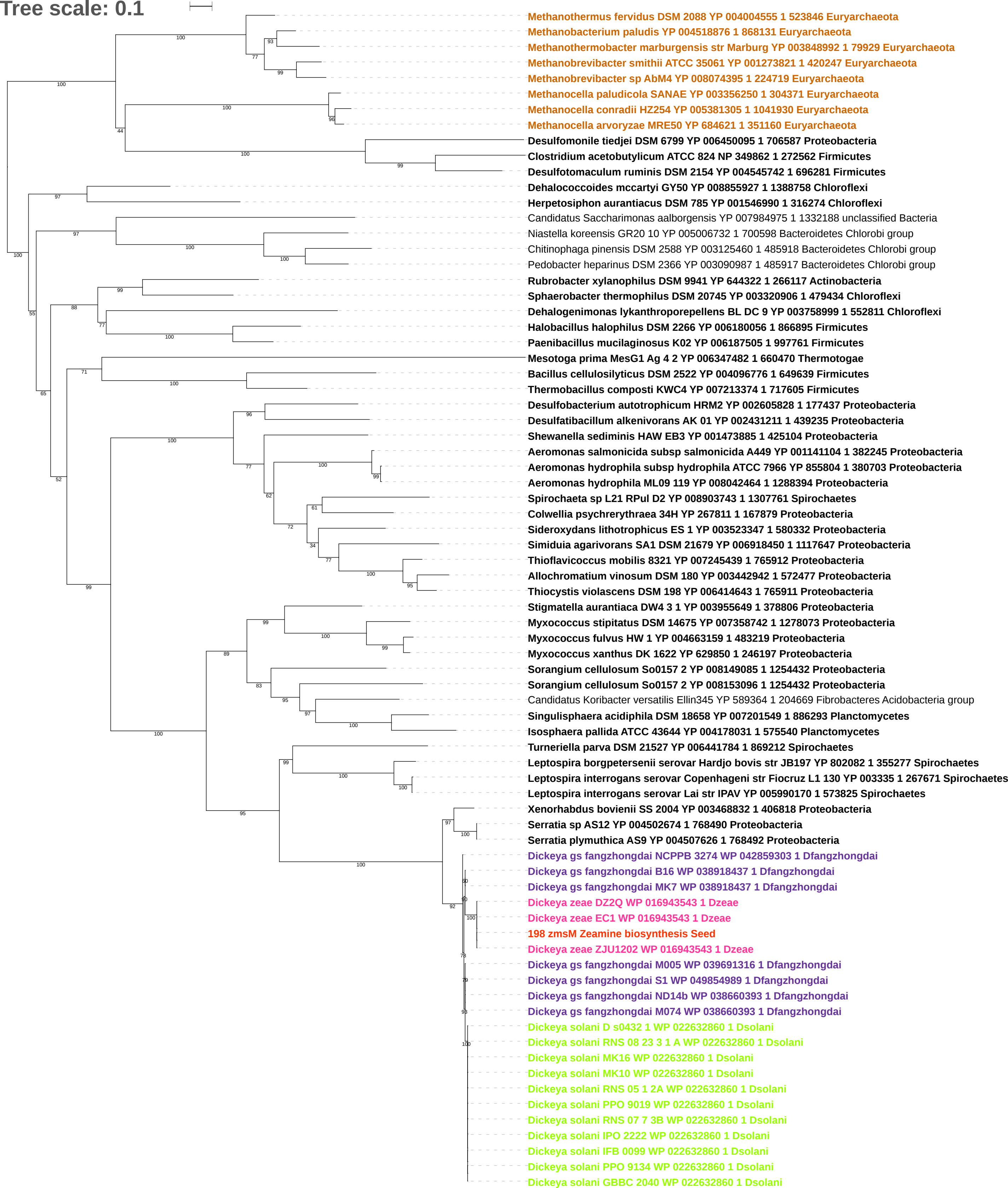
Tree scale: 0.1
Methanothermus fervidus DSM 2088 YP 004004555 1 523846 Euryarchaeota
Methanobacterium paludis YP 004518876 1 868131 Euryarchaeota
Methanothermobacter marburgensis str Marburg YP 003848992 1 79929 Euryarchaeota
Methanobrevibacter smithii ATCC 35061 YP 001273821 1 420247 Euryarchaeota
Methanobrevibacter sp AbM4 YP 008074395 1 224719 Euryarchaeota
Methanocella paludicola SANAE YP 003356250 1 304371 Euryarchaeota
Methanocella conradii HZ254 YP 005381305 1 1041930 Euryarchaeota
Methanocella arvoryzae MRE50 YP 684621 1 351160 Euryarchaeota
Desulfomonile tiedjei DSM 6799 YP 006450095 1 706587 Proteobacteria
Clostridium acetobutylicum ATCC 824 NP 349862 1 272562 Firmicutes
Desulfotomaculum ruminis DSM 2154 YP 004545742 1 696281 Firmicutes
Dehalococcoides mccartyi GY50 YP 008855927 1 1388758 Chloroflexi
Herpetosiphon aurantiacus DSM 785 YP 001546990 1 316274 Chloroflexi
Candidatus Saccharimonas aalborgensis YP 007984975 1 1332188 unclassified Bacteria
Niastella koreensis GR20 10 YP 005006732 1 700598 Bacteroidetes Chlorobi group
Chitinophaga pinensis DSM 2588 YP 003125460 1 485918 Bacteroidetes Chlorobi group
Pedobacter heparinus DSM 2366 YP 003090987 1 485917 Bacteroidetes Chlorobi group
Rubrobacter xylanophilus DSM 9941 YP 644322 1 266117 Actinobacteria
Sphaerobacter thermophilus DSM 20745 YP 003320906 1 479434 Chloroflexi
Dehalogenimonas lykanthroporepellens BL DC 9 YP 003758999 1 552811 Chloroflexi
Halobacillus halophilus DSM 2266 YP 006180056 1 866895 Firmicutes
Paenibacillus mucilaginosus K02 YP 006187505 1 997761 Firmicutes
Mesotoga prima MesG1 Ag 4 2 YP 006347482 1 660470 Thermotogae
Bacillus cellulosilyticus DSM 2522 YP 004096776 1 649639 Firmicutes
Thermobacillus composti KWC4 YP 007213374 1 717605 Firmicutes
Desulfobacterium autotrophicum HRM2 YP 002605828 1 177437 Proteobacteria
Desulfatibacillum alkenivorans AK 01 YP 002431211 1 439235 Proteobacteria
Shewanella sediminis HAW EB3 YP 001473885 1 425104 Proteobacteria
Aeromonas salmonicida subsp salmonicida A449 YP 001141104 1 382245 Proteobacteria
Aeromonas hydrophila subsp hydrophila ATCC 7966 YP 855804 1 380703 Proteobacteria
Aeromonas hydrophila ML09 119 YP 008042464 1 1288394 Proteobacteria
Spirochaeta sp L21 RPul D2 YP 008903743 1 1307761 Spirochaetes
Colwellia psychrerythraea 34H YP 267811 1 167879 Proteobacteria
Sideroxydans lithotrophicus ES 1 YP 003523347 1 580332 Proteobacteria
Simiduia agarivorans SA1 DSM 21679 YP 006918450 1 1117647 Proteobacteria
Thioflavicoccus mobilis 8321 YP 007245439 1 765912 Proteobacteria
Allochromatium vinosum DSM 180 YP 003442942 1 572477 Proteobacteria
Thiocystis violascens DSM 198 YP 006414643 1 765911 Proteobacteria
Stigmatella aurantiaca DW4 3 1 YP 003955649 1 378806 Proteobacteria
Myxococcus stipitatus DSM 14675 YP 007358742 1 1278073 Proteobacteria
Myxococcus fulvus HW 1 YP 004663159 1 483219 Proteobacteria
Myxococcus xanthus DK 1622 YP 629850 1 246197 Proteobacteria
Sorangium cellulosum So0157 2 YP 008149085 1 1254432 Proteobacteria
Sorangium cellulosum So0157 2 YP 008153096 1 1254432 Proteobacteria
Candidatus Koribacter versatilis Ellin345 YP 589364 1 204669 Fibrobacteres Acidobacteria group
Singulisphaera acidiphila DSM 18658 YP 007201549 1 886293 Planctomycetes
Isosphaera pallida ATCC 43644 YP 004178031 1 575540 Planctomycetes
Turneriella parva DSM 21527 YP 006441784 1 869212 Spirochaetes
Leptospira borgpetersenii serovar Hardjo bovis str JB197 YP 802082 1 355277 Spirochaetes
Leptospira interrogans serovar Copenhageni str Fiocruz L1 130 YP 003335 1 267671 Spirochaetes
Leptospira interrogans serovar Lai str IPAV YP 005990170 1 573825 Spirochaetes
Xenorhabdus bovienii SS 2004 YP 003468832 1 406818 Proteobacteria
Serratia sp AS12 YP 004502674 1 768490 Proteobacteria
Serratia plymuthica AS9 YP 004507626 1 768492 Proteobacteria
Dickeya gs fangzhongdai NCPPB 3274 WP 042859303 1 Dfangzhongdai
Dickeya gs fangzhongdai B16 WP 038918437 1 Dfangzhongdai
Dickeya gs fangzhongdai MK7 WP 038918437 1 Dfangzhongdai
Dickeya zeae DZ2Q WP 016943543 1 Dzeae
Dickeya zeae EC1 WP 016943543 1 Dzeae
198 zmsM Zeamine biosynthesis Seed
Dickeya zeae ZJU1202 WP 016943543 1 Dzeae
Dickeya gs fangzhongdai M005 WP 039691316 1 Dfangzhongdai
Dickeya gs fangzhongdai S1 WP 049854989 1 Dfangzhongdai
Dickeya gs fangzhongdai ND14b WP 038660393 1 Dfangzhongdai
Dickeya gs fangzhongdai M074 WP 038660393 1 Dfangzhongdai
Dickeya solani D s0432 1 WP 022632860 1 Dsolani
Dickeya solani RNS 08 23 3 1 A WP 022632860 1 Dsolani
Dickeya solani MK16 WP 022632860 1 Dsolani
Dickeya solani MK10 WP 022632860 1 Dsolani
Dickeya solani RNS 05 1 2A WP 022632860 1 Dsolani
Dickeya solani PPO 9019 WP 022632860 1 Dsolani
Dickeya solani RNS 07 7 3B WP 022632860 1 Dsolani
Dickeya solani IPO 2222 WP 022632860 1 Dsolani
Dickeya solani IFB 0099 WP 022632860 1 Dsolani
Dickeya solani PPO 9134 WP 022632860 1 Dsolani
Dickeya solani GBBC 2040 WP 022632860 1 Dsolani

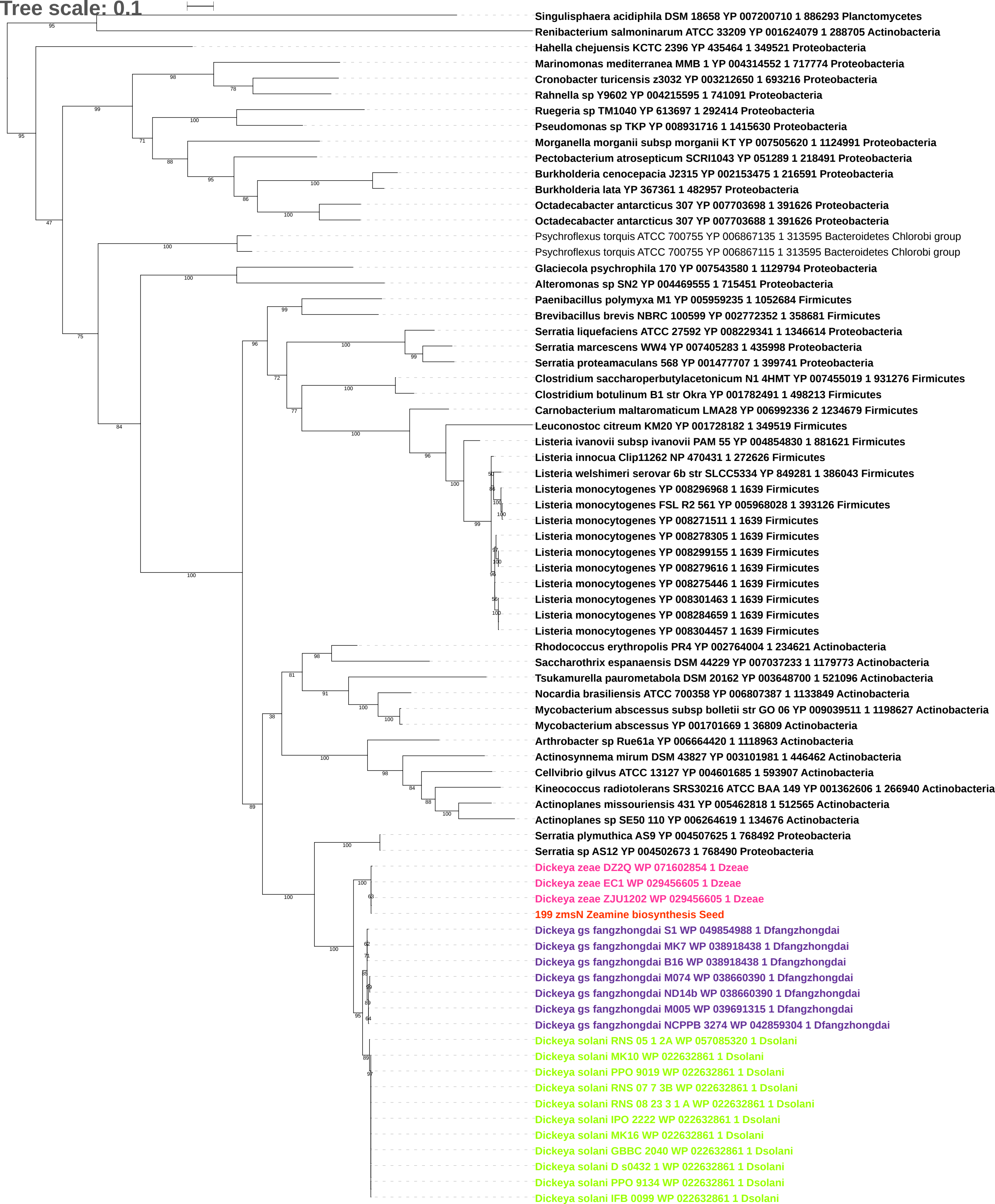
Tree scale: 0.1
Singulisphaera acidiphila DSM 18658 YP 007200710 1 886293 Planctomycetes
Renibacterium salmoninarum ATCC 33209 YP 001624079 1 288705 Actinobacteria
Hahella chejuensis KCTC 2396 YP 435464 1 349521 Proteobacteria
Marinomonas mediterranea MMB 1 YP 004314552 1 717774 Proteobacteria
Cronobacter turicensis z3032 YP 003212650 1 693216 Proteobacteria
Rahnella sp Y9602 YP 004215595 1 741091 Proteobacteria
Ruegeria sp TM1040 YP 613697 1 292414 Proteobacteria
Pseudomonas sp TKP YP 008931716 1 1415630 Proteobacteria
Morganella morganii subsp morganii KT YP 007505620 1 1124991 Proteobacteria
Pectobacterium atrosepticum SCRI1043 YP 051289 1 218491 Proteobacteria
Burkholderia cenocepacia J2315 YP 002153475 1 216591 Proteobacteria
Burkholderia lata YP 367361 1 482957 Proteobacteria
Octadecabacter antarcticus 307 YP 007703698 1 391626 Proteobacteria
Octadecabacter antarcticus 307 YP 007703688 1 391626 Proteobacteria
Psychroflexus torquis ATCC 700755 YP 006867135 1 313595 Bacteroidetes Chlorobi group
Psychroflexus torquis ATCC 700755 YP 006867115 1 313595 Bacteroidetes Chlorobi group
Glaciecola psychrophila 170 YP 007543580 1 1129794 Proteobacteria
Alteromonas sp SN2 YP 004469555 1 715451 Proteobacteria
Paenibacillus polymyxa M1 YP 005959235 1 1052684 Firmicutes
Brevibacillus brevis NBRC 100599 YP 002772352 1 358681 Firmicutes
Serratia liquefaciens ATCC 27592 YP 008229341 1 1346614 Proteobacteria
Serratia marcescens WW4 YP 007405283 1 435998 Proteobacteria
Serratia proteamaculans 568 YP 001477707 1 399741 Proteobacteria
Clostridium saccharoperbutylacetonicum N1 4HMT YP 007455019 1 931276 Firmicutes
Clostridium botulinum B1 str Okra YP 001782491 1 498213 Firmicutes
Carnobacterium maltaromaticum LMA28 YP 006992336 2 1234679 Firmicutes
Leuconostoc citreum KM20 YP 001728182 1 349519 Firmicutes
Listeria ivanovii subsp ivanovii PAM 55 YP 004854830 1 881621 Firmicutes
Listeria innocua Clip11262 NP 470431 1 272626 Firmicutes
Listeria welshimeri serovar 6b str SLCC5334 YP 849281 1 386043 Firmicutes
Listeria monocytogenes YP 008296968 1 1639 Firmicutes
Listeria monocytogenes FSL R2 561 YP 005968028 1 393126 Firmicutes
Listeria monocytogenes YP 008271511 1 1639 Firmicutes
Listeria monocytogenes YP 008278305 1 1639 Firmicutes
Listeria monocytogenes YP 008299155 1 1639 Firmicutes
Listeria monocytogenes YP 008279616 1 1639 Firmicutes
Listeria monocytogenes YP 008275446 1 1639 Firmicutes
Listeria monocytogenes YP 008301463 1 1639 Firmicutes
Listeria monocytogenes YP 008284659 1 1639 Firmicutes
Listeria monocytogenes YP 008304457 1 1639 Firmicutes
Rhodococcus erythropolis PR4 YP 002764004 1 234621 Actinobacteria
Saccharothrix espanaensis DSM 44229 YP 007037233 1 1179773 Actinobacteria
Tsukamurella paurometabola DSM 20162 YP 003648700 1 521096 Actinobacteria
Nocardia brasiliensis ATCC 700358 YP 006807387 1 1133849 Actinobacteria
Mycobacterium abscessus subsp bolletii str GO 06 YP 009039511 1 1198627 Actinobacteria
Mycobacterium abscessus YP 001701669 1 36809 Actinobacteria
Arthrobacter sp Rue61a YP 006664420 1 1118963 Actinobacteria
Actinosynnema mirum DSM 43827 YP 003101981 1 446462 Actinobacteria
Cellvibrio gilvus ATCC 13127 YP 004601685 1 593907 Actinobacteria
Kineococcus radiotolerans SRS30216 ATCC BAA 149 YP 001362606 1 266940 Actinobacteria
Actinoplanes missouriensis 431 YP 005462818 1 512565 Actinobacteria
Actinoplanes sp SE50 110 YP 006264619 1 134676 Actinobacteria
Serratia plymuthica AS9 YP 004507625 1 768492 Proteobacteria
Serratia sp AS12 YP 004502673 1 768490 Proteobacteria
Dickeya zeae DZ2Q WP 071602854 1 Dzeae
Dickeya zeae EC1 WP 029456605 1 Dzeae
Dickeya zeae ZJU1202 WP 029456605 1 Dzeae
199 zmsN Zeamine biosynthesis Seed
Dickeya gs fangzhongdai S1 WP 049854988 1 Dfangzhongdai
Dickeya gs fangzhongdai MK7 WP 038918438 1 Dfangzhongdai
Dickeya gs fangzhongdai B16 WP 038918438 1 Dfangzhongdai
Dickeya gs fangzhongdai M074 WP 038660390 1 Dfangzhongdai
Dickeya gs fangzhongdai ND14b WP 038660390 1 Dfangzhongdai
Dickeya gs fangzhongdai M005 WP 039691315 1 Dfangzhongdai
Dickeya gs fangzhongdai NCPPB 3274 WP 042859304 1 Dfangzhongdai
Dickeya solani RNS 05 1 2A WP 057085320 1 Dsolani
Dickeya solani MK10 WP 022632861 1 Dsolani
Dickeya solani PPO 9019 WP 022632861 1 Dsolani
Dickeya solani RNS 07 7 3B WP 022632861 1 Dsolani
Dickeya solani RNS 08 23 3 1 A WP 022632861 1 Dsolani
Dickeya solani IPO 2222 WP 022632861 1 Dsolani
Dickeya solani MK16 WP 022632861 1 Dsolani
Dickeya solani GBBC 2040 WP 022632861 1 Dsolani
Dickeya solani D s0432 1 WP 022632861 1 Dsolani
Dickeya solani PPO 9134 WP 022632861 1 Dsolani
Dickeya solani IFB 0099 WP 022632861 1 Dsolani

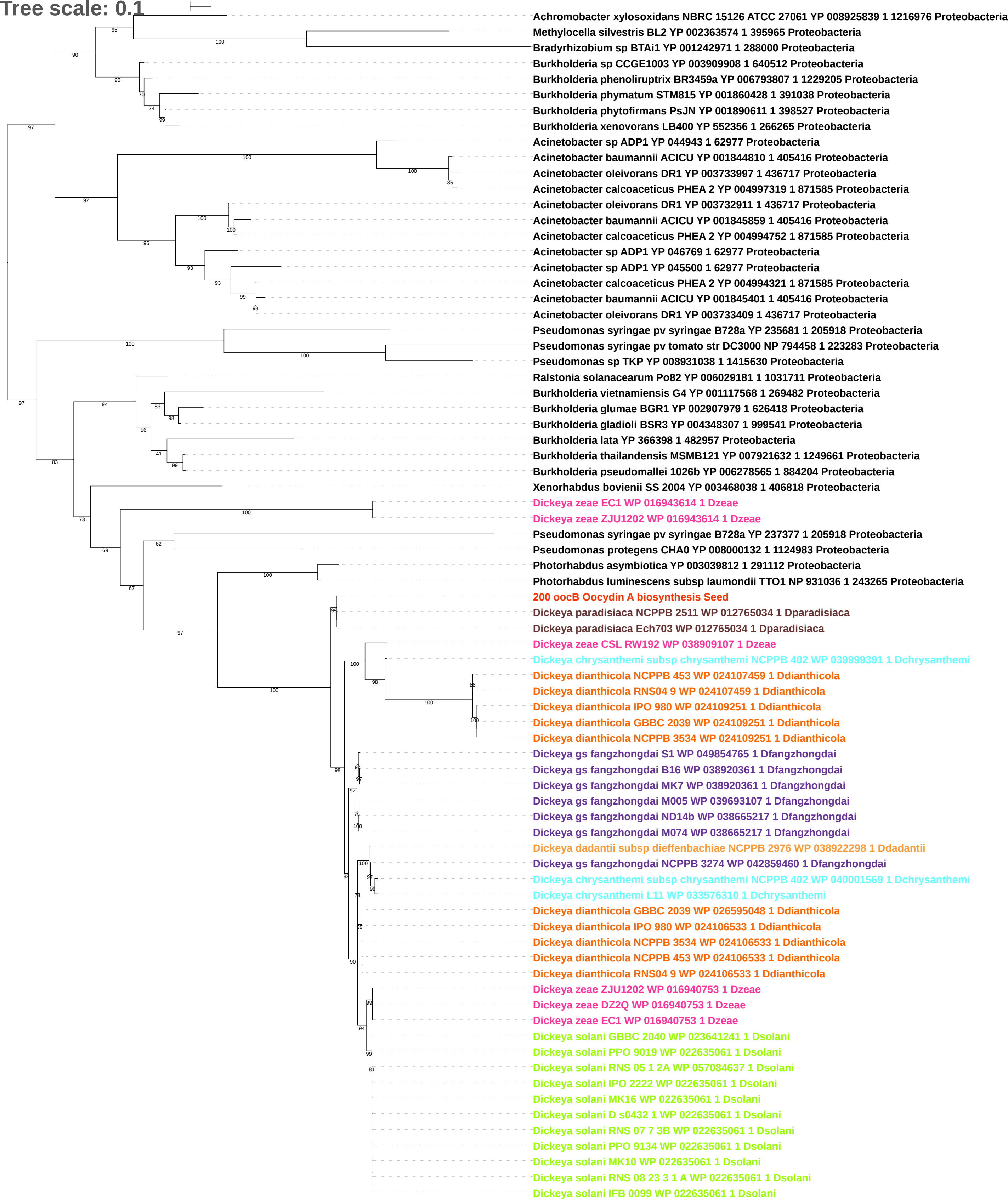

Tree scale: 0.1
Achromobacter xylosoxidans NBRC 15126 ATCC 27061 YP 008925839 1 1216976 Proteobacteria
Methylocella silvestris BL2 YP 002363574 1 395965 Proteobacteria
Bradyrhizobium sp BTAi1 YP 001242971 1 288000 Proteobacteria
Burkholderia sp CCGE1003 YP 003909908 1 640512 Proteobacteria
Burkholderia phenoliruptrix BR3459a YP 006793807 1 1229205 Proteobacteria
Burkholderia phymatum STM815 YP 001860428 1 391038 Proteobacteria
Burkholderia phytofirmans PsJN YP 001890611 1 398527 Proteobacteria
Burkholderia xenovorans LB400 YP 552356 1 266265 Proteobacteria
Acinetobacter sp ADP1 YP 044943 1 62977 Proteobacteria
Acinetobacter baumannii ACICU YP 001844810 1 405416 Proteobacteria
Acinetobacter oleivorans DR1 YP 003733997 1 436717 Proteobacteria
Acinetobacter calcoaceticus PHEA 2 YP 004997319 1 871585 Proteobacteria
Acinetobacter oleivorans DR1 YP 003732911 1 436717 Proteobacteria
Acinetobacter baumannii ACICU YP 001845859 1 405416 Proteobacteria
Acinetobacter calcoaceticus PHEA 2 YP 004994752 1 871585 Proteobacteria
Acinetobacter sp ADP1 YP 046769 1 62977 Proteobacteria
Acinetobacter sp ADP1 YP 045500 1 62977 Proteobacteria
Acinetobacter calcoaceticus PHEA 2 YP 004994321 1 871585 Proteobacteria
Acinetobacter baumannii ACICU YP 001845401 1 405416 Proteobacteria
Acinetobacter oleivorans DR1 YP 003733409 1 436717 Proteobacteria
Pseudomonas syringae pv syringae B728a YP 235681 1 205918 Proteobacteria
Pseudomonas syringae pv tomato str DC3000 NP 794458 1 223283 Proteobacteria
Pseudomonas sp TKP YP 008931038 1 1415630 Proteobacteria
Ralstonia solanacearum Po82 YP 006029181 1 1031711 Proteobacteria
Burkholderia vietnamiensis G4 YP 001117568 1 269482 Proteobacteria
Burkholderia glumae BGR1 YP 002907979 1 626418 Proteobacteria
Burkholderia gladioli BSR3 YP 004348307 1 999541 Proteobacteria
Burkholderia lata YP 366398 1 482957 Proteobacteria
Burkholderia thailandensis MSMB121 YP 007921632 1 1249661 Proteobacteria
Burkholderia pseudomallei 1026b YP 006278565 1 884204 Proteobacteria
Xenorhabdus bovienii SS 2004 YP 003468038 1 406818 Proteobacteria
Dickeya zeae EC1 WP 016943614 1 Dzeae
Dickeya zeae ZJU1202 WP 016943614 1 Dzeae
Pseudomonas syringae pv syringae B728a YP 237377 1 205918 Proteobacteria
Pseudomonas protegens CHA0 YP 008000132 1 1124983 Proteobacteria
Photorhabdus asymbiotica YP 003039812 1 291112 Proteobacteria
Photorhabdus luminescens subsp laumondii TTO1 NP 931036 1 243265 Proteobacteria
200 oocB Oocydin A biosynthesis Seed
Dickeya paradisiaca NCPPB 2511 WP 012765034 1 Dparadisiaca
Dickeya paradisiaca Ech703 WP 012765034 1 Dparadisiaca
Dickeya zeae CSL RW192 WP 038909107 1 Dzeae
Dickeya chrysanthemi subsp chrysanthemi NCPPB 402 WP 039999391 1 Dchrysanthemi
Dickeya dianthicola NCPPB 453 WP 024107459 1 Ddianthicola
Dickeya dianthicola RNS04 9 WP 024107459 1 Ddianthicola
Dickeya dianthicola IPO 980 WP 024109251 1 Ddianthicola
Dickeya dianthicola GBBC 2039 WP 024109251 1 Ddianthicola
Dickeya dianthicola NCPPB 3534 WP 024109251 1 Ddianthicola
Dickeya gs fangzhongdai S1 WP 049854765 1 Dfangzhongdai
Dickeya gs fangzhongdai B16 WP 038920361 1 Dfangzhongdai
Dickeya gs fangzhongdai MK7 WP 038920361 1 Dfangzhongdai
Dickeya gs fangzhongdai M005 WP 039693107 1 Dfangzhongdai
Dickeya gs fangzhongdai ND14b WP 038665217 1 Dfangzhongdai
Dickeya gs fangzhongdai M074 WP 038665217 1 Dfangzhongdai
Dickeya dadantii subsp dieffenbachiae NCPPB 2976 WP 038922298 1 Ddadantii
Dickeya gs fangzhongdai NCPPB 3274 WP 042859460 1 Dfangzhongdai
Dickeya chrysanthemi subsp chrysanthemi NCPPB 402 WP 040001569 1 Dchrysanthemi
Dickeya chrysanthemi L11 WP 033576310 1 Dchrysanthemi
Dickeya dianthicola GBBC 2039 WP 026595048 1 Ddianthicola
Dickeya dianthicola IPO 980 WP 024106533 1 Ddianthicola
Dickeya dianthicola NCPPB 3534 WP 024106533 1 Ddianthicola
Dickeya dianthicola NCPPB 453 WP 024106533 1 Ddianthicola
Dickeya dianthicola RNS04 9 WP 024106533 1 Ddianthicola
Dickeya zeae ZJU1202 WP 016940753 1 Dzeae
Dickeya zeae DZ2Q WP 016940753 1 Dzeae
Dickeya zeae EC1 WP 016940753 1 Dzeae
Dickeya solani GBBC 2040 WP 023641241 1 Dsolani
Dickeya solani PPO 9019 WP 022635061 1 Dsolani
Dickeya solani RNS 05 1 2A WP 057084637 1 Dsolani
Dickeya solani IPO 2222 WP 022635061 1 Dsolani
Dickeya solani MK16 WP 022635061 1 Dsolani
Dickeya solani D s0432 1 WP 022635061 1 Dsolani
Dickeya solani RNS 07 7 3B WP 022635061 1 Dsolani
Dickeya solani PPO 9134 WP 022635061 1 Dsolani
Dickeya solani MK10 WP 022635061 1 Dsolani
Dickeya solani RNS 08 23 3 1 A WP 022635061 1 Dsolani
Dickeya solani IFB 0099 WP 022635061 1 Dsolani

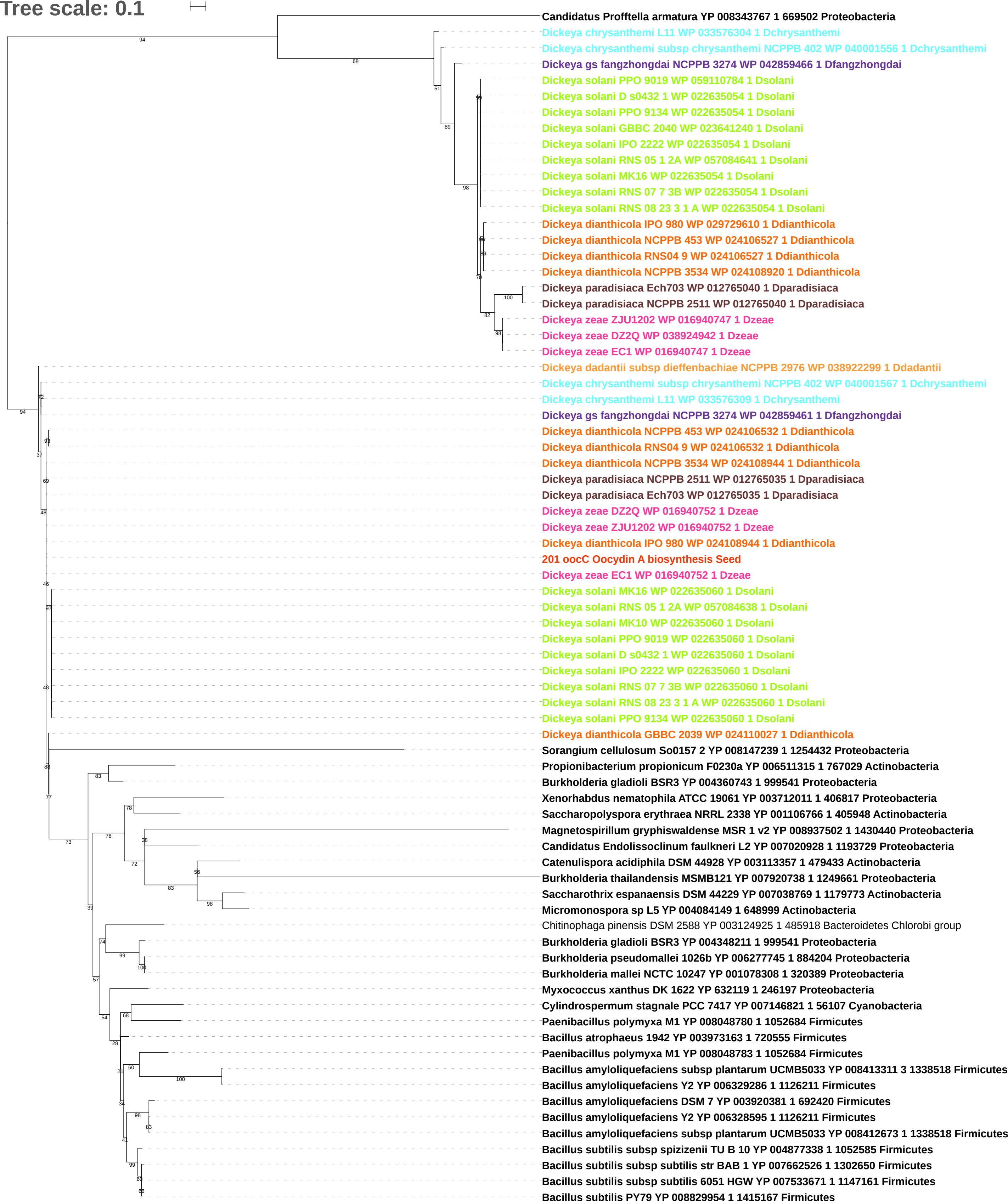
Tree scale: 0.1
Candidatus Profftella armatura YP 008343767 1 669502 Proteobacteria
Dickeya chrysanthemi L11 WP 033576304 1 Dchrysanthemi
Dickeya chrysanthemi subsp chrysanthemi NCPPB 402 WP 040001556 1 Dchrysanthemi
Dickeya gs fangzhongdai NCPPB 3274 WP 042859466 1 Dfangzhongdai
Dickeya solani PPO 9019 WP 059110784 1 Dsolani
Dickeya solani D s0432 1 WP 022635054 1 Dsolani
Dickeya solani PPO 9134 WP 022635054 1 Dsolani
Dickeya solani GBBC 2040 WP 023641240 1 Dsolani
Dickeya solani IPO 2222 WP 022635054 1 Dsolani
Dickeya solani RNS 05 1 2A WP 057084641 1 Dsolani
Dickeya solani MK16 WP 022635054 1 Dsolani
Dickeya solani RNS 07 7 3B WP 022635054 1 Dsolani
Dickeya solani RNS 08 23 3 1 A WP 022635054 1 Dsolani
Dickeya dianthicola IPO 980 WP 029729610 1 Ddianthicola
Dickeya dianthicola NCPPB 453 WP 024106527 1 Ddianthicola
Dickeya dianthicola RNS04 9 WP 024106527 1 Ddianthicola
Dickeya dianthicola NCPPB 3534 WP 024108920 1 Ddianthicola
Dickeya paradisiaca Ech703 WP 012765040 1 Dparadisiaca
Dickeya paradisiaca NCPPB 2511 WP 012765040 1 Dparadisiaca
Dickeya zeae ZJU1202 WP 016940747 1 Dzeae
Dickeya zeae DZ2Q WP 038924942 1 Dzeae
Dickeya zeae EC1 WP 016940747 1 Dzeae
Dickeya dadantii subsp dieffenbachiae NCPPB 2976 WP 038922299 1 Ddadantii
Dickeya chrysanthemi subsp chrysanthemi NCPPB 402 WP 040001567 1 Dchrysanthemi
Dickeya chrysanthemi L11 WP 033576309 1 Dchrysanthemi
Dickeya gs fangzhongdai NCPPB 3274 WP 042859461 1 Dfangzhongdai
Dickeya dianthicola NCPPB 453 WP 024106532 1 Ddianthicola
Dickeya dianthicola RNS04 9 WP 024106532 1 Ddianthicola
Dickeya dianthicola NCPPB 3534 WP 024108944 1 Ddianthicola
Dickeya paradisiaca NCPPB 2511 WP 012765035 1 Dparadisiaca
Dickeya paradisiaca Ech703 WP 012765035 1 Dparadisiaca
Dickeya zeae DZ2Q WP 016940752 1 Dzeae
Dickeya zeae ZJU1202 WP 016940752 1 Dzeae
Dickeya dianthicola IPO 980 WP 024108944 1 Ddianthicola
201 oocC Oocydin A biosynthesis Seed
Dickeya zeae EC1 WP 016940752 1 Dzeae
Dickeya solani MK16 WP 022635060 1 Dsolani
Dickeya solani RNS 05 1 2A WP 057084638 1 Dsolani
Dickeya solani MK10 WP 022635060 1 Dsolani
Dickeya solani PPO 9019 WP 022635060 1 Dsolani
Dickeya solani D s0432 1 WP 022635060 1 Dsolani
Dickeya solani IPO 2222 WP 022635060 1 Dsolani
Dickeya solani RNS 07 7 3B WP 022635060 1 Dsolani
Dickeya solani RNS 08 23 3 1 A WP 022635060 1 Dsolani
Dickeya solani PPO 9134 WP 022635060 1 Dsolani
Dickeya dianthicola GBBC 2039 WP 024110027 1 Ddianthicola
Sorangium cellulosum So0157 2 YP 008147239 1 1254432 Proteobacteria
Propionibacterium propionicum F0230a YP 006511315 1 767029 Actinobacteria
Burkholderia gladioli BSR3 YP 004360743 1 999541 Proteobacteria
Xenorhabdus nematophila ATCC 19061 YP 003712011 1 406817 Proteobacteria
Saccharopolyspora erythraea NRRL 2338 YP 001106766 1 405948 Actinobacteria
Magnetospirillum gryphiswaldense MSR 1 v2 YP 008937502 1 1430440 Proteobacteria
Candidatus Endolissoclinum faulkneri L2 YP 007020928 1 1193729 Proteobacteria
Catenulispora acidiphila DSM 44928 YP 003113357 1 479433 Actinobacteria
Burkholderia thailandensis MSMB121 YP 007920738 1 1249661 Proteobacteria
Saccharothrix espanaensis DSM 44229 YP 007038769 1 1179773 Actinobacteria
Micromonospora sp L5 YP 004084149 1 648999 Actinobacteria
Chitinophaga pinensis DSM 2588 YP 003124925 1 485918 Bacteroidetes Chlorobi group
Burkholderia gladioli BSR3 YP 004348211 1 999541 Proteobacteria
Burkholderia pseudomallei 1026b YP 006277745 1 884204 Proteobacteria
Burkholderia mallei NCTC 10247 YP 001078308 1 320389 Proteobacteria
Myxococcus xanthus DK 1622 YP 632119 1 246197 Proteobacteria
Cylindrospermum stagnale PCC 7417 YP 007146821 1 56107 Cyanobacteria
Paenibacillus polymyxa M1 YP 008048780 1 1052684 Firmicutes
Bacillus atrophaeus 1942 YP 003973163 1 720555 Firmicutes
Paenibacillus polymyxa M1 YP 008048783 1 1052684 Firmicutes
Bacillus amyloliquefaciens subsp plantarum UCMB5033 YP 008413311 3 1338518 Firmicutes
Bacillus amyloliquefaciens Y2 YP 006329286 1 1126211 Firmicutes
Bacillus amyloliquefaciens DSM 7 YP 003920381 1 692420 Firmicutes
Bacillus amyloliquefaciens Y2 YP 006328595 1 1126211 Firmicutes
Bacillus amyloliquefaciens subsp plantarum UCMB5033 YP 008412673 1 1338518 Firmicutes
Bacillus subtilis subsp spizizenii TU B 10 YP 004877338 1 1052585 Firmicutes
Bacillus subtilis subsp subtilis str BAB 1 YP 007662526 1 1302650 Firmicutes
Bacillus subtilis subsp subtilis 6051 HGW YP 007533671 1 1147161 Firmicutes
Bacillus subtilis PY79 YP 008829954 1 1415167 Firmicutes

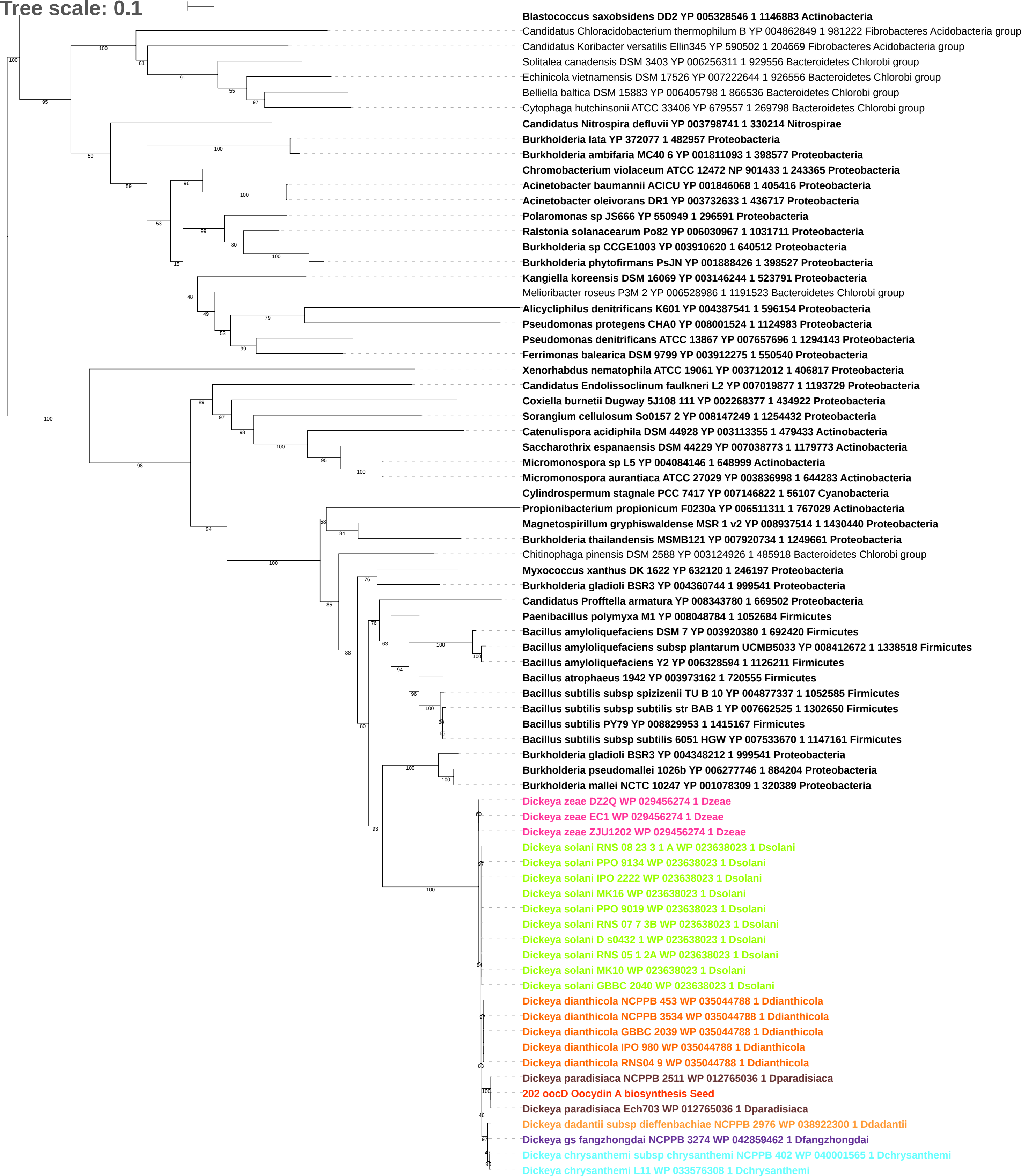
Tree scale: 0.1
Blastococcus saxobsidens DD2 YP 005328546 1 1146883 Actinobacteria
Candidatus Chloracidobacterium thermophilum B YP 004862849 1 981222 Fibrobacteres Acidobacteria group
Candidatus Koribacter versatilis Ellin345 YP 590502 1 204669 Fibrobacteres Acidobacteria group
Solitalea canadensis DSM 3403 YP 006256311 1 929556 Bacteroidetes Chlorobi group
Echinicola vietnamensis DSM 17526 YP 007222644 1 926556 Bacteroidetes Chlorobi group
Belliella baltica DSM 15883 YP 006405798 1 866536 Bacteroidetes Chlorobi group
Cytophaga hutchinsonii ATCC 33406 YP 679557 1 269798 Bacteroidetes Chlorobi group
Candidatus Nitrospira defluvii YP 003798741 1 330214 Nitrospirae
Burkholderia lata YP 372077 1 482957 Proteobacteria
Burkholderia ambifaria MC40 6 YP 001811093 1 398577 Proteobacteria
Chromobacterium violaceum ATCC 12472 NP 901433 1 243365 Proteobacteria
Acinetobacter baumannii ACICU YP 001846068 1 405416 Proteobacteria
Acinetobacter oleivorans DR1 YP 003732633 1 436717 Proteobacteria
Polaromonas sp JS666 YP 550949 1 296591 Proteobacteria
Ralstonia solanacearum Po82 YP 006030967 1 1031711 Proteobacteria
Burkholderia sp CCGE1003 YP 003910620 1 640512 Proteobacteria
Burkholderia phytofirmans PsJN YP 001888426 1 398527 Proteobacteria
Kangiella koreensis DSM 16069 YP 003146244 1 523791 Proteobacteria
Melioribacter roseus P3M 2 YP 006528986 1 1191523 Bacteroidetes Chlorobi group
Alicycliphilus denitrificans K601 YP 004387541 1 596154 Proteobacteria
Pseudomonas protegens CHA0 YP 008001524 1 1124983 Proteobacteria
Pseudomonas denitrificans ATCC 13867 YP 007657696 1 1294143 Proteobacteria
Ferrimonas balearica DSM 9799 YP 003912275 1 550540 Proteobacteria
Xenorhabdus nematophila ATCC 19061 YP 003712012 1 406817 Proteobacteria
Candidatus Endolissoclinum faulkneri L2 YP 007019877 1 1193729 Proteobacteria
Coxiella burnetii Dugway 5J108 111 YP 002268377 1 434922 Proteobacteria
Sorangium cellulosum So0157 2 YP 008147249 1 1254432 Proteobacteria
Catenulispora acidiphila DSM 44928 YP 003113355 1 479433 Actinobacteria
Saccharothrix espanaensis DSM 44229 YP 007038773 1 1179773 Actinobacteria
Micromonospora sp L5 YP 004084146 1 648999 Actinobacteria
Micromonospora aurantiaca ATCC 27029 YP 003836998 1 644283 Actinobacteria
Cylindrospermum stagnale PCC 7417 YP 007146822 1 56107 Cyanobacteria
Propionibacterium propionicum F0230a YP 006511311 1 767029 Actinobacteria
Magnetospirillum gryphiswaldense MSR 1 v2 YP 008937514 1 1430440 Proteobacteria
Burkholderia thailandensis MSMB121 YP 007920734 1 1249661 Proteobacteria
Chitinophaga pinensis DSM 2588 YP 003124926 1 485918 Bacteroidetes Chlorobi group
Myxococcus xanthus DK 1622 YP 632120 1 246197 Proteobacteria
Burkholderia gladioli BSR3 YP 004360744 1 999541 Proteobacteria
Candidatus Profftella armatura YP 008343780 1 669502 Proteobacteria
Paenibacillus polymyxa M1 YP 008048784 1 1052684 Firmicutes
Bacillus amyloliquefaciens DSM 7 YP 003920380 1 692420 Firmicutes
Bacillus amyloliquefaciens subsp plantarum UCMB5033 YP 008412672 1 1338518 Firmicutes
Bacillus amyloliquefaciens Y2 YP 006328594 1 1126211 Firmicutes
Bacillus atrophaeus 1942 YP 003973162 1 720555 Firmicutes
Bacillus subtilis subsp spizizenii TU B 10 YP 004877337 1 1052585 Firmicutes
Bacillus subtilis subsp subtilis str BAB 1 YP 007662525 1 1302650 Firmicutes
Bacillus subtilis PY79 YP 008829953 1 1415167 Firmicutes
Bacillus subtilis subsp subtilis 6051 HGW YP 007533670 1 1147161 Firmicutes
Burkholderia gladioli BSR3 YP 004348212 1 999541 Proteobacteria
Burkholderia pseudomallei 1026b YP 006277746 1 884204 Proteobacteria
Burkholderia mallei NCTC 10247 YP 001078309 1 320389 Proteobacteria
Dickeya zeae DZ2Q WP 029456274 1 Dzeae
Dickeya zeae EC1 WP 029456274 1 Dzeae
Dickeya zeae ZJU1202 WP 029456274 1 Dzeae
Dickeya solani RNS 08 23 3 1 A WP 023638023 1 Dsolani
Dickeya solani PPO 9134 WP 023638023 1 Dsolani
Dickeya solani IPO 2222 WP 023638023 1 Dsolani
Dickeya solani MK16 WP 023638023 1 Dsolani
Dickeya solani PPO 9019 WP 023638023 1 Dsolani
Dickeya solani RNS 07 7 3B WP 023638023 1 Dsolani
Dickeya solani D s0432 1 WP 023638023 1 Dsolani
Dickeya solani RNS 05 1 2A WP 023638023 1 Dsolani
Dickeya solani MK10 WP 023638023 1 Dsolani
Dickeya solani GBBC 2040 WP 023638023 1 Dsolani
Dickeya dianthicola NCPPB 453 WP 035044788 1 Ddianthicola
Dickeya dianthicola NCPPB 3534 WP 035044788 1 Ddianthicola
Dickeya dianthicola GBBC 2039 WP 035044788 1 Ddianthicola
Dickeya dianthicola IPO 980 WP 035044788 1 Ddianthicola
Dickeya dianthicola RNS04 9 WP 035044788 1 Ddianthicola
Dickeya paradisiaca NCPPB 2511 WP 012765036 1 Dparadisiaca
202 oocD Oocydin A biosynthesis Seed
Dickeya paradisiaca Ech703 WP 012765036 1 Dparadisiaca
Dickeya dadantii subsp dieffenbachiae NCPPB 2976 WP 038922300 1 Ddadantii
Dickeya gs fangzhongdai NCPPB 3274 WP 042859462 1 Dfangzhongdai
Dickeya chrysanthemi subsp chrysanthemi NCPPB 402 WP 040001565 1 Dchrysanthemi
Dickeya chrysanthemi L11 WP 033576308 1 Dchrysanthemi

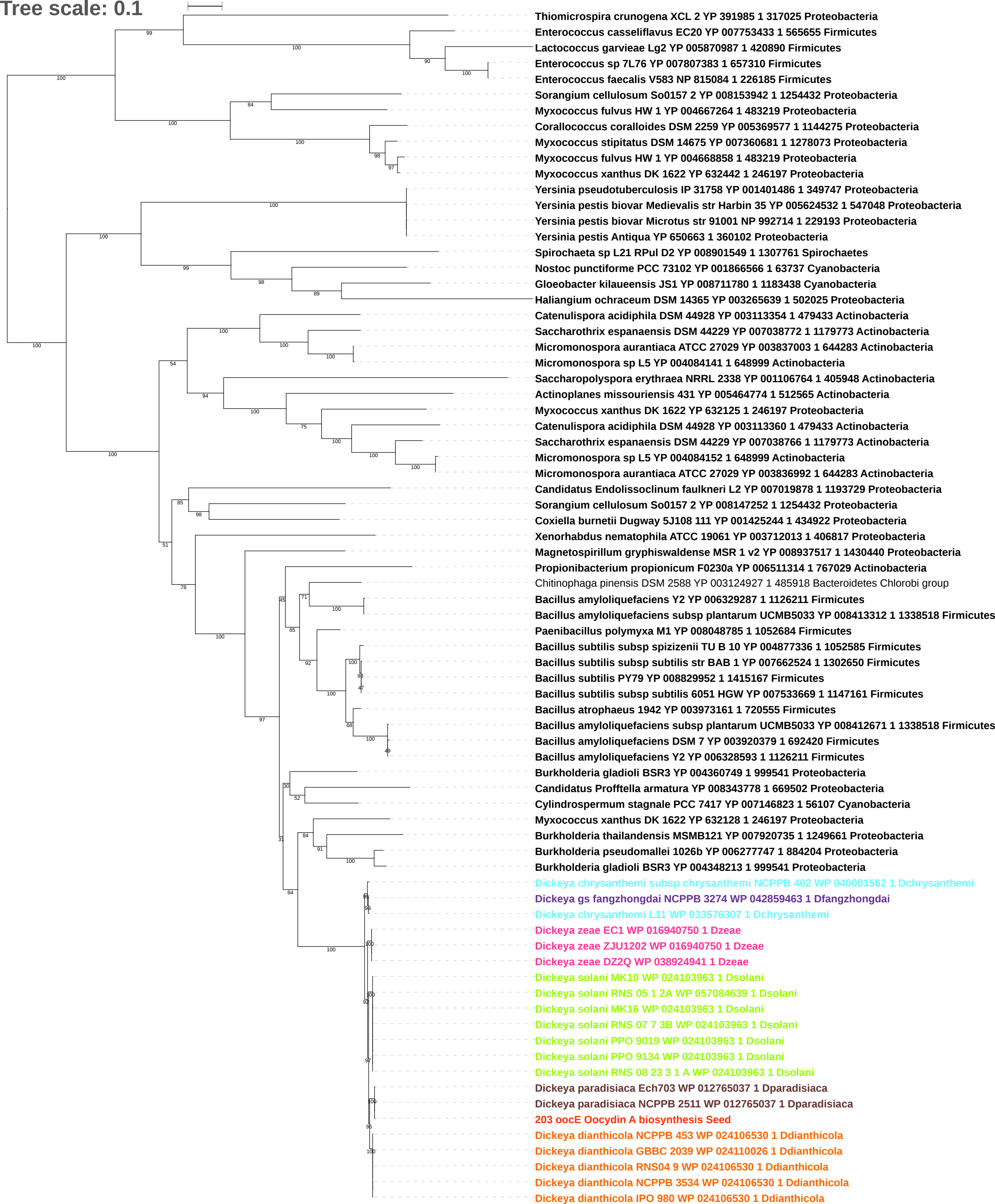

Tree scale: 0.1
Thiomicrospira crunogena XCL 2 YP 391985 1 317025 Proteobacteria
Enterococcus casseliflavus EC20 YP 007753433 1 565655 Firmicutes
Lactococcus garvieae Lg2 YP 005870987 1 420890 Firmicutes
Enterococcus sp 7L76 YP 007807383 1 657310 Firmicutes
Enterococcus faecalis V583 NP 815084 1 226185 Firmicutes
Sorangium cellulosum So0157 2 YP 008153942 1 1254432 Proteobacteria
Myxococcus fulvus HW 1 YP 004667264 1 483219 Proteobacteria
Corallococcus coralloides DSM 2259 YP 005369577 1 1144275 Proteobacteria
Myxococcus stipitatus DSM 14675 YP 007360681 1 1278073 Proteobacteria
Myxococcus fulvus HW 1 YP 004668858 1 483219 Proteobacteria
Myxococcus xanthus DK 1622 YP 632442 1 246197 Proteobacteria
Yersinia pseudotuberculosis IP 31758 YP 001401486 1 349747 Proteobacteria
Yersinia pestis biovar Medievalis str Harbin 35 YP 005624532 1 547048 Proteobacteria
Yersinia pestis biovar Microtus str 91001 NP 992714 1 229193 Proteobacteria
Yersinia pestis Antiqua YP 650663 1 360102 Proteobacteria
Spirochaeta sp L21 RPul D2 YP 008901549 1 1307761 Spirochaetes
Nostoc punctiforme PCC 73102 YP 001866566 1 63737 Cyanobacteria
Gloeobacter kilaueensis JS1 YP 008711780 1 1183438 Cyanobacteria
Haliangium ochraceum DSM 14365 YP 003265639 1 502025 Proteobacteria
Catenulispora acidiphila DSM 44928 YP 003113354 1 479433 Actinobacteria
Saccharothrix espanaensis DSM 44229 YP 007038772 1 1179773 Actinobacteria
Micromonospora aurantiaca ATCC 27029 YP 003837003 1 644283 Actinobacteria
Micromonospora sp L5 YP 004084141 1 648999 Actinobacteria
Saccharopolyspora erythraea NRRL 2338 YP 001106764 1 405948 Actinobacteria
Actinoplanes missouriensis 431 YP 005464774 1 512565 Actinobacteria
Myxococcus xanthus DK 1622 YP 632125 1 246197 Proteobacteria
Catenulispora acidiphila DSM 44928 YP 003113360 1 479433 Actinobacteria
Saccharothrix espanaensis DSM 44229 YP 007038766 1 1179773 Actinobacteria
Micromonospora sp L5 YP 004084152 1 648999 Actinobacteria
Micromonospora aurantiaca ATCC 27029 YP 003836992 1 644283 Actinobacteria
Candidatus Endolissoclinum faulkneri L2 YP 007019878 1 1193729 Proteobacteria
Sorangium cellulosum So0157 2 YP 008147252 1 1254432 Proteobacteria
Coxiella burnetii Dugway 5J108 111 YP 001425244 1 434922 Proteobacteria
Xenorhabdus nematophila ATCC 19061 YP 003712013 1 406817 Proteobacteria
Magnetospirillum gryphiswaldense MSR 1 v2 YP 008937517 1 1430440 Proteobacteria
Propionibacterium propionicum F0230a YP 006511314 1 767029 Actinobacteria
Chitinophaga pinensis DSM 2588 YP 003124927 1 485918 Bacteroidetes Chlorobi group
Bacillus amyloliquefaciens Y2 YP 006329287 1 1126211 Firmicutes
Bacillus amyloliquefaciens subsp plantarum UCMB5033 YP 008413312 1 1338518 Firmicutes
Paenibacillus polymyxa M1 YP 008048785 1 1052684 Firmicutes
Bacillus subtilis subsp spizizenii TU B 10 YP 004877336 1 1052585 Firmicutes
Bacillus subtilis subsp subtilis str BAB 1 YP 007662524 1 1302650 Firmicutes
Bacillus subtilis PY79 YP 008829952 1 1415167 Firmicutes
Bacillus subtilis subsp subtilis 6051 HGW YP 007533669 1 1147161 Firmicutes
Bacillus atrophaeus 1942 YP 003973161 1 720555 Firmicutes
Bacillus amyloliquefaciens subsp plantarum UCMB5033 YP 008412671 1 1338518 Firmicutes
Bacillus amyloliquefaciens DSM 7 YP 003920379 1 692420 Firmicutes
Bacillus amyloliquefaciens Y2 YP 006328593 1 1126211 Firmicutes
Burkholderia gladioli BSR3 YP 004360749 1 999541 Proteobacteria
Candidatus Profftella armatura YP 008343778 1 669502 Proteobacteria
Cylindrospermum stagnale PCC 7417 YP 007146823 1 56107 Cyanobacteria
Myxococcus xanthus DK 1622 YP 632128 1 246197 Proteobacteria
Burkholderia thailandensis MSMB121 YP 007920735 1 1249661 Proteobacteria
Burkholderia pseudomallei 1026b YP 006277747 1 884204 Proteobacteria
Burkholderia gladioli BSR3 YP 004348213 1 999541 Proteobacteria
Dickeya chrysanthemi subsp chrysanthemi NCPPB 402 WP 040001562 1 Dchrysanthemi
Dickeya gs fangzhongdai NCPPB 3274 WP 042859463 1 Dfangzhongdai
Dickeya chrysanthemi L11 WP 033576307 1 Dchrysanthemi
Dickeya zeae EC1 WP 016940750 1 Dzeae
Dickeya zeae ZJU1202 WP 016940750 1 Dzeae
Dickeya zeae DZ2Q WP 038924941 1 Dzeae
Dickeya solani MK10 WP 024103963 1 Dsolani
Dickeya solani RNS 05 1 2A WP 057084639 1 Dsolani
Dickeya solani MK16 WP 024103963 1 Dsolani
Dickeya solani RNS 07 7 3B WP 024103963 1 Dsolani
Dickeya solani PPO 9019 WP 024103963 1 Dsolani
Dickeya solani PPO 9134 WP 024103963 1 Dsolani
Dickeya solani RNS 08 23 3 1 A WP 024103963 1 Dsolani
Dickeya paradisiaca Ech703 WP 012765037 1 Dparadisiaca
Dickeya paradisiaca NCPPB 2511 WP 012765037 1 Dparadisiaca
203 oocE Oocydin A biosynthesis Seed
Dickeya dianthicola NCPPB 453 WP 024106530 1 Ddianthicola
Dickeya dianthicola GBBC 2039 WP 024110026 1 Ddianthicola
Dickeya dianthicola RNS04 9 WP 024106530 1 Ddianthicola
Dickeya dianthicola NCPPB 3534 WP 024106530 1 Ddianthicola
Dickeya dianthicola IPO 980 WP 024106530 1 Ddianthicola

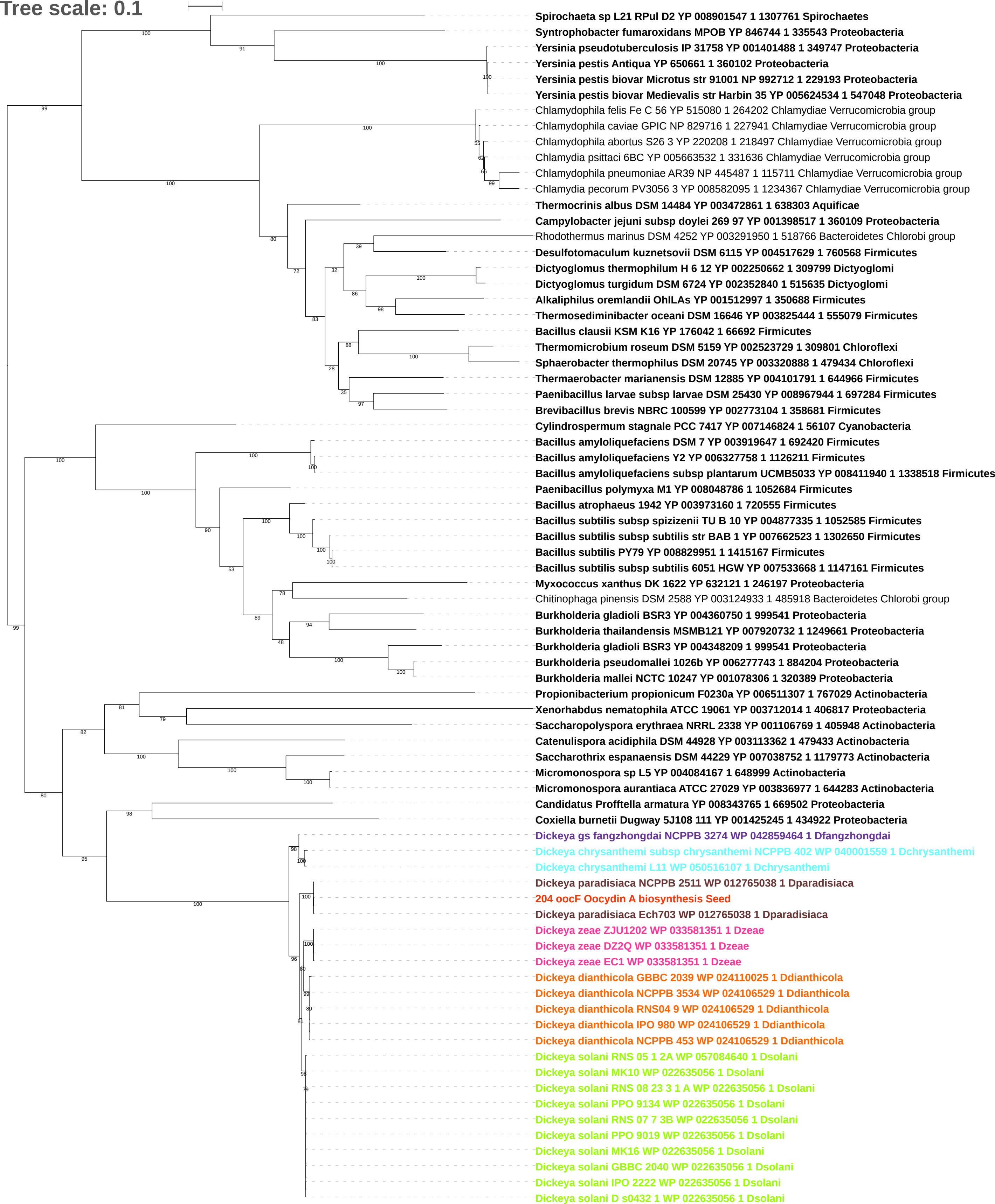
Tree scale: 0.1
Spirochaeta sp L21 RPul D2 YP 008901547 1 1307761 Spirochaetes
Syntrophobacter fumaroxidans MPOB YP 846744 1 335543 Proteobacteria
Yersinia pseudotuberculosis IP 31758 YP 001401488 1 349747 Proteobacteria
Yersinia pestis Antiqua YP 650661 1 360102 Proteobacteria
Yersinia pestis biovar Microtus str 91001 NP 992712 1 229193 Proteobacteria
Yersinia pestis biovar Medievalis str Harbin 35 YP 005624534 1 547048 Proteobacteria
Chlamydophila felis Fe C 56 YP 515080 1 264202 Chlamydiae Verrucomicrobia group
Chlamydophila caviae GPIC NP 829716 1 227941 Chlamydiae Verrucomicrobia group
Chlamydophila abortus S26 3 YP 220208 1 218497 Chlamydiae Verrucomicrobia group
Chlamydia psittaci 6BC YP 005663532 1 331636 Chlamydiae Verrucomicrobia group
Chlamydophila pneumoniae AR39 NP 445487 1 115711 Chlamydiae Verrucomicrobia group
Chlamydia pecorum PV3056 3 YP 008582095 1 1234367 Chlamydiae Verrucomicrobia group
Thermocrinis albus DSM 14484 YP 003472861 1 638303 Aquificae
Campylobacter jejuni subsp doylei 269 97 YP 001398517 1 360109 Proteobacteria
Rhodothermus marinus DSM 4252 YP 003291950 1 518766 Bacteroidetes Chlorobi group
Desulfotomaculum kuznetsovii DSM 6115 YP 004517629 1 760568 Firmicutes
Dictyoglomus thermophilum H 6 12 YP 002250662 1 309799 Dictyoglomi
Dictyoglomus turgidum DSM 6724 YP 002352840 1 515635 Dictyoglomi
Alkaliphilus oremlandii OhILAs YP 001512997 1 350688 Firmicutes
Thermosediminibacter oceani DSM 16646 YP 003825444 1 555079 Firmicutes
Bacillus clausii KSM K16 YP 176042 1 66692 Firmicutes
Thermomicrobium roseum DSM 5159 YP 002523729 1 309801 Chloroflexi
Sphaerobacter thermophilus DSM 20745 YP 003320888 1 479434 Chloroflexi
Thermaerobacter marianensis DSM 12885 YP 004101791 1 644966 Firmicutes
Paenibacillus larvae subsp larvae DSM 25430 YP 008967944 1 697284 Firmicutes
Brevibacillus brevis NBRC 100599 YP 002773104 1 358681 Firmicutes
Cylindrospermum stagnale PCC 7417 YP 007146824 1 56107 Cyanobacteria
Bacillus amyloliquefaciens DSM 7 YP 003919647 1 692420 Firmicutes
Bacillus amyloliquefaciens Y2 YP 006327758 1 1126211 Firmicutes
Bacillus amyloliquefaciens subsp plantarum UCMB5033 YP 008411940 1 1338518 Firmicutes
Paenibacillus polymyxa M1 YP 008048786 1 1052684 Firmicutes
Bacillus atrophaeus 1942 YP 003973160 1 720555 Firmicutes
Bacillus subtilis subsp spizizenii TU B 10 YP 004877335 1 1052585 Firmicutes
Bacillus subtilis subsp subtilis str BAB 1 YP 007662523 1 1302650 Firmicutes
Bacillus subtilis PY79 YP 008829951 1 1415167 Firmicutes
Bacillus subtilis subsp subtilis 6051 HGW YP 007533668 1 1147161 Firmicutes
Myxococcus xanthus DK 1622 YP 632121 1 246197 Proteobacteria
Chitinophaga pinensis DSM 2588 YP 003124933 1 485918 Bacteroidetes Chlorobi group
Burkholderia gladioli BSR3 YP 004360750 1 999541 Proteobacteria
Burkholderia thailandensis MSMB121 YP 007920732 1 1249661 Proteobacteria
Burkholderia gladioli BSR3 YP 004348209 1 999541 Proteobacteria
Burkholderia pseudomallei 1026b YP 006277743 1 884204 Proteobacteria
Burkholderia mallei NCTC 10247 YP 001078306 1 320389 Proteobacteria
Propionibacterium propionicum F0230a YP 006511307 1 767029 Actinobacteria
Xenorhabdus nematophila ATCC 19061 YP 003712014 1 406817 Proteobacteria
Saccharopolyspora erythraea NRRL 2338 YP 001106769 1 405948 Actinobacteria
Catenulispora acidiphila DSM 44928 YP 003113362 1 479433 Actinobacteria
Saccharothrix espanaensis DSM 44229 YP 007038752 1 1179773 Actinobacteria
Micromonospora sp L5 YP 004084167 1 648999 Actinobacteria
Micromonospora aurantiaca ATCC 27029 YP 003836977 1 644283 Actinobacteria
Candidatus Profftella armatura YP 008343765 1 669502 Proteobacteria
Coxiella burnetii Dugway 5J108 111 YP 001425245 1 434922 Proteobacteria
Dickeya gs fangzhongdai NCPPB 3274 WP 042859464 1 Dfangzhongdai
Dickeya chrysanthemi subsp chrysanthemi NCPPB 402 WP 040001559 1 Dchrysanthemi
Dickeya chrysanthemi L11 WP 050516107 1 Dchrysanthemi
Dickeya paradisiaca NCPPB 2511 WP 012765038 1 Dparadisiaca
204 oocF Oocydin A biosynthesis Seed
Dickeya paradisiaca Ech703 WP 012765038 1 Dparadisiaca
Dickeya zeae ZJU1202 WP 033581351 1 Dzeae
Dickeya zeae DZ2Q WP 033581351 1 Dzeae
Dickeya zeae EC1 WP 033581351 1 Dzeae
Dickeya dianthicola GBBC 2039 WP 024110025 1 Ddianthicola
Dickeya dianthicola NCPPB 3534 WP 024106529 1 Ddianthicola
Dickeya dianthicola RNS04 9 WP 024106529 1 Ddianthicola
Dickeya dianthicola IPO 980 WP 024106529 1 Ddianthicola
Dickeya dianthicola NCPPB 453 WP 024106529 1 Ddianthicola
Dickeya solani RNS 05 1 2A WP 057084640 1 Dsolani
Dickeya solani MK10 WP 022635056 1 Dsolani
Dickeya solani RNS 08 23 3 1 A WP 022635056 1 Dsolani
Dickeya solani PPO 9134 WP 022635056 1 Dsolani
Dickeya solani RNS 07 7 3B WP 022635056 1 Dsolani
Dickeya solani PPO 9019 WP 022635056 1 Dsolani
Dickeya solani MK16 WP 022635056 1 Dsolani
Dickeya solani GBBC 2040 WP 022635056 1 Dsolani
Dickeya solani IPO 2222 WP 022635056 1 Dsolani
Dickeya solani D s0432 1 WP 022635056 1 Dsolani

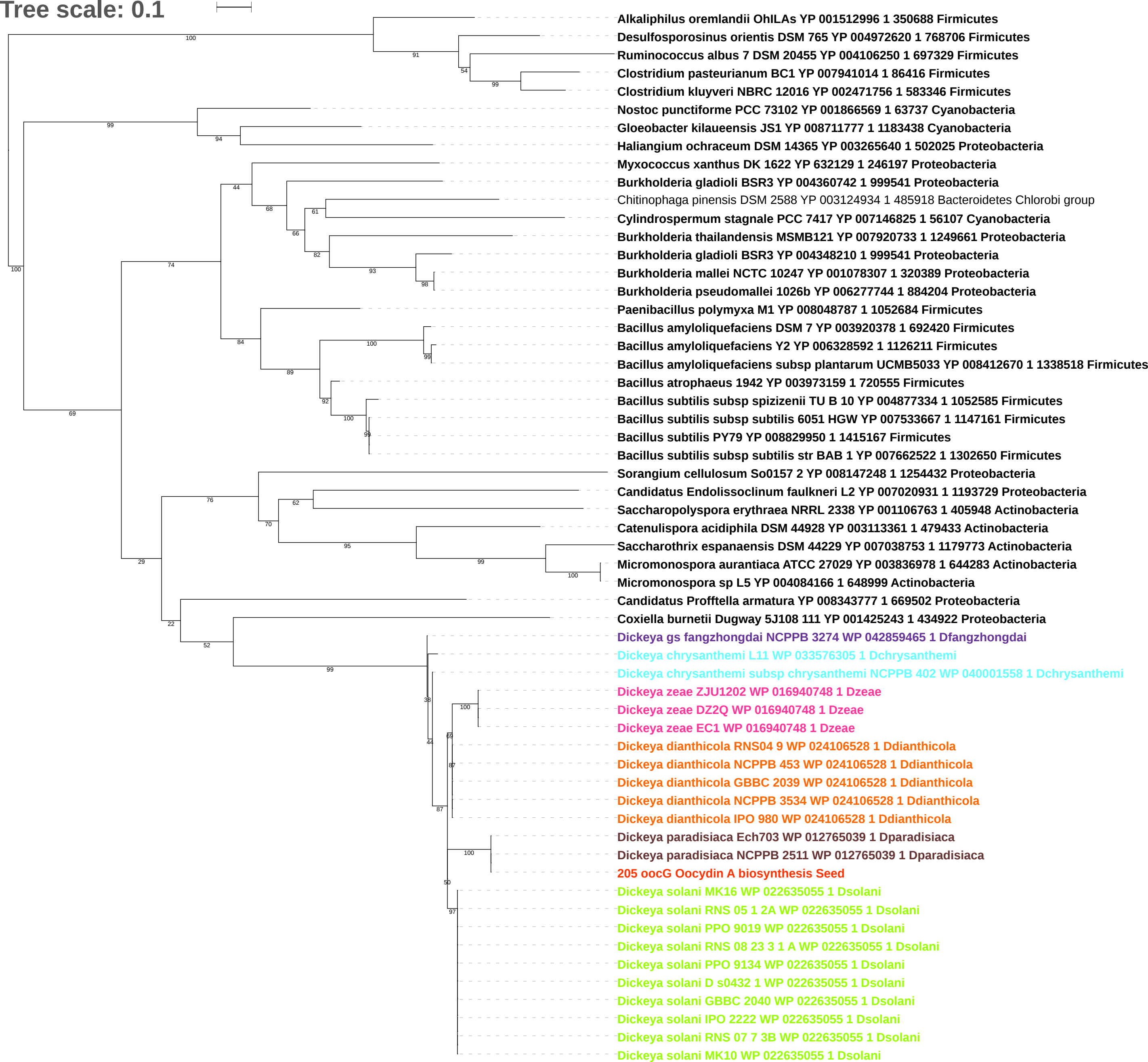

Tree scale: 0.1
Alkaliphilus oremlandii OhILAs YP 001512996 1 350688 Firmicutes
Desulfosporosinus orientis DSM 765 YP 004972620 1 768706 Firmicutes
Ruminococcus albus 7 DSM 20455 YP 004106250 1 697329 Firmicutes
Clostridium pasteurianum BC1 YP 007941014 1 86416 Firmicutes
Clostridium kluyveri NBRC 12016 YP 002471756 1 583346 Firmicutes
Nostoc punctiforme PCC 73102 YP 001866569 1 63737 Cyanobacteria
Gloeobacter kilaueensis JS1 YP 008711777 1 1183438 Cyanobacteria
Haliangium ochraceum DSM 14365 YP 003265640 1 502025 Proteobacteria
Myxococcus xanthus DK 1622 YP 632129 1 246197 Proteobacteria
Burkholderia gladioli BSR3 YP 004360742 1 999541 Proteobacteria
Chitinophaga pinensis DSM 2588 YP 003124934 1 485918 Bacteroidetes Chlorobi group
Cylindrospermum stagnale PCC 7417 YP 007146825 1 56107 Cyanobacteria
Burkholderia thailandensis MSMB121 YP 007920733 1 1249661 Proteobacteria
Burkholderia gladioli BSR3 YP 004348210 1 999541 Proteobacteria
Burkholderia mallei NCTC 10247 YP 001078307 1 320389 Proteobacteria
Burkholderia pseudomallei 1026b YP 006277744 1 884204 Proteobacteria
Paenibacillus polymyxa M1 YP 008048787 1 1052684 Firmicutes
Bacillus amyloliquefaciens DSM 7 YP 003920378 1 692420 Firmicutes
Bacillus amyloliquefaciens Y2 YP 006328592 1 1126211 Firmicutes
Bacillus amyloliquefaciens subsp plantarum UCMB5033 YP 008412670 1 1338518 Firmicutes
Bacillus atrophaeus 1942 YP 003973159 1 720555 Firmicutes
Bacillus subtilis subsp spizizenii TU B 10 YP 004877334 1 1052585 Firmicutes
Bacillus subtilis subsp subtilis 6051 HGW YP 007533667 1 1147161 Firmicutes
Bacillus subtilis PY79 YP 008829950 1 1415167 Firmicutes
Bacillus subtilis subsp subtilis str BAB 1 YP 007662522 1 1302650 Firmicutes
Sorangium cellulosum So0157 2 YP 008147248 1 1254432 Proteobacteria
Candidatus Endolissoclinum faulkneri L2 YP 007020931 1 1193729 Proteobacteria
Saccharopolyspora erythraea NRRL 2338 YP 001106763 1 405948 Actinobacteria
Catenulispora acidiphila DSM 44928 YP 003113361 1 479433 Actinobacteria
Saccharothrix espanaensis DSM 44229 YP 007038753 1 1179773 Actinobacteria
Micromonospora aurantiaca ATCC 27029 YP 003836978 1 644283 Actinobacteria
Micromonospora sp L5 YP 004084166 1 648999 Actinobacteria
Candidatus Profftella armatura YP 008343777 1 669502 Proteobacteria
Coxiella burnetii Dugway 5J108 111 YP 001425243 1 434922 Proteobacteria
Dickeya gs fangzhongdai NCPPB 3274 WP 042859465 1 Dfangzhongdai
Dickeya chrysanthemi L11 WP 033576305 1 Dchrysanthemi
Dickeya chrysanthemi subsp chrysanthemi NCPPB 402 WP 040001558 1 Dchrysanthemi
Dickeya zeae ZJU1202 WP 016940748 1 Dzeae
Dickeya zeae DZ2Q WP 016940748 1 Dzeae
Dickeya zeae EC1 WP 016940748 1 Dzeae
Dickeya dianthicola RNS04 9 WP 024106528 1 Ddianthicola
Dickeya dianthicola NCPPB 453 WP 024106528 1 Ddianthicola
Dickeya dianthicola GBBC 2039 WP 024106528 1 Ddianthicola
Dickeya dianthicola NCPPB 3534 WP 024106528 1 Ddianthicola
Dickeya dianthicola IPO 980 WP 024106528 1 Ddianthicola
Dickeya paradisiaca Ech703 WP 012765039 1 Dparadisiaca
Dickeya paradisiaca NCPPB 2511 WP 012765039 1 Dparadisiaca
205 oocG Oocydin A biosynthesis Seed
Dickeya solani MK16 WP 022635055 1 Dsolani
Dickeya solani RNS 05 1 2A WP 022635055 1 Dsolani
Dickeya solani PPO 9019 WP 022635055 1 Dsolani
Dickeya solani RNS 08 23 3 1 A WP 022635055 1 Dsolani
Dickeya solani PPO 9134 WP 022635055 1 Dsolani
Dickeya solani D s0432 1 WP 022635055 1 Dsolani
Dickeya solani GBBC 2040 WP 022635055 1 Dsolani
Dickeya solani IPO 2222 WP 022635055 1 Dsolani
Dickeya solani RNS 07 7 3B WP 022635055 1 Dsolani
Dickeya solani MK10 WP 022635055 1 Dsolani

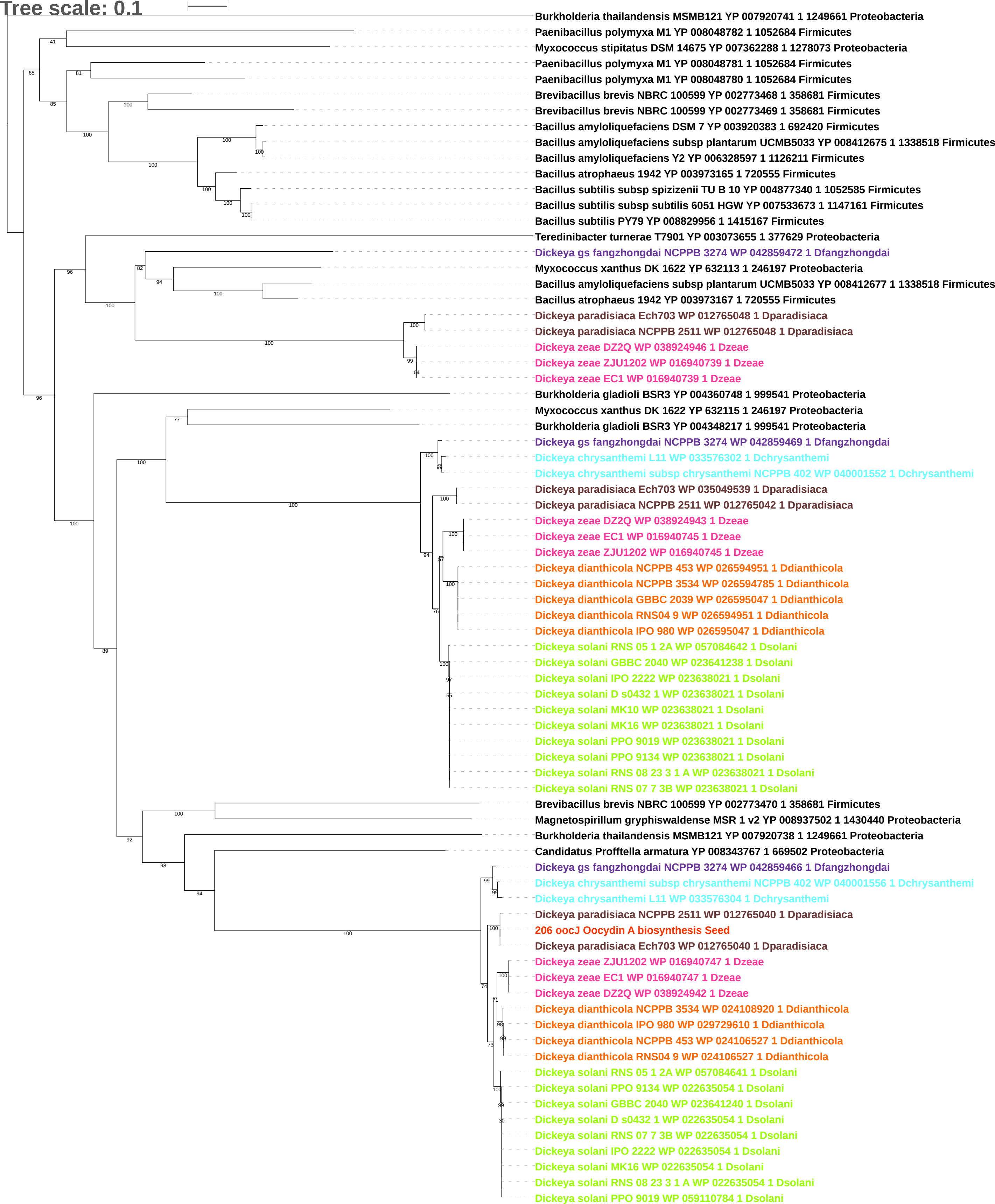
Tree scale: 0.1
Burkholderia thailandensis MSMB121 YP 007920741 1 1249661 Proteobacteria
Paenibacillus polymyxa M1 YP 008048782 1 1052684 Firmicutes
Myxococcus stipitatus DSM 14675 YP 007362288 1 1278073 Proteobacteria
Paenibacillus polymyxa M1 YP 008048781 1 1052684 Firmicutes
Paenibacillus polymyxa M1 YP 008048780 1 1052684 Firmicutes
Brevibacillus brevis NBRC 100599 YP 002773468 1 358681 Firmicutes
Brevibacillus brevis NBRC 100599 YP 002773469 1 358681 Firmicutes
Bacillus amyloliquefaciens DSM 7 YP 003920383 1 692420 Firmicutes
Bacillus amyloliquefaciens subsp plantarum UCMB5033 YP 008412675 1 1338518 Firmicutes
Bacillus amyloliquefaciens Y2 YP 006328597 1 1126211 Firmicutes
Bacillus atrophaeus 1942 YP 003973165 1 720555 Firmicutes
Bacillus subtilis subsp spizizenii TU B 10 YP 004877340 1 1052585 Firmicutes
Bacillus subtilis subsp subtilis 6051 HGW YP 007533673 1 1147161 Firmicutes
Bacillus subtilis PY79 YP 008829956 1 1415167 Firmicutes
Teredinibacter turnerae T7901 YP 003073655 1 377629 Proteobacteria
Dickeya gs fangzhongdai NCPPB 3274 WP 042859472 1 Dfangzhongdai
Myxococcus xanthus DK 1622 YP 632113 1 246197 Proteobacteria
Bacillus amyloliquefaciens subsp plantarum UCMB5033 YP 008412677 1 1338518 Firmicutes
Bacillus atrophaeus 1942 YP 003973167 1 720555 Firmicutes
Dickeya paradisiaca Ech703 WP 012765048 1 Dparadisiaca
Dickeya paradisiaca NCPPB 2511 WP 012765048 1 Dparadisiaca
Dickeya zeae DZ2Q WP 038924946 1 Dzeae
Dickeya zeae ZJU1202 WP 016940739 1 Dzeae
Dickeya zeae EC1 WP 016940739 1 Dzeae
Burkholderia gladioli BSR3 YP 004360748 1 999541 Proteobacteria
Myxococcus xanthus DK 1622 YP 632115 1 246197 Proteobacteria
Burkholderia gladioli BSR3 YP 004348217 1 999541 Proteobacteria
Dickeya gs fangzhongdai NCPPB 3274 WP 042859469 1 Dfangzhongdai
Dickeya chrysanthemi L11 WP 033576302 1 Dchrysanthemi
Dickeya chrysanthemi subsp chrysanthemi NCPPB 402 WP 040001552 1 Dchrysanthemi
Dickeya paradisiaca Ech703 WP 035049539 1 Dparadisiaca
Dickeya paradisiaca NCPPB 2511 WP 012765042 1 Dparadisiaca
Dickeya zeae DZ2Q WP 038924943 1 Dzeae
Dickeya zeae EC1 WP 016940745 1 Dzeae
Dickeya zeae ZJU1202 WP 016940745 1 Dzeae
Dickeya dianthicola NCPPB 453 WP 026594951 1 Ddianthicola
Dickeya dianthicola NCPPB 3534 WP 026594785 1 Ddianthicola
Dickeya dianthicola GBBC 2039 WP 026595047 1 Ddianthicola
Dickeya dianthicola RNS04 9 WP 026594951 1 Ddianthicola
Dickeya dianthicola IPO 980 WP 026595047 1 Ddianthicola
Dickeya solani RNS 05 1 2A WP 057084642 1 Dsolani
Dickeya solani GBBC 2040 WP 023641238 1 Dsolani
Dickeya solani IPO 2222 WP 023638021 1 Dsolani
Dickeya solani D s0432 1 WP 023638021 1 Dsolani
Dickeya solani MK10 WP 023638021 1 Dsolani
Dickeya solani MK16 WP 023638021 1 Dsolani
Dickeya solani PPO 9019 WP 023638021 1 Dsolani
Dickeya solani PPO 9134 WP 023638021 1 Dsolani
Dickeya solani RNS 08 23 3 1 A WP 023638021 1 Dsolani
Dickeya solani RNS 07 7 3B WP 023638021 1 Dsolani
Brevibacillus brevis NBRC 100599 YP 002773470 1 358681 Firmicutes
Magnetospirillum gryphiswaldense MSR 1 v2 YP 008937502 1 1430440 Proteobacteria
Burkholderia thailandensis MSMB121 YP 007920738 1 1249661 Proteobacteria
Candidatus Profftella armatura YP 008343767 1 669502 Proteobacteria
Dickeya gs fangzhongdai NCPPB 3274 WP 042859466 1 Dfangzhongdai
Dickeya chrysanthemi subsp chrysanthemi NCPPB 402 WP 040001556 1 Dchrysanthemi
Dickeya chrysanthemi L11 WP 033576304 1 Dchrysanthemi
Dickeya paradisiaca NCPPB 2511 WP 012765040 1 Dparadisiaca
206 oocJ Oocydin A biosynthesis Seed
Dickeya paradisiaca Ech703 WP 012765040 1 Dparadisiaca
Dickeya zeae ZJU1202 WP 016940747 1 Dzeae
Dickeya zeae EC1 WP 016940747 1 Dzeae
Dickeya zeae DZ2Q WP 038924942 1 Dzeae
Dickeya dianthicola NCPPB 3534 WP 024108920 1 Ddianthicola
Dickeya dianthicola IPO 980 WP 029729610 1 Ddianthicola
Dickeya dianthicola NCPPB 453 WP 024106527 1 Ddianthicola
Dickeya dianthicola RNS04 9 WP 024106527 1 Ddianthicola
Dickeya solani RNS 05 1 2A WP 057084641 1 Dsolani
Dickeya solani PPO 9134 WP 022635054 1 Dsolani
Dickeya solani GBBC 2040 WP 023641240 1 Dsolani
Dickeya solani D s0432 1 WP 022635054 1 Dsolani
Dickeya solani RNS 07 7 3B WP 022635054 1 Dsolani
Dickeya solani IPO 2222 WP 022635054 1 Dsolani
Dickeya solani MK16 WP 022635054 1 Dsolani
Dickeya solani RNS 08 23 3 1 A WP 022635054 1 Dsolani
Dickeya solani PPO 9019 WP 059110784 1 Dsolani

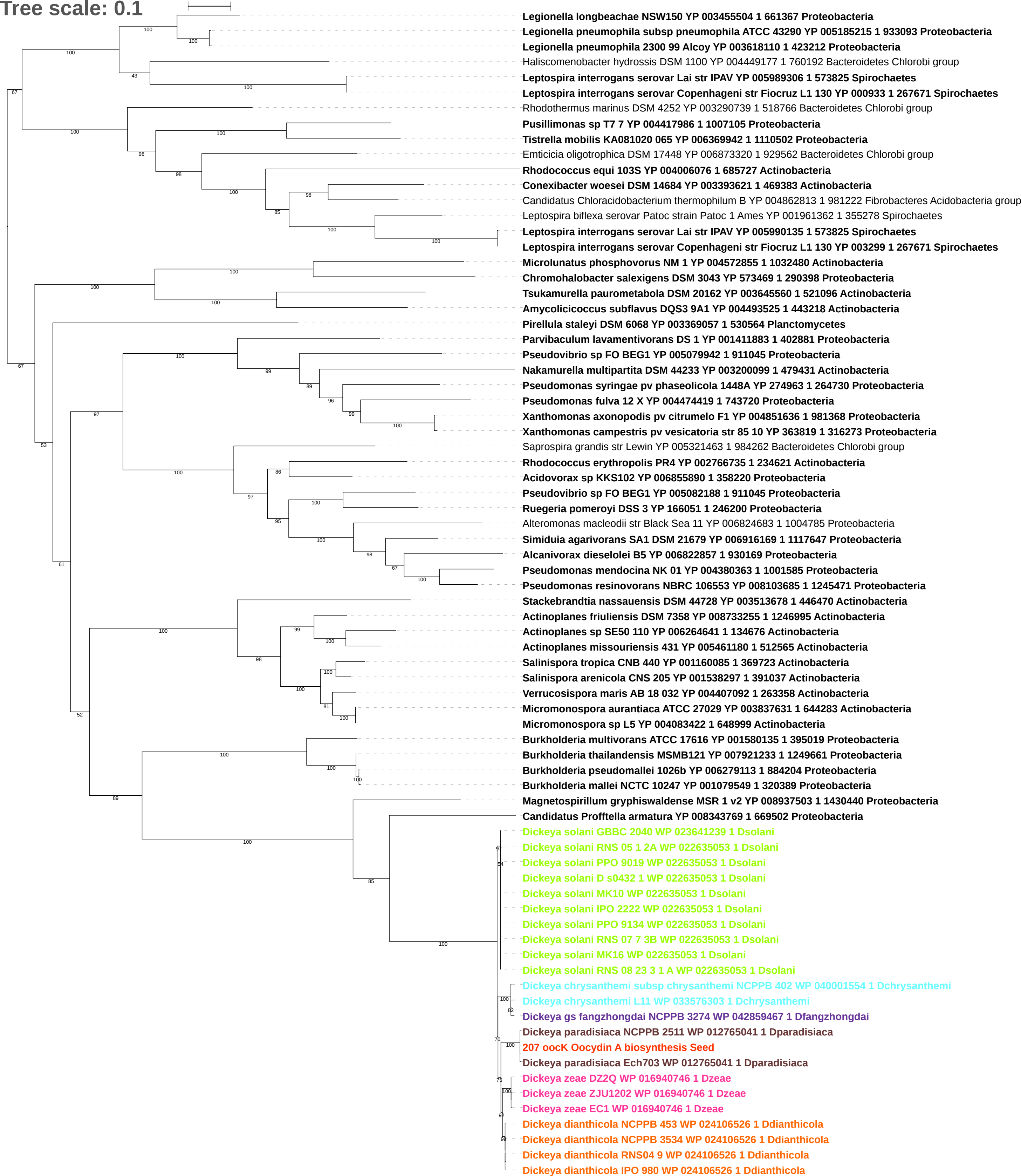
Tree scale: 0.1
Legionella longbeachae NSW150 YP 003455504 1 661367 Proteobacteria
Legionella pneumophila subsp pneumophila ATCC 43290 YP 005185215 1 933093 Proteobacteria
Legionella pneumophila 2300 99 Alcoy YP 003618110 1 423212 Proteobacteria
Haliscomenobacter hydrossis DSM 1100 YP 004449177 1 760192 Bacteroidetes Chlorobi group
Leptospira interrogans serovar Lai str IPAV YP 005989306 1 573825 Spirochaetes
Leptospira interrogans serovar Copenhageni str Fiocruz L1 130 YP 000933 1 267671 Spirochaetes
Rhodothermus marinus DSM 4252 YP 003290739 1 518766 Bacteroidetes Chlorobi group
Pusillimonas sp T7 7 YP 004417986 1 1007105 Proteobacteria
Tistrella mobilis KA081020 065 YP 006369942 1 1110502 Proteobacteria
Emticicia oligotrophica DSM 17448 YP 006873320 1 929562 Bacteroidetes Chlorobi group
Rhodococcus equi 103S YP 004006076 1 685727 Actinobacteria
Conexibacter woesei DSM 14684 YP 003393621 1 469383 Actinobacteria
Candidatus Chloracidobacterium thermophilum B YP 004862813 1 981222 Fibrobacteres Acidobacteria group
Leptospira biflexa serovar Patoc strain Patoc 1 Ames YP 001961362 1 355278 Spirochaetes
Leptospira interrogans serovar Lai str IPAV YP 005990135 1 573825 Spirochaetes
Leptospira interrogans serovar Copenhageni str Fiocruz L1 130 YP 003299 1 267671 Spirochaetes
Microlunatus phosphovorus NM 1 YP 004572855 1 1032480 Actinobacteria
Chromohalobacter salexigens DSM 3043 YP 573469 1 290398 Proteobacteria
Tsukamurella paurometabola DSM 20162 YP 003645560 1 521096 Actinobacteria
Amycolicicoccus subflavus DQS3 9A1 YP 004493525 1 443218 Actinobacteria
Pirellula staleyi DSM 6068 YP 003369057 1 530564 Planctomycetes
Parvibaculum lavamentivorans DS 1 YP 001411883 1 402881 Proteobacteria
Pseudovibrio sp FO BEG1 YP 005079942 1 911045 Proteobacteria
Nakamurella multipartita DSM 44233 YP 003200099 1 479431 Actinobacteria
Pseudomonas syringae pv phaseolicola 1448A YP 274963 1 264730 Proteobacteria
Pseudomonas fulva 12 X YP 004474419 1 743720 Proteobacteria
Xanthomonas axonopodis pv citrumelo F1 YP 004851636 1 981368 Proteobacteria
Xanthomonas campestris pv vesicatoria str 85 10 YP 363819 1 316273 Proteobacteria
Saprospira grandis str Lewin YP 005321463 1 984262 Bacteroidetes Chlorobi group
Rhodococcus erythropolis PR4 YP 002766735 1 234621 Actinobacteria
Acidovorax sp KKS102 YP 006855890 1 358220 Proteobacteria
Pseudovibrio sp FO BEG1 YP 005082188 1 911045 Proteobacteria
Ruegeria pomeroyi DSS 3 YP 166051 1 246200 Proteobacteria
Alteromonas macleodii str Black Sea 11 YP 006824683 1 1004785 Proteobacteria
Simiduia agarivorans SA1 DSM 21679 YP 006916169 1 1117647 Proteobacteria
Alcanivorax dieselolei B5 YP 006822857 1 930169 Proteobacteria
Pseudomonas mendocina NK 01 YP 004380363 1 1001585 Proteobacteria
Pseudomonas resinovorans NBRC 106553 YP 008103685 1 1245471 Proteobacteria
Stackebrandtia nassauensis DSM 44728 YP 003513678 1 446470 Actinobacteria
Actinoplanes friuliensis DSM 7358 YP 008733255 1 1246995 Actinobacteria
Actinoplanes sp SE50 110 YP 006264641 1 134676 Actinobacteria
Actinoplanes missouriensis 431 YP 005461180 1 512565 Actinobacteria
Salinispora tropica CNB 440 YP 001160085 1 369723 Actinobacteria
Salinispora arenicola CNS 205 YP 001538297 1 391037 Actinobacteria
Verrucosispora maris AB 18 032 YP 004407092 1 263358 Actinobacteria
Micromonospora aurantiaca ATCC 27029 YP 003837631 1 644283 Actinobacteria
Micromonospora sp L5 YP 004083422 1 648999 Actinobacteria
Burkholderia multivorans ATCC 17616 YP 001580135 1 395019 Proteobacteria
Burkholderia thailandensis MSMB121 YP 007921233 1 1249661 Proteobacteria
Burkholderia pseudomallei 1026b YP 006279113 1 884204 Proteobacteria
Burkholderia mallei NCTC 10247 YP 001079549 1 320389 Proteobacteria
Magnetospirillum gryphiswaldense MSR 1 v2 YP 008937503 1 1430440 Proteobacteria
Candidatus Profftella armatura YP 008343769 1 669502 Proteobacteria
Dickeya solani GBBC 2040 WP 023641239 1 Dsolani
Dickeya solani RNS 05 1 2A WP 022635053 1 Dsolani
Dickeya solani PPO 9019 WP 022635053 1 Dsolani
Dickeya solani D s0432 1 WP 022635053 1 Dsolani
Dickeya solani MK10 WP 022635053 1 Dsolani
Dickeya solani IPO 2222 WP 022635053 1 Dsolani
Dickeya solani PPO 9134 WP 022635053 1 Dsolani
Dickeya solani RNS 07 7 3B WP 022635053 1 Dsolani
Dickeya solani MK16 WP 022635053 1 Dsolani
Dickeya solani RNS 08 23 3 1 A WP 022635053 1 Dsolani
Dickeya chrysanthemi subsp chrysanthemi NCPPB 402 WP 040001554 1 Dchrysanthemi
Dickeya chrysanthemi L11 WP 033576303 1 Dchrysanthemi
Dickeya gs fangzhongdai NCPPB 3274 WP 042859467 1 Dfangzhongdai
Dickeya paradisiaca NCPPB 2511 WP 012765041 1 Dparadisiaca
207 oocK Oocydin A biosynthesis Seed
Dickeya paradisiaca Ech703 WP 012765041 1 Dparadisiaca
Dickeya zeae DZ2Q WP 016940746 1 Dzeae
Dickeya zeae ZJU1202 WP 016940746 1 Dzeae
Dickeya zeae EC1 WP 016940746 1 Dzeae
Dickeya dianthicola NCPPB 453 WP 024106526 1 Ddianthicola
Dickeya dianthicola NCPPB 3534 WP 024106526 1 Ddianthicola
Dickeya dianthicola RNS04 9 WP 024106526 1 Ddianthicola
Dickeya dianthicola IPO 980 WP 024106526 1 Ddianthicola

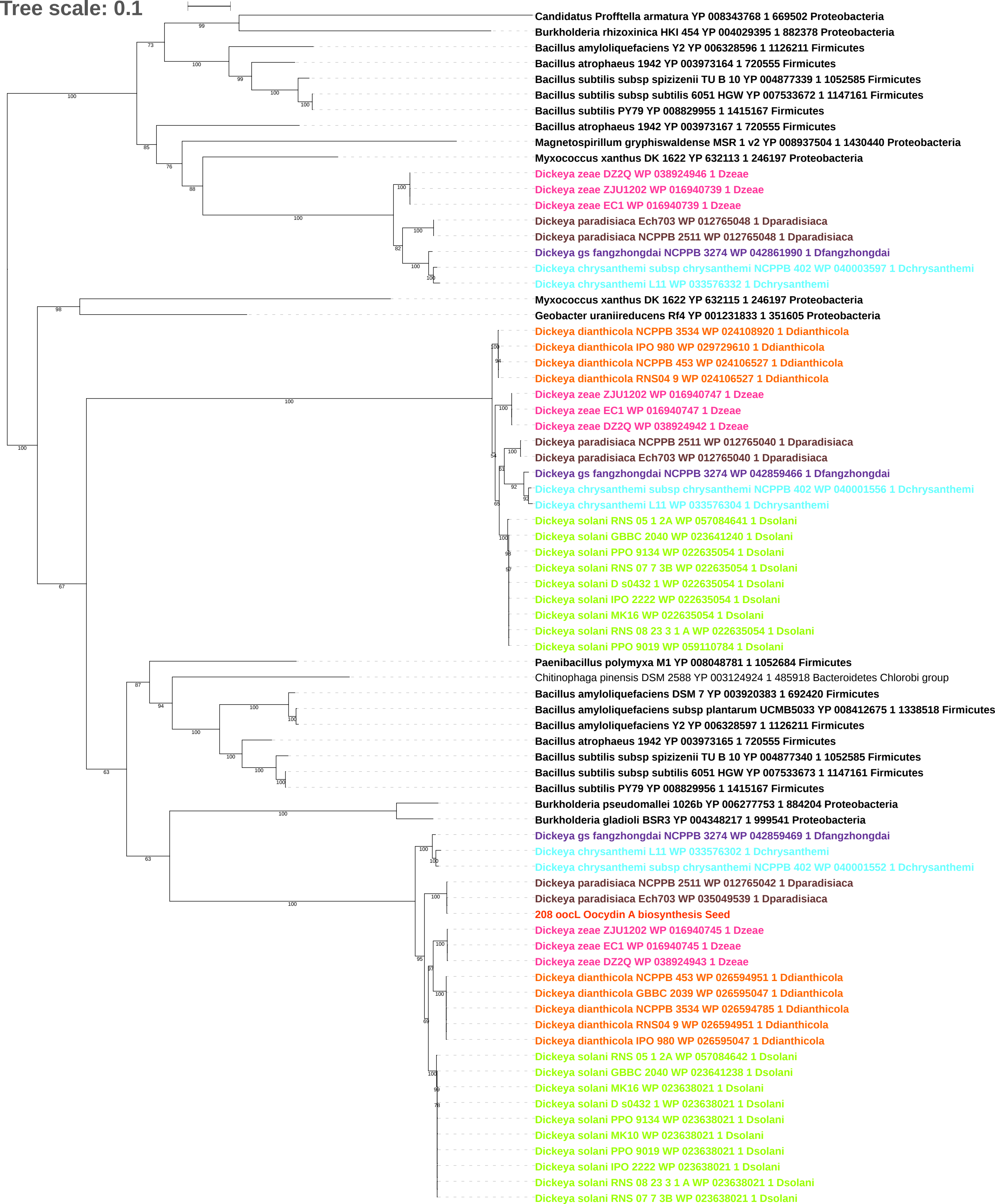
Tree scale: 0.1
Candidatus Profftella armatura YP 008343768 1 669502 Proteobacteria
Burkholderia rhizoxinica HKI 454 YP 004029395 1 882378 Proteobacteria
Bacillus amyloliquefaciens Y2 YP 006328596 1 1126211 Firmicutes
Bacillus atrophaeus 1942 YP 003973164 1 720555 Firmicutes
Bacillus subtilis subsp spizizenii TU B 10 YP 004877339 1 1052585 Firmicutes
Bacillus subtilis subsp subtilis 6051 HGW YP 007533672 1 1147161 Firmicutes
Bacillus subtilis PY79 YP 008829955 1 1415167 Firmicutes
Bacillus atrophaeus 1942 YP 003973167 1 720555 Firmicutes
Magnetospirillum gryphiswaldense MSR 1 v2 YP 008937504 1 1430440 Proteobacteria
Myxococcus xanthus DK 1622 YP 632113 1 246197 Proteobacteria
Dickeya zeae DZ2Q WP 038924946 1 Dzeae
Dickeya zeae ZJU1202 WP 016940739 1 Dzeae
Dickeya zeae EC1 WP 016940739 1 Dzeae
Dickeya paradisiaca Ech703 WP 012765048 1 Dparadisiaca
Dickeya paradisiaca NCPPB 2511 WP 012765048 1 Dparadisiaca
Dickeya gs fangzhongdai NCPPB 3274 WP 042861990 1 Dfangzhongdai
Dickeya chrysanthemi subsp chrysanthemi NCPPB 402 WP 040003597 1 Dchrysanthemi
Dickeya chrysanthemi L11 WP 033576332 1 Dchrysanthemi
Myxococcus xanthus DK 1622 YP 632115 1 246197 Proteobacteria
Geobacter uraniireducens Rf4 YP 001231833 1 351605 Proteobacteria
Dickeya dianthicola NCPPB 3534 WP 024108920 1 Ddianthicola
Dickeya dianthicola IPO 980 WP 029729610 1 Ddianthicola
Dickeya dianthicola NCPPB 453 WP 024106527 1 Ddianthicola
Dickeya dianthicola RNS04 9 WP 024106527 1 Ddianthicola
Dickeya zeae ZJU1202 WP 016940747 1 Dzeae
Dickeya zeae EC1 WP 016940747 1 Dzeae
Dickeya zeae DZ2Q WP 038924942 1 Dzeae
Dickeya paradisiaca NCPPB 2511 WP 012765040 1 Dparadisiaca
Dickeya paradisiaca Ech703 WP 012765040 1 Dparadisiaca
Dickeya gs fangzhongdai NCPPB 3274 WP 042859466 1 Dfangzhongdai
Dickeya chrysanthemi subsp chrysanthemi NCPPB 402 WP 040001556 1 Dchrysanthemi
Dickeya chrysanthemi L11 WP 033576304 1 Dchrysanthemi
Dickeya solani RNS 05 1 2A WP 057084641 1 Dsolani
Dickeya solani GBBC 2040 WP 023641240 1 Dsolani
Dickeya solani PPO 9134 WP 022635054 1 Dsolani
Dickeya solani RNS 07 7 3B WP 022635054 1 Dsolani
Dickeya solani D s0432 1 WP 022635054 1 Dsolani
Dickeya solani IPO 2222 WP 022635054 1 Dsolani
Dickeya solani MK16 WP 022635054 1 Dsolani
Dickeya solani RNS 08 23 3 1 A WP 022635054 1 Dsolani
Dickeya solani PPO 9019 WP 059110784 1 Dsolani
Paenibacillus polymyxa M1 YP 008048781 1 1052684 Firmicutes
Chitinophaga pinensis DSM 2588 YP 003124924 1 485918 Bacteroidetes Chlorobi group
Bacillus amyloliquefaciens DSM 7 YP 003920383 1 692420 Firmicutes
Bacillus amyloliquefaciens subsp plantarum UCMB5033 YP 008412675 1 1338518 Firmicutes
Bacillus amyloliquefaciens Y2 YP 006328597 1 1126211 Firmicutes
Bacillus atrophaeus 1942 YP 003973165 1 720555 Firmicutes
Bacillus subtilis subsp spizizenii TU B 10 YP 004877340 1 1052585 Firmicutes
Bacillus subtilis subsp subtilis 6051 HGW YP 007533673 1 1147161 Firmicutes
Bacillus subtilis PY79 YP 008829956 1 1415167 Firmicutes
Burkholderia pseudomallei 1026b YP 006277753 1 884204 Proteobacteria
Burkholderia gladioli BSR3 YP 004348217 1 999541 Proteobacteria
Dickeya gs fangzhongdai NCPPB 3274 WP 042859469 1 Dfangzhongdai
Dickeya chrysanthemi L11 WP 033576302 1 Dchrysanthemi
Dickeya chrysanthemi subsp chrysanthemi NCPPB 402 WP 040001552 1 Dchrysanthemi
Dickeya paradisiaca NCPPB 2511 WP 012765042 1 Dparadisiaca
Dickeya paradisiaca Ech703 WP 035049539 1 Dparadisiaca
208 oocL Oocydin A biosynthesis Seed
Dickeya zeae ZJU1202 WP 016940745 1 Dzeae
Dickeya zeae EC1 WP 016940745 1 Dzeae
Dickeya zeae DZ2Q WP 038924943 1 Dzeae
Dickeya dianthicola NCPPB 453 WP 026594951 1 Ddianthicola
Dickeya dianthicola GBBC 2039 WP 026595047 1 Ddianthicola
Dickeya dianthicola NCPPB 3534 WP 026594785 1 Ddianthicola
Dickeya dianthicola RNS04 9 WP 026594951 1 Ddianthicola
Dickeya dianthicola IPO 980 WP 026595047 1 Ddianthicola
Dickeya solani RNS 05 1 2A WP 057084642 1 Dsolani
Dickeya solani GBBC 2040 WP 023641238 1 Dsolani
Dickeya solani MK16 WP 023638021 1 Dsolani
Dickeya solani D s0432 1 WP 023638021 1 Dsolani
Dickeya solani PPO 9134 WP 023638021 1 Dsolani
Dickeya solani MK10 WP 023638021 1 Dsolani
Dickeya solani PPO 9019 WP 023638021 1 Dsolani
Dickeya solani IPO 2222 WP 023638021 1 Dsolani
Dickeya solani RNS 08 23 3 1 A WP 023638021 1 Dsolani
Dickeya solani RNS 07 7 3B WP 023638021 1 Dsolani

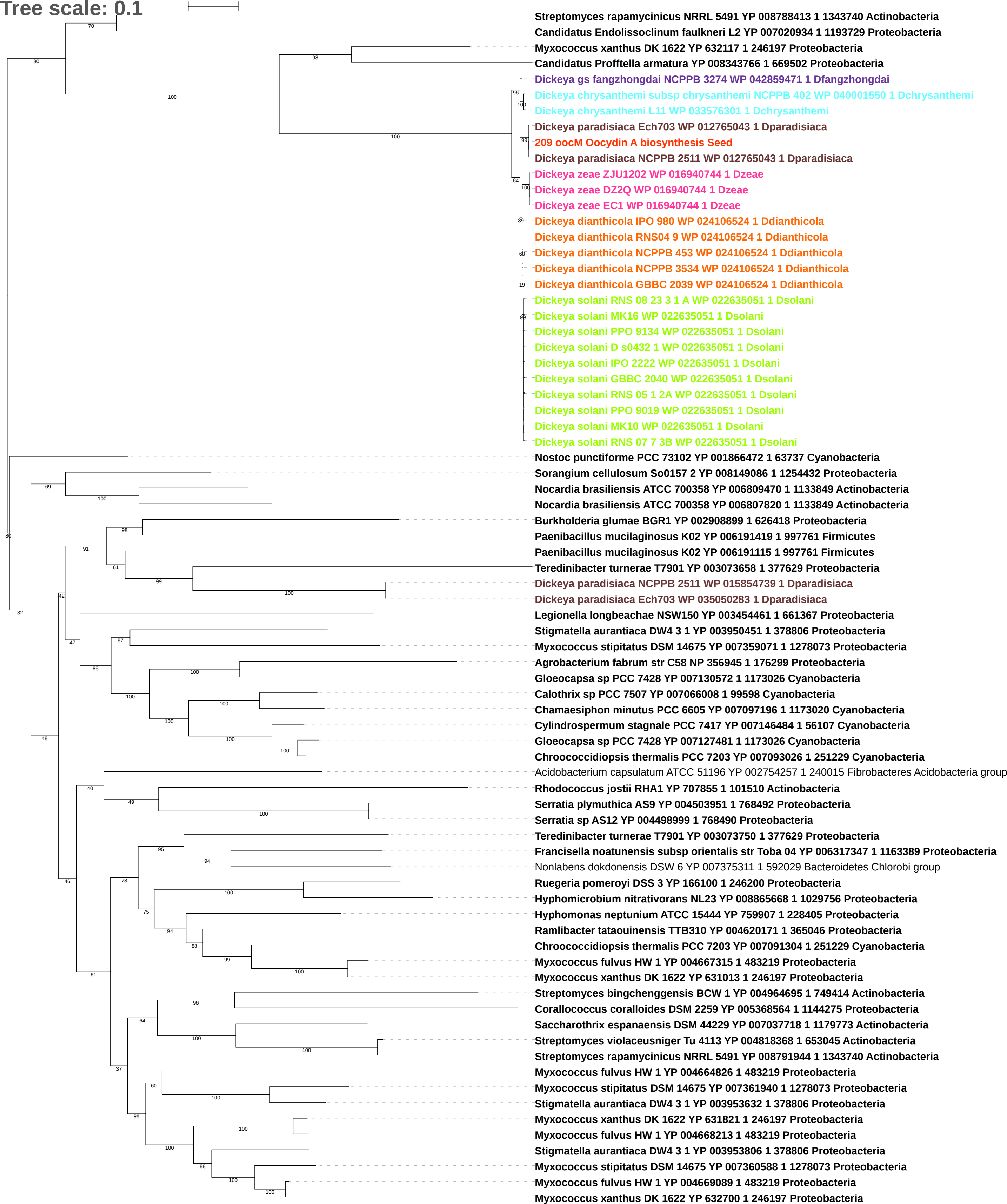
Tree scale: 0.1
Streptomyces rapamycinicus NRRL 5491 YP 008788413 1 1343740 Actinobacteria
Candidatus Endolissoclinum faulkneri L2 YP 007020934 1 1193729 Proteobacteria
Myxococcus xanthus DK 1622 YP 632117 1 246197 Proteobacteria
Candidatus Profftella armatura YP 008343766 1 669502 Proteobacteria
Dickeya gs fangzhongdai NCPPB 3274 WP 042859471 1 Dfangzhongdai
Dickeya chrysanthemi subsp chrysanthemi NCPPB 402 WP 040001550 1 Dchrysanthemi
Dickeya chrysanthemi L11 WP 033576301 1 Dchrysanthemi
Dickeya paradisiaca Ech703 WP 012765043 1 Dparadisiaca
209 oocM Oocydin A biosynthesis Seed
Dickeya paradisiaca NCPPB 2511 WP 012765043 1 Dparadisiaca
Dickeya zeae ZJU1202 WP 016940744 1 Dzeae
Dickeya zeae DZ2Q WP 016940744 1 Dzeae
Dickeya zeae EC1 WP 016940744 1 Dzeae
Dickeya dianthicola IPO 980 WP 024106524 1 Ddianthicola
Dickeya dianthicola RNS04 9 WP 024106524 1 Ddianthicola
Dickeya dianthicola NCPPB 453 WP 024106524 1 Ddianthicola
Dickeya dianthicola NCPPB 3534 WP 024106524 1 Ddianthicola
Dickeya dianthicola GBBC 2039 WP 024106524 1 Ddianthicola
Dickeya solani RNS 08 23 3 1 A WP 022635051 1 Dsolani
Dickeya solani MK16 WP 022635051 1 Dsolani
Dickeya solani PPO 9134 WP 022635051 1 Dsolani
Dickeya solani D s0432 1 WP 022635051 1 Dsolani
Dickeya solani IPO 2222 WP 022635051 1 Dsolani
Dickeya solani GBBC 2040 WP 022635051 1 Dsolani
Dickeya solani RNS 05 1 2A WP 022635051 1 Dsolani
Dickeya solani PPO 9019 WP 022635051 1 Dsolani
Dickeya solani MK10 WP 022635051 1 Dsolani
Dickeya solani RNS 07 7 3B WP 022635051 1 Dsolani
Nostoc punctiforme PCC 73102 YP 001866472 1 63737 Cyanobacteria
Sorangium cellulosum So0157 2 YP 008149086 1 1254432 Proteobacteria
Nocardia brasiliensis ATCC 700358 YP 006809470 1 1133849 Actinobacteria
Nocardia brasiliensis ATCC 700358 YP 006807820 1 1133849 Actinobacteria
Burkholderia glumae BGR1 YP 002908899 1 626418 Proteobacteria
Paenibacillus mucilaginosus K02 YP 006191419 1 997761 Firmicutes
Paenibacillus mucilaginosus K02 YP 006191115 1 997761 Firmicutes
Teredinibacter turnerae T7901 YP 003073658 1 377629 Proteobacteria
Dickeya paradisiaca NCPPB 2511 WP 015854739 1 Dparadisiaca
Dickeya paradisiaca Ech703 WP 035050283 1 Dparadisiaca
Legionella longbeachae NSW150 YP 003454461 1 661367 Proteobacteria
Stigmatella aurantiaca DW4 3 1 YP 003950451 1 378806 Proteobacteria
Myxococcus stipitatus DSM 14675 YP 007359071 1 1278073 Proteobacteria
Agrobacterium fabrum str C58 NP 356945 1 176299 Proteobacteria
Gloeocapsa sp PCC 7428 YP 007130572 1 1173026 Cyanobacteria
Calothrix sp PCC 7507 YP 007066008 1 99598 Cyanobacteria
Chamaesiphon minutus PCC 6605 YP 007097196 1 1173020 Cyanobacteria
Cylindrospermum stagnale PCC 7417 YP 007146484 1 56107 Cyanobacteria
Gloeocapsa sp PCC 7428 YP 007127481 1 1173026 Cyanobacteria
Chroococcidiopsis thermalis PCC 7203 YP 007093026 1 251229 Cyanobacteria
Acidobacterium capsulatum ATCC 51196 YP 002754257 1 240015 Fibrobacteres Acidobacteria group
Rhodococcus jostii RHA1 YP 707855 1 101510 Actinobacteria
Serratia plymuthica AS9 YP 004503951 1 768492 Proteobacteria
Serratia sp AS12 YP 004498999 1 768490 Proteobacteria
Teredinibacter turnerae T7901 YP 003073750 1 377629 Proteobacteria
Francisella noatunensis subsp orientalis str Toba 04 YP 006317347 1 1163389 Proteobacteria
Nonlabens dokdonensis DSW 6 YP 007375311 1 592029 Bacteroidetes Chlorobi group
Ruegeria pomeroyi DSS 3 YP 166100 1 246200 Proteobacteria
Hyphomicrobium nitrativorans NL23 YP 008865668 1 1029756 Proteobacteria
Hyphomonas neptunium ATCC 15444 YP 759907 1 228405 Proteobacteria
Ramlibacter tataouinensis TTB310 YP 004620171 1 365046 Proteobacteria
Chroococcidiopsis thermalis PCC 7203 YP 007091304 1 251229 Cyanobacteria
Myxococcus fulvus HW 1 YP 004667315 1 483219 Proteobacteria
Myxococcus xanthus DK 1622 YP 631013 1 246197 Proteobacteria
Streptomyces bingchenggensis BCW 1 YP 004964695 1 749414 Actinobacteria
Corallococcus coralloides DSM 2259 YP 005368564 1 1144275 Proteobacteria
Saccharothrix espanaensis DSM 44229 YP 007037718 1 1179773 Actinobacteria
Streptomyces violaceusniger Tu 4113 YP 004818368 1 653045 Actinobacteria
Streptomyces rapamycinicus NRRL 5491 YP 008791944 1 1343740 Actinobacteria
Myxococcus fulvus HW 1 YP 004664826 1 483219 Proteobacteria
Myxococcus stipitatus DSM 14675 YP 007361940 1 1278073 Proteobacteria
Stigmatella aurantiaca DW4 3 1 YP 003953632 1 378806 Proteobacteria
Myxococcus xanthus DK 1622 YP 631821 1 246197 Proteobacteria
Myxococcus fulvus HW 1 YP 004668213 1 483219 Proteobacteria
Stigmatella aurantiaca DW4 3 1 YP 003953806 1 378806 Proteobacteria
Myxococcus stipitatus DSM 14675 YP 007360588 1 1278073 Proteobacteria
Myxococcus fulvus HW 1 YP 004669089 1 483219 Proteobacteria
Myxococcus xanthus DK 1622 YP 632700 1 246197 Proteobacteria

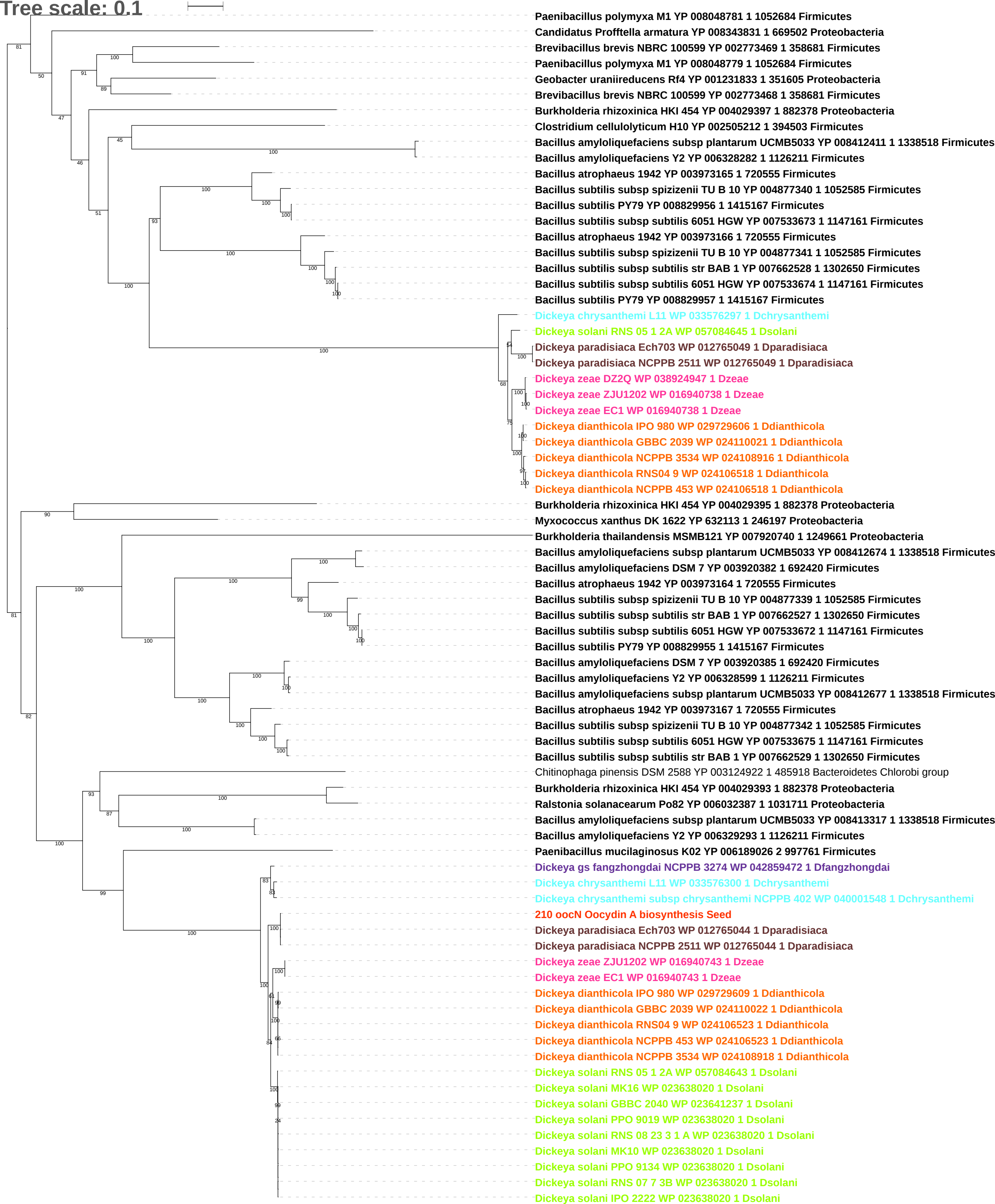
Tree scale: 0.1
Paenibacillus polymyxa M1 YP 008048781 1 1052684 Firmicutes
Candidatus Profftella armatura YP 008343831 1 669502 Proteobacteria
Brevibacillus brevis NBRC 100599 YP 002773469 1 358681 Firmicutes
Paenibacillus polymyxa M1 YP 008048779 1 1052684 Firmicutes
Geobacter uraniireducens Rf4 YP 001231833 1 351605 Proteobacteria
Brevibacillus brevis NBRC 100599 YP 002773468 1 358681 Firmicutes
Burkholderia rhizoxinica HKI 454 YP 004029397 1 882378 Proteobacteria
Clostridium cellulolyticum H10 YP 002505212 1 394503 Firmicutes
Bacillus amyloliquefaciens subsp plantarum UCMB5033 YP 008412411 1 1338518 Firmicutes
Bacillus amyloliquefaciens Y2 YP 006328282 1 1126211 Firmicutes
Bacillus atrophaeus 1942 YP 003973165 1 720555 Firmicutes
Bacillus subtilis subsp spizizenii TU B 10 YP 004877340 1 1052585 Firmicutes
Bacillus subtilis PY79 YP 008829956 1 1415167 Firmicutes
Bacillus subtilis subsp subtilis 6051 HGW YP 007533673 1 1147161 Firmicutes
Bacillus atrophaeus 1942 YP 003973166 1 720555 Firmicutes
Bacillus subtilis subsp spizizenii TU B 10 YP 004877341 1 1052585 Firmicutes
Bacillus subtilis subsp subtilis str BAB 1 YP 007662528 1 1302650 Firmicutes
Bacillus subtilis subsp subtilis 6051 HGW YP 007533674 1 1147161 Firmicutes
Bacillus subtilis PY79 YP 008829957 1 1415167 Firmicutes
Dickeya chrysanthemi L11 WP 033576297 1 Dchrysanthemi
Dickeya solani RNS 05 1 2A WP 057084645 1 Dsolani
Dickeya paradisiaca Ech703 WP 012765049 1 Dparadisiaca
Dickeya paradisiaca NCPPB 2511 WP 012765049 1 Dparadisiaca
Dickeya zeae DZ2Q WP 038924947 1 Dzeae
Dickeya zeae ZJU1202 WP 016940738 1 Dzeae
Dickeya zeae EC1 WP 016940738 1 Dzeae
Dickeya dianthicola IPO 980 WP 029729606 1 Ddianthicola
Dickeya dianthicola GBBC 2039 WP 024110021 1 Ddianthicola
Dickeya dianthicola NCPPB 3534 WP 024108916 1 Ddianthicola
Dickeya dianthicola RNS04 9 WP 024106518 1 Ddianthicola
Dickeya dianthicola NCPPB 453 WP 024106518 1 Ddianthicola
Burkholderia rhizoxinica HKI 454 YP 004029395 1 882378 Proteobacteria
Myxococcus xanthus DK 1622 YP 632113 1 246197 Proteobacteria
Burkholderia thailandensis MSMB121 YP 007920740 1 1249661 Proteobacteria
Bacillus amyloliquefaciens subsp plantarum UCMB5033 YP 008412674 1 1338518 Firmicutes
Bacillus amyloliquefaciens DSM 7 YP 003920382 1 692420 Firmicutes
Bacillus atrophaeus 1942 YP 003973164 1 720555 Firmicutes
Bacillus subtilis subsp spizizenii TU B 10 YP 004877339 1 1052585 Firmicutes
Bacillus subtilis subsp subtilis str BAB 1 YP 007662527 1 1302650 Firmicutes
Bacillus subtilis subsp subtilis 6051 HGW YP 007533672 1 1147161 Firmicutes
Bacillus subtilis PY79 YP 008829955 1 1415167 Firmicutes
Bacillus amyloliquefaciens DSM 7 YP 003920385 1 692420 Firmicutes
Bacillus amyloliquefaciens Y2 YP 006328599 1 1126211 Firmicutes
Bacillus amyloliquefaciens subsp plantarum UCMB5033 YP 008412677 1 1338518 Firmicutes
Bacillus atrophaeus 1942 YP 003973167 1 720555 Firmicutes
Bacillus subtilis subsp spizizenii TU B 10 YP 004877342 1 1052585 Firmicutes
Bacillus subtilis subsp subtilis 6051 HGW YP 007533675 1 1147161 Firmicutes
Bacillus subtilis subsp subtilis str BAB 1 YP 007662529 1 1302650 Firmicutes
Chitinophaga pinensis DSM 2588 YP 003124922 1 485918 Bacteroidetes Chlorobi group
Burkholderia rhizoxinica HKI 454 YP 004029393 1 882378 Proteobacteria
Ralstonia solanacearum Po82 YP 006032387 1 1031711 Proteobacteria
Bacillus amyloliquefaciens subsp plantarum UCMB5033 YP 008413317 1 1338518 Firmicutes
Bacillus amyloliquefaciens Y2 YP 006329293 1 1126211 Firmicutes
Paenibacillus mucilaginosus K02 YP 006189026 2 997761 Firmicutes
Dickeya gs fangzhongdai NCPPB 3274 WP 042859472 1 Dfangzhongdai
Dickeya chrysanthemi L11 WP 033576300 1 Dchrysanthemi
Dickeya chrysanthemi subsp chrysanthemi NCPPB 402 WP 040001548 1 Dchrysanthemi
210 oocN Oocydin A biosynthesis Seed
Dickeya paradisiaca Ech703 WP 012765044 1 Dparadisiaca
Dickeya paradisiaca NCPPB 2511 WP 012765044 1 Dparadisiaca
Dickeya zeae ZJU1202 WP 016940743 1 Dzeae
Dickeya zeae EC1 WP 016940743 1 Dzeae
Dickeya dianthicola IPO 980 WP 029729609 1 Ddianthicola
Dickeya dianthicola GBBC 2039 WP 024110022 1 Ddianthicola
Dickeya dianthicola RNS04 9 WP 024106523 1 Ddianthicola
Dickeya dianthicola NCPPB 453 WP 024106523 1 Ddianthicola
Dickeya dianthicola NCPPB 3534 WP 024108918 1 Ddianthicola
Dickeya solani RNS 05 1 2A WP 057084643 1 Dsolani
Dickeya solani MK16 WP 023638020 1 Dsolani
Dickeya solani GBBC 2040 WP 023641237 1 Dsolani
Dickeya solani PPO 9019 WP 023638020 1 Dsolani
Dickeya solani RNS 08 23 3 1 A WP 023638020 1 Dsolani
Dickeya solani MK10 WP 023638020 1 Dsolani
Dickeya solani PPO 9134 WP 023638020 1 Dsolani
Dickeya solani RNS 07 7 3B WP 023638020 1 Dsolani
Dickeya solani IPO 2222 WP 023638020 1 Dsolani

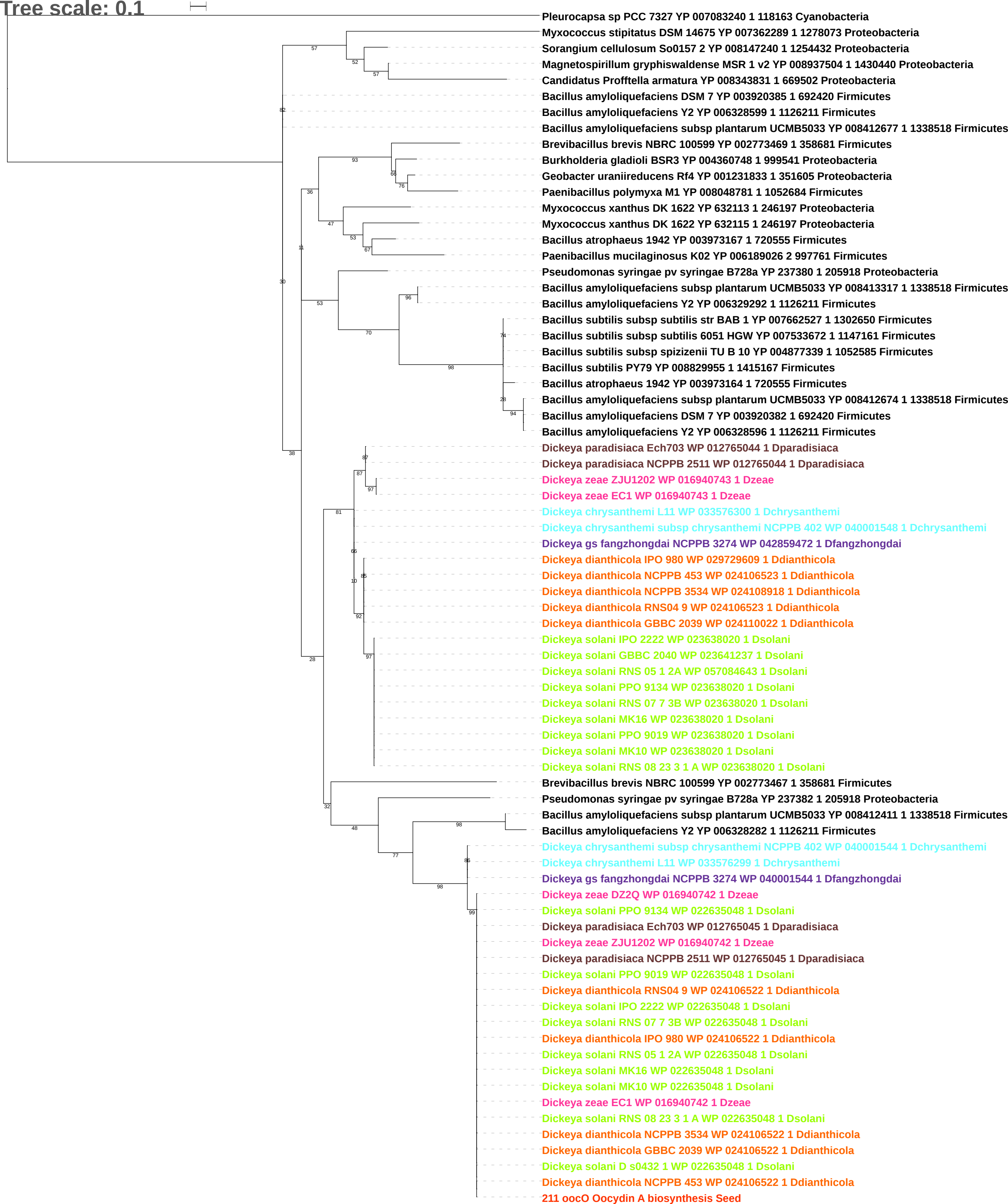
Tree scale: 0.1
Pleurocapsa sp PCC 7327 YP 007083240 1 118163 Cyanobacteria
Myxococcus stipitatus DSM 14675 YP 007362289 1 1278073 Proteobacteria
Sorangium cellulosum So0157 2 YP 008147240 1 1254432 Proteobacteria
Magnetospirillum gryphiswaldense MSR 1 v2 YP 008937504 1 1430440 Proteobacteria
Candidatus Profftella armatura YP 008343831 1 669502 Proteobacteria
Bacillus amyloliquefaciens DSM 7 YP 003920385 1 692420 Firmicutes
Bacillus amyloliquefaciens Y2 YP 006328599 1 1126211 Firmicutes
Bacillus amyloliquefaciens subsp plantarum UCMB5033 YP 008412677 1 1338518 Firmicutes
Brevibacillus brevis NBRC 100599 YP 002773469 1 358681 Firmicutes
Burkholderia gladioli BSR3 YP 004360748 1 999541 Proteobacteria
Geobacter uraniireducens Rf4 YP 001231833 1 351605 Proteobacteria
Paenibacillus polymyxa M1 YP 008048781 1 1052684 Firmicutes
Myxococcus xanthus DK 1622 YP 632113 1 246197 Proteobacteria
Myxococcus xanthus DK 1622 YP 632115 1 246197 Proteobacteria
Bacillus atrophaeus 1942 YP 003973167 1 720555 Firmicutes
Paenibacillus mucilaginosus K02 YP 006189026 2 997761 Firmicutes
Pseudomonas syringae pv syringae B728a YP 237380 1 205918 Proteobacteria
Bacillus amyloliquefaciens subsp plantarum UCMB5033 YP 008413317 1 1338518 Firmicutes
Bacillus amyloliquefaciens Y2 YP 006329292 1 1126211 Firmicutes
Bacillus subtilis subsp subtilis str BAB 1 YP 007662527 1 1302650 Firmicutes
Bacillus subtilis subsp subtilis 6051 HGW YP 007533672 1 1147161 Firmicutes
Bacillus subtilis subsp spizizenii TU B 10 YP 004877339 1 1052585 Firmicutes
Bacillus subtilis PY79 YP 008829955 1 1415167 Firmicutes
Bacillus atrophaeus 1942 YP 003973164 1 720555 Firmicutes
Bacillus amyloliquefaciens subsp plantarum UCMB5033 YP 008412674 1 1338518 Firmicutes
Bacillus amyloliquefaciens DSM 7 YP 003920382 1 692420 Firmicutes
Bacillus amyloliquefaciens Y2 YP 006328596 1 1126211 Firmicutes
Dickeya paradisiaca Ech703 WP 012765044 1 Dparadisiaca
Dickeya paradisiaca NCPPB 2511 WP 012765044 1 Dparadisiaca
Dickeya zeae ZJU1202 WP 016940743 1 Dzeae
Dickeya zeae EC1 WP 016940743 1 Dzeae
Dickeya chrysanthemi L11 WP 033576300 1 Dchrysanthemi
Dickeya chrysanthemi subsp chrysanthemi NCPPB 402 WP 040001548 1 Dchrysanthemi
Dickeya gs fangzhongdai NCPPB 3274 WP 042859472 1 Dfangzhongdai
Dickeya dianthicola IPO 980 WP 029729609 1 Ddianthicola
Dickeya dianthicola NCPPB 453 WP 024106523 1 Ddianthicola
Dickeya dianthicola NCPPB 3534 WP 024108918 1 Ddianthicola
Dickeya dianthicola RNS04 9 WP 024106523 1 Ddianthicola
Dickeya dianthicola GBBC 2039 WP 024110022 1 Ddianthicola
Dickeya solani IPO 2222 WP 023638020 1 Dsolani
Dickeya solani GBBC 2040 WP 023641237 1 Dsolani
Dickeya solani RNS 05 1 2A WP 057084643 1 Dsolani
Dickeya solani PPO 9134 WP 023638020 1 Dsolani
Dickeya solani RNS 07 7 3B WP 023638020 1 Dsolani
Dickeya solani MK16 WP 023638020 1 Dsolani
Dickeya solani PPO 9019 WP 023638020 1 Dsolani
Dickeya solani MK10 WP 023638020 1 Dsolani
Dickeya solani RNS 08 23 3 1 A WP 023638020 1 Dsolani
Brevibacillus brevis NBRC 100599 YP 002773467 1 358681 Firmicutes
Pseudomonas syringae pv syringae B728a YP 237382 1 205918 Proteobacteria
Bacillus amyloliquefaciens subsp plantarum UCMB5033 YP 008412411 1 1338518 Firmicutes
Bacillus amyloliquefaciens Y2 YP 006328282 1 1126211 Firmicutes
Dickeya chrysanthemi subsp chrysanthemi NCPPB 402 WP 040001544 1 Dchrysanthemi
Dickeya chrysanthemi L11 WP 033576299 1 Dchrysanthemi
Dickeya gs fangzhongdai NCPPB 3274 WP 040001544 1 Dfangzhongdai
Dickeya zeae DZ2Q WP 016940742 1 Dzeae
Dickeya solani PPO 9134 WP 022635048 1 Dsolani
Dickeya paradisiaca Ech703 WP 012765045 1 Dparadisiaca
Dickeya zeae ZJU1202 WP 016940742 1 Dzeae
Dickeya paradisiaca NCPPB 2511 WP 012765045 1 Dparadisiaca
Dickeya solani PPO 9019 WP 022635048 1 Dsolani
Dickeya dianthicola RNS04 9 WP 024106522 1 Ddianthicola
Dickeya solani IPO 2222 WP 022635048 1 Dsolani
Dickeya solani RNS 07 7 3B WP 022635048 1 Dsolani
Dickeya dianthicola IPO 980 WP 024106522 1 Ddianthicola
Dickeya solani RNS 05 1 2A WP 022635048 1 Dsolani
Dickeya solani MK16 WP 022635048 1 Dsolani
Dickeya solani MK10 WP 022635048 1 Dsolani
Dickeya zeae EC1 WP 016940742 1 Dzeae
Dickeya solani RNS 08 23 3 1 A WP 022635048 1 Dsolani
Dickeya dianthicola NCPPB 3534 WP 024106522 1 Ddianthicola
Dickeya dianthicola GBBC 2039 WP 024106522 1 Ddianthicola
Dickeya solani D s0432 1 WP 022635048 1 Dsolani
Dickeya dianthicola NCPPB 453 WP 024106522 1 Ddianthicola
211 oocO Oocydin A biosynthesis Seed

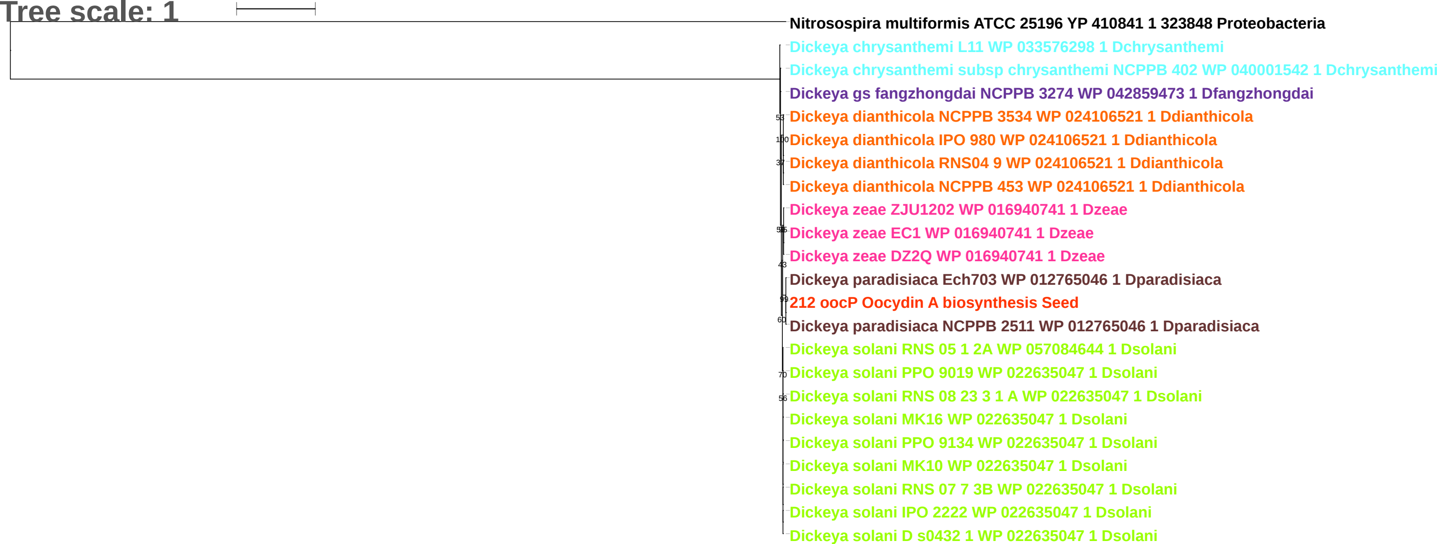
Tree scale: 1
Nitrosospira multiformis ATCC 25196 YP 410841 1 323848 Proteobacteria
Dickeya chrysanthemi L11 WP 033576298 1 Dchrysanthemi
Dickeya chrysanthemi subsp chrysanthemi NCPPB 402 WP 040001542 1 Dchrysanthemi
Dickeya gs fangzhongdai NCPPB 3274 WP 042859473 1 Dfangzhongdai
Dickeya dianthicola NCPPB 3534 WP 024106521 1 Ddianthicola
Dickeya dianthicola IPO 980 WP 024106521 1 Ddianthicola
Dickeya dianthicola RNS04 9 WP 024106521 1 Ddianthicola
Dickeya dianthicola NCPPB 453 WP 024106521 1 Ddianthicola
Dickeya zeae ZJU1202 WP 016940741 1 Dzeae
Dickeya zeae EC1 WP 016940741 1 Dzeae
Dickeya zeae DZ2Q WP 016940741 1 Dzeae
Dickeya paradisiaca Ech703 WP 012765046 1 Dparadisiaca
212 oocP Oocydin A biosynthesis Seed
Dickeya paradisiaca NCPPB 2511 WP 012765046 1 Dparadisiaca
Dickeya solani RNS 05 1 2A WP 057084644 1 Dsolani
Dickeya solani PPO 9019 WP 022635047 1 Dsolani
Dickeya solani RNS 08 23 3 1 A WP 022635047 1 Dsolani
Dickeya solani MK16 WP 022635047 1 Dsolani
Dickeya solani PPO 9134 WP 022635047 1 Dsolani
Dickeya solani MK10 WP 022635047 1 Dsolani
Dickeya solani RNS 07 7 3B WP 022635047 1 Dsolani
Dickeya solani IPO 2222 WP 022635047 1 Dsolani
Dickeya solani D s0432 1 WP 022635047 1 Dsolani

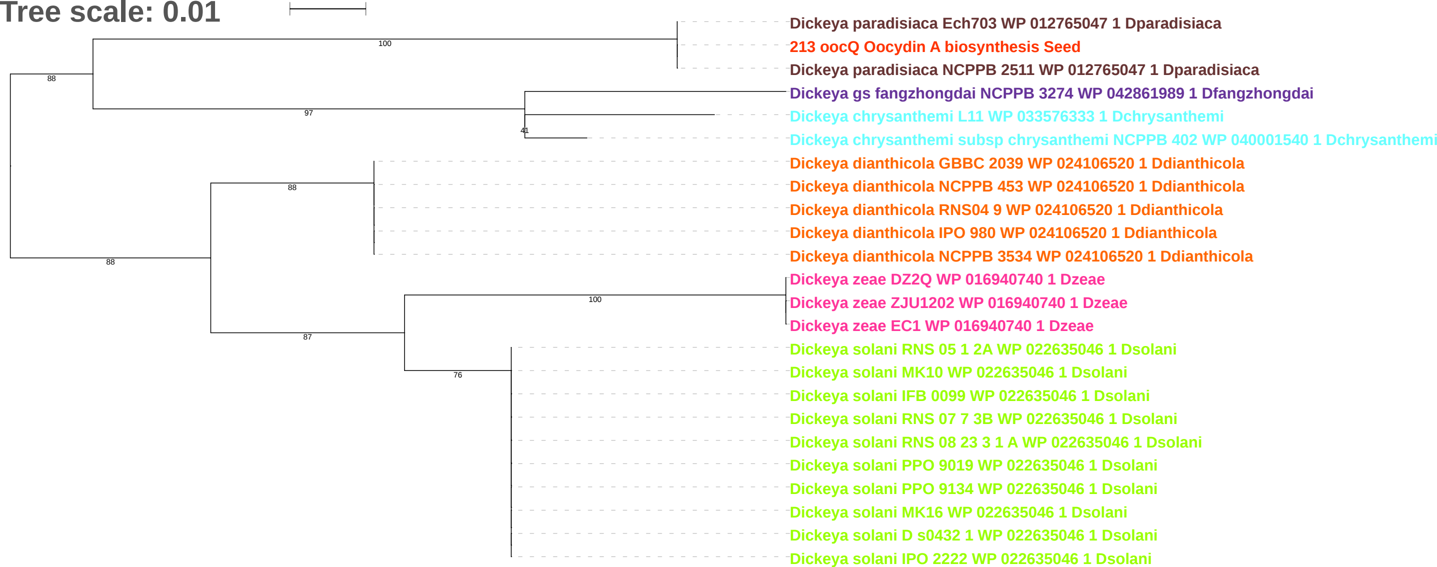

Tree scale: 0.01
100
88
97
41
88
88
100
87
76
Dickeya paradisiaca Ech703 WP 012765047 1 Dparadisiaca
213 oocQ Oocydin A biosynthesis Seed
Dickeya paradisiaca NCPPB 2511 WP 012765047 1 Dparadisiaca
Dickeya gs fangzhongdai NCPPB 3274 WP 042861989 1 Dfangzhongdai
Dickeya chrysanthemi L11 WP 033576333 1 Dchrysanthemi
Dickeya chrysanthemi subsp chrysanthemi NCPPB 402 WP 040001540 1 Dchrysanthemi
Dickeya dianthicola GBBC 2039 WP 024106520 1 Ddianthicola
Dickeya dianthicola NCPPB 453 WP 024106520 1 Ddianthicola
Dickeya dianthicola RNS04 9 WP 024106520 1 Ddianthicola
Dickeya dianthicola IPO 980 WP 024106520 1 Ddianthicola
Dickeya dianthicola NCPPB 3534 WP 024106520 1 Ddianthicola
Dickeya zeae DZ2Q WP 016940740 1 Dzeae
Dickeya zeae ZJU1202 WP 016940740 1 Dzeae
Dickeya zeae EC1 WP 016940740 1 Dzeae
Dickeya solani RNS 05 1 2A WP 022635046 1 Dsolani
Dickeya solani MK10 WP 022635046 1 Dsolani
Dickeya solani IFB 0099 WP 022635046 1 Dsolani
Dickeya solani RNS 07 7 3B WP 022635046 1 Dsolani
Dickeya solani RNS 08 23 3 1 A WP 022635046 1 Dsolani
Dickeya solani PPO 9019 WP 022635046 1 Dsolani
Dickeya solani PPO 9134 WP 022635046 1 Dsolani
Dickeya solani MK16 WP 022635046 1 Dsolani
Dickeya solani D s0432 1 WP 022635046 1 Dsolani
Dickeya solani IPO 2222 WP 022635046 1 Dsolani

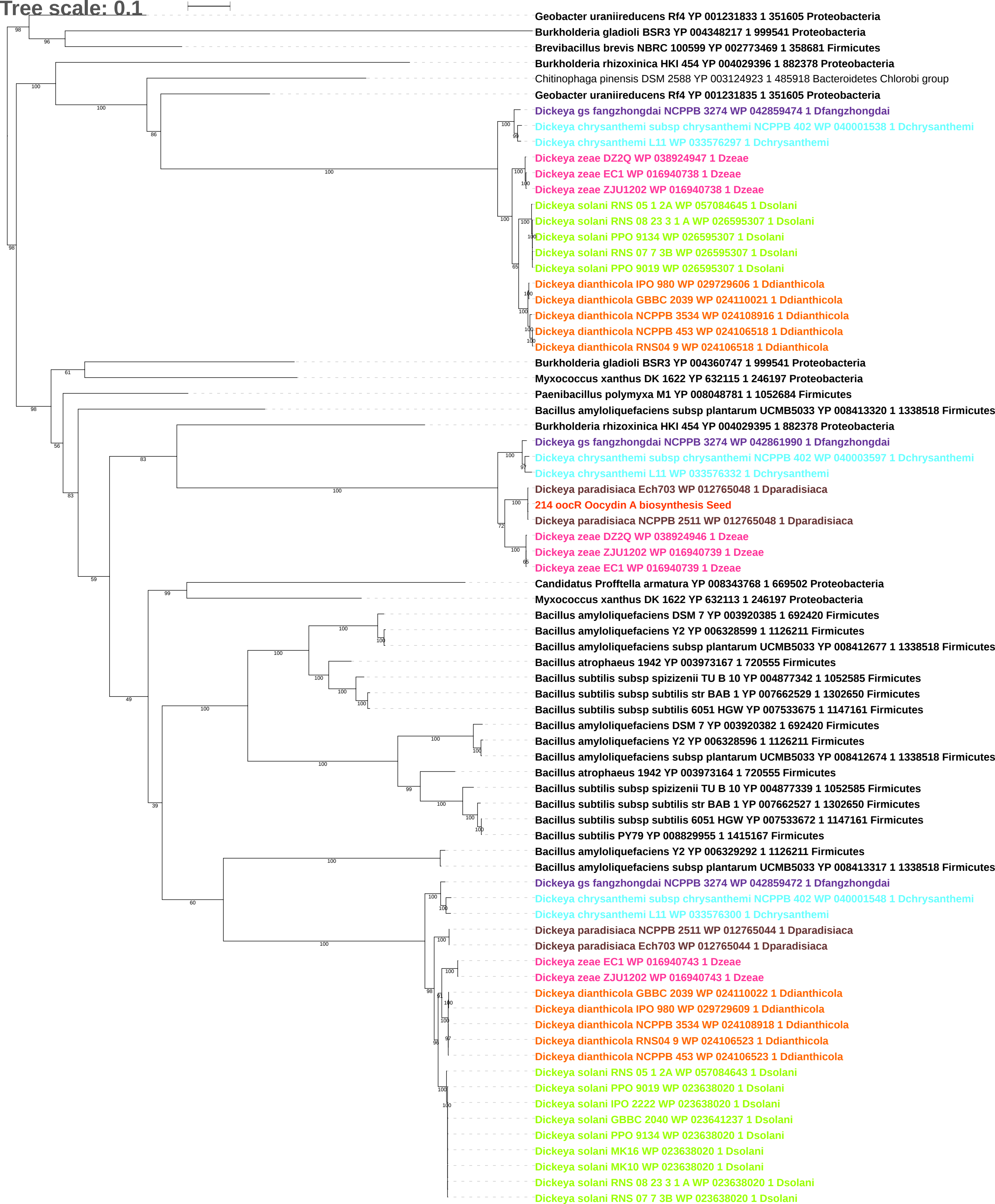
Tree scale: 0.1
Geobacter uraniireducens Rf4 YP 001231833 1 351605 Proteobacteria
Burkholderia gladioli BSR3 YP 004348217 1 999541 Proteobacteria
Brevibacillus brevis NBRC 100599 YP 002773469 1 358681 Firmicutes
Burkholderia rhizoxinica HKI 454 YP 004029396 1 882378 Proteobacteria
Chitinophaga pinensis DSM 2588 YP 003124923 1 485918 Bacteroidetes Chlorobi group
Geobacter uraniireducens Rf4 YP 001231835 1 351605 Proteobacteria
Dickeya gs fangzhongdai NCPPB 3274 WP 042859474 1 Dfangzhongdai
Dickeya chrysanthemi subsp chrysanthemi NCPPB 402 WP 040001538 1 Dchrysanthemi
Dickeya chrysanthemi L11 WP 033576297 1 Dchrysanthemi
Dickeya zeae DZ2Q WP 038924947 1 Dzeae
Dickeya zeae EC1 WP 016940738 1 Dzeae
Dickeya zeae ZJU1202 WP 016940738 1 Dzeae
Dickeya solani RNS 05 1 2A WP 057084645 1 Dsolani
Dickeya solani RNS 08 23 3 1 A WP 026595307 1 Dsolani
Dickeya solani PPO 9134 WP 026595307 1 Dsolani
Dickeya solani RNS 07 7 3B WP 026595307 1 Dsolani
Dickeya solani PPO 9019 WP 026595307 1 Dsolani
Dickeya dianthicola IPO 980 WP 029729606 1 Ddianthicola
Dickeya dianthicola GBBC 2039 WP 024110021 1 Ddianthicola
Dickeya dianthicola NCPPB 3534 WP 024108916 1 Ddianthicola
Dickeya dianthicola NCPPB 453 WP 024106518 1 Ddianthicola
Dickeya dianthicola RNS04 9 WP 024106518 1 Ddianthicola
Burkholderia gladioli BSR3 YP 004360747 1 999541 Proteobacteria
Myxococcus xanthus DK 1622 YP 632115 1 246197 Proteobacteria
Paenibacillus polymyxa M1 YP 008048781 1 1052684 Firmicutes
Bacillus amyloliquefaciens subsp plantarum UCMB5033 YP 008413320 1 1338518 Firmicutes
Burkholderia rhizoxinica HKI 454 YP 004029395 1 882378 Proteobacteria
Dickeya gs fangzhongdai NCPPB 3274 WP 042861990 1 Dfangzhongdai
Dickeya chrysanthemi subsp chrysanthemi NCPPB 402 WP 040003597 1 Dchrysanthemi
Dickeya chrysanthemi L11 WP 033576332 1 Dchrysanthemi
Dickeya paradisiaca Ech703 WP 012765048 1 Dparadisiaca
214 oocR Oocydin A biosynthesis Seed
Dickeya paradisiaca NCPPB 2511 WP 012765048 1 Dparadisiaca
Dickeya zeae DZ2Q WP 038924946 1 Dzeae
Dickeya zeae ZJU1202 WP 016940739 1 Dzeae
Dickeya zeae EC1 WP 016940739 1 Dzeae
Candidatus Profftella armatura YP 008343768 1 669502 Proteobacteria
Myxococcus xanthus DK 1622 YP 632113 1 246197 Proteobacteria
Bacillus amyloliquefaciens DSM 7 YP 003920385 1 692420 Firmicutes
Bacillus amyloliquefaciens Y2 YP 006328599 1 1126211 Firmicutes
Bacillus amyloliquefaciens subsp plantarum UCMB5033 YP 008412677 1 1338518 Firmicutes
Bacillus atrophaeus 1942 YP 003973167 1 720555 Firmicutes
Bacillus subtilis subsp spizizenii TU B 10 YP 004877342 1 1052585 Firmicutes
Bacillus subtilis subsp subtilis str BAB 1 YP 007662529 1 1302650 Firmicutes
Bacillus subtilis subsp subtilis 6051 HGW YP 007533675 1 1147161 Firmicutes
Bacillus amyloliquefaciens DSM 7 YP 003920382 1 692420 Firmicutes
Bacillus amyloliquefaciens Y2 YP 006328596 1 1126211 Firmicutes
Bacillus amyloliquefaciens subsp plantarum UCMB5033 YP 008412674 1 1338518 Firmicutes
Bacillus atrophaeus 1942 YP 003973164 1 720555 Firmicutes
Bacillus subtilis subsp spizizenii TU B 10 YP 004877339 1 1052585 Firmicutes
Bacillus subtilis subsp subtilis str BAB 1 YP 007662527 1 1302650 Firmicutes
Bacillus subtilis subsp subtilis 6051 HGW YP 007533672 1 1147161 Firmicutes
Bacillus subtilis PY79 YP 008829955 1 1415167 Firmicutes
Bacillus amyloliquefaciens Y2 YP 006329292 1 1126211 Firmicutes
Bacillus amyloliquefaciens subsp plantarum UCMB5033 YP 008413317 1 1338518 Firmicutes
Dickeya gs fangzhongdai NCPPB 3274 WP 042859472 1 Dfangzhongdai
Dickeya chrysanthemi subsp chrysanthemi NCPPB 402 WP 040001548 1 Dchrysanthemi
Dickeya chrysanthemi L11 WP 033576300 1 Dchrysanthemi
Dickeya paradisiaca NCPPB 2511 WP 012765044 1 Dparadisiaca
Dickeya paradisiaca Ech703 WP 012765044 1 Dparadisiaca
Dickeya zeae EC1 WP 016940743 1 Dzeae
Dickeya zeae ZJU1202 WP 016940743 1 Dzeae
Dickeya dianthicola GBBC 2039 WP 024110022 1 Ddianthicola
Dickeya dianthicola IPO 980 WP 029729609 1 Ddianthicola
Dickeya dianthicola NCPPB 3534 WP 024108918 1 Ddianthicola
Dickeya dianthicola RNS04 9 WP 024106523 1 Ddianthicola
Dickeya dianthicola NCPPB 453 WP 024106523 1 Ddianthicola
Dickeya solani RNS 05 1 2A WP 057084643 1 Dsolani
Dickeya solani PPO 9019 WP 023638020 1 Dsolani
Dickeya solani IPO 2222 WP 023638020 1 Dsolani
Dickeya solani GBBC 2040 WP 023641237 1 Dsolani
Dickeya solani PPO 9134 WP 023638020 1 Dsolani
Dickeya solani MK16 WP 023638020 1 Dsolani
Dickeya solani MK10 WP 023638020 1 Dsolani
Dickeya solani RNS 08 23 3 1 A WP 023638020 1 Dsolani
Dickeya solani RNS 07 7 3B WP 023638020 1 Dsolani

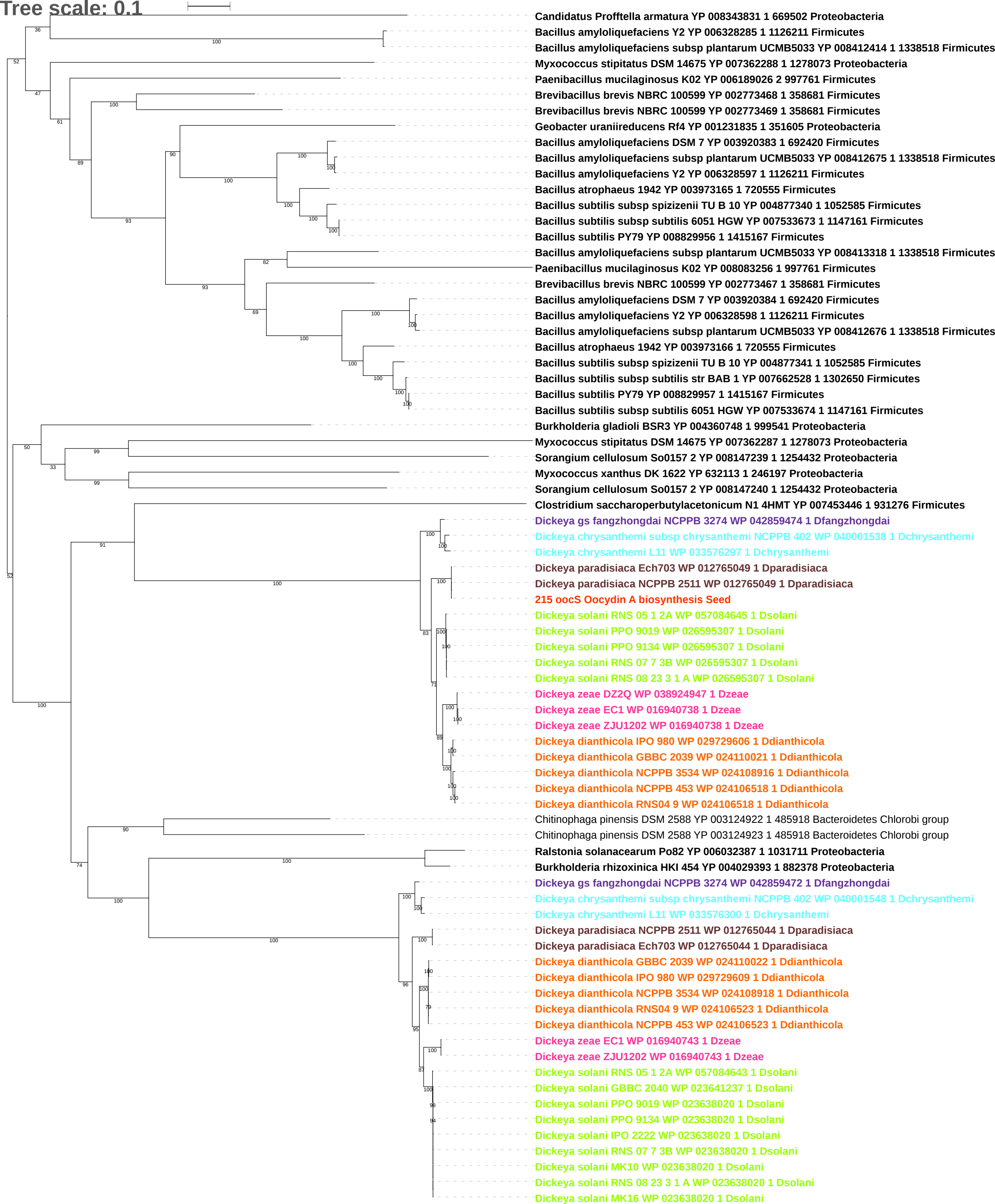
Tree scale: 0.1
Candidatus Profftella armatura YP 008343831 1 669502 Proteobacteria
Bacillus amyloliquefaciens Y2 YP 006328285 1 1126211 Firmicutes
Bacillus amyloliquefaciens subsp plantarum UCMB5033 YP 008412414 1 1338518 Firmicutes
Myxococcus stipitatus DSM 14675 YP 007362288 1 1278073 Proteobacteria
Paenibacillus mucilaginosus K02 YP 006189026 2 997761 Firmicutes
Brevibacillus brevis NBRC 100599 YP 002773468 1 358681 Firmicutes
Brevibacillus brevis NBRC 100599 YP 002773469 1 358681 Firmicutes
Geobacter uraniireducens Rf4 YP 001231835 1 351605 Proteobacteria
Bacillus amyloliquefaciens DSM 7 YP 003920383 1 692420 Firmicutes
Bacillus amyloliquefaciens subsp plantarum UCMB5033 YP 008412675 1 1338518 Firmicutes
Bacillus amyloliquefaciens Y2 YP 006328597 1 1126211 Firmicutes
Bacillus atrophaeus 1942 YP 003973165 1 720555 Firmicutes
Bacillus subtilis subsp spizizenii TU B 10 YP 004877340 1 1052585 Firmicutes
Bacillus subtilis subsp subtilis 6051 HGW YP 007533673 1 1147161 Firmicutes
Bacillus subtilis PY79 YP 008829956 1 1415167 Firmicutes
Bacillus amyloliquefaciens subsp plantarum UCMB5033 YP 008413318 1 1338518 Firmicutes
Paenibacillus mucilaginosus K02 YP 008083256 1 997761 Firmicutes
Brevibacillus brevis NBRC 100599 YP 002773467 1 358681 Firmicutes
Bacillus amyloliquefaciens DSM 7 YP 003920384 1 692420 Firmicutes
Bacillus amyloliquefaciens Y2 YP 006328598 1 1126211 Firmicutes
Bacillus amyloliquefaciens subsp plantarum UCMB5033 YP 008412676 1 1338518 Firmicutes
Bacillus atrophaeus 1942 YP 003973166 1 720555 Firmicutes
Bacillus subtilis subsp spizizenii TU B 10 YP 004877341 1 1052585 Firmicutes
Bacillus subtilis subsp subtilis str BAB 1 YP 007662528 1 1302650 Firmicutes
Bacillus subtilis PY79 YP 008829957 1 1415167 Firmicutes
Bacillus subtilis subsp subtilis 6051 HGW YP 007533674 1 1147161 Firmicutes
Burkholderia gladioli BSR3 YP 004360748 1 999541 Proteobacteria
Myxococcus stipitatus DSM 14675 YP 007362287 1 1278073 Proteobacteria
Sorangium cellulosum So0157 2 YP 008147239 1 1254432 Proteobacteria
Myxococcus xanthus DK 1622 YP 632113 1 246197 Proteobacteria
Sorangium cellulosum So0157 2 YP 008147240 1 1254432 Proteobacteria
Clostridium saccharoperbutylacetonicum N1 4HMT YP 007453446 1 931276 Firmicutes
Dickeya gs fangzhongdai NCPPB 3274 WP 042859474 1 Dfangzhongdai
Dickeya chrysanthemi subsp chrysanthemi NCPPB 402 WP 040001538 1 Dchrysanthemi
Dickeya chrysanthemi L11 WP 033576297 1 Dchrysanthemi
Dickeya paradisiaca Ech703 WP 012765049 1 Dparadisiaca
Dickeya paradisiaca NCPPB 2511 WP 012765049 1 Dparadisiaca
215 oocS Oocydin A biosynthesis Seed
Dickeya solani RNS 05 1 2A WP 057084645 1 Dsolani
Dickeya solani PPO 9019 WP 026595307 1 Dsolani
Dickeya solani PPO 9134 WP 026595307 1 Dsolani
Dickeya solani RNS 07 7 3B WP 026595307 1 Dsolani
Dickeya solani RNS 08 23 3 1 A WP 026595307 1 Dsolani
Dickeya zeae DZ2Q WP 038924947 1 Dzeae
Dickeya zeae EC1 WP 016940738 1 Dzeae
Dickeya zeae ZJU1202 WP 016940738 1 Dzeae
Dickeya dianthicola IPO 980 WP 029729606 1 Ddianthicola
Dickeya dianthicola GBBC 2039 WP 024110021 1 Ddianthicola
Dickeya dianthicola NCPPB 3534 WP 024108916 1 Ddianthicola
Dickeya dianthicola NCPPB 453 WP 024106518 1 Ddianthicola
Dickeya dianthicola RNS04 9 WP 024106518 1 Ddianthicola
Chitinophaga pinensis DSM 2588 YP 003124922 1 485918 Bacteroidetes Chlorobi group
Chitinophaga pinensis DSM 2588 YP 003124923 1 485918 Bacteroidetes Chlorobi group
Ralstonia solanacearum Po82 YP 006032387 1 1031711 Proteobacteria
Burkholderia rhizoxinica HKI 454 YP 004029393 1 882378 Proteobacteria
Dickeya gs fangzhongdai NCPPB 3274 WP 042859472 1 Dfangzhongdai
Dickeya chrysanthemi subsp chrysanthemi NCPPB 402 WP 040001548 1 Dchrysanthemi
Dickeya chrysanthemi L11 WP 033576300 1 Dchrysanthemi
Dickeya paradisiaca NCPPB 2511 WP 012765044 1 Dparadisiaca
Dickeya paradisiaca Ech703 WP 012765044 1 Dparadisiaca
Dickeya dianthicola GBBC 2039 WP 024110022 1 Ddianthicola
Dickeya dianthicola IPO 980 WP 029729609 1 Ddianthicola
Dickeya dianthicola NCPPB 3534 WP 024108918 1 Ddianthicola
Dickeya dianthicola RNS04 9 WP 024106523 1 Ddianthicola
Dickeya dianthicola NCPPB 453 WP 024106523 1 Ddianthicola
Dickeya zeae EC1 WP 016940743 1 Dzeae
Dickeya zeae ZJU1202 WP 016940743 1 Dzeae
Dickeya solani RNS 05 1 2A WP 057084643 1 Dsolani
Dickeya solani GBBC 2040 WP 023641237 1 Dsolani
Dickeya solani PPO 9019 WP 023638020 1 Dsolani
Dickeya solani PPO 9134 WP 023638020 1 Dsolani
Dickeya solani IPO 2222 WP 023638020 1 Dsolani
Dickeya solani RNS 07 7 3B WP 023638020 1 Dsolani
Dickeya solani MK10 WP 023638020 1 Dsolani
Dickeya solani RNS 08 23 3 1 A WP 023638020 1 Dsolani
Dickeya solani MK16 WP 023638020 1 Dsolani

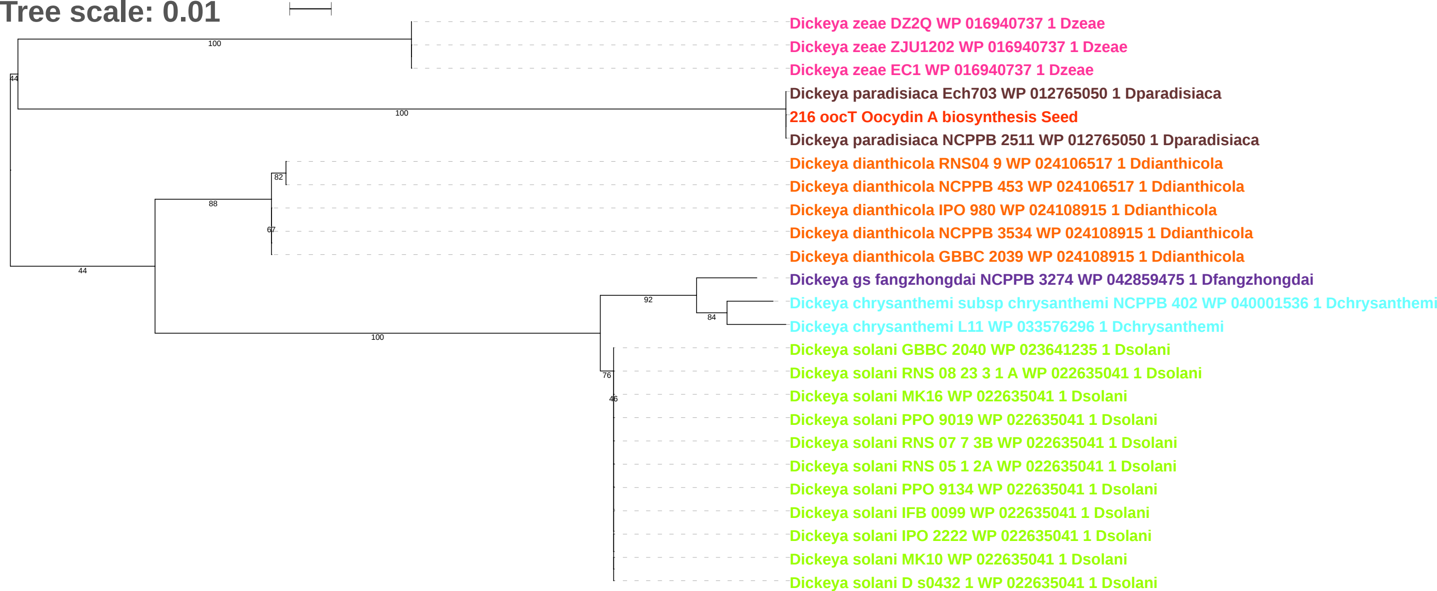
Tree scale: 0.01
100
44
100
82
88
67
44
92
84
100
76
46
Dickeya zeae DZ2Q WP 016940737 1 Dzeae
Dickeya zeae ZJU1202 WP 016940737 1 Dzeae
Dickeya zeae EC1 WP 016940737 1 Dzeae
Dickeya paradisiaca Ech703 WP 012765050 1 Dparadisiaca
216 oocT Oocydin A biosynthesis Seed
Dickeya paradisiaca NCPPB 2511 WP 012765050 1 Dparadisiaca
Dickeya dianthicola RNS04 9 WP 024106517 1 Ddianthicola
Dickeya dianthicola NCPPB 453 WP 024106517 1 Ddianthicola
Dickeya dianthicola IPO 980 WP 024108915 1 Ddianthicola
Dickeya dianthicola NCPPB 3534 WP 024108915 1 Ddianthicola
Dickeya dianthicola GBBC 2039 WP 024108915 1 Ddianthicola
Dickeya gs fangzhongdai NCPPB 3274 WP 042859475 1 Dfangzhongdai
Dickeya chrysanthemi subsp chrysanthemi NCPPB 402 WP 040001536 1 Dchrysanthemi
Dickeya chrysanthemi L11 WP 033576296 1 Dchrysanthemi
Dickeya solani GBBC 2040 WP 023641235 1 Dsolani
Dickeya solani RNS 08 23 3 1 A WP 022635041 1 Dsolani
Dickeya solani MK16 WP 022635041 1 Dsolani
Dickeya solani PPO 9019 WP 022635041 1 Dsolani
Dickeya solani RNS 07 7 3B WP 022635041 1 Dsolani
Dickeya solani RNS 05 1 2A WP 022635041 1 Dsolani
Dickeya solani PPO 9134 WP 022635041 1 Dsolani
Dickeya solani IFB 0099 WP 022635041 1 Dsolani
Dickeya solani IPO 2222 WP 022635041 1 Dsolani
Dickeya solani MK10 WP 022635041 1 Dsolani
Dickeya solani D s0432 1 WP 022635041 1 Dsolani

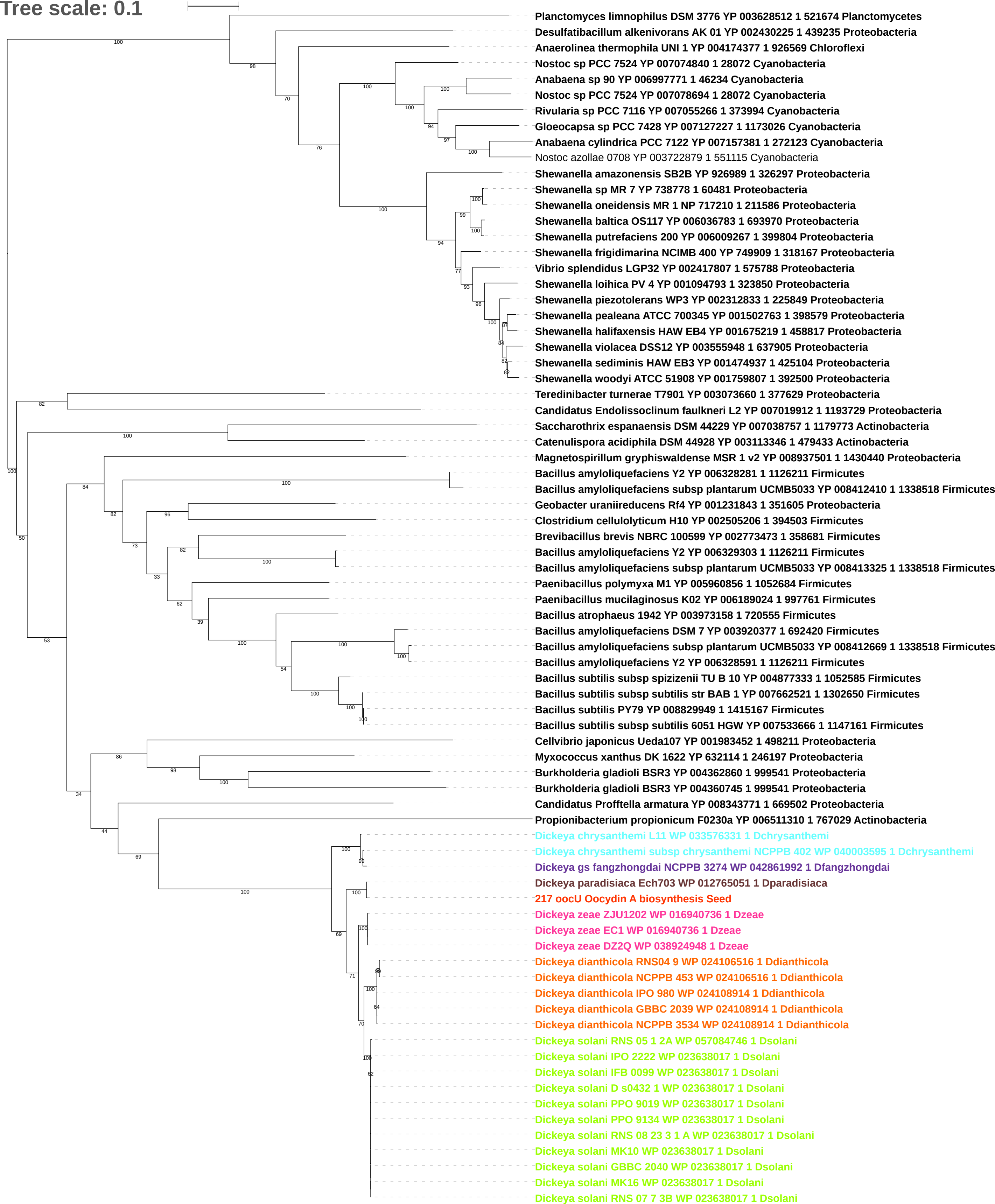
Tree scale: 0.1
Planctomyces limnophilus DSM 3776 YP 003628512 1 521674 Planctomycetes
Desulfatibacillum alkenivorans AK 01 YP 002430225 1 439235 Proteobacteria
Anaerolinea thermophila UNI 1 YP 004174377 1 926569 Chloroflexi
Nostoc sp PCC 7524 YP 007074840 1 28072 Cyanobacteria
Anabaena sp 90 YP 006997771 1 46234 Cyanobacteria
Nostoc sp PCC 7524 YP 007078694 1 28072 Cyanobacteria
Rivularia sp PCC 7116 YP 007055266 1 373994 Cyanobacteria
Gloeocapsa sp PCC 7428 YP 007127227 1 1173026 Cyanobacteria
Anabaena cylindrica PCC 7122 YP 007157381 1 272123 Cyanobacteria
Nostoc azollae 0708 YP 003722879 1 551115 Cyanobacteria
Shewanella amazonensis SB2B YP 926989 1 326297 Proteobacteria
Shewanella sp MR 7 YP 738778 1 60481 Proteobacteria
Shewanella oneidensis MR 1 NP 717210 1 211586 Proteobacteria
Shewanella baltica OS117 YP 006036783 1 693970 Proteobacteria
Shewanella putrefaciens 200 YP 006009267 1 399804 Proteobacteria
Shewanella frigidimarina NCIMB 400 YP 749909 1 318167 Proteobacteria
Vibrio splendidus LGP32 YP 002417807 1 575788 Proteobacteria
Shewanella loihica PV 4 YP 001094793 1 323850 Proteobacteria
Shewanella piezotolerans WP3 YP 002312833 1 225849 Proteobacteria
Shewanella pealeana ATCC 700345 YP 001502763 1 398579 Proteobacteria
Shewanella halifaxensis HAW EB4 YP 001675219 1 458817 Proteobacteria
Shewanella violacea DSS12 YP 003555948 1 637905 Proteobacteria
Shewanella sediminis HAW EB3 YP 001474937 1 425104 Proteobacteria
Shewanella woodyi ATCC 51908 YP 001759807 1 392500 Proteobacteria
Teredinibacter turnerae T7901 YP 003073660 1 377629 Proteobacteria
Candidatus Endolissoclinum faulkneri L2 YP 007019912 1 1193729 Proteobacteria
Saccharothrix espanaensis DSM 44229 YP 007038757 1 1179773 Actinobacteria
Catenulispora acidiphila DSM 44928 YP 003113346 1 479433 Actinobacteria
Magnetospirillum gryphiswaldense MSR 1 v2 YP 008937501 1 1430440 Proteobacteria
Bacillus amyloliquefaciens Y2 YP 006328281 1 1126211 Firmicutes
Bacillus amyloliquefaciens subsp plantarum UCMB5033 YP 008412410 1 1338518 Firmicutes
Geobacter uraniireducens Rf4 YP 001231843 1 351605 Proteobacteria
Clostridium cellulolyticum H10 YP 002505206 1 394503 Firmicutes
Brevibacillus brevis NBRC 100599 YP 002773473 1 358681 Firmicutes
Bacillus amyloliquefaciens Y2 YP 006329303 1 1126211 Firmicutes
Bacillus amyloliquefaciens subsp plantarum UCMB5033 YP 008413325 1 1338518 Firmicutes
Paenibacillus polymyxa M1 YP 005960856 1 1052684 Firmicutes
Paenibacillus mucilaginosus K02 YP 006189024 1 997761 Firmicutes
Bacillus atrophaeus 1942 YP 003973158 1 720555 Firmicutes
Bacillus amyloliquefaciens DSM 7 YP 003920377 1 692420 Firmicutes
Bacillus amyloliquefaciens subsp plantarum UCMB5033 YP 008412669 1 1338518 Firmicutes
Bacillus amyloliquefaciens Y2 YP 006328591 1 1126211 Firmicutes
Bacillus subtilis subsp spizizenii TU B 10 YP 004877333 1 1052585 Firmicutes
Bacillus subtilis subsp subtilis str BAB 1 YP 007662521 1 1302650 Firmicutes
Bacillus subtilis PY79 YP 008829949 1 1415167 Firmicutes
Bacillus subtilis subsp subtilis 6051 HGW YP 007533666 1 1147161 Firmicutes
Cellvibrio japonicus Ueda107 YP 001983452 1 498211 Proteobacteria
Myxococcus xanthus DK 1622 YP 632114 1 246197 Proteobacteria
Burkholderia gladioli BSR3 YP 004362860 1 999541 Proteobacteria
Burkholderia gladioli BSR3 YP 004360745 1 999541 Proteobacteria
Candidatus Profftella armatura YP 008343771 1 669502 Proteobacteria
Propionibacterium propionicum F0230a YP 006511310 1 767029 Actinobacteria
Dickeya chrysanthemi L11 WP 033576331 1 Dchrysanthemi
Dickeya chrysanthemi subsp chrysanthemi NCPPB 402 WP 040003595 1 Dchrysanthemi
Dickeya gs fangzhongdai NCPPB 3274 WP 042861992 1 Dfangzhongdai
Dickeya paradisiaca Ech703 WP 012765051 1 Dparadisiaca
217 oocU Oocydin A biosynthesis Seed
Dickeya zeae ZJU1202 WP 016940736 1 Dzeae
Dickeya zeae EC1 WP 016940736 1 Dzeae
Dickeya zeae DZ2Q WP 038924948 1 Dzeae
Dickeya dianthicola RNS04 9 WP 024106516 1 Ddianthicola
Dickeya dianthicola NCPPB 453 WP 024106516 1 Ddianthicola
Dickeya dianthicola IPO 980 WP 024108914 1 Ddianthicola
Dickeya dianthicola GBBC 2039 WP 024108914 1 Ddianthicola
Dickeya dianthicola NCPPB 3534 WP 024108914 1 Ddianthicola
Dickeya solani RNS 05 1 2A WP 057084746 1 Dsolani
Dickeya solani IPO 2222 WP 023638017 1 Dsolani
Dickeya solani IFB 0099 WP 023638017 1 Dsolani
Dickeya solani D s0432 1 WP 023638017 1 Dsolani
Dickeya solani PPO 9019 WP 023638017 1 Dsolani
Dickeya solani PPO 9134 WP 023638017 1 Dsolani
Dickeya solani RNS 08 23 3 1 A WP 023638017 1 Dsolani
Dickeya solani MK10 WP 023638017 1 Dsolani
Dickeya solani GBBC 2040 WP 023638017 1 Dsolani
Dickeya solani MK16 WP 023638017 1 Dsolani
Dickeya solani RNS 07 7 3B WP 023638017 1 Dsolani

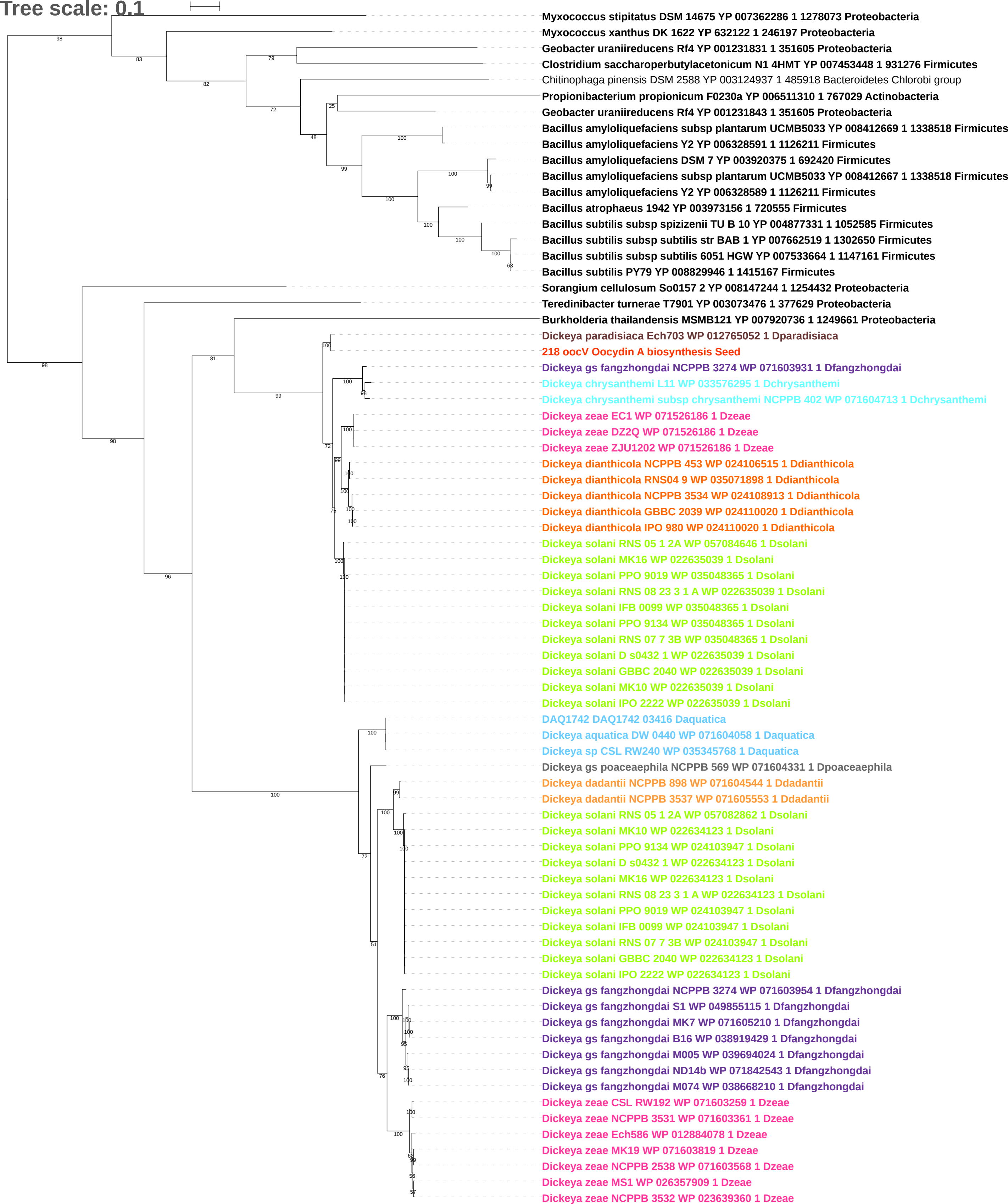
Tree scale: 0.1
Myxococcus stipitatus DSM 14675 YP 007362286 1 1278073 Proteobacteria
Myxococcus xanthus DK 1622 YP 632122 1 246197 Proteobacteria
Geobacter uraniireducens Rf4 YP 001231831 1 351605 Proteobacteria
Clostridium saccharoperbutylacetonicum N1 4HMT YP 007453448 1 931276 Firmicutes
Chitinophaga pinensis DSM 2588 YP 003124937 1 485918 Bacteroidetes Chlorobi group
Propionibacterium propionicum F0230a YP 006511310 1 767029 Actinobacteria
Geobacter uraniireducens Rf4 YP 001231843 1 351605 Proteobacteria
Bacillus amyloliquefaciens subsp plantarum UCMB5033 YP 008412669 1 1338518 Firmicutes
Bacillus amyloliquefaciens Y2 YP 006328591 1 1126211 Firmicutes
Bacillus amyloliquefaciens DSM 7 YP 003920375 1 692420 Firmicutes
Bacillus amyloliquefaciens subsp plantarum UCMB5033 YP 008412667 1 1338518 Firmicutes
Bacillus amyloliquefaciens Y2 YP 006328589 1 1126211 Firmicutes
Bacillus atrophaeus 1942 YP 003973156 1 720555 Firmicutes
Bacillus subtilis subsp spizizenii TU B 10 YP 004877331 1 1052585 Firmicutes
Bacillus subtilis subsp subtilis str BAB 1 YP 007662519 1 1302650 Firmicutes
Bacillus subtilis subsp subtilis 6051 HGW YP 007533664 1 1147161 Firmicutes
Bacillus subtilis PY79 YP 008829946 1 1415167 Firmicutes
Sorangium cellulosum So0157 2 YP 008147244 1 1254432 Proteobacteria
Teredinibacter turnerae T7901 YP 003073476 1 377629 Proteobacteria
Burkholderia thailandensis MSMB121 YP 007920736 1 1249661 Proteobacteria
Dickeya paradisiaca Ech703 WP 012765052 1 Dparadisiaca
218 oocV Oocydin A biosynthesis Seed
Dickeya gs fangzhongdai NCPPB 3274 WP 071603931 1 Dfangzhongdai
Dickeya chrysanthemi L11 WP 033576295 1 Dchrysanthemi
Dickeya chrysanthemi subsp chrysanthemi NCPPB 402 WP 071604713 1 Dchrysanthemi
Dickeya zeae EC1 WP 071526186 1 Dzeae
Dickeya zeae DZ2Q WP 071526186 1 Dzeae
Dickeya zeae ZJU1202 WP 071526186 1 Dzeae
Dickeya dianthicola NCPPB 453 WP 024106515 1 Ddianthicola
Dickeya dianthicola RNS04 9 WP 035071898 1 Ddianthicola
Dickeya dianthicola NCPPB 3534 WP 024108913 1 Ddianthicola
Dickeya dianthicola GBBC 2039 WP 024110020 1 Ddianthicola
Dickeya dianthicola IPO 980 WP 024110020 1 Ddianthicola
Dickeya solani RNS 05 1 2A WP 057084646 1 Dsolani
Dickeya solani MK16 WP 022635039 1 Dsolani
Dickeya solani PPO 9019 WP 035048365 1 Dsolani
Dickeya solani RNS 08 23 3 1 A WP 022635039 1 Dsolani
Dickeya solani IFB 0099 WP 035048365 1 Dsolani
Dickeya solani PPO 9134 WP 035048365 1 Dsolani
Dickeya solani RNS 07 7 3B WP 035048365 1 Dsolani
Dickeya solani D s0432 1 WP 022635039 1 Dsolani
Dickeya solani GBBC 2040 WP 022635039 1 Dsolani
Dickeya solani MK10 WP 022635039 1 Dsolani
Dickeya solani IPO 2222 WP 022635039 1 Dsolani
DAQ1742 DAQ1742 03416 Daquatica
Dickeya aquatica DW 0440 WP 071604058 1 Daquatica
Dickeya sp CSL RW240 WP 035345768 1 Daquatica
Dickeya gs poaceaephila NCPPB 569 WP 071604331 1 Dpoaceaephila
Dickeya dadantii NCPPB 898 WP 071604544 1 Ddadantii
Dickeya dadantii NCPPB 3537 WP 071605553 1 Ddadantii
Dickeya solani RNS 05 1 2A WP 057082862 1 Dsolani
Dickeya solani MK10 WP 022634123 1 Dsolani
Dickeya solani PPO 9134 WP 024103947 1 Dsolani
Dickeya solani D s0432 1 WP 022634123 1 Dsolani
Dickeya solani MK16 WP 022634123 1 Dsolani
Dickeya solani RNS 08 23 3 1 A WP 022634123 1 Dsolani
Dickeya solani PPO 9019 WP 024103947 1 Dsolani
Dickeya solani IFB 0099 WP 024103947 1 Dsolani
Dickeya solani RNS 07 7 3B WP 024103947 1 Dsolani
Dickeya solani GBBC 2040 WP 022634123 1 Dsolani
Dickeya solani IPO 2222 WP 022634123 1 Dsolani
Dickeya gs fangzhongdai NCPPB 3274 WP 071603954 1 Dfangzhongdai
Dickeya gs fangzhongdai S1 WP 049855115 1 Dfangzhongdai
Dickeya gs fangzhongdai MK7 WP 071605210 1 Dfangzhongdai
Dickeya gs fangzhongdai B16 WP 038919429 1 Dfangzhongdai
Dickeya gs fangzhongdai M005 WP 039694024 1 Dfangzhongdai
Dickeya gs fangzhongdai ND14b WP 071842543 1 Dfangzhongdai
Dickeya gs fangzhongdai M074 WP 038668210 1 Dfangzhongdai
Dickeya zeae CSL RW192 WP 071603259 1 Dzeae
Dickeya zeae NCPPB 3531 WP 071603361 1 Dzeae
Dickeya zeae Ech586 WP 012884078 1 Dzeae
Dickeya zeae MK19 WP 071603819 1 Dzeae
Dickeya zeae NCPPB 2538 WP 071603568 1 Dzeae
Dickeya zeae MS1 WP 026357909 1 Dzeae
Dickeya zeae NCPPB 3532 WP 023639360 1 Dzeae

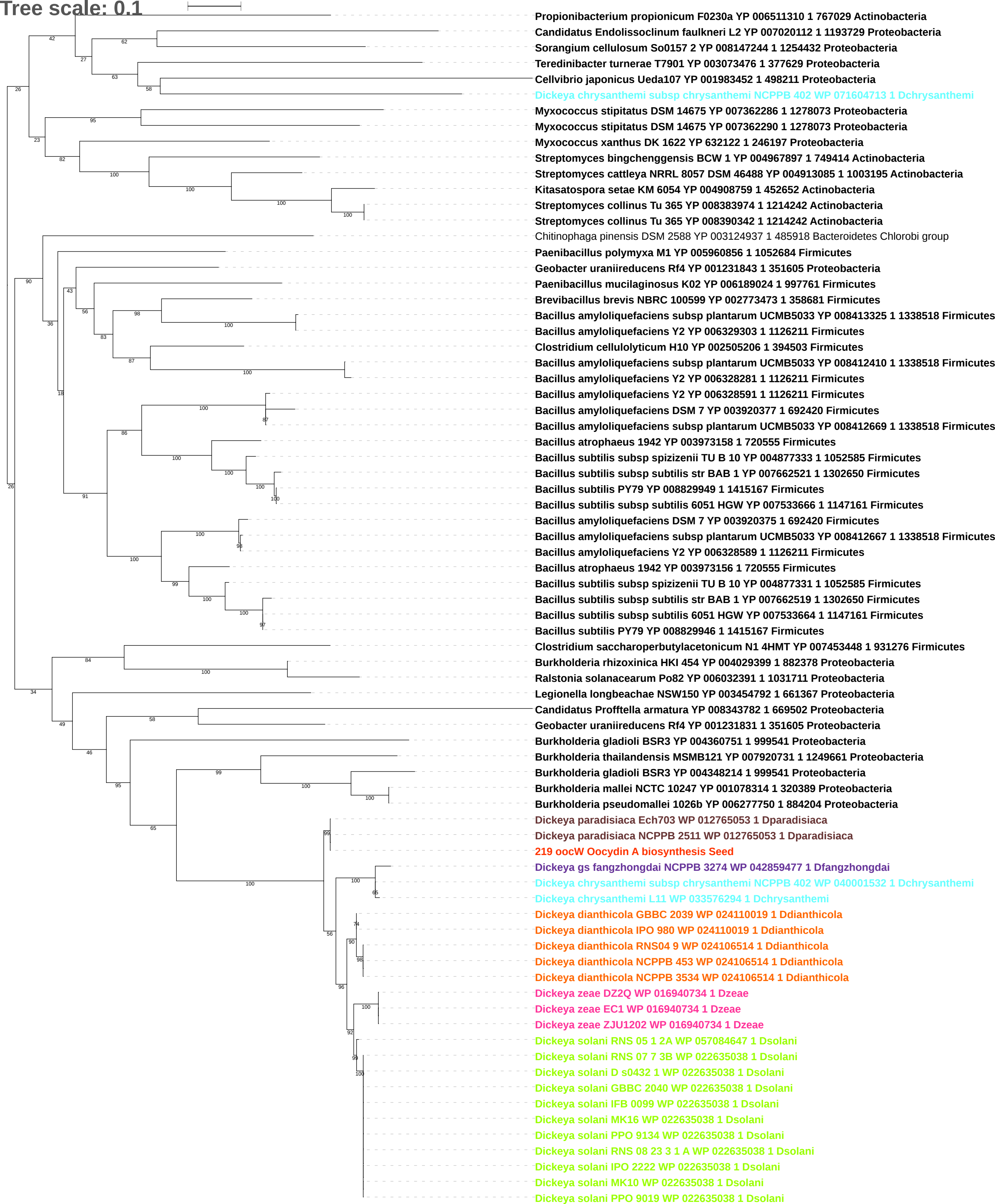

Tree scale: 0.1
Propionibacterium propionicum F0230a YP 006511310 1 767029 Actinobacteria
Candidatus Endolissoclinum faulkneri L2 YP 007020112 1 1193729 Proteobacteria
Sorangium cellulosum So0157 2 YP 008147244 1 1254432 Proteobacteria
Teredinibacter turnerae T7901 YP 003073476 1 377629 Proteobacteria
Cellvibrio japonicus Ueda107 YP 001983452 1 498211 Proteobacteria
Dickeya chrysanthemi subsp chrysanthemi NCPPB 402 WP 071604713 1 Dchrysanthemi
Myxococcus stipitatus DSM 14675 YP 007362286 1 1278073 Proteobacteria
Myxococcus stipitatus DSM 14675 YP 007362290 1 1278073 Proteobacteria
Myxococcus xanthus DK 1622 YP 632122 1 246197 Proteobacteria
Streptomyces bingchenggensis BCW 1 YP 004967897 1 749414 Actinobacteria
Streptomyces cattleya NRRL 8057 DSM 46488 YP 004913085 1 1003195 Actinobacteria
Kitasatospora setae KM 6054 YP 004908759 1 452652 Actinobacteria
Streptomyces collinus Tu 365 YP 008383974 1 1214242 Actinobacteria
Streptomyces collinus Tu 365 YP 008390342 1 1214242 Actinobacteria
Chitinophaga pinensis DSM 2588 YP 003124937 1 485918 Bacteroidetes Chlorobi group
Paenibacillus polymyxa M1 YP 005960856 1 1052684 Firmicutes
Geobacter uraniireducens Rf4 YP 001231843 1 351605 Proteobacteria
Paenibacillus mucilaginosus K02 YP 006189024 1 997761 Firmicutes
Brevibacillus brevis NBRC 100599 YP 002773473 1 358681 Firmicutes
Bacillus amyloliquefaciens subsp plantarum UCMB5033 YP 008413325 1 1338518 Firmicutes
Bacillus amyloliquefaciens Y2 YP 006329303 1 1126211 Firmicutes
Clostridium cellulolyticum H10 YP 002505206 1 394503 Firmicutes
Bacillus amyloliquefaciens subsp plantarum UCMB5033 YP 008412410 1 1338518 Firmicutes
Bacillus amyloliquefaciens Y2 YP 006328281 1 1126211 Firmicutes
Bacillus amyloliquefaciens Y2 YP 006328591 1 1126211 Firmicutes
Bacillus amyloliquefaciens DSM 7 YP 003920377 1 692420 Firmicutes
Bacillus amyloliquefaciens subsp plantarum UCMB5033 YP 008412669 1 1338518 Firmicutes
Bacillus atrophaeus 1942 YP 003973158 1 720555 Firmicutes
Bacillus subtilis subsp spizizenii TU B 10 YP 004877333 1 1052585 Firmicutes
Bacillus subtilis subsp subtilis str BAB 1 YP 007662521 1 1302650 Firmicutes
Bacillus subtilis PY79 YP 008829949 1 1415167 Firmicutes
Bacillus subtilis subsp subtilis 6051 HGW YP 007533666 1 1147161 Firmicutes
Bacillus amyloliquefaciens DSM 7 YP 003920375 1 692420 Firmicutes
Bacillus amyloliquefaciens subsp plantarum UCMB5033 YP 008412667 1 1338518 Firmicutes
Bacillus amyloliquefaciens Y2 YP 006328589 1 1126211 Firmicutes
Bacillus atrophaeus 1942 YP 003973156 1 720555 Firmicutes
Bacillus subtilis subsp spizizenii TU B 10 YP 004877331 1 1052585 Firmicutes
Bacillus subtilis subsp subtilis str BAB 1 YP 007662519 1 1302650 Firmicutes
Bacillus subtilis subsp subtilis 6051 HGW YP 007533664 1 1147161 Firmicutes
Bacillus subtilis PY79 YP 008829946 1 1415167 Firmicutes
Clostridium saccharoperbutylacetonicum N1 4HMT YP 007453448 1 931276 Firmicutes
Burkholderia rhizoxinica HKI 454 YP 004029399 1 882378 Proteobacteria
Ralstonia solanacearum Po82 YP 006032391 1 1031711 Proteobacteria
Legionella longbeachae NSW150 YP 003454792 1 661367 Proteobacteria
Candidatus Profftella armatura YP 008343782 1 669502 Proteobacteria
Geobacter uraniireducens Rf4 YP 001231831 1 351605 Proteobacteria
Burkholderia gladioli BSR3 YP 004360751 1 999541 Proteobacteria
Burkholderia thailandensis MSMB121 YP 007920731 1 1249661 Proteobacteria
Burkholderia gladioli BSR3 YP 004348214 1 999541 Proteobacteria
Burkholderia mallei NCTC 10247 YP 001078314 1 320389 Proteobacteria
Burkholderia pseudomallei 1026b YP 006277750 1 884204 Proteobacteria
Dickeya paradisiaca Ech703 WP 012765053 1 Dparadisiaca
Dickeya paradisiaca NCPPB 2511 WP 012765053 1 Dparadisiaca
219 oocW Oocydin A biosynthesis Seed
Dickeya gs fangzhongdai NCPPB 3274 WP 042859477 1 Dfangzhongdai
Dickeya chrysanthemi subsp chrysanthemi NCPPB 402 WP 040001532 1 Dchrysanthemi
Dickeya chrysanthemi L11 WP 033576294 1 Dchrysanthemi
Dickeya dianthicola GBBC 2039 WP 024110019 1 Ddianthicola
Dickeya dianthicola IPO 980 WP 024110019 1 Ddianthicola
Dickeya dianthicola RNS04 9 WP 024106514 1 Ddianthicola
Dickeya dianthicola NCPPB 453 WP 024106514 1 Ddianthicola
Dickeya dianthicola NCPPB 3534 WP 024106514 1 Ddianthicola
Dickeya zeae DZ2Q WP 016940734 1 Dzeae
Dickeya zeae EC1 WP 016940734 1 Dzeae
Dickeya zeae ZJU1202 WP 016940734 1 Dzeae
Dickeya solani RNS 05 1 2A WP 057084647 1 Dsolani
Dickeya solani RNS 07 7 3B WP 022635038 1 Dsolani
Dickeya solani D s0432 1 WP 022635038 1 Dsolani
Dickeya solani GBBC 2040 WP 022635038 1 Dsolani
Dickeya solani IFB 0099 WP 022635038 1 Dsolani
Dickeya solani MK16 WP 022635038 1 Dsolani
Dickeya solani PPO 9134 WP 022635038 1 Dsolani
Dickeya solani RNS 08 23 3 1 A WP 022635038 1 Dsolani
Dickeya solani IPO 2222 WP 022635038 1 Dsolani
Dickeya solani MK10 WP 022635038 1 Dsolani
Dickeya solani PPO 9019 WP 022635038 1 Dsolani

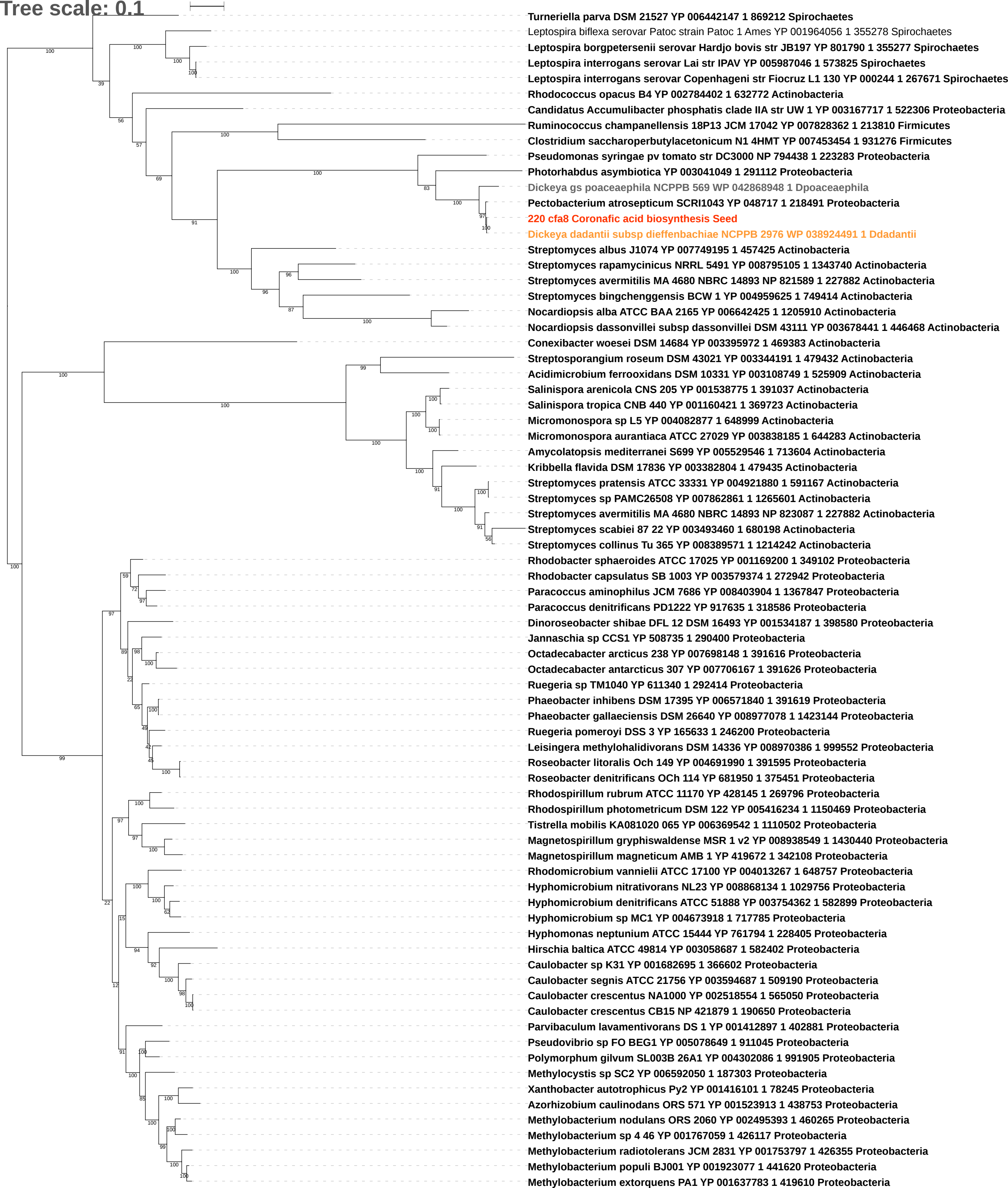
Tree scale: 0.1
Turneriella parva DSM 21527 YP 006442147 1 869212 Spirochaetes
Leptospira biflexa serovar Patoc strain Patoc 1 Ames YP 001964056 1 355278 Spirochaetes
Leptospira borgpetersenii serovar Hardjo bovis str JB197 YP 801790 1 355277 Spirochaetes
Leptospira interrogans serovar Lai str IPAV YP 005987046 1 573825 Spirochaetes
Leptospira interrogans serovar Copenhageni str Fiocruz L1 130 YP 000244 1 267671 Spirochaetes
Rhodococcus opacus B4 YP 002784402 1 632772 Actinobacteria
Candidatus Accumulibacter phosphatis clade IIA str UW 1 YP 003167717 1 522306 Proteobacteria
Ruminococcus champanellensis 18P13 JCM 17042 YP 007828362 1 213810 Firmicutes
Clostridium saccharoperbutylacetonicum N1 4HMT YP 007453454 1 931276 Firmicutes
Pseudomonas syringae pv tomato str DC3000 NP 794438 1 223283 Proteobacteria
Photorhabdus asymbiotica YP 003041049 1 291112 Proteobacteria
Dickeya gs poaceaephila NCPPB 569 WP 042868948 1 Dpoaceaephila
Pectobacterium atrosepticum SCRI1043 YP 048717 1 218491 Proteobacteria
220 cfa8 Coronafic acid biosynthesis Seed
Dickeya dadantii subsp dieffenbachiae NCPPB 2976 WP 038924491 1 Ddadantii
Streptomyces albus J1074 YP 007749195 1 457425 Actinobacteria
Streptomyces rapamycinicus NRRL 5491 YP 008795105 1 1343740 Actinobacteria
Streptomyces avermitilis MA 4680 NBRC 14893 NP 821589 1 227882 Actinobacteria
Streptomyces bingchenggensis BCW 1 YP 004959625 1 749414 Actinobacteria
Nocardiopsis alba ATCC BAA 2165 YP 006642425 1 1205910 Actinobacteria
Nocardiopsis dassonvillei subsp dassonvillei DSM 43111 YP 003678441 1 446468 Actinobacteria
Conexibacter woesei DSM 14684 YP 003395972 1 469383 Actinobacteria
Streptosporangium roseum DSM 43021 YP 003344191 1 479432 Actinobacteria
Acidimicrobium ferrooxidans DSM 10331 YP 003108749 1 525909 Actinobacteria
Salinispora arenicola CNS 205 YP 001538775 1 391037 Actinobacteria
Salinispora tropica CNB 440 YP 001160421 1 369723 Actinobacteria
Micromonospora sp L5 YP 004082877 1 648999 Actinobacteria
Micromonospora aurantiaca ATCC 27029 YP 003838185 1 644283 Actinobacteria
Amycolatopsis mediterranei S699 YP 005529546 1 713604 Actinobacteria
Kribbella flavida DSM 17836 YP 003382804 1 479435 Actinobacteria
Streptomyces pratensis ATCC 33331 YP 004921880 1 591167 Actinobacteria
Streptomyces sp PAMC26508 YP 007862861 1 1265601 Actinobacteria
Streptomyces avermitilis MA 4680 NBRC 14893 NP 823087 1 227882 Actinobacteria
Streptomyces scabiei 87 22 YP 003493460 1 680198 Actinobacteria
Streptomyces collinus Tu 365 YP 008389571 1 1214242 Actinobacteria
Rhodobacter sphaeroides ATCC 17025 YP 001169200 1 349102 Proteobacteria
Rhodobacter capsulatus SB 1003 YP 003579374 1 272942 Proteobacteria
Paracoccus aminophilus JCM 7686 YP 008403904 1 1367847 Proteobacteria
Paracoccus denitrificans PD1222 YP 917635 1 318586 Proteobacteria
Dinoroseobacter shibae DFL 12 DSM 16493 YP 001534187 1 398580 Proteobacteria
Jannaschia sp CCS1 YP 508735 1 290400 Proteobacteria
Octadecabacter arcticus 238 YP 007698148 1 391616 Proteobacteria
Octadecabacter antarcticus 307 YP 007706167 1 391626 Proteobacteria
Ruegeria sp TM1040 YP 611340 1 292414 Proteobacteria
Phaeobacter inhibens DSM 17395 YP 006571840 1 391619 Proteobacteria
Phaeobacter gallaeciensis DSM 26640 YP 008977078 1 1423144 Proteobacteria
Ruegeria pomeroyi DSS 3 YP 165633 1 246200 Proteobacteria
Leisingera methylohalidivorans DSM 14336 YP 008970386 1 999552 Proteobacteria
Roseobacter litoralis Och 149 YP 004691990 1 391595 Proteobacteria
Roseobacter denitrificans OCh 114 YP 681950 1 375451 Proteobacteria
Rhodospirillum rubrum ATCC 11170 YP 428145 1 269796 Proteobacteria
Rhodospirillum photometricum DSM 122 YP 005416234 1 1150469 Proteobacteria
Tistrella mobilis KA081020 065 YP 006369542 1 1110502 Proteobacteria
Magnetospirillum gryphiswaldense MSR 1 v2 YP 008938549 1 1430440 Proteobacteria
Magnetospirillum magneticum AMB 1 YP 419672 1 342108 Proteobacteria
Rhodomicrobium vannielii ATCC 17100 YP 004013267 1 648757 Proteobacteria
Hyphomicrobium nitrativorans NL23 YP 008868134 1 1029756 Proteobacteria
Hyphomicrobium denitrificans ATCC 51888 YP 003754362 1 582899 Proteobacteria
Hyphomicrobium sp MC1 YP 004673918 1 717785 Proteobacteria
Hyphomonas neptunium ATCC 15444 YP 761794 1 228405 Proteobacteria
Hirschia baltica ATCC 49814 YP 003058687 1 582402 Proteobacteria
Caulobacter sp K31 YP 001682695 1 366602 Proteobacteria
Caulobacter segnis ATCC 21756 YP 003594687 1 509190 Proteobacteria
Caulobacter crescentus NA1000 YP 002518554 1 565050 Proteobacteria
Caulobacter crescentus CB15 NP 421879 1 190650 Proteobacteria
Parvibaculum lavamentivorans DS 1 YP 001412897 1 402881 Proteobacteria
Pseudovibrio sp FO BEG1 YP 005078649 1 911045 Proteobacteria
Polymorphum gilvum SL003B 26A1 YP 004302086 1 991905 Proteobacteria
Methylocystis sp SC2 YP 006592050 1 187303 Proteobacteria
Xanthobacter autotrophicus Py2 YP 001416101 1 78245 Proteobacteria
Azorhizobium caulinodans ORS 571 YP 001523913 1 438753 Proteobacteria
Methylobacterium nodulans ORS 2060 YP 002495393 1 460265 Proteobacteria
Methylobacterium sp 4 46 YP 001767059 1 426117 Proteobacteria
Methylobacterium radiotolerans JCM 2831 YP 001753797 1 426355 Proteobacteria
Methylobacterium populi BJ001 YP 001923077 1 441620 Proteobacteria
Methylobacterium extorquens PA1 YP 001637783 1 419610 Proteobacteria

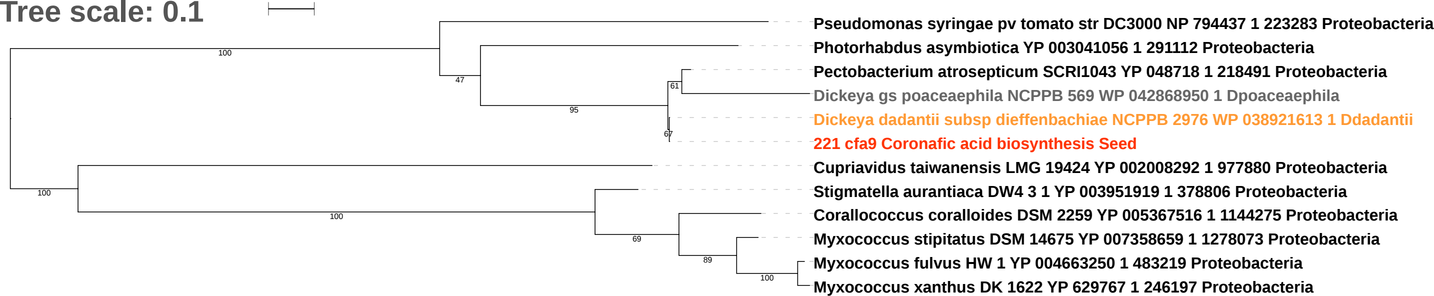
Tree scale: 0.1
Pseudomonas syringae pv tomato str DC3000 NP 794437 1 223283 Proteobacteria
Photorhabdus asymbiotica YP 003041056 1 291112 Proteobacteria
Pectobacterium atrosepticum SCRI1043 YP 048718 1 218491 Proteobacteria
Dickeya gs poaceaephila NCPPB 569 WP 042868950 1 Dpoaceaephila
Dickeya dadantii subsp dieffenbachiae NCPPB 2976 WP 038921613 1 Ddadantii
221 cfa9 Coronafic acid biosynthesis Seed
Cupriavidus taiwanensis LMG 19424 YP 002008292 1 977880 Proteobacteria
Stigmatella aurantiaca DW4 3 1 YP 003951919 1 378806 Proteobacteria
Corallococcus coralloides DSM 2259 YP 005367516 1 1144275 Proteobacteria
Myxococcus stipitatus DSM 14675 YP 007358659 1 1278073 Proteobacteria
Myxococcus fulvus HW 1 YP 004663250 1 483219 Proteobacteria
Myxococcus xanthus DK 1622 YP 629767 1 246197 Proteobacteria
100
47
61
95
67
100
100
69
89
100

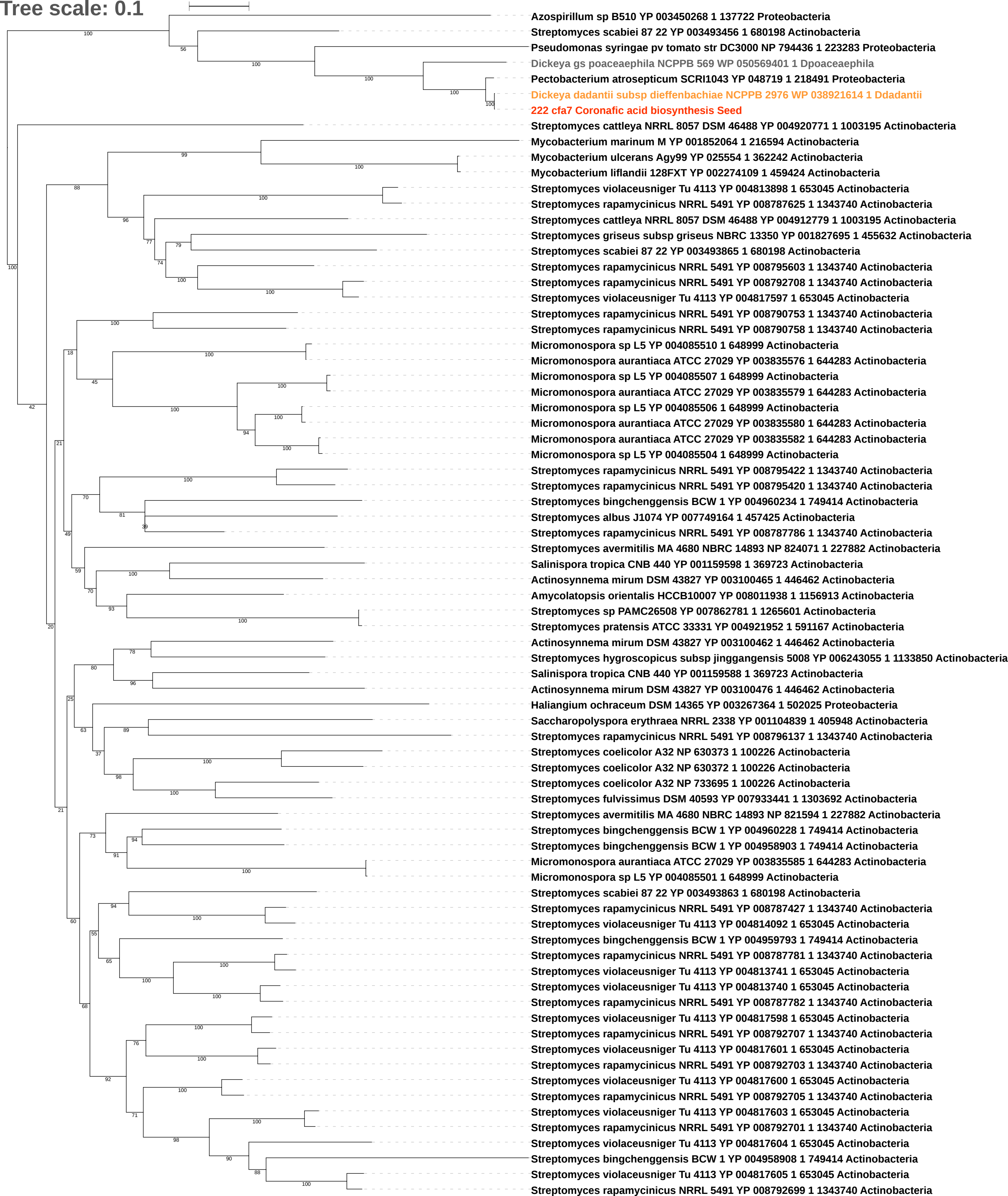

Tree scale: 0.1
Azospirillum sp B510 YP 003450268 1 137722 Proteobacteria
Streptomyces scabiei 87 22 YP 003493456 1 680198 Actinobacteria
Pseudomonas syringae pv tomato str DC3000 NP 794436 1 223283 Proteobacteria
Dickeya gs poaceaephila NCPPB 569 WP 050569401 1 Dpoaceaephila
Pectobacterium atrosepticum SCRI1043 YP 048719 1 218491 Proteobacteria
Dickeya dadantii subsp dieffenbachiae NCPPB 2976 WP 038921614 1 Ddadantii
222 cfa7 Coronafic acid biosynthesis Seed
Streptomyces cattleya NRRL 8057 DSM 46488 YP 004920771 1 1003195 Actinobacteria
Mycobacterium marinum M YP 001852064 1 216594 Actinobacteria
Mycobacterium ulcerans Agy99 YP 025554 1 362242 Actinobacteria
Mycobacterium liflandii 128FXT YP 002274109 1 459424 Actinobacteria
Streptomyces violaceusniger Tu 4113 YP 004813898 1 653045 Actinobacteria
Streptomyces rapamycinicus NRRL 5491 YP 008787625 1 1343740 Actinobacteria
Streptomyces cattleya NRRL 8057 DSM 46488 YP 004912779 1 1003195 Actinobacteria
Streptomyces griseus subsp griseus NBRC 13350 YP 001827695 1 455632 Actinobacteria
Streptomyces scabiei 87 22 YP 003493865 1 680198 Actinobacteria
Streptomyces rapamycinicus NRRL 5491 YP 008795603 1 1343740 Actinobacteria
Streptomyces rapamycinicus NRRL 5491 YP 008792708 1 1343740 Actinobacteria
Streptomyces violaceusniger Tu 4113 YP 004817597 1 653045 Actinobacteria
Streptomyces rapamycinicus NRRL 5491 YP 008790753 1 1343740 Actinobacteria
Streptomyces rapamycinicus NRRL 5491 YP 008790758 1 1343740 Actinobacteria
Micromonospora sp L5 YP 004085510 1 648999 Actinobacteria
Micromonospora aurantiaca ATCC 27029 YP 003835576 1 644283 Actinobacteria
Micromonospora sp L5 YP 004085507 1 648999 Actinobacteria
Micromonospora aurantiaca ATCC 27029 YP 003835579 1 644283 Actinobacteria
Micromonospora sp L5 YP 004085506 1 648999 Actinobacteria
Micromonospora aurantiaca ATCC 27029 YP 003835580 1 644283 Actinobacteria
Micromonospora aurantiaca ATCC 27029 YP 003835582 1 644283 Actinobacteria
Micromonospora sp L5 YP 004085504 1 648999 Actinobacteria
Streptomyces rapamycinicus NRRL 5491 YP 008795422 1 1343740 Actinobacteria
Streptomyces rapamycinicus NRRL 5491 YP 008795420 1 1343740 Actinobacteria
Streptomyces bingchenggensis BCW 1 YP 004960234 1 749414 Actinobacteria
Streptomyces albus J1074 YP 007749164 1 457425 Actinobacteria
Streptomyces rapamycinicus NRRL 5491 YP 008787786 1 1343740 Actinobacteria
Streptomyces avermitilis MA 4680 NBRC 14893 NP 824071 1 227882 Actinobacteria
Salinispora tropica CNB 440 YP 001159598 1 369723 Actinobacteria
Actinosynnema mirum DSM 43827 YP 003100465 1 446462 Actinobacteria
Amycolatopsis orientalis HCCB10007 YP 008011938 1 1156913 Actinobacteria
Streptomyces sp PAMC26508 YP 007862781 1 1265601 Actinobacteria
Streptomyces pratensis ATCC 33331 YP 004921952 1 591167 Actinobacteria
Actinosynnema mirum DSM 43827 YP 003100462 1 446462 Actinobacteria
Streptomyces hygroscopicus subsp jinggangensis 5008 YP 006243055 1 1133850 Actinobacteria
Salinispora tropica CNB 440 YP 001159588 1 369723 Actinobacteria
Actinosynnema mirum DSM 43827 YP 003100476 1 446462 Actinobacteria
Haliangium ochraceum DSM 14365 YP 003267364 1 502025 Proteobacteria
Saccharopolyspora erythraea NRRL 2338 YP 001104839 1 405948 Actinobacteria
Streptomyces rapamycinicus NRRL 5491 YP 008796137 1 1343740 Actinobacteria
Streptomyces coelicolor A32 NP 630373 1 100226 Actinobacteria
Streptomyces coelicolor A32 NP 630372 1 100226 Actinobacteria
Streptomyces coelicolor A32 NP 733695 1 100226 Actinobacteria
Streptomyces fulvissimus DSM 40593 YP 007933441 1 1303692 Actinobacteria
Streptomyces avermitilis MA 4680 NBRC 14893 NP 821594 1 227882 Actinobacteria
Streptomyces bingchenggensis BCW 1 YP 004960228 1 749414 Actinobacteria
Streptomyces bingchenggensis BCW 1 YP 004958903 1 749414 Actinobacteria
Micromonospora aurantiaca ATCC 27029 YP 003835585 1 644283 Actinobacteria
Micromonospora sp L5 YP 004085501 1 648999 Actinobacteria
Streptomyces scabiei 87 22 YP 003493863 1 680198 Actinobacteria
Streptomyces rapamycinicus NRRL 5491 YP 008787427 1 1343740 Actinobacteria
Streptomyces violaceusniger Tu 4113 YP 004814092 1 653045 Actinobacteria
Streptomyces bingchenggensis BCW 1 YP 004959793 1 749414 Actinobacteria
Streptomyces rapamycinicus NRRL 5491 YP 008787781 1 1343740 Actinobacteria
Streptomyces violaceusniger Tu 4113 YP 004813741 1 653045 Actinobacteria
Streptomyces violaceusniger Tu 4113 YP 004813740 1 653045 Actinobacteria
Streptomyces rapamycinicus NRRL 5491 YP 008787782 1 1343740 Actinobacteria
Streptomyces violaceusniger Tu 4113 YP 004817598 1 653045 Actinobacteria
Streptomyces rapamycinicus NRRL 5491 YP 008792707 1 1343740 Actinobacteria
Streptomyces violaceusniger Tu 4113 YP 004817601 1 653045 Actinobacteria
Streptomyces rapamycinicus NRRL 5491 YP 008792703 1 1343740 Actinobacteria
Streptomyces violaceusniger Tu 4113 YP 004817600 1 653045 Actinobacteria
Streptomyces rapamycinicus NRRL 5491 YP 008792705 1 1343740 Actinobacteria
Streptomyces violaceusniger Tu 4113 YP 004817603 1 653045 Actinobacteria
Streptomyces rapamycinicus NRRL 5491 YP 008792701 1 1343740 Actinobacteria
Streptomyces violaceusniger Tu 4113 YP 004817604 1 653045 Actinobacteria
Streptomyces bingchenggensis BCW 1 YP 004958908 1 749414 Actinobacteria
Streptomyces violaceusniger Tu 4113 YP 004817605 1 653045 Actinobacteria
Streptomyces rapamycinicus NRRL 5491 YP 008792699 1 1343740 Actinobacteria

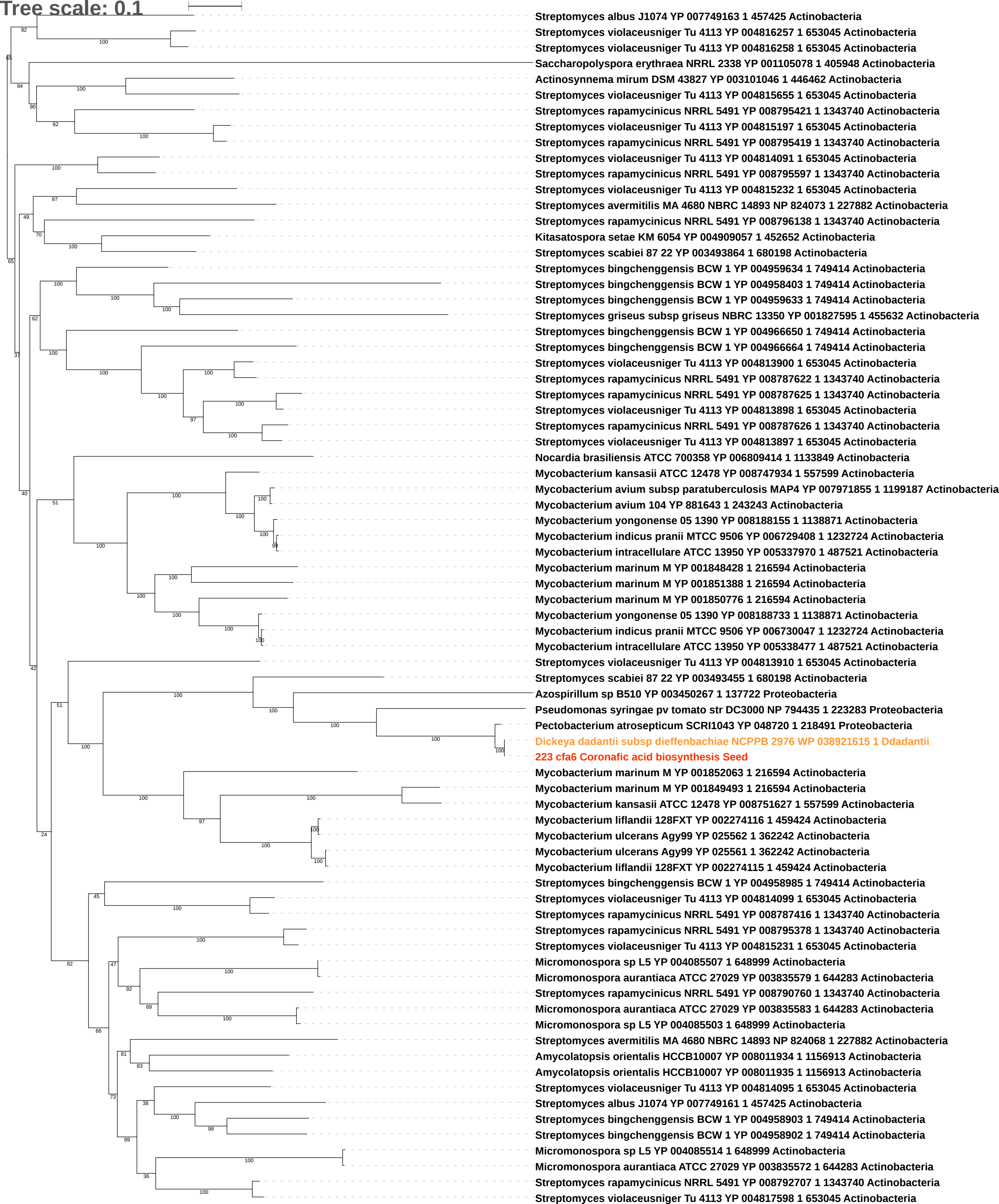
Tree scale: 0.1
Streptomyces albus J1074 YP 007749163 1 457425 Actinobacteria
Streptomyces violaceusniger Tu 4113 YP 004816257 1 653045 Actinobacteria
Streptomyces violaceusniger Tu 4113 YP 004816258 1 653045 Actinobacteria
Saccharopolyspora erythraea NRRL 2338 YP 001105078 1 405948 Actinobacteria
Actinosynnema mirum DSM 43827 YP 003101046 1 446462 Actinobacteria
Streptomyces violaceusniger Tu 4113 YP 004815655 1 653045 Actinobacteria
Streptomyces rapamycinicus NRRL 5491 YP 008795421 1 1343740 Actinobacteria
Streptomyces violaceusniger Tu 4113 YP 004815197 1 653045 Actinobacteria
Streptomyces rapamycinicus NRRL 5491 YP 008795419 1 1343740 Actinobacteria
Streptomyces violaceusniger Tu 4113 YP 004814091 1 653045 Actinobacteria
Streptomyces rapamycinicus NRRL 5491 YP 008795597 1 1343740 Actinobacteria
Streptomyces violaceusniger Tu 4113 YP 004815232 1 653045 Actinobacteria
Streptomyces avermitilis MA 4680 NBRC 14893 NP 824073 1 227882 Actinobacteria
Streptomyces rapamycinicus NRRL 5491 YP 008796138 1 1343740 Actinobacteria
Kitasatospora setae KM 6054 YP 004909057 1 452652 Actinobacteria
Streptomyces scabiei 87 22 YP 003493864 1 680198 Actinobacteria
Streptomyces bingchenggensis BCW 1 YP 004959634 1 749414 Actinobacteria
Streptomyces bingchenggensis BCW 1 YP 004958403 1 749414 Actinobacteria
Streptomyces bingchenggensis BCW 1 YP 004959633 1 749414 Actinobacteria
Streptomyces griseus subsp griseus NBRC 13350 YP 001827595 1 455632 Actinobacteria
Streptomyces bingchenggensis BCW 1 YP 004966650 1 749414 Actinobacteria
Streptomyces bingchenggensis BCW 1 YP 004966664 1 749414 Actinobacteria
Streptomyces violaceusniger Tu 4113 YP 004813900 1 653045 Actinobacteria
Streptomyces rapamycinicus NRRL 5491 YP 008787622 1 1343740 Actinobacteria
Streptomyces rapamycinicus NRRL 5491 YP 008787625 1 1343740 Actinobacteria
Streptomyces violaceusniger Tu 4113 YP 004813898 1 653045 Actinobacteria
Streptomyces rapamycinicus NRRL 5491 YP 008787626 1 1343740 Actinobacteria
Streptomyces violaceusniger Tu 4113 YP 004813897 1 653045 Actinobacteria
Nocardia brasiliensis ATCC 700358 YP 006809414 1 1133849 Actinobacteria
Mycobacterium kansasii ATCC 12478 YP 008747934 1 557599 Actinobacteria
Mycobacterium avium subsp paratuberculosis MAP4 YP 007971855 1 1199187 Actinobacteria
Mycobacterium avium 104 YP 881643 1 243243 Actinobacteria
Mycobacterium yongonense 05 1390 YP 008188155 1 1138871 Actinobacteria
Mycobacterium indicus pranii MTCC 9506 YP 006729408 1 1232724 Actinobacteria
Mycobacterium intracellulare ATCC 13950 YP 005337970 1 487521 Actinobacteria
Mycobacterium marinum M YP 001848428 1 216594 Actinobacteria
Mycobacterium marinum M YP 001851388 1 216594 Actinobacteria
Mycobacterium marinum M YP 001850776 1 216594 Actinobacteria
Mycobacterium yongonense 05 1390 YP 008188733 1 1138871 Actinobacteria
Mycobacterium indicus pranii MTCC 9506 YP 006730047 1 1232724 Actinobacteria
Mycobacterium intracellulare ATCC 13950 YP 005338477 1 487521 Actinobacteria
Streptomyces violaceusniger Tu 4113 YP 004813910 1 653045 Actinobacteria
Streptomyces scabiei 87 22 YP 003493455 1 680198 Actinobacteria
Azospirillum sp B510 YP 003450267 1 137722 Proteobacteria
Pseudomonas syringae pv tomato str DC3000 NP 794435 1 223283 Proteobacteria
Pectobacterium atrosepticum SCRI1043 YP 048720 1 218491 Proteobacteria
Dickeya dadantii subsp dieffenbachiae NCPPB 2976 WP 038921615 1 Ddadantii
223 cfa6 Coronafic acid biosynthesis Seed
Mycobacterium marinum M YP 001852063 1 216594 Actinobacteria
Mycobacterium marinum M YP 001849493 1 216594 Actinobacteria
Mycobacterium kansasii ATCC 12478 YP 008751627 1 557599 Actinobacteria
Mycobacterium liflandii 128FXT YP 002274116 1 459424 Actinobacteria
Mycobacterium ulcerans Agy99 YP 025562 1 362242 Actinobacteria
Mycobacterium ulcerans Agy99 YP 025561 1 362242 Actinobacteria
Mycobacterium liflandii 128FXT YP 002274115 1 459424 Actinobacteria
Streptomyces bingchenggensis BCW 1 YP 004958985 1 749414 Actinobacteria
Streptomyces violaceusniger Tu 4113 YP 004814099 1 653045 Actinobacteria
Streptomyces rapamycinicus NRRL 5491 YP 008787416 1 1343740 Actinobacteria
Streptomyces rapamycinicus NRRL 5491 YP 008795378 1 1343740 Actinobacteria
Streptomyces violaceusniger Tu 4113 YP 004815231 1 653045 Actinobacteria
Micromonospora sp L5 YP 004085507 1 648999 Actinobacteria
Micromonospora aurantiaca ATCC 27029 YP 003835579 1 644283 Actinobacteria
Streptomyces rapamycinicus NRRL 5491 YP 008790760 1 1343740 Actinobacteria
Micromonospora aurantiaca ATCC 27029 YP 003835583 1 644283 Actinobacteria
Micromonospora sp L5 YP 004085503 1 648999 Actinobacteria
Streptomyces avermitilis MA 4680 NBRC 14893 NP 824068 1 227882 Actinobacteria
Amycolatopsis orientalis HCCB10007 YP 008011934 1 1156913 Actinobacteria
Amycolatopsis orientalis HCCB10007 YP 008011935 1 1156913 Actinobacteria
Streptomyces violaceusniger Tu 4113 YP 004814095 1 653045 Actinobacteria
Streptomyces albus J1074 YP 007749161 1 457425 Actinobacteria
Streptomyces bingchenggensis BCW 1 YP 004958903 1 749414 Actinobacteria
Streptomyces bingchenggensis BCW 1 YP 004958902 1 749414 Actinobacteria
Micromonospora sp L5 YP 004085514 1 648999 Actinobacteria
Micromonospora aurantiaca ATCC 27029 YP 003835572 1 644283 Actinobacteria
Streptomyces rapamycinicus NRRL 5491 YP 008792707 1 1343740 Actinobacteria
Streptomyces violaceusniger Tu 4113 YP 004817598 1 653045 Actinobacteria

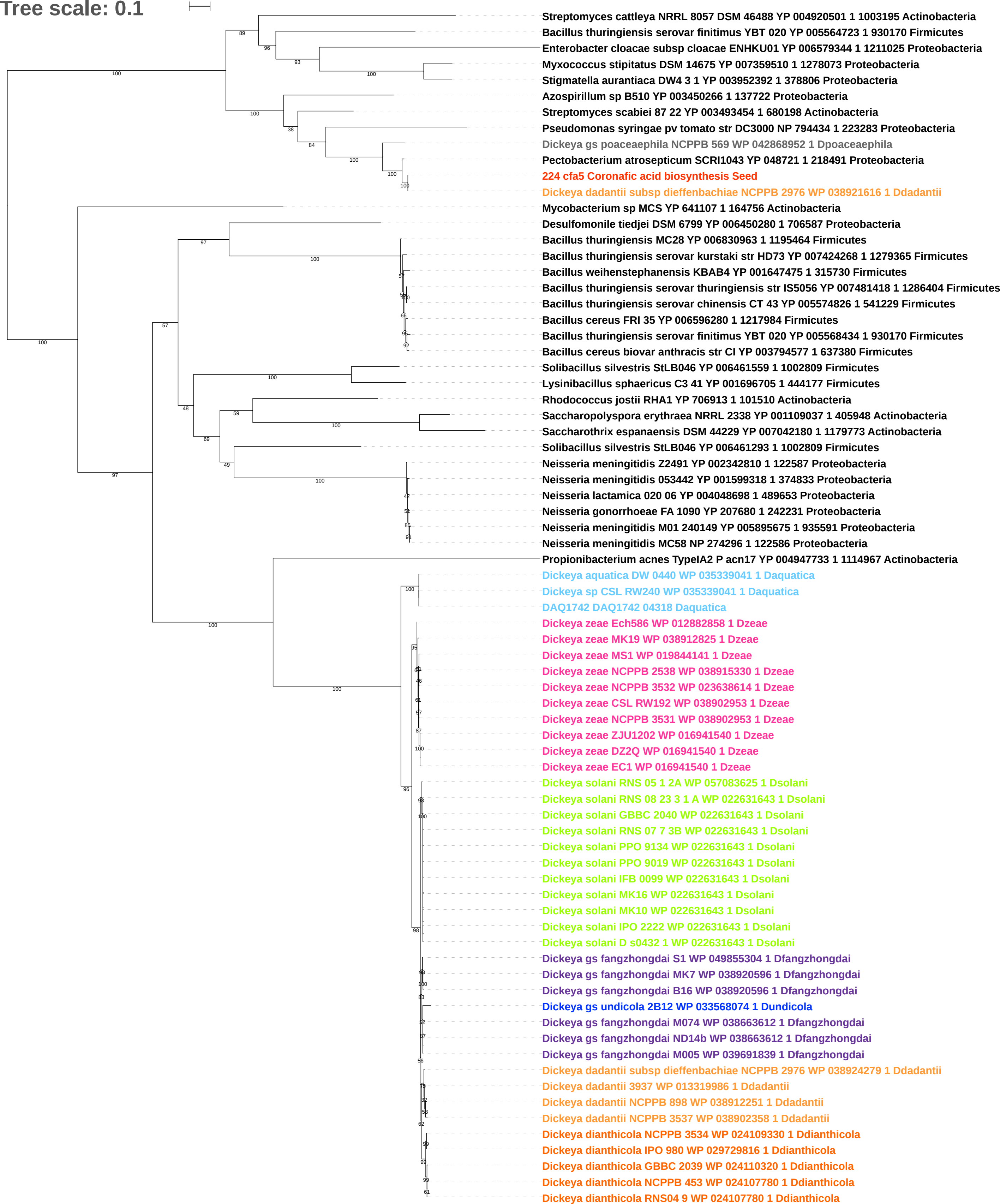
Tree scale: 0.1
Streptomyces cattleya NRRL 8057 DSM 46488 YP 004920501 1 1003195 Actinobacteria
Bacillus thuringiensis serovar finitimus YBT 020 YP 005564723 1 930170 Firmicutes
Enterobacter cloacae subsp cloacae ENHKU01 YP 006579344 1 1211025 Proteobacteria
Myxococcus stipitatus DSM 14675 YP 007359510 1 1278073 Proteobacteria
Stigmatella aurantiaca DW4 3 1 YP 003952392 1 378806 Proteobacteria
Azospirillum sp B510 YP 003450266 1 137722 Proteobacteria
Streptomyces scabiei 87 22 YP 003493454 1 680198 Actinobacteria
Pseudomonas syringae pv tomato str DC3000 NP 794434 1 223283 Proteobacteria
Dickeya gs poaceaephila NCPPB 569 WP 042868952 1 Dpoaceaephila
Pectobacterium atrosepticum SCRI1043 YP 048721 1 218491 Proteobacteria
224 cfa5 Coronafic acid biosynthesis Seed
Dickeya dadantii subsp dieffenbachiae NCPPB 2976 WP 038921616 1 Ddadantii
Mycobacterium sp MCS YP 641107 1 164756 Actinobacteria
Desulfomonile tiedjei DSM 6799 YP 006450280 1 706587 Proteobacteria
Bacillus thuringiensis MC28 YP 006830963 1 1195464 Firmicutes
Bacillus thuringiensis serovar kurstaki str HD73 YP 007424268 1 1279365 Firmicutes
Bacillus weihenstephanensis KBAB4 YP 001647475 1 315730 Firmicutes
Bacillus thuringiensis serovar thuringiensis str IS5056 YP 007481418 1 1286404 Firmicutes
Bacillus thuringiensis serovar chinensis CT 43 YP 005574826 1 541229 Firmicutes
Bacillus cereus FRI 35 YP 006596280 1 1217984 Firmicutes
Bacillus thuringiensis serovar finitimus YBT 020 YP 005568434 1 930170 Firmicutes
Bacillus cereus biovar anthracis str CI YP 003794577 1 637380 Firmicutes
Solibacillus silvestris StLB046 YP 006461559 1 1002809 Firmicutes
Lysinibacillus sphaericus C3 41 YP 001696705 1 444177 Firmicutes
Rhodococcus jostii RHA1 YP 706913 1 101510 Actinobacteria
Saccharopolyspora erythraea NRRL 2338 YP 001109037 1 405948 Actinobacteria
Saccharothrix espanaensis DSM 44229 YP 007042180 1 1179773 Actinobacteria
Solibacillus silvestris StLB046 YP 006461293 1 1002809 Firmicutes
Neisseria meningitidis Z2491 YP 002342810 1 122587 Proteobacteria
Neisseria meningitidis 053442 YP 001599318 1 374833 Proteobacteria
Neisseria lactamica 020 06 YP 004048698 1 489653 Proteobacteria
Neisseria gonorrhoeae FA 1090 YP 207680 1 242231 Proteobacteria
Neisseria meningitidis M01 240149 YP 005895675 1 935591 Proteobacteria
Neisseria meningitidis MC58 NP 274296 1 122586 Proteobacteria
Propionibacterium acnes TypeIA2 P acn17 YP 004947733 1 1114967 Actinobacteria
Dickeya aquatica DW 0440 WP 035339041 1 Daquatica
Dickeya sp CSL RW240 WP 035339041 1 Daquatica
DAQ1742 DAQ1742 04318 Daquatica
Dickeya zeae Ech586 WP 012882858 1 Dzeae
Dickeya zeae MK19 WP 038912825 1 Dzeae
Dickeya zeae MS1 WP 019844141 1 Dzeae
Dickeya zeae NCPPB 2538 WP 038915330 1 Dzeae
Dickeya zeae NCPPB 3532 WP 023638614 1 Dzeae
Dickeya zeae CSL RW192 WP 038902953 1 Dzeae
Dickeya zeae NCPPB 3531 WP 038902953 1 Dzeae
Dickeya zeae ZJU1202 WP 016941540 1 Dzeae
Dickeya zeae DZ2Q WP 016941540 1 Dzeae
Dickeya zeae EC1 WP 016941540 1 Dzeae
Dickeya solani RNS 05 1 2A WP 057083625 1 Dsolani
Dickeya solani RNS 08 23 3 1 A WP 022631643 1 Dsolani
Dickeya solani GBBC 2040 WP 022631643 1 Dsolani
Dickeya solani RNS 07 7 3B WP 022631643 1 Dsolani
Dickeya solani PPO 9134 WP 022631643 1 Dsolani
Dickeya solani PPO 9019 WP 022631643 1 Dsolani
Dickeya solani IFB 0099 WP 022631643 1 Dsolani
Dickeya solani MK16 WP 022631643 1 Dsolani
Dickeya solani MK10 WP 022631643 1 Dsolani
Dickeya solani IPO 2222 WP 022631643 1 Dsolani
Dickeya solani D s0432 1 WP 022631643 1 Dsolani
Dickeya gs fangzhongdai S1 WP 049855304 1 Dfangzhongdai
Dickeya gs fangzhongdai MK7 WP 038920596 1 Dfangzhongdai
Dickeya gs fangzhongdai B16 WP 038920596 1 Dfangzhongdai
Dickeya gs undicola 2B12 WP 033568074 1 Dundicola
Dickeya gs fangzhongdai M074 WP 038663612 1 Dfangzhongdai
Dickeya gs fangzhongdai ND14b WP 038663612 1 Dfangzhongdai
Dickeya gs fangzhongdai M005 WP 039691839 1 Dfangzhongdai
Dickeya dadantii subsp dieffenbachiae NCPPB 2976 WP 038924279 1 Ddadantii
Dickeya dadantii 3937 WP 013319986 1 Ddadantii
Dickeya dadantii NCPPB 898 WP 038912251 1 Ddadantii
Dickeya dadantii NCPPB 3537 WP 038902358 1 Ddadantii
Dickeya dianthicola NCPPB 3534 WP 024109330 1 Ddianthicola
Dickeya dianthicola IPO 980 WP 029729816 1 Ddianthicola
Dickeya dianthicola GBBC 2039 WP 024110320 1 Ddianthicola
Dickeya dianthicola NCPPB 453 WP 024107780 1 Ddianthicola
Dickeya dianthicola RNS04 9 WP 024107780 1 Ddianthicola

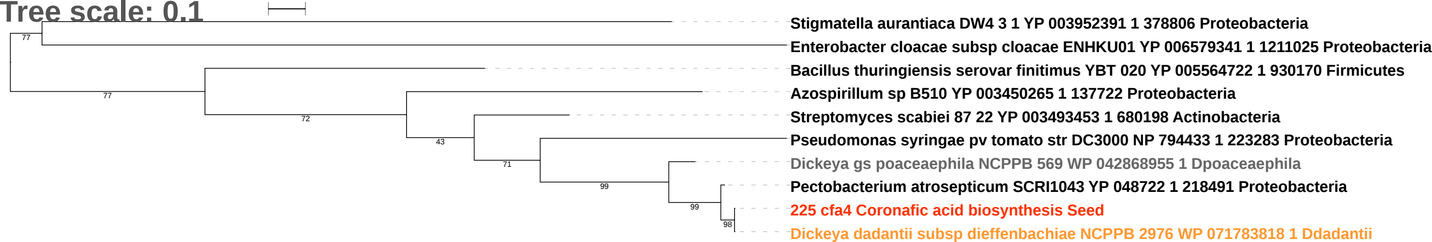
Tree scale: 0.1
Stigmatella aurantiaca DW4 3 1 YP 003952391 1 378806 Proteobacteria
Enterobacter cloacae subsp cloacae ENHKU01 YP 006579341 1 1211025 Proteobacteria
Bacillus thuringiensis serovar finitimus YBT 020 YP 005564722 1 930170 Firmicutes
Azospirillum sp B510 YP 003450265 1 137722 Proteobacteria
Streptomyces scabiei 87 22 YP 003493453 1 680198 Actinobacteria
Pseudomonas syringae pv tomato str DC3000 NP 794433 1 223283 Proteobacteria
Dickeya gs poaceaephila NCPPB 569 WP 042868955 1 Dpoaceaephila
Pectobacterium atrosepticum SCRI1043 YP 048722 1 218491 Proteobacteria
225 cfa4 Coronafic acid biosynthesis Seed
Dickeya dadantii subsp dieffenbachiae NCPPB 2976 WP 071783818 1 Ddadantii
77
77
72
43
71
99
99
98

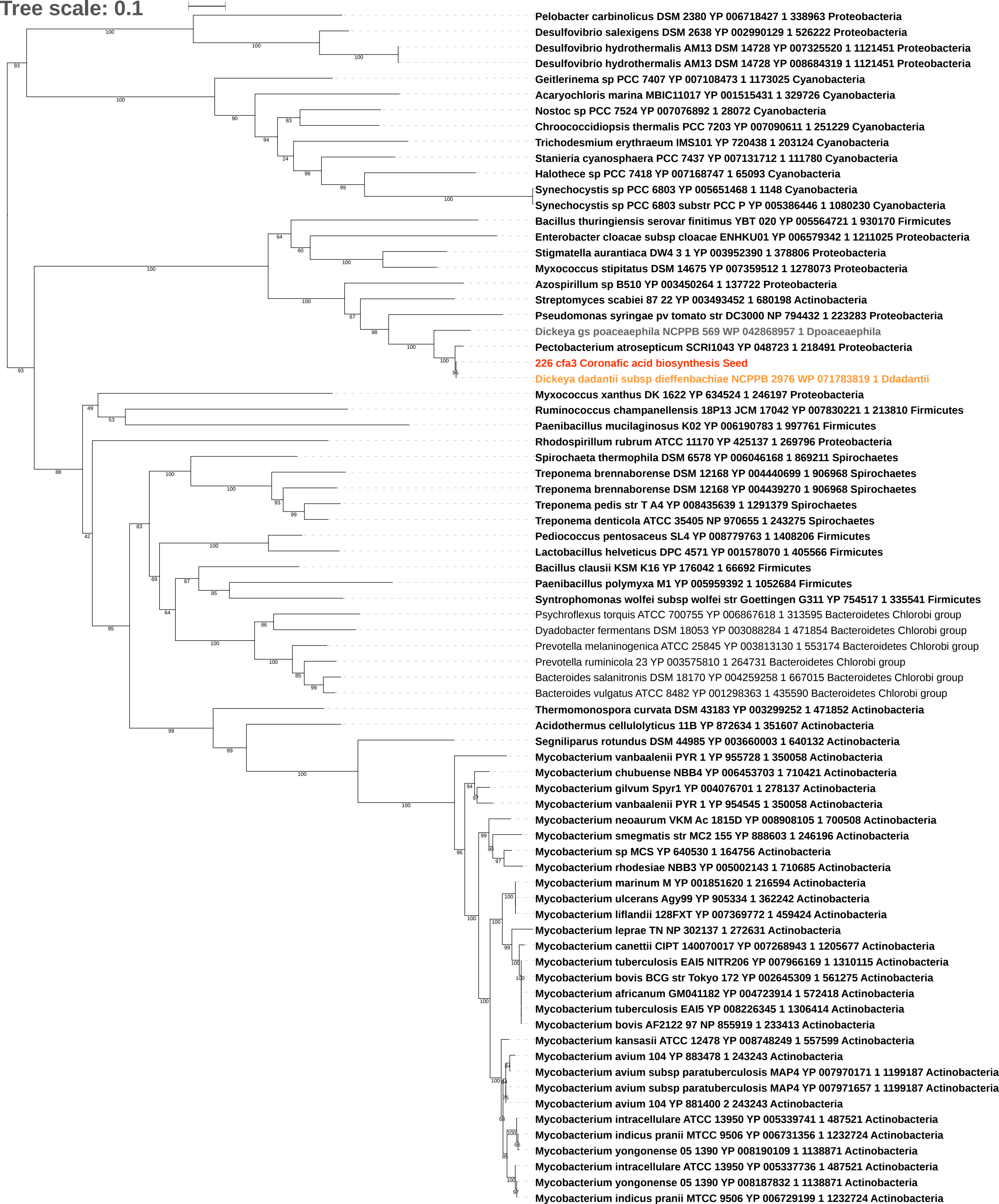

Tree scale: 0.1
Pelobacter carbinolicus DSM 2380 YP 006718427 1 338963 Proteobacteria
Desulfovibrio salexigens DSM 2638 YP 002990129 1 526222 Proteobacteria
Desulfovibrio hydrothermalis AM13 DSM 14728 YP 007325520 1 1121451 Proteobacteria
Desulfovibrio hydrothermalis AM13 DSM 14728 YP 008684319 1 1121451 Proteobacteria
Geitlerinema sp PCC 7407 YP 007108473 1 1173025 Cyanobacteria
Acaryochloris marina MBIC11017 YP 001515431 1 329726 Cyanobacteria
Nostoc sp PCC 7524 YP 007076892 1 28072 Cyanobacteria
Chroococcidiopsis thermalis PCC 7203 YP 007090611 1 251229 Cyanobacteria
Trichodesmium erythraeum IMS101 YP 720438 1 203124 Cyanobacteria
Stanieria cyanosphaera PCC 7437 YP 007131712 1 111780 Cyanobacteria
Halothece sp PCC 7418 YP 007168747 1 65093 Cyanobacteria
Synechocystis sp PCC 6803 YP 005651468 1 1148 Cyanobacteria
Synechocystis sp PCC 6803 substr PCC P YP 005386446 1 1080230 Cyanobacteria
Bacillus thuringiensis serovar finitimus YBT 020 YP 005564721 1 930170 Firmicutes
Enterobacter cloacae subsp cloacae ENHKU01 YP 006579342 1 1211025 Proteobacteria
Stigmatella aurantiaca DW4 3 1 YP 003952390 1 378806 Proteobacteria
Myxococcus stipitatus DSM 14675 YP 007359512 1 1278073 Proteobacteria
Azospirillum sp B510 YP 003450264 1 137722 Proteobacteria
Streptomyces scabiei 87 22 YP 003493452 1 680198 Actinobacteria
Pseudomonas syringae pv tomato str DC3000 NP 794432 1 223283 Proteobacteria
Dickeya gs poaceaephila NCPPB 569 WP 042868957 1 Dpoaceaephila
Pectobacterium atrosepticum SCRI1043 YP 048723 1 218491 Proteobacteria
226 cfa3 Coronafic acid biosynthesis Seed
Dickeya dadantii subsp dieffenbachiae NCPPB 2976 WP 071783819 1 Ddadantii
Myxococcus xanthus DK 1622 YP 634524 1 246197 Proteobacteria
Ruminococcus champanellensis 18P13 JCM 17042 YP 007830221 1 213810 Firmicutes
Paenibacillus mucilaginosus K02 YP 006190783 1 997761 Firmicutes
Rhodospirillum rubrum ATCC 11170 YP 425137 1 269796 Proteobacteria
Spirochaeta thermophila DSM 6578 YP 006046168 1 869211 Spirochaetes
Treponema brennaborense DSM 12168 YP 004440699 1 906968 Spirochaetes
Treponema brennaborense DSM 12168 YP 004439270 1 906968 Spirochaetes
Treponema pedis str T A4 YP 008435639 1 1291379 Spirochaetes
Treponema denticola ATCC 35405 NP 970655 1 243275 Spirochaetes
Pediococcus pentosaceus SL4 YP 008779763 1 1408206 Firmicutes
Lactobacillus helveticus DPC 4571 YP 001578070 1 405566 Firmicutes
Bacillus clausii KSM K16 YP 176042 1 66692 Firmicutes
Paenibacillus polymyxa M1 YP 005959392 1 1052684 Firmicutes
Syntrophomonas wolfei subsp wolfei str Goettingen G311 YP 754517 1 335541 Firmicutes
Psychroflexus torquis ATCC 700755 YP 006867618 1 313595 Bacteroidetes Chlorobi group
Dyadobacter fermentans DSM 18053 YP 003088284 1 471854 Bacteroidetes Chlorobi group
Prevotella melaninogenica ATCC 25845 YP 003813130 1 553174 Bacteroidetes Chlorobi group
Prevotella ruminicola 23 YP 003575810 1 264731 Bacteroidetes Chlorobi group
Bacteroides salanitronis DSM 18170 YP 004259258 1 667015 Bacteroidetes Chlorobi group
Bacteroides vulgatus ATCC 8482 YP 001298363 1 435590 Bacteroidetes Chlorobi group
Thermomonospora curvata DSM 43183 YP 003299252 1 471852 Actinobacteria
Acidothermus cellulolyticus 11B YP 872634 1 351607 Actinobacteria
Segniliparus rotundus DSM 44985 YP 003660003 1 640132 Actinobacteria
Mycobacterium vanbaalenii PYR 1 YP 955728 1 350058 Actinobacteria
Mycobacterium chubuense NBB4 YP 006453703 1 710421 Actinobacteria
Mycobacterium gilvum Spyr1 YP 004076701 1 278137 Actinobacteria
Mycobacterium vanbaalenii PYR 1 YP 954545 1 350058 Actinobacteria
Mycobacterium neoaurum VKM Ac 1815D YP 008908105 1 700508 Actinobacteria
Mycobacterium smegmatis str MC2 155 YP 888603 1 246196 Actinobacteria
Mycobacterium sp MCS YP 640530 1 164756 Actinobacteria
Mycobacterium rhodesiae NBB3 YP 005002143 1 710685 Actinobacteria
Mycobacterium marinum M YP 001851620 1 216594 Actinobacteria
Mycobacterium ulcerans Agy99 YP 905334 1 362242 Actinobacteria
Mycobacterium liflandii 128FXT YP 007369772 1 459424 Actinobacteria
Mycobacterium leprae TN NP 302137 1 272631 Actinobacteria
Mycobacterium canettii CIPT 140070017 YP 007268943 1 1205677 Actinobacteria
Mycobacterium tuberculosis EAI5 NITR206 YP 007966169 1 1310115 Actinobacteria
Mycobacterium bovis BCG str Tokyo 172 YP 002645309 1 561275 Actinobacteria
Mycobacterium africanum GM041182 YP 004723914 1 572418 Actinobacteria
Mycobacterium tuberculosis EAI5 YP 008226345 1 1306414 Actinobacteria
Mycobacterium bovis AF2122 97 NP 855919 1 233413 Actinobacteria
Mycobacterium kansasii ATCC 12478 YP 008748249 1 557599 Actinobacteria
Mycobacterium avium 104 YP 883478 1 243243 Actinobacteria
Mycobacterium avium subsp paratuberculosis MAP4 YP 007970171 1 1199187 Actinobacteria
Mycobacterium avium subsp paratuberculosis MAP4 YP 007971657 1 1199187 Actinobacteria
Mycobacterium avium 104 YP 881400 2 243243 Actinobacteria
Mycobacterium intracellulare ATCC 13950 YP 005339741 1 487521 Actinobacteria
Mycobacterium indicus pranii MTCC 9506 YP 006731356 1 1232724 Actinobacteria
Mycobacterium yongonense 05 1390 YP 008190109 1 1138871 Actinobacteria
Mycobacterium intracellulare ATCC 13950 YP 005337736 1 487521 Actinobacteria
Mycobacterium yongonense 05 1390 YP 008187832 1 1138871 Actinobacteria
Mycobacterium indicus pranii MTCC 9506 YP 006729199 1 1232724 Actinobacteria

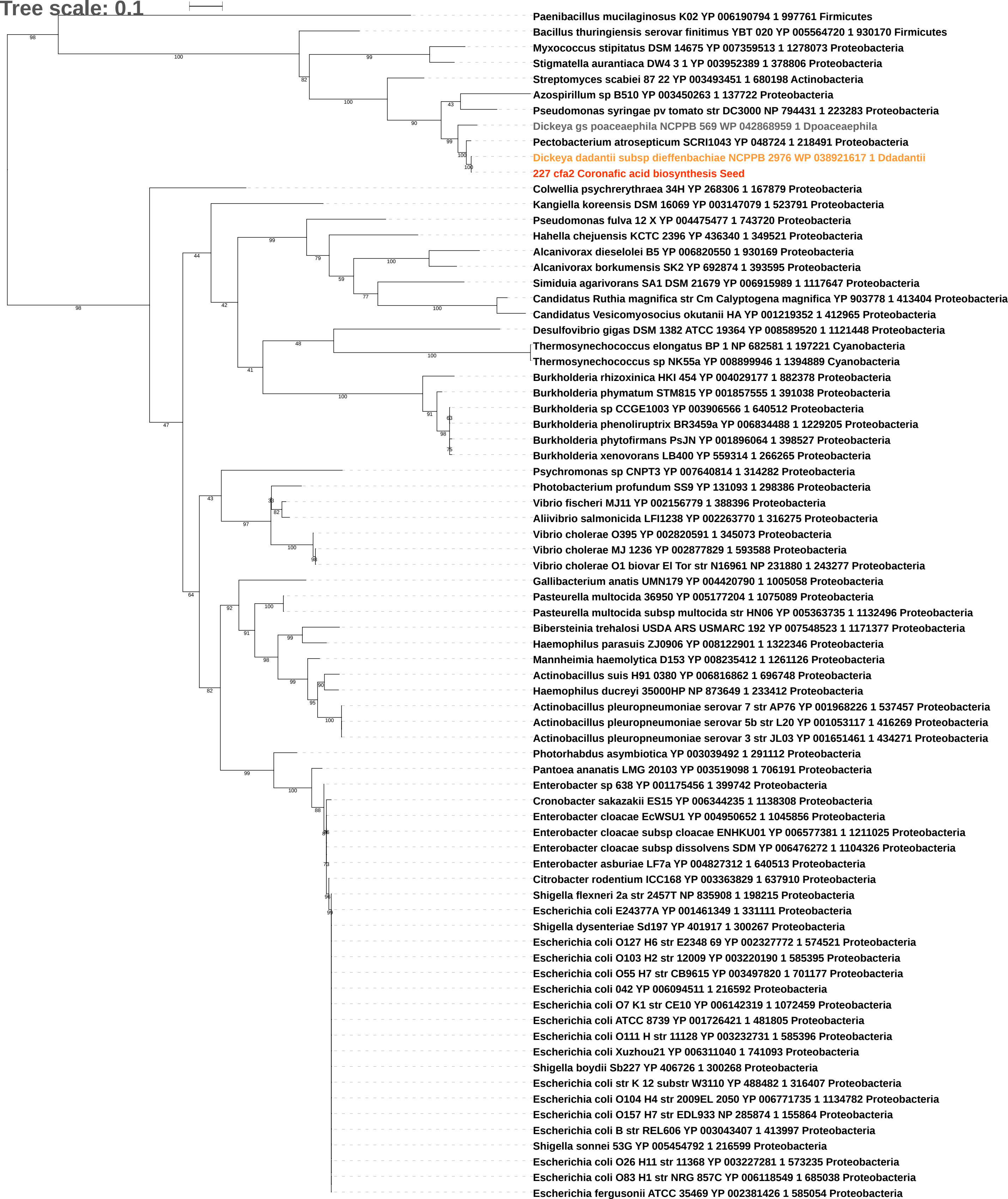
Tree scale: 0.1
Paenibacillus mucilaginosus K02 YP 006190794 1 997761 Firmicutes
Bacillus thuringiensis serovar finitimus YBT 020 YP 005564720 1 930170 Firmicutes
Myxococcus stipitatus DSM 14675 YP 007359513 1 1278073 Proteobacteria
Stigmatella aurantiaca DW4 3 1 YP 003952389 1 378806 Proteobacteria
Streptomyces scabiei 87 22 YP 003493451 1 680198 Actinobacteria
Azospirillum sp B510 YP 003450263 1 137722 Proteobacteria
Pseudomonas syringae pv tomato str DC3000 NP 794431 1 223283 Proteobacteria
Dickeya gs poaceaephila NCPPB 569 WP 042868959 1 Dpoaceaephila
Pectobacterium atrosepticum SCRI1043 YP 048724 1 218491 Proteobacteria
Dickeya dadantii subsp dieffenbachiae NCPPB 2976 WP 038921617 1 Ddadantii
227 cfa2 Coronafic acid biosynthesis Seed
Colwellia psychrerythraea 34H YP 268306 1 167879 Proteobacteria
Kangiella koreensis DSM 16069 YP 003147079 1 523791 Proteobacteria
Pseudomonas fulva 12 X YP 004475477 1 743720 Proteobacteria
Hahella chejuensis KCTC 2396 YP 436340 1 349521 Proteobacteria
Alcanivorax dieselolei B5 YP 006820550 1 930169 Proteobacteria
Alcanivorax borkumensis SK2 YP 692874 1 393595 Proteobacteria
Simiduia agarivorans SA1 DSM 21679 YP 006915989 1 1117647 Proteobacteria
Candidatus Ruthia magnifica str Cm Calyptogena magnifica YP 903778 1 413404 Proteobacteria
Candidatus Vesicomyosocius okutanii HA YP 001219352 1 412965 Proteobacteria
Desulfovibrio gigas DSM 1382 ATCC 19364 YP 008589520 1 1121448 Proteobacteria
Thermosynechococcus elongatus BP 1 NP 682581 1 197221 Cyanobacteria
Thermosynechococcus sp NK55a YP 008899946 1 1394889 Cyanobacteria
Burkholderia rhizoxinica HKI 454 YP 004029177 1 882378 Proteobacteria
Burkholderia phymatum STM815 YP 001857555 1 391038 Proteobacteria
Burkholderia sp CCGE1003 YP 003906566 1 640512 Proteobacteria
Burkholderia phenoliruptrix BR3459a YP 006834488 1 1229205 Proteobacteria
Burkholderia phytofirmans PsJN YP 001896064 1 398527 Proteobacteria
Burkholderia xenovorans LB400 YP 559314 1 266265 Proteobacteria
Psychromonas sp CNPT3 YP 007640814 1 314282 Proteobacteria
Photobacterium profundum SS9 YP 131093 1 298386 Proteobacteria
Vibrio fischeri MJ11 YP 002156779 1 388396 Proteobacteria
Aliivibrio salmonicida LFI1238 YP 002263770 1 316275 Proteobacteria
Vibrio cholerae O395 YP 002820591 1 345073 Proteobacteria
Vibrio cholerae MJ 1236 YP 002877829 1 593588 Proteobacteria
Vibrio cholerae O1 biovar El Tor str N16961 NP 231880 1 243277 Proteobacteria
Gallibacterium anatis UMN179 YP 004420790 1 1005058 Proteobacteria
Pasteurella multocida 36950 YP 005177204 1 1075089 Proteobacteria
Pasteurella multocida subsp multocida str HN06 YP 005363735 1 1132496 Proteobacteria
Bibersteinia trehalosi USDA ARS USMARC 192 YP 007548523 1 1171377 Proteobacteria
Haemophilus parasuis ZJ0906 YP 008122901 1 1322346 Proteobacteria
Mannheimia haemolytica D153 YP 008235412 1 1261126 Proteobacteria
Actinobacillus suis H91 0380 YP 006816862 1 696748 Proteobacteria
Haemophilus ducreyi 35000HP NP 873649 1 233412 Proteobacteria
Actinobacillus pleuropneumoniae serovar 7 str AP76 YP 001968226 1 537457 Proteobacteria
Actinobacillus pleuropneumoniae serovar 5b str L20 YP 001053117 1 416269 Proteobacteria
Actinobacillus pleuropneumoniae serovar 3 str JL03 YP 001651461 1 434271 Proteobacteria
Photorhabdus asymbiotica YP 003039492 1 291112 Proteobacteria
Pantoea ananatis LMG 20103 YP 003519098 1 706191 Proteobacteria
Enterobacter sp 638 YP 001175456 1 399742 Proteobacteria
Cronobacter sakazakii ES15 YP 006344235 1 1138308 Proteobacteria
Enterobacter cloacae EcWSU1 YP 004950652 1 1045856 Proteobacteria
Enterobacter cloacae subsp cloacae ENHKU01 YP 006577381 1 1211025 Proteobacteria
Enterobacter cloacae subsp dissolvens SDM YP 006476272 1 1104326 Proteobacteria
Enterobacter asburiae LF7a YP 004827312 1 640513 Proteobacteria
Citrobacter rodentium ICC168 YP 003363829 1 637910 Proteobacteria
Shigella flexneri 2a str 2457T NP 835908 1 198215 Proteobacteria
Escherichia coli E24377A YP 001461349 1 331111 Proteobacteria
Shigella dysenteriae Sd197 YP 401917 1 300267 Proteobacteria
Escherichia coli O127 H6 str E2348 69 YP 002327772 1 574521 Proteobacteria
Escherichia coli O103 H2 str 12009 YP 003220190 1 585395 Proteobacteria
Escherichia coli O55 H7 str CB9615 YP 003497820 1 701177 Proteobacteria
Escherichia coli 042 YP 006094511 1 216592 Proteobacteria
Escherichia coli O7 K1 str CE10 YP 006142319 1 1072459 Proteobacteria
Escherichia coli ATCC 8739 YP 001726421 1 481805 Proteobacteria
Escherichia coli O111 H str 11128 YP 003232731 1 585396 Proteobacteria
Escherichia coli Xuzhou21 YP 006311040 1 741093 Proteobacteria
Shigella boydii Sb227 YP 406726 1 300268 Proteobacteria
Escherichia coli str K 12 substr W3110 YP 488482 1 316407 Proteobacteria
Escherichia coli O104 H4 str 2009EL 2050 YP 006771735 1 1134782 Proteobacteria
Escherichia coli O157 H7 str EDL933 NP 285874 1 155864 Proteobacteria
Escherichia coli B str REL606 YP 003043407 1 413997 Proteobacteria
Shigella sonnei 53G YP 005454792 1 216599 Proteobacteria
Escherichia coli O26 H11 str 11368 YP 003227281 1 573235 Proteobacteria
Escherichia coli O83 H1 str NRG 857C YP 006118549 1 685038 Proteobacteria
Escherichia fergusonii ATCC 35469 YP 002381426 1 585054 Proteobacteria

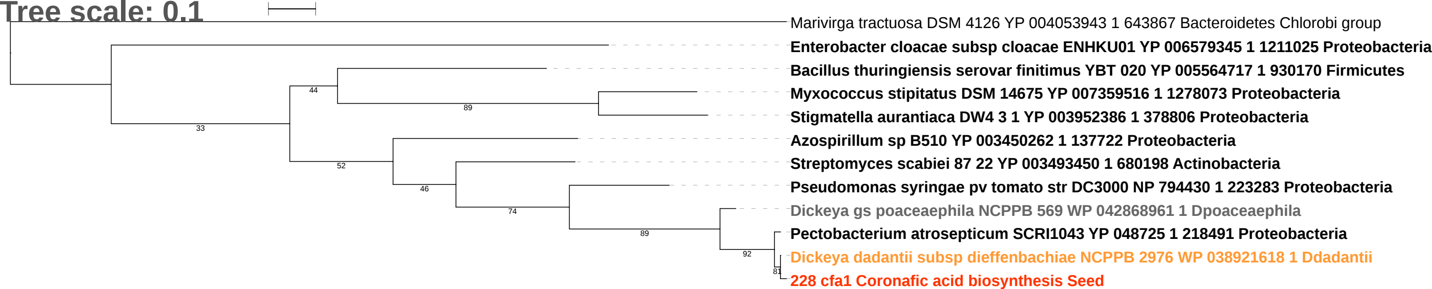
Tree scale: 0.1
Marivirga tractuosa DSM 4126 YP 004053943 1 643867 Bacteroidetes Chlorobi group
Enterobacter cloacae subsp cloacae ENHKU01 YP 006579345 1 1211025 Proteobacteria
Bacillus thuringiensis serovar finitimus YBT 020 YP 005564717 1 930170 Firmicutes
Myxococcus stipitatus DSM 14675 YP 007359516 1 1278073 Proteobacteria
Stigmatella aurantiaca DW4 3 1 YP 003952386 1 378806 Proteobacteria
Azospirillum sp B510 YP 003450262 1 137722 Proteobacteria
Streptomyces scabiei 87 22 YP 003493450 1 680198 Actinobacteria
Pseudomonas syringae pv tomato str DC3000 NP 794430 1 223283 Proteobacteria
Dickeya gs poaceaephila NCPPB 569 WP 042868961 1 Dpoaceaephila
Pectobacterium atrosepticum SCRI1043 YP 048725 1 218491 Proteobacteria
Dickeya dadantii subsp dieffenbachiae NCPPB 2976 WP 038921618 1 Ddadantii
228 cfa1 Coronafic acid biosynthesis Seed
44
89
33
52
46
74
89
92
81

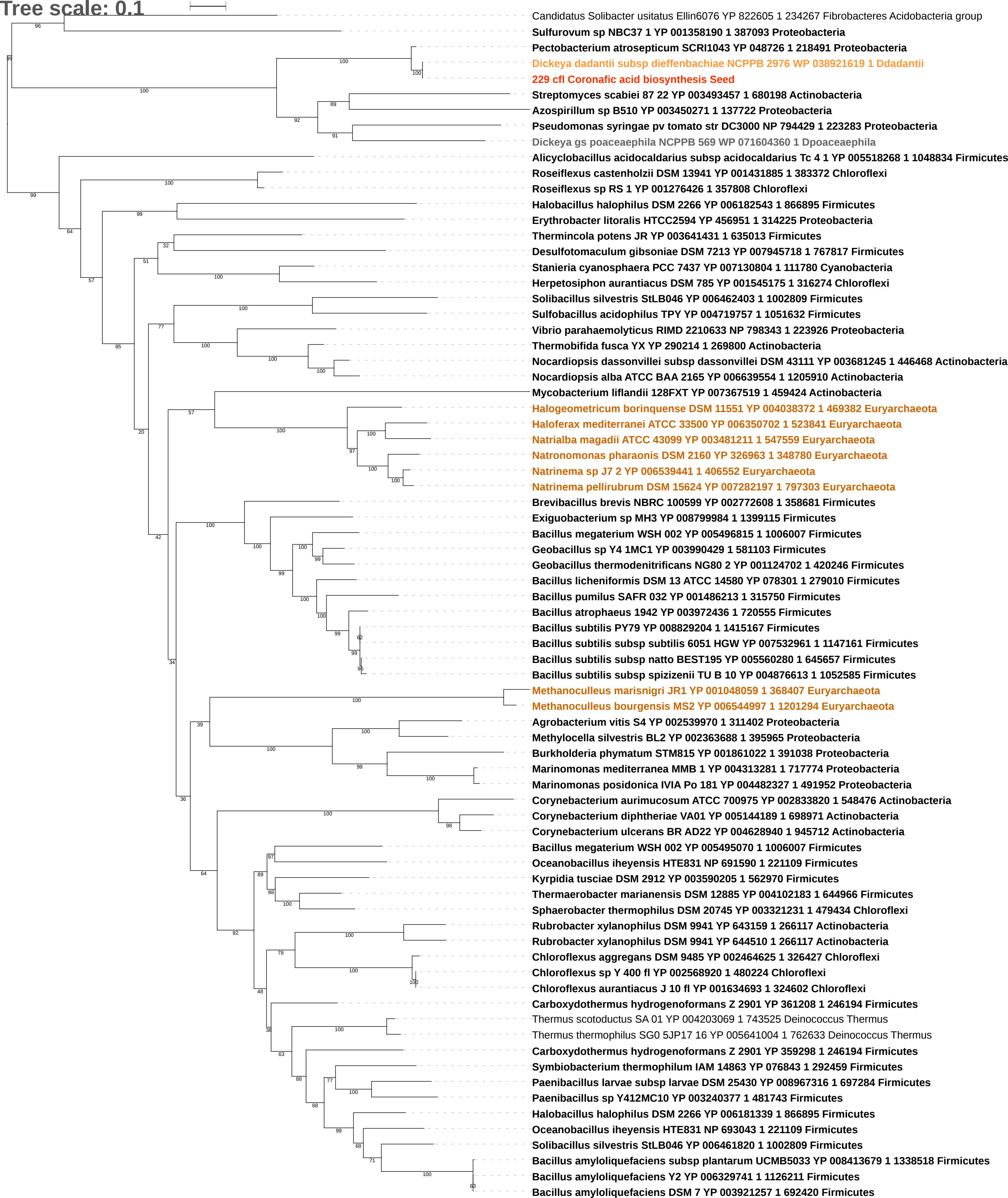
Tree scale: 0.1
Candidatus Solibacter usitatus Ellin6076 YP 822605 1 234267 Fibrobacteres Acidobacteria group
Sulfurovum sp NBC37 1 YP 001358190 1 387093 Proteobacteria
Pectobacterium atrosepticum SCRI1043 YP 048726 1 218491 Proteobacteria
Dickeya dadantii subsp dieffenbachiae NCPPB 2976 WP 038921619 1 Ddadantii
229 cfl Coronafic acid biosynthesis Seed
Streptomyces scabiei 87 22 YP 003493457 1 680198 Actinobacteria
Azospirillum sp B510 YP 003450271 1 137722 Proteobacteria
Pseudomonas syringae pv tomato str DC3000 NP 794429 1 223283 Proteobacteria
Dickeya gs poaceaephila NCPPB 569 WP 071604360 1 Dpoaceaephila
Alicyclobacillus acidocaldarius subsp acidocaldarius Tc 4 1 YP 005518268 1 1048834 Firmicutes
Roseiflexus castenholzii DSM 13941 YP 001431885 1 383372 Chloroflexi
Roseiflexus sp RS 1 YP 001276426 1 357808 Chloroflexi
Halobacillus halophilus DSM 2266 YP 006182543 1 866895 Firmicutes
Erythrobacter litoralis HTCC2594 YP 456951 1 314225 Proteobacteria
Thermincola potens JR YP 003641431 1 635013 Firmicutes
Desulfotomaculum gibsoniae DSM 7213 YP 007945718 1 767817 Firmicutes
Stanieria cyanosphaera PCC 7437 YP 007130804 1 111780 Cyanobacteria
Herpetosiphon aurantiacus DSM 785 YP 001545175 1 316274 Chloroflexi
Solibacillus silvestris StLB046 YP 006462403 1 1002809 Firmicutes
Sulfobacillus acidophilus TPY YP 004719757 1 1051632 Firmicutes
Vibrio parahaemolyticus RIMD 2210633 NP 798343 1 223926 Proteobacteria
Thermobifida fusca YX YP 290214 1 269800 Actinobacteria
Nocardiopsis dassonvillei subsp dassonvillei DSM 43111 YP 003681245 1 446468 Actinobacteria
Nocardiopsis alba ATCC BAA 2165 YP 006639554 1 1205910 Actinobacteria
Mycobacterium liflandii 128FXT YP 007367519 1 459424 Actinobacteria
Halogeometricum borinquense DSM 11551 YP 004038372 1 469382 Euryarchaeota
Haloferax mediterranei ATCC 33500 YP 006350702 1 523841 Euryarchaeota
Natrialba magadii ATCC 43099 YP 003481211 1 547559 Euryarchaeota
Natronomonas pharaonis DSM 2160 YP 326963 1 348780 Euryarchaeota
Natrinema sp J7 2 YP 006539441 1 406552 Euryarchaeota
Natrinema pellirubrum DSM 15624 YP 007282197 1 797303 Euryarchaeota
Brevibacillus brevis NBRC 100599 YP 002772608 1 358681 Firmicutes
Exiguobacterium sp MH3 YP 008799984 1 1399115 Firmicutes
Bacillus megaterium WSH 002 YP 005496815 1 1006007 Firmicutes
Geobacillus sp Y4 1MC1 YP 003990429 1 581103 Firmicutes
Geobacillus thermodenitrificans NG80 2 YP 001124702 1 420246 Firmicutes
Bacillus licheniformis DSM 13 ATCC 14580 YP 078301 1 279010 Firmicutes
Bacillus pumilus SAFR 032 YP 001486213 1 315750 Firmicutes
Bacillus atrophaeus 1942 YP 003972436 1 720555 Firmicutes
Bacillus subtilis PY79 YP 008829204 1 1415167 Firmicutes
Bacillus subtilis subsp subtilis 6051 HGW YP 007532961 1 1147161 Firmicutes
Bacillus subtilis subsp natto BEST195 YP 005560280 1 645657 Firmicutes
Bacillus subtilis subsp spizizenii TU B 10 YP 004876613 1 1052585 Firmicutes
Methanoculleus marisnigri JR1 YP 001048059 1 368407 Euryarchaeota
Methanoculleus bourgensis MS2 YP 006544997 1 1201294 Euryarchaeota
Agrobacterium vitis S4 YP 002539970 1 311402 Proteobacteria
Methylocella silvestris BL2 YP 002363688 1 395965 Proteobacteria
Burkholderia phymatum STM815 YP 001861022 1 391038 Proteobacteria
Marinomonas mediterranea MMB 1 YP 004313281 1 717774 Proteobacteria
Marinomonas posidonica IVIA Po 181 YP 004482327 1 491952 Proteobacteria
Corynebacterium aurimucosum ATCC 700975 YP 002833820 1 548476 Actinobacteria
Corynebacterium diphtheriae VA01 YP 005144189 1 698971 Actinobacteria
Corynebacterium ulcerans BR AD22 YP 004628940 1 945712 Actinobacteria
Bacillus megaterium WSH 002 YP 005495070 1 1006007 Firmicutes
Oceanobacillus iheyensis HTE831 NP 691590 1 221109 Firmicutes
Kyrpidia tusciae DSM 2912 YP 003590205 1 562970 Firmicutes
Thermaerobacter marianensis DSM 12885 YP 004102183 1 644966 Firmicutes
Sphaerobacter thermophilus DSM 20745 YP 003321231 1 479434 Chloroflexi
Rubrobacter xylanophilus DSM 9941 YP 643159 1 266117 Actinobacteria
Rubrobacter xylanophilus DSM 9941 YP 644510 1 266117 Actinobacteria
Chloroflexus aggregans DSM 9485 YP 002464625 1 326427 Chloroflexi
Chloroflexus sp Y 400 fl YP 002568920 1 480224 Chloroflexi
Chloroflexus aurantiacus J 10 fl YP 001634693 1 324602 Chloroflexi
Carboxydothermus hydrogenoformans Z 2901 YP 361208 1 246194 Firmicutes
Thermus scotoductus SA 01 YP 004203069 1 743525 Deinococcus Thermus
Thermus thermophilus SG0 5JP17 16 YP 005641004 1 762633 Deinococcus Thermus
Carboxydothermus hydrogenoformans Z 2901 YP 359298 1 246194 Firmicutes
Symbiobacterium thermophilum IAM 14863 YP 076843 1 292459 Firmicutes
Paenibacillus larvae subsp larvae DSM 25430 YP 008967316 1 697284 Firmicutes
Paenibacillus sp Y412MC10 YP 003240377 1 481743 Firmicutes
Halobacillus halophilus DSM 2266 YP 006181339 1 866895 Firmicutes
Oceanobacillus iheyensis HTE831 NP 693043 1 221109 Firmicutes
Solibacillus silvestris StLB046 YP 006461820 1 1002809 Firmicutes
Bacillus amyloliquefaciens subsp plantarum UCMB5033 YP 008413679 1 1338518 Firmicutes
Bacillus amyloliquefaciens Y2 YP 006329741 1 1126211 Firmicutes
Bacillus amyloliquefaciens DSM 7 YP 003921257 1 692420 Firmicutes

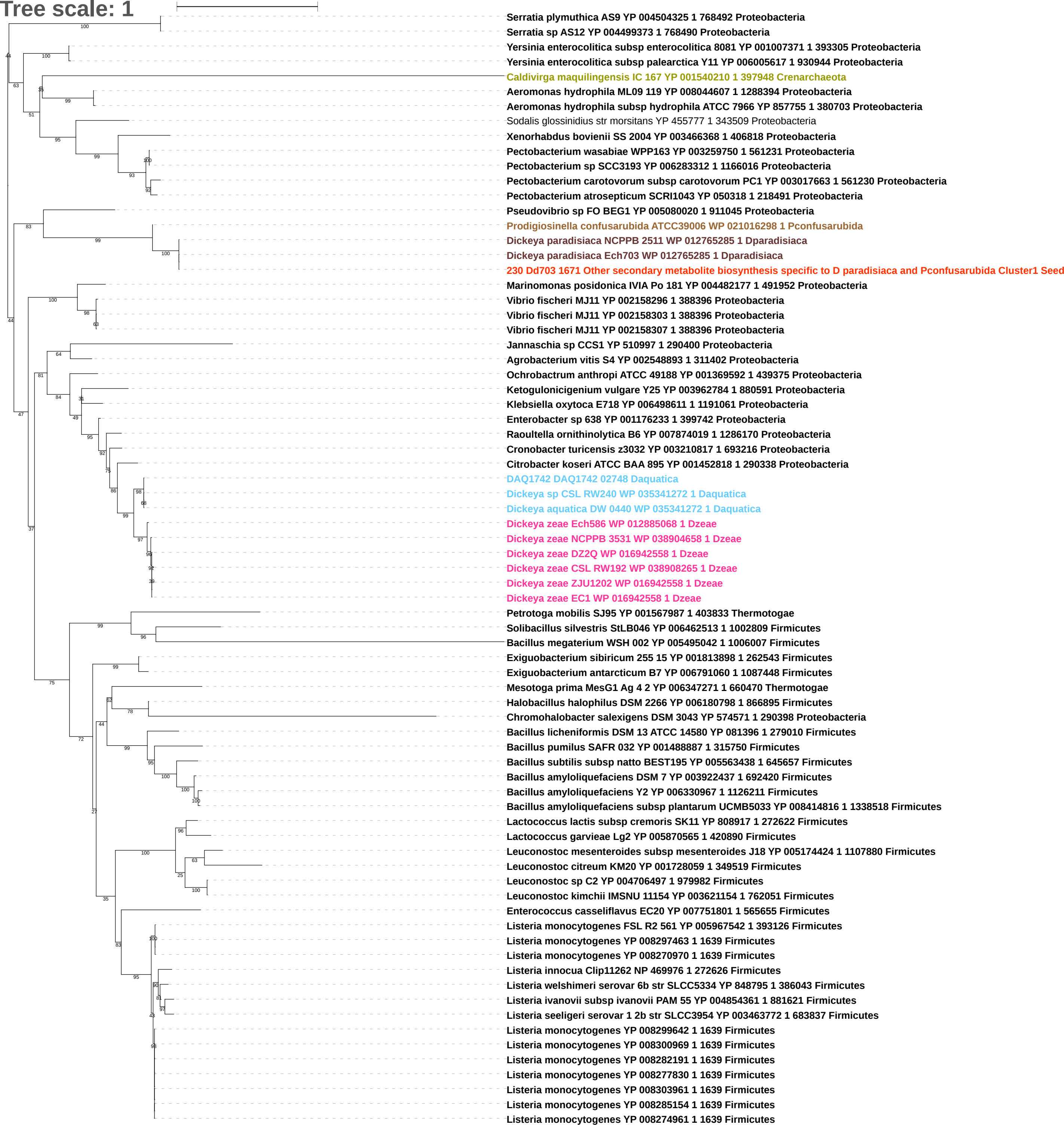
Tree scale: 1
Serratia plymuthica AS9 YP 004504325 1 768492 Proteobacteria
Serratia sp AS12 YP 004499373 1 768490 Proteobacteria
Yersinia enterocolitica subsp enterocolitica 8081 YP 001007371 1 393305 Proteobacteria
Yersinia enterocolitica subsp palearctica Y11 YP 006005617 1 930944 Proteobacteria
Caldivirga maquilingensis IC 167 YP 001540210 1 397948 Crenarchaeota
Aeromonas hydrophila ML09 119 YP 008044607 1 1288394 Proteobacteria
Aeromonas hydrophila subsp hydrophila ATCC 7966 YP 857755 1 380703 Proteobacteria
Sodalis glossinidius str morsitans YP 455777 1 343509 Proteobacteria
Xenorhabdus bovienii SS 2004 YP 003466368 1 406818 Proteobacteria
Pectobacterium wasabiae WPP163 YP 003259750 1 561231 Proteobacteria
Pectobacterium sp SCC3193 YP 006283312 1 1166016 Proteobacteria
Pectobacterium carotovorum subsp carotovorum PC1 YP 003017663 1 561230 Proteobacteria
Pectobacterium atrosepticum SCRI1043 YP 050318 1 218491 Proteobacteria
Pseudovibrio sp FO BEG1 YP 005080020 1 911045 Proteobacteria
Prodigiosinella confusarubida ATCC39006 WP 021016298 1 Pconfusarubida
Dickeya paradisiaca NCPPB 2511 WP 012765285 1 Dparadisiaca
Dickeya paradisiaca Ech703 WP 012765285 1 Dparadisiaca
230 Dd703 1671 Other secondary metabolite biosynthesis specific to D paradisiaca and Pconfusarubida Cluster1 Seed
Marinomonas posidonica IVIA Po 181 YP 004482177 1 491952 Proteobacteria
Vibrio fischeri MJ11 YP 002158296 1 388396 Proteobacteria
Vibrio fischeri MJ11 YP 002158303 1 388396 Proteobacteria
Vibrio fischeri MJ11 YP 002158307 1 388396 Proteobacteria
Jannaschia sp CCS1 YP 510997 1 290400 Proteobacteria
Agrobacterium vitis S4 YP 002548893 1 311402 Proteobacteria
Ochrobactrum anthropi ATCC 49188 YP 001369592 1 439375 Proteobacteria
Ketogulonicigenium vulgare Y25 YP 003962784 1 880591 Proteobacteria
Klebsiella oxytoca E718 YP 006498611 1 1191061 Proteobacteria
Enterobacter sp 638 YP 001176233 1 399742 Proteobacteria
Raoultella ornithinolytica B6 YP 007874019 1 1286170 Proteobacteria
Cronobacter turicensis z3032 YP 003210817 1 693216 Proteobacteria
Citrobacter koseri ATCC BAA 895 YP 001452818 1 290338 Proteobacteria
DAQ1742 DAQ1742 02748 Daquatica
Dickeya sp CSL RW240 WP 035341272 1 Daquatica
Dickeya aquatica DW 0440 WP 035341272 1 Daquatica
Dickeya zeae Ech586 WP 012885068 1 Dzeae
Dickeya zeae NCPPB 3531 WP 038904658 1 Dzeae
Dickeya zeae DZ2Q WP 016942558 1 Dzeae
Dickeya zeae CSL RW192 WP 038908265 1 Dzeae
Dickeya zeae ZJU1202 WP 016942558 1 Dzeae
Dickeya zeae EC1 WP 016942558 1 Dzeae
Petrotoga mobilis SJ95 YP 001567987 1 403833 Thermotogae
Solibacillus silvestris StLB046 YP 006462513 1 1002809 Firmicutes
Bacillus megaterium WSH 002 YP 005495042 1 1006007 Firmicutes
Exiguobacterium sibiricum 255 15 YP 001813898 1 262543 Firmicutes
Exiguobacterium antarcticum B7 YP 006791060 1 1087448 Firmicutes
Mesotoga prima MesG1 Ag 4 2 YP 006347271 1 660470 Thermotogae
Halobacillus halophilus DSM 2266 YP 006180798 1 866895 Firmicutes
Chromohalobacter salexigens DSM 3043 YP 574571 1 290398 Proteobacteria
Bacillus licheniformis DSM 13 ATCC 14580 YP 081396 1 279010 Firmicutes
Bacillus pumilus SAFR 032 YP 001488887 1 315750 Firmicutes
Bacillus subtilis subsp natto BEST195 YP 005563438 1 645657 Firmicutes
Bacillus amyloliquefaciens DSM 7 YP 003922437 1 692420 Firmicutes
Bacillus amyloliquefaciens Y2 YP 006330967 1 1126211 Firmicutes
Bacillus amyloliquefaciens subsp plantarum UCMB5033 YP 008414816 1 1338518 Firmicutes
Lactococcus lactis subsp cremoris SK11 YP 808917 1 272622 Firmicutes
Lactococcus garvieae Lg2 YP 005870565 1 420890 Firmicutes
Leuconostoc mesenteroides subsp mesenteroides J18 YP 005174424 1 1107880 Firmicutes
Leuconostoc citreum KM20 YP 001728059 1 349519 Firmicutes
Leuconostoc sp C2 YP 004706497 1 979982 Firmicutes
Leuconostoc kimchii IMSNU 11154 YP 003621154 1 762051 Firmicutes
Enterococcus casseliflavus EC20 YP 007751801 1 565655 Firmicutes
Listeria monocytogenes FSL R2 561 YP 005967542 1 393126 Firmicutes
Listeria monocytogenes YP 008297463 1 1639 Firmicutes
Listeria monocytogenes YP 008270970 1 1639 Firmicutes
Listeria innocua Clip11262 NP 469976 1 272626 Firmicutes
Listeria welshimeri serovar 6b str SLCC5334 YP 848795 1 386043 Firmicutes
Listeria ivanovii subsp ivanovii PAM 55 YP 004854361 1 881621 Firmicutes
Listeria seeligeri serovar 1 2b str SLCC3954 YP 003463772 1 683837 Firmicutes
Listeria monocytogenes YP 008299642 1 1639 Firmicutes
Listeria monocytogenes YP 008300969 1 1639 Firmicutes
Listeria monocytogenes YP 008282191 1 1639 Firmicutes
Listeria monocytogenes YP 008277830 1 1639 Firmicutes
Listeria monocytogenes YP 008303961 1 1639 Firmicutes
Listeria monocytogenes YP 008285154 1 1639 Firmicutes
Listeria monocytogenes YP 008274961 1 1639 Firmicutes

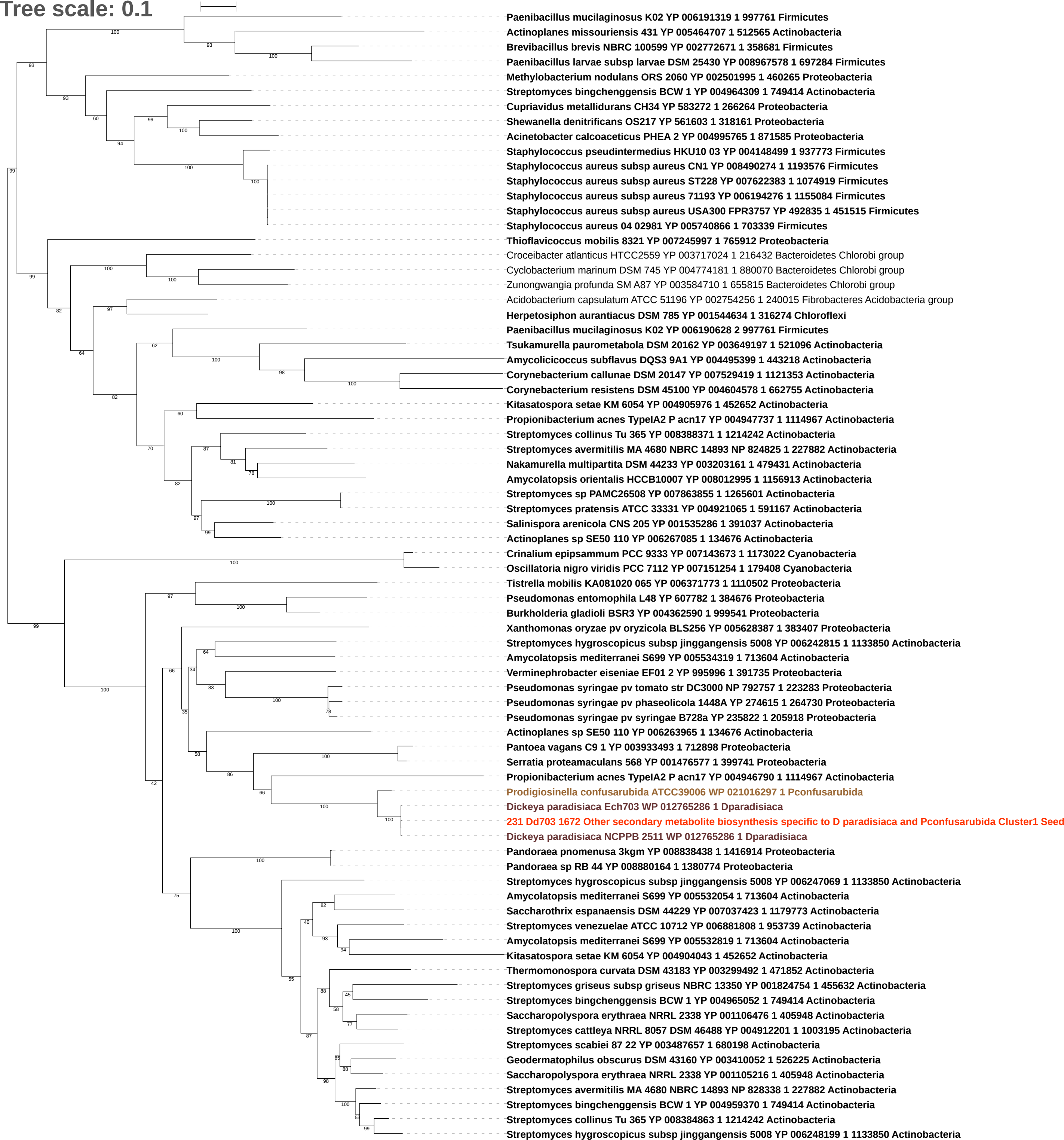
Tree scale: 0.1
Paenibacillus mucilaginosus K02 YP 006191319 1 997761 Firmicutes
Actinoplanes missouriensis 431 YP 005464707 1 512565 Actinobacteria
Brevibacillus brevis NBRC 100599 YP 002772671 1 358681 Firmicutes
Paenibacillus larvae subsp larvae DSM 25430 YP 008967578 1 697284 Firmicutes
Methylobacterium nodulans ORS 2060 YP 002501995 1 460265 Proteobacteria
Streptomyces bingchenggensis BCW 1 YP 004964309 1 749414 Actinobacteria
Cupriavidus metallidurans CH34 YP 583272 1 266264 Proteobacteria
Shewanella denitrificans OS217 YP 561603 1 318161 Proteobacteria
Acinetobacter calcoaceticus PHEA 2 YP 004995765 1 871585 Proteobacteria
Staphylococcus pseudintermedius HKU10 03 YP 004148499 1 937773 Firmicutes
Staphylococcus aureus subsp aureus CN1 YP 008490274 1 1193576 Firmicutes
Staphylococcus aureus subsp aureus ST228 YP 007622383 1 1074919 Firmicutes
Staphylococcus aureus subsp aureus 71193 YP 006194276 1 1155084 Firmicutes
Staphylococcus aureus subsp aureus USA300 FPR3757 YP 492835 1 451515 Firmicutes
Staphylococcus aureus 04 02981 YP 005740866 1 703339 Firmicutes
Thioflavicoccus mobilis 8321 YP 007245997 1 765912 Proteobacteria
Croceibacter atlanticus HTCC2559 YP 003717024 1 216432 Bacteroidetes Chlorobi group
Cyclobacterium marinum DSM 745 YP 004774181 1 880070 Bacteroidetes Chlorobi group
Zunongwangia profunda SM A87 YP 003584710 1 655815 Bacteroidetes Chlorobi group
Acidobacterium capsulatum ATCC 51196 YP 002754256 1 240015 Fibrobacteres Acidobacteria group
Herpetosiphon aurantiacus DSM 785 YP 001544634 1 316274 Chloroflexi
Paenibacillus mucilaginosus K02 YP 006190628 2 997761 Firmicutes
Tsukamurella paurometabola DSM 20162 YP 003649197 1 521096 Actinobacteria
Amycolicicoccus subflavus DQS3 9A1 YP 004495399 1 443218 Actinobacteria
Corynebacterium callunae DSM 20147 YP 007529419 1 1121353 Actinobacteria
Corynebacterium resistens DSM 45100 YP 004604578 1 662755 Actinobacteria
Kitasatospora setae KM 6054 YP 004905976 1 452652 Actinobacteria
Propionibacterium acnes TypeIA2 P acn17 YP 004947737 1 1114967 Actinobacteria
Streptomyces collinus Tu 365 YP 008388371 1 1214242 Actinobacteria
Streptomyces avermitilis MA 4680 NBRC 14893 NP 824825 1 227882 Actinobacteria
Nakamurella multipartita DSM 44233 YP 003203161 1 479431 Actinobacteria
Amycolatopsis orientalis HCCB10007 YP 008012995 1 1156913 Actinobacteria
Streptomyces sp PAMC26508 YP 007863855 1 1265601 Actinobacteria
Streptomyces pratensis ATCC 33331 YP 004921065 1 591167 Actinobacteria
Salinispora arenicola CNS 205 YP 001535286 1 391037 Actinobacteria
Actinoplanes sp SE50 110 YP 006267085 1 134676 Actinobacteria
Crinalium epipsammum PCC 9333 YP 007143673 1 1173022 Cyanobacteria
Oscillatoria nigro viridis PCC 7112 YP 007151254 1 179408 Cyanobacteria
Tistrella mobilis KA081020 065 YP 006371773 1 1110502 Proteobacteria
Pseudomonas entomophila L48 YP 607782 1 384676 Proteobacteria
Burkholderia gladioli BSR3 YP 004362590 1 999541 Proteobacteria
Xanthomonas oryzae pv oryzicola BLS256 YP 005628387 1 383407 Proteobacteria
Streptomyces hygroscopicus subsp jinggangensis 5008 YP 006242815 1 1133850 Actinobacteria
Amycolatopsis mediterranei S699 YP 005534319 1 713604 Actinobacteria
Verminephrobacter eiseniae EF01 2 YP 995996 1 391735 Proteobacteria
Pseudomonas syringae pv tomato str DC3000 NP 792757 1 223283 Proteobacteria
Pseudomonas syringae pv phaseolicola 1448A YP 274615 1 264730 Proteobacteria
Pseudomonas syringae pv syringae B728a YP 235822 1 205918 Proteobacteria
Actinoplanes sp SE50 110 YP 006263965 1 134676 Actinobacteria
Pantoea vagans C9 1 YP 003933493 1 712898 Proteobacteria
Serratia proteamaculans 568 YP 001476577 1 399741 Proteobacteria
Propionibacterium acnes TypeIA2 P acn17 YP 004946790 1 1114967 Actinobacteria
Prodigiosinella confusarubida ATCC39006 WP 021016297 1 Pconfusarubida
Dickeya paradisiaca Ech703 WP 012765286 1 Dparadisiaca
231 Dd703 1672 Other secondary metabolite biosynthesis specific to D paradisiaca and Pconfusarubida Cluster1 Seed
Dickeya paradisiaca NCPPB 2511 WP 012765286 1 Dparadisiaca
Pandoraea pnomenusa 3kgm YP 008838438 1 1416914 Proteobacteria
Pandoraea sp RB 44 YP 008880164 1 1380774 Proteobacteria
Streptomyces hygroscopicus subsp jinggangensis 5008 YP 006247069 1 1133850 Actinobacteria
Amycolatopsis mediterranei S699 YP 005532054 1 713604 Actinobacteria
Saccharothrix espanaensis DSM 44229 YP 007037423 1 1179773 Actinobacteria
Streptomyces venezuelae ATCC 10712 YP 006881808 1 953739 Actinobacteria
Amycolatopsis mediterranei S699 YP 005532819 1 713604 Actinobacteria
Kitasatospora setae KM 6054 YP 004904043 1 452652 Actinobacteria
Thermomonospora curvata DSM 43183 YP 003299492 1 471852 Actinobacteria
Streptomyces griseus subsp griseus NBRC 13350 YP 001824754 1 455632 Actinobacteria
Streptomyces bingchenggensis BCW 1 YP 004965052 1 749414 Actinobacteria
Saccharopolyspora erythraea NRRL 2338 YP 001106476 1 405948 Actinobacteria
Streptomyces cattleya NRRL 8057 DSM 46488 YP 004912201 1 1003195 Actinobacteria
Streptomyces scabiei 87 22 YP 003487657 1 680198 Actinobacteria
Geodermatophilus obscurus DSM 43160 YP 003410052 1 526225 Actinobacteria
Saccharopolyspora erythraea NRRL 2338 YP 001105216 1 405948 Actinobacteria
Streptomyces avermitilis MA 4680 NBRC 14893 NP 828338 1 227882 Actinobacteria
Streptomyces bingchenggensis BCW 1 YP 004959370 1 749414 Actinobacteria
Streptomyces collinus Tu 365 YP 008384863 1 1214242 Actinobacteria
Streptomyces hygroscopicus subsp jinggangensis 5008 YP 006248199 1 1133850 Actinobacteria

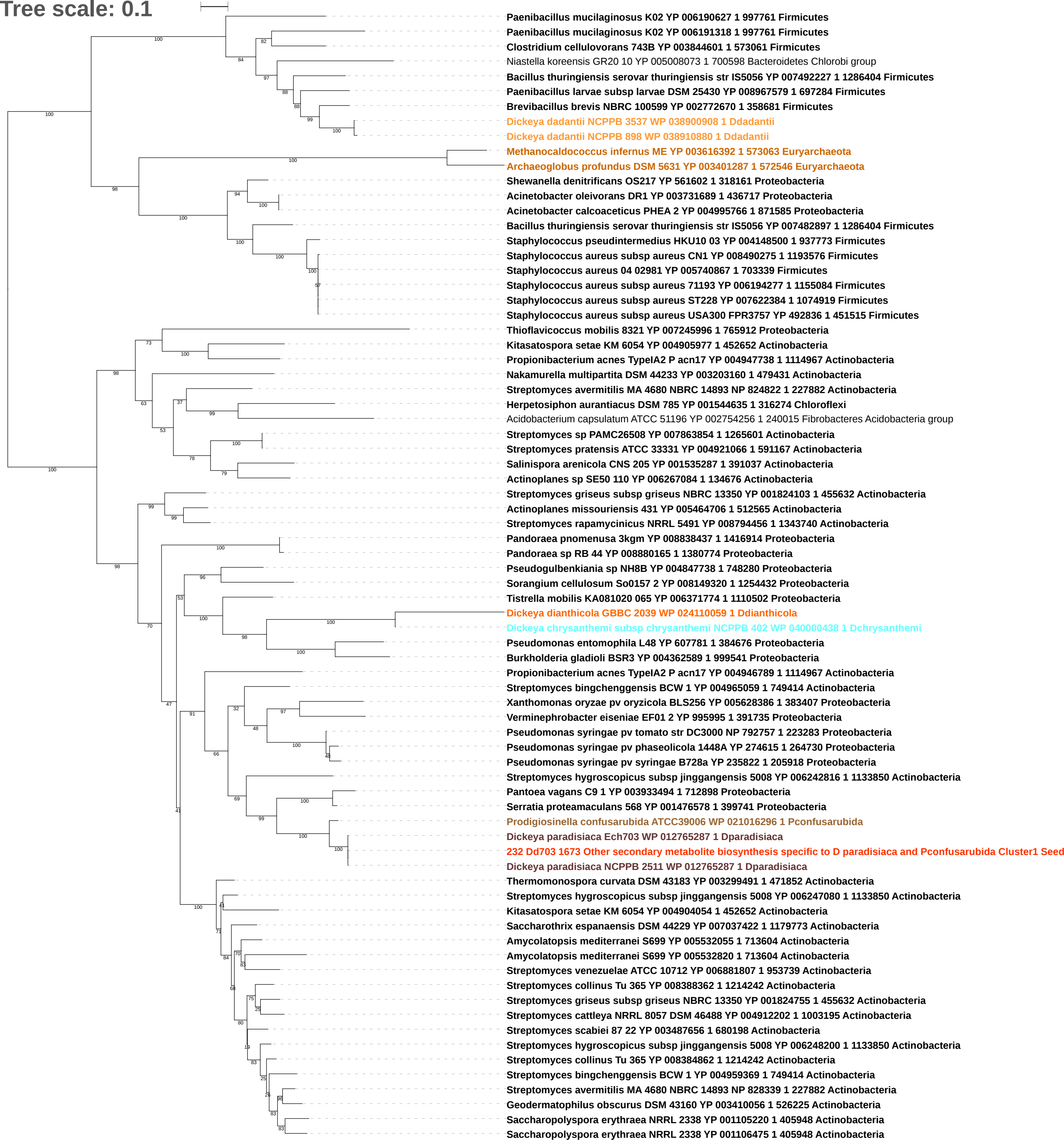
Tree scale: 0.1
Paenibacillus mucilaginosus K02 YP 006190627 1 997761 Firmicutes
Paenibacillus mucilaginosus K02 YP 006191318 1 997761 Firmicutes
Clostridium cellulovorans 743B YP 003844601 1 573061 Firmicutes
Niastella koreensis GR20 10 YP 005008073 1 700598 Bacteroidetes Chlorobi group
Bacillus thuringiensis serovar thuringiensis str IS5056 YP 007492227 1 1286404 Firmicutes
Paenibacillus larvae subsp larvae DSM 25430 YP 008967579 1 697284 Firmicutes
Brevibacillus brevis NBRC 100599 YP 002772670 1 358681 Firmicutes
Dickeya dadantii NCPPB 3537 WP 038900908 1 Ddadantii
Dickeya dadantii NCPPB 898 WP 038910880 1 Ddadantii
Methanocaldococcus infernus ME YP 003616392 1 573063 Euryarchaeota
Archaeoglobus profundus DSM 5631 YP 003401287 1 572546 Euryarchaeota
Shewanella denitrificans OS217 YP 561602 1 318161 Proteobacteria
Acinetobacter oleivorans DR1 YP 003731689 1 436717 Proteobacteria
Acinetobacter calcoaceticus PHEA 2 YP 004995766 1 871585 Proteobacteria
Bacillus thuringiensis serovar thuringiensis str IS5056 YP 007482897 1 1286404 Firmicutes
Staphylococcus pseudintermedius HKU10 03 YP 004148500 1 937773 Firmicutes
Staphylococcus aureus subsp aureus CN1 YP 008490275 1 1193576 Firmicutes
Staphylococcus aureus 04 02981 YP 005740867 1 703339 Firmicutes
Staphylococcus aureus subsp aureus 71193 YP 006194277 1 1155084 Firmicutes
Staphylococcus aureus subsp aureus ST228 YP 007622384 1 1074919 Firmicutes
Staphylococcus aureus subsp aureus USA300 FPR3757 YP 492836 1 451515 Firmicutes
Thioflavicoccus mobilis 8321 YP 007245996 1 765912 Proteobacteria
Kitasatospora setae KM 6054 YP 004905977 1 452652 Actinobacteria
Propionibacterium acnes TypeIA2 P acn17 YP 004947738 1 1114967 Actinobacteria
Nakamurella multipartita DSM 44233 YP 003203160 1 479431 Actinobacteria
Streptomyces avermitilis MA 4680 NBRC 14893 NP 824822 1 227882 Actinobacteria
Herpetosiphon aurantiacus DSM 785 YP 001544635 1 316274 Chloroflexi
Acidobacterium capsulatum ATCC 51196 YP 002754256 1 240015 Fibrobacteres Acidobacteria group
Streptomyces sp PAMC26508 YP 007863854 1 1265601 Actinobacteria
Streptomyces pratensis ATCC 33331 YP 004921066 1 591167 Actinobacteria
Salinispora arenicola CNS 205 YP 001535287 1 391037 Actinobacteria
Actinoplanes sp SE50 110 YP 006267084 1 134676 Actinobacteria
Streptomyces griseus subsp griseus NBRC 13350 YP 001824103 1 455632 Actinobacteria
Actinoplanes missouriensis 431 YP 005464706 1 512565 Actinobacteria
Streptomyces rapamycinicus NRRL 5491 YP 008794456 1 1343740 Actinobacteria
Pandoraea pnomenusa 3kgm YP 008838437 1 1416914 Proteobacteria
Pandoraea sp RB 44 YP 008880165 1 1380774 Proteobacteria
Pseudogulbenkiania sp NH8B YP 004847738 1 748280 Proteobacteria
Sorangium cellulosum So0157 2 YP 008149320 1 1254432 Proteobacteria
Tistrella mobilis KA081020 065 YP 006371774 1 1110502 Proteobacteria
Dickeya dianthicola GBBC 2039 WP 024110059 1 Ddianthicola
Dickeya chrysanthemi subsp chrysanthemi NCPPB 402 WP 040000438 1 Dchrysanthemi
Pseudomonas entomophila L48 YP 607781 1 384676 Proteobacteria
Burkholderia gladioli BSR3 YP 004362589 1 999541 Proteobacteria
Propionibacterium acnes TypeIA2 P acn17 YP 004946789 1 1114967 Actinobacteria
Streptomyces bingchenggensis BCW 1 YP 004965059 1 749414 Actinobacteria
Xanthomonas oryzae pv oryzicola BLS256 YP 005628386 1 383407 Proteobacteria
Verminephrobacter eiseniae EF01 2 YP 995995 1 391735 Proteobacteria
Pseudomonas syringae pv tomato str DC3000 NP 792757 1 223283 Proteobacteria
Pseudomonas syringae pv phaseolicola 1448A YP 274615 1 264730 Proteobacteria
Pseudomonas syringae pv syringae B728a YP 235822 1 205918 Proteobacteria
Streptomyces hygroscopicus subsp jinggangensis 5008 YP 006242816 1 1133850 Actinobacteria
Pantoea vagans C9 1 YP 003933494 1 712898 Proteobacteria
Serratia proteamaculans 568 YP 001476578 1 399741 Proteobacteria
Prodigiosinella confusarubida ATCC39006 WP 021016296 1 Pconfusarubida
Dickeya paradisiaca Ech703 WP 012765287 1 Dparadisiaca
232 Dd703 1673 Other secondary metabolite biosynthesis specific to D paradisiaca and Pconfusarubida Cluster1 Seed
Dickeya paradisiaca NCPPB 2511 WP 012765287 1 Dparadisiaca
Thermomonospora curvata DSM 43183 YP 003299491 1 471852 Actinobacteria
Streptomyces hygroscopicus subsp jinggangensis 5008 YP 006247080 1 1133850 Actinobacteria
Kitasatospora setae KM 6054 YP 004904054 1 452652 Actinobacteria
Saccharothrix espanaensis DSM 44229 YP 007037422 1 1179773 Actinobacteria
Amycolatopsis mediterranei S699 YP 005532055 1 713604 Actinobacteria
Amycolatopsis mediterranei S699 YP 005532820 1 713604 Actinobacteria
Streptomyces venezuelae ATCC 10712 YP 006881807 1 953739 Actinobacteria
Streptomyces collinus Tu 365 YP 008388362 1 1214242 Actinobacteria
Streptomyces griseus subsp griseus NBRC 13350 YP 001824755 1 455632 Actinobacteria
Streptomyces cattleya NRRL 8057 DSM 46488 YP 004912202 1 1003195 Actinobacteria
Streptomyces scabiei 87 22 YP 003487656 1 680198 Actinobacteria
Streptomyces hygroscopicus subsp jinggangensis 5008 YP 006248200 1 1133850 Actinobacteria
Streptomyces collinus Tu 365 YP 008384862 1 1214242 Actinobacteria
Streptomyces bingchenggensis BCW 1 YP 004959369 1 749414 Actinobacteria
Streptomyces avermitilis MA 4680 NBRC 14893 NP 828339 1 227882 Actinobacteria
Geodermatophilus obscurus DSM 43160 YP 003410056 1 526225 Actinobacteria
Saccharopolyspora erythraea NRRL 2338 YP 001105220 1 405948 Actinobacteria
Saccharopolyspora erythraea NRRL 2338 YP 001106475 1 405948 Actinobacteria

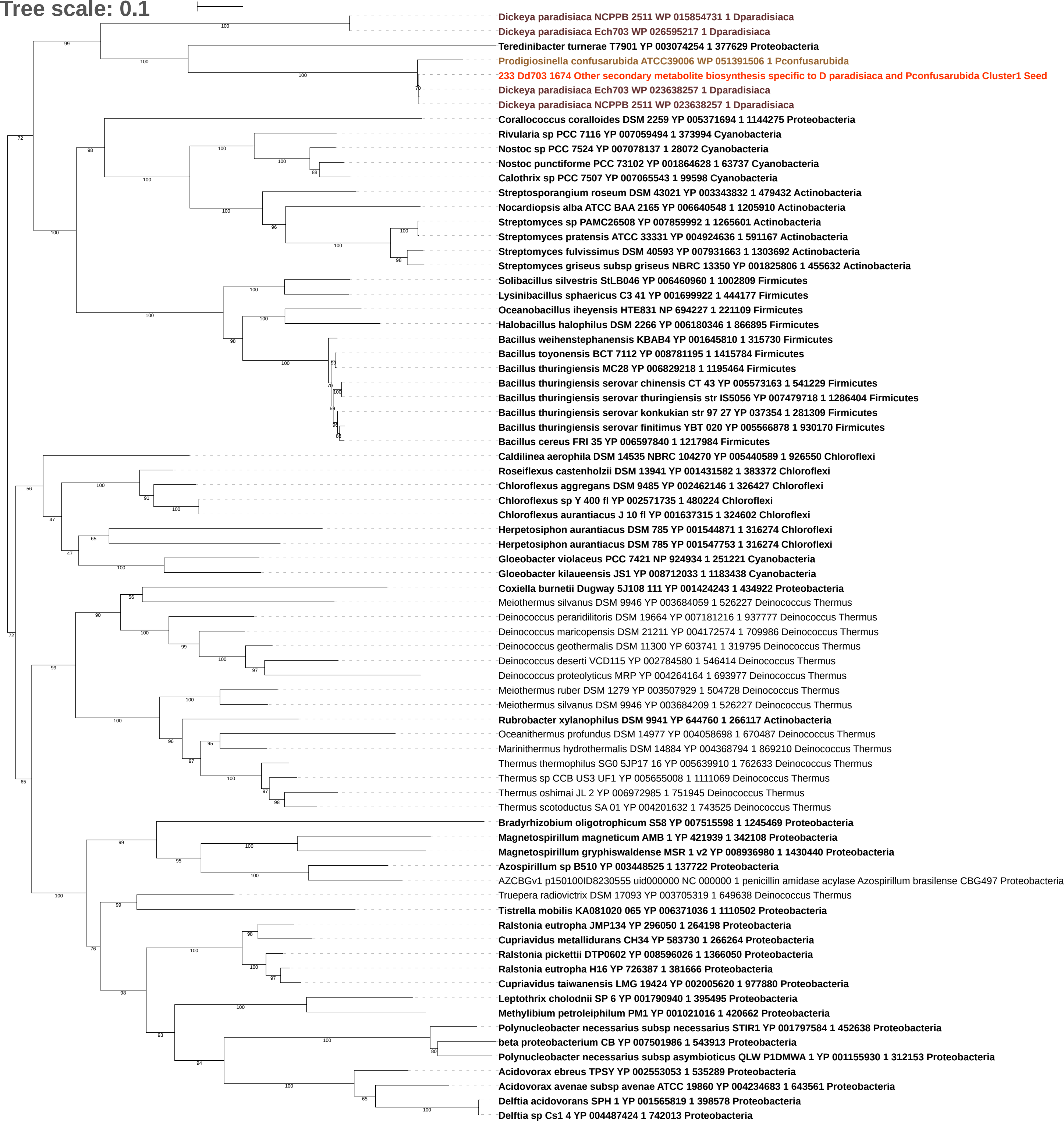
Tree scale: 0.1
Dickeya paradisiaca NCPPB 2511 WP 015854731 1 Dparadisiaca
Dickeya paradisiaca Ech703 WP 026595217 1 Dparadisiaca
Teredinibacter turnerae T7901 YP 003074254 1 377629 Proteobacteria
Prodigiosinella confusarubida ATCC39006 WP 051391506 1 Pconfusarubida
233 Dd703 1674 Other secondary metabolite biosynthesis specific to D paradisiaca and Pconfusarubida Cluster1 Seed
Dickeya paradisiaca Ech703 WP 023638257 1 Dparadisiaca
Dickeya paradisiaca NCPPB 2511 WP 023638257 1 Dparadisiaca
Corallococcus coralloides DSM 2259 YP 005371694 1 1144275 Proteobacteria
Rivularia sp PCC 7116 YP 007059494 1 373994 Cyanobacteria
Nostoc sp PCC 7524 YP 007078137 1 28072 Cyanobacteria
Nostoc punctiforme PCC 73102 YP 001864628 1 63737 Cyanobacteria
Calothrix sp PCC 7507 YP 007065543 1 99598 Cyanobacteria
Streptosporangium roseum DSM 43021 YP 003343832 1 479432 Actinobacteria
Nocardiopsis alba ATCC BAA 2165 YP 006640548 1 1205910 Actinobacteria
Streptomyces sp PAMC26508 YP 007859992 1 1265601 Actinobacteria
Streptomyces pratensis ATCC 33331 YP 004924636 1 591167 Actinobacteria
Streptomyces fulvissimus DSM 40593 YP 007931663 1 1303692 Actinobacteria
Streptomyces griseus subsp griseus NBRC 13350 YP 001825806 1 455632 Actinobacteria
Solibacillus silvestris StLB046 YP 006460960 1 1002809 Firmicutes
Lysinibacillus sphaericus C3 41 YP 001699922 1 444177 Firmicutes
Oceanobacillus iheyensis HTE831 NP 694227 1 221109 Firmicutes
Halobacillus halophilus DSM 2266 YP 006180346 1 866895 Firmicutes
Bacillus weihenstephanensis KBAB4 YP 001645810 1 315730 Firmicutes
Bacillus toyonensis BCT 7112 YP 008781195 1 1415784 Firmicutes
Bacillus thuringiensis MC28 YP 006829218 1 1195464 Firmicutes
Bacillus thuringiensis serovar chinensis CT 43 YP 005573163 1 541229 Firmicutes
Bacillus thuringiensis serovar thuringiensis str IS5056 YP 007479718 1 1286404 Firmicutes
Bacillus thuringiensis serovar konkukian str 97 27 YP 037354 1 281309 Firmicutes
Bacillus thuringiensis serovar finitimus YBT 020 YP 005566878 1 930170 Firmicutes
Bacillus cereus FRI 35 YP 006597840 1 1217984 Firmicutes
Caldilinea aerophila DSM 14535 NBRC 104270 YP 005440589 1 926550 Chloroflexi
Roseiflexus castenholzii DSM 13941 YP 001431582 1 383372 Chloroflexi
Chloroflexus aggregans DSM 9485 YP 002462146 1 326427 Chloroflexi
Chloroflexus sp Y 400 fl YP 002571735 1 480224 Chloroflexi
Chloroflexus aurantiacus J 10 fl YP 001637315 1 324602 Chloroflexi
Herpetosiphon aurantiacus DSM 785 YP 001544871 1 316274 Chloroflexi
Herpetosiphon aurantiacus DSM 785 YP 001547753 1 316274 Chloroflexi
Gloeobacter violaceus PCC 7421 NP 924934 1 251221 Cyanobacteria
Gloeobacter kilaueensis JS1 YP 008712033 1 1183438 Cyanobacteria
Coxiella burnetii Dugway 5J108 111 YP 001424243 1 434922 Proteobacteria
Meiothermus silvanus DSM 9946 YP 003684059 1 526227 Deinococcus Thermus
Deinococcus peraridilitoris DSM 19664 YP 007181216 1 937777 Deinococcus Thermus
Deinococcus maricopensis DSM 21211 YP 004172574 1 709986 Deinococcus Thermus
Deinococcus geothermalis DSM 11300 YP 603741 1 319795 Deinococcus Thermus
Deinococcus deserti VCD115 YP 002784580 1 546414 Deinococcus Thermus
Deinococcus proteolyticus MRP YP 004264164 1 693977 Deinococcus Thermus
Meiothermus ruber DSM 1279 YP 003507929 1 504728 Deinococcus Thermus
Meiothermus silvanus DSM 9946 YP 003684209 1 526227 Deinococcus Thermus
Rubrobacter xylanophilus DSM 9941 YP 644760 1 266117 Actinobacteria
Oceanithermus profundus DSM 14977 YP 004058698 1 670487 Deinococcus Thermus
Marinithermus hydrothermalis DSM 14884 YP 004368794 1 869210 Deinococcus Thermus
Thermus thermophilus SG0 5JP17 16 YP 005639910 1 762633 Deinococcus Thermus
Thermus sp CCB US3 UF1 YP 005655008 1 1111069 Deinococcus Thermus
Thermus oshimai JL 2 YP 006972985 1 751945 Deinococcus Thermus
Thermus scotoductus SA 01 YP 004201632 1 743525 Deinococcus Thermus
Bradyrhizobium oligotrophicum S58 YP 007515598 1 1245469 Proteobacteria
Magnetospirillum magneticum AMB 1 YP 421939 1 342108 Proteobacteria
Magnetospirillum gryphiswaldense MSR 1 v2 YP 008936980 1 1430440 Proteobacteria
Azospirillum sp B510 YP 003448525 1 137722 Proteobacteria
AZCBGv1 p150100ID8230555 uid000000 NC 000000 1 penicillin amidase acylase Azospirillum brasilense CBG497 Proteobacteria
Truepera radiovictrix DSM 17093 YP 003705319 1 649638 Deinococcus Thermus
Tistrella mobilis KA081020 065 YP 006371036 1 1110502 Proteobacteria
Ralstonia eutropha JMP134 YP 296050 1 264198 Proteobacteria
Cupriavidus metallidurans CH34 YP 583730 1 266264 Proteobacteria
Ralstonia pickettii DTP0602 YP 008596026 1 1366050 Proteobacteria
Ralstonia eutropha H16 YP 726387 1 381666 Proteobacteria
Cupriavidus taiwanensis LMG 19424 YP 002005620 1 977880 Proteobacteria
Leptothrix cholodnii SP 6 YP 001790940 1 395495 Proteobacteria
Methylibium petroleiphilum PM1 YP 001021016 1 420662 Proteobacteria
Polynucleobacter necessarius subsp necessarius STIR1 YP 001797584 1 452638 Proteobacteria
beta proteobacterium CB YP 007501986 1 543913 Proteobacteria
Polynucleobacter necessarius subsp asymbioticus QLW P1DMWA 1 YP 001155930 1 312153 Proteobacteria
Acidovorax ebreus TPSY YP 002553053 1 535289 Proteobacteria
Acidovorax avenae subsp avenae ATCC 19860 YP 004234683 1 643561 Proteobacteria
Delftia acidovorans SPH 1 YP 001565819 1 398578 Proteobacteria
Delftia sp Cs1 4 YP 004487424 1 742013 Proteobacteria

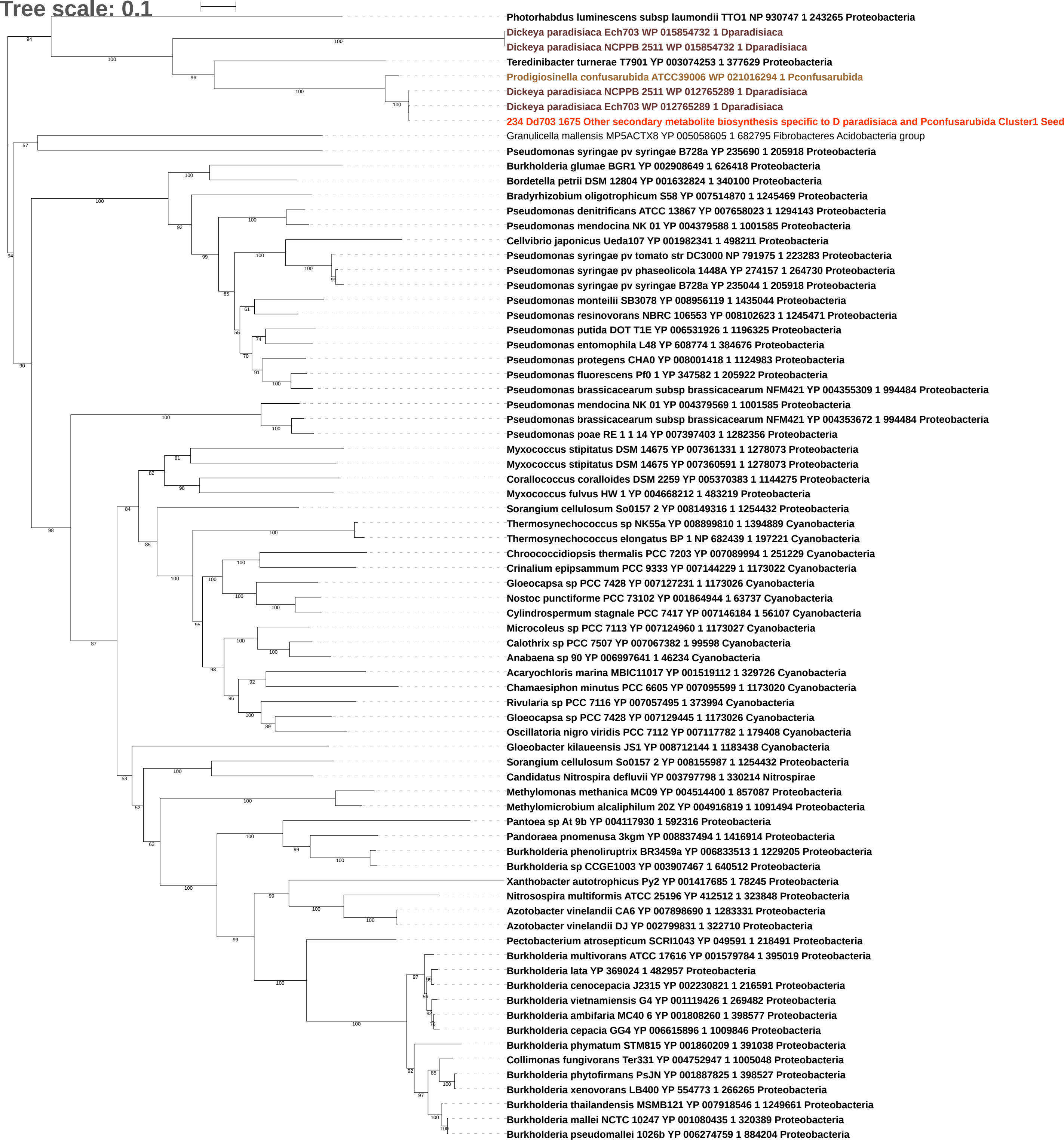
Tree scale: 0.1
Photorhabdus luminescens subsp laumondii TTO1 NP 930747 1 243265 Proteobacteria
Dickeya paradisiaca Ech703 WP 015854732 1 Dparadisiaca
Dickeya paradisiaca NCPPB 2511 WP 015854732 1 Dparadisiaca
Teredinibacter turnerae T7901 YP 003074253 1 377629 Proteobacteria
Prodigiosinella confusarubida ATCC39006 WP 021016294 1 Pconfusarubida
Dickeya paradisiaca NCPPB 2511 WP 012765289 1 Dparadisiaca
Dickeya paradisiaca Ech703 WP 012765289 1 Dparadisiaca
234 Dd703 1675 Other secondary metabolite biosynthesis specific to D paradisiaca and Pconfusarubida Cluster1 Seed
Granulicella mallensis MP5ACTX8 YP 005058605 1 682795 Fibrobacteres Acidobacteria group
Pseudomonas syringae pv syringae B728a YP 235690 1 205918 Proteobacteria
Burkholderia glumae BGR1 YP 002908649 1 626418 Proteobacteria
Bordetella petrii DSM 12804 YP 001632824 1 340100 Proteobacteria
Bradyrhizobium oligotrophicum S58 YP 007514870 1 1245469 Proteobacteria
Pseudomonas denitrificans ATCC 13867 YP 007658023 1 1294143 Proteobacteria
Pseudomonas mendocina NK 01 YP 004379588 1 1001585 Proteobacteria
Cellvibrio japonicus Ueda107 YP 001982341 1 498211 Proteobacteria
Pseudomonas syringae pv tomato str DC3000 NP 791975 1 223283 Proteobacteria
Pseudomonas syringae pv phaseolicola 1448A YP 274157 1 264730 Proteobacteria
Pseudomonas syringae pv syringae B728a YP 235044 1 205918 Proteobacteria
Pseudomonas monteilii SB3078 YP 008956119 1 1435044 Proteobacteria
Pseudomonas resinovorans NBRC 106553 YP 008102623 1 1245471 Proteobacteria
Pseudomonas putida DOT T1E YP 006531926 1 1196325 Proteobacteria
Pseudomonas entomophila L48 YP 608774 1 384676 Proteobacteria
Pseudomonas protegens CHA0 YP 008001418 1 1124983 Proteobacteria
Pseudomonas fluorescens Pf0 1 YP 347582 1 205922 Proteobacteria
Pseudomonas brassicacearum subsp brassicacearum NFM421 YP 004355309 1 994484 Proteobacteria
Pseudomonas mendocina NK 01 YP 004379569 1 1001585 Proteobacteria
Pseudomonas brassicacearum subsp brassicacearum NFM421 YP 004353672 1 994484 Proteobacteria
Pseudomonas poae RE 1 1 14 YP 007397403 1 1282356 Proteobacteria
Myxococcus stipitatus DSM 14675 YP 007361331 1 1278073 Proteobacteria
Myxococcus stipitatus DSM 14675 YP 007360591 1 1278073 Proteobacteria
Corallococcus coralloides DSM 2259 YP 005370383 1 1144275 Proteobacteria
Myxococcus fulvus HW 1 YP 004668212 1 483219 Proteobacteria
Sorangium cellulosum So0157 2 YP 008149316 1 1254432 Proteobacteria
Thermosynechococcus sp NK55a YP 008899810 1 1394889 Cyanobacteria
Thermosynechococcus elongatus BP 1 NP 682439 1 197221 Cyanobacteria
Chroococcidiopsis thermalis PCC 7203 YP 007089994 1 251229 Cyanobacteria
Crinalium epipsammum PCC 9333 YP 007144229 1 1173022 Cyanobacteria
Gloeocapsa sp PCC 7428 YP 007127231 1 1173026 Cyanobacteria
Nostoc punctiforme PCC 73102 YP 001864944 1 63737 Cyanobacteria
Cylindrospermum stagnale PCC 7417 YP 007146184 1 56107 Cyanobacteria
Microcoleus sp PCC 7113 YP 007124960 1 1173027 Cyanobacteria
Calothrix sp PCC 7507 YP 007067382 1 99598 Cyanobacteria
Anabaena sp 90 YP 006997641 1 46234 Cyanobacteria
Acaryochloris marina MBIC11017 YP 001519112 1 329726 Cyanobacteria
Chamaesiphon minutus PCC 6605 YP 007095599 1 1173020 Cyanobacteria
Rivularia sp PCC 7116 YP 007057495 1 373994 Cyanobacteria
Gloeocapsa sp PCC 7428 YP 007129445 1 1173026 Cyanobacteria
Oscillatoria nigro viridis PCC 7112 YP 007117782 1 179408 Cyanobacteria
Gloeobacter kilaueensis JS1 YP 008712144 1 1183438 Cyanobacteria
Sorangium cellulosum So0157 2 YP 008155987 1 1254432 Proteobacteria
Candidatus Nitrospira defluvii YP 003797798 1 330214 Nitrospirae
Methylomonas methanica MC09 YP 004514400 1 857087 Proteobacteria
Methylomicrobium alcaliphilum 20Z YP 004916819 1 1091494 Proteobacteria
Pantoea sp At 9b YP 004117930 1 592316 Proteobacteria
Pandoraea pnomenusa 3kgm YP 008837494 1 1416914 Proteobacteria
Burkholderia phenoliruptrix BR3459a YP 006833513 1 1229205 Proteobacteria
Burkholderia sp CCGE1003 YP 003907467 1 640512 Proteobacteria
Xanthobacter autotrophicus Py2 YP 001417685 1 78245 Proteobacteria
Nitrosospira multiformis ATCC 25196 YP 412512 1 323848 Proteobacteria
Azotobacter vinelandii CA6 YP 007898690 1 1283331 Proteobacteria
Azotobacter vinelandii DJ YP 002799831 1 322710 Proteobacteria
Pectobacterium atrosepticum SCRI1043 YP 049591 1 218491 Proteobacteria
Burkholderia multivorans ATCC 17616 YP 001579784 1 395019 Proteobacteria
Burkholderia lata YP 369024 1 482957 Proteobacteria
Burkholderia cenocepacia J2315 YP 002230821 1 216591 Proteobacteria
Burkholderia vietnamiensis G4 YP 001119426 1 269482 Proteobacteria
Burkholderia ambifaria MC40 6 YP 001808260 1 398577 Proteobacteria
Burkholderia cepacia GG4 YP 006615896 1 1009846 Proteobacteria
Burkholderia phymatum STM815 YP 001860209 1 391038 Proteobacteria
Collimonas fungivorans Ter331 YP 004752947 1 1005048 Proteobacteria
Burkholderia phytofirmans PsJN YP 001887825 1 398527 Proteobacteria
Burkholderia xenovorans LB400 YP 554773 1 266265 Proteobacteria
Burkholderia thailandensis MSMB121 YP 007918546 1 1249661 Proteobacteria
Burkholderia mallei NCTC 10247 YP 001080435 1 320389 Proteobacteria
Burkholderia pseudomallei 1026b YP 006274759 1 884204 Proteobacteria

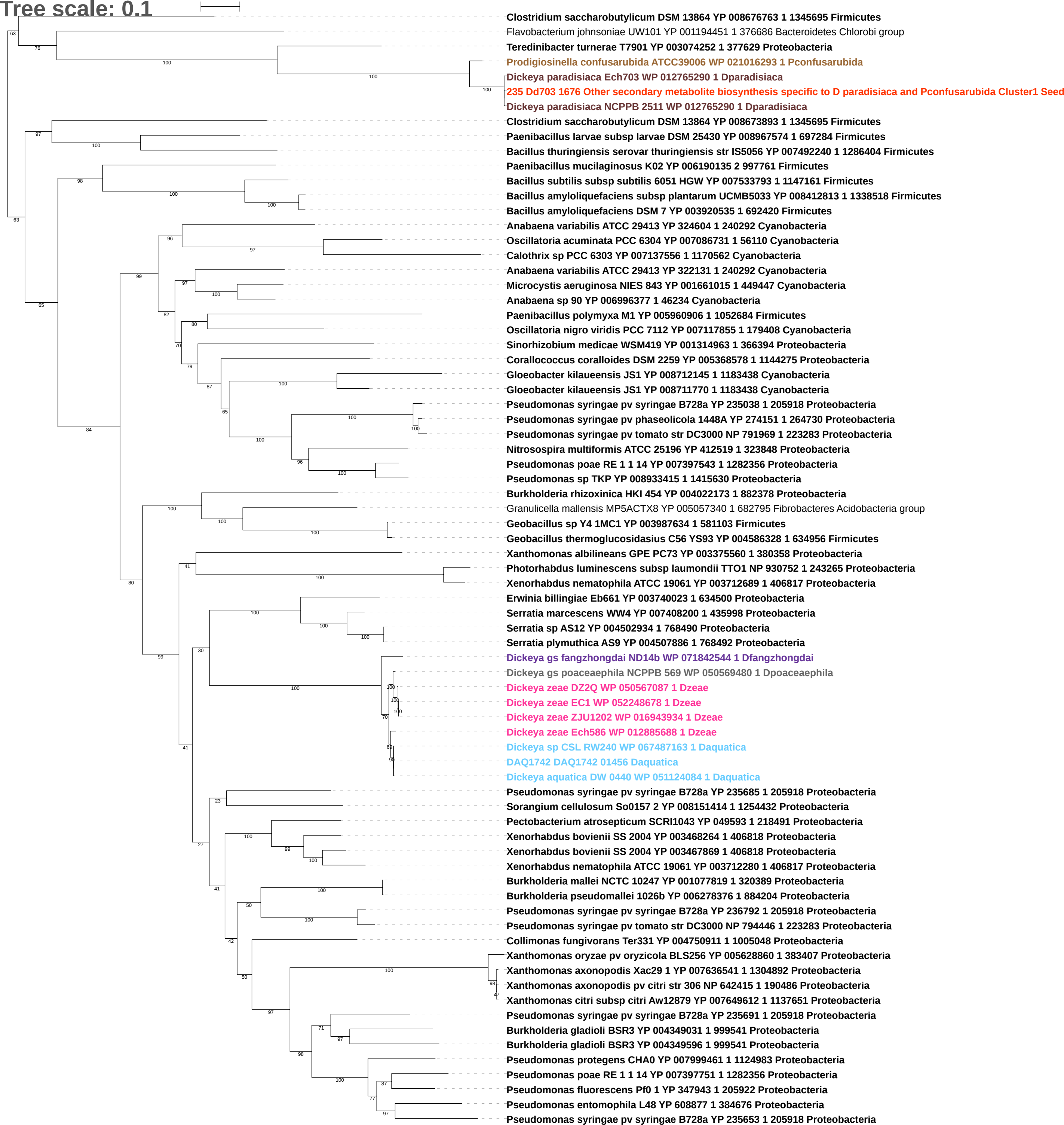
Tree scale: 0.1
Clostridium saccharobutylicum DSM 13864 YP 008676763 1 1345695 Firmicutes
Flavobacterium johnsoniae UW101 YP 001194451 1 376686 Bacteroidetes Chlorobi group
Teredinibacter turnerae T7901 YP 003074252 1 377629 Proteobacteria
Prodigiosinella confusarubida ATCC39006 WP 021016293 1 Pconfusarubida
Dickeya paradisiaca Ech703 WP 012765290 1 Dparadisiaca
235 Dd703 1676 Other secondary metabolite biosynthesis specific to D paradisiaca and Pconfusarubida Cluster1 Seed
Dickeya paradisiaca NCPPB 2511 WP 012765290 1 Dparadisiaca
Clostridium saccharobutylicum DSM 13864 YP 008673893 1 1345695 Firmicutes
Paenibacillus larvae subsp larvae DSM 25430 YP 008967574 1 697284 Firmicutes
Bacillus thuringiensis serovar thuringiensis str IS5056 YP 007492240 1 1286404 Firmicutes
Paenibacillus mucilaginosus K02 YP 006190135 2 997761 Firmicutes
Bacillus subtilis subsp subtilis 6051 HGW YP 007533793 1 1147161 Firmicutes
Bacillus amyloliquefaciens subsp plantarum UCMB5033 YP 008412813 1 1338518 Firmicutes
Bacillus amyloliquefaciens DSM 7 YP 003920535 1 692420 Firmicutes
Anabaena variabilis ATCC 29413 YP 324604 1 240292 Cyanobacteria
Oscillatoria acuminata PCC 6304 YP 007086731 1 56110 Cyanobacteria
Calothrix sp PCC 6303 YP 007137556 1 1170562 Cyanobacteria
Anabaena variabilis ATCC 29413 YP 322131 1 240292 Cyanobacteria
Microcystis aeruginosa NIES 843 YP 001661015 1 449447 Cyanobacteria
Anabaena sp 90 YP 006996377 1 46234 Cyanobacteria
Paenibacillus polymyxa M1 YP 005960906 1 1052684 Firmicutes
Oscillatoria nigro viridis PCC 7112 YP 007117855 1 179408 Cyanobacteria
Sinorhizobium medicae WSM419 YP 001314963 1 366394 Proteobacteria
Corallococcus coralloides DSM 2259 YP 005368578 1 1144275 Proteobacteria
Gloeobacter kilaueensis JS1 YP 008712145 1 1183438 Cyanobacteria
Gloeobacter kilaueensis JS1 YP 008711770 1 1183438 Cyanobacteria
Pseudomonas syringae pv syringae B728a YP 235038 1 205918 Proteobacteria
Pseudomonas syringae pv phaseolicola 1448A YP 274151 1 264730 Proteobacteria
Pseudomonas syringae pv tomato str DC3000 NP 791969 1 223283 Proteobacteria
Nitrosospira multiformis ATCC 25196 YP 412519 1 323848 Proteobacteria
Pseudomonas poae RE 1 1 14 YP 007397543 1 1282356 Proteobacteria
Pseudomonas sp TKP YP 008933415 1 1415630 Proteobacteria
Burkholderia rhizoxinica HKI 454 YP 004022173 1 882378 Proteobacteria
Granulicella mallensis MP5ACTX8 YP 005057340 1 682795 Fibrobacteres Acidobacteria group
Geobacillus sp Y4 1MC1 YP 003987634 1 581103 Firmicutes
Geobacillus thermoglucosidasius C56 YS93 YP 004586328 1 634956 Firmicutes
Xanthomonas albilineans GPE PC73 YP 003375560 1 380358 Proteobacteria
Photorhabdus luminescens subsp laumondii TTO1 NP 930752 1 243265 Proteobacteria
Xenorhabdus nematophila ATCC 19061 YP 003712689 1 406817 Proteobacteria
Erwinia billingiae Eb661 YP 003740023 1 634500 Proteobacteria
Serratia marcescens WW4 YP 007408200 1 435998 Proteobacteria
Serratia sp AS12 YP 004502934 1 768490 Proteobacteria
Serratia plymuthica AS9 YP 004507886 1 768492 Proteobacteria
Dickeya gs fangzhongdai ND14b WP 071842544 1 Dfangzhongdai
Dickeya gs poaceaephila NCPPB 569 WP 050569480 1 Dpoaceaephila
Dickeya zeae DZ2Q WP 050567087 1 Dzeae
Dickeya zeae EC1 WP 052248678 1 Dzeae
Dickeya zeae ZJU1202 WP 016943934 1 Dzeae
Dickeya zeae Ech586 WP 012885688 1 Dzeae
Dickeya sp CSL RW240 WP 067487163 1 Daquatica
DAQ1742 DAQ1742 01456 Daquatica
Dickeya aquatica DW 0440 WP 051124084 1 Daquatica
Pseudomonas syringae pv syringae B728a YP 235685 1 205918 Proteobacteria
Sorangium cellulosum So0157 2 YP 008151414 1 1254432 Proteobacteria
Pectobacterium atrosepticum SCRI1043 YP 049593 1 218491 Proteobacteria
Xenorhabdus bovienii SS 2004 YP 003468264 1 406818 Proteobacteria
Xenorhabdus bovienii SS 2004 YP 003467869 1 406818 Proteobacteria
Xenorhabdus nematophila ATCC 19061 YP 003712280 1 406817 Proteobacteria
Burkholderia mallei NCTC 10247 YP 001077819 1 320389 Proteobacteria
Burkholderia pseudomallei 1026b YP 006278376 1 884204 Proteobacteria
Pseudomonas syringae pv syringae B728a YP 236792 1 205918 Proteobacteria
Pseudomonas syringae pv tomato str DC3000 NP 794446 1 223283 Proteobacteria
Collimonas fungivorans Ter331 YP 004750911 1 1005048 Proteobacteria
Xanthomonas oryzae pv oryzicola BLS256 YP 005628860 1 383407 Proteobacteria
Xanthomonas axonopodis Xac29 1 YP 007636541 1 1304892 Proteobacteria
Xanthomonas axonopodis pv citri str 306 NP 642415 1 190486 Proteobacteria
Xanthomonas citri subsp citri Aw12879 YP 007649612 1 1137651 Proteobacteria
Pseudomonas syringae pv syringae B728a YP 235691 1 205918 Proteobacteria
Burkholderia gladioli BSR3 YP 004349031 1 999541 Proteobacteria
Burkholderia gladioli BSR3 YP 004349596 1 999541 Proteobacteria
Pseudomonas protegens CHA0 YP 007999461 1 1124983 Proteobacteria
Pseudomonas poae RE 1 1 14 YP 007397751 1 1282356 Proteobacteria
Pseudomonas fluorescens Pf0 1 YP 347943 1 205922 Proteobacteria
Pseudomonas entomophila L48 YP 608877 1 384676 Proteobacteria
Pseudomonas syringae pv syringae B728a YP 235653 1 205918 Proteobacteria

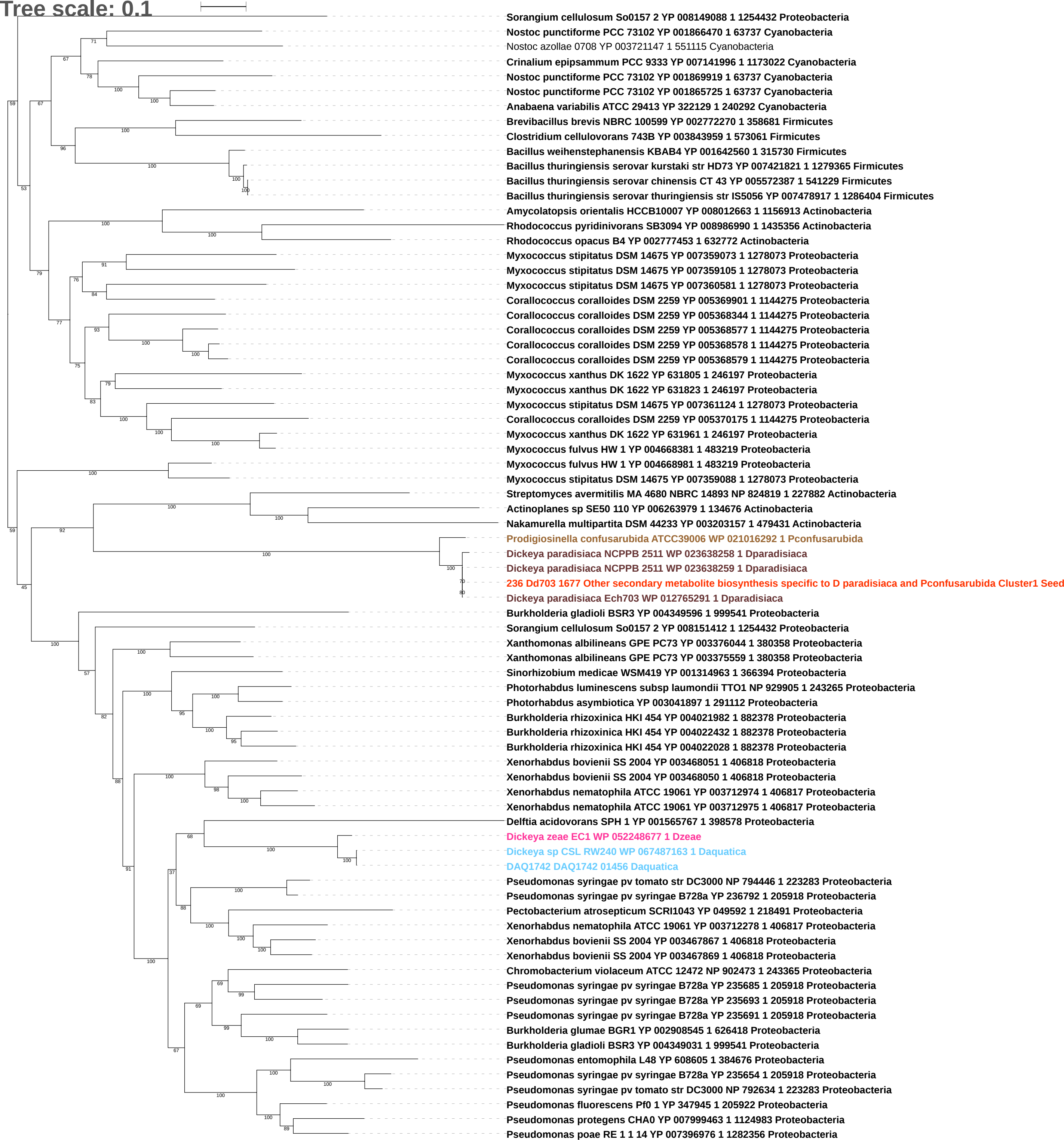
Tree scale: 0.1
Sorangium cellulosum So0157 2 YP 008149088 1 1254432 Proteobacteria
Nostoc punctiforme PCC 73102 YP 001866470 1 63737 Cyanobacteria
Nostoc azollae 0708 YP 003721147 1 551115 Cyanobacteria
Crinalium epipsammum PCC 9333 YP 007141996 1 1173022 Cyanobacteria
Nostoc punctiforme PCC 73102 YP 001869919 1 63737 Cyanobacteria
Nostoc punctiforme PCC 73102 YP 001865725 1 63737 Cyanobacteria
Anabaena variabilis ATCC 29413 YP 322129 1 240292 Cyanobacteria
Brevibacillus brevis NBRC 100599 YP 002772270 1 358681 Firmicutes
Clostridium cellulovorans 743B YP 003843959 1 573061 Firmicutes
Bacillus weihenstephanensis KBAB4 YP 001642560 1 315730 Firmicutes
Bacillus thuringiensis serovar kurstaki str HD73 YP 007421821 1 1279365 Firmicutes
Bacillus thuringiensis serovar chinensis CT 43 YP 005572387 1 541229 Firmicutes
Bacillus thuringiensis serovar thuringiensis str IS5056 YP 007478917 1 1286404 Firmicutes
Amycolatopsis orientalis HCCB10007 YP 008012663 1 1156913 Actinobacteria
Rhodococcus pyridinivorans SB3094 YP 008986990 1 1435356 Actinobacteria
Rhodococcus opacus B4 YP 002777453 1 632772 Actinobacteria
Myxococcus stipitatus DSM 14675 YP 007359073 1 1278073 Proteobacteria
Myxococcus stipitatus DSM 14675 YP 007359105 1 1278073 Proteobacteria
Myxococcus stipitatus DSM 14675 YP 007360581 1 1278073 Proteobacteria
Corallococcus coralloides DSM 2259 YP 005369901 1 1144275 Proteobacteria
Corallococcus coralloides DSM 2259 YP 005368344 1 1144275 Proteobacteria
Corallococcus coralloides DSM 2259 YP 005368577 1 1144275 Proteobacteria
Corallococcus coralloides DSM 2259 YP 005368578 1 1144275 Proteobacteria
Corallococcus coralloides DSM 2259 YP 005368579 1 1144275 Proteobacteria
Myxococcus xanthus DK 1622 YP 631805 1 246197 Proteobacteria
Myxococcus xanthus DK 1622 YP 631823 1 246197 Proteobacteria
Myxococcus stipitatus DSM 14675 YP 007361124 1 1278073 Proteobacteria
Corallococcus coralloides DSM 2259 YP 005370175 1 1144275 Proteobacteria
Myxococcus xanthus DK 1622 YP 631961 1 246197 Proteobacteria
Myxococcus fulvus HW 1 YP 004668381 1 483219 Proteobacteria
Myxococcus fulvus HW 1 YP 004668981 1 483219 Proteobacteria
Myxococcus stipitatus DSM 14675 YP 007359088 1 1278073 Proteobacteria
Streptomyces avermitilis MA 4680 NBRC 14893 NP 824819 1 227882 Actinobacteria
Actinoplanes sp SE50 110 YP 006263979 1 134676 Actinobacteria
Nakamurella multipartita DSM 44233 YP 003203157 1 479431 Actinobacteria
Prodigiosinella confusarubida ATCC39006 WP 021016292 1 Pconfusarubida
Dickeya paradisiaca NCPPB 2511 WP 023638258 1 Dparadisiaca
Dickeya paradisiaca NCPPB 2511 WP 023638259 1 Dparadisiaca
236 Dd703 1677 Other secondary metabolite biosynthesis specific to D paradisiaca and Pconfusarubida Cluster1 Seed
Dickeya paradisiaca Ech703 WP 012765291 1 Dparadisiaca
Burkholderia gladioli BSR3 YP 004349596 1 999541 Proteobacteria
Sorangium cellulosum So0157 2 YP 008151412 1 1254432 Proteobacteria
Xanthomonas albilineans GPE PC73 YP 003376044 1 380358 Proteobacteria
Xanthomonas albilineans GPE PC73 YP 003375559 1 380358 Proteobacteria
Sinorhizobium medicae WSM419 YP 001314963 1 366394 Proteobacteria
Photorhabdus luminescens subsp laumondii TTO1 NP 929905 1 243265 Proteobacteria
Photorhabdus asymbiotica YP 003041897 1 291112 Proteobacteria
Burkholderia rhizoxinica HKI 454 YP 004021982 1 882378 Proteobacteria
Burkholderia rhizoxinica HKI 454 YP 004022432 1 882378 Proteobacteria
Burkholderia rhizoxinica HKI 454 YP 004022028 1 882378 Proteobacteria
Xenorhabdus bovienii SS 2004 YP 003468051 1 406818 Proteobacteria
Xenorhabdus bovienii SS 2004 YP 003468050 1 406818 Proteobacteria
Xenorhabdus nematophila ATCC 19061 YP 003712974 1 406817 Proteobacteria
Xenorhabdus nematophila ATCC 19061 YP 003712975 1 406817 Proteobacteria
Delftia acidovorans SPH 1 YP 001565767 1 398578 Proteobacteria
Dickeya zeae EC1 WP 052248677 1 Dzeae
Dickeya sp CSL RW240 WP 067487163 1 Daquatica
DAQ1742 DAQ1742 01456 Daquatica
Pseudomonas syringae pv tomato str DC3000 NP 794446 1 223283 Proteobacteria
Pseudomonas syringae pv syringae B728a YP 236792 1 205918 Proteobacteria
Pectobacterium atrosepticum SCRI1043 YP 049592 1 218491 Proteobacteria
Xenorhabdus nematophila ATCC 19061 YP 003712278 1 406817 Proteobacteria
Xenorhabdus bovienii SS 2004 YP 003467867 1 406818 Proteobacteria
Xenorhabdus bovienii SS 2004 YP 003467869 1 406818 Proteobacteria
Chromobacterium violaceum ATCC 12472 NP 902473 1 243365 Proteobacteria
Pseudomonas syringae pv syringae B728a YP 235685 1 205918 Proteobacteria
Pseudomonas syringae pv syringae B728a YP 235693 1 205918 Proteobacteria
Pseudomonas syringae pv syringae B728a YP 235691 1 205918 Proteobacteria
Burkholderia glumae BGR1 YP 002908545 1 626418 Proteobacteria
Burkholderia gladioli BSR3 YP 004349031 1 999541 Proteobacteria
Pseudomonas entomophila L48 YP 608605 1 384676 Proteobacteria
Pseudomonas syringae pv syringae B728a YP 235654 1 205918 Proteobacteria
Pseudomonas syringae pv tomato str DC3000 NP 792634 1 223283 Proteobacteria
Pseudomonas fluorescens Pf0 1 YP 347945 1 205922 Proteobacteria
Pseudomonas protegens CHA0 YP 007999463 1 1124983 Proteobacteria
Pseudomonas poae RE 1 1 14 YP 007396976 1 1282356 Proteobacteria

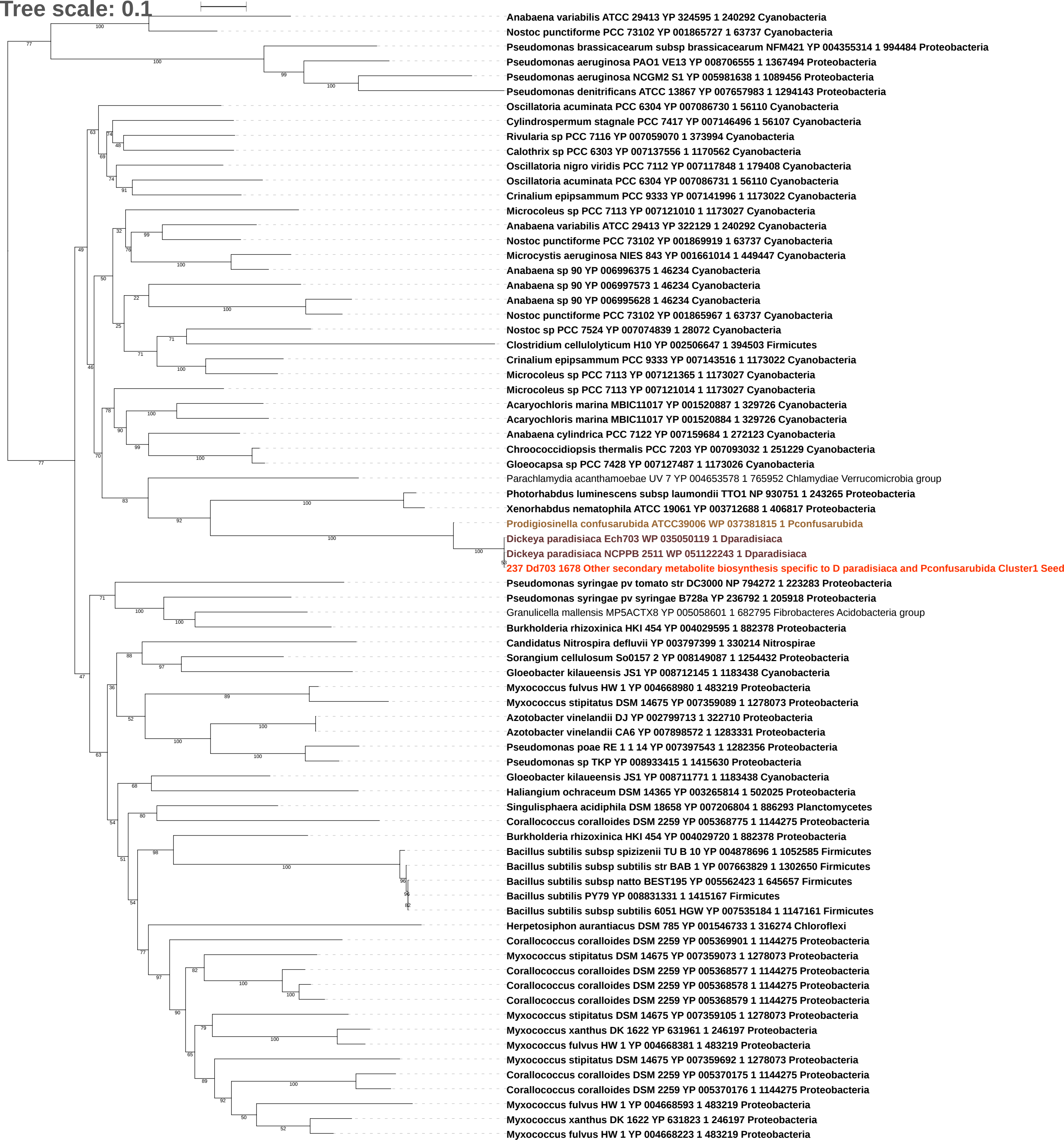
Tree scale: 0.1
Anabaena variabilis ATCC 29413 YP 324595 1 240292 Cyanobacteria
Nostoc punctiforme PCC 73102 YP 001865727 1 63737 Cyanobacteria
Pseudomonas brassicacearum subsp brassicacearum NFM421 YP 004355314 1 994484 Proteobacteria
Pseudomonas aeruginosa PAO1 VE13 YP 008706555 1 1367494 Proteobacteria
Pseudomonas aeruginosa NCGM2 S1 YP 005981638 1 1089456 Proteobacteria
Pseudomonas denitrificans ATCC 13867 YP 007657983 1 1294143 Proteobacteria
Oscillatoria acuminata PCC 6304 YP 007086730 1 56110 Cyanobacteria
Cylindrospermum stagnale PCC 7417 YP 007146496 1 56107 Cyanobacteria
Rivularia sp PCC 7116 YP 007059070 1 373994 Cyanobacteria
Calothrix sp PCC 6303 YP 007137556 1 1170562 Cyanobacteria
Oscillatoria nigro viridis PCC 7112 YP 007117848 1 179408 Cyanobacteria
Oscillatoria acuminata PCC 6304 YP 007086731 1 56110 Cyanobacteria
Crinalium epipsammum PCC 9333 YP 007141996 1 1173022 Cyanobacteria
Microcoleus sp PCC 7113 YP 007121010 1 1173027 Cyanobacteria
Anabaena variabilis ATCC 29413 YP 322129 1 240292 Cyanobacteria
Nostoc punctiforme PCC 73102 YP 001869919 1 63737 Cyanobacteria
Microcystis aeruginosa NIES 843 YP 001661014 1 449447 Cyanobacteria
Anabaena sp 90 YP 006996375 1 46234 Cyanobacteria
Anabaena sp 90 YP 006997573 1 46234 Cyanobacteria
Anabaena sp 90 YP 006995628 1 46234 Cyanobacteria
Nostoc punctiforme PCC 73102 YP 001865967 1 63737 Cyanobacteria
Nostoc sp PCC 7524 YP 007074839 1 28072 Cyanobacteria
Clostridium cellulolyticum H10 YP 002506647 1 394503 Firmicutes
Crinalium epipsammum PCC 9333 YP 007143516 1 1173022 Cyanobacteria
Microcoleus sp PCC 7113 YP 007121365 1 1173027 Cyanobacteria
Microcoleus sp PCC 7113 YP 007121014 1 1173027 Cyanobacteria
Acaryochloris marina MBIC11017 YP 001520887 1 329726 Cyanobacteria
Acaryochloris marina MBIC11017 YP 001520884 1 329726 Cyanobacteria
Anabaena cylindrica PCC 7122 YP 007159684 1 272123 Cyanobacteria
Chroococcidiopsis thermalis PCC 7203 YP 007093032 1 251229 Cyanobacteria
Gloeocapsa sp PCC 7428 YP 007127487 1 1173026 Cyanobacteria
Parachlamydia acanthamoebae UV 7 YP 004653578 1 765952 Chlamydiae Verrucomicrobia group
Photorhabdus luminescens subsp laumondii TTO1 NP 930751 1 243265 Proteobacteria
Xenorhabdus nematophila ATCC 19061 YP 003712688 1 406817 Proteobacteria
Prodigiosinella confusarubida ATCC39006 WP 037381815 1 Pconfusarubida
Dickeya paradisiaca Ech703 WP 035050119 1 Dparadisiaca
Dickeya paradisiaca NCPPB 2511 WP 051122243 1 Dparadisiaca
237 Dd703 1678 Other secondary metabolite biosynthesis specific to D paradisiaca and Pconfusarubida Cluster1 Seed
Pseudomonas syringae pv tomato str DC3000 NP 794272 1 223283 Proteobacteria
Pseudomonas syringae pv syringae B728a YP 236792 1 205918 Proteobacteria
Granulicella mallensis MP5ACTX8 YP 005058601 1 682795 Fibrobacteres Acidobacteria group
Burkholderia rhizoxinica HKI 454 YP 004029595 1 882378 Proteobacteria
Candidatus Nitrospira defluvii YP 003797399 1 330214 Nitrospirae
Sorangium cellulosum So0157 2 YP 008149087 1 1254432 Proteobacteria
Gloeobacter kilaueensis JS1 YP 008712145 1 1183438 Cyanobacteria
Myxococcus fulvus HW 1 YP 004668980 1 483219 Proteobacteria
Myxococcus stipitatus DSM 14675 YP 007359089 1 1278073 Proteobacteria
Azotobacter vinelandii DJ YP 002799713 1 322710 Proteobacteria
Azotobacter vinelandii CA6 YP 007898572 1 1283331 Proteobacteria
Pseudomonas poae RE 1 1 14 YP 007397543 1 1282356 Proteobacteria
Pseudomonas sp TKP YP 008933415 1 1415630 Proteobacteria
Gloeobacter kilaueensis JS1 YP 008711771 1 1183438 Cyanobacteria
Haliangium ochraceum DSM 14365 YP 003265814 1 502025 Proteobacteria
Singulisphaera acidiphila DSM 18658 YP 007206804 1 886293 Planctomycetes
Corallococcus coralloides DSM 2259 YP 005368775 1 1144275 Proteobacteria
Burkholderia rhizoxinica HKI 454 YP 004029720 1 882378 Proteobacteria
Bacillus subtilis subsp spizizenii TU B 10 YP 004878696 1 1052585 Firmicutes
Bacillus subtilis subsp subtilis str BAB 1 YP 007663829 1 1302650 Firmicutes
Bacillus subtilis subsp natto BEST195 YP 005562423 1 645657 Firmicutes
Bacillus subtilis PY79 YP 008831331 1 1415167 Firmicutes
Bacillus subtilis subsp subtilis 6051 HGW YP 007535184 1 1147161 Firmicutes
Herpetosiphon aurantiacus DSM 785 YP 001546733 1 316274 Chloroflexi
Corallococcus coralloides DSM 2259 YP 005369901 1 1144275 Proteobacteria
Myxococcus stipitatus DSM 14675 YP 007359073 1 1278073 Proteobacteria
Corallococcus coralloides DSM 2259 YP 005368577 1 1144275 Proteobacteria
Corallococcus coralloides DSM 2259 YP 005368578 1 1144275 Proteobacteria
Corallococcus coralloides DSM 2259 YP 005368579 1 1144275 Proteobacteria
Myxococcus stipitatus DSM 14675 YP 007359105 1 1278073 Proteobacteria
Myxococcus xanthus DK 1622 YP 631961 1 246197 Proteobacteria
Myxococcus fulvus HW 1 YP 004668381 1 483219 Proteobacteria
Myxococcus stipitatus DSM 14675 YP 007359692 1 1278073 Proteobacteria
Corallococcus coralloides DSM 2259 YP 005370175 1 1144275 Proteobacteria
Corallococcus coralloides DSM 2259 YP 005370176 1 1144275 Proteobacteria
Myxococcus fulvus HW 1 YP 004668593 1 483219 Proteobacteria
Myxococcus xanthus DK 1622 YP 631823 1 246197 Proteobacteria
Myxococcus fulvus HW 1 YP 004668223 1 483219 Proteobacteria

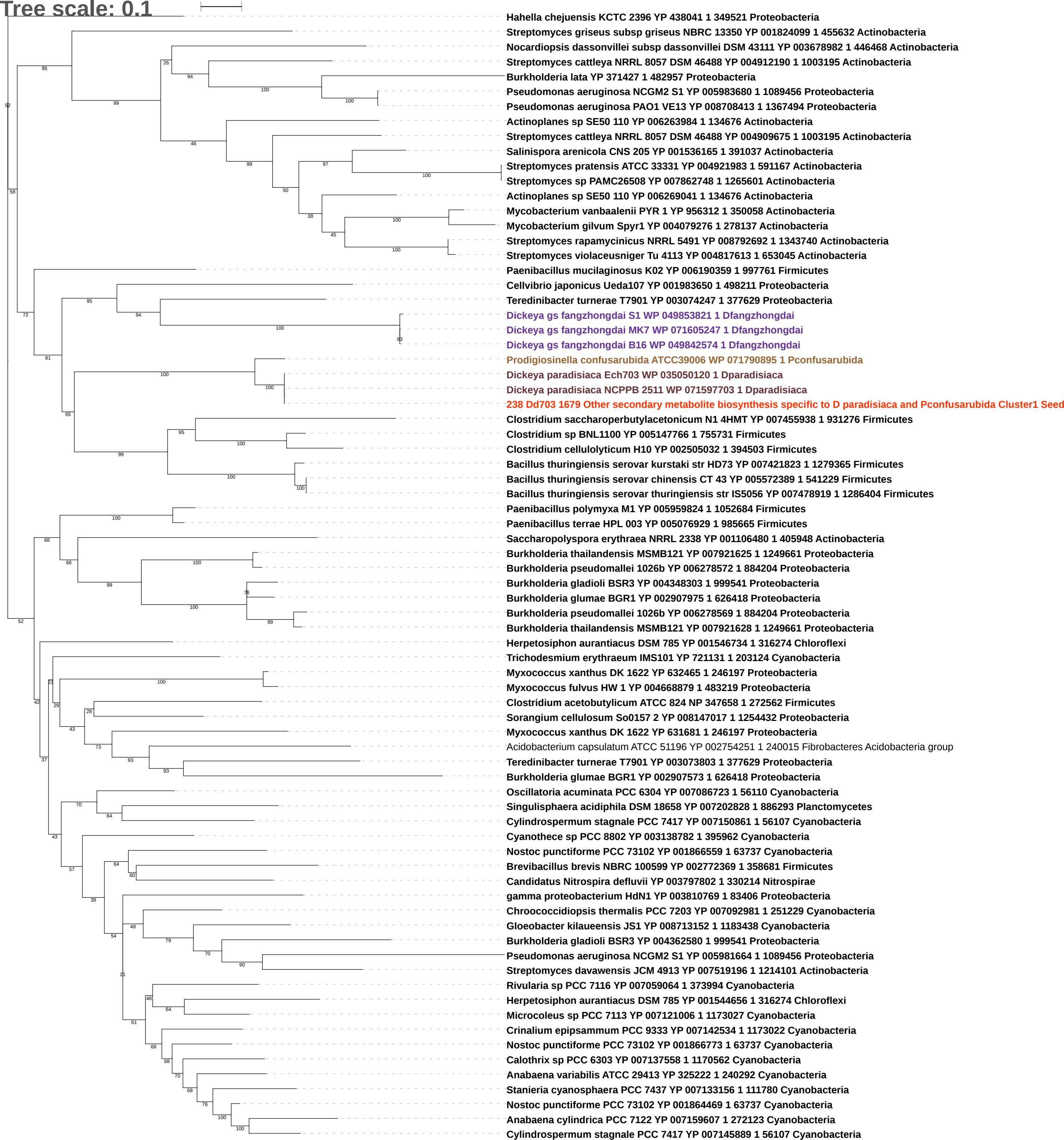
Tree scale: 0.1
Hahella chejuensis KCTC 2396 YP 438041 1 349521 Proteobacteria
Streptomyces griseus subsp griseus NBRC 13350 YP 001824099 1 455632 Actinobacteria
Nocardiopsis dassonvillei subsp dassonvillei DSM 43111 YP 003678982 1 446468 Actinobacteria
Streptomyces cattleya NRRL 8057 DSM 46488 YP 004912190 1 1003195 Actinobacteria
Burkholderia lata YP 371427 1 482957 Proteobacteria
Pseudomonas aeruginosa NCGM2 S1 YP 005983680 1 1089456 Proteobacteria
Pseudomonas aeruginosa PAO1 VE13 YP 008708413 1 1367494 Proteobacteria
Actinoplanes sp SE50 110 YP 006263984 1 134676 Actinobacteria
Streptomyces cattleya NRRL 8057 DSM 46488 YP 004909675 1 1003195 Actinobacteria
Salinispora arenicola CNS 205 YP 001536165 1 391037 Actinobacteria
Streptomyces pratensis ATCC 33331 YP 004921983 1 591167 Actinobacteria
Streptomyces sp PAMC26508 YP 007862748 1 1265601 Actinobacteria
Actinoplanes sp SE50 110 YP 006269041 1 134676 Actinobacteria
Mycobacterium vanbaalenii PYR 1 YP 956312 1 350058 Actinobacteria
Mycobacterium gilvum Spyr1 YP 004079276 1 278137 Actinobacteria
Streptomyces rapamycinicus NRRL 5491 YP 008792692 1 1343740 Actinobacteria
Streptomyces violaceusniger Tu 4113 YP 004817613 1 653045 Actinobacteria
Paenibacillus mucilaginosus K02 YP 006190359 1 997761 Firmicutes
Cellvibrio japonicus Ueda107 YP 001983650 1 498211 Proteobacteria
Teredinibacter turnerae T7901 YP 003074247 1 377629 Proteobacteria
Dickeya gs fangzhongdai S1 WP 049853821 1 Dfangzhongdai
Dickeya gs fangzhongdai MK7 WP 071605247 1 Dfangzhongdai
Dickeya gs fangzhongdai B16 WP 049842574 1 Dfangzhongdai
Prodigiosinella confusarubida ATCC39006 WP 071790895 1 Pconfusarubida
Dickeya paradisiaca Ech703 WP 035050120 1 Dparadisiaca
Dickeya paradisiaca NCPPB 2511 WP 071597703 1 Dparadisiaca
238 Dd703 1679 Other secondary metabolite biosynthesis specific to D paradisiaca and Pconfusarubida Cluster1 Seed
Clostridium saccharoperbutylacetonicum N1 4HMT YP 007455938 1 931276 Firmicutes
Clostridium sp BNL1100 YP 005147766 1 755731 Firmicutes
Clostridium cellulolyticum H10 YP 002505032 1 394503 Firmicutes
Bacillus thuringiensis serovar kurstaki str HD73 YP 007421823 1 1279365 Firmicutes
Bacillus thuringiensis serovar chinensis CT 43 YP 005572389 1 541229 Firmicutes
Bacillus thuringiensis serovar thuringiensis str IS5056 YP 007478919 1 1286404 Firmicutes
Paenibacillus polymyxa M1 YP 005959824 1 1052684 Firmicutes
Paenibacillus terrae HPL 003 YP 005076929 1 985665 Firmicutes
Saccharopolyspora erythraea NRRL 2338 YP 001106480 1 405948 Actinobacteria
Burkholderia thailandensis MSMB121 YP 007921625 1 1249661 Proteobacteria
Burkholderia pseudomallei 1026b YP 006278572 1 884204 Proteobacteria
Burkholderia gladioli BSR3 YP 004348303 1 999541 Proteobacteria
Burkholderia glumae BGR1 YP 002907975 1 626418 Proteobacteria
Burkholderia pseudomallei 1026b YP 006278569 1 884204 Proteobacteria
Burkholderia thailandensis MSMB121 YP 007921628 1 1249661 Proteobacteria
Herpetosiphon aurantiacus DSM 785 YP 001546734 1 316274 Chloroflexi
Trichodesmium erythraeum IMS101 YP 721131 1 203124 Cyanobacteria
Myxococcus xanthus DK 1622 YP 632465 1 246197 Proteobacteria
Myxococcus fulvus HW 1 YP 004668879 1 483219 Proteobacteria
Clostridium acetobutylicum ATCC 824 NP 347658 1 272562 Firmicutes
Sorangium cellulosum So0157 2 YP 008147017 1 1254432 Proteobacteria
Myxococcus xanthus DK 1622 YP 631681 1 246197 Proteobacteria
Acidobacterium capsulatum ATCC 51196 YP 002754251 1 240015 Fibrobacteres Acidobacteria group
Teredinibacter turnerae T7901 YP 003073803 1 377629 Proteobacteria
Burkholderia glumae BGR1 YP 002907573 1 626418 Proteobacteria
Oscillatoria acuminata PCC 6304 YP 007086723 1 56110 Cyanobacteria
Singulisphaera acidiphila DSM 18658 YP 007202828 1 886293 Planctomycetes
Cylindrospermum stagnale PCC 7417 YP 007150861 1 56107 Cyanobacteria
Cyanothece sp PCC 8802 YP 003138782 1 395962 Cyanobacteria
Nostoc punctiforme PCC 73102 YP 001866559 1 63737 Cyanobacteria
Brevibacillus brevis NBRC 100599 YP 002772369 1 358681 Firmicutes
Candidatus Nitrospira defluvii YP 003797802 1 330214 Nitrospirae
gamma proteobacterium HdN1 YP 003810769 1 83406 Proteobacteria
Chroococcidiopsis thermalis PCC 7203 YP 007092981 1 251229 Cyanobacteria
Gloeobacter kilaueensis JS1 YP 008713152 1 1183438 Cyanobacteria
Burkholderia gladioli BSR3 YP 004362580 1 999541 Proteobacteria
Pseudomonas aeruginosa NCGM2 S1 YP 005981664 1 1089456 Proteobacteria
Streptomyces davawensis JCM 4913 YP 007519196 1 1214101 Actinobacteria
Rivularia sp PCC 7116 YP 007059064 1 373994 Cyanobacteria
Herpetosiphon aurantiacus DSM 785 YP 001544656 1 316274 Chloroflexi
Microcoleus sp PCC 7113 YP 007121006 1 1173027 Cyanobacteria
Crinalium epipsammum PCC 9333 YP 007142534 1 1173022 Cyanobacteria
Nostoc punctiforme PCC 73102 YP 001866773 1 63737 Cyanobacteria
Calothrix sp PCC 6303 YP 007137558 1 1170562 Cyanobacteria
Anabaena variabilis ATCC 29413 YP 325222 1 240292 Cyanobacteria
Stanieria cyanosphaera PCC 7437 YP 007133156 1 111780 Cyanobacteria
Nostoc punctiforme PCC 73102 YP 001864469 1 63737 Cyanobacteria
Anabaena cylindrica PCC 7122 YP 007159607 1 272123 Cyanobacteria
Cylindrospermum stagnale PCC 7417 YP 007145889 1 56107 Cyanobacteria

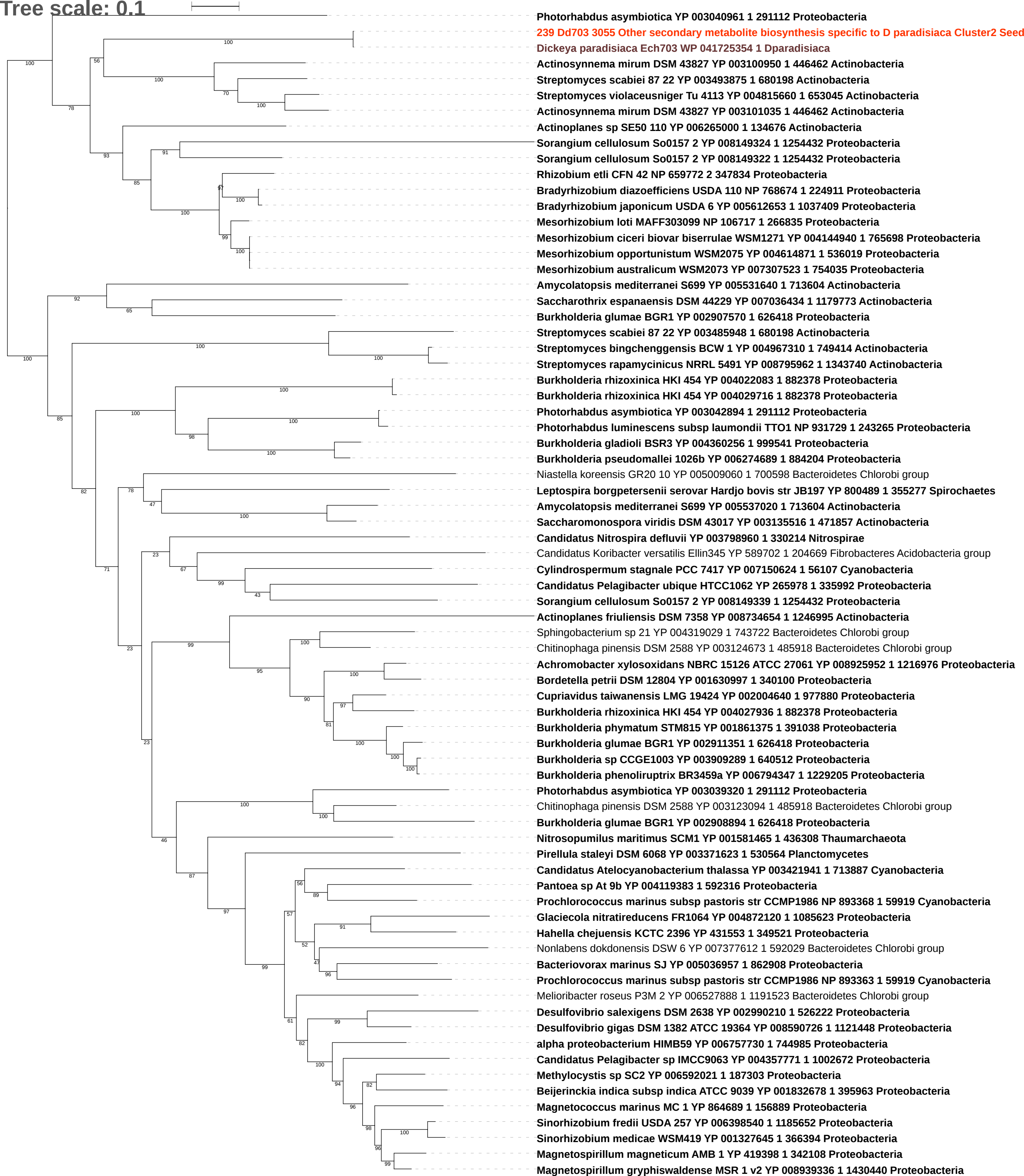
Tree scale: 0.1
Photorhabdus asymbiotica YP 003040961 1 291112 Proteobacteria
239 Dd703 3055 Other secondary metabolite biosynthesis specific to D paradisiaca Cluster2 Seed
Dickeya paradisiaca Ech703 WP 041725354 1 Dparadisiaca
Actinosynnema mirum DSM 43827 YP 003100950 1 446462 Actinobacteria
Streptomyces scabiei 87 22 YP 003493875 1 680198 Actinobacteria
Streptomyces violaceusniger Tu 4113 YP 004815660 1 653045 Actinobacteria
Actinosynnema mirum DSM 43827 YP 003101035 1 446462 Actinobacteria
Actinoplanes sp SE50 110 YP 006265000 1 134676 Actinobacteria
Sorangium cellulosum So0157 2 YP 008149324 1 1254432 Proteobacteria
Sorangium cellulosum So0157 2 YP 008149322 1 1254432 Proteobacteria
Rhizobium etli CFN 42 NP 659772 2 347834 Proteobacteria
Bradyrhizobium diazoefficiens USDA 110 NP 768674 1 224911 Proteobacteria
Bradyrhizobium japonicum USDA 6 YP 005612653 1 1037409 Proteobacteria
Mesorhizobium loti MAFF303099 NP 106717 1 266835 Proteobacteria
Mesorhizobium ciceri biovar biserrulae WSM1271 YP 004144940 1 765698 Proteobacteria
Mesorhizobium opportunistum WSM2075 YP 004614871 1 536019 Proteobacteria
Mesorhizobium australicum WSM2073 YP 007307523 1 754035 Proteobacteria
Amycolatopsis mediterranei S699 YP 005531640 1 713604 Actinobacteria
Saccharothrix espanaensis DSM 44229 YP 007036434 1 1179773 Actinobacteria
Burkholderia glumae BGR1 YP 002907570 1 626418 Proteobacteria
Streptomyces scabiei 87 22 YP 003485948 1 680198 Actinobacteria
Streptomyces bingchenggensis BCW 1 YP 004967310 1 749414 Actinobacteria
Streptomyces rapamycinicus NRRL 5491 YP 008795962 1 1343740 Actinobacteria
Burkholderia rhizoxinica HKI 454 YP 004022083 1 882378 Proteobacteria
Burkholderia rhizoxinica HKI 454 YP 004029716 1 882378 Proteobacteria
Photorhabdus asymbiotica YP 003042894 1 291112 Proteobacteria
Photorhabdus luminescens subsp laumondii TTO1 NP 931729 1 243265 Proteobacteria
Burkholderia gladioli BSR3 YP 004360256 1 999541 Proteobacteria
Burkholderia pseudomallei 1026b YP 006274689 1 884204 Proteobacteria
Niastella koreensis GR20 10 YP 005009060 1 700598 Bacteroidetes Chlorobi group
Leptospira borgpetersenii serovar Hardjo bovis str JB197 YP 800489 1 355277 Spirochaetes
Amycolatopsis mediterranei S699 YP 005537020 1 713604 Actinobacteria
Saccharomonospora viridis DSM 43017 YP 003135516 1 471857 Actinobacteria
Candidatus Nitrospira defluvii YP 003798960 1 330214 Nitrospirae
Candidatus Koribacter versatilis Ellin345 YP 589702 1 204669 Fibrobacteres Acidobacteria group
Cylindrospermum stagnale PCC 7417 YP 007150624 1 56107 Cyanobacteria
Candidatus Pelagibacter ubique HTCC1062 YP 265978 1 335992 Proteobacteria
Sorangium cellulosum So0157 2 YP 008149339 1 1254432 Proteobacteria
Actinoplanes friuliensis DSM 7358 YP 008734654 1 1246995 Actinobacteria
Sphingobacterium sp 21 YP 004319029 1 743722 Bacteroidetes Chlorobi group
Chitinophaga pinensis DSM 2588 YP 003124673 1 485918 Bacteroidetes Chlorobi group
Achromobacter xylosoxidans NBRC 15126 ATCC 27061 YP 008925952 1 1216976 Proteobacteria
Bordetella petrii DSM 12804 YP 001630997 1 340100 Proteobacteria
Cupriavidus taiwanensis LMG 19424 YP 002004640 1 977880 Proteobacteria
Burkholderia rhizoxinica HKI 454 YP 004027936 1 882378 Proteobacteria
Burkholderia phymatum STM815 YP 001861375 1 391038 Proteobacteria
Burkholderia glumae BGR1 YP 002911351 1 626418 Proteobacteria
Burkholderia sp CCGE1003 YP 003909289 1 640512 Proteobacteria
Burkholderia phenoliruptrix BR3459a YP 006794347 1 1229205 Proteobacteria
Photorhabdus asymbiotica YP 003039320 1 291112 Proteobacteria
Chitinophaga pinensis DSM 2588 YP 003123094 1 485918 Bacteroidetes Chlorobi group
Burkholderia glumae BGR1 YP 002908894 1 626418 Proteobacteria
Nitrosopumilus maritimus SCM1 YP 001581465 1 436308 Thaumarchaeota
Pirellula staleyi DSM 6068 YP 003371623 1 530564 Planctomycetes
Candidatus Atelocyanobacterium thalassa YP 003421941 1 713887 Cyanobacteria
Pantoea sp At 9b YP 004119383 1 592316 Proteobacteria
Prochlorococcus marinus subsp pastoris str CCMP1986 NP 893368 1 59919 Cyanobacteria
Glaciecola nitratireducens FR1064 YP 004872120 1 1085623 Proteobacteria
Hahella chejuensis KCTC 2396 YP 431553 1 349521 Proteobacteria
Nonlabens dokdonensis DSW 6 YP 007377612 1 592029 Bacteroidetes Chlorobi group
Bacteriovorax marinus SJ YP 005036957 1 862908 Proteobacteria
Prochlorococcus marinus subsp pastoris str CCMP1986 NP 893363 1 59919 Cyanobacteria
Melioribacter roseus P3M 2 YP 006527888 1 1191523 Bacteroidetes Chlorobi group
Desulfovibrio salexigens DSM 2638 YP 002990210 1 526222 Proteobacteria
Desulfovibrio gigas DSM 1382 ATCC 19364 YP 008590726 1 1121448 Proteobacteria
alpha proteobacterium HIMB59 YP 006757730 1 744985 Proteobacteria
Candidatus Pelagibacter sp IMCC9063 YP 004357771 1 1002672 Proteobacteria
Methylocystis sp SC2 YP 006592021 1 187303 Proteobacteria
Beijerinckia indica subsp indica ATCC 9039 YP 001832678 1 395963 Proteobacteria
Magnetococcus marinus MC 1 YP 864689 1 156889 Proteobacteria
Sinorhizobium fredii USDA 257 YP 006398540 1 1185652 Proteobacteria
Sinorhizobium medicae WSM419 YP 001327645 1 366394 Proteobacteria
Magnetospirillum magneticum AMB 1 YP 419398 1 342108 Proteobacteria
Magnetospirillum gryphiswaldense MSR 1 v2 YP 008939336 1 1430440 Proteobacteria

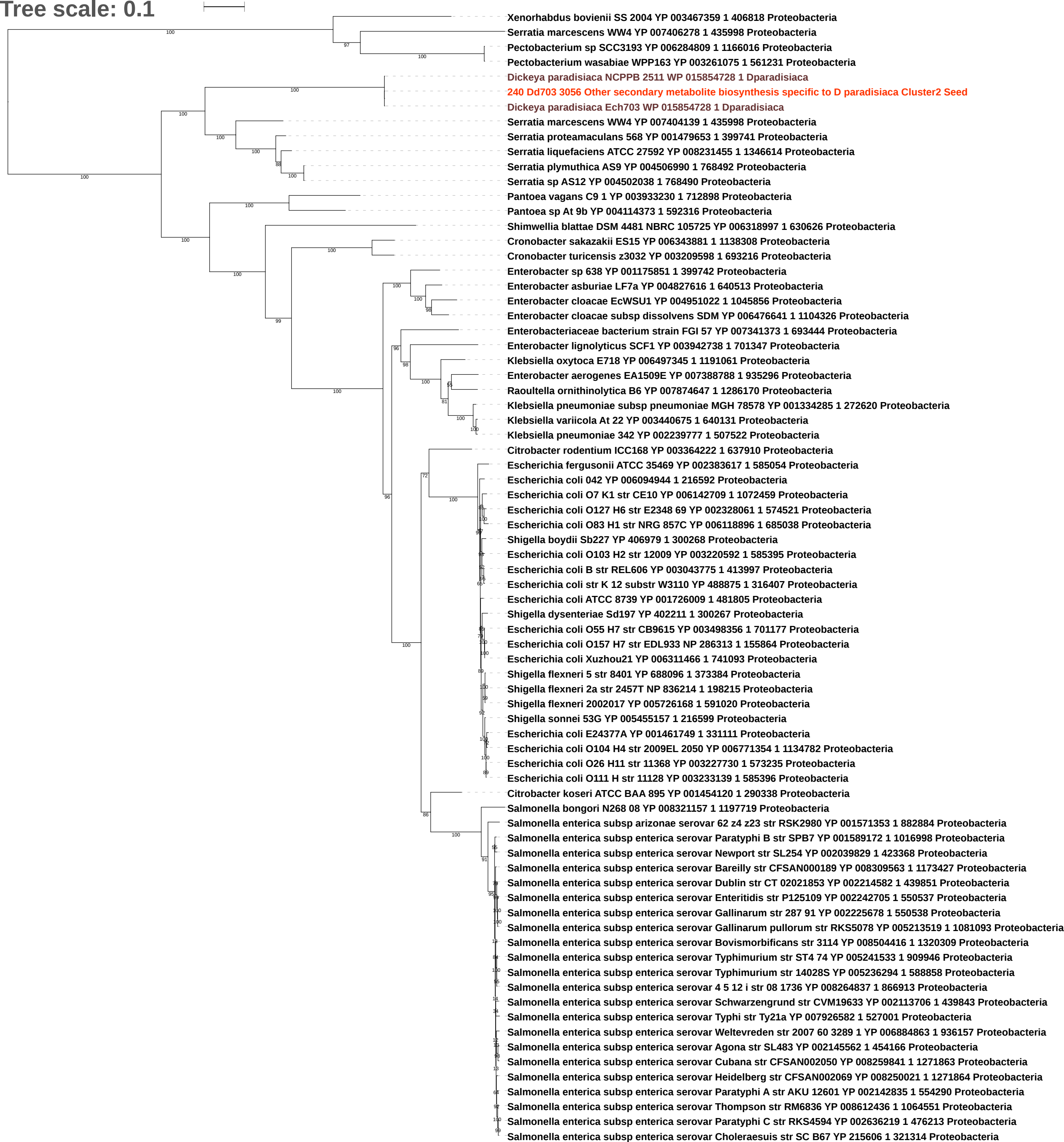

Tree scale: 0.1
Xenorhabdus bovienii SS 2004 YP 003467359 1 406818 Proteobacteria
Serratia marcescens WW4 YP 007406278 1 435998 Proteobacteria
Pectobacterium sp SCC3193 YP 006284809 1 1166016 Proteobacteria
Pectobacterium wasabiae WPP163 YP 003261075 1 561231 Proteobacteria
Dickeya paradisiaca NCPPB 2511 WP 015854728 1 Dparadisiaca
240 Dd703 3056 Other secondary metabolite biosynthesis specific to D paradisiaca Cluster2 Seed
Dickeya paradisiaca Ech703 WP 015854728 1 Dparadisiaca
Serratia marcescens WW4 YP 007404139 1 435998 Proteobacteria
Serratia proteamaculans 568 YP 001479653 1 399741 Proteobacteria
Serratia liquefaciens ATCC 27592 YP 008231455 1 1346614 Proteobacteria
Serratia plymuthica AS9 YP 004506990 1 768492 Proteobacteria
Serratia sp AS12 YP 004502038 1 768490 Proteobacteria
Pantoea vagans C9 1 YP 003933230 1 712898 Proteobacteria
Pantoea sp At 9b YP 004114373 1 592316 Proteobacteria
Shimwellia blattae DSM 4481 NBRC 105725 YP 006318997 1 630626 Proteobacteria
Cronobacter sakazakii ES15 YP 006343881 1 1138308 Proteobacteria
Cronobacter turicensis z3032 YP 003209598 1 693216 Proteobacteria
Enterobacter sp 638 YP 001175851 1 399742 Proteobacteria
Enterobacter asburiae LF7a YP 004827616 1 640513 Proteobacteria
Enterobacter cloacae EcWSU1 YP 004951022 1 1045856 Proteobacteria
Enterobacter cloacae subsp dissolvens SDM YP 006476641 1 1104326 Proteobacteria
Enterobacteriaceae bacterium strain FGI 57 YP 007341373 1 693444 Proteobacteria
Enterobacter lignolyticus SCF1 YP 003942738 1 701347 Proteobacteria
Klebsiella oxytoca E718 YP 006497345 1 1191061 Proteobacteria
Enterobacter aerogenes EA1509E YP 007388788 1 935296 Proteobacteria
Raoultella ornithinolytica B6 YP 007874647 1 1286170 Proteobacteria
Klebsiella pneumoniae subsp pneumoniae MGH 78578 YP 001334285 1 272620 Proteobacteria
Klebsiella variicola At 22 YP 003440675 1 640131 Proteobacteria
Klebsiella pneumoniae 342 YP 002239777 1 507522 Proteobacteria
Citrobacter rodentium ICC168 YP 003364222 1 637910 Proteobacteria
Escherichia fergusonii ATCC 35469 YP 002383617 1 585054 Proteobacteria
Escherichia coli 042 YP 006094944 1 216592 Proteobacteria
Escherichia coli O7 K1 str CE10 YP 006142709 1 1072459 Proteobacteria
Escherichia coli O127 H6 str E2348 69 YP 002328061 1 574521 Proteobacteria
Escherichia coli O83 H1 str NRG 857C YP 006118896 1 685038 Proteobacteria
Shigella boydii Sb227 YP 406979 1 300268 Proteobacteria
Escherichia coli O103 H2 str 12009 YP 003220592 1 585395 Proteobacteria
Escherichia coli B str REL606 YP 003043775 1 413997 Proteobacteria
Escherichia coli str K 12 substr W3110 YP 488875 1 316407 Proteobacteria
Escherichia coli ATCC 8739 YP 001726009 1 481805 Proteobacteria
Shigella dysenteriae Sd197 YP 402211 1 300267 Proteobacteria
Escherichia coli O55 H7 str CB9615 YP 003498356 1 701177 Proteobacteria
Escherichia coli O157 H7 str EDL933 NP 286313 1 155864 Proteobacteria
Escherichia coli Xuzhou21 YP 006311466 1 741093 Proteobacteria
Shigella flexneri 5 str 8401 YP 688096 1 373384 Proteobacteria
Shigella flexneri 2a str 2457T NP 836214 1 198215 Proteobacteria
Shigella flexneri 2002017 YP 005726168 1 591020 Proteobacteria
Shigella sonnei 53G YP 005455157 1 216599 Proteobacteria
Escherichia coli E24377A YP 001461749 1 331111 Proteobacteria
Escherichia coli O104 H4 str 2009EL 2050 YP 006771354 1 1134782 Proteobacteria
Escherichia coli O26 H11 str 11368 YP 003227730 1 573235 Proteobacteria
Escherichia coli O111 H str 11128 YP 003233139 1 585396 Proteobacteria
Citrobacter koseri ATCC BAA 895 YP 001454120 1 290338 Proteobacteria
Salmonella bongori N268 08 YP 008321157 1 1197719 Proteobacteria
Salmonella enterica subsp arizonae serovar 62 z4 z23 str RSK2980 YP 001571353 1 882884 Proteobacteria
Salmonella enterica subsp enterica serovar Paratyphi B str SPB7 YP 001589172 1 1016998 Proteobacteria
Salmonella enterica subsp enterica serovar Newport str SL254 YP 002039829 1 423368 Proteobacteria
Salmonella enterica subsp enterica serovar Bareilly str CFSAN000189 YP 008309563 1 1173427 Proteobacteria
Salmonella enterica subsp enterica serovar Dublin str CT 02021853 YP 002214582 1 439851 Proteobacteria
Salmonella enterica subsp enterica serovar Enteritidis str P125109 YP 002242705 1 550537 Proteobacteria
Salmonella enterica subsp enterica serovar Gallinarum str 287 91 YP 002225678 1 550538 Proteobacteria
Salmonella enterica subsp enterica serovar Gallinarum pullorum str RKS5078 YP 005213519 1 1081093 Proteobacteria
Salmonella enterica subsp enterica serovar Bovismorbificans str 3114 YP 008504416 1 1320309 Proteobacteria
Salmonella enterica subsp enterica serovar Typhimurium str ST4 74 YP 005241533 1 909946 Proteobacteria
Salmonella enterica subsp enterica serovar Typhimurium str 14028S YP 005236294 1 588858 Proteobacteria
Salmonella enterica subsp enterica serovar 4 5 12 i str 08 1736 YP 008264837 1 866913 Proteobacteria
Salmonella enterica subsp enterica serovar Schwarzengrund str CVM19633 YP 002113706 1 439843 Proteobacteria
Salmonella enterica subsp enterica serovar Typhi str Ty21a YP 007926582 1 527001 Proteobacteria
Salmonella enterica subsp enterica serovar Weltevreden str 2007 60 3289 1 YP 006884863 1 936157 Proteobacteria
Salmonella enterica subsp enterica serovar Agona str SL483 YP 002145562 1 454166 Proteobacteria
Salmonella enterica subsp enterica serovar Cubana str CFSAN002050 YP 008259841 1 1271863 Proteobacteria
Salmonella enterica subsp enterica serovar Heidelberg str CFSAN002069 YP 008250021 1 1271864 Proteobacteria
Salmonella enterica subsp enterica serovar Paratyphi A str AKU 12601 YP 002142835 1 554290 Proteobacteria
Salmonella enterica subsp enterica serovar Thompson str RM6836 YP 008612436 1 1064551 Proteobacteria
Salmonella enterica subsp enterica serovar Paratyphi C str RKS4594 YP 002636219 1 476213 Proteobacteria
Salmonella enterica subsp enterica serovar Choleraesuis str SC B67 YP 215606 1 321314 Proteobacteria

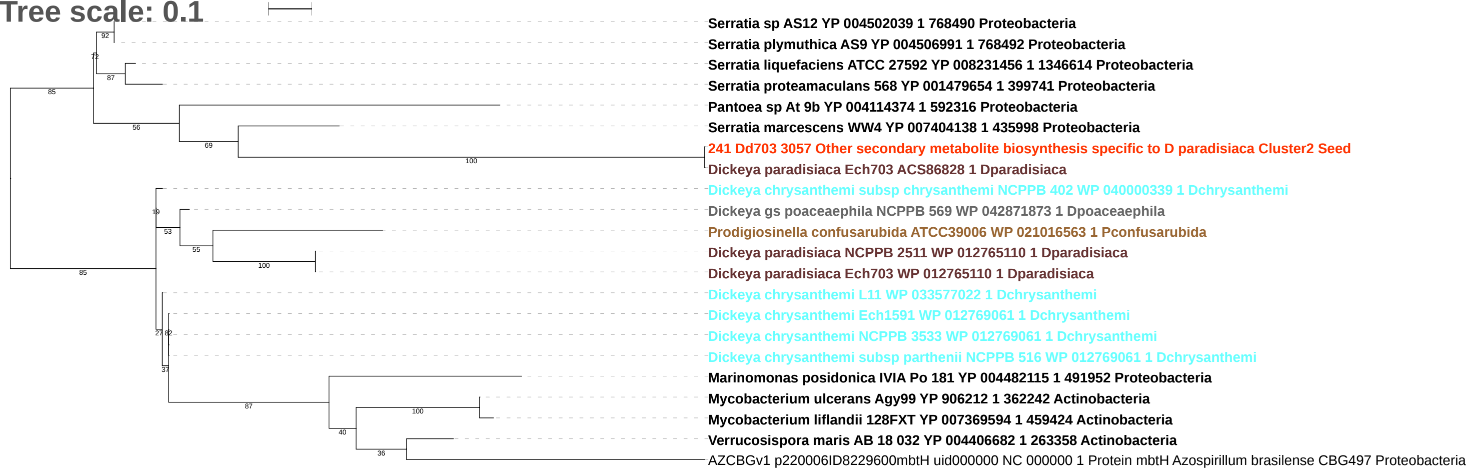
Tree scale: 0.1
Serratia sp AS12 YP 004502039 1 768490 Proteobacteria
Serratia plymuthica AS9 YP 004506991 1 768492 Proteobacteria
Serratia liquefaciens ATCC 27592 YP 008231456 1 1346614 Proteobacteria
Serratia proteamaculans 568 YP 001479654 1 399741 Proteobacteria
Pantoea sp At 9b YP 004114374 1 592316 Proteobacteria
Serratia marcescens WW4 YP 007404138 1 435998 Proteobacteria
241 Dd703 3057 Other secondary metabolite biosynthesis specific to D paradisiaca Cluster2 Seed
Dickeya paradisiaca Ech703 ACS86828 1 Dparadisiaca
Dickeya chrysanthemi subsp chrysanthemi NCPPB 402 WP 040000339 1 Dchrysanthemi
Dickeya gs poaceaephila NCPPB 569 WP 042871873 1 Dpoaceaephila
Prodigiosinella confusarubida ATCC39006 WP 021016563 1 Pconfusarubida
Dickeya paradisiaca NCPPB 2511 WP 012765110 1 Dparadisiaca
Dickeya paradisiaca Ech703 WP 012765110 1 Dparadisiaca
Dickeya chrysanthemi L11 WP 033577022 1 Dchrysanthemi
Dickeya chrysanthemi Ech1591 WP 012769061 1 Dchrysanthemi
Dickeya chrysanthemi NCPPB 3533 WP 012769061 1 Dchrysanthemi
Dickeya chrysanthemi subsp parthenii NCPPB 516 WP 012769061 1 Dchrysanthemi
Marinomonas posidonica IVIA Po 181 YP 004482115 1 491952 Proteobacteria
Mycobacterium ulcerans Agy99 YP 906212 1 362242 Actinobacteria
Mycobacterium liflandii 128FXT YP 007369594 1 459424 Actinobacteria
Verrucosispora maris AB 18 032 YP 004406682 1 263358 Actinobacteria
AZCBGv1 p220006ID8229600mbtH uid000000 NC 000000 1 Protein mbtH Azospirillum brasilense CBG497 Proteobacteria
92
72
87
85
56
69
100
19
53
55
100
85
27
82
37
87
100
40
36

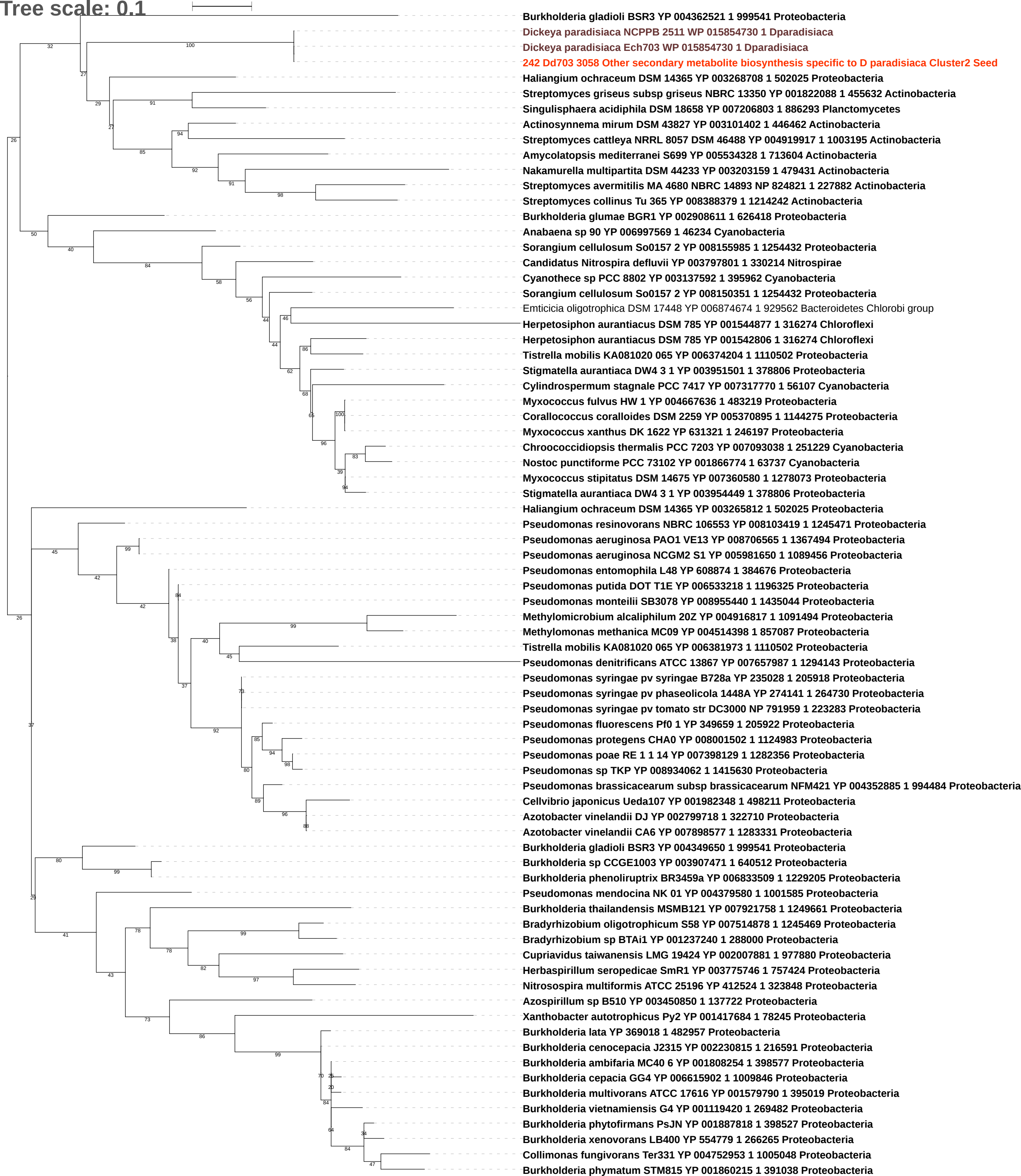
Tree scale: 0.1
Burkholderia gladioli BSR3 YP 004362521 1 999541 Proteobacteria
Dickeya paradisiaca NCPPB 2511 WP 015854730 1 Dparadisiaca
Dickeya paradisiaca Ech703 WP 015854730 1 Dparadisiaca
242 Dd703 3058 Other secondary metabolite biosynthesis specific to D paradisiaca Cluster2 Seed
Haliangium ochraceum DSM 14365 YP 003268708 1 502025 Proteobacteria
Streptomyces griseus subsp griseus NBRC 13350 YP 001822088 1 455632 Actinobacteria
Singulisphaera acidiphila DSM 18658 YP 007206803 1 886293 Planctomycetes
Actinosynnema mirum DSM 43827 YP 003101402 1 446462 Actinobacteria
Streptomyces cattleya NRRL 8057 DSM 46488 YP 004919917 1 1003195 Actinobacteria
Amycolatopsis mediterranei S699 YP 005534328 1 713604 Actinobacteria
Nakamurella multipartita DSM 44233 YP 003203159 1 479431 Actinobacteria
Streptomyces avermitilis MA 4680 NBRC 14893 NP 824821 1 227882 Actinobacteria
Streptomyces collinus Tu 365 YP 008388379 1 1214242 Actinobacteria
Burkholderia glumae BGR1 YP 002908611 1 626418 Proteobacteria
Anabaena sp 90 YP 006997569 1 46234 Cyanobacteria
Sorangium cellulosum So0157 2 YP 008155985 1 1254432 Proteobacteria
Candidatus Nitrospira defluvii YP 003797801 1 330214 Nitrospirae
Cyanothece sp PCC 8802 YP 003137592 1 395962 Cyanobacteria
Sorangium cellulosum So0157 2 YP 008150351 1 1254432 Proteobacteria
Emticicia oligotrophica DSM 17448 YP 006874674 1 929562 Bacteroidetes Chlorobi group
Herpetosiphon aurantiacus DSM 785 YP 001544877 1 316274 Chloroflexi
Herpetosiphon aurantiacus DSM 785 YP 001542806 1 316274 Chloroflexi
Tistrella mobilis KA081020 065 YP 006374204 1 1110502 Proteobacteria
Stigmatella aurantiaca DW4 3 1 YP 003951501 1 378806 Proteobacteria
Cylindrospermum stagnale PCC 7417 YP 007317770 1 56107 Cyanobacteria
Myxococcus fulvus HW 1 YP 004667636 1 483219 Proteobacteria
Corallococcus coralloides DSM 2259 YP 005370895 1 1144275 Proteobacteria
Myxococcus xanthus DK 1622 YP 631321 1 246197 Proteobacteria
Chroococcidiopsis thermalis PCC 7203 YP 007093038 1 251229 Cyanobacteria
Nostoc punctiforme PCC 73102 YP 001866774 1 63737 Cyanobacteria
Myxococcus stipitatus DSM 14675 YP 007360580 1 1278073 Proteobacteria
Stigmatella aurantiaca DW4 3 1 YP 003954449 1 378806 Proteobacteria
Haliangium ochraceum DSM 14365 YP 003265812 1 502025 Proteobacteria
Pseudomonas resinovorans NBRC 106553 YP 008103419 1 1245471 Proteobacteria
Pseudomonas aeruginosa PAO1 VE13 YP 008706565 1 1367494 Proteobacteria
Pseudomonas aeruginosa NCGM2 S1 YP 005981650 1 1089456 Proteobacteria
Pseudomonas entomophila L48 YP 608874 1 384676 Proteobacteria
Pseudomonas putida DOT T1E YP 006533218 1 1196325 Proteobacteria
Pseudomonas monteilii SB3078 YP 008955440 1 1435044 Proteobacteria
Methylomicrobium alcaliphilum 20Z YP 004916817 1 1091494 Proteobacteria
Methylomonas methanica MC09 YP 004514398 1 857087 Proteobacteria
Tistrella mobilis KA081020 065 YP 006381973 1 1110502 Proteobacteria
Pseudomonas denitrificans ATCC 13867 YP 007657987 1 1294143 Proteobacteria
Pseudomonas syringae pv syringae B728a YP 235028 1 205918 Proteobacteria
Pseudomonas syringae pv phaseolicola 1448A YP 274141 1 264730 Proteobacteria
Pseudomonas syringae pv tomato str DC3000 NP 791959 1 223283 Proteobacteria
Pseudomonas fluorescens Pf0 1 YP 349659 1 205922 Proteobacteria
Pseudomonas protegens CHA0 YP 008001502 1 1124983 Proteobacteria
Pseudomonas poae RE 1 1 14 YP 007398129 1 1282356 Proteobacteria
Pseudomonas sp TKP YP 008934062 1 1415630 Proteobacteria
Pseudomonas brassicacearum subsp brassicacearum NFM421 YP 004352885 1 994484 Proteobacteria
Cellvibrio japonicus Ueda107 YP 001982348 1 498211 Proteobacteria
Azotobacter vinelandii DJ YP 002799718 1 322710 Proteobacteria
Azotobacter vinelandii CA6 YP 007898577 1 1283331 Proteobacteria
Burkholderia gladioli BSR3 YP 004349650 1 999541 Proteobacteria
Burkholderia sp CCGE1003 YP 003907471 1 640512 Proteobacteria
Burkholderia phenoliruptrix BR3459a YP 006833509 1 1229205 Proteobacteria
Pseudomonas mendocina NK 01 YP 004379580 1 1001585 Proteobacteria
Burkholderia thailandensis MSMB121 YP 007921758 1 1249661 Proteobacteria
Bradyrhizobium oligotrophicum S58 YP 007514878 1 1245469 Proteobacteria
Bradyrhizobium sp BTAi1 YP 001237240 1 288000 Proteobacteria
Cupriavidus taiwanensis LMG 19424 YP 002007881 1 977880 Proteobacteria
Herbaspirillum seropedicae SmR1 YP 003775746 1 757424 Proteobacteria
Nitrosospira multiformis ATCC 25196 YP 412524 1 323848 Proteobacteria
Azospirillum sp B510 YP 003450850 1 137722 Proteobacteria
Xanthobacter autotrophicus Py2 YP 001417684 1 78245 Proteobacteria
Burkholderia lata YP 369018 1 482957 Proteobacteria
Burkholderia cenocepacia J2315 YP 002230815 1 216591 Proteobacteria
Burkholderia ambifaria MC40 6 YP 001808254 1 398577 Proteobacteria
Burkholderia cepacia GG4 YP 006615902 1 1009846 Proteobacteria
Burkholderia multivorans ATCC 17616 YP 001579790 1 395019 Proteobacteria
Burkholderia vietnamiensis G4 YP 001119420 1 269482 Proteobacteria
Burkholderia phytofirmans PsJN YP 001887818 1 398527 Proteobacteria
Burkholderia xenovorans LB400 YP 554779 1 266265 Proteobacteria
Collimonas fungivorans Ter331 YP 004752953 1 1005048 Proteobacteria
Burkholderia phymatum STM815 YP 001860215 1 391038 Proteobacteria

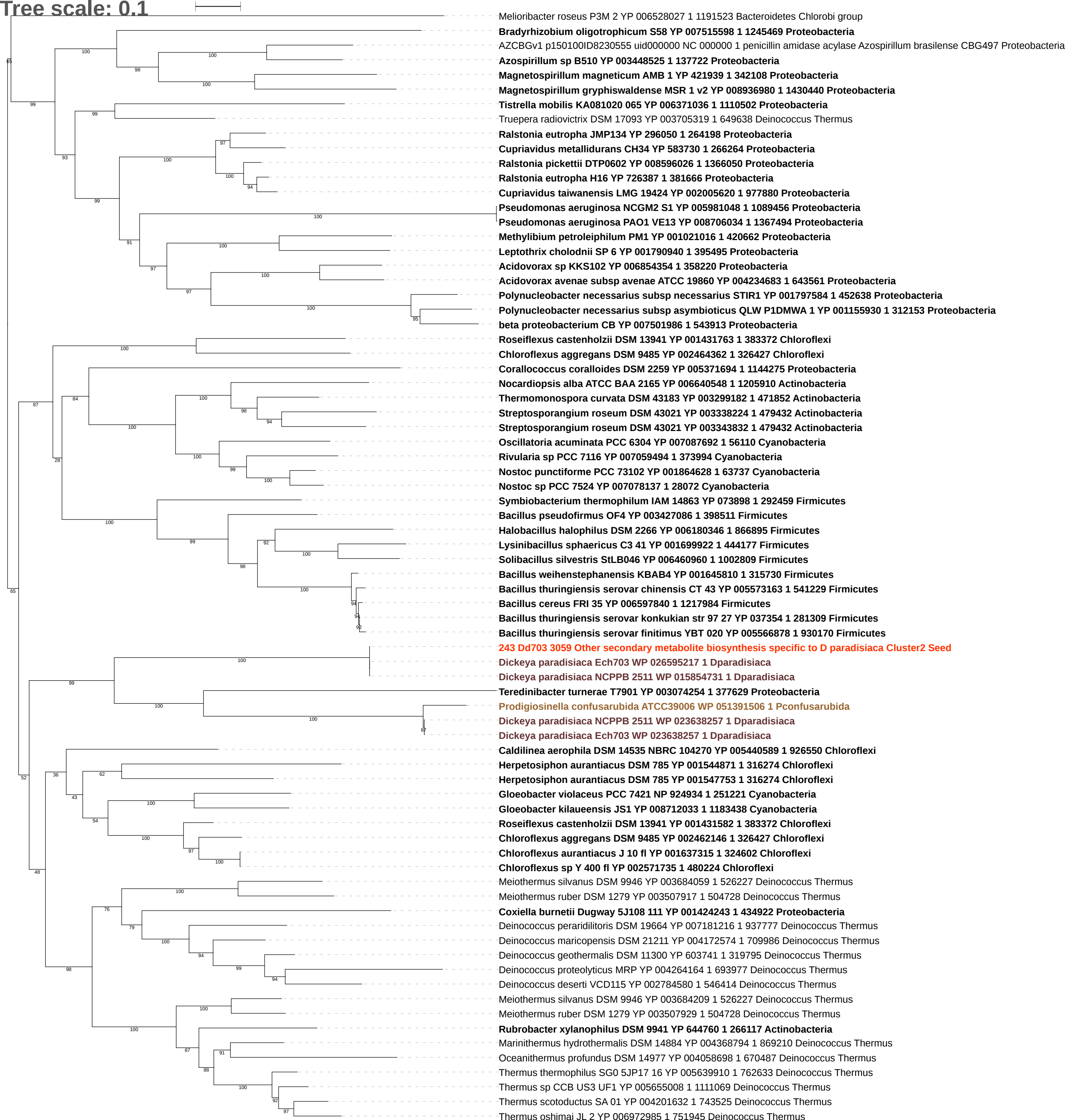
Tree scale: 0.1
Melioribacter roseus P3M 2 YP 006528027 1 1191523 Bacteroidetes Chlorobi group
Bradyrhizobium oligotrophicum S58 YP 007515598 1 1245469 Proteobacteria
AZCBGv1 p150100ID8230555 uid000000 NC 000000 1 penicillin amidase acylase Azospirillum brasilense CBG497 Proteobacteria
Azospirillum sp B510 YP 003448525 1 137722 Proteobacteria
Magnetospirillum magneticum AMB 1 YP 421939 1 342108 Proteobacteria
Magnetospirillum gryphiswaldense MSR 1 v2 YP 008936980 1 1430440 Proteobacteria
Tistrella mobilis KA081020 065 YP 006371036 1 1110502 Proteobacteria
Truepera radiovictrix DSM 17093 YP 003705319 1 649638 Deinococcus Thermus
Ralstonia eutropha JMP134 YP 296050 1 264198 Proteobacteria
Cupriavidus metallidurans CH34 YP 583730 1 266264 Proteobacteria
Ralstonia pickettii DTP0602 YP 008596026 1 1366050 Proteobacteria
Ralstonia eutropha H16 YP 726387 1 381666 Proteobacteria
Cupriavidus taiwanensis LMG 19424 YP 002005620 1 977880 Proteobacteria
Pseudomonas aeruginosa NCGM2 S1 YP 005981048 1 1089456 Proteobacteria
Pseudomonas aeruginosa PAO1 VE13 YP 008706034 1 1367494 Proteobacteria
Methylibium petroleiphilum PM1 YP 001021016 1 420662 Proteobacteria
Leptothrix cholodnii SP 6 YP 001790940 1 395495 Proteobacteria
Acidovorax sp KKS102 YP 006854354 1 358220 Proteobacteria
Acidovorax avenae subsp avenae ATCC 19860 YP 004234683 1 643561 Proteobacteria
Polynucleobacter necessarius subsp necessarius STIR1 YP 001797584 1 452638 Proteobacteria
Polynucleobacter necessarius subsp asymbioticus QLW P1DMWA 1 YP 001155930 1 312153 Proteobacteria
beta proteobacterium CB YP 007501986 1 543913 Proteobacteria
Roseiflexus castenholzii DSM 13941 YP 001431763 1 383372 Chloroflexi
Chloroflexus aggregans DSM 9485 YP 002464362 1 326427 Chloroflexi
Corallococcus coralloides DSM 2259 YP 005371694 1 1144275 Proteobacteria
Nocardiopsis alba ATCC BAA 2165 YP 006640548 1 1205910 Actinobacteria
Thermomonospora curvata DSM 43183 YP 003299182 1 471852 Actinobacteria
Streptosporangium roseum DSM 43021 YP 003338224 1 479432 Actinobacteria
Streptosporangium roseum DSM 43021 YP 003343832 1 479432 Actinobacteria
Oscillatoria acuminata PCC 6304 YP 007087692 1 56110 Cyanobacteria
Rivularia sp PCC 7116 YP 007059494 1 373994 Cyanobacteria
Nostoc punctiforme PCC 73102 YP 001864628 1 63737 Cyanobacteria
Nostoc sp PCC 7524 YP 007078137 1 28072 Cyanobacteria
Symbiobacterium thermophilum IAM 14863 YP 073898 1 292459 Firmicutes
Bacillus pseudofirmus OF4 YP 003427086 1 398511 Firmicutes
Halobacillus halophilus DSM 2266 YP 006180346 1 866895 Firmicutes
Lysinibacillus sphaericus C3 41 YP 001699922 1 444177 Firmicutes
Solibacillus silvestris StLB046 YP 006460960 1 1002809 Firmicutes
Bacillus weihenstephanensis KBAB4 YP 001645810 1 315730 Firmicutes
Bacillus thuringiensis serovar chinensis CT 43 YP 005573163 1 541229 Firmicutes
Bacillus cereus FRI 35 YP 006597840 1 1217984 Firmicutes
Bacillus thuringiensis serovar konkukian str 97 27 YP 037354 1 281309 Firmicutes
Bacillus thuringiensis serovar finitimus YBT 020 YP 005566878 1 930170 Firmicutes
243 Dd703 3059 Other secondary metabolite biosynthesis specific to D paradisiaca Cluster2 Seed
Dickeya paradisiaca Ech703 WP 026595217 1 Dparadisiaca
Dickeya paradisiaca NCPPB 2511 WP 015854731 1 Dparadisiaca
Teredinibacter turnerae T7901 YP 003074254 1 377629 Proteobacteria
Prodigiosinella confusarubida ATCC39006 WP 051391506 1 Pconfusarubida
Dickeya paradisiaca NCPPB 2511 WP 023638257 1 Dparadisiaca
Dickeya paradisiaca Ech703 WP 023638257 1 Dparadisiaca
Caldilinea aerophila DSM 14535 NBRC 104270 YP 005440589 1 926550 Chloroflexi
Herpetosiphon aurantiacus DSM 785 YP 001544871 1 316274 Chloroflexi
Herpetosiphon aurantiacus DSM 785 YP 001547753 1 316274 Chloroflexi
Gloeobacter violaceus PCC 7421 NP 924934 1 251221 Cyanobacteria
Gloeobacter kilaueensis JS1 YP 008712033 1 1183438 Cyanobacteria
Roseiflexus castenholzii DSM 13941 YP 001431582 1 383372 Chloroflexi
Chloroflexus aggregans DSM 9485 YP 002462146 1 326427 Chloroflexi
Chloroflexus aurantiacus J 10 fl YP 001637315 1 324602 Chloroflexi
Chloroflexus sp Y 400 fl YP 002571735 1 480224 Chloroflexi
Meiothermus silvanus DSM 9946 YP 003684059 1 526227 Deinococcus Thermus
Meiothermus ruber DSM 1279 YP 003507917 1 504728 Deinococcus Thermus
Coxiella burnetii Dugway 5J108 111 YP 001424243 1 434922 Proteobacteria
Deinococcus peraridilitoris DSM 19664 YP 007181216 1 937777 Deinococcus Thermus
Deinococcus maricopensis DSM 21211 YP 004172574 1 709986 Deinococcus Thermus
Deinococcus geothermalis DSM 11300 YP 603741 1 319795 Deinococcus Thermus
Deinococcus proteolyticus MRP YP 004264164 1 693977 Deinococcus Thermus
Deinococcus deserti VCD115 YP 002784580 1 546414 Deinococcus Thermus
Meiothermus silvanus DSM 9946 YP 003684209 1 526227 Deinococcus Thermus
Meiothermus ruber DSM 1279 YP 003507929 1 504728 Deinococcus Thermus
Rubrobacter xylanophilus DSM 9941 YP 644760 1 266117 Actinobacteria
Marinithermus hydrothermalis DSM 14884 YP 004368794 1 869210 Deinococcus Thermus
Oceanithermus profundus DSM 14977 YP 004058698 1 670487 Deinococcus Thermus
Thermus thermophilus SG0 5JP17 16 YP 005639910 1 762633 Deinococcus Thermus
Thermus sp CCB US3 UF1 YP 005655008 1 1111069 Deinococcus Thermus
Thermus scotoductus SA 01 YP 004201632 1 743525 Deinococcus Thermus
Thermus oshimai JL 2 YP 006972985 1 751945 Deinococcus Thermus

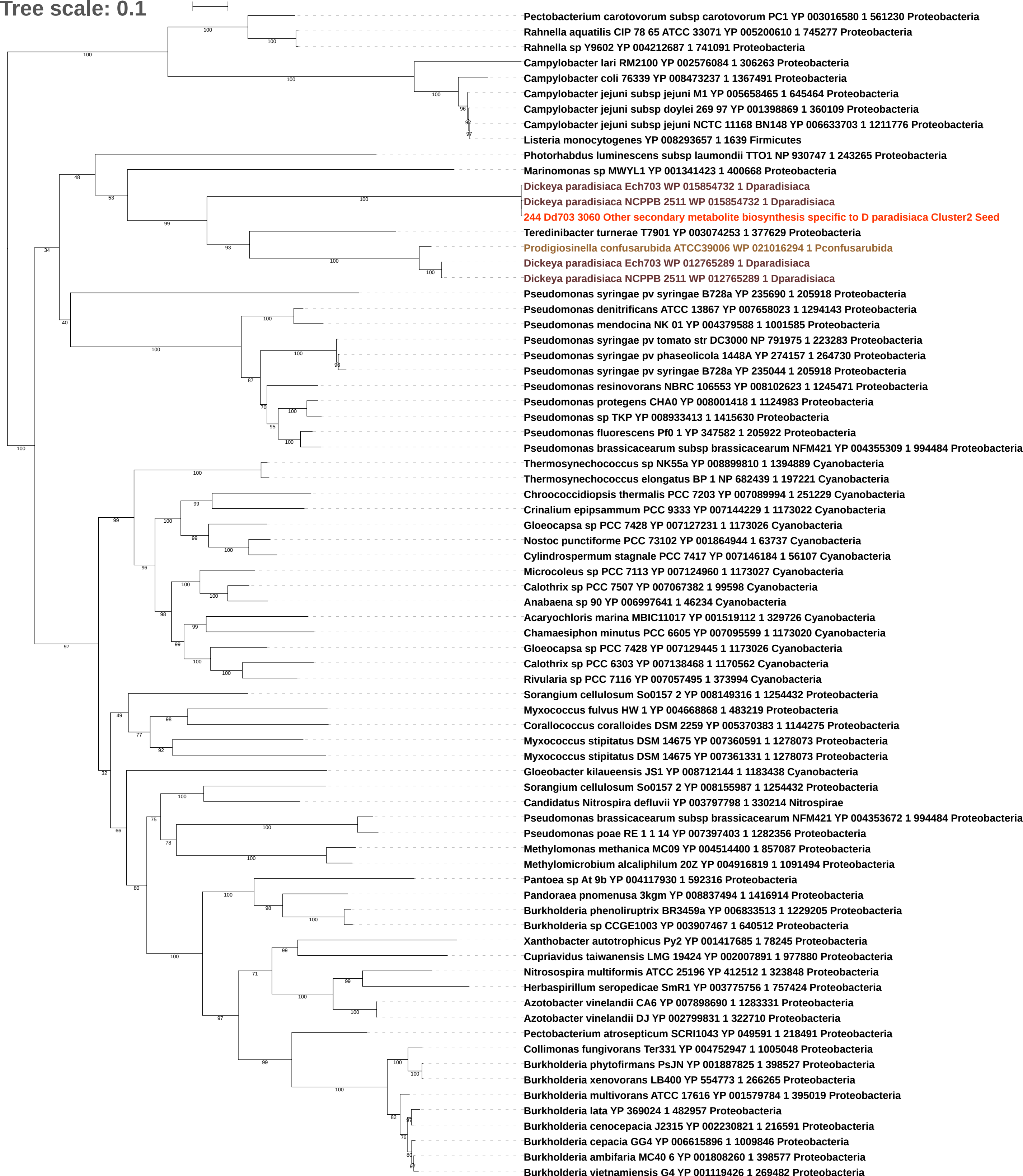
Tree scale: 0.1
Pectobacterium carotovorum subsp carotovorum PC1 YP 003016580 1 561230 Proteobacteria
Rahnella aquatilis CIP 78 65 ATCC 33071 YP 005200610 1 745277 Proteobacteria
Rahnella sp Y9602 YP 004212687 1 741091 Proteobacteria
Campylobacter lari RM2100 YP 002576084 1 306263 Proteobacteria
Campylobacter coli 76339 YP 008473237 1 1367491 Proteobacteria
Campylobacter jejuni subsp jejuni M1 YP 005658465 1 645464 Proteobacteria
Campylobacter jejuni subsp doylei 269 97 YP 001398869 1 360109 Proteobacteria
Campylobacter jejuni subsp jejuni NCTC 11168 BN148 YP 006633703 1 1211776 Proteobacteria
Listeria monocytogenes YP 008293657 1 1639 Firmicutes
Photorhabdus luminescens subsp laumondii TTO1 NP 930747 1 243265 Proteobacteria
Marinomonas sp MWYL1 YP 001341423 1 400668 Proteobacteria
Dickeya paradisiaca Ech703 WP 015854732 1 Dparadisiaca
Dickeya paradisiaca NCPPB 2511 WP 015854732 1 Dparadisiaca
244 Dd703 3060 Other secondary metabolite biosynthesis specific to D paradisiaca Cluster2 Seed
Teredinibacter turnerae T7901 YP 003074253 1 377629 Proteobacteria
Prodigiosinella confusarubida ATCC39006 WP 021016294 1 Pconfusarubida
Dickeya paradisiaca Ech703 WP 012765289 1 Dparadisiaca
Dickeya paradisiaca NCPPB 2511 WP 012765289 1 Dparadisiaca
Pseudomonas syringae pv syringae B728a YP 235690 1 205918 Proteobacteria
Pseudomonas denitrificans ATCC 13867 YP 007658023 1 1294143 Proteobacteria
Pseudomonas mendocina NK 01 YP 004379588 1 1001585 Proteobacteria
Pseudomonas syringae pv tomato str DC3000 NP 791975 1 223283 Proteobacteria
Pseudomonas syringae pv phaseolicola 1448A YP 274157 1 264730 Proteobacteria
Pseudomonas syringae pv syringae B728a YP 235044 1 205918 Proteobacteria
Pseudomonas resinovorans NBRC 106553 YP 008102623 1 1245471 Proteobacteria
Pseudomonas protegens CHA0 YP 008001418 1 1124983 Proteobacteria
Pseudomonas sp TKP YP 008933413 1 1415630 Proteobacteria
Pseudomonas fluorescens Pf0 1 YP 347582 1 205922 Proteobacteria
Pseudomonas brassicacearum subsp brassicacearum NFM421 YP 004355309 1 994484 Proteobacteria
Thermosynechococcus sp NK55a YP 008899810 1 1394889 Cyanobacteria
Thermosynechococcus elongatus BP 1 NP 682439 1 197221 Cyanobacteria
Chroococcidiopsis thermalis PCC 7203 YP 007089994 1 251229 Cyanobacteria
Crinalium epipsammum PCC 9333 YP 007144229 1 1173022 Cyanobacteria
Gloeocapsa sp PCC 7428 YP 007127231 1 1173026 Cyanobacteria
Nostoc punctiforme PCC 73102 YP 001864944 1 63737 Cyanobacteria
Cylindrospermum stagnale PCC 7417 YP 007146184 1 56107 Cyanobacteria
Microcoleus sp PCC 7113 YP 007124960 1 1173027 Cyanobacteria
Calothrix sp PCC 7507 YP 007067382 1 99598 Cyanobacteria
Anabaena sp 90 YP 006997641 1 46234 Cyanobacteria
Acaryochloris marina MBIC11017 YP 001519112 1 329726 Cyanobacteria
Chamaesiphon minutus PCC 6605 YP 007095599 1 1173020 Cyanobacteria
Gloeocapsa sp PCC 7428 YP 007129445 1 1173026 Cyanobacteria
Calothrix sp PCC 6303 YP 007138468 1 1170562 Cyanobacteria
Rivularia sp PCC 7116 YP 007057495 1 373994 Cyanobacteria
Sorangium cellulosum So0157 2 YP 008149316 1 1254432 Proteobacteria
Myxococcus fulvus HW 1 YP 004668868 1 483219 Proteobacteria
Corallococcus coralloides DSM 2259 YP 005370383 1 1144275 Proteobacteria
Myxococcus stipitatus DSM 14675 YP 007360591 1 1278073 Proteobacteria
Myxococcus stipitatus DSM 14675 YP 007361331 1 1278073 Proteobacteria
Gloeobacter kilaueensis JS1 YP 008712144 1 1183438 Cyanobacteria
Sorangium cellulosum So0157 2 YP 008155987 1 1254432 Proteobacteria
Candidatus Nitrospira defluvii YP 003797798 1 330214 Nitrospirae
Pseudomonas brassicacearum subsp brassicacearum NFM421 YP 004353672 1 994484 Proteobacteria
Pseudomonas poae RE 1 1 14 YP 007397403 1 1282356 Proteobacteria
Methylomonas methanica MC09 YP 004514400 1 857087 Proteobacteria
Methylomicrobium alcaliphilum 20Z YP 004916819 1 1091494 Proteobacteria
Pantoea sp At 9b YP 004117930 1 592316 Proteobacteria
Pandoraea pnomenusa 3kgm YP 008837494 1 1416914 Proteobacteria
Burkholderia phenoliruptrix BR3459a YP 006833513 1 1229205 Proteobacteria
Burkholderia sp CCGE1003 YP 003907467 1 640512 Proteobacteria
Xanthobacter autotrophicus Py2 YP 001417685 1 78245 Proteobacteria
Cupriavidus taiwanensis LMG 19424 YP 002007891 1 977880 Proteobacteria
Nitrosospira multiformis ATCC 25196 YP 412512 1 323848 Proteobacteria
Herbaspirillum seropedicae SmR1 YP 003775756 1 757424 Proteobacteria
Azotobacter vinelandii CA6 YP 007898690 1 1283331 Proteobacteria
Azotobacter vinelandii DJ YP 002799831 1 322710 Proteobacteria
Pectobacterium atrosepticum SCRI1043 YP 049591 1 218491 Proteobacteria
Collimonas fungivorans Ter331 YP 004752947 1 1005048 Proteobacteria
Burkholderia phytofirmans PsJN YP 001887825 1 398527 Proteobacteria
Burkholderia xenovorans LB400 YP 554773 1 266265 Proteobacteria
Burkholderia multivorans ATCC 17616 YP 001579784 1 395019 Proteobacteria
Burkholderia lata YP 369024 1 482957 Proteobacteria
Burkholderia cenocepacia J2315 YP 002230821 1 216591 Proteobacteria
Burkholderia cepacia GG4 YP 006615896 1 1009846 Proteobacteria
Burkholderia ambifaria MC40 6 YP 001808260 1 398577 Proteobacteria
Burkholderia vietnamiensis G4 YP 001119426 1 269482 Proteobacteria

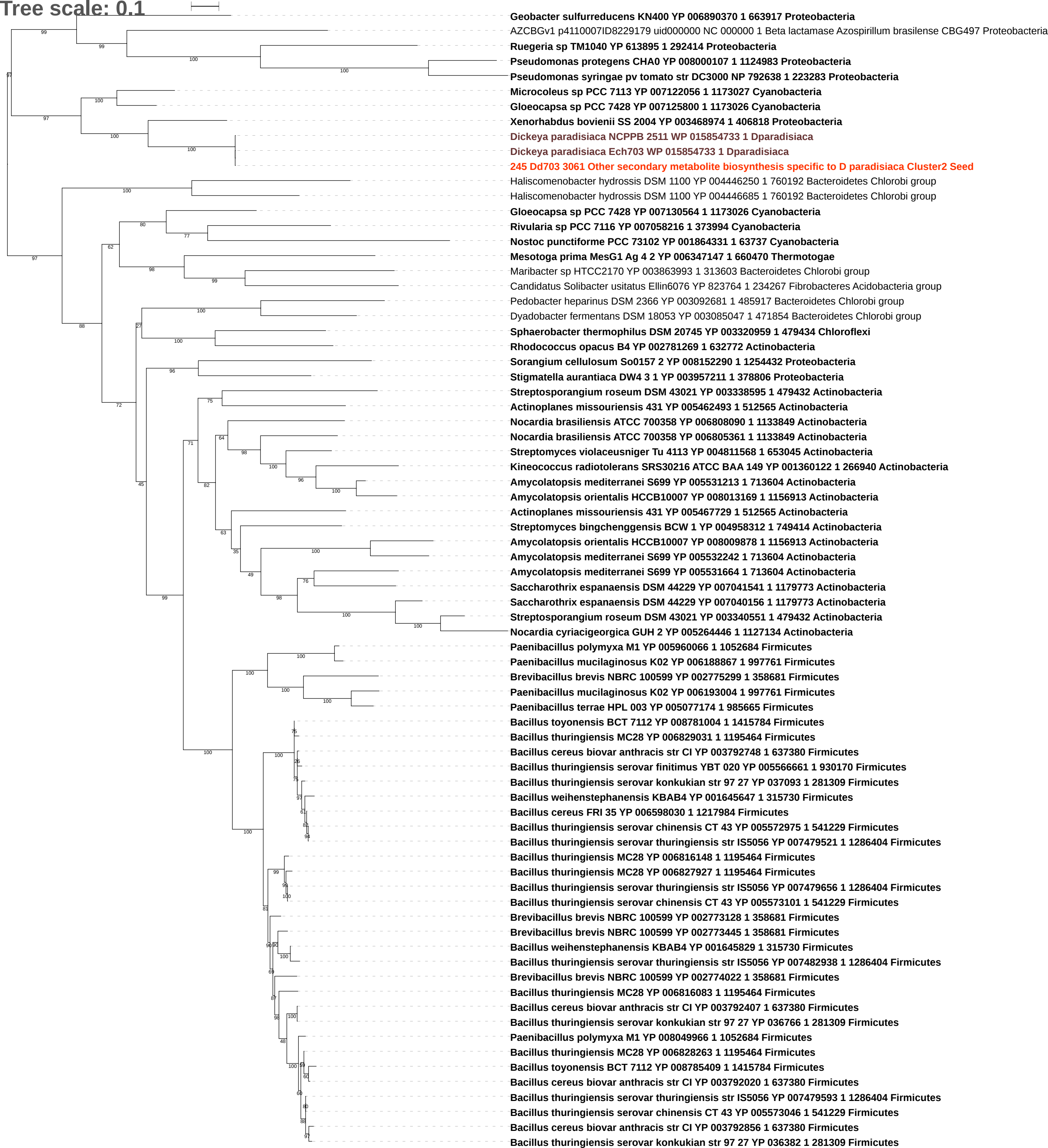

Tree scale: 0.1
Geobacter sulfurreducens KN400 YP 006890370 1 663917 Proteobacteria
AZCBGv1 p4110007ID8229179 uid000000 NC 000000 1 Beta lactamase Azospirillum brasilense CBG497 Proteobacteria
Ruegeria sp TM1040 YP 613895 1 292414 Proteobacteria
Pseudomonas protegens CHA0 YP 008000107 1 1124983 Proteobacteria
Pseudomonas syringae pv tomato str DC3000 NP 792638 1 223283 Proteobacteria
Microcoleus sp PCC 7113 YP 007122056 1 1173027 Cyanobacteria
Gloeocapsa sp PCC 7428 YP 007125800 1 1173026 Cyanobacteria
Xenorhabdus bovienii SS 2004 YP 003468974 1 406818 Proteobacteria
Dickeya paradisiaca NCPPB 2511 WP 015854733 1 Dparadisiaca
Dickeya paradisiaca Ech703 WP 015854733 1 Dparadisiaca
245 Dd703 3061 Other secondary metabolite biosynthesis specific to D paradisiaca Cluster2 Seed
Haliscomenobacter hydrossis DSM 1100 YP 004446250 1 760192 Bacteroidetes Chlorobi group
Haliscomenobacter hydrossis DSM 1100 YP 004446685 1 760192 Bacteroidetes Chlorobi group
Gloeocapsa sp PCC 7428 YP 007130564 1 1173026 Cyanobacteria
Rivularia sp PCC 7116 YP 007058216 1 373994 Cyanobacteria
Nostoc punctiforme PCC 73102 YP 001864331 1 63737 Cyanobacteria
Mesotoga prima MesG1 Ag 4 2 YP 006347147 1 660470 Thermotogae
Maribacter sp HTCC2170 YP 003863993 1 313603 Bacteroidetes Chlorobi group
Candidatus Solibacter usitatus Ellin6076 YP 823764 1 234267 Fibrobacteres Acidobacteria group
Pedobacter heparinus DSM 2366 YP 003092681 1 485917 Bacteroidetes Chlorobi group
Dyadobacter fermentans DSM 18053 YP 003085047 1 471854 Bacteroidetes Chlorobi group
Sphaerobacter thermophilus DSM 20745 YP 003320959 1 479434 Chloroflexi
Rhodococcus opacus B4 YP 002781269 1 632772 Actinobacteria
Sorangium cellulosum So0157 2 YP 008152290 1 1254432 Proteobacteria
Stigmatella aurantiaca DW4 3 1 YP 003957211 1 378806 Proteobacteria
Streptosporangium roseum DSM 43021 YP 003338595 1 479432 Actinobacteria
Actinoplanes missouriensis 431 YP 005462493 1 512565 Actinobacteria
Nocardia brasiliensis ATCC 700358 YP 006808090 1 1133849 Actinobacteria
Nocardia brasiliensis ATCC 700358 YP 006805361 1 1133849 Actinobacteria
Streptomyces violaceusniger Tu 4113 YP 004811568 1 653045 Actinobacteria
Kineococcus radiotolerans SRS30216 ATCC BAA 149 YP 001360122 1 266940 Actinobacteria
Amycolatopsis mediterranei S699 YP 005531213 1 713604 Actinobacteria
Amycolatopsis orientalis HCCB10007 YP 008013169 1 1156913 Actinobacteria
Actinoplanes missouriensis 431 YP 005467729 1 512565 Actinobacteria
Streptomyces bingchenggensis BCW 1 YP 004958312 1 749414 Actinobacteria
Amycolatopsis orientalis HCCB10007 YP 008009878 1 1156913 Actinobacteria
Amycolatopsis mediterranei S699 YP 005532242 1 713604 Actinobacteria
Amycolatopsis mediterranei S699 YP 005531664 1 713604 Actinobacteria
Saccharothrix espanaensis DSM 44229 YP 007041541 1 1179773 Actinobacteria
Saccharothrix espanaensis DSM 44229 YP 007040156 1 1179773 Actinobacteria
Streptosporangium roseum DSM 43021 YP 003340551 1 479432 Actinobacteria
Nocardia cyriacigeorgica GUH 2 YP 005264446 1 1127134 Actinobacteria
Paenibacillus polymyxa M1 YP 005960066 1 1052684 Firmicutes
Paenibacillus mucilaginosus K02 YP 006188867 1 997761 Firmicutes
Brevibacillus brevis NBRC 100599 YP 002775299 1 358681 Firmicutes
Paenibacillus mucilaginosus K02 YP 006193004 1 997761 Firmicutes
Paenibacillus terrae HPL 003 YP 005077174 1 985665 Firmicutes
Bacillus toyonensis BCT 7112 YP 008781004 1 1415784 Firmicutes
Bacillus thuringiensis MC28 YP 006829031 1 1195464 Firmicutes
Bacillus cereus biovar anthracis str CI YP 003792748 1 637380 Firmicutes
Bacillus thuringiensis serovar finitimus YBT 020 YP 005566661 1 930170 Firmicutes
Bacillus thuringiensis serovar konkukian str 97 27 YP 037093 1 281309 Firmicutes
Bacillus weihenstephanensis KBAB4 YP 001645647 1 315730 Firmicutes
Bacillus cereus FRI 35 YP 006598030 1 1217984 Firmicutes
Bacillus thuringiensis serovar chinensis CT 43 YP 005572975 1 541229 Firmicutes
Bacillus thuringiensis serovar thuringiensis str IS5056 YP 007479521 1 1286404 Firmicutes
Bacillus thuringiensis MC28 YP 006816148 1 1195464 Firmicutes
Bacillus thuringiensis MC28 YP 006827927 1 1195464 Firmicutes
Bacillus thuringiensis serovar thuringiensis str IS5056 YP 007479656 1 1286404 Firmicutes
Bacillus thuringiensis serovar chinensis CT 43 YP 005573101 1 541229 Firmicutes
Brevibacillus brevis NBRC 100599 YP 002773128 1 358681 Firmicutes
Brevibacillus brevis NBRC 100599 YP 002773445 1 358681 Firmicutes
Bacillus weihenstephanensis KBAB4 YP 001645829 1 315730 Firmicutes
Bacillus thuringiensis serovar thuringiensis str IS5056 YP 007482938 1 1286404 Firmicutes
Brevibacillus brevis NBRC 100599 YP 002774022 1 358681 Firmicutes
Bacillus thuringiensis MC28 YP 006816083 1 1195464 Firmicutes
Bacillus cereus biovar anthracis str CI YP 003792407 1 637380 Firmicutes
Bacillus thuringiensis serovar konkukian str 97 27 YP 036766 1 281309 Firmicutes
Paenibacillus polymyxa M1 YP 008049966 1 1052684 Firmicutes
Bacillus thuringiensis MC28 YP 006828263 1 1195464 Firmicutes
Bacillus toyonensis BCT 7112 YP 008785409 1 1415784 Firmicutes
Bacillus cereus biovar anthracis str CI YP 003792020 1 637380 Firmicutes
Bacillus thuringiensis serovar thuringiensis str IS5056 YP 007479593 1 1286404 Firmicutes
Bacillus thuringiensis serovar chinensis CT 43 YP 005573046 1 541229 Firmicutes
Bacillus cereus biovar anthracis str CI YP 003792856 1 637380 Firmicutes
Bacillus thuringiensis serovar konkukian str 97 27 YP 036382 1 281309 Firmicutes

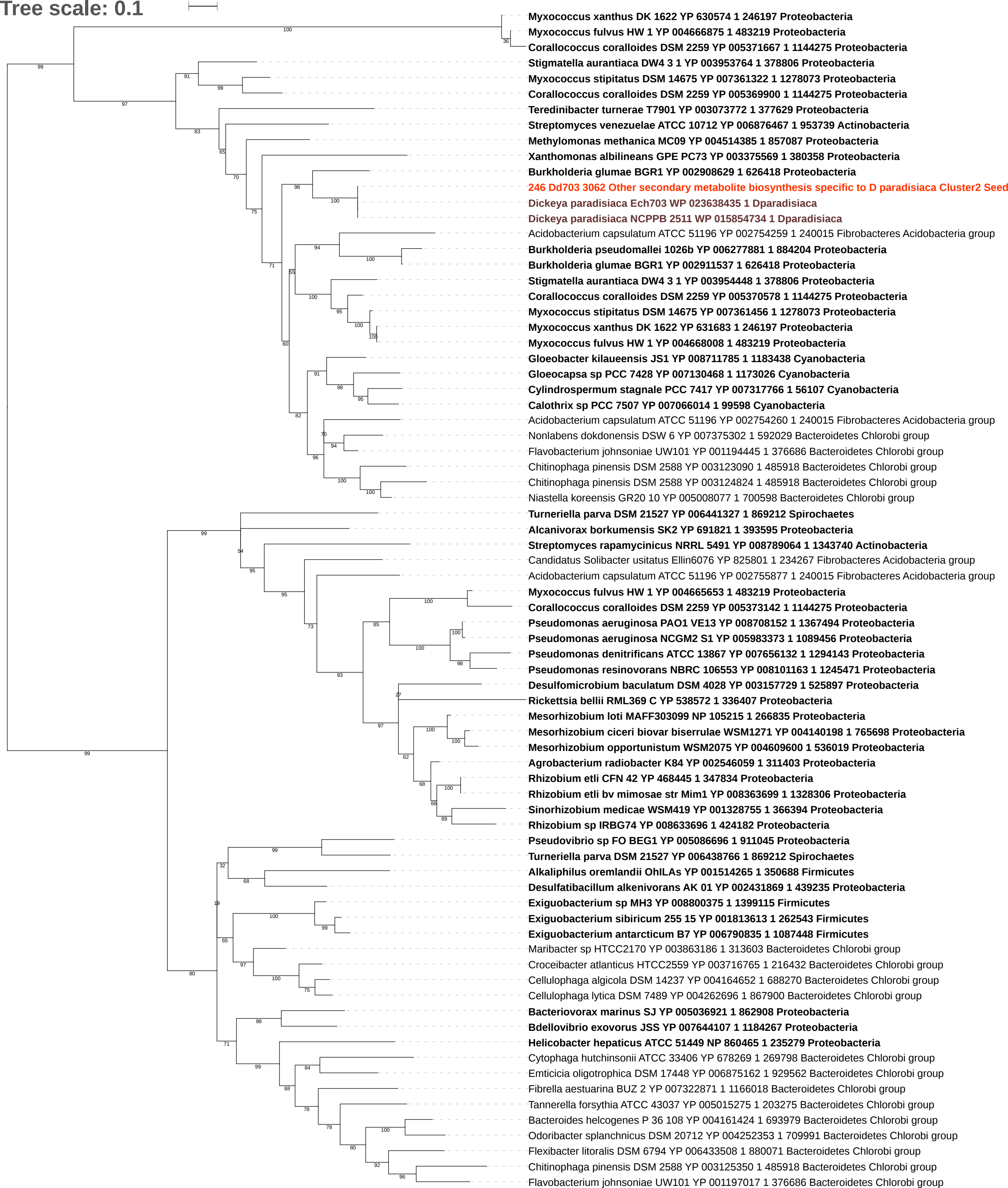

Tree scale: 0.1
Myxococcus xanthus DK 1622 YP 630574 1 246197 Proteobacteria
Myxococcus fulvus HW 1 YP 004666875 1 483219 Proteobacteria
Corallococcus coralloides DSM 2259 YP 005371667 1 1144275 Proteobacteria
Stigmatella aurantiaca DW4 3 1 YP 003953764 1 378806 Proteobacteria
Myxococcus stipitatus DSM 14675 YP 007361322 1 1278073 Proteobacteria
Corallococcus coralloides DSM 2259 YP 005369900 1 1144275 Proteobacteria
Teredinibacter turnerae T7901 YP 003073772 1 377629 Proteobacteria
Streptomyces venezuelae ATCC 10712 YP 006876467 1 953739 Actinobacteria
Methylomonas methanica MC09 YP 004514385 1 857087 Proteobacteria
Xanthomonas albilineans GPE PC73 YP 003375569 1 380358 Proteobacteria
Burkholderia glumae BGR1 YP 002908629 1 626418 Proteobacteria
246 Dd703 3062 Other secondary metabolite biosynthesis specific to D paradisiaca Cluster2 Seed
Dickeya paradisiaca Ech703 WP 023638435 1 Dparadisiaca
Dickeya paradisiaca NCPPB 2511 WP 015854734 1 Dparadisiaca
Acidobacterium capsulatum ATCC 51196 YP 002754259 1 240015 Fibrobacteres Acidobacteria group
Burkholderia pseudomallei 1026b YP 006277881 1 884204 Proteobacteria
Burkholderia glumae BGR1 YP 002911537 1 626418 Proteobacteria
Stigmatella aurantiaca DW4 3 1 YP 003954448 1 378806 Proteobacteria
Corallococcus coralloides DSM 2259 YP 005370578 1 1144275 Proteobacteria
Myxococcus stipitatus DSM 14675 YP 007361456 1 1278073 Proteobacteria
Myxococcus xanthus DK 1622 YP 631683 1 246197 Proteobacteria
Myxococcus fulvus HW 1 YP 004668008 1 483219 Proteobacteria
Gloeobacter kilaueensis JS1 YP 008711785 1 1183438 Cyanobacteria
Gloeocapsa sp PCC 7428 YP 007130468 1 1173026 Cyanobacteria
Cylindrospermum stagnale PCC 7417 YP 007317766 1 56107 Cyanobacteria
Calothrix sp PCC 7507 YP 007066014 1 99598 Cyanobacteria
Acidobacterium capsulatum ATCC 51196 YP 002754260 1 240015 Fibrobacteres Acidobacteria group
Nonlabens dokdonensis DSW 6 YP 007375302 1 592029 Bacteroidetes Chlorobi group
Flavobacterium johnsoniae UW101 YP 001194445 1 376686 Bacteroidetes Chlorobi group
Chitinophaga pinensis DSM 2588 YP 003123090 1 485918 Bacteroidetes Chlorobi group
Chitinophaga pinensis DSM 2588 YP 003124824 1 485918 Bacteroidetes Chlorobi group
Niastella koreensis GR20 10 YP 005008077 1 700598 Bacteroidetes Chlorobi group
Turneriella parva DSM 21527 YP 006441327 1 869212 Spirochaetes
Alcanivorax borkumensis SK2 YP 691821 1 393595 Proteobacteria
Streptomyces rapamycinicus NRRL 5491 YP 008789064 1 1343740 Actinobacteria
Candidatus Solibacter usitatus Ellin6076 YP 825801 1 234267 Fibrobacteres Acidobacteria group
Acidobacterium capsulatum ATCC 51196 YP 002755877 1 240015 Fibrobacteres Acidobacteria group
Myxococcus fulvus HW 1 YP 004665653 1 483219 Proteobacteria
Corallococcus coralloides DSM 2259 YP 005373142 1 1144275 Proteobacteria
Pseudomonas aeruginosa PAO1 VE13 YP 008708152 1 1367494 Proteobacteria
Pseudomonas aeruginosa NCGM2 S1 YP 005983373 1 1089456 Proteobacteria
Pseudomonas denitrificans ATCC 13867 YP 007656132 1 1294143 Proteobacteria
Pseudomonas resinovorans NBRC 106553 YP 008101163 1 1245471 Proteobacteria
Desulfomicrobium baculatum DSM 4028 YP 003157729 1 525897 Proteobacteria
Rickettsia bellii RML369 C YP 538572 1 336407 Proteobacteria
Mesorhizobium loti MAFF303099 NP 105215 1 266835 Proteobacteria
Mesorhizobium ciceri biovar biserrulae WSM1271 YP 004140198 1 765698 Proteobacteria
Mesorhizobium opportunistum WSM2075 YP 004609600 1 536019 Proteobacteria
Agrobacterium radiobacter K84 YP 002546059 1 311403 Proteobacteria
Rhizobium etli CFN 42 YP 468445 1 347834 Proteobacteria
Rhizobium etli bv mimosae str Mim1 YP 008363699 1 1328306 Proteobacteria
Sinorhizobium medicae WSM419 YP 001328755 1 366394 Proteobacteria
Rhizobium sp IRBG74 YP 008633696 1 424182 Proteobacteria
Pseudovibrio sp FO BEG1 YP 005086696 1 911045 Proteobacteria
Turneriella parva DSM 21527 YP 006438766 1 869212 Spirochaetes
Alkaliphilus oremlandii OhILAs YP 001514265 1 350688 Firmicutes
Desulfatibacillum alkenivorans AK 01 YP 002431869 1 439235 Proteobacteria
Exiguobacterium sp MH3 YP 008800375 1 1399115 Firmicutes
Exiguobacterium sibiricum 255 15 YP 001813613 1 262543 Firmicutes
Exiguobacterium antarcticum B7 YP 006790835 1 1087448 Firmicutes
Maribacter sp HTCC2170 YP 003863186 1 313603 Bacteroidetes Chlorobi group
Croceibacter atlanticus HTCC2559 YP 003716765 1 216432 Bacteroidetes Chlorobi group
Cellulophaga algicola DSM 14237 YP 004164652 1 688270 Bacteroidetes Chlorobi group
Cellulophaga lytica DSM 7489 YP 004262696 1 867900 Bacteroidetes Chlorobi group
Bacteriovorax marinus SJ YP 005036921 1 862908 Proteobacteria
Bdellovibrio exovorus JSS YP 007644107 1 1184267 Proteobacteria
Helicobacter hepaticus ATCC 51449 NP 860465 1 235279 Proteobacteria
Cytophaga hutchinsonii ATCC 33406 YP 678269 1 269798 Bacteroidetes Chlorobi group
Emticicia oligotrophica DSM 17448 YP 006875162 1 929562 Bacteroidetes Chlorobi group
Fibrella aestuarina BUZ 2 YP 007322871 1 1166018 Bacteroidetes Chlorobi group
Tannerella forsythia ATCC 43037 YP 005015275 1 203275 Bacteroidetes Chlorobi group
Bacteroides helcogenes P 36 108 YP 004161424 1 693979 Bacteroidetes Chlorobi group
Odoribacter splanchnicus DSM 20712 YP 004252353 1 709991 Bacteroidetes Chlorobi group
Flexibacter litoralis DSM 6794 YP 006433508 1 880071 Bacteroidetes Chlorobi group
Chitinophaga pinensis DSM 2588 YP 003125350 1 485918 Bacteroidetes Chlorobi group
Flavobacterium johnsoniae UW101 YP 001197017 1 376686 Bacteroidetes Chlorobi group

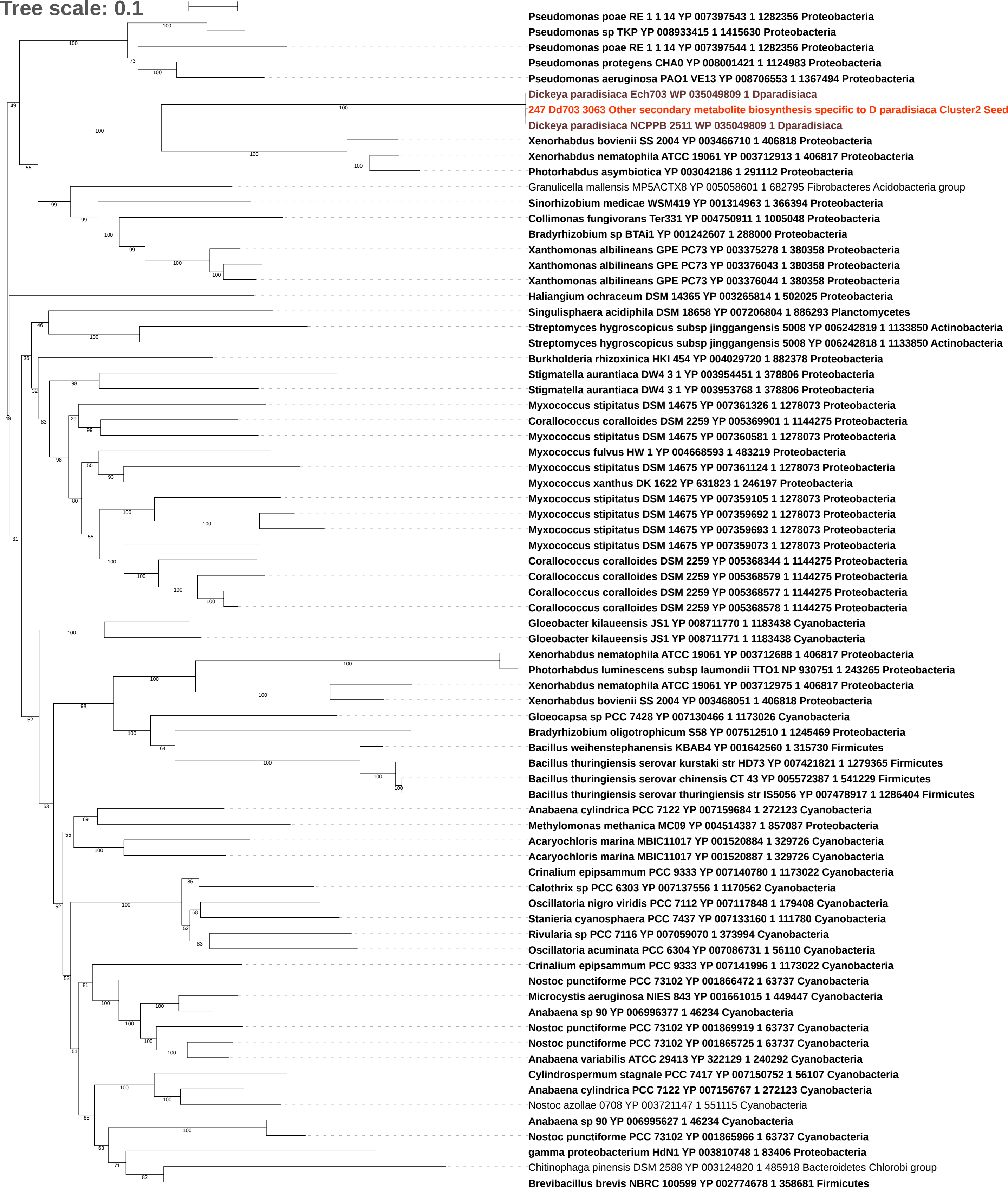

Tree scale: 0.1
Pseudomonas poae RE 1 1 14 YP 007397543 1 1282356 Proteobacteria
Pseudomonas sp TKP YP 008933415 1 1415630 Proteobacteria
Pseudomonas poae RE 1 1 14 YP 007397544 1 1282356 Proteobacteria
Pseudomonas protegens CHA0 YP 008001421 1 1124983 Proteobacteria
Pseudomonas aeruginosa PAO1 VE13 YP 008706553 1 1367494 Proteobacteria
Dickeya paradisiaca Ech703 WP 035049809 1 Dparadisiaca
247 Dd703 3063 Other secondary metabolite biosynthesis specific to D paradisiaca Cluster2 Seed
Dickeya paradisiaca NCPPB 2511 WP 035049809 1 Dparadisiaca
Xenorhabdus bovienii SS 2004 YP 003466710 1 406818 Proteobacteria
Xenorhabdus nematophila ATCC 19061 YP 003712913 1 406817 Proteobacteria
Photorhabdus asymbiotica YP 003042186 1 291112 Proteobacteria
Granulicella mallensis MP5ACTX8 YP 005058601 1 682795 Fibrobacteres Acidobacteria group
Sinorhizobium medicae WSM419 YP 001314963 1 366394 Proteobacteria
Collimonas fungivorans Ter331 YP 004750911 1 1005048 Proteobacteria
Bradyrhizobium sp BTAi1 YP 001242607 1 288000 Proteobacteria
Xanthomonas albilineans GPE PC73 YP 003375278 1 380358 Proteobacteria
Xanthomonas albilineans GPE PC73 YP 003376043 1 380358 Proteobacteria
Xanthomonas albilineans GPE PC73 YP 003376044 1 380358 Proteobacteria
Haliangium ochraceum DSM 14365 YP 003265814 1 502025 Proteobacteria
Singulisphaera acidiphila DSM 18658 YP 007206804 1 886293 Planctomycetes
Streptomyces hygroscopicus subsp jinggangensis 5008 YP 006242819 1 1133850 Actinobacteria
Streptomyces hygroscopicus subsp jinggangensis 5008 YP 006242818 1 1133850 Actinobacteria
Burkholderia rhizoxinica HKI 454 YP 004029720 1 882378 Proteobacteria
Stigmatella aurantiaca DW4 3 1 YP 003954451 1 378806 Proteobacteria
Stigmatella aurantiaca DW4 3 1 YP 003953768 1 378806 Proteobacteria
Myxococcus stipitatus DSM 14675 YP 007361326 1 1278073 Proteobacteria
Corallococcus coralloides DSM 2259 YP 005369901 1 1144275 Proteobacteria
Myxococcus stipitatus DSM 14675 YP 007360581 1 1278073 Proteobacteria
Myxococcus fulvus HW 1 YP 004668593 1 483219 Proteobacteria
Myxococcus stipitatus DSM 14675 YP 007361124 1 1278073 Proteobacteria
Myxococcus xanthus DK 1622 YP 631823 1 246197 Proteobacteria
Myxococcus stipitatus DSM 14675 YP 007359105 1 1278073 Proteobacteria
Myxococcus stipitatus DSM 14675 YP 007359692 1 1278073 Proteobacteria
Myxococcus stipitatus DSM 14675 YP 007359693 1 1278073 Proteobacteria
Myxococcus stipitatus DSM 14675 YP 007359073 1 1278073 Proteobacteria
Corallococcus coralloides DSM 2259 YP 005368344 1 1144275 Proteobacteria
Corallococcus coralloides DSM 2259 YP 005368579 1 1144275 Proteobacteria
Corallococcus coralloides DSM 2259 YP 005368577 1 1144275 Proteobacteria
Corallococcus coralloides DSM 2259 YP 005368578 1 1144275 Proteobacteria
Gloeobacter kilaueensis JS1 YP 008711770 1 1183438 Cyanobacteria
Gloeobacter kilaueensis JS1 YP 008711771 1 1183438 Cyanobacteria
Xenorhabdus nematophila ATCC 19061 YP 003712688 1 406817 Proteobacteria
Photorhabdus luminescens subsp laumondii TTO1 NP 930751 1 243265 Proteobacteria
Xenorhabdus nematophila ATCC 19061 YP 003712975 1 406817 Proteobacteria
Xenorhabdus bovienii SS 2004 YP 003468051 1 406818 Proteobacteria
Gloeocapsa sp PCC 7428 YP 007130466 1 1173026 Cyanobacteria
Bradyrhizobium oligotrophicum S58 YP 007512510 1 1245469 Proteobacteria
Bacillus weihenstephanensis KBAB4 YP 001642560 1 315730 Firmicutes
Bacillus thuringiensis serovar kurstaki str HD73 YP 007421821 1 1279365 Firmicutes
Bacillus thuringiensis serovar chinensis CT 43 YP 005572387 1 541229 Firmicutes
Bacillus thuringiensis serovar thuringiensis str IS5056 YP 007478917 1 1286404 Firmicutes
Anabaena cylindrica PCC 7122 YP 007159684 1 272123 Cyanobacteria
Methylomonas methanica MC09 YP 004514387 1 857087 Proteobacteria
Acaryochloris marina MBIC11017 YP 001520884 1 329726 Cyanobacteria
Acaryochloris marina MBIC11017 YP 001520887 1 329726 Cyanobacteria
Crinalium epipsammum PCC 9333 YP 007140780 1 1173022 Cyanobacteria
Calothrix sp PCC 6303 YP 007137556 1 1170562 Cyanobacteria
Oscillatoria nigro viridis PCC 7112 YP 007117848 1 179408 Cyanobacteria
Stanieria cyanosphaera PCC 7437 YP 007133160 1 111780 Cyanobacteria
Rivularia sp PCC 7116 YP 007059070 1 373994 Cyanobacteria
Oscillatoria acuminata PCC 6304 YP 007086731 1 56110 Cyanobacteria
Crinalium epipsammum PCC 9333 YP 007141996 1 1173022 Cyanobacteria
Nostoc punctiforme PCC 73102 YP 001866472 1 63737 Cyanobacteria
Microcystis aeruginosa NIES 843 YP 001661015 1 449447 Cyanobacteria
Anabaena sp 90 YP 006996377 1 46234 Cyanobacteria
Nostoc punctiforme PCC 73102 YP 001869919 1 63737 Cyanobacteria
Nostoc punctiforme PCC 73102 YP 001865725 1 63737 Cyanobacteria
Anabaena variabilis ATCC 29413 YP 322129 1 240292 Cyanobacteria
Cylindrospermum stagnale PCC 7417 YP 007150752 1 56107 Cyanobacteria
Anabaena cylindrica PCC 7122 YP 007156767 1 272123 Cyanobacteria
Nostoc azollae 0708 YP 003721147 1 551115 Cyanobacteria
Anabaena sp 90 YP 006995627 1 46234 Cyanobacteria
Nostoc punctiforme PCC 73102 YP 001865966 1 63737 Cyanobacteria
gamma proteobacterium HdN1 YP 003810748 1 83406 Proteobacteria
Chitinophaga pinensis DSM 2588 YP 003124820 1 485918 Bacteroidetes Chlorobi group
Brevibacillus brevis NBRC 100599 YP 002774678 1 358681 Firmicutes

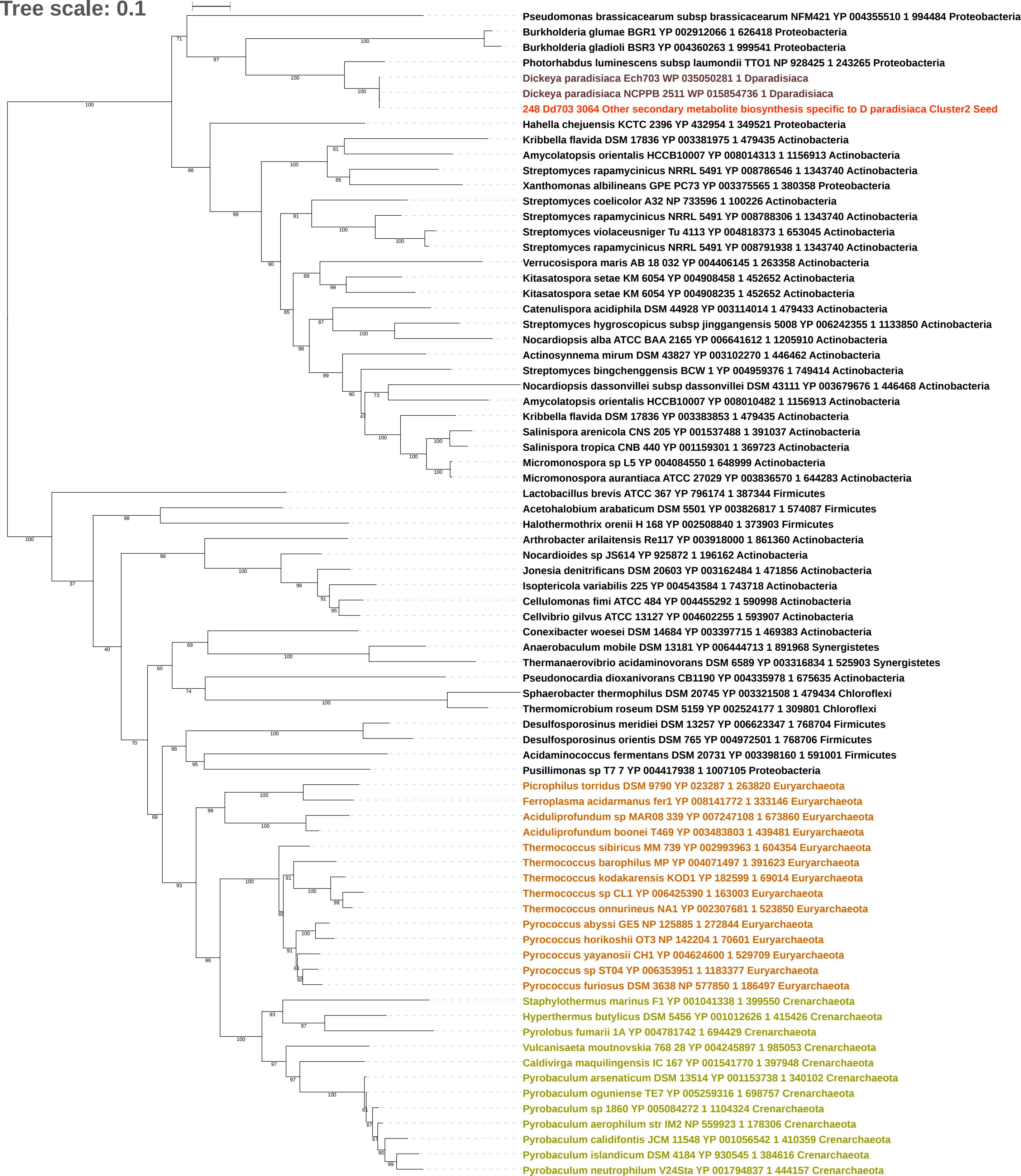
Tree scale: 0.1
Pseudomonas brassicacearum subsp brassicacearum NFM421 YP 004355510 1 994484 Proteobacteria
Burkholderia glumae BGR1 YP 002912066 1 626418 Proteobacteria
Burkholderia gladioli BSR3 YP 004360263 1 999541 Proteobacteria
Photorhabdus luminescens subsp laumondii TTO1 NP 928425 1 243265 Proteobacteria
Dickeya paradisiaca Ech703 WP 035050281 1 Dparadisiaca
Dickeya paradisiaca NCPPB 2511 WP 015854736 1 Dparadisiaca
248 Dd703 3064 Other secondary metabolite biosynthesis specific to D paradisiaca Cluster2 Seed
Hahella chejuensis KCTC 2396 YP 432954 1 349521 Proteobacteria
Kribbella flavida DSM 17836 YP 003381975 1 479435 Actinobacteria
Amycolatopsis orientalis HCCB10007 YP 008014313 1 1156913 Actinobacteria
Streptomyces rapamycinicus NRRL 5491 YP 008786546 1 1343740 Actinobacteria
Xanthomonas albilineans GPE PC73 YP 003375565 1 380358 Proteobacteria
Streptomyces coelicolor A32 NP 733596 1 100226 Actinobacteria
Streptomyces rapamycinicus NRRL 5491 YP 008788306 1 1343740 Actinobacteria
Streptomyces violaceusniger Tu 4113 YP 004818373 1 653045 Actinobacteria
Streptomyces rapamycinicus NRRL 5491 YP 008791938 1 1343740 Actinobacteria
Verrucosispora maris AB 18 032 YP 004406145 1 263358 Actinobacteria
Kitasatospora setae KM 6054 YP 004908458 1 452652 Actinobacteria
Kitasatospora setae KM 6054 YP 004908235 1 452652 Actinobacteria
Catenulispora acidiphila DSM 44928 YP 003114014 1 479433 Actinobacteria
Streptomyces hygroscopicus subsp jinggangensis 5008 YP 006242355 1 1133850 Actinobacteria
Nocardiopsis alba ATCC BAA 2165 YP 006641612 1 1205910 Actinobacteria
Actinosynnema mirum DSM 43827 YP 003102270 1 446462 Actinobacteria
Streptomyces bingchenggensis BCW 1 YP 004959376 1 749414 Actinobacteria
Nocardiopsis dassonvillei subsp dassonvillei DSM 43111 YP 003679676 1 446468 Actinobacteria
Amycolatopsis orientalis HCCB10007 YP 008010482 1 1156913 Actinobacteria
Kribbella flavida DSM 17836 YP 003383853 1 479435 Actinobacteria
Salinispora arenicola CNS 205 YP 001537488 1 391037 Actinobacteria
Salinispora tropica CNB 440 YP 001159301 1 369723 Actinobacteria
Micromonospora sp L5 YP 004084550 1 648999 Actinobacteria
Micromonospora aurantiaca ATCC 27029 YP 003836570 1 644283 Actinobacteria
Lactobacillus brevis ATCC 367 YP 796174 1 387344 Firmicutes
Acetohalobium arabaticum DSM 5501 YP 003826817 1 574087 Firmicutes
Halothermothrix orenii H 168 YP 002508840 1 373903 Firmicutes
Arthrobacter arilaitensis Re117 YP 003918000 1 861360 Actinobacteria
Nocardioides sp JS614 YP 925872 1 196162 Actinobacteria
Jonesia denitrificans DSM 20603 YP 003162484 1 471856 Actinobacteria
Isoptericola variabilis 225 YP 004543584 1 743718 Actinobacteria
Cellulomonas fimi ATCC 484 YP 004455292 1 590998 Actinobacteria
Cellvibrio gilvus ATCC 13127 YP 004602255 1 593907 Actinobacteria
Conexibacter woesei DSM 14684 YP 003397715 1 469383 Actinobacteria
Anaerobaculum mobile DSM 13181 YP 006444713 1 891968 Synergistetes
Thermanaerovibrio acidaminovorans DSM 6589 YP 003316834 1 525903 Synergistetes
Pseudonocardia dioxanivorans CB1190 YP 004335978 1 675635 Actinobacteria
Sphaerobacter thermophilus DSM 20745 YP 003321508 1 479434 Chloroflexi
Thermomicrobium roseum DSM 5159 YP 002524177 1 309801 Chloroflexi
Desulfosporosinus meridiei DSM 13257 YP 006623347 1 768704 Firmicutes
Desulfosporosinus orientis DSM 765 YP 004972501 1 768706 Firmicutes
Acidaminococcus fermentans DSM 20731 YP 003398160 1 591001 Firmicutes
Pusillimonas sp T7 7 YP 004417938 1 1007105 Proteobacteria
Picrophilus torridus DSM 9790 YP 023287 1 263820 Euryarchaeota
Ferroplasma acidarmanus fer1 YP 008141772 1 333146 Euryarchaeota
Aciduliprofundum sp MAR08 339 YP 007247108 1 673860 Euryarchaeota
Aciduliprofundum boonei T469 YP 003483803 1 439481 Euryarchaeota
Thermococcus sibiricus MM 739 YP 002993963 1 604354 Euryarchaeota
Thermococcus barophilus MP YP 004071497 1 391623 Euryarchaeota
Thermococcus kodakarensis KOD1 YP 182599 1 69014 Euryarchaeota
Thermococcus sp CL1 YP 006425390 1 163003 Euryarchaeota
Thermococcus onnurineus NA1 YP 002307681 1 523850 Euryarchaeota
Pyrococcus abyssi GE5 NP 125885 1 272844 Euryarchaeota
Pyrococcus horikoshii OT3 NP 142204 1 70601 Euryarchaeota
Pyrococcus yayanosii CH1 YP 004624600 1 529709 Euryarchaeota
Pyrococcus sp ST04 YP 006353951 1 1183377 Euryarchaeota
Pyrococcus furiosus DSM 3638 NP 577850 1 186497 Euryarchaeota
Staphylothermus marinus F1 YP 001041338 1 399550 Crenarchaeota
Hyperthermus butylicus DSM 5456 YP 001012626 1 415426 Crenarchaeota
Pyrolobus fumarii 1A YP 004781742 1 694429 Crenarchaeota
Vulcanisaeta moutnovskia 768 28 YP 004245897 1 985053 Crenarchaeota
Caldivirga maquilingensis IC 167 YP 001541770 1 397948 Crenarchaeota
Pyrobaculum arsenaticum DSM 13514 YP 001153738 1 340102 Crenarchaeota
Pyrobaculum oguniense TE7 YP 005259316 1 698757 Crenarchaeota
Pyrobaculum sp 1860 YP 005084272 1 1104324 Crenarchaeota
Pyrobaculum aerophilum str IM2 NP 559923 1 178306 Crenarchaeota
Pyrobaculum calidifontis JCM 11548 YP 001056542 1 410359 Crenarchaeota
Pyrobaculum islandicum DSM 4184 YP 930545 1 384616 Crenarchaeota
Pyrobaculum neutrophilum V24Sta YP 001794837 1 444157 Crenarchaeota

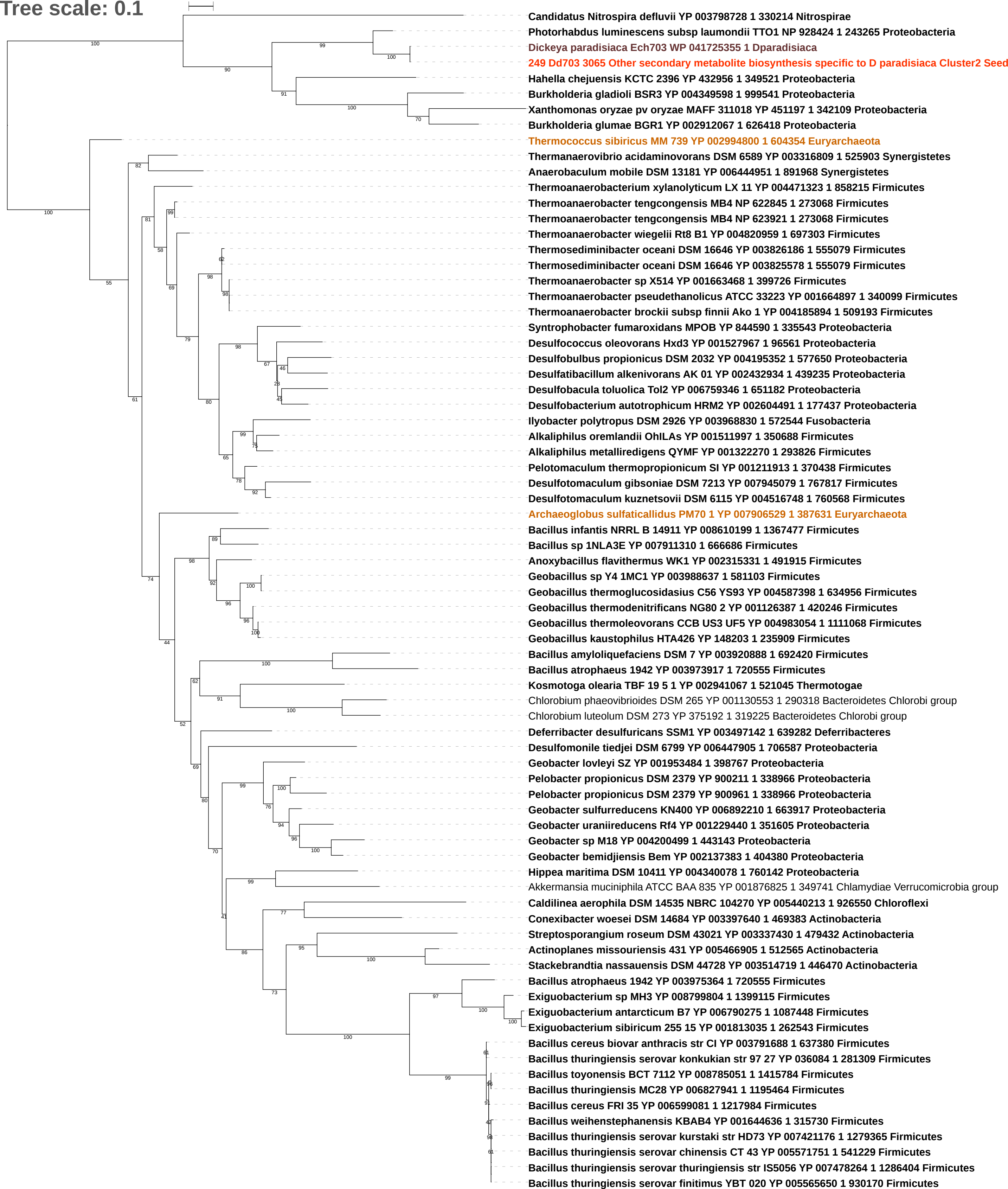
Tree scale: 0.1
Candidatus Nitrospira defluvii YP 003798728 1 330214 Nitrospirae
Photorhabdus luminescens subsp laumondii TTO1 NP 928424 1 243265 Proteobacteria
Dickeya paradisiaca Ech703 WP 041725355 1 Dparadisiaca
249 Dd703 3065 Other secondary metabolite biosynthesis specific to D paradisiaca Cluster2 Seed
Hahella chejuensis KCTC 2396 YP 432956 1 349521 Proteobacteria
Burkholderia gladioli BSR3 YP 004349598 1 999541 Proteobacteria
Xanthomonas oryzae pv oryzae MAFF 311018 YP 451197 1 342109 Proteobacteria
Burkholderia glumae BGR1 YP 002912067 1 626418 Proteobacteria
Thermococcus sibiricus MM 739 YP 002994800 1 604354 Euryarchaeota
Thermanaerovibrio acidaminovorans DSM 6589 YP 003316809 1 525903 Synergistetes
Anaerobaculum mobile DSM 13181 YP 006444951 1 891968 Synergistetes
Thermoanaerobacterium xylanolyticum LX 11 YP 004471323 1 858215 Firmicutes
Thermoanaerobacter tengcongensis MB4 NP 622845 1 273068 Firmicutes
Thermoanaerobacter tengcongensis MB4 NP 623921 1 273068 Firmicutes
Thermoanaerobacter wiegelii Rt8 B1 YP 004820959 1 697303 Firmicutes
Thermosediminibacter oceani DSM 16646 YP 003826186 1 555079 Firmicutes
Thermosediminibacter oceani DSM 16646 YP 003825578 1 555079 Firmicutes
Thermoanaerobacter sp X514 YP 001663468 1 399726 Firmicutes
Thermoanaerobacter pseudethanolicus ATCC 33223 YP 001664897 1 340099 Firmicutes
Thermoanaerobacter brockii subsp finnii Ako 1 YP 004185894 1 509193 Firmicutes
Syntrophobacter fumaroxidans MPOB YP 844590 1 335543 Proteobacteria
Desulfococcus oleovorans Hxd3 YP 001527967 1 96561 Proteobacteria
Desulfobulbus propionicus DSM 2032 YP 004195352 1 577650 Proteobacteria
Desulfatibacillum alkenivorans AK 01 YP 002432934 1 439235 Proteobacteria
Desulfobacula toluolica Tol2 YP 006759346 1 651182 Proteobacteria
Desulfobacterium autotrophicum HRM2 YP 002604491 1 177437 Proteobacteria
Ilyobacter polytropus DSM 2926 YP 003968830 1 572544 Fusobacteria
Alkaliphilus oremlandii OhILAs YP 001511997 1 350688 Firmicutes
Alkaliphilus metalliredigens QYMF YP 001322270 1 293826 Firmicutes
Pelotomaculum thermopropionicum SI YP 001211913 1 370438 Firmicutes
Desulfotomaculum gibsoniae DSM 7213 YP 007945079 1 767817 Firmicutes
Desulfotomaculum kuznetsovii DSM 6115 YP 004516748 1 760568 Firmicutes
Archaeoglobus sulfaticallidus PM70 1 YP 007906529 1 387631 Euryarchaeota
Bacillus infantis NRRL B 14911 YP 008610199 1 1367477 Firmicutes
Bacillus sp 1NLA3E YP 007911310 1 666686 Firmicutes
Anoxybacillus flavithermus WK1 YP 002315331 1 491915 Firmicutes
Geobacillus sp Y4 1MC1 YP 003988637 1 581103 Firmicutes
Geobacillus thermoglucosidasius C56 YS93 YP 004587398 1 634956 Firmicutes
Geobacillus thermodenitrificans NG80 2 YP 001126387 1 420246 Firmicutes
Geobacillus thermoleovorans CCB US3 UF5 YP 004983054 1 1111068 Firmicutes
Geobacillus kaustophilus HTA426 YP 148203 1 235909 Firmicutes
Bacillus amyloliquefaciens DSM 7 YP 003920888 1 692420 Firmicutes
Bacillus atrophaeus 1942 YP 003973917 1 720555 Firmicutes
Kosmotoga olearia TBF 19 5 1 YP 002941067 1 521045 Thermotogae
Chlorobium phaeovibrioides DSM 265 YP 001130553 1 290318 Bacteroidetes Chlorobi group
Chlorobium luteolum DSM 273 YP 375192 1 319225 Bacteroidetes Chlorobi group
Deferribacter desulfuricans SSM1 YP 003497142 1 639282 Deferribacteres
Desulfomonile tiedjei DSM 6799 YP 006447905 1 706587 Proteobacteria
Geobacter lovleyi SZ YP 001953484 1 398767 Proteobacteria
Pelobacter propionicus DSM 2379 YP 900211 1 338966 Proteobacteria
Pelobacter propionicus DSM 2379 YP 900961 1 338966 Proteobacteria
Geobacter sulfurreducens KN400 YP 006892210 1 663917 Proteobacteria
Geobacter uraniireducens Rf4 YP 001229440 1 351605 Proteobacteria
Geobacter sp M18 YP 004200499 1 443143 Proteobacteria
Geobacter bemidjiensis Bem YP 002137383 1 404380 Proteobacteria
Hippea maritima DSM 10411 YP 004340078 1 760142 Proteobacteria
Akkermansia muciniphila ATCC BAA 835 YP 001876825 1 349741 Chlamydiae Verrucomicrobia group
Caldilinea aerophila DSM 14535 NBRC 104270 YP 005440213 1 926550 Chloroflexi
Conexibacter woesei DSM 14684 YP 003397640 1 469383 Actinobacteria
Streptosporangium roseum DSM 43021 YP 003337430 1 479432 Actinobacteria
Actinoplanes missouriensis 431 YP 005466905 1 512565 Actinobacteria
Stackebrandtia nassauensis DSM 44728 YP 003514719 1 446470 Actinobacteria
Bacillus atrophaeus 1942 YP 003975364 1 720555 Firmicutes
Exiguobacterium sp MH3 YP 008799804 1 1399115 Firmicutes
Exiguobacterium antarcticum B7 YP 006790275 1 1087448 Firmicutes
Exiguobacterium sibiricum 255 15 YP 001813035 1 262543 Firmicutes
Bacillus cereus biovar anthracis str CI YP 003791688 1 637380 Firmicutes
Bacillus thuringiensis serovar konkukian str 97 27 YP 036084 1 281309 Firmicutes
Bacillus toyonensis BCT 7112 YP 008785051 1 1415784 Firmicutes
Bacillus thuringiensis MC28 YP 006827941 1 1195464 Firmicutes
Bacillus cereus FRI 35 YP 006599081 1 1217984 Firmicutes
Bacillus weihenstephanensis KBAB4 YP 001644636 1 315730 Firmicutes
Bacillus thuringiensis serovar kurstaki str HD73 YP 007421176 1 1279365 Firmicutes
Bacillus thuringiensis serovar chinensis CT 43 YP 005571751 1 541229 Firmicutes
Bacillus thuringiensis serovar thuringiensis str IS5056 YP 007478264 1 1286404 Firmicutes
Bacillus thuringiensis serovar finitimus YBT 020 YP 005565650 1 930170 Firmicutes

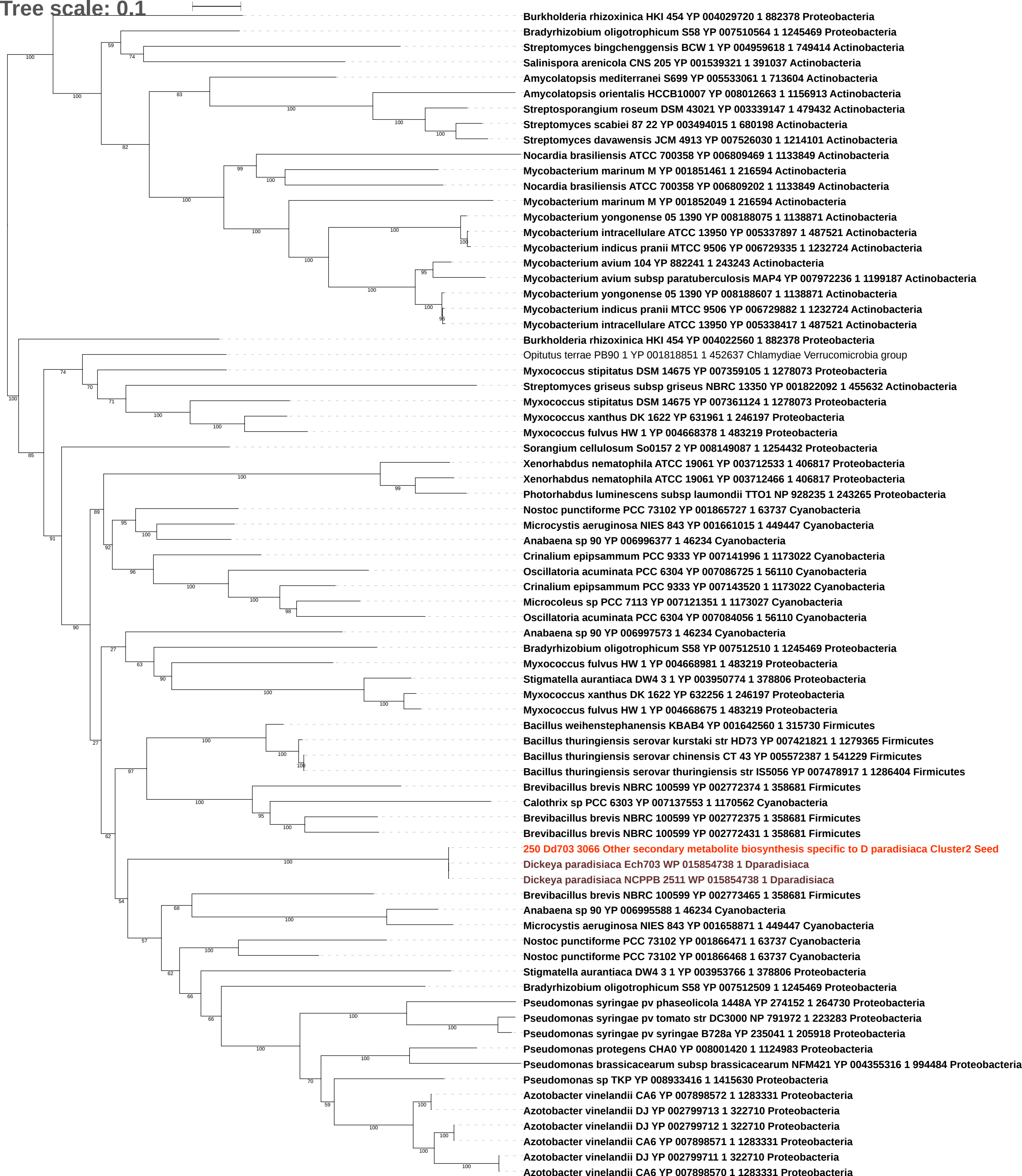

Tree scale: 0.1
Burkholderia rhizoxinica HKI 454 YP 004029720 1 882378 Proteobacteria
Bradyrhizobium oligotrophicum S58 YP 007510564 1 1245469 Proteobacteria
Streptomyces bingchenggensis BCW 1 YP 004959618 1 749414 Actinobacteria
Salinispora arenicola CNS 205 YP 001539321 1 391037 Actinobacteria
Amycolatopsis mediterranei S699 YP 005533061 1 713604 Actinobacteria
Amycolatopsis orientalis HCCB10007 YP 008012663 1 1156913 Actinobacteria
Streptosporangium roseum DSM 43021 YP 003339147 1 479432 Actinobacteria
Streptomyces scabiei 87 22 YP 003494015 1 680198 Actinobacteria
Streptomyces davawensis JCM 4913 YP 007526030 1 1214101 Actinobacteria
Nocardia brasiliensis ATCC 700358 YP 006809469 1 1133849 Actinobacteria
Mycobacterium marinum M YP 001851461 1 216594 Actinobacteria
Nocardia brasiliensis ATCC 700358 YP 006809202 1 1133849 Actinobacteria
Mycobacterium marinum M YP 001852049 1 216594 Actinobacteria
Mycobacterium yongonense 05 1390 YP 008188075 1 1138871 Actinobacteria
Mycobacterium intracellulare ATCC 13950 YP 005337897 1 487521 Actinobacteria
Mycobacterium indicus pranii MTCC 9506 YP 006729335 1 1232724 Actinobacteria
Mycobacterium avium 104 YP 882241 1 243243 Actinobacteria
Mycobacterium avium subsp paratuberculosis MAP4 YP 007972236 1 1199187 Actinobacteria
Mycobacterium yongonense 05 1390 YP 008188607 1 1138871 Actinobacteria
Mycobacterium indicus pranii MTCC 9506 YP 006729882 1 1232724 Actinobacteria
Mycobacterium intracellulare ATCC 13950 YP 005338417 1 487521 Actinobacteria
Burkholderia rhizoxinica HKI 454 YP 004022560 1 882378 Proteobacteria
Opitutus terrae PB90 1 YP 001818851 1 452637 Chlamydiae Verrucomicrobia group
Myxococcus stipitatus DSM 14675 YP 007359105 1 1278073 Proteobacteria
Streptomyces griseus subsp griseus NBRC 13350 YP 001822092 1 455632 Actinobacteria
Myxococcus stipitatus DSM 14675 YP 007361124 1 1278073 Proteobacteria
Myxococcus xanthus DK 1622 YP 631961 1 246197 Proteobacteria
Myxococcus fulvus HW 1 YP 004668378 1 483219 Proteobacteria
Sorangium cellulosum So0157 2 YP 008149087 1 1254432 Proteobacteria
Xenorhabdus nematophila ATCC 19061 YP 003712533 1 406817 Proteobacteria
Xenorhabdus nematophila ATCC 19061 YP 003712466 1 406817 Proteobacteria
Photorhabdus luminescens subsp laumondii TTO1 NP 928235 1 243265 Proteobacteria
Nostoc punctiforme PCC 73102 YP 001865727 1 63737 Cyanobacteria
Microcystis aeruginosa NIES 843 YP 001661015 1 449447 Cyanobacteria
Anabaena sp 90 YP 006996377 1 46234 Cyanobacteria
Crinalium epipsammum PCC 9333 YP 007141996 1 1173022 Cyanobacteria
Oscillatoria acuminata PCC 6304 YP 007086725 1 56110 Cyanobacteria
Crinalium epipsammum PCC 9333 YP 007143520 1 1173022 Cyanobacteria
Microcoleus sp PCC 7113 YP 007121351 1 1173027 Cyanobacteria
Oscillatoria acuminata PCC 6304 YP 007084056 1 56110 Cyanobacteria
Anabaena sp 90 YP 006997573 1 46234 Cyanobacteria
Bradyrhizobium oligotrophicum S58 YP 007512510 1 1245469 Proteobacteria
Myxococcus fulvus HW 1 YP 004668981 1 483219 Proteobacteria
Stigmatella aurantiaca DW4 3 1 YP 003950774 1 378806 Proteobacteria
Myxococcus xanthus DK 1622 YP 632256 1 246197 Proteobacteria
Myxococcus fulvus HW 1 YP 004668675 1 483219 Proteobacteria
Bacillus weihenstephanensis KBAB4 YP 001642560 1 315730 Firmicutes
Bacillus thuringiensis serovar kurstaki str HD73 YP 007421821 1 1279365 Firmicutes
Bacillus thuringiensis serovar chinensis CT 43 YP 005572387 1 541229 Firmicutes
Bacillus thuringiensis serovar thuringiensis str IS5056 YP 007478917 1 1286404 Firmicutes
Brevibacillus brevis NBRC 100599 YP 002772374 1 358681 Firmicutes
Calothrix sp PCC 6303 YP 007137553 1 1170562 Cyanobacteria
Brevibacillus brevis NBRC 100599 YP 002772375 1 358681 Firmicutes
Brevibacillus brevis NBRC 100599 YP 002772431 1 358681 Firmicutes
250 Dd703 3066 Other secondary metabolite biosynthesis specific to D paradisiaca Cluster2 Seed
Dickeya paradisiaca Ech703 WP 015854738 1 Dparadisiaca
Dickeya paradisiaca NCPPB 2511 WP 015854738 1 Dparadisiaca
Brevibacillus brevis NBRC 100599 YP 002773465 1 358681 Firmicutes
Anabaena sp 90 YP 006995588 1 46234 Cyanobacteria
Microcystis aeruginosa NIES 843 YP 001658871 1 449447 Cyanobacteria
Nostoc punctiforme PCC 73102 YP 001866471 1 63737 Cyanobacteria
Nostoc punctiforme PCC 73102 YP 001866468 1 63737 Cyanobacteria
Stigmatella aurantiaca DW4 3 1 YP 003953766 1 378806 Proteobacteria
Bradyrhizobium oligotrophicum S58 YP 007512509 1 1245469 Proteobacteria
Pseudomonas syringae pv phaseolicola 1448A YP 274152 1 264730 Proteobacteria
Pseudomonas syringae pv tomato str DC3000 NP 791972 1 223283 Proteobacteria
Pseudomonas syringae pv syringae B728a YP 235041 1 205918 Proteobacteria
Pseudomonas protegens CHA0 YP 008001420 1 1124983 Proteobacteria
Pseudomonas brassicacearum subsp brassicacearum NFM421 YP 004355316 1 994484 Proteobacteria
Pseudomonas sp TKP YP 008933416 1 1415630 Proteobacteria
Azotobacter vinelandii CA6 YP 007898572 1 1283331 Proteobacteria
Azotobacter vinelandii DJ YP 002799713 1 322710 Proteobacteria
Azotobacter vinelandii DJ YP 002799712 1 322710 Proteobacteria
Azotobacter vinelandii CA6 YP 007898571 1 1283331 Proteobacteria
Azotobacter vinelandii DJ YP 002799711 1 322710 Proteobacteria
Azotobacter vinelandii CA6 YP 007898570 1 1283331 Proteobacteria

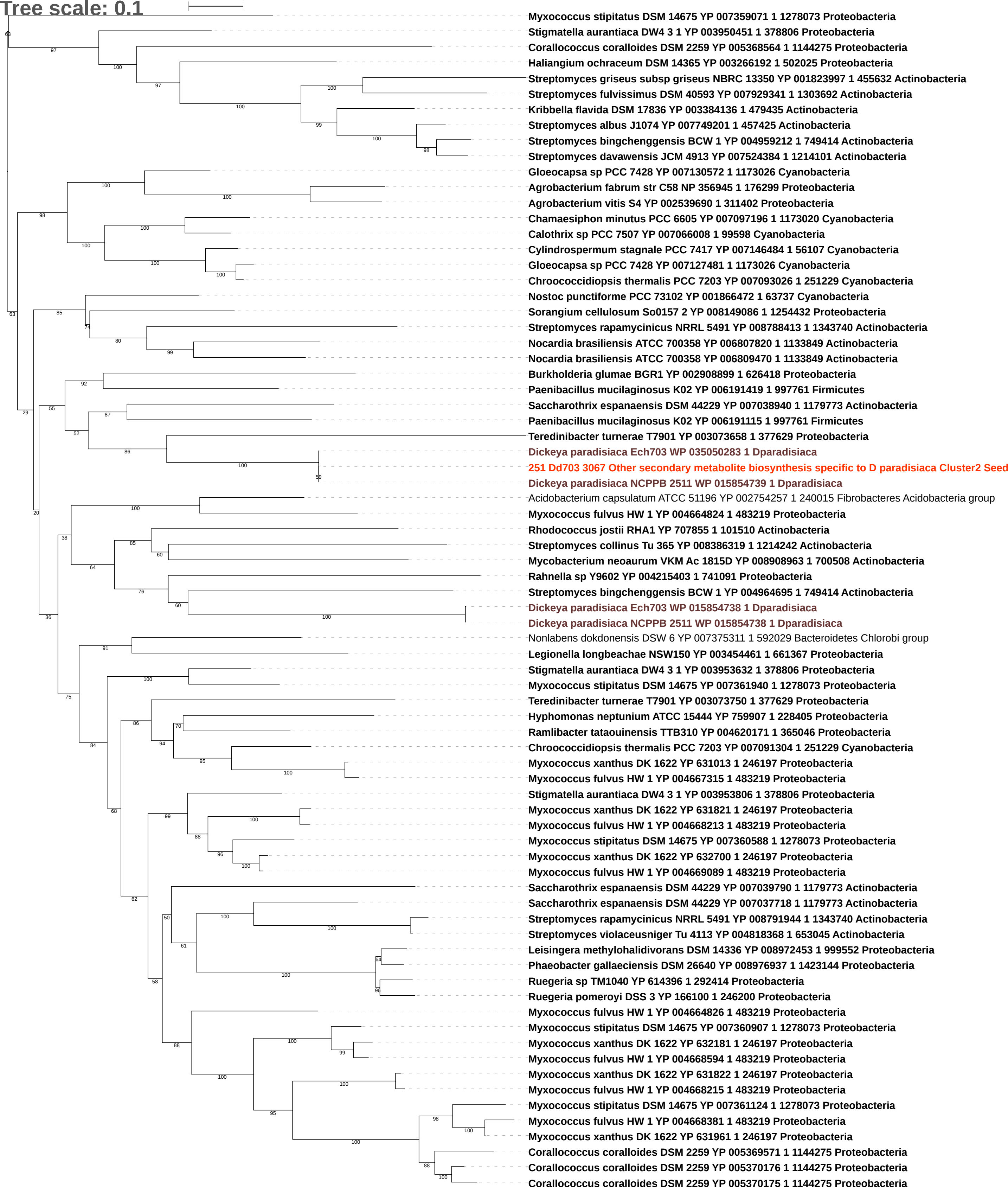

Tree scale: 0.1
Myxococcus stipitatus DSM 14675 YP 007359071 1 1278073 Proteobacteria
Stigmatella aurantiaca DW4 3 1 YP 003950451 1 378806 Proteobacteria
Corallococcus coralloides DSM 2259 YP 005368564 1 1144275 Proteobacteria
Haliangium ochraceum DSM 14365 YP 003266192 1 502025 Proteobacteria
Streptomyces griseus subsp griseus NBRC 13350 YP 001823997 1 455632 Actinobacteria
Streptomyces fulvissimus DSM 40593 YP 007929341 1 1303692 Actinobacteria
Kribbella flavida DSM 17836 YP 003384136 1 479435 Actinobacteria
Streptomyces albus J1074 YP 007749201 1 457425 Actinobacteria
Streptomyces bingchenggensis BCW 1 YP 004959212 1 749414 Actinobacteria
Streptomyces davawensis JCM 4913 YP 007524384 1 1214101 Actinobacteria
Gloeocapsa sp PCC 7428 YP 007130572 1 1173026 Cyanobacteria
Agrobacterium fabrum str C58 NP 356945 1 176299 Proteobacteria
Agrobacterium vitis S4 YP 002539690 1 311402 Proteobacteria
Chamaesiphon minutus PCC 6605 YP 007097196 1 1173020 Cyanobacteria
Calothrix sp PCC 7507 YP 007066008 1 99598 Cyanobacteria
Cylindrospermum stagnale PCC 7417 YP 007146484 1 56107 Cyanobacteria
Gloeocapsa sp PCC 7428 YP 007127481 1 1173026 Cyanobacteria
Chroococcidiopsis thermalis PCC 7203 YP 007093026 1 251229 Cyanobacteria
Nostoc punctiforme PCC 73102 YP 001866472 1 63737 Cyanobacteria
Sorangium cellulosum So0157 2 YP 008149086 1 1254432 Proteobacteria
Streptomyces rapamycinicus NRRL 5491 YP 008788413 1 1343740 Actinobacteria
Nocardia brasiliensis ATCC 700358 YP 006807820 1 1133849 Actinobacteria
Nocardia brasiliensis ATCC 700358 YP 006809470 1 1133849 Actinobacteria
Burkholderia glumae BGR1 YP 002908899 1 626418 Proteobacteria
Paenibacillus mucilaginosus K02 YP 006191419 1 997761 Firmicutes
Saccharothrix espanaensis DSM 44229 YP 007038940 1 1179773 Actinobacteria
Paenibacillus mucilaginosus K02 YP 006191115 1 997761 Firmicutes
Teredinibacter turnerae T7901 YP 003073658 1 377629 Proteobacteria
Dickeya paradisiaca Ech703 WP 035050283 1 Dparadisiaca
251 Dd703 3067 Other secondary metabolite biosynthesis specific to D paradisiaca Cluster2 Seed
Dickeya paradisiaca NCPPB 2511 WP 015854739 1 Dparadisiaca
Acidobacterium capsulatum ATCC 51196 YP 002754257 1 240015 Fibrobacteres Acidobacteria group
Myxococcus fulvus HW 1 YP 004664824 1 483219 Proteobacteria
Rhodococcus jostii RHA1 YP 707855 1 101510 Actinobacteria
Streptomyces collinus Tu 365 YP 008386319 1 1214242 Actinobacteria
Mycobacterium neoaurum VKM Ac 1815D YP 008908963 1 700508 Actinobacteria
Rahnella sp Y9602 YP 004215403 1 741091 Proteobacteria
Streptomyces bingchenggensis BCW 1 YP 004964695 1 749414 Actinobacteria
Dickeya paradisiaca Ech703 WP 015854738 1 Dparadisiaca
Dickeya paradisiaca NCPPB 2511 WP 015854738 1 Dparadisiaca
Nonlabens dokdonensis DSW 6 YP 007375311 1 592029 Bacteroidetes Chlorobi group
Legionella longbeachae NSW150 YP 003454461 1 661367 Proteobacteria
Stigmatella aurantiaca DW4 3 1 YP 003953632 1 378806 Proteobacteria
Myxococcus stipitatus DSM 14675 YP 007361940 1 1278073 Proteobacteria
Teredinibacter turnerae T7901 YP 003073750 1 377629 Proteobacteria
Hyphomonas neptunium ATCC 15444 YP 759907 1 228405 Proteobacteria
Ramlibacter tataouinensis TTB310 YP 004620171 1 365046 Proteobacteria
Chroococcidiopsis thermalis PCC 7203 YP 007091304 1 251229 Cyanobacteria
Myxococcus xanthus DK 1622 YP 631013 1 246197 Proteobacteria
Myxococcus fulvus HW 1 YP 004667315 1 483219 Proteobacteria
Stigmatella aurantiaca DW4 3 1 YP 003953806 1 378806 Proteobacteria
Myxococcus xanthus DK 1622 YP 631821 1 246197 Proteobacteria
Myxococcus fulvus HW 1 YP 004668213 1 483219 Proteobacteria
Myxococcus stipitatus DSM 14675 YP 007360588 1 1278073 Proteobacteria
Myxococcus xanthus DK 1622 YP 632700 1 246197 Proteobacteria
Myxococcus fulvus HW 1 YP 004669089 1 483219 Proteobacteria
Saccharothrix espanaensis DSM 44229 YP 007039790 1 1179773 Actinobacteria
Saccharothrix espanaensis DSM 44229 YP 007037718 1 1179773 Actinobacteria
Streptomyces rapamycinicus NRRL 5491 YP 008791944 1 1343740 Actinobacteria
Streptomyces violaceusniger Tu 4113 YP 004818368 1 653045 Actinobacteria
Leisingera methylohalidivorans DSM 14336 YP 008972453 1 999552 Proteobacteria
Phaeobacter gallaeciensis DSM 26640 YP 008976937 1 1423144 Proteobacteria
Ruegeria sp TM1040 YP 614396 1 292414 Proteobacteria
Ruegeria pomeroyi DSS 3 YP 166100 1 246200 Proteobacteria
Myxococcus fulvus HW 1 YP 004664826 1 483219 Proteobacteria
Myxococcus stipitatus DSM 14675 YP 007360907 1 1278073 Proteobacteria
Myxococcus xanthus DK 1622 YP 632181 1 246197 Proteobacteria
Myxococcus fulvus HW 1 YP 004668594 1 483219 Proteobacteria
Myxococcus xanthus DK 1622 YP 631822 1 246197 Proteobacteria
Myxococcus fulvus HW 1 YP 004668215 1 483219 Proteobacteria
Myxococcus stipitatus DSM 14675 YP 007361124 1 1278073 Proteobacteria
Myxococcus fulvus HW 1 YP 004668381 1 483219 Proteobacteria
Myxococcus xanthus DK 1622 YP 631961 1 246197 Proteobacteria
Corallococcus coralloides DSM 2259 YP 005369571 1 1144275 Proteobacteria
Corallococcus coralloides DSM 2259 YP 005370176 1 1144275 Proteobacteria
Corallococcus coralloides DSM 2259 YP 005370175 1 1144275 Proteobacteria

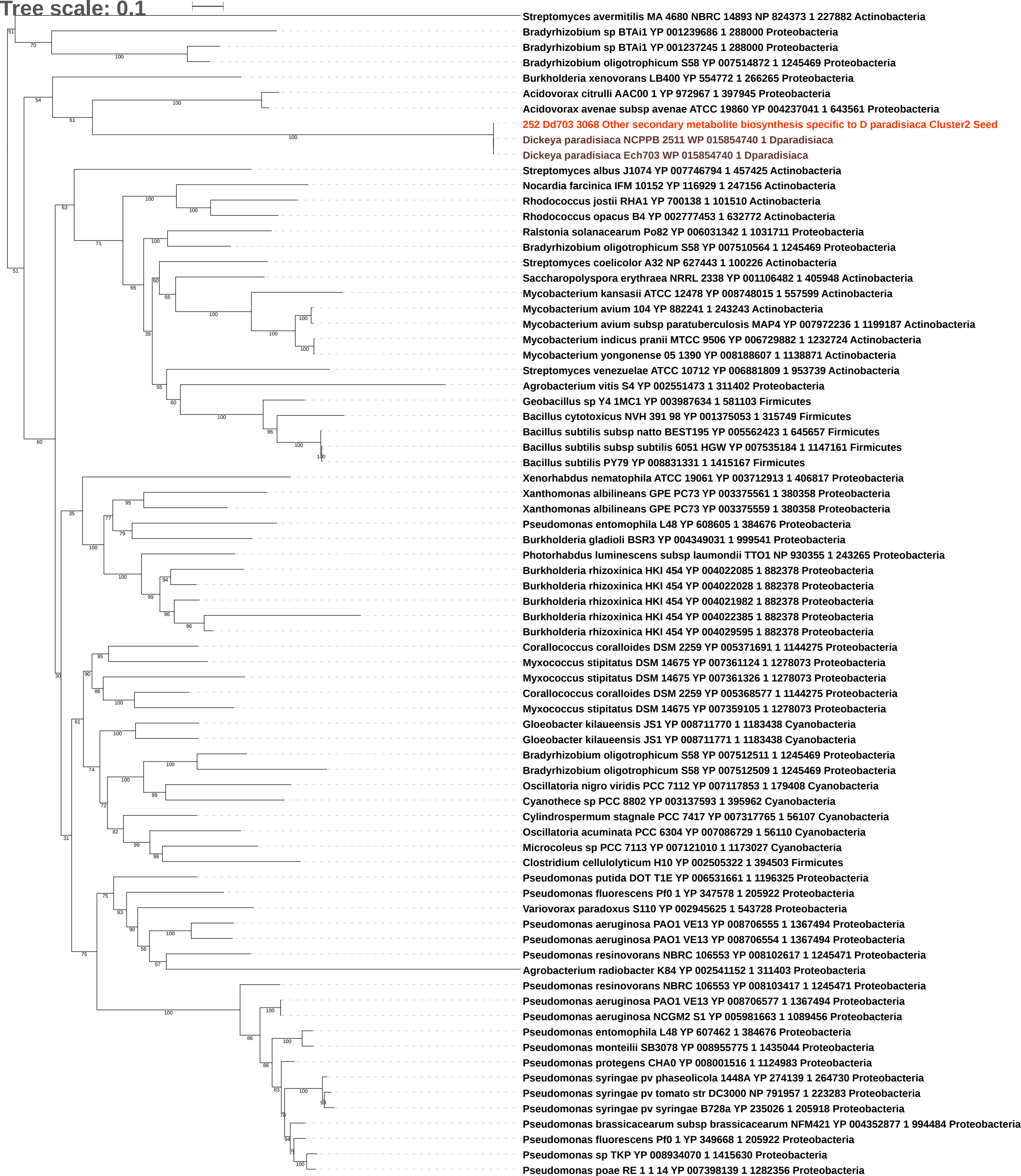

Tree scale: 0.1
Streptomyces avermitilis MA 4680 NBRC 14893 NP 824373 1 227882 Actinobacteria
Bradyrhizobium sp BTAi1 YP 001239686 1 288000 Proteobacteria
Bradyrhizobium sp BTAi1 YP 001237245 1 288000 Proteobacteria
Bradyrhizobium oligotrophicum S58 YP 007514872 1 1245469 Proteobacteria
Burkholderia xenovorans LB400 YP 554772 1 266265 Proteobacteria
Acidovorax citrulli AAC00 1 YP 972967 1 397945 Proteobacteria
Acidovorax avenae subsp avenae ATCC 19860 YP 004237041 1 643561 Proteobacteria
252 Dd703 3068 Other secondary metabolite biosynthesis specific to D paradisiaca Cluster2 Seed
Dickeya paradisiaca NCPPB 2511 WP 015854740 1 Dparadisiaca
Dickeya paradisiaca Ech703 WP 015854740 1 Dparadisiaca
Streptomyces albus J1074 YP 007746794 1 457425 Actinobacteria
Nocardia farcinica IFM 10152 YP 116929 1 247156 Actinobacteria
Rhodococcus jostii RHA1 YP 700138 1 101510 Actinobacteria
Rhodococcus opacus B4 YP 002777453 1 632772 Actinobacteria
Ralstonia solanacearum Po82 YP 006031342 1 1031711 Proteobacteria
Bradyrhizobium oligotrophicum S58 YP 007510564 1 1245469 Proteobacteria
Streptomyces coelicolor A32 NP 627443 1 100226 Actinobacteria
Saccharopolyspora erythraea NRRL 2338 YP 001106482 1 405948 Actinobacteria
Mycobacterium kansasii ATCC 12478 YP 008748015 1 557599 Actinobacteria
Mycobacterium avium 104 YP 882241 1 243243 Actinobacteria
Mycobacterium avium subsp paratuberculosis MAP4 YP 007972236 1 1199187 Actinobacteria
Mycobacterium indicus pranii MTCC 9506 YP 006729882 1 1232724 Actinobacteria
Mycobacterium yongonense 05 1390 YP 008188607 1 1138871 Actinobacteria
Streptomyces venezuelae ATCC 10712 YP 006881809 1 953739 Actinobacteria
Agrobacterium vitis S4 YP 002551473 1 311402 Proteobacteria
Geobacillus sp Y4 1MC1 YP 003987634 1 581103 Firmicutes
Bacillus cytotoxicus NVH 391 98 YP 001375053 1 315749 Firmicutes
Bacillus subtilis subsp natto BEST195 YP 005562423 1 645657 Firmicutes
Bacillus subtilis subsp subtilis 6051 HGW YP 007535184 1 1147161 Firmicutes
Bacillus subtilis PY79 YP 008831331 1 1415167 Firmicutes
Xenorhabdus nematophila ATCC 19061 YP 003712913 1 406817 Proteobacteria
Xanthomonas albilineans GPE PC73 YP 003375561 1 380358 Proteobacteria
Xanthomonas albilineans GPE PC73 YP 003375559 1 380358 Proteobacteria
Pseudomonas entomophila L48 YP 608605 1 384676 Proteobacteria
Burkholderia gladioli BSR3 YP 004349031 1 999541 Proteobacteria
Photorhabdus luminescens subsp laumondii TTO1 NP 930355 1 243265 Proteobacteria
Burkholderia rhizoxinica HKI 454 YP 004022085 1 882378 Proteobacteria
Burkholderia rhizoxinica HKI 454 YP 004022028 1 882378 Proteobacteria
Burkholderia rhizoxinica HKI 454 YP 004021982 1 882378 Proteobacteria
Burkholderia rhizoxinica HKI 454 YP 004022385 1 882378 Proteobacteria
Burkholderia rhizoxinica HKI 454 YP 004029595 1 882378 Proteobacteria
Corallococcus coralloides DSM 2259 YP 005371691 1 1144275 Proteobacteria
Myxococcus stipitatus DSM 14675 YP 007361124 1 1278073 Proteobacteria
Myxococcus stipitatus DSM 14675 YP 007361326 1 1278073 Proteobacteria
Corallococcus coralloides DSM 2259 YP 005368577 1 1144275 Proteobacteria
Myxococcus stipitatus DSM 14675 YP 007359105 1 1278073 Proteobacteria
Gloeobacter kilaueensis JS1 YP 008711770 1 1183438 Cyanobacteria
Gloeobacter kilaueensis JS1 YP 008711771 1 1183438 Cyanobacteria
Bradyrhizobium oligotrophicum S58 YP 007512511 1 1245469 Proteobacteria
Bradyrhizobium oligotrophicum S58 YP 007512509 1 1245469 Proteobacteria
Oscillatoria nigro viridis PCC 7112 YP 007117853 1 179408 Cyanobacteria
Cyanothece sp PCC 8802 YP 003137593 1 395962 Cyanobacteria
Cylindrospermum stagnale PCC 7417 YP 007317765 1 56107 Cyanobacteria
Oscillatoria acuminata PCC 6304 YP 007086729 1 56110 Cyanobacteria
Microcoleus sp PCC 7113 YP 007121010 1 1173027 Cyanobacteria
Clostridium cellulolyticum H10 YP 002505322 1 394503 Firmicutes
Pseudomonas putida DOT T1E YP 006531661 1 1196325 Proteobacteria
Pseudomonas fluorescens Pf0 1 YP 347578 1 205922 Proteobacteria
Variovorax paradoxus S110 YP 002945625 1 543728 Proteobacteria
Pseudomonas aeruginosa PAO1 VE13 YP 008706555 1 1367494 Proteobacteria
Pseudomonas aeruginosa PAO1 VE13 YP 008706554 1 1367494 Proteobacteria
Pseudomonas resinovorans NBRC 106553 YP 008102617 1 1245471 Proteobacteria
Agrobacterium radiobacter K84 YP 002541152 1 311403 Proteobacteria
Pseudomonas resinovorans NBRC 106553 YP 008103417 1 1245471 Proteobacteria
Pseudomonas aeruginosa PAO1 VE13 YP 008706577 1 1367494 Proteobacteria
Pseudomonas aeruginosa NCGM2 S1 YP 005981663 1 1089456 Proteobacteria
Pseudomonas entomophila L48 YP 607462 1 384676 Proteobacteria
Pseudomonas monteilii SB3078 YP 008955775 1 1435044 Proteobacteria
Pseudomonas protegens CHA0 YP 008001516 1 1124983 Proteobacteria
Pseudomonas syringae pv phaseolicola 1448A YP 274139 1 264730 Proteobacteria
Pseudomonas syringae pv tomato str DC3000 NP 791957 1 223283 Proteobacteria
Pseudomonas syringae pv syringae B728a YP 235026 1 205918 Proteobacteria
Pseudomonas brassicacearum subsp brassicacearum NFM421 YP 004352877 1 994484 Proteobacteria
Pseudomonas fluorescens Pf0 1 YP 349668 1 205922 Proteobacteria
Pseudomonas sp TKP YP 008934070 1 1415630 Proteobacteria
Pseudomonas poae RE 1 1 14 YP 007398139 1 1282356 Proteobacteria

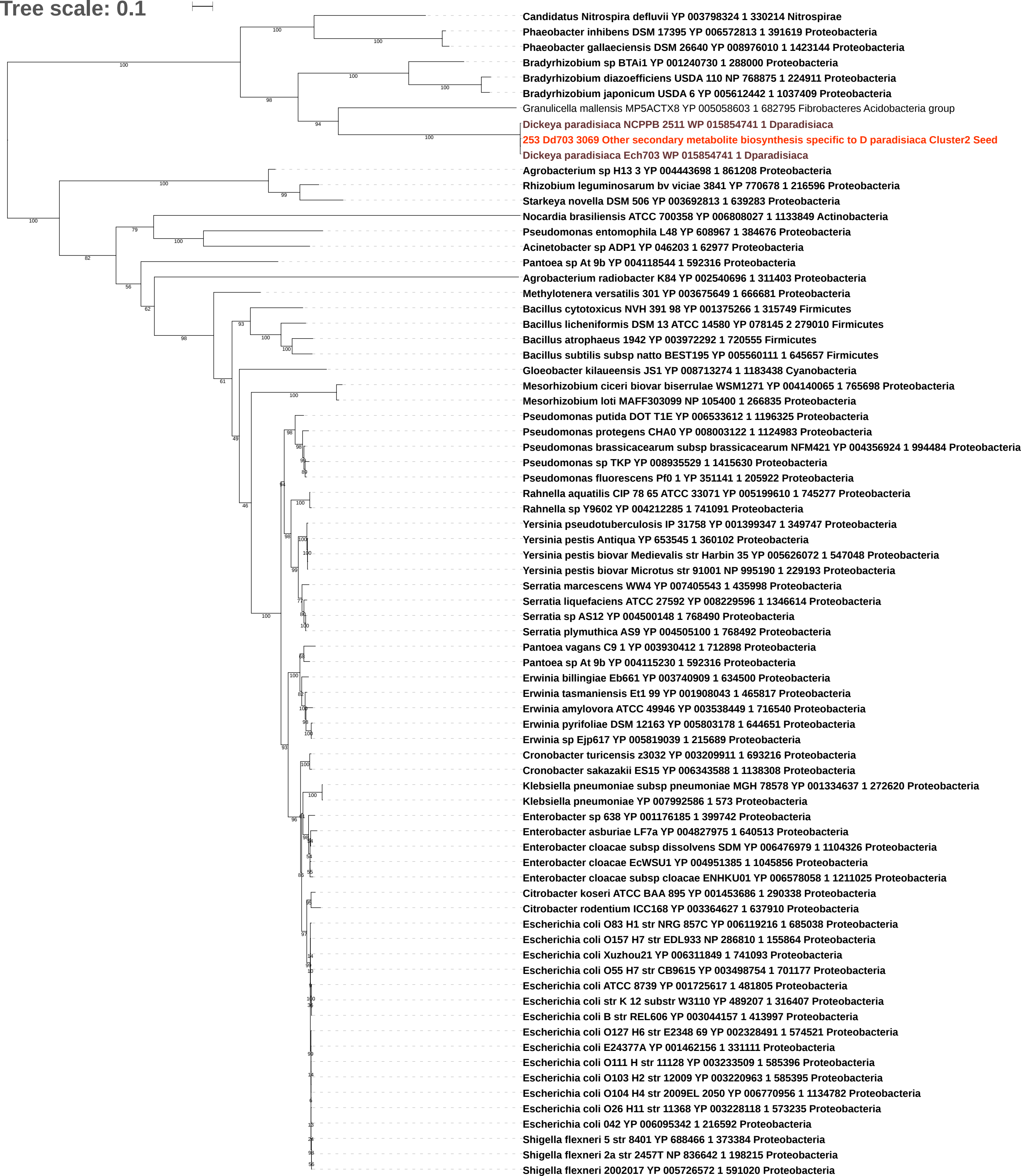
Tree scale: 0.1
Candidatus Nitrospira defluvii YP 003798324 1 330214 Nitrospirae
Phaeobacter inhibens DSM 17395 YP 006572813 1 391619 Proteobacteria
Phaeobacter gallaeciensis DSM 26640 YP 008976010 1 1423144 Proteobacteria
Bradyrhizobium sp BTAi1 YP 001240730 1 288000 Proteobacteria
Bradyrhizobium diazoefficiens USDA 110 NP 768875 1 224911 Proteobacteria
Bradyrhizobium japonicum USDA 6 YP 005612442 1 1037409 Proteobacteria
Granulicella mallensis MP5ACTX8 YP 005058603 1 682795 Fibrobacteres Acidobacteria group
Dickeya paradisiaca NCPPB 2511 WP 015854741 1 Dparadisiaca
253 Dd703 3069 Other secondary metabolite biosynthesis specific to D paradisiaca Cluster2 Seed
Dickeya paradisiaca Ech703 WP 015854741 1 Dparadisiaca
Agrobacterium sp H13 3 YP 004443698 1 861208 Proteobacteria
Rhizobium leguminosarum bv viciae 3841 YP 770678 1 216596 Proteobacteria
Starkeya novella DSM 506 YP 003692813 1 639283 Proteobacteria
Nocardia brasiliensis ATCC 700358 YP 006808027 1 1133849 Actinobacteria
Pseudomonas entomophila L48 YP 608967 1 384676 Proteobacteria
Acinetobacter sp ADP1 YP 046203 1 62977 Proteobacteria
Pantoea sp At 9b YP 004118544 1 592316 Proteobacteria
Agrobacterium radiobacter K84 YP 002540696 1 311403 Proteobacteria
Methylotenera versatilis 301 YP 003675649 1 666681 Proteobacteria
Bacillus cytotoxicus NVH 391 98 YP 001375266 1 315749 Firmicutes
Bacillus licheniformis DSM 13 ATCC 14580 YP 078145 2 279010 Firmicutes
Bacillus atrophaeus 1942 YP 003972292 1 720555 Firmicutes
Bacillus subtilis subsp natto BEST195 YP 005560111 1 645657 Firmicutes
Gloeobacter kilaueensis JS1 YP 008713274 1 1183438 Cyanobacteria
Mesorhizobium ciceri biovar biserrulae WSM1271 YP 004140065 1 765698 Proteobacteria
Mesorhizobium loti MAFF303099 NP 105400 1 266835 Proteobacteria
Pseudomonas putida DOT T1E YP 006533612 1 1196325 Proteobacteria
Pseudomonas protegens CHA0 YP 008003122 1 1124983 Proteobacteria
Pseudomonas brassicacearum subsp brassicacearum NFM421 YP 004356924 1 994484 Proteobacteria
Pseudomonas sp TKP YP 008935529 1 1415630 Proteobacteria
Pseudomonas fluorescens Pf0 1 YP 351141 1 205922 Proteobacteria
Rahnella aquatilis CIP 78 65 ATCC 33071 YP 005199610 1 745277 Proteobacteria
Rahnella sp Y9602 YP 004212285 1 741091 Proteobacteria
Yersinia pseudotuberculosis IP 31758 YP 001399347 1 349747 Proteobacteria
Yersinia pestis Antiqua YP 653545 1 360102 Proteobacteria
Yersinia pestis biovar Medievalis str Harbin 35 YP 005626072 1 547048 Proteobacteria
Yersinia pestis biovar Microtus str 91001 NP 995190 1 229193 Proteobacteria
Serratia marcescens WW4 YP 007405543 1 435998 Proteobacteria
Serratia liquefaciens ATCC 27592 YP 008229596 1 1346614 Proteobacteria
Serratia sp AS12 YP 004500148 1 768490 Proteobacteria
Serratia plymuthica AS9 YP 004505100 1 768492 Proteobacteria
Pantoea vagans C9 1 YP 003930412 1 712898 Proteobacteria
Pantoea sp At 9b YP 004115230 1 592316 Proteobacteria
Erwinia billingiae Eb661 YP 003740909 1 634500 Proteobacteria
Erwinia tasmaniensis Et1 99 YP 001908043 1 465817 Proteobacteria
Erwinia amylovora ATCC 49946 YP 003538449 1 716540 Proteobacteria
Erwinia pyrifoliae DSM 12163 YP 005803178 1 644651 Proteobacteria
Erwinia sp Ejp617 YP 005819039 1 215689 Proteobacteria
Cronobacter turicensis z3032 YP 003209911 1 693216 Proteobacteria
Cronobacter sakazakii ES15 YP 006343588 1 1138308 Proteobacteria
Klebsiella pneumoniae subsp pneumoniae MGH 78578 YP 001334637 1 272620 Proteobacteria
Klebsiella pneumoniae YP 007992586 1 573 Proteobacteria
Enterobacter sp 638 YP 001176185 1 399742 Proteobacteria
Enterobacter asburiae LF7a YP 004827975 1 640513 Proteobacteria
Enterobacter cloacae subsp dissolvens SDM YP 006476979 1 1104326 Proteobacteria
Enterobacter cloacae EcWSU1 YP 004951385 1 1045856 Proteobacteria
Enterobacter cloacae subsp cloacae ENHKU01 YP 006578058 1 1211025 Proteobacteria
Citrobacter koseri ATCC BAA 895 YP 001453686 1 290338 Proteobacteria
Citrobacter rodentium ICC168 YP 003364627 1 637910 Proteobacteria
Escherichia coli O83 H1 str NRG 857C YP 006119216 1 685038 Proteobacteria
Escherichia coli O157 H7 str EDL933 NP 286810 1 155864 Proteobacteria
Escherichia coli Xuzhou21 YP 006311849 1 741093 Proteobacteria
Escherichia coli O55 H7 str CB9615 YP 003498754 1 701177 Proteobacteria
Escherichia coli ATCC 8739 YP 001725617 1 481805 Proteobacteria
Escherichia coli str K 12 substr W3110 YP 489207 1 316407 Proteobacteria
Escherichia coli B str REL606 YP 003044157 1 413997 Proteobacteria
Escherichia coli O127 H6 str E2348 69 YP 002328491 1 574521 Proteobacteria
Escherichia coli E24377A YP 001462156 1 331111 Proteobacteria
Escherichia coli O111 H str 11128 YP 003233509 1 585396 Proteobacteria
Escherichia coli O103 H2 str 12009 YP 003220963 1 585395 Proteobacteria
Escherichia coli O104 H4 str 2009EL 2050 YP 006770956 1 1134782 Proteobacteria
Escherichia coli O26 H11 str 11368 YP 003228118 1 573235 Proteobacteria
Escherichia coli 042 YP 006095342 1 216592 Proteobacteria
Shigella flexneri 5 str 8401 YP 688466 1 373384 Proteobacteria
Shigella flexneri 2a str 2457T NP 836642 1 198215 Proteobacteria
Shigella flexneri 2002017 YP 005726572 1 591020 Proteobacteria

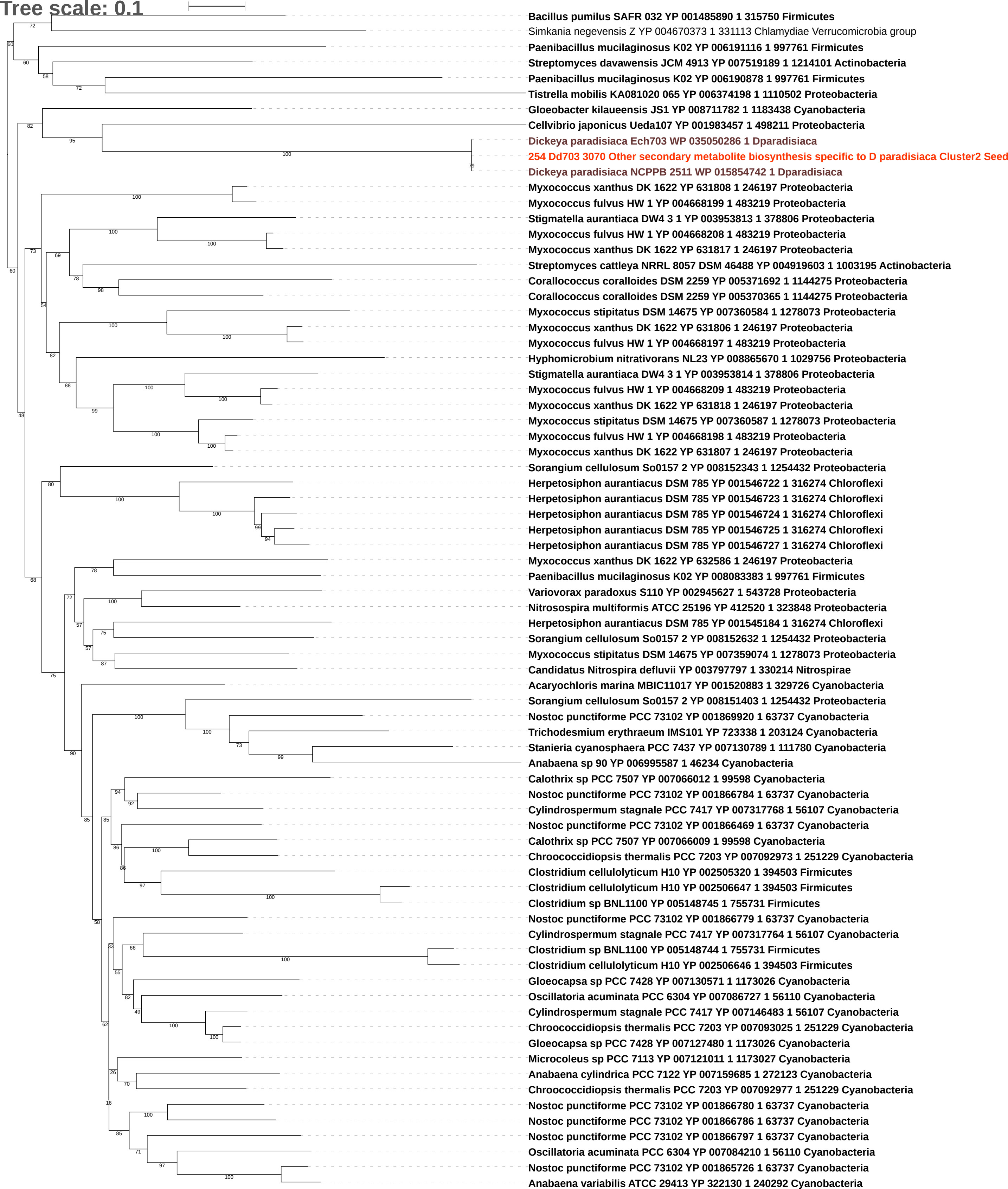
Tree scale: 0.1
Bacillus pumilus SAFR 032 YP 001485890 1 315750 Firmicutes
Simkania negevensis Z YP 004670373 1 331113 Chlamydiae Verrucomicrobia group
Paenibacillus mucilaginosus K02 YP 006191116 1 997761 Firmicutes
Streptomyces davawensis JCM 4913 YP 007519189 1 1214101 Actinobacteria
Paenibacillus mucilaginosus K02 YP 006190878 1 997761 Firmicutes
Tistrella mobilis KA081020 065 YP 006374198 1 1110502 Proteobacteria
Gloeobacter kilaueensis JS1 YP 008711782 1 1183438 Cyanobacteria
Cellvibrio japonicus Ueda107 YP 001983457 1 498211 Proteobacteria
Dickeya paradisiaca Ech703 WP 035050286 1 Dparadisiaca
254 Dd703 3070 Other secondary metabolite biosynthesis specific to D paradisiaca Cluster2 Seed
Dickeya paradisiaca NCPPB 2511 WP 015854742 1 Dparadisiaca
Myxococcus xanthus DK 1622 YP 631808 1 246197 Proteobacteria
Myxococcus fulvus HW 1 YP 004668199 1 483219 Proteobacteria
Stigmatella aurantiaca DW4 3 1 YP 003953813 1 378806 Proteobacteria
Myxococcus fulvus HW 1 YP 004668208 1 483219 Proteobacteria
Myxococcus xanthus DK 1622 YP 631817 1 246197 Proteobacteria
Streptomyces cattleya NRRL 8057 DSM 46488 YP 004919603 1 1003195 Actinobacteria
Corallococcus coralloides DSM 2259 YP 005371692 1 1144275 Proteobacteria
Corallococcus coralloides DSM 2259 YP 005370365 1 1144275 Proteobacteria
Myxococcus stipitatus DSM 14675 YP 007360584 1 1278073 Proteobacteria
Myxococcus xanthus DK 1622 YP 631806 1 246197 Proteobacteria
Myxococcus fulvus HW 1 YP 004668197 1 483219 Proteobacteria
Hyphomicrobium nitrativorans NL23 YP 008865670 1 1029756 Proteobacteria
Stigmatella aurantiaca DW4 3 1 YP 003953814 1 378806 Proteobacteria
Myxococcus fulvus HW 1 YP 004668209 1 483219 Proteobacteria
Myxococcus xanthus DK 1622 YP 631818 1 246197 Proteobacteria
Myxococcus stipitatus DSM 14675 YP 007360587 1 1278073 Proteobacteria
Myxococcus fulvus HW 1 YP 004668198 1 483219 Proteobacteria
Myxococcus xanthus DK 1622 YP 631807 1 246197 Proteobacteria
Sorangium cellulosum So0157 2 YP 008152343 1 1254432 Proteobacteria
Herpetosiphon aurantiacus DSM 785 YP 001546722 1 316274 Chloroflexi
Herpetosiphon aurantiacus DSM 785 YP 001546723 1 316274 Chloroflexi
Herpetosiphon aurantiacus DSM 785 YP 001546724 1 316274 Chloroflexi
Herpetosiphon aurantiacus DSM 785 YP 001546725 1 316274 Chloroflexi
Herpetosiphon aurantiacus DSM 785 YP 001546727 1 316274 Chloroflexi
Myxococcus xanthus DK 1622 YP 632586 1 246197 Proteobacteria
Paenibacillus mucilaginosus K02 YP 008083383 1 997761 Firmicutes
Variovorax paradoxus S110 YP 002945627 1 543728 Proteobacteria
Nitrosospira multiformis ATCC 25196 YP 412520 1 323848 Proteobacteria
Herpetosiphon aurantiacus DSM 785 YP 001545184 1 316274 Chloroflexi
Sorangium cellulosum So0157 2 YP 008152632 1 1254432 Proteobacteria
Myxococcus stipitatus DSM 14675 YP 007359074 1 1278073 Proteobacteria
Candidatus Nitrospira defluvii YP 003797797 1 330214 Nitrospirae
Acaryochloris marina MBIC11017 YP 001520883 1 329726 Cyanobacteria
Sorangium cellulosum So0157 2 YP 008151403 1 1254432 Proteobacteria
Nostoc punctiforme PCC 73102 YP 001869920 1 63737 Cyanobacteria
Trichodesmium erythraeum IMS101 YP 723338 1 203124 Cyanobacteria
Stanieria cyanosphaera PCC 7437 YP 007130789 1 111780 Cyanobacteria
Anabaena sp 90 YP 006995587 1 46234 Cyanobacteria
Calothrix sp PCC 7507 YP 007066012 1 99598 Cyanobacteria
Nostoc punctiforme PCC 73102 YP 001866784 1 63737 Cyanobacteria
Cylindrospermum stagnale PCC 7417 YP 007317768 1 56107 Cyanobacteria
Nostoc punctiforme PCC 73102 YP 001866469 1 63737 Cyanobacteria
Calothrix sp PCC 7507 YP 007066009 1 99598 Cyanobacteria
Chroococcidiopsis thermalis PCC 7203 YP 007092973 1 251229 Cyanobacteria
Clostridium cellulolyticum H10 YP 002505320 1 394503 Firmicutes
Clostridium cellulolyticum H10 YP 002506647 1 394503 Firmicutes
Clostridium sp BNL1100 YP 005148745 1 755731 Firmicutes
Nostoc punctiforme PCC 73102 YP 001866779 1 63737 Cyanobacteria
Cylindrospermum stagnale PCC 7417 YP 007317764 1 56107 Cyanobacteria
Clostridium sp BNL1100 YP 005148744 1 755731 Firmicutes
Clostridium cellulolyticum H10 YP 002506646 1 394503 Firmicutes
Gloeocapsa sp PCC 7428 YP 007130571 1 1173026 Cyanobacteria
Oscillatoria acuminata PCC 6304 YP 007086727 1 56110 Cyanobacteria
Cylindrospermum stagnale PCC 7417 YP 007146483 1 56107 Cyanobacteria
Chroococcidiopsis thermalis PCC 7203 YP 007093025 1 251229 Cyanobacteria
Gloeocapsa sp PCC 7428 YP 007127480 1 1173026 Cyanobacteria
Microcoleus sp PCC 7113 YP 007121011 1 1173027 Cyanobacteria
Anabaena cylindrica PCC 7122 YP 007159685 1 272123 Cyanobacteria
Chroococcidiopsis thermalis PCC 7203 YP 007092977 1 251229 Cyanobacteria
Nostoc punctiforme PCC 73102 YP 001866780 1 63737 Cyanobacteria
Nostoc punctiforme PCC 73102 YP 001866786 1 63737 Cyanobacteria
Nostoc punctiforme PCC 73102 YP 001866797 1 63737 Cyanobacteria
Oscillatoria acuminata PCC 6304 YP 007084210 1 56110 Cyanobacteria
Nostoc punctiforme PCC 73102 YP 001865726 1 63737 Cyanobacteria
Anabaena variabilis ATCC 29413 YP 322130 1 240292 Cyanobacteria

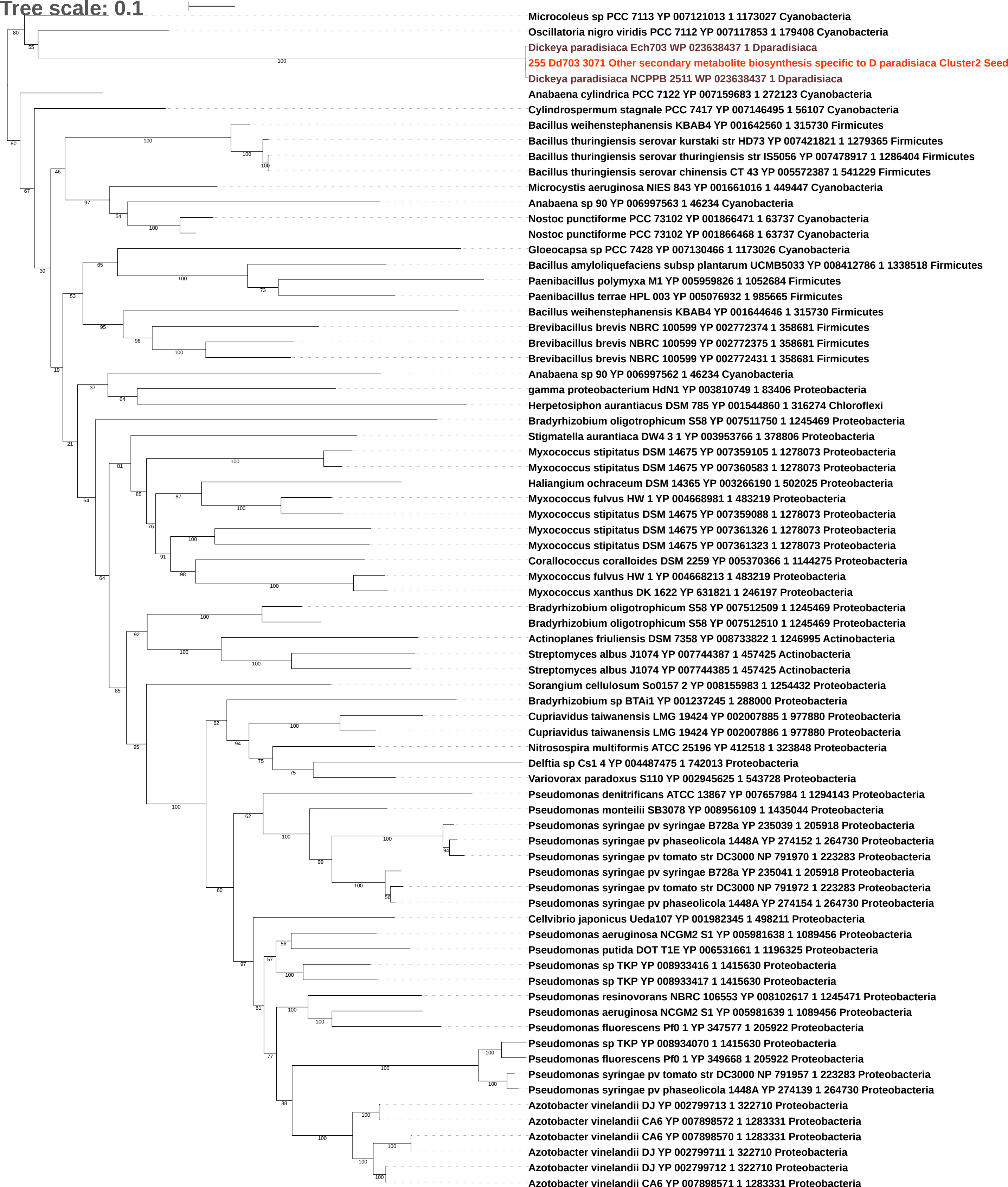
Tree scale: 0.1
Microcoleus sp PCC 7113 YP 007121013 1 1173027 Cyanobacteria
Oscillatoria nigro viridis PCC 7112 YP 007117853 1 179408 Cyanobacteria
Dickeya paradisiaca Ech703 WP 023638437 1 Dparadisiaca
255 Dd703 3071 Other secondary metabolite biosynthesis specific to D paradisiaca Cluster2 Seed
Dickeya paradisiaca NCPPB 2511 WP 023638437 1 Dparadisiaca
Anabaena cylindrica PCC 7122 YP 007159683 1 272123 Cyanobacteria
Cylindrospermum stagnale PCC 7417 YP 007146495 1 56107 Cyanobacteria
Bacillus weihenstephanensis KBAB4 YP 001642560 1 315730 Firmicutes
Bacillus thuringiensis serovar kurstaki str HD73 YP 007421821 1 1279365 Firmicutes
Bacillus thuringiensis serovar thuringiensis str IS5056 YP 007478917 1 1286404 Firmicutes
Bacillus thuringiensis serovar chinensis CT 43 YP 005572387 1 541229 Firmicutes
Microcystis aeruginosa NIES 843 YP 001661016 1 449447 Cyanobacteria
Anabaena sp 90 YP 006997563 1 46234 Cyanobacteria
Nostoc punctiforme PCC 73102 YP 001866471 1 63737 Cyanobacteria
Nostoc punctiforme PCC 73102 YP 001866468 1 63737 Cyanobacteria
Gloeocapsa sp PCC 7428 YP 007130466 1 1173026 Cyanobacteria
Bacillus amyloliquefaciens subsp plantarum UCMB5033 YP 008412786 1 1338518 Firmicutes
Paenibacillus polymyxa M1 YP 005959826 1 1052684 Firmicutes
Paenibacillus terrae HPL 003 YP 005076932 1 985665 Firmicutes
Bacillus weihenstephanensis KBAB4 YP 001644646 1 315730 Firmicutes
Brevibacillus brevis NBRC 100599 YP 002772374 1 358681 Firmicutes
Brevibacillus brevis NBRC 100599 YP 002772375 1 358681 Firmicutes
Brevibacillus brevis NBRC 100599 YP 002772431 1 358681 Firmicutes
Anabaena sp 90 YP 006997562 1 46234 Cyanobacteria
gamma proteobacterium HdN1 YP 003810749 1 83406 Proteobacteria
Herpetosiphon aurantiacus DSM 785 YP 001544860 1 316274 Chloroflexi
Bradyrhizobium oligotrophicum S58 YP 007511750 1 1245469 Proteobacteria
Stigmatella aurantiaca DW4 3 1 YP 003953766 1 378806 Proteobacteria
Myxococcus stipitatus DSM 14675 YP 007359105 1 1278073 Proteobacteria
Myxococcus stipitatus DSM 14675 YP 007360583 1 1278073 Proteobacteria
Haliangium ochraceum DSM 14365 YP 003266190 1 502025 Proteobacteria
Myxococcus fulvus HW 1 YP 004668981 1 483219 Proteobacteria
Myxococcus stipitatus DSM 14675 YP 007359088 1 1278073 Proteobacteria
Myxococcus stipitatus DSM 14675 YP 007361326 1 1278073 Proteobacteria
Myxococcus stipitatus DSM 14675 YP 007361323 1 1278073 Proteobacteria
Corallococcus coralloides DSM 2259 YP 005370366 1 1144275 Proteobacteria
Myxococcus fulvus HW 1 YP 004668213 1 483219 Proteobacteria
Myxococcus xanthus DK 1622 YP 631821 1 246197 Proteobacteria
Bradyrhizobium oligotrophicum S58 YP 007512509 1 1245469 Proteobacteria
Bradyrhizobium oligotrophicum S58 YP 007512510 1 1245469 Proteobacteria
Actinoplanes friuliensis DSM 7358 YP 008733822 1 1246995 Actinobacteria
Streptomyces albus J1074 YP 007744387 1 457425 Actinobacteria
Streptomyces albus J1074 YP 007744385 1 457425 Actinobacteria
Sorangium cellulosum So0157 2 YP 008155983 1 1254432 Proteobacteria
Bradyrhizobium sp BTAi1 YP 001237245 1 288000 Proteobacteria
Cupriavidus taiwanensis LMG 19424 YP 002007885 1 977880 Proteobacteria
Cupriavidus taiwanensis LMG 19424 YP 002007886 1 977880 Proteobacteria
Nitrosospira multiformis ATCC 25196 YP 412518 1 323848 Proteobacteria
Delftia sp Cs1 4 YP 004487475 1 742013 Proteobacteria
Variovorax paradoxus S110 YP 002945625 1 543728 Proteobacteria
Pseudomonas denitrificans ATCC 13867 YP 007657984 1 1294143 Proteobacteria
Pseudomonas monteilii SB3078 YP 008956109 1 1435044 Proteobacteria
Pseudomonas syringae pv syringae B728a YP 235039 1 205918 Proteobacteria
Pseudomonas syringae pv phaseolicola 1448A YP 274152 1 264730 Proteobacteria
Pseudomonas syringae pv tomato str DC3000 NP 791970 1 223283 Proteobacteria
Pseudomonas syringae pv syringae B728a YP 235041 1 205918 Proteobacteria
Pseudomonas syringae pv tomato str DC3000 NP 791972 1 223283 Proteobacteria
Pseudomonas syringae pv phaseolicola 1448A YP 274154 1 264730 Proteobacteria
Cellvibrio japonicus Ueda107 YP 001982345 1 498211 Proteobacteria
Pseudomonas aeruginosa NCGM2 S1 YP 005981638 1 1089456 Proteobacteria
Pseudomonas putida DOT T1E YP 006531661 1 1196325 Proteobacteria
Pseudomonas sp TKP YP 008933416 1 1415630 Proteobacteria
Pseudomonas sp TKP YP 008933417 1 1415630 Proteobacteria
Pseudomonas resinovorans NBRC 106553 YP 008102617 1 1245471 Proteobacteria
Pseudomonas aeruginosa NCGM2 S1 YP 005981639 1 1089456 Proteobacteria
Pseudomonas fluorescens Pf0 1 YP 347577 1 205922 Proteobacteria
Pseudomonas sp TKP YP 008934070 1 1415630 Proteobacteria
Pseudomonas fluorescens Pf0 1 YP 349668 1 205922 Proteobacteria
Pseudomonas syringae pv tomato str DC3000 NP 791957 1 223283 Proteobacteria
Pseudomonas syringae pv phaseolicola 1448A YP 274139 1 264730 Proteobacteria
Azotobacter vinelandii DJ YP 002799713 1 322710 Proteobacteria
Azotobacter vinelandii CA6 YP 007898572 1 1283331 Proteobacteria
Azotobacter vinelandii CA6 YP 007898570 1 1283331 Proteobacteria
Azotobacter vinelandii DJ YP 002799711 1 322710 Proteobacteria
Azotobacter vinelandii DJ YP 002799712 1 322710 Proteobacteria
Azotobacter vinelandii CA6 YP 007898571 1 1283331 Proteobacteria

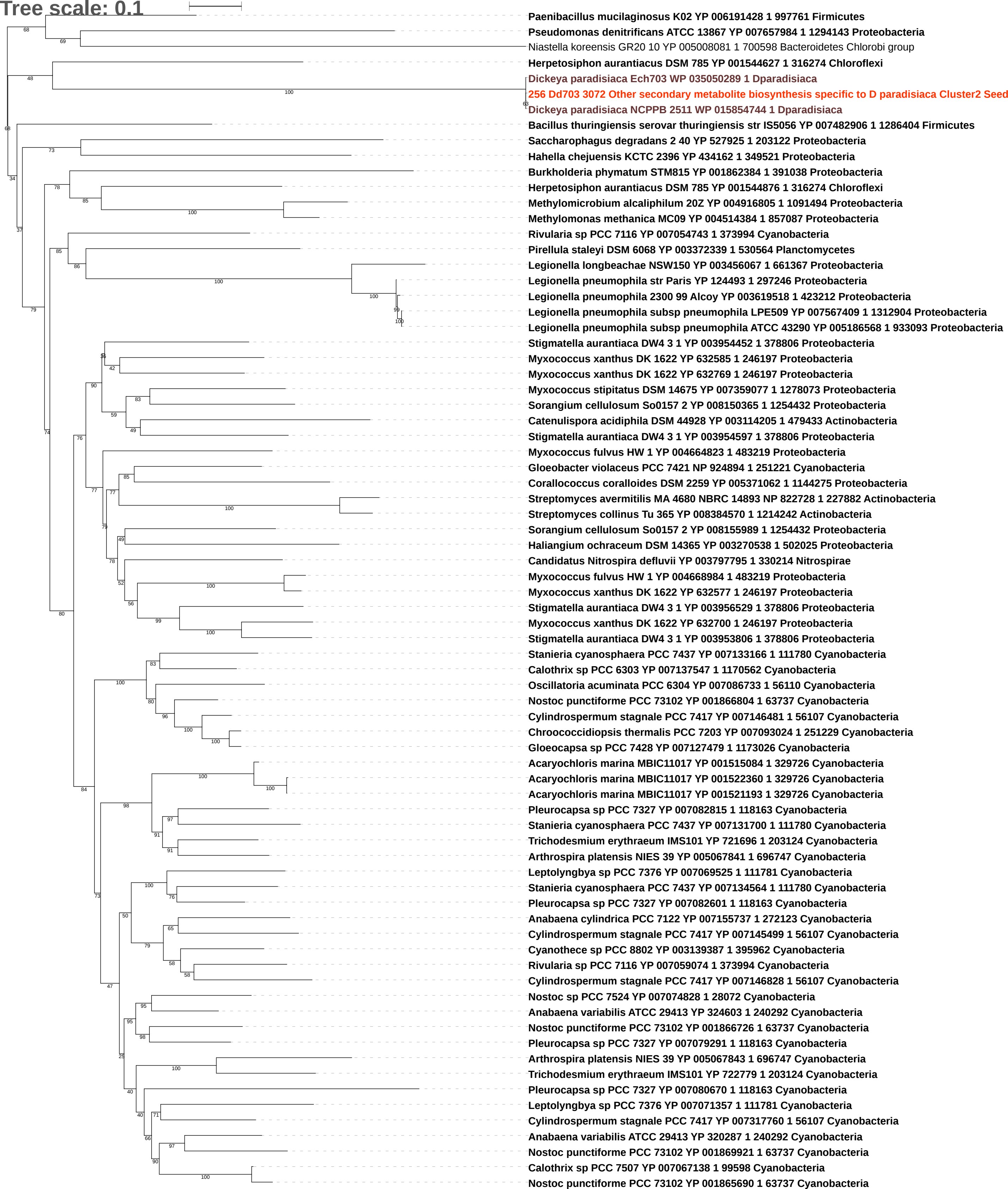
Tree scale: 0.1
Paenibacillus mucilaginosus K02 YP 006191428 1 997761 Firmicutes
Pseudomonas denitrificans ATCC 13867 YP 007657984 1 1294143 Proteobacteria
Niastella koreensis GR20 10 YP 005008081 1 700598 Bacteroidetes Chlorobi group
Herpetosiphon aurantiacus DSM 785 YP 001544627 1 316274 Chloroflexi
Dickeya paradisiaca Ech703 WP 035050289 1 Dparadisiaca
256 Dd703 3072 Other secondary metabolite biosynthesis specific to D paradisiaca Cluster2 Seed
Dickeya paradisiaca NCPPB 2511 WP 015854744 1 Dparadisiaca
Bacillus thuringiensis serovar thuringiensis str IS5056 YP 007482906 1 1286404 Firmicutes
Saccharophagus degradans 2 40 YP 527925 1 203122 Proteobacteria
Hahella chejuensis KCTC 2396 YP 434162 1 349521 Proteobacteria
Burkholderia phymatum STM815 YP 001862384 1 391038 Proteobacteria
Herpetosiphon aurantiacus DSM 785 YP 001544876 1 316274 Chloroflexi
Methylomicrobium alcaliphilum 20Z YP 004916805 1 1091494 Proteobacteria
Methylomonas methanica MC09 YP 004514384 1 857087 Proteobacteria
Rivularia sp PCC 7116 YP 007054743 1 373994 Cyanobacteria
Pirellula staleyi DSM 6068 YP 003372339 1 530564 Planctomycetes
Legionella longbeachae NSW150 YP 003456067 1 661367 Proteobacteria
Legionella pneumophila str Paris YP 124493 1 297246 Proteobacteria
Legionella pneumophila 2300 99 Alcoy YP 003619518 1 423212 Proteobacteria
Legionella pneumophila subsp pneumophila LPE509 YP 007567409 1 1312904 Proteobacteria
Legionella pneumophila subsp pneumophila ATCC 43290 YP 005186568 1 933093 Proteobacteria
Stigmatella aurantiaca DW4 3 1 YP 003954452 1 378806 Proteobacteria
Myxococcus xanthus DK 1622 YP 632585 1 246197 Proteobacteria
Myxococcus xanthus DK 1622 YP 632769 1 246197 Proteobacteria
Myxococcus stipitatus DSM 14675 YP 007359077 1 1278073 Proteobacteria
Sorangium cellulosum So0157 2 YP 008150365 1 1254432 Proteobacteria
Catenulispora acidiphila DSM 44928 YP 003114205 1 479433 Actinobacteria
Stigmatella aurantiaca DW4 3 1 YP 003954597 1 378806 Proteobacteria
Myxococcus fulvus HW 1 YP 004664823 1 483219 Proteobacteria
Gloeobacter violaceus PCC 7421 NP 924894 1 251221 Cyanobacteria
Corallococcus coralloides DSM 2259 YP 005371062 1 1144275 Proteobacteria
Streptomyces avermitilis MA 4680 NBRC 14893 NP 822728 1 227882 Actinobacteria
Streptomyces collinus Tu 365 YP 008384570 1 1214242 Actinobacteria
Sorangium cellulosum So0157 2 YP 008155989 1 1254432 Proteobacteria
Haliangium ochraceum DSM 14365 YP 003270538 1 502025 Proteobacteria
Candidatus Nitrospira defluvii YP 003797795 1 330214 Nitrospirae
Myxococcus fulvus HW 1 YP 004668984 1 483219 Proteobacteria
Myxococcus xanthus DK 1622 YP 632577 1 246197 Proteobacteria
Stigmatella aurantiaca DW4 3 1 YP 003956529 1 378806 Proteobacteria
Myxococcus xanthus DK 1622 YP 632700 1 246197 Proteobacteria
Stigmatella aurantiaca DW4 3 1 YP 003953806 1 378806 Proteobacteria
Stanieria cyanosphaera PCC 7437 YP 007133166 1 111780 Cyanobacteria
Calothrix sp PCC 6303 YP 007137547 1 1170562 Cyanobacteria
Oscillatoria acuminata PCC 6304 YP 007086733 1 56110 Cyanobacteria
Nostoc punctiforme PCC 73102 YP 001866804 1 63737 Cyanobacteria
Cylindrospermum stagnale PCC 7417 YP 007146481 1 56107 Cyanobacteria
Chroococcidiopsis thermalis PCC 7203 YP 007093024 1 251229 Cyanobacteria
Gloeocapsa sp PCC 7428 YP 007127479 1 1173026 Cyanobacteria
Acaryochloris marina MBIC11017 YP 001515084 1 329726 Cyanobacteria
Acaryochloris marina MBIC11017 YP 001522360 1 329726 Cyanobacteria
Acaryochloris marina MBIC11017 YP 001521193 1 329726 Cyanobacteria
Pleurocapsa sp PCC 7327 YP 007082815 1 118163 Cyanobacteria
Stanieria cyanosphaera PCC 7437 YP 007131700 1 111780 Cyanobacteria
Trichodesmium erythraeum IMS101 YP 721696 1 203124 Cyanobacteria
Arthrospira platensis NIES 39 YP 005067841 1 696747 Cyanobacteria
Leptolyngbya sp PCC 7376 YP 007069525 1 111781 Cyanobacteria
Stanieria cyanosphaera PCC 7437 YP 007134564 1 111780 Cyanobacteria
Pleurocapsa sp PCC 7327 YP 007082601 1 118163 Cyanobacteria
Anabaena cylindrica PCC 7122 YP 007155737 1 272123 Cyanobacteria
Cylindrospermum stagnale PCC 7417 YP 007145499 1 56107 Cyanobacteria
Cyanothece sp PCC 8802 YP 003139387 1 395962 Cyanobacteria
Rivularia sp PCC 7116 YP 007059074 1 373994 Cyanobacteria
Cylindrospermum stagnale PCC 7417 YP 007146828 1 56107 Cyanobacteria
Nostoc sp PCC 7524 YP 007074828 1 28072 Cyanobacteria
Anabaena variabilis ATCC 29413 YP 324603 1 240292 Cyanobacteria
Nostoc punctiforme PCC 73102 YP 001866726 1 63737 Cyanobacteria
Pleurocapsa sp PCC 7327 YP 007079291 1 118163 Cyanobacteria
Arthrospira platensis NIES 39 YP 005067843 1 696747 Cyanobacteria
Trichodesmium erythraeum IMS101 YP 722779 1 203124 Cyanobacteria
Pleurocapsa sp PCC 7327 YP 007080670 1 118163 Cyanobacteria
Leptolyngbya sp PCC 7376 YP 007071357 1 111781 Cyanobacteria
Cylindrospermum stagnale PCC 7417 YP 007317760 1 56107 Cyanobacteria
Anabaena variabilis ATCC 29413 YP 320287 1 240292 Cyanobacteria
Nostoc punctiforme PCC 73102 YP 001869921 1 63737 Cyanobacteria
Calothrix sp PCC 7507 YP 007067138 1 99598 Cyanobacteria
Nostoc punctiforme PCC 73102 YP 001865690 1 63737 Cyanobacteria

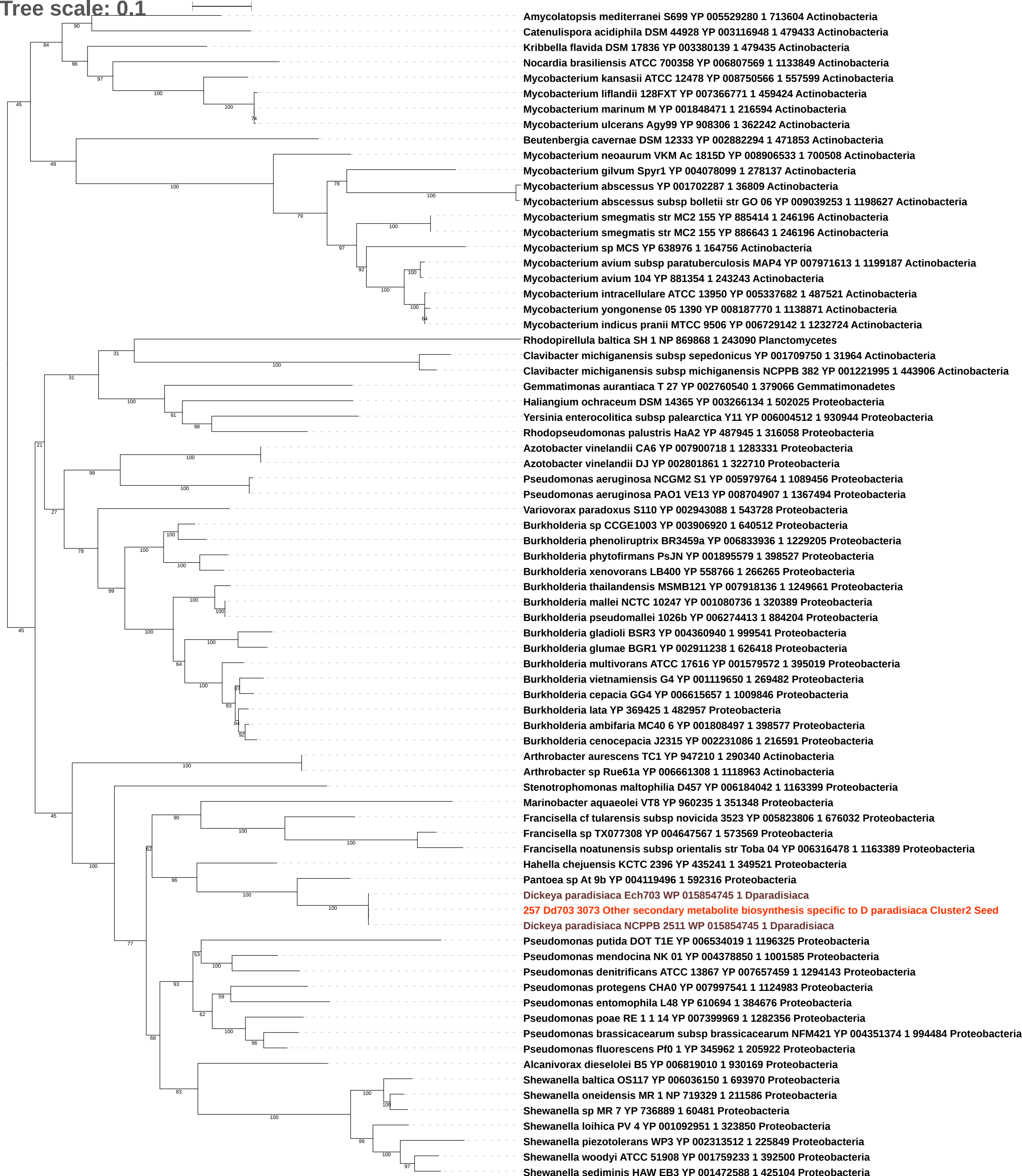
Tree scale: 0.1
Amycolatopsis mediterranei S699 YP 005529280 1 713604 Actinobacteria
Catenulispora acidiphila DSM 44928 YP 003116948 1 479433 Actinobacteria
Kribbella flavida DSM 17836 YP 003380139 1 479435 Actinobacteria
Nocardia brasiliensis ATCC 700358 YP 006807569 1 1133849 Actinobacteria
Mycobacterium kansasii ATCC 12478 YP 008750566 1 557599 Actinobacteria
Mycobacterium liflandii 128FXT YP 007366771 1 459424 Actinobacteria
Mycobacterium marinum M YP 001848471 1 216594 Actinobacteria
Mycobacterium ulcerans Agy99 YP 908306 1 362242 Actinobacteria
Beutenbergia cavernae DSM 12333 YP 002882294 1 471853 Actinobacteria
Mycobacterium neoaurum VKM Ac 1815D YP 008906533 1 700508 Actinobacteria
Mycobacterium gilvum Spyr1 YP 004078099 1 278137 Actinobacteria
Mycobacterium abscessus YP 001702287 1 36809 Actinobacteria
Mycobacterium abscessus subsp bolletii str GO 06 YP 009039253 1 1198627 Actinobacteria
Mycobacterium smegmatis str MC2 155 YP 885414 1 246196 Actinobacteria
Mycobacterium smegmatis str MC2 155 YP 886643 1 246196 Actinobacteria
Mycobacterium sp MCS YP 638976 1 164756 Actinobacteria
Mycobacterium avium subsp paratuberculosis MAP4 YP 007971613 1 1199187 Actinobacteria
Mycobacterium avium 104 YP 881354 1 243243 Actinobacteria
Mycobacterium intracellulare ATCC 13950 YP 005337682 1 487521 Actinobacteria
Mycobacterium yongonense 05 1390 YP 008187770 1 1138871 Actinobacteria
Mycobacterium indicus pranii MTCC 9506 YP 006729142 1 1232724 Actinobacteria
Rhodopirellula baltica SH 1 NP 869868 1 243090 Planctomycetes
Clavibacter michiganensis subsp sepedonicus YP 001709750 1 31964 Actinobacteria
Clavibacter michiganensis subsp michiganensis NCPPB 382 YP 001221995 1 443906 Actinobacteria
Gemmatimonas aurantiaca T 27 YP 002760540 1 379066 Gemmatimonadetes
Haliangium ochraceum DSM 14365 YP 003266134 1 502025 Proteobacteria
Yersinia enterocolitica subsp palearctica Y11 YP 006004512 1 930944 Proteobacteria
Rhodopseudomonas palustris HaA2 YP 487945 1 316058 Proteobacteria
Azotobacter vinelandii CA6 YP 007900718 1 1283331 Proteobacteria
Azotobacter vinelandii DJ YP 002801861 1 322710 Proteobacteria
Pseudomonas aeruginosa NCGM2 S1 YP 005979764 1 1089456 Proteobacteria
Pseudomonas aeruginosa PAO1 VE13 YP 008704907 1 1367494 Proteobacteria
Variovorax paradoxus S110 YP 002943088 1 543728 Proteobacteria
Burkholderia sp CCGE1003 YP 003906920 1 640512 Proteobacteria
Burkholderia phenoliruptrix BR3459a YP 006833936 1 1229205 Proteobacteria
Burkholderia phytofirmans PsJN YP 001895579 1 398527 Proteobacteria
Burkholderia xenovorans LB400 YP 558766 1 266265 Proteobacteria
Burkholderia thailandensis MSMB121 YP 007918136 1 1249661 Proteobacteria
Burkholderia mallei NCTC 10247 YP 001080736 1 320389 Proteobacteria
Burkholderia pseudomallei 1026b YP 006274413 1 884204 Proteobacteria
Burkholderia gladioli BSR3 YP 004360940 1 999541 Proteobacteria
Burkholderia glumae BGR1 YP 002911238 1 626418 Proteobacteria
Burkholderia multivorans ATCC 17616 YP 001579572 1 395019 Proteobacteria
Burkholderia vietnamiensis G4 YP 001119650 1 269482 Proteobacteria
Burkholderia cepacia GG4 YP 006615657 1 1009846 Proteobacteria
Burkholderia lata YP 369425 1 482957 Proteobacteria
Burkholderia ambifaria MC40 6 YP 001808497 1 398577 Proteobacteria
Burkholderia cenocepacia J2315 YP 002231086 1 216591 Proteobacteria
Arthrobacter aurescens TC1 YP 947210 1 290340 Actinobacteria
Arthrobacter sp Rue61a YP 006661308 1 1118963 Actinobacteria
Stenotrophomonas maltophilia D457 YP 006184042 1 1163399 Proteobacteria
Marinobacter aquaeolei VT8 YP 960235 1 351348 Proteobacteria
Francisella cf tularensis subsp novicida 3523 YP 005823806 1 676032 Proteobacteria
Francisella sp TX077308 YP 004647567 1 573569 Proteobacteria
Francisella noatunensis subsp orientalis str Toba 04 YP 006316478 1 1163389 Proteobacteria
Hahella chejuensis KCTC 2396 YP 435241 1 349521 Proteobacteria
Pantoea sp At 9b YP 004119496 1 592316 Proteobacteria
Dickeya paradisiaca Ech703 WP 015854745 1 Dparadisiaca
257 Dd703 3073 Other secondary metabolite biosynthesis specific to D paradisiaca Cluster2 Seed
Dickeya paradisiaca NCPPB 2511 WP 015854745 1 Dparadisiaca
Pseudomonas putida DOT T1E YP 006534019 1 1196325 Proteobacteria
Pseudomonas mendocina NK 01 YP 004378850 1 1001585 Proteobacteria
Pseudomonas denitrificans ATCC 13867 YP 007657459 1 1294143 Proteobacteria
Pseudomonas protegens CHA0 YP 007997541 1 1124983 Proteobacteria
Pseudomonas entomophila L48 YP 610694 1 384676 Proteobacteria
Pseudomonas poae RE 1 1 14 YP 007399969 1 1282356 Proteobacteria
Pseudomonas brassicacearum subsp brassicacearum NFM421 YP 004351374 1 994484 Proteobacteria
Pseudomonas fluorescens Pf0 1 YP 345962 1 205922 Proteobacteria
Alcanivorax dieselolei B5 YP 006819010 1 930169 Proteobacteria
Shewanella baltica OS117 YP 006036150 1 693970 Proteobacteria
Shewanella oneidensis MR 1 NP 719329 1 211586 Proteobacteria
Shewanella sp MR 7 YP 736889 1 60481 Proteobacteria
Shewanella loihica PV 4 YP 001092951 1 323850 Proteobacteria
Shewanella piezotolerans WP3 YP 002313512 1 225849 Proteobacteria
Shewanella woodyi ATCC 51908 YP 001759233 1 392500 Proteobacteria
Shewanella sediminis HAW EB3 YP 001472588 1 425104 Proteobacteria

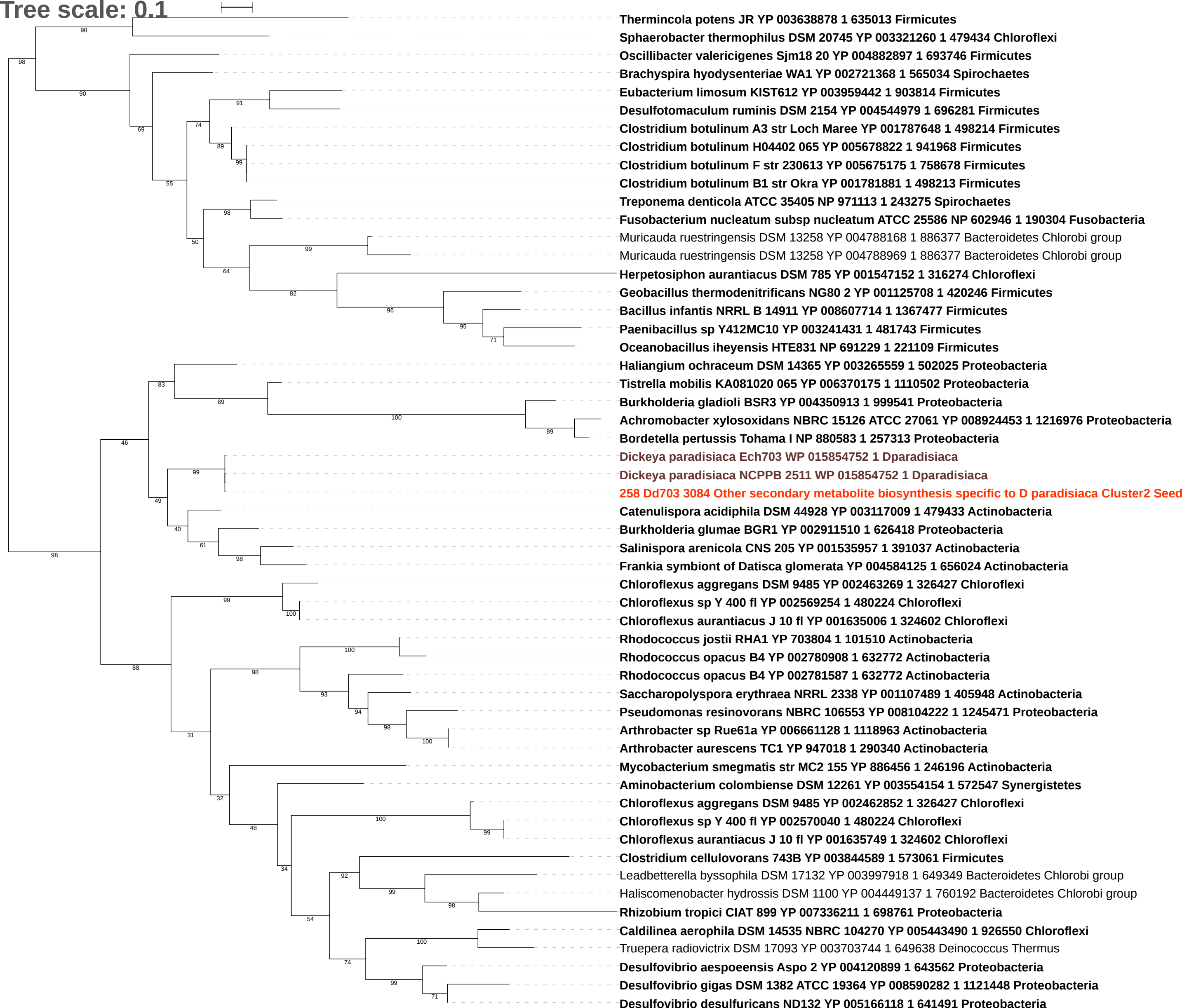
Tree scale: 0.1
Thermincola potens JR YP 003638878 1 635013 Firmicutes
Sphaerobacter thermophilus DSM 20745 YP 003321260 1 479434 Chloroflexi
Oscillibacter valericigenes Sjm18 20 YP 004882897 1 693746 Firmicutes
Brachyspira hyodysenteriae WA1 YP 002721368 1 565034 Spirochaetes
Eubacterium limosum KIST612 YP 003959442 1 903814 Firmicutes
Desulfotomaculum ruminis DSM 2154 YP 004544979 1 696281 Firmicutes
Clostridium botulinum A3 str Loch Maree YP 001787648 1 498214 Firmicutes
Clostridium botulinum H04402 065 YP 005678822 1 941968 Firmicutes
Clostridium botulinum F str 230613 YP 005675175 1 758678 Firmicutes
Clostridium botulinum B1 str Okra YP 001781881 1 498213 Firmicutes
Treponema denticola ATCC 35405 NP 971113 1 243275 Spirochaetes
Fusobacterium nucleatum subsp nucleatum ATCC 25586 NP 602946 1 190304 Fusobacteria
Muricauda ruestringensis DSM 13258 YP 004788168 1 886377 Bacteroidetes Chlorobi group
Muricauda ruestringensis DSM 13258 YP 004788969 1 886377 Bacteroidetes Chlorobi group
Herpetosiphon aurantiacus DSM 785 YP 001547152 1 316274 Chloroflexi
Geobacillus thermodenitrificans NG80 2 YP 001125708 1 420246 Firmicutes
Bacillus infantis NRRL B 14911 YP 008607714 1 1367477 Firmicutes
Paenibacillus sp Y412MC10 YP 003241431 1 481743 Firmicutes
Oceanobacillus iheyensis HTE831 NP 691229 1 221109 Firmicutes
Haliangium ochraceum DSM 14365 YP 003265559 1 502025 Proteobacteria
Tistrella mobilis KA081020 065 YP 006370175 1 1110502 Proteobacteria
Burkholderia gladioli BSR3 YP 004350913 1 999541 Proteobacteria
Achromobacter xylosoxidans NBRC 15126 ATCC 27061 YP 008924453 1 1216976 Proteobacteria
Bordetella pertussis Tohama I NP 880583 1 257313 Proteobacteria
Dickeya paradisiaca Ech703 WP 015854752 1 Dparadisiaca
Dickeya paradisiaca NCPPB 2511 WP 015854752 1 Dparadisiaca
258 Dd703 3084 Other secondary metabolite biosynthesis specific to D paradisiaca Cluster2 Seed
Catenulispora acidiphila DSM 44928 YP 003117009 1 479433 Actinobacteria
Burkholderia glumae BGR1 YP 002911510 1 626418 Proteobacteria
Salinispora arenicola CNS 205 YP 001535957 1 391037 Actinobacteria
Frankia symbiont of Datisca glomerata YP 004584125 1 656024 Actinobacteria
Chloroflexus aggregans DSM 9485 YP 002463269 1 326427 Chloroflexi
Chloroflexus sp Y 400 fl YP 002569254 1 480224 Chloroflexi
Chloroflexus aurantiacus J 10 fl YP 001635006 1 324602 Chloroflexi
Rhodococcus jostii RHA1 YP 703804 1 101510 Actinobacteria
Rhodococcus opacus B4 YP 002780908 1 632772 Actinobacteria
Rhodococcus opacus B4 YP 002781587 1 632772 Actinobacteria
Saccharopolyspora erythraea NRRL 2338 YP 001107489 1 405948 Actinobacteria
Pseudomonas resinovorans NBRC 106553 YP 008104222 1 1245471 Proteobacteria
Arthrobacter sp Rue61a YP 006661128 1 1118963 Actinobacteria
Arthrobacter aurescens TC1 YP 947018 1 290340 Actinobacteria
Mycobacterium smegmatis str MC2 155 YP 886456 1 246196 Actinobacteria
Aminobacterium colombiense DSM 12261 YP 003554154 1 572547 Synergistetes
Chloroflexus aggregans DSM 9485 YP 002462852 1 326427 Chloroflexi
Chloroflexus sp Y 400 fl YP 002570040 1 480224 Chloroflexi
Chloroflexus aurantiacus J 10 fl YP 001635749 1 324602 Chloroflexi
Clostridium cellulovorans 743B YP 003844589 1 573061 Firmicutes
Leadbetterella byssophila DSM 17132 YP 003997918 1 649349 Bacteroidetes Chlorobi group
Haliscomenobacter hydrossis DSM 1100 YP 004449137 1 760192 Bacteroidetes Chlorobi group
Rhizobium tropici CIAT 899 YP 007336211 1 698761 Proteobacteria
Caldilinea aerophila DSM 14535 NBRC 104270 YP 005443490 1 926550 Chloroflexi
Truepera radiovictrix DSM 17093 YP 003703744 1 649638 Deinococcus Thermus
Desulfovibrio aespoeensis Aspo 2 YP 004120899 1 643562 Proteobacteria
Desulfovibrio gigas DSM 1382 ATCC 19364 YP 008590282 1 1121448 Proteobacteria
Desulfovibrio desulfuricans ND132 YP 005166118 1 641491 Proteobacteria

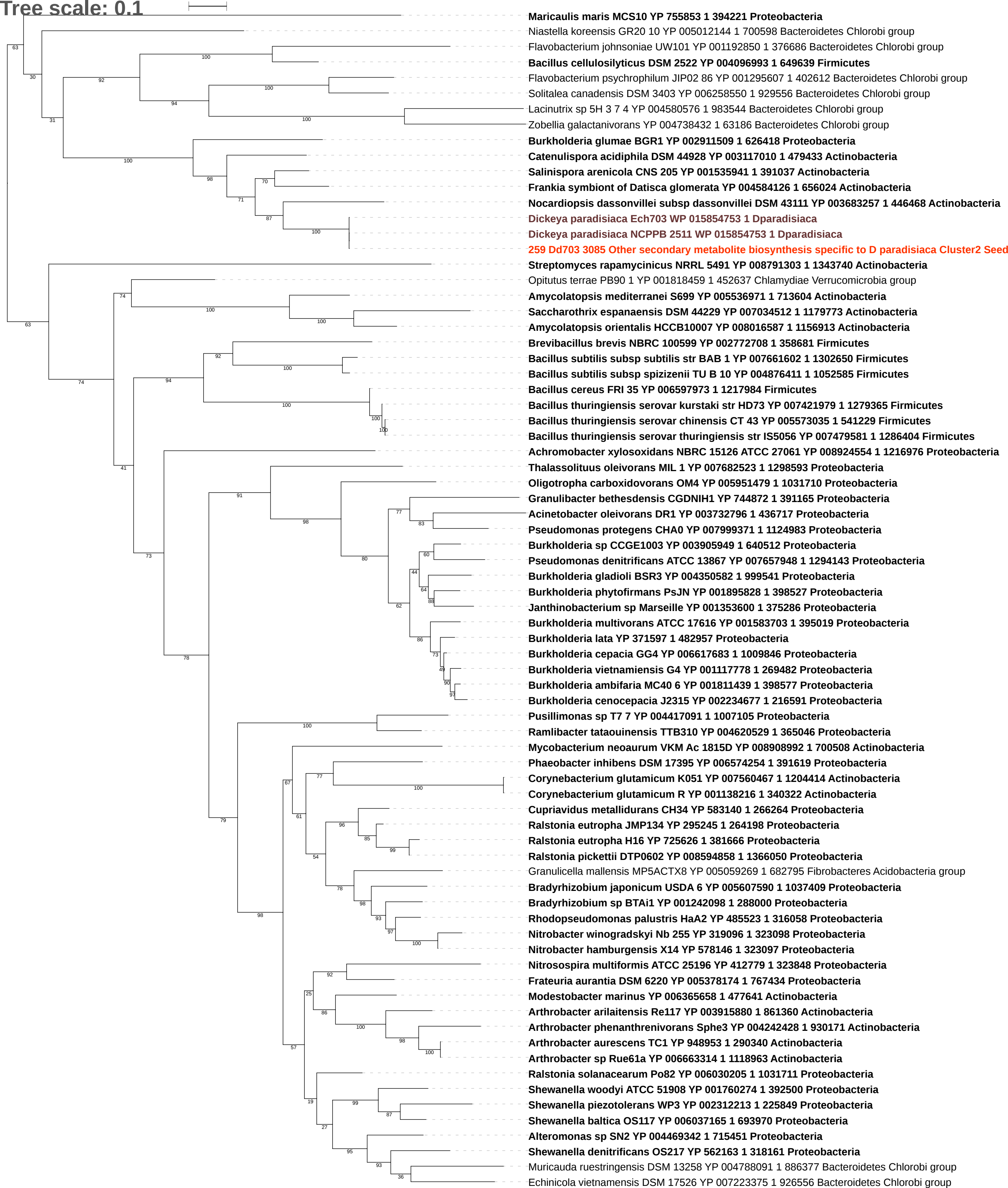
Tree scale: 0.1
Maricaulis maris MCS10 YP 755853 1 394221 Proteobacteria
Niastella koreensis GR20 10 YP 005012144 1 700598 Bacteroidetes Chlorobi group
Flavobacterium johnsoniae UW101 YP 001192850 1 376686 Bacteroidetes Chlorobi group
Bacillus cellulosilyticus DSM 2522 YP 004096993 1 649639 Firmicutes
Flavobacterium psychrophilum JIP02 86 YP 001295607 1 402612 Bacteroidetes Chlorobi group
Solitalea canadensis DSM 3403 YP 006258550 1 929556 Bacteroidetes Chlorobi group
Lacinutrix sp 5H 3 7 4 YP 004580576 1 983544 Bacteroidetes Chlorobi group
Zobellia galactanivorans YP 004738432 1 63186 Bacteroidetes Chlorobi group
Burkholderia glumae BGR1 YP 002911509 1 626418 Proteobacteria
Catenulispora acidiphila DSM 44928 YP 003117010 1 479433 Actinobacteria
Salinispora arenicola CNS 205 YP 001535941 1 391037 Actinobacteria
Frankia symbiont of Datisca glomerata YP 004584126 1 656024 Actinobacteria
Nocardiopsis dassonvillei subsp dassonvillei DSM 43111 YP 003683257 1 446468 Actinobacteria
Dickeya paradisiaca Ech703 WP 015854753 1 Dparadisiaca
Dickeya paradisiaca NCPPB 2511 WP 015854753 1 Dparadisiaca
259 Dd703 3085 Other secondary metabolite biosynthesis specific to D paradisiaca Cluster2 Seed
Streptomyces rapamycinicus NRRL 5491 YP 008791303 1 1343740 Actinobacteria
Opitutus terrae PB90 1 YP 001818459 1 452637 Chlamydiae Verrucomicrobia group
Amycolatopsis mediterranei S699 YP 005536971 1 713604 Actinobacteria
Saccharothrix espanaensis DSM 44229 YP 007034512 1 1179773 Actinobacteria
Amycolatopsis orientalis HCCB10007 YP 008016587 1 1156913 Actinobacteria
Brevibacillus brevis NBRC 100599 YP 002772708 1 358681 Firmicutes
Bacillus subtilis subsp subtilis str BAB 1 YP 007661602 1 1302650 Firmicutes
Bacillus subtilis subsp spizizenii TU B 10 YP 004876411 1 1052585 Firmicutes
Bacillus cereus FRI 35 YP 006597973 1 1217984 Firmicutes
Bacillus thuringiensis serovar kurstaki str HD73 YP 007421979 1 1279365 Firmicutes
Bacillus thuringiensis serovar chinensis CT 43 YP 005573035 1 541229 Firmicutes
Bacillus thuringiensis serovar thuringiensis str IS5056 YP 007479581 1 1286404 Firmicutes
Achromobacter xylosoxidans NBRC 15126 ATCC 27061 YP 008924554 1 1216976 Proteobacteria
Thalassolituus oleivorans MIL 1 YP 007682523 1 1298593 Proteobacteria
Oligotropha carboxidovorans OM4 YP 005951479 1 1031710 Proteobacteria
Granulibacter bethesdensis CGDNIH1 YP 744872 1 391165 Proteobacteria
Acinetobacter oleivorans DR1 YP 003732796 1 436717 Proteobacteria
Pseudomonas protegens CHA0 YP 007999371 1 1124983 Proteobacteria
Burkholderia sp CCGE1003 YP 003905949 1 640512 Proteobacteria
Pseudomonas denitrificans ATCC 13867 YP 007657948 1 1294143 Proteobacteria
Burkholderia gladioli BSR3 YP 004350582 1 999541 Proteobacteria
Burkholderia phytofirmans PsJN YP 001895828 1 398527 Proteobacteria
Janthinobacterium sp Marseille YP 001353600 1 375286 Proteobacteria
Burkholderia multivorans ATCC 17616 YP 001583703 1 395019 Proteobacteria
Burkholderia lata YP 371597 1 482957 Proteobacteria
Burkholderia cepacia GG4 YP 006617683 1 1009846 Proteobacteria
Burkholderia vietnamiensis G4 YP 001117778 1 269482 Proteobacteria
Burkholderia ambifaria MC40 6 YP 001811439 1 398577 Proteobacteria
Burkholderia cenocepacia J2315 YP 002234677 1 216591 Proteobacteria
Pusillimonas sp T7 7 YP 004417091 1 1007105 Proteobacteria
Ramlibacter tataouinensis TTB310 YP 004620529 1 365046 Proteobacteria
Mycobacterium neoaurum VKM Ac 1815D YP 008908992 1 700508 Actinobacteria
Phaeobacter inhibens DSM 17395 YP 006574254 1 391619 Proteobacteria
Corynebacterium glutamicum K051 YP 007560467 1 1204414 Actinobacteria
Corynebacterium glutamicum R YP 001138216 1 340322 Actinobacteria
Cupriavidus metallidurans CH34 YP 583140 1 266264 Proteobacteria
Ralstonia eutropha JMP134 YP 295245 1 264198 Proteobacteria
Ralstonia eutropha H16 YP 725626 1 381666 Proteobacteria
Ralstonia pickettii DTP0602 YP 008594858 1 1366050 Proteobacteria
Granulicella mallensis MP5ACTX8 YP 005059269 1 682795 Fibrobacteres Acidobacteria group
Bradyrhizobium japonicum USDA 6 YP 005607590 1 1037409 Proteobacteria
Bradyrhizobium sp BTAi1 YP 001242098 1 288000 Proteobacteria
Rhodopseudomonas palustris HaA2 YP 485523 1 316058 Proteobacteria
Nitrobacter winogradskyi Nb 255 YP 319096 1 323098 Proteobacteria
Nitrobacter hamburgensis X14 YP 578146 1 323097 Proteobacteria
Nitrosospira multiformis ATCC 25196 YP 412779 1 323848 Proteobacteria
Frateuria aurantia DSM 6220 YP 005378174 1 767434 Proteobacteria
Modestobacter marinus YP 006365658 1 477641 Actinobacteria
Arthrobacter arilaitensis Re117 YP 003915880 1 861360 Actinobacteria
Arthrobacter phenanthrenivorans Sphe3 YP 004242428 1 930171 Actinobacteria
Arthrobacter aurescens TC1 YP 948953 1 290340 Actinobacteria
Arthrobacter sp Rue61a YP 006663314 1 1118963 Actinobacteria
Ralstonia solanacearum Po82 YP 006030205 1 1031711 Proteobacteria
Shewanella woodyi ATCC 51908 YP 001760274 1 392500 Proteobacteria
Shewanella piezotolerans WP3 YP 002312213 1 225849 Proteobacteria
Shewanella baltica OS117 YP 006037165 1 693970 Proteobacteria
Alteromonas sp SN2 YP 004469342 1 715451 Proteobacteria
Shewanella denitrificans OS217 YP 562163 1 318161 Proteobacteria
Muricauda ruestringensis DSM 13258 YP 004788091 1 886377 Bacteroidetes Chlorobi group
Echinicola vietnamensis DSM 17526 YP 007223375 1 926556 Bacteroidetes Chlorobi group

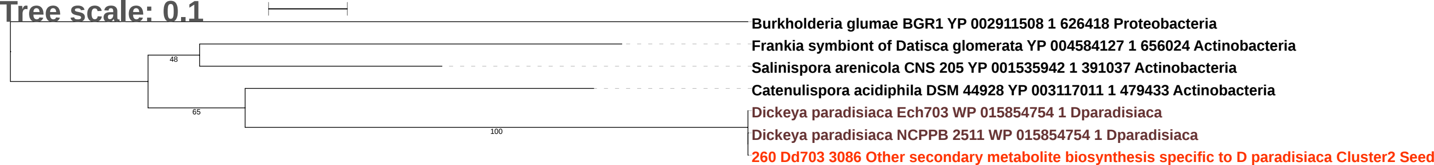
Tree scale: 0.1
Burkholderia glumae BGR1 YP 002911508 1 626418 Proteobacteria
Frankia symbiont of Datisca glomerata YP 004584127 1 656024 Actinobacteria
Salinispora arenicola CNS 205 YP 001535942 1 391037 Actinobacteria
Catenulispora acidiphila DSM 44928 YP 003117011 1 479433 Actinobacteria
Dickeya paradisiaca Ech703 WP 015854754 1 Dparadisiaca
Dickeya paradisiaca NCPPB 2511 WP 015854754 1 Dparadisiaca
260 Dd703 3086 Other secondary metabolite biosynthesis specific to D paradisiaca Cluster2 Seed
48
65
100

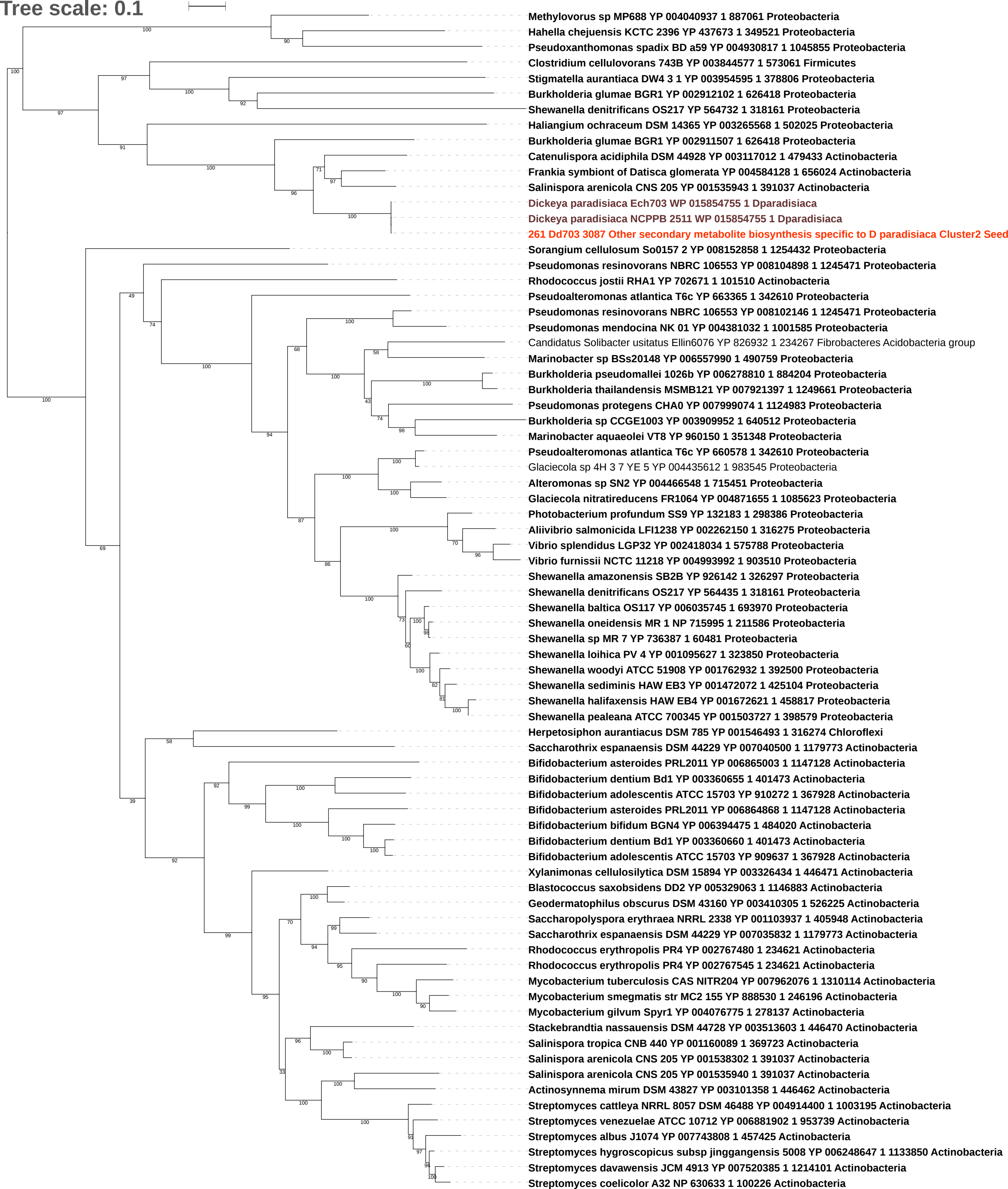
Tree scale: 0.1
Methylovorus sp MP688 YP 004040937 1 887061 Proteobacteria
Hahella chejuensis KCTC 2396 YP 437673 1 349521 Proteobacteria
Pseudoxanthomonas spadix BD a59 YP 004930817 1 1045855 Proteobacteria
Clostridium cellulovorans 743B YP 003844577 1 573061 Firmicutes
Stigmatella aurantiaca DW4 3 1 YP 003954595 1 378806 Proteobacteria
Burkholderia glumae BGR1 YP 002912102 1 626418 Proteobacteria
Shewanella denitrificans OS217 YP 564732 1 318161 Proteobacteria
Haliangium ochraceum DSM 14365 YP 003265568 1 502025 Proteobacteria
Burkholderia glumae BGR1 YP 002911507 1 626418 Proteobacteria
Catenulispora acidiphila DSM 44928 YP 003117012 1 479433 Actinobacteria
Frankia symbiont of Datisca glomerata YP 004584128 1 656024 Actinobacteria
Salinispora arenicola CNS 205 YP 001535943 1 391037 Actinobacteria
Dickeya paradisiaca Ech703 WP 015854755 1 Dparadisiaca
Dickeya paradisiaca NCPPB 2511 WP 015854755 1 Dparadisiaca
261 Dd703 3087 Other secondary metabolite biosynthesis specific to D paradisiaca Cluster2 Seed
Sorangium cellulosum So0157 2 YP 008152858 1 1254432 Proteobacteria
Pseudomonas resinovorans NBRC 106553 YP 008104898 1 1245471 Proteobacteria
Rhodococcus jostii RHA1 YP 702671 1 101510 Actinobacteria
Pseudoalteromonas atlantica T6c YP 663365 1 342610 Proteobacteria
Pseudomonas resinovorans NBRC 106553 YP 008102146 1 1245471 Proteobacteria
Pseudomonas mendocina NK 01 YP 004381032 1 1001585 Proteobacteria
Candidatus Solibacter usitatus Ellin6076 YP 826932 1 234267 Fibrobacteres Acidobacteria group
Marinobacter sp BSs20148 YP 006557990 1 490759 Proteobacteria
Burkholderia pseudomallei 1026b YP 006278810 1 884204 Proteobacteria
Burkholderia thailandensis MSMB121 YP 007921397 1 1249661 Proteobacteria
Pseudomonas protegens CHA0 YP 007999074 1 1124983 Proteobacteria
Burkholderia sp CCGE1003 YP 003909952 1 640512 Proteobacteria
Marinobacter aquaeolei VT8 YP 960150 1 351348 Proteobacteria
Pseudoalteromonas atlantica T6c YP 660578 1 342610 Proteobacteria
Glaciecola sp 4H 3 7 YE 5 YP 004435612 1 983545 Proteobacteria
Alteromonas sp SN2 YP 004466548 1 715451 Proteobacteria
Glaciecola nitratireducens FR1064 YP 004871655 1 1085623 Proteobacteria
Photobacterium profundum SS9 YP 132183 1 298386 Proteobacteria
Aliivibrio salmonicida LFI1238 YP 002262150 1 316275 Proteobacteria
Vibrio splendidus LGP32 YP 002418034 1 575788 Proteobacteria
Vibrio furnissii NCTC 11218 YP 004993992 1 903510 Proteobacteria
Shewanella amazonensis SB2B YP 926142 1 326297 Proteobacteria
Shewanella denitrificans OS217 YP 564435 1 318161 Proteobacteria
Shewanella baltica OS117 YP 006035745 1 693970 Proteobacteria
Shewanella oneidensis MR 1 NP 715995 1 211586 Proteobacteria
Shewanella sp MR 7 YP 736387 1 60481 Proteobacteria
Shewanella loihica PV 4 YP 001095627 1 323850 Proteobacteria
Shewanella woodyi ATCC 51908 YP 001762932 1 392500 Proteobacteria
Shewanella sediminis HAW EB3 YP 001472072 1 425104 Proteobacteria
Shewanella halifaxensis HAW EB4 YP 001672621 1 458817 Proteobacteria
Shewanella pealeana ATCC 700345 YP 001503727 1 398579 Proteobacteria
Herpetosiphon aurantiacus DSM 785 YP 001546493 1 316274 Chloroflexi
Saccharothrix espanaensis DSM 44229 YP 007040500 1 1179773 Actinobacteria
Bifidobacterium asteroides PRL2011 YP 006865003 1 1147128 Actinobacteria
Bifidobacterium dentium Bd1 YP 003360655 1 401473 Actinobacteria
Bifidobacterium adolescentis ATCC 15703 YP 910272 1 367928 Actinobacteria
Bifidobacterium asteroides PRL2011 YP 006864868 1 1147128 Actinobacteria
Bifidobacterium bifidum BGN4 YP 006394475 1 484020 Actinobacteria
Bifidobacterium dentium Bd1 YP 003360660 1 401473 Actinobacteria
Bifidobacterium adolescentis ATCC 15703 YP 909637 1 367928 Actinobacteria
Xylanimonas cellulosilytica DSM 15894 YP 003326434 1 446471 Actinobacteria
Blastococcus saxobsidens DD2 YP 005329063 1 1146883 Actinobacteria
Geodermatophilus obscurus DSM 43160 YP 003410305 1 526225 Actinobacteria
Saccharopolyspora erythraea NRRL 2338 YP 001103937 1 405948 Actinobacteria
Saccharothrix espanaensis DSM 44229 YP 007035832 1 1179773 Actinobacteria
Rhodococcus erythropolis PR4 YP 002767480 1 234621 Actinobacteria
Rhodococcus erythropolis PR4 YP 002767545 1 234621 Actinobacteria
Mycobacterium tuberculosis CAS NITR204 YP 007962076 1 1310114 Actinobacteria
Mycobacterium smegmatis str MC2 155 YP 888530 1 246196 Actinobacteria
Mycobacterium gilvum Spyr1 YP 004076775 1 278137 Actinobacteria
Stackebrandtia nassauensis DSM 44728 YP 003513603 1 446470 Actinobacteria
Salinispora tropica CNB 440 YP 001160089 1 369723 Actinobacteria
Salinispora arenicola CNS 205 YP 001538302 1 391037 Actinobacteria
Salinispora arenicola CNS 205 YP 001535940 1 391037 Actinobacteria
Actinosynnema mirum DSM 43827 YP 003101358 1 446462 Actinobacteria
Streptomyces cattleya NRRL 8057 DSM 46488 YP 004914400 1 1003195 Actinobacteria
Streptomyces venezuelae ATCC 10712 YP 006881902 1 953739 Actinobacteria
Streptomyces albus J1074 YP 007743808 1 457425 Actinobacteria
Streptomyces hygroscopicus subsp jinggangensis 5008 YP 006248647 1 1133850 Actinobacteria
Streptomyces davawensis JCM 4913 YP 007520385 1 1214101 Actinobacteria
Streptomyces coelicolor A32 NP 630633 1 100226 Actinobacteria

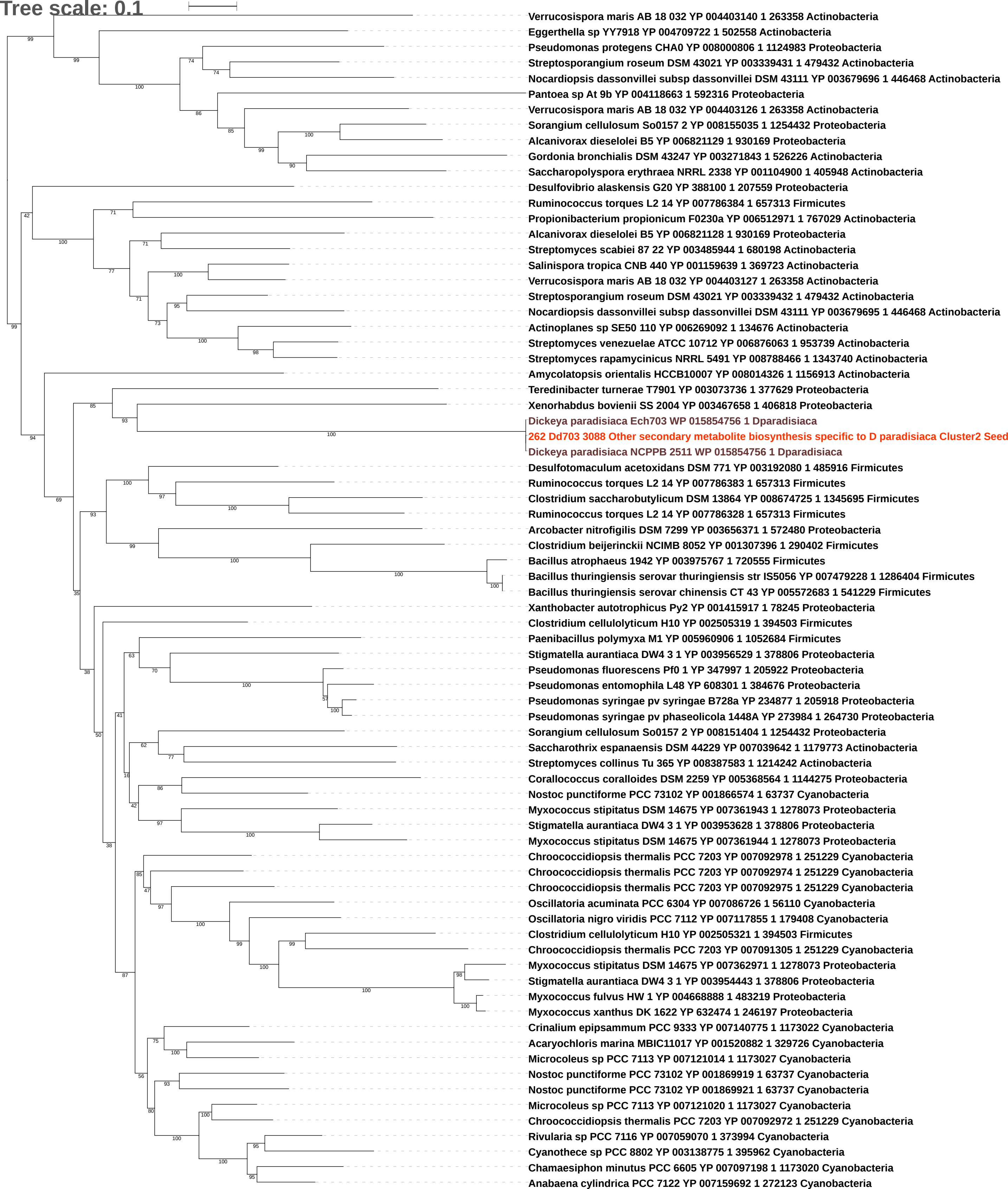

Tree scale: 0.1
Verrucosispora maris AB 18 032 YP 004403140 1 263358 Actinobacteria
Eggerthella sp YY7918 YP 004709722 1 502558 Actinobacteria
Pseudomonas protegens CHA0 YP 008000806 1 1124983 Proteobacteria
Streptosporangium roseum DSM 43021 YP 003339431 1 479432 Actinobacteria
Nocardiopsis dassonvillei subsp dassonvillei DSM 43111 YP 003679696 1 446468 Actinobacteria
Pantoea sp At 9b YP 004118663 1 592316 Proteobacteria
Verrucosispora maris AB 18 032 YP 004403126 1 263358 Actinobacteria
Sorangium cellulosum So0157 2 YP 008155035 1 1254432 Proteobacteria
Alcanivorax dieselolei B5 YP 006821129 1 930169 Proteobacteria
Gordonia bronchialis DSM 43247 YP 003271843 1 526226 Actinobacteria
Saccharopolyspora erythraea NRRL 2338 YP 001104900 1 405948 Actinobacteria
Desulfovibrio alaskensis G20 YP 388100 1 207559 Proteobacteria
Ruminococcus torques L2 14 YP 007786384 1 657313 Firmicutes
Propionibacterium propionicum F0230a YP 006512971 1 767029 Actinobacteria
Alcanivorax dieselolei B5 YP 006821128 1 930169 Proteobacteria
Streptomyces scabiei 87 22 YP 003485944 1 680198 Actinobacteria
Salinispora tropica CNB 440 YP 001159639 1 369723 Actinobacteria
Verrucosispora maris AB 18 032 YP 004403127 1 263358 Actinobacteria
Streptosporangium roseum DSM 43021 YP 003339432 1 479432 Actinobacteria
Nocardiopsis dassonvillei subsp dassonvillei DSM 43111 YP 003679695 1 446468 Actinobacteria
Actinoplanes sp SE50 110 YP 006269092 1 134676 Actinobacteria
Streptomyces venezuelae ATCC 10712 YP 006876063 1 953739 Actinobacteria
Streptomyces rapamycinicus NRRL 5491 YP 008788466 1 1343740 Actinobacteria
Amycolatopsis orientalis HCCB10007 YP 008014326 1 1156913 Actinobacteria
Teredinibacter turnerae T7901 YP 003073736 1 377629 Proteobacteria
Xenorhabdus bovienii SS 2004 YP 003467658 1 406818 Proteobacteria
Dickeya paradisiaca Ech703 WP 015854756 1 Dparadisiaca
262 Dd703 3088 Other secondary metabolite biosynthesis specific to D paradisiaca Cluster2 Seed
Dickeya paradisiaca NCPPB 2511 WP 015854756 1 Dparadisiaca
Desulfotomaculum acetoxidans DSM 771 YP 003192080 1 485916 Firmicutes
Ruminococcus torques L2 14 YP 007786383 1 657313 Firmicutes
Clostridium saccharobutylicum DSM 13864 YP 008674725 1 1345695 Firmicutes
Ruminococcus torques L2 14 YP 007786328 1 657313 Firmicutes
Arcobacter nitrofigilis DSM 7299 YP 003656371 1 572480 Proteobacteria
Clostridium beijerinckii NCIMB 8052 YP 001307396 1 290402 Firmicutes
Bacillus atrophaeus 1942 YP 003975767 1 720555 Firmicutes
Bacillus thuringiensis serovar thuringiensis str IS5056 YP 007479228 1 1286404 Firmicutes
Bacillus thuringiensis serovar chinensis CT 43 YP 005572683 1 541229 Firmicutes
Xanthobacter autotrophicus Py2 YP 001415917 1 78245 Proteobacteria
Clostridium cellulolyticum H10 YP 002505319 1 394503 Firmicutes
Paenibacillus polymyxa M1 YP 005960906 1 1052684 Firmicutes
Stigmatella aurantiaca DW4 3 1 YP 003956529 1 378806 Proteobacteria
Pseudomonas fluorescens Pf0 1 YP 347997 1 205922 Proteobacteria
Pseudomonas entomophila L48 YP 608301 1 384676 Proteobacteria
Pseudomonas syringae pv syringae B728a YP 234877 1 205918 Proteobacteria
Pseudomonas syringae pv phaseolicola 1448A YP 273984 1 264730 Proteobacteria
Sorangium cellulosum So0157 2 YP 008151404 1 1254432 Proteobacteria
Saccharothrix espanaensis DSM 44229 YP 007039642 1 1179773 Actinobacteria
Streptomyces collinus Tu 365 YP 008387583 1 1214242 Actinobacteria
Corallococcus coralloides DSM 2259 YP 005368564 1 1144275 Proteobacteria
Nostoc punctiforme PCC 73102 YP 001866574 1 63737 Cyanobacteria
Myxococcus stipitatus DSM 14675 YP 007361943 1 1278073 Proteobacteria
Stigmatella aurantiaca DW4 3 1 YP 003953628 1 378806 Proteobacteria
Myxococcus stipitatus DSM 14675 YP 007361944 1 1278073 Proteobacteria
Chroococcidiopsis thermalis PCC 7203 YP 007092978 1 251229 Cyanobacteria
Chroococcidiopsis thermalis PCC 7203 YP 007092974 1 251229 Cyanobacteria
Chroococcidiopsis thermalis PCC 7203 YP 007092975 1 251229 Cyanobacteria
Oscillatoria acuminata PCC 6304 YP 007086726 1 56110 Cyanobacteria
Oscillatoria nigro viridis PCC 7112 YP 007117855 1 179408 Cyanobacteria
Clostridium cellulolyticum H10 YP 002505321 1 394503 Firmicutes
Chroococcidiopsis thermalis PCC 7203 YP 007091305 1 251229 Cyanobacteria
Myxococcus stipitatus DSM 14675 YP 007362971 1 1278073 Proteobacteria
Stigmatella aurantiaca DW4 3 1 YP 003954443 1 378806 Proteobacteria
Myxococcus fulvus HW 1 YP 004668888 1 483219 Proteobacteria
Myxococcus xanthus DK 1622 YP 632474 1 246197 Proteobacteria
Crinalium epipsammum PCC 9333 YP 007140775 1 1173022 Cyanobacteria
Acaryochloris marina MBIC11017 YP 001520882 1 329726 Cyanobacteria
Microcoleus sp PCC 7113 YP 007121014 1 1173027 Cyanobacteria
Nostoc punctiforme PCC 73102 YP 001869919 1 63737 Cyanobacteria
Nostoc punctiforme PCC 73102 YP 001869921 1 63737 Cyanobacteria
Microcoleus sp PCC 7113 YP 007121020 1 1173027 Cyanobacteria
Chroococcidiopsis thermalis PCC 7203 YP 007092972 1 251229 Cyanobacteria
Rivularia sp PCC 7116 YP 007059070 1 373994 Cyanobacteria
Cyanothece sp PCC 8802 YP 003138775 1 395962 Cyanobacteria
Chamaesiphon minutus PCC 6605 YP 007097198 1 1173020 Cyanobacteria
Anabaena cylindrica PCC 7122 YP 007159692 1 272123 Cyanobacteria

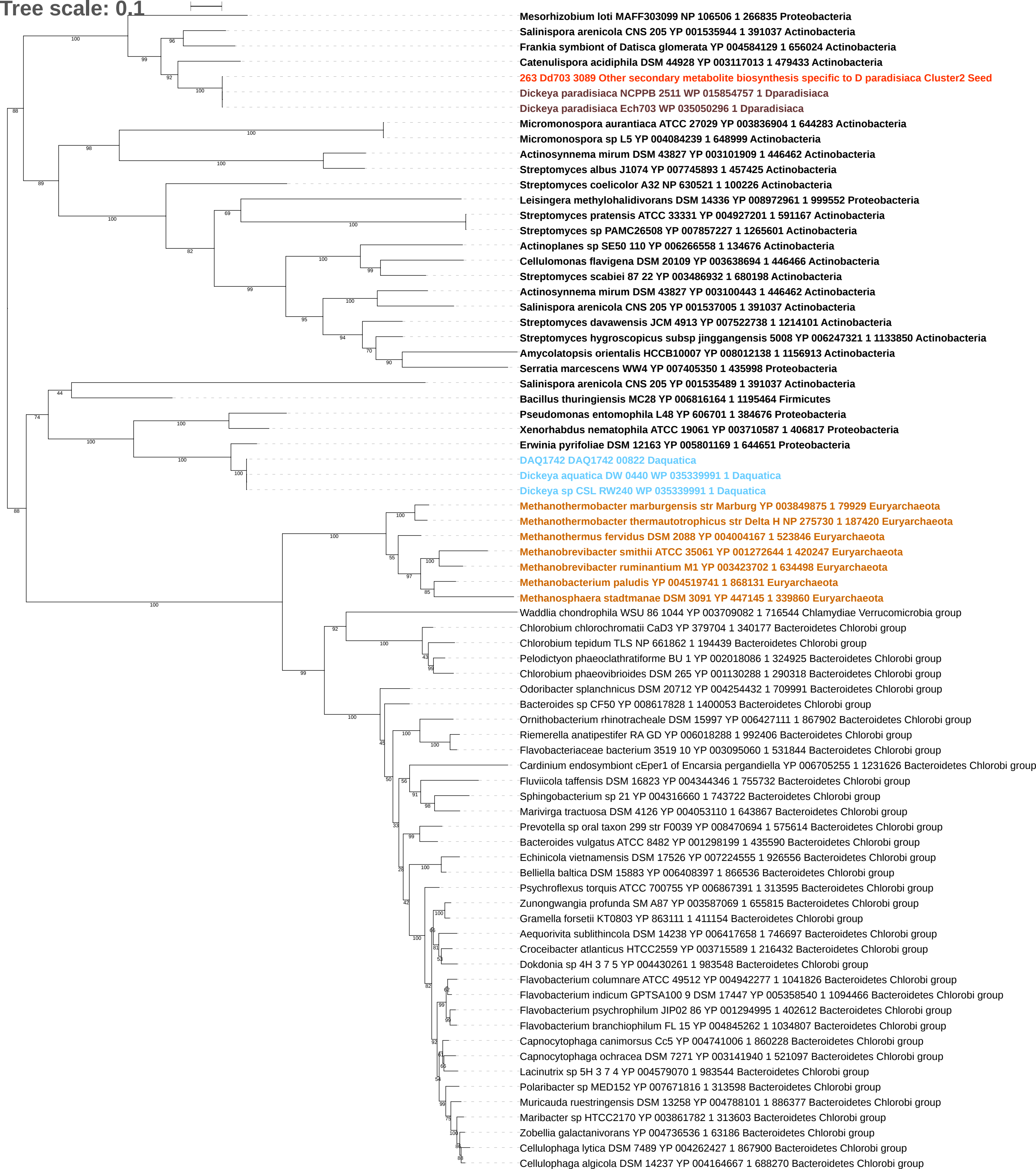
Tree scale: 0.1
Mesorhizobium loti MAFF303099 NP 106506 1 266835 Proteobacteria
Salinispora arenicola CNS 205 YP 001535944 1 391037 Actinobacteria
Frankia symbiont of Datisca glomerata YP 004584129 1 656024 Actinobacteria
Catenulispora acidiphila DSM 44928 YP 003117013 1 479433 Actinobacteria
263 Dd703 3089 Other secondary metabolite biosynthesis specific to D paradisiaca Cluster2 Seed
Dickeya paradisiaca NCPPB 2511 WP 015854757 1 Dparadisiaca
Dickeya paradisiaca Ech703 WP 035050296 1 Dparadisiaca
Micromonospora aurantiaca ATCC 27029 YP 003836904 1 644283 Actinobacteria
Micromonospora sp L5 YP 004084239 1 648999 Actinobacteria
Actinosynnema mirum DSM 43827 YP 003101909 1 446462 Actinobacteria
Streptomyces albus J1074 YP 007745893 1 457425 Actinobacteria
Streptomyces coelicolor A32 NP 630521 1 100226 Actinobacteria
Leisingera methylohalidivorans DSM 14336 YP 008972961 1 999552 Proteobacteria
Streptomyces pratensis ATCC 33331 YP 004927201 1 591167 Actinobacteria
Streptomyces sp PAMC26508 YP 007857227 1 1265601 Actinobacteria
Actinoplanes sp SE50 110 YP 006266558 1 134676 Actinobacteria
Cellulomonas flavigena DSM 20109 YP 003638694 1 446466 Actinobacteria
Streptomyces scabiei 87 22 YP 003486932 1 680198 Actinobacteria
Actinosynnema mirum DSM 43827 YP 003100443 1 446462 Actinobacteria
Salinispora arenicola CNS 205 YP 001537005 1 391037 Actinobacteria
Streptomyces davawensis JCM 4913 YP 007522738 1 1214101 Actinobacteria
Streptomyces hygroscopicus subsp jinggangensis 5008 YP 006247321 1 1133850 Actinobacteria
Amycolatopsis orientalis HCCB10007 YP 008012138 1 1156913 Actinobacteria
Serratia marcescens WW4 YP 007405350 1 435998 Proteobacteria
Salinispora arenicola CNS 205 YP 001535489 1 391037 Actinobacteria
Bacillus thuringiensis MC28 YP 006816164 1 1195464 Firmicutes
Pseudomonas entomophila L48 YP 606701 1 384676 Proteobacteria
Xenorhabdus nematophila ATCC 19061 YP 003710587 1 406817 Proteobacteria
Erwinia pyrifoliae DSM 12163 YP 005801169 1 644651 Proteobacteria
DAQ1742 DAQ1742 00822 Daquatica
Dickeya aquatica DW 0440 WP 035339991 1 Daquatica
Dickeya sp CSL RW240 WP 035339991 1 Daquatica
Methanothermobacter marburgensis str Marburg YP 003849875 1 79929 Euryarchaeota
Methanothermobacter thermautotrophicus str Delta H NP 275730 1 187420 Euryarchaeota
Methanothermus fervidus DSM 2088 YP 004004167 1 523846 Euryarchaeota
Methanobrevibacter smithii ATCC 35061 YP 001272644 1 420247 Euryarchaeota
Methanobrevibacter ruminantium M1 YP 003423702 1 634498 Euryarchaeota
Methanobacterium paludis YP 004519741 1 868131 Euryarchaeota
Methanosphaera stadtmanae DSM 3091 YP 447145 1 339860 Euryarchaeota
Waddlia chondrophila WSU 86 1044 YP 003709082 1 716544 Chlamydiae Verrucomicrobia group
Chlorobium chlorochromatii CaD3 YP 379704 1 340177 Bacteroidetes Chlorobi group
Chlorobium tepidum TLS NP 661862 1 194439 Bacteroidetes Chlorobi group
Pelodictyon phaeoclathratiforme BU 1 YP 002018086 1 324925 Bacteroidetes Chlorobi group
Chlorobium phaeovibrioides DSM 265 YP 001130288 1 290318 Bacteroidetes Chlorobi group
Odoribacter splanchnicus DSM 20712 YP 004254432 1 709991 Bacteroidetes Chlorobi group
Bacteroides sp CF50 YP 008617828 1 1400053 Bacteroidetes Chlorobi group
Ornithobacterium rhinotracheale DSM 15997 YP 006427111 1 867902 Bacteroidetes Chlorobi group
Riemerella anatipestifer RA GD YP 006018288 1 992406 Bacteroidetes Chlorobi group
Flavobacteriaceae bacterium 3519 10 YP 003095060 1 531844 Bacteroidetes Chlorobi group
Cardinium endosymbiont cEper1 of Encarsia pergandiella YP 006705255 1 1231626 Bacteroidetes Chlorobi group
Fluviicola taffensis DSM 16823 YP 004344346 1 755732 Bacteroidetes Chlorobi group
Sphingobacterium sp 21 YP 004316660 1 743722 Bacteroidetes Chlorobi group
Marivirga tractuosa DSM 4126 YP 004053110 1 643867 Bacteroidetes Chlorobi group
Prevotella sp oral taxon 299 str F0039 YP 008470694 1 575614 Bacteroidetes Chlorobi group
Bacteroides vulgatus ATCC 8482 YP 001298199 1 435590 Bacteroidetes Chlorobi group
Echinicola vietnamensis DSM 17526 YP 007224555 1 926556 Bacteroidetes Chlorobi group
Belliella baltica DSM 15883 YP 006408397 1 866536 Bacteroidetes Chlorobi group
Psychroflexus torquis ATCC 700755 YP 006867391 1 313595 Bacteroidetes Chlorobi group
Zunongwangia profunda SM A87 YP 003587069 1 655815 Bacteroidetes Chlorobi group
Gramella forsetii KT0803 YP 863111 1 411154 Bacteroidetes Chlorobi group
Aequorivita sublithincola DSM 14238 YP 006417658 1 746697 Bacteroidetes Chlorobi group
Croceibacter atlanticus HTCC2559 YP 003715589 1 216432 Bacteroidetes Chlorobi group
Dokdonia sp 4H 3 7 5 YP 004430261 1 983548 Bacteroidetes Chlorobi group
Flavobacterium columnare ATCC 49512 YP 004942277 1 1041826 Bacteroidetes Chlorobi group
Flavobacterium indicum GPTSA100 9 DSM 17447 YP 005358540 1 1094466 Bacteroidetes Chlorobi group
Flavobacterium psychrophilum JIP02 86 YP 001294995 1 402612 Bacteroidetes Chlorobi group
Flavobacterium branchiophilum FL 15 YP 004845262 1 1034807 Bacteroidetes Chlorobi group
Capnocytophaga canimorsus Cc5 YP 004741006 1 860228 Bacteroidetes Chlorobi group
Capnocytophaga ochracea DSM 7271 YP 003141940 1 521097 Bacteroidetes Chlorobi group
Lacinutrix sp 5H 3 7 4 YP 004579070 1 983544 Bacteroidetes Chlorobi group
Polaribacter sp MED152 YP 007671816 1 313598 Bacteroidetes Chlorobi group
Muricauda ruestringensis DSM 13258 YP 004788101 1 886377 Bacteroidetes Chlorobi group
Maribacter sp HTCC2170 YP 003861782 1 313603 Bacteroidetes Chlorobi group
Zobellia galactanivorans YP 004736536 1 63186 Bacteroidetes Chlorobi group
Cellulophaga lytica DSM 7489 YP 004262427 1 867900 Bacteroidetes Chlorobi group
Cellulophaga algicola DSM 14237 YP 004164667 1 688270 Bacteroidetes Chlorobi group

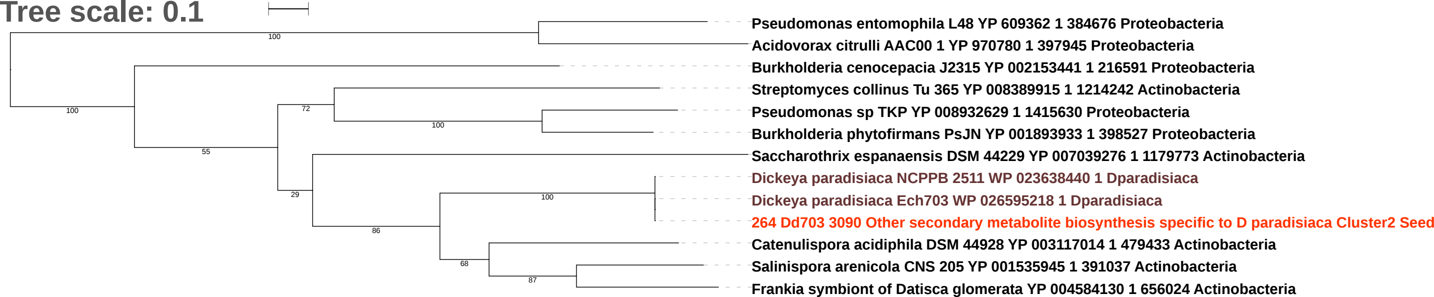
Tree scale: 0.1
Pseudomonas entomophila L48 YP 609362 1 384676 Proteobacteria
Acidovorax citrulli AAC00 1 YP 970780 1 397945 Proteobacteria
Burkholderia cenocepacia J2315 YP 002153441 1 216591 Proteobacteria
Streptomyces collinus Tu 365 YP 008389915 1 1214242 Actinobacteria
Pseudomonas sp TKP YP 008932629 1 1415630 Proteobacteria
Burkholderia phytofirmans PsJN YP 001893933 1 398527 Proteobacteria
Saccharothrix espanaensis DSM 44229 YP 007039276 1 1179773 Actinobacteria
Dickeya paradisiaca NCPPB 2511 WP 023638440 1 Dparadisiaca
Dickeya paradisiaca Ech703 WP 026595218 1 Dparadisiaca
264 Dd703 3090 Other secondary metabolite biosynthesis specific to D paradisiaca Cluster2 Seed
Catenulispora acidiphila DSM 44928 YP 003117014 1 479433 Actinobacteria
Salinispora arenicola CNS 205 YP 001535945 1 391037 Actinobacteria
Frankia symbiont of Datisca glomerata YP 004584130 1 656024 Actinobacteria
100
100
72
100
55
29
100
86
68
87

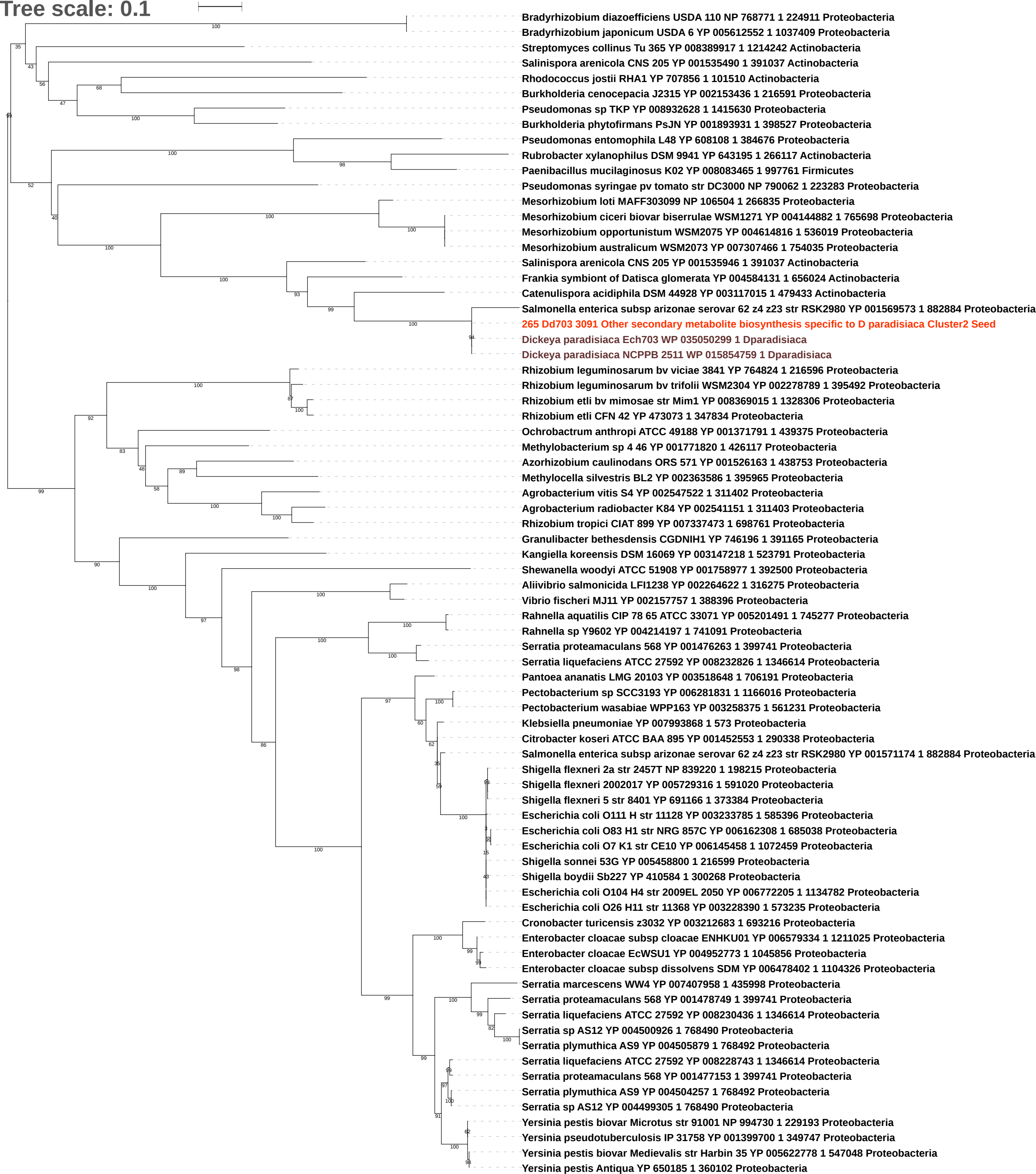
Tree scale: 0.1
Bradyrhizobium diazoefficiens USDA 110 NP 768771 1 224911 Proteobacteria
Bradyrhizobium japonicum USDA 6 YP 005612552 1 1037409 Proteobacteria
Streptomyces collinus Tu 365 YP 008389917 1 1214242 Actinobacteria
Salinispora arenicola CNS 205 YP 001535490 1 391037 Actinobacteria
Rhodococcus jostii RHA1 YP 707856 1 101510 Actinobacteria
Burkholderia cenocepacia J2315 YP 002153436 1 216591 Proteobacteria
Pseudomonas sp TKP YP 008932628 1 1415630 Proteobacteria
Burkholderia phytofirmans PsJN YP 001893931 1 398527 Proteobacteria
Pseudomonas entomophila L48 YP 608108 1 384676 Proteobacteria
Rubrobacter xylanophilus DSM 9941 YP 643195 1 266117 Actinobacteria
Paenibacillus mucilaginosus K02 YP 008083465 1 997761 Firmicutes
Pseudomonas syringae pv tomato str DC3000 NP 790062 1 223283 Proteobacteria
Mesorhizobium loti MAFF303099 NP 106504 1 266835 Proteobacteria
Mesorhizobium ciceri biovar biserrulae WSM1271 YP 004144882 1 765698 Proteobacteria
Mesorhizobium opportunistum WSM2075 YP 004614816 1 536019 Proteobacteria
Mesorhizobium australicum WSM2073 YP 007307466 1 754035 Proteobacteria
Salinispora arenicola CNS 205 YP 001535946 1 391037 Actinobacteria
Frankia symbiont of Datisca glomerata YP 004584131 1 656024 Actinobacteria
Catenulispora acidiphila DSM 44928 YP 003117015 1 479433 Actinobacteria
Salmonella enterica subsp arizonae serovar 62 z4 z23 str RSK2980 YP 001569573 1 882884 Proteobacteria
265 Dd703 3091 Other secondary metabolite biosynthesis specific to D paradisiaca Cluster2 Seed
Dickeya paradisiaca Ech703 WP 035050299 1 Dparadisiaca
Dickeya paradisiaca NCPPB 2511 WP 015854759 1 Dparadisiaca
Rhizobium leguminosarum bv viciae 3841 YP 764824 1 216596 Proteobacteria
Rhizobium leguminosarum bv trifolii WSM2304 YP 002278789 1 395492 Proteobacteria
Rhizobium etli bv mimosae str Mim1 YP 008369015 1 1328306 Proteobacteria
Rhizobium etli CFN 42 YP 473073 1 347834 Proteobacteria
Ochrobactrum anthropi ATCC 49188 YP 001371791 1 439375 Proteobacteria
Methylobacterium sp 4 46 YP 001771820 1 426117 Proteobacteria
Azorhizobium caulinodans ORS 571 YP 001526163 1 438753 Proteobacteria
Methylocella silvestris BL2 YP 002363586 1 395965 Proteobacteria
Agrobacterium vitis S4 YP 002547522 1 311402 Proteobacteria
Agrobacterium radiobacter K84 YP 002541151 1 311403 Proteobacteria
Rhizobium tropici CIAT 899 YP 007337473 1 698761 Proteobacteria
Granulibacter bethesdensis CGDNIH1 YP 746196 1 391165 Proteobacteria
Kangiella koreensis DSM 16069 YP 003147218 1 523791 Proteobacteria
Shewanella woodyi ATCC 51908 YP 001758977 1 392500 Proteobacteria
Aliivibrio salmonicida LFI1238 YP 002264622 1 316275 Proteobacteria
Vibrio fischeri MJ11 YP 002157757 1 388396 Proteobacteria
Rahnella aquatilis CIP 78 65 ATCC 33071 YP 005201491 1 745277 Proteobacteria
Rahnella sp Y9602 YP 004214197 1 741091 Proteobacteria
Serratia proteamaculans 568 YP 001476263 1 399741 Proteobacteria
Serratia liquefaciens ATCC 27592 YP 008232826 1 1346614 Proteobacteria
Pantoea ananatis LMG 20103 YP 003518648 1 706191 Proteobacteria
Pectobacterium sp SCC3193 YP 006281831 1 1166016 Proteobacteria
Pectobacterium wasabiae WPP163 YP 003258375 1 561231 Proteobacteria
Klebsiella pneumoniae YP 007993868 1 573 Proteobacteria
Citrobacter koseri ATCC BAA 895 YP 001452553 1 290338 Proteobacteria
Salmonella enterica subsp arizonae serovar 62 z4 z23 str RSK2980 YP 001571174 1 882884 Proteobacteria
Shigella flexneri 2a str 2457T NP 839220 1 198215 Proteobacteria
Shigella flexneri 2002017 YP 005729316 1 591020 Proteobacteria
Shigella flexneri 5 str 8401 YP 691166 1 373384 Proteobacteria
Escherichia coli O111 H str 11128 YP 003233785 1 585396 Proteobacteria
Escherichia coli O83 H1 str NRG 857C YP 006162308 1 685038 Proteobacteria
Escherichia coli O7 K1 str CE10 YP 006145458 1 1072459 Proteobacteria
Shigella sonnei 53G YP 005458800 1 216599 Proteobacteria
Shigella boydii Sb227 YP 410584 1 300268 Proteobacteria
Escherichia coli O104 H4 str 2009EL 2050 YP 006772205 1 1134782 Proteobacteria
Escherichia coli O26 H11 str 11368 YP 003228390 1 573235 Proteobacteria
Cronobacter turicensis z3032 YP 003212683 1 693216 Proteobacteria
Enterobacter cloacae subsp cloacae ENHKU01 YP 006579334 1 1211025 Proteobacteria
Enterobacter cloacae EcWSU1 YP 004952773 1 1045856 Proteobacteria
Enterobacter cloacae subsp dissolvens SDM YP 006478402 1 1104326 Proteobacteria
Serratia marcescens WW4 YP 007407958 1 435998 Proteobacteria
Serratia proteamaculans 568 YP 001478749 1 399741 Proteobacteria
Serratia liquefaciens ATCC 27592 YP 008230436 1 1346614 Proteobacteria
Serratia sp AS12 YP 004500926 1 768490 Proteobacteria
Serratia plymuthica AS9 YP 004505879 1 768492 Proteobacteria
Serratia liquefaciens ATCC 27592 YP 008228743 1 1346614 Proteobacteria
Serratia proteamaculans 568 YP 001477153 1 399741 Proteobacteria
Serratia plymuthica AS9 YP 004504257 1 768492 Proteobacteria
Serratia sp AS12 YP 004499305 1 768490 Proteobacteria
Yersinia pestis biovar Microtus str 91001 NP 994730 1 229193 Proteobacteria
Yersinia pseudotuberculosis IP 31758 YP 001399700 1 349747 Proteobacteria
Yersinia pestis biovar Medievalis str Harbin 35 YP 005622778 1 547048 Proteobacteria
Yersinia pestis Antiqua YP 650185 1 360102 Proteobacteria

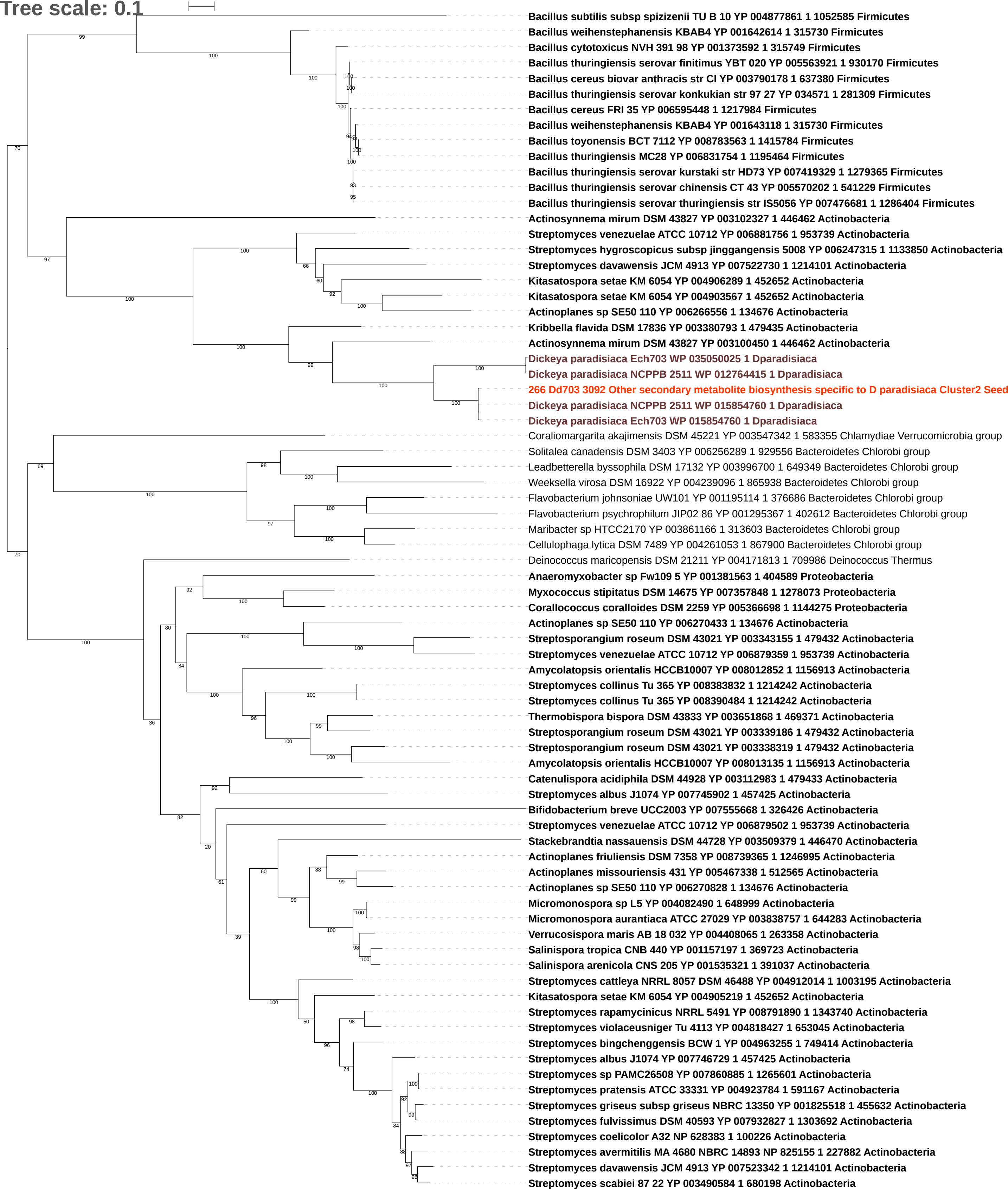
Tree scale: 0.1
Bacillus subtilis subsp spizizenii TU B 10 YP 004877861 1 1052585 Firmicutes
Bacillus weihenstephanensis KBAB4 YP 001642614 1 315730 Firmicutes
Bacillus cytotoxicus NVH 391 98 YP 001373592 1 315749 Firmicutes
Bacillus thuringiensis serovar finitimus YBT 020 YP 005563921 1 930170 Firmicutes
Bacillus cereus biovar anthracis str CI YP 003790178 1 637380 Firmicutes
Bacillus thuringiensis serovar konkukian str 97 27 YP 034571 1 281309 Firmicutes
Bacillus cereus FRI 35 YP 006595448 1 1217984 Firmicutes
Bacillus weihenstephanensis KBAB4 YP 001643118 1 315730 Firmicutes
Bacillus toyonensis BCT 7112 YP 008783563 1 1415784 Firmicutes
Bacillus thuringiensis MC28 YP 006831754 1 1195464 Firmicutes
Bacillus thuringiensis serovar kurstaki str HD73 YP 007419329 1 1279365 Firmicutes
Bacillus thuringiensis serovar chinensis CT 43 YP 005570202 1 541229 Firmicutes
Bacillus thuringiensis serovar thuringiensis str IS5056 YP 007476681 1 1286404 Firmicutes
Actinosynnema mirum DSM 43827 YP 003102327 1 446462 Actinobacteria
Streptomyces venezuelae ATCC 10712 YP 006881756 1 953739 Actinobacteria
Streptomyces hygroscopicus subsp jinggangensis 5008 YP 006247315 1 1133850 Actinobacteria
Streptomyces davawensis JCM 4913 YP 007522730 1 1214101 Actinobacteria
Kitasatospora setae KM 6054 YP 004906289 1 452652 Actinobacteria
Kitasatospora setae KM 6054 YP 004903567 1 452652 Actinobacteria
Actinoplanes sp SE50 110 YP 006266556 1 134676 Actinobacteria
Kribbella flavida DSM 17836 YP 003380793 1 479435 Actinobacteria
Actinosynnema mirum DSM 43827 YP 003100450 1 446462 Actinobacteria
Dickeya paradisiaca Ech703 WP 035050025 1 Dparadisiaca
Dickeya paradisiaca NCPPB 2511 WP 012764415 1 Dparadisiaca
266 Dd703 3092 Other secondary metabolite biosynthesis specific to D paradisiaca Cluster2 Seed
Dickeya paradisiaca NCPPB 2511 WP 015854760 1 Dparadisiaca
Dickeya paradisiaca Ech703 WP 015854760 1 Dparadisiaca
Coraliomargarita akajimensis DSM 45221 YP 003547342 1 583355 Chlamydiae Verrucomicrobia group
Solitalea canadensis DSM 3403 YP 006256289 1 929556 Bacteroidetes Chlorobi group
Leadbetterella byssophila DSM 17132 YP 003996700 1 649349 Bacteroidetes Chlorobi group
Weeksella virosa DSM 16922 YP 004239096 1 865938 Bacteroidetes Chlorobi group
Flavobacterium johnsoniae UW101 YP 001195114 1 376686 Bacteroidetes Chlorobi group
Flavobacterium psychrophilum JIP02 86 YP 001295367 1 402612 Bacteroidetes Chlorobi group
Maribacter sp HTCC2170 YP 003861166 1 313603 Bacteroidetes Chlorobi group
Cellulophaga lytica DSM 7489 YP 004261053 1 867900 Bacteroidetes Chlorobi group
Deinococcus maricopensis DSM 21211 YP 004171813 1 709986 Deinococcus Thermus
Anaeromyxobacter sp Fw109 5 YP 001381563 1 404589 Proteobacteria
Myxococcus stipitatus DSM 14675 YP 007357848 1 1278073 Proteobacteria
Corallococcus coralloides DSM 2259 YP 005366698 1 1144275 Proteobacteria
Actinoplanes sp SE50 110 YP 006270433 1 134676 Actinobacteria
Streptosporangium roseum DSM 43021 YP 003343155 1 479432 Actinobacteria
Streptomyces venezuelae ATCC 10712 YP 006879359 1 953739 Actinobacteria
Amycolatopsis orientalis HCCB10007 YP 008012852 1 1156913 Actinobacteria
Streptomyces collinus Tu 365 YP 008383832 1 1214242 Actinobacteria
Streptomyces collinus Tu 365 YP 008390484 1 1214242 Actinobacteria
Thermobispora bispora DSM 43833 YP 003651868 1 469371 Actinobacteria
Streptosporangium roseum DSM 43021 YP 003339186 1 479432 Actinobacteria
Streptosporangium roseum DSM 43021 YP 003338319 1 479432 Actinobacteria
Amycolatopsis orientalis HCCB10007 YP 008013135 1 1156913 Actinobacteria
Catenulispora acidiphila DSM 44928 YP 003112983 1 479433 Actinobacteria
Streptomyces albus J1074 YP 007745902 1 457425 Actinobacteria
Bifidobacterium breve UCC2003 YP 007555668 1 326426 Actinobacteria
Streptomyces venezuelae ATCC 10712 YP 006879502 1 953739 Actinobacteria
Stackebrandtia nassauensis DSM 44728 YP 003509379 1 446470 Actinobacteria
Actinoplanes friuliensis DSM 7358 YP 008739365 1 1246995 Actinobacteria
Actinoplanes missouriensis 431 YP 005467338 1 512565 Actinobacteria
Actinoplanes sp SE50 110 YP 006270828 1 134676 Actinobacteria
Micromonospora sp L5 YP 004082490 1 648999 Actinobacteria
Micromonospora aurantiaca ATCC 27029 YP 003838757 1 644283 Actinobacteria
Verrucosispora maris AB 18 032 YP 004408065 1 263358 Actinobacteria
Salinispora tropica CNB 440 YP 001157197 1 369723 Actinobacteria
Salinispora arenicola CNS 205 YP 001535321 1 391037 Actinobacteria
Streptomyces cattleya NRRL 8057 DSM 46488 YP 004912014 1 1003195 Actinobacteria
Kitasatospora setae KM 6054 YP 004905219 1 452652 Actinobacteria
Streptomyces rapamycinicus NRRL 5491 YP 008791890 1 1343740 Actinobacteria
Streptomyces violaceusniger Tu 4113 YP 004818427 1 653045 Actinobacteria
Streptomyces bingchenggensis BCW 1 YP 004963255 1 749414 Actinobacteria
Streptomyces albus J1074 YP 007746729 1 457425 Actinobacteria
Streptomyces sp PAMC26508 YP 007860885 1 1265601 Actinobacteria
Streptomyces pratensis ATCC 33331 YP 004923784 1 591167 Actinobacteria
Streptomyces griseus subsp griseus NBRC 13350 YP 001825518 1 455632 Actinobacteria
Streptomyces fulvissimus DSM 40593 YP 007932827 1 1303692 Actinobacteria
Streptomyces coelicolor A32 NP 628383 1 100226 Actinobacteria
Streptomyces avermitilis MA 4680 NBRC 14893 NP 825155 1 227882 Actinobacteria
Streptomyces davawensis JCM 4913 YP 007523342 1 1214101 Actinobacteria
Streptomyces scabiei 87 22 YP 003490584 1 680198 Actinobacteria

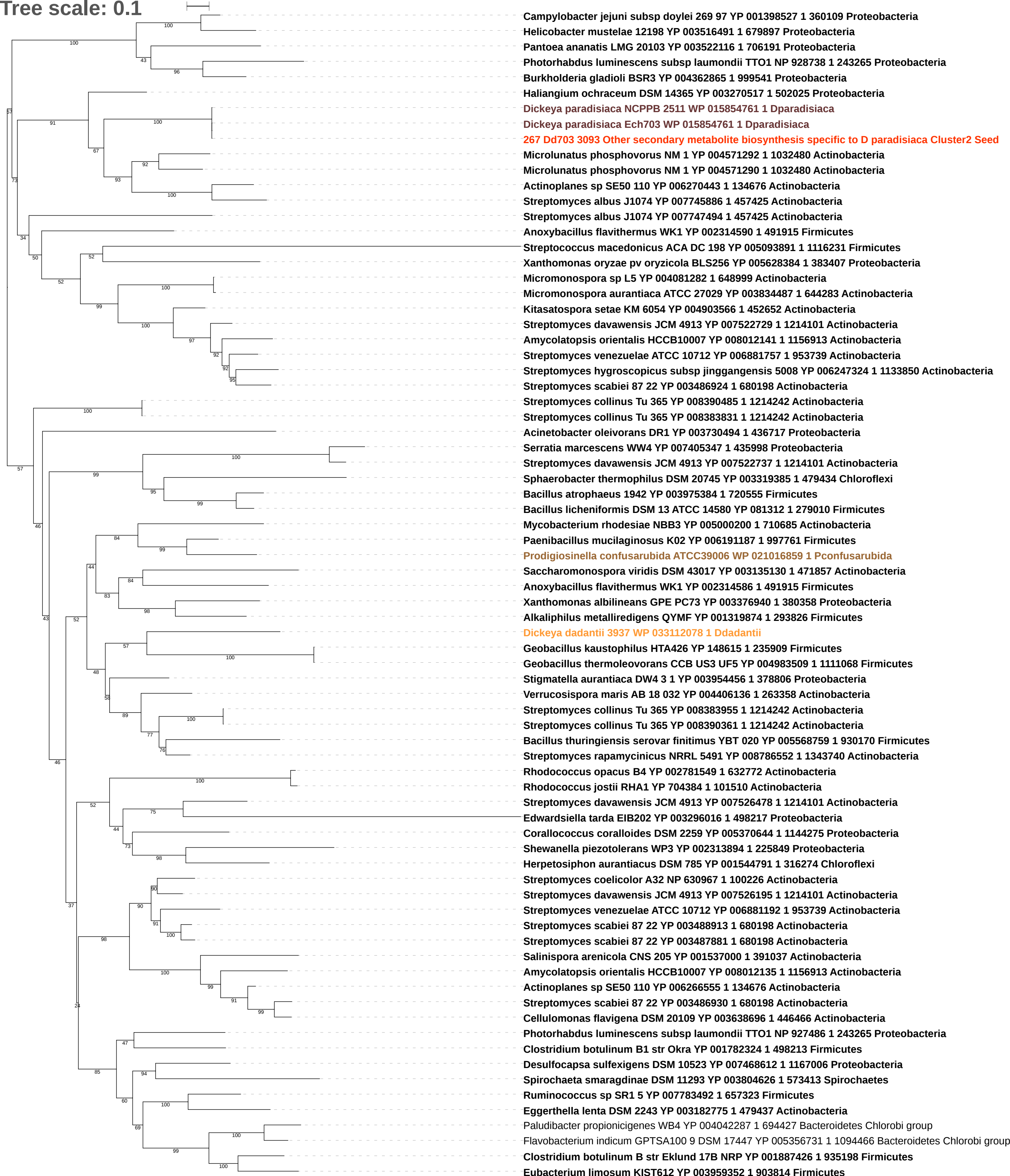
Tree scale: 0.1
Campylobacter jejuni subsp doylei 269 97 YP 001398527 1 360109 Proteobacteria
Helicobacter mustelae 12198 YP 003516491 1 679897 Proteobacteria
Pantoea ananatis LMG 20103 YP 003522116 1 706191 Proteobacteria
Photorhabdus luminescens subsp laumondii TTO1 NP 928738 1 243265 Proteobacteria
Burkholderia gladioli BSR3 YP 004362865 1 999541 Proteobacteria
Haliangium ochraceum DSM 14365 YP 003270517 1 502025 Proteobacteria
Dickeya paradisiaca NCPPB 2511 WP 015854761 1 Dparadisiaca
Dickeya paradisiaca Ech703 WP 015854761 1 Dparadisiaca
267 Dd703 3093 Other secondary metabolite biosynthesis specific to D paradisiaca Cluster2 Seed
Microlunatus phosphovorus NM 1 YP 004571292 1 1032480 Actinobacteria
Microlunatus phosphovorus NM 1 YP 004571290 1 1032480 Actinobacteria
Actinoplanes sp SE50 110 YP 006270443 1 134676 Actinobacteria
Streptomyces albus J1074 YP 007745886 1 457425 Actinobacteria
Streptomyces albus J1074 YP 007747494 1 457425 Actinobacteria
Anoxybacillus flavithermus WK1 YP 002314590 1 491915 Firmicutes
Streptococcus macedonicus ACA DC 198 YP 005093891 1 1116231 Firmicutes
Xanthomonas oryzae pv oryzicola BLS256 YP 005628384 1 383407 Proteobacteria
Micromonospora sp L5 YP 004081282 1 648999 Actinobacteria
Micromonospora aurantiaca ATCC 27029 YP 003834487 1 644283 Actinobacteria
Kitasatospora setae KM 6054 YP 004903566 1 452652 Actinobacteria
Streptomyces davawensis JCM 4913 YP 007522729 1 1214101 Actinobacteria
Amycolatopsis orientalis HCCB10007 YP 008012141 1 1156913 Actinobacteria
Streptomyces venezuelae ATCC 10712 YP 006881757 1 953739 Actinobacteria
Streptomyces hygroscopicus subsp jinggangensis 5008 YP 006247324 1 1133850 Actinobacteria
Streptomyces scabiei 87 22 YP 003486924 1 680198 Actinobacteria
Streptomyces collinus Tu 365 YP 008390485 1 1214242 Actinobacteria
Streptomyces collinus Tu 365 YP 008383831 1 1214242 Actinobacteria
Acinetobacter oleivorans DR1 YP 003730494 1 436717 Proteobacteria
Serratia marcescens WW4 YP 007405347 1 435998 Proteobacteria
Streptomyces davawensis JCM 4913 YP 007522737 1 1214101 Actinobacteria
Sphaerobacter thermophilus DSM 20745 YP 003319385 1 479434 Chloroflexi
Bacillus atrophaeus 1942 YP 003975384 1 720555 Firmicutes
Bacillus licheniformis DSM 13 ATCC 14580 YP 081312 1 279010 Firmicutes
Mycobacterium rhodesiae NBB3 YP 005000200 1 710685 Actinobacteria
Paenibacillus mucilaginosus K02 YP 006191187 1 997761 Firmicutes
Prodigiosinella confusarubida ATCC39006 WP 021016859 1 Pconfusarubida
Saccharomonospora viridis DSM 43017 YP 003135130 1 471857 Actinobacteria
Anoxybacillus flavithermus WK1 YP 002314586 1 491915 Firmicutes
Xanthomonas albilineans GPE PC73 YP 003376940 1 380358 Proteobacteria
Alkaliphilus metalliredigens QYMF YP 001319874 1 293826 Firmicutes
Dickeya dadantii 3937 WP 033112078 1 Ddadantii
Geobacillus kaustophilus HTA426 YP 148615 1 235909 Firmicutes
Geobacillus thermoleovorans CCB US3 UF5 YP 004983509 1 1111068 Firmicutes
Stigmatella aurantiaca DW4 3 1 YP 003954456 1 378806 Proteobacteria
Verrucosispora maris AB 18 032 YP 004406136 1 263358 Actinobacteria
Streptomyces collinus Tu 365 YP 008383955 1 1214242 Actinobacteria
Streptomyces collinus Tu 365 YP 008390361 1 1214242 Actinobacteria
Bacillus thuringiensis serovar finitimus YBT 020 YP 005568759 1 930170 Firmicutes
Streptomyces rapamycinicus NRRL 5491 YP 008786552 1 1343740 Actinobacteria
Rhodococcus opacus B4 YP 002781549 1 632772 Actinobacteria
Rhodococcus jostii RHA1 YP 704384 1 101510 Actinobacteria
Streptomyces davawensis JCM 4913 YP 007526478 1 1214101 Actinobacteria
Edwardsiella tarda EIB202 YP 003296016 1 498217 Proteobacteria
Corallococcus coralloides DSM 2259 YP 005370644 1 1144275 Proteobacteria
Shewanella piezotolerans WP3 YP 002313894 1 225849 Proteobacteria
Herpetosiphon aurantiacus DSM 785 YP 001544791 1 316274 Chloroflexi
Streptomyces coelicolor A32 NP 630967 1 100226 Actinobacteria
Streptomyces davawensis JCM 4913 YP 007526195 1 1214101 Actinobacteria
Streptomyces venezuelae ATCC 10712 YP 006881192 1 953739 Actinobacteria
Streptomyces scabiei 87 22 YP 003488913 1 680198 Actinobacteria
Streptomyces scabiei 87 22 YP 003487881 1 680198 Actinobacteria
Salinispora arenicola CNS 205 YP 001537000 1 391037 Actinobacteria
Amycolatopsis orientalis HCCB10007 YP 008012135 1 1156913 Actinobacteria
Actinoplanes sp SE50 110 YP 006266555 1 134676 Actinobacteria
Streptomyces scabiei 87 22 YP 003486930 1 680198 Actinobacteria
Cellulomonas flavigena DSM 20109 YP 003638696 1 446466 Actinobacteria
Photorhabdus luminescens subsp laumondii TTO1 NP 927486 1 243265 Proteobacteria
Clostridium botulinum B1 str Okra YP 001782324 1 498213 Firmicutes
Desulfocapsa sulfexigens DSM 10523 YP 007468612 1 1167006 Proteobacteria
Spirochaeta smaragdinae DSM 11293 YP 003804626 1 573413 Spirochaetes
Ruminococcus sp SR1 5 YP 007783492 1 657323 Firmicutes
Eggerthella lenta DSM 2243 YP 003182775 1 479437 Actinobacteria
Paludibacter propionicigenes WB4 YP 004042287 1 694427 Bacteroidetes Chlorobi group
Flavobacterium indicum GPTSA100 9 DSM 17447 YP 005356731 1 1094466 Bacteroidetes Chlorobi group
Clostridium botulinum B str Eklund 17B NRP YP 001887426 1 935198 Firmicutes
Eubacterium limosum KIST612 YP 003959352 1 903814 Firmicutes

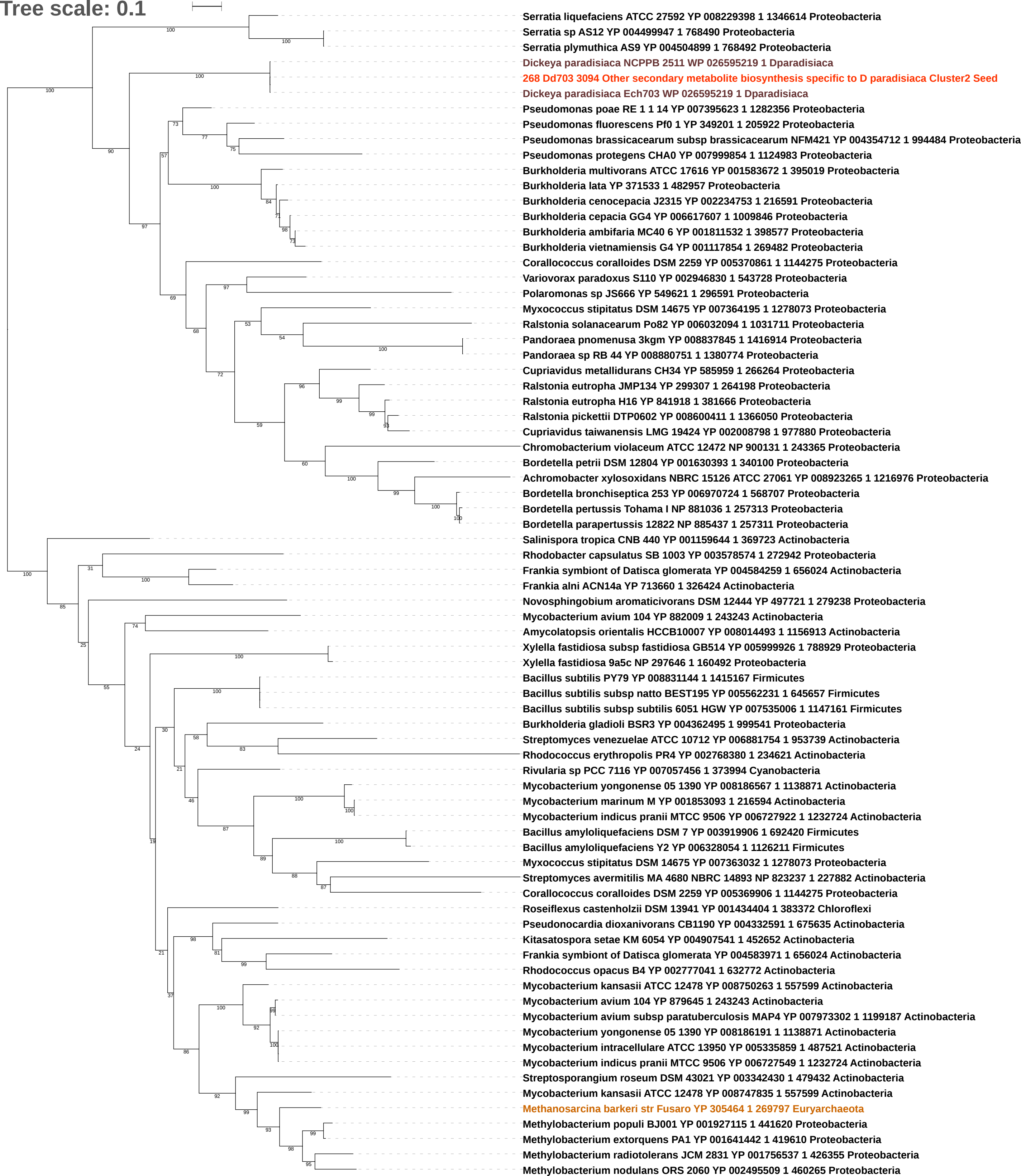

Tree scale: 0.1
Serratia liquefaciens ATCC 27592 YP 008229398 1 1346614 Proteobacteria
Serratia sp AS12 YP 004499947 1 768490 Proteobacteria
Serratia plymuthica AS9 YP 004504899 1 768492 Proteobacteria
Dickeya paradisiaca NCPPB 2511 WP 026595219 1 Dparadisiaca
268 Dd703 3094 Other secondary metabolite biosynthesis specific to D paradisiaca Cluster2 Seed
Dickeya paradisiaca Ech703 WP 026595219 1 Dparadisiaca
Pseudomonas poae RE 1 1 14 YP 007395623 1 1282356 Proteobacteria
Pseudomonas fluorescens Pf0 1 YP 349201 1 205922 Proteobacteria
Pseudomonas brassicacearum subsp brassicacearum NFM421 YP 004354712 1 994484 Proteobacteria
Pseudomonas protegens CHA0 YP 007999854 1 1124983 Proteobacteria
Burkholderia multivorans ATCC 17616 YP 001583672 1 395019 Proteobacteria
Burkholderia lata YP 371533 1 482957 Proteobacteria
Burkholderia cenocepacia J2315 YP 002234753 1 216591 Proteobacteria
Burkholderia cepacia GG4 YP 006617607 1 1009846 Proteobacteria
Burkholderia ambifaria MC40 6 YP 001811532 1 398577 Proteobacteria
Burkholderia vietnamiensis G4 YP 001117854 1 269482 Proteobacteria
Corallococcus coralloides DSM 2259 YP 005370861 1 1144275 Proteobacteria
Variovorax paradoxus S110 YP 002946830 1 543728 Proteobacteria
Polaromonas sp JS666 YP 549621 1 296591 Proteobacteria
Myxococcus stipitatus DSM 14675 YP 007364195 1 1278073 Proteobacteria
Ralstonia solanacearum Po82 YP 006032094 1 1031711 Proteobacteria
Pandoraea pnomenusa 3kgm YP 008837845 1 1416914 Proteobacteria
Pandoraea sp RB 44 YP 008880751 1 1380774 Proteobacteria
Cupriavidus metallidurans CH34 YP 585959 1 266264 Proteobacteria
Ralstonia eutropha JMP134 YP 299307 1 264198 Proteobacteria
Ralstonia eutropha H16 YP 841918 1 381666 Proteobacteria
Ralstonia pickettii DTP0602 YP 008600411 1 1366050 Proteobacteria
Cupriavidus taiwanensis LMG 19424 YP 002008798 1 977880 Proteobacteria
Chromobacterium violaceum ATCC 12472 NP 900131 1 243365 Proteobacteria
Bordetella petrii DSM 12804 YP 001630393 1 340100 Proteobacteria
Achromobacter xylosoxidans NBRC 15126 ATCC 27061 YP 008923265 1 1216976 Proteobacteria
Bordetella bronchiseptica 253 YP 006970724 1 568707 Proteobacteria
Bordetella pertussis Tohama I NP 881036 1 257313 Proteobacteria
Bordetella parapertussis 12822 NP 885437 1 257311 Proteobacteria
Salinispora tropica CNB 440 YP 001159644 1 369723 Actinobacteria
Rhodobacter capsulatus SB 1003 YP 003578574 1 272942 Proteobacteria
Frankia symbiont of Datisca glomerata YP 004584259 1 656024 Actinobacteria
Frankia alni ACN14a YP 713660 1 326424 Actinobacteria
Novosphingobium aromaticivorans DSM 12444 YP 497721 1 279238 Proteobacteria
Mycobacterium avium 104 YP 882009 1 243243 Actinobacteria
Amycolatopsis orientalis HCCB10007 YP 008014493 1 1156913 Actinobacteria
Xylella fastidiosa subsp fastidiosa GB514 YP 005999926 1 788929 Proteobacteria
Xylella fastidiosa 9a5c NP 297646 1 160492 Proteobacteria
Bacillus subtilis PY79 YP 008831144 1 1415167 Firmicutes
Bacillus subtilis subsp natto BEST195 YP 005562231 1 645657 Firmicutes
Bacillus subtilis subsp subtilis 6051 HGW YP 007535006 1 1147161 Firmicutes
Burkholderia gladioli BSR3 YP 004362495 1 999541 Proteobacteria
Streptomyces venezuelae ATCC 10712 YP 006881754 1 953739 Actinobacteria
Rhodococcus erythropolis PR4 YP 002768380 1 234621 Actinobacteria
Rivularia sp PCC 7116 YP 007057456 1 373994 Cyanobacteria
Mycobacterium yongonense 05 1390 YP 008186567 1 1138871 Actinobacteria
Mycobacterium marinum M YP 001853093 1 216594 Actinobacteria
Mycobacterium indicus pranii MTCC 9506 YP 006727922 1 1232724 Actinobacteria
Bacillus amyloliquefaciens DSM 7 YP 003919906 1 692420 Firmicutes
Bacillus amyloliquefaciens Y2 YP 006328054 1 1126211 Firmicutes
Myxococcus stipitatus DSM 14675 YP 007363032 1 1278073 Proteobacteria
Streptomyces avermitilis MA 4680 NBRC 14893 NP 823237 1 227882 Actinobacteria
Corallococcus coralloides DSM 2259 YP 005369906 1 1144275 Proteobacteria
Roseiflexus castenholzii DSM 13941 YP 001434404 1 383372 Chloroflexi
Pseudonocardia dioxanivorans CB1190 YP 004332591 1 675635 Actinobacteria
Kitasatospora setae KM 6054 YP 004907541 1 452652 Actinobacteria
Frankia symbiont of Datisca glomerata YP 004583971 1 656024 Actinobacteria
Rhodococcus opacus B4 YP 002777041 1 632772 Actinobacteria
Mycobacterium kansasii ATCC 12478 YP 008750263 1 557599 Actinobacteria
Mycobacterium avium 104 YP 879645 1 243243 Actinobacteria
Mycobacterium avium subsp paratuberculosis MAP4 YP 007973302 1 1199187 Actinobacteria
Mycobacterium yongonense 05 1390 YP 008186191 1 1138871 Actinobacteria
Mycobacterium intracellulare ATCC 13950 YP 005335859 1 487521 Actinobacteria
Mycobacterium indicus pranii MTCC 9506 YP 006727549 1 1232724 Actinobacteria
Streptosporangium roseum DSM 43021 YP 003342430 1 479432 Actinobacteria
Mycobacterium kansasii ATCC 12478 YP 008747835 1 557599 Actinobacteria
Methanosarcina barkeri str Fusaro YP 305464 1 269797 Euryarchaeota
Methylobacterium populi BJ001 YP 001927115 1 441620 Proteobacteria
Methylobacterium extorquens PA1 YP 001641442 1 419610 Proteobacteria
Methylobacterium radiotolerans JCM 2831 YP 001756537 1 426355 Proteobacteria
Methylobacterium nodulans ORS 2060 YP 002495509 1 460265 Proteobacteria

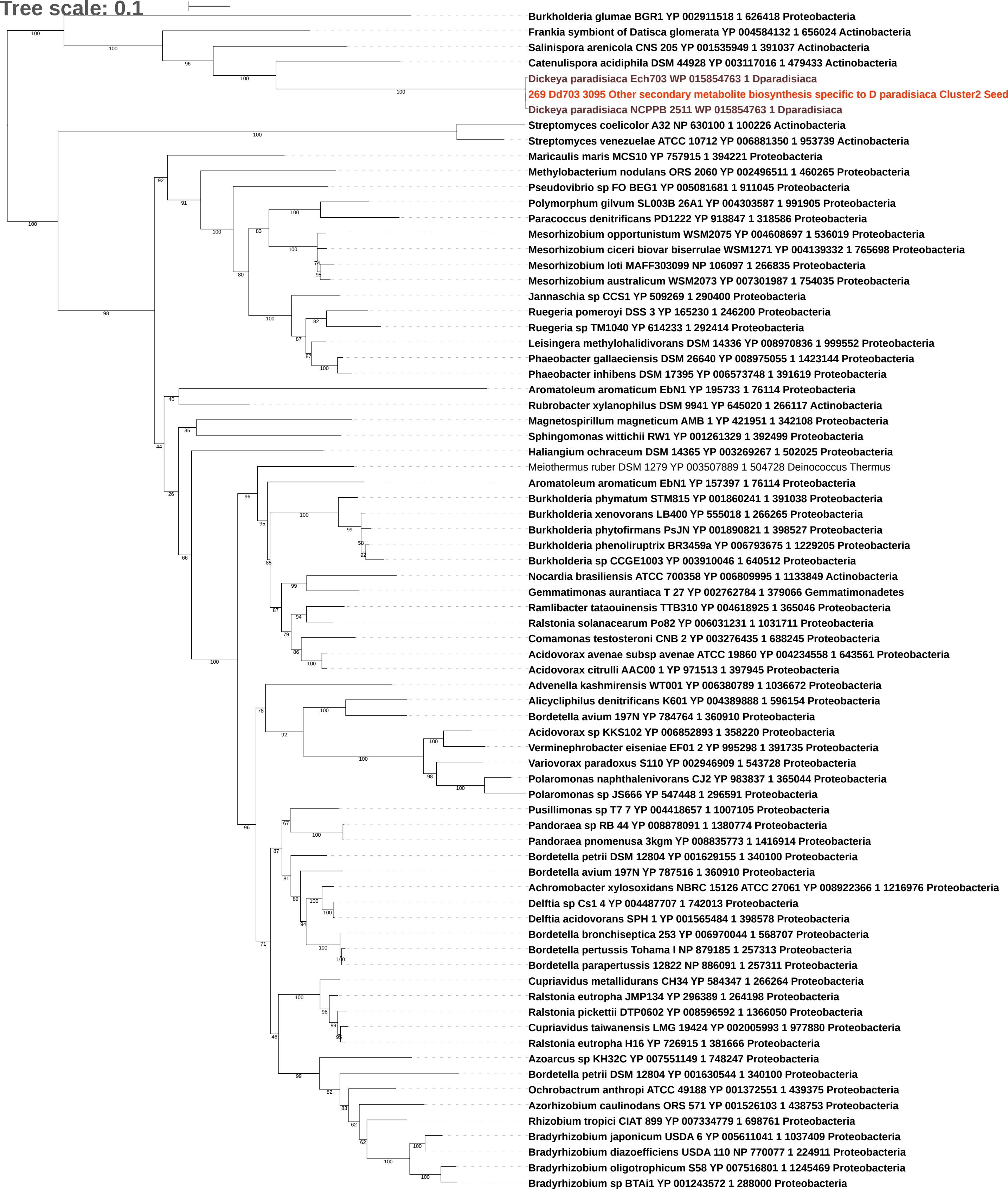
Tree scale: 0.1
Burkholderia glumae BGR1 YP 002911518 1 626418 Proteobacteria
Frankia symbiont of Datisca glomerata YP 004584132 1 656024 Actinobacteria
Salinispora arenicola CNS 205 YP 001535949 1 391037 Actinobacteria
Catenulispora acidiphila DSM 44928 YP 003117016 1 479433 Actinobacteria
Dickeya paradisiaca Ech703 WP 015854763 1 Dparadisiaca
269 Dd703 3095 Other secondary metabolite biosynthesis specific to D paradisiaca Cluster2 Seed
Dickeya paradisiaca NCPPB 2511 WP 015854763 1 Dparadisiaca
Streptomyces coelicolor A32 NP 630100 1 100226 Actinobacteria
Streptomyces venezuelae ATCC 10712 YP 006881350 1 953739 Actinobacteria
Maricaulis maris MCS10 YP 757915 1 394221 Proteobacteria
Methylobacterium nodulans ORS 2060 YP 002496511 1 460265 Proteobacteria
Pseudovibrio sp FO BEG1 YP 005081681 1 911045 Proteobacteria
Polymorphum gilvum SL003B 26A1 YP 004303587 1 991905 Proteobacteria
Paracoccus denitrificans PD1222 YP 918847 1 318586 Proteobacteria
Mesorhizobium opportunistum WSM2075 YP 004608697 1 536019 Proteobacteria
Mesorhizobium ciceri biovar biserrulae WSM1271 YP 004139332 1 765698 Proteobacteria
Mesorhizobium loti MAFF303099 NP 106097 1 266835 Proteobacteria
Mesorhizobium australicum WSM2073 YP 007301987 1 754035 Proteobacteria
Jannaschia sp CCS1 YP 509269 1 290400 Proteobacteria
Ruegeria pomeroyi DSS 3 YP 165230 1 246200 Proteobacteria
Ruegeria sp TM1040 YP 614233 1 292414 Proteobacteria
Leisingera methylohalidivorans DSM 14336 YP 008970836 1 999552 Proteobacteria
Phaeobacter gallaeciensis DSM 26640 YP 008975055 1 1423144 Proteobacteria
Phaeobacter inhibens DSM 17395 YP 006573748 1 391619 Proteobacteria
Aromatoleum aromaticum EbN1 YP 195733 1 76114 Proteobacteria
Rubrobacter xylanophilus DSM 9941 YP 645020 1 266117 Actinobacteria
Magnetospirillum magneticum AMB 1 YP 421951 1 342108 Proteobacteria
Sphingomonas wittichii RW1 YP 001261329 1 392499 Proteobacteria
Haliangium ochraceum DSM 14365 YP 003269267 1 502025 Proteobacteria
Meiothermus ruber DSM 1279 YP 003507889 1 504728 Deinococcus Thermus
Aromatoleum aromaticum EbN1 YP 157397 1 76114 Proteobacteria
Burkholderia phymatum STM815 YP 001860241 1 391038 Proteobacteria
Burkholderia xenovorans LB400 YP 555018 1 266265 Proteobacteria
Burkholderia phytofirmans PsJN YP 001890821 1 398527 Proteobacteria
Burkholderia phenoliruptrix BR3459a YP 006793675 1 1229205 Proteobacteria
Burkholderia sp CCGE1003 YP 003910046 1 640512 Proteobacteria
Nocardia brasiliensis ATCC 700358 YP 006809995 1 1133849 Actinobacteria
Gemmatimonas aurantiaca T 27 YP 002762784 1 379066 Gemmatimonadetes
Ramlibacter tataouinensis TTB310 YP 004618925 1 365046 Proteobacteria
Ralstonia solanacearum Po82 YP 006031231 1 1031711 Proteobacteria
Comamonas testosteroni CNB 2 YP 003276435 1 688245 Proteobacteria
Acidovorax avenae subsp avenae ATCC 19860 YP 004234558 1 643561 Proteobacteria
Acidovorax citrulli AAC00 1 YP 971513 1 397945 Proteobacteria
Advenella kashmirensis WT001 YP 006380789 1 1036672 Proteobacteria
Alicycliphilus denitrificans K601 YP 004389888 1 596154 Proteobacteria
Bordetella avium 197N YP 784764 1 360910 Proteobacteria
Acidovorax sp KKS102 YP 006852893 1 358220 Proteobacteria
Verminephrobacter eiseniae EF01 2 YP 995298 1 391735 Proteobacteria
Variovorax paradoxus S110 YP 002946909 1 543728 Proteobacteria
Polaromonas naphthalenivorans CJ2 YP 983837 1 365044 Proteobacteria
Polaromonas sp JS666 YP 547448 1 296591 Proteobacteria
Pusillimonas sp T7 7 YP 004418657 1 1007105 Proteobacteria
Pandoraea sp RB 44 YP 008878091 1 1380774 Proteobacteria
Pandoraea pnomenusa 3kgm YP 008835773 1 1416914 Proteobacteria
Bordetella petrii DSM 12804 YP 001629155 1 340100 Proteobacteria
Bordetella avium 197N YP 787516 1 360910 Proteobacteria
Achromobacter xylosoxidans NBRC 15126 ATCC 27061 YP 008922366 1 1216976 Proteobacteria
Delftia sp Cs1 4 YP 004487707 1 742013 Proteobacteria
Delftia acidovorans SPH 1 YP 001565484 1 398578 Proteobacteria
Bordetella bronchiseptica 253 YP 006970044 1 568707 Proteobacteria
Bordetella pertussis Tohama I NP 879185 1 257313 Proteobacteria
Bordetella parapertussis 12822 NP 886091 1 257311 Proteobacteria
Cupriavidus metallidurans CH34 YP 584347 1 266264 Proteobacteria
Ralstonia eutropha JMP134 YP 296389 1 264198 Proteobacteria
Ralstonia pickettii DTP0602 YP 008596592 1 1366050 Proteobacteria
Cupriavidus taiwanensis LMG 19424 YP 002005993 1 977880 Proteobacteria
Ralstonia eutropha H16 YP 726915 1 381666 Proteobacteria
Azoarcus sp KH32C YP 007551149 1 748247 Proteobacteria
Bordetella petrii DSM 12804 YP 001630544 1 340100 Proteobacteria
Ochrobactrum anthropi ATCC 49188 YP 001372551 1 439375 Proteobacteria
Azorhizobium caulinodans ORS 571 YP 001526103 1 438753 Proteobacteria
Rhizobium tropici CIAT 899 YP 007334779 1 698761 Proteobacteria
Bradyrhizobium japonicum USDA 6 YP 005611041 1 1037409 Proteobacteria
Bradyrhizobium diazoefficiens USDA 110 NP 770077 1 224911 Proteobacteria
Bradyrhizobium oligotrophicum S58 YP 007516801 1 1245469 Proteobacteria
Bradyrhizobium sp BTAi1 YP 001243572 1 288000 Proteobacteria

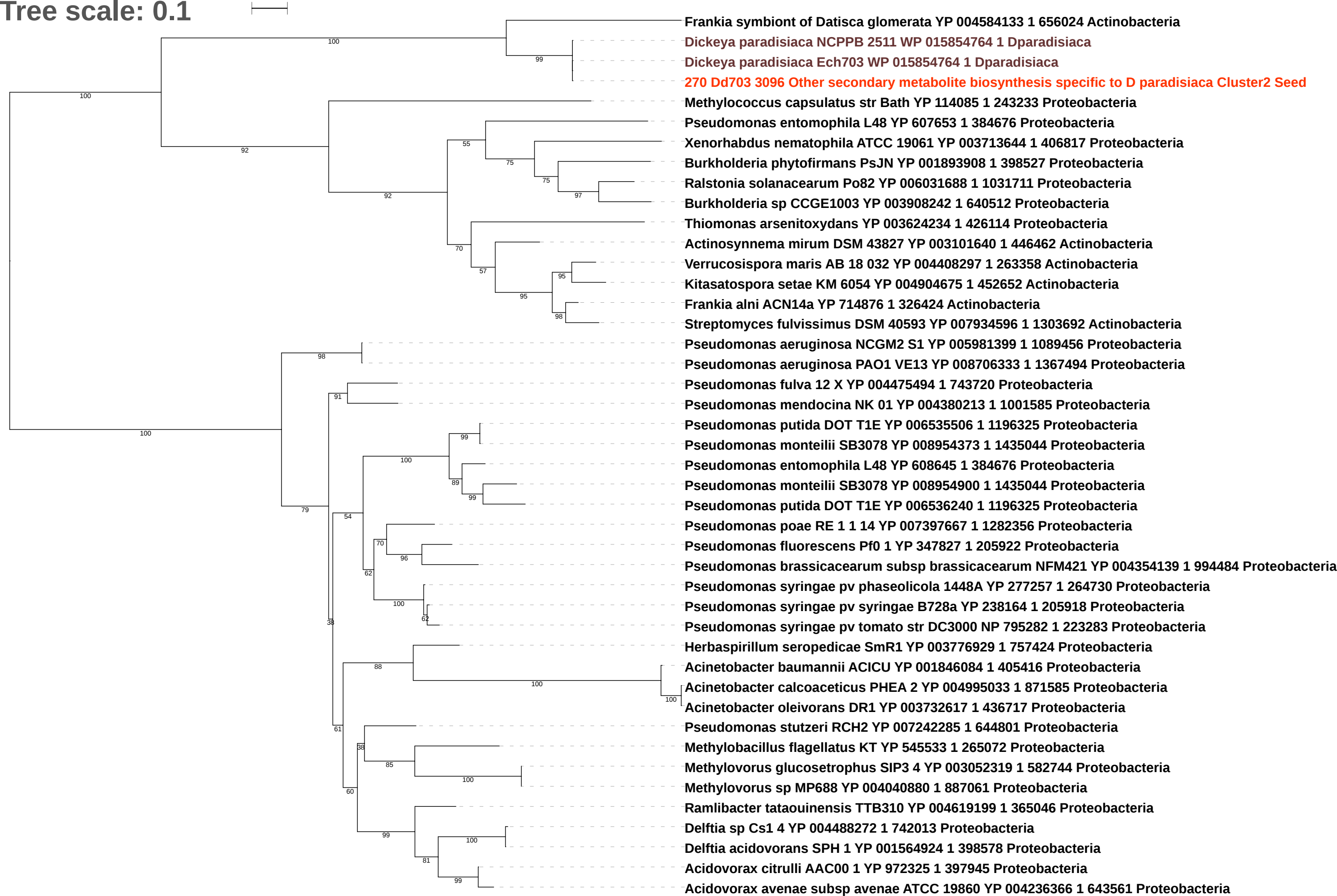
Tree scale: 0.1
Frankia symbiont of Datisca glomerata YP 004584133 1 656024 Actinobacteria
Dickeya paradisiaca NCPPB 2511 WP 015854764 1 Dparadisiaca
Dickeya paradisiaca Ech703 WP 015854764 1 Dparadisiaca
270 Dd703 3096 Other secondary metabolite biosynthesis specific to D paradisiaca Cluster2 Seed
Methylococcus capsulatus str Bath YP 114085 1 243233 Proteobacteria
Pseudomonas entomophila L48 YP 607653 1 384676 Proteobacteria
Xenorhabdus nematophila ATCC 19061 YP 003713644 1 406817 Proteobacteria
Burkholderia phytofirmans PsJN YP 001893908 1 398527 Proteobacteria
Ralstonia solanacearum Po82 YP 006031688 1 1031711 Proteobacteria
Burkholderia sp CCGE1003 YP 003908242 1 640512 Proteobacteria
Thiomonas arsenitoxydans YP 003624234 1 426114 Proteobacteria
Actinosynnema mirum DSM 43827 YP 003101640 1 446462 Actinobacteria
Verrucosispora maris AB 18 032 YP 004408297 1 263358 Actinobacteria
Kitasatospora setae KM 6054 YP 004904675 1 452652 Actinobacteria
Frankia alni ACN14a YP 714876 1 326424 Actinobacteria
Streptomyces fulvissimus DSM 40593 YP 007934596 1 1303692 Actinobacteria
Pseudomonas aeruginosa NCGM2 S1 YP 005981399 1 1089456 Proteobacteria
Pseudomonas aeruginosa PAO1 VE13 YP 008706333 1 1367494 Proteobacteria
Pseudomonas fulva 12 X YP 004475494 1 743720 Proteobacteria
Pseudomonas mendocina NK 01 YP 004380213 1 1001585 Proteobacteria
Pseudomonas putida DOT T1E YP 006535506 1 1196325 Proteobacteria
Pseudomonas monteilii SB3078 YP 008954373 1 1435044 Proteobacteria
Pseudomonas entomophila L48 YP 608645 1 384676 Proteobacteria
Pseudomonas monteilii SB3078 YP 008954900 1 1435044 Proteobacteria
Pseudomonas putida DOT T1E YP 006536240 1 1196325 Proteobacteria
Pseudomonas poae RE 1 1 14 YP 007397667 1 1282356 Proteobacteria
Pseudomonas fluorescens Pf0 1 YP 347827 1 205922 Proteobacteria
Pseudomonas brassicacearum subsp brassicacearum NFM421 YP 004354139 1 994484 Proteobacteria
Pseudomonas syringae pv phaseolicola 1448A YP 277257 1 264730 Proteobacteria
Pseudomonas syringae pv syringae B728a YP 238164 1 205918 Proteobacteria
Pseudomonas syringae pv tomato str DC3000 NP 795282 1 223283 Proteobacteria
Herbaspirillum seropedicae SmR1 YP 003776929 1 757424 Proteobacteria
Acinetobacter baumannii ACICU YP 001846084 1 405416 Proteobacteria
Acinetobacter calcoaceticus PHEA 2 YP 004995033 1 871585 Proteobacteria
Acinetobacter oleivorans DR1 YP 003732617 1 436717 Proteobacteria
Pseudomonas stutzeri RCH2 YP 007242285 1 644801 Proteobacteria
Methylobacillus flagellatus KT YP 545533 1 265072 Proteobacteria
Methylovorus glucosetrophus SIP3 4 YP 003052319 1 582744 Proteobacteria
Methylovorus sp MP688 YP 004040880 1 887061 Proteobacteria
Ramlibacter tataouinensis TTB310 YP 004619199 1 365046 Proteobacteria
Delftia sp Cs1 4 YP 004488272 1 742013 Proteobacteria
Delftia acidovorans SPH 1 YP 001564924 1 398578 Proteobacteria
Acidovorax citrulli AAC00 1 YP 972325 1 397945 Proteobacteria
Acidovorax avenae subsp avenae ATCC 19860 YP 004236366 1 643561 Proteobacteria

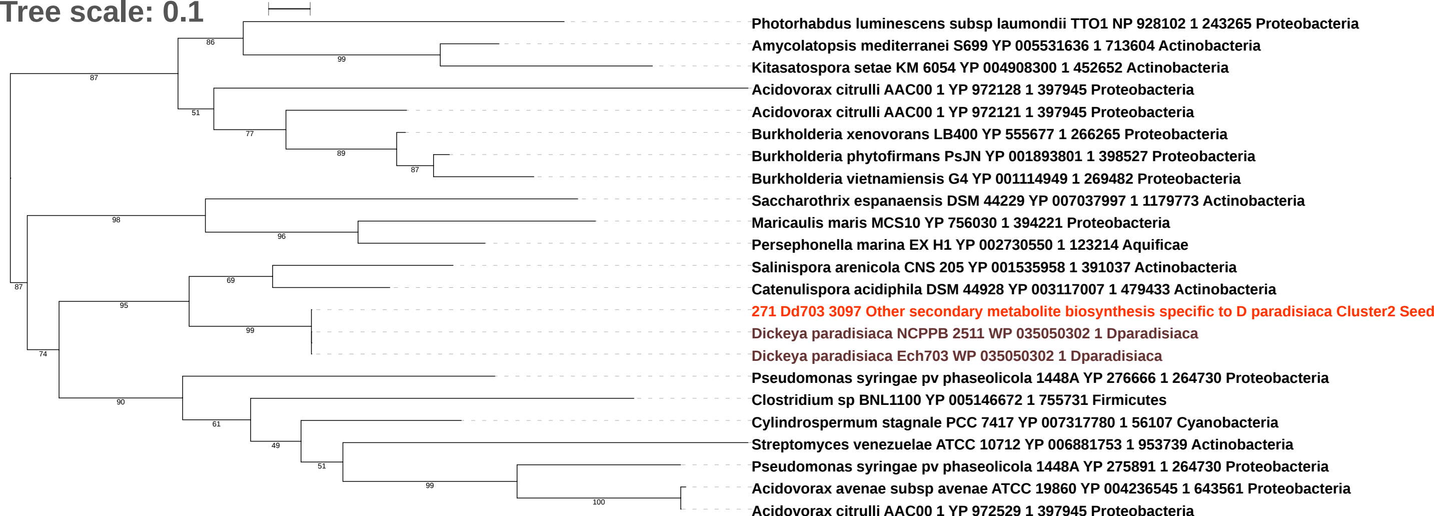
Tree scale: 0.1
Photorhabdus luminescens subsp laumondii TTO1 NP 928102 1 243265 Proteobacteria
Amycolatopsis mediterranei S699 YP 005531636 1 713604 Actinobacteria
Kitasatospora setae KM 6054 YP 004908300 1 452652 Actinobacteria
Acidovorax citrulli AAC00 1 YP 972128 1 397945 Proteobacteria
Acidovorax citrulli AAC00 1 YP 972121 1 397945 Proteobacteria
Burkholderia xenovorans LB400 YP 555677 1 266265 Proteobacteria
Burkholderia phytofirmans PsJN YP 001893801 1 398527 Proteobacteria
Burkholderia vietnamiensis G4 YP 001114949 1 269482 Proteobacteria
Saccharothrix espanaensis DSM 44229 YP 007037997 1 1179773 Actinobacteria
Maricaulis maris MCS10 YP 756030 1 394221 Proteobacteria
Persephonella marina EX H1 YP 002730550 1 123214 Aquificae
Salinispora arenicola CNS 205 YP 001535958 1 391037 Actinobacteria
Catenulispora acidiphila DSM 44928 YP 003117007 1 479433 Actinobacteria
271 Dd703 3097 Other secondary metabolite biosynthesis specific to D paradisiaca Cluster2 Seed
Dickeya paradisiaca NCPPB 2511 WP 035050302 1 Dparadisiaca
Dickeya paradisiaca Ech703 WP 035050302 1 Dparadisiaca
Pseudomonas syringae pv phaseolicola 1448A YP 276666 1 264730 Proteobacteria
Clostridium sp BNL1100 YP 005146672 1 755731 Firmicutes
Cylindrospermum stagnale PCC 7417 YP 007317780 1 56107 Cyanobacteria
Streptomyces venezuelae ATCC 10712 YP 006881753 1 953739 Actinobacteria
Pseudomonas syringae pv phaseolicola 1448A YP 275891 1 264730 Proteobacteria
Acidovorax avenae subsp avenae ATCC 19860 YP 004236545 1 643561 Proteobacteria
Acidovorax citrulli AAC00 1 YP 972529 1 397945 Proteobacteria
86
99
87
51
77
89
87
98
96
87
69
95
99
74
90
61
49
51
99
100

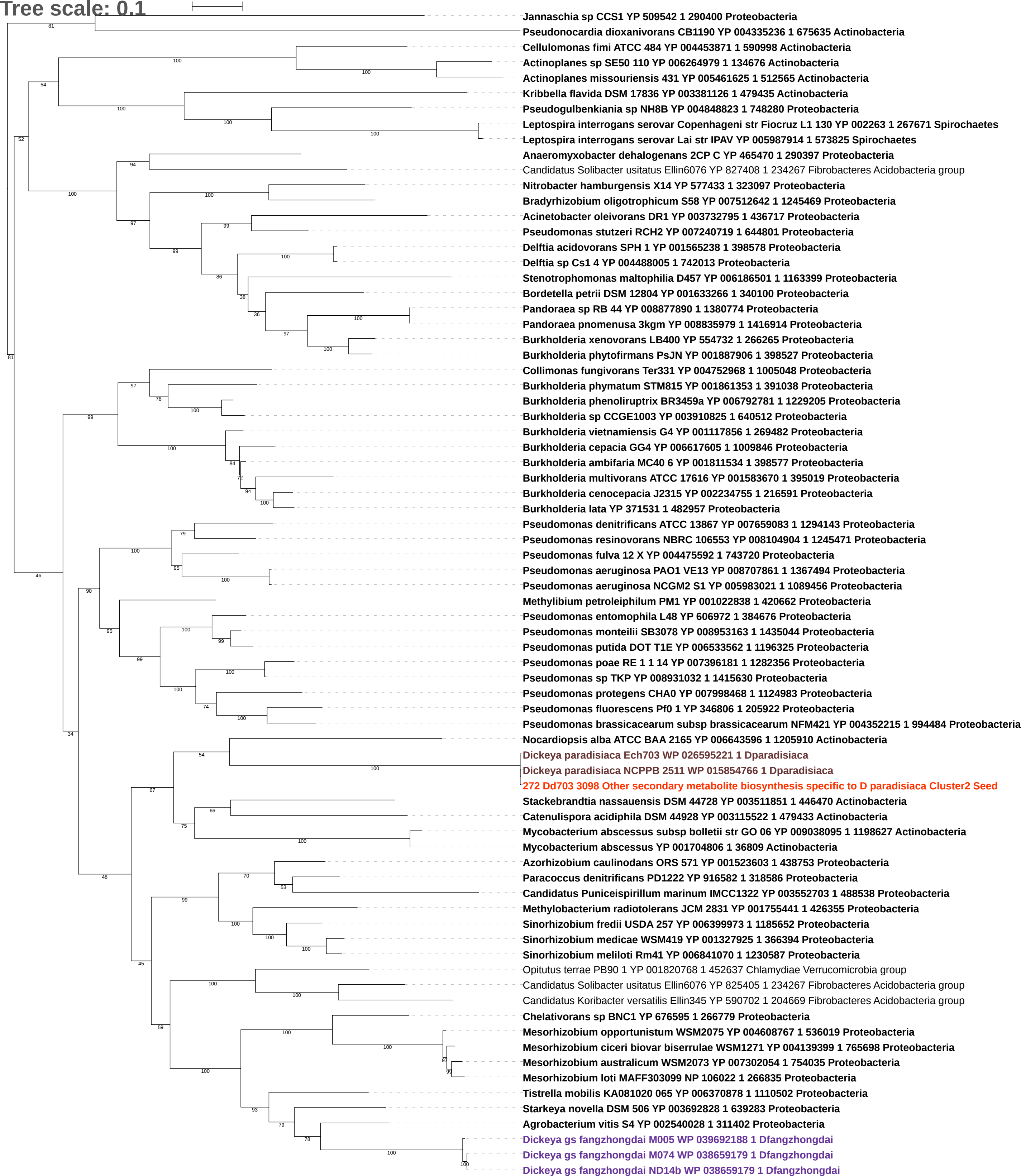
Tree scale: 0.1
Jannaschia sp CCS1 YP 509542 1 290400 Proteobacteria
Pseudonocardia dioxanivorans CB1190 YP 004335236 1 675635 Actinobacteria
Cellulomonas fimi ATCC 484 YP 004453871 1 590998 Actinobacteria
Actinoplanes sp SE50 110 YP 006264979 1 134676 Actinobacteria
Actinoplanes missouriensis 431 YP 005461625 1 512565 Actinobacteria
Kribbella flavida DSM 17836 YP 003381126 1 479435 Actinobacteria
Pseudogulbenkiania sp NH8B YP 004848823 1 748280 Proteobacteria
Leptospira interrogans serovar Copenhageni str Fiocruz L1 130 YP 002263 1 267671 Spirochaetes
Leptospira interrogans serovar Lai str IPAV YP 005987914 1 573825 Spirochaetes
Anaeromyxobacter dehalogenans 2CP C YP 465470 1 290397 Proteobacteria
Candidatus Solibacter usitatus Ellin6076 YP 827408 1 234267 Fibrobacteres Acidobacteria group
Nitrobacter hamburgensis X14 YP 577433 1 323097 Proteobacteria
Bradyrhizobium oligotrophicum S58 YP 007512642 1 1245469 Proteobacteria
Acinetobacter oleivorans DR1 YP 003732795 1 436717 Proteobacteria
Pseudomonas stutzeri RCH2 YP 007240719 1 644801 Proteobacteria
Delftia acidovorans SPH 1 YP 001565238 1 398578 Proteobacteria
Delftia sp Cs1 4 YP 004488005 1 742013 Proteobacteria
Stenotrophomonas maltophilia D457 YP 006186501 1 1163399 Proteobacteria
Bordetella petrii DSM 12804 YP 001633266 1 340100 Proteobacteria
Pandoraea sp RB 44 YP 008877890 1 1380774 Proteobacteria
Pandoraea pnomenusa 3kgm YP 008835979 1 1416914 Proteobacteria
Burkholderia xenovorans LB400 YP 554732 1 266265 Proteobacteria
Burkholderia phytofirmans PsJN YP 001887906 1 398527 Proteobacteria
Collimonas fungivorans Ter331 YP 004752968 1 1005048 Proteobacteria
Burkholderia phymatum STM815 YP 001861353 1 391038 Proteobacteria
Burkholderia phenoliruptrix BR3459a YP 006792781 1 1229205 Proteobacteria
Burkholderia sp CCGE1003 YP 003910825 1 640512 Proteobacteria
Burkholderia vietnamiensis G4 YP 001117856 1 269482 Proteobacteria
Burkholderia cepacia GG4 YP 006617605 1 1009846 Proteobacteria
Burkholderia ambifaria MC40 6 YP 001811534 1 398577 Proteobacteria
Burkholderia multivorans ATCC 17616 YP 001583670 1 395019 Proteobacteria
Burkholderia cenocepacia J2315 YP 002234755 1 216591 Proteobacteria
Burkholderia lata YP 371531 1 482957 Proteobacteria
Pseudomonas denitrificans ATCC 13867 YP 007659083 1 1294143 Proteobacteria
Pseudomonas resinovorans NBRC 106553 YP 008104904 1 1245471 Proteobacteria
Pseudomonas fulva 12 X YP 004475592 1 743720 Proteobacteria
Pseudomonas aeruginosa PAO1 VE13 YP 008707861 1 1367494 Proteobacteria
Pseudomonas aeruginosa NCGM2 S1 YP 005983021 1 1089456 Proteobacteria
Methylibium petroleiphilum PM1 YP 001022838 1 420662 Proteobacteria
Pseudomonas entomophila L48 YP 606972 1 384676 Proteobacteria
Pseudomonas monteilii SB3078 YP 008953163 1 1435044 Proteobacteria
Pseudomonas putida DOT T1E YP 006533562 1 1196325 Proteobacteria
Pseudomonas poae RE 1 1 14 YP 007396181 1 1282356 Proteobacteria
Pseudomonas sp TKP YP 008931032 1 1415630 Proteobacteria
Pseudomonas protegens CHA0 YP 007998468 1 1124983 Proteobacteria
Pseudomonas fluorescens Pf0 1 YP 346806 1 205922 Proteobacteria
Pseudomonas brassicacearum subsp brassicacearum NFM421 YP 004352215 1 994484 Proteobacteria
Nocardiopsis alba ATCC BAA 2165 YP 006643596 1 1205910 Actinobacteria
Dickeya paradisiaca Ech703 WP 026595221 1 Dparadisiaca
Dickeya paradisiaca NCPPB 2511 WP 015854766 1 Dparadisiaca
272 Dd703 3098 Other secondary metabolite biosynthesis specific to D paradisiaca Cluster2 Seed
Stackebrandtia nassauensis DSM 44728 YP 003511851 1 446470 Actinobacteria
Catenulispora acidiphila DSM 44928 YP 003115522 1 479433 Actinobacteria
Mycobacterium abscessus subsp bolletii str GO 06 YP 009038095 1 1198627 Actinobacteria
Mycobacterium abscessus YP 001704806 1 36809 Actinobacteria
Azorhizobium caulinodans ORS 571 YP 001523603 1 438753 Proteobacteria
Paracoccus denitrificans PD1222 YP 916582 1 318586 Proteobacteria
Candidatus Puniceispirillum marinum IMCC1322 YP 003552703 1 488538 Proteobacteria
Methylobacterium radiotolerans JCM 2831 YP 001755441 1 426355 Proteobacteria
Sinorhizobium fredii USDA 257 YP 006399973 1 1185652 Proteobacteria
Sinorhizobium medicae WSM419 YP 001327925 1 366394 Proteobacteria
Sinorhizobium meliloti Rm41 YP 006841070 1 1230587 Proteobacteria
Opitutus terrae PB90 1 YP 001820768 1 452637 Chlamydiae Verrucomicrobia group
Candidatus Solibacter usitatus Ellin6076 YP 825405 1 234267 Fibrobacteres Acidobacteria group
Candidatus Koribacter versatilis Ellin345 YP 590702 1 204669 Fibrobacteres Acidobacteria group
Chelativorans sp BNC1 YP 676595 1 266779 Proteobacteria
Mesorhizobium opportunistum WSM2075 YP 004608767 1 536019 Proteobacteria
Mesorhizobium ciceri biovar biserrulae WSM1271 YP 004139399 1 765698 Proteobacteria
Mesorhizobium australicum WSM2073 YP 007302054 1 754035 Proteobacteria
Mesorhizobium loti MAFF303099 NP 106022 1 266835 Proteobacteria
Tistrella mobilis KA081020 065 YP 006370878 1 1110502 Proteobacteria
Starkeya novella DSM 506 YP 003692828 1 639283 Proteobacteria
Agrobacterium vitis S4 YP 002540028 1 311402 Proteobacteria
Dickeya gs fangzhongdai M005 WP 039692188 1 Dfangzhongdai
Dickeya gs fangzhongdai M074 WP 038659179 1 Dfangzhongdai
Dickeya gs fangzhongdai ND14b WP 038659179 1 Dfangzhongdai

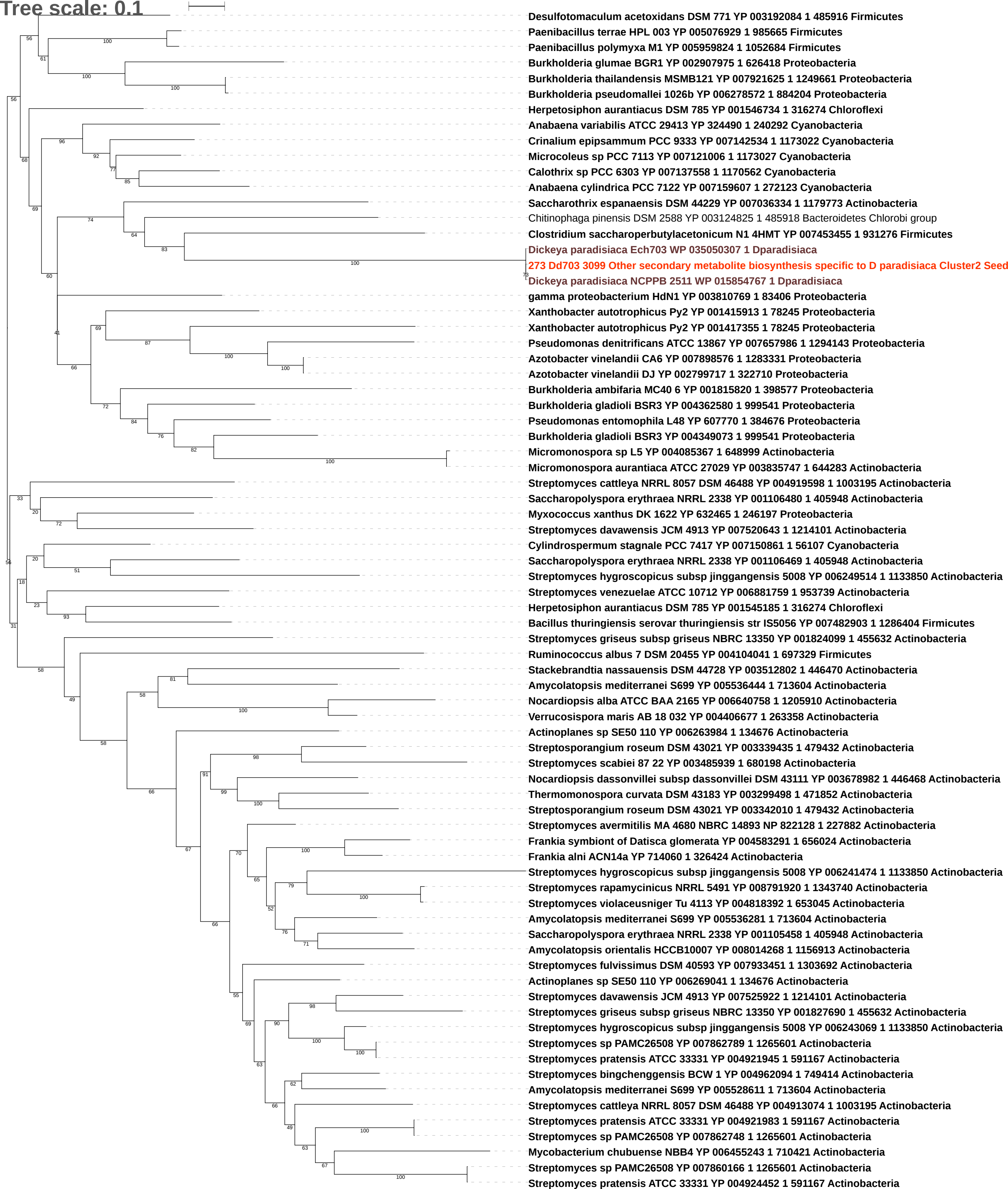

Tree scale: 0.1
Desulfotomaculum acetoxidans DSM 771 YP 003192084 1 485916 Firmicutes
Paenibacillus terrae HPL 003 YP 005076929 1 985665 Firmicutes
Paenibacillus polymyxa M1 YP 005959824 1 1052684 Firmicutes
Burkholderia glumae BGR1 YP 002907975 1 626418 Proteobacteria
Burkholderia thailandensis MSMB121 YP 007921625 1 1249661 Proteobacteria
Burkholderia pseudomallei 1026b YP 006278572 1 884204 Proteobacteria
Herpetosiphon aurantiacus DSM 785 YP 001546734 1 316274 Chloroflexi
Anabaena variabilis ATCC 29413 YP 324490 1 240292 Cyanobacteria
Crinalium epipsammum PCC 9333 YP 007142534 1 1173022 Cyanobacteria
Microcoleus sp PCC 7113 YP 007121006 1 1173027 Cyanobacteria
Calothrix sp PCC 6303 YP 007137558 1 1170562 Cyanobacteria
Anabaena cylindrica PCC 7122 YP 007159607 1 272123 Cyanobacteria
Saccharothrix espanaensis DSM 44229 YP 007036334 1 1179773 Actinobacteria
Chitinophaga pinensis DSM 2588 YP 003124825 1 485918 Bacteroidetes Chlorobi group
Clostridium saccharoperbutylacetonicum N1 4HMT YP 007453455 1 931276 Firmicutes
Dickeya paradisiaca Ech703 WP 035050307 1 Dparadisiaca
273 Dd703 3099 Other secondary metabolite biosynthesis specific to D paradisiaca Cluster2 Seed
Dickeya paradisiaca NCPPB 2511 WP 015854767 1 Dparadisiaca
gamma proteobacterium HdN1 YP 003810769 1 83406 Proteobacteria
Xanthobacter autotrophicus Py2 YP 001415913 1 78245 Proteobacteria
Xanthobacter autotrophicus Py2 YP 001417355 1 78245 Proteobacteria
Pseudomonas denitrificans ATCC 13867 YP 007657986 1 1294143 Proteobacteria
Azotobacter vinelandii CA6 YP 007898576 1 1283331 Proteobacteria
Azotobacter vinelandii DJ YP 002799717 1 322710 Proteobacteria
Burkholderia ambifaria MC40 6 YP 001815820 1 398577 Proteobacteria
Burkholderia gladioli BSR3 YP 004362580 1 999541 Proteobacteria
Pseudomonas entomophila L48 YP 607770 1 384676 Proteobacteria
Burkholderia gladioli BSR3 YP 004349073 1 999541 Proteobacteria
Micromonospora sp L5 YP 004085367 1 648999 Actinobacteria
Micromonospora aurantiaca ATCC 27029 YP 003835747 1 644283 Actinobacteria
Streptomyces cattleya NRRL 8057 DSM 46488 YP 004919598 1 1003195 Actinobacteria
Saccharopolyspora erythraea NRRL 2338 YP 001106480 1 405948 Actinobacteria
Myxococcus xanthus DK 1622 YP 632465 1 246197 Proteobacteria
Streptomyces davawensis JCM 4913 YP 007520643 1 1214101 Actinobacteria
Cylindrospermum stagnale PCC 7417 YP 007150861 1 56107 Cyanobacteria
Saccharopolyspora erythraea NRRL 2338 YP 001106469 1 405948 Actinobacteria
Streptomyces hygroscopicus subsp jinggangensis 5008 YP 006249514 1 1133850 Actinobacteria
Streptomyces venezuelae ATCC 10712 YP 006881759 1 953739 Actinobacteria
Herpetosiphon aurantiacus DSM 785 YP 001545185 1 316274 Chloroflexi
Bacillus thuringiensis serovar thuringiensis str IS5056 YP 007482903 1 1286404 Firmicutes
Streptomyces griseus subsp griseus NBRC 13350 YP 001824099 1 455632 Actinobacteria
Ruminococcus albus 7 DSM 20455 YP 004104041 1 697329 Firmicutes
Stackebrandtia nassauensis DSM 44728 YP 003512802 1 446470 Actinobacteria
Amycolatopsis mediterranei S699 YP 005536444 1 713604 Actinobacteria
Nocardiopsis alba ATCC BAA 2165 YP 006640758 1 1205910 Actinobacteria
Verrucosispora maris AB 18 032 YP 004406677 1 263358 Actinobacteria
Actinoplanes sp SE50 110 YP 006263984 1 134676 Actinobacteria
Streptosporangium roseum DSM 43021 YP 003339435 1 479432 Actinobacteria
Streptomyces scabiei 87 22 YP 003485939 1 680198 Actinobacteria
Nocardiopsis dassonvillei subsp dassonvillei DSM 43111 YP 003678982 1 446468 Actinobacteria
Thermomonospora curvata DSM 43183 YP 003299498 1 471852 Actinobacteria
Streptosporangium roseum DSM 43021 YP 003342010 1 479432 Actinobacteria
Streptomyces avermitilis MA 4680 NBRC 14893 NP 822128 1 227882 Actinobacteria
Frankia symbiont of Datisca glomerata YP 004583291 1 656024 Actinobacteria
Frankia alni ACN14a YP 714060 1 326424 Actinobacteria
Streptomyces hygroscopicus subsp jinggangensis 5008 YP 006241474 1 1133850 Actinobacteria
Streptomyces rapamycinicus NRRL 5491 YP 008791920 1 1343740 Actinobacteria
Streptomyces violaceusniger Tu 4113 YP 004818392 1 653045 Actinobacteria
Amycolatopsis mediterranei S699 YP 005536281 1 713604 Actinobacteria
Saccharopolyspora erythraea NRRL 2338 YP 001105458 1 405948 Actinobacteria
Amycolatopsis orientalis HCCB10007 YP 008014268 1 1156913 Actinobacteria
Streptomyces fulvissimus DSM 40593 YP 007933451 1 1303692 Actinobacteria
Actinoplanes sp SE50 110 YP 006269041 1 134676 Actinobacteria
Streptomyces davawensis JCM 4913 YP 007525922 1 1214101 Actinobacteria
Streptomyces griseus subsp griseus NBRC 13350 YP 001827690 1 455632 Actinobacteria
Streptomyces hygroscopicus subsp jinggangensis 5008 YP 006243069 1 1133850 Actinobacteria
Streptomyces sp PAMC26508 YP 007862789 1 1265601 Actinobacteria
Streptomyces pratensis ATCC 33331 YP 004921945 1 591167 Actinobacteria
Streptomyces bingchenggensis BCW 1 YP 004962094 1 749414 Actinobacteria
Amycolatopsis mediterranei S699 YP 005528611 1 713604 Actinobacteria
Streptomyces cattleya NRRL 8057 DSM 46488 YP 004913074 1 1003195 Actinobacteria
Streptomyces pratensis ATCC 33331 YP 004921983 1 591167 Actinobacteria
Streptomyces sp PAMC26508 YP 007862748 1 1265601 Actinobacteria
Mycobacterium chubuense NBB4 YP 006455243 1 710421 Actinobacteria
Streptomyces sp PAMC26508 YP 007860166 1 1265601 Actinobacteria
Streptomyces pratensis ATCC 33331 YP 004924452 1 591167 Actinobacteria

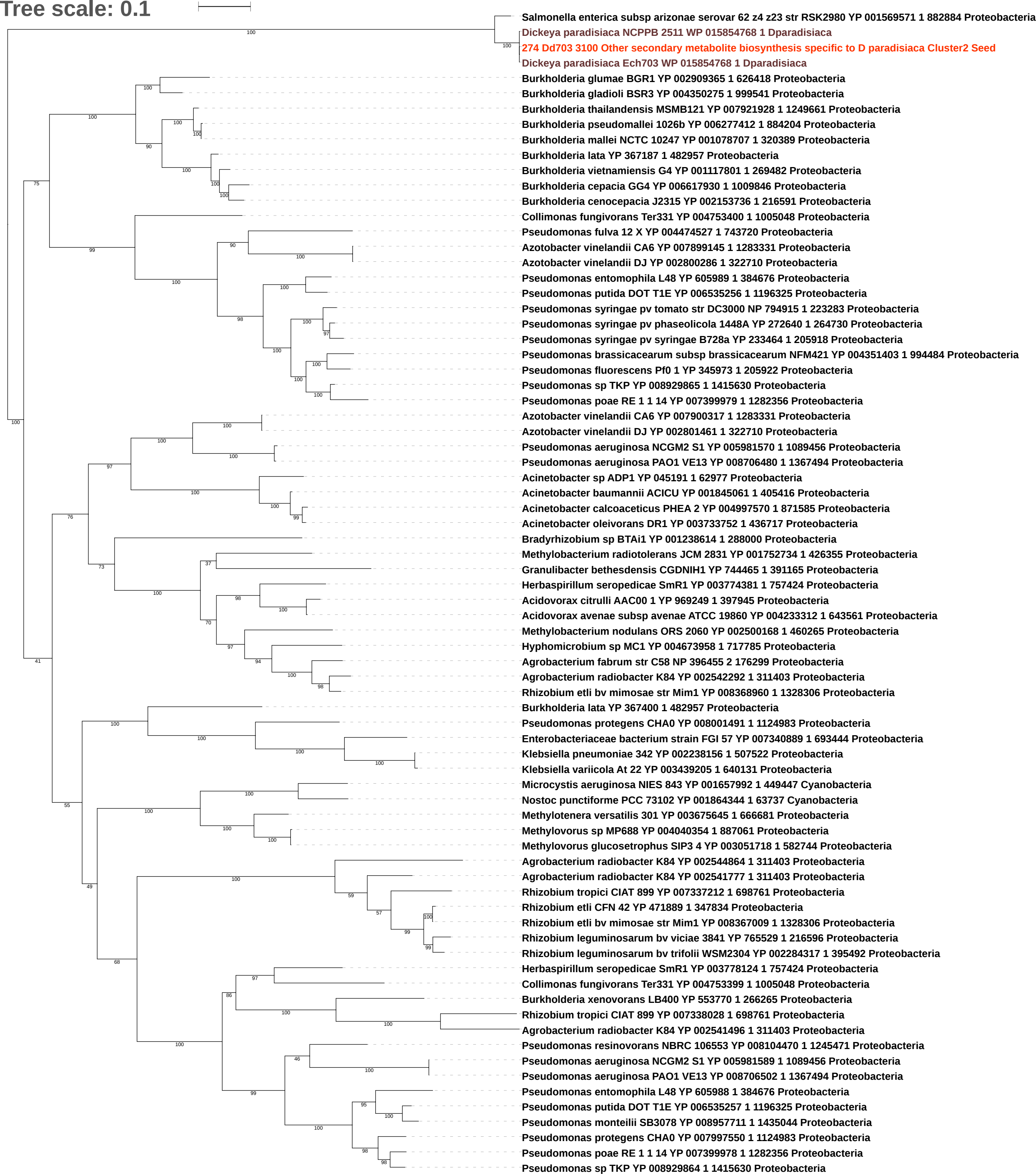
Tree scale: 0.1
Salmonella enterica subsp arizonae serovar 62 z4 z23 str RSK2980 YP 001569571 1 882884 Proteobacteria
Dickeya paradisiaca NCPPB 2511 WP 015854768 1 Dparadisiaca
274 Dd703 3100 Other secondary metabolite biosynthesis specific to D paradisiaca Cluster2 Seed
Dickeya paradisiaca Ech703 WP 015854768 1 Dparadisiaca
Burkholderia glumae BGR1 YP 002909365 1 626418 Proteobacteria
Burkholderia gladioli BSR3 YP 004350275 1 999541 Proteobacteria
Burkholderia thailandensis MSMB121 YP 007921928 1 1249661 Proteobacteria
Burkholderia pseudomallei 1026b YP 006277412 1 884204 Proteobacteria
Burkholderia mallei NCTC 10247 YP 001078707 1 320389 Proteobacteria
Burkholderia lata YP 367187 1 482957 Proteobacteria
Burkholderia vietnamiensis G4 YP 001117801 1 269482 Proteobacteria
Burkholderia cepacia GG4 YP 006617930 1 1009846 Proteobacteria
Burkholderia cenocepacia J2315 YP 002153736 1 216591 Proteobacteria
Collimonas fungivorans Ter331 YP 004753400 1 1005048 Proteobacteria
Pseudomonas fulva 12 X YP 004474527 1 743720 Proteobacteria
Azotobacter vinelandii CA6 YP 007899145 1 1283331 Proteobacteria
Azotobacter vinelandii DJ YP 002800286 1 322710 Proteobacteria
Pseudomonas entomophila L48 YP 605989 1 384676 Proteobacteria
Pseudomonas putida DOT T1E YP 006535256 1 1196325 Proteobacteria
Pseudomonas syringae pv tomato str DC3000 NP 794915 1 223283 Proteobacteria
Pseudomonas syringae pv phaseolicola 1448A YP 272640 1 264730 Proteobacteria
Pseudomonas syringae pv syringae B728a YP 233464 1 205918 Proteobacteria
Pseudomonas brassicacearum subsp brassicacearum NFM421 YP 004351403 1 994484 Proteobacteria
Pseudomonas fluorescens Pf0 1 YP 345973 1 205922 Proteobacteria
Pseudomonas sp TKP YP 008929865 1 1415630 Proteobacteria
Pseudomonas poae RE 1 1 14 YP 007399979 1 1282356 Proteobacteria
Azotobacter vinelandii CA6 YP 007900317 1 1283331 Proteobacteria
Azotobacter vinelandii DJ YP 002801461 1 322710 Proteobacteria
Pseudomonas aeruginosa NCGM2 S1 YP 005981570 1 1089456 Proteobacteria
Pseudomonas aeruginosa PAO1 VE13 YP 008706480 1 1367494 Proteobacteria
Acinetobacter sp ADP1 YP 045191 1 62977 Proteobacteria
Acinetobacter baumannii ACICU YP 001845061 1 405416 Proteobacteria
Acinetobacter calcoaceticus PHEA 2 YP 004997570 1 871585 Proteobacteria
Acinetobacter oleivorans DR1 YP 003733752 1 436717 Proteobacteria
Bradyrhizobium sp BTAi1 YP 001238614 1 288000 Proteobacteria
Methylobacterium radiotolerans JCM 2831 YP 001752734 1 426355 Proteobacteria
Granulibacter bethesdensis CGDNIH1 YP 744465 1 391165 Proteobacteria
Herbaspirillum seropedicae SmR1 YP 003774381 1 757424 Proteobacteria
Acidovorax citrulli AAC00 1 YP 969249 1 397945 Proteobacteria
Acidovorax avenae subsp avenae ATCC 19860 YP 004233312 1 643561 Proteobacteria
Methylobacterium nodulans ORS 2060 YP 002500168 1 460265 Proteobacteria
Hyphomicrobium sp MC1 YP 004673958 1 717785 Proteobacteria
Agrobacterium fabrum str C58 NP 396455 2 176299 Proteobacteria
Agrobacterium radiobacter K84 YP 002542292 1 311403 Proteobacteria
Rhizobium etli bv mimosae str Mim1 YP 008368960 1 1328306 Proteobacteria
Burkholderia lata YP 367400 1 482957 Proteobacteria
Pseudomonas protegens CHA0 YP 008001491 1 1124983 Proteobacteria
Enterobacteriaceae bacterium strain FGI 57 YP 007340889 1 693444 Proteobacteria
Klebsiella pneumoniae 342 YP 002238156 1 507522 Proteobacteria
Klebsiella variicola At 22 YP 003439205 1 640131 Proteobacteria
Microcystis aeruginosa NIES 843 YP 001657992 1 449447 Cyanobacteria
Nostoc punctiforme PCC 73102 YP 001864344 1 63737 Cyanobacteria
Methylotenera versatilis 301 YP 003675645 1 666681 Proteobacteria
Methylovorus sp MP688 YP 004040354 1 887061 Proteobacteria
Methylovorus glucosetrophus SIP3 4 YP 003051718 1 582744 Proteobacteria
Agrobacterium radiobacter K84 YP 002544864 1 311403 Proteobacteria
Agrobacterium radiobacter K84 YP 002541777 1 311403 Proteobacteria
Rhizobium tropici CIAT 899 YP 007337212 1 698761 Proteobacteria
Rhizobium etli CFN 42 YP 471889 1 347834 Proteobacteria
Rhizobium etli bv mimosae str Mim1 YP 008367009 1 1328306 Proteobacteria
Rhizobium leguminosarum bv viciae 3841 YP 765529 1 216596 Proteobacteria
Rhizobium leguminosarum bv trifolii WSM2304 YP 002284317 1 395492 Proteobacteria
Herbaspirillum seropedicae SmR1 YP 003778124 1 757424 Proteobacteria
Collimonas fungivorans Ter331 YP 004753399 1 1005048 Proteobacteria
Burkholderia xenovorans LB400 YP 553770 1 266265 Proteobacteria
Rhizobium tropici CIAT 899 YP 007338028 1 698761 Proteobacteria
Agrobacterium radiobacter K84 YP 002541496 1 311403 Proteobacteria
Pseudomonas resinovorans NBRC 106553 YP 008104470 1 1245471 Proteobacteria
Pseudomonas aeruginosa NCGM2 S1 YP 005981589 1 1089456 Proteobacteria
Pseudomonas aeruginosa PAO1 VE13 YP 008706502 1 1367494 Proteobacteria
Pseudomonas entomophila L48 YP 605988 1 384676 Proteobacteria
Pseudomonas putida DOT T1E YP 006535257 1 1196325 Proteobacteria
Pseudomonas monteilii SB3078 YP 008957711 1 1435044 Proteobacteria
Pseudomonas protegens CHA0 YP 007997550 1 1124983 Proteobacteria
Pseudomonas poae RE 1 1 14 YP 007399978 1 1282356 Proteobacteria
Pseudomonas sp TKP YP 008929864 1 1415630 Proteobacteria

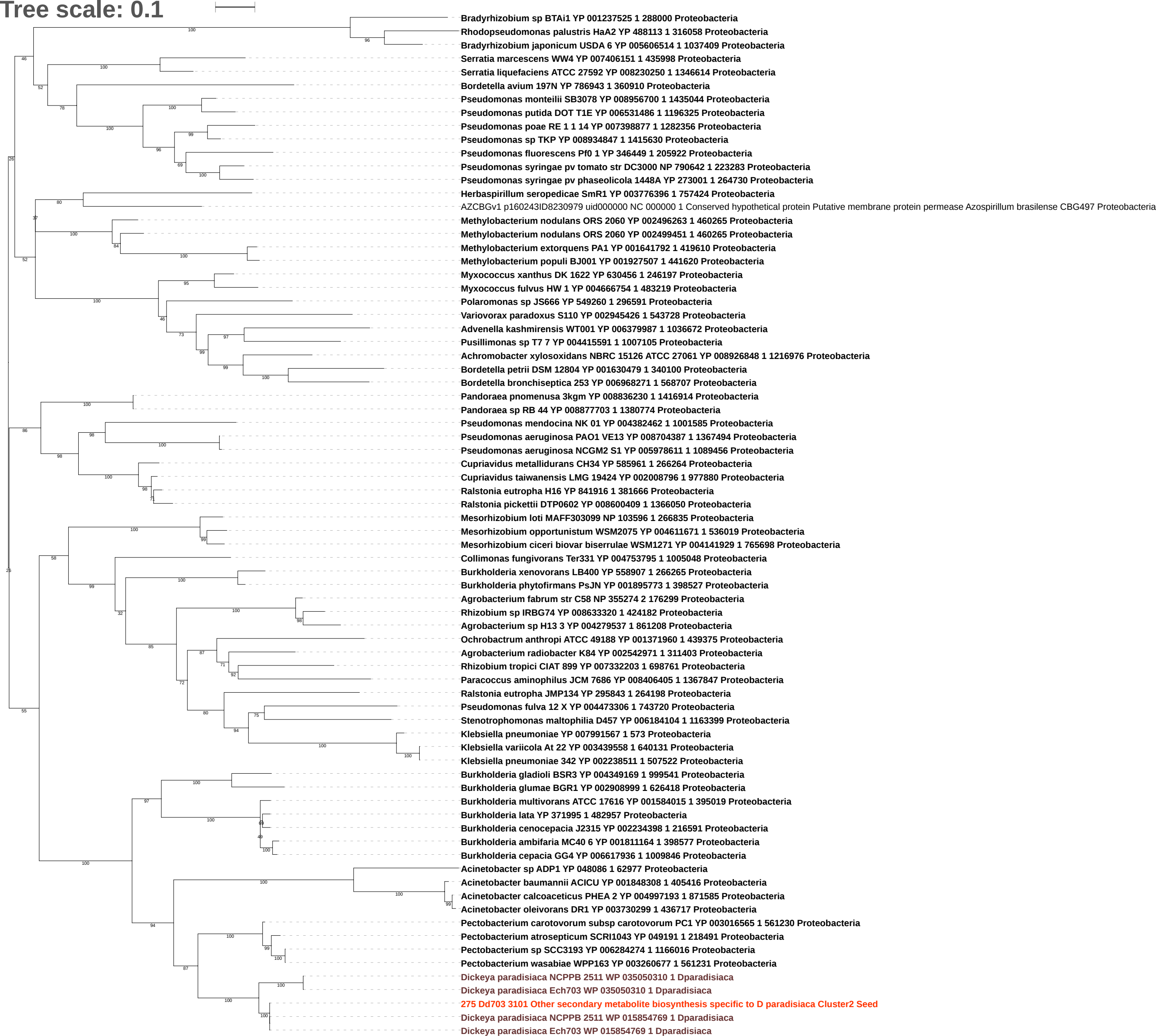
Tree scale: 0.1
Bradyrhizobium sp BTAi1 YP 001237525 1 288000 Proteobacteria
Rhodopseudomonas palustris HaA2 YP 488113 1 316058 Proteobacteria
Bradyrhizobium japonicum USDA 6 YP 005606514 1 1037409 Proteobacteria
Serratia marcescens WW4 YP 007406151 1 435998 Proteobacteria
Serratia liquefaciens ATCC 27592 YP 008230250 1 1346614 Proteobacteria
Bordetella avium 197N YP 786943 1 360910 Proteobacteria
Pseudomonas monteilii SB3078 YP 008956700 1 1435044 Proteobacteria
Pseudomonas putida DOT T1E YP 006531486 1 1196325 Proteobacteria
Pseudomonas poae RE 1 1 14 YP 007398877 1 1282356 Proteobacteria
Pseudomonas sp TKP YP 008934847 1 1415630 Proteobacteria
Pseudomonas fluorescens Pf0 1 YP 346449 1 205922 Proteobacteria
Pseudomonas syringae pv tomato str DC3000 NP 790642 1 223283 Proteobacteria
Pseudomonas syringae pv phaseolicola 1448A YP 273001 1 264730 Proteobacteria
Herbaspirillum seropedicae SmR1 YP 003776396 1 757424 Proteobacteria
AZCBGv1 p160243ID8230979 uid000000 NC 000000 1 Conserved hypothetical protein Putative membrane protein permease Azospirillum brasilense CBG497 Proteobacteria
Methylobacterium nodulans ORS 2060 YP 002496263 1 460265 Proteobacteria
Methylobacterium nodulans ORS 2060 YP 002499451 1 460265 Proteobacteria
Methylobacterium extorquens PA1 YP 001641792 1 419610 Proteobacteria
Methylobacterium populi BJ001 YP 001927507 1 441620 Proteobacteria
Myxococcus xanthus DK 1622 YP 630456 1 246197 Proteobacteria
Myxococcus fulvus HW 1 YP 004666754 1 483219 Proteobacteria
Polaromonas sp JS666 YP 549260 1 296591 Proteobacteria
Variovorax paradoxus S110 YP 002945426 1 543728 Proteobacteria
Advenella kashmirensis WT001 YP 006379987 1 1036672 Proteobacteria
Pusillimonas sp T7 7 YP 004415591 1 1007105 Proteobacteria
Achromobacter xylosoxidans NBRC 15126 ATCC 27061 YP 008926848 1 1216976 Proteobacteria
Bordetella petrii DSM 12804 YP 001630479 1 340100 Proteobacteria
Bordetella bronchiseptica 253 YP 006968271 1 568707 Proteobacteria
Pandoraea pnomenusa 3kgm YP 008836230 1 1416914 Proteobacteria
Pandoraea sp RB 44 YP 008877703 1 1380774 Proteobacteria
Pseudomonas mendocina NK 01 YP 004382462 1 1001585 Proteobacteria
Pseudomonas aeruginosa PAO1 VE13 YP 008704387 1 1367494 Proteobacteria
Pseudomonas aeruginosa NCGM2 S1 YP 005978611 1 1089456 Proteobacteria
Cupriavidus metallidurans CH34 YP 585961 1 266264 Proteobacteria
Cupriavidus taiwanensis LMG 19424 YP 002008796 1 977880 Proteobacteria
Ralstonia eutropha H16 YP 841916 1 381666 Proteobacteria
Ralstonia pickettii DTP0602 YP 008600409 1 1366050 Proteobacteria
Mesorhizobium loti MAFF303099 NP 103596 1 266835 Proteobacteria
Mesorhizobium opportunistum WSM2075 YP 004611671 1 536019 Proteobacteria
Mesorhizobium ciceri biovar biserrulae WSM1271 YP 004141929 1 765698 Proteobacteria
Collimonas fungivorans Ter331 YP 004753795 1 1005048 Proteobacteria
Burkholderia xenovorans LB400 YP 558907 1 266265 Proteobacteria
Burkholderia phytofirmans PsJN YP 001895773 1 398527 Proteobacteria
Agrobacterium fabrum str C58 NP 355274 2 176299 Proteobacteria
Rhizobium sp IRBG74 YP 008633320 1 424182 Proteobacteria
Agrobacterium sp H13 3 YP 004279537 1 861208 Proteobacteria
Ochrobactrum anthropi ATCC 49188 YP 001371960 1 439375 Proteobacteria
Agrobacterium radiobacter K84 YP 002542971 1 311403 Proteobacteria
Rhizobium tropici CIAT 899 YP 007332203 1 698761 Proteobacteria
Paracoccus aminophilus JCM 7686 YP 008406405 1 1367847 Proteobacteria
Ralstonia eutropha JMP134 YP 295843 1 264198 Proteobacteria
Pseudomonas fulva 12 X YP 004473306 1 743720 Proteobacteria
Stenotrophomonas maltophilia D457 YP 006184104 1 1163399 Proteobacteria
Klebsiella pneumoniae YP 007991567 1 573 Proteobacteria
Klebsiella variicola At 22 YP 003439558 1 640131 Proteobacteria
Klebsiella pneumoniae 342 YP 002238511 1 507522 Proteobacteria
Burkholderia gladioli BSR3 YP 004349169 1 999541 Proteobacteria
Burkholderia glumae BGR1 YP 002908999 1 626418 Proteobacteria
Burkholderia multivorans ATCC 17616 YP 001584015 1 395019 Proteobacteria
Burkholderia lata YP 371995 1 482957 Proteobacteria
Burkholderia cenocepacia J2315 YP 002234398 1 216591 Proteobacteria
Burkholderia ambifaria MC40 6 YP 001811164 1 398577 Proteobacteria
Burkholderia cepacia GG4 YP 006617936 1 1009846 Proteobacteria
Acinetobacter sp ADP1 YP 048086 1 62977 Proteobacteria
Acinetobacter baumannii ACICU YP 001848308 1 405416 Proteobacteria
Acinetobacter calcoaceticus PHEA 2 YP 004997193 1 871585 Proteobacteria
Acinetobacter oleivorans DR1 YP 003730299 1 436717 Proteobacteria
Pectobacterium carotovorum subsp carotovorum PC1 YP 003016565 1 561230 Proteobacteria
Pectobacterium atrosepticum SCRI1043 YP 049191 1 218491 Proteobacteria
Pectobacterium sp SCC3193 YP 006284274 1 1166016 Proteobacteria
Pectobacterium wasabiae WPP163 YP 003260677 1 561231 Proteobacteria
Dickeya paradisiaca NCPPB 2511 WP 035050310 1 Dparadisiaca
Dickeya paradisiaca Ech703 WP 035050310 1 Dparadisiaca
275 Dd703 3101 Other secondary metabolite biosynthesis specific to D paradisiaca Cluster2 Seed
Dickeya paradisiaca NCPPB 2511 WP 015854769 1 Dparadisiaca
Dickeya paradisiaca Ech703 WP 015854769 1 Dparadisiaca

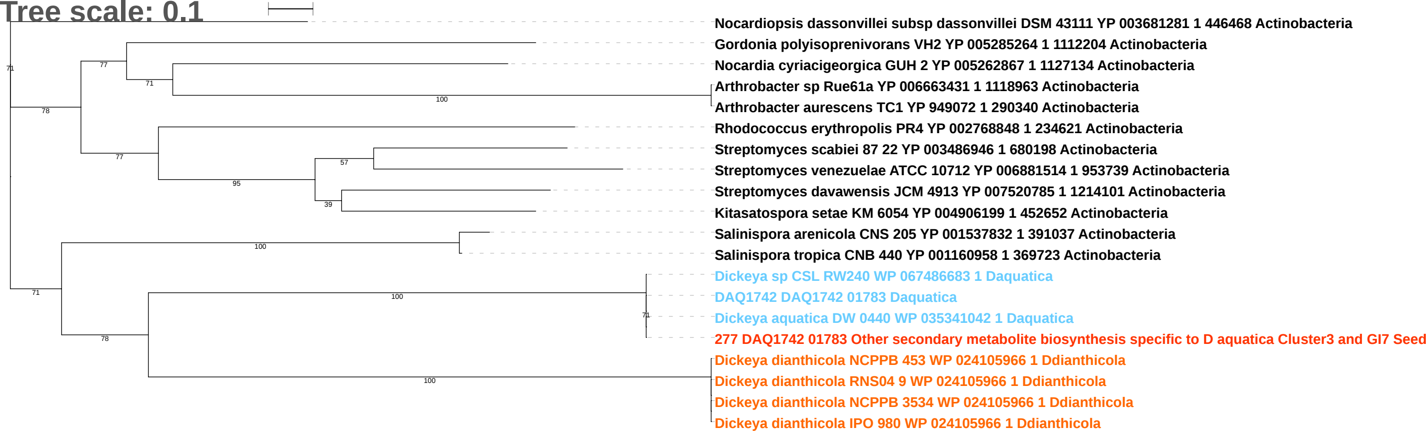
Tree scale: 0.1
Nocardiopsis dassonvillei subsp dassonvillei DSM 43111 YP 003681281 1 446468 Actinobacteria
Gordonia polyisoprenivorans VH2 YP 005285264 1 1112204 Actinobacteria
Nocardia cyriacigeorgica GUH 2 YP 005262867 1 1127134 Actinobacteria
Arthrobacter sp Rue61a YP 006663431 1 1118963 Actinobacteria
Arthrobacter aurescens TC1 YP 949072 1 290340 Actinobacteria
Rhodococcus erythropolis PR4 YP 002768848 1 234621 Actinobacteria
Streptomyces scabiei 87 22 YP 003486946 1 680198 Actinobacteria
Streptomyces venezuelae ATCC 10712 YP 006881514 1 953739 Actinobacteria
Streptomyces davawensis JCM 4913 YP 007520785 1 1214101 Actinobacteria
Kitasatospora setae KM 6054 YP 004906199 1 452652 Actinobacteria
Salinispora arenicola CNS 205 YP 001537832 1 391037 Actinobacteria
Salinispora tropica CNB 440 YP 001160958 1 369723 Actinobacteria
Dickeya sp CSL RW240 WP 067486683 1 Daquatica
DAQ1742 DAQ1742 01783 Daquatica
Dickeya aquatica DW 0440 WP 035341042 1 Daquatica
277 DAQ1742 01783 Other secondary metabolite biosynthesis specific to D aquatica Cluster3 and GI7 Seed
Dickeya dianthicola NCPPB 453 WP 024105966 1 Ddianthicola
Dickeya dianthicola RNS04 9 WP 024105966 1 Ddianthicola
Dickeya dianthicola NCPPB 3534 WP 024105966 1 Ddianthicola
Dickeya dianthicola IPO 980 WP 024105966 1 Ddianthicola
71
77
71
100
78
77
57
95
39
100
71
100
71
78
100

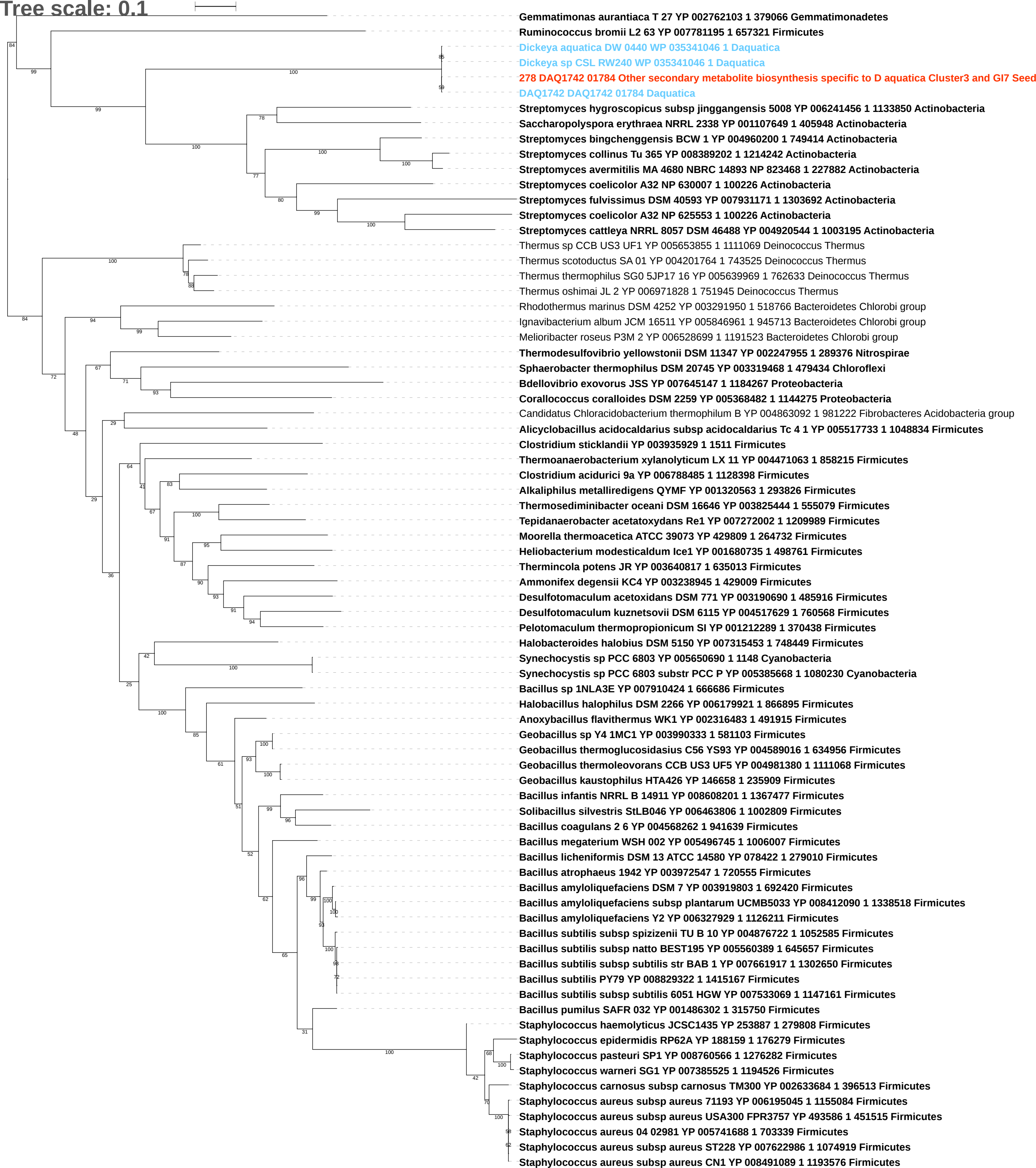
Tree scale: 0.1
Gemmatimonas aurantiaca T 27 YP 002762103 1 379066 Gemmatimonadetes
Ruminococcus bromii L2 63 YP 007781195 1 657321 Firmicutes
Dickeya aquatica DW 0440 WP 035341046 1 Daquatica
Dickeya sp CSL RW240 WP 035341046 1 Daquatica
278 DAQ1742 01784 Other secondary metabolite biosynthesis specific to D aquatica Cluster3 and GI7 Seed
DAQ1742 DAQ1742 01784 Daquatica
Streptomyces hygroscopicus subsp jinggangensis 5008 YP 006241456 1 1133850 Actinobacteria
Saccharopolyspora erythraea NRRL 2338 YP 001107649 1 405948 Actinobacteria
Streptomyces bingchenggensis BCW 1 YP 004960200 1 749414 Actinobacteria
Streptomyces collinus Tu 365 YP 008389202 1 1214242 Actinobacteria
Streptomyces avermitilis MA 4680 NBRC 14893 NP 823468 1 227882 Actinobacteria
Streptomyces coelicolor A32 NP 630007 1 100226 Actinobacteria
Streptomyces fulvissimus DSM 40593 YP 007931171 1 1303692 Actinobacteria
Streptomyces coelicolor A32 NP 625553 1 100226 Actinobacteria
Streptomyces cattleya NRRL 8057 DSM 46488 YP 004920544 1 1003195 Actinobacteria
Thermus sp CCB US3 UF1 YP 005653855 1 1111069 Deinococcus Thermus
Thermus scotoductus SA 01 YP 004201764 1 743525 Deinococcus Thermus
Thermus thermophilus SG0 5JP17 16 YP 005639969 1 762633 Deinococcus Thermus
Thermus oshimai JL 2 YP 006971828 1 751945 Deinococcus Thermus
Rhodothermus marinus DSM 4252 YP 003291950 1 518766 Bacteroidetes Chlorobi group
Ignavibacterium album JCM 16511 YP 005846961 1 945713 Bacteroidetes Chlorobi group
Melioribacter roseus P3M 2 YP 006528699 1 1191523 Bacteroidetes Chlorobi group
Thermodesulfovibrio yellowstonii DSM 11347 YP 002247955 1 289376 Nitrospirae
Sphaerobacter thermophilus DSM 20745 YP 003319468 1 479434 Chloroflexi
Bdellovibrio exovorus JSS YP 007645147 1 1184267 Proteobacteria
Corallococcus coralloides DSM 2259 YP 005368482 1 1144275 Proteobacteria
Candidatus Chloracidobacterium thermophilum B YP 004863092 1 981222 Fibrobacteres Acidobacteria group
Alicyclobacillus acidocaldarius subsp acidocaldarius Tc 4 1 YP 005517733 1 1048834 Firmicutes
Clostridium sticklandii YP 003935929 1 1511 Firmicutes
Thermoanaerobacterium xylanolyticum LX 11 YP 004471063 1 858215 Firmicutes
Clostridium acidurici 9a YP 006788485 1 1128398 Firmicutes
Alkaliphilus metalliredigens QYMF YP 001320563 1 293826 Firmicutes
Thermosediminibacter oceani DSM 16646 YP 003825444 1 555079 Firmicutes
Tepidanaerobacter acetatoxydans Re1 YP 007272002 1 1209989 Firmicutes
Moorella thermoacetica ATCC 39073 YP 429809 1 264732 Firmicutes
Heliobacterium modesticaldum Ice1 YP 001680735 1 498761 Firmicutes
Thermincola potens JR YP 003640817 1 635013 Firmicutes
Ammonifex degensii KC4 YP 003238945 1 429009 Firmicutes
Desulfotomaculum acetoxidans DSM 771 YP 003190690 1 485916 Firmicutes
Desulfotomaculum kuznetsovii DSM 6115 YP 004517629 1 760568 Firmicutes
Pelotomaculum thermopropionicum SI YP 001212289 1 370438 Firmicutes
Halobacteroides halobius DSM 5150 YP 007315453 1 748449 Firmicutes
Synechocystis sp PCC 6803 YP 005650690 1 1148 Cyanobacteria
Synechocystis sp PCC 6803 substr PCC P YP 005385668 1 1080230 Cyanobacteria
Bacillus sp 1NLA3E YP 007910424 1 666686 Firmicutes
Halobacillus halophilus DSM 2266 YP 006179921 1 866895 Firmicutes
Anoxybacillus flavithermus WK1 YP 002316483 1 491915 Firmicutes
Geobacillus sp Y4 1MC1 YP 003990333 1 581103 Firmicutes
Geobacillus thermoglucosidasius C56 YS93 YP 004589016 1 634956 Firmicutes
Geobacillus thermoleovorans CCB US3 UF5 YP 004981380 1 1111068 Firmicutes
Geobacillus kaustophilus HTA426 YP 146658 1 235909 Firmicutes
Bacillus infantis NRRL B 14911 YP 008608201 1 1367477 Firmicutes
Solibacillus silvestris StLB046 YP 006463806 1 1002809 Firmicutes
Bacillus coagulans 2 6 YP 004568262 1 941639 Firmicutes
Bacillus megaterium WSH 002 YP 005496745 1 1006007 Firmicutes
Bacillus licheniformis DSM 13 ATCC 14580 YP 078422 1 279010 Firmicutes
Bacillus atrophaeus 1942 YP 003972547 1 720555 Firmicutes
Bacillus amyloliquefaciens DSM 7 YP 003919803 1 692420 Firmicutes
Bacillus amyloliquefaciens subsp plantarum UCMB5033 YP 008412090 1 1338518 Firmicutes
Bacillus amyloliquefaciens Y2 YP 006327929 1 1126211 Firmicutes
Bacillus subtilis subsp spizizenii TU B 10 YP 004876722 1 1052585 Firmicutes
Bacillus subtilis subsp natto BEST195 YP 005560389 1 645657 Firmicutes
Bacillus subtilis subsp subtilis str BAB 1 YP 007661917 1 1302650 Firmicutes
Bacillus subtilis PY79 YP 008829322 1 1415167 Firmicutes
Bacillus subtilis subsp subtilis 6051 HGW YP 007533069 1 1147161 Firmicutes
Bacillus pumilus SAFR 032 YP 001486302 1 315750 Firmicutes
Staphylococcus haemolyticus JCSC1435 YP 253887 1 279808 Firmicutes
Staphylococcus epidermidis RP62A YP 188159 1 176279 Firmicutes
Staphylococcus pasteuri SP1 YP 008760566 1 1276282 Firmicutes
Staphylococcus warneri SG1 YP 007385525 1 1194526 Firmicutes
Staphylococcus carnosus subsp carnosus TM300 YP 002633684 1 396513 Firmicutes
Staphylococcus aureus subsp aureus 71193 YP 006195045 1 1155084 Firmicutes
Staphylococcus aureus subsp aureus USA300 FPR3757 YP 493586 1 451515 Firmicutes
Staphylococcus aureus 04 02981 YP 005741688 1 703339 Firmicutes
Staphylococcus aureus subsp aureus ST228 YP 007622986 1 1074919 Firmicutes
Staphylococcus aureus subsp aureus CN1 YP 008491089 1 1193576 Firmicutes

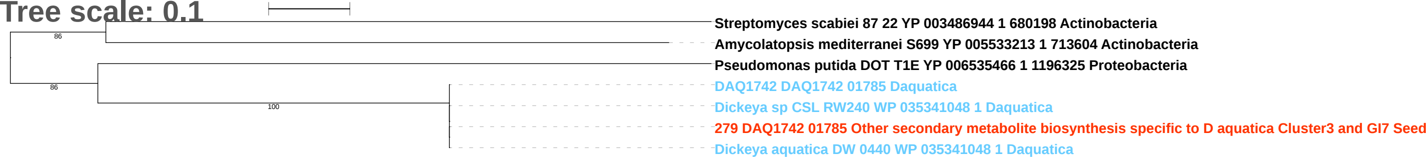
Tree scale: 0.1
86
86
100
Streptomyces scabiei 87 22 YP 003486944 1 680198 Actinobacteria
Amycolatopsis mediterranei S699 YP 005533213 1 713604 Actinobacteria
Pseudomonas putida DOT T1E YP 006535466 1 1196325 Proteobacteria
DAQ1742 DAQ1742 01785 Daquatica
Dickeya sp CSL RW240 WP 035341048 1 Daquatica
279 DAQ1742 01785 Other secondary metabolite biosynthesis specific to D aquatica Cluster3 and GI7 Seed
Dickeya aquatica DW 0440 WP 035341048 1 Daquatica

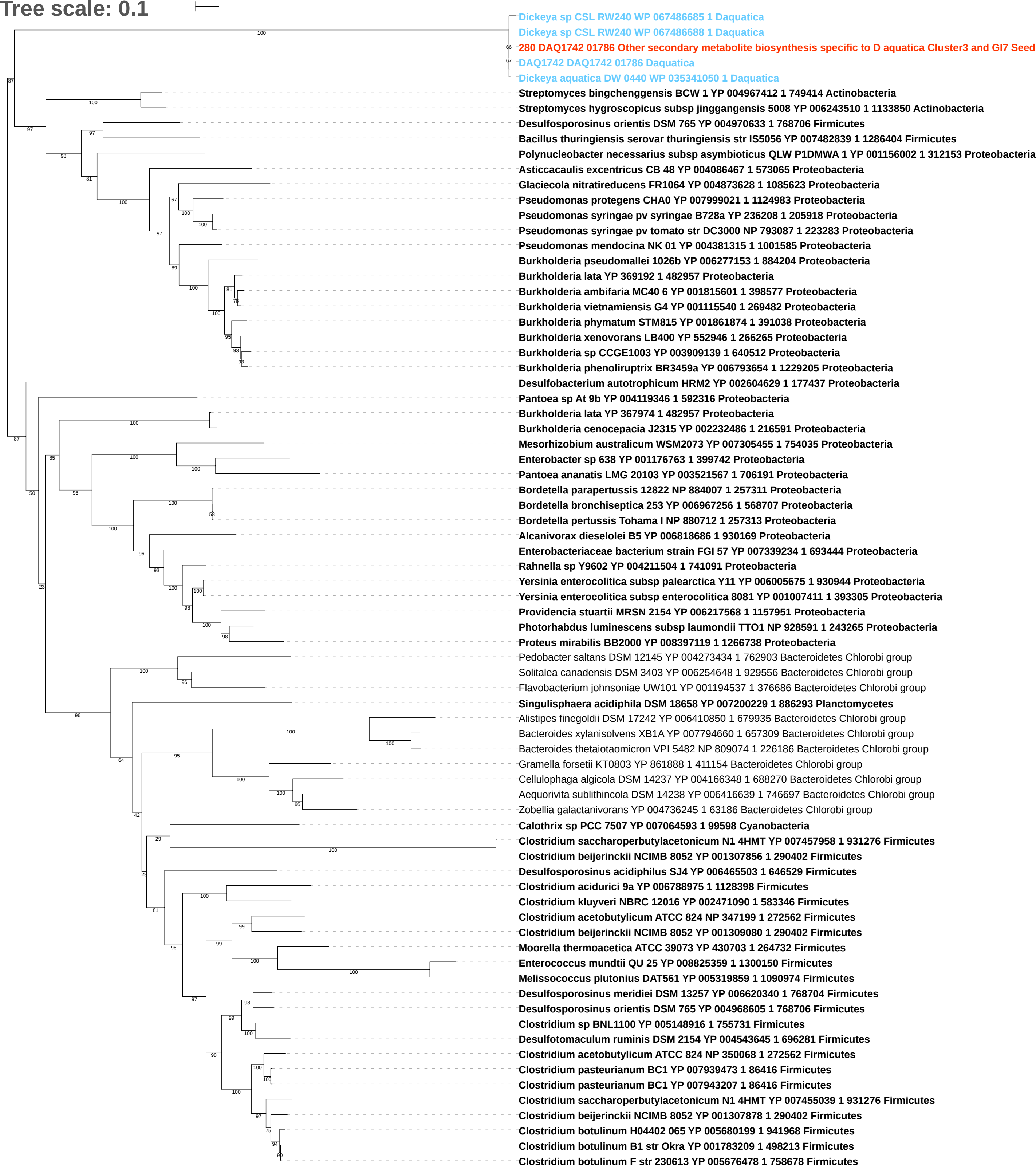

Tree scale: 0.1
Dickeya sp CSL RW240 WP 067486685 1 Daquatica
Dickeya sp CSL RW240 WP 067486688 1 Daquatica
280 DAQ1742 01786 Other secondary metabolite biosynthesis specific to D aquatica Cluster3 and GI7 Seed
DAQ1742 DAQ1742 01786 Daquatica
Dickeya aquatica DW 0440 WP 035341050 1 Daquatica
Streptomyces bingchenggensis BCW 1 YP 004967412 1 749414 Actinobacteria
Streptomyces hygroscopicus subsp jinggangensis 5008 YP 006243510 1 1133850 Actinobacteria
Desulfosporosinus orientis DSM 765 YP 004970633 1 768706 Firmicutes
Bacillus thuringiensis serovar thuringiensis str IS5056 YP 007482839 1 1286404 Firmicutes
Polynucleobacter necessarius subsp asymbioticus QLW P1DMWA 1 YP 001156002 1 312153 Proteobacteria
Asticcacaulis excentricus CB 48 YP 004086467 1 573065 Proteobacteria
Glaciecola nitratireducens FR1064 YP 004873628 1 1085623 Proteobacteria
Pseudomonas protegens CHA0 YP 007999021 1 1124983 Proteobacteria
Pseudomonas syringae pv syringae B728a YP 236208 1 205918 Proteobacteria
Pseudomonas syringae pv tomato str DC3000 NP 793087 1 223283 Proteobacteria
Pseudomonas mendocina NK 01 YP 004381315 1 1001585 Proteobacteria
Burkholderia pseudomallei 1026b YP 006277153 1 884204 Proteobacteria
Burkholderia lata YP 369192 1 482957 Proteobacteria
Burkholderia ambifaria MC40 6 YP 001815601 1 398577 Proteobacteria
Burkholderia vietnamiensis G4 YP 001115540 1 269482 Proteobacteria
Burkholderia phymatum STM815 YP 001861874 1 391038 Proteobacteria
Burkholderia xenovorans LB400 YP 552946 1 266265 Proteobacteria
Burkholderia sp CCGE1003 YP 003909139 1 640512 Proteobacteria
Burkholderia phenoliruptrix BR3459a YP 006793654 1 1229205 Proteobacteria
Desulfobacterium autotrophicum HRM2 YP 002604629 1 177437 Proteobacteria
Pantoea sp At 9b YP 004119346 1 592316 Proteobacteria
Burkholderia lata YP 367974 1 482957 Proteobacteria
Burkholderia cenocepacia J2315 YP 002232486 1 216591 Proteobacteria
Mesorhizobium australicum WSM2073 YP 007305455 1 754035 Proteobacteria
Enterobacter sp 638 YP 001176763 1 399742 Proteobacteria
Pantoea ananatis LMG 20103 YP 003521567 1 706191 Proteobacteria
Bordetella parapertussis 12822 NP 884007 1 257311 Proteobacteria
Bordetella bronchiseptica 253 YP 006967256 1 568707 Proteobacteria
Bordetella pertussis Tohama I NP 880712 1 257313 Proteobacteria
Alcanivorax dieselolei B5 YP 006818686 1 930169 Proteobacteria
Enterobacteriaceae bacterium strain FGI 57 YP 007339234 1 693444 Proteobacteria
Rahnella sp Y9602 YP 004211504 1 741091 Proteobacteria
Yersinia enterocolitica subsp palearctica Y11 YP 006005675 1 930944 Proteobacteria
Yersinia enterocolitica subsp enterocolitica 8081 YP 001007411 1 393305 Proteobacteria
Providencia stuartii MRSN 2154 YP 006217568 1 1157951 Proteobacteria
Photorhabdus luminescens subsp laumondii TTO1 NP 928591 1 243265 Proteobacteria
Proteus mirabilis BB2000 YP 008397119 1 1266738 Proteobacteria
Pedobacter saltans DSM 12145 YP 004273434 1 762903 Bacteroidetes Chlorobi group
Solitalea canadensis DSM 3403 YP 006254648 1 929556 Bacteroidetes Chlorobi group
Flavobacterium johnsoniae UW101 YP 001194537 1 376686 Bacteroidetes Chlorobi group
Singulisphaera acidiphila DSM 18658 YP 007200229 1 886293 Planctomycetes
Alistipes finegoldii DSM 17242 YP 006410850 1 679935 Bacteroidetes Chlorobi group
Bacteroides xylanisolvens XB1A YP 007794660 1 657309 Bacteroidetes Chlorobi group
Bacteroides thetaiotaomicron VPI 5482 NP 809074 1 226186 Bacteroidetes Chlorobi group
Gramella forsetii KT0803 YP 861888 1 411154 Bacteroidetes Chlorobi group
Cellulophaga algicola DSM 14237 YP 004166348 1 688270 Bacteroidetes Chlorobi group
Aequorivita sublithincola DSM 14238 YP 006416697 1 746697 Bacteroidetes Chlorobi group
Zobellia galactanivorans YP 004736245 1 63186 Bacteroidetes Chlorobi group
Calothrix sp PCC 7507 YP 007064593 1 99598 Cyanobacteria
Clostridium saccharoperbutylacetonicum N1 4HMT YP 007457958 1 931276 Firmicutes
Clostridium beijerinckii NCIMB 8052 YP 001307856 1 290402 Firmicutes
Desulfosporosinus acidiphilus SJ4 YP 006465503 1 646529 Firmicutes
Clostridium acidurici 9a YP 006788975 1 1128398 Firmicutes
Clostridium kluyveri NBRC 12016 YP 002471090 1 583346 Firmicutes
Clostridium acetobutylicum ATCC 824 NP 347199 1 272562 Firmicutes
Clostridium beijerinckii NCIMB 8052 YP 001309080 1 290402 Firmicutes
Moorella thermoacetica ATCC 39073 YP 430703 1 264732 Firmicutes
Enterococcus mundtii QU 25 YP 008825359 1 1300150 Firmicutes
Melissococcus plutonius DAT561 YP 005319859 1 1090974 Firmicutes
Desulfosporosinus meridiei DSM 13257 YP 006620340 1 768704 Firmicutes
Desulfosporosinus orientis DSM 765 YP 004968605 1 768706 Firmicutes
Clostridium sp BNL1100 YP 005148916 1 755731 Firmicutes
Desulfotomaculum ruminis DSM 2154 YP 004543645 1 696281 Firmicutes
Clostridium acetobutylicum ATCC 824 NP 350068 1 272562 Firmicutes
Clostridium pasteurianum BC1 YP 007939473 1 86416 Firmicutes
Clostridium pasteurianum BC1 YP 007943207 1 86416 Firmicutes
Clostridium saccharoperbutylacetonicum N1 4HMT YP 007455039 1 931276 Firmicutes
Clostridium beijerinckii NCIMB 8052 YP 001307878 1 290402 Firmicutes
Clostridium botulinum H04402 065 YP 005680199 1 941968 Firmicutes
Clostridium botulinum B1 str Okra YP 001783209 1 498213 Firmicutes
Clostridium botulinum F str 230613 YP 005676478 1 758678 Firmicutes

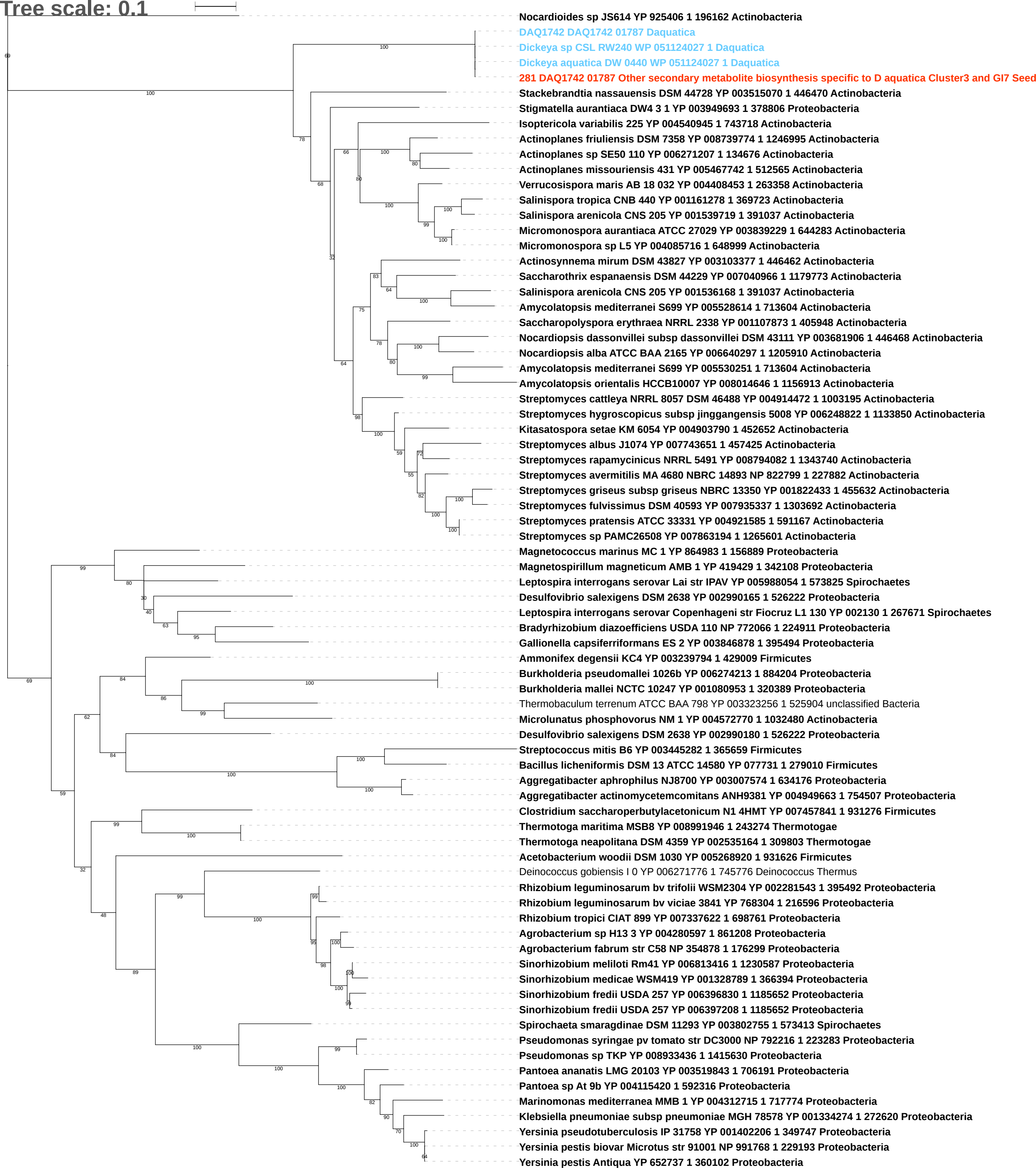
Tree scale: 0.1
Nocardioides sp JS614 YP 925406 1 196162 Actinobacteria
DAQ1742 DAQ1742 01787 Daquatica
Dickeya sp CSL RW240 WP 051124027 1 Daquatica
Dickeya aquatica DW 0440 WP 051124027 1 Daquatica
281 DAQ1742 01787 Other secondary metabolite biosynthesis specific to D aquatica Cluster3 and GI7 Seed
Stackebrandtia nassauensis DSM 44728 YP 003515070 1 446470 Actinobacteria
Stigmatella aurantiaca DW4 3 1 YP 003949693 1 378806 Proteobacteria
Isoptericola variabilis 225 YP 004540945 1 743718 Actinobacteria
Actinoplanes friuliensis DSM 7358 YP 008739774 1 1246995 Actinobacteria
Actinoplanes sp SE50 110 YP 006271207 1 134676 Actinobacteria
Actinoplanes missouriensis 431 YP 005467742 1 512565 Actinobacteria
Verrucosispora maris AB 18 032 YP 004408453 1 263358 Actinobacteria
Salinispora tropica CNB 440 YP 001161278 1 369723 Actinobacteria
Salinispora arenicola CNS 205 YP 001539719 1 391037 Actinobacteria
Micromonospora aurantiaca ATCC 27029 YP 003839229 1 644283 Actinobacteria
Micromonospora sp L5 YP 004085716 1 648999 Actinobacteria
Actinosynnema mirum DSM 43827 YP 003103377 1 446462 Actinobacteria
Saccharothrix espanaensis DSM 44229 YP 007040966 1 1179773 Actinobacteria
Salinispora arenicola CNS 205 YP 001536168 1 391037 Actinobacteria
Amycolatopsis mediterranei S699 YP 005528614 1 713604 Actinobacteria
Saccharopolyspora erythraea NRRL 2338 YP 001107873 1 405948 Actinobacteria
Nocardiopsis dassonvillei subsp dassonvillei DSM 43111 YP 003681906 1 446468 Actinobacteria
Nocardiopsis alba ATCC BAA 2165 YP 006640297 1 1205910 Actinobacteria
Amycolatopsis mediterranei S699 YP 005530251 1 713604 Actinobacteria
Amycolatopsis orientalis HCCB10007 YP 008014646 1 1156913 Actinobacteria
Streptomyces cattleya NRRL 8057 DSM 46488 YP 004914472 1 1003195 Actinobacteria
Streptomyces hygroscopicus subsp jinggangensis 5008 YP 006248822 1 1133850 Actinobacteria
Kitasatospora setae KM 6054 YP 004903790 1 452652 Actinobacteria
Streptomyces albus J1074 YP 007743651 1 457425 Actinobacteria
Streptomyces rapamycinicus NRRL 5491 YP 008794082 1 1343740 Actinobacteria
Streptomyces avermitilis MA 4680 NBRC 14893 NP 822799 1 227882 Actinobacteria
Streptomyces griseus subsp griseus NBRC 13350 YP 001822433 1 455632 Actinobacteria
Streptomyces fulvissimus DSM 40593 YP 007935337 1 1303692 Actinobacteria
Streptomyces pratensis ATCC 33331 YP 004921585 1 591167 Actinobacteria
Streptomyces sp PAMC26508 YP 007863194 1 1265601 Actinobacteria
Magnetococcus marinus MC 1 YP 864983 1 156889 Proteobacteria
Magnetospirillum magneticum AMB 1 YP 419429 1 342108 Proteobacteria
Leptospira interrogans serovar Lai str IPAV YP 005988054 1 573825 Spirochaetes
Desulfovibrio salexigens DSM 2638 YP 002990165 1 526222 Proteobacteria
Leptospira interrogans serovar Copenhageni str Fiocruz L1 130 YP 002130 1 267671 Spirochaetes
Bradyrhizobium diazoefficiens USDA 110 NP 772066 1 224911 Proteobacteria
Gallionella capsiferriformans ES 2 YP 003846878 1 395494 Proteobacteria
Ammonifex degensii KC4 YP 003239794 1 429009 Firmicutes
Burkholderia pseudomallei 1026b YP 006274213 1 884204 Proteobacteria
Burkholderia mallei NCTC 10247 YP 001080953 1 320389 Proteobacteria
Thermobaculum terrenum ATCC BAA 798 YP 003323256 1 525904 unclassified Bacteria
Microlunatus phosphovorus NM 1 YP 004572770 1 1032480 Actinobacteria
Desulfovibrio salexigens DSM 2638 YP 002990180 1 526222 Proteobacteria
Streptococcus mitis B6 YP 003445282 1 365659 Firmicutes
Bacillus licheniformis DSM 13 ATCC 14580 YP 077731 1 279010 Firmicutes
Aggregatibacter aphrophilus NJ8700 YP 003007574 1 634176 Proteobacteria
Aggregatibacter actinomycetemcomitans ANH9381 YP 004949663 1 754507 Proteobacteria
Clostridium saccharoperbutylacetonicum N1 4HMT YP 007457841 1 931276 Firmicutes
Thermotoga maritima MSB8 YP 008991946 1 243274 Thermotogae
Thermotoga neapolitana DSM 4359 YP 002535164 1 309803 Thermotogae
Acetobacterium woodii DSM 1030 YP 005268920 1 931626 Firmicutes
Deinococcus gobiensis I 0 YP 006271776 1 745776 Deinococcus Thermus
Rhizobium leguminosarum bv trifolii WSM2304 YP 002281543 1 395492 Proteobacteria
Rhizobium leguminosarum bv viciae 3841 YP 768304 1 216596 Proteobacteria
Rhizobium tropici CIAT 899 YP 007337622 1 698761 Proteobacteria
Agrobacterium sp H13 3 YP 004280597 1 861208 Proteobacteria
Agrobacterium fabrum str C58 NP 354878 1 176299 Proteobacteria
Sinorhizobium meliloti Rm41 YP 006813416 1 1230587 Proteobacteria
Sinorhizobium medicae WSM419 YP 001328789 1 366394 Proteobacteria
Sinorhizobium fredii USDA 257 YP 006396830 1 1185652 Proteobacteria
Sinorhizobium fredii USDA 257 YP 006397208 1 1185652 Proteobacteria
Spirochaeta smaragdinae DSM 11293 YP 003802755 1 573413 Spirochaetes
Pseudomonas syringae pv tomato str DC3000 NP 792216 1 223283 Proteobacteria
Pseudomonas sp TKP YP 008933436 1 1415630 Proteobacteria
Pantoea ananatis LMG 20103 YP 003519843 1 706191 Proteobacteria
Pantoea sp At 9b YP 004115420 1 592316 Proteobacteria
Marinomonas mediterranea MMB 1 YP 004312775 1 717774 Proteobacteria
Klebsiella pneumoniae subsp pneumoniae MGH 78578 YP 001334274 1 272620 Proteobacteria
Yersinia pseudotuberculosis IP 31758 YP 001402206 1 349747 Proteobacteria
Yersinia pestis biovar Microtus str 91001 NP 991768 1 229193 Proteobacteria
Yersinia pestis Antiqua YP 652737 1 360102 Proteobacteria

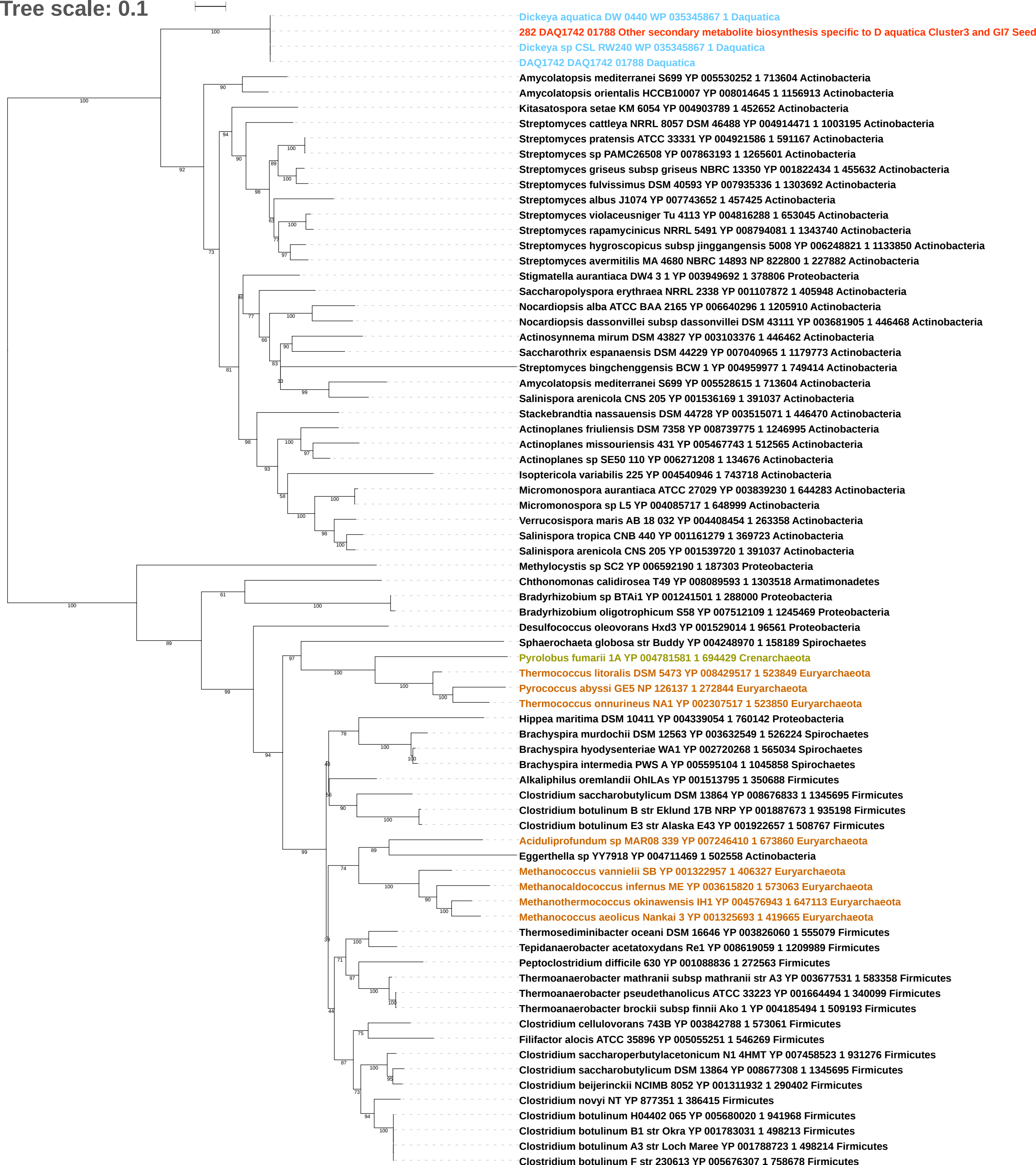
Tree scale: 0.1
Dickeya aquatica DW 0440 WP 035345867 1 Daquatica
282 DAQ1742 01788 Other secondary metabolite biosynthesis specific to D aquatica Cluster3 and GI7 Seed
Dickeya sp CSL RW240 WP 035345867 1 Daquatica
DAQ1742 DAQ1742 01788 Daquatica
Amycolatopsis mediterranei S699 YP 005530252 1 713604 Actinobacteria
Amycolatopsis orientalis HCCB10007 YP 008014645 1 1156913 Actinobacteria
Kitasatospora setae KM 6054 YP 004903789 1 452652 Actinobacteria
Streptomyces cattleya NRRL 8057 DSM 46488 YP 004914471 1 1003195 Actinobacteria
Streptomyces pratensis ATCC 33331 YP 004921586 1 591167 Actinobacteria
Streptomyces sp PAMC26508 YP 007863193 1 1265601 Actinobacteria
Streptomyces griseus subsp griseus NBRC 13350 YP 001822434 1 455632 Actinobacteria
Streptomyces fulvissimus DSM 40593 YP 007935336 1 1303692 Actinobacteria
Streptomyces albus J1074 YP 007743652 1 457425 Actinobacteria
Streptomyces violaceusniger Tu 4113 YP 004816288 1 653045 Actinobacteria
Streptomyces rapamycinicus NRRL 5491 YP 008794081 1 1343740 Actinobacteria
Streptomyces hygroscopicus subsp jinggangensis 5008 YP 006248821 1 1133850 Actinobacteria
Streptomyces avermitilis MA 4680 NBRC 14893 NP 822800 1 227882 Actinobacteria
Stigmatella aurantiaca DW4 3 1 YP 003949692 1 378806 Proteobacteria
Saccharopolyspora erythraea NRRL 2338 YP 001107872 1 405948 Actinobacteria
Nocardiopsis alba ATCC BAA 2165 YP 006640296 1 1205910 Actinobacteria
Nocardiopsis dassonvillei subsp dassonvillei DSM 43111 YP 003681905 1 446468 Actinobacteria
Actinosynnema mirum DSM 43827 YP 003103376 1 446462 Actinobacteria
Saccharothrix espanaensis DSM 44229 YP 007040965 1 1179773 Actinobacteria
Streptomyces bingchenggensis BCW 1 YP 004959977 1 749414 Actinobacteria
Amycolatopsis mediterranei S699 YP 005528615 1 713604 Actinobacteria
Salinispora arenicola CNS 205 YP 001536169 1 391037 Actinobacteria
Stackebrandtia nassauensis DSM 44728 YP 003515071 1 446470 Actinobacteria
Actinoplanes friuliensis DSM 7358 YP 008739775 1 1246995 Actinobacteria
Actinoplanes missouriensis 431 YP 005467743 1 512565 Actinobacteria
Actinoplanes sp SE50 110 YP 006271208 1 134676 Actinobacteria
Isoptericola variabilis 225 YP 004540946 1 743718 Actinobacteria
Micromonospora aurantiaca ATCC 27029 YP 003839230 1 644283 Actinobacteria
Micromonospora sp L5 YP 004085717 1 648999 Actinobacteria
Verrucosispora maris AB 18 032 YP 004408454 1 263358 Actinobacteria
Salinispora tropica CNB 440 YP 001161279 1 369723 Actinobacteria
Salinispora arenicola CNS 205 YP 001539720 1 391037 Actinobacteria
Methylocystis sp SC2 YP 006592190 1 187303 Proteobacteria
Chthonomonas calidirosea T49 YP 008089593 1 1303518 Armatimonadetes
Bradyrhizobium sp BTAi1 YP 001241501 1 288000 Proteobacteria
Bradyrhizobium oligotrophicum S58 YP 007512109 1 1245469 Proteobacteria
Desulfococcus oleovorans Hxd3 YP 001529014 1 96561 Proteobacteria
Sphaerochaeta globosa str Buddy YP 004248970 1 158189 Spirochaetes
Pyrolobus fumarii 1A YP 004781581 1 694429 Crenarchaeota
Thermococcus litoralis DSM 5473 YP 008429517 1 523849 Euryarchaeota
Pyrococcus abyssi GE5 NP 126137 1 272844 Euryarchaeota
Thermococcus onnurineus NA1 YP 002307517 1 523850 Euryarchaeota
Hippea maritima DSM 10411 YP 004339054 1 760142 Proteobacteria
Brachyspira murdochii DSM 12563 YP 003632549 1 526224 Spirochaetes
Brachyspira hyodysenteriae WA1 YP 002720268 1 565034 Spirochaetes
Brachyspira intermedia PWS A YP 005595104 1 1045858 Spirochaetes
Alkaliphilus oremlandii OhILAs YP 001513795 1 350688 Firmicutes
Clostridium saccharobutylicum DSM 13864 YP 008676833 1 1345695 Firmicutes
Clostridium botulinum B str Eklund 17B NRP YP 001887673 1 935198 Firmicutes
Clostridium botulinum E3 str Alaska E43 YP 001922657 1 508767 Firmicutes
Aciduliprofundum sp MAR08 339 YP 007246410 1 673860 Euryarchaeota
Eggerthella sp YY7918 YP 004711469 1 502558 Actinobacteria
Methanococcus vannielii SB YP 001322957 1 406327 Euryarchaeota
Methanocaldococcus infernus ME YP 003615820 1 573063 Euryarchaeota
Methanothermococcus okinawensis IH1 YP 004576943 1 647113 Euryarchaeota
Methanococcus aeolicus Nankai 3 YP 001325693 1 419665 Euryarchaeota
Thermosediminibacter oceani DSM 16646 YP 003826060 1 555079 Firmicutes
Tepidanaerobacter acetatoxydans Re1 YP 008619059 1 1209989 Firmicutes
Peptoclostridium difficile 630 YP 001088836 1 272563 Firmicutes
Thermoanaerobacter mathranii subsp mathranii str A3 YP 003677531 1 583358 Firmicutes
Thermoanaerobacter pseudethanolicus ATCC 33223 YP 001664494 1 340099 Firmicutes
Thermoanaerobacter brockii subsp finnii Ako 1 YP 004185494 1 509193 Firmicutes
Clostridium cellulovorans 743B YP 003842788 1 573061 Firmicutes
Filifactor alocis ATCC 35896 YP 005055251 1 546269 Firmicutes
Clostridium saccharoperbutylacetonicum N1 4HMT YP 007458523 1 931276 Firmicutes
Clostridium saccharobutylicum DSM 13864 YP 008677308 1 1345695 Firmicutes
Clostridium beijerinckii NCIMB 8052 YP 001311932 1 290402 Firmicutes
Clostridium novyi NT YP 877351 1 386415 Firmicutes
Clostridium botulinum H04402 065 YP 005680020 1 941968 Firmicutes
Clostridium botulinum B1 str Okra YP 001783031 1 498213 Firmicutes
Clostridium botulinum A3 str Loch Maree YP 001788723 1 498214 Firmicutes
Clostridium botulinum F str 230613 YP 005676307 1 758678 Firmicutes

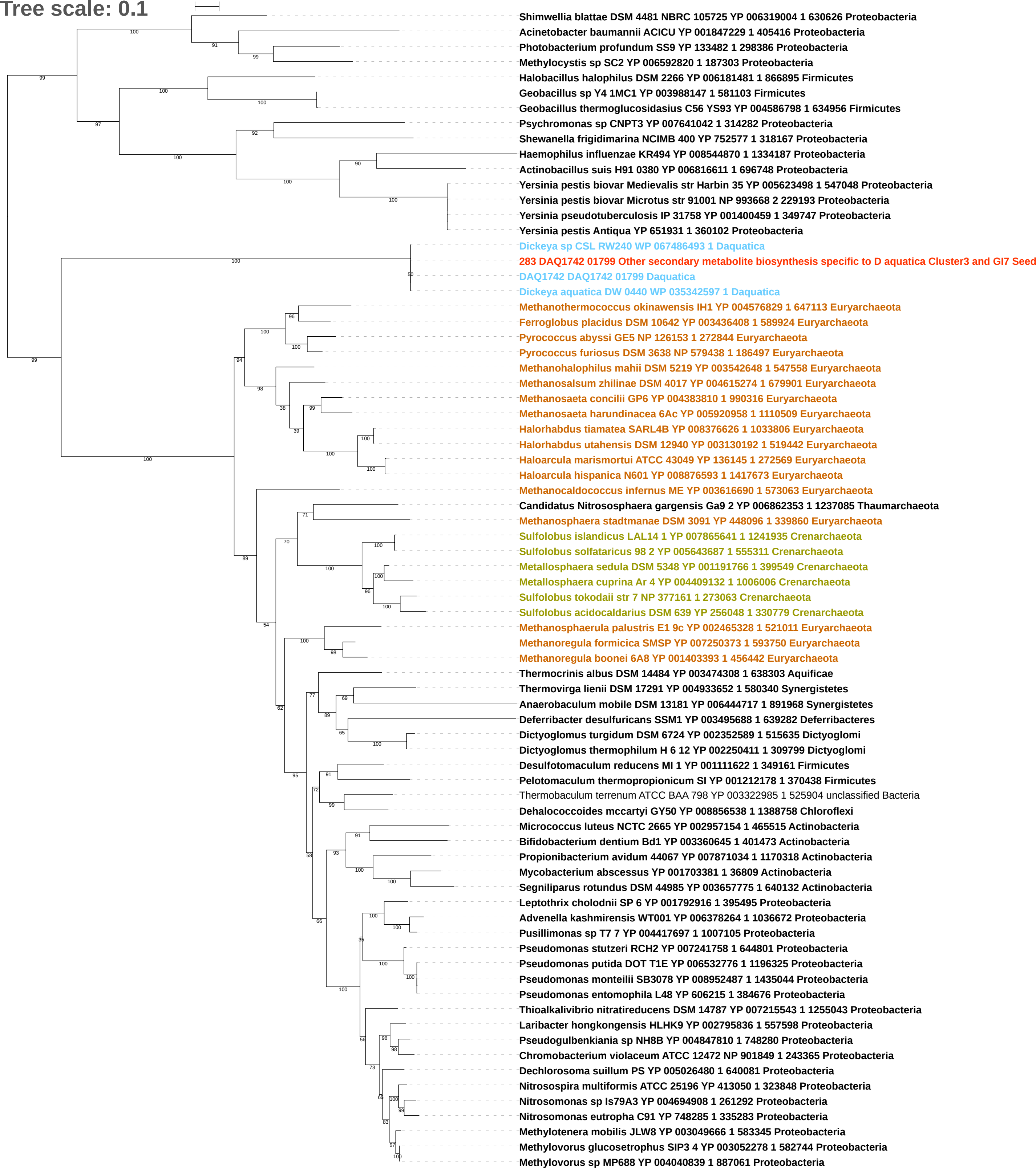

Tree scale: 0.1
Shimwellia blattae DSM 4481 NBRC 105725 YP 006319004 1 630626 Proteobacteria
Acinetobacter baumannii ACICU YP 001847229 1 405416 Proteobacteria
Photobacterium profundum SS9 YP 133482 1 298386 Proteobacteria
Methylocystis sp SC2 YP 006592820 1 187303 Proteobacteria
Halobacillus halophilus DSM 2266 YP 006181481 1 866895 Firmicutes
Geobacillus sp Y4 1MC1 YP 003988147 1 581103 Firmicutes
Geobacillus thermoglucosidasius C56 YS93 YP 004586798 1 634956 Firmicutes
Psychromonas sp CNPT3 YP 007641042 1 314282 Proteobacteria
Shewanella frigidimarina NCIMB 400 YP 752577 1 318167 Proteobacteria
Haemophilus influenzae KR494 YP 008544870 1 1334187 Proteobacteria
Actinobacillus suis H91 0380 YP 006816611 1 696748 Proteobacteria
Yersinia pestis biovar Medievalis str Harbin 35 YP 005623498 1 547048 Proteobacteria
Yersinia pestis biovar Microtus str 91001 NP 993668 2 229193 Proteobacteria
Yersinia pseudotuberculosis IP 31758 YP 001400459 1 349747 Proteobacteria
Yersinia pestis Antiqua YP 651931 1 360102 Proteobacteria
Dickeya sp CSL RW240 WP 067486493 1 Daquatica
283 DAQ1742 01799 Other secondary metabolite biosynthesis specific to D aquatica Cluster3 and GI7 Seed
DAQ1742 DAQ1742 01799 Daquatica
Dickeya aquatica DW 0440 WP 035342597 1 Daquatica
Methanothermococcus okinawensis IH1 YP 004576829 1 647113 Euryarchaeota
Ferroglobus placidus DSM 10642 YP 003436408 1 589924 Euryarchaeota
Pyrococcus abyssi GE5 NP 126153 1 272844 Euryarchaeota
Pyrococcus furiosus DSM 3638 NP 579438 1 186497 Euryarchaeota
Methanohalophilus mahii DSM 5219 YP 003542648 1 547558 Euryarchaeota
Methanosalsum zhilinae DSM 4017 YP 004615274 1 679901 Euryarchaeota
Methanosaeta concilii GP6 YP 004383810 1 990316 Euryarchaeota
Methanosaeta harundinacea 6Ac YP 005920958 1 1110509 Euryarchaeota
Halorhabdus tiamatea SARL4B YP 008376626 1 1033806 Euryarchaeota
Halorhabdus utahensis DSM 12940 YP 003130192 1 519442 Euryarchaeota
Haloarcula marismortui ATCC 43049 YP 136145 1 272569 Euryarchaeota
Haloarcula hispanica N601 YP 008876593 1 1417673 Euryarchaeota
Methanocaldococcus infernus ME YP 003616690 1 573063 Euryarchaeota
Candidatus Nitrososphaera gargensis Ga9 2 YP 006862353 1 1237085 Thaumarchaeota
Methanosphaera stadtmanae DSM 3091 YP 448096 1 339860 Euryarchaeota
Sulfolobus islandicus LAL14 1 YP 007865641 1 1241935 Crenarchaeota
Sulfolobus solfataricus 98 2 YP 005643687 1 555311 Crenarchaeota
Metallosphaera sedula DSM 5348 YP 001191766 1 399549 Crenarchaeota
Metallosphaera cuprina Ar 4 YP 004409132 1 1006006 Crenarchaeota
Sulfolobus tokodaii str 7 NP 377161 1 273063 Crenarchaeota
Sulfolobus acidocaldarius DSM 639 YP 256048 1 330779 Crenarchaeota
Methanosphaerula palustris E1 9c YP 002465328 1 521011 Euryarchaeota
Methanoregula formicica SMSP YP 007250373 1 593750 Euryarchaeota
Methanoregula boonei 6A8 YP 001403393 1 456442 Euryarchaeota
Thermocrinis albus DSM 14484 YP 003474308 1 638303 Aquificae
Thermovirga lienii DSM 17291 YP 004933652 1 580340 Synergistetes
Anaerobaculum mobile DSM 13181 YP 006444717 1 891968 Synergistetes
Deferribacter desulfuricans SSM1 YP 003495688 1 639282 Deferribacteres
Dictyoglomus turgidum DSM 6724 YP 002352589 1 515635 Dictyoglomi
Dictyoglomus thermophilum H 6 12 YP 002250411 1 309799 Dictyoglomi
Desulfotomaculum reducens MI 1 YP 001111622 1 349161 Firmicutes
Pelotomaculum thermopropionicum SI YP 001212178 1 370438 Firmicutes
Thermobaculum terrenum ATCC BAA 798 YP 003322985 1 525904 unclassified Bacteria
Dehalococcoides mccartyi GY50 YP 008856538 1 1388758 Chloroflexi
Micrococcus luteus NCTC 2665 YP 002957154 1 465515 Actinobacteria
Bifidobacterium dentium Bd1 YP 003360645 1 401473 Actinobacteria
Propionibacterium avidum 44067 YP 007871034 1 1170318 Actinobacteria
Mycobacterium abscessus YP 001703381 1 36809 Actinobacteria
Segniliparus rotundus DSM 44985 YP 003657775 1 640132 Actinobacteria
Leptothrix cholodnii SP 6 YP 001792916 1 395495 Proteobacteria
Advenella kashmirensis WT001 YP 006378264 1 1036672 Proteobacteria
Pusillimonas sp T7 7 YP 004417697 1 1007105 Proteobacteria
Pseudomonas stutzeri RCH2 YP 007241758 1 644801 Proteobacteria
Pseudomonas putida DOT T1E YP 006532776 1 1196325 Proteobacteria
Pseudomonas monteilii SB3078 YP 008952487 1 1435044 Proteobacteria
Pseudomonas entomophila L48 YP 606215 1 384676 Proteobacteria
Thioalkalivibrio nitratireducens DSM 14787 YP 007215543 1 1255043 Proteobacteria
Laribacter hongkongensis HLHK9 YP 002795836 1 557598 Proteobacteria
Pseudogulbenkiania sp NH8B YP 004847810 1 748280 Proteobacteria
Chromobacterium violaceum ATCC 12472 NP 901849 1 243365 Proteobacteria
Dechlorosoma suillum PS YP 005026480 1 640081 Proteobacteria
Nitrosospira multiformis ATCC 25196 YP 413050 1 323848 Proteobacteria
Nitrosomonas sp Is79A3 YP 004694908 1 261292 Proteobacteria
Nitrosomonas eutropha C91 YP 748285 1 335283 Proteobacteria
Methylotenera mobilis JLW8 YP 003049666 1 583345 Proteobacteria
Methylovorus glucosetrophus SIP3 4 YP 003052278 1 582744 Proteobacteria
Methylovorus sp MP688 YP 004040839 1 887061 Proteobacteria

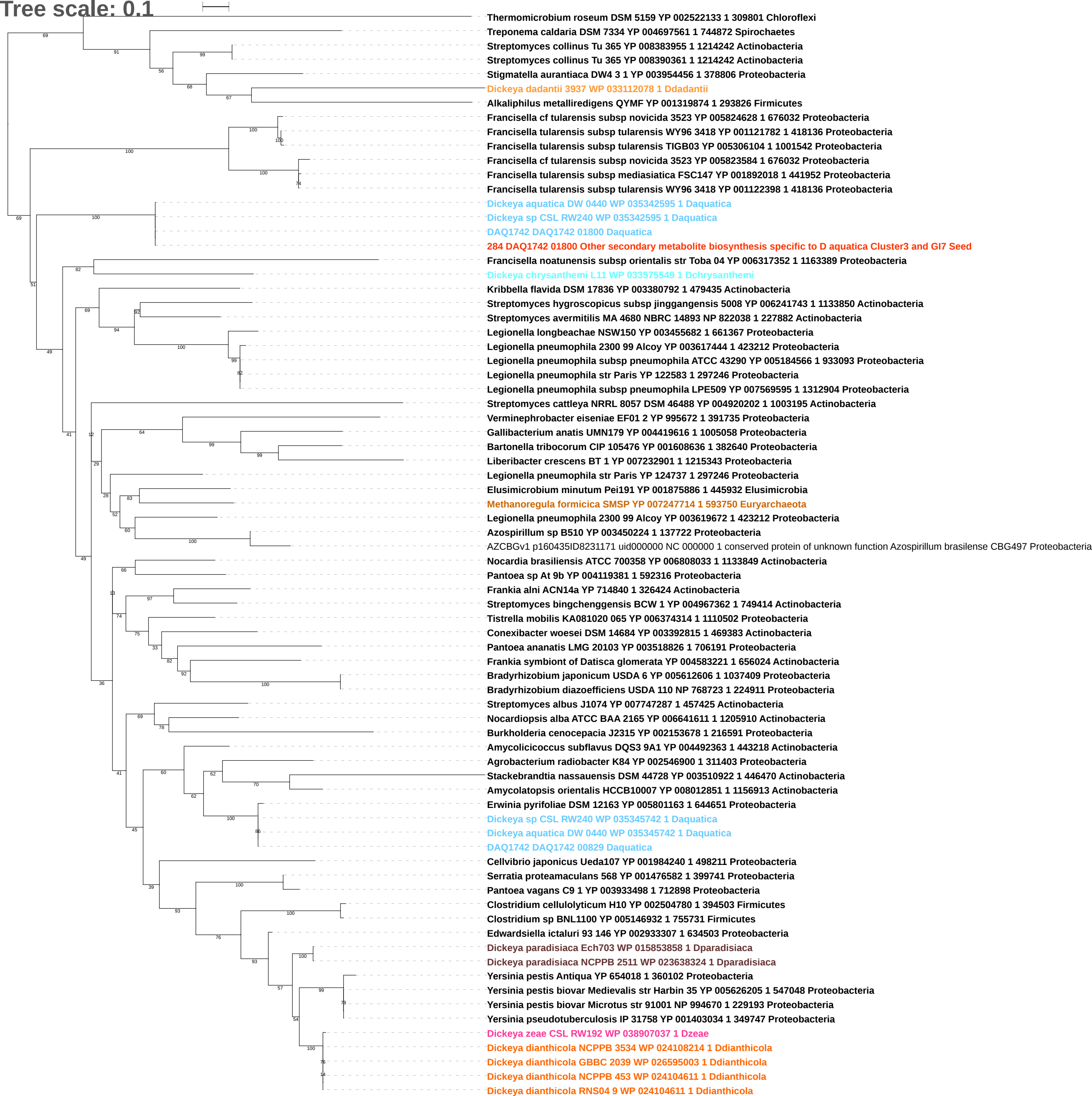
Tree scale: 0.1
Thermomicrobium roseum DSM 5159 YP 002522133 1 309801 Chloroflexi
Treponema caldaria DSM 7334 YP 004697561 1 744872 Spirochaetes
Streptomyces collinus Tu 365 YP 008383955 1 1214242 Actinobacteria
Streptomyces collinus Tu 365 YP 008390361 1 1214242 Actinobacteria
Stigmatella aurantiaca DW4 3 1 YP 003954456 1 378806 Proteobacteria
Dickeya dadantii 3937 WP 033112078 1 Ddadantii
Alkaliphilus metalliredigens QYMF YP 001319874 1 293826 Firmicutes
Francisella cf tularensis subsp novicida 3523 YP 005824628 1 676032 Proteobacteria
Francisella tularensis subsp tularensis WY96 3418 YP 001121782 1 418136 Proteobacteria
Francisella tularensis subsp tularensis TIGB03 YP 005306104 1 1001542 Proteobacteria
Francisella cf tularensis subsp novicida 3523 YP 005823584 1 676032 Proteobacteria
Francisella tularensis subsp mediasiatica FSC147 YP 001892018 1 441952 Proteobacteria
Francisella tularensis subsp tularensis WY96 3418 YP 001122398 1 418136 Proteobacteria
Dickeya aquatica DW 0440 WP 035342595 1 Daquatica
Dickeya sp CSL RW240 WP 035342595 1 Daquatica
DAQ1742 DAQ1742 01800 Daquatica
284 DAQ1742 01800 Other secondary metabolite biosynthesis specific to D aquatica Cluster3 and GI7 Seed
Francisella noatunensis subsp orientalis str Toba 04 YP 006317352 1 1163389 Proteobacteria
Dickeya chrysanthemi L11 WP 033575549 1 Dchrysanthemi
Kribbella flavida DSM 17836 YP 003380792 1 479435 Actinobacteria
Streptomyces hygroscopicus subsp jinggangensis 5008 YP 006241743 1 1133850 Actinobacteria
Streptomyces avermitilis MA 4680 NBRC 14893 NP 822038 1 227882 Actinobacteria
Legionella longbeachae NSW150 YP 003455682 1 661367 Proteobacteria
Legionella pneumophila 2300 99 Alcoy YP 003617444 1 423212 Proteobacteria
Legionella pneumophila subsp pneumophila ATCC 43290 YP 005184566 1 933093 Proteobacteria
Legionella pneumophila str Paris YP 122583 1 297246 Proteobacteria
Legionella pneumophila subsp pneumophila LPE509 YP 007569595 1 1312904 Proteobacteria
Streptomyces cattleya NRRL 8057 DSM 46488 YP 004920202 1 1003195 Actinobacteria
Verminephrobacter eiseniae EF01 2 YP 995672 1 391735 Proteobacteria
Gallibacterium anatis UMN179 YP 004419616 1 1005058 Proteobacteria
Bartonella tribocorum CIP 105476 YP 001608636 1 382640 Proteobacteria
Liberibacter crescens BT 1 YP 007232901 1 1215343 Proteobacteria
Legionella pneumophila str Paris YP 124737 1 297246 Proteobacteria
Elusimicrobium minutum Pei191 YP 001875886 1 445932 Elusimicrobia
Methanoregula formicica SMSP YP 007247714 1 593750 Euryarchaeota
Legionella pneumophila 2300 99 Alcoy YP 003619672 1 423212 Proteobacteria
Azospirillum sp B510 YP 003450224 1 137722 Proteobacteria
AZCBGv1 p160435ID8231171 uid000000 NC 000000 1 conserved protein of unknown function Azospirillum brasilense CBG497 Proteobacteria
Nocardia brasiliensis ATCC 700358 YP 006808033 1 1133849 Actinobacteria
Pantoea sp At 9b YP 004119381 1 592316 Proteobacteria
Frankia alni ACN14a YP 714840 1 326424 Actinobacteria
Streptomyces bingchenggensis BCW 1 YP 004967362 1 749414 Actinobacteria
Tistrella mobilis KA081020 065 YP 006374314 1 1110502 Proteobacteria
Conexibacter woesei DSM 14684 YP 003392815 1 469383 Actinobacteria
Pantoea ananatis LMG 20103 YP 003518826 1 706191 Proteobacteria
Frankia symbiont of Datisca glomerata YP 004583221 1 656024 Actinobacteria
Bradyrhizobium japonicum USDA 6 YP 005612606 1 1037409 Proteobacteria
Bradyrhizobium diazoefficiens USDA 110 NP 768723 1 224911 Proteobacteria
Streptomyces albus J1074 YP 007747287 1 457425 Actinobacteria
Nocardiopsis alba ATCC BAA 2165 YP 006641611 1 1205910 Actinobacteria
Burkholderia cenocepacia J2315 YP 002153678 1 216591 Proteobacteria
Amycolicicoccus subflavus DQS3 9A1 YP 004492363 1 443218 Actinobacteria
Agrobacterium radiobacter K84 YP 002546900 1 311403 Proteobacteria
Stackebrandtia nassauensis DSM 44728 YP 003510922 1 446470 Actinobacteria
Amycolatopsis orientalis HCCB10007 YP 008012851 1 1156913 Actinobacteria
Erwinia pyrifoliae DSM 12163 YP 005801163 1 644651 Proteobacteria
Dickeya sp CSL RW240 WP 035345742 1 Daquatica
Dickeya aquatica DW 0440 WP 035345742 1 Daquatica
DAQ1742 DAQ1742 00829 Daquatica
Cellvibrio japonicus Ueda107 YP 001984240 1 498211 Proteobacteria
Serratia proteamaculans 568 YP 001476582 1 399741 Proteobacteria
Pantoea vagans C9 1 YP 003933498 1 712898 Proteobacteria
Clostridium cellulolyticum H10 YP 002504780 1 394503 Firmicutes
Clostridium sp BNL1100 YP 005146932 1 755731 Firmicutes
Edwardsiella ictaluri 93 146 YP 002933307 1 634503 Proteobacteria
Dickeya paradisiaca Ech703 WP 015853858 1 Dparadisiaca
Dickeya paradisiaca NCPPB 2511 WP 023638324 1 Dparadisiaca
Yersinia pestis Antiqua YP 654018 1 360102 Proteobacteria
Yersinia pestis biovar Medievalis str Harbin 35 YP 005626205 1 547048 Proteobacteria
Yersinia pestis biovar Microtus str 91001 NP 994670 1 229193 Proteobacteria
Yersinia pseudotuberculosis IP 31758 YP 001403034 1 349747 Proteobacteria
Dickeya zeae CSL RW192 WP 038907037 1 Dzeae
Dickeya dianthicola NCPPB 3534 WP 024108214 1 Ddianthicola
Dickeya dianthicola GBBC 2039 WP 026595003 1 Ddianthicola
Dickeya dianthicola NCPPB 453 WP 024104611 1 Ddianthicola
Dickeya dianthicola RNS04 9 WP 024104611 1 Ddianthicola

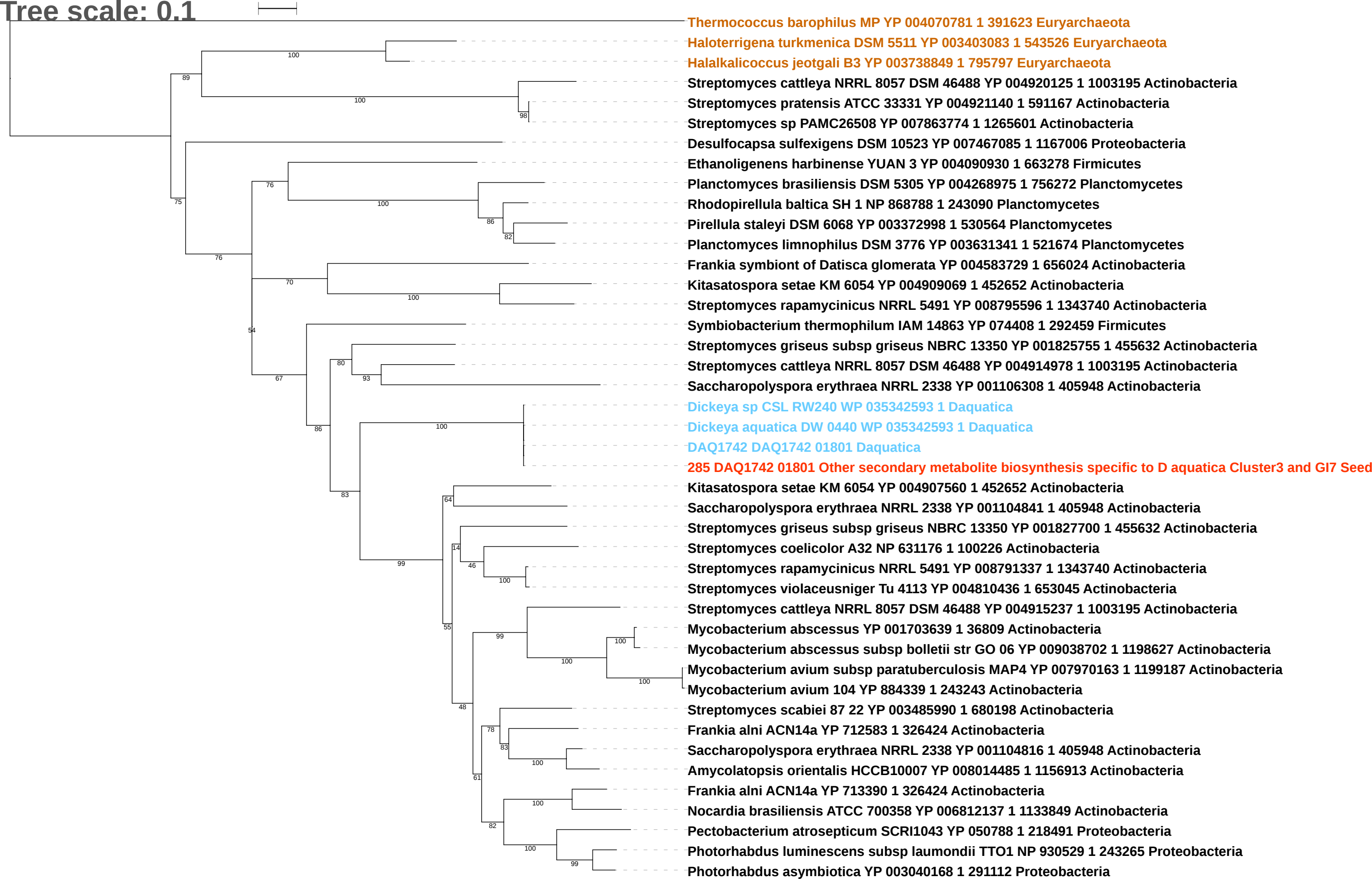
Tree scale: 0.1
Thermococcus barophilus MP YP 004070781 1 391623 Euryarchaeota
Haloterrigena turkmenica DSM 5511 YP 003403083 1 543526 Euryarchaeota
Halalkalicoccus jeotgali B3 YP 003738849 1 795797 Euryarchaeota
Streptomyces cattleya NRRL 8057 DSM 46488 YP 004920125 1 1003195 Actinobacteria
Streptomyces pratensis ATCC 33331 YP 004921140 1 591167 Actinobacteria
Streptomyces sp PAMC26508 YP 007863774 1 1265601 Actinobacteria
Desulfocapsa sulfexigens DSM 10523 YP 007467085 1 1167006 Proteobacteria
Ethanoligenens harbinense YUAN 3 YP 004090930 1 663278 Firmicutes
Planctomyces brasiliensis DSM 5305 YP 004268975 1 756272 Planctomycetes
Rhodopirellula baltica SH 1 NP 868788 1 243090 Planctomycetes
Pirellula staleyi DSM 6068 YP 003372998 1 530564 Planctomycetes
Planctomyces limnophilus DSM 3776 YP 003631341 1 521674 Planctomycetes
Frankia symbiont of Datisca glomerata YP 004583729 1 656024 Actinobacteria
Kitasatospora setae KM 6054 YP 004909069 1 452652 Actinobacteria
Streptomyces rapamycinicus NRRL 5491 YP 008795596 1 1343740 Actinobacteria
Symbiobacterium thermophilum IAM 14863 YP 074408 1 292459 Firmicutes
Streptomyces griseus subsp griseus NBRC 13350 YP 001825755 1 455632 Actinobacteria
Streptomyces cattleya NRRL 8057 DSM 46488 YP 004914978 1 1003195 Actinobacteria
Saccharopolyspora erythraea NRRL 2338 YP 001106308 1 405948 Actinobacteria
Dickeya sp CSL RW240 WP 035342593 1 Daquatica
Dickeya aquatica DW 0440 WP 035342593 1 Daquatica
DAQ1742 DAQ1742 01801 Daquatica
285 DAQ1742 01801 Other secondary metabolite biosynthesis specific to D aquatica Cluster3 and GI7 Seed
Kitasatospora setae KM 6054 YP 004907560 1 452652 Actinobacteria
Saccharopolyspora erythraea NRRL 2338 YP 001104841 1 405948 Actinobacteria
Streptomyces griseus subsp griseus NBRC 13350 YP 001827700 1 455632 Actinobacteria
Streptomyces coelicolor A32 NP 631176 1 100226 Actinobacteria
Streptomyces rapamycinicus NRRL 5491 YP 008791337 1 1343740 Actinobacteria
Streptomyces violaceusniger Tu 4113 YP 004810436 1 653045 Actinobacteria
Streptomyces cattleya NRRL 8057 DSM 46488 YP 004915237 1 1003195 Actinobacteria
Mycobacterium abscessus YP 001703639 1 36809 Actinobacteria
Mycobacterium abscessus subsp bolletii str GO 06 YP 009038702 1 1198627 Actinobacteria
Mycobacterium avium subsp paratuberculosis MAP4 YP 007970163 1 1199187 Actinobacteria
Mycobacterium avium 104 YP 884339 1 243243 Actinobacteria
Streptomyces scabiei 87 22 YP 003485990 1 680198 Actinobacteria
Frankia alni ACN14a YP 712583 1 326424 Actinobacteria
Saccharopolyspora erythraea NRRL 2338 YP 001104816 1 405948 Actinobacteria
Amycolatopsis orientalis HCCB10007 YP 008014485 1 1156913 Actinobacteria
Frankia alni ACN14a YP 713390 1 326424 Actinobacteria
Nocardia brasiliensis ATCC 700358 YP 006812137 1 1133849 Actinobacteria
Pectobacterium atrosepticum SCRI1043 YP 050788 1 218491 Proteobacteria
Photorhabdus luminescens subsp laumondii TTO1 NP 930529 1 243265 Proteobacteria
Photorhabdus asymbiotica YP 003040168 1 291112 Proteobacteria

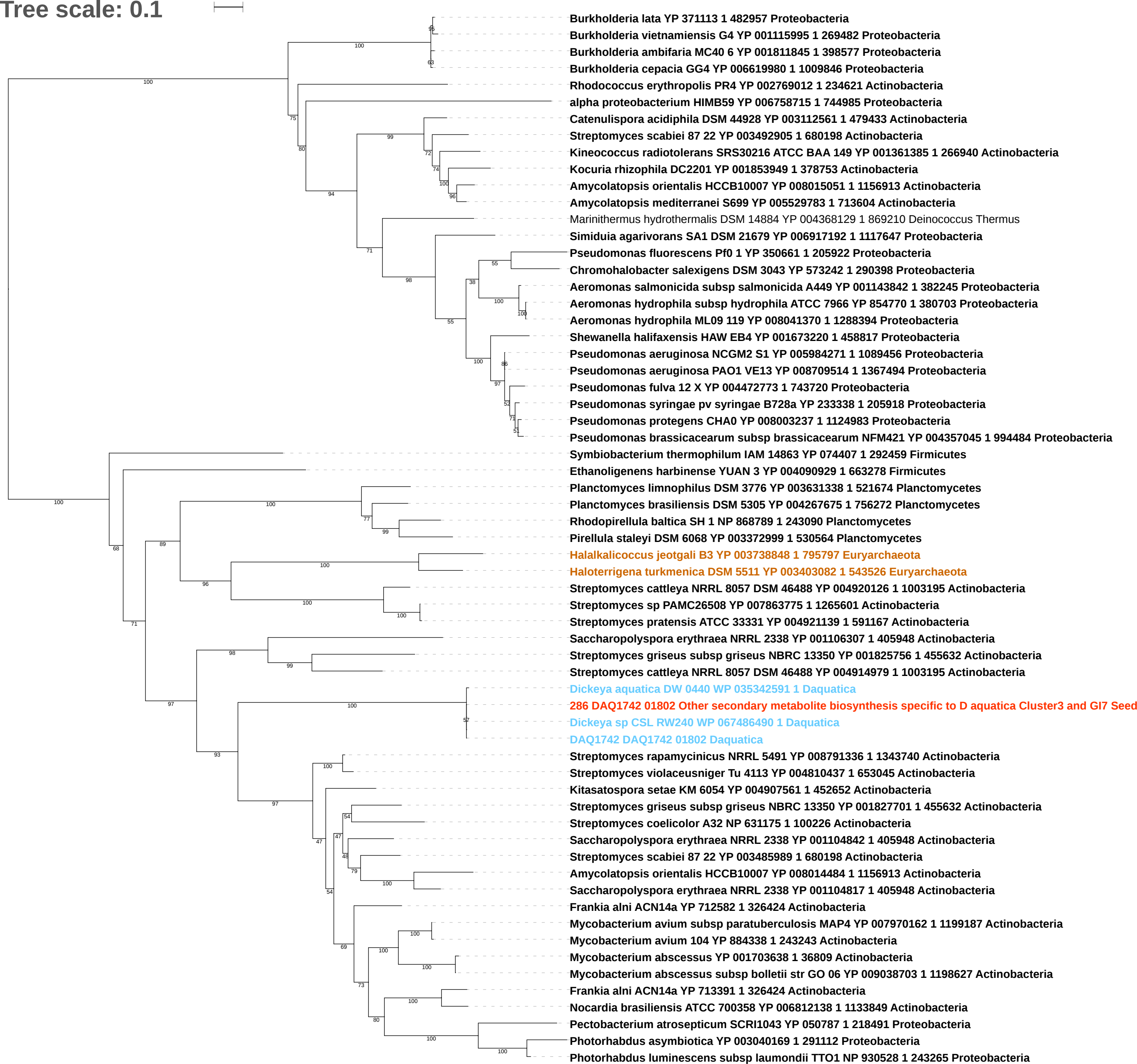

Tree scale: 0.1
Burkholderia lata YP 371113 1 482957 Proteobacteria
Burkholderia vietnamiensis G4 YP 001115995 1 269482 Proteobacteria
Burkholderia ambifaria MC40 6 YP 001811845 1 398577 Proteobacteria
Burkholderia cepacia GG4 YP 006619980 1 1009846 Proteobacteria
Rhodococcus erythropolis PR4 YP 002769012 1 234621 Actinobacteria
alpha proteobacterium HIMB59 YP 006758715 1 744985 Proteobacteria
Catenulispora acidiphila DSM 44928 YP 003112561 1 479433 Actinobacteria
Streptomyces scabiei 87 22 YP 003492905 1 680198 Actinobacteria
Kineococcus radiotolerans SRS30216 ATCC BAA 149 YP 001361385 1 266940 Actinobacteria
Kocuria rhizophila DC2201 YP 001853949 1 378753 Actinobacteria
Amycolatopsis orientalis HCCB10007 YP 008015051 1 1156913 Actinobacteria
Amycolatopsis mediterranei S699 YP 005529783 1 713604 Actinobacteria
Marinithermus hydrothermalis DSM 14884 YP 004368129 1 869210 Deinococcus Thermus
Simiduia agarivorans SA1 DSM 21679 YP 006917192 1 1117647 Proteobacteria
Pseudomonas fluorescens Pf0 1 YP 350661 1 205922 Proteobacteria
Chromohalobacter salexigens DSM 3043 YP 573242 1 290398 Proteobacteria
Aeromonas salmonicida subsp salmonicida A449 YP 001143842 1 382245 Proteobacteria
Aeromonas hydrophila subsp hydrophila ATCC 7966 YP 854770 1 380703 Proteobacteria
Aeromonas hydrophila ML09 119 YP 008041370 1 1288394 Proteobacteria
Shewanella halifaxensis HAW EB4 YP 001673220 1 458817 Proteobacteria
Pseudomonas aeruginosa NCGM2 S1 YP 005984271 1 1089456 Proteobacteria
Pseudomonas aeruginosa PAO1 VE13 YP 008709514 1 1367494 Proteobacteria
Pseudomonas fulva 12 X YP 004472773 1 743720 Proteobacteria
Pseudomonas syringae pv syringae B728a YP 233338 1 205918 Proteobacteria
Pseudomonas protegens CHA0 YP 008003237 1 1124983 Proteobacteria
Pseudomonas brassicacearum subsp brassicacearum NFM421 YP 004357045 1 994484 Proteobacteria
Symbiobacterium thermophilum IAM 14863 YP 074407 1 292459 Firmicutes
Ethanoligenens harbinense YUAN 3 YP 004090929 1 663278 Firmicutes
Planctomyces limnophilus DSM 3776 YP 003631338 1 521674 Planctomycetes
Planctomyces brasiliensis DSM 5305 YP 004267675 1 756272 Planctomycetes
Rhodopirellula baltica SH 1 NP 868789 1 243090 Planctomycetes
Pirellula staleyi DSM 6068 YP 003372999 1 530564 Planctomycetes
Halalkalicoccus jeotgali B3 YP 003738848 1 795797 Euryarchaeota
Haloterrigena turkmenica DSM 5511 YP 003403082 1 543526 Euryarchaeota
Streptomyces cattleya NRRL 8057 DSM 46488 YP 004920126 1 1003195 Actinobacteria
Streptomyces sp PAMC26508 YP 007863775 1 1265601 Actinobacteria
Streptomyces pratensis ATCC 33331 YP 004921139 1 591167 Actinobacteria
Saccharopolyspora erythraea NRRL 2338 YP 001106307 1 405948 Actinobacteria
Streptomyces griseus subsp griseus NBRC 13350 YP 001825756 1 455632 Actinobacteria
Streptomyces cattleya NRRL 8057 DSM 46488 YP 004914979 1 1003195 Actinobacteria
Dickeya aquatica DW 0440 WP 035342591 1 Daquatica
286 DAQ1742 01802 Other secondary metabolite biosynthesis specific to D aquatica Cluster3 and GI7 Seed
Dickeya sp CSL RW240 WP 067486490 1 Daquatica
DAQ1742 DAQ1742 01802 Daquatica
Streptomyces rapamycinicus NRRL 5491 YP 008791336 1 1343740 Actinobacteria
Streptomyces violaceusniger Tu 4113 YP 004810437 1 653045 Actinobacteria
Kitasatospora setae KM 6054 YP 004907561 1 452652 Actinobacteria
Streptomyces griseus subsp griseus NBRC 13350 YP 001827701 1 455632 Actinobacteria
Streptomyces coelicolor A32 NP 631175 1 100226 Actinobacteria
Saccharopolyspora erythraea NRRL 2338 YP 001104842 1 405948 Actinobacteria
Streptomyces scabiei 87 22 YP 003485989 1 680198 Actinobacteria
Amycolatopsis orientalis HCCB10007 YP 008014484 1 1156913 Actinobacteria
Saccharopolyspora erythraea NRRL 2338 YP 001104817 1 405948 Actinobacteria
Frankia alni ACN14a YP 712582 1 326424 Actinobacteria
Mycobacterium avium subsp paratuberculosis MAP4 YP 007970162 1 1199187 Actinobacteria
Mycobacterium avium 104 YP 884338 1 243243 Actinobacteria
Mycobacterium abscessus YP 001703638 1 36809 Actinobacteria
Mycobacterium abscessus subsp bolletii str GO 06 YP 009038703 1 1198627 Actinobacteria
Frankia alni ACN14a YP 713391 1 326424 Actinobacteria
Nocardia brasiliensis ATCC 700358 YP 006812138 1 1133849 Actinobacteria
Pectobacterium atrosepticum SCRI1043 YP 050787 1 218491 Proteobacteria
Photorhabdus asymbiotica YP 003040169 1 291112 Proteobacteria
Photorhabdus luminescens subsp laumondii TTO1 NP 930528 1 243265 Proteobacteria

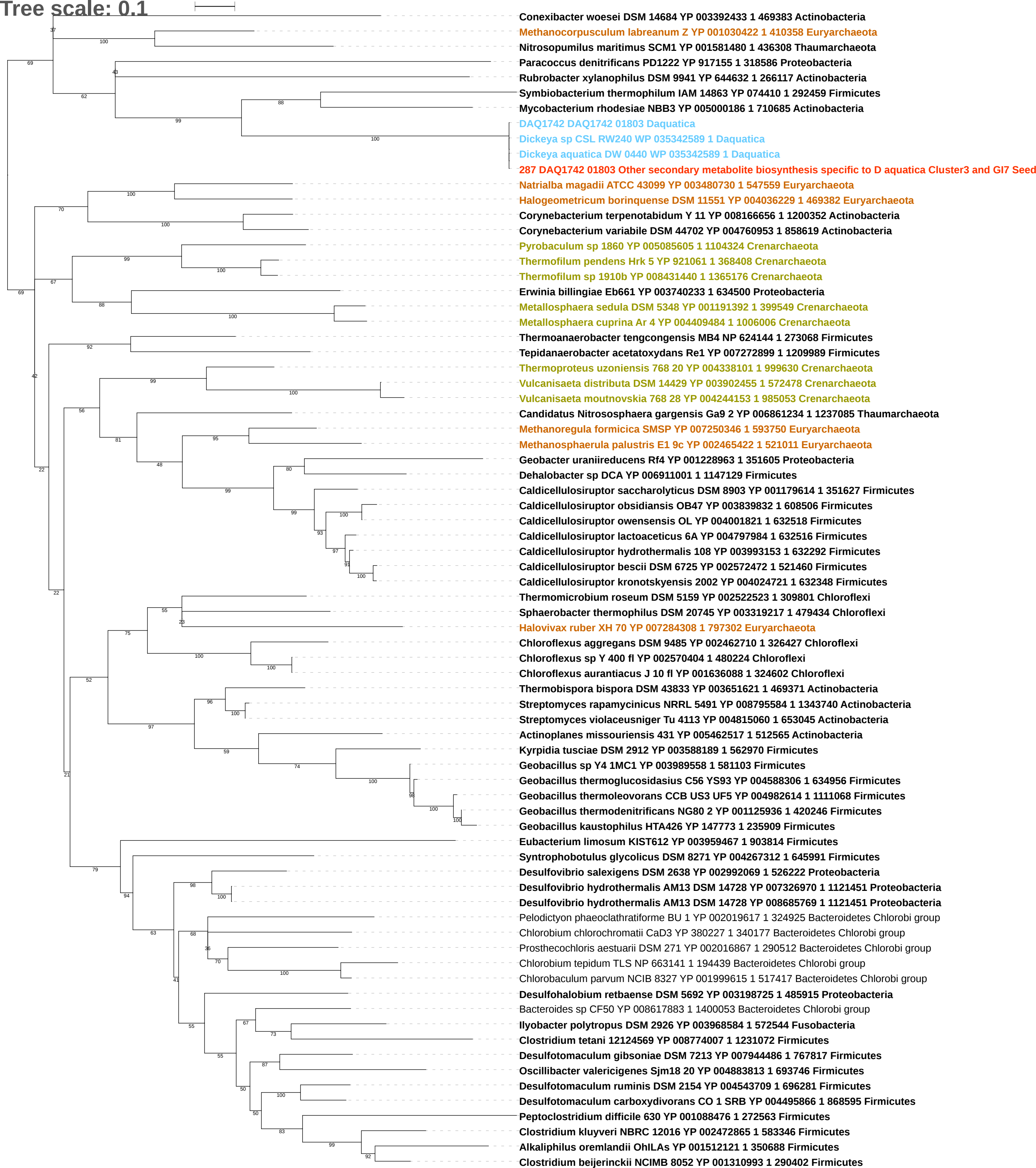
Tree scale: 0.1
Conexibacter woesei DSM 14684 YP 003392433 1 469383 Actinobacteria
Methanocorpusculum labreanum Z YP 001030422 1 410358 Euryarchaeota
Nitrosopumilus maritimus SCM1 YP 001581480 1 436308 Thaumarchaeota
Paracoccus denitrificans PD1222 YP 917155 1 318586 Proteobacteria
Rubrobacter xylanophilus DSM 9941 YP 644632 1 266117 Actinobacteria
Symbiobacterium thermophilum IAM 14863 YP 074410 1 292459 Firmicutes
Mycobacterium rhodesiae NBB3 YP 005000186 1 710685 Actinobacteria
DAQ1742 DAQ1742 01803 Daquatica
Dickeya sp CSL RW240 WP 035342589 1 Daquatica
Dickeya aquatica DW 0440 WP 035342589 1 Daquatica
287 DAQ1742 01803 Other secondary metabolite biosynthesis specific to D aquatica Cluster3 and GI7 Seed
Natrialba magadii ATCC 43099 YP 003480730 1 547559 Euryarchaeota
Halogeometricum borinquense DSM 11551 YP 004036229 1 469382 Euryarchaeota
Corynebacterium terpenotabidum Y 11 YP 008166656 1 1200352 Actinobacteria
Corynebacterium variabile DSM 44702 YP 004760953 1 858619 Actinobacteria
Pyrobaculum sp 1860 YP 005085605 1 1104324 Crenarchaeota
Thermofilum pendens Hrk 5 YP 921061 1 368408 Crenarchaeota
Thermofilum sp 1910b YP 008431440 1 1365176 Crenarchaeota
Erwinia billingiae Eb661 YP 003740233 1 634500 Proteobacteria
Metallosphaera sedula DSM 5348 YP 001191392 1 399549 Crenarchaeota
Metallosphaera cuprina Ar 4 YP 004409484 1 1006006 Crenarchaeota
Thermoanaerobacter tengcongensis MB4 NP 624144 1 273068 Firmicutes
Tepidanaerobacter acetatoxydans Re1 YP 007272899 1 1209989 Firmicutes
Thermoproteus uzoniensis 768 20 YP 004338101 1 999630 Crenarchaeota
Vulcanisaeta distributa DSM 14429 YP 003902455 1 572478 Crenarchaeota
Vulcanisaeta moutnovskia 768 28 YP 004244153 1 985053 Crenarchaeota
Candidatus Nitrososphaera gargensis Ga9 2 YP 006861234 1 1237085 Thaumarchaeota
Methanoregula formicica SMSP YP 007250346 1 593750 Euryarchaeota
Methanosphaerula palustris E1 9c YP 002465422 1 521011 Euryarchaeota
Geobacter uraniireducens Rf4 YP 001228963 1 351605 Proteobacteria
Dehalobacter sp DCA YP 006911001 1 1147129 Firmicutes
Caldicellulosiruptor saccharolyticus DSM 8903 YP 001179614 1 351627 Firmicutes
Caldicellulosiruptor obsidiansis OB47 YP 003839832 1 608506 Firmicutes
Caldicellulosiruptor owensensis OL YP 004001821 1 632518 Firmicutes
Caldicellulosiruptor lactoaceticus 6A YP 004797984 1 632516 Firmicutes
Caldicellulosiruptor hydrothermalis 108 YP 003993153 1 632292 Firmicutes
Caldicellulosiruptor bescii DSM 6725 YP 002572472 1 521460 Firmicutes
Caldicellulosiruptor kronotskyensis 2002 YP 004024721 1 632348 Firmicutes
Thermomicrobium roseum DSM 5159 YP 002522523 1 309801 Chloroflexi
Sphaerobacter thermophilus DSM 20745 YP 003319217 1 479434 Chloroflexi
Halovivax ruber XH 70 YP 007284308 1 797302 Euryarchaeota
Chloroflexus aggregans DSM 9485 YP 002462710 1 326427 Chloroflexi
Chloroflexus sp Y 400 fl YP 002570404 1 480224 Chloroflexi
Chloroflexus aurantiacus J 10 fl YP 001636088 1 324602 Chloroflexi
Thermobispora bispora DSM 43833 YP 003651621 1 469371 Actinobacteria
Streptomyces rapamycinicus NRRL 5491 YP 008795584 1 1343740 Actinobacteria
Streptomyces violaceusniger Tu 4113 YP 004815060 1 653045 Actinobacteria
Actinoplanes missouriensis 431 YP 005462517 1 512565 Actinobacteria
Kyrpidia tusciae DSM 2912 YP 003588189 1 562970 Firmicutes
Geobacillus sp Y4 1MC1 YP 003989558 1 581103 Firmicutes
Geobacillus thermoglucosidasius C56 YS93 YP 004588306 1 634956 Firmicutes
Geobacillus thermoleovorans CCB US3 UF5 YP 004982614 1 1111068 Firmicutes
Geobacillus thermodenitrificans NG80 2 YP 001125936 1 420246 Firmicutes
Geobacillus kaustophilus HTA426 YP 147773 1 235909 Firmicutes
Eubacterium limosum KIST612 YP 003959467 1 903814 Firmicutes
Syntrophobotulus glycolicus DSM 8271 YP 004267312 1 645991 Firmicutes
Desulfovibrio salexigens DSM 2638 YP 002992069 1 526222 Proteobacteria
Desulfovibrio hydrothermalis AM13 DSM 14728 YP 007326970 1 1121451 Proteobacteria
Desulfovibrio hydrothermalis AM13 DSM 14728 YP 008685769 1 1121451 Proteobacteria
Pelodictyon phaeoclathratiforme BU 1 YP 002019617 1 324925 Bacteroidetes Chlorobi group
Chlorobium chlorochromatii CaD3 YP 380227 1 340177 Bacteroidetes Chlorobi group
Prosthecochloris aestuarii DSM 271 YP 002016867 1 290512 Bacteroidetes Chlorobi group
Chlorobium tepidum TLS NP 663141 1 194439 Bacteroidetes Chlorobi group
Chlorobaculum parvum NCIB 8327 YP 001999615 1 517417 Bacteroidetes Chlorobi group
Desulfohalobium retbaense DSM 5692 YP 003198725 1 485915 Proteobacteria
Bacteroides sp CF50 YP 008617883 1 1400053 Bacteroidetes Chlorobi group
Ilyobacter polytropus DSM 2926 YP 003968584 1 572544 Fusobacteria
Clostridium tetani 12124569 YP 008774007 1 1231072 Firmicutes
Desulfotomaculum gibsoniae DSM 7213 YP 007944486 1 767817 Firmicutes
Oscillibacter valericigenes Sjm18 20 YP 004883813 1 693746 Firmicutes
Desulfotomaculum ruminis DSM 2154 YP 004543709 1 696281 Firmicutes
Desulfotomaculum carboxydivorans CO 1 SRB YP 004495866 1 868595 Firmicutes
Peptoclostridium difficile 630 YP 001088476 1 272563 Firmicutes
Clostridium kluyveri NBRC 12016 YP 002472865 1 583346 Firmicutes
Alkaliphilus oremlandii OhILAs YP 001512121 1 350688 Firmicutes
Clostridium beijerinckii NCIMB 8052 YP 001310993 1 290402 Firmicutes

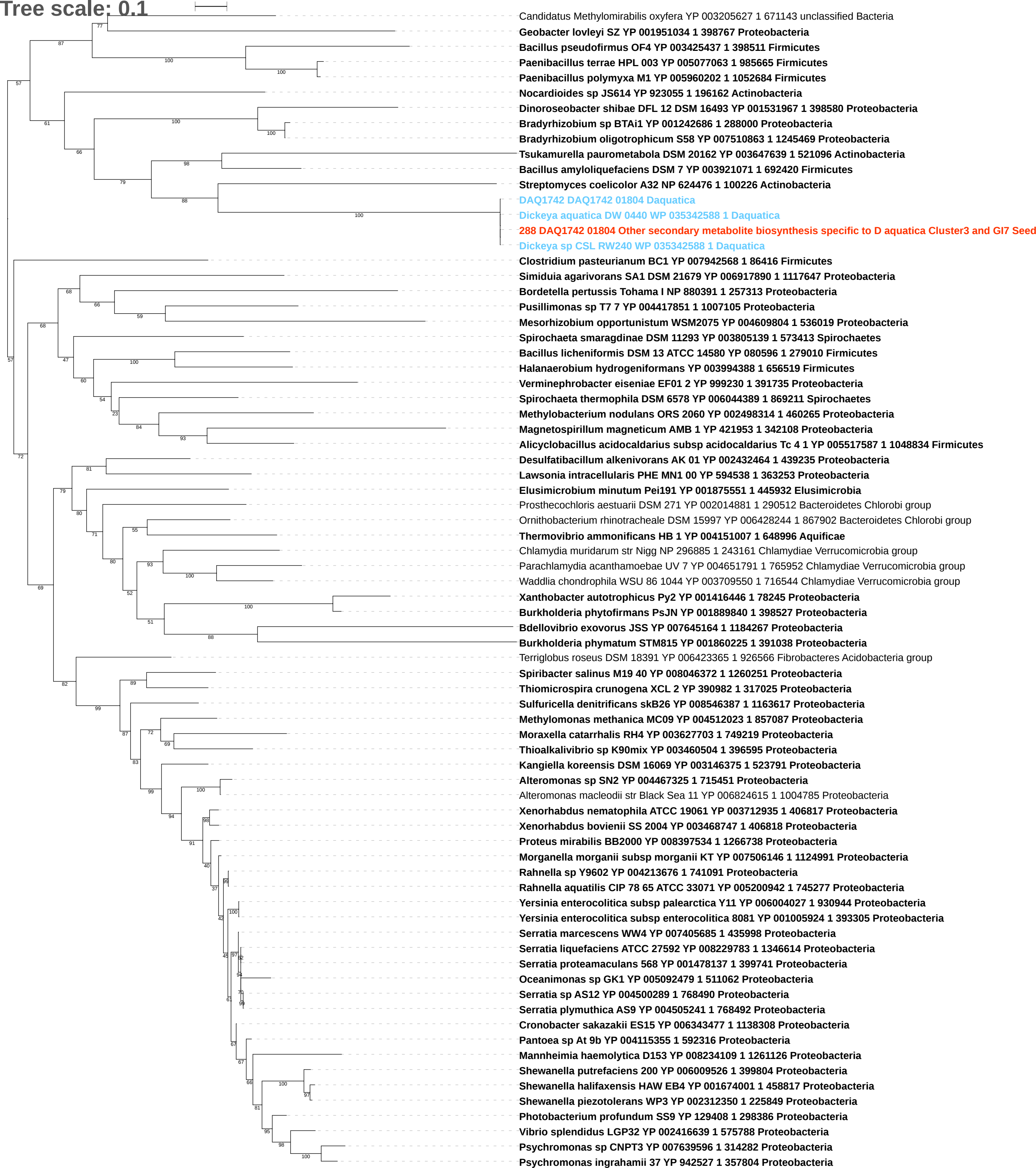
Tree scale: 0.1
Candidatus Methylomirabilis oxyfera YP 003205627 1 671143 unclassified Bacteria
Geobacter lovleyi SZ YP 001951034 1 398767 Proteobacteria
Bacillus pseudofirmus OF4 YP 003425437 1 398511 Firmicutes
Paenibacillus terrae HPL 003 YP 005077063 1 985665 Firmicutes
Paenibacillus polymyxa M1 YP 005960202 1 1052684 Firmicutes
Nocardioides sp JS614 YP 923055 1 196162 Actinobacteria
Dinoroseobacter shibae DFL 12 DSM 16493 YP 001531967 1 398580 Proteobacteria
Bradyrhizobium sp BTAi1 YP 001242686 1 288000 Proteobacteria
Bradyrhizobium oligotrophicum S58 YP 007510863 1 1245469 Proteobacteria
Tsukamurella paurometabola DSM 20162 YP 003647639 1 521096 Actinobacteria
Bacillus amyloliquefaciens DSM 7 YP 003921071 1 692420 Firmicutes
Streptomyces coelicolor A32 NP 624476 1 100226 Actinobacteria
DAQ1742 DAQ1742 01804 Daquatica
Dickeya aquatica DW 0440 WP 035342588 1 Daquatica
288 DAQ1742 01804 Other secondary metabolite biosynthesis specific to D aquatica Cluster3 and GI7 Seed
Dickeya sp CSL RW240 WP 035342588 1 Daquatica
Clostridium pasteurianum BC1 YP 007942568 1 86416 Firmicutes
Simiduia agarivorans SA1 DSM 21679 YP 006917890 1 1117647 Proteobacteria
Bordetella pertussis Tohama I NP 880391 1 257313 Proteobacteria
Pusillimonas sp T7 7 YP 004417851 1 1007105 Proteobacteria
Mesorhizobium opportunistum WSM2075 YP 004609804 1 536019 Proteobacteria
Spirochaeta smaragdinae DSM 11293 YP 003805139 1 573413 Spirochaetes
Bacillus licheniformis DSM 13 ATCC 14580 YP 080596 1 279010 Firmicutes
Halanaerobium hydrogeniformans YP 003994388 1 656519 Firmicutes
Verminephrobacter eiseniae EF01 2 YP 999230 1 391735 Proteobacteria
Spirochaeta thermophila DSM 6578 YP 006044389 1 869211 Spirochaetes
Methylobacterium nodulans ORS 2060 YP 002498314 1 460265 Proteobacteria
Magnetospirillum magneticum AMB 1 YP 421953 1 342108 Proteobacteria
Alicyclobacillus acidocaldarius subsp acidocaldarius Tc 4 1 YP 005517587 1 1048834 Firmicutes
Desulfatibacillum alkenivorans AK 01 YP 002432464 1 439235 Proteobacteria
Lawsonia intracellularis PHE MN1 00 YP 594538 1 363253 Proteobacteria
Elusimicrobium minutum Pei191 YP 001875551 1 445932 Elusimicrobia
Prosthecochloris aestuarii DSM 271 YP 002014881 1 290512 Bacteroidetes Chlorobi group
Ornithobacterium rhinotracheale DSM 15997 YP 006428244 1 867902 Bacteroidetes Chlorobi group
Thermovibrio ammonificans HB 1 YP 004151007 1 648996 Aquificae
Chlamydia muridarum str Nigg NP 296885 1 243161 Chlamydiae Verrucomicrobia group
Parachlamydia acanthamoebae UV 7 YP 004651791 1 765952 Chlamydiae Verrucomicrobia group
Waddlia chondrophila WSU 86 1044 YP 003709550 1 716544 Chlamydiae Verrucomicrobia group
Xanthobacter autotrophicus Py2 YP 001416446 1 78245 Proteobacteria
Burkholderia phytofirmans PsJN YP 001889840 1 398527 Proteobacteria
Bdellovibrio exovorus JSS YP 007645164 1 1184267 Proteobacteria
Burkholderia phymatum STM815 YP 001860225 1 391038 Proteobacteria
Terriglobus roseus DSM 18391 YP 006423365 1 926566 Fibrobacteres Acidobacteria group
Spiribacter salinus M19 40 YP 008046372 1 1260251 Proteobacteria
Thiomicrospira crunogena XCL 2 YP 390982 1 317025 Proteobacteria
Sulfuricella denitrificans skB26 YP 008546387 1 1163617 Proteobacteria
Methylomonas methanica MC09 YP 004512023 1 857087 Proteobacteria
Moraxella catarrhalis RH4 YP 003627703 1 749219 Proteobacteria
Thioalkalivibrio sp K90mix YP 003460504 1 396595 Proteobacteria
Kangiella koreensis DSM 16069 YP 003146375 1 523791 Proteobacteria
Alteromonas sp SN2 YP 004467325 1 715451 Proteobacteria
Alteromonas macleodii str Black Sea 11 YP 006824615 1 1004785 Proteobacteria
Xenorhabdus nematophila ATCC 19061 YP 003712935 1 406817 Proteobacteria
Xenorhabdus bovienii SS 2004 YP 003468747 1 406818 Proteobacteria
Proteus mirabilis BB2000 YP 008397534 1 1266738 Proteobacteria
Morganella morganii subsp morganii KT YP 007506146 1 1124991 Proteobacteria
Rahnella sp Y9602 YP 004213676 1 741091 Proteobacteria
Rahnella aquatilis CIP 78 65 ATCC 33071 YP 005200942 1 745277 Proteobacteria
Yersinia enterocolitica subsp palearctica Y11 YP 006004027 1 930944 Proteobacteria
Yersinia enterocolitica subsp enterocolitica 8081 YP 001005924 1 393305 Proteobacteria
Serratia marcescens WW4 YP 007405685 1 435998 Proteobacteria
Serratia liquefaciens ATCC 27592 YP 008229783 1 1346614 Proteobacteria
Serratia proteamaculans 568 YP 001478137 1 399741 Proteobacteria
Oceanimonas sp GK1 YP 005092479 1 511062 Proteobacteria
Serratia sp AS12 YP 004500289 1 768490 Proteobacteria
Serratia plymuthica AS9 YP 004505241 1 768492 Proteobacteria
Cronobacter sakazakii ES15 YP 006343477 1 1138308 Proteobacteria
Pantoea sp At 9b YP 004115355 1 592316 Proteobacteria
Mannheimia haemolytica D153 YP 008234109 1 1261126 Proteobacteria
Shewanella putrefaciens 200 YP 006009526 1 399804 Proteobacteria
Shewanella halifaxensis HAW EB4 YP 001674001 1 458817 Proteobacteria
Shewanella piezotolerans WP3 YP 002312350 1 225849 Proteobacteria
Photobacterium profundum SS9 YP 129408 1 298386 Proteobacteria
Vibrio splendidus LGP32 YP 002416639 1 575788 Proteobacteria
Psychromonas sp CNPT3 YP 007639596 1 314282 Proteobacteria
Psychromonas ingrahamii 37 YP 942527 1 357804 Proteobacteria

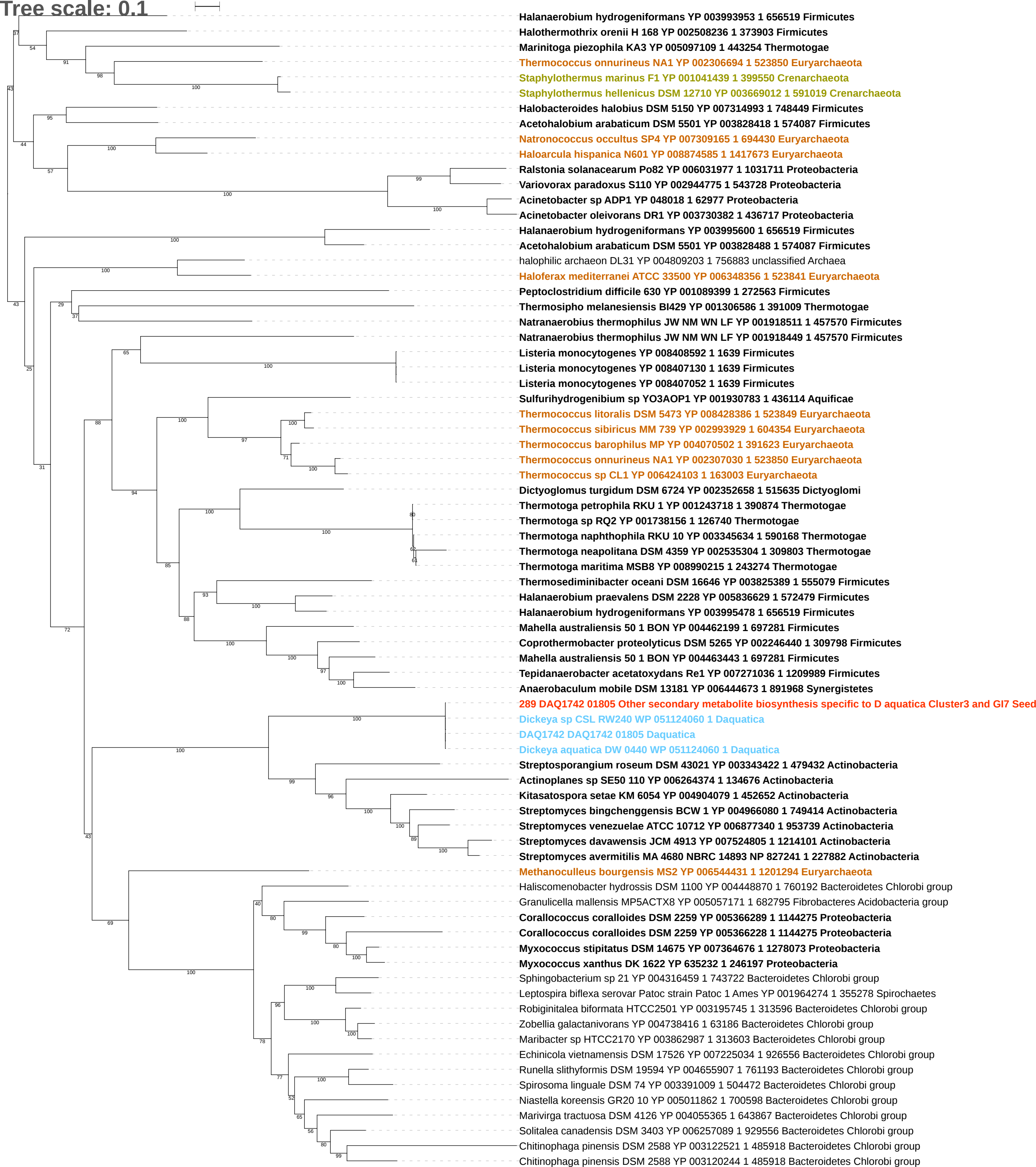
Tree scale: 0.1
Halanaerobium hydrogeniformans YP 003993953 1 656519 Firmicutes
Halothermothrix orenii H 168 YP 002508236 1 373903 Firmicutes
Marinitoga piezophila KA3 YP 005097109 1 443254 Thermotogae
Thermococcus onnurineus NA1 YP 002306694 1 523850 Euryarchaeota
Staphylothermus marinus F1 YP 001041439 1 399550 Crenarchaeota
Staphylothermus hellenicus DSM 12710 YP 003669012 1 591019 Crenarchaeota
Halobacteroides halobius DSM 5150 YP 007314993 1 748449 Firmicutes
Acetohalobium arabaticum DSM 5501 YP 003828418 1 574087 Firmicutes
Natronococcus occultus SP4 YP 007309165 1 694430 Euryarchaeota
Haloarcula hispanica N601 YP 008874585 1 1417673 Euryarchaeota
Ralstonia solanacearum Po82 YP 006031977 1 1031711 Proteobacteria
Variovorax paradoxus S110 YP 002944775 1 543728 Proteobacteria
Acinetobacter sp ADP1 YP 048018 1 62977 Proteobacteria
Acinetobacter oleivorans DR1 YP 003730382 1 436717 Proteobacteria
Halanaerobium hydrogeniformans YP 003995600 1 656519 Firmicutes
Acetohalobium arabaticum DSM 5501 YP 003828488 1 574087 Firmicutes
halophilic archaeon DL31 YP 004809203 1 756883 unclassified Archaea
Haloferax mediterranei ATCC 33500 YP 006348356 1 523841 Euryarchaeota
Peptoclostridium difficile 630 YP 001089399 1 272563 Firmicutes
Thermosipho melanesiensis BI429 YP 001306586 1 391009 Thermotogae
Natranaerobius thermophilus JW NM WN LF YP 001918511 1 457570 Firmicutes
Natranaerobius thermophilus JW NM WN LF YP 001918449 1 457570 Firmicutes
Listeria monocytogenes YP 008408592 1 1639 Firmicutes
Listeria monocytogenes YP 008407130 1 1639 Firmicutes
Listeria monocytogenes YP 008407052 1 1639 Firmicutes
Sulfurihydrogenibium sp YO3AOP1 YP 001930783 1 436114 Aquificae
Thermococcus litoralis DSM 5473 YP 008428386 1 523849 Euryarchaeota
Thermococcus sibiricus MM 739 YP 002993929 1 604354 Euryarchaeota
Thermococcus barophilus MP YP 004070502 1 391623 Euryarchaeota
Thermococcus onnurineus NA1 YP 002307030 1 523850 Euryarchaeota
Thermococcus sp CL1 YP 006424103 1 163003 Euryarchaeota
Dictyoglomus turgidum DSM 6724 YP 002352658 1 515635 Dictyoglomi
Thermotoga petrophila RKU 1 YP 001243718 1 390874 Thermotogae
Thermotoga sp RQ2 YP 001738156 1 126740 Thermotogae
Thermotoga naphthophila RKU 10 YP 003345634 1 590168 Thermotogae
Thermotoga neapolitana DSM 4359 YP 002535304 1 309803 Thermotogae
Thermotoga maritima MSB8 YP 008990215 1 243274 Thermotogae
Thermosediminibacter oceani DSM 16646 YP 003825389 1 555079 Firmicutes
Halanaerobium praevalens DSM 2228 YP 005836629 1 572479 Firmicutes
Halanaerobium hydrogeniformans YP 003995478 1 656519 Firmicutes
Mahella australiensis 50 1 BON YP 004462199 1 697281 Firmicutes
Coprothermobacter proteolyticus DSM 5265 YP 002246440 1 309798 Firmicutes
Mahella australiensis 50 1 BON YP 004463443 1 697281 Firmicutes
Tepidanaerobacter acetatoxydans Re1 YP 007271036 1 1209989 Firmicutes
Anaerobaculum mobile DSM 13181 YP 006444673 1 891968 Synergistetes
289 DAQ1742 01805 Other secondary metabolite biosynthesis specific to D aquatica Cluster3 and GI7 Seed
Dickeya sp CSL RW240 WP 051124060 1 Daquatica
DAQ1742 DAQ1742 01805 Daquatica
Dickeya aquatica DW 0440 WP 051124060 1 Daquatica
Streptosporangium roseum DSM 43021 YP 003343422 1 479432 Actinobacteria
Actinoplanes sp SE50 110 YP 006264374 1 134676 Actinobacteria
Kitasatospora setae KM 6054 YP 004904079 1 452652 Actinobacteria
Streptomyces bingchenggensis BCW 1 YP 004966080 1 749414 Actinobacteria
Streptomyces venezuelae ATCC 10712 YP 006877340 1 953739 Actinobacteria
Streptomyces davawensis JCM 4913 YP 007524805 1 1214101 Actinobacteria
Streptomyces avermitilis MA 4680 NBRC 14893 NP 827241 1 227882 Actinobacteria
Methanoculleus bourgensis MS2 YP 006544431 1 1201294 Euryarchaeota
Haliscomenobacter hydrossis DSM 1100 YP 004448870 1 760192 Bacteroidetes Chlorobi group
Granulicella mallensis MP5ACTX8 YP 005057171 1 682795 Fibrobacteres Acidobacteria group
Corallococcus coralloides DSM 2259 YP 005366289 1 1144275 Proteobacteria
Corallococcus coralloides DSM 2259 YP 005366228 1 1144275 Proteobacteria
Myxococcus stipitatus DSM 14675 YP 007364676 1 1278073 Proteobacteria
Myxococcus xanthus DK 1622 YP 635232 1 246197 Proteobacteria
Sphingobacterium sp 21 YP 004316459 1 743722 Bacteroidetes Chlorobi group
Leptospira biflexa serovar Patoc strain Patoc 1 Ames YP 001964274 1 355278 Spirochaetes
Robiginitalea biformata HTCC2501 YP 003195745 1 313596 Bacteroidetes Chlorobi group
Zobellia galactanivorans YP 004738416 1 63186 Bacteroidetes Chlorobi group
Maribacter sp HTCC2170 YP 003862987 1 313603 Bacteroidetes Chlorobi group
Echinicola vietnamensis DSM 17526 YP 007225034 1 926556 Bacteroidetes Chlorobi group
Runella slithyformis DSM 19594 YP 004655907 1 761193 Bacteroidetes Chlorobi group
Spirosoma linguale DSM 74 YP 003391009 1 504472 Bacteroidetes Chlorobi group
Niastella koreensis GR20 10 YP 005011862 1 700598 Bacteroidetes Chlorobi group
Marivirga tractuosa DSM 4126 YP 004055365 1 643867 Bacteroidetes Chlorobi group
Solitalea canadensis DSM 3403 YP 006257089 1 929556 Bacteroidetes Chlorobi group
Chitinophaga pinensis DSM 2588 YP 003122521 1 485918 Bacteroidetes Chlorobi group
Chitinophaga pinensis DSM 2588 YP 003120244 1 485918 Bacteroidetes Chlorobi group

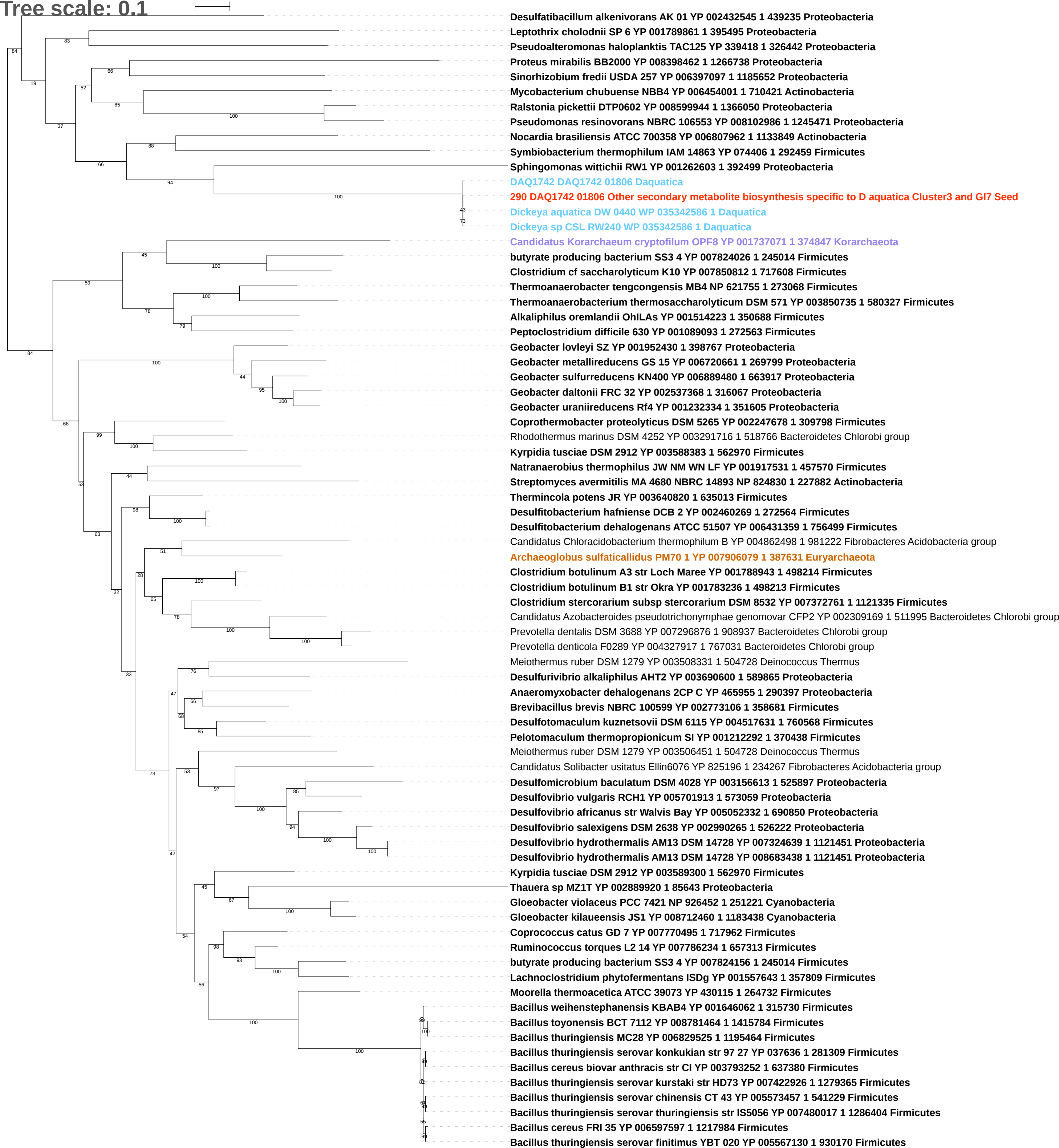
Tree scale: 0.1
Desulfatibacillum alkenivorans AK 01 YP 002432545 1 439235 Proteobacteria
Leptothrix cholodnii SP 6 YP 001789861 1 395495 Proteobacteria
Pseudoalteromonas haloplanktis TAC125 YP 339418 1 326442 Proteobacteria
Proteus mirabilis BB2000 YP 008398462 1 1266738 Proteobacteria
Sinorhizobium fredii USDA 257 YP 006397097 1 1185652 Proteobacteria
Mycobacterium chubuense NBB4 YP 006454001 1 710421 Actinobacteria
Ralstonia pickettii DTP0602 YP 008599944 1 1366050 Proteobacteria
Pseudomonas resinovorans NBRC 106553 YP 008102986 1 1245471 Proteobacteria
Nocardia brasiliensis ATCC 700358 YP 006807962 1 1133849 Actinobacteria
Symbiobacterium thermophilum IAM 14863 YP 074406 1 292459 Firmicutes
Sphingomonas wittichii RW1 YP 001262603 1 392499 Proteobacteria
DAQ1742 DAQ1742 01806 Daquatica
290 DAQ1742 01806 Other secondary metabolite biosynthesis specific to D aquatica Cluster3 and GI7 Seed
Dickeya aquatica DW 0440 WP 035342586 1 Daquatica
Dickeya sp CSL RW240 WP 035342586 1 Daquatica
Candidatus Korarchaeum cryptofilum OPF8 YP 001737071 1 374847 Korarchaeota
butyrate producing bacterium SS3 4 YP 007824026 1 245014 Firmicutes
Clostridium cf saccharolyticum K10 YP 007850812 1 717608 Firmicutes
Thermoanaerobacter tengcongensis MB4 NP 621755 1 273068 Firmicutes
Thermoanaerobacterium thermosaccharolyticum DSM 571 YP 003850735 1 580327 Firmicutes
Alkaliphilus oremlandii OhILAs YP 001514223 1 350688 Firmicutes
Peptoclostridium difficile 630 YP 001089093 1 272563 Firmicutes
Geobacter lovleyi SZ YP 001952430 1 398767 Proteobacteria
Geobacter metallireducens GS 15 YP 006720661 1 269799 Proteobacteria
Geobacter sulfurreducens KN400 YP 006889480 1 663917 Proteobacteria
Geobacter daltonii FRC 32 YP 002537368 1 316067 Proteobacteria
Geobacter uraniireducens Rf4 YP 001232334 1 351605 Proteobacteria
Coprothermobacter proteolyticus DSM 5265 YP 002247678 1 309798 Firmicutes
Rhodothermus marinus DSM 4252 YP 003291716 1 518766 Bacteroidetes Chlorobi group
Kyrpidia tusciae DSM 2912 YP 003588383 1 562970 Firmicutes
Natranaerobius thermophilus JW NM WN LF YP 001917531 1 457570 Firmicutes
Streptomyces avermitilis MA 4680 NBRC 14893 NP 824830 1 227882 Actinobacteria
Thermincola potens JR YP 003640820 1 635013 Firmicutes
Desulfitobacterium hafniense DCB 2 YP 002460269 1 272564 Firmicutes
Desulfitobacterium dehalogenans ATCC 51507 YP 006431359 1 756499 Firmicutes
Candidatus Chloracidobacterium thermophilum B YP 004862498 1 981222 Fibrobacteres Acidobacteria group
Archaeoglobus sulfaticallidus PM70 1 YP 007906079 1 387631 Euryarchaeota
Clostridium botulinum A3 str Loch Maree YP 001788943 1 498214 Firmicutes
Clostridium botulinum B1 str Okra YP 001783236 1 498213 Firmicutes
Clostridium stercorarium subsp stercorarium DSM 8532 YP 007372761 1 1121335 Firmicutes
Candidatus Azobacteroides pseudotrichonymphae genomovar CFP2 YP 002309169 1 511995 Bacteroidetes Chlorobi group
Prevotella dentalis DSM 3688 YP 007296876 1 908937 Bacteroidetes Chlorobi group
Prevotella denticola F0289 YP 004327917 1 767031 Bacteroidetes Chlorobi group
Meiothermus ruber DSM 1279 YP 003508331 1 504728 Deinococcus Thermus
Desulfurivibrio alkaliphilus AHT2 YP 003690600 1 589865 Proteobacteria
Anaeromyxobacter dehalogenans 2CP C YP 465955 1 290397 Proteobacteria
Brevibacillus brevis NBRC 100599 YP 002773106 1 358681 Firmicutes
Desulfotomaculum kuznetsovii DSM 6115 YP 004517631 1 760568 Firmicutes
Pelotomaculum thermopropionicum SI YP 001212292 1 370438 Firmicutes
Meiothermus ruber DSM 1279 YP 003506451 1 504728 Deinococcus Thermus
Candidatus Solibacter usitatus Ellin6076 YP 825196 1 234267 Fibrobacteres Acidobacteria group
Desulfomicrobium baculatum DSM 4028 YP 003156613 1 525897 Proteobacteria
Desulfovibrio vulgaris RCH1 YP 005701913 1 573059 Proteobacteria
Desulfovibrio africanus str Walvis Bay YP 005052332 1 690850 Proteobacteria
Desulfovibrio salexigens DSM 2638 YP 002990265 1 526222 Proteobacteria
Desulfovibrio hydrothermalis AM13 DSM 14728 YP 007324639 1 1121451 Proteobacteria
Desulfovibrio hydrothermalis AM13 DSM 14728 YP 008683438 1 1121451 Proteobacteria
Kyrpidia tusciae DSM 2912 YP 003589300 1 562970 Firmicutes
Thauera sp MZ1T YP 002889920 1 85643 Proteobacteria
Gloeobacter violaceus PCC 7421 NP 926452 1 251221 Cyanobacteria
Gloeobacter kilaueensis JS1 YP 008712460 1 1183438 Cyanobacteria
Coprococcus catus GD 7 YP 007770495 1 717962 Firmicutes
Ruminococcus torques L2 14 YP 007786234 1 657313 Firmicutes
butyrate producing bacterium SS3 4 YP 007824156 1 245014 Firmicutes
Lachnoclostridium phytofermentans ISDg YP 001557643 1 357809 Firmicutes
Moorella thermoacetica ATCC 39073 YP 430115 1 264732 Firmicutes
Bacillus weihenstephanensis KBAB4 YP 001646062 1 315730 Firmicutes
Bacillus toyonensis BCT 7112 YP 008781464 1 1415784 Firmicutes
Bacillus thuringiensis MC28 YP 006829525 1 1195464 Firmicutes
Bacillus thuringiensis serovar konkukian str 97 27 YP 037636 1 281309 Firmicutes
Bacillus cereus biovar anthracis str CI YP 003793252 1 637380 Firmicutes
Bacillus thuringiensis serovar kurstaki str HD73 YP 007422926 1 1279365 Firmicutes
Bacillus thuringiensis serovar chinensis CT 43 YP 005573457 1 541229 Firmicutes
Bacillus thuringiensis serovar thuringiensis str IS5056 YP 007480017 1 1286404 Firmicutes
Bacillus cereus FRI 35 YP 006597597 1 1217984 Firmicutes
Bacillus thuringiensis serovar finitimus YBT 020 YP 005567130 1 930170 Firmicutes

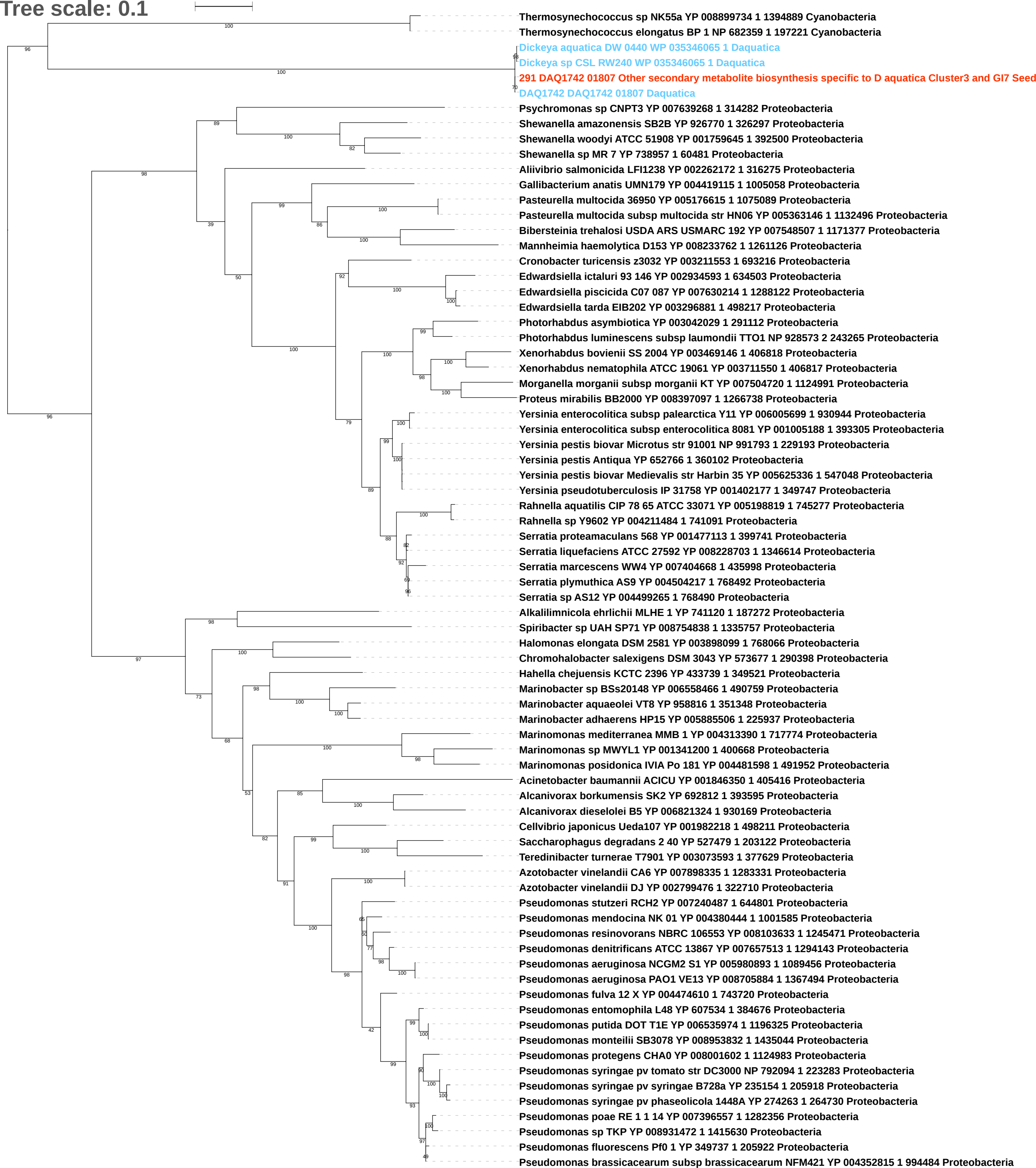
Tree scale: 0.1
Thermosynechococcus sp NK55a YP 008899734 1 1394889 Cyanobacteria
Thermosynechococcus elongatus BP 1 NP 682359 1 197221 Cyanobacteria
Dickeya aquatica DW 0440 WP 035346065 1 Daquatica
Dickeya sp CSL RW240 WP 035346065 1 Daquatica
291 DAQ1742 01807 Other secondary metabolite biosynthesis specific to D aquatica Cluster3 and GI7 Seed
DAQ1742 DAQ1742 01807 Daquatica
Psychromonas sp CNPT3 YP 007639268 1 314282 Proteobacteria
Shewanella amazonensis SB2B YP 926770 1 326297 Proteobacteria
Shewanella woodyi ATCC 51908 YP 001759645 1 392500 Proteobacteria
Shewanella sp MR 7 YP 738957 1 60481 Proteobacteria
Aliivibrio salmonicida LFI1238 YP 002262172 1 316275 Proteobacteria
Gallibacterium anatis UMN179 YP 004419115 1 1005058 Proteobacteria
Pasteurella multocida 36950 YP 005176615 1 1075089 Proteobacteria
Pasteurella multocida subsp multocida str HN06 YP 005363146 1 1132496 Proteobacteria
Bibersteinia trehalosi USDA ARS USMARC 192 YP 007548507 1 1171377 Proteobacteria
Mannheimia haemolytica D153 YP 008233762 1 1261126 Proteobacteria
Cronobacter turicensis z3032 YP 003211553 1 693216 Proteobacteria
Edwardsiella ictaluri 93 146 YP 002934593 1 634503 Proteobacteria
Edwardsiella piscicida C07 087 YP 007630214 1 1288122 Proteobacteria
Edwardsiella tarda EIB202 YP 003296881 1 498217 Proteobacteria
Photorhabdus asymbiotica YP 003042029 1 291112 Proteobacteria
Photorhabdus luminescens subsp laumondii TTO1 NP 928573 2 243265 Proteobacteria
Xenorhabdus bovienii SS 2004 YP 003469146 1 406818 Proteobacteria
Xenorhabdus nematophila ATCC 19061 YP 003711550 1 406817 Proteobacteria
Morganella morganii subsp morganii KT YP 007504720 1 1124991 Proteobacteria
Proteus mirabilis BB2000 YP 008397097 1 1266738 Proteobacteria
Yersinia enterocolitica subsp palearctica Y11 YP 006005699 1 930944 Proteobacteria
Yersinia enterocolitica subsp enterocolitica 8081 YP 001005188 1 393305 Proteobacteria
Yersinia pestis biovar Microtus str 91001 NP 991793 1 229193 Proteobacteria
Yersinia pestis Antiqua YP 652766 1 360102 Proteobacteria
Yersinia pestis biovar Medievalis str Harbin 35 YP 005625336 1 547048 Proteobacteria
Yersinia pseudotuberculosis IP 31758 YP 001402177 1 349747 Proteobacteria
Rahnella aquatilis CIP 78 65 ATCC 33071 YP 005198819 1 745277 Proteobacteria
Rahnella sp Y9602 YP 004211484 1 741091 Proteobacteria
Serratia proteamaculans 568 YP 001477113 1 399741 Proteobacteria
Serratia liquefaciens ATCC 27592 YP 008228703 1 1346614 Proteobacteria
Serratia marcescens WW4 YP 007404668 1 435998 Proteobacteria
Serratia plymuthica AS9 YP 004504217 1 768492 Proteobacteria
Serratia sp AS12 YP 004499265 1 768490 Proteobacteria
Alkalilimnicola ehrlichii MLHE 1 YP 741120 1 187272 Proteobacteria
Spiribacter sp UAH SP71 YP 008754838 1 1335757 Proteobacteria
Halomonas elongata DSM 2581 YP 003898099 1 768066 Proteobacteria
Chromohalobacter salexigens DSM 3043 YP 573677 1 290398 Proteobacteria
Hahella chejuensis KCTC 2396 YP 433739 1 349521 Proteobacteria
Marinobacter sp BSs20148 YP 006558466 1 490759 Proteobacteria
Marinobacter aquaeolei VT8 YP 958816 1 351348 Proteobacteria
Marinobacter adhaerens HP15 YP 005885506 1 225937 Proteobacteria
Marinomonas mediterranea MMB 1 YP 004313390 1 717774 Proteobacteria
Marinomonas sp MWYL1 YP 001341200 1 400668 Proteobacteria
Marinomonas posidonica IVIA Po 181 YP 004481598 1 491952 Proteobacteria
Acinetobacter baumannii ACICU YP 001846350 1 405416 Proteobacteria
Alcanivorax borkumensis SK2 YP 692812 1 393595 Proteobacteria
Alcanivorax dieselolei B5 YP 006821324 1 930169 Proteobacteria
Cellvibrio japonicus Ueda107 YP 001982218 1 498211 Proteobacteria
Saccharophagus degradans 2 40 YP 527479 1 203122 Proteobacteria
Teredinibacter turnerae T7901 YP 003073593 1 377629 Proteobacteria
Azotobacter vinelandii CA6 YP 007898335 1 1283331 Proteobacteria
Azotobacter vinelandii DJ YP 002799476 1 322710 Proteobacteria
Pseudomonas stutzeri RCH2 YP 007240487 1 644801 Proteobacteria
Pseudomonas mendocina NK 01 YP 004380444 1 1001585 Proteobacteria
Pseudomonas resinovorans NBRC 106553 YP 008103633 1 1245471 Proteobacteria
Pseudomonas denitrificans ATCC 13867 YP 007657513 1 1294143 Proteobacteria
Pseudomonas aeruginosa NCGM2 S1 YP 005980893 1 1089456 Proteobacteria
Pseudomonas aeruginosa PAO1 VE13 YP 008705884 1 1367494 Proteobacteria
Pseudomonas fulva 12 X YP 004474610 1 743720 Proteobacteria
Pseudomonas entomophila L48 YP 607534 1 384676 Proteobacteria
Pseudomonas putida DOT T1E YP 006535974 1 1196325 Proteobacteria
Pseudomonas monteilii SB3078 YP 008953832 1 1435044 Proteobacteria
Pseudomonas protegens CHA0 YP 008001602 1 1124983 Proteobacteria
Pseudomonas syringae pv tomato str DC3000 NP 792094 1 223283 Proteobacteria
Pseudomonas syringae pv syringae B728a YP 235154 1 205918 Proteobacteria
Pseudomonas syringae pv phaseolicola 1448A YP 274263 1 264730 Proteobacteria
Pseudomonas poae RE 1 1 14 YP 007396557 1 1282356 Proteobacteria
Pseudomonas sp TKP YP 008931472 1 1415630 Proteobacteria
Pseudomonas fluorescens Pf0 1 YP 349737 1 205922 Proteobacteria
Pseudomonas brassicacearum subsp brassicacearum NFM421 YP 004352815 1 994484 Proteobacteria

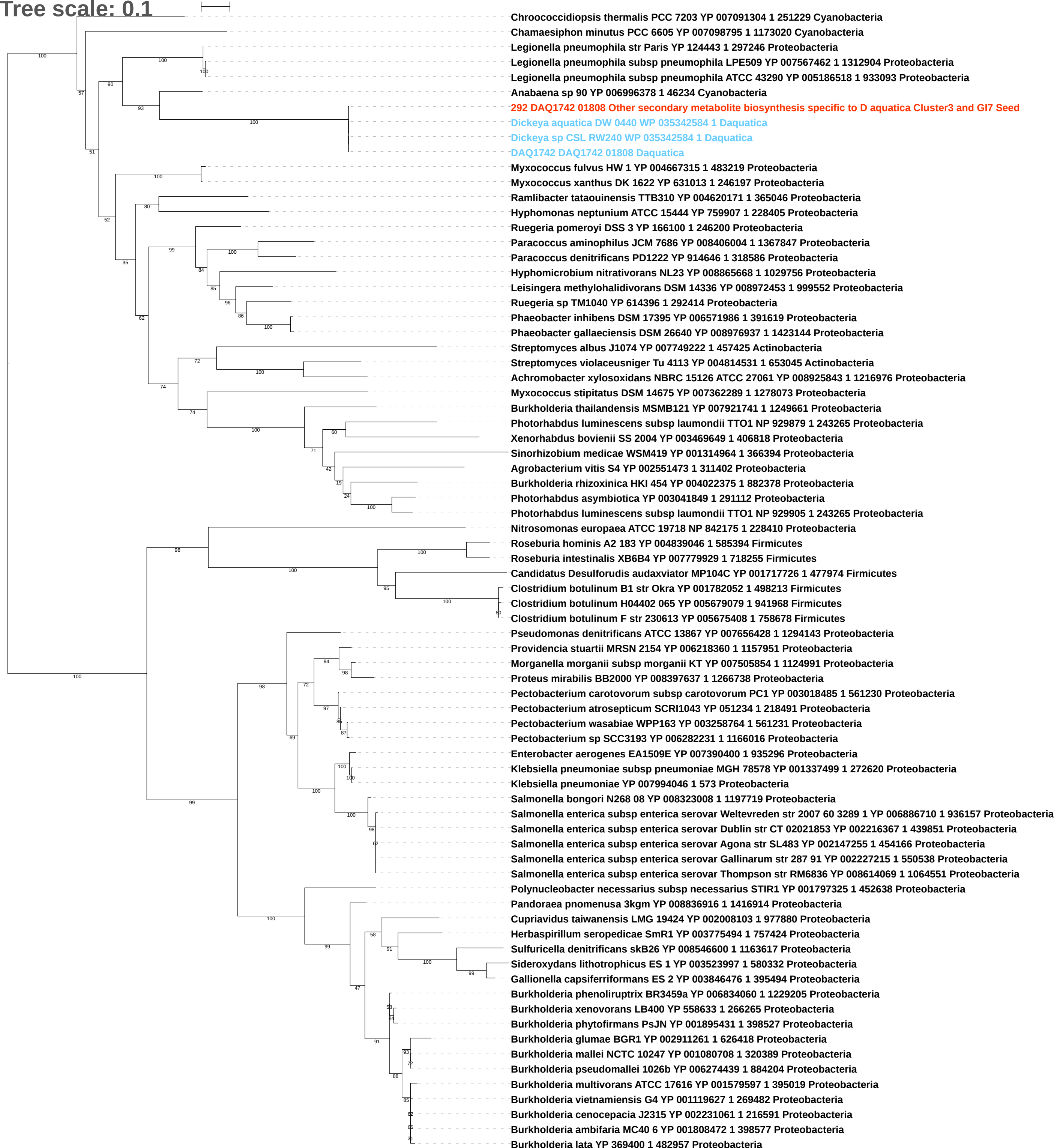
Tree scale: 0.1
Chroococcidiopsis thermalis PCC 7203 YP 007091304 1 251229 Cyanobacteria
Chamaesiphon minutus PCC 6605 YP 007098795 1 1173020 Cyanobacteria
Legionella pneumophila str Paris YP 124443 1 297246 Proteobacteria
Legionella pneumophila subsp pneumophila LPE509 YP 007567462 1 1312904 Proteobacteria
Legionella pneumophila subsp pneumophila ATCC 43290 YP 005186518 1 933093 Proteobacteria
Anabaena sp 90 YP 006996378 1 46234 Cyanobacteria
292 DAQ1742 01808 Other secondary metabolite biosynthesis specific to D aquatica Cluster3 and GI7 Seed
Dickeya aquatica DW 0440 WP 035342584 1 Daquatica
Dickeya sp CSL RW240 WP 035342584 1 Daquatica
DAQ1742 DAQ1742 01808 Daquatica
Myxococcus fulvus HW 1 YP 004667315 1 483219 Proteobacteria
Myxococcus xanthus DK 1622 YP 631013 1 246197 Proteobacteria
Ramlibacter tataouinensis TTB310 YP 004620171 1 365046 Proteobacteria
Hyphomonas neptunium ATCC 15444 YP 759907 1 228405 Proteobacteria
Ruegeria pomeroyi DSS 3 YP 166100 1 246200 Proteobacteria
Paracoccus aminophilus JCM 7686 YP 008406004 1 1367847 Proteobacteria
Paracoccus denitrificans PD1222 YP 914646 1 318586 Proteobacteria
Hyphomicrobium nitrativorans NL23 YP 008865668 1 1029756 Proteobacteria
Leisingera methylohalidivorans DSM 14336 YP 008972453 1 999552 Proteobacteria
Ruegeria sp TM1040 YP 614396 1 292414 Proteobacteria
Phaeobacter inhibens DSM 17395 YP 006571986 1 391619 Proteobacteria
Phaeobacter gallaeciensis DSM 26640 YP 008976937 1 1423144 Proteobacteria
Streptomyces albus J1074 YP 007749222 1 457425 Actinobacteria
Streptomyces violaceusniger Tu 4113 YP 004814531 1 653045 Actinobacteria
Achromobacter xylosoxidans NBRC 15126 ATCC 27061 YP 008925843 1 1216976 Proteobacteria
Myxococcus stipitatus DSM 14675 YP 007362289 1 1278073 Proteobacteria
Burkholderia thailandensis MSMB121 YP 007921741 1 1249661 Proteobacteria
Photorhabdus luminescens subsp laumondii TTO1 NP 929879 1 243265 Proteobacteria
Xenorhabdus bovienii SS 2004 YP 003469649 1 406818 Proteobacteria
Sinorhizobium medicae WSM419 YP 001314964 1 366394 Proteobacteria
Agrobacterium vitis S4 YP 002551473 1 311402 Proteobacteria
Burkholderia rhizoxinica HKI 454 YP 004022375 1 882378 Proteobacteria
Photorhabdus asymbiotica YP 003041849 1 291112 Proteobacteria
Photorhabdus luminescens subsp laumondii TTO1 NP 929905 1 243265 Proteobacteria
Nitrosomonas europaea ATCC 19718 NP 842175 1 228410 Proteobacteria
Roseburia hominis A2 183 YP 004839046 1 585394 Firmicutes
Roseburia intestinalis XB6B4 YP 007779929 1 718255 Firmicutes
Candidatus Desulforudis audaxviator MP104C YP 001717726 1 477974 Firmicutes
Clostridium botulinum B1 str Okra YP 001782052 1 498213 Firmicutes
Clostridium botulinum H04402 065 YP 005679079 1 941968 Firmicutes
Clostridium botulinum F str 230613 YP 005675408 1 758678 Firmicutes
Pseudomonas denitrificans ATCC 13867 YP 007656428 1 1294143 Proteobacteria
Providencia stuartii MRSN 2154 YP 006218360 1 1157951 Proteobacteria
Morganella morganii subsp morganii KT YP 007505854 1 1124991 Proteobacteria
Proteus mirabilis BB2000 YP 008397637 1 1266738 Proteobacteria
Pectobacterium carotovorum subsp carotovorum PC1 YP 003018485 1 561230 Proteobacteria
Pectobacterium atrosepticum SCRI1043 YP 051234 1 218491 Proteobacteria
Pectobacterium wasabiae WPP163 YP 003258764 1 561231 Proteobacteria
Pectobacterium sp SCC3193 YP 006282231 1 1166016 Proteobacteria
Enterobacter aerogenes EA1509E YP 007390400 1 935296 Proteobacteria
Klebsiella pneumoniae subsp pneumoniae MGH 78578 YP 001337499 1 272620 Proteobacteria
Klebsiella pneumoniae YP 007994046 1 573 Proteobacteria
Salmonella bongori N268 08 YP 008323008 1 1197719 Proteobacteria
Salmonella enterica subsp enterica serovar Weltevreden str 2007 60 3289 1 YP 006886710 1 936157 Proteobacteria
Salmonella enterica subsp enterica serovar Dublin str CT 02021853 YP 002216367 1 439851 Proteobacteria
Salmonella enterica subsp enterica serovar Agona str SL483 YP 002147255 1 454166 Proteobacteria
Salmonella enterica subsp enterica serovar Gallinarum str 287 91 YP 002227215 1 550538 Proteobacteria
Salmonella enterica subsp enterica serovar Thompson str RM6836 YP 008614069 1 1064551 Proteobacteria
Polynucleobacter necessarius subsp necessarius STIR1 YP 001797325 1 452638 Proteobacteria
Pandoraea pnomenusa 3kgm YP 008836916 1 1416914 Proteobacteria
Cupriavidus taiwanensis LMG 19424 YP 002008103 1 977880 Proteobacteria
Herbaspirillum seropedicae SmR1 YP 003775494 1 757424 Proteobacteria
Sulfuricella denitrificans skB26 YP 008546600 1 1163617 Proteobacteria
Sideroxydans lithotrophicus ES 1 YP 003523997 1 580332 Proteobacteria
Gallionella capsiferriformans ES 2 YP 003846476 1 395494 Proteobacteria
Burkholderia phenoliruptrix BR3459a YP 006834060 1 1229205 Proteobacteria
Burkholderia xenovorans LB400 YP 558633 1 266265 Proteobacteria
Burkholderia phytofirmans PsJN YP 001895431 1 398527 Proteobacteria
Burkholderia glumae BGR1 YP 002911261 1 626418 Proteobacteria
Burkholderia mallei NCTC 10247 YP 001080708 1 320389 Proteobacteria
Burkholderia pseudomallei 1026b YP 006274439 1 884204 Proteobacteria
Burkholderia multivorans ATCC 17616 YP 001579597 1 395019 Proteobacteria
Burkholderia vietnamiensis G4 YP 001119627 1 269482 Proteobacteria
Burkholderia cenocepacia J2315 YP 002231061 1 216591 Proteobacteria
Burkholderia ambifaria MC40 6 YP 001808472 1 398577 Proteobacteria
Burkholderia lata YP 369400 1 482957 Proteobacteria

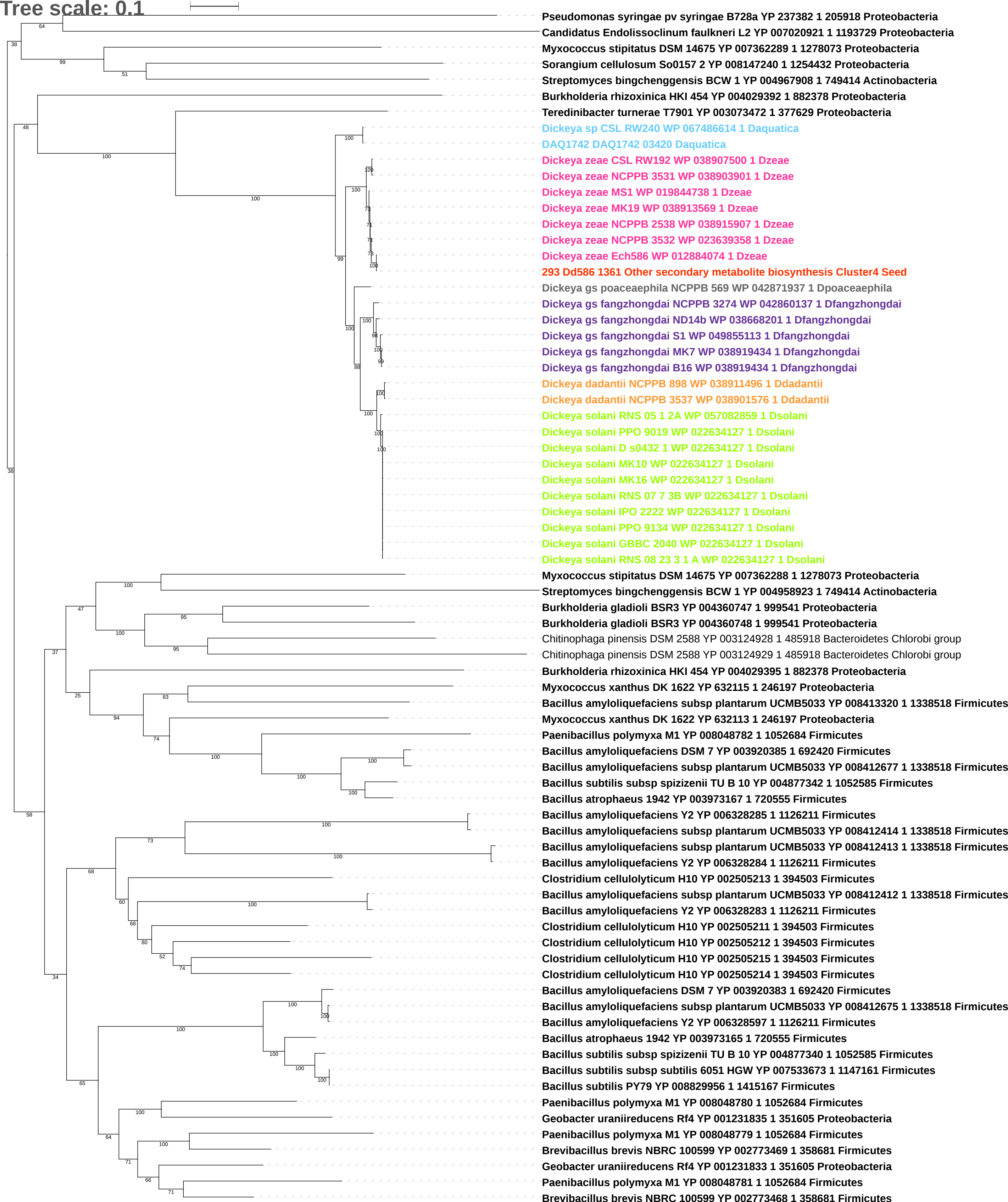

Tree scale: 0.1
Pseudomonas syringae pv syringae B728a YP 237382 1 205918 Proteobacteria
Candidatus Endolissoclinum faulkneri L2 YP 007020921 1 1193729 Proteobacteria
Myxococcus stipitatus DSM 14675 YP 007362289 1 1278073 Proteobacteria
Sorangium cellulosum So0157 2 YP 008147240 1 1254432 Proteobacteria
Streptomyces bingchenggensis BCW 1 YP 004967908 1 749414 Actinobacteria
Burkholderia rhizoxinica HKI 454 YP 004029392 1 882378 Proteobacteria
Teredinibacter turnerae T7901 YP 003073472 1 377629 Proteobacteria
Dickeya sp CSL RW240 WP 067486614 1 Daquatica
DAQ1742 DAQ1742 03420 Daquatica
Dickeya zeae CSL RW192 WP 038907500 1 Dzeae
Dickeya zeae NCPPB 3531 WP 038903901 1 Dzeae
Dickeya zeae MS1 WP 019844738 1 Dzeae
Dickeya zeae MK19 WP 038913569 1 Dzeae
Dickeya zeae NCPPB 2538 WP 038915907 1 Dzeae
Dickeya zeae NCPPB 3532 WP 023639358 1 Dzeae
Dickeya zeae Ech586 WP 012884074 1 Dzeae
293 Dd586 1361 Other secondary metabolite biosynthesis Cluster4 Seed
Dickeya gs poaceaephila NCPPB 569 WP 042871937 1 Dpoaceaephila
Dickeya gs fangzhongdai NCPPB 3274 WP 042860137 1 Dfangzhongdai
Dickeya gs fangzhongdai ND14b WP 038668201 1 Dfangzhongdai
Dickeya gs fangzhongdai S1 WP 049855113 1 Dfangzhongdai
Dickeya gs fangzhongdai MK7 WP 038919434 1 Dfangzhongdai
Dickeya gs fangzhongdai B16 WP 038919434 1 Dfangzhongdai
Dickeya dadantii NCPPB 898 WP 038911496 1 Ddadantii
Dickeya dadantii NCPPB 3537 WP 038901576 1 Ddadantii
Dickeya solani RNS 05 1 2A WP 057082859 1 Dsolani
Dickeya solani PPO 9019 WP 022634127 1 Dsolani
Dickeya solani D s0432 1 WP 022634127 1 Dsolani
Dickeya solani MK10 WP 022634127 1 Dsolani
Dickeya solani MK16 WP 022634127 1 Dsolani
Dickeya solani RNS 07 7 3B WP 022634127 1 Dsolani
Dickeya solani IPO 2222 WP 022634127 1 Dsolani
Dickeya solani PPO 9134 WP 022634127 1 Dsolani
Dickeya solani GBBC 2040 WP 022634127 1 Dsolani
Dickeya solani RNS 08 23 3 1 A WP 022634127 1 Dsolani
Myxococcus stipitatus DSM 14675 YP 007362288 1 1278073 Proteobacteria
Streptomyces bingchenggensis BCW 1 YP 004958923 1 749414 Actinobacteria
Burkholderia gladioli BSR3 YP 004360747 1 999541 Proteobacteria
Burkholderia gladioli BSR3 YP 004360748 1 999541 Proteobacteria
Chitinophaga pinensis DSM 2588 YP 003124928 1 485918 Bacteroidetes Chlorobi group
Chitinophaga pinensis DSM 2588 YP 003124929 1 485918 Bacteroidetes Chlorobi group
Burkholderia rhizoxinica HKI 454 YP 004029395 1 882378 Proteobacteria
Myxococcus xanthus DK 1622 YP 632115 1 246197 Proteobacteria
Bacillus amyloliquefaciens subsp plantarum UCMB5033 YP 008413320 1 1338518 Firmicutes
Myxococcus xanthus DK 1622 YP 632113 1 246197 Proteobacteria
Paenibacillus polymyxa M1 YP 008048782 1 1052684 Firmicutes
Bacillus amyloliquefaciens DSM 7 YP 003920385 1 692420 Firmicutes
Bacillus amyloliquefaciens subsp plantarum UCMB5033 YP 008412677 1 1338518 Firmicutes
Bacillus subtilis subsp spizizenii TU B 10 YP 004877342 1 1052585 Firmicutes
Bacillus atrophaeus 1942 YP 003973167 1 720555 Firmicutes
Bacillus amyloliquefaciens Y2 YP 006328285 1 1126211 Firmicutes
Bacillus amyloliquefaciens subsp plantarum UCMB5033 YP 008412414 1 1338518 Firmicutes
Bacillus amyloliquefaciens subsp plantarum UCMB5033 YP 008412413 1 1338518 Firmicutes
Bacillus amyloliquefaciens Y2 YP 006328284 1 1126211 Firmicutes
Clostridium cellulolyticum H10 YP 002505213 1 394503 Firmicutes
Bacillus amyloliquefaciens subsp plantarum UCMB5033 YP 008412412 1 1338518 Firmicutes
Bacillus amyloliquefaciens Y2 YP 006328283 1 1126211 Firmicutes
Clostridium cellulolyticum H10 YP 002505211 1 394503 Firmicutes
Clostridium cellulolyticum H10 YP 002505212 1 394503 Firmicutes
Clostridium cellulolyticum H10 YP 002505215 1 394503 Firmicutes
Clostridium cellulolyticum H10 YP 002505214 1 394503 Firmicutes
Bacillus amyloliquefaciens DSM 7 YP 003920383 1 692420 Firmicutes
Bacillus amyloliquefaciens subsp plantarum UCMB5033 YP 008412675 1 1338518 Firmicutes
Bacillus amyloliquefaciens Y2 YP 006328597 1 1126211 Firmicutes
Bacillus atrophaeus 1942 YP 003973165 1 720555 Firmicutes
Bacillus subtilis subsp spizizenii TU B 10 YP 004877340 1 1052585 Firmicutes
Bacillus subtilis subsp subtilis 6051 HGW YP 007533673 1 1147161 Firmicutes
Bacillus subtilis PY79 YP 008829956 1 1415167 Firmicutes
Paenibacillus polymyxa M1 YP 008048780 1 1052684 Firmicutes
Geobacter uraniireducens Rf4 YP 001231835 1 351605 Proteobacteria
Paenibacillus polymyxa M1 YP 008048779 1 1052684 Firmicutes
Brevibacillus brevis NBRC 100599 YP 002773469 1 358681 Firmicutes
Geobacter uraniireducens Rf4 YP 001231833 1 351605 Proteobacteria
Paenibacillus polymyxa M1 YP 008048781 1 1052684 Firmicutes
Brevibacillus brevis NBRC 100599 YP 002773468 1 358681 Firmicutes

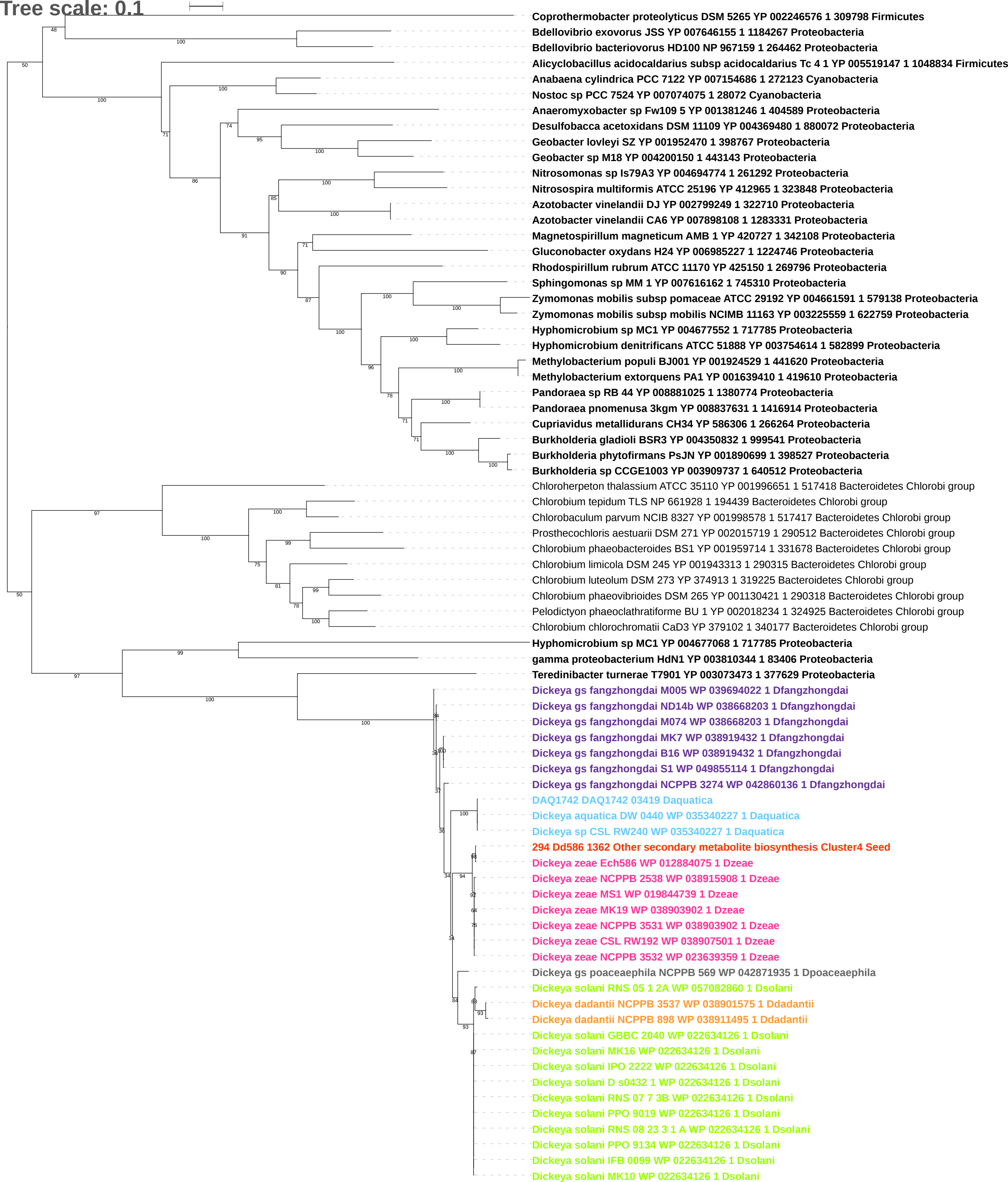
Tree scale: 0.1
Coprothermobacter proteolyticus DSM 5265 YP 002246576 1 309798 Firmicutes
Bdellovibrio exovorus JSS YP 007646155 1 1184267 Proteobacteria
Bdellovibrio bacteriovorus HD100 NP 967159 1 264462 Proteobacteria
Alicyclobacillus acidocaldarius subsp acidocaldarius Tc 4 1 YP 005519147 1 1048834 Firmicutes
Anabaena cylindrica PCC 7122 YP 007154686 1 272123 Cyanobacteria
Nostoc sp PCC 7524 YP 007074075 1 28072 Cyanobacteria
Anaeromyxobacter sp Fw109 5 YP 001381246 1 404589 Proteobacteria
Desulfobacca acetoxidans DSM 11109 YP 004369480 1 880072 Proteobacteria
Geobacter lovleyi SZ YP 001952470 1 398767 Proteobacteria
Geobacter sp M18 YP 004200150 1 443143 Proteobacteria
Nitrosomonas sp Is79A3 YP 004694774 1 261292 Proteobacteria
Nitrosospira multiformis ATCC 25196 YP 412965 1 323848 Proteobacteria
Azotobacter vinelandii DJ YP 002799249 1 322710 Proteobacteria
Azotobacter vinelandii CA6 YP 007898108 1 1283331 Proteobacteria
Magnetospirillum magneticum AMB 1 YP 420727 1 342108 Proteobacteria
Gluconobacter oxydans H24 YP 006985227 1 1224746 Proteobacteria
Rhodospirillum rubrum ATCC 11170 YP 425150 1 269796 Proteobacteria
Sphingomonas sp MM 1 YP 007616162 1 745310 Proteobacteria
Zymomonas mobilis subsp pomaceae ATCC 29192 YP 004661591 1 579138 Proteobacteria
Zymomonas mobilis subsp mobilis NCIMB 11163 YP 003225559 1 622759 Proteobacteria
Hyphomicrobium sp MC1 YP 004677552 1 717785 Proteobacteria
Hyphomicrobium denitrificans ATCC 51888 YP 003754614 1 582899 Proteobacteria
Methylobacterium populi BJ001 YP 001924529 1 441620 Proteobacteria
Methylobacterium extorquens PA1 YP 001639410 1 419610 Proteobacteria
Pandoraea sp RB 44 YP 008881025 1 1380774 Proteobacteria
Pandoraea pnomenusa 3kgm YP 008837631 1 1416914 Proteobacteria
Cupriavidus metallidurans CH34 YP 586306 1 266264 Proteobacteria
Burkholderia gladioli BSR3 YP 004350832 1 999541 Proteobacteria
Burkholderia phytofirmans PsJN YP 001890699 1 398527 Proteobacteria
Burkholderia sp CCGE1003 YP 003909737 1 640512 Proteobacteria
Chloroherpeton thalassium ATCC 35110 YP 001996651 1 517418 Bacteroidetes Chlorobi group
Chlorobium tepidum TLS NP 661928 1 194439 Bacteroidetes Chlorobi group
Chlorobaculum parvum NCIB 8327 YP 001998578 1 517417 Bacteroidetes Chlorobi group
Prosthecochloris aestuarii DSM 271 YP 002015719 1 290512 Bacteroidetes Chlorobi group
Chlorobium phaeobacteroides BS1 YP 001959714 1 331678 Bacteroidetes Chlorobi group
Chlorobium limicola DSM 245 YP 001943313 1 290315 Bacteroidetes Chlorobi group
Chlorobium luteolum DSM 273 YP 374913 1 319225 Bacteroidetes Chlorobi group
Chlorobium phaeovibrioides DSM 265 YP 001130421 1 290318 Bacteroidetes Chlorobi group
Pelodictyon phaeoclathratiforme BU 1 YP 002018234 1 324925 Bacteroidetes Chlorobi group
Chlorobium chlorochromatii CaD3 YP 379102 1 340177 Bacteroidetes Chlorobi group
Hyphomicrobium sp MC1 YP 004677068 1 717785 Proteobacteria
gamma proteobacterium HdN1 YP 003810344 1 83406 Proteobacteria
Teredinibacter turnerae T7901 YP 003073473 1 377629 Proteobacteria
Dickeya gs fangzhongdai M005 WP 039694022 1 Dfangzhongdai
Dickeya gs fangzhongdai ND14b WP 038668203 1 Dfangzhongdai
Dickeya gs fangzhongdai M074 WP 038668203 1 Dfangzhongdai
Dickeya gs fangzhongdai MK7 WP 038919432 1 Dfangzhongdai
Dickeya gs fangzhongdai B16 WP 038919432 1 Dfangzhongdai
Dickeya gs fangzhongdai S1 WP 049855114 1 Dfangzhongdai
Dickeya gs fangzhongdai NCPPB 3274 WP 042860136 1 Dfangzhongdai
DAQ1742 DAQ1742 03419 Daquatica
Dickeya aquatica DW 0440 WP 035340227 1 Daquatica
Dickeya sp CSL RW240 WP 035340227 1 Daquatica
294 Dd586 1362 Other secondary metabolite biosynthesis Cluster4 Seed
Dickeya zeae Ech586 WP 012884075 1 Dzeae
Dickeya zeae NCPPB 2538 WP 038915908 1 Dzeae
Dickeya zeae MS1 WP 019844739 1 Dzeae
Dickeya zeae MK19 WP 038903902 1 Dzeae
Dickeya zeae NCPPB 3531 WP 038903902 1 Dzeae
Dickeya zeae CSL RW192 WP 038907501 1 Dzeae
Dickeya zeae NCPPB 3532 WP 023639359 1 Dzeae
Dickeya gs poaceaephila NCPPB 569 WP 042871935 1 Dpoaceaephila
Dickeya solani RNS 05 1 2A WP 057082860 1 Dsolani
Dickeya dadantii NCPPB 3537 WP 038901575 1 Ddadantii
Dickeya dadantii NCPPB 898 WP 038911495 1 Ddadantii
Dickeya solani GBBC 2040 WP 022634126 1 Dsolani
Dickeya solani MK16 WP 022634126 1 Dsolani
Dickeya solani IPO 2222 WP 022634126 1 Dsolani
Dickeya solani D s0432 1 WP 022634126 1 Dsolani
Dickeya solani RNS 07 7 3B WP 022634126 1 Dsolani
Dickeya solani PPO 9019 WP 022634126 1 Dsolani
Dickeya solani RNS 08 23 3 1 A WP 022634126 1 Dsolani
Dickeya solani PPO 9134 WP 022634126 1 Dsolani
Dickeya solani IFB 0099 WP 022634126 1 Dsolani
Dickeya solani MK10 WP 022634126 1 Dsolani

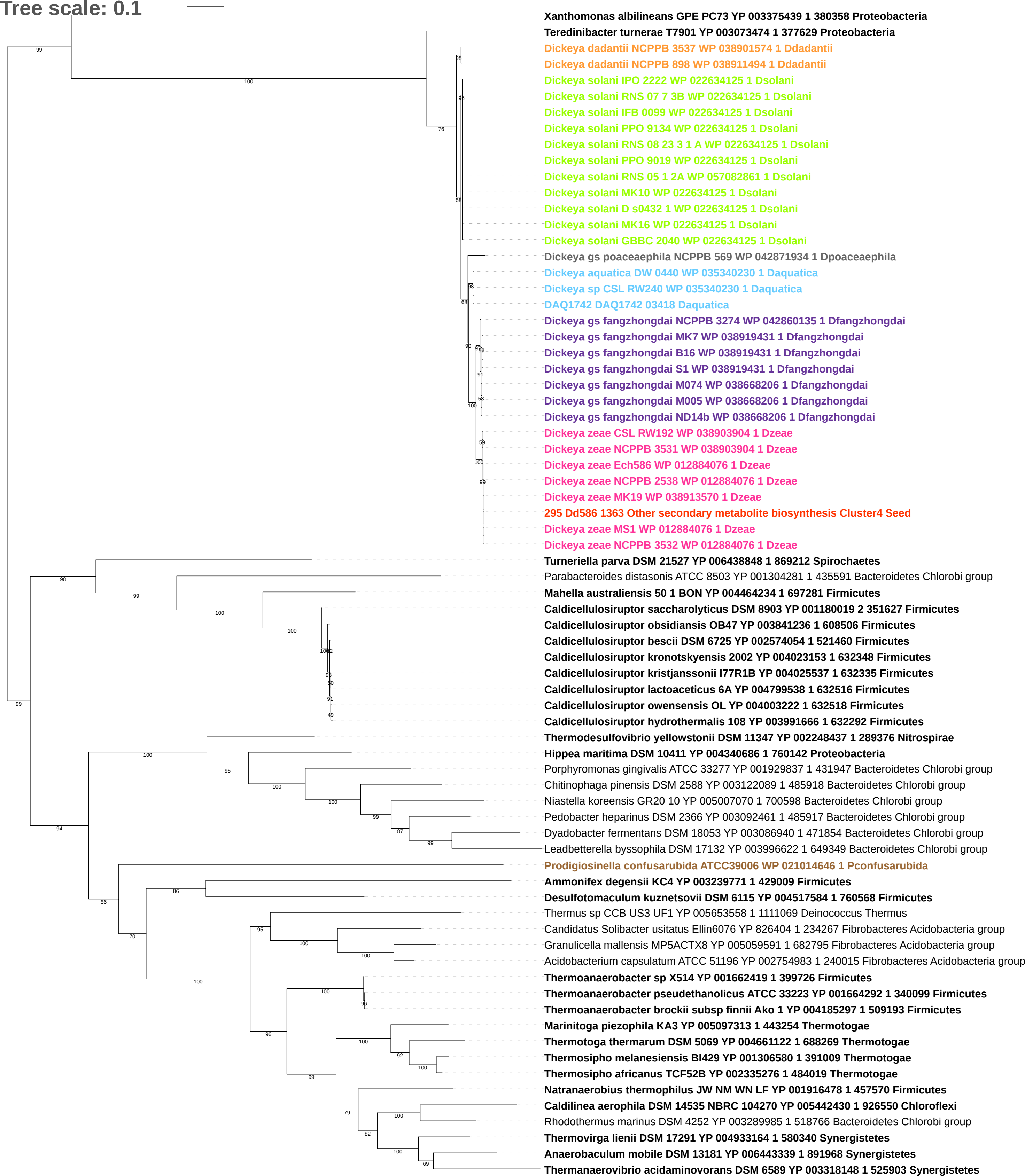
Tree scale: 0.1
Xanthomonas albilineans GPE PC73 YP 003375439 1 380358 Proteobacteria
Teredinibacter turnerae T7901 YP 003073474 1 377629 Proteobacteria
Dickeya dadantii NCPPB 3537 WP 038901574 1 Ddadantii
Dickeya dadantii NCPPB 898 WP 038911494 1 Ddadantii
Dickeya solani IPO 2222 WP 022634125 1 Dsolani
Dickeya solani RNS 07 7 3B WP 022634125 1 Dsolani
Dickeya solani IFB 0099 WP 022634125 1 Dsolani
Dickeya solani PPO 9134 WP 022634125 1 Dsolani
Dickeya solani RNS 08 23 3 1 A WP 022634125 1 Dsolani
Dickeya solani PPO 9019 WP 022634125 1 Dsolani
Dickeya solani RNS 05 1 2A WP 057082861 1 Dsolani
Dickeya solani MK10 WP 022634125 1 Dsolani
Dickeya solani D s0432 1 WP 022634125 1 Dsolani
Dickeya solani MK16 WP 022634125 1 Dsolani
Dickeya solani GBBC 2040 WP 022634125 1 Dsolani
Dickeya gs poaceaephila NCPPB 569 WP 042871934 1 Dpoaceaephila
Dickeya aquatica DW 0440 WP 035340230 1 Daquatica
Dickeya sp CSL RW240 WP 035340230 1 Daquatica
DAQ1742 DAQ1742 03418 Daquatica
Dickeya gs fangzhongdai NCPPB 3274 WP 042860135 1 Dfangzhongdai
Dickeya gs fangzhongdai MK7 WP 038919431 1 Dfangzhongdai
Dickeya gs fangzhongdai B16 WP 038919431 1 Dfangzhongdai
Dickeya gs fangzhongdai S1 WP 038919431 1 Dfangzhongdai
Dickeya gs fangzhongdai M074 WP 038668206 1 Dfangzhongdai
Dickeya gs fangzhongdai M005 WP 038668206 1 Dfangzhongdai
Dickeya gs fangzhongdai ND14b WP 038668206 1 Dfangzhongdai
Dickeya zeae CSL RW192 WP 038903904 1 Dzeae
Dickeya zeae NCPPB 3531 WP 038903904 1 Dzeae
Dickeya zeae Ech586 WP 012884076 1 Dzeae
Dickeya zeae NCPPB 2538 WP 012884076 1 Dzeae
Dickeya zeae MK19 WP 038913570 1 Dzeae
295 Dd586 1363 Other secondary metabolite biosynthesis Cluster4 Seed
Dickeya zeae MS1 WP 012884076 1 Dzeae
Dickeya zeae NCPPB 3532 WP 012884076 1 Dzeae
Turneriella parva DSM 21527 YP 006438848 1 869212 Spirochaetes
Parabacteroides distasonis ATCC 8503 YP 001304281 1 435591 Bacteroidetes Chlorobi group
Mahella australiensis 50 1 BON YP 004464234 1 697281 Firmicutes
Caldicellulosiruptor saccharolyticus DSM 8903 YP 001180019 2 351627 Firmicutes
Caldicellulosiruptor obsidiansis OB47 YP 003841236 1 608506 Firmicutes
Caldicellulosiruptor bescii DSM 6725 YP 002574054 1 521460 Firmicutes
Caldicellulosiruptor kronotskyensis 2002 YP 004023153 1 632348 Firmicutes
Caldicellulosiruptor kristjanssonii I77R1B YP 004025537 1 632335 Firmicutes
Caldicellulosiruptor lactoaceticus 6A YP 004799538 1 632516 Firmicutes
Caldicellulosiruptor owensensis OL YP 004003222 1 632518 Firmicutes
Caldicellulosiruptor hydrothermalis 108 YP 003991666 1 632292 Firmicutes
Thermodesulfovibrio yellowstonii DSM 11347 YP 002248437 1 289376 Nitrospirae
Hippea maritima DSM 10411 YP 004340686 1 760142 Proteobacteria
Porphyromonas gingivalis ATCC 33277 YP 001929837 1 431947 Bacteroidetes Chlorobi group
Chitinophaga pinensis DSM 2588 YP 003122089 1 485918 Bacteroidetes Chlorobi group
Niastella koreensis GR20 10 YP 005007070 1 700598 Bacteroidetes Chlorobi group
Pedobacter heparinus DSM 2366 YP 003092461 1 485917 Bacteroidetes Chlorobi group
Dyadobacter fermentans DSM 18053 YP 003086940 1 471854 Bacteroidetes Chlorobi group
Leadbetterella byssophila DSM 17132 YP 003996622 1 649349 Bacteroidetes Chlorobi group
Prodigiosinella confusarubida ATCC39006 WP 021014646 1 Pconfusarubida
Ammonifex degensii KC4 YP 003239771 1 429009 Firmicutes
Desulfotomaculum kuznetsovii DSM 6115 YP 004517584 1 760568 Firmicutes
Thermus sp CCB US3 UF1 YP 005653558 1 1111069 Deinococcus Thermus
Candidatus Solibacter usitatus Ellin6076 YP 826404 1 234267 Fibrobacteres Acidobacteria group
Granulicella mallensis MP5ACTX8 YP 005059591 1 682795 Fibrobacteres Acidobacteria group
Acidobacterium capsulatum ATCC 51196 YP 002754983 1 240015 Fibrobacteres Acidobacteria group
Thermoanaerobacter sp X514 YP 001662419 1 399726 Firmicutes
Thermoanaerobacter pseudethanolicus ATCC 33223 YP 001664292 1 340099 Firmicutes
Thermoanaerobacter brockii subsp finnii Ako 1 YP 004185297 1 509193 Firmicutes
Marinitoga piezophila KA3 YP 005097313 1 443254 Thermotogae
Thermotoga thermarum DSM 5069 YP 004661122 1 688269 Thermotogae
Thermosipho melanesiensis BI429 YP 001306580 1 391009 Thermotogae
Thermosipho africanus TCF52B YP 002335276 1 484019 Thermotogae
Natranaerobius thermophilus JW NM WN LF YP 001916478 1 457570 Firmicutes
Caldilinea aerophila DSM 14535 NBRC 104270 YP 005442430 1 926550 Chloroflexi
Rhodothermus marinus DSM 4252 YP 003289985 1 518766 Bacteroidetes Chlorobi group
Thermovirga lienii DSM 17291 YP 004933164 1 580340 Synergistetes
Anaerobaculum mobile DSM 13181 YP 006443339 1 891968 Synergistetes
Thermanaerovibrio acidaminovorans DSM 6589 YP 003318148 1 525903 Synergistetes

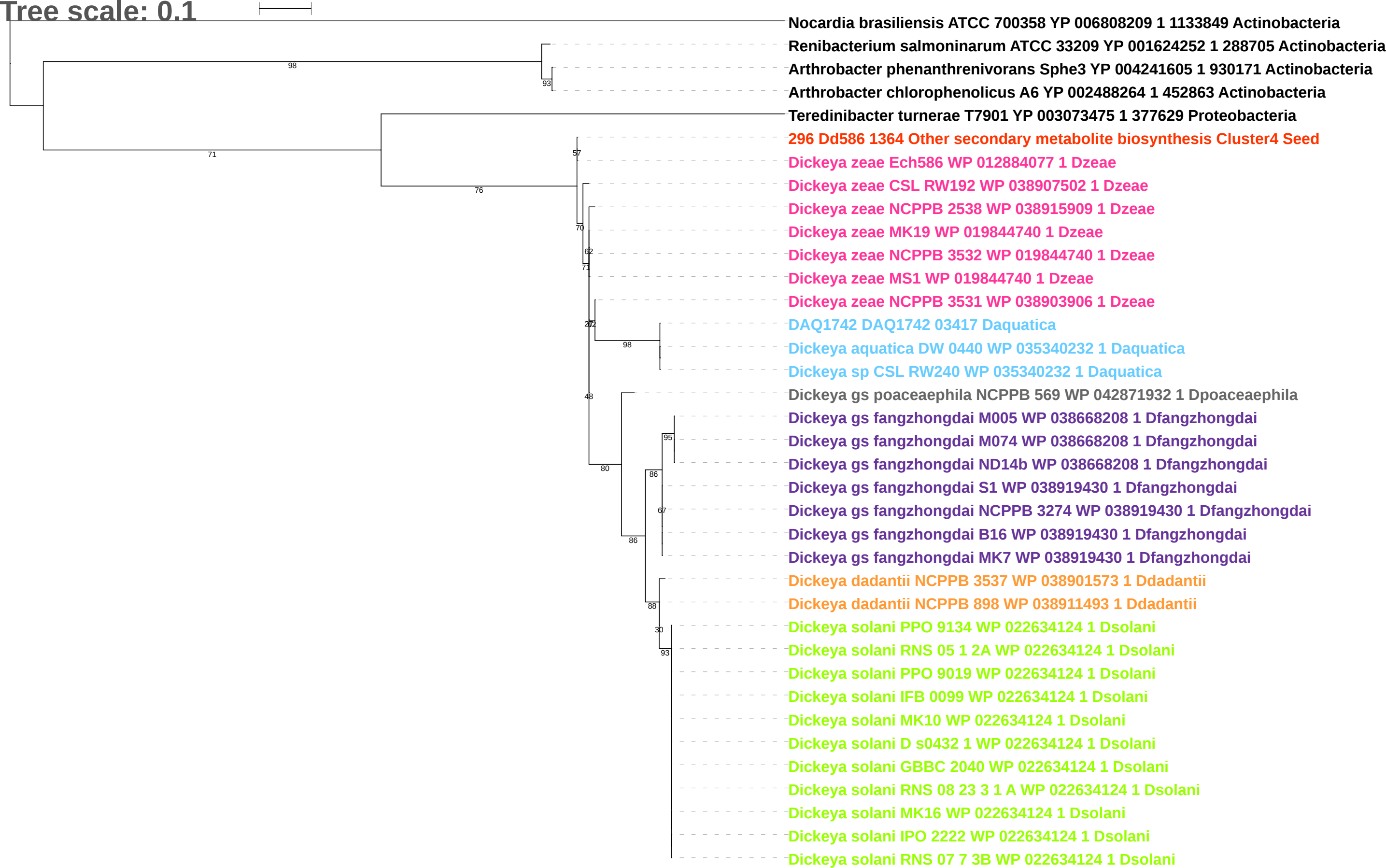

Tree scale: 0.1
Nocardia brasiliensis ATCC 700358 YP 006808209 1 1133849 Actinobacteria
Renibacterium salmoninarum ATCC 33209 YP 001624252 1 288705 Actinobacteria
Arthrobacter phenanthrenivorans Sphe3 YP 004241605 1 930171 Actinobacteria
Arthrobacter chlorophenolicus A6 YP 002488264 1 452863 Actinobacteria
Teredinibacter turnerae T7901 YP 003073475 1 377629 Proteobacteria
296 Dd586 1364 Other secondary metabolite biosynthesis Cluster4 Seed
Dickeya zeae Ech586 WP 012884077 1 Dzeae
Dickeya zeae CSL RW192 WP 038907502 1 Dzeae
Dickeya zeae NCPPB 2538 WP 038915909 1 Dzeae
Dickeya zeae MK19 WP 019844740 1 Dzeae
Dickeya zeae NCPPB 3532 WP 019844740 1 Dzeae
Dickeya zeae MS1 WP 019844740 1 Dzeae
Dickeya zeae NCPPB 3531 WP 038903906 1 Dzeae
DAQ1742 DAQ1742 03417 Daquatica
Dickeya aquatica DW 0440 WP 035340232 1 Daquatica
Dickeya sp CSL RW240 WP 035340232 1 Daquatica
Dickeya gs poaceaephila NCPPB 569 WP 042871932 1 Dpoaceaephila
Dickeya gs fangzhongdai M005 WP 038668208 1 Dfangzhongdai
Dickeya gs fangzhongdai M074 WP 038668208 1 Dfangzhongdai
Dickeya gs fangzhongdai ND14b WP 038668208 1 Dfangzhongdai
Dickeya gs fangzhongdai S1 WP 038919430 1 Dfangzhongdai
Dickeya gs fangzhongdai NCPPB 3274 WP 038919430 1 Dfangzhongdai
Dickeya gs fangzhongdai B16 WP 038919430 1 Dfangzhongdai
Dickeya gs fangzhongdai MK7 WP 038919430 1 Dfangzhongdai
Dickeya dadantii NCPPB 3537 WP 038901573 1 Ddadantii
Dickeya dadantii NCPPB 898 WP 038911493 1 Ddadantii
Dickeya solani PPO 9134 WP 022634124 1 Dsolani
Dickeya solani RNS 05 1 2A WP 022634124 1 Dsolani
Dickeya solani PPO 9019 WP 022634124 1 Dsolani
Dickeya solani IFB 0099 WP 022634124 1 Dsolani
Dickeya solani MK10 WP 022634124 1 Dsolani
Dickeya solani D s0432 1 WP 022634124 1 Dsolani
Dickeya solani GBBC 2040 WP 022634124 1 Dsolani
Dickeya solani RNS 08 23 3 1 A WP 022634124 1 Dsolani
Dickeya solani MK16 WP 022634124 1 Dsolani
Dickeya solani IPO 2222 WP 022634124 1 Dsolani
Dickeya solani RNS 07 7 3B WP 022634124 1 Dsolani

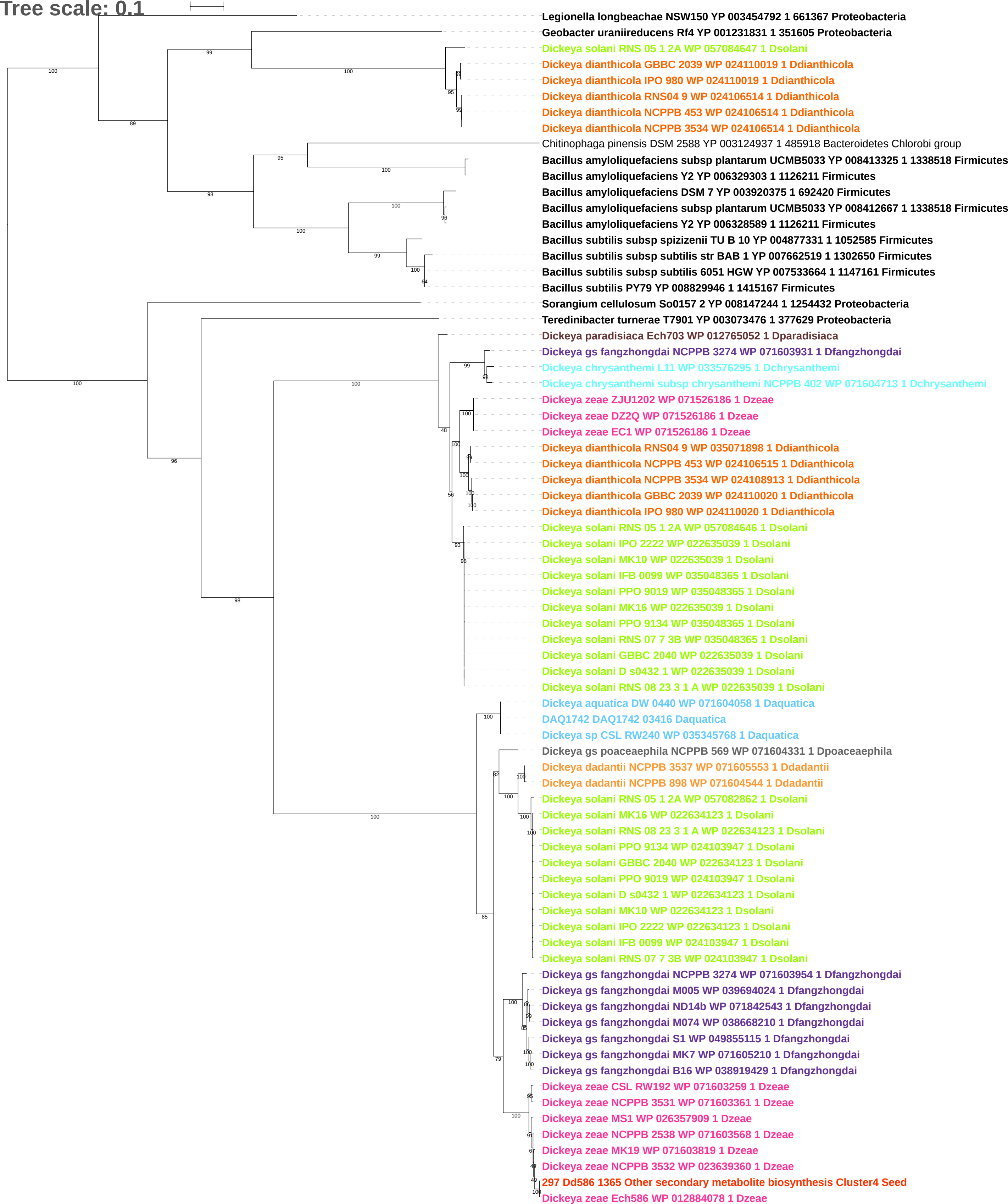
Tree scale: 0.1
Legionella longbeachae NSW150 YP 003454792 1 661367 Proteobacteria
Geobacter uraniireducens Rf4 YP 001231831 1 351605 Proteobacteria
Dickeya solani RNS 05 1 2A WP 057084647 1 Dsolani
Dickeya dianthicola GBBC 2039 WP 024110019 1 Ddianthicola
Dickeya dianthicola IPO 980 WP 024110019 1 Ddianthicola
Dickeya dianthicola RNS04 9 WP 024106514 1 Ddianthicola
Dickeya dianthicola NCPPB 453 WP 024106514 1 Ddianthicola
Dickeya dianthicola NCPPB 3534 WP 024106514 1 Ddianthicola
Chitinophaga pinensis DSM 2588 YP 003124937 1 485918 Bacteroidetes Chlorobi group
Bacillus amyloliquefaciens subsp plantarum UCMB5033 YP 008413325 1 1338518 Firmicutes
Bacillus amyloliquefaciens Y2 YP 006329303 1 1126211 Firmicutes
Bacillus amyloliquefaciens DSM 7 YP 003920375 1 692420 Firmicutes
Bacillus amyloliquefaciens subsp plantarum UCMB5033 YP 008412667 1 1338518 Firmicutes
Bacillus amyloliquefaciens Y2 YP 006328589 1 1126211 Firmicutes
Bacillus subtilis subsp spizizenii TU B 10 YP 004877331 1 1052585 Firmicutes
Bacillus subtilis subsp subtilis str BAB 1 YP 007662519 1 1302650 Firmicutes
Bacillus subtilis subsp subtilis 6051 HGW YP 007533664 1 1147161 Firmicutes
Bacillus subtilis PY79 YP 008829946 1 1415167 Firmicutes
Sorangium cellulosum So0157 2 YP 008147244 1 1254432 Proteobacteria
Teredinibacter turnerae T7901 YP 003073476 1 377629 Proteobacteria
Dickeya paradisiaca Ech703 WP 012765052 1 Dparadisiaca
Dickeya gs fangzhongdai NCPPB 3274 WP 071603931 1 Dfangzhongdai
Dickeya chrysanthemi L11 WP 033576295 1 Dchrysanthemi
Dickeya chrysanthemi subsp chrysanthemi NCPPB 402 WP 071604713 1 Dchrysanthemi
Dickeya zeae ZJU1202 WP 071526186 1 Dzeae
Dickeya zeae DZ2Q WP 071526186 1 Dzeae
Dickeya zeae EC1 WP 071526186 1 Dzeae
Dickeya dianthicola RNS04 9 WP 035071898 1 Ddianthicola
Dickeya dianthicola NCPPB 453 WP 024106515 1 Ddianthicola
Dickeya dianthicola NCPPB 3534 WP 024108913 1 Ddianthicola
Dickeya dianthicola GBBC 2039 WP 024110020 1 Ddianthicola
Dickeya dianthicola IPO 980 WP 024110020 1 Ddianthicola
Dickeya solani RNS 05 1 2A WP 057084646 1 Dsolani
Dickeya solani IPO 2222 WP 022635039 1 Dsolani
Dickeya solani MK10 WP 022635039 1 Dsolani
Dickeya solani IFB 0099 WP 035048365 1 Dsolani
Dickeya solani PPO 9019 WP 035048365 1 Dsolani
Dickeya solani MK16 WP 022635039 1 Dsolani
Dickeya solani PPO 9134 WP 035048365 1 Dsolani
Dickeya solani RNS 07 7 3B WP 035048365 1 Dsolani
Dickeya solani GBBC 2040 WP 022635039 1 Dsolani
Dickeya solani D s0432 1 WP 022635039 1 Dsolani
Dickeya solani RNS 08 23 3 1 A WP 022635039 1 Dsolani
Dickeya aquatica DW 0440 WP 071604058 1 Daquatica
DAQ1742 DAQ1742 03416 Daquatica
Dickeya sp CSL RW240 WP 035345768 1 Daquatica
Dickeya gs poaceaephila NCPPB 569 WP 071604331 1 Dpoaceaephila
Dickeya dadantii NCPPB 3537 WP 071605553 1 Ddadantii
Dickeya dadantii NCPPB 898 WP 071604544 1 Ddadantii
Dickeya solani RNS 05 1 2A WP 057082862 1 Dsolani
Dickeya solani MK16 WP 022634123 1 Dsolani
Dickeya solani RNS 08 23 3 1 A WP 022634123 1 Dsolani
Dickeya solani PPO 9134 WP 024103947 1 Dsolani
Dickeya solani GBBC 2040 WP 022634123 1 Dsolani
Dickeya solani PPO 9019 WP 024103947 1 Dsolani
Dickeya solani D s0432 1 WP 022634123 1 Dsolani
Dickeya solani MK10 WP 022634123 1 Dsolani
Dickeya solani IPO 2222 WP 022634123 1 Dsolani
Dickeya solani IFB 0099 WP 024103947 1 Dsolani
Dickeya solani RNS 07 7 3B WP 024103947 1 Dsolani
Dickeya gs fangzhongdai NCPPB 3274 WP 071603954 1 Dfangzhongdai
Dickeya gs fangzhongdai M005 WP 039694024 1 Dfangzhongdai
Dickeya gs fangzhongdai ND14b WP 071842543 1 Dfangzhongdai
Dickeya gs fangzhongdai M074 WP 038668210 1 Dfangzhongdai
Dickeya gs fangzhongdai S1 WP 049855115 1 Dfangzhongdai
Dickeya gs fangzhongdai MK7 WP 071605210 1 Dfangzhongdai
Dickeya gs fangzhongdai B16 WP 038919429 1 Dfangzhongdai
Dickeya zeae CSL RW192 WP 071603259 1 Dzeae
Dickeya zeae NCPPB 3531 WP 071603361 1 Dzeae
Dickeya zeae MS1 WP 026357909 1 Dzeae
Dickeya zeae NCPPB 2538 WP 071603568 1 Dzeae
Dickeya zeae MK19 WP 071603819 1 Dzeae
Dickeya zeae NCPPB 3532 WP 023639360 1 Dzeae
297 Dd586 1365 Other secondary metabolite biosynthesis Cluster4 Seed
Dickeya zeae Ech586 WP 012884078 1 Dzeae

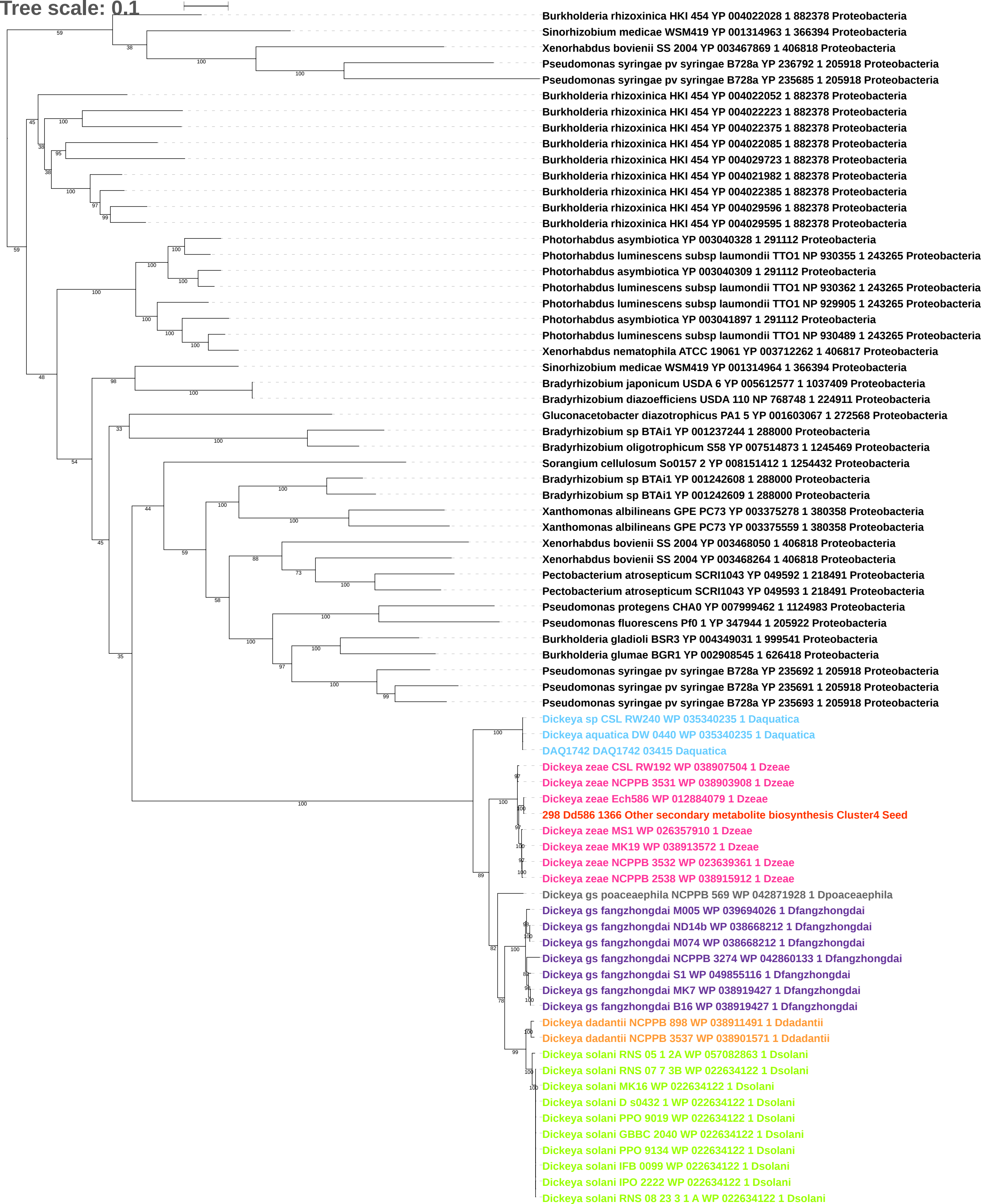
Tree scale: 0.1
Burkholderia rhizoxinica HKI 454 YP 004022028 1 882378 Proteobacteria
Sinorhizobium medicae WSM419 YP 001314963 1 366394 Proteobacteria
Xenorhabdus bovienii SS 2004 YP 003467869 1 406818 Proteobacteria
Pseudomonas syringae pv syringae B728a YP 236792 1 205918 Proteobacteria
Pseudomonas syringae pv syringae B728a YP 235685 1 205918 Proteobacteria
Burkholderia rhizoxinica HKI 454 YP 004022052 1 882378 Proteobacteria
Burkholderia rhizoxinica HKI 454 YP 004022223 1 882378 Proteobacteria
Burkholderia rhizoxinica HKI 454 YP 004022375 1 882378 Proteobacteria
Burkholderia rhizoxinica HKI 454 YP 004022085 1 882378 Proteobacteria
Burkholderia rhizoxinica HKI 454 YP 004029723 1 882378 Proteobacteria
Burkholderia rhizoxinica HKI 454 YP 004021982 1 882378 Proteobacteria
Burkholderia rhizoxinica HKI 454 YP 004022385 1 882378 Proteobacteria
Burkholderia rhizoxinica HKI 454 YP 004029596 1 882378 Proteobacteria
Burkholderia rhizoxinica HKI 454 YP 004029595 1 882378 Proteobacteria
Photorhabdus asymbiotica YP 003040328 1 291112 Proteobacteria
Photorhabdus luminescens subsp laumondii TTO1 NP 930355 1 243265 Proteobacteria
Photorhabdus asymbiotica YP 003040309 1 291112 Proteobacteria
Photorhabdus luminescens subsp laumondii TTO1 NP 930362 1 243265 Proteobacteria
Photorhabdus luminescens subsp laumondii TTO1 NP 929905 1 243265 Proteobacteria
Photorhabdus asymbiotica YP 003041897 1 291112 Proteobacteria
Photorhabdus luminescens subsp laumondii TTO1 NP 930489 1 243265 Proteobacteria
Xenorhabdus nematophila ATCC 19061 YP 003712262 1 406817 Proteobacteria
Sinorhizobium medicae WSM419 YP 001314964 1 366394 Proteobacteria
Bradyrhizobium japonicum USDA 6 YP 005612577 1 1037409 Proteobacteria
Bradyrhizobium diazoefficiens USDA 110 NP 768748 1 224911 Proteobacteria
Gluconacetobacter diazotrophicus PA1 5 YP 001603067 1 272568 Proteobacteria
Bradyrhizobium sp BTAi1 YP 001237244 1 288000 Proteobacteria
Bradyrhizobium oligotrophicum S58 YP 007514873 1 1245469 Proteobacteria
Sorangium cellulosum So0157 2 YP 008151412 1 1254432 Proteobacteria
Bradyrhizobium sp BTAi1 YP 001242608 1 288000 Proteobacteria
Bradyrhizobium sp BTAi1 YP 001242609 1 288000 Proteobacteria
Xanthomonas albilineans GPE PC73 YP 003375278 1 380358 Proteobacteria
Xanthomonas albilineans GPE PC73 YP 003375559 1 380358 Proteobacteria
Xenorhabdus bovienii SS 2004 YP 003468050 1 406818 Proteobacteria
Xenorhabdus bovienii SS 2004 YP 003468264 1 406818 Proteobacteria
Pectobacterium atrosepticum SCRI1043 YP 049592 1 218491 Proteobacteria
Pectobacterium atrosepticum SCRI1043 YP 049593 1 218491 Proteobacteria
Pseudomonas protegens CHA0 YP 007999462 1 1124983 Proteobacteria
Pseudomonas fluorescens Pf0 1 YP 347944 1 205922 Proteobacteria
Burkholderia gladioli BSR3 YP 004349031 1 999541 Proteobacteria
Burkholderia glumae BGR1 YP 002908545 1 626418 Proteobacteria
Pseudomonas syringae pv syringae B728a YP 235692 1 205918 Proteobacteria
Pseudomonas syringae pv syringae B728a YP 235691 1 205918 Proteobacteria
Pseudomonas syringae pv syringae B728a YP 235693 1 205918 Proteobacteria
Dickeya sp CSL RW240 WP 035340235 1 Daquatica
Dickeya aquatica DW 0440 WP 035340235 1 Daquatica
DAQ1742 DAQ1742 03415 Daquatica
Dickeya zeae CSL RW192 WP 038907504 1 Dzeae
Dickeya zeae NCPPB 3531 WP 038903908 1 Dzeae
Dickeya zeae Ech586 WP 012884079 1 Dzeae
298 Dd586 1366 Other secondary metabolite biosynthesis Cluster4 Seed
Dickeya zeae MS1 WP 026357910 1 Dzeae
Dickeya zeae MK19 WP 038913572 1 Dzeae
Dickeya zeae NCPPB 3532 WP 023639361 1 Dzeae
Dickeya zeae NCPPB 2538 WP 038915912 1 Dzeae
Dickeya gs poaceaephila NCPPB 569 WP 042871928 1 Dpoaceaephila
Dickeya gs fangzhongdai M005 WP 039694026 1 Dfangzhongdai
Dickeya gs fangzhongdai ND14b WP 038668212 1 Dfangzhongdai
Dickeya gs fangzhongdai M074 WP 038668212 1 Dfangzhongdai
Dickeya gs fangzhongdai NCPPB 3274 WP 042860133 1 Dfangzhongdai
Dickeya gs fangzhongdai S1 WP 049855116 1 Dfangzhongdai
Dickeya gs fangzhongdai MK7 WP 038919427 1 Dfangzhongdai
Dickeya gs fangzhongdai B16 WP 038919427 1 Dfangzhongdai
Dickeya dadantii NCPPB 898 WP 038911491 1 Ddadantii
Dickeya dadantii NCPPB 3537 WP 038901571 1 Ddadantii
Dickeya solani RNS 05 1 2A WP 057082863 1 Dsolani
Dickeya solani RNS 07 7 3B WP 022634122 1 Dsolani
Dickeya solani MK16 WP 022634122 1 Dsolani
Dickeya solani D s0432 1 WP 022634122 1 Dsolani
Dickeya solani PPO 9019 WP 022634122 1 Dsolani
Dickeya solani GBBC 2040 WP 022634122 1 Dsolani
Dickeya solani PPO 9134 WP 022634122 1 Dsolani
Dickeya solani IFB 0099 WP 022634122 1 Dsolani
Dickeya solani IPO 2222 WP 022634122 1 Dsolani
Dickeya solani RNS 08 23 3 1 A WP 022634122 1 Dsolani

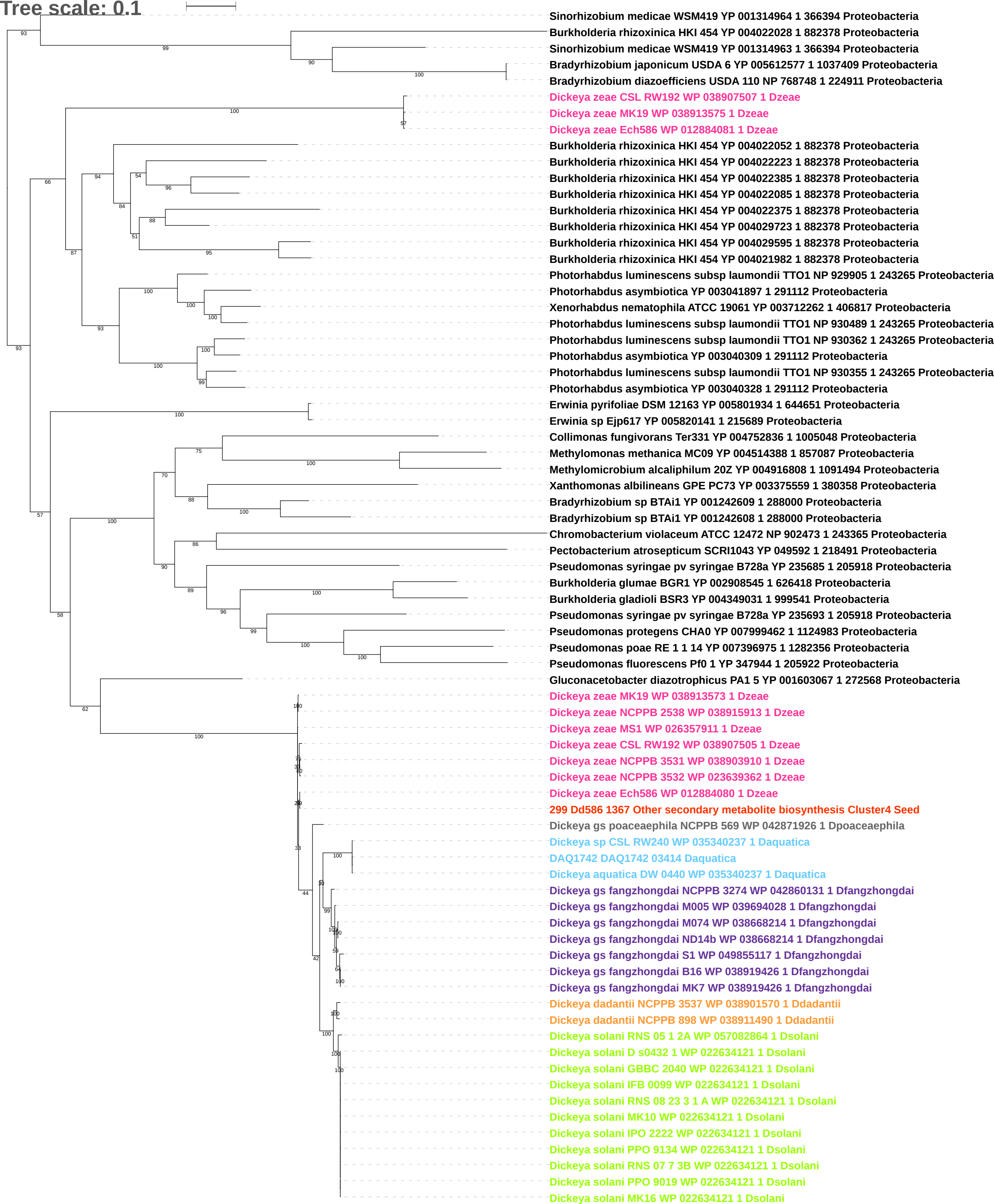
Tree scale: 0.1
Sinorhizobium medicae WSM419 YP 001314964 1 366394 Proteobacteria
Burkholderia rhizoxinica HKI 454 YP 004022028 1 882378 Proteobacteria
Sinorhizobium medicae WSM419 YP 001314963 1 366394 Proteobacteria
Bradyrhizobium japonicum USDA 6 YP 005612577 1 1037409 Proteobacteria
Bradyrhizobium diazoefficiens USDA 110 NP 768748 1 224911 Proteobacteria
Dickeya zeae CSL RW192 WP 038907507 1 Dzeae
Dickeya zeae MK19 WP 038913575 1 Dzeae
Dickeya zeae Ech586 WP 012884081 1 Dzeae
Burkholderia rhizoxinica HKI 454 YP 004022052 1 882378 Proteobacteria
Burkholderia rhizoxinica HKI 454 YP 004022223 1 882378 Proteobacteria
Burkholderia rhizoxinica HKI 454 YP 004022385 1 882378 Proteobacteria
Burkholderia rhizoxinica HKI 454 YP 004022085 1 882378 Proteobacteria
Burkholderia rhizoxinica HKI 454 YP 004022375 1 882378 Proteobacteria
Burkholderia rhizoxinica HKI 454 YP 004029723 1 882378 Proteobacteria
Burkholderia rhizoxinica HKI 454 YP 004029595 1 882378 Proteobacteria
Burkholderia rhizoxinica HKI 454 YP 004021982 1 882378 Proteobacteria
Photorhabdus luminescens subsp laumondii TTO1 NP 929905 1 243265 Proteobacteria
Photorhabdus asymbiotica YP 003041897 1 291112 Proteobacteria
Xenorhabdus nematophila ATCC 19061 YP 003712262 1 406817 Proteobacteria
Photorhabdus luminescens subsp laumondii TTO1 NP 930489 1 243265 Proteobacteria
Photorhabdus luminescens subsp laumondii TTO1 NP 930362 1 243265 Proteobacteria
Photorhabdus asymbiotica YP 003040309 1 291112 Proteobacteria
Photorhabdus luminescens subsp laumondii TTO1 NP 930355 1 243265 Proteobacteria
Photorhabdus asymbiotica YP 003040328 1 291112 Proteobacteria
Erwinia pyrifoliae DSM 12163 YP 005801934 1 644651 Proteobacteria
Erwinia sp Ejp617 YP 005820141 1 215689 Proteobacteria
Collimonas fungivorans Ter331 YP 004752836 1 1005048 Proteobacteria
Methylomonas methanica MC09 YP 004514388 1 857087 Proteobacteria
Methylomicrobium alcaliphilum 20Z YP 004916808 1 1091494 Proteobacteria
Xanthomonas albilineans GPE PC73 YP 003375559 1 380358 Proteobacteria
Bradyrhizobium sp BTAi1 YP 001242609 1 288000 Proteobacteria
Bradyrhizobium sp BTAi1 YP 001242608 1 288000 Proteobacteria
Chromobacterium violaceum ATCC 12472 NP 902473 1 243365 Proteobacteria
Pectobacterium atrosepticum SCRI1043 YP 049592 1 218491 Proteobacteria
Pseudomonas syringae pv syringae B728a YP 235685 1 205918 Proteobacteria
Burkholderia glumae BGR1 YP 002908545 1 626418 Proteobacteria
Burkholderia gladioli BSR3 YP 004349031 1 999541 Proteobacteria
Pseudomonas syringae pv syringae B728a YP 235693 1 205918 Proteobacteria
Pseudomonas protegens CHA0 YP 007999462 1 1124983 Proteobacteria
Pseudomonas poae RE 1 1 14 YP 007396975 1 1282356 Proteobacteria
Pseudomonas fluorescens Pf0 1 YP 347944 1 205922 Proteobacteria
Gluconacetobacter diazotrophicus PA1 5 YP 001603067 1 272568 Proteobacteria
Dickeya zeae MK19 WP 038913573 1 Dzeae
Dickeya zeae NCPPB 2538 WP 038915913 1 Dzeae
Dickeya zeae MS1 WP 026357911 1 Dzeae
Dickeya zeae CSL RW192 WP 038907505 1 Dzeae
Dickeya zeae NCPPB 3531 WP 038903910 1 Dzeae
Dickeya zeae NCPPB 3532 WP 023639362 1 Dzeae
Dickeya zeae Ech586 WP 012884080 1 Dzeae
299 Dd586 1367 Other secondary metabolite biosynthesis Cluster4 Seed
Dickeya gs poaceaephila NCPPB 569 WP 042871926 1 Dpoaceaephila
Dickeya sp CSL RW240 WP 035340237 1 Daquatica
DAQ1742 DAQ1742 03414 Daquatica
Dickeya aquatica DW 0440 WP 035340237 1 Daquatica
Dickeya gs fangzhongdai NCPPB 3274 WP 042860131 1 Dfangzhongdai
Dickeya gs fangzhongdai M005 WP 039694028 1 Dfangzhongdai
Dickeya gs fangzhongdai M074 WP 038668214 1 Dfangzhongdai
Dickeya gs fangzhongdai ND14b WP 038668214 1 Dfangzhongdai
Dickeya gs fangzhongdai S1 WP 049855117 1 Dfangzhongdai
Dickeya gs fangzhongdai B16 WP 038919426 1 Dfangzhongdai
Dickeya gs fangzhongdai MK7 WP 038919426 1 Dfangzhongdai
Dickeya dadantii NCPPB 3537 WP 038901570 1 Ddadantii
Dickeya dadantii NCPPB 898 WP 038911490 1 Ddadantii
Dickeya solani RNS 05 1 2A WP 057082864 1 Dsolani
Dickeya solani D s0432 1 WP 022634121 1 Dsolani
Dickeya solani GBBC 2040 WP 022634121 1 Dsolani
Dickeya solani IFB 0099 WP 022634121 1 Dsolani
Dickeya solani RNS 08 23 3 1 A WP 022634121 1 Dsolani
Dickeya solani MK10 WP 022634121 1 Dsolani
Dickeya solani IPO 2222 WP 022634121 1 Dsolani
Dickeya solani PPO 9134 WP 022634121 1 Dsolani
Dickeya solani RNS 07 7 3B WP 022634121 1 Dsolani
Dickeya solani PPO 9019 WP 022634121 1 Dsolani
Dickeya solani MK16 WP 022634121 1 Dsolani

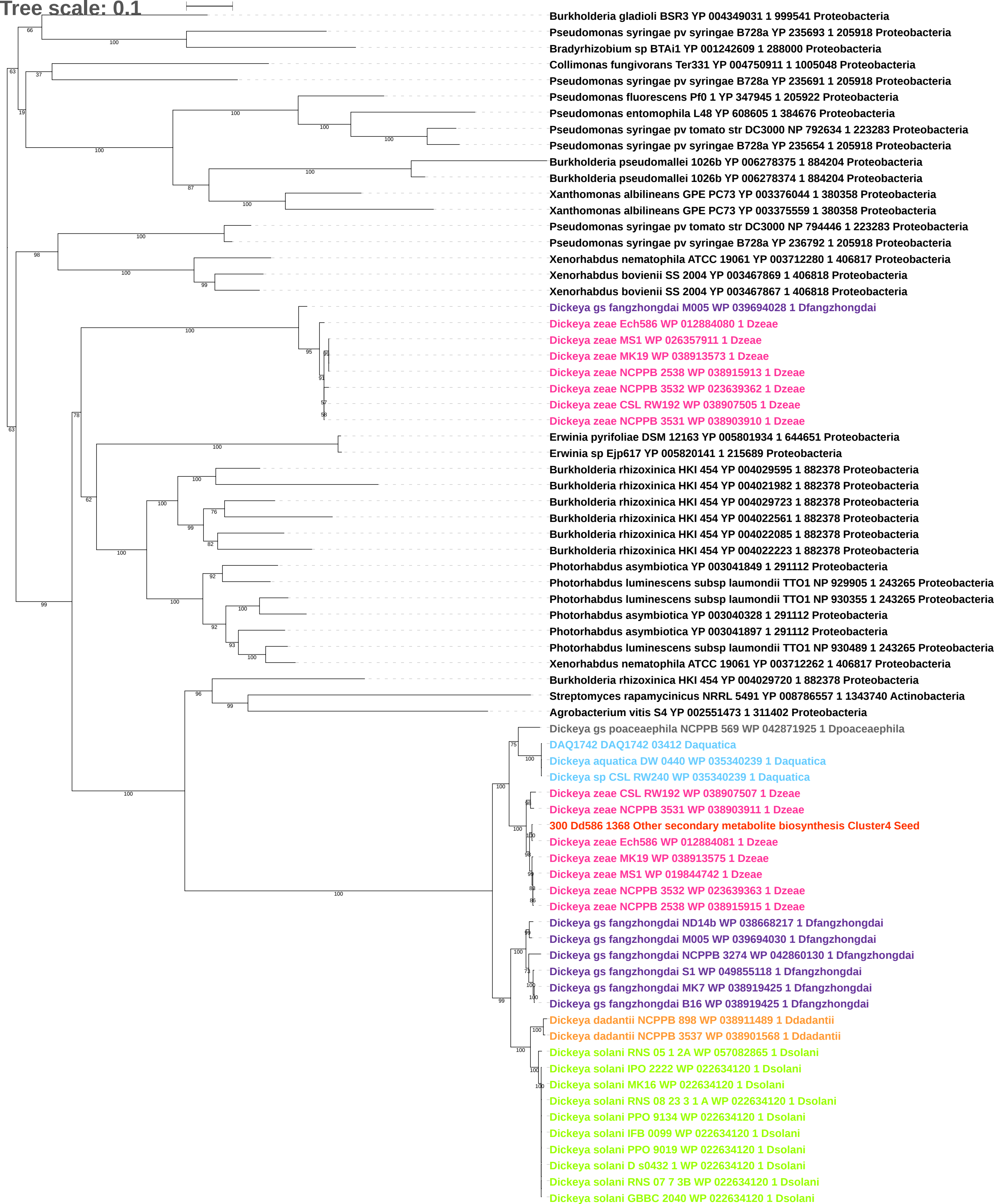

Tree scale: 0.1
Burkholderia gladioli BSR3 YP 004349031 1 999541 Proteobacteria
Pseudomonas syringae pv syringae B728a YP 235693 1 205918 Proteobacteria
Bradyrhizobium sp BTAi1 YP 001242609 1 288000 Proteobacteria
Collimonas fungivorans Ter331 YP 004750911 1 1005048 Proteobacteria
Pseudomonas syringae pv syringae B728a YP 235691 1 205918 Proteobacteria
Pseudomonas fluorescens Pf0 1 YP 347945 1 205922 Proteobacteria
Pseudomonas entomophila L48 YP 608605 1 384676 Proteobacteria
Pseudomonas syringae pv tomato str DC3000 NP 792634 1 223283 Proteobacteria
Pseudomonas syringae pv syringae B728a YP 235654 1 205918 Proteobacteria
Burkholderia pseudomallei 1026b YP 006278375 1 884204 Proteobacteria
Burkholderia pseudomallei 1026b YP 006278374 1 884204 Proteobacteria
Xanthomonas albilineans GPE PC73 YP 003376044 1 380358 Proteobacteria
Xanthomonas albilineans GPE PC73 YP 003375559 1 380358 Proteobacteria
Pseudomonas syringae pv tomato str DC3000 NP 794446 1 223283 Proteobacteria
Pseudomonas syringae pv syringae B728a YP 236792 1 205918 Proteobacteria
Xenorhabdus nematophila ATCC 19061 YP 003712280 1 406817 Proteobacteria
Xenorhabdus bovienii SS 2004 YP 003467869 1 406818 Proteobacteria
Xenorhabdus bovienii SS 2004 YP 003467867 1 406818 Proteobacteria
Dickeya gs fangzhongdai M005 WP 039694028 1 Dfangzhongdai
Dickeya zeae Ech586 WP 012884080 1 Dzeae
Dickeya zeae MS1 WP 026357911 1 Dzeae
Dickeya zeae MK19 WP 038913573 1 Dzeae
Dickeya zeae NCPPB 2538 WP 038915913 1 Dzeae
Dickeya zeae NCPPB 3532 WP 023639362 1 Dzeae
Dickeya zeae CSL RW192 WP 038907505 1 Dzeae
Dickeya zeae NCPPB 3531 WP 038903910 1 Dzeae
Erwinia pyrifoliae DSM 12163 YP 005801934 1 644651 Proteobacteria
Erwinia sp Ejp617 YP 005820141 1 215689 Proteobacteria
Burkholderia rhizoxinica HKI 454 YP 004029595 1 882378 Proteobacteria
Burkholderia rhizoxinica HKI 454 YP 004021982 1 882378 Proteobacteria
Burkholderia rhizoxinica HKI 454 YP 004029723 1 882378 Proteobacteria
Burkholderia rhizoxinica HKI 454 YP 004022561 1 882378 Proteobacteria
Burkholderia rhizoxinica HKI 454 YP 004022085 1 882378 Proteobacteria
Burkholderia rhizoxinica HKI 454 YP 004022223 1 882378 Proteobacteria
Photorhabdus asymbiotica YP 003041849 1 291112 Proteobacteria
Photorhabdus luminescens subsp laumondii TTO1 NP 929905 1 243265 Proteobacteria
Photorhabdus luminescens subsp laumondii TTO1 NP 930355 1 243265 Proteobacteria
Photorhabdus asymbiotica YP 003040328 1 291112 Proteobacteria
Photorhabdus asymbiotica YP 003041897 1 291112 Proteobacteria
Photorhabdus luminescens subsp laumondii TTO1 NP 930489 1 243265 Proteobacteria
Xenorhabdus nematophila ATCC 19061 YP 003712262 1 406817 Proteobacteria
Burkholderia rhizoxinica HKI 454 YP 004029720 1 882378 Proteobacteria
Streptomyces rapamycinicus NRRL 5491 YP 008786557 1 1343740 Actinobacteria
Agrobacterium vitis S4 YP 002551473 1 311402 Proteobacteria
Dickeya gs poaceaephila NCPPB 569 WP 042871925 1 Dpoaceaephila
DAQ1742 DAQ1742 03412 Daquatica
Dickeya aquatica DW 0440 WP 035340239 1 Daquatica
Dickeya sp CSL RW240 WP 035340239 1 Daquatica
Dickeya zeae CSL RW192 WP 038907507 1 Dzeae
Dickeya zeae NCPPB 3531 WP 038903911 1 Dzeae
300 Dd586 1368 Other secondary metabolite biosynthesis Cluster4 Seed
Dickeya zeae Ech586 WP 012884081 1 Dzeae
Dickeya zeae MK19 WP 038913575 1 Dzeae
Dickeya zeae MS1 WP 019844742 1 Dzeae
Dickeya zeae NCPPB 3532 WP 023639363 1 Dzeae
Dickeya zeae NCPPB 2538 WP 038915915 1 Dzeae
Dickeya gs fangzhongdai ND14b WP 038668217 1 Dfangzhongdai
Dickeya gs fangzhongdai M005 WP 039694030 1 Dfangzhongdai
Dickeya gs fangzhongdai NCPPB 3274 WP 042860130 1 Dfangzhongdai
Dickeya gs fangzhongdai S1 WP 049855118 1 Dfangzhongdai
Dickeya gs fangzhongdai MK7 WP 038919425 1 Dfangzhongdai
Dickeya gs fangzhongdai B16 WP 038919425 1 Dfangzhongdai
Dickeya dadantii NCPPB 898 WP 038911489 1 Ddadantii
Dickeya dadantii NCPPB 3537 WP 038901568 1 Ddadantii
Dickeya solani RNS 05 1 2A WP 057082865 1 Dsolani
Dickeya solani IPO 2222 WP 022634120 1 Dsolani
Dickeya solani MK16 WP 022634120 1 Dsolani
Dickeya solani RNS 08 23 3 1 A WP 022634120 1 Dsolani
Dickeya solani PPO 9134 WP 022634120 1 Dsolani
Dickeya solani IFB 0099 WP 022634120 1 Dsolani
Dickeya solani PPO 9019 WP 022634120 1 Dsolani
Dickeya solani D s0432 1 WP 022634120 1 Dsolani
Dickeya solani RNS 07 7 3B WP 022634120 1 Dsolani
Dickeya solani GBBC 2040 WP 022634120 1 Dsolani

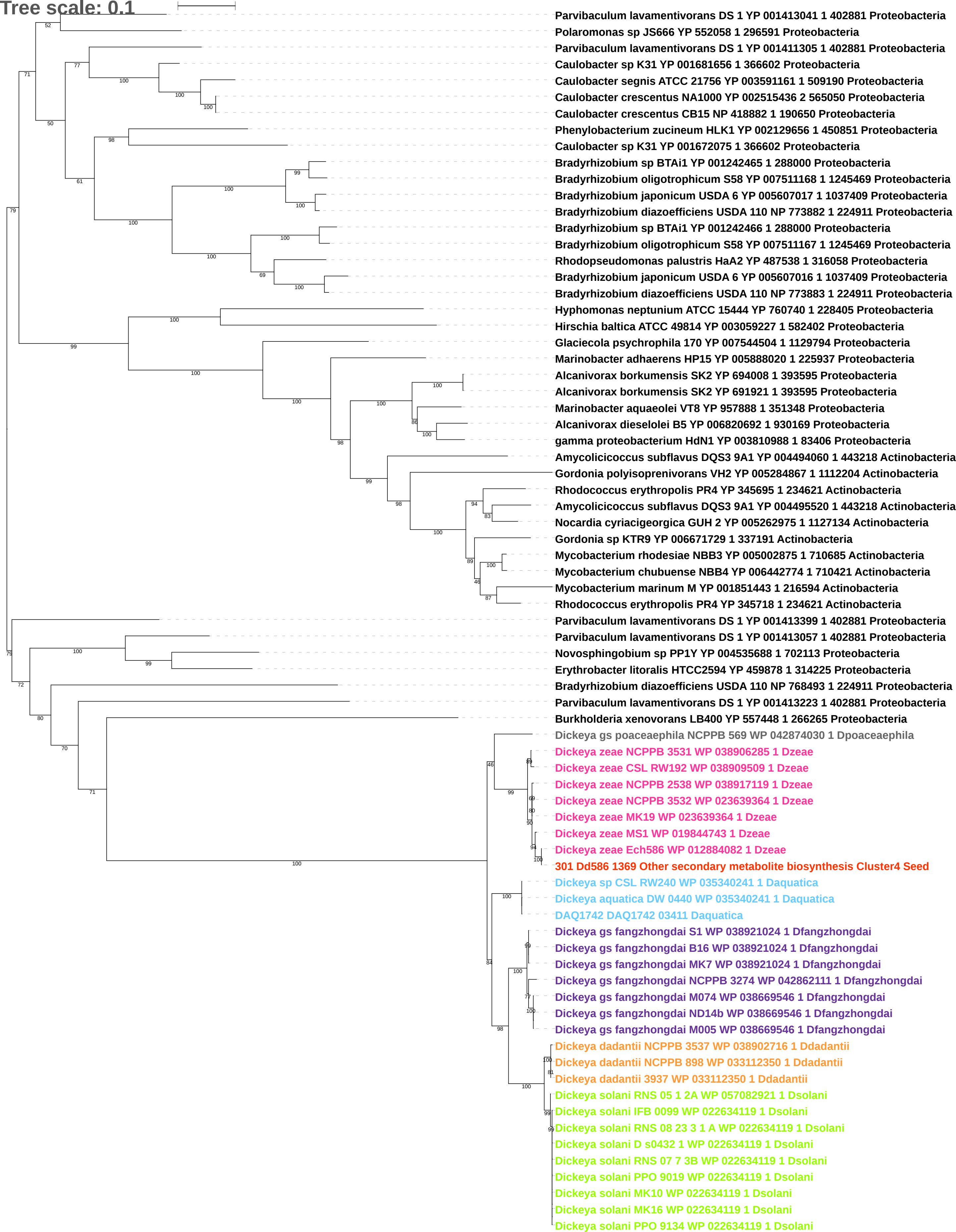
Tree scale: 0.1
Parvibaculum lavamentivorans DS 1 YP 001413041 1 402881 Proteobacteria
Polaromonas sp JS666 YP 552058 1 296591 Proteobacteria
Parvibaculum lavamentivorans DS 1 YP 001411305 1 402881 Proteobacteria
Caulobacter sp K31 YP 001681656 1 366602 Proteobacteria
Caulobacter segnis ATCC 21756 YP 003591161 1 509190 Proteobacteria
Caulobacter crescentus NA1000 YP 002515436 2 565050 Proteobacteria
Caulobacter crescentus CB15 NP 418882 1 190650 Proteobacteria
Phenylobacterium zucineum HLK1 YP 002129656 1 450851 Proteobacteria
Caulobacter sp K31 YP 001672075 1 366602 Proteobacteria
Bradyrhizobium sp BTAi1 YP 001242465 1 288000 Proteobacteria
Bradyrhizobium oligotrophicum S58 YP 007511168 1 1245469 Proteobacteria
Bradyrhizobium japonicum USDA 6 YP 005607017 1 1037409 Proteobacteria
Bradyrhizobium diazoefficiens USDA 110 NP 773882 1 224911 Proteobacteria
Bradyrhizobium sp BTAi1 YP 001242466 1 288000 Proteobacteria
Bradyrhizobium oligotrophicum S58 YP 007511167 1 1245469 Proteobacteria
Rhodopseudomonas palustris HaA2 YP 487538 1 316058 Proteobacteria
Bradyrhizobium japonicum USDA 6 YP 005607016 1 1037409 Proteobacteria
Bradyrhizobium diazoefficiens USDA 110 NP 773883 1 224911 Proteobacteria
Hyphomonas neptunium ATCC 15444 YP 760740 1 228405 Proteobacteria
Hirschia baltica ATCC 49814 YP 003059227 1 582402 Proteobacteria
Glaciecola psychrophila 170 YP 007544504 1 1129794 Proteobacteria
Marinobacter adhaerens HP15 YP 005888020 1 225937 Proteobacteria
Alcanivorax borkumensis SK2 YP 694008 1 393595 Proteobacteria
Alcanivorax borkumensis SK2 YP 691921 1 393595 Proteobacteria
Marinobacter aquaeolei VT8 YP 957888 1 351348 Proteobacteria
Alcanivorax dieselolei B5 YP 006820692 1 930169 Proteobacteria
gamma proteobacterium HdN1 YP 003810988 1 83406 Proteobacteria
Amycolicicoccus subflavus DQS3 9A1 YP 004494060 1 443218 Actinobacteria
Gordonia polyisoprenivorans VH2 YP 005284867 1 1112204 Actinobacteria
Rhodococcus erythropolis PR4 YP 345695 1 234621 Actinobacteria
Amycolicicoccus subflavus DQS3 9A1 YP 004495520 1 443218 Actinobacteria
Nocardia cyriacigeorgica GUH 2 YP 005262975 1 1127134 Actinobacteria
Gordonia sp KTR9 YP 006671729 1 337191 Actinobacteria
Mycobacterium rhodesiae NBB3 YP 005002875 1 710685 Actinobacteria
Mycobacterium chubuense NBB4 YP 006442774 1 710421 Actinobacteria
Mycobacterium marinum M YP 001851443 1 216594 Actinobacteria
Rhodococcus erythropolis PR4 YP 345718 1 234621 Actinobacteria
Parvibaculum lavamentivorans DS 1 YP 001413399 1 402881 Proteobacteria
Parvibaculum lavamentivorans DS 1 YP 001413057 1 402881 Proteobacteria
Novosphingobium sp PP1Y YP 004535688 1 702113 Proteobacteria
Erythrobacter litoralis HTCC2594 YP 459878 1 314225 Proteobacteria
Bradyrhizobium diazoefficiens USDA 110 NP 768493 1 224911 Proteobacteria
Parvibaculum lavamentivorans DS 1 YP 001413223 1 402881 Proteobacteria
Burkholderia xenovorans LB400 YP 557448 1 266265 Proteobacteria
Dickeya gs poaceaephila NCPPB 569 WP 042874030 1 Dpoaceaephila
Dickeya zeae NCPPB 3531 WP 038906285 1 Dzeae
Dickeya zeae CSL RW192 WP 038909509 1 Dzeae
Dickeya zeae NCPPB 2538 WP 038917119 1 Dzeae
Dickeya zeae NCPPB 3532 WP 023639364 1 Dzeae
Dickeya zeae MK19 WP 023639364 1 Dzeae
Dickeya zeae MS1 WP 019844743 1 Dzeae
Dickeya zeae Ech586 WP 012884082 1 Dzeae
301 Dd586 1369 Other secondary metabolite biosynthesis Cluster4 Seed
Dickeya sp CSL RW240 WP 035340241 1 Daquatica
Dickeya aquatica DW 0440 WP 035340241 1 Daquatica
DAQ1742 DAQ1742 03411 Daquatica
Dickeya gs fangzhongdai S1 WP 038921024 1 Dfangzhongdai
Dickeya gs fangzhongdai B16 WP 038921024 1 Dfangzhongdai
Dickeya gs fangzhongdai MK7 WP 038921024 1 Dfangzhongdai
Dickeya gs fangzhongdai NCPPB 3274 WP 042862111 1 Dfangzhongdai
Dickeya gs fangzhongdai M074 WP 038669546 1 Dfangzhongdai
Dickeya gs fangzhongdai ND14b WP 038669546 1 Dfangzhongdai
Dickeya gs fangzhongdai M005 WP 038669546 1 Dfangzhongdai
Dickeya dadantii NCPPB 3537 WP 038902716 1 Ddadantii
Dickeya dadantii NCPPB 898 WP 033112350 1 Ddadantii
Dickeya dadantii 3937 WP 033112350 1 Ddadantii
Dickeya solani RNS 05 1 2A WP 057082921 1 Dsolani
Dickeya solani IFB 0099 WP 022634119 1 Dsolani
Dickeya solani RNS 08 23 3 1 A WP 022634119 1 Dsolani
Dickeya solani D s0432 1 WP 022634119 1 Dsolani
Dickeya solani RNS 07 7 3B WP 022634119 1 Dsolani
Dickeya solani PPO 9019 WP 022634119 1 Dsolani
Dickeya solani MK10 WP 022634119 1 Dsolani
Dickeya solani MK16 WP 022634119 1 Dsolani
Dickeya solani PPO 9134 WP 022634119 1 Dsolani

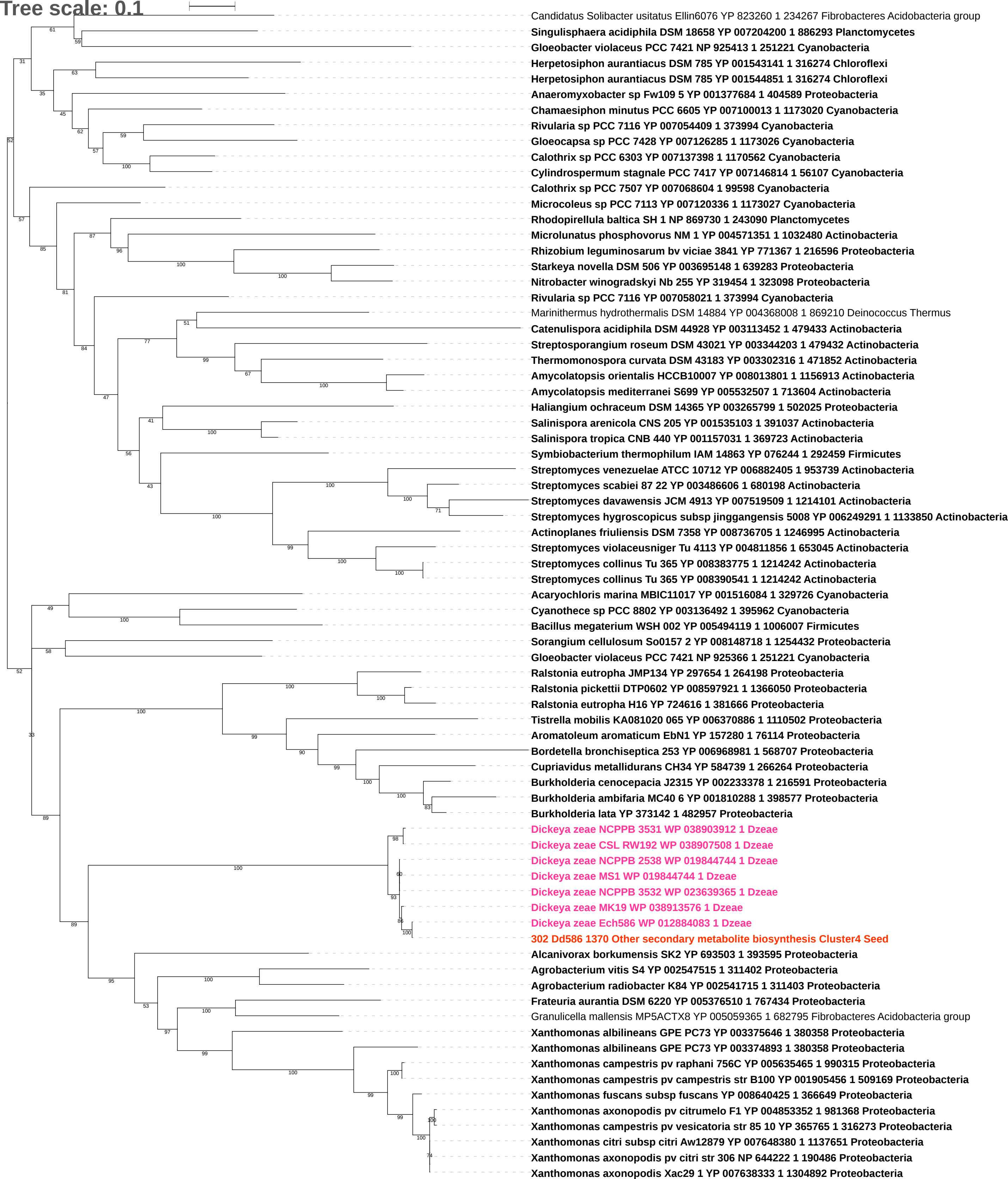
Tree scale: 0.1
Candidatus Solibacter usitatus Ellin6076 YP 823260 1 234267 Fibrobacteres Acidobacteria group
Singulisphaera acidiphila DSM 18658 YP 007204200 1 886293 Planctomycetes
Gloeobacter violaceus PCC 7421 NP 925413 1 251221 Cyanobacteria
Herpetosiphon aurantiacus DSM 785 YP 001543141 1 316274 Chloroflexi
Herpetosiphon aurantiacus DSM 785 YP 001544851 1 316274 Chloroflexi
Anaeromyxobacter sp Fw109 5 YP 001377684 1 404589 Proteobacteria
Chamaesiphon minutus PCC 6605 YP 007100013 1 1173020 Cyanobacteria
Rivularia sp PCC 7116 YP 007054409 1 373994 Cyanobacteria
Gloeocapsa sp PCC 7428 YP 007126285 1 1173026 Cyanobacteria
Calothrix sp PCC 6303 YP 007137398 1 1170562 Cyanobacteria
Cylindrospermum stagnale PCC 7417 YP 007146814 1 56107 Cyanobacteria
Calothrix sp PCC 7507 YP 007068604 1 99598 Cyanobacteria
Microcoleus sp PCC 7113 YP 007120336 1 1173027 Cyanobacteria
Rhodopirellula baltica SH 1 NP 869730 1 243090 Planctomycetes
Microlunatus phosphovorus NM 1 YP 004571351 1 1032480 Actinobacteria
Rhizobium leguminosarum bv viciae 3841 YP 771367 1 216596 Proteobacteria
Starkeya novella DSM 506 YP 003695148 1 639283 Proteobacteria
Nitrobacter winogradskyi Nb 255 YP 319454 1 323098 Proteobacteria
Rivularia sp PCC 7116 YP 007058021 1 373994 Cyanobacteria
Marinithermus hydrothermalis DSM 14884 YP 004368008 1 869210 Deinococcus Thermus
Catenulispora acidiphila DSM 44928 YP 003113452 1 479433 Actinobacteria
Streptosporangium roseum DSM 43021 YP 003344203 1 479432 Actinobacteria
Thermomonospora curvata DSM 43183 YP 003302316 1 471852 Actinobacteria
Amycolatopsis orientalis HCCB10007 YP 008013801 1 1156913 Actinobacteria
Amycolatopsis mediterranei S699 YP 005532507 1 713604 Actinobacteria
Haliangium ochraceum DSM 14365 YP 003265799 1 502025 Proteobacteria
Salinispora arenicola CNS 205 YP 001535103 1 391037 Actinobacteria
Salinispora tropica CNB 440 YP 001157031 1 369723 Actinobacteria
Symbiobacterium thermophilum IAM 14863 YP 076244 1 292459 Firmicutes
Streptomyces venezuelae ATCC 10712 YP 006882405 1 953739 Actinobacteria
Streptomyces scabiei 87 22 YP 003486606 1 680198 Actinobacteria
Streptomyces davawensis JCM 4913 YP 007519509 1 1214101 Actinobacteria
Streptomyces hygroscopicus subsp jinggangensis 5008 YP 006249291 1 1133850 Actinobacteria
Actinoplanes friuliensis DSM 7358 YP 008736705 1 1246995 Actinobacteria
Streptomyces violaceusniger Tu 4113 YP 004811856 1 653045 Actinobacteria
Streptomyces collinus Tu 365 YP 008383775 1 1214242 Actinobacteria
Streptomyces collinus Tu 365 YP 008390541 1 1214242 Actinobacteria
Acaryochloris marina MBIC11017 YP 001516084 1 329726 Cyanobacteria
Cyanothece sp PCC 8802 YP 003136492 1 395962 Cyanobacteria
Bacillus megaterium WSH 002 YP 005494119 1 1006007 Firmicutes
Sorangium cellulosum So0157 2 YP 008148718 1 1254432 Proteobacteria
Gloeobacter violaceus PCC 7421 NP 925366 1 251221 Cyanobacteria
Ralstonia eutropha JMP134 YP 297654 1 264198 Proteobacteria
Ralstonia pickettii DTP0602 YP 008597921 1 1366050 Proteobacteria
Ralstonia eutropha H16 YP 724616 1 381666 Proteobacteria
Tistrella mobilis KA081020 065 YP 006370886 1 1110502 Proteobacteria
Aromatoleum aromaticum EbN1 YP 157280 1 76114 Proteobacteria
Bordetella bronchiseptica 253 YP 006968981 1 568707 Proteobacteria
Cupriavidus metallidurans CH34 YP 584739 1 266264 Proteobacteria
Burkholderia cenocepacia J2315 YP 002233378 1 216591 Proteobacteria
Burkholderia ambifaria MC40 6 YP 001810288 1 398577 Proteobacteria
Burkholderia lata YP 373142 1 482957 Proteobacteria
Dickeya zeae NCPPB 3531 WP 038903912 1 Dzeae
Dickeya zeae CSL RW192 WP 038907508 1 Dzeae
Dickeya zeae NCPPB 2538 WP 019844744 1 Dzeae
Dickeya zeae MS1 WP 019844744 1 Dzeae
Dickeya zeae NCPPB 3532 WP 023639365 1 Dzeae
Dickeya zeae MK19 WP 038913576 1 Dzeae
Dickeya zeae Ech586 WP 012884083 1 Dzeae
302 Dd586 1370 Other secondary metabolite biosynthesis Cluster4 Seed
Alcanivorax borkumensis SK2 YP 693503 1 393595 Proteobacteria
Agrobacterium vitis S4 YP 002547515 1 311402 Proteobacteria
Agrobacterium radiobacter K84 YP 002541715 1 311403 Proteobacteria
Frateuria aurantia DSM 6220 YP 005376510 1 767434 Proteobacteria
Granulicella mallensis MP5ACTX8 YP 005059365 1 682795 Fibrobacteres Acidobacteria group
Xanthomonas albilineans GPE PC73 YP 003375646 1 380358 Proteobacteria
Xanthomonas albilineans GPE PC73 YP 003374893 1 380358 Proteobacteria
Xanthomonas campestris pv raphani 756C YP 005635465 1 990315 Proteobacteria
Xanthomonas campestris pv campestris str B100 YP 001905456 1 509169 Proteobacteria
Xanthomonas fuscans subsp fuscans YP 008640425 1 366649 Proteobacteria
Xanthomonas axonopodis pv citrumelo F1 YP 004853352 1 981368 Proteobacteria
Xanthomonas campestris pv vesicatoria str 85 10 YP 365765 1 316273 Proteobacteria
Xanthomonas citri subsp citri Aw12879 YP 007648380 1 1137651 Proteobacteria
Xanthomonas axonopodis pv citri str 306 NP 644222 1 190486 Proteobacteria
Xanthomonas axonopodis Xac29 1 YP 007638333 1 1304892 Proteobacteria

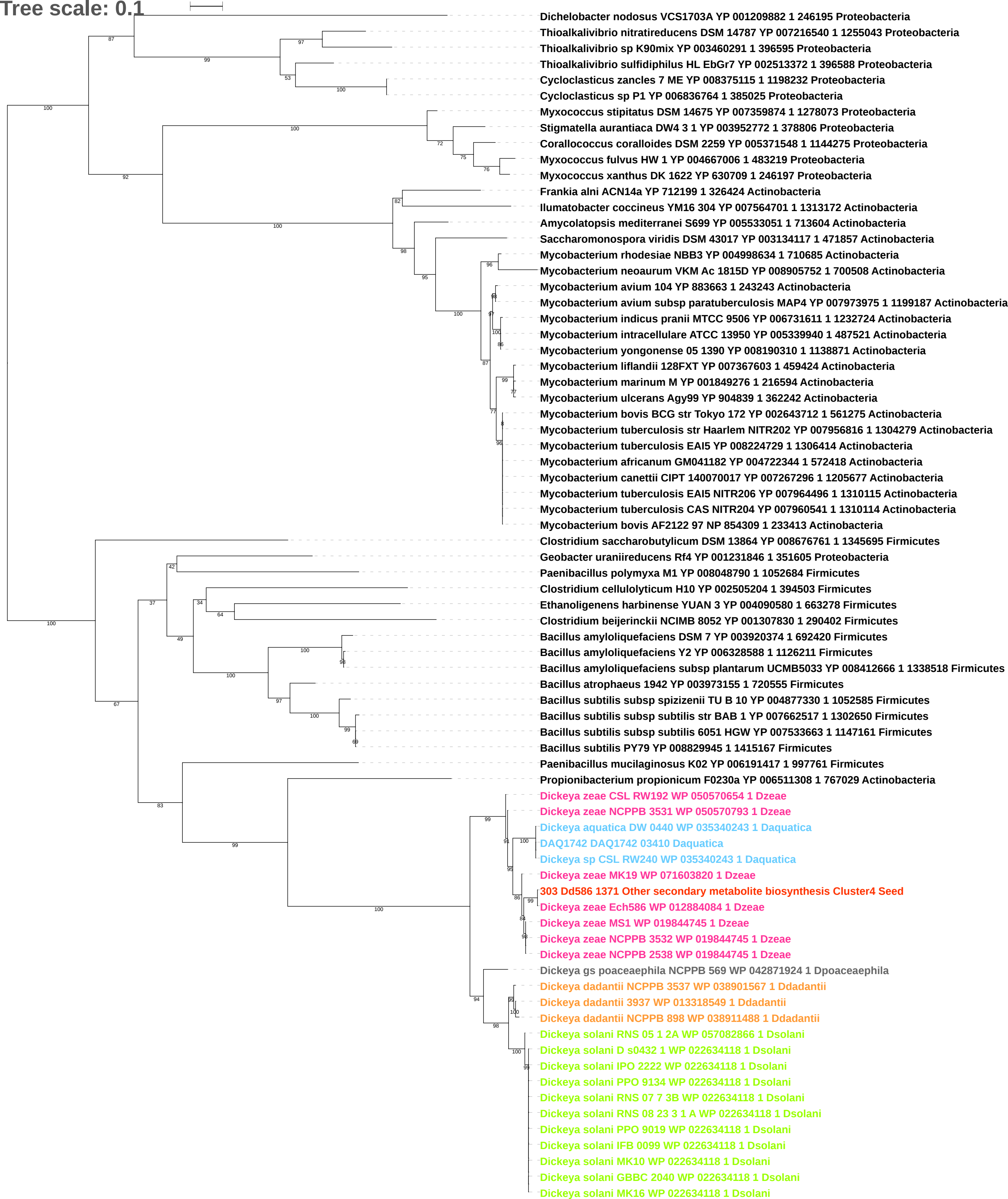
Tree scale: 0.1
Dichelobacter nodosus VCS1703A YP 001209882 1 246195 Proteobacteria
Thioalkalivibrio nitratireducens DSM 14787 YP 007216540 1 1255043 Proteobacteria
Thioalkalivibrio sp K90mix YP 003460291 1 396595 Proteobacteria
Thioalkalivibrio sulfidiphilus HL EbGr7 YP 002513372 1 396588 Proteobacteria
Cycloclasticus zancles 7 ME YP 008375115 1 1198232 Proteobacteria
Cycloclasticus sp P1 YP 006836764 1 385025 Proteobacteria
Myxococcus stipitatus DSM 14675 YP 007359874 1 1278073 Proteobacteria
Stigmatella aurantiaca DW4 3 1 YP 003952772 1 378806 Proteobacteria
Corallococcus coralloides DSM 2259 YP 005371548 1 1144275 Proteobacteria
Myxococcus fulvus HW 1 YP 004667006 1 483219 Proteobacteria
Myxococcus xanthus DK 1622 YP 630709 1 246197 Proteobacteria
Frankia alni ACN14a YP 712199 1 326424 Actinobacteria
Ilumatobacter coccineus YM16 304 YP 007564701 1 1313172 Actinobacteria
Amycolatopsis mediterranei S699 YP 005533051 1 713604 Actinobacteria
Saccharomonospora viridis DSM 43017 YP 003134117 1 471857 Actinobacteria
Mycobacterium rhodesiae NBB3 YP 004998634 1 710685 Actinobacteria
Mycobacterium neoaurum VKM Ac 1815D YP 008905752 1 700508 Actinobacteria
Mycobacterium avium 104 YP 883663 1 243243 Actinobacteria
Mycobacterium avium subsp paratuberculosis MAP4 YP 007973975 1 1199187 Actinobacteria
Mycobacterium indicus pranii MTCC 9506 YP 006731611 1 1232724 Actinobacteria
Mycobacterium intracellulare ATCC 13950 YP 005339940 1 487521 Actinobacteria
Mycobacterium yongonense 05 1390 YP 008190310 1 1138871 Actinobacteria
Mycobacterium liflandii 128FXT YP 007367603 1 459424 Actinobacteria
Mycobacterium marinum M YP 001849276 1 216594 Actinobacteria
Mycobacterium ulcerans Agy99 YP 904839 1 362242 Actinobacteria
Mycobacterium bovis BCG str Tokyo 172 YP 002643712 1 561275 Actinobacteria
Mycobacterium tuberculosis str Haarlem NITR202 YP 007956816 1 1304279 Actinobacteria
Mycobacterium tuberculosis EAI5 YP 008224729 1 1306414 Actinobacteria
Mycobacterium africanum GM041182 YP 004722344 1 572418 Actinobacteria
Mycobacterium canettii CIPT 140070017 YP 007267296 1 1205677 Actinobacteria
Mycobacterium tuberculosis EAI5 NITR206 YP 007964496 1 1310115 Actinobacteria
Mycobacterium tuberculosis CAS NITR204 YP 007960541 1 1310114 Actinobacteria
Mycobacterium bovis AF2122 97 NP 854309 1 233413 Actinobacteria
Clostridium saccharobutylicum DSM 13864 YP 008676761 1 1345695 Firmicutes
Geobacter uraniireducens Rf4 YP 001231846 1 351605 Proteobacteria
Paenibacillus polymyxa M1 YP 008048790 1 1052684 Firmicutes
Clostridium cellulolyticum H10 YP 002505204 1 394503 Firmicutes
Ethanoligenens harbinense YUAN 3 YP 004090580 1 663278 Firmicutes
Clostridium beijerinckii NCIMB 8052 YP 001307830 1 290402 Firmicutes
Bacillus amyloliquefaciens DSM 7 YP 003920374 1 692420 Firmicutes
Bacillus amyloliquefaciens Y2 YP 006328588 1 1126211 Firmicutes
Bacillus amyloliquefaciens subsp plantarum UCMB5033 YP 008412666 1 1338518 Firmicutes
Bacillus atrophaeus 1942 YP 003973155 1 720555 Firmicutes
Bacillus subtilis subsp spizizenii TU B 10 YP 004877330 1 1052585 Firmicutes
Bacillus subtilis subsp subtilis str BAB 1 YP 007662517 1 1302650 Firmicutes
Bacillus subtilis subsp subtilis 6051 HGW YP 007533663 1 1147161 Firmicutes
Bacillus subtilis PY79 YP 008829945 1 1415167 Firmicutes
Paenibacillus mucilaginosus K02 YP 006191417 1 997761 Firmicutes
Propionibacterium propionicum F0230a YP 006511308 1 767029 Actinobacteria
Dickeya zeae CSL RW192 WP 050570654 1 Dzeae
Dickeya zeae NCPPB 3531 WP 050570793 1 Dzeae
Dickeya aquatica DW 0440 WP 035340243 1 Daquatica
DAQ1742 DAQ1742 03410 Daquatica
Dickeya sp CSL RW240 WP 035340243 1 Daquatica
Dickeya zeae MK19 WP 071603820 1 Dzeae
303 Dd586 1371 Other secondary metabolite biosynthesis Cluster4 Seed
Dickeya zeae Ech586 WP 012884084 1 Dzeae
Dickeya zeae MS1 WP 019844745 1 Dzeae
Dickeya zeae NCPPB 3532 WP 019844745 1 Dzeae
Dickeya zeae NCPPB 2538 WP 019844745 1 Dzeae
Dickeya gs poaceaephila NCPPB 569 WP 042871924 1 Dpoaceaephila
Dickeya dadantii NCPPB 3537 WP 038901567 1 Ddadantii
Dickeya dadantii 3937 WP 013318549 1 Ddadantii
Dickeya dadantii NCPPB 898 WP 038911488 1 Ddadantii
Dickeya solani RNS 05 1 2A WP 057082866 1 Dsolani
Dickeya solani D s0432 1 WP 022634118 1 Dsolani
Dickeya solani IPO 2222 WP 022634118 1 Dsolani
Dickeya solani PPO 9134 WP 022634118 1 Dsolani
Dickeya solani RNS 07 7 3B WP 022634118 1 Dsolani
Dickeya solani RNS 08 23 3 1 A WP 022634118 1 Dsolani
Dickeya solani PPO 9019 WP 022634118 1 Dsolani
Dickeya solani IFB 0099 WP 022634118 1 Dsolani
Dickeya solani MK10 WP 022634118 1 Dsolani
Dickeya solani GBBC 2040 WP 022634118 1 Dsolani
Dickeya solani MK16 WP 022634118 1 Dsolani

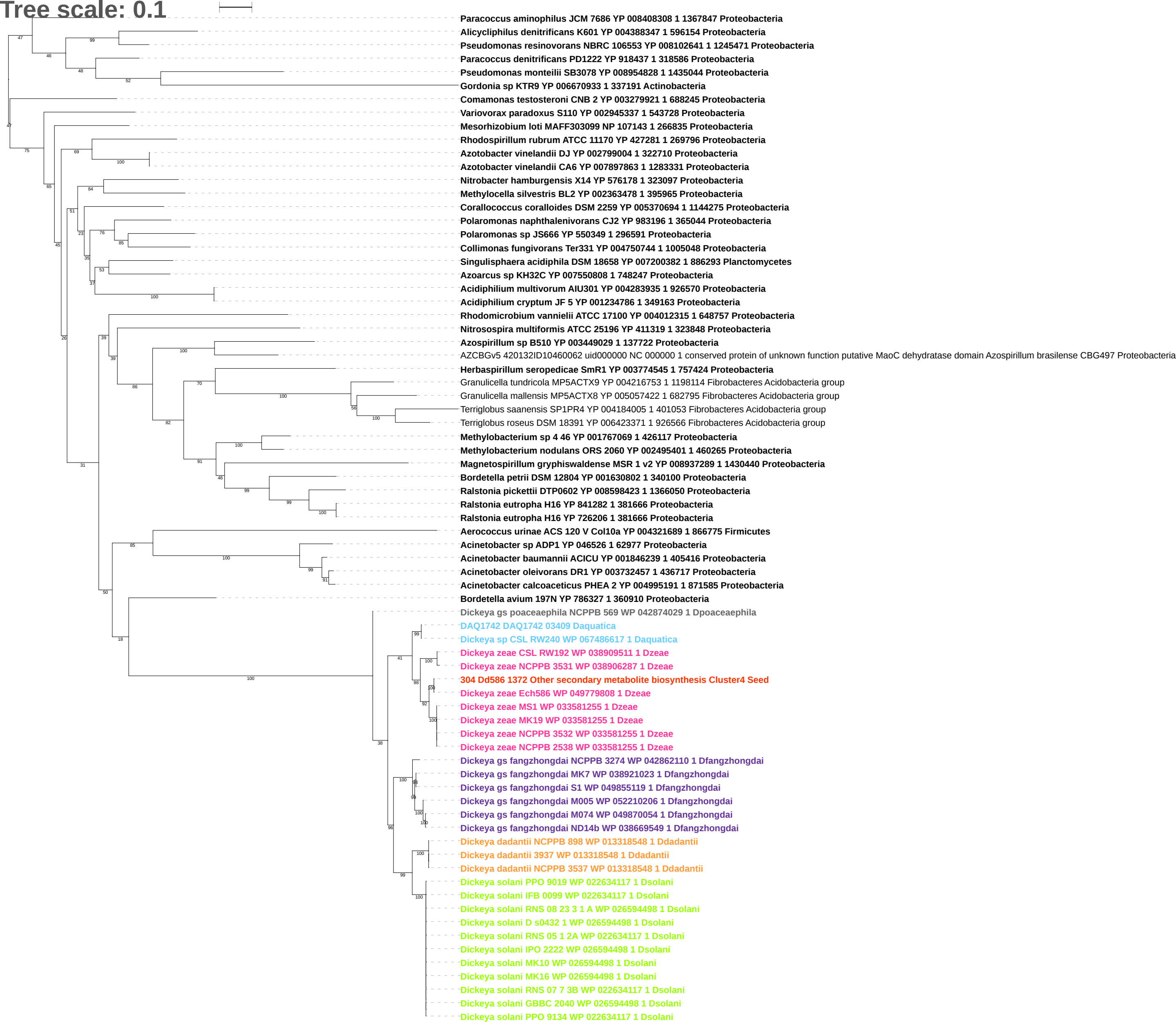
Tree scale: 0.1
Paracoccus aminophilus JCM 7686 YP 008408308 1 1367847 Proteobacteria
Alicycliphilus denitrificans K601 YP 004388347 1 596154 Proteobacteria
Pseudomonas resinovorans NBRC 106553 YP 008102641 1 1245471 Proteobacteria
Paracoccus denitrificans PD1222 YP 918437 1 318586 Proteobacteria
Pseudomonas monteilii SB3078 YP 008954828 1 1435044 Proteobacteria
Gordonia sp KTR9 YP 006670933 1 337191 Actinobacteria
Comamonas testosteroni CNB 2 YP 003279921 1 688245 Proteobacteria
Variovorax paradoxus S110 YP 002945337 1 543728 Proteobacteria
Mesorhizobium loti MAFF303099 NP 107143 1 266835 Proteobacteria
Rhodospirillum rubrum ATCC 11170 YP 427281 1 269796 Proteobacteria
Azotobacter vinelandii DJ YP 002799004 1 322710 Proteobacteria
Azotobacter vinelandii CA6 YP 007897863 1 1283331 Proteobacteria
Nitrobacter hamburgensis X14 YP 576178 1 323097 Proteobacteria
Methylocella silvestris BL2 YP 002363478 1 395965 Proteobacteria
Corallococcus coralloides DSM 2259 YP 005370694 1 1144275 Proteobacteria
Polaromonas naphthalenivorans CJ2 YP 983196 1 365044 Proteobacteria
Polaromonas sp JS666 YP 550349 1 296591 Proteobacteria
Collimonas fungivorans Ter331 YP 004750744 1 1005048 Proteobacteria
Singulisphaera acidiphila DSM 18658 YP 007200382 1 886293 Planctomycetes
Azoarcus sp KH32C YP 007550808 1 748247 Proteobacteria
Acidiphilium multivorum AIU301 YP 004283935 1 926570 Proteobacteria
Acidiphilium cryptum JF 5 YP 001234786 1 349163 Proteobacteria
Rhodomicrobium vannielii ATCC 17100 YP 004012315 1 648757 Proteobacteria
Nitrosospira multiformis ATCC 25196 YP 411319 1 323848 Proteobacteria
Azospirillum sp B510 YP 003449029 1 137722 Proteobacteria
AZCBGv5 420132ID10460062 uid000000 NC 000000 1 conserved protein of unknown function putative MaoC dehydratase domain Azospirillum brasilense CBG497 Proteobacteria
Herbaspirillum seropedicae SmR1 YP 003774545 1 757424 Proteobacteria
Granulicella tundricola MP5ACTX9 YP 004216753 1 1198114 Fibrobacteres Acidobacteria group
Granulicella mallensis MP5ACTX8 YP 005057422 1 682795 Fibrobacteres Acidobacteria group
Terriglobus saanensis SP1PR4 YP 004184005 1 401053 Fibrobacteres Acidobacteria group
Terriglobus roseus DSM 18391 YP 006423371 1 926566 Fibrobacteres Acidobacteria group
Methylobacterium sp 4 46 YP 001767069 1 426117 Proteobacteria
Methylobacterium nodulans ORS 2060 YP 002495401 1 460265 Proteobacteria
Magnetospirillum gryphiswaldense MSR 1 v2 YP 008937289 1 1430440 Proteobacteria
Bordetella petrii DSM 12804 YP 001630802 1 340100 Proteobacteria
Ralstonia pickettii DTP0602 YP 008598423 1 1366050 Proteobacteria
Ralstonia eutropha H16 YP 841282 1 381666 Proteobacteria
Ralstonia eutropha H16 YP 726206 1 381666 Proteobacteria
Aerococcus urinae ACS 120 V Col10a YP 004321689 1 866775 Firmicutes
Acinetobacter sp ADP1 YP 046526 1 62977 Proteobacteria
Acinetobacter baumannii ACICU YP 001846239 1 405416 Proteobacteria
Acinetobacter oleivorans DR1 YP 003732457 1 436717 Proteobacteria
Acinetobacter calcoaceticus PHEA 2 YP 004995191 1 871585 Proteobacteria
Bordetella avium 197N YP 786327 1 360910 Proteobacteria
Dickeya gs poaceaephila NCPPB 569 WP 042874029 1 Dpoaceaephila
DAQ1742 DAQ1742 03409 Daquatica
Dickeya sp CSL RW240 WP 067486617 1 Daquatica
Dickeya zeae CSL RW192 WP 038909511 1 Dzeae
Dickeya zeae NCPPB 3531 WP 038906287 1 Dzeae
304 Dd586 1372 Other secondary metabolite biosynthesis Cluster4 Seed
Dickeya zeae Ech586 WP 049779808 1 Dzeae
Dickeya zeae MS1 WP 033581255 1 Dzeae
Dickeya zeae MK19 WP 033581255 1 Dzeae
Dickeya zeae NCPPB 3532 WP 033581255 1 Dzeae
Dickeya zeae NCPPB 2538 WP 033581255 1 Dzeae
Dickeya gs fangzhongdai NCPPB 3274 WP 042862110 1 Dfangzhongdai
Dickeya gs fangzhongdai MK7 WP 038921023 1 Dfangzhongdai
Dickeya gs fangzhongdai S1 WP 049855119 1 Dfangzhongdai
Dickeya gs fangzhongdai M005 WP 052210206 1 Dfangzhongdai
Dickeya gs fangzhongdai M074 WP 049870054 1 Dfangzhongdai
Dickeya gs fangzhongdai ND14b WP 038669549 1 Dfangzhongdai
Dickeya dadantii NCPPB 898 WP 013318548 1 Ddadantii
Dickeya dadantii 3937 WP 013318548 1 Ddadantii
Dickeya dadantii NCPPB 3537 WP 013318548 1 Ddadantii
Dickeya solani PPO 9019 WP 022634117 1 Dsolani
Dickeya solani IFB 0099 WP 022634117 1 Dsolani
Dickeya solani RNS 08 23 3 1 A WP 026594498 1 Dsolani
Dickeya solani D s0432 1 WP 026594498 1 Dsolani
Dickeya solani RNS 05 1 2A WP 022634117 1 Dsolani
Dickeya solani IPO 2222 WP 026594498 1 Dsolani
Dickeya solani MK10 WP 026594498 1 Dsolani
Dickeya solani MK16 WP 026594498 1 Dsolani
Dickeya solani RNS 07 7 3B WP 022634117 1 Dsolani
Dickeya solani GBBC 2040 WP 026594498 1 Dsolani
Dickeya solani PPO 9134 WP 022634117 1 Dsolani

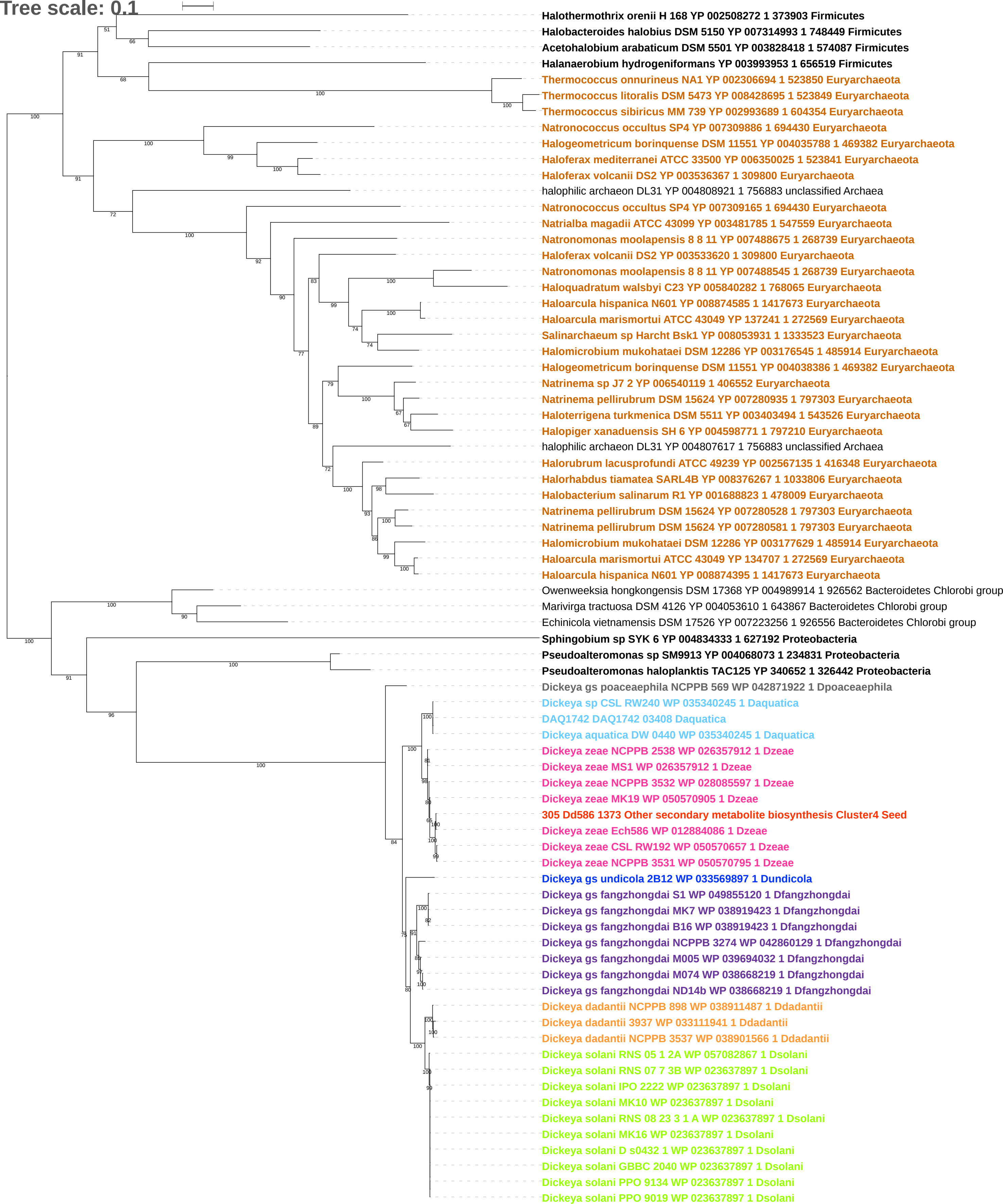
Tree scale: 0.1
Halothermothrix orenii H 168 YP 002508272 1 373903 Firmicutes
Halobacteroides halobius DSM 5150 YP 007314993 1 748449 Firmicutes
Acetohalobium arabaticum DSM 5501 YP 003828418 1 574087 Firmicutes
Halanaerobium hydrogeniformans YP 003993953 1 656519 Firmicutes
Thermococcus onnurineus NA1 YP 002306694 1 523850 Euryarchaeota
Thermococcus litoralis DSM 5473 YP 008428695 1 523849 Euryarchaeota
Thermococcus sibiricus MM 739 YP 002993689 1 604354 Euryarchaeota
Natronococcus occultus SP4 YP 007309886 1 694430 Euryarchaeota
Halogeometricum borinquense DSM 11551 YP 004035788 1 469382 Euryarchaeota
Haloferax mediterranei ATCC 33500 YP 006350025 1 523841 Euryarchaeota
Haloferax volcanii DS2 YP 003536367 1 309800 Euryarchaeota
halophilic archaeon DL31 YP 004808921 1 756883 unclassified Archaea
Natronococcus occultus SP4 YP 007309165 1 694430 Euryarchaeota
Natrialba magadii ATCC 43099 YP 003481785 1 547559 Euryarchaeota
Natronomonas moolapensis 8 8 11 YP 007488675 1 268739 Euryarchaeota
Haloferax volcanii DS2 YP 003533620 1 309800 Euryarchaeota
Natronomonas moolapensis 8 8 11 YP 007488545 1 268739 Euryarchaeota
Haloquadratum walsbyi C23 YP 005840282 1 768065 Euryarchaeota
Haloarcula hispanica N601 YP 008874585 1 1417673 Euryarchaeota
Haloarcula marismortui ATCC 43049 YP 137241 1 272569 Euryarchaeota
Salinarchaeum sp Harcht Bsk1 YP 008053931 1 1333523 Euryarchaeota
Halomicrobium mukohataei DSM 12286 YP 003176545 1 485914 Euryarchaeota
Halogeometricum borinquense DSM 11551 YP 004038386 1 469382 Euryarchaeota
Natrinema sp J7 2 YP 006540119 1 406552 Euryarchaeota
Natrinema pellirubrum DSM 15624 YP 007280935 1 797303 Euryarchaeota
Haloterrigena turkmenica DSM 5511 YP 003403494 1 543526 Euryarchaeota
Halopiger xanaduensis SH 6 YP 004598771 1 797210 Euryarchaeota
halophilic archaeon DL31 YP 004807617 1 756883 unclassified Archaea
Halorubrum lacusprofundi ATCC 49239 YP 002567135 1 416348 Euryarchaeota
Halorhabdus tiamatea SARL4B YP 008376267 1 1033806 Euryarchaeota
Halobacterium salinarum R1 YP 001688823 1 478009 Euryarchaeota
Natrinema pellirubrum DSM 15624 YP 007280528 1 797303 Euryarchaeota
Natrinema pellirubrum DSM 15624 YP 007280581 1 797303 Euryarchaeota
Halomicrobium mukohataei DSM 12286 YP 003177629 1 485914 Euryarchaeota
Haloarcula marismortui ATCC 43049 YP 134707 1 272569 Euryarchaeota
Haloarcula hispanica N601 YP 008874395 1 1417673 Euryarchaeota
Owenweeksia hongkongensis DSM 17368 YP 004989914 1 926562 Bacteroidetes Chlorobi group
Marivirga tractuosa DSM 4126 YP 004053610 1 643867 Bacteroidetes Chlorobi group
Echinicola vietnamensis DSM 17526 YP 007223256 1 926556 Bacteroidetes Chlorobi group
Sphingobium sp SYK 6 YP 004834333 1 627192 Proteobacteria
Pseudoalteromonas sp SM9913 YP 004068073 1 234831 Proteobacteria
Pseudoalteromonas haloplanktis TAC125 YP 340652 1 326442 Proteobacteria
Dickeya gs poaceaephila NCPPB 569 WP 042871922 1 Dpoaceaephila
Dickeya sp CSL RW240 WP 035340245 1 Daquatica
DAQ1742 DAQ1742 03408 Daquatica
Dickeya aquatica DW 0440 WP 035340245 1 Daquatica
Dickeya zeae NCPPB 2538 WP 026357912 1 Dzeae
Dickeya zeae MS1 WP 026357912 1 Dzeae
Dickeya zeae NCPPB 3532 WP 028085597 1 Dzeae
Dickeya zeae MK19 WP 050570905 1 Dzeae
305 Dd586 1373 Other secondary metabolite biosynthesis Cluster4 Seed
Dickeya zeae Ech586 WP 012884086 1 Dzeae
Dickeya zeae CSL RW192 WP 050570657 1 Dzeae
Dickeya zeae NCPPB 3531 WP 050570795 1 Dzeae
Dickeya gs undicola 2B12 WP 033569897 1 Dundicola
Dickeya gs fangzhongdai S1 WP 049855120 1 Dfangzhongdai
Dickeya gs fangzhongdai MK7 WP 038919423 1 Dfangzhongdai
Dickeya gs fangzhongdai B16 WP 038919423 1 Dfangzhongdai
Dickeya gs fangzhongdai NCPPB 3274 WP 042860129 1 Dfangzhongdai
Dickeya gs fangzhongdai M005 WP 039694032 1 Dfangzhongdai
Dickeya gs fangzhongdai M074 WP 038668219 1 Dfangzhongdai
Dickeya gs fangzhongdai ND14b WP 038668219 1 Dfangzhongdai
Dickeya dadantii NCPPB 898 WP 038911487 1 Ddadantii
Dickeya dadantii 3937 WP 033111941 1 Ddadantii
Dickeya dadantii NCPPB 3537 WP 038901566 1 Ddadantii
Dickeya solani RNS 05 1 2A WP 057082867 1 Dsolani
Dickeya solani RNS 07 7 3B WP 023637897 1 Dsolani
Dickeya solani IPO 2222 WP 023637897 1 Dsolani
Dickeya solani MK10 WP 023637897 1 Dsolani
Dickeya solani RNS 08 23 3 1 A WP 023637897 1 Dsolani
Dickeya solani MK16 WP 023637897 1 Dsolani
Dickeya solani D s0432 1 WP 023637897 1 Dsolani
Dickeya solani GBBC 2040 WP 023637897 1 Dsolani
Dickeya solani PPO 9134 WP 023637897 1 Dsolani
Dickeya solani PPO 9019 WP 023637897 1 Dsolani

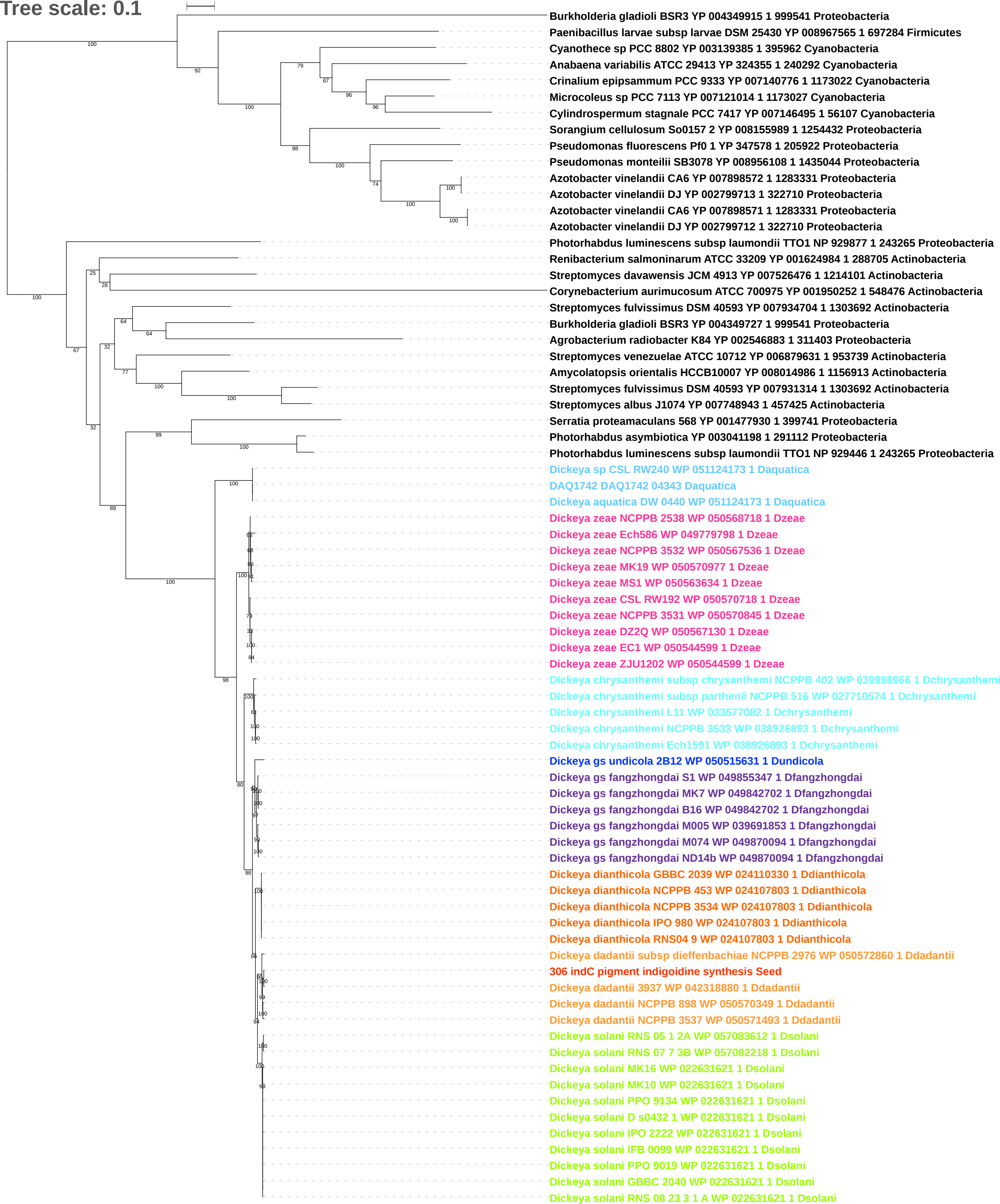
Tree scale: 0.1
Burkholderia gladioli BSR3 YP 004349915 1 999541 Proteobacteria
Paenibacillus larvae subsp larvae DSM 25430 YP 008967565 1 697284 Firmicutes
Cyanothece sp PCC 8802 YP 003139385 1 395962 Cyanobacteria
Anabaena variabilis ATCC 29413 YP 324355 1 240292 Cyanobacteria
Crinalium epipsammum PCC 9333 YP 007140776 1 1173022 Cyanobacteria
Microcoleus sp PCC 7113 YP 007121014 1 1173027 Cyanobacteria
Cylindrospermum stagnale PCC 7417 YP 007146495 1 56107 Cyanobacteria
Sorangium cellulosum So0157 2 YP 008155989 1 1254432 Proteobacteria
Pseudomonas fluorescens Pf0 1 YP 347578 1 205922 Proteobacteria
Pseudomonas monteilii SB3078 YP 008956108 1 1435044 Proteobacteria
Azotobacter vinelandii CA6 YP 007898572 1 1283331 Proteobacteria
Azotobacter vinelandii DJ YP 002799713 1 322710 Proteobacteria
Azotobacter vinelandii CA6 YP 007898571 1 1283331 Proteobacteria
Azotobacter vinelandii DJ YP 002799712 1 322710 Proteobacteria
Photorhabdus luminescens subsp laumondii TTO1 NP 929877 1 243265 Proteobacteria
Renibacterium salmoninarum ATCC 33209 YP 001624984 1 288705 Actinobacteria
Streptomyces davawensis JCM 4913 YP 007526476 1 1214101 Actinobacteria
Corynebacterium aurimucosum ATCC 700975 YP 001950252 1 548476 Actinobacteria
Streptomyces fulvissimus DSM 40593 YP 007934704 1 1303692 Actinobacteria
Burkholderia gladioli BSR3 YP 004349727 1 999541 Proteobacteria
Agrobacterium radiobacter K84 YP 002546883 1 311403 Proteobacteria
Streptomyces venezuelae ATCC 10712 YP 006879631 1 953739 Actinobacteria
Amycolatopsis orientalis HCCB10007 YP 008014986 1 1156913 Actinobacteria
Streptomyces fulvissimus DSM 40593 YP 007931314 1 1303692 Actinobacteria
Streptomyces albus J1074 YP 007748943 1 457425 Actinobacteria
Serratia proteamaculans 568 YP 001477930 1 399741 Proteobacteria
Photorhabdus asymbiotica YP 003041198 1 291112 Proteobacteria
Photorhabdus luminescens subsp laumondii TTO1 NP 929446 1 243265 Proteobacteria
Dickeya sp CSL RW240 WP 051124173 1 Daquatica
DAQ1742 DAQ1742 04343 Daquatica
Dickeya aquatica DW 0440 WP 051124173 1 Daquatica
Dickeya zeae NCPPB 2538 WP 050568718 1 Dzeae
Dickeya zeae Ech586 WP 049779798 1 Dzeae
Dickeya zeae NCPPB 3532 WP 050567536 1 Dzeae
Dickeya zeae MK19 WP 050570977 1 Dzeae
Dickeya zeae MS1 WP 050563634 1 Dzeae
Dickeya zeae CSL RW192 WP 050570718 1 Dzeae
Dickeya zeae NCPPB 3531 WP 050570845 1 Dzeae
Dickeya zeae DZ2Q WP 050567130 1 Dzeae
Dickeya zeae EC1 WP 050544599 1 Dzeae
Dickeya zeae ZJU1202 WP 050544599 1 Dzeae
Dickeya chrysanthemi subsp chrysanthemi NCPPB 402 WP 039998966 1 Dchrysanthemi
Dickeya chrysanthemi subsp parthenii NCPPB 516 WP 027710574 1 Dchrysanthemi
Dickeya chrysanthemi L11 WP 033577082 1 Dchrysanthemi
Dickeya chrysanthemi NCPPB 3533 WP 038926893 1 Dchrysanthemi
Dickeya chrysanthemi Ech1591 WP 038926893 1 Dchrysanthemi
Dickeya gs undicola 2B12 WP 050515631 1 Dundicola
Dickeya gs fangzhongdai S1 WP 049855347 1 Dfangzhongdai
Dickeya gs fangzhongdai MK7 WP 049842702 1 Dfangzhongdai
Dickeya gs fangzhongdai B16 WP 049842702 1 Dfangzhongdai
Dickeya gs fangzhongdai M005 WP 039691853 1 Dfangzhongdai
Dickeya gs fangzhongdai M074 WP 049870094 1 Dfangzhongdai
Dickeya gs fangzhongdai ND14b WP 049870094 1 Dfangzhongdai
Dickeya dianthicola GBBC 2039 WP 024110330 1 Ddianthicola
Dickeya dianthicola NCPPB 453 WP 024107803 1 Ddianthicola
Dickeya dianthicola NCPPB 3534 WP 024107803 1 Ddianthicola
Dickeya dianthicola IPO 980 WP 024107803 1 Ddianthicola
Dickeya dianthicola RNS04 9 WP 024107803 1 Ddianthicola
Dickeya dadantii subsp dieffenbachiae NCPPB 2976 WP 050572860 1 Ddadantii
306 indC pigment indigoidine synthesis Seed
Dickeya dadantii 3937 WP 042318880 1 Ddadantii
Dickeya dadantii NCPPB 898 WP 050570349 1 Ddadantii
Dickeya dadantii NCPPB 3537 WP 050571493 1 Ddadantii
Dickeya solani RNS 05 1 2A WP 057083612 1 Dsolani
Dickeya solani RNS 07 7 3B WP 057082218 1 Dsolani
Dickeya solani MK16 WP 022631621 1 Dsolani
Dickeya solani MK10 WP 022631621 1 Dsolani
Dickeya solani PPO 9134 WP 022631621 1 Dsolani
Dickeya solani D s0432 1 WP 022631621 1 Dsolani
Dickeya solani IPO 2222 WP 022631621 1 Dsolani
Dickeya solani IFB 0099 WP 022631621 1 Dsolani
Dickeya solani PPO 9019 WP 022631621 1 Dsolani
Dickeya solani GBBC 2040 WP 022631621 1 Dsolani
Dickeya solani RNS 08 23 3 1 A WP 022631621 1 Dsolani

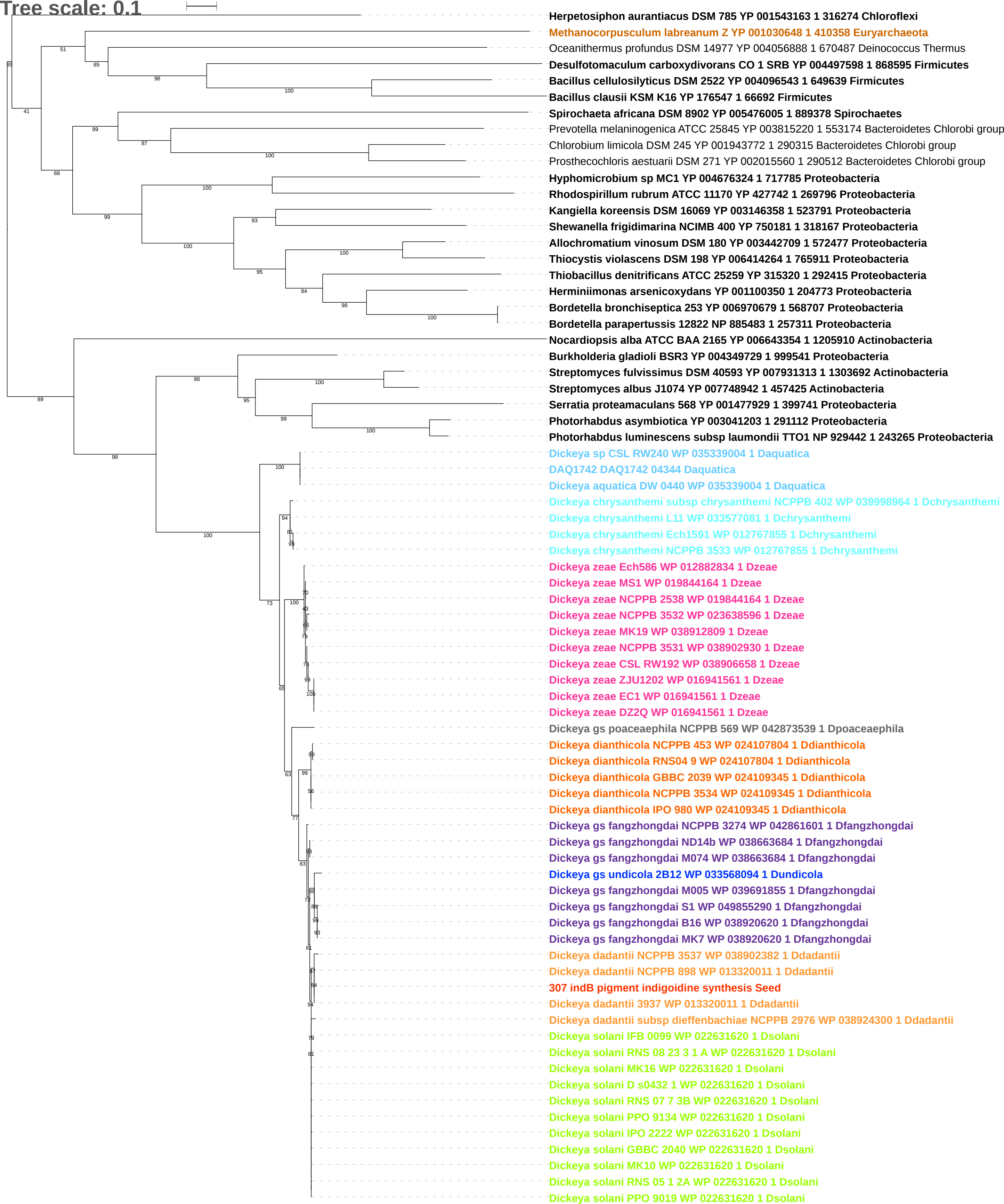
Tree scale: 0.1
Herpetosiphon aurantiacus DSM 785 YP 001543163 1 316274 Chloroflexi
Methanocorpusculum labreanum Z YP 001030648 1 410358 Euryarchaeota
Oceanithermus profundus DSM 14977 YP 004056888 1 670487 Deinococcus Thermus
Desulfotomaculum carboxydivorans CO 1 SRB YP 004497598 1 868595 Firmicutes
Bacillus cellulosilyticus DSM 2522 YP 004096543 1 649639 Firmicutes
Bacillus clausii KSM K16 YP 176547 1 66692 Firmicutes
Spirochaeta africana DSM 8902 YP 005476005 1 889378 Spirochaetes
Prevotella melaninogenica ATCC 25845 YP 003815220 1 553174 Bacteroidetes Chlorobi group
Chlorobium limicola DSM 245 YP 001943772 1 290315 Bacteroidetes Chlorobi group
Prosthecochloris aestuarii DSM 271 YP 002015560 1 290512 Bacteroidetes Chlorobi group
Hyphomicrobium sp MC1 YP 004676324 1 717785 Proteobacteria
Rhodospirillum rubrum ATCC 11170 YP 427742 1 269796 Proteobacteria
Kangiella koreensis DSM 16069 YP 003146358 1 523791 Proteobacteria
Shewanella frigidimarina NCIMB 400 YP 750181 1 318167 Proteobacteria
Allochromatium vinosum DSM 180 YP 003442709 1 572477 Proteobacteria
Thiocystis violascens DSM 198 YP 006414264 1 765911 Proteobacteria
Thiobacillus denitrificans ATCC 25259 YP 315320 1 292415 Proteobacteria
Herminiimonas arsenicoxydans YP 001100350 1 204773 Proteobacteria
Bordetella bronchiseptica 253 YP 006970679 1 568707 Proteobacteria
Bordetella parapertussis 12822 NP 885483 1 257311 Proteobacteria
Nocardiopsis alba ATCC BAA 2165 YP 006643354 1 1205910 Actinobacteria
Burkholderia gladioli BSR3 YP 004349729 1 999541 Proteobacteria
Streptomyces fulvissimus DSM 40593 YP 007931313 1 1303692 Actinobacteria
Streptomyces albus J1074 YP 007748942 1 457425 Actinobacteria
Serratia proteamaculans 568 YP 001477929 1 399741 Proteobacteria
Photorhabdus asymbiotica YP 003041203 1 291112 Proteobacteria
Photorhabdus luminescens subsp laumondii TTO1 NP 929442 1 243265 Proteobacteria
Dickeya sp CSL RW240 WP 035339004 1 Daquatica
DAQ1742 DAQ1742 04344 Daquatica
Dickeya aquatica DW 0440 WP 035339004 1 Daquatica
Dickeya chrysanthemi subsp chrysanthemi NCPPB 402 WP 039998964 1 Dchrysanthemi
Dickeya chrysanthemi L11 WP 033577081 1 Dchrysanthemi
Dickeya chrysanthemi Ech1591 WP 012767855 1 Dchrysanthemi
Dickeya chrysanthemi NCPPB 3533 WP 012767855 1 Dchrysanthemi
Dickeya zeae Ech586 WP 012882834 1 Dzeae
Dickeya zeae MS1 WP 019844164 1 Dzeae
Dickeya zeae NCPPB 2538 WP 019844164 1 Dzeae
Dickeya zeae NCPPB 3532 WP 023638596 1 Dzeae
Dickeya zeae MK19 WP 038912809 1 Dzeae
Dickeya zeae NCPPB 3531 WP 038902930 1 Dzeae
Dickeya zeae CSL RW192 WP 038906658 1 Dzeae
Dickeya zeae ZJU1202 WP 016941561 1 Dzeae
Dickeya zeae EC1 WP 016941561 1 Dzeae
Dickeya zeae DZ2Q WP 016941561 1 Dzeae
Dickeya gs poaceaephila NCPPB 569 WP 042873539 1 Dpoaceaephila
Dickeya dianthicola NCPPB 453 WP 024107804 1 Ddianthicola
Dickeya dianthicola RNS04 9 WP 024107804 1 Ddianthicola
Dickeya dianthicola GBBC 2039 WP 024109345 1 Ddianthicola
Dickeya dianthicola NCPPB 3534 WP 024109345 1 Ddianthicola
Dickeya dianthicola IPO 980 WP 024109345 1 Ddianthicola
Dickeya gs fangzhongdai NCPPB 3274 WP 042861601 1 Dfangzhongdai
Dickeya gs fangzhongdai ND14b WP 038663684 1 Dfangzhongdai
Dickeya gs fangzhongdai M074 WP 038663684 1 Dfangzhongdai
Dickeya gs undicola 2B12 WP 033568094 1 Dundicola
Dickeya gs fangzhongdai M005 WP 039691855 1 Dfangzhongdai
Dickeya gs fangzhongdai S1 WP 049855290 1 Dfangzhongdai
Dickeya gs fangzhongdai B16 WP 038920620 1 Dfangzhongdai
Dickeya gs fangzhongdai MK7 WP 038920620 1 Dfangzhongdai
Dickeya dadantii NCPPB 3537 WP 038902382 1 Ddadantii
Dickeya dadantii NCPPB 898 WP 013320011 1 Ddadantii
307 indB pigment indigoidine synthesis Seed
Dickeya dadantii 3937 WP 013320011 1 Ddadantii
Dickeya dadantii subsp dieffenbachiae NCPPB 2976 WP 038924300 1 Ddadantii
Dickeya solani IFB 0099 WP 022631620 1 Dsolani
Dickeya solani RNS 08 23 3 1 A WP 022631620 1 Dsolani
Dickeya solani MK16 WP 022631620 1 Dsolani
Dickeya solani D s0432 1 WP 022631620 1 Dsolani
Dickeya solani RNS 07 7 3B WP 022631620 1 Dsolani
Dickeya solani PPO 9134 WP 022631620 1 Dsolani
Dickeya solani IPO 2222 WP 022631620 1 Dsolani
Dickeya solani GBBC 2040 WP 022631620 1 Dsolani
Dickeya solani MK10 WP 022631620 1 Dsolani
Dickeya solani RNS 05 1 2A WP 022631620 1 Dsolani
Dickeya solani PPO 9019 WP 022631620 1 Dsolani

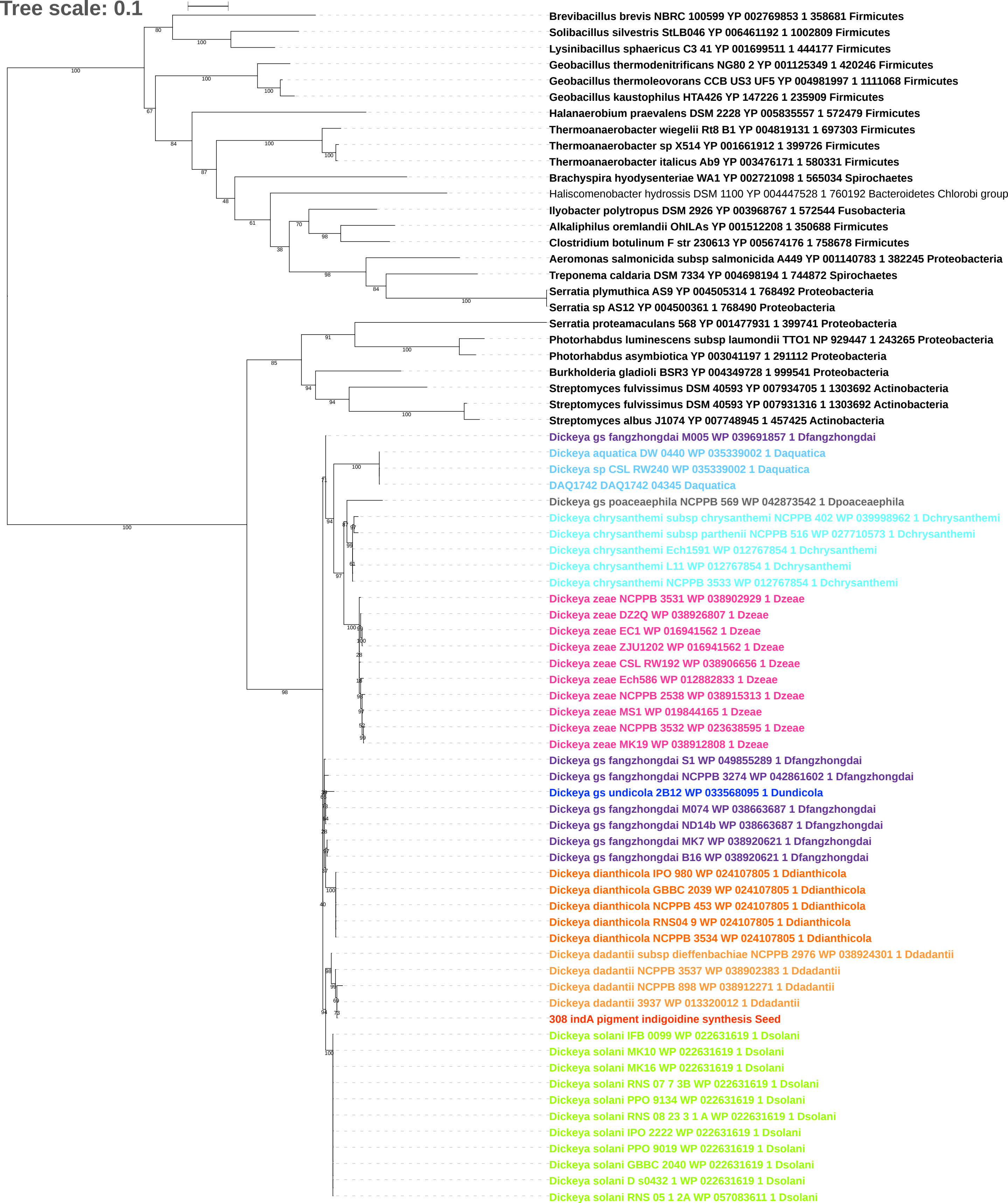
Tree scale: 0.1
Brevibacillus brevis NBRC 100599 YP 002769853 1 358681 Firmicutes
Solibacillus silvestris StLB046 YP 006461192 1 1002809 Firmicutes
Lysinibacillus sphaericus C3 41 YP 001699511 1 444177 Firmicutes
Geobacillus thermodenitrificans NG80 2 YP 001125349 1 420246 Firmicutes
Geobacillus thermoleovorans CCB US3 UF5 YP 004981997 1 1111068 Firmicutes
Geobacillus kaustophilus HTA426 YP 147226 1 235909 Firmicutes
Halanaerobium praevalens DSM 2228 YP 005835557 1 572479 Firmicutes
Thermoanaerobacter wiegelii Rt8 B1 YP 004819131 1 697303 Firmicutes
Thermoanaerobacter sp X514 YP 001661912 1 399726 Firmicutes
Thermoanaerobacter italicus Ab9 YP 003476171 1 580331 Firmicutes
Brachyspira hyodysenteriae WA1 YP 002721098 1 565034 Spirochaetes
Haliscomenobacter hydrossis DSM 1100 YP 004447528 1 760192 Bacteroidetes Chlorobi group
Ilyobacter polytropus DSM 2926 YP 003968767 1 572544 Fusobacteria
Alkaliphilus oremlandii OhILAs YP 001512208 1 350688 Firmicutes
Clostridium botulinum F str 230613 YP 005674176 1 758678 Firmicutes
Aeromonas salmonicida subsp salmonicida A449 YP 001140783 1 382245 Proteobacteria
Treponema caldaria DSM 7334 YP 004698194 1 744872 Spirochaetes
Serratia plymuthica AS9 YP 004505314 1 768492 Proteobacteria
Serratia sp AS12 YP 004500361 1 768490 Proteobacteria
Serratia proteamaculans 568 YP 001477931 1 399741 Proteobacteria
Photorhabdus luminescens subsp laumondii TTO1 NP 929447 1 243265 Proteobacteria
Photorhabdus asymbiotica YP 003041197 1 291112 Proteobacteria
Burkholderia gladioli BSR3 YP 004349728 1 999541 Proteobacteria
Streptomyces fulvissimus DSM 40593 YP 007934705 1 1303692 Actinobacteria
Streptomyces fulvissimus DSM 40593 YP 007931316 1 1303692 Actinobacteria
Streptomyces albus J1074 YP 007748945 1 457425 Actinobacteria
Dickeya gs fangzhongdai M005 WP 039691857 1 Dfangzhongdai
Dickeya aquatica DW 0440 WP 035339002 1 Daquatica
Dickeya sp CSL RW240 WP 035339002 1 Daquatica
DAQ1742 DAQ1742 04345 Daquatica
Dickeya gs poaceaephila NCPPB 569 WP 042873542 1 Dpoaceaephila
Dickeya chrysanthemi subsp chrysanthemi NCPPB 402 WP 039998962 1 Dchrysanthemi
Dickeya chrysanthemi subsp parthenii NCPPB 516 WP 027710573 1 Dchrysanthemi
Dickeya chrysanthemi Ech1591 WP 012767854 1 Dchrysanthemi
Dickeya chrysanthemi L11 WP 012767854 1 Dchrysanthemi
Dickeya chrysanthemi NCPPB 3533 WP 012767854 1 Dchrysanthemi
Dickeya zeae NCPPB 3531 WP 038902929 1 Dzeae
Dickeya zeae DZ2Q WP 038926807 1 Dzeae
Dickeya zeae EC1 WP 016941562 1 Dzeae
Dickeya zeae ZJU1202 WP 016941562 1 Dzeae
Dickeya zeae CSL RW192 WP 038906656 1 Dzeae
Dickeya zeae Ech586 WP 012882833 1 Dzeae
Dickeya zeae NCPPB 2538 WP 038915313 1 Dzeae
Dickeya zeae MS1 WP 019844165 1 Dzeae
Dickeya zeae NCPPB 3532 WP 023638595 1 Dzeae
Dickeya zeae MK19 WP 038912808 1 Dzeae
Dickeya gs fangzhongdai S1 WP 049855289 1 Dfangzhongdai
Dickeya gs fangzhongdai NCPPB 3274 WP 042861602 1 Dfangzhongdai
Dickeya gs undicola 2B12 WP 033568095 1 Dundicola
Dickeya gs fangzhongdai M074 WP 038663687 1 Dfangzhongdai
Dickeya gs fangzhongdai ND14b WP 038663687 1 Dfangzhongdai
Dickeya gs fangzhongdai MK7 WP 038920621 1 Dfangzhongdai
Dickeya gs fangzhongdai B16 WP 038920621 1 Dfangzhongdai
Dickeya dianthicola IPO 980 WP 024107805 1 Ddianthicola
Dickeya dianthicola GBBC 2039 WP 024107805 1 Ddianthicola
Dickeya dianthicola NCPPB 453 WP 024107805 1 Ddianthicola
Dickeya dianthicola RNS04 9 WP 024107805 1 Ddianthicola
Dickeya dianthicola NCPPB 3534 WP 024107805 1 Ddianthicola
Dickeya dadantii subsp dieffenbachiae NCPPB 2976 WP 038924301 1 Ddadantii
Dickeya dadantii NCPPB 3537 WP 038902383 1 Ddadantii
Dickeya dadantii NCPPB 898 WP 038912271 1 Ddadantii
Dickeya dadantii 3937 WP 013320012 1 Ddadantii
308 indA pigment indigoidine synthesis Seed
Dickeya solani IFB 0099 WP 022631619 1 Dsolani
Dickeya solani MK10 WP 022631619 1 Dsolani
Dickeya solani MK16 WP 022631619 1 Dsolani
Dickeya solani RNS 07 7 3B WP 022631619 1 Dsolani
Dickeya solani PPO 9134 WP 022631619 1 Dsolani
Dickeya solani RNS 08 23 3 1 A WP 022631619 1 Dsolani
Dickeya solani IPO 2222 WP 022631619 1 Dsolani
Dickeya solani PPO 9019 WP 022631619 1 Dsolani
Dickeya solani GBBC 2040 WP 022631619 1 Dsolani
Dickeya solani D s0432 1 WP 022631619 1 Dsolani
Dickeya solani RNS 05 1 2A WP 057083611 1 Dsolani

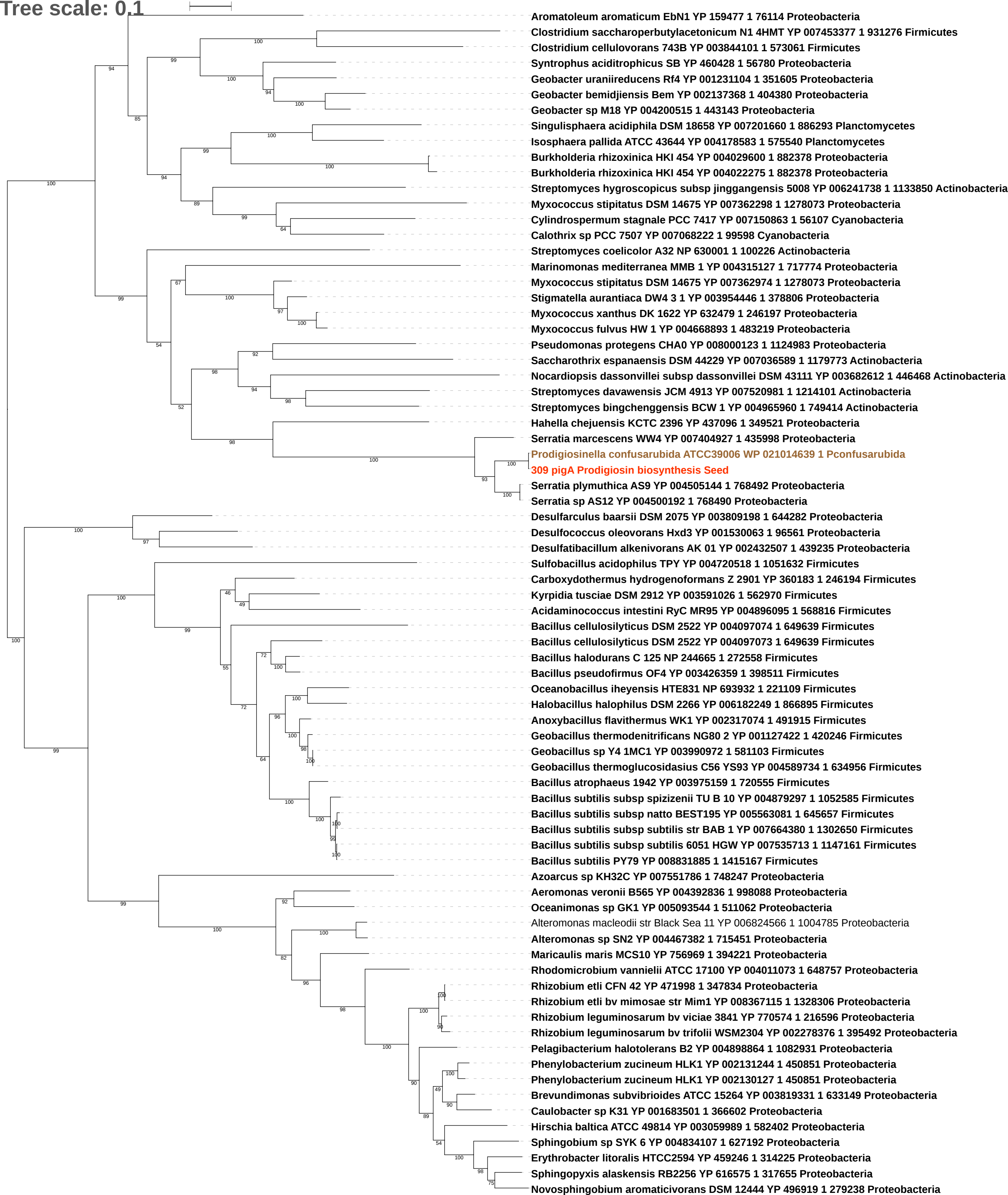

Tree scale: 0.1
Aromatoleum aromaticum EbN1 YP 159477 1 76114 Proteobacteria
Clostridium saccharoperbutylacetonicum N1 4HMT YP 007453377 1 931276 Firmicutes
Clostridium cellulovorans 743B YP 003844101 1 573061 Firmicutes
Syntrophus aciditrophicus SB YP 460428 1 56780 Proteobacteria
Geobacter uraniireducens Rf4 YP 001231104 1 351605 Proteobacteria
Geobacter bemidjiensis Bem YP 002137368 1 404380 Proteobacteria
Geobacter sp M18 YP 004200515 1 443143 Proteobacteria
Singulisphaera acidiphila DSM 18658 YP 007201660 1 886293 Planctomycetes
Isosphaera pallida ATCC 43644 YP 004178583 1 575540 Planctomycetes
Burkholderia rhizoxinica HKI 454 YP 004029600 1 882378 Proteobacteria
Burkholderia rhizoxinica HKI 454 YP 004022275 1 882378 Proteobacteria
Streptomyces hygroscopicus subsp jinggangensis 5008 YP 006241738 1 1133850 Actinobacteria
Myxococcus stipitatus DSM 14675 YP 007362298 1 1278073 Proteobacteria
Cylindrospermum stagnale PCC 7417 YP 007150863 1 56107 Cyanobacteria
Calothrix sp PCC 7507 YP 007068222 1 99598 Cyanobacteria
Streptomyces coelicolor A32 NP 630001 1 100226 Actinobacteria
Marinomonas mediterranea MMB 1 YP 004315127 1 717774 Proteobacteria
Myxococcus stipitatus DSM 14675 YP 007362974 1 1278073 Proteobacteria
Stigmatella aurantiaca DW4 3 1 YP 003954446 1 378806 Proteobacteria
Myxococcus xanthus DK 1622 YP 632479 1 246197 Proteobacteria
Myxococcus fulvus HW 1 YP 004668893 1 483219 Proteobacteria
Pseudomonas protegens CHA0 YP 008000123 1 1124983 Proteobacteria
Saccharothrix espanaensis DSM 44229 YP 007036589 1 1179773 Actinobacteria
Nocardiopsis dassonvillei subsp dassonvillei DSM 43111 YP 003682612 1 446468 Actinobacteria
Streptomyces davawensis JCM 4913 YP 007520981 1 1214101 Actinobacteria
Streptomyces bingchenggensis BCW 1 YP 004965960 1 749414 Actinobacteria
Hahella chejuensis KCTC 2396 YP 437096 1 349521 Proteobacteria
Serratia marcescens WW4 YP 007404927 1 435998 Proteobacteria
Prodigiosinella confusarubida ATCC39006 WP 021014639 1 Pconfusarubida
309 pigA Prodigiosin biosynthesis Seed
Serratia plymuthica AS9 YP 004505144 1 768492 Proteobacteria
Serratia sp AS12 YP 004500192 1 768490 Proteobacteria
Desulfarculus baarsii DSM 2075 YP 003809198 1 644282 Proteobacteria
Desulfococcus oleovorans Hxd3 YP 001530063 1 96561 Proteobacteria
Desulfatibacillum alkenivorans AK 01 YP 002432507 1 439235 Proteobacteria
Sulfobacillus acidophilus TPY YP 004720518 1 1051632 Firmicutes
Carboxydothermus hydrogenoformans Z 2901 YP 360183 1 246194 Firmicutes
Kyrpidia tusciae DSM 2912 YP 003591026 1 562970 Firmicutes
Acidaminococcus intestini RyC MR95 YP 004896095 1 568816 Firmicutes
Bacillus cellulosilyticus DSM 2522 YP 004097074 1 649639 Firmicutes
Bacillus cellulosilyticus DSM 2522 YP 004097073 1 649639 Firmicutes
Bacillus halodurans C 125 NP 244665 1 272558 Firmicutes
Bacillus pseudofirmus OF4 YP 003426359 1 398511 Firmicutes
Oceanobacillus iheyensis HTE831 NP 693932 1 221109 Firmicutes
Halobacillus halophilus DSM 2266 YP 006182249 1 866895 Firmicutes
Anoxybacillus flavithermus WK1 YP 002317074 1 491915 Firmicutes
Geobacillus thermodenitrificans NG80 2 YP 001127422 1 420246 Firmicutes
Geobacillus sp Y4 1MC1 YP 003990972 1 581103 Firmicutes
Geobacillus thermoglucosidasius C56 YS93 YP 004589734 1 634956 Firmicutes
Bacillus atrophaeus 1942 YP 003975159 1 720555 Firmicutes
Bacillus subtilis subsp spizizenii TU B 10 YP 004879297 1 1052585 Firmicutes
Bacillus subtilis subsp natto BEST195 YP 005563081 1 645657 Firmicutes
Bacillus subtilis subsp subtilis str BAB 1 YP 007664380 1 1302650 Firmicutes
Bacillus subtilis subsp subtilis 6051 HGW YP 007535713 1 1147161 Firmicutes
Bacillus subtilis PY79 YP 008831885 1 1415167 Firmicutes
Azoarcus sp KH32C YP 007551786 1 748247 Proteobacteria
Aeromonas veronii B565 YP 004392836 1 998088 Proteobacteria
Oceanimonas sp GK1 YP 005093544 1 511062 Proteobacteria
Alteromonas macleodii str Black Sea 11 YP 006824566 1 1004785 Proteobacteria
Alteromonas sp SN2 YP 004467382 1 715451 Proteobacteria
Maricaulis maris MCS10 YP 756969 1 394221 Proteobacteria
Rhodomicrobium vannielii ATCC 17100 YP 004011073 1 648757 Proteobacteria
Rhizobium etli CFN 42 YP 471998 1 347834 Proteobacteria
Rhizobium etli bv mimosae str Mim1 YP 008367115 1 1328306 Proteobacteria
Rhizobium leguminosarum bv viciae 3841 YP 770574 1 216596 Proteobacteria
Rhizobium leguminosarum bv trifolii WSM2304 YP 002278376 1 395492 Proteobacteria
Pelagibacterium halotolerans B2 YP 004898864 1 1082931 Proteobacteria
Phenylobacterium zucineum HLK1 YP 002131244 1 450851 Proteobacteria
Phenylobacterium zucineum HLK1 YP 002130127 1 450851 Proteobacteria
Brevundimonas subvibrioides ATCC 15264 YP 003819331 1 633149 Proteobacteria
Caulobacter sp K31 YP 001683501 1 366602 Proteobacteria
Hirschia baltica ATCC 49814 YP 003059989 1 582402 Proteobacteria
Sphingobium sp SYK 6 YP 004834107 1 627192 Proteobacteria
Erythrobacter litoralis HTCC2594 YP 459246 1 314225 Proteobacteria
Sphingopyxis alaskensis RB2256 YP 616575 1 317655 Proteobacteria
Novosphingobium aromaticivorans DSM 12444 YP 496919 1 279238 Proteobacteria

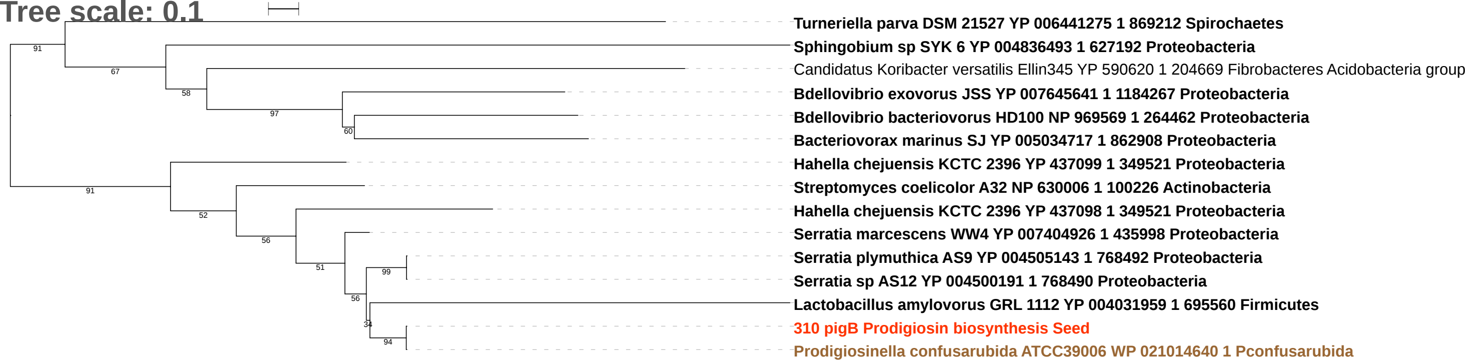
Tree scale: 0.1
Turneriella parva DSM 21527 YP 006441275 1 869212 Spirochaetes
Sphingobium sp SYK 6 YP 004836493 1 627192 Proteobacteria
Candidatus Koribacter versatilis Ellin345 YP 590620 1 204669 Fibrobacteres Acidobacteria group
Bdellovibrio exovorus JSS YP 007645641 1 1184267 Proteobacteria
Bdellovibrio bacteriovorus HD100 NP 969569 1 264462 Proteobacteria
Bacteriovorax marinus SJ YP 005034717 1 862908 Proteobacteria
Hahella chejuensis KCTC 2396 YP 437099 1 349521 Proteobacteria
Streptomyces coelicolor A32 NP 630006 1 100226 Actinobacteria
Hahella chejuensis KCTC 2396 YP 437098 1 349521 Proteobacteria
Serratia marcescens WW4 YP 007404926 1 435998 Proteobacteria
Serratia plymuthica AS9 YP 004505143 1 768492 Proteobacteria
Serratia sp AS12 YP 004500191 1 768490 Proteobacteria
Lactobacillus amylovorus GRL 1112 YP 004031959 1 695560 Firmicutes
310 pigB Prodigiosin biosynthesis Seed
Prodigiosinella confusarubida ATCC39006 WP 021014640 1 Pconfusarubida
91
67
58
97
60
91
52
56
51
99
56
34
94

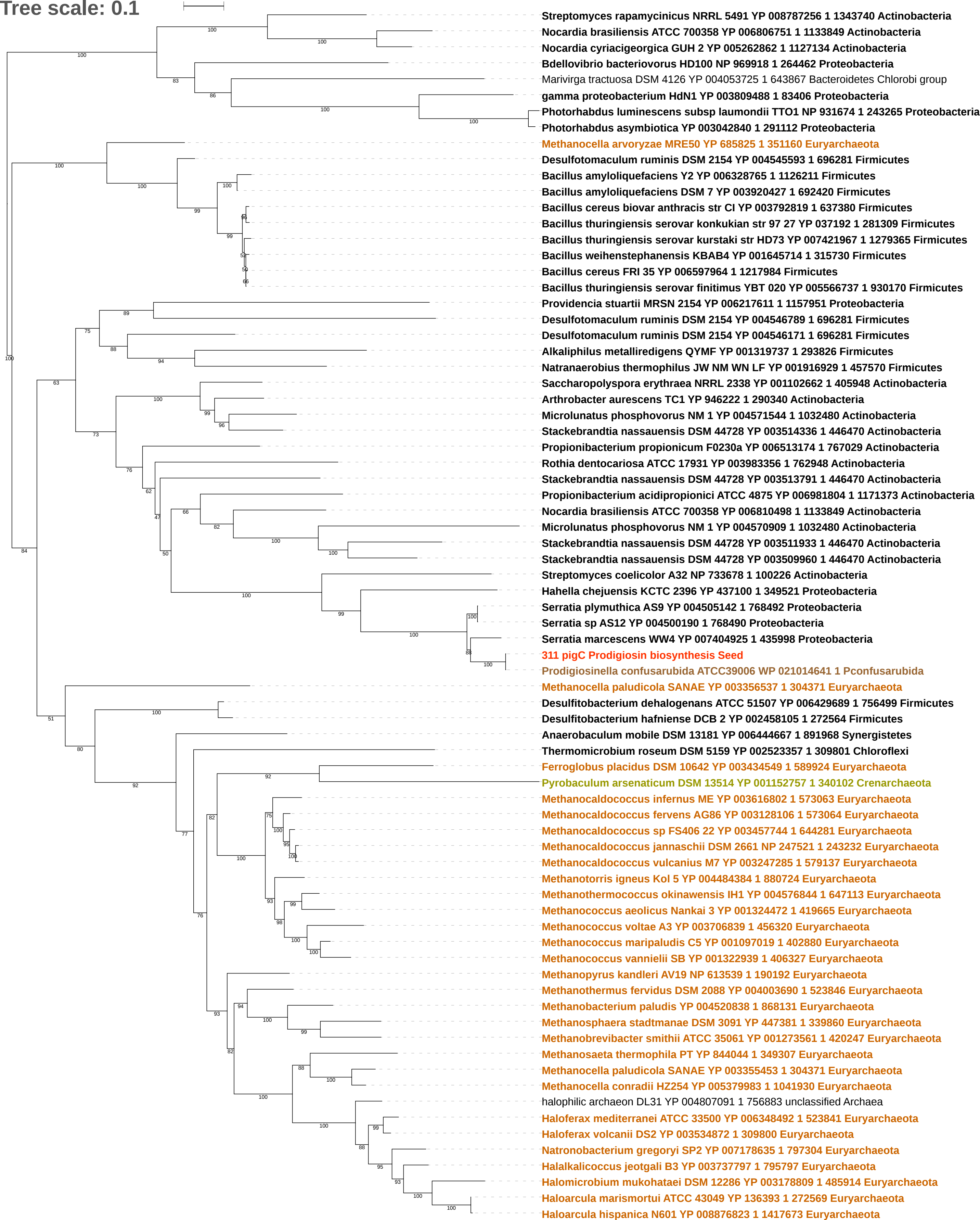
Tree scale: 0.1
Streptomyces rapamycinicus NRRL 5491 YP 008787256 1 1343740 Actinobacteria
Nocardia brasiliensis ATCC 700358 YP 006806751 1 1133849 Actinobacteria
Nocardia cyriacigeorgica GUH 2 YP 005262862 1 1127134 Actinobacteria
Bdellovibrio bacteriovorus HD100 NP 969918 1 264462 Proteobacteria
Marivirga tractuosa DSM 4126 YP 004053725 1 643867 Bacteroidetes Chlorobi group
gamma proteobacterium HdN1 YP 003809488 1 83406 Proteobacteria
Photorhabdus luminescens subsp laumondii TTO1 NP 931674 1 243265 Proteobacteria
Photorhabdus asymbiotica YP 003042840 1 291112 Proteobacteria
Methanocella arvoryzae MRE50 YP 685825 1 351160 Euryarchaeota
Desulfotomaculum ruminis DSM 2154 YP 004545593 1 696281 Firmicutes
Bacillus amyloliquefaciens Y2 YP 006328765 1 1126211 Firmicutes
Bacillus amyloliquefaciens DSM 7 YP 003920427 1 692420 Firmicutes
Bacillus cereus biovar anthracis str CI YP 003792819 1 637380 Firmicutes
Bacillus thuringiensis serovar konkukian str 97 27 YP 037192 1 281309 Firmicutes
Bacillus thuringiensis serovar kurstaki str HD73 YP 007421967 1 1279365 Firmicutes
Bacillus weihenstephanensis KBAB4 YP 001645714 1 315730 Firmicutes
Bacillus cereus FRI 35 YP 006597964 1 1217984 Firmicutes
Bacillus thuringiensis serovar finitimus YBT 020 YP 005566737 1 930170 Firmicutes
Providencia stuartii MRSN 2154 YP 006217611 1 1157951 Proteobacteria
Desulfotomaculum ruminis DSM 2154 YP 004546789 1 696281 Firmicutes
Desulfotomaculum ruminis DSM 2154 YP 004546171 1 696281 Firmicutes
Alkaliphilus metalliredigens QYMF YP 001319737 1 293826 Firmicutes
Natranaerobius thermophilus JW NM WN LF YP 001916929 1 457570 Firmicutes
Saccharopolyspora erythraea NRRL 2338 YP 001102662 1 405948 Actinobacteria
Arthrobacter aurescens TC1 YP 946222 1 290340 Actinobacteria
Microlunatus phosphovorus NM 1 YP 004571544 1 1032480 Actinobacteria
Stackebrandtia nassauensis DSM 44728 YP 003514336 1 446470 Actinobacteria
Propionibacterium propionicum F0230a YP 006513174 1 767029 Actinobacteria
Rothia dentocariosa ATCC 17931 YP 003983356 1 762948 Actinobacteria
Stackebrandtia nassauensis DSM 44728 YP 003513791 1 446470 Actinobacteria
Propionibacterium acidipropionici ATCC 4875 YP 006981804 1 1171373 Actinobacteria
Nocardia brasiliensis ATCC 700358 YP 006810498 1 1133849 Actinobacteria
Microlunatus phosphovorus NM 1 YP 004570909 1 1032480 Actinobacteria
Stackebrandtia nassauensis DSM 44728 YP 003511933 1 446470 Actinobacteria
Stackebrandtia nassauensis DSM 44728 YP 003509960 1 446470 Actinobacteria
Streptomyces coelicolor A32 NP 733678 1 100226 Actinobacteria
Hahella chejuensis KCTC 2396 YP 437100 1 349521 Proteobacteria
Serratia plymuthica AS9 YP 004505142 1 768492 Proteobacteria
Serratia sp AS12 YP 004500190 1 768490 Proteobacteria
Serratia marcescens WW4 YP 007404925 1 435998 Proteobacteria
311 pigC Prodigiosin biosynthesis Seed
Prodigiosinella confusarubida ATCC39006 WP 021014641 1 Pconfusarubida
Methanocella paludicola SANAE YP 003356537 1 304371 Euryarchaeota
Desulfitobacterium dehalogenans ATCC 51507 YP 006429689 1 756499 Firmicutes
Desulfitobacterium hafniense DCB 2 YP 002458105 1 272564 Firmicutes
Anaerobaculum mobile DSM 13181 YP 006444667 1 891968 Synergistetes
Thermomicrobium roseum DSM 5159 YP 002523357 1 309801 Chloroflexi
Ferroglobus placidus DSM 10642 YP 003434549 1 589924 Euryarchaeota
Pyrobaculum arsenaticum DSM 13514 YP 001152757 1 340102 Crenarchaeota
Methanocaldococcus infernus ME YP 003616802 1 573063 Euryarchaeota
Methanocaldococcus fervens AG86 YP 003128106 1 573064 Euryarchaeota
Methanocaldococcus sp FS406 22 YP 003457744 1 644281 Euryarchaeota
Methanocaldococcus jannaschii DSM 2661 NP 247521 1 243232 Euryarchaeota
Methanocaldococcus vulcanius M7 YP 003247285 1 579137 Euryarchaeota
Methanotorris igneus Kol 5 YP 004484384 1 880724 Euryarchaeota
Methanothermococcus okinawensis IH1 YP 004576844 1 647113 Euryarchaeota
Methanococcus aeolicus Nankai 3 YP 001324472 1 419665 Euryarchaeota
Methanococcus voltae A3 YP 003706839 1 456320 Euryarchaeota
Methanococcus maripaludis C5 YP 001097019 1 402880 Euryarchaeota
Methanococcus vannielii SB YP 001322939 1 406327 Euryarchaeota
Methanopyrus kandleri AV19 NP 613539 1 190192 Euryarchaeota
Methanothermus fervidus DSM 2088 YP 004003690 1 523846 Euryarchaeota
Methanobacterium paludis YP 004520838 1 868131 Euryarchaeota
Methanosphaera stadtmanae DSM 3091 YP 447381 1 339860 Euryarchaeota
Methanobrevibacter smithii ATCC 35061 YP 001273561 1 420247 Euryarchaeota
Methanosaeta thermophila PT YP 844044 1 349307 Euryarchaeota
Methanocella paludicola SANAE YP 003355453 1 304371 Euryarchaeota
Methanocella conradii HZ254 YP 005379983 1 1041930 Euryarchaeota
halophilic archaeon DL31 YP 004807091 1 756883 unclassified Archaea
Haloferax mediterranei ATCC 33500 YP 006348492 1 523841 Euryarchaeota
Haloferax volcanii DS2 YP 003534872 1 309800 Euryarchaeota
Natronobacterium gregoryi SP2 YP 007178635 1 797304 Euryarchaeota
Halalkalicoccus jeotgali B3 YP 003737797 1 795797 Euryarchaeota
Halomicrobium mukohataei DSM 12286 YP 003178809 1 485914 Euryarchaeota
Haloarcula marismortui ATCC 43049 YP 136393 1 272569 Euryarchaeota
Haloarcula hispanica N601 YP 008876823 1 1417673 Euryarchaeota

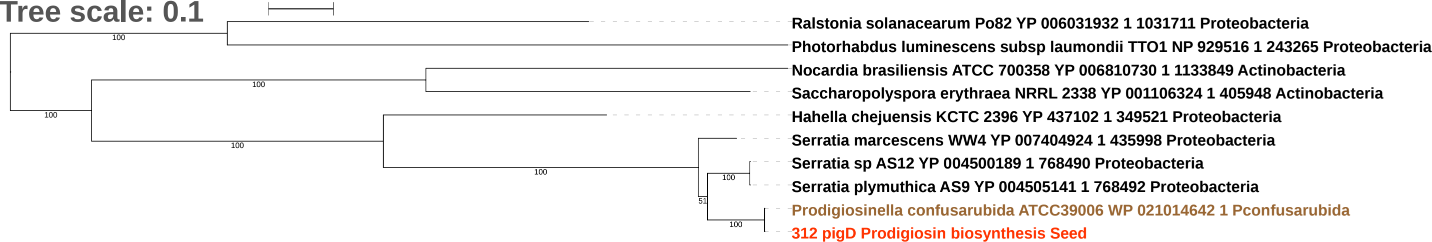

Tree scale: 0.1
Ralstonia solanacearum Po82 YP 006031932 1 1031711 Proteobacteria
Photorhabdus luminescens subsp laumondii TTO1 NP 929516 1 243265 Proteobacteria
Nocardia brasiliensis ATCC 700358 YP 006810730 1 1133849 Actinobacteria
Saccharopolyspora erythraea NRRL 2338 YP 001106324 1 405948 Actinobacteria
Hahella chejuensis KCTC 2396 YP 437102 1 349521 Proteobacteria
Serratia marcescens WW4 YP 007404924 1 435998 Proteobacteria
Serratia sp AS12 YP 004500189 1 768490 Proteobacteria
Serratia plymuthica AS9 YP 004505141 1 768492 Proteobacteria
Prodigiosinella confusarubida ATCC39006 WP 021014642 1 Pconfusarubida
312 pigD Prodigiosin biosynthesis Seed
100
100
100
100
100
100
100
51
100

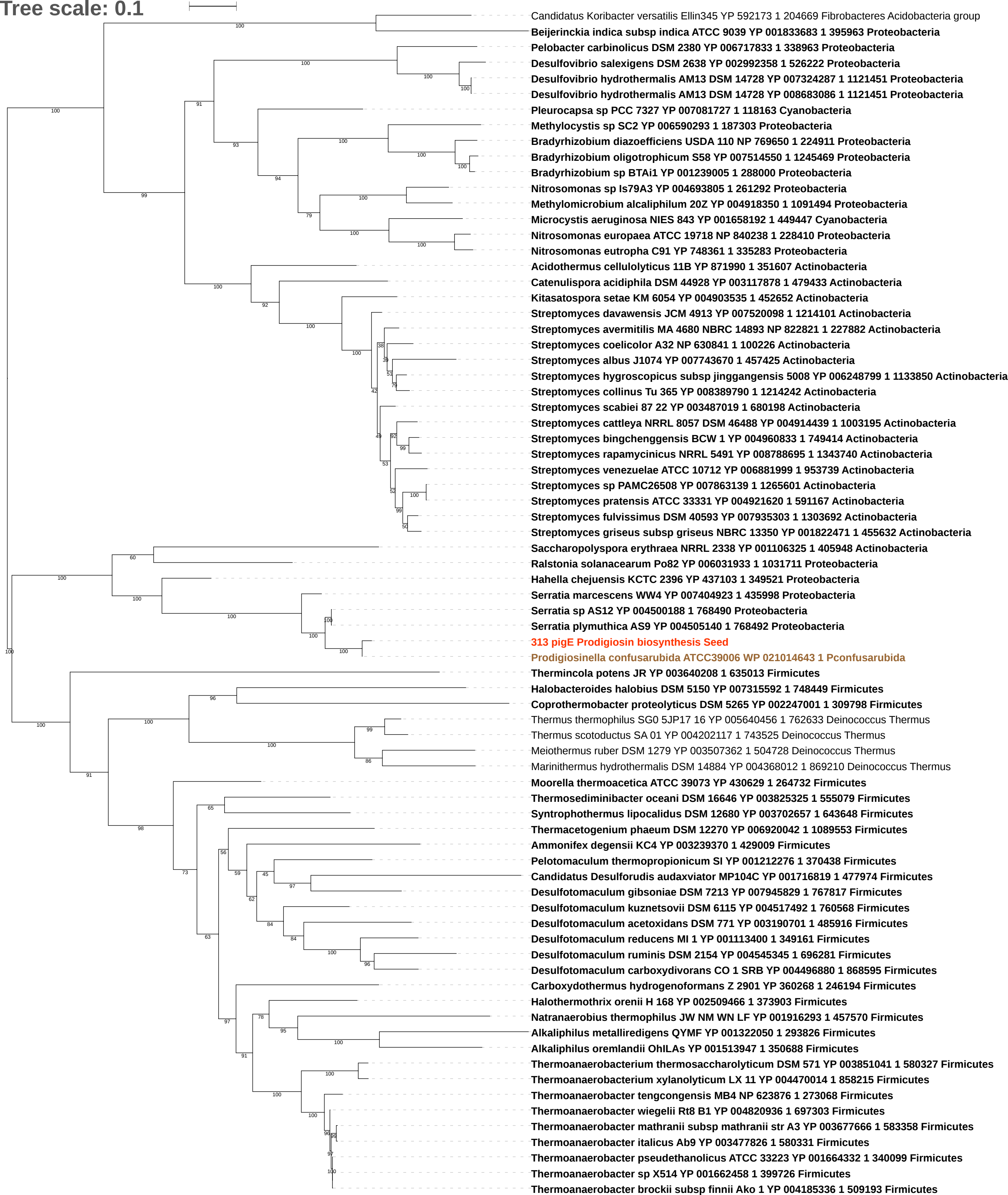
Tree scale: 0.1
Candidatus Koribacter versatilis Ellin345 YP 592173 1 204669 Fibrobacteres Acidobacteria group
Beijerinckia indica subsp indica ATCC 9039 YP 001833683 1 395963 Proteobacteria
Pelobacter carbinolicus DSM 2380 YP 006717833 1 338963 Proteobacteria
Desulfovibrio salexigens DSM 2638 YP 002992358 1 526222 Proteobacteria
Desulfovibrio hydrothermalis AM13 DSM 14728 YP 007324287 1 1121451 Proteobacteria
Desulfovibrio hydrothermalis AM13 DSM 14728 YP 008683086 1 1121451 Proteobacteria
Pleurocapsa sp PCC 7327 YP 007081727 1 118163 Cyanobacteria
Methylocystis sp SC2 YP 006590293 1 187303 Proteobacteria
Bradyrhizobium diazoefficiens USDA 110 NP 769650 1 224911 Proteobacteria
Bradyrhizobium oligotrophicum S58 YP 007514550 1 1245469 Proteobacteria
Bradyrhizobium sp BTAi1 YP 001239005 1 288000 Proteobacteria
Nitrosomonas sp Is79A3 YP 004693805 1 261292 Proteobacteria
Methylomicrobium alcaliphilum 20Z YP 004918350 1 1091494 Proteobacteria
Microcystis aeruginosa NIES 843 YP 001658192 1 449447 Cyanobacteria
Nitrosomonas europaea ATCC 19718 NP 840238 1 228410 Proteobacteria
Nitrosomonas eutropha C91 YP 748361 1 335283 Proteobacteria
Acidothermus cellulolyticus 11B YP 871990 1 351607 Actinobacteria
Catenulispora acidiphila DSM 44928 YP 003117878 1 479433 Actinobacteria
Kitasatospora setae KM 6054 YP 004903535 1 452652 Actinobacteria
Streptomyces davawensis JCM 4913 YP 007520098 1 1214101 Actinobacteria
Streptomyces avermitilis MA 4680 NBRC 14893 NP 822821 1 227882 Actinobacteria
Streptomyces coelicolor A32 NP 630841 1 100226 Actinobacteria
Streptomyces albus J1074 YP 007743670 1 457425 Actinobacteria
Streptomyces hygroscopicus subsp jinggangensis 5008 YP 006248799 1 1133850 Actinobacteria
Streptomyces collinus Tu 365 YP 008389790 1 1214242 Actinobacteria
Streptomyces scabiei 87 22 YP 003487019 1 680198 Actinobacteria
Streptomyces cattleya NRRL 8057 DSM 46488 YP 004914439 1 1003195 Actinobacteria
Streptomyces bingchenggensis BCW 1 YP 004960833 1 749414 Actinobacteria
Streptomyces rapamycinicus NRRL 5491 YP 008788695 1 1343740 Actinobacteria
Streptomyces venezuelae ATCC 10712 YP 006881999 1 953739 Actinobacteria
Streptomyces sp PAMC26508 YP 007863139 1 1265601 Actinobacteria
Streptomyces pratensis ATCC 33331 YP 004921620 1 591167 Actinobacteria
Streptomyces fulvissimus DSM 40593 YP 007935303 1 1303692 Actinobacteria
Streptomyces griseus subsp griseus NBRC 13350 YP 001822471 1 455632 Actinobacteria
Saccharopolyspora erythraea NRRL 2338 YP 001106325 1 405948 Actinobacteria
Ralstonia solanacearum Po82 YP 006031933 1 1031711 Proteobacteria
Hahella chejuensis KCTC 2396 YP 437103 1 349521 Proteobacteria
Serratia marcescens WW4 YP 007404923 1 435998 Proteobacteria
Serratia sp AS12 YP 004500188 1 768490 Proteobacteria
Serratia plymuthica AS9 YP 004505140 1 768492 Proteobacteria
313 pigE Prodigiosin biosynthesis Seed
Prodigiosinella confusarubida ATCC39006 WP 021014643 1 Pconfusarubida
Thermincola potens JR YP 003640208 1 635013 Firmicutes
Halobacteroides halobius DSM 5150 YP 007315592 1 748449 Firmicutes
Coprothermobacter proteolyticus DSM 5265 YP 002247001 1 309798 Firmicutes
Thermus thermophilus SG0 5JP17 16 YP 005640456 1 762633 Deinococcus Thermus
Thermus scotoductus SA 01 YP 004202117 1 743525 Deinococcus Thermus
Meiothermus ruber DSM 1279 YP 003507362 1 504728 Deinococcus Thermus
Marinithermus hydrothermalis DSM 14884 YP 004368012 1 869210 Deinococcus Thermus
Moorella thermoacetica ATCC 39073 YP 430629 1 264732 Firmicutes
Thermosediminibacter oceani DSM 16646 YP 003825325 1 555079 Firmicutes
Syntrophothermus lipocalidus DSM 12680 YP 003702657 1 643648 Firmicutes
Thermacetogenium phaeum DSM 12270 YP 006920042 1 1089553 Firmicutes
Ammonifex degensii KC4 YP 003239370 1 429009 Firmicutes
Pelotomaculum thermopropionicum SI YP 001212276 1 370438 Firmicutes
Candidatus Desulforudis audaxviator MP104C YP 001716819 1 477974 Firmicutes
Desulfotomaculum gibsoniae DSM 7213 YP 007945829 1 767817 Firmicutes
Desulfotomaculum kuznetsovii DSM 6115 YP 004517492 1 760568 Firmicutes
Desulfotomaculum acetoxidans DSM 771 YP 003190701 1 485916 Firmicutes
Desulfotomaculum reducens MI 1 YP 001113400 1 349161 Firmicutes
Desulfotomaculum ruminis DSM 2154 YP 004545345 1 696281 Firmicutes
Desulfotomaculum carboxydivorans CO 1 SRB YP 004496880 1 868595 Firmicutes
Carboxydothermus hydrogenoformans Z 2901 YP 360268 1 246194 Firmicutes
Halothermothrix orenii H 168 YP 002509466 1 373903 Firmicutes
Natranaerobius thermophilus JW NM WN LF YP 001916293 1 457570 Firmicutes
Alkaliphilus metalliredigens QYMF YP 001322050 1 293826 Firmicutes
Alkaliphilus oremlandii OhILAs YP 001513947 1 350688 Firmicutes
Thermoanaerobacterium thermosaccharolyticum DSM 571 YP 003851041 1 580327 Firmicutes
Thermoanaerobacterium xylanolyticum LX 11 YP 004470014 1 858215 Firmicutes
Thermoanaerobacter tengcongensis MB4 NP 623876 1 273068 Firmicutes
Thermoanaerobacter wiegelii Rt8 B1 YP 004820936 1 697303 Firmicutes
Thermoanaerobacter mathranii subsp mathranii str A3 YP 003677666 1 583358 Firmicutes
Thermoanaerobacter italicus Ab9 YP 003477826 1 580331 Firmicutes
Thermoanaerobacter pseudethanolicus ATCC 33223 YP 001664332 1 340099 Firmicutes
Thermoanaerobacter sp X514 YP 001662458 1 399726 Firmicutes
Thermoanaerobacter brockii subsp finnii Ako 1 YP 004185336 1 509193 Firmicutes

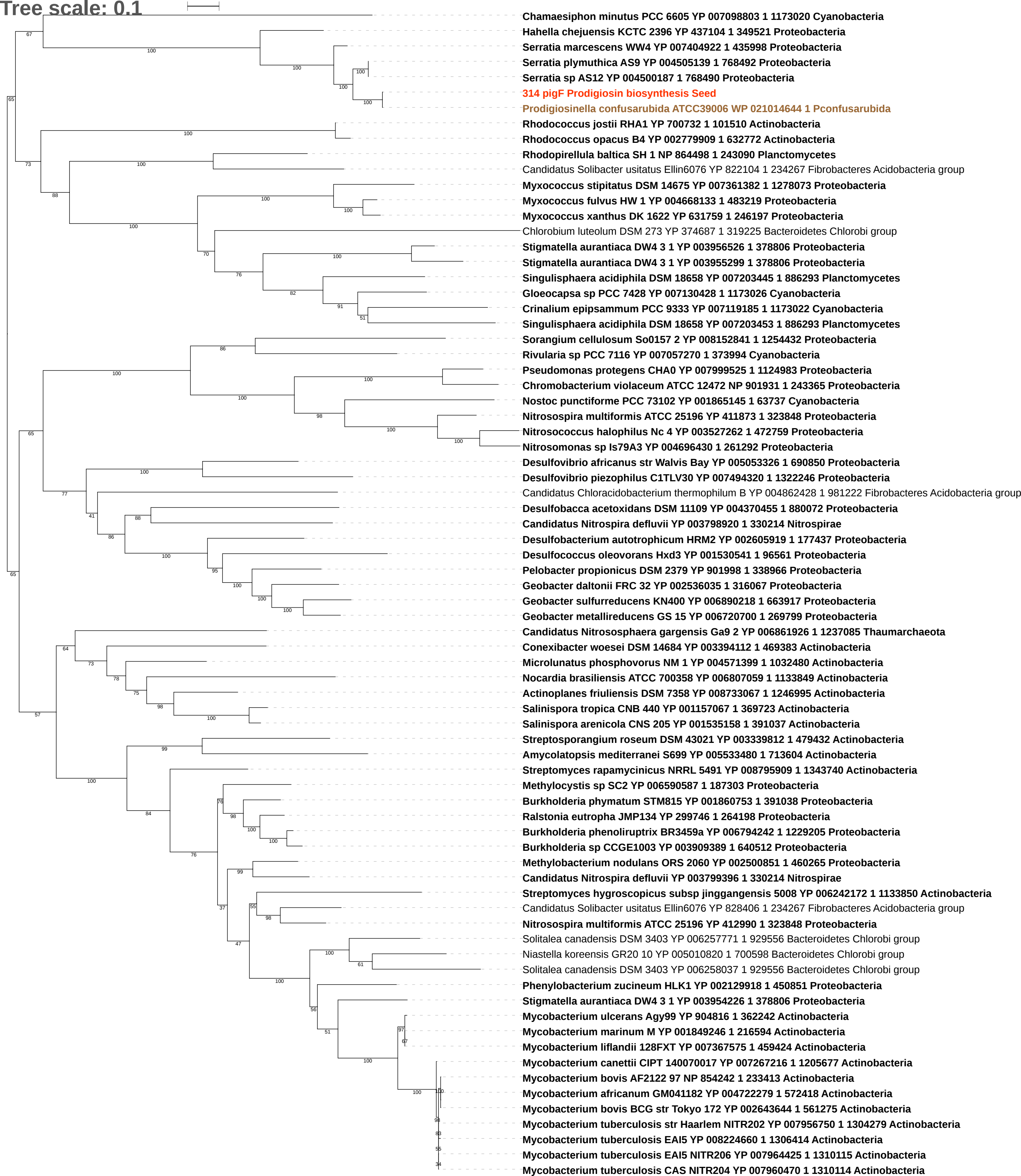
Tree scale: 0.1
Chamaesiphon minutus PCC 6605 YP 007098803 1 1173020 Cyanobacteria
Hahella chejuensis KCTC 2396 YP 437104 1 349521 Proteobacteria
Serratia marcescens WW4 YP 007404922 1 435998 Proteobacteria
Serratia plymuthica AS9 YP 004505139 1 768492 Proteobacteria
Serratia sp AS12 YP 004500187 1 768490 Proteobacteria
314 pigF Prodigiosin biosynthesis Seed
Prodigiosinella confusarubida ATCC39006 WP 021014644 1 Pconfusarubida
Rhodococcus jostii RHA1 YP 700732 1 101510 Actinobacteria
Rhodococcus opacus B4 YP 002779909 1 632772 Actinobacteria
Rhodopirellula baltica SH 1 NP 864498 1 243090 Planctomycetes
Candidatus Solibacter usitatus Ellin6076 YP 822104 1 234267 Fibrobacteres Acidobacteria group
Myxococcus stipitatus DSM 14675 YP 007361382 1 1278073 Proteobacteria
Myxococcus fulvus HW 1 YP 004668133 1 483219 Proteobacteria
Myxococcus xanthus DK 1622 YP 631759 1 246197 Proteobacteria
Chlorobium luteolum DSM 273 YP 374687 1 319225 Bacteroidetes Chlorobi group
Stigmatella aurantiaca DW4 3 1 YP 003956526 1 378806 Proteobacteria
Stigmatella aurantiaca DW4 3 1 YP 003955299 1 378806 Proteobacteria
Singulisphaera acidiphila DSM 18658 YP 007203445 1 886293 Planctomycetes
Gloeocapsa sp PCC 7428 YP 007130428 1 1173026 Cyanobacteria
Crinalium epipsammum PCC 9333 YP 007119185 1 1173022 Cyanobacteria
Singulisphaera acidiphila DSM 18658 YP 007203453 1 886293 Planctomycetes
Sorangium cellulosum So0157 2 YP 008152841 1 1254432 Proteobacteria
Rivularia sp PCC 7116 YP 007057270 1 373994 Cyanobacteria
Pseudomonas protegens CHA0 YP 007999525 1 1124983 Proteobacteria
Chromobacterium violaceum ATCC 12472 NP 901931 1 243365 Proteobacteria
Nostoc punctiforme PCC 73102 YP 001865145 1 63737 Cyanobacteria
Nitrosospira multiformis ATCC 25196 YP 411873 1 323848 Proteobacteria
Nitrosococcus halophilus Nc 4 YP 003527262 1 472759 Proteobacteria
Nitrosomonas sp Is79A3 YP 004696430 1 261292 Proteobacteria
Desulfovibrio africanus str Walvis Bay YP 005053326 1 690850 Proteobacteria
Desulfovibrio piezophilus C1TLV30 YP 007494320 1 1322246 Proteobacteria
Candidatus Chloracidobacterium thermophilum B YP 004862428 1 981222 Fibrobacteres Acidobacteria group
Desulfobacca acetoxidans DSM 11109 YP 004370455 1 880072 Proteobacteria
Candidatus Nitrospira defluvii YP 003798920 1 330214 Nitrospirae
Desulfobacterium autotrophicum HRM2 YP 002605919 1 177437 Proteobacteria
Desulfococcus oleovorans Hxd3 YP 001530541 1 96561 Proteobacteria
Pelobacter propionicus DSM 2379 YP 901998 1 338966 Proteobacteria
Geobacter daltonii FRC 32 YP 002536035 1 316067 Proteobacteria
Geobacter sulfurreducens KN400 YP 006890218 1 663917 Proteobacteria
Geobacter metallireducens GS 15 YP 006720700 1 269799 Proteobacteria
Candidatus Nitrososphaera gargensis Ga9 2 YP 006861926 1 1237085 Thaumarchaeota
Conexibacter woesei DSM 14684 YP 003394112 1 469383 Actinobacteria
Microlunatus phosphovorus NM 1 YP 004571399 1 1032480 Actinobacteria
Nocardia brasiliensis ATCC 700358 YP 006807059 1 1133849 Actinobacteria
Actinoplanes friuliensis DSM 7358 YP 008733067 1 1246995 Actinobacteria
Salinispora tropica CNB 440 YP 001157067 1 369723 Actinobacteria
Salinispora arenicola CNS 205 YP 001535158 1 391037 Actinobacteria
Streptosporangium roseum DSM 43021 YP 003339812 1 479432 Actinobacteria
Amycolatopsis mediterranei S699 YP 005533480 1 713604 Actinobacteria
Streptomyces rapamycinicus NRRL 5491 YP 008795909 1 1343740 Actinobacteria
Methylocystis sp SC2 YP 006590587 1 187303 Proteobacteria
Burkholderia phymatum STM815 YP 001860753 1 391038 Proteobacteria
Ralstonia eutropha JMP134 YP 299746 1 264198 Proteobacteria
Burkholderia phenoliruptrix BR3459a YP 006794242 1 1229205 Proteobacteria
Burkholderia sp CCGE1003 YP 003909389 1 640512 Proteobacteria
Methylobacterium nodulans ORS 2060 YP 002500851 1 460265 Proteobacteria
Candidatus Nitrospira defluvii YP 003799396 1 330214 Nitrospirae
Streptomyces hygroscopicus subsp jinggangensis 5008 YP 006242172 1 1133850 Actinobacteria
Candidatus Solibacter usitatus Ellin6076 YP 828406 1 234267 Fibrobacteres Acidobacteria group
Nitrosospira multiformis ATCC 25196 YP 412990 1 323848 Proteobacteria
Solitalea canadensis DSM 3403 YP 006257771 1 929556 Bacteroidetes Chlorobi group
Niastella koreensis GR20 10 YP 005010820 1 700598 Bacteroidetes Chlorobi group
Solitalea canadensis DSM 3403 YP 006258037 1 929556 Bacteroidetes Chlorobi group
Phenylobacterium zucineum HLK1 YP 002129918 1 450851 Proteobacteria
Stigmatella aurantiaca DW4 3 1 YP 003954226 1 378806 Proteobacteria
Mycobacterium ulcerans Agy99 YP 904816 1 362242 Actinobacteria
Mycobacterium marinum M YP 001849246 1 216594 Actinobacteria
Mycobacterium liflandii 128FXT YP 007367575 1 459424 Actinobacteria
Mycobacterium canettii CIPT 140070017 YP 007267216 1 1205677 Actinobacteria
Mycobacterium bovis AF2122 97 NP 854242 1 233413 Actinobacteria
Mycobacterium africanum GM041182 YP 004722279 1 572418 Actinobacteria
Mycobacterium bovis BCG str Tokyo 172 YP 002643644 1 561275 Actinobacteria
Mycobacterium tuberculosis str Haarlem NITR202 YP 007956750 1 1304279 Actinobacteria
Mycobacterium tuberculosis EAI5 YP 008224660 1 1306414 Actinobacteria
Mycobacterium tuberculosis EAI5 NITR206 YP 007964425 1 1310115 Actinobacteria
Mycobacterium tuberculosis CAS NITR204 YP 007960470 1 1310114 Actinobacteria

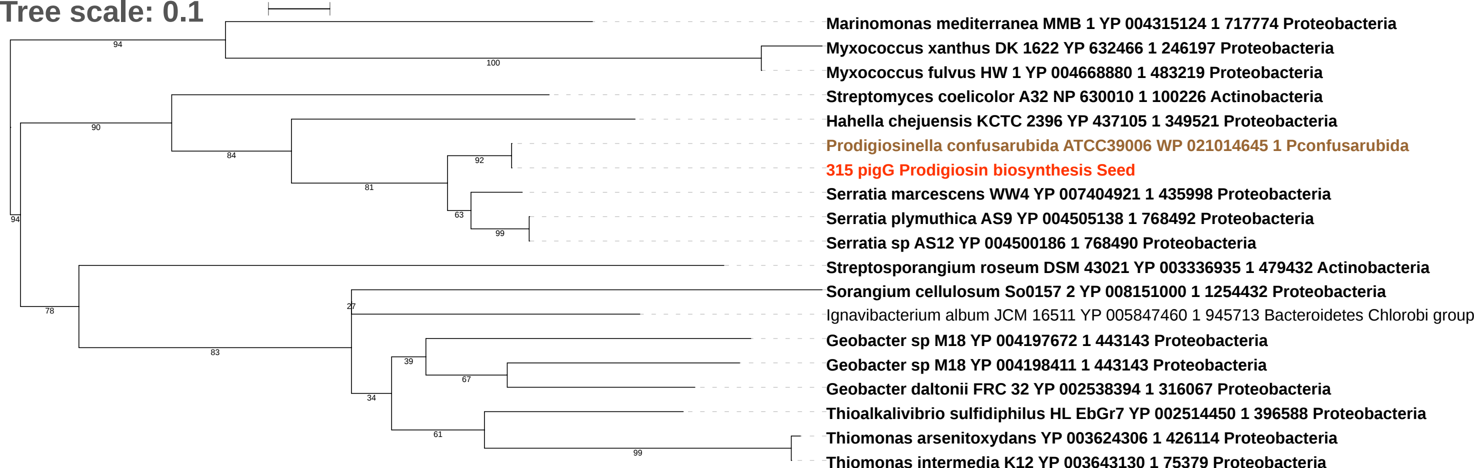

Tree scale: 0.1
Marinomonas mediterranea MMB 1 YP 004315124 1 717774 Proteobacteria
Myxococcus xanthus DK 1622 YP 632466 1 246197 Proteobacteria
Myxococcus fulvus HW 1 YP 004668880 1 483219 Proteobacteria
Streptomyces coelicolor A32 NP 630010 1 100226 Actinobacteria
Hahella chejuensis KCTC 2396 YP 437105 1 349521 Proteobacteria
Prodigiosinella confusarubida ATCC39006 WP 021014645 1 Pconfusarubida
315 pigG Prodigiosin biosynthesis Seed
Serratia marcescens WW4 YP 007404921 1 435998 Proteobacteria
Serratia plymuthica AS9 YP 004505138 1 768492 Proteobacteria
Serratia sp AS12 YP 004500186 1 768490 Proteobacteria
Streptosporangium roseum DSM 43021 YP 003336935 1 479432 Actinobacteria
Sorangium cellulosum So0157 2 YP 008151000 1 1254432 Proteobacteria
Ignavibacterium album JCM 16511 YP 005847460 1 945713 Bacteroidetes Chlorobi group
Geobacter sp M18 YP 004197672 1 443143 Proteobacteria
Geobacter sp M18 YP 004198411 1 443143 Proteobacteria
Geobacter daltonii FRC 32 YP 002538394 1 316067 Proteobacteria
Thioalkalivibrio sulfidiphilus HL EbGr7 YP 002514450 1 396588 Proteobacteria
Thiomonas arsenitoxydans YP 003624306 1 426114 Proteobacteria
Thiomonas intermedia K12 YP 003643130 1 75379 Proteobacteria
94
100
90
84
92
81
63
99
94
78
27
83
39
67
34
61
99

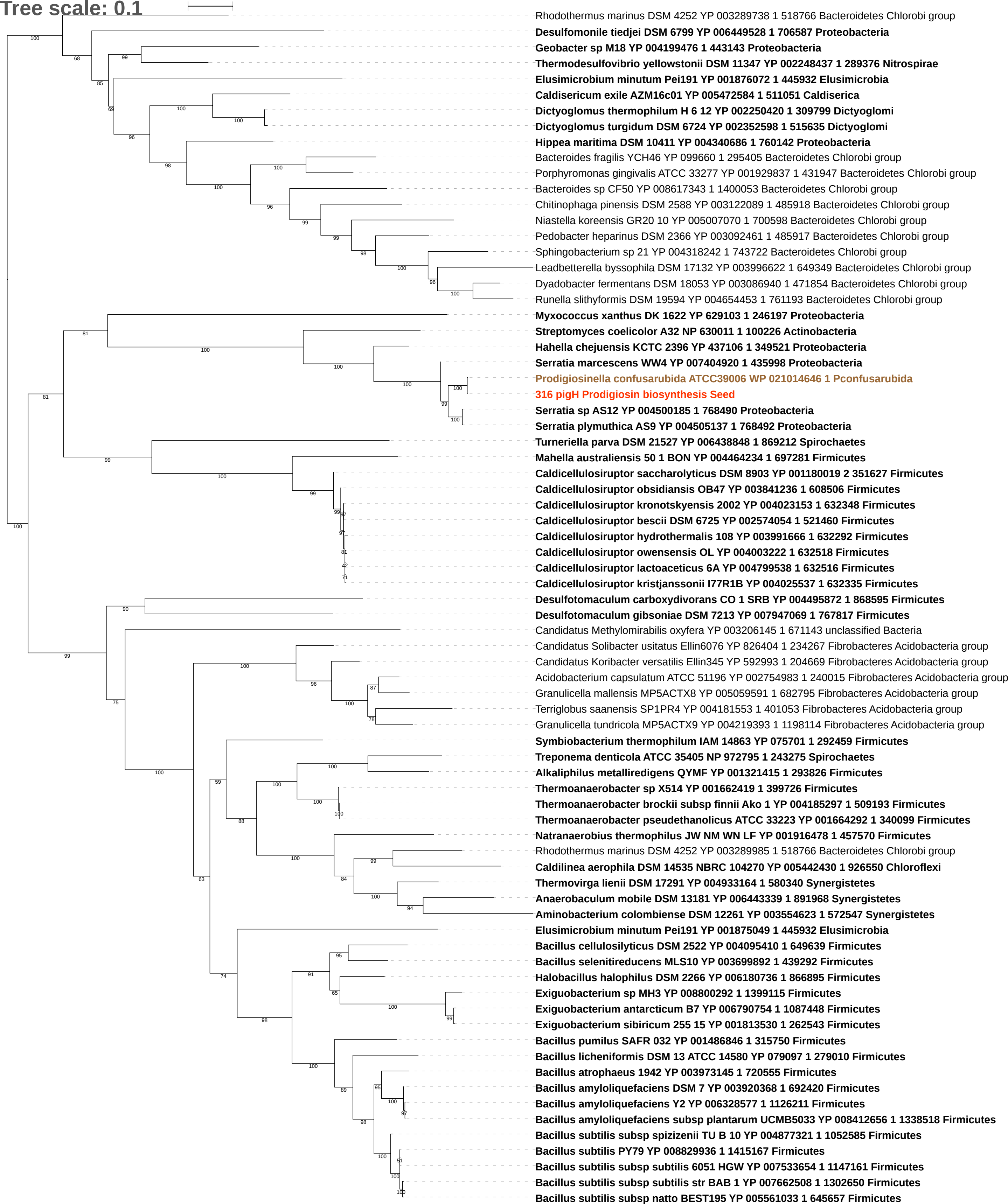
Tree scale: 0.1
Rhodothermus marinus DSM 4252 YP 003289738 1 518766 Bacteroidetes Chlorobi group
Desulfomonile tiedjei DSM 6799 YP 006449528 1 706587 Proteobacteria
Geobacter sp M18 YP 004199476 1 443143 Proteobacteria
Thermodesulfovibrio yellowstonii DSM 11347 YP 002248437 1 289376 Nitrospirae
Elusimicrobium minutum Pei191 YP 001876072 1 445932 Elusimicrobia
Caldisericum exile AZM16c01 YP 005472584 1 511051 Caldiserica
Dictyoglomus thermophilum H 6 12 YP 002250420 1 309799 Dictyoglomi
Dictyoglomus turgidum DSM 6724 YP 002352598 1 515635 Dictyoglomi
Hippea maritima DSM 10411 YP 004340686 1 760142 Proteobacteria
Bacteroides fragilis YCH46 YP 099660 1 295405 Bacteroidetes Chlorobi group
Porphyromonas gingivalis ATCC 33277 YP 001929837 1 431947 Bacteroidetes Chlorobi group
Bacteroides sp CF50 YP 008617343 1 1400053 Bacteroidetes Chlorobi group
Chitinophaga pinensis DSM 2588 YP 003122089 1 485918 Bacteroidetes Chlorobi group
Niastella koreensis GR20 10 YP 005007070 1 700598 Bacteroidetes Chlorobi group
Pedobacter heparinus DSM 2366 YP 003092461 1 485917 Bacteroidetes Chlorobi group
Sphingobacterium sp 21 YP 004318242 1 743722 Bacteroidetes Chlorobi group
Leadbetterella byssophila DSM 17132 YP 003996622 1 649349 Bacteroidetes Chlorobi group
Dyadobacter fermentans DSM 18053 YP 003086940 1 471854 Bacteroidetes Chlorobi group
Runella slithyformis DSM 19594 YP 004654453 1 761193 Bacteroidetes Chlorobi group
Myxococcus xanthus DK 1622 YP 629103 1 246197 Proteobacteria
Streptomyces coelicolor A32 NP 630011 1 100226 Actinobacteria
Hahella chejuensis KCTC 2396 YP 437106 1 349521 Proteobacteria
Serratia marcescens WW4 YP 007404920 1 435998 Proteobacteria
Prodigiosinella confusarubida ATCC39006 WP 021014646 1 Pconfusarubida
316 pigH Prodigiosin biosynthesis Seed
Serratia sp AS12 YP 004500185 1 768490 Proteobacteria
Serratia plymuthica AS9 YP 004505137 1 768492 Proteobacteria
Turneriella parva DSM 21527 YP 006438848 1 869212 Spirochaetes
Mahella australiensis 50 1 BON YP 004464234 1 697281 Firmicutes
Caldicellulosiruptor saccharolyticus DSM 8903 YP 001180019 2 351627 Firmicutes
Caldicellulosiruptor obsidiansis OB47 YP 003841236 1 608506 Firmicutes
Caldicellulosiruptor kronotskyensis 2002 YP 004023153 1 632348 Firmicutes
Caldicellulosiruptor bescii DSM 6725 YP 002574054 1 521460 Firmicutes
Caldicellulosiruptor hydrothermalis 108 YP 003991666 1 632292 Firmicutes
Caldicellulosiruptor owensensis OL YP 004003222 1 632518 Firmicutes
Caldicellulosiruptor lactoaceticus 6A YP 004799538 1 632516 Firmicutes
Caldicellulosiruptor kristjanssonii I77R1B YP 004025537 1 632335 Firmicutes
Desulfotomaculum carboxydivorans CO 1 SRB YP 004495872 1 868595 Firmicutes
Desulfotomaculum gibsoniae DSM 7213 YP 007947069 1 767817 Firmicutes
Candidatus Methylomirabilis oxyfera YP 003206145 1 671143 unclassified Bacteria
Candidatus Solibacter usitatus Ellin6076 YP 826404 1 234267 Fibrobacteres Acidobacteria group
Candidatus Koribacter versatilis Ellin345 YP 592993 1 204669 Fibrobacteres Acidobacteria group
Acidobacterium capsulatum ATCC 51196 YP 002754983 1 240015 Fibrobacteres Acidobacteria group
Granulicella mallensis MP5ACTX8 YP 005059591 1 682795 Fibrobacteres Acidobacteria group
Terriglobus saanensis SP1PR4 YP 004181553 1 401053 Fibrobacteres Acidobacteria group
Granulicella tundricola MP5ACTX9 YP 004219393 1 1198114 Fibrobacteres Acidobacteria group
Symbiobacterium thermophilum IAM 14863 YP 075701 1 292459 Firmicutes
Treponema denticola ATCC 35405 NP 972795 1 243275 Spirochaetes
Alkaliphilus metalliredigens QYMF YP 001321415 1 293826 Firmicutes
Thermoanaerobacter sp X514 YP 001662419 1 399726 Firmicutes
Thermoanaerobacter brockii subsp finnii Ako 1 YP 004185297 1 509193 Firmicutes
Thermoanaerobacter pseudethanolicus ATCC 33223 YP 001664292 1 340099 Firmicutes
Natranaerobius thermophilus JW NM WN LF YP 001916478 1 457570 Firmicutes
Rhodothermus marinus DSM 4252 YP 003289985 1 518766 Bacteroidetes Chlorobi group
Caldilinea aerophila DSM 14535 NBRC 104270 YP 005442430 1 926550 Chloroflexi
Thermovirga lienii DSM 17291 YP 004933164 1 580340 Synergistetes
Anaerobaculum mobile DSM 13181 YP 006443339 1 891968 Synergistetes
Aminobacterium colombiense DSM 12261 YP 003554623 1 572547 Synergistetes
Elusimicrobium minutum Pei191 YP 001875049 1 445932 Elusimicrobia
Bacillus cellulosilyticus DSM 2522 YP 004095410 1 649639 Firmicutes
Bacillus selenitireducens MLS10 YP 003699892 1 439292 Firmicutes
Halobacillus halophilus DSM 2266 YP 006180736 1 866895 Firmicutes
Exiguobacterium sp MH3 YP 008800292 1 1399115 Firmicutes
Exiguobacterium antarcticum B7 YP 006790754 1 1087448 Firmicutes
Exiguobacterium sibiricum 255 15 YP 001813530 1 262543 Firmicutes
Bacillus pumilus SAFR 032 YP 001486846 1 315750 Firmicutes
Bacillus licheniformis DSM 13 ATCC 14580 YP 079097 1 279010 Firmicutes
Bacillus atrophaeus 1942 YP 003973145 1 720555 Firmicutes
Bacillus amyloliquefaciens DSM 7 YP 003920368 1 692420 Firmicutes
Bacillus amyloliquefaciens Y2 YP 006328577 1 1126211 Firmicutes
Bacillus amyloliquefaciens subsp plantarum UCMB5033 YP 008412656 1 1338518 Firmicutes
Bacillus subtilis subsp spizizenii TU B 10 YP 004877321 1 1052585 Firmicutes
Bacillus subtilis PY79 YP 008829936 1 1415167 Firmicutes
Bacillus subtilis subsp subtilis 6051 HGW YP 007533654 1 1147161 Firmicutes
Bacillus subtilis subsp subtilis str BAB 1 YP 007662508 1 1302650 Firmicutes
Bacillus subtilis subsp natto BEST195 YP 005561033 1 645657 Firmicutes

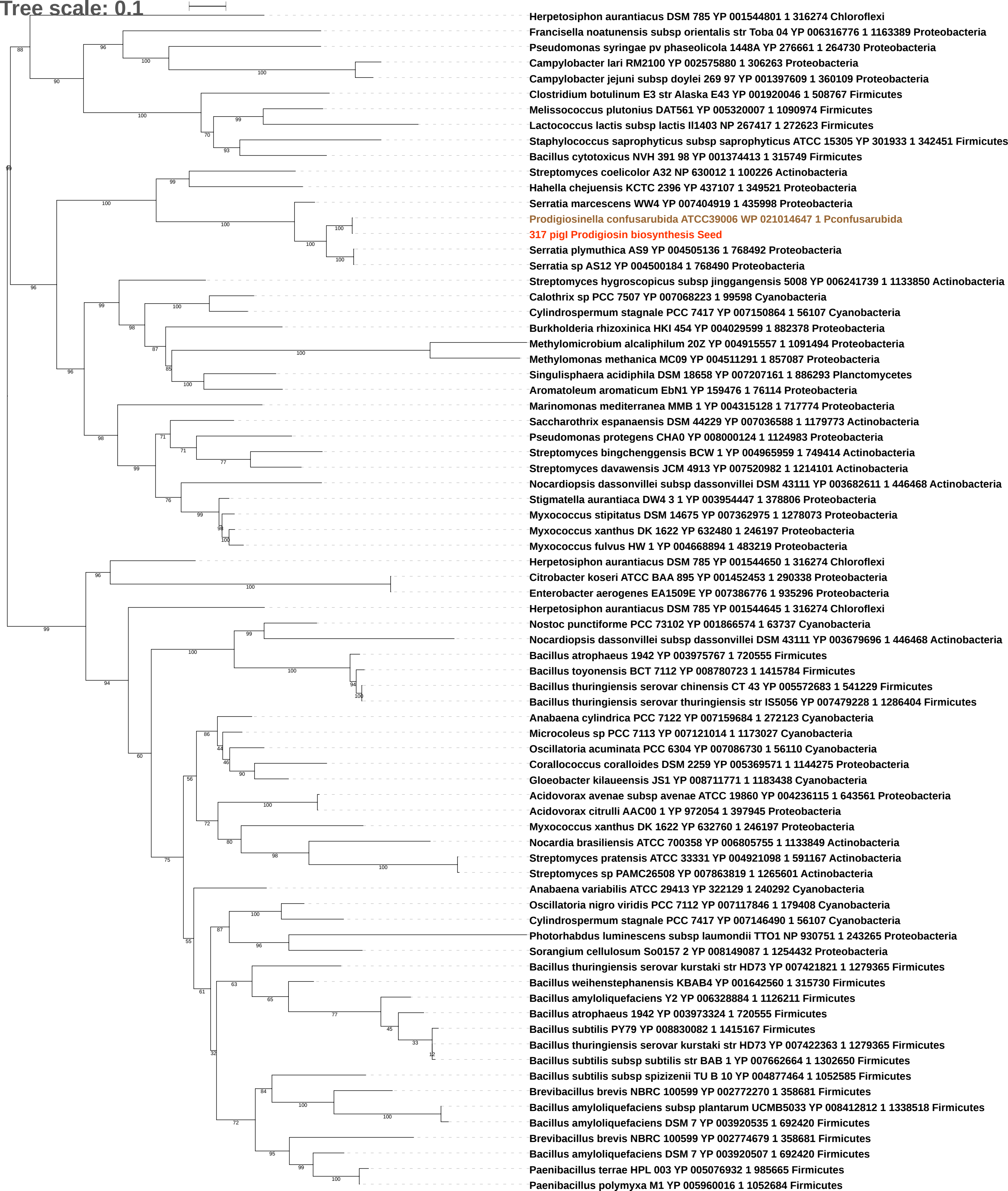
Tree scale: 0.1
Herpetosiphon aurantiacus DSM 785 YP 001544801 1 316274 Chloroflexi
Francisella noatunensis subsp orientalis str Toba 04 YP 006316776 1 1163389 Proteobacteria
Pseudomonas syringae pv phaseolicola 1448A YP 276661 1 264730 Proteobacteria
Campylobacter lari RM2100 YP 002575880 1 306263 Proteobacteria
Campylobacter jejuni subsp doylei 269 97 YP 001397609 1 360109 Proteobacteria
Clostridium botulinum E3 str Alaska E43 YP 001920046 1 508767 Firmicutes
Melissococcus plutonius DAT561 YP 005320007 1 1090974 Firmicutes
Lactococcus lactis subsp lactis Il1403 NP 267417 1 272623 Firmicutes
Staphylococcus saprophyticus subsp saprophyticus ATCC 15305 YP 301933 1 342451 Firmicutes
Bacillus cytotoxicus NVH 391 98 YP 001374413 1 315749 Firmicutes
Streptomyces coelicolor A32 NP 630012 1 100226 Actinobacteria
Hahella chejuensis KCTC 2396 YP 437107 1 349521 Proteobacteria
Serratia marcescens WW4 YP 007404919 1 435998 Proteobacteria
Prodigiosinella confusarubida ATCC39006 WP 021014647 1 Pconfusarubida
317 pigI Prodigiosin biosynthesis Seed
Serratia plymuthica AS9 YP 004505136 1 768492 Proteobacteria
Serratia sp AS12 YP 004500184 1 768490 Proteobacteria
Streptomyces hygroscopicus subsp jinggangensis 5008 YP 006241739 1 1133850 Actinobacteria
Calothrix sp PCC 7507 YP 007068223 1 99598 Cyanobacteria
Cylindrospermum stagnale PCC 7417 YP 007150864 1 56107 Cyanobacteria
Burkholderia rhizoxinica HKI 454 YP 004029599 1 882378 Proteobacteria
Methylomicrobium alcaliphilum 20Z YP 004915557 1 1091494 Proteobacteria
Methylomonas methanica MC09 YP 004511291 1 857087 Proteobacteria
Singulisphaera acidiphila DSM 18658 YP 007207161 1 886293 Planctomycetes
Aromatoleum aromaticum EbN1 YP 159476 1 76114 Proteobacteria
Marinomonas mediterranea MMB 1 YP 004315128 1 717774 Proteobacteria
Saccharothrix espanaensis DSM 44229 YP 007036588 1 1179773 Actinobacteria
Pseudomonas protegens CHA0 YP 008000124 1 1124983 Proteobacteria
Streptomyces bingchenggensis BCW 1 YP 004965959 1 749414 Actinobacteria
Streptomyces davawensis JCM 4913 YP 007520982 1 1214101 Actinobacteria
Nocardiopsis dassonvillei subsp dassonvillei DSM 43111 YP 003682611 1 446468 Actinobacteria
Stigmatella aurantiaca DW4 3 1 YP 003954447 1 378806 Proteobacteria
Myxococcus stipitatus DSM 14675 YP 007362975 1 1278073 Proteobacteria
Myxococcus xanthus DK 1622 YP 632480 1 246197 Proteobacteria
Myxococcus fulvus HW 1 YP 004668894 1 483219 Proteobacteria
Herpetosiphon aurantiacus DSM 785 YP 001544650 1 316274 Chloroflexi
Citrobacter koseri ATCC BAA 895 YP 001452453 1 290338 Proteobacteria
Enterobacter aerogenes EA1509E YP 007386776 1 935296 Proteobacteria
Herpetosiphon aurantiacus DSM 785 YP 001544645 1 316274 Chloroflexi
Nostoc punctiforme PCC 73102 YP 001866574 1 63737 Cyanobacteria
Nocardiopsis dassonvillei subsp dassonvillei DSM 43111 YP 003679696 1 446468 Actinobacteria
Bacillus atrophaeus 1942 YP 003975767 1 720555 Firmicutes
Bacillus toyonensis BCT 7112 YP 008780723 1 1415784 Firmicutes
Bacillus thuringiensis serovar chinensis CT 43 YP 005572683 1 541229 Firmicutes
Bacillus thuringiensis serovar thuringiensis str IS5056 YP 007479228 1 1286404 Firmicutes
Anabaena cylindrica PCC 7122 YP 007159684 1 272123 Cyanobacteria
Microcoleus sp PCC 7113 YP 007121014 1 1173027 Cyanobacteria
Oscillatoria acuminata PCC 6304 YP 007086730 1 56110 Cyanobacteria
Corallococcus coralloides DSM 2259 YP 005369571 1 1144275 Proteobacteria
Gloeobacter kilaueensis JS1 YP 008711771 1 1183438 Cyanobacteria
Acidovorax avenae subsp avenae ATCC 19860 YP 004236115 1 643561 Proteobacteria
Acidovorax citrulli AAC00 1 YP 972054 1 397945 Proteobacteria
Myxococcus xanthus DK 1622 YP 632760 1 246197 Proteobacteria
Nocardia brasiliensis ATCC 700358 YP 006805755 1 1133849 Actinobacteria
Streptomyces pratensis ATCC 33331 YP 004921098 1 591167 Actinobacteria
Streptomyces sp PAMC26508 YP 007863819 1 1265601 Actinobacteria
Anabaena variabilis ATCC 29413 YP 322129 1 240292 Cyanobacteria
Oscillatoria nigro viridis PCC 7112 YP 007117846 1 179408 Cyanobacteria
Cylindrospermum stagnale PCC 7417 YP 007146490 1 56107 Cyanobacteria
Photorhabdus luminescens subsp laumondii TTO1 NP 930751 1 243265 Proteobacteria
Sorangium cellulosum So0157 2 YP 008149087 1 1254432 Proteobacteria
Bacillus thuringiensis serovar kurstaki str HD73 YP 007421821 1 1279365 Firmicutes
Bacillus weihenstephanensis KBAB4 YP 001642560 1 315730 Firmicutes
Bacillus amyloliquefaciens Y2 YP 006328884 1 1126211 Firmicutes
Bacillus atrophaeus 1942 YP 003973324 1 720555 Firmicutes
Bacillus subtilis PY79 YP 008830082 1 1415167 Firmicutes
Bacillus thuringiensis serovar kurstaki str HD73 YP 007422363 1 1279365 Firmicutes
Bacillus subtilis subsp subtilis str BAB 1 YP 007662664 1 1302650 Firmicutes
Bacillus subtilis subsp spizizenii TU B 10 YP 004877464 1 1052585 Firmicutes
Brevibacillus brevis NBRC 100599 YP 002772270 1 358681 Firmicutes
Bacillus amyloliquefaciens subsp plantarum UCMB5033 YP 008412812 1 1338518 Firmicutes
Bacillus amyloliquefaciens DSM 7 YP 003920535 1 692420 Firmicutes
Brevibacillus brevis NBRC 100599 YP 002774679 1 358681 Firmicutes
Bacillus amyloliquefaciens DSM 7 YP 003920507 1 692420 Firmicutes
Paenibacillus terrae HPL 003 YP 005076932 1 985665 Firmicutes
Paenibacillus polymyxa M1 YP 005960016 1 1052684 Firmicutes

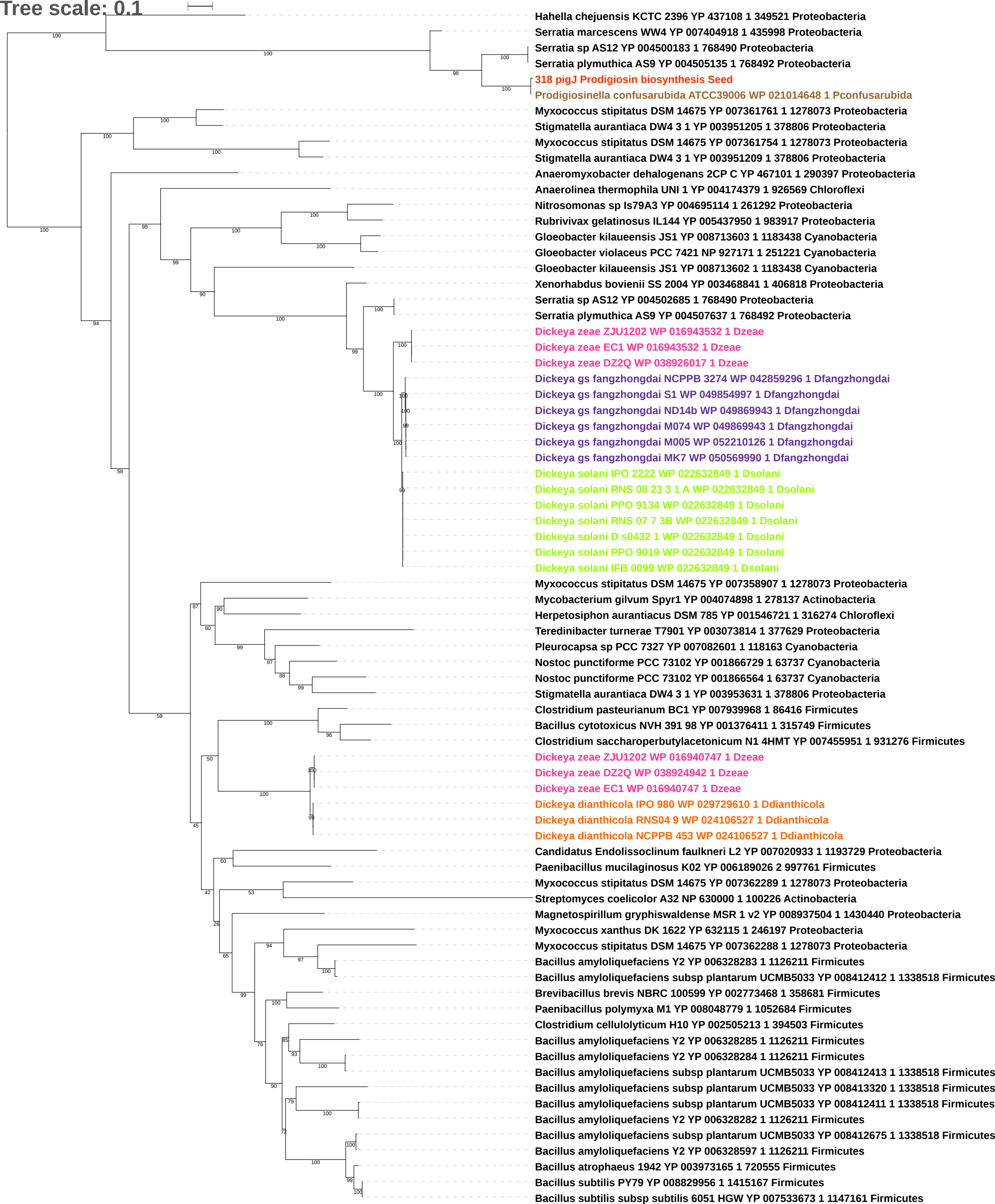
Tree scale: 0.1
Hahella chejuensis KCTC 2396 YP 437108 1 349521 Proteobacteria
Serratia marcescens WW4 YP 007404918 1 435998 Proteobacteria
Serratia sp AS12 YP 004500183 1 768490 Proteobacteria
Serratia plymuthica AS9 YP 004505135 1 768492 Proteobacteria
318 pigJ Prodigiosin biosynthesis Seed
Prodigiosinella confusarubida ATCC39006 WP 021014648 1 Pconfusarubida
Myxococcus stipitatus DSM 14675 YP 007361761 1 1278073 Proteobacteria
Stigmatella aurantiaca DW4 3 1 YP 003951205 1 378806 Proteobacteria
Myxococcus stipitatus DSM 14675 YP 007361754 1 1278073 Proteobacteria
Stigmatella aurantiaca DW4 3 1 YP 003951209 1 378806 Proteobacteria
Anaeromyxobacter dehalogenans 2CP C YP 467101 1 290397 Proteobacteria
Anaerolinea thermophila UNI 1 YP 004174379 1 926569 Chloroflexi
Nitrosomonas sp Is79A3 YP 004695114 1 261292 Proteobacteria
Rubrivivax gelatinosus IL144 YP 005437950 1 983917 Proteobacteria
Gloeobacter kilaueensis JS1 YP 008713603 1 1183438 Cyanobacteria
Gloeobacter violaceus PCC 7421 NP 927171 1 251221 Cyanobacteria
Gloeobacter kilaueensis JS1 YP 008713602 1 1183438 Cyanobacteria
Xenorhabdus bovienii SS 2004 YP 003468841 1 406818 Proteobacteria
Serratia sp AS12 YP 004502685 1 768490 Proteobacteria
Serratia plymuthica AS9 YP 004507637 1 768492 Proteobacteria
Dickeya zeae ZJU1202 WP 016943532 1 Dzeae
Dickeya zeae EC1 WP 016943532 1 Dzeae
Dickeya zeae DZ2Q WP 038926017 1 Dzeae
Dickeya gs fangzhongdai NCPPB 3274 WP 042859296 1 Dfangzhongdai
Dickeya gs fangzhongdai S1 WP 049854997 1 Dfangzhongdai
Dickeya gs fangzhongdai ND14b WP 049869943 1 Dfangzhongdai
Dickeya gs fangzhongdai M074 WP 049869943 1 Dfangzhongdai
Dickeya gs fangzhongdai M005 WP 052210126 1 Dfangzhongdai
Dickeya gs fangzhongdai MK7 WP 050569990 1 Dfangzhongdai
Dickeya solani IPO 2222 WP 022632849 1 Dsolani
Dickeya solani RNS 08 23 3 1 A WP 022632849 1 Dsolani
Dickeya solani PPO 9134 WP 022632849 1 Dsolani
Dickeya solani RNS 07 7 3B WP 022632849 1 Dsolani
Dickeya solani D s0432 1 WP 022632849 1 Dsolani
Dickeya solani PPO 9019 WP 022632849 1 Dsolani
Dickeya solani IFB 0099 WP 022632849 1 Dsolani
Myxococcus stipitatus DSM 14675 YP 007358907 1 1278073 Proteobacteria
Mycobacterium gilvum Spyr1 YP 004074898 1 278137 Actinobacteria
Herpetosiphon aurantiacus DSM 785 YP 001546721 1 316274 Chloroflexi
Teredinibacter turnerae T7901 YP 003073814 1 377629 Proteobacteria
Pleurocapsa sp PCC 7327 YP 007082601 1 118163 Cyanobacteria
Nostoc punctiforme PCC 73102 YP 001866729 1 63737 Cyanobacteria
Nostoc punctiforme PCC 73102 YP 001866564 1 63737 Cyanobacteria
Stigmatella aurantiaca DW4 3 1 YP 003953631 1 378806 Proteobacteria
Clostridium pasteurianum BC1 YP 007939968 1 86416 Firmicutes
Bacillus cytotoxicus NVH 391 98 YP 001376411 1 315749 Firmicutes
Clostridium saccharoperbutylacetonicum N1 4HMT YP 007455951 1 931276 Firmicutes
Dickeya zeae ZJU1202 WP 016940747 1 Dzeae
Dickeya zeae DZ2Q WP 038924942 1 Dzeae
Dickeya zeae EC1 WP 016940747 1 Dzeae
Dickeya dianthicola IPO 980 WP 029729610 1 Ddianthicola
Dickeya dianthicola RNS04 9 WP 024106527 1 Ddianthicola
Dickeya dianthicola NCPPB 453 WP 024106527 1 Ddianthicola
Candidatus Endolissoclinum faulkneri L2 YP 007020933 1 1193729 Proteobacteria
Paenibacillus mucilaginosus K02 YP 006189026 2 997761 Firmicutes
Myxococcus stipitatus DSM 14675 YP 007362289 1 1278073 Proteobacteria
Streptomyces coelicolor A32 NP 630000 1 100226 Actinobacteria
Magnetospirillum gryphiswaldense MSR 1 v2 YP 008937504 1 1430440 Proteobacteria
Myxococcus xanthus DK 1622 YP 632115 1 246197 Proteobacteria
Myxococcus stipitatus DSM 14675 YP 007362288 1 1278073 Proteobacteria
Bacillus amyloliquefaciens Y2 YP 006328283 1 1126211 Firmicutes
Bacillus amyloliquefaciens subsp plantarum UCMB5033 YP 008412412 1 1338518 Firmicutes
Brevibacillus brevis NBRC 100599 YP 002773468 1 358681 Firmicutes
Paenibacillus polymyxa M1 YP 008048779 1 1052684 Firmicutes
Clostridium cellulolyticum H10 YP 002505213 1 394503 Firmicutes
Bacillus amyloliquefaciens Y2 YP 006328285 1 1126211 Firmicutes
Bacillus amyloliquefaciens Y2 YP 006328284 1 1126211 Firmicutes
Bacillus amyloliquefaciens subsp plantarum UCMB5033 YP 008412413 1 1338518 Firmicutes
Bacillus amyloliquefaciens subsp plantarum UCMB5033 YP 008413320 1 1338518 Firmicutes
Bacillus amyloliquefaciens subsp plantarum UCMB5033 YP 008412411 1 1338518 Firmicutes
Bacillus amyloliquefaciens Y2 YP 006328282 1 1126211 Firmicutes
Bacillus amyloliquefaciens subsp plantarum UCMB5033 YP 008412675 1 1338518 Firmicutes
Bacillus amyloliquefaciens Y2 YP 006328597 1 1126211 Firmicutes
Bacillus atrophaeus 1942 YP 003973165 1 720555 Firmicutes
Bacillus subtilis PY79 YP 008829956 1 1415167 Firmicutes
Bacillus subtilis subsp subtilis 6051 HGW YP 007533673 1 1147161 Firmicutes

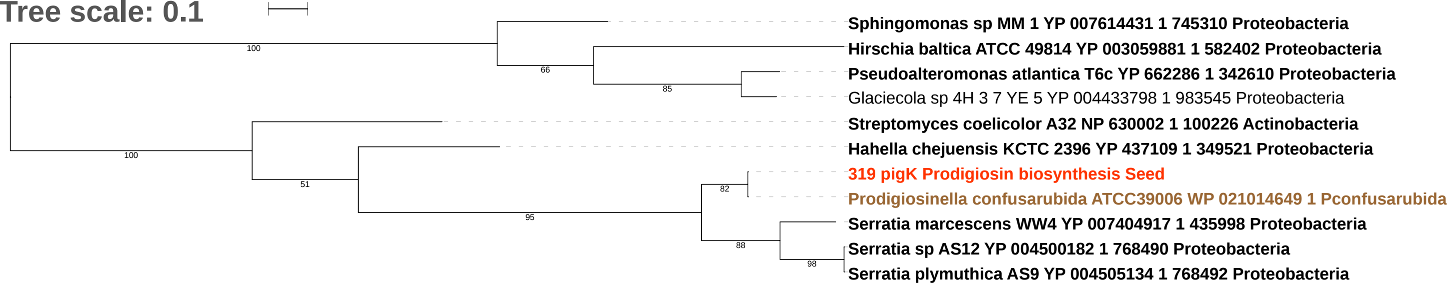

Tree scale: 0.1
Sphingomonas sp MM 1 YP 007614431 1 745310 Proteobacteria
Hirschia baltica ATCC 49814 YP 003059881 1 582402 Proteobacteria
Pseudoalteromonas atlantica T6c YP 662286 1 342610 Proteobacteria
Glaciecola sp 4H 3 7 YE 5 YP 004433798 1 983545 Proteobacteria
Streptomyces coelicolor A32 NP 630002 1 100226 Actinobacteria
Hahella chejuensis KCTC 2396 YP 437109 1 349521 Proteobacteria
319 pigK Prodigiosin biosynthesis Seed
Prodigiosinella confusarubida ATCC39006 WP 021014649 1 Pconfusarubida
Serratia marcescens WW4 YP 007404917 1 435998 Proteobacteria
Serratia sp AS12 YP 004500182 1 768490 Proteobacteria
Serratia plymuthica AS9 YP 004505134 1 768492 Proteobacteria
100
66
85
100
51
82
95
88
98

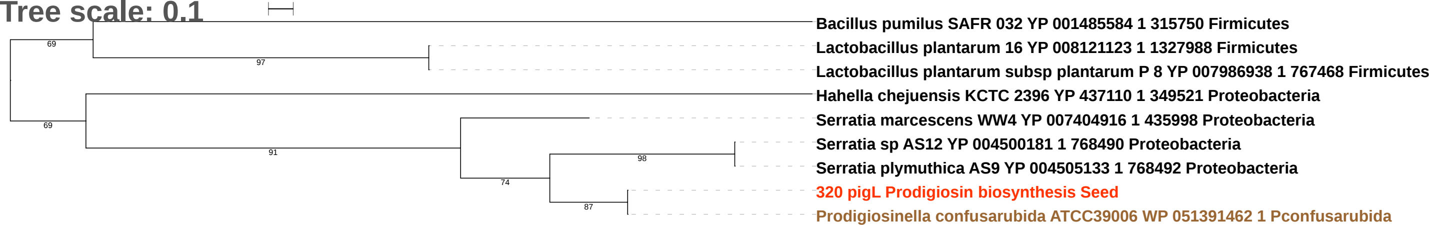

Tree scale: 0.1
Bacillus pumilus SAFR 032 YP 001485584 1 315750 Firmicutes
Lactobacillus plantarum 16 YP 008121123 1 1327988 Firmicutes
Lactobacillus plantarum subsp plantarum P 8 YP 007986938 1 767468 Firmicutes
Hahella chejuensis KCTC 2396 YP 437110 1 349521 Proteobacteria
Serratia marcescens WW4 YP 007404916 1 435998 Proteobacteria
Serratia sp AS12 YP 004500181 1 768490 Proteobacteria
Serratia plymuthica AS9 YP 004505133 1 768492 Proteobacteria
320 pigL Prodigiosin biosynthesis Seed
Prodigiosinella confusarubida ATCC39006 WP 051391462 1 Pconfusarubida
69
97
69
91
98
74
87

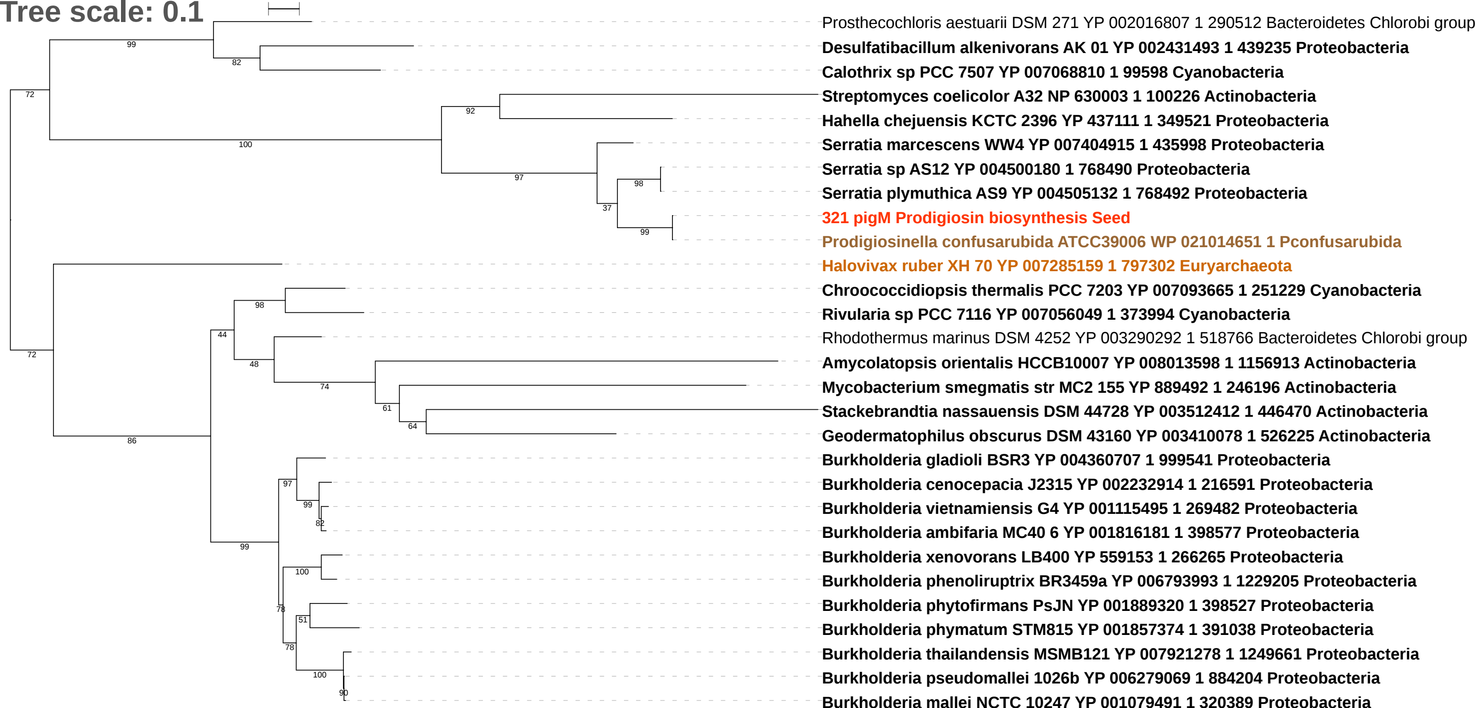
Tree scale: 0.1
Prosthecochloris aestuarii DSM 271 YP 002016807 1 290512 Bacteroidetes Chlorobi group
Desulfatibacillum alkenivorans AK 01 YP 002431493 1 439235 Proteobacteria
Calothrix sp PCC 7507 YP 007068810 1 99598 Cyanobacteria
Streptomyces coelicolor A32 NP 630003 1 100226 Actinobacteria
Hahella chejuensis KCTC 2396 YP 437111 1 349521 Proteobacteria
Serratia marcescens WW4 YP 007404915 1 435998 Proteobacteria
Serratia sp AS12 YP 004500180 1 768490 Proteobacteria
Serratia plymuthica AS9 YP 004505132 1 768492 Proteobacteria
321 pigM Prodigiosin biosynthesis Seed
Prodigiosinella confusarubida ATCC39006 WP 021014651 1 Pconfusarubida
Halovivax ruber XH 70 YP 007285159 1 797302 Euryarchaeota
Chroococcidiopsis thermalis PCC 7203 YP 007093665 1 251229 Cyanobacteria
Rivularia sp PCC 7116 YP 007056049 1 373994 Cyanobacteria
Rhodothermus marinus DSM 4252 YP 003290292 1 518766 Bacteroidetes Chlorobi group
Amycolatopsis orientalis HCCB10007 YP 008013598 1 1156913 Actinobacteria
Mycobacterium smegmatis str MC2 155 YP 889492 1 246196 Actinobacteria
Stackebrandtia nassauensis DSM 44728 YP 003512412 1 446470 Actinobacteria
Geodermatophilus obscurus DSM 43160 YP 003410078 1 526225 Actinobacteria
Burkholderia gladioli BSR3 YP 004360707 1 999541 Proteobacteria
Burkholderia cenocepacia J2315 YP 002232914 1 216591 Proteobacteria
Burkholderia vietnamiensis G4 YP 001115495 1 269482 Proteobacteria
Burkholderia ambifaria MC40 6 YP 001816181 1 398577 Proteobacteria
Burkholderia xenovorans LB400 YP 559153 1 266265 Proteobacteria
Burkholderia phenoliruptrix BR3459a YP 006793993 1 1229205 Proteobacteria
Burkholderia phytofirmans PsJN YP 001889320 1 398527 Proteobacteria
Burkholderia phymatum STM815 YP 001857374 1 391038 Proteobacteria
Burkholderia thailandensis MSMB121 YP 007921278 1 1249661 Proteobacteria
Burkholderia pseudomallei 1026b YP 006279069 1 884204 Proteobacteria
Burkholderia mallei NCTC 10247 YP 001079491 1 320389 Proteobacteria

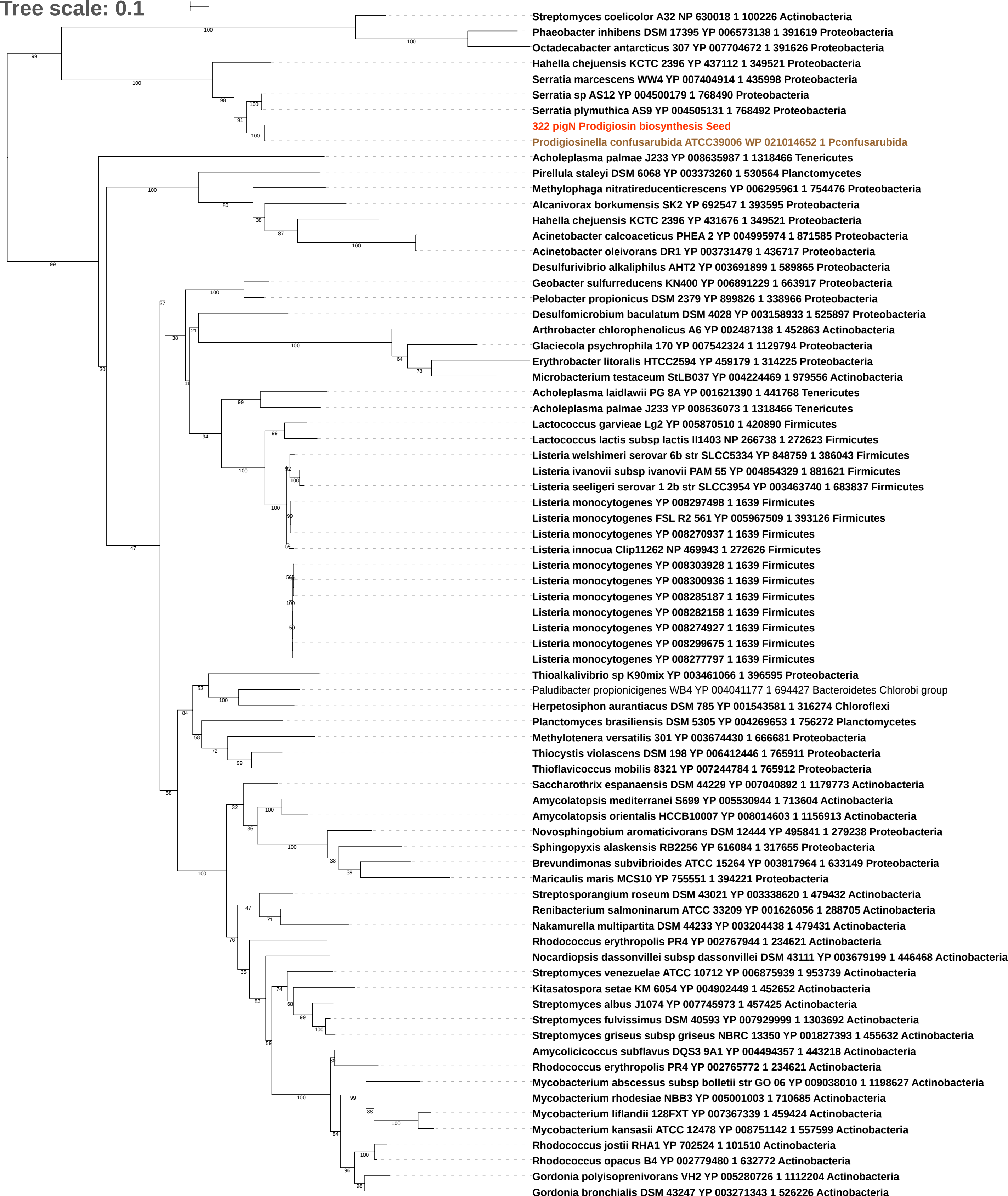
Tree scale: 0.1
Streptomyces coelicolor A32 NP 630018 1 100226 Actinobacteria
Phaeobacter inhibens DSM 17395 YP 006573138 1 391619 Proteobacteria
Octadecabacter antarcticus 307 YP 007704672 1 391626 Proteobacteria
Hahella chejuensis KCTC 2396 YP 437112 1 349521 Proteobacteria
Serratia marcescens WW4 YP 007404914 1 435998 Proteobacteria
Serratia sp AS12 YP 004500179 1 768490 Proteobacteria
Serratia plymuthica AS9 YP 004505131 1 768492 Proteobacteria
322 pigN Prodigiosin biosynthesis Seed
Prodigiosinella confusarubida ATCC39006 WP 021014652 1 Pconfusarubida
Acholeplasma palmae J233 YP 008635987 1 1318466 Tenericutes
Pirellula staleyi DSM 6068 YP 003373260 1 530564 Planctomycetes
Methylophaga nitratireducenticrescens YP 006295961 1 754476 Proteobacteria
Alcanivorax borkumensis SK2 YP 692547 1 393595 Proteobacteria
Hahella chejuensis KCTC 2396 YP 431676 1 349521 Proteobacteria
Acinetobacter calcoaceticus PHEA 2 YP 004995974 1 871585 Proteobacteria
Acinetobacter oleivorans DR1 YP 003731479 1 436717 Proteobacteria
Desulfurivibrio alkaliphilus AHT2 YP 003691899 1 589865 Proteobacteria
Geobacter sulfurreducens KN400 YP 006891229 1 663917 Proteobacteria
Pelobacter propionicus DSM 2379 YP 899826 1 338966 Proteobacteria
Desulfomicrobium baculatum DSM 4028 YP 003158933 1 525897 Proteobacteria
Arthrobacter chlorophenolicus A6 YP 002487138 1 452863 Actinobacteria
Glaciecola psychrophila 170 YP 007542324 1 1129794 Proteobacteria
Erythrobacter litoralis HTCC2594 YP 459179 1 314225 Proteobacteria
Microbacterium testaceum StLB037 YP 004224469 1 979556 Actinobacteria
Acholeplasma laidlawii PG 8A YP 001621390 1 441768 Tenericutes
Acholeplasma palmae J233 YP 008636073 1 1318466 Tenericutes
Lactococcus garvieae Lg2 YP 005870510 1 420890 Firmicutes
Lactococcus lactis subsp lactis Il1403 NP 266738 1 272623 Firmicutes
Listeria welshimeri serovar 6b str SLCC5334 YP 848759 1 386043 Firmicutes
Listeria ivanovii subsp ivanovii PAM 55 YP 004854329 1 881621 Firmicutes
Listeria seeligeri serovar 1 2b str SLCC3954 YP 003463740 1 683837 Firmicutes
Listeria monocytogenes YP 008297498 1 1639 Firmicutes
Listeria monocytogenes FSL R2 561 YP 005967509 1 393126 Firmicutes
Listeria monocytogenes YP 008270937 1 1639 Firmicutes
Listeria innocua Clip11262 NP 469943 1 272626 Firmicutes
Listeria monocytogenes YP 008303928 1 1639 Firmicutes
Listeria monocytogenes YP 008300936 1 1639 Firmicutes
Listeria monocytogenes YP 008285187 1 1639 Firmicutes
Listeria monocytogenes YP 008282158 1 1639 Firmicutes
Listeria monocytogenes YP 008274927 1 1639 Firmicutes
Listeria monocytogenes YP 008299675 1 1639 Firmicutes
Listeria monocytogenes YP 008277797 1 1639 Firmicutes
Thioalkalivibrio sp K90mix YP 003461066 1 396595 Proteobacteria
Paludibacter propionicigenes WB4 YP 004041177 1 694427 Bacteroidetes Chlorobi group
Herpetosiphon aurantiacus DSM 785 YP 001543581 1 316274 Chloroflexi
Planctomyces brasiliensis DSM 5305 YP 004269653 1 756272 Planctomycetes
Methylotenera versatilis 301 YP 003674430 1 666681 Proteobacteria
Thiocystis violascens DSM 198 YP 006412446 1 765911 Proteobacteria
Thioflavicoccus mobilis 8321 YP 007244784 1 765912 Proteobacteria
Saccharothrix espanaensis DSM 44229 YP 007040892 1 1179773 Actinobacteria
Amycolatopsis mediterranei S699 YP 005530944 1 713604 Actinobacteria
Amycolatopsis orientalis HCCB10007 YP 008014603 1 1156913 Actinobacteria
Novosphingobium aromaticivorans DSM 12444 YP 495841 1 279238 Proteobacteria
Sphingopyxis alaskensis RB2256 YP 616084 1 317655 Proteobacteria
Brevundimonas subvibrioides ATCC 15264 YP 003817964 1 633149 Proteobacteria
Maricaulis maris MCS10 YP 755551 1 394221 Proteobacteria
Streptosporangium roseum DSM 43021 YP 003338620 1 479432 Actinobacteria
Renibacterium salmoninarum ATCC 33209 YP 001626056 1 288705 Actinobacteria
Nakamurella multipartita DSM 44233 YP 003204438 1 479431 Actinobacteria
Rhodococcus erythropolis PR4 YP 002767944 1 234621 Actinobacteria
Nocardiopsis dassonvillei subsp dassonvillei DSM 43111 YP 003679199 1 446468 Actinobacteria
Streptomyces venezuelae ATCC 10712 YP 006875939 1 953739 Actinobacteria
Kitasatospora setae KM 6054 YP 004902449 1 452652 Actinobacteria
Streptomyces albus J1074 YP 007745973 1 457425 Actinobacteria
Streptomyces fulvissimus DSM 40593 YP 007929999 1 1303692 Actinobacteria
Streptomyces griseus subsp griseus NBRC 13350 YP 001827393 1 455632 Actinobacteria
Amycolicicoccus subflavus DQS3 9A1 YP 004494357 1 443218 Actinobacteria
Rhodococcus erythropolis PR4 YP 002765772 1 234621 Actinobacteria
Mycobacterium abscessus subsp bolletii str GO 06 YP 009038010 1 1198627 Actinobacteria
Mycobacterium rhodesiae NBB3 YP 005001003 1 710685 Actinobacteria
Mycobacterium liflandii 128FXT YP 007367339 1 459424 Actinobacteria
Mycobacterium kansasii ATCC 12478 YP 008751142 1 557599 Actinobacteria
Rhodococcus jostii RHA1 YP 702524 1 101510 Actinobacteria
Rhodococcus opacus B4 YP 002779480 1 632772 Actinobacteria
Gordonia polyisoprenivorans VH2 YP 005280726 1 1112204 Actinobacteria
Gordonia bronchialis DSM 43247 YP 003271343 1 526226 Actinobacteria

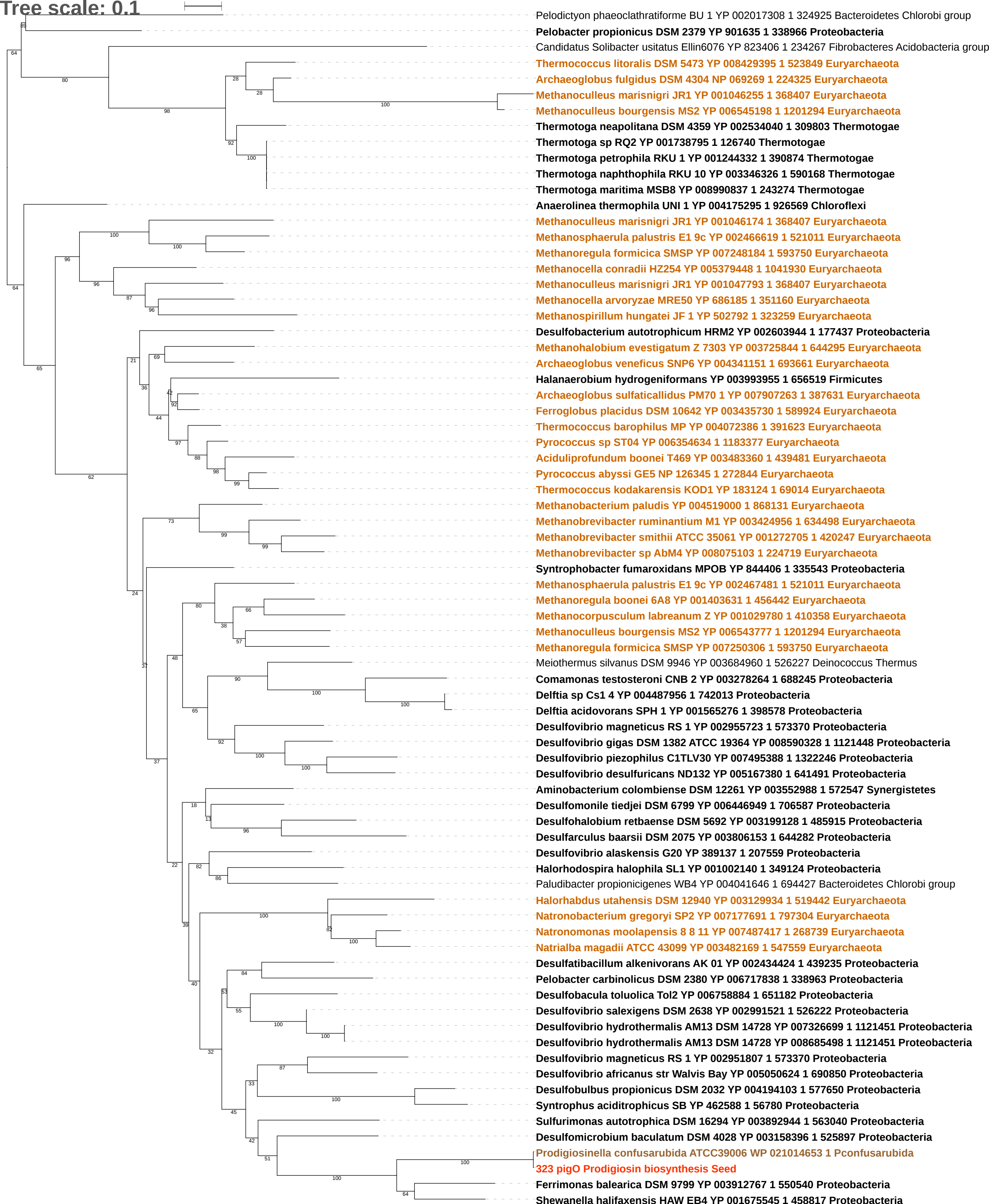
Tree scale: 0.1
Pelodictyon phaeoclathratiforme BU 1 YP 002017308 1 324925 Bacteroidetes Chlorobi group
Pelobacter propionicus DSM 2379 YP 901635 1 338966 Proteobacteria
Candidatus Solibacter usitatus Ellin6076 YP 823406 1 234267 Fibrobacteres Acidobacteria group
Thermococcus litoralis DSM 5473 YP 008429395 1 523849 Euryarchaeota
Archaeoglobus fulgidus DSM 4304 NP 069269 1 224325 Euryarchaeota
Methanoculleus marisnigri JR1 YP 001046255 1 368407 Euryarchaeota
Methanoculleus bourgensis MS2 YP 006545198 1 1201294 Euryarchaeota
Thermotoga neapolitana DSM 4359 YP 002534040 1 309803 Thermotogae
Thermotoga sp RQ2 YP 001738795 1 126740 Thermotogae
Thermotoga petrophila RKU 1 YP 001244332 1 390874 Thermotogae
Thermotoga naphthophila RKU 10 YP 003346326 1 590168 Thermotogae
Thermotoga maritima MSB8 YP 008990837 1 243274 Thermotogae
Anaerolinea thermophila UNI 1 YP 004175295 1 926569 Chloroflexi
Methanoculleus marisnigri JR1 YP 001046174 1 368407 Euryarchaeota
Methanosphaerula palustris E1 9c YP 002466619 1 521011 Euryarchaeota
Methanoregula formicica SMSP YP 007248184 1 593750 Euryarchaeota
Methanocella conradii HZ254 YP 005379448 1 1041930 Euryarchaeota
Methanoculleus marisnigri JR1 YP 001047793 1 368407 Euryarchaeota
Methanocella arvoryzae MRE50 YP 686185 1 351160 Euryarchaeota
Methanospirillum hungatei JF 1 YP 502792 1 323259 Euryarchaeota
Desulfobacterium autotrophicum HRM2 YP 002603944 1 177437 Proteobacteria
Methanohalobium evestigatum Z 7303 YP 003725844 1 644295 Euryarchaeota
Archaeoglobus veneficus SNP6 YP 004341151 1 693661 Euryarchaeota
Halanaerobium hydrogeniformans YP 003993955 1 656519 Firmicutes
Archaeoglobus sulfaticallidus PM70 1 YP 007907263 1 387631 Euryarchaeota
Ferroglobus placidus DSM 10642 YP 003435730 1 589924 Euryarchaeota
Thermococcus barophilus MP YP 004072386 1 391623 Euryarchaeota
Pyrococcus sp ST04 YP 006354634 1 1183377 Euryarchaeota
Aciduliprofundum boonei T469 YP 003483360 1 439481 Euryarchaeota
Pyrococcus abyssi GE5 NP 126345 1 272844 Euryarchaeota
Thermococcus kodakarensis KOD1 YP 183124 1 69014 Euryarchaeota
Methanobacterium paludis YP 004519000 1 868131 Euryarchaeota
Methanobrevibacter ruminantium M1 YP 003424956 1 634498 Euryarchaeota
Methanobrevibacter smithii ATCC 35061 YP 001272705 1 420247 Euryarchaeota
Methanobrevibacter sp AbM4 YP 008075103 1 224719 Euryarchaeota
Syntrophobacter fumaroxidans MPOB YP 844406 1 335543 Proteobacteria
Methanosphaerula palustris E1 9c YP 002467481 1 521011 Euryarchaeota
Methanoregula boonei 6A8 YP 001403631 1 456442 Euryarchaeota
Methanocorpusculum labreanum Z YP 001029780 1 410358 Euryarchaeota
Methanoculleus bourgensis MS2 YP 006543777 1 1201294 Euryarchaeota
Methanoregula formicica SMSP YP 007250306 1 593750 Euryarchaeota
Meiothermus silvanus DSM 9946 YP 003684960 1 526227 Deinococcus Thermus
Comamonas testosteroni CNB 2 YP 003278264 1 688245 Proteobacteria
Delftia sp Cs1 4 YP 004487956 1 742013 Proteobacteria
Delftia acidovorans SPH 1 YP 001565276 1 398578 Proteobacteria
Desulfovibrio magneticus RS 1 YP 002955723 1 573370 Proteobacteria
Desulfovibrio gigas DSM 1382 ATCC 19364 YP 008590328 1 1121448 Proteobacteria
Desulfovibrio piezophilus C1TLV30 YP 007495388 1 1322246 Proteobacteria
Desulfovibrio desulfuricans ND132 YP 005167380 1 641491 Proteobacteria
Aminobacterium colombiense DSM 12261 YP 003552988 1 572547 Synergistetes
Desulfomonile tiedjei DSM 6799 YP 006446949 1 706587 Proteobacteria
Desulfohalobium retbaense DSM 5692 YP 003199128 1 485915 Proteobacteria
Desulfarculus baarsii DSM 2075 YP 003806153 1 644282 Proteobacteria
Desulfovibrio alaskensis G20 YP 389137 1 207559 Proteobacteria
Halorhodospira halophila SL1 YP 001002140 1 349124 Proteobacteria
Paludibacter propionicigenes WB4 YP 004041646 1 694427 Bacteroidetes Chlorobi group
Halorhabdus utahensis DSM 12940 YP 003129934 1 519442 Euryarchaeota
Natronobacterium gregoryi SP2 YP 007177691 1 797304 Euryarchaeota
Natronomonas moolapensis 8 8 11 YP 007487417 1 268739 Euryarchaeota
Natrialba magadii ATCC 43099 YP 003482169 1 547559 Euryarchaeota
Desulfatibacillum alkenivorans AK 01 YP 002434424 1 439235 Proteobacteria
Pelobacter carbinolicus DSM 2380 YP 006717838 1 338963 Proteobacteria
Desulfobacula toluolica Tol2 YP 006758884 1 651182 Proteobacteria
Desulfovibrio salexigens DSM 2638 YP 002991521 1 526222 Proteobacteria
Desulfovibrio hydrothermalis AM13 DSM 14728 YP 007326699 1 1121451 Proteobacteria
Desulfovibrio hydrothermalis AM13 DSM 14728 YP 008685498 1 1121451 Proteobacteria
Desulfovibrio magneticus RS 1 YP 002951807 1 573370 Proteobacteria
Desulfovibrio africanus str Walvis Bay YP 005050624 1 690850 Proteobacteria
Desulfobulbus propionicus DSM 2032 YP 004194103 1 577650 Proteobacteria
Syntrophus aciditrophicus SB YP 462588 1 56780 Proteobacteria
Sulfurimonas autotrophica DSM 16294 YP 003892944 1 563040 Proteobacteria
Desulfomicrobium baculatum DSM 4028 YP 003158396 1 525897 Proteobacteria
Prodigiosinella confusarubida ATCC39006 WP 021014653 1 Pconfusarubida
323 pigO Prodigiosin biosynthesis Seed
Ferrimonas balearica DSM 9799 YP 003912767 1 550540 Proteobacteria
Shewanella halifaxensis HAW EB4 YP 001675545 1 458817 Proteobacteria

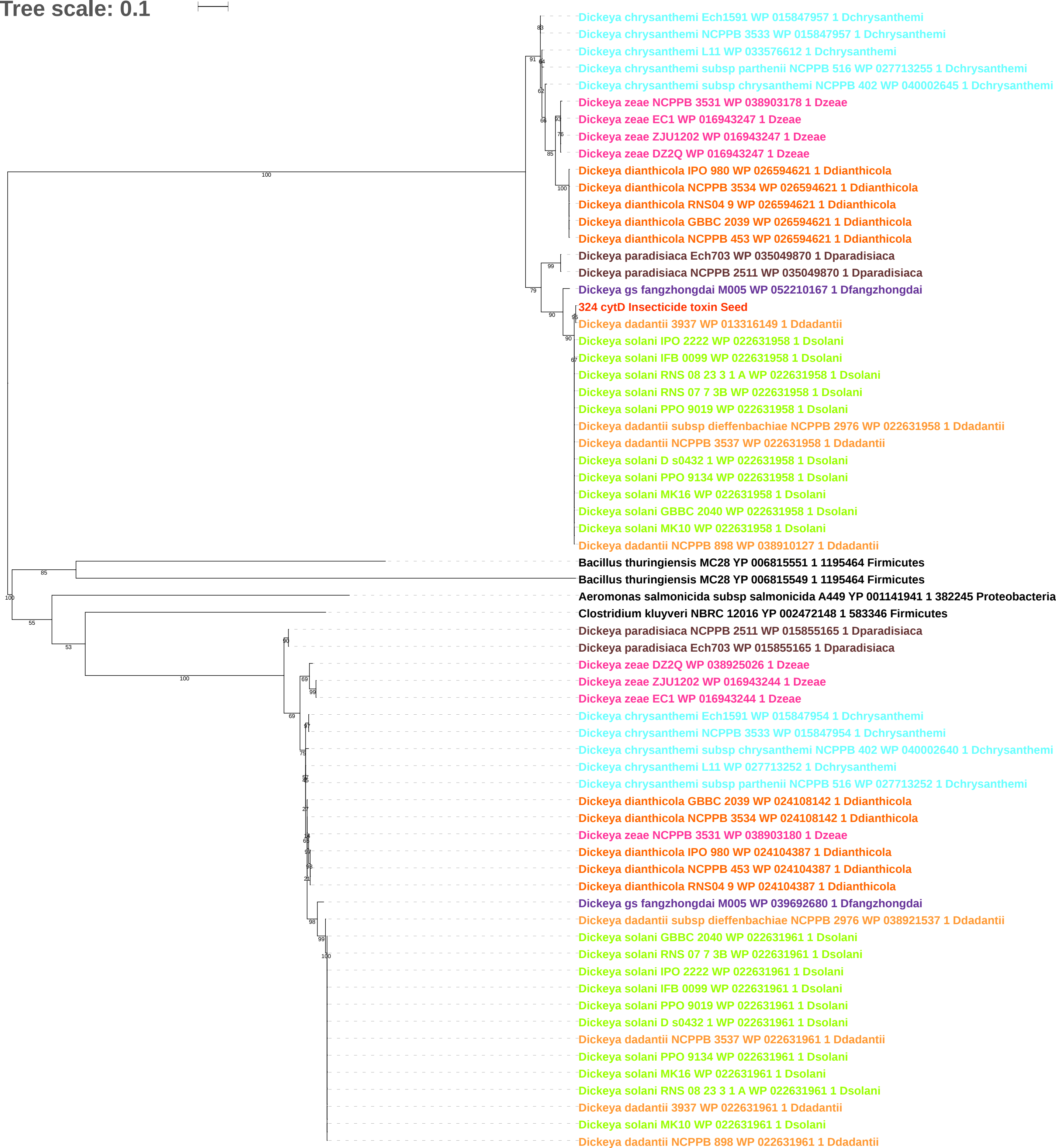
Tree scale: 0.1
Dickeya chrysanthemi Ech1591 WP 015847957 1 Dchrysanthemi
Dickeya chrysanthemi NCPPB 3533 WP 015847957 1 Dchrysanthemi
Dickeya chrysanthemi L11 WP 033576612 1 Dchrysanthemi
Dickeya chrysanthemi subsp parthenii NCPPB 516 WP 027713255 1 Dchrysanthemi
Dickeya chrysanthemi subsp chrysanthemi NCPPB 402 WP 040002645 1 Dchrysanthemi
Dickeya zeae NCPPB 3531 WP 038903178 1 Dzeae
Dickeya zeae EC1 WP 016943247 1 Dzeae
Dickeya zeae ZJU1202 WP 016943247 1 Dzeae
Dickeya zeae DZ2Q WP 016943247 1 Dzeae
Dickeya dianthicola IPO 980 WP 026594621 1 Ddianthicola
Dickeya dianthicola NCPPB 3534 WP 026594621 1 Ddianthicola
Dickeya dianthicola RNS04 9 WP 026594621 1 Ddianthicola
Dickeya dianthicola GBBC 2039 WP 026594621 1 Ddianthicola
Dickeya dianthicola NCPPB 453 WP 026594621 1 Ddianthicola
Dickeya paradisiaca Ech703 WP 035049870 1 Dparadisiaca
Dickeya paradisiaca NCPPB 2511 WP 035049870 1 Dparadisiaca
Dickeya gs fangzhongdai M005 WP 052210167 1 Dfangzhongdai
324 cytD Insecticide toxin Seed
Dickeya dadantii 3937 WP 013316149 1 Ddadantii
Dickeya solani IPO 2222 WP 022631958 1 Dsolani
Dickeya solani IFB 0099 WP 022631958 1 Dsolani
Dickeya solani RNS 08 23 3 1 A WP 022631958 1 Dsolani
Dickeya solani RNS 07 7 3B WP 022631958 1 Dsolani
Dickeya solani PPO 9019 WP 022631958 1 Dsolani
Dickeya dadantii subsp dieffenbachiae NCPPB 2976 WP 022631958 1 Ddadantii
Dickeya dadantii NCPPB 3537 WP 022631958 1 Ddadantii
Dickeya solani D s0432 1 WP 022631958 1 Dsolani
Dickeya solani PPO 9134 WP 022631958 1 Dsolani
Dickeya solani MK16 WP 022631958 1 Dsolani
Dickeya solani GBBC 2040 WP 022631958 1 Dsolani
Dickeya solani MK10 WP 022631958 1 Dsolani
Dickeya dadantii NCPPB 898 WP 038910127 1 Ddadantii
Bacillus thuringiensis MC28 YP 006815551 1 1195464 Firmicutes
Bacillus thuringiensis MC28 YP 006815549 1 1195464 Firmicutes
Aeromonas salmonicida subsp salmonicida A449 YP 001141941 1 382245 Proteobacteria
Clostridium kluyveri NBRC 12016 YP 002472148 1 583346 Firmicutes
Dickeya paradisiaca NCPPB 2511 WP 015855165 1 Dparadisiaca
Dickeya paradisiaca Ech703 WP 015855165 1 Dparadisiaca
Dickeya zeae DZ2Q WP 038925026 1 Dzeae
Dickeya zeae ZJU1202 WP 016943244 1 Dzeae
Dickeya zeae EC1 WP 016943244 1 Dzeae
Dickeya chrysanthemi Ech1591 WP 015847954 1 Dchrysanthemi
Dickeya chrysanthemi NCPPB 3533 WP 015847954 1 Dchrysanthemi
Dickeya chrysanthemi subsp chrysanthemi NCPPB 402 WP 040002640 1 Dchrysanthemi
Dickeya chrysanthemi L11 WP 027713252 1 Dchrysanthemi
Dickeya chrysanthemi subsp parthenii NCPPB 516 WP 027713252 1 Dchrysanthemi
Dickeya dianthicola GBBC 2039 WP 024108142 1 Ddianthicola
Dickeya dianthicola NCPPB 3534 WP 024108142 1 Ddianthicola
Dickeya zeae NCPPB 3531 WP 038903180 1 Dzeae
Dickeya dianthicola IPO 980 WP 024104387 1 Ddianthicola
Dickeya dianthicola NCPPB 453 WP 024104387 1 Ddianthicola
Dickeya dianthicola RNS04 9 WP 024104387 1 Ddianthicola
Dickeya gs fangzhongdai M005 WP 039692680 1 Dfangzhongdai
Dickeya dadantii subsp dieffenbachiae NCPPB 2976 WP 038921537 1 Ddadantii
Dickeya solani GBBC 2040 WP 022631961 1 Dsolani
Dickeya solani RNS 07 7 3B WP 022631961 1 Dsolani
Dickeya solani IPO 2222 WP 022631961 1 Dsolani
Dickeya solani IFB 0099 WP 022631961 1 Dsolani
Dickeya solani PPO 9019 WP 022631961 1 Dsolani
Dickeya solani D s0432 1 WP 022631961 1 Dsolani
Dickeya dadantii NCPPB 3537 WP 022631961 1 Ddadantii
Dickeya solani PPO 9134 WP 022631961 1 Dsolani
Dickeya solani MK16 WP 022631961 1 Dsolani
Dickeya solani RNS 08 23 3 1 A WP 022631961 1 Dsolani
Dickeya dadantii 3937 WP 022631961 1 Ddadantii
Dickeya solani MK10 WP 022631961 1 Dsolani
Dickeya dadantii NCPPB 898 WP 022631961 1 Ddadantii

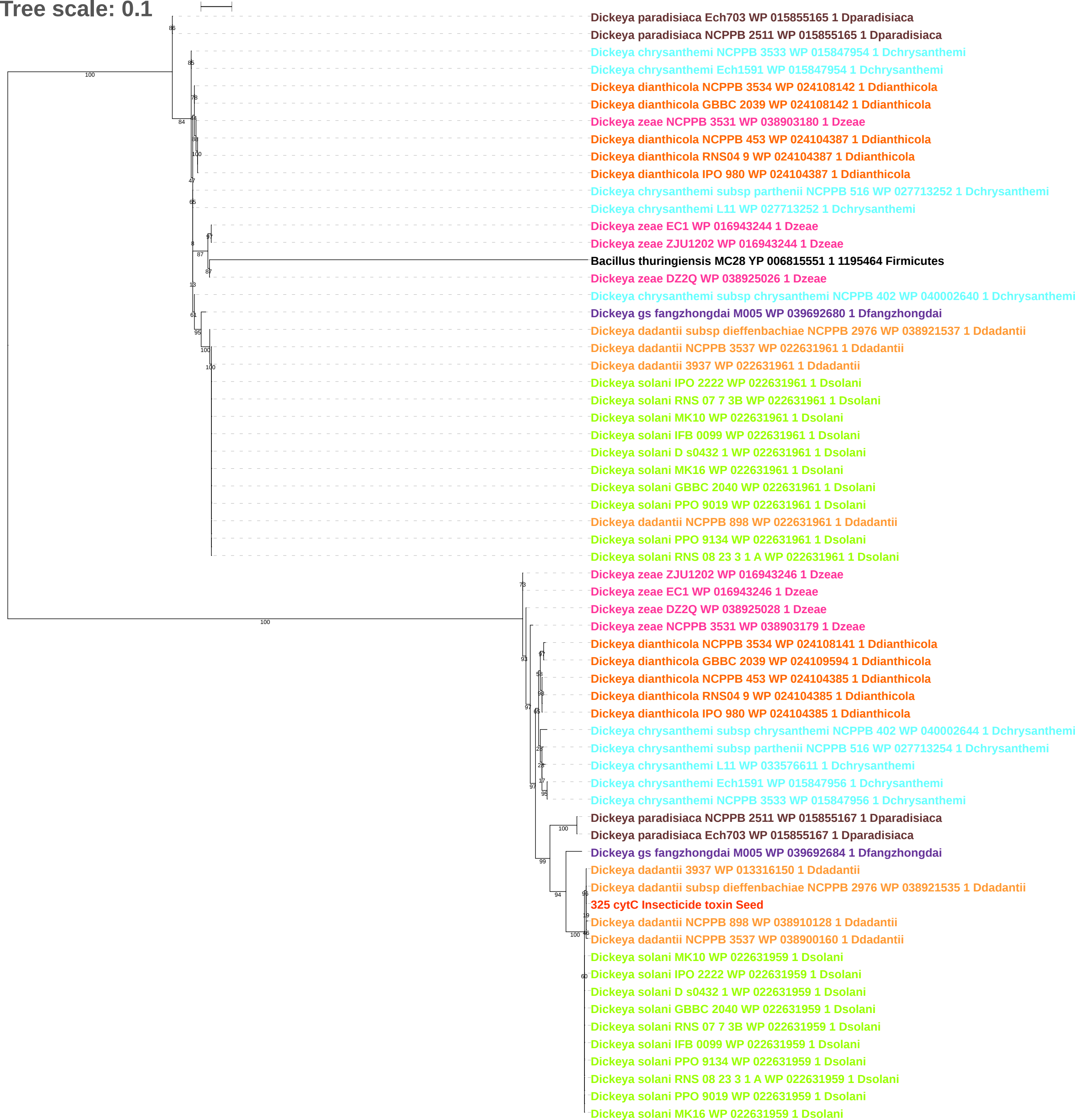

Tree scale: 0.1
Dickeya paradisiaca Ech703 WP 015855165 1 Dparadisiaca
Dickeya paradisiaca NCPPB 2511 WP 015855165 1 Dparadisiaca
Dickeya chrysanthemi NCPPB 3533 WP 015847954 1 Dchrysanthemi
Dickeya chrysanthemi Ech1591 WP 015847954 1 Dchrysanthemi
Dickeya dianthicola NCPPB 3534 WP 024108142 1 Ddianthicola
Dickeya dianthicola GBBC 2039 WP 024108142 1 Ddianthicola
Dickeya zeae NCPPB 3531 WP 038903180 1 Dzeae
Dickeya dianthicola NCPPB 453 WP 024104387 1 Ddianthicola
Dickeya dianthicola RNS04 9 WP 024104387 1 Ddianthicola
Dickeya dianthicola IPO 980 WP 024104387 1 Ddianthicola
Dickeya chrysanthemi subsp parthenii NCPPB 516 WP 027713252 1 Dchrysanthemi
Dickeya chrysanthemi L11 WP 027713252 1 Dchrysanthemi
Dickeya zeae EC1 WP 016943244 1 Dzeae
Dickeya zeae ZJU1202 WP 016943244 1 Dzeae
Bacillus thuringiensis MC28 YP 006815551 1 1195464 Firmicutes
Dickeya zeae DZ2Q WP 038925026 1 Dzeae
Dickeya chrysanthemi subsp chrysanthemi NCPPB 402 WP 040002640 1 Dchrysanthemi
Dickeya gs fangzhongdai M005 WP 039692680 1 Dfangzhongdai
Dickeya dadantii subsp dieffenbachiae NCPPB 2976 WP 038921537 1 Ddadantii
Dickeya dadantii NCPPB 3537 WP 022631961 1 Ddadantii
Dickeya dadantii 3937 WP 022631961 1 Ddadantii
Dickeya solani IPO 2222 WP 022631961 1 Dsolani
Dickeya solani RNS 07 7 3B WP 022631961 1 Dsolani
Dickeya solani MK10 WP 022631961 1 Dsolani
Dickeya solani IFB 0099 WP 022631961 1 Dsolani
Dickeya solani D s0432 1 WP 022631961 1 Dsolani
Dickeya solani MK16 WP 022631961 1 Dsolani
Dickeya solani GBBC 2040 WP 022631961 1 Dsolani
Dickeya solani PPO 9019 WP 022631961 1 Dsolani
Dickeya dadantii NCPPB 898 WP 022631961 1 Ddadantii
Dickeya solani PPO 9134 WP 022631961 1 Dsolani
Dickeya solani RNS 08 23 3 1 A WP 022631961 1 Dsolani
Dickeya zeae ZJU1202 WP 016943246 1 Dzeae
Dickeya zeae EC1 WP 016943246 1 Dzeae
Dickeya zeae DZ2Q WP 038925028 1 Dzeae
Dickeya zeae NCPPB 3531 WP 038903179 1 Dzeae
Dickeya dianthicola NCPPB 3534 WP 024108141 1 Ddianthicola
Dickeya dianthicola GBBC 2039 WP 024109594 1 Ddianthicola
Dickeya dianthicola NCPPB 453 WP 024104385 1 Ddianthicola
Dickeya dianthicola RNS04 9 WP 024104385 1 Ddianthicola
Dickeya dianthicola IPO 980 WP 024104385 1 Ddianthicola
Dickeya chrysanthemi subsp chrysanthemi NCPPB 402 WP 040002644 1 Dchrysanthemi
Dickeya chrysanthemi subsp parthenii NCPPB 516 WP 027713254 1 Dchrysanthemi
Dickeya chrysanthemi L11 WP 033576611 1 Dchrysanthemi
Dickeya chrysanthemi Ech1591 WP 015847956 1 Dchrysanthemi
Dickeya chrysanthemi NCPPB 3533 WP 015847956 1 Dchrysanthemi
Dickeya paradisiaca NCPPB 2511 WP 015855167 1 Dparadisiaca
Dickeya paradisiaca Ech703 WP 015855167 1 Dparadisiaca
Dickeya gs fangzhongdai M005 WP 039692684 1 Dfangzhongdai
Dickeya dadantii 3937 WP 013316150 1 Ddadantii
Dickeya dadantii subsp dieffenbachiae NCPPB 2976 WP 038921535 1 Ddadantii
325 cytC Insecticide toxin Seed
Dickeya dadantii NCPPB 898 WP 038910128 1 Ddadantii
Dickeya dadantii NCPPB 3537 WP 038900160 1 Ddadantii
Dickeya solani MK10 WP 022631959 1 Dsolani
Dickeya solani IPO 2222 WP 022631959 1 Dsolani
Dickeya solani D s0432 1 WP 022631959 1 Dsolani
Dickeya solani GBBC 2040 WP 022631959 1 Dsolani
Dickeya solani RNS 07 7 3B WP 022631959 1 Dsolani
Dickeya solani IFB 0099 WP 022631959 1 Dsolani
Dickeya solani PPO 9134 WP 022631959 1 Dsolani
Dickeya solani RNS 08 23 3 1 A WP 022631959 1 Dsolani
Dickeya solani PPO 9019 WP 022631959 1 Dsolani
Dickeya solani MK16 WP 022631959 1 Dsolani

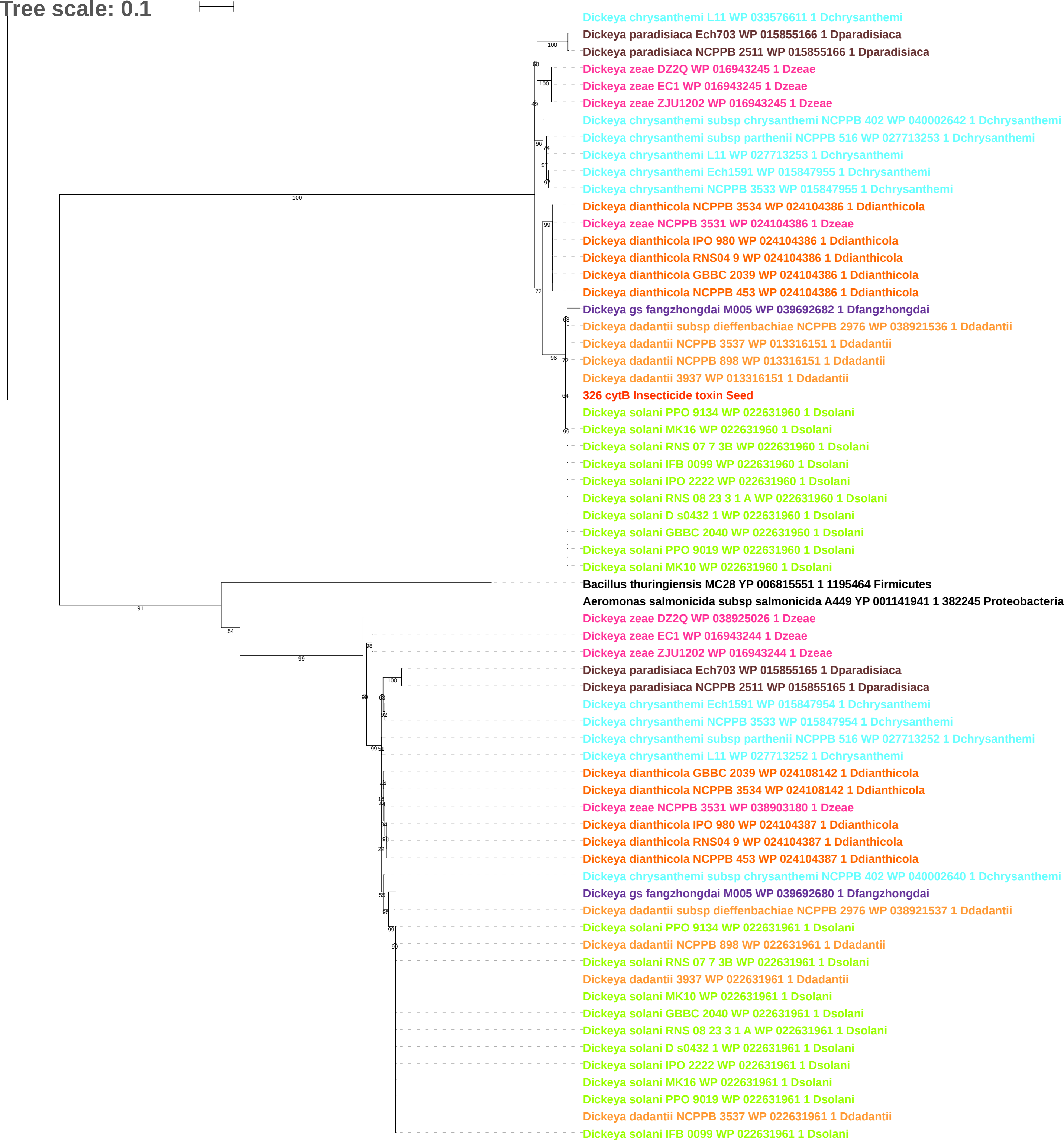
Tree scale: 0.1
Dickeya chrysanthemi L11 WP 033576611 1 Dchrysanthemi
Dickeya paradisiaca Ech703 WP 015855166 1 Dparadisiaca
Dickeya paradisiaca NCPPB 2511 WP 015855166 1 Dparadisiaca
Dickeya zeae DZ2Q WP 016943245 1 Dzeae
Dickeya zeae EC1 WP 016943245 1 Dzeae
Dickeya zeae ZJU1202 WP 016943245 1 Dzeae
Dickeya chrysanthemi subsp chrysanthemi NCPPB 402 WP 040002642 1 Dchrysanthemi
Dickeya chrysanthemi subsp parthenii NCPPB 516 WP 027713253 1 Dchrysanthemi
Dickeya chrysanthemi L11 WP 027713253 1 Dchrysanthemi
Dickeya chrysanthemi Ech1591 WP 015847955 1 Dchrysanthemi
Dickeya chrysanthemi NCPPB 3533 WP 015847955 1 Dchrysanthemi
Dickeya dianthicola NCPPB 3534 WP 024104386 1 Ddianthicola
Dickeya zeae NCPPB 3531 WP 024104386 1 Dzeae
Dickeya dianthicola IPO 980 WP 024104386 1 Ddianthicola
Dickeya dianthicola RNS04 9 WP 024104386 1 Ddianthicola
Dickeya dianthicola GBBC 2039 WP 024104386 1 Ddianthicola
Dickeya dianthicola NCPPB 453 WP 024104386 1 Ddianthicola
Dickeya gs fangzhongdai M005 WP 039692682 1 Dfangzhongdai
Dickeya dadantii subsp dieffenbachiae NCPPB 2976 WP 038921536 1 Ddadantii
Dickeya dadantii NCPPB 3537 WP 013316151 1 Ddadantii
Dickeya dadantii NCPPB 898 WP 013316151 1 Ddadantii
Dickeya dadantii 3937 WP 013316151 1 Ddadantii
326 cytB Insecticide toxin Seed
Dickeya solani PPO 9134 WP 022631960 1 Dsolani
Dickeya solani MK16 WP 022631960 1 Dsolani
Dickeya solani RNS 07 7 3B WP 022631960 1 Dsolani
Dickeya solani IFB 0099 WP 022631960 1 Dsolani
Dickeya solani IPO 2222 WP 022631960 1 Dsolani
Dickeya solani RNS 08 23 3 1 A WP 022631960 1 Dsolani
Dickeya solani D s0432 1 WP 022631960 1 Dsolani
Dickeya solani GBBC 2040 WP 022631960 1 Dsolani
Dickeya solani PPO 9019 WP 022631960 1 Dsolani
Dickeya solani MK10 WP 022631960 1 Dsolani
Bacillus thuringiensis MC28 YP 006815551 1 1195464 Firmicutes
Aeromonas salmonicida subsp salmonicida A449 YP 001141941 1 382245 Proteobacteria
Dickeya zeae DZ2Q WP 038925026 1 Dzeae
Dickeya zeae EC1 WP 016943244 1 Dzeae
Dickeya zeae ZJU1202 WP 016943244 1 Dzeae
Dickeya paradisiaca Ech703 WP 015855165 1 Dparadisiaca
Dickeya paradisiaca NCPPB 2511 WP 015855165 1 Dparadisiaca
Dickeya chrysanthemi Ech1591 WP 015847954 1 Dchrysanthemi
Dickeya chrysanthemi NCPPB 3533 WP 015847954 1 Dchrysanthemi
Dickeya chrysanthemi subsp parthenii NCPPB 516 WP 027713252 1 Dchrysanthemi
Dickeya chrysanthemi L11 WP 027713252 1 Dchrysanthemi
Dickeya dianthicola GBBC 2039 WP 024108142 1 Ddianthicola
Dickeya dianthicola NCPPB 3534 WP 024108142 1 Ddianthicola
Dickeya zeae NCPPB 3531 WP 038903180 1 Dzeae
Dickeya dianthicola IPO 980 WP 024104387 1 Ddianthicola
Dickeya dianthicola RNS04 9 WP 024104387 1 Ddianthicola
Dickeya dianthicola NCPPB 453 WP 024104387 1 Ddianthicola
Dickeya chrysanthemi subsp chrysanthemi NCPPB 402 WP 040002640 1 Dchrysanthemi
Dickeya gs fangzhongdai M005 WP 039692680 1 Dfangzhongdai
Dickeya dadantii subsp dieffenbachiae NCPPB 2976 WP 038921537 1 Ddadantii
Dickeya solani PPO 9134 WP 022631961 1 Dsolani
Dickeya dadantii NCPPB 898 WP 022631961 1 Ddadantii
Dickeya solani RNS 07 7 3B WP 022631961 1 Dsolani
Dickeya dadantii 3937 WP 022631961 1 Ddadantii
Dickeya solani MK10 WP 022631961 1 Dsolani
Dickeya solani GBBC 2040 WP 022631961 1 Dsolani
Dickeya solani RNS 08 23 3 1 A WP 022631961 1 Dsolani
Dickeya solani D s0432 1 WP 022631961 1 Dsolani
Dickeya solani IPO 2222 WP 022631961 1 Dsolani
Dickeya solani MK16 WP 022631961 1 Dsolani
Dickeya solani PPO 9019 WP 022631961 1 Dsolani
Dickeya dadantii NCPPB 3537 WP 022631961 1 Ddadantii
Dickeya solani IFB 0099 WP 022631961 1 Dsolani

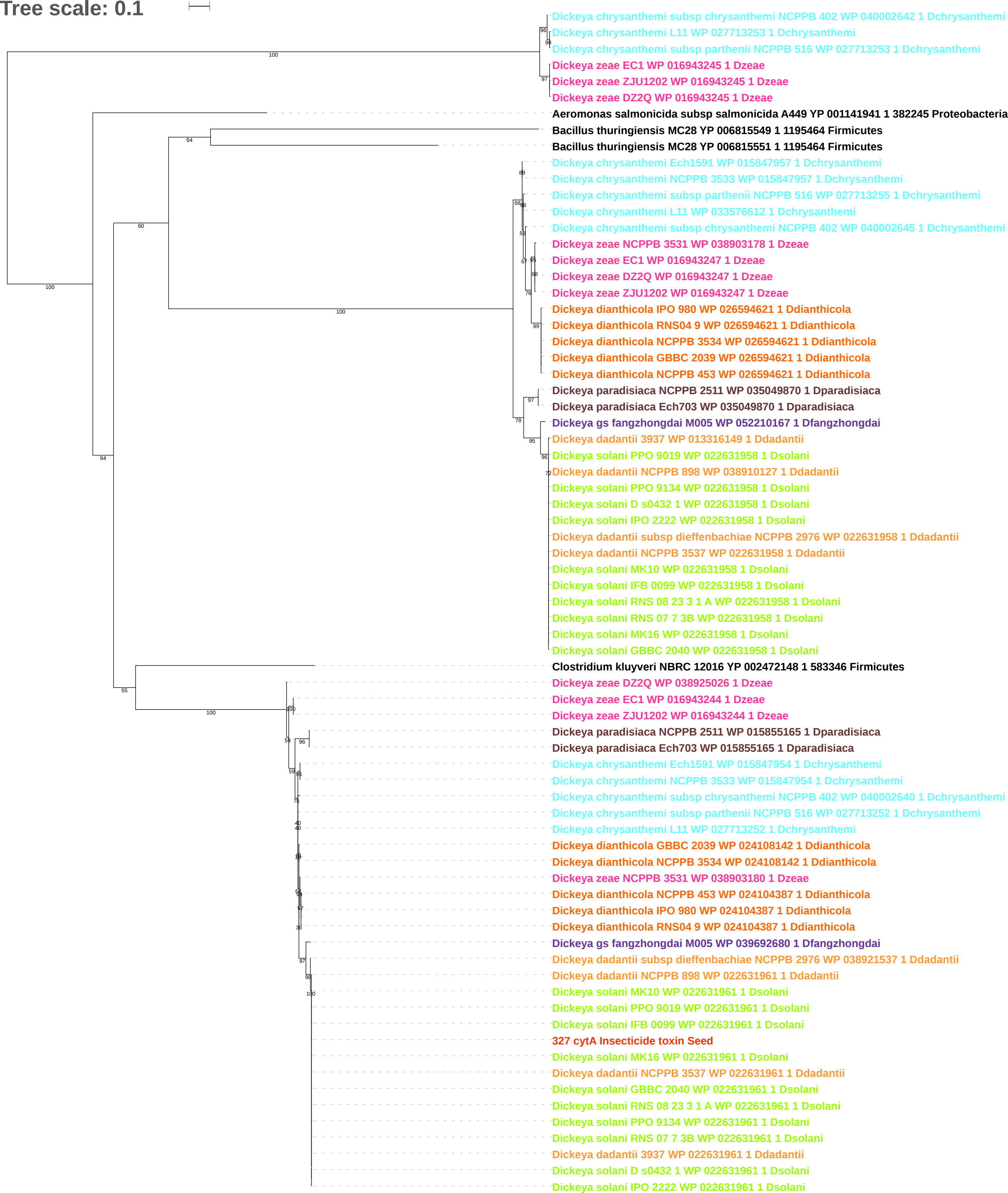
Tree scale: 0.1
Dickeya chrysanthemi subsp chrysanthemi NCPPB 402 WP 040002642 1 Dchrysanthemi
Dickeya chrysanthemi L11 WP 027713253 1 Dchrysanthemi
Dickeya chrysanthemi subsp parthenii NCPPB 516 WP 027713253 1 Dchrysanthemi
Dickeya zeae EC1 WP 016943245 1 Dzeae
Dickeya zeae ZJU1202 WP 016943245 1 Dzeae
Dickeya zeae DZ2Q WP 016943245 1 Dzeae
Aeromonas salmonicida subsp salmonicida A449 YP 001141941 1 382245 Proteobacteria
Bacillus thuringiensis MC28 YP 006815549 1 1195464 Firmicutes
Bacillus thuringiensis MC28 YP 006815551 1 1195464 Firmicutes
Dickeya chrysanthemi Ech1591 WP 015847957 1 Dchrysanthemi
Dickeya chrysanthemi NCPPB 3533 WP 015847957 1 Dchrysanthemi
Dickeya chrysanthemi subsp parthenii NCPPB 516 WP 027713255 1 Dchrysanthemi
Dickeya chrysanthemi L11 WP 033576612 1 Dchrysanthemi
Dickeya chrysanthemi subsp chrysanthemi NCPPB 402 WP 040002645 1 Dchrysanthemi
Dickeya zeae NCPPB 3531 WP 038903178 1 Dzeae
Dickeya zeae EC1 WP 016943247 1 Dzeae
Dickeya zeae DZ2Q WP 016943247 1 Dzeae
Dickeya zeae ZJU1202 WP 016943247 1 Dzeae
Dickeya dianthicola IPO 980 WP 026594621 1 Ddianthicola
Dickeya dianthicola RNS04 9 WP 026594621 1 Ddianthicola
Dickeya dianthicola NCPPB 3534 WP 026594621 1 Ddianthicola
Dickeya dianthicola GBBC 2039 WP 026594621 1 Ddianthicola
Dickeya dianthicola NCPPB 453 WP 026594621 1 Ddianthicola
Dickeya paradisiaca NCPPB 2511 WP 035049870 1 Dparadisiaca
Dickeya paradisiaca Ech703 WP 035049870 1 Dparadisiaca
Dickeya gs fangzhongdai M005 WP 052210167 1 Dfangzhongdai
Dickeya dadantii 3937 WP 013316149 1 Ddadantii
Dickeya solani PPO 9019 WP 022631958 1 Dsolani
Dickeya dadantii NCPPB 898 WP 038910127 1 Ddadantii
Dickeya solani PPO 9134 WP 022631958 1 Dsolani
Dickeya solani D s0432 1 WP 022631958 1 Dsolani
Dickeya solani IPO 2222 WP 022631958 1 Dsolani
Dickeya dadantii subsp dieffenbachiae NCPPB 2976 WP 022631958 1 Ddadantii
Dickeya dadantii NCPPB 3537 WP 022631958 1 Ddadantii
Dickeya solani MK10 WP 022631958 1 Dsolani
Dickeya solani IFB 0099 WP 022631958 1 Dsolani
Dickeya solani RNS 08 23 3 1 A WP 022631958 1 Dsolani
Dickeya solani RNS 07 7 3B WP 022631958 1 Dsolani
Dickeya solani MK16 WP 022631958 1 Dsolani
Dickeya solani GBBC 2040 WP 022631958 1 Dsolani
Clostridium kluyveri NBRC 12016 YP 002472148 1 583346 Firmicutes
Dickeya zeae DZ2Q WP 038925026 1 Dzeae
Dickeya zeae EC1 WP 016943244 1 Dzeae
Dickeya zeae ZJU1202 WP 016943244 1 Dzeae
Dickeya paradisiaca NCPPB 2511 WP 015855165 1 Dparadisiaca
Dickeya paradisiaca Ech703 WP 015855165 1 Dparadisiaca
Dickeya chrysanthemi Ech1591 WP 015847954 1 Dchrysanthemi
Dickeya chrysanthemi NCPPB 3533 WP 015847954 1 Dchrysanthemi
Dickeya chrysanthemi subsp chrysanthemi NCPPB 402 WP 040002640 1 Dchrysanthemi
Dickeya chrysanthemi subsp parthenii NCPPB 516 WP 027713252 1 Dchrysanthemi
Dickeya chrysanthemi L11 WP 027713252 1 Dchrysanthemi
Dickeya dianthicola GBBC 2039 WP 024108142 1 Ddianthicola
Dickeya dianthicola NCPPB 3534 WP 024108142 1 Ddianthicola
Dickeya zeae NCPPB 3531 WP 038903180 1 Dzeae
Dickeya dianthicola NCPPB 453 WP 024104387 1 Ddianthicola
Dickeya dianthicola IPO 980 WP 024104387 1 Ddianthicola
Dickeya dianthicola RNS04 9 WP 024104387 1 Ddianthicola
Dickeya gs fangzhongdai M005 WP 039692680 1 Dfangzhongdai
Dickeya dadantii subsp dieffenbachiae NCPPB 2976 WP 038921537 1 Ddadantii
Dickeya dadantii NCPPB 898 WP 022631961 1 Ddadantii
Dickeya solani MK10 WP 022631961 1 Dsolani
Dickeya solani PPO 9019 WP 022631961 1 Dsolani
Dickeya solani IFB 0099 WP 022631961 1 Dsolani
327 cytA Insecticide toxin Seed
Dickeya solani MK16 WP 022631961 1 Dsolani
Dickeya dadantii NCPPB 3537 WP 022631961 1 Ddadantii
Dickeya solani GBBC 2040 WP 022631961 1 Dsolani
Dickeya solani RNS 08 23 3 1 A WP 022631961 1 Dsolani
Dickeya solani PPO 9134 WP 022631961 1 Dsolani
Dickeya solani RNS 07 7 3B WP 022631961 1 Dsolani
Dickeya dadantii 3937 WP 022631961 1 Ddadantii
Dickeya solani D s0432 1 WP 022631961 1 Dsolani
Dickeya solani IPO 2222 WP 022631961 1 Dsolani

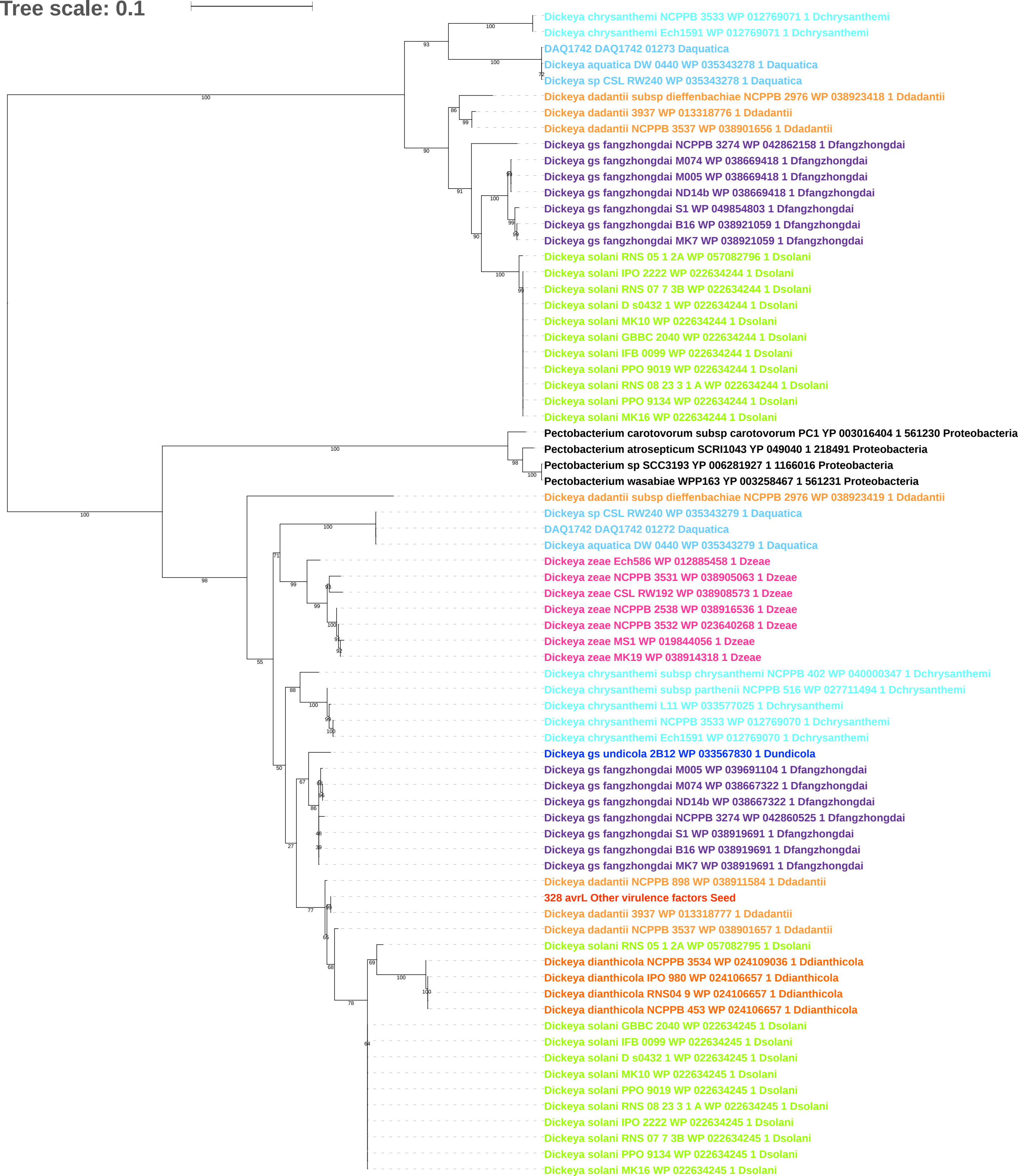

Tree scale: 0.1
Dickeya chrysanthemi NCPPB 3533 WP 012769071 1 Dchrysanthemi
Dickeya chrysanthemi Ech1591 WP 012769071 1 Dchrysanthemi
DAQ1742 DAQ1742 01273 Daquatica
Dickeya aquatica DW 0440 WP 035343278 1 Daquatica
Dickeya sp CSL RW240 WP 035343278 1 Daquatica
Dickeya dadantii subsp dieffenbachiae NCPPB 2976 WP 038923418 1 Ddadantii
Dickeya dadantii 3937 WP 013318776 1 Ddadantii
Dickeya dadantii NCPPB 3537 WP 038901656 1 Ddadantii
Dickeya gs fangzhongdai NCPPB 3274 WP 042862158 1 Dfangzhongdai
Dickeya gs fangzhongdai M074 WP 038669418 1 Dfangzhongdai
Dickeya gs fangzhongdai M005 WP 038669418 1 Dfangzhongdai
Dickeya gs fangzhongdai ND14b WP 038669418 1 Dfangzhongdai
Dickeya gs fangzhongdai S1 WP 049854803 1 Dfangzhongdai
Dickeya gs fangzhongdai B16 WP 038921059 1 Dfangzhongdai
Dickeya gs fangzhongdai MK7 WP 038921059 1 Dfangzhongdai
Dickeya solani RNS 05 1 2A WP 057082796 1 Dsolani
Dickeya solani IPO 2222 WP 022634244 1 Dsolani
Dickeya solani RNS 07 7 3B WP 022634244 1 Dsolani
Dickeya solani D s0432 1 WP 022634244 1 Dsolani
Dickeya solani MK10 WP 022634244 1 Dsolani
Dickeya solani GBBC 2040 WP 022634244 1 Dsolani
Dickeya solani IFB 0099 WP 022634244 1 Dsolani
Dickeya solani PPO 9019 WP 022634244 1 Dsolani
Dickeya solani RNS 08 23 3 1 A WP 022634244 1 Dsolani
Dickeya solani PPO 9134 WP 022634244 1 Dsolani
Dickeya solani MK16 WP 022634244 1 Dsolani
Pectobacterium carotovorum subsp carotovorum PC1 YP 003016404 1 561230 Proteobacteria
Pectobacterium atrosepticum SCRI1043 YP 049040 1 218491 Proteobacteria
Pectobacterium sp SCC3193 YP 006281927 1 1166016 Proteobacteria
Pectobacterium wasabiae WPP163 YP 003258467 1 561231 Proteobacteria
Dickeya dadantii subsp dieffenbachiae NCPPB 2976 WP 038923419 1 Ddadantii
Dickeya sp CSL RW240 WP 035343279 1 Daquatica
DAQ1742 DAQ1742 01272 Daquatica
Dickeya aquatica DW 0440 WP 035343279 1 Daquatica
Dickeya zeae Ech586 WP 012885458 1 Dzeae
Dickeya zeae NCPPB 3531 WP 038905063 1 Dzeae
Dickeya zeae CSL RW192 WP 038908573 1 Dzeae
Dickeya zeae NCPPB 2538 WP 038916536 1 Dzeae
Dickeya zeae NCPPB 3532 WP 023640268 1 Dzeae
Dickeya zeae MS1 WP 019844056 1 Dzeae
Dickeya zeae MK19 WP 038914318 1 Dzeae
Dickeya chrysanthemi subsp chrysanthemi NCPPB 402 WP 040000347 1 Dchrysanthemi
Dickeya chrysanthemi subsp parthenii NCPPB 516 WP 027711494 1 Dchrysanthemi
Dickeya chrysanthemi L11 WP 033577025 1 Dchrysanthemi
Dickeya chrysanthemi NCPPB 3533 WP 012769070 1 Dchrysanthemi
Dickeya chrysanthemi Ech1591 WP 012769070 1 Dchrysanthemi
Dickeya gs undicola 2B12 WP 033567830 1 Dundicola
Dickeya gs fangzhongdai M005 WP 039691104 1 Dfangzhongdai
Dickeya gs fangzhongdai M074 WP 038667322 1 Dfangzhongdai
Dickeya gs fangzhongdai ND14b WP 038667322 1 Dfangzhongdai
Dickeya gs fangzhongdai NCPPB 3274 WP 042860525 1 Dfangzhongdai
Dickeya gs fangzhongdai S1 WP 038919691 1 Dfangzhongdai
Dickeya gs fangzhongdai B16 WP 038919691 1 Dfangzhongdai
Dickeya gs fangzhongdai MK7 WP 038919691 1 Dfangzhongdai
Dickeya dadantii NCPPB 898 WP 038911584 1 Ddadantii
328 avrL Other virulence factors Seed
Dickeya dadantii 3937 WP 013318777 1 Ddadantii
Dickeya dadantii NCPPB 3537 WP 038901657 1 Ddadantii
Dickeya solani RNS 05 1 2A WP 057082795 1 Dsolani
Dickeya dianthicola NCPPB 3534 WP 024109036 1 Ddianthicola
Dickeya dianthicola IPO 980 WP 024106657 1 Ddianthicola
Dickeya dianthicola RNS04 9 WP 024106657 1 Ddianthicola
Dickeya dianthicola NCPPB 453 WP 024106657 1 Ddianthicola
Dickeya solani GBBC 2040 WP 022634245 1 Dsolani
Dickeya solani IFB 0099 WP 022634245 1 Dsolani
Dickeya solani D s0432 1 WP 022634245 1 Dsolani
Dickeya solani MK10 WP 022634245 1 Dsolani
Dickeya solani PPO 9019 WP 022634245 1 Dsolani
Dickeya solani RNS 08 23 3 1 A WP 022634245 1 Dsolani
Dickeya solani IPO 2222 WP 022634245 1 Dsolani
Dickeya solani RNS 07 7 3B WP 022634245 1 Dsolani
Dickeya solani PPO 9134 WP 022634245 1 Dsolani
Dickeya solani MK16 WP 022634245 1 Dsolani

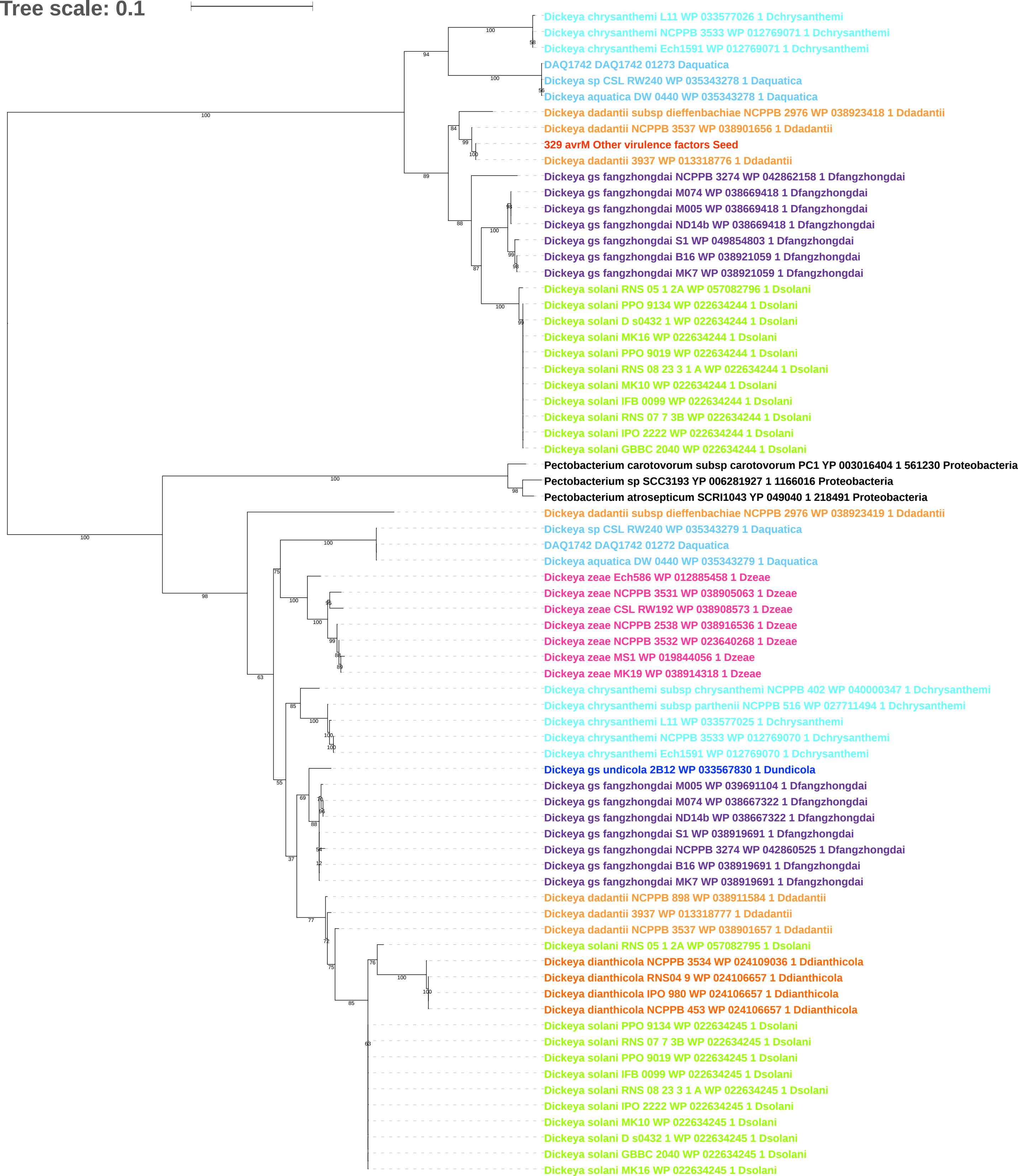
Tree scale: 0.1
Dickeya chrysanthemi L11 WP 033577026 1 Dchrysanthemi
Dickeya chrysanthemi NCPPB 3533 WP 012769071 1 Dchrysanthemi
Dickeya chrysanthemi Ech1591 WP 012769071 1 Dchrysanthemi
DAQ1742 DAQ1742 01273 Daquatica
Dickeya sp CSL RW240 WP 035343278 1 Daquatica
Dickeya aquatica DW 0440 WP 035343278 1 Daquatica
Dickeya dadantii subsp dieffenbachiae NCPPB 2976 WP 038923418 1 Ddadantii
Dickeya dadantii NCPPB 3537 WP 038901656 1 Ddadantii
329 avrM Other virulence factors Seed
Dickeya dadantii 3937 WP 013318776 1 Ddadantii
Dickeya gs fangzhongdai NCPPB 3274 WP 042862158 1 Dfangzhongdai
Dickeya gs fangzhongdai M074 WP 038669418 1 Dfangzhongdai
Dickeya gs fangzhongdai M005 WP 038669418 1 Dfangzhongdai
Dickeya gs fangzhongdai ND14b WP 038669418 1 Dfangzhongdai
Dickeya gs fangzhongdai S1 WP 049854803 1 Dfangzhongdai
Dickeya gs fangzhongdai B16 WP 038921059 1 Dfangzhongdai
Dickeya gs fangzhongdai MK7 WP 038921059 1 Dfangzhongdai
Dickeya solani RNS 05 1 2A WP 057082796 1 Dsolani
Dickeya solani PPO 9134 WP 022634244 1 Dsolani
Dickeya solani D s0432 1 WP 022634244 1 Dsolani
Dickeya solani MK16 WP 022634244 1 Dsolani
Dickeya solani PPO 9019 WP 022634244 1 Dsolani
Dickeya solani RNS 08 23 3 1 A WP 022634244 1 Dsolani
Dickeya solani MK10 WP 022634244 1 Dsolani
Dickeya solani IFB 0099 WP 022634244 1 Dsolani
Dickeya solani RNS 07 7 3B WP 022634244 1 Dsolani
Dickeya solani IPO 2222 WP 022634244 1 Dsolani
Dickeya solani GBBC 2040 WP 022634244 1 Dsolani
Pectobacterium carotovorum subsp carotovorum PC1 YP 003016404 1 561230 Proteobacteria
Pectobacterium sp SCC3193 YP 006281927 1 1166016 Proteobacteria
Pectobacterium atrosepticum SCRI1043 YP 049040 1 218491 Proteobacteria
Dickeya dadantii subsp dieffenbachiae NCPPB 2976 WP 038923419 1 Ddadantii
Dickeya sp CSL RW240 WP 035343279 1 Daquatica
DAQ1742 DAQ1742 01272 Daquatica
Dickeya aquatica DW 0440 WP 035343279 1 Daquatica
Dickeya zeae Ech586 WP 012885458 1 Dzeae
Dickeya zeae NCPPB 3531 WP 038905063 1 Dzeae
Dickeya zeae CSL RW192 WP 038908573 1 Dzeae
Dickeya zeae NCPPB 2538 WP 038916536 1 Dzeae
Dickeya zeae NCPPB 3532 WP 023640268 1 Dzeae
Dickeya zeae MS1 WP 019844056 1 Dzeae
Dickeya zeae MK19 WP 038914318 1 Dzeae
Dickeya chrysanthemi subsp chrysanthemi NCPPB 402 WP 040000347 1 Dchrysanthemi
Dickeya chrysanthemi subsp parthenii NCPPB 516 WP 027711494 1 Dchrysanthemi
Dickeya chrysanthemi L11 WP 033577025 1 Dchrysanthemi
Dickeya chrysanthemi NCPPB 3533 WP 012769070 1 Dchrysanthemi
Dickeya chrysanthemi Ech1591 WP 012769070 1 Dchrysanthemi
Dickeya gs undicola 2B12 WP 033567830 1 Dundicola
Dickeya gs fangzhongdai M005 WP 039691104 1 Dfangzhongdai
Dickeya gs fangzhongdai M074 WP 038667322 1 Dfangzhongdai
Dickeya gs fangzhongdai ND14b WP 038667322 1 Dfangzhongdai
Dickeya gs fangzhongdai S1 WP 038919691 1 Dfangzhongdai
Dickeya gs fangzhongdai NCPPB 3274 WP 042860525 1 Dfangzhongdai
Dickeya gs fangzhongdai B16 WP 038919691 1 Dfangzhongdai
Dickeya gs fangzhongdai MK7 WP 038919691 1 Dfangzhongdai
Dickeya dadantii NCPPB 898 WP 038911584 1 Ddadantii
Dickeya dadantii 3937 WP 013318777 1 Ddadantii
Dickeya dadantii NCPPB 3537 WP 038901657 1 Ddadantii
Dickeya solani RNS 05 1 2A WP 057082795 1 Dsolani
Dickeya dianthicola NCPPB 3534 WP 024109036 1 Ddianthicola
Dickeya dianthicola RNS04 9 WP 024106657 1 Ddianthicola
Dickeya dianthicola IPO 980 WP 024106657 1 Ddianthicola
Dickeya dianthicola NCPPB 453 WP 024106657 1 Ddianthicola
Dickeya solani PPO 9134 WP 022634245 1 Dsolani
Dickeya solani RNS 07 7 3B WP 022634245 1 Dsolani
Dickeya solani PPO 9019 WP 022634245 1 Dsolani
Dickeya solani IFB 0099 WP 022634245 1 Dsolani
Dickeya solani RNS 08 23 3 1 A WP 022634245 1 Dsolani
Dickeya solani IPO 2222 WP 022634245 1 Dsolani
Dickeya solani MK10 WP 022634245 1 Dsolani
Dickeya solani D s0432 1 WP 022634245 1 Dsolani
Dickeya solani GBBC 2040 WP 022634245 1 Dsolani
Dickeya solani MK16 WP 022634245 1 Dsolani

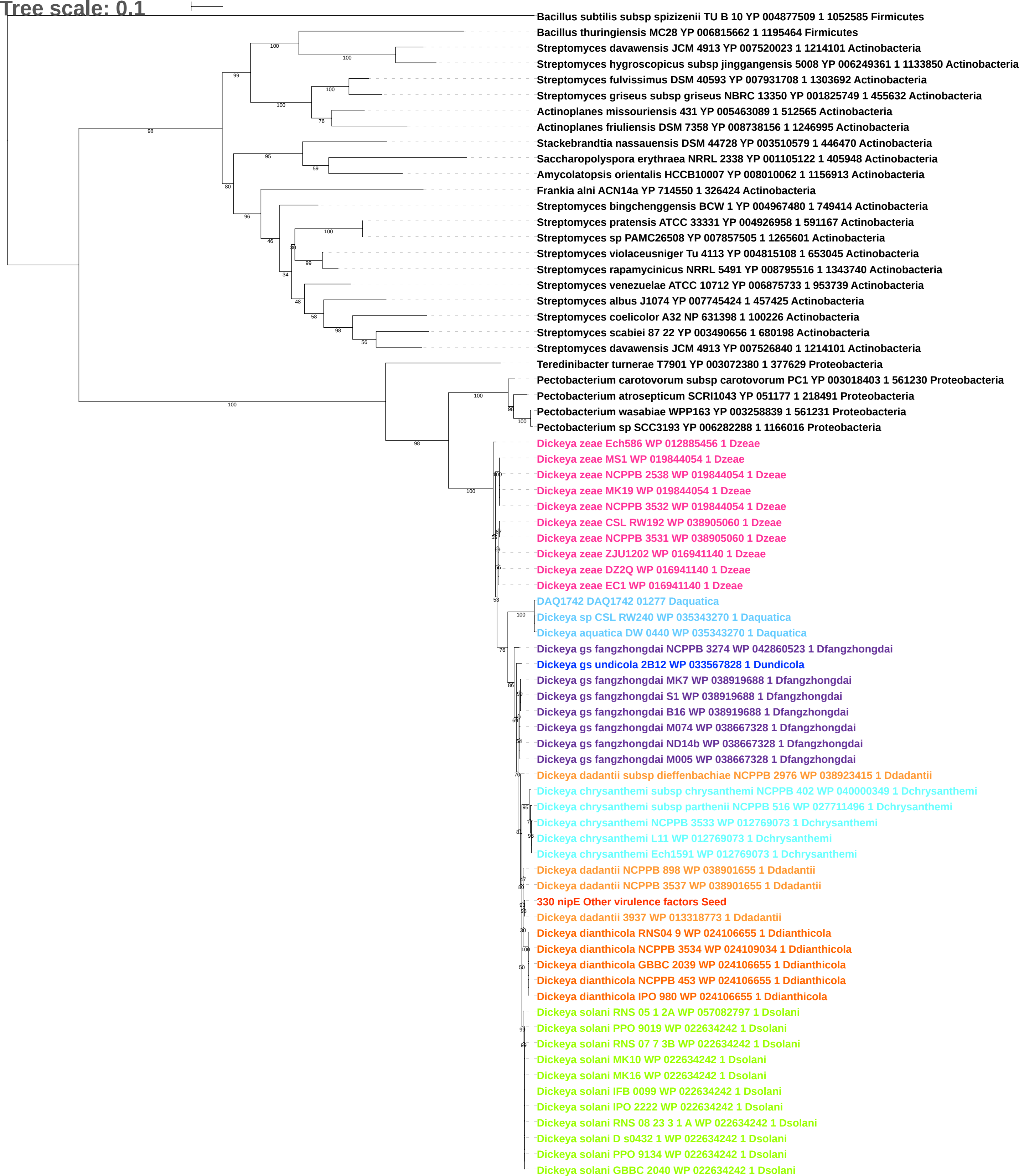

Tree scale: 0.1
Bacillus subtilis subsp spizizenii TU B 10 YP 004877509 1 1052585 Firmicutes
Bacillus thuringiensis MC28 YP 006815662 1 1195464 Firmicutes
Streptomyces davawensis JCM 4913 YP 007520023 1 1214101 Actinobacteria
Streptomyces hygroscopicus subsp jinggangensis 5008 YP 006249361 1 1133850 Actinobacteria
Streptomyces fulvissimus DSM 40593 YP 007931708 1 1303692 Actinobacteria
Streptomyces griseus subsp griseus NBRC 13350 YP 001825749 1 455632 Actinobacteria
Actinoplanes missouriensis 431 YP 005463089 1 512565 Actinobacteria
Actinoplanes friuliensis DSM 7358 YP 008738156 1 1246995 Actinobacteria
Stackebrandtia nassauensis DSM 44728 YP 003510579 1 446470 Actinobacteria
Saccharopolyspora erythraea NRRL 2338 YP 001105122 1 405948 Actinobacteria
Amycolatopsis orientalis HCCB10007 YP 008010062 1 1156913 Actinobacteria
Frankia alni ACN14a YP 714550 1 326424 Actinobacteria
Streptomyces bingchenggensis BCW 1 YP 004967480 1 749414 Actinobacteria
Streptomyces pratensis ATCC 33331 YP 004926958 1 591167 Actinobacteria
Streptomyces sp PAMC26508 YP 007857505 1 1265601 Actinobacteria
Streptomyces violaceusniger Tu 4113 YP 004815108 1 653045 Actinobacteria
Streptomyces rapamycinicus NRRL 5491 YP 008795516 1 1343740 Actinobacteria
Streptomyces venezuelae ATCC 10712 YP 006875733 1 953739 Actinobacteria
Streptomyces albus J1074 YP 007745424 1 457425 Actinobacteria
Streptomyces coelicolor A32 NP 631398 1 100226 Actinobacteria
Streptomyces scabiei 87 22 YP 003490656 1 680198 Actinobacteria
Streptomyces davawensis JCM 4913 YP 007526840 1 1214101 Actinobacteria
Teredinibacter turnerae T7901 YP 003072380 1 377629 Proteobacteria
Pectobacterium carotovorum subsp carotovorum PC1 YP 003018403 1 561230 Proteobacteria
Pectobacterium atrosepticum SCRI1043 YP 051177 1 218491 Proteobacteria
Pectobacterium wasabiae WPP163 YP 003258839 1 561231 Proteobacteria
Pectobacterium sp SCC3193 YP 006282288 1 1166016 Proteobacteria
Dickeya zeae Ech586 WP 012885456 1 Dzeae
Dickeya zeae MS1 WP 019844054 1 Dzeae
Dickeya zeae NCPPB 2538 WP 019844054 1 Dzeae
Dickeya zeae MK19 WP 019844054 1 Dzeae
Dickeya zeae NCPPB 3532 WP 019844054 1 Dzeae
Dickeya zeae CSL RW192 WP 038905060 1 Dzeae
Dickeya zeae NCPPB 3531 WP 038905060 1 Dzeae
Dickeya zeae ZJU1202 WP 016941140 1 Dzeae
Dickeya zeae DZ2Q WP 016941140 1 Dzeae
Dickeya zeae EC1 WP 016941140 1 Dzeae
DAQ1742 DAQ1742 01277 Daquatica
Dickeya sp CSL RW240 WP 035343270 1 Daquatica
Dickeya aquatica DW 0440 WP 035343270 1 Daquatica
Dickeya gs fangzhongdai NCPPB 3274 WP 042860523 1 Dfangzhongdai
Dickeya gs undicola 2B12 WP 033567828 1 Dundicola
Dickeya gs fangzhongdai MK7 WP 038919688 1 Dfangzhongdai
Dickeya gs fangzhongdai S1 WP 038919688 1 Dfangzhongdai
Dickeya gs fangzhongdai B16 WP 038919688 1 Dfangzhongdai
Dickeya gs fangzhongdai M074 WP 038667328 1 Dfangzhongdai
Dickeya gs fangzhongdai ND14b WP 038667328 1 Dfangzhongdai
Dickeya gs fangzhongdai M005 WP 038667328 1 Dfangzhongdai
Dickeya dadantii subsp dieffenbachiae NCPPB 2976 WP 038923415 1 Ddadantii
Dickeya chrysanthemi subsp chrysanthemi NCPPB 402 WP 040000349 1 Dchrysanthemi
Dickeya chrysanthemi subsp parthenii NCPPB 516 WP 027711496 1 Dchrysanthemi
Dickeya chrysanthemi NCPPB 3533 WP 012769073 1 Dchrysanthemi
Dickeya chrysanthemi L11 WP 012769073 1 Dchrysanthemi
Dickeya chrysanthemi Ech1591 WP 012769073 1 Dchrysanthemi
Dickeya dadantii NCPPB 898 WP 038901655 1 Ddadantii
Dickeya dadantii NCPPB 3537 WP 038901655 1 Ddadantii
330 nipE Other virulence factors Seed
Dickeya dadantii 3937 WP 013318773 1 Ddadantii
Dickeya dianthicola RNS04 9 WP 024106655 1 Ddianthicola
Dickeya dianthicola NCPPB 3534 WP 024109034 1 Ddianthicola
Dickeya dianthicola GBBC 2039 WP 024106655 1 Ddianthicola
Dickeya dianthicola NCPPB 453 WP 024106655 1 Ddianthicola
Dickeya dianthicola IPO 980 WP 024106655 1 Ddianthicola
Dickeya solani RNS 05 1 2A WP 057082797 1 Dsolani
Dickeya solani PPO 9019 WP 022634242 1 Dsolani
Dickeya solani RNS 07 7 3B WP 022634242 1 Dsolani
Dickeya solani MK10 WP 022634242 1 Dsolani
Dickeya solani MK16 WP 022634242 1 Dsolani
Dickeya solani IFB 0099 WP 022634242 1 Dsolani
Dickeya solani IPO 2222 WP 022634242 1 Dsolani
Dickeya solani RNS 08 23 3 1 A WP 022634242 1 Dsolani
Dickeya solani D s0432 1 WP 022634242 1 Dsolani
Dickeya solani PPO 9134 WP 022634242 1 Dsolani
Dickeya solani GBBC 2040 WP 022634242 1 Dsolani

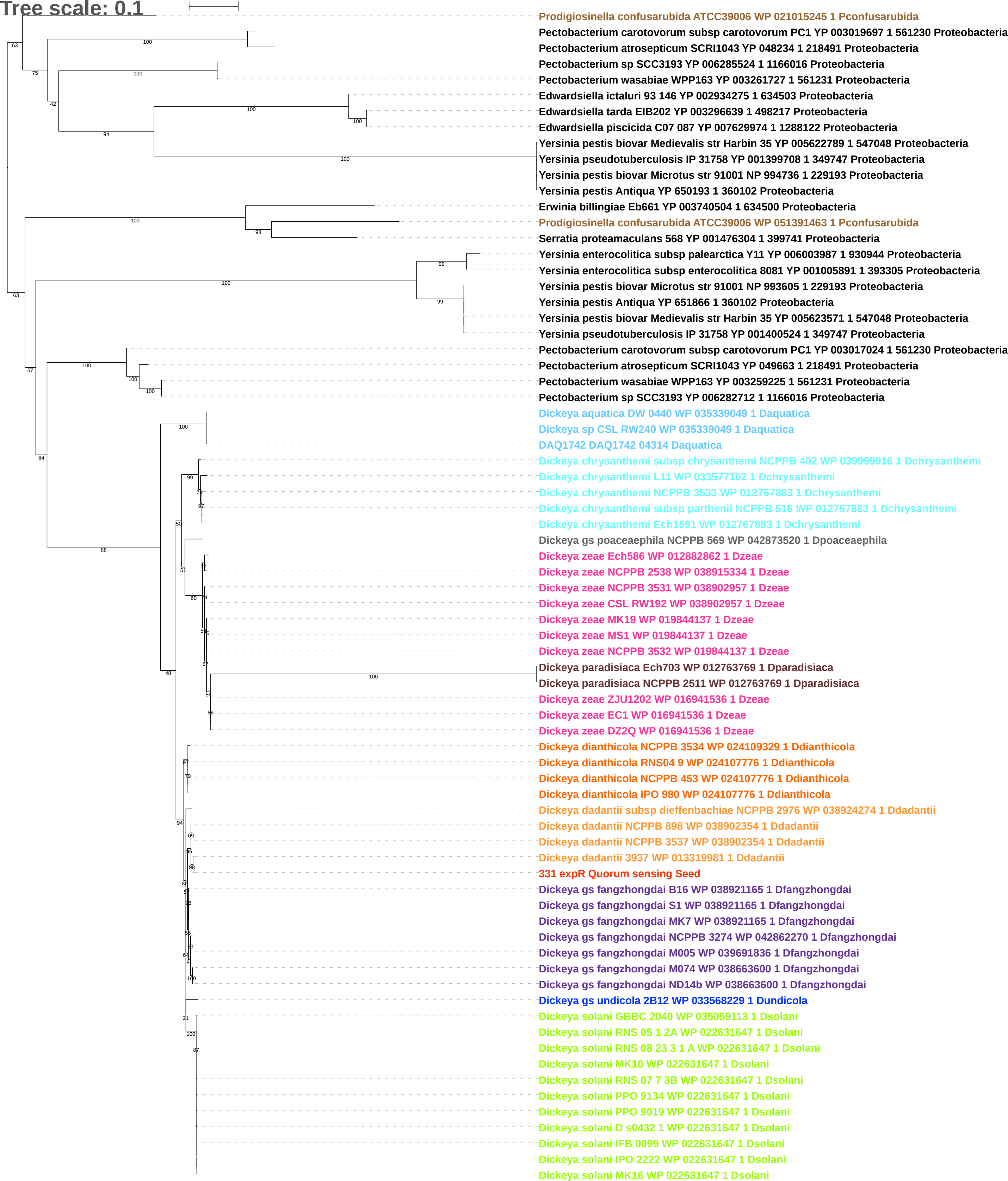
Tree scale: 0.1
Prodigiosinella confusarubida ATCC39006 WP 021015245 1 Pconfusarubida
Pectobacterium carotovorum subsp carotovorum PC1 YP 003019697 1 561230 Proteobacteria
Pectobacterium atrosepticum SCRI1043 YP 048234 1 218491 Proteobacteria
Pectobacterium sp SCC3193 YP 006285524 1 1166016 Proteobacteria
Pectobacterium wasabiae WPP163 YP 003261727 1 561231 Proteobacteria
Edwardsiella ictaluri 93 146 YP 002934275 1 634503 Proteobacteria
Edwardsiella tarda EIB202 YP 003296639 1 498217 Proteobacteria
Edwardsiella piscicida C07 087 YP 007629974 1 1288122 Proteobacteria
Yersinia pestis biovar Medievalis str Harbin 35 YP 005622789 1 547048 Proteobacteria
Yersinia pseudotuberculosis IP 31758 YP 001399708 1 349747 Proteobacteria
Yersinia pestis biovar Microtus str 91001 NP 994736 1 229193 Proteobacteria
Yersinia pestis Antiqua YP 650193 1 360102 Proteobacteria
Erwinia billingiae Eb661 YP 003740504 1 634500 Proteobacteria
Prodigiosinella confusarubida ATCC39006 WP 051391463 1 Pconfusarubida
Serratia proteamaculans 568 YP 001476304 1 399741 Proteobacteria
Yersinia enterocolitica subsp palearctica Y11 YP 006003987 1 930944 Proteobacteria
Yersinia enterocolitica subsp enterocolitica 8081 YP 001005891 1 393305 Proteobacteria
Yersinia pestis biovar Microtus str 91001 NP 993605 1 229193 Proteobacteria
Yersinia pestis Antiqua YP 651866 1 360102 Proteobacteria
Yersinia pestis biovar Medievalis str Harbin 35 YP 005623571 1 547048 Proteobacteria
Yersinia pseudotuberculosis IP 31758 YP 001400524 1 349747 Proteobacteria
Pectobacterium carotovorum subsp carotovorum PC1 YP 003017024 1 561230 Proteobacteria
Pectobacterium atrosepticum SCRI1043 YP 049663 1 218491 Proteobacteria
Pectobacterium wasabiae WPP163 YP 003259225 1 561231 Proteobacteria
Pectobacterium sp SCC3193 YP 006282712 1 1166016 Proteobacteria
Dickeya aquatica DW 0440 WP 035339049 1 Daquatica
Dickeya sp CSL RW240 WP 035339049 1 Daquatica
DAQ1742 DAQ1742 04314 Daquatica
Dickeya chrysanthemi subsp chrysanthemi NCPPB 402 WP 039999016 1 Dchrysanthemi
Dickeya chrysanthemi L11 WP 033577102 1 Dchrysanthemi
Dickeya chrysanthemi NCPPB 3533 WP 012767883 1 Dchrysanthemi
Dickeya chrysanthemi subsp parthenii NCPPB 516 WP 012767883 1 Dchrysanthemi
Dickeya chrysanthemi Ech1591 WP 012767883 1 Dchrysanthemi
Dickeya gs poaceaephila NCPPB 569 WP 042873520 1 Dpoaceaephila
Dickeya zeae Ech586 WP 012882862 1 Dzeae
Dickeya zeae NCPPB 2538 WP 038915334 1 Dzeae
Dickeya zeae NCPPB 3531 WP 038902957 1 Dzeae
Dickeya zeae CSL RW192 WP 038902957 1 Dzeae
Dickeya zeae MK19 WP 019844137 1 Dzeae
Dickeya zeae MS1 WP 019844137 1 Dzeae
Dickeya zeae NCPPB 3532 WP 019844137 1 Dzeae
Dickeya paradisiaca Ech703 WP 012763769 1 Dparadisiaca
Dickeya paradisiaca NCPPB 2511 WP 012763769 1 Dparadisiaca
Dickeya zeae ZJU1202 WP 016941536 1 Dzeae
Dickeya zeae EC1 WP 016941536 1 Dzeae
Dickeya zeae DZ2Q WP 016941536 1 Dzeae
Dickeya dianthicola NCPPB 3534 WP 024109329 1 Ddianthicola
Dickeya dianthicola RNS04 9 WP 024107776 1 Ddianthicola
Dickeya dianthicola NCPPB 453 WP 024107776 1 Ddianthicola
Dickeya dianthicola IPO 980 WP 024107776 1 Ddianthicola
Dickeya dadantii subsp dieffenbachiae NCPPB 2976 WP 038924274 1 Ddadantii
Dickeya dadantii NCPPB 898 WP 038902354 1 Ddadantii
Dickeya dadantii NCPPB 3537 WP 038902354 1 Ddadantii
Dickeya dadantii 3937 WP 013319981 1 Ddadantii
331 expR Quorum sensing Seed
Dickeya gs fangzhongdai B16 WP 038921165 1 Dfangzhongdai
Dickeya gs fangzhongdai S1 WP 038921165 1 Dfangzhongdai
Dickeya gs fangzhongdai MK7 WP 038921165 1 Dfangzhongdai
Dickeya gs fangzhongdai NCPPB 3274 WP 042862270 1 Dfangzhongdai
Dickeya gs fangzhongdai M005 WP 039691836 1 Dfangzhongdai
Dickeya gs fangzhongdai M074 WP 038663600 1 Dfangzhongdai
Dickeya gs fangzhongdai ND14b WP 038663600 1 Dfangzhongdai
Dickeya gs undicola 2B12 WP 033568229 1 Dundicola
Dickeya solani GBBC 2040 WP 035059113 1 Dsolani
Dickeya solani RNS 05 1 2A WP 022631647 1 Dsolani
Dickeya solani RNS 08 23 3 1 A WP 022631647 1 Dsolani
Dickeya solani MK10 WP 022631647 1 Dsolani
Dickeya solani RNS 07 7 3B WP 022631647 1 Dsolani
Dickeya solani PPO 9134 WP 022631647 1 Dsolani
Dickeya solani PPO 9019 WP 022631647 1 Dsolani
Dickeya solani D s0432 1 WP 022631647 1 Dsolani
Dickeya solani IFB 0099 WP 022631647 1 Dsolani
Dickeya solani IPO 2222 WP 022631647 1 Dsolani
Dickeya solani MK16 WP 022631647 1 Dsolani

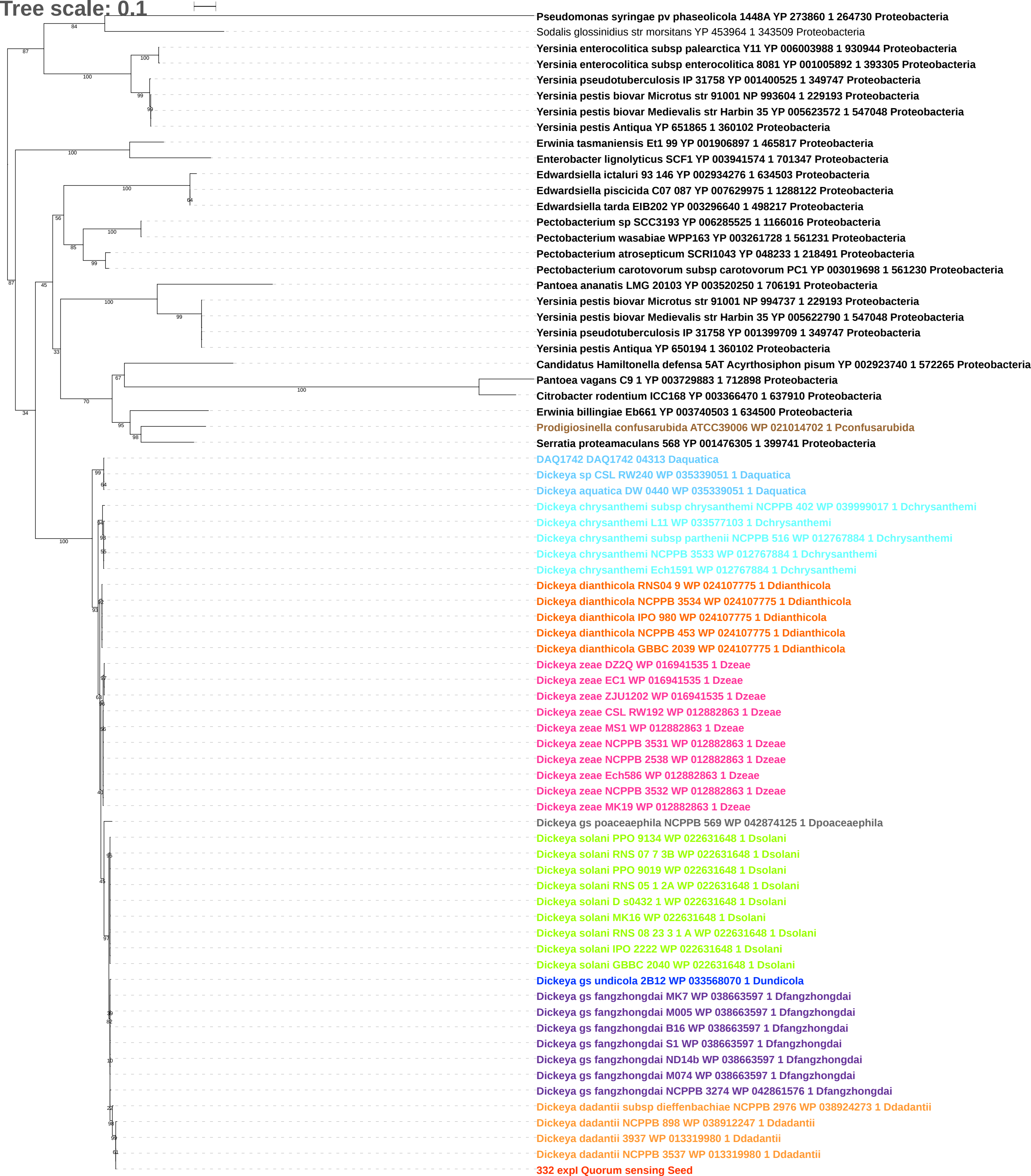

Tree scale: 0.1
Pseudomonas syringae pv phaseolicola 1448A YP 273860 1 264730 Proteobacteria
Sodalis glossinidius str morsitans YP 453964 1 343509 Proteobacteria
Yersinia enterocolitica subsp palearctica Y11 YP 006003988 1 930944 Proteobacteria
Yersinia enterocolitica subsp enterocolitica 8081 YP 001005892 1 393305 Proteobacteria
Yersinia pseudotuberculosis IP 31758 YP 001400525 1 349747 Proteobacteria
Yersinia pestis biovar Microtus str 91001 NP 993604 1 229193 Proteobacteria
Yersinia pestis biovar Medievalis str Harbin 35 YP 005623572 1 547048 Proteobacteria
Yersinia pestis Antiqua YP 651865 1 360102 Proteobacteria
Erwinia tasmaniensis Et1 99 YP 001906897 1 465817 Proteobacteria
Enterobacter lignolyticus SCF1 YP 003941574 1 701347 Proteobacteria
Edwardsiella ictaluri 93 146 YP 002934276 1 634503 Proteobacteria
Edwardsiella piscicida C07 087 YP 007629975 1 1288122 Proteobacteria
Edwardsiella tarda EIB202 YP 003296640 1 498217 Proteobacteria
Pectobacterium sp SCC3193 YP 006285525 1 1166016 Proteobacteria
Pectobacterium wasabiae WPP163 YP 003261728 1 561231 Proteobacteria
Pectobacterium atrosepticum SCRI1043 YP 048233 1 218491 Proteobacteria
Pectobacterium carotovorum subsp carotovorum PC1 YP 003019698 1 561230 Proteobacteria
Pantoea ananatis LMG 20103 YP 003520250 1 706191 Proteobacteria
Yersinia pestis biovar Microtus str 91001 NP 994737 1 229193 Proteobacteria
Yersinia pestis biovar Medievalis str Harbin 35 YP 005622790 1 547048 Proteobacteria
Yersinia pseudotuberculosis IP 31758 YP 001399709 1 349747 Proteobacteria
Yersinia pestis Antiqua YP 650194 1 360102 Proteobacteria
Candidatus Hamiltonella defensa 5AT Acyrthosiphon pisum YP 002923740 1 572265 Proteobacteria
Pantoea vagans C9 1 YP 003729883 1 712898 Proteobacteria
Citrobacter rodentium ICC168 YP 003366470 1 637910 Proteobacteria
Erwinia billingiae Eb661 YP 003740503 1 634500 Proteobacteria
Prodigiosinella confusarubida ATCC39006 WP 021014702 1 Pconfusarubida
Serratia proteamaculans 568 YP 001476305 1 399741 Proteobacteria
DAQ1742 DAQ1742 04313 Daquatica
Dickeya sp CSL RW240 WP 035339051 1 Daquatica
Dickeya aquatica DW 0440 WP 035339051 1 Daquatica
Dickeya chrysanthemi subsp chrysanthemi NCPPB 402 WP 039999017 1 Dchrysanthemi
Dickeya chrysanthemi L11 WP 033577103 1 Dchrysanthemi
Dickeya chrysanthemi subsp parthenii NCPPB 516 WP 012767884 1 Dchrysanthemi
Dickeya chrysanthemi NCPPB 3533 WP 012767884 1 Dchrysanthemi
Dickeya chrysanthemi Ech1591 WP 012767884 1 Dchrysanthemi
Dickeya dianthicola RNS04 9 WP 024107775 1 Ddianthicola
Dickeya dianthicola NCPPB 3534 WP 024107775 1 Ddianthicola
Dickeya dianthicola IPO 980 WP 024107775 1 Ddianthicola
Dickeya dianthicola NCPPB 453 WP 024107775 1 Ddianthicola
Dickeya dianthicola GBBC 2039 WP 024107775 1 Ddianthicola
Dickeya zeae DZ2Q WP 016941535 1 Dzeae
Dickeya zeae EC1 WP 016941535 1 Dzeae
Dickeya zeae ZJU1202 WP 016941535 1 Dzeae
Dickeya zeae CSL RW192 WP 012882863 1 Dzeae
Dickeya zeae MS1 WP 012882863 1 Dzeae
Dickeya zeae NCPPB 3531 WP 012882863 1 Dzeae
Dickeya zeae NCPPB 2538 WP 012882863 1 Dzeae
Dickeya zeae Ech586 WP 012882863 1 Dzeae
Dickeya zeae NCPPB 3532 WP 012882863 1 Dzeae
Dickeya zeae MK19 WP 012882863 1 Dzeae
Dickeya gs poaceaephila NCPPB 569 WP 042874125 1 Dpoaceaephila
Dickeya solani PPO 9134 WP 022631648 1 Dsolani
Dickeya solani RNS 07 7 3B WP 022631648 1 Dsolani
Dickeya solani PPO 9019 WP 022631648 1 Dsolani
Dickeya solani RNS 05 1 2A WP 022631648 1 Dsolani
Dickeya solani D s0432 1 WP 022631648 1 Dsolani
Dickeya solani MK16 WP 022631648 1 Dsolani
Dickeya solani RNS 08 23 3 1 A WP 022631648 1 Dsolani
Dickeya solani IPO 2222 WP 022631648 1 Dsolani
Dickeya solani GBBC 2040 WP 022631648 1 Dsolani
Dickeya gs undicola 2B12 WP 033568070 1 Dundicola
Dickeya gs fangzhongdai MK7 WP 038663597 1 Dfangzhongdai
Dickeya gs fangzhongdai M005 WP 038663597 1 Dfangzhongdai
Dickeya gs fangzhongdai B16 WP 038663597 1 Dfangzhongdai
Dickeya gs fangzhongdai S1 WP 038663597 1 Dfangzhongdai
Dickeya gs fangzhongdai ND14b WP 038663597 1 Dfangzhongdai
Dickeya gs fangzhongdai M074 WP 038663597 1 Dfangzhongdai
Dickeya gs fangzhongdai NCPPB 3274 WP 042861576 1 Dfangzhongdai
Dickeya dadantii subsp dieffenbachiae NCPPB 2976 WP 038924273 1 Ddadantii
Dickeya dadantii NCPPB 898 WP 038912247 1 Ddadantii
Dickeya dadantii 3937 WP 013319980 1 Ddadantii
Dickeya dadantii NCPPB 3537 WP 013319980 1 Ddadantii
332 expl Quorum sensing Seed

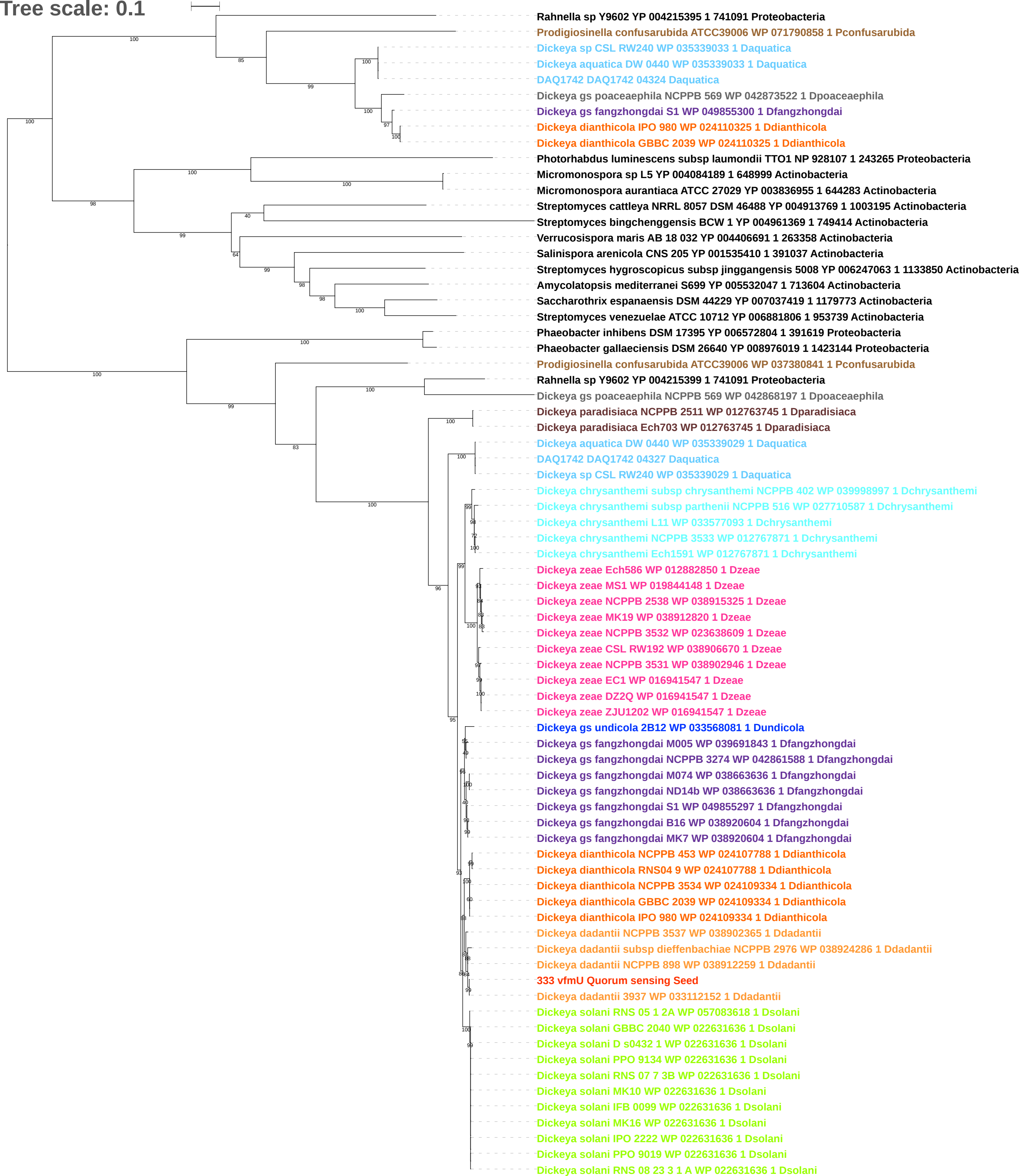
Tree scale: 0.1
Rahnella sp Y9602 YP 004215395 1 741091 Proteobacteria
Prodigiosinella confusarubida ATCC39006 WP 071790858 1 Pconfusarubida
Dickeya sp CSL RW240 WP 035339033 1 Daquatica
Dickeya aquatica DW 0440 WP 035339033 1 Daquatica
DAQ1742 DAQ1742 04324 Daquatica
Dickeya gs poaceaephila NCPPB 569 WP 042873522 1 Dpoaceaephila
Dickeya gs fangzhongdai S1 WP 049855300 1 Dfangzhongdai
Dickeya dianthicola IPO 980 WP 024110325 1 Ddianthicola
Dickeya dianthicola GBBC 2039 WP 024110325 1 Ddianthicola
Photorhabdus luminescens subsp laumondii TTO1 NP 928107 1 243265 Proteobacteria
Micromonospora sp L5 YP 004084189 1 648999 Actinobacteria
Micromonospora aurantiaca ATCC 27029 YP 003836955 1 644283 Actinobacteria
Streptomyces cattleya NRRL 8057 DSM 46488 YP 004913769 1 1003195 Actinobacteria
Streptomyces bingchenggensis BCW 1 YP 004961369 1 749414 Actinobacteria
Verrucosispora maris AB 18 032 YP 004406691 1 263358 Actinobacteria
Salinispora arenicola CNS 205 YP 001535410 1 391037 Actinobacteria
Streptomyces hygroscopicus subsp jinggangensis 5008 YP 006247063 1 1133850 Actinobacteria
Amycolatopsis mediterranei S699 YP 005532047 1 713604 Actinobacteria
Saccharothrix espanaensis DSM 44229 YP 007037419 1 1179773 Actinobacteria
Streptomyces venezuelae ATCC 10712 YP 006881806 1 953739 Actinobacteria
Phaeobacter inhibens DSM 17395 YP 006572804 1 391619 Proteobacteria
Phaeobacter gallaeciensis DSM 26640 YP 008976019 1 1423144 Proteobacteria
Prodigiosinella confusarubida ATCC39006 WP 037380841 1 Pconfusarubida
Rahnella sp Y9602 YP 004215399 1 741091 Proteobacteria
Dickeya gs poaceaephila NCPPB 569 WP 042868197 1 Dpoaceaephila
Dickeya paradisiaca NCPPB 2511 WP 012763745 1 Dparadisiaca
Dickeya paradisiaca Ech703 WP 012763745 1 Dparadisiaca
Dickeya aquatica DW 0440 WP 035339029 1 Daquatica
DAQ1742 DAQ1742 04327 Daquatica
Dickeya sp CSL RW240 WP 035339029 1 Daquatica
Dickeya chrysanthemi subsp chrysanthemi NCPPB 402 WP 039998997 1 Dchrysanthemi
Dickeya chrysanthemi subsp parthenii NCPPB 516 WP 027710587 1 Dchrysanthemi
Dickeya chrysanthemi L11 WP 033577093 1 Dchrysanthemi
Dickeya chrysanthemi NCPPB 3533 WP 012767871 1 Dchrysanthemi
Dickeya chrysanthemi Ech1591 WP 012767871 1 Dchrysanthemi
Dickeya zeae Ech586 WP 012882850 1 Dzeae
Dickeya zeae MS1 WP 019844148 1 Dzeae
Dickeya zeae NCPPB 2538 WP 038915325 1 Dzeae
Dickeya zeae MK19 WP 038912820 1 Dzeae
Dickeya zeae NCPPB 3532 WP 023638609 1 Dzeae
Dickeya zeae CSL RW192 WP 038906670 1 Dzeae
Dickeya zeae NCPPB 3531 WP 038902946 1 Dzeae
Dickeya zeae EC1 WP 016941547 1 Dzeae
Dickeya zeae DZ2Q WP 016941547 1 Dzeae
Dickeya zeae ZJU1202 WP 016941547 1 Dzeae
Dickeya gs undicola 2B12 WP 033568081 1 Dundicola
Dickeya gs fangzhongdai M005 WP 039691843 1 Dfangzhongdai
Dickeya gs fangzhongdai NCPPB 3274 WP 042861588 1 Dfangzhongdai
Dickeya gs fangzhongdai M074 WP 038663636 1 Dfangzhongdai
Dickeya gs fangzhongdai ND14b WP 038663636 1 Dfangzhongdai
Dickeya gs fangzhongdai S1 WP 049855297 1 Dfangzhongdai
Dickeya gs fangzhongdai B16 WP 038920604 1 Dfangzhongdai
Dickeya gs fangzhongdai MK7 WP 038920604 1 Dfangzhongdai
Dickeya dianthicola NCPPB 453 WP 024107788 1 Ddianthicola
Dickeya dianthicola RNS04 9 WP 024107788 1 Ddianthicola
Dickeya dianthicola NCPPB 3534 WP 024109334 1 Ddianthicola
Dickeya dianthicola GBBC 2039 WP 024109334 1 Ddianthicola
Dickeya dianthicola IPO 980 WP 024109334 1 Ddianthicola
Dickeya dadantii NCPPB 3537 WP 038902365 1 Ddadantii
Dickeya dadantii subsp dieffenbachiae NCPPB 2976 WP 038924286 1 Ddadantii
Dickeya dadantii NCPPB 898 WP 038912259 1 Ddadantii
333 vfmU Quorum sensing Seed
Dickeya dadantii 3937 WP 033112152 1 Ddadantii
Dickeya solani RNS 05 1 2A WP 057083618 1 Dsolani
Dickeya solani GBBC 2040 WP 022631636 1 Dsolani
Dickeya solani D s0432 1 WP 022631636 1 Dsolani
Dickeya solani PPO 9134 WP 022631636 1 Dsolani
Dickeya solani RNS 07 7 3B WP 022631636 1 Dsolani
Dickeya solani MK10 WP 022631636 1 Dsolani
Dickeya solani IFB 0099 WP 022631636 1 Dsolani
Dickeya solani MK16 WP 022631636 1 Dsolani
Dickeya solani IPO 2222 WP 022631636 1 Dsolani
Dickeya solani PPO 9019 WP 022631636 1 Dsolani
Dickeya solani RNS 08 23 3 1 A WP 022631636 1 Dsolani

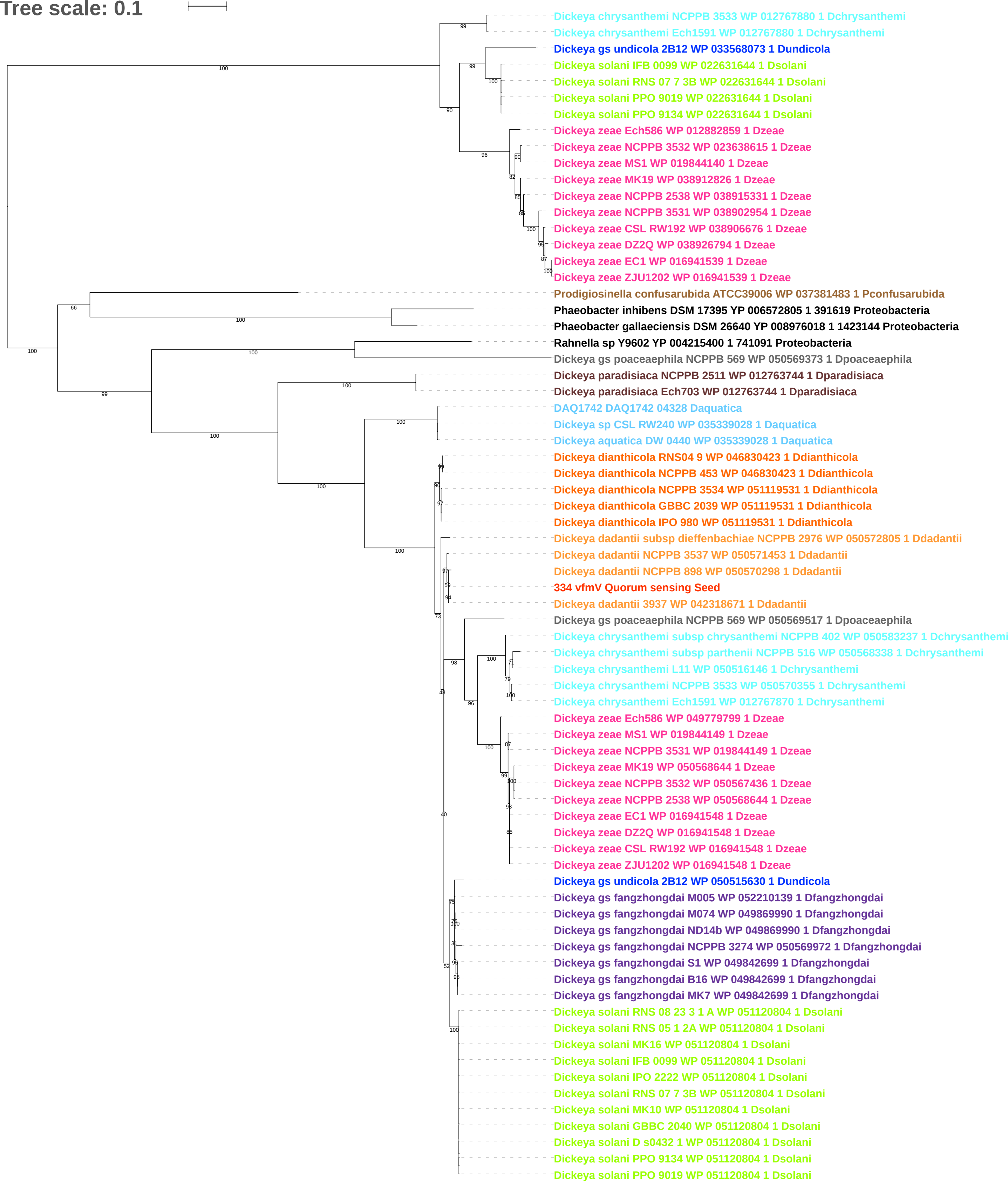
Tree scale: 0.1
Dickeya chrysanthemi NCPPB 3533 WP 012767880 1 Dchrysanthemi
Dickeya chrysanthemi Ech1591 WP 012767880 1 Dchrysanthemi
Dickeya gs undicola 2B12 WP 033568073 1 Dundicola
Dickeya solani IFB 0099 WP 022631644 1 Dsolani
Dickeya solani RNS 07 7 3B WP 022631644 1 Dsolani
Dickeya solani PPO 9019 WP 022631644 1 Dsolani
Dickeya solani PPO 9134 WP 022631644 1 Dsolani
Dickeya zeae Ech586 WP 012882859 1 Dzeae
Dickeya zeae NCPPB 3532 WP 023638615 1 Dzeae
Dickeya zeae MS1 WP 019844140 1 Dzeae
Dickeya zeae MK19 WP 038912826 1 Dzeae
Dickeya zeae NCPPB 2538 WP 038915331 1 Dzeae
Dickeya zeae NCPPB 3531 WP 038902954 1 Dzeae
Dickeya zeae CSL RW192 WP 038906676 1 Dzeae
Dickeya zeae DZ2Q WP 038926794 1 Dzeae
Dickeya zeae EC1 WP 016941539 1 Dzeae
Dickeya zeae ZJU1202 WP 016941539 1 Dzeae
Prodigiosinella confusarubida ATCC39006 WP 037381483 1 Pconfusarubida
Phaeobacter inhibens DSM 17395 YP 006572805 1 391619 Proteobacteria
Phaeobacter gallaeciensis DSM 26640 YP 008976018 1 1423144 Proteobacteria
Rahnella sp Y9602 YP 004215400 1 741091 Proteobacteria
Dickeya gs poaceaephila NCPPB 569 WP 050569373 1 Dpoaceaephila
Dickeya paradisiaca NCPPB 2511 WP 012763744 1 Dparadisiaca
Dickeya paradisiaca Ech703 WP 012763744 1 Dparadisiaca
DAQ1742 DAQ1742 04328 Daquatica
Dickeya sp CSL RW240 WP 035339028 1 Daquatica
Dickeya aquatica DW 0440 WP 035339028 1 Daquatica
Dickeya dianthicola RNS04 9 WP 046830423 1 Ddianthicola
Dickeya dianthicola NCPPB 453 WP 046830423 1 Ddianthicola
Dickeya dianthicola NCPPB 3534 WP 051119531 1 Ddianthicola
Dickeya dianthicola GBBC 2039 WP 051119531 1 Ddianthicola
Dickeya dianthicola IPO 980 WP 051119531 1 Ddianthicola
Dickeya dadantii subsp dieffenbachiae NCPPB 2976 WP 050572805 1 Ddadantii
Dickeya dadantii NCPPB 3537 WP 050571453 1 Ddadantii
Dickeya dadantii NCPPB 898 WP 050570298 1 Ddadantii
334 vfmV Quorum sensing Seed
Dickeya dadantii 3937 WP 042318671 1 Ddadantii
Dickeya gs poaceaephila NCPPB 569 WP 050569517 1 Dpoaceaephila
Dickeya chrysanthemi subsp chrysanthemi NCPPB 402 WP 050583237 1 Dchrysanthemi
Dickeya chrysanthemi subsp parthenii NCPPB 516 WP 050568338 1 Dchrysanthemi
Dickeya chrysanthemi L11 WP 050516146 1 Dchrysanthemi
Dickeya chrysanthemi NCPPB 3533 WP 050570355 1 Dchrysanthemi
Dickeya chrysanthemi Ech1591 WP 012767870 1 Dchrysanthemi
Dickeya zeae Ech586 WP 049779799 1 Dzeae
Dickeya zeae MS1 WP 019844149 1 Dzeae
Dickeya zeae NCPPB 3531 WP 019844149 1 Dzeae
Dickeya zeae MK19 WP 050568644 1 Dzeae
Dickeya zeae NCPPB 3532 WP 050567436 1 Dzeae
Dickeya zeae NCPPB 2538 WP 050568644 1 Dzeae
Dickeya zeae EC1 WP 016941548 1 Dzeae
Dickeya zeae DZ2Q WP 016941548 1 Dzeae
Dickeya zeae CSL RW192 WP 016941548 1 Dzeae
Dickeya zeae ZJU1202 WP 016941548 1 Dzeae
Dickeya gs undicola 2B12 WP 050515630 1 Dundicola
Dickeya gs fangzhongdai M005 WP 052210139 1 Dfangzhongdai
Dickeya gs fangzhongdai M074 WP 049869990 1 Dfangzhongdai
Dickeya gs fangzhongdai ND14b WP 049869990 1 Dfangzhongdai
Dickeya gs fangzhongdai NCPPB 3274 WP 050569972 1 Dfangzhongdai
Dickeya gs fangzhongdai S1 WP 049842699 1 Dfangzhongdai
Dickeya gs fangzhongdai B16 WP 049842699 1 Dfangzhongdai
Dickeya gs fangzhongdai MK7 WP 049842699 1 Dfangzhongdai
Dickeya solani RNS 08 23 3 1 A WP 051120804 1 Dsolani
Dickeya solani RNS 05 1 2A WP 051120804 1 Dsolani
Dickeya solani MK16 WP 051120804 1 Dsolani
Dickeya solani IFB 0099 WP 051120804 1 Dsolani
Dickeya solani IPO 2222 WP 051120804 1 Dsolani
Dickeya solani RNS 07 7 3B WP 051120804 1 Dsolani
Dickeya solani MK10 WP 051120804 1 Dsolani
Dickeya solani GBBC 2040 WP 051120804 1 Dsolani
Dickeya solani D s0432 1 WP 051120804 1 Dsolani
Dickeya solani PPO 9134 WP 051120804 1 Dsolani
Dickeya solani PPO 9019 WP 051120804 1 Dsolani

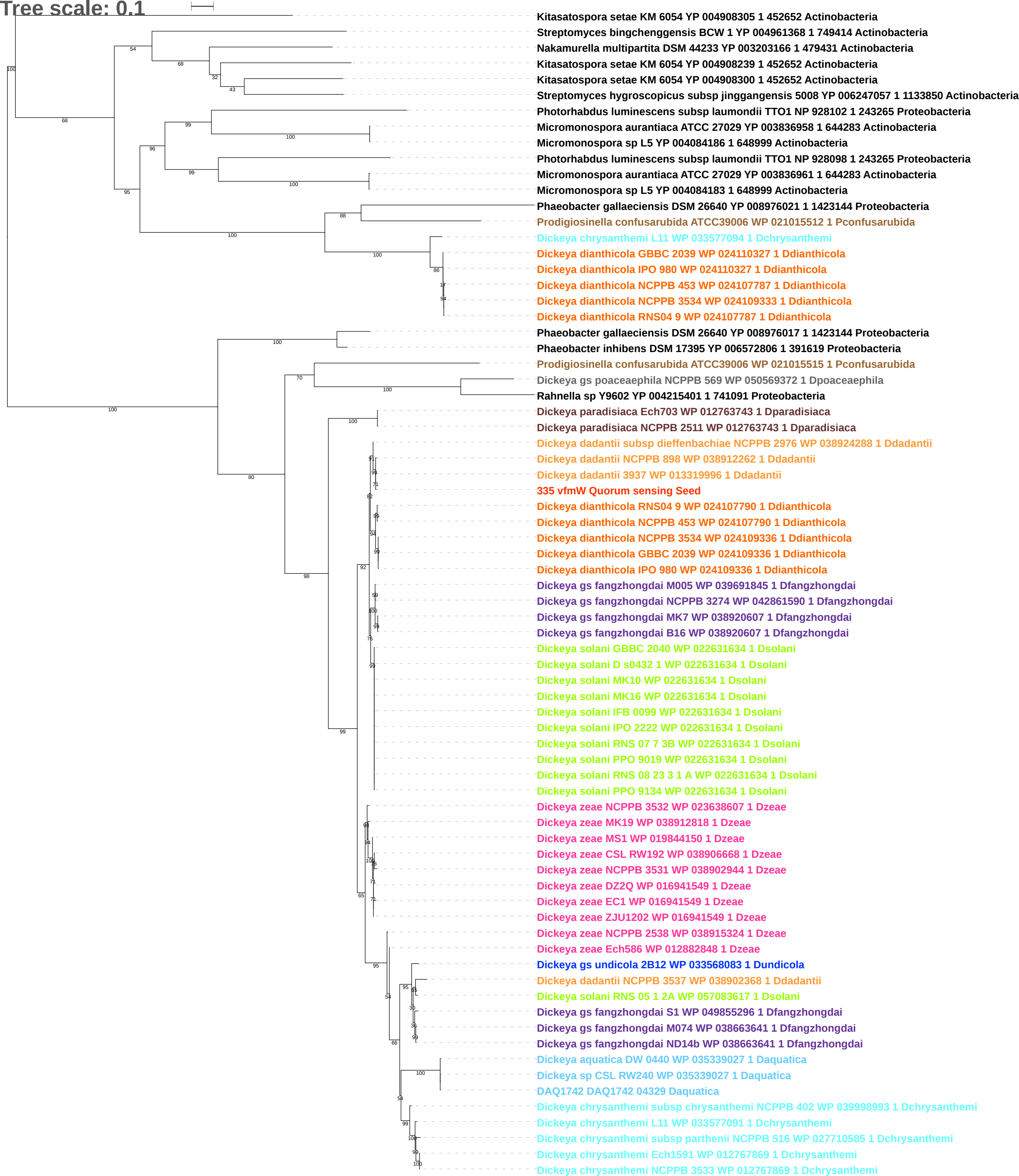
Tree scale: 0.1
Kitasatospora setae KM 6054 YP 004908305 1 452652 Actinobacteria
Streptomyces bingchenggensis BCW 1 YP 004961368 1 749414 Actinobacteria
Nakamurella multipartita DSM 44233 YP 003203166 1 479431 Actinobacteria
Kitasatospora setae KM 6054 YP 004908239 1 452652 Actinobacteria
Kitasatospora setae KM 6054 YP 004908300 1 452652 Actinobacteria
Streptomyces hygroscopicus subsp jinggangensis 5008 YP 006247057 1 1133850 Actinobacteria
Photorhabdus luminescens subsp laumondii TTO1 NP 928102 1 243265 Proteobacteria
Micromonospora aurantiaca ATCC 27029 YP 003836958 1 644283 Actinobacteria
Micromonospora sp L5 YP 004084186 1 648999 Actinobacteria
Photorhabdus luminescens subsp laumondii TTO1 NP 928098 1 243265 Proteobacteria
Micromonospora aurantiaca ATCC 27029 YP 003836961 1 644283 Actinobacteria
Micromonospora sp L5 YP 004084183 1 648999 Actinobacteria
Phaeobacter gallaeciensis DSM 26640 YP 008976021 1 1423144 Proteobacteria
Prodigiosinella confusarubida ATCC39006 WP 021015512 1 Pconfusarubida
Dickeya chrysanthemi L11 WP 033577094 1 Dchrysanthemi
Dickeya dianthicola GBBC 2039 WP 024110327 1 Ddianthicola
Dickeya dianthicola IPO 980 WP 024110327 1 Ddianthicola
Dickeya dianthicola NCPPB 453 WP 024107787 1 Ddianthicola
Dickeya dianthicola NCPPB 3534 WP 024109333 1 Ddianthicola
Dickeya dianthicola RNS04 9 WP 024107787 1 Ddianthicola
Phaeobacter gallaeciensis DSM 26640 YP 008976017 1 1423144 Proteobacteria
Phaeobacter inhibens DSM 17395 YP 006572806 1 391619 Proteobacteria
Prodigiosinella confusarubida ATCC39006 WP 021015515 1 Pconfusarubida
Dickeya gs poaceaephila NCPPB 569 WP 050569372 1 Dpoaceaephila
Rahnella sp Y9602 YP 004215401 1 741091 Proteobacteria
Dickeya paradisiaca Ech703 WP 012763743 1 Dparadisiaca
Dickeya paradisiaca NCPPB 2511 WP 012763743 1 Dparadisiaca
Dickeya dadantii subsp dieffenbachiae NCPPB 2976 WP 038924288 1 Ddadantii
Dickeya dadantii NCPPB 898 WP 038912262 1 Ddadantii
Dickeya dadantii 3937 WP 013319996 1 Ddadantii
335 vfmW Quorum sensing Seed
Dickeya dianthicola RNS04 9 WP 024107790 1 Ddianthicola
Dickeya dianthicola NCPPB 453 WP 024107790 1 Ddianthicola
Dickeya dianthicola NCPPB 3534 WP 024109336 1 Ddianthicola
Dickeya dianthicola GBBC 2039 WP 024109336 1 Ddianthicola
Dickeya dianthicola IPO 980 WP 024109336 1 Ddianthicola
Dickeya gs fangzhongdai M005 WP 039691845 1 Dfangzhongdai
Dickeya gs fangzhongdai NCPPB 3274 WP 042861590 1 Dfangzhongdai
Dickeya gs fangzhongdai MK7 WP 038920607 1 Dfangzhongdai
Dickeya gs fangzhongdai B16 WP 038920607 1 Dfangzhongdai
Dickeya solani GBBC 2040 WP 022631634 1 Dsolani
Dickeya solani D s0432 1 WP 022631634 1 Dsolani
Dickeya solani MK10 WP 022631634 1 Dsolani
Dickeya solani MK16 WP 022631634 1 Dsolani
Dickeya solani IFB 0099 WP 022631634 1 Dsolani
Dickeya solani IPO 2222 WP 022631634 1 Dsolani
Dickeya solani RNS 07 7 3B WP 022631634 1 Dsolani
Dickeya solani PPO 9019 WP 022631634 1 Dsolani
Dickeya solani RNS 08 23 3 1 A WP 022631634 1 Dsolani
Dickeya solani PPO 9134 WP 022631634 1 Dsolani
Dickeya zeae NCPPB 3532 WP 023638607 1 Dzeae
Dickeya zeae MK19 WP 038912818 1 Dzeae
Dickeya zeae MS1 WP 019844150 1 Dzeae
Dickeya zeae CSL RW192 WP 038906668 1 Dzeae
Dickeya zeae NCPPB 3531 WP 038902944 1 Dzeae
Dickeya zeae DZ2Q WP 016941549 1 Dzeae
Dickeya zeae EC1 WP 016941549 1 Dzeae
Dickeya zeae ZJU1202 WP 016941549 1 Dzeae
Dickeya zeae NCPPB 2538 WP 038915324 1 Dzeae
Dickeya zeae Ech586 WP 012882848 1 Dzeae
Dickeya gs undicola 2B12 WP 033568083 1 Dundicola
Dickeya dadantii NCPPB 3537 WP 038902368 1 Ddadantii
Dickeya solani RNS 05 1 2A WP 057083617 1 Dsolani
Dickeya gs fangzhongdai S1 WP 049855296 1 Dfangzhongdai
Dickeya gs fangzhongdai M074 WP 038663641 1 Dfangzhongdai
Dickeya gs fangzhongdai ND14b WP 038663641 1 Dfangzhongdai
Dickeya aquatica DW 0440 WP 035339027 1 Daquatica
Dickeya sp CSL RW240 WP 035339027 1 Daquatica
DAQ1742 DAQ1742 04329 Daquatica
Dickeya chrysanthemi subsp chrysanthemi NCPPB 402 WP 039998993 1 Dchrysanthemi
Dickeya chrysanthemi L11 WP 033577091 1 Dchrysanthemi
Dickeya chrysanthemi subsp parthenii NCPPB 516 WP 027710585 1 Dchrysanthemi
Dickeya chrysanthemi Ech1591 WP 012767869 1 Dchrysanthemi
Dickeya chrysanthemi NCPPB 3533 WP 012767869 1 Dchrysanthemi

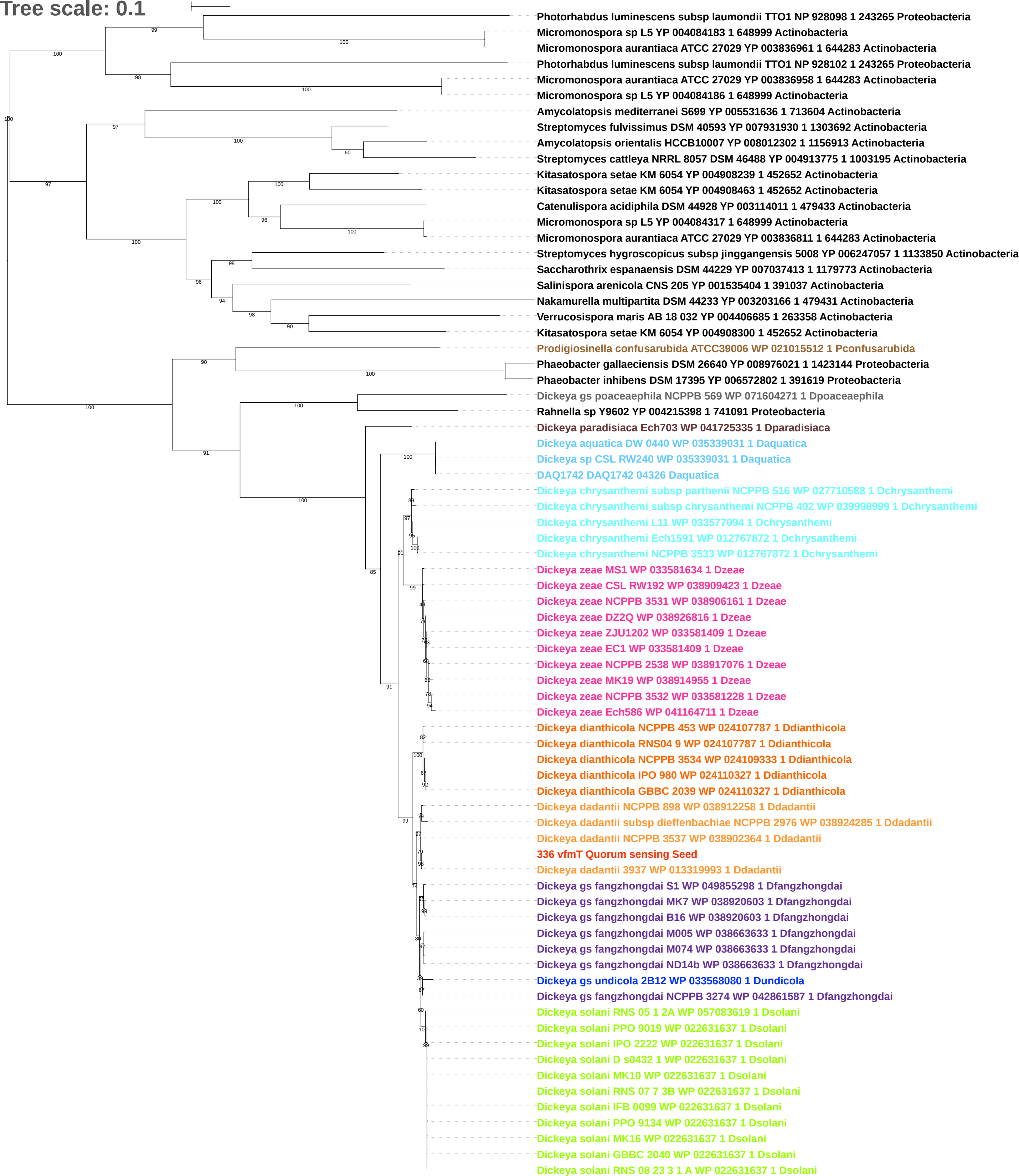

Tree scale: 0.1
Photorhabdus luminescens subsp laumondii TTO1 NP 928098 1 243265 Proteobacteria
Micromonospora sp L5 YP 004084183 1 648999 Actinobacteria
Micromonospora aurantiaca ATCC 27029 YP 003836961 1 644283 Actinobacteria
Photorhabdus luminescens subsp laumondii TTO1 NP 928102 1 243265 Proteobacteria
Micromonospora aurantiaca ATCC 27029 YP 003836958 1 644283 Actinobacteria
Micromonospora sp L5 YP 004084186 1 648999 Actinobacteria
Amycolatopsis mediterranei S699 YP 005531636 1 713604 Actinobacteria
Streptomyces fulvissimus DSM 40593 YP 007931930 1 1303692 Actinobacteria
Amycolatopsis orientalis HCCB10007 YP 008012302 1 1156913 Actinobacteria
Streptomyces cattleya NRRL 8057 DSM 46488 YP 004913775 1 1003195 Actinobacteria
Kitasatospora setae KM 6054 YP 004908239 1 452652 Actinobacteria
Kitasatospora setae KM 6054 YP 004908463 1 452652 Actinobacteria
Catenulispora acidiphila DSM 44928 YP 003114011 1 479433 Actinobacteria
Micromonospora sp L5 YP 004084317 1 648999 Actinobacteria
Micromonospora aurantiaca ATCC 27029 YP 003836811 1 644283 Actinobacteria
Streptomyces hygroscopicus subsp jinggangensis 5008 YP 006247057 1 1133850 Actinobacteria
Saccharothrix espanaensis DSM 44229 YP 007037413 1 1179773 Actinobacteria
Salinispora arenicola CNS 205 YP 001535404 1 391037 Actinobacteria
Nakamurella multipartita DSM 44233 YP 003203166 1 479431 Actinobacteria
Verrucosispora maris AB 18 032 YP 004406685 1 263358 Actinobacteria
Kitasatospora setae KM 6054 YP 004908300 1 452652 Actinobacteria
Prodigiosinella confusarubida ATCC39006 WP 021015512 1 Pconfusarubida
Phaeobacter gallaeciensis DSM 26640 YP 008976021 1 1423144 Proteobacteria
Phaeobacter inhibens DSM 17395 YP 006572802 1 391619 Proteobacteria
Dickeya gs poaceaephila NCPPB 569 WP 071604271 1 Dpoaceaephila
Rahnella sp Y9602 YP 004215398 1 741091 Proteobacteria
Dickeya paradisiaca Ech703 WP 041725335 1 Dparadisiaca
Dickeya aquatica DW 0440 WP 035339031 1 Daquatica
Dickeya sp CSL RW240 WP 035339031 1 Daquatica
DAQ1742 DAQ1742 04326 Daquatica
Dickeya chrysanthemi subsp parthenii NCPPB 516 WP 027710588 1 Dchrysanthemi
Dickeya chrysanthemi subsp chrysanthemi NCPPB 402 WP 039998999 1 Dchrysanthemi
Dickeya chrysanthemi L11 WP 033577094 1 Dchrysanthemi
Dickeya chrysanthemi Ech1591 WP 012767872 1 Dchrysanthemi
Dickeya chrysanthemi NCPPB 3533 WP 012767872 1 Dchrysanthemi
Dickeya zeae MS1 WP 033581634 1 Dzeae
Dickeya zeae CSL RW192 WP 038909423 1 Dzeae
Dickeya zeae NCPPB 3531 WP 038906161 1 Dzeae
Dickeya zeae DZ2Q WP 038926816 1 Dzeae
Dickeya zeae ZJU1202 WP 033581409 1 Dzeae
Dickeya zeae EC1 WP 033581409 1 Dzeae
Dickeya zeae NCPPB 2538 WP 038917076 1 Dzeae
Dickeya zeae MK19 WP 038914955 1 Dzeae
Dickeya zeae NCPPB 3532 WP 033581228 1 Dzeae
Dickeya zeae Ech586 WP 041164711 1 Dzeae
Dickeya dianthicola NCPPB 453 WP 024107787 1 Ddianthicola
Dickeya dianthicola RNS04 9 WP 024107787 1 Ddianthicola
Dickeya dianthicola NCPPB 3534 WP 024109333 1 Ddianthicola
Dickeya dianthicola IPO 980 WP 024110327 1 Ddianthicola
Dickeya dianthicola GBBC 2039 WP 024110327 1 Ddianthicola
Dickeya dadantii NCPPB 898 WP 038912258 1 Ddadantii
Dickeya dadantii subsp dieffenbachiae NCPPB 2976 WP 038924285 1 Ddadantii
Dickeya dadantii NCPPB 3537 WP 038902364 1 Ddadantii
336 vfmT Quorum sensing Seed
Dickeya dadantii 3937 WP 013319993 1 Ddadantii
Dickeya gs fangzhongdai S1 WP 049855298 1 Dfangzhongdai
Dickeya gs fangzhongdai MK7 WP 038920603 1 Dfangzhongdai
Dickeya gs fangzhongdai B16 WP 038920603 1 Dfangzhongdai
Dickeya gs fangzhongdai M005 WP 038663633 1 Dfangzhongdai
Dickeya gs fangzhongdai M074 WP 038663633 1 Dfangzhongdai
Dickeya gs fangzhongdai ND14b WP 038663633 1 Dfangzhongdai
Dickeya gs undicola 2B12 WP 033568080 1 Dundicola
Dickeya gs fangzhongdai NCPPB 3274 WP 042861587 1 Dfangzhongdai
Dickeya solani RNS 05 1 2A WP 057083619 1 Dsolani
Dickeya solani PPO 9019 WP 022631637 1 Dsolani
Dickeya solani IPO 2222 WP 022631637 1 Dsolani
Dickeya solani D s0432 1 WP 022631637 1 Dsolani
Dickeya solani MK10 WP 022631637 1 Dsolani
Dickeya solani RNS 07 7 3B WP 022631637 1 Dsolani
Dickeya solani IFB 0099 WP 022631637 1 Dsolani
Dickeya solani PPO 9134 WP 022631637 1 Dsolani
Dickeya solani MK16 WP 022631637 1 Dsolani
Dickeya solani GBBC 2040 WP 022631637 1 Dsolani
Dickeya solani RNS 08 23 3 1 A WP 022631637 1 Dsolani

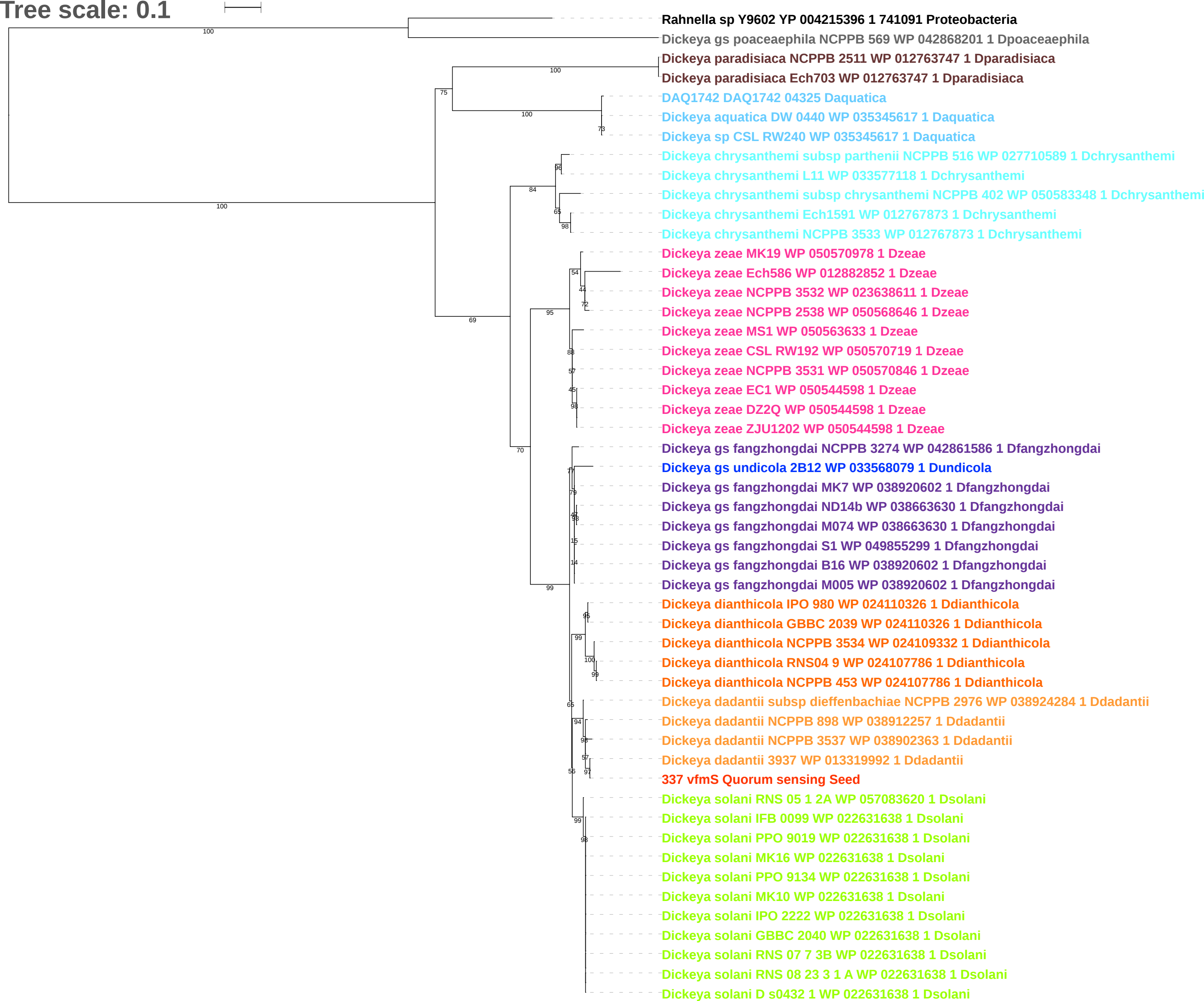
Tree scale: 0.1
Rahnella sp Y9602 YP 004215396 1 741091 Proteobacteria
Dickeya gs poaceaephila NCPPB 569 WP 042868201 1 Dpoaceaephila
Dickeya paradisiaca NCPPB 2511 WP 012763747 1 Dparadisiaca
Dickeya paradisiaca Ech703 WP 012763747 1 Dparadisiaca
DAQ1742 DAQ1742 04325 Daquatica
Dickeya aquatica DW 0440 WP 035345617 1 Daquatica
Dickeya sp CSL RW240 WP 035345617 1 Daquatica
Dickeya chrysanthemi subsp parthenii NCPPB 516 WP 027710589 1 Dchrysanthemi
Dickeya chrysanthemi L11 WP 033577118 1 Dchrysanthemi
Dickeya chrysanthemi subsp chrysanthemi NCPPB 402 WP 050583348 1 Dchrysanthemi
Dickeya chrysanthemi Ech1591 WP 012767873 1 Dchrysanthemi
Dickeya chrysanthemi NCPPB 3533 WP 012767873 1 Dchrysanthemi
Dickeya zeae MK19 WP 050570978 1 Dzeae
Dickeya zeae Ech586 WP 012882852 1 Dzeae
Dickeya zeae NCPPB 3532 WP 023638611 1 Dzeae
Dickeya zeae NCPPB 2538 WP 050568646 1 Dzeae
Dickeya zeae MS1 WP 050563633 1 Dzeae
Dickeya zeae CSL RW192 WP 050570719 1 Dzeae
Dickeya zeae NCPPB 3531 WP 050570846 1 Dzeae
Dickeya zeae EC1 WP 050544598 1 Dzeae
Dickeya zeae DZ2Q WP 050544598 1 Dzeae
Dickeya zeae ZJU1202 WP 050544598 1 Dzeae
Dickeya gs fangzhongdai NCPPB 3274 WP 042861586 1 Dfangzhongdai
Dickeya gs undicola 2B12 WP 033568079 1 Dundicola
Dickeya gs fangzhongdai MK7 WP 038920602 1 Dfangzhongdai
Dickeya gs fangzhongdai ND14b WP 038663630 1 Dfangzhongdai
Dickeya gs fangzhongdai M074 WP 038663630 1 Dfangzhongdai
Dickeya gs fangzhongdai S1 WP 049855299 1 Dfangzhongdai
Dickeya gs fangzhongdai B16 WP 038920602 1 Dfangzhongdai
Dickeya gs fangzhongdai M005 WP 038920602 1 Dfangzhongdai
Dickeya dianthicola IPO 980 WP 024110326 1 Ddianthicola
Dickeya dianthicola GBBC 2039 WP 024110326 1 Ddianthicola
Dickeya dianthicola NCPPB 3534 WP 024109332 1 Ddianthicola
Dickeya dianthicola RNS04 9 WP 024107786 1 Ddianthicola
Dickeya dianthicola NCPPB 453 WP 024107786 1 Ddianthicola
Dickeya dadantii subsp dieffenbachiae NCPPB 2976 WP 038924284 1 Ddadantii
Dickeya dadantii NCPPB 898 WP 038912257 1 Ddadantii
Dickeya dadantii NCPPB 3537 WP 038902363 1 Ddadantii
Dickeya dadantii 3937 WP 013319992 1 Ddadantii
337 vfmS Quorum sensing Seed
Dickeya solani RNS 05 1 2A WP 057083620 1 Dsolani
Dickeya solani IFB 0099 WP 022631638 1 Dsolani
Dickeya solani PPO 9019 WP 022631638 1 Dsolani
Dickeya solani MK16 WP 022631638 1 Dsolani
Dickeya solani PPO 9134 WP 022631638 1 Dsolani
Dickeya solani MK10 WP 022631638 1 Dsolani
Dickeya solani IPO 2222 WP 022631638 1 Dsolani
Dickeya solani GBBC 2040 WP 022631638 1 Dsolani
Dickeya solani RNS 07 7 3B WP 022631638 1 Dsolani
Dickeya solani RNS 08 23 3 1 A WP 022631638 1 Dsolani
Dickeya solani D s0432 1 WP 022631638 1 Dsolani

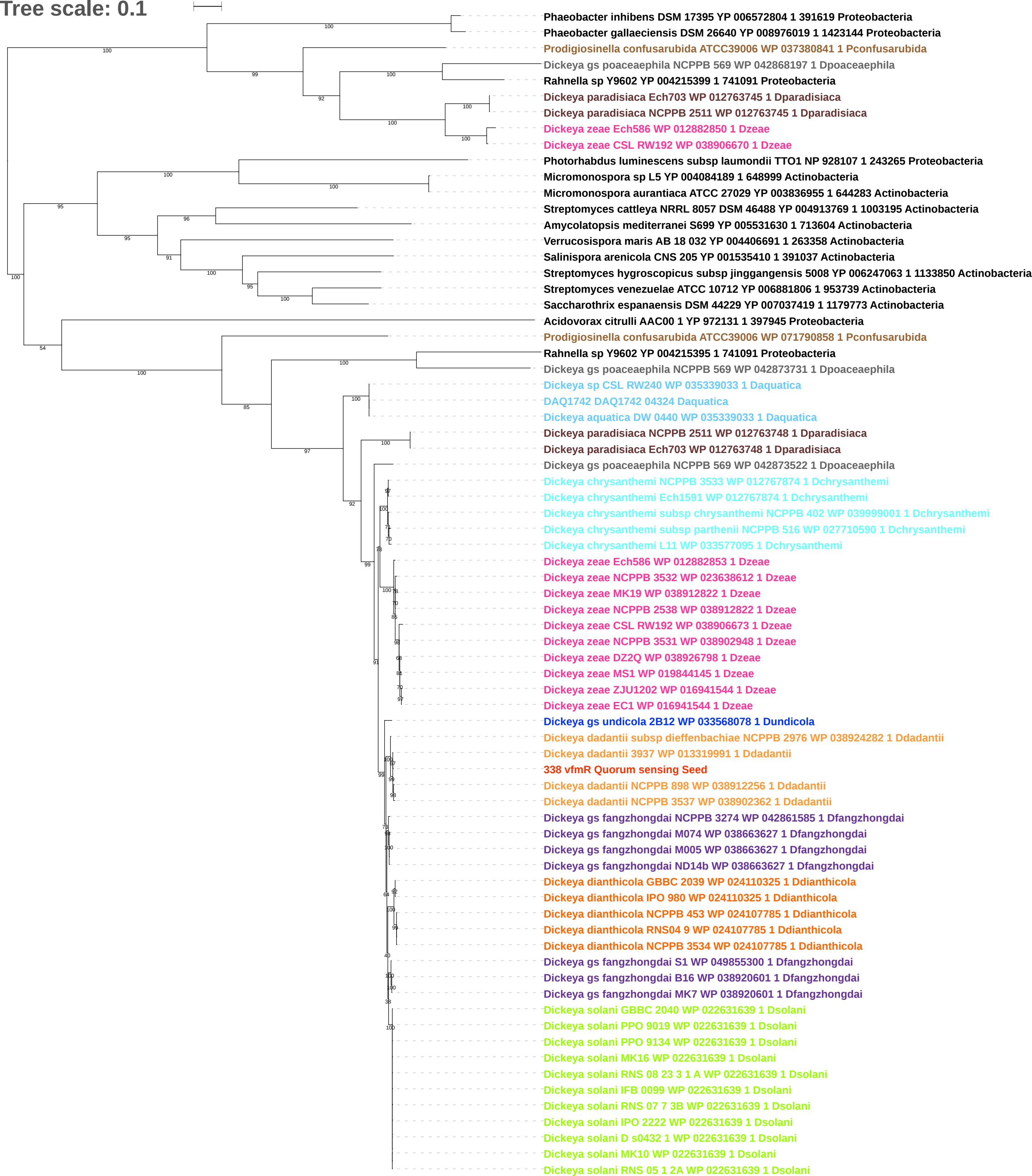
Tree scale: 0.1
Phaeobacter inhibens DSM 17395 YP 006572804 1 391619 Proteobacteria
Phaeobacter gallaeciensis DSM 26640 YP 008976019 1 1423144 Proteobacteria
Prodigiosinella confusarubida ATCC39006 WP 037380841 1 Pconfusarubida
Dickeya gs poaceaephila NCPPB 569 WP 042868197 1 Dpoaceaephila
Rahnella sp Y9602 YP 004215399 1 741091 Proteobacteria
Dickeya paradisiaca Ech703 WP 012763745 1 Dparadisiaca
Dickeya paradisiaca NCPPB 2511 WP 012763745 1 Dparadisiaca
Dickeya zeae Ech586 WP 012882850 1 Dzeae
Dickeya zeae CSL RW192 WP 038906670 1 Dzeae
Photorhabdus luminescens subsp laumondii TTO1 NP 928107 1 243265 Proteobacteria
Micromonospora sp L5 YP 004084189 1 648999 Actinobacteria
Micromonospora aurantiaca ATCC 27029 YP 003836955 1 644283 Actinobacteria
Streptomyces cattleya NRRL 8057 DSM 46488 YP 004913769 1 1003195 Actinobacteria
Amycolatopsis mediterranei S699 YP 005531630 1 713604 Actinobacteria
Verrucosispora maris AB 18 032 YP 004406691 1 263358 Actinobacteria
Salinispora arenicola CNS 205 YP 001535410 1 391037 Actinobacteria
Streptomyces hygroscopicus subsp jinggangensis 5008 YP 006247063 1 1133850 Actinobacteria
Streptomyces venezuelae ATCC 10712 YP 006881806 1 953739 Actinobacteria
Saccharothrix espanaensis DSM 44229 YP 007037419 1 1179773 Actinobacteria
Acidovorax citrulli AAC00 1 YP 972131 1 397945 Proteobacteria
Prodigiosinella confusarubida ATCC39006 WP 071790858 1 Pconfusarubida
Rahnella sp Y9602 YP 004215395 1 741091 Proteobacteria
Dickeya gs poaceaephila NCPPB 569 WP 042873731 1 Dpoaceaephila
Dickeya sp CSL RW240 WP 035339033 1 Daquatica
DAQ1742 DAQ1742 04324 Daquatica
Dickeya aquatica DW 0440 WP 035339033 1 Daquatica
Dickeya paradisiaca NCPPB 2511 WP 012763748 1 Dparadisiaca
Dickeya paradisiaca Ech703 WP 012763748 1 Dparadisiaca
Dickeya gs poaceaephila NCPPB 569 WP 042873522 1 Dpoaceaephila
Dickeya chrysanthemi NCPPB 3533 WP 012767874 1 Dchrysanthemi
Dickeya chrysanthemi Ech1591 WP 012767874 1 Dchrysanthemi
Dickeya chrysanthemi subsp chrysanthemi NCPPB 402 WP 039999001 1 Dchrysanthemi
Dickeya chrysanthemi subsp parthenii NCPPB 516 WP 027710590 1 Dchrysanthemi
Dickeya chrysanthemi L11 WP 033577095 1 Dchrysanthemi
Dickeya zeae Ech586 WP 012882853 1 Dzeae
Dickeya zeae NCPPB 3532 WP 023638612 1 Dzeae
Dickeya zeae MK19 WP 038912822 1 Dzeae
Dickeya zeae NCPPB 2538 WP 038912822 1 Dzeae
Dickeya zeae CSL RW192 WP 038906673 1 Dzeae
Dickeya zeae NCPPB 3531 WP 038902948 1 Dzeae
Dickeya zeae DZ2Q WP 038926798 1 Dzeae
Dickeya zeae MS1 WP 019844145 1 Dzeae
Dickeya zeae ZJU1202 WP 016941544 1 Dzeae
Dickeya zeae EC1 WP 016941544 1 Dzeae
Dickeya gs undicola 2B12 WP 033568078 1 Dundicola
Dickeya dadantii subsp dieffenbachiae NCPPB 2976 WP 038924282 1 Ddadantii
Dickeya dadantii 3937 WP 013319991 1 Ddadantii
338 vfmR Quorum sensing Seed
Dickeya dadantii NCPPB 898 WP 038912256 1 Ddadantii
Dickeya dadantii NCPPB 3537 WP 038902362 1 Ddadantii
Dickeya gs fangzhongdai NCPPB 3274 WP 042861585 1 Dfangzhongdai
Dickeya gs fangzhongdai M074 WP 038663627 1 Dfangzhongdai
Dickeya gs fangzhongdai M005 WP 038663627 1 Dfangzhongdai
Dickeya gs fangzhongdai ND14b WP 038663627 1 Dfangzhongdai
Dickeya dianthicola GBBC 2039 WP 024110325 1 Ddianthicola
Dickeya dianthicola IPO 980 WP 024110325 1 Ddianthicola
Dickeya dianthicola NCPPB 453 WP 024107785 1 Ddianthicola
Dickeya dianthicola RNS04 9 WP 024107785 1 Ddianthicola
Dickeya dianthicola NCPPB 3534 WP 024107785 1 Ddianthicola
Dickeya gs fangzhongdai S1 WP 049855300 1 Dfangzhongdai
Dickeya gs fangzhongdai B16 WP 038920601 1 Dfangzhongdai
Dickeya gs fangzhongdai MK7 WP 038920601 1 Dfangzhongdai
Dickeya solani GBBC 2040 WP 022631639 1 Dsolani
Dickeya solani PPO 9019 WP 022631639 1 Dsolani
Dickeya solani PPO 9134 WP 022631639 1 Dsolani
Dickeya solani MK16 WP 022631639 1 Dsolani
Dickeya solani RNS 08 23 3 1 A WP 022631639 1 Dsolani
Dickeya solani IFB 0099 WP 022631639 1 Dsolani
Dickeya solani RNS 07 7 3B WP 022631639 1 Dsolani
Dickeya solani IPO 2222 WP 022631639 1 Dsolani
Dickeya solani D s0432 1 WP 022631639 1 Dsolani
Dickeya solani MK10 WP 022631639 1 Dsolani
Dickeya solani RNS 05 1 2A WP 022631639 1 Dsolani

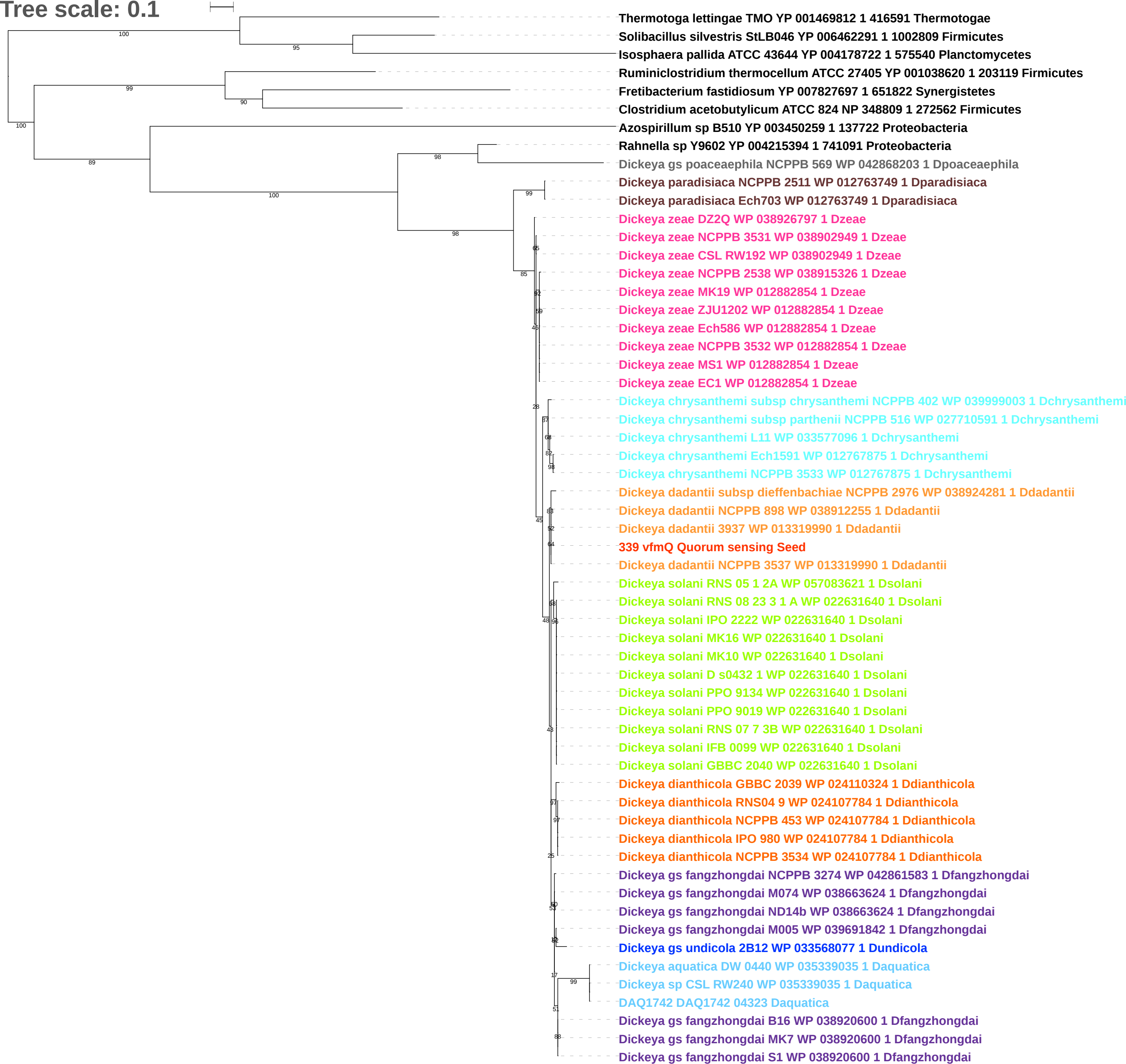
Tree scale: 0.1
Thermotoga lettingae TMO YP 001469812 1 416591 Thermotogae
Solibacillus silvestris StLB046 YP 006462291 1 1002809 Firmicutes
Isosphaera pallida ATCC 43644 YP 004178722 1 575540 Planctomycetes
Ruminiclostridium thermocellum ATCC 27405 YP 001038620 1 203119 Firmicutes
Fretibacterium fastidiosum YP 007827697 1 651822 Synergistetes
Clostridium acetobutylicum ATCC 824 NP 348809 1 272562 Firmicutes
Azospirillum sp B510 YP 003450259 1 137722 Proteobacteria
Rahnella sp Y9602 YP 004215394 1 741091 Proteobacteria
Dickeya gs poaceaephila NCPPB 569 WP 042868203 1 Dpoaceaephila
Dickeya paradisiaca NCPPB 2511 WP 012763749 1 Dparadisiaca
Dickeya paradisiaca Ech703 WP 012763749 1 Dparadisiaca
Dickeya zeae DZ2Q WP 038926797 1 Dzeae
Dickeya zeae NCPPB 3531 WP 038902949 1 Dzeae
Dickeya zeae CSL RW192 WP 038902949 1 Dzeae
Dickeya zeae NCPPB 2538 WP 038915326 1 Dzeae
Dickeya zeae MK19 WP 012882854 1 Dzeae
Dickeya zeae ZJU1202 WP 012882854 1 Dzeae
Dickeya zeae Ech586 WP 012882854 1 Dzeae
Dickeya zeae NCPPB 3532 WP 012882854 1 Dzeae
Dickeya zeae MS1 WP 012882854 1 Dzeae
Dickeya zeae EC1 WP 012882854 1 Dzeae
Dickeya chrysanthemi subsp chrysanthemi NCPPB 402 WP 039999003 1 Dchrysanthemi
Dickeya chrysanthemi subsp parthenii NCPPB 516 WP 027710591 1 Dchrysanthemi
Dickeya chrysanthemi L11 WP 033577096 1 Dchrysanthemi
Dickeya chrysanthemi Ech1591 WP 012767875 1 Dchrysanthemi
Dickeya chrysanthemi NCPPB 3533 WP 012767875 1 Dchrysanthemi
Dickeya dadantii subsp dieffenbachiae NCPPB 2976 WP 038924281 1 Ddadantii
Dickeya dadantii NCPPB 898 WP 038912255 1 Ddadantii
Dickeya dadantii 3937 WP 013319990 1 Ddadantii
339 vfmQ Quorum sensing Seed
Dickeya dadantii NCPPB 3537 WP 013319990 1 Ddadantii
Dickeya solani RNS 05 1 2A WP 057083621 1 Dsolani
Dickeya solani RNS 08 23 3 1 A WP 022631640 1 Dsolani
Dickeya solani IPO 2222 WP 022631640 1 Dsolani
Dickeya solani MK16 WP 022631640 1 Dsolani
Dickeya solani MK10 WP 022631640 1 Dsolani
Dickeya solani D s0432 1 WP 022631640 1 Dsolani
Dickeya solani PPO 9134 WP 022631640 1 Dsolani
Dickeya solani PPO 9019 WP 022631640 1 Dsolani
Dickeya solani RNS 07 7 3B WP 022631640 1 Dsolani
Dickeya solani IFB 0099 WP 022631640 1 Dsolani
Dickeya solani GBBC 2040 WP 022631640 1 Dsolani
Dickeya dianthicola GBBC 2039 WP 024110324 1 Ddianthicola
Dickeya dianthicola RNS04 9 WP 024107784 1 Ddianthicola
Dickeya dianthicola NCPPB 453 WP 024107784 1 Ddianthicola
Dickeya dianthicola IPO 980 WP 024107784 1 Ddianthicola
Dickeya dianthicola NCPPB 3534 WP 024107784 1 Ddianthicola
Dickeya gs fangzhongdai NCPPB 3274 WP 042861583 1 Dfangzhongdai
Dickeya gs fangzhongdai M074 WP 038663624 1 Dfangzhongdai
Dickeya gs fangzhongdai ND14b WP 038663624 1 Dfangzhongdai
Dickeya gs fangzhongdai M005 WP 039691842 1 Dfangzhongdai
Dickeya gs undicola 2B12 WP 033568077 1 Dundicola
Dickeya aquatica DW 0440 WP 035339035 1 Daquatica
Dickeya sp CSL RW240 WP 035339035 1 Daquatica
DAQ1742 DAQ1742 04323 Daquatica
Dickeya gs fangzhongdai B16 WP 038920600 1 Dfangzhongdai
Dickeya gs fangzhongdai MK7 WP 038920600 1 Dfangzhongdai
Dickeya gs fangzhongdai S1 WP 038920600 1 Dfangzhongdai

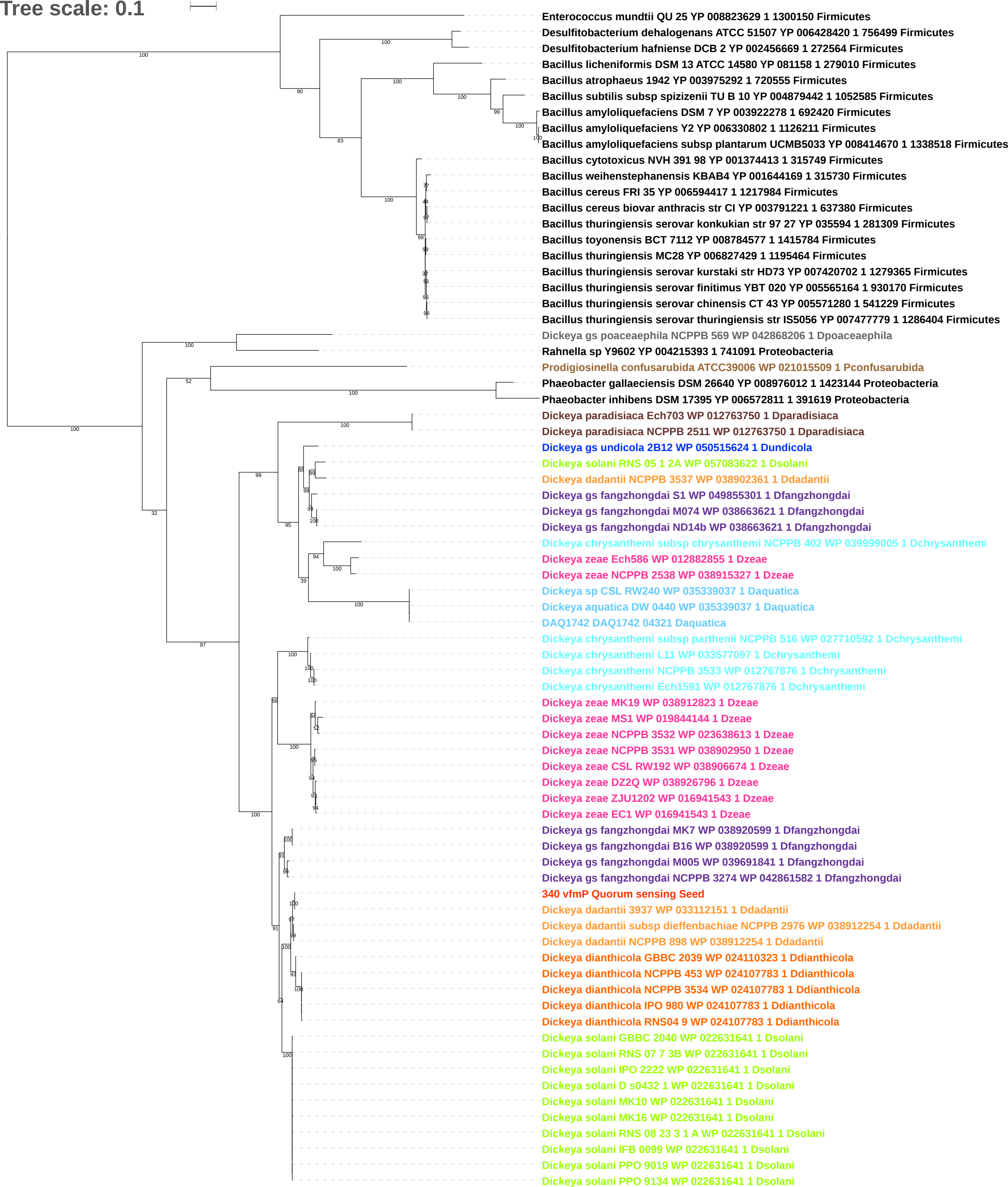
Tree scale: 0.1
Enterococcus mundtii QU 25 YP 008823629 1 1300150 Firmicutes
Desulfitobacterium dehalogenans ATCC 51507 YP 006428420 1 756499 Firmicutes
Desulfitobacterium hafniense DCB 2 YP 002456669 1 272564 Firmicutes
Bacillus licheniformis DSM 13 ATCC 14580 YP 081158 1 279010 Firmicutes
Bacillus atrophaeus 1942 YP 003975292 1 720555 Firmicutes
Bacillus subtilis subsp spizizenii TU B 10 YP 004879442 1 1052585 Firmicutes
Bacillus amyloliquefaciens DSM 7 YP 003922278 1 692420 Firmicutes
Bacillus amyloliquefaciens Y2 YP 006330802 1 1126211 Firmicutes
Bacillus amyloliquefaciens subsp plantarum UCMB5033 YP 008414670 1 1338518 Firmicutes
Bacillus cytotoxicus NVH 391 98 YP 001374413 1 315749 Firmicutes
Bacillus weihenstephanensis KBAB4 YP 001644169 1 315730 Firmicutes
Bacillus cereus FRI 35 YP 006594417 1 1217984 Firmicutes
Bacillus cereus biovar anthracis str CI YP 003791221 1 637380 Firmicutes
Bacillus thuringiensis serovar konkukian str 97 27 YP 035594 1 281309 Firmicutes
Bacillus toyonensis BCT 7112 YP 008784577 1 1415784 Firmicutes
Bacillus thuringiensis MC28 YP 006827429 1 1195464 Firmicutes
Bacillus thuringiensis serovar kurstaki str HD73 YP 007420702 1 1279365 Firmicutes
Bacillus thuringiensis serovar finitimus YBT 020 YP 005565164 1 930170 Firmicutes
Bacillus thuringiensis serovar chinensis CT 43 YP 005571280 1 541229 Firmicutes
Bacillus thuringiensis serovar thuringiensis str IS5056 YP 007477779 1 1286404 Firmicutes
Dickeya gs poaceaephila NCPPB 569 WP 042868206 1 Dpoaceaephila
Rahnella sp Y9602 YP 004215393 1 741091 Proteobacteria
Prodigiosinella confusarubida ATCC39006 WP 021015509 1 Pconfusarubida
Phaeobacter gallaeciensis DSM 26640 YP 008976012 1 1423144 Proteobacteria
Phaeobacter inhibens DSM 17395 YP 006572811 1 391619 Proteobacteria
Dickeya paradisiaca Ech703 WP 012763750 1 Dparadisiaca
Dickeya paradisiaca NCPPB 2511 WP 012763750 1 Dparadisiaca
Dickeya gs undicola 2B12 WP 050515624 1 Dundicola
Dickeya solani RNS 05 1 2A WP 057083622 1 Dsolani
Dickeya dadantii NCPPB 3537 WP 038902361 1 Ddadantii
Dickeya gs fangzhongdai S1 WP 049855301 1 Dfangzhongdai
Dickeya gs fangzhongdai M074 WP 038663621 1 Dfangzhongdai
Dickeya gs fangzhongdai ND14b WP 038663621 1 Dfangzhongdai
Dickeya chrysanthemi subsp chrysanthemi NCPPB 402 WP 039999005 1 Dchrysanthemi
Dickeya zeae Ech586 WP 012882855 1 Dzeae
Dickeya zeae NCPPB 2538 WP 038915327 1 Dzeae
Dickeya sp CSL RW240 WP 035339037 1 Daquatica
Dickeya aquatica DW 0440 WP 035339037 1 Daquatica
DAQ1742 DAQ1742 04321 Daquatica
Dickeya chrysanthemi subsp parthenii NCPPB 516 WP 027710592 1 Dchrysanthemi
Dickeya chrysanthemi L11 WP 033577097 1 Dchrysanthemi
Dickeya chrysanthemi NCPPB 3533 WP 012767876 1 Dchrysanthemi
Dickeya chrysanthemi Ech1591 WP 012767876 1 Dchrysanthemi
Dickeya zeae MK19 WP 038912823 1 Dzeae
Dickeya zeae MS1 WP 019844144 1 Dzeae
Dickeya zeae NCPPB 3532 WP 023638613 1 Dzeae
Dickeya zeae NCPPB 3531 WP 038902950 1 Dzeae
Dickeya zeae CSL RW192 WP 038906674 1 Dzeae
Dickeya zeae DZ2Q WP 038926796 1 Dzeae
Dickeya zeae ZJU1202 WP 016941543 1 Dzeae
Dickeya zeae EC1 WP 016941543 1 Dzeae
Dickeya gs fangzhongdai MK7 WP 038920599 1 Dfangzhongdai
Dickeya gs fangzhongdai B16 WP 038920599 1 Dfangzhongdai
Dickeya gs fangzhongdai M005 WP 039691841 1 Dfangzhongdai
Dickeya gs fangzhongdai NCPPB 3274 WP 042861582 1 Dfangzhongdai
340 vfmP Quorum sensing Seed
Dickeya dadantii 3937 WP 033112151 1 Ddadantii
Dickeya dadantii subsp dieffenbachiae NCPPB 2976 WP 038912254 1 Ddadantii
Dickeya dadantii NCPPB 898 WP 038912254 1 Ddadantii
Dickeya dianthicola GBBC 2039 WP 024110323 1 Ddianthicola
Dickeya dianthicola NCPPB 453 WP 024107783 1 Ddianthicola
Dickeya dianthicola NCPPB 3534 WP 024107783 1 Ddianthicola
Dickeya dianthicola IPO 980 WP 024107783 1 Ddianthicola
Dickeya dianthicola RNS04 9 WP 024107783 1 Ddianthicola
Dickeya solani GBBC 2040 WP 022631641 1 Dsolani
Dickeya solani RNS 07 7 3B WP 022631641 1 Dsolani
Dickeya solani IPO 2222 WP 022631641 1 Dsolani
Dickeya solani D s0432 1 WP 022631641 1 Dsolani
Dickeya solani MK10 WP 022631641 1 Dsolani
Dickeya solani MK16 WP 022631641 1 Dsolani
Dickeya solani RNS 08 23 3 1 A WP 022631641 1 Dsolani
Dickeya solani IFB 0099 WP 022631641 1 Dsolani
Dickeya solani PPO 9019 WP 022631641 1 Dsolani
Dickeya solani PPO 9134 WP 022631641 1 Dsolani

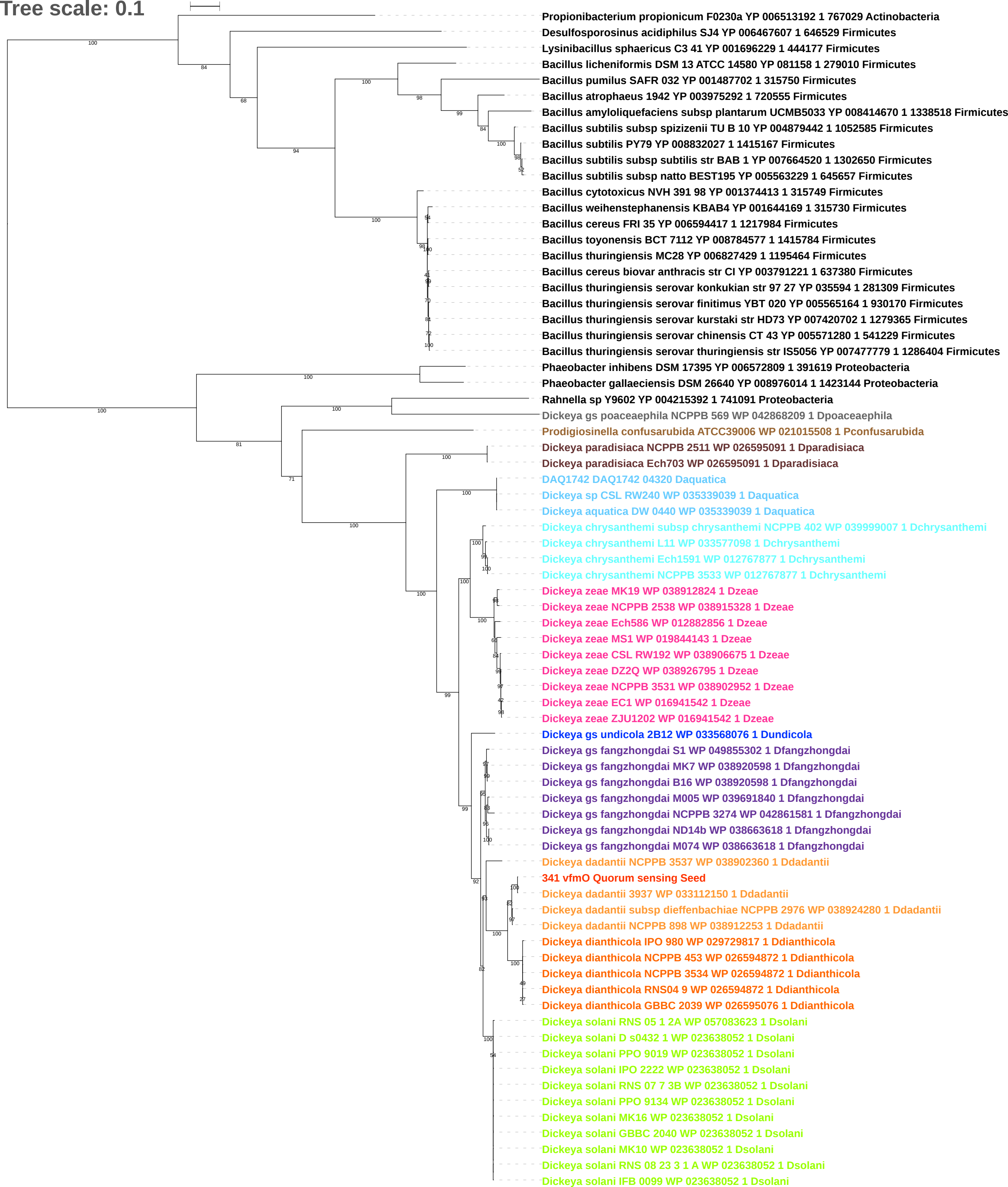

Tree scale: 0.1
Propionibacterium propionicum F0230a YP 006513192 1 767029 Actinobacteria
Desulfosporosinus acidiphilus SJ4 YP 006467607 1 646529 Firmicutes
Lysinibacillus sphaericus C3 41 YP 001696229 1 444177 Firmicutes
Bacillus licheniformis DSM 13 ATCC 14580 YP 081158 1 279010 Firmicutes
Bacillus pumilus SAFR 032 YP 001487702 1 315750 Firmicutes
Bacillus atrophaeus 1942 YP 003975292 1 720555 Firmicutes
Bacillus amyloliquefaciens subsp plantarum UCMB5033 YP 008414670 1 1338518 Firmicutes
Bacillus subtilis subsp spizizenii TU B 10 YP 004879442 1 1052585 Firmicutes
Bacillus subtilis PY79 YP 008832027 1 1415167 Firmicutes
Bacillus subtilis subsp subtilis str BAB 1 YP 007664520 1 1302650 Firmicutes
Bacillus subtilis subsp natto BEST195 YP 005563229 1 645657 Firmicutes
Bacillus cytotoxicus NVH 391 98 YP 001374413 1 315749 Firmicutes
Bacillus weihenstephanensis KBAB4 YP 001644169 1 315730 Firmicutes
Bacillus cereus FRI 35 YP 006594417 1 1217984 Firmicutes
Bacillus toyonensis BCT 7112 YP 008784577 1 1415784 Firmicutes
Bacillus thuringiensis MC28 YP 006827429 1 1195464 Firmicutes
Bacillus cereus biovar anthracis str CI YP 003791221 1 637380 Firmicutes
Bacillus thuringiensis serovar konkukian str 97 27 YP 035594 1 281309 Firmicutes
Bacillus thuringiensis serovar finitimus YBT 020 YP 005565164 1 930170 Firmicutes
Bacillus thuringiensis serovar kurstaki str HD73 YP 007420702 1 1279365 Firmicutes
Bacillus thuringiensis serovar chinensis CT 43 YP 005571280 1 541229 Firmicutes
Bacillus thuringiensis serovar thuringiensis str IS5056 YP 007477779 1 1286404 Firmicutes
Phaeobacter inhibens DSM 17395 YP 006572809 1 391619 Proteobacteria
Phaeobacter gallaeciensis DSM 26640 YP 008976014 1 1423144 Proteobacteria
Rahnella sp Y9602 YP 004215392 1 741091 Proteobacteria
Dickeya gs poaceaephila NCPPB 569 WP 042868209 1 Dpoaceaephila
Prodigiosinella confusarubida ATCC39006 WP 021015508 1 Pconfusarubida
Dickeya paradisiaca NCPPB 2511 WP 026595091 1 Dparadisiaca
Dickeya paradisiaca Ech703 WP 026595091 1 Dparadisiaca
DAQ1742 DAQ1742 04320 Daquatica
Dickeya sp CSL RW240 WP 035339039 1 Daquatica
Dickeya aquatica DW 0440 WP 035339039 1 Daquatica
Dickeya chrysanthemi subsp chrysanthemi NCPPB 402 WP 039999007 1 Dchrysanthemi
Dickeya chrysanthemi L11 WP 033577098 1 Dchrysanthemi
Dickeya chrysanthemi Ech1591 WP 012767877 1 Dchrysanthemi
Dickeya chrysanthemi NCPPB 3533 WP 012767877 1 Dchrysanthemi
Dickeya zeae MK19 WP 038912824 1 Dzeae
Dickeya zeae NCPPB 2538 WP 038915328 1 Dzeae
Dickeya zeae Ech586 WP 012882856 1 Dzeae
Dickeya zeae MS1 WP 019844143 1 Dzeae
Dickeya zeae CSL RW192 WP 038906675 1 Dzeae
Dickeya zeae DZ2Q WP 038926795 1 Dzeae
Dickeya zeae NCPPB 3531 WP 038902952 1 Dzeae
Dickeya zeae EC1 WP 016941542 1 Dzeae
Dickeya zeae ZJU1202 WP 016941542 1 Dzeae
Dickeya gs undicola 2B12 WP 033568076 1 Dundicola
Dickeya gs fangzhongdai S1 WP 049855302 1 Dfangzhongdai
Dickeya gs fangzhongdai MK7 WP 038920598 1 Dfangzhongdai
Dickeya gs fangzhongdai B16 WP 038920598 1 Dfangzhongdai
Dickeya gs fangzhongdai M005 WP 039691840 1 Dfangzhongdai
Dickeya gs fangzhongdai NCPPB 3274 WP 042861581 1 Dfangzhongdai
Dickeya gs fangzhongdai ND14b WP 038663618 1 Dfangzhongdai
Dickeya gs fangzhongdai M074 WP 038663618 1 Dfangzhongdai
Dickeya dadantii NCPPB 3537 WP 038902360 1 Ddadantii
341 vfmO Quorum sensing Seed
Dickeya dadantii 3937 WP 033112150 1 Ddadantii
Dickeya dadantii subsp dieffenbachiae NCPPB 2976 WP 038924280 1 Ddadantii
Dickeya dadantii NCPPB 898 WP 038912253 1 Ddadantii
Dickeya dianthicola IPO 980 WP 029729817 1 Ddianthicola
Dickeya dianthicola NCPPB 453 WP 026594872 1 Ddianthicola
Dickeya dianthicola NCPPB 3534 WP 026594872 1 Ddianthicola
Dickeya dianthicola RNS04 9 WP 026594872 1 Ddianthicola
Dickeya dianthicola GBBC 2039 WP 026595076 1 Ddianthicola
Dickeya solani RNS 05 1 2A WP 057083623 1 Dsolani
Dickeya solani D s0432 1 WP 023638052 1 Dsolani
Dickeya solani PPO 9019 WP 023638052 1 Dsolani
Dickeya solani IPO 2222 WP 023638052 1 Dsolani
Dickeya solani RNS 07 7 3B WP 023638052 1 Dsolani
Dickeya solani PPO 9134 WP 023638052 1 Dsolani
Dickeya solani MK16 WP 023638052 1 Dsolani
Dickeya solani GBBC 2040 WP 023638052 1 Dsolani
Dickeya solani MK10 WP 023638052 1 Dsolani
Dickeya solani RNS 08 23 3 1 A WP 023638052 1 Dsolani
Dickeya solani IFB 0099 WP 023638052 1 Dsolani

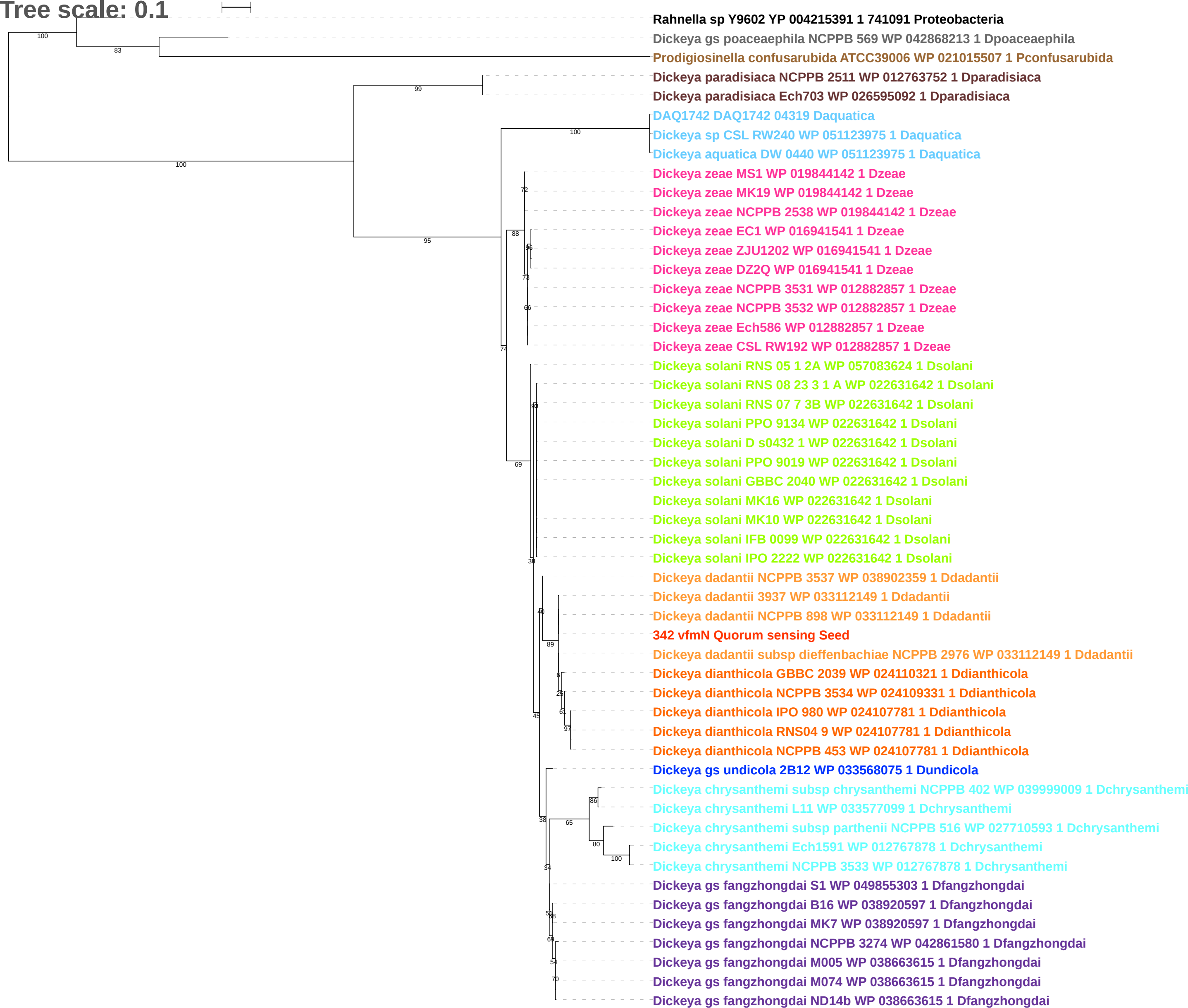
Tree scale: 0.1
Rahnella sp Y9602 YP 004215391 1 741091 Proteobacteria
Dickeya gs poaceaephila NCPPB 569 WP 042868213 1 Dpoaceaephila
Prodigiosinella confusarubida ATCC39006 WP 021015507 1 Pconfusarubida
Dickeya paradisiaca NCPPB 2511 WP 012763752 1 Dparadisiaca
Dickeya paradisiaca Ech703 WP 026595092 1 Dparadisiaca
DAQ1742 DAQ1742 04319 Daquatica
Dickeya sp CSL RW240 WP 051123975 1 Daquatica
Dickeya aquatica DW 0440 WP 051123975 1 Daquatica
Dickeya zeae MS1 WP 019844142 1 Dzeae
Dickeya zeae MK19 WP 019844142 1 Dzeae
Dickeya zeae NCPPB 2538 WP 019844142 1 Dzeae
Dickeya zeae EC1 WP 016941541 1 Dzeae
Dickeya zeae ZJU1202 WP 016941541 1 Dzeae
Dickeya zeae DZ2Q WP 016941541 1 Dzeae
Dickeya zeae NCPPB 3531 WP 012882857 1 Dzeae
Dickeya zeae NCPPB 3532 WP 012882857 1 Dzeae
Dickeya zeae Ech586 WP 012882857 1 Dzeae
Dickeya zeae CSL RW192 WP 012882857 1 Dzeae
Dickeya solani RNS 05 1 2A WP 057083624 1 Dsolani
Dickeya solani RNS 08 23 3 1 A WP 022631642 1 Dsolani
Dickeya solani RNS 07 7 3B WP 022631642 1 Dsolani
Dickeya solani PPO 9134 WP 022631642 1 Dsolani
Dickeya solani D s0432 1 WP 022631642 1 Dsolani
Dickeya solani PPO 9019 WP 022631642 1 Dsolani
Dickeya solani GBBC 2040 WP 022631642 1 Dsolani
Dickeya solani MK16 WP 022631642 1 Dsolani
Dickeya solani MK10 WP 022631642 1 Dsolani
Dickeya solani IFB 0099 WP 022631642 1 Dsolani
Dickeya solani IPO 2222 WP 022631642 1 Dsolani
Dickeya dadantii NCPPB 3537 WP 038902359 1 Ddadantii
Dickeya dadantii 3937 WP 033112149 1 Ddadantii
Dickeya dadantii NCPPB 898 WP 033112149 1 Ddadantii
342 vfmN Quorum sensing Seed
Dickeya dadantii subsp dieffenbachiae NCPPB 2976 WP 033112149 1 Ddadantii
Dickeya dianthicola GBBC 2039 WP 024110321 1 Ddianthicola
Dickeya dianthicola NCPPB 3534 WP 024109331 1 Ddianthicola
Dickeya dianthicola IPO 980 WP 024107781 1 Ddianthicola
Dickeya dianthicola RNS04 9 WP 024107781 1 Ddianthicola
Dickeya dianthicola NCPPB 453 WP 024107781 1 Ddianthicola
Dickeya gs undicola 2B12 WP 033568075 1 Dundicola
Dickeya chrysanthemi subsp chrysanthemi NCPPB 402 WP 039999009 1 Dchrysanthemi
Dickeya chrysanthemi L11 WP 033577099 1 Dchrysanthemi
Dickeya chrysanthemi subsp parthenii NCPPB 516 WP 027710593 1 Dchrysanthemi
Dickeya chrysanthemi Ech1591 WP 012767878 1 Dchrysanthemi
Dickeya chrysanthemi NCPPB 3533 WP 012767878 1 Dchrysanthemi
Dickeya gs fangzhongdai S1 WP 049855303 1 Dfangzhongdai
Dickeya gs fangzhongdai B16 WP 038920597 1 Dfangzhongdai
Dickeya gs fangzhongdai MK7 WP 038920597 1 Dfangzhongdai
Dickeya gs fangzhongdai NCPPB 3274 WP 042861580 1 Dfangzhongdai
Dickeya gs fangzhongdai M005 WP 038663615 1 Dfangzhongdai
Dickeya gs fangzhongdai M074 WP 038663615 1 Dfangzhongdai
Dickeya gs fangzhongdai ND14b WP 038663615 1 Dfangzhongdai

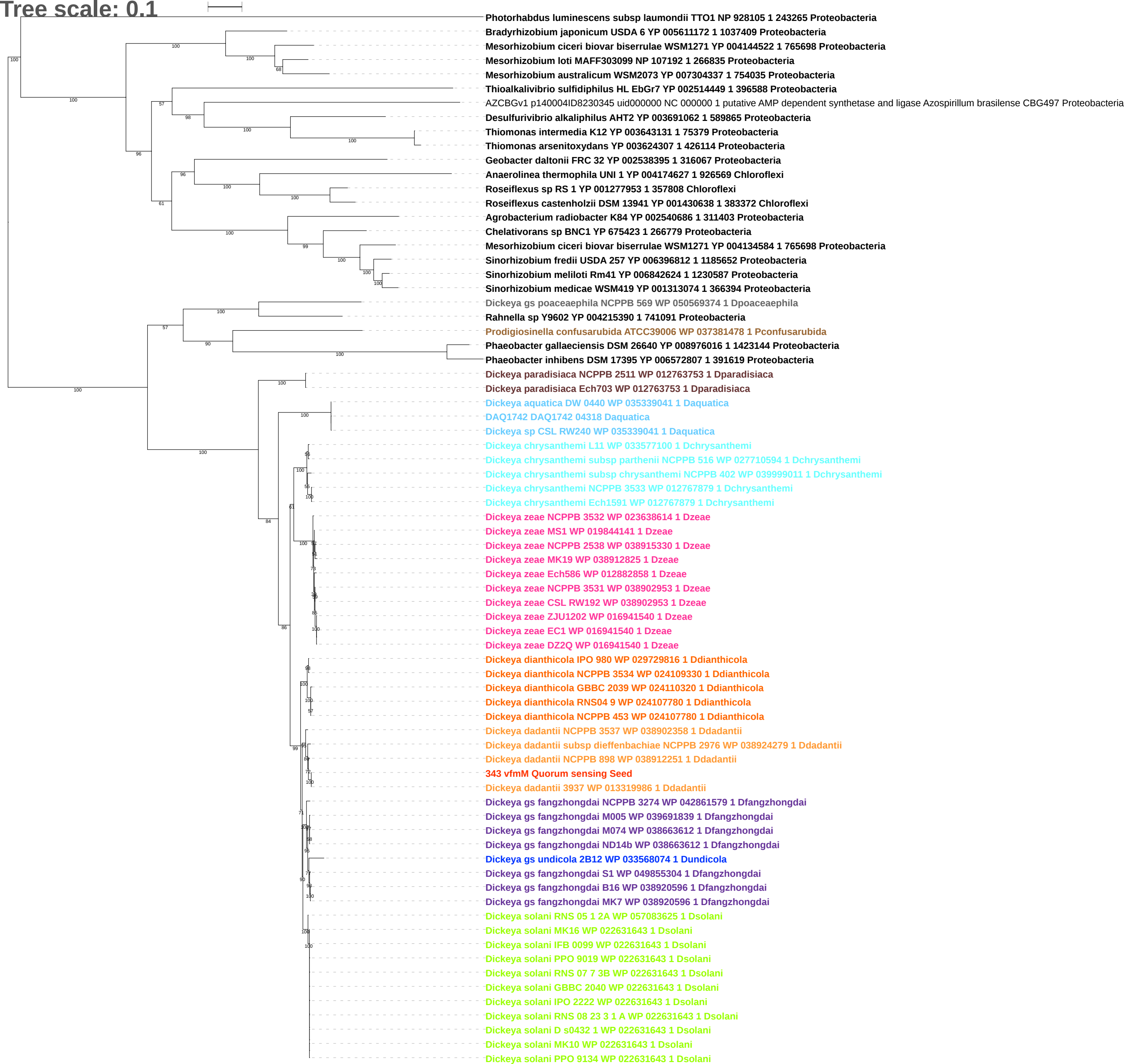
Tree scale: 0.1
Photorhabdus luminescens subsp laumondii TTO1 NP 928105 1 243265 Proteobacteria
Bradyrhizobium japonicum USDA 6 YP 005611172 1 1037409 Proteobacteria
Mesorhizobium ciceri biovar biserrulae WSM1271 YP 004144522 1 765698 Proteobacteria
Mesorhizobium loti MAFF303099 NP 107192 1 266835 Proteobacteria
Mesorhizobium australicum WSM2073 YP 007304337 1 754035 Proteobacteria
Thioalkalivibrio sulfidiphilus HL EbGr7 YP 002514449 1 396588 Proteobacteria
AZCBGv1 p140004ID8230345 uid000000 NC 000000 1 putative AMP dependent synthetase and ligase Azospirillum brasilense CBG497 Proteobacteria
Desulfurivibrio alkaliphilus AHT2 YP 003691062 1 589865 Proteobacteria
Thiomonas intermedia K12 YP 003643131 1 75379 Proteobacteria
Thiomonas arsenitoxydans YP 003624307 1 426114 Proteobacteria
Geobacter daltonii FRC 32 YP 002538395 1 316067 Proteobacteria
Anaerolinea thermophila UNI 1 YP 004174627 1 926569 Chloroflexi
Roseiflexus sp RS 1 YP 001277953 1 357808 Chloroflexi
Roseiflexus castenholzii DSM 13941 YP 001430638 1 383372 Chloroflexi
Agrobacterium radiobacter K84 YP 002540686 1 311403 Proteobacteria
Chelativorans sp BNC1 YP 675423 1 266779 Proteobacteria
Mesorhizobium ciceri biovar biserrulae WSM1271 YP 004134584 1 765698 Proteobacteria
Sinorhizobium fredii USDA 257 YP 006396812 1 1185652 Proteobacteria
Sinorhizobium meliloti Rm41 YP 006842624 1 1230587 Proteobacteria
Sinorhizobium medicae WSM419 YP 001313074 1 366394 Proteobacteria
Dickeya gs poaceaephila NCPPB 569 WP 050569374 1 Dpoaceaephila
Rahnella sp Y9602 YP 004215390 1 741091 Proteobacteria
Prodigiosinella confusarubida ATCC39006 WP 037381478 1 Pconfusarubida
Phaeobacter gallaeciensis DSM 26640 YP 008976016 1 1423144 Proteobacteria
Phaeobacter inhibens DSM 17395 YP 006572807 1 391619 Proteobacteria
Dickeya paradisiaca NCPPB 2511 WP 012763753 1 Dparadisiaca
Dickeya paradisiaca Ech703 WP 012763753 1 Dparadisiaca
Dickeya aquatica DW 0440 WP 035339041 1 Daquatica
DAQ1742 DAQ1742 04318 Daquatica
Dickeya sp CSL RW240 WP 035339041 1 Daquatica
Dickeya chrysanthemi L11 WP 033577100 1 Dchrysanthemi
Dickeya chrysanthemi subsp parthenii NCPPB 516 WP 027710594 1 Dchrysanthemi
Dickeya chrysanthemi subsp chrysanthemi NCPPB 402 WP 039999011 1 Dchrysanthemi
Dickeya chrysanthemi NCPPB 3533 WP 012767879 1 Dchrysanthemi
Dickeya chrysanthemi Ech1591 WP 012767879 1 Dchrysanthemi
Dickeya zeae NCPPB 3532 WP 023638614 1 Dzeae
Dickeya zeae MS1 WP 019844141 1 Dzeae
Dickeya zeae NCPPB 2538 WP 038915330 1 Dzeae
Dickeya zeae MK19 WP 038912825 1 Dzeae
Dickeya zeae Ech586 WP 012882858 1 Dzeae
Dickeya zeae NCPPB 3531 WP 038902953 1 Dzeae
Dickeya zeae CSL RW192 WP 038902953 1 Dzeae
Dickeya zeae ZJU1202 WP 016941540 1 Dzeae
Dickeya zeae EC1 WP 016941540 1 Dzeae
Dickeya zeae DZ2Q WP 016941540 1 Dzeae
Dickeya dianthicola IPO 980 WP 029729816 1 Ddianthicola
Dickeya dianthicola NCPPB 3534 WP 024109330 1 Ddianthicola
Dickeya dianthicola GBBC 2039 WP 024110320 1 Ddianthicola
Dickeya dianthicola RNS04 9 WP 024107780 1 Ddianthicola
Dickeya dianthicola NCPPB 453 WP 024107780 1 Ddianthicola
Dickeya dadantii NCPPB 3537 WP 038902358 1 Ddadantii
Dickeya dadantii subsp dieffenbachiae NCPPB 2976 WP 038924279 1 Ddadantii
Dickeya dadantii NCPPB 898 WP 038912251 1 Ddadantii
343 vfmM Quorum sensing Seed
Dickeya dadantii 3937 WP 013319986 1 Ddadantii
Dickeya gs fangzhongdai NCPPB 3274 WP 042861579 1 Dfangzhongdai
Dickeya gs fangzhongdai M005 WP 039691839 1 Dfangzhongdai
Dickeya gs fangzhongdai M074 WP 038663612 1 Dfangzhongdai
Dickeya gs fangzhongdai ND14b WP 038663612 1 Dfangzhongdai
Dickeya gs undicola 2B12 WP 033568074 1 Dundicola
Dickeya gs fangzhongdai S1 WP 049855304 1 Dfangzhongdai
Dickeya gs fangzhongdai B16 WP 038920596 1 Dfangzhongdai
Dickeya gs fangzhongdai MK7 WP 038920596 1 Dfangzhongdai
Dickeya solani RNS 05 1 2A WP 057083625 1 Dsolani
Dickeya solani MK16 WP 022631643 1 Dsolani
Dickeya solani IFB 0099 WP 022631643 1 Dsolani
Dickeya solani PPO 9019 WP 022631643 1 Dsolani
Dickeya solani RNS 07 7 3B WP 022631643 1 Dsolani
Dickeya solani GBBC 2040 WP 022631643 1 Dsolani
Dickeya solani IPO 2222 WP 022631643 1 Dsolani
Dickeya solani RNS 08 23 3 1 A WP 022631643 1 Dsolani
Dickeya solani D s0432 1 WP 022631643 1 Dsolani
Dickeya solani MK10 WP 022631643 1 Dsolani
Dickeya solani PPO 9134 WP 022631643 1 Dsolani

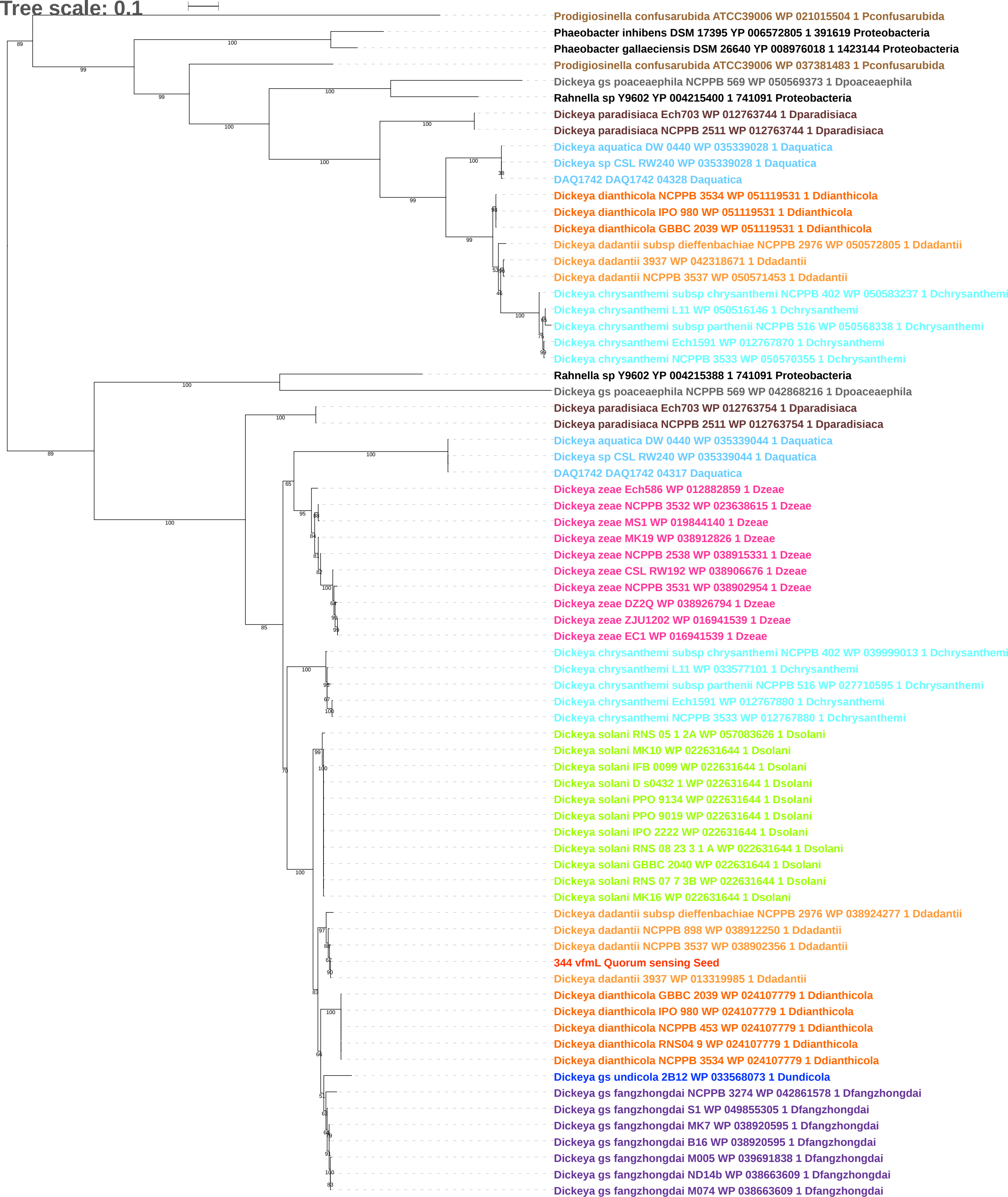
Tree scale: 0.1
Prodigiosinella confusarubida ATCC39006 WP 021015504 1 Pconfusarubida
Phaeobacter inhibens DSM 17395 YP 006572805 1 391619 Proteobacteria
Phaeobacter gallaeciensis DSM 26640 YP 008976018 1 1423144 Proteobacteria
Prodigiosinella confusarubida ATCC39006 WP 037381483 1 Pconfusarubida
Dickeya gs poaceaephila NCPPB 569 WP 050569373 1 Dpoaceaephila
Rahnella sp Y9602 YP 004215400 1 741091 Proteobacteria
Dickeya paradisiaca Ech703 WP 012763744 1 Dparadisiaca
Dickeya paradisiaca NCPPB 2511 WP 012763744 1 Dparadisiaca
Dickeya aquatica DW 0440 WP 035339028 1 Daquatica
Dickeya sp CSL RW240 WP 035339028 1 Daquatica
DAQ1742 DAQ1742 04328 Daquatica
Dickeya dianthicola NCPPB 3534 WP 051119531 1 Ddianthicola
Dickeya dianthicola IPO 980 WP 051119531 1 Ddianthicola
Dickeya dianthicola GBBC 2039 WP 051119531 1 Ddianthicola
Dickeya dadantii subsp dieffenbachiae NCPPB 2976 WP 050572805 1 Ddadantii
Dickeya dadantii 3937 WP 042318671 1 Ddadantii
Dickeya dadantii NCPPB 3537 WP 050571453 1 Ddadantii
Dickeya chrysanthemi subsp chrysanthemi NCPPB 402 WP 050583237 1 Dchrysanthemi
Dickeya chrysanthemi L11 WP 050516146 1 Dchrysanthemi
Dickeya chrysanthemi subsp parthenii NCPPB 516 WP 050568338 1 Dchrysanthemi
Dickeya chrysanthemi Ech1591 WP 012767870 1 Dchrysanthemi
Dickeya chrysanthemi NCPPB 3533 WP 050570355 1 Dchrysanthemi
Rahnella sp Y9602 YP 004215388 1 741091 Proteobacteria
Dickeya gs poaceaephila NCPPB 569 WP 042868216 1 Dpoaceaephila
Dickeya paradisiaca Ech703 WP 012763754 1 Dparadisiaca
Dickeya paradisiaca NCPPB 2511 WP 012763754 1 Dparadisiaca
Dickeya aquatica DW 0440 WP 035339044 1 Daquatica
Dickeya sp CSL RW240 WP 035339044 1 Daquatica
DAQ1742 DAQ1742 04317 Daquatica
Dickeya zeae Ech586 WP 012882859 1 Dzeae
Dickeya zeae NCPPB 3532 WP 023638615 1 Dzeae
Dickeya zeae MS1 WP 019844140 1 Dzeae
Dickeya zeae MK19 WP 038912826 1 Dzeae
Dickeya zeae NCPPB 2538 WP 038915331 1 Dzeae
Dickeya zeae CSL RW192 WP 038906676 1 Dzeae
Dickeya zeae NCPPB 3531 WP 038902954 1 Dzeae
Dickeya zeae DZ2Q WP 038926794 1 Dzeae
Dickeya zeae ZJU1202 WP 016941539 1 Dzeae
Dickeya zeae EC1 WP 016941539 1 Dzeae
Dickeya chrysanthemi subsp chrysanthemi NCPPB 402 WP 039999013 1 Dchrysanthemi
Dickeya chrysanthemi L11 WP 033577101 1 Dchrysanthemi
Dickeya chrysanthemi subsp parthenii NCPPB 516 WP 027710595 1 Dchrysanthemi
Dickeya chrysanthemi Ech1591 WP 012767880 1 Dchrysanthemi
Dickeya chrysanthemi NCPPB 3533 WP 012767880 1 Dchrysanthemi
Dickeya solani RNS 05 1 2A WP 057083626 1 Dsolani
Dickeya solani MK10 WP 022631644 1 Dsolani
Dickeya solani IFB 0099 WP 022631644 1 Dsolani
Dickeya solani D s0432 1 WP 022631644 1 Dsolani
Dickeya solani PPO 9134 WP 022631644 1 Dsolani
Dickeya solani PPO 9019 WP 022631644 1 Dsolani
Dickeya solani IPO 2222 WP 022631644 1 Dsolani
Dickeya solani RNS 08 23 3 1 A WP 022631644 1 Dsolani
Dickeya solani GBBC 2040 WP 022631644 1 Dsolani
Dickeya solani RNS 07 7 3B WP 022631644 1 Dsolani
Dickeya solani MK16 WP 022631644 1 Dsolani
Dickeya dadantii subsp dieffenbachiae NCPPB 2976 WP 038924277 1 Ddadantii
Dickeya dadantii NCPPB 898 WP 038912250 1 Ddadantii
Dickeya dadantii NCPPB 3537 WP 038902356 1 Ddadantii
344 vfmL Quorum sensing Seed
Dickeya dadantii 3937 WP 013319985 1 Ddadantii
Dickeya dianthicola GBBC 2039 WP 024107779 1 Ddianthicola
Dickeya dianthicola IPO 980 WP 024107779 1 Ddianthicola
Dickeya dianthicola NCPPB 453 WP 024107779 1 Ddianthicola
Dickeya dianthicola RNS04 9 WP 024107779 1 Ddianthicola
Dickeya dianthicola NCPPB 3534 WP 024107779 1 Ddianthicola
Dickeya gs undicola 2B12 WP 033568073 1 Dundicola
Dickeya gs fangzhongdai NCPPB 3274 WP 042861578 1 Dfangzhongdai
Dickeya gs fangzhongdai S1 WP 049855305 1 Dfangzhongdai
Dickeya gs fangzhongdai MK7 WP 038920595 1 Dfangzhongdai
Dickeya gs fangzhongdai B16 WP 038920595 1 Dfangzhongdai
Dickeya gs fangzhongdai M005 WP 039691838 1 Dfangzhongdai
Dickeya gs fangzhongdai ND14b WP 038663609 1 Dfangzhongdai
Dickeya gs fangzhongdai M074 WP 038663609 1 Dfangzhongdai

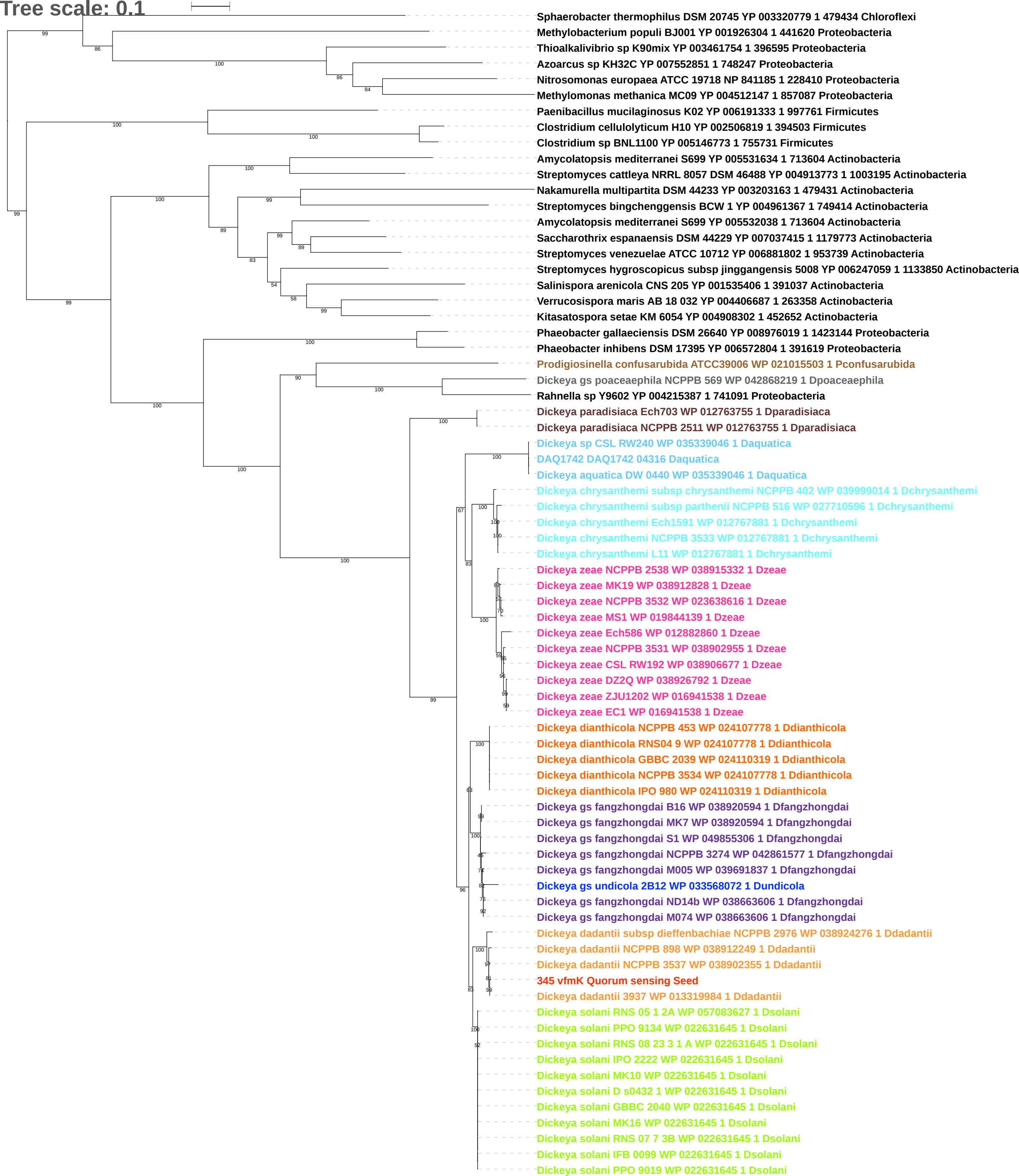
Tree scale: 0.1
Sphaerobacter thermophilus DSM 20745 YP 003320779 1 479434 Chloroflexi
Methylobacterium populi BJ001 YP 001926304 1 441620 Proteobacteria
Thioalkalivibrio sp K90mix YP 003461754 1 396595 Proteobacteria
Azoarcus sp KH32C YP 007552851 1 748247 Proteobacteria
Nitrosomonas europaea ATCC 19718 NP 841185 1 228410 Proteobacteria
Methylomonas methanica MC09 YP 004512147 1 857087 Proteobacteria
Paenibacillus mucilaginosus K02 YP 006191333 1 997761 Firmicutes
Clostridium cellulolyticum H10 YP 002506819 1 394503 Firmicutes
Clostridium sp BNL1100 YP 005146773 1 755731 Firmicutes
Amycolatopsis mediterranei S699 YP 005531634 1 713604 Actinobacteria
Streptomyces cattleya NRRL 8057 DSM 46488 YP 004913773 1 1003195 Actinobacteria
Nakamurella multipartita DSM 44233 YP 003203163 1 479431 Actinobacteria
Streptomyces bingchenggensis BCW 1 YP 004961367 1 749414 Actinobacteria
Amycolatopsis mediterranei S699 YP 005532038 1 713604 Actinobacteria
Saccharothrix espanaensis DSM 44229 YP 007037415 1 1179773 Actinobacteria
Streptomyces venezuelae ATCC 10712 YP 006881802 1 953739 Actinobacteria
Streptomyces hygroscopicus subsp jinggangensis 5008 YP 006247059 1 1133850 Actinobacteria
Salinispora arenicola CNS 205 YP 001535406 1 391037 Actinobacteria
Verrucosispora maris AB 18 032 YP 004406687 1 263358 Actinobacteria
Kitasatospora setae KM 6054 YP 004908302 1 452652 Actinobacteria
Phaeobacter gallaeciensis DSM 26640 YP 008976019 1 1423144 Proteobacteria
Phaeobacter inhibens DSM 17395 YP 006572804 1 391619 Proteobacteria
Prodigiosinella confusarubida ATCC39006 WP 021015503 1 Pconfusarubida
Dickeya gs poaceaephila NCPPB 569 WP 042868219 1 Dpoaceaephila
Rahnella sp Y9602 YP 004215387 1 741091 Proteobacteria
Dickeya paradisiaca Ech703 WP 012763755 1 Dparadisiaca
Dickeya paradisiaca NCPPB 2511 WP 012763755 1 Dparadisiaca
Dickeya sp CSL RW240 WP 035339046 1 Daquatica
DAQ1742 DAQ1742 04316 Daquatica
Dickeya aquatica DW 0440 WP 035339046 1 Daquatica
Dickeya chrysanthemi subsp chrysanthemi NCPPB 402 WP 039999014 1 Dchrysanthemi
Dickeya chrysanthemi subsp parthenii NCPPB 516 WP 027710596 1 Dchrysanthemi
Dickeya chrysanthemi Ech1591 WP 012767881 1 Dchrysanthemi
Dickeya chrysanthemi NCPPB 3533 WP 012767881 1 Dchrysanthemi
Dickeya chrysanthemi L11 WP 012767881 1 Dchrysanthemi
Dickeya zeae NCPPB 2538 WP 038915332 1 Dzeae
Dickeya zeae MK19 WP 038912828 1 Dzeae
Dickeya zeae NCPPB 3532 WP 023638616 1 Dzeae
Dickeya zeae MS1 WP 019844139 1 Dzeae
Dickeya zeae Ech586 WP 012882860 1 Dzeae
Dickeya zeae NCPPB 3531 WP 038902955 1 Dzeae
Dickeya zeae CSL RW192 WP 038906677 1 Dzeae
Dickeya zeae DZ2Q WP 038926792 1 Dzeae
Dickeya zeae ZJU1202 WP 016941538 1 Dzeae
Dickeya zeae EC1 WP 016941538 1 Dzeae
Dickeya dianthicola NCPPB 453 WP 024107778 1 Ddianthicola
Dickeya dianthicola RNS04 9 WP 024107778 1 Ddianthicola
Dickeya dianthicola GBBC 2039 WP 024110319 1 Ddianthicola
Dickeya dianthicola NCPPB 3534 WP 024107778 1 Ddianthicola
Dickeya dianthicola IPO 980 WP 024110319 1 Ddianthicola
Dickeya gs fangzhongdai B16 WP 038920594 1 Dfangzhongdai
Dickeya gs fangzhongdai MK7 WP 038920594 1 Dfangzhongdai
Dickeya gs fangzhongdai S1 WP 049855306 1 Dfangzhongdai
Dickeya gs fangzhongdai NCPPB 3274 WP 042861577 1 Dfangzhongdai
Dickeya gs fangzhongdai M005 WP 039691837 1 Dfangzhongdai
Dickeya gs undicola 2B12 WP 033568072 1 Dundicola
Dickeya gs fangzhongdai ND14b WP 038663606 1 Dfangzhongdai
Dickeya gs fangzhongdai M074 WP 038663606 1 Dfangzhongdai
Dickeya dadantii subsp dieffenbachiae NCPPB 2976 WP 038924276 1 Ddadantii
Dickeya dadantii NCPPB 898 WP 038912249 1 Ddadantii
Dickeya dadantii NCPPB 3537 WP 038902355 1 Ddadantii
345 vfmK Quorum sensing Seed
Dickeya dadantii 3937 WP 013319984 1 Ddadantii
Dickeya solani RNS 05 1 2A WP 057083627 1 Dsolani
Dickeya solani PPO 9134 WP 022631645 1 Dsolani
Dickeya solani RNS 08 23 3 1 A WP 022631645 1 Dsolani
Dickeya solani IPO 2222 WP 022631645 1 Dsolani
Dickeya solani MK10 WP 022631645 1 Dsolani
Dickeya solani D s0432 1 WP 022631645 1 Dsolani
Dickeya solani GBBC 2040 WP 022631645 1 Dsolani
Dickeya solani MK16 WP 022631645 1 Dsolani
Dickeya solani RNS 07 7 3B WP 022631645 1 Dsolani
Dickeya solani IFB 0099 WP 022631645 1 Dsolani
Dickeya solani PPO 9019 WP 022631645 1 Dsolani

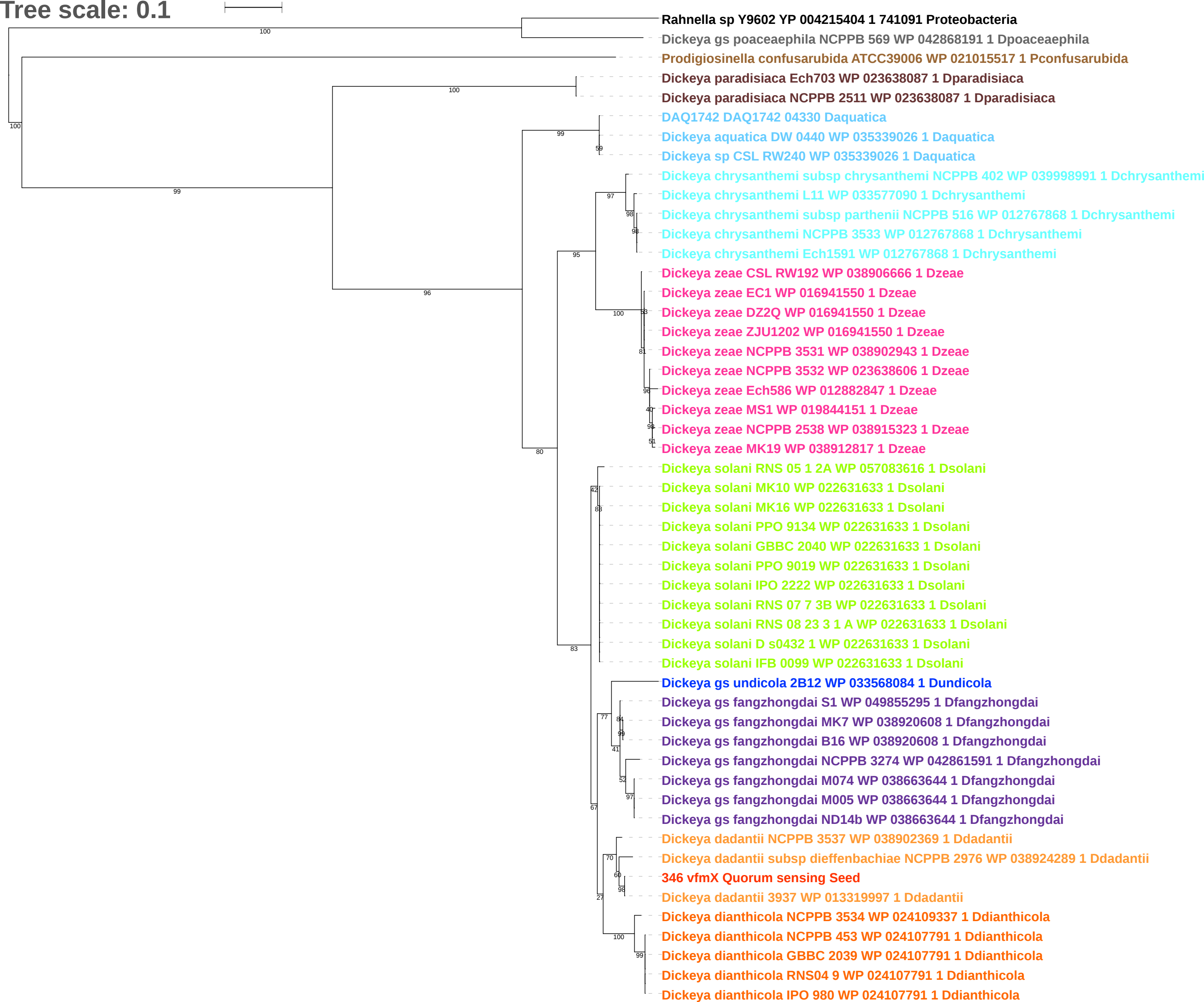
Tree scale: 0.1
Rahnella sp Y9602 YP 004215404 1 741091 Proteobacteria
Dickeya gs poaceaephila NCPPB 569 WP 042868191 1 Dpoaceaephila
Prodigiosinella confusarubida ATCC39006 WP 021015517 1 Pconfusarubida
Dickeya paradisiaca Ech703 WP 023638087 1 Dparadisiaca
Dickeya paradisiaca NCPPB 2511 WP 023638087 1 Dparadisiaca
DAQ1742 DAQ1742 04330 Daquatica
Dickeya aquatica DW 0440 WP 035339026 1 Daquatica
Dickeya sp CSL RW240 WP 035339026 1 Daquatica
Dickeya chrysanthemi subsp chrysanthemi NCPPB 402 WP 039998991 1 Dchrysanthemi
Dickeya chrysanthemi L11 WP 033577090 1 Dchrysanthemi
Dickeya chrysanthemi subsp parthenii NCPPB 516 WP 012767868 1 Dchrysanthemi
Dickeya chrysanthemi NCPPB 3533 WP 012767868 1 Dchrysanthemi
Dickeya chrysanthemi Ech1591 WP 012767868 1 Dchrysanthemi
Dickeya zeae CSL RW192 WP 038906666 1 Dzeae
Dickeya zeae EC1 WP 016941550 1 Dzeae
Dickeya zeae DZ2Q WP 016941550 1 Dzeae
Dickeya zeae ZJU1202 WP 016941550 1 Dzeae
Dickeya zeae NCPPB 3531 WP 038902943 1 Dzeae
Dickeya zeae NCPPB 3532 WP 023638606 1 Dzeae
Dickeya zeae Ech586 WP 012882847 1 Dzeae
Dickeya zeae MS1 WP 019844151 1 Dzeae
Dickeya zeae NCPPB 2538 WP 038915323 1 Dzeae
Dickeya zeae MK19 WP 038912817 1 Dzeae
Dickeya solani RNS 05 1 2A WP 057083616 1 Dsolani
Dickeya solani MK10 WP 022631633 1 Dsolani
Dickeya solani MK16 WP 022631633 1 Dsolani
Dickeya solani PPO 9134 WP 022631633 1 Dsolani
Dickeya solani GBBC 2040 WP 022631633 1 Dsolani
Dickeya solani PPO 9019 WP 022631633 1 Dsolani
Dickeya solani IPO 2222 WP 022631633 1 Dsolani
Dickeya solani RNS 07 7 3B WP 022631633 1 Dsolani
Dickeya solani RNS 08 23 3 1 A WP 022631633 1 Dsolani
Dickeya solani D s0432 1 WP 022631633 1 Dsolani
Dickeya solani IFB 0099 WP 022631633 1 Dsolani
Dickeya gs undicola 2B12 WP 033568084 1 Dundicola
Dickeya gs fangzhongdai S1 WP 049855295 1 Dfangzhongdai
Dickeya gs fangzhongdai MK7 WP 038920608 1 Dfangzhongdai
Dickeya gs fangzhongdai B16 WP 038920608 1 Dfangzhongdai
Dickeya gs fangzhongdai NCPPB 3274 WP 042861591 1 Dfangzhongdai
Dickeya gs fangzhongdai M074 WP 038663644 1 Dfangzhongdai
Dickeya gs fangzhongdai M005 WP 038663644 1 Dfangzhongdai
Dickeya gs fangzhongdai ND14b WP 038663644 1 Dfangzhongdai
Dickeya dadantii NCPPB 3537 WP 038902369 1 Ddadantii
Dickeya dadantii subsp dieffenbachiae NCPPB 2976 WP 038924289 1 Ddadantii
346 vfmX Quorum sensing Seed
Dickeya dadantii 3937 WP 013319997 1 Ddadantii
Dickeya dianthicola NCPPB 3534 WP 024109337 1 Ddianthicola
Dickeya dianthicola NCPPB 453 WP 024107791 1 Ddianthicola
Dickeya dianthicola GBBC 2039 WP 024107791 1 Ddianthicola
Dickeya dianthicola RNS04 9 WP 024107791 1 Ddianthicola
Dickeya dianthicola IPO 980 WP 024107791 1 Ddianthicola

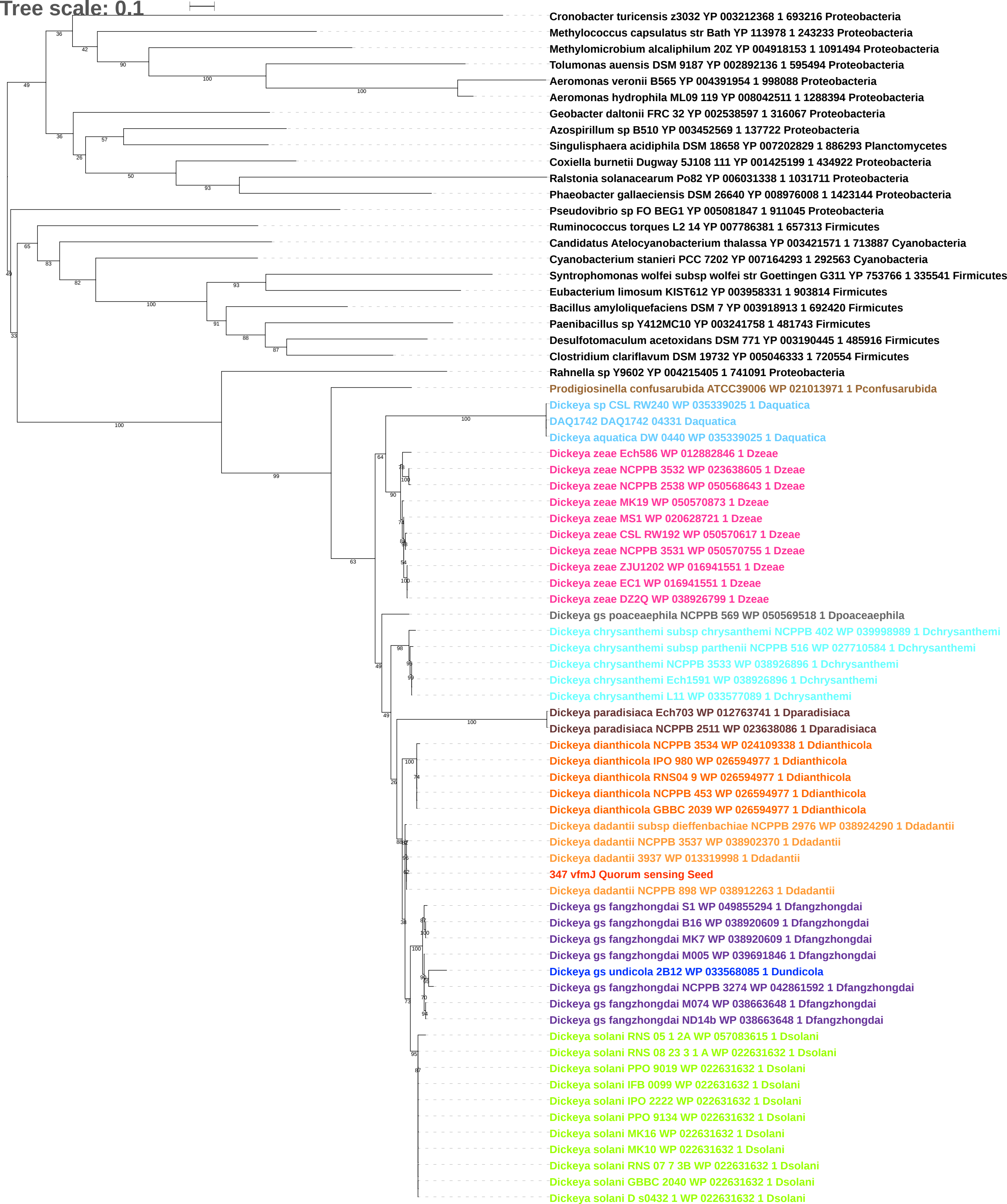
Tree scale: 0.1
Cronobacter turicensis z3032 YP 003212368 1 693216 Proteobacteria
Methylococcus capsulatus str Bath YP 113978 1 243233 Proteobacteria
Methylomicrobium alcaliphilum 20Z YP 004918153 1 1091494 Proteobacteria
Tolumonas auensis DSM 9187 YP 002892136 1 595494 Proteobacteria
Aeromonas veronii B565 YP 004391954 1 998088 Proteobacteria
Aeromonas hydrophila ML09 119 YP 008042511 1 1288394 Proteobacteria
Geobacter daltonii FRC 32 YP 002538597 1 316067 Proteobacteria
Azospirillum sp B510 YP 003452569 1 137722 Proteobacteria
Singulisphaera acidiphila DSM 18658 YP 007202829 1 886293 Planctomycetes
Coxiella burnetii Dugway 5J108 111 YP 001425199 1 434922 Proteobacteria
Ralstonia solanacearum Po82 YP 006031338 1 1031711 Proteobacteria
Phaeobacter gallaeciensis DSM 26640 YP 008976008 1 1423144 Proteobacteria
Pseudovibrio sp FO BEG1 YP 005081847 1 911045 Proteobacteria
Ruminococcus torques L2 14 YP 007786381 1 657313 Firmicutes
Candidatus Atelocyanobacterium thalassa YP 003421571 1 713887 Cyanobacteria
Cyanobacterium stanieri PCC 7202 YP 007164293 1 292563 Cyanobacteria
Syntrophomonas wolfei subsp wolfei str Goettingen G311 YP 753766 1 335541 Firmicutes
Eubacterium limosum KIST612 YP 003958331 1 903814 Firmicutes
Bacillus amyloliquefaciens DSM 7 YP 003918913 1 692420 Firmicutes
Paenibacillus sp Y412MC10 YP 003241758 1 481743 Firmicutes
Desulfotomaculum acetoxidans DSM 771 YP 003190445 1 485916 Firmicutes
Clostridium clariflavum DSM 19732 YP 005046333 1 720554 Firmicutes
Rahnella sp Y9602 YP 004215405 1 741091 Proteobacteria
Prodigiosinella confusarubida ATCC39006 WP 021013971 1 Pconfusarubida
Dickeya sp CSL RW240 WP 035339025 1 Daquatica
DAQ1742 DAQ1742 04331 Daquatica
Dickeya aquatica DW 0440 WP 035339025 1 Daquatica
Dickeya zeae Ech586 WP 012882846 1 Dzeae
Dickeya zeae NCPPB 3532 WP 023638605 1 Dzeae
Dickeya zeae NCPPB 2538 WP 050568643 1 Dzeae
Dickeya zeae MK19 WP 050570873 1 Dzeae
Dickeya zeae MS1 WP 020628721 1 Dzeae
Dickeya zeae CSL RW192 WP 050570617 1 Dzeae
Dickeya zeae NCPPB 3531 WP 050570755 1 Dzeae
Dickeya zeae ZJU1202 WP 016941551 1 Dzeae
Dickeya zeae EC1 WP 016941551 1 Dzeae
Dickeya zeae DZ2Q WP 038926799 1 Dzeae
Dickeya gs poaceaephila NCPPB 569 WP 050569518 1 Dpoaceaephila
Dickeya chrysanthemi subsp chrysanthemi NCPPB 402 WP 039998989 1 Dchrysanthemi
Dickeya chrysanthemi subsp parthenii NCPPB 516 WP 027710584 1 Dchrysanthemi
Dickeya chrysanthemi NCPPB 3533 WP 038926896 1 Dchrysanthemi
Dickeya chrysanthemi Ech1591 WP 038926896 1 Dchrysanthemi
Dickeya chrysanthemi L11 WP 033577089 1 Dchrysanthemi
Dickeya paradisiaca Ech703 WP 012763741 1 Dparadisiaca
Dickeya paradisiaca NCPPB 2511 WP 023638086 1 Dparadisiaca
Dickeya dianthicola NCPPB 3534 WP 024109338 1 Ddianthicola
Dickeya dianthicola IPO 980 WP 026594977 1 Ddianthicola
Dickeya dianthicola RNS04 9 WP 026594977 1 Ddianthicola
Dickeya dianthicola NCPPB 453 WP 026594977 1 Ddianthicola
Dickeya dianthicola GBBC 2039 WP 026594977 1 Ddianthicola
Dickeya dadantii subsp dieffenbachiae NCPPB 2976 WP 038924290 1 Ddadantii
Dickeya dadantii NCPPB 3537 WP 038902370 1 Ddadantii
Dickeya dadantii 3937 WP 013319998 1 Ddadantii
347 vfmJ Quorum sensing Seed
Dickeya dadantii NCPPB 898 WP 038912263 1 Ddadantii
Dickeya gs fangzhongdai S1 WP 049855294 1 Dfangzhongdai
Dickeya gs fangzhongdai B16 WP 038920609 1 Dfangzhongdai
Dickeya gs fangzhongdai MK7 WP 038920609 1 Dfangzhongdai
Dickeya gs fangzhongdai M005 WP 039691846 1 Dfangzhongdai
Dickeya gs undicola 2B12 WP 033568085 1 Dundicola
Dickeya gs fangzhongdai NCPPB 3274 WP 042861592 1 Dfangzhongdai
Dickeya gs fangzhongdai M074 WP 038663648 1 Dfangzhongdai
Dickeya gs fangzhongdai ND14b WP 038663648 1 Dfangzhongdai
Dickeya solani RNS 05 1 2A WP 057083615 1 Dsolani
Dickeya solani RNS 08 23 3 1 A WP 022631632 1 Dsolani
Dickeya solani PPO 9019 WP 022631632 1 Dsolani
Dickeya solani IFB 0099 WP 022631632 1 Dsolani
Dickeya solani IPO 2222 WP 022631632 1 Dsolani
Dickeya solani PPO 9134 WP 022631632 1 Dsolani
Dickeya solani MK16 WP 022631632 1 Dsolani
Dickeya solani MK10 WP 022631632 1 Dsolani
Dickeya solani RNS 07 7 3B WP 022631632 1 Dsolani
Dickeya solani GBBC 2040 WP 022631632 1 Dsolani
Dickeya solani D s0432 1 WP 022631632 1 Dsolani

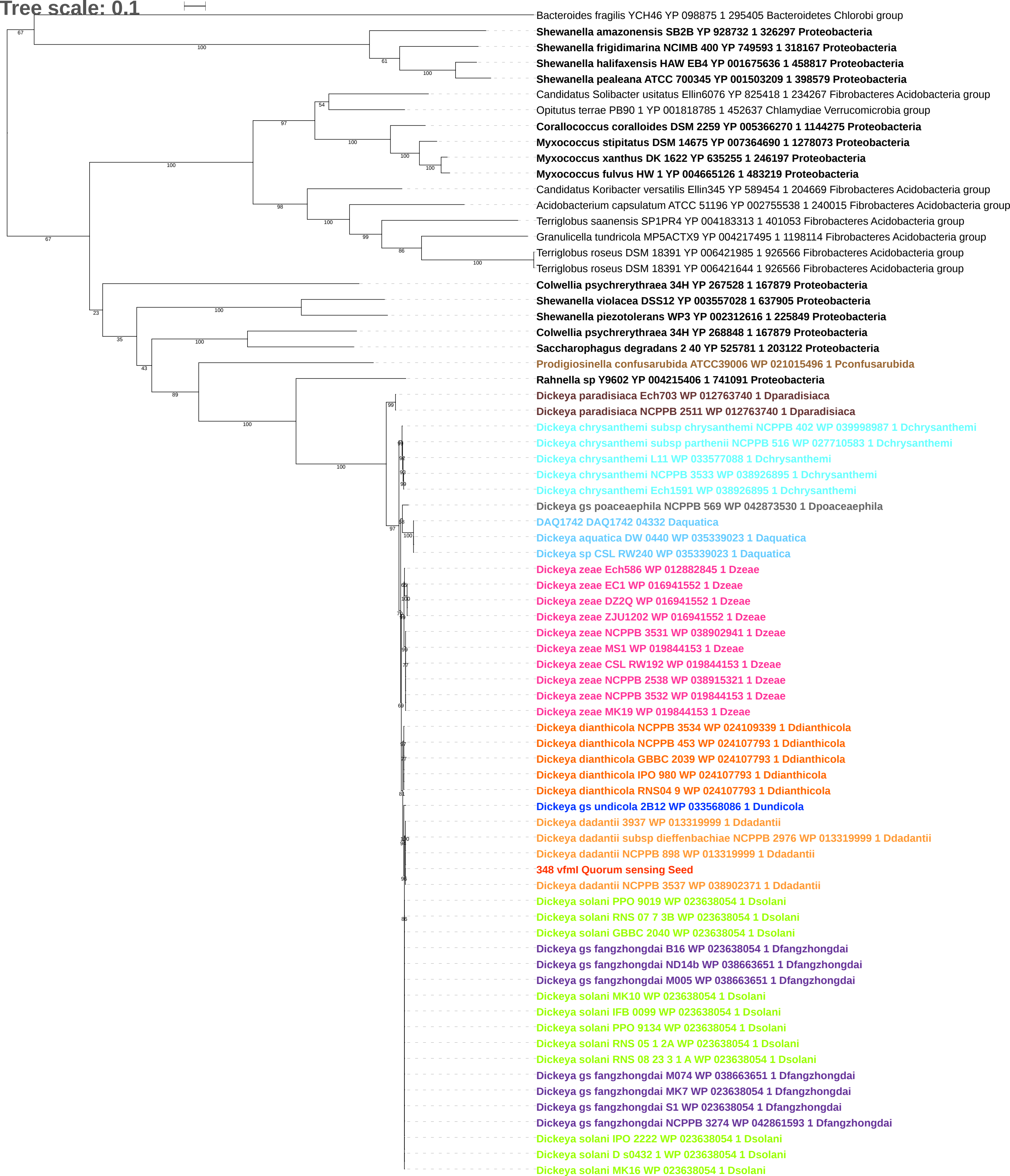
Tree scale: 0.1
Bacteroides fragilis YCH46 YP 098875 1 295405 Bacteroidetes Chlorobi group
Shewanella amazonensis SB2B YP 928732 1 326297 Proteobacteria
Shewanella frigidimarina NCIMB 400 YP 749593 1 318167 Proteobacteria
Shewanella halifaxensis HAW EB4 YP 001675636 1 458817 Proteobacteria
Shewanella pealeana ATCC 700345 YP 001503209 1 398579 Proteobacteria
Candidatus Solibacter usitatus Ellin6076 YP 825418 1 234267 Fibrobacteres Acidobacteria group
Opitutus terrae PB90 1 YP 001818785 1 452637 Chlamydiae Verrucomicrobia group
Corallococcus coralloides DSM 2259 YP 005366270 1 1144275 Proteobacteria
Myxococcus stipitatus DSM 14675 YP 007364690 1 1278073 Proteobacteria
Myxococcus xanthus DK 1622 YP 635255 1 246197 Proteobacteria
Myxococcus fulvus HW 1 YP 004665126 1 483219 Proteobacteria
Candidatus Koribacter versatilis Ellin345 YP 589454 1 204669 Fibrobacteres Acidobacteria group
Acidobacterium capsulatum ATCC 51196 YP 002755538 1 240015 Fibrobacteres Acidobacteria group
Terriglobus saanensis SP1PR4 YP 004183313 1 401053 Fibrobacteres Acidobacteria group
Granulicella tundricola MP5ACTX9 YP 004217495 1 1198114 Fibrobacteres Acidobacteria group
Terriglobus roseus DSM 18391 YP 006421985 1 926566 Fibrobacteres Acidobacteria group
Terriglobus roseus DSM 18391 YP 006421644 1 926566 Fibrobacteres Acidobacteria group
Colwellia psychrerythraea 34H YP 267528 1 167879 Proteobacteria
Shewanella violacea DSS12 YP 003557028 1 637905 Proteobacteria
Shewanella piezotolerans WP3 YP 002312616 1 225849 Proteobacteria
Colwellia psychrerythraea 34H YP 268848 1 167879 Proteobacteria
Saccharophagus degradans 2 40 YP 525781 1 203122 Proteobacteria
Prodigiosinella confusarubida ATCC39006 WP 021015496 1 Pconfusarubida
Rahnella sp Y9602 YP 004215406 1 741091 Proteobacteria
Dickeya paradisiaca Ech703 WP 012763740 1 Dparadisiaca
Dickeya paradisiaca NCPPB 2511 WP 012763740 1 Dparadisiaca
Dickeya chrysanthemi subsp chrysanthemi NCPPB 402 WP 039998987 1 Dchrysanthemi
Dickeya chrysanthemi subsp parthenii NCPPB 516 WP 027710583 1 Dchrysanthemi
Dickeya chrysanthemi L11 WP 033577088 1 Dchrysanthemi
Dickeya chrysanthemi NCPPB 3533 WP 038926895 1 Dchrysanthemi
Dickeya chrysanthemi Ech1591 WP 038926895 1 Dchrysanthemi
Dickeya gs poaceaephila NCPPB 569 WP 042873530 1 Dpoaceaephila
DAQ1742 DAQ1742 04332 Daquatica
Dickeya aquatica DW 0440 WP 035339023 1 Daquatica
Dickeya sp CSL RW240 WP 035339023 1 Daquatica
Dickeya zeae Ech586 WP 012882845 1 Dzeae
Dickeya zeae EC1 WP 016941552 1 Dzeae
Dickeya zeae DZ2Q WP 016941552 1 Dzeae
Dickeya zeae ZJU1202 WP 016941552 1 Dzeae
Dickeya zeae NCPPB 3531 WP 038902941 1 Dzeae
Dickeya zeae MS1 WP 019844153 1 Dzeae
Dickeya zeae CSL RW192 WP 019844153 1 Dzeae
Dickeya zeae NCPPB 2538 WP 038915321 1 Dzeae
Dickeya zeae NCPPB 3532 WP 019844153 1 Dzeae
Dickeya zeae MK19 WP 019844153 1 Dzeae
Dickeya dianthicola NCPPB 3534 WP 024109339 1 Ddianthicola
Dickeya dianthicola NCPPB 453 WP 024107793 1 Ddianthicola
Dickeya dianthicola GBBC 2039 WP 024107793 1 Ddianthicola
Dickeya dianthicola IPO 980 WP 024107793 1 Ddianthicola
Dickeya dianthicola RNS04 9 WP 024107793 1 Ddianthicola
Dickeya gs undicola 2B12 WP 033568086 1 Dundicola
Dickeya dadantii 3937 WP 013319999 1 Ddadantii
Dickeya dadantii subsp dieffenbachiae NCPPB 2976 WP 013319999 1 Ddadantii
Dickeya dadantii NCPPB 898 WP 013319999 1 Ddadantii
348 vfmI Quorum sensing Seed
Dickeya dadantii NCPPB 3537 WP 038902371 1 Ddadantii
Dickeya solani PPO 9019 WP 023638054 1 Dsolani
Dickeya solani RNS 07 7 3B WP 023638054 1 Dsolani
Dickeya solani GBBC 2040 WP 023638054 1 Dsolani
Dickeya gs fangzhongdai B16 WP 023638054 1 Dfangzhongdai
Dickeya gs fangzhongdai ND14b WP 038663651 1 Dfangzhongdai
Dickeya gs fangzhongdai M005 WP 038663651 1 Dfangzhongdai
Dickeya solani MK10 WP 023638054 1 Dsolani
Dickeya solani IFB 0099 WP 023638054 1 Dsolani
Dickeya solani PPO 9134 WP 023638054 1 Dsolani
Dickeya solani RNS 05 1 2A WP 023638054 1 Dsolani
Dickeya solani RNS 08 23 3 1 A WP 023638054 1 Dsolani
Dickeya gs fangzhongdai M074 WP 038663651 1 Dfangzhongdai
Dickeya gs fangzhongdai MK7 WP 023638054 1 Dfangzhongdai
Dickeya gs fangzhongdai S1 WP 023638054 1 Dfangzhongdai
Dickeya gs fangzhongdai NCPPB 3274 WP 042861593 1 Dfangzhongdai
Dickeya solani IPO 2222 WP 023638054 1 Dsolani
Dickeya solani D s0432 1 WP 023638054 1 Dsolani
Dickeya solani MK16 WP 023638054 1 Dsolani

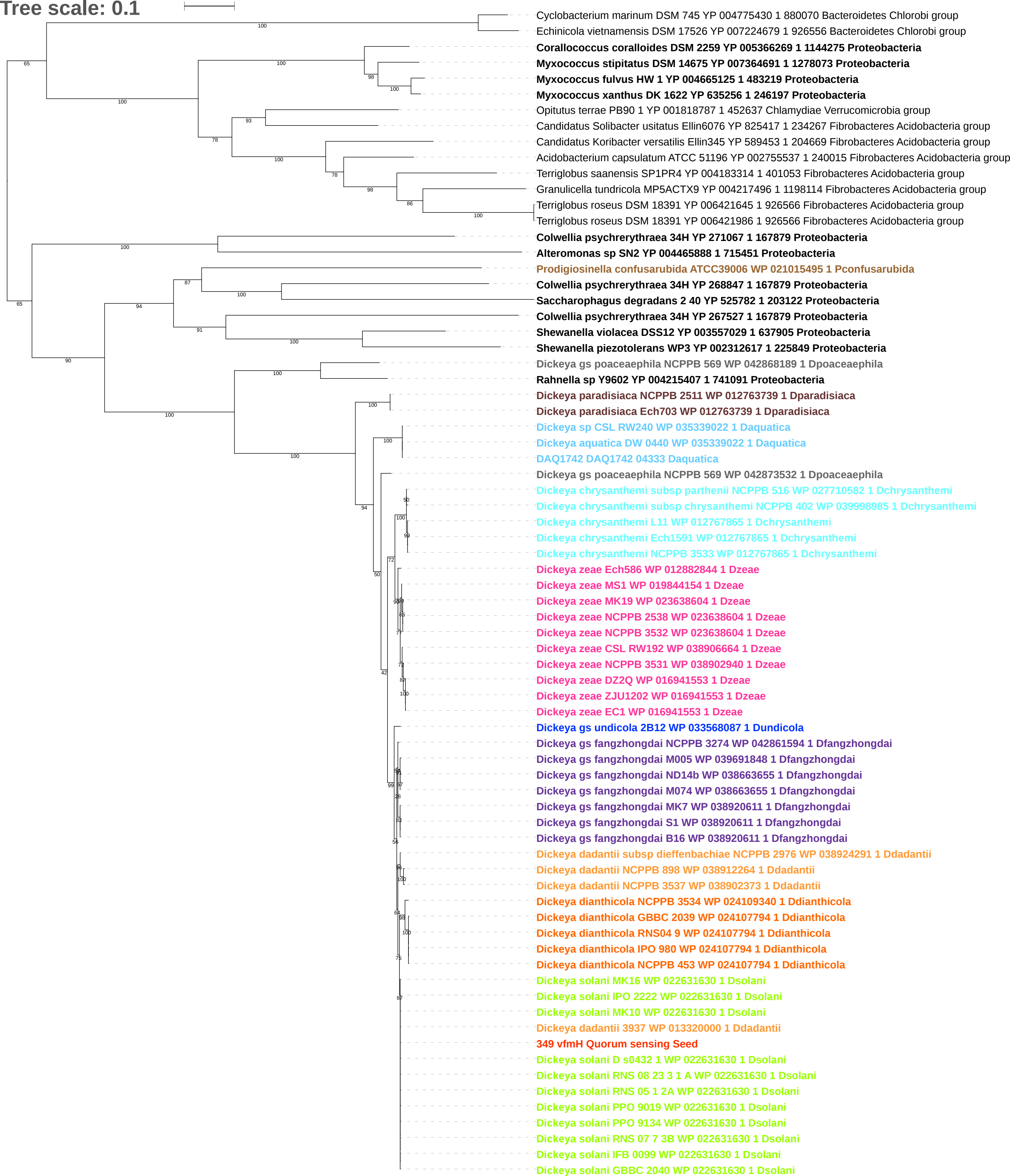
Tree scale: 0.1
Cyclobacterium marinum DSM 745 YP 004775430 1 880070 Bacteroidetes Chlorobi group
Echinicola vietnamensis DSM 17526 YP 007224679 1 926556 Bacteroidetes Chlorobi group
Corallococcus coralloides DSM 2259 YP 005366269 1 1144275 Proteobacteria
Myxococcus stipitatus DSM 14675 YP 007364691 1 1278073 Proteobacteria
Myxococcus fulvus HW 1 YP 004665125 1 483219 Proteobacteria
Myxococcus xanthus DK 1622 YP 635256 1 246197 Proteobacteria
Opitutus terrae PB90 1 YP 001818787 1 452637 Chlamydiae Verrucomicrobia group
Candidatus Solibacter usitatus Ellin6076 YP 825417 1 234267 Fibrobacteres Acidobacteria group
Candidatus Koribacter versatilis Ellin345 YP 589453 1 204669 Fibrobacteres Acidobacteria group
Acidobacterium capsulatum ATCC 51196 YP 002755537 1 240015 Fibrobacteres Acidobacteria group
Terriglobus saanensis SP1PR4 YP 004183314 1 401053 Fibrobacteres Acidobacteria group
Granulicella tundricola MP5ACTX9 YP 004217496 1 1198114 Fibrobacteres Acidobacteria group
Terriglobus roseus DSM 18391 YP 006421645 1 926566 Fibrobacteres Acidobacteria group
Terriglobus roseus DSM 18391 YP 006421986 1 926566 Fibrobacteres Acidobacteria group
Colwellia psychrerythraea 34H YP 271067 1 167879 Proteobacteria
Alteromonas sp SN2 YP 004465888 1 715451 Proteobacteria
Prodigiosinella confusarubida ATCC39006 WP 021015495 1 Pconfusarubida
Colwellia psychrerythraea 34H YP 268847 1 167879 Proteobacteria
Saccharophagus degradans 2 40 YP 525782 1 203122 Proteobacteria
Colwellia psychrerythraea 34H YP 267527 1 167879 Proteobacteria
Shewanella violacea DSS12 YP 003557029 1 637905 Proteobacteria
Shewanella piezotolerans WP3 YP 002312617 1 225849 Proteobacteria
Dickeya gs poaceaephila NCPPB 569 WP 042868189 1 Dpoaceaephila
Rahnella sp Y9602 YP 004215407 1 741091 Proteobacteria
Dickeya paradisiaca NCPPB 2511 WP 012763739 1 Dparadisiaca
Dickeya paradisiaca Ech703 WP 012763739 1 Dparadisiaca
Dickeya sp CSL RW240 WP 035339022 1 Daquatica
Dickeya aquatica DW 0440 WP 035339022 1 Daquatica
DAQ1742 DAQ1742 04333 Daquatica
Dickeya gs poaceaephila NCPPB 569 WP 042873532 1 Dpoaceaephila
Dickeya chrysanthemi subsp parthenii NCPPB 516 WP 027710582 1 Dchrysanthemi
Dickeya chrysanthemi subsp chrysanthemi NCPPB 402 WP 039998985 1 Dchrysanthemi
Dickeya chrysanthemi L11 WP 012767865 1 Dchrysanthemi
Dickeya chrysanthemi Ech1591 WP 012767865 1 Dchrysanthemi
Dickeya chrysanthemi NCPPB 3533 WP 012767865 1 Dchrysanthemi
Dickeya zeae Ech586 WP 012882844 1 Dzeae
Dickeya zeae MS1 WP 019844154 1 Dzeae
Dickeya zeae MK19 WP 023638604 1 Dzeae
Dickeya zeae NCPPB 2538 WP 023638604 1 Dzeae
Dickeya zeae NCPPB 3532 WP 023638604 1 Dzeae
Dickeya zeae CSL RW192 WP 038906664 1 Dzeae
Dickeya zeae NCPPB 3531 WP 038902940 1 Dzeae
Dickeya zeae DZ2Q WP 016941553 1 Dzeae
Dickeya zeae ZJU1202 WP 016941553 1 Dzeae
Dickeya zeae EC1 WP 016941553 1 Dzeae
Dickeya gs undicola 2B12 WP 033568087 1 Dundicola
Dickeya gs fangzhongdai NCPPB 3274 WP 042861594 1 Dfangzhongdai
Dickeya gs fangzhongdai M005 WP 039691848 1 Dfangzhongdai
Dickeya gs fangzhongdai ND14b WP 038663655 1 Dfangzhongdai
Dickeya gs fangzhongdai M074 WP 038663655 1 Dfangzhongdai
Dickeya gs fangzhongdai MK7 WP 038920611 1 Dfangzhongdai
Dickeya gs fangzhongdai S1 WP 038920611 1 Dfangzhongdai
Dickeya gs fangzhongdai B16 WP 038920611 1 Dfangzhongdai
Dickeya dadantii subsp dieffenbachiae NCPPB 2976 WP 038924291 1 Ddadantii
Dickeya dadantii NCPPB 898 WP 038912264 1 Ddadantii
Dickeya dadantii NCPPB 3537 WP 038902373 1 Ddadantii
Dickeya dianthicola NCPPB 3534 WP 024109340 1 Ddianthicola
Dickeya dianthicola GBBC 2039 WP 024107794 1 Ddianthicola
Dickeya dianthicola RNS04 9 WP 024107794 1 Ddianthicola
Dickeya dianthicola IPO 980 WP 024107794 1 Ddianthicola
Dickeya dianthicola NCPPB 453 WP 024107794 1 Ddianthicola
Dickeya solani MK16 WP 022631630 1 Dsolani
Dickeya solani IPO 2222 WP 022631630 1 Dsolani
Dickeya solani MK10 WP 022631630 1 Dsolani
Dickeya dadantii 3937 WP 013320000 1 Ddadantii
349 vfmH Quorum sensing Seed
Dickeya solani D s0432 1 WP 022631630 1 Dsolani
Dickeya solani RNS 08 23 3 1 A WP 022631630 1 Dsolani
Dickeya solani RNS 05 1 2A WP 022631630 1 Dsolani
Dickeya solani PPO 9019 WP 022631630 1 Dsolani
Dickeya solani PPO 9134 WP 022631630 1 Dsolani
Dickeya solani RNS 07 7 3B WP 022631630 1 Dsolani
Dickeya solani IFB 0099 WP 022631630 1 Dsolani
Dickeya solani GBBC 2040 WP 022631630 1 Dsolani

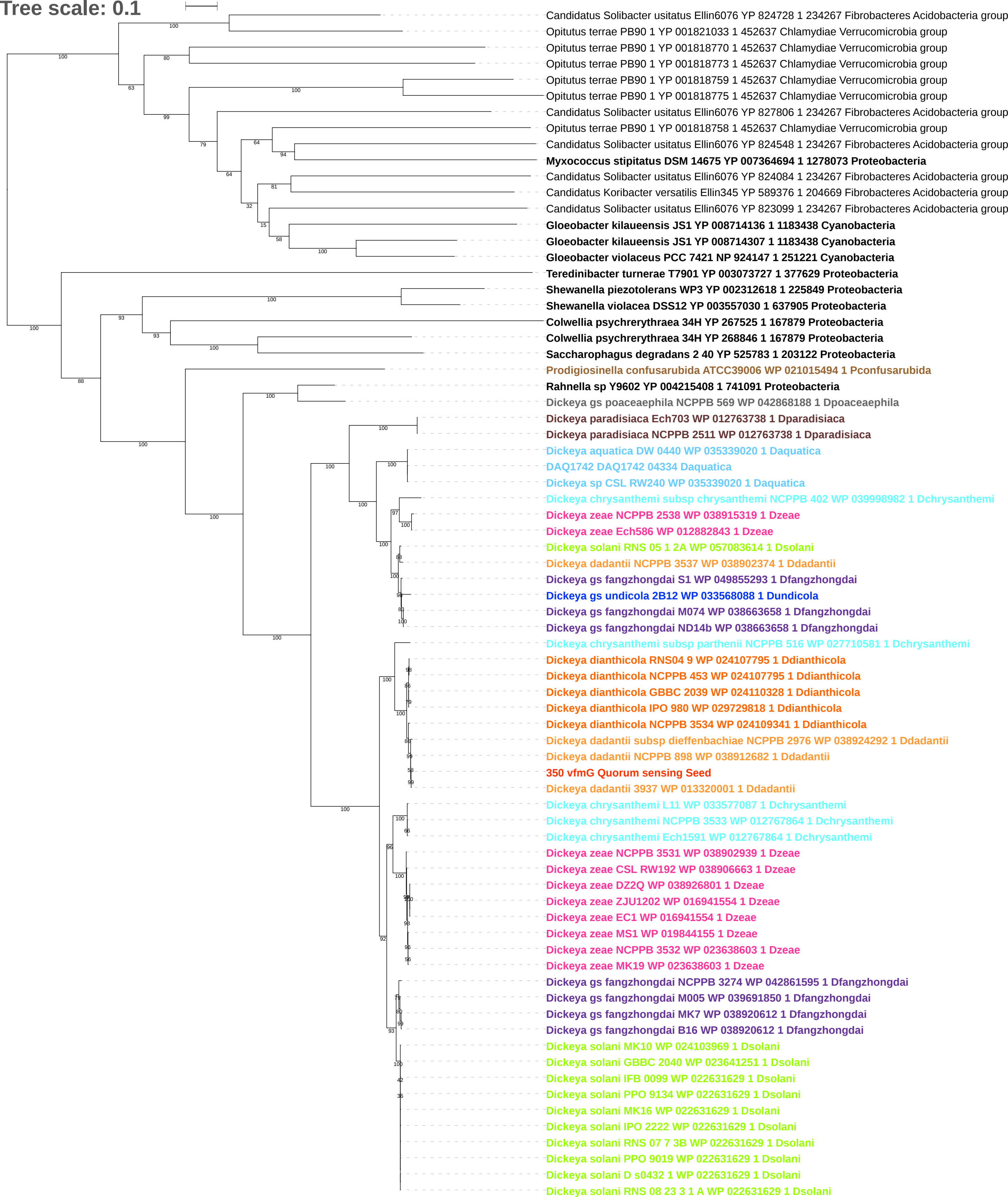

Tree scale: 0.1
Candidatus Solibacter usitatus Ellin6076 YP 824728 1 234267 Fibrobacteres Acidobacteria group
Opitutus terrae PB90 1 YP 001821033 1 452637 Chlamydiae Verrucomicrobia group
Opitutus terrae PB90 1 YP 001818770 1 452637 Chlamydiae Verrucomicrobia group
Opitutus terrae PB90 1 YP 001818773 1 452637 Chlamydiae Verrucomicrobia group
Opitutus terrae PB90 1 YP 001818759 1 452637 Chlamydiae Verrucomicrobia group
Opitutus terrae PB90 1 YP 001818775 1 452637 Chlamydiae Verrucomicrobia group
Candidatus Solibacter usitatus Ellin6076 YP 827806 1 234267 Fibrobacteres Acidobacteria group
Opitutus terrae PB90 1 YP 001818758 1 452637 Chlamydiae Verrucomicrobia group
Candidatus Solibacter usitatus Ellin6076 YP 824548 1 234267 Fibrobacteres Acidobacteria group
Myxococcus stipitatus DSM 14675 YP 007364694 1 1278073 Proteobacteria
Candidatus Solibacter usitatus Ellin6076 YP 824084 1 234267 Fibrobacteres Acidobacteria group
Candidatus Koribacter versatilis Ellin345 YP 589376 1 204669 Fibrobacteres Acidobacteria group
Candidatus Solibacter usitatus Ellin6076 YP 823099 1 234267 Fibrobacteres Acidobacteria group
Gloeobacter kilaueensis JS1 YP 008714136 1 1183438 Cyanobacteria
Gloeobacter kilaueensis JS1 YP 008714307 1 1183438 Cyanobacteria
Gloeobacter violaceus PCC 7421 NP 924147 1 251221 Cyanobacteria
Teredinibacter turnerae T7901 YP 003073727 1 377629 Proteobacteria
Shewanella piezotolerans WP3 YP 002312618 1 225849 Proteobacteria
Shewanella violacea DSS12 YP 003557030 1 637905 Proteobacteria
Colwellia psychrerythraea 34H YP 267525 1 167879 Proteobacteria
Colwellia psychrerythraea 34H YP 268846 1 167879 Proteobacteria
Saccharophagus degradans 2 40 YP 525783 1 203122 Proteobacteria
Prodigiosinella confusarubida ATCC39006 WP 021015494 1 Pconfusarubida
Rahnella sp Y9602 YP 004215408 1 741091 Proteobacteria
Dickeya gs poaceaephila NCPPB 569 WP 042868188 1 Dpoaceaephila
Dickeya paradisiaca Ech703 WP 012763738 1 Dparadisiaca
Dickeya paradisiaca NCPPB 2511 WP 012763738 1 Dparadisiaca
Dickeya aquatica DW 0440 WP 035339020 1 Daquatica
DAQ1742 DAQ1742 04334 Daquatica
Dickeya sp CSL RW240 WP 035339020 1 Daquatica
Dickeya chrysanthemi subsp chrysanthemi NCPPB 402 WP 039998982 1 Dchrysanthemi
Dickeya zeae NCPPB 2538 WP 038915319 1 Dzeae
Dickeya zeae Ech586 WP 012882843 1 Dzeae
Dickeya solani RNS 05 1 2A WP 057083614 1 Dsolani
Dickeya dadantii NCPPB 3537 WP 038902374 1 Ddadantii
Dickeya gs fangzhongdai S1 WP 049855293 1 Dfangzhongdai
Dickeya gs undicola 2B12 WP 033568088 1 Dundicola
Dickeya gs fangzhongdai M074 WP 038663658 1 Dfangzhongdai
Dickeya gs fangzhongdai ND14b WP 038663658 1 Dfangzhongdai
Dickeya chrysanthemi subsp parthenii NCPPB 516 WP 027710581 1 Dchrysanthemi
Dickeya dianthicola RNS04 9 WP 024107795 1 Ddianthicola
Dickeya dianthicola NCPPB 453 WP 024107795 1 Ddianthicola
Dickeya dianthicola GBBC 2039 WP 024110328 1 Ddianthicola
Dickeya dianthicola IPO 980 WP 029729818 1 Ddianthicola
Dickeya dianthicola NCPPB 3534 WP 024109341 1 Ddianthicola
Dickeya dadantii subsp dieffenbachiae NCPPB 2976 WP 038924292 1 Ddadantii
Dickeya dadantii NCPPB 898 WP 038912682 1 Ddadantii
350 vfmG Quorum sensing Seed
Dickeya dadantii 3937 WP 013320001 1 Ddadantii
Dickeya chrysanthemi L11 WP 033577087 1 Dchrysanthemi
Dickeya chrysanthemi NCPPB 3533 WP 012767864 1 Dchrysanthemi
Dickeya chrysanthemi Ech1591 WP 012767864 1 Dchrysanthemi
Dickeya zeae NCPPB 3531 WP 038902939 1 Dzeae
Dickeya zeae CSL RW192 WP 038906663 1 Dzeae
Dickeya zeae DZ2Q WP 038926801 1 Dzeae
Dickeya zeae ZJU1202 WP 016941554 1 Dzeae
Dickeya zeae EC1 WP 016941554 1 Dzeae
Dickeya zeae MS1 WP 019844155 1 Dzeae
Dickeya zeae NCPPB 3532 WP 023638603 1 Dzeae
Dickeya zeae MK19 WP 023638603 1 Dzeae
Dickeya gs fangzhongdai NCPPB 3274 WP 042861595 1 Dfangzhongdai
Dickeya gs fangzhongdai M005 WP 039691850 1 Dfangzhongdai
Dickeya gs fangzhongdai MK7 WP 038920612 1 Dfangzhongdai
Dickeya gs fangzhongdai B16 WP 038920612 1 Dfangzhongdai
Dickeya solani MK10 WP 024103969 1 Dsolani
Dickeya solani GBBC 2040 WP 023641251 1 Dsolani
Dickeya solani IFB 0099 WP 022631629 1 Dsolani
Dickeya solani PPO 9134 WP 022631629 1 Dsolani
Dickeya solani MK16 WP 022631629 1 Dsolani
Dickeya solani IPO 2222 WP 022631629 1 Dsolani
Dickeya solani RNS 07 7 3B WP 022631629 1 Dsolani
Dickeya solani PPO 9019 WP 022631629 1 Dsolani
Dickeya solani D s0432 1 WP 022631629 1 Dsolani
Dickeya solani RNS 08 23 3 1 A WP 022631629 1 Dsolani

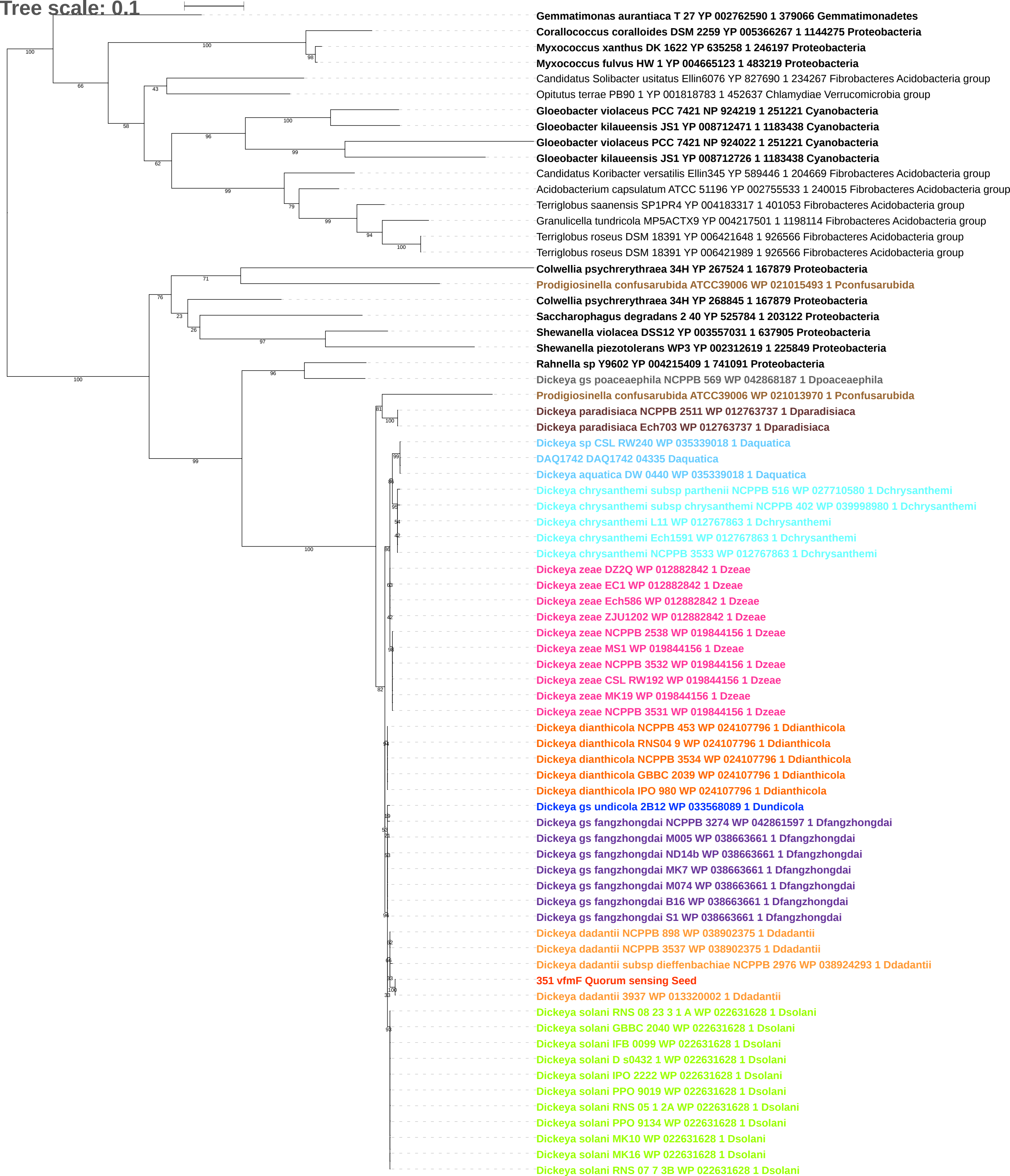
Tree scale: 0.1
Gemmatimonas aurantiaca T 27 YP 002762590 1 379066 Gemmatimonadetes
Corallococcus coralloides DSM 2259 YP 005366267 1 1144275 Proteobacteria
Myxococcus xanthus DK 1622 YP 635258 1 246197 Proteobacteria
Myxococcus fulvus HW 1 YP 004665123 1 483219 Proteobacteria
Candidatus Solibacter usitatus Ellin6076 YP 827690 1 234267 Fibrobacteres Acidobacteria group
Opitutus terrae PB90 1 YP 001818783 1 452637 Chlamydiae Verrucomicrobia group
Gloeobacter violaceus PCC 7421 NP 924219 1 251221 Cyanobacteria
Gloeobacter kilaueensis JS1 YP 008712471 1 1183438 Cyanobacteria
Gloeobacter violaceus PCC 7421 NP 924022 1 251221 Cyanobacteria
Gloeobacter kilaueensis JS1 YP 008712726 1 1183438 Cyanobacteria
Candidatus Koribacter versatilis Ellin345 YP 589446 1 204669 Fibrobacteres Acidobacteria group
Acidobacterium capsulatum ATCC 51196 YP 002755533 1 240015 Fibrobacteres Acidobacteria group
Terriglobus saanensis SP1PR4 YP 004183317 1 401053 Fibrobacteres Acidobacteria group
Granulicella tundricola MP5ACTX9 YP 004217501 1 1198114 Fibrobacteres Acidobacteria group
Terriglobus roseus DSM 18391 YP 006421648 1 926566 Fibrobacteres Acidobacteria group
Terriglobus roseus DSM 18391 YP 006421989 1 926566 Fibrobacteres Acidobacteria group
Colwellia psychrerythraea 34H YP 267524 1 167879 Proteobacteria
Prodigiosinella confusarubida ATCC39006 WP 021015493 1 Pconfusarubida
Colwellia psychrerythraea 34H YP 268845 1 167879 Proteobacteria
Saccharophagus degradans 2 40 YP 525784 1 203122 Proteobacteria
Shewanella violacea DSS12 YP 003557031 1 637905 Proteobacteria
Shewanella piezotolerans WP3 YP 002312619 1 225849 Proteobacteria
Rahnella sp Y9602 YP 004215409 1 741091 Proteobacteria
Dickeya gs poaceaephila NCPPB 569 WP 042868187 1 Dpoaceaephila
Prodigiosinella confusarubida ATCC39006 WP 021013970 1 Pconfusarubida
Dickeya paradisiaca NCPPB 2511 WP 012763737 1 Dparadisiaca
Dickeya paradisiaca Ech703 WP 012763737 1 Dparadisiaca
Dickeya sp CSL RW240 WP 035339018 1 Daquatica
DAQ1742 DAQ1742 04335 Daquatica
Dickeya aquatica DW 0440 WP 035339018 1 Daquatica
Dickeya chrysanthemi subsp parthenii NCPPB 516 WP 027710580 1 Dchrysanthemi
Dickeya chrysanthemi subsp chrysanthemi NCPPB 402 WP 039998980 1 Dchrysanthemi
Dickeya chrysanthemi L11 WP 012767863 1 Dchrysanthemi
Dickeya chrysanthemi Ech1591 WP 012767863 1 Dchrysanthemi
Dickeya chrysanthemi NCPPB 3533 WP 012767863 1 Dchrysanthemi
Dickeya zeae DZ2Q WP 012882842 1 Dzeae
Dickeya zeae EC1 WP 012882842 1 Dzeae
Dickeya zeae Ech586 WP 012882842 1 Dzeae
Dickeya zeae ZJU1202 WP 012882842 1 Dzeae
Dickeya zeae NCPPB 2538 WP 019844156 1 Dzeae
Dickeya zeae MS1 WP 019844156 1 Dzeae
Dickeya zeae NCPPB 3532 WP 019844156 1 Dzeae
Dickeya zeae CSL RW192 WP 019844156 1 Dzeae
Dickeya zeae MK19 WP 019844156 1 Dzeae
Dickeya zeae NCPPB 3531 WP 019844156 1 Dzeae
Dickeya dianthicola NCPPB 453 WP 024107796 1 Ddianthicola
Dickeya dianthicola RNS04 9 WP 024107796 1 Ddianthicola
Dickeya dianthicola NCPPB 3534 WP 024107796 1 Ddianthicola
Dickeya dianthicola GBBC 2039 WP 024107796 1 Ddianthicola
Dickeya dianthicola IPO 980 WP 024107796 1 Ddianthicola
Dickeya gs undicola 2B12 WP 033568089 1 Dundicola
Dickeya gs fangzhongdai NCPPB 3274 WP 042861597 1 Dfangzhongdai
Dickeya gs fangzhongdai M005 WP 038663661 1 Dfangzhongdai
Dickeya gs fangzhongdai ND14b WP 038663661 1 Dfangzhongdai
Dickeya gs fangzhongdai MK7 WP 038663661 1 Dfangzhongdai
Dickeya gs fangzhongdai M074 WP 038663661 1 Dfangzhongdai
Dickeya gs fangzhongdai B16 WP 038663661 1 Dfangzhongdai
Dickeya gs fangzhongdai S1 WP 038663661 1 Dfangzhongdai
Dickeya dadantii NCPPB 898 WP 038902375 1 Ddadantii
Dickeya dadantii NCPPB 3537 WP 038902375 1 Ddadantii
Dickeya dadantii subsp dieffenbachiae NCPPB 2976 WP 038924293 1 Ddadantii
351 vfmF Quorum sensing Seed
Dickeya dadantii 3937 WP 013320002 1 Ddadantii
Dickeya solani RNS 08 23 3 1 A WP 022631628 1 Dsolani
Dickeya solani GBBC 2040 WP 022631628 1 Dsolani
Dickeya solani IFB 0099 WP 022631628 1 Dsolani
Dickeya solani D s0432 1 WP 022631628 1 Dsolani
Dickeya solani IPO 2222 WP 022631628 1 Dsolani
Dickeya solani PPO 9019 WP 022631628 1 Dsolani
Dickeya solani RNS 05 1 2A WP 022631628 1 Dsolani
Dickeya solani PPO 9134 WP 022631628 1 Dsolani
Dickeya solani MK10 WP 022631628 1 Dsolani
Dickeya solani MK16 WP 022631628 1 Dsolani
Dickeya solani RNS 07 7 3B WP 022631628 1 Dsolani

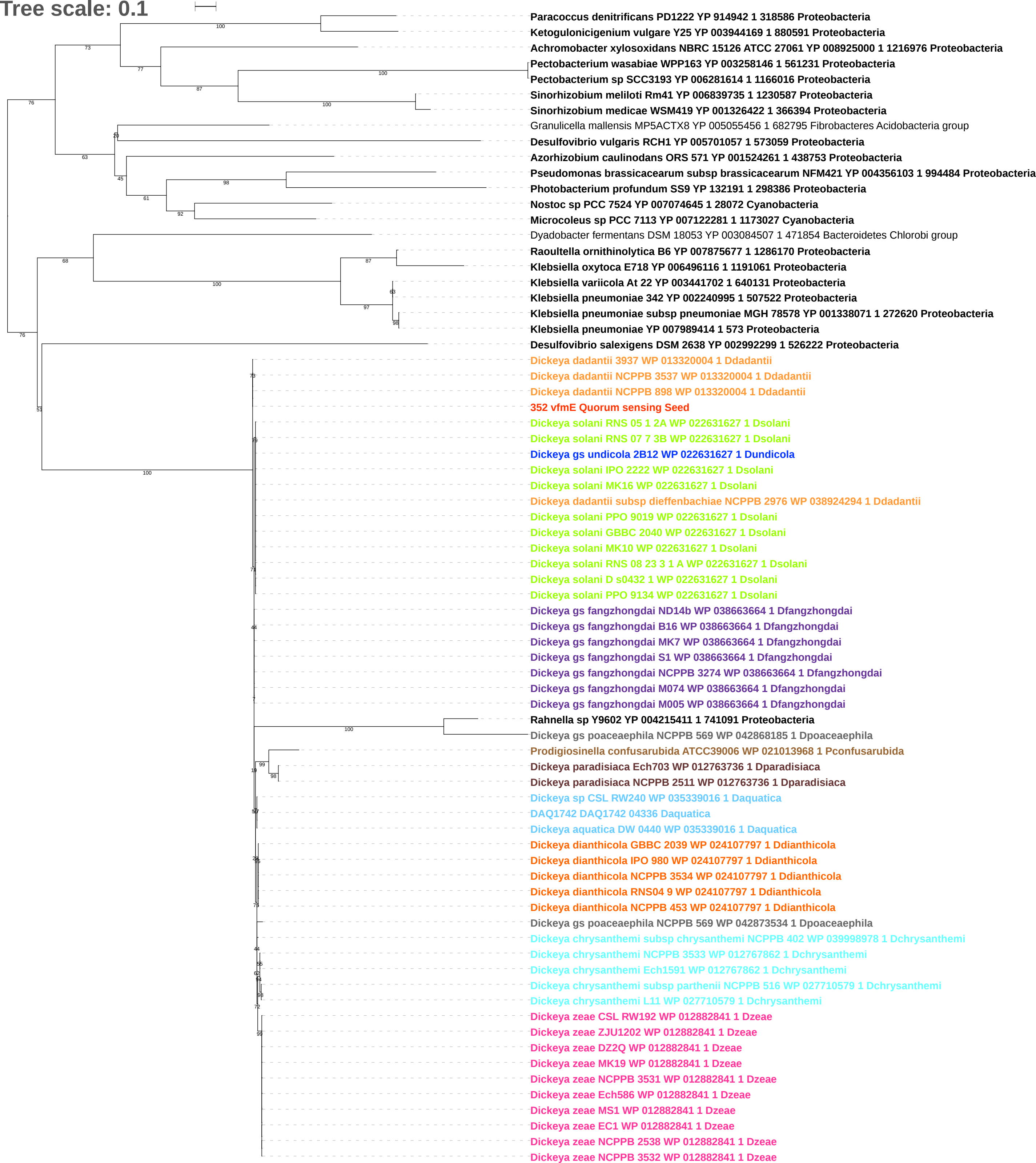

Tree scale: 0.1
Paracoccus denitrificans PD1222 YP 914942 1 318586 Proteobacteria
Ketogulonicigenium vulgare Y25 YP 003944169 1 880591 Proteobacteria
Achromobacter xylosoxidans NBRC 15126 ATCC 27061 YP 008925000 1 1216976 Proteobacteria
Pectobacterium wasabiae WPP163 YP 003258146 1 561231 Proteobacteria
Pectobacterium sp SCC3193 YP 006281614 1 1166016 Proteobacteria
Sinorhizobium meliloti Rm41 YP 006839735 1 1230587 Proteobacteria
Sinorhizobium medicae WSM419 YP 001326422 1 366394 Proteobacteria
Granulicella mallensis MP5ACTX8 YP 005055456 1 682795 Fibrobacteres Acidobacteria group
Desulfovibrio vulgaris RCH1 YP 005701057 1 573059 Proteobacteria
Azorhizobium caulinodans ORS 571 YP 001524261 1 438753 Proteobacteria
Pseudomonas brassicacearum subsp brassicacearum NFM421 YP 004356103 1 994484 Proteobacteria
Photobacterium profundum SS9 YP 132191 1 298386 Proteobacteria
Nostoc sp PCC 7524 YP 007074645 1 28072 Cyanobacteria
Microcoleus sp PCC 7113 YP 007122281 1 1173027 Cyanobacteria
Dyadobacter fermentans DSM 18053 YP 003084507 1 471854 Bacteroidetes Chlorobi group
Raoultella ornithinolytica B6 YP 007875677 1 1286170 Proteobacteria
Klebsiella oxytoca E718 YP 006496116 1 1191061 Proteobacteria
Klebsiella variicola At 22 YP 003441702 1 640131 Proteobacteria
Klebsiella pneumoniae 342 YP 002240995 1 507522 Proteobacteria
Klebsiella pneumoniae subsp pneumoniae MGH 78578 YP 001338071 1 272620 Proteobacteria
Klebsiella pneumoniae YP 007989414 1 573 Proteobacteria
Desulfovibrio salexigens DSM 2638 YP 002992299 1 526222 Proteobacteria
Dickeya dadantii 3937 WP 013320004 1 Ddadantii
Dickeya dadantii NCPPB 3537 WP 013320004 1 Ddadantii
Dickeya dadantii NCPPB 898 WP 013320004 1 Ddadantii
352 vfmE Quorum sensing Seed
Dickeya solani RNS 05 1 2A WP 022631627 1 Dsolani
Dickeya solani RNS 07 7 3B WP 022631627 1 Dsolani
Dickeya gs undicola 2B12 WP 022631627 1 Dundicola
Dickeya solani IPO 2222 WP 022631627 1 Dsolani
Dickeya solani MK16 WP 022631627 1 Dsolani
Dickeya dadantii subsp dieffenbachiae NCPPB 2976 WP 038924294 1 Ddadantii
Dickeya solani PPO 9019 WP 022631627 1 Dsolani
Dickeya solani GBBC 2040 WP 022631627 1 Dsolani
Dickeya solani MK10 WP 022631627 1 Dsolani
Dickeya solani RNS 08 23 3 1 A WP 022631627 1 Dsolani
Dickeya solani D s0432 1 WP 022631627 1 Dsolani
Dickeya solani PPO 9134 WP 022631627 1 Dsolani
Dickeya gs fangzhongdai ND14b WP 038663664 1 Dfangzhongdai
Dickeya gs fangzhongdai B16 WP 038663664 1 Dfangzhongdai
Dickeya gs fangzhongdai MK7 WP 038663664 1 Dfangzhongdai
Dickeya gs fangzhongdai S1 WP 038663664 1 Dfangzhongdai
Dickeya gs fangzhongdai NCPPB 3274 WP 038663664 1 Dfangzhongdai
Dickeya gs fangzhongdai M074 WP 038663664 1 Dfangzhongdai
Dickeya gs fangzhongdai M005 WP 038663664 1 Dfangzhongdai
Rahnella sp Y9602 YP 004215411 1 741091 Proteobacteria
Dickeya gs poaceaephila NCPPB 569 WP 042868185 1 Dpoaceaephila
Prodigiosinella confusarubida ATCC39006 WP 021013968 1 Pconfusarubida
Dickeya paradisiaca Ech703 WP 012763736 1 Dparadisiaca
Dickeya paradisiaca NCPPB 2511 WP 012763736 1 Dparadisiaca
Dickeya sp CSL RW240 WP 035339016 1 Daquatica
DAQ1742 DAQ1742 04336 Daquatica
Dickeya aquatica DW 0440 WP 035339016 1 Daquatica
Dickeya dianthicola GBBC 2039 WP 024107797 1 Ddianthicola
Dickeya dianthicola IPO 980 WP 024107797 1 Ddianthicola
Dickeya dianthicola NCPPB 3534 WP 024107797 1 Ddianthicola
Dickeya dianthicola RNS04 9 WP 024107797 1 Ddianthicola
Dickeya dianthicola NCPPB 453 WP 024107797 1 Ddianthicola
Dickeya gs poaceaephila NCPPB 569 WP 042873534 1 Dpoaceaephila
Dickeya chrysanthemi subsp chrysanthemi NCPPB 402 WP 039998978 1 Dchrysanthemi
Dickeya chrysanthemi NCPPB 3533 WP 012767862 1 Dchrysanthemi
Dickeya chrysanthemi Ech1591 WP 012767862 1 Dchrysanthemi
Dickeya chrysanthemi subsp parthenii NCPPB 516 WP 027710579 1 Dchrysanthemi
Dickeya chrysanthemi L11 WP 027710579 1 Dchrysanthemi
Dickeya zeae CSL RW192 WP 012882841 1 Dzeae
Dickeya zeae ZJU1202 WP 012882841 1 Dzeae
Dickeya zeae DZ2Q WP 012882841 1 Dzeae
Dickeya zeae MK19 WP 012882841 1 Dzeae
Dickeya zeae NCPPB 3531 WP 012882841 1 Dzeae
Dickeya zeae Ech586 WP 012882841 1 Dzeae
Dickeya zeae MS1 WP 012882841 1 Dzeae
Dickeya zeae EC1 WP 012882841 1 Dzeae
Dickeya zeae NCPPB 2538 WP 012882841 1 Dzeae
Dickeya zeae NCPPB 3532 WP 012882841 1 Dzeae

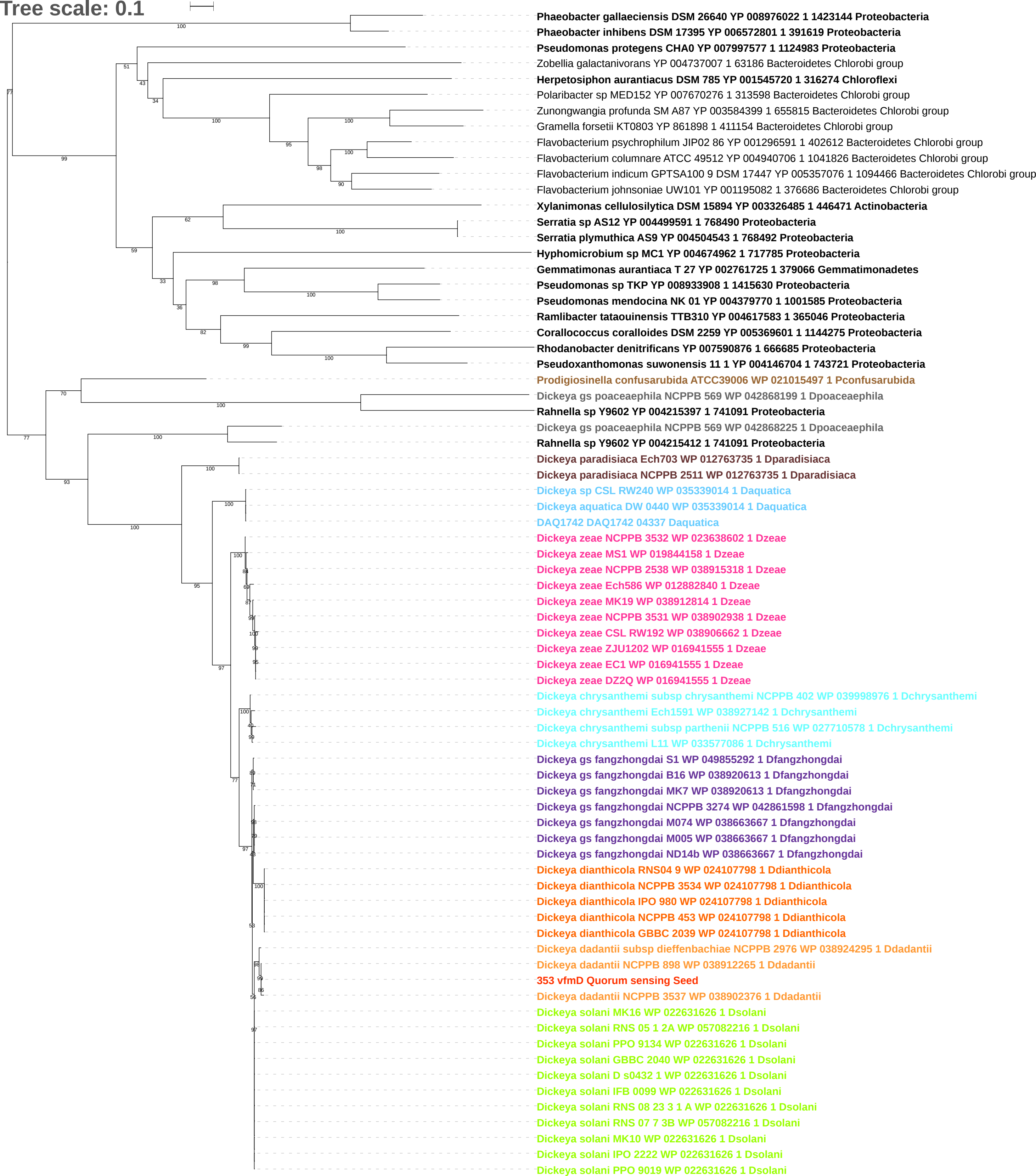
Tree scale: 0.1
Phaeobacter gallaeciensis DSM 26640 YP 008976022 1 1423144 Proteobacteria
Phaeobacter inhibens DSM 17395 YP 006572801 1 391619 Proteobacteria
Pseudomonas protegens CHA0 YP 007997577 1 1124983 Proteobacteria
Zobellia galactanivorans YP 004737007 1 63186 Bacteroidetes Chlorobi group
Herpetosiphon aurantiacus DSM 785 YP 001545720 1 316274 Chloroflexi
Polaribacter sp MED152 YP 007670276 1 313598 Bacteroidetes Chlorobi group
Zunongwangia profunda SM A87 YP 003584399 1 655815 Bacteroidetes Chlorobi group
Gramella forsetii KT0803 YP 861898 1 411154 Bacteroidetes Chlorobi group
Flavobacterium psychrophilum JIP02 86 YP 001296591 1 402612 Bacteroidetes Chlorobi group
Flavobacterium columnare ATCC 49512 YP 004940706 1 1041826 Bacteroidetes Chlorobi group
Flavobacterium indicum GPTSA100 9 DSM 17447 YP 005357076 1 1094466 Bacteroidetes Chlorobi group
Flavobacterium johnsoniae UW101 YP 001195082 1 376686 Bacteroidetes Chlorobi group
Xylanimonas cellulosilytica DSM 15894 YP 003326485 1 446471 Actinobacteria
Serratia sp AS12 YP 004499591 1 768490 Proteobacteria
Serratia plymuthica AS9 YP 004504543 1 768492 Proteobacteria
Hyphomicrobium sp MC1 YP 004674962 1 717785 Proteobacteria
Gemmatimonas aurantiaca T 27 YP 002761725 1 379066 Gemmatimonadetes
Pseudomonas sp TKP YP 008933908 1 1415630 Proteobacteria
Pseudomonas mendocina NK 01 YP 004379770 1 1001585 Proteobacteria
Ramlibacter tataouinensis TTB310 YP 004617583 1 365046 Proteobacteria
Corallococcus coralloides DSM 2259 YP 005369601 1 1144275 Proteobacteria
Rhodanobacter denitrificans YP 007590876 1 666685 Proteobacteria
Pseudoxanthomonas suwonensis 11 1 YP 004146704 1 743721 Proteobacteria
Prodigiosinella confusarubida ATCC39006 WP 021015497 1 Pconfusarubida
Dickeya gs poaceaephila NCPPB 569 WP 042868199 1 Dpoaceaephila
Rahnella sp Y9602 YP 004215397 1 741091 Proteobacteria
Dickeya gs poaceaephila NCPPB 569 WP 042868225 1 Dpoaceaephila
Rahnella sp Y9602 YP 004215412 1 741091 Proteobacteria
Dickeya paradisiaca Ech703 WP 012763735 1 Dparadisiaca
Dickeya paradisiaca NCPPB 2511 WP 012763735 1 Dparadisiaca
Dickeya sp CSL RW240 WP 035339014 1 Daquatica
Dickeya aquatica DW 0440 WP 035339014 1 Daquatica
DAQ1742 DAQ1742 04337 Daquatica
Dickeya zeae NCPPB 3532 WP 023638602 1 Dzeae
Dickeya zeae MS1 WP 019844158 1 Dzeae
Dickeya zeae NCPPB 2538 WP 038915318 1 Dzeae
Dickeya zeae Ech586 WP 012882840 1 Dzeae
Dickeya zeae MK19 WP 038912814 1 Dzeae
Dickeya zeae NCPPB 3531 WP 038902938 1 Dzeae
Dickeya zeae CSL RW192 WP 038906662 1 Dzeae
Dickeya zeae ZJU1202 WP 016941555 1 Dzeae
Dickeya zeae EC1 WP 016941555 1 Dzeae
Dickeya zeae DZ2Q WP 016941555 1 Dzeae
Dickeya chrysanthemi subsp chrysanthemi NCPPB 402 WP 039998976 1 Dchrysanthemi
Dickeya chrysanthemi Ech1591 WP 038927142 1 Dchrysanthemi
Dickeya chrysanthemi subsp parthenii NCPPB 516 WP 027710578 1 Dchrysanthemi
Dickeya chrysanthemi L11 WP 033577086 1 Dchrysanthemi
Dickeya gs fangzhongdai S1 WP 049855292 1 Dfangzhongdai
Dickeya gs fangzhongdai B16 WP 038920613 1 Dfangzhongdai
Dickeya gs fangzhongdai MK7 WP 038920613 1 Dfangzhongdai
Dickeya gs fangzhongdai NCPPB 3274 WP 042861598 1 Dfangzhongdai
Dickeya gs fangzhongdai M074 WP 038663667 1 Dfangzhongdai
Dickeya gs fangzhongdai M005 WP 038663667 1 Dfangzhongdai
Dickeya gs fangzhongdai ND14b WP 038663667 1 Dfangzhongdai
Dickeya dianthicola RNS04 9 WP 024107798 1 Ddianthicola
Dickeya dianthicola NCPPB 3534 WP 024107798 1 Ddianthicola
Dickeya dianthicola IPO 980 WP 024107798 1 Ddianthicola
Dickeya dianthicola NCPPB 453 WP 024107798 1 Ddianthicola
Dickeya dianthicola GBBC 2039 WP 024107798 1 Ddianthicola
Dickeya dadantii subsp dieffenbachiae NCPPB 2976 WP 038924295 1 Ddadantii
Dickeya dadantii NCPPB 898 WP 038912265 1 Ddadantii
353 vfmD Quorum sensing Seed
Dickeya dadantii NCPPB 3537 WP 038902376 1 Ddadantii
Dickeya solani MK16 WP 022631626 1 Dsolani
Dickeya solani RNS 05 1 2A WP 057082216 1 Dsolani
Dickeya solani PPO 9134 WP 022631626 1 Dsolani
Dickeya solani GBBC 2040 WP 022631626 1 Dsolani
Dickeya solani D s0432 1 WP 022631626 1 Dsolani
Dickeya solani IFB 0099 WP 022631626 1 Dsolani
Dickeya solani RNS 08 23 3 1 A WP 022631626 1 Dsolani
Dickeya solani RNS 07 7 3B WP 057082216 1 Dsolani
Dickeya solani MK10 WP 022631626 1 Dsolani
Dickeya solani IPO 2222 WP 022631626 1 Dsolani
Dickeya solani PPO 9019 WP 022631626 1 Dsolani

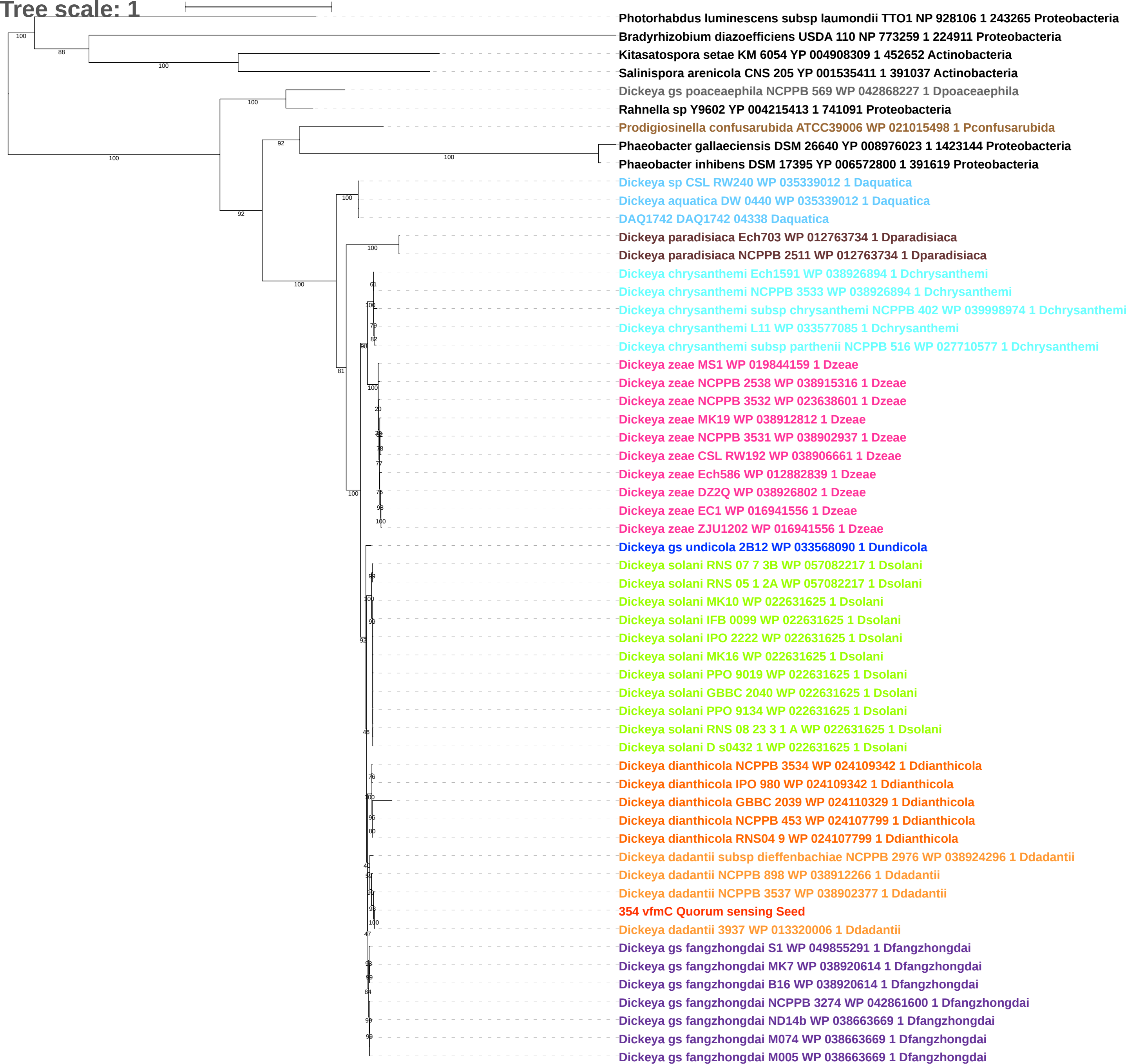
Tree scale: 1
Photorhabdus luminescens subsp laumondii TTO1 NP 928106 1 243265 Proteobacteria
Bradyrhizobium diazoefficiens USDA 110 NP 773259 1 224911 Proteobacteria
Kitasatospora setae KM 6054 YP 004908309 1 452652 Actinobacteria
Salinispora arenicola CNS 205 YP 001535411 1 391037 Actinobacteria
Dickeya gs poaceaephila NCPPB 569 WP 042868227 1 Dpoaceaephila
Rahnella sp Y9602 YP 004215413 1 741091 Proteobacteria
Prodigiosinella confusarubida ATCC39006 WP 021015498 1 Pconfusarubida
Phaeobacter gallaeciensis DSM 26640 YP 008976023 1 1423144 Proteobacteria
Phaeobacter inhibens DSM 17395 YP 006572800 1 391619 Proteobacteria
Dickeya sp CSL RW240 WP 035339012 1 Daquatica
Dickeya aquatica DW 0440 WP 035339012 1 Daquatica
DAQ1742 DAQ1742 04338 Daquatica
Dickeya paradisiaca Ech703 WP 012763734 1 Dparadisiaca
Dickeya paradisiaca NCPPB 2511 WP 012763734 1 Dparadisiaca
Dickeya chrysanthemi Ech1591 WP 038926894 1 Dchrysanthemi
Dickeya chrysanthemi NCPPB 3533 WP 038926894 1 Dchrysanthemi
Dickeya chrysanthemi subsp chrysanthemi NCPPB 402 WP 039998974 1 Dchrysanthemi
Dickeya chrysanthemi L11 WP 033577085 1 Dchrysanthemi
Dickeya chrysanthemi subsp parthenii NCPPB 516 WP 027710577 1 Dchrysanthemi
Dickeya zeae MS1 WP 019844159 1 Dzeae
Dickeya zeae NCPPB 2538 WP 038915316 1 Dzeae
Dickeya zeae NCPPB 3532 WP 023638601 1 Dzeae
Dickeya zeae MK19 WP 038912812 1 Dzeae
Dickeya zeae NCPPB 3531 WP 038902937 1 Dzeae
Dickeya zeae CSL RW192 WP 038906661 1 Dzeae
Dickeya zeae Ech586 WP 012882839 1 Dzeae
Dickeya zeae DZ2Q WP 038926802 1 Dzeae
Dickeya zeae EC1 WP 016941556 1 Dzeae
Dickeya zeae ZJU1202 WP 016941556 1 Dzeae
Dickeya gs undicola 2B12 WP 033568090 1 Dundicola
Dickeya solani RNS 07 7 3B WP 057082217 1 Dsolani
Dickeya solani RNS 05 1 2A WP 057082217 1 Dsolani
Dickeya solani MK10 WP 022631625 1 Dsolani
Dickeya solani IFB 0099 WP 022631625 1 Dsolani
Dickeya solani IPO 2222 WP 022631625 1 Dsolani
Dickeya solani MK16 WP 022631625 1 Dsolani
Dickeya solani PPO 9019 WP 022631625 1 Dsolani
Dickeya solani GBBC 2040 WP 022631625 1 Dsolani
Dickeya solani PPO 9134 WP 022631625 1 Dsolani
Dickeya solani RNS 08 23 3 1 A WP 022631625 1 Dsolani
Dickeya solani D s0432 1 WP 022631625 1 Dsolani
Dickeya dianthicola NCPPB 3534 WP 024109342 1 Ddianthicola
Dickeya dianthicola IPO 980 WP 024109342 1 Ddianthicola
Dickeya dianthicola GBBC 2039 WP 024110329 1 Ddianthicola
Dickeya dianthicola NCPPB 453 WP 024107799 1 Ddianthicola
Dickeya dianthicola RNS04 9 WP 024107799 1 Ddianthicola
Dickeya dadantii subsp dieffenbachiae NCPPB 2976 WP 038924296 1 Ddadantii
Dickeya dadantii NCPPB 898 WP 038912266 1 Ddadantii
Dickeya dadantii NCPPB 3537 WP 038902377 1 Ddadantii
354 vfmC Quorum sensing Seed
Dickeya dadantii 3937 WP 013320006 1 Ddadantii
Dickeya gs fangzhongdai S1 WP 049855291 1 Dfangzhongdai
Dickeya gs fangzhongdai MK7 WP 038920614 1 Dfangzhongdai
Dickeya gs fangzhongdai B16 WP 038920614 1 Dfangzhongdai
Dickeya gs fangzhongdai NCPPB 3274 WP 042861600 1 Dfangzhongdai
Dickeya gs fangzhongdai ND14b WP 038663669 1 Dfangzhongdai
Dickeya gs fangzhongdai M074 WP 038663669 1 Dfangzhongdai
Dickeya gs fangzhongdai M005 WP 038663669 1 Dfangzhongdai

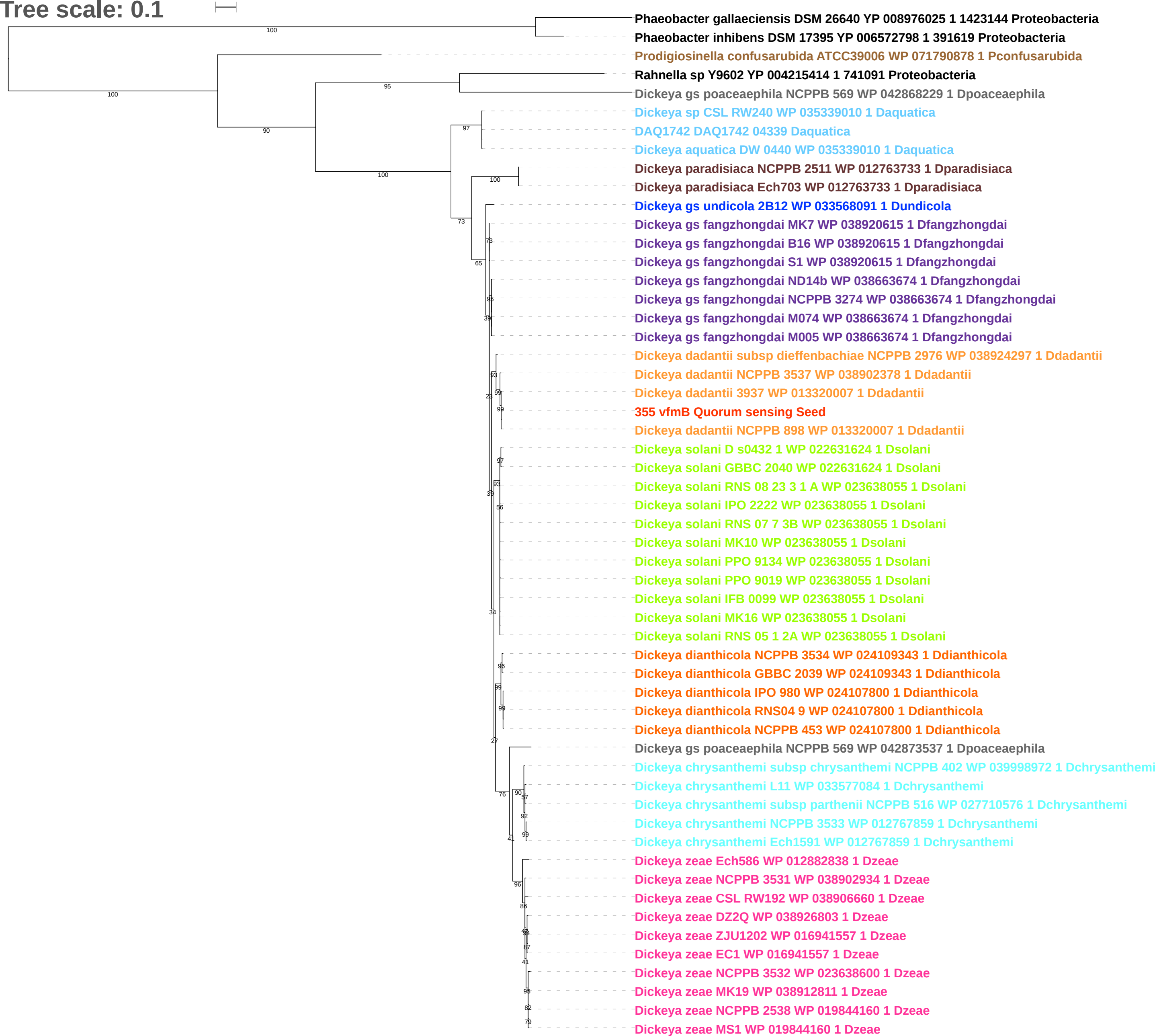

Tree scale: 0.1
Phaeobacter gallaeciensis DSM 26640 YP 008976025 1 1423144 Proteobacteria
Phaeobacter inhibens DSM 17395 YP 006572798 1 391619 Proteobacteria
Prodigiosinella confusarubida ATCC39006 WP 071790878 1 Pconfusarubida
Rahnella sp Y9602 YP 004215414 1 741091 Proteobacteria
Dickeya gs poaceaephila NCPPB 569 WP 042868229 1 Dpoaceaephila
Dickeya sp CSL RW240 WP 035339010 1 Daquatica
DAQ1742 DAQ1742 04339 Daquatica
Dickeya aquatica DW 0440 WP 035339010 1 Daquatica
Dickeya paradisiaca NCPPB 2511 WP 012763733 1 Dparadisiaca
Dickeya paradisiaca Ech703 WP 012763733 1 Dparadisiaca
Dickeya gs undicola 2B12 WP 033568091 1 Dundicola
Dickeya gs fangzhongdai MK7 WP 038920615 1 Dfangzhongdai
Dickeya gs fangzhongdai B16 WP 038920615 1 Dfangzhongdai
Dickeya gs fangzhongdai S1 WP 038920615 1 Dfangzhongdai
Dickeya gs fangzhongdai ND14b WP 038663674 1 Dfangzhongdai
Dickeya gs fangzhongdai NCPPB 3274 WP 038663674 1 Dfangzhongdai
Dickeya gs fangzhongdai M074 WP 038663674 1 Dfangzhongdai
Dickeya gs fangzhongdai M005 WP 038663674 1 Dfangzhongdai
Dickeya dadantii subsp dieffenbachiae NCPPB 2976 WP 038924297 1 Ddadantii
Dickeya dadantii NCPPB 3537 WP 038902378 1 Ddadantii
Dickeya dadantii 3937 WP 013320007 1 Ddadantii
355 vfmB Quorum sensing Seed
Dickeya dadantii NCPPB 898 WP 013320007 1 Ddadantii
Dickeya solani D s0432 1 WP 022631624 1 Dsolani
Dickeya solani GBBC 2040 WP 022631624 1 Dsolani
Dickeya solani RNS 08 23 3 1 A WP 023638055 1 Dsolani
Dickeya solani IPO 2222 WP 023638055 1 Dsolani
Dickeya solani RNS 07 7 3B WP 023638055 1 Dsolani
Dickeya solani MK10 WP 023638055 1 Dsolani
Dickeya solani PPO 9134 WP 023638055 1 Dsolani
Dickeya solani PPO 9019 WP 023638055 1 Dsolani
Dickeya solani IFB 0099 WP 023638055 1 Dsolani
Dickeya solani MK16 WP 023638055 1 Dsolani
Dickeya solani RNS 05 1 2A WP 023638055 1 Dsolani
Dickeya dianthicola NCPPB 3534 WP 024109343 1 Ddianthicola
Dickeya dianthicola GBBC 2039 WP 024109343 1 Ddianthicola
Dickeya dianthicola IPO 980 WP 024107800 1 Ddianthicola
Dickeya dianthicola RNS04 9 WP 024107800 1 Ddianthicola
Dickeya dianthicola NCPPB 453 WP 024107800 1 Ddianthicola
Dickeya gs poaceaephila NCPPB 569 WP 042873537 1 Dpoaceaephila
Dickeya chrysanthemi subsp chrysanthemi NCPPB 402 WP 039998972 1 Dchrysanthemi
Dickeya chrysanthemi L11 WP 033577084 1 Dchrysanthemi
Dickeya chrysanthemi subsp parthenii NCPPB 516 WP 027710576 1 Dchrysanthemi
Dickeya chrysanthemi NCPPB 3533 WP 012767859 1 Dchrysanthemi
Dickeya chrysanthemi Ech1591 WP 012767859 1 Dchrysanthemi
Dickeya zeae Ech586 WP 012882838 1 Dzeae
Dickeya zeae NCPPB 3531 WP 038902934 1 Dzeae
Dickeya zeae CSL RW192 WP 038906660 1 Dzeae
Dickeya zeae DZ2Q WP 038926803 1 Dzeae
Dickeya zeae ZJU1202 WP 016941557 1 Dzeae
Dickeya zeae EC1 WP 016941557 1 Dzeae
Dickeya zeae NCPPB 3532 WP 023638600 1 Dzeae
Dickeya zeae MK19 WP 038912811 1 Dzeae
Dickeya zeae NCPPB 2538 WP 019844160 1 Dzeae
Dickeya zeae MS1 WP 019844160 1 Dzeae

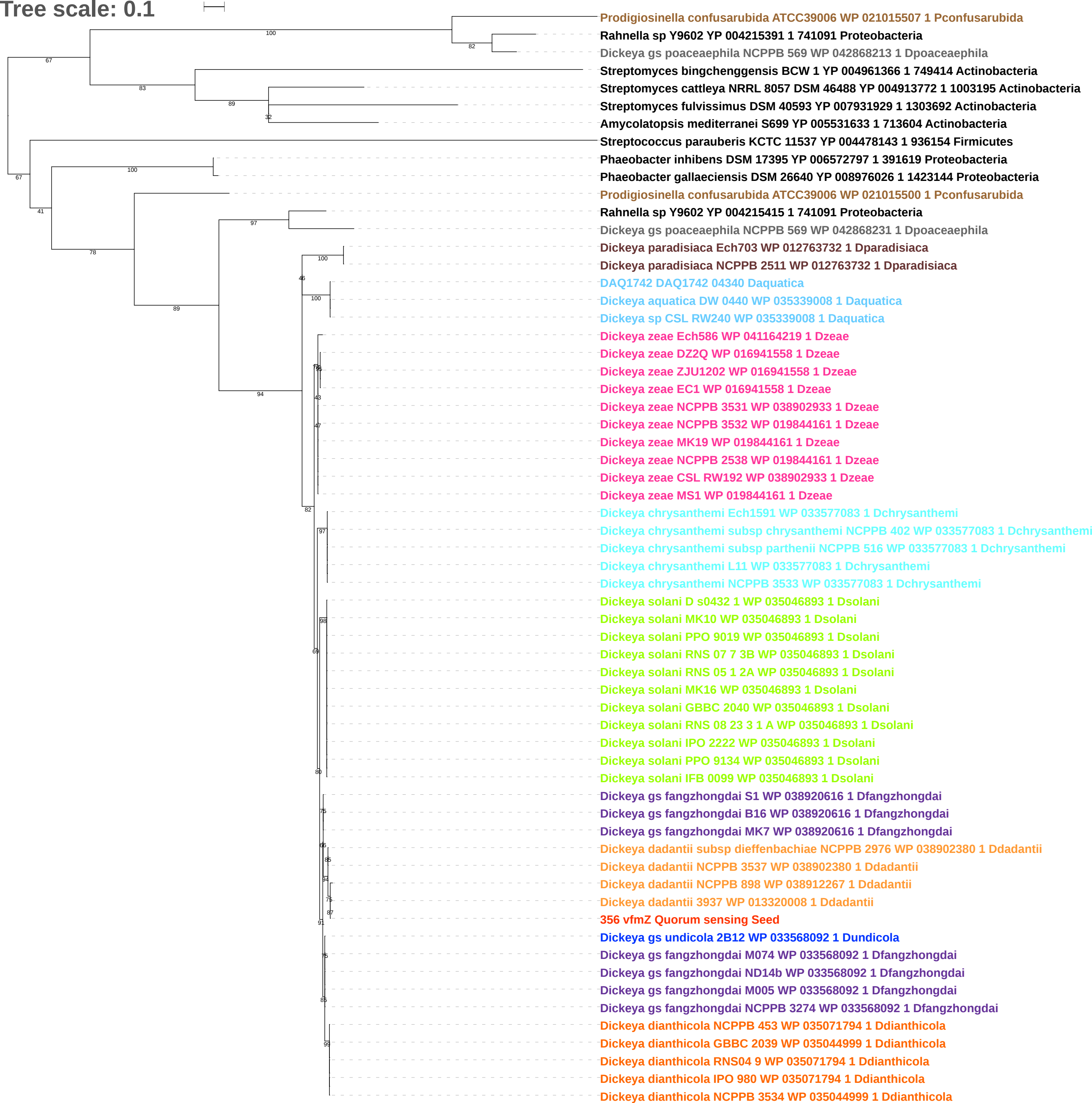

Tree scale: 0.1
Prodigiosinella confusarubida ATCC39006 WP 021015507 1 Pconfusarubida
Rahnella sp Y9602 YP 004215391 1 741091 Proteobacteria
Dickeya gs poaceaephila NCPPB 569 WP 042868213 1 Dpoaceaephila
Streptomyces bingchenggensis BCW 1 YP 004961366 1 749414 Actinobacteria
Streptomyces cattleya NRRL 8057 DSM 46488 YP 004913772 1 1003195 Actinobacteria
Streptomyces fulvissimus DSM 40593 YP 007931929 1 1303692 Actinobacteria
Amycolatopsis mediterranei S699 YP 005531633 1 713604 Actinobacteria
Streptococcus parauberis KCTC 11537 YP 004478143 1 936154 Firmicutes
Phaeobacter inhibens DSM 17395 YP 006572797 1 391619 Proteobacteria
Phaeobacter gallaeciensis DSM 26640 YP 008976026 1 1423144 Proteobacteria
Prodigiosinella confusarubida ATCC39006 WP 021015500 1 Pconfusarubida
Rahnella sp Y9602 YP 004215415 1 741091 Proteobacteria
Dickeya gs poaceaephila NCPPB 569 WP 042868231 1 Dpoaceaephila
Dickeya paradisiaca Ech703 WP 012763732 1 Dparadisiaca
Dickeya paradisiaca NCPPB 2511 WP 012763732 1 Dparadisiaca
DAQ1742 DAQ1742 04340 Daquatica
Dickeya aquatica DW 0440 WP 035339008 1 Daquatica
Dickeya sp CSL RW240 WP 035339008 1 Daquatica
Dickeya zeae Ech586 WP 041164219 1 Dzeae
Dickeya zeae DZ2Q WP 016941558 1 Dzeae
Dickeya zeae ZJU1202 WP 016941558 1 Dzeae
Dickeya zeae EC1 WP 016941558 1 Dzeae
Dickeya zeae NCPPB 3531 WP 038902933 1 Dzeae
Dickeya zeae NCPPB 3532 WP 019844161 1 Dzeae
Dickeya zeae MK19 WP 019844161 1 Dzeae
Dickeya zeae NCPPB 2538 WP 019844161 1 Dzeae
Dickeya zeae CSL RW192 WP 038902933 1 Dzeae
Dickeya zeae MS1 WP 019844161 1 Dzeae
Dickeya chrysanthemi Ech1591 WP 033577083 1 Dchrysanthemi
Dickeya chrysanthemi subsp chrysanthemi NCPPB 402 WP 033577083 1 Dchrysanthemi
Dickeya chrysanthemi subsp parthenii NCPPB 516 WP 033577083 1 Dchrysanthemi
Dickeya chrysanthemi L11 WP 033577083 1 Dchrysanthemi
Dickeya chrysanthemi NCPPB 3533 WP 033577083 1 Dchrysanthemi
Dickeya solani D s0432 1 WP 035046893 1 Dsolani
Dickeya solani MK10 WP 035046893 1 Dsolani
Dickeya solani PPO 9019 WP 035046893 1 Dsolani
Dickeya solani RNS 07 7 3B WP 035046893 1 Dsolani
Dickeya solani RNS 05 1 2A WP 035046893 1 Dsolani
Dickeya solani MK16 WP 035046893 1 Dsolani
Dickeya solani GBBC 2040 WP 035046893 1 Dsolani
Dickeya solani RNS 08 23 3 1 A WP 035046893 1 Dsolani
Dickeya solani IPO 2222 WP 035046893 1 Dsolani
Dickeya solani PPO 9134 WP 035046893 1 Dsolani
Dickeya solani IFB 0099 WP 035046893 1 Dsolani
Dickeya gs fangzhongdai S1 WP 038920616 1 Dfangzhongdai
Dickeya gs fangzhongdai B16 WP 038920616 1 Dfangzhongdai
Dickeya gs fangzhongdai MK7 WP 038920616 1 Dfangzhongdai
Dickeya dadantii subsp dieffenbachiae NCPPB 2976 WP 038902380 1 Ddadantii
Dickeya dadantii NCPPB 3537 WP 038902380 1 Ddadantii
Dickeya dadantii NCPPB 898 WP 038912267 1 Ddadantii
Dickeya dadantii 3937 WP 013320008 1 Ddadantii
356 vfmZ Quorum sensing Seed
Dickeya gs undicola 2B12 WP 033568092 1 Dundicola
Dickeya gs fangzhongdai M074 WP 033568092 1 Dfangzhongdai
Dickeya gs fangzhongdai ND14b WP 033568092 1 Dfangzhongdai
Dickeya gs fangzhongdai M005 WP 033568092 1 Dfangzhongdai
Dickeya gs fangzhongdai NCPPB 3274 WP 033568092 1 Dfangzhongdai
Dickeya dianthicola NCPPB 453 WP 035071794 1 Ddianthicola
Dickeya dianthicola GBBC 2039 WP 035044999 1 Ddianthicola
Dickeya dianthicola RNS04 9 WP 035071794 1 Ddianthicola
Dickeya dianthicola IPO 980 WP 035071794 1 Ddianthicola
Dickeya dianthicola NCPPB 3534 WP 035044999 1 Ddianthicola

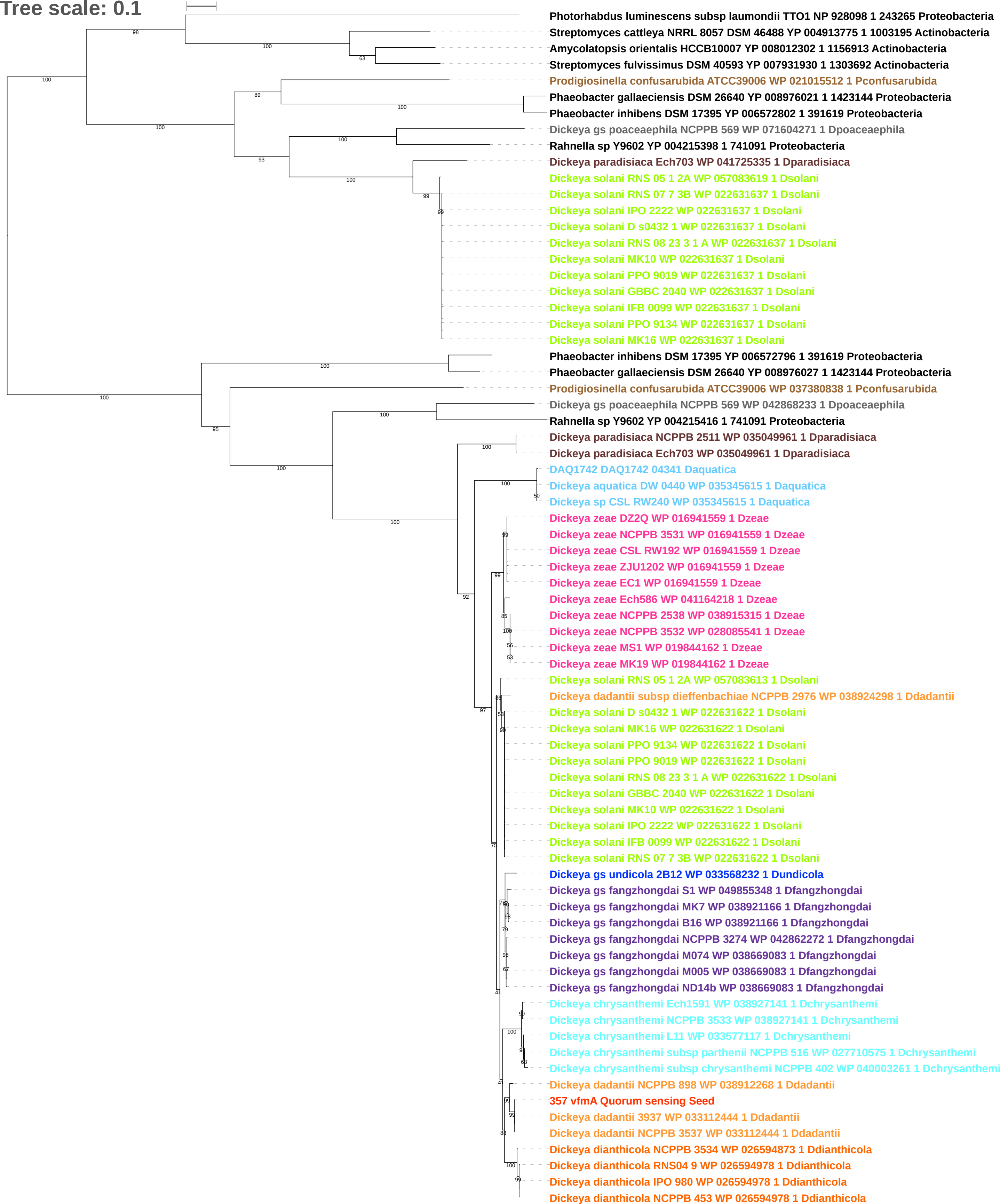

Tree scale: 0.1
Photorhabdus luminescens subsp laumondii TTO1 NP 928098 1 243265 Proteobacteria
Streptomyces cattleya NRRL 8057 DSM 46488 YP 004913775 1 1003195 Actinobacteria
Amycolatopsis orientalis HCCB10007 YP 008012302 1 1156913 Actinobacteria
Streptomyces fulvissimus DSM 40593 YP 007931930 1 1303692 Actinobacteria
Prodigiosinella confusarubida ATCC39006 WP 021015512 1 Pconfusarubida
Phaeobacter gallaeciensis DSM 26640 YP 008976021 1 1423144 Proteobacteria
Phaeobacter inhibens DSM 17395 YP 006572802 1 391619 Proteobacteria
Dickeya gs poaceaephila NCPPB 569 WP 071604271 1 Dpoaceaephila
Rahnella sp Y9602 YP 004215398 1 741091 Proteobacteria
Dickeya paradisiaca Ech703 WP 041725335 1 Dparadisiaca
Dickeya solani RNS 05 1 2A WP 057083619 1 Dsolani
Dickeya solani RNS 07 7 3B WP 022631637 1 Dsolani
Dickeya solani IPO 2222 WP 022631637 1 Dsolani
Dickeya solani D s0432 1 WP 022631637 1 Dsolani
Dickeya solani RNS 08 23 3 1 A WP 022631637 1 Dsolani
Dickeya solani MK10 WP 022631637 1 Dsolani
Dickeya solani PPO 9019 WP 022631637 1 Dsolani
Dickeya solani GBBC 2040 WP 022631637 1 Dsolani
Dickeya solani IFB 0099 WP 022631637 1 Dsolani
Dickeya solani PPO 9134 WP 022631637 1 Dsolani
Dickeya solani MK16 WP 022631637 1 Dsolani
Phaeobacter inhibens DSM 17395 YP 006572796 1 391619 Proteobacteria
Phaeobacter gallaeciensis DSM 26640 YP 008976027 1 1423144 Proteobacteria
Prodigiosinella confusarubida ATCC39006 WP 037380838 1 Pconfusarubida
Dickeya gs poaceaephila NCPPB 569 WP 042868233 1 Dpoaceaephila
Rahnella sp Y9602 YP 004215416 1 741091 Proteobacteria
Dickeya paradisiaca NCPPB 2511 WP 035049961 1 Dparadisiaca
Dickeya paradisiaca Ech703 WP 035049961 1 Dparadisiaca
DAQ1742 DAQ1742 04341 Daquatica
Dickeya aquatica DW 0440 WP 035345615 1 Daquatica
Dickeya sp CSL RW240 WP 035345615 1 Daquatica
Dickeya zeae DZ2Q WP 016941559 1 Dzeae
Dickeya zeae NCPPB 3531 WP 016941559 1 Dzeae
Dickeya zeae CSL RW192 WP 016941559 1 Dzeae
Dickeya zeae ZJU1202 WP 016941559 1 Dzeae
Dickeya zeae EC1 WP 016941559 1 Dzeae
Dickeya zeae Ech586 WP 041164218 1 Dzeae
Dickeya zeae NCPPB 2538 WP 038915315 1 Dzeae
Dickeya zeae NCPPB 3532 WP 028085541 1 Dzeae
Dickeya zeae MS1 WP 019844162 1 Dzeae
Dickeya zeae MK19 WP 019844162 1 Dzeae
Dickeya solani RNS 05 1 2A WP 057083613 1 Dsolani
Dickeya dadantii subsp dieffenbachiae NCPPB 2976 WP 038924298 1 Ddadantii
Dickeya solani D s0432 1 WP 022631622 1 Dsolani
Dickeya solani MK16 WP 022631622 1 Dsolani
Dickeya solani PPO 9134 WP 022631622 1 Dsolani
Dickeya solani PPO 9019 WP 022631622 1 Dsolani
Dickeya solani RNS 08 23 3 1 A WP 022631622 1 Dsolani
Dickeya solani GBBC 2040 WP 022631622 1 Dsolani
Dickeya solani MK10 WP 022631622 1 Dsolani
Dickeya solani IPO 2222 WP 022631622 1 Dsolani
Dickeya solani IFB 0099 WP 022631622 1 Dsolani
Dickeya solani RNS 07 7 3B WP 022631622 1 Dsolani
Dickeya gs undicola 2B12 WP 033568232 1 Dundicola
Dickeya gs fangzhongdai S1 WP 049855348 1 Dfangzhongdai
Dickeya gs fangzhongdai MK7 WP 038921166 1 Dfangzhongdai
Dickeya gs fangzhongdai B16 WP 038921166 1 Dfangzhongdai
Dickeya gs fangzhongdai NCPPB 3274 WP 042862272 1 Dfangzhongdai
Dickeya gs fangzhongdai M074 WP 038669083 1 Dfangzhongdai
Dickeya gs fangzhongdai M005 WP 038669083 1 Dfangzhongdai
Dickeya gs fangzhongdai ND14b WP 038669083 1 Dfangzhongdai
Dickeya chrysanthemi Ech1591 WP 038927141 1 Dchrysanthemi
Dickeya chrysanthemi NCPPB 3533 WP 038927141 1 Dchrysanthemi
Dickeya chrysanthemi L11 WP 033577117 1 Dchrysanthemi
Dickeya chrysanthemi subsp parthenii NCPPB 516 WP 027710575 1 Dchrysanthemi
Dickeya chrysanthemi subsp chrysanthemi NCPPB 402 WP 040003261 1 Dchrysanthemi
Dickeya dadantii NCPPB 898 WP 038912268 1 Ddadantii
357 vfmA Quorum sensing Seed
Dickeya dadantii 3937 WP 033112444 1 Ddadantii
Dickeya dadantii NCPPB 3537 WP 033112444 1 Ddadantii
Dickeya dianthicola NCPPB 3534 WP 026594873 1 Ddianthicola
Dickeya dianthicola RNS04 9 WP 026594978 1 Ddianthicola
Dickeya dianthicola IPO 980 WP 026594978 1 Ddianthicola
Dickeya dianthicola NCPPB 453 WP 026594978 1 Ddianthicola

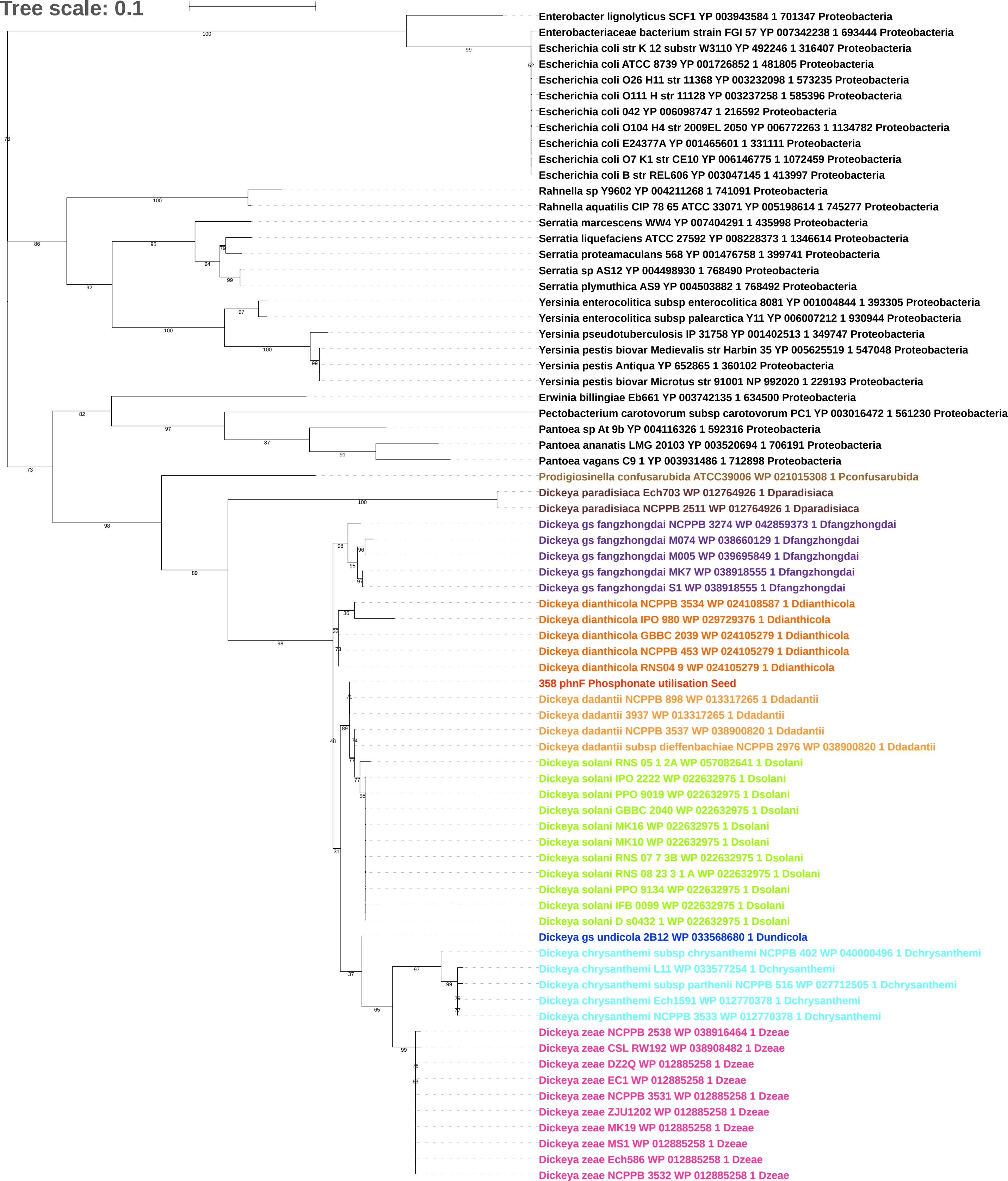
Tree scale: 0.1
Enterobacter lignolyticus SCF1 YP 003943584 1 701347 Proteobacteria
Enterobacteriaceae bacterium strain FGI 57 YP 007342238 1 693444 Proteobacteria
Escherichia coli str K 12 substr W3110 YP 492246 1 316407 Proteobacteria
Escherichia coli ATCC 8739 YP 001726852 1 481805 Proteobacteria
Escherichia coli O26 H11 str 11368 YP 003232098 1 573235 Proteobacteria
Escherichia coli O111 H str 11128 YP 003237258 1 585396 Proteobacteria
Escherichia coli 042 YP 006098747 1 216592 Proteobacteria
Escherichia coli O104 H4 str 2009EL 2050 YP 006772263 1 1134782 Proteobacteria
Escherichia coli E24377A YP 001465601 1 331111 Proteobacteria
Escherichia coli O7 K1 str CE10 YP 006146775 1 1072459 Proteobacteria
Escherichia coli B str REL606 YP 003047145 1 413997 Proteobacteria
Rahnella sp Y9602 YP 004211268 1 741091 Proteobacteria
Rahnella aquatilis CIP 78 65 ATCC 33071 YP 005198614 1 745277 Proteobacteria
Serratia marcescens WW4 YP 007404291 1 435998 Proteobacteria
Serratia liquefaciens ATCC 27592 YP 008228373 1 1346614 Proteobacteria
Serratia proteamaculans 568 YP 001476758 1 399741 Proteobacteria
Serratia sp AS12 YP 004498930 1 768490 Proteobacteria
Serratia plymuthica AS9 YP 004503882 1 768492 Proteobacteria
Yersinia enterocolitica subsp enterocolitica 8081 YP 001004844 1 393305 Proteobacteria
Yersinia enterocolitica subsp palearctica Y11 YP 006007212 1 930944 Proteobacteria
Yersinia pseudotuberculosis IP 31758 YP 001402513 1 349747 Proteobacteria
Yersinia pestis biovar Medievalis str Harbin 35 YP 005625519 1 547048 Proteobacteria
Yersinia pestis Antiqua YP 652865 1 360102 Proteobacteria
Yersinia pestis biovar Microtus str 91001 NP 992020 1 229193 Proteobacteria
Erwinia billingiae Eb661 YP 003742135 1 634500 Proteobacteria
Pectobacterium carotovorum subsp carotovorum PC1 YP 003016472 1 561230 Proteobacteria
Pantoea sp At 9b YP 004116326 1 592316 Proteobacteria
Pantoea ananatis LMG 20103 YP 003520694 1 706191 Proteobacteria
Pantoea vagans C9 1 YP 003931486 1 712898 Proteobacteria
Prodigiosinella confusarubida ATCC39006 WP 021015308 1 Pconfusarubida
Dickeya paradisiaca Ech703 WP 012764926 1 Dparadisiaca
Dickeya paradisiaca NCPPB 2511 WP 012764926 1 Dparadisiaca
Dickeya gs fangzhongdai NCPPB 3274 WP 042859373 1 Dfangzhongdai
Dickeya gs fangzhongdai M074 WP 038660129 1 Dfangzhongdai
Dickeya gs fangzhongdai M005 WP 039695849 1 Dfangzhongdai
Dickeya gs fangzhongdai MK7 WP 038918555 1 Dfangzhongdai
Dickeya gs fangzhongdai S1 WP 038918555 1 Dfangzhongdai
Dickeya dianthicola NCPPB 3534 WP 024108587 1 Ddianthicola
Dickeya dianthicola IPO 980 WP 029729376 1 Ddianthicola
Dickeya dianthicola GBBC 2039 WP 024105279 1 Ddianthicola
Dickeya dianthicola NCPPB 453 WP 024105279 1 Ddianthicola
Dickeya dianthicola RNS04 9 WP 024105279 1 Ddianthicola
358 phnF Phosphonate utilisation Seed
Dickeya dadantii NCPPB 898 WP 013317265 1 Ddadantii
Dickeya dadantii 3937 WP 013317265 1 Ddadantii
Dickeya dadantii NCPPB 3537 WP 038900820 1 Ddadantii
Dickeya dadantii subsp dieffenbachiae NCPPB 2976 WP 038900820 1 Ddadantii
Dickeya solani RNS 05 1 2A WP 057082641 1 Dsolani
Dickeya solani IPO 2222 WP 022632975 1 Dsolani
Dickeya solani PPO 9019 WP 022632975 1 Dsolani
Dickeya solani GBBC 2040 WP 022632975 1 Dsolani
Dickeya solani MK16 WP 022632975 1 Dsolani
Dickeya solani MK10 WP 022632975 1 Dsolani
Dickeya solani RNS 07 7 3B WP 022632975 1 Dsolani
Dickeya solani RNS 08 23 3 1 A WP 022632975 1 Dsolani
Dickeya solani PPO 9134 WP 022632975 1 Dsolani
Dickeya solani IFB 0099 WP 022632975 1 Dsolani
Dickeya solani D s0432 1 WP 022632975 1 Dsolani
Dickeya gs undicola 2B12 WP 033568680 1 Dundicola
Dickeya chrysanthemi subsp chrysanthemi NCPPB 402 WP 040000496 1 Dchrysanthemi
Dickeya chrysanthemi L11 WP 033577254 1 Dchrysanthemi
Dickeya chrysanthemi subsp parthenii NCPPB 516 WP 027712505 1 Dchrysanthemi
Dickeya chrysanthemi Ech1591 WP 012770378 1 Dchrysanthemi
Dickeya chrysanthemi NCPPB 3533 WP 012770378 1 Dchrysanthemi
Dickeya zeae NCPPB 2538 WP 038916464 1 Dzeae
Dickeya zeae CSL RW192 WP 038908482 1 Dzeae
Dickeya zeae DZ2Q WP 012885258 1 Dzeae
Dickeya zeae EC1 WP 012885258 1 Dzeae
Dickeya zeae NCPPB 3531 WP 012885258 1 Dzeae
Dickeya zeae ZJU1202 WP 012885258 1 Dzeae
Dickeya zeae MK19 WP 012885258 1 Dzeae
Dickeya zeae MS1 WP 012885258 1 Dzeae
Dickeya zeae Ech586 WP 012885258 1 Dzeae
Dickeya zeae NCPPB 3532 WP 012885258 1 Dzeae

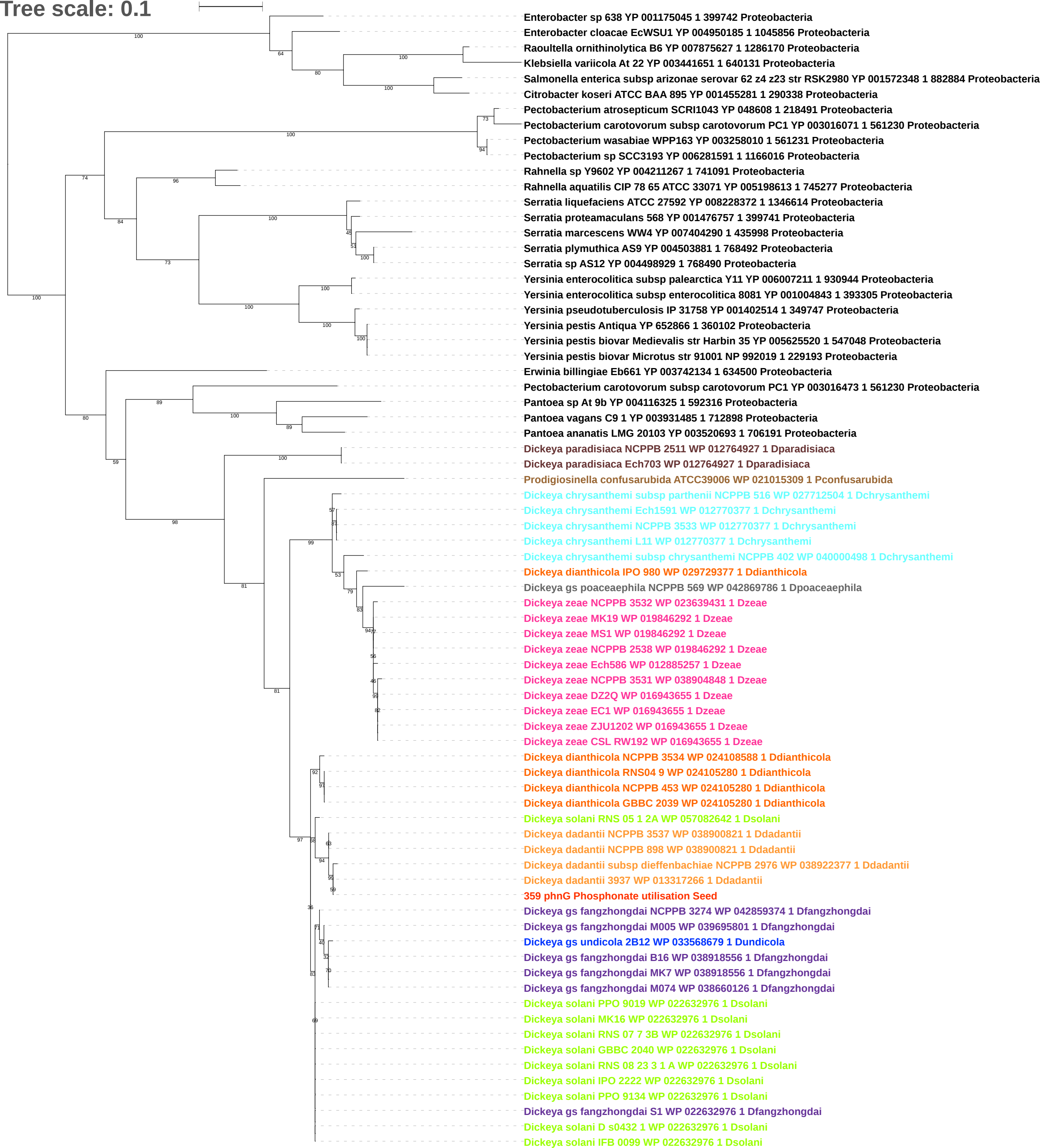
Tree scale: 0.1
Enterobacter sp 638 YP 001175045 1 399742 Proteobacteria
Enterobacter cloacae EcWSU1 YP 004950185 1 1045856 Proteobacteria
Raoultella ornithinolytica B6 YP 007875627 1 1286170 Proteobacteria
Klebsiella variicola At 22 YP 003441651 1 640131 Proteobacteria
Salmonella enterica subsp arizonae serovar 62 z4 z23 str RSK2980 YP 001572348 1 882884 Proteobacteria
Citrobacter koseri ATCC BAA 895 YP 001455281 1 290338 Proteobacteria
Pectobacterium atrosepticum SCRI1043 YP 048608 1 218491 Proteobacteria
Pectobacterium carotovorum subsp carotovorum PC1 YP 003016071 1 561230 Proteobacteria
Pectobacterium wasabiae WPP163 YP 003258010 1 561231 Proteobacteria
Pectobacterium sp SCC3193 YP 006281591 1 1166016 Proteobacteria
Rahnella sp Y9602 YP 004211267 1 741091 Proteobacteria
Rahnella aquatilis CIP 78 65 ATCC 33071 YP 005198613 1 745277 Proteobacteria
Serratia liquefaciens ATCC 27592 YP 008228372 1 1346614 Proteobacteria
Serratia proteamaculans 568 YP 001476757 1 399741 Proteobacteria
Serratia marcescens WW4 YP 007404290 1 435998 Proteobacteria
Serratia plymuthica AS9 YP 004503881 1 768492 Proteobacteria
Serratia sp AS12 YP 004498929 1 768490 Proteobacteria
Yersinia enterocolitica subsp palearctica Y11 YP 006007211 1 930944 Proteobacteria
Yersinia enterocolitica subsp enterocolitica 8081 YP 001004843 1 393305 Proteobacteria
Yersinia pseudotuberculosis IP 31758 YP 001402514 1 349747 Proteobacteria
Yersinia pestis Antiqua YP 652866 1 360102 Proteobacteria
Yersinia pestis biovar Medievalis str Harbin 35 YP 005625520 1 547048 Proteobacteria
Yersinia pestis biovar Microtus str 91001 NP 992019 1 229193 Proteobacteria
Erwinia billingiae Eb661 YP 003742134 1 634500 Proteobacteria
Pectobacterium carotovorum subsp carotovorum PC1 YP 003016473 1 561230 Proteobacteria
Pantoea sp At 9b YP 004116325 1 592316 Proteobacteria
Pantoea vagans C9 1 YP 003931485 1 712898 Proteobacteria
Pantoea ananatis LMG 20103 YP 003520693 1 706191 Proteobacteria
Dickeya paradisiaca NCPPB 2511 WP 012764927 1 Dparadisiaca
Dickeya paradisiaca Ech703 WP 012764927 1 Dparadisiaca
Prodigiosinella confusarubida ATCC39006 WP 021015309 1 Pconfusarubida
Dickeya chrysanthemi subsp parthenii NCPPB 516 WP 027712504 1 Dchrysanthemi
Dickeya chrysanthemi Ech1591 WP 012770377 1 Dchrysanthemi
Dickeya chrysanthemi NCPPB 3533 WP 012770377 1 Dchrysanthemi
Dickeya chrysanthemi L11 WP 012770377 1 Dchrysanthemi
Dickeya chrysanthemi subsp chrysanthemi NCPPB 402 WP 040000498 1 Dchrysanthemi
Dickeya dianthicola IPO 980 WP 029729377 1 Ddianthicola
Dickeya gs poaceaephila NCPPB 569 WP 042869786 1 Dpoaceaephila
Dickeya zeae NCPPB 3532 WP 023639431 1 Dzeae
Dickeya zeae MK19 WP 019846292 1 Dzeae
Dickeya zeae MS1 WP 019846292 1 Dzeae
Dickeya zeae NCPPB 2538 WP 019846292 1 Dzeae
Dickeya zeae Ech586 WP 012885257 1 Dzeae
Dickeya zeae NCPPB 3531 WP 038904848 1 Dzeae
Dickeya zeae DZ2Q WP 016943655 1 Dzeae
Dickeya zeae EC1 WP 016943655 1 Dzeae
Dickeya zeae ZJU1202 WP 016943655 1 Dzeae
Dickeya zeae CSL RW192 WP 016943655 1 Dzeae
Dickeya dianthicola NCPPB 3534 WP 024108588 1 Ddianthicola
Dickeya dianthicola RNS04 9 WP 024105280 1 Ddianthicola
Dickeya dianthicola NCPPB 453 WP 024105280 1 Ddianthicola
Dickeya dianthicola GBBC 2039 WP 024105280 1 Ddianthicola
Dickeya solani RNS 05 1 2A WP 057082642 1 Dsolani
Dickeya dadantii NCPPB 3537 WP 038900821 1 Ddadantii
Dickeya dadantii NCPPB 898 WP 038900821 1 Ddadantii
Dickeya dadantii subsp dieffenbachiae NCPPB 2976 WP 038922377 1 Ddadantii
Dickeya dadantii 3937 WP 013317266 1 Ddadantii
359 phnG Phosphonate utilisation Seed
Dickeya gs fangzhongdai NCPPB 3274 WP 042859374 1 Dfangzhongdai
Dickeya gs fangzhongdai M005 WP 039695801 1 Dfangzhongdai
Dickeya gs undicola 2B12 WP 033568679 1 Dundicola
Dickeya gs fangzhongdai B16 WP 038918556 1 Dfangzhongdai
Dickeya gs fangzhongdai MK7 WP 038918556 1 Dfangzhongdai
Dickeya gs fangzhongdai M074 WP 038660126 1 Dfangzhongdai
Dickeya solani PPO 9019 WP 022632976 1 Dsolani
Dickeya solani MK16 WP 022632976 1 Dsolani
Dickeya solani RNS 07 7 3B WP 022632976 1 Dsolani
Dickeya solani GBBC 2040 WP 022632976 1 Dsolani
Dickeya solani RNS 08 23 3 1 A WP 022632976 1 Dsolani
Dickeya solani IPO 2222 WP 022632976 1 Dsolani
Dickeya solani PPO 9134 WP 022632976 1 Dsolani
Dickeya gs fangzhongdai S1 WP 022632976 1 Dfangzhongdai
Dickeya solani D s0432 1 WP 022632976 1 Dsolani
Dickeya solani IFB 0099 WP 022632976 1 Dsolani

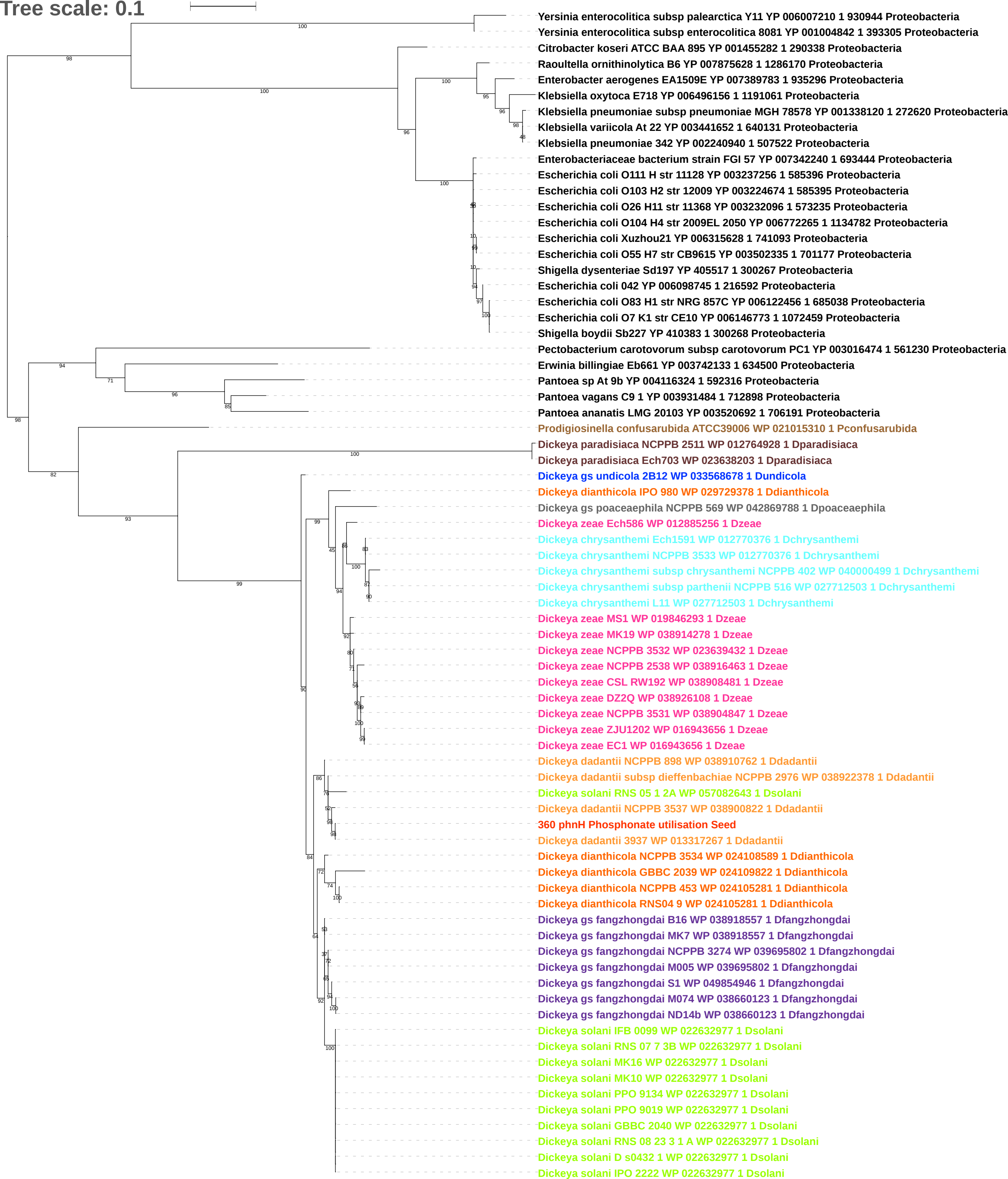
Tree scale: 0.1
Yersinia enterocolitica subsp palearctica Y11 YP 006007210 1 930944 Proteobacteria
Yersinia enterocolitica subsp enterocolitica 8081 YP 001004842 1 393305 Proteobacteria
Citrobacter koseri ATCC BAA 895 YP 001455282 1 290338 Proteobacteria
Raoultella ornithinolytica B6 YP 007875628 1 1286170 Proteobacteria
Enterobacter aerogenes EA1509E YP 007389783 1 935296 Proteobacteria
Klebsiella oxytoca E718 YP 006496156 1 1191061 Proteobacteria
Klebsiella pneumoniae subsp pneumoniae MGH 78578 YP 001338120 1 272620 Proteobacteria
Klebsiella variicola At 22 YP 003441652 1 640131 Proteobacteria
Klebsiella pneumoniae 342 YP 002240940 1 507522 Proteobacteria
Enterobacteriaceae bacterium strain FGI 57 YP 007342240 1 693444 Proteobacteria
Escherichia coli O111 H str 11128 YP 003237256 1 585396 Proteobacteria
Escherichia coli O103 H2 str 12009 YP 003224674 1 585395 Proteobacteria
Escherichia coli O26 H11 str 11368 YP 003232096 1 573235 Proteobacteria
Escherichia coli O104 H4 str 2009EL 2050 YP 006772265 1 1134782 Proteobacteria
Escherichia coli Xuzhou21 YP 006315628 1 741093 Proteobacteria
Escherichia coli O55 H7 str CB9615 YP 003502335 1 701177 Proteobacteria
Shigella dysenteriae Sd197 YP 405517 1 300267 Proteobacteria
Escherichia coli 042 YP 006098745 1 216592 Proteobacteria
Escherichia coli O83 H1 str NRG 857C YP 006122456 1 685038 Proteobacteria
Escherichia coli O7 K1 str CE10 YP 006146773 1 1072459 Proteobacteria
Shigella boydii Sb227 YP 410383 1 300268 Proteobacteria
Pectobacterium carotovorum subsp carotovorum PC1 YP 003016474 1 561230 Proteobacteria
Erwinia billingiae Eb661 YP 003742133 1 634500 Proteobacteria
Pantoea sp At 9b YP 004116324 1 592316 Proteobacteria
Pantoea vagans C9 1 YP 003931484 1 712898 Proteobacteria
Pantoea ananatis LMG 20103 YP 003520692 1 706191 Proteobacteria
Prodigiosinella confusarubida ATCC39006 WP 021015310 1 Pconfusarubida
Dickeya paradisiaca NCPPB 2511 WP 012764928 1 Dparadisiaca
Dickeya paradisiaca Ech703 WP 023638203 1 Dparadisiaca
Dickeya gs undicola 2B12 WP 033568678 1 Dundicola
Dickeya dianthicola IPO 980 WP 029729378 1 Ddianthicola
Dickeya gs poaceaephila NCPPB 569 WP 042869788 1 Dpoaceaephila
Dickeya zeae Ech586 WP 012885256 1 Dzeae
Dickeya chrysanthemi Ech1591 WP 012770376 1 Dchrysanthemi
Dickeya chrysanthemi NCPPB 3533 WP 012770376 1 Dchrysanthemi
Dickeya chrysanthemi subsp chrysanthemi NCPPB 402 WP 040000499 1 Dchrysanthemi
Dickeya chrysanthemi subsp parthenii NCPPB 516 WP 027712503 1 Dchrysanthemi
Dickeya chrysanthemi L11 WP 027712503 1 Dchrysanthemi
Dickeya zeae MS1 WP 019846293 1 Dzeae
Dickeya zeae MK19 WP 038914278 1 Dzeae
Dickeya zeae NCPPB 3532 WP 023639432 1 Dzeae
Dickeya zeae NCPPB 2538 WP 038916463 1 Dzeae
Dickeya zeae CSL RW192 WP 038908481 1 Dzeae
Dickeya zeae DZ2Q WP 038926108 1 Dzeae
Dickeya zeae NCPPB 3531 WP 038904847 1 Dzeae
Dickeya zeae ZJU1202 WP 016943656 1 Dzeae
Dickeya zeae EC1 WP 016943656 1 Dzeae
Dickeya dadantii NCPPB 898 WP 038910762 1 Ddadantii
Dickeya dadantii subsp dieffenbachiae NCPPB 2976 WP 038922378 1 Ddadantii
Dickeya solani RNS 05 1 2A WP 057082643 1 Dsolani
Dickeya dadantii NCPPB 3537 WP 038900822 1 Ddadantii
360 phnH Phosphonate utilisation Seed
Dickeya dadantii 3937 WP 013317267 1 Ddadantii
Dickeya dianthicola NCPPB 3534 WP 024108589 1 Ddianthicola
Dickeya dianthicola GBBC 2039 WP 024109822 1 Ddianthicola
Dickeya dianthicola NCPPB 453 WP 024105281 1 Ddianthicola
Dickeya dianthicola RNS04 9 WP 024105281 1 Ddianthicola
Dickeya gs fangzhongdai B16 WP 038918557 1 Dfangzhongdai
Dickeya gs fangzhongdai MK7 WP 038918557 1 Dfangzhongdai
Dickeya gs fangzhongdai NCPPB 3274 WP 039695802 1 Dfangzhongdai
Dickeya gs fangzhongdai M005 WP 039695802 1 Dfangzhongdai
Dickeya gs fangzhongdai S1 WP 049854946 1 Dfangzhongdai
Dickeya gs fangzhongdai M074 WP 038660123 1 Dfangzhongdai
Dickeya gs fangzhongdai ND14b WP 038660123 1 Dfangzhongdai
Dickeya solani IFB 0099 WP 022632977 1 Dsolani
Dickeya solani RNS 07 7 3B WP 022632977 1 Dsolani
Dickeya solani MK16 WP 022632977 1 Dsolani
Dickeya solani MK10 WP 022632977 1 Dsolani
Dickeya solani PPO 9134 WP 022632977 1 Dsolani
Dickeya solani PPO 9019 WP 022632977 1 Dsolani
Dickeya solani GBBC 2040 WP 022632977 1 Dsolani
Dickeya solani RNS 08 23 3 1 A WP 022632977 1 Dsolani
Dickeya solani D s0432 1 WP 022632977 1 Dsolani
Dickeya solani IPO 2222 WP 022632977 1 Dsolani

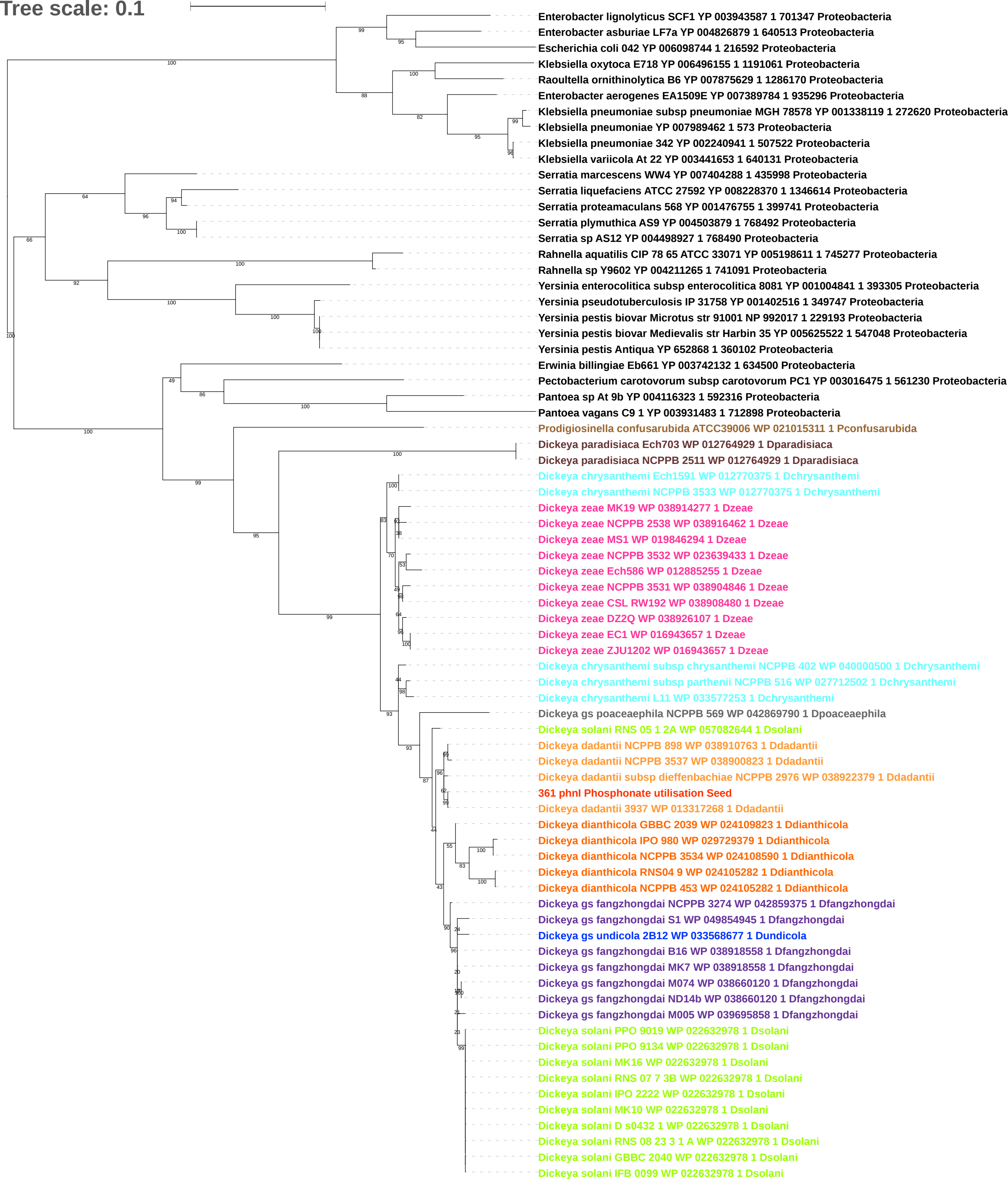
Tree scale: 0.1
Enterobacter lignolyticus SCF1 YP 003943587 1 701347 Proteobacteria
Enterobacter asburiae LF7a YP 004826879 1 640513 Proteobacteria
Escherichia coli 042 YP 006098744 1 216592 Proteobacteria
Klebsiella oxytoca E718 YP 006496155 1 1191061 Proteobacteria
Raoultella ornithinolytica B6 YP 007875629 1 1286170 Proteobacteria
Enterobacter aerogenes EA1509E YP 007389784 1 935296 Proteobacteria
Klebsiella pneumoniae subsp pneumoniae MGH 78578 YP 001338119 1 272620 Proteobacteria
Klebsiella pneumoniae YP 007989462 1 573 Proteobacteria
Klebsiella pneumoniae 342 YP 002240941 1 507522 Proteobacteria
Klebsiella variicola At 22 YP 003441653 1 640131 Proteobacteria
Serratia marcescens WW4 YP 007404288 1 435998 Proteobacteria
Serratia liquefaciens ATCC 27592 YP 008228370 1 1346614 Proteobacteria
Serratia proteamaculans 568 YP 001476755 1 399741 Proteobacteria
Serratia plymuthica AS9 YP 004503879 1 768492 Proteobacteria
Serratia sp AS12 YP 004498927 1 768490 Proteobacteria
Rahnella aquatilis CIP 78 65 ATCC 33071 YP 005198611 1 745277 Proteobacteria
Rahnella sp Y9602 YP 004211265 1 741091 Proteobacteria
Yersinia enterocolitica subsp enterocolitica 8081 YP 001004841 1 393305 Proteobacteria
Yersinia pseudotuberculosis IP 31758 YP 001402516 1 349747 Proteobacteria
Yersinia pestis biovar Microtus str 91001 NP 992017 1 229193 Proteobacteria
Yersinia pestis biovar Medievalis str Harbin 35 YP 005625522 1 547048 Proteobacteria
Yersinia pestis Antiqua YP 652868 1 360102 Proteobacteria
Erwinia billingiae Eb661 YP 003742132 1 634500 Proteobacteria
Pectobacterium carotovorum subsp carotovorum PC1 YP 003016475 1 561230 Proteobacteria
Pantoea sp At 9b YP 004116323 1 592316 Proteobacteria
Pantoea vagans C9 1 YP 003931483 1 712898 Proteobacteria
Prodigiosinella confusarubida ATCC39006 WP 021015311 1 Pconfusarubida
Dickeya paradisiaca Ech703 WP 012764929 1 Dparadisiaca
Dickeya paradisiaca NCPPB 2511 WP 012764929 1 Dparadisiaca
Dickeya chrysanthemi Ech1591 WP 012770375 1 Dchrysanthemi
Dickeya chrysanthemi NCPPB 3533 WP 012770375 1 Dchrysanthemi
Dickeya zeae MK19 WP 038914277 1 Dzeae
Dickeya zeae NCPPB 2538 WP 038916462 1 Dzeae
Dickeya zeae MS1 WP 019846294 1 Dzeae
Dickeya zeae NCPPB 3532 WP 023639433 1 Dzeae
Dickeya zeae Ech586 WP 012885255 1 Dzeae
Dickeya zeae NCPPB 3531 WP 038904846 1 Dzeae
Dickeya zeae CSL RW192 WP 038908480 1 Dzeae
Dickeya zeae DZ2Q WP 038926107 1 Dzeae
Dickeya zeae EC1 WP 016943657 1 Dzeae
Dickeya zeae ZJU1202 WP 016943657 1 Dzeae
Dickeya chrysanthemi subsp chrysanthemi NCPPB 402 WP 040000500 1 Dchrysanthemi
Dickeya chrysanthemi subsp parthenii NCPPB 516 WP 027712502 1 Dchrysanthemi
Dickeya chrysanthemi L11 WP 033577253 1 Dchrysanthemi
Dickeya gs poaceaephila NCPPB 569 WP 042869790 1 Dpoaceaephila
Dickeya solani RNS 05 1 2A WP 057082644 1 Dsolani
Dickeya dadantii NCPPB 898 WP 038910763 1 Ddadantii
Dickeya dadantii NCPPB 3537 WP 038900823 1 Ddadantii
Dickeya dadantii subsp dieffenbachiae NCPPB 2976 WP 038922379 1 Ddadantii
361 phnI Phosphonate utilisation Seed
Dickeya dadantii 3937 WP 013317268 1 Ddadantii
Dickeya dianthicola GBBC 2039 WP 024109823 1 Ddianthicola
Dickeya dianthicola IPO 980 WP 029729379 1 Ddianthicola
Dickeya dianthicola NCPPB 3534 WP 024108590 1 Ddianthicola
Dickeya dianthicola RNS04 9 WP 024105282 1 Ddianthicola
Dickeya dianthicola NCPPB 453 WP 024105282 1 Ddianthicola
Dickeya gs fangzhongdai NCPPB 3274 WP 042859375 1 Dfangzhongdai
Dickeya gs fangzhongdai S1 WP 049854945 1 Dfangzhongdai
Dickeya gs undicola 2B12 WP 033568677 1 Dundicola
Dickeya gs fangzhongdai B16 WP 038918558 1 Dfangzhongdai
Dickeya gs fangzhongdai MK7 WP 038918558 1 Dfangzhongdai
Dickeya gs fangzhongdai M074 WP 038660120 1 Dfangzhongdai
Dickeya gs fangzhongdai ND14b WP 038660120 1 Dfangzhongdai
Dickeya gs fangzhongdai M005 WP 039695858 1 Dfangzhongdai
Dickeya solani PPO 9019 WP 022632978 1 Dsolani
Dickeya solani PPO 9134 WP 022632978 1 Dsolani
Dickeya solani MK16 WP 022632978 1 Dsolani
Dickeya solani RNS 07 7 3B WP 022632978 1 Dsolani
Dickeya solani IPO 2222 WP 022632978 1 Dsolani
Dickeya solani MK10 WP 022632978 1 Dsolani
Dickeya solani D s0432 1 WP 022632978 1 Dsolani
Dickeya solani RNS 08 23 3 1 A WP 022632978 1 Dsolani
Dickeya solani GBBC 2040 WP 022632978 1 Dsolani
Dickeya solani IFB 0099 WP 022632978 1 Dsolani

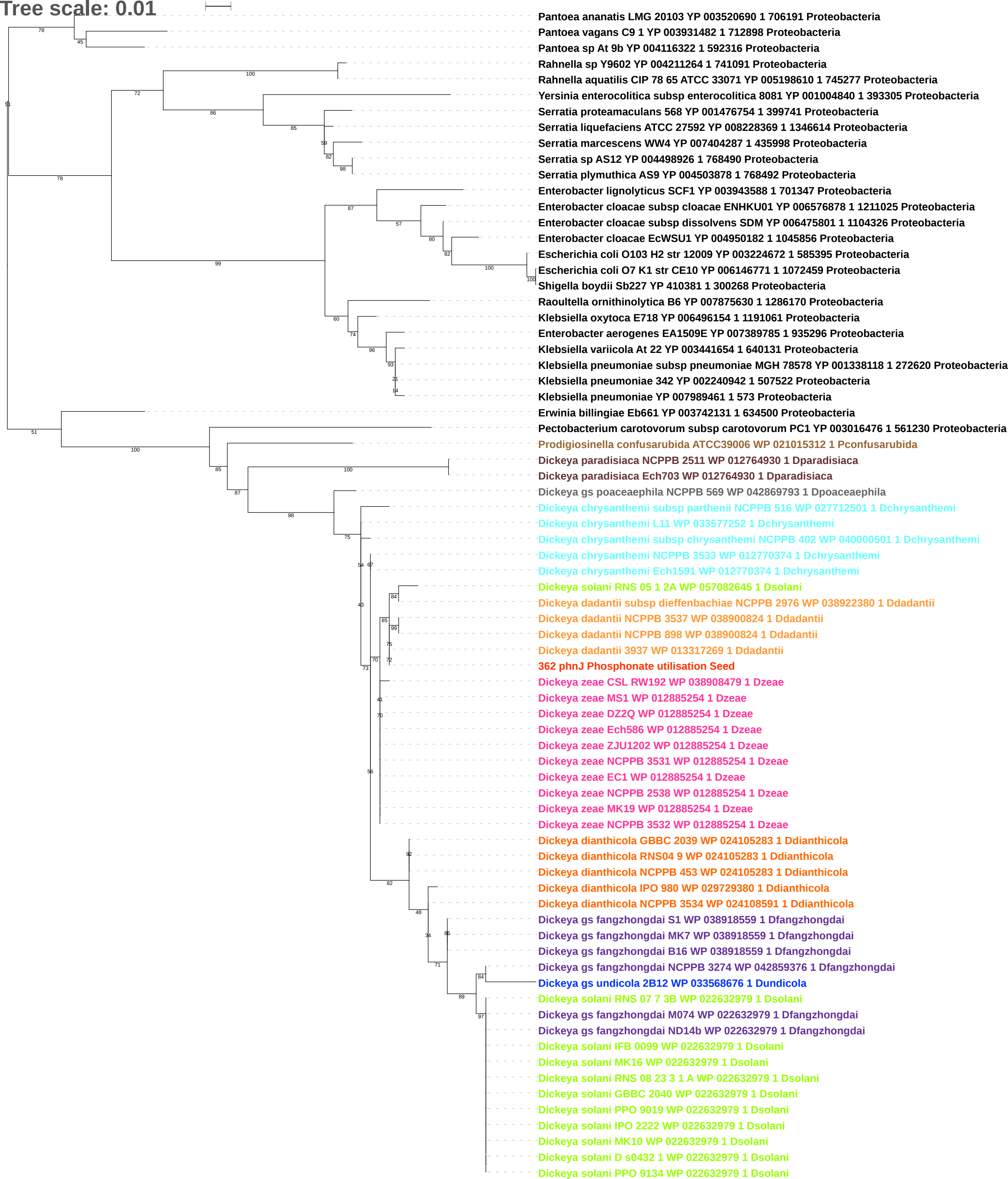
Tree scale: 0.01
Pantoea ananatis LMG 20103 YP 003520690 1 706191 Proteobacteria
Pantoea vagans C9 1 YP 003931482 1 712898 Proteobacteria
Pantoea sp At 9b YP 004116322 1 592316 Proteobacteria
Rahnella sp Y9602 YP 004211264 1 741091 Proteobacteria
Rahnella aquatilis CIP 78 65 ATCC 33071 YP 005198610 1 745277 Proteobacteria
Yersinia enterocolitica subsp enterocolitica 8081 YP 001004840 1 393305 Proteobacteria
Serratia proteamaculans 568 YP 001476754 1 399741 Proteobacteria
Serratia liquefaciens ATCC 27592 YP 008228369 1 1346614 Proteobacteria
Serratia marcescens WW4 YP 007404287 1 435998 Proteobacteria
Serratia sp AS12 YP 004498926 1 768490 Proteobacteria
Serratia plymuthica AS9 YP 004503878 1 768492 Proteobacteria
Enterobacter lignolyticus SCF1 YP 003943588 1 701347 Proteobacteria
Enterobacter cloacae subsp cloacae ENHKU01 YP 006576878 1 1211025 Proteobacteria
Enterobacter cloacae subsp dissolvens SDM YP 006475801 1 1104326 Proteobacteria
Enterobacter cloacae EcWSU1 YP 004950182 1 1045856 Proteobacteria
Escherichia coli O103 H2 str 12009 YP 003224672 1 585395 Proteobacteria
Escherichia coli O7 K1 str CE10 YP 006146771 1 1072459 Proteobacteria
Shigella boydii Sb227 YP 410381 1 300268 Proteobacteria
Raoultella ornithinolytica B6 YP 007875630 1 1286170 Proteobacteria
Klebsiella oxytoca E718 YP 006496154 1 1191061 Proteobacteria
Enterobacter aerogenes EA1509E YP 007389785 1 935296 Proteobacteria
Klebsiella variicola At 22 YP 003441654 1 640131 Proteobacteria
Klebsiella pneumoniae subsp pneumoniae MGH 78578 YP 001338118 1 272620 Proteobacteria
Klebsiella pneumoniae 342 YP 002240942 1 507522 Proteobacteria
Klebsiella pneumoniae YP 007989461 1 573 Proteobacteria
Erwinia billingiae Eb661 YP 003742131 1 634500 Proteobacteria
Pectobacterium carotovorum subsp carotovorum PC1 YP 003016476 1 561230 Proteobacteria
Prodigiosinella confusarubida ATCC39006 WP 021015312 1 Pconfusarubida
Dickeya paradisiaca NCPPB 2511 WP 012764930 1 Dparadisiaca
Dickeya paradisiaca Ech703 WP 012764930 1 Dparadisiaca
Dickeya gs poaceaephila NCPPB 569 WP 042869793 1 Dpoaceaephila
Dickeya chrysanthemi subsp parthenii NCPPB 516 WP 027712501 1 Dchrysanthemi
Dickeya chrysanthemi L11 WP 033577252 1 Dchrysanthemi
Dickeya chrysanthemi subsp chrysanthemi NCPPB 402 WP 040000501 1 Dchrysanthemi
Dickeya chrysanthemi NCPPB 3533 WP 012770374 1 Dchrysanthemi
Dickeya chrysanthemi Ech1591 WP 012770374 1 Dchrysanthemi
Dickeya solani RNS 05 1 2A WP 057082645 1 Dsolani
Dickeya dadantii subsp dieffenbachiae NCPPB 2976 WP 038922380 1 Ddadantii
Dickeya dadantii NCPPB 3537 WP 038900824 1 Ddadantii
Dickeya dadantii NCPPB 898 WP 038900824 1 Ddadantii
Dickeya dadantii 3937 WP 013317269 1 Ddadantii
362 phnJ Phosphonate utilisation Seed
Dickeya zeae CSL RW192 WP 038908479 1 Dzeae
Dickeya zeae MS1 WP 012885254 1 Dzeae
Dickeya zeae DZ2Q WP 012885254 1 Dzeae
Dickeya zeae Ech586 WP 012885254 1 Dzeae
Dickeya zeae ZJU1202 WP 012885254 1 Dzeae
Dickeya zeae NCPPB 3531 WP 012885254 1 Dzeae
Dickeya zeae EC1 WP 012885254 1 Dzeae
Dickeya zeae NCPPB 2538 WP 012885254 1 Dzeae
Dickeya zeae MK19 WP 012885254 1 Dzeae
Dickeya zeae NCPPB 3532 WP 012885254 1 Dzeae
Dickeya dianthicola GBBC 2039 WP 024105283 1 Ddianthicola
Dickeya dianthicola RNS04 9 WP 024105283 1 Ddianthicola
Dickeya dianthicola NCPPB 453 WP 024105283 1 Ddianthicola
Dickeya dianthicola IPO 980 WP 029729380 1 Ddianthicola
Dickeya dianthicola NCPPB 3534 WP 024108591 1 Ddianthicola
Dickeya gs fangzhongdai S1 WP 038918559 1 Dfangzhongdai
Dickeya gs fangzhongdai MK7 WP 038918559 1 Dfangzhongdai
Dickeya gs fangzhongdai B16 WP 038918559 1 Dfangzhongdai
Dickeya gs fangzhongdai NCPPB 3274 WP 042859376 1 Dfangzhongdai
Dickeya gs undicola 2B12 WP 033568676 1 Dundicola
Dickeya solani RNS 07 7 3B WP 022632979 1 Dsolani
Dickeya gs fangzhongdai M074 WP 022632979 1 Dfangzhongdai
Dickeya gs fangzhongdai ND14b WP 022632979 1 Dfangzhongdai
Dickeya solani IFB 0099 WP 022632979 1 Dsolani
Dickeya solani MK16 WP 022632979 1 Dsolani
Dickeya solani RNS 08 23 3 1 A WP 022632979 1 Dsolani
Dickeya solani GBBC 2040 WP 022632979 1 Dsolani
Dickeya solani PPO 9019 WP 022632979 1 Dsolani
Dickeya solani IPO 2222 WP 022632979 1 Dsolani
Dickeya solani MK10 WP 022632979 1 Dsolani
Dickeya solani D s0432 1 WP 022632979 1 Dsolani
Dickeya solani PPO 9134 WP 022632979 1 Dsolani

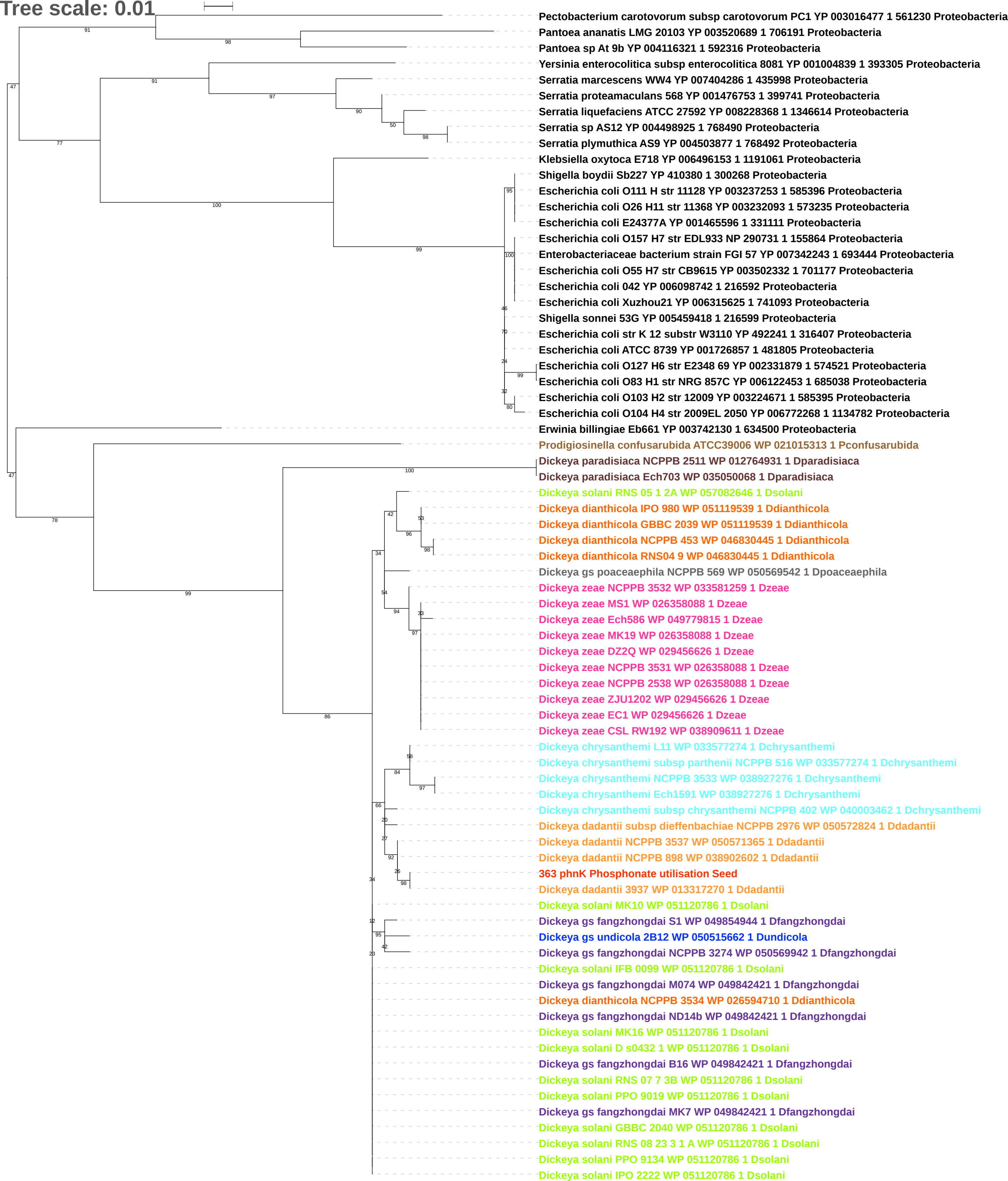
Tree scale: 0.01
Pectobacterium carotovorum subsp carotovorum PC1 YP 003016477 1 561230 Proteobacteria
Pantoea ananatis LMG 20103 YP 003520689 1 706191 Proteobacteria
Pantoea sp At 9b YP 004116321 1 592316 Proteobacteria
Yersinia enterocolitica subsp enterocolitica 8081 YP 001004839 1 393305 Proteobacteria
Serratia marcescens WW4 YP 007404286 1 435998 Proteobacteria
Serratia proteamaculans 568 YP 001476753 1 399741 Proteobacteria
Serratia liquefaciens ATCC 27592 YP 008228368 1 1346614 Proteobacteria
Serratia sp AS12 YP 004498925 1 768490 Proteobacteria
Serratia plymuthica AS9 YP 004503877 1 768492 Proteobacteria
Klebsiella oxytoca E718 YP 006496153 1 1191061 Proteobacteria
Shigella boydii Sb227 YP 410380 1 300268 Proteobacteria
Escherichia coli O111 H str 11128 YP 003237253 1 585396 Proteobacteria
Escherichia coli O26 H11 str 11368 YP 003232093 1 573235 Proteobacteria
Escherichia coli E24377A YP 001465596 1 331111 Proteobacteria
Escherichia coli O157 H7 str EDL933 NP 290731 1 155864 Proteobacteria
Enterobacteriaceae bacterium strain FGI 57 YP 007342243 1 693444 Proteobacteria
Escherichia coli O55 H7 str CB9615 YP 003502332 1 701177 Proteobacteria
Escherichia coli 042 YP 006098742 1 216592 Proteobacteria
Escherichia coli Xuzhou21 YP 006315625 1 741093 Proteobacteria
Shigella sonnei 53G YP 005459418 1 216599 Proteobacteria
Escherichia coli str K 12 substr W3110 YP 492241 1 316407 Proteobacteria
Escherichia coli ATCC 8739 YP 001726857 1 481805 Proteobacteria
Escherichia coli O127 H6 str E2348 69 YP 002331879 1 574521 Proteobacteria
Escherichia coli O83 H1 str NRG 857C YP 006122453 1 685038 Proteobacteria
Escherichia coli O103 H2 str 12009 YP 003224671 1 585395 Proteobacteria
Escherichia coli O104 H4 str 2009EL 2050 YP 006772268 1 1134782 Proteobacteria
Erwinia billingiae Eb661 YP 003742130 1 634500 Proteobacteria
Prodigiosinella confusarubida ATCC39006 WP 021015313 1 Pconfusarubida
Dickeya paradisiaca NCPPB 2511 WP 012764931 1 Dparadisiaca
Dickeya paradisiaca Ech703 WP 035050068 1 Dparadisiaca
Dickeya solani RNS 05 1 2A WP 057082646 1 Dsolani
Dickeya dianthicola IPO 980 WP 051119539 1 Ddianthicola
Dickeya dianthicola GBBC 2039 WP 051119539 1 Ddianthicola
Dickeya dianthicola NCPPB 453 WP 046830445 1 Ddianthicola
Dickeya dianthicola RNS04 9 WP 046830445 1 Ddianthicola
Dickeya gs poaceaephila NCPPB 569 WP 050569542 1 Dpoaceaephila
Dickeya zeae NCPPB 3532 WP 033581259 1 Dzeae
Dickeya zeae MS1 WP 026358088 1 Dzeae
Dickeya zeae Ech586 WP 049779815 1 Dzeae
Dickeya zeae MK19 WP 026358088 1 Dzeae
Dickeya zeae DZ2Q WP 029456626 1 Dzeae
Dickeya zeae NCPPB 3531 WP 026358088 1 Dzeae
Dickeya zeae NCPPB 2538 WP 026358088 1 Dzeae
Dickeya zeae ZJU1202 WP 029456626 1 Dzeae
Dickeya zeae EC1 WP 029456626 1 Dzeae
Dickeya zeae CSL RW192 WP 038909611 1 Dzeae
Dickeya chrysanthemi L11 WP 033577274 1 Dchrysanthemi
Dickeya chrysanthemi subsp parthenii NCPPB 516 WP 033577274 1 Dchrysanthemi
Dickeya chrysanthemi NCPPB 3533 WP 038927276 1 Dchrysanthemi
Dickeya chrysanthemi Ech1591 WP 038927276 1 Dchrysanthemi
Dickeya chrysanthemi subsp chrysanthemi NCPPB 402 WP 040003462 1 Dchrysanthemi
Dickeya dadantii subsp dieffenbachiae NCPPB 2976 WP 050572824 1 Ddadantii
Dickeya dadantii NCPPB 3537 WP 050571365 1 Ddadantii
Dickeya dadantii NCPPB 898 WP 038902602 1 Ddadantii
363 phnK Phosphonate utilisation Seed
Dickeya dadantii 3937 WP 013317270 1 Ddadantii
Dickeya solani MK10 WP 051120786 1 Dsolani
Dickeya gs fangzhongdai S1 WP 049854944 1 Dfangzhongdai
Dickeya gs undicola 2B12 WP 050515662 1 Dundicola
Dickeya gs fangzhongdai NCPPB 3274 WP 050569942 1 Dfangzhongdai
Dickeya solani IFB 0099 WP 051120786 1 Dsolani
Dickeya gs fangzhongdai M074 WP 049842421 1 Dfangzhongdai
Dickeya dianthicola NCPPB 3534 WP 026594710 1 Ddianthicola
Dickeya gs fangzhongdai ND14b WP 049842421 1 Dfangzhongdai
Dickeya solani MK16 WP 051120786 1 Dsolani
Dickeya solani D s0432 1 WP 051120786 1 Dsolani
Dickeya gs fangzhongdai B16 WP 049842421 1 Dfangzhongdai
Dickeya solani RNS 07 7 3B WP 051120786 1 Dsolani
Dickeya solani PPO 9019 WP 051120786 1 Dsolani
Dickeya gs fangzhongdai MK7 WP 049842421 1 Dfangzhongdai
Dickeya solani GBBC 2040 WP 051120786 1 Dsolani
Dickeya solani RNS 08 23 3 1 A WP 051120786 1 Dsolani
Dickeya solani PPO 9134 WP 051120786 1 Dsolani
Dickeya solani IPO 2222 WP 051120786 1 Dsolani

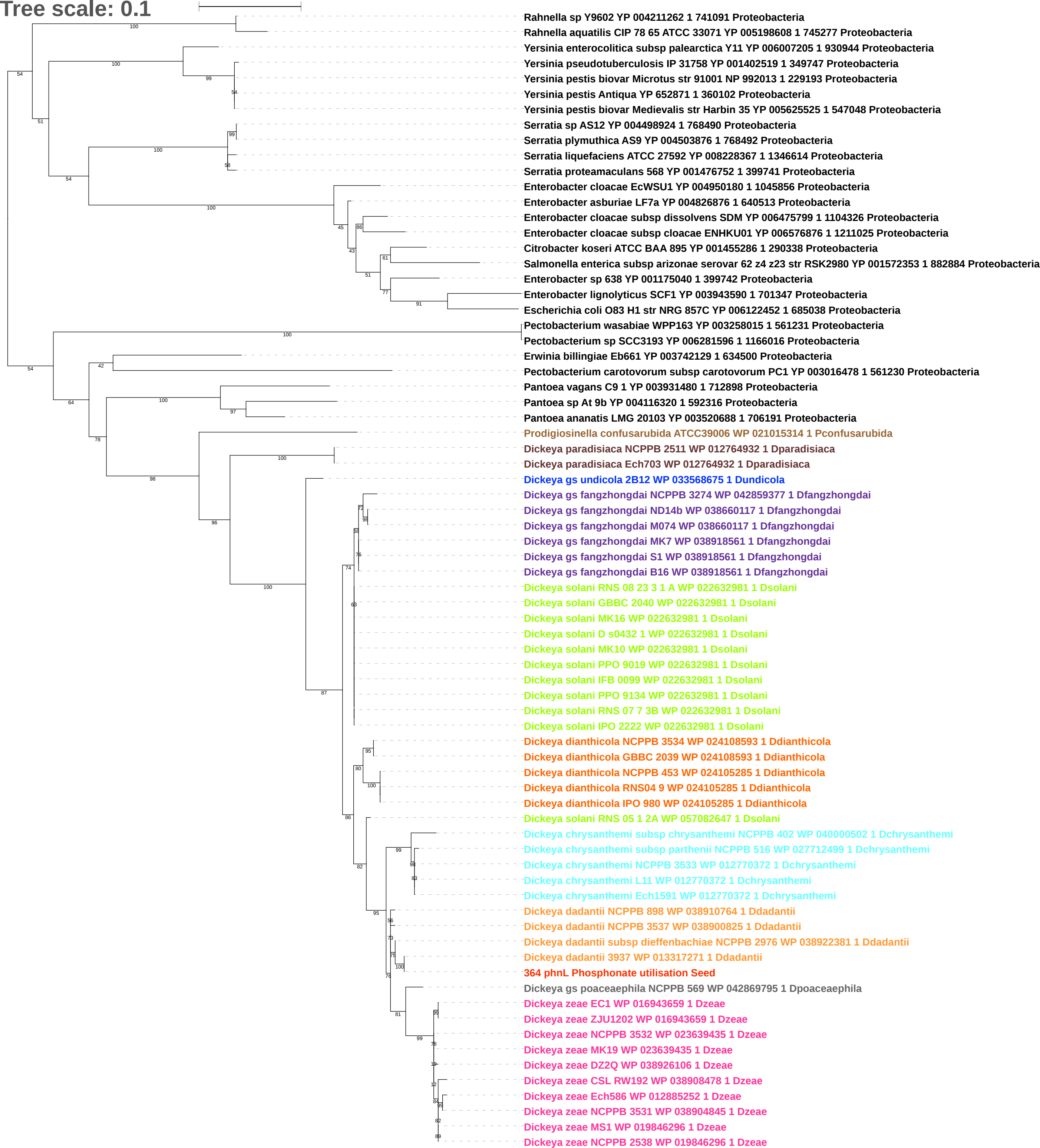
Tree scale: 0.1
Rahnella sp Y9602 YP 004211262 1 741091 Proteobacteria
Rahnella aquatilis CIP 78 65 ATCC 33071 YP 005198608 1 745277 Proteobacteria
Yersinia enterocolitica subsp palearctica Y11 YP 006007205 1 930944 Proteobacteria
Yersinia pseudotuberculosis IP 31758 YP 001402519 1 349747 Proteobacteria
Yersinia pestis biovar Microtus str 91001 NP 992013 1 229193 Proteobacteria
Yersinia pestis Antiqua YP 652871 1 360102 Proteobacteria
Yersinia pestis biovar Medievalis str Harbin 35 YP 005625525 1 547048 Proteobacteria
Serratia sp AS12 YP 004498924 1 768490 Proteobacteria
Serratia plymuthica AS9 YP 004503876 1 768492 Proteobacteria
Serratia liquefaciens ATCC 27592 YP 008228367 1 1346614 Proteobacteria
Serratia proteamaculans 568 YP 001476752 1 399741 Proteobacteria
Enterobacter cloacae EcWSU1 YP 004950180 1 1045856 Proteobacteria
Enterobacter asburiae LF7a YP 004826876 1 640513 Proteobacteria
Enterobacter cloacae subsp dissolvens SDM YP 006475799 1 1104326 Proteobacteria
Enterobacter cloacae subsp cloacae ENHKU01 YP 006576876 1 1211025 Proteobacteria
Citrobacter koseri ATCC BAA 895 YP 001455286 1 290338 Proteobacteria
Salmonella enterica subsp arizonae serovar 62 z4 z23 str RSK2980 YP 001572353 1 882884 Proteobacteria
Enterobacter sp 638 YP 001175040 1 399742 Proteobacteria
Enterobacter lignolyticus SCF1 YP 003943590 1 701347 Proteobacteria
Escherichia coli O83 H1 str NRG 857C YP 006122452 1 685038 Proteobacteria
Pectobacterium wasabiae WPP163 YP 003258015 1 561231 Proteobacteria
Pectobacterium sp SCC3193 YP 006281596 1 1166016 Proteobacteria
Erwinia billingiae Eb661 YP 003742129 1 634500 Proteobacteria
Pectobacterium carotovorum subsp carotovorum PC1 YP 003016478 1 561230 Proteobacteria
Pantoea vagans C9 1 YP 003931480 1 712898 Proteobacteria
Pantoea sp At 9b YP 004116320 1 592316 Proteobacteria
Pantoea ananatis LMG 20103 YP 003520688 1 706191 Proteobacteria
Prodigiosinella confusarubida ATCC39006 WP 021015314 1 Pconfusarubida
Dickeya paradisiaca NCPPB 2511 WP 012764932 1 Dparadisiaca
Dickeya paradisiaca Ech703 WP 012764932 1 Dparadisiaca
Dickeya gs undicola 2B12 WP 033568675 1 Dundicola
Dickeya gs fangzhongdai NCPPB 3274 WP 042859377 1 Dfangzhongdai
Dickeya gs fangzhongdai ND14b WP 038660117 1 Dfangzhongdai
Dickeya gs fangzhongdai M074 WP 038660117 1 Dfangzhongdai
Dickeya gs fangzhongdai MK7 WP 038918561 1 Dfangzhongdai
Dickeya gs fangzhongdai S1 WP 038918561 1 Dfangzhongdai
Dickeya gs fangzhongdai B16 WP 038918561 1 Dfangzhongdai
Dickeya solani RNS 08 23 3 1 A WP 022632981 1 Dsolani
Dickeya solani GBBC 2040 WP 022632981 1 Dsolani
Dickeya solani MK16 WP 022632981 1 Dsolani
Dickeya solani D s0432 1 WP 022632981 1 Dsolani
Dickeya solani MK10 WP 022632981 1 Dsolani
Dickeya solani PPO 9019 WP 022632981 1 Dsolani
Dickeya solani IFB 0099 WP 022632981 1 Dsolani
Dickeya solani PPO 9134 WP 022632981 1 Dsolani
Dickeya solani RNS 07 7 3B WP 022632981 1 Dsolani
Dickeya solani IPO 2222 WP 022632981 1 Dsolani
Dickeya dianthicola NCPPB 3534 WP 024108593 1 Ddianthicola
Dickeya dianthicola GBBC 2039 WP 024108593 1 Ddianthicola
Dickeya dianthicola NCPPB 453 WP 024105285 1 Ddianthicola
Dickeya dianthicola RNS04 9 WP 024105285 1 Ddianthicola
Dickeya dianthicola IPO 980 WP 024105285 1 Ddianthicola
Dickeya solani RNS 05 1 2A WP 057082647 1 Dsolani
Dickeya chrysanthemi subsp chrysanthemi NCPPB 402 WP 040000502 1 Dchrysanthemi
Dickeya chrysanthemi subsp parthenii NCPPB 516 WP 027712499 1 Dchrysanthemi
Dickeya chrysanthemi NCPPB 3533 WP 012770372 1 Dchrysanthemi
Dickeya chrysanthemi L11 WP 012770372 1 Dchrysanthemi
Dickeya chrysanthemi Ech1591 WP 012770372 1 Dchrysanthemi
Dickeya dadantii NCPPB 898 WP 038910764 1 Ddadantii
Dickeya dadantii NCPPB 3537 WP 038900825 1 Ddadantii
Dickeya dadantii subsp dieffenbachiae NCPPB 2976 WP 038922381 1 Ddadantii
Dickeya dadantii 3937 WP 013317271 1 Ddadantii
364 phnL Phosphonate utilisation Seed
Dickeya gs poaceaephila NCPPB 569 WP 042869795 1 Dpoaceaephila
Dickeya zeae EC1 WP 016943659 1 Dzeae
Dickeya zeae ZJU1202 WP 016943659 1 Dzeae
Dickeya zeae NCPPB 3532 WP 023639435 1 Dzeae
Dickeya zeae MK19 WP 023639435 1 Dzeae
Dickeya zeae DZ2Q WP 038926106 1 Dzeae
Dickeya zeae CSL RW192 WP 038908478 1 Dzeae
Dickeya zeae Ech586 WP 012885252 1 Dzeae
Dickeya zeae NCPPB 3531 WP 038904845 1 Dzeae
Dickeya zeae MS1 WP 019846296 1 Dzeae
Dickeya zeae NCPPB 2538 WP 019846296 1 Dzeae

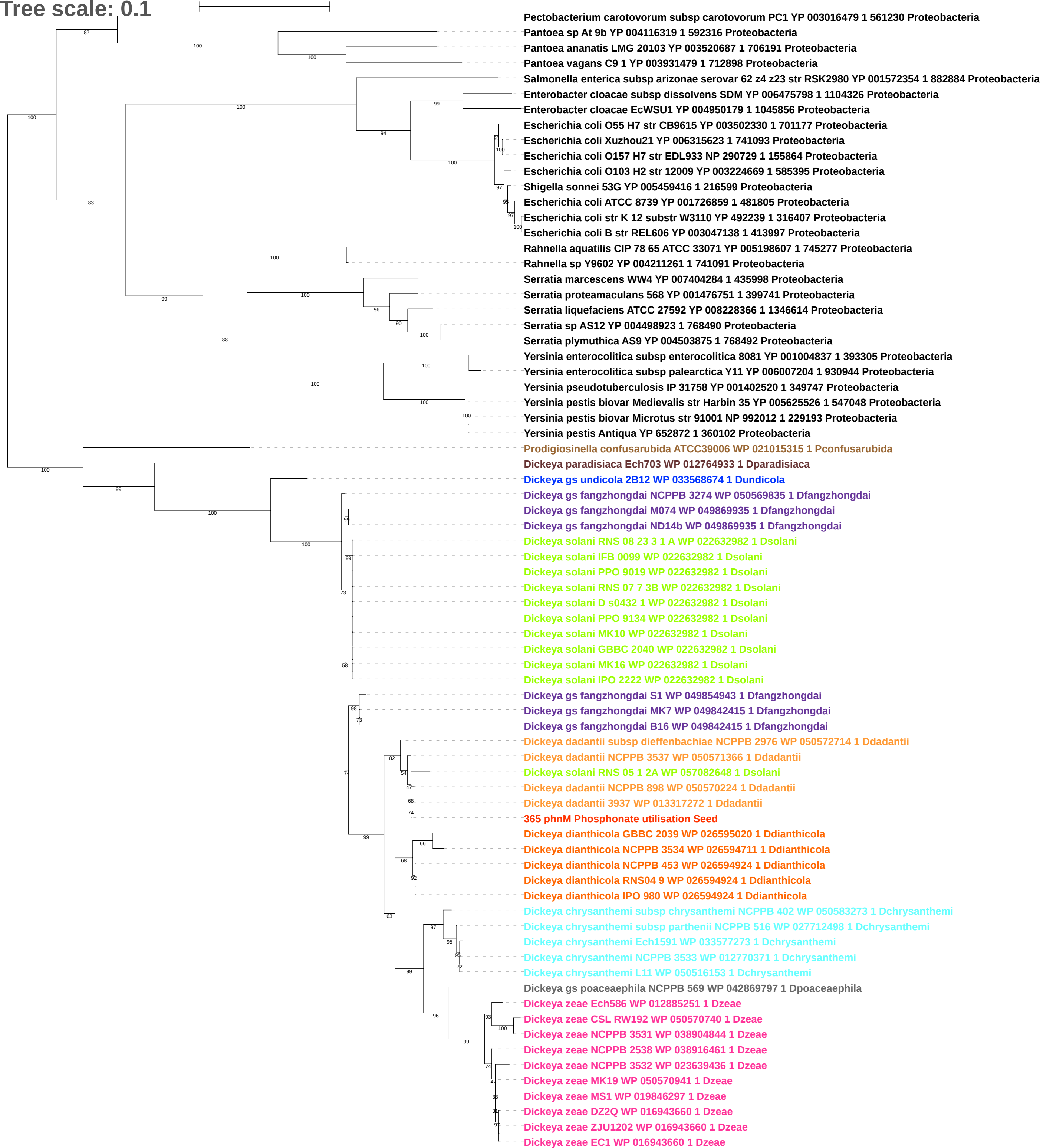
Tree scale: 0.1
Pectobacterium carotovorum subsp carotovorum PC1 YP 003016479 1 561230 Proteobacteria
Pantoea sp At 9b YP 004116319 1 592316 Proteobacteria
Pantoea ananatis LMG 20103 YP 003520687 1 706191 Proteobacteria
Pantoea vagans C9 1 YP 003931479 1 712898 Proteobacteria
Salmonella enterica subsp arizonae serovar 62 z4 z23 str RSK2980 YP 001572354 1 882884 Proteobacteria
Enterobacter cloacae subsp dissolvens SDM YP 006475798 1 1104326 Proteobacteria
Enterobacter cloacae EcWSU1 YP 004950179 1 1045856 Proteobacteria
Escherichia coli O55 H7 str CB9615 YP 003502330 1 701177 Proteobacteria
Escherichia coli Xuzhou21 YP 006315623 1 741093 Proteobacteria
Escherichia coli O157 H7 str EDL933 NP 290729 1 155864 Proteobacteria
Escherichia coli O103 H2 str 12009 YP 003224669 1 585395 Proteobacteria
Shigella sonnei 53G YP 005459416 1 216599 Proteobacteria
Escherichia coli ATCC 8739 YP 001726859 1 481805 Proteobacteria
Escherichia coli str K 12 substr W3110 YP 492239 1 316407 Proteobacteria
Escherichia coli B str REL606 YP 003047138 1 413997 Proteobacteria
Rahnella aquatilis CIP 78 65 ATCC 33071 YP 005198607 1 745277 Proteobacteria
Rahnella sp Y9602 YP 004211261 1 741091 Proteobacteria
Serratia marcescens WW4 YP 007404284 1 435998 Proteobacteria
Serratia proteamaculans 568 YP 001476751 1 399741 Proteobacteria
Serratia liquefaciens ATCC 27592 YP 008228366 1 1346614 Proteobacteria
Serratia sp AS12 YP 004498923 1 768490 Proteobacteria
Serratia plymuthica AS9 YP 004503875 1 768492 Proteobacteria
Yersinia enterocolitica subsp enterocolitica 8081 YP 001004837 1 393305 Proteobacteria
Yersinia enterocolitica subsp palearctica Y11 YP 006007204 1 930944 Proteobacteria
Yersinia pseudotuberculosis IP 31758 YP 001402520 1 349747 Proteobacteria
Yersinia pestis biovar Medievalis str Harbin 35 YP 005625526 1 547048 Proteobacteria
Yersinia pestis biovar Microtus str 91001 NP 992012 1 229193 Proteobacteria
Yersinia pestis Antiqua YP 652872 1 360102 Proteobacteria
Prodigiosinella confusarubida ATCC39006 WP 021015315 1 Pconfusarubida
Dickeya paradisiaca Ech703 WP 012764933 1 Dparadisiaca
Dickeya gs undicola 2B12 WP 033568674 1 Dundicola
Dickeya gs fangzhongdai NCPPB 3274 WP 050569835 1 Dfangzhongdai
Dickeya gs fangzhongdai M074 WP 049869935 1 Dfangzhongdai
Dickeya gs fangzhongdai ND14b WP 049869935 1 Dfangzhongdai
Dickeya solani RNS 08 23 3 1 A WP 022632982 1 Dsolani
Dickeya solani IFB 0099 WP 022632982 1 Dsolani
Dickeya solani PPO 9019 WP 022632982 1 Dsolani
Dickeya solani RNS 07 7 3B WP 022632982 1 Dsolani
Dickeya solani D s0432 1 WP 022632982 1 Dsolani
Dickeya solani PPO 9134 WP 022632982 1 Dsolani
Dickeya solani MK10 WP 022632982 1 Dsolani
Dickeya solani GBBC 2040 WP 022632982 1 Dsolani
Dickeya solani MK16 WP 022632982 1 Dsolani
Dickeya solani IPO 2222 WP 022632982 1 Dsolani
Dickeya gs fangzhongdai S1 WP 049854943 1 Dfangzhongdai
Dickeya gs fangzhongdai MK7 WP 049842415 1 Dfangzhongdai
Dickeya gs fangzhongdai B16 WP 049842415 1 Dfangzhongdai
Dickeya dadantii subsp dieffenbachiae NCPPB 2976 WP 050572714 1 Ddadantii
Dickeya dadantii NCPPB 3537 WP 050571366 1 Ddadantii
Dickeya solani RNS 05 1 2A WP 057082648 1 Dsolani
Dickeya dadantii NCPPB 898 WP 050570224 1 Ddadantii
Dickeya dadantii 3937 WP 013317272 1 Ddadantii
365 phnM Phosphonate utilisation Seed
Dickeya dianthicola GBBC 2039 WP 026595020 1 Ddianthicola
Dickeya dianthicola NCPPB 3534 WP 026594711 1 Ddianthicola
Dickeya dianthicola NCPPB 453 WP 026594924 1 Ddianthicola
Dickeya dianthicola RNS04 9 WP 026594924 1 Ddianthicola
Dickeya dianthicola IPO 980 WP 026594924 1 Ddianthicola
Dickeya chrysanthemi subsp chrysanthemi NCPPB 402 WP 050583273 1 Dchrysanthemi
Dickeya chrysanthemi subsp parthenii NCPPB 516 WP 027712498 1 Dchrysanthemi
Dickeya chrysanthemi Ech1591 WP 033577273 1 Dchrysanthemi
Dickeya chrysanthemi NCPPB 3533 WP 012770371 1 Dchrysanthemi
Dickeya chrysanthemi L11 WP 050516153 1 Dchrysanthemi
Dickeya gs poaceaephila NCPPB 569 WP 042869797 1 Dpoaceaephila
Dickeya zeae Ech586 WP 012885251 1 Dzeae
Dickeya zeae CSL RW192 WP 050570740 1 Dzeae
Dickeya zeae NCPPB 3531 WP 038904844 1 Dzeae
Dickeya zeae NCPPB 2538 WP 038916461 1 Dzeae
Dickeya zeae NCPPB 3532 WP 023639436 1 Dzeae
Dickeya zeae MK19 WP 050570941 1 Dzeae
Dickeya zeae MS1 WP 019846297 1 Dzeae
Dickeya zeae DZ2Q WP 016943660 1 Dzeae
Dickeya zeae ZJU1202 WP 016943660 1 Dzeae
Dickeya zeae EC1 WP 016943660 1 Dzeae

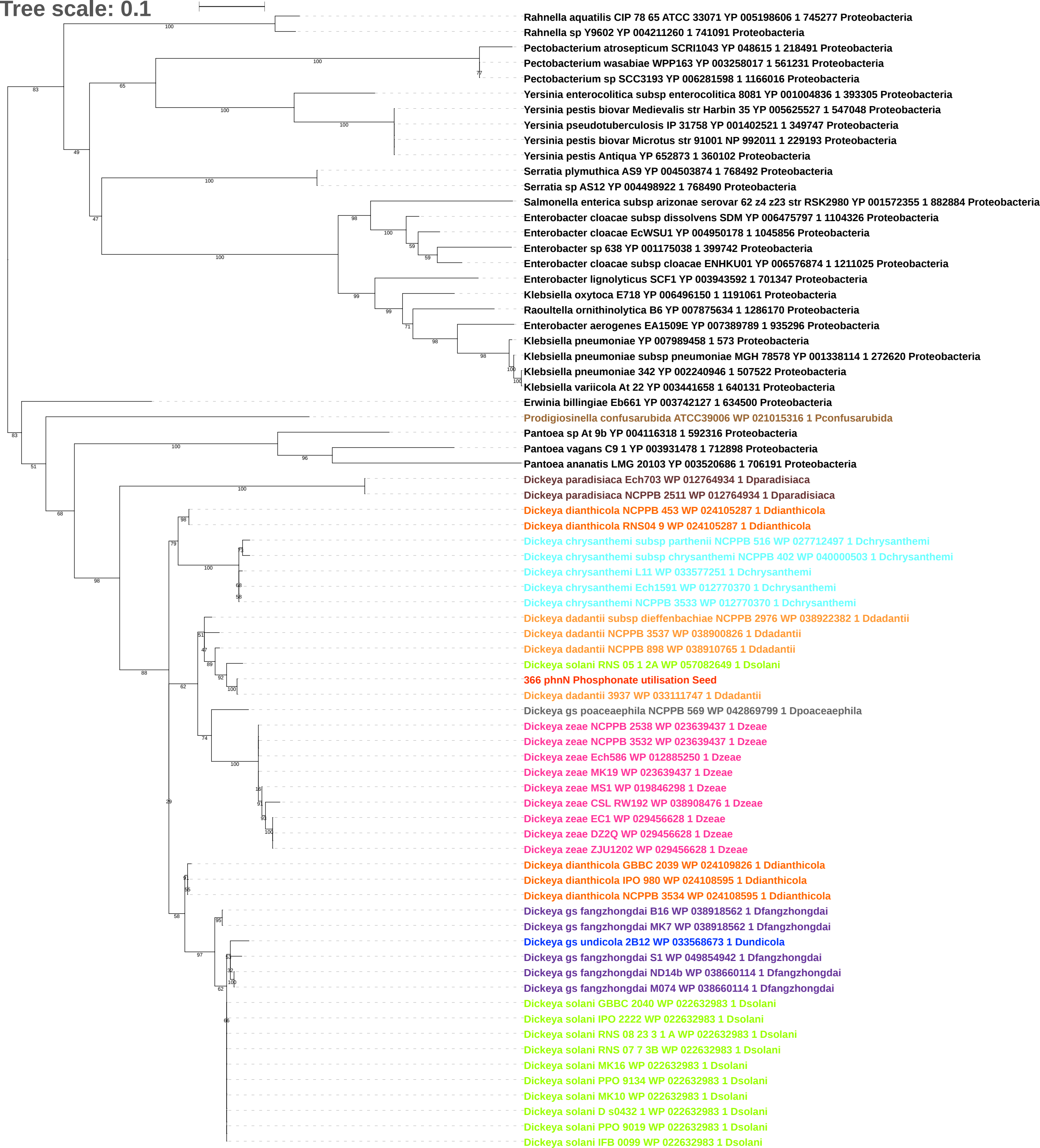
Tree scale: 0.1
Rahnella aquatilis CIP 78 65 ATCC 33071 YP 005198606 1 745277 Proteobacteria
Rahnella sp Y9602 YP 004211260 1 741091 Proteobacteria
Pectobacterium atrosepticum SCRI1043 YP 048615 1 218491 Proteobacteria
Pectobacterium wasabiae WPP163 YP 003258017 1 561231 Proteobacteria
Pectobacterium sp SCC3193 YP 006281598 1 1166016 Proteobacteria
Yersinia enterocolitica subsp enterocolitica 8081 YP 001004836 1 393305 Proteobacteria
Yersinia pestis biovar Medievalis str Harbin 35 YP 005625527 1 547048 Proteobacteria
Yersinia pseudotuberculosis IP 31758 YP 001402521 1 349747 Proteobacteria
Yersinia pestis biovar Microtus str 91001 NP 992011 1 229193 Proteobacteria
Yersinia pestis Antiqua YP 652873 1 360102 Proteobacteria
Serratia plymuthica AS9 YP 004503874 1 768492 Proteobacteria
Serratia sp AS12 YP 004498922 1 768490 Proteobacteria
Salmonella enterica subsp arizonae serovar 62 z4 z23 str RSK2980 YP 001572355 1 882884 Proteobacteria
Enterobacter cloacae subsp dissolvens SDM YP 006475797 1 1104326 Proteobacteria
Enterobacter cloacae EcWSU1 YP 004950178 1 1045856 Proteobacteria
Enterobacter sp 638 YP 001175038 1 399742 Proteobacteria
Enterobacter cloacae subsp cloacae ENHKU01 YP 006576874 1 1211025 Proteobacteria
Enterobacter lignolyticus SCF1 YP 003943592 1 701347 Proteobacteria
Klebsiella oxytoca E718 YP 006496150 1 1191061 Proteobacteria
Raoultella ornithinolytica B6 YP 007875634 1 1286170 Proteobacteria
Enterobacter aerogenes EA1509E YP 007389789 1 935296 Proteobacteria
Klebsiella pneumoniae YP 007989458 1 573 Proteobacteria
Klebsiella pneumoniae subsp pneumoniae MGH 78578 YP 001338114 1 272620 Proteobacteria
Klebsiella pneumoniae 342 YP 002240946 1 507522 Proteobacteria
Klebsiella variicola At 22 YP 003441658 1 640131 Proteobacteria
Erwinia billingiae Eb661 YP 003742127 1 634500 Proteobacteria
Prodigiosinella confusarubida ATCC39006 WP 021015316 1 Pconfusarubida
Pantoea sp At 9b YP 004116318 1 592316 Proteobacteria
Pantoea vagans C9 1 YP 003931478 1 712898 Proteobacteria
Pantoea ananatis LMG 20103 YP 003520686 1 706191 Proteobacteria
Dickeya paradisiaca Ech703 WP 012764934 1 Dparadisiaca
Dickeya paradisiaca NCPPB 2511 WP 012764934 1 Dparadisiaca
Dickeya dianthicola NCPPB 453 WP 024105287 1 Ddianthicola
Dickeya dianthicola RNS04 9 WP 024105287 1 Ddianthicola
Dickeya chrysanthemi subsp parthenii NCPPB 516 WP 027712497 1 Dchrysanthemi
Dickeya chrysanthemi subsp chrysanthemi NCPPB 402 WP 040000503 1 Dchrysanthemi
Dickeya chrysanthemi L11 WP 033577251 1 Dchrysanthemi
Dickeya chrysanthemi Ech1591 WP 012770370 1 Dchrysanthemi
Dickeya chrysanthemi NCPPB 3533 WP 012770370 1 Dchrysanthemi
Dickeya dadantii subsp dieffenbachiae NCPPB 2976 WP 038922382 1 Ddadantii
Dickeya dadantii NCPPB 3537 WP 038900826 1 Ddadantii
Dickeya dadantii NCPPB 898 WP 038910765 1 Ddadantii
Dickeya solani RNS 05 1 2A WP 057082649 1 Dsolani
366 phnN Phosphonate utilisation Seed
Dickeya dadantii 3937 WP 033111747 1 Ddadantii
Dickeya gs poaceaephila NCPPB 569 WP 042869799 1 Dpoaceaephila
Dickeya zeae NCPPB 2538 WP 023639437 1 Dzeae
Dickeya zeae NCPPB 3532 WP 023639437 1 Dzeae
Dickeya zeae Ech586 WP 012885250 1 Dzeae
Dickeya zeae MK19 WP 023639437 1 Dzeae
Dickeya zeae MS1 WP 019846298 1 Dzeae
Dickeya zeae CSL RW192 WP 038908476 1 Dzeae
Dickeya zeae EC1 WP 029456628 1 Dzeae
Dickeya zeae DZ2Q WP 029456628 1 Dzeae
Dickeya zeae ZJU1202 WP 029456628 1 Dzeae
Dickeya dianthicola GBBC 2039 WP 024109826 1 Ddianthicola
Dickeya dianthicola IPO 980 WP 024108595 1 Ddianthicola
Dickeya dianthicola NCPPB 3534 WP 024108595 1 Ddianthicola
Dickeya gs fangzhongdai B16 WP 038918562 1 Dfangzhongdai
Dickeya gs fangzhongdai MK7 WP 038918562 1 Dfangzhongdai
Dickeya gs undicola 2B12 WP 033568673 1 Dundicola
Dickeya gs fangzhongdai S1 WP 049854942 1 Dfangzhongdai
Dickeya gs fangzhongdai ND14b WP 038660114 1 Dfangzhongdai
Dickeya gs fangzhongdai M074 WP 038660114 1 Dfangzhongdai
Dickeya solani GBBC 2040 WP 022632983 1 Dsolani
Dickeya solani IPO 2222 WP 022632983 1 Dsolani
Dickeya solani RNS 08 23 3 1 A WP 022632983 1 Dsolani
Dickeya solani RNS 07 7 3B WP 022632983 1 Dsolani
Dickeya solani MK16 WP 022632983 1 Dsolani
Dickeya solani PPO 9134 WP 022632983 1 Dsolani
Dickeya solani MK10 WP 022632983 1 Dsolani
Dickeya solani D s0432 1 WP 022632983 1 Dsolani
Dickeya solani PPO 9019 WP 022632983 1 Dsolani
Dickeya solani IFB 0099 WP 022632983 1 Dsolani

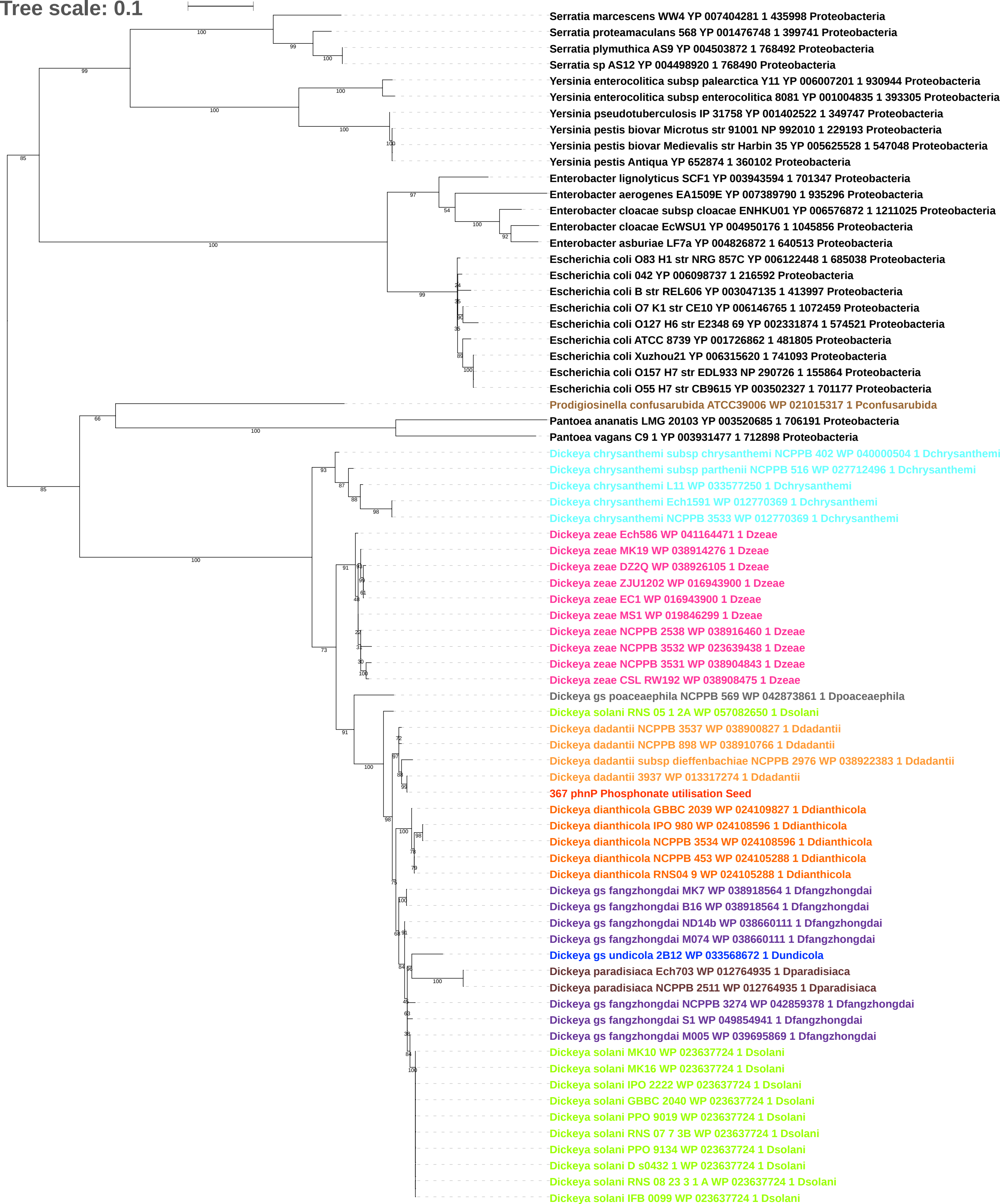

Tree scale: 0.1
Serratia marcescens WW4 YP 007404281 1 435998 Proteobacteria
Serratia proteamaculans 568 YP 001476748 1 399741 Proteobacteria
Serratia plymuthica AS9 YP 004503872 1 768492 Proteobacteria
Serratia sp AS12 YP 004498920 1 768490 Proteobacteria
Yersinia enterocolitica subsp palearctica Y11 YP 006007201 1 930944 Proteobacteria
Yersinia enterocolitica subsp enterocolitica 8081 YP 001004835 1 393305 Proteobacteria
Yersinia pseudotuberculosis IP 31758 YP 001402522 1 349747 Proteobacteria
Yersinia pestis biovar Microtus str 91001 NP 992010 1 229193 Proteobacteria
Yersinia pestis biovar Medievalis str Harbin 35 YP 005625528 1 547048 Proteobacteria
Yersinia pestis Antiqua YP 652874 1 360102 Proteobacteria
Enterobacter lignolyticus SCF1 YP 003943594 1 701347 Proteobacteria
Enterobacter aerogenes EA1509E YP 007389790 1 935296 Proteobacteria
Enterobacter cloacae subsp cloacae ENHKU01 YP 006576872 1 1211025 Proteobacteria
Enterobacter cloacae EcWSU1 YP 004950176 1 1045856 Proteobacteria
Enterobacter asburiae LF7a YP 004826872 1 640513 Proteobacteria
Escherichia coli O83 H1 str NRG 857C YP 006122448 1 685038 Proteobacteria
Escherichia coli 042 YP 006098737 1 216592 Proteobacteria
Escherichia coli B str REL606 YP 003047135 1 413997 Proteobacteria
Escherichia coli O7 K1 str CE10 YP 006146765 1 1072459 Proteobacteria
Escherichia coli O127 H6 str E2348 69 YP 002331874 1 574521 Proteobacteria
Escherichia coli ATCC 8739 YP 001726862 1 481805 Proteobacteria
Escherichia coli Xuzhou21 YP 006315620 1 741093 Proteobacteria
Escherichia coli O157 H7 str EDL933 NP 290726 1 155864 Proteobacteria
Escherichia coli O55 H7 str CB9615 YP 003502327 1 701177 Proteobacteria
Prodigiosinella confusarubida ATCC39006 WP 021015317 1 Pconfusarubida
Pantoea ananatis LMG 20103 YP 003520685 1 706191 Proteobacteria
Pantoea vagans C9 1 YP 003931477 1 712898 Proteobacteria
Dickeya chrysanthemi subsp chrysanthemi NCPPB 402 WP 040000504 1 Dchrysanthemi
Dickeya chrysanthemi subsp parthenii NCPPB 516 WP 027712496 1 Dchrysanthemi
Dickeya chrysanthemi L11 WP 033577250 1 Dchrysanthemi
Dickeya chrysanthemi Ech1591 WP 012770369 1 Dchrysanthemi
Dickeya chrysanthemi NCPPB 3533 WP 012770369 1 Dchrysanthemi
Dickeya zeae Ech586 WP 041164471 1 Dzeae
Dickeya zeae MK19 WP 038914276 1 Dzeae
Dickeya zeae DZ2Q WP 038926105 1 Dzeae
Dickeya zeae ZJU1202 WP 016943900 1 Dzeae
Dickeya zeae EC1 WP 016943900 1 Dzeae
Dickeya zeae MS1 WP 019846299 1 Dzeae
Dickeya zeae NCPPB 2538 WP 038916460 1 Dzeae
Dickeya zeae NCPPB 3532 WP 023639438 1 Dzeae
Dickeya zeae NCPPB 3531 WP 038904843 1 Dzeae
Dickeya zeae CSL RW192 WP 038908475 1 Dzeae
Dickeya gs poaceaephila NCPPB 569 WP 042873861 1 Dpoaceaephila
Dickeya solani RNS 05 1 2A WP 057082650 1 Dsolani
Dickeya dadantii NCPPB 3537 WP 038900827 1 Ddadantii
Dickeya dadantii NCPPB 898 WP 038910766 1 Ddadantii
Dickeya dadantii subsp dieffenbachiae NCPPB 2976 WP 038922383 1 Ddadantii
Dickeya dadantii 3937 WP 013317274 1 Ddadantii
367 phnP Phosphonate utilisation Seed
Dickeya dianthicola GBBC 2039 WP 024109827 1 Ddianthicola
Dickeya dianthicola IPO 980 WP 024108596 1 Ddianthicola
Dickeya dianthicola NCPPB 3534 WP 024108596 1 Ddianthicola
Dickeya dianthicola NCPPB 453 WP 024105288 1 Ddianthicola
Dickeya dianthicola RNS04 9 WP 024105288 1 Ddianthicola
Dickeya gs fangzhongdai MK7 WP 038918564 1 Dfangzhongdai
Dickeya gs fangzhongdai B16 WP 038918564 1 Dfangzhongdai
Dickeya gs fangzhongdai ND14b WP 038660111 1 Dfangzhongdai
Dickeya gs fangzhongdai M074 WP 038660111 1 Dfangzhongdai
Dickeya gs undicola 2B12 WP 033568672 1 Dundicola
Dickeya paradisiaca Ech703 WP 012764935 1 Dparadisiaca
Dickeya paradisiaca NCPPB 2511 WP 012764935 1 Dparadisiaca
Dickeya gs fangzhongdai NCPPB 3274 WP 042859378 1 Dfangzhongdai
Dickeya gs fangzhongdai S1 WP 049854941 1 Dfangzhongdai
Dickeya gs fangzhongdai M005 WP 039695869 1 Dfangzhongdai
Dickeya solani MK10 WP 023637724 1 Dsolani
Dickeya solani MK16 WP 023637724 1 Dsolani
Dickeya solani IPO 2222 WP 023637724 1 Dsolani
Dickeya solani GBBC 2040 WP 023637724 1 Dsolani
Dickeya solani PPO 9019 WP 023637724 1 Dsolani
Dickeya solani RNS 07 7 3B WP 023637724 1 Dsolani
Dickeya solani PPO 9134 WP 023637724 1 Dsolani
Dickeya solani D s0432 1 WP 023637724 1 Dsolani
Dickeya solani RNS 08 23 3 1 A WP 023637724 1 Dsolani
Dickeya solani IFB 0099 WP 023637724 1 Dsolani

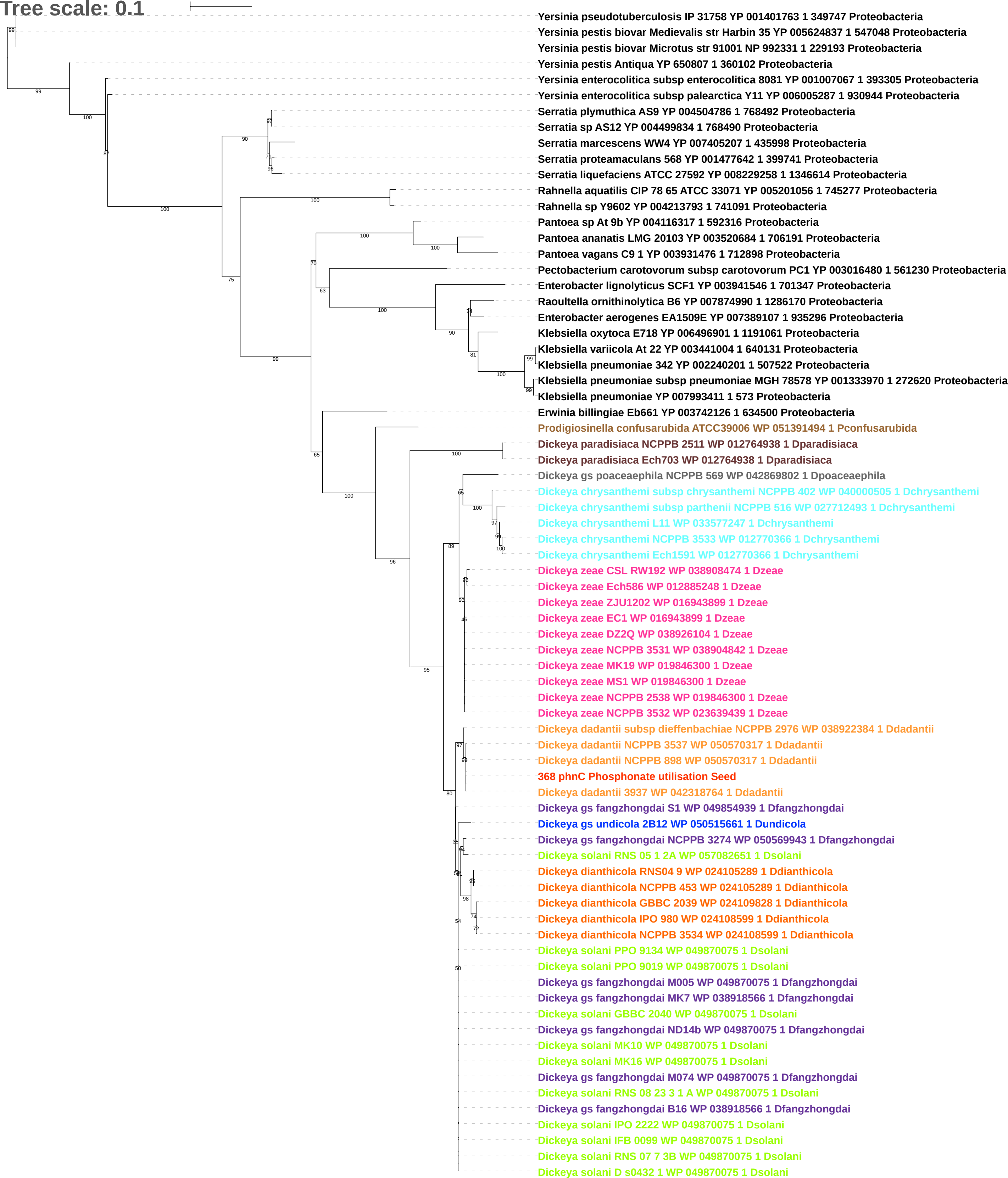
Tree scale: 0.1
Yersinia pseudotuberculosis IP 31758 YP 001401763 1 349747 Proteobacteria
Yersinia pestis biovar Medievalis str Harbin 35 YP 005624837 1 547048 Proteobacteria
Yersinia pestis biovar Microtus str 91001 NP 992331 1 229193 Proteobacteria
Yersinia pestis Antiqua YP 650807 1 360102 Proteobacteria
Yersinia enterocolitica subsp enterocolitica 8081 YP 001007067 1 393305 Proteobacteria
Yersinia enterocolitica subsp palearctica Y11 YP 006005287 1 930944 Proteobacteria
Serratia plymuthica AS9 YP 004504786 1 768492 Proteobacteria
Serratia sp AS12 YP 004499834 1 768490 Proteobacteria
Serratia marcescens WW4 YP 007405207 1 435998 Proteobacteria
Serratia proteamaculans 568 YP 001477642 1 399741 Proteobacteria
Serratia liquefaciens ATCC 27592 YP 008229258 1 1346614 Proteobacteria
Rahnella aquatilis CIP 78 65 ATCC 33071 YP 005201056 1 745277 Proteobacteria
Rahnella sp Y9602 YP 004213793 1 741091 Proteobacteria
Pantoea sp At 9b YP 004116317 1 592316 Proteobacteria
Pantoea ananatis LMG 20103 YP 003520684 1 706191 Proteobacteria
Pantoea vagans C9 1 YP 003931476 1 712898 Proteobacteria
Pectobacterium carotovorum subsp carotovorum PC1 YP 003016480 1 561230 Proteobacteria
Enterobacter lignolyticus SCF1 YP 003941546 1 701347 Proteobacteria
Raoultella ornithinolytica B6 YP 007874990 1 1286170 Proteobacteria
Enterobacter aerogenes EA1509E YP 007389107 1 935296 Proteobacteria
Klebsiella oxytoca E718 YP 006496901 1 1191061 Proteobacteria
Klebsiella variicola At 22 YP 003441004 1 640131 Proteobacteria
Klebsiella pneumoniae 342 YP 002240201 1 507522 Proteobacteria
Klebsiella pneumoniae subsp pneumoniae MGH 78578 YP 001333970 1 272620 Proteobacteria
Klebsiella pneumoniae YP 007993411 1 573 Proteobacteria
Erwinia billingiae Eb661 YP 003742126 1 634500 Proteobacteria
Prodigiosinella confusarubida ATCC39006 WP 051391494 1 Pconfusarubida
Dickeya paradisiaca NCPPB 2511 WP 012764938 1 Dparadisiaca
Dickeya paradisiaca Ech703 WP 012764938 1 Dparadisiaca
Dickeya gs poaceaephila NCPPB 569 WP 042869802 1 Dpoaceaephila
Dickeya chrysanthemi subsp chrysanthemi NCPPB 402 WP 040000505 1 Dchrysanthemi
Dickeya chrysanthemi subsp parthenii NCPPB 516 WP 027712493 1 Dchrysanthemi
Dickeya chrysanthemi L11 WP 033577247 1 Dchrysanthemi
Dickeya chrysanthemi NCPPB 3533 WP 012770366 1 Dchrysanthemi
Dickeya chrysanthemi Ech1591 WP 012770366 1 Dchrysanthemi
Dickeya zeae CSL RW192 WP 038908474 1 Dzeae
Dickeya zeae Ech586 WP 012885248 1 Dzeae
Dickeya zeae ZJU1202 WP 016943899 1 Dzeae
Dickeya zeae EC1 WP 016943899 1 Dzeae
Dickeya zeae DZ2Q WP 038926104 1 Dzeae
Dickeya zeae NCPPB 3531 WP 038904842 1 Dzeae
Dickeya zeae MK19 WP 019846300 1 Dzeae
Dickeya zeae MS1 WP 019846300 1 Dzeae
Dickeya zeae NCPPB 2538 WP 019846300 1 Dzeae
Dickeya zeae NCPPB 3532 WP 023639439 1 Dzeae
Dickeya dadantii subsp dieffenbachiae NCPPB 2976 WP 038922384 1 Ddadantii
Dickeya dadantii NCPPB 3537 WP 050570317 1 Ddadantii
Dickeya dadantii NCPPB 898 WP 050570317 1 Ddadantii
368 phnC Phosphonate utilisation Seed
Dickeya dadantii 3937 WP 042318764 1 Ddadantii
Dickeya gs fangzhongdai S1 WP 049854939 1 Dfangzhongdai
Dickeya gs undicola 2B12 WP 050515661 1 Dundicola
Dickeya gs fangzhongdai NCPPB 3274 WP 050569943 1 Dfangzhongdai
Dickeya solani RNS 05 1 2A WP 057082651 1 Dsolani
Dickeya dianthicola RNS04 9 WP 024105289 1 Ddianthicola
Dickeya dianthicola NCPPB 453 WP 024105289 1 Ddianthicola
Dickeya dianthicola GBBC 2039 WP 024109828 1 Ddianthicola
Dickeya dianthicola IPO 980 WP 024108599 1 Ddianthicola
Dickeya dianthicola NCPPB 3534 WP 024108599 1 Ddianthicola
Dickeya solani PPO 9134 WP 049870075 1 Dsolani
Dickeya solani PPO 9019 WP 049870075 1 Dsolani
Dickeya gs fangzhongdai M005 WP 049870075 1 Dfangzhongdai
Dickeya gs fangzhongdai MK7 WP 038918566 1 Dfangzhongdai
Dickeya solani GBBC 2040 WP 049870075 1 Dsolani
Dickeya gs fangzhongdai ND14b WP 049870075 1 Dfangzhongdai
Dickeya solani MK10 WP 049870075 1 Dsolani
Dickeya solani MK16 WP 049870075 1 Dsolani
Dickeya gs fangzhongdai M074 WP 049870075 1 Dfangzhongdai
Dickeya solani RNS 08 23 3 1 A WP 049870075 1 Dsolani
Dickeya gs fangzhongdai B16 WP 038918566 1 Dfangzhongdai
Dickeya solani IPO 2222 WP 049870075 1 Dsolani
Dickeya solani IFB 0099 WP 049870075 1 Dsolani
Dickeya solani RNS 07 7 3B WP 049870075 1 Dsolani
Dickeya solani D s0432 1 WP 049870075 1 Dsolani

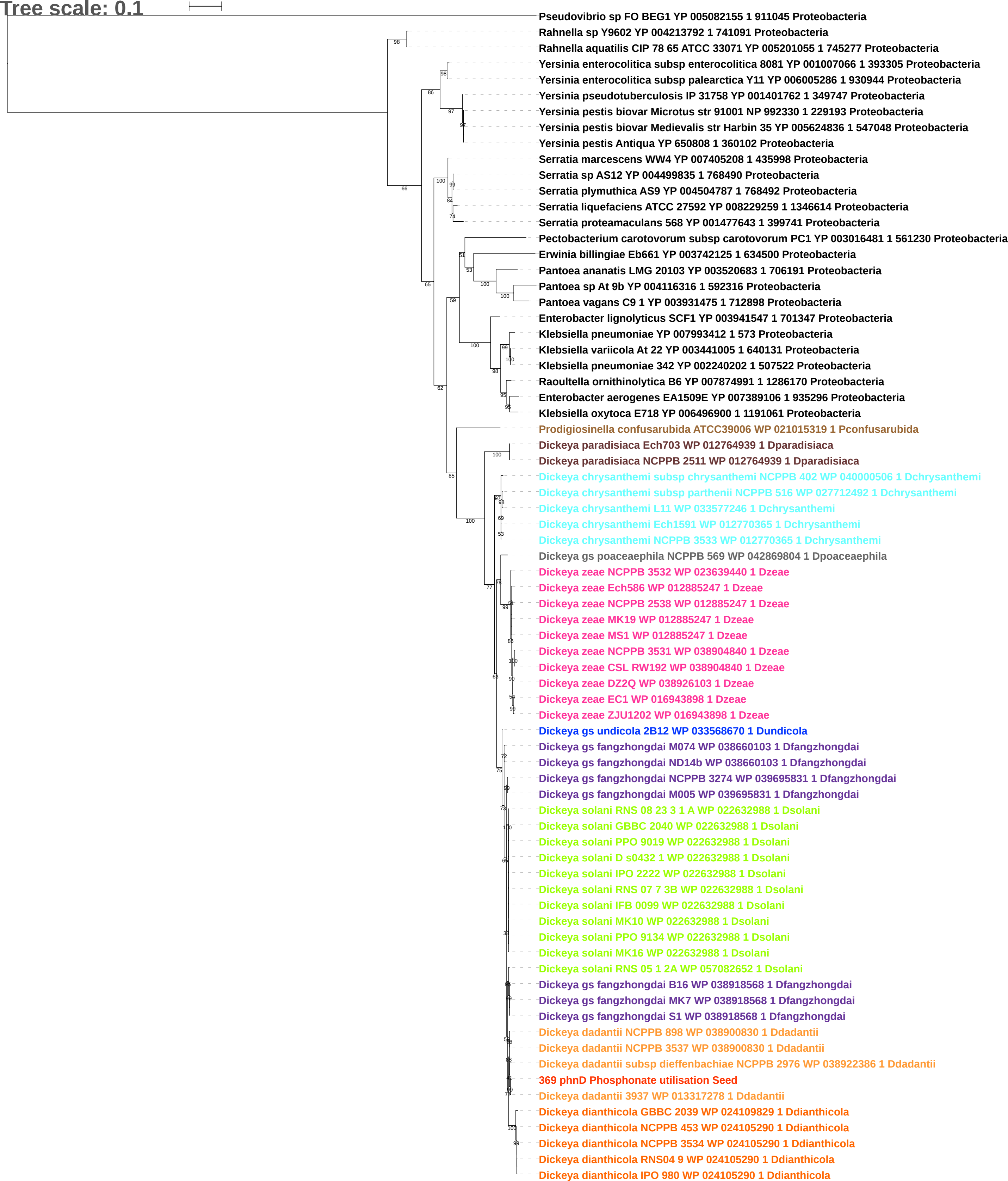
Tree scale: 0.1
Pseudovibrio sp FO BEG1 YP 005082155 1 911045 Proteobacteria
Rahnella sp Y9602 YP 004213792 1 741091 Proteobacteria
Rahnella aquatilis CIP 78 65 ATCC 33071 YP 005201055 1 745277 Proteobacteria
Yersinia enterocolitica subsp enterocolitica 8081 YP 001007066 1 393305 Proteobacteria
Yersinia enterocolitica subsp palearctica Y11 YP 006005286 1 930944 Proteobacteria
Yersinia pseudotuberculosis IP 31758 YP 001401762 1 349747 Proteobacteria
Yersinia pestis biovar Microtus str 91001 NP 992330 1 229193 Proteobacteria
Yersinia pestis biovar Medievalis str Harbin 35 YP 005624836 1 547048 Proteobacteria
Yersinia pestis Antiqua YP 650808 1 360102 Proteobacteria
Serratia marcescens WW4 YP 007405208 1 435998 Proteobacteria
Serratia sp AS12 YP 004499835 1 768490 Proteobacteria
Serratia plymuthica AS9 YP 004504787 1 768492 Proteobacteria
Serratia liquefaciens ATCC 27592 YP 008229259 1 1346614 Proteobacteria
Serratia proteamaculans 568 YP 001477643 1 399741 Proteobacteria
Pectobacterium carotovorum subsp carotovorum PC1 YP 003016481 1 561230 Proteobacteria
Erwinia billingiae Eb661 YP 003742125 1 634500 Proteobacteria
Pantoea ananatis LMG 20103 YP 003520683 1 706191 Proteobacteria
Pantoea sp At 9b YP 004116316 1 592316 Proteobacteria
Pantoea vagans C9 1 YP 003931475 1 712898 Proteobacteria
Enterobacter lignolyticus SCF1 YP 003941547 1 701347 Proteobacteria
Klebsiella pneumoniae YP 007993412 1 573 Proteobacteria
Klebsiella variicola At 22 YP 003441005 1 640131 Proteobacteria
Klebsiella pneumoniae 342 YP 002240202 1 507522 Proteobacteria
Raoultella ornithinolytica B6 YP 007874991 1 1286170 Proteobacteria
Enterobacter aerogenes EA1509E YP 007389106 1 935296 Proteobacteria
Klebsiella oxytoca E718 YP 006496900 1 1191061 Proteobacteria
Prodigiosinella confusarubida ATCC39006 WP 021015319 1 Pconfusarubida
Dickeya paradisiaca Ech703 WP 012764939 1 Dparadisiaca
Dickeya paradisiaca NCPPB 2511 WP 012764939 1 Dparadisiaca
Dickeya chrysanthemi subsp chrysanthemi NCPPB 402 WP 040000506 1 Dchrysanthemi
Dickeya chrysanthemi subsp parthenii NCPPB 516 WP 027712492 1 Dchrysanthemi
Dickeya chrysanthemi L11 WP 033577246 1 Dchrysanthemi
Dickeya chrysanthemi Ech1591 WP 012770365 1 Dchrysanthemi
Dickeya chrysanthemi NCPPB 3533 WP 012770365 1 Dchrysanthemi
Dickeya gs poaceaephila NCPPB 569 WP 042869804 1 Dpoaceaephila
Dickeya zeae NCPPB 3532 WP 023639440 1 Dzeae
Dickeya zeae Ech586 WP 012885247 1 Dzeae
Dickeya zeae NCPPB 2538 WP 012885247 1 Dzeae
Dickeya zeae MK19 WP 012885247 1 Dzeae
Dickeya zeae MS1 WP 012885247 1 Dzeae
Dickeya zeae NCPPB 3531 WP 038904840 1 Dzeae
Dickeya zeae CSL RW192 WP 038904840 1 Dzeae
Dickeya zeae DZ2Q WP 038926103 1 Dzeae
Dickeya zeae EC1 WP 016943898 1 Dzeae
Dickeya zeae ZJU1202 WP 016943898 1 Dzeae
Dickeya gs undicola 2B12 WP 033568670 1 Dundicola
Dickeya gs fangzhongdai M074 WP 038660103 1 Dfangzhongdai
Dickeya gs fangzhongdai ND14b WP 038660103 1 Dfangzhongdai
Dickeya gs fangzhongdai NCPPB 3274 WP 039695831 1 Dfangzhongdai
Dickeya gs fangzhongdai M005 WP 039695831 1 Dfangzhongdai
Dickeya solani RNS 08 23 3 1 A WP 022632988 1 Dsolani
Dickeya solani GBBC 2040 WP 022632988 1 Dsolani
Dickeya solani PPO 9019 WP 022632988 1 Dsolani
Dickeya solani D s0432 1 WP 022632988 1 Dsolani
Dickeya solani IPO 2222 WP 022632988 1 Dsolani
Dickeya solani RNS 07 7 3B WP 022632988 1 Dsolani
Dickeya solani IFB 0099 WP 022632988 1 Dsolani
Dickeya solani MK10 WP 022632988 1 Dsolani
Dickeya solani PPO 9134 WP 022632988 1 Dsolani
Dickeya solani MK16 WP 022632988 1 Dsolani
Dickeya solani RNS 05 1 2A WP 057082652 1 Dsolani
Dickeya gs fangzhongdai B16 WP 038918568 1 Dfangzhongdai
Dickeya gs fangzhongdai MK7 WP 038918568 1 Dfangzhongdai
Dickeya gs fangzhongdai S1 WP 038918568 1 Dfangzhongdai
Dickeya dadantii NCPPB 898 WP 038900830 1 Ddadantii
Dickeya dadantii NCPPB 3537 WP 038900830 1 Ddadantii
Dickeya dadantii subsp dieffenbachiae NCPPB 2976 WP 038922386 1 Ddadantii
369 phnD Phosphonate utilisation Seed
Dickeya dadantii 3937 WP 013317278 1 Ddadantii
Dickeya dianthicola GBBC 2039 WP 024109829 1 Ddianthicola
Dickeya dianthicola NCPPB 453 WP 024105290 1 Ddianthicola
Dickeya dianthicola NCPPB 3534 WP 024105290 1 Ddianthicola
Dickeya dianthicola RNS04 9 WP 024105290 1 Ddianthicola
Dickeya dianthicola IPO 980 WP 024105290 1 Ddianthicola

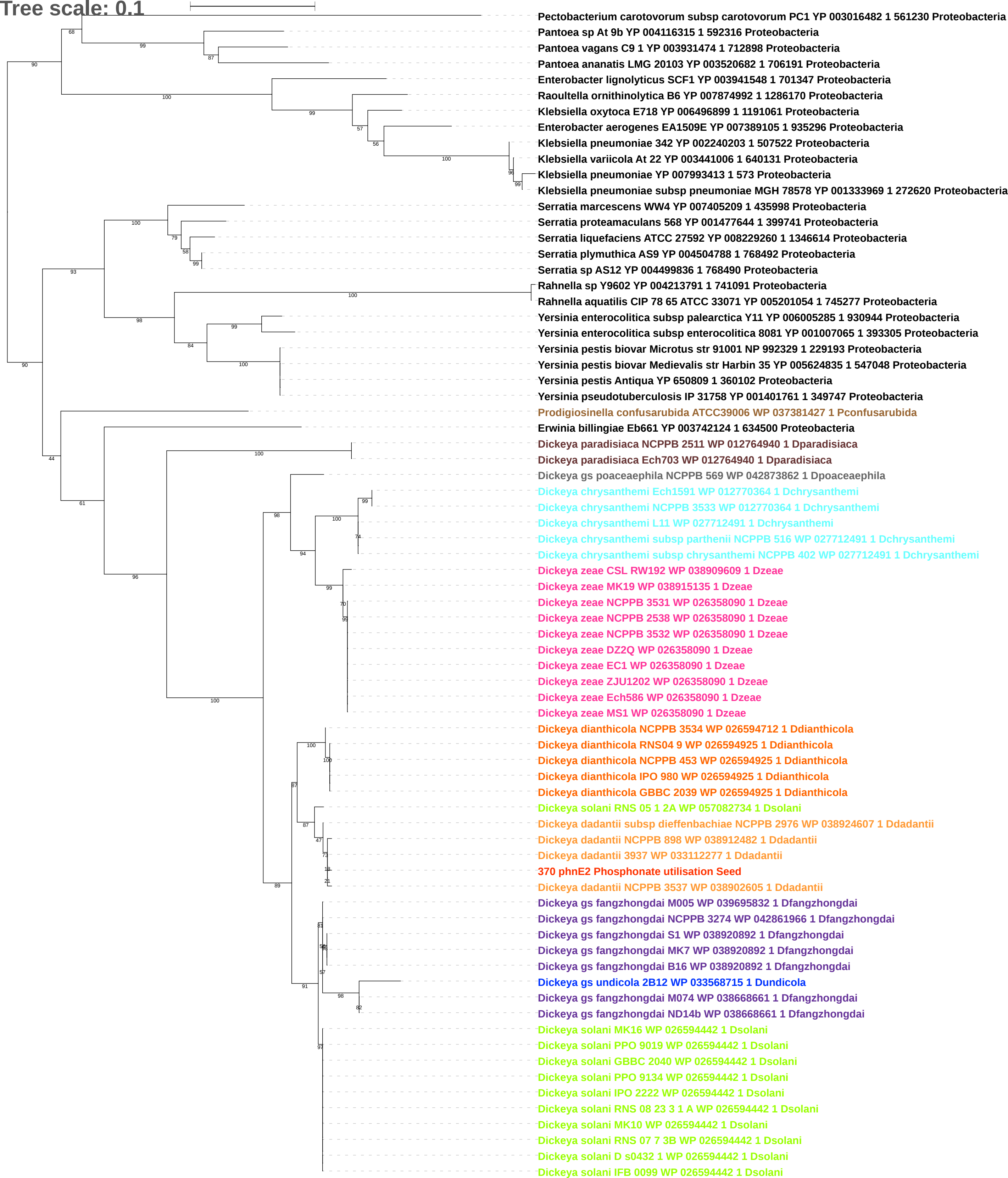
Tree scale: 0.1
Pectobacterium carotovorum subsp carotovorum PC1 YP 003016482 1 561230 Proteobacteria
Pantoea sp At 9b YP 004116315 1 592316 Proteobacteria
Pantoea vagans C9 1 YP 003931474 1 712898 Proteobacteria
Pantoea ananatis LMG 20103 YP 003520682 1 706191 Proteobacteria
Enterobacter lignolyticus SCF1 YP 003941548 1 701347 Proteobacteria
Raoultella ornithinolytica B6 YP 007874992 1 1286170 Proteobacteria
Klebsiella oxytoca E718 YP 006496899 1 1191061 Proteobacteria
Enterobacter aerogenes EA1509E YP 007389105 1 935296 Proteobacteria
Klebsiella pneumoniae 342 YP 002240203 1 507522 Proteobacteria
Klebsiella variicola At 22 YP 003441006 1 640131 Proteobacteria
Klebsiella pneumoniae YP 007993413 1 573 Proteobacteria
Klebsiella pneumoniae subsp pneumoniae MGH 78578 YP 001333969 1 272620 Proteobacteria
Serratia marcescens WW4 YP 007405209 1 435998 Proteobacteria
Serratia proteamaculans 568 YP 001477644 1 399741 Proteobacteria
Serratia liquefaciens ATCC 27592 YP 008229260 1 1346614 Proteobacteria
Serratia plymuthica AS9 YP 004504788 1 768492 Proteobacteria
Serratia sp AS12 YP 004499836 1 768490 Proteobacteria
Rahnella sp Y9602 YP 004213791 1 741091 Proteobacteria
Rahnella aquatilis CIP 78 65 ATCC 33071 YP 005201054 1 745277 Proteobacteria
Yersinia enterocolitica subsp palearctica Y11 YP 006005285 1 930944 Proteobacteria
Yersinia enterocolitica subsp enterocolitica 8081 YP 001007065 1 393305 Proteobacteria
Yersinia pestis biovar Microtus str 91001 NP 992329 1 229193 Proteobacteria
Yersinia pestis biovar Medievalis str Harbin 35 YP 005624835 1 547048 Proteobacteria
Yersinia pestis Antiqua YP 650809 1 360102 Proteobacteria
Yersinia pseudotuberculosis IP 31758 YP 001401761 1 349747 Proteobacteria
Prodigiosinella confusarubida ATCC39006 WP 037381427 1 Pconfusarubida
Erwinia billingiae Eb661 YP 003742124 1 634500 Proteobacteria
Dickeya paradisiaca NCPPB 2511 WP 012764940 1 Dparadisiaca
Dickeya paradisiaca Ech703 WP 012764940 1 Dparadisiaca
Dickeya gs poaceaephila NCPPB 569 WP 042873862 1 Dpoaceaephila
Dickeya chrysanthemi Ech1591 WP 012770364 1 Dchrysanthemi
Dickeya chrysanthemi NCPPB 3533 WP 012770364 1 Dchrysanthemi
Dickeya chrysanthemi L11 WP 027712491 1 Dchrysanthemi
Dickeya chrysanthemi subsp parthenii NCPPB 516 WP 027712491 1 Dchrysanthemi
Dickeya chrysanthemi subsp chrysanthemi NCPPB 402 WP 027712491 1 Dchrysanthemi
Dickeya zeae CSL RW192 WP 038909609 1 Dzeae
Dickeya zeae MK19 WP 038915135 1 Dzeae
Dickeya zeae NCPPB 3531 WP 026358090 1 Dzeae
Dickeya zeae NCPPB 2538 WP 026358090 1 Dzeae
Dickeya zeae NCPPB 3532 WP 026358090 1 Dzeae
Dickeya zeae DZ2Q WP 026358090 1 Dzeae
Dickeya zeae EC1 WP 026358090 1 Dzeae
Dickeya zeae ZJU1202 WP 026358090 1 Dzeae
Dickeya zeae Ech586 WP 026358090 1 Dzeae
Dickeya zeae MS1 WP 026358090 1 Dzeae
Dickeya dianthicola NCPPB 3534 WP 026594712 1 Ddianthicola
Dickeya dianthicola RNS04 9 WP 026594925 1 Ddianthicola
Dickeya dianthicola NCPPB 453 WP 026594925 1 Ddianthicola
Dickeya dianthicola IPO 980 WP 026594925 1 Ddianthicola
Dickeya dianthicola GBBC 2039 WP 026594925 1 Ddianthicola
Dickeya solani RNS 05 1 2A WP 057082734 1 Dsolani
Dickeya dadantii subsp dieffenbachiae NCPPB 2976 WP 038924607 1 Ddadantii
Dickeya dadantii NCPPB 898 WP 038912482 1 Ddadantii
Dickeya dadantii 3937 WP 033112277 1 Ddadantii
370 phnE2 Phosphonate utilisation Seed
Dickeya dadantii NCPPB 3537 WP 038902605 1 Ddadantii
Dickeya gs fangzhongdai M005 WP 039695832 1 Dfangzhongdai
Dickeya gs fangzhongdai NCPPB 3274 WP 042861966 1 Dfangzhongdai
Dickeya gs fangzhongdai S1 WP 038920892 1 Dfangzhongdai
Dickeya gs fangzhongdai MK7 WP 038920892 1 Dfangzhongdai
Dickeya gs fangzhongdai B16 WP 038920892 1 Dfangzhongdai
Dickeya gs undicola 2B12 WP 033568715 1 Dundicola
Dickeya gs fangzhongdai M074 WP 038668661 1 Dfangzhongdai
Dickeya gs fangzhongdai ND14b WP 038668661 1 Dfangzhongdai
Dickeya solani MK16 WP 026594442 1 Dsolani
Dickeya solani PPO 9019 WP 026594442 1 Dsolani
Dickeya solani GBBC 2040 WP 026594442 1 Dsolani
Dickeya solani PPO 9134 WP 026594442 1 Dsolani
Dickeya solani IPO 2222 WP 026594442 1 Dsolani
Dickeya solani RNS 08 23 3 1 A WP 026594442 1 Dsolani
Dickeya solani MK10 WP 026594442 1 Dsolani
Dickeya solani RNS 07 7 3B WP 026594442 1 Dsolani
Dickeya solani D s0432 1 WP 026594442 1 Dsolani
Dickeya solani IFB 0099 WP 026594442 1 Dsolani

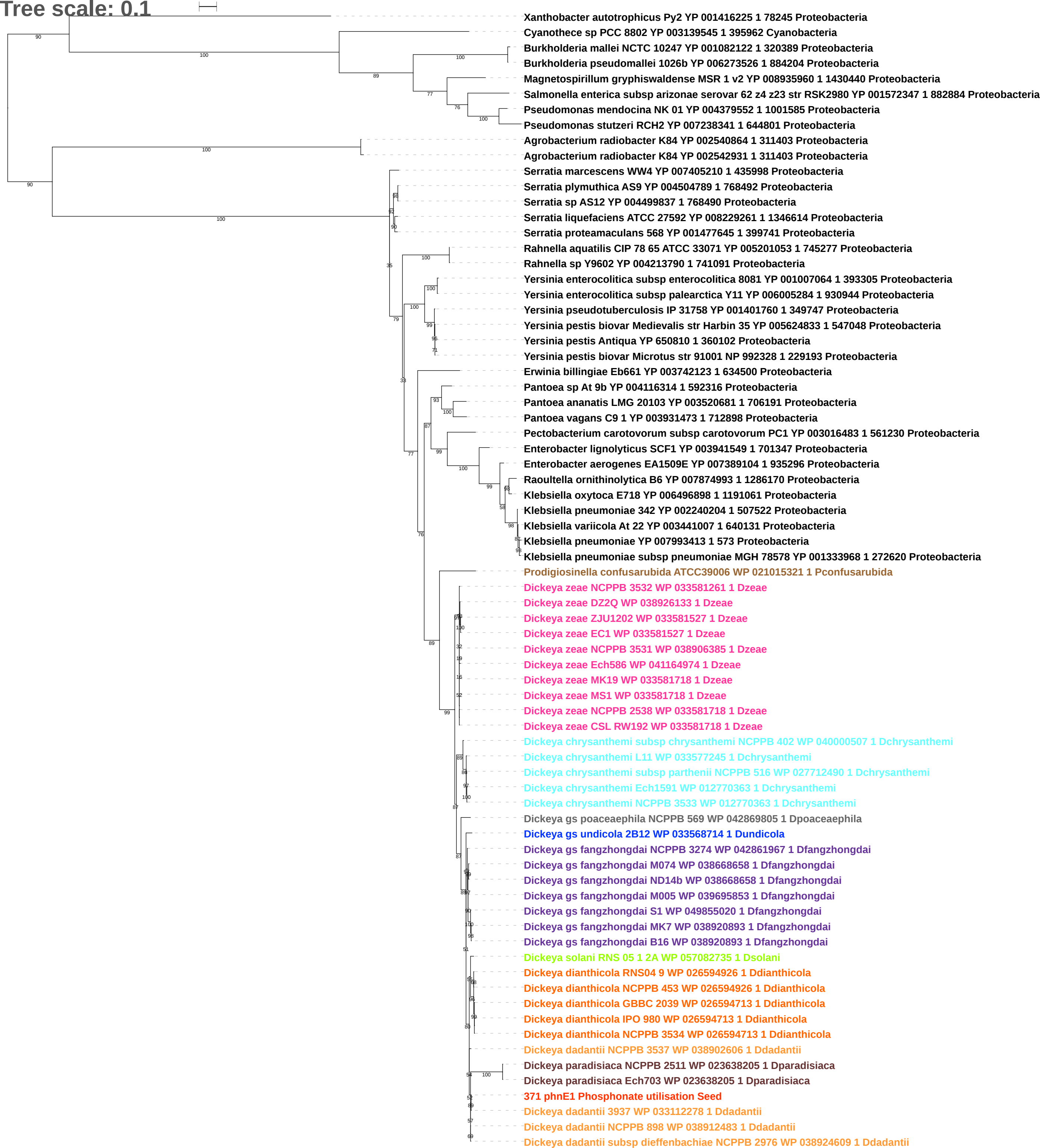
Tree scale: 0.1
Xanthobacter autotrophicus Py2 YP 001416225 1 78245 Proteobacteria
Cyanothece sp PCC 8802 YP 003139545 1 395962 Cyanobacteria
Burkholderia mallei NCTC 10247 YP 001082122 1 320389 Proteobacteria
Burkholderia pseudomallei 1026b YP 006273526 1 884204 Proteobacteria
Magnetospirillum gryphiswaldense MSR 1 v2 YP 008935960 1 1430440 Proteobacteria
Salmonella enterica subsp arizonae serovar 62 z4 z23 str RSK2980 YP 001572347 1 882884 Proteobacteria
Pseudomonas mendocina NK 01 YP 004379552 1 1001585 Proteobacteria
Pseudomonas stutzeri RCH2 YP 007238341 1 644801 Proteobacteria
Agrobacterium radiobacter K84 YP 002540864 1 311403 Proteobacteria
Agrobacterium radiobacter K84 YP 002542931 1 311403 Proteobacteria
Serratia marcescens WW4 YP 007405210 1 435998 Proteobacteria
Serratia plymuthica AS9 YP 004504789 1 768492 Proteobacteria
Serratia sp AS12 YP 004499837 1 768490 Proteobacteria
Serratia liquefaciens ATCC 27592 YP 008229261 1 1346614 Proteobacteria
Serratia proteamaculans 568 YP 001477645 1 399741 Proteobacteria
Rahnella aquatilis CIP 78 65 ATCC 33071 YP 005201053 1 745277 Proteobacteria
Rahnella sp Y9602 YP 004213790 1 741091 Proteobacteria
Yersinia enterocolitica subsp enterocolitica 8081 YP 001007064 1 393305 Proteobacteria
Yersinia enterocolitica subsp palearctica Y11 YP 006005284 1 930944 Proteobacteria
Yersinia pseudotuberculosis IP 31758 YP 001401760 1 349747 Proteobacteria
Yersinia pestis biovar Medievalis str Harbin 35 YP 005624833 1 547048 Proteobacteria
Yersinia pestis Antiqua YP 650810 1 360102 Proteobacteria
Yersinia pestis biovar Microtus str 91001 NP 992328 1 229193 Proteobacteria
Erwinia billingiae Eb661 YP 003742123 1 634500 Proteobacteria
Pantoea sp At 9b YP 004116314 1 592316 Proteobacteria
Pantoea ananatis LMG 20103 YP 003520681 1 706191 Proteobacteria
Pantoea vagans C9 1 YP 003931473 1 712898 Proteobacteria
Pectobacterium carotovorum subsp carotovorum PC1 YP 003016483 1 561230 Proteobacteria
Enterobacter lignolyticus SCF1 YP 003941549 1 701347 Proteobacteria
Enterobacter aerogenes EA1509E YP 007389104 1 935296 Proteobacteria
Raoultella ornithinolytica B6 YP 007874993 1 1286170 Proteobacteria
Klebsiella oxytoca E718 YP 006496898 1 1191061 Proteobacteria
Klebsiella pneumoniae 342 YP 002240204 1 507522 Proteobacteria
Klebsiella variicola At 22 YP 003441007 1 640131 Proteobacteria
Klebsiella pneumoniae YP 007993413 1 573 Proteobacteria
Klebsiella pneumoniae subsp pneumoniae MGH 78578 YP 001333968 1 272620 Proteobacteria
Prodigiosinella confusarubida ATCC39006 WP 021015321 1 Pconfusarubida
Dickeya zeae NCPPB 3532 WP 033581261 1 Dzeae
Dickeya zeae DZ2Q WP 038926133 1 Dzeae
Dickeya zeae ZJU1202 WP 033581527 1 Dzeae
Dickeya zeae EC1 WP 033581527 1 Dzeae
Dickeya zeae NCPPB 3531 WP 038906385 1 Dzeae
Dickeya zeae Ech586 WP 041164974 1 Dzeae
Dickeya zeae MK19 WP 033581718 1 Dzeae
Dickeya zeae MS1 WP 033581718 1 Dzeae
Dickeya zeae NCPPB 2538 WP 033581718 1 Dzeae
Dickeya zeae CSL RW192 WP 033581718 1 Dzeae
Dickeya chrysanthemi subsp chrysanthemi NCPPB 402 WP 040000507 1 Dchrysanthemi
Dickeya chrysanthemi L11 WP 033577245 1 Dchrysanthemi
Dickeya chrysanthemi subsp parthenii NCPPB 516 WP 027712490 1 Dchrysanthemi
Dickeya chrysanthemi Ech1591 WP 012770363 1 Dchrysanthemi
Dickeya chrysanthemi NCPPB 3533 WP 012770363 1 Dchrysanthemi
Dickeya gs poaceaephila NCPPB 569 WP 042869805 1 Dpoaceaephila
Dickeya gs undicola 2B12 WP 033568714 1 Dundicola
Dickeya gs fangzhongdai NCPPB 3274 WP 042861967 1 Dfangzhongdai
Dickeya gs fangzhongdai M074 WP 038668658 1 Dfangzhongdai
Dickeya gs fangzhongdai ND14b WP 038668658 1 Dfangzhongdai
Dickeya gs fangzhongdai M005 WP 039695853 1 Dfangzhongdai
Dickeya gs fangzhongdai S1 WP 049855020 1 Dfangzhongdai
Dickeya gs fangzhongdai MK7 WP 038920893 1 Dfangzhongdai
Dickeya gs fangzhongdai B16 WP 038920893 1 Dfangzhongdai
Dickeya solani RNS 05 1 2A WP 057082735 1 Dsolani
Dickeya dianthicola RNS04 9 WP 026594926 1 Ddianthicola
Dickeya dianthicola NCPPB 453 WP 026594926 1 Ddianthicola
Dickeya dianthicola GBBC 2039 WP 026594713 1 Ddianthicola
Dickeya dianthicola IPO 980 WP 026594713 1 Ddianthicola
Dickeya dianthicola NCPPB 3534 WP 026594713 1 Ddianthicola
Dickeya dadantii NCPPB 3537 WP 038902606 1 Ddadantii
Dickeya paradisiaca NCPPB 2511 WP 023638205 1 Dparadisiaca
Dickeya paradisiaca Ech703 WP 023638205 1 Dparadisiaca
371 phnE1 Phosphonate utilisation Seed
Dickeya dadantii 3937 WP 033112278 1 Ddadantii
Dickeya dadantii NCPPB 898 WP 038912483 1 Ddadantii
Dickeya dadantii subsp dieffenbachiae NCPPB 2976 WP 038924609 1 Ddadantii

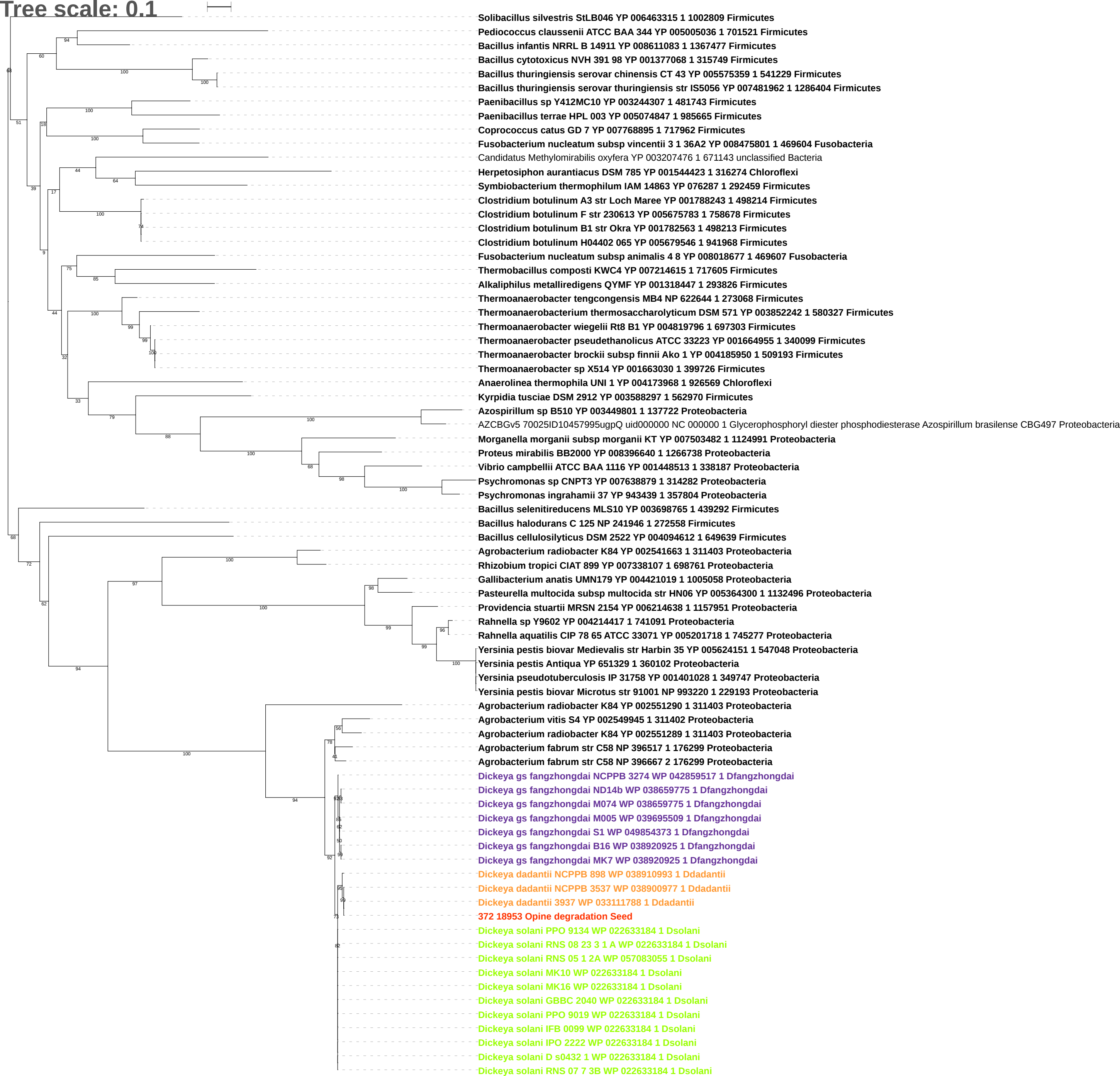
Tree scale: 0.1
Solibacillus silvestris StLB046 YP 006463315 1 1002809 Firmicutes
Pediococcus claussenii ATCC BAA 344 YP 005005036 1 701521 Firmicutes
Bacillus infantis NRRL B 14911 YP 008611083 1 1367477 Firmicutes
Bacillus cytotoxicus NVH 391 98 YP 001377068 1 315749 Firmicutes
Bacillus thuringiensis serovar chinensis CT 43 YP 005575359 1 541229 Firmicutes
Bacillus thuringiensis serovar thuringiensis str IS5056 YP 007481962 1 1286404 Firmicutes
Paenibacillus sp Y412MC10 YP 003244307 1 481743 Firmicutes
Paenibacillus terrae HPL 003 YP 005074847 1 985665 Firmicutes
Coprococcus catus GD 7 YP 007768895 1 717962 Firmicutes
Fusobacterium nucleatum subsp vincentii 3 1 36A2 YP 008475801 1 469604 Fusobacteria
Candidatus Methylomirabilis oxyfera YP 003207476 1 671143 unclassified Bacteria
Herpetosiphon aurantiacus DSM 785 YP 001544423 1 316274 Chloroflexi
Symbiobacterium thermophilum IAM 14863 YP 076287 1 292459 Firmicutes
Clostridium botulinum A3 str Loch Maree YP 001788243 1 498214 Firmicutes
Clostridium botulinum F str 230613 YP 005675783 1 758678 Firmicutes
Clostridium botulinum B1 str Okra YP 001782563 1 498213 Firmicutes
Clostridium botulinum H04402 065 YP 005679546 1 941968 Firmicutes
Fusobacterium nucleatum subsp animalis 4 8 YP 008018677 1 469607 Fusobacteria
Thermobacillus composti KWC4 YP 007214615 1 717605 Firmicutes
Alkaliphilus metalliredigens QYMF YP 001318447 1 293826 Firmicutes
Thermoanaerobacter tengcongensis MB4 NP 622644 1 273068 Firmicutes
Thermoanaerobacterium thermosaccharolyticum DSM 571 YP 003852242 1 580327 Firmicutes
Thermoanaerobacter wiegelii Rt8 B1 YP 004819796 1 697303 Firmicutes
Thermoanaerobacter pseudethanolicus ATCC 33223 YP 001664955 1 340099 Firmicutes
Thermoanaerobacter brockii subsp finnii Ako 1 YP 004185950 1 509193 Firmicutes
Thermoanaerobacter sp X514 YP 001663030 1 399726 Firmicutes
Anaerolinea thermophila UNI 1 YP 004173968 1 926569 Chloroflexi
Kyrpidia tusciae DSM 2912 YP 003588297 1 562970 Firmicutes
Azospirillum sp B510 YP 003449801 1 137722 Proteobacteria
AZCBGv5 70025ID10457995ugpQ uid000000 NC 000000 1 Glycerophosphoryl diester phosphodiesterase Azospirillum brasilense CBG497 Proteobacteria
Morganella morganii subsp morganii KT YP 007503482 1 1124991 Proteobacteria
Proteus mirabilis BB2000 YP 008396640 1 1266738 Proteobacteria
Vibrio campbellii ATCC BAA 1116 YP 001448513 1 338187 Proteobacteria
Psychromonas sp CNPT3 YP 007638879 1 314282 Proteobacteria
Psychromonas ingrahamii 37 YP 943439 1 357804 Proteobacteria
Bacillus selenitireducens MLS10 YP 003698765 1 439292 Firmicutes
Bacillus halodurans C 125 NP 241946 1 272558 Firmicutes
Bacillus cellulosilyticus DSM 2522 YP 004094612 1 649639 Firmicutes
Agrobacterium radiobacter K84 YP 002541663 1 311403 Proteobacteria
Rhizobium tropici CIAT 899 YP 007338107 1 698761 Proteobacteria
Gallibacterium anatis UMN179 YP 004421019 1 1005058 Proteobacteria
Pasteurella multocida subsp multocida str HN06 YP 005364300 1 1132496 Proteobacteria
Providencia stuartii MRSN 2154 YP 006214638 1 1157951 Proteobacteria
Rahnella sp Y9602 YP 004214417 1 741091 Proteobacteria
Rahnella aquatilis CIP 78 65 ATCC 33071 YP 005201718 1 745277 Proteobacteria
Yersinia pestis biovar Medievalis str Harbin 35 YP 005624151 1 547048 Proteobacteria
Yersinia pestis Antiqua YP 651329 1 360102 Proteobacteria
Yersinia pseudotuberculosis IP 31758 YP 001401028 1 349747 Proteobacteria
Yersinia pestis biovar Microtus str 91001 NP 993220 1 229193 Proteobacteria
Agrobacterium radiobacter K84 YP 002551290 1 311403 Proteobacteria
Agrobacterium vitis S4 YP 002549945 1 311402 Proteobacteria
Agrobacterium radiobacter K84 YP 002551289 1 311403 Proteobacteria
Agrobacterium fabrum str C58 NP 396517 1 176299 Proteobacteria
Agrobacterium fabrum str C58 NP 396667 2 176299 Proteobacteria
Dickeya gs fangzhongdai NCPPB 3274 WP 042859517 1 Dfangzhongdai
Dickeya gs fangzhongdai ND14b WP 038659775 1 Dfangzhongdai
Dickeya gs fangzhongdai M074 WP 038659775 1 Dfangzhongdai
Dickeya gs fangzhongdai M005 WP 039695509 1 Dfangzhongdai
Dickeya gs fangzhongdai S1 WP 049854373 1 Dfangzhongdai
Dickeya gs fangzhongdai B16 WP 038920925 1 Dfangzhongdai
Dickeya gs fangzhongdai MK7 WP 038920925 1 Dfangzhongdai
Dickeya dadantii NCPPB 898 WP 038910993 1 Ddadantii
Dickeya dadantii NCPPB 3537 WP 038900977 1 Ddadantii
Dickeya dadantii 3937 WP 033111788 1 Ddadantii
372 18953 Opine degradation Seed
Dickeya solani PPO 9134 WP 022633184 1 Dsolani
Dickeya solani RNS 08 23 3 1 A WP 022633184 1 Dsolani
Dickeya solani RNS 05 1 2A WP 057083055 1 Dsolani
Dickeya solani MK10 WP 022633184 1 Dsolani
Dickeya solani MK16 WP 022633184 1 Dsolani
Dickeya solani GBBC 2040 WP 022633184 1 Dsolani
Dickeya solani PPO 9019 WP 022633184 1 Dsolani
Dickeya solani IFB 0099 WP 022633184 1 Dsolani
Dickeya solani IPO 2222 WP 022633184 1 Dsolani
Dickeya solani D s0432 1 WP 022633184 1 Dsolani
Dickeya solani RNS 07 7 3B WP 022633184 1 Dsolani

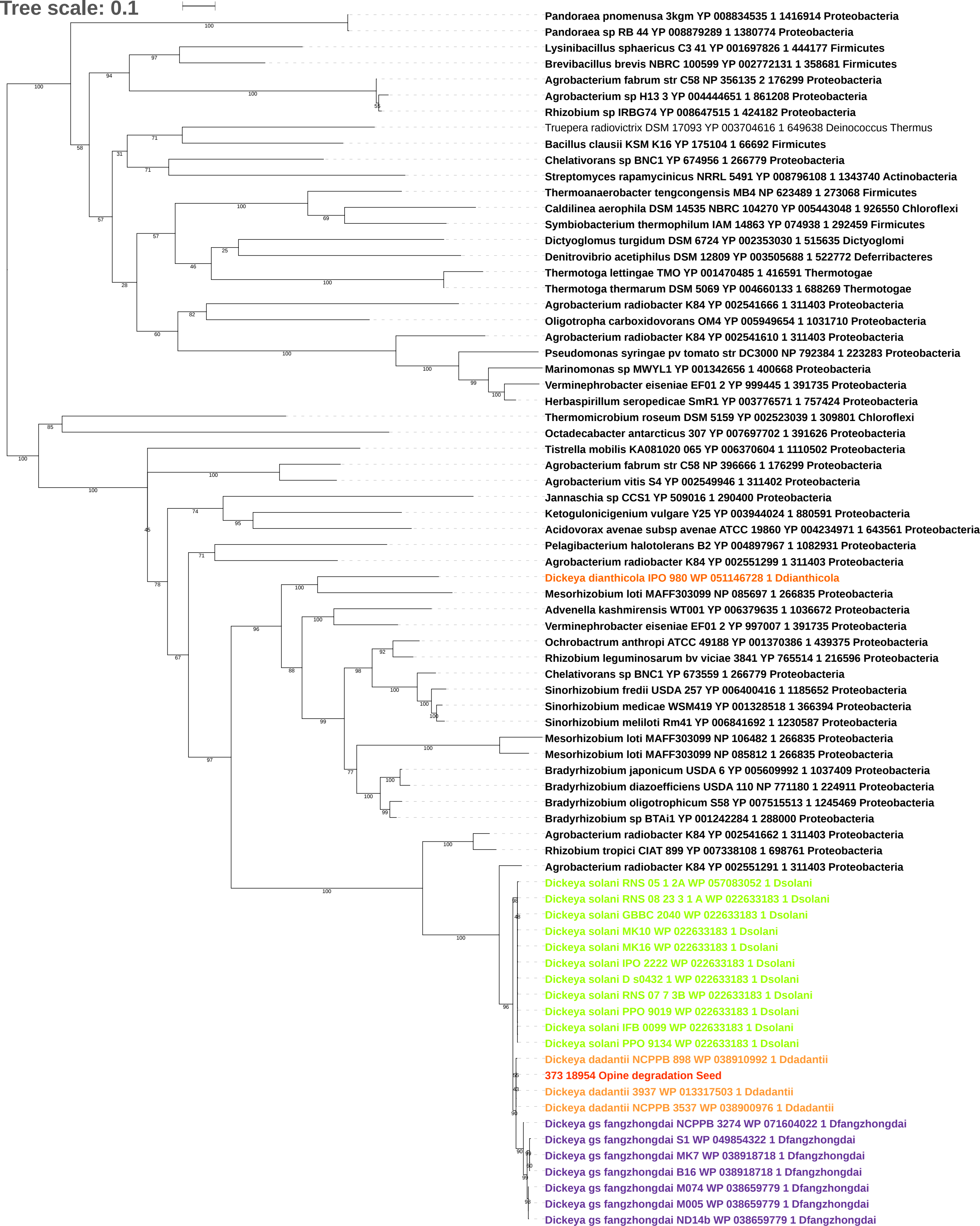
Tree scale: 0.1
Pandoraea pnomenusa 3kgm YP 008834535 1 1416914 Proteobacteria
Pandoraea sp RB 44 YP 008879289 1 1380774 Proteobacteria
Lysinibacillus sphaericus C3 41 YP 001697826 1 444177 Firmicutes
Brevibacillus brevis NBRC 100599 YP 002772131 1 358681 Firmicutes
Agrobacterium fabrum str C58 NP 356135 2 176299 Proteobacteria
Agrobacterium sp H13 3 YP 004444651 1 861208 Proteobacteria
Rhizobium sp IRBG74 YP 008647515 1 424182 Proteobacteria
Truepera radiovictrix DSM 17093 YP 003704616 1 649638 Deinococcus Thermus
Bacillus clausii KSM K16 YP 175104 1 66692 Firmicutes
Chelativorans sp BNC1 YP 674956 1 266779 Proteobacteria
Streptomyces rapamycinicus NRRL 5491 YP 008796108 1 1343740 Actinobacteria
Thermoanaerobacter tengcongensis MB4 NP 623489 1 273068 Firmicutes
Caldilinea aerophila DSM 14535 NBRC 104270 YP 005443048 1 926550 Chloroflexi
Symbiobacterium thermophilum IAM 14863 YP 074938 1 292459 Firmicutes
Dictyoglomus turgidum DSM 6724 YP 002353030 1 515635 Dictyoglomi
Denitrovibrio acetiphilus DSM 12809 YP 003505688 1 522772 Deferribacteres
Thermotoga lettingae TMO YP 001470485 1 416591 Thermotogae
Thermotoga thermarum DSM 5069 YP 004660133 1 688269 Thermotogae
Agrobacterium radiobacter K84 YP 002541666 1 311403 Proteobacteria
Oligotropha carboxidovorans OM4 YP 005949654 1 1031710 Proteobacteria
Agrobacterium radiobacter K84 YP 002541610 1 311403 Proteobacteria
Pseudomonas syringae pv tomato str DC3000 NP 792384 1 223283 Proteobacteria
Marinomonas sp MWYL1 YP 001342656 1 400668 Proteobacteria
Verminephrobacter eiseniae EF01 2 YP 999445 1 391735 Proteobacteria
Herbaspirillum seropedicae SmR1 YP 003776571 1 757424 Proteobacteria
Thermomicrobium roseum DSM 5159 YP 002523039 1 309801 Chloroflexi
Octadecabacter antarcticus 307 YP 007697702 1 391626 Proteobacteria
Tistrella mobilis KA081020 065 YP 006370604 1 1110502 Proteobacteria
Agrobacterium fabrum str C58 NP 396666 1 176299 Proteobacteria
Agrobacterium vitis S4 YP 002549946 1 311402 Proteobacteria
Jannaschia sp CCS1 YP 509016 1 290400 Proteobacteria
Ketogulonicigenium vulgare Y25 YP 003944024 1 880591 Proteobacteria
Acidovorax avenae subsp avenae ATCC 19860 YP 004234971 1 643561 Proteobacteria
Pelagibacterium halotolerans B2 YP 004897967 1 1082931 Proteobacteria
Agrobacterium radiobacter K84 YP 002551299 1 311403 Proteobacteria
Dickeya dianthicola IPO 980 WP 051146728 1 Ddianthicola
Mesorhizobium loti MAFF303099 NP 085697 1 266835 Proteobacteria
Advenella kashmirensis WT001 YP 006379635 1 1036672 Proteobacteria
Verminephrobacter eiseniae EF01 2 YP 997007 1 391735 Proteobacteria
Ochrobactrum anthropi ATCC 49188 YP 001370386 1 439375 Proteobacteria
Rhizobium leguminosarum bv viciae 3841 YP 765514 1 216596 Proteobacteria
Chelativorans sp BNC1 YP 673559 1 266779 Proteobacteria
Sinorhizobium fredii USDA 257 YP 006400416 1 1185652 Proteobacteria
Sinorhizobium medicae WSM419 YP 001328518 1 366394 Proteobacteria
Sinorhizobium meliloti Rm41 YP 006841692 1 1230587 Proteobacteria
Mesorhizobium loti MAFF303099 NP 106482 1 266835 Proteobacteria
Mesorhizobium loti MAFF303099 NP 085812 1 266835 Proteobacteria
Bradyrhizobium japonicum USDA 6 YP 005609992 1 1037409 Proteobacteria
Bradyrhizobium diazoefficiens USDA 110 NP 771180 1 224911 Proteobacteria
Bradyrhizobium oligotrophicum S58 YP 007515513 1 1245469 Proteobacteria
Bradyrhizobium sp BTAi1 YP 001242284 1 288000 Proteobacteria
Agrobacterium radiobacter K84 YP 002541662 1 311403 Proteobacteria
Rhizobium tropici CIAT 899 YP 007338108 1 698761 Proteobacteria
Agrobacterium radiobacter K84 YP 002551291 1 311403 Proteobacteria
Dickeya solani RNS 05 1 2A WP 057083052 1 Dsolani
Dickeya solani RNS 08 23 3 1 A WP 022633183 1 Dsolani
Dickeya solani GBBC 2040 WP 022633183 1 Dsolani
Dickeya solani MK10 WP 022633183 1 Dsolani
Dickeya solani MK16 WP 022633183 1 Dsolani
Dickeya solani IPO 2222 WP 022633183 1 Dsolani
Dickeya solani D s0432 1 WP 022633183 1 Dsolani
Dickeya solani RNS 07 7 3B WP 022633183 1 Dsolani
Dickeya solani PPO 9019 WP 022633183 1 Dsolani
Dickeya solani IFB 0099 WP 022633183 1 Dsolani
Dickeya solani PPO 9134 WP 022633183 1 Dsolani
Dickeya dadantii NCPPB 898 WP 038910992 1 Ddadantii
373 18954 Opine degradation Seed
Dickeya dadantii 3937 WP 013317503 1 Ddadantii
Dickeya dadantii NCPPB 3537 WP 038900976 1 Ddadantii
Dickeya gs fangzhongdai NCPPB 3274 WP 071604022 1 Dfangzhongdai
Dickeya gs fangzhongdai S1 WP 049854322 1 Dfangzhongdai
Dickeya gs fangzhongdai MK7 WP 038918718 1 Dfangzhongdai
Dickeya gs fangzhongdai B16 WP 038918718 1 Dfangzhongdai
Dickeya gs fangzhongdai M074 WP 038659779 1 Dfangzhongdai
Dickeya gs fangzhongdai M005 WP 038659779 1 Dfangzhongdai
Dickeya gs fangzhongdai ND14b WP 038659779 1 Dfangzhongdai

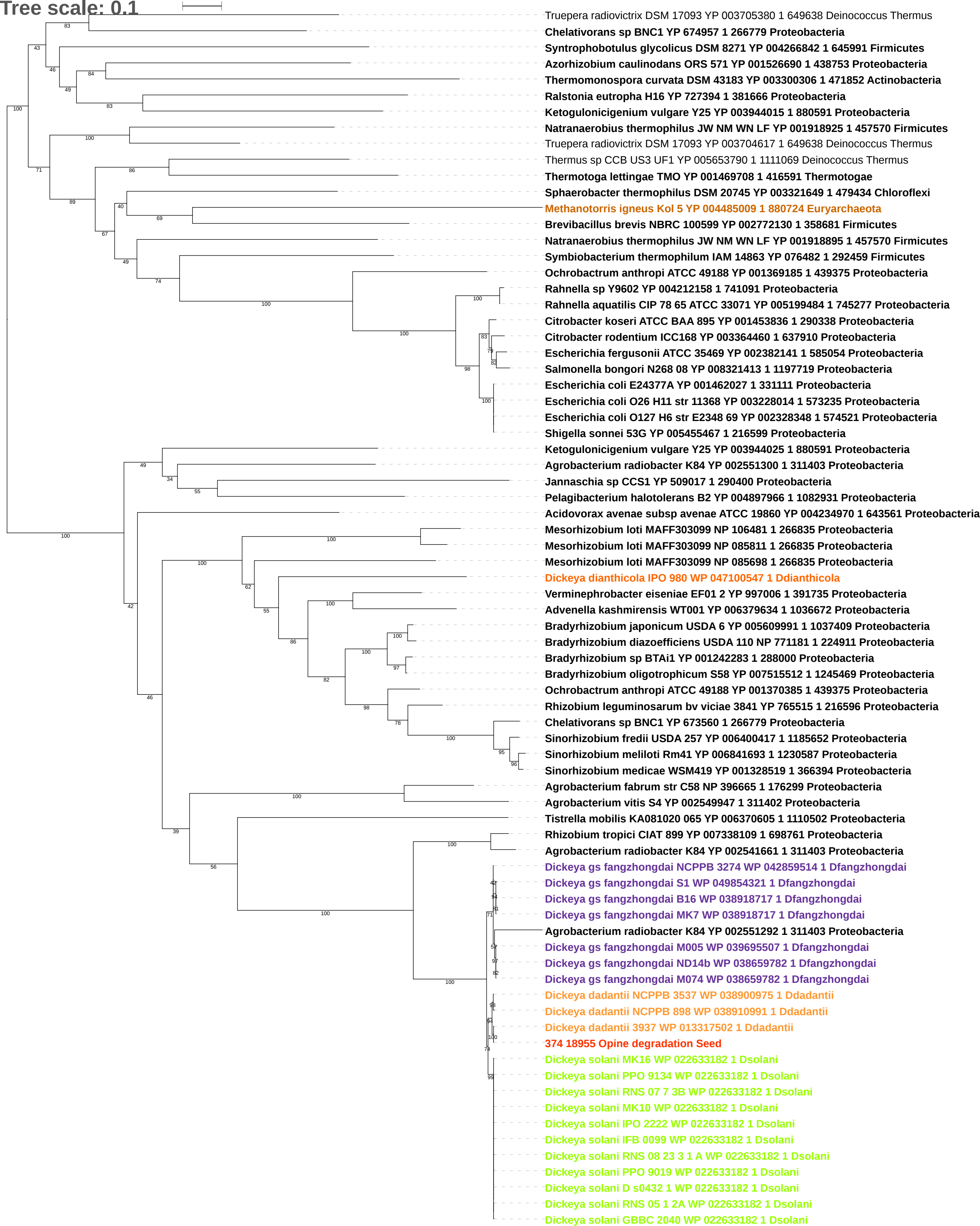
Tree scale: 0.1
Truepera radiovictrix DSM 17093 YP 003705380 1 649638 Deinococcus Thermus
Chelativorans sp BNC1 YP 674957 1 266779 Proteobacteria
Syntrophobotulus glycolicus DSM 8271 YP 004266842 1 645991 Firmicutes
Azorhizobium caulinodans ORS 571 YP 001526690 1 438753 Proteobacteria
Thermomonospora curvata DSM 43183 YP 003300306 1 471852 Actinobacteria
Ralstonia eutropha H16 YP 727394 1 381666 Proteobacteria
Ketogulonicigenium vulgare Y25 YP 003944015 1 880591 Proteobacteria
Natranaerobius thermophilus JW NM WN LF YP 001918925 1 457570 Firmicutes
Truepera radiovictrix DSM 17093 YP 003704617 1 649638 Deinococcus Thermus
Thermus sp CCB US3 UF1 YP 005653790 1 1111069 Deinococcus Thermus
Thermotoga lettingae TMO YP 001469708 1 416591 Thermotogae
Sphaerobacter thermophilus DSM 20745 YP 003321649 1 479434 Chloroflexi
Methanotorris igneus Kol 5 YP 004485009 1 880724 Euryarchaeota
Brevibacillus brevis NBRC 100599 YP 002772130 1 358681 Firmicutes
Natranaerobius thermophilus JW NM WN LF YP 001918895 1 457570 Firmicutes
Symbiobacterium thermophilum IAM 14863 YP 076482 1 292459 Firmicutes
Ochrobactrum anthropi ATCC 49188 YP 001369185 1 439375 Proteobacteria
Rahnella sp Y9602 YP 004212158 1 741091 Proteobacteria
Rahnella aquatilis CIP 78 65 ATCC 33071 YP 005199484 1 745277 Proteobacteria
Citrobacter koseri ATCC BAA 895 YP 001453836 1 290338 Proteobacteria
Citrobacter rodentium ICC168 YP 003364460 1 637910 Proteobacteria
Escherichia fergusonii ATCC 35469 YP 002382141 1 585054 Proteobacteria
Salmonella bongori N268 08 YP 008321413 1 1197719 Proteobacteria
Escherichia coli E24377A YP 001462027 1 331111 Proteobacteria
Escherichia coli O26 H11 str 11368 YP 003228014 1 573235 Proteobacteria
Escherichia coli O127 H6 str E2348 69 YP 002328348 1 574521 Proteobacteria
Shigella sonnei 53G YP 005455467 1 216599 Proteobacteria
Ketogulonicigenium vulgare Y25 YP 003944025 1 880591 Proteobacteria
Agrobacterium radiobacter K84 YP 002551300 1 311403 Proteobacteria
Jannaschia sp CCS1 YP 509017 1 290400 Proteobacteria
Pelagibacterium halotolerans B2 YP 004897966 1 1082931 Proteobacteria
Acidovorax avenae subsp avenae ATCC 19860 YP 004234970 1 643561 Proteobacteria
Mesorhizobium loti MAFF303099 NP 106481 1 266835 Proteobacteria
Mesorhizobium loti MAFF303099 NP 085811 1 266835 Proteobacteria
Mesorhizobium loti MAFF303099 NP 085698 1 266835 Proteobacteria
Dickeya dianthicola IPO 980 WP 047100547 1 Ddianthicola
Verminephrobacter eiseniae EF01 2 YP 997006 1 391735 Proteobacteria
Advenella kashmirensis WT001 YP 006379634 1 1036672 Proteobacteria
Bradyrhizobium japonicum USDA 6 YP 005609991 1 1037409 Proteobacteria
Bradyrhizobium diazoefficiens USDA 110 NP 771181 1 224911 Proteobacteria
Bradyrhizobium sp BTAi1 YP 001242283 1 288000 Proteobacteria
Bradyrhizobium oligotrophicum S58 YP 007515512 1 1245469 Proteobacteria
Ochrobactrum anthropi ATCC 49188 YP 001370385 1 439375 Proteobacteria
Rhizobium leguminosarum bv viciae 3841 YP 765515 1 216596 Proteobacteria
Chelativorans sp BNC1 YP 673560 1 266779 Proteobacteria
Sinorhizobium fredii USDA 257 YP 006400417 1 1185652 Proteobacteria
Sinorhizobium meliloti Rm41 YP 006841693 1 1230587 Proteobacteria
Sinorhizobium medicae WSM419 YP 001328519 1 366394 Proteobacteria
Agrobacterium fabrum str C58 NP 396665 1 176299 Proteobacteria
Agrobacterium vitis S4 YP 002549947 1 311402 Proteobacteria
Tistrella mobilis KA081020 065 YP 006370605 1 1110502 Proteobacteria
Rhizobium tropici CIAT 899 YP 007338109 1 698761 Proteobacteria
Agrobacterium radiobacter K84 YP 002541661 1 311403 Proteobacteria
Dickeya gs fangzhongdai NCPPB 3274 WP 042859514 1 Dfangzhongdai
Dickeya gs fangzhongdai S1 WP 049854321 1 Dfangzhongdai
Dickeya gs fangzhongdai B16 WP 038918717 1 Dfangzhongdai
Dickeya gs fangzhongdai MK7 WP 038918717 1 Dfangzhongdai
Agrobacterium radiobacter K84 YP 002551292 1 311403 Proteobacteria
Dickeya gs fangzhongdai M005 WP 039695507 1 Dfangzhongdai
Dickeya gs fangzhongdai ND14b WP 038659782 1 Dfangzhongdai
Dickeya gs fangzhongdai M074 WP 038659782 1 Dfangzhongdai
Dickeya dadantii NCPPB 3537 WP 038900975 1 Ddadantii
Dickeya dadantii NCPPB 898 WP 038910991 1 Ddadantii
Dickeya dadantii 3937 WP 013317502 1 Ddadantii
374 18955 Opine degradation Seed
Dickeya solani MK16 WP 022633182 1 Dsolani
Dickeya solani PPO 9134 WP 022633182 1 Dsolani
Dickeya solani RNS 07 7 3B WP 022633182 1 Dsolani
Dickeya solani MK10 WP 022633182 1 Dsolani
Dickeya solani IPO 2222 WP 022633182 1 Dsolani
Dickeya solani IFB 0099 WP 022633182 1 Dsolani
Dickeya solani RNS 08 23 3 1 A WP 022633182 1 Dsolani
Dickeya solani PPO 9019 WP 022633182 1 Dsolani
Dickeya solani D s0432 1 WP 022633182 1 Dsolani
Dickeya solani RNS 05 1 2A WP 022633182 1 Dsolani
Dickeya solani GBBC 2040 WP 022633182 1 Dsolani

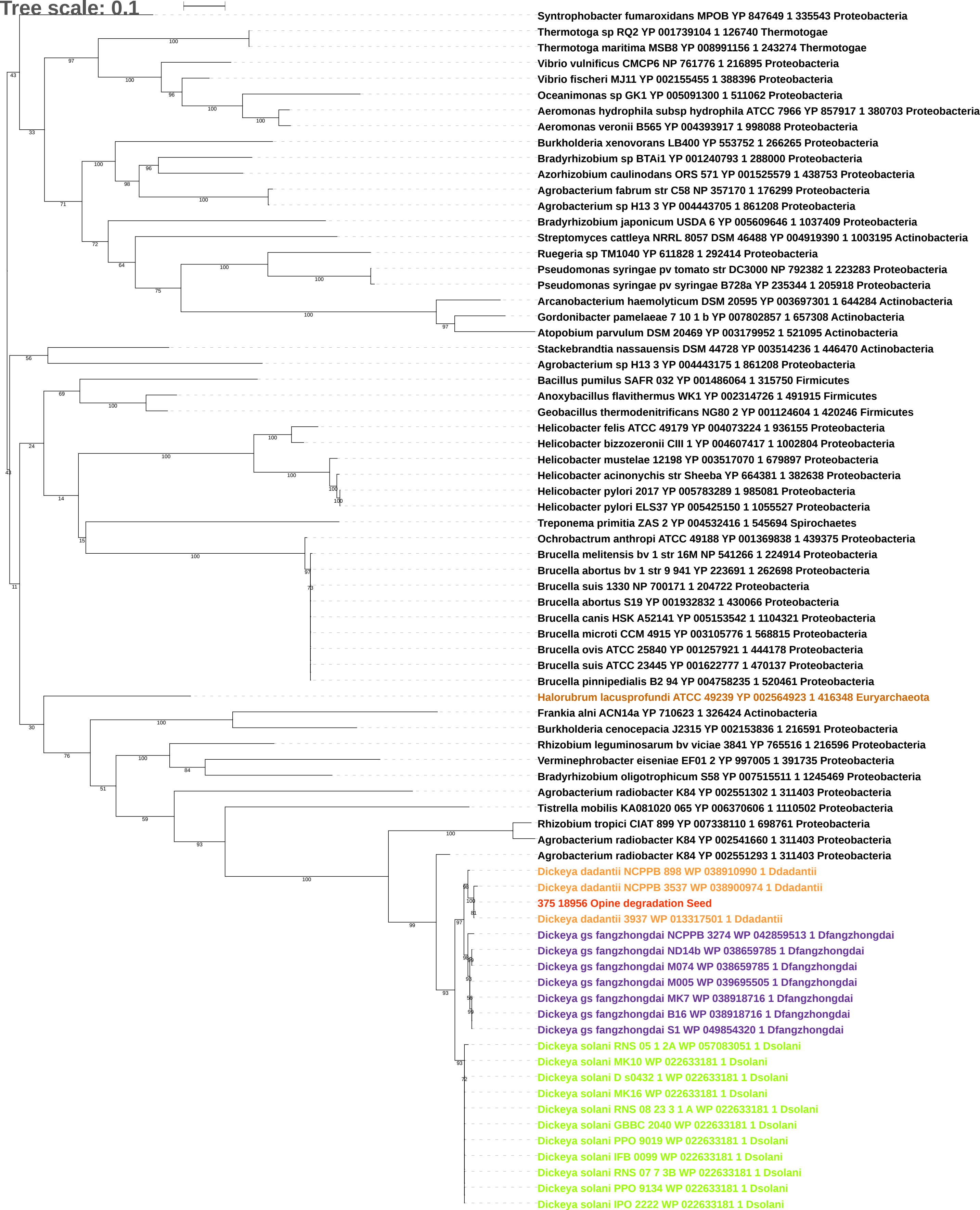

Tree scale: 0.1
Syntrophobacter fumaroxidans MPOB YP 847649 1 335543 Proteobacteria
Thermotoga sp RQ2 YP 001739104 1 126740 Thermotogae
Thermotoga maritima MSB8 YP 008991156 1 243274 Thermotogae
Vibrio vulnificus CMCP6 NP 761776 1 216895 Proteobacteria
Vibrio fischeri MJ11 YP 002155455 1 388396 Proteobacteria
Oceanimonas sp GK1 YP 005091300 1 511062 Proteobacteria
Aeromonas hydrophila subsp hydrophila ATCC 7966 YP 857917 1 380703 Proteobacteria
Aeromonas veronii B565 YP 004393917 1 998088 Proteobacteria
Burkholderia xenovorans LB400 YP 553752 1 266265 Proteobacteria
Bradyrhizobium sp BTAi1 YP 001240793 1 288000 Proteobacteria
Azorhizobium caulinodans ORS 571 YP 001525579 1 438753 Proteobacteria
Agrobacterium fabrum str C58 NP 357170 1 176299 Proteobacteria
Agrobacterium sp H13 3 YP 004443705 1 861208 Proteobacteria
Bradyrhizobium japonicum USDA 6 YP 005609646 1 1037409 Proteobacteria
Streptomyces cattleya NRRL 8057 DSM 46488 YP 004919390 1 1003195 Actinobacteria
Ruegeria sp TM1040 YP 611828 1 292414 Proteobacteria
Pseudomonas syringae pv tomato str DC3000 NP 792382 1 223283 Proteobacteria
Pseudomonas syringae pv syringae B728a YP 235344 1 205918 Proteobacteria
Arcanobacterium haemolyticum DSM 20595 YP 003697301 1 644284 Actinobacteria
Gordonibacter pamelaeae 7 10 1 b YP 007802857 1 657308 Actinobacteria
Atopobium parvulum DSM 20469 YP 003179952 1 521095 Actinobacteria
Stackebrandtia nassauensis DSM 44728 YP 003514236 1 446470 Actinobacteria
Agrobacterium sp H13 3 YP 004443175 1 861208 Proteobacteria
Bacillus pumilus SAFR 032 YP 001486064 1 315750 Firmicutes
Anoxybacillus flavithermus WK1 YP 002314726 1 491915 Firmicutes
Geobacillus thermodenitrificans NG80 2 YP 001124604 1 420246 Firmicutes
Helicobacter felis ATCC 49179 YP 004073224 1 936155 Proteobacteria
Helicobacter bizzozeronii CIII 1 YP 004607417 1 1002804 Proteobacteria
Helicobacter mustelae 12198 YP 003517070 1 679897 Proteobacteria
Helicobacter acinonychis str Sheeba YP 664381 1 382638 Proteobacteria
Helicobacter pylori 2017 YP 005783289 1 985081 Proteobacteria
Helicobacter pylori ELS37 YP 005425150 1 1055527 Proteobacteria
Treponema primitia ZAS 2 YP 004532416 1 545694 Spirochaetes
Ochrobactrum anthropi ATCC 49188 YP 001369838 1 439375 Proteobacteria
Brucella melitensis bv 1 str 16M NP 541266 1 224914 Proteobacteria
Brucella abortus bv 1 str 9 941 YP 223691 1 262698 Proteobacteria
Brucella suis 1330 NP 700171 1 204722 Proteobacteria
Brucella abortus S19 YP 001932832 1 430066 Proteobacteria
Brucella canis HSK A52141 YP 005153542 1 1104321 Proteobacteria
Brucella microti CCM 4915 YP 003105776 1 568815 Proteobacteria
Brucella ovis ATCC 25840 YP 001257921 1 444178 Proteobacteria
Brucella suis ATCC 23445 YP 001622777 1 470137 Proteobacteria
Brucella pinnipedialis B2 94 YP 004758235 1 520461 Proteobacteria
Halorubrum lacusprofundi ATCC 49239 YP 002564923 1 416348 Euryarchaeota
Frankia alni ACN14a YP 710623 1 326424 Actinobacteria
Burkholderia cenocepacia J2315 YP 002153836 1 216591 Proteobacteria
Rhizobium leguminosarum bv viciae 3841 YP 765516 1 216596 Proteobacteria
Verminephrobacter eiseniae EF01 2 YP 997005 1 391735 Proteobacteria
Bradyrhizobium oligotrophicum S58 YP 007515511 1 1245469 Proteobacteria
Agrobacterium radiobacter K84 YP 002551302 1 311403 Proteobacteria
Tistrella mobilis KA081020 065 YP 006370606 1 1110502 Proteobacteria
Rhizobium tropici CIAT 899 YP 007338110 1 698761 Proteobacteria
Agrobacterium radiobacter K84 YP 002541660 1 311403 Proteobacteria
Agrobacterium radiobacter K84 YP 002551293 1 311403 Proteobacteria
Dickeya dadantii NCPPB 898 WP 038910990 1 Ddadantii
Dickeya dadantii NCPPB 3537 WP 038900974 1 Ddadantii
375 18956 Opine degradation Seed
Dickeya dadantii 3937 WP 013317501 1 Ddadantii
Dickeya gs fangzhongdai NCPPB 3274 WP 042859513 1 Dfangzhongdai
Dickeya gs fangzhongdai ND14b WP 038659785 1 Dfangzhongdai
Dickeya gs fangzhongdai M074 WP 038659785 1 Dfangzhongdai
Dickeya gs fangzhongdai M005 WP 039695505 1 Dfangzhongdai
Dickeya gs fangzhongdai MK7 WP 038918716 1 Dfangzhongdai
Dickeya gs fangzhongdai B16 WP 038918716 1 Dfangzhongdai
Dickeya gs fangzhongdai S1 WP 049854320 1 Dfangzhongdai
Dickeya solani RNS 05 1 2A WP 057083051 1 Dsolani
Dickeya solani MK10 WP 022633181 1 Dsolani
Dickeya solani D s0432 1 WP 022633181 1 Dsolani
Dickeya solani MK16 WP 022633181 1 Dsolani
Dickeya solani RNS 08 23 3 1 A WP 022633181 1 Dsolani
Dickeya solani GBBC 2040 WP 022633181 1 Dsolani
Dickeya solani PPO 9019 WP 022633181 1 Dsolani
Dickeya solani IFB 0099 WP 022633181 1 Dsolani
Dickeya solani RNS 07 7 3B WP 022633181 1 Dsolani
Dickeya solani PPO 9134 WP 022633181 1 Dsolani
Dickeya solani IPO 2222 WP 022633181 1 Dsolani

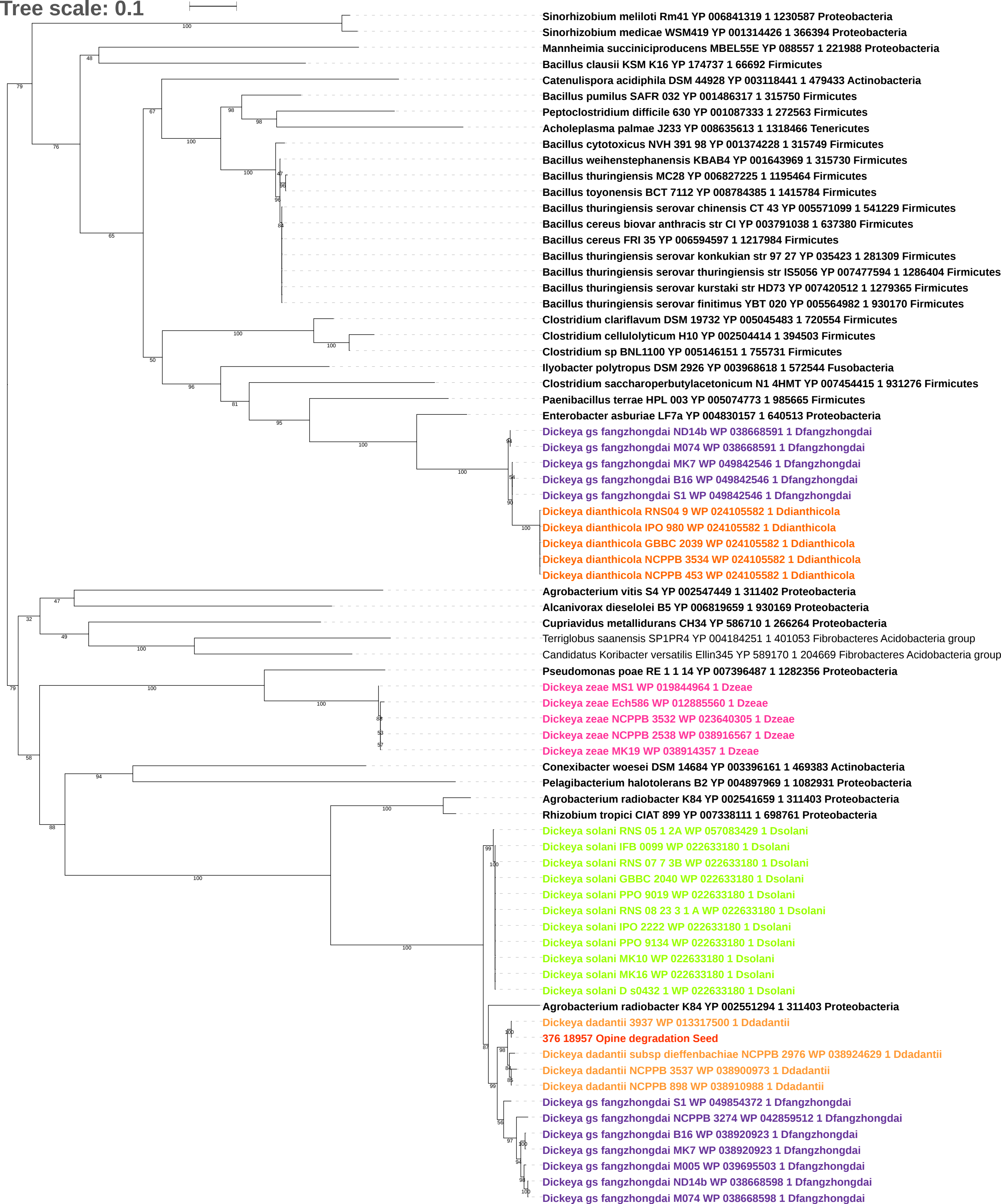
Tree scale: 0.1
Sinorhizobium meliloti Rm41 YP 006841319 1 1230587 Proteobacteria
Sinorhizobium medicae WSM419 YP 001314426 1 366394 Proteobacteria
Mannheimia succiniciproducens MBEL55E YP 088557 1 221988 Proteobacteria
Bacillus clausii KSM K16 YP 174737 1 66692 Firmicutes
Catenulispora acidiphila DSM 44928 YP 003118441 1 479433 Actinobacteria
Bacillus pumilus SAFR 032 YP 001486317 1 315750 Firmicutes
Peptoclostridium difficile 630 YP 001087333 1 272563 Firmicutes
Acholeplasma palmae J233 YP 008635613 1 1318466 Tenericutes
Bacillus cytotoxicus NVH 391 98 YP 001374228 1 315749 Firmicutes
Bacillus weihenstephanensis KBAB4 YP 001643969 1 315730 Firmicutes
Bacillus thuringiensis MC28 YP 006827225 1 1195464 Firmicutes
Bacillus toyonensis BCT 7112 YP 008784385 1 1415784 Firmicutes
Bacillus thuringiensis serovar chinensis CT 43 YP 005571099 1 541229 Firmicutes
Bacillus cereus biovar anthracis str CI YP 003791038 1 637380 Firmicutes
Bacillus cereus FRI 35 YP 006594597 1 1217984 Firmicutes
Bacillus thuringiensis serovar konkukian str 97 27 YP 035423 1 281309 Firmicutes
Bacillus thuringiensis serovar thuringiensis str IS5056 YP 007477594 1 1286404 Firmicutes
Bacillus thuringiensis serovar kurstaki str HD73 YP 007420512 1 1279365 Firmicutes
Bacillus thuringiensis serovar finitimus YBT 020 YP 005564982 1 930170 Firmicutes
Clostridium clariflavum DSM 19732 YP 005045483 1 720554 Firmicutes
Clostridium cellulolyticum H10 YP 002504414 1 394503 Firmicutes
Clostridium sp BNL1100 YP 005146151 1 755731 Firmicutes
Ilyobacter polytropus DSM 2926 YP 003968618 1 572544 Fusobacteria
Clostridium saccharoperbutylacetonicum N1 4HMT YP 007454415 1 931276 Firmicutes
Paenibacillus terrae HPL 003 YP 005074773 1 985665 Firmicutes
Enterobacter asburiae LF7a YP 004830157 1 640513 Proteobacteria
Dickeya gs fangzhongdai ND14b WP 038668591 1 Dfangzhongdai
Dickeya gs fangzhongdai M074 WP 038668591 1 Dfangzhongdai
Dickeya gs fangzhongdai MK7 WP 049842546 1 Dfangzhongdai
Dickeya gs fangzhongdai B16 WP 049842546 1 Dfangzhongdai
Dickeya gs fangzhongdai S1 WP 049842546 1 Dfangzhongdai
Dickeya dianthicola RNS04 9 WP 024105582 1 Ddianthicola
Dickeya dianthicola IPO 980 WP 024105582 1 Ddianthicola
Dickeya dianthicola GBBC 2039 WP 024105582 1 Ddianthicola
Dickeya dianthicola NCPPB 3534 WP 024105582 1 Ddianthicola
Dickeya dianthicola NCPPB 453 WP 024105582 1 Ddianthicola
Agrobacterium vitis S4 YP 002547449 1 311402 Proteobacteria
Alcanivorax dieselolei B5 YP 006819659 1 930169 Proteobacteria
Cupriavidus metallidurans CH34 YP 586710 1 266264 Proteobacteria
Terriglobus saanensis SP1PR4 YP 004184251 1 401053 Fibrobacteres Acidobacteria group
Candidatus Koribacter versatilis Ellin345 YP 589170 1 204669 Fibrobacteres Acidobacteria group
Pseudomonas poae RE 1 1 14 YP 007396487 1 1282356 Proteobacteria
Dickeya zeae MS1 WP 019844964 1 Dzeae
Dickeya zeae Ech586 WP 012885560 1 Dzeae
Dickeya zeae NCPPB 3532 WP 023640305 1 Dzeae
Dickeya zeae NCPPB 2538 WP 038916567 1 Dzeae
Dickeya zeae MK19 WP 038914357 1 Dzeae
Conexibacter woesei DSM 14684 YP 003396161 1 469383 Actinobacteria
Pelagibacterium halotolerans B2 YP 004897969 1 1082931 Proteobacteria
Agrobacterium radiobacter K84 YP 002541659 1 311403 Proteobacteria
Rhizobium tropici CIAT 899 YP 007338111 1 698761 Proteobacteria
Dickeya solani RNS 05 1 2A WP 057083429 1 Dsolani
Dickeya solani IFB 0099 WP 022633180 1 Dsolani
Dickeya solani RNS 07 7 3B WP 022633180 1 Dsolani
Dickeya solani GBBC 2040 WP 022633180 1 Dsolani
Dickeya solani PPO 9019 WP 022633180 1 Dsolani
Dickeya solani RNS 08 23 3 1 A WP 022633180 1 Dsolani
Dickeya solani IPO 2222 WP 022633180 1 Dsolani
Dickeya solani PPO 9134 WP 022633180 1 Dsolani
Dickeya solani MK10 WP 022633180 1 Dsolani
Dickeya solani MK16 WP 022633180 1 Dsolani
Dickeya solani D s0432 1 WP 022633180 1 Dsolani
Agrobacterium radiobacter K84 YP 002551294 1 311403 Proteobacteria
Dickeya dadantii 3937 WP 013317500 1 Ddadantii
376 18957 Opine degradation Seed
Dickeya dadantii subsp dieffenbachiae NCPPB 2976 WP 038924629 1 Ddadantii
Dickeya dadantii NCPPB 3537 WP 038900973 1 Ddadantii
Dickeya dadantii NCPPB 898 WP 038910988 1 Ddadantii
Dickeya gs fangzhongdai S1 WP 049854372 1 Dfangzhongdai
Dickeya gs fangzhongdai NCPPB 3274 WP 042859512 1 Dfangzhongdai
Dickeya gs fangzhongdai B16 WP 038920923 1 Dfangzhongdai
Dickeya gs fangzhongdai MK7 WP 038920923 1 Dfangzhongdai
Dickeya gs fangzhongdai M005 WP 039695503 1 Dfangzhongdai
Dickeya gs fangzhongdai ND14b WP 038668598 1 Dfangzhongdai
Dickeya gs fangzhongdai M074 WP 038668598 1 Dfangzhongdai

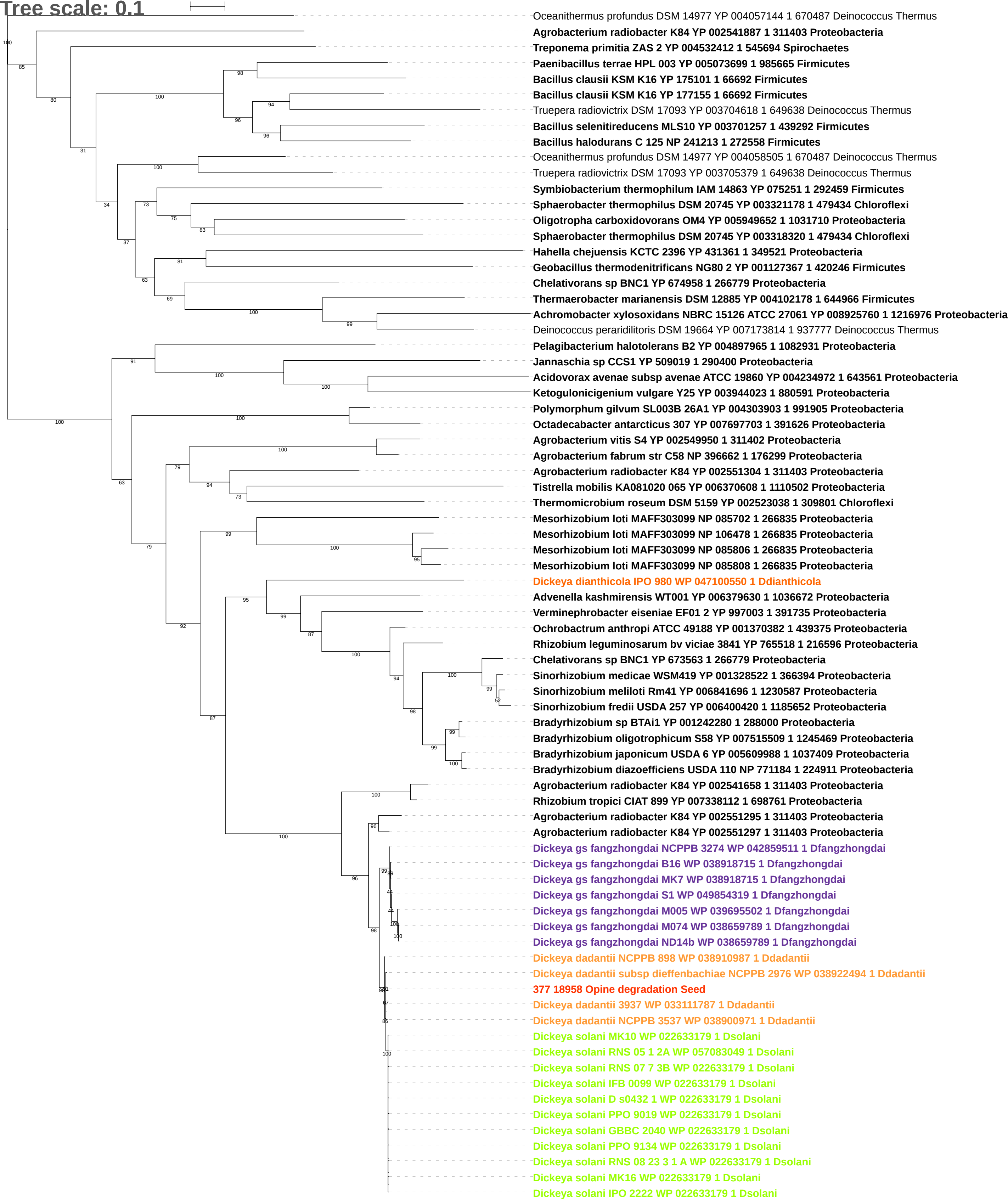
Tree scale: 0.1
Oceanithermus profundus DSM 14977 YP 004057144 1 670487 Deinococcus Thermus
Agrobacterium radiobacter K84 YP 002541887 1 311403 Proteobacteria
Treponema primitia ZAS 2 YP 004532412 1 545694 Spirochaetes
Paenibacillus terrae HPL 003 YP 005073699 1 985665 Firmicutes
Bacillus clausii KSM K16 YP 175101 1 66692 Firmicutes
Bacillus clausii KSM K16 YP 177155 1 66692 Firmicutes
Truepera radiovictrix DSM 17093 YP 003704618 1 649638 Deinococcus Thermus
Bacillus selenitireducens MLS10 YP 003701257 1 439292 Firmicutes
Bacillus halodurans C 125 NP 241213 1 272558 Firmicutes
Oceanithermus profundus DSM 14977 YP 004058505 1 670487 Deinococcus Thermus
Truepera radiovictrix DSM 17093 YP 003705379 1 649638 Deinococcus Thermus
Symbiobacterium thermophilum IAM 14863 YP 075251 1 292459 Firmicutes
Sphaerobacter thermophilus DSM 20745 YP 003321178 1 479434 Chloroflexi
Oligotropha carboxidovorans OM4 YP 005949652 1 1031710 Proteobacteria
Sphaerobacter thermophilus DSM 20745 YP 003318320 1 479434 Chloroflexi
Hahella chejuensis KCTC 2396 YP 431361 1 349521 Proteobacteria
Geobacillus thermodenitrificans NG80 2 YP 001127367 1 420246 Firmicutes
Chelativorans sp BNC1 YP 674958 1 266779 Proteobacteria
Thermaerobacter marianensis DSM 12885 YP 004102178 1 644966 Firmicutes
Achromobacter xylosoxidans NBRC 15126 ATCC 27061 YP 008925760 1 1216976 Proteobacteria
Deinococcus peraridilitoris DSM 19664 YP 007173814 1 937777 Deinococcus Thermus
Pelagibacterium halotolerans B2 YP 004897965 1 1082931 Proteobacteria
Jannaschia sp CCS1 YP 509019 1 290400 Proteobacteria
Acidovorax avenae subsp avenae ATCC 19860 YP 004234972 1 643561 Proteobacteria
Ketogulonicigenium vulgare Y25 YP 003944023 1 880591 Proteobacteria
Polymorphum gilvum SL003B 26A1 YP 004303903 1 991905 Proteobacteria
Octadecabacter antarcticus 307 YP 007697703 1 391626 Proteobacteria
Agrobacterium vitis S4 YP 002549950 1 311402 Proteobacteria
Agrobacterium fabrum str C58 NP 396662 1 176299 Proteobacteria
Agrobacterium radiobacter K84 YP 002551304 1 311403 Proteobacteria
Tistrella mobilis KA081020 065 YP 006370608 1 1110502 Proteobacteria
Thermomicrobium roseum DSM 5159 YP 002523038 1 309801 Chloroflexi
Mesorhizobium loti MAFF303099 NP 085702 1 266835 Proteobacteria
Mesorhizobium loti MAFF303099 NP 106478 1 266835 Proteobacteria
Mesorhizobium loti MAFF303099 NP 085806 1 266835 Proteobacteria
Mesorhizobium loti MAFF303099 NP 085808 1 266835 Proteobacteria
Dickeya dianthicola IPO 980 WP 047100550 1 Ddianthicola
Advenella kashmirensis WT001 YP 006379630 1 1036672 Proteobacteria
Verminephrobacter eiseniae EF01 2 YP 997003 1 391735 Proteobacteria
Ochrobactrum anthropi ATCC 49188 YP 001370382 1 439375 Proteobacteria
Rhizobium leguminosarum bv viciae 3841 YP 765518 1 216596 Proteobacteria
Chelativorans sp BNC1 YP 673563 1 266779 Proteobacteria
Sinorhizobium medicae WSM419 YP 001328522 1 366394 Proteobacteria
Sinorhizobium meliloti Rm41 YP 006841696 1 1230587 Proteobacteria
Sinorhizobium fredii USDA 257 YP 006400420 1 1185652 Proteobacteria
Bradyrhizobium sp BTAi1 YP 001242280 1 288000 Proteobacteria
Bradyrhizobium oligotrophicum S58 YP 007515509 1 1245469 Proteobacteria
Bradyrhizobium japonicum USDA 6 YP 005609988 1 1037409 Proteobacteria
Bradyrhizobium diazoefficiens USDA 110 NP 771184 1 224911 Proteobacteria
Agrobacterium radiobacter K84 YP 002541658 1 311403 Proteobacteria
Rhizobium tropici CIAT 899 YP 007338112 1 698761 Proteobacteria
Agrobacterium radiobacter K84 YP 002551295 1 311403 Proteobacteria
Agrobacterium radiobacter K84 YP 002551297 1 311403 Proteobacteria
Dickeya gs fangzhongdai NCPPB 3274 WP 042859511 1 Dfangzhongdai
Dickeya gs fangzhongdai B16 WP 038918715 1 Dfangzhongdai
Dickeya gs fangzhongdai MK7 WP 038918715 1 Dfangzhongdai
Dickeya gs fangzhongdai S1 WP 049854319 1 Dfangzhongdai
Dickeya gs fangzhongdai M005 WP 039695502 1 Dfangzhongdai
Dickeya gs fangzhongdai M074 WP 038659789 1 Dfangzhongdai
Dickeya gs fangzhongdai ND14b WP 038659789 1 Dfangzhongdai
Dickeya dadantii NCPPB 898 WP 038910987 1 Ddadantii
Dickeya dadantii subsp dieffenbachiae NCPPB 2976 WP 038922494 1 Ddadantii
377 18958 Opine degradation Seed
Dickeya dadantii 3937 WP 033111787 1 Ddadantii
Dickeya dadantii NCPPB 3537 WP 038900971 1 Ddadantii
Dickeya solani MK10 WP 022633179 1 Dsolani
Dickeya solani RNS 05 1 2A WP 057083049 1 Dsolani
Dickeya solani RNS 07 7 3B WP 022633179 1 Dsolani
Dickeya solani IFB 0099 WP 022633179 1 Dsolani
Dickeya solani D s0432 1 WP 022633179 1 Dsolani
Dickeya solani PPO 9019 WP 022633179 1 Dsolani
Dickeya solani GBBC 2040 WP 022633179 1 Dsolani
Dickeya solani PPO 9134 WP 022633179 1 Dsolani
Dickeya solani RNS 08 23 3 1 A WP 022633179 1 Dsolani
Dickeya solani MK16 WP 022633179 1 Dsolani
Dickeya solani IPO 2222 WP 022633179 1 Dsolani

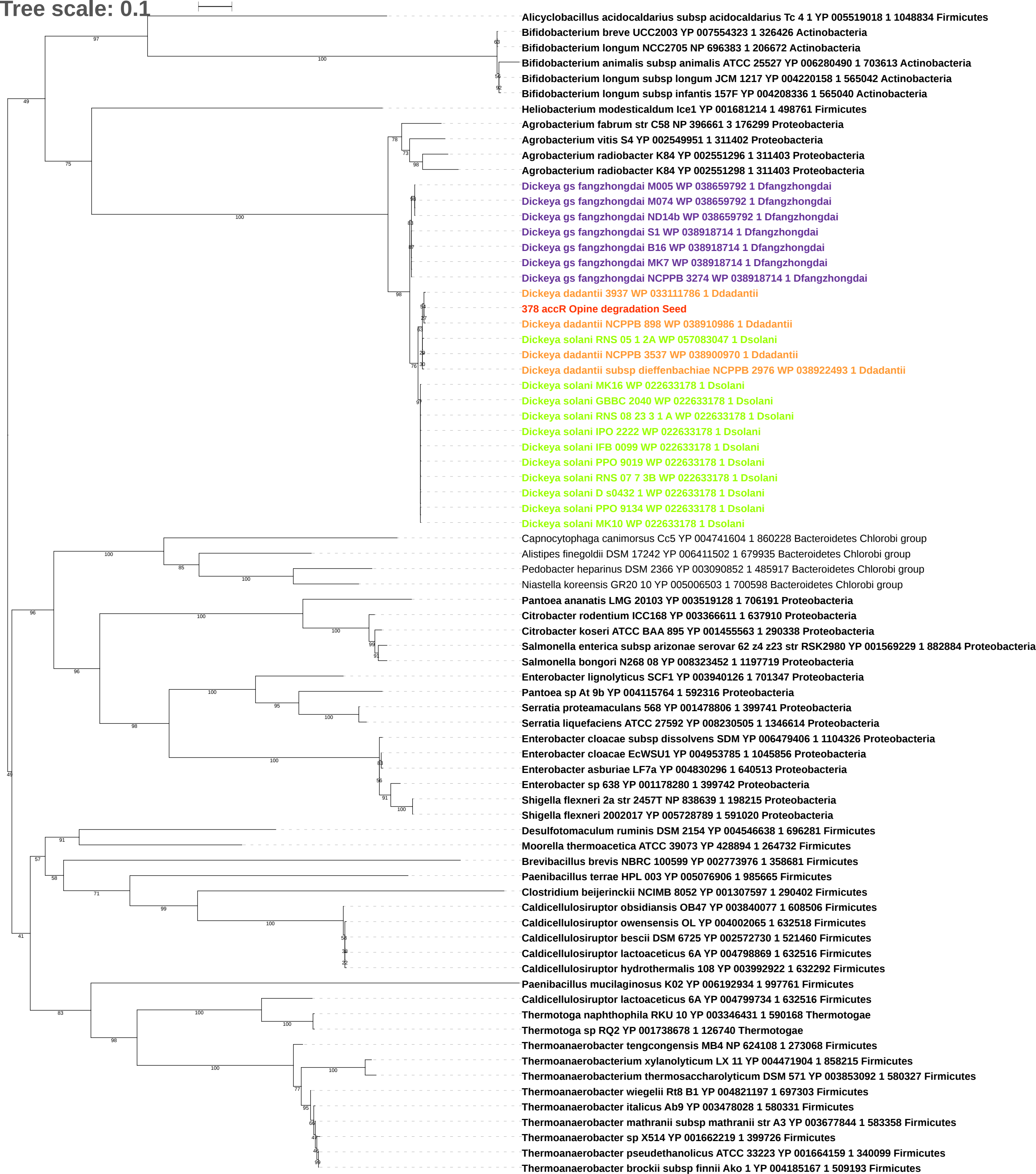
Tree scale: 0.1
Alicyclobacillus acidocaldarius subsp acidocaldarius Tc 4 1 YP 005519018 1 1048834 Firmicutes
Bifidobacterium breve UCC2003 YP 007554323 1 326426 Actinobacteria
Bifidobacterium longum NCC2705 NP 696383 1 206672 Actinobacteria
Bifidobacterium animalis subsp animalis ATCC 25527 YP 006280490 1 703613 Actinobacteria
Bifidobacterium longum subsp longum JCM 1217 YP 004220158 1 565042 Actinobacteria
Bifidobacterium longum subsp infantis 157F YP 004208336 1 565040 Actinobacteria
Heliobacterium modesticaldum Ice1 YP 001681214 1 498761 Firmicutes
Agrobacterium fabrum str C58 NP 396661 3 176299 Proteobacteria
Agrobacterium vitis S4 YP 002549951 1 311402 Proteobacteria
Agrobacterium radiobacter K84 YP 002551296 1 311403 Proteobacteria
Agrobacterium radiobacter K84 YP 002551298 1 311403 Proteobacteria
Dickeya gs fangzhongdai M005 WP 038659792 1 Dfangzhongdai
Dickeya gs fangzhongdai M074 WP 038659792 1 Dfangzhongdai
Dickeya gs fangzhongdai ND14b WP 038659792 1 Dfangzhongdai
Dickeya gs fangzhongdai S1 WP 038918714 1 Dfangzhongdai
Dickeya gs fangzhongdai B16 WP 038918714 1 Dfangzhongdai
Dickeya gs fangzhongdai MK7 WP 038918714 1 Dfangzhongdai
Dickeya gs fangzhongdai NCPPB 3274 WP 038918714 1 Dfangzhongdai
Dickeya dadantii 3937 WP 033111786 1 Ddadantii
378 accR Opine degradation Seed
Dickeya dadantii NCPPB 898 WP 038910986 1 Ddadantii
Dickeya solani RNS 05 1 2A WP 057083047 1 Dsolani
Dickeya dadantii NCPPB 3537 WP 038900970 1 Ddadantii
Dickeya dadantii subsp dieffenbachiae NCPPB 2976 WP 038922493 1 Ddadantii
Dickeya solani MK16 WP 022633178 1 Dsolani
Dickeya solani GBBC 2040 WP 022633178 1 Dsolani
Dickeya solani RNS 08 23 3 1 A WP 022633178 1 Dsolani
Dickeya solani IPO 2222 WP 022633178 1 Dsolani
Dickeya solani IFB 0099 WP 022633178 1 Dsolani
Dickeya solani PPO 9019 WP 022633178 1 Dsolani
Dickeya solani RNS 07 7 3B WP 022633178 1 Dsolani
Dickeya solani D s0432 1 WP 022633178 1 Dsolani
Dickeya solani PPO 9134 WP 022633178 1 Dsolani
Dickeya solani MK10 WP 022633178 1 Dsolani
Capnocytophaga canimorsus Cc5 YP 004741604 1 860228 Bacteroidetes Chlorobi group
Alistipes finegoldii DSM 17242 YP 006411502 1 679935 Bacteroidetes Chlorobi group
Pedobacter heparinus DSM 2366 YP 003090852 1 485917 Bacteroidetes Chlorobi group
Niastella koreensis GR20 10 YP 005006503 1 700598 Bacteroidetes Chlorobi group
Pantoea ananatis LMG 20103 YP 003519128 1 706191 Proteobacteria
Citrobacter rodentium ICC168 YP 003366611 1 637910 Proteobacteria
Citrobacter koseri ATCC BAA 895 YP 001455563 1 290338 Proteobacteria
Salmonella enterica subsp arizonae serovar 62 z4 z23 str RSK2980 YP 001569229 1 882884 Proteobacteria
Salmonella bongori N268 08 YP 008323452 1 1197719 Proteobacteria
Enterobacter lignolyticus SCF1 YP 003940126 1 701347 Proteobacteria
Pantoea sp At 9b YP 004115764 1 592316 Proteobacteria
Serratia proteamaculans 568 YP 001478806 1 399741 Proteobacteria
Serratia liquefaciens ATCC 27592 YP 008230505 1 1346614 Proteobacteria
Enterobacter cloacae subsp dissolvens SDM YP 006479406 1 1104326 Proteobacteria
Enterobacter cloacae EcWSU1 YP 004953785 1 1045856 Proteobacteria
Enterobacter asburiae LF7a YP 004830296 1 640513 Proteobacteria
Enterobacter sp 638 YP 001178280 1 399742 Proteobacteria
Shigella flexneri 2a str 2457T NP 838639 1 198215 Proteobacteria
Shigella flexneri 2002017 YP 005728789 1 591020 Proteobacteria
Desulfotomaculum ruminis DSM 2154 YP 004546638 1 696281 Firmicutes
Moorella thermoacetica ATCC 39073 YP 428894 1 264732 Firmicutes
Brevibacillus brevis NBRC 100599 YP 002773976 1 358681 Firmicutes
Paenibacillus terrae HPL 003 YP 005076906 1 985665 Firmicutes
Clostridium beijerinckii NCIMB 8052 YP 001307597 1 290402 Firmicutes
Caldicellulosiruptor obsidiansis OB47 YP 003840077 1 608506 Firmicutes
Caldicellulosiruptor owensensis OL YP 004002065 1 632518 Firmicutes
Caldicellulosiruptor bescii DSM 6725 YP 002572730 1 521460 Firmicutes
Caldicellulosiruptor lactoaceticus 6A YP 004798869 1 632516 Firmicutes
Caldicellulosiruptor hydrothermalis 108 YP 003992922 1 632292 Firmicutes
Paenibacillus mucilaginosus K02 YP 006192934 1 997761 Firmicutes
Caldicellulosiruptor lactoaceticus 6A YP 004799734 1 632516 Firmicutes
Thermotoga naphthophila RKU 10 YP 003346431 1 590168 Thermotogae
Thermotoga sp RQ2 YP 001738678 1 126740 Thermotogae
Thermoanaerobacter tengcongensis MB4 NP 624108 1 273068 Firmicutes
Thermoanaerobacterium xylanolyticum LX 11 YP 004471904 1 858215 Firmicutes
Thermoanaerobacterium thermosaccharolyticum DSM 571 YP 003853092 1 580327 Firmicutes
Thermoanaerobacter wiegelii Rt8 B1 YP 004821197 1 697303 Firmicutes
Thermoanaerobacter italicus Ab9 YP 003478028 1 580331 Firmicutes
Thermoanaerobacter mathranii subsp mathranii str A3 YP 003677844 1 583358 Firmicutes
Thermoanaerobacter sp X514 YP 001662219 1 399726 Firmicutes
Thermoanaerobacter pseudethanolicus ATCC 33223 YP 001664159 1 340099 Firmicutes
Thermoanaerobacter brockii subsp finnii Ako 1 YP 004185167 1 509193 Firmicutes

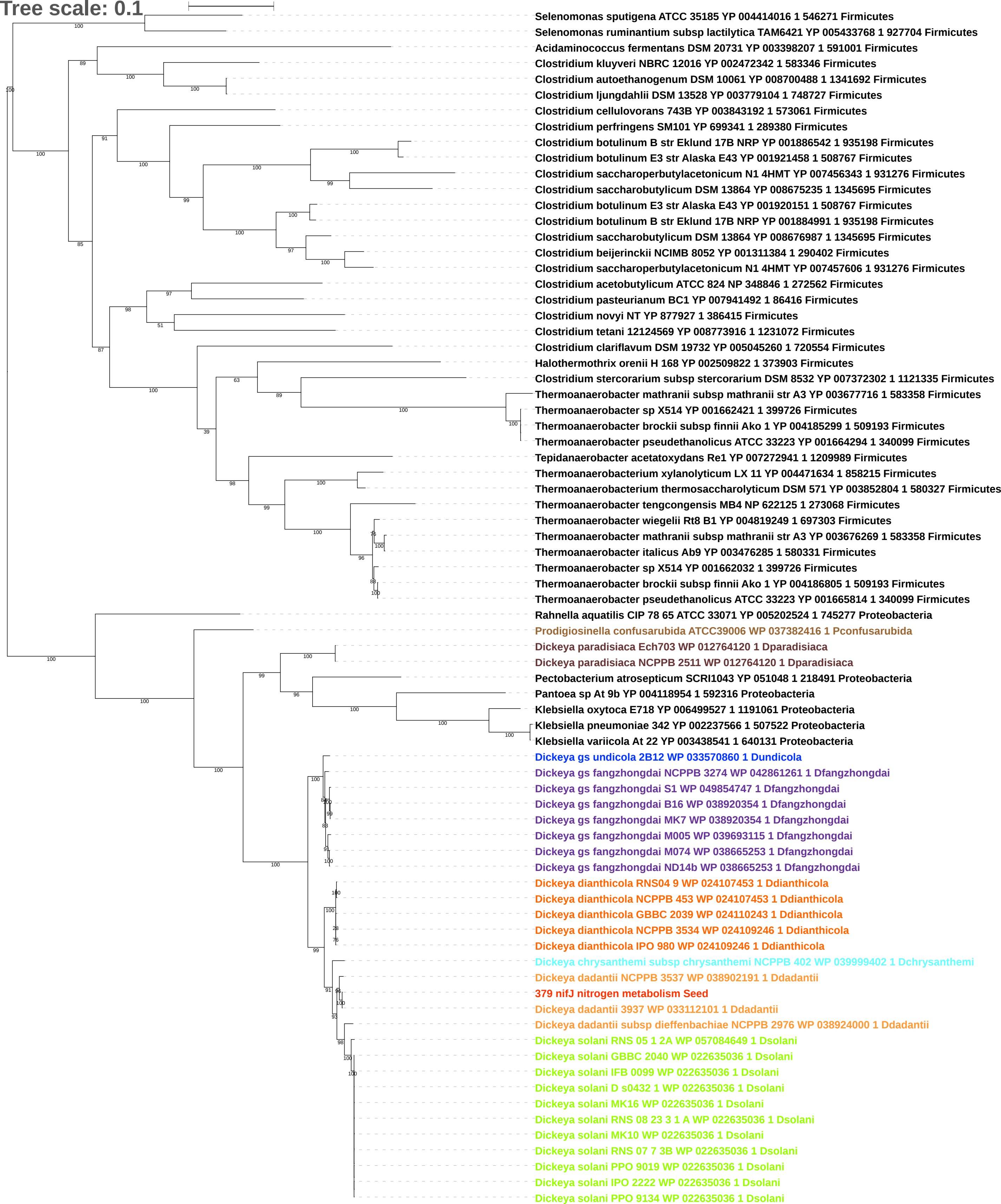
Tree scale: 0.1
Selenomonas sputigena ATCC 35185 YP 004414016 1 546271 Firmicutes
Selenomonas ruminantium subsp lactilytica TAM6421 YP 005433768 1 927704 Firmicutes
Acidaminococcus fermentans DSM 20731 YP 003398207 1 591001 Firmicutes
Clostridium kluyveri NBRC 12016 YP 002472342 1 583346 Firmicutes
Clostridium autoethanogenum DSM 10061 YP 008700488 1 1341692 Firmicutes
Clostridium ljungdahlii DSM 13528 YP 003779104 1 748727 Firmicutes
Clostridium cellulovorans 743B YP 003843192 1 573061 Firmicutes
Clostridium perfringens SM101 YP 699341 1 289380 Firmicutes
Clostridium botulinum B str Eklund 17B NRP YP 001886542 1 935198 Firmicutes
Clostridium botulinum E3 str Alaska E43 YP 001921458 1 508767 Firmicutes
Clostridium saccharoperbutylacetonicum N1 4HMT YP 007456343 1 931276 Firmicutes
Clostridium saccharobutylicum DSM 13864 YP 008675235 1 1345695 Firmicutes
Clostridium botulinum E3 str Alaska E43 YP 001920151 1 508767 Firmicutes
Clostridium botulinum B str Eklund 17B NRP YP 001884991 1 935198 Firmicutes
Clostridium saccharobutylicum DSM 13864 YP 008676987 1 1345695 Firmicutes
Clostridium beijerinckii NCIMB 8052 YP 001311384 1 290402 Firmicutes
Clostridium saccharoperbutylacetonicum N1 4HMT YP 007457606 1 931276 Firmicutes
Clostridium acetobutylicum ATCC 824 NP 348846 1 272562 Firmicutes
Clostridium pasteurianum BC1 YP 007941492 1 86416 Firmicutes
Clostridium novyi NT YP 877927 1 386415 Firmicutes
Clostridium tetani 12124569 YP 008773916 1 1231072 Firmicutes
Clostridium clariflavum DSM 19732 YP 005045260 1 720554 Firmicutes
Halothermothrix orenii H 168 YP 002509822 1 373903 Firmicutes
Clostridium stercorarium subsp stercorarium DSM 8532 YP 007372302 1 1121335 Firmicutes
Thermoanaerobacter mathranii subsp mathranii str A3 YP 003677716 1 583358 Firmicutes
Thermoanaerobacter sp X514 YP 001662421 1 399726 Firmicutes
Thermoanaerobacter brockii subsp finnii Ako 1 YP 004185299 1 509193 Firmicutes
Thermoanaerobacter pseudethanolicus ATCC 33223 YP 001664294 1 340099 Firmicutes
Tepidanaerobacter acetatoxydans Re1 YP 007272941 1 1209989 Firmicutes
Thermoanaerobacterium xylanolyticum LX 11 YP 004471634 1 858215 Firmicutes
Thermoanaerobacterium thermosaccharolyticum DSM 571 YP 003852804 1 580327 Firmicutes
Thermoanaerobacter tengcongensis MB4 NP 622125 1 273068 Firmicutes
Thermoanaerobacter wiegelii Rt8 B1 YP 004819249 1 697303 Firmicutes
Thermoanaerobacter mathranii subsp mathranii str A3 YP 003676269 1 583358 Firmicutes
Thermoanaerobacter italicus Ab9 YP 003476285 1 580331 Firmicutes
Thermoanaerobacter sp X514 YP 001662032 1 399726 Firmicutes
Thermoanaerobacter brockii subsp finnii Ako 1 YP 004186805 1 509193 Firmicutes
Thermoanaerobacter pseudethanolicus ATCC 33223 YP 001665814 1 340099 Firmicutes
Rahnella aquatilis CIP 78 65 ATCC 33071 YP 005202524 1 745277 Proteobacteria
Prodigiosinella confusarubida ATCC39006 WP 037382416 1 Pconfusarubida
Dickeya paradisiaca Ech703 WP 012764120 1 Dparadisiaca
Dickeya paradisiaca NCPPB 2511 WP 012764120 1 Dparadisiaca
Pectobacterium atrosepticum SCRI1043 YP 051048 1 218491 Proteobacteria
Pantoea sp At 9b YP 004118954 1 592316 Proteobacteria
Klebsiella oxytoca E718 YP 006499527 1 1191061 Proteobacteria
Klebsiella pneumoniae 342 YP 002237566 1 507522 Proteobacteria
Klebsiella variicola At 22 YP 003438541 1 640131 Proteobacteria
Dickeya gs undicola 2B12 WP 033570860 1 Dundicola
Dickeya gs fangzhongdai NCPPB 3274 WP 042861261 1 Dfangzhongdai
Dickeya gs fangzhongdai S1 WP 049854747 1 Dfangzhongdai
Dickeya gs fangzhongdai B16 WP 038920354 1 Dfangzhongdai
Dickeya gs fangzhongdai MK7 WP 038920354 1 Dfangzhongdai
Dickeya gs fangzhongdai M005 WP 039693115 1 Dfangzhongdai
Dickeya gs fangzhongdai M074 WP 038665253 1 Dfangzhongdai
Dickeya gs fangzhongdai ND14b WP 038665253 1 Dfangzhongdai
Dickeya dianthicola RNS04 9 WP 024107453 1 Ddianthicola
Dickeya dianthicola NCPPB 453 WP 024107453 1 Ddianthicola
Dickeya dianthicola GBBC 2039 WP 024110243 1 Ddianthicola
Dickeya dianthicola NCPPB 3534 WP 024109246 1 Ddianthicola
Dickeya dianthicola IPO 980 WP 024109246 1 Ddianthicola
Dickeya chrysanthemi subsp chrysanthemi NCPPB 402 WP 039999402 1 Dchrysanthemi
Dickeya dadantii NCPPB 3537 WP 038902191 1 Ddadantii
379 nifJ nitrogen metabolism Seed
Dickeya dadantii 3937 WP 033112101 1 Ddadantii
Dickeya dadantii subsp dieffenbachiae NCPPB 2976 WP 038924000 1 Ddadantii
Dickeya solani RNS 05 1 2A WP 057084649 1 Dsolani
Dickeya solani GBBC 2040 WP 022635036 1 Dsolani
Dickeya solani IFB 0099 WP 022635036 1 Dsolani
Dickeya solani D s0432 1 WP 022635036 1 Dsolani
Dickeya solani MK16 WP 022635036 1 Dsolani
Dickeya solani RNS 08 23 3 1 A WP 022635036 1 Dsolani
Dickeya solani MK10 WP 022635036 1 Dsolani
Dickeya solani RNS 07 7 3B WP 022635036 1 Dsolani
Dickeya solani PPO 9019 WP 022635036 1 Dsolani
Dickeya solani IPO 2222 WP 022635036 1 Dsolani
Dickeya solani PPO 9134 WP 022635036 1 Dsolani

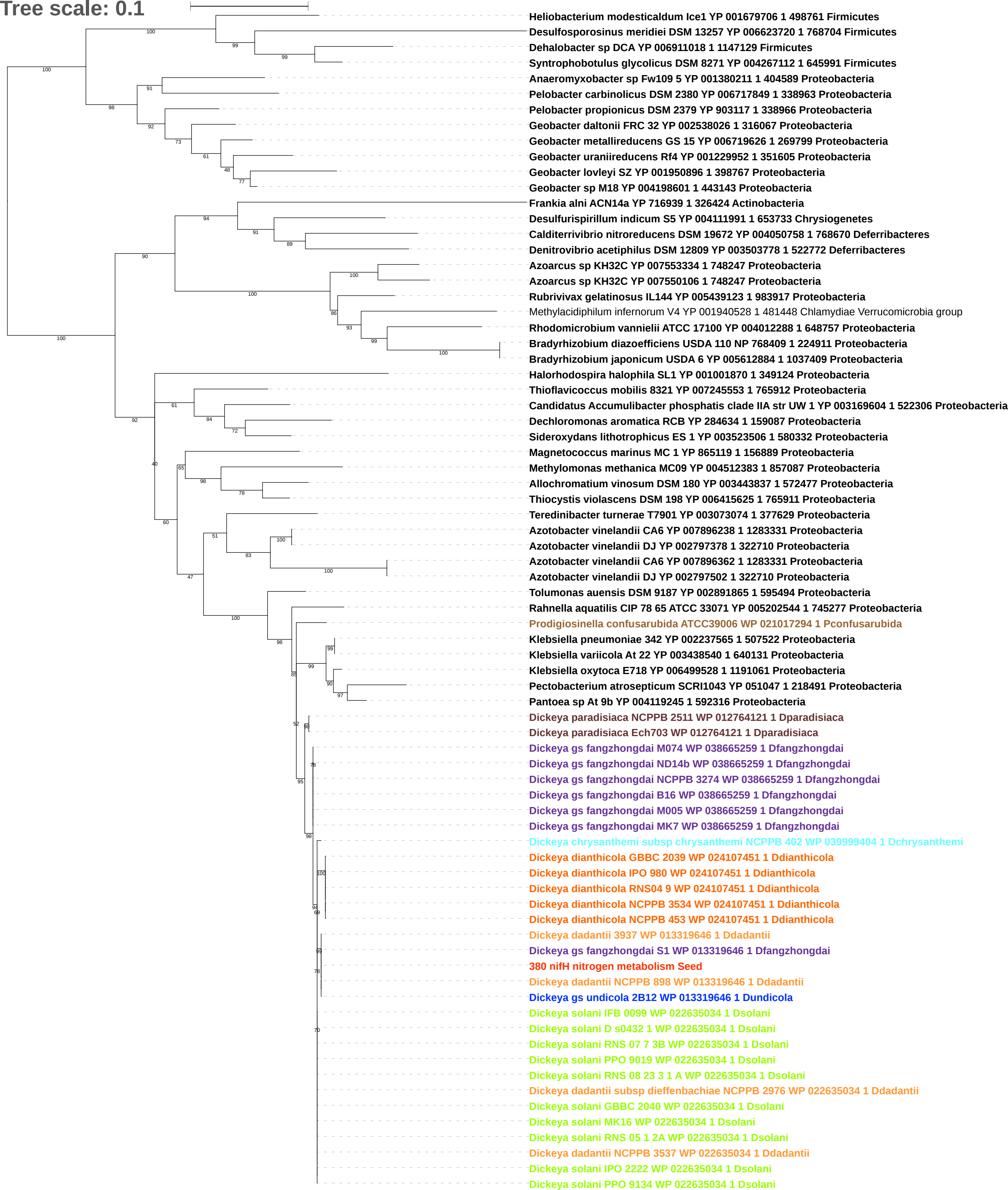
Tree scale: 0.1
Heliobacterium modesticaldum Ice1 YP 001679706 1 498761 Firmicutes
Desulfosporosinus meridiei DSM 13257 YP 006623720 1 768704 Firmicutes
Dehalobacter sp DCA YP 006911018 1 1147129 Firmicutes
Syntrophobotulus glycolicus DSM 8271 YP 004267112 1 645991 Firmicutes
Anaeromyxobacter sp Fw109 5 YP 001380211 1 404589 Proteobacteria
Pelobacter carbinolicus DSM 2380 YP 006717849 1 338963 Proteobacteria
Pelobacter propionicus DSM 2379 YP 903117 1 338966 Proteobacteria
Geobacter daltonii FRC 32 YP 002538026 1 316067 Proteobacteria
Geobacter metallireducens GS 15 YP 006719626 1 269799 Proteobacteria
Geobacter uraniireducens Rf4 YP 001229952 1 351605 Proteobacteria
Geobacter lovleyi SZ YP 001950896 1 398767 Proteobacteria
Geobacter sp M18 YP 004198601 1 443143 Proteobacteria
Frankia alni ACN14a YP 716939 1 326424 Actinobacteria
Desulfurispirillum indicum S5 YP 004111991 1 653733 Chrysiogenetes
Calditerrivibrio nitroreducens DSM 19672 YP 004050758 1 768670 Deferribacteres
Denitrovibrio acetiphilus DSM 12809 YP 003503778 1 522772 Deferribacteres
Azoarcus sp KH32C YP 007553334 1 748247 Proteobacteria
Azoarcus sp KH32C YP 007550106 1 748247 Proteobacteria
Rubrivivax gelatinosus IL144 YP 005439123 1 983917 Proteobacteria
Methylacidiphilum infernorum V4 YP 001940528 1 481448 Chlamydiae Verrucomicrobia group
Rhodomicrobium vannielii ATCC 17100 YP 004012288 1 648757 Proteobacteria
Bradyrhizobium diazoefficiens USDA 110 NP 768409 1 224911 Proteobacteria
Bradyrhizobium japonicum USDA 6 YP 005612884 1 1037409 Proteobacteria
Halorhodospira halophila SL1 YP 001001870 1 349124 Proteobacteria
Thioflavicoccus mobilis 8321 YP 007245553 1 765912 Proteobacteria
Candidatus Accumulibacter phosphatis clade IIA str UW 1 YP 003169604 1 522306 Proteobacteria
Dechloromonas aromatica RCB YP 284634 1 159087 Proteobacteria
Sideroxydans lithotrophicus ES 1 YP 003523506 1 580332 Proteobacteria
Magnetococcus marinus MC 1 YP 865119 1 156889 Proteobacteria
Methylomonas methanica MC09 YP 004512383 1 857087 Proteobacteria
Allochromatium vinosum DSM 180 YP 003443837 1 572477 Proteobacteria
Thiocystis violascens DSM 198 YP 006415625 1 765911 Proteobacteria
Teredinibacter turnerae T7901 YP 003073074 1 377629 Proteobacteria
Azotobacter vinelandii CA6 YP 007896238 1 1283331 Proteobacteria
Azotobacter vinelandii DJ YP 002797378 1 322710 Proteobacteria
Azotobacter vinelandii CA6 YP 007896362 1 1283331 Proteobacteria
Azotobacter vinelandii DJ YP 002797502 1 322710 Proteobacteria
Tolumonas auensis DSM 9187 YP 002891865 1 595494 Proteobacteria
Rahnella aquatilis CIP 78 65 ATCC 33071 YP 005202544 1 745277 Proteobacteria
Prodigiosinella confusarubida ATCC39006 WP 021017294 1 Pconfusarubida
Klebsiella pneumoniae 342 YP 002237565 1 507522 Proteobacteria
Klebsiella variicola At 22 YP 003438540 1 640131 Proteobacteria
Klebsiella oxytoca E718 YP 006499528 1 1191061 Proteobacteria
Pectobacterium atrosepticum SCRI1043 YP 051047 1 218491 Proteobacteria
Pantoea sp At 9b YP 004119245 1 592316 Proteobacteria
Dickeya paradisiaca NCPPB 2511 WP 012764121 1 Dparadisiaca
Dickeya paradisiaca Ech703 WP 012764121 1 Dparadisiaca
Dickeya gs fangzhongdai M074 WP 038665259 1 Dfangzhongdai
Dickeya gs fangzhongdai ND14b WP 038665259 1 Dfangzhongdai
Dickeya gs fangzhongdai NCPPB 3274 WP 038665259 1 Dfangzhongdai
Dickeya gs fangzhongdai B16 WP 038665259 1 Dfangzhongdai
Dickeya gs fangzhongdai M005 WP 038665259 1 Dfangzhongdai
Dickeya gs fangzhongdai MK7 WP 038665259 1 Dfangzhongdai
Dickeya chrysanthemi subsp chrysanthemi NCPPB 402 WP 039999404 1 Dchrysanthemi
Dickeya dianthicola GBBC 2039 WP 024107451 1 Ddianthicola
Dickeya dianthicola IPO 980 WP 024107451 1 Ddianthicola
Dickeya dianthicola RNS04 9 WP 024107451 1 Ddianthicola
Dickeya dianthicola NCPPB 3534 WP 024107451 1 Ddianthicola
Dickeya dianthicola NCPPB 453 WP 024107451 1 Ddianthicola
Dickeya dadantii 3937 WP 013319646 1 Ddadantii
Dickeya gs fangzhongdai S1 WP 013319646 1 Dfangzhongdai
380 nifH nitrogen metabolism Seed
Dickeya dadantii NCPPB 898 WP 013319646 1 Ddadantii
Dickeya gs undicola 2B12 WP 013319646 1 Dundicola
Dickeya solani IFB 0099 WP 022635034 1 Dsolani
Dickeya solani D s0432 1 WP 022635034 1 Dsolani
Dickeya solani RNS 07 7 3B WP 022635034 1 Dsolani
Dickeya solani PPO 9019 WP 022635034 1 Dsolani
Dickeya solani RNS 08 23 3 1 A WP 022635034 1 Dsolani
Dickeya dadantii subsp dieffenbachiae NCPPB 2976 WP 022635034 1 Ddadantii
Dickeya solani GBBC 2040 WP 022635034 1 Dsolani
Dickeya solani MK16 WP 022635034 1 Dsolani
Dickeya solani RNS 05 1 2A WP 022635034 1 Dsolani
Dickeya dadantii NCPPB 3537 WP 022635034 1 Ddadantii
Dickeya solani IPO 2222 WP 022635034 1 Dsolani
Dickeya solani PPO 9134 WP 022635034 1 Dsolani

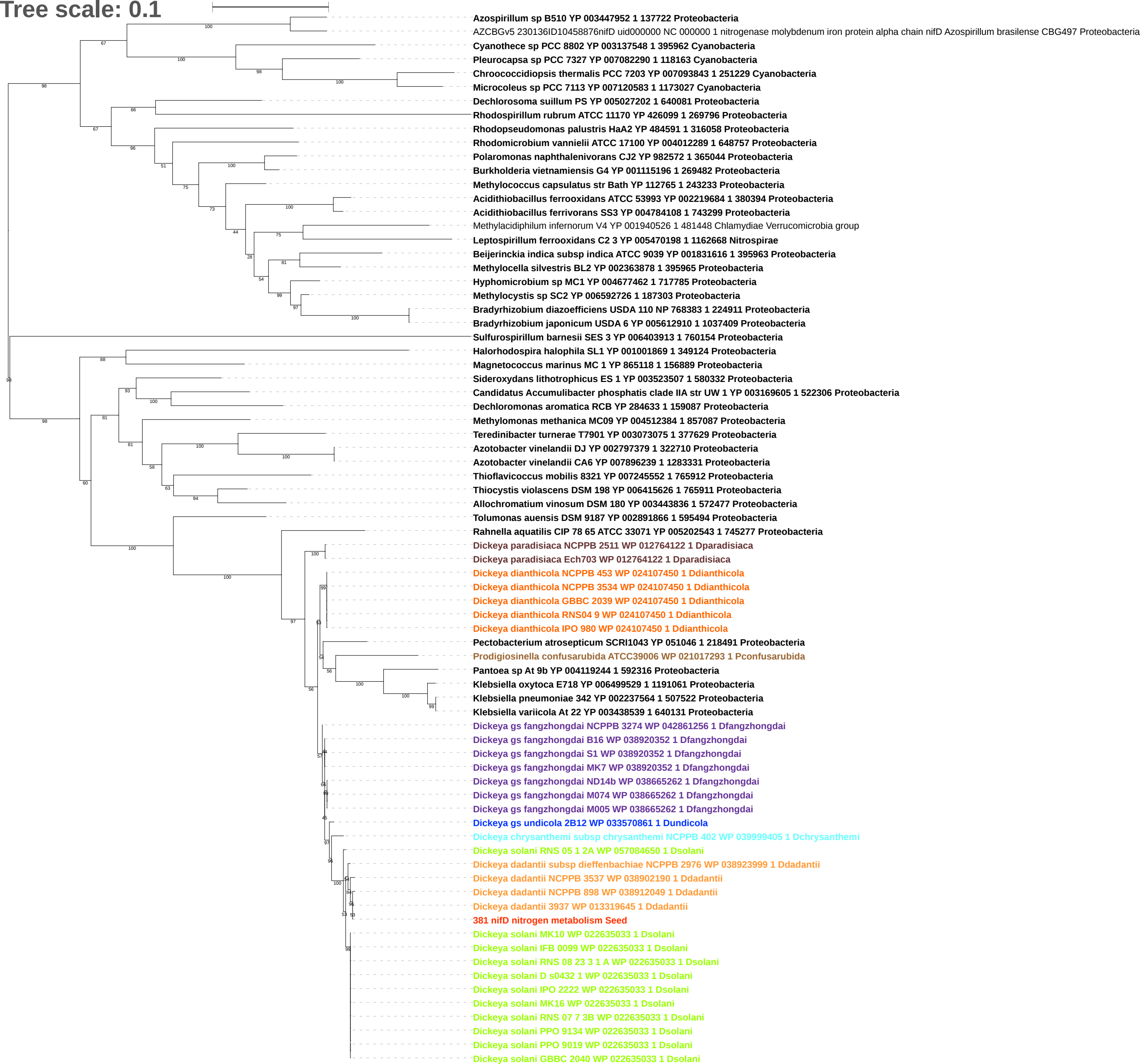
Tree scale: 0.1
Azospirillum sp B510 YP 003447952 1 137722 Proteobacteria
AZCBGv5 230136ID10458876nifD uid000000 NC 000000 1 nitrogenase molybdenum iron protein alpha chain nifD Azospirillum brasilense CBG497 Proteobacteria
Cyanothece sp PCC 8802 YP 003137548 1 395962 Cyanobacteria
Pleurocapsa sp PCC 7327 YP 007082290 1 118163 Cyanobacteria
Chroococcidiopsis thermalis PCC 7203 YP 007093843 1 251229 Cyanobacteria
Microcoleus sp PCC 7113 YP 007120583 1 1173027 Cyanobacteria
Dechlorosoma suillum PS YP 005027202 1 640081 Proteobacteria
Rhodospirillum rubrum ATCC 11170 YP 426099 1 269796 Proteobacteria
Rhodopseudomonas palustris HaA2 YP 484591 1 316058 Proteobacteria
Rhodomicrobium vannielii ATCC 17100 YP 004012289 1 648757 Proteobacteria
Polaromonas naphthalenivorans CJ2 YP 982572 1 365044 Proteobacteria
Burkholderia vietnamiensis G4 YP 001115196 1 269482 Proteobacteria
Methylococcus capsulatus str Bath YP 112765 1 243233 Proteobacteria
Acidithiobacillus ferrooxidans ATCC 53993 YP 002219684 1 380394 Proteobacteria
Acidithiobacillus ferrivorans SS3 YP 004784108 1 743299 Proteobacteria
Methylacidiphilum infernorum V4 YP 001940526 1 481448 Chlamydiae Verrucomicrobia group
Leptospirillum ferrooxidans C2 3 YP 005470198 1 1162668 Nitrospirae
Beijerinckia indica subsp indica ATCC 9039 YP 001831616 1 395963 Proteobacteria
Methylocella silvestris BL2 YP 002363878 1 395965 Proteobacteria
Hyphomicrobium sp MC1 YP 004677462 1 717785 Proteobacteria
Methylocystis sp SC2 YP 006592726 1 187303 Proteobacteria
Bradyrhizobium diazoefficiens USDA 110 NP 768383 1 224911 Proteobacteria
Bradyrhizobium japonicum USDA 6 YP 005612910 1 1037409 Proteobacteria
Sulfurospirillum barnesii SES 3 YP 006403913 1 760154 Proteobacteria
Halorhodospira halophila SL1 YP 001001869 1 349124 Proteobacteria
Magnetococcus marinus MC 1 YP 865118 1 156889 Proteobacteria
Sideroxydans lithotrophicus ES 1 YP 003523507 1 580332 Proteobacteria
Candidatus Accumulibacter phosphatis clade IIA str UW 1 YP 003169605 1 522306 Proteobacteria
Dechloromonas aromatica RCB YP 284633 1 159087 Proteobacteria
Methylomonas methanica MC09 YP 004512384 1 857087 Proteobacteria
Teredinibacter turnerae T7901 YP 003073075 1 377629 Proteobacteria
Azotobacter vinelandii DJ YP 002797379 1 322710 Proteobacteria
Azotobacter vinelandii CA6 YP 007896239 1 1283331 Proteobacteria
Thioflavicoccus mobilis 8321 YP 007245552 1 765912 Proteobacteria
Thiocystis violascens DSM 198 YP 006415626 1 765911 Proteobacteria
Allochromatium vinosum DSM 180 YP 003443836 1 572477 Proteobacteria
Tolumonas auensis DSM 9187 YP 002891866 1 595494 Proteobacteria
Rahnella aquatilis CIP 78 65 ATCC 33071 YP 005202543 1 745277 Proteobacteria
Dickeya paradisiaca NCPPB 2511 WP 012764122 1 Dparadisiaca
Dickeya paradisiaca Ech703 WP 012764122 1 Dparadisiaca
Dickeya dianthicola NCPPB 453 WP 024107450 1 Ddianthicola
Dickeya dianthicola NCPPB 3534 WP 024107450 1 Ddianthicola
Dickeya dianthicola GBBC 2039 WP 024107450 1 Ddianthicola
Dickeya dianthicola RNS04 9 WP 024107450 1 Ddianthicola
Dickeya dianthicola IPO 980 WP 024107450 1 Ddianthicola
Pectobacterium atrosepticum SCRI1043 YP 051046 1 218491 Proteobacteria
Prodigiosinella confusarubida ATCC39006 WP 021017293 1 Pconfusarubida
Pantoea sp At 9b YP 004119244 1 592316 Proteobacteria
Klebsiella oxytoca E718 YP 006499529 1 1191061 Proteobacteria
Klebsiella pneumoniae 342 YP 002237564 1 507522 Proteobacteria
Klebsiella variicola At 22 YP 003438539 1 640131 Proteobacteria
Dickeya gs fangzhongdai NCPPB 3274 WP 042861256 1 Dfangzhongdai
Dickeya gs fangzhongdai B16 WP 038920352 1 Dfangzhongdai
Dickeya gs fangzhongdai S1 WP 038920352 1 Dfangzhongdai
Dickeya gs fangzhongdai MK7 WP 038920352 1 Dfangzhongdai
Dickeya gs fangzhongdai ND14b WP 038665262 1 Dfangzhongdai
Dickeya gs fangzhongdai M074 WP 038665262 1 Dfangzhongdai
Dickeya gs fangzhongdai M005 WP 038665262 1 Dfangzhongdai
Dickeya gs undicola 2B12 WP 033570861 1 Dundicola
Dickeya chrysanthemi subsp chrysanthemi NCPPB 402 WP 039999405 1 Dchrysanthemi
Dickeya solani RNS 05 1 2A WP 057084650 1 Dsolani
Dickeya dadantii subsp dieffenbachiae NCPPB 2976 WP 038923999 1 Ddadantii
Dickeya dadantii NCPPB 3537 WP 038902190 1 Ddadantii
Dickeya dadantii NCPPB 898 WP 038912049 1 Ddadantii
Dickeya dadantii 3937 WP 013319645 1 Ddadantii
381 nifD nitrogen metabolism Seed
Dickeya solani MK10 WP 022635033 1 Dsolani
Dickeya solani IFB 0099 WP 022635033 1 Dsolani
Dickeya solani RNS 08 23 3 1 A WP 022635033 1 Dsolani
Dickeya solani D s0432 1 WP 022635033 1 Dsolani
Dickeya solani IPO 2222 WP 022635033 1 Dsolani
Dickeya solani MK16 WP 022635033 1 Dsolani
Dickeya solani RNS 07 7 3B WP 022635033 1 Dsolani
Dickeya solani PPO 9134 WP 022635033 1 Dsolani
Dickeya solani PPO 9019 WP 022635033 1 Dsolani
Dickeya solani GBBC 2040 WP 022635033 1 Dsolani

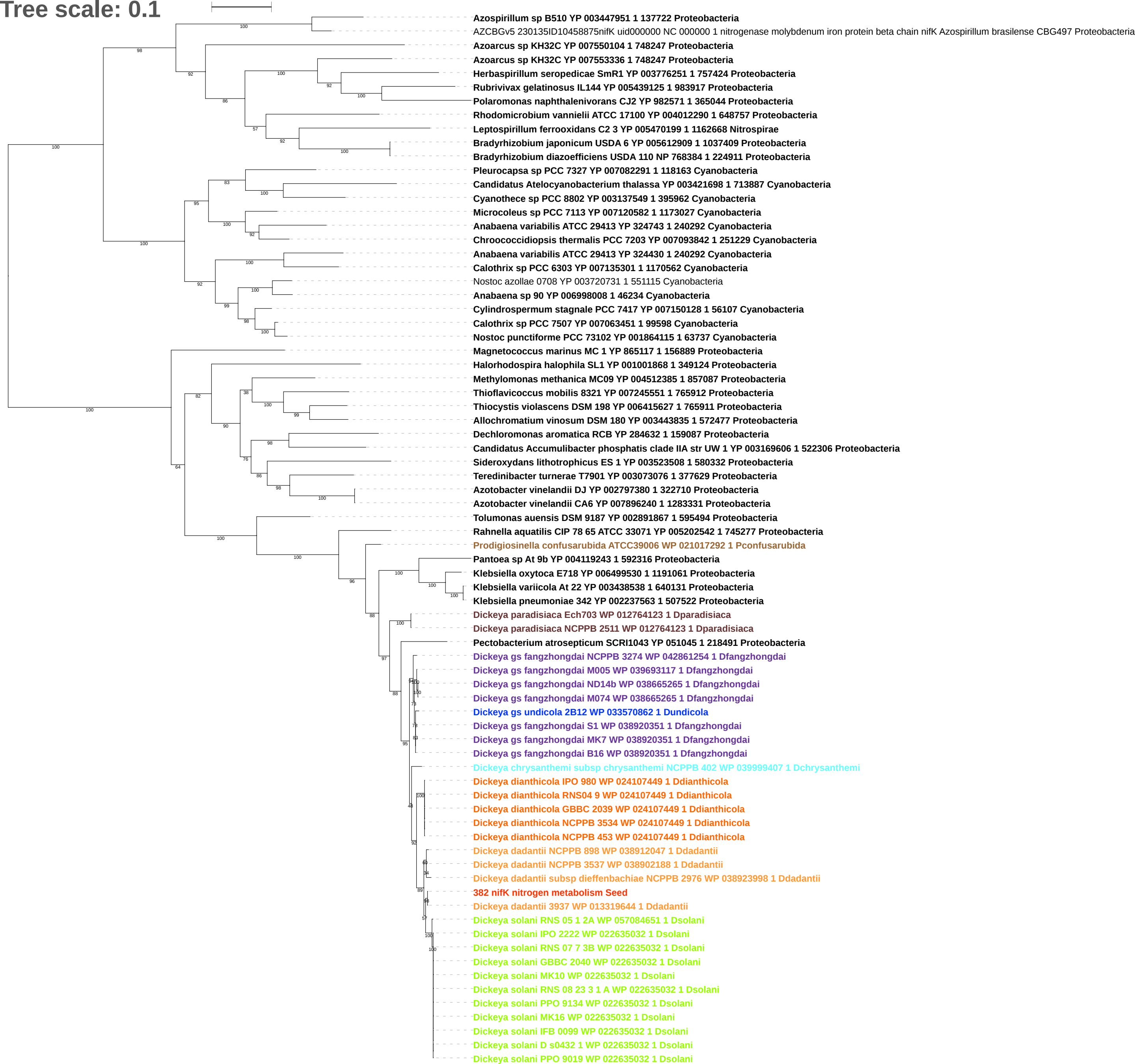
Tree scale: 0.1
Azospirillum sp B510 YP 003447951 1 137722 Proteobacteria
AZCBGv5 230135ID10458875nifK uid000000 NC 000000 1 nitrogenase molybdenum iron protein beta chain nifK Azospirillum brasilense CBG497 Proteobacteria
Azoarcus sp KH32C YP 007550104 1 748247 Proteobacteria
Azoarcus sp KH32C YP 007553336 1 748247 Proteobacteria
Herbaspirillum seropedicae SmR1 YP 003776251 1 757424 Proteobacteria
Rubrivivax gelatinosus IL144 YP 005439125 1 983917 Proteobacteria
Polaromonas naphthalenivorans CJ2 YP 982571 1 365044 Proteobacteria
Rhodomicrobium vannielii ATCC 17100 YP 004012290 1 648757 Proteobacteria
Leptospirillum ferrooxidans C2 3 YP 005470199 1 1162668 Nitrospirae
Bradyrhizobium japonicum USDA 6 YP 005612909 1 1037409 Proteobacteria
Bradyrhizobium diazoefficiens USDA 110 NP 768384 1 224911 Proteobacteria
Pleurocapsa sp PCC 7327 YP 007082291 1 118163 Cyanobacteria
Candidatus Atelocyanobacterium thalassa YP 003421698 1 713887 Cyanobacteria
Cyanothece sp PCC 8802 YP 003137549 1 395962 Cyanobacteria
Microcoleus sp PCC 7113 YP 007120582 1 1173027 Cyanobacteria
Anabaena variabilis ATCC 29413 YP 324743 1 240292 Cyanobacteria
Chroococcidiopsis thermalis PCC 7203 YP 007093842 1 251229 Cyanobacteria
Anabaena variabilis ATCC 29413 YP 324430 1 240292 Cyanobacteria
Calothrix sp PCC 6303 YP 007135301 1 1170562 Cyanobacteria
Nostoc azollae 0708 YP 003720731 1 551115 Cyanobacteria
Anabaena sp 90 YP 006998008 1 46234 Cyanobacteria
Cylindrospermum stagnale PCC 7417 YP 007150128 1 56107 Cyanobacteria
Calothrix sp PCC 7507 YP 007063451 1 99598 Cyanobacteria
Nostoc punctiforme PCC 73102 YP 001864115 1 63737 Cyanobacteria
Magnetococcus marinus MC 1 YP 865117 1 156889 Proteobacteria
Halorhodospira halophila SL1 YP 001001868 1 349124 Proteobacteria
Methylomonas methanica MC09 YP 004512385 1 857087 Proteobacteria
Thioflavicoccus mobilis 8321 YP 007245551 1 765912 Proteobacteria
Thiocystis violascens DSM 198 YP 006415627 1 765911 Proteobacteria
Allochromatium vinosum DSM 180 YP 003443835 1 572477 Proteobacteria
Dechloromonas aromatica RCB YP 284632 1 159087 Proteobacteria
Candidatus Accumulibacter phosphatis clade IIA str UW 1 YP 003169606 1 522306 Proteobacteria
Sideroxydans lithotrophicus ES 1 YP 003523508 1 580332 Proteobacteria
Teredinibacter turnerae T7901 YP 003073076 1 377629 Proteobacteria
Azotobacter vinelandii DJ YP 002797380 1 322710 Proteobacteria
Azotobacter vinelandii CA6 YP 007896240 1 1283331 Proteobacteria
Tolumonas auensis DSM 9187 YP 002891867 1 595494 Proteobacteria
Rahnella aquatilis CIP 78 65 ATCC 33071 YP 005202542 1 745277 Proteobacteria
Prodigiosinella confusarubida ATCC39006 WP 021017292 1 Pconfusarubida
Pantoea sp At 9b YP 004119243 1 592316 Proteobacteria
Klebsiella oxytoca E718 YP 006499530 1 1191061 Proteobacteria
Klebsiella variicola At 22 YP 003438538 1 640131 Proteobacteria
Klebsiella pneumoniae 342 YP 002237563 1 507522 Proteobacteria
Dickeya paradisiaca Ech703 WP 012764123 1 Dparadisiaca
Dickeya paradisiaca NCPPB 2511 WP 012764123 1 Dparadisiaca
Pectobacterium atrosepticum SCRI1043 YP 051045 1 218491 Proteobacteria
Dickeya gs fangzhongdai NCPPB 3274 WP 042861254 1 Dfangzhongdai
Dickeya gs fangzhongdai M005 WP 039693117 1 Dfangzhongdai
Dickeya gs fangzhongdai ND14b WP 038665265 1 Dfangzhongdai
Dickeya gs fangzhongdai M074 WP 038665265 1 Dfangzhongdai
Dickeya gs undicola 2B12 WP 033570862 1 Dundicola
Dickeya gs fangzhongdai S1 WP 038920351 1 Dfangzhongdai
Dickeya gs fangzhongdai MK7 WP 038920351 1 Dfangzhongdai
Dickeya gs fangzhongdai B16 WP 038920351 1 Dfangzhongdai
Dickeya chrysanthemi subsp chrysanthemi NCPPB 402 WP 039999407 1 Dchrysanthemi
Dickeya dianthicola IPO 980 WP 024107449 1 Ddianthicola
Dickeya dianthicola RNS04 9 WP 024107449 1 Ddianthicola
Dickeya dianthicola GBBC 2039 WP 024107449 1 Ddianthicola
Dickeya dianthicola NCPPB 3534 WP 024107449 1 Ddianthicola
Dickeya dianthicola NCPPB 453 WP 024107449 1 Ddianthicola
Dickeya dadantii NCPPB 898 WP 038912047 1 Ddadantii
Dickeya dadantii NCPPB 3537 WP 038902188 1 Ddadantii
Dickeya dadantii subsp dieffenbachiae NCPPB 2976 WP 038923998 1 Ddadantii
382 nifK nitrogen metabolism Seed
Dickeya dadantii 3937 WP 013319644 1 Ddadantii
Dickeya solani RNS 05 1 2A WP 057084651 1 Dsolani
Dickeya solani IPO 2222 WP 022635032 1 Dsolani
Dickeya solani RNS 07 7 3B WP 022635032 1 Dsolani
Dickeya solani GBBC 2040 WP 022635032 1 Dsolani
Dickeya solani MK10 WP 022635032 1 Dsolani
Dickeya solani RNS 08 23 3 1 A WP 022635032 1 Dsolani
Dickeya solani PPO 9134 WP 022635032 1 Dsolani
Dickeya solani MK16 WP 022635032 1 Dsolani
Dickeya solani IFB 0099 WP 022635032 1 Dsolani
Dickeya solani D s0432 1 WP 022635032 1 Dsolani
Dickeya solani PPO 9019 WP 022635032 1 Dsolani

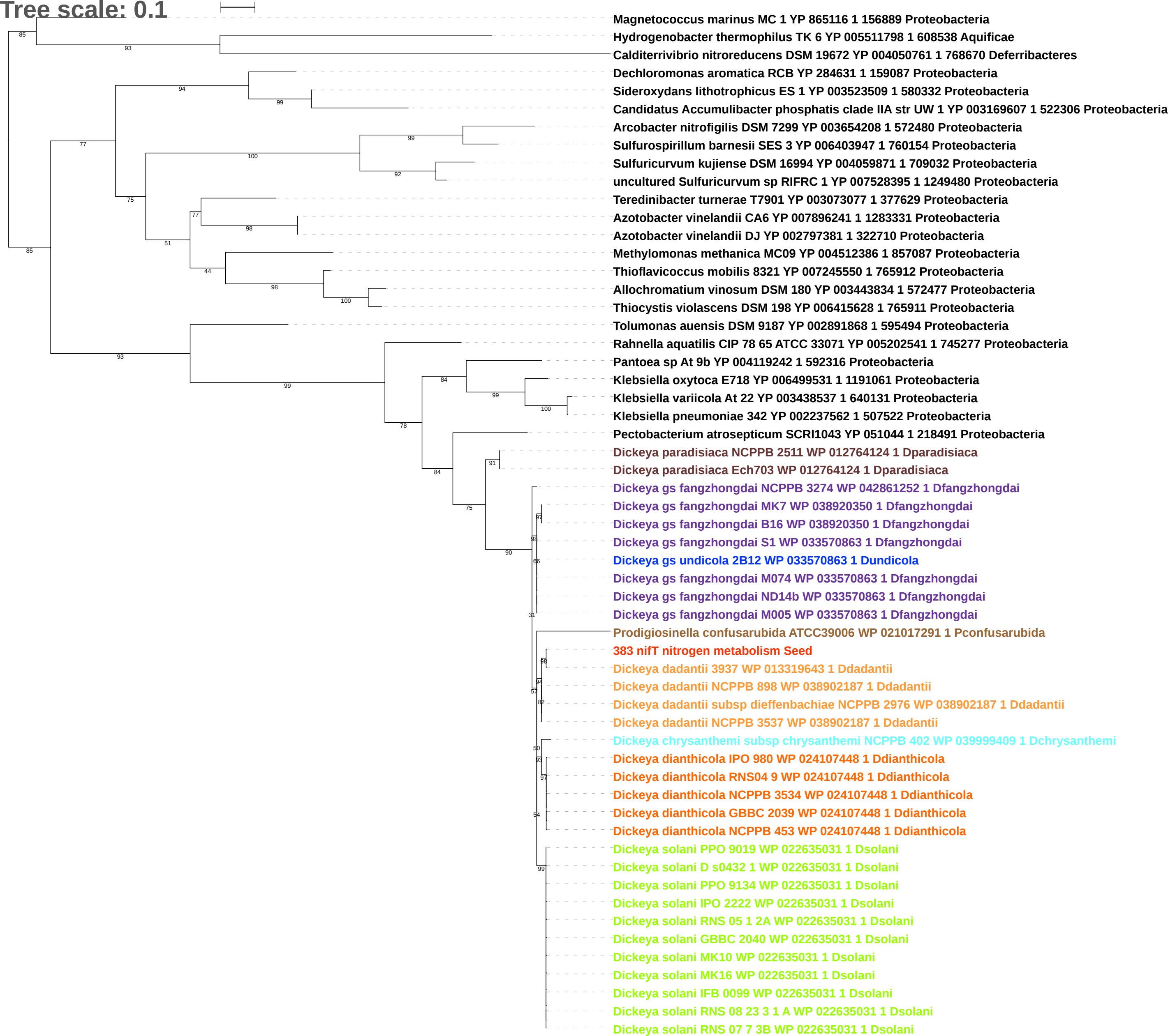
Tree scale: 0.1
Magnetococcus marinus MC 1 YP 865116 1 156889 Proteobacteria
Hydrogenobacter thermophilus TK 6 YP 005511798 1 608538 Aquificae
Calditerrivibrio nitroreducens DSM 19672 YP 004050761 1 768670 Deferribacteres
Dechloromonas aromatica RCB YP 284631 1 159087 Proteobacteria
Sideroxydans lithotrophicus ES 1 YP 003523509 1 580332 Proteobacteria
Candidatus Accumulibacter phosphatis clade IIA str UW 1 YP 003169607 1 522306 Proteobacteria
Arcobacter nitrofigilis DSM 7299 YP 003654208 1 572480 Proteobacteria
Sulfurospirillum barnesii SES 3 YP 006403947 1 760154 Proteobacteria
Sulfuricurvum kujiense DSM 16994 YP 004059871 1 709032 Proteobacteria
uncultured Sulfuricurvum sp RIFRC 1 YP 007528395 1 1249480 Proteobacteria
Teredinibacter turnerae T7901 YP 003073077 1 377629 Proteobacteria
Azotobacter vinelandii CA6 YP 007896241 1 1283331 Proteobacteria
Azotobacter vinelandii DJ YP 002797381 1 322710 Proteobacteria
Methylomonas methanica MC09 YP 004512386 1 857087 Proteobacteria
Thioflavicoccus mobilis 8321 YP 007245550 1 765912 Proteobacteria
Allochromatium vinosum DSM 180 YP 003443834 1 572477 Proteobacteria
Thiocystis violascens DSM 198 YP 006415628 1 765911 Proteobacteria
Tolumonas auensis DSM 9187 YP 002891868 1 595494 Proteobacteria
Rahnella aquatilis CIP 78 65 ATCC 33071 YP 005202541 1 745277 Proteobacteria
Pantoea sp At 9b YP 004119242 1 592316 Proteobacteria
Klebsiella oxytoca E718 YP 006499531 1 1191061 Proteobacteria
Klebsiella variicola At 22 YP 003438537 1 640131 Proteobacteria
Klebsiella pneumoniae 342 YP 002237562 1 507522 Proteobacteria
Pectobacterium atrosepticum SCRI1043 YP 051044 1 218491 Proteobacteria
Dickeya paradisiaca NCPPB 2511 WP 012764124 1 Dparadisiaca
Dickeya paradisiaca Ech703 WP 012764124 1 Dparadisiaca
Dickeya gs fangzhongdai NCPPB 3274 WP 042861252 1 Dfangzhongdai
Dickeya gs fangzhongdai MK7 WP 038920350 1 Dfangzhongdai
Dickeya gs fangzhongdai B16 WP 038920350 1 Dfangzhongdai
Dickeya gs fangzhongdai S1 WP 033570863 1 Dfangzhongdai
Dickeya gs undicola 2B12 WP 033570863 1 Dundicola
Dickeya gs fangzhongdai M074 WP 033570863 1 Dfangzhongdai
Dickeya gs fangzhongdai ND14b WP 033570863 1 Dfangzhongdai
Dickeya gs fangzhongdai M005 WP 033570863 1 Dfangzhongdai
Prodigiosinella confusarubida ATCC39006 WP 021017291 1 Pconfusarubida
383 nifT nitrogen metabolism Seed
Dickeya dadantii 3937 WP 013319643 1 Ddadantii
Dickeya dadantii NCPPB 898 WP 038902187 1 Ddadantii
Dickeya dadantii subsp dieffenbachiae NCPPB 2976 WP 038902187 1 Ddadantii
Dickeya dadantii NCPPB 3537 WP 038902187 1 Ddadantii
Dickeya chrysanthemi subsp chrysanthemi NCPPB 402 WP 039999409 1 Dchrysanthemi
Dickeya dianthicola IPO 980 WP 024107448 1 Ddianthicola
Dickeya dianthicola RNS04 9 WP 024107448 1 Ddianthicola
Dickeya dianthicola NCPPB 3534 WP 024107448 1 Ddianthicola
Dickeya dianthicola GBBC 2039 WP 024107448 1 Ddianthicola
Dickeya dianthicola NCPPB 453 WP 024107448 1 Ddianthicola
Dickeya solani PPO 9019 WP 022635031 1 Dsolani
Dickeya solani D s0432 1 WP 022635031 1 Dsolani
Dickeya solani PPO 9134 WP 022635031 1 Dsolani
Dickeya solani IPO 2222 WP 022635031 1 Dsolani
Dickeya solani RNS 05 1 2A WP 022635031 1 Dsolani
Dickeya solani GBBC 2040 WP 022635031 1 Dsolani
Dickeya solani MK10 WP 022635031 1 Dsolani
Dickeya solani MK16 WP 022635031 1 Dsolani
Dickeya solani IFB 0099 WP 022635031 1 Dsolani
Dickeya solani RNS 08 23 3 1 A WP 022635031 1 Dsolani
Dickeya solani RNS 07 7 3B WP 022635031 1 Dsolani

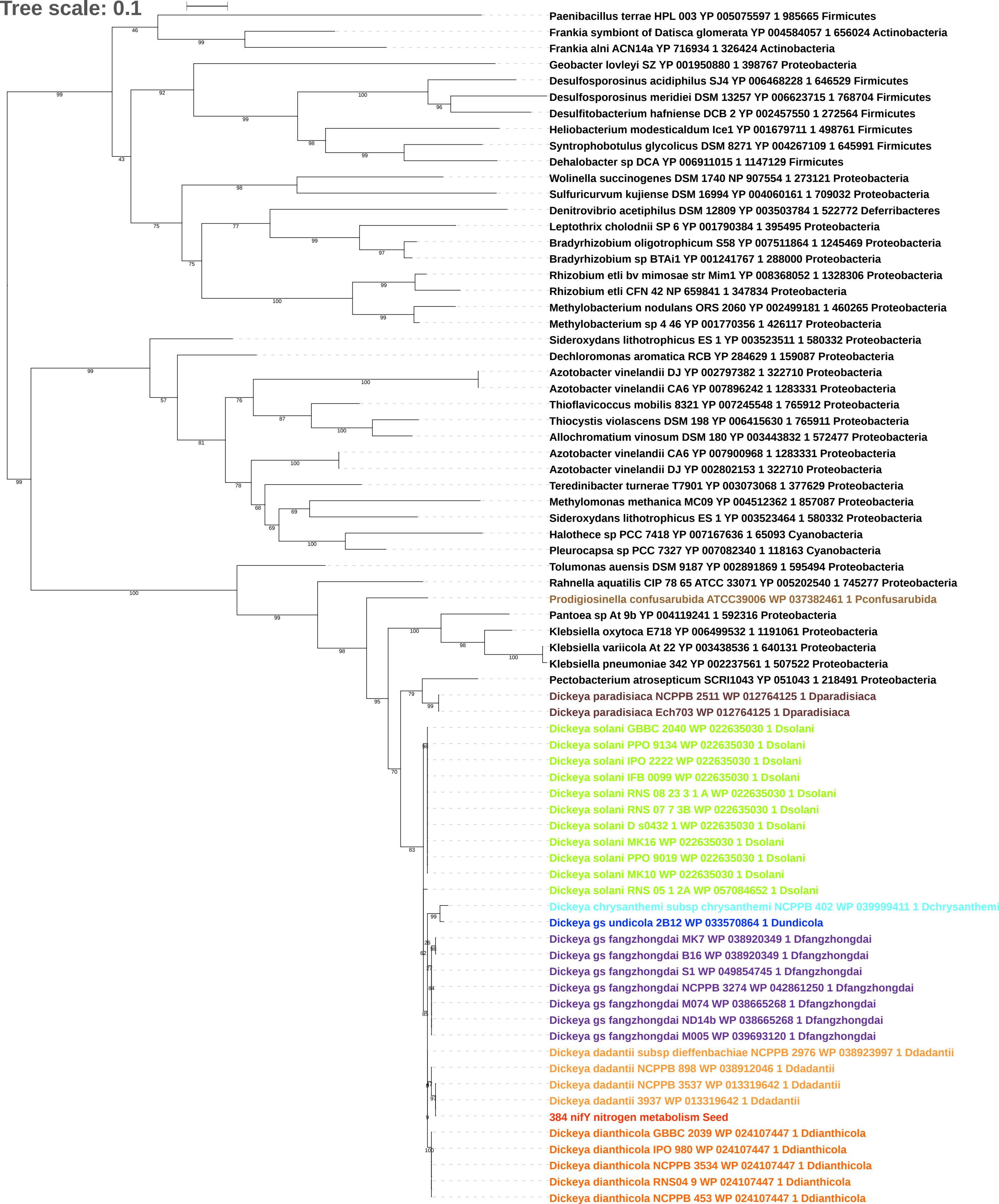
Tree scale: 0.1
Paenibacillus terrae HPL 003 YP 005075597 1 985665 Firmicutes
Frankia symbiont of Datisca glomerata YP 004584057 1 656024 Actinobacteria
Frankia alni ACN14a YP 716934 1 326424 Actinobacteria
Geobacter lovleyi SZ YP 001950880 1 398767 Proteobacteria
Desulfosporosinus acidiphilus SJ4 YP 006468228 1 646529 Firmicutes
Desulfosporosinus meridiei DSM 13257 YP 006623715 1 768704 Firmicutes
Desulfitobacterium hafniense DCB 2 YP 002457550 1 272564 Firmicutes
Heliobacterium modesticaldum Ice1 YP 001679711 1 498761 Firmicutes
Syntrophobotulus glycolicus DSM 8271 YP 004267109 1 645991 Firmicutes
Dehalobacter sp DCA YP 006911015 1 1147129 Firmicutes
Wolinella succinogenes DSM 1740 NP 907554 1 273121 Proteobacteria
Sulfuricurvum kujiense DSM 16994 YP 004060161 1 709032 Proteobacteria
Denitrovibrio acetiphilus DSM 12809 YP 003503784 1 522772 Deferribacteres
Leptothrix cholodnii SP 6 YP 001790384 1 395495 Proteobacteria
Bradyrhizobium oligotrophicum S58 YP 007511864 1 1245469 Proteobacteria
Bradyrhizobium sp BTAi1 YP 001241767 1 288000 Proteobacteria
Rhizobium etli bv mimosae str Mim1 YP 008368052 1 1328306 Proteobacteria
Rhizobium etli CFN 42 NP 659841 1 347834 Proteobacteria
Methylobacterium nodulans ORS 2060 YP 002499181 1 460265 Proteobacteria
Methylobacterium sp 4 46 YP 001770356 1 426117 Proteobacteria
Sideroxydans lithotrophicus ES 1 YP 003523511 1 580332 Proteobacteria
Dechloromonas aromatica RCB YP 284629 1 159087 Proteobacteria
Azotobacter vinelandii DJ YP 002797382 1 322710 Proteobacteria
Azotobacter vinelandii CA6 YP 007896242 1 1283331 Proteobacteria
Thioflavicoccus mobilis 8321 YP 007245548 1 765912 Proteobacteria
Thiocystis violascens DSM 198 YP 006415630 1 765911 Proteobacteria
Allochromatium vinosum DSM 180 YP 003443832 1 572477 Proteobacteria
Azotobacter vinelandii CA6 YP 007900968 1 1283331 Proteobacteria
Azotobacter vinelandii DJ YP 002802153 1 322710 Proteobacteria
Teredinibacter turnerae T7901 YP 003073068 1 377629 Proteobacteria
Methylomonas methanica MC09 YP 004512362 1 857087 Proteobacteria
Sideroxydans lithotrophicus ES 1 YP 003523464 1 580332 Proteobacteria
Halothece sp PCC 7418 YP 007167636 1 65093 Cyanobacteria
Pleurocapsa sp PCC 7327 YP 007082340 1 118163 Cyanobacteria
Tolumonas auensis DSM 9187 YP 002891869 1 595494 Proteobacteria
Rahnella aquatilis CIP 78 65 ATCC 33071 YP 005202540 1 745277 Proteobacteria
Prodigiosinella confusarubida ATCC39006 WP 037382461 1 Pconfusarubida
Pantoea sp At 9b YP 004119241 1 592316 Proteobacteria
Klebsiella oxytoca E718 YP 006499532 1 1191061 Proteobacteria
Klebsiella variicola At 22 YP 003438536 1 640131 Proteobacteria
Klebsiella pneumoniae 342 YP 002237561 1 507522 Proteobacteria
Pectobacterium atrosepticum SCRI1043 YP 051043 1 218491 Proteobacteria
Dickeya paradisiaca NCPPB 2511 WP 012764125 1 Dparadisiaca
Dickeya paradisiaca Ech703 WP 012764125 1 Dparadisiaca
Dickeya solani GBBC 2040 WP 022635030 1 Dsolani
Dickeya solani PPO 9134 WP 022635030 1 Dsolani
Dickeya solani IPO 2222 WP 022635030 1 Dsolani
Dickeya solani IFB 0099 WP 022635030 1 Dsolani
Dickeya solani RNS 08 23 3 1 A WP 022635030 1 Dsolani
Dickeya solani RNS 07 7 3B WP 022635030 1 Dsolani
Dickeya solani D s0432 1 WP 022635030 1 Dsolani
Dickeya solani MK16 WP 022635030 1 Dsolani
Dickeya solani PPO 9019 WP 022635030 1 Dsolani
Dickeya solani MK10 WP 022635030 1 Dsolani
Dickeya solani RNS 05 1 2A WP 057084652 1 Dsolani
Dickeya chrysanthemi subsp chrysanthemi NCPPB 402 WP 039999411 1 Dchrysanthemi
Dickeya gs undicola 2B12 WP 033570864 1 Dundicola
Dickeya gs fangzhongdai MK7 WP 038920349 1 Dfangzhongdai
Dickeya gs fangzhongdai B16 WP 038920349 1 Dfangzhongdai
Dickeya gs fangzhongdai S1 WP 049854745 1 Dfangzhongdai
Dickeya gs fangzhongdai NCPPB 3274 WP 042861250 1 Dfangzhongdai
Dickeya gs fangzhongdai M074 WP 038665268 1 Dfangzhongdai
Dickeya gs fangzhongdai ND14b WP 038665268 1 Dfangzhongdai
Dickeya gs fangzhongdai M005 WP 039693120 1 Dfangzhongdai
Dickeya dadantii subsp dieffenbachiae NCPPB 2976 WP 038923997 1 Ddadantii
Dickeya dadantii NCPPB 898 WP 038912046 1 Ddadantii
Dickeya dadantii NCPPB 3537 WP 013319642 1 Ddadantii
Dickeya dadantii 3937 WP 013319642 1 Ddadantii
384 nifY nitrogen metabolism Seed
Dickeya dianthicola GBBC 2039 WP 024107447 1 Ddianthicola
Dickeya dianthicola IPO 980 WP 024107447 1 Ddianthicola
Dickeya dianthicola NCPPB 3534 WP 024107447 1 Ddianthicola
Dickeya dianthicola RNS04 9 WP 024107447 1 Ddianthicola
Dickeya dianthicola NCPPB 453 WP 024107447 1 Ddianthicola

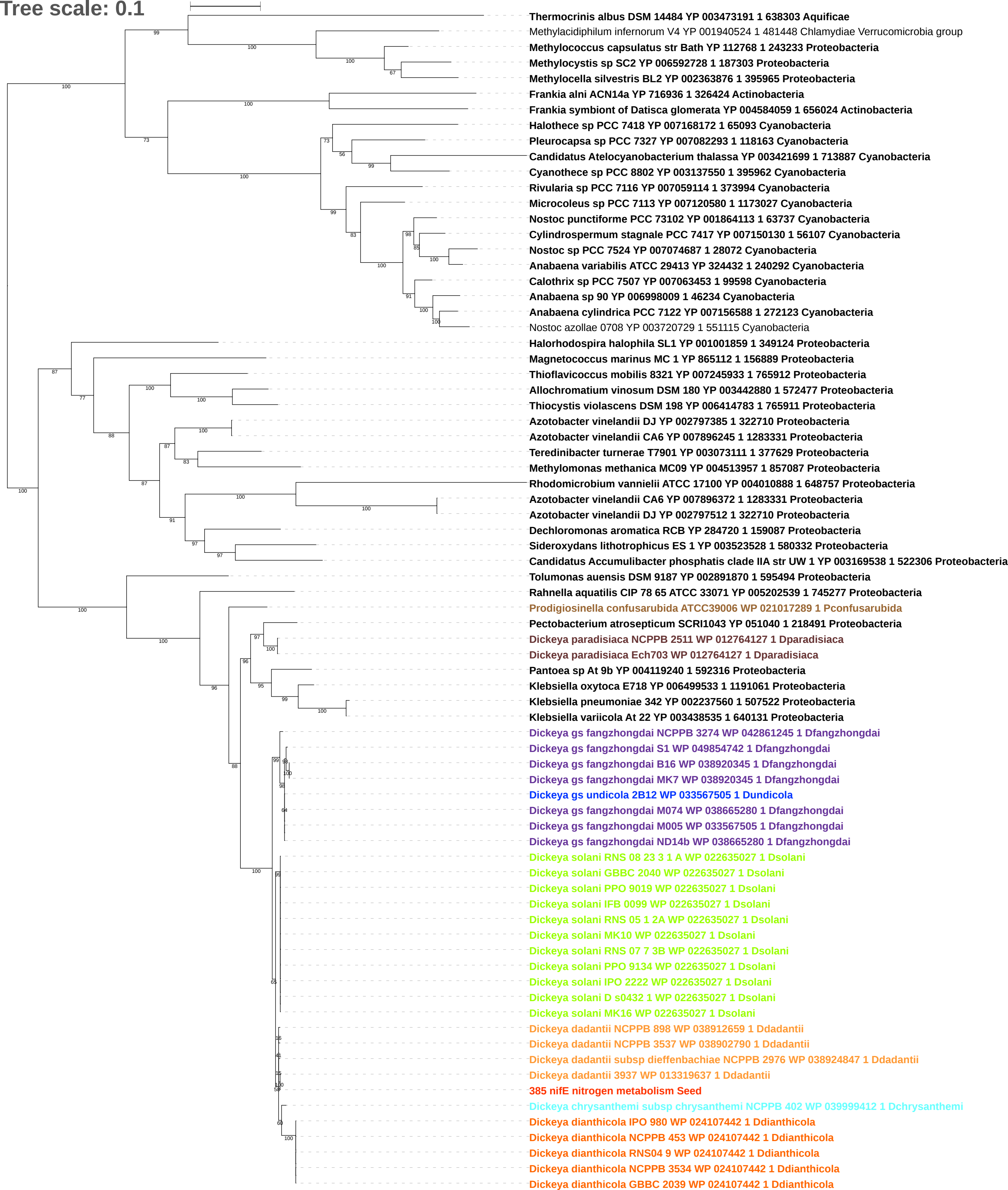
Tree scale: 0.1
Thermocrinis albus DSM 14484 YP 003473191 1 638303 Aquificae
Methylacidiphilum infernorum V4 YP 001940524 1 481448 Chlamydiae Verrucomicrobia group
Methylococcus capsulatus str Bath YP 112768 1 243233 Proteobacteria
Methylocystis sp SC2 YP 006592728 1 187303 Proteobacteria
Methylocella silvestris BL2 YP 002363876 1 395965 Proteobacteria
Frankia alni ACN14a YP 716936 1 326424 Actinobacteria
Frankia symbiont of Datisca glomerata YP 004584059 1 656024 Actinobacteria
Halothece sp PCC 7418 YP 007168172 1 65093 Cyanobacteria
Pleurocapsa sp PCC 7327 YP 007082293 1 118163 Cyanobacteria
Candidatus Atelocyanobacterium thalassa YP 003421699 1 713887 Cyanobacteria
Cyanothece sp PCC 8802 YP 003137550 1 395962 Cyanobacteria
Rivularia sp PCC 7116 YP 007059114 1 373994 Cyanobacteria
Microcoleus sp PCC 7113 YP 007120580 1 1173027 Cyanobacteria
Nostoc punctiforme PCC 73102 YP 001864113 1 63737 Cyanobacteria
Cylindrospermum stagnale PCC 7417 YP 007150130 1 56107 Cyanobacteria
Nostoc sp PCC 7524 YP 007074687 1 28072 Cyanobacteria
Anabaena variabilis ATCC 29413 YP 324432 1 240292 Cyanobacteria
Calothrix sp PCC 7507 YP 007063453 1 99598 Cyanobacteria
Anabaena sp 90 YP 006998009 1 46234 Cyanobacteria
Anabaena cylindrica PCC 7122 YP 007156588 1 272123 Cyanobacteria
Nostoc azollae 0708 YP 003720729 1 551115 Cyanobacteria
Halorhodospira halophila SL1 YP 001001859 1 349124 Proteobacteria
Magnetococcus marinus MC 1 YP 865112 1 156889 Proteobacteria
Thioflavicoccus mobilis 8321 YP 007245933 1 765912 Proteobacteria
Allochromatium vinosum DSM 180 YP 003442880 1 572477 Proteobacteria
Thiocystis violascens DSM 198 YP 006414783 1 765911 Proteobacteria
Azotobacter vinelandii DJ YP 002797385 1 322710 Proteobacteria
Azotobacter vinelandii CA6 YP 007896245 1 1283331 Proteobacteria
Teredinibacter turnerae T7901 YP 003073111 1 377629 Proteobacteria
Methylomonas methanica MC09 YP 004513957 1 857087 Proteobacteria
Rhodomicrobium vannielii ATCC 17100 YP 004010888 1 648757 Proteobacteria
Azotobacter vinelandii CA6 YP 007896372 1 1283331 Proteobacteria
Azotobacter vinelandii DJ YP 002797512 1 322710 Proteobacteria
Dechloromonas aromatica RCB YP 284720 1 159087 Proteobacteria
Sideroxydans lithotrophicus ES 1 YP 003523528 1 580332 Proteobacteria
Candidatus Accumulibacter phosphatis clade IIA str UW 1 YP 003169538 1 522306 Proteobacteria
Tolumonas auensis DSM 9187 YP 002891870 1 595494 Proteobacteria
Rahnella aquatilis CIP 78 65 ATCC 33071 YP 005202539 1 745277 Proteobacteria
Prodigiosinella confusarubida ATCC39006 WP 021017289 1 Pconfusarubida
Pectobacterium atrosepticum SCRI1043 YP 051040 1 218491 Proteobacteria
Dickeya paradisiaca NCPPB 2511 WP 012764127 1 Dparadisiaca
Dickeya paradisiaca Ech703 WP 012764127 1 Dparadisiaca
Pantoea sp At 9b YP 004119240 1 592316 Proteobacteria
Klebsiella oxytoca E718 YP 006499533 1 1191061 Proteobacteria
Klebsiella pneumoniae 342 YP 002237560 1 507522 Proteobacteria
Klebsiella variicola At 22 YP 003438535 1 640131 Proteobacteria
Dickeya gs fangzhongdai NCPPB 3274 WP 042861245 1 Dfangzhongdai
Dickeya gs fangzhongdai S1 WP 049854742 1 Dfangzhongdai
Dickeya gs fangzhongdai B16 WP 038920345 1 Dfangzhongdai
Dickeya gs fangzhongdai MK7 WP 038920345 1 Dfangzhongdai
Dickeya gs undicola 2B12 WP 033567505 1 Dundicola
Dickeya gs fangzhongdai M074 WP 038665280 1 Dfangzhongdai
Dickeya gs fangzhongdai M005 WP 033567505 1 Dfangzhongdai
Dickeya gs fangzhongdai ND14b WP 038665280 1 Dfangzhongdai
Dickeya solani RNS 08 23 3 1 A WP 022635027 1 Dsolani
Dickeya solani GBBC 2040 WP 022635027 1 Dsolani
Dickeya solani PPO 9019 WP 022635027 1 Dsolani
Dickeya solani IFB 0099 WP 022635027 1 Dsolani
Dickeya solani RNS 05 1 2A WP 022635027 1 Dsolani
Dickeya solani MK10 WP 022635027 1 Dsolani
Dickeya solani RNS 07 7 3B WP 022635027 1 Dsolani
Dickeya solani PPO 9134 WP 022635027 1 Dsolani
Dickeya solani IPO 2222 WP 022635027 1 Dsolani
Dickeya solani D s0432 1 WP 022635027 1 Dsolani
Dickeya solani MK16 WP 022635027 1 Dsolani
Dickeya dadantii NCPPB 898 WP 038912659 1 Ddadantii
Dickeya dadantii NCPPB 3537 WP 038902790 1 Ddadantii
Dickeya dadantii subsp dieffenbachiae NCPPB 2976 WP 038924847 1 Ddadantii
Dickeya dadantii 3937 WP 013319637 1 Ddadantii
385 nifE nitrogen metabolism Seed
Dickeya chrysanthemi subsp chrysanthemi NCPPB 402 WP 039999412 1 Dchrysanthemi
Dickeya dianthicola IPO 980 WP 024107442 1 Ddianthicola
Dickeya dianthicola NCPPB 453 WP 024107442 1 Ddianthicola
Dickeya dianthicola RNS04 9 WP 024107442 1 Ddianthicola
Dickeya dianthicola NCPPB 3534 WP 024107442 1 Ddianthicola
Dickeya dianthicola GBBC 2039 WP 024107442 1 Ddianthicola

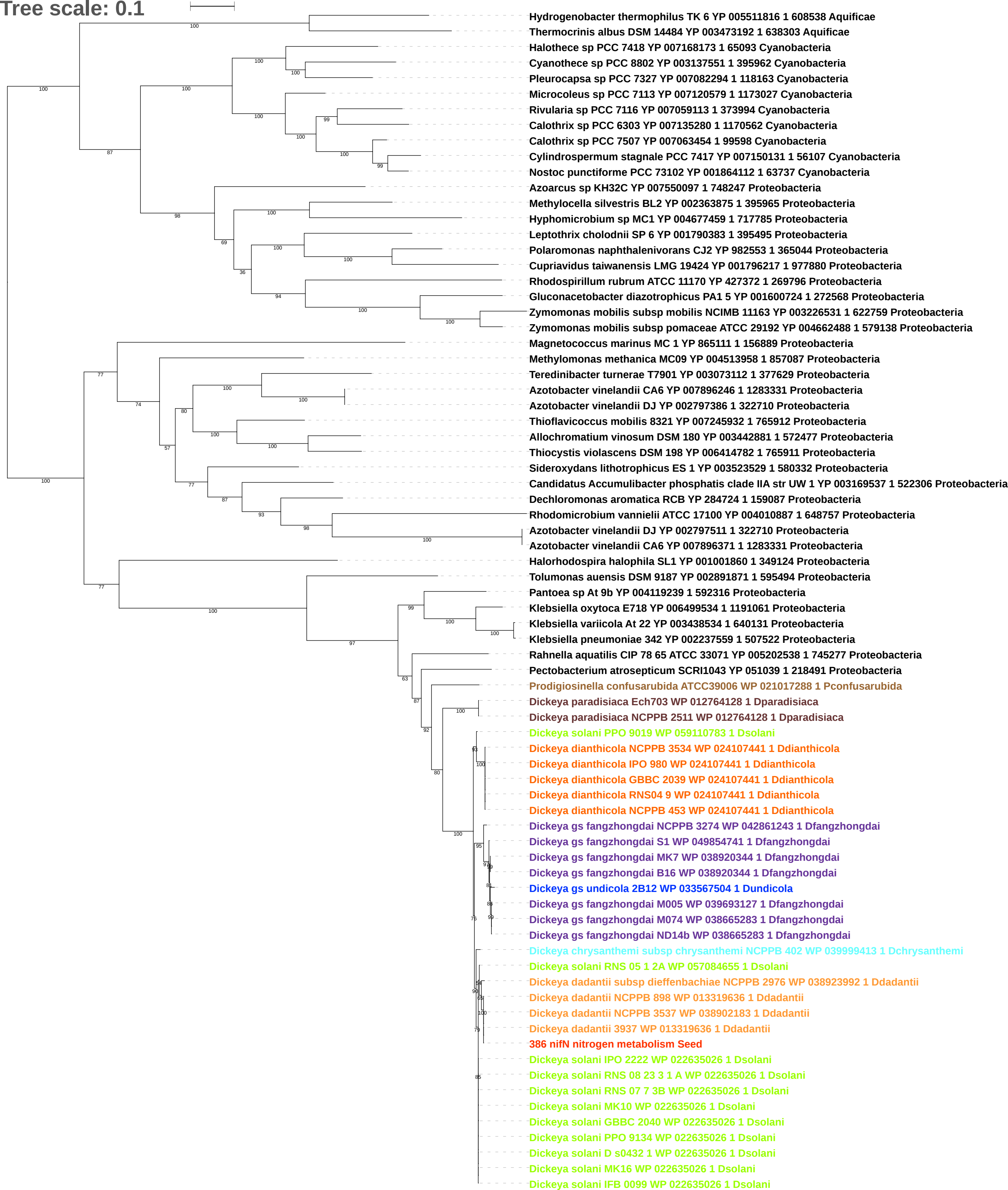

Tree scale: 0.1
Hydrogenobacter thermophilus TK 6 YP 005511816 1 608538 Aquificae
Thermocrinis albus DSM 14484 YP 003473192 1 638303 Aquificae
Halothece sp PCC 7418 YP 007168173 1 65093 Cyanobacteria
Cyanothece sp PCC 8802 YP 003137551 1 395962 Cyanobacteria
Pleurocapsa sp PCC 7327 YP 007082294 1 118163 Cyanobacteria
Microcoleus sp PCC 7113 YP 007120579 1 1173027 Cyanobacteria
Rivularia sp PCC 7116 YP 007059113 1 373994 Cyanobacteria
Calothrix sp PCC 6303 YP 007135280 1 1170562 Cyanobacteria
Calothrix sp PCC 7507 YP 007063454 1 99598 Cyanobacteria
Cylindrospermum stagnale PCC 7417 YP 007150131 1 56107 Cyanobacteria
Nostoc punctiforme PCC 73102 YP 001864112 1 63737 Cyanobacteria
Azoarcus sp KH32C YP 007550097 1 748247 Proteobacteria
Methylocella silvestris BL2 YP 002363875 1 395965 Proteobacteria
Hyphomicrobium sp MC1 YP 004677459 1 717785 Proteobacteria
Leptothrix cholodnii SP 6 YP 001790383 1 395495 Proteobacteria
Polaromonas naphthalenivorans CJ2 YP 982553 1 365044 Proteobacteria
Cupriavidus taiwanensis LMG 19424 YP 001796217 1 977880 Proteobacteria
Rhodospirillum rubrum ATCC 11170 YP 427372 1 269796 Proteobacteria
Gluconacetobacter diazotrophicus PA1 5 YP 001600724 1 272568 Proteobacteria
Zymomonas mobilis subsp mobilis NCIMB 11163 YP 003226531 1 622759 Proteobacteria
Zymomonas mobilis subsp pomaceae ATCC 29192 YP 004662488 1 579138 Proteobacteria
Magnetococcus marinus MC 1 YP 865111 1 156889 Proteobacteria
Methylomonas methanica MC09 YP 004513958 1 857087 Proteobacteria
Teredinibacter turnerae T7901 YP 003073112 1 377629 Proteobacteria
Azotobacter vinelandii CA6 YP 007896246 1 1283331 Proteobacteria
Azotobacter vinelandii DJ YP 002797386 1 322710 Proteobacteria
Thioflavicoccus mobilis 8321 YP 007245932 1 765912 Proteobacteria
Allochromatium vinosum DSM 180 YP 003442881 1 572477 Proteobacteria
Thiocystis violascens DSM 198 YP 006414782 1 765911 Proteobacteria
Sideroxydans lithotrophicus ES 1 YP 003523529 1 580332 Proteobacteria
Candidatus Accumulibacter phosphatis clade IIA str UW 1 YP 003169537 1 522306 Proteobacteria
Dechloromonas aromatica RCB YP 284724 1 159087 Proteobacteria
Rhodomicrobium vannielii ATCC 17100 YP 004010887 1 648757 Proteobacteria
Azotobacter vinelandii DJ YP 002797511 1 322710 Proteobacteria
Azotobacter vinelandii CA6 YP 007896371 1 1283331 Proteobacteria
Halorhodospira halophila SL1 YP 001001860 1 349124 Proteobacteria
Tolumonas auensis DSM 9187 YP 002891871 1 595494 Proteobacteria
Pantoea sp At 9b YP 004119239 1 592316 Proteobacteria
Klebsiella oxytoca E718 YP 006499534 1 1191061 Proteobacteria
Klebsiella variicola At 22 YP 003438534 1 640131 Proteobacteria
Klebsiella pneumoniae 342 YP 002237559 1 507522 Proteobacteria
Rahnella aquatilis CIP 78 65 ATCC 33071 YP 005202538 1 745277 Proteobacteria
Pectobacterium atrosepticum SCRI1043 YP 051039 1 218491 Proteobacteria
Prodigiosinella confusarubida ATCC39006 WP 021017288 1 Pconfusarubida
Dickeya paradisiaca Ech703 WP 012764128 1 Dparadisiaca
Dickeya paradisiaca NCPPB 2511 WP 012764128 1 Dparadisiaca
Dickeya solani PPO 9019 WP 059110783 1 Dsolani
Dickeya dianthicola NCPPB 3534 WP 024107441 1 Ddianthicola
Dickeya dianthicola IPO 980 WP 024107441 1 Ddianthicola
Dickeya dianthicola GBBC 2039 WP 024107441 1 Ddianthicola
Dickeya dianthicola RNS04 9 WP 024107441 1 Ddianthicola
Dickeya dianthicola NCPPB 453 WP 024107441 1 Ddianthicola
Dickeya gs fangzhongdai NCPPB 3274 WP 042861243 1 Dfangzhongdai
Dickeya gs fangzhongdai S1 WP 049854741 1 Dfangzhongdai
Dickeya gs fangzhongdai MK7 WP 038920344 1 Dfangzhongdai
Dickeya gs fangzhongdai B16 WP 038920344 1 Dfangzhongdai
Dickeya gs undicola 2B12 WP 033567504 1 Dundicola
Dickeya gs fangzhongdai M005 WP 039693127 1 Dfangzhongdai
Dickeya gs fangzhongdai M074 WP 038665283 1 Dfangzhongdai
Dickeya gs fangzhongdai ND14b WP 038665283 1 Dfangzhongdai
Dickeya chrysanthemi subsp chrysanthemi NCPPB 402 WP 039999413 1 Dchrysanthemi
Dickeya solani RNS 05 1 2A WP 057084655 1 Dsolani
Dickeya dadantii subsp dieffenbachiae NCPPB 2976 WP 038923992 1 Ddadantii
Dickeya dadantii NCPPB 898 WP 013319636 1 Ddadantii
Dickeya dadantii NCPPB 3537 WP 038902183 1 Ddadantii
Dickeya dadantii 3937 WP 013319636 1 Ddadantii
386 nifN nitrogen metabolism Seed
Dickeya solani IPO 2222 WP 022635026 1 Dsolani
Dickeya solani RNS 08 23 3 1 A WP 022635026 1 Dsolani
Dickeya solani RNS 07 7 3B WP 022635026 1 Dsolani
Dickeya solani MK10 WP 022635026 1 Dsolani
Dickeya solani GBBC 2040 WP 022635026 1 Dsolani
Dickeya solani PPO 9134 WP 022635026 1 Dsolani
Dickeya solani D s0432 1 WP 022635026 1 Dsolani
Dickeya solani MK16 WP 022635026 1 Dsolani
Dickeya solani IFB 0099 WP 022635026 1 Dsolani

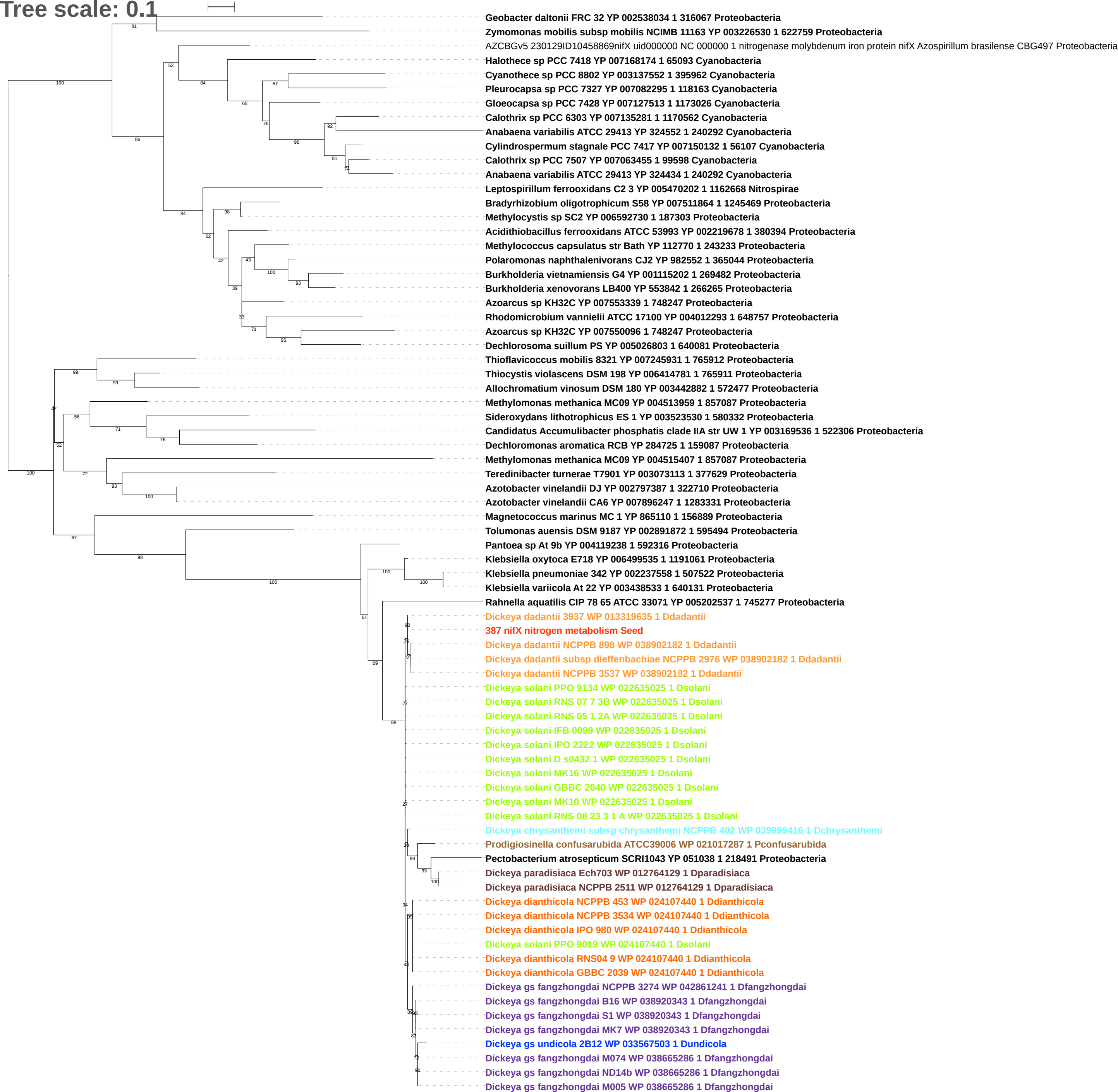
Tree scale: 0.1
Geobacter daltonii FRC 32 YP 002538034 1 316067 Proteobacteria
Zymomonas mobilis subsp mobilis NCIMB 11163 YP 003226530 1 622759 Proteobacteria
AZCBGv5 230129ID10458869nifX uid000000 NC 000000 1 nitrogenase molybdenum iron protein nifX Azospirillum brasilense CBG497 Proteobacteria
Halothece sp PCC 7418 YP 007168174 1 65093 Cyanobacteria
Cyanothece sp PCC 8802 YP 003137552 1 395962 Cyanobacteria
Pleurocapsa sp PCC 7327 YP 007082295 1 118163 Cyanobacteria
Gloeocapsa sp PCC 7428 YP 007127513 1 1173026 Cyanobacteria
Calothrix sp PCC 6303 YP 007135281 1 1170562 Cyanobacteria
Anabaena variabilis ATCC 29413 YP 324552 1 240292 Cyanobacteria
Cylindrospermum stagnale PCC 7417 YP 007150132 1 56107 Cyanobacteria
Calothrix sp PCC 7507 YP 007063455 1 99598 Cyanobacteria
Anabaena variabilis ATCC 29413 YP 324434 1 240292 Cyanobacteria
Leptospirillum ferrooxidans C2 3 YP 005470202 1 1162668 Nitrospirae
Bradyrhizobium oligotrophicum S58 YP 007511864 1 1245469 Proteobacteria
Methylocystis sp SC2 YP 006592730 1 187303 Proteobacteria
Acidithiobacillus ferrooxidans ATCC 53993 YP 002219678 1 380394 Proteobacteria
Methylococcus capsulatus str Bath YP 112770 1 243233 Proteobacteria
Polaromonas naphthalenivorans CJ2 YP 982552 1 365044 Proteobacteria
Burkholderia vietnamiensis G4 YP 001115202 1 269482 Proteobacteria
Burkholderia xenovorans LB400 YP 553842 1 266265 Proteobacteria
Azoarcus sp KH32C YP 007553339 1 748247 Proteobacteria
Rhodomicrobium vannielii ATCC 17100 YP 004012293 1 648757 Proteobacteria
Azoarcus sp KH32C YP 007550096 1 748247 Proteobacteria
Dechlorosoma suillum PS YP 005026803 1 640081 Proteobacteria
Thioflavicoccus mobilis 8321 YP 007245931 1 765912 Proteobacteria
Thiocystis violascens DSM 198 YP 006414781 1 765911 Proteobacteria
Allochromatium vinosum DSM 180 YP 003442882 1 572477 Proteobacteria
Methylomonas methanica MC09 YP 004513959 1 857087 Proteobacteria
Sideroxydans lithotrophicus ES 1 YP 003523530 1 580332 Proteobacteria
Candidatus Accumulibacter phosphatis clade IIA str UW 1 YP 003169536 1 522306 Proteobacteria
Dechloromonas aromatica RCB YP 284725 1 159087 Proteobacteria
Methylomonas methanica MC09 YP 004515407 1 857087 Proteobacteria
Teredinibacter turnerae T7901 YP 003073113 1 377629 Proteobacteria
Azotobacter vinelandii DJ YP 002797387 1 322710 Proteobacteria
Azotobacter vinelandii CA6 YP 007896247 1 1283331 Proteobacteria
Magnetococcus marinus MC 1 YP 865110 1 156889 Proteobacteria
Tolumonas auensis DSM 9187 YP 002891872 1 595494 Proteobacteria
Pantoea sp At 9b YP 004119238 1 592316 Proteobacteria
Klebsiella oxytoca E718 YP 006499535 1 1191061 Proteobacteria
Klebsiella pneumoniae 342 YP 002237558 1 507522 Proteobacteria
Klebsiella variicola At 22 YP 003438533 1 640131 Proteobacteria
Rahnella aquatilis CIP 78 65 ATCC 33071 YP 005202537 1 745277 Proteobacteria
Dickeya dadantii 3937 WP 013319635 1 Ddadantii
387 nifX nitrogen metabolism Seed
Dickeya dadantii NCPPB 898 WP 038902182 1 Ddadantii
Dickeya dadantii subsp dieffenbachiae NCPPB 2976 WP 038902182 1 Ddadantii
Dickeya dadantii NCPPB 3537 WP 038902182 1 Ddadantii
Dickeya solani PPO 9134 WP 022635025 1 Dsolani
Dickeya solani RNS 07 7 3B WP 022635025 1 Dsolani
Dickeya solani RNS 05 1 2A WP 022635025 1 Dsolani
Dickeya solani IFB 0099 WP 022635025 1 Dsolani
Dickeya solani IPO 2222 WP 022635025 1 Dsolani
Dickeya solani D s0432 1 WP 022635025 1 Dsolani
Dickeya solani MK16 WP 022635025 1 Dsolani
Dickeya solani GBBC 2040 WP 022635025 1 Dsolani
Dickeya solani MK10 WP 022635025 1 Dsolani
Dickeya solani RNS 08 23 3 1 A WP 022635025 1 Dsolani
Dickeya chrysanthemi subsp chrysanthemi NCPPB 402 WP 039999416 1 Dchrysanthemi
Prodigiosinella confusarubida ATCC39006 WP 021017287 1 Pconfusarubida
Pectobacterium atrosepticum SCRI1043 YP 051038 1 218491 Proteobacteria
Dickeya paradisiaca Ech703 WP 012764129 1 Dparadisiaca
Dickeya paradisiaca NCPPB 2511 WP 012764129 1 Dparadisiaca
Dickeya dianthicola NCPPB 453 WP 024107440 1 Ddianthicola
Dickeya dianthicola NCPPB 3534 WP 024107440 1 Ddianthicola
Dickeya dianthicola IPO 980 WP 024107440 1 Ddianthicola
Dickeya solani PPO 9019 WP 024107440 1 Dsolani
Dickeya dianthicola RNS04 9 WP 024107440 1 Ddianthicola
Dickeya dianthicola GBBC 2039 WP 024107440 1 Ddianthicola
Dickeya gs fangzhongdai NCPPB 3274 WP 042861241 1 Dfangzhongdai
Dickeya gs fangzhongdai B16 WP 038920343 1 Dfangzhongdai
Dickeya gs fangzhongdai S1 WP 038920343 1 Dfangzhongdai
Dickeya gs fangzhongdai MK7 WP 038920343 1 Dfangzhongdai
Dickeya gs undicola 2B12 WP 033567503 1 Dundicola
Dickeya gs fangzhongdai M074 WP 038665286 1 Dfangzhongdai
Dickeya gs fangzhongdai ND14b WP 038665286 1 Dfangzhongdai
Dickeya gs fangzhongdai M005 WP 038665286 1 Dfangzhongdai

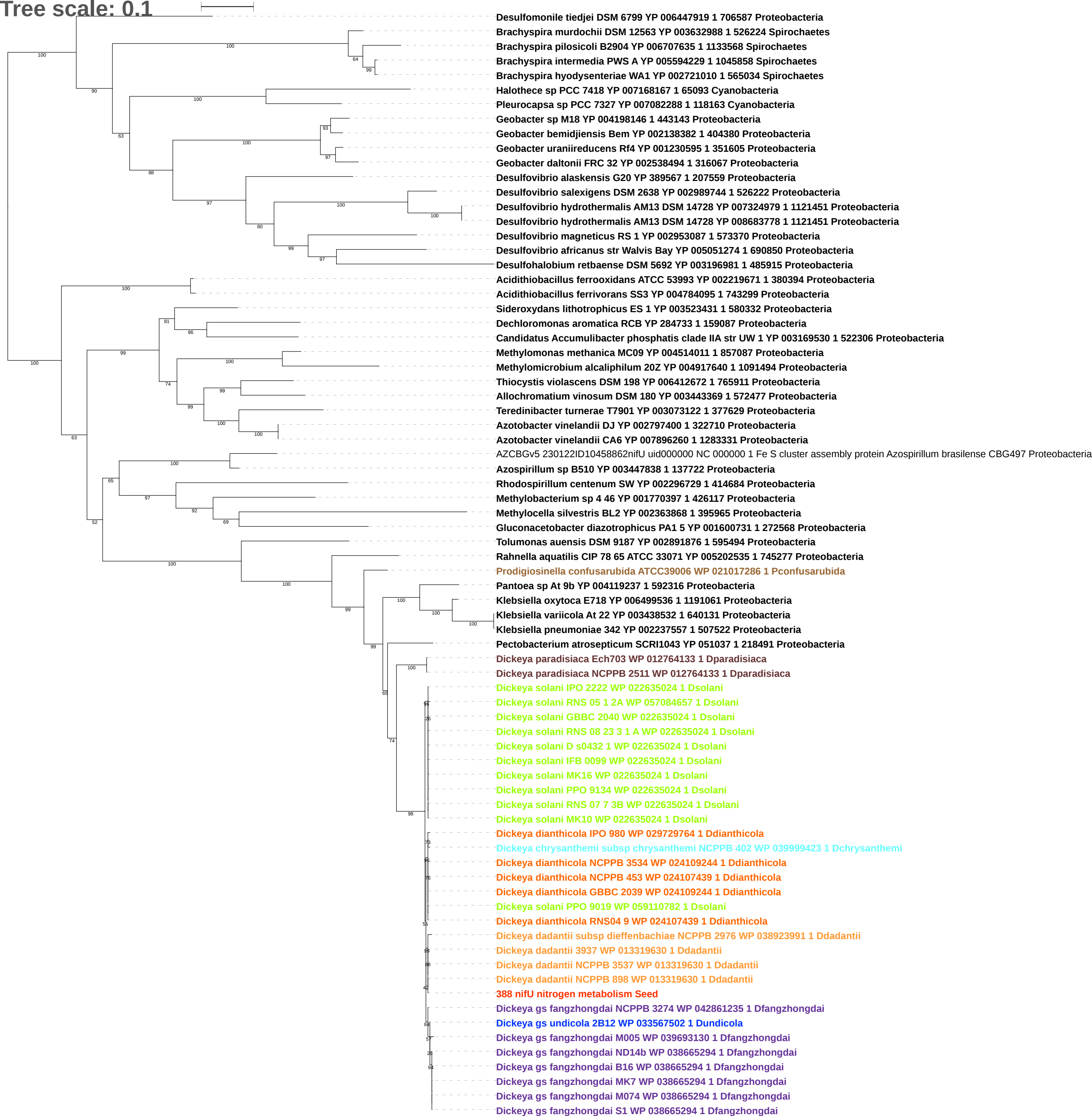
Tree scale: 0.1
Desulfomonile tiedjei DSM 6799 YP 006447919 1 706587 Proteobacteria
Brachyspira murdochii DSM 12563 YP 003632988 1 526224 Spirochaetes
Brachyspira pilosicoli B2904 YP 006707635 1 1133568 Spirochaetes
Brachyspira intermedia PWS A YP 005594229 1 1045858 Spirochaetes
Brachyspira hyodysenteriae WA1 YP 002721010 1 565034 Spirochaetes
Halothece sp PCC 7418 YP 007168167 1 65093 Cyanobacteria
Pleurocapsa sp PCC 7327 YP 007082288 1 118163 Cyanobacteria
Geobacter sp M18 YP 004198146 1 443143 Proteobacteria
Geobacter bemidjiensis Bem YP 002138382 1 404380 Proteobacteria
Geobacter uraniireducens Rf4 YP 001230595 1 351605 Proteobacteria
Geobacter daltonii FRC 32 YP 002538494 1 316067 Proteobacteria
Desulfovibrio alaskensis G20 YP 389567 1 207559 Proteobacteria
Desulfovibrio salexigens DSM 2638 YP 002989744 1 526222 Proteobacteria
Desulfovibrio hydrothermalis AM13 DSM 14728 YP 007324979 1 1121451 Proteobacteria
Desulfovibrio hydrothermalis AM13 DSM 14728 YP 008683778 1 1121451 Proteobacteria
Desulfovibrio magneticus RS 1 YP 002953087 1 573370 Proteobacteria
Desulfovibrio africanus str Walvis Bay YP 005051274 1 690850 Proteobacteria
Desulfohalobium retbaense DSM 5692 YP 003196981 1 485915 Proteobacteria
Acidithiobacillus ferrooxidans ATCC 53993 YP 002219671 1 380394 Proteobacteria
Acidithiobacillus ferrivorans SS3 YP 004784095 1 743299 Proteobacteria
Sideroxydans lithotrophicus ES 1 YP 003523431 1 580332 Proteobacteria
Dechloromonas aromatica RCB YP 284733 1 159087 Proteobacteria
Candidatus Accumulibacter phosphatis clade IIA str UW 1 YP 003169530 1 522306 Proteobacteria
Methylomonas methanica MC09 YP 004514011 1 857087 Proteobacteria
Methylomicrobium alcaliphilum 20Z YP 004917640 1 1091494 Proteobacteria
Thiocystis violascens DSM 198 YP 006412672 1 765911 Proteobacteria
Allochromatium vinosum DSM 180 YP 003443369 1 572477 Proteobacteria
Teredinibacter turnerae T7901 YP 003073122 1 377629 Proteobacteria
Azotobacter vinelandii DJ YP 002797400 1 322710 Proteobacteria
Azotobacter vinelandii CA6 YP 007896260 1 1283331 Proteobacteria
AZCBGv5 230122ID10458862nifU uid000000 NC 000000 1 Fe S cluster assembly protein Azospirillum brasilense CBG497 Proteobacteria
Azospirillum sp B510 YP 003447838 1 137722 Proteobacteria
Rhodospirillum centenum SW YP 002296729 1 414684 Proteobacteria
Methylobacterium sp 4 46 YP 001770397 1 426117 Proteobacteria
Methylocella silvestris BL2 YP 002363868 1 395965 Proteobacteria
Gluconacetobacter diazotrophicus PA1 5 YP 001600731 1 272568 Proteobacteria
Tolumonas auensis DSM 9187 YP 002891876 1 595494 Proteobacteria
Rahnella aquatilis CIP 78 65 ATCC 33071 YP 005202535 1 745277 Proteobacteria
Prodigiosinella confusarubida ATCC39006 WP 021017286 1 Pconfusarubida
Pantoea sp At 9b YP 004119237 1 592316 Proteobacteria
Klebsiella oxytoca E718 YP 006499536 1 1191061 Proteobacteria
Klebsiella variicola At 22 YP 003438532 1 640131 Proteobacteria
Klebsiella pneumoniae 342 YP 002237557 1 507522 Proteobacteria
Pectobacterium atrosepticum SCRI1043 YP 051037 1 218491 Proteobacteria
Dickeya paradisiaca Ech703 WP 012764133 1 Dparadisiaca
Dickeya paradisiaca NCPPB 2511 WP 012764133 1 Dparadisiaca
Dickeya solani IPO 2222 WP 022635024 1 Dsolani
Dickeya solani RNS 05 1 2A WP 057084657 1 Dsolani
Dickeya solani GBBC 2040 WP 022635024 1 Dsolani
Dickeya solani RNS 08 23 3 1 A WP 022635024 1 Dsolani
Dickeya solani D s0432 1 WP 022635024 1 Dsolani
Dickeya solani IFB 0099 WP 022635024 1 Dsolani
Dickeya solani MK16 WP 022635024 1 Dsolani
Dickeya solani PPO 9134 WP 022635024 1 Dsolani
Dickeya solani RNS 07 7 3B WP 022635024 1 Dsolani
Dickeya solani MK10 WP 022635024 1 Dsolani
Dickeya dianthicola IPO 980 WP 029729764 1 Ddianthicola
Dickeya chrysanthemi subsp chrysanthemi NCPPB 402 WP 039999423 1 Dchrysanthemi
Dickeya dianthicola NCPPB 3534 WP 024109244 1 Ddianthicola
Dickeya dianthicola NCPPB 453 WP 024107439 1 Ddianthicola
Dickeya dianthicola GBBC 2039 WP 024109244 1 Ddianthicola
Dickeya solani PPO 9019 WP 059110782 1 Dsolani
Dickeya dianthicola RNS04 9 WP 024107439 1 Ddianthicola
Dickeya dadantii subsp dieffenbachiae NCPPB 2976 WP 038923991 1 Ddadantii
Dickeya dadantii 3937 WP 013319630 1 Ddadantii
Dickeya dadantii NCPPB 3537 WP 013319630 1 Ddadantii
Dickeya dadantii NCPPB 898 WP 013319630 1 Ddadantii
388 nifU nitrogen metabolism Seed
Dickeya gs fangzhongdai NCPPB 3274 WP 042861235 1 Dfangzhongdai
Dickeya gs undicola 2B12 WP 033567502 1 Dundicola
Dickeya gs fangzhongdai M005 WP 039693130 1 Dfangzhongdai
Dickeya gs fangzhongdai ND14b WP 038665294 1 Dfangzhongdai
Dickeya gs fangzhongdai B16 WP 038665294 1 Dfangzhongdai
Dickeya gs fangzhongdai MK7 WP 038665294 1 Dfangzhongdai
Dickeya gs fangzhongdai M074 WP 038665294 1 Dfangzhongdai
Dickeya gs fangzhongdai S1 WP 038665294 1 Dfangzhongdai

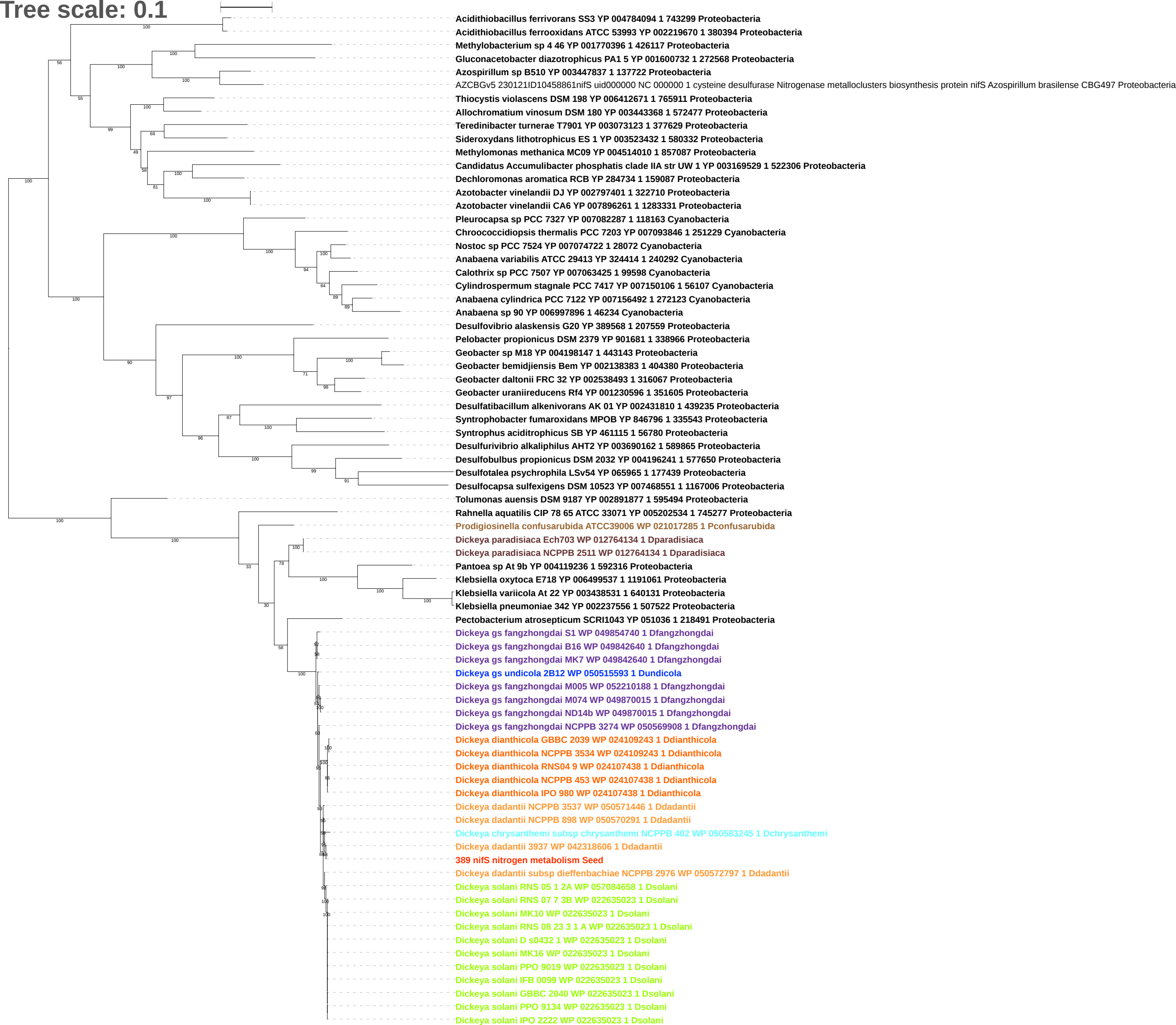
Tree scale: 0.1
Acidithiobacillus ferrivorans SS3 YP 004784094 1 743299 Proteobacteria
Acidithiobacillus ferrooxidans ATCC 53993 YP 002219670 1 380394 Proteobacteria
Methylobacterium sp 4 46 YP 001770396 1 426117 Proteobacteria
Gluconacetobacter diazotrophicus PA1 5 YP 001600732 1 272568 Proteobacteria
Azospirillum sp B510 YP 003447837 1 137722 Proteobacteria
AZCBGv5 230121ID10458861nifS uid000000 NC 000000 1 cysteine desulfurase Nitrogenase metalloclusters biosynthesis protein nifS Azospirillum brasilense CBG497 Proteobacteria
Thiocystis violascens DSM 198 YP 006412671 1 765911 Proteobacteria
Allochromatium vinosum DSM 180 YP 003443368 1 572477 Proteobacteria
Teredinibacter turnerae T7901 YP 003073123 1 377629 Proteobacteria
Sideroxydans lithotrophicus ES 1 YP 003523432 1 580332 Proteobacteria
Methylomonas methanica MC09 YP 004514010 1 857087 Proteobacteria
Candidatus Accumulibacter phosphatis clade IIA str UW 1 YP 003169529 1 522306 Proteobacteria
Dechloromonas aromatica RCB YP 284734 1 159087 Proteobacteria
Azotobacter vinelandii DJ YP 002797401 1 322710 Proteobacteria
Azotobacter vinelandii CA6 YP 007896261 1 1283331 Proteobacteria
Pleurocapsa sp PCC 7327 YP 007082287 1 118163 Cyanobacteria
Chroococcidiopsis thermalis PCC 7203 YP 007093846 1 251229 Cyanobacteria
Nostoc sp PCC 7524 YP 007074722 1 28072 Cyanobacteria
Anabaena variabilis ATCC 29413 YP 324414 1 240292 Cyanobacteria
Calothrix sp PCC 7507 YP 007063425 1 99598 Cyanobacteria
Cylindrospermum stagnale PCC 7417 YP 007150106 1 56107 Cyanobacteria
Anabaena cylindrica PCC 7122 YP 007156492 1 272123 Cyanobacteria
Anabaena sp 90 YP 006997896 1 46234 Cyanobacteria
Desulfovibrio alaskensis G20 YP 389568 1 207559 Proteobacteria
Pelobacter propionicus DSM 2379 YP 901681 1 338966 Proteobacteria
Geobacter sp M18 YP 004198147 1 443143 Proteobacteria
Geobacter bemidjiensis Bem YP 002138383 1 404380 Proteobacteria
Geobacter daltonii FRC 32 YP 002538493 1 316067 Proteobacteria
Geobacter uraniireducens Rf4 YP 001230596 1 351605 Proteobacteria
Desulfatibacillum alkenivorans AK 01 YP 002431810 1 439235 Proteobacteria
Syntrophobacter fumaroxidans MPOB YP 846796 1 335543 Proteobacteria
Syntrophus aciditrophicus SB YP 461115 1 56780 Proteobacteria
Desulfurivibrio alkaliphilus AHT2 YP 003690162 1 589865 Proteobacteria
Desulfobulbus propionicus DSM 2032 YP 004196241 1 577650 Proteobacteria
Desulfotalea psychrophila LSv54 YP 065965 1 177439 Proteobacteria
Desulfocapsa sulfexigens DSM 10523 YP 007468551 1 1167006 Proteobacteria
Tolumonas auensis DSM 9187 YP 002891877 1 595494 Proteobacteria
Rahnella aquatilis CIP 78 65 ATCC 33071 YP 005202534 1 745277 Proteobacteria
Prodigiosinella confusarubida ATCC39006 WP 021017285 1 Pconfusarubida
Dickeya paradisiaca Ech703 WP 012764134 1 Dparadisiaca
Dickeya paradisiaca NCPPB 2511 WP 012764134 1 Dparadisiaca
Pantoea sp At 9b YP 004119236 1 592316 Proteobacteria
Klebsiella oxytoca E718 YP 006499537 1 1191061 Proteobacteria
Klebsiella variicola At 22 YP 003438531 1 640131 Proteobacteria
Klebsiella pneumoniae 342 YP 002237556 1 507522 Proteobacteria
Pectobacterium atrosepticum SCRI1043 YP 051036 1 218491 Proteobacteria
Dickeya gs fangzhongdai S1 WP 049854740 1 Dfangzhongdai
Dickeya gs fangzhongdai B16 WP 049842640 1 Dfangzhongdai
Dickeya gs fangzhongdai MK7 WP 049842640 1 Dfangzhongdai
Dickeya gs undicola 2B12 WP 050515593 1 Dundicola
Dickeya gs fangzhongdai M005 WP 052210188 1 Dfangzhongdai
Dickeya gs fangzhongdai M074 WP 049870015 1 Dfangzhongdai
Dickeya gs fangzhongdai ND14b WP 049870015 1 Dfangzhongdai
Dickeya gs fangzhongdai NCPPB 3274 WP 050569908 1 Dfangzhongdai
Dickeya dianthicola GBBC 2039 WP 024109243 1 Ddianthicola
Dickeya dianthicola NCPPB 3534 WP 024109243 1 Ddianthicola
Dickeya dianthicola RNS04 9 WP 024107438 1 Ddianthicola
Dickeya dianthicola NCPPB 453 WP 024107438 1 Ddianthicola
Dickeya dianthicola IPO 980 WP 024107438 1 Ddianthicola
Dickeya dadantii NCPPB 3537 WP 050571446 1 Ddadantii
Dickeya dadantii NCPPB 898 WP 050570291 1 Ddadantii
Dickeya chrysanthemi subsp chrysanthemi NCPPB 402 WP 050583245 1 Dchrysanthemi
Dickeya dadantii 3937 WP 042318606 1 Ddadantii
389 nifS nitrogen metabolism Seed
Dickeya dadantii subsp dieffenbachiae NCPPB 2976 WP 050572797 1 Ddadantii
Dickeya solani RNS 05 1 2A WP 057084658 1 Dsolani
Dickeya solani RNS 07 7 3B WP 022635023 1 Dsolani
Dickeya solani MK10 WP 022635023 1 Dsolani
Dickeya solani RNS 08 23 3 1 A WP 022635023 1 Dsolani
Dickeya solani D s0432 1 WP 022635023 1 Dsolani
Dickeya solani MK16 WP 022635023 1 Dsolani
Dickeya solani PPO 9019 WP 022635023 1 Dsolani
Dickeya solani IFB 0099 WP 022635023 1 Dsolani
Dickeya solani GBBC 2040 WP 022635023 1 Dsolani
Dickeya solani PPO 9134 WP 022635023 1 Dsolani
Dickeya solani IPO 2222 WP 022635023 1 Dsolani

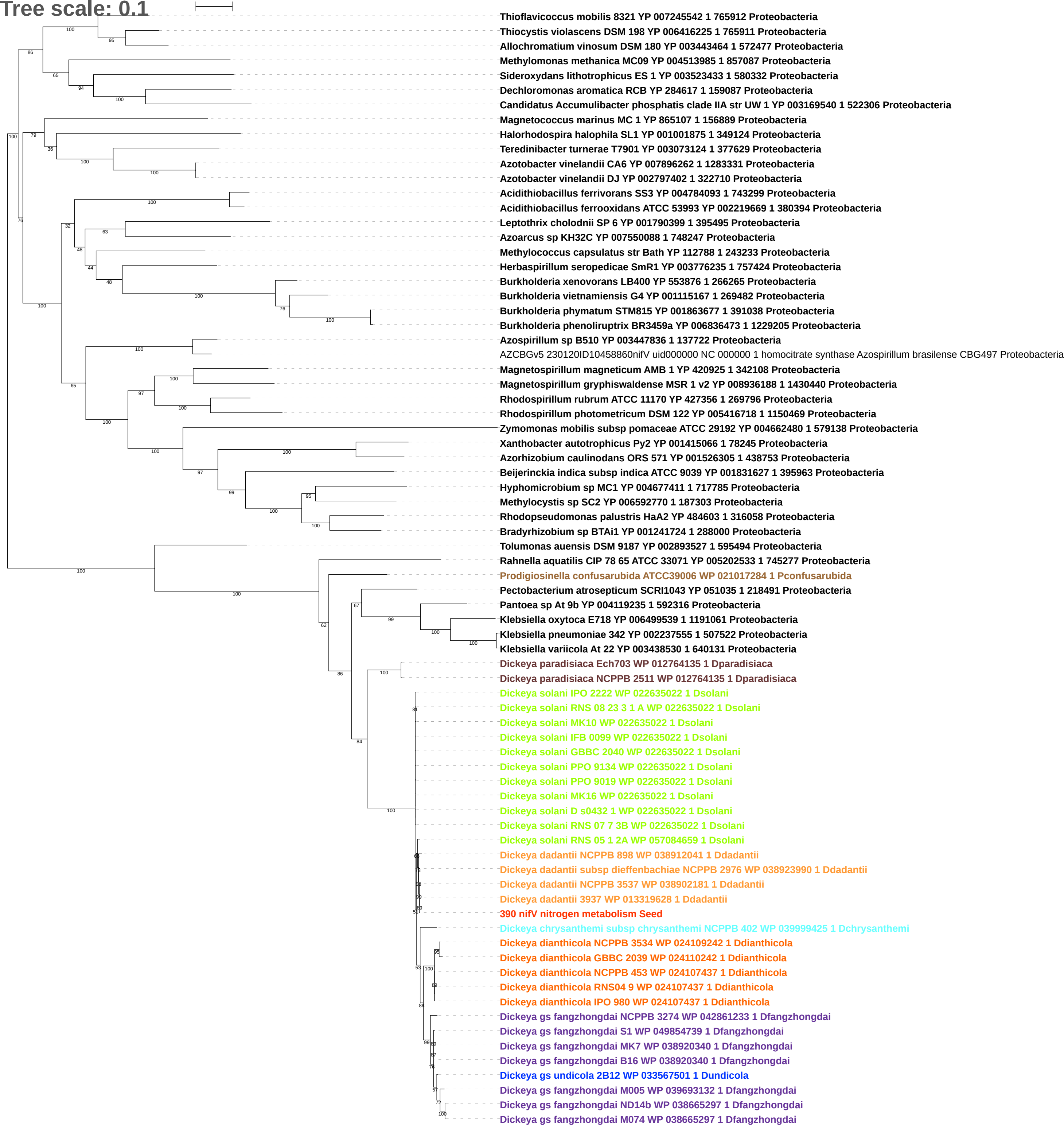
Tree scale: 0.1
Thioflavicoccus mobilis 8321 YP 007245542 1 765912 Proteobacteria
Thiocystis violascens DSM 198 YP 006416225 1 765911 Proteobacteria
Allochromatium vinosum DSM 180 YP 003443464 1 572477 Proteobacteria
Methylomonas methanica MC09 YP 004513985 1 857087 Proteobacteria
Sideroxydans lithotrophicus ES 1 YP 003523433 1 580332 Proteobacteria
Dechloromonas aromatica RCB YP 284617 1 159087 Proteobacteria
Candidatus Accumulibacter phosphatis clade IIA str UW 1 YP 003169540 1 522306 Proteobacteria
Magnetococcus marinus MC 1 YP 865107 1 156889 Proteobacteria
Halorhodospira halophila SL1 YP 001001875 1 349124 Proteobacteria
Teredinibacter turnerae T7901 YP 003073124 1 377629 Proteobacteria
Azotobacter vinelandii CA6 YP 007896262 1 1283331 Proteobacteria
Azotobacter vinelandii DJ YP 002797402 1 322710 Proteobacteria
Acidithiobacillus ferrivorans SS3 YP 004784093 1 743299 Proteobacteria
Acidithiobacillus ferrooxidans ATCC 53993 YP 002219669 1 380394 Proteobacteria
Leptothrix cholodnii SP 6 YP 001790399 1 395495 Proteobacteria
Azoarcus sp KH32C YP 007550088 1 748247 Proteobacteria
Methylococcus capsulatus str Bath YP 112788 1 243233 Proteobacteria
Herbaspirillum seropedicae SmR1 YP 003776235 1 757424 Proteobacteria
Burkholderia xenovorans LB400 YP 553876 1 266265 Proteobacteria
Burkholderia vietnamiensis G4 YP 001115167 1 269482 Proteobacteria
Burkholderia phymatum STM815 YP 001863677 1 391038 Proteobacteria
Burkholderia phenoliruptrix BR3459a YP 006836473 1 1229205 Proteobacteria
Azospirillum sp B510 YP 003447836 1 137722 Proteobacteria
AZCBGv5 230120ID10458860nifV uid000000 NC 000000 1 homocitrate synthase Azospirillum brasilense CBG497 Proteobacteria
Magnetospirillum magneticum AMB 1 YP 420925 1 342108 Proteobacteria
Magnetospirillum gryphiswaldense MSR 1 v2 YP 008936188 1 1430440 Proteobacteria
Rhodospirillum rubrum ATCC 11170 YP 427356 1 269796 Proteobacteria
Rhodospirillum photometricum DSM 122 YP 005416718 1 1150469 Proteobacteria
Zymomonas mobilis subsp pomaceae ATCC 29192 YP 004662480 1 579138 Proteobacteria
Xanthobacter autotrophicus Py2 YP 001415066 1 78245 Proteobacteria
Azorhizobium caulinodans ORS 571 YP 001526305 1 438753 Proteobacteria
Beijerinckia indica subsp indica ATCC 9039 YP 001831627 1 395963 Proteobacteria
Hyphomicrobium sp MC1 YP 004677411 1 717785 Proteobacteria
Methylocystis sp SC2 YP 006592770 1 187303 Proteobacteria
Rhodopseudomonas palustris HaA2 YP 484603 1 316058 Proteobacteria
Bradyrhizobium sp BTAi1 YP 001241724 1 288000 Proteobacteria
Tolumonas auensis DSM 9187 YP 002893527 1 595494 Proteobacteria
Rahnella aquatilis CIP 78 65 ATCC 33071 YP 005202533 1 745277 Proteobacteria
Prodigiosinella confusarubida ATCC39006 WP 021017284 1 Pconfusarubida
Pectobacterium atrosepticum SCRI1043 YP 051035 1 218491 Proteobacteria
Pantoea sp At 9b YP 004119235 1 592316 Proteobacteria
Klebsiella oxytoca E718 YP 006499539 1 1191061 Proteobacteria
Klebsiella pneumoniae 342 YP 002237555 1 507522 Proteobacteria
Klebsiella variicola At 22 YP 003438530 1 640131 Proteobacteria
Dickeya paradisiaca Ech703 WP 012764135 1 Dparadisiaca
Dickeya paradisiaca NCPPB 2511 WP 012764135 1 Dparadisiaca
Dickeya solani IPO 2222 WP 022635022 1 Dsolani
Dickeya solani RNS 08 23 3 1 A WP 022635022 1 Dsolani
Dickeya solani MK10 WP 022635022 1 Dsolani
Dickeya solani IFB 0099 WP 022635022 1 Dsolani
Dickeya solani GBBC 2040 WP 022635022 1 Dsolani
Dickeya solani PPO 9134 WP 022635022 1 Dsolani
Dickeya solani PPO 9019 WP 022635022 1 Dsolani
Dickeya solani MK16 WP 022635022 1 Dsolani
Dickeya solani D s0432 1 WP 022635022 1 Dsolani
Dickeya solani RNS 07 7 3B WP 022635022 1 Dsolani
Dickeya solani RNS 05 1 2A WP 057084659 1 Dsolani
Dickeya dadantii NCPPB 898 WP 038912041 1 Ddadantii
Dickeya dadantii subsp dieffenbachiae NCPPB 2976 WP 038923990 1 Ddadantii
Dickeya dadantii NCPPB 3537 WP 038902181 1 Ddadantii
Dickeya dadantii 3937 WP 013319628 1 Ddadantii
390 nifV nitrogen metabolism Seed
Dickeya chrysanthemi subsp chrysanthemi NCPPB 402 WP 039999425 1 Dchrysanthemi
Dickeya dianthicola NCPPB 3534 WP 024109242 1 Ddianthicola
Dickeya dianthicola GBBC 2039 WP 024110242 1 Ddianthicola
Dickeya dianthicola NCPPB 453 WP 024107437 1 Ddianthicola
Dickeya dianthicola RNS04 9 WP 024107437 1 Ddianthicola
Dickeya dianthicola IPO 980 WP 024107437 1 Ddianthicola
Dickeya gs fangzhongdai NCPPB 3274 WP 042861233 1 Dfangzhongdai
Dickeya gs fangzhongdai S1 WP 049854739 1 Dfangzhongdai
Dickeya gs fangzhongdai MK7 WP 038920340 1 Dfangzhongdai
Dickeya gs fangzhongdai B16 WP 038920340 1 Dfangzhongdai
Dickeya gs undicola 2B12 WP 033567501 1 Dundicola
Dickeya gs fangzhongdai M005 WP 039693132 1 Dfangzhongdai
Dickeya gs fangzhongdai ND14b WP 038665297 1 Dfangzhongdai
Dickeya gs fangzhongdai M074 WP 038665297 1 Dfangzhongdai

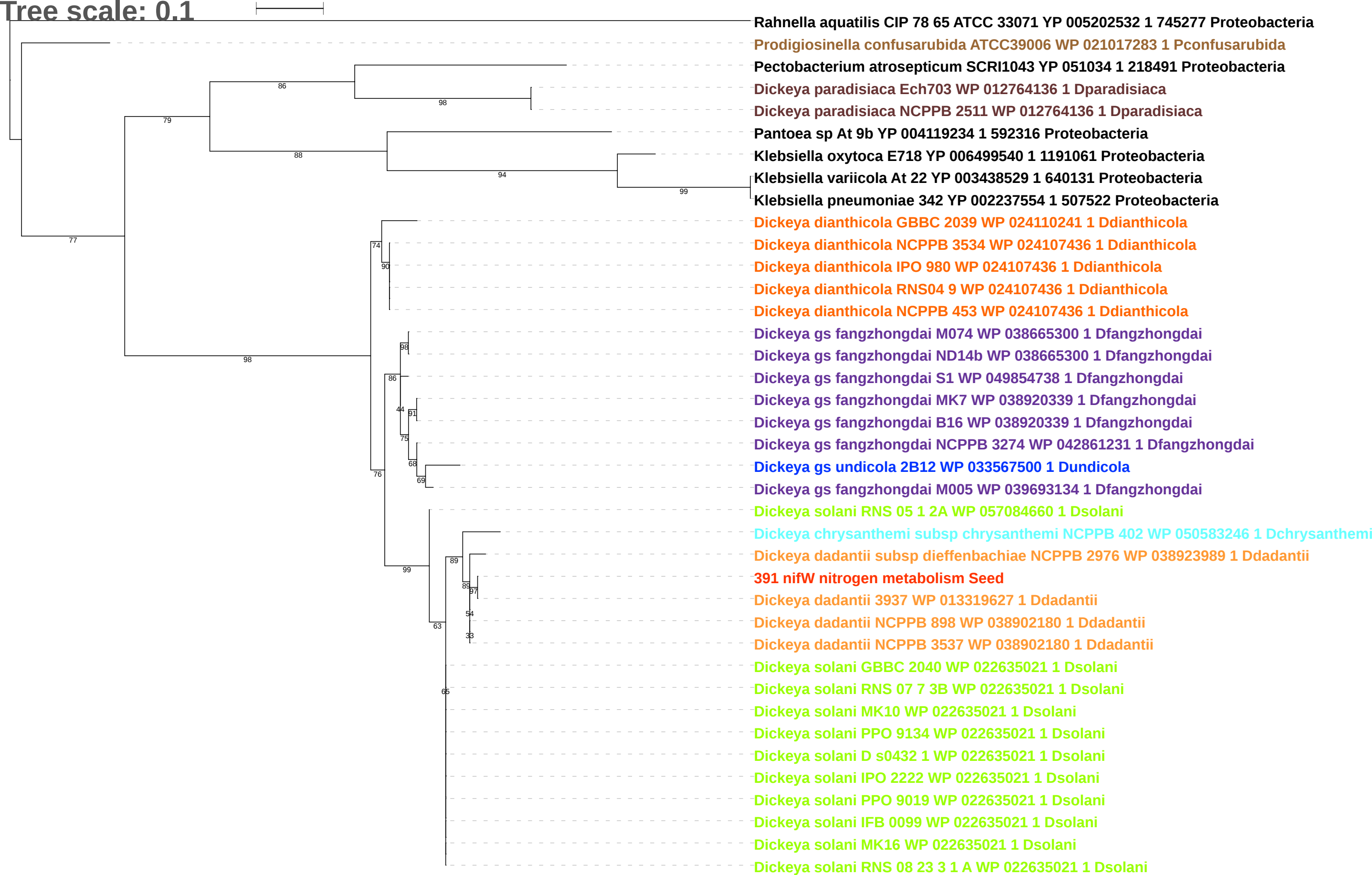
Tree scale: 0.1
Rahnella aquatilis CIP 78 65 ATCC 33071 YP 005202532 1 745277 Proteobacteria
Prodigiosinella confusarubida ATCC39006 WP 021017283 1 Pconfusarubida
Pectobacterium atrosepticum SCRI1043 YP 051034 1 218491 Proteobacteria
Dickeya paradisiaca Ech703 WP 012764136 1 Dparadisiaca
Dickeya paradisiaca NCPPB 2511 WP 012764136 1 Dparadisiaca
Pantoea sp At 9b YP 004119234 1 592316 Proteobacteria
Klebsiella oxytoca E718 YP 006499540 1 1191061 Proteobacteria
Klebsiella variicola At 22 YP 003438529 1 640131 Proteobacteria
Klebsiella pneumoniae 342 YP 002237554 1 507522 Proteobacteria
Dickeya dianthicola GBBC 2039 WP 024110241 1 Ddianthicola
Dickeya dianthicola NCPPB 3534 WP 024107436 1 Ddianthicola
Dickeya dianthicola IPO 980 WP 024107436 1 Ddianthicola
Dickeya dianthicola RNS04 9 WP 024107436 1 Ddianthicola
Dickeya dianthicola NCPPB 453 WP 024107436 1 Ddianthicola
Dickeya gs fangzhongdai M074 WP 038665300 1 Dfangzhongdai
Dickeya gs fangzhongdai ND14b WP 038665300 1 Dfangzhongdai
Dickeya gs fangzhongdai S1 WP 049854738 1 Dfangzhongdai
Dickeya gs fangzhongdai MK7 WP 038920339 1 Dfangzhongdai
Dickeya gs fangzhongdai B16 WP 038920339 1 Dfangzhongdai
Dickeya gs fangzhongdai NCPPB 3274 WP 042861231 1 Dfangzhongdai
Dickeya gs undicola 2B12 WP 033567500 1 Dundicola
Dickeya gs fangzhongdai M005 WP 039693134 1 Dfangzhongdai
Dickeya solani RNS 05 1 2A WP 057084660 1 Dsolani
Dickeya chrysanthemi subsp chrysanthemi NCPPB 402 WP 050583246 1 Dchrysanthemi
Dickeya dadantii subsp dieffenbachiae NCPPB 2976 WP 038923989 1 Ddadantii
391 nifW nitrogen metabolism Seed
Dickeya dadantii 3937 WP 013319627 1 Ddadantii
Dickeya dadantii NCPPB 898 WP 038902180 1 Ddadantii
Dickeya dadantii NCPPB 3537 WP 038902180 1 Ddadantii
Dickeya solani GBBC 2040 WP 022635021 1 Dsolani
Dickeya solani RNS 07 7 3B WP 022635021 1 Dsolani
Dickeya solani MK10 WP 022635021 1 Dsolani
Dickeya solani PPO 9134 WP 022635021 1 Dsolani
Dickeya solani D s0432 1 WP 022635021 1 Dsolani
Dickeya solani IPO 2222 WP 022635021 1 Dsolani
Dickeya solani PPO 9019 WP 022635021 1 Dsolani
Dickeya solani IFB 0099 WP 022635021 1 Dsolani
Dickeya solani MK16 WP 022635021 1 Dsolani
Dickeya solani RNS 08 23 3 1 A WP 022635021 1 Dsolani

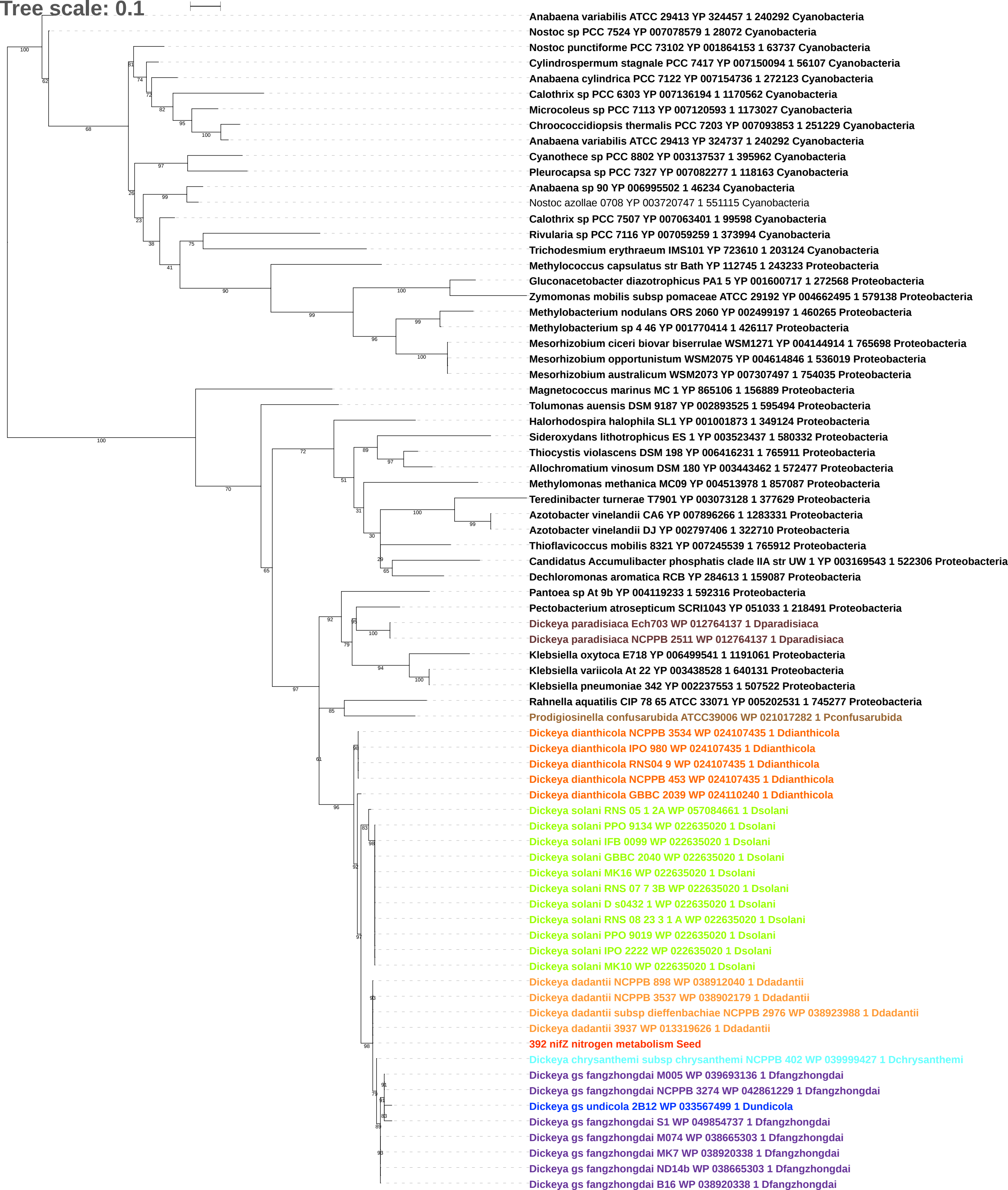

Tree scale: 0.1
Anabaena variabilis ATCC 29413 YP 324457 1 240292 Cyanobacteria
Nostoc sp PCC 7524 YP 007078579 1 28072 Cyanobacteria
Nostoc punctiforme PCC 73102 YP 001864153 1 63737 Cyanobacteria
Cylindrospermum stagnale PCC 7417 YP 007150094 1 56107 Cyanobacteria
Anabaena cylindrica PCC 7122 YP 007154736 1 272123 Cyanobacteria
Calothrix sp PCC 6303 YP 007136194 1 1170562 Cyanobacteria
Microcoleus sp PCC 7113 YP 007120593 1 1173027 Cyanobacteria
Chroococcidiopsis thermalis PCC 7203 YP 007093853 1 251229 Cyanobacteria
Anabaena variabilis ATCC 29413 YP 324737 1 240292 Cyanobacteria
Cyanothece sp PCC 8802 YP 003137537 1 395962 Cyanobacteria
Pleurocapsa sp PCC 7327 YP 007082277 1 118163 Cyanobacteria
Anabaena sp 90 YP 006995502 1 46234 Cyanobacteria
Nostoc azollae 0708 YP 003720747 1 551115 Cyanobacteria
Calothrix sp PCC 7507 YP 007063401 1 99598 Cyanobacteria
Rivularia sp PCC 7116 YP 007059259 1 373994 Cyanobacteria
Trichodesmium erythraeum IMS101 YP 723610 1 203124 Cyanobacteria
Methylococcus capsulatus str Bath YP 112745 1 243233 Proteobacteria
Gluconacetobacter diazotrophicus PA1 5 YP 001600717 1 272568 Proteobacteria
Zymomonas mobilis subsp pomaceae ATCC 29192 YP 004662495 1 579138 Proteobacteria
Methylobacterium nodulans ORS 2060 YP 002499197 1 460265 Proteobacteria
Methylobacterium sp 4 46 YP 001770414 1 426117 Proteobacteria
Mesorhizobium ciceri biovar biserrulae WSM1271 YP 004144914 1 765698 Proteobacteria
Mesorhizobium opportunistum WSM2075 YP 004614846 1 536019 Proteobacteria
Mesorhizobium australicum WSM2073 YP 007307497 1 754035 Proteobacteria
Magnetococcus marinus MC 1 YP 865106 1 156889 Proteobacteria
Tolumonas auensis DSM 9187 YP 002893525 1 595494 Proteobacteria
Halorhodospira halophila SL1 YP 001001873 1 349124 Proteobacteria
Sideroxydans lithotrophicus ES 1 YP 003523437 1 580332 Proteobacteria
Thiocystis violascens DSM 198 YP 006416231 1 765911 Proteobacteria
Allochromatium vinosum DSM 180 YP 003443462 1 572477 Proteobacteria
Methylomonas methanica MC09 YP 004513978 1 857087 Proteobacteria
Teredinibacter turnerae T7901 YP 003073128 1 377629 Proteobacteria
Azotobacter vinelandii CA6 YP 007896266 1 1283331 Proteobacteria
Azotobacter vinelandii DJ YP 002797406 1 322710 Proteobacteria
Thioflavicoccus mobilis 8321 YP 007245539 1 765912 Proteobacteria
Candidatus Accumulibacter phosphatis clade IIA str UW 1 YP 003169543 1 522306 Proteobacteria
Dechloromonas aromatica RCB YP 284613 1 159087 Proteobacteria
Pantoea sp At 9b YP 004119233 1 592316 Proteobacteria
Pectobacterium atrosepticum SCRI1043 YP 051033 1 218491 Proteobacteria
Dickeya paradisiaca Ech703 WP 012764137 1 Dparadisiaca
Dickeya paradisiaca NCPPB 2511 WP 012764137 1 Dparadisiaca
Klebsiella oxytoca E718 YP 006499541 1 1191061 Proteobacteria
Klebsiella variicola At 22 YP 003438528 1 640131 Proteobacteria
Klebsiella pneumoniae 342 YP 002237553 1 507522 Proteobacteria
Rahnella aquatilis CIP 78 65 ATCC 33071 YP 005202531 1 745277 Proteobacteria
Prodigiosinella confusarubida ATCC39006 WP 021017282 1 Pconfusarubida
Dickeya dianthicola NCPPB 3534 WP 024107435 1 Ddianthicola
Dickeya dianthicola IPO 980 WP 024107435 1 Ddianthicola
Dickeya dianthicola RNS04 9 WP 024107435 1 Ddianthicola
Dickeya dianthicola NCPPB 453 WP 024107435 1 Ddianthicola
Dickeya dianthicola GBBC 2039 WP 024110240 1 Ddianthicola
Dickeya solani RNS 05 1 2A WP 057084661 1 Dsolani
Dickeya solani PPO 9134 WP 022635020 1 Dsolani
Dickeya solani IFB 0099 WP 022635020 1 Dsolani
Dickeya solani GBBC 2040 WP 022635020 1 Dsolani
Dickeya solani MK16 WP 022635020 1 Dsolani
Dickeya solani RNS 07 7 3B WP 022635020 1 Dsolani
Dickeya solani D s0432 1 WP 022635020 1 Dsolani
Dickeya solani RNS 08 23 3 1 A WP 022635020 1 Dsolani
Dickeya solani PPO 9019 WP 022635020 1 Dsolani
Dickeya solani IPO 2222 WP 022635020 1 Dsolani
Dickeya solani MK10 WP 022635020 1 Dsolani
Dickeya dadantii NCPPB 898 WP 038912040 1 Ddadantii
Dickeya dadantii NCPPB 3537 WP 038902179 1 Ddadantii
Dickeya dadantii subsp dieffenbachiae NCPPB 2976 WP 038923988 1 Ddadantii
Dickeya dadantii 3937 WP 013319626 1 Ddadantii
392 nifZ nitrogen metabolism Seed
Dickeya chrysanthemi subsp chrysanthemi NCPPB 402 WP 039999427 1 Dchrysanthemi
Dickeya gs fangzhongdai M005 WP 039693136 1 Dfangzhongdai
Dickeya gs fangzhongdai NCPPB 3274 WP 042861229 1 Dfangzhongdai
Dickeya gs undicola 2B12 WP 033567499 1 Dundicola
Dickeya gs fangzhongdai S1 WP 049854737 1 Dfangzhongdai
Dickeya gs fangzhongdai M074 WP 038665303 1 Dfangzhongdai
Dickeya gs fangzhongdai MK7 WP 038920338 1 Dfangzhongdai
Dickeya gs fangzhongdai ND14b WP 038665303 1 Dfangzhongdai
Dickeya gs fangzhongdai B16 WP 038920338 1 Dfangzhongdai

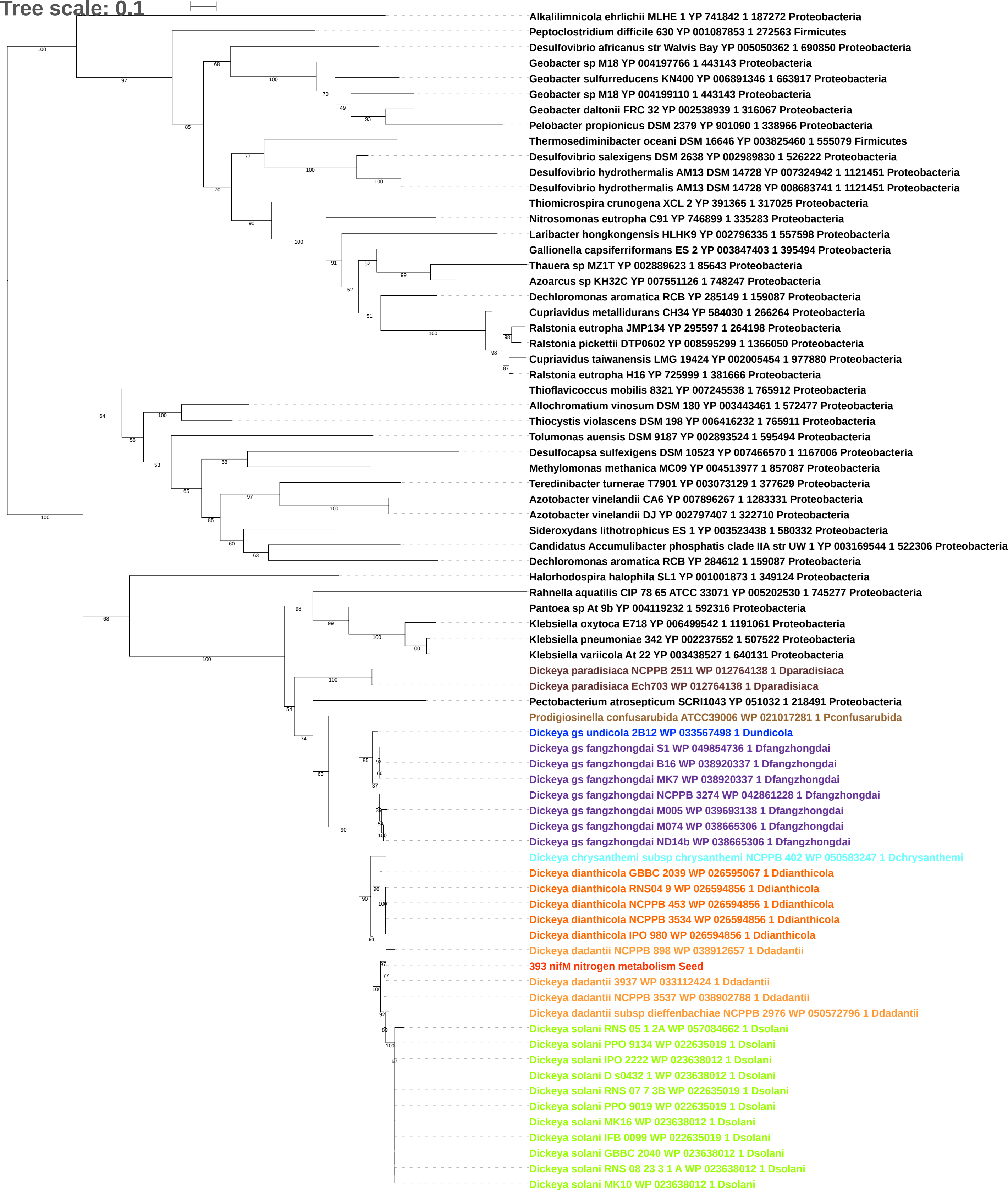
Tree scale: 0.1
Alkalilimnicola ehrlichii MLHE 1 YP 741842 1 187272 Proteobacteria
Peptoclostridium difficile 630 YP 001087853 1 272563 Firmicutes
Desulfovibrio africanus str Walvis Bay YP 005050362 1 690850 Proteobacteria
Geobacter sp M18 YP 004197766 1 443143 Proteobacteria
Geobacter sulfurreducens KN400 YP 006891346 1 663917 Proteobacteria
Geobacter sp M18 YP 004199110 1 443143 Proteobacteria
Geobacter daltonii FRC 32 YP 002538939 1 316067 Proteobacteria
Pelobacter propionicus DSM 2379 YP 901090 1 338966 Proteobacteria
Thermosediminibacter oceani DSM 16646 YP 003825460 1 555079 Firmicutes
Desulfovibrio salexigens DSM 2638 YP 002989830 1 526222 Proteobacteria
Desulfovibrio hydrothermalis AM13 DSM 14728 YP 007324942 1 1121451 Proteobacteria
Desulfovibrio hydrothermalis AM13 DSM 14728 YP 008683741 1 1121451 Proteobacteria
Thiomicrospira crunogena XCL 2 YP 391365 1 317025 Proteobacteria
Nitrosomonas eutropha C91 YP 746899 1 335283 Proteobacteria
Laribacter hongkongensis HLHK9 YP 002796335 1 557598 Proteobacteria
Gallionella capsiferriformans ES 2 YP 003847403 1 395494 Proteobacteria
Thauera sp MZ1T YP 002889623 1 85643 Proteobacteria
Azoarcus sp KH32C YP 007551126 1 748247 Proteobacteria
Dechloromonas aromatica RCB YP 285149 1 159087 Proteobacteria
Cupriavidus metallidurans CH34 YP 584030 1 266264 Proteobacteria
Ralstonia eutropha JMP134 YP 295597 1 264198 Proteobacteria
Ralstonia pickettii DTP0602 YP 008595299 1 1366050 Proteobacteria
Cupriavidus taiwanensis LMG 19424 YP 002005454 1 977880 Proteobacteria
Ralstonia eutropha H16 YP 725999 1 381666 Proteobacteria
Thioflavicoccus mobilis 8321 YP 007245538 1 765912 Proteobacteria
Allochromatium vinosum DSM 180 YP 003443461 1 572477 Proteobacteria
Thiocystis violascens DSM 198 YP 006416232 1 765911 Proteobacteria
Tolumonas auensis DSM 9187 YP 002893524 1 595494 Proteobacteria
Desulfocapsa sulfexigens DSM 10523 YP 007466570 1 1167006 Proteobacteria
Methylomonas methanica MC09 YP 004513977 1 857087 Proteobacteria
Teredinibacter turnerae T7901 YP 003073129 1 377629 Proteobacteria
Azotobacter vinelandii CA6 YP 007896267 1 1283331 Proteobacteria
Azotobacter vinelandii DJ YP 002797407 1 322710 Proteobacteria
Sideroxydans lithotrophicus ES 1 YP 003523438 1 580332 Proteobacteria
Candidatus Accumulibacter phosphatis clade IIA str UW 1 YP 003169544 1 522306 Proteobacteria
Dechloromonas aromatica RCB YP 284612 1 159087 Proteobacteria
Halorhodospira halophila SL1 YP 001001873 1 349124 Proteobacteria
Rahnella aquatilis CIP 78 65 ATCC 33071 YP 005202530 1 745277 Proteobacteria
Pantoea sp At 9b YP 004119232 1 592316 Proteobacteria
Klebsiella oxytoca E718 YP 006499542 1 1191061 Proteobacteria
Klebsiella pneumoniae 342 YP 002237552 1 507522 Proteobacteria
Klebsiella variicola At 22 YP 003438527 1 640131 Proteobacteria
Dickeya paradisiaca NCPPB 2511 WP 012764138 1 Dparadisiaca
Dickeya paradisiaca Ech703 WP 012764138 1 Dparadisiaca
Pectobacterium atrosepticum SCRI1043 YP 051032 1 218491 Proteobacteria
Prodigiosinella confusarubida ATCC39006 WP 021017281 1 Pconfusarubida
Dickeya gs undicola 2B12 WP 033567498 1 Dundicola
Dickeya gs fangzhongdai S1 WP 049854736 1 Dfangzhongdai
Dickeya gs fangzhongdai B16 WP 038920337 1 Dfangzhongdai
Dickeya gs fangzhongdai MK7 WP 038920337 1 Dfangzhongdai
Dickeya gs fangzhongdai NCPPB 3274 WP 042861228 1 Dfangzhongdai
Dickeya gs fangzhongdai M005 WP 039693138 1 Dfangzhongdai
Dickeya gs fangzhongdai M074 WP 038665306 1 Dfangzhongdai
Dickeya gs fangzhongdai ND14b WP 038665306 1 Dfangzhongdai
Dickeya chrysanthemi subsp chrysanthemi NCPPB 402 WP 050583247 1 Dchrysanthemi
Dickeya dianthicola GBBC 2039 WP 026595067 1 Ddianthicola
Dickeya dianthicola RNS04 9 WP 026594856 1 Ddianthicola
Dickeya dianthicola NCPPB 453 WP 026594856 1 Ddianthicola
Dickeya dianthicola NCPPB 3534 WP 026594856 1 Ddianthicola
Dickeya dianthicola IPO 980 WP 026594856 1 Ddianthicola
Dickeya dadantii NCPPB 898 WP 038912657 1 Ddadantii
393 nifM nitrogen metabolism Seed
Dickeya dadantii 3937 WP 033112424 1 Ddadantii
Dickeya dadantii NCPPB 3537 WP 038902788 1 Ddadantii
Dickeya dadantii subsp dieffenbachiae NCPPB 2976 WP 050572796 1 Ddadantii
Dickeya solani RNS 05 1 2A WP 057084662 1 Dsolani
Dickeya solani PPO 9134 WP 022635019 1 Dsolani
Dickeya solani IPO 2222 WP 023638012 1 Dsolani
Dickeya solani D s0432 1 WP 023638012 1 Dsolani
Dickeya solani RNS 07 7 3B WP 022635019 1 Dsolani
Dickeya solani PPO 9019 WP 022635019 1 Dsolani
Dickeya solani MK16 WP 023638012 1 Dsolani
Dickeya solani IFB 0099 WP 022635019 1 Dsolani
Dickeya solani GBBC 2040 WP 023638012 1 Dsolani
Dickeya solani RNS 08 23 3 1 A WP 023638012 1 Dsolani
Dickeya solani MK10 WP 023638012 1 Dsolani

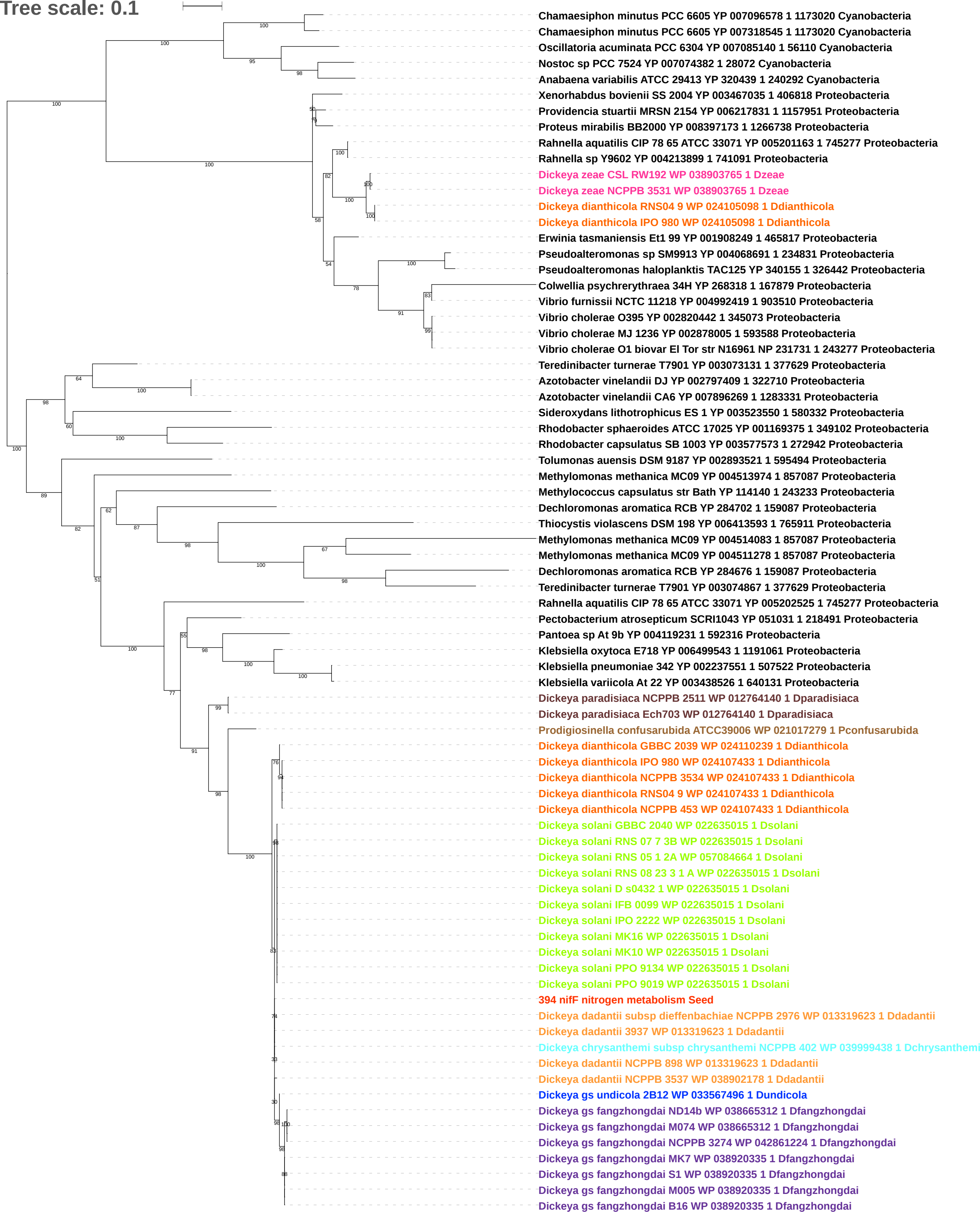
Tree scale: 0.1
Chamaesiphon minutus PCC 6605 YP 007096578 1 1173020 Cyanobacteria
Chamaesiphon minutus PCC 6605 YP 007318545 1 1173020 Cyanobacteria
Oscillatoria acuminata PCC 6304 YP 007085140 1 56110 Cyanobacteria
Nostoc sp PCC 7524 YP 007074382 1 28072 Cyanobacteria
Anabaena variabilis ATCC 29413 YP 320439 1 240292 Cyanobacteria
Xenorhabdus bovienii SS 2004 YP 003467035 1 406818 Proteobacteria
Providencia stuartii MRSN 2154 YP 006217831 1 1157951 Proteobacteria
Proteus mirabilis BB2000 YP 008397173 1 1266738 Proteobacteria
Rahnella aquatilis CIP 78 65 ATCC 33071 YP 005201163 1 745277 Proteobacteria
Rahnella sp Y9602 YP 004213899 1 741091 Proteobacteria
Dickeya zeae CSL RW192 WP 038903765 1 Dzeae
Dickeya zeae NCPPB 3531 WP 038903765 1 Dzeae
Dickeya dianthicola RNS04 9 WP 024105098 1 Ddianthicola
Dickeya dianthicola IPO 980 WP 024105098 1 Ddianthicola
Erwinia tasmaniensis Et1 99 YP 001908249 1 465817 Proteobacteria
Pseudoalteromonas sp SM9913 YP 004068691 1 234831 Proteobacteria
Pseudoalteromonas haloplanktis TAC125 YP 340155 1 326442 Proteobacteria
Colwellia psychrerythraea 34H YP 268318 1 167879 Proteobacteria
Vibrio furnissii NCTC 11218 YP 004992419 1 903510 Proteobacteria
Vibrio cholerae O395 YP 002820442 1 345073 Proteobacteria
Vibrio cholerae MJ 1236 YP 002878005 1 593588 Proteobacteria
Vibrio cholerae O1 biovar El Tor str N16961 NP 231731 1 243277 Proteobacteria
Teredinibacter turnerae T7901 YP 003073131 1 377629 Proteobacteria
Azotobacter vinelandii DJ YP 002797409 1 322710 Proteobacteria
Azotobacter vinelandii CA6 YP 007896269 1 1283331 Proteobacteria
Sideroxydans lithotrophicus ES 1 YP 003523550 1 580332 Proteobacteria
Rhodobacter sphaeroides ATCC 17025 YP 001169375 1 349102 Proteobacteria
Rhodobacter capsulatus SB 1003 YP 003577573 1 272942 Proteobacteria
Tolumonas auensis DSM 9187 YP 002893521 1 595494 Proteobacteria
Methylomonas methanica MC09 YP 004513974 1 857087 Proteobacteria
Methylococcus capsulatus str Bath YP 114140 1 243233 Proteobacteria
Dechloromonas aromatica RCB YP 284702 1 159087 Proteobacteria
Thiocystis violascens DSM 198 YP 006413593 1 765911 Proteobacteria
Methylomonas methanica MC09 YP 004514083 1 857087 Proteobacteria
Methylomonas methanica MC09 YP 004511278 1 857087 Proteobacteria
Dechloromonas aromatica RCB YP 284676 1 159087 Proteobacteria
Teredinibacter turnerae T7901 YP 003074867 1 377629 Proteobacteria
Rahnella aquatilis CIP 78 65 ATCC 33071 YP 005202525 1 745277 Proteobacteria
Pectobacterium atrosepticum SCRI1043 YP 051031 1 218491 Proteobacteria
Pantoea sp At 9b YP 004119231 1 592316 Proteobacteria
Klebsiella oxytoca E718 YP 006499543 1 1191061 Proteobacteria
Klebsiella pneumoniae 342 YP 002237551 1 507522 Proteobacteria
Klebsiella variicola At 22 YP 003438526 1 640131 Proteobacteria
Dickeya paradisiaca NCPPB 2511 WP 012764140 1 Dparadisiaca
Dickeya paradisiaca Ech703 WP 012764140 1 Dparadisiaca
Prodigiosinella confusarubida ATCC39006 WP 021017279 1 Pconfusarubida
Dickeya dianthicola GBBC 2039 WP 024110239 1 Ddianthicola
Dickeya dianthicola IPO 980 WP 024107433 1 Ddianthicola
Dickeya dianthicola NCPPB 3534 WP 024107433 1 Ddianthicola
Dickeya dianthicola RNS04 9 WP 024107433 1 Ddianthicola
Dickeya dianthicola NCPPB 453 WP 024107433 1 Ddianthicola
Dickeya solani GBBC 2040 WP 022635015 1 Dsolani
Dickeya solani RNS 07 7 3B WP 022635015 1 Dsolani
Dickeya solani RNS 05 1 2A WP 057084664 1 Dsolani
Dickeya solani RNS 08 23 3 1 A WP 022635015 1 Dsolani
Dickeya solani D s0432 1 WP 022635015 1 Dsolani
Dickeya solani IFB 0099 WP 022635015 1 Dsolani
Dickeya solani IPO 2222 WP 022635015 1 Dsolani
Dickeya solani MK16 WP 022635015 1 Dsolani
Dickeya solani MK10 WP 022635015 1 Dsolani
Dickeya solani PPO 9134 WP 022635015 1 Dsolani
Dickeya solani PPO 9019 WP 022635015 1 Dsolani
394 nifF nitrogen metabolism Seed
Dickeya dadantii subsp dieffenbachiae NCPPB 2976 WP 013319623 1 Ddadantii
Dickeya dadantii 3937 WP 013319623 1 Ddadantii
Dickeya chrysanthemi subsp chrysanthemi NCPPB 402 WP 039999438 1 Dchrysanthemi
Dickeya dadantii NCPPB 898 WP 013319623 1 Ddadantii
Dickeya dadantii NCPPB 3537 WP 038902178 1 Ddadantii
Dickeya gs undicola 2B12 WP 033567496 1 Dundicola
Dickeya gs fangzhongdai ND14b WP 038665312 1 Dfangzhongdai
Dickeya gs fangzhongdai M074 WP 038665312 1 Dfangzhongdai
Dickeya gs fangzhongdai NCPPB 3274 WP 042861224 1 Dfangzhongdai
Dickeya gs fangzhongdai MK7 WP 038920335 1 Dfangzhongdai
Dickeya gs fangzhongdai S1 WP 038920335 1 Dfangzhongdai
Dickeya gs fangzhongdai M005 WP 038920335 1 Dfangzhongdai
Dickeya gs fangzhongdai B16 WP 038920335 1 Dfangzhongdai

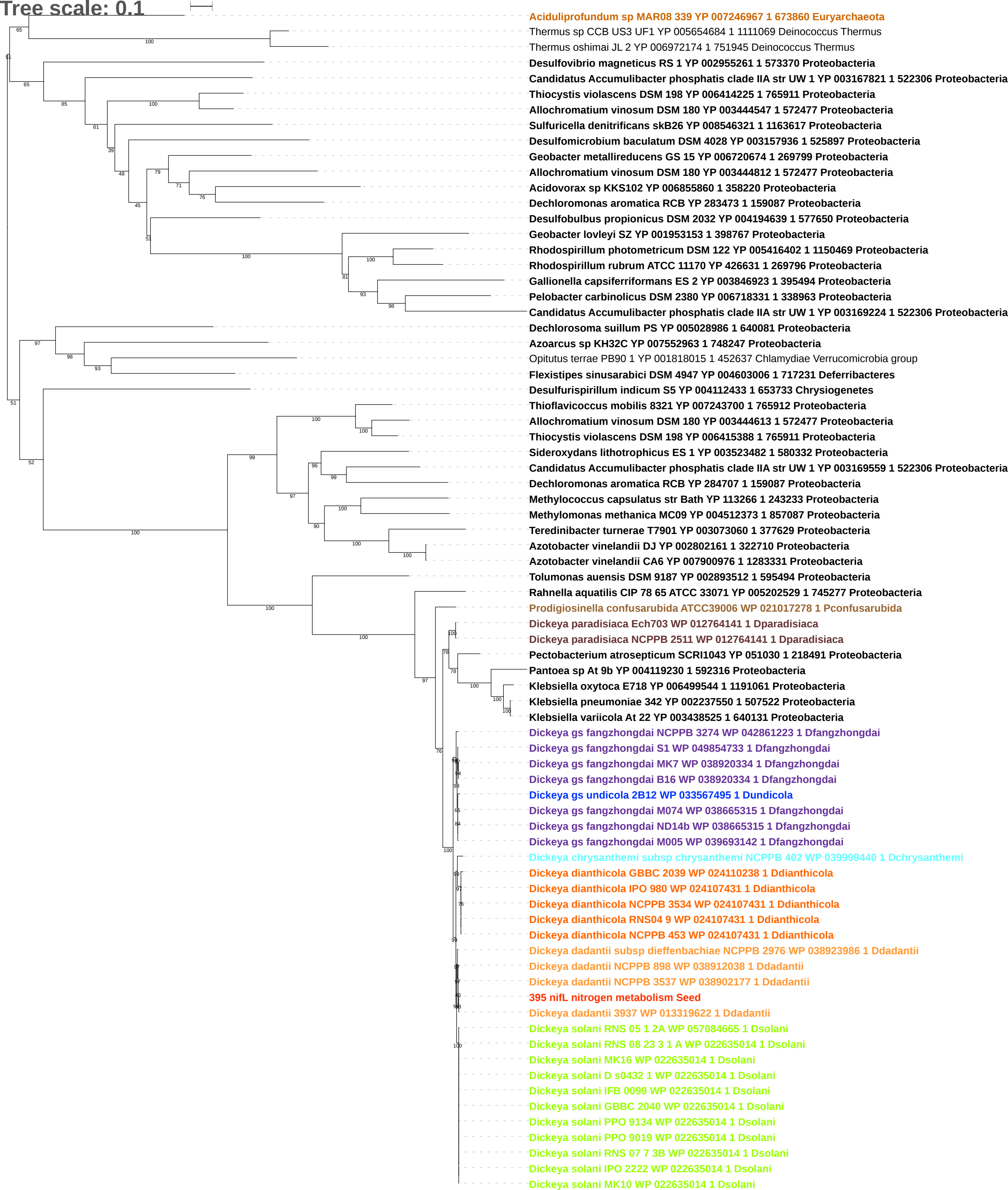

Tree scale: 0.1
Aciduliprofundum sp MAR08 339 YP 007246967 1 673860 Euryarchaeota
Thermus sp CCB US3 UF1 YP 005654684 1 1111069 Deinococcus Thermus
Thermus oshimai JL 2 YP 006972174 1 751945 Deinococcus Thermus
Desulfovibrio magneticus RS 1 YP 002955261 1 573370 Proteobacteria
Candidatus Accumulibacter phosphatis clade IIA str UW 1 YP 003167821 1 522306 Proteobacteria
Thiocystis violascens DSM 198 YP 006414225 1 765911 Proteobacteria
Allochromatium vinosum DSM 180 YP 003444547 1 572477 Proteobacteria
Sulfuricella denitrificans skB26 YP 008546321 1 1163617 Proteobacteria
Desulfomicrobium baculatum DSM 4028 YP 003157936 1 525897 Proteobacteria
Geobacter metallireducens GS 15 YP 006720674 1 269799 Proteobacteria
Allochromatium vinosum DSM 180 YP 003444812 1 572477 Proteobacteria
Acidovorax sp KKS102 YP 006855860 1 358220 Proteobacteria
Dechloromonas aromatica RCB YP 283473 1 159087 Proteobacteria
Desulfobulbus propionicus DSM 2032 YP 004194639 1 577650 Proteobacteria
Geobacter lovleyi SZ YP 001953153 1 398767 Proteobacteria
Rhodospirillum photometricum DSM 122 YP 005416402 1 1150469 Proteobacteria
Rhodospirillum rubrum ATCC 11170 YP 426631 1 269796 Proteobacteria
Gallionella capsiferriformans ES 2 YP 003846923 1 395494 Proteobacteria
Pelobacter carbinolicus DSM 2380 YP 006718331 1 338963 Proteobacteria
Candidatus Accumulibacter phosphatis clade IIA str UW 1 YP 003169224 1 522306 Proteobacteria
Dechlorosoma suillum PS YP 005028986 1 640081 Proteobacteria
Azoarcus sp KH32C YP 007552963 1 748247 Proteobacteria
Opitutus terrae PB90 1 YP 001818015 1 452637 Chlamydiae Verrucomicrobia group
Flexistipes sinusarabici DSM 4947 YP 004603006 1 717231 Deferribacteres
Desulfurispirillum indicum S5 YP 004112433 1 653733 Chrysiogenetes
Thioflavicoccus mobilis 8321 YP 007243700 1 765912 Proteobacteria
Allochromatium vinosum DSM 180 YP 003444613 1 572477 Proteobacteria
Thiocystis violascens DSM 198 YP 006415388 1 765911 Proteobacteria
Sideroxydans lithotrophicus ES 1 YP 003523482 1 580332 Proteobacteria
Candidatus Accumulibacter phosphatis clade IIA str UW 1 YP 003169559 1 522306 Proteobacteria
Dechloromonas aromatica RCB YP 284707 1 159087 Proteobacteria
Methylococcus capsulatus str Bath YP 113266 1 243233 Proteobacteria
Methylomonas methanica MC09 YP 004512373 1 857087 Proteobacteria
Teredinibacter turnerae T7901 YP 003073060 1 377629 Proteobacteria
Azotobacter vinelandii DJ YP 002802161 1 322710 Proteobacteria
Azotobacter vinelandii CA6 YP 007900976 1 1283331 Proteobacteria
Tolumonas auensis DSM 9187 YP 002893512 1 595494 Proteobacteria
Rahnella aquatilis CIP 78 65 ATCC 33071 YP 005202529 1 745277 Proteobacteria
Prodigiosinella confusarubida ATCC39006 WP 021017278 1 Pconfusarubida
Dickeya paradisiaca Ech703 WP 012764141 1 Dparadisiaca
Dickeya paradisiaca NCPPB 2511 WP 012764141 1 Dparadisiaca
Pectobacterium atrosepticum SCRI1043 YP 051030 1 218491 Proteobacteria
Pantoea sp At 9b YP 004119230 1 592316 Proteobacteria
Klebsiella oxytoca E718 YP 006499544 1 1191061 Proteobacteria
Klebsiella pneumoniae 342 YP 002237550 1 507522 Proteobacteria
Klebsiella variicola At 22 YP 003438525 1 640131 Proteobacteria
Dickeya gs fangzhongdai NCPPB 3274 WP 042861223 1 Dfangzhongdai
Dickeya gs fangzhongdai S1 WP 049854733 1 Dfangzhongdai
Dickeya gs fangzhongdai MK7 WP 038920334 1 Dfangzhongdai
Dickeya gs fangzhongdai B16 WP 038920334 1 Dfangzhongdai
Dickeya gs undicola 2B12 WP 033567495 1 Dundicola
Dickeya gs fangzhongdai M074 WP 038665315 1 Dfangzhongdai
Dickeya gs fangzhongdai ND14b WP 038665315 1 Dfangzhongdai
Dickeya gs fangzhongdai M005 WP 039693142 1 Dfangzhongdai
Dickeya chrysanthemi subsp chrysanthemi NCPPB 402 WP 039999440 1 Dchrysanthemi
Dickeya dianthicola GBBC 2039 WP 024110238 1 Ddianthicola
Dickeya dianthicola IPO 980 WP 024107431 1 Ddianthicola
Dickeya dianthicola NCPPB 3534 WP 024107431 1 Ddianthicola
Dickeya dianthicola RNS04 9 WP 024107431 1 Ddianthicola
Dickeya dianthicola NCPPB 453 WP 024107431 1 Ddianthicola
Dickeya dadantii subsp dieffenbachiae NCPPB 2976 WP 038923986 1 Ddadantii
Dickeya dadantii NCPPB 898 WP 038912038 1 Ddadantii
Dickeya dadantii NCPPB 3537 WP 038902177 1 Ddadantii
395 nifL nitrogen metabolism Seed
Dickeya dadantii 3937 WP 013319622 1 Ddadantii
Dickeya solani RNS 05 1 2A WP 057084665 1 Dsolani
Dickeya solani RNS 08 23 3 1 A WP 022635014 1 Dsolani
Dickeya solani MK16 WP 022635014 1 Dsolani
Dickeya solani D s0432 1 WP 022635014 1 Dsolani
Dickeya solani IFB 0099 WP 022635014 1 Dsolani
Dickeya solani GBBC 2040 WP 022635014 1 Dsolani
Dickeya solani PPO 9134 WP 022635014 1 Dsolani
Dickeya solani PPO 9019 WP 022635014 1 Dsolani
Dickeya solani RNS 07 7 3B WP 022635014 1 Dsolani
Dickeya solani IPO 2222 WP 022635014 1 Dsolani
Dickeya solani MK10 WP 022635014 1 Dsolani

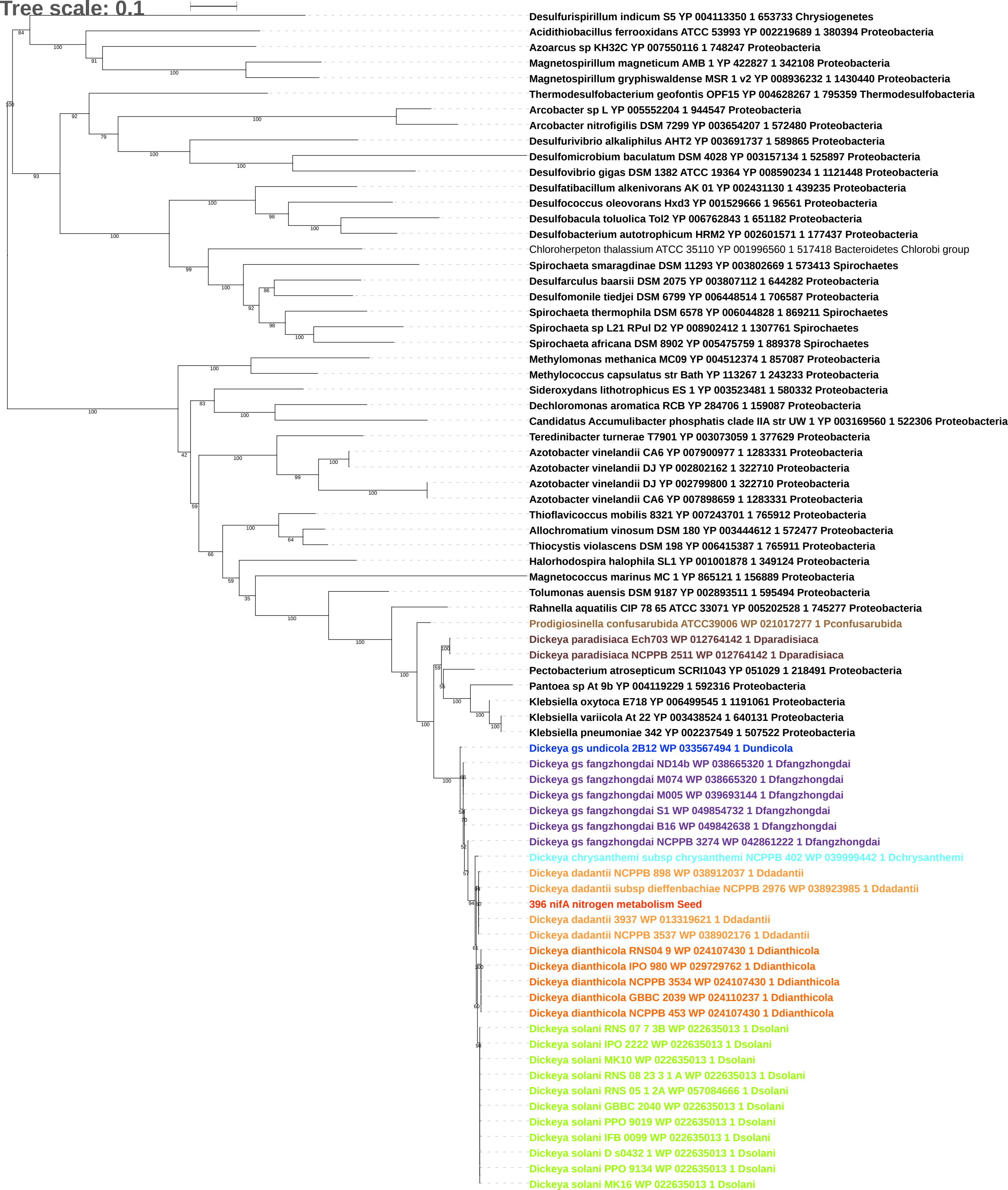
Tree scale: 0.1
Desulfurispirillum indicum S5 YP 004113350 1 653733 Chrysiogenetes
Acidithiobacillus ferrooxidans ATCC 53993 YP 002219689 1 380394 Proteobacteria
Azoarcus sp KH32C YP 007550116 1 748247 Proteobacteria
Magnetospirillum magneticum AMB 1 YP 422827 1 342108 Proteobacteria
Magnetospirillum gryphiswaldense MSR 1 v2 YP 008936232 1 1430440 Proteobacteria
Thermodesulfobacterium geofontis OPF15 YP 004628267 1 795359 Thermodesulfobacteria
Arcobacter sp L YP 005552204 1 944547 Proteobacteria
Arcobacter nitrofigilis DSM 7299 YP 003654207 1 572480 Proteobacteria
Desulfurivibrio alkaliphilus AHT2 YP 003691737 1 589865 Proteobacteria
Desulfomicrobium baculatum DSM 4028 YP 003157134 1 525897 Proteobacteria
Desulfovibrio gigas DSM 1382 ATCC 19364 YP 008590234 1 1121448 Proteobacteria
Desulfatibacillum alkenivorans AK 01 YP 002431130 1 439235 Proteobacteria
Desulfococcus oleovorans Hxd3 YP 001529666 1 96561 Proteobacteria
Desulfobacula toluolica Tol2 YP 006762843 1 651182 Proteobacteria
Desulfobacterium autotrophicum HRM2 YP 002601571 1 177437 Proteobacteria
Chloroherpeton thalassium ATCC 35110 YP 001996560 1 517418 Bacteroidetes Chlorobi group
Spirochaeta smaragdinae DSM 11293 YP 003802669 1 573413 Spirochaetes
Desulfarculus baarsii DSM 2075 YP 003807112 1 644282 Proteobacteria
Desulfomonile tiedjei DSM 6799 YP 006448514 1 706587 Proteobacteria
Spirochaeta thermophila DSM 6578 YP 006044828 1 869211 Spirochaetes
Spirochaeta sp L21 RPul D2 YP 008902412 1 1307761 Spirochaetes
Spirochaeta africana DSM 8902 YP 005475759 1 889378 Spirochaetes
Methylomonas methanica MC09 YP 004512374 1 857087 Proteobacteria
Methylococcus capsulatus str Bath YP 113267 1 243233 Proteobacteria
Sideroxydans lithotrophicus ES 1 YP 003523481 1 580332 Proteobacteria
Dechloromonas aromatica RCB YP 284706 1 159087 Proteobacteria
Candidatus Accumulibacter phosphatis clade IIA str UW 1 YP 003169560 1 522306 Proteobacteria
Teredinibacter turnerae T7901 YP 003073059 1 377629 Proteobacteria
Azotobacter vinelandii CA6 YP 007900977 1 1283331 Proteobacteria
Azotobacter vinelandii DJ YP 002802162 1 322710 Proteobacteria
Azotobacter vinelandii DJ YP 002799800 1 322710 Proteobacteria
Azotobacter vinelandii CA6 YP 007898659 1 1283331 Proteobacteria
Thioflavicoccus mobilis 8321 YP 007243701 1 765912 Proteobacteria
Allochromatium vinosum DSM 180 YP 003444612 1 572477 Proteobacteria
Thiocystis violascens DSM 198 YP 006415387 1 765911 Proteobacteria
Halorhodospira halophila SL1 YP 001001878 1 349124 Proteobacteria
Magnetococcus marinus MC 1 YP 865121 1 156889 Proteobacteria
Tolumonas auensis DSM 9187 YP 002893511 1 595494 Proteobacteria
Rahnella aquatilis CIP 78 65 ATCC 33071 YP 005202528 1 745277 Proteobacteria
Prodigiosinella confusarubida ATCC39006 WP 021017277 1 Pconfusarubida
Dickeya paradisiaca Ech703 WP 012764142 1 Dparadisiaca
Dickeya paradisiaca NCPPB 2511 WP 012764142 1 Dparadisiaca
Pectobacterium atrosepticum SCRI1043 YP 051029 1 218491 Proteobacteria
Pantoea sp At 9b YP 004119229 1 592316 Proteobacteria
Klebsiella oxytoca E718 YP 006499545 1 1191061 Proteobacteria
Klebsiella variicola At 22 YP 003438524 1 640131 Proteobacteria
Klebsiella pneumoniae 342 YP 002237549 1 507522 Proteobacteria
Dickeya gs undicola 2B12 WP 033567494 1 Dundicola
Dickeya gs fangzhongdai ND14b WP 038665320 1 Dfangzhongdai
Dickeya gs fangzhongdai M074 WP 038665320 1 Dfangzhongdai
Dickeya gs fangzhongdai M005 WP 039693144 1 Dfangzhongdai
Dickeya gs fangzhongdai S1 WP 049854732 1 Dfangzhongdai
Dickeya gs fangzhongdai B16 WP 049842638 1 Dfangzhongdai
Dickeya gs fangzhongdai NCPPB 3274 WP 042861222 1 Dfangzhongdai
Dickeya chrysanthemi subsp chrysanthemi NCPPB 402 WP 039999442 1 Dchrysanthemi
Dickeya dadantii NCPPB 898 WP 038912037 1 Ddadantii
Dickeya dadantii subsp dieffenbachiae NCPPB 2976 WP 038923985 1 Ddadantii
396 nifA nitrogen metabolism Seed
Dickeya dadantii 3937 WP 013319621 1 Ddadantii
Dickeya dadantii NCPPB 3537 WP 038902176 1 Ddadantii
Dickeya dianthicola RNS04 9 WP 024107430 1 Ddianthicola
Dickeya dianthicola IPO 980 WP 029729762 1 Ddianthicola
Dickeya dianthicola NCPPB 3534 WP 024107430 1 Ddianthicola
Dickeya dianthicola GBBC 2039 WP 024110237 1 Ddianthicola
Dickeya dianthicola NCPPB 453 WP 024107430 1 Ddianthicola
Dickeya solani RNS 07 7 3B WP 022635013 1 Dsolani
Dickeya solani IPO 2222 WP 022635013 1 Dsolani
Dickeya solani MK10 WP 022635013 1 Dsolani
Dickeya solani RNS 08 23 3 1 A WP 022635013 1 Dsolani
Dickeya solani RNS 05 1 2A WP 057084666 1 Dsolani
Dickeya solani GBBC 2040 WP 022635013 1 Dsolani
Dickeya solani PPO 9019 WP 022635013 1 Dsolani
Dickeya solani IFB 0099 WP 022635013 1 Dsolani
Dickeya solani D s0432 1 WP 022635013 1 Dsolani
Dickeya solani PPO 9134 WP 022635013 1 Dsolani
Dickeya solani MK16 WP 022635013 1 Dsolani

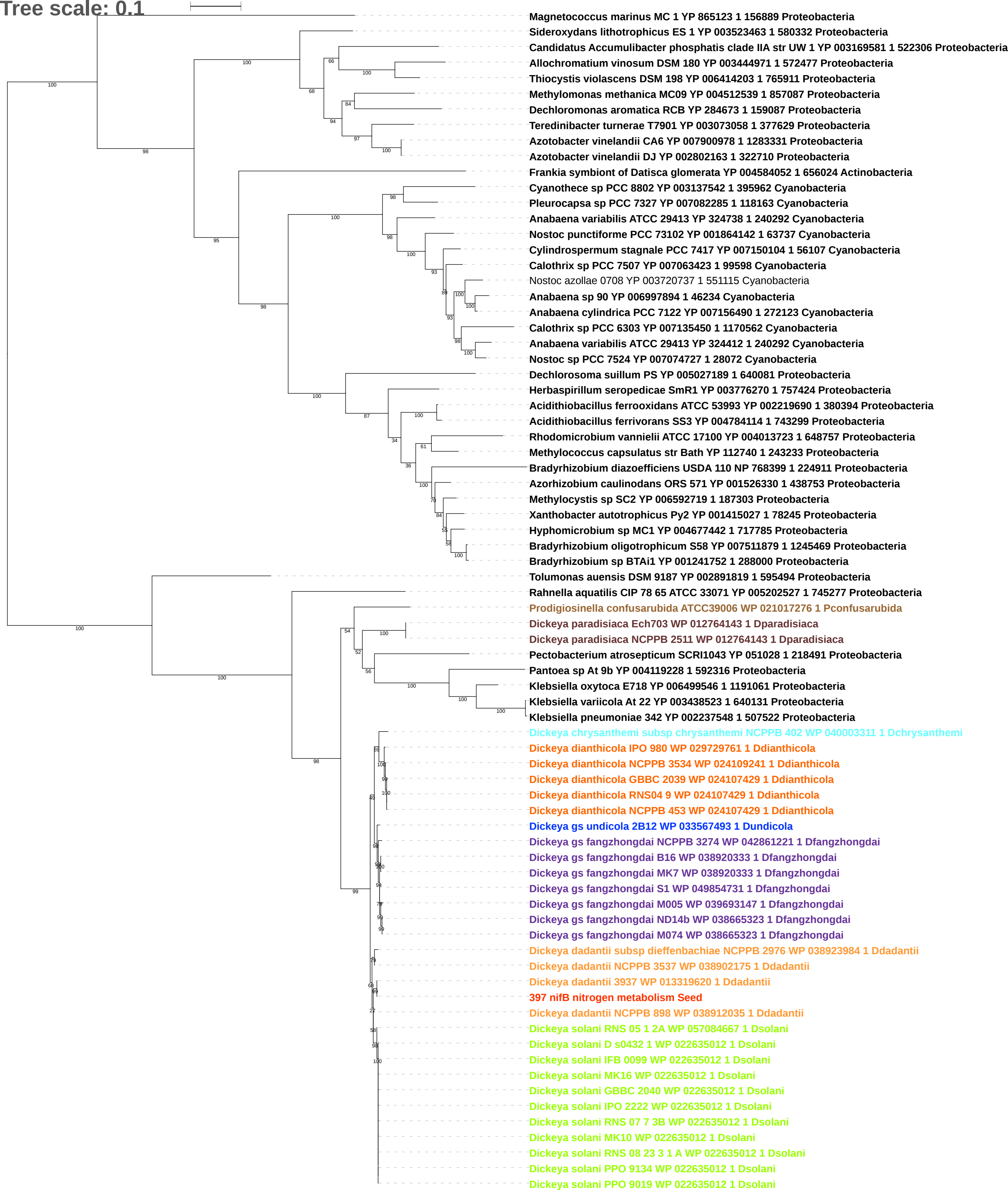
Tree scale: 0.1
Magnetococcus marinus MC 1 YP 865123 1 156889 Proteobacteria
Sideroxydans lithotrophicus ES 1 YP 003523463 1 580332 Proteobacteria
Candidatus Accumulibacter phosphatis clade IIA str UW 1 YP 003169581 1 522306 Proteobacteria
Allochromatium vinosum DSM 180 YP 003444971 1 572477 Proteobacteria
Thiocystis violascens DSM 198 YP 006414203 1 765911 Proteobacteria
Methylomonas methanica MC09 YP 004512539 1 857087 Proteobacteria
Dechloromonas aromatica RCB YP 284673 1 159087 Proteobacteria
Teredinibacter turnerae T7901 YP 003073058 1 377629 Proteobacteria
Azotobacter vinelandii CA6 YP 007900978 1 1283331 Proteobacteria
Azotobacter vinelandii DJ YP 002802163 1 322710 Proteobacteria
Frankia symbiont of Datisca glomerata YP 004584052 1 656024 Actinobacteria
Cyanothece sp PCC 8802 YP 003137542 1 395962 Cyanobacteria
Pleurocapsa sp PCC 7327 YP 007082285 1 118163 Cyanobacteria
Anabaena variabilis ATCC 29413 YP 324738 1 240292 Cyanobacteria
Nostoc punctiforme PCC 73102 YP 001864142 1 63737 Cyanobacteria
Cylindrospermum stagnale PCC 7417 YP 007150104 1 56107 Cyanobacteria
Calothrix sp PCC 7507 YP 007063423 1 99598 Cyanobacteria
Nostoc azollae 0708 YP 003720737 1 551115 Cyanobacteria
Anabaena sp 90 YP 006997894 1 46234 Cyanobacteria
Anabaena cylindrica PCC 7122 YP 007156490 1 272123 Cyanobacteria
Calothrix sp PCC 6303 YP 007135450 1 1170562 Cyanobacteria
Anabaena variabilis ATCC 29413 YP 324412 1 240292 Cyanobacteria
Nostoc sp PCC 7524 YP 007074727 1 28072 Cyanobacteria
Dechlorosoma suillum PS YP 005027189 1 640081 Proteobacteria
Herbaspirillum seropedicae SmR1 YP 003776270 1 757424 Proteobacteria
Acidithiobacillus ferrooxidans ATCC 53993 YP 002219690 1 380394 Proteobacteria
Acidithiobacillus ferrivorans SS3 YP 004784114 1 743299 Proteobacteria
Rhodomicrobium vannielii ATCC 17100 YP 004013723 1 648757 Proteobacteria
Methylococcus capsulatus str Bath YP 112740 1 243233 Proteobacteria
Bradyrhizobium diazoefficiens USDA 110 NP 768399 1 224911 Proteobacteria
Azorhizobium caulinodans ORS 571 YP 001526330 1 438753 Proteobacteria
Methylocystis sp SC2 YP 006592719 1 187303 Proteobacteria
Xanthobacter autotrophicus Py2 YP 001415027 1 78245 Proteobacteria
Hyphomicrobium sp MC1 YP 004677442 1 717785 Proteobacteria
Bradyrhizobium oligotrophicum S58 YP 007511879 1 1245469 Proteobacteria
Bradyrhizobium sp BTAi1 YP 001241752 1 288000 Proteobacteria
Tolumonas auensis DSM 9187 YP 002891819 1 595494 Proteobacteria
Rahnella aquatilis CIP 78 65 ATCC 33071 YP 005202527 1 745277 Proteobacteria
Prodigiosinella confusarubida ATCC39006 WP 021017276 1 Pconfusarubida
Dickeya paradisiaca Ech703 WP 012764143 1 Dparadisiaca
Dickeya paradisiaca NCPPB 2511 WP 012764143 1 Dparadisiaca
Pectobacterium atrosepticum SCRI1043 YP 051028 1 218491 Proteobacteria
Pantoea sp At 9b YP 004119228 1 592316 Proteobacteria
Klebsiella oxytoca E718 YP 006499546 1 1191061 Proteobacteria
Klebsiella variicola At 22 YP 003438523 1 640131 Proteobacteria
Klebsiella pneumoniae 342 YP 002237548 1 507522 Proteobacteria
Dickeya chrysanthemi subsp chrysanthemi NCPPB 402 WP 040003311 1 Dchrysanthemi
Dickeya dianthicola IPO 980 WP 029729761 1 Ddianthicola
Dickeya dianthicola NCPPB 3534 WP 024109241 1 Ddianthicola
Dickeya dianthicola GBBC 2039 WP 024107429 1 Ddianthicola
Dickeya dianthicola RNS04 9 WP 024107429 1 Ddianthicola
Dickeya dianthicola NCPPB 453 WP 024107429 1 Ddianthicola
Dickeya gs undicola 2B12 WP 033567493 1 Dundicola
Dickeya gs fangzhongdai NCPPB 3274 WP 042861221 1 Dfangzhongdai
Dickeya gs fangzhongdai B16 WP 038920333 1 Dfangzhongdai
Dickeya gs fangzhongdai MK7 WP 038920333 1 Dfangzhongdai
Dickeya gs fangzhongdai S1 WP 049854731 1 Dfangzhongdai
Dickeya gs fangzhongdai M005 WP 039693147 1 Dfangzhongdai
Dickeya gs fangzhongdai ND14b WP 038665323 1 Dfangzhongdai
Dickeya gs fangzhongdai M074 WP 038665323 1 Dfangzhongdai
Dickeya dadantii subsp dieffenbachiae NCPPB 2976 WP 038923984 1 Ddadantii
Dickeya dadantii NCPPB 3537 WP 038902175 1 Ddadantii
Dickeya dadantii 3937 WP 013319620 1 Ddadantii
397 nifB nitrogen metabolism Seed
Dickeya dadantii NCPPB 898 WP 038912035 1 Ddadantii
Dickeya solani RNS 05 1 2A WP 057084667 1 Dsolani
Dickeya solani D s0432 1 WP 022635012 1 Dsolani
Dickeya solani IFB 0099 WP 022635012 1 Dsolani
Dickeya solani MK16 WP 022635012 1 Dsolani
Dickeya solani GBBC 2040 WP 022635012 1 Dsolani
Dickeya solani IPO 2222 WP 022635012 1 Dsolani
Dickeya solani RNS 07 7 3B WP 022635012 1 Dsolani
Dickeya solani MK10 WP 022635012 1 Dsolani
Dickeya solani RNS 08 23 3 1 A WP 022635012 1 Dsolani
Dickeya solani PPO 9134 WP 022635012 1 Dsolani
Dickeya solani PPO 9019 WP 022635012 1 Dsolani

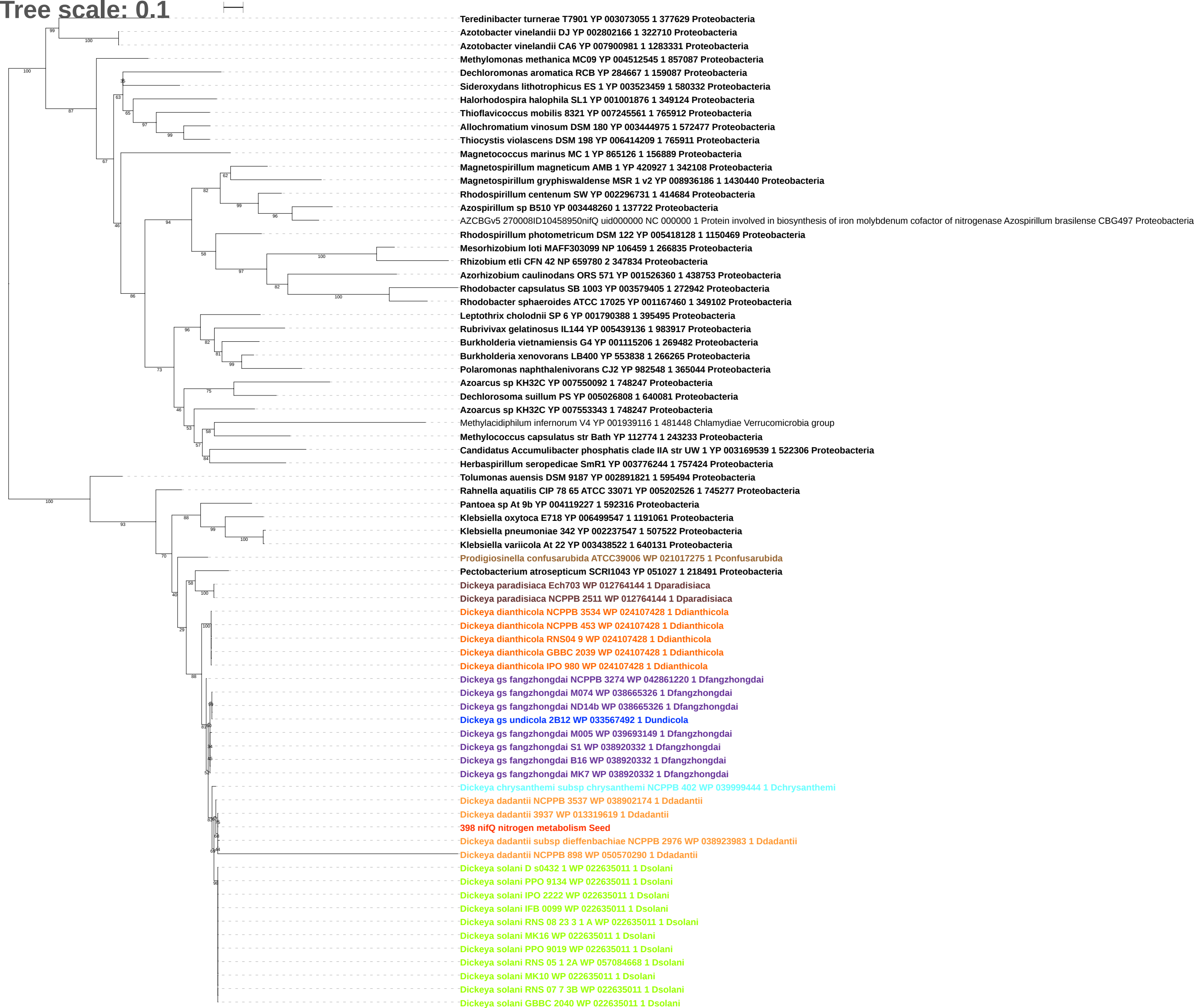
Tree scale: 0.1
Teredinibacter turnerae T7901 YP 003073055 1 377629 Proteobacteria
Azotobacter vinelandii DJ YP 002802166 1 322710 Proteobacteria
Azotobacter vinelandii CA6 YP 007900981 1 1283331 Proteobacteria
Methylomonas methanica MC09 YP 004512545 1 857087 Proteobacteria
Dechloromonas aromatica RCB YP 284667 1 159087 Proteobacteria
Sideroxydans lithotrophicus ES 1 YP 003523459 1 580332 Proteobacteria
Halorhodospira halophila SL1 YP 001001876 1 349124 Proteobacteria
Thioflavicoccus mobilis 8321 YP 007245561 1 765912 Proteobacteria
Allochromatium vinosum DSM 180 YP 003444975 1 572477 Proteobacteria
Thiocystis violascens DSM 198 YP 006414209 1 765911 Proteobacteria
Magnetococcus marinus MC 1 YP 865126 1 156889 Proteobacteria
Magnetospirillum magneticum AMB 1 YP 420927 1 342108 Proteobacteria
Magnetospirillum gryphiswaldense MSR 1 v2 YP 008936186 1 1430440 Proteobacteria
Rhodospirillum centenum SW YP 002296731 1 414684 Proteobacteria
Azospirillum sp B510 YP 003448260 1 137722 Proteobacteria
AZCBGv5 270008ID10458950nifQ uid000000 NC 000000 1 Protein involved in biosynthesis of iron molybdenum cofactor of nitrogenase Azospirillum brasilense CBG497 Proteobacteria
Rhodospirillum photometricum DSM 122 YP 005418128 1 1150469 Proteobacteria
Mesorhizobium loti MAFF303099 NP 106459 1 266835 Proteobacteria
Rhizobium etli CFN 42 NP 659780 2 347834 Proteobacteria
Azorhizobium caulinodans ORS 571 YP 001526360 1 438753 Proteobacteria
Rhodobacter capsulatus SB 1003 YP 003579405 1 272942 Proteobacteria
Rhodobacter sphaeroides ATCC 17025 YP 001167460 1 349102 Proteobacteria
Leptothrix cholodnii SP 6 YP 001790388 1 395495 Proteobacteria
Rubrivivax gelatinosus IL144 YP 005439136 1 983917 Proteobacteria
Burkholderia vietnamiensis G4 YP 001115206 1 269482 Proteobacteria
Burkholderia xenovorans LB400 YP 553838 1 266265 Proteobacteria
Polaromonas naphthalenivorans CJ2 YP 982548 1 365044 Proteobacteria
Azoarcus sp KH32C YP 007550092 1 748247 Proteobacteria
Dechlorosoma suillum PS YP 005026808 1 640081 Proteobacteria
Azoarcus sp KH32C YP 007553343 1 748247 Proteobacteria
Methylacidiphilum infernorum V4 YP 001939116 1 481448 Chlamydiae Verrucomicrobia group
Methylococcus capsulatus str Bath YP 112774 1 243233 Proteobacteria
Candidatus Accumulibacter phosphatis clade IIA str UW 1 YP 003169539 1 522306 Proteobacteria
Herbaspirillum seropedicae SmR1 YP 003776244 1 757424 Proteobacteria
Tolumonas auensis DSM 9187 YP 002891821 1 595494 Proteobacteria
Rahnella aquatilis CIP 78 65 ATCC 33071 YP 005202526 1 745277 Proteobacteria
Pantoea sp At 9b YP 004119227 1 592316 Proteobacteria
Klebsiella oxytoca E718 YP 006499547 1 1191061 Proteobacteria
Klebsiella pneumoniae 342 YP 002237547 1 507522 Proteobacteria
Klebsiella variicola At 22 YP 003438522 1 640131 Proteobacteria
Prodigiosinella confusarubida ATCC39006 WP 021017275 1 Pconfusarubida
Pectobacterium atrosepticum SCRI1043 YP 051027 1 218491 Proteobacteria
Dickeya paradisiaca Ech703 WP 012764144 1 Dparadisiaca
Dickeya paradisiaca NCPPB 2511 WP 012764144 1 Dparadisiaca
Dickeya dianthicola NCPPB 3534 WP 024107428 1 Ddianthicola
Dickeya dianthicola NCPPB 453 WP 024107428 1 Ddianthicola
Dickeya dianthicola RNS04 9 WP 024107428 1 Ddianthicola
Dickeya dianthicola GBBC 2039 WP 024107428 1 Ddianthicola
Dickeya dianthicola IPO 980 WP 024107428 1 Ddianthicola
Dickeya gs fangzhongdai NCPPB 3274 WP 042861220 1 Dfangzhongdai
Dickeya gs fangzhongdai M074 WP 038665326 1 Dfangzhongdai
Dickeya gs fangzhongdai ND14b WP 038665326 1 Dfangzhongdai
Dickeya gs undicola 2B12 WP 033567492 1 Dundicola
Dickeya gs fangzhongdai M005 WP 039693149 1 Dfangzhongdai
Dickeya gs fangzhongdai S1 WP 038920332 1 Dfangzhongdai
Dickeya gs fangzhongdai B16 WP 038920332 1 Dfangzhongdai
Dickeya gs fangzhongdai MK7 WP 038920332 1 Dfangzhongdai
Dickeya chrysanthemi subsp chrysanthemi NCPPB 402 WP 039999444 1 Dchrysanthemi
Dickeya dadantii NCPPB 3537 WP 038902174 1 Ddadantii
Dickeya dadantii 3937 WP 013319619 1 Ddadantii
398 nifQ nitrogen metabolism Seed
Dickeya dadantii subsp dieffenbachiae NCPPB 2976 WP 038923983 1 Ddadantii
Dickeya dadantii NCPPB 898 WP 050570290 1 Ddadantii
Dickeya solani D s0432 1 WP 022635011 1 Dsolani
Dickeya solani PPO 9134 WP 022635011 1 Dsolani
Dickeya solani IPO 2222 WP 022635011 1 Dsolani
Dickeya solani IFB 0099 WP 022635011 1 Dsolani
Dickeya solani RNS 08 23 3 1 A WP 022635011 1 Dsolani
Dickeya solani MK16 WP 022635011 1 Dsolani
Dickeya solani PPO 9019 WP 022635011 1 Dsolani
Dickeya solani RNS 05 1 2A WP 057084668 1 Dsolani
Dickeya solani MK10 WP 022635011 1 Dsolani
Dickeya solani RNS 07 7 3B WP 022635011 1 Dsolani
Dickeya solani GBBC 2040 WP 022635011 1 Dsolani

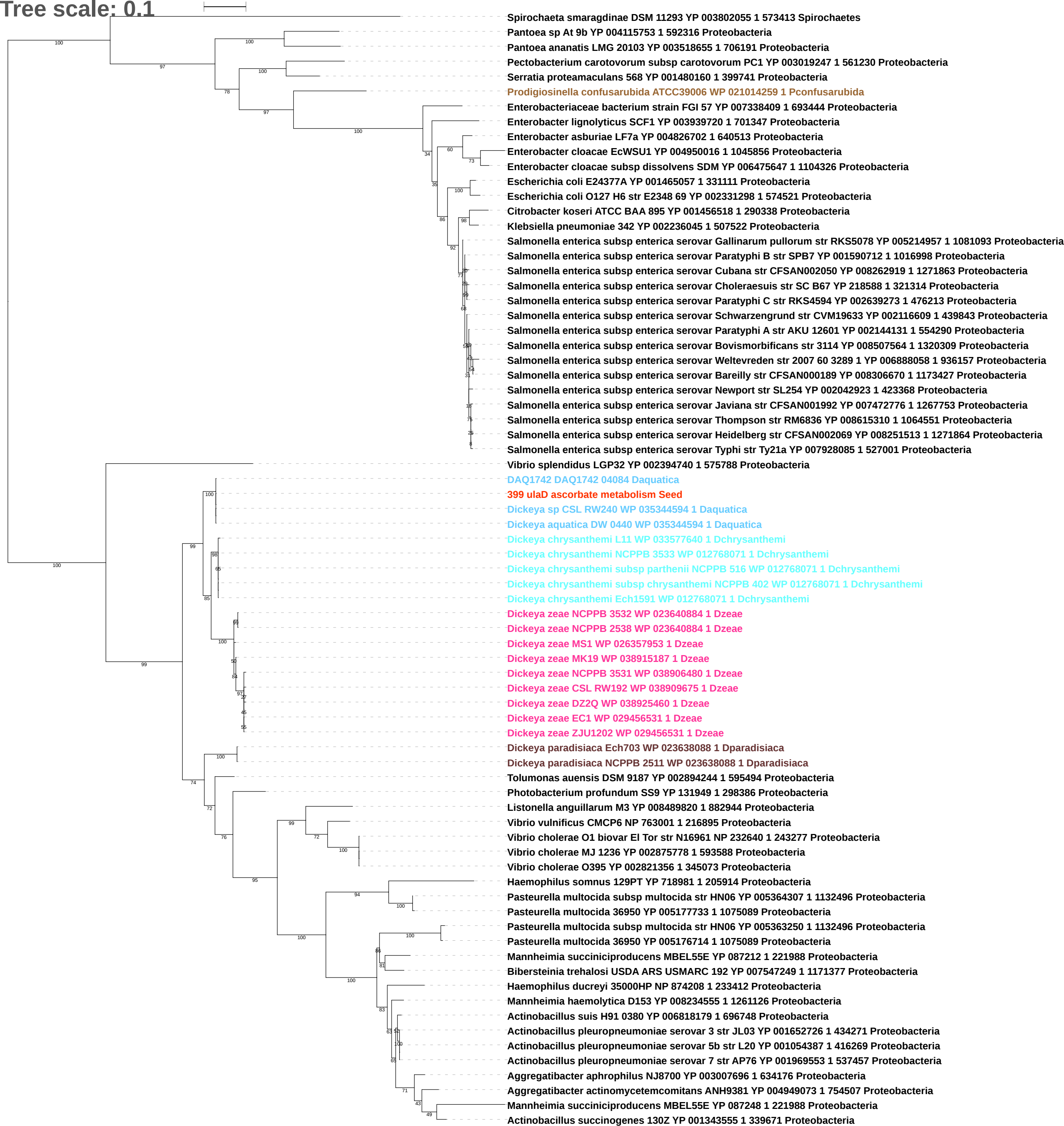
Tree scale: 0.1
Spirochaeta smaragdinae DSM 11293 YP 003802055 1 573413 Spirochaetes
Pantoea sp At 9b YP 004115753 1 592316 Proteobacteria
Pantoea ananatis LMG 20103 YP 003518655 1 706191 Proteobacteria
Pectobacterium carotovorum subsp carotovorum PC1 YP 003019247 1 561230 Proteobacteria
Serratia proteamaculans 568 YP 001480160 1 399741 Proteobacteria
Prodigiosinella confusarubida ATCC39006 WP 021014259 1 Pconfusarubida
Enterobacteriaceae bacterium strain FGI 57 YP 007338409 1 693444 Proteobacteria
Enterobacter lignolyticus SCF1 YP 003939720 1 701347 Proteobacteria
Enterobacter asburiae LF7a YP 004826702 1 640513 Proteobacteria
Enterobacter cloacae EcWSU1 YP 004950016 1 1045856 Proteobacteria
Enterobacter cloacae subsp dissolvens SDM YP 006475647 1 1104326 Proteobacteria
Escherichia coli E24377A YP 001465057 1 331111 Proteobacteria
Escherichia coli O127 H6 str E2348 69 YP 002331298 1 574521 Proteobacteria
Citrobacter koseri ATCC BAA 895 YP 001456518 1 290338 Proteobacteria
Klebsiella pneumoniae 342 YP 002236045 1 507522 Proteobacteria
Salmonella enterica subsp enterica serovar Gallinarum pullorum str RKS5078 YP 005214957 1 1081093 Proteobacteria
Salmonella enterica subsp enterica serovar Paratyphi B str SPB7 YP 001590712 1 1016998 Proteobacteria
Salmonella enterica subsp enterica serovar Cubana str CFSAN002050 YP 008262919 1 1271863 Proteobacteria
Salmonella enterica subsp enterica serovar Choleraesuis str SC B67 YP 218588 1 321314 Proteobacteria
Salmonella enterica subsp enterica serovar Paratyphi C str RKS4594 YP 002639273 1 476213 Proteobacteria
Salmonella enterica subsp enterica serovar Schwarzengrund str CVM19633 YP 002116609 1 439843 Proteobacteria
Salmonella enterica subsp enterica serovar Paratyphi A str AKU 12601 YP 002144131 1 554290 Proteobacteria
Salmonella enterica subsp enterica serovar Bovismorbificans str 3114 YP 008507564 1 1320309 Proteobacteria
Salmonella enterica subsp enterica serovar Weltevreden str 2007 60 3289 1 YP 006888058 1 936157 Proteobacteria
Salmonella enterica subsp enterica serovar Bareilly str CFSAN000189 YP 008306670 1 1173427 Proteobacteria
Salmonella enterica subsp enterica serovar Newport str SL254 YP 002042923 1 423368 Proteobacteria
Salmonella enterica subsp enterica serovar Javiana str CFSAN001992 YP 007472776 1 1267753 Proteobacteria
Salmonella enterica subsp enterica serovar Thompson str RM6836 YP 008615310 1 1064551 Proteobacteria
Salmonella enterica subsp enterica serovar Heidelberg str CFSAN002069 YP 008251513 1 1271864 Proteobacteria
Salmonella enterica subsp enterica serovar Typhi str Ty21a YP 007928085 1 527001 Proteobacteria
Vibrio splendidus LGP32 YP 002394740 1 575788 Proteobacteria
DAQ1742 DAQ1742 04084 Daquatica
399 ulaD ascorbate metabolism Seed
Dickeya sp CSL RW240 WP 035344594 1 Daquatica
Dickeya aquatica DW 0440 WP 035344594 1 Daquatica
Dickeya chrysanthemi L11 WP 033577640 1 Dchrysanthemi
Dickeya chrysanthemi NCPPB 3533 WP 012768071 1 Dchrysanthemi
Dickeya chrysanthemi subsp parthenii NCPPB 516 WP 012768071 1 Dchrysanthemi
Dickeya chrysanthemi subsp chrysanthemi NCPPB 402 WP 012768071 1 Dchrysanthemi
Dickeya chrysanthemi Ech1591 WP 012768071 1 Dchrysanthemi
Dickeya zeae NCPPB 3532 WP 023640884 1 Dzeae
Dickeya zeae NCPPB 2538 WP 023640884 1 Dzeae
Dickeya zeae MS1 WP 026357953 1 Dzeae
Dickeya zeae MK19 WP 038915187 1 Dzeae
Dickeya zeae NCPPB 3531 WP 038906480 1 Dzeae
Dickeya zeae CSL RW192 WP 038909675 1 Dzeae
Dickeya zeae DZ2Q WP 038925460 1 Dzeae
Dickeya zeae EC1 WP 029456531 1 Dzeae
Dickeya zeae ZJU1202 WP 029456531 1 Dzeae
Dickeya paradisiaca Ech703 WP 023638088 1 Dparadisiaca
Dickeya paradisiaca NCPPB 2511 WP 023638088 1 Dparadisiaca
Tolumonas auensis DSM 9187 YP 002894244 1 595494 Proteobacteria
Photobacterium profundum SS9 YP 131949 1 298386 Proteobacteria
Listonella anguillarum M3 YP 008489820 1 882944 Proteobacteria
Vibrio vulnificus CMCP6 NP 763001 1 216895 Proteobacteria
Vibrio cholerae O1 biovar El Tor str N16961 NP 232640 1 243277 Proteobacteria
Vibrio cholerae MJ 1236 YP 002875778 1 593588 Proteobacteria
Vibrio cholerae O395 YP 002821356 1 345073 Proteobacteria
Haemophilus somnus 129PT YP 718981 1 205914 Proteobacteria
Pasteurella multocida subsp multocida str HN06 YP 005364307 1 1132496 Proteobacteria
Pasteurella multocida 36950 YP 005177733 1 1075089 Proteobacteria
Pasteurella multocida subsp multocida str HN06 YP 005363250 1 1132496 Proteobacteria
Pasteurella multocida 36950 YP 005176714 1 1075089 Proteobacteria
Mannheimia succiniciproducens MBEL55E YP 087212 1 221988 Proteobacteria
Bibersteinia trehalosi USDA ARS USMARC 192 YP 007547249 1 1171377 Proteobacteria
Haemophilus ducreyi 35000HP NP 874208 1 233412 Proteobacteria
Mannheimia haemolytica D153 YP 008234555 1 1261126 Proteobacteria
Actinobacillus suis H91 0380 YP 006818179 1 696748 Proteobacteria
Actinobacillus pleuropneumoniae serovar 3 str JL03 YP 001652726 1 434271 Proteobacteria
Actinobacillus pleuropneumoniae serovar 5b str L20 YP 001054387 1 416269 Proteobacteria
Actinobacillus pleuropneumoniae serovar 7 str AP76 YP 001969553 1 537457 Proteobacteria
Aggregatibacter aphrophilus NJ8700 YP 003007696 1 634176 Proteobacteria
Aggregatibacter actinomycetemcomitans ANH9381 YP 004949073 1 754507 Proteobacteria
Mannheimia succiniciproducens MBEL55E YP 087248 1 221988 Proteobacteria
Actinobacillus succinogenes 130Z YP 001343555 1 339671 Proteobacteria

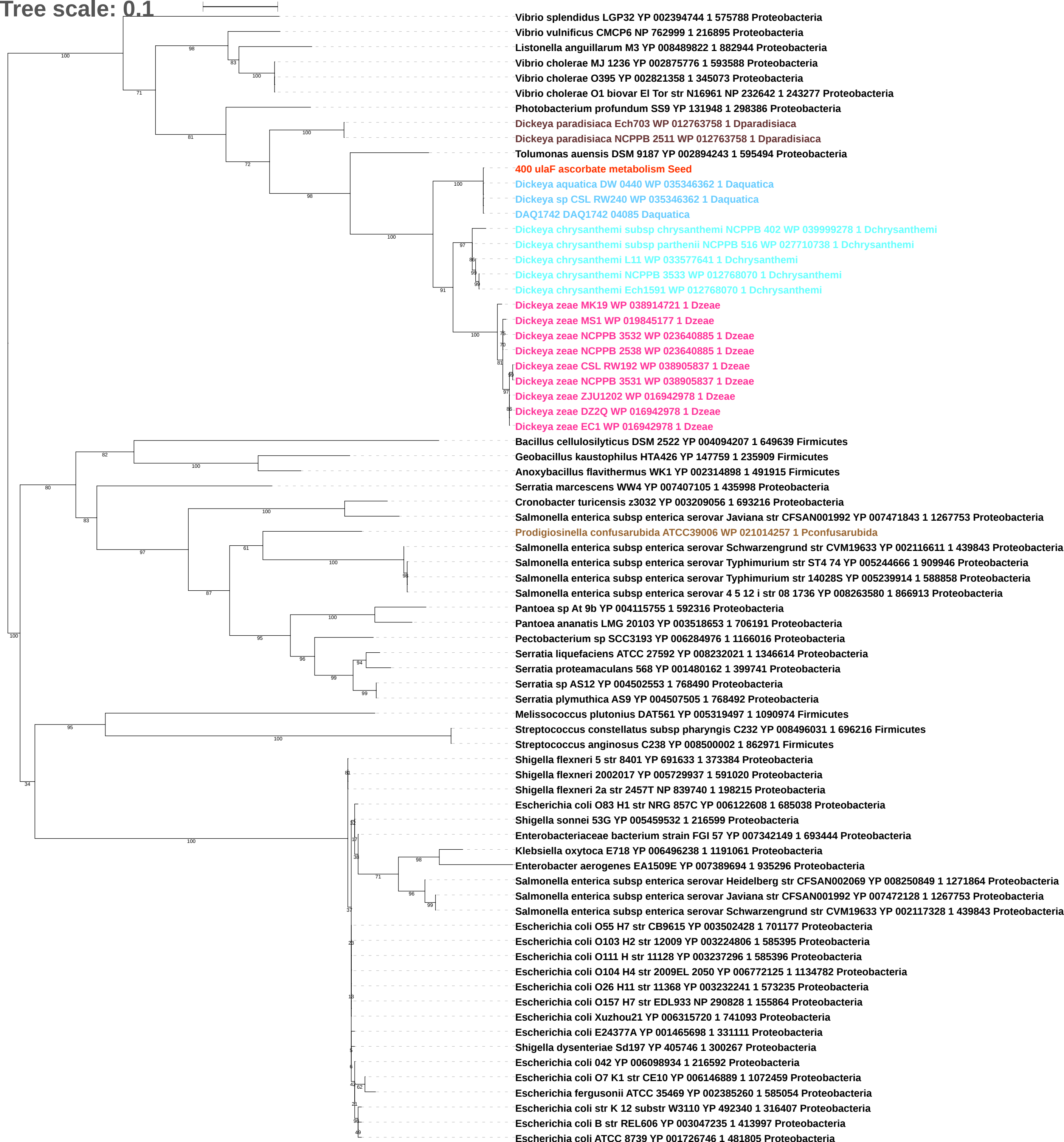

Tree scale: 0.1
Vibrio splendidus LGP32 YP 002394744 1 575788 Proteobacteria
Vibrio vulnificus CMCP6 NP 762999 1 216895 Proteobacteria
Listonella anguillarum M3 YP 008489822 1 882944 Proteobacteria
Vibrio cholerae MJ 1236 YP 002875776 1 593588 Proteobacteria
Vibrio cholerae O395 YP 002821358 1 345073 Proteobacteria
Vibrio cholerae O1 biovar El Tor str N16961 NP 232642 1 243277 Proteobacteria
Photobacterium profundum SS9 YP 131948 1 298386 Proteobacteria
Dickeya paradisiaca Ech703 WP 012763758 1 Dparadisiaca
Dickeya paradisiaca NCPPB 2511 WP 012763758 1 Dparadisiaca
Tolumonas auensis DSM 9187 YP 002894243 1 595494 Proteobacteria
400 ulaF ascorbate metabolism Seed
Dickeya aquatica DW 0440 WP 035346362 1 Daquatica
Dickeya sp CSL RW240 WP 035346362 1 Daquatica
DAQ1742 DAQ1742 04085 Daquatica
Dickeya chrysanthemi subsp chrysanthemi NCPPB 402 WP 039999278 1 Dchrysanthemi
Dickeya chrysanthemi subsp parthenii NCPPB 516 WP 027710738 1 Dchrysanthemi
Dickeya chrysanthemi L11 WP 033577641 1 Dchrysanthemi
Dickeya chrysanthemi NCPPB 3533 WP 012768070 1 Dchrysanthemi
Dickeya chrysanthemi Ech1591 WP 012768070 1 Dchrysanthemi
Dickeya zeae MK19 WP 038914721 1 Dzeae
Dickeya zeae MS1 WP 019845177 1 Dzeae
Dickeya zeae NCPPB 3532 WP 023640885 1 Dzeae
Dickeya zeae NCPPB 2538 WP 023640885 1 Dzeae
Dickeya zeae CSL RW192 WP 038905837 1 Dzeae
Dickeya zeae NCPPB 3531 WP 038905837 1 Dzeae
Dickeya zeae ZJU1202 WP 016942978 1 Dzeae
Dickeya zeae DZ2Q WP 016942978 1 Dzeae
Dickeya zeae EC1 WP 016942978 1 Dzeae
Bacillus cellulosilyticus DSM 2522 YP 004094207 1 649639 Firmicutes
Geobacillus kaustophilus HTA426 YP 147759 1 235909 Firmicutes
Anoxybacillus flavithermus WK1 YP 002314898 1 491915 Firmicutes
Serratia marcescens WW4 YP 007407105 1 435998 Proteobacteria
Cronobacter turicensis z3032 YP 003209056 1 693216 Proteobacteria
Salmonella enterica subsp enterica serovar Javiana str CFSAN001992 YP 007471843 1 1267753 Proteobacteria
Prodigiosinella confusarubida ATCC39006 WP 021014257 1 Pconfusarubida
Salmonella enterica subsp enterica serovar Schwarzengrund str CVM19633 YP 002116611 1 439843 Proteobacteria
Salmonella enterica subsp enterica serovar Typhimurium str ST4 74 YP 005244666 1 909946 Proteobacteria
Salmonella enterica subsp enterica serovar Typhimurium str 14028S YP 005239914 1 588858 Proteobacteria
Salmonella enterica subsp enterica serovar 4 5 12 i str 08 1736 YP 008263580 1 866913 Proteobacteria
Pantoea sp At 9b YP 004115755 1 592316 Proteobacteria
Pantoea ananatis LMG 20103 YP 003518653 1 706191 Proteobacteria
Pectobacterium sp SCC3193 YP 006284976 1 1166016 Proteobacteria
Serratia liquefaciens ATCC 27592 YP 008232021 1 1346614 Proteobacteria
Serratia proteamaculans 568 YP 001480162 1 399741 Proteobacteria
Serratia sp AS12 YP 004502553 1 768490 Proteobacteria
Serratia plymuthica AS9 YP 004507505 1 768492 Proteobacteria
Melissococcus plutonius DAT561 YP 005319497 1 1090974 Firmicutes
Streptococcus constellatus subsp pharyngis C232 YP 008496031 1 696216 Firmicutes
Streptococcus anginosus C238 YP 008500002 1 862971 Firmicutes
Shigella flexneri 5 str 8401 YP 691633 1 373384 Proteobacteria
Shigella flexneri 2002017 YP 005729937 1 591020 Proteobacteria
Shigella flexneri 2a str 2457T NP 839740 1 198215 Proteobacteria
Escherichia coli O83 H1 str NRG 857C YP 006122608 1 685038 Proteobacteria
Shigella sonnei 53G YP 005459532 1 216599 Proteobacteria
Enterobacteriaceae bacterium strain FGI 57 YP 007342149 1 693444 Proteobacteria
Klebsiella oxytoca E718 YP 006496238 1 1191061 Proteobacteria
Enterobacter aerogenes EA1509E YP 007389694 1 935296 Proteobacteria
Salmonella enterica subsp enterica serovar Heidelberg str CFSAN002069 YP 008250849 1 1271864 Proteobacteria
Salmonella enterica subsp enterica serovar Javiana str CFSAN001992 YP 007472128 1 1267753 Proteobacteria
Salmonella enterica subsp enterica serovar Schwarzengrund str CVM19633 YP 002117328 1 439843 Proteobacteria
Escherichia coli O55 H7 str CB9615 YP 003502428 1 701177 Proteobacteria
Escherichia coli O103 H2 str 12009 YP 003224806 1 585395 Proteobacteria
Escherichia coli O111 H str 11128 YP 003237296 1 585396 Proteobacteria
Escherichia coli O104 H4 str 2009EL 2050 YP 006772125 1 1134782 Proteobacteria
Escherichia coli O26 H11 str 11368 YP 003232241 1 573235 Proteobacteria
Escherichia coli O157 H7 str EDL933 NP 290828 1 155864 Proteobacteria
Escherichia coli Xuzhou21 YP 006315720 1 741093 Proteobacteria
Escherichia coli E24377A YP 001465698 1 331111 Proteobacteria
Shigella dysenteriae Sd197 YP 405746 1 300267 Proteobacteria
Escherichia coli 042 YP 006098934 1 216592 Proteobacteria
Escherichia coli O7 K1 str CE10 YP 006146889 1 1072459 Proteobacteria
Escherichia fergusonii ATCC 35469 YP 002385260 1 585054 Proteobacteria
Escherichia coli str K 12 substr W3110 YP 492340 1 316407 Proteobacteria
Escherichia coli B str REL606 YP 003047235 1 413997 Proteobacteria
Escherichia coli ATCC 8739 YP 001726746 1 481805 Proteobacteria

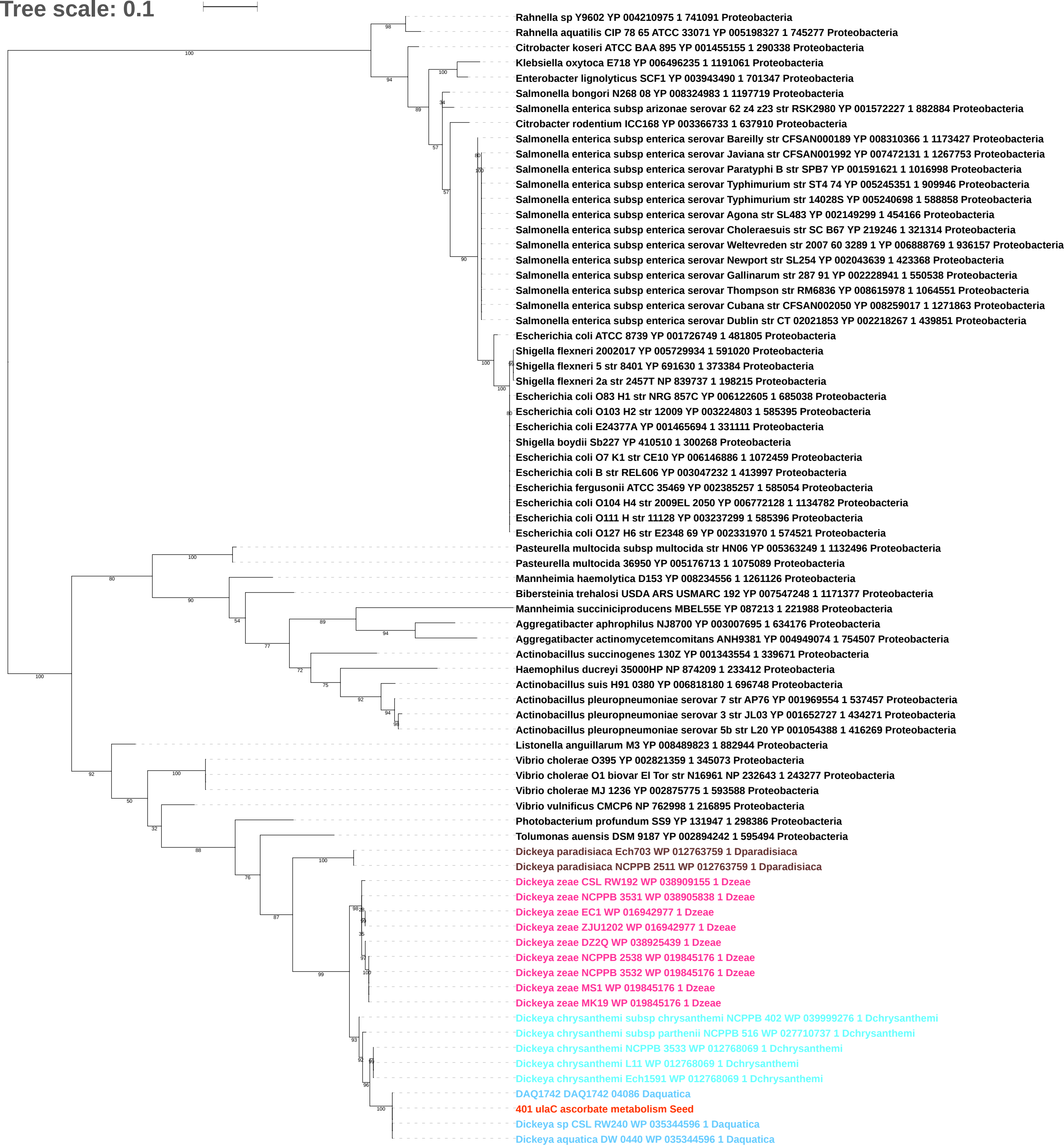
Tree scale: 0.1
Rahnella sp Y9602 YP 004210975 1 741091 Proteobacteria
Rahnella aquatilis CIP 78 65 ATCC 33071 YP 005198327 1 745277 Proteobacteria
Citrobacter koseri ATCC BAA 895 YP 001455155 1 290338 Proteobacteria
Klebsiella oxytoca E718 YP 006496235 1 1191061 Proteobacteria
Enterobacter lignolyticus SCF1 YP 003943490 1 701347 Proteobacteria
Salmonella bongori N268 08 YP 008324983 1 1197719 Proteobacteria
Salmonella enterica subsp arizonae serovar 62 z4 z23 str RSK2980 YP 001572227 1 882884 Proteobacteria
Citrobacter rodentium ICC168 YP 003366733 1 637910 Proteobacteria
Salmonella enterica subsp enterica serovar Bareilly str CFSAN000189 YP 008310366 1 1173427 Proteobacteria
Salmonella enterica subsp enterica serovar Javiana str CFSAN001992 YP 007472131 1 1267753 Proteobacteria
Salmonella enterica subsp enterica serovar Paratyphi B str SPB7 YP 001591621 1 1016998 Proteobacteria
Salmonella enterica subsp enterica serovar Typhimurium str ST4 74 YP 005245351 1 909946 Proteobacteria
Salmonella enterica subsp enterica serovar Typhimurium str 14028S YP 005240698 1 588858 Proteobacteria
Salmonella enterica subsp enterica serovar Agona str SL483 YP 002149299 1 454166 Proteobacteria
Salmonella enterica subsp enterica serovar Choleraesuis str SC B67 YP 219246 1 321314 Proteobacteria
Salmonella enterica subsp enterica serovar Weltevreden str 2007 60 3289 1 YP 006888769 1 936157 Proteobacteria
Salmonella enterica subsp enterica serovar Newport str SL254 YP 002043639 1 423368 Proteobacteria
Salmonella enterica subsp enterica serovar Gallinarum str 287 91 YP 002228941 1 550538 Proteobacteria
Salmonella enterica subsp enterica serovar Thompson str RM6836 YP 008615978 1 1064551 Proteobacteria
Salmonella enterica subsp enterica serovar Cubana str CFSAN002050 YP 008259017 1 1271863 Proteobacteria
Salmonella enterica subsp enterica serovar Dublin str CT 02021853 YP 002218267 1 439851 Proteobacteria
Escherichia coli ATCC 8739 YP 001726749 1 481805 Proteobacteria
Shigella flexneri 2002017 YP 005729934 1 591020 Proteobacteria
Shigella flexneri 5 str 8401 YP 691630 1 373384 Proteobacteria
Shigella flexneri 2a str 2457T NP 839737 1 198215 Proteobacteria
Escherichia coli O83 H1 str NRG 857C YP 006122605 1 685038 Proteobacteria
Escherichia coli O103 H2 str 12009 YP 003224803 1 585395 Proteobacteria
Escherichia coli E24377A YP 001465694 1 331111 Proteobacteria
Shigella boydii Sb227 YP 410510 1 300268 Proteobacteria
Escherichia coli O7 K1 str CE10 YP 006146886 1 1072459 Proteobacteria
Escherichia coli B str REL606 YP 003047232 1 413997 Proteobacteria
Escherichia fergusonii ATCC 35469 YP 002385257 1 585054 Proteobacteria
Escherichia coli O104 H4 str 2009EL 2050 YP 006772128 1 1134782 Proteobacteria
Escherichia coli O111 H str 11128 YP 003237299 1 585396 Proteobacteria
Escherichia coli O127 H6 str E2348 69 YP 002331970 1 574521 Proteobacteria
Pasteurella multocida subsp multocida str HN06 YP 005363249 1 1132496 Proteobacteria
Pasteurella multocida 36950 YP 005176713 1 1075089 Proteobacteria
Mannheimia haemolytica D153 YP 008234556 1 1261126 Proteobacteria
Bibersteinia trehalosi USDA ARS USMARC 192 YP 007547248 1 1171377 Proteobacteria
Mannheimia succiniciproducens MBEL55E YP 087213 1 221988 Proteobacteria
Aggregatibacter aphrophilus NJ8700 YP 003007695 1 634176 Proteobacteria
Aggregatibacter actinomycetemcomitans ANH9381 YP 004949074 1 754507 Proteobacteria
Actinobacillus succinogenes 130Z YP 001343554 1 339671 Proteobacteria
Haemophilus ducreyi 35000HP NP 874209 1 233412 Proteobacteria
Actinobacillus suis H91 0380 YP 006818180 1 696748 Proteobacteria
Actinobacillus pleuropneumoniae serovar 7 str AP76 YP 001969554 1 537457 Proteobacteria
Actinobacillus pleuropneumoniae serovar 3 str JL03 YP 001652727 1 434271 Proteobacteria
Actinobacillus pleuropneumoniae serovar 5b str L20 YP 001054388 1 416269 Proteobacteria
Listonella anguillarum M3 YP 008489823 1 882944 Proteobacteria
Vibrio cholerae O395 YP 002821359 1 345073 Proteobacteria
Vibrio cholerae O1 biovar El Tor str N16961 NP 232643 1 243277 Proteobacteria
Vibrio cholerae MJ 1236 YP 002875775 1 593588 Proteobacteria
Vibrio vulnificus CMCP6 NP 762998 1 216895 Proteobacteria
Photobacterium profundum SS9 YP 131947 1 298386 Proteobacteria
Tolumonas auensis DSM 9187 YP 002894242 1 595494 Proteobacteria
Dickeya paradisiaca Ech703 WP 012763759 1 Dparadisiaca
Dickeya paradisiaca NCPPB 2511 WP 012763759 1 Dparadisiaca
Dickeya zeae CSL RW192 WP 038909155 1 Dzeae
Dickeya zeae NCPPB 3531 WP 038905838 1 Dzeae
Dickeya zeae EC1 WP 016942977 1 Dzeae
Dickeya zeae ZJU1202 WP 016942977 1 Dzeae
Dickeya zeae DZ2Q WP 038925439 1 Dzeae
Dickeya zeae NCPPB 2538 WP 019845176 1 Dzeae
Dickeya zeae NCPPB 3532 WP 019845176 1 Dzeae
Dickeya zeae MS1 WP 019845176 1 Dzeae
Dickeya zeae MK19 WP 019845176 1 Dzeae
Dickeya chrysanthemi subsp chrysanthemi NCPPB 402 WP 039999276 1 Dchrysanthemi
Dickeya chrysanthemi subsp parthenii NCPPB 516 WP 027710737 1 Dchrysanthemi
Dickeya chrysanthemi NCPPB 3533 WP 012768069 1 Dchrysanthemi
Dickeya chrysanthemi L11 WP 012768069 1 Dchrysanthemi
Dickeya chrysanthemi Ech1591 WP 012768069 1 Dchrysanthemi
DAQ1742 DAQ1742 04086 Daquatica
401 ulaC ascorbate metabolism Seed
Dickeya sp CSL RW240 WP 035344596 1 Daquatica
Dickeya aquatica DW 0440 WP 035344596 1 Daquatica

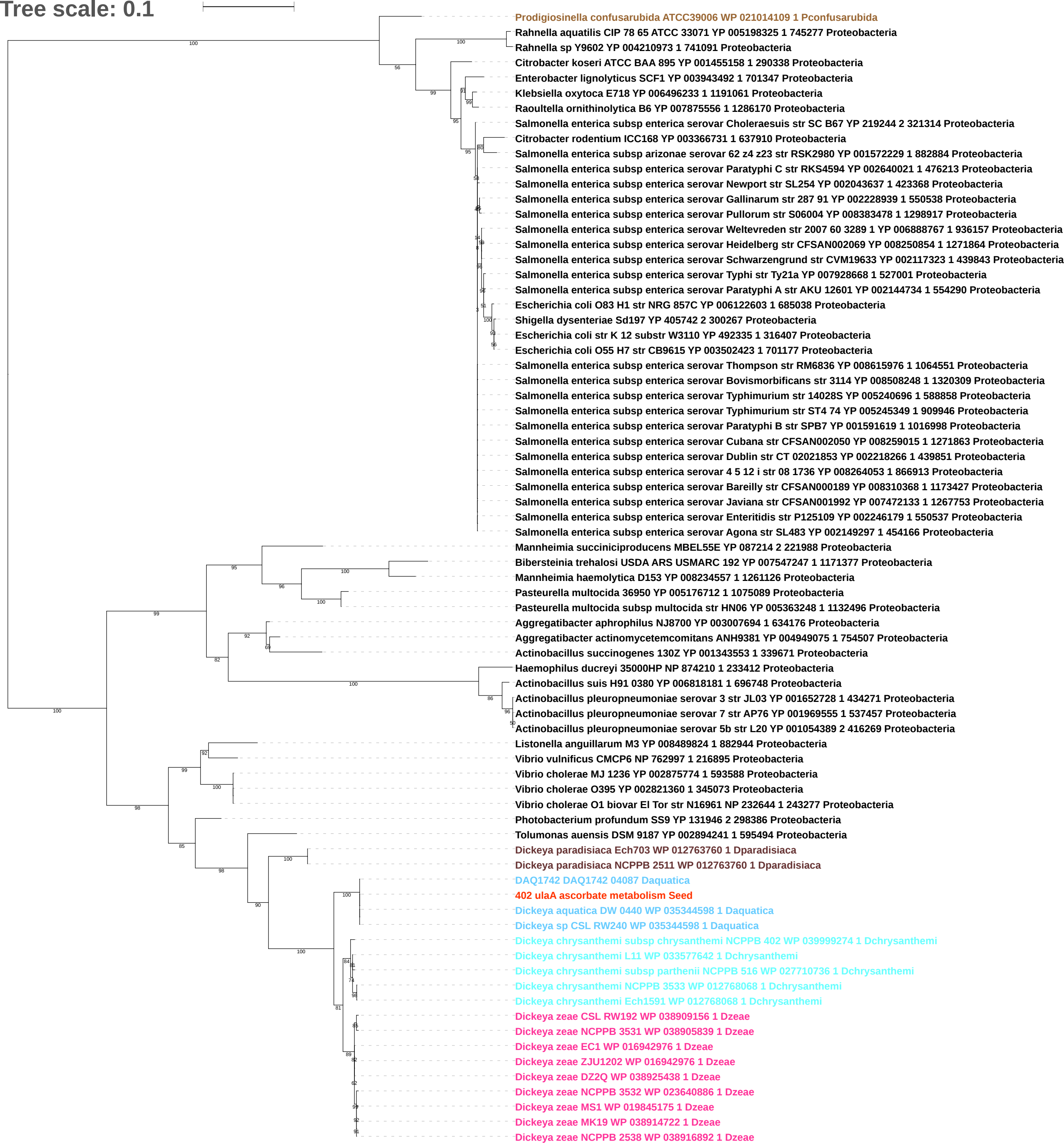
Tree scale: 0.1
Prodigiosinella confusarubida ATCC39006 WP 021014109 1 Pconfusarubida
Rahnella aquatilis CIP 78 65 ATCC 33071 YP 005198325 1 745277 Proteobacteria
Rahnella sp Y9602 YP 004210973 1 741091 Proteobacteria
Citrobacter koseri ATCC BAA 895 YP 001455158 1 290338 Proteobacteria
Enterobacter lignolyticus SCF1 YP 003943492 1 701347 Proteobacteria
Klebsiella oxytoca E718 YP 006496233 1 1191061 Proteobacteria
Raoultella ornithinolytica B6 YP 007875556 1 1286170 Proteobacteria
Salmonella enterica subsp enterica serovar Choleraesuis str SC B67 YP 219244 2 321314 Proteobacteria
Citrobacter rodentium ICC168 YP 003366731 1 637910 Proteobacteria
Salmonella enterica subsp arizonae serovar 62 z4 z23 str RSK2980 YP 001572229 1 882884 Proteobacteria
Salmonella enterica subsp enterica serovar Paratyphi C str RKS4594 YP 002640021 1 476213 Proteobacteria
Salmonella enterica subsp enterica serovar Newport str SL254 YP 002043637 1 423368 Proteobacteria
Salmonella enterica subsp enterica serovar Gallinarum str 287 91 YP 002228939 1 550538 Proteobacteria
Salmonella enterica subsp enterica serovar Pullorum str S06004 YP 008383478 1 1298917 Proteobacteria
Salmonella enterica subsp enterica serovar Weltevreden str 2007 60 3289 1 YP 006888767 1 936157 Proteobacteria
Salmonella enterica subsp enterica serovar Heidelberg str CFSAN002069 YP 008250854 1 1271864 Proteobacteria
Salmonella enterica subsp enterica serovar Schwarzengrund str CVM19633 YP 002117323 1 439843 Proteobacteria
Salmonella enterica subsp enterica serovar Typhi str Ty21a YP 007928668 1 527001 Proteobacteria
Salmonella enterica subsp enterica serovar Paratyphi A str AKU 12601 YP 002144734 1 554290 Proteobacteria
Escherichia coli O83 H1 str NRG 857C YP 006122603 1 685038 Proteobacteria
Shigella dysenteriae Sd197 YP 405742 2 300267 Proteobacteria
Escherichia coli str K 12 substr W3110 YP 492335 1 316407 Proteobacteria
Escherichia coli O55 H7 str CB9615 YP 003502423 1 701177 Proteobacteria
Salmonella enterica subsp enterica serovar Thompson str RM6836 YP 008615976 1 1064551 Proteobacteria
Salmonella enterica subsp enterica serovar Bovismorbificans str 3114 YP 008508248 1 1320309 Proteobacteria
Salmonella enterica subsp enterica serovar Typhimurium str 14028S YP 005240696 1 588858 Proteobacteria
Salmonella enterica subsp enterica serovar Typhimurium str ST4 74 YP 005245349 1 909946 Proteobacteria
Salmonella enterica subsp enterica serovar Paratyphi B str SPB7 YP 001591619 1 1016998 Proteobacteria
Salmonella enterica subsp enterica serovar Cubana str CFSAN002050 YP 008259015 1 1271863 Proteobacteria
Salmonella enterica subsp enterica serovar Dublin str CT 02021853 YP 002218266 1 439851 Proteobacteria
Salmonella enterica subsp enterica serovar 4 5 12 i str 08 1736 YP 008264053 1 866913 Proteobacteria
Salmonella enterica subsp enterica serovar Bareilly str CFSAN000189 YP 008310368 1 1173427 Proteobacteria
Salmonella enterica subsp enterica serovar Javiana str CFSAN001992 YP 007472133 1 1267753 Proteobacteria
Salmonella enterica subsp enterica serovar Enteritidis str P125109 YP 002246179 1 550537 Proteobacteria
Salmonella enterica subsp enterica serovar Agona str SL483 YP 002149297 1 454166 Proteobacteria
Mannheimia succiniciproducens MBEL55E YP 087214 2 221988 Proteobacteria
Bibersteinia trehalosi USDA ARS USMARC 192 YP 007547247 1 1171377 Proteobacteria
Mannheimia haemolytica D153 YP 008234557 1 1261126 Proteobacteria
Pasteurella multocida 36950 YP 005176712 1 1075089 Proteobacteria
Pasteurella multocida subsp multocida str HN06 YP 005363248 1 1132496 Proteobacteria
Aggregatibacter aphrophilus NJ8700 YP 003007694 1 634176 Proteobacteria
Aggregatibacter actinomycetemcomitans ANH9381 YP 004949075 1 754507 Proteobacteria
Actinobacillus succinogenes 130Z YP 001343553 1 339671 Proteobacteria
Haemophilus ducreyi 35000HP NP 874210 1 233412 Proteobacteria
Actinobacillus suis H91 0380 YP 006818181 1 696748 Proteobacteria
Actinobacillus pleuropneumoniae serovar 3 str JL03 YP 001652728 1 434271 Proteobacteria
Actinobacillus pleuropneumoniae serovar 7 str AP76 YP 001969555 1 537457 Proteobacteria
Actinobacillus pleuropneumoniae serovar 5b str L20 YP 001054389 2 416269 Proteobacteria
Listonella anguillarum M3 YP 008489824 1 882944 Proteobacteria
Vibrio vulnificus CMCP6 NP 762997 1 216895 Proteobacteria
Vibrio cholerae MJ 1236 YP 002875774 1 593588 Proteobacteria
Vibrio cholerae O395 YP 002821360 1 345073 Proteobacteria
Vibrio cholerae O1 biovar El Tor str N16961 NP 232644 1 243277 Proteobacteria
Photobacterium profundum SS9 YP 131946 2 298386 Proteobacteria
Tolumonas auensis DSM 9187 YP 002894241 1 595494 Proteobacteria
Dickeya paradisiaca Ech703 WP 012763760 1 Dparadisiaca
Dickeya paradisiaca NCPPB 2511 WP 012763760 1 Dparadisiaca
DAQ1742 DAQ1742 04087 Daquatica
402 ulaA ascorbate metabolism Seed
Dickeya aquatica DW 0440 WP 035344598 1 Daquatica
Dickeya sp CSL RW240 WP 035344598 1 Daquatica
Dickeya chrysanthemi subsp chrysanthemi NCPPB 402 WP 039999274 1 Dchrysanthemi
Dickeya chrysanthemi L11 WP 033577642 1 Dchrysanthemi
Dickeya chrysanthemi subsp parthenii NCPPB 516 WP 027710736 1 Dchrysanthemi
Dickeya chrysanthemi NCPPB 3533 WP 012768068 1 Dchrysanthemi
Dickeya chrysanthemi Ech1591 WP 012768068 1 Dchrysanthemi
Dickeya zeae CSL RW192 WP 038909156 1 Dzeae
Dickeya zeae NCPPB 3531 WP 038905839 1 Dzeae
Dickeya zeae EC1 WP 016942976 1 Dzeae
Dickeya zeae ZJU1202 WP 016942976 1 Dzeae
Dickeya zeae DZ2Q WP 038925438 1 Dzeae
Dickeya zeae NCPPB 3532 WP 023640886 1 Dzeae
Dickeya zeae MS1 WP 019845175 1 Dzeae
Dickeya zeae MK19 WP 038914722 1 Dzeae
Dickeya zeae NCPPB 2538 WP 038916892 1 Dzeae

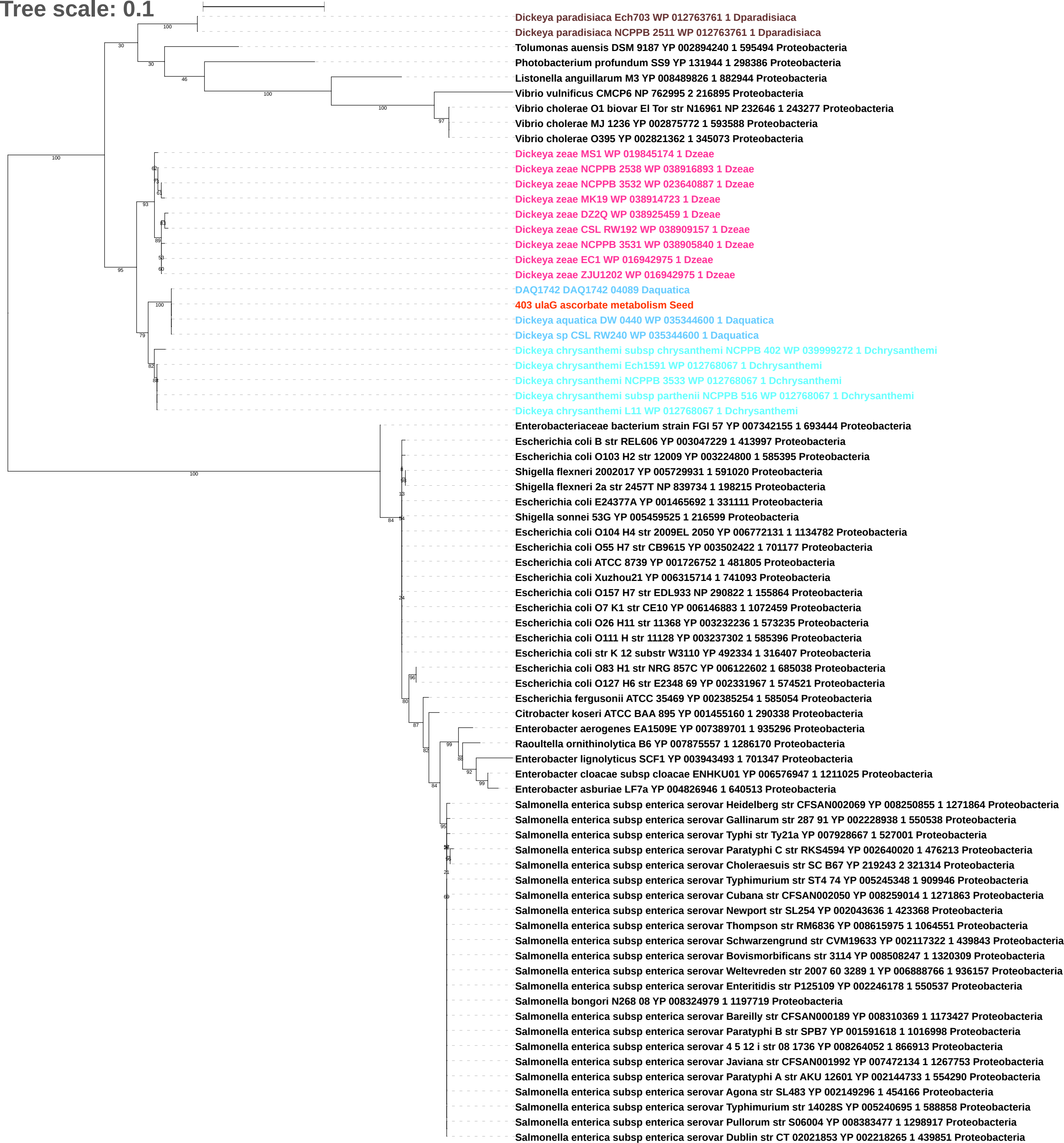
Tree scale: 0.1
Dickeya paradisiaca Ech703 WP 012763761 1 Dparadisiaca
Dickeya paradisiaca NCPPB 2511 WP 012763761 1 Dparadisiaca
Tolumonas auensis DSM 9187 YP 002894240 1 595494 Proteobacteria
Photobacterium profundum SS9 YP 131944 1 298386 Proteobacteria
Listonella anguillarum M3 YP 008489826 1 882944 Proteobacteria
Vibrio vulnificus CMCP6 NP 762995 2 216895 Proteobacteria
Vibrio cholerae O1 biovar El Tor str N16961 NP 232646 1 243277 Proteobacteria
Vibrio cholerae MJ 1236 YP 002875772 1 593588 Proteobacteria
Vibrio cholerae O395 YP 002821362 1 345073 Proteobacteria
Dickeya zeae MS1 WP 019845174 1 Dzeae
Dickeya zeae NCPPB 2538 WP 038916893 1 Dzeae
Dickeya zeae NCPPB 3532 WP 023640887 1 Dzeae
Dickeya zeae MK19 WP 038914723 1 Dzeae
Dickeya zeae DZ2Q WP 038925459 1 Dzeae
Dickeya zeae CSL RW192 WP 038909157 1 Dzeae
Dickeya zeae NCPPB 3531 WP 038905840 1 Dzeae
Dickeya zeae EC1 WP 016942975 1 Dzeae
Dickeya zeae ZJU1202 WP 016942975 1 Dzeae
DAQ1742 DAQ1742 04089 Daquatica
403 ulaG ascorbate metabolism Seed
Dickeya aquatica DW 0440 WP 035344600 1 Daquatica
Dickeya sp CSL RW240 WP 035344600 1 Daquatica
Dickeya chrysanthemi subsp chrysanthemi NCPPB 402 WP 039999272 1 Dchrysanthemi
Dickeya chrysanthemi Ech1591 WP 012768067 1 Dchrysanthemi
Dickeya chrysanthemi NCPPB 3533 WP 012768067 1 Dchrysanthemi
Dickeya chrysanthemi subsp parthenii NCPPB 516 WP 012768067 1 Dchrysanthemi
Dickeya chrysanthemi L11 WP 012768067 1 Dchrysanthemi
Enterobacteriaceae bacterium strain FGI 57 YP 007342155 1 693444 Proteobacteria
Escherichia coli B str REL606 YP 003047229 1 413997 Proteobacteria
Escherichia coli O103 H2 str 12009 YP 003224800 1 585395 Proteobacteria
Shigella flexneri 2002017 YP 005729931 1 591020 Proteobacteria
Shigella flexneri 2a str 2457T NP 839734 1 198215 Proteobacteria
Escherichia coli E24377A YP 001465692 1 331111 Proteobacteria
Shigella sonnei 53G YP 005459525 1 216599 Proteobacteria
Escherichia coli O104 H4 str 2009EL 2050 YP 006772131 1 1134782 Proteobacteria
Escherichia coli O55 H7 str CB9615 YP 003502422 1 701177 Proteobacteria
Escherichia coli ATCC 8739 YP 001726752 1 481805 Proteobacteria
Escherichia coli Xuzhou21 YP 006315714 1 741093 Proteobacteria
Escherichia coli O157 H7 str EDL933 NP 290822 1 155864 Proteobacteria
Escherichia coli O7 K1 str CE10 YP 006146883 1 1072459 Proteobacteria
Escherichia coli O26 H11 str 11368 YP 003232236 1 573235 Proteobacteria
Escherichia coli O111 H str 11128 YP 003237302 1 585396 Proteobacteria
Escherichia coli str K 12 substr W3110 YP 492334 1 316407 Proteobacteria
Escherichia coli O83 H1 str NRG 857C YP 006122602 1 685038 Proteobacteria
Escherichia coli O127 H6 str E2348 69 YP 002331967 1 574521 Proteobacteria
Escherichia fergusonii ATCC 35469 YP 002385254 1 585054 Proteobacteria
Citrobacter koseri ATCC BAA 895 YP 001455160 1 290338 Proteobacteria
Enterobacter aerogenes EA1509E YP 007389701 1 935296 Proteobacteria
Raoultella ornithinolytica B6 YP 007875557 1 1286170 Proteobacteria
Enterobacter lignolyticus SCF1 YP 003943493 1 701347 Proteobacteria
Enterobacter cloacae subsp cloacae ENHKU01 YP 006576947 1 1211025 Proteobacteria
Enterobacter asburiae LF7a YP 004826946 1 640513 Proteobacteria
Salmonella enterica subsp enterica serovar Heidelberg str CFSAN002069 YP 008250855 1 1271864 Proteobacteria
Salmonella enterica subsp enterica serovar Gallinarum str 287 91 YP 002228938 1 550538 Proteobacteria
Salmonella enterica subsp enterica serovar Typhi str Ty21a YP 007928667 1 527001 Proteobacteria
Salmonella enterica subsp enterica serovar Paratyphi C str RKS4594 YP 002640020 1 476213 Proteobacteria
Salmonella enterica subsp enterica serovar Choleraesuis str SC B67 YP 219243 2 321314 Proteobacteria
Salmonella enterica subsp enterica serovar Typhimurium str ST4 74 YP 005245348 1 909946 Proteobacteria
Salmonella enterica subsp enterica serovar Cubana str CFSAN002050 YP 008259014 1 1271863 Proteobacteria
Salmonella enterica subsp enterica serovar Newport str SL254 YP 002043636 1 423368 Proteobacteria
Salmonella enterica subsp enterica serovar Thompson str RM6836 YP 008615975 1 1064551 Proteobacteria
Salmonella enterica subsp enterica serovar Schwarzengrund str CVM19633 YP 002117322 1 439843 Proteobacteria
Salmonella enterica subsp enterica serovar Bovismorbificans str 3114 YP 008508247 1 1320309 Proteobacteria
Salmonella enterica subsp enterica serovar Weltevreden str 2007 60 3289 1 YP 006888766 1 936157 Proteobacteria
Salmonella enterica subsp enterica serovar Enteritidis str P125109 YP 002246178 1 550537 Proteobacteria
Salmonella bongori N268 08 YP 008324979 1 1197719 Proteobacteria
Salmonella enterica subsp enterica serovar Bareilly str CFSAN000189 YP 008310369 1 1173427 Proteobacteria
Salmonella enterica subsp enterica serovar Paratyphi B str SPB7 YP 001591618 1 1016998 Proteobacteria
Salmonella enterica subsp enterica serovar 4 5 12 i str 08 1736 YP 008264052 1 866913 Proteobacteria
Salmonella enterica subsp enterica serovar Javiana str CFSAN001992 YP 007472134 1 1267753 Proteobacteria
Salmonella enterica subsp enterica serovar Paratyphi A str AKU 12601 YP 002144733 1 554290 Proteobacteria
Salmonella enterica subsp enterica serovar Agona str SL483 YP 002149296 1 454166 Proteobacteria
Salmonella enterica subsp enterica serovar Typhimurium str 14028S YP 005240695 1 588858 Proteobacteria
Salmonella enterica subsp enterica serovar Pullorum str S06004 YP 008383477 1 1298917 Proteobacteria
Salmonella enterica subsp enterica serovar Dublin str CT 02021853 YP 002218265 1 439851 Proteobacteria

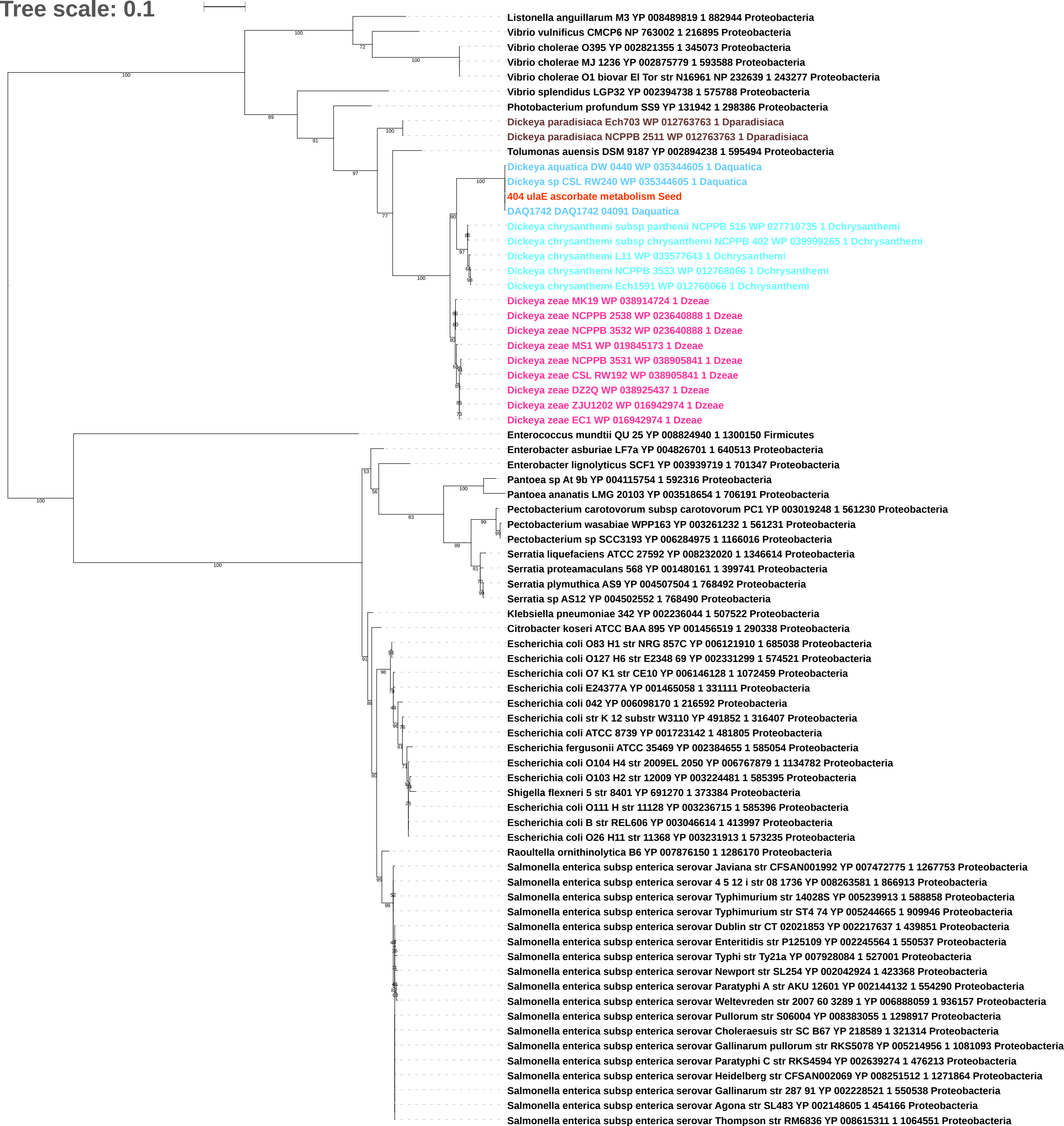
Tree scale: 0.1
Listonella anguillarum M3 YP 008489819 1 882944 Proteobacteria
Vibrio vulnificus CMCP6 NP 763002 1 216895 Proteobacteria
Vibrio cholerae O395 YP 002821355 1 345073 Proteobacteria
Vibrio cholerae MJ 1236 YP 002875779 1 593588 Proteobacteria
Vibrio cholerae O1 biovar El Tor str N16961 NP 232639 1 243277 Proteobacteria
Vibrio splendidus LGP32 YP 002394738 1 575788 Proteobacteria
Photobacterium profundum SS9 YP 131942 1 298386 Proteobacteria
Dickeya paradisiaca Ech703 WP 012763763 1 Dparadisiaca
Dickeya paradisiaca NCPPB 2511 WP 012763763 1 Dparadisiaca
Tolumonas auensis DSM 9187 YP 002894238 1 595494 Proteobacteria
Dickeya aquatica DW 0440 WP 035344605 1 Daquatica
Dickeya sp CSL RW240 WP 035344605 1 Daquatica
404 ulaE ascorbate metabolism Seed
DAQ1742 DAQ1742 04091 Daquatica
Dickeya chrysanthemi subsp parthenii NCPPB 516 WP 027710735 1 Dchrysanthemi
Dickeya chrysanthemi subsp chrysanthemi NCPPB 402 WP 039999265 1 Dchrysanthemi
Dickeya chrysanthemi L11 WP 033577643 1 Dchrysanthemi
Dickeya chrysanthemi NCPPB 3533 WP 012768066 1 Dchrysanthemi
Dickeya chrysanthemi Ech1591 WP 012768066 1 Dchrysanthemi
Dickeya zeae MK19 WP 038914724 1 Dzeae
Dickeya zeae NCPPB 2538 WP 023640888 1 Dzeae
Dickeya zeae NCPPB 3532 WP 023640888 1 Dzeae
Dickeya zeae MS1 WP 019845173 1 Dzeae
Dickeya zeae NCPPB 3531 WP 038905841 1 Dzeae
Dickeya zeae CSL RW192 WP 038905841 1 Dzeae
Dickeya zeae DZ2Q WP 038925437 1 Dzeae
Dickeya zeae ZJU1202 WP 016942974 1 Dzeae
Dickeya zeae EC1 WP 016942974 1 Dzeae
Enterococcus mundtii QU 25 YP 008824940 1 1300150 Firmicutes
Enterobacter asburiae LF7a YP 004826701 1 640513 Proteobacteria
Enterobacter lignolyticus SCF1 YP 003939719 1 701347 Proteobacteria
Pantoea sp At 9b YP 004115754 1 592316 Proteobacteria
Pantoea ananatis LMG 20103 YP 003518654 1 706191 Proteobacteria
Pectobacterium carotovorum subsp carotovorum PC1 YP 003019248 1 561230 Proteobacteria
Pectobacterium wasabiae WPP163 YP 003261232 1 561231 Proteobacteria
Pectobacterium sp SCC3193 YP 006284975 1 1166016 Proteobacteria
Serratia liquefaciens ATCC 27592 YP 008232020 1 1346614 Proteobacteria
Serratia proteamaculans 568 YP 001480161 1 399741 Proteobacteria
Serratia plymuthica AS9 YP 004507504 1 768492 Proteobacteria
Serratia sp AS12 YP 004502552 1 768490 Proteobacteria
Klebsiella pneumoniae 342 YP 002236044 1 507522 Proteobacteria
Citrobacter koseri ATCC BAA 895 YP 001456519 1 290338 Proteobacteria
Escherichia coli O83 H1 str NRG 857C YP 006121910 1 685038 Proteobacteria
Escherichia coli O127 H6 str E2348 69 YP 002331299 1 574521 Proteobacteria
Escherichia coli O7 K1 str CE10 YP 006146128 1 1072459 Proteobacteria
Escherichia coli E24377A YP 001465058 1 331111 Proteobacteria
Escherichia coli 042 YP 006098170 1 216592 Proteobacteria
Escherichia coli str K 12 substr W3110 YP 491852 1 316407 Proteobacteria
Escherichia coli ATCC 8739 YP 001723142 1 481805 Proteobacteria
Escherichia fergusonii ATCC 35469 YP 002384655 1 585054 Proteobacteria
Escherichia coli O104 H4 str 2009EL 2050 YP 006767879 1 1134782 Proteobacteria
Escherichia coli O103 H2 str 12009 YP 003224481 1 585395 Proteobacteria
Shigella flexneri 5 str 8401 YP 691270 1 373384 Proteobacteria
Escherichia coli O111 H str 11128 YP 003236715 1 585396 Proteobacteria
Escherichia coli B str REL606 YP 003046614 1 413997 Proteobacteria
Escherichia coli O26 H11 str 11368 YP 003231913 1 573235 Proteobacteria
Raoultella ornithinolytica B6 YP 007876150 1 1286170 Proteobacteria
Salmonella enterica subsp enterica serovar Javiana str CFSAN001992 YP 007472775 1 1267753 Proteobacteria
Salmonella enterica subsp enterica serovar 4 5 12 i str 08 1736 YP 008263581 1 866913 Proteobacteria
Salmonella enterica subsp enterica serovar Typhimurium str 14028S YP 005239913 1 588858 Proteobacteria
Salmonella enterica subsp enterica serovar Typhimurium str ST4 74 YP 005244665 1 909946 Proteobacteria
Salmonella enterica subsp enterica serovar Dublin str CT 02021853 YP 002217637 1 439851 Proteobacteria
Salmonella enterica subsp enterica serovar Enteritidis str P125109 YP 002245564 1 550537 Proteobacteria
Salmonella enterica subsp enterica serovar Typhi str Ty21a YP 007928084 1 527001 Proteobacteria
Salmonella enterica subsp enterica serovar Newport str SL254 YP 002042924 1 423368 Proteobacteria
Salmonella enterica subsp enterica serovar Paratyphi A str AKU 12601 YP 002144132 1 554290 Proteobacteria
Salmonella enterica subsp enterica serovar Weltevreden str 2007 60 3289 1 YP 006888059 1 936157 Proteobacteria
Salmonella enterica subsp enterica serovar Pullorum str S06004 YP 008383055 1 1298917 Proteobacteria
Salmonella enterica subsp enterica serovar Choleraesuis str SC B67 YP 218589 1 321314 Proteobacteria
Salmonella enterica subsp enterica serovar Gallinarum pullorum str RKS5078 YP 005214956 1 1081093 Proteobacteria
Salmonella enterica subsp enterica serovar Paratyphi C str RKS4594 YP 002639274 1 476213 Proteobacteria
Salmonella enterica subsp enterica serovar Heidelberg str CFSAN002069 YP 008251512 1 1271864 Proteobacteria
Salmonella enterica subsp enterica serovar Gallinarum str 287 91 YP 002228521 1 550538 Proteobacteria
Salmonella enterica subsp enterica serovar Agona str SL483 YP 002148605 1 454166 Proteobacteria
Salmonella enterica subsp enterica serovar Thompson str RM6836 YP 008615311 1 1064551 Proteobacteria

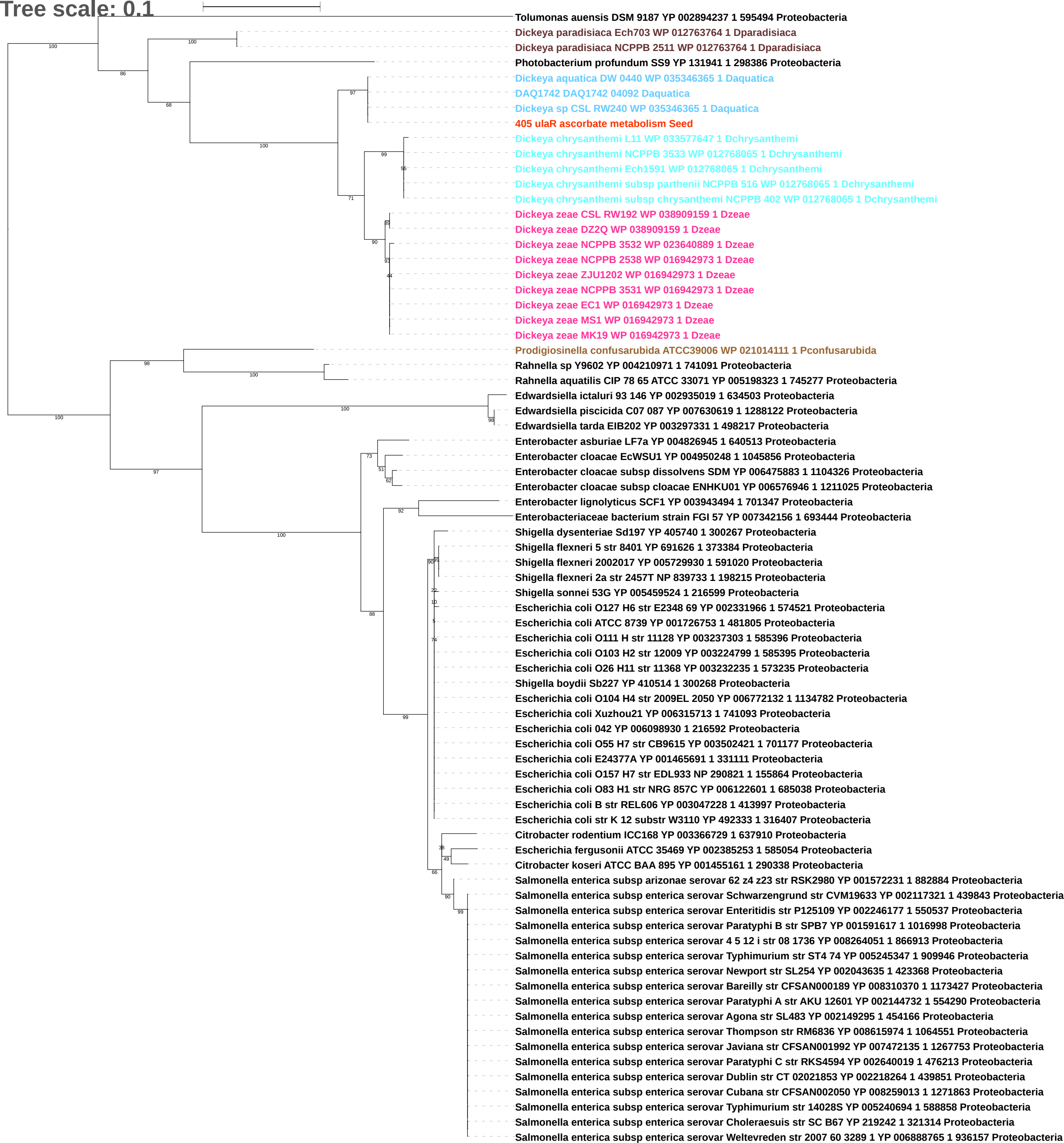
Tree scale: 0.1
Tolumonas auensis DSM 9187 YP 002894237 1 595494 Proteobacteria
Dickeya paradisiaca Ech703 WP 012763764 1 Dparadisiaca
Dickeya paradisiaca NCPPB 2511 WP 012763764 1 Dparadisiaca
Photobacterium profundum SS9 YP 131941 1 298386 Proteobacteria
Dickeya aquatica DW 0440 WP 035346365 1 Daquatica
DAQ1742 DAQ1742 04092 Daquatica
Dickeya sp CSL RW240 WP 035346365 1 Daquatica
405 ulaR ascorbate metabolism Seed
Dickeya chrysanthemi L11 WP 033577647 1 Dchrysanthemi
Dickeya chrysanthemi NCPPB 3533 WP 012768065 1 Dchrysanthemi
Dickeya chrysanthemi Ech1591 WP 012768065 1 Dchrysanthemi
Dickeya chrysanthemi subsp parthenii NCPPB 516 WP 012768065 1 Dchrysanthemi
Dickeya chrysanthemi subsp chrysanthemi NCPPB 402 WP 012768065 1 Dchrysanthemi
Dickeya zeae CSL RW192 WP 038909159 1 Dzeae
Dickeya zeae DZ2Q WP 038909159 1 Dzeae
Dickeya zeae NCPPB 3532 WP 023640889 1 Dzeae
Dickeya zeae NCPPB 2538 WP 016942973 1 Dzeae
Dickeya zeae ZJU1202 WP 016942973 1 Dzeae
Dickeya zeae NCPPB 3531 WP 016942973 1 Dzeae
Dickeya zeae EC1 WP 016942973 1 Dzeae
Dickeya zeae MS1 WP 016942973 1 Dzeae
Dickeya zeae MK19 WP 016942973 1 Dzeae
Prodigiosinella confusarubida ATCC39006 WP 021014111 1 Pconfusarubida
Rahnella sp Y9602 YP 004210971 1 741091 Proteobacteria
Rahnella aquatilis CIP 78 65 ATCC 33071 YP 005198323 1 745277 Proteobacteria
Edwardsiella ictaluri 93 146 YP 002935019 1 634503 Proteobacteria
Edwardsiella piscicida C07 087 YP 007630619 1 1288122 Proteobacteria
Edwardsiella tarda EIB202 YP 003297331 1 498217 Proteobacteria
Enterobacter asburiae LF7a YP 004826945 1 640513 Proteobacteria
Enterobacter cloacae EcWSU1 YP 004950248 1 1045856 Proteobacteria
Enterobacter cloacae subsp dissolvens SDM YP 006475883 1 1104326 Proteobacteria
Enterobacter cloacae subsp cloacae ENHKU01 YP 006576946 1 1211025 Proteobacteria
Enterobacter lignolyticus SCF1 YP 003943494 1 701347 Proteobacteria
Enterobacteriaceae bacterium strain FGI 57 YP 007342156 1 693444 Proteobacteria
Shigella dysenteriae Sd197 YP 405740 1 300267 Proteobacteria
Shigella flexneri 5 str 8401 YP 691626 1 373384 Proteobacteria
Shigella flexneri 2002017 YP 005729930 1 591020 Proteobacteria
Shigella flexneri 2a str 2457T NP 839733 1 198215 Proteobacteria
Shigella sonnei 53G YP 005459524 1 216599 Proteobacteria
Escherichia coli O127 H6 str E2348 69 YP 002331966 1 574521 Proteobacteria
Escherichia coli ATCC 8739 YP 001726753 1 481805 Proteobacteria
Escherichia coli O111 H str 11128 YP 003237303 1 585396 Proteobacteria
Escherichia coli O103 H2 str 12009 YP 003224799 1 585395 Proteobacteria
Escherichia coli O26 H11 str 11368 YP 003232235 1 573235 Proteobacteria
Shigella boydii Sb227 YP 410514 1 300268 Proteobacteria
Escherichia coli O104 H4 str 2009EL 2050 YP 006772132 1 1134782 Proteobacteria
Escherichia coli Xuzhou21 YP 006315713 1 741093 Proteobacteria
Escherichia coli 042 YP 006098930 1 216592 Proteobacteria
Escherichia coli O55 H7 str CB9615 YP 003502421 1 701177 Proteobacteria
Escherichia coli E24377A YP 001465691 1 331111 Proteobacteria
Escherichia coli O157 H7 str EDL933 NP 290821 1 155864 Proteobacteria
Escherichia coli O83 H1 str NRG 857C YP 006122601 1 685038 Proteobacteria
Escherichia coli B str REL606 YP 003047228 1 413997 Proteobacteria
Escherichia coli str K 12 substr W3110 YP 492333 1 316407 Proteobacteria
Citrobacter rodentium ICC168 YP 003366729 1 637910 Proteobacteria
Escherichia fergusonii ATCC 35469 YP 002385253 1 585054 Proteobacteria
Citrobacter koseri ATCC BAA 895 YP 001455161 1 290338 Proteobacteria
Salmonella enterica subsp arizonae serovar 62 z4 z23 str RSK2980 YP 001572231 1 882884 Proteobacteria
Salmonella enterica subsp enterica serovar Schwarzengrund str CVM19633 YP 002117321 1 439843 Proteobacteria
Salmonella enterica subsp enterica serovar Enteritidis str P125109 YP 002246177 1 550537 Proteobacteria
Salmonella enterica subsp enterica serovar Paratyphi B str SPB7 YP 001591617 1 1016998 Proteobacteria
Salmonella enterica subsp enterica serovar 4 5 12 i str 08 1736 YP 008264051 1 866913 Proteobacteria
Salmonella enterica subsp enterica serovar Typhimurium str ST4 74 YP 005245347 1 909946 Proteobacteria
Salmonella enterica subsp enterica serovar Newport str SL254 YP 002043635 1 423368 Proteobacteria
Salmonella enterica subsp enterica serovar Bareilly str CFSAN000189 YP 008310370 1 1173427 Proteobacteria
Salmonella enterica subsp enterica serovar Paratyphi A str AKU 12601 YP 002144732 1 554290 Proteobacteria
Salmonella enterica subsp enterica serovar Agona str SL483 YP 002149295 1 454166 Proteobacteria
Salmonella enterica subsp enterica serovar Thompson str RM6836 YP 008615974 1 1064551 Proteobacteria
Salmonella enterica subsp enterica serovar Javiana str CFSAN001992 YP 007472135 1 1267753 Proteobacteria
Salmonella enterica subsp enterica serovar Paratyphi C str RKS4594 YP 002640019 1 476213 Proteobacteria
Salmonella enterica subsp enterica serovar Dublin str CT 02021853 YP 002218264 1 439851 Proteobacteria
Salmonella enterica subsp enterica serovar Cubana str CFSAN002050 YP 008259013 1 1271863 Proteobacteria
Salmonella enterica subsp enterica serovar Typhimurium str 14028S YP 005240694 1 588858 Proteobacteria
Salmonella enterica subsp enterica serovar Choleraesuis str SC B67 YP 219242 1 321314 Proteobacteria
Salmonella enterica subsp enterica serovar Weltevreden str 2007 60 3289 1 YP 006888765 1 936157 Proteobacteria

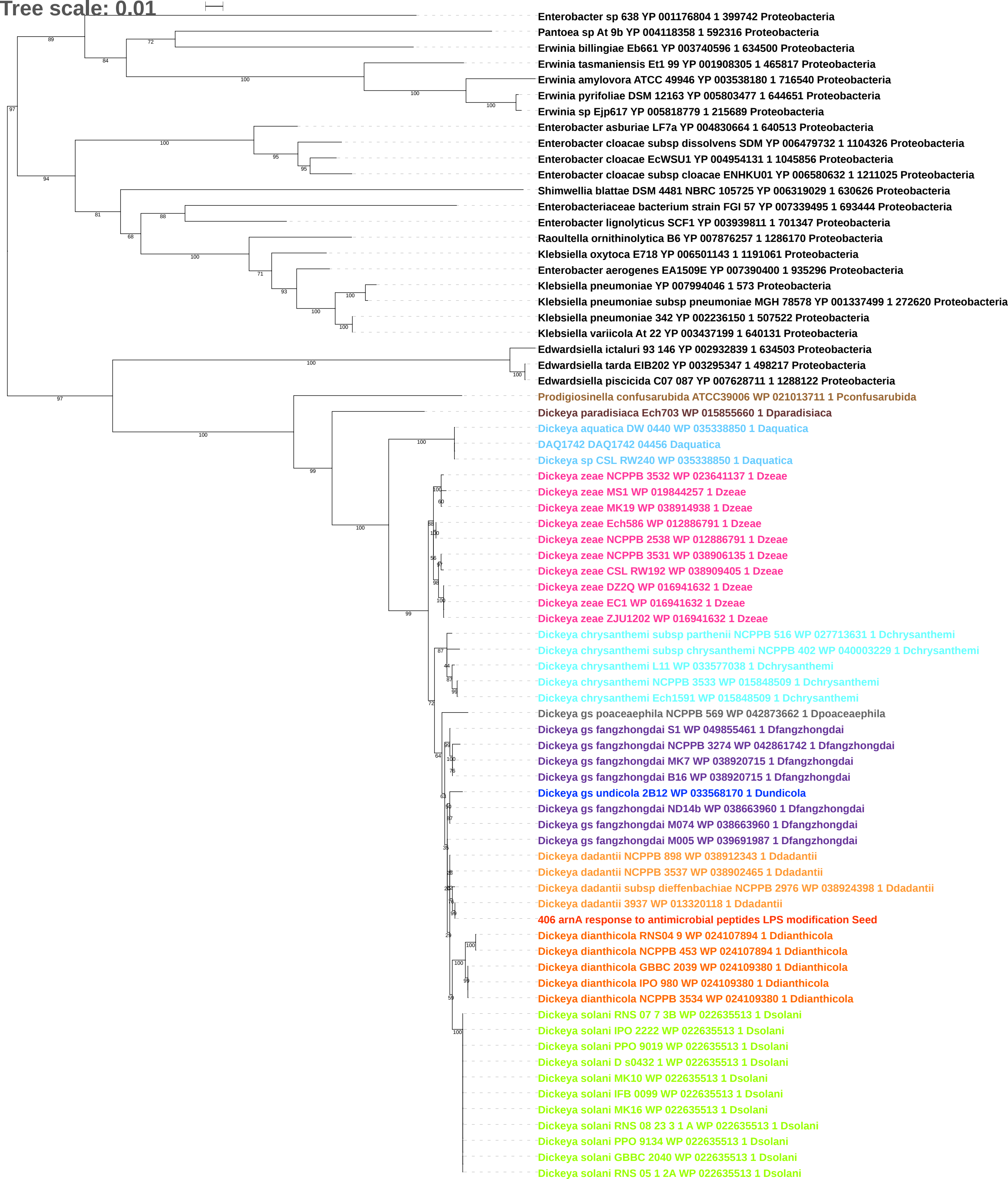
Tree scale: 0.01
Enterobacter sp 638 YP 001176804 1 399742 Proteobacteria
Pantoea sp At 9b YP 004118358 1 592316 Proteobacteria
Erwinia billingiae Eb661 YP 003740596 1 634500 Proteobacteria
Erwinia tasmaniensis Et1 99 YP 001908305 1 465817 Proteobacteria
Erwinia amylovora ATCC 49946 YP 003538180 1 716540 Proteobacteria
Erwinia pyrifoliae DSM 12163 YP 005803477 1 644651 Proteobacteria
Erwinia sp Ejp617 YP 005818779 1 215689 Proteobacteria
Enterobacter asburiae LF7a YP 004830664 1 640513 Proteobacteria
Enterobacter cloacae subsp dissolvens SDM YP 006479732 1 1104326 Proteobacteria
Enterobacter cloacae EcWSU1 YP 004954131 1 1045856 Proteobacteria
Enterobacter cloacae subsp cloacae ENHKU01 YP 006580632 1 1211025 Proteobacteria
Shimwellia blattae DSM 4481 NBRC 105725 YP 006319029 1 630626 Proteobacteria
Enterobacteriaceae bacterium strain FGI 57 YP 007339495 1 693444 Proteobacteria
Enterobacter lignolyticus SCF1 YP 003939811 1 701347 Proteobacteria
Raoultella ornithinolytica B6 YP 007876257 1 1286170 Proteobacteria
Klebsiella oxytoca E718 YP 006501143 1 1191061 Proteobacteria
Enterobacter aerogenes EA1509E YP 007390400 1 935296 Proteobacteria
Klebsiella pneumoniae YP 007994046 1 573 Proteobacteria
Klebsiella pneumoniae subsp pneumoniae MGH 78578 YP 001337499 1 272620 Proteobacteria
Klebsiella pneumoniae 342 YP 002236150 1 507522 Proteobacteria
Klebsiella variicola At 22 YP 003437199 1 640131 Proteobacteria
Edwardsiella ictaluri 93 146 YP 002932839 1 634503 Proteobacteria
Edwardsiella tarda EIB202 YP 003295347 1 498217 Proteobacteria
Edwardsiella piscicida C07 087 YP 007628711 1 1288122 Proteobacteria
Prodigiosinella confusarubida ATCC39006 WP 021013711 1 Pconfusarubida
Dickeya paradisiaca Ech703 WP 015855660 1 Dparadisiaca
Dickeya aquatica DW 0440 WP 035338850 1 Daquatica
DAQ1742 DAQ1742 04456 Daquatica
Dickeya sp CSL RW240 WP 035338850 1 Daquatica
Dickeya zeae NCPPB 3532 WP 023641137 1 Dzeae
Dickeya zeae MS1 WP 019844257 1 Dzeae
Dickeya zeae MK19 WP 038914938 1 Dzeae
Dickeya zeae Ech586 WP 012886791 1 Dzeae
Dickeya zeae NCPPB 2538 WP 012886791 1 Dzeae
Dickeya zeae NCPPB 3531 WP 038906135 1 Dzeae
Dickeya zeae CSL RW192 WP 038909405 1 Dzeae
Dickeya zeae DZ2Q WP 016941632 1 Dzeae
Dickeya zeae EC1 WP 016941632 1 Dzeae
Dickeya zeae ZJU1202 WP 016941632 1 Dzeae
Dickeya chrysanthemi subsp parthenii NCPPB 516 WP 027713631 1 Dchrysanthemi
Dickeya chrysanthemi subsp chrysanthemi NCPPB 402 WP 040003229 1 Dchrysanthemi
Dickeya chrysanthemi L11 WP 033577038 1 Dchrysanthemi
Dickeya chrysanthemi NCPPB 3533 WP 015848509 1 Dchrysanthemi
Dickeya chrysanthemi Ech1591 WP 015848509 1 Dchrysanthemi
Dickeya gs poaceaephila NCPPB 569 WP 042873662 1 Dpoaceaephila
Dickeya gs fangzhongdai S1 WP 049855461 1 Dfangzhongdai
Dickeya gs fangzhongdai NCPPB 3274 WP 042861742 1 Dfangzhongdai
Dickeya gs fangzhongdai MK7 WP 038920715 1 Dfangzhongdai
Dickeya gs fangzhongdai B16 WP 038920715 1 Dfangzhongdai
Dickeya gs undicola 2B12 WP 033568170 1 Dundicola
Dickeya gs fangzhongdai ND14b WP 038663960 1 Dfangzhongdai
Dickeya gs fangzhongdai M074 WP 038663960 1 Dfangzhongdai
Dickeya gs fangzhongdai M005 WP 039691987 1 Dfangzhongdai
Dickeya dadantii NCPPB 898 WP 038912343 1 Ddadantii
Dickeya dadantii NCPPB 3537 WP 038902465 1 Ddadantii
Dickeya dadantii subsp dieffenbachiae NCPPB 2976 WP 038924398 1 Ddadantii
Dickeya dadantii 3937 WP 013320118 1 Ddadantii
406 arnA response to antimicrobial peptides LPS modification Seed
Dickeya dianthicola RNS04 9 WP 024107894 1 Ddianthicola
Dickeya dianthicola NCPPB 453 WP 024107894 1 Ddianthicola
Dickeya dianthicola GBBC 2039 WP 024109380 1 Ddianthicola
Dickeya dianthicola IPO 980 WP 024109380 1 Ddianthicola
Dickeya dianthicola NCPPB 3534 WP 024109380 1 Ddianthicola
Dickeya solani RNS 07 7 3B WP 022635513 1 Dsolani
Dickeya solani IPO 2222 WP 022635513 1 Dsolani
Dickeya solani PPO 9019 WP 022635513 1 Dsolani
Dickeya solani D s0432 1 WP 022635513 1 Dsolani
Dickeya solani MK10 WP 022635513 1 Dsolani
Dickeya solani IFB 0099 WP 022635513 1 Dsolani
Dickeya solani MK16 WP 022635513 1 Dsolani
Dickeya solani RNS 08 23 3 1 A WP 022635513 1 Dsolani
Dickeya solani PPO 9134 WP 022635513 1 Dsolani
Dickeya solani GBBC 2040 WP 022635513 1 Dsolani
Dickeya solani RNS 05 1 2A WP 022635513 1 Dsolani

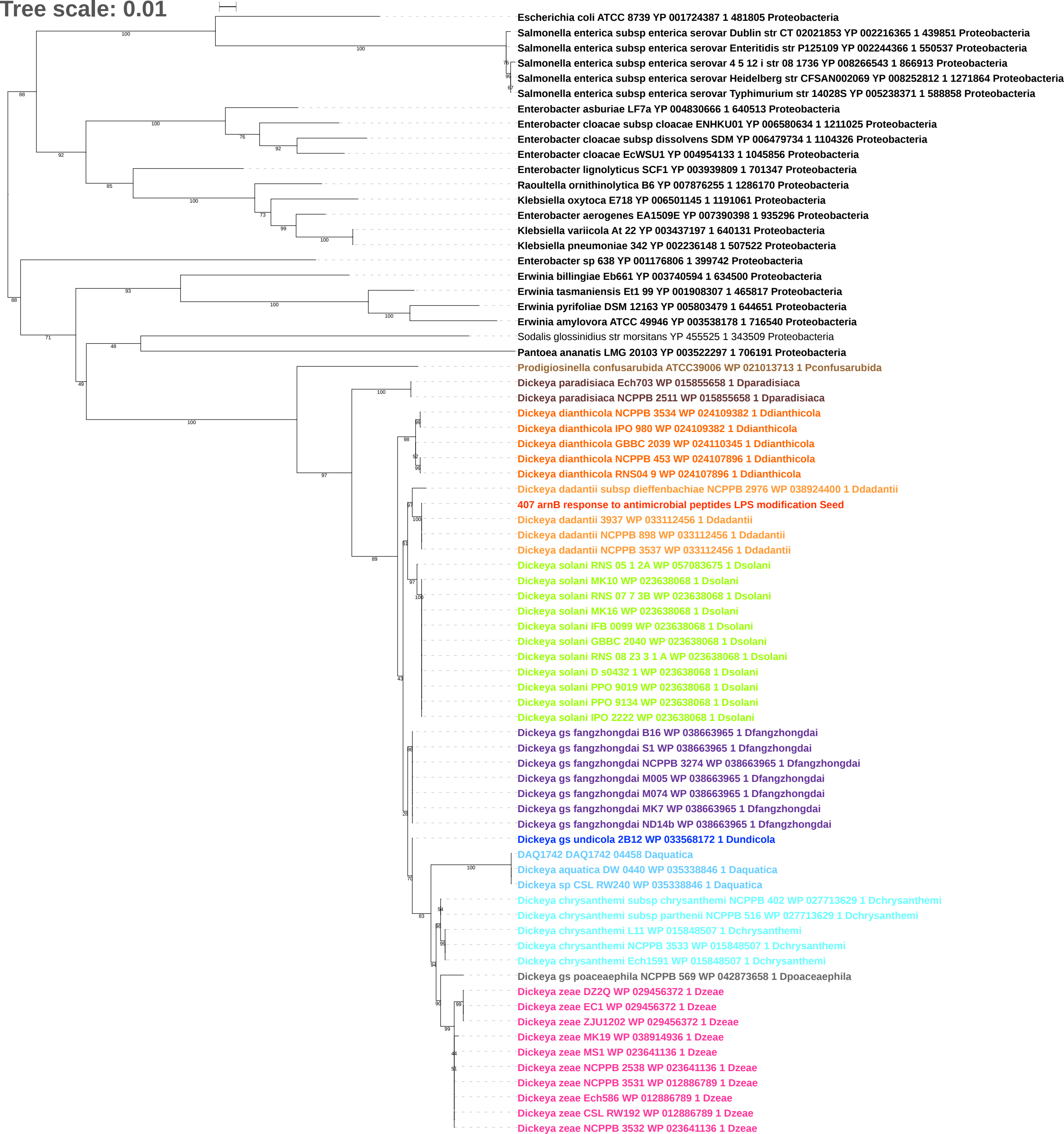
Tree scale: 0.01
Escherichia coli ATCC 8739 YP 001724387 1 481805 Proteobacteria
Salmonella enterica subsp enterica serovar Dublin str CT 02021853 YP 002216365 1 439851 Proteobacteria
Salmonella enterica subsp enterica serovar Enteritidis str P125109 YP 002244366 1 550537 Proteobacteria
Salmonella enterica subsp enterica serovar 4 5 12 i str 08 1736 YP 008266543 1 866913 Proteobacteria
Salmonella enterica subsp enterica serovar Heidelberg str CFSAN002069 YP 008252812 1 1271864 Proteobacteria
Salmonella enterica subsp enterica serovar Typhimurium str 14028S YP 005238371 1 588858 Proteobacteria
Enterobacter asburiae LF7a YP 004830666 1 640513 Proteobacteria
Enterobacter cloacae subsp cloacae ENHKU01 YP 006580634 1 1211025 Proteobacteria
Enterobacter cloacae subsp dissolvens SDM YP 006479734 1 1104326 Proteobacteria
Enterobacter cloacae EcWSU1 YP 004954133 1 1045856 Proteobacteria
Enterobacter lignolyticus SCF1 YP 003939809 1 701347 Proteobacteria
Raoultella ornithinolytica B6 YP 007876255 1 1286170 Proteobacteria
Klebsiella oxytoca E718 YP 006501145 1 1191061 Proteobacteria
Enterobacter aerogenes EA1509E YP 007390398 1 935296 Proteobacteria
Klebsiella variicola At 22 YP 003437197 1 640131 Proteobacteria
Klebsiella pneumoniae 342 YP 002236148 1 507522 Proteobacteria
Enterobacter sp 638 YP 001176806 1 399742 Proteobacteria
Erwinia billingiae Eb661 YP 003740594 1 634500 Proteobacteria
Erwinia tasmaniensis Et1 99 YP 001908307 1 465817 Proteobacteria
Erwinia pyrifoliae DSM 12163 YP 005803479 1 644651 Proteobacteria
Erwinia amylovora ATCC 49946 YP 003538178 1 716540 Proteobacteria
Sodalis glossinidius str morsitans YP 455525 1 343509 Proteobacteria
Pantoea ananatis LMG 20103 YP 003522297 1 706191 Proteobacteria
Prodigiosinella confusarubida ATCC39006 WP 021013713 1 Pconfusarubida
Dickeya paradisiaca Ech703 WP 015855658 1 Dparadisiaca
Dickeya paradisiaca NCPPB 2511 WP 015855658 1 Dparadisiaca
Dickeya dianthicola NCPPB 3534 WP 024109382 1 Ddianthicola
Dickeya dianthicola IPO 980 WP 024109382 1 Ddianthicola
Dickeya dianthicola GBBC 2039 WP 024110345 1 Ddianthicola
Dickeya dianthicola NCPPB 453 WP 024107896 1 Ddianthicola
Dickeya dianthicola RNS04 9 WP 024107896 1 Ddianthicola
Dickeya dadantii subsp dieffenbachiae NCPPB 2976 WP 038924400 1 Ddadantii
407 arnB response to antimicrobial peptides LPS modification Seed
Dickeya dadantii 3937 WP 033112456 1 Ddadantii
Dickeya dadantii NCPPB 898 WP 033112456 1 Ddadantii
Dickeya dadantii NCPPB 3537 WP 033112456 1 Ddadantii
Dickeya solani RNS 05 1 2A WP 057083675 1 Dsolani
Dickeya solani MK10 WP 023638068 1 Dsolani
Dickeya solani RNS 07 7 3B WP 023638068 1 Dsolani
Dickeya solani MK16 WP 023638068 1 Dsolani
Dickeya solani IFB 0099 WP 023638068 1 Dsolani
Dickeya solani GBBC 2040 WP 023638068 1 Dsolani
Dickeya solani RNS 08 23 3 1 A WP 023638068 1 Dsolani
Dickeya solani D s0432 1 WP 023638068 1 Dsolani
Dickeya solani PPO 9019 WP 023638068 1 Dsolani
Dickeya solani PPO 9134 WP 023638068 1 Dsolani
Dickeya solani IPO 2222 WP 023638068 1 Dsolani
Dickeya gs fangzhongdai B16 WP 038663965 1 Dfangzhongdai
Dickeya gs fangzhongdai S1 WP 038663965 1 Dfangzhongdai
Dickeya gs fangzhongdai NCPPB 3274 WP 038663965 1 Dfangzhongdai
Dickeya gs fangzhongdai M005 WP 038663965 1 Dfangzhongdai
Dickeya gs fangzhongdai M074 WP 038663965 1 Dfangzhongdai
Dickeya gs fangzhongdai MK7 WP 038663965 1 Dfangzhongdai
Dickeya gs fangzhongdai ND14b WP 038663965 1 Dfangzhongdai
Dickeya gs undicola 2B12 WP 033568172 1 Dundicola
DAQ1742 DAQ1742 04458 Daquatica
Dickeya aquatica DW 0440 WP 035338846 1 Daquatica
Dickeya sp CSL RW240 WP 035338846 1 Daquatica
Dickeya chrysanthemi subsp chrysanthemi NCPPB 402 WP 027713629 1 Dchrysanthemi
Dickeya chrysanthemi subsp parthenii NCPPB 516 WP 027713629 1 Dchrysanthemi
Dickeya chrysanthemi L11 WP 015848507 1 Dchrysanthemi
Dickeya chrysanthemi NCPPB 3533 WP 015848507 1 Dchrysanthemi
Dickeya chrysanthemi Ech1591 WP 015848507 1 Dchrysanthemi
Dickeya gs poaceaephila NCPPB 569 WP 042873658 1 Dpoaceaephila
Dickeya zeae DZ2Q WP 029456372 1 Dzeae
Dickeya zeae EC1 WP 029456372 1 Dzeae
Dickeya zeae ZJU1202 WP 029456372 1 Dzeae
Dickeya zeae MK19 WP 038914936 1 Dzeae
Dickeya zeae MS1 WP 023641136 1 Dzeae
Dickeya zeae NCPPB 2538 WP 023641136 1 Dzeae
Dickeya zeae NCPPB 3531 WP 012886789 1 Dzeae
Dickeya zeae Ech586 WP 012886789 1 Dzeae
Dickeya zeae CSL RW192 WP 012886789 1 Dzeae
Dickeya zeae NCPPB 3532 WP 023641136 1 Dzeae

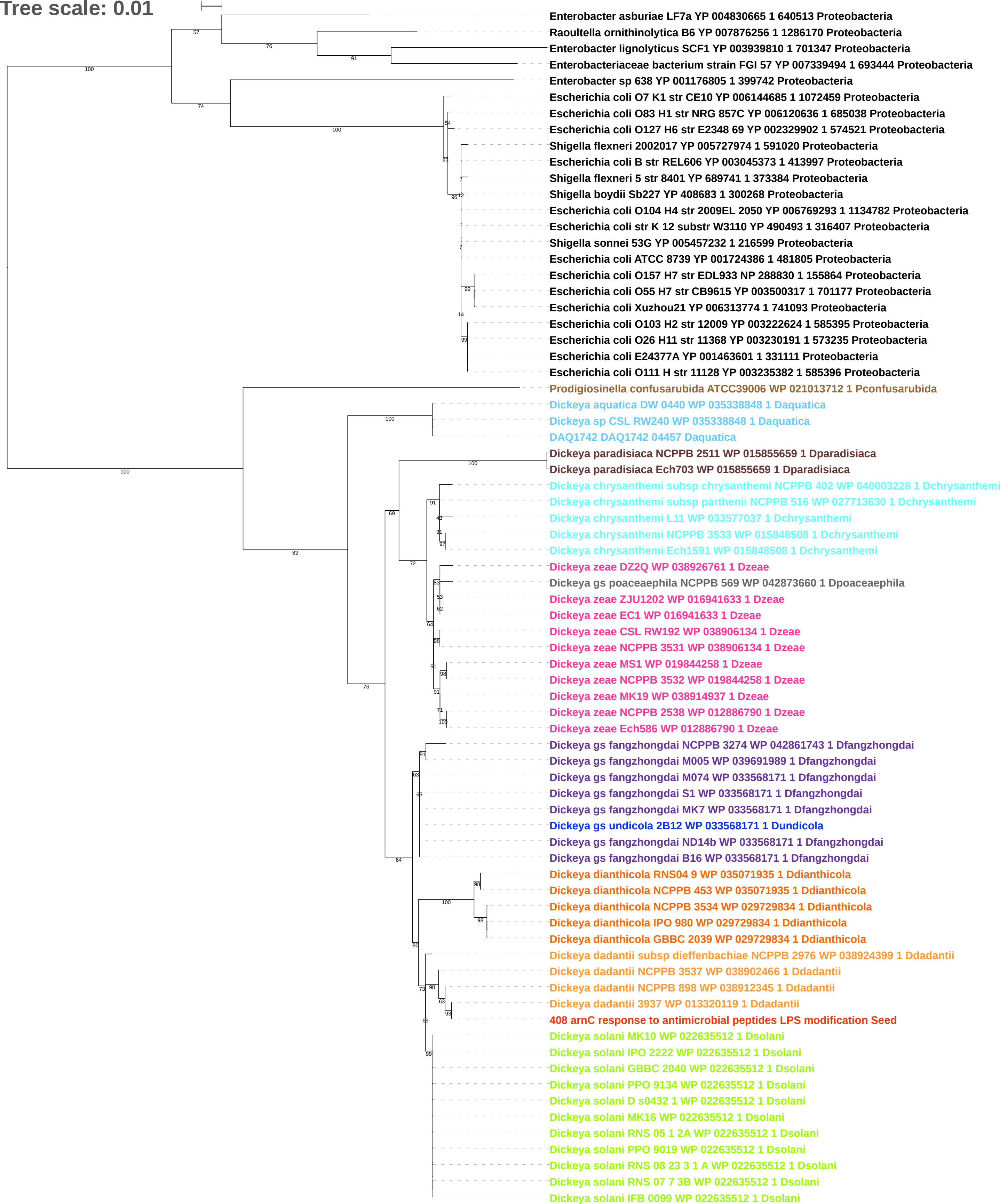
Tree scale: 0.01
Enterobacter asburiae LF7a YP 004830665 1 640513 Proteobacteria
Raoultella ornithinolytica B6 YP 007876256 1 1286170 Proteobacteria
Enterobacter lignolyticus SCF1 YP 003939810 1 701347 Proteobacteria
Enterobacteriaceae bacterium strain FGI 57 YP 007339494 1 693444 Proteobacteria
Enterobacter sp 638 YP 001176805 1 399742 Proteobacteria
Escherichia coli O7 K1 str CE10 YP 006144685 1 1072459 Proteobacteria
Escherichia coli O83 H1 str NRG 857C YP 006120636 1 685038 Proteobacteria
Escherichia coli O127 H6 str E2348 69 YP 002329902 1 574521 Proteobacteria
Shigella flexneri 2002017 YP 005727974 1 591020 Proteobacteria
Escherichia coli B str REL606 YP 003045373 1 413997 Proteobacteria
Shigella flexneri 5 str 8401 YP 689741 1 373384 Proteobacteria
Shigella boydii Sb227 YP 408683 1 300268 Proteobacteria
Escherichia coli O104 H4 str 2009EL 2050 YP 006769293 1 1134782 Proteobacteria
Escherichia coli str K 12 substr W3110 YP 490493 1 316407 Proteobacteria
Shigella sonnei 53G YP 005457232 1 216599 Proteobacteria
Escherichia coli ATCC 8739 YP 001724386 1 481805 Proteobacteria
Escherichia coli O157 H7 str EDL933 NP 288830 1 155864 Proteobacteria
Escherichia coli O55 H7 str CB9615 YP 003500317 1 701177 Proteobacteria
Escherichia coli Xuzhou21 YP 006313774 1 741093 Proteobacteria
Escherichia coli O103 H2 str 12009 YP 003222624 1 585395 Proteobacteria
Escherichia coli O26 H11 str 11368 YP 003230191 1 573235 Proteobacteria
Escherichia coli E24377A YP 001463601 1 331111 Proteobacteria
Escherichia coli O111 H str 11128 YP 003235382 1 585396 Proteobacteria
Prodigiosinella confusarubida ATCC39006 WP 021013712 1 Pconfusarubida
Dickeya aquatica DW 0440 WP 035338848 1 Daquatica
Dickeya sp CSL RW240 WP 035338848 1 Daquatica
DAQ1742 DAQ1742 04457 Daquatica
Dickeya paradisiaca NCPPB 2511 WP 015855659 1 Dparadisiaca
Dickeya paradisiaca Ech703 WP 015855659 1 Dparadisiaca
Dickeya chrysanthemi subsp chrysanthemi NCPPB 402 WP 040003228 1 Dchrysanthemi
Dickeya chrysanthemi subsp parthenii NCPPB 516 WP 027713630 1 Dchrysanthemi
Dickeya chrysanthemi L11 WP 033577037 1 Dchrysanthemi
Dickeya chrysanthemi NCPPB 3533 WP 015848508 1 Dchrysanthemi
Dickeya chrysanthemi Ech1591 WP 015848508 1 Dchrysanthemi
Dickeya zeae DZ2Q WP 038926761 1 Dzeae
Dickeya gs poaceaephila NCPPB 569 WP 042873660 1 Dpoaceaephila
Dickeya zeae ZJU1202 WP 016941633 1 Dzeae
Dickeya zeae EC1 WP 016941633 1 Dzeae
Dickeya zeae CSL RW192 WP 038906134 1 Dzeae
Dickeya zeae NCPPB 3531 WP 038906134 1 Dzeae
Dickeya zeae MS1 WP 019844258 1 Dzeae
Dickeya zeae NCPPB 3532 WP 019844258 1 Dzeae
Dickeya zeae MK19 WP 038914937 1 Dzeae
Dickeya zeae NCPPB 2538 WP 012886790 1 Dzeae
Dickeya zeae Ech586 WP 012886790 1 Dzeae
Dickeya gs fangzhongdai NCPPB 3274 WP 042861743 1 Dfangzhongdai
Dickeya gs fangzhongdai M005 WP 039691989 1 Dfangzhongdai
Dickeya gs fangzhongdai M074 WP 033568171 1 Dfangzhongdai
Dickeya gs fangzhongdai S1 WP 033568171 1 Dfangzhongdai
Dickeya gs fangzhongdai MK7 WP 033568171 1 Dfangzhongdai
Dickeya gs undicola 2B12 WP 033568171 1 Dundicola
Dickeya gs fangzhongdai ND14b WP 033568171 1 Dfangzhongdai
Dickeya gs fangzhongdai B16 WP 033568171 1 Dfangzhongdai
Dickeya dianthicola RNS04 9 WP 035071935 1 Ddianthicola
Dickeya dianthicola NCPPB 453 WP 035071935 1 Ddianthicola
Dickeya dianthicola NCPPB 3534 WP 029729834 1 Ddianthicola
Dickeya dianthicola IPO 980 WP 029729834 1 Ddianthicola
Dickeya dianthicola GBBC 2039 WP 029729834 1 Ddianthicola
Dickeya dadantii subsp dieffenbachiae NCPPB 2976 WP 038924399 1 Ddadantii
Dickeya dadantii NCPPB 3537 WP 038902466 1 Ddadantii
Dickeya dadantii NCPPB 898 WP 038912345 1 Ddadantii
Dickeya dadantii 3937 WP 013320119 1 Ddadantii
408 arnC response to antimicrobial peptides LPS modification Seed
Dickeya solani MK10 WP 022635512 1 Dsolani
Dickeya solani IPO 2222 WP 022635512 1 Dsolani
Dickeya solani GBBC 2040 WP 022635512 1 Dsolani
Dickeya solani PPO 9134 WP 022635512 1 Dsolani
Dickeya solani D s0432 1 WP 022635512 1 Dsolani
Dickeya solani MK16 WP 022635512 1 Dsolani
Dickeya solani RNS 05 1 2A WP 022635512 1 Dsolani
Dickeya solani PPO 9019 WP 022635512 1 Dsolani
Dickeya solani RNS 08 23 3 1 A WP 022635512 1 Dsolani
Dickeya solani RNS 07 7 3B WP 022635512 1 Dsolani
Dickeya solani IFB 0099 WP 022635512 1 Dsolani

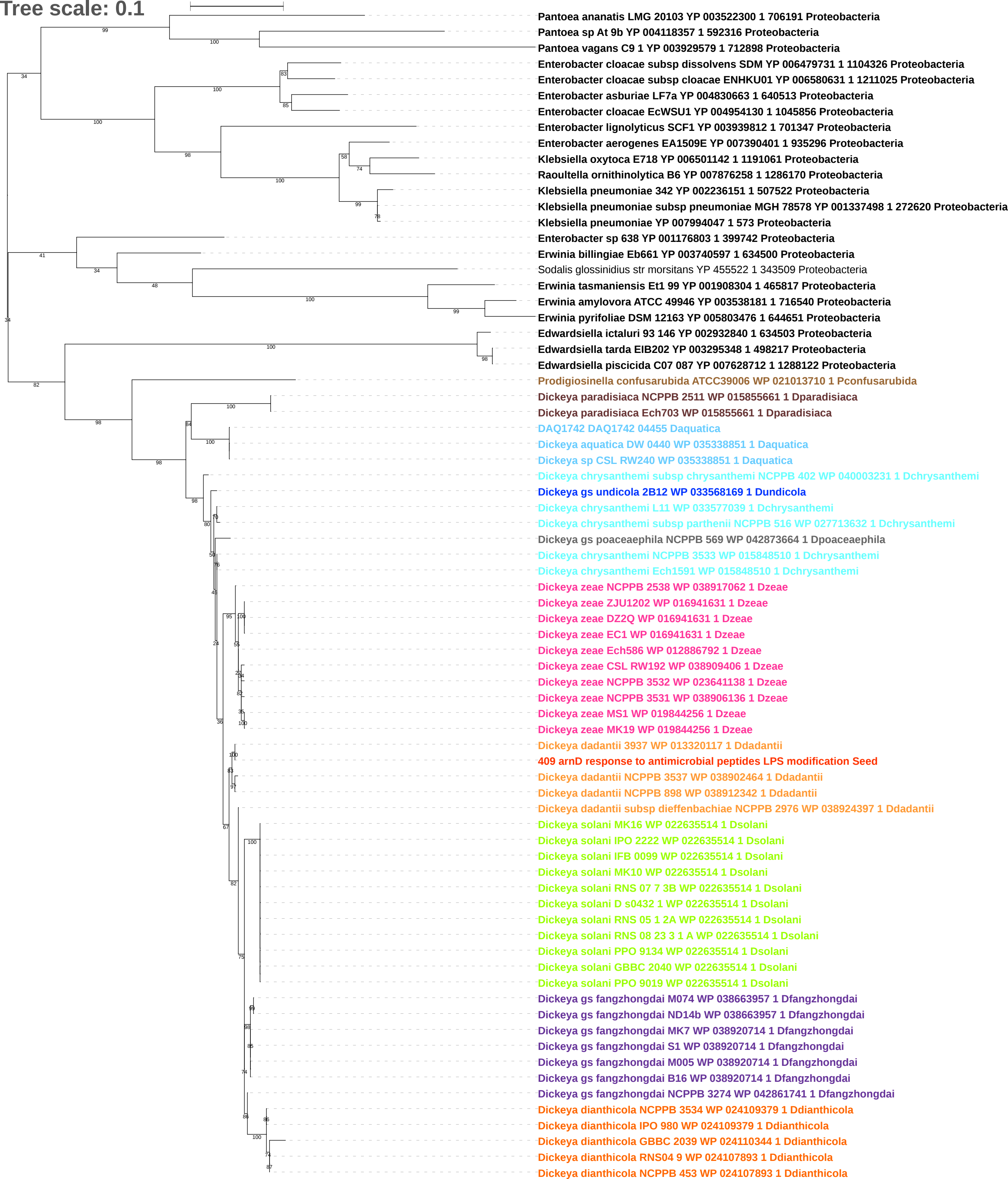
Tree scale: 0.1
Pantoea ananatis LMG 20103 YP 003522300 1 706191 Proteobacteria
Pantoea sp At 9b YP 004118357 1 592316 Proteobacteria
Pantoea vagans C9 1 YP 003929579 1 712898 Proteobacteria
Enterobacter cloacae subsp dissolvens SDM YP 006479731 1 1104326 Proteobacteria
Enterobacter cloacae subsp cloacae ENHKU01 YP 006580631 1 1211025 Proteobacteria
Enterobacter asburiae LF7a YP 004830663 1 640513 Proteobacteria
Enterobacter cloacae EcWSU1 YP 004954130 1 1045856 Proteobacteria
Enterobacter lignolyticus SCF1 YP 003939812 1 701347 Proteobacteria
Enterobacter aerogenes EA1509E YP 007390401 1 935296 Proteobacteria
Klebsiella oxytoca E718 YP 006501142 1 1191061 Proteobacteria
Raoultella ornithinolytica B6 YP 007876258 1 1286170 Proteobacteria
Klebsiella pneumoniae 342 YP 002236151 1 507522 Proteobacteria
Klebsiella pneumoniae subsp pneumoniae MGH 78578 YP 001337498 1 272620 Proteobacteria
Klebsiella pneumoniae YP 007994047 1 573 Proteobacteria
Enterobacter sp 638 YP 001176803 1 399742 Proteobacteria
Erwinia billingiae Eb661 YP 003740597 1 634500 Proteobacteria
Sodalis glossinidius str morsitans YP 455522 1 343509 Proteobacteria
Erwinia tasmaniensis Et1 99 YP 001908304 1 465817 Proteobacteria
Erwinia amylovora ATCC 49946 YP 003538181 1 716540 Proteobacteria
Erwinia pyrifoliae DSM 12163 YP 005803476 1 644651 Proteobacteria
Edwardsiella ictaluri 93 146 YP 002932840 1 634503 Proteobacteria
Edwardsiella tarda EIB202 YP 003295348 1 498217 Proteobacteria
Edwardsiella piscicida C07 087 YP 007628712 1 1288122 Proteobacteria
Prodigiosinella confusarubida ATCC39006 WP 021013710 1 Pconfusarubida
Dickeya paradisiaca NCPPB 2511 WP 015855661 1 Dparadisiaca
Dickeya paradisiaca Ech703 WP 015855661 1 Dparadisiaca
DAQ1742 DAQ1742 04455 Daquatica
Dickeya aquatica DW 0440 WP 035338851 1 Daquatica
Dickeya sp CSL RW240 WP 035338851 1 Daquatica
Dickeya chrysanthemi subsp chrysanthemi NCPPB 402 WP 040003231 1 Dchrysanthemi
Dickeya gs undicola 2B12 WP 033568169 1 Dundicola
Dickeya chrysanthemi L11 WP 033577039 1 Dchrysanthemi
Dickeya chrysanthemi subsp parthenii NCPPB 516 WP 027713632 1 Dchrysanthemi
Dickeya gs poaceaephila NCPPB 569 WP 042873664 1 Dpoaceaephila
Dickeya chrysanthemi NCPPB 3533 WP 015848510 1 Dchrysanthemi
Dickeya chrysanthemi Ech1591 WP 015848510 1 Dchrysanthemi
Dickeya zeae NCPPB 2538 WP 038917062 1 Dzeae
Dickeya zeae ZJU1202 WP 016941631 1 Dzeae
Dickeya zeae DZ2Q WP 016941631 1 Dzeae
Dickeya zeae EC1 WP 016941631 1 Dzeae
Dickeya zeae Ech586 WP 012886792 1 Dzeae
Dickeya zeae CSL RW192 WP 038909406 1 Dzeae
Dickeya zeae NCPPB 3532 WP 023641138 1 Dzeae
Dickeya zeae NCPPB 3531 WP 038906136 1 Dzeae
Dickeya zeae MS1 WP 019844256 1 Dzeae
Dickeya zeae MK19 WP 019844256 1 Dzeae
Dickeya dadantii 3937 WP 013320117 1 Ddadantii
409 arnD response to antimicrobial peptides LPS modification Seed
Dickeya dadantii NCPPB 3537 WP 038902464 1 Ddadantii
Dickeya dadantii NCPPB 898 WP 038912342 1 Ddadantii
Dickeya dadantii subsp dieffenbachiae NCPPB 2976 WP 038924397 1 Ddadantii
Dickeya solani MK16 WP 022635514 1 Dsolani
Dickeya solani IPO 2222 WP 022635514 1 Dsolani
Dickeya solani IFB 0099 WP 022635514 1 Dsolani
Dickeya solani MK10 WP 022635514 1 Dsolani
Dickeya solani RNS 07 7 3B WP 022635514 1 Dsolani
Dickeya solani D s0432 1 WP 022635514 1 Dsolani
Dickeya solani RNS 05 1 2A WP 022635514 1 Dsolani
Dickeya solani RNS 08 23 3 1 A WP 022635514 1 Dsolani
Dickeya solani PPO 9134 WP 022635514 1 Dsolani
Dickeya solani GBBC 2040 WP 022635514 1 Dsolani
Dickeya solani PPO 9019 WP 022635514 1 Dsolani
Dickeya gs fangzhongdai M074 WP 038663957 1 Dfangzhongdai
Dickeya gs fangzhongdai ND14b WP 038663957 1 Dfangzhongdai
Dickeya gs fangzhongdai MK7 WP 038920714 1 Dfangzhongdai
Dickeya gs fangzhongdai S1 WP 038920714 1 Dfangzhongdai
Dickeya gs fangzhongdai M005 WP 038920714 1 Dfangzhongdai
Dickeya gs fangzhongdai B16 WP 038920714 1 Dfangzhongdai
Dickeya gs fangzhongdai NCPPB 3274 WP 042861741 1 Dfangzhongdai
Dickeya dianthicola NCPPB 3534 WP 024109379 1 Ddianthicola
Dickeya dianthicola IPO 980 WP 024109379 1 Ddianthicola
Dickeya dianthicola GBBC 2039 WP 024110344 1 Ddianthicola
Dickeya dianthicola RNS04 9 WP 024107893 1 Ddianthicola
Dickeya dianthicola NCPPB 453 WP 024107893 1 Ddianthicola

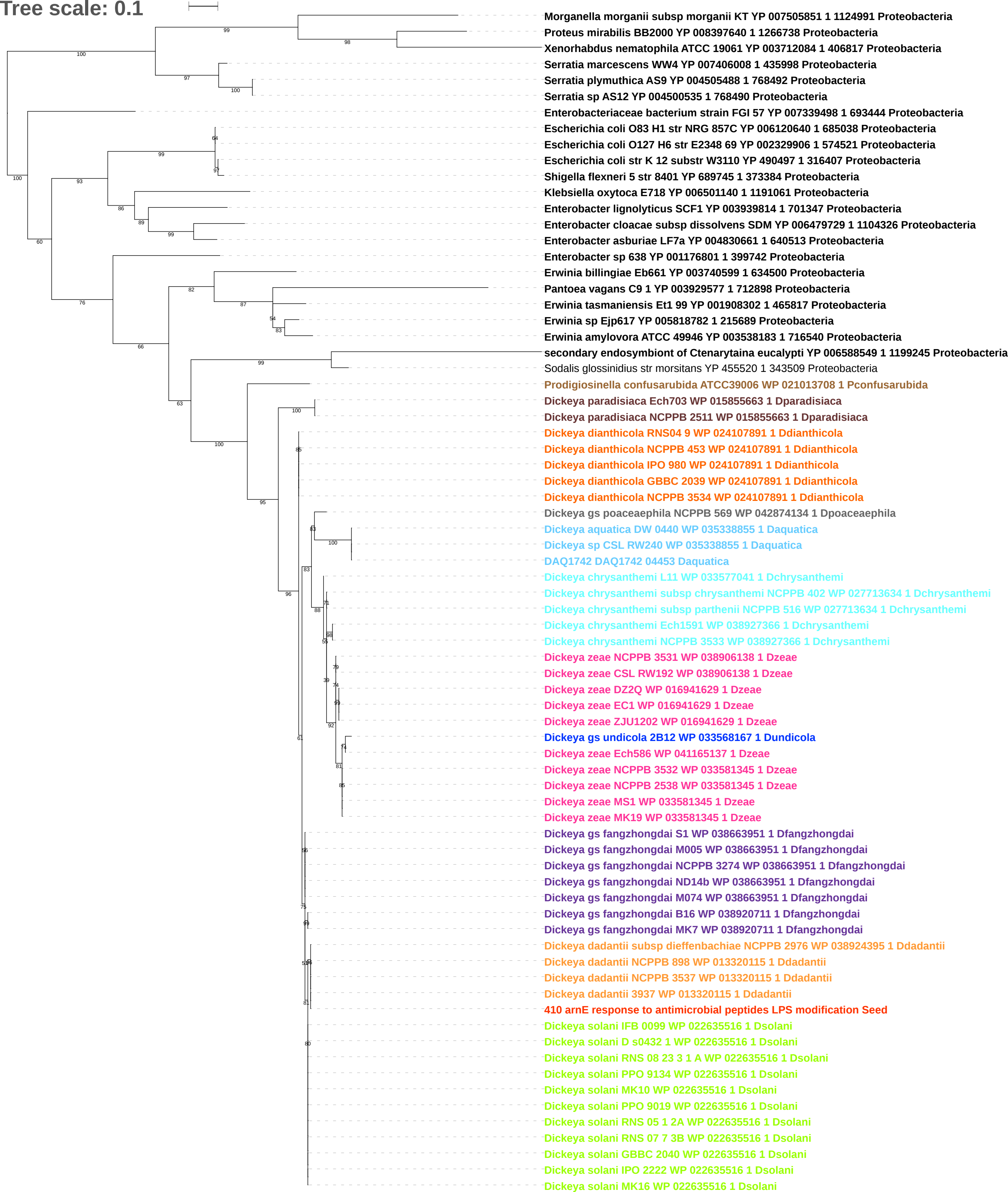
Tree scale: 0.1
Morganella morganii subsp morganii KT YP 007505851 1 1124991 Proteobacteria
Proteus mirabilis BB2000 YP 008397640 1 1266738 Proteobacteria
Xenorhabdus nematophila ATCC 19061 YP 003712084 1 406817 Proteobacteria
Serratia marcescens WW4 YP 007406008 1 435998 Proteobacteria
Serratia plymuthica AS9 YP 004505488 1 768492 Proteobacteria
Serratia sp AS12 YP 004500535 1 768490 Proteobacteria
Enterobacteriaceae bacterium strain FGI 57 YP 007339498 1 693444 Proteobacteria
Escherichia coli O83 H1 str NRG 857C YP 006120640 1 685038 Proteobacteria
Escherichia coli O127 H6 str E2348 69 YP 002329906 1 574521 Proteobacteria
Escherichia coli str K 12 substr W3110 YP 490497 1 316407 Proteobacteria
Shigella flexneri 5 str 8401 YP 689745 1 373384 Proteobacteria
Klebsiella oxytoca E718 YP 006501140 1 1191061 Proteobacteria
Enterobacter lignolyticus SCF1 YP 003939814 1 701347 Proteobacteria
Enterobacter cloacae subsp dissolvens SDM YP 006479729 1 1104326 Proteobacteria
Enterobacter asburiae LF7a YP 004830661 1 640513 Proteobacteria
Enterobacter sp 638 YP 001176801 1 399742 Proteobacteria
Erwinia billingiae Eb661 YP 003740599 1 634500 Proteobacteria
Pantoea vagans C9 1 YP 003929577 1 712898 Proteobacteria
Erwinia tasmaniensis Et1 99 YP 001908302 1 465817 Proteobacteria
Erwinia sp Ejp617 YP 005818782 1 215689 Proteobacteria
Erwinia amylovora ATCC 49946 YP 003538183 1 716540 Proteobacteria
secondary endosymbiont of Ctenarytaina eucalypti YP 006588549 1 1199245 Proteobacteria
Sodalis glossinidius str morsitans YP 455520 1 343509 Proteobacteria
Prodigiosinella confusarubida ATCC39006 WP 021013708 1 Pconfusarubida
Dickeya paradisiaca Ech703 WP 015855663 1 Dparadisiaca
Dickeya paradisiaca NCPPB 2511 WP 015855663 1 Dparadisiaca
Dickeya dianthicola RNS04 9 WP 024107891 1 Ddianthicola
Dickeya dianthicola NCPPB 453 WP 024107891 1 Ddianthicola
Dickeya dianthicola IPO 980 WP 024107891 1 Ddianthicola
Dickeya dianthicola GBBC 2039 WP 024107891 1 Ddianthicola
Dickeya dianthicola NCPPB 3534 WP 024107891 1 Ddianthicola
Dickeya gs poaceaephila NCPPB 569 WP 042874134 1 Dpoaceaephila
Dickeya aquatica DW 0440 WP 035338855 1 Daquatica
Dickeya sp CSL RW240 WP 035338855 1 Daquatica
DAQ1742 DAQ1742 04453 Daquatica
Dickeya chrysanthemi L11 WP 033577041 1 Dchrysanthemi
Dickeya chrysanthemi subsp chrysanthemi NCPPB 402 WP 027713634 1 Dchrysanthemi
Dickeya chrysanthemi subsp parthenii NCPPB 516 WP 027713634 1 Dchrysanthemi
Dickeya chrysanthemi Ech1591 WP 038927366 1 Dchrysanthemi
Dickeya chrysanthemi NCPPB 3533 WP 038927366 1 Dchrysanthemi
Dickeya zeae NCPPB 3531 WP 038906138 1 Dzeae
Dickeya zeae CSL RW192 WP 038906138 1 Dzeae
Dickeya zeae DZ2Q WP 016941629 1 Dzeae
Dickeya zeae EC1 WP 016941629 1 Dzeae
Dickeya zeae ZJU1202 WP 016941629 1 Dzeae
Dickeya gs undicola 2B12 WP 033568167 1 Dundicola
Dickeya zeae Ech586 WP 041165137 1 Dzeae
Dickeya zeae NCPPB 3532 WP 033581345 1 Dzeae
Dickeya zeae NCPPB 2538 WP 033581345 1 Dzeae
Dickeya zeae MS1 WP 033581345 1 Dzeae
Dickeya zeae MK19 WP 033581345 1 Dzeae
Dickeya gs fangzhongdai S1 WP 038663951 1 Dfangzhongdai
Dickeya gs fangzhongdai M005 WP 038663951 1 Dfangzhongdai
Dickeya gs fangzhongdai NCPPB 3274 WP 038663951 1 Dfangzhongdai
Dickeya gs fangzhongdai ND14b WP 038663951 1 Dfangzhongdai
Dickeya gs fangzhongdai M074 WP 038663951 1 Dfangzhongdai
Dickeya gs fangzhongdai B16 WP 038920711 1 Dfangzhongdai
Dickeya gs fangzhongdai MK7 WP 038920711 1 Dfangzhongdai
Dickeya dadantii subsp dieffenbachiae NCPPB 2976 WP 038924395 1 Ddadantii
Dickeya dadantii NCPPB 898 WP 013320115 1 Ddadantii
Dickeya dadantii NCPPB 3537 WP 013320115 1 Ddadantii
Dickeya dadantii 3937 WP 013320115 1 Ddadantii
410 arnE response to antimicrobial peptides LPS modification Seed
Dickeya solani IFB 0099 WP 022635516 1 Dsolani
Dickeya solani D s0432 1 WP 022635516 1 Dsolani
Dickeya solani RNS 08 23 3 1 A WP 022635516 1 Dsolani
Dickeya solani PPO 9134 WP 022635516 1 Dsolani
Dickeya solani MK10 WP 022635516 1 Dsolani
Dickeya solani PPO 9019 WP 022635516 1 Dsolani
Dickeya solani RNS 05 1 2A WP 022635516 1 Dsolani
Dickeya solani RNS 07 7 3B WP 022635516 1 Dsolani
Dickeya solani GBBC 2040 WP 022635516 1 Dsolani
Dickeya solani IPO 2222 WP 022635516 1 Dsolani
Dickeya solani MK16 WP 022635516 1 Dsolani

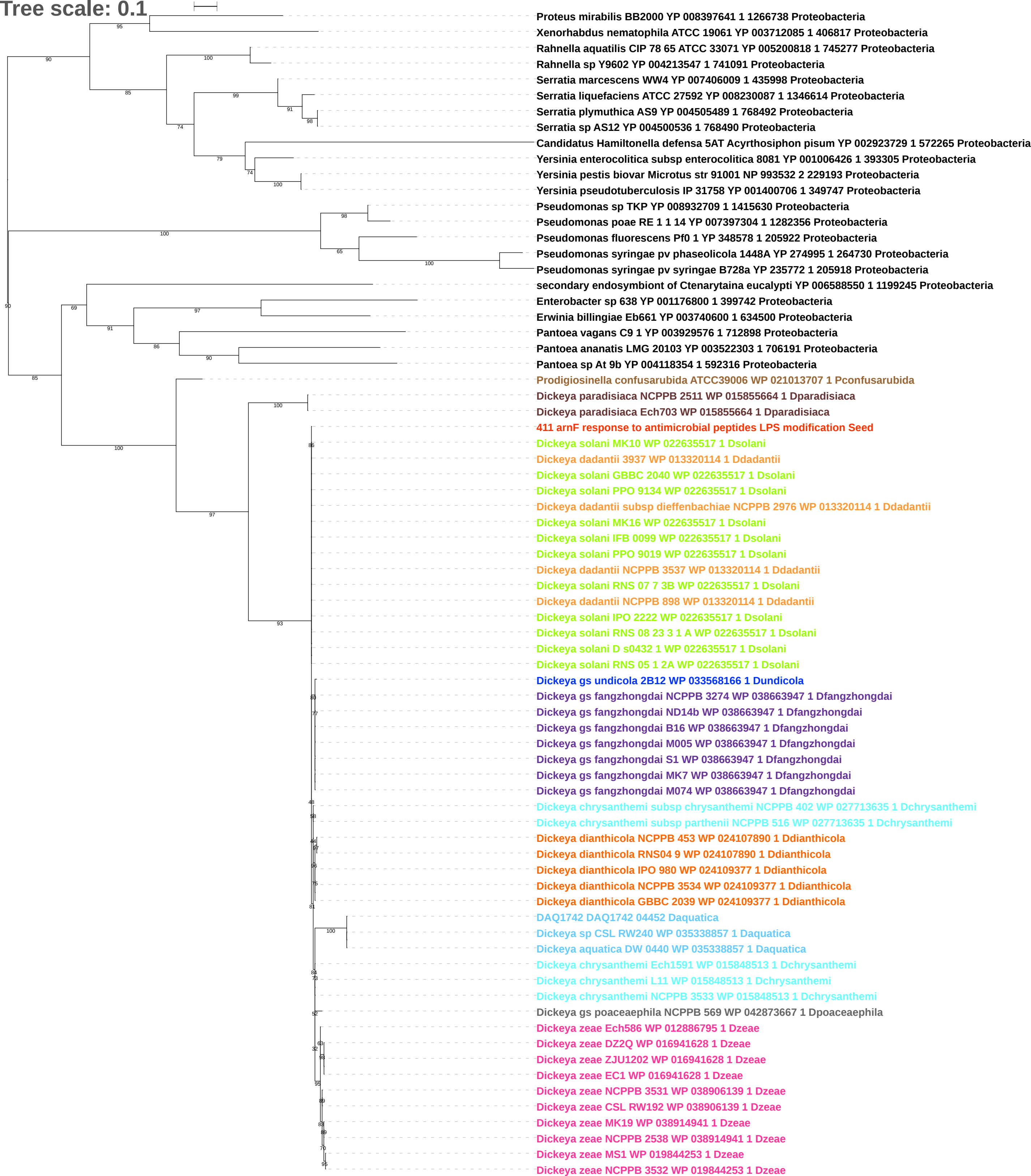
Tree scale: 0.1
Proteus mirabilis BB2000 YP 008397641 1 1266738 Proteobacteria
Xenorhabdus nematophila ATCC 19061 YP 003712085 1 406817 Proteobacteria
Rahnella aquatilis CIP 78 65 ATCC 33071 YP 005200818 1 745277 Proteobacteria
Rahnella sp Y9602 YP 004213547 1 741091 Proteobacteria
Serratia marcescens WW4 YP 007406009 1 435998 Proteobacteria
Serratia liquefaciens ATCC 27592 YP 008230087 1 1346614 Proteobacteria
Serratia plymuthica AS9 YP 004505489 1 768492 Proteobacteria
Serratia sp AS12 YP 004500536 1 768490 Proteobacteria
Candidatus Hamiltonella defensa 5AT Acyrthosiphon pisum YP 002923729 1 572265 Proteobacteria
Yersinia enterocolitica subsp enterocolitica 8081 YP 001006426 1 393305 Proteobacteria
Yersinia pestis biovar Microtus str 91001 NP 993532 2 229193 Proteobacteria
Yersinia pseudotuberculosis IP 31758 YP 001400706 1 349747 Proteobacteria
Pseudomonas sp TKP YP 008932709 1 1415630 Proteobacteria
Pseudomonas poae RE 1 1 14 YP 007397304 1 1282356 Proteobacteria
Pseudomonas fluorescens Pf0 1 YP 348578 1 205922 Proteobacteria
Pseudomonas syringae pv phaseolicola 1448A YP 274995 1 264730 Proteobacteria
Pseudomonas syringae pv syringae B728a YP 235772 1 205918 Proteobacteria
secondary endosymbiont of Ctenarytaina eucalypti YP 006588550 1 1199245 Proteobacteria
Enterobacter sp 638 YP 001176800 1 399742 Proteobacteria
Erwinia billingiae Eb661 YP 003740600 1 634500 Proteobacteria
Pantoea vagans C9 1 YP 003929576 1 712898 Proteobacteria
Pantoea ananatis LMG 20103 YP 003522303 1 706191 Proteobacteria
Pantoea sp At 9b YP 004118354 1 592316 Proteobacteria
Prodigiosinella confusarubida ATCC39006 WP 021013707 1 Pconfusarubida
Dickeya paradisiaca NCPPB 2511 WP 015855664 1 Dparadisiaca
Dickeya paradisiaca Ech703 WP 015855664 1 Dparadisiaca
411 arnF response to antimicrobial peptides LPS modification Seed
Dickeya solani MK10 WP 022635517 1 Dsolani
Dickeya dadantii 3937 WP 013320114 1 Ddadantii
Dickeya solani GBBC 2040 WP 022635517 1 Dsolani
Dickeya solani PPO 9134 WP 022635517 1 Dsolani
Dickeya dadantii subsp dieffenbachiae NCPPB 2976 WP 013320114 1 Ddadantii
Dickeya solani MK16 WP 022635517 1 Dsolani
Dickeya solani IFB 0099 WP 022635517 1 Dsolani
Dickeya solani PPO 9019 WP 022635517 1 Dsolani
Dickeya dadantii NCPPB 3537 WP 013320114 1 Ddadantii
Dickeya solani RNS 07 7 3B WP 022635517 1 Dsolani
Dickeya dadantii NCPPB 898 WP 013320114 1 Ddadantii
Dickeya solani IPO 2222 WP 022635517 1 Dsolani
Dickeya solani RNS 08 23 3 1 A WP 022635517 1 Dsolani
Dickeya solani D s0432 1 WP 022635517 1 Dsolani
Dickeya solani RNS 05 1 2A WP 022635517 1 Dsolani
Dickeya gs undicola 2B12 WP 033568166 1 Dundicola
Dickeya gs fangzhongdai NCPPB 3274 WP 038663947 1 Dfangzhongdai
Dickeya gs fangzhongdai ND14b WP 038663947 1 Dfangzhongdai
Dickeya gs fangzhongdai B16 WP 038663947 1 Dfangzhongdai
Dickeya gs fangzhongdai M005 WP 038663947 1 Dfangzhongdai
Dickeya gs fangzhongdai S1 WP 038663947 1 Dfangzhongdai
Dickeya gs fangzhongdai MK7 WP 038663947 1 Dfangzhongdai
Dickeya gs fangzhongdai M074 WP 038663947 1 Dfangzhongdai
Dickeya chrysanthemi subsp chrysanthemi NCPPB 402 WP 027713635 1 Dchrysanthemi
Dickeya chrysanthemi subsp parthenii NCPPB 516 WP 027713635 1 Dchrysanthemi
Dickeya dianthicola NCPPB 453 WP 024107890 1 Ddianthicola
Dickeya dianthicola RNS04 9 WP 024107890 1 Ddianthicola
Dickeya dianthicola IPO 980 WP 024109377 1 Ddianthicola
Dickeya dianthicola NCPPB 3534 WP 024109377 1 Ddianthicola
Dickeya dianthicola GBBC 2039 WP 024109377 1 Ddianthicola
DAQ1742 DAQ1742 04452 Daquatica
Dickeya sp CSL RW240 WP 035338857 1 Daquatica
Dickeya aquatica DW 0440 WP 035338857 1 Daquatica
Dickeya chrysanthemi Ech1591 WP 015848513 1 Dchrysanthemi
Dickeya chrysanthemi L11 WP 015848513 1 Dchrysanthemi
Dickeya chrysanthemi NCPPB 3533 WP 015848513 1 Dchrysanthemi
Dickeya gs poaceaephila NCPPB 569 WP 042873667 1 Dpoaceaephila
Dickeya zeae Ech586 WP 012886795 1 Dzeae
Dickeya zeae DZ2Q WP 016941628 1 Dzeae
Dickeya zeae ZJU1202 WP 016941628 1 Dzeae
Dickeya zeae EC1 WP 016941628 1 Dzeae
Dickeya zeae NCPPB 3531 WP 038906139 1 Dzeae
Dickeya zeae CSL RW192 WP 038906139 1 Dzeae
Dickeya zeae MK19 WP 038914941 1 Dzeae
Dickeya zeae NCPPB 2538 WP 038914941 1 Dzeae
Dickeya zeae MS1 WP 019844253 1 Dzeae
Dickeya zeae NCPPB 3532 WP 019844253 1 Dzeae

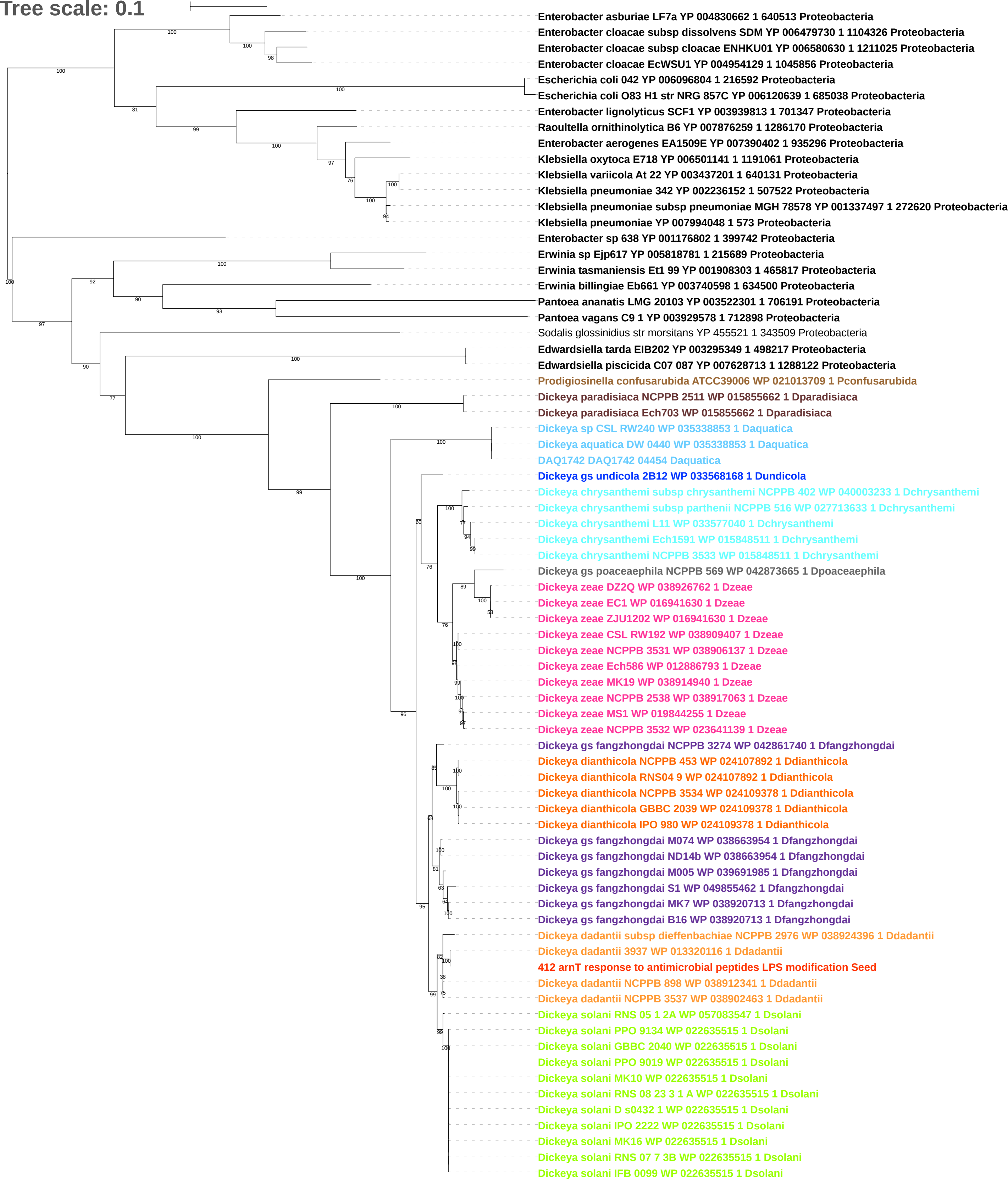
Tree scale: 0.1
Enterobacter asburiae LF7a YP 004830662 1 640513 Proteobacteria
Enterobacter cloacae subsp dissolvens SDM YP 006479730 1 1104326 Proteobacteria
Enterobacter cloacae subsp cloacae ENHKU01 YP 006580630 1 1211025 Proteobacteria
Enterobacter cloacae EcWSU1 YP 004954129 1 1045856 Proteobacteria
Escherichia coli 042 YP 006096804 1 216592 Proteobacteria
Escherichia coli O83 H1 str NRG 857C YP 006120639 1 685038 Proteobacteria
Enterobacter lignolyticus SCF1 YP 003939813 1 701347 Proteobacteria
Raoultella ornithinolytica B6 YP 007876259 1 1286170 Proteobacteria
Enterobacter aerogenes EA1509E YP 007390402 1 935296 Proteobacteria
Klebsiella oxytoca E718 YP 006501141 1 1191061 Proteobacteria
Klebsiella variicola At 22 YP 003437201 1 640131 Proteobacteria
Klebsiella pneumoniae 342 YP 002236152 1 507522 Proteobacteria
Klebsiella pneumoniae subsp pneumoniae MGH 78578 YP 001337497 1 272620 Proteobacteria
Klebsiella pneumoniae YP 007994048 1 573 Proteobacteria
Enterobacter sp 638 YP 001176802 1 399742 Proteobacteria
Erwinia sp Ejp617 YP 005818781 1 215689 Proteobacteria
Erwinia tasmaniensis Et1 99 YP 001908303 1 465817 Proteobacteria
Erwinia billingiae Eb661 YP 003740598 1 634500 Proteobacteria
Pantoea ananatis LMG 20103 YP 003522301 1 706191 Proteobacteria
Pantoea vagans C9 1 YP 003929578 1 712898 Proteobacteria
Sodalis glossinidius str morsitans YP 455521 1 343509 Proteobacteria
Edwardsiella tarda EIB202 YP 003295349 1 498217 Proteobacteria
Edwardsiella piscicida C07 087 YP 007628713 1 1288122 Proteobacteria
Prodigiosinella confusarubida ATCC39006 WP 021013709 1 Pconfusarubida
Dickeya paradisiaca NCPPB 2511 WP 015855662 1 Dparadisiaca
Dickeya paradisiaca Ech703 WP 015855662 1 Dparadisiaca
Dickeya sp CSL RW240 WP 035338853 1 Daquatica
Dickeya aquatica DW 0440 WP 035338853 1 Daquatica
DAQ1742 DAQ1742 04454 Daquatica
Dickeya gs undicola 2B12 WP 033568168 1 Dundicola
Dickeya chrysanthemi subsp chrysanthemi NCPPB 402 WP 040003233 1 Dchrysanthemi
Dickeya chrysanthemi subsp parthenii NCPPB 516 WP 027713633 1 Dchrysanthemi
Dickeya chrysanthemi L11 WP 033577040 1 Dchrysanthemi
Dickeya chrysanthemi Ech1591 WP 015848511 1 Dchrysanthemi
Dickeya chrysanthemi NCPPB 3533 WP 015848511 1 Dchrysanthemi
Dickeya gs poaceaephila NCPPB 569 WP 042873665 1 Dpoaceaephila
Dickeya zeae DZ2Q WP 038926762 1 Dzeae
Dickeya zeae EC1 WP 016941630 1 Dzeae
Dickeya zeae ZJU1202 WP 016941630 1 Dzeae
Dickeya zeae CSL RW192 WP 038909407 1 Dzeae
Dickeya zeae NCPPB 3531 WP 038906137 1 Dzeae
Dickeya zeae Ech586 WP 012886793 1 Dzeae
Dickeya zeae MK19 WP 038914940 1 Dzeae
Dickeya zeae NCPPB 2538 WP 038917063 1 Dzeae
Dickeya zeae MS1 WP 019844255 1 Dzeae
Dickeya zeae NCPPB 3532 WP 023641139 1 Dzeae
Dickeya gs fangzhongdai NCPPB 3274 WP 042861740 1 Dfangzhongdai
Dickeya dianthicola NCPPB 453 WP 024107892 1 Ddianthicola
Dickeya dianthicola RNS04 9 WP 024107892 1 Ddianthicola
Dickeya dianthicola NCPPB 3534 WP 024109378 1 Ddianthicola
Dickeya dianthicola GBBC 2039 WP 024109378 1 Ddianthicola
Dickeya dianthicola IPO 980 WP 024109378 1 Ddianthicola
Dickeya gs fangzhongdai M074 WP 038663954 1 Dfangzhongdai
Dickeya gs fangzhongdai ND14b WP 038663954 1 Dfangzhongdai
Dickeya gs fangzhongdai M005 WP 039691985 1 Dfangzhongdai
Dickeya gs fangzhongdai S1 WP 049855462 1 Dfangzhongdai
Dickeya gs fangzhongdai MK7 WP 038920713 1 Dfangzhongdai
Dickeya gs fangzhongdai B16 WP 038920713 1 Dfangzhongdai
Dickeya dadantii subsp dieffenbachiae NCPPB 2976 WP 038924396 1 Ddadantii
Dickeya dadantii 3937 WP 013320116 1 Ddadantii
412 arnT response to antimicrobial peptides LPS modification Seed
Dickeya dadantii NCPPB 898 WP 038912341 1 Ddadantii
Dickeya dadantii NCPPB 3537 WP 038902463 1 Ddadantii
Dickeya solani RNS 05 1 2A WP 057083547 1 Dsolani
Dickeya solani PPO 9134 WP 022635515 1 Dsolani
Dickeya solani GBBC 2040 WP 022635515 1 Dsolani
Dickeya solani PPO 9019 WP 022635515 1 Dsolani
Dickeya solani MK10 WP 022635515 1 Dsolani
Dickeya solani RNS 08 23 3 1 A WP 022635515 1 Dsolani
Dickeya solani D s0432 1 WP 022635515 1 Dsolani
Dickeya solani IPO 2222 WP 022635515 1 Dsolani
Dickeya solani MK16 WP 022635515 1 Dsolani
Dickeya solani RNS 07 7 3B WP 022635515 1 Dsolani
Dickeya solani IFB 0099 WP 022635515 1 Dsolani

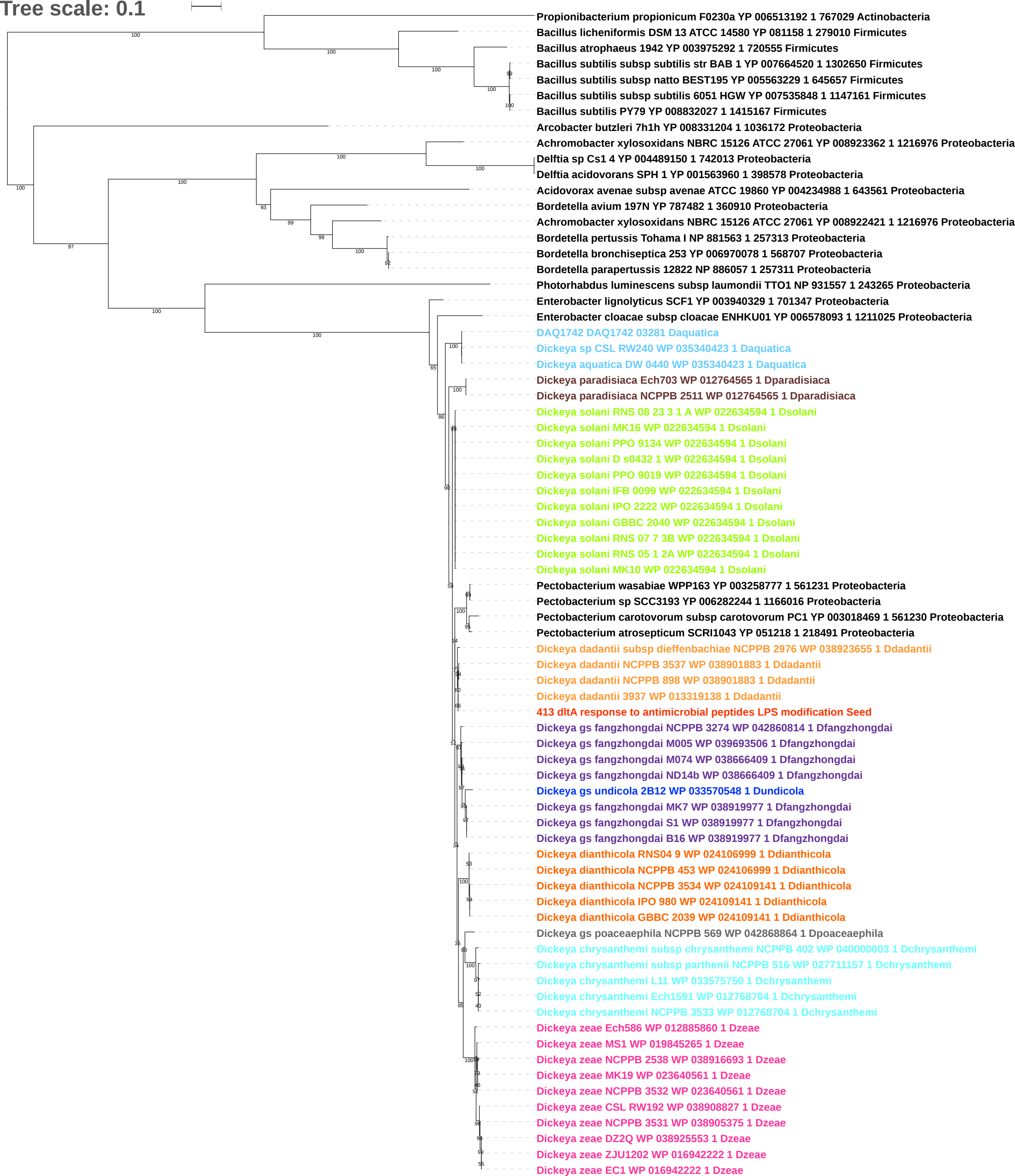

Tree scale: 0.1
Propionibacterium propionicum F0230a YP 006513192 1 767029 Actinobacteria
Bacillus licheniformis DSM 13 ATCC 14580 YP 081158 1 279010 Firmicutes
Bacillus atrophaeus 1942 YP 003975292 1 720555 Firmicutes
Bacillus subtilis subsp subtilis str BAB 1 YP 007664520 1 1302650 Firmicutes
Bacillus subtilis subsp natto BEST195 YP 005563229 1 645657 Firmicutes
Bacillus subtilis subsp subtilis 6051 HGW YP 007535848 1 1147161 Firmicutes
Bacillus subtilis PY79 YP 008832027 1 1415167 Firmicutes
Arcobacter butzleri 7h1h YP 008331204 1 1036172 Proteobacteria
Achromobacter xylosoxidans NBRC 15126 ATCC 27061 YP 008923362 1 1216976 Proteobacteria
Delftia sp Cs1 4 YP 004489150 1 742013 Proteobacteria
Delftia acidovorans SPH 1 YP 001563960 1 398578 Proteobacteria
Acidovorax avenae subsp avenae ATCC 19860 YP 004234988 1 643561 Proteobacteria
Bordetella avium 197N YP 787482 1 360910 Proteobacteria
Achromobacter xylosoxidans NBRC 15126 ATCC 27061 YP 008922421 1 1216976 Proteobacteria
Bordetella pertussis Tohama I NP 881563 1 257313 Proteobacteria
Bordetella bronchiseptica 253 YP 006970078 1 568707 Proteobacteria
Bordetella parapertussis 12822 NP 886057 1 257311 Proteobacteria
Photorhabdus luminescens subsp laumondii TTO1 NP 931557 1 243265 Proteobacteria
Enterobacter lignolyticus SCF1 YP 003940329 1 701347 Proteobacteria
Enterobacter cloacae subsp cloacae ENHKU01 YP 006578093 1 1211025 Proteobacteria
DAQ1742 DAQ1742 03281 Daquatica
Dickeya sp CSL RW240 WP 035340423 1 Daquatica
Dickeya aquatica DW 0440 WP 035340423 1 Daquatica
Dickeya paradisiaca Ech703 WP 012764565 1 Dparadisiaca
Dickeya paradisiaca NCPPB 2511 WP 012764565 1 Dparadisiaca
Dickeya solani RNS 08 23 3 1 A WP 022634594 1 Dsolani
Dickeya solani MK16 WP 022634594 1 Dsolani
Dickeya solani PPO 9134 WP 022634594 1 Dsolani
Dickeya solani D s0432 1 WP 022634594 1 Dsolani
Dickeya solani PPO 9019 WP 022634594 1 Dsolani
Dickeya solani IFB 0099 WP 022634594 1 Dsolani
Dickeya solani IPO 2222 WP 022634594 1 Dsolani
Dickeya solani GBBC 2040 WP 022634594 1 Dsolani
Dickeya solani RNS 07 7 3B WP 022634594 1 Dsolani
Dickeya solani RNS 05 1 2A WP 022634594 1 Dsolani
Dickeya solani MK10 WP 022634594 1 Dsolani
Pectobacterium wasabiae WPP163 YP 003258777 1 561231 Proteobacteria
Pectobacterium sp SCC3193 YP 006282244 1 1166016 Proteobacteria
Pectobacterium carotovorum subsp carotovorum PC1 YP 003018469 1 561230 Proteobacteria
Pectobacterium atrosepticum SCRI1043 YP 051218 1 218491 Proteobacteria
Dickeya dadantii subsp dieffenbachiae NCPPB 2976 WP 038923655 1 Ddadantii
Dickeya dadantii NCPPB 3537 WP 038901883 1 Ddadantii
Dickeya dadantii NCPPB 898 WP 038901883 1 Ddadantii
Dickeya dadantii 3937 WP 013319138 1 Ddadantii
413 dltA response to antimicrobial peptides LPS modification Seed
Dickeya gs fangzhongdai NCPPB 3274 WP 042860814 1 Dfangzhongdai
Dickeya gs fangzhongdai M005 WP 039693506 1 Dfangzhongdai
Dickeya gs fangzhongdai M074 WP 038666409 1 Dfangzhongdai
Dickeya gs fangzhongdai ND14b WP 038666409 1 Dfangzhongdai
Dickeya gs undicola 2B12 WP 033570548 1 Dundicola
Dickeya gs fangzhongdai MK7 WP 038919977 1 Dfangzhongdai
Dickeya gs fangzhongdai S1 WP 038919977 1 Dfangzhongdai
Dickeya gs fangzhongdai B16 WP 038919977 1 Dfangzhongdai
Dickeya dianthicola RNS04 9 WP 024106999 1 Ddianthicola
Dickeya dianthicola NCPPB 453 WP 024106999 1 Ddianthicola
Dickeya dianthicola NCPPB 3534 WP 024109141 1 Ddianthicola
Dickeya dianthicola IPO 980 WP 024109141 1 Ddianthicola
Dickeya dianthicola GBBC 2039 WP 024109141 1 Ddianthicola
Dickeya gs poaceaephila NCPPB 569 WP 042868864 1 Dpoaceaephila
Dickeya chrysanthemi subsp chrysanthemi NCPPB 402 WP 040000003 1 Dchrysanthemi
Dickeya chrysanthemi subsp parthenii NCPPB 516 WP 027711157 1 Dchrysanthemi
Dickeya chrysanthemi L11 WP 033575750 1 Dchrysanthemi
Dickeya chrysanthemi Ech1591 WP 012768704 1 Dchrysanthemi
Dickeya chrysanthemi NCPPB 3533 WP 012768704 1 Dchrysanthemi
Dickeya zeae Ech586 WP 012885860 1 Dzeae
Dickeya zeae MS1 WP 019845265 1 Dzeae
Dickeya zeae NCPPB 2538 WP 038916693 1 Dzeae
Dickeya zeae MK19 WP 023640561 1 Dzeae
Dickeya zeae NCPPB 3532 WP 023640561 1 Dzeae
Dickeya zeae CSL RW192 WP 038908827 1 Dzeae
Dickeya zeae NCPPB 3531 WP 038905375 1 Dzeae
Dickeya zeae DZ2Q WP 038925553 1 Dzeae
Dickeya zeae ZJU1202 WP 016942222 1 Dzeae
Dickeya zeae EC1 WP 016942222 1 Dzeae

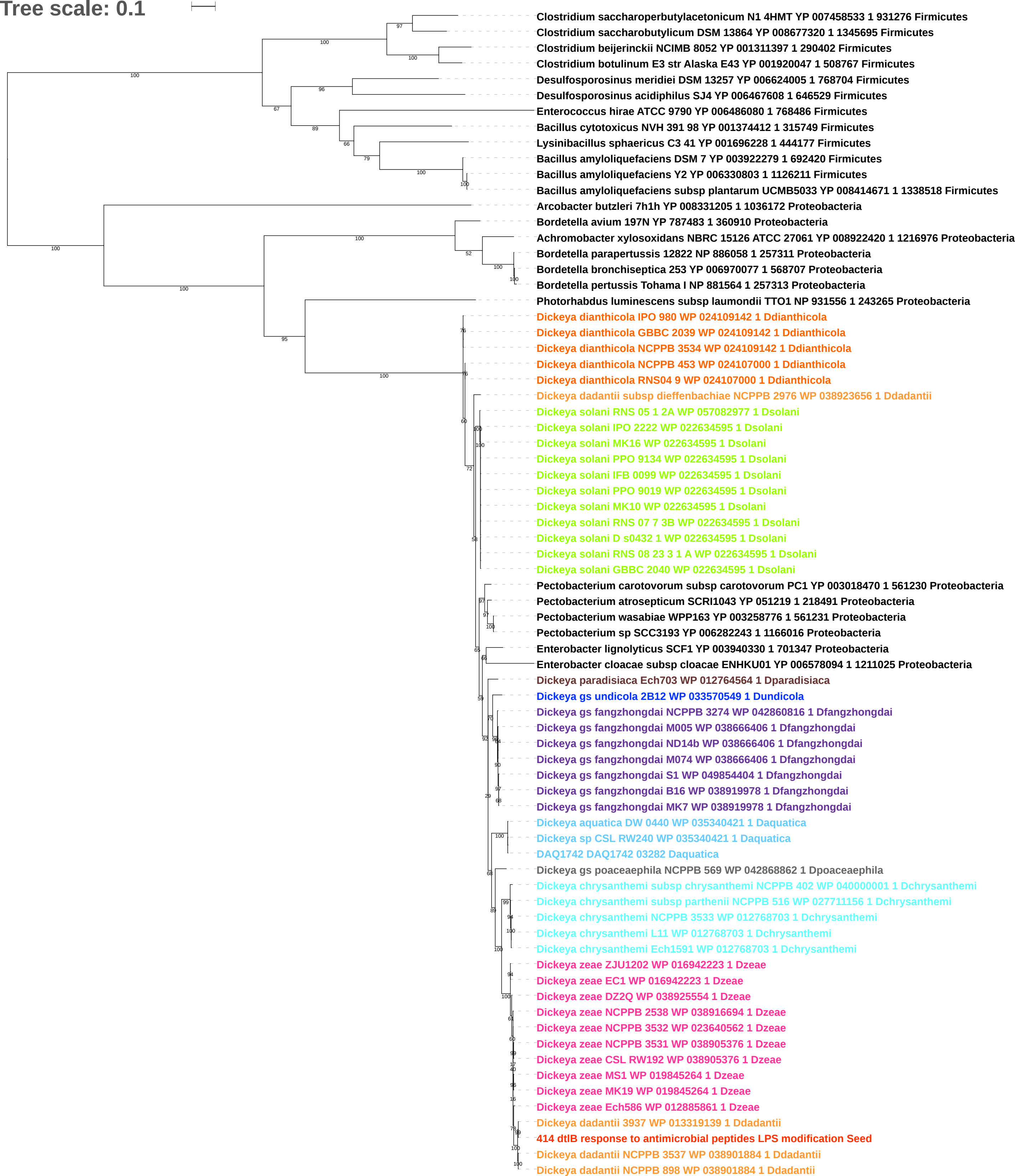
Tree scale: 0.1
Clostridium saccharoperbutylacetonicum N1 4HMT YP 007458533 1 931276 Firmicutes
Clostridium saccharobutylicum DSM 13864 YP 008677320 1 1345695 Firmicutes
Clostridium beijerinckii NCIMB 8052 YP 001311397 1 290402 Firmicutes
Clostridium botulinum E3 str Alaska E43 YP 001920047 1 508767 Firmicutes
Desulfosporosinus meridiei DSM 13257 YP 006624005 1 768704 Firmicutes
Desulfosporosinus acidiphilus SJ4 YP 006467608 1 646529 Firmicutes
Enterococcus hirae ATCC 9790 YP 006486080 1 768486 Firmicutes
Bacillus cytotoxicus NVH 391 98 YP 001374412 1 315749 Firmicutes
Lysinibacillus sphaericus C3 41 YP 001696228 1 444177 Firmicutes
Bacillus amyloliquefaciens DSM 7 YP 003922279 1 692420 Firmicutes
Bacillus amyloliquefaciens Y2 YP 006330803 1 1126211 Firmicutes
Bacillus amyloliquefaciens subsp plantarum UCMB5033 YP 008414671 1 1338518 Firmicutes
Arcobacter butzleri 7h1h YP 008331205 1 1036172 Proteobacteria
Bordetella avium 197N YP 787483 1 360910 Proteobacteria
Achromobacter xylosoxidans NBRC 15126 ATCC 27061 YP 008922420 1 1216976 Proteobacteria
Bordetella parapertussis 12822 NP 886058 1 257311 Proteobacteria
Bordetella bronchiseptica 253 YP 006970077 1 568707 Proteobacteria
Bordetella pertussis Tohama I NP 881564 1 257313 Proteobacteria
Photorhabdus luminescens subsp laumondii TTO1 NP 931556 1 243265 Proteobacteria
Dickeya dianthicola IPO 980 WP 024109142 1 Ddianthicola
Dickeya dianthicola GBBC 2039 WP 024109142 1 Ddianthicola
Dickeya dianthicola NCPPB 3534 WP 024109142 1 Ddianthicola
Dickeya dianthicola NCPPB 453 WP 024107000 1 Ddianthicola
Dickeya dianthicola RNS04 9 WP 024107000 1 Ddianthicola
Dickeya dadantii subsp dieffenbachiae NCPPB 2976 WP 038923656 1 Ddadantii
Dickeya solani RNS 05 1 2A WP 057082977 1 Dsolani
Dickeya solani IPO 2222 WP 022634595 1 Dsolani
Dickeya solani MK16 WP 022634595 1 Dsolani
Dickeya solani PPO 9134 WP 022634595 1 Dsolani
Dickeya solani IFB 0099 WP 022634595 1 Dsolani
Dickeya solani PPO 9019 WP 022634595 1 Dsolani
Dickeya solani MK10 WP 022634595 1 Dsolani
Dickeya solani RNS 07 7 3B WP 022634595 1 Dsolani
Dickeya solani D s0432 1 WP 022634595 1 Dsolani
Dickeya solani RNS 08 23 3 1 A WP 022634595 1 Dsolani
Dickeya solani GBBC 2040 WP 022634595 1 Dsolani
Pectobacterium carotovorum subsp carotovorum PC1 YP 003018470 1 561230 Proteobacteria
Pectobacterium atrosepticum SCRI1043 YP 051219 1 218491 Proteobacteria
Pectobacterium wasabiae WPP163 YP 003258776 1 561231 Proteobacteria
Pectobacterium sp SCC3193 YP 006282243 1 1166016 Proteobacteria
Enterobacter lignolyticus SCF1 YP 003940330 1 701347 Proteobacteria
Enterobacter cloacae subsp cloacae ENHKU01 YP 006578094 1 1211025 Proteobacteria
Dickeya paradisiaca Ech703 WP 012764564 1 Dparadisiaca
Dickeya gs undicola 2B12 WP 033570549 1 Dundicola
Dickeya gs fangzhongdai NCPPB 3274 WP 042860816 1 Dfangzhongdai
Dickeya gs fangzhongdai M005 WP 038666406 1 Dfangzhongdai
Dickeya gs fangzhongdai ND14b WP 038666406 1 Dfangzhongdai
Dickeya gs fangzhongdai M074 WP 038666406 1 Dfangzhongdai
Dickeya gs fangzhongdai S1 WP 049854404 1 Dfangzhongdai
Dickeya gs fangzhongdai B16 WP 038919978 1 Dfangzhongdai
Dickeya gs fangzhongdai MK7 WP 038919978 1 Dfangzhongdai
Dickeya aquatica DW 0440 WP 035340421 1 Daquatica
Dickeya sp CSL RW240 WP 035340421 1 Daquatica
DAQ1742 DAQ1742 03282 Daquatica
Dickeya gs poaceaephila NCPPB 569 WP 042868862 1 Dpoaceaephila
Dickeya chrysanthemi subsp chrysanthemi NCPPB 402 WP 040000001 1 Dchrysanthemi
Dickeya chrysanthemi subsp parthenii NCPPB 516 WP 027711156 1 Dchrysanthemi
Dickeya chrysanthemi NCPPB 3533 WP 012768703 1 Dchrysanthemi
Dickeya chrysanthemi L11 WP 012768703 1 Dchrysanthemi
Dickeya chrysanthemi Ech1591 WP 012768703 1 Dchrysanthemi
Dickeya zeae ZJU1202 WP 016942223 1 Dzeae
Dickeya zeae EC1 WP 016942223 1 Dzeae
Dickeya zeae DZ2Q WP 038925554 1 Dzeae
Dickeya zeae NCPPB 2538 WP 038916694 1 Dzeae
Dickeya zeae NCPPB 3532 WP 023640562 1 Dzeae
Dickeya zeae NCPPB 3531 WP 038905376 1 Dzeae
Dickeya zeae CSL RW192 WP 038905376 1 Dzeae
Dickeya zeae MS1 WP 019845264 1 Dzeae
Dickeya zeae MK19 WP 019845264 1 Dzeae
Dickeya zeae Ech586 WP 012885861 1 Dzeae
Dickeya dadantii 3937 WP 013319139 1 Ddadantii
414 dtlB response to antimicrobial peptides LPS modification Seed
Dickeya dadantii NCPPB 3537 WP 038901884 1 Ddadantii
Dickeya dadantii NCPPB 898 WP 038901884 1 Ddadantii

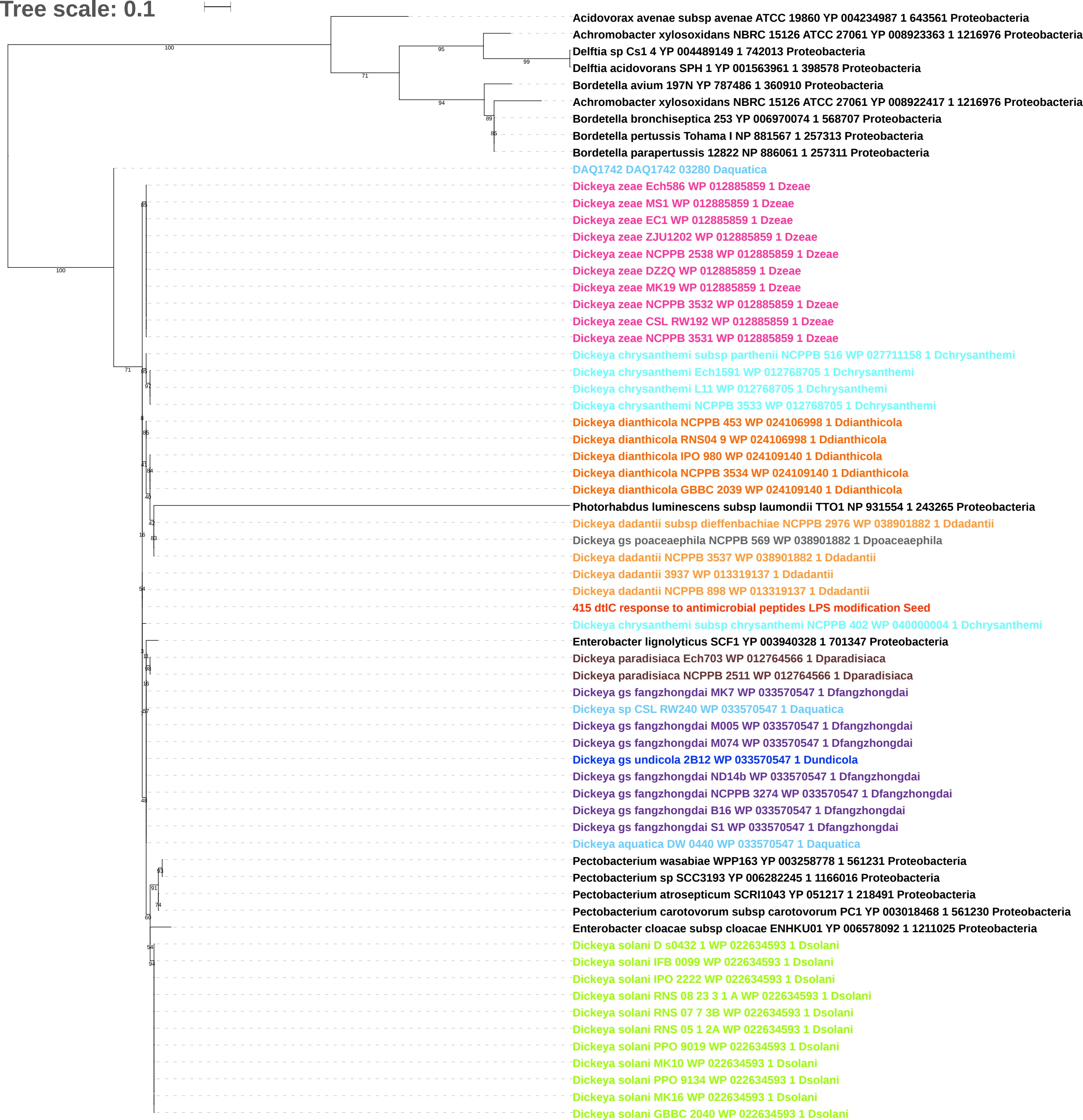
Tree scale: 0.1
Acidovorax avenae subsp avenae ATCC 19860 YP 004234987 1 643561 Proteobacteria
Achromobacter xylosoxidans NBRC 15126 ATCC 27061 YP 008923363 1 1216976 Proteobacteria
Delftia sp Cs1 4 YP 004489149 1 742013 Proteobacteria
Delftia acidovorans SPH 1 YP 001563961 1 398578 Proteobacteria
Bordetella avium 197N YP 787486 1 360910 Proteobacteria
Achromobacter xylosoxidans NBRC 15126 ATCC 27061 YP 008922417 1 1216976 Proteobacteria
Bordetella bronchiseptica 253 YP 006970074 1 568707 Proteobacteria
Bordetella pertussis Tohama I NP 881567 1 257313 Proteobacteria
Bordetella parapertussis 12822 NP 886061 1 257311 Proteobacteria
DAQ1742 DAQ1742 03280 Daquatica
Dickeya zeae Ech586 WP 012885859 1 Dzeae
Dickeya zeae MS1 WP 012885859 1 Dzeae
Dickeya zeae EC1 WP 012885859 1 Dzeae
Dickeya zeae ZJU1202 WP 012885859 1 Dzeae
Dickeya zeae NCPPB 2538 WP 012885859 1 Dzeae
Dickeya zeae DZ2Q WP 012885859 1 Dzeae
Dickeya zeae MK19 WP 012885859 1 Dzeae
Dickeya zeae NCPPB 3532 WP 012885859 1 Dzeae
Dickeya zeae CSL RW192 WP 012885859 1 Dzeae
Dickeya zeae NCPPB 3531 WP 012885859 1 Dzeae
Dickeya chrysanthemi subsp parthenii NCPPB 516 WP 027711158 1 Dchrysanthemi
Dickeya chrysanthemi Ech1591 WP 012768705 1 Dchrysanthemi
Dickeya chrysanthemi L11 WP 012768705 1 Dchrysanthemi
Dickeya chrysanthemi NCPPB 3533 WP 012768705 1 Dchrysanthemi
Dickeya dianthicola NCPPB 453 WP 024106998 1 Ddianthicola
Dickeya dianthicola RNS04 9 WP 024106998 1 Ddianthicola
Dickeya dianthicola IPO 980 WP 024109140 1 Ddianthicola
Dickeya dianthicola NCPPB 3534 WP 024109140 1 Ddianthicola
Dickeya dianthicola GBBC 2039 WP 024109140 1 Ddianthicola
Photorhabdus luminescens subsp laumondii TTO1 NP 931554 1 243265 Proteobacteria
Dickeya dadantii subsp dieffenbachiae NCPPB 2976 WP 038901882 1 Ddadantii
Dickeya gs poaceaephila NCPPB 569 WP 038901882 1 Dpoaceaephila
Dickeya dadantii NCPPB 3537 WP 038901882 1 Ddadantii
Dickeya dadantii 3937 WP 013319137 1 Ddadantii
Dickeya dadantii NCPPB 898 WP 013319137 1 Ddadantii
415 dtlC response to antimicrobial peptides LPS modification Seed
Dickeya chrysanthemi subsp chrysanthemi NCPPB 402 WP 040000004 1 Dchrysanthemi
Enterobacter lignolyticus SCF1 YP 003940328 1 701347 Proteobacteria
Dickeya paradisiaca Ech703 WP 012764566 1 Dparadisiaca
Dickeya paradisiaca NCPPB 2511 WP 012764566 1 Dparadisiaca
Dickeya gs fangzhongdai MK7 WP 033570547 1 Dfangzhongdai
Dickeya sp CSL RW240 WP 033570547 1 Daquatica
Dickeya gs fangzhongdai M005 WP 033570547 1 Dfangzhongdai
Dickeya gs fangzhongdai M074 WP 033570547 1 Dfangzhongdai
Dickeya gs undicola 2B12 WP 033570547 1 Dundicola
Dickeya gs fangzhongdai ND14b WP 033570547 1 Dfangzhongdai
Dickeya gs fangzhongdai NCPPB 3274 WP 033570547 1 Dfangzhongdai
Dickeya gs fangzhongdai B16 WP 033570547 1 Dfangzhongdai
Dickeya gs fangzhongdai S1 WP 033570547 1 Dfangzhongdai
Dickeya aquatica DW 0440 WP 033570547 1 Daquatica
Pectobacterium wasabiae WPP163 YP 003258778 1 561231 Proteobacteria
Pectobacterium sp SCC3193 YP 006282245 1 1166016 Proteobacteria
Pectobacterium atrosepticum SCRI1043 YP 051217 1 218491 Proteobacteria
Pectobacterium carotovorum subsp carotovorum PC1 YP 003018468 1 561230 Proteobacteria
Enterobacter cloacae subsp cloacae ENHKU01 YP 006578092 1 1211025 Proteobacteria
Dickeya solani D s0432 1 WP 022634593 1 Dsolani
Dickeya solani IFB 0099 WP 022634593 1 Dsolani
Dickeya solani IPO 2222 WP 022634593 1 Dsolani
Dickeya solani RNS 08 23 3 1 A WP 022634593 1 Dsolani
Dickeya solani RNS 07 7 3B WP 022634593 1 Dsolani
Dickeya solani RNS 05 1 2A WP 022634593 1 Dsolani
Dickeya solani PPO 9019 WP 022634593 1 Dsolani
Dickeya solani MK10 WP 022634593 1 Dsolani
Dickeya solani PPO 9134 WP 022634593 1 Dsolani
Dickeya solani MK16 WP 022634593 1 Dsolani
Dickeya solani GBBC 2040 WP 022634593 1 Dsolani

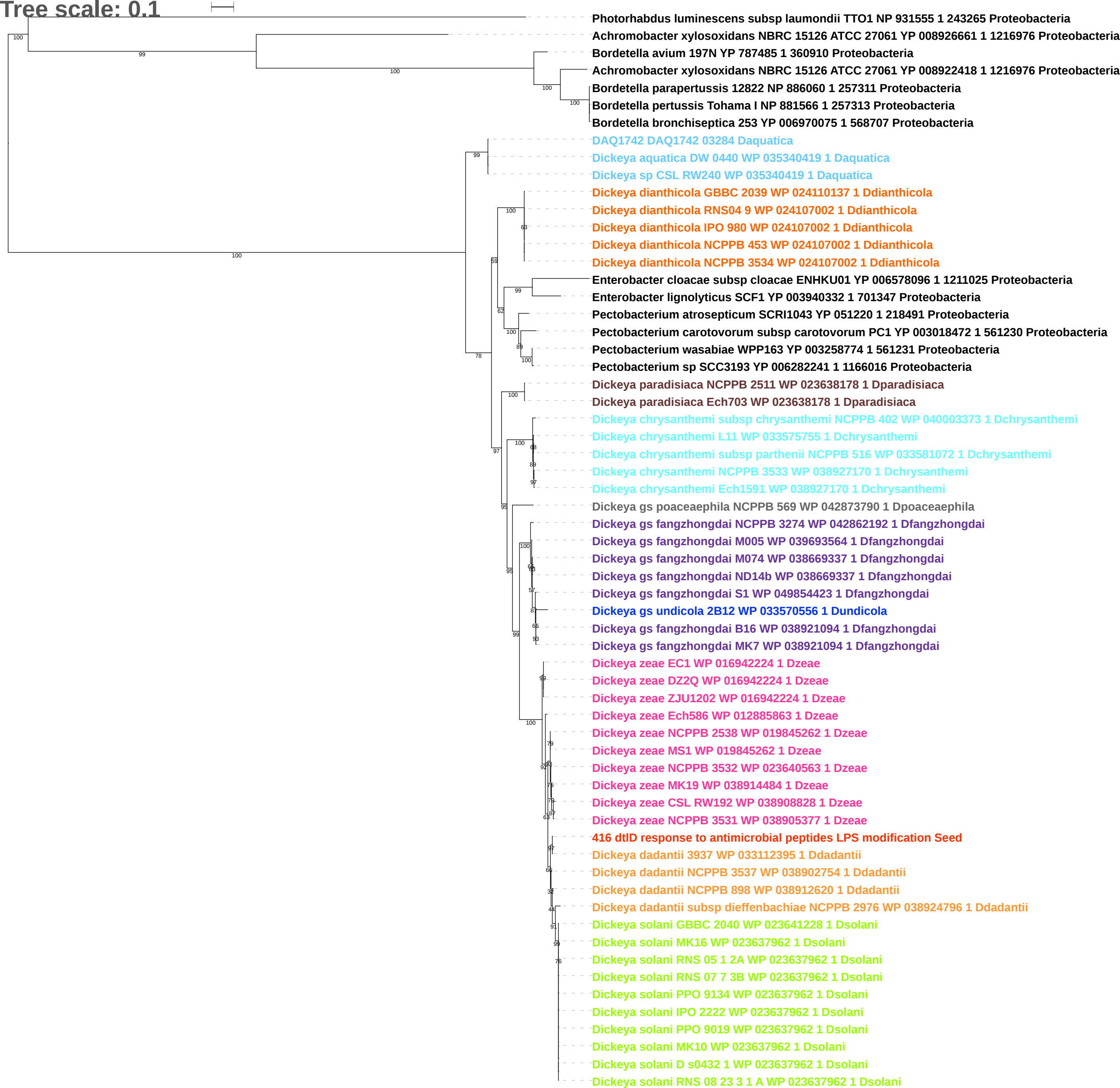
Tree scale: 0.1
Photorhabdus luminescens subsp laumondii TTO1 NP 931555 1 243265 Proteobacteria
Achromobacter xylosoxidans NBRC 15126 ATCC 27061 YP 008926661 1 1216976 Proteobacteria
Bordetella avium 197N YP 787485 1 360910 Proteobacteria
Achromobacter xylosoxidans NBRC 15126 ATCC 27061 YP 008922418 1 1216976 Proteobacteria
Bordetella parapertussis 12822 NP 886060 1 257311 Proteobacteria
Bordetella pertussis Tohama I NP 881566 1 257313 Proteobacteria
Bordetella bronchiseptica 253 YP 006970075 1 568707 Proteobacteria
DAQ1742 DAQ1742 03284 Daquatica
Dickeya aquatica DW 0440 WP 035340419 1 Daquatica
Dickeya sp CSL RW240 WP 035340419 1 Daquatica
Dickeya dianthicola GBBC 2039 WP 024110137 1 Ddianthicola
Dickeya dianthicola RNS04 9 WP 024107002 1 Ddianthicola
Dickeya dianthicola IPO 980 WP 024107002 1 Ddianthicola
Dickeya dianthicola NCPPB 453 WP 024107002 1 Ddianthicola
Dickeya dianthicola NCPPB 3534 WP 024107002 1 Ddianthicola
Enterobacter cloacae subsp cloacae ENHKU01 YP 006578096 1 1211025 Proteobacteria
Enterobacter lignolyticus SCF1 YP 003940332 1 701347 Proteobacteria
Pectobacterium atrosepticum SCRI1043 YP 051220 1 218491 Proteobacteria
Pectobacterium carotovorum subsp carotovorum PC1 YP 003018472 1 561230 Proteobacteria
Pectobacterium wasabiae WPP163 YP 003258774 1 561231 Proteobacteria
Pectobacterium sp SCC3193 YP 006282241 1 1166016 Proteobacteria
Dickeya paradisiaca NCPPB 2511 WP 023638178 1 Dparadisiaca
Dickeya paradisiaca Ech703 WP 023638178 1 Dparadisiaca
Dickeya chrysanthemi subsp chrysanthemi NCPPB 402 WP 040003373 1 Dchrysanthemi
Dickeya chrysanthemi L11 WP 033575755 1 Dchrysanthemi
Dickeya chrysanthemi subsp parthenii NCPPB 516 WP 033581072 1 Dchrysanthemi
Dickeya chrysanthemi NCPPB 3533 WP 038927170 1 Dchrysanthemi
Dickeya chrysanthemi Ech1591 WP 038927170 1 Dchrysanthemi
Dickeya gs poaceaephila NCPPB 569 WP 042873790 1 Dpoaceaephila
Dickeya gs fangzhongdai NCPPB 3274 WP 042862192 1 Dfangzhongdai
Dickeya gs fangzhongdai M005 WP 039693564 1 Dfangzhongdai
Dickeya gs fangzhongdai M074 WP 038669337 1 Dfangzhongdai
Dickeya gs fangzhongdai ND14b WP 038669337 1 Dfangzhongdai
Dickeya gs fangzhongdai S1 WP 049854423 1 Dfangzhongdai
Dickeya gs undicola 2B12 WP 033570556 1 Dundicola
Dickeya gs fangzhongdai B16 WP 038921094 1 Dfangzhongdai
Dickeya gs fangzhongdai MK7 WP 038921094 1 Dfangzhongdai
Dickeya zeae EC1 WP 016942224 1 Dzeae
Dickeya zeae DZ2Q WP 016942224 1 Dzeae
Dickeya zeae ZJU1202 WP 016942224 1 Dzeae
Dickeya zeae Ech586 WP 012885863 1 Dzeae
Dickeya zeae NCPPB 2538 WP 019845262 1 Dzeae
Dickeya zeae MS1 WP 019845262 1 Dzeae
Dickeya zeae NCPPB 3532 WP 023640563 1 Dzeae
Dickeya zeae MK19 WP 038914484 1 Dzeae
Dickeya zeae CSL RW192 WP 038908828 1 Dzeae
Dickeya zeae NCPPB 3531 WP 038905377 1 Dzeae
416 dtlD response to antimicrobial peptides LPS modification Seed
Dickeya dadantii 3937 WP 033112395 1 Ddadantii
Dickeya dadantii NCPPB 3537 WP 038902754 1 Ddadantii
Dickeya dadantii NCPPB 898 WP 038912620 1 Ddadantii
Dickeya dadantii subsp dieffenbachiae NCPPB 2976 WP 038924796 1 Ddadantii
Dickeya solani GBBC 2040 WP 023641228 1 Dsolani
Dickeya solani MK16 WP 023637962 1 Dsolani
Dickeya solani RNS 05 1 2A WP 023637962 1 Dsolani
Dickeya solani RNS 07 7 3B WP 023637962 1 Dsolani
Dickeya solani PPO 9134 WP 023637962 1 Dsolani
Dickeya solani IPO 2222 WP 023637962 1 Dsolani
Dickeya solani PPO 9019 WP 023637962 1 Dsolani
Dickeya solani MK10 WP 023637962 1 Dsolani
Dickeya solani D s0432 1 WP 023637962 1 Dsolani
Dickeya solani RNS 08 23 3 1 A WP 023637962 1 Dsolani

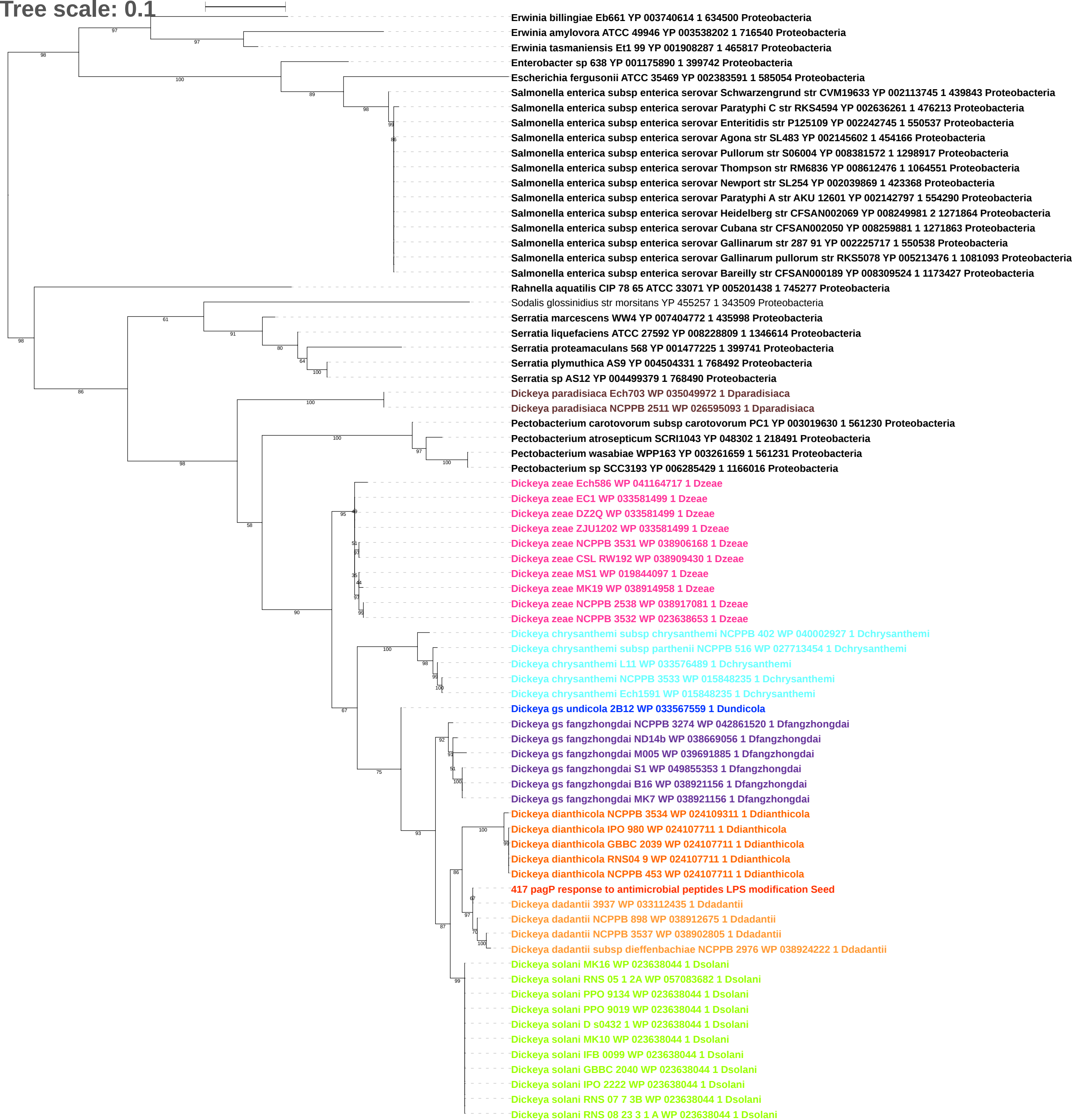
Tree scale: 0.1
Erwinia billingiae Eb661 YP 003740614 1 634500 Proteobacteria
Erwinia amylovora ATCC 49946 YP 003538202 1 716540 Proteobacteria
Erwinia tasmaniensis Et1 99 YP 001908287 1 465817 Proteobacteria
Enterobacter sp 638 YP 001175890 1 399742 Proteobacteria
Escherichia fergusonii ATCC 35469 YP 002383591 1 585054 Proteobacteria
Salmonella enterica subsp enterica serovar Schwarzengrund str CVM19633 YP 002113745 1 439843 Proteobacteria
Salmonella enterica subsp enterica serovar Paratyphi C str RKS4594 YP 002636261 1 476213 Proteobacteria
Salmonella enterica subsp enterica serovar Enteritidis str P125109 YP 002242745 1 550537 Proteobacteria
Salmonella enterica subsp enterica serovar Agona str SL483 YP 002145602 1 454166 Proteobacteria
Salmonella enterica subsp enterica serovar Pullorum str S06004 YP 008381572 1 1298917 Proteobacteria
Salmonella enterica subsp enterica serovar Thompson str RM6836 YP 008612476 1 1064551 Proteobacteria
Salmonella enterica subsp enterica serovar Newport str SL254 YP 002039869 1 423368 Proteobacteria
Salmonella enterica subsp enterica serovar Paratyphi A str AKU 12601 YP 002142797 1 554290 Proteobacteria
Salmonella enterica subsp enterica serovar Heidelberg str CFSAN002069 YP 008249981 2 1271864 Proteobacteria
Salmonella enterica subsp enterica serovar Cubana str CFSAN002050 YP 008259881 1 1271863 Proteobacteria
Salmonella enterica subsp enterica serovar Gallinarum str 287 91 YP 002225717 1 550538 Proteobacteria
Salmonella enterica subsp enterica serovar Gallinarum pullorum str RKS5078 YP 005213476 1 1081093 Proteobacteria
Salmonella enterica subsp enterica serovar Bareilly str CFSAN000189 YP 008309524 1 1173427 Proteobacteria
Rahnella aquatilis CIP 78 65 ATCC 33071 YP 005201438 1 745277 Proteobacteria
Sodalis glossinidius str morsitans YP 455257 1 343509 Proteobacteria
Serratia marcescens WW4 YP 007404772 1 435998 Proteobacteria
Serratia liquefaciens ATCC 27592 YP 008228809 1 1346614 Proteobacteria
Serratia proteamaculans 568 YP 001477225 1 399741 Proteobacteria
Serratia plymuthica AS9 YP 004504331 1 768492 Proteobacteria
Serratia sp AS12 YP 004499379 1 768490 Proteobacteria
Dickeya paradisiaca Ech703 WP 035049972 1 Dparadisiaca
Dickeya paradisiaca NCPPB 2511 WP 026595093 1 Dparadisiaca
Pectobacterium carotovorum subsp carotovorum PC1 YP 003019630 1 561230 Proteobacteria
Pectobacterium atrosepticum SCRI1043 YP 048302 1 218491 Proteobacteria
Pectobacterium wasabiae WPP163 YP 003261659 1 561231 Proteobacteria
Pectobacterium sp SCC3193 YP 006285429 1 1166016 Proteobacteria
Dickeya zeae Ech586 WP 041164717 1 Dzeae
Dickeya zeae EC1 WP 033581499 1 Dzeae
Dickeya zeae DZ2Q WP 033581499 1 Dzeae
Dickeya zeae ZJU1202 WP 033581499 1 Dzeae
Dickeya zeae NCPPB 3531 WP 038906168 1 Dzeae
Dickeya zeae CSL RW192 WP 038909430 1 Dzeae
Dickeya zeae MS1 WP 019844097 1 Dzeae
Dickeya zeae MK19 WP 038914958 1 Dzeae
Dickeya zeae NCPPB 2538 WP 038917081 1 Dzeae
Dickeya zeae NCPPB 3532 WP 023638653 1 Dzeae
Dickeya chrysanthemi subsp chrysanthemi NCPPB 402 WP 040002927 1 Dchrysanthemi
Dickeya chrysanthemi subsp parthenii NCPPB 516 WP 027713454 1 Dchrysanthemi
Dickeya chrysanthemi L11 WP 033576489 1 Dchrysanthemi
Dickeya chrysanthemi NCPPB 3533 WP 015848235 1 Dchrysanthemi
Dickeya chrysanthemi Ech1591 WP 015848235 1 Dchrysanthemi
Dickeya gs undicola 2B12 WP 033567559 1 Dundicola
Dickeya gs fangzhongdai NCPPB 3274 WP 042861520 1 Dfangzhongdai
Dickeya gs fangzhongdai ND14b WP 038669056 1 Dfangzhongdai
Dickeya gs fangzhongdai M005 WP 039691885 1 Dfangzhongdai
Dickeya gs fangzhongdai S1 WP 049855353 1 Dfangzhongdai
Dickeya gs fangzhongdai B16 WP 038921156 1 Dfangzhongdai
Dickeya gs fangzhongdai MK7 WP 038921156 1 Dfangzhongdai
Dickeya dianthicola NCPPB 3534 WP 024109311 1 Ddianthicola
Dickeya dianthicola IPO 980 WP 024107711 1 Ddianthicola
Dickeya dianthicola GBBC 2039 WP 024107711 1 Ddianthicola
Dickeya dianthicola RNS04 9 WP 024107711 1 Ddianthicola
Dickeya dianthicola NCPPB 453 WP 024107711 1 Ddianthicola
417 pagP response to antimicrobial peptides LPS modification Seed
Dickeya dadantii 3937 WP 033112435 1 Ddadantii
Dickeya dadantii NCPPB 898 WP 038912675 1 Ddadantii
Dickeya dadantii NCPPB 3537 WP 038902805 1 Ddadantii
Dickeya dadantii subsp dieffenbachiae NCPPB 2976 WP 038924222 1 Ddadantii
Dickeya solani MK16 WP 023638044 1 Dsolani
Dickeya solani RNS 05 1 2A WP 057083682 1 Dsolani
Dickeya solani PPO 9134 WP 023638044 1 Dsolani
Dickeya solani PPO 9019 WP 023638044 1 Dsolani
Dickeya solani D s0432 1 WP 023638044 1 Dsolani
Dickeya solani MK10 WP 023638044 1 Dsolani
Dickeya solani IFB 0099 WP 023638044 1 Dsolani
Dickeya solani GBBC 2040 WP 023638044 1 Dsolani
Dickeya solani IPO 2222 WP 023638044 1 Dsolani
Dickeya solani RNS 07 7 3B WP 023638044 1 Dsolani
Dickeya solani RNS 08 23 3 1 A WP 023638044 1 Dsolani

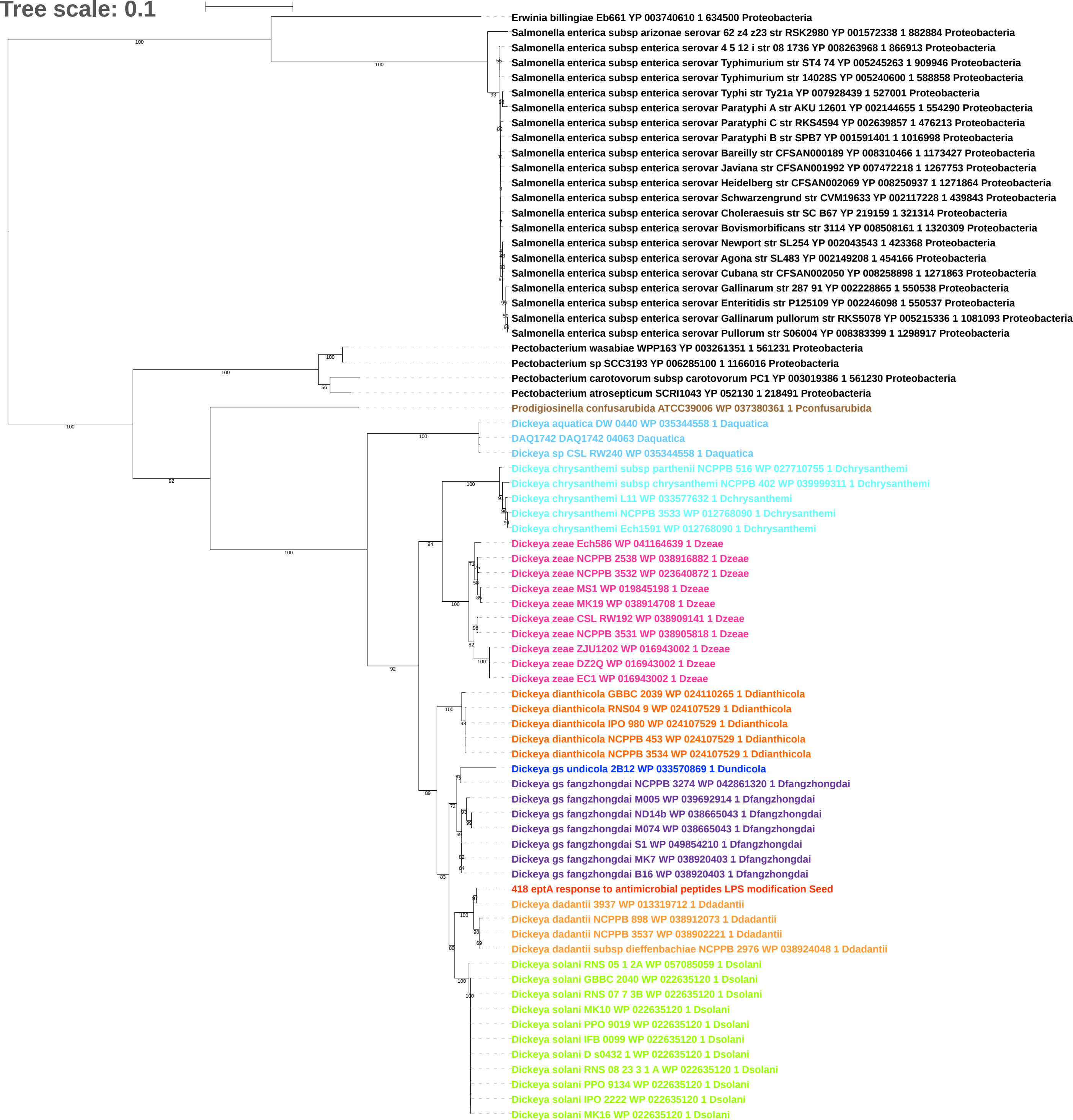
Tree scale: 0.1
Erwinia billingiae Eb661 YP 003740610 1 634500 Proteobacteria
Salmonella enterica subsp arizonae serovar 62 z4 z23 str RSK2980 YP 001572338 1 882884 Proteobacteria
Salmonella enterica subsp enterica serovar 4 5 12 i str 08 1736 YP 008263968 1 866913 Proteobacteria
Salmonella enterica subsp enterica serovar Typhimurium str ST4 74 YP 005245263 1 909946 Proteobacteria
Salmonella enterica subsp enterica serovar Typhimurium str 14028S YP 005240600 1 588858 Proteobacteria
Salmonella enterica subsp enterica serovar Typhi str Ty21a YP 007928439 1 527001 Proteobacteria
Salmonella enterica subsp enterica serovar Paratyphi A str AKU 12601 YP 002144655 1 554290 Proteobacteria
Salmonella enterica subsp enterica serovar Paratyphi C str RKS4594 YP 002639857 1 476213 Proteobacteria
Salmonella enterica subsp enterica serovar Paratyphi B str SPB7 YP 001591401 1 1016998 Proteobacteria
Salmonella enterica subsp enterica serovar Bareilly str CFSAN000189 YP 008310466 1 1173427 Proteobacteria
Salmonella enterica subsp enterica serovar Javiana str CFSAN001992 YP 007472218 1 1267753 Proteobacteria
Salmonella enterica subsp enterica serovar Heidelberg str CFSAN002069 YP 008250937 1 1271864 Proteobacteria
Salmonella enterica subsp enterica serovar Schwarzengrund str CVM19633 YP 002117228 1 439843 Proteobacteria
Salmonella enterica subsp enterica serovar Choleraesuis str SC B67 YP 219159 1 321314 Proteobacteria
Salmonella enterica subsp enterica serovar Bovismorbificans str 3114 YP 008508161 1 1320309 Proteobacteria
Salmonella enterica subsp enterica serovar Newport str SL254 YP 002043543 1 423368 Proteobacteria
Salmonella enterica subsp enterica serovar Agona str SL483 YP 002149208 1 454166 Proteobacteria
Salmonella enterica subsp enterica serovar Cubana str CFSAN002050 YP 008258898 1 1271863 Proteobacteria
Salmonella enterica subsp enterica serovar Gallinarum str 287 91 YP 002228865 1 550538 Proteobacteria
Salmonella enterica subsp enterica serovar Enteritidis str P125109 YP 002246098 1 550537 Proteobacteria
Salmonella enterica subsp enterica serovar Gallinarum pullorum str RKS5078 YP 005215336 1 1081093 Proteobacteria
Salmonella enterica subsp enterica serovar Pullorum str S06004 YP 008383399 1 1298917 Proteobacteria
Pectobacterium wasabiae WPP163 YP 003261351 1 561231 Proteobacteria
Pectobacterium sp SCC3193 YP 006285100 1 1166016 Proteobacteria
Pectobacterium carotovorum subsp carotovorum PC1 YP 003019386 1 561230 Proteobacteria
Pectobacterium atrosepticum SCRI1043 YP 052130 1 218491 Proteobacteria
Prodigiosinella confusarubida ATCC39006 WP 037380361 1 Pconfusarubida
Dickeya aquatica DW 0440 WP 035344558 1 Daquatica
DAQ1742 DAQ1742 04063 Daquatica
Dickeya sp CSL RW240 WP 035344558 1 Daquatica
Dickeya chrysanthemi subsp parthenii NCPPB 516 WP 027710755 1 Dchrysanthemi
Dickeya chrysanthemi subsp chrysanthemi NCPPB 402 WP 039999311 1 Dchrysanthemi
Dickeya chrysanthemi L11 WP 033577632 1 Dchrysanthemi
Dickeya chrysanthemi NCPPB 3533 WP 012768090 1 Dchrysanthemi
Dickeya chrysanthemi Ech1591 WP 012768090 1 Dchrysanthemi
Dickeya zeae Ech586 WP 041164639 1 Dzeae
Dickeya zeae NCPPB 2538 WP 038916882 1 Dzeae
Dickeya zeae NCPPB 3532 WP 023640872 1 Dzeae
Dickeya zeae MS1 WP 019845198 1 Dzeae
Dickeya zeae MK19 WP 038914708 1 Dzeae
Dickeya zeae CSL RW192 WP 038909141 1 Dzeae
Dickeya zeae NCPPB 3531 WP 038905818 1 Dzeae
Dickeya zeae ZJU1202 WP 016943002 1 Dzeae
Dickeya zeae DZ2Q WP 016943002 1 Dzeae
Dickeya zeae EC1 WP 016943002 1 Dzeae
Dickeya dianthicola GBBC 2039 WP 024110265 1 Ddianthicola
Dickeya dianthicola RNS04 9 WP 024107529 1 Ddianthicola
Dickeya dianthicola IPO 980 WP 024107529 1 Ddianthicola
Dickeya dianthicola NCPPB 453 WP 024107529 1 Ddianthicola
Dickeya dianthicola NCPPB 3534 WP 024107529 1 Ddianthicola
Dickeya gs undicola 2B12 WP 033570869 1 Dundicola
Dickeya gs fangzhongdai NCPPB 3274 WP 042861320 1 Dfangzhongdai
Dickeya gs fangzhongdai M005 WP 039692914 1 Dfangzhongdai
Dickeya gs fangzhongdai ND14b WP 038665043 1 Dfangzhongdai
Dickeya gs fangzhongdai M074 WP 038665043 1 Dfangzhongdai
Dickeya gs fangzhongdai S1 WP 049854210 1 Dfangzhongdai
Dickeya gs fangzhongdai MK7 WP 038920403 1 Dfangzhongdai
Dickeya gs fangzhongdai B16 WP 038920403 1 Dfangzhongdai
418 eptA response to antimicrobial peptides LPS modification Seed
Dickeya dadantii 3937 WP 013319712 1 Ddadantii
Dickeya dadantii NCPPB 898 WP 038912073 1 Ddadantii
Dickeya dadantii NCPPB 3537 WP 038902221 1 Ddadantii
Dickeya dadantii subsp dieffenbachiae NCPPB 2976 WP 038924048 1 Ddadantii
Dickeya solani RNS 05 1 2A WP 057085059 1 Dsolani
Dickeya solani GBBC 2040 WP 022635120 1 Dsolani
Dickeya solani RNS 07 7 3B WP 022635120 1 Dsolani
Dickeya solani MK10 WP 022635120 1 Dsolani
Dickeya solani PPO 9019 WP 022635120 1 Dsolani
Dickeya solani IFB 0099 WP 022635120 1 Dsolani
Dickeya solani D s0432 1 WP 022635120 1 Dsolani
Dickeya solani RNS 08 23 3 1 A WP 022635120 1 Dsolani
Dickeya solani PPO 9134 WP 022635120 1 Dsolani
Dickeya solani IPO 2222 WP 022635120 1 Dsolani
Dickeya solani MK16 WP 022635120 1 Dsolani

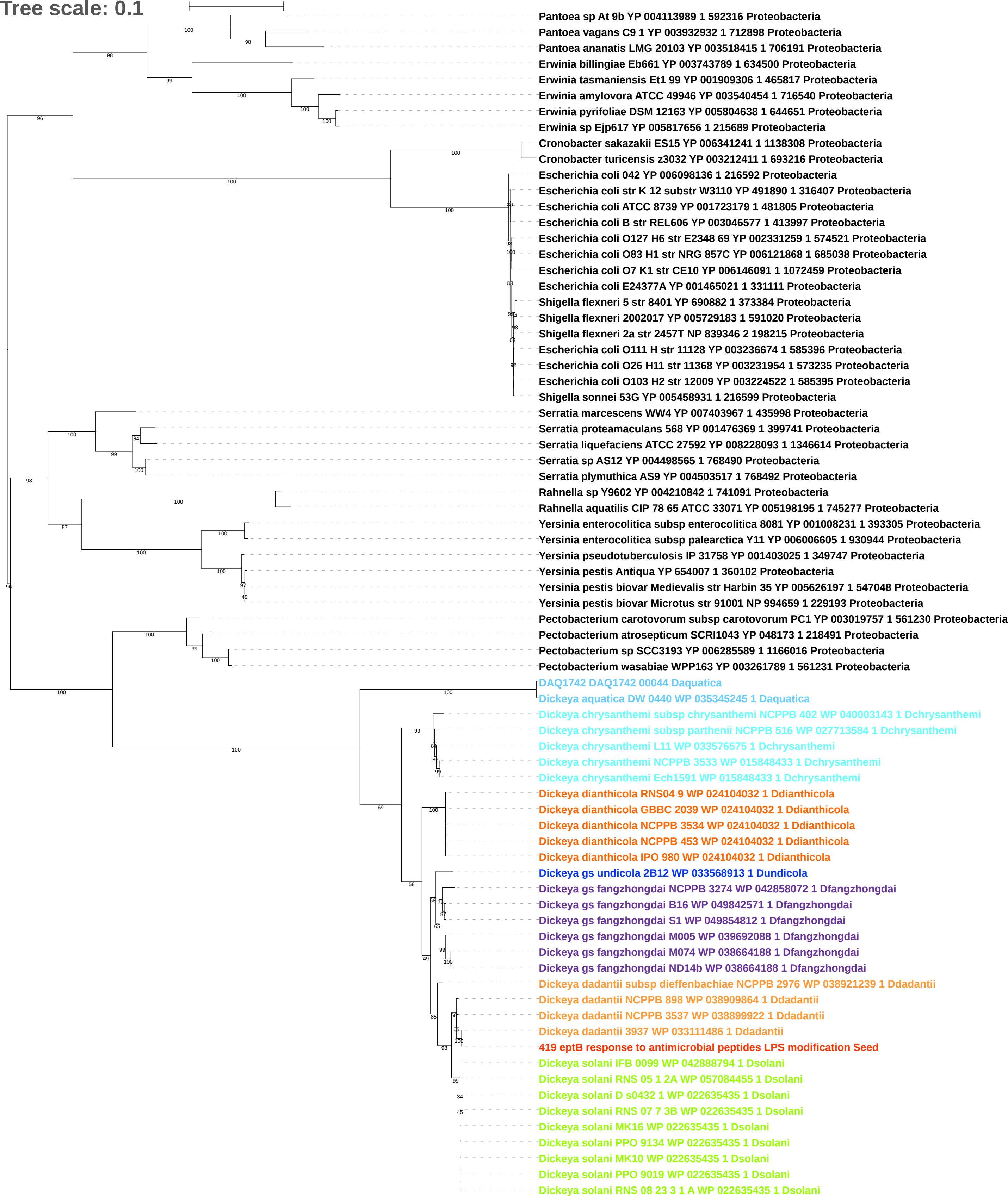
Tree scale: 0.1
Pantoea sp At 9b YP 004113989 1 592316 Proteobacteria
Pantoea vagans C9 1 YP 003932932 1 712898 Proteobacteria
Pantoea ananatis LMG 20103 YP 003518415 1 706191 Proteobacteria
Erwinia billingiae Eb661 YP 003743789 1 634500 Proteobacteria
Erwinia tasmaniensis Et1 99 YP 001909306 1 465817 Proteobacteria
Erwinia amylovora ATCC 49946 YP 003540454 1 716540 Proteobacteria
Erwinia pyrifoliae DSM 12163 YP 005804638 1 644651 Proteobacteria
Erwinia sp Ejp617 YP 005817656 1 215689 Proteobacteria
Cronobacter sakazakii ES15 YP 006341241 1 1138308 Proteobacteria
Cronobacter turicensis z3032 YP 003212411 1 693216 Proteobacteria
Escherichia coli 042 YP 006098136 1 216592 Proteobacteria
Escherichia coli str K 12 substr W3110 YP 491890 1 316407 Proteobacteria
Escherichia coli ATCC 8739 YP 001723179 1 481805 Proteobacteria
Escherichia coli B str REL606 YP 003046577 1 413997 Proteobacteria
Escherichia coli O127 H6 str E2348 69 YP 002331259 1 574521 Proteobacteria
Escherichia coli O83 H1 str NRG 857C YP 006121868 1 685038 Proteobacteria
Escherichia coli O7 K1 str CE10 YP 006146091 1 1072459 Proteobacteria
Escherichia coli E24377A YP 001465021 1 331111 Proteobacteria
Shigella flexneri 5 str 8401 YP 690882 1 373384 Proteobacteria
Shigella flexneri 2002017 YP 005729183 1 591020 Proteobacteria
Shigella flexneri 2a str 2457T NP 839346 2 198215 Proteobacteria
Escherichia coli O111 H str 11128 YP 003236674 1 585396 Proteobacteria
Escherichia coli O26 H11 str 11368 YP 003231954 1 573235 Proteobacteria
Escherichia coli O103 H2 str 12009 YP 003224522 1 585395 Proteobacteria
Shigella sonnei 53G YP 005458931 1 216599 Proteobacteria
Serratia marcescens WW4 YP 007403967 1 435998 Proteobacteria
Serratia proteamaculans 568 YP 001476369 1 399741 Proteobacteria
Serratia liquefaciens ATCC 27592 YP 008228093 1 1346614 Proteobacteria
Serratia sp AS12 YP 004498565 1 768490 Proteobacteria
Serratia plymuthica AS9 YP 004503517 1 768492 Proteobacteria
Rahnella sp Y9602 YP 004210842 1 741091 Proteobacteria
Rahnella aquatilis CIP 78 65 ATCC 33071 YP 005198195 1 745277 Proteobacteria
Yersinia enterocolitica subsp enterocolitica 8081 YP 001008231 1 393305 Proteobacteria
Yersinia enterocolitica subsp palearctica Y11 YP 006006605 1 930944 Proteobacteria
Yersinia pseudotuberculosis IP 31758 YP 001403025 1 349747 Proteobacteria
Yersinia pestis Antiqua YP 654007 1 360102 Proteobacteria
Yersinia pestis biovar Medievalis str Harbin 35 YP 005626197 1 547048 Proteobacteria
Yersinia pestis biovar Microtus str 91001 NP 994659 1 229193 Proteobacteria
Pectobacterium carotovorum subsp carotovorum PC1 YP 003019757 1 561230 Proteobacteria
Pectobacterium atrosepticum SCRI1043 YP 048173 1 218491 Proteobacteria
Pectobacterium sp SCC3193 YP 006285589 1 1166016 Proteobacteria
Pectobacterium wasabiae WPP163 YP 003261789 1 561231 Proteobacteria
DAQ1742 DAQ1742 00044 Daquatica
Dickeya aquatica DW 0440 WP 035345245 1 Daquatica
Dickeya chrysanthemi subsp chrysanthemi NCPPB 402 WP 040003143 1 Dchrysanthemi
Dickeya chrysanthemi subsp parthenii NCPPB 516 WP 027713584 1 Dchrysanthemi
Dickeya chrysanthemi L11 WP 033576575 1 Dchrysanthemi
Dickeya chrysanthemi NCPPB 3533 WP 015848433 1 Dchrysanthemi
Dickeya chrysanthemi Ech1591 WP 015848433 1 Dchrysanthemi
Dickeya dianthicola RNS04 9 WP 024104032 1 Ddianthicola
Dickeya dianthicola GBBC 2039 WP 024104032 1 Ddianthicola
Dickeya dianthicola NCPPB 3534 WP 024104032 1 Ddianthicola
Dickeya dianthicola NCPPB 453 WP 024104032 1 Ddianthicola
Dickeya dianthicola IPO 980 WP 024104032 1 Ddianthicola
Dickeya gs undicola 2B12 WP 033568913 1 Dundicola
Dickeya gs fangzhongdai NCPPB 3274 WP 042858072 1 Dfangzhongdai
Dickeya gs fangzhongdai B16 WP 049842571 1 Dfangzhongdai
Dickeya gs fangzhongdai S1 WP 049854812 1 Dfangzhongdai
Dickeya gs fangzhongdai M005 WP 039692088 1 Dfangzhongdai
Dickeya gs fangzhongdai M074 WP 038664188 1 Dfangzhongdai
Dickeya gs fangzhongdai ND14b WP 038664188 1 Dfangzhongdai
Dickeya dadantii subsp dieffenbachiae NCPPB 2976 WP 038921239 1 Ddadantii
Dickeya dadantii NCPPB 898 WP 038909864 1 Ddadantii
Dickeya dadantii NCPPB 3537 WP 038899922 1 Ddadantii
Dickeya dadantii 3937 WP 033111486 1 Ddadantii
419 eptB response to antimicrobial peptides LPS modification Seed
Dickeya solani IFB 0099 WP 042888794 1 Dsolani
Dickeya solani RNS 05 1 2A WP 057084455 1 Dsolani
Dickeya solani D s0432 1 WP 022635435 1 Dsolani
Dickeya solani RNS 07 7 3B WP 022635435 1 Dsolani
Dickeya solani MK16 WP 022635435 1 Dsolani
Dickeya solani PPO 9134 WP 022635435 1 Dsolani
Dickeya solani MK10 WP 022635435 1 Dsolani
Dickeya solani PPO 9019 WP 022635435 1 Dsolani
Dickeya solani RNS 08 23 3 1 A WP 022635435 1 Dsolani

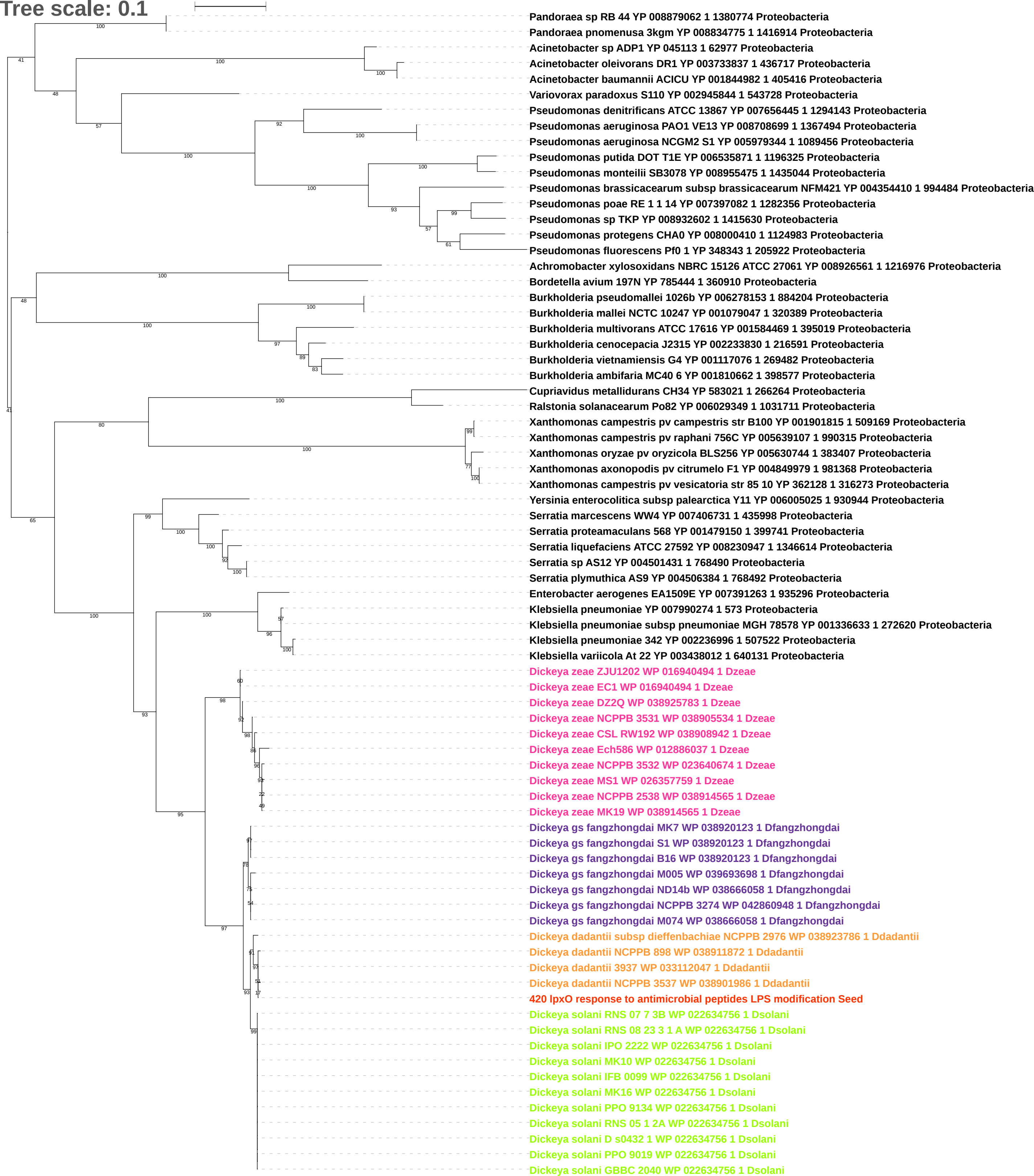

Tree scale: 0.1
Pandoraea sp RB 44 YP 008879062 1 1380774 Proteobacteria
Pandoraea pnomenusa 3kgm YP 008834775 1 1416914 Proteobacteria
Acinetobacter sp ADP1 YP 045113 1 62977 Proteobacteria
Acinetobacter oleivorans DR1 YP 003733837 1 436717 Proteobacteria
Acinetobacter baumannii ACICU YP 001844982 1 405416 Proteobacteria
Variovorax paradoxus S110 YP 002945844 1 543728 Proteobacteria
Pseudomonas denitrificans ATCC 13867 YP 007656445 1 1294143 Proteobacteria
Pseudomonas aeruginosa PAO1 VE13 YP 008708699 1 1367494 Proteobacteria
Pseudomonas aeruginosa NCGM2 S1 YP 005979344 1 1089456 Proteobacteria
Pseudomonas putida DOT T1E YP 006535871 1 1196325 Proteobacteria
Pseudomonas monteilii SB3078 YP 008955475 1 1435044 Proteobacteria
Pseudomonas brassicacearum subsp brassicacearum NFM421 YP 004354410 1 994484 Proteobacteria
Pseudomonas poae RE 1 1 14 YP 007397082 1 1282356 Proteobacteria
Pseudomonas sp TKP YP 008932602 1 1415630 Proteobacteria
Pseudomonas protegens CHA0 YP 008000410 1 1124983 Proteobacteria
Pseudomonas fluorescens Pf0 1 YP 348343 1 205922 Proteobacteria
Achromobacter xylosoxidans NBRC 15126 ATCC 27061 YP 008926561 1 1216976 Proteobacteria
Bordetella avium 197N YP 785444 1 360910 Proteobacteria
Burkholderia pseudomallei 1026b YP 006278153 1 884204 Proteobacteria
Burkholderia mallei NCTC 10247 YP 001079047 1 320389 Proteobacteria
Burkholderia multivorans ATCC 17616 YP 001584469 1 395019 Proteobacteria
Burkholderia cenocepacia J2315 YP 002233830 1 216591 Proteobacteria
Burkholderia vietnamiensis G4 YP 001117076 1 269482 Proteobacteria
Burkholderia ambifaria MC40 6 YP 001810662 1 398577 Proteobacteria
Cupriavidus metallidurans CH34 YP 583021 1 266264 Proteobacteria
Ralstonia solanacearum Po82 YP 006029349 1 1031711 Proteobacteria
Xanthomonas campestris pv campestris str B100 YP 001901815 1 509169 Proteobacteria
Xanthomonas campestris pv raphani 756C YP 005639107 1 990315 Proteobacteria
Xanthomonas oryzae pv oryzicola BLS256 YP 005630744 1 383407 Proteobacteria
Xanthomonas axonopodis pv citrumelo F1 YP 004849979 1 981368 Proteobacteria
Xanthomonas campestris pv vesicatoria str 85 10 YP 362128 1 316273 Proteobacteria
Yersinia enterocolitica subsp palearctica Y11 YP 006005025 1 930944 Proteobacteria
Serratia marcescens WW4 YP 007406731 1 435998 Proteobacteria
Serratia proteamaculans 568 YP 001479150 1 399741 Proteobacteria
Serratia liquefaciens ATCC 27592 YP 008230947 1 1346614 Proteobacteria
Serratia sp AS12 YP 004501431 1 768490 Proteobacteria
Serratia plymuthica AS9 YP 004506384 1 768492 Proteobacteria
Enterobacter aerogenes EA1509E YP 007391263 1 935296 Proteobacteria
Klebsiella pneumoniae YP 007990274 1 573 Proteobacteria
Klebsiella pneumoniae subsp pneumoniae MGH 78578 YP 001336633 1 272620 Proteobacteria
Klebsiella pneumoniae 342 YP 002236996 1 507522 Proteobacteria
Klebsiella variicola At 22 YP 003438012 1 640131 Proteobacteria
Dickeya zeae ZJU1202 WP 016940494 1 Dzeae
Dickeya zeae EC1 WP 016940494 1 Dzeae
Dickeya zeae DZ2Q WP 038925783 1 Dzeae
Dickeya zeae NCPPB 3531 WP 038905534 1 Dzeae
Dickeya zeae CSL RW192 WP 038908942 1 Dzeae
Dickeya zeae Ech586 WP 012886037 1 Dzeae
Dickeya zeae NCPPB 3532 WP 023640674 1 Dzeae
Dickeya zeae MS1 WP 026357759 1 Dzeae
Dickeya zeae NCPPB 2538 WP 038914565 1 Dzeae
Dickeya zeae MK19 WP 038914565 1 Dzeae
Dickeya gs fangzhongdai MK7 WP 038920123 1 Dfangzhongdai
Dickeya gs fangzhongdai S1 WP 038920123 1 Dfangzhongdai
Dickeya gs fangzhongdai B16 WP 038920123 1 Dfangzhongdai
Dickeya gs fangzhongdai M005 WP 039693698 1 Dfangzhongdai
Dickeya gs fangzhongdai ND14b WP 038666058 1 Dfangzhongdai
Dickeya gs fangzhongdai NCPPB 3274 WP 042860948 1 Dfangzhongdai
Dickeya gs fangzhongdai M074 WP 038666058 1 Dfangzhongdai
Dickeya dadantii subsp dieffenbachiae NCPPB 2976 WP 038923786 1 Ddadantii
Dickeya dadantii NCPPB 898 WP 038911872 1 Ddadantii
Dickeya dadantii 3937 WP 033112047 1 Ddadantii
Dickeya dadantii NCPPB 3537 WP 038901986 1 Ddadantii
420 lpxO response to antimicrobial peptides LPS modification Seed
Dickeya solani RNS 07 7 3B WP 022634756 1 Dsolani
Dickeya solani RNS 08 23 3 1 A WP 022634756 1 Dsolani
Dickeya solani IPO 2222 WP 022634756 1 Dsolani
Dickeya solani MK10 WP 022634756 1 Dsolani
Dickeya solani IFB 0099 WP 022634756 1 Dsolani
Dickeya solani MK16 WP 022634756 1 Dsolani
Dickeya solani PPO 9134 WP 022634756 1 Dsolani
Dickeya solani RNS 05 1 2A WP 022634756 1 Dsolani
Dickeya solani D s0432 1 WP 022634756 1 Dsolani
Dickeya solani PPO 9019 WP 022634756 1 Dsolani
Dickeya solani GBBC 2040 WP 022634756 1 Dsolani

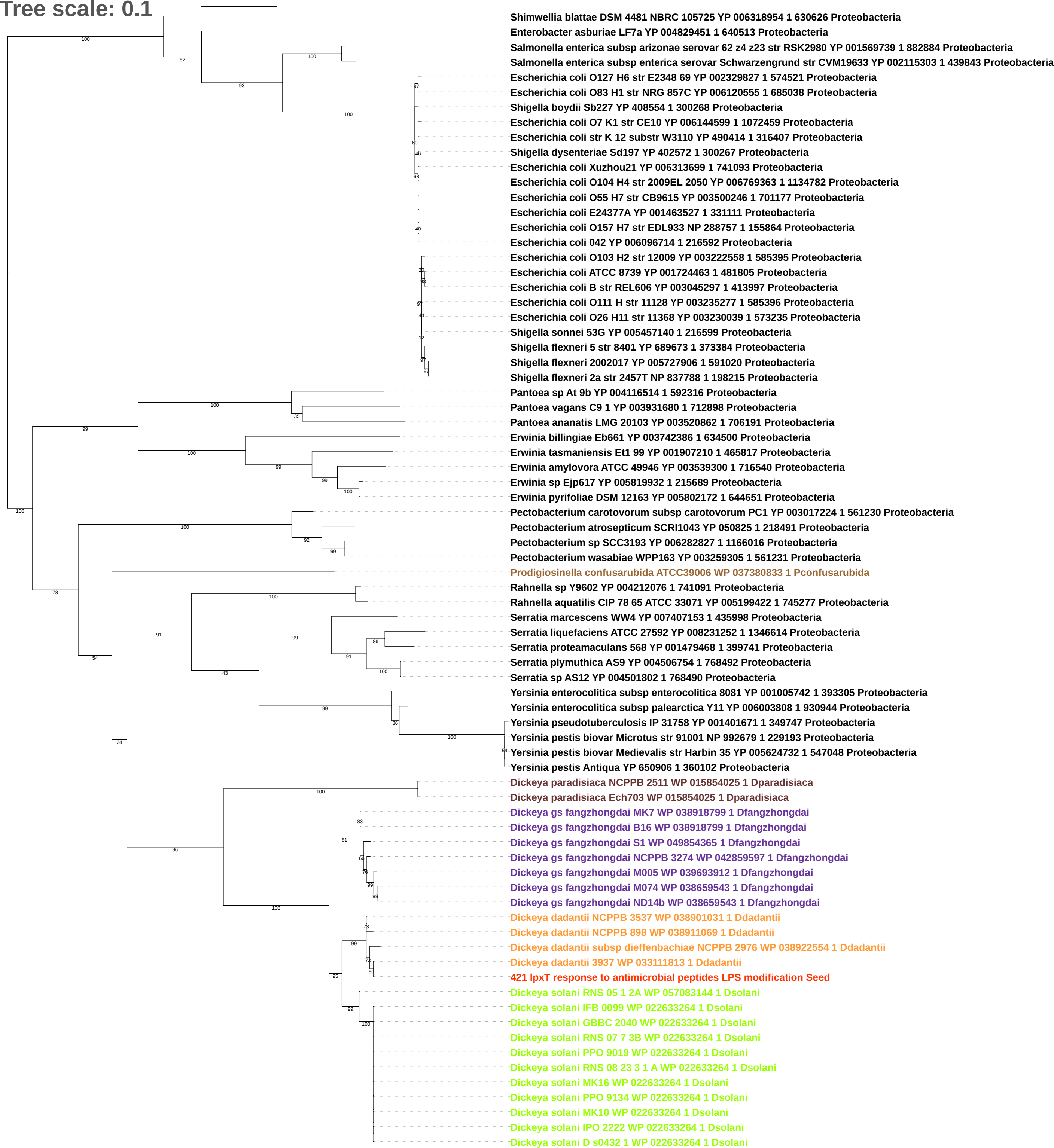
Tree scale: 0.1
Shimwellia blattae DSM 4481 NBRC 105725 YP 006318954 1 630626 Proteobacteria
Enterobacter asburiae LF7a YP 004829451 1 640513 Proteobacteria
Salmonella enterica subsp arizonae serovar 62 z4 z23 str RSK2980 YP 001569739 1 882884 Proteobacteria
Salmonella enterica subsp enterica serovar Schwarzengrund str CVM19633 YP 002115303 1 439843 Proteobacteria
Escherichia coli O127 H6 str E2348 69 YP 002329827 1 574521 Proteobacteria
Escherichia coli O83 H1 str NRG 857C YP 006120555 1 685038 Proteobacteria
Shigella boydii Sb227 YP 408554 1 300268 Proteobacteria
Escherichia coli O7 K1 str CE10 YP 006144599 1 1072459 Proteobacteria
Escherichia coli str K 12 substr W3110 YP 490414 1 316407 Proteobacteria
Shigella dysenteriae Sd197 YP 402572 1 300267 Proteobacteria
Escherichia coli Xuzhou21 YP 006313699 1 741093 Proteobacteria
Escherichia coli O104 H4 str 2009EL 2050 YP 006769363 1 1134782 Proteobacteria
Escherichia coli O55 H7 str CB9615 YP 003500246 1 701177 Proteobacteria
Escherichia coli E24377A YP 001463527 1 331111 Proteobacteria
Escherichia coli O157 H7 str EDL933 NP 288757 1 155864 Proteobacteria
Escherichia coli 042 YP 006096714 1 216592 Proteobacteria
Escherichia coli O103 H2 str 12009 YP 003222558 1 585395 Proteobacteria
Escherichia coli ATCC 8739 YP 001724463 1 481805 Proteobacteria
Escherichia coli B str REL606 YP 003045297 1 413997 Proteobacteria
Escherichia coli O111 H str 11128 YP 003235277 1 585396 Proteobacteria
Escherichia coli O26 H11 str 11368 YP 003230039 1 573235 Proteobacteria
Shigella sonnei 53G YP 005457140 1 216599 Proteobacteria
Shigella flexneri 5 str 8401 YP 689673 1 373384 Proteobacteria
Shigella flexneri 2002017 YP 005727906 1 591020 Proteobacteria
Shigella flexneri 2a str 2457T NP 837788 1 198215 Proteobacteria
Pantoea sp At 9b YP 004116514 1 592316 Proteobacteria
Pantoea vagans C9 1 YP 003931680 1 712898 Proteobacteria
Pantoea ananatis LMG 20103 YP 003520862 1 706191 Proteobacteria
Erwinia billingiae Eb661 YP 003742386 1 634500 Proteobacteria
Erwinia tasmaniensis Et1 99 YP 001907210 1 465817 Proteobacteria
Erwinia amylovora ATCC 49946 YP 003539300 1 716540 Proteobacteria
Erwinia sp Ejp617 YP 005819932 1 215689 Proteobacteria
Erwinia pyrifoliae DSM 12163 YP 005802172 1 644651 Proteobacteria
Pectobacterium carotovorum subsp carotovorum PC1 YP 003017224 1 561230 Proteobacteria
Pectobacterium atrosepticum SCRI1043 YP 050825 1 218491 Proteobacteria
Pectobacterium sp SCC3193 YP 006282827 1 1166016 Proteobacteria
Pectobacterium wasabiae WPP163 YP 003259305 1 561231 Proteobacteria
Prodigiosinella confusarubida ATCC39006 WP 037380833 1 Pconfusarubida
Rahnella sp Y9602 YP 004212076 1 741091 Proteobacteria
Rahnella aquatilis CIP 78 65 ATCC 33071 YP 005199422 1 745277 Proteobacteria
Serratia marcescens WW4 YP 007407153 1 435998 Proteobacteria
Serratia liquefaciens ATCC 27592 YP 008231252 1 1346614 Proteobacteria
Serratia proteamaculans 568 YP 001479468 1 399741 Proteobacteria
Serratia plymuthica AS9 YP 004506754 1 768492 Proteobacteria
Serratia sp AS12 YP 004501802 1 768490 Proteobacteria
Yersinia enterocolitica subsp enterocolitica 8081 YP 001005742 1 393305 Proteobacteria
Yersinia enterocolitica subsp palearctica Y11 YP 006003808 1 930944 Proteobacteria
Yersinia pseudotuberculosis IP 31758 YP 001401671 1 349747 Proteobacteria
Yersinia pestis biovar Microtus str 91001 NP 992679 1 229193 Proteobacteria
Yersinia pestis biovar Medievalis str Harbin 35 YP 005624732 1 547048 Proteobacteria
Yersinia pestis Antiqua YP 650906 1 360102 Proteobacteria
Dickeya paradisiaca NCPPB 2511 WP 015854025 1 Dparadisiaca
Dickeya paradisiaca Ech703 WP 015854025 1 Dparadisiaca
Dickeya gs fangzhongdai MK7 WP 038918799 1 Dfangzhongdai
Dickeya gs fangzhongdai B16 WP 038918799 1 Dfangzhongdai
Dickeya gs fangzhongdai S1 WP 049854365 1 Dfangzhongdai
Dickeya gs fangzhongdai NCPPB 3274 WP 042859597 1 Dfangzhongdai
Dickeya gs fangzhongdai M005 WP 039693912 1 Dfangzhongdai
Dickeya gs fangzhongdai M074 WP 038659543 1 Dfangzhongdai
Dickeya gs fangzhongdai ND14b WP 038659543 1 Dfangzhongdai
Dickeya dadantii NCPPB 3537 WP 038901031 1 Ddadantii
Dickeya dadantii NCPPB 898 WP 038911069 1 Ddadantii
Dickeya dadantii subsp dieffenbachiae NCPPB 2976 WP 038922554 1 Ddadantii
Dickeya dadantii 3937 WP 033111813 1 Ddadantii
421 lpxT response to antimicrobial peptides LPS modification Seed
Dickeya solani RNS 05 1 2A WP 057083144 1 Dsolani
Dickeya solani IFB 0099 WP 022633264 1 Dsolani
Dickeya solani GBBC 2040 WP 022633264 1 Dsolani
Dickeya solani RNS 07 7 3B WP 022633264 1 Dsolani
Dickeya solani PPO 9019 WP 022633264 1 Dsolani
Dickeya solani RNS 08 23 3 1 A WP 022633264 1 Dsolani
Dickeya solani MK16 WP 022633264 1 Dsolani
Dickeya solani PPO 9134 WP 022633264 1 Dsolani
Dickeya solani MK10 WP 022633264 1 Dsolani
Dickeya solani IPO 2222 WP 022633264 1 Dsolani
Dickeya solani D s0432 1 WP 022633264 1 Dsolani

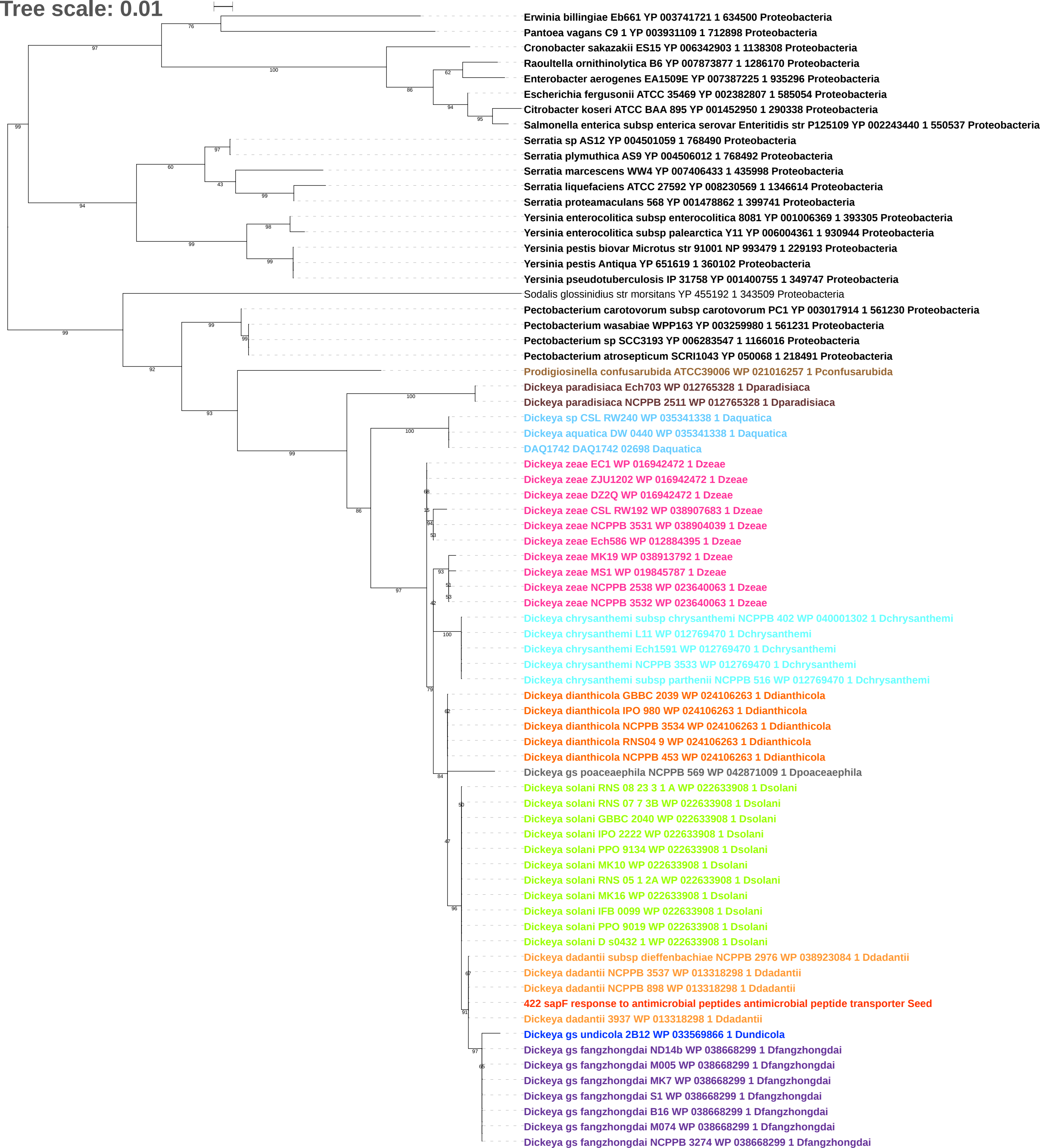

Tree scale: 0.01
Erwinia billingiae Eb661 YP 003741721 1 634500 Proteobacteria
Pantoea vagans C9 1 YP 003931109 1 712898 Proteobacteria
Cronobacter sakazakii ES15 YP 006342903 1 1138308 Proteobacteria
Raoultella ornithinolytica B6 YP 007873877 1 1286170 Proteobacteria
Enterobacter aerogenes EA1509E YP 007387225 1 935296 Proteobacteria
Escherichia fergusonii ATCC 35469 YP 002382807 1 585054 Proteobacteria
Citrobacter koseri ATCC BAA 895 YP 001452950 1 290338 Proteobacteria
Salmonella enterica subsp enterica serovar Enteritidis str P125109 YP 002243440 1 550537 Proteobacteria
Serratia sp AS12 YP 004501059 1 768490 Proteobacteria
Serratia plymuthica AS9 YP 004506012 1 768492 Proteobacteria
Serratia marcescens WW4 YP 007406433 1 435998 Proteobacteria
Serratia liquefaciens ATCC 27592 YP 008230569 1 1346614 Proteobacteria
Serratia proteamaculans 568 YP 001478862 1 399741 Proteobacteria
Yersinia enterocolitica subsp enterocolitica 8081 YP 001006369 1 393305 Proteobacteria
Yersinia enterocolitica subsp palearctica Y11 YP 006004361 1 930944 Proteobacteria
Yersinia pestis biovar Microtus str 91001 NP 993479 1 229193 Proteobacteria
Yersinia pestis Antiqua YP 651619 1 360102 Proteobacteria
Yersinia pseudotuberculosis IP 31758 YP 001400755 1 349747 Proteobacteria
Sodalis glossinidius str morsitans YP 455192 1 343509 Proteobacteria
Pectobacterium carotovorum subsp carotovorum PC1 YP 003017914 1 561230 Proteobacteria
Pectobacterium wasabiae WPP163 YP 003259980 1 561231 Proteobacteria
Pectobacterium sp SCC3193 YP 006283547 1 1166016 Proteobacteria
Pectobacterium atrosepticum SCRI1043 YP 050068 1 218491 Proteobacteria
Prodigiosinella confusarubida ATCC39006 WP 021016257 1 Pconfusarubida
Dickeya paradisiaca Ech703 WP 012765328 1 Dparadisiaca
Dickeya paradisiaca NCPPB 2511 WP 012765328 1 Dparadisiaca
Dickeya sp CSL RW240 WP 035341338 1 Daquatica
Dickeya aquatica DW 0440 WP 035341338 1 Daquatica
DAQ1742 DAQ1742 02698 Daquatica
Dickeya zeae EC1 WP 016942472 1 Dzeae
Dickeya zeae ZJU1202 WP 016942472 1 Dzeae
Dickeya zeae DZ2Q WP 016942472 1 Dzeae
Dickeya zeae CSL RW192 WP 038907683 1 Dzeae
Dickeya zeae NCPPB 3531 WP 038904039 1 Dzeae
Dickeya zeae Ech586 WP 012884395 1 Dzeae
Dickeya zeae MK19 WP 038913792 1 Dzeae
Dickeya zeae MS1 WP 019845787 1 Dzeae
Dickeya zeae NCPPB 2538 WP 023640063 1 Dzeae
Dickeya zeae NCPPB 3532 WP 023640063 1 Dzeae
Dickeya chrysanthemi subsp chrysanthemi NCPPB 402 WP 040001302 1 Dchrysanthemi
Dickeya chrysanthemi L11 WP 012769470 1 Dchrysanthemi
Dickeya chrysanthemi Ech1591 WP 012769470 1 Dchrysanthemi
Dickeya chrysanthemi NCPPB 3533 WP 012769470 1 Dchrysanthemi
Dickeya chrysanthemi subsp parthenii NCPPB 516 WP 012769470 1 Dchrysanthemi
Dickeya dianthicola GBBC 2039 WP 024106263 1 Ddianthicola
Dickeya dianthicola IPO 980 WP 024106263 1 Ddianthicola
Dickeya dianthicola NCPPB 3534 WP 024106263 1 Ddianthicola
Dickeya dianthicola RNS04 9 WP 024106263 1 Ddianthicola
Dickeya dianthicola NCPPB 453 WP 024106263 1 Ddianthicola
Dickeya gs poaceaephila NCPPB 569 WP 042871009 1 Dpoaceaephila
Dickeya solani RNS 08 23 3 1 A WP 022633908 1 Dsolani
Dickeya solani RNS 07 7 3B WP 022633908 1 Dsolani
Dickeya solani GBBC 2040 WP 022633908 1 Dsolani
Dickeya solani IPO 2222 WP 022633908 1 Dsolani
Dickeya solani PPO 9134 WP 022633908 1 Dsolani
Dickeya solani MK10 WP 022633908 1 Dsolani
Dickeya solani RNS 05 1 2A WP 022633908 1 Dsolani
Dickeya solani MK16 WP 022633908 1 Dsolani
Dickeya solani IFB 0099 WP 022633908 1 Dsolani
Dickeya solani PPO 9019 WP 022633908 1 Dsolani
Dickeya solani D s0432 1 WP 022633908 1 Dsolani
Dickeya dadantii subsp dieffenbachiae NCPPB 2976 WP 038923084 1 Ddadantii
Dickeya dadantii NCPPB 3537 WP 013318298 1 Ddadantii
Dickeya dadantii NCPPB 898 WP 013318298 1 Ddadantii
422 sapF response to antimicrobial peptides antimicrobial peptide transporter Seed
Dickeya dadantii 3937 WP 013318298 1 Ddadantii
Dickeya gs undicola 2B12 WP 033569866 1 Dundicola
Dickeya gs fangzhongdai ND14b WP 038668299 1 Dfangzhongdai
Dickeya gs fangzhongdai M005 WP 038668299 1 Dfangzhongdai
Dickeya gs fangzhongdai MK7 WP 038668299 1 Dfangzhongdai
Dickeya gs fangzhongdai S1 WP 038668299 1 Dfangzhongdai
Dickeya gs fangzhongdai B16 WP 038668299 1 Dfangzhongdai
Dickeya gs fangzhongdai M074 WP 038668299 1 Dfangzhongdai
Dickeya gs fangzhongdai NCPPB 3274 WP 038668299 1 Dfangzhongdai

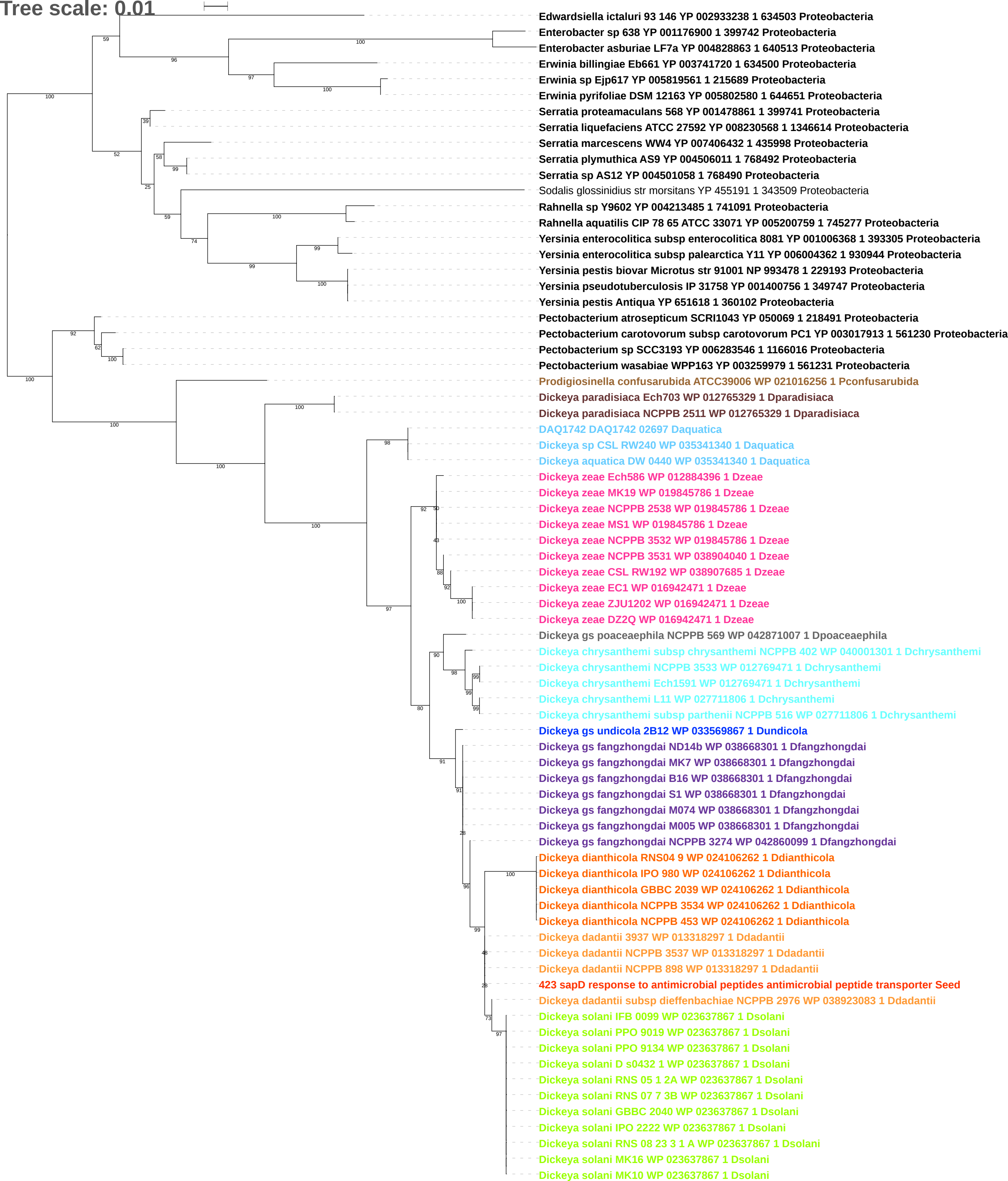
Tree scale: 0.01
Edwardsiella ictaluri 93 146 YP 002933238 1 634503 Proteobacteria
Enterobacter sp 638 YP 001176900 1 399742 Proteobacteria
Enterobacter asburiae LF7a YP 004828863 1 640513 Proteobacteria
Erwinia billingiae Eb661 YP 003741720 1 634500 Proteobacteria
Erwinia sp Ejp617 YP 005819561 1 215689 Proteobacteria
Erwinia pyrifoliae DSM 12163 YP 005802580 1 644651 Proteobacteria
Serratia proteamaculans 568 YP 001478861 1 399741 Proteobacteria
Serratia liquefaciens ATCC 27592 YP 008230568 1 1346614 Proteobacteria
Serratia marcescens WW4 YP 007406432 1 435998 Proteobacteria
Serratia plymuthica AS9 YP 004506011 1 768492 Proteobacteria
Serratia sp AS12 YP 004501058 1 768490 Proteobacteria
Sodalis glossinidius str morsitans YP 455191 1 343509 Proteobacteria
Rahnella sp Y9602 YP 004213485 1 741091 Proteobacteria
Rahnella aquatilis CIP 78 65 ATCC 33071 YP 005200759 1 745277 Proteobacteria
Yersinia enterocolitica subsp enterocolitica 8081 YP 001006368 1 393305 Proteobacteria
Yersinia enterocolitica subsp palearctica Y11 YP 006004362 1 930944 Proteobacteria
Yersinia pestis biovar Microtus str 91001 NP 993478 1 229193 Proteobacteria
Yersinia pseudotuberculosis IP 31758 YP 001400756 1 349747 Proteobacteria
Yersinia pestis Antiqua YP 651618 1 360102 Proteobacteria
Pectobacterium atrosepticum SCRI1043 YP 050069 1 218491 Proteobacteria
Pectobacterium carotovorum subsp carotovorum PC1 YP 003017913 1 561230 Proteobacteria
Pectobacterium sp SCC3193 YP 006283546 1 1166016 Proteobacteria
Pectobacterium wasabiae WPP163 YP 003259979 1 561231 Proteobacteria
Prodigiosinella confusarubida ATCC39006 WP 021016256 1 Pconfusarubida
Dickeya paradisiaca Ech703 WP 012765329 1 Dparadisiaca
Dickeya paradisiaca NCPPB 2511 WP 012765329 1 Dparadisiaca
DAQ1742 DAQ1742 02697 Daquatica
Dickeya sp CSL RW240 WP 035341340 1 Daquatica
Dickeya aquatica DW 0440 WP 035341340 1 Daquatica
Dickeya zeae Ech586 WP 012884396 1 Dzeae
Dickeya zeae MK19 WP 019845786 1 Dzeae
Dickeya zeae NCPPB 2538 WP 019845786 1 Dzeae
Dickeya zeae MS1 WP 019845786 1 Dzeae
Dickeya zeae NCPPB 3532 WP 019845786 1 Dzeae
Dickeya zeae NCPPB 3531 WP 038904040 1 Dzeae
Dickeya zeae CSL RW192 WP 038907685 1 Dzeae
Dickeya zeae EC1 WP 016942471 1 Dzeae
Dickeya zeae ZJU1202 WP 016942471 1 Dzeae
Dickeya zeae DZ2Q WP 016942471 1 Dzeae
Dickeya gs poaceaephila NCPPB 569 WP 042871007 1 Dpoaceaephila
Dickeya chrysanthemi subsp chrysanthemi NCPPB 402 WP 040001301 1 Dchrysanthemi
Dickeya chrysanthemi NCPPB 3533 WP 012769471 1 Dchrysanthemi
Dickeya chrysanthemi Ech1591 WP 012769471 1 Dchrysanthemi
Dickeya chrysanthemi L11 WP 027711806 1 Dchrysanthemi
Dickeya chrysanthemi subsp parthenii NCPPB 516 WP 027711806 1 Dchrysanthemi
Dickeya gs undicola 2B12 WP 033569867 1 Dundicola
Dickeya gs fangzhongdai ND14b WP 038668301 1 Dfangzhongdai
Dickeya gs fangzhongdai MK7 WP 038668301 1 Dfangzhongdai
Dickeya gs fangzhongdai B16 WP 038668301 1 Dfangzhongdai
Dickeya gs fangzhongdai S1 WP 038668301 1 Dfangzhongdai
Dickeya gs fangzhongdai M074 WP 038668301 1 Dfangzhongdai
Dickeya gs fangzhongdai M005 WP 038668301 1 Dfangzhongdai
Dickeya gs fangzhongdai NCPPB 3274 WP 042860099 1 Dfangzhongdai
Dickeya dianthicola RNS04 9 WP 024106262 1 Ddianthicola
Dickeya dianthicola IPO 980 WP 024106262 1 Ddianthicola
Dickeya dianthicola GBBC 2039 WP 024106262 1 Ddianthicola
Dickeya dianthicola NCPPB 3534 WP 024106262 1 Ddianthicola
Dickeya dianthicola NCPPB 453 WP 024106262 1 Ddianthicola
Dickeya dadantii 3937 WP 013318297 1 Ddadantii
Dickeya dadantii NCPPB 3537 WP 013318297 1 Ddadantii
Dickeya dadantii NCPPB 898 WP 013318297 1 Ddadantii
423 sapD response to antimicrobial peptides antimicrobial peptide transporter Seed
Dickeya dadantii subsp dieffenbachiae NCPPB 2976 WP 038923083 1 Ddadantii
Dickeya solani IFB 0099 WP 023637867 1 Dsolani
Dickeya solani PPO 9019 WP 023637867 1 Dsolani
Dickeya solani PPO 9134 WP 023637867 1 Dsolani
Dickeya solani D s0432 1 WP 023637867 1 Dsolani
Dickeya solani RNS 05 1 2A WP 023637867 1 Dsolani
Dickeya solani RNS 07 7 3B WP 023637867 1 Dsolani
Dickeya solani GBBC 2040 WP 023637867 1 Dsolani
Dickeya solani IPO 2222 WP 023637867 1 Dsolani
Dickeya solani RNS 08 23 3 1 A WP 023637867 1 Dsolani
Dickeya solani MK16 WP 023637867 1 Dsolani
Dickeya solani MK10 WP 023637867 1 Dsolani

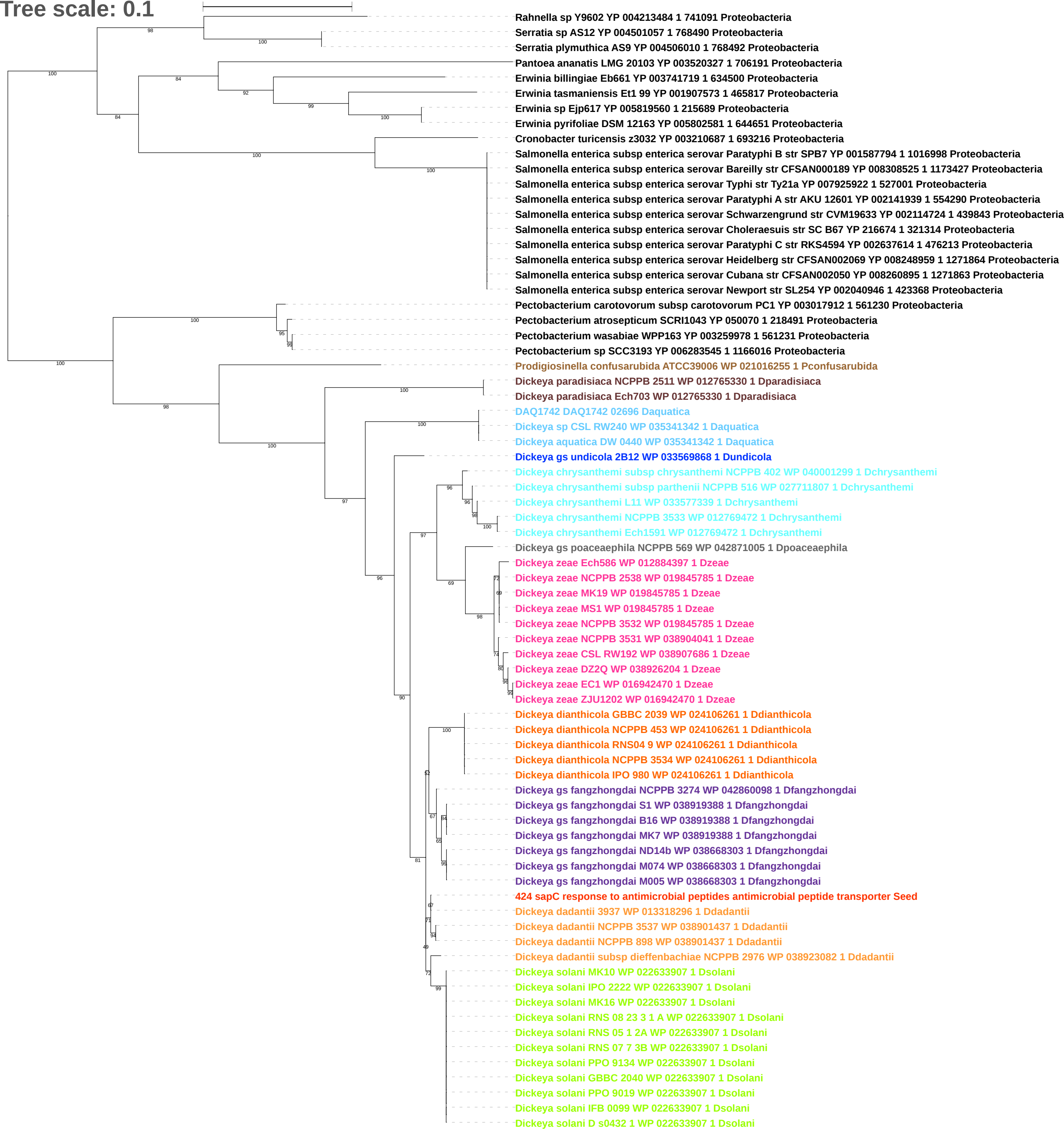
Tree scale: 0.1
Rahnella sp Y9602 YP 004213484 1 741091 Proteobacteria
Serratia sp AS12 YP 004501057 1 768490 Proteobacteria
Serratia plymuthica AS9 YP 004506010 1 768492 Proteobacteria
Pantoea ananatis LMG 20103 YP 003520327 1 706191 Proteobacteria
Erwinia billingiae Eb661 YP 003741719 1 634500 Proteobacteria
Erwinia tasmaniensis Et1 99 YP 001907573 1 465817 Proteobacteria
Erwinia sp Ejp617 YP 005819560 1 215689 Proteobacteria
Erwinia pyrifoliae DSM 12163 YP 005802581 1 644651 Proteobacteria
Cronobacter turicensis z3032 YP 003210687 1 693216 Proteobacteria
Salmonella enterica subsp enterica serovar Paratyphi B str SPB7 YP 001587794 1 1016998 Proteobacteria
Salmonella enterica subsp enterica serovar Bareilly str CFSAN000189 YP 008308525 1 1173427 Proteobacteria
Salmonella enterica subsp enterica serovar Typhi str Ty21a YP 007925922 1 527001 Proteobacteria
Salmonella enterica subsp enterica serovar Paratyphi A str AKU 12601 YP 002141939 1 554290 Proteobacteria
Salmonella enterica subsp enterica serovar Schwarzengrund str CVM19633 YP 002114724 1 439843 Proteobacteria
Salmonella enterica subsp enterica serovar Choleraesuis str SC B67 YP 216674 1 321314 Proteobacteria
Salmonella enterica subsp enterica serovar Paratyphi C str RKS4594 YP 002637614 1 476213 Proteobacteria
Salmonella enterica subsp enterica serovar Heidelberg str CFSAN002069 YP 008248959 1 1271864 Proteobacteria
Salmonella enterica subsp enterica serovar Cubana str CFSAN002050 YP 008260895 1 1271863 Proteobacteria
Salmonella enterica subsp enterica serovar Newport str SL254 YP 002040946 1 423368 Proteobacteria
Pectobacterium carotovorum subsp carotovorum PC1 YP 003017912 1 561230 Proteobacteria
Pectobacterium atrosepticum SCRI1043 YP 050070 1 218491 Proteobacteria
Pectobacterium wasabiae WPP163 YP 003259978 1 561231 Proteobacteria
Pectobacterium sp SCC3193 YP 006283545 1 1166016 Proteobacteria
Prodigiosinella confusarubida ATCC39006 WP 021016255 1 Pconfusarubida
Dickeya paradisiaca NCPPB 2511 WP 012765330 1 Dparadisiaca
Dickeya paradisiaca Ech703 WP 012765330 1 Dparadisiaca
DAQ1742 DAQ1742 02696 Daquatica
Dickeya sp CSL RW240 WP 035341342 1 Daquatica
Dickeya aquatica DW 0440 WP 035341342 1 Daquatica
Dickeya gs undicola 2B12 WP 033569868 1 Dundicola
Dickeya chrysanthemi subsp chrysanthemi NCPPB 402 WP 040001299 1 Dchrysanthemi
Dickeya chrysanthemi subsp parthenii NCPPB 516 WP 027711807 1 Dchrysanthemi
Dickeya chrysanthemi L11 WP 033577339 1 Dchrysanthemi
Dickeya chrysanthemi NCPPB 3533 WP 012769472 1 Dchrysanthemi
Dickeya chrysanthemi Ech1591 WP 012769472 1 Dchrysanthemi
Dickeya gs poaceaephila NCPPB 569 WP 042871005 1 Dpoaceaephila
Dickeya zeae Ech586 WP 012884397 1 Dzeae
Dickeya zeae NCPPB 2538 WP 019845785 1 Dzeae
Dickeya zeae MK19 WP 019845785 1 Dzeae
Dickeya zeae MS1 WP 019845785 1 Dzeae
Dickeya zeae NCPPB 3532 WP 019845785 1 Dzeae
Dickeya zeae NCPPB 3531 WP 038904041 1 Dzeae
Dickeya zeae CSL RW192 WP 038907686 1 Dzeae
Dickeya zeae DZ2Q WP 038926204 1 Dzeae
Dickeya zeae EC1 WP 016942470 1 Dzeae
Dickeya zeae ZJU1202 WP 016942470 1 Dzeae
Dickeya dianthicola GBBC 2039 WP 024106261 1 Ddianthicola
Dickeya dianthicola NCPPB 453 WP 024106261 1 Ddianthicola
Dickeya dianthicola RNS04 9 WP 024106261 1 Ddianthicola
Dickeya dianthicola NCPPB 3534 WP 024106261 1 Ddianthicola
Dickeya dianthicola IPO 980 WP 024106261 1 Ddianthicola
Dickeya gs fangzhongdai NCPPB 3274 WP 042860098 1 Dfangzhongdai
Dickeya gs fangzhongdai S1 WP 038919388 1 Dfangzhongdai
Dickeya gs fangzhongdai B16 WP 038919388 1 Dfangzhongdai
Dickeya gs fangzhongdai MK7 WP 038919388 1 Dfangzhongdai
Dickeya gs fangzhongdai ND14b WP 038668303 1 Dfangzhongdai
Dickeya gs fangzhongdai M074 WP 038668303 1 Dfangzhongdai
Dickeya gs fangzhongdai M005 WP 038668303 1 Dfangzhongdai
424 sapC response to antimicrobial peptides antimicrobial peptide transporter Seed
Dickeya dadantii 3937 WP 013318296 1 Ddadantii
Dickeya dadantii NCPPB 3537 WP 038901437 1 Ddadantii
Dickeya dadantii NCPPB 898 WP 038901437 1 Ddadantii
Dickeya dadantii subsp dieffenbachiae NCPPB 2976 WP 038923082 1 Ddadantii
Dickeya solani MK10 WP 022633907 1 Dsolani
Dickeya solani IPO 2222 WP 022633907 1 Dsolani
Dickeya solani MK16 WP 022633907 1 Dsolani
Dickeya solani RNS 08 23 3 1 A WP 022633907 1 Dsolani
Dickeya solani RNS 05 1 2A WP 022633907 1 Dsolani
Dickeya solani RNS 07 7 3B WP 022633907 1 Dsolani
Dickeya solani PPO 9134 WP 022633907 1 Dsolani
Dickeya solani GBBC 2040 WP 022633907 1 Dsolani
Dickeya solani PPO 9019 WP 022633907 1 Dsolani
Dickeya solani IFB 0099 WP 022633907 1 Dsolani
Dickeya solani D s0432 1 WP 022633907 1 Dsolani

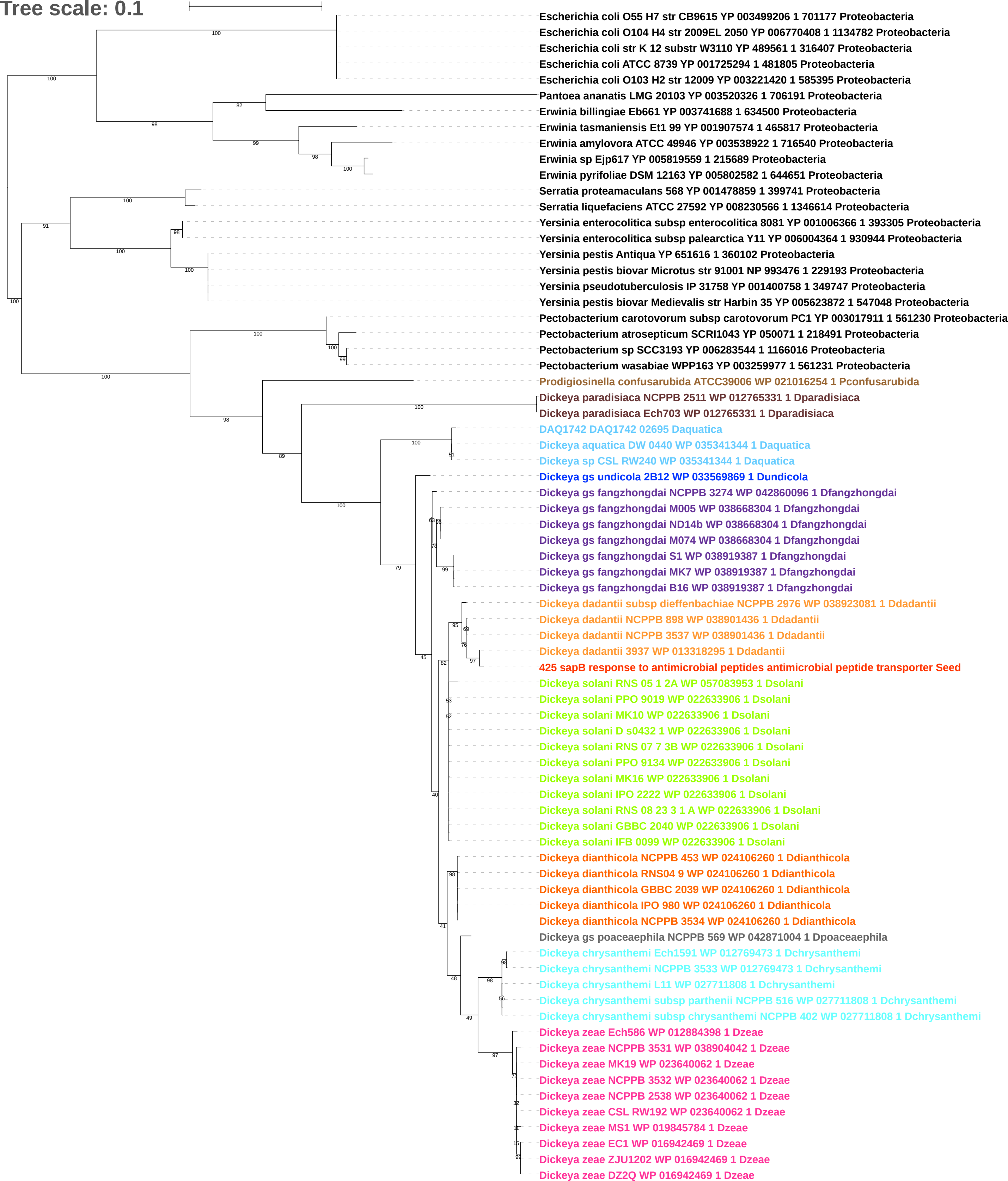
Tree scale: 0.1
Escherichia coli O55 H7 str CB9615 YP 003499206 1 701177 Proteobacteria
Escherichia coli O104 H4 str 2009EL 2050 YP 006770408 1 1134782 Proteobacteria
Escherichia coli str K 12 substr W3110 YP 489561 1 316407 Proteobacteria
Escherichia coli ATCC 8739 YP 001725294 1 481805 Proteobacteria
Escherichia coli O103 H2 str 12009 YP 003221420 1 585395 Proteobacteria
Pantoea ananatis LMG 20103 YP 003520326 1 706191 Proteobacteria
Erwinia billingiae Eb661 YP 003741688 1 634500 Proteobacteria
Erwinia tasmaniensis Et1 99 YP 001907574 1 465817 Proteobacteria
Erwinia amylovora ATCC 49946 YP 003538922 1 716540 Proteobacteria
Erwinia sp Ejp617 YP 005819559 1 215689 Proteobacteria
Erwinia pyrifoliae DSM 12163 YP 005802582 1 644651 Proteobacteria
Serratia proteamaculans 568 YP 001478859 1 399741 Proteobacteria
Serratia liquefaciens ATCC 27592 YP 008230566 1 1346614 Proteobacteria
Yersinia enterocolitica subsp enterocolitica 8081 YP 001006366 1 393305 Proteobacteria
Yersinia enterocolitica subsp palearctica Y11 YP 006004364 1 930944 Proteobacteria
Yersinia pestis Antiqua YP 651616 1 360102 Proteobacteria
Yersinia pestis biovar Microtus str 91001 NP 993476 1 229193 Proteobacteria
Yersinia pseudotuberculosis IP 31758 YP 001400758 1 349747 Proteobacteria
Yersinia pestis biovar Medievalis str Harbin 35 YP 005623872 1 547048 Proteobacteria
Pectobacterium carotovorum subsp carotovorum PC1 YP 003017911 1 561230 Proteobacteria
Pectobacterium atrosepticum SCRI1043 YP 050071 1 218491 Proteobacteria
Pectobacterium sp SCC3193 YP 006283544 1 1166016 Proteobacteria
Pectobacterium wasabiae WPP163 YP 003259977 1 561231 Proteobacteria
Prodigiosinella confusarubida ATCC39006 WP 021016254 1 Pconfusarubida
Dickeya paradisiaca NCPPB 2511 WP 012765331 1 Dparadisiaca
Dickeya paradisiaca Ech703 WP 012765331 1 Dparadisiaca
DAQ1742 DAQ1742 02695 Daquatica
Dickeya aquatica DW 0440 WP 035341344 1 Daquatica
Dickeya sp CSL RW240 WP 035341344 1 Daquatica
Dickeya gs undicola 2B12 WP 033569869 1 Dundicola
Dickeya gs fangzhongdai NCPPB 3274 WP 042860096 1 Dfangzhongdai
Dickeya gs fangzhongdai M005 WP 038668304 1 Dfangzhongdai
Dickeya gs fangzhongdai ND14b WP 038668304 1 Dfangzhongdai
Dickeya gs fangzhongdai M074 WP 038668304 1 Dfangzhongdai
Dickeya gs fangzhongdai S1 WP 038919387 1 Dfangzhongdai
Dickeya gs fangzhongdai MK7 WP 038919387 1 Dfangzhongdai
Dickeya gs fangzhongdai B16 WP 038919387 1 Dfangzhongdai
Dickeya dadantii subsp dieffenbachiae NCPPB 2976 WP 038923081 1 Ddadantii
Dickeya dadantii NCPPB 898 WP 038901436 1 Ddadantii
Dickeya dadantii NCPPB 3537 WP 038901436 1 Ddadantii
Dickeya dadantii 3937 WP 013318295 1 Ddadantii
425 sapB response to antimicrobial peptides antimicrobial peptide transporter Seed
Dickeya solani RNS 05 1 2A WP 057083953 1 Dsolani
Dickeya solani PPO 9019 WP 022633906 1 Dsolani
Dickeya solani MK10 WP 022633906 1 Dsolani
Dickeya solani D s0432 1 WP 022633906 1 Dsolani
Dickeya solani RNS 07 7 3B WP 022633906 1 Dsolani
Dickeya solani PPO 9134 WP 022633906 1 Dsolani
Dickeya solani MK16 WP 022633906 1 Dsolani
Dickeya solani IPO 2222 WP 022633906 1 Dsolani
Dickeya solani RNS 08 23 3 1 A WP 022633906 1 Dsolani
Dickeya solani GBBC 2040 WP 022633906 1 Dsolani
Dickeya solani IFB 0099 WP 022633906 1 Dsolani
Dickeya dianthicola NCPPB 453 WP 024106260 1 Ddianthicola
Dickeya dianthicola RNS04 9 WP 024106260 1 Ddianthicola
Dickeya dianthicola GBBC 2039 WP 024106260 1 Ddianthicola
Dickeya dianthicola IPO 980 WP 024106260 1 Ddianthicola
Dickeya dianthicola NCPPB 3534 WP 024106260 1 Ddianthicola
Dickeya gs poaceaephila NCPPB 569 WP 042871004 1 Dpoaceaephila
Dickeya chrysanthemi Ech1591 WP 012769473 1 Dchrysanthemi
Dickeya chrysanthemi NCPPB 3533 WP 012769473 1 Dchrysanthemi
Dickeya chrysanthemi L11 WP 027711808 1 Dchrysanthemi
Dickeya chrysanthemi subsp parthenii NCPPB 516 WP 027711808 1 Dchrysanthemi
Dickeya chrysanthemi subsp chrysanthemi NCPPB 402 WP 027711808 1 Dchrysanthemi
Dickeya zeae Ech586 WP 012884398 1 Dzeae
Dickeya zeae NCPPB 3531 WP 038904042 1 Dzeae
Dickeya zeae MK19 WP 023640062 1 Dzeae
Dickeya zeae NCPPB 3532 WP 023640062 1 Dzeae
Dickeya zeae NCPPB 2538 WP 023640062 1 Dzeae
Dickeya zeae CSL RW192 WP 023640062 1 Dzeae
Dickeya zeae MS1 WP 019845784 1 Dzeae
Dickeya zeae EC1 WP 016942469 1 Dzeae
Dickeya zeae ZJU1202 WP 016942469 1 Dzeae
Dickeya zeae DZ2Q WP 016942469 1 Dzeae

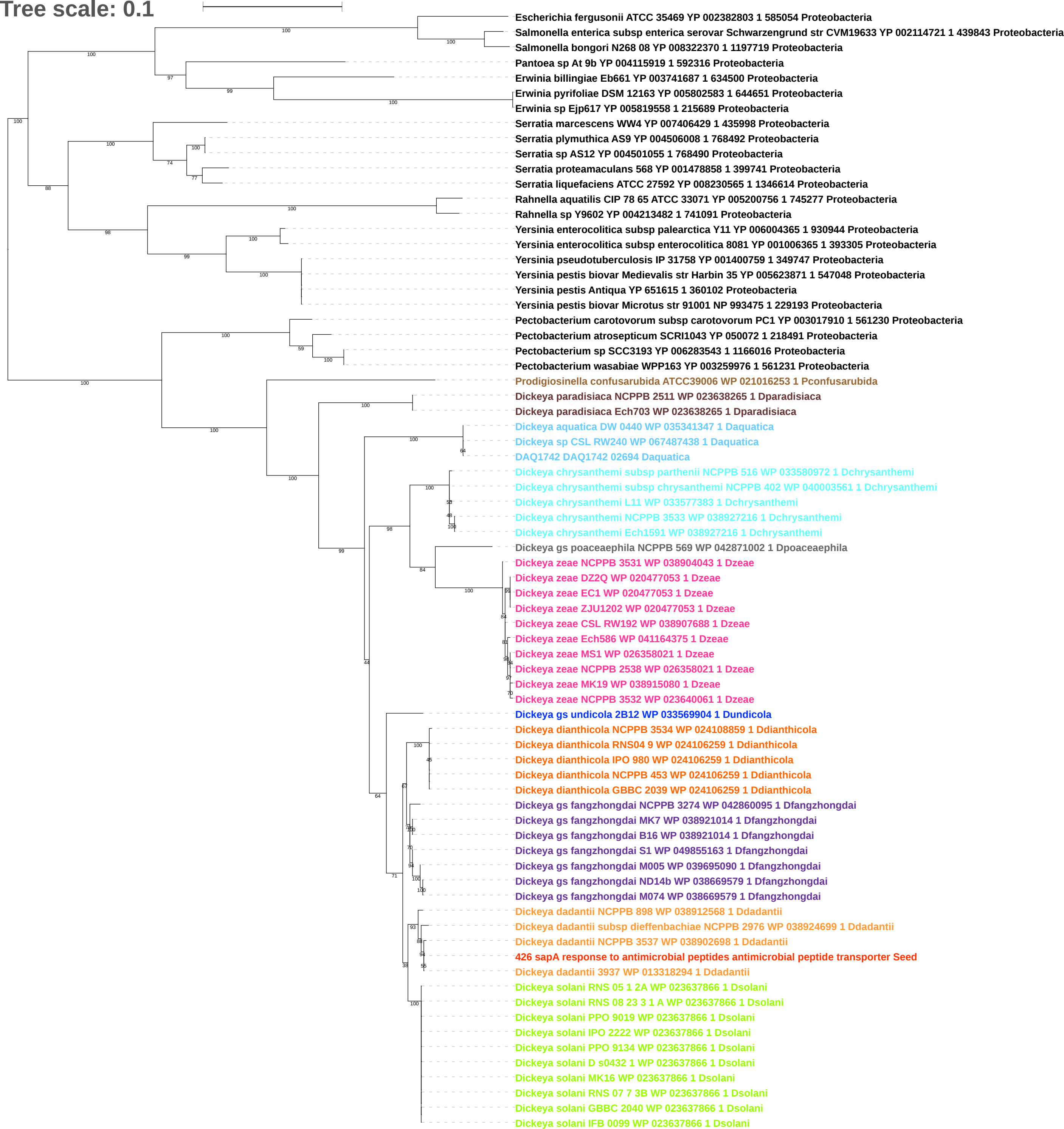
Tree scale: 0.1
Escherichia fergusonii ATCC 35469 YP 002382803 1 585054 Proteobacteria
Salmonella enterica subsp enterica serovar Schwarzengrund str CVM19633 YP 002114721 1 439843 Proteobacteria
Salmonella bongori N268 08 YP 008322370 1 1197719 Proteobacteria
Pantoea sp At 9b YP 004115919 1 592316 Proteobacteria
Erwinia billingiae Eb661 YP 003741687 1 634500 Proteobacteria
Erwinia pyrifoliae DSM 12163 YP 005802583 1 644651 Proteobacteria
Erwinia sp Ejp617 YP 005819558 1 215689 Proteobacteria
Serratia marcescens WW4 YP 007406429 1 435998 Proteobacteria
Serratia plymuthica AS9 YP 004506008 1 768492 Proteobacteria
Serratia sp AS12 YP 004501055 1 768490 Proteobacteria
Serratia proteamaculans 568 YP 001478858 1 399741 Proteobacteria
Serratia liquefaciens ATCC 27592 YP 008230565 1 1346614 Proteobacteria
Rahnella aquatilis CIP 78 65 ATCC 33071 YP 005200756 1 745277 Proteobacteria
Rahnella sp Y9602 YP 004213482 1 741091 Proteobacteria
Yersinia enterocolitica subsp palearctica Y11 YP 006004365 1 930944 Proteobacteria
Yersinia enterocolitica subsp enterocolitica 8081 YP 001006365 1 393305 Proteobacteria
Yersinia pseudotuberculosis IP 31758 YP 001400759 1 349747 Proteobacteria
Yersinia pestis biovar Medievalis str Harbin 35 YP 005623871 1 547048 Proteobacteria
Yersinia pestis Antiqua YP 651615 1 360102 Proteobacteria
Yersinia pestis biovar Microtus str 91001 NP 993475 1 229193 Proteobacteria
Pectobacterium carotovorum subsp carotovorum PC1 YP 003017910 1 561230 Proteobacteria
Pectobacterium atrosepticum SCRI1043 YP 050072 1 218491 Proteobacteria
Pectobacterium sp SCC3193 YP 006283543 1 1166016 Proteobacteria
Pectobacterium wasabiae WPP163 YP 003259976 1 561231 Proteobacteria
Prodigiosinella confusarubida ATCC39006 WP 021016253 1 Pconfusarubida
Dickeya paradisiaca NCPPB 2511 WP 023638265 1 Dparadisiaca
Dickeya paradisiaca Ech703 WP 023638265 1 Dparadisiaca
Dickeya aquatica DW 0440 WP 035341347 1 Daquatica
Dickeya sp CSL RW240 WP 067487438 1 Daquatica
DAQ1742 DAQ1742 02694 Daquatica
Dickeya chrysanthemi subsp parthenii NCPPB 516 WP 033580972 1 Dchrysanthemi
Dickeya chrysanthemi subsp chrysanthemi NCPPB 402 WP 040003561 1 Dchrysanthemi
Dickeya chrysanthemi L11 WP 033577383 1 Dchrysanthemi
Dickeya chrysanthemi NCPPB 3533 WP 038927216 1 Dchrysanthemi
Dickeya chrysanthemi Ech1591 WP 038927216 1 Dchrysanthemi
Dickeya gs poaceaephila NCPPB 569 WP 042871002 1 Dpoaceaephila
Dickeya zeae NCPPB 3531 WP 038904043 1 Dzeae
Dickeya zeae DZ2Q WP 020477053 1 Dzeae
Dickeya zeae EC1 WP 020477053 1 Dzeae
Dickeya zeae ZJU1202 WP 020477053 1 Dzeae
Dickeya zeae CSL RW192 WP 038907688 1 Dzeae
Dickeya zeae Ech586 WP 041164375 1 Dzeae
Dickeya zeae MS1 WP 026358021 1 Dzeae
Dickeya zeae NCPPB 2538 WP 026358021 1 Dzeae
Dickeya zeae MK19 WP 038915080 1 Dzeae
Dickeya zeae NCPPB 3532 WP 023640061 1 Dzeae
Dickeya gs undicola 2B12 WP 033569904 1 Dundicola
Dickeya dianthicola NCPPB 3534 WP 024108859 1 Ddianthicola
Dickeya dianthicola RNS04 9 WP 024106259 1 Ddianthicola
Dickeya dianthicola IPO 980 WP 024106259 1 Ddianthicola
Dickeya dianthicola NCPPB 453 WP 024106259 1 Ddianthicola
Dickeya dianthicola GBBC 2039 WP 024106259 1 Ddianthicola
Dickeya gs fangzhongdai NCPPB 3274 WP 042860095 1 Dfangzhongdai
Dickeya gs fangzhongdai MK7 WP 038921014 1 Dfangzhongdai
Dickeya gs fangzhongdai B16 WP 038921014 1 Dfangzhongdai
Dickeya gs fangzhongdai S1 WP 049855163 1 Dfangzhongdai
Dickeya gs fangzhongdai M005 WP 039695090 1 Dfangzhongdai
Dickeya gs fangzhongdai ND14b WP 038669579 1 Dfangzhongdai
Dickeya gs fangzhongdai M074 WP 038669579 1 Dfangzhongdai
Dickeya dadantii NCPPB 898 WP 038912568 1 Ddadantii
Dickeya dadantii subsp dieffenbachiae NCPPB 2976 WP 038924699 1 Ddadantii
Dickeya dadantii NCPPB 3537 WP 038902698 1 Ddadantii
426 sapA response to antimicrobial peptides antimicrobial peptide transporter Seed
Dickeya dadantii 3937 WP 013318294 1 Ddadantii
Dickeya solani RNS 05 1 2A WP 023637866 1 Dsolani
Dickeya solani RNS 08 23 3 1 A WP 023637866 1 Dsolani
Dickeya solani PPO 9019 WP 023637866 1 Dsolani
Dickeya solani IPO 2222 WP 023637866 1 Dsolani
Dickeya solani PPO 9134 WP 023637866 1 Dsolani
Dickeya solani D s0432 1 WP 023637866 1 Dsolani
Dickeya solani MK16 WP 023637866 1 Dsolani
Dickeya solani RNS 07 7 3B WP 023637866 1 Dsolani
Dickeya solani GBBC 2040 WP 023637866 1 Dsolani
Dickeya solani IFB 0099 WP 023637866 1 Dsolani

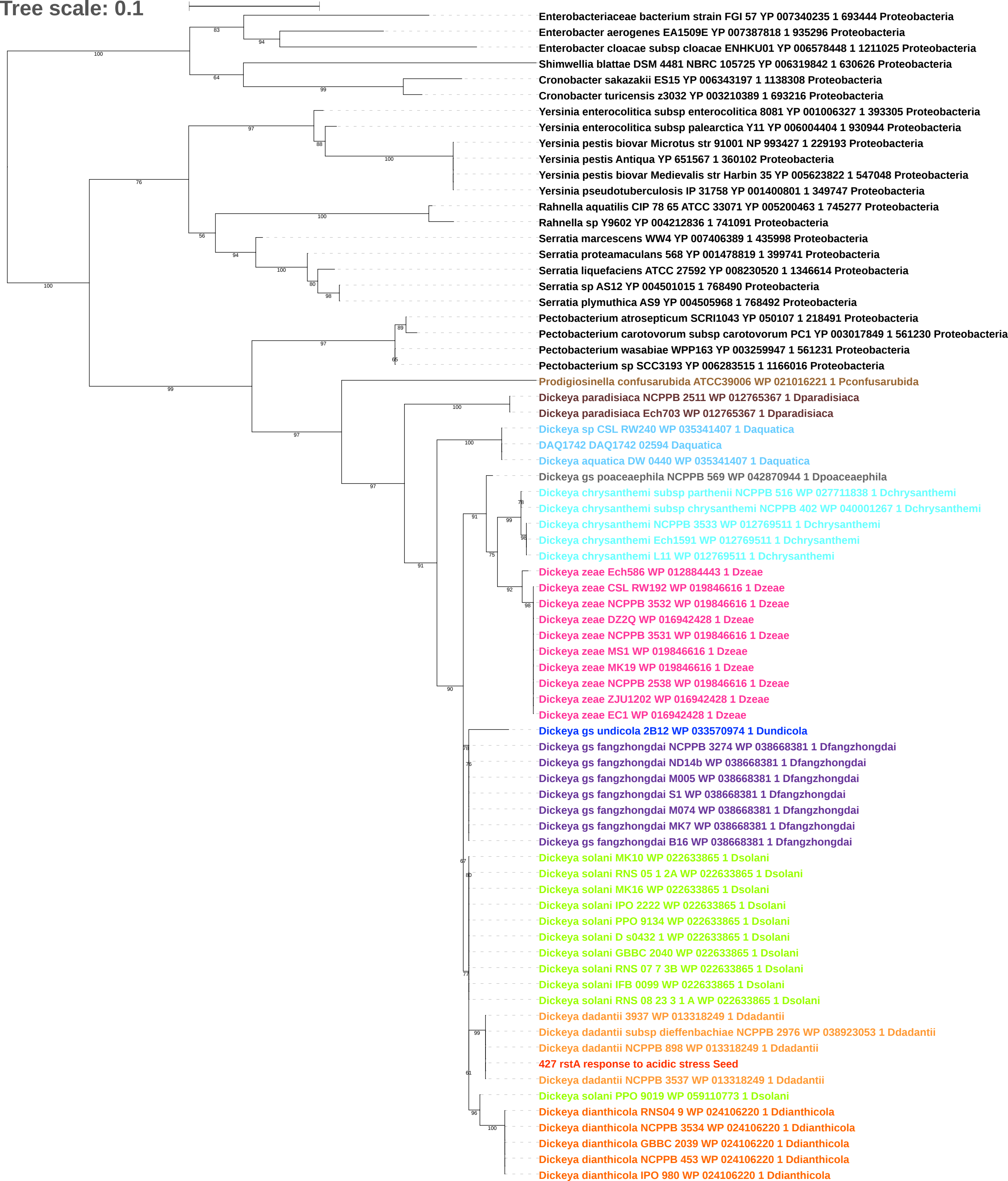
Tree scale: 0.1
Enterobacteriaceae bacterium strain FGI 57 YP 007340235 1 693444 Proteobacteria
Enterobacter aerogenes EA1509E YP 007387818 1 935296 Proteobacteria
Enterobacter cloacae subsp cloacae ENHKU01 YP 006578448 1 1211025 Proteobacteria
Shimwellia blattae DSM 4481 NBRC 105725 YP 006319842 1 630626 Proteobacteria
Cronobacter sakazakii ES15 YP 006343197 1 1138308 Proteobacteria
Cronobacter turicensis z3032 YP 003210389 1 693216 Proteobacteria
Yersinia enterocolitica subsp enterocolitica 8081 YP 001006327 1 393305 Proteobacteria
Yersinia enterocolitica subsp palearctica Y11 YP 006004404 1 930944 Proteobacteria
Yersinia pestis biovar Microtus str 91001 NP 993427 1 229193 Proteobacteria
Yersinia pestis Antiqua YP 651567 1 360102 Proteobacteria
Yersinia pestis biovar Medievalis str Harbin 35 YP 005623822 1 547048 Proteobacteria
Yersinia pseudotuberculosis IP 31758 YP 001400801 1 349747 Proteobacteria
Rahnella aquatilis CIP 78 65 ATCC 33071 YP 005200463 1 745277 Proteobacteria
Rahnella sp Y9602 YP 004212836 1 741091 Proteobacteria
Serratia marcescens WW4 YP 007406389 1 435998 Proteobacteria
Serratia proteamaculans 568 YP 001478819 1 399741 Proteobacteria
Serratia liquefaciens ATCC 27592 YP 008230520 1 1346614 Proteobacteria
Serratia sp AS12 YP 004501015 1 768490 Proteobacteria
Serratia plymuthica AS9 YP 004505968 1 768492 Proteobacteria
Pectobacterium atrosepticum SCRI1043 YP 050107 1 218491 Proteobacteria
Pectobacterium carotovorum subsp carotovorum PC1 YP 003017849 1 561230 Proteobacteria
Pectobacterium wasabiae WPP163 YP 003259947 1 561231 Proteobacteria
Pectobacterium sp SCC3193 YP 006283515 1 1166016 Proteobacteria
Prodigiosinella confusarubida ATCC39006 WP 021016221 1 Pconfusarubida
Dickeya paradisiaca NCPPB 2511 WP 012765367 1 Dparadisiaca
Dickeya paradisiaca Ech703 WP 012765367 1 Dparadisiaca
Dickeya sp CSL RW240 WP 035341407 1 Daquatica
DAQ1742 DAQ1742 02594 Daquatica
Dickeya aquatica DW 0440 WP 035341407 1 Daquatica
Dickeya gs poaceaephila NCPPB 569 WP 042870944 1 Dpoaceaephila
Dickeya chrysanthemi subsp parthenii NCPPB 516 WP 027711838 1 Dchrysanthemi
Dickeya chrysanthemi subsp chrysanthemi NCPPB 402 WP 040001267 1 Dchrysanthemi
Dickeya chrysanthemi NCPPB 3533 WP 012769511 1 Dchrysanthemi
Dickeya chrysanthemi Ech1591 WP 012769511 1 Dchrysanthemi
Dickeya chrysanthemi L11 WP 012769511 1 Dchrysanthemi
Dickeya zeae Ech586 WP 012884443 1 Dzeae
Dickeya zeae CSL RW192 WP 019846616 1 Dzeae
Dickeya zeae NCPPB 3532 WP 019846616 1 Dzeae
Dickeya zeae DZ2Q WP 016942428 1 Dzeae
Dickeya zeae NCPPB 3531 WP 019846616 1 Dzeae
Dickeya zeae MS1 WP 019846616 1 Dzeae
Dickeya zeae MK19 WP 019846616 1 Dzeae
Dickeya zeae NCPPB 2538 WP 019846616 1 Dzeae
Dickeya zeae ZJU1202 WP 016942428 1 Dzeae
Dickeya zeae EC1 WP 016942428 1 Dzeae
Dickeya gs undicola 2B12 WP 033570974 1 Dundicola
Dickeya gs fangzhongdai NCPPB 3274 WP 038668381 1 Dfangzhongdai
Dickeya gs fangzhongdai ND14b WP 038668381 1 Dfangzhongdai
Dickeya gs fangzhongdai M005 WP 038668381 1 Dfangzhongdai
Dickeya gs fangzhongdai S1 WP 038668381 1 Dfangzhongdai
Dickeya gs fangzhongdai M074 WP 038668381 1 Dfangzhongdai
Dickeya gs fangzhongdai MK7 WP 038668381 1 Dfangzhongdai
Dickeya gs fangzhongdai B16 WP 038668381 1 Dfangzhongdai
Dickeya solani MK10 WP 022633865 1 Dsolani
Dickeya solani RNS 05 1 2A WP 022633865 1 Dsolani
Dickeya solani MK16 WP 022633865 1 Dsolani
Dickeya solani IPO 2222 WP 022633865 1 Dsolani
Dickeya solani PPO 9134 WP 022633865 1 Dsolani
Dickeya solani D s0432 1 WP 022633865 1 Dsolani
Dickeya solani GBBC 2040 WP 022633865 1 Dsolani
Dickeya solani RNS 07 7 3B WP 022633865 1 Dsolani
Dickeya solani IFB 0099 WP 022633865 1 Dsolani
Dickeya solani RNS 08 23 3 1 A WP 022633865 1 Dsolani
Dickeya dadantii 3937 WP 013318249 1 Ddadantii
Dickeya dadantii subsp dieffenbachiae NCPPB 2976 WP 038923053 1 Ddadantii
Dickeya dadantii NCPPB 898 WP 013318249 1 Ddadantii
427 rstA response to acidic stress Seed
Dickeya dadantii NCPPB 3537 WP 013318249 1 Ddadantii
Dickeya solani PPO 9019 WP 059110773 1 Dsolani
Dickeya dianthicola RNS04 9 WP 024106220 1 Ddianthicola
Dickeya dianthicola NCPPB 3534 WP 024106220 1 Ddianthicola
Dickeya dianthicola GBBC 2039 WP 024106220 1 Ddianthicola
Dickeya dianthicola NCPPB 453 WP 024106220 1 Ddianthicola
Dickeya dianthicola IPO 980 WP 024106220 1 Ddianthicola

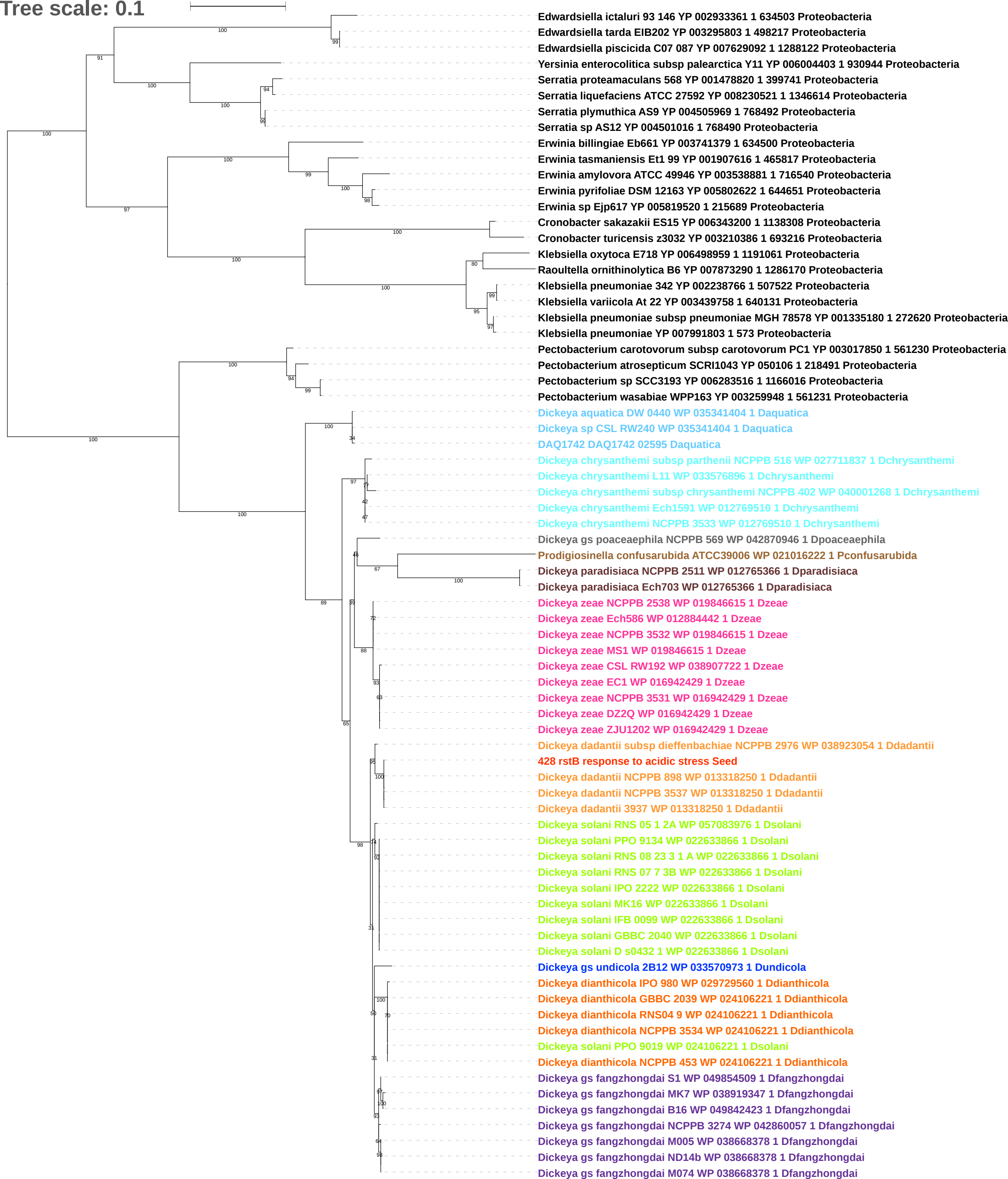
Tree scale: 0.1
Edwardsiella ictaluri 93 146 YP 002933361 1 634503 Proteobacteria
Edwardsiella tarda EIB202 YP 003295803 1 498217 Proteobacteria
Edwardsiella piscicida C07 087 YP 007629092 1 1288122 Proteobacteria
Yersinia enterocolitica subsp palearctica Y11 YP 006004403 1 930944 Proteobacteria
Serratia proteamaculans 568 YP 001478820 1 399741 Proteobacteria
Serratia liquefaciens ATCC 27592 YP 008230521 1 1346614 Proteobacteria
Serratia plymuthica AS9 YP 004505969 1 768492 Proteobacteria
Serratia sp AS12 YP 004501016 1 768490 Proteobacteria
Erwinia billingiae Eb661 YP 003741379 1 634500 Proteobacteria
Erwinia tasmaniensis Et1 99 YP 001907616 1 465817 Proteobacteria
Erwinia amylovora ATCC 49946 YP 003538881 1 716540 Proteobacteria
Erwinia pyrifoliae DSM 12163 YP 005802622 1 644651 Proteobacteria
Erwinia sp Ejp617 YP 005819520 1 215689 Proteobacteria
Cronobacter sakazakii ES15 YP 006343200 1 1138308 Proteobacteria
Cronobacter turicensis z3032 YP 003210386 1 693216 Proteobacteria
Klebsiella oxytoca E718 YP 006498959 1 1191061 Proteobacteria
Raoultella ornithinolytica B6 YP 007873290 1 1286170 Proteobacteria
Klebsiella pneumoniae 342 YP 002238766 1 507522 Proteobacteria
Klebsiella variicola At 22 YP 003439758 1 640131 Proteobacteria
Klebsiella pneumoniae subsp pneumoniae MGH 78578 YP 001335180 1 272620 Proteobacteria
Klebsiella pneumoniae YP 007991803 1 573 Proteobacteria
Pectobacterium carotovorum subsp carotovorum PC1 YP 003017850 1 561230 Proteobacteria
Pectobacterium atrosepticum SCRI1043 YP 050106 1 218491 Proteobacteria
Pectobacterium sp SCC3193 YP 006283516 1 1166016 Proteobacteria
Pectobacterium wasabiae WPP163 YP 003259948 1 561231 Proteobacteria
Dickeya aquatica DW 0440 WP 035341404 1 Daquatica
Dickeya sp CSL RW240 WP 035341404 1 Daquatica
DAQ1742 DAQ1742 02595 Daquatica
Dickeya chrysanthemi subsp parthenii NCPPB 516 WP 027711837 1 Dchrysanthemi
Dickeya chrysanthemi L11 WP 033576896 1 Dchrysanthemi
Dickeya chrysanthemi subsp chrysanthemi NCPPB 402 WP 040001268 1 Dchrysanthemi
Dickeya chrysanthemi Ech1591 WP 012769510 1 Dchrysanthemi
Dickeya chrysanthemi NCPPB 3533 WP 012769510 1 Dchrysanthemi
Dickeya gs poaceaephila NCPPB 569 WP 042870946 1 Dpoaceaephila
Prodigiosinella confusarubida ATCC39006 WP 021016222 1 Pconfusarubida
Dickeya paradisiaca NCPPB 2511 WP 012765366 1 Dparadisiaca
Dickeya paradisiaca Ech703 WP 012765366 1 Dparadisiaca
Dickeya zeae NCPPB 2538 WP 019846615 1 Dzeae
Dickeya zeae Ech586 WP 012884442 1 Dzeae
Dickeya zeae NCPPB 3532 WP 019846615 1 Dzeae
Dickeya zeae MS1 WP 019846615 1 Dzeae
Dickeya zeae CSL RW192 WP 038907722 1 Dzeae
Dickeya zeae EC1 WP 016942429 1 Dzeae
Dickeya zeae NCPPB 3531 WP 016942429 1 Dzeae
Dickeya zeae DZ2Q WP 016942429 1 Dzeae
Dickeya zeae ZJU1202 WP 016942429 1 Dzeae
Dickeya dadantii subsp dieffenbachiae NCPPB 2976 WP 038923054 1 Ddadantii
428 rstB response to acidic stress Seed
Dickeya dadantii NCPPB 898 WP 013318250 1 Ddadantii
Dickeya dadantii NCPPB 3537 WP 013318250 1 Ddadantii
Dickeya dadantii 3937 WP 013318250 1 Ddadantii
Dickeya solani RNS 05 1 2A WP 057083976 1 Dsolani
Dickeya solani PPO 9134 WP 022633866 1 Dsolani
Dickeya solani RNS 08 23 3 1 A WP 022633866 1 Dsolani
Dickeya solani RNS 07 7 3B WP 022633866 1 Dsolani
Dickeya solani IPO 2222 WP 022633866 1 Dsolani
Dickeya solani MK16 WP 022633866 1 Dsolani
Dickeya solani IFB 0099 WP 022633866 1 Dsolani
Dickeya solani GBBC 2040 WP 022633866 1 Dsolani
Dickeya solani D s0432 1 WP 022633866 1 Dsolani
Dickeya gs undicola 2B12 WP 033570973 1 Dundicola
Dickeya dianthicola IPO 980 WP 029729560 1 Ddianthicola
Dickeya dianthicola GBBC 2039 WP 024106221 1 Ddianthicola
Dickeya dianthicola RNS04 9 WP 024106221 1 Ddianthicola
Dickeya dianthicola NCPPB 3534 WP 024106221 1 Ddianthicola
Dickeya solani PPO 9019 WP 024106221 1 Dsolani
Dickeya dianthicola NCPPB 453 WP 024106221 1 Ddianthicola
Dickeya gs fangzhongdai S1 WP 049854509 1 Dfangzhongdai
Dickeya gs fangzhongdai MK7 WP 038919347 1 Dfangzhongdai
Dickeya gs fangzhongdai B16 WP 049842423 1 Dfangzhongdai
Dickeya gs fangzhongdai NCPPB 3274 WP 042860057 1 Dfangzhongdai
Dickeya gs fangzhongdai M005 WP 038668378 1 Dfangzhongdai
Dickeya gs fangzhongdai ND14b WP 038668378 1 Dfangzhongdai
Dickeya gs fangzhongdai M074 WP 038668378 1 Dfangzhongdai

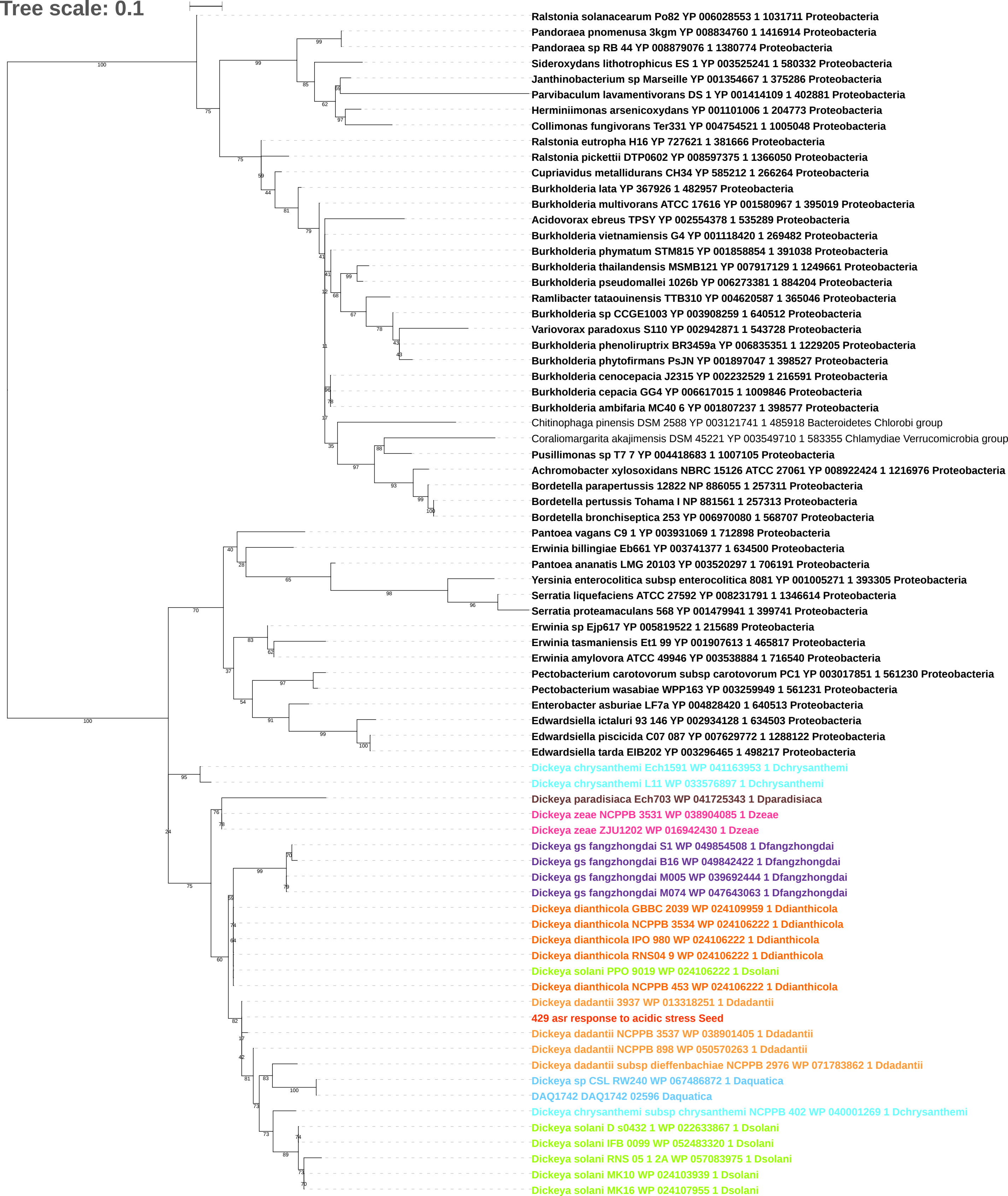
Tree scale: 0.1
Ralstonia solanacearum Po82 YP 006028553 1 1031711 Proteobacteria
Pandoraea pnomenusa 3kgm YP 008834760 1 1416914 Proteobacteria
Pandoraea sp RB 44 YP 008879076 1 1380774 Proteobacteria
Sideroxydans lithotrophicus ES 1 YP 003525241 1 580332 Proteobacteria
Janthinobacterium sp Marseille YP 001354667 1 375286 Proteobacteria
Parvibaculum lavamentivorans DS 1 YP 001414109 1 402881 Proteobacteria
Herminiimonas arsenicoxydans YP 001101006 1 204773 Proteobacteria
Collimonas fungivorans Ter331 YP 004754521 1 1005048 Proteobacteria
Ralstonia eutropha H16 YP 727621 1 381666 Proteobacteria
Ralstonia pickettii DTP0602 YP 008597375 1 1366050 Proteobacteria
Cupriavidus metallidurans CH34 YP 585212 1 266264 Proteobacteria
Burkholderia lata YP 367926 1 482957 Proteobacteria
Burkholderia multivorans ATCC 17616 YP 001580967 1 395019 Proteobacteria
Acidovorax ebreus TPSY YP 002554378 1 535289 Proteobacteria
Burkholderia vietnamiensis G4 YP 001118420 1 269482 Proteobacteria
Burkholderia phymatum STM815 YP 001858854 1 391038 Proteobacteria
Burkholderia thailandensis MSMB121 YP 007917129 1 1249661 Proteobacteria
Burkholderia pseudomallei 1026b YP 006273381 1 884204 Proteobacteria
Ramlibacter tataouinensis TTB310 YP 004620587 1 365046 Proteobacteria
Burkholderia sp CCGE1003 YP 003908259 1 640512 Proteobacteria
Variovorax paradoxus S110 YP 002942871 1 543728 Proteobacteria
Burkholderia phenoliruptrix BR3459a YP 006835351 1 1229205 Proteobacteria
Burkholderia phytofirmans PsJN YP 001897047 1 398527 Proteobacteria
Burkholderia cenocepacia J2315 YP 002232529 1 216591 Proteobacteria
Burkholderia cepacia GG4 YP 006617015 1 1009846 Proteobacteria
Burkholderia ambifaria MC40 6 YP 001807237 1 398577 Proteobacteria
Chitinophaga pinensis DSM 2588 YP 003121741 1 485918 Bacteroidetes Chlorobi group
Coraliomargarita akajimensis DSM 45221 YP 003549710 1 583355 Chlamydiae Verrucomicrobia group
Pusillimonas sp T7 7 YP 004418683 1 1007105 Proteobacteria
Achromobacter xylosoxidans NBRC 15126 ATCC 27061 YP 008922424 1 1216976 Proteobacteria
Bordetella parapertussis 12822 NP 886055 1 257311 Proteobacteria
Bordetella pertussis Tohama I NP 881561 1 257313 Proteobacteria
Bordetella bronchiseptica 253 YP 006970080 1 568707 Proteobacteria
Pantoea vagans C9 1 YP 003931069 1 712898 Proteobacteria
Erwinia billingiae Eb661 YP 003741377 1 634500 Proteobacteria
Pantoea ananatis LMG 20103 YP 003520297 1 706191 Proteobacteria
Yersinia enterocolitica subsp enterocolitica 8081 YP 001005271 1 393305 Proteobacteria
Serratia liquefaciens ATCC 27592 YP 008231791 1 1346614 Proteobacteria
Serratia proteamaculans 568 YP 001479941 1 399741 Proteobacteria
Erwinia sp Ejp617 YP 005819522 1 215689 Proteobacteria
Erwinia tasmaniensis Et1 99 YP 001907613 1 465817 Proteobacteria
Erwinia amylovora ATCC 49946 YP 003538884 1 716540 Proteobacteria
Pectobacterium carotovorum subsp carotovorum PC1 YP 003017851 1 561230 Proteobacteria
Pectobacterium wasabiae WPP163 YP 003259949 1 561231 Proteobacteria
Enterobacter asburiae LF7a YP 004828420 1 640513 Proteobacteria
Edwardsiella ictaluri 93 146 YP 002934128 1 634503 Proteobacteria
Edwardsiella piscicida C07 087 YP 007629772 1 1288122 Proteobacteria
Edwardsiella tarda EIB202 YP 003296465 1 498217 Proteobacteria
Dickeya chrysanthemi Ech1591 WP 041163953 1 Dchrysanthemi
Dickeya chrysanthemi L11 WP 033576897 1 Dchrysanthemi
Dickeya paradisiaca Ech703 WP 041725343 1 Dparadisiaca
Dickeya zeae NCPPB 3531 WP 038904085 1 Dzeae
Dickeya zeae ZJU1202 WP 016942430 1 Dzeae
Dickeya gs fangzhongdai S1 WP 049854508 1 Dfangzhongdai
Dickeya gs fangzhongdai B16 WP 049842422 1 Dfangzhongdai
Dickeya gs fangzhongdai M005 WP 039692444 1 Dfangzhongdai
Dickeya gs fangzhongdai M074 WP 047643063 1 Dfangzhongdai
Dickeya dianthicola GBBC 2039 WP 024109959 1 Ddianthicola
Dickeya dianthicola NCPPB 3534 WP 024106222 1 Ddianthicola
Dickeya dianthicola IPO 980 WP 024106222 1 Ddianthicola
Dickeya dianthicola RNS04 9 WP 024106222 1 Ddianthicola
Dickeya solani PPO 9019 WP 024106222 1 Dsolani
Dickeya dianthicola NCPPB 453 WP 024106222 1 Ddianthicola
Dickeya dadantii 3937 WP 013318251 1 Ddadantii
429 asr response to acidic stress Seed
Dickeya dadantii NCPPB 3537 WP 038901405 1 Ddadantii
Dickeya dadantii NCPPB 898 WP 050570263 1 Ddadantii
Dickeya dadantii subsp dieffenbachiae NCPPB 2976 WP 071783862 1 Ddadantii
Dickeya sp CSL RW240 WP 067486872 1 Daquatica
DAQ1742 DAQ1742 02596 Daquatica
Dickeya chrysanthemi subsp chrysanthemi NCPPB 402 WP 040001269 1 Dchrysanthemi
Dickeya solani D s0432 1 WP 022633867 1 Dsolani
Dickeya solani IFB 0099 WP 052483320 1 Dsolani
Dickeya solani RNS 05 1 2A WP 057083975 1 Dsolani
Dickeya solani MK10 WP 024103939 1 Dsolani
Dickeya solani MK16 WP 024107955 1 Dsolani

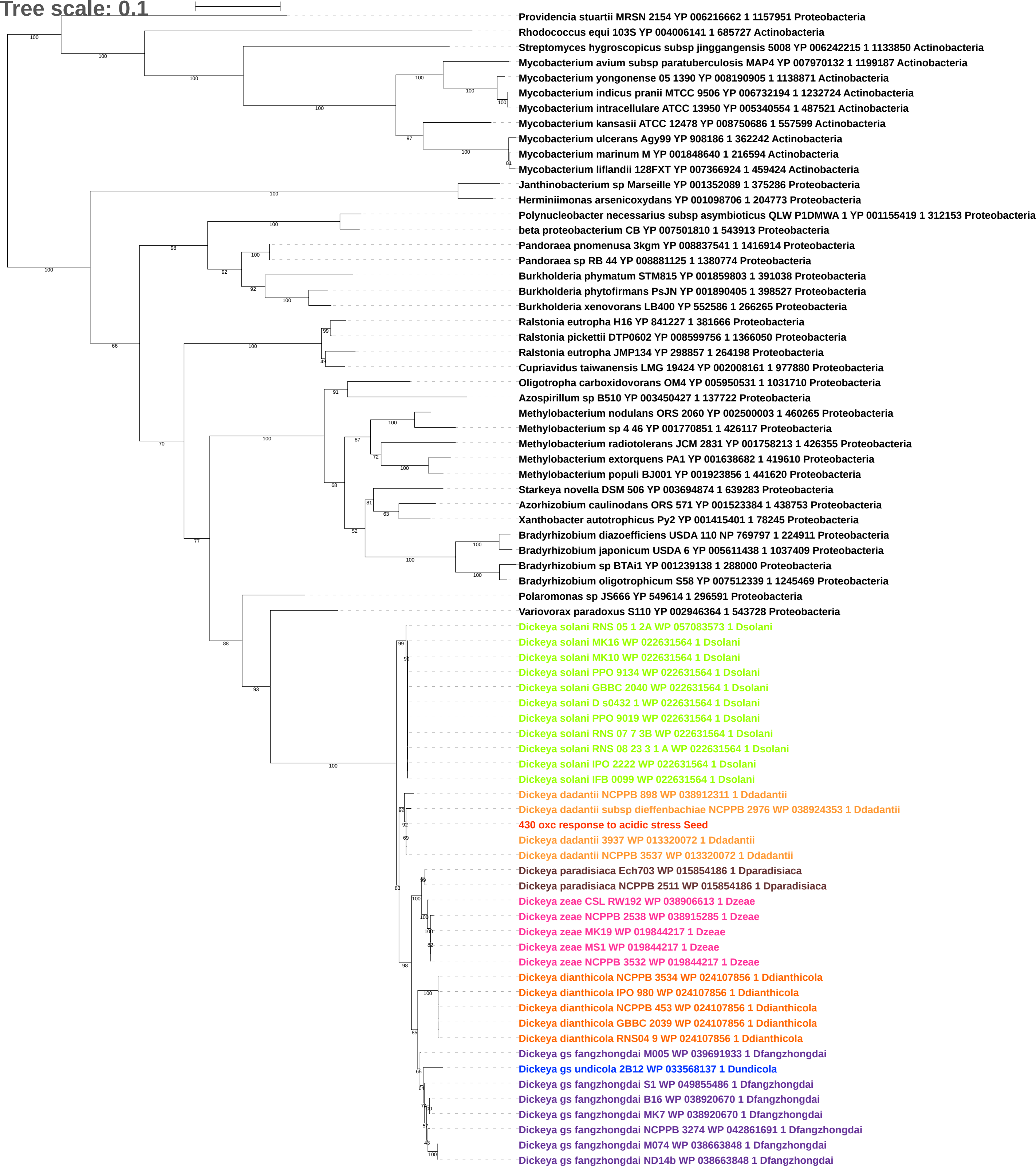
Tree scale: 0.1
Providencia stuartii MRSN 2154 YP 006216662 1 1157951 Proteobacteria
Rhodococcus equi 103S YP 004006141 1 685727 Actinobacteria
Streptomyces hygroscopicus subsp jinggangensis 5008 YP 006242215 1 1133850 Actinobacteria
Mycobacterium avium subsp paratuberculosis MAP4 YP 007970132 1 1199187 Actinobacteria
Mycobacterium yongonense 05 1390 YP 008190905 1 1138871 Actinobacteria
Mycobacterium indicus pranii MTCC 9506 YP 006732194 1 1232724 Actinobacteria
Mycobacterium intracellulare ATCC 13950 YP 005340554 1 487521 Actinobacteria
Mycobacterium kansasii ATCC 12478 YP 008750686 1 557599 Actinobacteria
Mycobacterium ulcerans Agy99 YP 908186 1 362242 Actinobacteria
Mycobacterium marinum M YP 001848640 1 216594 Actinobacteria
Mycobacterium liflandii 128FXT YP 007366924 1 459424 Actinobacteria
Janthinobacterium sp Marseille YP 001352089 1 375286 Proteobacteria
Herminiimonas arsenicoxydans YP 001098706 1 204773 Proteobacteria
Polynucleobacter necessarius subsp asymbioticus QLW P1DMWA 1 YP 001155419 1 312153 Proteobacteria
beta proteobacterium CB YP 007501810 1 543913 Proteobacteria
Pandoraea pnomenusa 3kgm YP 008837541 1 1416914 Proteobacteria
Pandoraea sp RB 44 YP 008881125 1 1380774 Proteobacteria
Burkholderia phymatum STM815 YP 001859803 1 391038 Proteobacteria
Burkholderia phytofirmans PsJN YP 001890405 1 398527 Proteobacteria
Burkholderia xenovorans LB400 YP 552586 1 266265 Proteobacteria
Ralstonia eutropha H16 YP 841227 1 381666 Proteobacteria
Ralstonia pickettii DTP0602 YP 008599756 1 1366050 Proteobacteria
Ralstonia eutropha JMP134 YP 298857 1 264198 Proteobacteria
Cupriavidus taiwanensis LMG 19424 YP 002008161 1 977880 Proteobacteria
Oligotropha carboxidovorans OM4 YP 005950531 1 1031710 Proteobacteria
Azospirillum sp B510 YP 003450427 1 137722 Proteobacteria
Methylobacterium nodulans ORS 2060 YP 002500003 1 460265 Proteobacteria
Methylobacterium sp 4 46 YP 001770851 1 426117 Proteobacteria
Methylobacterium radiotolerans JCM 2831 YP 001758213 1 426355 Proteobacteria
Methylobacterium extorquens PA1 YP 001638682 1 419610 Proteobacteria
Methylobacterium populi BJ001 YP 001923856 1 441620 Proteobacteria
Starkeya novella DSM 506 YP 003694874 1 639283 Proteobacteria
Azorhizobium caulinodans ORS 571 YP 001523384 1 438753 Proteobacteria
Xanthobacter autotrophicus Py2 YP 001415401 1 78245 Proteobacteria
Bradyrhizobium diazoefficiens USDA 110 NP 769797 1 224911 Proteobacteria
Bradyrhizobium japonicum USDA 6 YP 005611438 1 1037409 Proteobacteria
Bradyrhizobium sp BTAi1 YP 001239138 1 288000 Proteobacteria
Bradyrhizobium oligotrophicum S58 YP 007512339 1 1245469 Proteobacteria
Polaromonas sp JS666 YP 549614 1 296591 Proteobacteria
Variovorax paradoxus S110 YP 002946364 1 543728 Proteobacteria
Dickeya solani RNS 05 1 2A WP 057083573 1 Dsolani
Dickeya solani MK16 WP 022631564 1 Dsolani
Dickeya solani MK10 WP 022631564 1 Dsolani
Dickeya solani PPO 9134 WP 022631564 1 Dsolani
Dickeya solani GBBC 2040 WP 022631564 1 Dsolani
Dickeya solani D s0432 1 WP 022631564 1 Dsolani
Dickeya solani PPO 9019 WP 022631564 1 Dsolani
Dickeya solani RNS 07 7 3B WP 022631564 1 Dsolani
Dickeya solani RNS 08 23 3 1 A WP 022631564 1 Dsolani
Dickeya solani IPO 2222 WP 022631564 1 Dsolani
Dickeya solani IFB 0099 WP 022631564 1 Dsolani
Dickeya dadantii NCPPB 898 WP 038912311 1 Ddadantii
Dickeya dadantii subsp dieffenbachiae NCPPB 2976 WP 038924353 1 Ddadantii
430 oxc response to acidic stress Seed
Dickeya dadantii 3937 WP 013320072 1 Ddadantii
Dickeya dadantii NCPPB 3537 WP 013320072 1 Ddadantii
Dickeya paradisiaca Ech703 WP 015854186 1 Dparadisiaca
Dickeya paradisiaca NCPPB 2511 WP 015854186 1 Dparadisiaca
Dickeya zeae CSL RW192 WP 038906613 1 Dzeae
Dickeya zeae NCPPB 2538 WP 038915285 1 Dzeae
Dickeya zeae MK19 WP 019844217 1 Dzeae
Dickeya zeae MS1 WP 019844217 1 Dzeae
Dickeya zeae NCPPB 3532 WP 019844217 1 Dzeae
Dickeya dianthicola NCPPB 3534 WP 024107856 1 Ddianthicola
Dickeya dianthicola IPO 980 WP 024107856 1 Ddianthicola
Dickeya dianthicola NCPPB 453 WP 024107856 1 Ddianthicola
Dickeya dianthicola GBBC 2039 WP 024107856 1 Ddianthicola
Dickeya dianthicola RNS04 9 WP 024107856 1 Ddianthicola
Dickeya gs fangzhongdai M005 WP 039691933 1 Dfangzhongdai
Dickeya gs undicola 2B12 WP 033568137 1 Dundicola
Dickeya gs fangzhongdai S1 WP 049855486 1 Dfangzhongdai
Dickeya gs fangzhongdai B16 WP 038920670 1 Dfangzhongdai
Dickeya gs fangzhongdai MK7 WP 038920670 1 Dfangzhongdai
Dickeya gs fangzhongdai NCPPB 3274 WP 042861691 1 Dfangzhongdai
Dickeya gs fangzhongdai M074 WP 038663848 1 Dfangzhongdai
Dickeya gs fangzhongdai ND14b WP 038663848 1 Dfangzhongdai

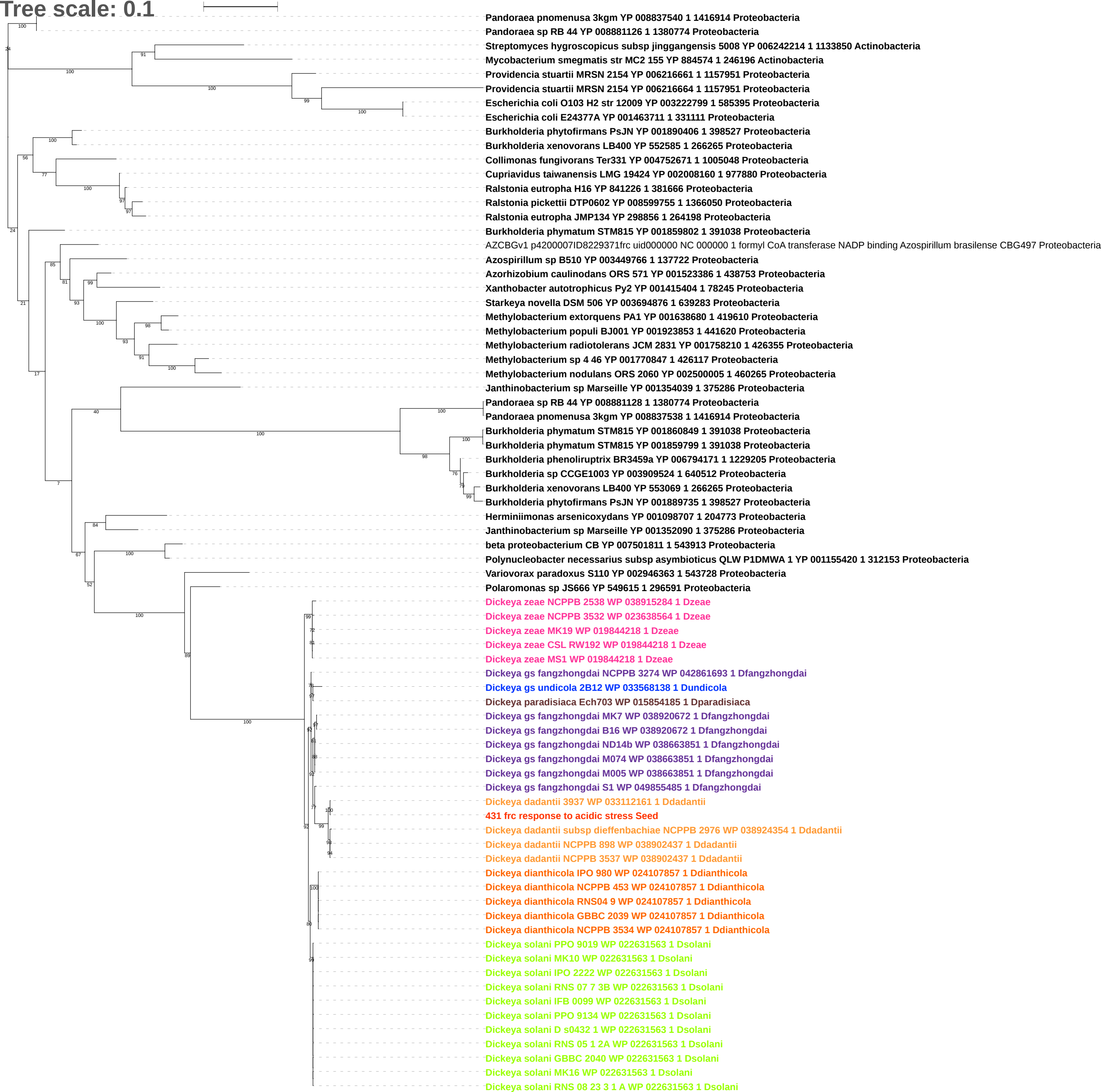
Tree scale: 0.1
Pandoraea pnomenusa 3kgm YP 008837540 1 1416914 Proteobacteria
Pandoraea sp RB 44 YP 008881126 1 1380774 Proteobacteria
Streptomyces hygroscopicus subsp jinggangensis 5008 YP 006242214 1 1133850 Actinobacteria
Mycobacterium smegmatis str MC2 155 YP 884574 1 246196 Actinobacteria
Providencia stuartii MRSN 2154 YP 006216661 1 1157951 Proteobacteria
Providencia stuartii MRSN 2154 YP 006216664 1 1157951 Proteobacteria
Escherichia coli O103 H2 str 12009 YP 003222799 1 585395 Proteobacteria
Escherichia coli E24377A YP 001463711 1 331111 Proteobacteria
Burkholderia phytofirmans PsJN YP 001890406 1 398527 Proteobacteria
Burkholderia xenovorans LB400 YP 552585 1 266265 Proteobacteria
Collimonas fungivorans Ter331 YP 004752671 1 1005048 Proteobacteria
Cupriavidus taiwanensis LMG 19424 YP 002008160 1 977880 Proteobacteria
Ralstonia eutropha H16 YP 841226 1 381666 Proteobacteria
Ralstonia pickettii DTP0602 YP 008599755 1 1366050 Proteobacteria
Ralstonia eutropha JMP134 YP 298856 1 264198 Proteobacteria
Burkholderia phymatum STM815 YP 001859802 1 391038 Proteobacteria
AZCBGv1 p4200007ID8229371frc uid000000 NC 000000 1 formyl CoA transferase NADP binding Azospirillum brasilense CBG497 Proteobacteria
Azospirillum sp B510 YP 003449766 1 137722 Proteobacteria
Azorhizobium caulinodans ORS 571 YP 001523386 1 438753 Proteobacteria
Xanthobacter autotrophicus Py2 YP 001415404 1 78245 Proteobacteria
Starkeya novella DSM 506 YP 003694876 1 639283 Proteobacteria
Methylobacterium extorquens PA1 YP 001638680 1 419610 Proteobacteria
Methylobacterium populi BJ001 YP 001923853 1 441620 Proteobacteria
Methylobacterium radiotolerans JCM 2831 YP 001758210 1 426355 Proteobacteria
Methylobacterium sp 4 46 YP 001770847 1 426117 Proteobacteria
Methylobacterium nodulans ORS 2060 YP 002500005 1 460265 Proteobacteria
Janthinobacterium sp Marseille YP 001354039 1 375286 Proteobacteria
Pandoraea sp RB 44 YP 008881128 1 1380774 Proteobacteria
Pandoraea pnomenusa 3kgm YP 008837538 1 1416914 Proteobacteria
Burkholderia phymatum STM815 YP 001860849 1 391038 Proteobacteria
Burkholderia phymatum STM815 YP 001859799 1 391038 Proteobacteria
Burkholderia phenoliruptrix BR3459a YP 006794171 1 1229205 Proteobacteria
Burkholderia sp CCGE1003 YP 003909524 1 640512 Proteobacteria
Burkholderia xenovorans LB400 YP 553069 1 266265 Proteobacteria
Burkholderia phytofirmans PsJN YP 001889735 1 398527 Proteobacteria
Herminiimonas arsenicoxydans YP 001098707 1 204773 Proteobacteria
Janthinobacterium sp Marseille YP 001352090 1 375286 Proteobacteria
beta proteobacterium CB YP 007501811 1 543913 Proteobacteria
Polynucleobacter necessarius subsp asymbioticus QLW P1DMWA 1 YP 001155420 1 312153 Proteobacteria
Variovorax paradoxus S110 YP 002946363 1 543728 Proteobacteria
Polaromonas sp JS666 YP 549615 1 296591 Proteobacteria
Dickeya zeae NCPPB 2538 WP 038915284 1 Dzeae
Dickeya zeae NCPPB 3532 WP 023638564 1 Dzeae
Dickeya zeae MK19 WP 019844218 1 Dzeae
Dickeya zeae CSL RW192 WP 019844218 1 Dzeae
Dickeya zeae MS1 WP 019844218 1 Dzeae
Dickeya gs fangzhongdai NCPPB 3274 WP 042861693 1 Dfangzhongdai
Dickeya gs undicola 2B12 WP 033568138 1 Dundicola
Dickeya paradisiaca Ech703 WP 015854185 1 Dparadisiaca
Dickeya gs fangzhongdai MK7 WP 038920672 1 Dfangzhongdai
Dickeya gs fangzhongdai B16 WP 038920672 1 Dfangzhongdai
Dickeya gs fangzhongdai ND14b WP 038663851 1 Dfangzhongdai
Dickeya gs fangzhongdai M074 WP 038663851 1 Dfangzhongdai
Dickeya gs fangzhongdai M005 WP 038663851 1 Dfangzhongdai
Dickeya gs fangzhongdai S1 WP 049855485 1 Dfangzhongdai
Dickeya dadantii 3937 WP 033112161 1 Ddadantii
431 frc response to acidic stress Seed
Dickeya dadantii subsp dieffenbachiae NCPPB 2976 WP 038924354 1 Ddadantii
Dickeya dadantii NCPPB 898 WP 038902437 1 Ddadantii
Dickeya dadantii NCPPB 3537 WP 038902437 1 Ddadantii
Dickeya dianthicola IPO 980 WP 024107857 1 Ddianthicola
Dickeya dianthicola NCPPB 453 WP 024107857 1 Ddianthicola
Dickeya dianthicola RNS04 9 WP 024107857 1 Ddianthicola
Dickeya dianthicola GBBC 2039 WP 024107857 1 Ddianthicola
Dickeya dianthicola NCPPB 3534 WP 024107857 1 Ddianthicola
Dickeya solani PPO 9019 WP 022631563 1 Dsolani
Dickeya solani MK10 WP 022631563 1 Dsolani
Dickeya solani IPO 2222 WP 022631563 1 Dsolani
Dickeya solani RNS 07 7 3B WP 022631563 1 Dsolani
Dickeya solani IFB 0099 WP 022631563 1 Dsolani
Dickeya solani PPO 9134 WP 022631563 1 Dsolani
Dickeya solani D s0432 1 WP 022631563 1 Dsolani
Dickeya solani RNS 05 1 2A WP 022631563 1 Dsolani
Dickeya solani GBBC 2040 WP 022631563 1 Dsolani
Dickeya solani MK16 WP 022631563 1 Dsolani
Dickeya solani RNS 08 23 3 1 A WP 022631563 1 Dsolani

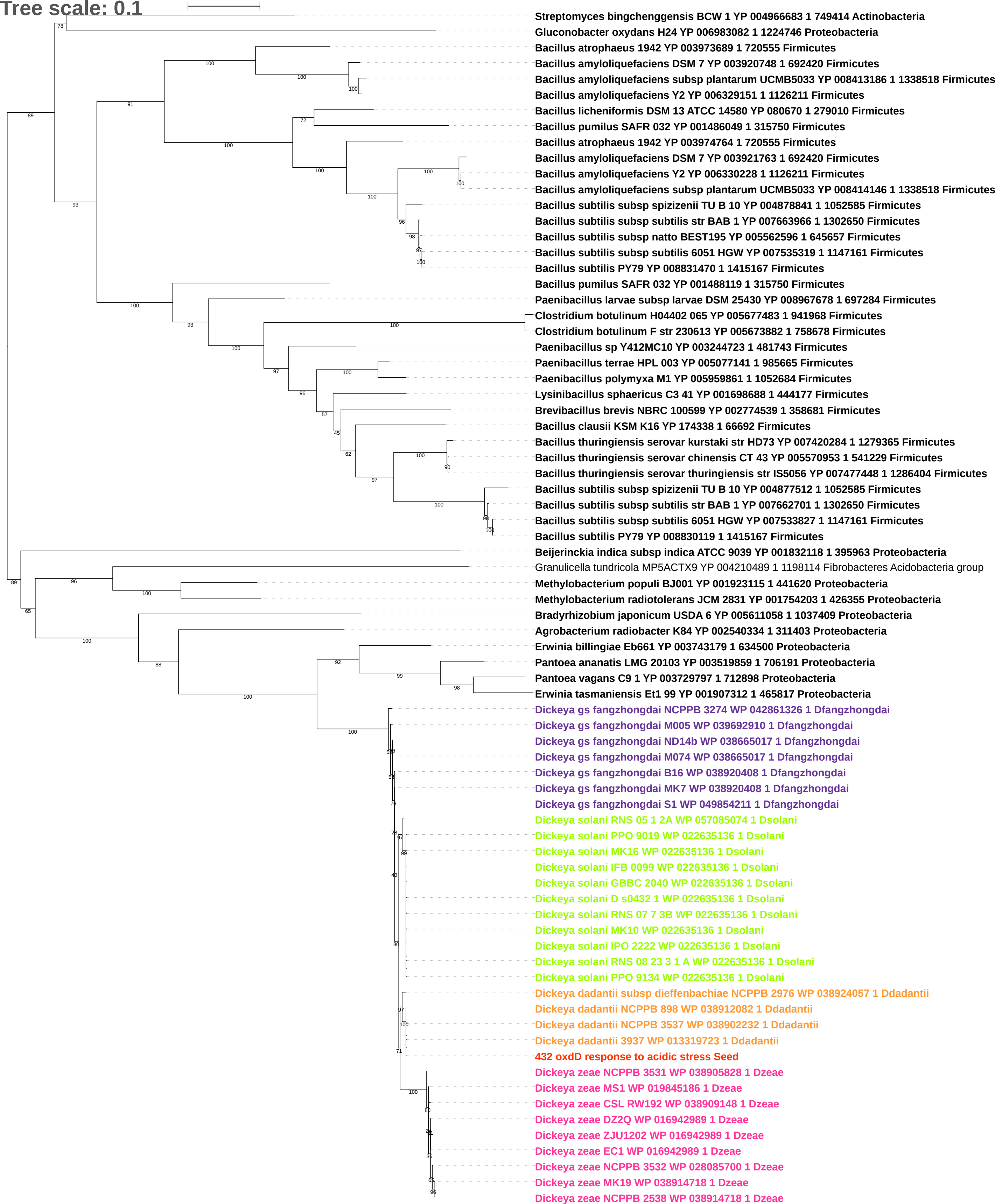
Tree scale: 0.1
Streptomyces bingchenggensis BCW 1 YP 004966683 1 749414 Actinobacteria
Gluconobacter oxydans H24 YP 006983082 1 1224746 Proteobacteria
Bacillus atrophaeus 1942 YP 003973689 1 720555 Firmicutes
Bacillus amyloliquefaciens DSM 7 YP 003920748 1 692420 Firmicutes
Bacillus amyloliquefaciens subsp plantarum UCMB5033 YP 008413186 1 1338518 Firmicutes
Bacillus amyloliquefaciens Y2 YP 006329151 1 1126211 Firmicutes
Bacillus licheniformis DSM 13 ATCC 14580 YP 080670 1 279010 Firmicutes
Bacillus pumilus SAFR 032 YP 001486049 1 315750 Firmicutes
Bacillus atrophaeus 1942 YP 003974764 1 720555 Firmicutes
Bacillus amyloliquefaciens DSM 7 YP 003921763 1 692420 Firmicutes
Bacillus amyloliquefaciens Y2 YP 006330228 1 1126211 Firmicutes
Bacillus amyloliquefaciens subsp plantarum UCMB5033 YP 008414146 1 1338518 Firmicutes
Bacillus subtilis subsp spizizenii TU B 10 YP 004878841 1 1052585 Firmicutes
Bacillus subtilis subsp subtilis str BAB 1 YP 007663966 1 1302650 Firmicutes
Bacillus subtilis subsp natto BEST195 YP 005562596 1 645657 Firmicutes
Bacillus subtilis subsp subtilis 6051 HGW YP 007535319 1 1147161 Firmicutes
Bacillus subtilis PY79 YP 008831470 1 1415167 Firmicutes
Bacillus pumilus SAFR 032 YP 001488119 1 315750 Firmicutes
Paenibacillus larvae subsp larvae DSM 25430 YP 008967678 1 697284 Firmicutes
Clostridium botulinum H04402 065 YP 005677483 1 941968 Firmicutes
Clostridium botulinum F str 230613 YP 005673882 1 758678 Firmicutes
Paenibacillus sp Y412MC10 YP 003244723 1 481743 Firmicutes
Paenibacillus terrae HPL 003 YP 005077141 1 985665 Firmicutes
Paenibacillus polymyxa M1 YP 005959861 1 1052684 Firmicutes
Lysinibacillus sphaericus C3 41 YP 001698688 1 444177 Firmicutes
Brevibacillus brevis NBRC 100599 YP 002774539 1 358681 Firmicutes
Bacillus clausii KSM K16 YP 174338 1 66692 Firmicutes
Bacillus thuringiensis serovar kurstaki str HD73 YP 007420284 1 1279365 Firmicutes
Bacillus thuringiensis serovar chinensis CT 43 YP 005570953 1 541229 Firmicutes
Bacillus thuringiensis serovar thuringiensis str IS5056 YP 007477448 1 1286404 Firmicutes
Bacillus subtilis subsp spizizenii TU B 10 YP 004877512 1 1052585 Firmicutes
Bacillus subtilis subsp subtilis str BAB 1 YP 007662701 1 1302650 Firmicutes
Bacillus subtilis subsp subtilis 6051 HGW YP 007533827 1 1147161 Firmicutes
Bacillus subtilis PY79 YP 008830119 1 1415167 Firmicutes
Beijerinckia indica subsp indica ATCC 9039 YP 001832118 1 395963 Proteobacteria
Granulicella tundricola MP5ACTX9 YP 004210489 1 1198114 Fibrobacteres Acidobacteria group
Methylobacterium populi BJ001 YP 001923115 1 441620 Proteobacteria
Methylobacterium radiotolerans JCM 2831 YP 001754203 1 426355 Proteobacteria
Bradyrhizobium japonicum USDA 6 YP 005611058 1 1037409 Proteobacteria
Agrobacterium radiobacter K84 YP 002540334 1 311403 Proteobacteria
Erwinia billingiae Eb661 YP 003743179 1 634500 Proteobacteria
Pantoea ananatis LMG 20103 YP 003519859 1 706191 Proteobacteria
Pantoea vagans C9 1 YP 003729797 1 712898 Proteobacteria
Erwinia tasmaniensis Et1 99 YP 001907312 1 465817 Proteobacteria
Dickeya gs fangzhongdai NCPPB 3274 WP 042861326 1 Dfangzhongdai
Dickeya gs fangzhongdai M005 WP 039692910 1 Dfangzhongdai
Dickeya gs fangzhongdai ND14b WP 038665017 1 Dfangzhongdai
Dickeya gs fangzhongdai M074 WP 038665017 1 Dfangzhongdai
Dickeya gs fangzhongdai B16 WP 038920408 1 Dfangzhongdai
Dickeya gs fangzhongdai MK7 WP 038920408 1 Dfangzhongdai
Dickeya gs fangzhongdai S1 WP 049854211 1 Dfangzhongdai
Dickeya solani RNS 05 1 2A WP 057085074 1 Dsolani
Dickeya solani PPO 9019 WP 022635136 1 Dsolani
Dickeya solani MK16 WP 022635136 1 Dsolani
Dickeya solani IFB 0099 WP 022635136 1 Dsolani
Dickeya solani GBBC 2040 WP 022635136 1 Dsolani
Dickeya solani D s0432 1 WP 022635136 1 Dsolani
Dickeya solani RNS 07 7 3B WP 022635136 1 Dsolani
Dickeya solani MK10 WP 022635136 1 Dsolani
Dickeya solani IPO 2222 WP 022635136 1 Dsolani
Dickeya solani RNS 08 23 3 1 A WP 022635136 1 Dsolani
Dickeya solani PPO 9134 WP 022635136 1 Dsolani
Dickeya dadantii subsp dieffenbachiae NCPPB 2976 WP 038924057 1 Ddadantii
Dickeya dadantii NCPPB 898 WP 038912082 1 Ddadantii
Dickeya dadantii NCPPB 3537 WP 038902232 1 Ddadantii
Dickeya dadantii 3937 WP 013319723 1 Ddadantii
432 oxdD response to acidic stress Seed
Dickeya zeae NCPPB 3531 WP 038905828 1 Dzeae
Dickeya zeae MS1 WP 019845186 1 Dzeae
Dickeya zeae CSL RW192 WP 038909148 1 Dzeae
Dickeya zeae DZ2Q WP 016942989 1 Dzeae
Dickeya zeae ZJU1202 WP 016942989 1 Dzeae
Dickeya zeae EC1 WP 016942989 1 Dzeae
Dickeya zeae NCPPB 3532 WP 028085700 1 Dzeae
Dickeya zeae MK19 WP 038914718 1 Dzeae
Dickeya zeae NCPPB 2538 WP 038914718 1 Dzeae

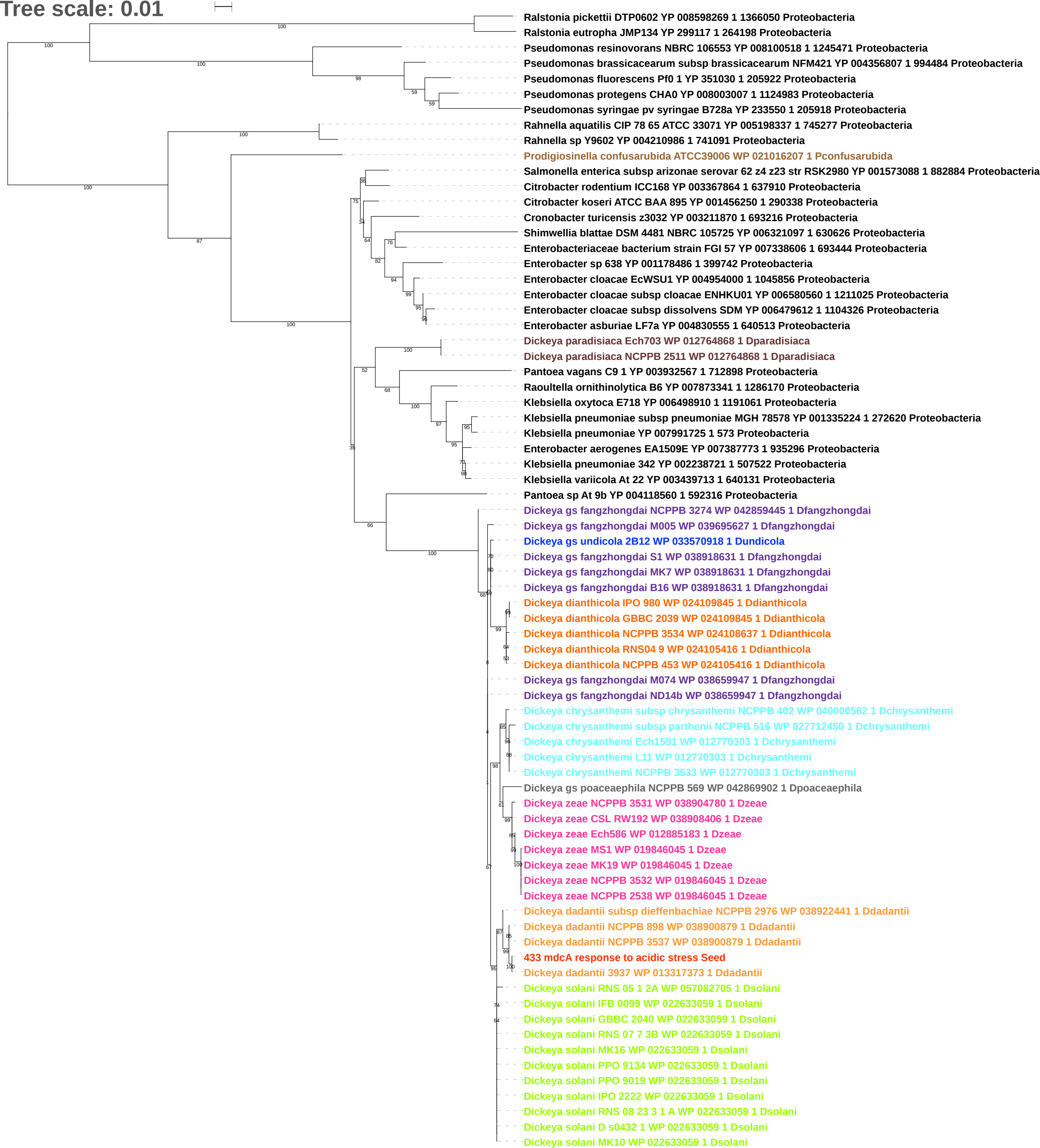
Tree scale: 0.01
Ralstonia pickettii DTP0602 YP 008598269 1 1366050 Proteobacteria
Ralstonia eutropha JMP134 YP 299117 1 264198 Proteobacteria
Pseudomonas resinovorans NBRC 106553 YP 008100518 1 1245471 Proteobacteria
Pseudomonas brassicacearum subsp brassicacearum NFM421 YP 004356807 1 994484 Proteobacteria
Pseudomonas fluorescens Pf0 1 YP 351030 1 205922 Proteobacteria
Pseudomonas protegens CHA0 YP 008003007 1 1124983 Proteobacteria
Pseudomonas syringae pv syringae B728a YP 233550 1 205918 Proteobacteria
Rahnella aquatilis CIP 78 65 ATCC 33071 YP 005198337 1 745277 Proteobacteria
Rahnella sp Y9602 YP 004210986 1 741091 Proteobacteria
Prodigiosinella confusarubida ATCC39006 WP 021016207 1 Pconfusarubida
Salmonella enterica subsp arizonae serovar 62 z4 z23 str RSK2980 YP 001573088 1 882884 Proteobacteria
Citrobacter rodentium ICC168 YP 003367864 1 637910 Proteobacteria
Citrobacter koseri ATCC BAA 895 YP 001456250 1 290338 Proteobacteria
Cronobacter turicensis z3032 YP 003211870 1 693216 Proteobacteria
Shimwellia blattae DSM 4481 NBRC 105725 YP 006321097 1 630626 Proteobacteria
Enterobacteriaceae bacterium strain FGI 57 YP 007338606 1 693444 Proteobacteria
Enterobacter sp 638 YP 001178486 1 399742 Proteobacteria
Enterobacter cloacae EcWSU1 YP 004954000 1 1045856 Proteobacteria
Enterobacter cloacae subsp cloacae ENHKU01 YP 006580560 1 1211025 Proteobacteria
Enterobacter cloacae subsp dissolvens SDM YP 006479612 1 1104326 Proteobacteria
Enterobacter asburiae LF7a YP 004830555 1 640513 Proteobacteria
Dickeya paradisiaca Ech703 WP 012764868 1 Dparadisiaca
Dickeya paradisiaca NCPPB 2511 WP 012764868 1 Dparadisiaca
Pantoea vagans C9 1 YP 003932567 1 712898 Proteobacteria
Raoultella ornithinolytica B6 YP 007873341 1 1286170 Proteobacteria
Klebsiella oxytoca E718 YP 006498910 1 1191061 Proteobacteria
Klebsiella pneumoniae subsp pneumoniae MGH 78578 YP 001335224 1 272620 Proteobacteria
Klebsiella pneumoniae YP 007991725 1 573 Proteobacteria
Enterobacter aerogenes EA1509E YP 007387773 1 935296 Proteobacteria
Klebsiella pneumoniae 342 YP 002238721 1 507522 Proteobacteria
Klebsiella variicola At 22 YP 003439713 1 640131 Proteobacteria
Pantoea sp At 9b YP 004118560 1 592316 Proteobacteria
Dickeya gs fangzhongdai NCPPB 3274 WP 042859445 1 Dfangzhongdai
Dickeya gs fangzhongdai M005 WP 039695627 1 Dfangzhongdai
Dickeya gs undicola 2B12 WP 033570918 1 Dundicola
Dickeya gs fangzhongdai S1 WP 038918631 1 Dfangzhongdai
Dickeya gs fangzhongdai MK7 WP 038918631 1 Dfangzhongdai
Dickeya gs fangzhongdai B16 WP 038918631 1 Dfangzhongdai
Dickeya dianthicola IPO 980 WP 024109845 1 Ddianthicola
Dickeya dianthicola GBBC 2039 WP 024109845 1 Ddianthicola
Dickeya dianthicola NCPPB 3534 WP 024108637 1 Ddianthicola
Dickeya dianthicola RNS04 9 WP 024105416 1 Ddianthicola
Dickeya dianthicola NCPPB 453 WP 024105416 1 Ddianthicola
Dickeya gs fangzhongdai M074 WP 038659947 1 Dfangzhongdai
Dickeya gs fangzhongdai ND14b WP 038659947 1 Dfangzhongdai
Dickeya chrysanthemi subsp chrysanthemi NCPPB 402 WP 040000562 1 Dchrysanthemi
Dickeya chrysanthemi subsp parthenii NCPPB 516 WP 027712450 1 Dchrysanthemi
Dickeya chrysanthemi Ech1591 WP 012770303 1 Dchrysanthemi
Dickeya chrysanthemi L11 WP 012770303 1 Dchrysanthemi
Dickeya chrysanthemi NCPPB 3533 WP 012770303 1 Dchrysanthemi
Dickeya gs poaceaephila NCPPB 569 WP 042869902 1 Dpoaceaephila
Dickeya zeae NCPPB 3531 WP 038904780 1 Dzeae
Dickeya zeae CSL RW192 WP 038908406 1 Dzeae
Dickeya zeae Ech586 WP 012885183 1 Dzeae
Dickeya zeae MS1 WP 019846045 1 Dzeae
Dickeya zeae MK19 WP 019846045 1 Dzeae
Dickeya zeae NCPPB 3532 WP 019846045 1 Dzeae
Dickeya zeae NCPPB 2538 WP 019846045 1 Dzeae
Dickeya dadantii subsp dieffenbachiae NCPPB 2976 WP 038922441 1 Ddadantii
Dickeya dadantii NCPPB 898 WP 038900879 1 Ddadantii
Dickeya dadantii NCPPB 3537 WP 038900879 1 Ddadantii
433 mdcA response to acidic stress Seed
Dickeya dadantii 3937 WP 013317373 1 Ddadantii
Dickeya solani RNS 05 1 2A WP 057082705 1 Dsolani
Dickeya solani IFB 0099 WP 022633059 1 Dsolani
Dickeya solani GBBC 2040 WP 022633059 1 Dsolani
Dickeya solani RNS 07 7 3B WP 022633059 1 Dsolani
Dickeya solani MK16 WP 022633059 1 Dsolani
Dickeya solani PPO 9134 WP 022633059 1 Dsolani
Dickeya solani PPO 9019 WP 022633059 1 Dsolani
Dickeya solani IPO 2222 WP 022633059 1 Dsolani
Dickeya solani RNS 08 23 3 1 A WP 022633059 1 Dsolani
Dickeya solani D s0432 1 WP 022633059 1 Dsolani
Dickeya solani MK10 WP 022633059 1 Dsolani

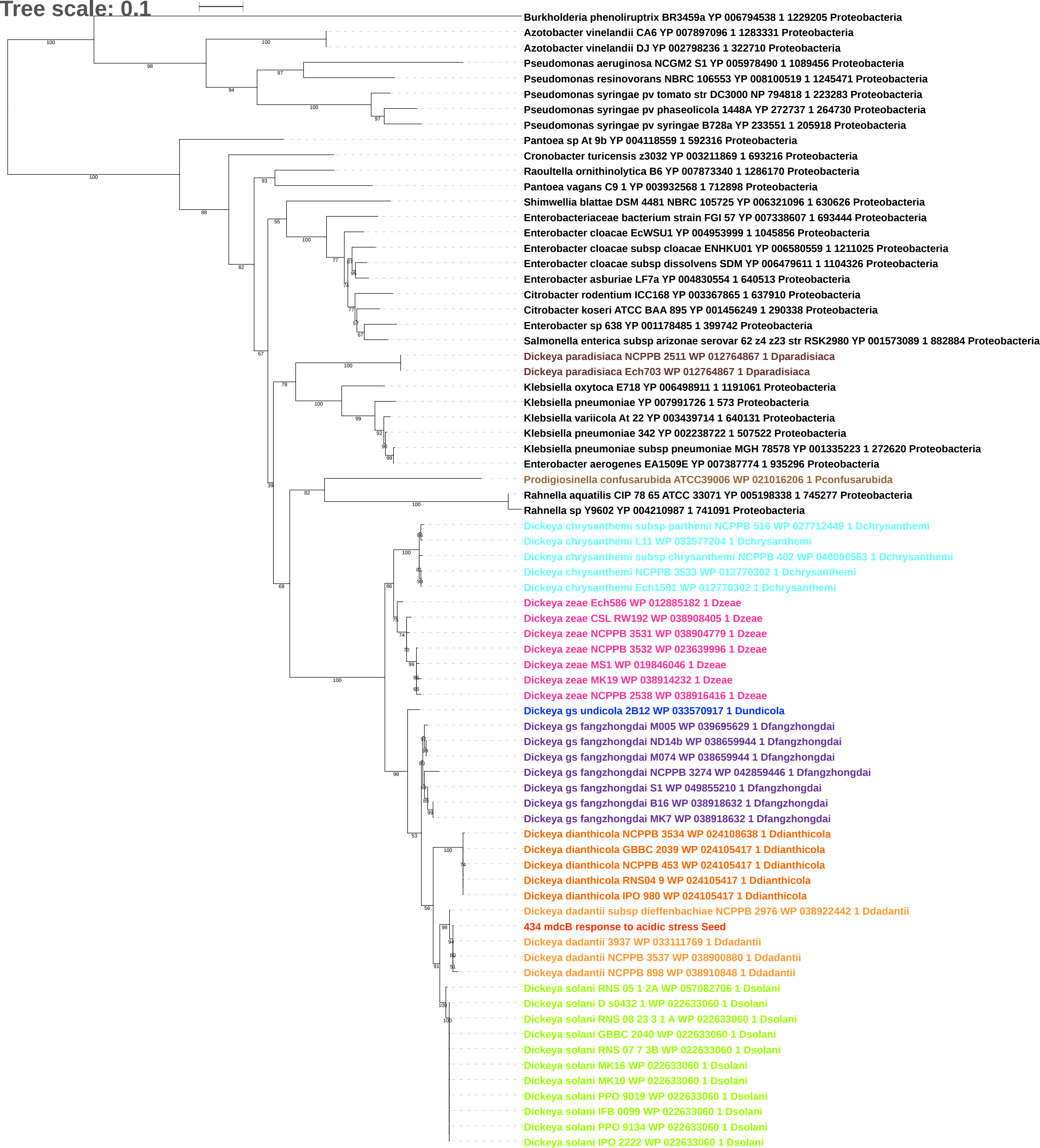
Tree scale: 0.1
Burkholderia phenoliruptrix BR3459a YP 006794538 1 1229205 Proteobacteria
Azotobacter vinelandii CA6 YP 007897096 1 1283331 Proteobacteria
Azotobacter vinelandii DJ YP 002798236 1 322710 Proteobacteria
Pseudomonas aeruginosa NCGM2 S1 YP 005978490 1 1089456 Proteobacteria
Pseudomonas resinovorans NBRC 106553 YP 008100519 1 1245471 Proteobacteria
Pseudomonas syringae pv tomato str DC3000 NP 794818 1 223283 Proteobacteria
Pseudomonas syringae pv phaseolicola 1448A YP 272737 1 264730 Proteobacteria
Pseudomonas syringae pv syringae B728a YP 233551 1 205918 Proteobacteria
Pantoea sp At 9b YP 004118559 1 592316 Proteobacteria
Cronobacter turicensis z3032 YP 003211869 1 693216 Proteobacteria
Raoultella ornithinolytica B6 YP 007873340 1 1286170 Proteobacteria
Pantoea vagans C9 1 YP 003932568 1 712898 Proteobacteria
Shimwellia blattae DSM 4481 NBRC 105725 YP 006321096 1 630626 Proteobacteria
Enterobacteriaceae bacterium strain FGI 57 YP 007338607 1 693444 Proteobacteria
Enterobacter cloacae EcWSU1 YP 004953999 1 1045856 Proteobacteria
Enterobacter cloacae subsp cloacae ENHKU01 YP 006580559 1 1211025 Proteobacteria
Enterobacter cloacae subsp dissolvens SDM YP 006479611 1 1104326 Proteobacteria
Enterobacter asburiae LF7a YP 004830554 1 640513 Proteobacteria
Citrobacter rodentium ICC168 YP 003367865 1 637910 Proteobacteria
Citrobacter koseri ATCC BAA 895 YP 001456249 1 290338 Proteobacteria
Enterobacter sp 638 YP 001178485 1 399742 Proteobacteria
Salmonella enterica subsp arizonae serovar 62 z4 z23 str RSK2980 YP 001573089 1 882884 Proteobacteria
Dickeya paradisiaca NCPPB 2511 WP 012764867 1 Dparadisiaca
Dickeya paradisiaca Ech703 WP 012764867 1 Dparadisiaca
Klebsiella oxytoca E718 YP 006498911 1 1191061 Proteobacteria
Klebsiella pneumoniae YP 007991726 1 573 Proteobacteria
Klebsiella variicola At 22 YP 003439714 1 640131 Proteobacteria
Klebsiella pneumoniae 342 YP 002238722 1 507522 Proteobacteria
Klebsiella pneumoniae subsp pneumoniae MGH 78578 YP 001335223 1 272620 Proteobacteria
Enterobacter aerogenes EA1509E YP 007387774 1 935296 Proteobacteria
Prodigiosinella confusarubida ATCC39006 WP 021016206 1 Pconfusarubida
Rahnella aquatilis CIP 78 65 ATCC 33071 YP 005198338 1 745277 Proteobacteria
Rahnella sp Y9602 YP 004210987 1 741091 Proteobacteria
Dickeya chrysanthemi subsp parthenii NCPPB 516 WP 027712449 1 Dchrysanthemi
Dickeya chrysanthemi L11 WP 033577204 1 Dchrysanthemi
Dickeya chrysanthemi subsp chrysanthemi NCPPB 402 WP 040000563 1 Dchrysanthemi
Dickeya chrysanthemi NCPPB 3533 WP 012770302 1 Dchrysanthemi
Dickeya chrysanthemi Ech1591 WP 012770302 1 Dchrysanthemi
Dickeya zeae Ech586 WP 012885182 1 Dzeae
Dickeya zeae CSL RW192 WP 038908405 1 Dzeae
Dickeya zeae NCPPB 3531 WP 038904779 1 Dzeae
Dickeya zeae NCPPB 3532 WP 023639996 1 Dzeae
Dickeya zeae MS1 WP 019846046 1 Dzeae
Dickeya zeae MK19 WP 038914232 1 Dzeae
Dickeya zeae NCPPB 2538 WP 038916416 1 Dzeae
Dickeya gs undicola 2B12 WP 033570917 1 Dundicola
Dickeya gs fangzhongdai M005 WP 039695629 1 Dfangzhongdai
Dickeya gs fangzhongdai ND14b WP 038659944 1 Dfangzhongdai
Dickeya gs fangzhongdai M074 WP 038659944 1 Dfangzhongdai
Dickeya gs fangzhongdai NCPPB 3274 WP 042859446 1 Dfangzhongdai
Dickeya gs fangzhongdai S1 WP 049855210 1 Dfangzhongdai
Dickeya gs fangzhongdai B16 WP 038918632 1 Dfangzhongdai
Dickeya gs fangzhongdai MK7 WP 038918632 1 Dfangzhongdai
Dickeya dianthicola NCPPB 3534 WP 024108638 1 Ddianthicola
Dickeya dianthicola GBBC 2039 WP 024105417 1 Ddianthicola
Dickeya dianthicola NCPPB 453 WP 024105417 1 Ddianthicola
Dickeya dianthicola RNS04 9 WP 024105417 1 Ddianthicola
Dickeya dianthicola IPO 980 WP 024105417 1 Ddianthicola
Dickeya dadantii subsp dieffenbachiae NCPPB 2976 WP 038922442 1 Ddadantii
434 mdcB response to acidic stress Seed
Dickeya dadantii 3937 WP 033111769 1 Ddadantii
Dickeya dadantii NCPPB 3537 WP 038900880 1 Ddadantii
Dickeya dadantii NCPPB 898 WP 038910848 1 Ddadantii
Dickeya solani RNS 05 1 2A WP 057082706 1 Dsolani
Dickeya solani D s0432 1 WP 022633060 1 Dsolani
Dickeya solani RNS 08 23 3 1 A WP 022633060 1 Dsolani
Dickeya solani GBBC 2040 WP 022633060 1 Dsolani
Dickeya solani RNS 07 7 3B WP 022633060 1 Dsolani
Dickeya solani MK16 WP 022633060 1 Dsolani
Dickeya solani MK10 WP 022633060 1 Dsolani
Dickeya solani PPO 9019 WP 022633060 1 Dsolani
Dickeya solani IFB 0099 WP 022633060 1 Dsolani
Dickeya solani PPO 9134 WP 022633060 1 Dsolani
Dickeya solani IPO 2222 WP 022633060 1 Dsolani

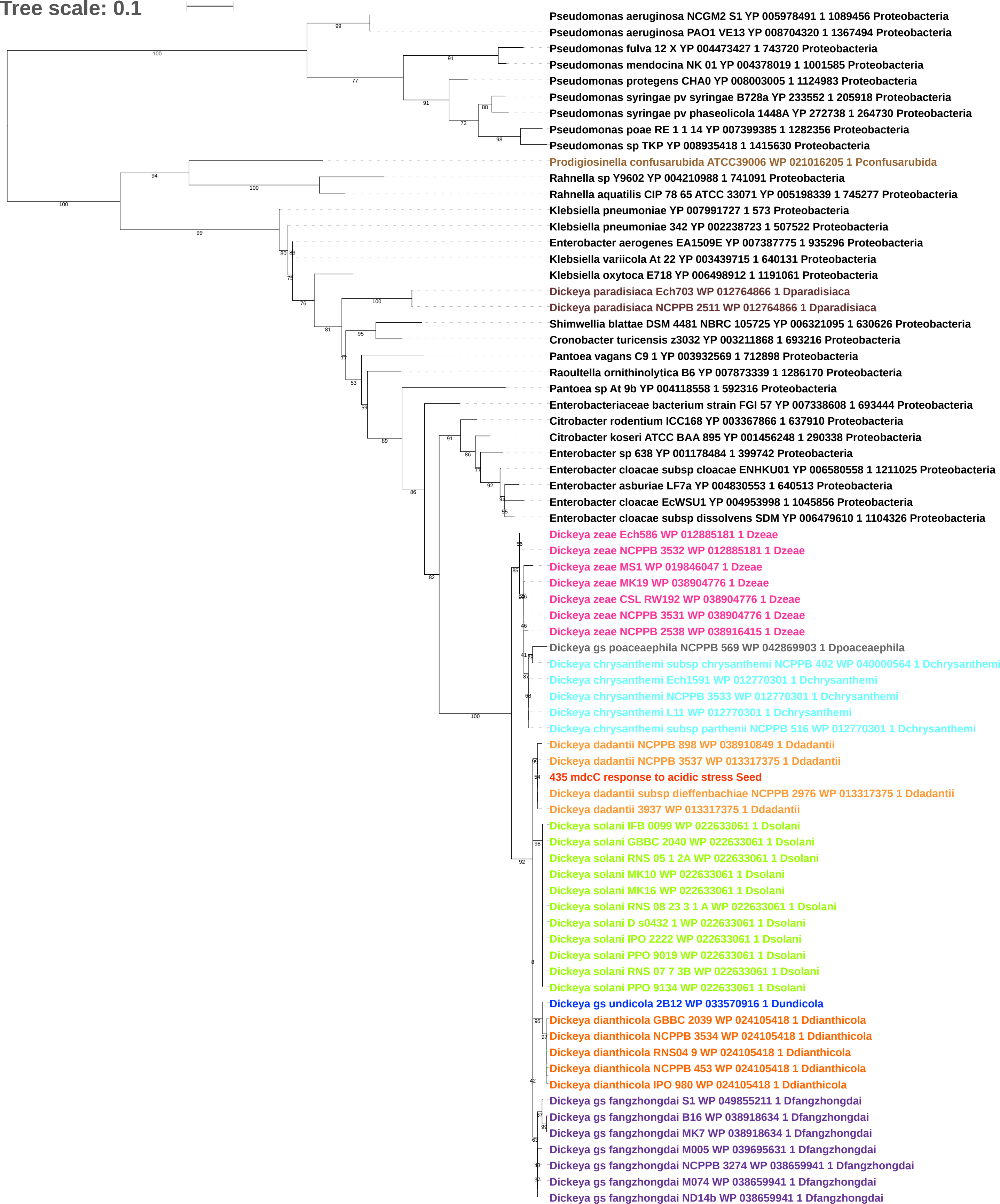
Tree scale: 0.1
Pseudomonas aeruginosa NCGM2 S1 YP 005978491 1 1089456 Proteobacteria
Pseudomonas aeruginosa PAO1 VE13 YP 008704320 1 1367494 Proteobacteria
Pseudomonas fulva 12 X YP 004473427 1 743720 Proteobacteria
Pseudomonas mendocina NK 01 YP 004378019 1 1001585 Proteobacteria
Pseudomonas protegens CHA0 YP 008003005 1 1124983 Proteobacteria
Pseudomonas syringae pv syringae B728a YP 233552 1 205918 Proteobacteria
Pseudomonas syringae pv phaseolicola 1448A YP 272738 1 264730 Proteobacteria
Pseudomonas poae RE 1 1 14 YP 007399385 1 1282356 Proteobacteria
Pseudomonas sp TKP YP 008935418 1 1415630 Proteobacteria
Prodigiosinella confusarubida ATCC39006 WP 021016205 1 Pconfusarubida
Rahnella sp Y9602 YP 004210988 1 741091 Proteobacteria
Rahnella aquatilis CIP 78 65 ATCC 33071 YP 005198339 1 745277 Proteobacteria
Klebsiella pneumoniae YP 007991727 1 573 Proteobacteria
Klebsiella pneumoniae 342 YP 002238723 1 507522 Proteobacteria
Enterobacter aerogenes EA1509E YP 007387775 1 935296 Proteobacteria
Klebsiella variicola At 22 YP 003439715 1 640131 Proteobacteria
Klebsiella oxytoca E718 YP 006498912 1 1191061 Proteobacteria
Dickeya paradisiaca Ech703 WP 012764866 1 Dparadisiaca
Dickeya paradisiaca NCPPB 2511 WP 012764866 1 Dparadisiaca
Shimwellia blattae DSM 4481 NBRC 105725 YP 006321095 1 630626 Proteobacteria
Cronobacter turicensis z3032 YP 003211868 1 693216 Proteobacteria
Pantoea vagans C9 1 YP 003932569 1 712898 Proteobacteria
Raoultella ornithinolytica B6 YP 007873339 1 1286170 Proteobacteria
Pantoea sp At 9b YP 004118558 1 592316 Proteobacteria
Enterobacteriaceae bacterium strain FGI 57 YP 007338608 1 693444 Proteobacteria
Citrobacter rodentium ICC168 YP 003367866 1 637910 Proteobacteria
Citrobacter koseri ATCC BAA 895 YP 001456248 1 290338 Proteobacteria
Enterobacter sp 638 YP 001178484 1 399742 Proteobacteria
Enterobacter cloacae subsp cloacae ENHKU01 YP 006580553 1 1211025 Proteobacteria
Enterobacter asburiae LF7a YP 004830553 1 640513 Proteobacteria
Enterobacter cloacae EcWSU1 YP 004953998 1 1045856 Proteobacteria
Enterobacter cloacae subsp dissolvens SDM YP 006479610 1 1104326 Proteobacteria
Dickeya zeae Ech586 WP 012885181 1 Dzeae
Dickeya zeae NCPPB 3532 WP 012885181 1 Dzeae
Dickeya zeae MS1 WP 019846047 1 Dzeae
Dickeya zeae MK19 WP 038904776 1 Dzeae
Dickeya zeae CSL RW192 WP 038904776 1 Dzeae
Dickeya zeae NCPPB 3531 WP 038904776 1 Dzeae
Dickeya zeae NCPPB 2538 WP 038916415 1 Dzeae
Dickeya gs poaceaephila NCPPB 569 WP 042869903 1 Dpoaceaephila
Dickeya chrysanthemi subsp chrysanthemi NCPPB 402 WP 040000564 1 Dchrysanthemi
Dickeya chrysanthemi Ech1591 WP 012770301 1 Dchrysanthemi
Dickeya chrysanthemi NCPPB 3533 WP 012770301 1 Dchrysanthemi
Dickeya chrysanthemi L11 WP 012770301 1 Dchrysanthemi
Dickeya chrysanthemi subsp parthenii NCPPB 516 WP 012770301 1 Dchrysanthemi
Dickeya dadantii NCPPB 898 WP 038910849 1 Ddadantii
Dickeya dadantii NCPPB 3537 WP 013317375 1 Ddadantii
435 mdcC response to acidic stress Seed
Dickeya dadantii subsp dieffenbachiae NCPPB 2976 WP 013317375 1 Ddadantii
Dickeya dadantii 3937 WP 013317375 1 Ddadantii
Dickeya solani IFB 0099 WP 022633061 1 Dsolani
Dickeya solani GBBC 2040 WP 022633061 1 Dsolani
Dickeya solani RNS 05 1 2A WP 022633061 1 Dsolani
Dickeya solani MK10 WP 022633061 1 Dsolani
Dickeya solani MK16 WP 022633061 1 Dsolani
Dickeya solani RNS 08 23 3 1 A WP 022633061 1 Dsolani
Dickeya solani D s0432 1 WP 022633061 1 Dsolani
Dickeya solani IPO 2222 WP 022633061 1 Dsolani
Dickeya solani PPO 9019 WP 022633061 1 Dsolani
Dickeya solani RNS 07 7 3B WP 022633061 1 Dsolani
Dickeya solani PPO 9134 WP 022633061 1 Dsolani
Dickeya gs undicola 2B12 WP 033570916 1 Dundicola
Dickeya dianthicola GBBC 2039 WP 024105418 1 Ddianthicola
Dickeya dianthicola NCPPB 3534 WP 024105418 1 Ddianthicola
Dickeya dianthicola RNS04 9 WP 024105418 1 Ddianthicola
Dickeya dianthicola NCPPB 453 WP 024105418 1 Ddianthicola
Dickeya dianthicola IPO 980 WP 024105418 1 Ddianthicola
Dickeya gs fangzhongdai S1 WP 049855211 1 Dfangzhongdai
Dickeya gs fangzhongdai B16 WP 038918634 1 Dfangzhongdai
Dickeya gs fangzhongdai MK7 WP 038918634 1 Dfangzhongdai
Dickeya gs fangzhongdai M005 WP 039695631 1 Dfangzhongdai
Dickeya gs fangzhongdai NCPPB 3274 WP 038659941 1 Dfangzhongdai
Dickeya gs fangzhongdai M074 WP 038659941 1 Dfangzhongdai
Dickeya gs fangzhongdai ND14b WP 038659941 1 Dfangzhongdai

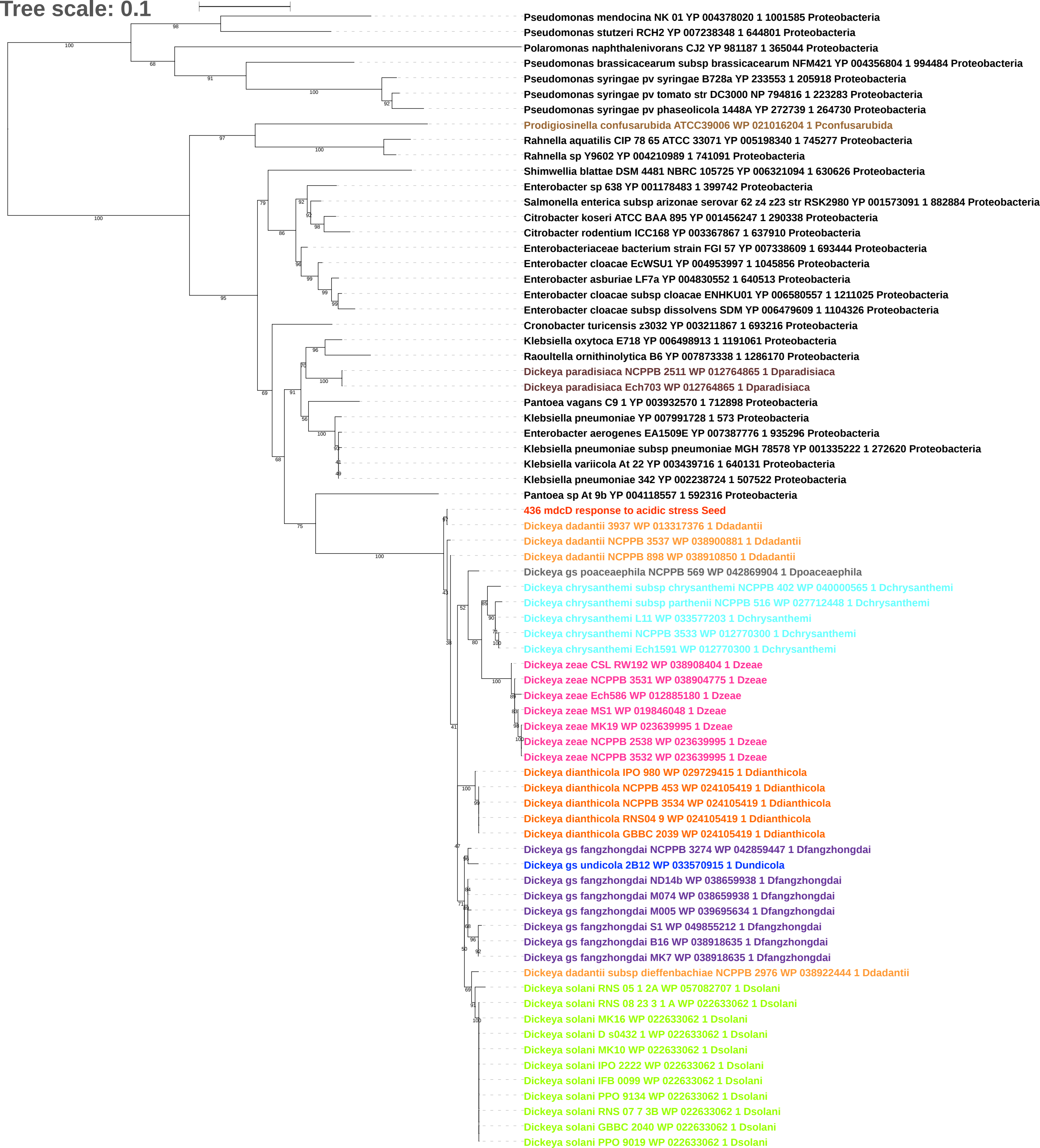
Tree scale: 0.1
Pseudomonas mendocina NK 01 YP 004378020 1 1001585 Proteobacteria
Pseudomonas stutzeri RCH2 YP 007238348 1 644801 Proteobacteria
Polaromonas naphthalenivorans CJ2 YP 981187 1 365044 Proteobacteria
Pseudomonas brassicacearum subsp brassicacearum NFM421 YP 004356804 1 994484 Proteobacteria
Pseudomonas syringae pv syringae B728a YP 233553 1 205918 Proteobacteria
Pseudomonas syringae pv tomato str DC3000 NP 794816 1 223283 Proteobacteria
Pseudomonas syringae pv phaseolicola 1448A YP 272739 1 264730 Proteobacteria
Prodigiosinella confusarubida ATCC39006 WP 021016204 1 Pconfusarubida
Rahnella aquatilis CIP 78 65 ATCC 33071 YP 005198340 1 745277 Proteobacteria
Rahnella sp Y9602 YP 004210989 1 741091 Proteobacteria
Shimwellia blattae DSM 4481 NBRC 105725 YP 006321094 1 630626 Proteobacteria
Enterobacter sp 638 YP 001178483 1 399742 Proteobacteria
Salmonella enterica subsp arizonae serovar 62 z4 z23 str RSK2980 YP 001573091 1 882884 Proteobacteria
Citrobacter koseri ATCC BAA 895 YP 001456247 1 290338 Proteobacteria
Citrobacter rodentium ICC168 YP 003367867 1 637910 Proteobacteria
Enterobacteriaceae bacterium strain FGI 57 YP 007338609 1 693444 Proteobacteria
Enterobacter cloacae EcWSU1 YP 004953997 1 1045856 Proteobacteria
Enterobacter asburiae LF7a YP 004830552 1 640513 Proteobacteria
Enterobacter cloacae subsp cloacae ENHKU01 YP 006580557 1 1211025 Proteobacteria
Enterobacter cloacae subsp dissolvens SDM YP 006479609 1 1104326 Proteobacteria
Cronobacter turicensis z3032 YP 003211867 1 693216 Proteobacteria
Klebsiella oxytoca E718 YP 006498913 1 1191061 Proteobacteria
Raoultella ornithinolytica B6 YP 007873338 1 1286170 Proteobacteria
Dickeya paradisiaca NCPPB 2511 WP 012764865 1 Dparadisiaca
Dickeya paradisiaca Ech703 WP 012764865 1 Dparadisiaca
Pantoea vagans C9 1 YP 003932570 1 712898 Proteobacteria
Klebsiella pneumoniae YP 007991728 1 573 Proteobacteria
Enterobacter aerogenes EA1509E YP 007387776 1 935296 Proteobacteria
Klebsiella pneumoniae subsp pneumoniae MGH 78578 YP 001335222 1 272620 Proteobacteria
Klebsiella variicola At 22 YP 003439716 1 640131 Proteobacteria
Klebsiella pneumoniae 342 YP 002238724 1 507522 Proteobacteria
Pantoea sp At 9b YP 004118557 1 592316 Proteobacteria
436 mdcD response to acidic stress Seed
Dickeya dadantii 3937 WP 013317376 1 Ddadantii
Dickeya dadantii NCPPB 3537 WP 038900881 1 Ddadantii
Dickeya dadantii NCPPB 898 WP 038910850 1 Ddadantii
Dickeya gs poaceaephila NCPPB 569 WP 042869904 1 Dpoaceaephila
Dickeya chrysanthemi subsp chrysanthemi NCPPB 402 WP 040000565 1 Dchrysanthemi
Dickeya chrysanthemi subsp parthenii NCPPB 516 WP 027712448 1 Dchrysanthemi
Dickeya chrysanthemi L11 WP 033577203 1 Dchrysanthemi
Dickeya chrysanthemi NCPPB 3533 WP 012770300 1 Dchrysanthemi
Dickeya chrysanthemi Ech1591 WP 012770300 1 Dchrysanthemi
Dickeya zeae CSL RW192 WP 038908404 1 Dzeae
Dickeya zeae NCPPB 3531 WP 038904775 1 Dzeae
Dickeya zeae Ech586 WP 012885180 1 Dzeae
Dickeya zeae MS1 WP 019846048 1 Dzeae
Dickeya zeae MK19 WP 023639995 1 Dzeae
Dickeya zeae NCPPB 2538 WP 023639995 1 Dzeae
Dickeya zeae NCPPB 3532 WP 023639995 1 Dzeae
Dickeya dianthicola IPO 980 WP 029729415 1 Ddianthicola
Dickeya dianthicola NCPPB 453 WP 024105419 1 Ddianthicola
Dickeya dianthicola NCPPB 3534 WP 024105419 1 Ddianthicola
Dickeya dianthicola RNS04 9 WP 024105419 1 Ddianthicola
Dickeya dianthicola GBBC 2039 WP 024105419 1 Ddianthicola
Dickeya gs fangzhongdai NCPPB 3274 WP 042859447 1 Dfangzhongdai
Dickeya gs undicola 2B12 WP 033570915 1 Dundicola
Dickeya gs fangzhongdai ND14b WP 038659938 1 Dfangzhongdai
Dickeya gs fangzhongdai M074 WP 038659938 1 Dfangzhongdai
Dickeya gs fangzhongdai M005 WP 039695634 1 Dfangzhongdai
Dickeya gs fangzhongdai S1 WP 049855212 1 Dfangzhongdai
Dickeya gs fangzhongdai B16 WP 038918635 1 Dfangzhongdai
Dickeya gs fangzhongdai MK7 WP 038918635 1 Dfangzhongdai
Dickeya dadantii subsp dieffenbachiae NCPPB 2976 WP 038922444 1 Ddadantii
Dickeya solani RNS 05 1 2A WP 057082707 1 Dsolani
Dickeya solani RNS 08 23 3 1 A WP 022633062 1 Dsolani
Dickeya solani MK16 WP 022633062 1 Dsolani
Dickeya solani D s0432 1 WP 022633062 1 Dsolani
Dickeya solani MK10 WP 022633062 1 Dsolani
Dickeya solani IPO 2222 WP 022633062 1 Dsolani
Dickeya solani IFB 0099 WP 022633062 1 Dsolani
Dickeya solani PPO 9134 WP 022633062 1 Dsolani
Dickeya solani RNS 07 7 3B WP 022633062 1 Dsolani
Dickeya solani GBBC 2040 WP 022633062 1 Dsolani
Dickeya solani PPO 9019 WP 022633062 1 Dsolani

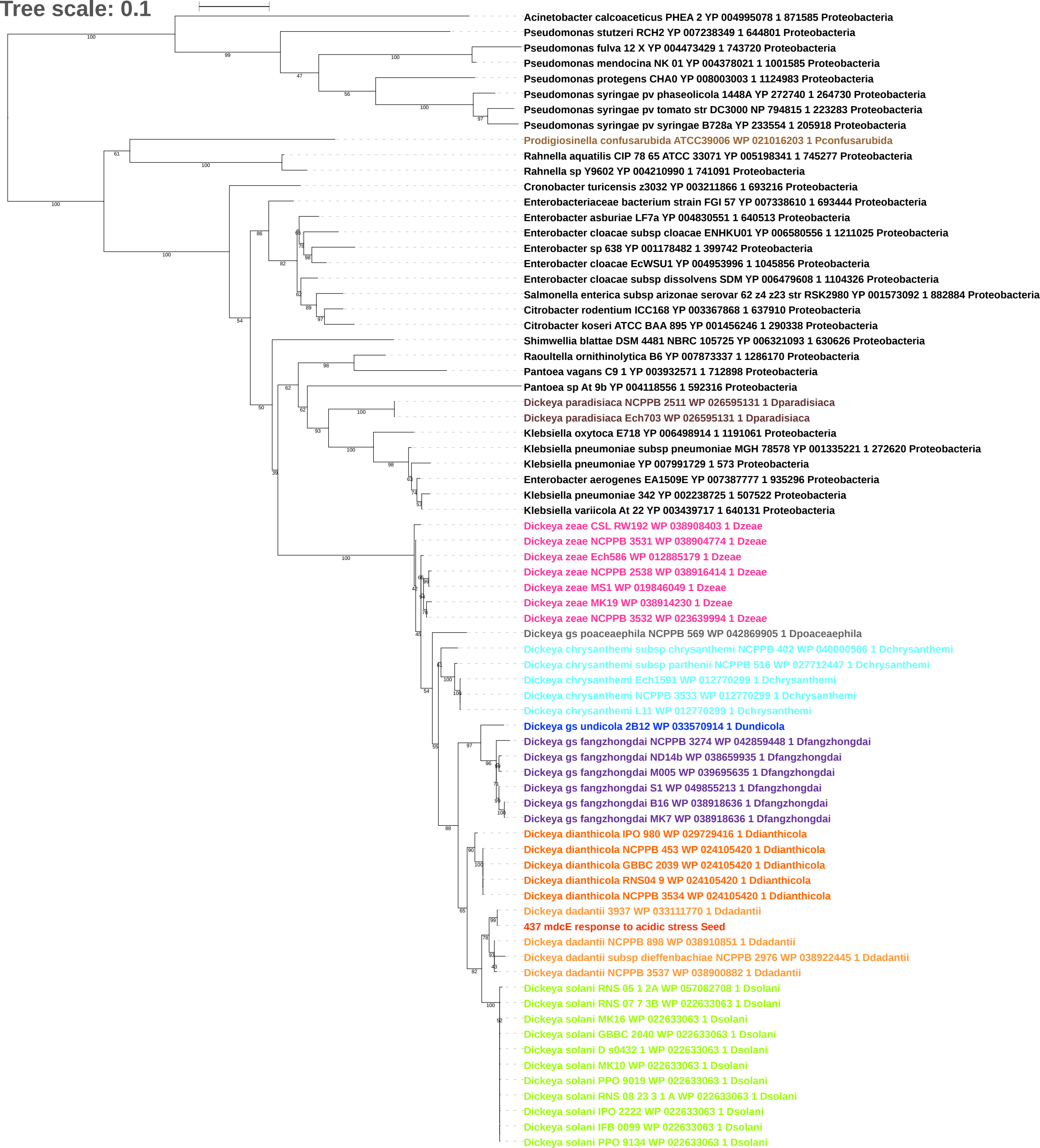
Tree scale: 0.1
Acinetobacter calcoaceticus PHEA 2 YP 004995078 1 871585 Proteobacteria
Pseudomonas stutzeri RCH2 YP 007238349 1 644801 Proteobacteria
Pseudomonas fulva 12 X YP 004473429 1 743720 Proteobacteria
Pseudomonas mendocina NK 01 YP 004378021 1 1001585 Proteobacteria
Pseudomonas protegens CHA0 YP 008003003 1 1124983 Proteobacteria
Pseudomonas syringae pv phaseolicola 1448A YP 272740 1 264730 Proteobacteria
Pseudomonas syringae pv tomato str DC3000 NP 794815 1 223283 Proteobacteria
Pseudomonas syringae pv syringae B728a YP 233554 1 205918 Proteobacteria
Prodigiosinella confusarubida ATCC39006 WP 021016203 1 Pconfusarubida
Rahnella aquatilis CIP 78 65 ATCC 33071 YP 005198341 1 745277 Proteobacteria
Rahnella sp Y9602 YP 004210990 1 741091 Proteobacteria
Cronobacter turicensis z3032 YP 003211866 1 693216 Proteobacteria
Enterobacteriaceae bacterium strain FGI 57 YP 007338610 1 693444 Proteobacteria
Enterobacter asburiae LF7a YP 004830551 1 640513 Proteobacteria
Enterobacter cloacae subsp cloacae ENHKU01 YP 006580556 1 1211025 Proteobacteria
Enterobacter sp 638 YP 001178482 1 399742 Proteobacteria
Enterobacter cloacae EcWSU1 YP 004953996 1 1045856 Proteobacteria
Enterobacter cloacae subsp dissolvens SDM YP 006479608 1 1104326 Proteobacteria
Salmonella enterica subsp arizonae serovar 62 z4 z23 str RSK2980 YP 001573092 1 882884 Proteobacteria
Citrobacter rodentium ICC168 YP 003367868 1 637910 Proteobacteria
Citrobacter koseri ATCC BAA 895 YP 001456246 1 290338 Proteobacteria
Shimwellia blattae DSM 4481 NBRC 105725 YP 006321093 1 630626 Proteobacteria
Raoultella ornithinolytica B6 YP 007873337 1 1286170 Proteobacteria
Pantoea vagans C9 1 YP 003932571 1 712898 Proteobacteria
Pantoea sp At 9b YP 004118556 1 592316 Proteobacteria
Dickeya paradisiaca NCPPB 2511 WP 026595131 1 Dparadisiaca
Dickeya paradisiaca Ech703 WP 026595131 1 Dparadisiaca
Klebsiella oxytoca E718 YP 006498914 1 1191061 Proteobacteria
Klebsiella pneumoniae subsp pneumoniae MGH 78578 YP 001335221 1 272620 Proteobacteria
Klebsiella pneumoniae YP 007991729 1 573 Proteobacteria
Enterobacter aerogenes EA1509E YP 007387777 1 935296 Proteobacteria
Klebsiella pneumoniae 342 YP 002238725 1 507522 Proteobacteria
Klebsiella variicola At 22 YP 003439717 1 640131 Proteobacteria
Dickeya zeae CSL RW192 WP 038908403 1 Dzeae
Dickeya zeae NCPPB 3531 WP 038904774 1 Dzeae
Dickeya zeae Ech586 WP 012885179 1 Dzeae
Dickeya zeae NCPPB 2538 WP 038916414 1 Dzeae
Dickeya zeae MS1 WP 019846049 1 Dzeae
Dickeya zeae MK19 WP 038914230 1 Dzeae
Dickeya zeae NCPPB 3532 WP 023639994 1 Dzeae
Dickeya gs poaceaephila NCPPB 569 WP 042869905 1 Dpoaceaephila
Dickeya chrysanthemi subsp chrysanthemi NCPPB 402 WP 040000566 1 Dchrysanthemi
Dickeya chrysanthemi subsp parthenii NCPPB 516 WP 027712447 1 Dchrysanthemi
Dickeya chrysanthemi Ech1591 WP 012770299 1 Dchrysanthemi
Dickeya chrysanthemi NCPPB 3533 WP 012770299 1 Dchrysanthemi
Dickeya chrysanthemi L11 WP 012770299 1 Dchrysanthemi
Dickeya gs undicola 2B12 WP 033570914 1 Dundicola
Dickeya gs fangzhongdai NCPPB 3274 WP 042859448 1 Dfangzhongdai
Dickeya gs fangzhongdai ND14b WP 038659935 1 Dfangzhongdai
Dickeya gs fangzhongdai M005 WP 039695635 1 Dfangzhongdai
Dickeya gs fangzhongdai S1 WP 049855213 1 Dfangzhongdai
Dickeya gs fangzhongdai B16 WP 038918636 1 Dfangzhongdai
Dickeya gs fangzhongdai MK7 WP 038918636 1 Dfangzhongdai
Dickeya dianthicola IPO 980 WP 029729416 1 Ddianthicola
Dickeya dianthicola NCPPB 453 WP 024105420 1 Ddianthicola
Dickeya dianthicola GBBC 2039 WP 024105420 1 Ddianthicola
Dickeya dianthicola RNS04 9 WP 024105420 1 Ddianthicola
Dickeya dianthicola NCPPB 3534 WP 024105420 1 Ddianthicola
Dickeya dadantii 3937 WP 033111770 1 Ddadantii
437 mdcE response to acidic stress Seed
Dickeya dadantii NCPPB 898 WP 038910851 1 Ddadantii
Dickeya dadantii subsp dieffenbachiae NCPPB 2976 WP 038922445 1 Ddadantii
Dickeya dadantii NCPPB 3537 WP 038900882 1 Ddadantii
Dickeya solani RNS 05 1 2A WP 057082708 1 Dsolani
Dickeya solani RNS 07 7 3B WP 022633063 1 Dsolani
Dickeya solani MK16 WP 022633063 1 Dsolani
Dickeya solani GBBC 2040 WP 022633063 1 Dsolani
Dickeya solani D s0432 1 WP 022633063 1 Dsolani
Dickeya solani MK10 WP 022633063 1 Dsolani
Dickeya solani PPO 9019 WP 022633063 1 Dsolani
Dickeya solani RNS 08 23 3 1 A WP 022633063 1 Dsolani
Dickeya solani IPO 2222 WP 022633063 1 Dsolani
Dickeya solani IFB 0099 WP 022633063 1 Dsolani
Dickeya solani PPO 9134 WP 022633063 1 Dsolani

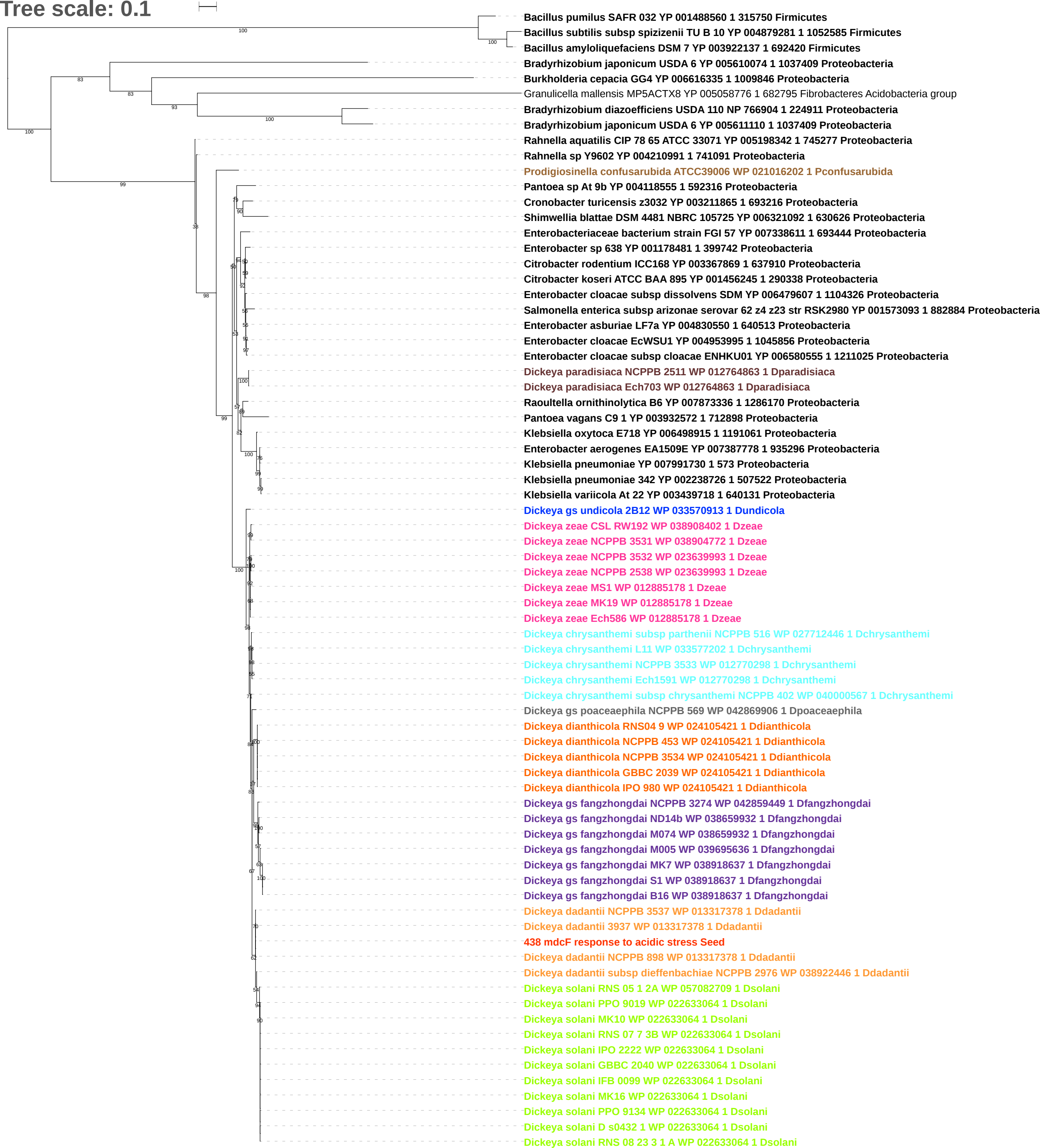
Tree scale: 0.1
Bacillus pumilus SAFR 032 YP 001488560 1 315750 Firmicutes
Bacillus subtilis subsp spizizenii TU B 10 YP 004879281 1 1052585 Firmicutes
Bacillus amyloliquefaciens DSM 7 YP 003922137 1 692420 Firmicutes
Bradyrhizobium japonicum USDA 6 YP 005610074 1 1037409 Proteobacteria
Burkholderia cepacia GG4 YP 006616335 1 1009846 Proteobacteria
Granulicella mallensis MP5ACTX8 YP 005058776 1 682795 Fibrobacteres Acidobacteria group
Bradyrhizobium diazoefficiens USDA 110 NP 766904 1 224911 Proteobacteria
Bradyrhizobium japonicum USDA 6 YP 005611110 1 1037409 Proteobacteria
Rahnella aquatilis CIP 78 65 ATCC 33071 YP 005198342 1 745277 Proteobacteria
Rahnella sp Y9602 YP 004210991 1 741091 Proteobacteria
Prodigiosinella confusarubida ATCC39006 WP 021016202 1 Pconfusarubida
Pantoea sp At 9b YP 004118555 1 592316 Proteobacteria
Cronobacter turicensis z3032 YP 003211865 1 693216 Proteobacteria
Shimwellia blattae DSM 4481 NBRC 105725 YP 006321092 1 630626 Proteobacteria
Enterobacteriaceae bacterium strain FGI 57 YP 007338611 1 693444 Proteobacteria
Enterobacter sp 638 YP 001178481 1 399742 Proteobacteria
Citrobacter rodentium ICC168 YP 003367869 1 637910 Proteobacteria
Citrobacter koseri ATCC BAA 895 YP 001456245 1 290338 Proteobacteria
Enterobacter cloacae subsp dissolvens SDM YP 006479607 1 1104326 Proteobacteria
Salmonella enterica subsp arizonae serovar 62 z4 z23 str RSK2980 YP 001573093 1 882884 Proteobacteria
Enterobacter asburiae LF7a YP 004830550 1 640513 Proteobacteria
Enterobacter cloacae EcWSU1 YP 004953995 1 1045856 Proteobacteria
Enterobacter cloacae subsp cloacae ENHKU01 YP 006580555 1 1211025 Proteobacteria
Dickeya paradisiaca NCPPB 2511 WP 012764863 1 Dparadisiaca
Dickeya paradisiaca Ech703 WP 012764863 1 Dparadisiaca
Raoultella ornithinolytica B6 YP 007873336 1 1286170 Proteobacteria
Pantoea vagans C9 1 YP 003932572 1 712898 Proteobacteria
Klebsiella oxytoca E718 YP 006498915 1 1191061 Proteobacteria
Enterobacter aerogenes EA1509E YP 007387778 1 935296 Proteobacteria
Klebsiella pneumoniae YP 007991730 1 573 Proteobacteria
Klebsiella pneumoniae 342 YP 002238726 1 507522 Proteobacteria
Klebsiella variicola At 22 YP 003439718 1 640131 Proteobacteria
Dickeya gs undicola 2B12 WP 033570913 1 Dundicola
Dickeya zeae CSL RW192 WP 038908402 1 Dzeae
Dickeya zeae NCPPB 3531 WP 038904772 1 Dzeae
Dickeya zeae NCPPB 3532 WP 023639993 1 Dzeae
Dickeya zeae NCPPB 2538 WP 023639993 1 Dzeae
Dickeya zeae MS1 WP 012885178 1 Dzeae
Dickeya zeae MK19 WP 012885178 1 Dzeae
Dickeya zeae Ech586 WP 012885178 1 Dzeae
Dickeya chrysanthemi subsp parthenii NCPPB 516 WP 027712446 1 Dchrysanthemi
Dickeya chrysanthemi L11 WP 033577202 1 Dchrysanthemi
Dickeya chrysanthemi NCPPB 3533 WP 012770298 1 Dchrysanthemi
Dickeya chrysanthemi Ech1591 WP 012770298 1 Dchrysanthemi
Dickeya chrysanthemi subsp chrysanthemi NCPPB 402 WP 040000567 1 Dchrysanthemi
Dickeya gs poaceaephila NCPPB 569 WP 042869906 1 Dpoaceaephila
Dickeya dianthicola RNS04 9 WP 024105421 1 Ddianthicola
Dickeya dianthicola NCPPB 453 WP 024105421 1 Ddianthicola
Dickeya dianthicola NCPPB 3534 WP 024105421 1 Ddianthicola
Dickeya dianthicola GBBC 2039 WP 024105421 1 Ddianthicola
Dickeya dianthicola IPO 980 WP 024105421 1 Ddianthicola
Dickeya gs fangzhongdai NCPPB 3274 WP 042859449 1 Dfangzhongdai
Dickeya gs fangzhongdai ND14b WP 038659932 1 Dfangzhongdai
Dickeya gs fangzhongdai M074 WP 038659932 1 Dfangzhongdai
Dickeya gs fangzhongdai M005 WP 039695636 1 Dfangzhongdai
Dickeya gs fangzhongdai MK7 WP 038918637 1 Dfangzhongdai
Dickeya gs fangzhongdai S1 WP 038918637 1 Dfangzhongdai
Dickeya gs fangzhongdai B16 WP 038918637 1 Dfangzhongdai
Dickeya dadantii NCPPB 3537 WP 013317378 1 Ddadantii
Dickeya dadantii 3937 WP 013317378 1 Ddadantii
438 mdcF response to acidic stress Seed
Dickeya dadantii NCPPB 898 WP 013317378 1 Ddadantii
Dickeya dadantii subsp dieffenbachiae NCPPB 2976 WP 038922446 1 Ddadantii
Dickeya solani RNS 05 1 2A WP 057082709 1 Dsolani
Dickeya solani PPO 9019 WP 022633064 1 Dsolani
Dickeya solani MK10 WP 022633064 1 Dsolani
Dickeya solani RNS 07 7 3B WP 022633064 1 Dsolani
Dickeya solani IPO 2222 WP 022633064 1 Dsolani
Dickeya solani GBBC 2040 WP 022633064 1 Dsolani
Dickeya solani IFB 0099 WP 022633064 1 Dsolani
Dickeya solani MK16 WP 022633064 1 Dsolani
Dickeya solani PPO 9134 WP 022633064 1 Dsolani
Dickeya solani D s0432 1 WP 022633064 1 Dsolani
Dickeya solani RNS 08 23 3 1 A WP 022633064 1 Dsolani

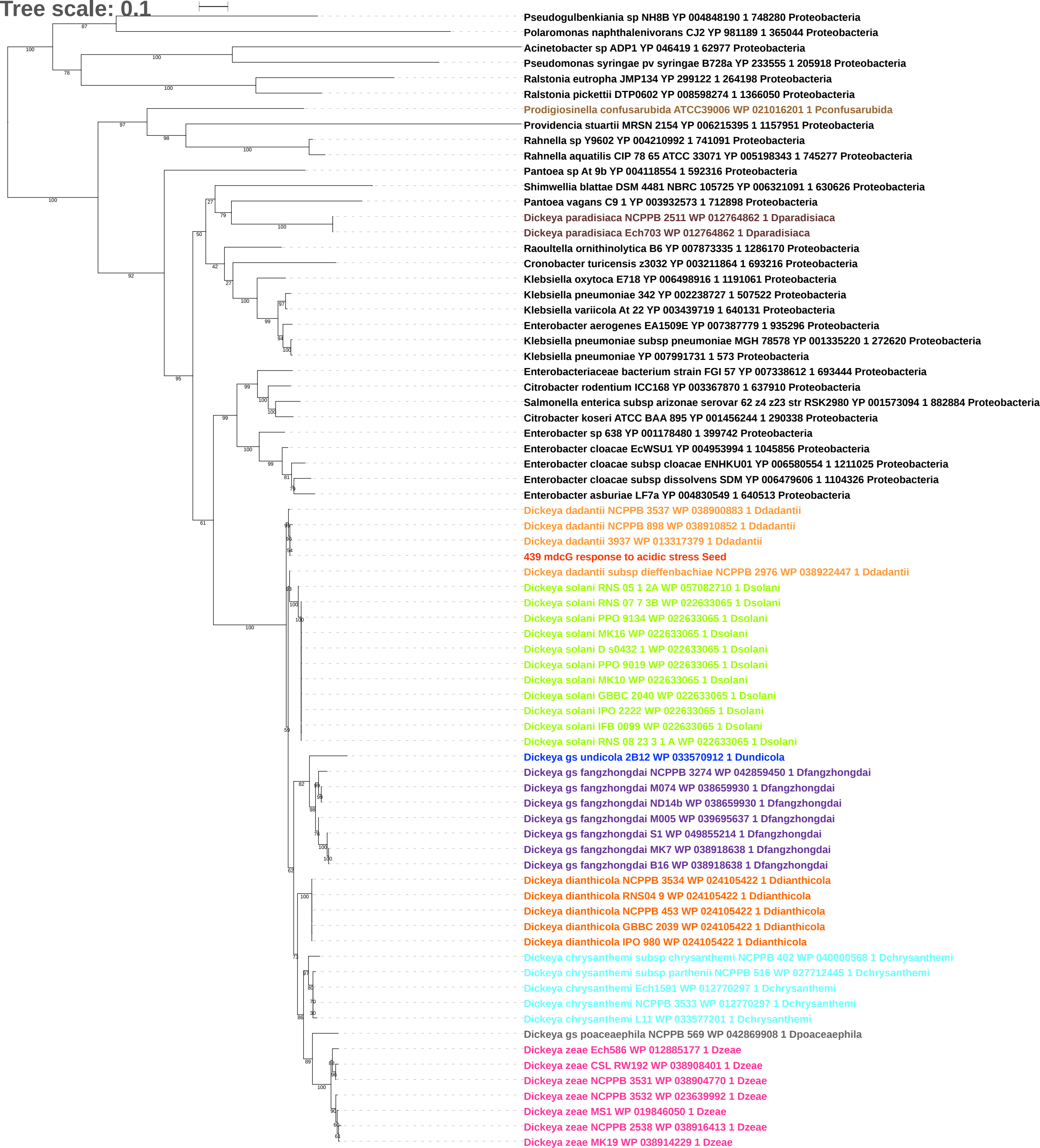
Tree scale: 0.1
Pseudogulbenkiania sp NH8B YP 004848190 1 748280 Proteobacteria
Polaromonas naphthalenivorans CJ2 YP 981189 1 365044 Proteobacteria
Acinetobacter sp ADP1 YP 046419 1 62977 Proteobacteria
Pseudomonas syringae pv syringae B728a YP 233555 1 205918 Proteobacteria
Ralstonia eutropha JMP134 YP 299122 1 264198 Proteobacteria
Ralstonia pickettii DTP0602 YP 008598274 1 1366050 Proteobacteria
Prodigiosinella confusarubida ATCC39006 WP 021016201 1 Pconfusarubida
Providencia stuartii MRSN 2154 YP 006215395 1 1157951 Proteobacteria
Rahnella sp Y9602 YP 004210992 1 741091 Proteobacteria
Rahnella aquatilis CIP 78 65 ATCC 33071 YP 005198343 1 745277 Proteobacteria
Pantoea sp At 9b YP 004118554 1 592316 Proteobacteria
Shimwellia blattae DSM 4481 NBRC 105725 YP 006321091 1 630626 Proteobacteria
Pantoea vagans C9 1 YP 003932573 1 712898 Proteobacteria
Dickeya paradisiaca NCPPB 2511 WP 012764862 1 Dparadisiaca
Dickeya paradisiaca Ech703 WP 012764862 1 Dparadisiaca
Raoultella ornithinolytica B6 YP 007873335 1 1286170 Proteobacteria
Cronobacter turicensis z3032 YP 003211864 1 693216 Proteobacteria
Klebsiella oxytoca E718 YP 006498916 1 1191061 Proteobacteria
Klebsiella pneumoniae 342 YP 002238727 1 507522 Proteobacteria
Klebsiella variicola At 22 YP 003439719 1 640131 Proteobacteria
Enterobacter aerogenes EA1509E YP 007387779 1 935296 Proteobacteria
Klebsiella pneumoniae subsp pneumoniae MGH 78578 YP 001335220 1 272620 Proteobacteria
Klebsiella pneumoniae YP 007991731 1 573 Proteobacteria
Enterobacteriaceae bacterium strain FGI 57 YP 007338612 1 693444 Proteobacteria
Citrobacter rodentium ICC168 YP 003367870 1 637910 Proteobacteria
Salmonella enterica subsp arizonae serovar 62 z4 z23 str RSK2980 YP 001573094 1 882884 Proteobacteria
Citrobacter koseri ATCC BAA 895 YP 001456244 1 290338 Proteobacteria
Enterobacter sp 638 YP 001178480 1 399742 Proteobacteria
Enterobacter cloacae EcWSU1 YP 004953994 1 1045856 Proteobacteria
Enterobacter cloacae subsp cloacae ENHKU01 YP 006580554 1 1211025 Proteobacteria
Enterobacter cloacae subsp dissolvens SDM YP 006479606 1 1104326 Proteobacteria
Enterobacter asburiae LF7a YP 004830549 1 640513 Proteobacteria
Dickeya dadantii NCPPB 3537 WP 038900883 1 Ddadantii
Dickeya dadantii NCPPB 898 WP 038910852 1 Ddadantii
Dickeya dadantii 3937 WP 013317379 1 Ddadantii
439 mdcG response to acidic stress Seed
Dickeya dadantii subsp dieffenbachiae NCPPB 2976 WP 038922447 1 Ddadantii
Dickeya solani RNS 05 1 2A WP 057082710 1 Dsolani
Dickeya solani RNS 07 7 3B WP 022633065 1 Dsolani
Dickeya solani PPO 9134 WP 022633065 1 Dsolani
Dickeya solani MK16 WP 022633065 1 Dsolani
Dickeya solani D s0432 1 WP 022633065 1 Dsolani
Dickeya solani PPO 9019 WP 022633065 1 Dsolani
Dickeya solani MK10 WP 022633065 1 Dsolani
Dickeya solani GBBC 2040 WP 022633065 1 Dsolani
Dickeya solani IPO 2222 WP 022633065 1 Dsolani
Dickeya solani IFB 0099 WP 022633065 1 Dsolani
Dickeya solani RNS 08 23 3 1 A WP 022633065 1 Dsolani
Dickeya gs undicola 2B12 WP 033570912 1 Dundicola
Dickeya gs fangzhongdai NCPPB 3274 WP 042859450 1 Dfangzhongdai
Dickeya gs fangzhongdai M074 WP 038659930 1 Dfangzhongdai
Dickeya gs fangzhongdai ND14b WP 038659930 1 Dfangzhongdai
Dickeya gs fangzhongdai M005 WP 039695637 1 Dfangzhongdai
Dickeya gs fangzhongdai S1 WP 049855214 1 Dfangzhongdai
Dickeya gs fangzhongdai MK7 WP 038918638 1 Dfangzhongdai
Dickeya gs fangzhongdai B16 WP 038918638 1 Dfangzhongdai
Dickeya dianthicola NCPPB 3534 WP 024105422 1 Ddianthicola
Dickeya dianthicola RNS04 9 WP 024105422 1 Ddianthicola
Dickeya dianthicola NCPPB 453 WP 024105422 1 Ddianthicola
Dickeya dianthicola GBBC 2039 WP 024105422 1 Ddianthicola
Dickeya dianthicola IPO 980 WP 024105422 1 Ddianthicola
Dickeya chrysanthemi subsp chrysanthemi NCPPB 402 WP 040000568 1 Dchrysanthemi
Dickeya chrysanthemi subsp parthenii NCPPB 516 WP 027712445 1 Dchrysanthemi
Dickeya chrysanthemi Ech1591 WP 012770297 1 Dchrysanthemi
Dickeya chrysanthemi NCPPB 3533 WP 012770297 1 Dchrysanthemi
Dickeya chrysanthemi L11 WP 033577201 1 Dchrysanthemi
Dickeya gs poaceaephila NCPPB 569 WP 042869908 1 Dpoaceaephila
Dickeya zeae Ech586 WP 012885177 1 Dzeae
Dickeya zeae CSL RW192 WP 038908401 1 Dzeae
Dickeya zeae NCPPB 3531 WP 038904770 1 Dzeae
Dickeya zeae NCPPB 3532 WP 023639992 1 Dzeae
Dickeya zeae MS1 WP 019846050 1 Dzeae
Dickeya zeae NCPPB 2538 WP 038916413 1 Dzeae
Dickeya zeae MK19 WP 038914229 1 Dzeae

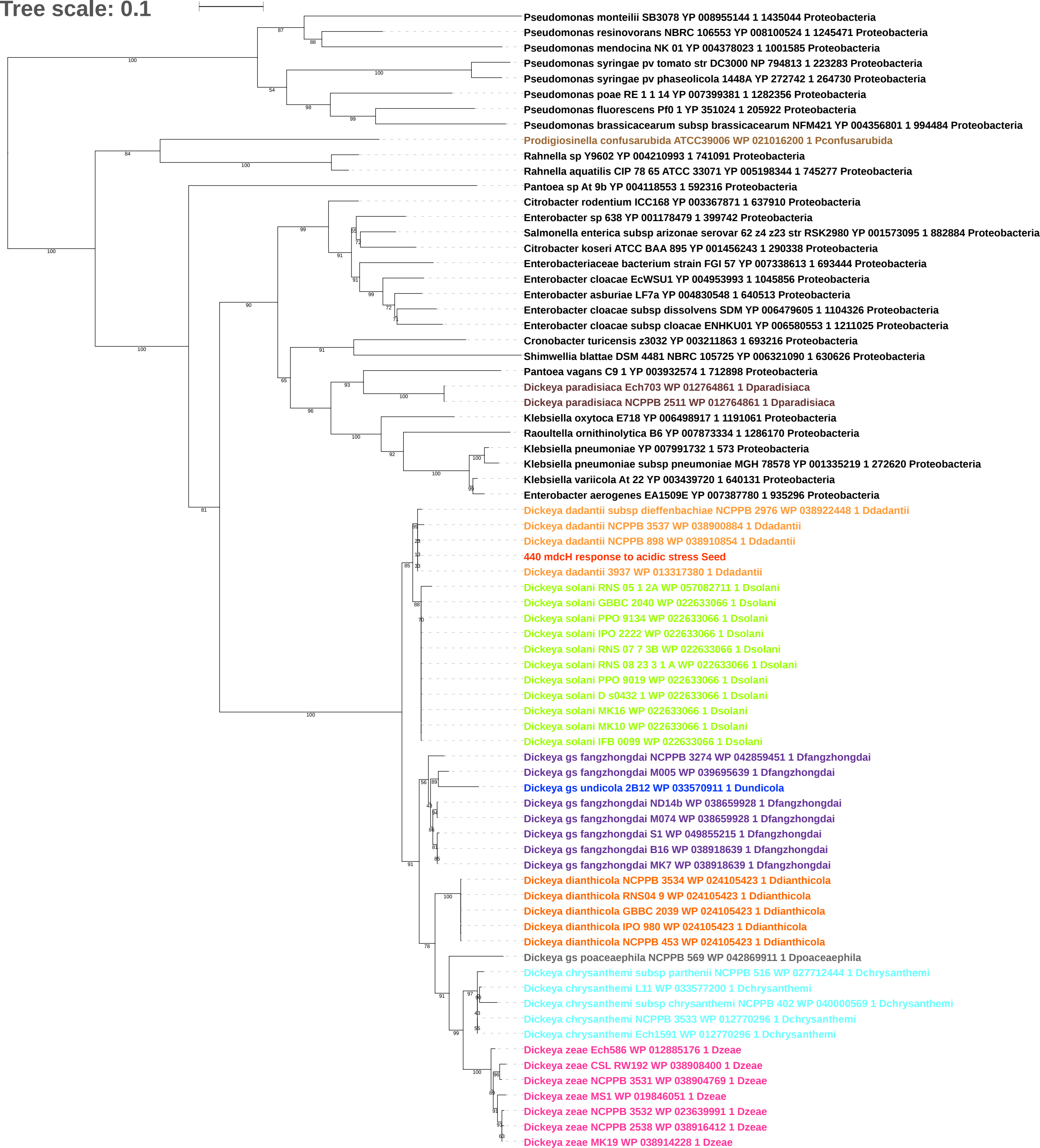
Tree scale: 0.1
Pseudomonas monteilii SB3078 YP 008955144 1 1435044 Proteobacteria
Pseudomonas resinovorans NBRC 106553 YP 008100524 1 1245471 Proteobacteria
Pseudomonas mendocina NK 01 YP 004378023 1 1001585 Proteobacteria
Pseudomonas syringae pv tomato str DC3000 NP 794813 1 223283 Proteobacteria
Pseudomonas syringae pv phaseolicola 1448A YP 272742 1 264730 Proteobacteria
Pseudomonas poae RE 1 1 14 YP 007399381 1 1282356 Proteobacteria
Pseudomonas fluorescens Pf0 1 YP 351024 1 205922 Proteobacteria
Pseudomonas brassicacearum subsp brassicacearum NFM421 YP 004356801 1 994484 Proteobacteria
Prodigiosinella confusarubida ATCC39006 WP 021016200 1 Pconfusarubida
Rahnella sp Y9602 YP 004210993 1 741091 Proteobacteria
Rahnella aquatilis CIP 78 65 ATCC 33071 YP 005198344 1 745277 Proteobacteria
Pantoea sp At 9b YP 004118553 1 592316 Proteobacteria
Citrobacter rodentium ICC168 YP 003367871 1 637910 Proteobacteria
Enterobacter sp 638 YP 001178479 1 399742 Proteobacteria
Salmonella enterica subsp arizonae serovar 62 z4 z23 str RSK2980 YP 001573095 1 882884 Proteobacteria
Citrobacter koseri ATCC BAA 895 YP 001456243 1 290338 Proteobacteria
Enterobacteriaceae bacterium strain FGI 57 YP 007338613 1 693444 Proteobacteria
Enterobacter cloacae EcWSU1 YP 004953993 1 1045856 Proteobacteria
Enterobacter asburiae LF7a YP 004830548 1 640513 Proteobacteria
Enterobacter cloacae subsp dissolvens SDM YP 006479605 1 1104326 Proteobacteria
Enterobacter cloacae subsp cloacae ENHKU01 YP 006580553 1 1211025 Proteobacteria
Cronobacter turicensis z3032 YP 003211863 1 693216 Proteobacteria
Shimwellia blattae DSM 4481 NBRC 105725 YP 006321090 1 630626 Proteobacteria
Pantoea vagans C9 1 YP 003932574 1 712898 Proteobacteria
Dickeya paradisiaca Ech703 WP 012764861 1 Dparadisiaca
Dickeya paradisiaca NCPPB 2511 WP 012764861 1 Dparadisiaca
Klebsiella oxytoca E718 YP 006498917 1 1191061 Proteobacteria
Raoultella ornithinolytica B6 YP 007873334 1 1286170 Proteobacteria
Klebsiella pneumoniae YP 007991732 1 573 Proteobacteria
Klebsiella pneumoniae subsp pneumoniae MGH 78578 YP 001335219 1 272620 Proteobacteria
Klebsiella variicola At 22 YP 003439720 1 640131 Proteobacteria
Enterobacter aerogenes EA1509E YP 007387780 1 935296 Proteobacteria
Dickeya dadantii subsp dieffenbachiae NCPPB 2976 WP 038922448 1 Ddadantii
Dickeya dadantii NCPPB 3537 WP 038900884 1 Ddadantii
Dickeya dadantii NCPPB 898 WP 038910854 1 Ddadantii
440 mdcH response to acidic stress Seed
Dickeya dadantii 3937 WP 013317380 1 Ddadantii
Dickeya solani RNS 05 1 2A WP 057082711 1 Dsolani
Dickeya solani GBBC 2040 WP 022633066 1 Dsolani
Dickeya solani PPO 9134 WP 022633066 1 Dsolani
Dickeya solani IPO 2222 WP 022633066 1 Dsolani
Dickeya solani RNS 07 7 3B WP 022633066 1 Dsolani
Dickeya solani RNS 08 23 3 1 A WP 022633066 1 Dsolani
Dickeya solani PPO 9019 WP 022633066 1 Dsolani
Dickeya solani D s0432 1 WP 022633066 1 Dsolani
Dickeya solani MK16 WP 022633066 1 Dsolani
Dickeya solani MK10 WP 022633066 1 Dsolani
Dickeya solani IFB 0099 WP 022633066 1 Dsolani
Dickeya gs fangzhongdai NCPPB 3274 WP 042859451 1 Dfangzhongdai
Dickeya gs fangzhongdai M005 WP 039695639 1 Dfangzhongdai
Dickeya gs undicola 2B12 WP 033570911 1 Dundicola
Dickeya gs fangzhongdai ND14b WP 038659928 1 Dfangzhongdai
Dickeya gs fangzhongdai M074 WP 038659928 1 Dfangzhongdai
Dickeya gs fangzhongdai S1 WP 049855215 1 Dfangzhongdai
Dickeya gs fangzhongdai B16 WP 038918639 1 Dfangzhongdai
Dickeya gs fangzhongdai MK7 WP 038918639 1 Dfangzhongdai
Dickeya dianthicola NCPPB 3534 WP 024105423 1 Ddianthicola
Dickeya dianthicola RNS04 9 WP 024105423 1 Ddianthicola
Dickeya dianthicola GBBC 2039 WP 024105423 1 Ddianthicola
Dickeya dianthicola IPO 980 WP 024105423 1 Ddianthicola
Dickeya dianthicola NCPPB 453 WP 024105423 1 Ddianthicola
Dickeya gs poaceaephila NCPPB 569 WP 042869911 1 Dpoaceaephila
Dickeya chrysanthemi subsp parthenii NCPPB 516 WP 027712444 1 Dchrysanthemi
Dickeya chrysanthemi L11 WP 033577200 1 Dchrysanthemi
Dickeya chrysanthemi subsp chrysanthemi NCPPB 402 WP 040000569 1 Dchrysanthemi
Dickeya chrysanthemi NCPPB 3533 WP 012770296 1 Dchrysanthemi
Dickeya chrysanthemi Ech1591 WP 012770296 1 Dchrysanthemi
Dickeya zeae Ech586 WP 012885176 1 Dzeae
Dickeya zeae CSL RW192 WP 038908400 1 Dzeae
Dickeya zeae NCPPB 3531 WP 038904769 1 Dzeae
Dickeya zeae MS1 WP 019846051 1 Dzeae
Dickeya zeae NCPPB 3532 WP 023639991 1 Dzeae
Dickeya zeae NCPPB 2538 WP 038916412 1 Dzeae
Dickeya zeae MK19 WP 038914228 1 Dzeae

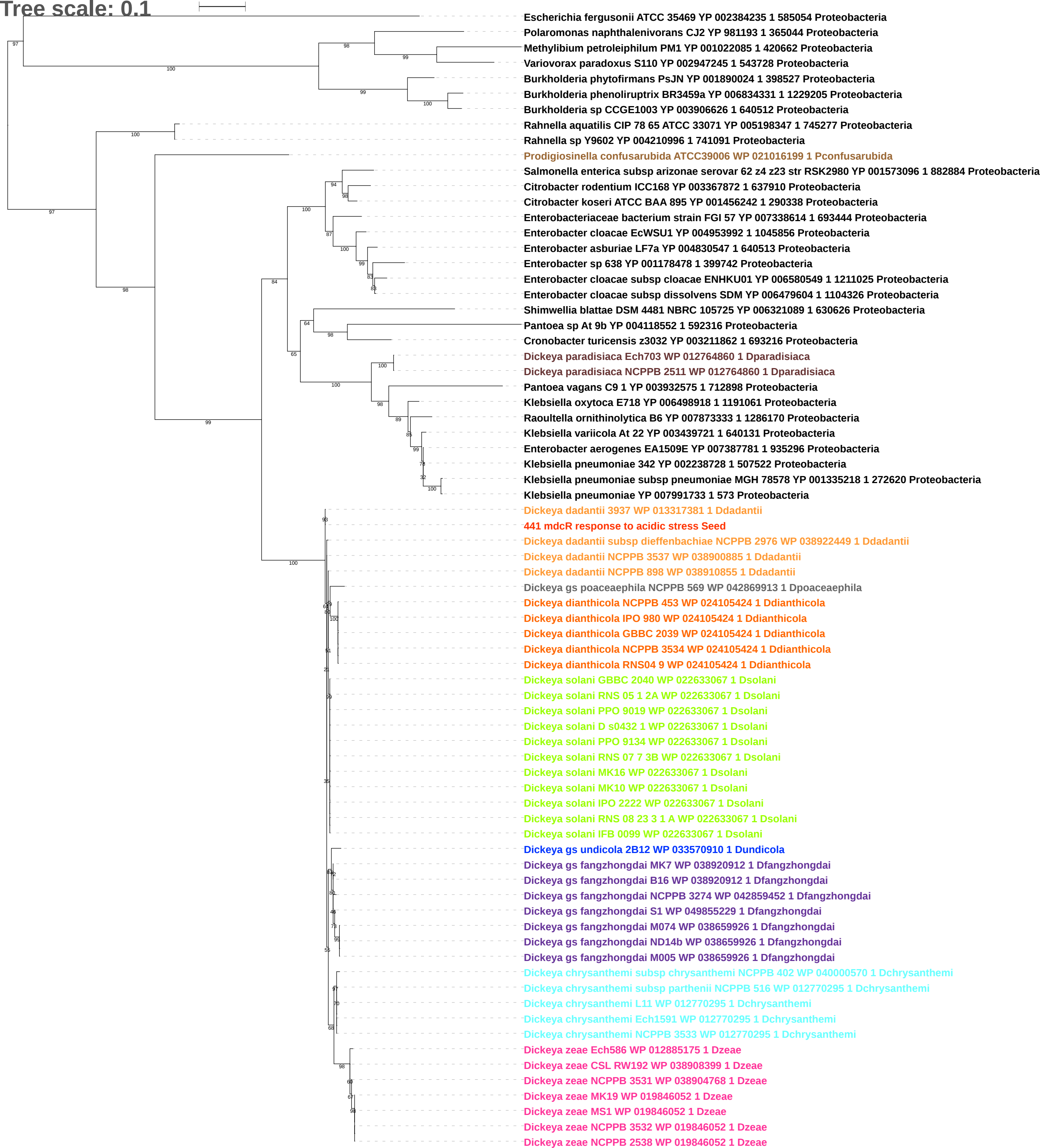
Tree scale: 0.1
Escherichia fergusonii ATCC 35469 YP 002384235 1 585054 Proteobacteria
Polaromonas naphthalenivorans CJ2 YP 981193 1 365044 Proteobacteria
Methylibium petroleiphilum PM1 YP 001022085 1 420662 Proteobacteria
Variovorax paradoxus S110 YP 002947245 1 543728 Proteobacteria
Burkholderia phytofirmans PsJN YP 001890024 1 398527 Proteobacteria
Burkholderia phenoliruptrix BR3459a YP 006834331 1 1229205 Proteobacteria
Burkholderia sp CCGE1003 YP 003906626 1 640512 Proteobacteria
Rahnella aquatilis CIP 78 65 ATCC 33071 YP 005198347 1 745277 Proteobacteria
Rahnella sp Y9602 YP 004210996 1 741091 Proteobacteria
Prodigiosinella confusarubida ATCC39006 WP 021016199 1 Pconfusarubida
Salmonella enterica subsp arizonae serovar 62 z4 z23 str RSK2980 YP 001573096 1 882884 Proteobacteria
Citrobacter rodentium ICC168 YP 003367872 1 637910 Proteobacteria
Citrobacter koseri ATCC BAA 895 YP 001456242 1 290338 Proteobacteria
Enterobacteriaceae bacterium strain FGI 57 YP 007338614 1 693444 Proteobacteria
Enterobacter cloacae EcWSU1 YP 004953992 1 1045856 Proteobacteria
Enterobacter asburiae LF7a YP 004830547 1 640513 Proteobacteria
Enterobacter sp 638 YP 001178478 1 399742 Proteobacteria
Enterobacter cloacae subsp cloacae ENHKU01 YP 006580549 1 1211025 Proteobacteria
Enterobacter cloacae subsp dissolvens SDM YP 006479604 1 1104326 Proteobacteria
Shimwellia blattae DSM 4481 NBRC 105725 YP 006321089 1 630626 Proteobacteria
Pantoea sp At 9b YP 004118552 1 592316 Proteobacteria
Cronobacter turicensis z3032 YP 003211862 1 693216 Proteobacteria
Dickeya paradisiaca Ech703 WP 012764860 1 Dparadisiaca
Dickeya paradisiaca NCPPB 2511 WP 012764860 1 Dparadisiaca
Pantoea vagans C9 1 YP 003932575 1 712898 Proteobacteria
Klebsiella oxytoca E718 YP 006498918 1 1191061 Proteobacteria
Raoultella ornithinolytica B6 YP 007873333 1 1286170 Proteobacteria
Klebsiella variicola At 22 YP 003439721 1 640131 Proteobacteria
Enterobacter aerogenes EA1509E YP 007387781 1 935296 Proteobacteria
Klebsiella pneumoniae 342 YP 002238728 1 507522 Proteobacteria
Klebsiella pneumoniae subsp pneumoniae MGH 78578 YP 001335218 1 272620 Proteobacteria
Klebsiella pneumoniae YP 007991733 1 573 Proteobacteria
Dickeya dadantii 3937 WP 013317381 1 Ddadantii
441 mdcR response to acidic stress Seed
Dickeya dadantii subsp dieffenbachiae NCPPB 2976 WP 038922449 1 Ddadantii
Dickeya dadantii NCPPB 3537 WP 038900885 1 Ddadantii
Dickeya dadantii NCPPB 898 WP 038910855 1 Ddadantii
Dickeya gs poaceaephila NCPPB 569 WP 042869913 1 Dpoaceaephila
Dickeya dianthicola NCPPB 453 WP 024105424 1 Ddianthicola
Dickeya dianthicola IPO 980 WP 024105424 1 Ddianthicola
Dickeya dianthicola GBBC 2039 WP 024105424 1 Ddianthicola
Dickeya dianthicola NCPPB 3534 WP 024105424 1 Ddianthicola
Dickeya dianthicola RNS04 9 WP 024105424 1 Ddianthicola
Dickeya solani GBBC 2040 WP 022633067 1 Dsolani
Dickeya solani RNS 05 1 2A WP 022633067 1 Dsolani
Dickeya solani PPO 9019 WP 022633067 1 Dsolani
Dickeya solani D s0432 1 WP 022633067 1 Dsolani
Dickeya solani PPO 9134 WP 022633067 1 Dsolani
Dickeya solani RNS 07 7 3B WP 022633067 1 Dsolani
Dickeya solani MK16 WP 022633067 1 Dsolani
Dickeya solani MK10 WP 022633067 1 Dsolani
Dickeya solani IPO 2222 WP 022633067 1 Dsolani
Dickeya solani RNS 08 23 3 1 A WP 022633067 1 Dsolani
Dickeya solani IFB 0099 WP 022633067 1 Dsolani
Dickeya gs undicola 2B12 WP 033570910 1 Dundicola
Dickeya gs fangzhongdai MK7 WP 038920912 1 Dfangzhongdai
Dickeya gs fangzhongdai B16 WP 038920912 1 Dfangzhongdai
Dickeya gs fangzhongdai NCPPB 3274 WP 042859452 1 Dfangzhongdai
Dickeya gs fangzhongdai S1 WP 049855229 1 Dfangzhongdai
Dickeya gs fangzhongdai M074 WP 038659926 1 Dfangzhongdai
Dickeya gs fangzhongdai ND14b WP 038659926 1 Dfangzhongdai
Dickeya gs fangzhongdai M005 WP 038659926 1 Dfangzhongdai
Dickeya chrysanthemi subsp chrysanthemi NCPPB 402 WP 040000570 1 Dchrysanthemi
Dickeya chrysanthemi subsp parthenii NCPPB 516 WP 012770295 1 Dchrysanthemi
Dickeya chrysanthemi L11 WP 012770295 1 Dchrysanthemi
Dickeya chrysanthemi Ech1591 WP 012770295 1 Dchrysanthemi
Dickeya chrysanthemi NCPPB 3533 WP 012770295 1 Dchrysanthemi
Dickeya zeae Ech586 WP 012885175 1 Dzeae
Dickeya zeae CSL RW192 WP 038908399 1 Dzeae
Dickeya zeae NCPPB 3531 WP 038904768 1 Dzeae
Dickeya zeae MK19 WP 019846052 1 Dzeae
Dickeya zeae MS1 WP 019846052 1 Dzeae
Dickeya zeae NCPPB 3532 WP 019846052 1 Dzeae
Dickeya zeae NCPPB 2538 WP 019846052 1 Dzeae

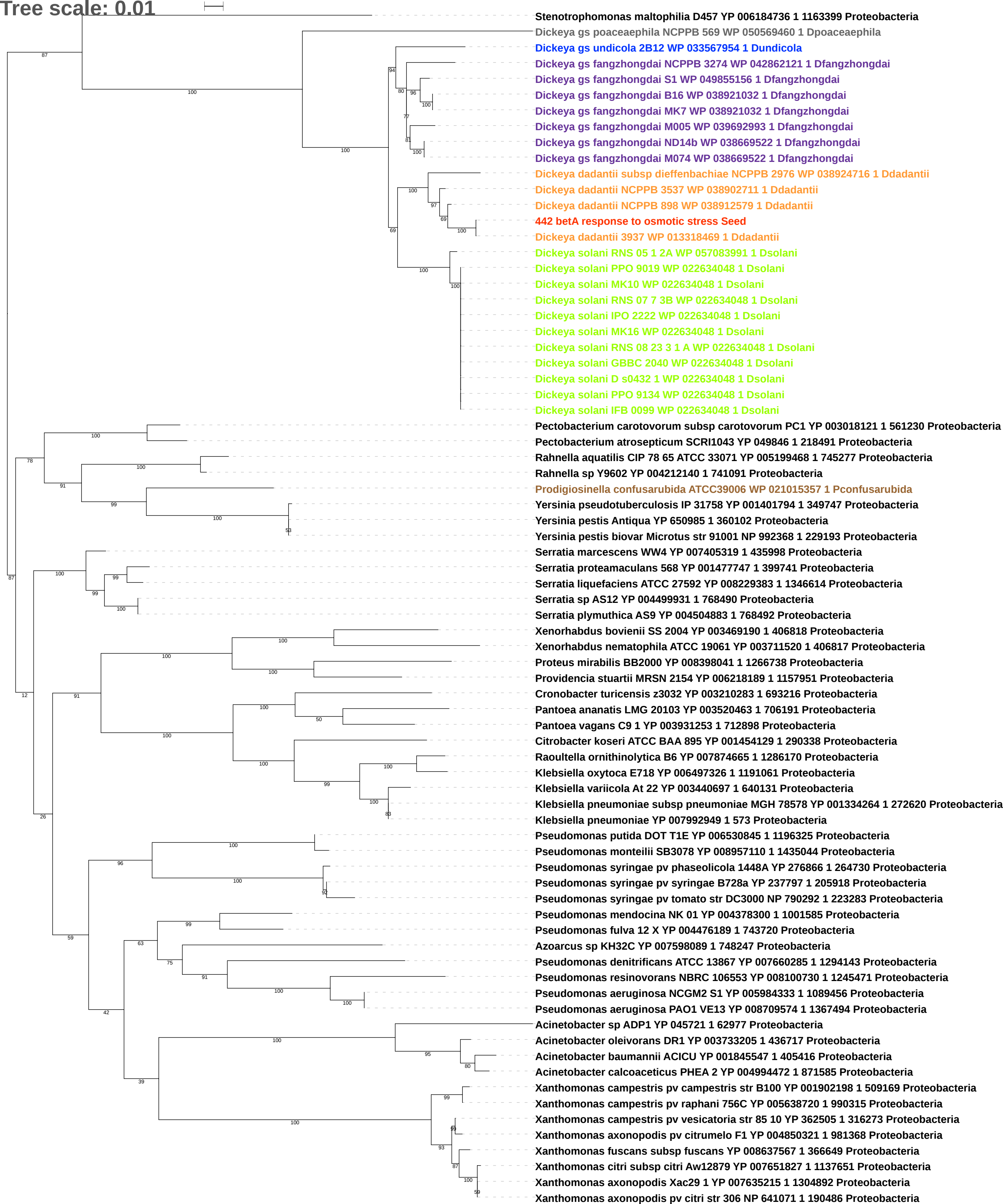
Tree scale: 0.01
Stenotrophomonas maltophilia D457 YP 006184736 1 1163399 Proteobacteria
Dickeya gs poaceaephila NCPPB 569 WP 050569460 1 Dpoaceaephila
Dickeya gs undicola 2B12 WP 033567954 1 Dundicola
Dickeya gs fangzhongdai NCPPB 3274 WP 042862121 1 Dfangzhongdai
Dickeya gs fangzhongdai S1 WP 049855156 1 Dfangzhongdai
Dickeya gs fangzhongdai B16 WP 038921032 1 Dfangzhongdai
Dickeya gs fangzhongdai MK7 WP 038921032 1 Dfangzhongdai
Dickeya gs fangzhongdai M005 WP 039692993 1 Dfangzhongdai
Dickeya gs fangzhongdai ND14b WP 038669522 1 Dfangzhongdai
Dickeya gs fangzhongdai M074 WP 038669522 1 Dfangzhongdai
Dickeya dadantii subsp dieffenbachiae NCPPB 2976 WP 038924716 1 Ddadantii
Dickeya dadantii NCPPB 3537 WP 038902711 1 Ddadantii
Dickeya dadantii NCPPB 898 WP 038912579 1 Ddadantii
442 betA response to osmotic stress Seed
Dickeya dadantii 3937 WP 013318469 1 Ddadantii
Dickeya solani RNS 05 1 2A WP 057083991 1 Dsolani
Dickeya solani PPO 9019 WP 022634048 1 Dsolani
Dickeya solani MK10 WP 022634048 1 Dsolani
Dickeya solani RNS 07 7 3B WP 022634048 1 Dsolani
Dickeya solani IPO 2222 WP 022634048 1 Dsolani
Dickeya solani MK16 WP 022634048 1 Dsolani
Dickeya solani RNS 08 23 3 1 A WP 022634048 1 Dsolani
Dickeya solani GBBC 2040 WP 022634048 1 Dsolani
Dickeya solani D s0432 1 WP 022634048 1 Dsolani
Dickeya solani PPO 9134 WP 022634048 1 Dsolani
Dickeya solani IFB 0099 WP 022634048 1 Dsolani
Pectobacterium carotovorum subsp carotovorum PC1 YP 003018121 1 561230 Proteobacteria
Pectobacterium atrosepticum SCRI1043 YP 049846 1 218491 Proteobacteria
Rahnella aquatilis CIP 78 65 ATCC 33071 YP 005199468 1 745277 Proteobacteria
Rahnella sp Y9602 YP 004212140 1 741091 Proteobacteria
Prodigiosinella confusarubida ATCC39006 WP 021015357 1 Pconfusarubida
Yersinia pseudotuberculosis IP 31758 YP 001401794 1 349747 Proteobacteria
Yersinia pestis Antiqua YP 650985 1 360102 Proteobacteria
Yersinia pestis biovar Microtus str 91001 NP 992368 1 229193 Proteobacteria
Serratia marcescens WW4 YP 007405319 1 435998 Proteobacteria
Serratia proteamaculans 568 YP 001477747 1 399741 Proteobacteria
Serratia liquefaciens ATCC 27592 YP 008229383 1 1346614 Proteobacteria
Serratia sp AS12 YP 004499931 1 768490 Proteobacteria
Serratia plymuthica AS9 YP 004504883 1 768492 Proteobacteria
Xenorhabdus bovienii SS 2004 YP 003469190 1 406818 Proteobacteria
Xenorhabdus nematophila ATCC 19061 YP 003711520 1 406817 Proteobacteria
Proteus mirabilis BB2000 YP 008398041 1 1266738 Proteobacteria
Providencia stuartii MRSN 2154 YP 006218189 1 1157951 Proteobacteria
Cronobacter turicensis z3032 YP 003210283 1 693216 Proteobacteria
Pantoea ananatis LMG 20103 YP 003520463 1 706191 Proteobacteria
Pantoea vagans C9 1 YP 003931253 1 712898 Proteobacteria
Citrobacter koseri ATCC BAA 895 YP 001454129 1 290338 Proteobacteria
Raoultella ornithinolytica B6 YP 007874665 1 1286170 Proteobacteria
Klebsiella oxytoca E718 YP 006497326 1 1191061 Proteobacteria
Klebsiella variicola At 22 YP 003440697 1 640131 Proteobacteria
Klebsiella pneumoniae subsp pneumoniae MGH 78578 YP 001334264 1 272620 Proteobacteria
Klebsiella pneumoniae YP 007992949 1 573 Proteobacteria
Pseudomonas putida DOT T1E YP 006530845 1 1196325 Proteobacteria
Pseudomonas monteilii SB3078 YP 008957110 1 1435044 Proteobacteria
Pseudomonas syringae pv phaseolicola 1448A YP 276866 1 264730 Proteobacteria
Pseudomonas syringae pv syringae B728a YP 237797 1 205918 Proteobacteria
Pseudomonas syringae pv tomato str DC3000 NP 790292 1 223283 Proteobacteria
Pseudomonas mendocina NK 01 YP 004378300 1 1001585 Proteobacteria
Pseudomonas fulva 12 X YP 004476189 1 743720 Proteobacteria
Azoarcus sp KH32C YP 007598089 1 748247 Proteobacteria
Pseudomonas denitrificans ATCC 13867 YP 007660285 1 1294143 Proteobacteria
Pseudomonas resinovorans NBRC 106553 YP 008100730 1 1245471 Proteobacteria
Pseudomonas aeruginosa NCGM2 S1 YP 005984333 1 1089456 Proteobacteria
Pseudomonas aeruginosa PAO1 VE13 YP 008709574 1 1367494 Proteobacteria
Acinetobacter sp ADP1 YP 045721 1 62977 Proteobacteria
Acinetobacter oleivorans DR1 YP 003733205 1 436717 Proteobacteria
Acinetobacter baumannii ACICU YP 001845547 1 405416 Proteobacteria
Acinetobacter calcoaceticus PHEA 2 YP 004994472 1 871585 Proteobacteria
Xanthomonas campestris pv campestris str B100 YP 001902198 1 509169 Proteobacteria
Xanthomonas campestris pv raphani 756C YP 005638720 1 990315 Proteobacteria
Xanthomonas campestris pv vesicatoria str 85 10 YP 362505 1 316273 Proteobacteria
Xanthomonas axonopodis pv citrumelo F1 YP 004850321 1 981368 Proteobacteria
Xanthomonas fuscans subsp fuscans YP 008637567 1 366649 Proteobacteria
Xanthomonas citri subsp citri Aw12879 YP 007651827 1 1137651 Proteobacteria
Xanthomonas axonopodis Xac29 1 YP 007635215 1 1304892 Proteobacteria
Xanthomonas axonopodis pv citri str 306 NP 641071 1 190486 Proteobacteria

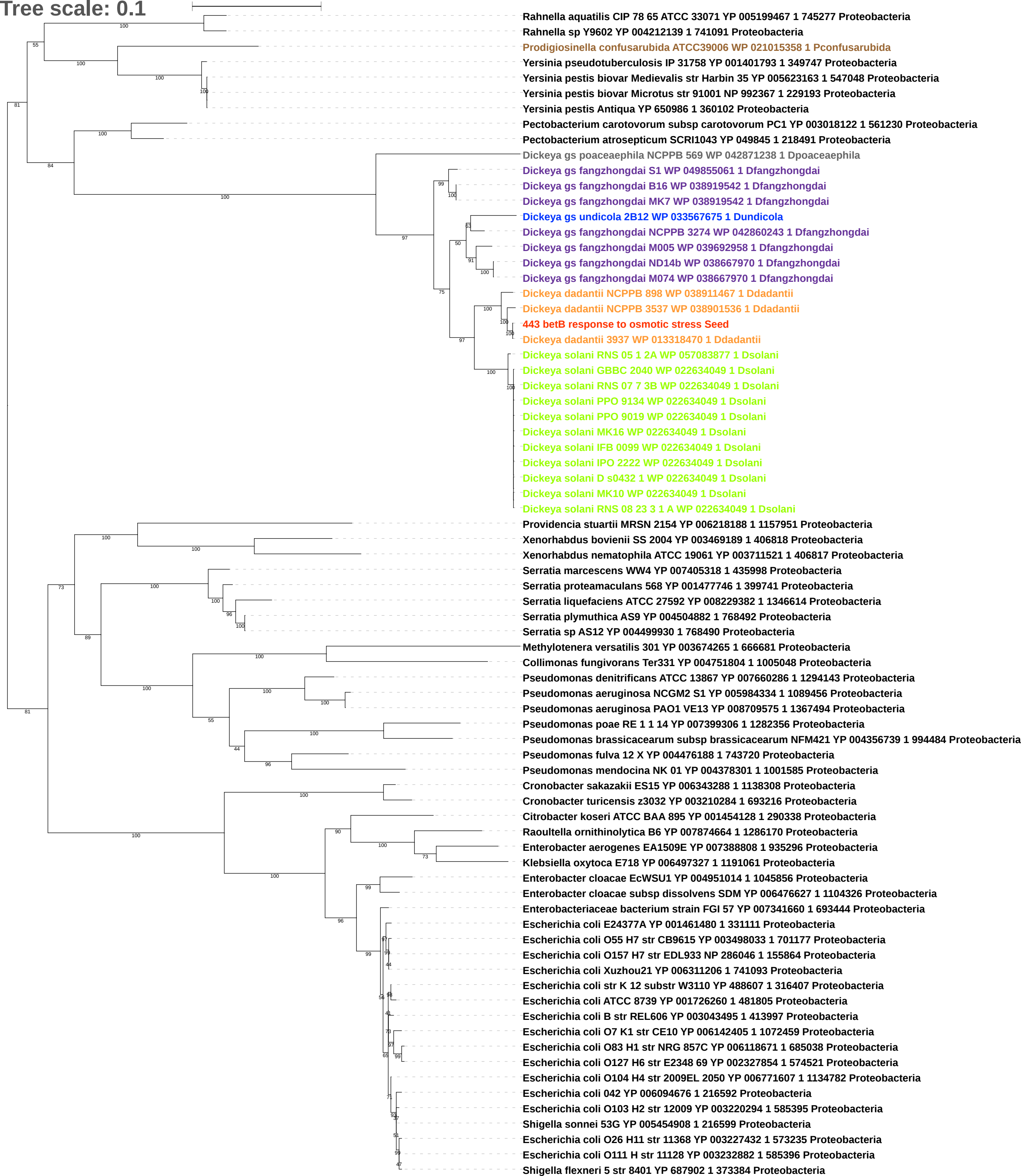
Tree scale: 0.1
Rahnella aquatilis CIP 78 65 ATCC 33071 YP 005199467 1 745277 Proteobacteria
Rahnella sp Y9602 YP 004212139 1 741091 Proteobacteria
Prodigiosinella confusarubida ATCC39006 WP 021015358 1 Pconfusarubida
Yersinia pseudotuberculosis IP 31758 YP 001401793 1 349747 Proteobacteria
Yersinia pestis biovar Medievalis str Harbin 35 YP 005623163 1 547048 Proteobacteria
Yersinia pestis biovar Microtus str 91001 NP 992367 1 229193 Proteobacteria
Yersinia pestis Antiqua YP 650986 1 360102 Proteobacteria
Pectobacterium carotovorum subsp carotovorum PC1 YP 003018122 1 561230 Proteobacteria
Pectobacterium atrosepticum SCRI1043 YP 049845 1 218491 Proteobacteria
Dickeya gs poaceaephila NCPPB 569 WP 042871238 1 Dpoaceaephila
Dickeya gs fangzhongdai S1 WP 049855061 1 Dfangzhongdai
Dickeya gs fangzhongdai B16 WP 038919542 1 Dfangzhongdai
Dickeya gs fangzhongdai MK7 WP 038919542 1 Dfangzhongdai
Dickeya gs undicola 2B12 WP 033567675 1 Dundicola
Dickeya gs fangzhongdai NCPPB 3274 WP 042860243 1 Dfangzhongdai
Dickeya gs fangzhongdai M005 WP 039692958 1 Dfangzhongdai
Dickeya gs fangzhongdai ND14b WP 038667970 1 Dfangzhongdai
Dickeya gs fangzhongdai M074 WP 038667970 1 Dfangzhongdai
Dickeya dadantii NCPPB 898 WP 038911467 1 Ddadantii
Dickeya dadantii NCPPB 3537 WP 038901536 1 Ddadantii
443 betB response to osmotic stress Seed
Dickeya dadantii 3937 WP 013318470 1 Ddadantii
Dickeya solani RNS 05 1 2A WP 057083877 1 Dsolani
Dickeya solani GBBC 2040 WP 022634049 1 Dsolani
Dickeya solani RNS 07 7 3B WP 022634049 1 Dsolani
Dickeya solani PPO 9134 WP 022634049 1 Dsolani
Dickeya solani PPO 9019 WP 022634049 1 Dsolani
Dickeya solani MK16 WP 022634049 1 Dsolani
Dickeya solani IFB 0099 WP 022634049 1 Dsolani
Dickeya solani IPO 2222 WP 022634049 1 Dsolani
Dickeya solani D s0432 1 WP 022634049 1 Dsolani
Dickeya solani MK10 WP 022634049 1 Dsolani
Dickeya solani RNS 08 23 3 1 A WP 022634049 1 Dsolani
Providencia stuartii MRSN 2154 YP 006218188 1 1157951 Proteobacteria
Xenorhabdus bovienii SS 2004 YP 003469189 1 406818 Proteobacteria
Xenorhabdus nematophila ATCC 19061 YP 003711521 1 406817 Proteobacteria
Serratia marcescens WW4 YP 007405318 1 435998 Proteobacteria
Serratia proteamaculans 568 YP 001477746 1 399741 Proteobacteria
Serratia liquefaciens ATCC 27592 YP 008229382 1 1346614 Proteobacteria
Serratia plymuthica AS9 YP 004504882 1 768492 Proteobacteria
Serratia sp AS12 YP 004499930 1 768490 Proteobacteria
Methylotenera versatilis 301 YP 003674265 1 666681 Proteobacteria
Collimonas fungivorans Ter331 YP 004751804 1 1005048 Proteobacteria
Pseudomonas denitrificans ATCC 13867 YP 007660286 1 1294143 Proteobacteria
Pseudomonas aeruginosa NCGM2 S1 YP 005984334 1 1089456 Proteobacteria
Pseudomonas aeruginosa PAO1 VE13 YP 008709575 1 1367494 Proteobacteria
Pseudomonas poae RE 1 1 14 YP 007399306 1 1282356 Proteobacteria
Pseudomonas brassicacearum subsp brassicacearum NFM421 YP 004356739 1 994484 Proteobacteria
Pseudomonas fulva 12 X YP 004476188 1 743720 Proteobacteria
Pseudomonas mendocina NK 01 YP 004378301 1 1001585 Proteobacteria
Cronobacter sakazakii ES15 YP 006343288 1 1138308 Proteobacteria
Cronobacter turicensis z3032 YP 003210284 1 693216 Proteobacteria
Citrobacter koseri ATCC BAA 895 YP 001454128 1 290338 Proteobacteria
Raoultella ornithinolytica B6 YP 007874664 1 1286170 Proteobacteria
Enterobacter aerogenes EA1509E YP 007388808 1 935296 Proteobacteria
Klebsiella oxytoca E718 YP 006497327 1 1191061 Proteobacteria
Enterobacter cloacae EcWSU1 YP 004951014 1 1045856 Proteobacteria
Enterobacter cloacae subsp dissolvens SDM YP 006476627 1 1104326 Proteobacteria
Enterobacteriaceae bacterium strain FGI 57 YP 007341660 1 693444 Proteobacteria
Escherichia coli E24377A YP 001461480 1 331111 Proteobacteria
Escherichia coli O55 H7 str CB9615 YP 003498033 1 701177 Proteobacteria
Escherichia coli O157 H7 str EDL933 NP 286046 1 155864 Proteobacteria
Escherichia coli Xuzhou21 YP 006311206 1 741093 Proteobacteria
Escherichia coli str K 12 substr W3110 YP 488607 1 316407 Proteobacteria
Escherichia coli ATCC 8739 YP 001726260 1 481805 Proteobacteria
Escherichia coli B str REL606 YP 003043495 1 413997 Proteobacteria
Escherichia coli O7 K1 str CE10 YP 006142405 1 1072459 Proteobacteria
Escherichia coli O83 H1 str NRG 857C YP 006118671 1 685038 Proteobacteria
Escherichia coli O127 H6 str E2348 69 YP 002327854 1 574521 Proteobacteria
Escherichia coli O104 H4 str 2009EL 2050 YP 006771607 1 1134782 Proteobacteria
Escherichia coli 042 YP 006094676 1 216592 Proteobacteria
Escherichia coli O103 H2 str 12009 YP 003220294 1 585395 Proteobacteria
Shigella sonnei 53G YP 005454908 1 216599 Proteobacteria
Escherichia coli O26 H11 str 11368 YP 003227432 1 573235 Proteobacteria
Escherichia coli O111 H str 11128 YP 003232882 1 585396 Proteobacteria
Shigella flexneri 5 str 8401 YP 687902 1 373384 Proteobacteria

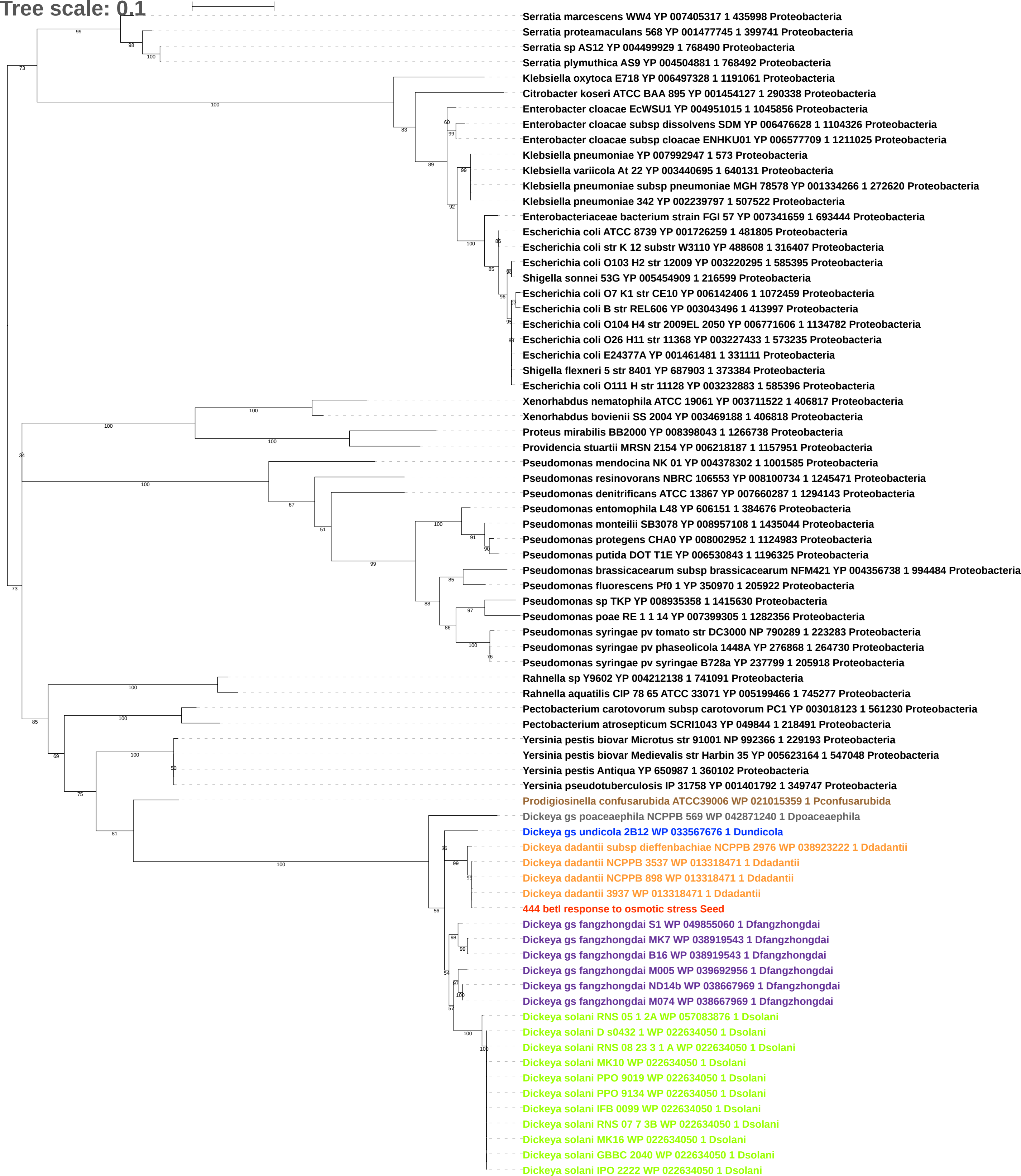

Tree scale: 0.1
Serratia marcescens WW4 YP 007405317 1 435998 Proteobacteria
Serratia proteamaculans 568 YP 001477745 1 399741 Proteobacteria
Serratia sp AS12 YP 004499929 1 768490 Proteobacteria
Serratia plymuthica AS9 YP 004504881 1 768492 Proteobacteria
Klebsiella oxytoca E718 YP 006497328 1 1191061 Proteobacteria
Citrobacter koseri ATCC BAA 895 YP 001454127 1 290338 Proteobacteria
Enterobacter cloacae EcWSU1 YP 004951015 1 1045856 Proteobacteria
Enterobacter cloacae subsp dissolvens SDM YP 006476628 1 1104326 Proteobacteria
Enterobacter cloacae subsp cloacae ENHKU01 YP 006577709 1 1211025 Proteobacteria
Klebsiella pneumoniae YP 007992947 1 573 Proteobacteria
Klebsiella variicola At 22 YP 003440695 1 640131 Proteobacteria
Klebsiella pneumoniae subsp pneumoniae MGH 78578 YP 001334266 1 272620 Proteobacteria
Klebsiella pneumoniae 342 YP 002239797 1 507522 Proteobacteria
Enterobacteriaceae bacterium strain FGI 57 YP 007341659 1 693444 Proteobacteria
Escherichia coli ATCC 8739 YP 001726259 1 481805 Proteobacteria
Escherichia coli str K 12 substr W3110 YP 488608 1 316407 Proteobacteria
Escherichia coli O103 H2 str 12009 YP 003220295 1 585395 Proteobacteria
Shigella sonnei 53G YP 005454909 1 216599 Proteobacteria
Escherichia coli O7 K1 str CE10 YP 006142406 1 1072459 Proteobacteria
Escherichia coli B str REL606 YP 003043496 1 413997 Proteobacteria
Escherichia coli O104 H4 str 2009EL 2050 YP 006771606 1 1134782 Proteobacteria
Escherichia coli O26 H11 str 11368 YP 003227433 1 573235 Proteobacteria
Escherichia coli E24377A YP 001461481 1 331111 Proteobacteria
Shigella flexneri 5 str 8401 YP 687903 1 373384 Proteobacteria
Escherichia coli O111 H str 11128 YP 003232883 1 585396 Proteobacteria
Xenorhabdus nematophila ATCC 19061 YP 003711522 1 406817 Proteobacteria
Xenorhabdus bovienii SS 2004 YP 003469188 1 406818 Proteobacteria
Proteus mirabilis BB2000 YP 008398043 1 1266738 Proteobacteria
Providencia stuartii MRSN 2154 YP 006218187 1 1157951 Proteobacteria
Pseudomonas mendocina NK 01 YP 004378302 1 1001585 Proteobacteria
Pseudomonas resinovorans NBRC 106553 YP 008100734 1 1245471 Proteobacteria
Pseudomonas denitrificans ATCC 13867 YP 007660287 1 1294143 Proteobacteria
Pseudomonas entomophila L48 YP 606151 1 384676 Proteobacteria
Pseudomonas monteilii SB3078 YP 008957108 1 1435044 Proteobacteria
Pseudomonas protegens CHA0 YP 008002952 1 1124983 Proteobacteria
Pseudomonas putida DOT T1E YP 006530843 1 1196325 Proteobacteria
Pseudomonas brassicacearum subsp brassicacearum NFM421 YP 004356738 1 994484 Proteobacteria
Pseudomonas fluorescens Pf0 1 YP 350970 1 205922 Proteobacteria
Pseudomonas sp TKP YP 008935358 1 1415630 Proteobacteria
Pseudomonas poae RE 1 1 14 YP 007399305 1 1282356 Proteobacteria
Pseudomonas syringae pv tomato str DC3000 NP 790289 1 223283 Proteobacteria
Pseudomonas syringae pv phaseolicola 1448A YP 276868 1 264730 Proteobacteria
Pseudomonas syringae pv syringae B728a YP 237799 1 205918 Proteobacteria
Rahnella sp Y9602 YP 004212138 1 741091 Proteobacteria
Rahnella aquatilis CIP 78 65 ATCC 33071 YP 005199466 1 745277 Proteobacteria
Pectobacterium carotovorum subsp carotovorum PC1 YP 003018123 1 561230 Proteobacteria
Pectobacterium atrosepticum SCRI1043 YP 049844 1 218491 Proteobacteria
Yersinia pestis biovar Microtus str 91001 NP 992366 1 229193 Proteobacteria
Yersinia pestis biovar Medievalis str Harbin 35 YP 005623164 1 547048 Proteobacteria
Yersinia pestis Antiqua YP 650987 1 360102 Proteobacteria
Yersinia pseudotuberculosis IP 31758 YP 001401792 1 349747 Proteobacteria
Prodigiosinella confusarubida ATCC39006 WP 021015359 1 Pconfusarubida
Dickeya gs poaceaephila NCPPB 569 WP 042871240 1 Dpoaceaephila
Dickeya gs undicola 2B12 WP 033567676 1 Dundicola
Dickeya dadantii subsp dieffenbachiae NCPPB 2976 WP 038923222 1 Ddadantii
Dickeya dadantii NCPPB 3537 WP 013318471 1 Ddadantii
Dickeya dadantii NCPPB 898 WP 013318471 1 Ddadantii
Dickeya dadantii 3937 WP 013318471 1 Ddadantii
444 betI response to osmotic stress Seed
Dickeya gs fangzhongdai S1 WP 049855060 1 Dfangzhongdai
Dickeya gs fangzhongdai MK7 WP 038919543 1 Dfangzhongdai
Dickeya gs fangzhongdai B16 WP 038919543 1 Dfangzhongdai
Dickeya gs fangzhongdai M005 WP 039692956 1 Dfangzhongdai
Dickeya gs fangzhongdai ND14b WP 038667969 1 Dfangzhongdai
Dickeya gs fangzhongdai M074 WP 038667969 1 Dfangzhongdai
Dickeya solani RNS 05 1 2A WP 057083876 1 Dsolani
Dickeya solani D s0432 1 WP 022634050 1 Dsolani
Dickeya solani RNS 08 23 3 1 A WP 022634050 1 Dsolani
Dickeya solani MK10 WP 022634050 1 Dsolani
Dickeya solani PPO 9019 WP 022634050 1 Dsolani
Dickeya solani PPO 9134 WP 022634050 1 Dsolani
Dickeya solani IFB 0099 WP 022634050 1 Dsolani
Dickeya solani RNS 07 7 3B WP 022634050 1 Dsolani
Dickeya solani MK16 WP 022634050 1 Dsolani
Dickeya solani GBBC 2040 WP 022634050 1 Dsolani
Dickeya solani IPO 2222 WP 022634050 1 Dsolani

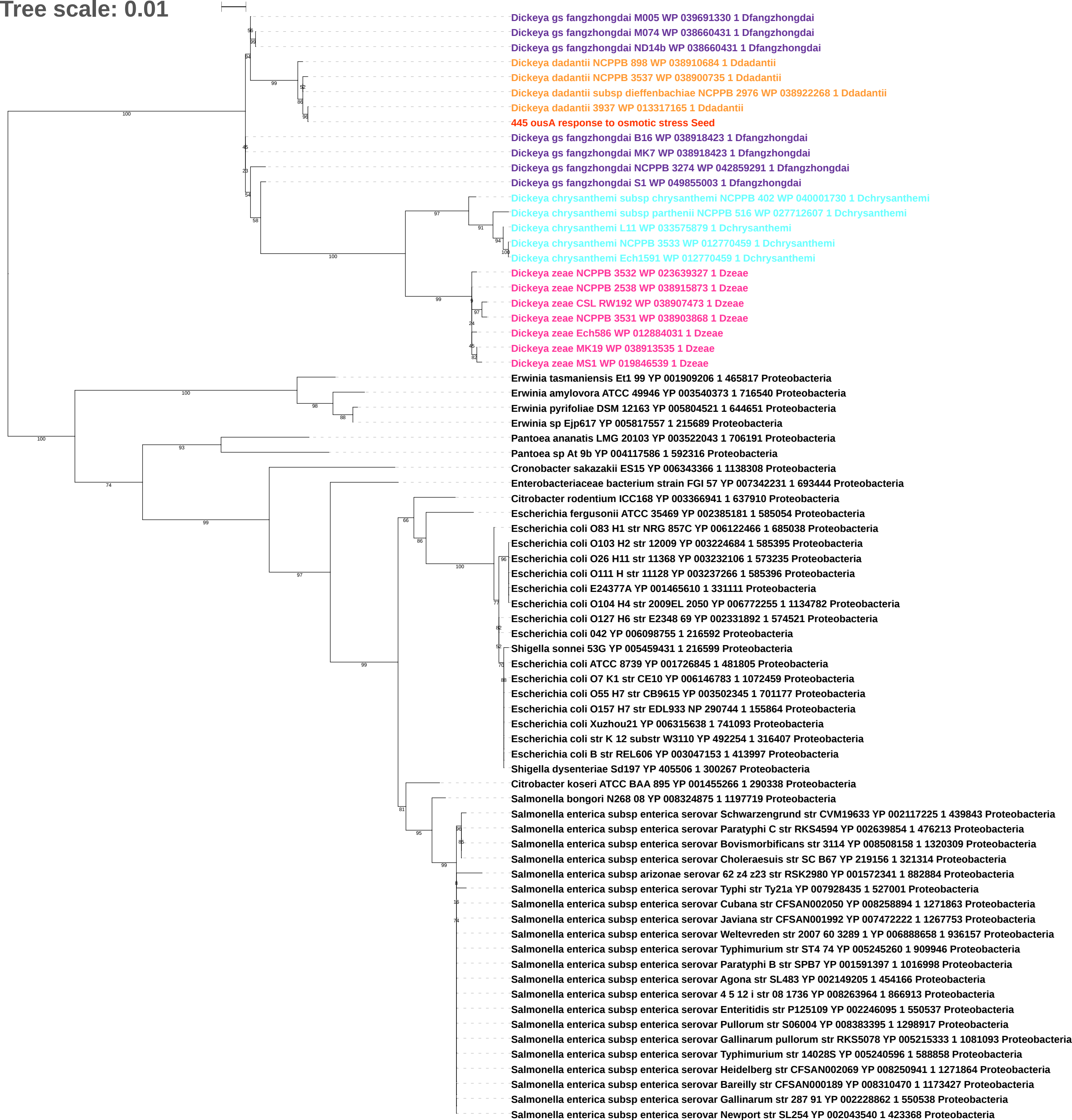
Tree scale: 0.01
Dickeya gs fangzhongdai M005 WP 039691330 1 Dfangzhongdai
Dickeya gs fangzhongdai M074 WP 038660431 1 Dfangzhongdai
Dickeya gs fangzhongdai ND14b WP 038660431 1 Dfangzhongdai
Dickeya dadantii NCPPB 898 WP 038910684 1 Ddadantii
Dickeya dadantii NCPPB 3537 WP 038900735 1 Ddadantii
Dickeya dadantii subsp dieffenbachiae NCPPB 2976 WP 038922268 1 Ddadantii
Dickeya dadantii 3937 WP 013317165 1 Ddadantii
445 ousA response to osmotic stress Seed
Dickeya gs fangzhongdai B16 WP 038918423 1 Dfangzhongdai
Dickeya gs fangzhongdai MK7 WP 038918423 1 Dfangzhongdai
Dickeya gs fangzhongdai NCPPB 3274 WP 042859291 1 Dfangzhongdai
Dickeya gs fangzhongdai S1 WP 049855003 1 Dfangzhongdai
Dickeya chrysanthemi subsp chrysanthemi NCPPB 402 WP 040001730 1 Dchrysanthemi
Dickeya chrysanthemi subsp parthenii NCPPB 516 WP 027712607 1 Dchrysanthemi
Dickeya chrysanthemi L11 WP 033575879 1 Dchrysanthemi
Dickeya chrysanthemi NCPPB 3533 WP 012770459 1 Dchrysanthemi
Dickeya chrysanthemi Ech1591 WP 012770459 1 Dchrysanthemi
Dickeya zeae NCPPB 3532 WP 023639327 1 Dzeae
Dickeya zeae NCPPB 2538 WP 038915873 1 Dzeae
Dickeya zeae CSL RW192 WP 038907473 1 Dzeae
Dickeya zeae NCPPB 3531 WP 038903868 1 Dzeae
Dickeya zeae Ech586 WP 012884031 1 Dzeae
Dickeya zeae MK19 WP 038913535 1 Dzeae
Dickeya zeae MS1 WP 019846539 1 Dzeae
Erwinia tasmaniensis Et1 99 YP 001909206 1 465817 Proteobacteria
Erwinia amylovora ATCC 49946 YP 003540373 1 716540 Proteobacteria
Erwinia pyrifoliae DSM 12163 YP 005804521 1 644651 Proteobacteria
Erwinia sp Ejp617 YP 005817557 1 215689 Proteobacteria
Pantoea ananatis LMG 20103 YP 003522043 1 706191 Proteobacteria
Pantoea sp At 9b YP 004117586 1 592316 Proteobacteria
Cronobacter sakazakii ES15 YP 006343366 1 1138308 Proteobacteria
Enterobacteriaceae bacterium strain FGI 57 YP 007342231 1 693444 Proteobacteria
Citrobacter rodentium ICC168 YP 003366941 1 637910 Proteobacteria
Escherichia fergusonii ATCC 35469 YP 002385181 1 585054 Proteobacteria
Escherichia coli O83 H1 str NRG 857C YP 006122466 1 685038 Proteobacteria
Escherichia coli O103 H2 str 12009 YP 003224684 1 585395 Proteobacteria
Escherichia coli O26 H11 str 11368 YP 003232106 1 573235 Proteobacteria
Escherichia coli O111 H str 11128 YP 003237266 1 585396 Proteobacteria
Escherichia coli E24377A YP 001465610 1 331111 Proteobacteria
Escherichia coli O104 H4 str 2009EL 2050 YP 006772255 1 1134782 Proteobacteria
Escherichia coli O127 H6 str E2348 69 YP 002331892 1 574521 Proteobacteria
Escherichia coli 042 YP 006098755 1 216592 Proteobacteria
Shigella sonnei 53G YP 005459431 1 216599 Proteobacteria
Escherichia coli ATCC 8739 YP 001726845 1 481805 Proteobacteria
Escherichia coli O7 K1 str CE10 YP 006146783 1 1072459 Proteobacteria
Escherichia coli O55 H7 str CB9615 YP 003502345 1 701177 Proteobacteria
Escherichia coli O157 H7 str EDL933 NP 290744 1 155864 Proteobacteria
Escherichia coli Xuzhou21 YP 006315638 1 741093 Proteobacteria
Escherichia coli str K 12 substr W3110 YP 492254 1 316407 Proteobacteria
Escherichia coli B str REL606 YP 003047153 1 413997 Proteobacteria
Shigella dysenteriae Sd197 YP 405506 1 300267 Proteobacteria
Citrobacter koseri ATCC BAA 895 YP 001455266 1 290338 Proteobacteria
Salmonella bongori N268 08 YP 008324875 1 1197719 Proteobacteria
Salmonella enterica subsp enterica serovar Schwarzengrund str CVM19633 YP 002117225 1 439843 Proteobacteria
Salmonella enterica subsp enterica serovar Paratyphi C str RKS4594 YP 002639854 1 476213 Proteobacteria
Salmonella enterica subsp enterica serovar Bovismorbificans str 3114 YP 008508158 1 1320309 Proteobacteria
Salmonella enterica subsp enterica serovar Choleraesuis str SC B67 YP 219156 1 321314 Proteobacteria
Salmonella enterica subsp arizonae serovar 62 z4 z23 str RSK2980 YP 001572341 1 882884 Proteobacteria
Salmonella enterica subsp enterica serovar Typhi str Ty21a YP 007928435 1 527001 Proteobacteria
Salmonella enterica subsp enterica serovar Cubana str CFSAN002050 YP 008258894 1 1271863 Proteobacteria
Salmonella enterica subsp enterica serovar Javiana str CFSAN001992 YP 007472222 1 1267753 Proteobacteria
Salmonella enterica subsp enterica serovar Weltevreden str 2007 60 3289 1 YP 006888658 1 936157 Proteobacteria
Salmonella enterica subsp enterica serovar Typhimurium str ST4 74 YP 005245260 1 909946 Proteobacteria
Salmonella enterica subsp enterica serovar Paratyphi B str SPB7 YP 001591397 1 1016998 Proteobacteria
Salmonella enterica subsp enterica serovar Agona str SL483 YP 002149205 1 454166 Proteobacteria
Salmonella enterica subsp enterica serovar 4 5 12 i str 08 1736 YP 008263964 1 866913 Proteobacteria
Salmonella enterica subsp enterica serovar Enteritidis str P125109 YP 002246095 1 550537 Proteobacteria
Salmonella enterica subsp enterica serovar Pullorum str S06004 YP 008383395 1 1298917 Proteobacteria
Salmonella enterica subsp enterica serovar Gallinarum pullorum str RKS5078 YP 005215333 1 1081093 Proteobacteria
Salmonella enterica subsp enterica serovar Typhimurium str 14028S YP 005240596 1 588858 Proteobacteria
Salmonella enterica subsp enterica serovar Heidelberg str CFSAN002069 YP 008250941 1 1271864 Proteobacteria
Salmonella enterica subsp enterica serovar Bareilly str CFSAN000189 YP 008310470 1 1173427 Proteobacteria
Salmonella enterica subsp enterica serovar Gallinarum str 287 91 YP 002228862 1 550538 Proteobacteria
Salmonella enterica subsp enterica serovar Newport str SL254 YP 002043540 1 423368 Proteobacteria

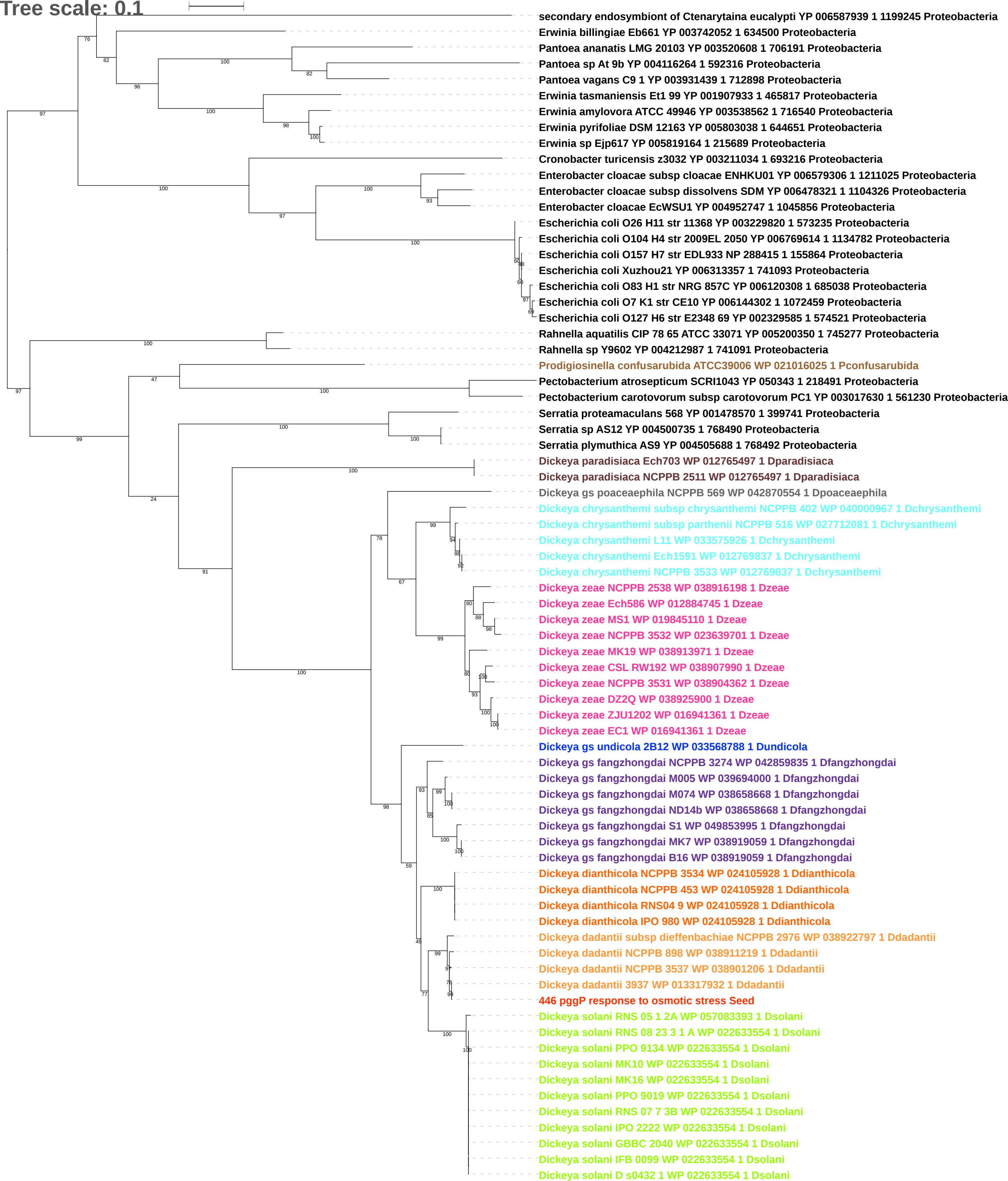
Tree scale: 0.1
secondary endosymbiont of Ctenarytaina eucalypti YP 006587939 1 1199245 Proteobacteria
Erwinia billingiae Eb661 YP 003742052 1 634500 Proteobacteria
Pantoea ananatis LMG 20103 YP 003520608 1 706191 Proteobacteria
Pantoea sp At 9b YP 004116264 1 592316 Proteobacteria
Pantoea vagans C9 1 YP 003931439 1 712898 Proteobacteria
Erwinia tasmaniensis Et1 99 YP 001907933 1 465817 Proteobacteria
Erwinia amylovora ATCC 49946 YP 003538562 1 716540 Proteobacteria
Erwinia pyrifoliae DSM 12163 YP 005803038 1 644651 Proteobacteria
Erwinia sp Ejp617 YP 005819164 1 215689 Proteobacteria
Cronobacter turicensis z3032 YP 003211034 1 693216 Proteobacteria
Enterobacter cloacae subsp cloacae ENHKU01 YP 006579306 1 1211025 Proteobacteria
Enterobacter cloacae subsp dissolvens SDM YP 006478321 1 1104326 Proteobacteria
Enterobacter cloacae EcWSU1 YP 004952747 1 1045856 Proteobacteria
Escherichia coli O26 H11 str 11368 YP 003229820 1 573235 Proteobacteria
Escherichia coli O104 H4 str 2009EL 2050 YP 006769614 1 1134782 Proteobacteria
Escherichia coli O157 H7 str EDL933 NP 288415 1 155864 Proteobacteria
Escherichia coli Xuzhou21 YP 006313357 1 741093 Proteobacteria
Escherichia coli O83 H1 str NRG 857C YP 006120308 1 685038 Proteobacteria
Escherichia coli O7 K1 str CE10 YP 006144302 1 1072459 Proteobacteria
Escherichia coli O127 H6 str E2348 69 YP 002329585 1 574521 Proteobacteria
Rahnella aquatilis CIP 78 65 ATCC 33071 YP 005200350 1 745277 Proteobacteria
Rahnella sp Y9602 YP 004212987 1 741091 Proteobacteria
Prodigiosinella confusarubida ATCC39006 WP 021016025 1 Pconfusarubida
Pectobacterium atrosepticum SCRI1043 YP 050343 1 218491 Proteobacteria
Pectobacterium carotovorum subsp carotovorum PC1 YP 003017630 1 561230 Proteobacteria
Serratia proteamaculans 568 YP 001478570 1 399741 Proteobacteria
Serratia sp AS12 YP 004500735 1 768490 Proteobacteria
Serratia plymuthica AS9 YP 004505688 1 768492 Proteobacteria
Dickeya paradisiaca Ech703 WP 012765497 1 Dparadisiaca
Dickeya paradisiaca NCPPB 2511 WP 012765497 1 Dparadisiaca
Dickeya gs poaceaephila NCPPB 569 WP 042870554 1 Dpoaceaephila
Dickeya chrysanthemi subsp chrysanthemi NCPPB 402 WP 040000967 1 Dchrysanthemi
Dickeya chrysanthemi subsp parthenii NCPPB 516 WP 027712081 1 Dchrysanthemi
Dickeya chrysanthemi L11 WP 033575926 1 Dchrysanthemi
Dickeya chrysanthemi Ech1591 WP 012769837 1 Dchrysanthemi
Dickeya chrysanthemi NCPPB 3533 WP 012769837 1 Dchrysanthemi
Dickeya zeae NCPPB 2538 WP 038916198 1 Dzeae
Dickeya zeae Ech586 WP 012884745 1 Dzeae
Dickeya zeae MS1 WP 019845110 1 Dzeae
Dickeya zeae NCPPB 3532 WP 023639701 1 Dzeae
Dickeya zeae MK19 WP 038913971 1 Dzeae
Dickeya zeae CSL RW192 WP 038907990 1 Dzeae
Dickeya zeae NCPPB 3531 WP 038904362 1 Dzeae
Dickeya zeae DZ2Q WP 038925900 1 Dzeae
Dickeya zeae ZJU1202 WP 016941361 1 Dzeae
Dickeya zeae EC1 WP 016941361 1 Dzeae
Dickeya gs undicola 2B12 WP 033568788 1 Dundicola
Dickeya gs fangzhongdai NCPPB 3274 WP 042859835 1 Dfangzhongdai
Dickeya gs fangzhongdai M005 WP 039694000 1 Dfangzhongdai
Dickeya gs fangzhongdai M074 WP 038658668 1 Dfangzhongdai
Dickeya gs fangzhongdai ND14b WP 038658668 1 Dfangzhongdai
Dickeya gs fangzhongdai S1 WP 049853995 1 Dfangzhongdai
Dickeya gs fangzhongdai MK7 WP 038919059 1 Dfangzhongdai
Dickeya gs fangzhongdai B16 WP 038919059 1 Dfangzhongdai
Dickeya dianthicola NCPPB 3534 WP 024105928 1 Ddianthicola
Dickeya dianthicola NCPPB 453 WP 024105928 1 Ddianthicola
Dickeya dianthicola RNS04 9 WP 024105928 1 Ddianthicola
Dickeya dianthicola IPO 980 WP 024105928 1 Ddianthicola
Dickeya dadantii subsp dieffenbachiae NCPPB 2976 WP 038922797 1 Ddadantii
Dickeya dadantii NCPPB 898 WP 038911219 1 Ddadantii
Dickeya dadantii NCPPB 3537 WP 038901206 1 Ddadantii
Dickeya dadantii 3937 WP 013317932 1 Ddadantii
446 pggP response to osmotic stress Seed
Dickeya solani RNS 05 1 2A WP 057083393 1 Dsolani
Dickeya solani RNS 08 23 3 1 A WP 022633554 1 Dsolani
Dickeya solani PPO 9134 WP 022633554 1 Dsolani
Dickeya solani MK10 WP 022633554 1 Dsolani
Dickeya solani MK16 WP 022633554 1 Dsolani
Dickeya solani PPO 9019 WP 022633554 1 Dsolani
Dickeya solani RNS 07 7 3B WP 022633554 1 Dsolani
Dickeya solani IPO 2222 WP 022633554 1 Dsolani
Dickeya solani GBBC 2040 WP 022633554 1 Dsolani
Dickeya solani IFB 0099 WP 022633554 1 Dsolani
Dickeya solani D s0432 1 WP 022633554 1 Dsolani

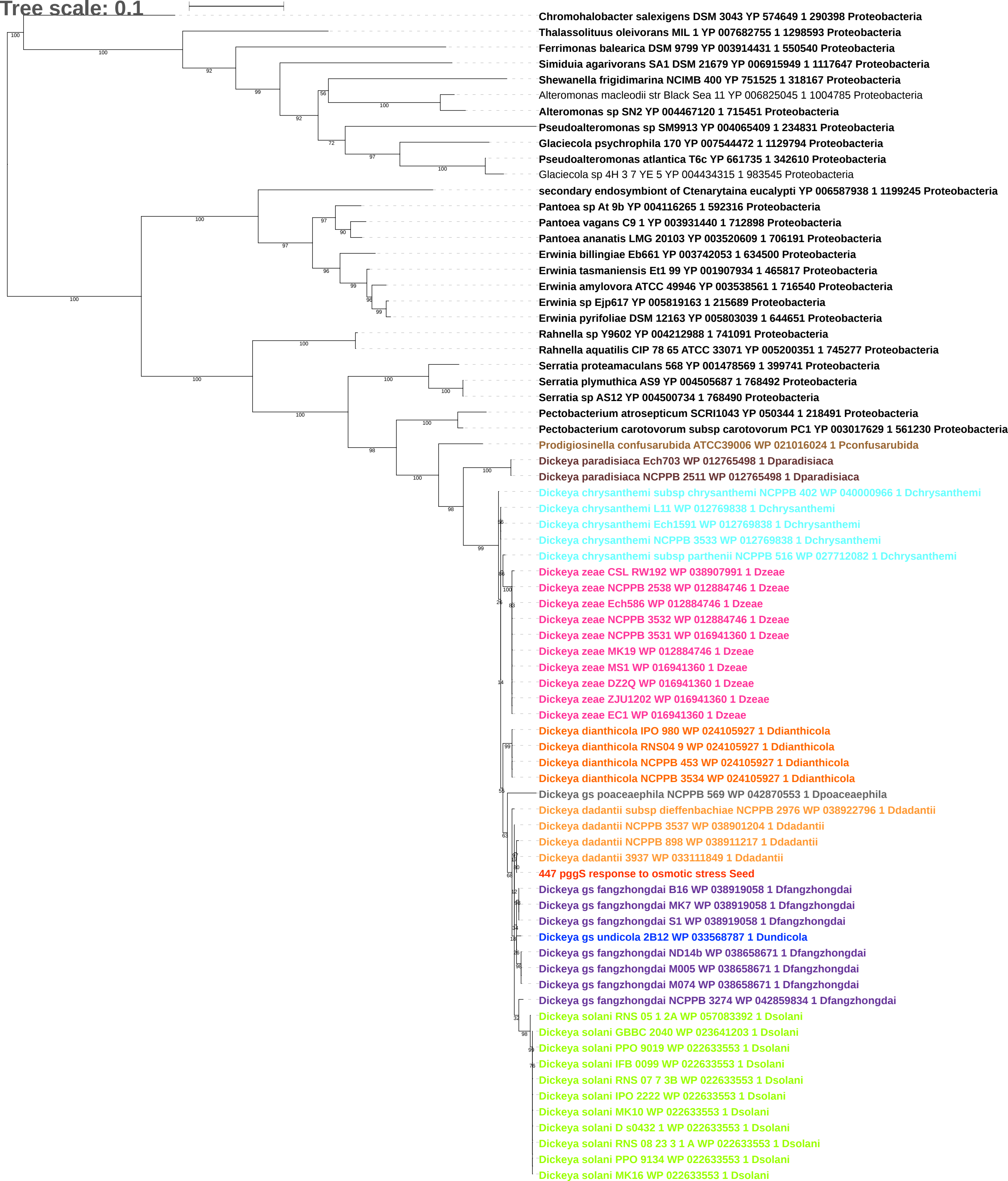
Tree scale: 0.1
Chromohalobacter salexigens DSM 3043 YP 574649 1 290398 Proteobacteria
Thalassolituus oleivorans MIL 1 YP 007682755 1 1298593 Proteobacteria
Ferrimonas balearica DSM 9799 YP 003914431 1 550540 Proteobacteria
Simiduia agarivorans SA1 DSM 21679 YP 006915949 1 1117647 Proteobacteria
Shewanella frigidimarina NCIMB 400 YP 751525 1 318167 Proteobacteria
Alteromonas macleodii str Black Sea 11 YP 006825045 1 1004785 Proteobacteria
Alteromonas sp SN2 YP 004467120 1 715451 Proteobacteria
Pseudoalteromonas sp SM9913 YP 004065409 1 234831 Proteobacteria
Glaciecola psychrophila 170 YP 007544472 1 1129794 Proteobacteria
Pseudoalteromonas atlantica T6c YP 661735 1 342610 Proteobacteria
Glaciecola sp 4H 3 7 YE 5 YP 004434315 1 983545 Proteobacteria
secondary endosymbiont of Ctenarytaina eucalypti YP 006587938 1 1199245 Proteobacteria
Pantoea sp At 9b YP 004116265 1 592316 Proteobacteria
Pantoea vagans C9 1 YP 003931440 1 712898 Proteobacteria
Pantoea ananatis LMG 20103 YP 003520609 1 706191 Proteobacteria
Erwinia billingiae Eb661 YP 003742053 1 634500 Proteobacteria
Erwinia tasmaniensis Et1 99 YP 001907934 1 465817 Proteobacteria
Erwinia amylovora ATCC 49946 YP 003538561 1 716540 Proteobacteria
Erwinia sp Ejp617 YP 005819163 1 215689 Proteobacteria
Erwinia pyrifoliae DSM 12163 YP 005803039 1 644651 Proteobacteria
Rahnella sp Y9602 YP 004212988 1 741091 Proteobacteria
Rahnella aquatilis CIP 78 65 ATCC 33071 YP 005200351 1 745277 Proteobacteria
Serratia proteamaculans 568 YP 001478569 1 399741 Proteobacteria
Serratia plymuthica AS9 YP 004505687 1 768492 Proteobacteria
Serratia sp AS12 YP 004500734 1 768490 Proteobacteria
Pectobacterium atrosepticum SCRI1043 YP 050344 1 218491 Proteobacteria
Pectobacterium carotovorum subsp carotovorum PC1 YP 003017629 1 561230 Proteobacteria
Prodigiosinella confusarubida ATCC39006 WP 021016024 1 Pconfusarubida
Dickeya paradisiaca Ech703 WP 012765498 1 Dparadisiaca
Dickeya paradisiaca NCPPB 2511 WP 012765498 1 Dparadisiaca
Dickeya chrysanthemi subsp chrysanthemi NCPPB 402 WP 040000966 1 Dchrysanthemi
Dickeya chrysanthemi L11 WP 012769838 1 Dchrysanthemi
Dickeya chrysanthemi Ech1591 WP 012769838 1 Dchrysanthemi
Dickeya chrysanthemi NCPPB 3533 WP 012769838 1 Dchrysanthemi
Dickeya chrysanthemi subsp parthenii NCPPB 516 WP 027712082 1 Dchrysanthemi
Dickeya zeae CSL RW192 WP 038907991 1 Dzeae
Dickeya zeae NCPPB 2538 WP 012884746 1 Dzeae
Dickeya zeae Ech586 WP 012884746 1 Dzeae
Dickeya zeae NCPPB 3532 WP 012884746 1 Dzeae
Dickeya zeae NCPPB 3531 WP 016941360 1 Dzeae
Dickeya zeae MK19 WP 012884746 1 Dzeae
Dickeya zeae MS1 WP 016941360 1 Dzeae
Dickeya zeae DZ2Q WP 016941360 1 Dzeae
Dickeya zeae ZJU1202 WP 016941360 1 Dzeae
Dickeya zeae EC1 WP 016941360 1 Dzeae
Dickeya dianthicola IPO 980 WP 024105927 1 Ddianthicola
Dickeya dianthicola RNS04 9 WP 024105927 1 Ddianthicola
Dickeya dianthicola NCPPB 453 WP 024105927 1 Ddianthicola
Dickeya dianthicola NCPPB 3534 WP 024105927 1 Ddianthicola
Dickeya gs poaceaephila NCPPB 569 WP 042870553 1 Dpoaceaephila
Dickeya dadantii subsp dieffenbachiae NCPPB 2976 WP 038922796 1 Ddadantii
Dickeya dadantii NCPPB 3537 WP 038901204 1 Ddadantii
Dickeya dadantii NCPPB 898 WP 038911217 1 Ddadantii
Dickeya dadantii 3937 WP 033111849 1 Ddadantii
447 pggS response to osmotic stress Seed
Dickeya gs fangzhongdai B16 WP 038919058 1 Dfangzhongdai
Dickeya gs fangzhongdai MK7 WP 038919058 1 Dfangzhongdai
Dickeya gs fangzhongdai S1 WP 038919058 1 Dfangzhongdai
Dickeya gs undicola 2B12 WP 033568787 1 Dundicola
Dickeya gs fangzhongdai ND14b WP 038658671 1 Dfangzhongdai
Dickeya gs fangzhongdai M005 WP 038658671 1 Dfangzhongdai
Dickeya gs fangzhongdai M074 WP 038658671 1 Dfangzhongdai
Dickeya gs fangzhongdai NCPPB 3274 WP 042859834 1 Dfangzhongdai
Dickeya solani RNS 05 1 2A WP 057083392 1 Dsolani
Dickeya solani GBBC 2040 WP 023641203 1 Dsolani
Dickeya solani PPO 9019 WP 022633553 1 Dsolani
Dickeya solani IFB 0099 WP 022633553 1 Dsolani
Dickeya solani RNS 07 7 3B WP 022633553 1 Dsolani
Dickeya solani IPO 2222 WP 022633553 1 Dsolani
Dickeya solani MK10 WP 022633553 1 Dsolani
Dickeya solani D s0432 1 WP 022633553 1 Dsolani
Dickeya solani RNS 08 23 3 1 A WP 022633553 1 Dsolani
Dickeya solani PPO 9134 WP 022633553 1 Dsolani
Dickeya solani MK16 WP 022633553 1 Dsolani

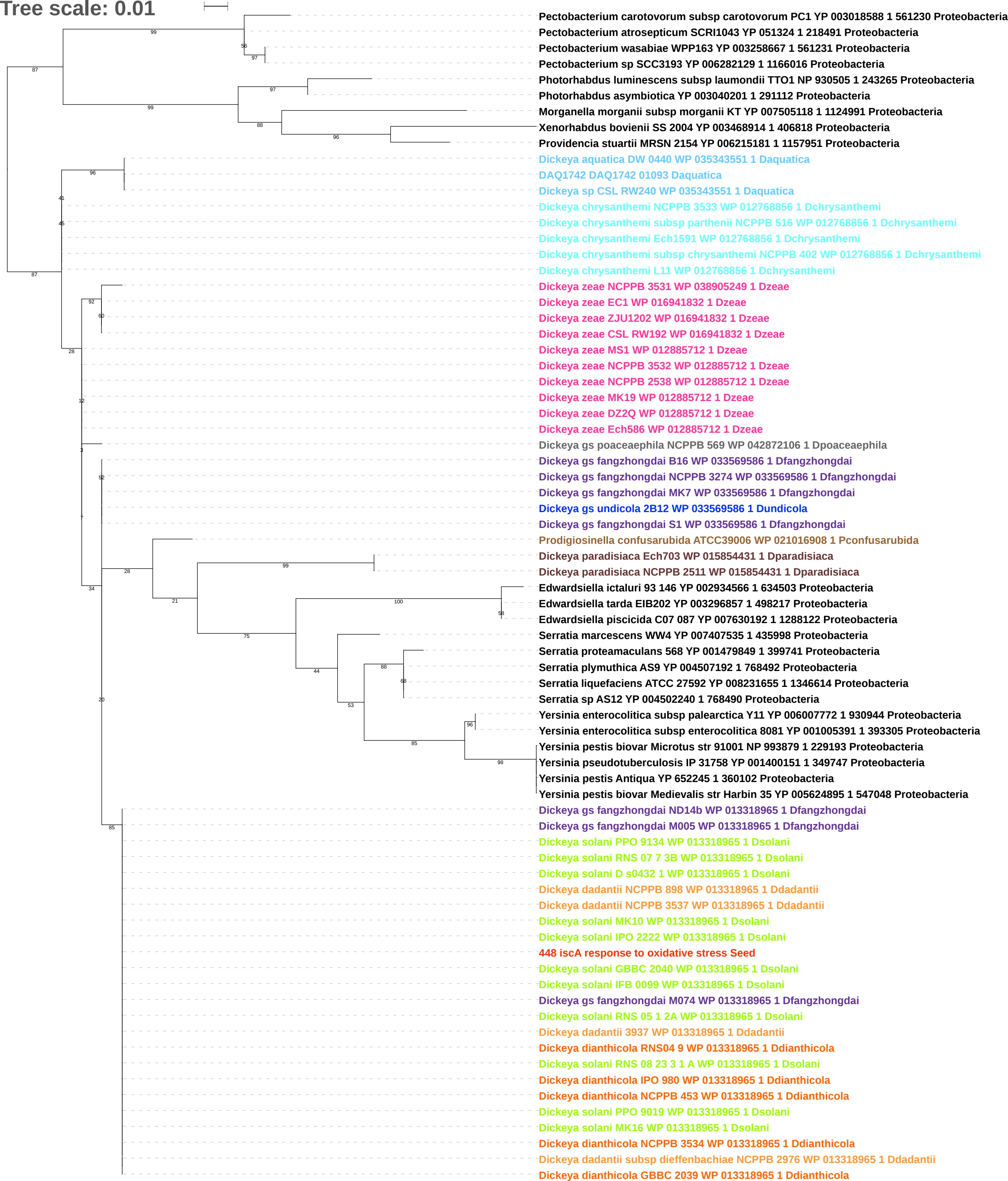
Tree scale: 0.01
Pectobacterium carotovorum subsp carotovorum PC1 YP 003018588 1 561230 Proteobacteria
Pectobacterium atrosepticum SCRI1043 YP 051324 1 218491 Proteobacteria
Pectobacterium wasabiae WPP163 YP 003258667 1 561231 Proteobacteria
Pectobacterium sp SCC3193 YP 006282129 1 1166016 Proteobacteria
Photorhabdus luminescens subsp laumondii TTO1 NP 930505 1 243265 Proteobacteria
Photorhabdus asymbiotica YP 003040201 1 291112 Proteobacteria
Morganella morganii subsp morganii KT YP 007505118 1 1124991 Proteobacteria
Xenorhabdus bovienii SS 2004 YP 003468914 1 406818 Proteobacteria
Providencia stuartii MRSN 2154 YP 006215181 1 1157951 Proteobacteria
Dickeya aquatica DW 0440 WP 035343551 1 Daquatica
DAQ1742 DAQ1742 01093 Daquatica
Dickeya sp CSL RW240 WP 035343551 1 Daquatica
Dickeya chrysanthemi NCPPB 3533 WP 012768856 1 Dchrysanthemi
Dickeya chrysanthemi subsp parthenii NCPPB 516 WP 012768856 1 Dchrysanthemi
Dickeya chrysanthemi Ech1591 WP 012768856 1 Dchrysanthemi
Dickeya chrysanthemi subsp chrysanthemi NCPPB 402 WP 012768856 1 Dchrysanthemi
Dickeya chrysanthemi L11 WP 012768856 1 Dchrysanthemi
Dickeya zeae NCPPB 3531 WP 038905249 1 Dzeae
Dickeya zeae EC1 WP 016941832 1 Dzeae
Dickeya zeae ZJU1202 WP 016941832 1 Dzeae
Dickeya zeae CSL RW192 WP 016941832 1 Dzeae
Dickeya zeae MS1 WP 012885712 1 Dzeae
Dickeya zeae NCPPB 3532 WP 012885712 1 Dzeae
Dickeya zeae NCPPB 2538 WP 012885712 1 Dzeae
Dickeya zeae MK19 WP 012885712 1 Dzeae
Dickeya zeae DZ2Q WP 012885712 1 Dzeae
Dickeya zeae Ech586 WP 012885712 1 Dzeae
Dickeya gs poaceaephila NCPPB 569 WP 042872106 1 Dpoaceaephila
Dickeya gs fangzhongdai B16 WP 033569586 1 Dfangzhongdai
Dickeya gs fangzhongdai NCPPB 3274 WP 033569586 1 Dfangzhongdai
Dickeya gs fangzhongdai MK7 WP 033569586 1 Dfangzhongdai
Dickeya gs undicola 2B12 WP 033569586 1 Dundicola
Dickeya gs fangzhongdai S1 WP 033569586 1 Dfangzhongdai
Prodigiosinella confusarubida ATCC39006 WP 021016908 1 Pconfusarubida
Dickeya paradisiaca Ech703 WP 015854431 1 Dparadisiaca
Dickeya paradisiaca NCPPB 2511 WP 015854431 1 Dparadisiaca
Edwardsiella ictaluri 93 146 YP 002934566 1 634503 Proteobacteria
Edwardsiella tarda EIB202 YP 003296857 1 498217 Proteobacteria
Edwardsiella piscicida C07 087 YP 007630192 1 1288122 Proteobacteria
Serratia marcescens WW4 YP 007407535 1 435998 Proteobacteria
Serratia proteamaculans 568 YP 001479849 1 399741 Proteobacteria
Serratia plymuthica AS9 YP 004507192 1 768492 Proteobacteria
Serratia liquefaciens ATCC 27592 YP 008231655 1 1346614 Proteobacteria
Serratia sp AS12 YP 004502240 1 768490 Proteobacteria
Yersinia enterocolitica subsp palearctica Y11 YP 006007772 1 930944 Proteobacteria
Yersinia enterocolitica subsp enterocolitica 8081 YP 001005391 1 393305 Proteobacteria
Yersinia pestis biovar Microtus str 91001 NP 993879 1 229193 Proteobacteria
Yersinia pseudotuberculosis IP 31758 YP 001400151 1 349747 Proteobacteria
Yersinia pestis Antiqua YP 652245 1 360102 Proteobacteria
Yersinia pestis biovar Medievalis str Harbin 35 YP 005624895 1 547048 Proteobacteria
Dickeya gs fangzhongdai ND14b WP 013318965 1 Dfangzhongdai
Dickeya gs fangzhongdai M005 WP 013318965 1 Dfangzhongdai
Dickeya solani PPO 9134 WP 013318965 1 Dsolani
Dickeya solani RNS 07 7 3B WP 013318965 1 Dsolani
Dickeya solani D s0432 1 WP 013318965 1 Dsolani
Dickeya dadantii NCPPB 898 WP 013318965 1 Ddadantii
Dickeya dadantii NCPPB 3537 WP 013318965 1 Ddadantii
Dickeya solani MK10 WP 013318965 1 Dsolani
Dickeya solani IPO 2222 WP 013318965 1 Dsolani
448 iscA response to oxidative stress Seed
Dickeya solani GBBC 2040 WP 013318965 1 Dsolani
Dickeya solani IFB 0099 WP 013318965 1 Dsolani
Dickeya gs fangzhongdai M074 WP 013318965 1 Dfangzhongdai
Dickeya solani RNS 05 1 2A WP 013318965 1 Dsolani
Dickeya dadantii 3937 WP 013318965 1 Ddadantii
Dickeya dianthicola RNS04 9 WP 013318965 1 Ddianthicola
Dickeya solani RNS 08 23 3 1 A WP 013318965 1 Dsolani
Dickeya dianthicola IPO 980 WP 013318965 1 Ddianthicola
Dickeya dianthicola NCPPB 453 WP 013318965 1 Ddianthicola
Dickeya solani PPO 9019 WP 013318965 1 Dsolani
Dickeya solani MK16 WP 013318965 1 Dsolani
Dickeya dianthicola NCPPB 3534 WP 013318965 1 Ddianthicola
Dickeya dadantii subsp dieffenbachiae NCPPB 2976 WP 013318965 1 Ddadantii
Dickeya dianthicola GBBC 2039 WP 013318965 1 Ddianthicola

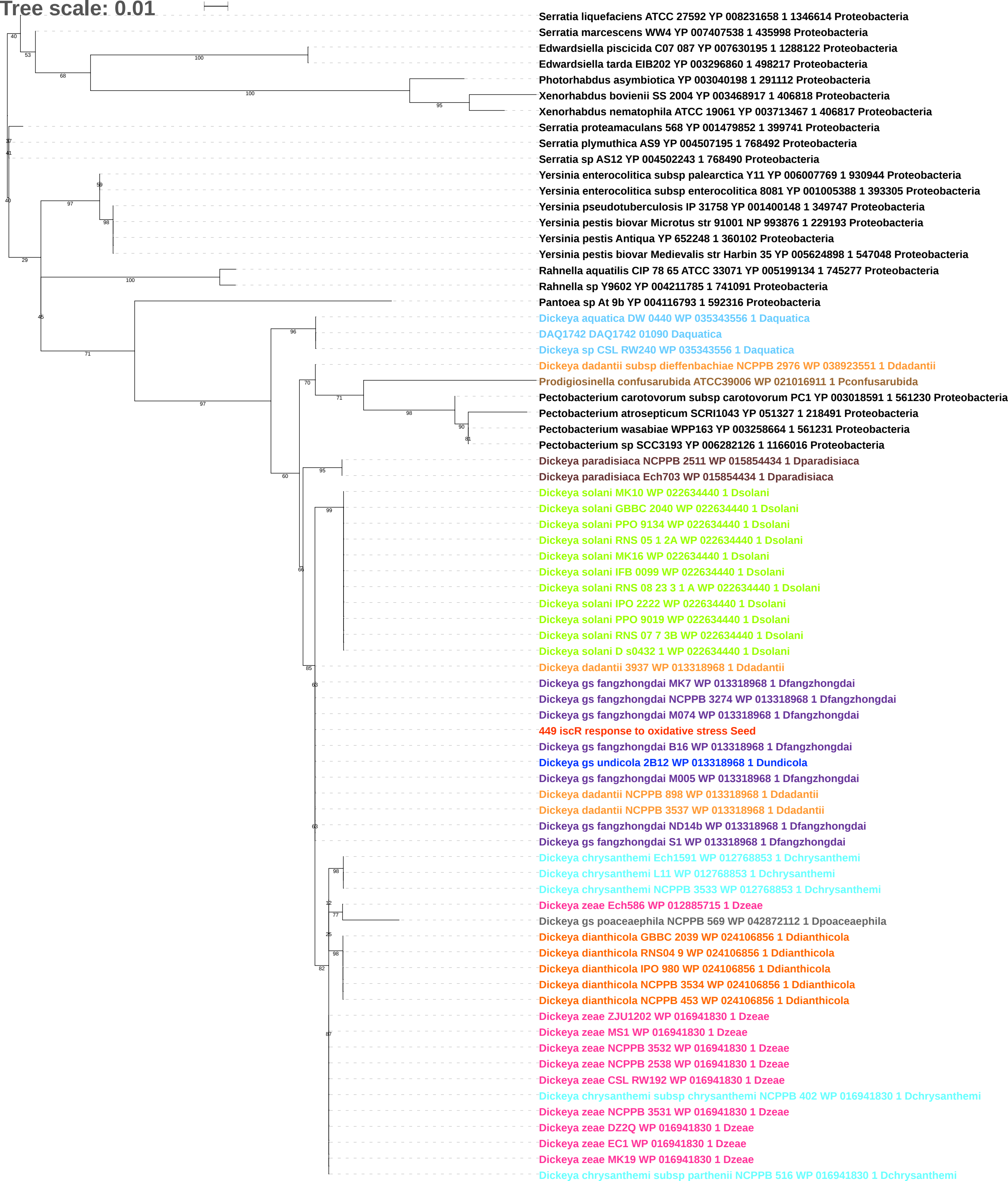
Tree scale: 0.01
Serratia liquefaciens ATCC 27592 YP 008231658 1 1346614 Proteobacteria
Serratia marcescens WW4 YP 007407538 1 435998 Proteobacteria
Edwardsiella piscicida C07 087 YP 007630195 1 1288122 Proteobacteria
Edwardsiella tarda EIB202 YP 003296860 1 498217 Proteobacteria
Photorhabdus asymbiotica YP 003040198 1 291112 Proteobacteria
Xenorhabdus bovienii SS 2004 YP 003468917 1 406818 Proteobacteria
Xenorhabdus nematophila ATCC 19061 YP 003713467 1 406817 Proteobacteria
Serratia proteamaculans 568 YP 001479852 1 399741 Proteobacteria
Serratia plymuthica AS9 YP 004507195 1 768492 Proteobacteria
Serratia sp AS12 YP 004502243 1 768490 Proteobacteria
Yersinia enterocolitica subsp palearctica Y11 YP 006007769 1 930944 Proteobacteria
Yersinia enterocolitica subsp enterocolitica 8081 YP 001005388 1 393305 Proteobacteria
Yersinia pseudotuberculosis IP 31758 YP 001400148 1 349747 Proteobacteria
Yersinia pestis biovar Microtus str 91001 NP 993876 1 229193 Proteobacteria
Yersinia pestis Antiqua YP 652248 1 360102 Proteobacteria
Yersinia pestis biovar Medievalis str Harbin 35 YP 005624898 1 547048 Proteobacteria
Rahnella aquatilis CIP 78 65 ATCC 33071 YP 005199134 1 745277 Proteobacteria
Rahnella sp Y9602 YP 004211785 1 741091 Proteobacteria
Pantoea sp At 9b YP 004116793 1 592316 Proteobacteria
Dickeya aquatica DW 0440 WP 035343556 1 Daquatica
DAQ1742 DAQ1742 01090 Daquatica
Dickeya sp CSL RW240 WP 035343556 1 Daquatica
Dickeya dadantii subsp dieffenbachiae NCPPB 2976 WP 038923551 1 Ddadantii
Prodigiosinella confusarubida ATCC39006 WP 021016911 1 Pconfusarubida
Pectobacterium carotovorum subsp carotovorum PC1 YP 003018591 1 561230 Proteobacteria
Pectobacterium atrosepticum SCRI1043 YP 051327 1 218491 Proteobacteria
Pectobacterium wasabiae WPP163 YP 003258664 1 561231 Proteobacteria
Pectobacterium sp SCC3193 YP 006282126 1 1166016 Proteobacteria
Dickeya paradisiaca NCPPB 2511 WP 015854434 1 Dparadisiaca
Dickeya paradisiaca Ech703 WP 015854434 1 Dparadisiaca
Dickeya solani MK10 WP 022634440 1 Dsolani
Dickeya solani GBBC 2040 WP 022634440 1 Dsolani
Dickeya solani PPO 9134 WP 022634440 1 Dsolani
Dickeya solani RNS 05 1 2A WP 022634440 1 Dsolani
Dickeya solani MK16 WP 022634440 1 Dsolani
Dickeya solani IFB 0099 WP 022634440 1 Dsolani
Dickeya solani RNS 08 23 3 1 A WP 022634440 1 Dsolani
Dickeya solani IPO 2222 WP 022634440 1 Dsolani
Dickeya solani PPO 9019 WP 022634440 1 Dsolani
Dickeya solani RNS 07 7 3B WP 022634440 1 Dsolani
Dickeya solani D s0432 1 WP 022634440 1 Dsolani
Dickeya dadantii 3937 WP 013318968 1 Ddadantii
Dickeya gs fangzhongdai MK7 WP 013318968 1 Dfangzhongdai
Dickeya gs fangzhongdai NCPPB 3274 WP 013318968 1 Dfangzhongdai
Dickeya gs fangzhongdai M074 WP 013318968 1 Dfangzhongdai
449 iscR response to oxidative stress Seed
Dickeya gs fangzhongdai B16 WP 013318968 1 Dfangzhongdai
Dickeya gs undicola 2B12 WP 013318968 1 Dundicola
Dickeya gs fangzhongdai M005 WP 013318968 1 Dfangzhongdai
Dickeya dadantii NCPPB 898 WP 013318968 1 Ddadantii
Dickeya dadantii NCPPB 3537 WP 013318968 1 Ddadantii
Dickeya gs fangzhongdai ND14b WP 013318968 1 Dfangzhongdai
Dickeya gs fangzhongdai S1 WP 013318968 1 Dfangzhongdai
Dickeya chrysanthemi Ech1591 WP 012768853 1 Dchrysanthemi
Dickeya chrysanthemi L11 WP 012768853 1 Dchrysanthemi
Dickeya chrysanthemi NCPPB 3533 WP 012768853 1 Dchrysanthemi
Dickeya zeae Ech586 WP 012885715 1 Dzeae
Dickeya gs poaceaephila NCPPB 569 WP 042872112 1 Dpoaceaephila
Dickeya dianthicola GBBC 2039 WP 024106856 1 Ddianthicola
Dickeya dianthicola RNS04 9 WP 024106856 1 Ddianthicola
Dickeya dianthicola IPO 980 WP 024106856 1 Ddianthicola
Dickeya dianthicola NCPPB 3534 WP 024106856 1 Ddianthicola
Dickeya dianthicola NCPPB 453 WP 024106856 1 Ddianthicola
Dickeya zeae ZJU1202 WP 016941830 1 Dzeae
Dickeya zeae MS1 WP 016941830 1 Dzeae
Dickeya zeae NCPPB 3532 WP 016941830 1 Dzeae
Dickeya zeae NCPPB 2538 WP 016941830 1 Dzeae
Dickeya zeae CSL RW192 WP 016941830 1 Dzeae
Dickeya chrysanthemi subsp chrysanthemi NCPPB 402 WP 016941830 1 Dchrysanthemi
Dickeya zeae NCPPB 3531 WP 016941830 1 Dzeae
Dickeya zeae DZ2Q WP 016941830 1 Dzeae
Dickeya zeae EC1 WP 016941830 1 Dzeae
Dickeya zeae MK19 WP 016941830 1 Dzeae
Dickeya chrysanthemi subsp parthenii NCPPB 516 WP 016941830 1 Dchrysanthemi

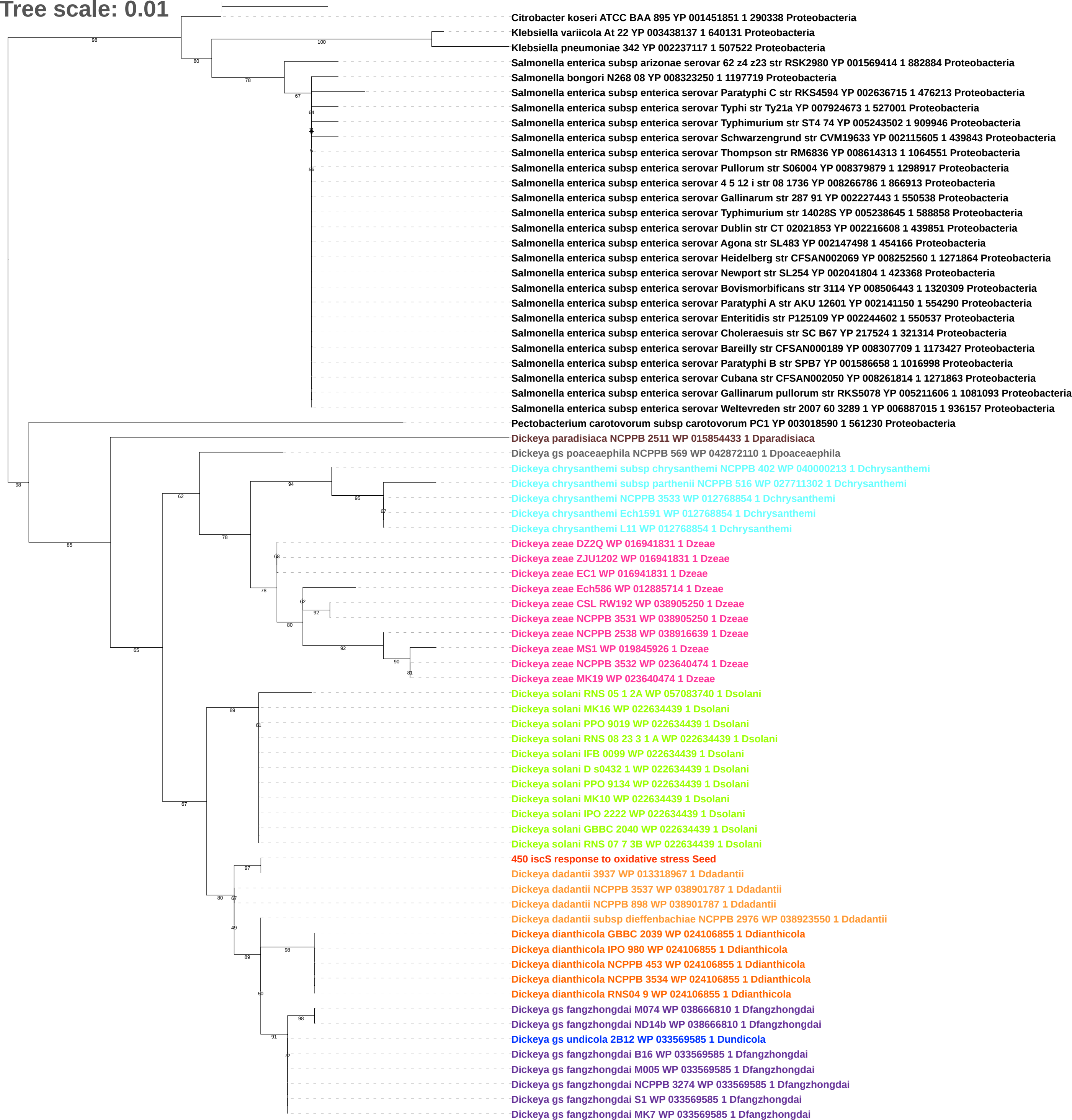

Tree scale: 0.01
Citrobacter koseri ATCC BAA 895 YP 001451851 1 290338 Proteobacteria
Klebsiella variicola At 22 YP 003438137 1 640131 Proteobacteria
Klebsiella pneumoniae 342 YP 002237117 1 507522 Proteobacteria
Salmonella enterica subsp arizonae serovar 62 z4 z23 str RSK2980 YP 001569414 1 882884 Proteobacteria
Salmonella bongori N268 08 YP 008323250 1 1197719 Proteobacteria
Salmonella enterica subsp enterica serovar Paratyphi C str RKS4594 YP 002636715 1 476213 Proteobacteria
Salmonella enterica subsp enterica serovar Typhi str Ty21a YP 007924673 1 527001 Proteobacteria
Salmonella enterica subsp enterica serovar Typhimurium str ST4 74 YP 005243502 1 909946 Proteobacteria
Salmonella enterica subsp enterica serovar Schwarzengrund str CVM19633 YP 002115605 1 439843 Proteobacteria
Salmonella enterica subsp enterica serovar Thompson str RM6836 YP 008614313 1 1064551 Proteobacteria
Salmonella enterica subsp enterica serovar Pullorum str S06004 YP 008379879 1 1298917 Proteobacteria
Salmonella enterica subsp enterica serovar 4 5 12 i str 08 1736 YP 008266786 1 866913 Proteobacteria
Salmonella enterica subsp enterica serovar Gallinarum str 287 91 YP 002227443 1 550538 Proteobacteria
Salmonella enterica subsp enterica serovar Typhimurium str 14028S YP 005238645 1 588858 Proteobacteria
Salmonella enterica subsp enterica serovar Dublin str CT 02021853 YP 002216608 1 439851 Proteobacteria
Salmonella enterica subsp enterica serovar Agona str SL483 YP 002147498 1 454166 Proteobacteria
Salmonella enterica subsp enterica serovar Heidelberg str CFSAN002069 YP 008252560 1 1271864 Proteobacteria
Salmonella enterica subsp enterica serovar Newport str SL254 YP 002041804 1 423368 Proteobacteria
Salmonella enterica subsp enterica serovar Bovismorbificans str 3114 YP 008506443 1 1320309 Proteobacteria
Salmonella enterica subsp enterica serovar Paratyphi A str AKU 12601 YP 002141150 1 554290 Proteobacteria
Salmonella enterica subsp enterica serovar Enteritidis str P125109 YP 002244602 1 550537 Proteobacteria
Salmonella enterica subsp enterica serovar Choleraesuis str SC B67 YP 217524 1 321314 Proteobacteria
Salmonella enterica subsp enterica serovar Bareilly str CFSAN000189 YP 008307709 1 1173427 Proteobacteria
Salmonella enterica subsp enterica serovar Paratyphi B str SPB7 YP 001586658 1 1016998 Proteobacteria
Salmonella enterica subsp enterica serovar Cubana str CFSAN002050 YP 008261814 1 1271863 Proteobacteria
Salmonella enterica subsp enterica serovar Gallinarum pullorum str RKS5078 YP 005211606 1 1081093 Proteobacteria
Salmonella enterica subsp enterica serovar Weltevreden str 2007 60 3289 1 YP 006887015 1 936157 Proteobacteria
Pectobacterium carotovorum subsp carotovorum PC1 YP 003018590 1 561230 Proteobacteria
Dickeya paradisiaca NCPPB 2511 WP 015854433 1 Dparadisiaca
Dickeya gs poaceaephila NCPPB 569 WP 042872110 1 Dpoaceaephila
Dickeya chrysanthemi subsp chrysanthemi NCPPB 402 WP 040000213 1 Dchrysanthemi
Dickeya chrysanthemi subsp parthenii NCPPB 516 WP 027711302 1 Dchrysanthemi
Dickeya chrysanthemi NCPPB 3533 WP 012768854 1 Dchrysanthemi
Dickeya chrysanthemi Ech1591 WP 012768854 1 Dchrysanthemi
Dickeya chrysanthemi L11 WP 012768854 1 Dchrysanthemi
Dickeya zeae DZ2Q WP 016941831 1 Dzeae
Dickeya zeae ZJU1202 WP 016941831 1 Dzeae
Dickeya zeae EC1 WP 016941831 1 Dzeae
Dickeya zeae Ech586 WP 012885714 1 Dzeae
Dickeya zeae CSL RW192 WP 038905250 1 Dzeae
Dickeya zeae NCPPB 3531 WP 038905250 1 Dzeae
Dickeya zeae NCPPB 2538 WP 038916639 1 Dzeae
Dickeya zeae MS1 WP 019845926 1 Dzeae
Dickeya zeae NCPPB 3532 WP 023640474 1 Dzeae
Dickeya zeae MK19 WP 023640474 1 Dzeae
Dickeya solani RNS 05 1 2A WP 057083740 1 Dsolani
Dickeya solani MK16 WP 022634439 1 Dsolani
Dickeya solani PPO 9019 WP 022634439 1 Dsolani
Dickeya solani RNS 08 23 3 1 A WP 022634439 1 Dsolani
Dickeya solani IFB 0099 WP 022634439 1 Dsolani
Dickeya solani D s0432 1 WP 022634439 1 Dsolani
Dickeya solani PPO 9134 WP 022634439 1 Dsolani
Dickeya solani MK10 WP 022634439 1 Dsolani
Dickeya solani IPO 2222 WP 022634439 1 Dsolani
Dickeya solani GBBC 2040 WP 022634439 1 Dsolani
Dickeya solani RNS 07 7 3B WP 022634439 1 Dsolani
450 iscS response to oxidative stress Seed
Dickeya dadantii 3937 WP 013318967 1 Ddadantii
Dickeya dadantii NCPPB 3537 WP 038901787 1 Ddadantii
Dickeya dadantii NCPPB 898 WP 038901787 1 Ddadantii
Dickeya dadantii subsp dieffenbachiae NCPPB 2976 WP 038923550 1 Ddadantii
Dickeya dianthicola GBBC 2039 WP 024106855 1 Ddianthicola
Dickeya dianthicola IPO 980 WP 024106855 1 Ddianthicola
Dickeya dianthicola NCPPB 453 WP 024106855 1 Ddianthicola
Dickeya dianthicola NCPPB 3534 WP 024106855 1 Ddianthicola
Dickeya dianthicola RNS04 9 WP 024106855 1 Ddianthicola
Dickeya gs fangzhongdai M074 WP 038666810 1 Dfangzhongdai
Dickeya gs fangzhongdai ND14b WP 038666810 1 Dfangzhongdai
Dickeya gs undicola 2B12 WP 033569585 1 Dundicola
Dickeya gs fangzhongdai B16 WP 033569585 1 Dfangzhongdai
Dickeya gs fangzhongdai M005 WP 033569585 1 Dfangzhongdai
Dickeya gs fangzhongdai NCPPB 3274 WP 033569585 1 Dfangzhongdai
Dickeya gs fangzhongdai S1 WP 033569585 1 Dfangzhongdai
Dickeya gs fangzhongdai MK7 WP 033569585 1 Dfangzhongdai

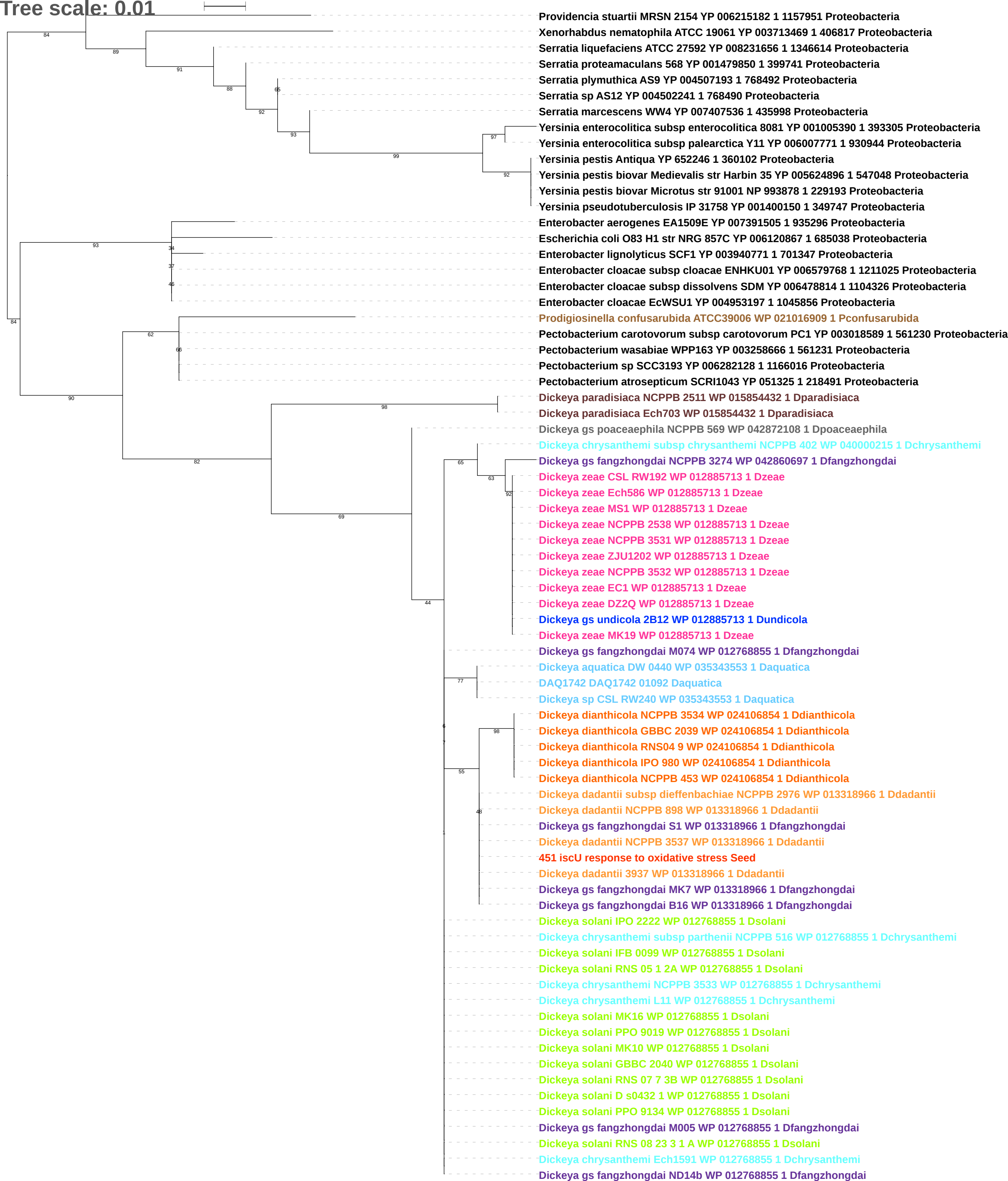
Tree scale: 0.01
Providencia stuartii MRSN 2154 YP 006215182 1 1157951 Proteobacteria
Xenorhabdus nematophila ATCC 19061 YP 003713469 1 406817 Proteobacteria
Serratia liquefaciens ATCC 27592 YP 008231656 1 1346614 Proteobacteria
Serratia proteamaculans 568 YP 001479850 1 399741 Proteobacteria
Serratia plymuthica AS9 YP 004507193 1 768492 Proteobacteria
Serratia sp AS12 YP 004502241 1 768490 Proteobacteria
Serratia marcescens WW4 YP 007407536 1 435998 Proteobacteria
Yersinia enterocolitica subsp enterocolitica 8081 YP 001005390 1 393305 Proteobacteria
Yersinia enterocolitica subsp palearctica Y11 YP 006007771 1 930944 Proteobacteria
Yersinia pestis Antiqua YP 652246 1 360102 Proteobacteria
Yersinia pestis biovar Medievalis str Harbin 35 YP 005624896 1 547048 Proteobacteria
Yersinia pestis biovar Microtus str 91001 NP 993878 1 229193 Proteobacteria
Yersinia pseudotuberculosis IP 31758 YP 001400150 1 349747 Proteobacteria
Enterobacter aerogenes EA1509E YP 007391505 1 935296 Proteobacteria
Escherichia coli O83 H1 str NRG 857C YP 006120867 1 685038 Proteobacteria
Enterobacter lignolyticus SCF1 YP 003940771 1 701347 Proteobacteria
Enterobacter cloacae subsp cloacae ENHKU01 YP 006579768 1 1211025 Proteobacteria
Enterobacter cloacae subsp dissolvens SDM YP 006478814 1 1104326 Proteobacteria
Enterobacter cloacae EcWSU1 YP 004953197 1 1045856 Proteobacteria
Prodigiosinella confusarubida ATCC39006 WP 021016909 1 Pconfusarubida
Pectobacterium carotovorum subsp carotovorum PC1 YP 003018589 1 561230 Proteobacteria
Pectobacterium wasabiae WPP163 YP 003258666 1 561231 Proteobacteria
Pectobacterium sp SCC3193 YP 006282128 1 1166016 Proteobacteria
Pectobacterium atrosepticum SCRI1043 YP 051325 1 218491 Proteobacteria
Dickeya paradisiaca NCPPB 2511 WP 015854432 1 Dparadisiaca
Dickeya paradisiaca Ech703 WP 015854432 1 Dparadisiaca
Dickeya gs poaceaephila NCPPB 569 WP 042872108 1 Dpoaceaephila
Dickeya chrysanthemi subsp chrysanthemi NCPPB 402 WP 040000215 1 Dchrysanthemi
Dickeya gs fangzhongdai NCPPB 3274 WP 042860697 1 Dfangzhongdai
Dickeya zeae CSL RW192 WP 012885713 1 Dzeae
Dickeya zeae Ech586 WP 012885713 1 Dzeae
Dickeya zeae MS1 WP 012885713 1 Dzeae
Dickeya zeae NCPPB 2538 WP 012885713 1 Dzeae
Dickeya zeae NCPPB 3531 WP 012885713 1 Dzeae
Dickeya zeae ZJU1202 WP 012885713 1 Dzeae
Dickeya zeae NCPPB 3532 WP 012885713 1 Dzeae
Dickeya zeae EC1 WP 012885713 1 Dzeae
Dickeya zeae DZ2Q WP 012885713 1 Dzeae
Dickeya gs undicola 2B12 WP 012885713 1 Dundicola
Dickeya zeae MK19 WP 012885713 1 Dzeae
Dickeya gs fangzhongdai M074 WP 012768855 1 Dfangzhongdai
Dickeya aquatica DW 0440 WP 035343553 1 Daquatica
DAQ1742 DAQ1742 01092 Daquatica
Dickeya sp CSL RW240 WP 035343553 1 Daquatica
Dickeya dianthicola NCPPB 3534 WP 024106854 1 Ddianthicola
Dickeya dianthicola GBBC 2039 WP 024106854 1 Ddianthicola
Dickeya dianthicola RNS04 9 WP 024106854 1 Ddianthicola
Dickeya dianthicola IPO 980 WP 024106854 1 Ddianthicola
Dickeya dianthicola NCPPB 453 WP 024106854 1 Ddianthicola
Dickeya dadantii subsp dieffenbachiae NCPPB 2976 WP 013318966 1 Ddadantii
Dickeya dadantii NCPPB 898 WP 013318966 1 Ddadantii
Dickeya gs fangzhongdai S1 WP 013318966 1 Dfangzhongdai
Dickeya dadantii NCPPB 3537 WP 013318966 1 Ddadantii
451 iscU response to oxidative stress Seed
Dickeya dadantii 3937 WP 013318966 1 Ddadantii
Dickeya gs fangzhongdai MK7 WP 013318966 1 Dfangzhongdai
Dickeya gs fangzhongdai B16 WP 013318966 1 Dfangzhongdai
Dickeya solani IPO 2222 WP 012768855 1 Dsolani
Dickeya chrysanthemi subsp parthenii NCPPB 516 WP 012768855 1 Dchrysanthemi
Dickeya solani IFB 0099 WP 012768855 1 Dsolani
Dickeya solani RNS 05 1 2A WP 012768855 1 Dsolani
Dickeya chrysanthemi NCPPB 3533 WP 012768855 1 Dchrysanthemi
Dickeya chrysanthemi L11 WP 012768855 1 Dchrysanthemi
Dickeya solani MK16 WP 012768855 1 Dsolani
Dickeya solani PPO 9019 WP 012768855 1 Dsolani
Dickeya solani MK10 WP 012768855 1 Dsolani
Dickeya solani GBBC 2040 WP 012768855 1 Dsolani
Dickeya solani RNS 07 7 3B WP 012768855 1 Dsolani
Dickeya solani D s0432 1 WP 012768855 1 Dsolani
Dickeya solani PPO 9134 WP 012768855 1 Dsolani
Dickeya gs fangzhongdai M005 WP 012768855 1 Dfangzhongdai
Dickeya solani RNS 08 23 3 1 A WP 012768855 1 Dsolani
Dickeya chrysanthemi Ech1591 WP 012768855 1 Dchrysanthemi
Dickeya gs fangzhongdai ND14b WP 012768855 1 Dfangzhongdai

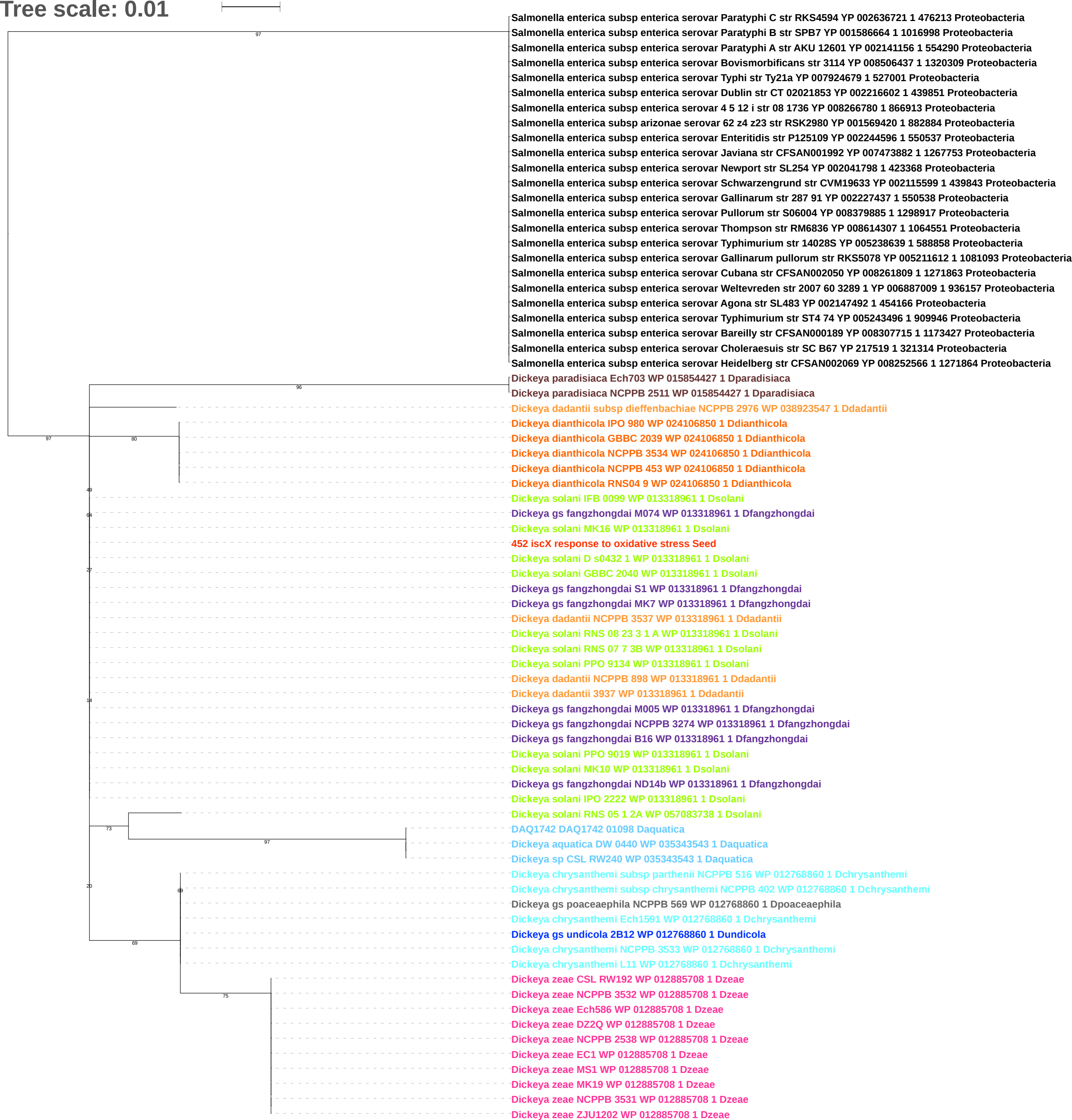
Tree scale: 0.01
Salmonella enterica subsp enterica serovar Paratyphi C str RKS4594 YP 002636721 1 476213 Proteobacteria
Salmonella enterica subsp enterica serovar Paratyphi B str SPB7 YP 001586664 1 1016998 Proteobacteria
Salmonella enterica subsp enterica serovar Paratyphi A str AKU 12601 YP 002141156 1 554290 Proteobacteria
Salmonella enterica subsp enterica serovar Bovismorbificans str 3114 YP 008506437 1 1320309 Proteobacteria
Salmonella enterica subsp enterica serovar Typhi str Ty21a YP 007924679 1 527001 Proteobacteria
Salmonella enterica subsp enterica serovar Dublin str CT 02021853 YP 002216602 1 439851 Proteobacteria
Salmonella enterica subsp enterica serovar 4 5 12 i str 08 1736 YP 008266780 1 866913 Proteobacteria
Salmonella enterica subsp arizonae serovar 62 z4 z23 str RSK2980 YP 001569420 1 882884 Proteobacteria
Salmonella enterica subsp enterica serovar Enteritidis str P125109 YP 002244596 1 550537 Proteobacteria
Salmonella enterica subsp enterica serovar Javiana str CFSAN001992 YP 007473882 1 1267753 Proteobacteria
Salmonella enterica subsp enterica serovar Newport str SL254 YP 002041798 1 423368 Proteobacteria
Salmonella enterica subsp enterica serovar Schwarzengrund str CVM19633 YP 002115599 1 439843 Proteobacteria
Salmonella enterica subsp enterica serovar Gallinarum str 287 91 YP 002227437 1 550538 Proteobacteria
Salmonella enterica subsp enterica serovar Pullorum str S06004 YP 008379885 1 1298917 Proteobacteria
Salmonella enterica subsp enterica serovar Thompson str RM6836 YP 008614307 1 1064551 Proteobacteria
Salmonella enterica subsp enterica serovar Typhimurium str 14028S YP 005238639 1 588858 Proteobacteria
Salmonella enterica subsp enterica serovar Gallinarum pullorum str RKS5078 YP 005211612 1 1081093 Proteobacteria
Salmonella enterica subsp enterica serovar Cubana str CFSAN002050 YP 008261809 1 1271863 Proteobacteria
Salmonella enterica subsp enterica serovar Weltevreden str 2007 60 3289 1 YP 006887009 1 936157 Proteobacteria
Salmonella enterica subsp enterica serovar Agona str SL483 YP 002147492 1 454166 Proteobacteria
Salmonella enterica subsp enterica serovar Typhimurium str ST4 74 YP 005243496 1 909946 Proteobacteria
Salmonella enterica subsp enterica serovar Bareilly str CFSAN000189 YP 008307715 1 1173427 Proteobacteria
Salmonella enterica subsp enterica serovar Choleraesuis str SC B67 YP 217519 1 321314 Proteobacteria
Salmonella enterica subsp enterica serovar Heidelberg str CFSAN002069 YP 008252566 1 1271864 Proteobacteria
Dickeya paradisiaca Ech703 WP 015854427 1 Dparadisiaca
Dickeya paradisiaca NCPPB 2511 WP 015854427 1 Dparadisiaca
Dickeya dadantii subsp dieffenbachiae NCPPB 2976 WP 038923547 1 Ddadantii
Dickeya dianthicola IPO 980 WP 024106850 1 Ddianthicola
Dickeya dianthicola GBBC 2039 WP 024106850 1 Ddianthicola
Dickeya dianthicola NCPPB 3534 WP 024106850 1 Ddianthicola
Dickeya dianthicola NCPPB 453 WP 024106850 1 Ddianthicola
Dickeya dianthicola RNS04 9 WP 024106850 1 Ddianthicola
Dickeya solani IFB 0099 WP 013318961 1 Dsolani
Dickeya gs fangzhongdai M074 WP 013318961 1 Dfangzhongdai
Dickeya solani MK16 WP 013318961 1 Dsolani
452 iscX response to oxidative stress Seed
Dickeya solani D s0432 1 WP 013318961 1 Dsolani
Dickeya solani GBBC 2040 WP 013318961 1 Dsolani
Dickeya gs fangzhongdai S1 WP 013318961 1 Dfangzhongdai
Dickeya gs fangzhongdai MK7 WP 013318961 1 Dfangzhongdai
Dickeya dadantii NCPPB 3537 WP 013318961 1 Ddadantii
Dickeya solani RNS 08 23 3 1 A WP 013318961 1 Dsolani
Dickeya solani RNS 07 7 3B WP 013318961 1 Dsolani
Dickeya solani PPO 9134 WP 013318961 1 Dsolani
Dickeya dadantii NCPPB 898 WP 013318961 1 Ddadantii
Dickeya dadantii 3937 WP 013318961 1 Ddadantii
Dickeya gs fangzhongdai M005 WP 013318961 1 Dfangzhongdai
Dickeya gs fangzhongdai NCPPB 3274 WP 013318961 1 Dfangzhongdai
Dickeya gs fangzhongdai B16 WP 013318961 1 Dfangzhongdai
Dickeya solani PPO 9019 WP 013318961 1 Dsolani
Dickeya solani MK10 WP 013318961 1 Dsolani
Dickeya gs fangzhongdai ND14b WP 013318961 1 Dfangzhongdai
Dickeya solani IPO 2222 WP 013318961 1 Dsolani
Dickeya solani RNS 05 1 2A WP 057083738 1 Dsolani
DAQ1742 DAQ1742 01098 Daquatica
Dickeya aquatica DW 0440 WP 035343543 1 Daquatica
Dickeya sp CSL RW240 WP 035343543 1 Daquatica
Dickeya chrysanthemi subsp parthenii NCPPB 516 WP 012768860 1 Dchrysanthemi
Dickeya chrysanthemi subsp chrysanthemi NCPPB 402 WP 012768860 1 Dchrysanthemi
Dickeya gs poaceaephila NCPPB 569 WP 012768860 1 Dpoaceaephila
Dickeya chrysanthemi Ech1591 WP 012768860 1 Dchrysanthemi
Dickeya gs undicola 2B12 WP 012768860 1 Dundicola
Dickeya chrysanthemi NCPPB 3533 WP 012768860 1 Dchrysanthemi
Dickeya chrysanthemi L11 WP 012768860 1 Dchrysanthemi
Dickeya zeae CSL RW192 WP 012885708 1 Dzeae
Dickeya zeae NCPPB 3532 WP 012885708 1 Dzeae
Dickeya zeae Ech586 WP 012885708 1 Dzeae
Dickeya zeae DZ2Q WP 012885708 1 Dzeae
Dickeya zeae NCPPB 2538 WP 012885708 1 Dzeae
Dickeya zeae EC1 WP 012885708 1 Dzeae
Dickeya zeae MS1 WP 012885708 1 Dzeae
Dickeya zeae MK19 WP 012885708 1 Dzeae
Dickeya zeae NCPPB 3531 WP 012885708 1 Dzeae
Dickeya zeae ZJU1202 WP 012885708 1 Dzeae

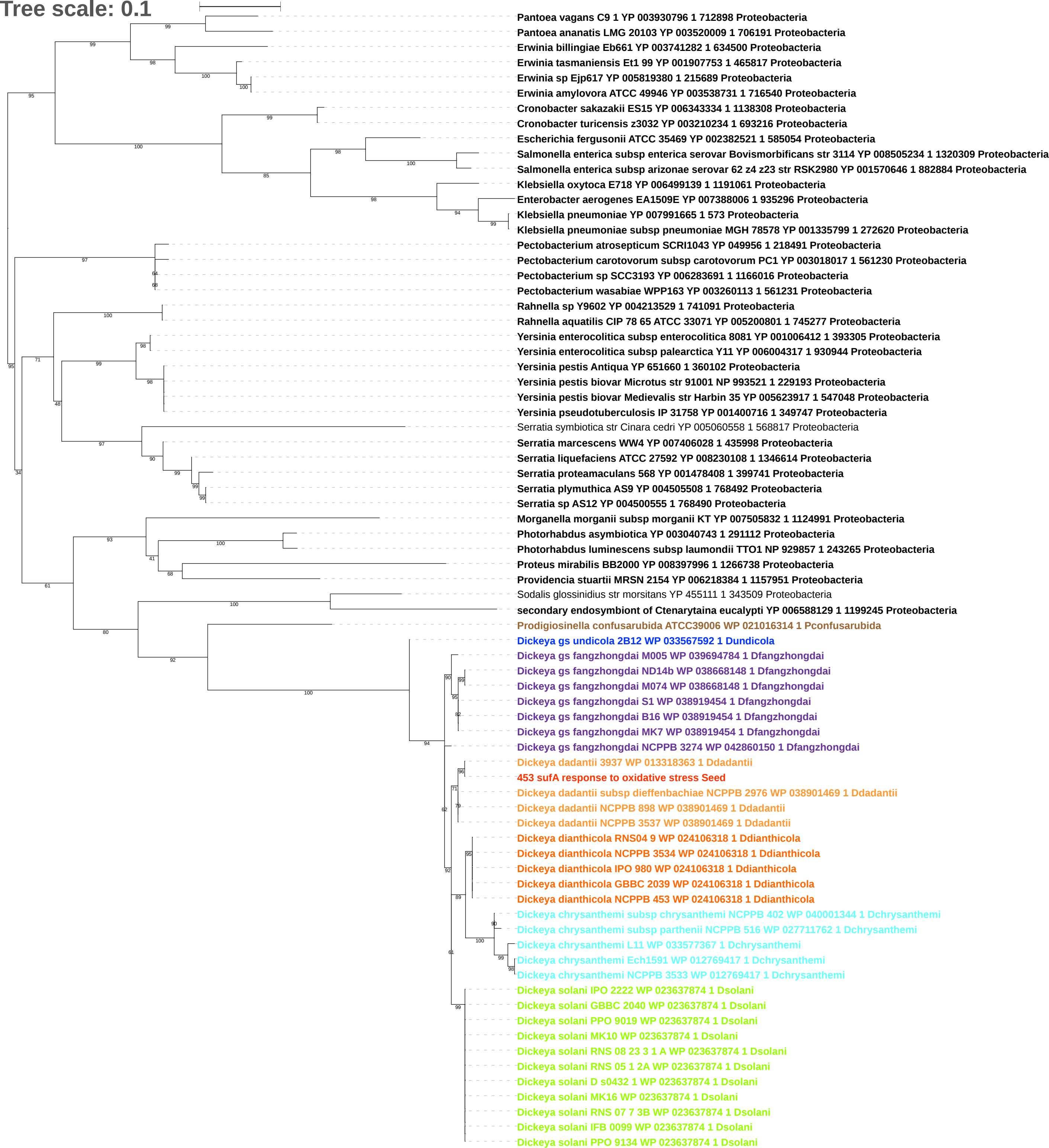
Tree scale: 0.1
Pantoea vagans C9 1 YP 003930796 1 712898 Proteobacteria
Pantoea ananatis LMG 20103 YP 003520009 1 706191 Proteobacteria
Erwinia billingiae Eb661 YP 003741282 1 634500 Proteobacteria
Erwinia tasmaniensis Et1 99 YP 001907753 1 465817 Proteobacteria
Erwinia sp Ejp617 YP 005819380 1 215689 Proteobacteria
Erwinia amylovora ATCC 49946 YP 003538731 1 716540 Proteobacteria
Cronobacter sakazakii ES15 YP 006343334 1 1138308 Proteobacteria
Cronobacter turicensis z3032 YP 003210234 1 693216 Proteobacteria
Escherichia fergusonii ATCC 35469 YP 002382521 1 585054 Proteobacteria
Salmonella enterica subsp enterica serovar Bovismorbificans str 3114 YP 008505234 1 1320309 Proteobacteria
Salmonella enterica subsp arizonae serovar 62 z4 z23 str RSK2980 YP 001570646 1 882884 Proteobacteria
Klebsiella oxytoca E718 YP 006499139 1 1191061 Proteobacteria
Enterobacter aerogenes EA1509E YP 007388006 1 935296 Proteobacteria
Klebsiella pneumoniae YP 007991665 1 573 Proteobacteria
Klebsiella pneumoniae subsp pneumoniae MGH 78578 YP 001335799 1 272620 Proteobacteria
Pectobacterium atrosepticum SCRI1043 YP 049956 1 218491 Proteobacteria
Pectobacterium carotovorum subsp carotovorum PC1 YP 003018017 1 561230 Proteobacteria
Pectobacterium sp SCC3193 YP 006283691 1 1166016 Proteobacteria
Pectobacterium wasabiae WPP163 YP 003260113 1 561231 Proteobacteria
Rahnella sp Y9602 YP 004213529 1 741091 Proteobacteria
Rahnella aquatilis CIP 78 65 ATCC 33071 YP 005200801 1 745277 Proteobacteria
Yersinia enterocolitica subsp enterocolitica 8081 YP 001006412 1 393305 Proteobacteria
Yersinia enterocolitica subsp palearctica Y11 YP 006004317 1 930944 Proteobacteria
Yersinia pestis Antiqua YP 651660 1 360102 Proteobacteria
Yersinia pestis biovar Microtus str 91001 NP 993521 1 229193 Proteobacteria
Yersinia pestis biovar Medievalis str Harbin 35 YP 005623917 1 547048 Proteobacteria
Yersinia pseudotuberculosis IP 31758 YP 001400716 1 349747 Proteobacteria
Serratia symbiotica str Cinara cedri YP 005060558 1 568817 Proteobacteria
Serratia marcescens WW4 YP 007406028 1 435998 Proteobacteria
Serratia liquefaciens ATCC 27592 YP 008230108 1 1346614 Proteobacteria
Serratia proteamaculans 568 YP 001478408 1 399741 Proteobacteria
Serratia plymuthica AS9 YP 004505508 1 768492 Proteobacteria
Serratia sp AS12 YP 004500555 1 768490 Proteobacteria
Morganella morganii subsp morganii KT YP 007505832 1 1124991 Proteobacteria
Photorhabdus asymbiotica YP 003040743 1 291112 Proteobacteria
Photorhabdus luminescens subsp laumondii TTO1 NP 929857 1 243265 Proteobacteria
Proteus mirabilis BB2000 YP 008397996 1 1266738 Proteobacteria
Providencia stuartii MRSN 2154 YP 006218384 1 1157951 Proteobacteria
Sodalis glossinidius str morsitans YP 455111 1 343509 Proteobacteria
secondary endosymbiont of Ctenarytaina eucalypti YP 006588129 1 1199245 Proteobacteria
Prodigiosinella confusarubida ATCC39006 WP 021016314 1 Pconfusarubida
Dickeya gs undicola 2B12 WP 033567592 1 Dundicola
Dickeya gs fangzhongdai M005 WP 039694784 1 Dfangzhongdai
Dickeya gs fangzhongdai ND14b WP 038668148 1 Dfangzhongdai
Dickeya gs fangzhongdai M074 WP 038668148 1 Dfangzhongdai
Dickeya gs fangzhongdai S1 WP 038919454 1 Dfangzhongdai
Dickeya gs fangzhongdai B16 WP 038919454 1 Dfangzhongdai
Dickeya gs fangzhongdai MK7 WP 038919454 1 Dfangzhongdai
Dickeya gs fangzhongdai NCPPB 3274 WP 042860150 1 Dfangzhongdai
Dickeya dadantii 3937 WP 013318363 1 Ddadantii
453 sufA response to oxidative stress Seed
Dickeya dadantii subsp dieffenbachiae NCPPB 2976 WP 038901469 1 Ddadantii
Dickeya dadantii NCPPB 898 WP 038901469 1 Ddadantii
Dickeya dadantii NCPPB 3537 WP 038901469 1 Ddadantii
Dickeya dianthicola RNS04 9 WP 024106318 1 Ddianthicola
Dickeya dianthicola NCPPB 3534 WP 024106318 1 Ddianthicola
Dickeya dianthicola IPO 980 WP 024106318 1 Ddianthicola
Dickeya dianthicola GBBC 2039 WP 024106318 1 Ddianthicola
Dickeya dianthicola NCPPB 453 WP 024106318 1 Ddianthicola
Dickeya chrysanthemi subsp chrysanthemi NCPPB 402 WP 040001344 1 Dchrysanthemi
Dickeya chrysanthemi subsp parthenii NCPPB 516 WP 027711762 1 Dchrysanthemi
Dickeya chrysanthemi L11 WP 033577367 1 Dchrysanthemi
Dickeya chrysanthemi Ech1591 WP 012769417 1 Dchrysanthemi
Dickeya chrysanthemi NCPPB 3533 WP 012769417 1 Dchrysanthemi
Dickeya solani IPO 2222 WP 023637874 1 Dsolani
Dickeya solani GBBC 2040 WP 023637874 1 Dsolani
Dickeya solani PPO 9019 WP 023637874 1 Dsolani
Dickeya solani MK10 WP 023637874 1 Dsolani
Dickeya solani RNS 08 23 3 1 A WP 023637874 1 Dsolani
Dickeya solani RNS 05 1 2A WP 023637874 1 Dsolani
Dickeya solani D s0432 1 WP 023637874 1 Dsolani
Dickeya solani MK16 WP 023637874 1 Dsolani
Dickeya solani RNS 07 7 3B WP 023637874 1 Dsolani
Dickeya solani IFB 0099 WP 023637874 1 Dsolani
Dickeya solani PPO 9134 WP 023637874 1 Dsolani

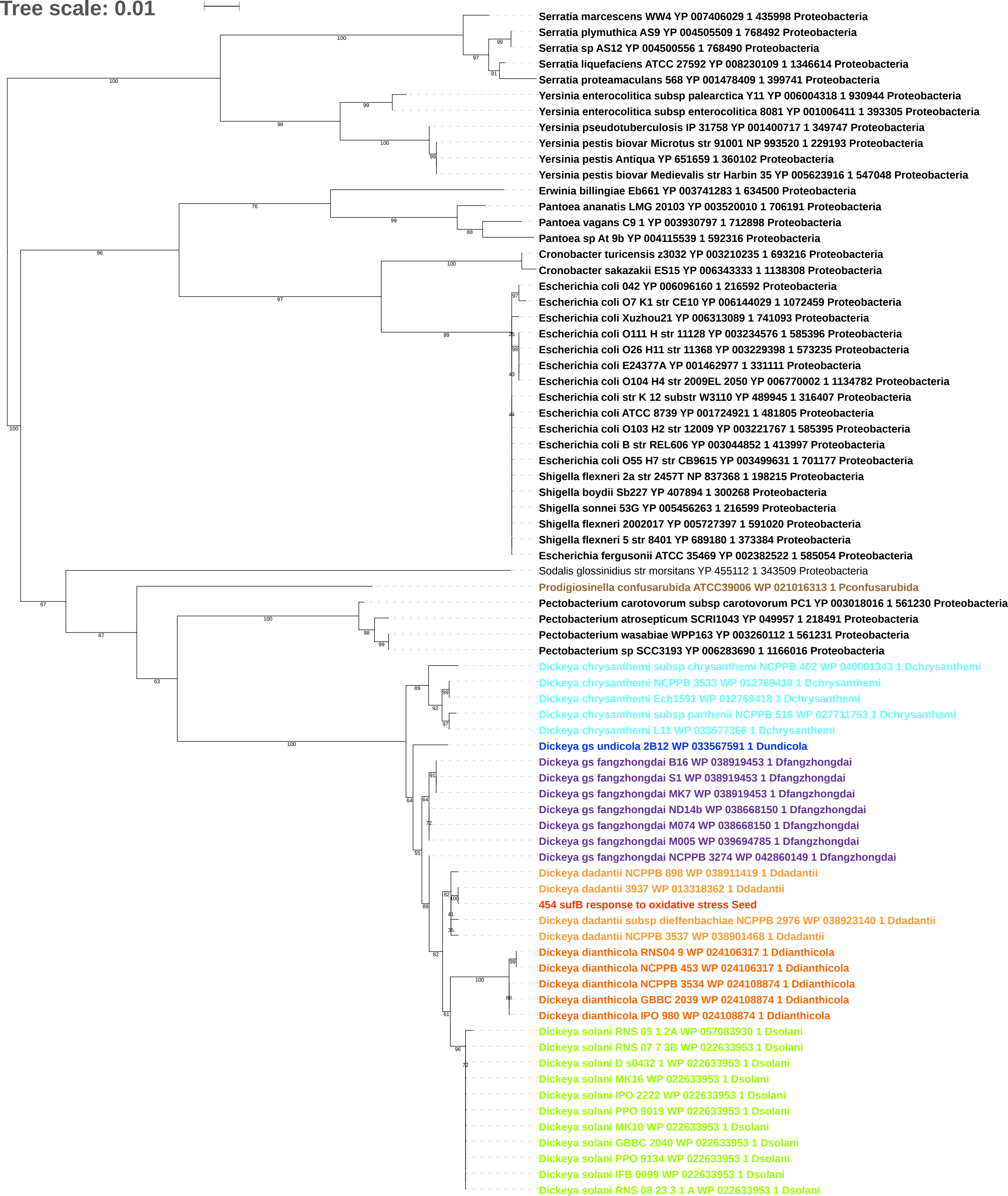
Tree scale: 0.01
Serratia marcescens WW4 YP 007406029 1 435998 Proteobacteria
Serratia plymuthica AS9 YP 004505509 1 768492 Proteobacteria
Serratia sp AS12 YP 004500556 1 768490 Proteobacteria
Serratia liquefaciens ATCC 27592 YP 008230109 1 1346614 Proteobacteria
Serratia proteamaculans 568 YP 001478409 1 399741 Proteobacteria
Yersinia enterocolitica subsp palearctica Y11 YP 006004318 1 930944 Proteobacteria
Yersinia enterocolitica subsp enterocolitica 8081 YP 001006411 1 393305 Proteobacteria
Yersinia pseudotuberculosis IP 31758 YP 001400717 1 349747 Proteobacteria
Yersinia pestis biovar Microtus str 91001 NP 993520 1 229193 Proteobacteria
Yersinia pestis Antiqua YP 651659 1 360102 Proteobacteria
Yersinia pestis biovar Medievalis str Harbin 35 YP 005623916 1 547048 Proteobacteria
Erwinia billingiae Eb661 YP 003741283 1 634500 Proteobacteria
Pantoea ananatis LMG 20103 YP 003520010 1 706191 Proteobacteria
Pantoea vagans C9 1 YP 003930797 1 712898 Proteobacteria
Pantoea sp At 9b YP 004115539 1 592316 Proteobacteria
Cronobacter turicensis z3032 YP 003210235 1 693216 Proteobacteria
Cronobacter sakazakii ES15 YP 006343333 1 1138308 Proteobacteria
Escherichia coli 042 YP 006096160 1 216592 Proteobacteria
Escherichia coli O7 K1 str CE10 YP 006144029 1 1072459 Proteobacteria
Escherichia coli Xuzhou21 YP 006313089 1 741093 Proteobacteria
Escherichia coli O111 H str 11128 YP 003234576 1 585396 Proteobacteria
Escherichia coli O26 H11 str 11368 YP 003229398 1 573235 Proteobacteria
Escherichia coli E24377A YP 001462977 1 331111 Proteobacteria
Escherichia coli O104 H4 str 2009EL 2050 YP 006770002 1 1134782 Proteobacteria
Escherichia coli str K 12 substr W3110 YP 489945 1 316407 Proteobacteria
Escherichia coli ATCC 8739 YP 001724921 1 481805 Proteobacteria
Escherichia coli O103 H2 str 12009 YP 003221767 1 585395 Proteobacteria
Escherichia coli B str REL606 YP 003044852 1 413997 Proteobacteria
Escherichia coli O55 H7 str CB9615 YP 003499631 1 701177 Proteobacteria
Shigella flexneri 2a str 2457T NP 837368 1 198215 Proteobacteria
Shigella boydii Sb227 YP 407894 1 300268 Proteobacteria
Shigella sonnei 53G YP 005456263 1 216599 Proteobacteria
Shigella flexneri 2002017 YP 005727397 1 591020 Proteobacteria
Shigella flexneri 5 str 8401 YP 689180 1 373384 Proteobacteria
Escherichia fergusonii ATCC 35469 YP 002382522 1 585054 Proteobacteria
Sodalis glossinidius str morsitans YP 455112 1 343509 Proteobacteria
Prodigiosinella confusarubida ATCC39006 WP 021016313 1 Pconfusarubida
Pectobacterium carotovorum subsp carotovorum PC1 YP 003018016 1 561230 Proteobacteria
Pectobacterium atrosepticum SCRI1043 YP 049957 1 218491 Proteobacteria
Pectobacterium wasabiae WPP163 YP 003260112 1 561231 Proteobacteria
Pectobacterium sp SCC3193 YP 006283690 1 1166016 Proteobacteria
Dickeya chrysanthemi subsp chrysanthemi NCPPB 402 WP 040001343 1 Dchrysanthemi
Dickeya chrysanthemi NCPPB 3533 WP 012769418 1 Dchrysanthemi
Dickeya chrysanthemi Ech1591 WP 012769418 1 Dchrysanthemi
Dickeya chrysanthemi subsp parthenii NCPPB 516 WP 027711763 1 Dchrysanthemi
Dickeya chrysanthemi L11 WP 033577366 1 Dchrysanthemi
Dickeya gs undicola 2B12 WP 033567591 1 Dundicola
Dickeya gs fangzhongdai B16 WP 038919453 1 Dfangzhongdai
Dickeya gs fangzhongdai S1 WP 038919453 1 Dfangzhongdai
Dickeya gs fangzhongdai MK7 WP 038919453 1 Dfangzhongdai
Dickeya gs fangzhongdai ND14b WP 038668150 1 Dfangzhongdai
Dickeya gs fangzhongdai M074 WP 038668150 1 Dfangzhongdai
Dickeya gs fangzhongdai M005 WP 039694785 1 Dfangzhongdai
Dickeya gs fangzhongdai NCPPB 3274 WP 042860149 1 Dfangzhongdai
Dickeya dadantii NCPPB 898 WP 038911419 1 Ddadantii
Dickeya dadantii 3937 WP 013318362 1 Ddadantii
454 sufB response to oxidative stress Seed
Dickeya dadantii subsp dieffenbachiae NCPPB 2976 WP 038923140 1 Ddadantii
Dickeya dadantii NCPPB 3537 WP 038901468 1 Ddadantii
Dickeya dianthicola RNS04 9 WP 024106317 1 Ddianthicola
Dickeya dianthicola NCPPB 453 WP 024106317 1 Ddianthicola
Dickeya dianthicola NCPPB 3534 WP 024108874 1 Ddianthicola
Dickeya dianthicola GBBC 2039 WP 024108874 1 Ddianthicola
Dickeya dianthicola IPO 980 WP 024108874 1 Ddianthicola
Dickeya solani RNS 05 1 2A WP 057083930 1 Dsolani
Dickeya solani RNS 07 7 3B WP 022633953 1 Dsolani
Dickeya solani D s0432 1 WP 022633953 1 Dsolani
Dickeya solani MK16 WP 022633953 1 Dsolani
Dickeya solani IPO 2222 WP 022633953 1 Dsolani
Dickeya solani PPO 9019 WP 022633953 1 Dsolani
Dickeya solani MK10 WP 022633953 1 Dsolani
Dickeya solani GBBC 2040 WP 022633953 1 Dsolani
Dickeya solani PPO 9134 WP 022633953 1 Dsolani
Dickeya solani IFB 0099 WP 022633953 1 Dsolani
Dickeya solani RNS 08 23 3 1 A WP 022633953 1 Dsolani

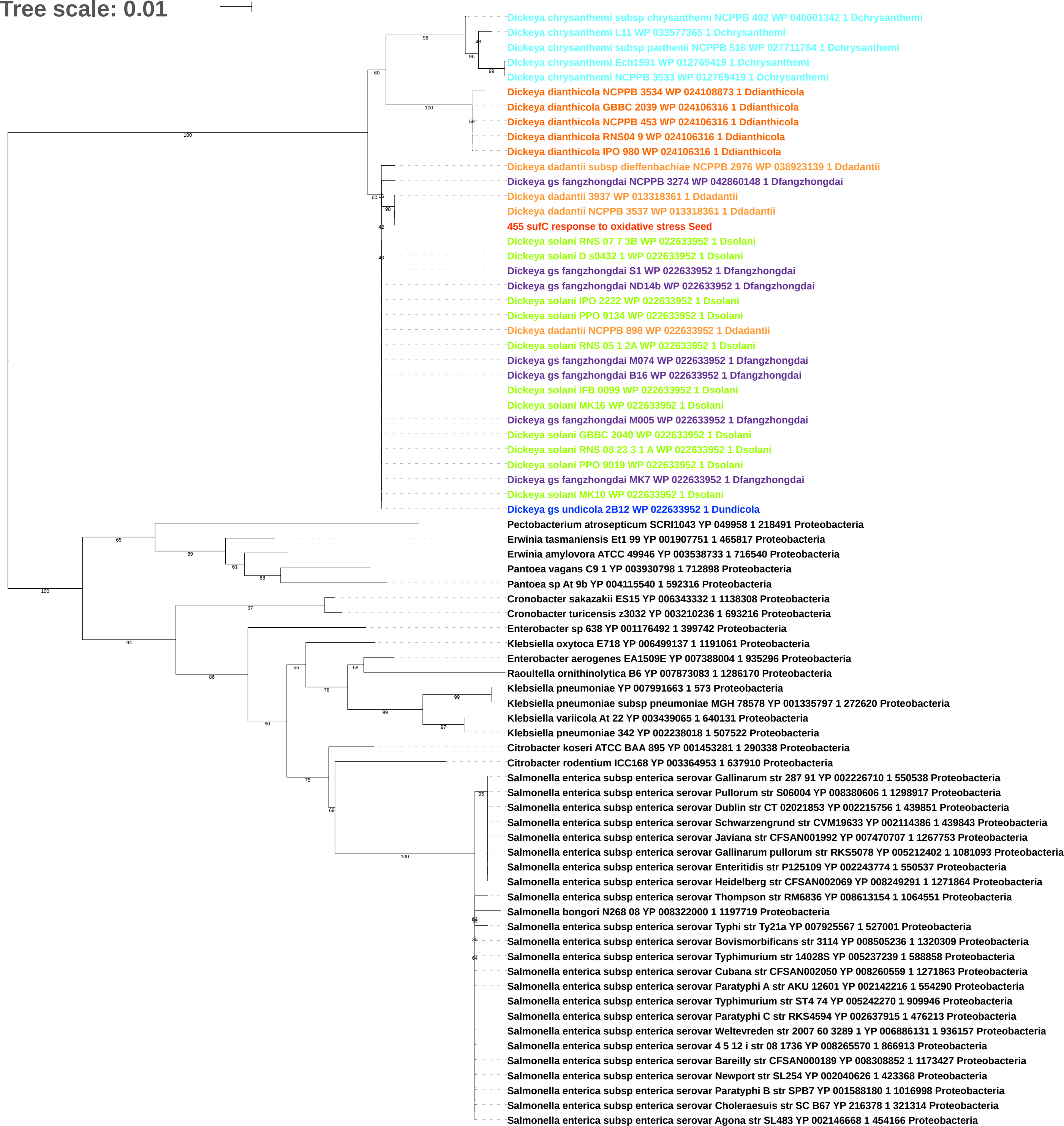
Tree scale: 0.01
Dickeya chrysanthemi subsp chrysanthemi NCPPB 402 WP 040001342 1 Dchrysanthemi
Dickeya chrysanthemi L11 WP 033577365 1 Dchrysanthemi
Dickeya chrysanthemi subsp parthenii NCPPB 516 WP 027711764 1 Dchrysanthemi
Dickeya chrysanthemi Ech1591 WP 012769419 1 Dchrysanthemi
Dickeya chrysanthemi NCPPB 3533 WP 012769419 1 Dchrysanthemi
Dickeya dianthicola NCPPB 3534 WP 024108873 1 Ddianthicola
Dickeya dianthicola GBBC 2039 WP 024106316 1 Ddianthicola
Dickeya dianthicola NCPPB 453 WP 024106316 1 Ddianthicola
Dickeya dianthicola RNS04 9 WP 024106316 1 Ddianthicola
Dickeya dianthicola IPO 980 WP 024106316 1 Ddianthicola
Dickeya dadantii subsp dieffenbachiae NCPPB 2976 WP 038923139 1 Ddadantii
Dickeya gs fangzhongdai NCPPB 3274 WP 042860148 1 Dfangzhongdai
Dickeya dadantii 3937 WP 013318361 1 Ddadantii
Dickeya dadantii NCPPB 3537 WP 013318361 1 Ddadantii
455 sufC response to oxidative stress Seed
Dickeya solani RNS 07 7 3B WP 022633952 1 Dsolani
Dickeya solani D s0432 1 WP 022633952 1 Dsolani
Dickeya gs fangzhongdai S1 WP 022633952 1 Dfangzhongdai
Dickeya gs fangzhongdai ND14b WP 022633952 1 Dfangzhongdai
Dickeya solani IPO 2222 WP 022633952 1 Dsolani
Dickeya solani PPO 9134 WP 022633952 1 Dsolani
Dickeya dadantii NCPPB 898 WP 022633952 1 Ddadantii
Dickeya solani RNS 05 1 2A WP 022633952 1 Dsolani
Dickeya gs fangzhongdai M074 WP 022633952 1 Dfangzhongdai
Dickeya gs fangzhongdai B16 WP 022633952 1 Dfangzhongdai
Dickeya solani IFB 0099 WP 022633952 1 Dsolani
Dickeya solani MK16 WP 022633952 1 Dsolani
Dickeya gs fangzhongdai M005 WP 022633952 1 Dfangzhongdai
Dickeya solani GBBC 2040 WP 022633952 1 Dsolani
Dickeya solani RNS 08 23 3 1 A WP 022633952 1 Dsolani
Dickeya solani PPO 9019 WP 022633952 1 Dsolani
Dickeya gs fangzhongdai MK7 WP 022633952 1 Dfangzhongdai
Dickeya solani MK10 WP 022633952 1 Dsolani
Dickeya gs undicola 2B12 WP 022633952 1 Dundicola
Pectobacterium atrosepticum SCRI1043 YP 049958 1 218491 Proteobacteria
Erwinia tasmaniensis Et1 99 YP 001907751 1 465817 Proteobacteria
Erwinia amylovora ATCC 49946 YP 003538733 1 716540 Proteobacteria
Pantoea vagans C9 1 YP 003930798 1 712898 Proteobacteria
Pantoea sp At 9b YP 004115540 1 592316 Proteobacteria
Cronobacter sakazakii ES15 YP 006343332 1 1138308 Proteobacteria
Cronobacter turicensis z3032 YP 003210236 1 693216 Proteobacteria
Enterobacter sp 638 YP 001176492 1 399742 Proteobacteria
Klebsiella oxytoca E718 YP 006499137 1 1191061 Proteobacteria
Enterobacter aerogenes EA1509E YP 007388004 1 935296 Proteobacteria
Raoultella ornithinolytica B6 YP 007873083 1 1286170 Proteobacteria
Klebsiella pneumoniae YP 007991663 1 573 Proteobacteria
Klebsiella pneumoniae subsp pneumoniae MGH 78578 YP 001335797 1 272620 Proteobacteria
Klebsiella variicola At 22 YP 003439065 1 640131 Proteobacteria
Klebsiella pneumoniae 342 YP 002238018 1 507522 Proteobacteria
Citrobacter koseri ATCC BAA 895 YP 001453281 1 290338 Proteobacteria
Citrobacter rodentium ICC168 YP 003364953 1 637910 Proteobacteria
Salmonella enterica subsp enterica serovar Gallinarum str 287 91 YP 002226710 1 550538 Proteobacteria
Salmonella enterica subsp enterica serovar Pullorum str S06004 YP 008380606 1 1298917 Proteobacteria
Salmonella enterica subsp enterica serovar Dublin str CT 02021853 YP 002215756 1 439851 Proteobacteria
Salmonella enterica subsp enterica serovar Schwarzengrund str CVM19633 YP 002114386 1 439843 Proteobacteria
Salmonella enterica subsp enterica serovar Javiana str CFSAN001992 YP 007470707 1 1267753 Proteobacteria
Salmonella enterica subsp enterica serovar Gallinarum pullorum str RKS5078 YP 005212402 1 1081093 Proteobacteria
Salmonella enterica subsp enterica serovar Enteritidis str P125109 YP 002243774 1 550537 Proteobacteria
Salmonella enterica subsp enterica serovar Heidelberg str CFSAN002069 YP 008249291 1 1271864 Proteobacteria
Salmonella enterica subsp enterica serovar Thompson str RM6836 YP 008613154 1 1064551 Proteobacteria
Salmonella bongori N268 08 YP 008322000 1 1197719 Proteobacteria
Salmonella enterica subsp enterica serovar Typhi str Ty21a YP 007925567 1 527001 Proteobacteria
Salmonella enterica subsp enterica serovar Bovismorbificans str 3114 YP 008505236 1 1320309 Proteobacteria
Salmonella enterica subsp enterica serovar Typhimurium str 14028S YP 005237239 1 588858 Proteobacteria
Salmonella enterica subsp enterica serovar Cubana str CFSAN002050 YP 008260559 1 1271863 Proteobacteria
Salmonella enterica subsp enterica serovar Paratyphi A str AKU 12601 YP 002142216 1 554290 Proteobacteria
Salmonella enterica subsp enterica serovar Typhimurium str ST4 74 YP 005242270 1 909946 Proteobacteria
Salmonella enterica subsp enterica serovar Paratyphi C str RKS4594 YP 002637915 1 476213 Proteobacteria
Salmonella enterica subsp enterica serovar Weltevreden str 2007 60 3289 1 YP 006886131 1 936157 Proteobacteria
Salmonella enterica subsp enterica serovar 4 5 12 i str 08 1736 YP 008265570 1 866913 Proteobacteria
Salmonella enterica subsp enterica serovar Bareilly str CFSAN000189 YP 008308852 1 1173427 Proteobacteria
Salmonella enterica subsp enterica serovar Newport str SL254 YP 002040626 1 423368 Proteobacteria
Salmonella enterica subsp enterica serovar Paratyphi B str SPB7 YP 001588180 1 1016998 Proteobacteria
Salmonella enterica subsp enterica serovar Choleraesuis str SC B67 YP 216378 1 321314 Proteobacteria
Salmonella enterica subsp enterica serovar Agona str SL483 YP 002146668 1 454166 Proteobacteria

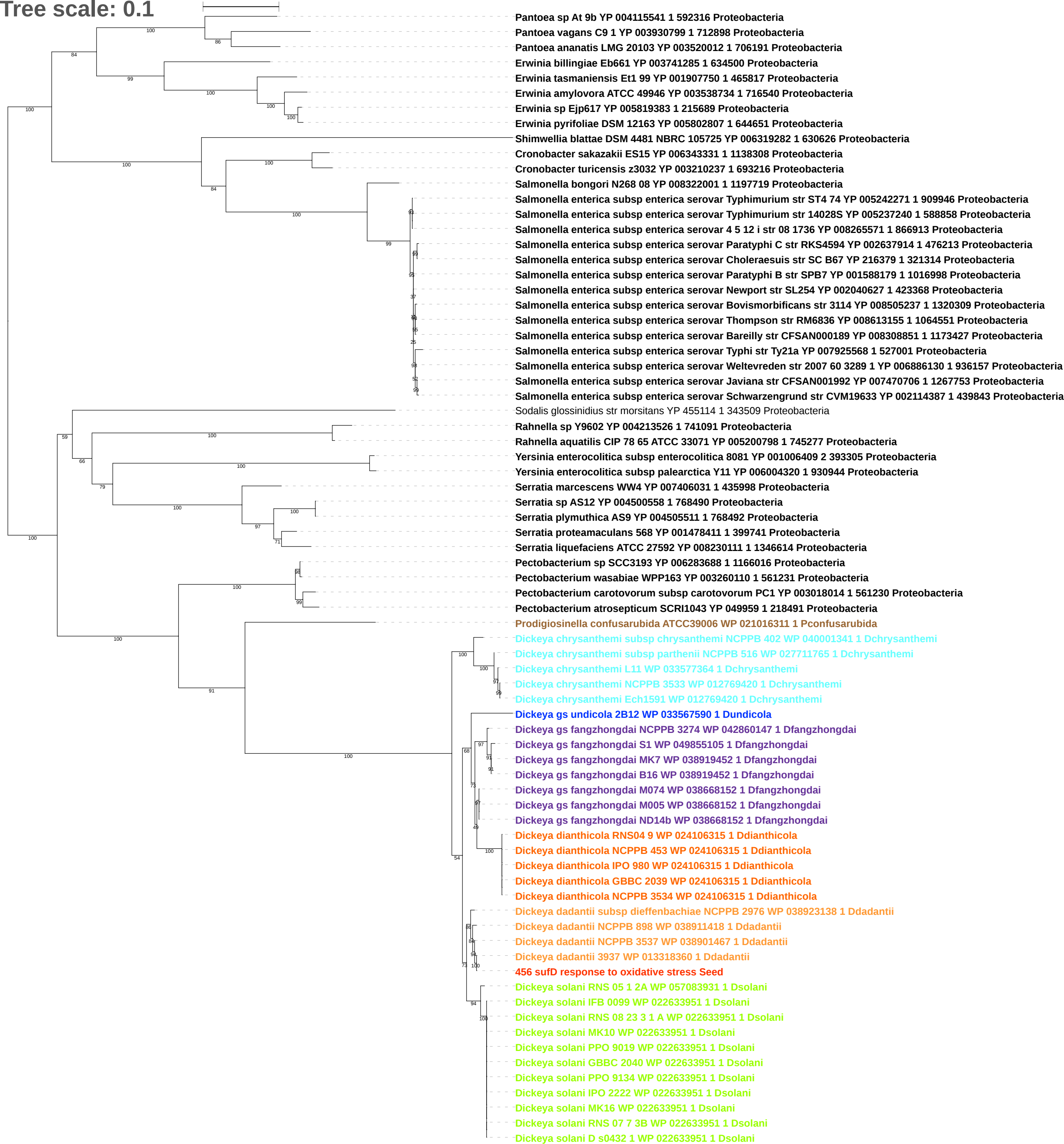

Tree scale: 0.1
Pantoea sp At 9b YP 004115541 1 592316 Proteobacteria
Pantoea vagans C9 1 YP 003930799 1 712898 Proteobacteria
Pantoea ananatis LMG 20103 YP 003520012 1 706191 Proteobacteria
Erwinia billingiae Eb661 YP 003741285 1 634500 Proteobacteria
Erwinia tasmaniensis Et1 99 YP 001907750 1 465817 Proteobacteria
Erwinia amylovora ATCC 49946 YP 003538734 1 716540 Proteobacteria
Erwinia sp Ejp617 YP 005819383 1 215689 Proteobacteria
Erwinia pyrifoliae DSM 12163 YP 005802807 1 644651 Proteobacteria
Shimwellia blattae DSM 4481 NBRC 105725 YP 006319282 1 630626 Proteobacteria
Cronobacter sakazakii ES15 YP 006343331 1 1138308 Proteobacteria
Cronobacter turicensis z3032 YP 003210237 1 693216 Proteobacteria
Salmonella bongori N268 08 YP 008322001 1 1197719 Proteobacteria
Salmonella enterica subsp enterica serovar Typhimurium str ST4 74 YP 005242271 1 909946 Proteobacteria
Salmonella enterica subsp enterica serovar Typhimurium str 14028S YP 005237240 1 588858 Proteobacteria
Salmonella enterica subsp enterica serovar 4 5 12 i str 08 1736 YP 008265571 1 866913 Proteobacteria
Salmonella enterica subsp enterica serovar Paratyphi C str RKS4594 YP 002637914 1 476213 Proteobacteria
Salmonella enterica subsp enterica serovar Choleraesuis str SC B67 YP 216379 1 321314 Proteobacteria
Salmonella enterica subsp enterica serovar Paratyphi B str SPB7 YP 001588179 1 1016998 Proteobacteria
Salmonella enterica subsp enterica serovar Newport str SL254 YP 002040627 1 423368 Proteobacteria
Salmonella enterica subsp enterica serovar Bovismorbificans str 3114 YP 008505237 1 1320309 Proteobacteria
Salmonella enterica subsp enterica serovar Thompson str RM6836 YP 008613155 1 1064551 Proteobacteria
Salmonella enterica subsp enterica serovar Bareilly str CFSAN000189 YP 008308851 1 1173427 Proteobacteria
Salmonella enterica subsp enterica serovar Typhi str Ty21a YP 007925568 1 527001 Proteobacteria
Salmonella enterica subsp enterica serovar Weltevreden str 2007 60 3289 1 YP 006886130 1 936157 Proteobacteria
Salmonella enterica subsp enterica serovar Javiana str CFSAN001992 YP 007470706 1 1267753 Proteobacteria
Salmonella enterica subsp enterica serovar Schwarzengrund str CVM19633 YP 002114387 1 439843 Proteobacteria
Sodalis glossinidius str morsitans YP 455114 1 343509 Proteobacteria
Rahnella sp Y9602 YP 004213526 1 741091 Proteobacteria
Rahnella aquatilis CIP 78 65 ATCC 33071 YP 005200798 1 745277 Proteobacteria
Yersinia enterocolitica subsp enterocolitica 8081 YP 001006409 2 393305 Proteobacteria
Yersinia enterocolitica subsp palearctica Y11 YP 006004320 1 930944 Proteobacteria
Serratia marcescens WW4 YP 007406031 1 435998 Proteobacteria
Serratia sp AS12 YP 004500558 1 768490 Proteobacteria
Serratia plymuthica AS9 YP 004505511 1 768492 Proteobacteria
Serratia proteamaculans 568 YP 001478411 1 399741 Proteobacteria
Serratia liquefaciens ATCC 27592 YP 008230111 1 1346614 Proteobacteria
Pectobacterium sp SCC3193 YP 006283688 1 1166016 Proteobacteria
Pectobacterium wasabiae WPP163 YP 003260110 1 561231 Proteobacteria
Pectobacterium carotovorum subsp carotovorum PC1 YP 003018014 1 561230 Proteobacteria
Pectobacterium atrosepticum SCRI1043 YP 049959 1 218491 Proteobacteria
Prodigiosinella confusarubida ATCC39006 WP 021016311 1 Pconfusarubida
Dickeya chrysanthemi subsp chrysanthemi NCPPB 402 WP 040001341 1 Dchrysanthemi
Dickeya chrysanthemi subsp parthenii NCPPB 516 WP 027711765 1 Dchrysanthemi
Dickeya chrysanthemi L11 WP 033577364 1 Dchrysanthemi
Dickeya chrysanthemi NCPPB 3533 WP 012769420 1 Dchrysanthemi
Dickeya chrysanthemi Ech1591 WP 012769420 1 Dchrysanthemi
Dickeya gs undicola 2B12 WP 033567590 1 Dundicola
Dickeya gs fangzhongdai NCPPB 3274 WP 042860147 1 Dfangzhongdai
Dickeya gs fangzhongdai S1 WP 049855105 1 Dfangzhongdai
Dickeya gs fangzhongdai MK7 WP 038919452 1 Dfangzhongdai
Dickeya gs fangzhongdai B16 WP 038919452 1 Dfangzhongdai
Dickeya gs fangzhongdai M074 WP 038668152 1 Dfangzhongdai
Dickeya gs fangzhongdai M005 WP 038668152 1 Dfangzhongdai
Dickeya gs fangzhongdai ND14b WP 038668152 1 Dfangzhongdai
Dickeya dianthicola RNS04 9 WP 024106315 1 Ddianthicola
Dickeya dianthicola NCPPB 453 WP 024106315 1 Ddianthicola
Dickeya dianthicola IPO 980 WP 024106315 1 Ddianthicola
Dickeya dianthicola GBBC 2039 WP 024106315 1 Ddianthicola
Dickeya dianthicola NCPPB 3534 WP 024106315 1 Ddianthicola
Dickeya dadantii subsp dieffenbachiae NCPPB 2976 WP 038923138 1 Ddadantii
Dickeya dadantii NCPPB 898 WP 038911418 1 Ddadantii
Dickeya dadantii NCPPB 3537 WP 038901467 1 Ddadantii
Dickeya dadantii 3937 WP 013318360 1 Ddadantii
456 sufD response to oxidative stress Seed
Dickeya solani RNS 05 1 2A WP 057083931 1 Dsolani
Dickeya solani IFB 0099 WP 022633951 1 Dsolani
Dickeya solani RNS 08 23 3 1 A WP 022633951 1 Dsolani
Dickeya solani MK10 WP 022633951 1 Dsolani
Dickeya solani PPO 9019 WP 022633951 1 Dsolani
Dickeya solani GBBC 2040 WP 022633951 1 Dsolani
Dickeya solani PPO 9134 WP 022633951 1 Dsolani
Dickeya solani IPO 2222 WP 022633951 1 Dsolani
Dickeya solani MK16 WP 022633951 1 Dsolani
Dickeya solani RNS 07 7 3B WP 022633951 1 Dsolani
Dickeya solani D s0432 1 WP 022633951 1 Dsolani

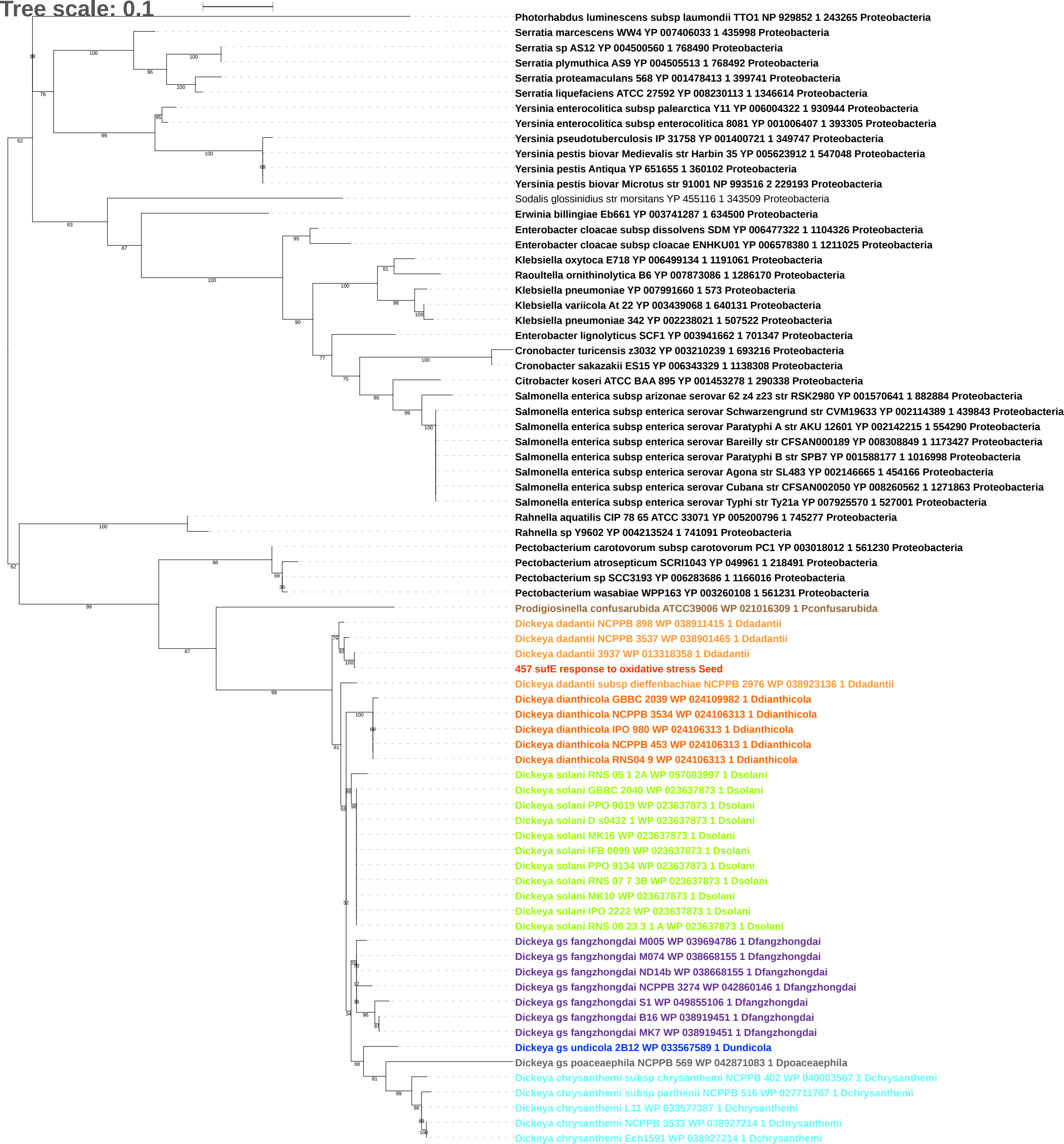
Tree scale: 0.1
Photorhabdus luminescens subsp laumondii TTO1 NP 929852 1 243265 Proteobacteria
Serratia marcescens WW4 YP 007406033 1 435998 Proteobacteria
Serratia sp AS12 YP 004500560 1 768490 Proteobacteria
Serratia plymuthica AS9 YP 004505513 1 768492 Proteobacteria
Serratia proteamaculans 568 YP 001478413 1 399741 Proteobacteria
Serratia liquefaciens ATCC 27592 YP 008230113 1 1346614 Proteobacteria
Yersinia enterocolitica subsp palearctica Y11 YP 006004322 1 930944 Proteobacteria
Yersinia enterocolitica subsp enterocolitica 8081 YP 001006407 1 393305 Proteobacteria
Yersinia pseudotuberculosis IP 31758 YP 001400721 1 349747 Proteobacteria
Yersinia pestis biovar Medievalis str Harbin 35 YP 005623912 1 547048 Proteobacteria
Yersinia pestis Antiqua YP 651655 1 360102 Proteobacteria
Yersinia pestis biovar Microtus str 91001 NP 993516 2 229193 Proteobacteria
Sodalis glossinidius str morsitans YP 455116 1 343509 Proteobacteria
Erwinia billingiae Eb661 YP 003741287 1 634500 Proteobacteria
Enterobacter cloacae subsp dissolvens SDM YP 006477322 1 1104326 Proteobacteria
Enterobacter cloacae subsp cloacae ENHKU01 YP 006578380 1 1211025 Proteobacteria
Klebsiella oxytoca E718 YP 006499134 1 1191061 Proteobacteria
Raoultella ornithinolytica B6 YP 007873086 1 1286170 Proteobacteria
Klebsiella pneumoniae YP 007991660 1 573 Proteobacteria
Klebsiella variicola At 22 YP 003439068 1 640131 Proteobacteria
Klebsiella pneumoniae 342 YP 002238021 1 507522 Proteobacteria
Enterobacter lignolyticus SCF1 YP 003941662 1 701347 Proteobacteria
Cronobacter turicensis z3032 YP 003210239 1 693216 Proteobacteria
Cronobacter sakazakii ES15 YP 006343329 1 1138308 Proteobacteria
Citrobacter koseri ATCC BAA 895 YP 001453278 1 290338 Proteobacteria
Salmonella enterica subsp arizonae serovar 62 z4 z23 str RSK2980 YP 001570641 1 882884 Proteobacteria
Salmonella enterica subsp enterica serovar Schwarzengrund str CVM19633 YP 002114389 1 439843 Proteobacteria
Salmonella enterica subsp enterica serovar Paratyphi A str AKU 12601 YP 002142215 1 554290 Proteobacteria
Salmonella enterica subsp enterica serovar Bareilly str CFSAN000189 YP 008308849 1 1173427 Proteobacteria
Salmonella enterica subsp enterica serovar Paratyphi B str SPB7 YP 001588177 1 1016998 Proteobacteria
Salmonella enterica subsp enterica serovar Agona str SL483 YP 002146665 1 454166 Proteobacteria
Salmonella enterica subsp enterica serovar Cubana str CFSAN002050 YP 008260562 1 1271863 Proteobacteria
Salmonella enterica subsp enterica serovar Typhi str Ty21a YP 007925570 1 527001 Proteobacteria
Rahnella aquatilis CIP 78 65 ATCC 33071 YP 005200796 1 745277 Proteobacteria
Rahnella sp Y9602 YP 004213524 1 741091 Proteobacteria
Pectobacterium carotovorum subsp carotovorum PC1 YP 003018012 1 561230 Proteobacteria
Pectobacterium atrosepticum SCRI1043 YP 049961 1 218491 Proteobacteria
Pectobacterium sp SCC3193 YP 006283686 1 1166016 Proteobacteria
Pectobacterium wasabiae WPP163 YP 003260108 1 561231 Proteobacteria
Prodigiosinella confusarubida ATCC39006 WP 021016309 1 Pconfusarubida
Dickeya dadantii NCPPB 898 WP 038911415 1 Ddadantii
Dickeya dadantii NCPPB 3537 WP 038901465 1 Ddadantii
Dickeya dadantii 3937 WP 013318358 1 Ddadantii
457 sufE response to oxidative stress Seed
Dickeya dadantii subsp dieffenbachiae NCPPB 2976 WP 038923136 1 Ddadantii
Dickeya dianthicola GBBC 2039 WP 024109982 1 Ddianthicola
Dickeya dianthicola NCPPB 3534 WP 024106313 1 Ddianthicola
Dickeya dianthicola IPO 980 WP 024106313 1 Ddianthicola
Dickeya dianthicola NCPPB 453 WP 024106313 1 Ddianthicola
Dickeya dianthicola RNS04 9 WP 024106313 1 Ddianthicola
Dickeya solani RNS 05 1 2A WP 057083997 1 Dsolani
Dickeya solani GBBC 2040 WP 023637873 1 Dsolani
Dickeya solani PPO 9019 WP 023637873 1 Dsolani
Dickeya solani D s0432 1 WP 023637873 1 Dsolani
Dickeya solani MK16 WP 023637873 1 Dsolani
Dickeya solani IFB 0099 WP 023637873 1 Dsolani
Dickeya solani PPO 9134 WP 023637873 1 Dsolani
Dickeya solani RNS 07 7 3B WP 023637873 1 Dsolani
Dickeya solani MK10 WP 023637873 1 Dsolani
Dickeya solani IPO 2222 WP 023637873 1 Dsolani
Dickeya solani RNS 08 23 3 1 A WP 023637873 1 Dsolani
Dickeya gs fangzhongdai M005 WP 039694786 1 Dfangzhongdai
Dickeya gs fangzhongdai M074 WP 038668155 1 Dfangzhongdai
Dickeya gs fangzhongdai ND14b WP 038668155 1 Dfangzhongdai
Dickeya gs fangzhongdai NCPPB 3274 WP 042860146 1 Dfangzhongdai
Dickeya gs fangzhongdai S1 WP 049855106 1 Dfangzhongdai
Dickeya gs fangzhongdai B16 WP 038919451 1 Dfangzhongdai
Dickeya gs fangzhongdai MK7 WP 038919451 1 Dfangzhongdai
Dickeya gs undicola 2B12 WP 033567589 1 Dundicola
Dickeya gs poaceaephila NCPPB 569 WP 042871083 1 Dpoaceaephila
Dickeya chrysanthemi subsp chrysanthemi NCPPB 402 WP 040003567 1 Dchrysanthemi
Dickeya chrysanthemi subsp parthenii NCPPB 516 WP 027711767 1 Dchrysanthemi
Dickeya chrysanthemi L11 WP 033577387 1 Dchrysanthemi
Dickeya chrysanthemi NCPPB 3533 WP 038927214 1 Dchrysanthemi
Dickeya chrysanthemi Ech1591 WP 038927214 1 Dchrysanthemi

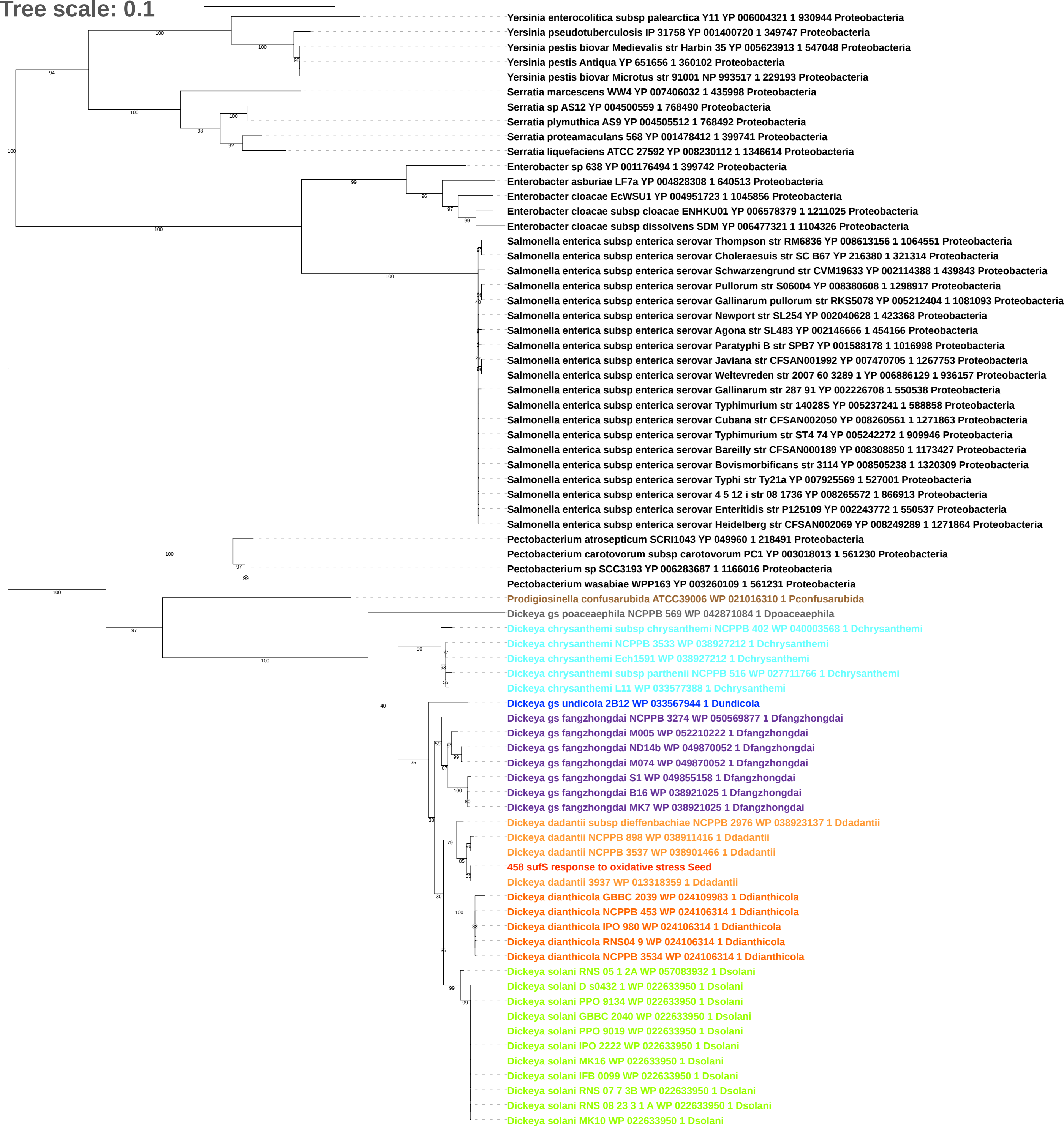

Tree scale: 0.1
Yersinia enterocolitica subsp palearctica Y11 YP 006004321 1 930944 Proteobacteria
Yersinia pseudotuberculosis IP 31758 YP 001400720 1 349747 Proteobacteria
Yersinia pestis biovar Medievalis str Harbin 35 YP 005623913 1 547048 Proteobacteria
Yersinia pestis Antiqua YP 651656 1 360102 Proteobacteria
Yersinia pestis biovar Microtus str 91001 NP 993517 1 229193 Proteobacteria
Serratia marcescens WW4 YP 007406032 1 435998 Proteobacteria
Serratia sp AS12 YP 004500559 1 768490 Proteobacteria
Serratia plymuthica AS9 YP 004505512 1 768492 Proteobacteria
Serratia proteamaculans 568 YP 001478412 1 399741 Proteobacteria
Serratia liquefaciens ATCC 27592 YP 008230112 1 1346614 Proteobacteria
Enterobacter sp 638 YP 001176494 1 399742 Proteobacteria
Enterobacter asburiae LF7a YP 004828308 1 640513 Proteobacteria
Enterobacter cloacae EcWSU1 YP 004951723 1 1045856 Proteobacteria
Enterobacter cloacae subsp cloacae ENHKU01 YP 006578379 1 1211025 Proteobacteria
Enterobacter cloacae subsp dissolvens SDM YP 006477321 1 1104326 Proteobacteria
Salmonella enterica subsp enterica serovar Thompson str RM6836 YP 008613156 1 1064551 Proteobacteria
Salmonella enterica subsp enterica serovar Choleraesuis str SC B67 YP 216380 1 321314 Proteobacteria
Salmonella enterica subsp enterica serovar Schwarzengrund str CVM19633 YP 002114388 1 439843 Proteobacteria
Salmonella enterica subsp enterica serovar Pullorum str S06004 YP 008380608 1 1298917 Proteobacteria
Salmonella enterica subsp enterica serovar Gallinarum pullorum str RKS5078 YP 005212404 1 1081093 Proteobacteria
Salmonella enterica subsp enterica serovar Newport str SL254 YP 002040628 1 423368 Proteobacteria
Salmonella enterica subsp enterica serovar Agona str SL483 YP 002146666 1 454166 Proteobacteria
Salmonella enterica subsp enterica serovar Paratyphi B str SPB7 YP 001588178 1 1016998 Proteobacteria
Salmonella enterica subsp enterica serovar Javiana str CFSAN001992 YP 007470705 1 1267753 Proteobacteria
Salmonella enterica subsp enterica serovar Weltevreden str 2007 60 3289 1 YP 006886129 1 936157 Proteobacteria
Salmonella enterica subsp enterica serovar Gallinarum str 287 91 YP 002226708 1 550538 Proteobacteria
Salmonella enterica subsp enterica serovar Typhimurium str 14028S YP 005237241 1 588858 Proteobacteria
Salmonella enterica subsp enterica serovar Cubana str CFSAN002050 YP 008260561 1 1271863 Proteobacteria
Salmonella enterica subsp enterica serovar Typhimurium str ST4 74 YP 005242272 1 909946 Proteobacteria
Salmonella enterica subsp enterica serovar Bareilly str CFSAN000189 YP 008308850 1 1173427 Proteobacteria
Salmonella enterica subsp enterica serovar Bovismorbificans str 3114 YP 008505238 1 1320309 Proteobacteria
Salmonella enterica subsp enterica serovar Typhi str Ty21a YP 007925569 1 527001 Proteobacteria
Salmonella enterica subsp enterica serovar 4 5 12 i str 08 1736 YP 008265572 1 866913 Proteobacteria
Salmonella enterica subsp enterica serovar Enteritidis str P125109 YP 002243772 1 550537 Proteobacteria
Salmonella enterica subsp enterica serovar Heidelberg str CFSAN002069 YP 008249289 1 1271864 Proteobacteria
Pectobacterium atrosepticum SCRI1043 YP 049960 1 218491 Proteobacteria
Pectobacterium carotovorum subsp carotovorum PC1 YP 003018013 1 561230 Proteobacteria
Pectobacterium sp SCC3193 YP 006283687 1 1166016 Proteobacteria
Pectobacterium wasabiae WPP163 YP 003260109 1 561231 Proteobacteria
Prodigiosinella confusarubida ATCC39006 WP 021016310 1 Pconfusarubida
Dickeya gs poaceaephila NCPPB 569 WP 042871084 1 Dpoaceaephila
Dickeya chrysanthemi subsp chrysanthemi NCPPB 402 WP 040003568 1 Dchrysanthemi
Dickeya chrysanthemi NCPPB 3533 WP 038927212 1 Dchrysanthemi
Dickeya chrysanthemi Ech1591 WP 038927212 1 Dchrysanthemi
Dickeya chrysanthemi subsp parthenii NCPPB 516 WP 027711766 1 Dchrysanthemi
Dickeya chrysanthemi L11 WP 033577388 1 Dchrysanthemi
Dickeya gs undicola 2B12 WP 033567944 1 Dundicola
Dickeya gs fangzhongdai NCPPB 3274 WP 050569877 1 Dfangzhongdai
Dickeya gs fangzhongdai M005 WP 052210222 1 Dfangzhongdai
Dickeya gs fangzhongdai ND14b WP 049870052 1 Dfangzhongdai
Dickeya gs fangzhongdai M074 WP 049870052 1 Dfangzhongdai
Dickeya gs fangzhongdai S1 WP 049855158 1 Dfangzhongdai
Dickeya gs fangzhongdai B16 WP 038921025 1 Dfangzhongdai
Dickeya gs fangzhongdai MK7 WP 038921025 1 Dfangzhongdai
Dickeya dadantii subsp dieffenbachiae NCPPB 2976 WP 038923137 1 Ddadantii
Dickeya dadantii NCPPB 898 WP 038911416 1 Ddadantii
Dickeya dadantii NCPPB 3537 WP 038901466 1 Ddadantii
458 sufS response to oxidative stress Seed
Dickeya dadantii 3937 WP 013318359 1 Ddadantii
Dickeya dianthicola GBBC 2039 WP 024109983 1 Ddianthicola
Dickeya dianthicola NCPPB 453 WP 024106314 1 Ddianthicola
Dickeya dianthicola IPO 980 WP 024106314 1 Ddianthicola
Dickeya dianthicola RNS04 9 WP 024106314 1 Ddianthicola
Dickeya dianthicola NCPPB 3534 WP 024106314 1 Ddianthicola
Dickeya solani RNS 05 1 2A WP 057083932 1 Dsolani
Dickeya solani D s0432 1 WP 022633950 1 Dsolani
Dickeya solani PPO 9134 WP 022633950 1 Dsolani
Dickeya solani GBBC 2040 WP 022633950 1 Dsolani
Dickeya solani PPO 9019 WP 022633950 1 Dsolani
Dickeya solani IPO 2222 WP 022633950 1 Dsolani
Dickeya solani MK16 WP 022633950 1 Dsolani
Dickeya solani IFB 0099 WP 022633950 1 Dsolani
Dickeya solani RNS 07 7 3B WP 022633950 1 Dsolani
Dickeya solani RNS 08 23 3 1 A WP 022633950 1 Dsolani
Dickeya solani MK10 WP 022633950 1 Dsolani

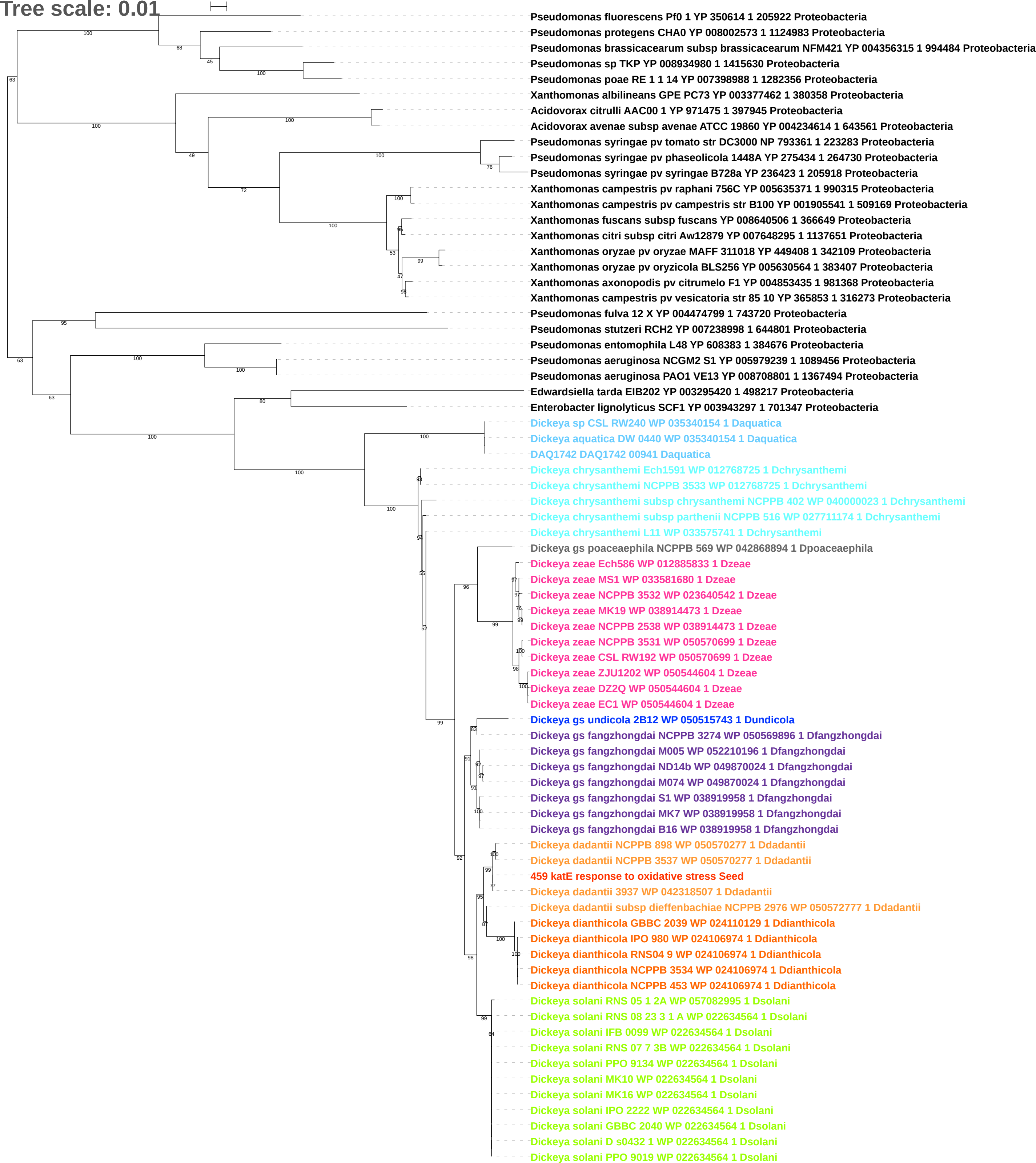
Tree scale: 0.01
Pseudomonas fluorescens Pf0 1 YP 350614 1 205922 Proteobacteria
Pseudomonas protegens CHA0 YP 008002573 1 1124983 Proteobacteria
Pseudomonas brassicacearum subsp brassicacearum NFM421 YP 004356315 1 994484 Proteobacteria
Pseudomonas sp TKP YP 008934980 1 1415630 Proteobacteria
Pseudomonas poae RE 1 1 14 YP 007398988 1 1282356 Proteobacteria
Xanthomonas albilineans GPE PC73 YP 003377462 1 380358 Proteobacteria
Acidovorax citrulli AAC00 1 YP 971475 1 397945 Proteobacteria
Acidovorax avenae subsp avenae ATCC 19860 YP 004234614 1 643561 Proteobacteria
Pseudomonas syringae pv tomato str DC3000 NP 793361 1 223283 Proteobacteria
Pseudomonas syringae pv phaseolicola 1448A YP 275434 1 264730 Proteobacteria
Pseudomonas syringae pv syringae B728a YP 236423 1 205918 Proteobacteria
Xanthomonas campestris pv raphani 756C YP 005635371 1 990315 Proteobacteria
Xanthomonas campestris pv campestris str B100 YP 001905541 1 509169 Proteobacteria
Xanthomonas fuscans subsp fuscans YP 008640506 1 366649 Proteobacteria
Xanthomonas citri subsp citri Aw12879 YP 007648295 1 1137651 Proteobacteria
Xanthomonas oryzae pv oryzae MAFF 311018 YP 449408 1 342109 Proteobacteria
Xanthomonas oryzae pv oryzicola BLS256 YP 005630564 1 383407 Proteobacteria
Xanthomonas axonopodis pv citrumelo F1 YP 004853435 1 981368 Proteobacteria
Xanthomonas campestris pv vesicatoria str 85 10 YP 365853 1 316273 Proteobacteria
Pseudomonas fulva 12 X YP 004474799 1 743720 Proteobacteria
Pseudomonas stutzeri RCH2 YP 007238998 1 644801 Proteobacteria
Pseudomonas entomophila L48 YP 608383 1 384676 Proteobacteria
Pseudomonas aeruginosa NCGM2 S1 YP 005979239 1 1089456 Proteobacteria
Pseudomonas aeruginosa PAO1 VE13 YP 008708801 1 1367494 Proteobacteria
Edwardsiella tarda EIB202 YP 003295420 1 498217 Proteobacteria
Enterobacter lignolyticus SCF1 YP 003943297 1 701347 Proteobacteria
Dickeya sp CSL RW240 WP 035340154 1 Daquatica
Dickeya aquatica DW 0440 WP 035340154 1 Daquatica
DAQ1742 DAQ1742 00941 Daquatica
Dickeya chrysanthemi Ech1591 WP 012768725 1 Dchrysanthemi
Dickeya chrysanthemi NCPPB 3533 WP 012768725 1 Dchrysanthemi
Dickeya chrysanthemi subsp chrysanthemi NCPPB 402 WP 040000023 1 Dchrysanthemi
Dickeya chrysanthemi subsp parthenii NCPPB 516 WP 027711174 1 Dchrysanthemi
Dickeya chrysanthemi L11 WP 033575741 1 Dchrysanthemi
Dickeya gs poaceaephila NCPPB 569 WP 042868894 1 Dpoaceaephila
Dickeya zeae Ech586 WP 012885833 1 Dzeae
Dickeya zeae MS1 WP 033581680 1 Dzeae
Dickeya zeae NCPPB 3532 WP 023640542 1 Dzeae
Dickeya zeae MK19 WP 038914473 1 Dzeae
Dickeya zeae NCPPB 2538 WP 038914473 1 Dzeae
Dickeya zeae NCPPB 3531 WP 050570699 1 Dzeae
Dickeya zeae CSL RW192 WP 050570699 1 Dzeae
Dickeya zeae ZJU1202 WP 050544604 1 Dzeae
Dickeya zeae DZ2Q WP 050544604 1 Dzeae
Dickeya zeae EC1 WP 050544604 1 Dzeae
Dickeya gs undicola 2B12 WP 050515743 1 Dundicola
Dickeya gs fangzhongdai NCPPB 3274 WP 050569896 1 Dfangzhongdai
Dickeya gs fangzhongdai M005 WP 052210196 1 Dfangzhongdai
Dickeya gs fangzhongdai ND14b WP 049870024 1 Dfangzhongdai
Dickeya gs fangzhongdai M074 WP 049870024 1 Dfangzhongdai
Dickeya gs fangzhongdai S1 WP 038919958 1 Dfangzhongdai
Dickeya gs fangzhongdai MK7 WP 038919958 1 Dfangzhongdai
Dickeya gs fangzhongdai B16 WP 038919958 1 Dfangzhongdai
Dickeya dadantii NCPPB 898 WP 050570277 1 Ddadantii
Dickeya dadantii NCPPB 3537 WP 050570277 1 Ddadantii
459 katE response to oxidative stress Seed
Dickeya dadantii 3937 WP 042318507 1 Ddadantii
Dickeya dadantii subsp dieffenbachiae NCPPB 2976 WP 050572777 1 Ddadantii
Dickeya dianthicola GBBC 2039 WP 024110129 1 Ddianthicola
Dickeya dianthicola IPO 980 WP 024106974 1 Ddianthicola
Dickeya dianthicola RNS04 9 WP 024106974 1 Ddianthicola
Dickeya dianthicola NCPPB 3534 WP 024106974 1 Ddianthicola
Dickeya dianthicola NCPPB 453 WP 024106974 1 Ddianthicola
Dickeya solani RNS 05 1 2A WP 057082995 1 Dsolani
Dickeya solani RNS 08 23 3 1 A WP 022634564 1 Dsolani
Dickeya solani IFB 0099 WP 022634564 1 Dsolani
Dickeya solani RNS 07 7 3B WP 022634564 1 Dsolani
Dickeya solani PPO 9134 WP 022634564 1 Dsolani
Dickeya solani MK10 WP 022634564 1 Dsolani
Dickeya solani MK16 WP 022634564 1 Dsolani
Dickeya solani IPO 2222 WP 022634564 1 Dsolani
Dickeya solani GBBC 2040 WP 022634564 1 Dsolani
Dickeya solani D s0432 1 WP 022634564 1 Dsolani
Dickeya solani PPO 9019 WP 022634564 1 Dsolani

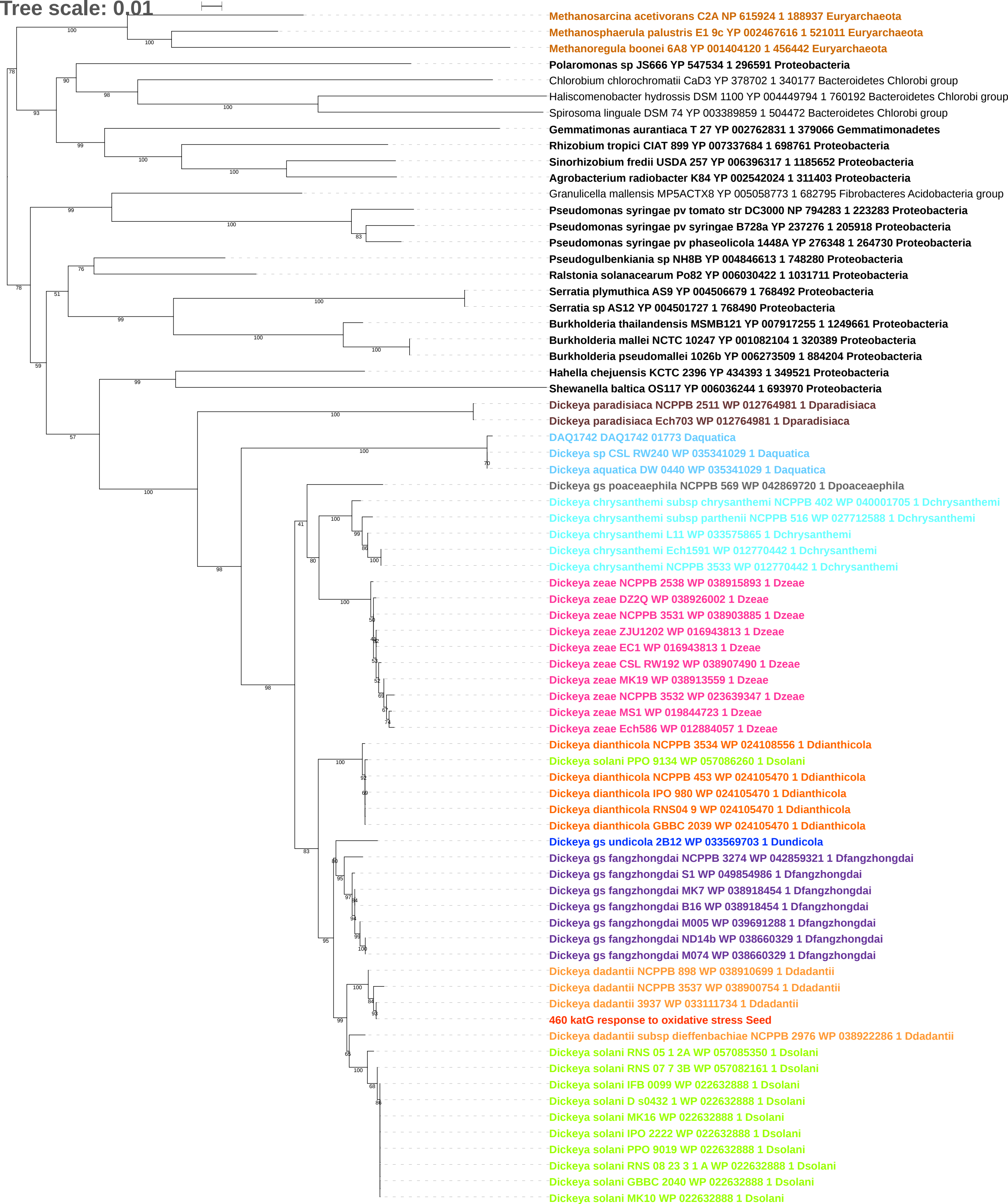
Tree scale: 0.01
Methanosarcina acetivorans C2A NP 615924 1 188937 Euryarchaeota
Methanosphaerula palustris E1 9c YP 002467616 1 521011 Euryarchaeota
Methanoregula boonei 6A8 YP 001404120 1 456442 Euryarchaeota
Polaromonas sp JS666 YP 547534 1 296591 Proteobacteria
Chlorobium chlorochromatii CaD3 YP 378702 1 340177 Bacteroidetes Chlorobi group
Haliscomenobacter hydrossis DSM 1100 YP 004449794 1 760192 Bacteroidetes Chlorobi group
Spirosoma linguale DSM 74 YP 003389859 1 504472 Bacteroidetes Chlorobi group
Gemmatimonas aurantiaca T 27 YP 002762831 1 379066 Gemmatimonadetes
Rhizobium tropici CIAT 899 YP 007337684 1 698761 Proteobacteria
Sinorhizobium fredii USDA 257 YP 006396317 1 1185652 Proteobacteria
Agrobacterium radiobacter K84 YP 002542024 1 311403 Proteobacteria
Granulicella mallensis MP5ACTX8 YP 005058773 1 682795 Fibrobacteres Acidobacteria group
Pseudomonas syringae pv tomato str DC3000 NP 794283 1 223283 Proteobacteria
Pseudomonas syringae pv syringae B728a YP 237276 1 205918 Proteobacteria
Pseudomonas syringae pv phaseolicola 1448A YP 276348 1 264730 Proteobacteria
Pseudogulbenkiania sp NH8B YP 004846613 1 748280 Proteobacteria
Ralstonia solanacearum Po82 YP 006030422 1 1031711 Proteobacteria
Serratia plymuthica AS9 YP 004506679 1 768492 Proteobacteria
Serratia sp AS12 YP 004501727 1 768490 Proteobacteria
Burkholderia thailandensis MSMB121 YP 007917255 1 1249661 Proteobacteria
Burkholderia mallei NCTC 10247 YP 001082104 1 320389 Proteobacteria
Burkholderia pseudomallei 1026b YP 006273509 1 884204 Proteobacteria
Hahella chejuensis KCTC 2396 YP 434393 1 349521 Proteobacteria
Shewanella baltica OS117 YP 006036244 1 693970 Proteobacteria
Dickeya paradisiaca NCPPB 2511 WP 012764981 1 Dparadisiaca
Dickeya paradisiaca Ech703 WP 012764981 1 Dparadisiaca
DAQ1742 DAQ1742 01773 Daquatica
Dickeya sp CSL RW240 WP 035341029 1 Daquatica
Dickeya aquatica DW 0440 WP 035341029 1 Daquatica
Dickeya gs poaceaephila NCPPB 569 WP 042869720 1 Dpoaceaephila
Dickeya chrysanthemi subsp chrysanthemi NCPPB 402 WP 040001705 1 Dchrysanthemi
Dickeya chrysanthemi subsp parthenii NCPPB 516 WP 027712588 1 Dchrysanthemi
Dickeya chrysanthemi L11 WP 033575865 1 Dchrysanthemi
Dickeya chrysanthemi Ech1591 WP 012770442 1 Dchrysanthemi
Dickeya chrysanthemi NCPPB 3533 WP 012770442 1 Dchrysanthemi
Dickeya zeae NCPPB 2538 WP 038915893 1 Dzeae
Dickeya zeae DZ2Q WP 038926002 1 Dzeae
Dickeya zeae NCPPB 3531 WP 038903885 1 Dzeae
Dickeya zeae ZJU1202 WP 016943813 1 Dzeae
Dickeya zeae EC1 WP 016943813 1 Dzeae
Dickeya zeae CSL RW192 WP 038907490 1 Dzeae
Dickeya zeae MK19 WP 038913559 1 Dzeae
Dickeya zeae NCPPB 3532 WP 023639347 1 Dzeae
Dickeya zeae MS1 WP 019844723 1 Dzeae
Dickeya zeae Ech586 WP 012884057 1 Dzeae
Dickeya dianthicola NCPPB 3534 WP 024108556 1 Ddianthicola
Dickeya solani PPO 9134 WP 057086260 1 Dsolani
Dickeya dianthicola NCPPB 453 WP 024105470 1 Ddianthicola
Dickeya dianthicola IPO 980 WP 024105470 1 Ddianthicola
Dickeya dianthicola RNS04 9 WP 024105470 1 Ddianthicola
Dickeya dianthicola GBBC 2039 WP 024105470 1 Ddianthicola
Dickeya gs undicola 2B12 WP 033569703 1 Dundicola
Dickeya gs fangzhongdai NCPPB 3274 WP 042859321 1 Dfangzhongdai
Dickeya gs fangzhongdai S1 WP 049854986 1 Dfangzhongdai
Dickeya gs fangzhongdai MK7 WP 038918454 1 Dfangzhongdai
Dickeya gs fangzhongdai B16 WP 038918454 1 Dfangzhongdai
Dickeya gs fangzhongdai M005 WP 039691288 1 Dfangzhongdai
Dickeya gs fangzhongdai ND14b WP 038660329 1 Dfangzhongdai
Dickeya gs fangzhongdai M074 WP 038660329 1 Dfangzhongdai
Dickeya dadantii NCPPB 898 WP 038910699 1 Ddadantii
Dickeya dadantii NCPPB 3537 WP 038900754 1 Ddadantii
Dickeya dadantii 3937 WP 033111734 1 Ddadantii
460 katG response to oxidative stress Seed
Dickeya dadantii subsp dieffenbachiae NCPPB 2976 WP 038922286 1 Ddadantii
Dickeya solani RNS 05 1 2A WP 057085350 1 Dsolani
Dickeya solani RNS 07 7 3B WP 057082161 1 Dsolani
Dickeya solani IFB 0099 WP 022632888 1 Dsolani
Dickeya solani D s0432 1 WP 022632888 1 Dsolani
Dickeya solani MK16 WP 022632888 1 Dsolani
Dickeya solani IPO 2222 WP 022632888 1 Dsolani
Dickeya solani PPO 9019 WP 022632888 1 Dsolani
Dickeya solani RNS 08 23 3 1 A WP 022632888 1 Dsolani
Dickeya solani GBBC 2040 WP 022632888 1 Dsolani
Dickeya solani MK10 WP 022632888 1 Dsolani

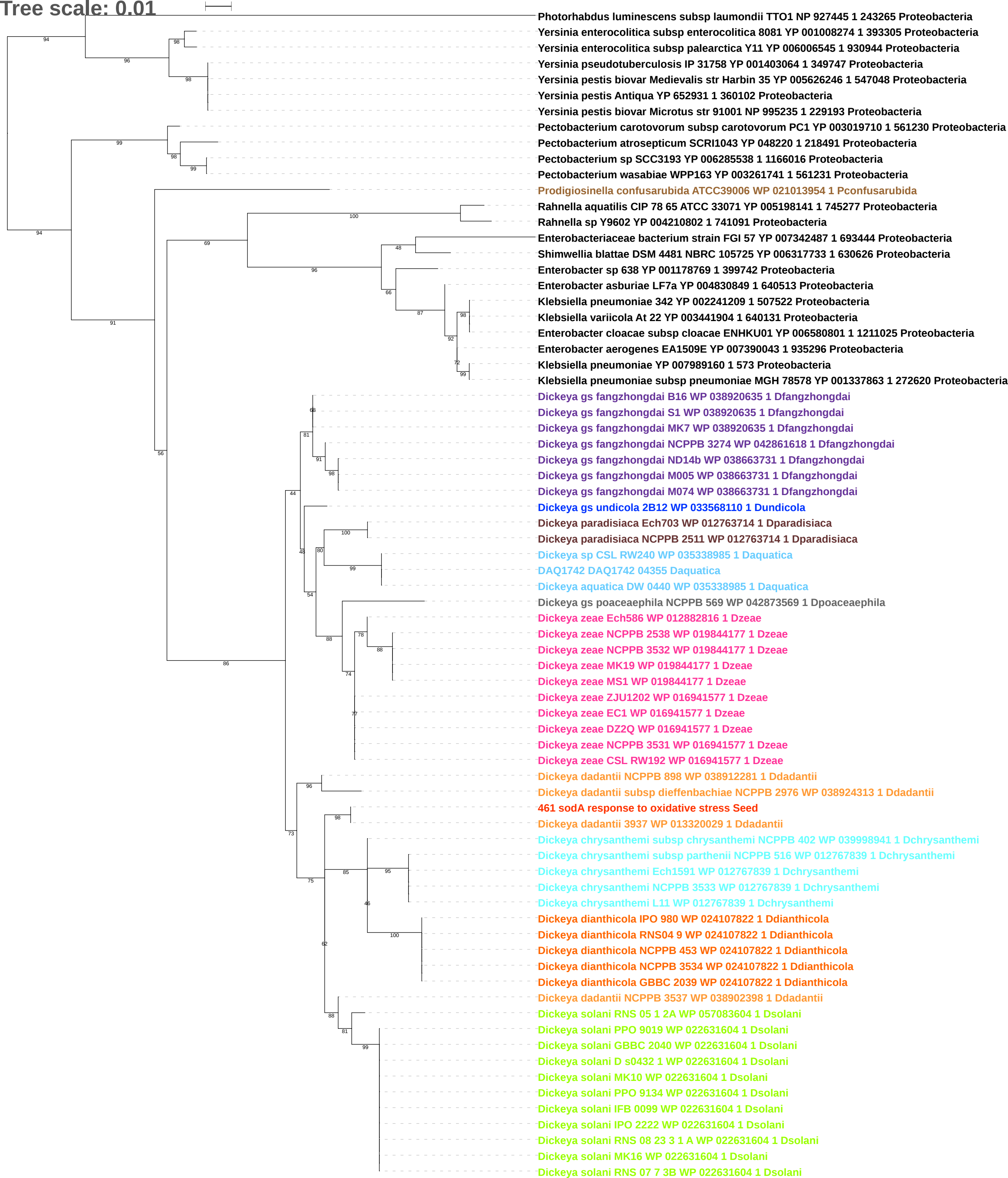
Tree scale: 0.01
Photorhabdus luminescens subsp laumondii TTO1 NP 927445 1 243265 Proteobacteria
Yersinia enterocolitica subsp enterocolitica 8081 YP 001008274 1 393305 Proteobacteria
Yersinia enterocolitica subsp palearctica Y11 YP 006006545 1 930944 Proteobacteria
Yersinia pseudotuberculosis IP 31758 YP 001403064 1 349747 Proteobacteria
Yersinia pestis biovar Medievalis str Harbin 35 YP 005626246 1 547048 Proteobacteria
Yersinia pestis Antiqua YP 652931 1 360102 Proteobacteria
Yersinia pestis biovar Microtus str 91001 NP 995235 1 229193 Proteobacteria
Pectobacterium carotovorum subsp carotovorum PC1 YP 003019710 1 561230 Proteobacteria
Pectobacterium atrosepticum SCRI1043 YP 048220 1 218491 Proteobacteria
Pectobacterium sp SCC3193 YP 006285538 1 1166016 Proteobacteria
Pectobacterium wasabiae WPP163 YP 003261741 1 561231 Proteobacteria
Prodigiosinella confusarubida ATCC39006 WP 021013954 1 Pconfusarubida
Rahnella aquatilis CIP 78 65 ATCC 33071 YP 005198141 1 745277 Proteobacteria
Rahnella sp Y9602 YP 004210802 1 741091 Proteobacteria
Enterobacteriaceae bacterium strain FGI 57 YP 007342487 1 693444 Proteobacteria
Shimwellia blattae DSM 4481 NBRC 105725 YP 006317733 1 630626 Proteobacteria
Enterobacter sp 638 YP 001178769 1 399742 Proteobacteria
Enterobacter asburiae LF7a YP 004830849 1 640513 Proteobacteria
Klebsiella pneumoniae 342 YP 002241209 1 507522 Proteobacteria
Klebsiella variicola At 22 YP 003441904 1 640131 Proteobacteria
Enterobacter cloacae subsp cloacae ENHKU01 YP 006580801 1 1211025 Proteobacteria
Enterobacter aerogenes EA1509E YP 007390043 1 935296 Proteobacteria
Klebsiella pneumoniae YP 007989160 1 573 Proteobacteria
Klebsiella pneumoniae subsp pneumoniae MGH 78578 YP 001337863 1 272620 Proteobacteria
Dickeya gs fangzhongdai B16 WP 038920635 1 Dfangzhongdai
Dickeya gs fangzhongdai S1 WP 038920635 1 Dfangzhongdai
Dickeya gs fangzhongdai MK7 WP 038920635 1 Dfangzhongdai
Dickeya gs fangzhongdai NCPPB 3274 WP 042861618 1 Dfangzhongdai
Dickeya gs fangzhongdai ND14b WP 038663731 1 Dfangzhongdai
Dickeya gs fangzhongdai M005 WP 038663731 1 Dfangzhongdai
Dickeya gs fangzhongdai M074 WP 038663731 1 Dfangzhongdai
Dickeya gs undicola 2B12 WP 033568110 1 Dundicola
Dickeya paradisiaca Ech703 WP 012763714 1 Dparadisiaca
Dickeya paradisiaca NCPPB 2511 WP 012763714 1 Dparadisiaca
Dickeya sp CSL RW240 WP 035338985 1 Daquatica
DAQ1742 DAQ1742 04355 Daquatica
Dickeya aquatica DW 0440 WP 035338985 1 Daquatica
Dickeya gs poaceaephila NCPPB 569 WP 042873569 1 Dpoaceaephila
Dickeya zeae Ech586 WP 012882816 1 Dzeae
Dickeya zeae NCPPB 2538 WP 019844177 1 Dzeae
Dickeya zeae NCPPB 3532 WP 019844177 1 Dzeae
Dickeya zeae MK19 WP 019844177 1 Dzeae
Dickeya zeae MS1 WP 019844177 1 Dzeae
Dickeya zeae ZJU1202 WP 016941577 1 Dzeae
Dickeya zeae EC1 WP 016941577 1 Dzeae
Dickeya zeae DZ2Q WP 016941577 1 Dzeae
Dickeya zeae NCPPB 3531 WP 016941577 1 Dzeae
Dickeya zeae CSL RW192 WP 016941577 1 Dzeae
Dickeya dadantii NCPPB 898 WP 038912281 1 Ddadantii
Dickeya dadantii subsp dieffenbachiae NCPPB 2976 WP 038924313 1 Ddadantii
461 sodA response to oxidative stress Seed
Dickeya dadantii 3937 WP 013320029 1 Ddadantii
Dickeya chrysanthemi subsp chrysanthemi NCPPB 402 WP 039998941 1 Dchrysanthemi
Dickeya chrysanthemi subsp parthenii NCPPB 516 WP 012767839 1 Dchrysanthemi
Dickeya chrysanthemi Ech1591 WP 012767839 1 Dchrysanthemi
Dickeya chrysanthemi NCPPB 3533 WP 012767839 1 Dchrysanthemi
Dickeya chrysanthemi L11 WP 012767839 1 Dchrysanthemi
Dickeya dianthicola IPO 980 WP 024107822 1 Ddianthicola
Dickeya dianthicola RNS04 9 WP 024107822 1 Ddianthicola
Dickeya dianthicola NCPPB 453 WP 024107822 1 Ddianthicola
Dickeya dianthicola NCPPB 3534 WP 024107822 1 Ddianthicola
Dickeya dianthicola GBBC 2039 WP 024107822 1 Ddianthicola
Dickeya dadantii NCPPB 3537 WP 038902398 1 Ddadantii
Dickeya solani RNS 05 1 2A WP 057083604 1 Dsolani
Dickeya solani PPO 9019 WP 022631604 1 Dsolani
Dickeya solani GBBC 2040 WP 022631604 1 Dsolani
Dickeya solani D s0432 1 WP 022631604 1 Dsolani
Dickeya solani MK10 WP 022631604 1 Dsolani
Dickeya solani PPO 9134 WP 022631604 1 Dsolani
Dickeya solani IFB 0099 WP 022631604 1 Dsolani
Dickeya solani IPO 2222 WP 022631604 1 Dsolani
Dickeya solani RNS 08 23 3 1 A WP 022631604 1 Dsolani
Dickeya solani MK16 WP 022631604 1 Dsolani
Dickeya solani RNS 07 7 3B WP 022631604 1 Dsolani

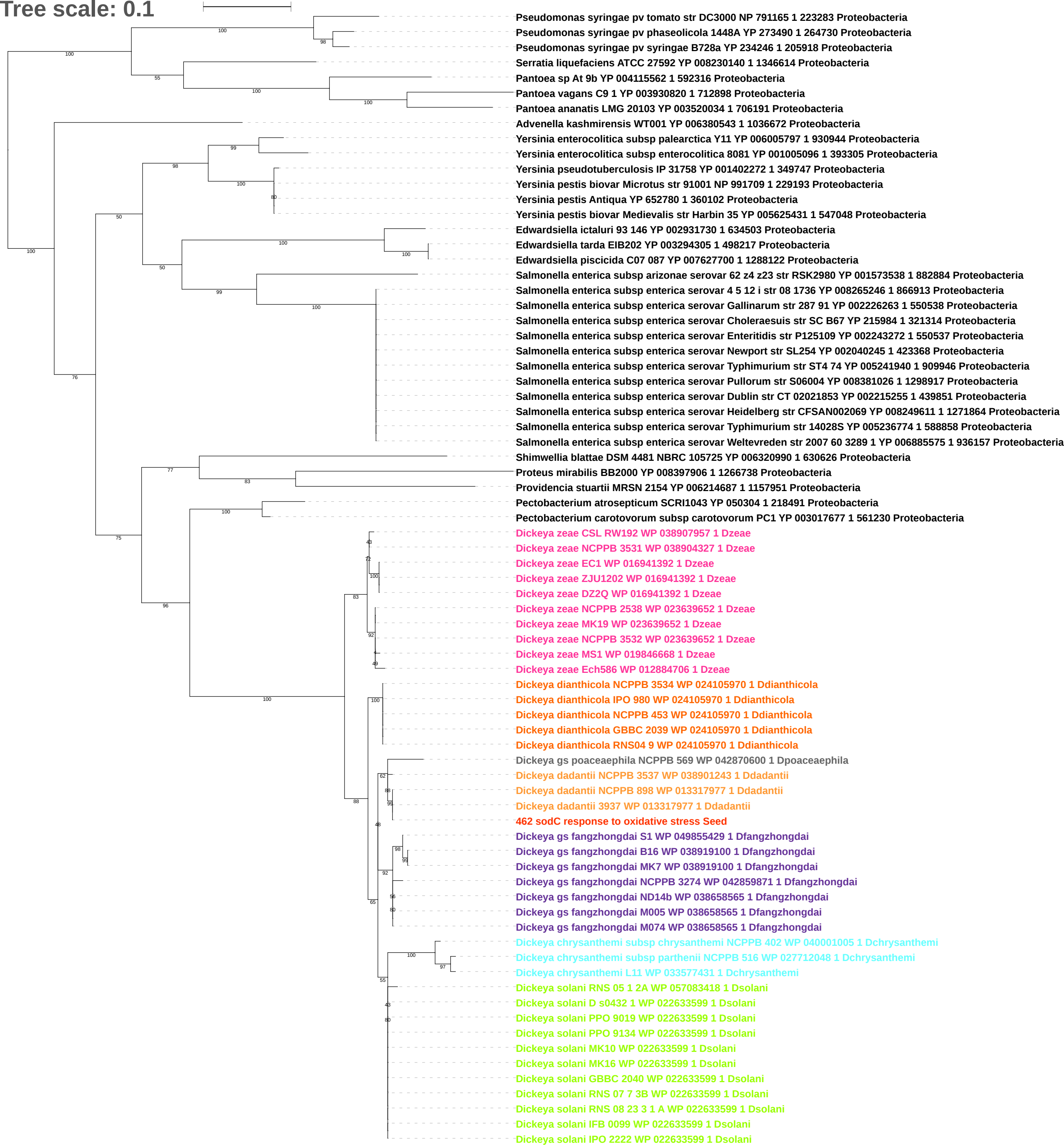

Tree scale: 0.1
Pseudomonas syringae pv tomato str DC3000 NP 791165 1 223283 Proteobacteria
Pseudomonas syringae pv phaseolicola 1448A YP 273490 1 264730 Proteobacteria
Pseudomonas syringae pv syringae B728a YP 234246 1 205918 Proteobacteria
Serratia liquefaciens ATCC 27592 YP 008230140 1 1346614 Proteobacteria
Pantoea sp At 9b YP 004115562 1 592316 Proteobacteria
Pantoea vagans C9 1 YP 003930820 1 712898 Proteobacteria
Pantoea ananatis LMG 20103 YP 003520034 1 706191 Proteobacteria
Advenella kashmirensis WT001 YP 006380543 1 1036672 Proteobacteria
Yersinia enterocolitica subsp palearctica Y11 YP 006005797 1 930944 Proteobacteria
Yersinia enterocolitica subsp enterocolitica 8081 YP 001005096 1 393305 Proteobacteria
Yersinia pseudotuberculosis IP 31758 YP 001402272 1 349747 Proteobacteria
Yersinia pestis biovar Microtus str 91001 NP 991709 1 229193 Proteobacteria
Yersinia pestis Antiqua YP 652780 1 360102 Proteobacteria
Yersinia pestis biovar Medievalis str Harbin 35 YP 005625431 1 547048 Proteobacteria
Edwardsiella ictaluri 93 146 YP 002931730 1 634503 Proteobacteria
Edwardsiella tarda EIB202 YP 003294305 1 498217 Proteobacteria
Edwardsiella piscicida C07 087 YP 007627700 1 1288122 Proteobacteria
Salmonella enterica subsp arizonae serovar 62 z4 z23 str RSK2980 YP 001573538 1 882884 Proteobacteria
Salmonella enterica subsp enterica serovar 4 5 12 i str 08 1736 YP 008265246 1 866913 Proteobacteria
Salmonella enterica subsp enterica serovar Gallinarum str 287 91 YP 002226263 1 550538 Proteobacteria
Salmonella enterica subsp enterica serovar Choleraesuis str SC B67 YP 215984 1 321314 Proteobacteria
Salmonella enterica subsp enterica serovar Enteritidis str P125109 YP 002243272 1 550537 Proteobacteria
Salmonella enterica subsp enterica serovar Newport str SL254 YP 002040245 1 423368 Proteobacteria
Salmonella enterica subsp enterica serovar Typhimurium str ST4 74 YP 005241940 1 909946 Proteobacteria
Salmonella enterica subsp enterica serovar Pullorum str S06004 YP 008381026 1 1298917 Proteobacteria
Salmonella enterica subsp enterica serovar Dublin str CT 02021853 YP 002215255 1 439851 Proteobacteria
Salmonella enterica subsp enterica serovar Heidelberg str CFSAN002069 YP 008249611 1 1271864 Proteobacteria
Salmonella enterica subsp enterica serovar Typhimurium str 14028S YP 005236774 1 588858 Proteobacteria
Salmonella enterica subsp enterica serovar Weltevreden str 2007 60 3289 1 YP 006885575 1 936157 Proteobacteria
Shimwellia blattae DSM 4481 NBRC 105725 YP 006320990 1 630626 Proteobacteria
Proteus mirabilis BB2000 YP 008397906 1 1266738 Proteobacteria
Providencia stuartii MRSN 2154 YP 006214687 1 1157951 Proteobacteria
Pectobacterium atrosepticum SCRI1043 YP 050304 1 218491 Proteobacteria
Pectobacterium carotovorum subsp carotovorum PC1 YP 003017677 1 561230 Proteobacteria
Dickeya zeae CSL RW192 WP 038907957 1 Dzeae
Dickeya zeae NCPPB 3531 WP 038904327 1 Dzeae
Dickeya zeae EC1 WP 016941392 1 Dzeae
Dickeya zeae ZJU1202 WP 016941392 1 Dzeae
Dickeya zeae DZ2Q WP 016941392 1 Dzeae
Dickeya zeae NCPPB 2538 WP 023639652 1 Dzeae
Dickeya zeae MK19 WP 023639652 1 Dzeae
Dickeya zeae NCPPB 3532 WP 023639652 1 Dzeae
Dickeya zeae MS1 WP 019846668 1 Dzeae
Dickeya zeae Ech586 WP 012884706 1 Dzeae
Dickeya dianthicola NCPPB 3534 WP 024105970 1 Ddianthicola
Dickeya dianthicola IPO 980 WP 024105970 1 Ddianthicola
Dickeya dianthicola NCPPB 453 WP 024105970 1 Ddianthicola
Dickeya dianthicola GBBC 2039 WP 024105970 1 Ddianthicola
Dickeya dianthicola RNS04 9 WP 024105970 1 Ddianthicola
Dickeya gs poaceaephila NCPPB 569 WP 042870600 1 Dpoaceaephila
Dickeya dadantii NCPPB 3537 WP 038901243 1 Ddadantii
Dickeya dadantii NCPPB 898 WP 013317977 1 Ddadantii
Dickeya dadantii 3937 WP 013317977 1 Ddadantii
462 sodC response to oxidative stress Seed
Dickeya gs fangzhongdai S1 WP 049855429 1 Dfangzhongdai
Dickeya gs fangzhongdai B16 WP 038919100 1 Dfangzhongdai
Dickeya gs fangzhongdai MK7 WP 038919100 1 Dfangzhongdai
Dickeya gs fangzhongdai NCPPB 3274 WP 042859871 1 Dfangzhongdai
Dickeya gs fangzhongdai ND14b WP 038658565 1 Dfangzhongdai
Dickeya gs fangzhongdai M005 WP 038658565 1 Dfangzhongdai
Dickeya gs fangzhongdai M074 WP 038658565 1 Dfangzhongdai
Dickeya chrysanthemi subsp chrysanthemi NCPPB 402 WP 040001005 1 Dchrysanthemi
Dickeya chrysanthemi subsp parthenii NCPPB 516 WP 027712048 1 Dchrysanthemi
Dickeya chrysanthemi L11 WP 033577431 1 Dchrysanthemi
Dickeya solani RNS 05 1 2A WP 057083418 1 Dsolani
Dickeya solani D s0432 1 WP 022633599 1 Dsolani
Dickeya solani PPO 9019 WP 022633599 1 Dsolani
Dickeya solani PPO 9134 WP 022633599 1 Dsolani
Dickeya solani MK10 WP 022633599 1 Dsolani
Dickeya solani MK16 WP 022633599 1 Dsolani
Dickeya solani GBBC 2040 WP 022633599 1 Dsolani
Dickeya solani RNS 07 7 3B WP 022633599 1 Dsolani
Dickeya solani RNS 08 23 3 1 A WP 022633599 1 Dsolani
Dickeya solani IFB 0099 WP 022633599 1 Dsolani
Dickeya solani IPO 2222 WP 022633599 1 Dsolani

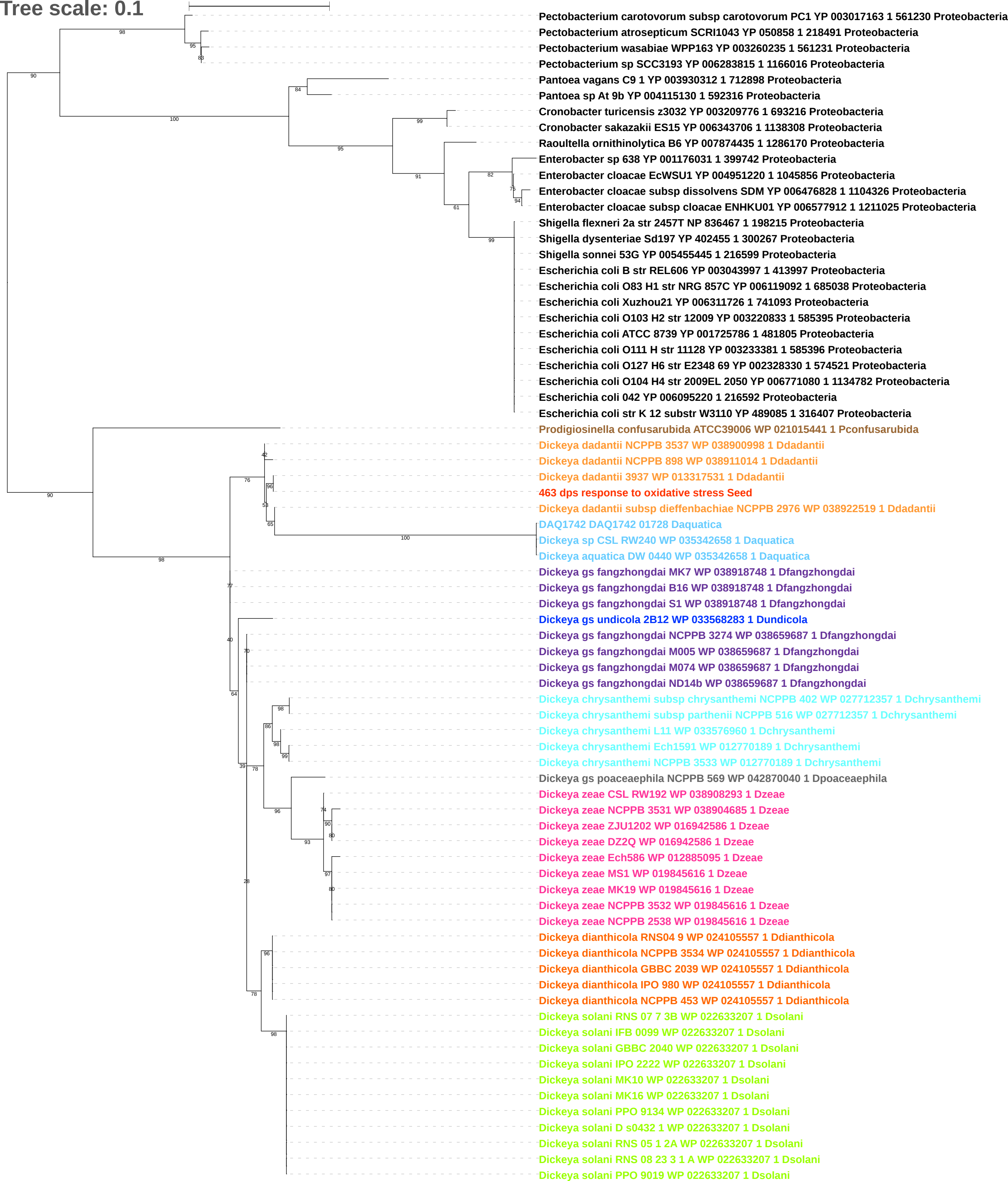
Tree scale: 0.1
Pectobacterium carotovorum subsp carotovorum PC1 YP 003017163 1 561230 Proteobacteria
Pectobacterium atrosepticum SCRI1043 YP 050858 1 218491 Proteobacteria
Pectobacterium wasabiae WPP163 YP 003260235 1 561231 Proteobacteria
Pectobacterium sp SCC3193 YP 006283815 1 1166016 Proteobacteria
Pantoea vagans C9 1 YP 003930312 1 712898 Proteobacteria
Pantoea sp At 9b YP 004115130 1 592316 Proteobacteria
Cronobacter turicensis z3032 YP 003209776 1 693216 Proteobacteria
Cronobacter sakazakii ES15 YP 006343706 1 1138308 Proteobacteria
Raoultella ornithinolytica B6 YP 007874435 1 1286170 Proteobacteria
Enterobacter sp 638 YP 001176031 1 399742 Proteobacteria
Enterobacter cloacae EcWSU1 YP 004951220 1 1045856 Proteobacteria
Enterobacter cloacae subsp dissolvens SDM YP 006476828 1 1104326 Proteobacteria
Enterobacter cloacae subsp cloacae ENHKU01 YP 006577912 1 1211025 Proteobacteria
Shigella flexneri 2a str 2457T NP 836467 1 198215 Proteobacteria
Shigella dysenteriae Sd197 YP 402455 1 300267 Proteobacteria
Shigella sonnei 53G YP 005455445 1 216599 Proteobacteria
Escherichia coli B str REL606 YP 003043997 1 413997 Proteobacteria
Escherichia coli O83 H1 str NRG 857C YP 006119092 1 685038 Proteobacteria
Escherichia coli Xuzhou21 YP 006311726 1 741093 Proteobacteria
Escherichia coli O103 H2 str 12009 YP 003220833 1 585395 Proteobacteria
Escherichia coli ATCC 8739 YP 001725786 1 481805 Proteobacteria
Escherichia coli O111 H str 11128 YP 003233381 1 585396 Proteobacteria
Escherichia coli O127 H6 str E2348 69 YP 002328330 1 574521 Proteobacteria
Escherichia coli O104 H4 str 2009EL 2050 YP 006771080 1 1134782 Proteobacteria
Escherichia coli 042 YP 006095220 1 216592 Proteobacteria
Escherichia coli str K 12 substr W3110 YP 489085 1 316407 Proteobacteria
Prodigiosinella confusarubida ATCC39006 WP 021015441 1 Pconfusarubida
Dickeya dadantii NCPPB 3537 WP 038900998 1 Ddadantii
Dickeya dadantii NCPPB 898 WP 038911014 1 Ddadantii
Dickeya dadantii 3937 WP 013317531 1 Ddadantii
463 dps response to oxidative stress Seed
Dickeya dadantii subsp dieffenbachiae NCPPB 2976 WP 038922519 1 Ddadantii
DAQ1742 DAQ1742 01728 Daquatica
Dickeya sp CSL RW240 WP 035342658 1 Daquatica
Dickeya aquatica DW 0440 WP 035342658 1 Daquatica
Dickeya gs fangzhongdai MK7 WP 038918748 1 Dfangzhongdai
Dickeya gs fangzhongdai B16 WP 038918748 1 Dfangzhongdai
Dickeya gs fangzhongdai S1 WP 038918748 1 Dfangzhongdai
Dickeya gs undicola 2B12 WP 033568283 1 Dundicola
Dickeya gs fangzhongdai NCPPB 3274 WP 038659687 1 Dfangzhongdai
Dickeya gs fangzhongdai M005 WP 038659687 1 Dfangzhongdai
Dickeya gs fangzhongdai M074 WP 038659687 1 Dfangzhongdai
Dickeya gs fangzhongdai ND14b WP 038659687 1 Dfangzhongdai
Dickeya chrysanthemi subsp chrysanthemi NCPPB 402 WP 027712357 1 Dchrysanthemi
Dickeya chrysanthemi subsp parthenii NCPPB 516 WP 027712357 1 Dchrysanthemi
Dickeya chrysanthemi L11 WP 033576960 1 Dchrysanthemi
Dickeya chrysanthemi Ech1591 WP 012770189 1 Dchrysanthemi
Dickeya chrysanthemi NCPPB 3533 WP 012770189 1 Dchrysanthemi
Dickeya gs poaceaephila NCPPB 569 WP 042870040 1 Dpoaceaephila
Dickeya zeae CSL RW192 WP 038908293 1 Dzeae
Dickeya zeae NCPPB 3531 WP 038904685 1 Dzeae
Dickeya zeae ZJU1202 WP 016942586 1 Dzeae
Dickeya zeae DZ2Q WP 016942586 1 Dzeae
Dickeya zeae Ech586 WP 012885095 1 Dzeae
Dickeya zeae MS1 WP 019845616 1 Dzeae
Dickeya zeae MK19 WP 019845616 1 Dzeae
Dickeya zeae NCPPB 3532 WP 019845616 1 Dzeae
Dickeya zeae NCPPB 2538 WP 019845616 1 Dzeae
Dickeya dianthicola RNS04 9 WP 024105557 1 Ddianthicola
Dickeya dianthicola NCPPB 3534 WP 024105557 1 Ddianthicola
Dickeya dianthicola GBBC 2039 WP 024105557 1 Ddianthicola
Dickeya dianthicola IPO 980 WP 024105557 1 Ddianthicola
Dickeya dianthicola NCPPB 453 WP 024105557 1 Ddianthicola
Dickeya solani RNS 07 7 3B WP 022633207 1 Dsolani
Dickeya solani IFB 0099 WP 022633207 1 Dsolani
Dickeya solani GBBC 2040 WP 022633207 1 Dsolani
Dickeya solani IPO 2222 WP 022633207 1 Dsolani
Dickeya solani MK10 WP 022633207 1 Dsolani
Dickeya solani MK16 WP 022633207 1 Dsolani
Dickeya solani PPO 9134 WP 022633207 1 Dsolani
Dickeya solani D s0432 1 WP 022633207 1 Dsolani
Dickeya solani RNS 05 1 2A WP 022633207 1 Dsolani
Dickeya solani RNS 08 23 3 1 A WP 022633207 1 Dsolani
Dickeya solani PPO 9019 WP 022633207 1 Dsolani

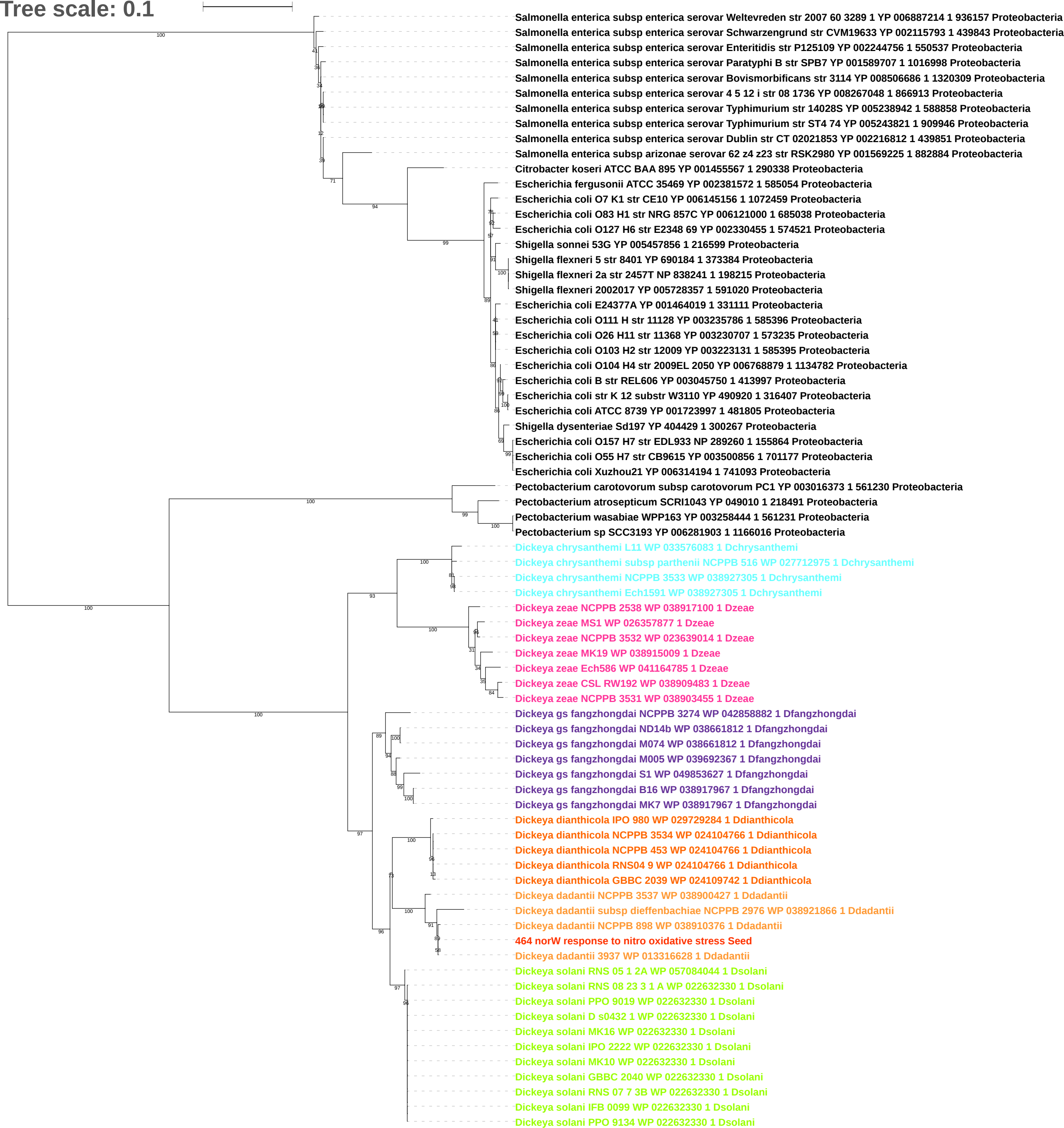
Tree scale: 0.1
Salmonella enterica subsp enterica serovar Weltevreden str 2007 60 3289 1 YP 006887214 1 936157 Proteobacteria
Salmonella enterica subsp enterica serovar Schwarzengrund str CVM19633 YP 002115793 1 439843 Proteobacteria
Salmonella enterica subsp enterica serovar Enteritidis str P125109 YP 002244756 1 550537 Proteobacteria
Salmonella enterica subsp enterica serovar Paratyphi B str SPB7 YP 001589707 1 1016998 Proteobacteria
Salmonella enterica subsp enterica serovar Bovismorbificans str 3114 YP 008506686 1 1320309 Proteobacteria
Salmonella enterica subsp enterica serovar 4 5 12 i str 08 1736 YP 008267048 1 866913 Proteobacteria
Salmonella enterica subsp enterica serovar Typhimurium str 14028S YP 005238942 1 588858 Proteobacteria
Salmonella enterica subsp enterica serovar Typhimurium str ST4 74 YP 005243821 1 909946 Proteobacteria
Salmonella enterica subsp enterica serovar Dublin str CT 02021853 YP 002216812 1 439851 Proteobacteria
Salmonella enterica subsp arizonae serovar 62 z4 z23 str RSK2980 YP 001569225 1 882884 Proteobacteria
Citrobacter koseri ATCC BAA 895 YP 001455567 1 290338 Proteobacteria
Escherichia fergusonii ATCC 35469 YP 002381572 1 585054 Proteobacteria
Escherichia coli O7 K1 str CE10 YP 006145156 1 1072459 Proteobacteria
Escherichia coli O83 H1 str NRG 857C YP 006121000 1 685038 Proteobacteria
Escherichia coli O127 H6 str E2348 69 YP 002330455 1 574521 Proteobacteria
Shigella sonnei 53G YP 005457856 1 216599 Proteobacteria
Shigella flexneri 5 str 8401 YP 690184 1 373384 Proteobacteria
Shigella flexneri 2a str 2457T NP 838241 1 198215 Proteobacteria
Shigella flexneri 2002017 YP 005728357 1 591020 Proteobacteria
Escherichia coli E24377A YP 001464019 1 331111 Proteobacteria
Escherichia coli O111 H str 11128 YP 003235786 1 585396 Proteobacteria
Escherichia coli O26 H11 str 11368 YP 003230707 1 573235 Proteobacteria
Escherichia coli O103 H2 str 12009 YP 003223131 1 585395 Proteobacteria
Escherichia coli O104 H4 str 2009EL 2050 YP 006768879 1 1134782 Proteobacteria
Escherichia coli B str REL606 YP 003045750 1 413997 Proteobacteria
Escherichia coli str K 12 substr W3110 YP 490920 1 316407 Proteobacteria
Escherichia coli ATCC 8739 YP 001723997 1 481805 Proteobacteria
Shigella dysenteriae Sd197 YP 404429 1 300267 Proteobacteria
Escherichia coli O157 H7 str EDL933 NP 289260 1 155864 Proteobacteria
Escherichia coli O55 H7 str CB9615 YP 003500856 1 701177 Proteobacteria
Escherichia coli Xuzhou21 YP 006314194 1 741093 Proteobacteria
Pectobacterium carotovorum subsp carotovorum PC1 YP 003016373 1 561230 Proteobacteria
Pectobacterium atrosepticum SCRI1043 YP 049010 1 218491 Proteobacteria
Pectobacterium wasabiae WPP163 YP 003258444 1 561231 Proteobacteria
Pectobacterium sp SCC3193 YP 006281903 1 1166016 Proteobacteria
Dickeya chrysanthemi L11 WP 033576083 1 Dchrysanthemi
Dickeya chrysanthemi subsp parthenii NCPPB 516 WP 027712975 1 Dchrysanthemi
Dickeya chrysanthemi NCPPB 3533 WP 038927305 1 Dchrysanthemi
Dickeya chrysanthemi Ech1591 WP 038927305 1 Dchrysanthemi
Dickeya zeae NCPPB 2538 WP 038917100 1 Dzeae
Dickeya zeae MS1 WP 026357877 1 Dzeae
Dickeya zeae NCPPB 3532 WP 023639014 1 Dzeae
Dickeya zeae MK19 WP 038915009 1 Dzeae
Dickeya zeae Ech586 WP 041164785 1 Dzeae
Dickeya zeae CSL RW192 WP 038909483 1 Dzeae
Dickeya zeae NCPPB 3531 WP 038903455 1 Dzeae
Dickeya gs fangzhongdai NCPPB 3274 WP 042858882 1 Dfangzhongdai
Dickeya gs fangzhongdai ND14b WP 038661812 1 Dfangzhongdai
Dickeya gs fangzhongdai M074 WP 038661812 1 Dfangzhongdai
Dickeya gs fangzhongdai M005 WP 039692367 1 Dfangzhongdai
Dickeya gs fangzhongdai S1 WP 049853627 1 Dfangzhongdai
Dickeya gs fangzhongdai B16 WP 038917967 1 Dfangzhongdai
Dickeya gs fangzhongdai MK7 WP 038917967 1 Dfangzhongdai
Dickeya dianthicola IPO 980 WP 029729284 1 Ddianthicola
Dickeya dianthicola NCPPB 3534 WP 024104766 1 Ddianthicola
Dickeya dianthicola NCPPB 453 WP 024104766 1 Ddianthicola
Dickeya dianthicola RNS04 9 WP 024104766 1 Ddianthicola
Dickeya dianthicola GBBC 2039 WP 024109742 1 Ddianthicola
Dickeya dadantii NCPPB 3537 WP 038900427 1 Ddadantii
Dickeya dadantii subsp dieffenbachiae NCPPB 2976 WP 038921866 1 Ddadantii
Dickeya dadantii NCPPB 898 WP 038910376 1 Ddadantii
464 norW response to nitro oxidative stress Seed
Dickeya dadantii 3937 WP 013316628 1 Ddadantii
Dickeya solani RNS 05 1 2A WP 057084044 1 Dsolani
Dickeya solani RNS 08 23 3 1 A WP 022632330 1 Dsolani
Dickeya solani PPO 9019 WP 022632330 1 Dsolani
Dickeya solani D s0432 1 WP 022632330 1 Dsolani
Dickeya solani MK16 WP 022632330 1 Dsolani
Dickeya solani IPO 2222 WP 022632330 1 Dsolani
Dickeya solani MK10 WP 022632330 1 Dsolani
Dickeya solani GBBC 2040 WP 022632330 1 Dsolani
Dickeya solani RNS 07 7 3B WP 022632330 1 Dsolani
Dickeya solani IFB 0099 WP 022632330 1 Dsolani
Dickeya solani PPO 9134 WP 022632330 1 Dsolani

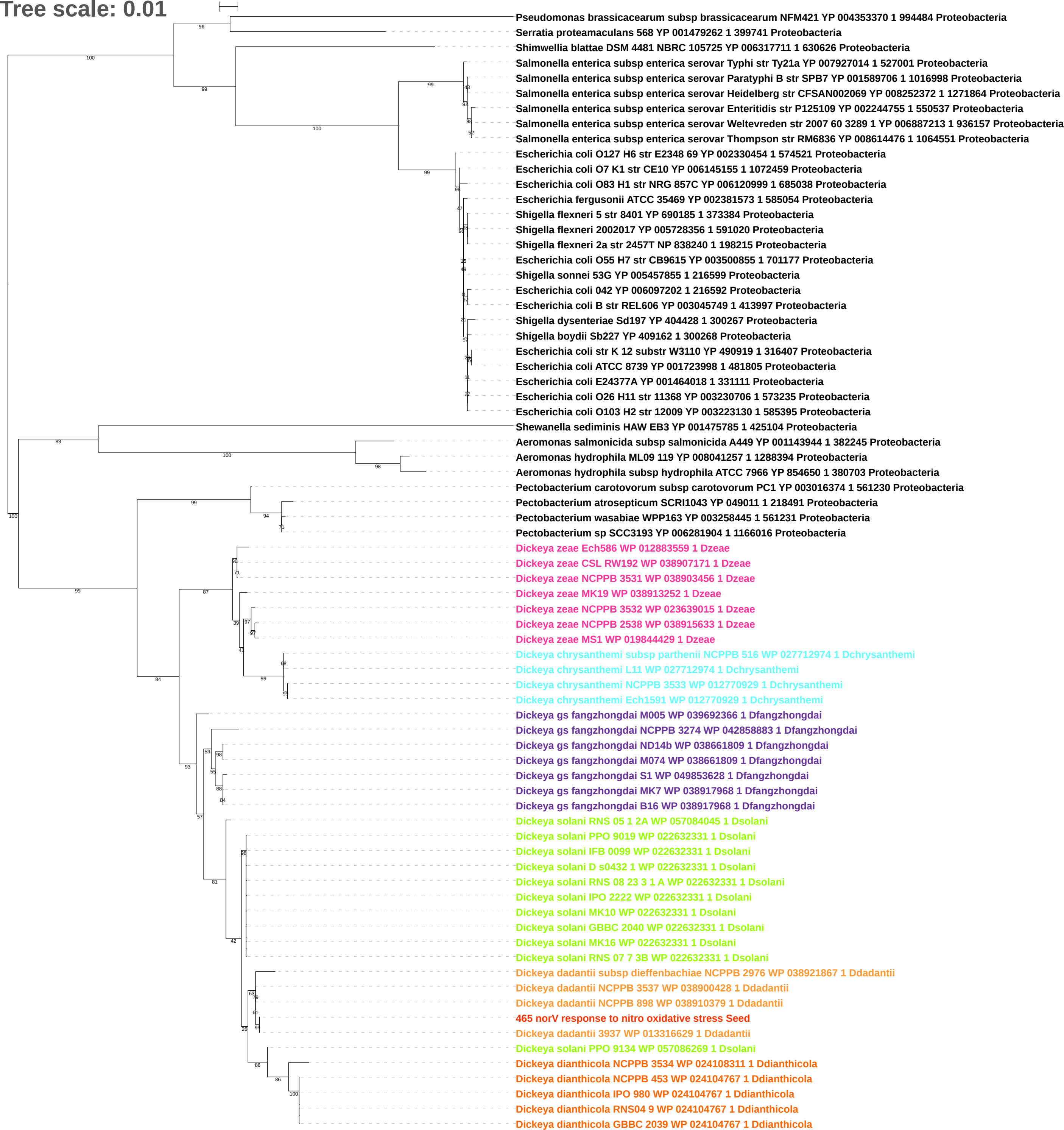
Tree scale: 0.01
Pseudomonas brassicacearum subsp brassicacearum NFM421 YP 004353370 1 994484 Proteobacteria
Serratia proteamaculans 568 YP 001479262 1 399741 Proteobacteria
Shimwellia blattae DSM 4481 NBRC 105725 YP 006317711 1 630626 Proteobacteria
Salmonella enterica subsp enterica serovar Typhi str Ty21a YP 007927014 1 527001 Proteobacteria
Salmonella enterica subsp enterica serovar Paratyphi B str SPB7 YP 001589706 1 1016998 Proteobacteria
Salmonella enterica subsp enterica serovar Heidelberg str CFSAN002069 YP 008252372 1 1271864 Proteobacteria
Salmonella enterica subsp enterica serovar Enteritidis str P125109 YP 002244755 1 550537 Proteobacteria
Salmonella enterica subsp enterica serovar Weltevreden str 2007 60 3289 1 YP 006887213 1 936157 Proteobacteria
Salmonella enterica subsp enterica serovar Thompson str RM6836 YP 008614476 1 1064551 Proteobacteria
Escherichia coli O127 H6 str E2348 69 YP 002330454 1 574521 Proteobacteria
Escherichia coli O7 K1 str CE10 YP 006145155 1 1072459 Proteobacteria
Escherichia coli O83 H1 str NRG 857C YP 006120999 1 685038 Proteobacteria
Escherichia fergusonii ATCC 35469 YP 002381573 1 585054 Proteobacteria
Shigella flexneri 5 str 8401 YP 690185 1 373384 Proteobacteria
Shigella flexneri 2002017 YP 005728356 1 591020 Proteobacteria
Shigella flexneri 2a str 2457T NP 838240 1 198215 Proteobacteria
Escherichia coli O55 H7 str CB9615 YP 003500855 1 701177 Proteobacteria
Shigella sonnei 53G YP 005457855 1 216599 Proteobacteria
Escherichia coli 042 YP 006097202 1 216592 Proteobacteria
Escherichia coli B str REL606 YP 003045749 1 413997 Proteobacteria
Shigella dysenteriae Sd197 YP 404428 1 300267 Proteobacteria
Shigella boydii Sb227 YP 409162 1 300268 Proteobacteria
Escherichia coli str K 12 substr W3110 YP 490919 1 316407 Proteobacteria
Escherichia coli ATCC 8739 YP 001723998 1 481805 Proteobacteria
Escherichia coli E24377A YP 001464018 1 331111 Proteobacteria
Escherichia coli O26 H11 str 11368 YP 003230706 1 573235 Proteobacteria
Escherichia coli O103 H2 str 12009 YP 003223130 1 585395 Proteobacteria
Shewanella sediminis HAW EB3 YP 001475785 1 425104 Proteobacteria
Aeromonas salmonicida subsp salmonicida A449 YP 001143944 1 382245 Proteobacteria
Aeromonas hydrophila ML09 119 YP 008041257 1 1288394 Proteobacteria
Aeromonas hydrophila subsp hydrophila ATCC 7966 YP 854650 1 380703 Proteobacteria
Pectobacterium carotovorum subsp carotovorum PC1 YP 003016374 1 561230 Proteobacteria
Pectobacterium atrosepticum SCRI1043 YP 049011 1 218491 Proteobacteria
Pectobacterium wasabiae WPP163 YP 003258445 1 561231 Proteobacteria
Pectobacterium sp SCC3193 YP 006281904 1 1166016 Proteobacteria
Dickeya zeae Ech586 WP 012883559 1 Dzeae
Dickeya zeae CSL RW192 WP 038907171 1 Dzeae
Dickeya zeae NCPPB 3531 WP 038903456 1 Dzeae
Dickeya zeae MK19 WP 038913252 1 Dzeae
Dickeya zeae NCPPB 3532 WP 023639015 1 Dzeae
Dickeya zeae NCPPB 2538 WP 038915633 1 Dzeae
Dickeya zeae MS1 WP 019844429 1 Dzeae
Dickeya chrysanthemi subsp parthenii NCPPB 516 WP 027712974 1 Dchrysanthemi
Dickeya chrysanthemi L11 WP 027712974 1 Dchrysanthemi
Dickeya chrysanthemi NCPPB 3533 WP 012770929 1 Dchrysanthemi
Dickeya chrysanthemi Ech1591 WP 012770929 1 Dchrysanthemi
Dickeya gs fangzhongdai M005 WP 039692366 1 Dfangzhongdai
Dickeya gs fangzhongdai NCPPB 3274 WP 042858883 1 Dfangzhongdai
Dickeya gs fangzhongdai ND14b WP 038661809 1 Dfangzhongdai
Dickeya gs fangzhongdai M074 WP 038661809 1 Dfangzhongdai
Dickeya gs fangzhongdai S1 WP 049853628 1 Dfangzhongdai
Dickeya gs fangzhongdai MK7 WP 038917968 1 Dfangzhongdai
Dickeya gs fangzhongdai B16 WP 038917968 1 Dfangzhongdai
Dickeya solani RNS 05 1 2A WP 057084045 1 Dsolani
Dickeya solani PPO 9019 WP 022632331 1 Dsolani
Dickeya solani IFB 0099 WP 022632331 1 Dsolani
Dickeya solani D s0432 1 WP 022632331 1 Dsolani
Dickeya solani RNS 08 23 3 1 A WP 022632331 1 Dsolani
Dickeya solani IPO 2222 WP 022632331 1 Dsolani
Dickeya solani MK10 WP 022632331 1 Dsolani
Dickeya solani GBBC 2040 WP 022632331 1 Dsolani
Dickeya solani MK16 WP 022632331 1 Dsolani
Dickeya solani RNS 07 7 3B WP 022632331 1 Dsolani
Dickeya dadantii subsp dieffenbachiae NCPPB 2976 WP 038921867 1 Ddadantii
Dickeya dadantii NCPPB 3537 WP 038900428 1 Ddadantii
Dickeya dadantii NCPPB 898 WP 038910379 1 Ddadantii
465 norV response to nitro oxidative stress Seed
Dickeya dadantii 3937 WP 013316629 1 Ddadantii
Dickeya solani PPO 9134 WP 057086269 1 Dsolani
Dickeya dianthicola NCPPB 3534 WP 024108311 1 Ddianthicola
Dickeya dianthicola NCPPB 453 WP 024104767 1 Ddianthicola
Dickeya dianthicola IPO 980 WP 024104767 1 Ddianthicola
Dickeya dianthicola RNS04 9 WP 024104767 1 Ddianthicola
Dickeya dianthicola GBBC 2039 WP 024104767 1 Ddianthicola

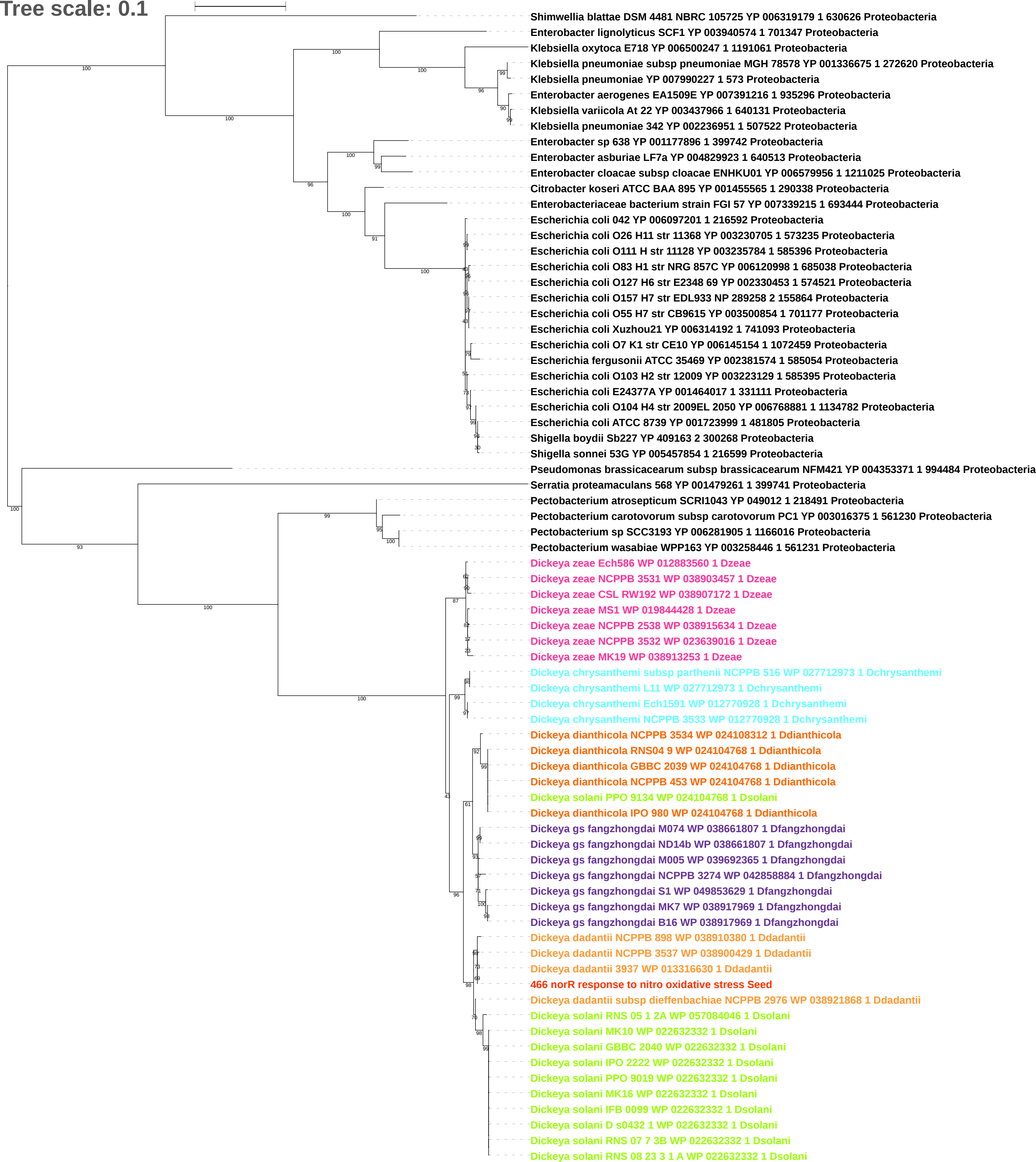
Tree scale: 0.1
Shimwellia blattae DSM 4481 NBRC 105725 YP 006319179 1 630626 Proteobacteria
Enterobacter lignolyticus SCF1 YP 003940574 1 701347 Proteobacteria
Klebsiella oxytoca E718 YP 006500247 1 1191061 Proteobacteria
Klebsiella pneumoniae subsp pneumoniae MGH 78578 YP 001336675 1 272620 Proteobacteria
Klebsiella pneumoniae YP 007990227 1 573 Proteobacteria
Enterobacter aerogenes EA1509E YP 007391216 1 935296 Proteobacteria
Klebsiella variicola At 22 YP 003437966 1 640131 Proteobacteria
Klebsiella pneumoniae 342 YP 002236951 1 507522 Proteobacteria
Enterobacter sp 638 YP 001177896 1 399742 Proteobacteria
Enterobacter asburiae LF7a YP 004829923 1 640513 Proteobacteria
Enterobacter cloacae subsp cloacae ENHKU01 YP 006579956 1 1211025 Proteobacteria
Citrobacter koseri ATCC BAA 895 YP 001455565 1 290338 Proteobacteria
Enterobacteriaceae bacterium strain FGI 57 YP 007339215 1 693444 Proteobacteria
Escherichia coli 042 YP 006097201 1 216592 Proteobacteria
Escherichia coli O26 H11 str 11368 YP 003230705 1 573235 Proteobacteria
Escherichia coli O111 H str 11128 YP 003235784 1 585396 Proteobacteria
Escherichia coli O83 H1 str NRG 857C YP 006120998 1 685038 Proteobacteria
Escherichia coli O127 H6 str E2348 69 YP 002330453 1 574521 Proteobacteria
Escherichia coli O157 H7 str EDL933 NP 289258 2 155864 Proteobacteria
Escherichia coli O55 H7 str CB9615 YP 003500854 1 701177 Proteobacteria
Escherichia coli Xuzhou21 YP 006314192 1 741093 Proteobacteria
Escherichia coli O7 K1 str CE10 YP 006145154 1 1072459 Proteobacteria
Escherichia fergusonii ATCC 35469 YP 002381574 1 585054 Proteobacteria
Escherichia coli O103 H2 str 12009 YP 003223129 1 585395 Proteobacteria
Escherichia coli E24377A YP 001464017 1 331111 Proteobacteria
Escherichia coli O104 H4 str 2009EL 2050 YP 006768881 1 1134782 Proteobacteria
Escherichia coli ATCC 8739 YP 001723999 1 481805 Proteobacteria
Shigella boydii Sb227 YP 409163 2 300268 Proteobacteria
Shigella sonnei 53G YP 005457854 1 216599 Proteobacteria
Pseudomonas brassicacearum subsp brassicacearum NFM421 YP 004353371 1 994484 Proteobacteria
Serratia proteamaculans 568 YP 001479261 1 399741 Proteobacteria
Pectobacterium atrosepticum SCRI1043 YP 049012 1 218491 Proteobacteria
Pectobacterium carotovorum subsp carotovorum PC1 YP 003016375 1 561230 Proteobacteria
Pectobacterium sp SCC3193 YP 006281905 1 1166016 Proteobacteria
Pectobacterium wasabiae WPP163 YP 003258446 1 561231 Proteobacteria
Dickeya zeae Ech586 WP 012883560 1 Dzeae
Dickeya zeae NCPPB 3531 WP 038903457 1 Dzeae
Dickeya zeae CSL RW192 WP 038907172 1 Dzeae
Dickeya zeae MS1 WP 019844428 1 Dzeae
Dickeya zeae NCPPB 2538 WP 038915634 1 Dzeae
Dickeya zeae NCPPB 3532 WP 023639016 1 Dzeae
Dickeya zeae MK19 WP 038913253 1 Dzeae
Dickeya chrysanthemi subsp parthenii NCPPB 516 WP 027712973 1 Dchrysanthemi
Dickeya chrysanthemi L11 WP 027712973 1 Dchrysanthemi
Dickeya chrysanthemi Ech1591 WP 012770928 1 Dchrysanthemi
Dickeya chrysanthemi NCPPB 3533 WP 012770928 1 Dchrysanthemi
Dickeya dianthicola NCPPB 3534 WP 024108312 1 Ddianthicola
Dickeya dianthicola RNS04 9 WP 024104768 1 Ddianthicola
Dickeya dianthicola GBBC 2039 WP 024104768 1 Ddianthicola
Dickeya dianthicola NCPPB 453 WP 024104768 1 Ddianthicola
Dickeya solani PPO 9134 WP 024104768 1 Dsolani
Dickeya dianthicola IPO 980 WP 024104768 1 Ddianthicola
Dickeya gs fangzhongdai M074 WP 038661807 1 Dfangzhongdai
Dickeya gs fangzhongdai ND14b WP 038661807 1 Dfangzhongdai
Dickeya gs fangzhongdai M005 WP 039692365 1 Dfangzhongdai
Dickeya gs fangzhongdai NCPPB 3274 WP 042858884 1 Dfangzhongdai
Dickeya gs fangzhongdai S1 WP 049853629 1 Dfangzhongdai
Dickeya gs fangzhongdai MK7 WP 038917969 1 Dfangzhongdai
Dickeya gs fangzhongdai B16 WP 038917969 1 Dfangzhongdai
Dickeya dadantii NCPPB 898 WP 038910380 1 Ddadantii
Dickeya dadantii NCPPB 3537 WP 038900429 1 Ddadantii
Dickeya dadantii 3937 WP 013316630 1 Ddadantii
466 norR response to nitro oxidative stress Seed
Dickeya dadantii subsp dieffenbachiae NCPPB 2976 WP 038921868 1 Ddadantii
Dickeya solani RNS 05 1 2A WP 057084046 1 Dsolani
Dickeya solani MK10 WP 022632332 1 Dsolani
Dickeya solani GBBC 2040 WP 022632332 1 Dsolani
Dickeya solani IPO 2222 WP 022632332 1 Dsolani
Dickeya solani PPO 9019 WP 022632332 1 Dsolani
Dickeya solani MK16 WP 022632332 1 Dsolani
Dickeya solani IFB 0099 WP 022632332 1 Dsolani
Dickeya solani D s0432 1 WP 022632332 1 Dsolani
Dickeya solani RNS 07 7 3B WP 022632332 1 Dsolani
Dickeya solani RNS 08 23 3 1 A WP 022632332 1 Dsolani

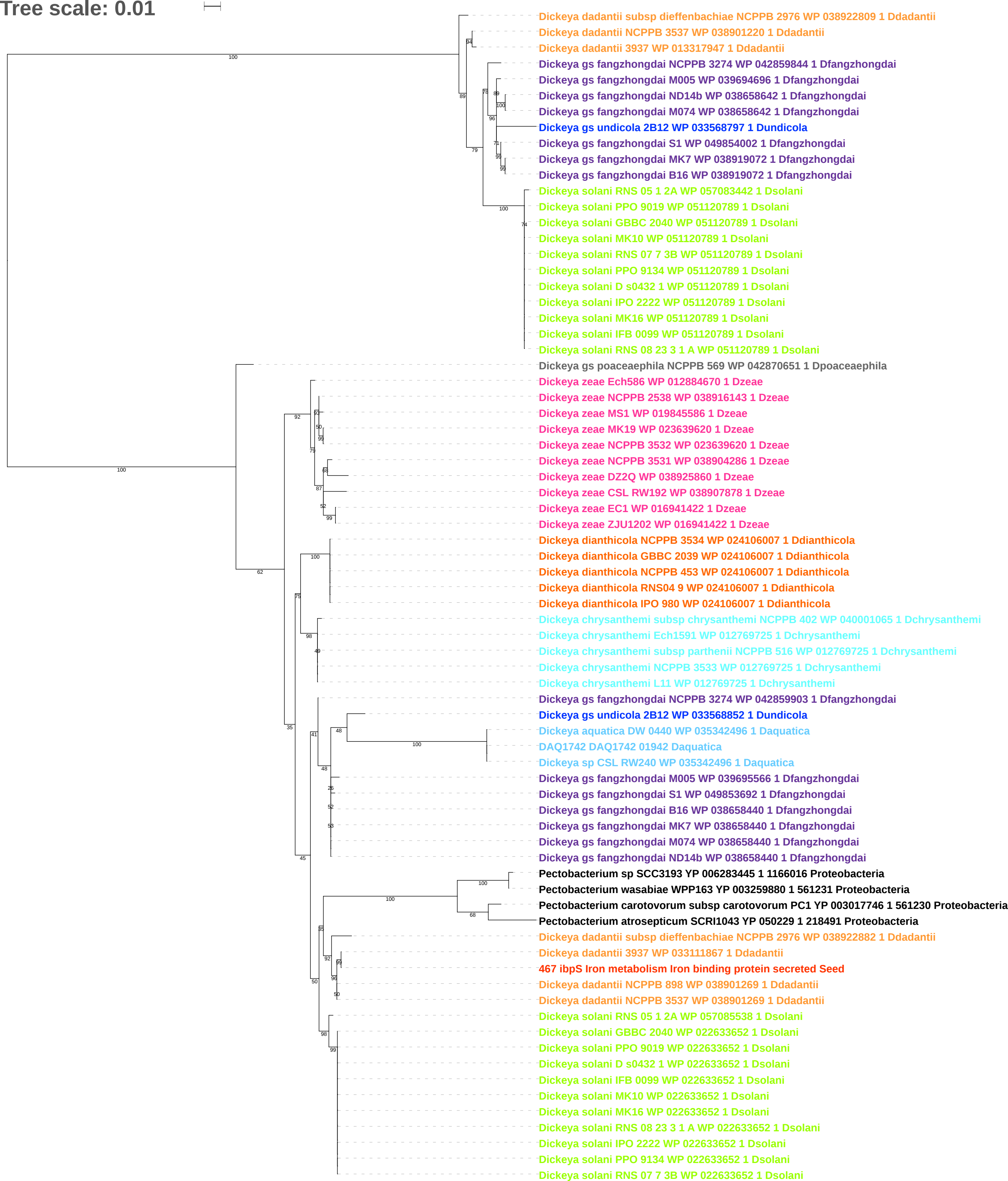
Tree scale: 0.01
Dickeya dadantii subsp dieffenbachiae NCPPB 2976 WP 038922809 1 Ddadantii
Dickeya dadantii NCPPB 3537 WP 038901220 1 Ddadantii
Dickeya dadantii 3937 WP 013317947 1 Ddadantii
Dickeya gs fangzhongdai NCPPB 3274 WP 042859844 1 Dfangzhongdai
Dickeya gs fangzhongdai M005 WP 039694696 1 Dfangzhongdai
Dickeya gs fangzhongdai ND14b WP 038658642 1 Dfangzhongdai
Dickeya gs fangzhongdai M074 WP 038658642 1 Dfangzhongdai
Dickeya gs undicola 2B12 WP 033568797 1 Dundicola
Dickeya gs fangzhongdai S1 WP 049854002 1 Dfangzhongdai
Dickeya gs fangzhongdai MK7 WP 038919072 1 Dfangzhongdai
Dickeya gs fangzhongdai B16 WP 038919072 1 Dfangzhongdai
Dickeya solani RNS 05 1 2A WP 057083442 1 Dsolani
Dickeya solani PPO 9019 WP 051120789 1 Dsolani
Dickeya solani GBBC 2040 WP 051120789 1 Dsolani
Dickeya solani MK10 WP 051120789 1 Dsolani
Dickeya solani RNS 07 7 3B WP 051120789 1 Dsolani
Dickeya solani PPO 9134 WP 051120789 1 Dsolani
Dickeya solani D s0432 1 WP 051120789 1 Dsolani
Dickeya solani IPO 2222 WP 051120789 1 Dsolani
Dickeya solani MK16 WP 051120789 1 Dsolani
Dickeya solani IFB 0099 WP 051120789 1 Dsolani
Dickeya solani RNS 08 23 3 1 A WP 051120789 1 Dsolani
Dickeya gs poaceaephila NCPPB 569 WP 042870651 1 Dpoaceaephila
Dickeya zeae Ech586 WP 012884670 1 Dzeae
Dickeya zeae NCPPB 2538 WP 038916143 1 Dzeae
Dickeya zeae MS1 WP 019845586 1 Dzeae
Dickeya zeae MK19 WP 023639620 1 Dzeae
Dickeya zeae NCPPB 3532 WP 023639620 1 Dzeae
Dickeya zeae NCPPB 3531 WP 038904286 1 Dzeae
Dickeya zeae DZ2Q WP 038925860 1 Dzeae
Dickeya zeae CSL RW192 WP 038907878 1 Dzeae
Dickeya zeae EC1 WP 016941422 1 Dzeae
Dickeya zeae ZJU1202 WP 016941422 1 Dzeae
Dickeya dianthicola NCPPB 3534 WP 024106007 1 Ddianthicola
Dickeya dianthicola GBBC 2039 WP 024106007 1 Ddianthicola
Dickeya dianthicola NCPPB 453 WP 024106007 1 Ddianthicola
Dickeya dianthicola RNS04 9 WP 024106007 1 Ddianthicola
Dickeya dianthicola IPO 980 WP 024106007 1 Ddianthicola
Dickeya chrysanthemi subsp chrysanthemi NCPPB 402 WP 040001065 1 Dchrysanthemi
Dickeya chrysanthemi Ech1591 WP 012769725 1 Dchrysanthemi
Dickeya chrysanthemi subsp parthenii NCPPB 516 WP 012769725 1 Dchrysanthemi
Dickeya chrysanthemi NCPPB 3533 WP 012769725 1 Dchrysanthemi
Dickeya chrysanthemi L11 WP 012769725 1 Dchrysanthemi
Dickeya gs fangzhongdai NCPPB 3274 WP 042859903 1 Dfangzhongdai
Dickeya gs undicola 2B12 WP 033568852 1 Dundicola
Dickeya aquatica DW 0440 WP 035342496 1 Daquatica
DAQ1742 DAQ1742 01942 Daquatica
Dickeya sp CSL RW240 WP 035342496 1 Daquatica
Dickeya gs fangzhongdai M005 WP 039695566 1 Dfangzhongdai
Dickeya gs fangzhongdai S1 WP 049853692 1 Dfangzhongdai
Dickeya gs fangzhongdai B16 WP 038658440 1 Dfangzhongdai
Dickeya gs fangzhongdai MK7 WP 038658440 1 Dfangzhongdai
Dickeya gs fangzhongdai M074 WP 038658440 1 Dfangzhongdai
Dickeya gs fangzhongdai ND14b WP 038658440 1 Dfangzhongdai
Pectobacterium sp SCC3193 YP 006283445 1 1166016 Proteobacteria
Pectobacterium wasabiae WPP163 YP 003259880 1 561231 Proteobacteria
Pectobacterium carotovorum subsp carotovorum PC1 YP 003017746 1 561230 Proteobacteria
Pectobacterium atrosepticum SCRI1043 YP 050229 1 218491 Proteobacteria
Dickeya dadantii subsp dieffenbachiae NCPPB 2976 WP 038922882 1 Ddadantii
Dickeya dadantii 3937 WP 033111867 1 Ddadantii
467 ibpS Iron metabolism Iron binding protein secreted Seed
Dickeya dadantii NCPPB 898 WP 038901269 1 Ddadantii
Dickeya dadantii NCPPB 3537 WP 038901269 1 Ddadantii
Dickeya solani RNS 05 1 2A WP 057085538 1 Dsolani
Dickeya solani GBBC 2040 WP 022633652 1 Dsolani
Dickeya solani PPO 9019 WP 022633652 1 Dsolani
Dickeya solani D s0432 1 WP 022633652 1 Dsolani
Dickeya solani IFB 0099 WP 022633652 1 Dsolani
Dickeya solani MK10 WP 022633652 1 Dsolani
Dickeya solani MK16 WP 022633652 1 Dsolani
Dickeya solani RNS 08 23 3 1 A WP 022633652 1 Dsolani
Dickeya solani IPO 2222 WP 022633652 1 Dsolani
Dickeya solani PPO 9134 WP 022633652 1 Dsolani
Dickeya solani RNS 07 7 3B WP 022633652 1 Dsolani

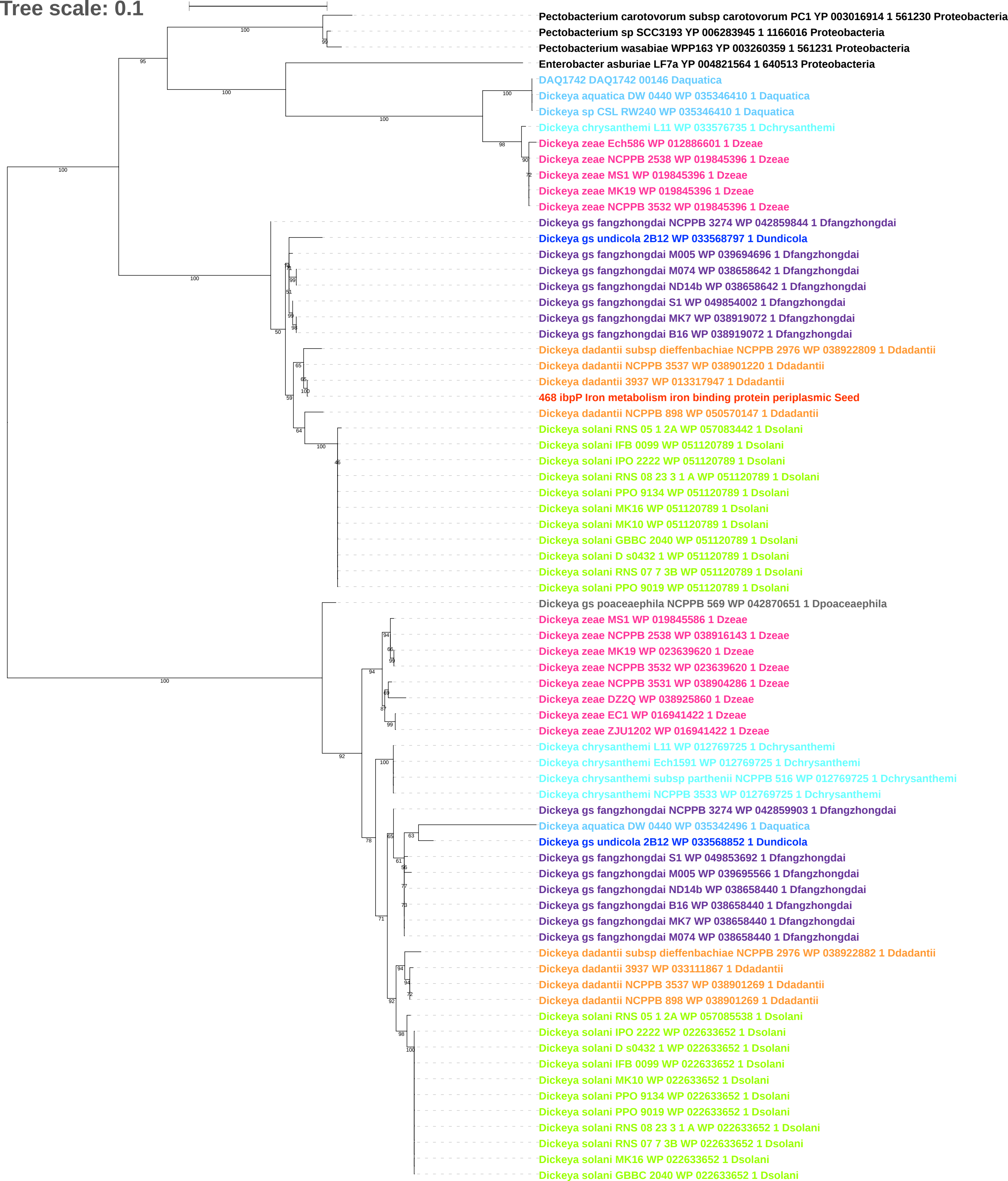
Tree scale: 0.1
Pectobacterium carotovorum subsp carotovorum PC1 YP 003016914 1 561230 Proteobacteria
Pectobacterium sp SCC3193 YP 006283945 1 1166016 Proteobacteria
Pectobacterium wasabiae WPP163 YP 003260359 1 561231 Proteobacteria
Enterobacter asburiae LF7a YP 004821564 1 640513 Proteobacteria
DAQ1742 DAQ1742 00146 Daquatica
Dickeya aquatica DW 0440 WP 035346410 1 Daquatica
Dickeya sp CSL RW240 WP 035346410 1 Daquatica
Dickeya chrysanthemi L11 WP 033576735 1 Dchrysanthemi
Dickeya zeae Ech586 WP 012886601 1 Dzeae
Dickeya zeae NCPPB 2538 WP 019845396 1 Dzeae
Dickeya zeae MS1 WP 019845396 1 Dzeae
Dickeya zeae MK19 WP 019845396 1 Dzeae
Dickeya zeae NCPPB 3532 WP 019845396 1 Dzeae
Dickeya gs fangzhongdai NCPPB 3274 WP 042859844 1 Dfangzhongdai
Dickeya gs undicola 2B12 WP 033568797 1 Dundicola
Dickeya gs fangzhongdai M005 WP 039694696 1 Dfangzhongdai
Dickeya gs fangzhongdai M074 WP 038658642 1 Dfangzhongdai
Dickeya gs fangzhongdai ND14b WP 038658642 1 Dfangzhongdai
Dickeya gs fangzhongdai S1 WP 049854002 1 Dfangzhongdai
Dickeya gs fangzhongdai MK7 WP 038919072 1 Dfangzhongdai
Dickeya gs fangzhongdai B16 WP 038919072 1 Dfangzhongdai
Dickeya dadantii subsp dieffenbachiae NCPPB 2976 WP 038922809 1 Ddadantii
Dickeya dadantii NCPPB 3537 WP 038901220 1 Ddadantii
Dickeya dadantii 3937 WP 013317947 1 Ddadantii
468 ibpP Iron metabolism iron binding protein periplasmic Seed
Dickeya dadantii NCPPB 898 WP 050570147 1 Ddadantii
Dickeya solani RNS 05 1 2A WP 057083442 1 Dsolani
Dickeya solani IFB 0099 WP 051120789 1 Dsolani
Dickeya solani IPO 2222 WP 051120789 1 Dsolani
Dickeya solani RNS 08 23 3 1 A WP 051120789 1 Dsolani
Dickeya solani PPO 9134 WP 051120789 1 Dsolani
Dickeya solani MK16 WP 051120789 1 Dsolani
Dickeya solani MK10 WP 051120789 1 Dsolani
Dickeya solani GBBC 2040 WP 051120789 1 Dsolani
Dickeya solani D s0432 1 WP 051120789 1 Dsolani
Dickeya solani RNS 07 7 3B WP 051120789 1 Dsolani
Dickeya solani PPO 9019 WP 051120789 1 Dsolani
Dickeya gs poaceaephila NCPPB 569 WP 042870651 1 Dpoaceaephila
Dickeya zeae MS1 WP 019845586 1 Dzeae
Dickeya zeae NCPPB 2538 WP 038916143 1 Dzeae
Dickeya zeae MK19 WP 023639620 1 Dzeae
Dickeya zeae NCPPB 3532 WP 023639620 1 Dzeae
Dickeya zeae NCPPB 3531 WP 038904286 1 Dzeae
Dickeya zeae DZ2Q WP 038925860 1 Dzeae
Dickeya zeae EC1 WP 016941422 1 Dzeae
Dickeya zeae ZJU1202 WP 016941422 1 Dzeae
Dickeya chrysanthemi L11 WP 012769725 1 Dchrysanthemi
Dickeya chrysanthemi Ech1591 WP 012769725 1 Dchrysanthemi
Dickeya chrysanthemi subsp parthenii NCPPB 516 WP 012769725 1 Dchrysanthemi
Dickeya chrysanthemi NCPPB 3533 WP 012769725 1 Dchrysanthemi
Dickeya gs fangzhongdai NCPPB 3274 WP 042859903 1 Dfangzhongdai
Dickeya aquatica DW 0440 WP 035342496 1 Daquatica
Dickeya gs undicola 2B12 WP 033568852 1 Dundicola
Dickeya gs fangzhongdai S1 WP 049853692 1 Dfangzhongdai
Dickeya gs fangzhongdai M005 WP 039695566 1 Dfangzhongdai
Dickeya gs fangzhongdai ND14b WP 038658440 1 Dfangzhongdai
Dickeya gs fangzhongdai B16 WP 038658440 1 Dfangzhongdai
Dickeya gs fangzhongdai MK7 WP 038658440 1 Dfangzhongdai
Dickeya gs fangzhongdai M074 WP 038658440 1 Dfangzhongdai
Dickeya dadantii subsp dieffenbachiae NCPPB 2976 WP 038922882 1 Ddadantii
Dickeya dadantii 3937 WP 033111867 1 Ddadantii
Dickeya dadantii NCPPB 3537 WP 038901269 1 Ddadantii
Dickeya dadantii NCPPB 898 WP 038901269 1 Ddadantii
Dickeya solani RNS 05 1 2A WP 057085538 1 Dsolani
Dickeya solani IPO 2222 WP 022633652 1 Dsolani
Dickeya solani D s0432 1 WP 022633652 1 Dsolani
Dickeya solani IFB 0099 WP 022633652 1 Dsolani
Dickeya solani MK10 WP 022633652 1 Dsolani
Dickeya solani PPO 9134 WP 022633652 1 Dsolani
Dickeya solani PPO 9019 WP 022633652 1 Dsolani
Dickeya solani RNS 08 23 3 1 A WP 022633652 1 Dsolani
Dickeya solani RNS 07 7 3B WP 022633652 1 Dsolani
Dickeya solani MK16 WP 022633652 1 Dsolani
Dickeya solani GBBC 2040 WP 022633652 1 Dsolani

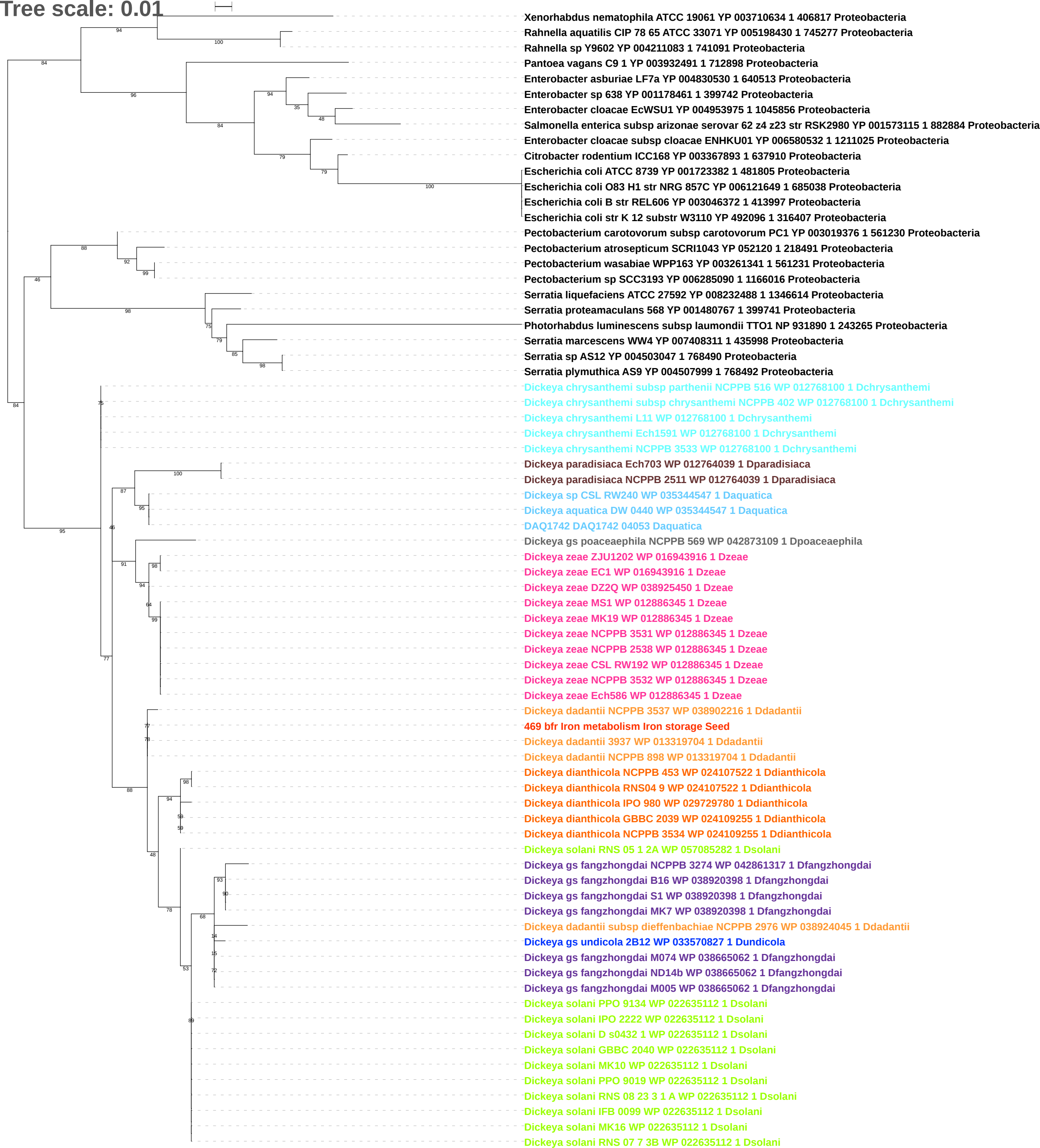
Tree scale: 0.01
Xenorhabdus nematophila ATCC 19061 YP 003710634 1 406817 Proteobacteria
Rahnella aquatilis CIP 78 65 ATCC 33071 YP 005198430 1 745277 Proteobacteria
Rahnella sp Y9602 YP 004211083 1 741091 Proteobacteria
Pantoea vagans C9 1 YP 003932491 1 712898 Proteobacteria
Enterobacter asburiae LF7a YP 004830530 1 640513 Proteobacteria
Enterobacter sp 638 YP 001178461 1 399742 Proteobacteria
Enterobacter cloacae EcWSU1 YP 004953975 1 1045856 Proteobacteria
Salmonella enterica subsp arizonae serovar 62 z4 z23 str RSK2980 YP 001573115 1 882884 Proteobacteria
Enterobacter cloacae subsp cloacae ENHKU01 YP 006580532 1 1211025 Proteobacteria
Citrobacter rodentium ICC168 YP 003367893 1 637910 Proteobacteria
Escherichia coli ATCC 8739 YP 001723382 1 481805 Proteobacteria
Escherichia coli O83 H1 str NRG 857C YP 006121649 1 685038 Proteobacteria
Escherichia coli B str REL606 YP 003046372 1 413997 Proteobacteria
Escherichia coli str K 12 substr W3110 YP 492096 1 316407 Proteobacteria
Pectobacterium carotovorum subsp carotovorum PC1 YP 003019376 1 561230 Proteobacteria
Pectobacterium atrosepticum SCRI1043 YP 052120 1 218491 Proteobacteria
Pectobacterium wasabiae WPP163 YP 003261341 1 561231 Proteobacteria
Pectobacterium sp SCC3193 YP 006285090 1 1166016 Proteobacteria
Serratia liquefaciens ATCC 27592 YP 008232488 1 1346614 Proteobacteria
Serratia proteamaculans 568 YP 001480767 1 399741 Proteobacteria
Photorhabdus luminescens subsp laumondii TTO1 NP 931890 1 243265 Proteobacteria
Serratia marcescens WW4 YP 007408311 1 435998 Proteobacteria
Serratia sp AS12 YP 004503047 1 768490 Proteobacteria
Serratia plymuthica AS9 YP 004507999 1 768492 Proteobacteria
Dickeya chrysanthemi subsp parthenii NCPPB 516 WP 012768100 1 Dchrysanthemi
Dickeya chrysanthemi subsp chrysanthemi NCPPB 402 WP 012768100 1 Dchrysanthemi
Dickeya chrysanthemi L11 WP 012768100 1 Dchrysanthemi
Dickeya chrysanthemi Ech1591 WP 012768100 1 Dchrysanthemi
Dickeya chrysanthemi NCPPB 3533 WP 012768100 1 Dchrysanthemi
Dickeya paradisiaca Ech703 WP 012764039 1 Dparadisiaca
Dickeya paradisiaca NCPPB 2511 WP 012764039 1 Dparadisiaca
Dickeya sp CSL RW240 WP 035344547 1 Daquatica
Dickeya aquatica DW 0440 WP 035344547 1 Daquatica
DAQ1742 DAQ1742 04053 Daquatica
Dickeya gs poaceaephila NCPPB 569 WP 042873109 1 Dpoaceaephila
Dickeya zeae ZJU1202 WP 016943916 1 Dzeae
Dickeya zeae EC1 WP 016943916 1 Dzeae
Dickeya zeae DZ2Q WP 038925450 1 Dzeae
Dickeya zeae MS1 WP 012886345 1 Dzeae
Dickeya zeae MK19 WP 012886345 1 Dzeae
Dickeya zeae NCPPB 3531 WP 012886345 1 Dzeae
Dickeya zeae NCPPB 2538 WP 012886345 1 Dzeae
Dickeya zeae CSL RW192 WP 012886345 1 Dzeae
Dickeya zeae NCPPB 3532 WP 012886345 1 Dzeae
Dickeya zeae Ech586 WP 012886345 1 Dzeae
Dickeya dadantii NCPPB 3537 WP 038902216 1 Ddadantii
469 bfr Iron metabolism Iron storage Seed
Dickeya dadantii 3937 WP 013319704 1 Ddadantii
Dickeya dadantii NCPPB 898 WP 013319704 1 Ddadantii
Dickeya dianthicola NCPPB 453 WP 024107522 1 Ddianthicola
Dickeya dianthicola RNS04 9 WP 024107522 1 Ddianthicola
Dickeya dianthicola IPO 980 WP 029729780 1 Ddianthicola
Dickeya dianthicola GBBC 2039 WP 024109255 1 Ddianthicola
Dickeya dianthicola NCPPB 3534 WP 024109255 1 Ddianthicola
Dickeya solani RNS 05 1 2A WP 057085282 1 Dsolani
Dickeya gs fangzhongdai NCPPB 3274 WP 042861317 1 Dfangzhongdai
Dickeya gs fangzhongdai B16 WP 038920398 1 Dfangzhongdai
Dickeya gs fangzhongdai S1 WP 038920398 1 Dfangzhongdai
Dickeya gs fangzhongdai MK7 WP 038920398 1 Dfangzhongdai
Dickeya dadantii subsp dieffenbachiae NCPPB 2976 WP 038924045 1 Ddadantii
Dickeya gs undicola 2B12 WP 033570827 1 Dundicola
Dickeya gs fangzhongdai M074 WP 038665062 1 Dfangzhongdai
Dickeya gs fangzhongdai ND14b WP 038665062 1 Dfangzhongdai
Dickeya gs fangzhongdai M005 WP 038665062 1 Dfangzhongdai
Dickeya solani PPO 9134 WP 022635112 1 Dsolani
Dickeya solani IPO 2222 WP 022635112 1 Dsolani
Dickeya solani D s0432 1 WP 022635112 1 Dsolani
Dickeya solani GBBC 2040 WP 022635112 1 Dsolani
Dickeya solani MK10 WP 022635112 1 Dsolani
Dickeya solani PPO 9019 WP 022635112 1 Dsolani
Dickeya solani RNS 08 23 3 1 A WP 022635112 1 Dsolani
Dickeya solani IFB 0099 WP 022635112 1 Dsolani
Dickeya solani MK16 WP 022635112 1 Dsolani
Dickeya solani RNS 07 7 3B WP 022635112 1 Dsolani

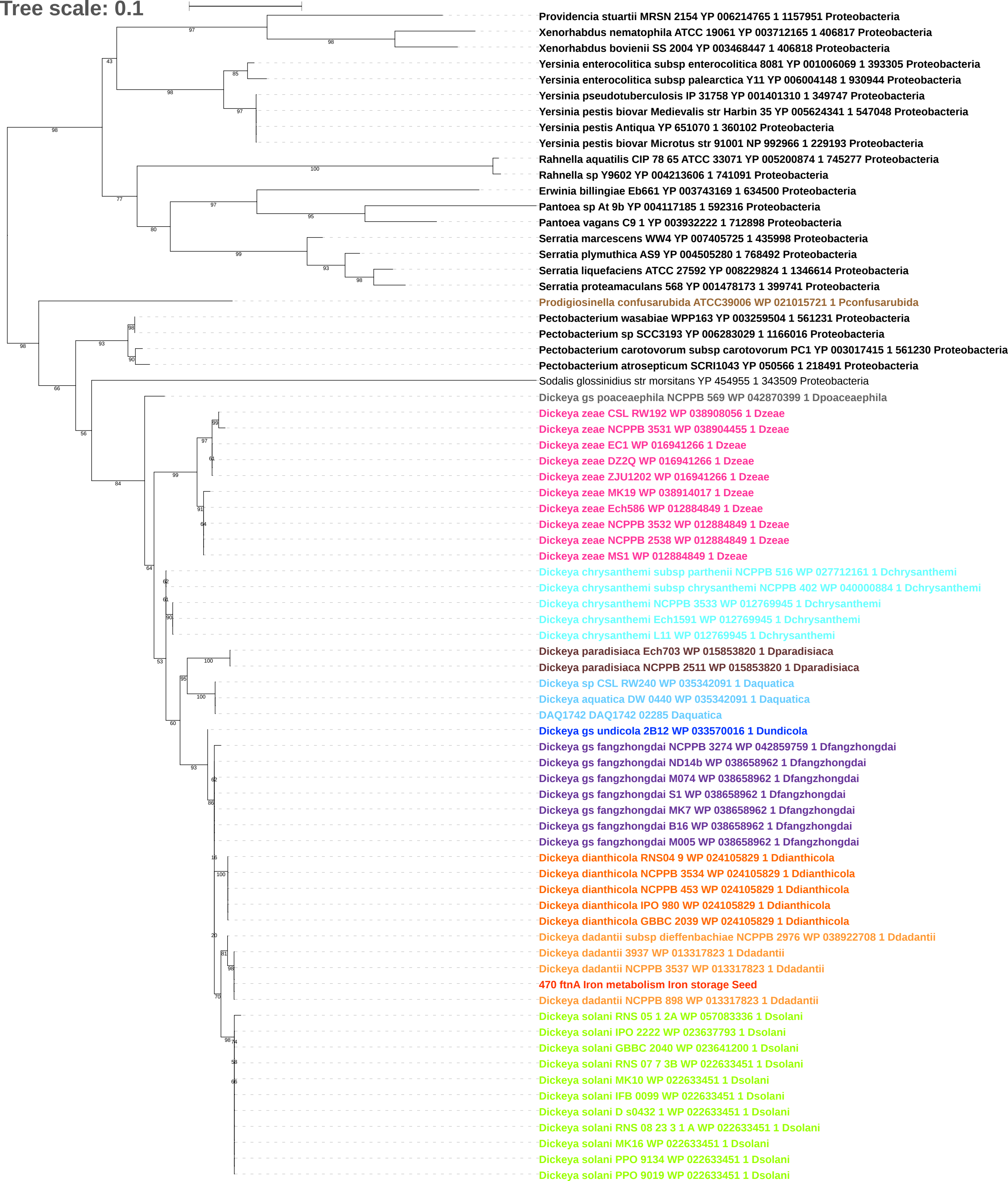
Tree scale: 0.1
Providencia stuartii MRSN 2154 YP 006214765 1 1157951 Proteobacteria
Xenorhabdus nematophila ATCC 19061 YP 003712165 1 406817 Proteobacteria
Xenorhabdus bovienii SS 2004 YP 003468447 1 406818 Proteobacteria
Yersinia enterocolitica subsp enterocolitica 8081 YP 001006069 1 393305 Proteobacteria
Yersinia enterocolitica subsp palearctica Y11 YP 006004148 1 930944 Proteobacteria
Yersinia pseudotuberculosis IP 31758 YP 001401310 1 349747 Proteobacteria
Yersinia pestis biovar Medievalis str Harbin 35 YP 005624341 1 547048 Proteobacteria
Yersinia pestis Antiqua YP 651070 1 360102 Proteobacteria
Yersinia pestis biovar Microtus str 91001 NP 992966 1 229193 Proteobacteria
Rahnella aquatilis CIP 78 65 ATCC 33071 YP 005200874 1 745277 Proteobacteria
Rahnella sp Y9602 YP 004213606 1 741091 Proteobacteria
Erwinia billingiae Eb661 YP 003743169 1 634500 Proteobacteria
Pantoea sp At 9b YP 004117185 1 592316 Proteobacteria
Pantoea vagans C9 1 YP 003932222 1 712898 Proteobacteria
Serratia marcescens WW4 YP 007405725 1 435998 Proteobacteria
Serratia plymuthica AS9 YP 004505280 1 768492 Proteobacteria
Serratia liquefaciens ATCC 27592 YP 008229824 1 1346614 Proteobacteria
Serratia proteamaculans 568 YP 001478173 1 399741 Proteobacteria
Prodigiosinella confusarubida ATCC39006 WP 021015721 1 Pconfusarubida
Pectobacterium wasabiae WPP163 YP 003259504 1 561231 Proteobacteria
Pectobacterium sp SCC3193 YP 006283029 1 1166016 Proteobacteria
Pectobacterium carotovorum subsp carotovorum PC1 YP 003017415 1 561230 Proteobacteria
Pectobacterium atrosepticum SCRI1043 YP 050566 1 218491 Proteobacteria
Sodalis glossinidius str morsitans YP 454955 1 343509 Proteobacteria
Dickeya gs poaceaephila NCPPB 569 WP 042870399 1 Dpoaceaephila
Dickeya zeae CSL RW192 WP 038908056 1 Dzeae
Dickeya zeae NCPPB 3531 WP 038904455 1 Dzeae
Dickeya zeae EC1 WP 016941266 1 Dzeae
Dickeya zeae DZ2Q WP 016941266 1 Dzeae
Dickeya zeae ZJU1202 WP 016941266 1 Dzeae
Dickeya zeae MK19 WP 038914017 1 Dzeae
Dickeya zeae Ech586 WP 012884849 1 Dzeae
Dickeya zeae NCPPB 3532 WP 012884849 1 Dzeae
Dickeya zeae NCPPB 2538 WP 012884849 1 Dzeae
Dickeya zeae MS1 WP 012884849 1 Dzeae
Dickeya chrysanthemi subsp parthenii NCPPB 516 WP 027712161 1 Dchrysanthemi
Dickeya chrysanthemi subsp chrysanthemi NCPPB 402 WP 040000884 1 Dchrysanthemi
Dickeya chrysanthemi NCPPB 3533 WP 012769945 1 Dchrysanthemi
Dickeya chrysanthemi Ech1591 WP 012769945 1 Dchrysanthemi
Dickeya chrysanthemi L11 WP 012769945 1 Dchrysanthemi
Dickeya paradisiaca Ech703 WP 015853820 1 Dparadisiaca
Dickeya paradisiaca NCPPB 2511 WP 015853820 1 Dparadisiaca
Dickeya sp CSL RW240 WP 035342091 1 Daquatica
Dickeya aquatica DW 0440 WP 035342091 1 Daquatica
DAQ1742 DAQ1742 02285 Daquatica
Dickeya gs undicola 2B12 WP 033570016 1 Dundicola
Dickeya gs fangzhongdai NCPPB 3274 WP 042859759 1 Dfangzhongdai
Dickeya gs fangzhongdai ND14b WP 038658962 1 Dfangzhongdai
Dickeya gs fangzhongdai M074 WP 038658962 1 Dfangzhongdai
Dickeya gs fangzhongdai S1 WP 038658962 1 Dfangzhongdai
Dickeya gs fangzhongdai MK7 WP 038658962 1 Dfangzhongdai
Dickeya gs fangzhongdai B16 WP 038658962 1 Dfangzhongdai
Dickeya gs fangzhongdai M005 WP 038658962 1 Dfangzhongdai
Dickeya dianthicola RNS04 9 WP 024105829 1 Ddianthicola
Dickeya dianthicola NCPPB 3534 WP 024105829 1 Ddianthicola
Dickeya dianthicola NCPPB 453 WP 024105829 1 Ddianthicola
Dickeya dianthicola IPO 980 WP 024105829 1 Ddianthicola
Dickeya dianthicola GBBC 2039 WP 024105829 1 Ddianthicola
Dickeya dadantii subsp dieffenbachiae NCPPB 2976 WP 038922708 1 Ddadantii
Dickeya dadantii 3937 WP 013317823 1 Ddadantii
Dickeya dadantii NCPPB 3537 WP 013317823 1 Ddadantii
470 ftnA Iron metabolism Iron storage Seed
Dickeya dadantii NCPPB 898 WP 013317823 1 Ddadantii
Dickeya solani RNS 05 1 2A WP 057083336 1 Dsolani
Dickeya solani IPO 2222 WP 023637793 1 Dsolani
Dickeya solani GBBC 2040 WP 023641200 1 Dsolani
Dickeya solani RNS 07 7 3B WP 022633451 1 Dsolani
Dickeya solani MK10 WP 022633451 1 Dsolani
Dickeya solani IFB 0099 WP 022633451 1 Dsolani
Dickeya solani D s0432 1 WP 022633451 1 Dsolani
Dickeya solani RNS 08 23 3 1 A WP 022633451 1 Dsolani
Dickeya solani MK16 WP 022633451 1 Dsolani
Dickeya solani PPO 9134 WP 022633451 1 Dsolani
Dickeya solani PPO 9019 WP 022633451 1 Dsolani

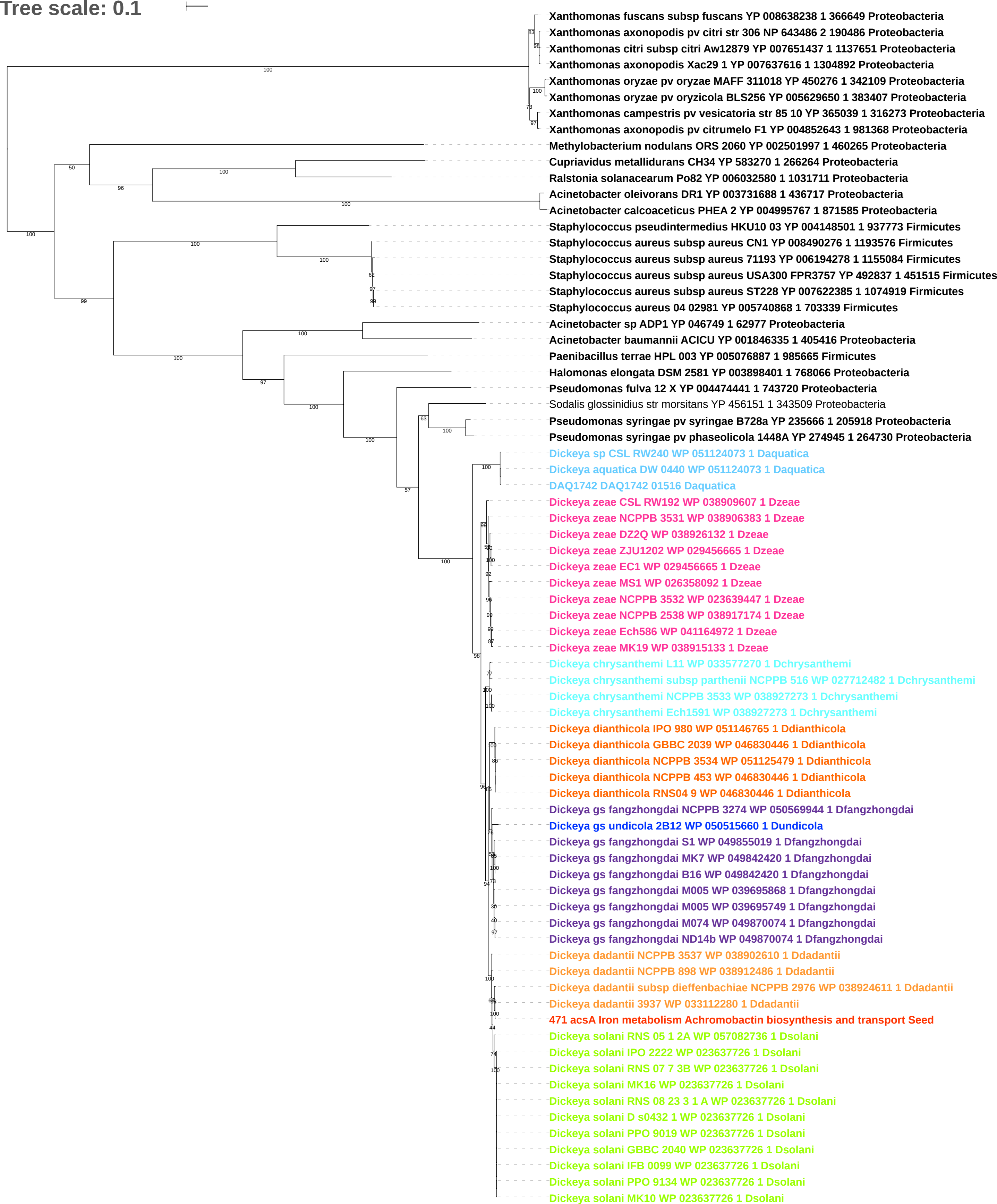
Tree scale: 0.1
Xanthomonas fuscans subsp fuscans YP 008638238 1 366649 Proteobacteria
Xanthomonas axonopodis pv citri str 306 NP 643486 2 190486 Proteobacteria
Xanthomonas citri subsp citri Aw12879 YP 007651437 1 1137651 Proteobacteria
Xanthomonas axonopodis Xac29 1 YP 007637616 1 1304892 Proteobacteria
Xanthomonas oryzae pv oryzae MAFF 311018 YP 450276 1 342109 Proteobacteria
Xanthomonas oryzae pv oryzicola BLS256 YP 005629650 1 383407 Proteobacteria
Xanthomonas campestris pv vesicatoria str 85 10 YP 365039 1 316273 Proteobacteria
Xanthomonas axonopodis pv citrumelo F1 YP 004852643 1 981368 Proteobacteria
Methylobacterium nodulans ORS 2060 YP 002501997 1 460265 Proteobacteria
Cupriavidus metallidurans CH34 YP 583270 1 266264 Proteobacteria
Ralstonia solanacearum Po82 YP 006032580 1 1031711 Proteobacteria
Acinetobacter oleivorans DR1 YP 003731688 1 436717 Proteobacteria
Acinetobacter calcoaceticus PHEA 2 YP 004995767 1 871585 Proteobacteria
Staphylococcus pseudintermedius HKU10 03 YP 004148501 1 937773 Firmicutes
Staphylococcus aureus subsp aureus CN1 YP 008490276 1 1193576 Firmicutes
Staphylococcus aureus subsp aureus 71193 YP 006194278 1 1155084 Firmicutes
Staphylococcus aureus subsp aureus USA300 FPR3757 YP 492837 1 451515 Firmicutes
Staphylococcus aureus subsp aureus ST228 YP 007622385 1 1074919 Firmicutes
Staphylococcus aureus 04 02981 YP 005740868 1 703339 Firmicutes
Acinetobacter sp ADP1 YP 046749 1 62977 Proteobacteria
Acinetobacter baumannii ACICU YP 001846335 1 405416 Proteobacteria
Paenibacillus terrae HPL 003 YP 005076887 1 985665 Firmicutes
Halomonas elongata DSM 2581 YP 003898401 1 768066 Proteobacteria
Pseudomonas fulva 12 X YP 004474441 1 743720 Proteobacteria
Sodalis glossinidius str morsitans YP 456151 1 343509 Proteobacteria
Pseudomonas syringae pv syringae B728a YP 235666 1 205918 Proteobacteria
Pseudomonas syringae pv phaseolicola 1448A YP 274945 1 264730 Proteobacteria
Dickeya sp CSL RW240 WP 051124073 1 Daquatica
Dickeya aquatica DW 0440 WP 051124073 1 Daquatica
DAQ1742 DAQ1742 01516 Daquatica
Dickeya zeae CSL RW192 WP 038909607 1 Dzeae
Dickeya zeae NCPPB 3531 WP 038906383 1 Dzeae
Dickeya zeae DZ2Q WP 038926132 1 Dzeae
Dickeya zeae ZJU1202 WP 029456665 1 Dzeae
Dickeya zeae EC1 WP 029456665 1 Dzeae
Dickeya zeae MS1 WP 026358092 1 Dzeae
Dickeya zeae NCPPB 3532 WP 023639447 1 Dzeae
Dickeya zeae NCPPB 2538 WP 038917174 1 Dzeae
Dickeya zeae Ech586 WP 041164972 1 Dzeae
Dickeya zeae MK19 WP 038915133 1 Dzeae
Dickeya chrysanthemi L11 WP 033577270 1 Dchrysanthemi
Dickeya chrysanthemi subsp parthenii NCPPB 516 WP 027712482 1 Dchrysanthemi
Dickeya chrysanthemi NCPPB 3533 WP 038927273 1 Dchrysanthemi
Dickeya chrysanthemi Ech1591 WP 038927273 1 Dchrysanthemi
Dickeya dianthicola IPO 980 WP 051146765 1 Ddianthicola
Dickeya dianthicola GBBC 2039 WP 046830446 1 Ddianthicola
Dickeya dianthicola NCPPB 3534 WP 051125479 1 Ddianthicola
Dickeya dianthicola NCPPB 453 WP 046830446 1 Ddianthicola
Dickeya dianthicola RNS04 9 WP 046830446 1 Ddianthicola
Dickeya gs fangzhongdai NCPPB 3274 WP 050569944 1 Dfangzhongdai
Dickeya gs undicola 2B12 WP 050515660 1 Dundicola
Dickeya gs fangzhongdai S1 WP 049855019 1 Dfangzhongdai
Dickeya gs fangzhongdai MK7 WP 049842420 1 Dfangzhongdai
Dickeya gs fangzhongdai B16 WP 049842420 1 Dfangzhongdai
Dickeya gs fangzhongdai M005 WP 039695868 1 Dfangzhongdai
Dickeya gs fangzhongdai M005 WP 039695749 1 Dfangzhongdai
Dickeya gs fangzhongdai M074 WP 049870074 1 Dfangzhongdai
Dickeya gs fangzhongdai ND14b WP 049870074 1 Dfangzhongdai
Dickeya dadantii NCPPB 3537 WP 038902610 1 Ddadantii
Dickeya dadantii NCPPB 898 WP 038912486 1 Ddadantii
Dickeya dadantii subsp dieffenbachiae NCPPB 2976 WP 038924611 1 Ddadantii
Dickeya dadantii 3937 WP 033112280 1 Ddadantii
471 acsA Iron metabolism Achromobactin biosynthesis and transport Seed
Dickeya solani RNS 05 1 2A WP 057082736 1 Dsolani
Dickeya solani IPO 2222 WP 023637726 1 Dsolani
Dickeya solani RNS 07 7 3B WP 023637726 1 Dsolani
Dickeya solani MK16 WP 023637726 1 Dsolani
Dickeya solani RNS 08 23 3 1 A WP 023637726 1 Dsolani
Dickeya solani D s0432 1 WP 023637726 1 Dsolani
Dickeya solani PPO 9019 WP 023637726 1 Dsolani
Dickeya solani GBBC 2040 WP 023637726 1 Dsolani
Dickeya solani IFB 0099 WP 023637726 1 Dsolani
Dickeya solani PPO 9134 WP 023637726 1 Dsolani
Dickeya solani MK10 WP 023637726 1 Dsolani

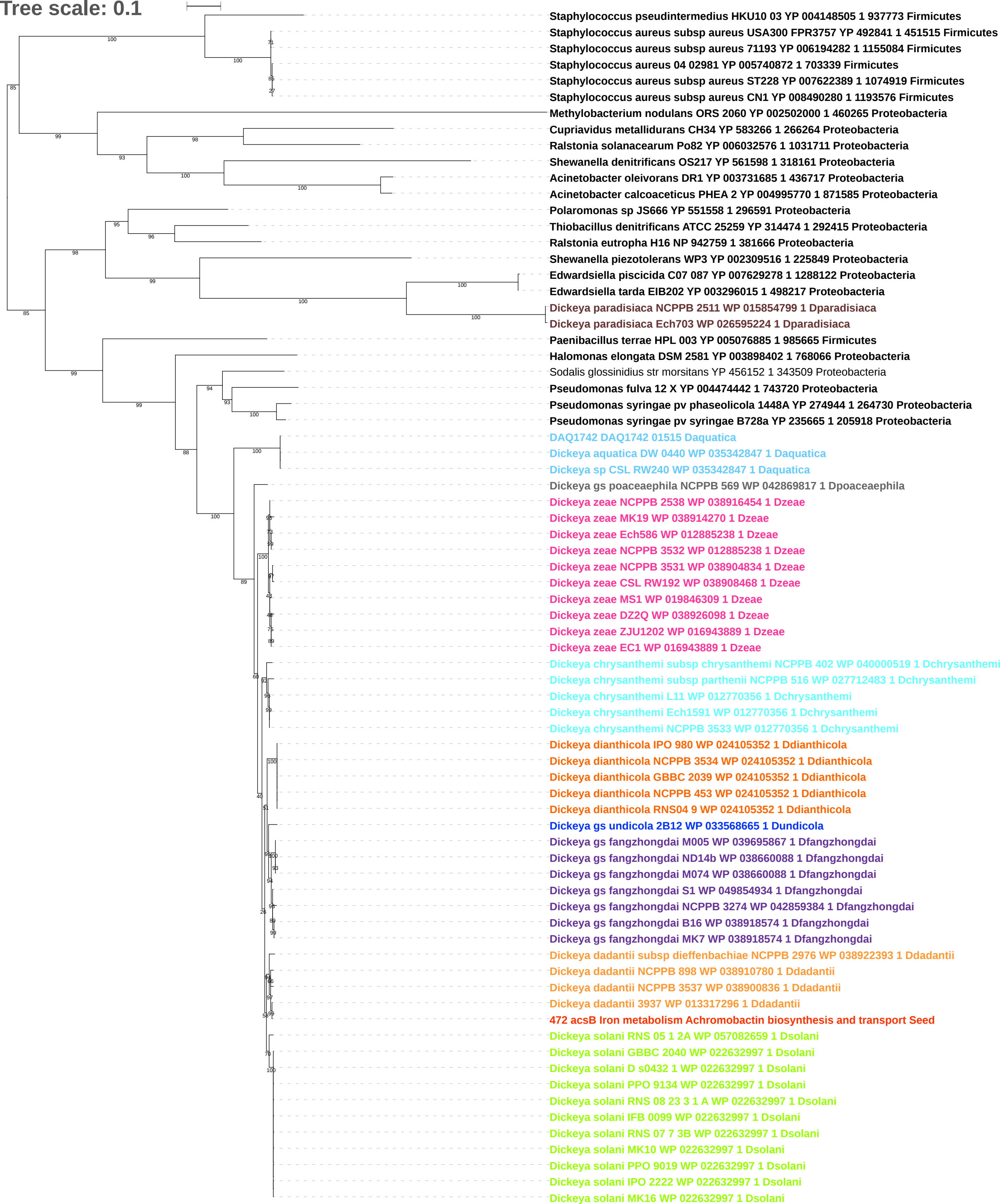
Tree scale: 0.1
Staphylococcus pseudintermedius HKU10 03 YP 004148505 1 937773 Firmicutes
Staphylococcus aureus subsp aureus USA300 FPR3757 YP 492841 1 451515 Firmicutes
Staphylococcus aureus subsp aureus 71193 YP 006194282 1 1155084 Firmicutes
Staphylococcus aureus 04 02981 YP 005740872 1 703339 Firmicutes
Staphylococcus aureus subsp aureus ST228 YP 007622389 1 1074919 Firmicutes
Staphylococcus aureus subsp aureus CN1 YP 008490280 1 1193576 Firmicutes
Methylobacterium nodulans ORS 2060 YP 002502000 1 460265 Proteobacteria
Cupriavidus metallidurans CH34 YP 583266 1 266264 Proteobacteria
Ralstonia solanacearum Po82 YP 006032576 1 1031711 Proteobacteria
Shewanella denitrificans OS217 YP 561598 1 318161 Proteobacteria
Acinetobacter oleivorans DR1 YP 003731685 1 436717 Proteobacteria
Acinetobacter calcoaceticus PHEA 2 YP 004995770 1 871585 Proteobacteria
Polaromonas sp JS666 YP 551558 1 296591 Proteobacteria
Thiobacillus denitrificans ATCC 25259 YP 314474 1 292415 Proteobacteria
Ralstonia eutropha H16 NP 942759 1 381666 Proteobacteria
Shewanella piezotolerans WP3 YP 002309516 1 225849 Proteobacteria
Edwardsiella piscicida C07 087 YP 007629278 1 1288122 Proteobacteria
Edwardsiella tarda EIB202 YP 003296015 1 498217 Proteobacteria
Dickeya paradisiaca NCPPB 2511 WP 015854799 1 Dparadisiaca
Dickeya paradisiaca Ech703 WP 026595224 1 Dparadisiaca
Paenibacillus terrae HPL 003 YP 005076885 1 985665 Firmicutes
Halomonas elongata DSM 2581 YP 003898402 1 768066 Proteobacteria
Sodalis glossinidius str morsitans YP 456152 1 343509 Proteobacteria
Pseudomonas fulva 12 X YP 004474442 1 743720 Proteobacteria
Pseudomonas syringae pv phaseolicola 1448A YP 274944 1 264730 Proteobacteria
Pseudomonas syringae pv syringae B728a YP 235665 1 205918 Proteobacteria
DAQ1742 DAQ1742 01515 Daquatica
Dickeya aquatica DW 0440 WP 035342847 1 Daquatica
Dickeya sp CSL RW240 WP 035342847 1 Daquatica
Dickeya gs poaceaephila NCPPB 569 WP 042869817 1 Dpoaceaephila
Dickeya zeae NCPPB 2538 WP 038916454 1 Dzeae
Dickeya zeae MK19 WP 038914270 1 Dzeae
Dickeya zeae Ech586 WP 012885238 1 Dzeae
Dickeya zeae NCPPB 3532 WP 012885238 1 Dzeae
Dickeya zeae NCPPB 3531 WP 038904834 1 Dzeae
Dickeya zeae CSL RW192 WP 038908468 1 Dzeae
Dickeya zeae MS1 WP 019846309 1 Dzeae
Dickeya zeae DZ2Q WP 038926098 1 Dzeae
Dickeya zeae ZJU1202 WP 016943889 1 Dzeae
Dickeya zeae EC1 WP 016943889 1 Dzeae
Dickeya chrysanthemi subsp chrysanthemi NCPPB 402 WP 040000519 1 Dchrysanthemi
Dickeya chrysanthemi subsp parthenii NCPPB 516 WP 027712483 1 Dchrysanthemi
Dickeya chrysanthemi L11 WP 012770356 1 Dchrysanthemi
Dickeya chrysanthemi Ech1591 WP 012770356 1 Dchrysanthemi
Dickeya chrysanthemi NCPPB 3533 WP 012770356 1 Dchrysanthemi
Dickeya dianthicola IPO 980 WP 024105352 1 Ddianthicola
Dickeya dianthicola NCPPB 3534 WP 024105352 1 Ddianthicola
Dickeya dianthicola GBBC 2039 WP 024105352 1 Ddianthicola
Dickeya dianthicola NCPPB 453 WP 024105352 1 Ddianthicola
Dickeya dianthicola RNS04 9 WP 024105352 1 Ddianthicola
Dickeya gs undicola 2B12 WP 033568665 1 Dundicola
Dickeya gs fangzhongdai M005 WP 039695867 1 Dfangzhongdai
Dickeya gs fangzhongdai ND14b WP 038660088 1 Dfangzhongdai
Dickeya gs fangzhongdai M074 WP 038660088 1 Dfangzhongdai
Dickeya gs fangzhongdai S1 WP 049854934 1 Dfangzhongdai
Dickeya gs fangzhongdai NCPPB 3274 WP 042859384 1 Dfangzhongdai
Dickeya gs fangzhongdai B16 WP 038918574 1 Dfangzhongdai
Dickeya gs fangzhongdai MK7 WP 038918574 1 Dfangzhongdai
Dickeya dadantii subsp dieffenbachiae NCPPB 2976 WP 038922393 1 Ddadantii
Dickeya dadantii NCPPB 898 WP 038910780 1 Ddadantii
Dickeya dadantii NCPPB 3537 WP 038900836 1 Ddadantii
Dickeya dadantii 3937 WP 013317296 1 Ddadantii
472 acsB Iron metabolism Achromobactin biosynthesis and transport Seed
Dickeya solani RNS 05 1 2A WP 057082659 1 Dsolani
Dickeya solani GBBC 2040 WP 022632997 1 Dsolani
Dickeya solani D s0432 1 WP 022632997 1 Dsolani
Dickeya solani PPO 9134 WP 022632997 1 Dsolani
Dickeya solani RNS 08 23 3 1 A WP 022632997 1 Dsolani
Dickeya solani IFB 0099 WP 022632997 1 Dsolani
Dickeya solani RNS 07 7 3B WP 022632997 1 Dsolani
Dickeya solani MK10 WP 022632997 1 Dsolani
Dickeya solani PPO 9019 WP 022632997 1 Dsolani
Dickeya solani IPO 2222 WP 022632997 1 Dsolani
Dickeya solani MK16 WP 022632997 1 Dsolani

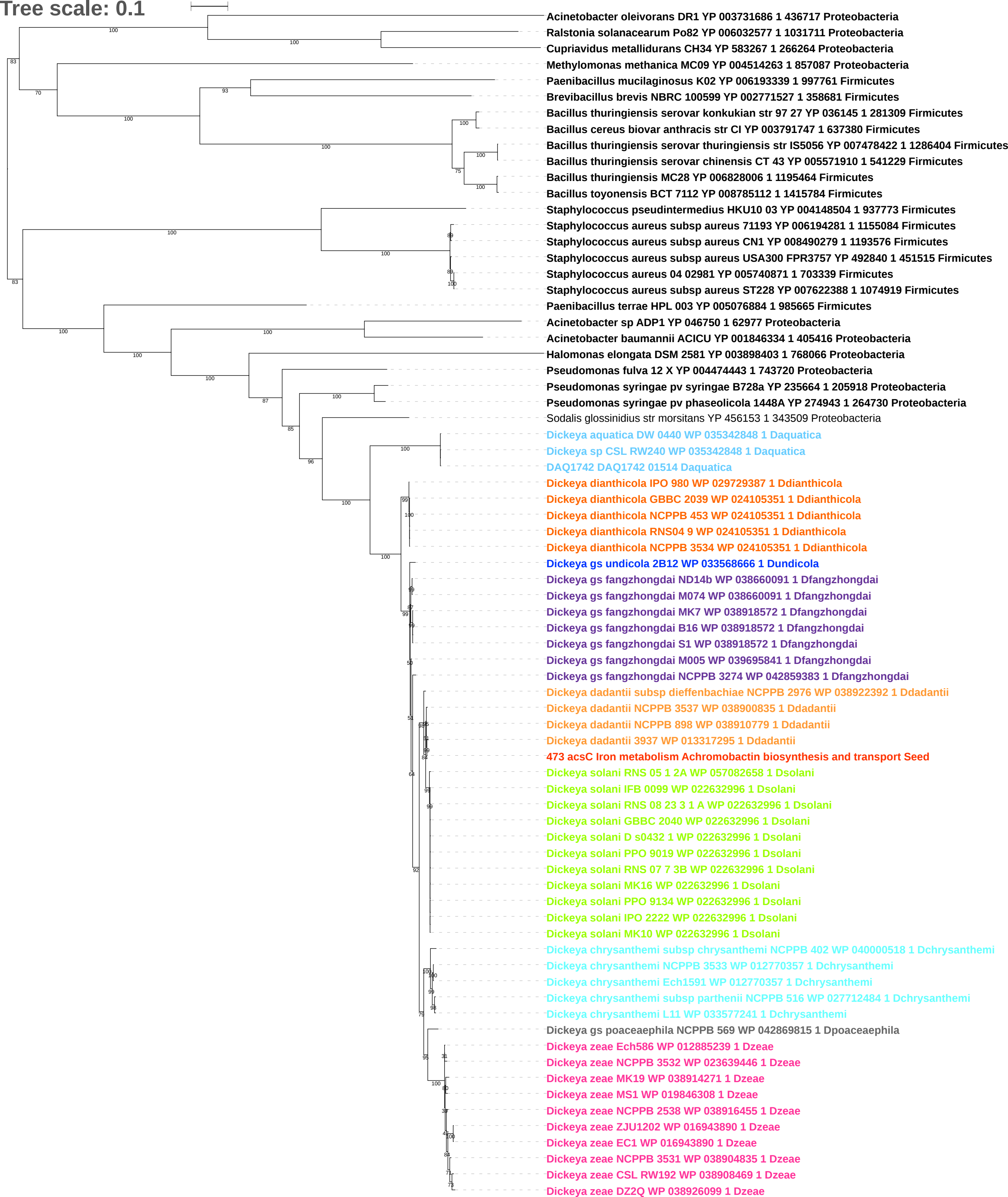

Tree scale: 0.1
Acinetobacter oleivorans DR1 YP 003731686 1 436717 Proteobacteria
Ralstonia solanacearum Po82 YP 006032577 1 1031711 Proteobacteria
Cupriavidus metallidurans CH34 YP 583267 1 266264 Proteobacteria
Methylomonas methanica MC09 YP 004514263 1 857087 Proteobacteria
Paenibacillus mucilaginosus K02 YP 006193339 1 997761 Firmicutes
Brevibacillus brevis NBRC 100599 YP 002771527 1 358681 Firmicutes
Bacillus thuringiensis serovar konkukian str 97 27 YP 036145 1 281309 Firmicutes
Bacillus cereus biovar anthracis str CI YP 003791747 1 637380 Firmicutes
Bacillus thuringiensis serovar thuringiensis str IS5056 YP 007478422 1 1286404 Firmicutes
Bacillus thuringiensis serovar chinensis CT 43 YP 005571910 1 541229 Firmicutes
Bacillus thuringiensis MC28 YP 006828006 1 1195464 Firmicutes
Bacillus toyonensis BCT 7112 YP 008785112 1 1415784 Firmicutes
Staphylococcus pseudintermedius HKU10 03 YP 004148504 1 937773 Firmicutes
Staphylococcus aureus subsp aureus 71193 YP 006194281 1 1155084 Firmicutes
Staphylococcus aureus subsp aureus CN1 YP 008490279 1 1193576 Firmicutes
Staphylococcus aureus subsp aureus USA300 FPR3757 YP 492840 1 451515 Firmicutes
Staphylococcus aureus 04 02981 YP 005740871 1 703339 Firmicutes
Staphylococcus aureus subsp aureus ST228 YP 007622388 1 1074919 Firmicutes
Paenibacillus terrae HPL 003 YP 005076884 1 985665 Firmicutes
Acinetobacter sp ADP1 YP 046750 1 62977 Proteobacteria
Acinetobacter baumannii ACICU YP 001846334 1 405416 Proteobacteria
Halomonas elongata DSM 2581 YP 003898403 1 768066 Proteobacteria
Pseudomonas fulva 12 X YP 004474443 1 743720 Proteobacteria
Pseudomonas syringae pv syringae B728a YP 235664 1 205918 Proteobacteria
Pseudomonas syringae pv phaseolicola 1448A YP 274943 1 264730 Proteobacteria
Sodalis glossinidius str morsitans YP 456153 1 343509 Proteobacteria
Dickeya aquatica DW 0440 WP 035342848 1 Daquatica
Dickeya sp CSL RW240 WP 035342848 1 Daquatica
DAQ1742 DAQ1742 01514 Daquatica
Dickeya dianthicola IPO 980 WP 029729387 1 Ddianthicola
Dickeya dianthicola GBBC 2039 WP 024105351 1 Ddianthicola
Dickeya dianthicola NCPPB 453 WP 024105351 1 Ddianthicola
Dickeya dianthicola RNS04 9 WP 024105351 1 Ddianthicola
Dickeya dianthicola NCPPB 3534 WP 024105351 1 Ddianthicola
Dickeya gs undicola 2B12 WP 033568666 1 Dundicola
Dickeya gs fangzhongdai ND14b WP 038660091 1 Dfangzhongdai
Dickeya gs fangzhongdai M074 WP 038660091 1 Dfangzhongdai
Dickeya gs fangzhongdai MK7 WP 038918572 1 Dfangzhongdai
Dickeya gs fangzhongdai B16 WP 038918572 1 Dfangzhongdai
Dickeya gs fangzhongdai S1 WP 038918572 1 Dfangzhongdai
Dickeya gs fangzhongdai M005 WP 039695841 1 Dfangzhongdai
Dickeya gs fangzhongdai NCPPB 3274 WP 042859383 1 Dfangzhongdai
Dickeya dadantii subsp dieffenbachiae NCPPB 2976 WP 038922392 1 Ddadantii
Dickeya dadantii NCPPB 3537 WP 038900835 1 Ddadantii
Dickeya dadantii NCPPB 898 WP 038910779 1 Ddadantii
Dickeya dadantii 3937 WP 013317295 1 Ddadantii
473 acsC Iron metabolism Achromobactin biosynthesis and transport Seed
Dickeya solani RNS 05 1 2A WP 057082658 1 Dsolani
Dickeya solani IFB 0099 WP 022632996 1 Dsolani
Dickeya solani RNS 08 23 3 1 A WP 022632996 1 Dsolani
Dickeya solani GBBC 2040 WP 022632996 1 Dsolani
Dickeya solani D s0432 1 WP 022632996 1 Dsolani
Dickeya solani PPO 9019 WP 022632996 1 Dsolani
Dickeya solani RNS 07 7 3B WP 022632996 1 Dsolani
Dickeya solani MK16 WP 022632996 1 Dsolani
Dickeya solani PPO 9134 WP 022632996 1 Dsolani
Dickeya solani IPO 2222 WP 022632996 1 Dsolani
Dickeya solani MK10 WP 022632996 1 Dsolani
Dickeya chrysanthemi subsp chrysanthemi NCPPB 402 WP 040000518 1 Dchrysanthemi
Dickeya chrysanthemi NCPPB 3533 WP 012770357 1 Dchrysanthemi
Dickeya chrysanthemi Ech1591 WP 012770357 1 Dchrysanthemi
Dickeya chrysanthemi subsp parthenii NCPPB 516 WP 027712484 1 Dchrysanthemi
Dickeya chrysanthemi L11 WP 033577241 1 Dchrysanthemi
Dickeya gs poaceaephila NCPPB 569 WP 042869815 1 Dpoaceaephila
Dickeya zeae Ech586 WP 012885239 1 Dzeae
Dickeya zeae NCPPB 3532 WP 023639446 1 Dzeae
Dickeya zeae MK19 WP 038914271 1 Dzeae
Dickeya zeae MS1 WP 019846308 1 Dzeae
Dickeya zeae NCPPB 2538 WP 038916455 1 Dzeae
Dickeya zeae ZJU1202 WP 016943890 1 Dzeae
Dickeya zeae EC1 WP 016943890 1 Dzeae
Dickeya zeae NCPPB 3531 WP 038904835 1 Dzeae
Dickeya zeae CSL RW192 WP 038908469 1 Dzeae
Dickeya zeae DZ2Q WP 038926099 1 Dzeae

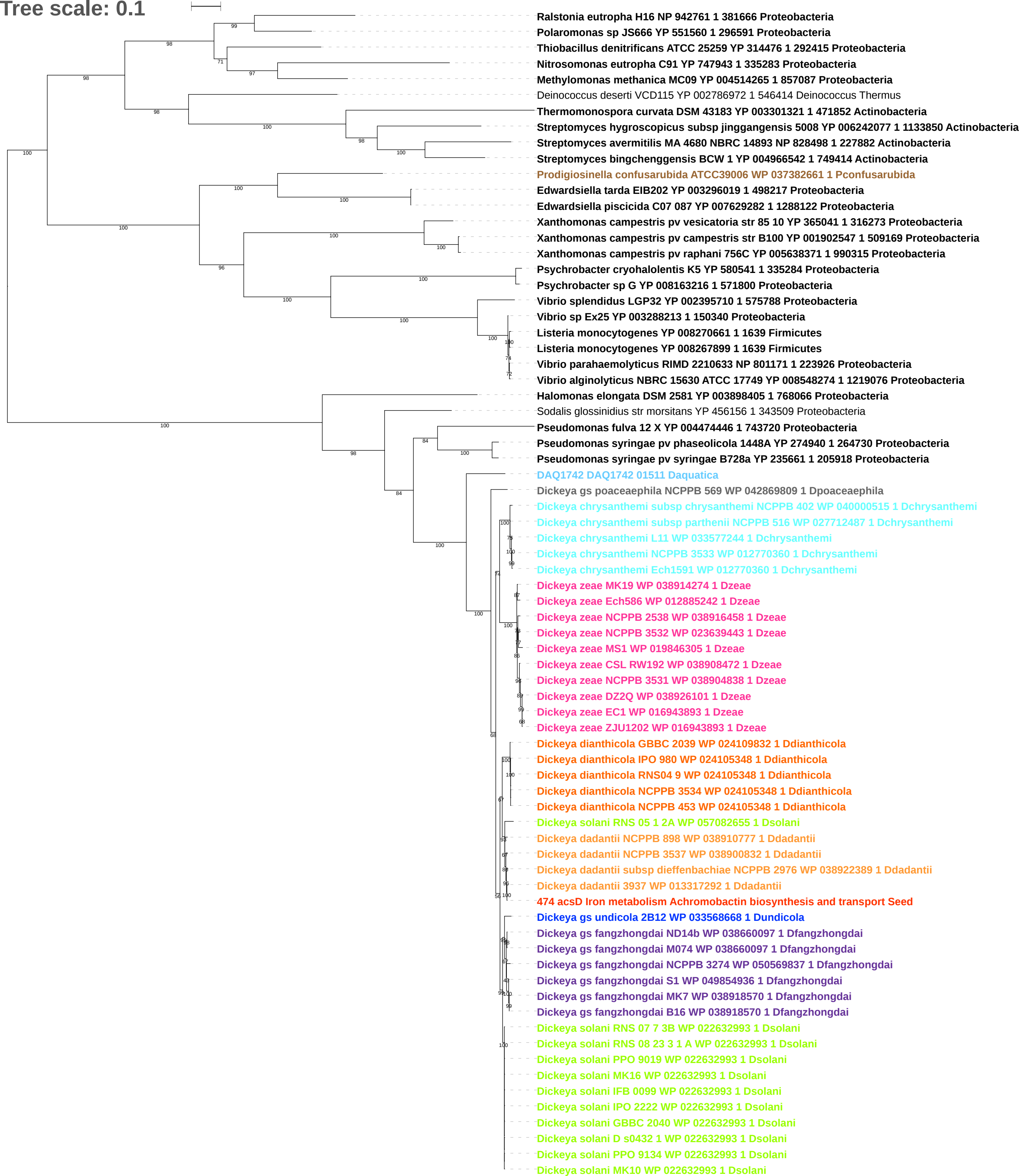
Tree scale: 0.1
Ralstonia eutropha H16 NP 942761 1 381666 Proteobacteria
Polaromonas sp JS666 YP 551560 1 296591 Proteobacteria
Thiobacillus denitrificans ATCC 25259 YP 314476 1 292415 Proteobacteria
Nitrosomonas eutropha C91 YP 747943 1 335283 Proteobacteria
Methylomonas methanica MC09 YP 004514265 1 857087 Proteobacteria
Deinococcus deserti VCD115 YP 002786972 1 546414 Deinococcus Thermus
Thermomonospora curvata DSM 43183 YP 003301321 1 471852 Actinobacteria
Streptomyces hygroscopicus subsp jinggangensis 5008 YP 006242077 1 1133850 Actinobacteria
Streptomyces avermitilis MA 4680 NBRC 14893 NP 828498 1 227882 Actinobacteria
Streptomyces bingchenggensis BCW 1 YP 004966542 1 749414 Actinobacteria
Prodigiosinella confusarubida ATCC39006 WP 037382661 1 Pconfusarubida
Edwardsiella tarda EIB202 YP 003296019 1 498217 Proteobacteria
Edwardsiella piscicida C07 087 YP 007629282 1 1288122 Proteobacteria
Xanthomonas campestris pv vesicatoria str 85 10 YP 365041 1 316273 Proteobacteria
Xanthomonas campestris pv campestris str B100 YP 001902547 1 509169 Proteobacteria
Xanthomonas campestris pv raphani 756C YP 005638371 1 990315 Proteobacteria
Psychrobacter cryohalolentis K5 YP 580541 1 335284 Proteobacteria
Psychrobacter sp G YP 008163216 1 571800 Proteobacteria
Vibrio splendidus LGP32 YP 002395710 1 575788 Proteobacteria
Vibrio sp Ex25 YP 003288213 1 150340 Proteobacteria
Listeria monocytogenes YP 008270661 1 1639 Firmicutes
Listeria monocytogenes YP 008267899 1 1639 Firmicutes
Vibrio parahaemolyticus RIMD 2210633 NP 801171 1 223926 Proteobacteria
Vibrio alginolyticus NBRC 15630 ATCC 17749 YP 008548274 1 1219076 Proteobacteria
Halomonas elongata DSM 2581 YP 003898405 1 768066 Proteobacteria
Sodalis glossinidius str morsitans YP 456156 1 343509 Proteobacteria
Pseudomonas fulva 12 X YP 004474446 1 743720 Proteobacteria
Pseudomonas syringae pv phaseolicola 1448A YP 274940 1 264730 Proteobacteria
Pseudomonas syringae pv syringae B728a YP 235661 1 205918 Proteobacteria
DAQ1742 DAQ1742 01511 Daquatica
Dickeya gs poaceaephila NCPPB 569 WP 042869809 1 Dpoaceaephila
Dickeya chrysanthemi subsp chrysanthemi NCPPB 402 WP 040000515 1 Dchrysanthemi
Dickeya chrysanthemi subsp parthenii NCPPB 516 WP 027712487 1 Dchrysanthemi
Dickeya chrysanthemi L11 WP 033577244 1 Dchrysanthemi
Dickeya chrysanthemi NCPPB 3533 WP 012770360 1 Dchrysanthemi
Dickeya chrysanthemi Ech1591 WP 012770360 1 Dchrysanthemi
Dickeya zeae MK19 WP 038914274 1 Dzeae
Dickeya zeae Ech586 WP 012885242 1 Dzeae
Dickeya zeae NCPPB 2538 WP 038916458 1 Dzeae
Dickeya zeae NCPPB 3532 WP 023639443 1 Dzeae
Dickeya zeae MS1 WP 019846305 1 Dzeae
Dickeya zeae CSL RW192 WP 038908472 1 Dzeae
Dickeya zeae NCPPB 3531 WP 038904838 1 Dzeae
Dickeya zeae DZ2Q WP 038926101 1 Dzeae
Dickeya zeae EC1 WP 016943893 1 Dzeae
Dickeya zeae ZJU1202 WP 016943893 1 Dzeae
Dickeya dianthicola GBBC 2039 WP 024109832 1 Ddianthicola
Dickeya dianthicola IPO 980 WP 024105348 1 Ddianthicola
Dickeya dianthicola RNS04 9 WP 024105348 1 Ddianthicola
Dickeya dianthicola NCPPB 3534 WP 024105348 1 Ddianthicola
Dickeya dianthicola NCPPB 453 WP 024105348 1 Ddianthicola
Dickeya solani RNS 05 1 2A WP 057082655 1 Dsolani
Dickeya dadantii NCPPB 898 WP 038910777 1 Ddadantii
Dickeya dadantii NCPPB 3537 WP 038900832 1 Ddadantii
Dickeya dadantii subsp dieffenbachiae NCPPB 2976 WP 038922389 1 Ddadantii
Dickeya dadantii 3937 WP 013317292 1 Ddadantii
474 acsD Iron metabolism Achromobactin biosynthesis and transport Seed
Dickeya gs undicola 2B12 WP 033568668 1 Dundicola
Dickeya gs fangzhongdai ND14b WP 038660097 1 Dfangzhongdai
Dickeya gs fangzhongdai M074 WP 038660097 1 Dfangzhongdai
Dickeya gs fangzhongdai NCPPB 3274 WP 050569837 1 Dfangzhongdai
Dickeya gs fangzhongdai S1 WP 049854936 1 Dfangzhongdai
Dickeya gs fangzhongdai MK7 WP 038918570 1 Dfangzhongdai
Dickeya gs fangzhongdai B16 WP 038918570 1 Dfangzhongdai
Dickeya solani RNS 07 7 3B WP 022632993 1 Dsolani
Dickeya solani RNS 08 23 3 1 A WP 022632993 1 Dsolani
Dickeya solani PPO 9019 WP 022632993 1 Dsolani
Dickeya solani MK16 WP 022632993 1 Dsolani
Dickeya solani IFB 0099 WP 022632993 1 Dsolani
Dickeya solani IPO 2222 WP 022632993 1 Dsolani
Dickeya solani GBBC 2040 WP 022632993 1 Dsolani
Dickeya solani D s0432 1 WP 022632993 1 Dsolani
Dickeya solani PPO 9134 WP 022632993 1 Dsolani
Dickeya solani MK10 WP 022632993 1 Dsolani

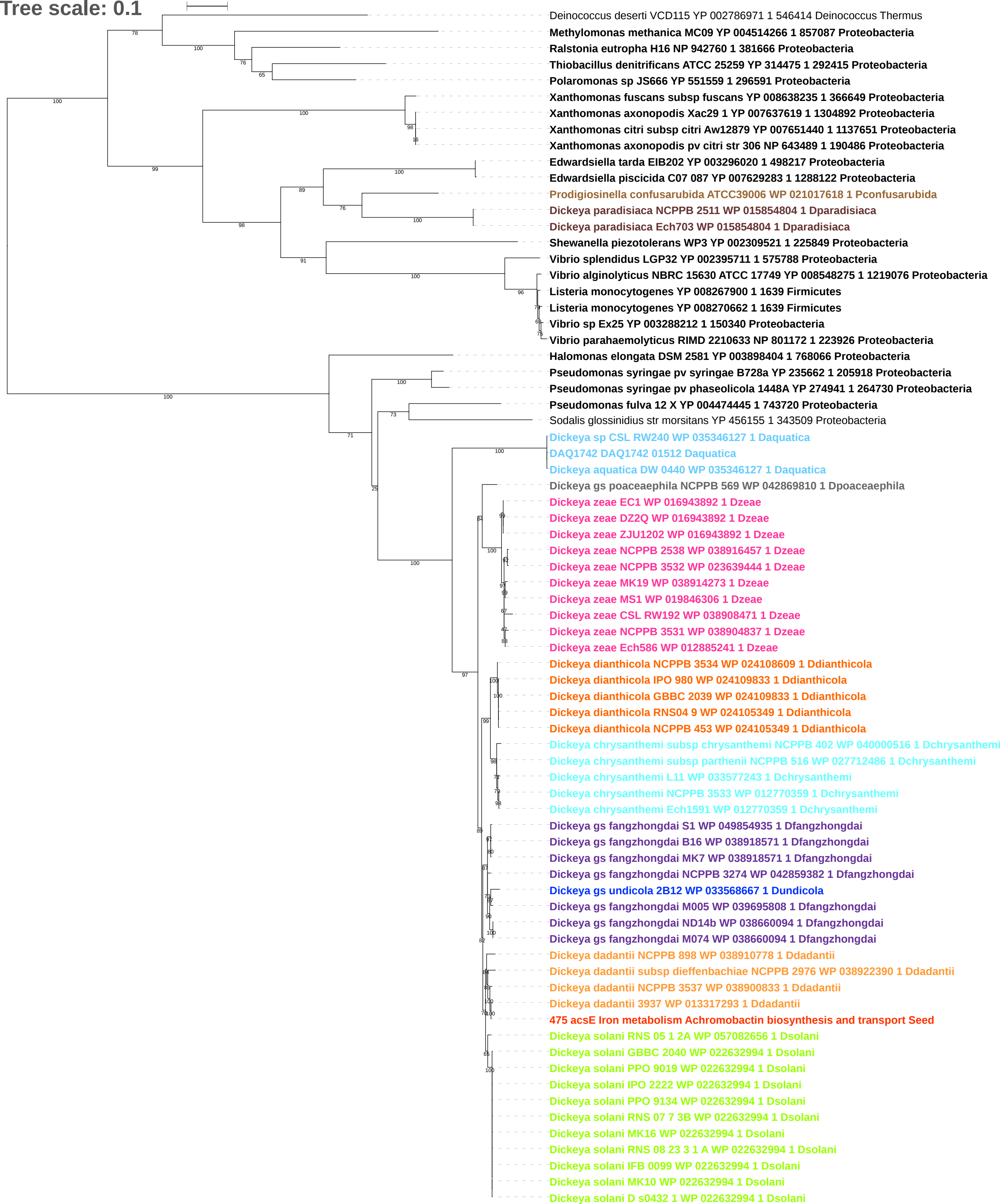
Tree scale: 0.1
Deinococcus deserti VCD115 YP 002786971 1 546414 Deinococcus Thermus
Methylomonas methanica MC09 YP 004514266 1 857087 Proteobacteria
Ralstonia eutropha H16 NP 942760 1 381666 Proteobacteria
Thiobacillus denitrificans ATCC 25259 YP 314475 1 292415 Proteobacteria
Polaromonas sp JS666 YP 551559 1 296591 Proteobacteria
Xanthomonas fuscans subsp fuscans YP 008638235 1 366649 Proteobacteria
Xanthomonas axonopodis Xac29 1 YP 007637619 1 1304892 Proteobacteria
Xanthomonas citri subsp citri Aw12879 YP 007651440 1 1137651 Proteobacteria
Xanthomonas axonopodis pv citri str 306 NP 643489 1 190486 Proteobacteria
Edwardsiella tarda EIB202 YP 003296020 1 498217 Proteobacteria
Edwardsiella piscicida C07 087 YP 007629283 1 1288122 Proteobacteria
Prodigiosinella confusarubida ATCC39006 WP 021017618 1 Pconfusarubida
Dickeya paradisiaca NCPPB 2511 WP 015854804 1 Dparadisiaca
Dickeya paradisiaca Ech703 WP 015854804 1 Dparadisiaca
Shewanella piezotolerans WP3 YP 002309521 1 225849 Proteobacteria
Vibrio splendidus LGP32 YP 002395711 1 575788 Proteobacteria
Vibrio alginolyticus NBRC 15630 ATCC 17749 YP 008548275 1 1219076 Proteobacteria
Listeria monocytogenes YP 008267900 1 1639 Firmicutes
Listeria monocytogenes YP 008270662 1 1639 Firmicutes
Vibrio sp Ex25 YP 003288212 1 150340 Proteobacteria
Vibrio parahaemolyticus RIMD 2210633 NP 801172 1 223926 Proteobacteria
Halomonas elongata DSM 2581 YP 003898404 1 768066 Proteobacteria
Pseudomonas syringae pv syringae B728a YP 235662 1 205918 Proteobacteria
Pseudomonas syringae pv phaseolicola 1448A YP 274941 1 264730 Proteobacteria
Pseudomonas fulva 12 X YP 004474445 1 743720 Proteobacteria
Sodalis glossinidius str morsitans YP 456155 1 343509 Proteobacteria
Dickeya sp CSL RW240 WP 035346127 1 Daquatica
DAQ1742 DAQ1742 01512 Daquatica
Dickeya aquatica DW 0440 WP 035346127 1 Daquatica
Dickeya gs poaceaephila NCPPB 569 WP 042869810 1 Dpoaceaephila
Dickeya zeae EC1 WP 016943892 1 Dzeae
Dickeya zeae DZ2Q WP 016943892 1 Dzeae
Dickeya zeae ZJU1202 WP 016943892 1 Dzeae
Dickeya zeae NCPPB 2538 WP 038916457 1 Dzeae
Dickeya zeae NCPPB 3532 WP 023639444 1 Dzeae
Dickeya zeae MK19 WP 038914273 1 Dzeae
Dickeya zeae MS1 WP 019846306 1 Dzeae
Dickeya zeae CSL RW192 WP 038908471 1 Dzeae
Dickeya zeae NCPPB 3531 WP 038904837 1 Dzeae
Dickeya zeae Ech586 WP 012885241 1 Dzeae
Dickeya dianthicola NCPPB 3534 WP 024108609 1 Ddianthicola
Dickeya dianthicola IPO 980 WP 024109833 1 Ddianthicola
Dickeya dianthicola GBBC 2039 WP 024109833 1 Ddianthicola
Dickeya dianthicola RNS04 9 WP 024105349 1 Ddianthicola
Dickeya dianthicola NCPPB 453 WP 024105349 1 Ddianthicola
Dickeya chrysanthemi subsp chrysanthemi NCPPB 402 WP 040000516 1 Dchrysanthemi
Dickeya chrysanthemi subsp parthenii NCPPB 516 WP 027712486 1 Dchrysanthemi
Dickeya chrysanthemi L11 WP 033577243 1 Dchrysanthemi
Dickeya chrysanthemi NCPPB 3533 WP 012770359 1 Dchrysanthemi
Dickeya chrysanthemi Ech1591 WP 012770359 1 Dchrysanthemi
Dickeya gs fangzhongdai S1 WP 049854935 1 Dfangzhongdai
Dickeya gs fangzhongdai B16 WP 038918571 1 Dfangzhongdai
Dickeya gs fangzhongdai MK7 WP 038918571 1 Dfangzhongdai
Dickeya gs fangzhongdai NCPPB 3274 WP 042859382 1 Dfangzhongdai
Dickeya gs undicola 2B12 WP 033568667 1 Dundicola
Dickeya gs fangzhongdai M005 WP 039695808 1 Dfangzhongdai
Dickeya gs fangzhongdai ND14b WP 038660094 1 Dfangzhongdai
Dickeya gs fangzhongdai M074 WP 038660094 1 Dfangzhongdai
Dickeya dadantii NCPPB 898 WP 038910778 1 Ddadantii
Dickeya dadantii subsp dieffenbachiae NCPPB 2976 WP 038922390 1 Ddadantii
Dickeya dadantii NCPPB 3537 WP 038900833 1 Ddadantii
Dickeya dadantii 3937 WP 013317293 1 Ddadantii
475 acsE Iron metabolism Achromobactin biosynthesis and transport Seed
Dickeya solani RNS 05 1 2A WP 057082656 1 Dsolani
Dickeya solani GBBC 2040 WP 022632994 1 Dsolani
Dickeya solani PPO 9019 WP 022632994 1 Dsolani
Dickeya solani IPO 2222 WP 022632994 1 Dsolani
Dickeya solani PPO 9134 WP 022632994 1 Dsolani
Dickeya solani RNS 07 7 3B WP 022632994 1 Dsolani
Dickeya solani MK16 WP 022632994 1 Dsolani
Dickeya solani RNS 08 23 3 1 A WP 022632994 1 Dsolani
Dickeya solani IFB 0099 WP 022632994 1 Dsolani
Dickeya solani MK10 WP 022632994 1 Dsolani
Dickeya solani D s0432 1 WP 022632994 1 Dsolani

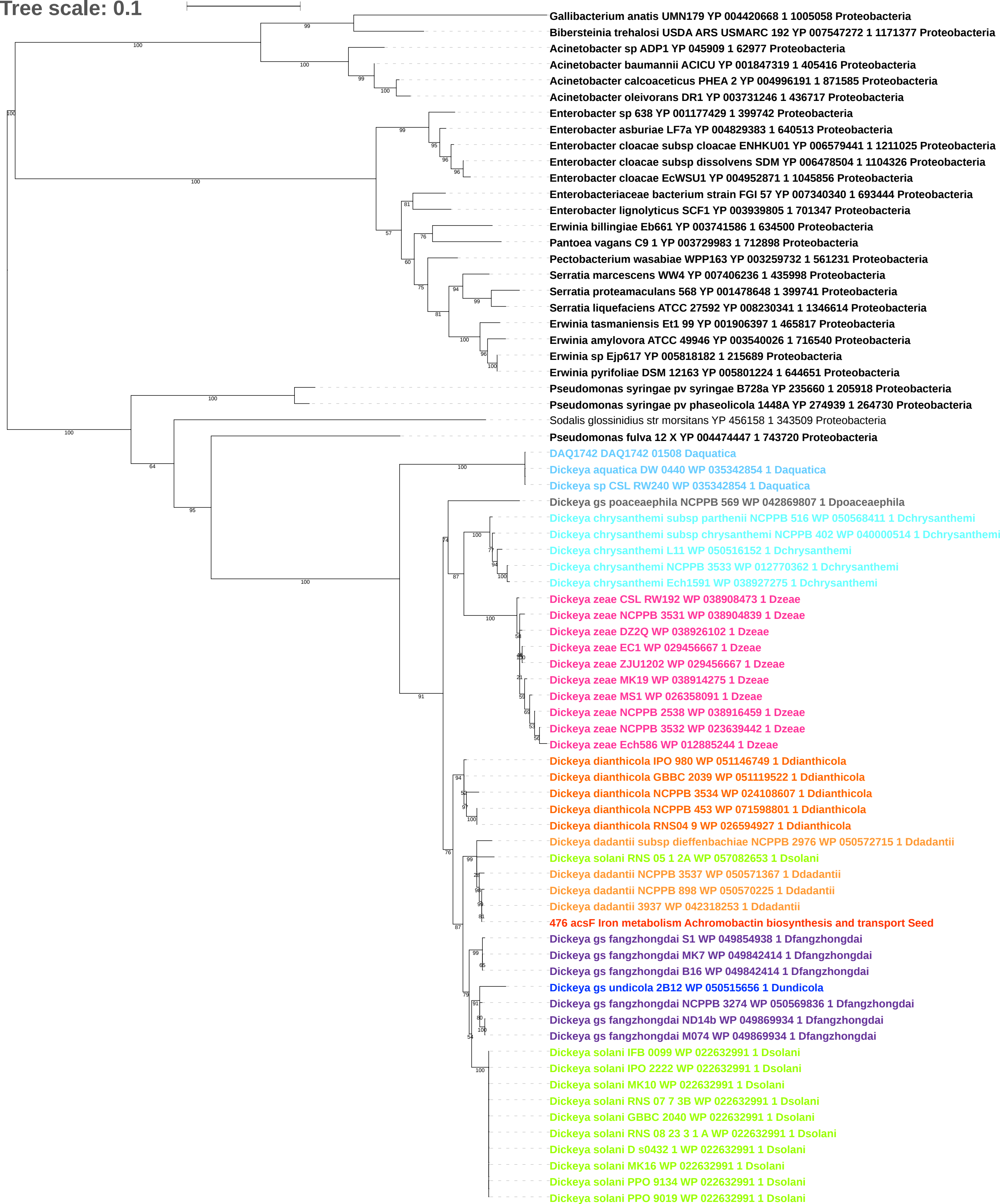
Tree scale: 0.1
Gallibacterium anatis UMN179 YP 004420668 1 1005058 Proteobacteria
Bibersteinia trehalosi USDA ARS USMARC 192 YP 007547272 1 1171377 Proteobacteria
Acinetobacter sp ADP1 YP 045909 1 62977 Proteobacteria
Acinetobacter baumannii ACICU YP 001847319 1 405416 Proteobacteria
Acinetobacter calcoaceticus PHEA 2 YP 004996191 1 871585 Proteobacteria
Acinetobacter oleivorans DR1 YP 003731246 1 436717 Proteobacteria
Enterobacter sp 638 YP 001177429 1 399742 Proteobacteria
Enterobacter asburiae LF7a YP 004829383 1 640513 Proteobacteria
Enterobacter cloacae subsp cloacae ENHKU01 YP 006579441 1 1211025 Proteobacteria
Enterobacter cloacae subsp dissolvens SDM YP 006478504 1 1104326 Proteobacteria
Enterobacter cloacae EcWSU1 YP 004952871 1 1045856 Proteobacteria
Enterobacteriaceae bacterium strain FGI 57 YP 007340340 1 693444 Proteobacteria
Enterobacter lignolyticus SCF1 YP 003939805 1 701347 Proteobacteria
Erwinia billingiae Eb661 YP 003741586 1 634500 Proteobacteria
Pantoea vagans C9 1 YP 003729983 1 712898 Proteobacteria
Pectobacterium wasabiae WPP163 YP 003259732 1 561231 Proteobacteria
Serratia marcescens WW4 YP 007406236 1 435998 Proteobacteria
Serratia proteamaculans 568 YP 001478648 1 399741 Proteobacteria
Serratia liquefaciens ATCC 27592 YP 008230341 1 1346614 Proteobacteria
Erwinia tasmaniensis Et1 99 YP 001906397 1 465817 Proteobacteria
Erwinia amylovora ATCC 49946 YP 003540026 1 716540 Proteobacteria
Erwinia sp Ejp617 YP 005818182 1 215689 Proteobacteria
Erwinia pyrifoliae DSM 12163 YP 005801224 1 644651 Proteobacteria
Pseudomonas syringae pv syringae B728a YP 235660 1 205918 Proteobacteria
Pseudomonas syringae pv phaseolicola 1448A YP 274939 1 264730 Proteobacteria
Sodalis glossinidius str morsitans YP 456158 1 343509 Proteobacteria
Pseudomonas fulva 12 X YP 004474447 1 743720 Proteobacteria
DAQ1742 DAQ1742 01508 Daquatica
Dickeya aquatica DW 0440 WP 035342854 1 Daquatica
Dickeya sp CSL RW240 WP 035342854 1 Daquatica
Dickeya gs poaceaephila NCPPB 569 WP 042869807 1 Dpoaceaephila
Dickeya chrysanthemi subsp parthenii NCPPB 516 WP 050568411 1 Dchrysanthemi
Dickeya chrysanthemi subsp chrysanthemi NCPPB 402 WP 040000514 1 Dchrysanthemi
Dickeya chrysanthemi L11 WP 050516152 1 Dchrysanthemi
Dickeya chrysanthemi NCPPB 3533 WP 012770362 1 Dchrysanthemi
Dickeya chrysanthemi Ech1591 WP 038927275 1 Dchrysanthemi
Dickeya zeae CSL RW192 WP 038908473 1 Dzeae
Dickeya zeae NCPPB 3531 WP 038904839 1 Dzeae
Dickeya zeae DZ2Q WP 038926102 1 Dzeae
Dickeya zeae EC1 WP 029456667 1 Dzeae
Dickeya zeae ZJU1202 WP 029456667 1 Dzeae
Dickeya zeae MK19 WP 038914275 1 Dzeae
Dickeya zeae MS1 WP 026358091 1 Dzeae
Dickeya zeae NCPPB 2538 WP 038916459 1 Dzeae
Dickeya zeae NCPPB 3532 WP 023639442 1 Dzeae
Dickeya zeae Ech586 WP 012885244 1 Dzeae
Dickeya dianthicola IPO 980 WP 051146749 1 Ddianthicola
Dickeya dianthicola GBBC 2039 WP 051119522 1 Ddianthicola
Dickeya dianthicola NCPPB 3534 WP 024108607 1 Ddianthicola
Dickeya dianthicola NCPPB 453 WP 071598801 1 Ddianthicola
Dickeya dianthicola RNS04 9 WP 026594927 1 Ddianthicola
Dickeya dadantii subsp dieffenbachiae NCPPB 2976 WP 050572715 1 Ddadantii
Dickeya solani RNS 05 1 2A WP 057082653 1 Dsolani
Dickeya dadantii NCPPB 3537 WP 050571367 1 Ddadantii
Dickeya dadantii NCPPB 898 WP 050570225 1 Ddadantii
Dickeya dadantii 3937 WP 042318253 1 Ddadantii
476 acsF Iron metabolism Achromobactin biosynthesis and transport Seed
Dickeya gs fangzhongdai S1 WP 049854938 1 Dfangzhongdai
Dickeya gs fangzhongdai MK7 WP 049842414 1 Dfangzhongdai
Dickeya gs fangzhongdai B16 WP 049842414 1 Dfangzhongdai
Dickeya gs undicola 2B12 WP 050515656 1 Dundicola
Dickeya gs fangzhongdai NCPPB 3274 WP 050569836 1 Dfangzhongdai
Dickeya gs fangzhongdai ND14b WP 049869934 1 Dfangzhongdai
Dickeya gs fangzhongdai M074 WP 049869934 1 Dfangzhongdai
Dickeya solani IFB 0099 WP 022632991 1 Dsolani
Dickeya solani IPO 2222 WP 022632991 1 Dsolani
Dickeya solani MK10 WP 022632991 1 Dsolani
Dickeya solani RNS 07 7 3B WP 022632991 1 Dsolani
Dickeya solani GBBC 2040 WP 022632991 1 Dsolani
Dickeya solani RNS 08 23 3 1 A WP 022632991 1 Dsolani
Dickeya solani D s0432 1 WP 022632991 1 Dsolani
Dickeya solani MK16 WP 022632991 1 Dsolani
Dickeya solani PPO 9134 WP 022632991 1 Dsolani
Dickeya solani PPO 9019 WP 022632991 1 Dsolani

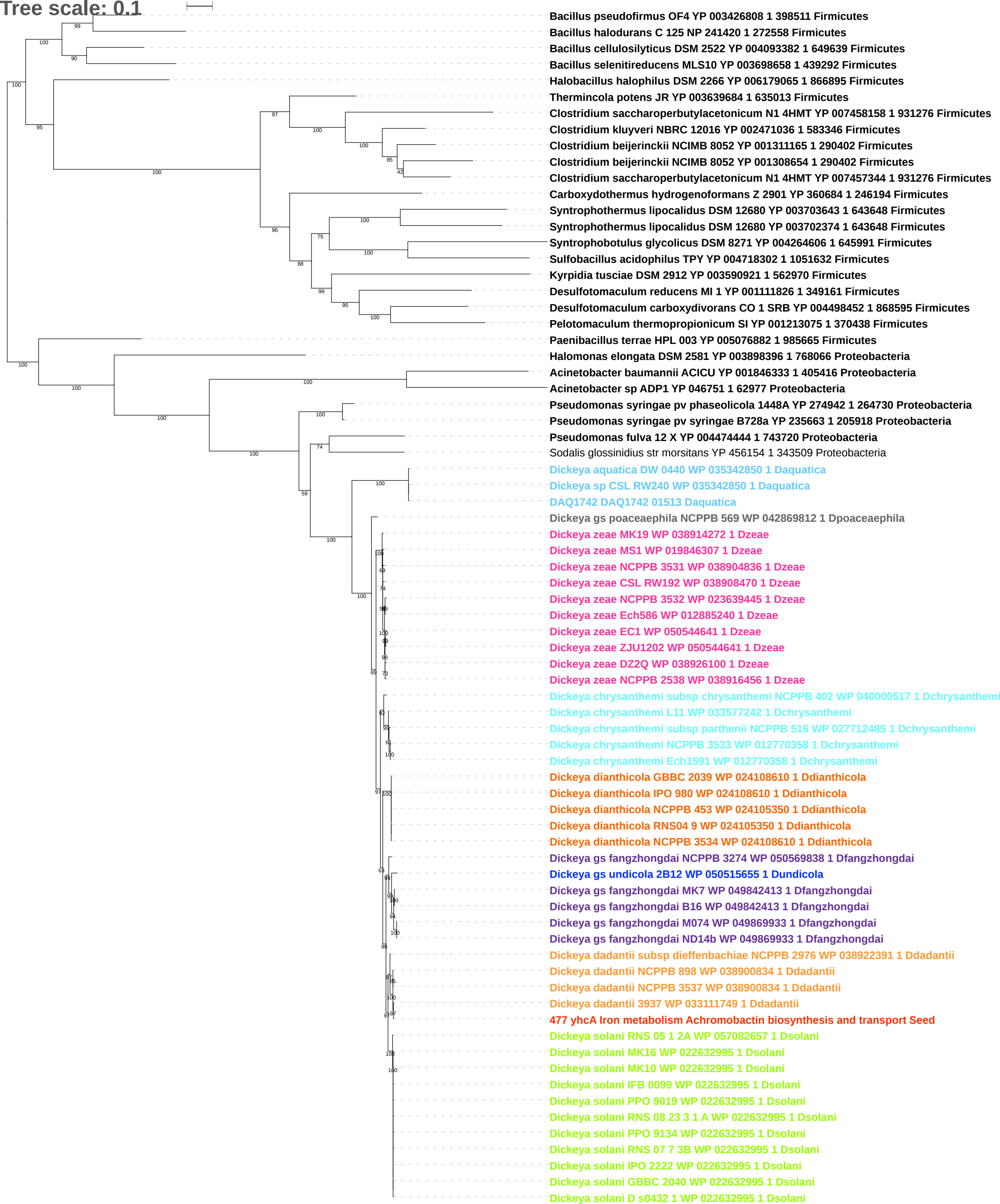

Tree scale: 0.1
Bacillus pseudofirmus OF4 YP 003426808 1 398511 Firmicutes
Bacillus halodurans C 125 NP 241420 1 272558 Firmicutes
Bacillus cellulosilyticus DSM 2522 YP 004093382 1 649639 Firmicutes
Bacillus selenitireducens MLS10 YP 003698658 1 439292 Firmicutes
Halobacillus halophilus DSM 2266 YP 006179065 1 866895 Firmicutes
Thermincola potens JR YP 003639684 1 635013 Firmicutes
Clostridium saccharoperbutylacetonicum N1 4HMT YP 007458158 1 931276 Firmicutes
Clostridium kluyveri NBRC 12016 YP 002471036 1 583346 Firmicutes
Clostridium beijerinckii NCIMB 8052 YP 001311165 1 290402 Firmicutes
Clostridium beijerinckii NCIMB 8052 YP 001308654 1 290402 Firmicutes
Clostridium saccharoperbutylacetonicum N1 4HMT YP 007457344 1 931276 Firmicutes
Carboxydothermus hydrogenoformans Z 2901 YP 360684 1 246194 Firmicutes
Syntrophothermus lipocalidus DSM 12680 YP 003703643 1 643648 Firmicutes
Syntrophothermus lipocalidus DSM 12680 YP 003702374 1 643648 Firmicutes
Syntrophobotulus glycolicus DSM 8271 YP 004264606 1 645991 Firmicutes
Sulfobacillus acidophilus TPY YP 004718302 1 1051632 Firmicutes
Kyrpidia tusciae DSM 2912 YP 003590921 1 562970 Firmicutes
Desulfotomaculum reducens MI 1 YP 001111826 1 349161 Firmicutes
Desulfotomaculum carboxydivorans CO 1 SRB YP 004498452 1 868595 Firmicutes
Pelotomaculum thermopropionicum SI YP 001213075 1 370438 Firmicutes
Paenibacillus terrae HPL 003 YP 005076882 1 985665 Firmicutes
Halomonas elongata DSM 2581 YP 003898396 1 768066 Proteobacteria
Acinetobacter baumannii ACICU YP 001846333 1 405416 Proteobacteria
Acinetobacter sp ADP1 YP 046751 1 62977 Proteobacteria
Pseudomonas syringae pv phaseolicola 1448A YP 274942 1 264730 Proteobacteria
Pseudomonas syringae pv syringae B728a YP 235663 1 205918 Proteobacteria
Pseudomonas fulva 12 X YP 004474444 1 743720 Proteobacteria
Sodalis glossinidius str morsitans YP 456154 1 343509 Proteobacteria
Dickeya aquatica DW 0440 WP 035342850 1 Daquatica
Dickeya sp CSL RW240 WP 035342850 1 Daquatica
DAQ1742 DAQ1742 01513 Daquatica
Dickeya gs poaceaephila NCPPB 569 WP 042869812 1 Dpoaceaephila
Dickeya zeae MK19 WP 038914272 1 Dzeae
Dickeya zeae MS1 WP 019846307 1 Dzeae
Dickeya zeae NCPPB 3531 WP 038904836 1 Dzeae
Dickeya zeae CSL RW192 WP 038908470 1 Dzeae
Dickeya zeae NCPPB 3532 WP 023639445 1 Dzeae
Dickeya zeae Ech586 WP 012885240 1 Dzeae
Dickeya zeae EC1 WP 050544641 1 Dzeae
Dickeya zeae ZJU1202 WP 050544641 1 Dzeae
Dickeya zeae DZ2Q WP 038926100 1 Dzeae
Dickeya zeae NCPPB 2538 WP 038916456 1 Dzeae
Dickeya chrysanthemi subsp chrysanthemi NCPPB 402 WP 040000517 1 Dchrysanthemi
Dickeya chrysanthemi L11 WP 033577242 1 Dchrysanthemi
Dickeya chrysanthemi subsp parthenii NCPPB 516 WP 027712485 1 Dchrysanthemi
Dickeya chrysanthemi NCPPB 3533 WP 012770358 1 Dchrysanthemi
Dickeya chrysanthemi Ech1591 WP 012770358 1 Dchrysanthemi
Dickeya dianthicola GBBC 2039 WP 024108610 1 Ddianthicola
Dickeya dianthicola IPO 980 WP 024108610 1 Ddianthicola
Dickeya dianthicola NCPPB 453 WP 024105350 1 Ddianthicola
Dickeya dianthicola RNS04 9 WP 024105350 1 Ddianthicola
Dickeya dianthicola NCPPB 3534 WP 024108610 1 Ddianthicola
Dickeya gs fangzhongdai NCPPB 3274 WP 050569838 1 Dfangzhongdai
Dickeya gs undicola 2B12 WP 050515655 1 Dundicola
Dickeya gs fangzhongdai MK7 WP 049842413 1 Dfangzhongdai
Dickeya gs fangzhongdai B16 WP 049842413 1 Dfangzhongdai
Dickeya gs fangzhongdai M074 WP 049869933 1 Dfangzhongdai
Dickeya gs fangzhongdai ND14b WP 049869933 1 Dfangzhongdai
Dickeya dadantii subsp dieffenbachiae NCPPB 2976 WP 038922391 1 Ddadantii
Dickeya dadantii NCPPB 898 WP 038900834 1 Ddadantii
Dickeya dadantii NCPPB 3537 WP 038900834 1 Ddadantii
Dickeya dadantii 3937 WP 033111749 1 Ddadantii
477 yhcA Iron metabolism Achromobactin biosynthesis and transport Seed
Dickeya solani RNS 05 1 2A WP 057082657 1 Dsolani
Dickeya solani MK16 WP 022632995 1 Dsolani
Dickeya solani MK10 WP 022632995 1 Dsolani
Dickeya solani IFB 0099 WP 022632995 1 Dsolani
Dickeya solani PPO 9019 WP 022632995 1 Dsolani
Dickeya solani RNS 08 23 3 1 A WP 022632995 1 Dsolani
Dickeya solani PPO 9134 WP 022632995 1 Dsolani
Dickeya solani RNS 07 7 3B WP 022632995 1 Dsolani
Dickeya solani IPO 2222 WP 022632995 1 Dsolani
Dickeya solani GBBC 2040 WP 022632995 1 Dsolani
Dickeya solani D s0432 1 WP 022632995 1 Dsolani

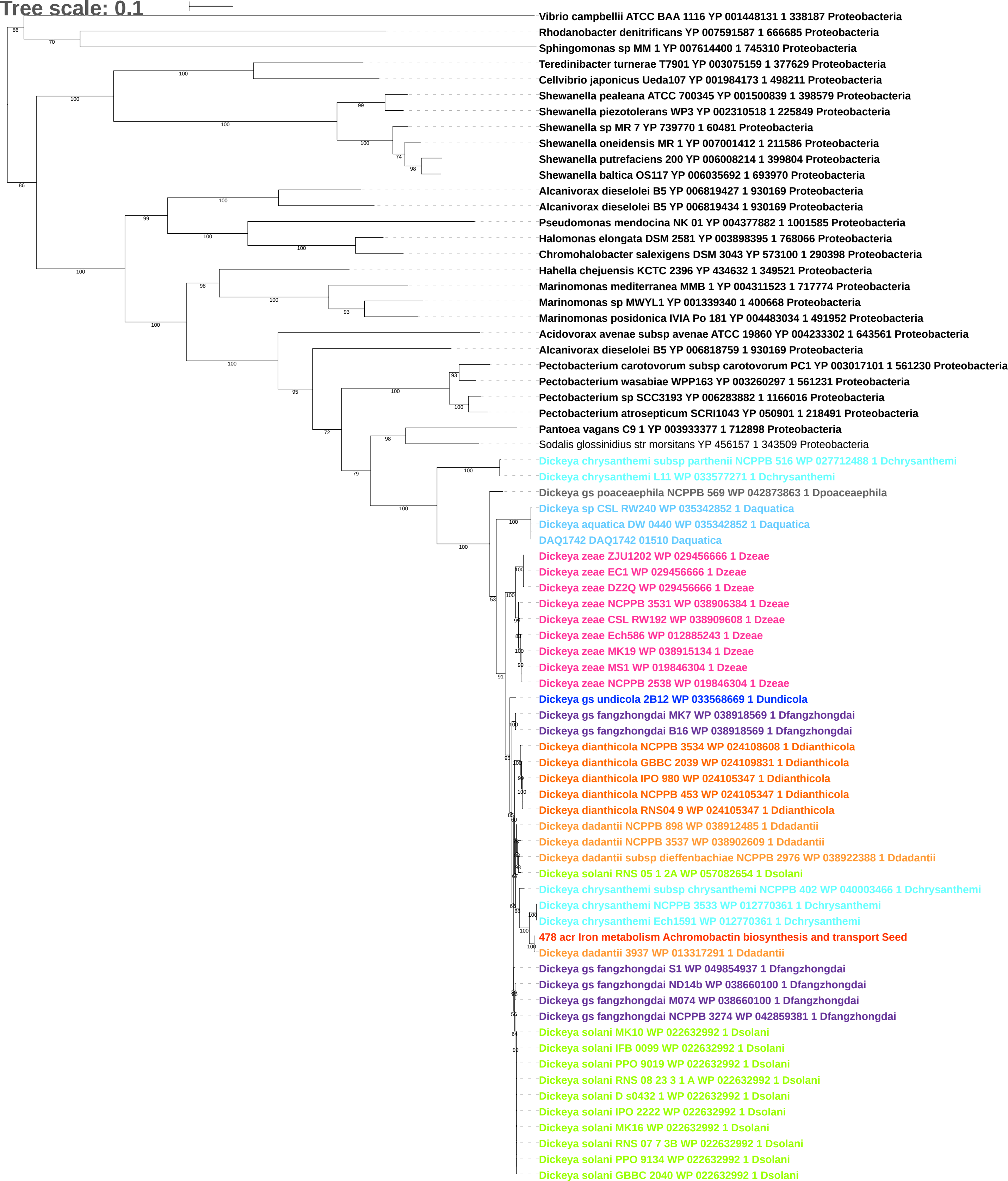
Tree scale: 0.1
Vibrio campbellii ATCC BAA 1116 YP 001448131 1 338187 Proteobacteria
Rhodanobacter denitrificans YP 007591587 1 666685 Proteobacteria
Sphingomonas sp MM 1 YP 007614400 1 745310 Proteobacteria
Teredinibacter turnerae T7901 YP 003075159 1 377629 Proteobacteria
Cellvibrio japonicus Ueda107 YP 001984173 1 498211 Proteobacteria
Shewanella pealeana ATCC 700345 YP 001500839 1 398579 Proteobacteria
Shewanella piezotolerans WP3 YP 002310518 1 225849 Proteobacteria
Shewanella sp MR 7 YP 739770 1 60481 Proteobacteria
Shewanella oneidensis MR 1 YP 007001412 1 211586 Proteobacteria
Shewanella putrefaciens 200 YP 006008214 1 399804 Proteobacteria
Shewanella baltica OS117 YP 006035692 1 693970 Proteobacteria
Alcanivorax dieselolei B5 YP 006819427 1 930169 Proteobacteria
Alcanivorax dieselolei B5 YP 006819434 1 930169 Proteobacteria
Pseudomonas mendocina NK 01 YP 004377882 1 1001585 Proteobacteria
Halomonas elongata DSM 2581 YP 003898395 1 768066 Proteobacteria
Chromohalobacter salexigens DSM 3043 YP 573100 1 290398 Proteobacteria
Hahella chejuensis KCTC 2396 YP 434632 1 349521 Proteobacteria
Marinomonas mediterranea MMB 1 YP 004311523 1 717774 Proteobacteria
Marinomonas sp MWYL1 YP 001339340 1 400668 Proteobacteria
Marinomonas posidonica IVIA Po 181 YP 004483034 1 491952 Proteobacteria
Acidovorax avenae subsp avenae ATCC 19860 YP 004233302 1 643561 Proteobacteria
Alcanivorax dieselolei B5 YP 006818759 1 930169 Proteobacteria
Pectobacterium carotovorum subsp carotovorum PC1 YP 003017101 1 561230 Proteobacteria
Pectobacterium wasabiae WPP163 YP 003260297 1 561231 Proteobacteria
Pectobacterium sp SCC3193 YP 006283882 1 1166016 Proteobacteria
Pectobacterium atrosepticum SCRI1043 YP 050901 1 218491 Proteobacteria
Pantoea vagans C9 1 YP 003933377 1 712898 Proteobacteria
Sodalis glossinidius str morsitans YP 456157 1 343509 Proteobacteria
Dickeya chrysanthemi subsp parthenii NCPPB 516 WP 027712488 1 Dchrysanthemi
Dickeya chrysanthemi L11 WP 033577271 1 Dchrysanthemi
Dickeya gs poaceaephila NCPPB 569 WP 042873863 1 Dpoaceaephila
Dickeya sp CSL RW240 WP 035342852 1 Daquatica
Dickeya aquatica DW 0440 WP 035342852 1 Daquatica
DAQ1742 DAQ1742 01510 Daquatica
Dickeya zeae ZJU1202 WP 029456666 1 Dzeae
Dickeya zeae EC1 WP 029456666 1 Dzeae
Dickeya zeae DZ2Q WP 029456666 1 Dzeae
Dickeya zeae NCPPB 3531 WP 038906384 1 Dzeae
Dickeya zeae CSL RW192 WP 038909608 1 Dzeae
Dickeya zeae Ech586 WP 012885243 1 Dzeae
Dickeya zeae MK19 WP 038915134 1 Dzeae
Dickeya zeae MS1 WP 019846304 1 Dzeae
Dickeya zeae NCPPB 2538 WP 019846304 1 Dzeae
Dickeya gs undicola 2B12 WP 033568669 1 Dundicola
Dickeya gs fangzhongdai MK7 WP 038918569 1 Dfangzhongdai
Dickeya gs fangzhongdai B16 WP 038918569 1 Dfangzhongdai
Dickeya dianthicola NCPPB 3534 WP 024108608 1 Ddianthicola
Dickeya dianthicola GBBC 2039 WP 024109831 1 Ddianthicola
Dickeya dianthicola IPO 980 WP 024105347 1 Ddianthicola
Dickeya dianthicola NCPPB 453 WP 024105347 1 Ddianthicola
Dickeya dianthicola RNS04 9 WP 024105347 1 Ddianthicola
Dickeya dadantii NCPPB 898 WP 038912485 1 Ddadantii
Dickeya dadantii NCPPB 3537 WP 038902609 1 Ddadantii
Dickeya dadantii subsp dieffenbachiae NCPPB 2976 WP 038922388 1 Ddadantii
Dickeya solani RNS 05 1 2A WP 057082654 1 Dsolani
Dickeya chrysanthemi subsp chrysanthemi NCPPB 402 WP 040003466 1 Dchrysanthemi
Dickeya chrysanthemi NCPPB 3533 WP 012770361 1 Dchrysanthemi
Dickeya chrysanthemi Ech1591 WP 012770361 1 Dchrysanthemi
478 acr Iron metabolism Achromobactin biosynthesis and transport Seed
Dickeya dadantii 3937 WP 013317291 1 Ddadantii
Dickeya gs fangzhongdai S1 WP 049854937 1 Dfangzhongdai
Dickeya gs fangzhongdai ND14b WP 038660100 1 Dfangzhongdai
Dickeya gs fangzhongdai M074 WP 038660100 1 Dfangzhongdai
Dickeya gs fangzhongdai NCPPB 3274 WP 042859381 1 Dfangzhongdai
Dickeya solani MK10 WP 022632992 1 Dsolani
Dickeya solani IFB 0099 WP 022632992 1 Dsolani
Dickeya solani PPO 9019 WP 022632992 1 Dsolani
Dickeya solani RNS 08 23 3 1 A WP 022632992 1 Dsolani
Dickeya solani D s0432 1 WP 022632992 1 Dsolani
Dickeya solani IPO 2222 WP 022632992 1 Dsolani
Dickeya solani MK16 WP 022632992 1 Dsolani
Dickeya solani RNS 07 7 3B WP 022632992 1 Dsolani
Dickeya solani PPO 9134 WP 022632992 1 Dsolani
Dickeya solani GBBC 2040 WP 022632992 1 Dsolani

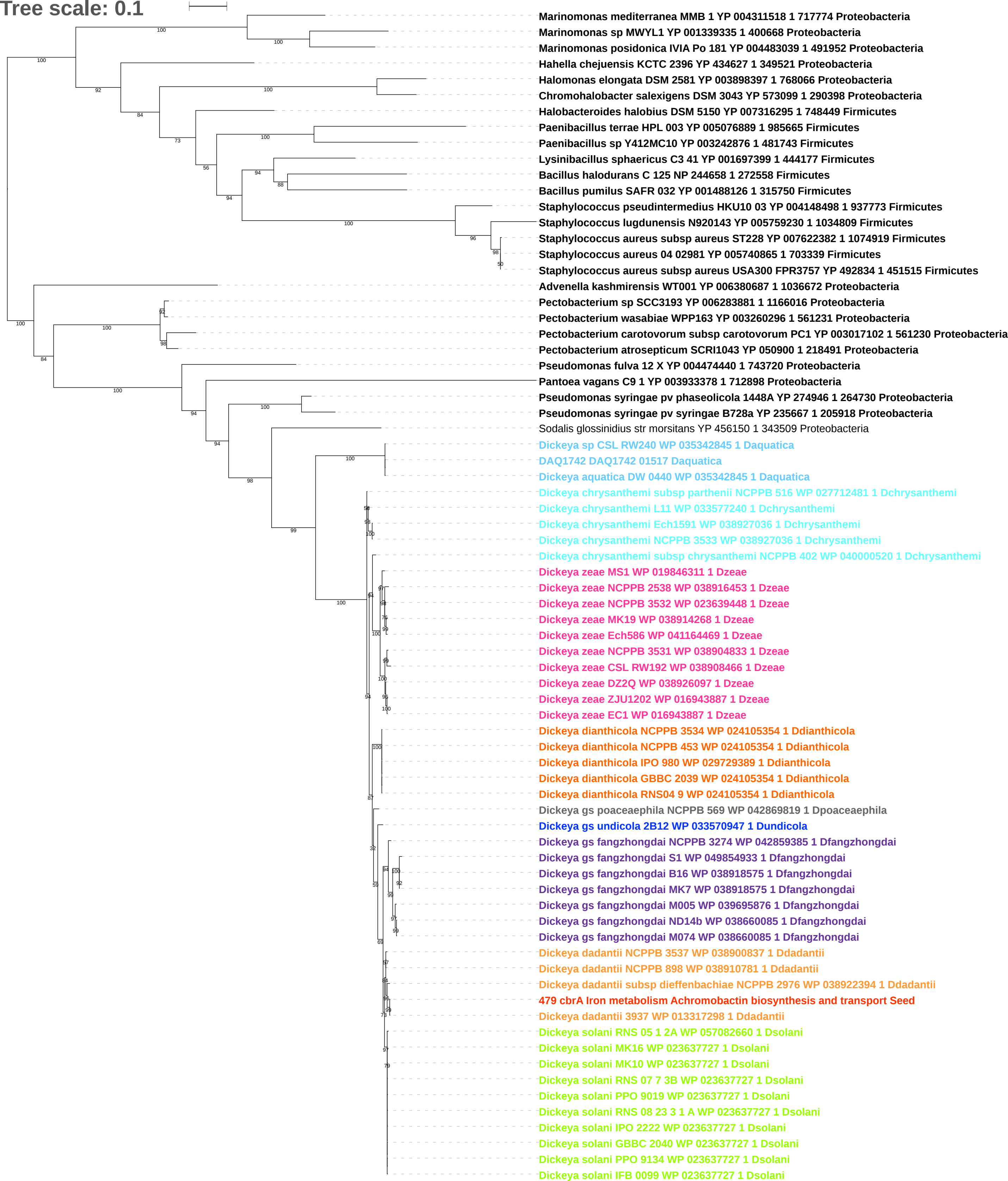
Tree scale: 0.1
Marinomonas mediterranea MMB 1 YP 004311518 1 717774 Proteobacteria
Marinomonas sp MWYL1 YP 001339335 1 400668 Proteobacteria
Marinomonas posidonica IVIA Po 181 YP 004483039 1 491952 Proteobacteria
Hahella chejuensis KCTC 2396 YP 434627 1 349521 Proteobacteria
Halomonas elongata DSM 2581 YP 003898397 1 768066 Proteobacteria
Chromohalobacter salexigens DSM 3043 YP 573099 1 290398 Proteobacteria
Halobacteroides halobius DSM 5150 YP 007316295 1 748449 Firmicutes
Paenibacillus terrae HPL 003 YP 005076889 1 985665 Firmicutes
Paenibacillus sp Y412MC10 YP 003242876 1 481743 Firmicutes
Lysinibacillus sphaericus C3 41 YP 001697399 1 444177 Firmicutes
Bacillus halodurans C 125 NP 244658 1 272558 Firmicutes
Bacillus pumilus SAFR 032 YP 001488126 1 315750 Firmicutes
Staphylococcus pseudintermedius HKU10 03 YP 004148498 1 937773 Firmicutes
Staphylococcus lugdunensis N920143 YP 005759230 1 1034809 Firmicutes
Staphylococcus aureus subsp aureus ST228 YP 007622382 1 1074919 Firmicutes
Staphylococcus aureus 04 02981 YP 005740865 1 703339 Firmicutes
Staphylococcus aureus subsp aureus USA300 FPR3757 YP 492834 1 451515 Firmicutes
Advenella kashmirensis WT001 YP 006380687 1 1036672 Proteobacteria
Pectobacterium sp SCC3193 YP 006283881 1 1166016 Proteobacteria
Pectobacterium wasabiae WPP163 YP 003260296 1 561231 Proteobacteria
Pectobacterium carotovorum subsp carotovorum PC1 YP 003017102 1 561230 Proteobacteria
Pectobacterium atrosepticum SCRI1043 YP 050900 1 218491 Proteobacteria
Pseudomonas fulva 12 X YP 004474440 1 743720 Proteobacteria
Pantoea vagans C9 1 YP 003933378 1 712898 Proteobacteria
Pseudomonas syringae pv phaseolicola 1448A YP 274946 1 264730 Proteobacteria
Pseudomonas syringae pv syringae B728a YP 235667 1 205918 Proteobacteria
Sodalis glossinidius str morsitans YP 456150 1 343509 Proteobacteria
Dickeya sp CSL RW240 WP 035342845 1 Daquatica
DAQ1742 DAQ1742 01517 Daquatica
Dickeya aquatica DW 0440 WP 035342845 1 Daquatica
Dickeya chrysanthemi subsp parthenii NCPPB 516 WP 027712481 1 Dchrysanthemi
Dickeya chrysanthemi L11 WP 033577240 1 Dchrysanthemi
Dickeya chrysanthemi Ech1591 WP 038927036 1 Dchrysanthemi
Dickeya chrysanthemi NCPPB 3533 WP 038927036 1 Dchrysanthemi
Dickeya chrysanthemi subsp chrysanthemi NCPPB 402 WP 040000520 1 Dchrysanthemi
Dickeya zeae MS1 WP 019846311 1 Dzeae
Dickeya zeae NCPPB 2538 WP 038916453 1 Dzeae
Dickeya zeae NCPPB 3532 WP 023639448 1 Dzeae
Dickeya zeae MK19 WP 038914268 1 Dzeae
Dickeya zeae Ech586 WP 041164469 1 Dzeae
Dickeya zeae NCPPB 3531 WP 038904833 1 Dzeae
Dickeya zeae CSL RW192 WP 038908466 1 Dzeae
Dickeya zeae DZ2Q WP 038926097 1 Dzeae
Dickeya zeae ZJU1202 WP 016943887 1 Dzeae
Dickeya zeae EC1 WP 016943887 1 Dzeae
Dickeya dianthicola NCPPB 3534 WP 024105354 1 Ddianthicola
Dickeya dianthicola NCPPB 453 WP 024105354 1 Ddianthicola
Dickeya dianthicola IPO 980 WP 029729389 1 Ddianthicola
Dickeya dianthicola GBBC 2039 WP 024105354 1 Ddianthicola
Dickeya dianthicola RNS04 9 WP 024105354 1 Ddianthicola
Dickeya gs poaceaephila NCPPB 569 WP 042869819 1 Dpoaceaephila
Dickeya gs undicola 2B12 WP 033570947 1 Dundicola
Dickeya gs fangzhongdai NCPPB 3274 WP 042859385 1 Dfangzhongdai
Dickeya gs fangzhongdai S1 WP 049854933 1 Dfangzhongdai
Dickeya gs fangzhongdai B16 WP 038918575 1 Dfangzhongdai
Dickeya gs fangzhongdai MK7 WP 038918575 1 Dfangzhongdai
Dickeya gs fangzhongdai M005 WP 039695876 1 Dfangzhongdai
Dickeya gs fangzhongdai ND14b WP 038660085 1 Dfangzhongdai
Dickeya gs fangzhongdai M074 WP 038660085 1 Dfangzhongdai
Dickeya dadantii NCPPB 3537 WP 038900837 1 Ddadantii
Dickeya dadantii NCPPB 898 WP 038910781 1 Ddadantii
Dickeya dadantii subsp dieffenbachiae NCPPB 2976 WP 038922394 1 Ddadantii
479 cbrA Iron metabolism Achromobactin biosynthesis and transport Seed
Dickeya dadantii 3937 WP 013317298 1 Ddadantii
Dickeya solani RNS 05 1 2A WP 057082660 1 Dsolani
Dickeya solani MK16 WP 023637727 1 Dsolani
Dickeya solani MK10 WP 023637727 1 Dsolani
Dickeya solani RNS 07 7 3B WP 023637727 1 Dsolani
Dickeya solani PPO 9019 WP 023637727 1 Dsolani
Dickeya solani RNS 08 23 3 1 A WP 023637727 1 Dsolani
Dickeya solani IPO 2222 WP 023637727 1 Dsolani
Dickeya solani GBBC 2040 WP 023637727 1 Dsolani
Dickeya solani PPO 9134 WP 023637727 1 Dsolani
Dickeya solani IFB 0099 WP 023637727 1 Dsolani

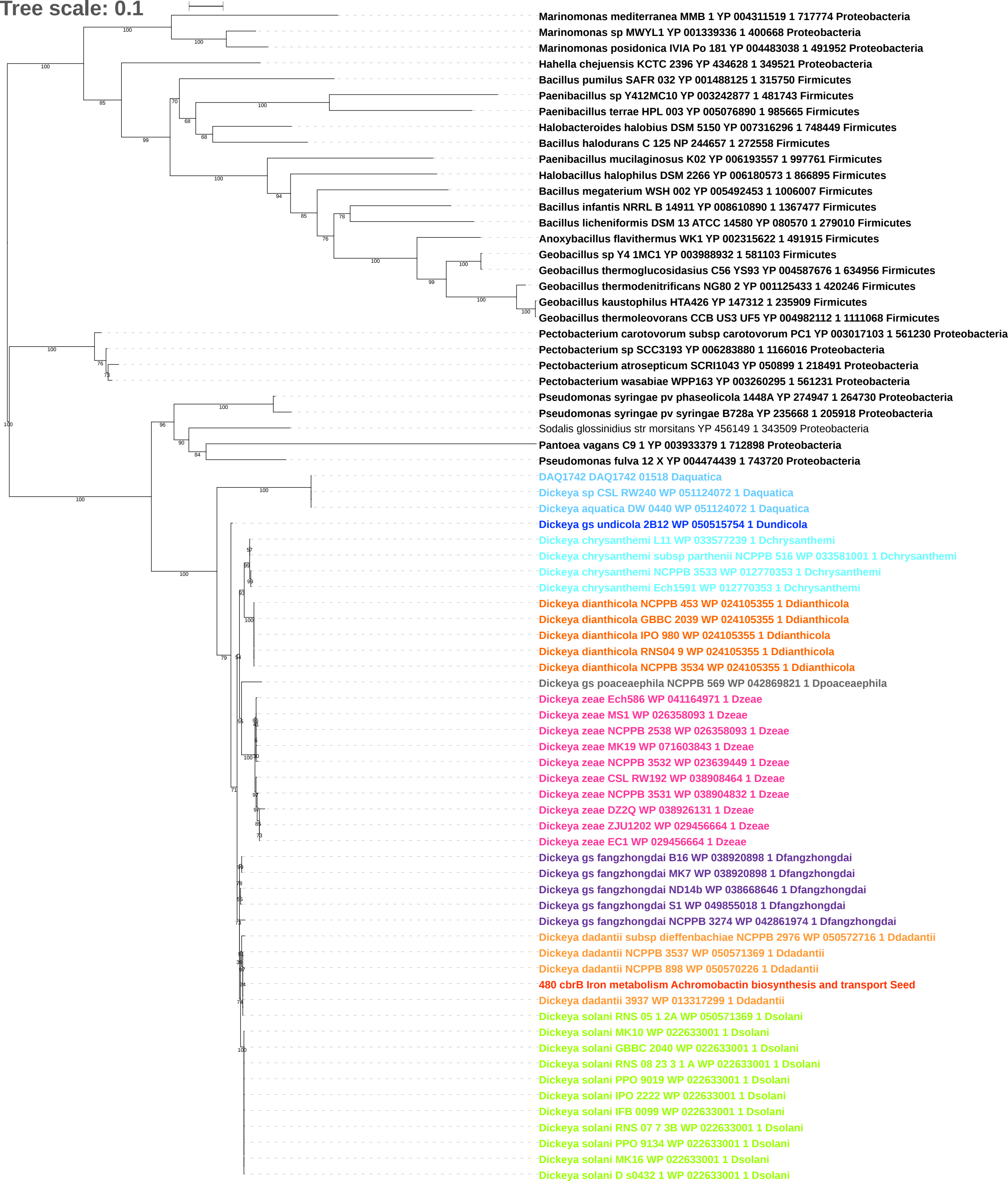
Tree scale: 0.1
Marinomonas mediterranea MMB 1 YP 004311519 1 717774 Proteobacteria
Marinomonas sp MWYL1 YP 001339336 1 400668 Proteobacteria
Marinomonas posidonica IVIA Po 181 YP 004483038 1 491952 Proteobacteria
Hahella chejuensis KCTC 2396 YP 434628 1 349521 Proteobacteria
Bacillus pumilus SAFR 032 YP 001488125 1 315750 Firmicutes
Paenibacillus sp Y412MC10 YP 003242877 1 481743 Firmicutes
Paenibacillus terrae HPL 003 YP 005076890 1 985665 Firmicutes
Halobacteroides halobius DSM 5150 YP 007316296 1 748449 Firmicutes
Bacillus halodurans C 125 NP 244657 1 272558 Firmicutes
Paenibacillus mucilaginosus K02 YP 006193557 1 997761 Firmicutes
Halobacillus halophilus DSM 2266 YP 006180573 1 866895 Firmicutes
Bacillus megaterium WSH 002 YP 005492453 1 1006007 Firmicutes
Bacillus infantis NRRL B 14911 YP 008610890 1 1367477 Firmicutes
Bacillus licheniformis DSM 13 ATCC 14580 YP 080570 1 279010 Firmicutes
Anoxybacillus flavithermus WK1 YP 002315622 1 491915 Firmicutes
Geobacillus sp Y4 1MC1 YP 003988932 1 581103 Firmicutes
Geobacillus thermoglucosidasius C56 YS93 YP 004587676 1 634956 Firmicutes
Geobacillus thermodenitrificans NG80 2 YP 001125433 1 420246 Firmicutes
Geobacillus kaustophilus HTA426 YP 147312 1 235909 Firmicutes
Geobacillus thermoleovorans CCB US3 UF5 YP 004982112 1 1111068 Firmicutes
Pectobacterium carotovorum subsp carotovorum PC1 YP 003017103 1 561230 Proteobacteria
Pectobacterium sp SCC3193 YP 006283880 1 1166016 Proteobacteria
Pectobacterium atrosepticum SCRI1043 YP 050899 1 218491 Proteobacteria
Pectobacterium wasabiae WPP163 YP 003260295 1 561231 Proteobacteria
Pseudomonas syringae pv phaseolicola 1448A YP 274947 1 264730 Proteobacteria
Pseudomonas syringae pv syringae B728a YP 235668 1 205918 Proteobacteria
Sodalis glossinidius str morsitans YP 456149 1 343509 Proteobacteria
Pantoea vagans C9 1 YP 003933379 1 712898 Proteobacteria
Pseudomonas fulva 12 X YP 004474439 1 743720 Proteobacteria
DAQ1742 DAQ1742 01518 Daquatica
Dickeya sp CSL RW240 WP 051124072 1 Daquatica
Dickeya aquatica DW 0440 WP 051124072 1 Daquatica
Dickeya gs undicola 2B12 WP 050515754 1 Dundicola
Dickeya chrysanthemi L11 WP 033577239 1 Dchrysanthemi
Dickeya chrysanthemi subsp parthenii NCPPB 516 WP 033581001 1 Dchrysanthemi
Dickeya chrysanthemi NCPPB 3533 WP 012770353 1 Dchrysanthemi
Dickeya chrysanthemi Ech1591 WP 012770353 1 Dchrysanthemi
Dickeya dianthicola NCPPB 453 WP 024105355 1 Ddianthicola
Dickeya dianthicola GBBC 2039 WP 024105355 1 Ddianthicola
Dickeya dianthicola IPO 980 WP 024105355 1 Ddianthicola
Dickeya dianthicola RNS04 9 WP 024105355 1 Ddianthicola
Dickeya dianthicola NCPPB 3534 WP 024105355 1 Ddianthicola
Dickeya gs poaceaephila NCPPB 569 WP 042869821 1 Dpoaceaephila
Dickeya zeae Ech586 WP 041164971 1 Dzeae
Dickeya zeae MS1 WP 026358093 1 Dzeae
Dickeya zeae NCPPB 2538 WP 026358093 1 Dzeae
Dickeya zeae MK19 WP 071603843 1 Dzeae
Dickeya zeae NCPPB 3532 WP 023639449 1 Dzeae
Dickeya zeae CSL RW192 WP 038908464 1 Dzeae
Dickeya zeae NCPPB 3531 WP 038904832 1 Dzeae
Dickeya zeae DZ2Q WP 038926131 1 Dzeae
Dickeya zeae ZJU1202 WP 029456664 1 Dzeae
Dickeya zeae EC1 WP 029456664 1 Dzeae
Dickeya gs fangzhongdai B16 WP 038920898 1 Dfangzhongdai
Dickeya gs fangzhongdai MK7 WP 038920898 1 Dfangzhongdai
Dickeya gs fangzhongdai ND14b WP 038668646 1 Dfangzhongdai
Dickeya gs fangzhongdai S1 WP 049855018 1 Dfangzhongdai
Dickeya gs fangzhongdai NCPPB 3274 WP 042861974 1 Dfangzhongdai
Dickeya dadantii subsp dieffenbachiae NCPPB 2976 WP 050572716 1 Ddadantii
Dickeya dadantii NCPPB 3537 WP 050571369 1 Ddadantii
Dickeya dadantii NCPPB 898 WP 050570226 1 Ddadantii
480 cbrB Iron metabolism Achromobactin biosynthesis and transport Seed
Dickeya dadantii 3937 WP 013317299 1 Ddadantii
Dickeya solani RNS 05 1 2A WP 050571369 1 Dsolani
Dickeya solani MK10 WP 022633001 1 Dsolani
Dickeya solani GBBC 2040 WP 022633001 1 Dsolani
Dickeya solani RNS 08 23 3 1 A WP 022633001 1 Dsolani
Dickeya solani PPO 9019 WP 022633001 1 Dsolani
Dickeya solani IPO 2222 WP 022633001 1 Dsolani
Dickeya solani IFB 0099 WP 022633001 1 Dsolani
Dickeya solani RNS 07 7 3B WP 022633001 1 Dsolani
Dickeya solani PPO 9134 WP 022633001 1 Dsolani
Dickeya solani MK16 WP 022633001 1 Dsolani
Dickeya solani D s0432 1 WP 022633001 1 Dsolani

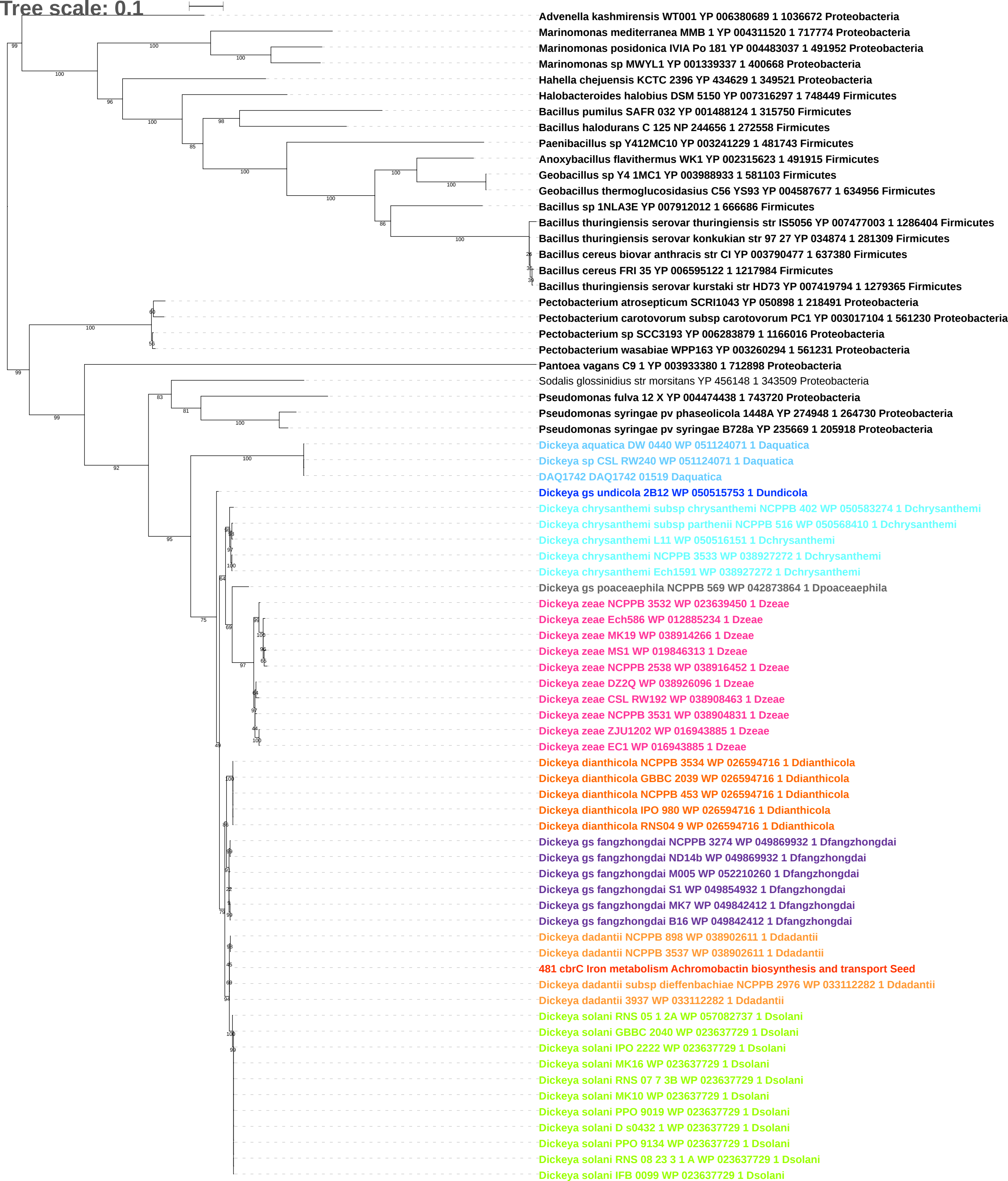
Tree scale: 0.1
Advenella kashmirensis WT001 YP 006380689 1 1036672 Proteobacteria
Marinomonas mediterranea MMB 1 YP 004311520 1 717774 Proteobacteria
Marinomonas posidonica IVIA Po 181 YP 004483037 1 491952 Proteobacteria
Marinomonas sp MWYL1 YP 001339337 1 400668 Proteobacteria
Hahella chejuensis KCTC 2396 YP 434629 1 349521 Proteobacteria
Halobacteroides halobius DSM 5150 YP 007316297 1 748449 Firmicutes
Bacillus pumilus SAFR 032 YP 001488124 1 315750 Firmicutes
Bacillus halodurans C 125 NP 244656 1 272558 Firmicutes
Paenibacillus sp Y412MC10 YP 003241229 1 481743 Firmicutes
Anoxybacillus flavithermus WK1 YP 002315623 1 491915 Firmicutes
Geobacillus sp Y4 1MC1 YP 003988933 1 581103 Firmicutes
Geobacillus thermoglucosidasius C56 YS93 YP 004587677 1 634956 Firmicutes
Bacillus sp 1NLA3E YP 007912012 1 666686 Firmicutes
Bacillus thuringiensis serovar thuringiensis str IS5056 YP 007477003 1 1286404 Firmicutes
Bacillus thuringiensis serovar konkukian str 97 27 YP 034874 1 281309 Firmicutes
Bacillus cereus biovar anthracis str CI YP 003790477 1 637380 Firmicutes
Bacillus cereus FRI 35 YP 006595122 1 1217984 Firmicutes
Bacillus thuringiensis serovar kurstaki str HD73 YP 007419794 1 1279365 Firmicutes
Pectobacterium atrosepticum SCRI1043 YP 050898 1 218491 Proteobacteria
Pectobacterium carotovorum subsp carotovorum PC1 YP 003017104 1 561230 Proteobacteria
Pectobacterium sp SCC3193 YP 006283879 1 1166016 Proteobacteria
Pectobacterium wasabiae WPP163 YP 003260294 1 561231 Proteobacteria
Pantoea vagans C9 1 YP 003933380 1 712898 Proteobacteria
Sodalis glossinidius str morsitans YP 456148 1 343509 Proteobacteria
Pseudomonas fulva 12 X YP 004474438 1 743720 Proteobacteria
Pseudomonas syringae pv phaseolicola 1448A YP 274948 1 264730 Proteobacteria
Pseudomonas syringae pv syringae B728a YP 235669 1 205918 Proteobacteria
Dickeya aquatica DW 0440 WP 051124071 1 Daquatica
Dickeya sp CSL RW240 WP 051124071 1 Daquatica
DAQ1742 DAQ1742 01519 Daquatica
Dickeya gs undicola 2B12 WP 050515753 1 Dundicola
Dickeya chrysanthemi subsp chrysanthemi NCPPB 402 WP 050583274 1 Dchrysanthemi
Dickeya chrysanthemi subsp parthenii NCPPB 516 WP 050568410 1 Dchrysanthemi
Dickeya chrysanthemi L11 WP 050516151 1 Dchrysanthemi
Dickeya chrysanthemi NCPPB 3533 WP 038927272 1 Dchrysanthemi
Dickeya chrysanthemi Ech1591 WP 038927272 1 Dchrysanthemi
Dickeya gs poaceaephila NCPPB 569 WP 042873864 1 Dpoaceaephila
Dickeya zeae NCPPB 3532 WP 023639450 1 Dzeae
Dickeya zeae Ech586 WP 012885234 1 Dzeae
Dickeya zeae MK19 WP 038914266 1 Dzeae
Dickeya zeae MS1 WP 019846313 1 Dzeae
Dickeya zeae NCPPB 2538 WP 038916452 1 Dzeae
Dickeya zeae DZ2Q WP 038926096 1 Dzeae
Dickeya zeae CSL RW192 WP 038908463 1 Dzeae
Dickeya zeae NCPPB 3531 WP 038904831 1 Dzeae
Dickeya zeae ZJU1202 WP 016943885 1 Dzeae
Dickeya zeae EC1 WP 016943885 1 Dzeae
Dickeya dianthicola NCPPB 3534 WP 026594716 1 Ddianthicola
Dickeya dianthicola GBBC 2039 WP 026594716 1 Ddianthicola
Dickeya dianthicola NCPPB 453 WP 026594716 1 Ddianthicola
Dickeya dianthicola IPO 980 WP 026594716 1 Ddianthicola
Dickeya dianthicola RNS04 9 WP 026594716 1 Ddianthicola
Dickeya gs fangzhongdai NCPPB 3274 WP 049869932 1 Dfangzhongdai
Dickeya gs fangzhongdai ND14b WP 049869932 1 Dfangzhongdai
Dickeya gs fangzhongdai M005 WP 052210260 1 Dfangzhongdai
Dickeya gs fangzhongdai S1 WP 049854932 1 Dfangzhongdai
Dickeya gs fangzhongdai MK7 WP 049842412 1 Dfangzhongdai
Dickeya gs fangzhongdai B16 WP 049842412 1 Dfangzhongdai
Dickeya dadantii NCPPB 898 WP 038902611 1 Ddadantii
Dickeya dadantii NCPPB 3537 WP 038902611 1 Ddadantii
481 cbrC Iron metabolism Achromobactin biosynthesis and transport Seed
Dickeya dadantii subsp dieffenbachiae NCPPB 2976 WP 033112282 1 Ddadantii
Dickeya dadantii 3937 WP 033112282 1 Ddadantii
Dickeya solani RNS 05 1 2A WP 057082737 1 Dsolani
Dickeya solani GBBC 2040 WP 023637729 1 Dsolani
Dickeya solani IPO 2222 WP 023637729 1 Dsolani
Dickeya solani MK16 WP 023637729 1 Dsolani
Dickeya solani RNS 07 7 3B WP 023637729 1 Dsolani
Dickeya solani MK10 WP 023637729 1 Dsolani
Dickeya solani PPO 9019 WP 023637729 1 Dsolani
Dickeya solani D s0432 1 WP 023637729 1 Dsolani
Dickeya solani PPO 9134 WP 023637729 1 Dsolani
Dickeya solani RNS 08 23 3 1 A WP 023637729 1 Dsolani
Dickeya solani IFB 0099 WP 023637729 1 Dsolani

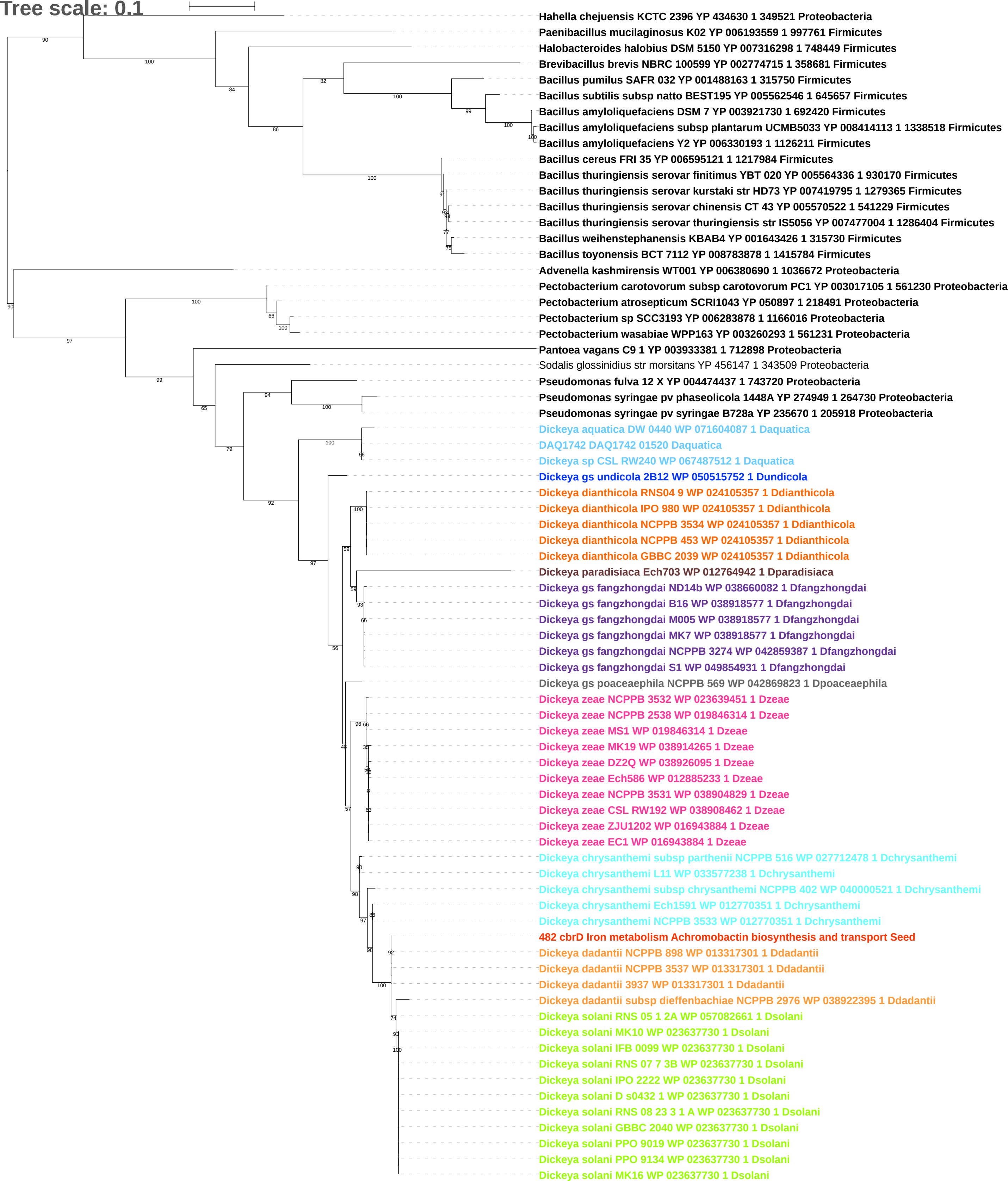
Tree scale: 0.1
Hahella chejuensis KCTC 2396 YP 434630 1 349521 Proteobacteria
Paenibacillus mucilaginosus K02 YP 006193559 1 997761 Firmicutes
Halobacteroides halobius DSM 5150 YP 007316298 1 748449 Firmicutes
Brevibacillus brevis NBRC 100599 YP 002774715 1 358681 Firmicutes
Bacillus pumilus SAFR 032 YP 001488163 1 315750 Firmicutes
Bacillus subtilis subsp natto BEST195 YP 005562546 1 645657 Firmicutes
Bacillus amyloliquefaciens DSM 7 YP 003921730 1 692420 Firmicutes
Bacillus amyloliquefaciens subsp plantarum UCMB5033 YP 008414113 1 1338518 Firmicutes
Bacillus amyloliquefaciens Y2 YP 006330193 1 1126211 Firmicutes
Bacillus cereus FRI 35 YP 006595121 1 1217984 Firmicutes
Bacillus thuringiensis serovar finitimus YBT 020 YP 005564336 1 930170 Firmicutes
Bacillus thuringiensis serovar kurstaki str HD73 YP 007419795 1 1279365 Firmicutes
Bacillus thuringiensis serovar chinensis CT 43 YP 005570522 1 541229 Firmicutes
Bacillus thuringiensis serovar thuringiensis str IS5056 YP 007477004 1 1286404 Firmicutes
Bacillus weihenstephanensis KBAB4 YP 001643426 1 315730 Firmicutes
Bacillus toyonensis BCT 7112 YP 008783878 1 1415784 Firmicutes
Advenella kashmirensis WT001 YP 006380690 1 1036672 Proteobacteria
Pectobacterium carotovorum subsp carotovorum PC1 YP 003017105 1 561230 Proteobacteria
Pectobacterium atrosepticum SCRI1043 YP 050897 1 218491 Proteobacteria
Pectobacterium sp SCC3193 YP 006283878 1 1166016 Proteobacteria
Pectobacterium wasabiae WPP163 YP 003260293 1 561231 Proteobacteria
Pantoea vagans C9 1 YP 003933381 1 712898 Proteobacteria
Sodalis glossinidius str morsitans YP 456147 1 343509 Proteobacteria
Pseudomonas fulva 12 X YP 004474437 1 743720 Proteobacteria
Pseudomonas syringae pv phaseolicola 1448A YP 274949 1 264730 Proteobacteria
Pseudomonas syringae pv syringae B728a YP 235670 1 205918 Proteobacteria
Dickeya aquatica DW 0440 WP 071604087 1 Daquatica
DAQ1742 DAQ1742 01520 Daquatica
Dickeya sp CSL RW240 WP 067487512 1 Daquatica
Dickeya gs undicola 2B12 WP 050515752 1 Dundicola
Dickeya dianthicola RNS04 9 WP 024105357 1 Ddianthicola
Dickeya dianthicola IPO 980 WP 024105357 1 Ddianthicola
Dickeya dianthicola NCPPB 3534 WP 024105357 1 Ddianthicola
Dickeya dianthicola NCPPB 453 WP 024105357 1 Ddianthicola
Dickeya dianthicola GBBC 2039 WP 024105357 1 Ddianthicola
Dickeya paradisiaca Ech703 WP 012764942 1 Dparadisiaca
Dickeya gs fangzhongdai ND14b WP 038660082 1 Dfangzhongdai
Dickeya gs fangzhongdai B16 WP 038918577 1 Dfangzhongdai
Dickeya gs fangzhongdai M005 WP 038918577 1 Dfangzhongdai
Dickeya gs fangzhongdai MK7 WP 038918577 1 Dfangzhongdai
Dickeya gs fangzhongdai NCPPB 3274 WP 042859387 1 Dfangzhongdai
Dickeya gs fangzhongdai S1 WP 049854931 1 Dfangzhongdai
Dickeya gs poaceaephila NCPPB 569 WP 042869823 1 Dpoaceaephila
Dickeya zeae NCPPB 3532 WP 023639451 1 Dzeae
Dickeya zeae NCPPB 2538 WP 019846314 1 Dzeae
Dickeya zeae MS1 WP 019846314 1 Dzeae
Dickeya zeae MK19 WP 038914265 1 Dzeae
Dickeya zeae DZ2Q WP 038926095 1 Dzeae
Dickeya zeae Ech586 WP 012885233 1 Dzeae
Dickeya zeae NCPPB 3531 WP 038904829 1 Dzeae
Dickeya zeae CSL RW192 WP 038908462 1 Dzeae
Dickeya zeae ZJU1202 WP 016943884 1 Dzeae
Dickeya zeae EC1 WP 016943884 1 Dzeae
Dickeya chrysanthemi subsp parthenii NCPPB 516 WP 027712478 1 Dchrysanthemi
Dickeya chrysanthemi L11 WP 033577238 1 Dchrysanthemi
Dickeya chrysanthemi subsp chrysanthemi NCPPB 402 WP 040000521 1 Dchrysanthemi
Dickeya chrysanthemi Ech1591 WP 012770351 1 Dchrysanthemi
Dickeya chrysanthemi NCPPB 3533 WP 012770351 1 Dchrysanthemi
482 cbrD Iron metabolism Achromobactin biosynthesis and transport Seed
Dickeya dadantii NCPPB 898 WP 013317301 1 Ddadantii
Dickeya dadantii NCPPB 3537 WP 013317301 1 Ddadantii
Dickeya dadantii 3937 WP 013317301 1 Ddadantii
Dickeya dadantii subsp dieffenbachiae NCPPB 2976 WP 038922395 1 Ddadantii
Dickeya solani RNS 05 1 2A WP 057082661 1 Dsolani
Dickeya solani MK10 WP 023637730 1 Dsolani
Dickeya solani IFB 0099 WP 023637730 1 Dsolani
Dickeya solani RNS 07 7 3B WP 023637730 1 Dsolani
Dickeya solani IPO 2222 WP 023637730 1 Dsolani
Dickeya solani D s0432 1 WP 023637730 1 Dsolani
Dickeya solani RNS 08 23 3 1 A WP 023637730 1 Dsolani
Dickeya solani GBBC 2040 WP 023637730 1 Dsolani
Dickeya solani PPO 9019 WP 023637730 1 Dsolani
Dickeya solani PPO 9134 WP 023637730 1 Dsolani
Dickeya solani MK16 WP 023637730 1 Dsolani

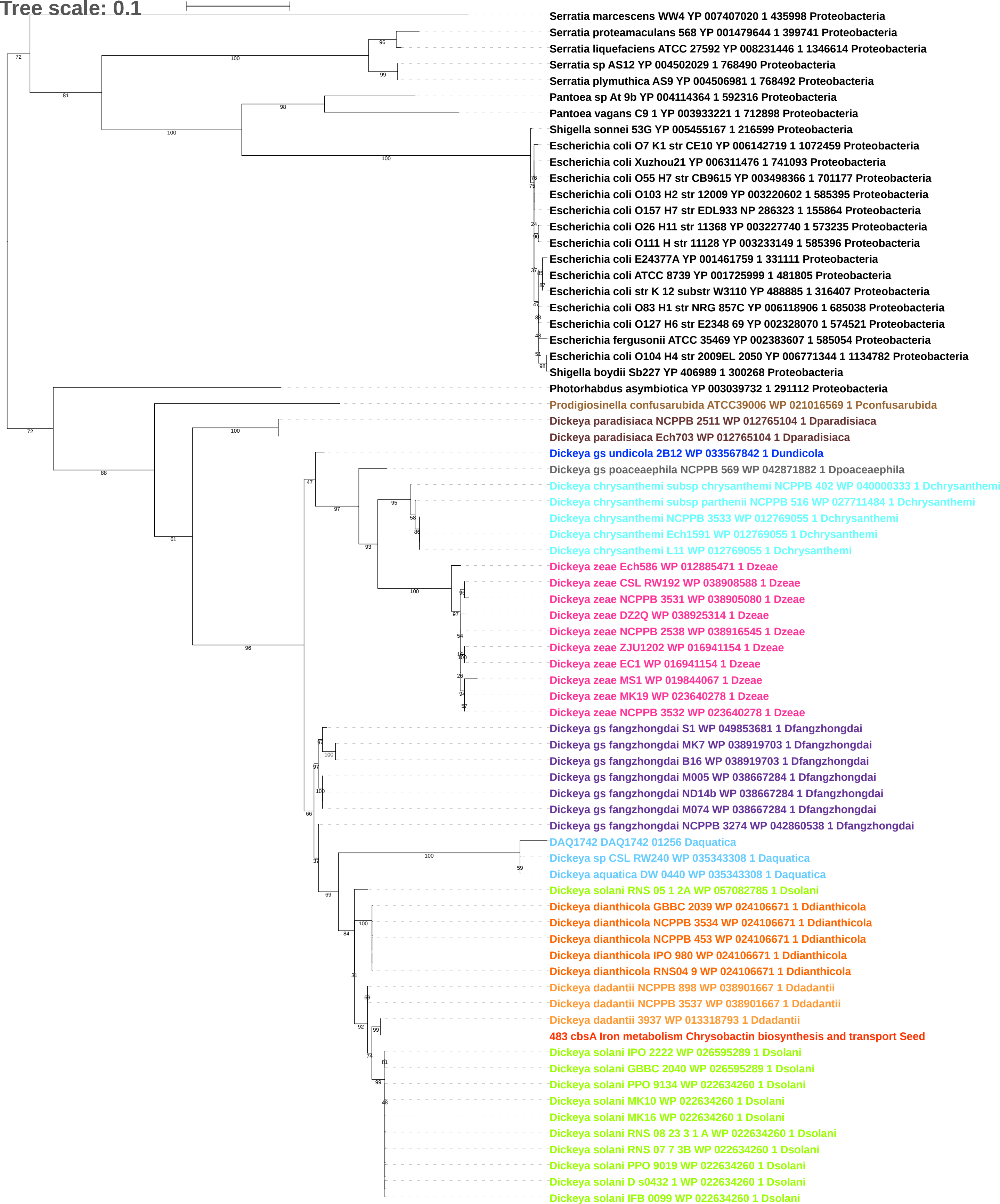

Tree scale: 0.1
Serratia marcescens WW4 YP 007407020 1 435998 Proteobacteria
Serratia proteamaculans 568 YP 001479644 1 399741 Proteobacteria
Serratia liquefaciens ATCC 27592 YP 008231446 1 1346614 Proteobacteria
Serratia sp AS12 YP 004502029 1 768490 Proteobacteria
Serratia plymuthica AS9 YP 004506981 1 768492 Proteobacteria
Pantoea sp At 9b YP 004114364 1 592316 Proteobacteria
Pantoea vagans C9 1 YP 003933221 1 712898 Proteobacteria
Shigella sonnei 53G YP 005455167 1 216599 Proteobacteria
Escherichia coli O7 K1 str CE10 YP 006142719 1 1072459 Proteobacteria
Escherichia coli Xuzhou21 YP 006311476 1 741093 Proteobacteria
Escherichia coli O55 H7 str CB9615 YP 003498366 1 701177 Proteobacteria
Escherichia coli O103 H2 str 12009 YP 003220602 1 585395 Proteobacteria
Escherichia coli O157 H7 str EDL933 NP 286323 1 155864 Proteobacteria
Escherichia coli O26 H11 str 11368 YP 003227740 1 573235 Proteobacteria
Escherichia coli O111 H str 11128 YP 003233149 1 585396 Proteobacteria
Escherichia coli E24377A YP 001461759 1 331111 Proteobacteria
Escherichia coli ATCC 8739 YP 001725999 1 481805 Proteobacteria
Escherichia coli str K 12 substr W3110 YP 488885 1 316407 Proteobacteria
Escherichia coli O83 H1 str NRG 857C YP 006118906 1 685038 Proteobacteria
Escherichia coli O127 H6 str E2348 69 YP 002328070 1 574521 Proteobacteria
Escherichia fergusonii ATCC 35469 YP 002383607 1 585054 Proteobacteria
Escherichia coli O104 H4 str 2009EL 2050 YP 006771344 1 1134782 Proteobacteria
Shigella boydii Sb227 YP 406989 1 300268 Proteobacteria
Photorhabdus asymbiotica YP 003039732 1 291112 Proteobacteria
Prodigiosinella confusarubida ATCC39006 WP 021016569 1 Pconfusarubida
Dickeya paradisiaca NCPPB 2511 WP 012765104 1 Dparadisiaca
Dickeya paradisiaca Ech703 WP 012765104 1 Dparadisiaca
Dickeya gs undicola 2B12 WP 033567842 1 Dundicola
Dickeya gs poaceaephila NCPPB 569 WP 042871882 1 Dpoaceaephila
Dickeya chrysanthemi subsp chrysanthemi NCPPB 402 WP 040000333 1 Dchrysanthemi
Dickeya chrysanthemi subsp parthenii NCPPB 516 WP 027711484 1 Dchrysanthemi
Dickeya chrysanthemi NCPPB 3533 WP 012769055 1 Dchrysanthemi
Dickeya chrysanthemi Ech1591 WP 012769055 1 Dchrysanthemi
Dickeya chrysanthemi L11 WP 012769055 1 Dchrysanthemi
Dickeya zeae Ech586 WP 012885471 1 Dzeae
Dickeya zeae CSL RW192 WP 038908588 1 Dzeae
Dickeya zeae NCPPB 3531 WP 038905080 1 Dzeae
Dickeya zeae DZ2Q WP 038925314 1 Dzeae
Dickeya zeae NCPPB 2538 WP 038916545 1 Dzeae
Dickeya zeae ZJU1202 WP 016941154 1 Dzeae
Dickeya zeae EC1 WP 016941154 1 Dzeae
Dickeya zeae MS1 WP 019844067 1 Dzeae
Dickeya zeae MK19 WP 023640278 1 Dzeae
Dickeya zeae NCPPB 3532 WP 023640278 1 Dzeae
Dickeya gs fangzhongdai S1 WP 049853681 1 Dfangzhongdai
Dickeya gs fangzhongdai MK7 WP 038919703 1 Dfangzhongdai
Dickeya gs fangzhongdai B16 WP 038919703 1 Dfangzhongdai
Dickeya gs fangzhongdai M005 WP 038667284 1 Dfangzhongdai
Dickeya gs fangzhongdai ND14b WP 038667284 1 Dfangzhongdai
Dickeya gs fangzhongdai M074 WP 038667284 1 Dfangzhongdai
Dickeya gs fangzhongdai NCPPB 3274 WP 042860538 1 Dfangzhongdai
DAQ1742 DAQ1742 01256 Daquatica
Dickeya sp CSL RW240 WP 035343308 1 Daquatica
Dickeya aquatica DW 0440 WP 035343308 1 Daquatica
Dickeya solani RNS 05 1 2A WP 057082785 1 Dsolani
Dickeya dianthicola GBBC 2039 WP 024106671 1 Ddianthicola
Dickeya dianthicola NCPPB 3534 WP 024106671 1 Ddianthicola
Dickeya dianthicola NCPPB 453 WP 024106671 1 Ddianthicola
Dickeya dianthicola IPO 980 WP 024106671 1 Ddianthicola
Dickeya dianthicola RNS04 9 WP 024106671 1 Ddianthicola
Dickeya dadantii NCPPB 898 WP 038901667 1 Ddadantii
Dickeya dadantii NCPPB 3537 WP 038901667 1 Ddadantii
Dickeya dadantii 3937 WP 013318793 1 Ddadantii
483 cbsA Iron metabolism Chrysobactin biosynthesis and transport Seed
Dickeya solani IPO 2222 WP 026595289 1 Dsolani
Dickeya solani GBBC 2040 WP 026595289 1 Dsolani
Dickeya solani PPO 9134 WP 022634260 1 Dsolani
Dickeya solani MK10 WP 022634260 1 Dsolani
Dickeya solani MK16 WP 022634260 1 Dsolani
Dickeya solani RNS 08 23 3 1 A WP 022634260 1 Dsolani
Dickeya solani RNS 07 7 3B WP 022634260 1 Dsolani
Dickeya solani PPO 9019 WP 022634260 1 Dsolani
Dickeya solani D s0432 1 WP 022634260 1 Dsolani
Dickeya solani IFB 0099 WP 022634260 1 Dsolani

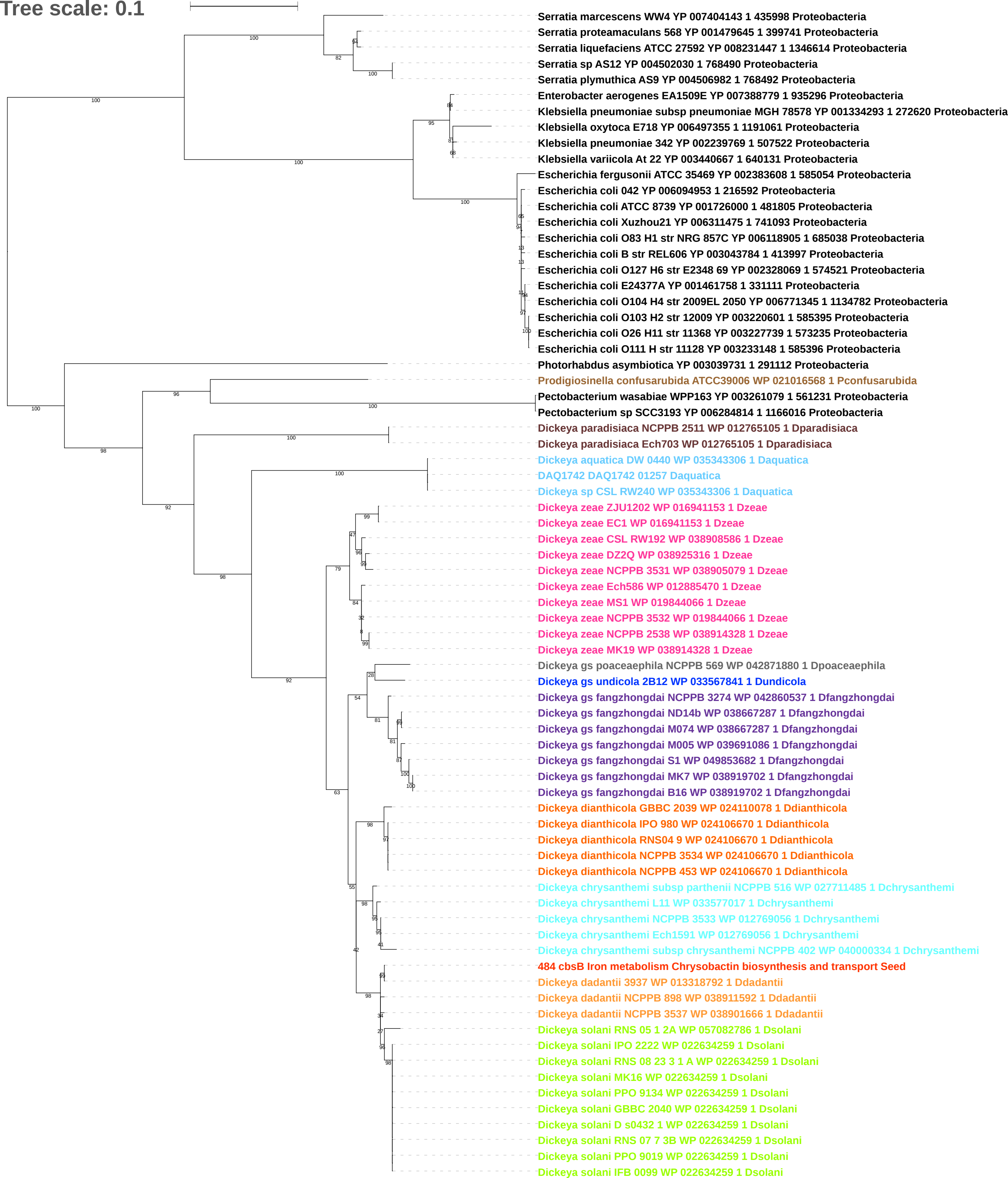

Tree scale: 0.1
Serratia marcescens WW4 YP 007404143 1 435998 Proteobacteria
Serratia proteamaculans 568 YP 001479645 1 399741 Proteobacteria
Serratia liquefaciens ATCC 27592 YP 008231447 1 1346614 Proteobacteria
Serratia sp AS12 YP 004502030 1 768490 Proteobacteria
Serratia plymuthica AS9 YP 004506982 1 768492 Proteobacteria
Enterobacter aerogenes EA1509E YP 007388779 1 935296 Proteobacteria
Klebsiella pneumoniae subsp pneumoniae MGH 78578 YP 001334293 1 272620 Proteobacteria
Klebsiella oxytoca E718 YP 006497355 1 1191061 Proteobacteria
Klebsiella pneumoniae 342 YP 002239769 1 507522 Proteobacteria
Klebsiella variicola At 22 YP 003440667 1 640131 Proteobacteria
Escherichia fergusonii ATCC 35469 YP 002383608 1 585054 Proteobacteria
Escherichia coli 042 YP 006094953 1 216592 Proteobacteria
Escherichia coli ATCC 8739 YP 001726000 1 481805 Proteobacteria
Escherichia coli Xuzhou21 YP 006311475 1 741093 Proteobacteria
Escherichia coli O83 H1 str NRG 857C YP 006118905 1 685038 Proteobacteria
Escherichia coli B str REL606 YP 003043784 1 413997 Proteobacteria
Escherichia coli O127 H6 str E2348 69 YP 002328069 1 574521 Proteobacteria
Escherichia coli E24377A YP 001461758 1 331111 Proteobacteria
Escherichia coli O104 H4 str 2009EL 2050 YP 006771345 1 1134782 Proteobacteria
Escherichia coli O103 H2 str 12009 YP 003220601 1 585395 Proteobacteria
Escherichia coli O26 H11 str 11368 YP 003227739 1 573235 Proteobacteria
Escherichia coli O111 H str 11128 YP 003233148 1 585396 Proteobacteria
Photorhabdus asymbiotica YP 003039731 1 291112 Proteobacteria
Prodigiosinella confusarubida ATCC39006 WP 021016568 1 Pconfusarubida
Pectobacterium wasabiae WPP163 YP 003261079 1 561231 Proteobacteria
Pectobacterium sp SCC3193 YP 006284814 1 1166016 Proteobacteria
Dickeya paradisiaca NCPPB 2511 WP 012765105 1 Dparadisiaca
Dickeya paradisiaca Ech703 WP 012765105 1 Dparadisiaca
Dickeya aquatica DW 0440 WP 035343306 1 Daquatica
DAQ1742 DAQ1742 01257 Daquatica
Dickeya sp CSL RW240 WP 035343306 1 Daquatica
Dickeya zeae ZJU1202 WP 016941153 1 Dzeae
Dickeya zeae EC1 WP 016941153 1 Dzeae
Dickeya zeae CSL RW192 WP 038908586 1 Dzeae
Dickeya zeae DZ2Q WP 038925316 1 Dzeae
Dickeya zeae NCPPB 3531 WP 038905079 1 Dzeae
Dickeya zeae Ech586 WP 012885470 1 Dzeae
Dickeya zeae MS1 WP 019844066 1 Dzeae
Dickeya zeae NCPPB 3532 WP 019844066 1 Dzeae
Dickeya zeae NCPPB 2538 WP 038914328 1 Dzeae
Dickeya zeae MK19 WP 038914328 1 Dzeae
Dickeya gs poaceaephila NCPPB 569 WP 042871880 1 Dpoaceaephila
Dickeya gs undicola 2B12 WP 033567841 1 Dundicola
Dickeya gs fangzhongdai NCPPB 3274 WP 042860537 1 Dfangzhongdai
Dickeya gs fangzhongdai ND14b WP 038667287 1 Dfangzhongdai
Dickeya gs fangzhongdai M074 WP 038667287 1 Dfangzhongdai
Dickeya gs fangzhongdai M005 WP 039691086 1 Dfangzhongdai
Dickeya gs fangzhongdai S1 WP 049853682 1 Dfangzhongdai
Dickeya gs fangzhongdai MK7 WP 038919702 1 Dfangzhongdai
Dickeya gs fangzhongdai B16 WP 038919702 1 Dfangzhongdai
Dickeya dianthicola GBBC 2039 WP 024110078 1 Ddianthicola
Dickeya dianthicola IPO 980 WP 024106670 1 Ddianthicola
Dickeya dianthicola RNS04 9 WP 024106670 1 Ddianthicola
Dickeya dianthicola NCPPB 3534 WP 024106670 1 Ddianthicola
Dickeya dianthicola NCPPB 453 WP 024106670 1 Ddianthicola
Dickeya chrysanthemi subsp parthenii NCPPB 516 WP 027711485 1 Dchrysanthemi
Dickeya chrysanthemi L11 WP 033577017 1 Dchrysanthemi
Dickeya chrysanthemi NCPPB 3533 WP 012769056 1 Dchrysanthemi
Dickeya chrysanthemi Ech1591 WP 012769056 1 Dchrysanthemi
Dickeya chrysanthemi subsp chrysanthemi NCPPB 402 WP 040000334 1 Dchrysanthemi
484 cbsB Iron metabolism Chrysobactin biosynthesis and transport Seed
Dickeya dadantii 3937 WP 013318792 1 Ddadantii
Dickeya dadantii NCPPB 898 WP 038911592 1 Ddadantii
Dickeya dadantii NCPPB 3537 WP 038901666 1 Ddadantii
Dickeya solani RNS 05 1 2A WP 057082786 1 Dsolani
Dickeya solani IPO 2222 WP 022634259 1 Dsolani
Dickeya solani RNS 08 23 3 1 A WP 022634259 1 Dsolani
Dickeya solani MK16 WP 022634259 1 Dsolani
Dickeya solani PPO 9134 WP 022634259 1 Dsolani
Dickeya solani GBBC 2040 WP 022634259 1 Dsolani
Dickeya solani D s0432 1 WP 022634259 1 Dsolani
Dickeya solani RNS 07 7 3B WP 022634259 1 Dsolani
Dickeya solani PPO 9019 WP 022634259 1 Dsolani
Dickeya solani IFB 0099 WP 022634259 1 Dsolani

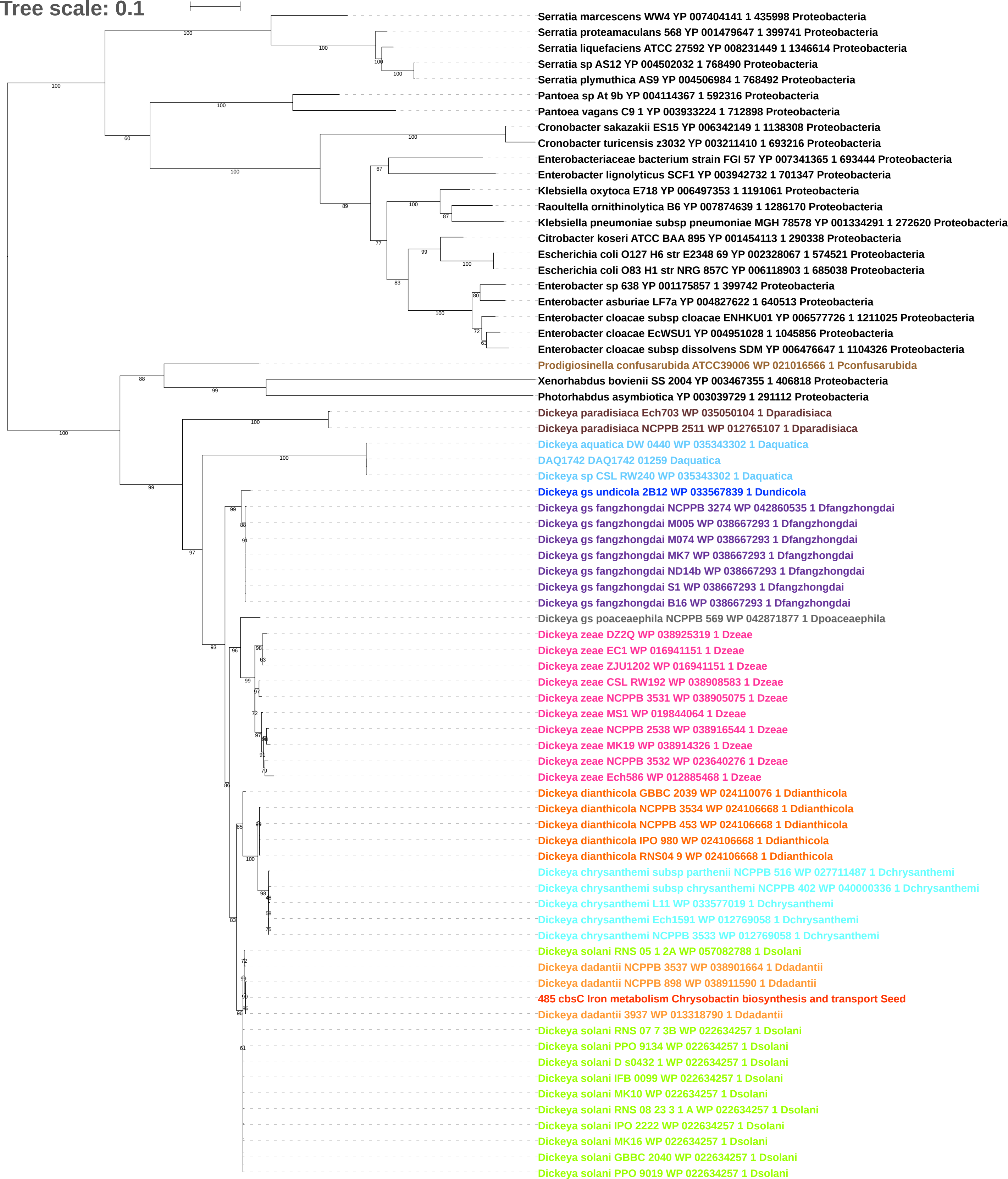

Tree scale: 0.1
Serratia marcescens WW4 YP 007404141 1 435998 Proteobacteria
Serratia proteamaculans 568 YP 001479647 1 399741 Proteobacteria
Serratia liquefaciens ATCC 27592 YP 008231449 1 1346614 Proteobacteria
Serratia sp AS12 YP 004502032 1 768490 Proteobacteria
Serratia plymuthica AS9 YP 004506984 1 768492 Proteobacteria
Pantoea sp At 9b YP 004114367 1 592316 Proteobacteria
Pantoea vagans C9 1 YP 003933224 1 712898 Proteobacteria
Cronobacter sakazakii ES15 YP 006342149 1 1138308 Proteobacteria
Cronobacter turicensis z3032 YP 003211410 1 693216 Proteobacteria
Enterobacteriaceae bacterium strain FGI 57 YP 007341365 1 693444 Proteobacteria
Enterobacter lignolyticus SCF1 YP 003942732 1 701347 Proteobacteria
Klebsiella oxytoca E718 YP 006497353 1 1191061 Proteobacteria
Raoultella ornithinolytica B6 YP 007874639 1 1286170 Proteobacteria
Klebsiella pneumoniae subsp pneumoniae MGH 78578 YP 001334291 1 272620 Proteobacteria
Citrobacter koseri ATCC BAA 895 YP 001454113 1 290338 Proteobacteria
Escherichia coli O127 H6 str E2348 69 YP 002328067 1 574521 Proteobacteria
Escherichia coli O83 H1 str NRG 857C YP 006118903 1 685038 Proteobacteria
Enterobacter sp 638 YP 001175857 1 399742 Proteobacteria
Enterobacter asburiae LF7a YP 004827622 1 640513 Proteobacteria
Enterobacter cloacae subsp cloacae ENHKU01 YP 006577726 1 1211025 Proteobacteria
Enterobacter cloacae EcWSU1 YP 004951028 1 1045856 Proteobacteria
Enterobacter cloacae subsp dissolvens SDM YP 006476647 1 1104326 Proteobacteria
Prodigiosinella confusarubida ATCC39006 WP 021016566 1 Pconfusarubida
Xenorhabdus bovienii SS 2004 YP 003467355 1 406818 Proteobacteria
Photorhabdus asymbiotica YP 003039729 1 291112 Proteobacteria
Dickeya paradisiaca Ech703 WP 035050104 1 Dparadisiaca
Dickeya paradisiaca NCPPB 2511 WP 012765107 1 Dparadisiaca
Dickeya aquatica DW 0440 WP 035343302 1 Daquatica
DAQ1742 DAQ1742 01259 Daquatica
Dickeya sp CSL RW240 WP 035343302 1 Daquatica
Dickeya gs undicola 2B12 WP 033567839 1 Dundicola
Dickeya gs fangzhongdai NCPPB 3274 WP 042860535 1 Dfangzhongdai
Dickeya gs fangzhongdai M005 WP 038667293 1 Dfangzhongdai
Dickeya gs fangzhongdai M074 WP 038667293 1 Dfangzhongdai
Dickeya gs fangzhongdai MK7 WP 038667293 1 Dfangzhongdai
Dickeya gs fangzhongdai ND14b WP 038667293 1 Dfangzhongdai
Dickeya gs fangzhongdai S1 WP 038667293 1 Dfangzhongdai
Dickeya gs fangzhongdai B16 WP 038667293 1 Dfangzhongdai
Dickeya gs poaceaephila NCPPB 569 WP 042871877 1 Dpoaceaephila
Dickeya zeae DZ2Q WP 038925319 1 Dzeae
Dickeya zeae EC1 WP 016941151 1 Dzeae
Dickeya zeae ZJU1202 WP 016941151 1 Dzeae
Dickeya zeae CSL RW192 WP 038908583 1 Dzeae
Dickeya zeae NCPPB 3531 WP 038905075 1 Dzeae
Dickeya zeae MS1 WP 019844064 1 Dzeae
Dickeya zeae NCPPB 2538 WP 038916544 1 Dzeae
Dickeya zeae MK19 WP 038914326 1 Dzeae
Dickeya zeae NCPPB 3532 WP 023640276 1 Dzeae
Dickeya zeae Ech586 WP 012885468 1 Dzeae
Dickeya dianthicola GBBC 2039 WP 024110076 1 Ddianthicola
Dickeya dianthicola NCPPB 3534 WP 024106668 1 Ddianthicola
Dickeya dianthicola NCPPB 453 WP 024106668 1 Ddianthicola
Dickeya dianthicola IPO 980 WP 024106668 1 Ddianthicola
Dickeya dianthicola RNS04 9 WP 024106668 1 Ddianthicola
Dickeya chrysanthemi subsp parthenii NCPPB 516 WP 027711487 1 Dchrysanthemi
Dickeya chrysanthemi subsp chrysanthemi NCPPB 402 WP 040000336 1 Dchrysanthemi
Dickeya chrysanthemi L11 WP 033577019 1 Dchrysanthemi
Dickeya chrysanthemi Ech1591 WP 012769058 1 Dchrysanthemi
Dickeya chrysanthemi NCPPB 3533 WP 012769058 1 Dchrysanthemi
Dickeya solani RNS 05 1 2A WP 057082788 1 Dsolani
Dickeya dadantii NCPPB 3537 WP 038901664 1 Ddadantii
Dickeya dadantii NCPPB 898 WP 038911590 1 Ddadantii
485 cbsC Iron metabolism Chrysobactin biosynthesis and transport Seed
Dickeya dadantii 3937 WP 013318790 1 Ddadantii
Dickeya solani RNS 07 7 3B WP 022634257 1 Dsolani
Dickeya solani PPO 9134 WP 022634257 1 Dsolani
Dickeya solani D s0432 1 WP 022634257 1 Dsolani
Dickeya solani IFB 0099 WP 022634257 1 Dsolani
Dickeya solani MK10 WP 022634257 1 Dsolani
Dickeya solani RNS 08 23 3 1 A WP 022634257 1 Dsolani
Dickeya solani IPO 2222 WP 022634257 1 Dsolani
Dickeya solani MK16 WP 022634257 1 Dsolani
Dickeya solani GBBC 2040 WP 022634257 1 Dsolani
Dickeya solani PPO 9019 WP 022634257 1 Dsolani

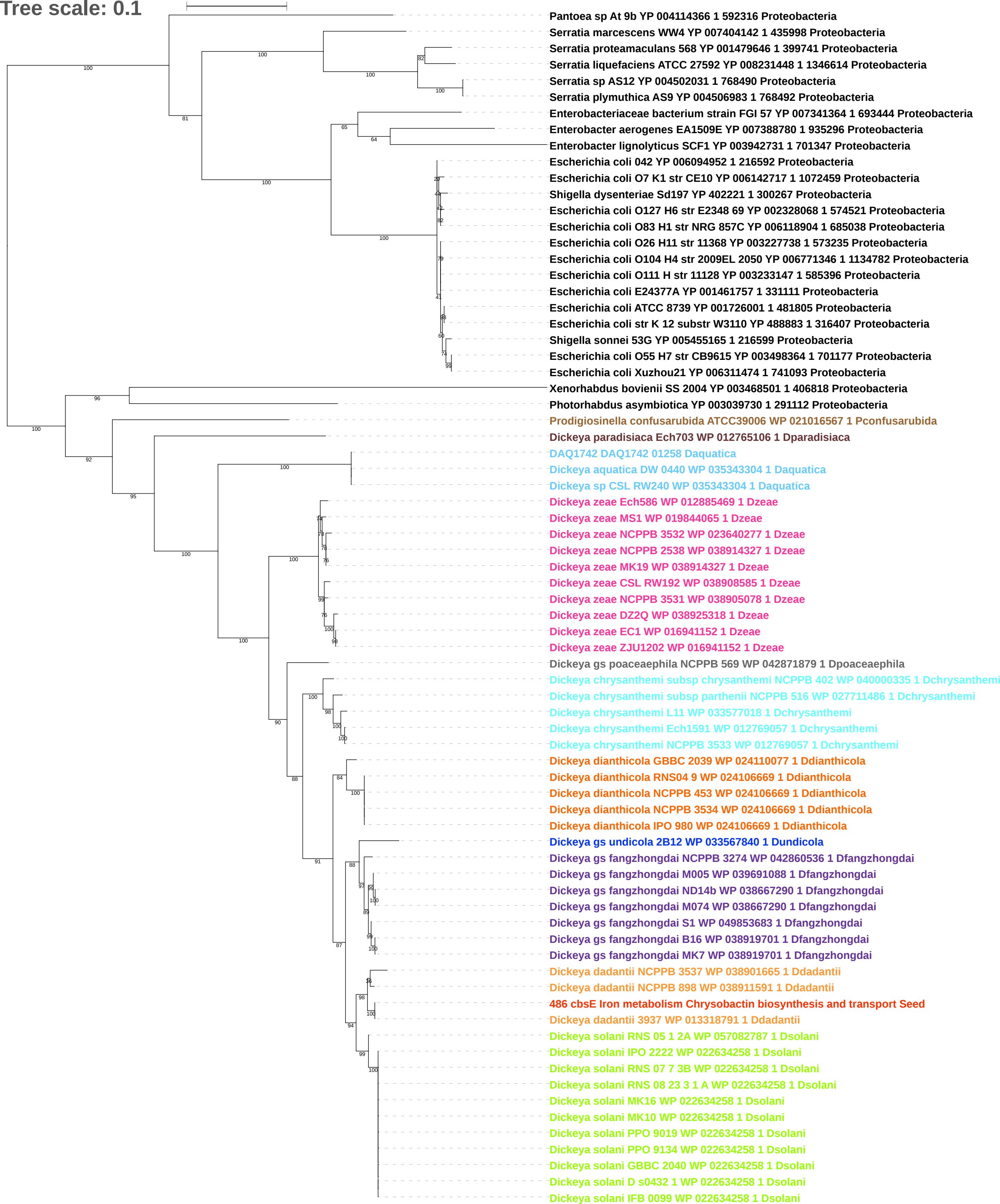
Tree scale: 0.1
Pantoea sp At 9b YP 004114366 1 592316 Proteobacteria
Serratia marcescens WW4 YP 007404142 1 435998 Proteobacteria
Serratia proteamaculans 568 YP 001479646 1 399741 Proteobacteria
Serratia liquefaciens ATCC 27592 YP 008231448 1 1346614 Proteobacteria
Serratia sp AS12 YP 004502031 1 768490 Proteobacteria
Serratia plymuthica AS9 YP 004506983 1 768492 Proteobacteria
Enterobacteriaceae bacterium strain FGI 57 YP 007341364 1 693444 Proteobacteria
Enterobacter aerogenes EA1509E YP 007388780 1 935296 Proteobacteria
Enterobacter lignolyticus SCF1 YP 003942731 1 701347 Proteobacteria
Escherichia coli 042 YP 006094952 1 216592 Proteobacteria
Escherichia coli O7 K1 str CE10 YP 006142717 1 1072459 Proteobacteria
Shigella dysenteriae Sd197 YP 402221 1 300267 Proteobacteria
Escherichia coli O127 H6 str E2348 69 YP 002328068 1 574521 Proteobacteria
Escherichia coli O83 H1 str NRG 857C YP 006118904 1 685038 Proteobacteria
Escherichia coli O26 H11 str 11368 YP 003227738 1 573235 Proteobacteria
Escherichia coli O104 H4 str 2009EL 2050 YP 006771346 1 1134782 Proteobacteria
Escherichia coli O111 H str 11128 YP 003233147 1 585396 Proteobacteria
Escherichia coli E24377A YP 001461757 1 331111 Proteobacteria
Escherichia coli ATCC 8739 YP 001726001 1 481805 Proteobacteria
Escherichia coli str K 12 substr W3110 YP 488883 1 316407 Proteobacteria
Shigella sonnei 53G YP 005455165 1 216599 Proteobacteria
Escherichia coli O55 H7 str CB9615 YP 003498364 1 701177 Proteobacteria
Escherichia coli Xuzhou21 YP 006311474 1 741093 Proteobacteria
Xenorhabdus bovienii SS 2004 YP 003468501 1 406818 Proteobacteria
Photorhabdus asymbiotica YP 003039730 1 291112 Proteobacteria
Prodigiosinella confusarubida ATCC39006 WP 021016567 1 Pconfusarubida
Dickeya paradisiaca Ech703 WP 012765106 1 Dparadisiaca
DAQ1742 DAQ1742 01258 Daquatica
Dickeya aquatica DW 0440 WP 035343304 1 Daquatica
Dickeya sp CSL RW240 WP 035343304 1 Daquatica
Dickeya zeae Ech586 WP 012885469 1 Dzeae
Dickeya zeae MS1 WP 019844065 1 Dzeae
Dickeya zeae NCPPB 3532 WP 023640277 1 Dzeae
Dickeya zeae NCPPB 2538 WP 038914327 1 Dzeae
Dickeya zeae MK19 WP 038914327 1 Dzeae
Dickeya zeae CSL RW192 WP 038908585 1 Dzeae
Dickeya zeae NCPPB 3531 WP 038905078 1 Dzeae
Dickeya zeae DZ2Q WP 038925318 1 Dzeae
Dickeya zeae EC1 WP 016941152 1 Dzeae
Dickeya zeae ZJU1202 WP 016941152 1 Dzeae
Dickeya gs poaceaephila NCPPB 569 WP 042871879 1 Dpoaceaephila
Dickeya chrysanthemi subsp chrysanthemi NCPPB 402 WP 040000335 1 Dchrysanthemi
Dickeya chrysanthemi subsp parthenii NCPPB 516 WP 027711486 1 Dchrysanthemi
Dickeya chrysanthemi L11 WP 033577018 1 Dchrysanthemi
Dickeya chrysanthemi Ech1591 WP 012769057 1 Dchrysanthemi
Dickeya chrysanthemi NCPPB 3533 WP 012769057 1 Dchrysanthemi
Dickeya dianthicola GBBC 2039 WP 024110077 1 Ddianthicola
Dickeya dianthicola RNS04 9 WP 024106669 1 Ddianthicola
Dickeya dianthicola NCPPB 453 WP 024106669 1 Ddianthicola
Dickeya dianthicola NCPPB 3534 WP 024106669 1 Ddianthicola
Dickeya dianthicola IPO 980 WP 024106669 1 Ddianthicola
Dickeya gs undicola 2B12 WP 033567840 1 Dundicola
Dickeya gs fangzhongdai NCPPB 3274 WP 042860536 1 Dfangzhongdai
Dickeya gs fangzhongdai M005 WP 039691088 1 Dfangzhongdai
Dickeya gs fangzhongdai ND14b WP 038667290 1 Dfangzhongdai
Dickeya gs fangzhongdai M074 WP 038667290 1 Dfangzhongdai
Dickeya gs fangzhongdai S1 WP 049853683 1 Dfangzhongdai
Dickeya gs fangzhongdai B16 WP 038919701 1 Dfangzhongdai
Dickeya gs fangzhongdai MK7 WP 038919701 1 Dfangzhongdai
Dickeya dadantii NCPPB 3537 WP 038901665 1 Ddadantii
Dickeya dadantii NCPPB 898 WP 038911591 1 Ddadantii
486 cbsE Iron metabolism Chrysobactin biosynthesis and transport Seed
Dickeya dadantii 3937 WP 013318791 1 Ddadantii
Dickeya solani RNS 05 1 2A WP 057082787 1 Dsolani
Dickeya solani IPO 2222 WP 022634258 1 Dsolani
Dickeya solani RNS 07 7 3B WP 022634258 1 Dsolani
Dickeya solani RNS 08 23 3 1 A WP 022634258 1 Dsolani
Dickeya solani MK16 WP 022634258 1 Dsolani
Dickeya solani MK10 WP 022634258 1 Dsolani
Dickeya solani PPO 9019 WP 022634258 1 Dsolani
Dickeya solani PPO 9134 WP 022634258 1 Dsolani
Dickeya solani GBBC 2040 WP 022634258 1 Dsolani
Dickeya solani D s0432 1 WP 022634258 1 Dsolani
Dickeya solani IFB 0099 WP 022634258 1 Dsolani

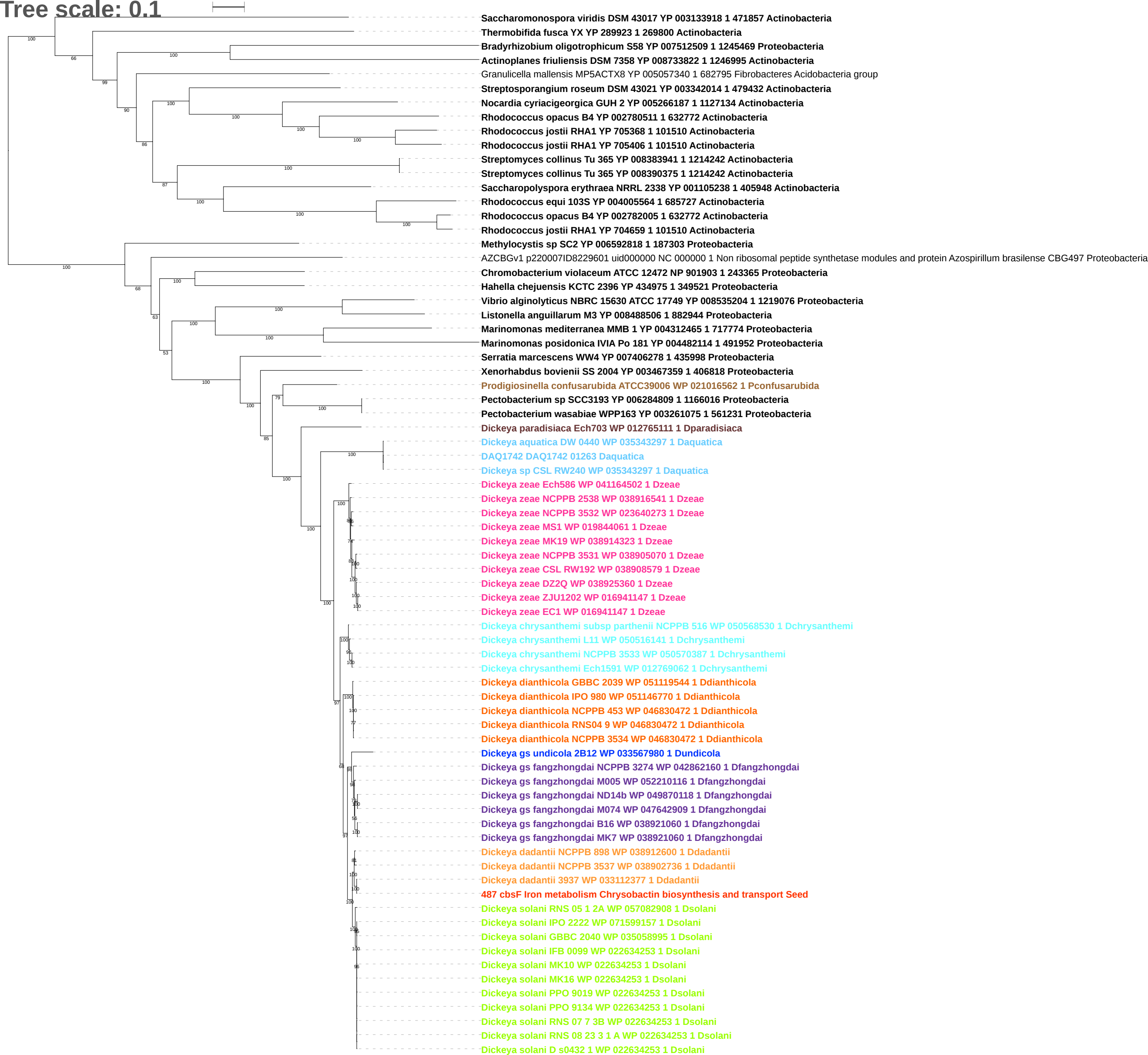
Tree scale: 0.1
Saccharomonospora viridis DSM 43017 YP 003133918 1 471857 Actinobacteria
Thermobifida fusca YX YP 289923 1 269800 Actinobacteria
Bradyrhizobium oligotrophicum S58 YP 007512509 1 1245469 Proteobacteria
Actinoplanes friuliensis DSM 7358 YP 008733822 1 1246995 Actinobacteria
Granulicella mallensis MP5ACTX8 YP 005057340 1 682795 Fibrobacteres Acidobacteria group
Streptosporangium roseum DSM 43021 YP 003342014 1 479432 Actinobacteria
Nocardia cyriacigeorgica GUH 2 YP 005266187 1 1127134 Actinobacteria
Rhodococcus opacus B4 YP 002780511 1 632772 Actinobacteria
Rhodococcus jostii RHA1 YP 705368 1 101510 Actinobacteria
Rhodococcus jostii RHA1 YP 705406 1 101510 Actinobacteria
Streptomyces collinus Tu 365 YP 008383941 1 1214242 Actinobacteria
Streptomyces collinus Tu 365 YP 008390375 1 1214242 Actinobacteria
Saccharopolyspora erythraea NRRL 2338 YP 001105238 1 405948 Actinobacteria
Rhodococcus equi 103S YP 004005564 1 685727 Actinobacteria
Rhodococcus opacus B4 YP 002782005 1 632772 Actinobacteria
Rhodococcus jostii RHA1 YP 704659 1 101510 Actinobacteria
Methylocystis sp SC2 YP 006592818 1 187303 Proteobacteria
AZCBGv1 p220007ID8229601 uid000000 NC 000000 1 Non ribosomal peptide synthetase modules and protein Azospirillum brasilense CBG497 Proteobacteria
Chromobacterium violaceum ATCC 12472 NP 901903 1 243365 Proteobacteria
Hahella chejuensis KCTC 2396 YP 434975 1 349521 Proteobacteria
Vibrio alginolyticus NBRC 15630 ATCC 17749 YP 008535204 1 1219076 Proteobacteria
Listonella anguillarum M3 YP 008488506 1 882944 Proteobacteria
Marinomonas mediterranea MMB 1 YP 004312465 1 717774 Proteobacteria
Marinomonas posidonica IVIA Po 181 YP 004482114 1 491952 Proteobacteria
Serratia marcescens WW4 YP 007406278 1 435998 Proteobacteria
Xenorhabdus bovienii SS 2004 YP 003467359 1 406818 Proteobacteria
Prodigiosinella confusarubida ATCC39006 WP 021016562 1 Pconfusarubida
Pectobacterium sp SCC3193 YP 006284809 1 1166016 Proteobacteria
Pectobacterium wasabiae WPP163 YP 003261075 1 561231 Proteobacteria
Dickeya paradisiaca Ech703 WP 012765111 1 Dparadisiaca
Dickeya aquatica DW 0440 WP 035343297 1 Daquatica
DAQ1742 DAQ1742 01263 Daquatica
Dickeya sp CSL RW240 WP 035343297 1 Daquatica
Dickeya zeae Ech586 WP 041164502 1 Dzeae
Dickeya zeae NCPPB 2538 WP 038916541 1 Dzeae
Dickeya zeae NCPPB 3532 WP 023640273 1 Dzeae
Dickeya zeae MS1 WP 019844061 1 Dzeae
Dickeya zeae MK19 WP 038914323 1 Dzeae
Dickeya zeae NCPPB 3531 WP 038905070 1 Dzeae
Dickeya zeae CSL RW192 WP 038908579 1 Dzeae
Dickeya zeae DZ2Q WP 038925360 1 Dzeae
Dickeya zeae ZJU1202 WP 016941147 1 Dzeae
Dickeya zeae EC1 WP 016941147 1 Dzeae
Dickeya chrysanthemi subsp parthenii NCPPB 516 WP 050568530 1 Dchrysanthemi
Dickeya chrysanthemi L11 WP 050516141 1 Dchrysanthemi
Dickeya chrysanthemi NCPPB 3533 WP 050570387 1 Dchrysanthemi
Dickeya chrysanthemi Ech1591 WP 012769062 1 Dchrysanthemi
Dickeya dianthicola GBBC 2039 WP 051119544 1 Ddianthicola
Dickeya dianthicola IPO 980 WP 051146770 1 Ddianthicola
Dickeya dianthicola NCPPB 453 WP 046830472 1 Ddianthicola
Dickeya dianthicola RNS04 9 WP 046830472 1 Ddianthicola
Dickeya dianthicola NCPPB 3534 WP 046830472 1 Ddianthicola
Dickeya gs undicola 2B12 WP 033567980 1 Dundicola
Dickeya gs fangzhongdai NCPPB 3274 WP 042862160 1 Dfangzhongdai
Dickeya gs fangzhongdai M005 WP 052210116 1 Dfangzhongdai
Dickeya gs fangzhongdai ND14b WP 049870118 1 Dfangzhongdai
Dickeya gs fangzhongdai M074 WP 047642909 1 Dfangzhongdai
Dickeya gs fangzhongdai B16 WP 038921060 1 Dfangzhongdai
Dickeya gs fangzhongdai MK7 WP 038921060 1 Dfangzhongdai
Dickeya dadantii NCPPB 898 WP 038912600 1 Ddadantii
Dickeya dadantii NCPPB 3537 WP 038902736 1 Ddadantii
Dickeya dadantii 3937 WP 033112377 1 Ddadantii
487 cbsF Iron metabolism Chrysobactin biosynthesis and transport Seed
Dickeya solani RNS 05 1 2A WP 057082908 1 Dsolani
Dickeya solani IPO 2222 WP 071599157 1 Dsolani
Dickeya solani GBBC 2040 WP 035058995 1 Dsolani
Dickeya solani IFB 0099 WP 022634253 1 Dsolani
Dickeya solani MK10 WP 022634253 1 Dsolani
Dickeya solani MK16 WP 022634253 1 Dsolani
Dickeya solani PPO 9019 WP 022634253 1 Dsolani
Dickeya solani PPO 9134 WP 022634253 1 Dsolani
Dickeya solani RNS 07 7 3B WP 022634253 1 Dsolani
Dickeya solani RNS 08 23 3 1 A WP 022634253 1 Dsolani
Dickeya solani D s0432 1 WP 022634253 1 Dsolani

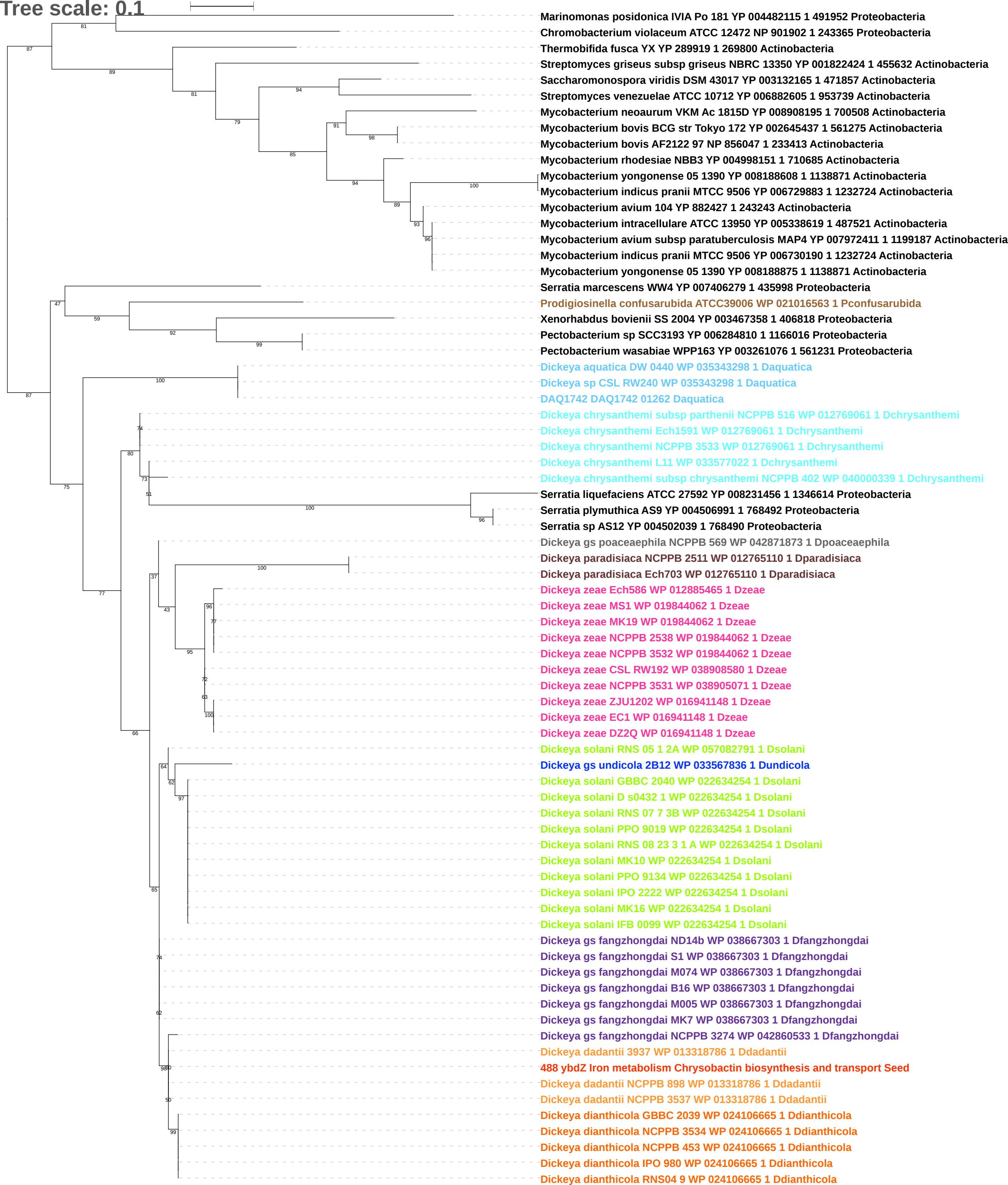
Tree scale: 0.1
Marinomonas posidonica IVIA Po 181 YP 004482115 1 491952 Proteobacteria
Chromobacterium violaceum ATCC 12472 NP 901902 1 243365 Proteobacteria
Thermobifida fusca YX YP 289919 1 269800 Actinobacteria
Streptomyces griseus subsp griseus NBRC 13350 YP 001822424 1 455632 Actinobacteria
Saccharomonospora viridis DSM 43017 YP 003132165 1 471857 Actinobacteria
Streptomyces venezuelae ATCC 10712 YP 006882605 1 953739 Actinobacteria
Mycobacterium neoaurum VKM Ac 1815D YP 008908195 1 700508 Actinobacteria
Mycobacterium bovis BCG str Tokyo 172 YP 002645437 1 561275 Actinobacteria
Mycobacterium bovis AF2122 97 NP 856047 1 233413 Actinobacteria
Mycobacterium rhodesiae NBB3 YP 004998151 1 710685 Actinobacteria
Mycobacterium yongonense 05 1390 YP 008188608 1 1138871 Actinobacteria
Mycobacterium indicus pranii MTCC 9506 YP 006729883 1 1232724 Actinobacteria
Mycobacterium avium 104 YP 882427 1 243243 Actinobacteria
Mycobacterium intracellulare ATCC 13950 YP 005338619 1 487521 Actinobacteria
Mycobacterium avium subsp paratuberculosis MAP4 YP 007972411 1 1199187 Actinobacteria
Mycobacterium indicus pranii MTCC 9506 YP 006730190 1 1232724 Actinobacteria
Mycobacterium yongonense 05 1390 YP 008188875 1 1138871 Actinobacteria
Serratia marcescens WW4 YP 007406279 1 435998 Proteobacteria
Prodigiosinella confusarubida ATCC39006 WP 021016563 1 Pconfusarubida
Xenorhabdus bovienii SS 2004 YP 003467358 1 406818 Proteobacteria
Pectobacterium sp SCC3193 YP 006284810 1 1166016 Proteobacteria
Pectobacterium wasabiae WPP163 YP 003261076 1 561231 Proteobacteria
Dickeya aquatica DW 0440 WP 035343298 1 Daquatica
Dickeya sp CSL RW240 WP 035343298 1 Daquatica
DAQ1742 DAQ1742 01262 Daquatica
Dickeya chrysanthemi subsp parthenii NCPPB 516 WP 012769061 1 Dchrysanthemi
Dickeya chrysanthemi Ech1591 WP 012769061 1 Dchrysanthemi
Dickeya chrysanthemi NCPPB 3533 WP 012769061 1 Dchrysanthemi
Dickeya chrysanthemi L11 WP 033577022 1 Dchrysanthemi
Dickeya chrysanthemi subsp chrysanthemi NCPPB 402 WP 040000339 1 Dchrysanthemi
Serratia liquefaciens ATCC 27592 YP 008231456 1 1346614 Proteobacteria
Serratia plymuthica AS9 YP 004506991 1 768492 Proteobacteria
Serratia sp AS12 YP 004502039 1 768490 Proteobacteria
Dickeya gs poaceaephila NCPPB 569 WP 042871873 1 Dpoaceaephila
Dickeya paradisiaca NCPPB 2511 WP 012765110 1 Dparadisiaca
Dickeya paradisiaca Ech703 WP 012765110 1 Dparadisiaca
Dickeya zeae Ech586 WP 012885465 1 Dzeae
Dickeya zeae MS1 WP 019844062 1 Dzeae
Dickeya zeae MK19 WP 019844062 1 Dzeae
Dickeya zeae NCPPB 2538 WP 019844062 1 Dzeae
Dickeya zeae NCPPB 3532 WP 019844062 1 Dzeae
Dickeya zeae CSL RW192 WP 038908580 1 Dzeae
Dickeya zeae NCPPB 3531 WP 038905071 1 Dzeae
Dickeya zeae ZJU1202 WP 016941148 1 Dzeae
Dickeya zeae EC1 WP 016941148 1 Dzeae
Dickeya zeae DZ2Q WP 016941148 1 Dzeae
Dickeya solani RNS 05 1 2A WP 057082791 1 Dsolani
Dickeya gs undicola 2B12 WP 033567836 1 Dundicola
Dickeya solani GBBC 2040 WP 022634254 1 Dsolani
Dickeya solani D s0432 1 WP 022634254 1 Dsolani
Dickeya solani RNS 07 7 3B WP 022634254 1 Dsolani
Dickeya solani PPO 9019 WP 022634254 1 Dsolani
Dickeya solani RNS 08 23 3 1 A WP 022634254 1 Dsolani
Dickeya solani MK10 WP 022634254 1 Dsolani
Dickeya solani PPO 9134 WP 022634254 1 Dsolani
Dickeya solani IPO 2222 WP 022634254 1 Dsolani
Dickeya solani MK16 WP 022634254 1 Dsolani
Dickeya solani IFB 0099 WP 022634254 1 Dsolani
Dickeya gs fangzhongdai ND14b WP 038667303 1 Dfangzhongdai
Dickeya gs fangzhongdai S1 WP 038667303 1 Dfangzhongdai
Dickeya gs fangzhongdai M074 WP 038667303 1 Dfangzhongdai
Dickeya gs fangzhongdai B16 WP 038667303 1 Dfangzhongdai
Dickeya gs fangzhongdai M005 WP 038667303 1 Dfangzhongdai
Dickeya gs fangzhongdai MK7 WP 038667303 1 Dfangzhongdai
Dickeya gs fangzhongdai NCPPB 3274 WP 042860533 1 Dfangzhongdai
Dickeya dadantii 3937 WP 013318786 1 Ddadantii
488 ybdZ Iron metabolism Chrysobactin biosynthesis and transport Seed
Dickeya dadantii NCPPB 898 WP 013318786 1 Ddadantii
Dickeya dadantii NCPPB 3537 WP 013318786 1 Ddadantii
Dickeya dianthicola GBBC 2039 WP 024106665 1 Ddianthicola
Dickeya dianthicola NCPPB 3534 WP 024106665 1 Ddianthicola
Dickeya dianthicola NCPPB 453 WP 024106665 1 Ddianthicola
Dickeya dianthicola IPO 980 WP 024106665 1 Ddianthicola
Dickeya dianthicola RNS04 9 WP 024106665 1 Ddianthicola

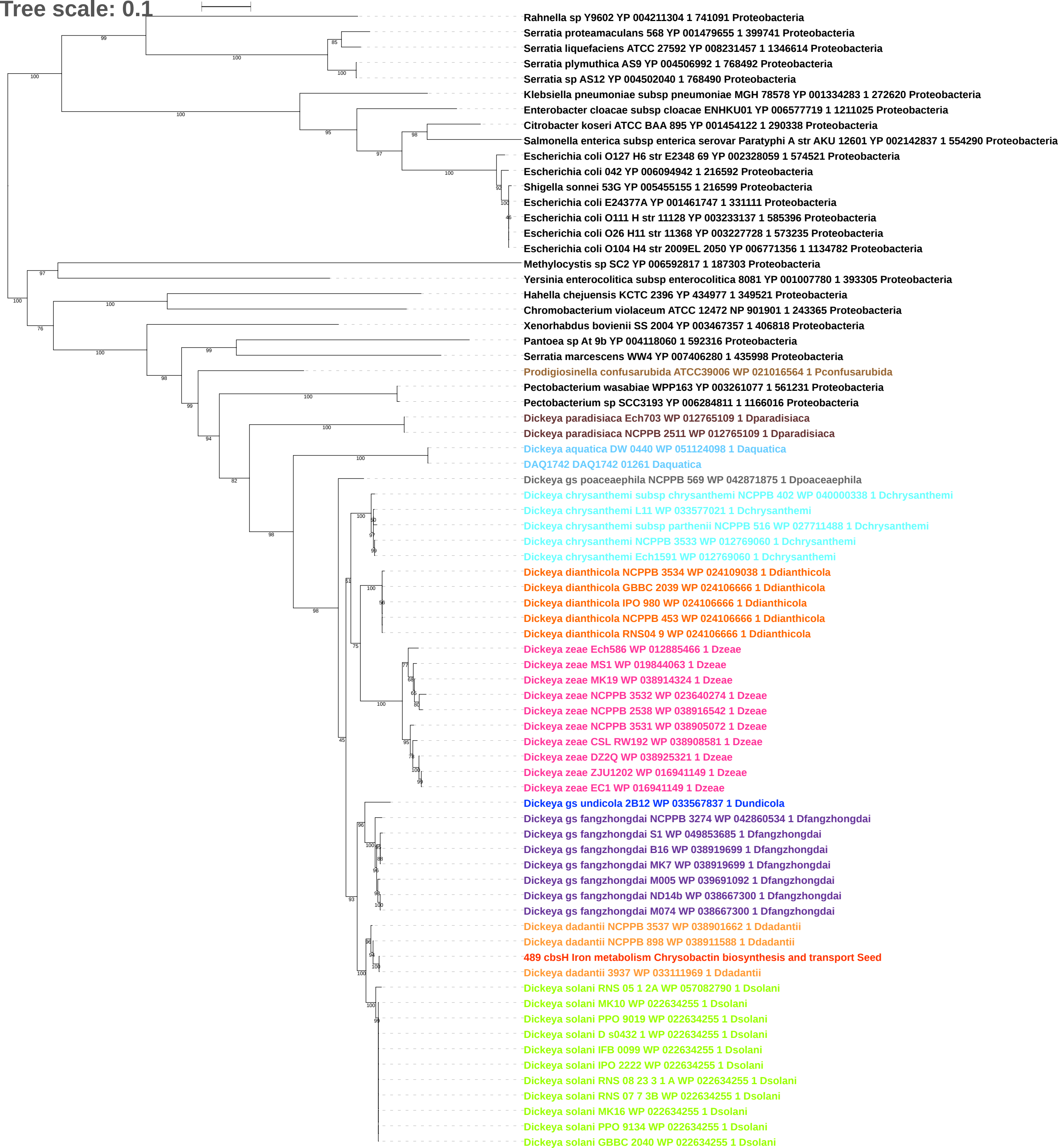
Tree scale: 0.1
Rahnella sp Y9602 YP 004211304 1 741091 Proteobacteria
Serratia proteamaculans 568 YP 001479655 1 399741 Proteobacteria
Serratia liquefaciens ATCC 27592 YP 008231457 1 1346614 Proteobacteria
Serratia plymuthica AS9 YP 004506992 1 768492 Proteobacteria
Serratia sp AS12 YP 004502040 1 768490 Proteobacteria
Klebsiella pneumoniae subsp pneumoniae MGH 78578 YP 001334283 1 272620 Proteobacteria
Enterobacter cloacae subsp cloacae ENHKU01 YP 006577719 1 1211025 Proteobacteria
Citrobacter koseri ATCC BAA 895 YP 001454122 1 290338 Proteobacteria
Salmonella enterica subsp enterica serovar Paratyphi A str AKU 12601 YP 002142837 1 554290 Proteobacteria
Escherichia coli O127 H6 str E2348 69 YP 002328059 1 574521 Proteobacteria
Escherichia coli 042 YP 006094942 1 216592 Proteobacteria
Shigella sonnei 53G YP 005455155 1 216599 Proteobacteria
Escherichia coli E24377A YP 001461747 1 331111 Proteobacteria
Escherichia coli O111 H str 11128 YP 003233137 1 585396 Proteobacteria
Escherichia coli O26 H11 str 11368 YP 003227728 1 573235 Proteobacteria
Escherichia coli O104 H4 str 2009EL 2050 YP 006771356 1 1134782 Proteobacteria
Methylocystis sp SC2 YP 006592817 1 187303 Proteobacteria
Yersinia enterocolitica subsp enterocolitica 8081 YP 001007780 1 393305 Proteobacteria
Hahella chejuensis KCTC 2396 YP 434977 1 349521 Proteobacteria
Chromobacterium violaceum ATCC 12472 NP 901901 1 243365 Proteobacteria
Xenorhabdus bovienii SS 2004 YP 003467357 1 406818 Proteobacteria
Pantoea sp At 9b YP 004118060 1 592316 Proteobacteria
Serratia marcescens WW4 YP 007406280 1 435998 Proteobacteria
Prodigiosinella confusarubida ATCC39006 WP 021016564 1 Pconfusarubida
Pectobacterium wasabiae WPP163 YP 003261077 1 561231 Proteobacteria
Pectobacterium sp SCC3193 YP 006284811 1 1166016 Proteobacteria
Dickeya paradisiaca Ech703 WP 012765109 1 Dparadisiaca
Dickeya paradisiaca NCPPB 2511 WP 012765109 1 Dparadisiaca
Dickeya aquatica DW 0440 WP 051124098 1 Daquatica
DAQ1742 DAQ1742 01261 Daquatica
Dickeya gs poaceaephila NCPPB 569 WP 042871875 1 Dpoaceaephila
Dickeya chrysanthemi subsp chrysanthemi NCPPB 402 WP 040000338 1 Dchrysanthemi
Dickeya chrysanthemi L11 WP 033577021 1 Dchrysanthemi
Dickeya chrysanthemi subsp parthenii NCPPB 516 WP 027711488 1 Dchrysanthemi
Dickeya chrysanthemi NCPPB 3533 WP 012769060 1 Dchrysanthemi
Dickeya chrysanthemi Ech1591 WP 012769060 1 Dchrysanthemi
Dickeya dianthicola NCPPB 3534 WP 024109038 1 Ddianthicola
Dickeya dianthicola GBBC 2039 WP 024106666 1 Ddianthicola
Dickeya dianthicola IPO 980 WP 024106666 1 Ddianthicola
Dickeya dianthicola NCPPB 453 WP 024106666 1 Ddianthicola
Dickeya dianthicola RNS04 9 WP 024106666 1 Ddianthicola
Dickeya zeae Ech586 WP 012885466 1 Dzeae
Dickeya zeae MS1 WP 019844063 1 Dzeae
Dickeya zeae MK19 WP 038914324 1 Dzeae
Dickeya zeae NCPPB 3532 WP 023640274 1 Dzeae
Dickeya zeae NCPPB 2538 WP 038916542 1 Dzeae
Dickeya zeae NCPPB 3531 WP 038905072 1 Dzeae
Dickeya zeae CSL RW192 WP 038908581 1 Dzeae
Dickeya zeae DZ2Q WP 038925321 1 Dzeae
Dickeya zeae ZJU1202 WP 016941149 1 Dzeae
Dickeya zeae EC1 WP 016941149 1 Dzeae
Dickeya gs undicola 2B12 WP 033567837 1 Dundicola
Dickeya gs fangzhongdai NCPPB 3274 WP 042860534 1 Dfangzhongdai
Dickeya gs fangzhongdai S1 WP 049853685 1 Dfangzhongdai
Dickeya gs fangzhongdai B16 WP 038919699 1 Dfangzhongdai
Dickeya gs fangzhongdai MK7 WP 038919699 1 Dfangzhongdai
Dickeya gs fangzhongdai M005 WP 039691092 1 Dfangzhongdai
Dickeya gs fangzhongdai ND14b WP 038667300 1 Dfangzhongdai
Dickeya gs fangzhongdai M074 WP 038667300 1 Dfangzhongdai
Dickeya dadantii NCPPB 3537 WP 038901662 1 Ddadantii
Dickeya dadantii NCPPB 898 WP 038911588 1 Ddadantii
489 cbsH Iron metabolism Chrysobactin biosynthesis and transport Seed
Dickeya dadantii 3937 WP 033111969 1 Ddadantii
Dickeya solani RNS 05 1 2A WP 057082790 1 Dsolani
Dickeya solani MK10 WP 022634255 1 Dsolani
Dickeya solani PPO 9019 WP 022634255 1 Dsolani
Dickeya solani D s0432 1 WP 022634255 1 Dsolani
Dickeya solani IFB 0099 WP 022634255 1 Dsolani
Dickeya solani IPO 2222 WP 022634255 1 Dsolani
Dickeya solani RNS 08 23 3 1 A WP 022634255 1 Dsolani
Dickeya solani RNS 07 7 3B WP 022634255 1 Dsolani
Dickeya solani MK16 WP 022634255 1 Dsolani
Dickeya solani PPO 9134 WP 022634255 1 Dsolani
Dickeya solani GBBC 2040 WP 022634255 1 Dsolani

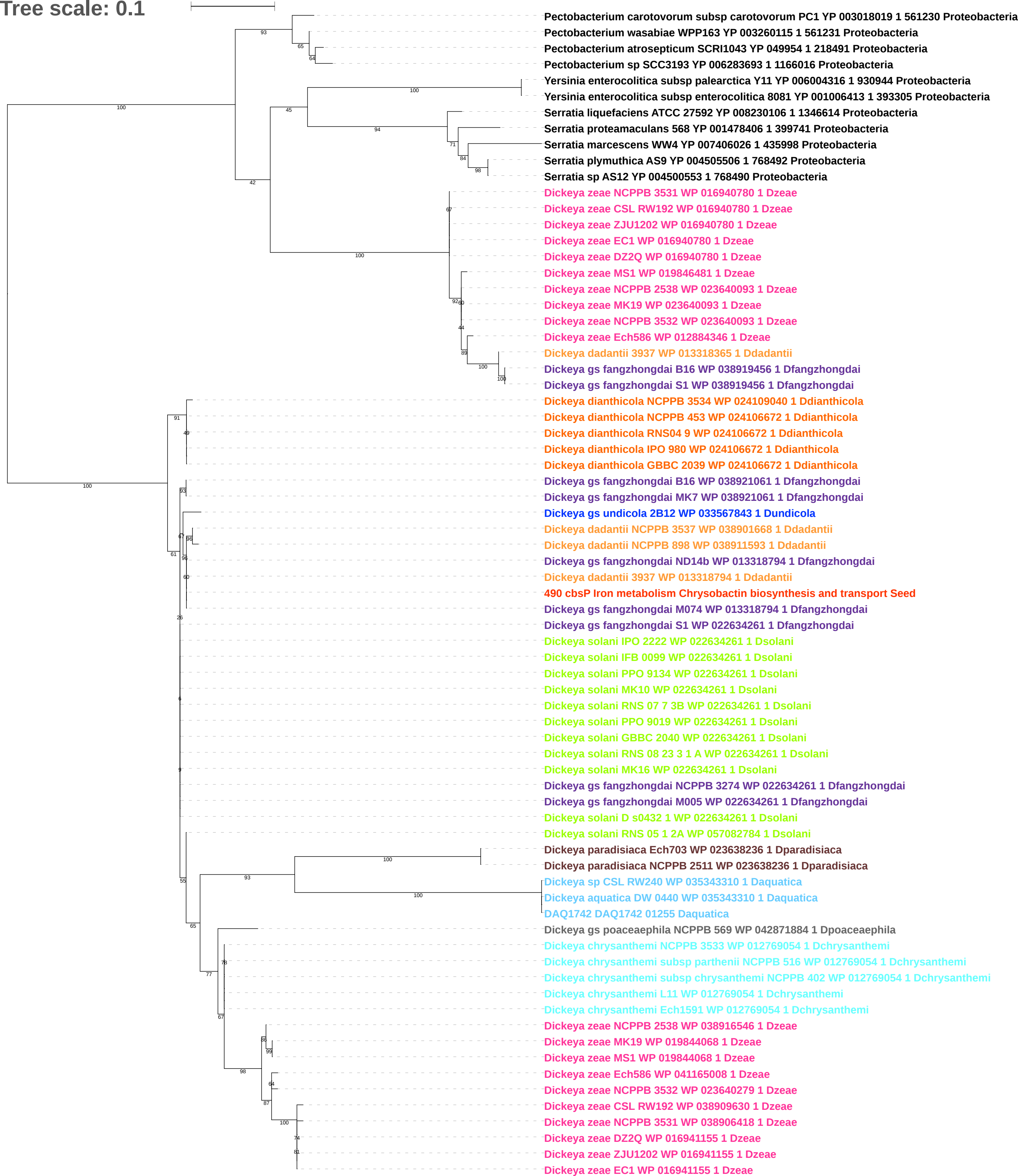
Tree scale: 0.1
Pectobacterium carotovorum subsp carotovorum PC1 YP 003018019 1 561230 Proteobacteria
Pectobacterium wasabiae WPP163 YP 003260115 1 561231 Proteobacteria
Pectobacterium atrosepticum SCRI1043 YP 049954 1 218491 Proteobacteria
Pectobacterium sp SCC3193 YP 006283693 1 1166016 Proteobacteria
Yersinia enterocolitica subsp palearctica Y11 YP 006004316 1 930944 Proteobacteria
Yersinia enterocolitica subsp enterocolitica 8081 YP 001006413 1 393305 Proteobacteria
Serratia liquefaciens ATCC 27592 YP 008230106 1 1346614 Proteobacteria
Serratia proteamaculans 568 YP 001478406 1 399741 Proteobacteria
Serratia marcescens WW4 YP 007406026 1 435998 Proteobacteria
Serratia plymuthica AS9 YP 004505506 1 768492 Proteobacteria
Serratia sp AS12 YP 004500553 1 768490 Proteobacteria
Dickeya zeae NCPPB 3531 WP 016940780 1 Dzeae
Dickeya zeae CSL RW192 WP 016940780 1 Dzeae
Dickeya zeae ZJU1202 WP 016940780 1 Dzeae
Dickeya zeae EC1 WP 016940780 1 Dzeae
Dickeya zeae DZ2Q WP 016940780 1 Dzeae
Dickeya zeae MS1 WP 019846481 1 Dzeae
Dickeya zeae NCPPB 2538 WP 023640093 1 Dzeae
Dickeya zeae MK19 WP 023640093 1 Dzeae
Dickeya zeae NCPPB 3532 WP 023640093 1 Dzeae
Dickeya zeae Ech586 WP 012884346 1 Dzeae
Dickeya dadantii 3937 WP 013318365 1 Ddadantii
Dickeya gs fangzhongdai B16 WP 038919456 1 Dfangzhongdai
Dickeya gs fangzhongdai S1 WP 038919456 1 Dfangzhongdai
Dickeya dianthicola NCPPB 3534 WP 024109040 1 Ddianthicola
Dickeya dianthicola NCPPB 453 WP 024106672 1 Ddianthicola
Dickeya dianthicola RNS04 9 WP 024106672 1 Ddianthicola
Dickeya dianthicola IPO 980 WP 024106672 1 Ddianthicola
Dickeya dianthicola GBBC 2039 WP 024106672 1 Ddianthicola
Dickeya gs fangzhongdai B16 WP 038921061 1 Dfangzhongdai
Dickeya gs fangzhongdai MK7 WP 038921061 1 Dfangzhongdai
Dickeya gs undicola 2B12 WP 033567843 1 Dundicola
Dickeya dadantii NCPPB 3537 WP 038901668 1 Ddadantii
Dickeya dadantii NCPPB 898 WP 038911593 1 Ddadantii
Dickeya gs fangzhongdai ND14b WP 013318794 1 Dfangzhongdai
Dickeya dadantii 3937 WP 013318794 1 Ddadantii
490 cbsP Iron metabolism Chrysobactin biosynthesis and transport Seed
Dickeya gs fangzhongdai M074 WP 013318794 1 Dfangzhongdai
Dickeya gs fangzhongdai S1 WP 022634261 1 Dfangzhongdai
Dickeya solani IPO 2222 WP 022634261 1 Dsolani
Dickeya solani IFB 0099 WP 022634261 1 Dsolani
Dickeya solani PPO 9134 WP 022634261 1 Dsolani
Dickeya solani MK10 WP 022634261 1 Dsolani
Dickeya solani RNS 07 7 3B WP 022634261 1 Dsolani
Dickeya solani PPO 9019 WP 022634261 1 Dsolani
Dickeya solani GBBC 2040 WP 022634261 1 Dsolani
Dickeya solani RNS 08 23 3 1 A WP 022634261 1 Dsolani
Dickeya solani MK16 WP 022634261 1 Dsolani
Dickeya gs fangzhongdai NCPPB 3274 WP 022634261 1 Dfangzhongdai
Dickeya gs fangzhongdai M005 WP 022634261 1 Dfangzhongdai
Dickeya solani D s0432 1 WP 022634261 1 Dsolani
Dickeya solani RNS 05 1 2A WP 057082784 1 Dsolani
Dickeya paradisiaca Ech703 WP 023638236 1 Dparadisiaca
Dickeya paradisiaca NCPPB 2511 WP 023638236 1 Dparadisiaca
Dickeya sp CSL RW240 WP 035343310 1 Daquatica
Dickeya aquatica DW 0440 WP 035343310 1 Daquatica
DAQ1742 DAQ1742 01255 Daquatica
Dickeya gs poaceaephila NCPPB 569 WP 042871884 1 Dpoaceaephila
Dickeya chrysanthemi NCPPB 3533 WP 012769054 1 Dchrysanthemi
Dickeya chrysanthemi subsp parthenii NCPPB 516 WP 012769054 1 Dchrysanthemi
Dickeya chrysanthemi subsp chrysanthemi NCPPB 402 WP 012769054 1 Dchrysanthemi
Dickeya chrysanthemi L11 WP 012769054 1 Dchrysanthemi
Dickeya chrysanthemi Ech1591 WP 012769054 1 Dchrysanthemi
Dickeya zeae NCPPB 2538 WP 038916546 1 Dzeae
Dickeya zeae MK19 WP 019844068 1 Dzeae
Dickeya zeae MS1 WP 019844068 1 Dzeae
Dickeya zeae Ech586 WP 041165008 1 Dzeae
Dickeya zeae NCPPB 3532 WP 023640279 1 Dzeae
Dickeya zeae CSL RW192 WP 038909630 1 Dzeae
Dickeya zeae NCPPB 3531 WP 038906418 1 Dzeae
Dickeya zeae DZ2Q WP 016941155 1 Dzeae
Dickeya zeae ZJU1202 WP 016941155 1 Dzeae
Dickeya zeae EC1 WP 016941155 1 Dzeae

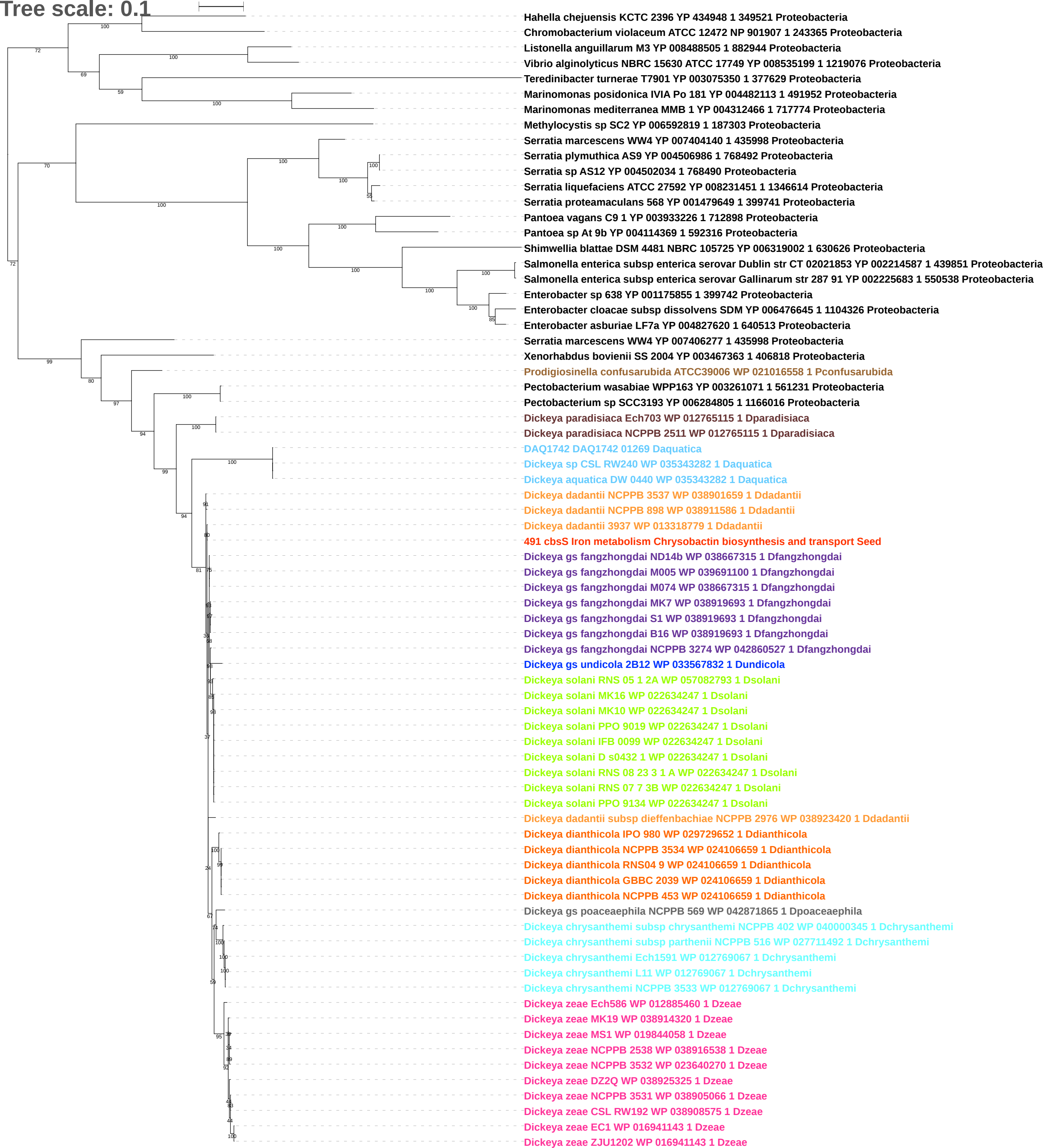
Tree scale: 0.1
Hahella chejuensis KCTC 2396 YP 434948 1 349521 Proteobacteria
Chromobacterium violaceum ATCC 12472 NP 901907 1 243365 Proteobacteria
Listonella anguillarum M3 YP 008488505 1 882944 Proteobacteria
Vibrio alginolyticus NBRC 15630 ATCC 17749 YP 008535199 1 1219076 Proteobacteria
Teredinibacter turnerae T7901 YP 003075350 1 377629 Proteobacteria
Marinomonas posidonica IVIA Po 181 YP 004482113 1 491952 Proteobacteria
Marinomonas mediterranea MMB 1 YP 004312466 1 717774 Proteobacteria
Methylocystis sp SC2 YP 006592819 1 187303 Proteobacteria
Serratia marcescens WW4 YP 007404140 1 435998 Proteobacteria
Serratia plymuthica AS9 YP 004506986 1 768492 Proteobacteria
Serratia sp AS12 YP 004502034 1 768490 Proteobacteria
Serratia liquefaciens ATCC 27592 YP 008231451 1 1346614 Proteobacteria
Serratia proteamaculans 568 YP 001479649 1 399741 Proteobacteria
Pantoea vagans C9 1 YP 003933226 1 712898 Proteobacteria
Pantoea sp At 9b YP 004114369 1 592316 Proteobacteria
Shimwellia blattae DSM 4481 NBRC 105725 YP 006319002 1 630626 Proteobacteria
Salmonella enterica subsp enterica serovar Dublin str CT 02021853 YP 002214587 1 439851 Proteobacteria
Salmonella enterica subsp enterica serovar Gallinarum str 287 91 YP 002225683 1 550538 Proteobacteria
Enterobacter sp 638 YP 001175855 1 399742 Proteobacteria
Enterobacter cloacae subsp dissolvens SDM YP 006476645 1 1104326 Proteobacteria
Enterobacter asburiae LF7a YP 004827620 1 640513 Proteobacteria
Serratia marcescens WW4 YP 007406277 1 435998 Proteobacteria
Xenorhabdus bovienii SS 2004 YP 003467363 1 406818 Proteobacteria
Prodigiosinella confusarubida ATCC39006 WP 021016558 1 Pconfusarubida
Pectobacterium wasabiae WPP163 YP 003261071 1 561231 Proteobacteria
Pectobacterium sp SCC3193 YP 006284805 1 1166016 Proteobacteria
Dickeya paradisiaca Ech703 WP 012765115 1 Dparadisiaca
Dickeya paradisiaca NCPPB 2511 WP 012765115 1 Dparadisiaca
DAQ1742 DAQ1742 01269 Daquatica
Dickeya sp CSL RW240 WP 035343282 1 Daquatica
Dickeya aquatica DW 0440 WP 035343282 1 Daquatica
Dickeya dadantii NCPPB 3537 WP 038901659 1 Ddadantii
Dickeya dadantii NCPPB 898 WP 038911586 1 Ddadantii
Dickeya dadantii 3937 WP 013318779 1 Ddadantii
491 cbsS Iron metabolism Chrysobactin biosynthesis and transport Seed
Dickeya gs fangzhongdai ND14b WP 038667315 1 Dfangzhongdai
Dickeya gs fangzhongdai M005 WP 039691100 1 Dfangzhongdai
Dickeya gs fangzhongdai M074 WP 038667315 1 Dfangzhongdai
Dickeya gs fangzhongdai MK7 WP 038919693 1 Dfangzhongdai
Dickeya gs fangzhongdai S1 WP 038919693 1 Dfangzhongdai
Dickeya gs fangzhongdai B16 WP 038919693 1 Dfangzhongdai
Dickeya gs fangzhongdai NCPPB 3274 WP 042860527 1 Dfangzhongdai
Dickeya gs undicola 2B12 WP 033567832 1 Dundicola
Dickeya solani RNS 05 1 2A WP 057082793 1 Dsolani
Dickeya solani MK16 WP 022634247 1 Dsolani
Dickeya solani MK10 WP 022634247 1 Dsolani
Dickeya solani PPO 9019 WP 022634247 1 Dsolani
Dickeya solani IFB 0099 WP 022634247 1 Dsolani
Dickeya solani D s0432 1 WP 022634247 1 Dsolani
Dickeya solani RNS 08 23 3 1 A WP 022634247 1 Dsolani
Dickeya solani RNS 07 7 3B WP 022634247 1 Dsolani
Dickeya solani PPO 9134 WP 022634247 1 Dsolani
Dickeya dadantii subsp dieffenbachiae NCPPB 2976 WP 038923420 1 Ddadantii
Dickeya dianthicola IPO 980 WP 029729652 1 Ddianthicola
Dickeya dianthicola NCPPB 3534 WP 024106659 1 Ddianthicola
Dickeya dianthicola RNS04 9 WP 024106659 1 Ddianthicola
Dickeya dianthicola GBBC 2039 WP 024106659 1 Ddianthicola
Dickeya dianthicola NCPPB 453 WP 024106659 1 Ddianthicola
Dickeya gs poaceaephila NCPPB 569 WP 042871865 1 Dpoaceaephila
Dickeya chrysanthemi subsp chrysanthemi NCPPB 402 WP 040000345 1 Dchrysanthemi
Dickeya chrysanthemi subsp parthenii NCPPB 516 WP 027711492 1 Dchrysanthemi
Dickeya chrysanthemi Ech1591 WP 012769067 1 Dchrysanthemi
Dickeya chrysanthemi L11 WP 012769067 1 Dchrysanthemi
Dickeya chrysanthemi NCPPB 3533 WP 012769067 1 Dchrysanthemi
Dickeya zeae Ech586 WP 012885460 1 Dzeae
Dickeya zeae MK19 WP 038914320 1 Dzeae
Dickeya zeae MS1 WP 019844058 1 Dzeae
Dickeya zeae NCPPB 2538 WP 038916538 1 Dzeae
Dickeya zeae NCPPB 3532 WP 023640270 1 Dzeae
Dickeya zeae DZ2Q WP 038925325 1 Dzeae
Dickeya zeae NCPPB 3531 WP 038905066 1 Dzeae
Dickeya zeae CSL RW192 WP 038908575 1 Dzeae
Dickeya zeae EC1 WP 016941143 1 Dzeae
Dickeya zeae ZJU1202 WP 016941143 1 Dzeae

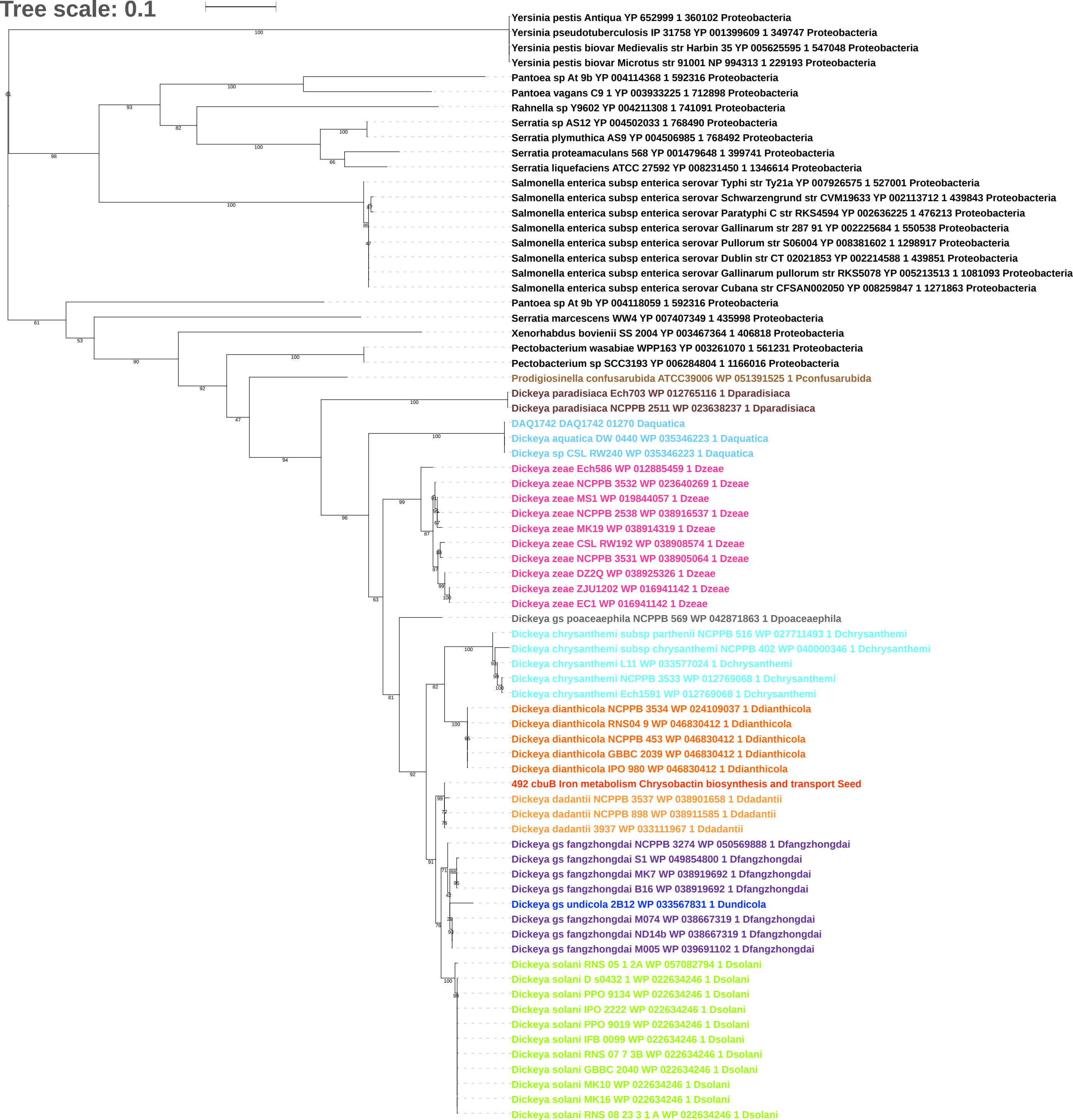
Tree scale: 0.1
Yersinia pestis Antiqua YP 652999 1 360102 Proteobacteria
Yersinia pseudotuberculosis IP 31758 YP 001399609 1 349747 Proteobacteria
Yersinia pestis biovar Medievalis str Harbin 35 YP 005625595 1 547048 Proteobacteria
Yersinia pestis biovar Microtus str 91001 NP 994313 1 229193 Proteobacteria
Pantoea sp At 9b YP 004114368 1 592316 Proteobacteria
Pantoea vagans C9 1 YP 003933225 1 712898 Proteobacteria
Rahnella sp Y9602 YP 004211308 1 741091 Proteobacteria
Serratia sp AS12 YP 004502033 1 768490 Proteobacteria
Serratia plymuthica AS9 YP 004506985 1 768492 Proteobacteria
Serratia proteamaculans 568 YP 001479648 1 399741 Proteobacteria
Serratia liquefaciens ATCC 27592 YP 008231450 1 1346614 Proteobacteria
Salmonella enterica subsp enterica serovar Typhi str Ty21a YP 007926575 1 527001 Proteobacteria
Salmonella enterica subsp enterica serovar Schwarzengrund str CVM19633 YP 002113712 1 439843 Proteobacteria
Salmonella enterica subsp enterica serovar Paratyphi C str RKS4594 YP 002636225 1 476213 Proteobacteria
Salmonella enterica subsp enterica serovar Gallinarum str 287 91 YP 002225684 1 550538 Proteobacteria
Salmonella enterica subsp enterica serovar Pullorum str S06004 YP 008381602 1 1298917 Proteobacteria
Salmonella enterica subsp enterica serovar Dublin str CT 02021853 YP 002214588 1 439851 Proteobacteria
Salmonella enterica subsp enterica serovar Gallinarum pullorum str RKS5078 YP 005213513 1 1081093 Proteobacteria
Salmonella enterica subsp enterica serovar Cubana str CFSAN002050 YP 008259847 1 1271863 Proteobacteria
Pantoea sp At 9b YP 004118059 1 592316 Proteobacteria
Serratia marcescens WW4 YP 007407349 1 435998 Proteobacteria
Xenorhabdus bovienii SS 2004 YP 003467364 1 406818 Proteobacteria
Pectobacterium wasabiae WPP163 YP 003261070 1 561231 Proteobacteria
Pectobacterium sp SCC3193 YP 006284804 1 1166016 Proteobacteria
Prodigiosinella confusarubida ATCC39006 WP 051391525 1 Pconfusarubida
Dickeya paradisiaca Ech703 WP 012765116 1 Dparadisiaca
Dickeya paradisiaca NCPPB 2511 WP 023638237 1 Dparadisiaca
DAQ1742 DAQ1742 01270 Daquatica
Dickeya aquatica DW 0440 WP 035346223 1 Daquatica
Dickeya sp CSL RW240 WP 035346223 1 Daquatica
Dickeya zeae Ech586 WP 012885459 1 Dzeae
Dickeya zeae NCPPB 3532 WP 023640269 1 Dzeae
Dickeya zeae MS1 WP 019844057 1 Dzeae
Dickeya zeae NCPPB 2538 WP 038916537 1 Dzeae
Dickeya zeae MK19 WP 038914319 1 Dzeae
Dickeya zeae CSL RW192 WP 038908574 1 Dzeae
Dickeya zeae NCPPB 3531 WP 038905064 1 Dzeae
Dickeya zeae DZ2Q WP 038925326 1 Dzeae
Dickeya zeae ZJU1202 WP 016941142 1 Dzeae
Dickeya zeae EC1 WP 016941142 1 Dzeae
Dickeya gs poaceaephila NCPPB 569 WP 042871863 1 Dpoaceaephila
Dickeya chrysanthemi subsp parthenii NCPPB 516 WP 027711493 1 Dchrysanthemi
Dickeya chrysanthemi subsp chrysanthemi NCPPB 402 WP 040000346 1 Dchrysanthemi
Dickeya chrysanthemi L11 WP 033577024 1 Dchrysanthemi
Dickeya chrysanthemi NCPPB 3533 WP 012769068 1 Dchrysanthemi
Dickeya chrysanthemi Ech1591 WP 012769068 1 Dchrysanthemi
Dickeya dianthicola NCPPB 3534 WP 024109037 1 Ddianthicola
Dickeya dianthicola RNS04 9 WP 046830412 1 Ddianthicola
Dickeya dianthicola NCPPB 453 WP 046830412 1 Ddianthicola
Dickeya dianthicola GBBC 2039 WP 046830412 1 Ddianthicola
Dickeya dianthicola IPO 980 WP 046830412 1 Ddianthicola
492 cbuB Iron metabolism Chrysobactin biosynthesis and transport Seed
Dickeya dadantii NCPPB 3537 WP 038901658 1 Ddadantii
Dickeya dadantii NCPPB 898 WP 038911585 1 Ddadantii
Dickeya dadantii 3937 WP 033111967 1 Ddadantii
Dickeya gs fangzhongdai NCPPB 3274 WP 050569888 1 Dfangzhongdai
Dickeya gs fangzhongdai S1 WP 049854800 1 Dfangzhongdai
Dickeya gs fangzhongdai MK7 WP 038919692 1 Dfangzhongdai
Dickeya gs fangzhongdai B16 WP 038919692 1 Dfangzhongdai
Dickeya gs undicola 2B12 WP 033567831 1 Dundicola
Dickeya gs fangzhongdai M074 WP 038667319 1 Dfangzhongdai
Dickeya gs fangzhongdai ND14b WP 038667319 1 Dfangzhongdai
Dickeya gs fangzhongdai M005 WP 039691102 1 Dfangzhongdai
Dickeya solani RNS 05 1 2A WP 057082794 1 Dsolani
Dickeya solani D s0432 1 WP 022634246 1 Dsolani
Dickeya solani PPO 9134 WP 022634246 1 Dsolani
Dickeya solani IPO 2222 WP 022634246 1 Dsolani
Dickeya solani PPO 9019 WP 022634246 1 Dsolani
Dickeya solani IFB 0099 WP 022634246 1 Dsolani
Dickeya solani RNS 07 7 3B WP 022634246 1 Dsolani
Dickeya solani GBBC 2040 WP 022634246 1 Dsolani
Dickeya solani MK10 WP 022634246 1 Dsolani
Dickeya solani MK16 WP 022634246 1 Dsolani
Dickeya solani RNS 08 23 3 1 A WP 022634246 1 Dsolani

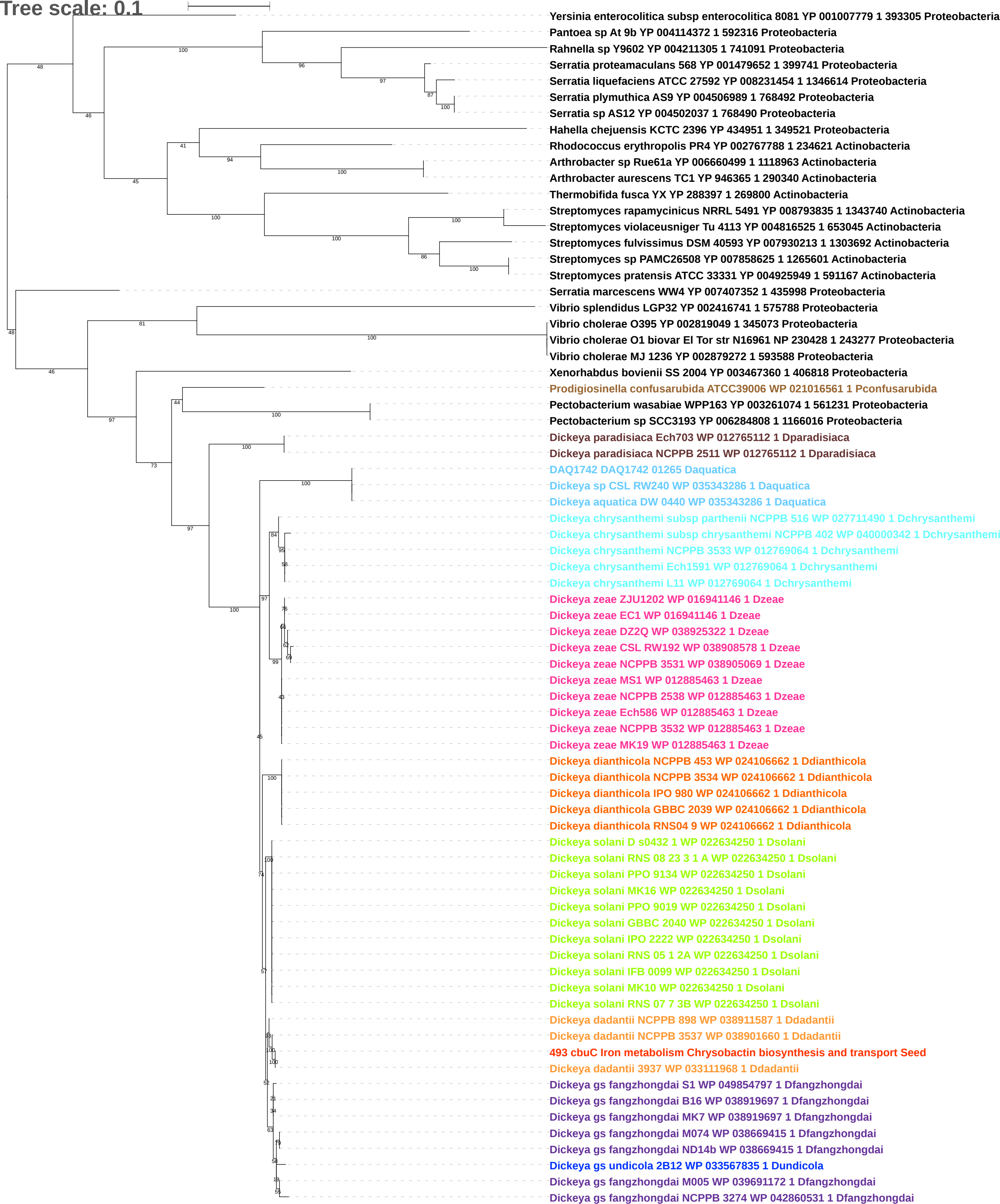

Tree scale: 0.1
Yersinia enterocolitica subsp enterocolitica 8081 YP 001007779 1 393305 Proteobacteria
Pantoea sp At 9b YP 004114372 1 592316 Proteobacteria
Rahnella sp Y9602 YP 004211305 1 741091 Proteobacteria
Serratia proteamaculans 568 YP 001479652 1 399741 Proteobacteria
Serratia liquefaciens ATCC 27592 YP 008231454 1 1346614 Proteobacteria
Serratia plymuthica AS9 YP 004506989 1 768492 Proteobacteria
Serratia sp AS12 YP 004502037 1 768490 Proteobacteria
Hahella chejuensis KCTC 2396 YP 434951 1 349521 Proteobacteria
Rhodococcus erythropolis PR4 YP 002767788 1 234621 Actinobacteria
Arthrobacter sp Rue61a YP 006660499 1 1118963 Actinobacteria
Arthrobacter aurescens TC1 YP 946365 1 290340 Actinobacteria
Thermobifida fusca YX YP 288397 1 269800 Actinobacteria
Streptomyces rapamycinicus NRRL 5491 YP 008793835 1 1343740 Actinobacteria
Streptomyces violaceusniger Tu 4113 YP 004816525 1 653045 Actinobacteria
Streptomyces fulvissimus DSM 40593 YP 007930213 1 1303692 Actinobacteria
Streptomyces sp PAMC26508 YP 007858625 1 1265601 Actinobacteria
Streptomyces pratensis ATCC 33331 YP 004925949 1 591167 Actinobacteria
Serratia marcescens WW4 YP 007407352 1 435998 Proteobacteria
Vibrio splendidus LGP32 YP 002416741 1 575788 Proteobacteria
Vibrio cholerae O395 YP 002819049 1 345073 Proteobacteria
Vibrio cholerae O1 biovar El Tor str N16961 NP 230428 1 243277 Proteobacteria
Vibrio cholerae MJ 1236 YP 002879272 1 593588 Proteobacteria
Xenorhabdus bovienii SS 2004 YP 003467360 1 406818 Proteobacteria
Prodigiosinella confusarubida ATCC39006 WP 021016561 1 Pconfusarubida
Pectobacterium wasabiae WPP163 YP 003261074 1 561231 Proteobacteria
Pectobacterium sp SCC3193 YP 006284808 1 1166016 Proteobacteria
Dickeya paradisiaca Ech703 WP 012765112 1 Dparadisiaca
Dickeya paradisiaca NCPPB 2511 WP 012765112 1 Dparadisiaca
DAQ1742 DAQ1742 01265 Daquatica
Dickeya sp CSL RW240 WP 035343286 1 Daquatica
Dickeya aquatica DW 0440 WP 035343286 1 Daquatica
Dickeya chrysanthemi subsp parthenii NCPPB 516 WP 027711490 1 Dchrysanthemi
Dickeya chrysanthemi subsp chrysanthemi NCPPB 402 WP 040000342 1 Dchrysanthemi
Dickeya chrysanthemi NCPPB 3533 WP 012769064 1 Dchrysanthemi
Dickeya chrysanthemi Ech1591 WP 012769064 1 Dchrysanthemi
Dickeya chrysanthemi L11 WP 012769064 1 Dchrysanthemi
Dickeya zeae ZJU1202 WP 016941146 1 Dzeae
Dickeya zeae EC1 WP 016941146 1 Dzeae
Dickeya zeae DZ2Q WP 038925322 1 Dzeae
Dickeya zeae CSL RW192 WP 038908578 1 Dzeae
Dickeya zeae NCPPB 3531 WP 038905069 1 Dzeae
Dickeya zeae MS1 WP 012885463 1 Dzeae
Dickeya zeae NCPPB 2538 WP 012885463 1 Dzeae
Dickeya zeae Ech586 WP 012885463 1 Dzeae
Dickeya zeae NCPPB 3532 WP 012885463 1 Dzeae
Dickeya zeae MK19 WP 012885463 1 Dzeae
Dickeya dianthicola NCPPB 453 WP 024106662 1 Ddianthicola
Dickeya dianthicola NCPPB 3534 WP 024106662 1 Ddianthicola
Dickeya dianthicola IPO 980 WP 024106662 1 Ddianthicola
Dickeya dianthicola GBBC 2039 WP 024106662 1 Ddianthicola
Dickeya dianthicola RNS04 9 WP 024106662 1 Ddianthicola
Dickeya solani D s0432 1 WP 022634250 1 Dsolani
Dickeya solani RNS 08 23 3 1 A WP 022634250 1 Dsolani
Dickeya solani PPO 9134 WP 022634250 1 Dsolani
Dickeya solani MK16 WP 022634250 1 Dsolani
Dickeya solani PPO 9019 WP 022634250 1 Dsolani
Dickeya solani GBBC 2040 WP 022634250 1 Dsolani
Dickeya solani IPO 2222 WP 022634250 1 Dsolani
Dickeya solani RNS 05 1 2A WP 022634250 1 Dsolani
Dickeya solani IFB 0099 WP 022634250 1 Dsolani
Dickeya solani MK10 WP 022634250 1 Dsolani
Dickeya solani RNS 07 7 3B WP 022634250 1 Dsolani
Dickeya dadantii NCPPB 898 WP 038911587 1 Ddadantii
Dickeya dadantii NCPPB 3537 WP 038901660 1 Ddadantii
493 cbuC Iron metabolism Chrysobactin biosynthesis and transport Seed
Dickeya dadantii 3937 WP 033111968 1 Ddadantii
Dickeya gs fangzhongdai S1 WP 049854797 1 Dfangzhongdai
Dickeya gs fangzhongdai B16 WP 038919697 1 Dfangzhongdai
Dickeya gs fangzhongdai MK7 WP 038919697 1 Dfangzhongdai
Dickeya gs fangzhongdai M074 WP 038669415 1 Dfangzhongdai
Dickeya gs fangzhongdai ND14b WP 038669415 1 Dfangzhongdai
Dickeya gs undicola 2B12 WP 033567835 1 Dundicola
Dickeya gs fangzhongdai M005 WP 039691172 1 Dfangzhongdai
Dickeya gs fangzhongdai NCPPB 3274 WP 042860531 1 Dfangzhongdai

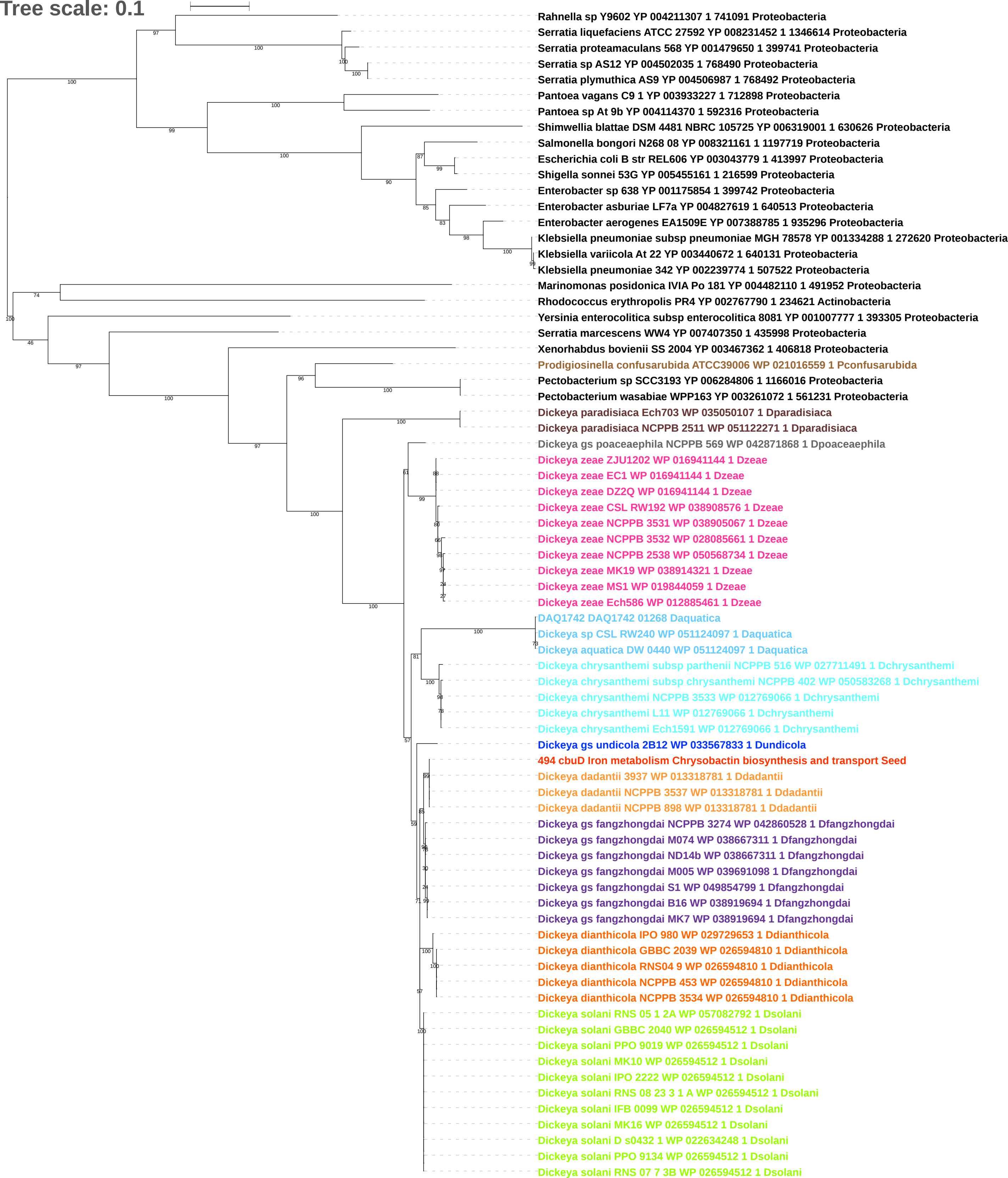
Tree scale: 0.1
Rahnella sp Y9602 YP 004211307 1 741091 Proteobacteria
Serratia liquefaciens ATCC 27592 YP 008231452 1 1346614 Proteobacteria
Serratia proteamaculans 568 YP 001479650 1 399741 Proteobacteria
Serratia sp AS12 YP 004502035 1 768490 Proteobacteria
Serratia plymuthica AS9 YP 004506987 1 768492 Proteobacteria
Pantoea vagans C9 1 YP 003933227 1 712898 Proteobacteria
Pantoea sp At 9b YP 004114370 1 592316 Proteobacteria
Shimwellia blattae DSM 4481 NBRC 105725 YP 006319001 1 630626 Proteobacteria
Salmonella bongori N268 08 YP 008321161 1 1197719 Proteobacteria
Escherichia coli B str REL606 YP 003043779 1 413997 Proteobacteria
Shigella sonnei 53G YP 005455161 1 216599 Proteobacteria
Enterobacter sp 638 YP 001175854 1 399742 Proteobacteria
Enterobacter asburiae LF7a YP 004827619 1 640513 Proteobacteria
Enterobacter aerogenes EA1509E YP 007388785 1 935296 Proteobacteria
Klebsiella pneumoniae subsp pneumoniae MGH 78578 YP 001334288 1 272620 Proteobacteria
Klebsiella variicola At 22 YP 003440672 1 640131 Proteobacteria
Klebsiella pneumoniae 342 YP 002239774 1 507522 Proteobacteria
Marinomonas posidonica IVIA Po 181 YP 004482110 1 491952 Proteobacteria
Rhodococcus erythropolis PR4 YP 002767790 1 234621 Actinobacteria
Yersinia enterocolitica subsp enterocolitica 8081 YP 001007777 1 393305 Proteobacteria
Serratia marcescens WW4 YP 007407350 1 435998 Proteobacteria
Xenorhabdus bovienii SS 2004 YP 003467362 1 406818 Proteobacteria
Prodigiosinella confusarubida ATCC39006 WP 021016559 1 Pconfusarubida
Pectobacterium sp SCC3193 YP 006284806 1 1166016 Proteobacteria
Pectobacterium wasabiae WPP163 YP 003261072 1 561231 Proteobacteria
Dickeya paradisiaca Ech703 WP 035050107 1 Dparadisiaca
Dickeya paradisiaca NCPPB 2511 WP 051122271 1 Dparadisiaca
Dickeya gs poaceaephila NCPPB 569 WP 042871868 1 Dpoaceaephila
Dickeya zeae ZJU1202 WP 016941144 1 Dzeae
Dickeya zeae EC1 WP 016941144 1 Dzeae
Dickeya zeae DZ2Q WP 016941144 1 Dzeae
Dickeya zeae CSL RW192 WP 038908576 1 Dzeae
Dickeya zeae NCPPB 3531 WP 038905067 1 Dzeae
Dickeya zeae NCPPB 3532 WP 028085661 1 Dzeae
Dickeya zeae NCPPB 2538 WP 050568734 1 Dzeae
Dickeya zeae MK19 WP 038914321 1 Dzeae
Dickeya zeae MS1 WP 019844059 1 Dzeae
Dickeya zeae Ech586 WP 012885461 1 Dzeae
DAQ1742 DAQ1742 01268 Daquatica
Dickeya sp CSL RW240 WP 051124097 1 Daquatica
Dickeya aquatica DW 0440 WP 051124097 1 Daquatica
Dickeya chrysanthemi subsp parthenii NCPPB 516 WP 027711491 1 Dchrysanthemi
Dickeya chrysanthemi subsp chrysanthemi NCPPB 402 WP 050583268 1 Dchrysanthemi
Dickeya chrysanthemi NCPPB 3533 WP 012769066 1 Dchrysanthemi
Dickeya chrysanthemi L11 WP 012769066 1 Dchrysanthemi
Dickeya chrysanthemi Ech1591 WP 012769066 1 Dchrysanthemi
Dickeya gs undicola 2B12 WP 033567833 1 Dundicola
494 cbuD Iron metabolism Chrysobactin biosynthesis and transport Seed
Dickeya dadantii 3937 WP 013318781 1 Ddadantii
Dickeya dadantii NCPPB 3537 WP 013318781 1 Ddadantii
Dickeya dadantii NCPPB 898 WP 013318781 1 Ddadantii
Dickeya gs fangzhongdai NCPPB 3274 WP 042860528 1 Dfangzhongdai
Dickeya gs fangzhongdai M074 WP 038667311 1 Dfangzhongdai
Dickeya gs fangzhongdai ND14b WP 038667311 1 Dfangzhongdai
Dickeya gs fangzhongdai M005 WP 039691098 1 Dfangzhongdai
Dickeya gs fangzhongdai S1 WP 049854799 1 Dfangzhongdai
Dickeya gs fangzhongdai B16 WP 038919694 1 Dfangzhongdai
Dickeya gs fangzhongdai MK7 WP 038919694 1 Dfangzhongdai
Dickeya dianthicola IPO 980 WP 029729653 1 Ddianthicola
Dickeya dianthicola GBBC 2039 WP 026594810 1 Ddianthicola
Dickeya dianthicola RNS04 9 WP 026594810 1 Ddianthicola
Dickeya dianthicola NCPPB 453 WP 026594810 1 Ddianthicola
Dickeya dianthicola NCPPB 3534 WP 026594810 1 Ddianthicola
Dickeya solani RNS 05 1 2A WP 057082792 1 Dsolani
Dickeya solani GBBC 2040 WP 026594512 1 Dsolani
Dickeya solani PPO 9019 WP 026594512 1 Dsolani
Dickeya solani MK10 WP 026594512 1 Dsolani
Dickeya solani IPO 2222 WP 026594512 1 Dsolani
Dickeya solani RNS 08 23 3 1 A WP 026594512 1 Dsolani
Dickeya solani IFB 0099 WP 026594512 1 Dsolani
Dickeya solani MK16 WP 026594512 1 Dsolani
Dickeya solani D s0432 1 WP 022634248 1 Dsolani
Dickeya solani PPO 9134 WP 026594512 1 Dsolani
Dickeya solani RNS 07 7 3B WP 026594512 1 Dsolani

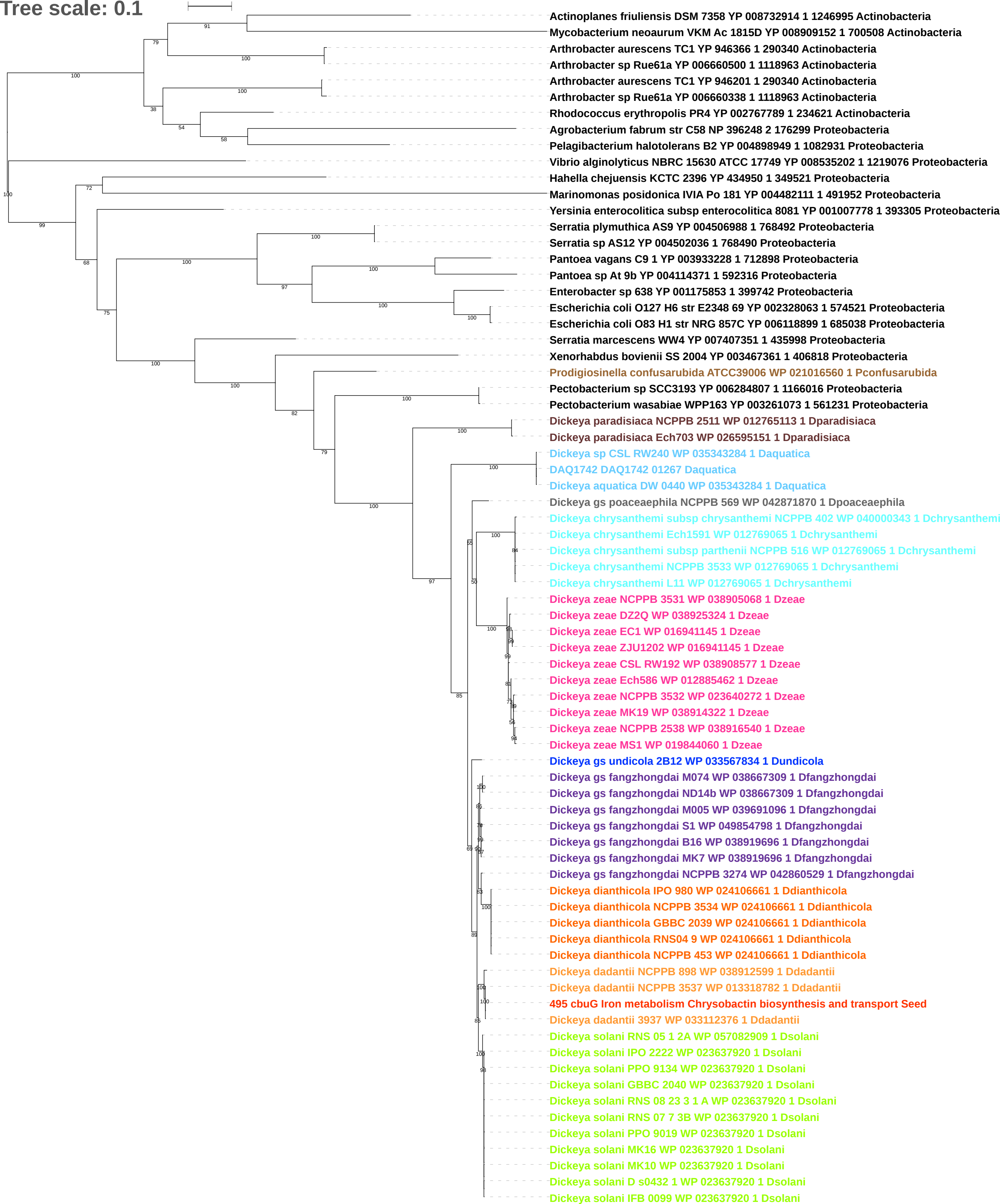

Tree scale: 0.1
Actinoplanes friuliensis DSM 7358 YP 008732914 1 1246995 Actinobacteria
Mycobacterium neoaurum VKM Ac 1815D YP 008909152 1 700508 Actinobacteria
Arthrobacter aurescens TC1 YP 946366 1 290340 Actinobacteria
Arthrobacter sp Rue61a YP 006660500 1 1118963 Actinobacteria
Arthrobacter aurescens TC1 YP 946201 1 290340 Actinobacteria
Arthrobacter sp Rue61a YP 006660338 1 1118963 Actinobacteria
Rhodococcus erythropolis PR4 YP 002767789 1 234621 Actinobacteria
Agrobacterium fabrum str C58 NP 396248 2 176299 Proteobacteria
Pelagibacterium halotolerans B2 YP 004898949 1 1082931 Proteobacteria
Vibrio alginolyticus NBRC 15630 ATCC 17749 YP 008535202 1 1219076 Proteobacteria
Hahella chejuensis KCTC 2396 YP 434950 1 349521 Proteobacteria
Marinomonas posidonica IVIA Po 181 YP 004482111 1 491952 Proteobacteria
Yersinia enterocolitica subsp enterocolitica 8081 YP 001007778 1 393305 Proteobacteria
Serratia plymuthica AS9 YP 004506988 1 768492 Proteobacteria
Serratia sp AS12 YP 004502036 1 768490 Proteobacteria
Pantoea vagans C9 1 YP 003933228 1 712898 Proteobacteria
Pantoea sp At 9b YP 004114371 1 592316 Proteobacteria
Enterobacter sp 638 YP 001175853 1 399742 Proteobacteria
Escherichia coli O127 H6 str E2348 69 YP 002328063 1 574521 Proteobacteria
Escherichia coli O83 H1 str NRG 857C YP 006118899 1 685038 Proteobacteria
Serratia marcescens WW4 YP 007407351 1 435998 Proteobacteria
Xenorhabdus bovienii SS 2004 YP 003467361 1 406818 Proteobacteria
Prodigiosinella confusarubida ATCC39006 WP 021016560 1 Pconfusarubida
Pectobacterium sp SCC3193 YP 006284807 1 1166016 Proteobacteria
Pectobacterium wasabiae WPP163 YP 003261073 1 561231 Proteobacteria
Dickeya paradisiaca NCPPB 2511 WP 012765113 1 Dparadisiaca
Dickeya paradisiaca Ech703 WP 026595151 1 Dparadisiaca
Dickeya sp CSL RW240 WP 035343284 1 Daquatica
DAQ1742 DAQ1742 01267 Daquatica
Dickeya aquatica DW 0440 WP 035343284 1 Daquatica
Dickeya gs poaceaephila NCPPB 569 WP 042871870 1 Dpoaceaephila
Dickeya chrysanthemi subsp chrysanthemi NCPPB 402 WP 040000343 1 Dchrysanthemi
Dickeya chrysanthemi Ech1591 WP 012769065 1 Dchrysanthemi
Dickeya chrysanthemi subsp parthenii NCPPB 516 WP 012769065 1 Dchrysanthemi
Dickeya chrysanthemi NCPPB 3533 WP 012769065 1 Dchrysanthemi
Dickeya chrysanthemi L11 WP 012769065 1 Dchrysanthemi
Dickeya zeae NCPPB 3531 WP 038905068 1 Dzeae
Dickeya zeae DZ2Q WP 038925324 1 Dzeae
Dickeya zeae EC1 WP 016941145 1 Dzeae
Dickeya zeae ZJU1202 WP 016941145 1 Dzeae
Dickeya zeae CSL RW192 WP 038908577 1 Dzeae
Dickeya zeae Ech586 WP 012885462 1 Dzeae
Dickeya zeae NCPPB 3532 WP 023640272 1 Dzeae
Dickeya zeae MK19 WP 038914322 1 Dzeae
Dickeya zeae NCPPB 2538 WP 038916540 1 Dzeae
Dickeya zeae MS1 WP 019844060 1 Dzeae
Dickeya gs undicola 2B12 WP 033567834 1 Dundicola
Dickeya gs fangzhongdai M074 WP 038667309 1 Dfangzhongdai
Dickeya gs fangzhongdai ND14b WP 038667309 1 Dfangzhongdai
Dickeya gs fangzhongdai M005 WP 039691096 1 Dfangzhongdai
Dickeya gs fangzhongdai S1 WP 049854798 1 Dfangzhongdai
Dickeya gs fangzhongdai B16 WP 038919696 1 Dfangzhongdai
Dickeya gs fangzhongdai MK7 WP 038919696 1 Dfangzhongdai
Dickeya gs fangzhongdai NCPPB 3274 WP 042860529 1 Dfangzhongdai
Dickeya dianthicola IPO 980 WP 024106661 1 Ddianthicola
Dickeya dianthicola NCPPB 3534 WP 024106661 1 Ddianthicola
Dickeya dianthicola GBBC 2039 WP 024106661 1 Ddianthicola
Dickeya dianthicola RNS04 9 WP 024106661 1 Ddianthicola
Dickeya dianthicola NCPPB 453 WP 024106661 1 Ddianthicola
Dickeya dadantii NCPPB 898 WP 038912599 1 Ddadantii
Dickeya dadantii NCPPB 3537 WP 013318782 1 Ddadantii
495 cbuG Iron metabolism Chrysobactin biosynthesis and transport Seed
Dickeya dadantii 3937 WP 033112376 1 Ddadantii
Dickeya solani RNS 05 1 2A WP 057082909 1 Dsolani
Dickeya solani IPO 2222 WP 023637920 1 Dsolani
Dickeya solani PPO 9134 WP 023637920 1 Dsolani
Dickeya solani GBBC 2040 WP 023637920 1 Dsolani
Dickeya solani RNS 08 23 3 1 A WP 023637920 1 Dsolani
Dickeya solani RNS 07 7 3B WP 023637920 1 Dsolani
Dickeya solani PPO 9019 WP 023637920 1 Dsolani
Dickeya solani MK16 WP 023637920 1 Dsolani
Dickeya solani MK10 WP 023637920 1 Dsolani
Dickeya solani D s0432 1 WP 023637920 1 Dsolani
Dickeya solani IFB 0099 WP 023637920 1 Dsolani

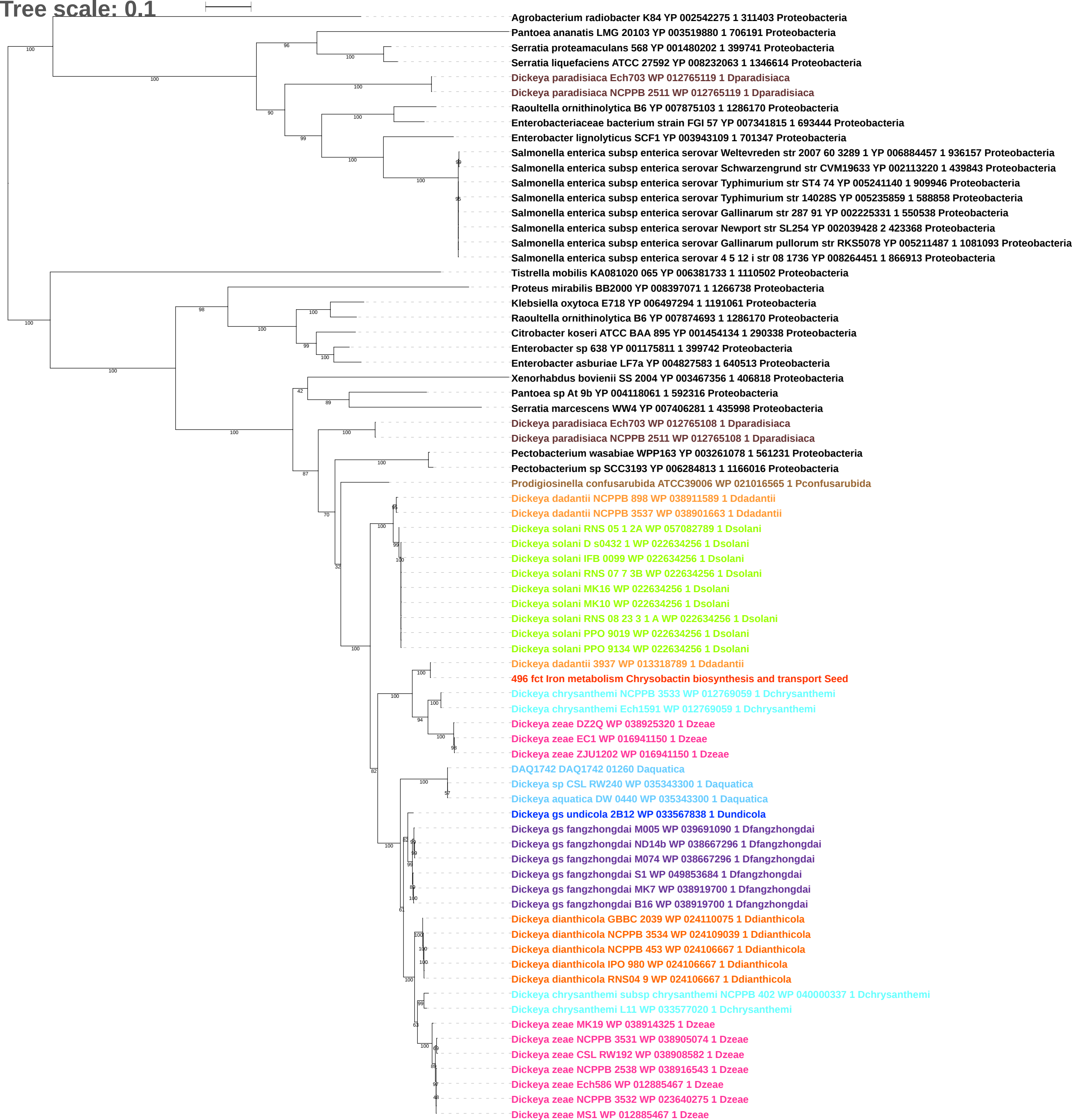
Tree scale: 0.1
Agrobacterium radiobacter K84 YP 002542275 1 311403 Proteobacteria
Pantoea ananatis LMG 20103 YP 003519880 1 706191 Proteobacteria
Serratia proteamaculans 568 YP 001480202 1 399741 Proteobacteria
Serratia liquefaciens ATCC 27592 YP 008232063 1 1346614 Proteobacteria
Dickeya paradisiaca Ech703 WP 012765119 1 Dparadisiaca
Dickeya paradisiaca NCPPB 2511 WP 012765119 1 Dparadisiaca
Raoultella ornithinolytica B6 YP 007875103 1 1286170 Proteobacteria
Enterobacteriaceae bacterium strain FGI 57 YP 007341815 1 693444 Proteobacteria
Enterobacter lignolyticus SCF1 YP 003943109 1 701347 Proteobacteria
Salmonella enterica subsp enterica serovar Weltevreden str 2007 60 3289 1 YP 006884457 1 936157 Proteobacteria
Salmonella enterica subsp enterica serovar Schwarzengrund str CVM19633 YP 002113220 1 439843 Proteobacteria
Salmonella enterica subsp enterica serovar Typhimurium str ST4 74 YP 005241140 1 909946 Proteobacteria
Salmonella enterica subsp enterica serovar Typhimurium str 14028S YP 005235859 1 588858 Proteobacteria
Salmonella enterica subsp enterica serovar Gallinarum str 287 91 YP 002225331 1 550538 Proteobacteria
Salmonella enterica subsp enterica serovar Newport str SL254 YP 002039428 2 423368 Proteobacteria
Salmonella enterica subsp enterica serovar Gallinarum pullorum str RKS5078 YP 005211487 1 1081093 Proteobacteria
Salmonella enterica subsp enterica serovar 4 5 12 i str 08 1736 YP 008264451 1 866913 Proteobacteria
Tistrella mobilis KA081020 065 YP 006381733 1 1110502 Proteobacteria
Proteus mirabilis BB2000 YP 008397071 1 1266738 Proteobacteria
Klebsiella oxytoca E718 YP 006497294 1 1191061 Proteobacteria
Raoultella ornithinolytica B6 YP 007874693 1 1286170 Proteobacteria
Citrobacter koseri ATCC BAA 895 YP 001454134 1 290338 Proteobacteria
Enterobacter sp 638 YP 001175811 1 399742 Proteobacteria
Enterobacter asburiae LF7a YP 004827583 1 640513 Proteobacteria
Xenorhabdus bovienii SS 2004 YP 003467356 1 406818 Proteobacteria
Pantoea sp At 9b YP 004118061 1 592316 Proteobacteria
Serratia marcescens WW4 YP 007406281 1 435998 Proteobacteria
Dickeya paradisiaca Ech703 WP 012765108 1 Dparadisiaca
Dickeya paradisiaca NCPPB 2511 WP 012765108 1 Dparadisiaca
Pectobacterium wasabiae WPP163 YP 003261078 1 561231 Proteobacteria
Pectobacterium sp SCC3193 YP 006284813 1 1166016 Proteobacteria
Prodigiosinella confusarubida ATCC39006 WP 021016565 1 Pconfusarubida
Dickeya dadantii NCPPB 898 WP 038911589 1 Ddadantii
Dickeya dadantii NCPPB 3537 WP 038901663 1 Ddadantii
Dickeya solani RNS 05 1 2A WP 057082789 1 Dsolani
Dickeya solani D s0432 1 WP 022634256 1 Dsolani
Dickeya solani IFB 0099 WP 022634256 1 Dsolani
Dickeya solani RNS 07 7 3B WP 022634256 1 Dsolani
Dickeya solani MK16 WP 022634256 1 Dsolani
Dickeya solani MK10 WP 022634256 1 Dsolani
Dickeya solani RNS 08 23 3 1 A WP 022634256 1 Dsolani
Dickeya solani PPO 9019 WP 022634256 1 Dsolani
Dickeya solani PPO 9134 WP 022634256 1 Dsolani
Dickeya dadantii 3937 WP 013318789 1 Ddadantii
496 fct Iron metabolism Chrysobactin biosynthesis and transport Seed
Dickeya chrysanthemi NCPPB 3533 WP 012769059 1 Dchrysanthemi
Dickeya chrysanthemi Ech1591 WP 012769059 1 Dchrysanthemi
Dickeya zeae DZ2Q WP 038925320 1 Dzeae
Dickeya zeae EC1 WP 016941150 1 Dzeae
Dickeya zeae ZJU1202 WP 016941150 1 Dzeae
DAQ1742 DAQ1742 01260 Daquatica
Dickeya sp CSL RW240 WP 035343300 1 Daquatica
Dickeya aquatica DW 0440 WP 035343300 1 Daquatica
Dickeya gs undicola 2B12 WP 033567838 1 Dundicola
Dickeya gs fangzhongdai M005 WP 039691090 1 Dfangzhongdai
Dickeya gs fangzhongdai ND14b WP 038667296 1 Dfangzhongdai
Dickeya gs fangzhongdai M074 WP 038667296 1 Dfangzhongdai
Dickeya gs fangzhongdai S1 WP 049853684 1 Dfangzhongdai
Dickeya gs fangzhongdai MK7 WP 038919700 1 Dfangzhongdai
Dickeya gs fangzhongdai B16 WP 038919700 1 Dfangzhongdai
Dickeya dianthicola GBBC 2039 WP 024110075 1 Ddianthicola
Dickeya dianthicola NCPPB 3534 WP 024109039 1 Ddianthicola
Dickeya dianthicola NCPPB 453 WP 024106667 1 Ddianthicola
Dickeya dianthicola IPO 980 WP 024106667 1 Ddianthicola
Dickeya dianthicola RNS04 9 WP 024106667 1 Ddianthicola
Dickeya chrysanthemi subsp chrysanthemi NCPPB 402 WP 040000337 1 Dchrysanthemi
Dickeya chrysanthemi L11 WP 033577020 1 Dchrysanthemi
Dickeya zeae MK19 WP 038914325 1 Dzeae
Dickeya zeae NCPPB 3531 WP 038905074 1 Dzeae
Dickeya zeae CSL RW192 WP 038908582 1 Dzeae
Dickeya zeae NCPPB 2538 WP 038916543 1 Dzeae
Dickeya zeae Ech586 WP 012885467 1 Dzeae
Dickeya zeae NCPPB 3532 WP 023640275 1 Dzeae
Dickeya zeae MS1 WP 012885467 1 Dzeae

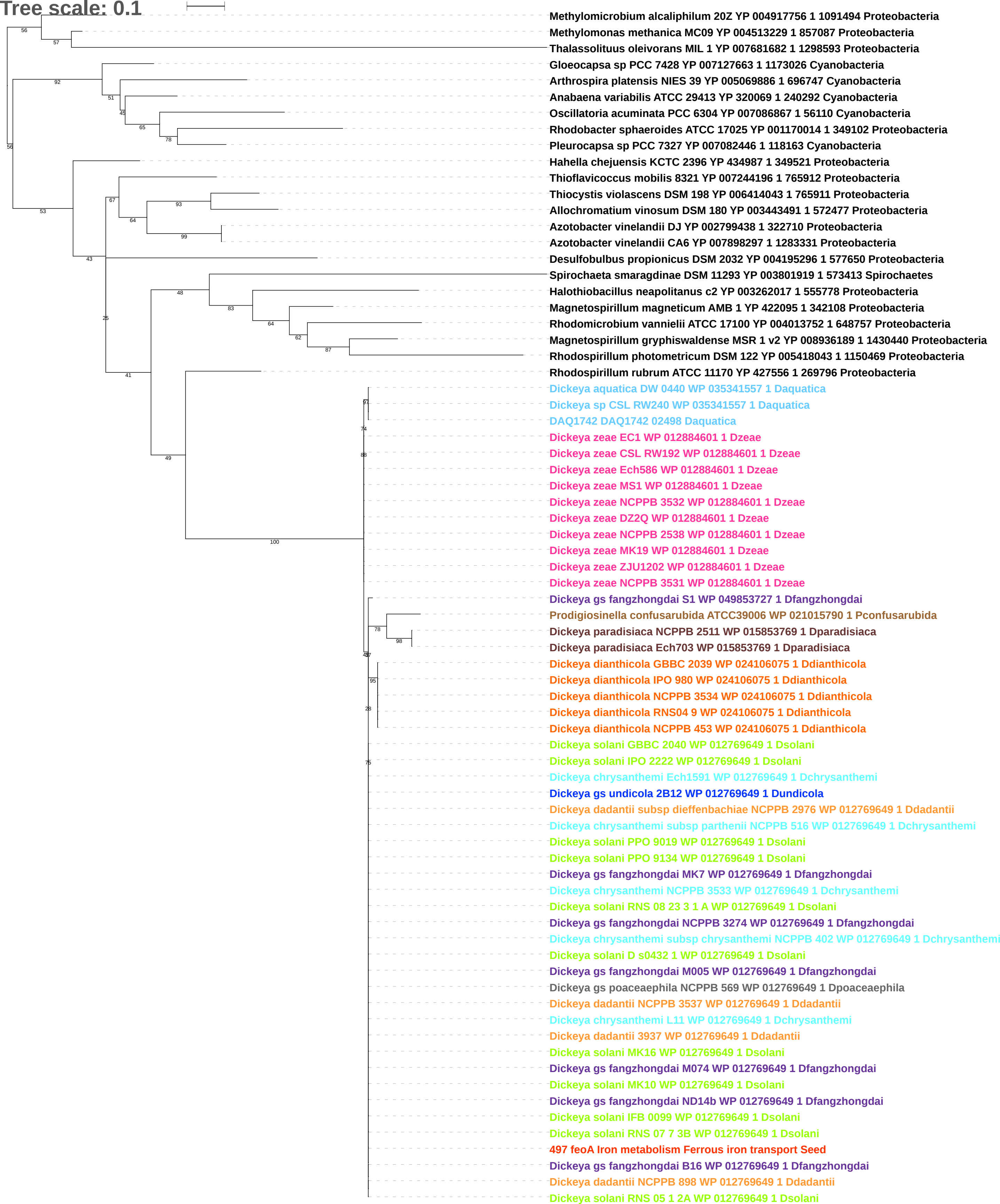
Tree scale: 0.1
Methylomicrobium alcaliphilum 20Z YP 004917756 1 1091494 Proteobacteria
Methylomonas methanica MC09 YP 004513229 1 857087 Proteobacteria
Thalassolituus oleivorans MIL 1 YP 007681682 1 1298593 Proteobacteria
Gloeocapsa sp PCC 7428 YP 007127663 1 1173026 Cyanobacteria
Arthrospira platensis NIES 39 YP 005069886 1 696747 Cyanobacteria
Anabaena variabilis ATCC 29413 YP 320069 1 240292 Cyanobacteria
Oscillatoria acuminata PCC 6304 YP 007086867 1 56110 Cyanobacteria
Rhodobacter sphaeroides ATCC 17025 YP 001170014 1 349102 Proteobacteria
Pleurocapsa sp PCC 7327 YP 007082446 1 118163 Cyanobacteria
Hahella chejuensis KCTC 2396 YP 434987 1 349521 Proteobacteria
Thioflavicoccus mobilis 8321 YP 007244196 1 765912 Proteobacteria
Thiocystis violascens DSM 198 YP 006414043 1 765911 Proteobacteria
Allochromatium vinosum DSM 180 YP 003443491 1 572477 Proteobacteria
Azotobacter vinelandii DJ YP 002799438 1 322710 Proteobacteria
Azotobacter vinelandii CA6 YP 007898297 1 1283331 Proteobacteria
Desulfobulbus propionicus DSM 2032 YP 004195296 1 577650 Proteobacteria
Spirochaeta smaragdinae DSM 11293 YP 003801919 1 573413 Spirochaetes
Halothiobacillus neapolitanus c2 YP 003262017 1 555778 Proteobacteria
Magnetospirillum magneticum AMB 1 YP 422095 1 342108 Proteobacteria
Rhodomicrobium vannielii ATCC 17100 YP 004013752 1 648757 Proteobacteria
Magnetospirillum gryphiswaldense MSR 1 v2 YP 008936189 1 1430440 Proteobacteria
Rhodospirillum photometricum DSM 122 YP 005418043 1 1150469 Proteobacteria
Rhodospirillum rubrum ATCC 11170 YP 427556 1 269796 Proteobacteria
Dickeya aquatica DW 0440 WP 035341557 1 Daquatica
Dickeya sp CSL RW240 WP 035341557 1 Daquatica
DAQ1742 DAQ1742 02498 Daquatica
Dickeya zeae EC1 WP 012884601 1 Dzeae
Dickeya zeae CSL RW192 WP 012884601 1 Dzeae
Dickeya zeae Ech586 WP 012884601 1 Dzeae
Dickeya zeae MS1 WP 012884601 1 Dzeae
Dickeya zeae NCPPB 3532 WP 012884601 1 Dzeae
Dickeya zeae DZ2Q WP 012884601 1 Dzeae
Dickeya zeae NCPPB 2538 WP 012884601 1 Dzeae
Dickeya zeae MK19 WP 012884601 1 Dzeae
Dickeya zeae ZJU1202 WP 012884601 1 Dzeae
Dickeya zeae NCPPB 3531 WP 012884601 1 Dzeae
Dickeya gs fangzhongdai S1 WP 049853727 1 Dfangzhongdai
Prodigiosinella confusarubida ATCC39006 WP 021015790 1 Pconfusarubida
Dickeya paradisiaca NCPPB 2511 WP 015853769 1 Dparadisiaca
Dickeya paradisiaca Ech703 WP 015853769 1 Dparadisiaca
Dickeya dianthicola GBBC 2039 WP 024106075 1 Ddianthicola
Dickeya dianthicola IPO 980 WP 024106075 1 Ddianthicola
Dickeya dianthicola NCPPB 3534 WP 024106075 1 Ddianthicola
Dickeya dianthicola RNS04 9 WP 024106075 1 Ddianthicola
Dickeya dianthicola NCPPB 453 WP 024106075 1 Ddianthicola
Dickeya solani GBBC 2040 WP 012769649 1 Dsolani
Dickeya solani IPO 2222 WP 012769649 1 Dsolani
Dickeya chrysanthemi Ech1591 WP 012769649 1 Dchrysanthemi
Dickeya gs undicola 2B12 WP 012769649 1 Dundicola
Dickeya dadantii subsp dieffenbachiae NCPPB 2976 WP 012769649 1 Ddadantii
Dickeya chrysanthemi subsp parthenii NCPPB 516 WP 012769649 1 Dchrysanthemi
Dickeya solani PPO 9019 WP 012769649 1 Dsolani
Dickeya solani PPO 9134 WP 012769649 1 Dsolani
Dickeya gs fangzhongdai MK7 WP 012769649 1 Dfangzhongdai
Dickeya chrysanthemi NCPPB 3533 WP 012769649 1 Dchrysanthemi
Dickeya solani RNS 08 23 3 1 A WP 012769649 1 Dsolani
Dickeya gs fangzhongdai NCPPB 3274 WP 012769649 1 Dfangzhongdai
Dickeya chrysanthemi subsp chrysanthemi NCPPB 402 WP 012769649 1 Dchrysanthemi
Dickeya solani D s0432 1 WP 012769649 1 Dsolani
Dickeya gs fangzhongdai M005 WP 012769649 1 Dfangzhongdai
Dickeya gs poaceaephila NCPPB 569 WP 012769649 1 Dpoaceaephila
Dickeya dadantii NCPPB 3537 WP 012769649 1 Ddadantii
Dickeya chrysanthemi L11 WP 012769649 1 Dchrysanthemi
Dickeya dadantii 3937 WP 012769649 1 Ddadantii
Dickeya solani MK16 WP 012769649 1 Dsolani
Dickeya gs fangzhongdai M074 WP 012769649 1 Dfangzhongdai
Dickeya solani MK10 WP 012769649 1 Dsolani
Dickeya gs fangzhongdai ND14b WP 012769649 1 Dfangzhongdai
Dickeya solani IFB 0099 WP 012769649 1 Dsolani
Dickeya solani RNS 07 7 3B WP 012769649 1 Dsolani
497 feoA Iron metabolism Ferrous iron transport Seed
Dickeya gs fangzhongdai B16 WP 012769649 1 Dfangzhongdai
Dickeya dadantii NCPPB 898 WP 012769649 1 Ddadantii
Dickeya solani RNS 05 1 2A WP 012769649 1 Dsolani

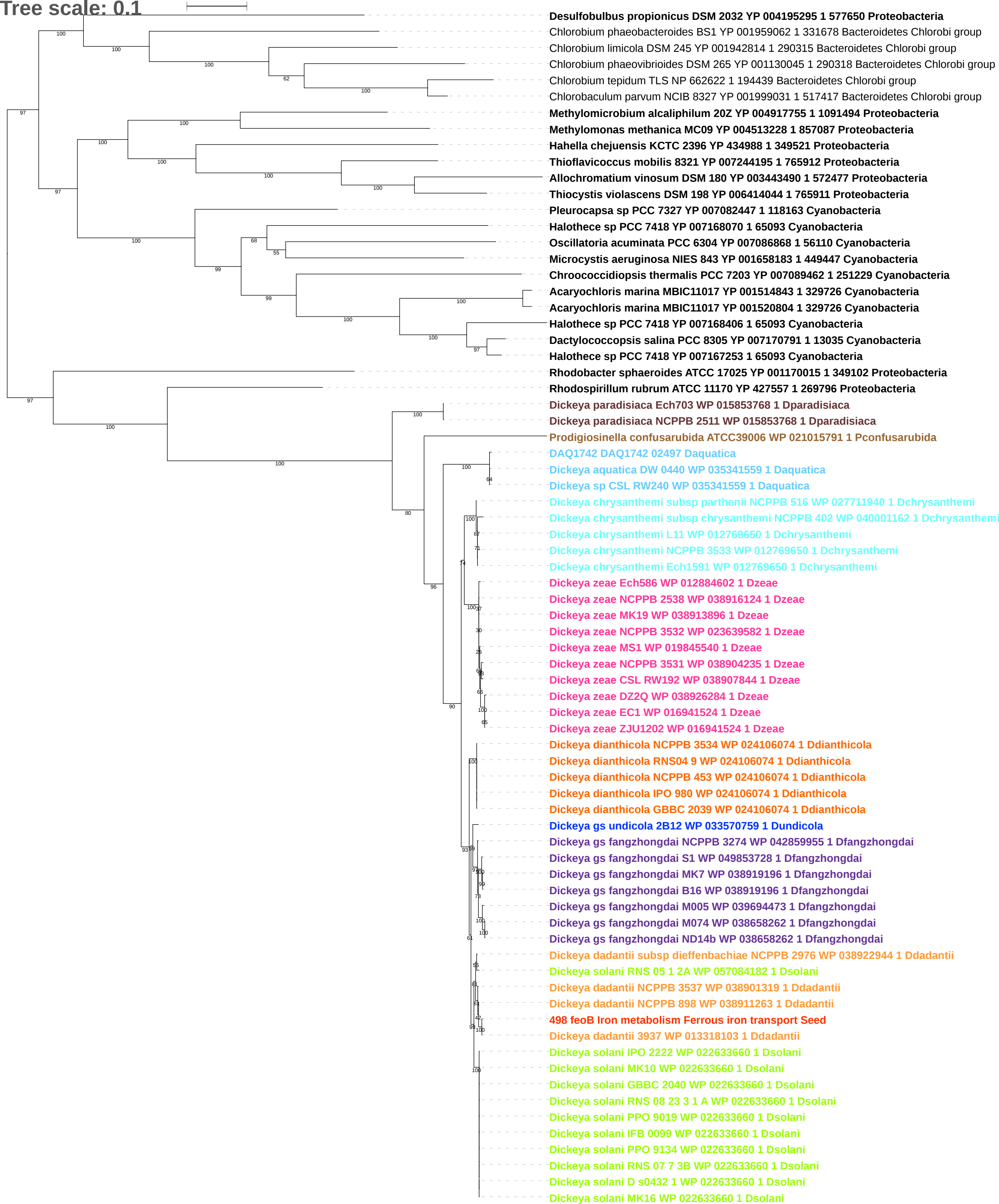
Tree scale: 0.1
Desulfobulbus propionicus DSM 2032 YP 004195295 1 577650 Proteobacteria
Chlorobium phaeobacteroides BS1 YP 001959062 1 331678 Bacteroidetes Chlorobi group
Chlorobium limicola DSM 245 YP 001942814 1 290315 Bacteroidetes Chlorobi group
Chlorobium phaeovibrioides DSM 265 YP 001130045 1 290318 Bacteroidetes Chlorobi group
Chlorobium tepidum TLS NP 662622 1 194439 Bacteroidetes Chlorobi group
Chlorobaculum parvum NCIB 8327 YP 001999031 1 517417 Bacteroidetes Chlorobi group
Methylomicrobium alcaliphilum 20Z YP 004917755 1 1091494 Proteobacteria
Methylomonas methanica MC09 YP 004513228 1 857087 Proteobacteria
Hahella chejuensis KCTC 2396 YP 434988 1 349521 Proteobacteria
Thioflavicoccus mobilis 8321 YP 007244195 1 765912 Proteobacteria
Allochromatium vinosum DSM 180 YP 003443490 1 572477 Proteobacteria
Thiocystis violascens DSM 198 YP 006414044 1 765911 Proteobacteria
Pleurocapsa sp PCC 7327 YP 007082447 1 118163 Cyanobacteria
Halothece sp PCC 7418 YP 007168070 1 65093 Cyanobacteria
Oscillatoria acuminata PCC 6304 YP 007086868 1 56110 Cyanobacteria
Microcystis aeruginosa NIES 843 YP 001658183 1 449447 Cyanobacteria
Chroococcidiopsis thermalis PCC 7203 YP 007089462 1 251229 Cyanobacteria
Acaryochloris marina MBIC11017 YP 001514843 1 329726 Cyanobacteria
Acaryochloris marina MBIC11017 YP 001520804 1 329726 Cyanobacteria
Halothece sp PCC 7418 YP 007168406 1 65093 Cyanobacteria
Dactylococcopsis salina PCC 8305 YP 007170791 1 13035 Cyanobacteria
Halothece sp PCC 7418 YP 007167253 1 65093 Cyanobacteria
Rhodobacter sphaeroides ATCC 17025 YP 001170015 1 349102 Proteobacteria
Rhodospirillum rubrum ATCC 11170 YP 427557 1 269796 Proteobacteria
Dickeya paradisiaca Ech703 WP 015853768 1 Dparadisiaca
Dickeya paradisiaca NCPPB 2511 WP 015853768 1 Dparadisiaca
Prodigiosinella confusarubida ATCC39006 WP 021015791 1 Pconfusarubida
DAQ1742 DAQ1742 02497 Daquatica
Dickeya aquatica DW 0440 WP 035341559 1 Daquatica
Dickeya sp CSL RW240 WP 035341559 1 Daquatica
Dickeya chrysanthemi subsp parthenii NCPPB 516 WP 027711940 1 Dchrysanthemi
Dickeya chrysanthemi subsp chrysanthemi NCPPB 402 WP 040001162 1 Dchrysanthemi
Dickeya chrysanthemi L11 WP 012769650 1 Dchrysanthemi
Dickeya chrysanthemi NCPPB 3533 WP 012769650 1 Dchrysanthemi
Dickeya chrysanthemi Ech1591 WP 012769650 1 Dchrysanthemi
Dickeya zeae Ech586 WP 012884602 1 Dzeae
Dickeya zeae NCPPB 2538 WP 038916124 1 Dzeae
Dickeya zeae MK19 WP 038913896 1 Dzeae
Dickeya zeae NCPPB 3532 WP 023639582 1 Dzeae
Dickeya zeae MS1 WP 019845540 1 Dzeae
Dickeya zeae NCPPB 3531 WP 038904235 1 Dzeae
Dickeya zeae CSL RW192 WP 038907844 1 Dzeae
Dickeya zeae DZ2Q WP 038926284 1 Dzeae
Dickeya zeae EC1 WP 016941524 1 Dzeae
Dickeya zeae ZJU1202 WP 016941524 1 Dzeae
Dickeya dianthicola NCPPB 3534 WP 024106074 1 Ddianthicola
Dickeya dianthicola RNS04 9 WP 024106074 1 Ddianthicola
Dickeya dianthicola NCPPB 453 WP 024106074 1 Ddianthicola
Dickeya dianthicola IPO 980 WP 024106074 1 Ddianthicola
Dickeya dianthicola GBBC 2039 WP 024106074 1 Ddianthicola
Dickeya gs undicola 2B12 WP 033570759 1 Dundicola
Dickeya gs fangzhongdai NCPPB 3274 WP 042859955 1 Dfangzhongdai
Dickeya gs fangzhongdai S1 WP 049853728 1 Dfangzhongdai
Dickeya gs fangzhongdai MK7 WP 038919196 1 Dfangzhongdai
Dickeya gs fangzhongdai B16 WP 038919196 1 Dfangzhongdai
Dickeya gs fangzhongdai M005 WP 039694473 1 Dfangzhongdai
Dickeya gs fangzhongdai M074 WP 038658262 1 Dfangzhongdai
Dickeya gs fangzhongdai ND14b WP 038658262 1 Dfangzhongdai
Dickeya dadantii subsp dieffenbachiae NCPPB 2976 WP 038922944 1 Ddadantii
Dickeya solani RNS 05 1 2A WP 057084182 1 Dsolani
Dickeya dadantii NCPPB 3537 WP 038901319 1 Ddadantii
Dickeya dadantii NCPPB 898 WP 038911263 1 Ddadantii
498 feoB Iron metabolism Ferrous iron transport Seed
Dickeya dadantii 3937 WP 013318103 1 Ddadantii
Dickeya solani IPO 2222 WP 022633660 1 Dsolani
Dickeya solani MK10 WP 022633660 1 Dsolani
Dickeya solani GBBC 2040 WP 022633660 1 Dsolani
Dickeya solani RNS 08 23 3 1 A WP 022633660 1 Dsolani
Dickeya solani PPO 9019 WP 022633660 1 Dsolani
Dickeya solani IFB 0099 WP 022633660 1 Dsolani
Dickeya solani PPO 9134 WP 022633660 1 Dsolani
Dickeya solani RNS 07 7 3B WP 022633660 1 Dsolani
Dickeya solani D s0432 1 WP 022633660 1 Dsolani
Dickeya solani MK16 WP 022633660 1 Dsolani

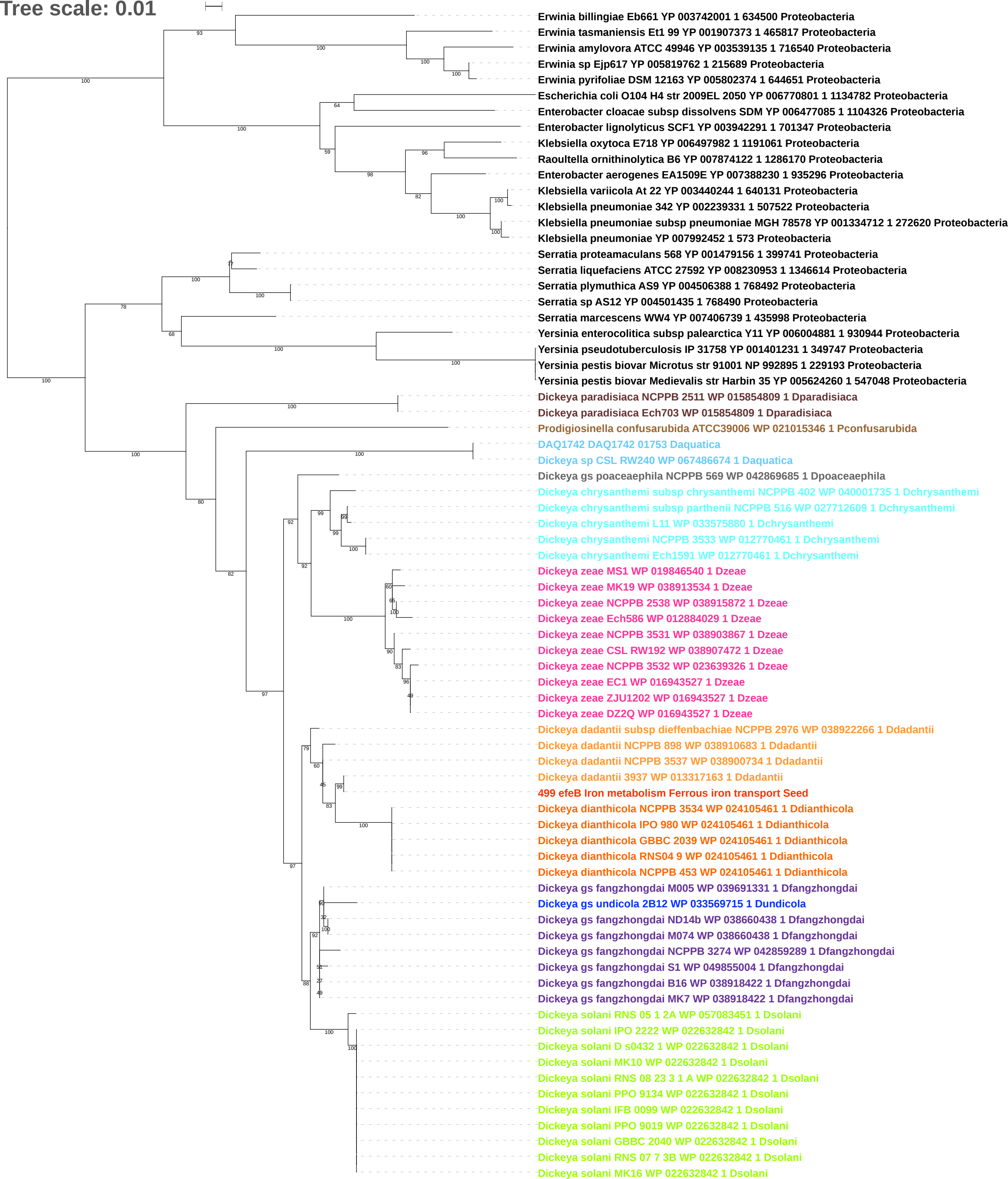
Tree scale: 0.01
Erwinia billingiae Eb661 YP 003742001 1 634500 Proteobacteria
Erwinia tasmaniensis Et1 99 YP 001907373 1 465817 Proteobacteria
Erwinia amylovora ATCC 49946 YP 003539135 1 716540 Proteobacteria
Erwinia sp Ejp617 YP 005819762 1 215689 Proteobacteria
Erwinia pyrifoliae DSM 12163 YP 005802374 1 644651 Proteobacteria
Escherichia coli O104 H4 str 2009EL 2050 YP 006770801 1 1134782 Proteobacteria
Enterobacter cloacae subsp dissolvens SDM YP 006477085 1 1104326 Proteobacteria
Enterobacter lignolyticus SCF1 YP 003942291 1 701347 Proteobacteria
Klebsiella oxytoca E718 YP 006497982 1 1191061 Proteobacteria
Raoultella ornithinolytica B6 YP 007874122 1 1286170 Proteobacteria
Enterobacter aerogenes EA1509E YP 007388230 1 935296 Proteobacteria
Klebsiella variicola At 22 YP 003440244 1 640131 Proteobacteria
Klebsiella pneumoniae 342 YP 002239331 1 507522 Proteobacteria
Klebsiella pneumoniae subsp pneumoniae MGH 78578 YP 001334712 1 272620 Proteobacteria
Klebsiella pneumoniae YP 007992452 1 573 Proteobacteria
Serratia proteamaculans 568 YP 001479156 1 399741 Proteobacteria
Serratia liquefaciens ATCC 27592 YP 008230953 1 1346614 Proteobacteria
Serratia plymuthica AS9 YP 004506388 1 768492 Proteobacteria
Serratia sp AS12 YP 004501435 1 768490 Proteobacteria
Serratia marcescens WW4 YP 007406739 1 435998 Proteobacteria
Yersinia enterocolitica subsp palearctica Y11 YP 006004881 1 930944 Proteobacteria
Yersinia pseudotuberculosis IP 31758 YP 001401231 1 349747 Proteobacteria
Yersinia pestis biovar Microtus str 91001 NP 992895 1 229193 Proteobacteria
Yersinia pestis biovar Medievalis str Harbin 35 YP 005624260 1 547048 Proteobacteria
Dickeya paradisiaca NCPPB 2511 WP 015854809 1 Dparadisiaca
Dickeya paradisiaca Ech703 WP 015854809 1 Dparadisiaca
Prodigiosinella confusarubida ATCC39006 WP 021015346 1 Pconfusarubida
DAQ1742 DAQ1742 01753 Daquatica
Dickeya sp CSL RW240 WP 067486674 1 Daquatica
Dickeya gs poaceaephila NCPPB 569 WP 042869685 1 Dpoaceaephila
Dickeya chrysanthemi subsp chrysanthemi NCPPB 402 WP 040001735 1 Dchrysanthemi
Dickeya chrysanthemi subsp parthenii NCPPB 516 WP 027712609 1 Dchrysanthemi
Dickeya chrysanthemi L11 WP 033575880 1 Dchrysanthemi
Dickeya chrysanthemi NCPPB 3533 WP 012770461 1 Dchrysanthemi
Dickeya chrysanthemi Ech1591 WP 012770461 1 Dchrysanthemi
Dickeya zeae MS1 WP 019846540 1 Dzeae
Dickeya zeae MK19 WP 038913534 1 Dzeae
Dickeya zeae NCPPB 2538 WP 038915872 1 Dzeae
Dickeya zeae Ech586 WP 012884029 1 Dzeae
Dickeya zeae NCPPB 3531 WP 038903867 1 Dzeae
Dickeya zeae CSL RW192 WP 038907472 1 Dzeae
Dickeya zeae NCPPB 3532 WP 023639326 1 Dzeae
Dickeya zeae EC1 WP 016943527 1 Dzeae
Dickeya zeae ZJU1202 WP 016943527 1 Dzeae
Dickeya zeae DZ2Q WP 016943527 1 Dzeae
Dickeya dadantii subsp dieffenbachiae NCPPB 2976 WP 038922266 1 Ddadantii
Dickeya dadantii NCPPB 898 WP 038910683 1 Ddadantii
Dickeya dadantii NCPPB 3537 WP 038900734 1 Ddadantii
Dickeya dadantii 3937 WP 013317163 1 Ddadantii
499 efeB Iron metabolism Ferrous iron transport Seed
Dickeya dianthicola NCPPB 3534 WP 024105461 1 Ddianthicola
Dickeya dianthicola IPO 980 WP 024105461 1 Ddianthicola
Dickeya dianthicola GBBC 2039 WP 024105461 1 Ddianthicola
Dickeya dianthicola RNS04 9 WP 024105461 1 Ddianthicola
Dickeya dianthicola NCPPB 453 WP 024105461 1 Ddianthicola
Dickeya gs fangzhongdai M005 WP 039691331 1 Dfangzhongdai
Dickeya gs undicola 2B12 WP 033569715 1 Dundicola
Dickeya gs fangzhongdai ND14b WP 038660438 1 Dfangzhongdai
Dickeya gs fangzhongdai M074 WP 038660438 1 Dfangzhongdai
Dickeya gs fangzhongdai NCPPB 3274 WP 042859289 1 Dfangzhongdai
Dickeya gs fangzhongdai S1 WP 049855004 1 Dfangzhongdai
Dickeya gs fangzhongdai B16 WP 038918422 1 Dfangzhongdai
Dickeya gs fangzhongdai MK7 WP 038918422 1 Dfangzhongdai
Dickeya solani RNS 05 1 2A WP 057083451 1 Dsolani
Dickeya solani IPO 2222 WP 022632842 1 Dsolani
Dickeya solani D s0432 1 WP 022632842 1 Dsolani
Dickeya solani MK10 WP 022632842 1 Dsolani
Dickeya solani RNS 08 23 3 1 A WP 022632842 1 Dsolani
Dickeya solani PPO 9134 WP 022632842 1 Dsolani
Dickeya solani IFB 0099 WP 022632842 1 Dsolani
Dickeya solani PPO 9019 WP 022632842 1 Dsolani
Dickeya solani GBBC 2040 WP 022632842 1 Dsolani
Dickeya solani RNS 07 7 3B WP 022632842 1 Dsolani
Dickeya solani MK16 WP 022632842 1 Dsolani

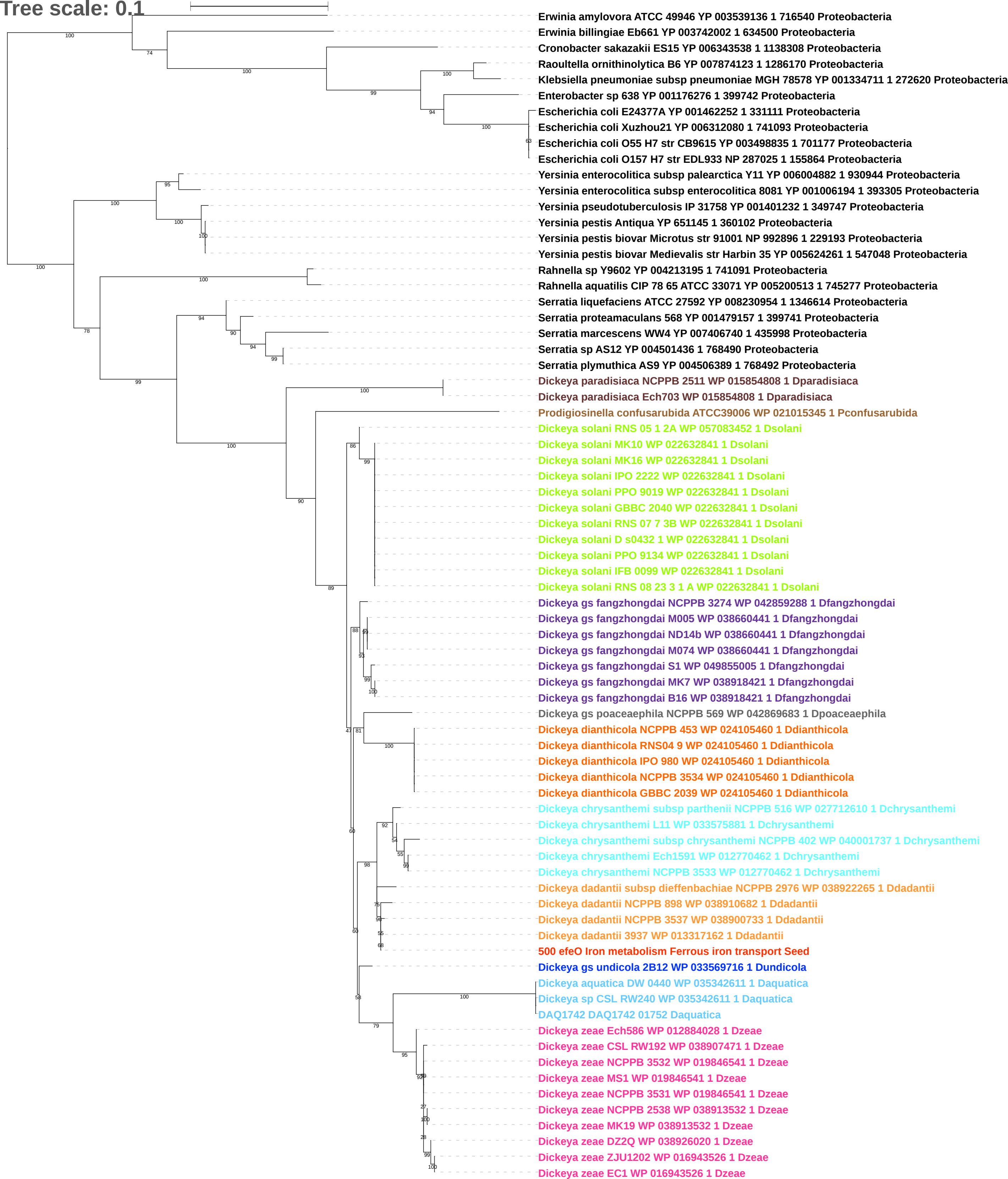
Tree scale: 0.1
Erwinia amylovora ATCC 49946 YP 003539136 1 716540 Proteobacteria
Erwinia billingiae Eb661 YP 003742002 1 634500 Proteobacteria
Cronobacter sakazakii ES15 YP 006343538 1 1138308 Proteobacteria
Raoultella ornithinolytica B6 YP 007874123 1 1286170 Proteobacteria
Klebsiella pneumoniae subsp pneumoniae MGH 78578 YP 001334711 1 272620 Proteobacteria
Enterobacter sp 638 YP 001176276 1 399742 Proteobacteria
Escherichia coli E24377A YP 001462252 1 331111 Proteobacteria
Escherichia coli Xuzhou21 YP 006312080 1 741093 Proteobacteria
Escherichia coli O55 H7 str CB9615 YP 003498835 1 701177 Proteobacteria
Escherichia coli O157 H7 str EDL933 NP 287025 1 155864 Proteobacteria
Yersinia enterocolitica subsp palearctica Y11 YP 006004882 1 930944 Proteobacteria
Yersinia enterocolitica subsp enterocolitica 8081 YP 001006194 1 393305 Proteobacteria
Yersinia pseudotuberculosis IP 31758 YP 001401232 1 349747 Proteobacteria
Yersinia pestis Antiqua YP 651145 1 360102 Proteobacteria
Yersinia pestis biovar Microtus str 91001 NP 992896 1 229193 Proteobacteria
Yersinia pestis biovar Medievalis str Harbin 35 YP 005624261 1 547048 Proteobacteria
Rahnella sp Y9602 YP 004213195 1 741091 Proteobacteria
Rahnella aquatilis CIP 78 65 ATCC 33071 YP 005200513 1 745277 Proteobacteria
Serratia liquefaciens ATCC 27592 YP 008230954 1 1346614 Proteobacteria
Serratia proteamaculans 568 YP 001479157 1 399741 Proteobacteria
Serratia marcescens WW4 YP 007406740 1 435998 Proteobacteria
Serratia sp AS12 YP 004501436 1 768490 Proteobacteria
Serratia plymuthica AS9 YP 004506389 1 768492 Proteobacteria
Dickeya paradisiaca NCPPB 2511 WP 015854808 1 Dparadisiaca
Dickeya paradisiaca Ech703 WP 015854808 1 Dparadisiaca
Prodigiosinella confusarubida ATCC39006 WP 021015345 1 Pconfusarubida
Dickeya solani RNS 05 1 2A WP 057083452 1 Dsolani
Dickeya solani MK10 WP 022632841 1 Dsolani
Dickeya solani MK16 WP 022632841 1 Dsolani
Dickeya solani IPO 2222 WP 022632841 1 Dsolani
Dickeya solani PPO 9019 WP 022632841 1 Dsolani
Dickeya solani GBBC 2040 WP 022632841 1 Dsolani
Dickeya solani RNS 07 7 3B WP 022632841 1 Dsolani
Dickeya solani D s0432 1 WP 022632841 1 Dsolani
Dickeya solani PPO 9134 WP 022632841 1 Dsolani
Dickeya solani IFB 0099 WP 022632841 1 Dsolani
Dickeya solani RNS 08 23 3 1 A WP 022632841 1 Dsolani
Dickeya gs fangzhongdai NCPPB 3274 WP 042859288 1 Dfangzhongdai
Dickeya gs fangzhongdai M005 WP 038660441 1 Dfangzhongdai
Dickeya gs fangzhongdai ND14b WP 038660441 1 Dfangzhongdai
Dickeya gs fangzhongdai M074 WP 038660441 1 Dfangzhongdai
Dickeya gs fangzhongdai S1 WP 049855005 1 Dfangzhongdai
Dickeya gs fangzhongdai MK7 WP 038918421 1 Dfangzhongdai
Dickeya gs fangzhongdai B16 WP 038918421 1 Dfangzhongdai
Dickeya gs poaceaephila NCPPB 569 WP 042869683 1 Dpoaceaephila
Dickeya dianthicola NCPPB 453 WP 024105460 1 Ddianthicola
Dickeya dianthicola RNS04 9 WP 024105460 1 Ddianthicola
Dickeya dianthicola IPO 980 WP 024105460 1 Ddianthicola
Dickeya dianthicola NCPPB 3534 WP 024105460 1 Ddianthicola
Dickeya dianthicola GBBC 2039 WP 024105460 1 Ddianthicola
Dickeya chrysanthemi subsp parthenii NCPPB 516 WP 027712610 1 Dchrysanthemi
Dickeya chrysanthemi L11 WP 033575881 1 Dchrysanthemi
Dickeya chrysanthemi subsp chrysanthemi NCPPB 402 WP 040001737 1 Dchrysanthemi
Dickeya chrysanthemi Ech1591 WP 012770462 1 Dchrysanthemi
Dickeya chrysanthemi NCPPB 3533 WP 012770462 1 Dchrysanthemi
Dickeya dadantii subsp dieffenbachiae NCPPB 2976 WP 038922265 1 Ddadantii
Dickeya dadantii NCPPB 898 WP 038910682 1 Ddadantii
Dickeya dadantii NCPPB 3537 WP 038900733 1 Ddadantii
Dickeya dadantii 3937 WP 013317162 1 Ddadantii
500 efeO Iron metabolism Ferrous iron transport Seed
Dickeya gs undicola 2B12 WP 033569716 1 Dundicola
Dickeya aquatica DW 0440 WP 035342611 1 Daquatica
Dickeya sp CSL RW240 WP 035342611 1 Daquatica
DAQ1742 DAQ1742 01752 Daquatica
Dickeya zeae Ech586 WP 012884028 1 Dzeae
Dickeya zeae CSL RW192 WP 038907471 1 Dzeae
Dickeya zeae NCPPB 3532 WP 019846541 1 Dzeae
Dickeya zeae MS1 WP 019846541 1 Dzeae
Dickeya zeae NCPPB 3531 WP 019846541 1 Dzeae
Dickeya zeae NCPPB 2538 WP 038913532 1 Dzeae
Dickeya zeae MK19 WP 038913532 1 Dzeae
Dickeya zeae DZ2Q WP 038926020 1 Dzeae
Dickeya zeae ZJU1202 WP 016943526 1 Dzeae
Dickeya zeae EC1 WP 016943526 1 Dzeae

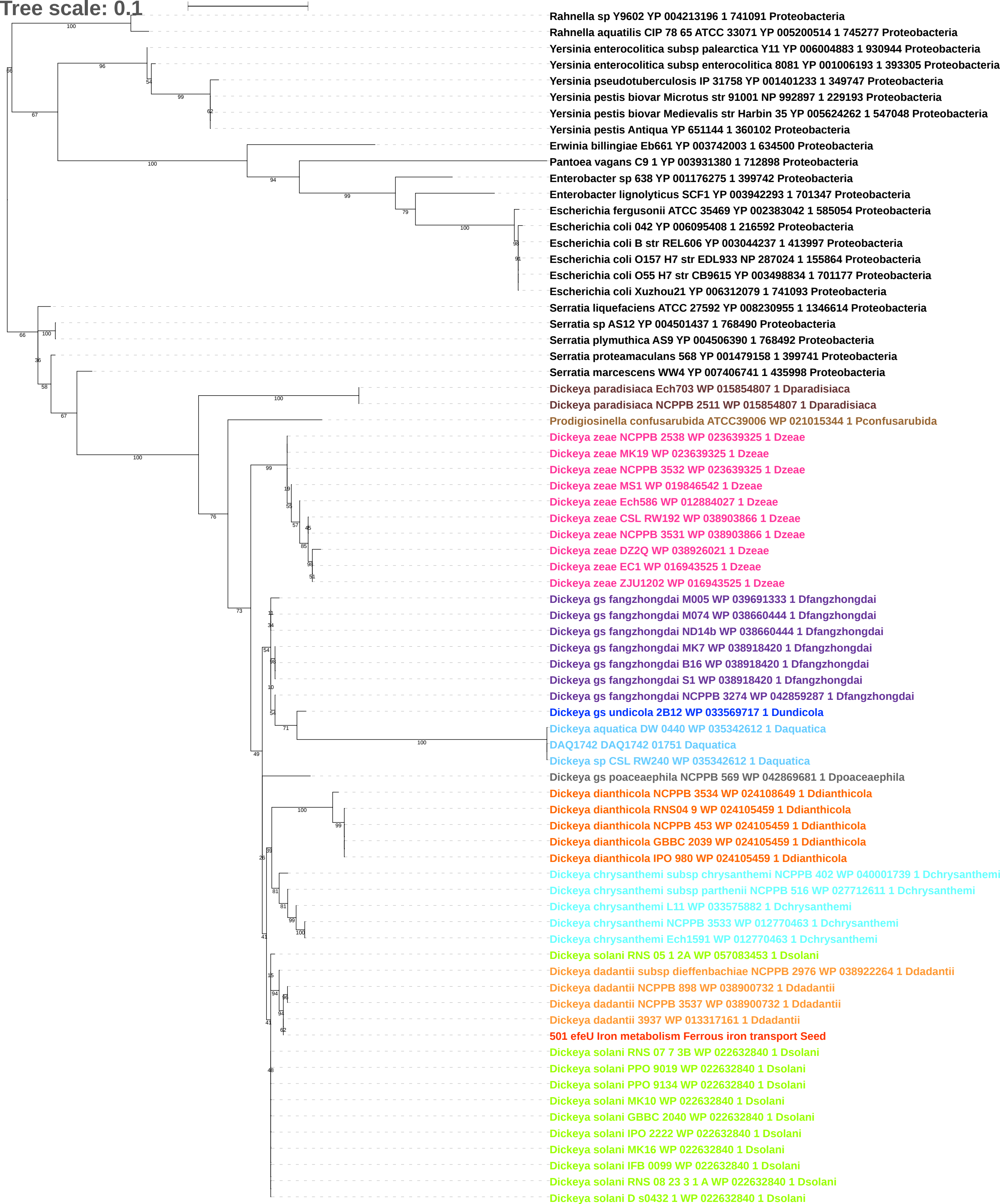

Tree scale: 0.1
Rahnella sp Y9602 YP 004213196 1 741091 Proteobacteria
Rahnella aquatilis CIP 78 65 ATCC 33071 YP 005200514 1 745277 Proteobacteria
Yersinia enterocolitica subsp palearctica Y11 YP 006004883 1 930944 Proteobacteria
Yersinia enterocolitica subsp enterocolitica 8081 YP 001006193 1 393305 Proteobacteria
Yersinia pseudotuberculosis IP 31758 YP 001401233 1 349747 Proteobacteria
Yersinia pestis biovar Microtus str 91001 NP 992897 1 229193 Proteobacteria
Yersinia pestis biovar Medievalis str Harbin 35 YP 005624262 1 547048 Proteobacteria
Yersinia pestis Antiqua YP 651144 1 360102 Proteobacteria
Erwinia billingiae Eb661 YP 003742003 1 634500 Proteobacteria
Pantoea vagans C9 1 YP 003931380 1 712898 Proteobacteria
Enterobacter sp 638 YP 001176275 1 399742 Proteobacteria
Enterobacter lignolyticus SCF1 YP 003942293 1 701347 Proteobacteria
Escherichia fergusonii ATCC 35469 YP 002383042 1 585054 Proteobacteria
Escherichia coli 042 YP 006095408 1 216592 Proteobacteria
Escherichia coli B str REL606 YP 003044237 1 413997 Proteobacteria
Escherichia coli O157 H7 str EDL933 NP 287024 1 155864 Proteobacteria
Escherichia coli O55 H7 str CB9615 YP 003498834 1 701177 Proteobacteria
Escherichia coli Xuzhou21 YP 006312079 1 741093 Proteobacteria
Serratia liquefaciens ATCC 27592 YP 008230955 1 1346614 Proteobacteria
Serratia sp AS12 YP 004501437 1 768490 Proteobacteria
Serratia plymuthica AS9 YP 004506390 1 768492 Proteobacteria
Serratia proteamaculans 568 YP 001479158 1 399741 Proteobacteria
Serratia marcescens WW4 YP 007406741 1 435998 Proteobacteria
Dickeya paradisiaca Ech703 WP 015854807 1 Dparadisiaca
Dickeya paradisiaca NCPPB 2511 WP 015854807 1 Dparadisiaca
Prodigiosinella confusarubida ATCC39006 WP 021015344 1 Pconfusarubida
Dickeya zeae NCPPB 2538 WP 023639325 1 Dzeae
Dickeya zeae MK19 WP 023639325 1 Dzeae
Dickeya zeae NCPPB 3532 WP 023639325 1 Dzeae
Dickeya zeae MS1 WP 019846542 1 Dzeae
Dickeya zeae Ech586 WP 012884027 1 Dzeae
Dickeya zeae CSL RW192 WP 038903866 1 Dzeae
Dickeya zeae NCPPB 3531 WP 038903866 1 Dzeae
Dickeya zeae DZ2Q WP 038926021 1 Dzeae
Dickeya zeae EC1 WP 016943525 1 Dzeae
Dickeya zeae ZJU1202 WP 016943525 1 Dzeae
Dickeya gs fangzhongdai M005 WP 039691333 1 Dfangzhongdai
Dickeya gs fangzhongdai M074 WP 038660444 1 Dfangzhongdai
Dickeya gs fangzhongdai ND14b WP 038660444 1 Dfangzhongdai
Dickeya gs fangzhongdai MK7 WP 038918420 1 Dfangzhongdai
Dickeya gs fangzhongdai B16 WP 038918420 1 Dfangzhongdai
Dickeya gs fangzhongdai S1 WP 038918420 1 Dfangzhongdai
Dickeya gs fangzhongdai NCPPB 3274 WP 042859287 1 Dfangzhongdai
Dickeya gs undicola 2B12 WP 033569717 1 Dundicola
Dickeya aquatica DW 0440 WP 035342612 1 Daquatica
DAQ1742 DAQ1742 01751 Daquatica
Dickeya sp CSL RW240 WP 035342612 1 Daquatica
Dickeya gs poaceaephila NCPPB 569 WP 042869681 1 Dpoaceaephila
Dickeya dianthicola NCPPB 3534 WP 024108649 1 Ddianthicola
Dickeya dianthicola RNS04 9 WP 024105459 1 Ddianthicola
Dickeya dianthicola NCPPB 453 WP 024105459 1 Ddianthicola
Dickeya dianthicola GBBC 2039 WP 024105459 1 Ddianthicola
Dickeya dianthicola IPO 980 WP 024105459 1 Ddianthicola
Dickeya chrysanthemi subsp chrysanthemi NCPPB 402 WP 040001739 1 Dchrysanthemi
Dickeya chrysanthemi subsp parthenii NCPPB 516 WP 027712611 1 Dchrysanthemi
Dickeya chrysanthemi L11 WP 033575882 1 Dchrysanthemi
Dickeya chrysanthemi NCPPB 3533 WP 012770463 1 Dchrysanthemi
Dickeya chrysanthemi Ech1591 WP 012770463 1 Dchrysanthemi
Dickeya solani RNS 05 1 2A WP 057083453 1 Dsolani
Dickeya dadantii subsp dieffenbachiae NCPPB 2976 WP 038922264 1 Ddadantii
Dickeya dadantii NCPPB 898 WP 038900732 1 Ddadantii
Dickeya dadantii NCPPB 3537 WP 038900732 1 Ddadantii
Dickeya dadantii 3937 WP 013317161 1 Ddadantii
501 efeU Iron metabolism Ferrous iron transport Seed
Dickeya solani RNS 07 7 3B WP 022632840 1 Dsolani
Dickeya solani PPO 9019 WP 022632840 1 Dsolani
Dickeya solani PPO 9134 WP 022632840 1 Dsolani
Dickeya solani MK10 WP 022632840 1 Dsolani
Dickeya solani GBBC 2040 WP 022632840 1 Dsolani
Dickeya solani IPO 2222 WP 022632840 1 Dsolani
Dickeya solani MK16 WP 022632840 1 Dsolani
Dickeya solani IFB 0099 WP 022632840 1 Dsolani
Dickeya solani RNS 08 23 3 1 A WP 022632840 1 Dsolani
Dickeya solani D s0432 1 WP 022632840 1 Dsolani

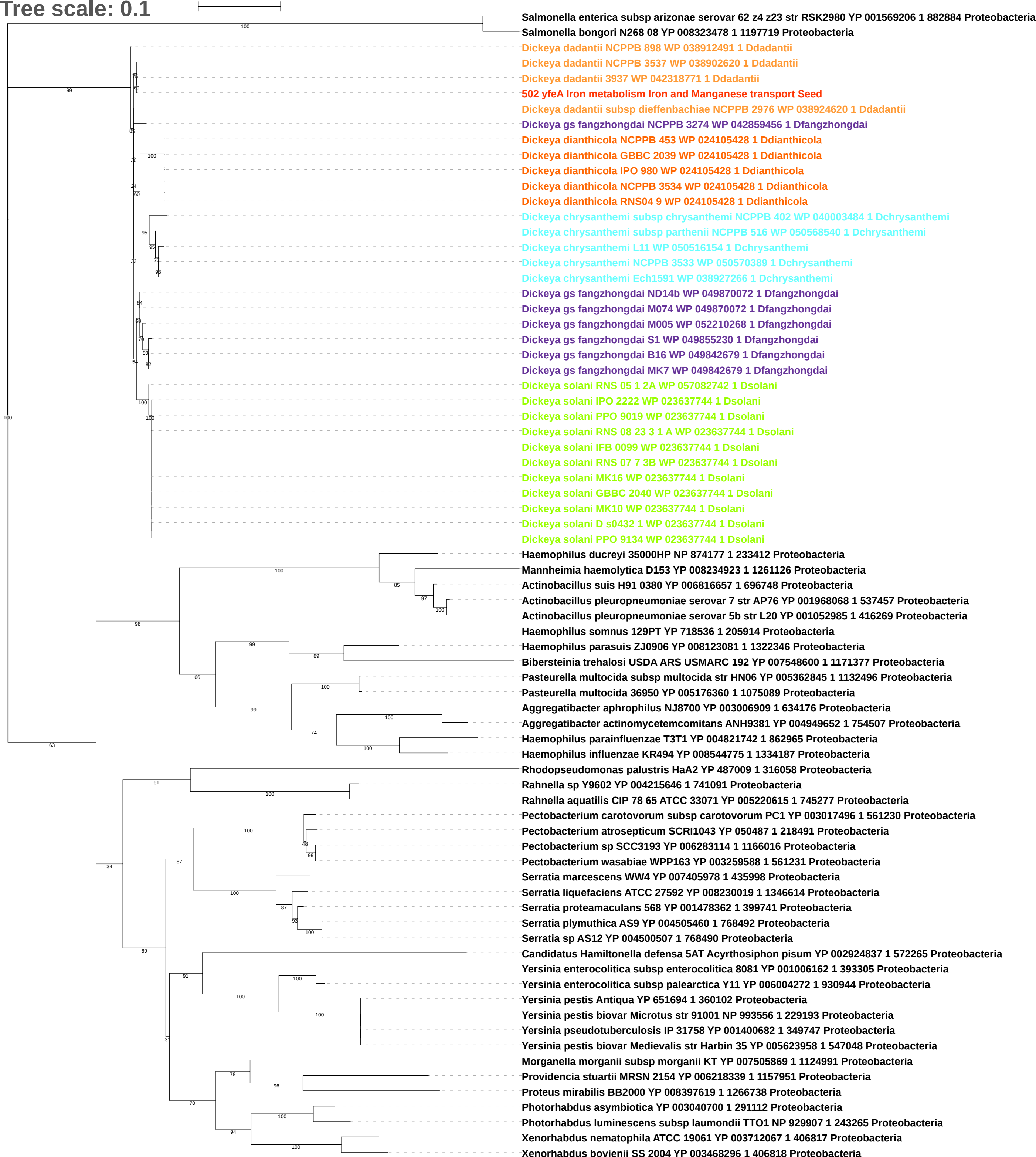
Tree scale: 0.1
Salmonella enterica subsp arizonae serovar 62 z4 z23 str RSK2980 YP 001569206 1 882884 Proteobacteria
Salmonella bongori N268 08 YP 008323478 1 1197719 Proteobacteria
Dickeya dadantii NCPPB 898 WP 038912491 1 Ddadantii
Dickeya dadantii NCPPB 3537 WP 038902620 1 Ddadantii
Dickeya dadantii 3937 WP 042318771 1 Ddadantii
502 yfeA Iron metabolism Iron and Manganese transport Seed
Dickeya dadantii subsp dieffenbachiae NCPPB 2976 WP 038924620 1 Ddadantii
Dickeya gs fangzhongdai NCPPB 3274 WP 042859456 1 Dfangzhongdai
Dickeya dianthicola NCPPB 453 WP 024105428 1 Ddianthicola
Dickeya dianthicola GBBC 2039 WP 024105428 1 Ddianthicola
Dickeya dianthicola IPO 980 WP 024105428 1 Ddianthicola
Dickeya dianthicola NCPPB 3534 WP 024105428 1 Ddianthicola
Dickeya dianthicola RNS04 9 WP 024105428 1 Ddianthicola
Dickeya chrysanthemi subsp chrysanthemi NCPPB 402 WP 040003484 1 Dchrysanthemi
Dickeya chrysanthemi subsp parthenii NCPPB 516 WP 050568540 1 Dchrysanthemi
Dickeya chrysanthemi L11 WP 050516154 1 Dchrysanthemi
Dickeya chrysanthemi NCPPB 3533 WP 050570389 1 Dchrysanthemi
Dickeya chrysanthemi Ech1591 WP 038927266 1 Dchrysanthemi
Dickeya gs fangzhongdai ND14b WP 049870072 1 Dfangzhongdai
Dickeya gs fangzhongdai M074 WP 049870072 1 Dfangzhongdai
Dickeya gs fangzhongdai M005 WP 052210268 1 Dfangzhongdai
Dickeya gs fangzhongdai S1 WP 049855230 1 Dfangzhongdai
Dickeya gs fangzhongdai B16 WP 049842679 1 Dfangzhongdai
Dickeya gs fangzhongdai MK7 WP 049842679 1 Dfangzhongdai
Dickeya solani RNS 05 1 2A WP 057082742 1 Dsolani
Dickeya solani IPO 2222 WP 023637744 1 Dsolani
Dickeya solani PPO 9019 WP 023637744 1 Dsolani
Dickeya solani RNS 08 23 3 1 A WP 023637744 1 Dsolani
Dickeya solani IFB 0099 WP 023637744 1 Dsolani
Dickeya solani RNS 07 7 3B WP 023637744 1 Dsolani
Dickeya solani MK16 WP 023637744 1 Dsolani
Dickeya solani GBBC 2040 WP 023637744 1 Dsolani
Dickeya solani MK10 WP 023637744 1 Dsolani
Dickeya solani D s0432 1 WP 023637744 1 Dsolani
Dickeya solani PPO 9134 WP 023637744 1 Dsolani
Haemophilus ducreyi 35000HP NP 874177 1 233412 Proteobacteria
Mannheimia haemolytica D153 YP 008234923 1 1261126 Proteobacteria
Actinobacillus suis H91 0380 YP 006816657 1 696748 Proteobacteria
Actinobacillus pleuropneumoniae serovar 7 str AP76 YP 001968068 1 537457 Proteobacteria
Actinobacillus pleuropneumoniae serovar 5b str L20 YP 001052985 1 416269 Proteobacteria
Haemophilus somnus 129PT YP 718536 1 205914 Proteobacteria
Haemophilus parasuis ZJ0906 YP 008123081 1 1322346 Proteobacteria
Bibersteinia trehalosi USDA ARS USMARC 192 YP 007548600 1 1171377 Proteobacteria
Pasteurella multocida subsp multocida str HN06 YP 005362845 1 1132496 Proteobacteria
Pasteurella multocida 36950 YP 005176360 1 1075089 Proteobacteria
Aggregatibacter aphrophilus NJ8700 YP 003006909 1 634176 Proteobacteria
Aggregatibacter actinomycetemcomitans ANH9381 YP 004949652 1 754507 Proteobacteria
Haemophilus parainfluenzae T3T1 YP 004821742 1 862965 Proteobacteria
Haemophilus influenzae KR494 YP 008544775 1 1334187 Proteobacteria
Rhodopseudomonas palustris HaA2 YP 487009 1 316058 Proteobacteria
Rahnella sp Y9602 YP 004215646 1 741091 Proteobacteria
Rahnella aquatilis CIP 78 65 ATCC 33071 YP 005220615 1 745277 Proteobacteria
Pectobacterium carotovorum subsp carotovorum PC1 YP 003017496 1 561230 Proteobacteria
Pectobacterium atrosepticum SCRI1043 YP 050487 1 218491 Proteobacteria
Pectobacterium sp SCC3193 YP 006283114 1 1166016 Proteobacteria
Pectobacterium wasabiae WPP163 YP 003259588 1 561231 Proteobacteria
Serratia marcescens WW4 YP 007405978 1 435998 Proteobacteria
Serratia liquefaciens ATCC 27592 YP 008230019 1 1346614 Proteobacteria
Serratia proteamaculans 568 YP 001478362 1 399741 Proteobacteria
Serratia plymuthica AS9 YP 004505460 1 768492 Proteobacteria
Serratia sp AS12 YP 004500507 1 768490 Proteobacteria
Candidatus Hamiltonella defensa 5AT Acyrthosiphon pisum YP 002924837 1 572265 Proteobacteria
Yersinia enterocolitica subsp enterocolitica 8081 YP 001006162 1 393305 Proteobacteria
Yersinia enterocolitica subsp palearctica Y11 YP 006004272 1 930944 Proteobacteria
Yersinia pestis Antiqua YP 651694 1 360102 Proteobacteria
Yersinia pestis biovar Microtus str 91001 NP 993556 1 229193 Proteobacteria
Yersinia pseudotuberculosis IP 31758 YP 001400682 1 349747 Proteobacteria
Yersinia pestis biovar Medievalis str Harbin 35 YP 005623958 1 547048 Proteobacteria
Morganella morganii subsp morganii KT YP 007505869 1 1124991 Proteobacteria
Providencia stuartii MRSN 2154 YP 006218339 1 1157951 Proteobacteria
Proteus mirabilis BB2000 YP 008397619 1 1266738 Proteobacteria
Photorhabdus asymbiotica YP 003040700 1 291112 Proteobacteria
Photorhabdus luminescens subsp laumondii TTO1 NP 929907 1 243265 Proteobacteria
Xenorhabdus nematophila ATCC 19061 YP 003712067 1 406817 Proteobacteria
Xenorhabdus bovienii SS 2004 YP 003468296 1 406818 Proteobacteria

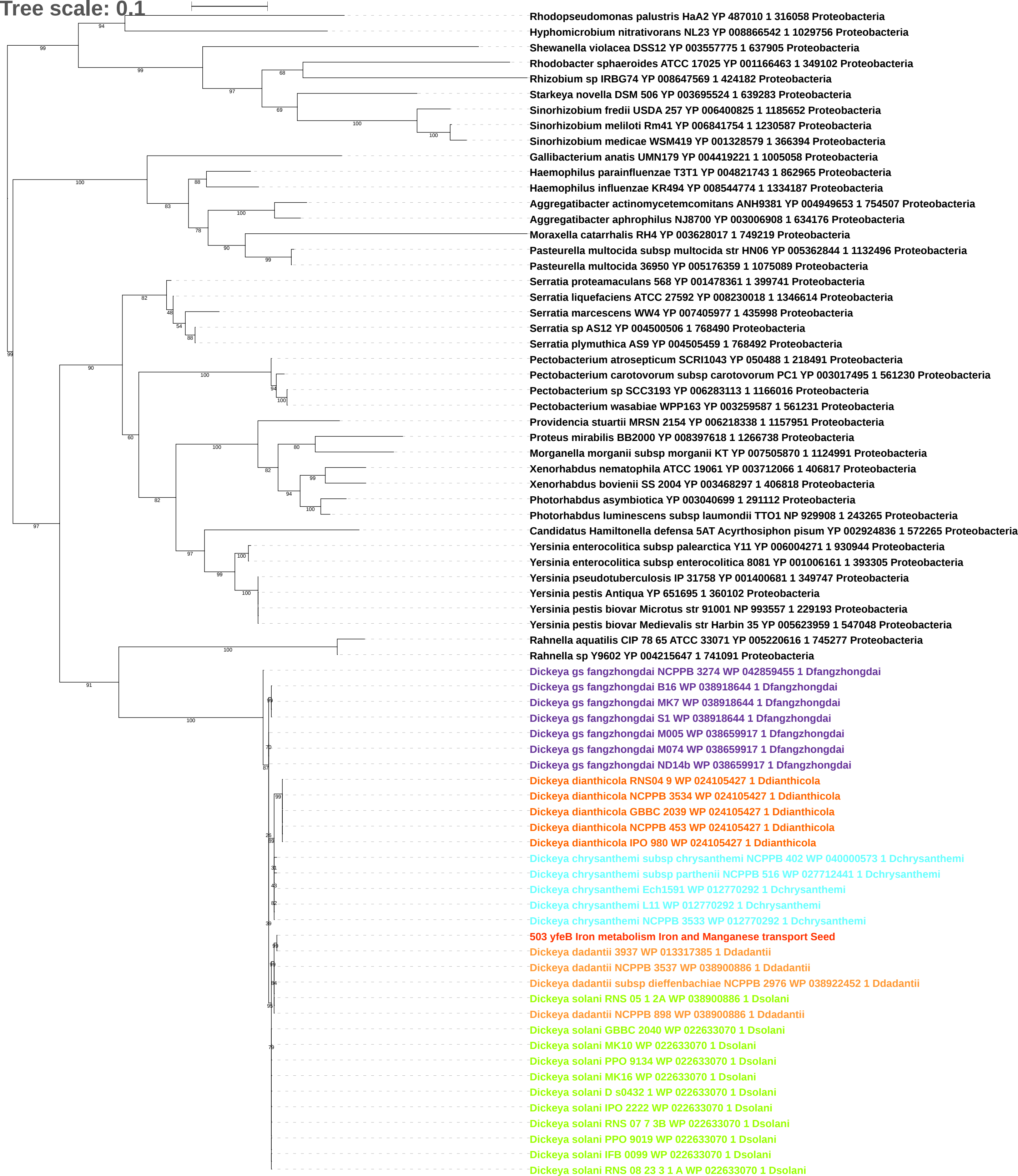
Tree scale: 0.1
Rhodopseudomonas palustris HaA2 YP 487010 1 316058 Proteobacteria
Hyphomicrobium nitrativorans NL23 YP 008866542 1 1029756 Proteobacteria
Shewanella violacea DSS12 YP 003557775 1 637905 Proteobacteria
Rhodobacter sphaeroides ATCC 17025 YP 001166463 1 349102 Proteobacteria
Rhizobium sp IRBG74 YP 008647569 1 424182 Proteobacteria
Starkeya novella DSM 506 YP 003695524 1 639283 Proteobacteria
Sinorhizobium fredii USDA 257 YP 006400825 1 1185652 Proteobacteria
Sinorhizobium meliloti Rm41 YP 006841754 1 1230587 Proteobacteria
Sinorhizobium medicae WSM419 YP 001328579 1 366394 Proteobacteria
Gallibacterium anatis UMN179 YP 004419221 1 1005058 Proteobacteria
Haemophilus parainfluenzae T3T1 YP 004821743 1 862965 Proteobacteria
Haemophilus influenzae KR494 YP 008544774 1 1334187 Proteobacteria
Aggregatibacter actinomycetemcomitans ANH9381 YP 004949653 1 754507 Proteobacteria
Aggregatibacter aphrophilus NJ8700 YP 003006908 1 634176 Proteobacteria
Moraxella catarrhalis RH4 YP 003628017 1 749219 Proteobacteria
Pasteurella multocida subsp multocida str HN06 YP 005362844 1 1132496 Proteobacteria
Pasteurella multocida 36950 YP 005176359 1 1075089 Proteobacteria
Serratia proteamaculans 568 YP 001478361 1 399741 Proteobacteria
Serratia liquefaciens ATCC 27592 YP 008230018 1 1346614 Proteobacteria
Serratia marcescens WW4 YP 007405977 1 435998 Proteobacteria
Serratia sp AS12 YP 004500506 1 768490 Proteobacteria
Serratia plymuthica AS9 YP 004505459 1 768492 Proteobacteria
Pectobacterium atrosepticum SCRI1043 YP 050488 1 218491 Proteobacteria
Pectobacterium carotovorum subsp carotovorum PC1 YP 003017495 1 561230 Proteobacteria
Pectobacterium sp SCC3193 YP 006283113 1 1166016 Proteobacteria
Pectobacterium wasabiae WPP163 YP 003259587 1 561231 Proteobacteria
Providencia stuartii MRSN 2154 YP 006218338 1 1157951 Proteobacteria
Proteus mirabilis BB2000 YP 008397618 1 1266738 Proteobacteria
Morganella morganii subsp morganii KT YP 007505870 1 1124991 Proteobacteria
Xenorhabdus nematophila ATCC 19061 YP 003712066 1 406817 Proteobacteria
Xenorhabdus bovienii SS 2004 YP 003468297 1 406818 Proteobacteria
Photorhabdus asymbiotica YP 003040699 1 291112 Proteobacteria
Photorhabdus luminescens subsp laumondii TTO1 NP 929908 1 243265 Proteobacteria
Candidatus Hamiltonella defensa 5AT Acyrthosiphon pisum YP 002924836 1 572265 Proteobacteria
Yersinia enterocolitica subsp palearctica Y11 YP 006004271 1 930944 Proteobacteria
Yersinia enterocolitica subsp enterocolitica 8081 YP 001006161 1 393305 Proteobacteria
Yersinia pseudotuberculosis IP 31758 YP 001400681 1 349747 Proteobacteria
Yersinia pestis Antiqua YP 651695 1 360102 Proteobacteria
Yersinia pestis biovar Microtus str 91001 NP 993557 1 229193 Proteobacteria
Yersinia pestis biovar Medievalis str Harbin 35 YP 005623959 1 547048 Proteobacteria
Rahnella aquatilis CIP 78 65 ATCC 33071 YP 005220616 1 745277 Proteobacteria
Rahnella sp Y9602 YP 004215647 1 741091 Proteobacteria
Dickeya gs fangzhongdai NCPPB 3274 WP 042859455 1 Dfangzhongdai
Dickeya gs fangzhongdai B16 WP 038918644 1 Dfangzhongdai
Dickeya gs fangzhongdai MK7 WP 038918644 1 Dfangzhongdai
Dickeya gs fangzhongdai S1 WP 038918644 1 Dfangzhongdai
Dickeya gs fangzhongdai M005 WP 038659917 1 Dfangzhongdai
Dickeya gs fangzhongdai M074 WP 038659917 1 Dfangzhongdai
Dickeya gs fangzhongdai ND14b WP 038659917 1 Dfangzhongdai
Dickeya dianthicola RNS04 9 WP 024105427 1 Ddianthicola
Dickeya dianthicola NCPPB 3534 WP 024105427 1 Ddianthicola
Dickeya dianthicola GBBC 2039 WP 024105427 1 Ddianthicola
Dickeya dianthicola NCPPB 453 WP 024105427 1 Ddianthicola
Dickeya dianthicola IPO 980 WP 024105427 1 Ddianthicola
Dickeya chrysanthemi subsp chrysanthemi NCPPB 402 WP 040000573 1 Dchrysanthemi
Dickeya chrysanthemi subsp parthenii NCPPB 516 WP 027712441 1 Dchrysanthemi
Dickeya chrysanthemi Ech1591 WP 012770292 1 Dchrysanthemi
Dickeya chrysanthemi L11 WP 012770292 1 Dchrysanthemi
Dickeya chrysanthemi NCPPB 3533 WP 012770292 1 Dchrysanthemi
503 yfeB Iron metabolism Iron and Manganese transport Seed
Dickeya dadantii 3937 WP 013317385 1 Ddadantii
Dickeya dadantii NCPPB 3537 WP 038900886 1 Ddadantii
Dickeya dadantii subsp dieffenbachiae NCPPB 2976 WP 038922452 1 Ddadantii
Dickeya solani RNS 05 1 2A WP 038900886 1 Dsolani
Dickeya dadantii NCPPB 898 WP 038900886 1 Ddadantii
Dickeya solani GBBC 2040 WP 022633070 1 Dsolani
Dickeya solani MK10 WP 022633070 1 Dsolani
Dickeya solani PPO 9134 WP 022633070 1 Dsolani
Dickeya solani MK16 WP 022633070 1 Dsolani
Dickeya solani D s0432 1 WP 022633070 1 Dsolani
Dickeya solani IPO 2222 WP 022633070 1 Dsolani
Dickeya solani RNS 07 7 3B WP 022633070 1 Dsolani
Dickeya solani PPO 9019 WP 022633070 1 Dsolani
Dickeya solani IFB 0099 WP 022633070 1 Dsolani
Dickeya solani RNS 08 23 3 1 A WP 022633070 1 Dsolani

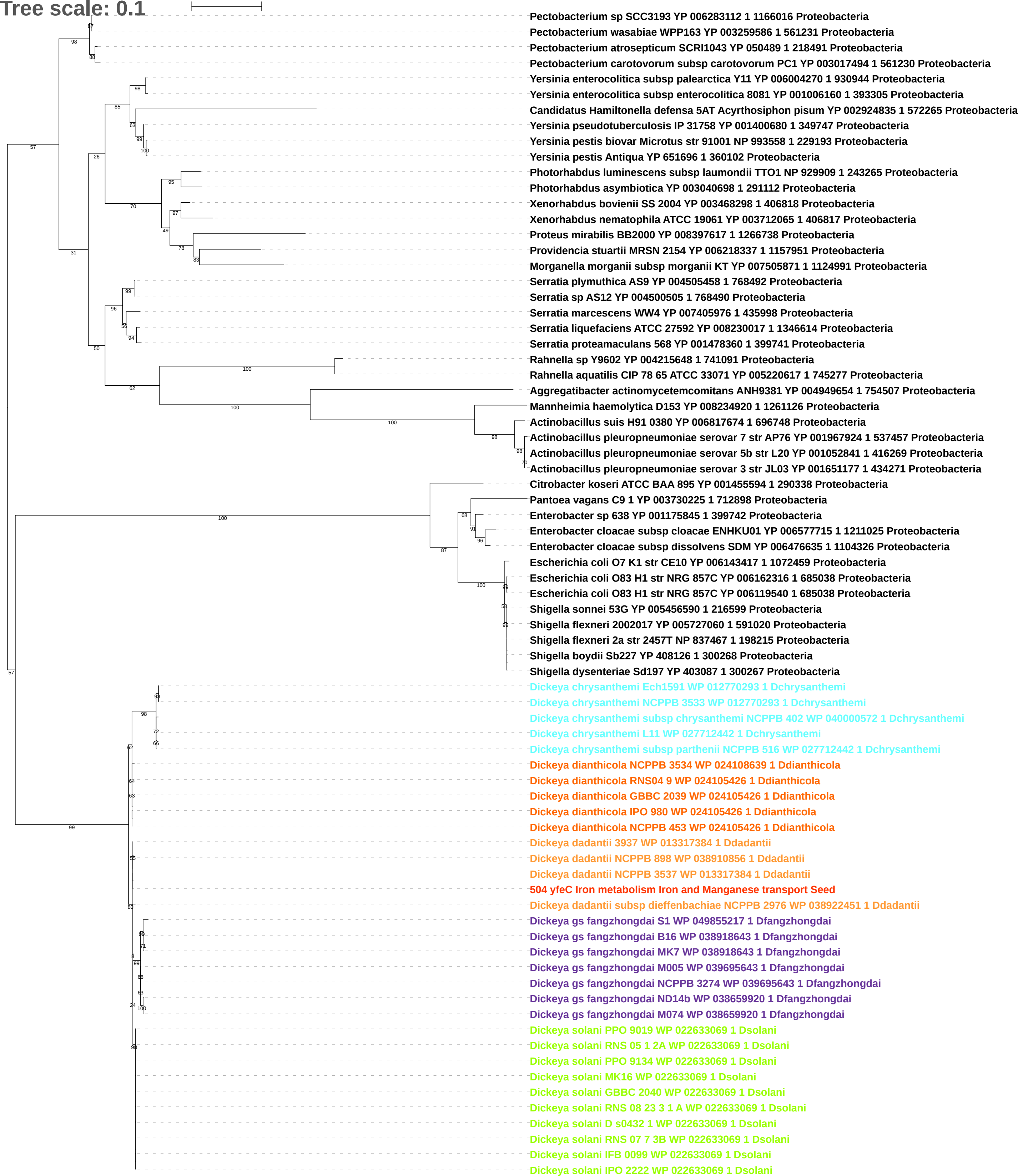
Tree scale: 0.1
Pectobacterium sp SCC3193 YP 006283112 1 1166016 Proteobacteria
Pectobacterium wasabiae WPP163 YP 003259586 1 561231 Proteobacteria
Pectobacterium atrosepticum SCRI1043 YP 050489 1 218491 Proteobacteria
Pectobacterium carotovorum subsp carotovorum PC1 YP 003017494 1 561230 Proteobacteria
Yersinia enterocolitica subsp palearctica Y11 YP 006004270 1 930944 Proteobacteria
Yersinia enterocolitica subsp enterocolitica 8081 YP 001006160 1 393305 Proteobacteria
Candidatus Hamiltonella defensa 5AT Acyrthosiphon pisum YP 002924835 1 572265 Proteobacteria
Yersinia pseudotuberculosis IP 31758 YP 001400680 1 349747 Proteobacteria
Yersinia pestis biovar Microtus str 91001 NP 993558 1 229193 Proteobacteria
Yersinia pestis Antiqua YP 651696 1 360102 Proteobacteria
Photorhabdus luminescens subsp laumondii TTO1 NP 929909 1 243265 Proteobacteria
Photorhabdus asymbiotica YP 003040698 1 291112 Proteobacteria
Xenorhabdus bovienii SS 2004 YP 003468298 1 406818 Proteobacteria
Xenorhabdus nematophila ATCC 19061 YP 003712065 1 406817 Proteobacteria
Proteus mirabilis BB2000 YP 008397617 1 1266738 Proteobacteria
Providencia stuartii MRSN 2154 YP 006218337 1 1157951 Proteobacteria
Morganella morganii subsp morganii KT YP 007505871 1 1124991 Proteobacteria
Serratia plymuthica AS9 YP 004505458 1 768492 Proteobacteria
Serratia sp AS12 YP 004500505 1 768490 Proteobacteria
Serratia marcescens WW4 YP 007405976 1 435998 Proteobacteria
Serratia liquefaciens ATCC 27592 YP 008230017 1 1346614 Proteobacteria
Serratia proteamaculans 568 YP 001478360 1 399741 Proteobacteria
Rahnella sp Y9602 YP 004215648 1 741091 Proteobacteria
Rahnella aquatilis CIP 78 65 ATCC 33071 YP 005220617 1 745277 Proteobacteria
Aggregatibacter actinomycetemcomitans ANH9381 YP 004949654 1 754507 Proteobacteria
Mannheimia haemolytica D153 YP 008234920 1 1261126 Proteobacteria
Actinobacillus suis H91 0380 YP 006817674 1 696748 Proteobacteria
Actinobacillus pleuropneumoniae serovar 7 str AP76 YP 001967924 1 537457 Proteobacteria
Actinobacillus pleuropneumoniae serovar 5b str L20 YP 001052841 1 416269 Proteobacteria
Actinobacillus pleuropneumoniae serovar 3 str JL03 YP 001651177 1 434271 Proteobacteria
Citrobacter koseri ATCC BAA 895 YP 001455594 1 290338 Proteobacteria
Pantoea vagans C9 1 YP 003730225 1 712898 Proteobacteria
Enterobacter sp 638 YP 001175845 1 399742 Proteobacteria
Enterobacter cloacae subsp cloacae ENHKU01 YP 006577715 1 1211025 Proteobacteria
Enterobacter cloacae subsp dissolvens SDM YP 006476635 1 1104326 Proteobacteria
Escherichia coli O7 K1 str CE10 YP 006143417 1 1072459 Proteobacteria
Escherichia coli O83 H1 str NRG 857C YP 006162316 1 685038 Proteobacteria
Escherichia coli O83 H1 str NRG 857C YP 006119540 1 685038 Proteobacteria
Shigella sonnei 53G YP 005456590 1 216599 Proteobacteria
Shigella flexneri 2002017 YP 005727060 1 591020 Proteobacteria
Shigella flexneri 2a str 2457T NP 837467 1 198215 Proteobacteria
Shigella boydii Sb227 YP 408126 1 300268 Proteobacteria
Shigella dysenteriae Sd197 YP 403087 1 300267 Proteobacteria
Dickeya chrysanthemi Ech1591 WP 012770293 1 Dchrysanthemi
Dickeya chrysanthemi NCPPB 3533 WP 012770293 1 Dchrysanthemi
Dickeya chrysanthemi subsp chrysanthemi NCPPB 402 WP 040000572 1 Dchrysanthemi
Dickeya chrysanthemi L11 WP 027712442 1 Dchrysanthemi
Dickeya chrysanthemi subsp parthenii NCPPB 516 WP 027712442 1 Dchrysanthemi
Dickeya dianthicola NCPPB 3534 WP 024108639 1 Ddianthicola
Dickeya dianthicola RNS04 9 WP 024105426 1 Ddianthicola
Dickeya dianthicola GBBC 2039 WP 024105426 1 Ddianthicola
Dickeya dianthicola IPO 980 WP 024105426 1 Ddianthicola
Dickeya dianthicola NCPPB 453 WP 024105426 1 Ddianthicola
Dickeya dadantii 3937 WP 013317384 1 Ddadantii
Dickeya dadantii NCPPB 898 WP 038910856 1 Ddadantii
Dickeya dadantii NCPPB 3537 WP 013317384 1 Ddadantii
504 yfeC Iron metabolism Iron and Manganese transport Seed
Dickeya dadantii subsp dieffenbachiae NCPPB 2976 WP 038922451 1 Ddadantii
Dickeya gs fangzhongdai S1 WP 049855217 1 Dfangzhongdai
Dickeya gs fangzhongdai B16 WP 038918643 1 Dfangzhongdai
Dickeya gs fangzhongdai MK7 WP 038918643 1 Dfangzhongdai
Dickeya gs fangzhongdai M005 WP 039695643 1 Dfangzhongdai
Dickeya gs fangzhongdai NCPPB 3274 WP 039695643 1 Dfangzhongdai
Dickeya gs fangzhongdai ND14b WP 038659920 1 Dfangzhongdai
Dickeya gs fangzhongdai M074 WP 038659920 1 Dfangzhongdai
Dickeya solani PPO 9019 WP 022633069 1 Dsolani
Dickeya solani RNS 05 1 2A WP 022633069 1 Dsolani
Dickeya solani PPO 9134 WP 022633069 1 Dsolani
Dickeya solani MK16 WP 022633069 1 Dsolani
Dickeya solani GBBC 2040 WP 022633069 1 Dsolani
Dickeya solani RNS 08 23 3 1 A WP 022633069 1 Dsolani
Dickeya solani D s0432 1 WP 022633069 1 Dsolani
Dickeya solani RNS 07 7 3B WP 022633069 1 Dsolani
Dickeya solani IFB 0099 WP 022633069 1 Dsolani
Dickeya solani IPO 2222 WP 022633069 1 Dsolani

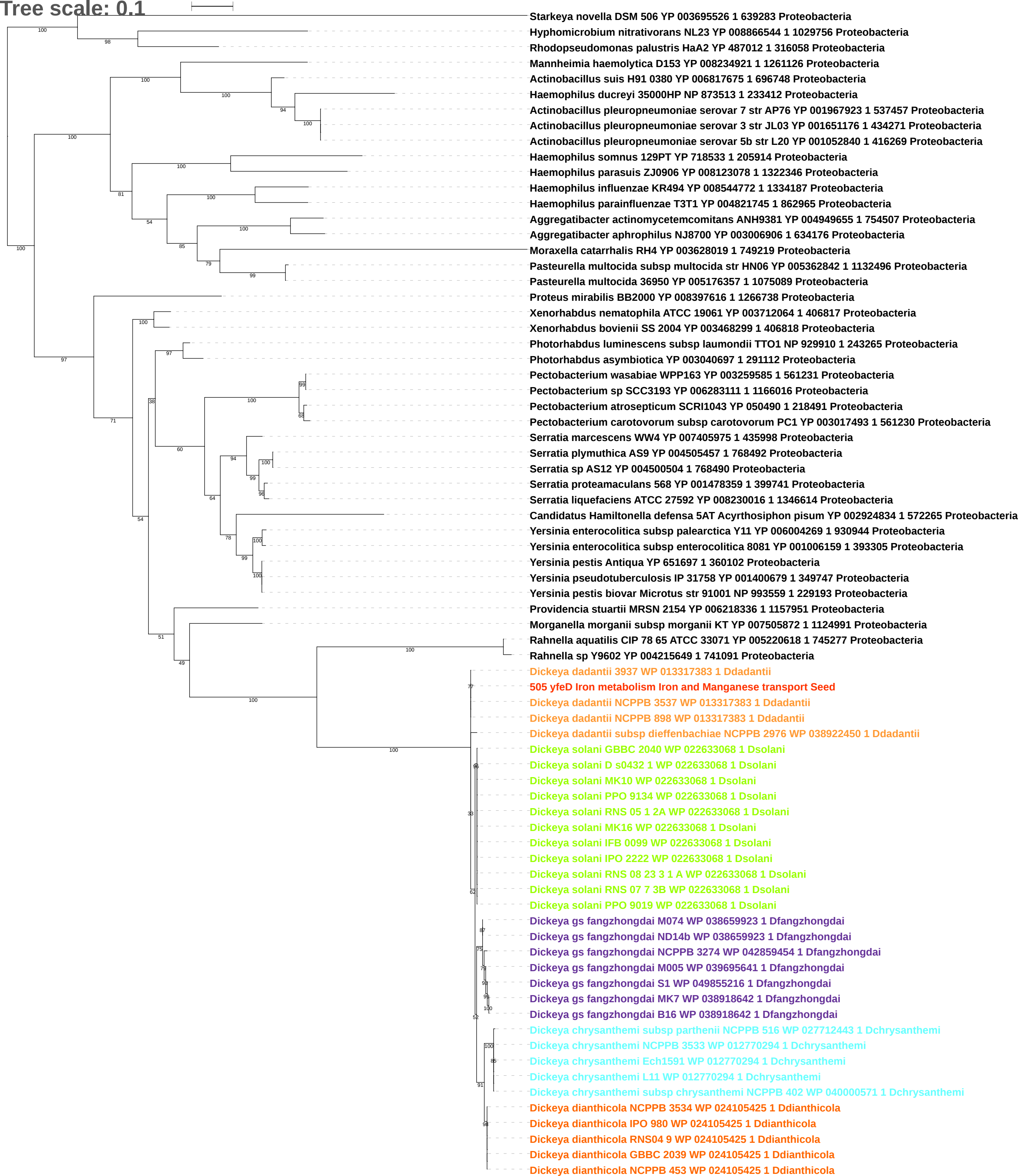
Tree scale: 0.1
Starkeya novella DSM 506 YP 003695526 1 639283 Proteobacteria
Hyphomicrobium nitrativorans NL23 YP 008866544 1 1029756 Proteobacteria
Rhodopseudomonas palustris HaA2 YP 487012 1 316058 Proteobacteria
Mannheimia haemolytica D153 YP 008234921 1 1261126 Proteobacteria
Actinobacillus suis H91 0380 YP 006817675 1 696748 Proteobacteria
Haemophilus ducreyi 35000HP NP 873513 1 233412 Proteobacteria
Actinobacillus pleuropneumoniae serovar 7 str AP76 YP 001967923 1 537457 Proteobacteria
Actinobacillus pleuropneumoniae serovar 3 str JL03 YP 001651176 1 434271 Proteobacteria
Actinobacillus pleuropneumoniae serovar 5b str L20 YP 001052840 1 416269 Proteobacteria
Haemophilus somnus 129PT YP 718533 1 205914 Proteobacteria
Haemophilus parasuis ZJ0906 YP 008123078 1 1322346 Proteobacteria
Haemophilus influenzae KR494 YP 008544772 1 1334187 Proteobacteria
Haemophilus parainfluenzae T3T1 YP 004821745 1 862965 Proteobacteria
Aggregatibacter actinomycetemcomitans ANH9381 YP 004949655 1 754507 Proteobacteria
Aggregatibacter aphrophilus NJ8700 YP 003006906 1 634176 Proteobacteria
Moraxella catarrhalis RH4 YP 003628019 1 749219 Proteobacteria
Pasteurella multocida subsp multocida str HN06 YP 005362842 1 1132496 Proteobacteria
Pasteurella multocida 36950 YP 005176357 1 1075089 Proteobacteria
Proteus mirabilis BB2000 YP 008397616 1 1266738 Proteobacteria
Xenorhabdus nematophila ATCC 19061 YP 003712064 1 406817 Proteobacteria
Xenorhabdus bovienii SS 2004 YP 003468299 1 406818 Proteobacteria
Photorhabdus luminescens subsp laumondii TTO1 NP 929910 1 243265 Proteobacteria
Photorhabdus asymbiotica YP 003040697 1 291112 Proteobacteria
Pectobacterium wasabiae WPP163 YP 003259585 1 561231 Proteobacteria
Pectobacterium sp SCC3193 YP 006283111 1 1166016 Proteobacteria
Pectobacterium atrosepticum SCRI1043 YP 050490 1 218491 Proteobacteria
Pectobacterium carotovorum subsp carotovorum PC1 YP 003017493 1 561230 Proteobacteria
Serratia marcescens WW4 YP 007405975 1 435998 Proteobacteria
Serratia plymuthica AS9 YP 004505457 1 768492 Proteobacteria
Serratia sp AS12 YP 004500504 1 768490 Proteobacteria
Serratia proteamaculans 568 YP 001478359 1 399741 Proteobacteria
Serratia liquefaciens ATCC 27592 YP 008230016 1 1346614 Proteobacteria
Candidatus Hamiltonella defensa 5AT Acyrthosiphon pisum YP 002924834 1 572265 Proteobacteria
Yersinia enterocolitica subsp palearctica Y11 YP 006004269 1 930944 Proteobacteria
Yersinia enterocolitica subsp enterocolitica 8081 YP 001006159 1 393305 Proteobacteria
Yersinia pestis Antiqua YP 651697 1 360102 Proteobacteria
Yersinia pseudotuberculosis IP 31758 YP 001400679 1 349747 Proteobacteria
Yersinia pestis biovar Microtus str 91001 NP 993559 1 229193 Proteobacteria
Providencia stuartii MRSN 2154 YP 006218336 1 1157951 Proteobacteria
Morganella morganii subsp morganii KT YP 007505872 1 1124991 Proteobacteria
Rahnella aquatilis CIP 78 65 ATCC 33071 YP 005220618 1 745277 Proteobacteria
Rahnella sp Y9602 YP 004215649 1 741091 Proteobacteria
Dickeya dadantii 3937 WP 013317383 1 Ddadantii
505 yfeD Iron metabolism Iron and Manganese transport Seed
Dickeya dadantii NCPPB 3537 WP 013317383 1 Ddadantii
Dickeya dadantii NCPPB 898 WP 013317383 1 Ddadantii
Dickeya dadantii subsp dieffenbachiae NCPPB 2976 WP 038922450 1 Ddadantii
Dickeya solani GBBC 2040 WP 022633068 1 Dsolani
Dickeya solani D s0432 1 WP 022633068 1 Dsolani
Dickeya solani MK10 WP 022633068 1 Dsolani
Dickeya solani PPO 9134 WP 022633068 1 Dsolani
Dickeya solani RNS 05 1 2A WP 022633068 1 Dsolani
Dickeya solani MK16 WP 022633068 1 Dsolani
Dickeya solani IFB 0099 WP 022633068 1 Dsolani
Dickeya solani IPO 2222 WP 022633068 1 Dsolani
Dickeya solani RNS 08 23 3 1 A WP 022633068 1 Dsolani
Dickeya solani RNS 07 7 3B WP 022633068 1 Dsolani
Dickeya solani PPO 9019 WP 022633068 1 Dsolani
Dickeya gs fangzhongdai M074 WP 038659923 1 Dfangzhongdai
Dickeya gs fangzhongdai ND14b WP 038659923 1 Dfangzhongdai
Dickeya gs fangzhongdai NCPPB 3274 WP 042859454 1 Dfangzhongdai
Dickeya gs fangzhongdai M005 WP 039695641 1 Dfangzhongdai
Dickeya gs fangzhongdai S1 WP 049855216 1 Dfangzhongdai
Dickeya gs fangzhongdai MK7 WP 038918642 1 Dfangzhongdai
Dickeya gs fangzhongdai B16 WP 038918642 1 Dfangzhongdai
Dickeya chrysanthemi subsp parthenii NCPPB 516 WP 027712443 1 Dchrysanthemi
Dickeya chrysanthemi NCPPB 3533 WP 012770294 1 Dchrysanthemi
Dickeya chrysanthemi Ech1591 WP 012770294 1 Dchrysanthemi
Dickeya chrysanthemi L11 WP 012770294 1 Dchrysanthemi
Dickeya chrysanthemi subsp chrysanthemi NCPPB 402 WP 040000571 1 Dchrysanthemi
Dickeya dianthicola NCPPB 3534 WP 024105425 1 Ddianthicola
Dickeya dianthicola IPO 980 WP 024105425 1 Ddianthicola
Dickeya dianthicola RNS04 9 WP 024105425 1 Ddianthicola
Dickeya dianthicola GBBC 2039 WP 024105425 1 Ddianthicola
Dickeya dianthicola NCPPB 453 WP 024105425 1 Ddianthicola

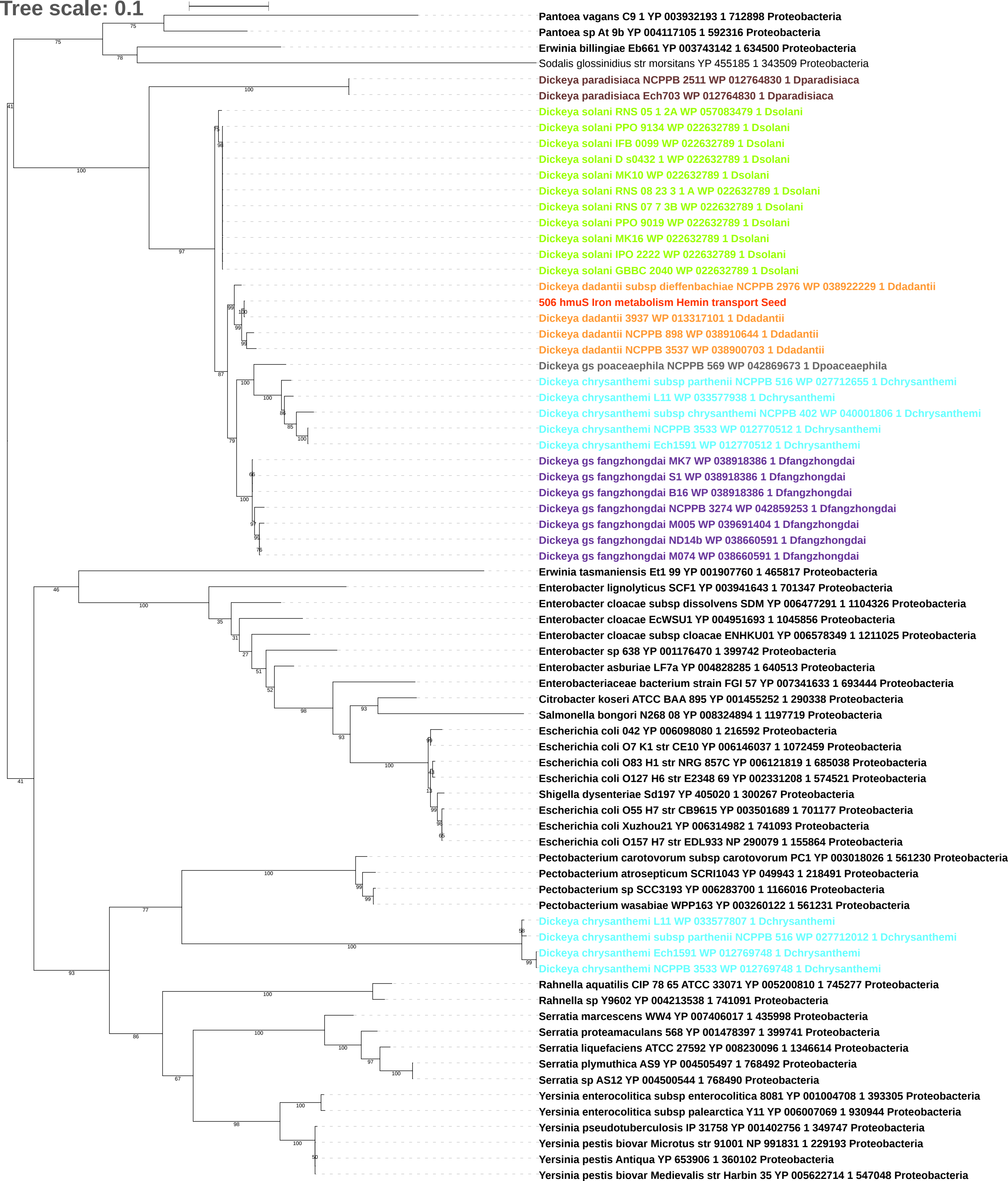
Tree scale: 0.1
Pantoea vagans C9 1 YP 003932193 1 712898 Proteobacteria
Pantoea sp At 9b YP 004117105 1 592316 Proteobacteria
Erwinia billingiae Eb661 YP 003743142 1 634500 Proteobacteria
Sodalis glossinidius str morsitans YP 455185 1 343509 Proteobacteria
Dickeya paradisiaca NCPPB 2511 WP 012764830 1 Dparadisiaca
Dickeya paradisiaca Ech703 WP 012764830 1 Dparadisiaca
Dickeya solani RNS 05 1 2A WP 057083479 1 Dsolani
Dickeya solani PPO 9134 WP 022632789 1 Dsolani
Dickeya solani IFB 0099 WP 022632789 1 Dsolani
Dickeya solani D s0432 1 WP 022632789 1 Dsolani
Dickeya solani MK10 WP 022632789 1 Dsolani
Dickeya solani RNS 08 23 3 1 A WP 022632789 1 Dsolani
Dickeya solani RNS 07 7 3B WP 022632789 1 Dsolani
Dickeya solani PPO 9019 WP 022632789 1 Dsolani
Dickeya solani MK16 WP 022632789 1 Dsolani
Dickeya solani IPO 2222 WP 022632789 1 Dsolani
Dickeya solani GBBC 2040 WP 022632789 1 Dsolani
Dickeya dadantii subsp dieffenbachiae NCPPB 2976 WP 038922229 1 Ddadantii
506 hmuS Iron metabolism Hemin transport Seed
Dickeya dadantii 3937 WP 013317101 1 Ddadantii
Dickeya dadantii NCPPB 898 WP 038910644 1 Ddadantii
Dickeya dadantii NCPPB 3537 WP 038900703 1 Ddadantii
Dickeya gs poaceaephila NCPPB 569 WP 042869673 1 Dpoaceaephila
Dickeya chrysanthemi subsp parthenii NCPPB 516 WP 027712655 1 Dchrysanthemi
Dickeya chrysanthemi L11 WP 033577938 1 Dchrysanthemi
Dickeya chrysanthemi subsp chrysanthemi NCPPB 402 WP 040001806 1 Dchrysanthemi
Dickeya chrysanthemi NCPPB 3533 WP 012770512 1 Dchrysanthemi
Dickeya chrysanthemi Ech1591 WP 012770512 1 Dchrysanthemi
Dickeya gs fangzhongdai MK7 WP 038918386 1 Dfangzhongdai
Dickeya gs fangzhongdai S1 WP 038918386 1 Dfangzhongdai
Dickeya gs fangzhongdai B16 WP 038918386 1 Dfangzhongdai
Dickeya gs fangzhongdai NCPPB 3274 WP 042859253 1 Dfangzhongdai
Dickeya gs fangzhongdai M005 WP 039691404 1 Dfangzhongdai
Dickeya gs fangzhongdai ND14b WP 038660591 1 Dfangzhongdai
Dickeya gs fangzhongdai M074 WP 038660591 1 Dfangzhongdai
Erwinia tasmaniensis Et1 99 YP 001907760 1 465817 Proteobacteria
Enterobacter lignolyticus SCF1 YP 003941643 1 701347 Proteobacteria
Enterobacter cloacae subsp dissolvens SDM YP 006477291 1 1104326 Proteobacteria
Enterobacter cloacae EcWSU1 YP 004951693 1 1045856 Proteobacteria
Enterobacter cloacae subsp cloacae ENHKU01 YP 006578349 1 1211025 Proteobacteria
Enterobacter sp 638 YP 001176470 1 399742 Proteobacteria
Enterobacter asburiae LF7a YP 004828285 1 640513 Proteobacteria
Enterobacteriaceae bacterium strain FGI 57 YP 007341633 1 693444 Proteobacteria
Citrobacter koseri ATCC BAA 895 YP 001455252 1 290338 Proteobacteria
Salmonella bongori N268 08 YP 008324894 1 1197719 Proteobacteria
Escherichia coli 042 YP 006098080 1 216592 Proteobacteria
Escherichia coli O7 K1 str CE10 YP 006146037 1 1072459 Proteobacteria
Escherichia coli O83 H1 str NRG 857C YP 006121819 1 685038 Proteobacteria
Escherichia coli O127 H6 str E2348 69 YP 002331208 1 574521 Proteobacteria
Shigella dysenteriae Sd197 YP 405020 1 300267 Proteobacteria
Escherichia coli O55 H7 str CB9615 YP 003501689 1 701177 Proteobacteria
Escherichia coli Xuzhou21 YP 006314982 1 741093 Proteobacteria
Escherichia coli O157 H7 str EDL933 NP 290079 1 155864 Proteobacteria
Pectobacterium carotovorum subsp carotovorum PC1 YP 003018026 1 561230 Proteobacteria
Pectobacterium atrosepticum SCRI1043 YP 049943 1 218491 Proteobacteria
Pectobacterium sp SCC3193 YP 006283700 1 1166016 Proteobacteria
Pectobacterium wasabiae WPP163 YP 003260122 1 561231 Proteobacteria
Dickeya chrysanthemi L11 WP 033577807 1 Dchrysanthemi
Dickeya chrysanthemi subsp parthenii NCPPB 516 WP 027712012 1 Dchrysanthemi
Dickeya chrysanthemi Ech1591 WP 012769748 1 Dchrysanthemi
Dickeya chrysanthemi NCPPB 3533 WP 012769748 1 Dchrysanthemi
Rahnella aquatilis CIP 78 65 ATCC 33071 YP 005200810 1 745277 Proteobacteria
Rahnella sp Y9602 YP 004213538 1 741091 Proteobacteria
Serratia marcescens WW4 YP 007406017 1 435998 Proteobacteria
Serratia proteamaculans 568 YP 001478397 1 399741 Proteobacteria
Serratia liquefaciens ATCC 27592 YP 008230096 1 1346614 Proteobacteria
Serratia plymuthica AS9 YP 004505497 1 768492 Proteobacteria
Serratia sp AS12 YP 004500544 1 768490 Proteobacteria
Yersinia enterocolitica subsp enterocolitica 8081 YP 001004708 1 393305 Proteobacteria
Yersinia enterocolitica subsp palearctica Y11 YP 006007069 1 930944 Proteobacteria
Yersinia pseudotuberculosis IP 31758 YP 001402756 1 349747 Proteobacteria
Yersinia pestis biovar Microtus str 91001 NP 991831 1 229193 Proteobacteria
Yersinia pestis Antiqua YP 653906 1 360102 Proteobacteria
Yersinia pestis biovar Medievalis str Harbin 35 YP 005622714 1 547048 Proteobacteria

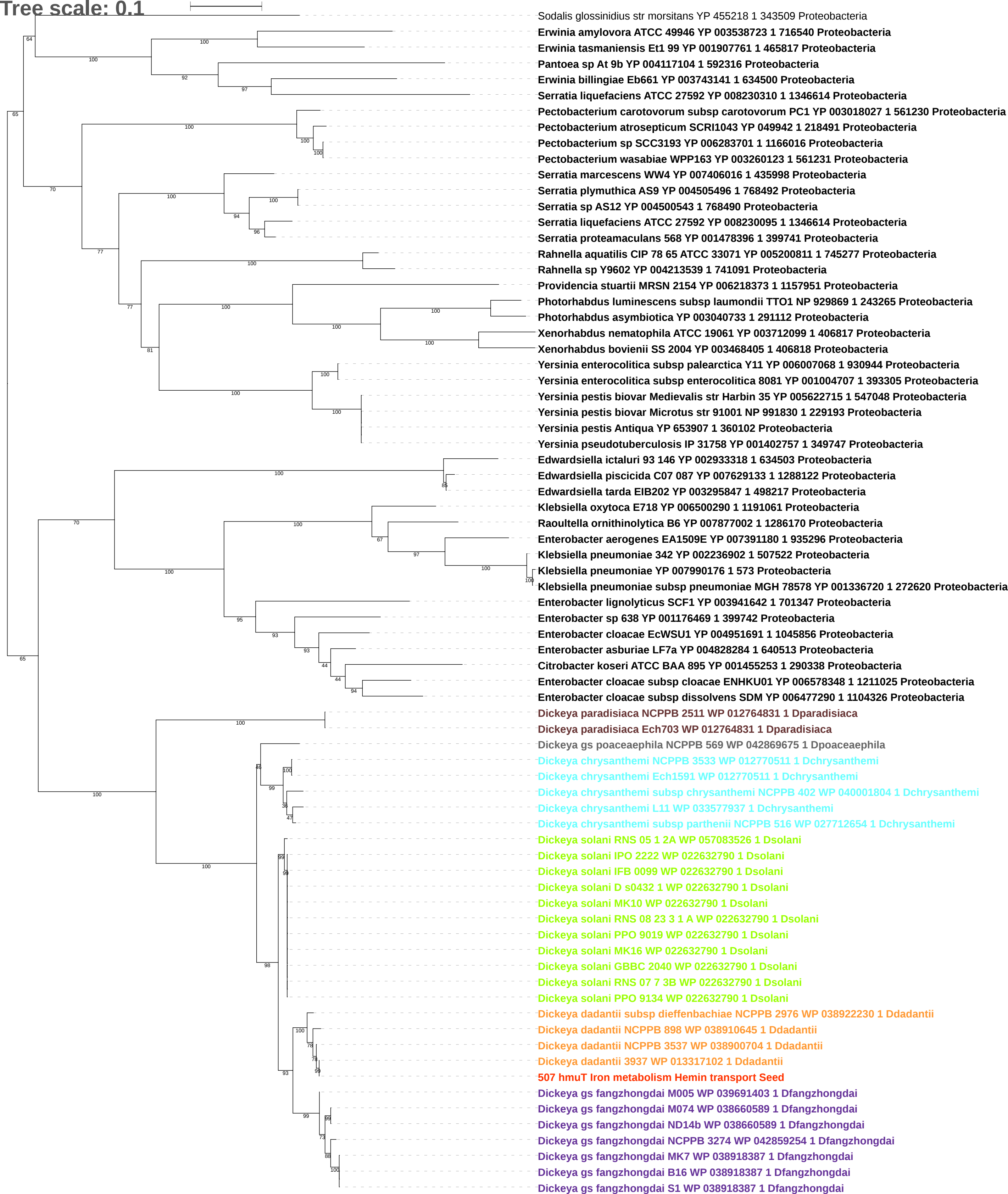

Tree scale: 0.1
Sodalis glossinidius str morsitans YP 455218 1 343509 Proteobacteria
Erwinia amylovora ATCC 49946 YP 003538723 1 716540 Proteobacteria
Erwinia tasmaniensis Et1 99 YP 001907761 1 465817 Proteobacteria
Pantoea sp At 9b YP 004117104 1 592316 Proteobacteria
Erwinia billingiae Eb661 YP 003743141 1 634500 Proteobacteria
Serratia liquefaciens ATCC 27592 YP 008230310 1 1346614 Proteobacteria
Pectobacterium carotovorum subsp carotovorum PC1 YP 003018027 1 561230 Proteobacteria
Pectobacterium atrosepticum SCRI1043 YP 049942 1 218491 Proteobacteria
Pectobacterium sp SCC3193 YP 006283701 1 1166016 Proteobacteria
Pectobacterium wasabiae WPP163 YP 003260123 1 561231 Proteobacteria
Serratia marcescens WW4 YP 007406016 1 435998 Proteobacteria
Serratia plymuthica AS9 YP 004505496 1 768492 Proteobacteria
Serratia sp AS12 YP 004500543 1 768490 Proteobacteria
Serratia liquefaciens ATCC 27592 YP 008230095 1 1346614 Proteobacteria
Serratia proteamaculans 568 YP 001478396 1 399741 Proteobacteria
Rahnella aquatilis CIP 78 65 ATCC 33071 YP 005200811 1 745277 Proteobacteria
Rahnella sp Y9602 YP 004213539 1 741091 Proteobacteria
Providencia stuartii MRSN 2154 YP 006218373 1 1157951 Proteobacteria
Photorhabdus luminescens subsp laumondii TTO1 NP 929869 1 243265 Proteobacteria
Photorhabdus asymbiotica YP 003040733 1 291112 Proteobacteria
Xenorhabdus nematophila ATCC 19061 YP 003712099 1 406817 Proteobacteria
Xenorhabdus bovienii SS 2004 YP 003468405 1 406818 Proteobacteria
Yersinia enterocolitica subsp palearctica Y11 YP 006007068 1 930944 Proteobacteria
Yersinia enterocolitica subsp enterocolitica 8081 YP 001004707 1 393305 Proteobacteria
Yersinia pestis biovar Medievalis str Harbin 35 YP 005622715 1 547048 Proteobacteria
Yersinia pestis biovar Microtus str 91001 NP 991830 1 229193 Proteobacteria
Yersinia pestis Antiqua YP 653907 1 360102 Proteobacteria
Yersinia pseudotuberculosis IP 31758 YP 001402757 1 349747 Proteobacteria
Edwardsiella ictaluri 93 146 YP 002933318 1 634503 Proteobacteria
Edwardsiella piscicida C07 087 YP 007629133 1 1288122 Proteobacteria
Edwardsiella tarda EIB202 YP 003295847 1 498217 Proteobacteria
Klebsiella oxytoca E718 YP 006500290 1 1191061 Proteobacteria
Raoultella ornithinolytica B6 YP 007877002 1 1286170 Proteobacteria
Enterobacter aerogenes EA1509E YP 007391180 1 935296 Proteobacteria
Klebsiella pneumoniae 342 YP 002236902 1 507522 Proteobacteria
Klebsiella pneumoniae YP 007990176 1 573 Proteobacteria
Klebsiella pneumoniae subsp pneumoniae MGH 78578 YP 001336720 1 272620 Proteobacteria
Enterobacter lignolyticus SCF1 YP 003941642 1 701347 Proteobacteria
Enterobacter sp 638 YP 001176469 1 399742 Proteobacteria
Enterobacter cloacae EcWSU1 YP 004951691 1 1045856 Proteobacteria
Enterobacter asburiae LF7a YP 004828284 1 640513 Proteobacteria
Citrobacter koseri ATCC BAA 895 YP 001455253 1 290338 Proteobacteria
Enterobacter cloacae subsp cloacae ENHKU01 YP 006578348 1 1211025 Proteobacteria
Enterobacter cloacae subsp dissolvens SDM YP 006477290 1 1104326 Proteobacteria
Dickeya paradisiaca NCPPB 2511 WP 012764831 1 Dparadisiaca
Dickeya paradisiaca Ech703 WP 012764831 1 Dparadisiaca
Dickeya gs poaceaephila NCPPB 569 WP 042869675 1 Dpoaceaephila
Dickeya chrysanthemi NCPPB 3533 WP 012770511 1 Dchrysanthemi
Dickeya chrysanthemi Ech1591 WP 012770511 1 Dchrysanthemi
Dickeya chrysanthemi subsp chrysanthemi NCPPB 402 WP 040001804 1 Dchrysanthemi
Dickeya chrysanthemi L11 WP 033577937 1 Dchrysanthemi
Dickeya chrysanthemi subsp parthenii NCPPB 516 WP 027712654 1 Dchrysanthemi
Dickeya solani RNS 05 1 2A WP 057083526 1 Dsolani
Dickeya solani IPO 2222 WP 022632790 1 Dsolani
Dickeya solani IFB 0099 WP 022632790 1 Dsolani
Dickeya solani D s0432 1 WP 022632790 1 Dsolani
Dickeya solani MK10 WP 022632790 1 Dsolani
Dickeya solani RNS 08 23 3 1 A WP 022632790 1 Dsolani
Dickeya solani PPO 9019 WP 022632790 1 Dsolani
Dickeya solani MK16 WP 022632790 1 Dsolani
Dickeya solani GBBC 2040 WP 022632790 1 Dsolani
Dickeya solani RNS 07 7 3B WP 022632790 1 Dsolani
Dickeya solani PPO 9134 WP 022632790 1 Dsolani
Dickeya dadantii subsp dieffenbachiae NCPPB 2976 WP 038922230 1 Ddadantii
Dickeya dadantii NCPPB 898 WP 038910645 1 Ddadantii
Dickeya dadantii NCPPB 3537 WP 038900704 1 Ddadantii
Dickeya dadantii 3937 WP 013317102 1 Ddadantii
507 hmuT Iron metabolism Hemin transport Seed
Dickeya gs fangzhongdai M005 WP 039691403 1 Dfangzhongdai
Dickeya gs fangzhongdai M074 WP 038660589 1 Dfangzhongdai
Dickeya gs fangzhongdai ND14b WP 038660589 1 Dfangzhongdai
Dickeya gs fangzhongdai NCPPB 3274 WP 042859254 1 Dfangzhongdai
Dickeya gs fangzhongdai MK7 WP 038918387 1 Dfangzhongdai
Dickeya gs fangzhongdai B16 WP 038918387 1 Dfangzhongdai
Dickeya gs fangzhongdai S1 WP 038918387 1 Dfangzhongdai

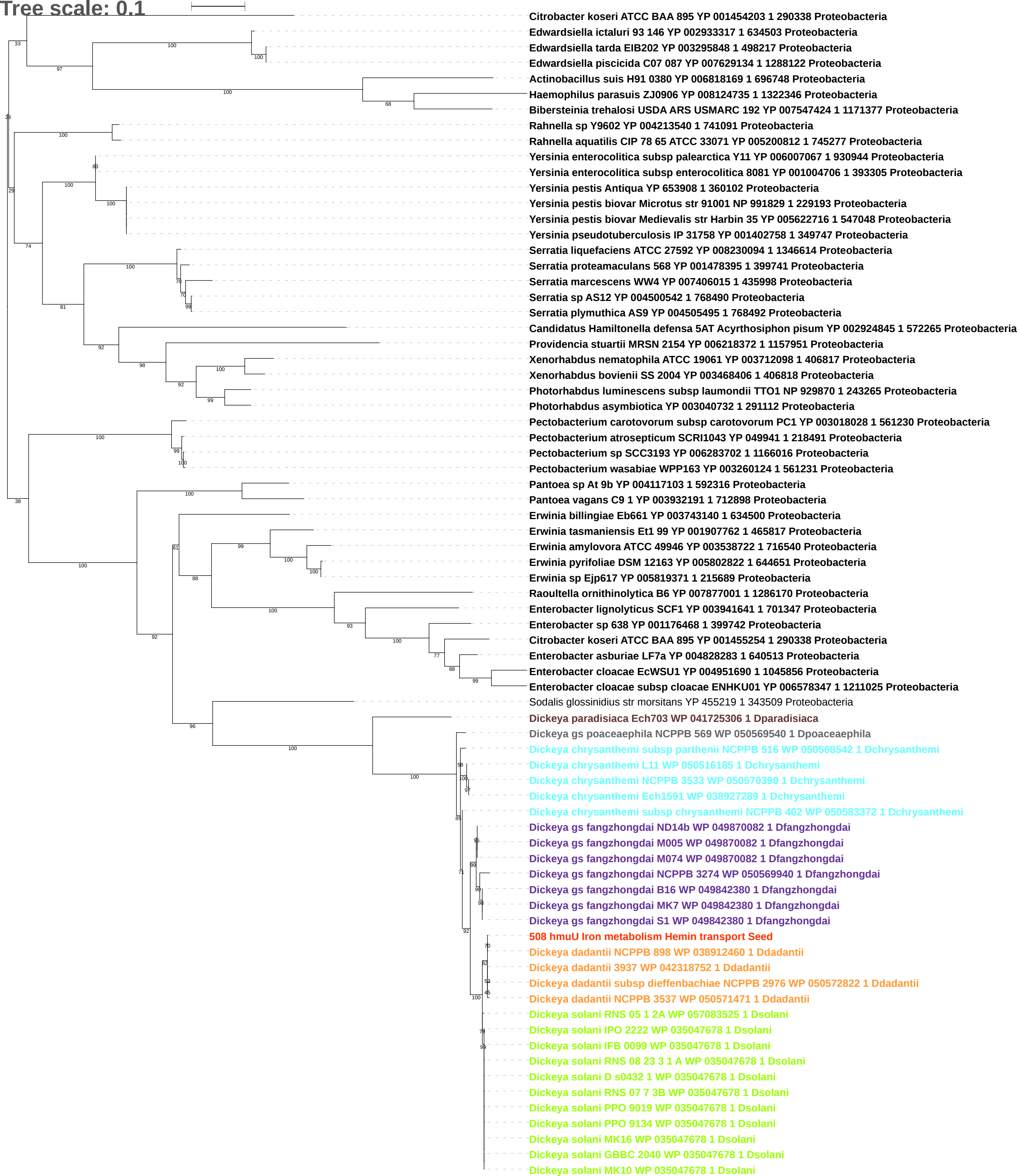
Tree scale: 0.1
Citrobacter koseri ATCC BAA 895 YP 001454203 1 290338 Proteobacteria
Edwardsiella ictaluri 93 146 YP 002933317 1 634503 Proteobacteria
Edwardsiella tarda EIB202 YP 003295848 1 498217 Proteobacteria
Edwardsiella piscicida C07 087 YP 007629134 1 1288122 Proteobacteria
Actinobacillus suis H91 0380 YP 006818169 1 696748 Proteobacteria
Haemophilus parasuis ZJ0906 YP 008124735 1 1322346 Proteobacteria
Bibersteinia trehalosi USDA ARS USMARC 192 YP 007547424 1 1171377 Proteobacteria
Rahnella sp Y9602 YP 004213540 1 741091 Proteobacteria
Rahnella aquatilis CIP 78 65 ATCC 33071 YP 005200812 1 745277 Proteobacteria
Yersinia enterocolitica subsp palearctica Y11 YP 006007067 1 930944 Proteobacteria
Yersinia enterocolitica subsp enterocolitica 8081 YP 001004706 1 393305 Proteobacteria
Yersinia pestis Antiqua YP 653908 1 360102 Proteobacteria
Yersinia pestis biovar Microtus str 91001 NP 991829 1 229193 Proteobacteria
Yersinia pestis biovar Medievalis str Harbin 35 YP 005622716 1 547048 Proteobacteria
Yersinia pseudotuberculosis IP 31758 YP 001402758 1 349747 Proteobacteria
Serratia liquefaciens ATCC 27592 YP 008230094 1 1346614 Proteobacteria
Serratia proteamaculans 568 YP 001478395 1 399741 Proteobacteria
Serratia marcescens WW4 YP 007406015 1 435998 Proteobacteria
Serratia sp AS12 YP 004500542 1 768490 Proteobacteria
Serratia plymuthica AS9 YP 004505495 1 768492 Proteobacteria
Candidatus Hamiltonella defensa 5AT Acyrthosiphon pisum YP 002924845 1 572265 Proteobacteria
Providencia stuartii MRSN 2154 YP 006218372 1 1157951 Proteobacteria
Xenorhabdus nematophila ATCC 19061 YP 003712098 1 406817 Proteobacteria
Xenorhabdus bovienii SS 2004 YP 003468406 1 406818 Proteobacteria
Photorhabdus luminescens subsp laumondii TTO1 NP 929870 1 243265 Proteobacteria
Photorhabdus asymbiotica YP 003040732 1 291112 Proteobacteria
Pectobacterium carotovorum subsp carotovorum PC1 YP 003018028 1 561230 Proteobacteria
Pectobacterium atrosepticum SCRI1043 YP 049941 1 218491 Proteobacteria
Pectobacterium sp SCC3193 YP 006283702 1 1166016 Proteobacteria
Pectobacterium wasabiae WPP163 YP 003260124 1 561231 Proteobacteria
Pantoea sp At 9b YP 004117103 1 592316 Proteobacteria
Pantoea vagans C9 1 YP 003932191 1 712898 Proteobacteria
Erwinia billingiae Eb661 YP 003743140 1 634500 Proteobacteria
Erwinia tasmaniensis Et1 99 YP 001907762 1 465817 Proteobacteria
Erwinia amylovora ATCC 49946 YP 003538722 1 716540 Proteobacteria
Erwinia pyrifoliae DSM 12163 YP 005802822 1 644651 Proteobacteria
Erwinia sp Ejp617 YP 005819371 1 215689 Proteobacteria
Raoultella ornithinolytica B6 YP 007877001 1 1286170 Proteobacteria
Enterobacter lignolyticus SCF1 YP 003941641 1 701347 Proteobacteria
Enterobacter sp 638 YP 001176468 1 399742 Proteobacteria
Citrobacter koseri ATCC BAA 895 YP 001455254 1 290338 Proteobacteria
Enterobacter asburiae LF7a YP 004828283 1 640513 Proteobacteria
Enterobacter cloacae EcWSU1 YP 004951690 1 1045856 Proteobacteria
Enterobacter cloacae subsp cloacae ENHKU01 YP 006578347 1 1211025 Proteobacteria
Sodalis glossinidius str morsitans YP 455219 1 343509 Proteobacteria
Dickeya paradisiaca Ech703 WP 041725306 1 Dparadisiaca
Dickeya gs poaceaephila NCPPB 569 WP 050569540 1 Dpoaceaephila
Dickeya chrysanthemi subsp parthenii NCPPB 516 WP 050568542 1 Dchrysanthemi
Dickeya chrysanthemi L11 WP 050516185 1 Dchrysanthemi
Dickeya chrysanthemi NCPPB 3533 WP 050570390 1 Dchrysanthemi
Dickeya chrysanthemi Ech1591 WP 038927289 1 Dchrysanthemi
Dickeya chrysanthemi subsp chrysanthemi NCPPB 402 WP 050583372 1 Dchrysanthemi
Dickeya gs fangzhongdai ND14b WP 049870082 1 Dfangzhongdai
Dickeya gs fangzhongdai M005 WP 049870082 1 Dfangzhongdai
Dickeya gs fangzhongdai M074 WP 049870082 1 Dfangzhongdai
Dickeya gs fangzhongdai NCPPB 3274 WP 050569940 1 Dfangzhongdai
Dickeya gs fangzhongdai B16 WP 049842380 1 Dfangzhongdai
Dickeya gs fangzhongdai MK7 WP 049842380 1 Dfangzhongdai
Dickeya gs fangzhongdai S1 WP 049842380 1 Dfangzhongdai
508 hmuU Iron metabolism Hemin transport Seed
Dickeya dadantii NCPPB 898 WP 038912460 1 Ddadantii
Dickeya dadantii 3937 WP 042318752 1 Ddadantii
Dickeya dadantii subsp dieffenbachiae NCPPB 2976 WP 050572822 1 Ddadantii
Dickeya dadantii NCPPB 3537 WP 050571471 1 Ddadantii
Dickeya solani RNS 05 1 2A WP 057083525 1 Dsolani
Dickeya solani IPO 2222 WP 035047678 1 Dsolani
Dickeya solani IFB 0099 WP 035047678 1 Dsolani
Dickeya solani RNS 08 23 3 1 A WP 035047678 1 Dsolani
Dickeya solani D s0432 1 WP 035047678 1 Dsolani
Dickeya solani RNS 07 7 3B WP 035047678 1 Dsolani
Dickeya solani PPO 9019 WP 035047678 1 Dsolani
Dickeya solani PPO 9134 WP 035047678 1 Dsolani
Dickeya solani MK16 WP 035047678 1 Dsolani
Dickeya solani GBBC 2040 WP 035047678 1 Dsolani
Dickeya solani MK10 WP 035047678 1 Dsolani

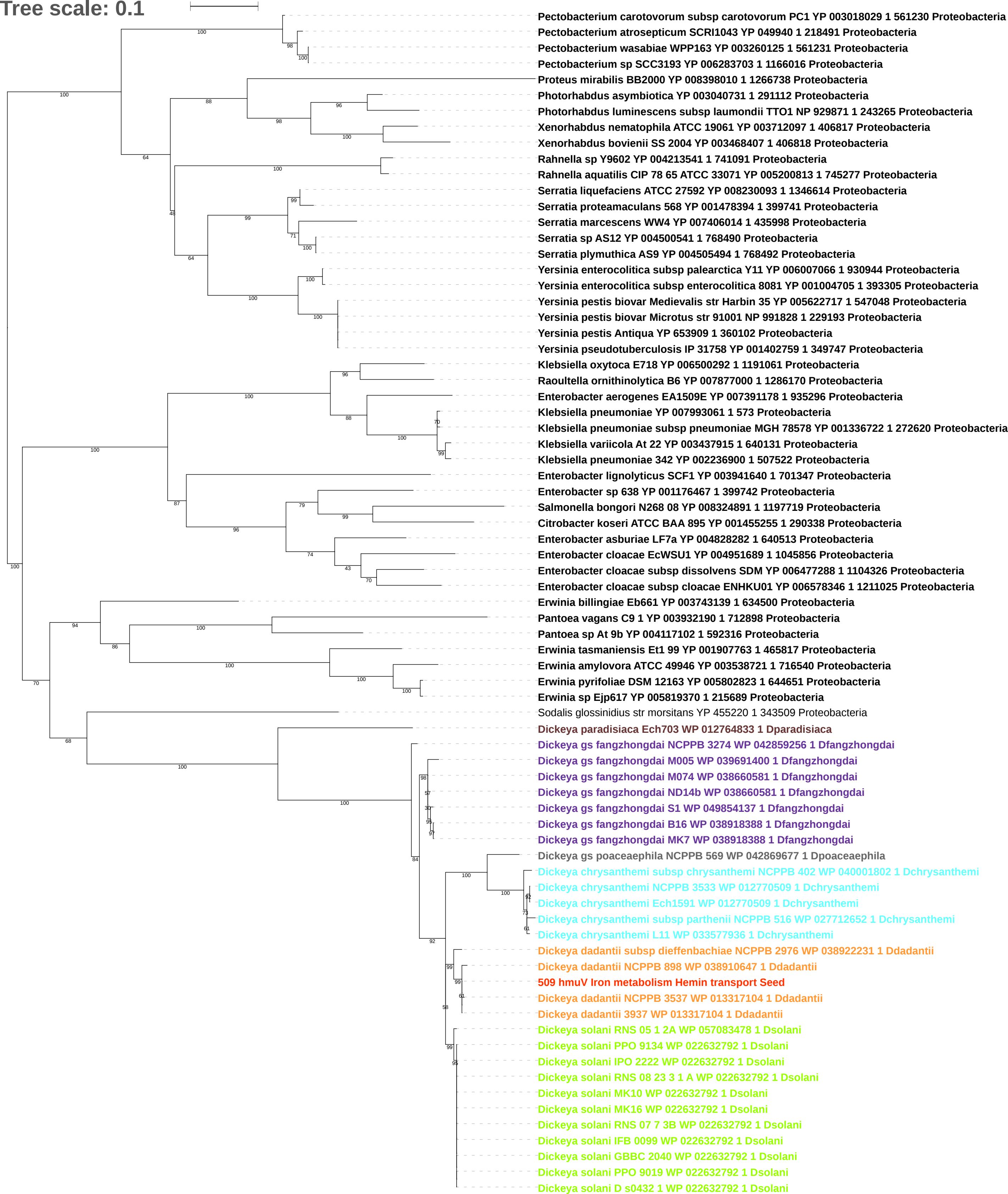
Tree scale: 0.1
Pectobacterium carotovorum subsp carotovorum PC1 YP 003018029 1 561230 Proteobacteria
Pectobacterium atrosepticum SCRI1043 YP 049940 1 218491 Proteobacteria
Pectobacterium wasabiae WPP163 YP 003260125 1 561231 Proteobacteria
Pectobacterium sp SCC3193 YP 006283703 1 1166016 Proteobacteria
Proteus mirabilis BB2000 YP 008398010 1 1266738 Proteobacteria
Photorhabdus asymbiotica YP 003040731 1 291112 Proteobacteria
Photorhabdus luminescens subsp laumondii TTO1 NP 929871 1 243265 Proteobacteria
Xenorhabdus nematophila ATCC 19061 YP 003712097 1 406817 Proteobacteria
Xenorhabdus bovienii SS 2004 YP 003468407 1 406818 Proteobacteria
Rahnella sp Y9602 YP 004213541 1 741091 Proteobacteria
Rahnella aquatilis CIP 78 65 ATCC 33071 YP 005200813 1 745277 Proteobacteria
Serratia liquefaciens ATCC 27592 YP 008230093 1 1346614 Proteobacteria
Serratia proteamaculans 568 YP 001478394 1 399741 Proteobacteria
Serratia marcescens WW4 YP 007406014 1 435998 Proteobacteria
Serratia sp AS12 YP 004500541 1 768490 Proteobacteria
Serratia plymuthica AS9 YP 004505494 1 768492 Proteobacteria
Yersinia enterocolitica subsp palearctica Y11 YP 006007066 1 930944 Proteobacteria
Yersinia enterocolitica subsp enterocolitica 8081 YP 001004705 1 393305 Proteobacteria
Yersinia pestis biovar Medievalis str Harbin 35 YP 005622717 1 547048 Proteobacteria
Yersinia pestis biovar Microtus str 91001 NP 991828 1 229193 Proteobacteria
Yersinia pestis Antiqua YP 653909 1 360102 Proteobacteria
Yersinia pseudotuberculosis IP 31758 YP 001402759 1 349747 Proteobacteria
Klebsiella oxytoca E718 YP 006500292 1 1191061 Proteobacteria
Raoultella ornithinolytica B6 YP 007877000 1 1286170 Proteobacteria
Enterobacter aerogenes EA1509E YP 007391178 1 935296 Proteobacteria
Klebsiella pneumoniae YP 007993061 1 573 Proteobacteria
Klebsiella pneumoniae subsp pneumoniae MGH 78578 YP 001336722 1 272620 Proteobacteria
Klebsiella variicola At 22 YP 003437915 1 640131 Proteobacteria
Klebsiella pneumoniae 342 YP 002236900 1 507522 Proteobacteria
Enterobacter lignolyticus SCF1 YP 003941640 1 701347 Proteobacteria
Enterobacter sp 638 YP 001176467 1 399742 Proteobacteria
Salmonella bongori N268 08 YP 008324891 1 1197719 Proteobacteria
Citrobacter koseri ATCC BAA 895 YP 001455255 1 290338 Proteobacteria
Enterobacter asburiae LF7a YP 004828282 1 640513 Proteobacteria
Enterobacter cloacae EcWSU1 YP 004951689 1 1045856 Proteobacteria
Enterobacter cloacae subsp dissolvens SDM YP 006477288 1 1104326 Proteobacteria
Enterobacter cloacae subsp cloacae ENHKU01 YP 006578346 1 1211025 Proteobacteria
Erwinia billingiae Eb661 YP 003743139 1 634500 Proteobacteria
Pantoea vagans C9 1 YP 003932190 1 712898 Proteobacteria
Pantoea sp At 9b YP 004117102 1 592316 Proteobacteria
Erwinia tasmaniensis Et1 99 YP 001907763 1 465817 Proteobacteria
Erwinia amylovora ATCC 49946 YP 003538721 1 716540 Proteobacteria
Erwinia pyrifoliae DSM 12163 YP 005802823 1 644651 Proteobacteria
Erwinia sp Ejp617 YP 005819370 1 215689 Proteobacteria
Sodalis glossinidius str morsitans YP 455220 1 343509 Proteobacteria
Dickeya paradisiaca Ech703 WP 012764833 1 Dparadisiaca
Dickeya gs fangzhongdai NCPPB 3274 WP 042859256 1 Dfangzhongdai
Dickeya gs fangzhongdai M005 WP 039691400 1 Dfangzhongdai
Dickeya gs fangzhongdai M074 WP 038660581 1 Dfangzhongdai
Dickeya gs fangzhongdai ND14b WP 038660581 1 Dfangzhongdai
Dickeya gs fangzhongdai S1 WP 049854137 1 Dfangzhongdai
Dickeya gs fangzhongdai B16 WP 038918388 1 Dfangzhongdai
Dickeya gs fangzhongdai MK7 WP 038918388 1 Dfangzhongdai
Dickeya gs poaceaephila NCPPB 569 WP 042869677 1 Dpoaceaephila
Dickeya chrysanthemi subsp chrysanthemi NCPPB 402 WP 040001802 1 Dchrysanthemi
Dickeya chrysanthemi NCPPB 3533 WP 012770509 1 Dchrysanthemi
Dickeya chrysanthemi Ech1591 WP 012770509 1 Dchrysanthemi
Dickeya chrysanthemi subsp parthenii NCPPB 516 WP 027712652 1 Dchrysanthemi
Dickeya chrysanthemi L11 WP 033577936 1 Dchrysanthemi
Dickeya dadantii subsp dieffenbachiae NCPPB 2976 WP 038922231 1 Ddadantii
Dickeya dadantii NCPPB 898 WP 038910647 1 Ddadantii
509 hmuV Iron metabolism Hemin transport Seed
Dickeya dadantii NCPPB 3537 WP 013317104 1 Ddadantii
Dickeya dadantii 3937 WP 013317104 1 Ddadantii
Dickeya solani RNS 05 1 2A WP 057083478 1 Dsolani
Dickeya solani PPO 9134 WP 022632792 1 Dsolani
Dickeya solani IPO 2222 WP 022632792 1 Dsolani
Dickeya solani RNS 08 23 3 1 A WP 022632792 1 Dsolani
Dickeya solani MK10 WP 022632792 1 Dsolani
Dickeya solani MK16 WP 022632792 1 Dsolani
Dickeya solani RNS 07 7 3B WP 022632792 1 Dsolani
Dickeya solani IFB 0099 WP 022632792 1 Dsolani
Dickeya solani GBBC 2040 WP 022632792 1 Dsolani
Dickeya solani PPO 9019 WP 022632792 1 Dsolani
Dickeya solani D s0432 1 WP 022632792 1 Dsolani

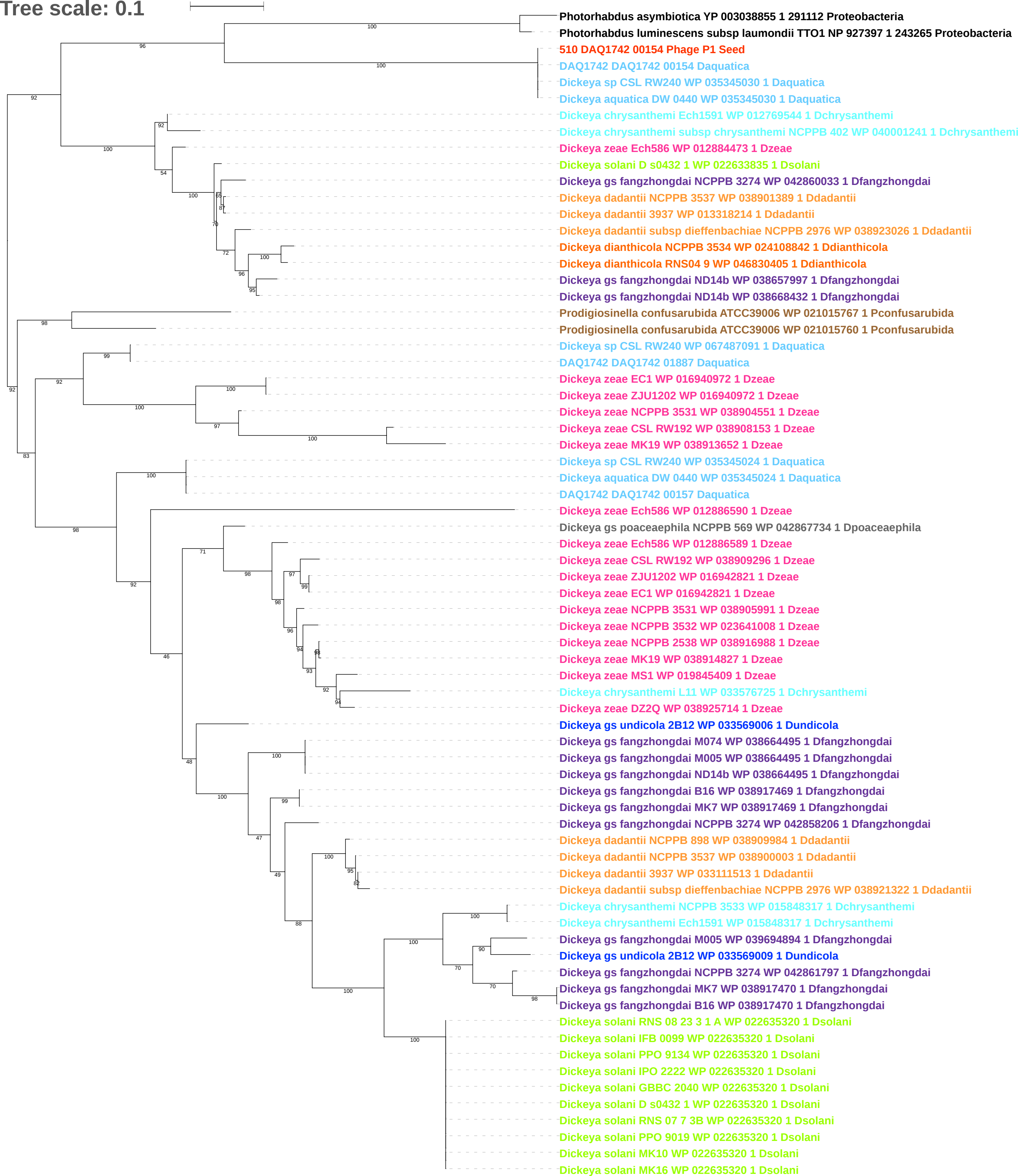
Tree scale: 0.1
Photorhabdus asymbiotica YP 003038855 1 291112 Proteobacteria
Photorhabdus luminescens subsp laumondii TTO1 NP 927397 1 243265 Proteobacteria
510 DAQ1742 00154 Phage P1 Seed
DAQ1742 DAQ1742 00154 Daquatica
Dickeya sp CSL RW240 WP 035345030 1 Daquatica
Dickeya aquatica DW 0440 WP 035345030 1 Daquatica
Dickeya chrysanthemi Ech1591 WP 012769544 1 Dchrysanthemi
Dickeya chrysanthemi subsp chrysanthemi NCPPB 402 WP 040001241 1 Dchrysanthemi
Dickeya zeae Ech586 WP 012884473 1 Dzeae
Dickeya solani D s0432 1 WP 022633835 1 Dsolani
Dickeya gs fangzhongdai NCPPB 3274 WP 042860033 1 Dfangzhongdai
Dickeya dadantii NCPPB 3537 WP 038901389 1 Ddadantii
Dickeya dadantii 3937 WP 013318214 1 Ddadantii
Dickeya dadantii subsp dieffenbachiae NCPPB 2976 WP 038923026 1 Ddadantii
Dickeya dianthicola NCPPB 3534 WP 024108842 1 Ddianthicola
Dickeya dianthicola RNS04 9 WP 046830405 1 Ddianthicola
Dickeya gs fangzhongdai ND14b WP 038657997 1 Dfangzhongdai
Dickeya gs fangzhongdai ND14b WP 038668432 1 Dfangzhongdai
Prodigiosinella confusarubida ATCC39006 WP 021015767 1 Pconfusarubida
Prodigiosinella confusarubida ATCC39006 WP 021015760 1 Pconfusarubida
Dickeya sp CSL RW240 WP 067487091 1 Daquatica
DAQ1742 DAQ1742 01887 Daquatica
Dickeya zeae EC1 WP 016940972 1 Dzeae
Dickeya zeae ZJU1202 WP 016940972 1 Dzeae
Dickeya zeae NCPPB 3531 WP 038904551 1 Dzeae
Dickeya zeae CSL RW192 WP 038908153 1 Dzeae
Dickeya zeae MK19 WP 038913652 1 Dzeae
Dickeya sp CSL RW240 WP 035345024 1 Daquatica
Dickeya aquatica DW 0440 WP 035345024 1 Daquatica
DAQ1742 DAQ1742 00157 Daquatica
Dickeya zeae Ech586 WP 012886590 1 Dzeae
Dickeya gs poaceaephila NCPPB 569 WP 042867734 1 Dpoaceaephila
Dickeya zeae Ech586 WP 012886589 1 Dzeae
Dickeya zeae CSL RW192 WP 038909296 1 Dzeae
Dickeya zeae ZJU1202 WP 016942821 1 Dzeae
Dickeya zeae EC1 WP 016942821 1 Dzeae
Dickeya zeae NCPPB 3531 WP 038905991 1 Dzeae
Dickeya zeae NCPPB 3532 WP 023641008 1 Dzeae
Dickeya zeae NCPPB 2538 WP 038916988 1 Dzeae
Dickeya zeae MK19 WP 038914827 1 Dzeae
Dickeya zeae MS1 WP 019845409 1 Dzeae
Dickeya chrysanthemi L11 WP 033576725 1 Dchrysanthemi
Dickeya zeae DZ2Q WP 038925714 1 Dzeae
Dickeya gs undicola 2B12 WP 033569006 1 Dundicola
Dickeya gs fangzhongdai M074 WP 038664495 1 Dfangzhongdai
Dickeya gs fangzhongdai M005 WP 038664495 1 Dfangzhongdai
Dickeya gs fangzhongdai ND14b WP 038664495 1 Dfangzhongdai
Dickeya gs fangzhongdai B16 WP 038917469 1 Dfangzhongdai
Dickeya gs fangzhongdai MK7 WP 038917469 1 Dfangzhongdai
Dickeya gs fangzhongdai NCPPB 3274 WP 042858206 1 Dfangzhongdai
Dickeya dadantii NCPPB 898 WP 038909984 1 Ddadantii
Dickeya dadantii NCPPB 3537 WP 038900003 1 Ddadantii
Dickeya dadantii 3937 WP 033111513 1 Ddadantii
Dickeya dadantii subsp dieffenbachiae NCPPB 2976 WP 038921322 1 Ddadantii
Dickeya chrysanthemi NCPPB 3533 WP 015848317 1 Dchrysanthemi
Dickeya chrysanthemi Ech1591 WP 015848317 1 Dchrysanthemi
Dickeya gs fangzhongdai M005 WP 039694894 1 Dfangzhongdai
Dickeya gs undicola 2B12 WP 033569009 1 Dundicola
Dickeya gs fangzhongdai NCPPB 3274 WP 042861797 1 Dfangzhongdai
Dickeya gs fangzhongdai MK7 WP 038917470 1 Dfangzhongdai
Dickeya gs fangzhongdai B16 WP 038917470 1 Dfangzhongdai
Dickeya solani RNS 08 23 3 1 A WP 022635320 1 Dsolani
Dickeya solani IFB 0099 WP 022635320 1 Dsolani
Dickeya solani PPO 9134 WP 022635320 1 Dsolani
Dickeya solani IPO 2222 WP 022635320 1 Dsolani
Dickeya solani GBBC 2040 WP 022635320 1 Dsolani
Dickeya solani D s0432 1 WP 022635320 1 Dsolani
Dickeya solani RNS 07 7 3B WP 022635320 1 Dsolani
Dickeya solani PPO 9019 WP 022635320 1 Dsolani
Dickeya solani MK10 WP 022635320 1 Dsolani
Dickeya solani MK16 WP 022635320 1 Dsolani

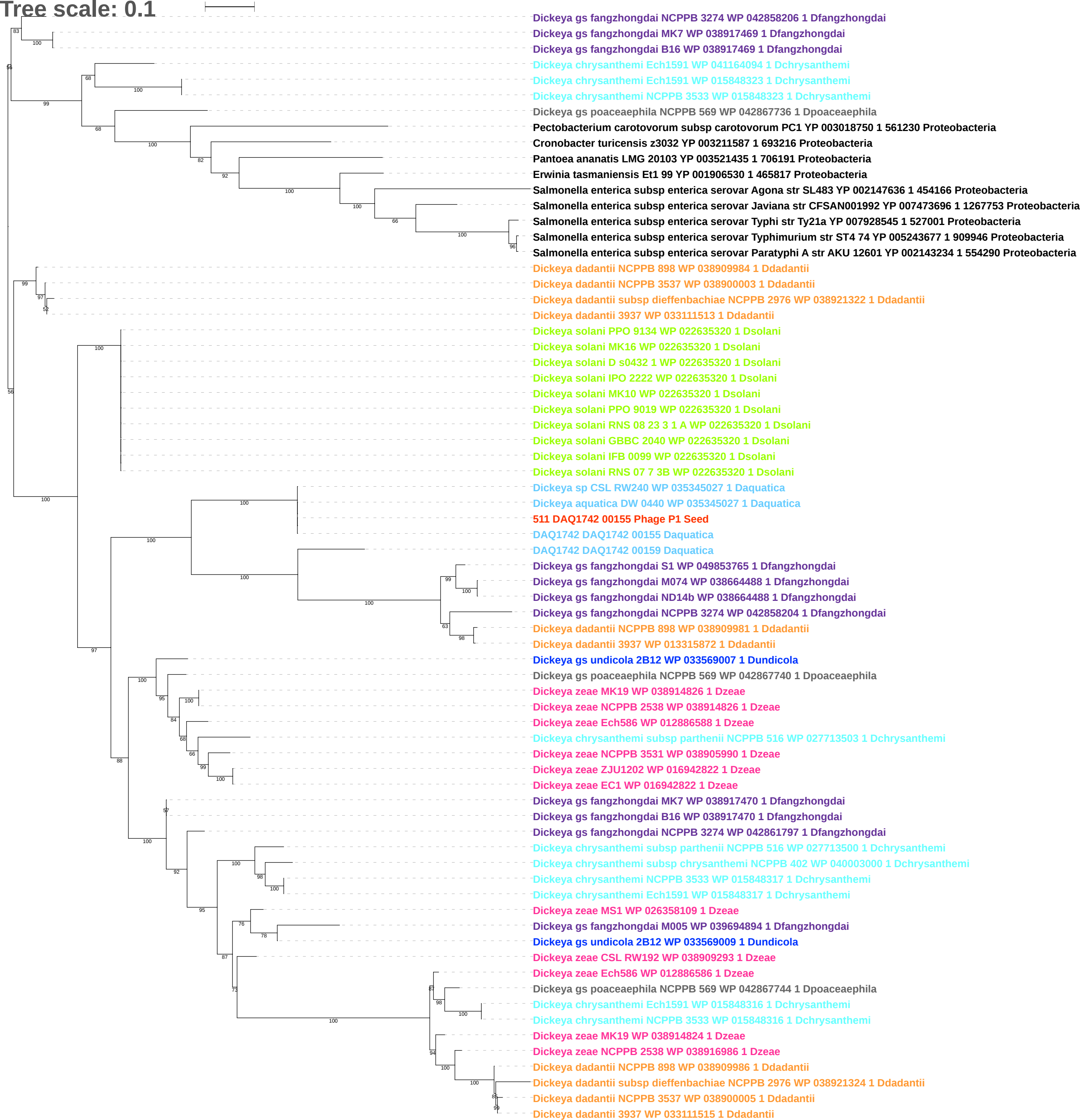
Tree scale: 0.1
Dickeya gs fangzhongdai NCPPB 3274 WP 042858206 1 Dfangzhongdai
Dickeya gs fangzhongdai MK7 WP 038917469 1 Dfangzhongdai
Dickeya gs fangzhongdai B16 WP 038917469 1 Dfangzhongdai
Dickeya chrysanthemi Ech1591 WP 041164094 1 Dchrysanthemi
Dickeya chrysanthemi Ech1591 WP 015848323 1 Dchrysanthemi
Dickeya chrysanthemi NCPPB 3533 WP 015848323 1 Dchrysanthemi
Dickeya gs poaceaephila NCPPB 569 WP 042867736 1 Dpoaceaephila
Pectobacterium carotovorum subsp carotovorum PC1 YP 003018750 1 561230 Proteobacteria
Cronobacter turicensis z3032 YP 003211587 1 693216 Proteobacteria
Pantoea ananatis LMG 20103 YP 003521435 1 706191 Proteobacteria
Erwinia tasmaniensis Et1 99 YP 001906530 1 465817 Proteobacteria
Salmonella enterica subsp enterica serovar Agona str SL483 YP 002147636 1 454166 Proteobacteria
Salmonella enterica subsp enterica serovar Javiana str CFSAN001992 YP 007473696 1 1267753 Proteobacteria
Salmonella enterica subsp enterica serovar Typhi str Ty21a YP 007928545 1 527001 Proteobacteria
Salmonella enterica subsp enterica serovar Typhimurium str ST4 74 YP 005243677 1 909946 Proteobacteria
Salmonella enterica subsp enterica serovar Paratyphi A str AKU 12601 YP 002143234 1 554290 Proteobacteria
Dickeya dadantii NCPPB 898 WP 038909984 1 Ddadantii
Dickeya dadantii NCPPB 3537 WP 038900003 1 Ddadantii
Dickeya dadantii subsp dieffenbachiae NCPPB 2976 WP 038921322 1 Ddadantii
Dickeya dadantii 3937 WP 033111513 1 Ddadantii
Dickeya solani PPO 9134 WP 022635320 1 Dsolani
Dickeya solani MK16 WP 022635320 1 Dsolani
Dickeya solani D s0432 1 WP 022635320 1 Dsolani
Dickeya solani IPO 2222 WP 022635320 1 Dsolani
Dickeya solani MK10 WP 022635320 1 Dsolani
Dickeya solani PPO 9019 WP 022635320 1 Dsolani
Dickeya solani RNS 08 23 3 1 A WP 022635320 1 Dsolani
Dickeya solani GBBC 2040 WP 022635320 1 Dsolani
Dickeya solani IFB 0099 WP 022635320 1 Dsolani
Dickeya solani RNS 07 7 3B WP 022635320 1 Dsolani
Dickeya sp CSL RW240 WP 035345027 1 Daquatica
Dickeya aquatica DW 0440 WP 035345027 1 Daquatica
511 DAQ1742 00155 Phage P1 Seed
DAQ1742 DAQ1742 00155 Daquatica
DAQ1742 DAQ1742 00159 Daquatica
Dickeya gs fangzhongdai S1 WP 049853765 1 Dfangzhongdai
Dickeya gs fangzhongdai M074 WP 038664488 1 Dfangzhongdai
Dickeya gs fangzhongdai ND14b WP 038664488 1 Dfangzhongdai
Dickeya gs fangzhongdai NCPPB 3274 WP 042858204 1 Dfangzhongdai
Dickeya dadantii NCPPB 898 WP 038909981 1 Ddadantii
Dickeya dadantii 3937 WP 013315872 1 Ddadantii
Dickeya gs undicola 2B12 WP 033569007 1 Dundicola
Dickeya gs poaceaephila NCPPB 569 WP 042867740 1 Dpoaceaephila
Dickeya zeae MK19 WP 038914826 1 Dzeae
Dickeya zeae NCPPB 2538 WP 038914826 1 Dzeae
Dickeya zeae Ech586 WP 012886588 1 Dzeae
Dickeya chrysanthemi subsp parthenii NCPPB 516 WP 027713503 1 Dchrysanthemi
Dickeya zeae NCPPB 3531 WP 038905990 1 Dzeae
Dickeya zeae ZJU1202 WP 016942822 1 Dzeae
Dickeya zeae EC1 WP 016942822 1 Dzeae
Dickeya gs fangzhongdai MK7 WP 038917470 1 Dfangzhongdai
Dickeya gs fangzhongdai B16 WP 038917470 1 Dfangzhongdai
Dickeya gs fangzhongdai NCPPB 3274 WP 042861797 1 Dfangzhongdai
Dickeya chrysanthemi subsp parthenii NCPPB 516 WP 027713500 1 Dchrysanthemi
Dickeya chrysanthemi subsp chrysanthemi NCPPB 402 WP 040003000 1 Dchrysanthemi
Dickeya chrysanthemi NCPPB 3533 WP 015848317 1 Dchrysanthemi
Dickeya chrysanthemi Ech1591 WP 015848317 1 Dchrysanthemi
Dickeya zeae MS1 WP 026358109 1 Dzeae
Dickeya gs fangzhongdai M005 WP 039694894 1 Dfangzhongdai
Dickeya gs undicola 2B12 WP 033569009 1 Dundicola
Dickeya zeae CSL RW192 WP 038909293 1 Dzeae
Dickeya zeae Ech586 WP 012886586 1 Dzeae
Dickeya gs poaceaephila NCPPB 569 WP 042867744 1 Dpoaceaephila
Dickeya chrysanthemi Ech1591 WP 015848316 1 Dchrysanthemi
Dickeya chrysanthemi NCPPB 3533 WP 015848316 1 Dchrysanthemi
Dickeya zeae MK19 WP 038914824 1 Dzeae
Dickeya zeae NCPPB 2538 WP 038916986 1 Dzeae
Dickeya dadantii NCPPB 898 WP 038909986 1 Ddadantii
Dickeya dadantii subsp dieffenbachiae NCPPB 2976 WP 038921324 1 Ddadantii
Dickeya dadantii NCPPB 3537 WP 038900005 1 Ddadantii
Dickeya dadantii 3937 WP 033111515 1 Ddadantii

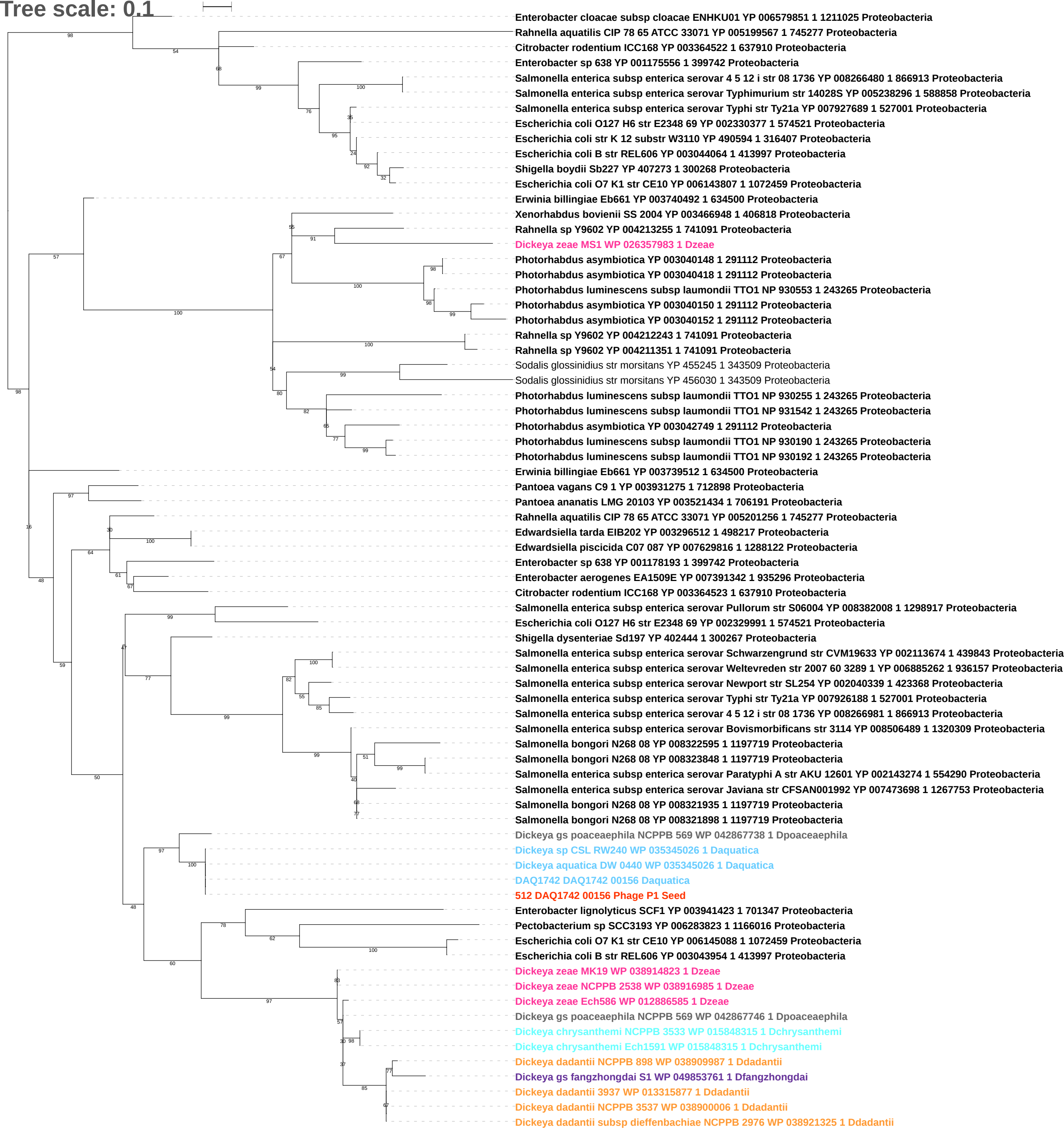
Tree scale: 0.1
Enterobacter cloacae subsp cloacae ENHKU01 YP 006579851 1 1211025 Proteobacteria
Rahnella aquatilis CIP 78 65 ATCC 33071 YP 005199567 1 745277 Proteobacteria
Citrobacter rodentium ICC168 YP 003364522 1 637910 Proteobacteria
Enterobacter sp 638 YP 001175556 1 399742 Proteobacteria
Salmonella enterica subsp enterica serovar 4 5 12 i str 08 1736 YP 008266480 1 866913 Proteobacteria
Salmonella enterica subsp enterica serovar Typhimurium str 14028S YP 005238296 1 588858 Proteobacteria
Salmonella enterica subsp enterica serovar Typhi str Ty21a YP 007927689 1 527001 Proteobacteria
Escherichia coli O127 H6 str E2348 69 YP 002330377 1 574521 Proteobacteria
Escherichia coli str K 12 substr W3110 YP 490594 1 316407 Proteobacteria
Escherichia coli B str REL606 YP 003044064 1 413997 Proteobacteria
Shigella boydii Sb227 YP 407273 1 300268 Proteobacteria
Escherichia coli O7 K1 str CE10 YP 006143807 1 1072459 Proteobacteria
Erwinia billingiae Eb661 YP 003740492 1 634500 Proteobacteria
Xenorhabdus bovienii SS 2004 YP 003466948 1 406818 Proteobacteria
Rahnella sp Y9602 YP 004213255 1 741091 Proteobacteria
Dickeya zeae MS1 WP 026357983 1 Dzeae
Photorhabdus asymbiotica YP 003040148 1 291112 Proteobacteria
Photorhabdus asymbiotica YP 003040418 1 291112 Proteobacteria
Photorhabdus luminescens subsp laumondii TTO1 NP 930553 1 243265 Proteobacteria
Photorhabdus asymbiotica YP 003040150 1 291112 Proteobacteria
Photorhabdus asymbiotica YP 003040152 1 291112 Proteobacteria
Rahnella sp Y9602 YP 004212243 1 741091 Proteobacteria
Rahnella sp Y9602 YP 004211351 1 741091 Proteobacteria
Sodalis glossinidius str morsitans YP 455245 1 343509 Proteobacteria
Sodalis glossinidius str morsitans YP 456030 1 343509 Proteobacteria
Photorhabdus luminescens subsp laumondii TTO1 NP 930255 1 243265 Proteobacteria
Photorhabdus luminescens subsp laumondii TTO1 NP 931542 1 243265 Proteobacteria
Photorhabdus asymbiotica YP 003042749 1 291112 Proteobacteria
Photorhabdus luminescens subsp laumondii TTO1 NP 930190 1 243265 Proteobacteria
Photorhabdus luminescens subsp laumondii TTO1 NP 930192 1 243265 Proteobacteria
Erwinia billingiae Eb661 YP 003739512 1 634500 Proteobacteria
Pantoea vagans C9 1 YP 003931275 1 712898 Proteobacteria
Pantoea ananatis LMG 20103 YP 003521434 1 706191 Proteobacteria
Rahnella aquatilis CIP 78 65 ATCC 33071 YP 005201256 1 745277 Proteobacteria
Edwardsiella tarda EIB202 YP 003296512 1 498217 Proteobacteria
Edwardsiella piscicida C07 087 YP 007629816 1 1288122 Proteobacteria
Enterobacter sp 638 YP 001178193 1 399742 Proteobacteria
Enterobacter aerogenes EA1509E YP 007391342 1 935296 Proteobacteria
Citrobacter rodentium ICC168 YP 003364523 1 637910 Proteobacteria
Salmonella enterica subsp enterica serovar Pullorum str S06004 YP 008382008 1 1298917 Proteobacteria
Escherichia coli O127 H6 str E2348 69 YP 002329991 1 574521 Proteobacteria
Shigella dysenteriae Sd197 YP 402444 1 300267 Proteobacteria
Salmonella enterica subsp enterica serovar Schwarzengrund str CVM19633 YP 002113674 1 439843 Proteobacteria
Salmonella enterica subsp enterica serovar Weltevreden str 2007 60 3289 1 YP 006885262 1 936157 Proteobacteria
Salmonella enterica subsp enterica serovar Newport str SL254 YP 002040339 1 423368 Proteobacteria
Salmonella enterica subsp enterica serovar Typhi str Ty21a YP 007926188 1 527001 Proteobacteria
Salmonella enterica subsp enterica serovar 4 5 12 i str 08 1736 YP 008266981 1 866913 Proteobacteria
Salmonella enterica subsp enterica serovar Bovismorbificans str 3114 YP 008506489 1 1320309 Proteobacteria
Salmonella bongori N268 08 YP 008322595 1 1197719 Proteobacteria
Salmonella bongori N268 08 YP 008323848 1 1197719 Proteobacteria
Salmonella enterica subsp enterica serovar Paratyphi A str AKU 12601 YP 002143274 1 554290 Proteobacteria
Salmonella enterica subsp enterica serovar Javiana str CFSAN001992 YP 007473698 1 1267753 Proteobacteria
Salmonella bongori N268 08 YP 008321935 1 1197719 Proteobacteria
Salmonella bongori N268 08 YP 008321898 1 1197719 Proteobacteria
Dickeya gs poaceaephila NCPPB 569 WP 042867738 1 Dpoaceaephila
Dickeya sp CSL RW240 WP 035345026 1 Daquatica
Dickeya aquatica DW 0440 WP 035345026 1 Daquatica
DAQ1742 DAQ1742 00156 Daquatica
512 DAQ1742 00156 Phage P1 Seed
Enterobacter lignolyticus SCF1 YP 003941423 1 701347 Proteobacteria
Pectobacterium sp SCC3193 YP 006283823 1 1166016 Proteobacteria
Escherichia coli O7 K1 str CE10 YP 006145088 1 1072459 Proteobacteria
Escherichia coli B str REL606 YP 003043954 1 413997 Proteobacteria
Dickeya zeae MK19 WP 038914823 1 Dzeae
Dickeya zeae NCPPB 2538 WP 038916985 1 Dzeae
Dickeya zeae Ech586 WP 012886585 1 Dzeae
Dickeya gs poaceaephila NCPPB 569 WP 042867746 1 Dpoaceaephila
Dickeya chrysanthemi NCPPB 3533 WP 015848315 1 Dchrysanthemi
Dickeya chrysanthemi Ech1591 WP 015848315 1 Dchrysanthemi
Dickeya dadantii NCPPB 898 WP 038909987 1 Ddadantii
Dickeya gs fangzhongdai S1 WP 049853761 1 Dfangzhongdai
Dickeya dadantii 3937 WP 013315877 1 Ddadantii
Dickeya dadantii NCPPB 3537 WP 038900006 1 Ddadantii
Dickeya dadantii subsp dieffenbachiae NCPPB 2976 WP 038921325 1 Ddadantii

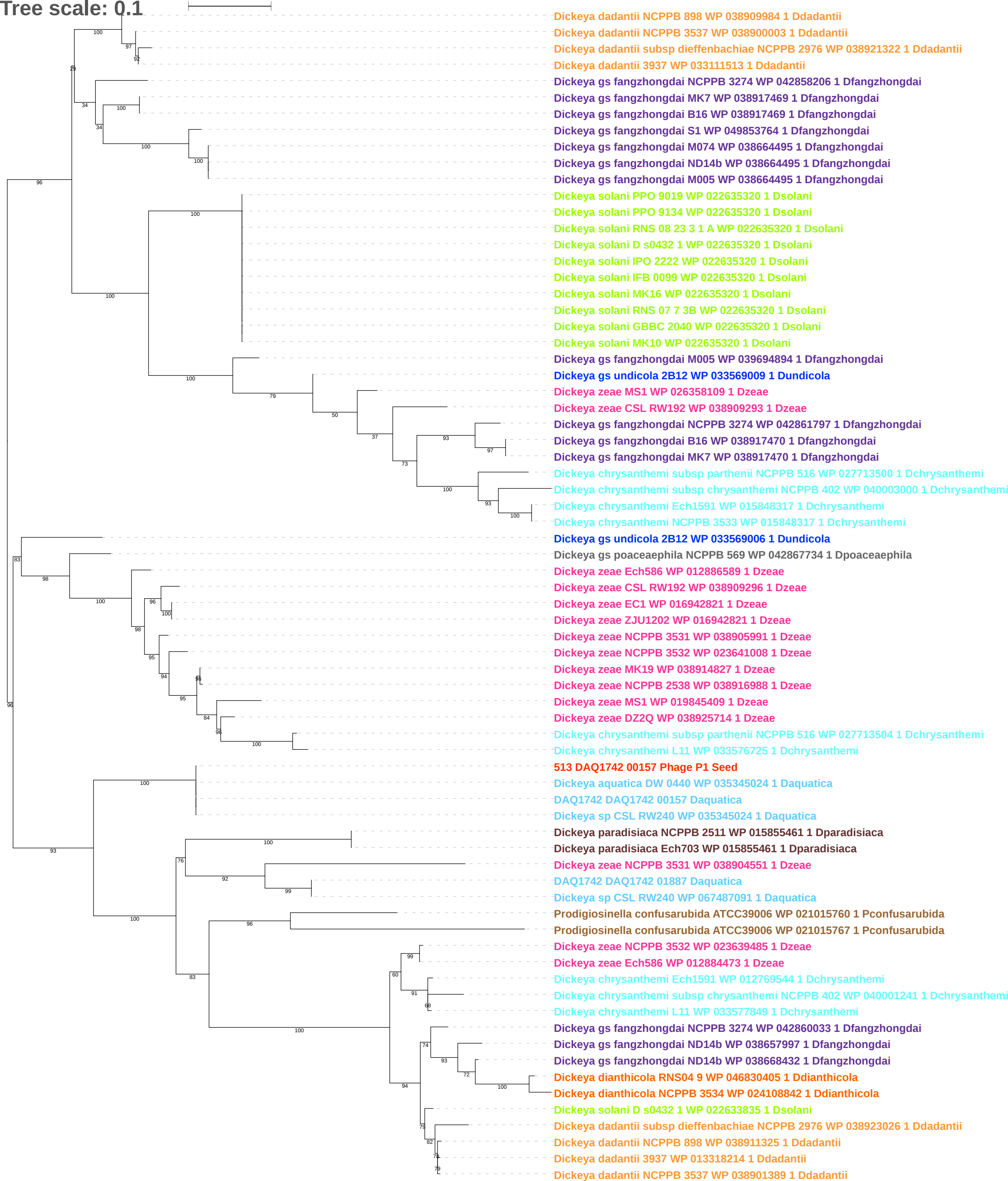
Tree scale: 0.1
Dickeya dadantii NCPPB 898 WP 038909984 1 Ddadantii
Dickeya dadantii NCPPB 3537 WP 038900003 1 Ddadantii
Dickeya dadantii subsp dieffenbachiae NCPPB 2976 WP 038921322 1 Ddadantii
Dickeya dadantii 3937 WP 033111513 1 Ddadantii
Dickeya gs fangzhongdai NCPPB 3274 WP 042858206 1 Dfangzhongdai
Dickeya gs fangzhongdai MK7 WP 038917469 1 Dfangzhongdai
Dickeya gs fangzhongdai B16 WP 038917469 1 Dfangzhongdai
Dickeya gs fangzhongdai S1 WP 049853764 1 Dfangzhongdai
Dickeya gs fangzhongdai M074 WP 038664495 1 Dfangzhongdai
Dickeya gs fangzhongdai ND14b WP 038664495 1 Dfangzhongdai
Dickeya gs fangzhongdai M005 WP 038664495 1 Dfangzhongdai
Dickeya solani PPO 9019 WP 022635320 1 Dsolani
Dickeya solani PPO 9134 WP 022635320 1 Dsolani
Dickeya solani RNS 08 23 3 1 A WP 022635320 1 Dsolani
Dickeya solani D s0432 1 WP 022635320 1 Dsolani
Dickeya solani IPO 2222 WP 022635320 1 Dsolani
Dickeya solani IFB 0099 WP 022635320 1 Dsolani
Dickeya solani MK16 WP 022635320 1 Dsolani
Dickeya solani RNS 07 7 3B WP 022635320 1 Dsolani
Dickeya solani GBBC 2040 WP 022635320 1 Dsolani
Dickeya solani MK10 WP 022635320 1 Dsolani
Dickeya gs fangzhongdai M005 WP 039694894 1 Dfangzhongdai
Dickeya gs undicola 2B12 WP 033569009 1 Dundicola
Dickeya zeae MS1 WP 026358109 1 Dzeae
Dickeya zeae CSL RW192 WP 038909293 1 Dzeae
Dickeya gs fangzhongdai NCPPB 3274 WP 042861797 1 Dfangzhongdai
Dickeya gs fangzhongdai B16 WP 038917470 1 Dfangzhongdai
Dickeya gs fangzhongdai MK7 WP 038917470 1 Dfangzhongdai
Dickeya chrysanthemi subsp parthenii NCPPB 516 WP 027713500 1 Dchrysanthemi
Dickeya chrysanthemi subsp chrysanthemi NCPPB 402 WP 040003000 1 Dchrysanthemi
Dickeya chrysanthemi Ech1591 WP 015848317 1 Dchrysanthemi
Dickeya chrysanthemi NCPPB 3533 WP 015848317 1 Dchrysanthemi
Dickeya gs undicola 2B12 WP 033569006 1 Dundicola
Dickeya gs poaceaephila NCPPB 569 WP 042867734 1 Dpoaceaephila
Dickeya zeae Ech586 WP 012886589 1 Dzeae
Dickeya zeae CSL RW192 WP 038909296 1 Dzeae
Dickeya zeae EC1 WP 016942821 1 Dzeae
Dickeya zeae ZJU1202 WP 016942821 1 Dzeae
Dickeya zeae NCPPB 3531 WP 038905991 1 Dzeae
Dickeya zeae NCPPB 3532 WP 023641008 1 Dzeae
Dickeya zeae MK19 WP 038914827 1 Dzeae
Dickeya zeae NCPPB 2538 WP 038916988 1 Dzeae
Dickeya zeae MS1 WP 019845409 1 Dzeae
Dickeya zeae DZ2Q WP 038925714 1 Dzeae
Dickeya chrysanthemi subsp parthenii NCPPB 516 WP 027713504 1 Dchrysanthemi
Dickeya chrysanthemi L11 WP 033576725 1 Dchrysanthemi
513 DAQ1742 00157 Phage P1 Seed
Dickeya aquatica DW 0440 WP 035345024 1 Daquatica
DAQ1742 DAQ1742 00157 Daquatica
Dickeya sp CSL RW240 WP 035345024 1 Daquatica
Dickeya paradisiaca NCPPB 2511 WP 015855461 1 Dparadisiaca
Dickeya paradisiaca Ech703 WP 015855461 1 Dparadisiaca
Dickeya zeae NCPPB 3531 WP 038904551 1 Dzeae
DAQ1742 DAQ1742 01887 Daquatica
Dickeya sp CSL RW240 WP 067487091 1 Daquatica
Prodigiosinella confusarubida ATCC39006 WP 021015760 1 Pconfusarubida
Prodigiosinella confusarubida ATCC39006 WP 021015767 1 Pconfusarubida
Dickeya zeae NCPPB 3532 WP 023639485 1 Dzeae
Dickeya zeae Ech586 WP 012884473 1 Dzeae
Dickeya chrysanthemi Ech1591 WP 012769544 1 Dchrysanthemi
Dickeya chrysanthemi subsp chrysanthemi NCPPB 402 WP 040001241 1 Dchrysanthemi
Dickeya chrysanthemi L11 WP 033577849 1 Dchrysanthemi
Dickeya gs fangzhongdai NCPPB 3274 WP 042860033 1 Dfangzhongdai
Dickeya gs fangzhongdai ND14b WP 038657997 1 Dfangzhongdai
Dickeya gs fangzhongdai ND14b WP 038668432 1 Dfangzhongdai
Dickeya dianthicola RNS04 9 WP 046830405 1 Ddianthicola
Dickeya dianthicola NCPPB 3534 WP 024108842 1 Ddianthicola
Dickeya solani D s0432 1 WP 022633835 1 Dsolani
Dickeya dadantii subsp dieffenbachiae NCPPB 2976 WP 038923026 1 Ddadantii
Dickeya dadantii NCPPB 898 WP 038911325 1 Ddadantii
Dickeya dadantii 3937 WP 013318214 1 Ddadantii
Dickeya dadantii NCPPB 3537 WP 038901389 1 Ddadantii

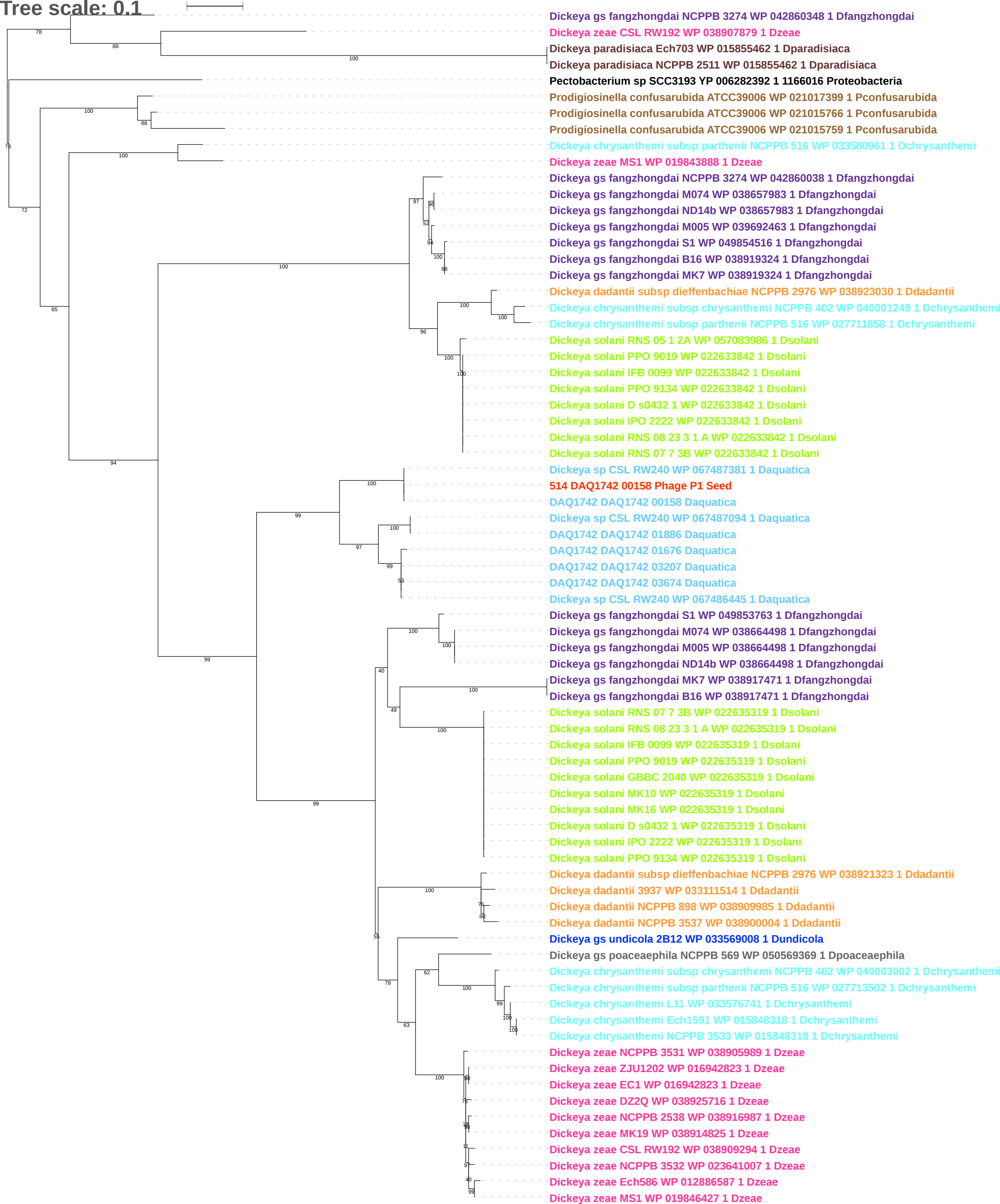
Tree scale: 0.1
Dickeya gs fangzhongdai NCPPB 3274 WP 042860348 1 Dfangzhongdai
Dickeya zeae CSL RW192 WP 038907879 1 Dzeae
Dickeya paradisiaca Ech703 WP 015855462 1 Dparadisiaca
Dickeya paradisiaca NCPPB 2511 WP 015855462 1 Dparadisiaca
Pectobacterium sp SCC3193 YP 006282392 1 1166016 Proteobacteria
Prodigiosinella confusarubida ATCC39006 WP 021017399 1 Pconfusarubida
Prodigiosinella confusarubida ATCC39006 WP 021015766 1 Pconfusarubida
Prodigiosinella confusarubida ATCC39006 WP 021015759 1 Pconfusarubida
Dickeya chrysanthemi subsp parthenii NCPPB 516 WP 033580961 1 Dchrysanthemi
Dickeya zeae MS1 WP 019843888 1 Dzeae
Dickeya gs fangzhongdai NCPPB 3274 WP 042860038 1 Dfangzhongdai
Dickeya gs fangzhongdai M074 WP 038657983 1 Dfangzhongdai
Dickeya gs fangzhongdai ND14b WP 038657983 1 Dfangzhongdai
Dickeya gs fangzhongdai M005 WP 039692463 1 Dfangzhongdai
Dickeya gs fangzhongdai S1 WP 049854516 1 Dfangzhongdai
Dickeya gs fangzhongdai B16 WP 038919324 1 Dfangzhongdai
Dickeya gs fangzhongdai MK7 WP 038919324 1 Dfangzhongdai
Dickeya dadantii subsp dieffenbachiae NCPPB 2976 WP 038923030 1 Ddadantii
Dickeya chrysanthemi subsp chrysanthemi NCPPB 402 WP 040001249 1 Dchrysanthemi
Dickeya chrysanthemi subsp parthenii NCPPB 516 WP 027711858 1 Dchrysanthemi
Dickeya solani RNS 05 1 2A WP 057083986 1 Dsolani
Dickeya solani PPO 9019 WP 022633842 1 Dsolani
Dickeya solani IFB 0099 WP 022633842 1 Dsolani
Dickeya solani PPO 9134 WP 022633842 1 Dsolani
Dickeya solani D s0432 1 WP 022633842 1 Dsolani
Dickeya solani IPO 2222 WP 022633842 1 Dsolani
Dickeya solani RNS 08 23 3 1 A WP 022633842 1 Dsolani
Dickeya solani RNS 07 7 3B WP 022633842 1 Dsolani
Dickeya sp CSL RW240 WP 067487381 1 Daquatica
514 DAQ1742 00158 Phage P1 Seed
DAQ1742 DAQ1742 00158 Daquatica
Dickeya sp CSL RW240 WP 067487094 1 Daquatica
DAQ1742 DAQ1742 01886 Daquatica
DAQ1742 DAQ1742 01676 Daquatica
DAQ1742 DAQ1742 03207 Daquatica
DAQ1742 DAQ1742 03674 Daquatica
Dickeya sp CSL RW240 WP 067486445 1 Daquatica
Dickeya gs fangzhongdai S1 WP 049853763 1 Dfangzhongdai
Dickeya gs fangzhongdai M074 WP 038664498 1 Dfangzhongdai
Dickeya gs fangzhongdai M005 WP 038664498 1 Dfangzhongdai
Dickeya gs fangzhongdai ND14b WP 038664498 1 Dfangzhongdai
Dickeya gs fangzhongdai MK7 WP 038917471 1 Dfangzhongdai
Dickeya gs fangzhongdai B16 WP 038917471 1 Dfangzhongdai
Dickeya solani RNS 07 7 3B WP 022635319 1 Dsolani
Dickeya solani RNS 08 23 3 1 A WP 022635319 1 Dsolani
Dickeya solani IFB 0099 WP 022635319 1 Dsolani
Dickeya solani PPO 9019 WP 022635319 1 Dsolani
Dickeya solani GBBC 2040 WP 022635319 1 Dsolani
Dickeya solani MK10 WP 022635319 1 Dsolani
Dickeya solani MK16 WP 022635319 1 Dsolani
Dickeya solani D s0432 1 WP 022635319 1 Dsolani
Dickeya solani IPO 2222 WP 022635319 1 Dsolani
Dickeya solani PPO 9134 WP 022635319 1 Dsolani
Dickeya dadantii subsp dieffenbachiae NCPPB 2976 WP 038921323 1 Ddadantii
Dickeya dadantii 3937 WP 033111514 1 Ddadantii
Dickeya dadantii NCPPB 898 WP 038909985 1 Ddadantii
Dickeya dadantii NCPPB 3537 WP 038900004 1 Ddadantii
Dickeya gs undicola 2B12 WP 033569008 1 Dundicola
Dickeya gs poaceaephila NCPPB 569 WP 050569369 1 Dpoaceaephila
Dickeya chrysanthemi subsp chrysanthemi NCPPB 402 WP 040003002 1 Dchrysanthemi
Dickeya chrysanthemi subsp parthenii NCPPB 516 WP 027713502 1 Dchrysanthemi
Dickeya chrysanthemi L11 WP 033576741 1 Dchrysanthemi
Dickeya chrysanthemi Ech1591 WP 015848318 1 Dchrysanthemi
Dickeya chrysanthemi NCPPB 3533 WP 015848318 1 Dchrysanthemi
Dickeya zeae NCPPB 3531 WP 038905989 1 Dzeae
Dickeya zeae ZJU1202 WP 016942823 1 Dzeae
Dickeya zeae EC1 WP 016942823 1 Dzeae
Dickeya zeae DZ2Q WP 038925716 1 Dzeae
Dickeya zeae NCPPB 2538 WP 038916987 1 Dzeae
Dickeya zeae MK19 WP 038914825 1 Dzeae
Dickeya zeae CSL RW192 WP 038909294 1 Dzeae
Dickeya zeae NCPPB 3532 WP 023641007 1 Dzeae
Dickeya zeae Ech586 WP 012886587 1 Dzeae
Dickeya zeae MS1 WP 019846427 1 Dzeae

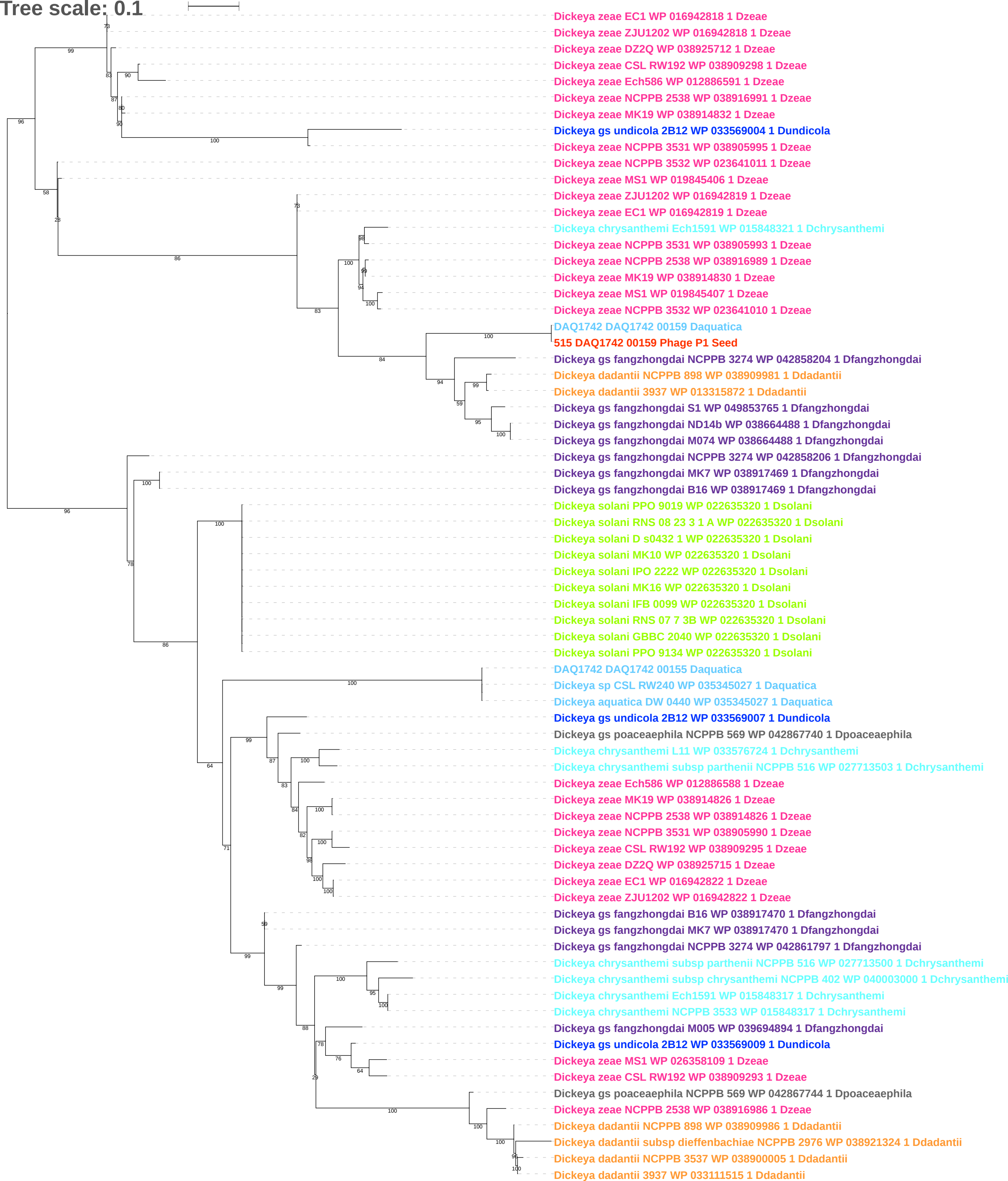
Tree scale: 0.1
Dickeya zeae EC1 WP 016942818 1 Dzeae
Dickeya zeae ZJU1202 WP 016942818 1 Dzeae
Dickeya zeae DZ2Q WP 038925712 1 Dzeae
Dickeya zeae CSL RW192 WP 038909298 1 Dzeae
Dickeya zeae Ech586 WP 012886591 1 Dzeae
Dickeya zeae NCPPB 2538 WP 038916991 1 Dzeae
Dickeya zeae MK19 WP 038914832 1 Dzeae
Dickeya gs undicola 2B12 WP 033569004 1 Dundicola
Dickeya zeae NCPPB 3531 WP 038905995 1 Dzeae
Dickeya zeae NCPPB 3532 WP 023641011 1 Dzeae
Dickeya zeae MS1 WP 019845406 1 Dzeae
Dickeya zeae ZJU1202 WP 016942819 1 Dzeae
Dickeya zeae EC1 WP 016942819 1 Dzeae
Dickeya chrysanthemi Ech1591 WP 015848321 1 Dchrysanthemi
Dickeya zeae NCPPB 3531 WP 038905993 1 Dzeae
Dickeya zeae NCPPB 2538 WP 038916989 1 Dzeae
Dickeya zeae MK19 WP 038914830 1 Dzeae
Dickeya zeae MS1 WP 019845407 1 Dzeae
Dickeya zeae NCPPB 3532 WP 023641010 1 Dzeae
DAQ1742 DAQ1742 00159 Daquatica
515 DAQ1742 00159 Phage P1 Seed
Dickeya gs fangzhongdai NCPPB 3274 WP 042858204 1 Dfangzhongdai
Dickeya dadantii NCPPB 898 WP 038909981 1 Ddadantii
Dickeya dadantii 3937 WP 013315872 1 Ddadantii
Dickeya gs fangzhongdai S1 WP 049853765 1 Dfangzhongdai
Dickeya gs fangzhongdai ND14b WP 038664488 1 Dfangzhongdai
Dickeya gs fangzhongdai M074 WP 038664488 1 Dfangzhongdai
Dickeya gs fangzhongdai NCPPB 3274 WP 042858206 1 Dfangzhongdai
Dickeya gs fangzhongdai MK7 WP 038917469 1 Dfangzhongdai
Dickeya gs fangzhongdai B16 WP 038917469 1 Dfangzhongdai
Dickeya solani PPO 9019 WP 022635320 1 Dsolani
Dickeya solani RNS 08 23 3 1 A WP 022635320 1 Dsolani
Dickeya solani D s0432 1 WP 022635320 1 Dsolani
Dickeya solani MK10 WP 022635320 1 Dsolani
Dickeya solani IPO 2222 WP 022635320 1 Dsolani
Dickeya solani MK16 WP 022635320 1 Dsolani
Dickeya solani IFB 0099 WP 022635320 1 Dsolani
Dickeya solani RNS 07 7 3B WP 022635320 1 Dsolani
Dickeya solani GBBC 2040 WP 022635320 1 Dsolani
Dickeya solani PPO 9134 WP 022635320 1 Dsolani
DAQ1742 DAQ1742 00155 Daquatica
Dickeya sp CSL RW240 WP 035345027 1 Daquatica
Dickeya aquatica DW 0440 WP 035345027 1 Daquatica
Dickeya gs undicola 2B12 WP 033569007 1 Dundicola
Dickeya gs poaceaephila NCPPB 569 WP 042867740 1 Dpoaceaephila
Dickeya chrysanthemi L11 WP 033576724 1 Dchrysanthemi
Dickeya chrysanthemi subsp parthenii NCPPB 516 WP 027713503 1 Dchrysanthemi
Dickeya zeae Ech586 WP 012886588 1 Dzeae
Dickeya zeae MK19 WP 038914826 1 Dzeae
Dickeya zeae NCPPB 2538 WP 038914826 1 Dzeae
Dickeya zeae NCPPB 3531 WP 038905990 1 Dzeae
Dickeya zeae CSL RW192 WP 038909295 1 Dzeae
Dickeya zeae DZ2Q WP 038925715 1 Dzeae
Dickeya zeae EC1 WP 016942822 1 Dzeae
Dickeya zeae ZJU1202 WP 016942822 1 Dzeae
Dickeya gs fangzhongdai B16 WP 038917470 1 Dfangzhongdai
Dickeya gs fangzhongdai MK7 WP 038917470 1 Dfangzhongdai
Dickeya gs fangzhongdai NCPPB 3274 WP 042861797 1 Dfangzhongdai
Dickeya chrysanthemi subsp parthenii NCPPB 516 WP 027713500 1 Dchrysanthemi
Dickeya chrysanthemi subsp chrysanthemi NCPPB 402 WP 040003000 1 Dchrysanthemi
Dickeya chrysanthemi Ech1591 WP 015848317 1 Dchrysanthemi
Dickeya chrysanthemi NCPPB 3533 WP 015848317 1 Dchrysanthemi
Dickeya gs fangzhongdai M005 WP 039694894 1 Dfangzhongdai
Dickeya gs undicola 2B12 WP 033569009 1 Dundicola
Dickeya zeae MS1 WP 026358109 1 Dzeae
Dickeya zeae CSL RW192 WP 038909293 1 Dzeae
Dickeya gs poaceaephila NCPPB 569 WP 042867744 1 Dpoaceaephila
Dickeya zeae NCPPB 2538 WP 038916986 1 Dzeae
Dickeya dadantii NCPPB 898 WP 038909986 1 Ddadantii
Dickeya dadantii subsp dieffenbachiae NCPPB 2976 WP 038921324 1 Ddadantii
Dickeya dadantii NCPPB 3537 WP 038900005 1 Ddadantii
Dickeya dadantii 3937 WP 033111515 1 Ddadantii

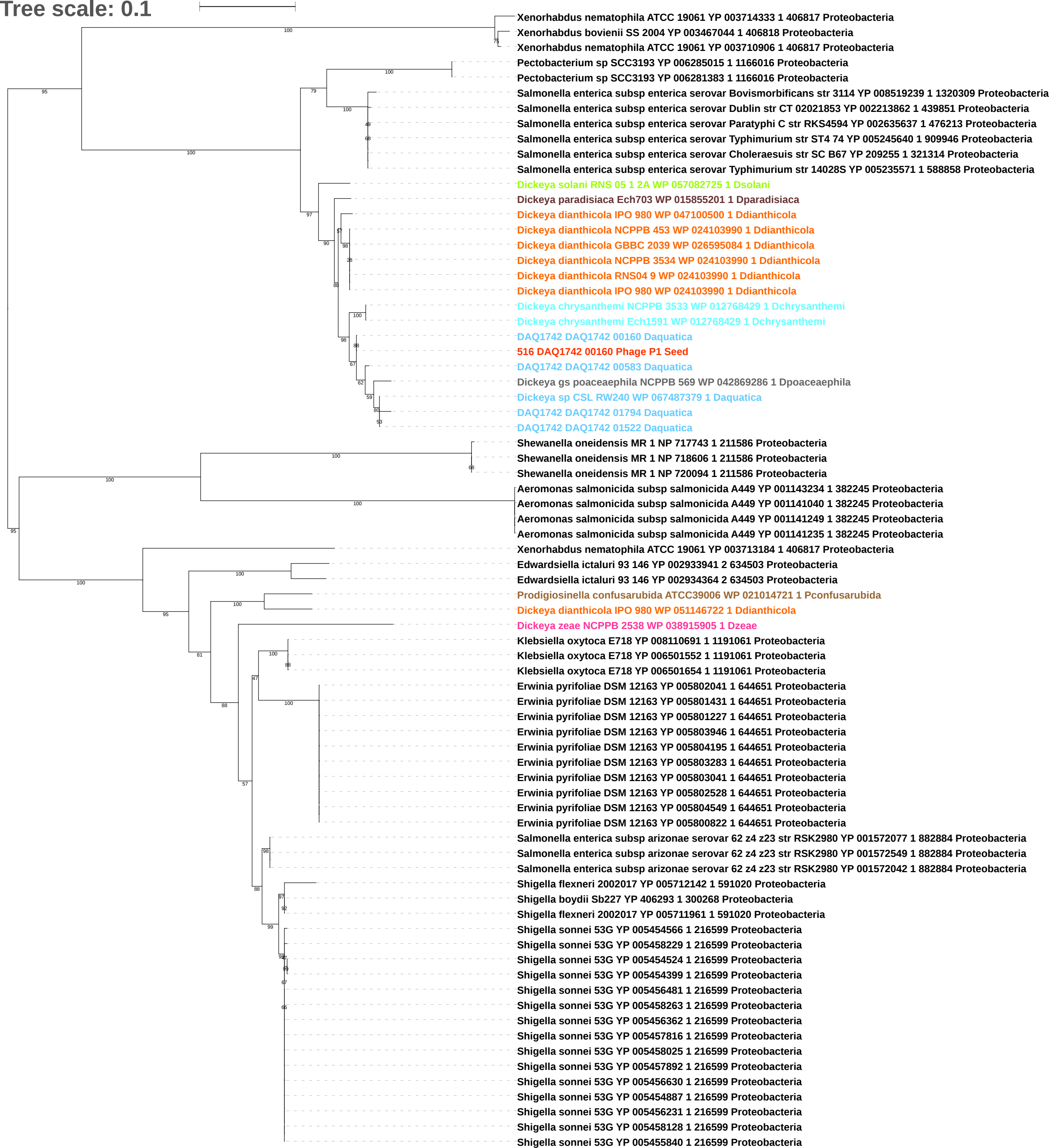
Tree scale: 0.1
Xenorhabdus nematophila ATCC 19061 YP 003714333 1 406817 Proteobacteria
Xenorhabdus bovienii SS 2004 YP 003467044 1 406818 Proteobacteria
Xenorhabdus nematophila ATCC 19061 YP 003710906 1 406817 Proteobacteria
Pectobacterium sp SCC3193 YP 006285015 1 1166016 Proteobacteria
Pectobacterium sp SCC3193 YP 006281383 1 1166016 Proteobacteria
Salmonella enterica subsp enterica serovar Bovismorbificans str 3114 YP 008519239 1 1320309 Proteobacteria
Salmonella enterica subsp enterica serovar Dublin str CT 02021853 YP 002213862 1 439851 Proteobacteria
Salmonella enterica subsp enterica serovar Paratyphi C str RKS4594 YP 002635637 1 476213 Proteobacteria
Salmonella enterica subsp enterica serovar Typhimurium str ST4 74 YP 005245640 1 909946 Proteobacteria
Salmonella enterica subsp enterica serovar Choleraesuis str SC B67 YP 209255 1 321314 Proteobacteria
Salmonella enterica subsp enterica serovar Typhimurium str 14028S YP 005235571 1 588858 Proteobacteria
Dickeya solani RNS 05 1 2A WP 057082725 1 Dsolani
Dickeya paradisiaca Ech703 WP 015855201 1 Dparadisiaca
Dickeya dianthicola IPO 980 WP 047100500 1 Ddianthicola
Dickeya dianthicola NCPPB 453 WP 024103990 1 Ddianthicola
Dickeya dianthicola GBBC 2039 WP 026595084 1 Ddianthicola
Dickeya dianthicola NCPPB 3534 WP 024103990 1 Ddianthicola
Dickeya dianthicola RNS04 9 WP 024103990 1 Ddianthicola
Dickeya dianthicola IPO 980 WP 024103990 1 Ddianthicola
Dickeya chrysanthemi NCPPB 3533 WP 012768429 1 Dchrysanthemi
Dickeya chrysanthemi Ech1591 WP 012768429 1 Dchrysanthemi
DAQ1742 DAQ1742 00160 Daquatica
516 DAQ1742 00160 Phage P1 Seed
DAQ1742 DAQ1742 00583 Daquatica
Dickeya gs poaceaephila NCPPB 569 WP 042869286 1 Dpoaceaephila
Dickeya sp CSL RW240 WP 067487379 1 Daquatica
DAQ1742 DAQ1742 01794 Daquatica
DAQ1742 DAQ1742 01522 Daquatica
Shewanella oneidensis MR 1 NP 717743 1 211586 Proteobacteria
Shewanella oneidensis MR 1 NP 718606 1 211586 Proteobacteria
Shewanella oneidensis MR 1 NP 720094 1 211586 Proteobacteria
Aeromonas salmonicida subsp salmonicida A449 YP 001143234 1 382245 Proteobacteria
Aeromonas salmonicida subsp salmonicida A449 YP 001141040 1 382245 Proteobacteria
Aeromonas salmonicida subsp salmonicida A449 YP 001141249 1 382245 Proteobacteria
Aeromonas salmonicida subsp salmonicida A449 YP 001141235 1 382245 Proteobacteria
Xenorhabdus nematophila ATCC 19061 YP 003713184 1 406817 Proteobacteria
Edwardsiella ictaluri 93 146 YP 002933941 2 634503 Proteobacteria
Edwardsiella ictaluri 93 146 YP 002934364 2 634503 Proteobacteria
Prodigiosinella confusarubida ATCC39006 WP 021014721 1 Pconfusarubida
Dickeya dianthicola IPO 980 WP 051146722 1 Ddianthicola
Dickeya zeae NCPPB 2538 WP 038915905 1 Dzeae
Klebsiella oxytoca E718 YP 008110691 1 1191061 Proteobacteria
Klebsiella oxytoca E718 YP 006501552 1 1191061 Proteobacteria
Klebsiella oxytoca E718 YP 006501654 1 1191061 Proteobacteria
Erwinia pyrifoliae DSM 12163 YP 005802041 1 644651 Proteobacteria
Erwinia pyrifoliae DSM 12163 YP 005801431 1 644651 Proteobacteria
Erwinia pyrifoliae DSM 12163 YP 005801227 1 644651 Proteobacteria
Erwinia pyrifoliae DSM 12163 YP 005803946 1 644651 Proteobacteria
Erwinia pyrifoliae DSM 12163 YP 005804195 1 644651 Proteobacteria
Erwinia pyrifoliae DSM 12163 YP 005803283 1 644651 Proteobacteria
Erwinia pyrifoliae DSM 12163 YP 005803041 1 644651 Proteobacteria
Erwinia pyrifoliae DSM 12163 YP 005802528 1 644651 Proteobacteria
Erwinia pyrifoliae DSM 12163 YP 005804549 1 644651 Proteobacteria
Erwinia pyrifoliae DSM 12163 YP 005800822 1 644651 Proteobacteria
Salmonella enterica subsp arizonae serovar 62 z4 z23 str RSK2980 YP 001572077 1 882884 Proteobacteria
Salmonella enterica subsp arizonae serovar 62 z4 z23 str RSK2980 YP 001572549 1 882884 Proteobacteria
Salmonella enterica subsp arizonae serovar 62 z4 z23 str RSK2980 YP 001572042 1 882884 Proteobacteria
Shigella flexneri 2002017 YP 005712142 1 591020 Proteobacteria
Shigella boydii Sb227 YP 406293 1 300268 Proteobacteria
Shigella flexneri 2002017 YP 005711961 1 591020 Proteobacteria
Shigella sonnei 53G YP 005454566 1 216599 Proteobacteria
Shigella sonnei 53G YP 005458229 1 216599 Proteobacteria
Shigella sonnei 53G YP 005454524 1 216599 Proteobacteria
Shigella sonnei 53G YP 005454399 1 216599 Proteobacteria
Shigella sonnei 53G YP 005456481 1 216599 Proteobacteria
Shigella sonnei 53G YP 005458263 1 216599 Proteobacteria
Shigella sonnei 53G YP 005456362 1 216599 Proteobacteria
Shigella sonnei 53G YP 005457816 1 216599 Proteobacteria
Shigella sonnei 53G YP 005458025 1 216599 Proteobacteria
Shigella sonnei 53G YP 005457892 1 216599 Proteobacteria
Shigella sonnei 53G YP 005456630 1 216599 Proteobacteria
Shigella sonnei 53G YP 005454887 1 216599 Proteobacteria
Shigella sonnei 53G YP 005456231 1 216599 Proteobacteria
Shigella sonnei 53G YP 005458128 1 216599 Proteobacteria
Shigella sonnei 53G YP 005455840 1 216599 Proteobacteria

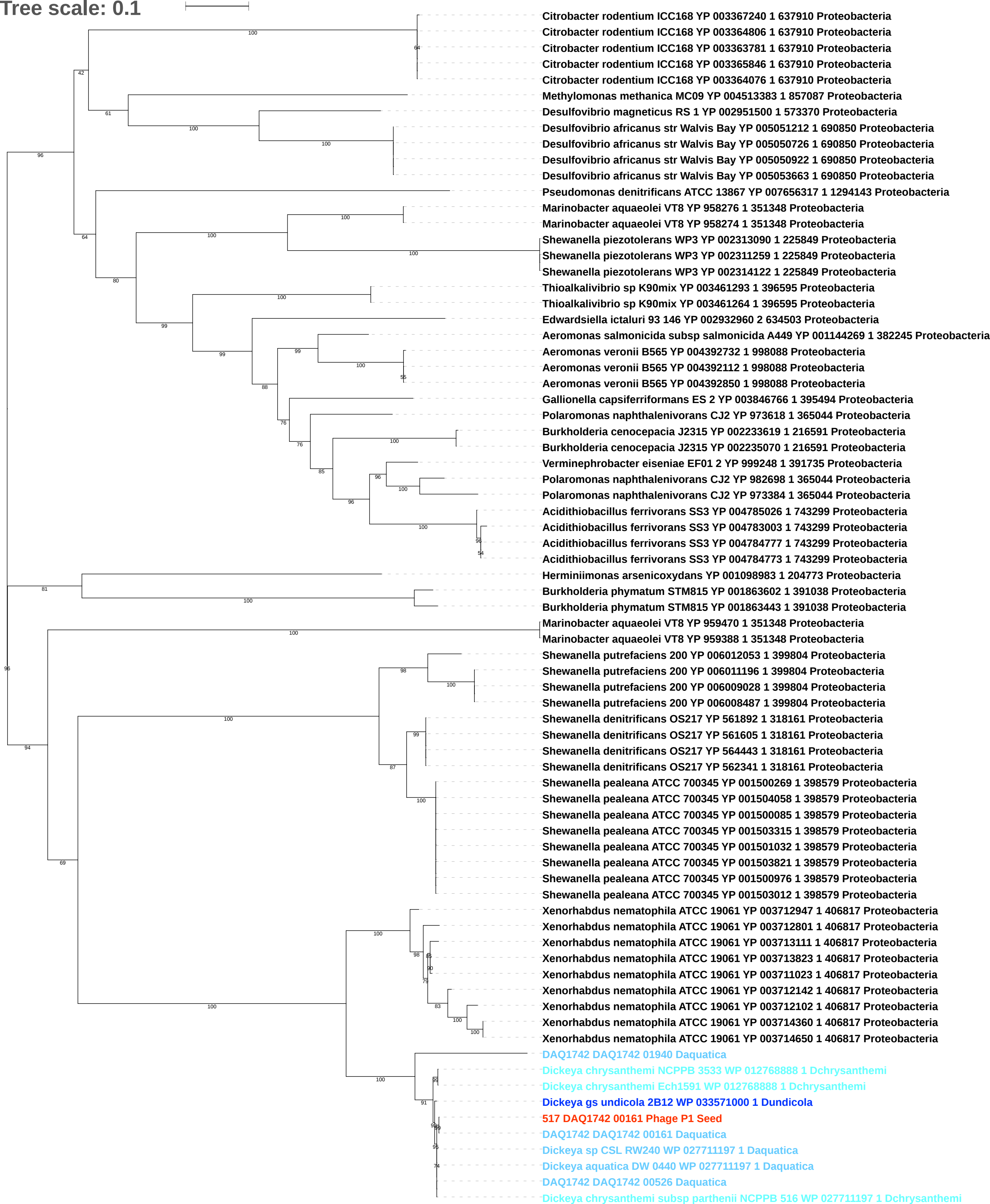
Tree scale: 0.1
Citrobacter rodentium ICC168 YP 003367240 1 637910 Proteobacteria
Citrobacter rodentium ICC168 YP 003364806 1 637910 Proteobacteria
Citrobacter rodentium ICC168 YP 003363781 1 637910 Proteobacteria
Citrobacter rodentium ICC168 YP 003365846 1 637910 Proteobacteria
Citrobacter rodentium ICC168 YP 003364076 1 637910 Proteobacteria
Methylomonas methanica MC09 YP 004513383 1 857087 Proteobacteria
Desulfovibrio magneticus RS 1 YP 002951500 1 573370 Proteobacteria
Desulfovibrio africanus str Walvis Bay YP 005051212 1 690850 Proteobacteria
Desulfovibrio africanus str Walvis Bay YP 005050726 1 690850 Proteobacteria
Desulfovibrio africanus str Walvis Bay YP 005050922 1 690850 Proteobacteria
Desulfovibrio africanus str Walvis Bay YP 005053663 1 690850 Proteobacteria
Pseudomonas denitrificans ATCC 13867 YP 007656317 1 1294143 Proteobacteria
Marinobacter aquaeolei VT8 YP 958276 1 351348 Proteobacteria
Marinobacter aquaeolei VT8 YP 958274 1 351348 Proteobacteria
Shewanella piezotolerans WP3 YP 002313090 1 225849 Proteobacteria
Shewanella piezotolerans WP3 YP 002311259 1 225849 Proteobacteria
Shewanella piezotolerans WP3 YP 002314122 1 225849 Proteobacteria
Thioalkalivibrio sp K90mix YP 003461293 1 396595 Proteobacteria
Thioalkalivibrio sp K90mix YP 003461264 1 396595 Proteobacteria
Edwardsiella ictaluri 93 146 YP 002932960 2 634503 Proteobacteria
Aeromonas salmonicida subsp salmonicida A449 YP 001144269 1 382245 Proteobacteria
Aeromonas veronii B565 YP 004392732 1 998088 Proteobacteria
Aeromonas veronii B565 YP 004392112 1 998088 Proteobacteria
Aeromonas veronii B565 YP 004392850 1 998088 Proteobacteria
Gallionella capsiferriformans ES 2 YP 003846766 1 395494 Proteobacteria
Polaromonas naphthalenivorans CJ2 YP 973618 1 365044 Proteobacteria
Burkholderia cenocepacia J2315 YP 002233619 1 216591 Proteobacteria
Burkholderia cenocepacia J2315 YP 002235070 1 216591 Proteobacteria
Verminephrobacter eiseniae EF01 2 YP 999248 1 391735 Proteobacteria
Polaromonas naphthalenivorans CJ2 YP 982698 1 365044 Proteobacteria
Polaromonas naphthalenivorans CJ2 YP 973384 1 365044 Proteobacteria
Acidithiobacillus ferrivorans SS3 YP 004785026 1 743299 Proteobacteria
Acidithiobacillus ferrivorans SS3 YP 004783003 1 743299 Proteobacteria
Acidithiobacillus ferrivorans SS3 YP 004784777 1 743299 Proteobacteria
Acidithiobacillus ferrivorans SS3 YP 004784773 1 743299 Proteobacteria
Herminiimonas arsenicoxydans YP 001098983 1 204773 Proteobacteria
Burkholderia phymatum STM815 YP 001863602 1 391038 Proteobacteria
Burkholderia phymatum STM815 YP 001863443 1 391038 Proteobacteria
Marinobacter aquaeolei VT8 YP 959470 1 351348 Proteobacteria
Marinobacter aquaeolei VT8 YP 959388 1 351348 Proteobacteria
Shewanella putrefaciens 200 YP 006012053 1 399804 Proteobacteria
Shewanella putrefaciens 200 YP 006011196 1 399804 Proteobacteria
Shewanella putrefaciens 200 YP 006009028 1 399804 Proteobacteria
Shewanella putrefaciens 200 YP 006008487 1 399804 Proteobacteria
Shewanella denitrificans OS217 YP 561892 1 318161 Proteobacteria
Shewanella denitrificans OS217 YP 561605 1 318161 Proteobacteria
Shewanella denitrificans OS217 YP 564443 1 318161 Proteobacteria
Shewanella denitrificans OS217 YP 562341 1 318161 Proteobacteria
Shewanella pealeana ATCC 700345 YP 001500269 1 398579 Proteobacteria
Shewanella pealeana ATCC 700345 YP 001504058 1 398579 Proteobacteria
Shewanella pealeana ATCC 700345 YP 001500085 1 398579 Proteobacteria
Shewanella pealeana ATCC 700345 YP 001503315 1 398579 Proteobacteria
Shewanella pealeana ATCC 700345 YP 001501032 1 398579 Proteobacteria
Shewanella pealeana ATCC 700345 YP 001503821 1 398579 Proteobacteria
Shewanella pealeana ATCC 700345 YP 001500976 1 398579 Proteobacteria
Shewanella pealeana ATCC 700345 YP 001503012 1 398579 Proteobacteria
Xenorhabdus nematophila ATCC 19061 YP 003712947 1 406817 Proteobacteria
Xenorhabdus nematophila ATCC 19061 YP 003712801 1 406817 Proteobacteria
Xenorhabdus nematophila ATCC 19061 YP 003713111 1 406817 Proteobacteria
Xenorhabdus nematophila ATCC 19061 YP 003713823 1 406817 Proteobacteria
Xenorhabdus nematophila ATCC 19061 YP 003711023 1 406817 Proteobacteria
Xenorhabdus nematophila ATCC 19061 YP 003712142 1 406817 Proteobacteria
Xenorhabdus nematophila ATCC 19061 YP 003712102 1 406817 Proteobacteria
Xenorhabdus nematophila ATCC 19061 YP 003714360 1 406817 Proteobacteria
Xenorhabdus nematophila ATCC 19061 YP 003714650 1 406817 Proteobacteria
DAQ1742 DAQ1742 01940 Daquatica
Dickeya chrysanthemi NCPPB 3533 WP 012768888 1 Dchrysanthemi
Dickeya chrysanthemi Ech1591 WP 012768888 1 Dchrysanthemi
Dickeya gs undicola 2B12 WP 033571000 1 Dundicola
517 DAQ1742 00161 Phage P1 Seed
DAQ1742 DAQ1742 00161 Daquatica
Dickeya sp CSL RW240 WP 027711197 1 Daquatica
Dickeya aquatica DW 0440 WP 027711197 1 Daquatica
DAQ1742 DAQ1742 00526 Daquatica
Dickeya chrysanthemi subsp parthenii NCPPB 516 WP 027711197 1 Dchrysanthemi

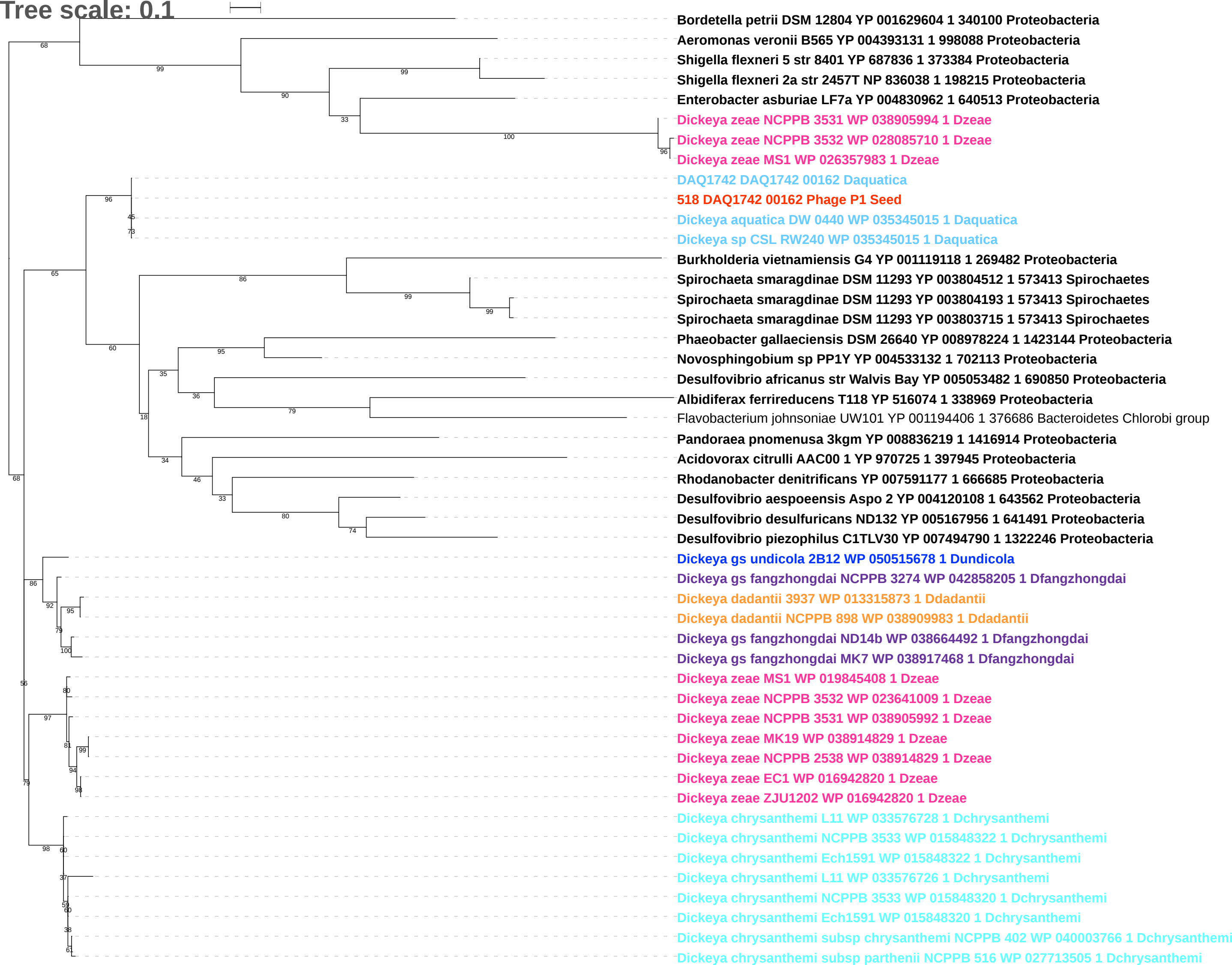
Tree scale: 0.1
Bordetella petrii DSM 12804 YP 001629604 1 340100 Proteobacteria
Aeromonas veronii B565 YP 004393131 1 998088 Proteobacteria
Shigella flexneri 5 str 8401 YP 687836 1 373384 Proteobacteria
Shigella flexneri 2a str 2457T NP 836038 1 198215 Proteobacteria
Enterobacter asburiae LF7a YP 004830962 1 640513 Proteobacteria
Dickeya zeae NCPPB 3531 WP 038905994 1 Dzeae
Dickeya zeae NCPPB 3532 WP 028085710 1 Dzeae
Dickeya zeae MS1 WP 026357983 1 Dzeae
DAQ1742 DAQ1742 00162 Daquatica
518 DAQ1742 00162 Phage P1 Seed
Dickeya aquatica DW 0440 WP 035345015 1 Daquatica
Dickeya sp CSL RW240 WP 035345015 1 Daquatica
Burkholderia vietnamiensis G4 YP 001119118 1 269482 Proteobacteria
Spirochaeta smaragdinae DSM 11293 YP 003804512 1 573413 Spirochaetes
Spirochaeta smaragdinae DSM 11293 YP 003804193 1 573413 Spirochaetes
Spirochaeta smaragdinae DSM 11293 YP 003803715 1 573413 Spirochaetes
Phaeobacter gallaeciensis DSM 26640 YP 008978224 1 1423144 Proteobacteria
Novosphingobium sp PP1Y YP 004533132 1 702113 Proteobacteria
Desulfovibrio africanus str Walvis Bay YP 005053482 1 690850 Proteobacteria
Albidiferax ferrireducens T118 YP 516074 1 338969 Proteobacteria
Flavobacterium johnsoniae UW101 YP 001194406 1 376686 Bacteroidetes Chlorobi group
Pandoraea pnomenusa 3kgm YP 008836219 1 1416914 Proteobacteria
Acidovorax citrulli AAC00 1 YP 970725 1 397945 Proteobacteria
Rhodanobacter denitrificans YP 007591177 1 666685 Proteobacteria
Desulfovibrio aespoeensis Aspo 2 YP 004120108 1 643562 Proteobacteria
Desulfovibrio desulfuricans ND132 YP 005167956 1 641491 Proteobacteria
Desulfovibrio piezophilus C1TLV30 YP 007494790 1 1322246 Proteobacteria
Dickeya gs undicola 2B12 WP 050515678 1 Dundicola
Dickeya gs fangzhongdai NCPPB 3274 WP 042858205 1 Dfangzhongdai
Dickeya dadantii 3937 WP 013315873 1 Ddadantii
Dickeya dadantii NCPPB 898 WP 038909983 1 Ddadantii
Dickeya gs fangzhongdai ND14b WP 038664492 1 Dfangzhongdai
Dickeya gs fangzhongdai MK7 WP 038917468 1 Dfangzhongdai
Dickeya zeae MS1 WP 019845408 1 Dzeae
Dickeya zeae NCPPB 3532 WP 023641009 1 Dzeae
Dickeya zeae NCPPB 3531 WP 038905992 1 Dzeae
Dickeya zeae MK19 WP 038914829 1 Dzeae
Dickeya zeae NCPPB 2538 WP 038914829 1 Dzeae
Dickeya zeae EC1 WP 016942820 1 Dzeae
Dickeya zeae ZJU1202 WP 016942820 1 Dzeae
Dickeya chrysanthemi L11 WP 033576728 1 Dchrysanthemi
Dickeya chrysanthemi NCPPB 3533 WP 015848322 1 Dchrysanthemi
Dickeya chrysanthemi Ech1591 WP 015848322 1 Dchrysanthemi
Dickeya chrysanthemi L11 WP 033576726 1 Dchrysanthemi
Dickeya chrysanthemi NCPPB 3533 WP 015848320 1 Dchrysanthemi
Dickeya chrysanthemi Ech1591 WP 015848320 1 Dchrysanthemi
Dickeya chrysanthemi subsp chrysanthemi NCPPB 402 WP 040003766 1 Dchrysanthemi
Dickeya chrysanthemi subsp parthenii NCPPB 516 WP 027713505 1 Dchrysanthemi

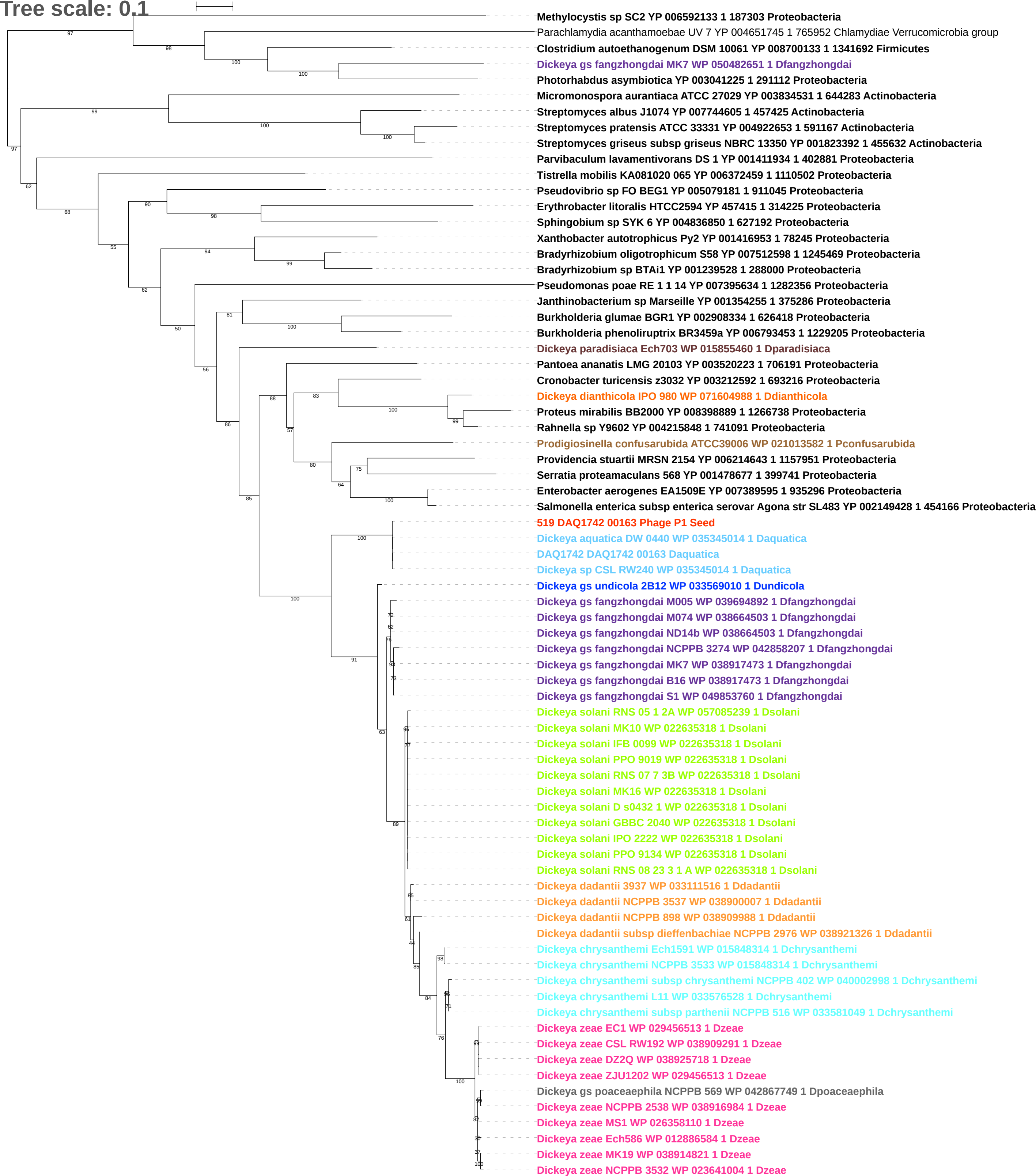
Tree scale: 0.1
Methylocystis sp SC2 YP 006592133 1 187303 Proteobacteria
Parachlamydia acanthamoebae UV 7 YP 004651745 1 765952 Chlamydiae Verrucomicrobia group
Clostridium autoethanogenum DSM 10061 YP 008700133 1 1341692 Firmicutes
Dickeya gs fangzhongdai MK7 WP 050482651 1 Dfangzhongdai
Photorhabdus asymbiotica YP 003041225 1 291112 Proteobacteria
Micromonospora aurantiaca ATCC 27029 YP 003834531 1 644283 Actinobacteria
Streptomyces albus J1074 YP 007744605 1 457425 Actinobacteria
Streptomyces pratensis ATCC 33331 YP 004922653 1 591167 Actinobacteria
Streptomyces griseus subsp griseus NBRC 13350 YP 001823392 1 455632 Actinobacteria
Parvibaculum lavamentivorans DS 1 YP 001411934 1 402881 Proteobacteria
Tistrella mobilis KA081020 065 YP 006372459 1 1110502 Proteobacteria
Pseudovibrio sp FO BEG1 YP 005079181 1 911045 Proteobacteria
Erythrobacter litoralis HTCC2594 YP 457415 1 314225 Proteobacteria
Sphingobium sp SYK 6 YP 004836850 1 627192 Proteobacteria
Xanthobacter autotrophicus Py2 YP 001416953 1 78245 Proteobacteria
Bradyrhizobium oligotrophicum S58 YP 007512598 1 1245469 Proteobacteria
Bradyrhizobium sp BTAi1 YP 001239528 1 288000 Proteobacteria
Pseudomonas poae RE 1 1 14 YP 007395634 1 1282356 Proteobacteria
Janthinobacterium sp Marseille YP 001354255 1 375286 Proteobacteria
Burkholderia glumae BGR1 YP 002908334 1 626418 Proteobacteria
Burkholderia phenoliruptrix BR3459a YP 006793453 1 1229205 Proteobacteria
Dickeya paradisiaca Ech703 WP 015855460 1 Dparadisiaca
Pantoea ananatis LMG 20103 YP 003520223 1 706191 Proteobacteria
Cronobacter turicensis z3032 YP 003212592 1 693216 Proteobacteria
Dickeya dianthicola IPO 980 WP 071604988 1 Ddianthicola
Proteus mirabilis BB2000 YP 008398889 1 1266738 Proteobacteria
Rahnella sp Y9602 YP 004215848 1 741091 Proteobacteria
Prodigiosinella confusarubida ATCC39006 WP 021013582 1 Pconfusarubida
Providencia stuartii MRSN 2154 YP 006214643 1 1157951 Proteobacteria
Serratia proteamaculans 568 YP 001478677 1 399741 Proteobacteria
Enterobacter aerogenes EA1509E YP 007389595 1 935296 Proteobacteria
Salmonella enterica subsp enterica serovar Agona str SL483 YP 002149428 1 454166 Proteobacteria
519 DAQ1742 00163 Phage P1 Seed
Dickeya aquatica DW 0440 WP 035345014 1 Daquatica
DAQ1742 DAQ1742 00163 Daquatica
Dickeya sp CSL RW240 WP 035345014 1 Daquatica
Dickeya gs undicola 2B12 WP 033569010 1 Dundicola
Dickeya gs fangzhongdai M005 WP 039694892 1 Dfangzhongdai
Dickeya gs fangzhongdai M074 WP 038664503 1 Dfangzhongdai
Dickeya gs fangzhongdai ND14b WP 038664503 1 Dfangzhongdai
Dickeya gs fangzhongdai NCPPB 3274 WP 042858207 1 Dfangzhongdai
Dickeya gs fangzhongdai MK7 WP 038917473 1 Dfangzhongdai
Dickeya gs fangzhongdai B16 WP 038917473 1 Dfangzhongdai
Dickeya gs fangzhongdai S1 WP 049853760 1 Dfangzhongdai
Dickeya solani RNS 05 1 2A WP 057085239 1 Dsolani
Dickeya solani MK10 WP 022635318 1 Dsolani
Dickeya solani IFB 0099 WP 022635318 1 Dsolani
Dickeya solani PPO 9019 WP 022635318 1 Dsolani
Dickeya solani RNS 07 7 3B WP 022635318 1 Dsolani
Dickeya solani MK16 WP 022635318 1 Dsolani
Dickeya solani D s0432 1 WP 022635318 1 Dsolani
Dickeya solani GBBC 2040 WP 022635318 1 Dsolani
Dickeya solani IPO 2222 WP 022635318 1 Dsolani
Dickeya solani PPO 9134 WP 022635318 1 Dsolani
Dickeya solani RNS 08 23 3 1 A WP 022635318 1 Dsolani
Dickeya dadantii 3937 WP 033111516 1 Ddadantii
Dickeya dadantii NCPPB 3537 WP 038900007 1 Ddadantii
Dickeya dadantii NCPPB 898 WP 038909988 1 Ddadantii
Dickeya dadantii subsp dieffenbachiae NCPPB 2976 WP 038921326 1 Ddadantii
Dickeya chrysanthemi Ech1591 WP 015848314 1 Dchrysanthemi
Dickeya chrysanthemi NCPPB 3533 WP 015848314 1 Dchrysanthemi
Dickeya chrysanthemi subsp chrysanthemi NCPPB 402 WP 040002998 1 Dchrysanthemi
Dickeya chrysanthemi L11 WP 033576528 1 Dchrysanthemi
Dickeya chrysanthemi subsp parthenii NCPPB 516 WP 033581049 1 Dchrysanthemi
Dickeya zeae EC1 WP 029456513 1 Dzeae
Dickeya zeae CSL RW192 WP 038909291 1 Dzeae
Dickeya zeae DZ2Q WP 038925718 1 Dzeae
Dickeya zeae ZJU1202 WP 029456513 1 Dzeae
Dickeya gs poaceaephila NCPPB 569 WP 042867749 1 Dpoaceaephila
Dickeya zeae NCPPB 2538 WP 038916984 1 Dzeae
Dickeya zeae MS1 WP 026358110 1 Dzeae
Dickeya zeae Ech586 WP 012886584 1 Dzeae
Dickeya zeae MK19 WP 038914821 1 Dzeae
Dickeya zeae NCPPB 3532 WP 023641004 1 Dzeae

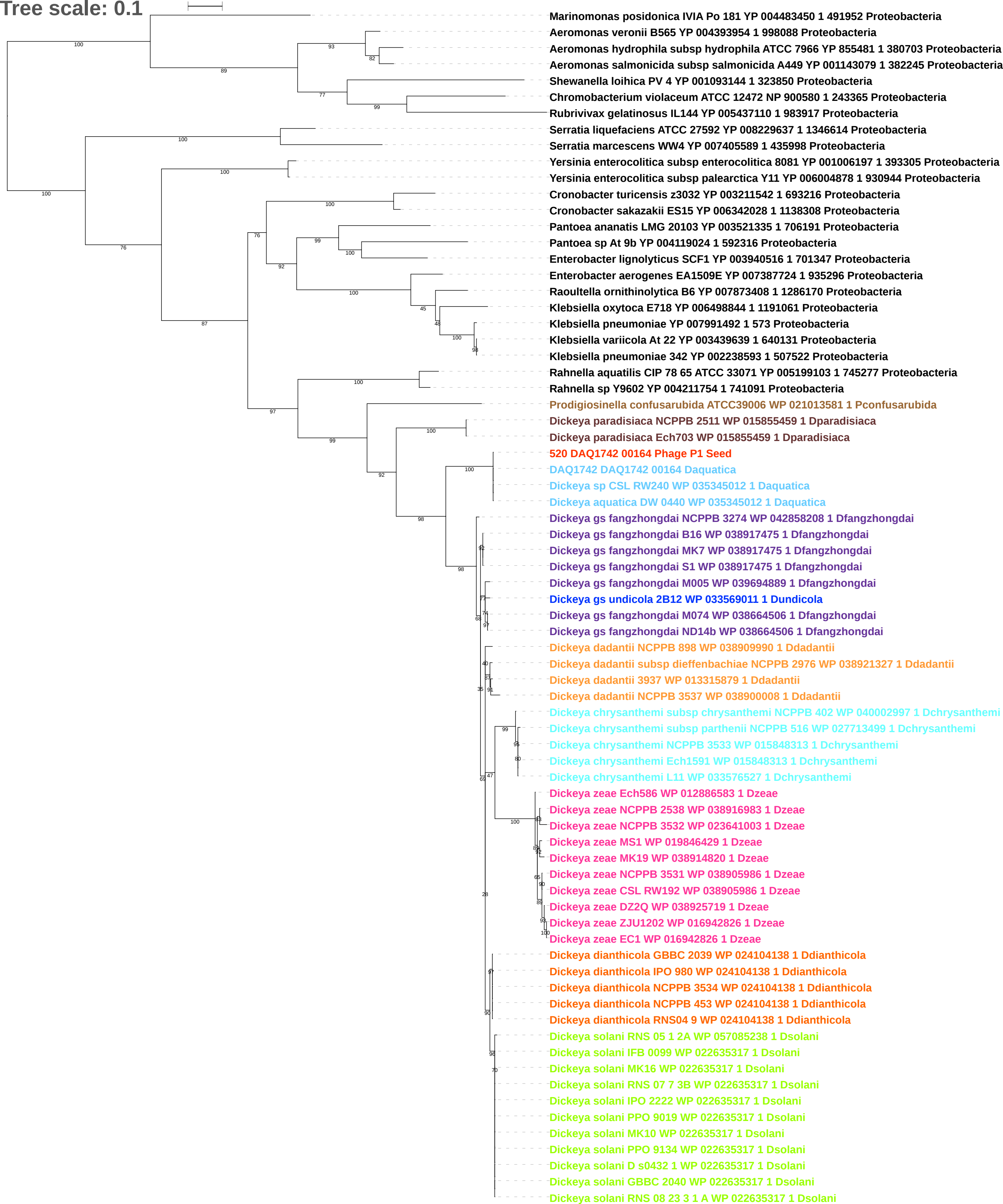

Tree scale: 0.1
Marinomonas posidonica IVIA Po 181 YP 004483450 1 491952 Proteobacteria
Aeromonas veronii B565 YP 004393954 1 998088 Proteobacteria
Aeromonas hydrophila subsp hydrophila ATCC 7966 YP 855481 1 380703 Proteobacteria
Aeromonas salmonicida subsp salmonicida A449 YP 001143079 1 382245 Proteobacteria
Shewanella loihica PV 4 YP 001093144 1 323850 Proteobacteria
Chromobacterium violaceum ATCC 12472 NP 900580 1 243365 Proteobacteria
Rubrivivax gelatinosus IL144 YP 005437110 1 983917 Proteobacteria
Serratia liquefaciens ATCC 27592 YP 008229637 1 1346614 Proteobacteria
Serratia marcescens WW4 YP 007405589 1 435998 Proteobacteria
Yersinia enterocolitica subsp enterocolitica 8081 YP 001006197 1 393305 Proteobacteria
Yersinia enterocolitica subsp palearctica Y11 YP 006004878 1 930944 Proteobacteria
Cronobacter turicensis z3032 YP 003211542 1 693216 Proteobacteria
Cronobacter sakazakii ES15 YP 006342028 1 1138308 Proteobacteria
Pantoea ananatis LMG 20103 YP 003521335 1 706191 Proteobacteria
Pantoea sp At 9b YP 004119024 1 592316 Proteobacteria
Enterobacter lignolyticus SCF1 YP 003940516 1 701347 Proteobacteria
Enterobacter aerogenes EA1509E YP 007387724 1 935296 Proteobacteria
Raoultella ornithinolytica B6 YP 007873408 1 1286170 Proteobacteria
Klebsiella oxytoca E718 YP 006498844 1 1191061 Proteobacteria
Klebsiella pneumoniae YP 007991492 1 573 Proteobacteria
Klebsiella variicola At 22 YP 003439639 1 640131 Proteobacteria
Klebsiella pneumoniae 342 YP 002238593 1 507522 Proteobacteria
Rahnella aquatilis CIP 78 65 ATCC 33071 YP 005199103 1 745277 Proteobacteria
Rahnella sp Y9602 YP 004211754 1 741091 Proteobacteria
Prodigiosinella confusarubida ATCC39006 WP 021013581 1 Pconfusarubida
Dickeya paradisiaca NCPPB 2511 WP 015855459 1 Dparadisiaca
Dickeya paradisiaca Ech703 WP 015855459 1 Dparadisiaca
520 DAQ1742 00164 Phage P1 Seed
DAQ1742 DAQ1742 00164 Daquatica
Dickeya sp CSL RW240 WP 035345012 1 Daquatica
Dickeya aquatica DW 0440 WP 035345012 1 Daquatica
Dickeya gs fangzhongdai NCPPB 3274 WP 042858208 1 Dfangzhongdai
Dickeya gs fangzhongdai B16 WP 038917475 1 Dfangzhongdai
Dickeya gs fangzhongdai MK7 WP 038917475 1 Dfangzhongdai
Dickeya gs fangzhongdai S1 WP 038917475 1 Dfangzhongdai
Dickeya gs fangzhongdai M005 WP 039694889 1 Dfangzhongdai
Dickeya gs undicola 2B12 WP 033569011 1 Dundicola
Dickeya gs fangzhongdai M074 WP 038664506 1 Dfangzhongdai
Dickeya gs fangzhongdai ND14b WP 038664506 1 Dfangzhongdai
Dickeya dadantii NCPPB 898 WP 038909990 1 Ddadantii
Dickeya dadantii subsp dieffenbachiae NCPPB 2976 WP 038921327 1 Ddadantii
Dickeya dadantii 3937 WP 013315879 1 Ddadantii
Dickeya dadantii NCPPB 3537 WP 038900008 1 Ddadantii
Dickeya chrysanthemi subsp chrysanthemi NCPPB 402 WP 040002997 1 Dchrysanthemi
Dickeya chrysanthemi subsp parthenii NCPPB 516 WP 027713499 1 Dchrysanthemi
Dickeya chrysanthemi NCPPB 3533 WP 015848313 1 Dchrysanthemi
Dickeya chrysanthemi Ech1591 WP 015848313 1 Dchrysanthemi
Dickeya chrysanthemi L11 WP 033576527 1 Dchrysanthemi
Dickeya zeae Ech586 WP 012886583 1 Dzeae
Dickeya zeae NCPPB 2538 WP 038916983 1 Dzeae
Dickeya zeae NCPPB 3532 WP 023641003 1 Dzeae
Dickeya zeae MS1 WP 019846429 1 Dzeae
Dickeya zeae MK19 WP 038914820 1 Dzeae
Dickeya zeae NCPPB 3531 WP 038905986 1 Dzeae
Dickeya zeae CSL RW192 WP 038905986 1 Dzeae
Dickeya zeae DZ2Q WP 038925719 1 Dzeae
Dickeya zeae ZJU1202 WP 016942826 1 Dzeae
Dickeya zeae EC1 WP 016942826 1 Dzeae
Dickeya dianthicola GBBC 2039 WP 024104138 1 Ddianthicola
Dickeya dianthicola IPO 980 WP 024104138 1 Ddianthicola
Dickeya dianthicola NCPPB 3534 WP 024104138 1 Ddianthicola
Dickeya dianthicola NCPPB 453 WP 024104138 1 Ddianthicola
Dickeya dianthicola RNS04 9 WP 024104138 1 Ddianthicola
Dickeya solani RNS 05 1 2A WP 057085238 1 Dsolani
Dickeya solani IFB 0099 WP 022635317 1 Dsolani
Dickeya solani MK16 WP 022635317 1 Dsolani
Dickeya solani RNS 07 7 3B WP 022635317 1 Dsolani
Dickeya solani IPO 2222 WP 022635317 1 Dsolani
Dickeya solani PPO 9019 WP 022635317 1 Dsolani
Dickeya solani MK10 WP 022635317 1 Dsolani
Dickeya solani PPO 9134 WP 022635317 1 Dsolani
Dickeya solani D s0432 1 WP 022635317 1 Dsolani
Dickeya solani GBBC 2040 WP 022635317 1 Dsolani
Dickeya solani RNS 08 23 3 1 A WP 022635317 1 Dsolani

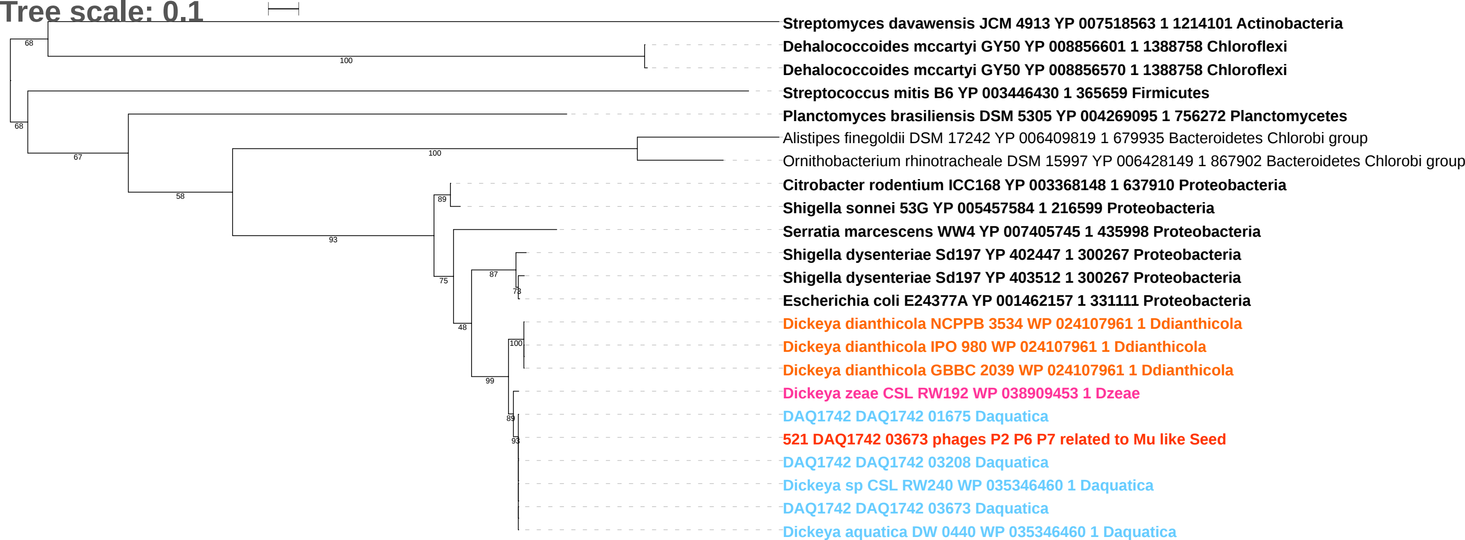
Tree scale: 0.1
Streptomyces davawensis JCM 4913 YP 007518563 1 1214101 Actinobacteria
Dehalococcoides mccartyi GY50 YP 008856601 1 1388758 Chloroflexi
Dehalococcoides mccartyi GY50 YP 008856570 1 1388758 Chloroflexi
Streptococcus mitis B6 YP 003446430 1 365659 Firmicutes
Planctomyces brasiliensis DSM 5305 YP 004269095 1 756272 Planctomycetes
Alistipes finegoldii DSM 17242 YP 006409819 1 679935 Bacteroidetes Chlorobi group
Ornithobacterium rhinotracheale DSM 15997 YP 006428149 1 867902 Bacteroidetes Chlorobi group
Citrobacter rodentium ICC168 YP 003368148 1 637910 Proteobacteria
Shigella sonnei 53G YP 005457584 1 216599 Proteobacteria
Serratia marcescens WW4 YP 007405745 1 435998 Proteobacteria
Shigella dysenteriae Sd197 YP 402447 1 300267 Proteobacteria
Shigella dysenteriae Sd197 YP 403512 1 300267 Proteobacteria
Escherichia coli E24377A YP 001462157 1 331111 Proteobacteria
Dickeya dianthicola NCPPB 3534 WP 024107961 1 Ddianthicola
Dickeya dianthicola IPO 980 WP 024107961 1 Ddianthicola
Dickeya dianthicola GBBC 2039 WP 024107961 1 Ddianthicola
Dickeya zeae CSL RW192 WP 038909453 1 Dzeae
DAQ1742 DAQ1742 01675 Daquatica
521 DAQ1742 03673 phages P2 P6 P7 related to Mu like Seed
DAQ1742 DAQ1742 03208 Daquatica
Dickeya sp CSL RW240 WP 035346460 1 Daquatica
DAQ1742 DAQ1742 03673 Daquatica
Dickeya aquatica DW 0440 WP 035346460 1 Daquatica
68
100
68
67
100
58
89
93
75
87
78
48
100
99
89
93

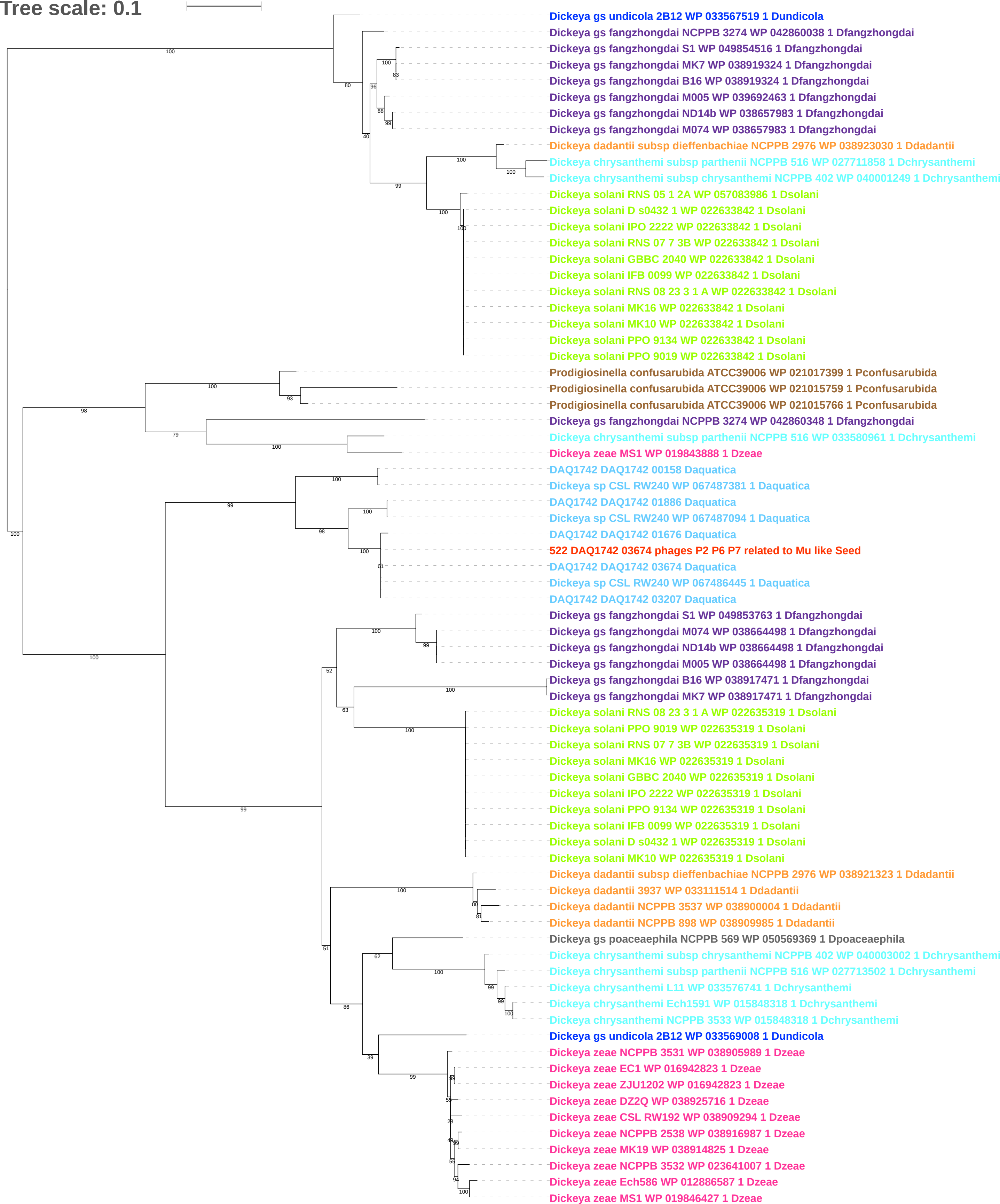
Tree scale: 0.1
Dickeya gs undicola 2B12 WP 033567519 1 Dundicola
Dickeya gs fangzhongdai NCPPB 3274 WP 042860038 1 Dfangzhongdai
Dickeya gs fangzhongdai S1 WP 049854516 1 Dfangzhongdai
Dickeya gs fangzhongdai MK7 WP 038919324 1 Dfangzhongdai
Dickeya gs fangzhongdai B16 WP 038919324 1 Dfangzhongdai
Dickeya gs fangzhongdai M005 WP 039692463 1 Dfangzhongdai
Dickeya gs fangzhongdai ND14b WP 038657983 1 Dfangzhongdai
Dickeya gs fangzhongdai M074 WP 038657983 1 Dfangzhongdai
Dickeya dadantii subsp dieffenbachiae NCPPB 2976 WP 038923030 1 Ddadantii
Dickeya chrysanthemi subsp parthenii NCPPB 516 WP 027711858 1 Dchrysanthemi
Dickeya chrysanthemi subsp chrysanthemi NCPPB 402 WP 040001249 1 Dchrysanthemi
Dickeya solani RNS 05 1 2A WP 057083986 1 Dsolani
Dickeya solani D s0432 1 WP 022633842 1 Dsolani
Dickeya solani IPO 2222 WP 022633842 1 Dsolani
Dickeya solani RNS 07 7 3B WP 022633842 1 Dsolani
Dickeya solani GBBC 2040 WP 022633842 1 Dsolani
Dickeya solani IFB 0099 WP 022633842 1 Dsolani
Dickeya solani RNS 08 23 3 1 A WP 022633842 1 Dsolani
Dickeya solani MK16 WP 022633842 1 Dsolani
Dickeya solani MK10 WP 022633842 1 Dsolani
Dickeya solani PPO 9134 WP 022633842 1 Dsolani
Dickeya solani PPO 9019 WP 022633842 1 Dsolani
Prodigiosinella confusarubida ATCC39006 WP 021017399 1 Pconfusarubida
Prodigiosinella confusarubida ATCC39006 WP 021015759 1 Pconfusarubida
Prodigiosinella confusarubida ATCC39006 WP 021015766 1 Pconfusarubida
Dickeya gs fangzhongdai NCPPB 3274 WP 042860348 1 Dfangzhongdai
Dickeya chrysanthemi subsp parthenii NCPPB 516 WP 033580961 1 Dchrysanthemi
Dickeya zeae MS1 WP 019843888 1 Dzeae
DAQ1742 DAQ1742 00158 Daquatica
Dickeya sp CSL RW240 WP 067487381 1 Daquatica
DAQ1742 DAQ1742 01886 Daquatica
Dickeya sp CSL RW240 WP 067487094 1 Daquatica
DAQ1742 DAQ1742 01676 Daquatica
522 DAQ1742 03674 phages P2 P6 P7 related to Mu like Seed
DAQ1742 DAQ1742 03674 Daquatica
Dickeya sp CSL RW240 WP 067486445 1 Daquatica
DAQ1742 DAQ1742 03207 Daquatica
Dickeya gs fangzhongdai S1 WP 049853763 1 Dfangzhongdai
Dickeya gs fangzhongdai M074 WP 038664498 1 Dfangzhongdai
Dickeya gs fangzhongdai ND14b WP 038664498 1 Dfangzhongdai
Dickeya gs fangzhongdai M005 WP 038664498 1 Dfangzhongdai
Dickeya gs fangzhongdai B16 WP 038917471 1 Dfangzhongdai
Dickeya gs fangzhongdai MK7 WP 038917471 1 Dfangzhongdai
Dickeya solani RNS 08 23 3 1 A WP 022635319 1 Dsolani
Dickeya solani PPO 9019 WP 022635319 1 Dsolani
Dickeya solani RNS 07 7 3B WP 022635319 1 Dsolani
Dickeya solani MK16 WP 022635319 1 Dsolani
Dickeya solani GBBC 2040 WP 022635319 1 Dsolani
Dickeya solani IPO 2222 WP 022635319 1 Dsolani
Dickeya solani PPO 9134 WP 022635319 1 Dsolani
Dickeya solani IFB 0099 WP 022635319 1 Dsolani
Dickeya solani D s0432 1 WP 022635319 1 Dsolani
Dickeya solani MK10 WP 022635319 1 Dsolani
Dickeya dadantii subsp dieffenbachiae NCPPB 2976 WP 038921323 1 Ddadantii
Dickeya dadantii 3937 WP 033111514 1 Ddadantii
Dickeya dadantii NCPPB 3537 WP 038900004 1 Ddadantii
Dickeya dadantii NCPPB 898 WP 038909985 1 Ddadantii
Dickeya gs poaceaephila NCPPB 569 WP 050569369 1 Dpoaceaephila
Dickeya chrysanthemi subsp chrysanthemi NCPPB 402 WP 040003002 1 Dchrysanthemi
Dickeya chrysanthemi subsp parthenii NCPPB 516 WP 027713502 1 Dchrysanthemi
Dickeya chrysanthemi L11 WP 033576741 1 Dchrysanthemi
Dickeya chrysanthemi Ech1591 WP 015848318 1 Dchrysanthemi
Dickeya chrysanthemi NCPPB 3533 WP 015848318 1 Dchrysanthemi
Dickeya gs undicola 2B12 WP 033569008 1 Dundicola
Dickeya zeae NCPPB 3531 WP 038905989 1 Dzeae
Dickeya zeae EC1 WP 016942823 1 Dzeae
Dickeya zeae ZJU1202 WP 016942823 1 Dzeae
Dickeya zeae DZ2Q WP 038925716 1 Dzeae
Dickeya zeae CSL RW192 WP 038909294 1 Dzeae
Dickeya zeae NCPPB 2538 WP 038916987 1 Dzeae
Dickeya zeae MK19 WP 038914825 1 Dzeae
Dickeya zeae NCPPB 3532 WP 023641007 1 Dzeae
Dickeya zeae Ech586 WP 012886587 1 Dzeae
Dickeya zeae MS1 WP 019846427 1 Dzeae

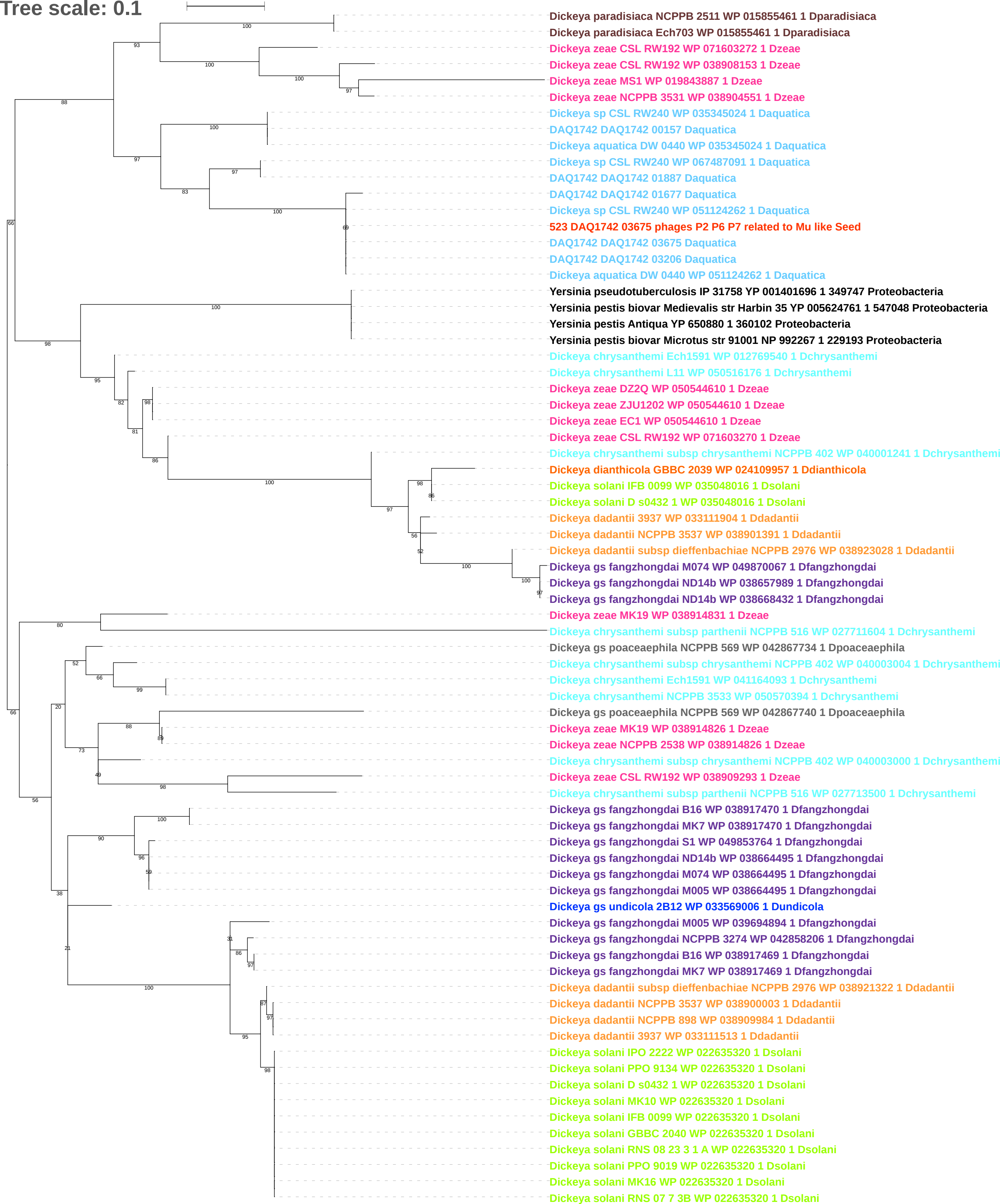
Tree scale: 0.1
Dickeya paradisiaca NCPPB 2511 WP 015855461 1 Dparadisiaca
Dickeya paradisiaca Ech703 WP 015855461 1 Dparadisiaca
Dickeya zeae CSL RW192 WP 071603272 1 Dzeae
Dickeya zeae CSL RW192 WP 038908153 1 Dzeae
Dickeya zeae MS1 WP 019843887 1 Dzeae
Dickeya zeae NCPPB 3531 WP 038904551 1 Dzeae
Dickeya sp CSL RW240 WP 035345024 1 Daquatica
DAQ1742 DAQ1742 00157 Daquatica
Dickeya aquatica DW 0440 WP 035345024 1 Daquatica
Dickeya sp CSL RW240 WP 067487091 1 Daquatica
DAQ1742 DAQ1742 01887 Daquatica
DAQ1742 DAQ1742 01677 Daquatica
Dickeya sp CSL RW240 WP 051124262 1 Daquatica
523 DAQ1742 03675 phages P2 P6 P7 related to Mu like Seed
DAQ1742 DAQ1742 03675 Daquatica
DAQ1742 DAQ1742 03206 Daquatica
Dickeya aquatica DW 0440 WP 051124262 1 Daquatica
Yersinia pseudotuberculosis IP 31758 YP 001401696 1 349747 Proteobacteria
Yersinia pestis biovar Medievalis str Harbin 35 YP 005624761 1 547048 Proteobacteria
Yersinia pestis Antiqua YP 650880 1 360102 Proteobacteria
Yersinia pestis biovar Microtus str 91001 NP 992267 1 229193 Proteobacteria
Dickeya chrysanthemi Ech1591 WP 012769540 1 Dchrysanthemi
Dickeya chrysanthemi L11 WP 050516176 1 Dchrysanthemi
Dickeya zeae DZ2Q WP 050544610 1 Dzeae
Dickeya zeae ZJU1202 WP 050544610 1 Dzeae
Dickeya zeae EC1 WP 050544610 1 Dzeae
Dickeya zeae CSL RW192 WP 071603270 1 Dzeae
Dickeya chrysanthemi subsp chrysanthemi NCPPB 402 WP 040001241 1 Dchrysanthemi
Dickeya dianthicola GBBC 2039 WP 024109957 1 Ddianthicola
Dickeya solani IFB 0099 WP 035048016 1 Dsolani
Dickeya solani D s0432 1 WP 035048016 1 Dsolani
Dickeya dadantii 3937 WP 033111904 1 Ddadantii
Dickeya dadantii NCPPB 3537 WP 038901391 1 Ddadantii
Dickeya dadantii subsp dieffenbachiae NCPPB 2976 WP 038923028 1 Ddadantii
Dickeya gs fangzhongdai M074 WP 049870067 1 Dfangzhongdai
Dickeya gs fangzhongdai ND14b WP 038657989 1 Dfangzhongdai
Dickeya gs fangzhongdai ND14b WP 038668432 1 Dfangzhongdai
Dickeya zeae MK19 WP 038914831 1 Dzeae
Dickeya chrysanthemi subsp parthenii NCPPB 516 WP 027711604 1 Dchrysanthemi
Dickeya gs poaceaephila NCPPB 569 WP 042867734 1 Dpoaceaephila
Dickeya chrysanthemi subsp chrysanthemi NCPPB 402 WP 040003004 1 Dchrysanthemi
Dickeya chrysanthemi Ech1591 WP 041164093 1 Dchrysanthemi
Dickeya chrysanthemi NCPPB 3533 WP 050570394 1 Dchrysanthemi
Dickeya gs poaceaephila NCPPB 569 WP 042867740 1 Dpoaceaephila
Dickeya zeae MK19 WP 038914826 1 Dzeae
Dickeya zeae NCPPB 2538 WP 038914826 1 Dzeae
Dickeya chrysanthemi subsp chrysanthemi NCPPB 402 WP 040003000 1 Dchrysanthemi
Dickeya zeae CSL RW192 WP 038909293 1 Dzeae
Dickeya chrysanthemi subsp parthenii NCPPB 516 WP 027713500 1 Dchrysanthemi
Dickeya gs fangzhongdai B16 WP 038917470 1 Dfangzhongdai
Dickeya gs fangzhongdai MK7 WP 038917470 1 Dfangzhongdai
Dickeya gs fangzhongdai S1 WP 049853764 1 Dfangzhongdai
Dickeya gs fangzhongdai ND14b WP 038664495 1 Dfangzhongdai
Dickeya gs fangzhongdai M074 WP 038664495 1 Dfangzhongdai
Dickeya gs fangzhongdai M005 WP 038664495 1 Dfangzhongdai
Dickeya gs undicola 2B12 WP 033569006 1 Dundicola
Dickeya gs fangzhongdai M005 WP 039694894 1 Dfangzhongdai
Dickeya gs fangzhongdai NCPPB 3274 WP 042858206 1 Dfangzhongdai
Dickeya gs fangzhongdai B16 WP 038917469 1 Dfangzhongdai
Dickeya gs fangzhongdai MK7 WP 038917469 1 Dfangzhongdai
Dickeya dadantii subsp dieffenbachiae NCPPB 2976 WP 038921322 1 Ddadantii
Dickeya dadantii NCPPB 3537 WP 038900003 1 Ddadantii
Dickeya dadantii NCPPB 898 WP 038909984 1 Ddadantii
Dickeya dadantii 3937 WP 033111513 1 Ddadantii
Dickeya solani IPO 2222 WP 022635320 1 Dsolani
Dickeya solani PPO 9134 WP 022635320 1 Dsolani
Dickeya solani D s0432 1 WP 022635320 1 Dsolani
Dickeya solani MK10 WP 022635320 1 Dsolani
Dickeya solani IFB 0099 WP 022635320 1 Dsolani
Dickeya solani GBBC 2040 WP 022635320 1 Dsolani
Dickeya solani RNS 08 23 3 1 A WP 022635320 1 Dsolani
Dickeya solani PPO 9019 WP 022635320 1 Dsolani
Dickeya solani MK16 WP 022635320 1 Dsolani
Dickeya solani RNS 07 7 3B WP 022635320 1 Dsolani

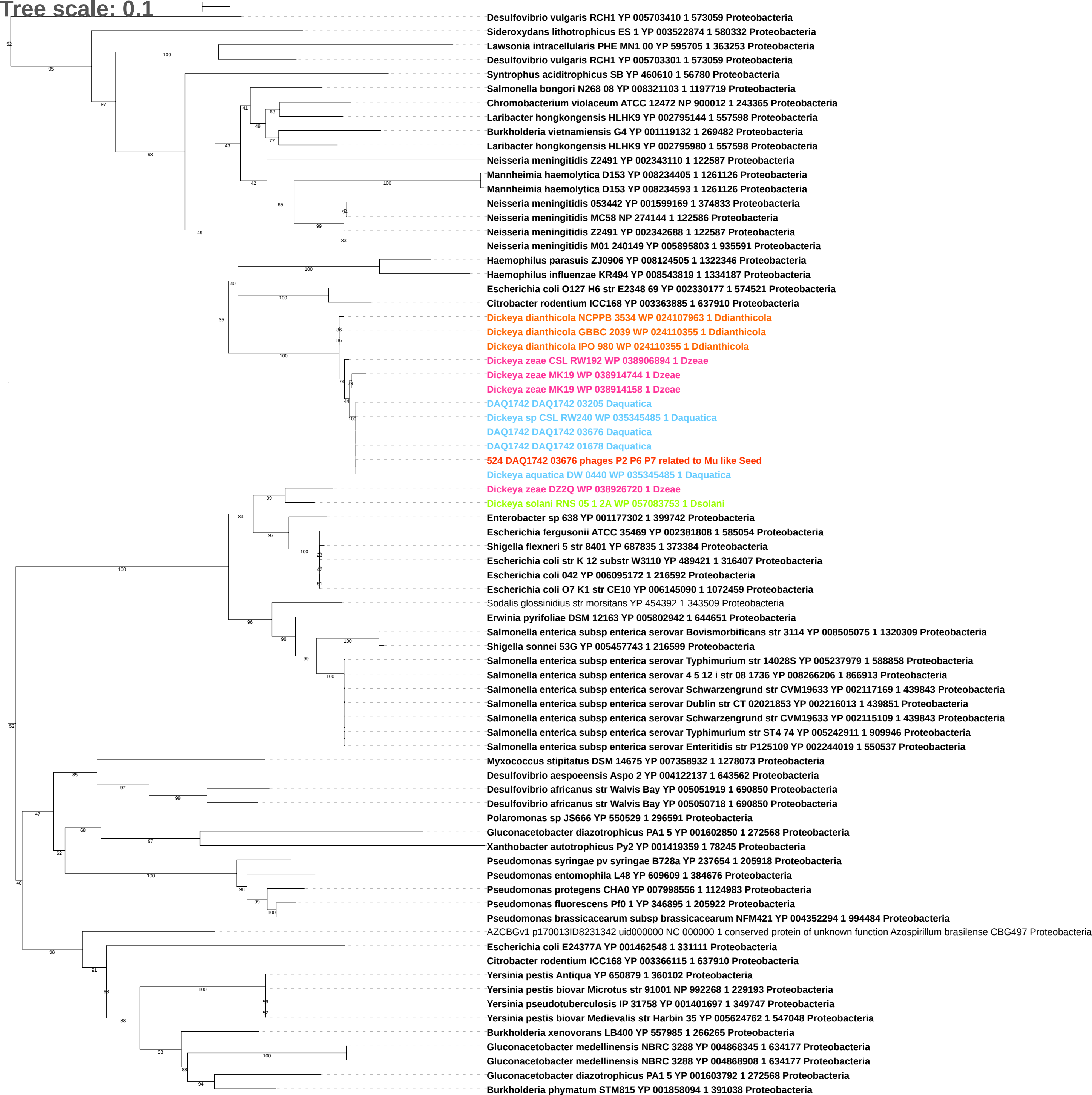
Tree scale: 0.1
Desulfovibrio vulgaris RCH1 YP 005703410 1 573059 Proteobacteria
Sideroxydans lithotrophicus ES 1 YP 003522874 1 580332 Proteobacteria
Lawsonia intracellularis PHE MN1 00 YP 595705 1 363253 Proteobacteria
Desulfovibrio vulgaris RCH1 YP 005703301 1 573059 Proteobacteria
Syntrophus aciditrophicus SB YP 460610 1 56780 Proteobacteria
Salmonella bongori N268 08 YP 008321103 1 1197719 Proteobacteria
Chromobacterium violaceum ATCC 12472 NP 900012 1 243365 Proteobacteria
Laribacter hongkongensis HLHK9 YP 002795144 1 557598 Proteobacteria
Burkholderia vietnamiensis G4 YP 001119132 1 269482 Proteobacteria
Laribacter hongkongensis HLHK9 YP 002795980 1 557598 Proteobacteria
Neisseria meningitidis Z2491 YP 002343110 1 122587 Proteobacteria
Mannheimia haemolytica D153 YP 008234405 1 1261126 Proteobacteria
Mannheimia haemolytica D153 YP 008234593 1 1261126 Proteobacteria
Neisseria meningitidis 053442 YP 001599169 1 374833 Proteobacteria
Neisseria meningitidis MC58 NP 274144 1 122586 Proteobacteria
Neisseria meningitidis Z2491 YP 002342688 1 122587 Proteobacteria
Neisseria meningitidis M01 240149 YP 005895803 1 935591 Proteobacteria
Haemophilus parasuis ZJ0906 YP 008124505 1 1322346 Proteobacteria
Haemophilus influenzae KR494 YP 008543819 1 1334187 Proteobacteria
Escherichia coli O127 H6 str E2348 69 YP 002330177 1 574521 Proteobacteria
Citrobacter rodentium ICC168 YP 003363885 1 637910 Proteobacteria
Dickeya dianthicola NCPPB 3534 WP 024107963 1 Ddianthicola
Dickeya dianthicola GBBC 2039 WP 024110355 1 Ddianthicola
Dickeya dianthicola IPO 980 WP 024110355 1 Ddianthicola
Dickeya zeae CSL RW192 WP 038906894 1 Dzeae
Dickeya zeae MK19 WP 038914744 1 Dzeae
Dickeya zeae MK19 WP 038914158 1 Dzeae
DAQ1742 DAQ1742 03205 Daquatica
Dickeya sp CSL RW240 WP 035345485 1 Daquatica
DAQ1742 DAQ1742 03676 Daquatica
DAQ1742 DAQ1742 01678 Daquatica
524 DAQ1742 03676 phages P2 P6 P7 related to Mu like Seed
Dickeya aquatica DW 0440 WP 035345485 1 Daquatica
Dickeya zeae DZ2Q WP 038926720 1 Dzeae
Dickeya solani RNS 05 1 2A WP 057083753 1 Dsolani
Enterobacter sp 638 YP 001177302 1 399742 Proteobacteria
Escherichia fergusonii ATCC 35469 YP 002381808 1 585054 Proteobacteria
Shigella flexneri 5 str 8401 YP 687835 1 373384 Proteobacteria
Escherichia coli str K 12 substr W3110 YP 489421 1 316407 Proteobacteria
Escherichia coli 042 YP 006095172 1 216592 Proteobacteria
Escherichia coli O7 K1 str CE10 YP 006145090 1 1072459 Proteobacteria
Sodalis glossinidius str morsitans YP 454392 1 343509 Proteobacteria
Erwinia pyrifoliae DSM 12163 YP 005802942 1 644651 Proteobacteria
Salmonella enterica subsp enterica serovar Bovismorbificans str 3114 YP 008505075 1 1320309 Proteobacteria
Shigella sonnei 53G YP 005457743 1 216599 Proteobacteria
Salmonella enterica subsp enterica serovar Typhimurium str 14028S YP 005237979 1 588858 Proteobacteria
Salmonella enterica subsp enterica serovar 4 5 12 i str 08 1736 YP 008266206 1 866913 Proteobacteria
Salmonella enterica subsp enterica serovar Schwarzengrund str CVM19633 YP 002117169 1 439843 Proteobacteria
Salmonella enterica subsp enterica serovar Dublin str CT 02021853 YP 002216013 1 439851 Proteobacteria
Salmonella enterica subsp enterica serovar Schwarzengrund str CVM19633 YP 002115109 1 439843 Proteobacteria
Salmonella enterica subsp enterica serovar Typhimurium str ST4 74 YP 005242911 1 909946 Proteobacteria
Salmonella enterica subsp enterica serovar Enteritidis str P125109 YP 002244019 1 550537 Proteobacteria
Myxococcus stipitatus DSM 14675 YP 007358932 1 1278073 Proteobacteria
Desulfovibrio aespoeensis Aspo 2 YP 004122137 1 643562 Proteobacteria
Desulfovibrio africanus str Walvis Bay YP 005051919 1 690850 Proteobacteria
Desulfovibrio africanus str Walvis Bay YP 005050718 1 690850 Proteobacteria
Polaromonas sp JS666 YP 550529 1 296591 Proteobacteria
Gluconacetobacter diazotrophicus PA1 5 YP 001602850 1 272568 Proteobacteria
Xanthobacter autotrophicus Py2 YP 001419359 1 78245 Proteobacteria
Pseudomonas syringae pv syringae B728a YP 237654 1 205918 Proteobacteria
Pseudomonas entomophila L48 YP 609609 1 384676 Proteobacteria
Pseudomonas protegens CHA0 YP 007998556 1 1124983 Proteobacteria
Pseudomonas fluorescens Pf0 1 YP 346895 1 205922 Proteobacteria
Pseudomonas brassicacearum subsp brassicacearum NFM421 YP 004352294 1 994484 Proteobacteria
AZCBGv1 p170013ID8231342 uid000000 NC 000000 1 conserved protein of unknown function Azospirillum brasilense CBG497 Proteobacteria
Escherichia coli E24377A YP 001462548 1 331111 Proteobacteria
Citrobacter rodentium ICC168 YP 003366115 1 637910 Proteobacteria
Yersinia pestis Antiqua YP 650879 1 360102 Proteobacteria
Yersinia pestis biovar Microtus str 91001 NP 992268 1 229193 Proteobacteria
Yersinia pseudotuberculosis IP 31758 YP 001401697 1 349747 Proteobacteria
Yersinia pestis biovar Medievalis str Harbin 35 YP 005624762 1 547048 Proteobacteria
Burkholderia xenovorans LB400 YP 557985 1 266265 Proteobacteria
Gluconacetobacter medellinensis NBRC 3288 YP 004868345 1 634177 Proteobacteria
Gluconacetobacter medellinensis NBRC 3288 YP 004868908 1 634177 Proteobacteria
Gluconacetobacter diazotrophicus PA1 5 YP 001603792 1 272568 Proteobacteria
Burkholderia phymatum STM815 YP 001858094 1 391038 Proteobacteria

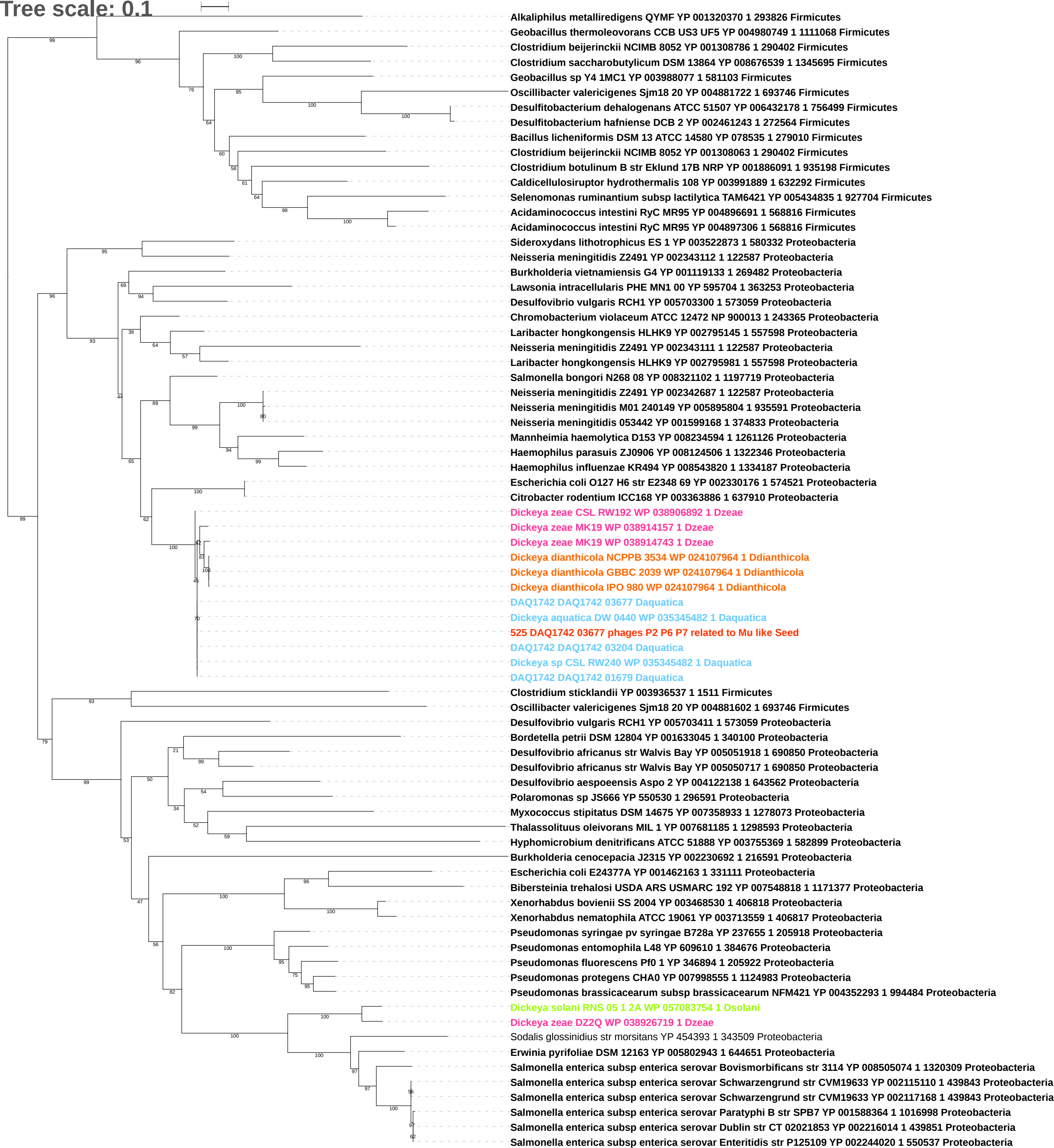
Tree scale: 0.1
Alkaliphilus metalliredigens QYMF YP 001320370 1 293826 Firmicutes
Geobacillus thermoleovorans CCB US3 UF5 YP 004980749 1 1111068 Firmicutes
Clostridium beijerinckii NCIMB 8052 YP 001308786 1 290402 Firmicutes
Clostridium saccharobutylicum DSM 13864 YP 008676539 1 1345695 Firmicutes
Geobacillus sp Y4 1MC1 YP 003988077 1 581103 Firmicutes
Oscillibacter valericigenes Sjm18 20 YP 004881722 1 693746 Firmicutes
Desulfitobacterium dehalogenans ATCC 51507 YP 006432178 1 756499 Firmicutes
Desulfitobacterium hafniense DCB 2 YP 002461243 1 272564 Firmicutes
Bacillus licheniformis DSM 13 ATCC 14580 YP 078535 1 279010 Firmicutes
Clostridium beijerinckii NCIMB 8052 YP 001308063 1 290402 Firmicutes
Clostridium botulinum B str Eklund 17B NRP YP 001886091 1 935198 Firmicutes
Caldicellulosiruptor hydrothermalis 108 YP 003991889 1 632292 Firmicutes
Selenomonas ruminantium subsp lactilytica TAM6421 YP 005434835 1 927704 Firmicutes
Acidaminococcus intestini RyC MR95 YP 004896691 1 568816 Firmicutes
Acidaminococcus intestini RyC MR95 YP 004897306 1 568816 Firmicutes
Sideroxydans lithotrophicus ES 1 YP 003522873 1 580332 Proteobacteria
Neisseria meningitidis Z2491 YP 002343112 1 122587 Proteobacteria
Burkholderia vietnamiensis G4 YP 001119133 1 269482 Proteobacteria
Lawsonia intracellularis PHE MN1 00 YP 595704 1 363253 Proteobacteria
Desulfovibrio vulgaris RCH1 YP 005703300 1 573059 Proteobacteria
Chromobacterium violaceum ATCC 12472 NP 900013 1 243365 Proteobacteria
Laribacter hongkongensis HLHK9 YP 002795145 1 557598 Proteobacteria
Neisseria meningitidis Z2491 YP 002343111 1 122587 Proteobacteria
Laribacter hongkongensis HLHK9 YP 002795981 1 557598 Proteobacteria
Salmonella bongori N268 08 YP 008321102 1 1197719 Proteobacteria
Neisseria meningitidis Z2491 YP 002342687 1 122587 Proteobacteria
Neisseria meningitidis M01 240149 YP 005895804 1 935591 Proteobacteria
Neisseria meningitidis 053442 YP 001599168 1 374833 Proteobacteria
Mannheimia haemolytica D153 YP 008234594 1 1261126 Proteobacteria
Haemophilus parasuis ZJ0906 YP 008124506 1 1322346 Proteobacteria
Haemophilus influenzae KR494 YP 008543820 1 1334187 Proteobacteria
Escherichia coli O127 H6 str E2348 69 YP 002330176 1 574521 Proteobacteria
Citrobacter rodentium ICC168 YP 003363886 1 637910 Proteobacteria
Dickeya zeae CSL RW192 WP 038906892 1 Dzeae
Dickeya zeae MK19 WP 038914157 1 Dzeae
Dickeya zeae MK19 WP 038914743 1 Dzeae
Dickeya dianthicola NCPPB 3534 WP 024107964 1 Ddianthicola
Dickeya dianthicola GBBC 2039 WP 024107964 1 Ddianthicola
Dickeya dianthicola IPO 980 WP 024107964 1 Ddianthicola
DAQ1742 DAQ1742 03677 Daquatica
Dickeya aquatica DW 0440 WP 035345482 1 Daquatica
525 DAQ1742 03677 phages P2 P6 P7 related to Mu like Seed
DAQ1742 DAQ1742 03204 Daquatica
Dickeya sp CSL RW240 WP 035345482 1 Daquatica
DAQ1742 DAQ1742 01679 Daquatica
Clostridium sticklandii YP 003936537 1 1511 Firmicutes
Oscillibacter valericigenes Sjm18 20 YP 004881602 1 693746 Firmicutes
Desulfovibrio vulgaris RCH1 YP 005703411 1 573059 Proteobacteria
Bordetella petrii DSM 12804 YP 001633045 1 340100 Proteobacteria
Desulfovibrio africanus str Walvis Bay YP 005051918 1 690850 Proteobacteria
Desulfovibrio africanus str Walvis Bay YP 005050717 1 690850 Proteobacteria
Desulfovibrio aespoeensis Aspo 2 YP 004122138 1 643562 Proteobacteria
Polaromonas sp JS666 YP 550530 1 296591 Proteobacteria
Myxococcus stipitatus DSM 14675 YP 007358933 1 1278073 Proteobacteria
Thalassolituus oleivorans MIL 1 YP 007681185 1 1298593 Proteobacteria
Hyphomicrobium denitrificans ATCC 51888 YP 003755369 1 582899 Proteobacteria
Burkholderia cenocepacia J2315 YP 002230692 1 216591 Proteobacteria
Escherichia coli E24377A YP 001462163 1 331111 Proteobacteria
Bibersteinia trehalosi USDA ARS USMARC 192 YP 007548818 1 1171377 Proteobacteria
Xenorhabdus bovienii SS 2004 YP 003468530 1 406818 Proteobacteria
Xenorhabdus nematophila ATCC 19061 YP 003713559 1 406817 Proteobacteria
Pseudomonas syringae pv syringae B728a YP 237655 1 205918 Proteobacteria
Pseudomonas entomophila L48 YP 609610 1 384676 Proteobacteria
Pseudomonas fluorescens Pf0 1 YP 346894 1 205922 Proteobacteria
Pseudomonas protegens CHA0 YP 007998555 1 1124983 Proteobacteria
Pseudomonas brassicacearum subsp brassicacearum NFM421 YP 004352293 1 994484 Proteobacteria
Dickeya solani RNS 05 1 2A WP 057083754 1 Dsolani
Dickeya zeae DZ2Q WP 038926719 1 Dzeae
Sodalis glossinidius str morsitans YP 454393 1 343509 Proteobacteria
Erwinia pyrifoliae DSM 12163 YP 005802943 1 644651 Proteobacteria
Salmonella enterica subsp enterica serovar Bovismorbificans str 3114 YP 008505074 1 1320309 Proteobacteria
Salmonella enterica subsp enterica serovar Schwarzengrund str CVM19633 YP 002115110 1 439843 Proteobacteria
Salmonella enterica subsp enterica serovar Schwarzengrund str CVM19633 YP 002117168 1 439843 Proteobacteria
Salmonella enterica subsp enterica serovar Paratyphi B str SPB7 YP 001588364 1 1016998 Proteobacteria
Salmonella enterica subsp enterica serovar Dublin str CT 02021853 YP 002216014 1 439851 Proteobacteria
Salmonella enterica subsp enterica serovar Enteritidis str P125109 YP 002244020 1 550537 Proteobacteria

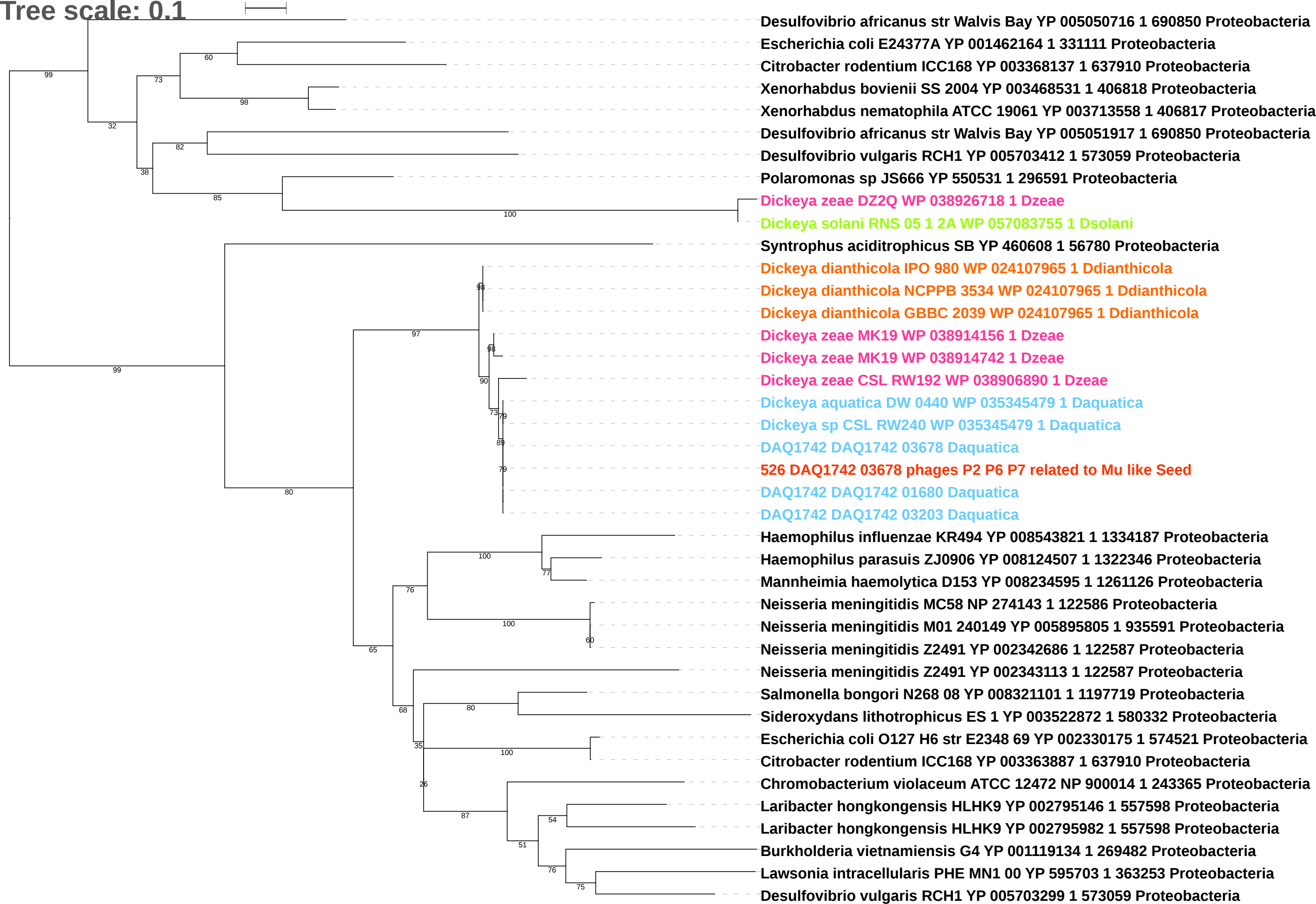

Tree scale: 0.1
Desulfovibrio africanus str Walvis Bay YP 005050716 1 690850 Proteobacteria
Escherichia coli E24377A YP 001462164 1 331111 Proteobacteria
Citrobacter rodentium ICC168 YP 003368137 1 637910 Proteobacteria
Xenorhabdus bovienii SS 2004 YP 003468531 1 406818 Proteobacteria
Xenorhabdus nematophila ATCC 19061 YP 003713558 1 406817 Proteobacteria
Desulfovibrio africanus str Walvis Bay YP 005051917 1 690850 Proteobacteria
Desulfovibrio vulgaris RCH1 YP 005703412 1 573059 Proteobacteria
Polaromonas sp JS666 YP 550531 1 296591 Proteobacteria
Dickeya zeae DZ2Q WP 038926718 1 Dzeae
Dickeya solani RNS 05 1 2A WP 057083755 1 Dsolani
Syntrophus aciditrophicus SB YP 460608 1 56780 Proteobacteria
Dickeya dianthicola IPO 980 WP 024107965 1 Ddianthicola
Dickeya dianthicola NCPPB 3534 WP 024107965 1 Ddianthicola
Dickeya dianthicola GBBC 2039 WP 024107965 1 Ddianthicola
Dickeya zeae MK19 WP 038914156 1 Dzeae
Dickeya zeae MK19 WP 038914742 1 Dzeae
Dickeya zeae CSL RW192 WP 038906890 1 Dzeae
Dickeya aquatica DW 0440 WP 035345479 1 Daquatica
Dickeya sp CSL RW240 WP 035345479 1 Daquatica
DAQ1742 DAQ1742 03678 Daquatica
526 DAQ1742 03678 phages P2 P6 P7 related to Mu like Seed
DAQ1742 DAQ1742 01680 Daquatica
DAQ1742 DAQ1742 03203 Daquatica
Haemophilus influenzae KR494 YP 008543821 1 1334187 Proteobacteria
Haemophilus parasuis ZJ0906 YP 008124507 1 1322346 Proteobacteria
Mannheimia haemolytica D153 YP 008234595 1 1261126 Proteobacteria
Neisseria meningitidis MC58 NP 274143 1 122586 Proteobacteria
Neisseria meningitidis M01 240149 YP 005895805 1 935591 Proteobacteria
Neisseria meningitidis Z2491 YP 002342686 1 122587 Proteobacteria
Neisseria meningitidis Z2491 YP 002343113 1 122587 Proteobacteria
Salmonella bongori N268 08 YP 008321101 1 1197719 Proteobacteria
Sideroxydans lithotrophicus ES 1 YP 003522872 1 580332 Proteobacteria
Escherichia coli O127 H6 str E2348 69 YP 002330175 1 574521 Proteobacteria
Citrobacter rodentium ICC168 YP 003363887 1 637910 Proteobacteria
Chromobacterium violaceum ATCC 12472 NP 900014 1 243365 Proteobacteria
Laribacter hongkongensis HLHK9 YP 002795146 1 557598 Proteobacteria
Laribacter hongkongensis HLHK9 YP 002795982 1 557598 Proteobacteria
Burkholderia vietnamiensis G4 YP 001119134 1 269482 Proteobacteria
Lawsonia intracellularis PHE MN1 00 YP 595703 1 363253 Proteobacteria
Desulfovibrio vulgaris RCH1 YP 005703299 1 573059 Proteobacteria

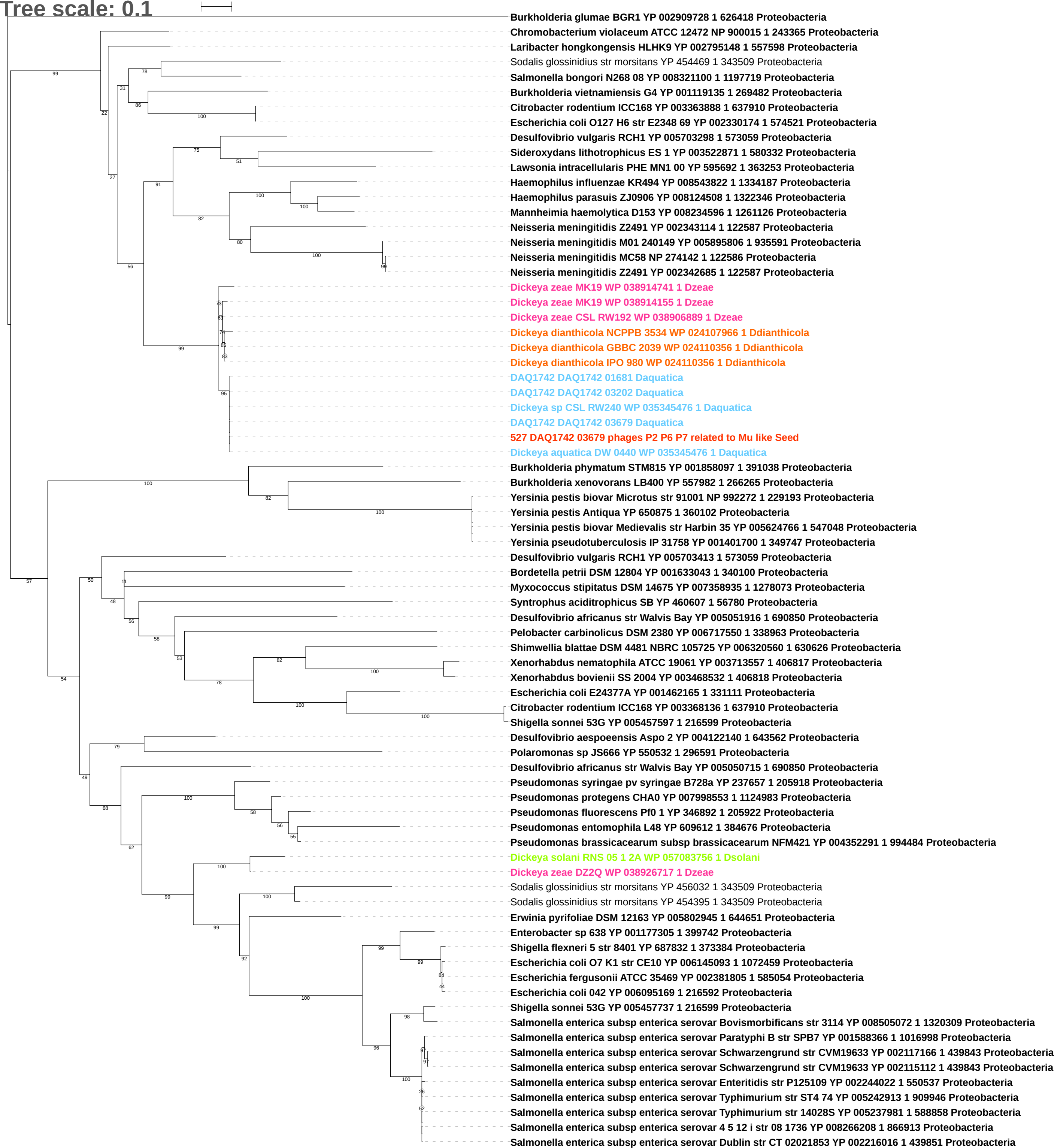

Tree scale: 0.1
Burkholderia glumae BGR1 YP 002909728 1 626418 Proteobacteria
Chromobacterium violaceum ATCC 12472 NP 900015 1 243365 Proteobacteria
Laribacter hongkongensis HLHK9 YP 002795148 1 557598 Proteobacteria
Sodalis glossinidius str morsitans YP 454469 1 343509 Proteobacteria
Salmonella bongori N268 08 YP 008321100 1 1197719 Proteobacteria
Burkholderia vietnamiensis G4 YP 001119135 1 269482 Proteobacteria
Citrobacter rodentium ICC168 YP 003363888 1 637910 Proteobacteria
Escherichia coli O127 H6 str E2348 69 YP 002330174 1 574521 Proteobacteria
Desulfovibrio vulgaris RCH1 YP 005703298 1 573059 Proteobacteria
Sideroxydans lithotrophicus ES 1 YP 003522871 1 580332 Proteobacteria
Lawsonia intracellularis PHE MN1 00 YP 595692 1 363253 Proteobacteria
Haemophilus influenzae KR494 YP 008543822 1 1334187 Proteobacteria
Haemophilus parasuis ZJ0906 YP 008124508 1 1322346 Proteobacteria
Mannheimia haemolytica D153 YP 008234596 1 1261126 Proteobacteria
Neisseria meningitidis Z2491 YP 002343114 1 122587 Proteobacteria
Neisseria meningitidis M01 240149 YP 005895806 1 935591 Proteobacteria
Neisseria meningitidis MC58 NP 274142 1 122586 Proteobacteria
Neisseria meningitidis Z2491 YP 002342685 1 122587 Proteobacteria
Dickeya zeae MK19 WP 038914741 1 Dzeae
Dickeya zeae MK19 WP 038914155 1 Dzeae
Dickeya zeae CSL RW192 WP 038906889 1 Dzeae
Dickeya dianthicola NCPPB 3534 WP 024107966 1 Ddianthicola
Dickeya dianthicola GBBC 2039 WP 024110356 1 Ddianthicola
Dickeya dianthicola IPO 980 WP 024110356 1 Ddianthicola
DAQ1742 DAQ1742 01681 Daquatica
DAQ1742 DAQ1742 03202 Daquatica
Dickeya sp CSL RW240 WP 035345476 1 Daquatica
DAQ1742 DAQ1742 03679 Daquatica
527 DAQ1742 03679 phages P2 P6 P7 related to Mu like Seed
Dickeya aquatica DW 0440 WP 035345476 1 Daquatica
Burkholderia phymatum STM815 YP 001858097 1 391038 Proteobacteria
Burkholderia xenovorans LB400 YP 557982 1 266265 Proteobacteria
Yersinia pestis biovar Microtus str 91001 NP 992272 1 229193 Proteobacteria
Yersinia pestis Antiqua YP 650875 1 360102 Proteobacteria
Yersinia pestis biovar Medievalis str Harbin 35 YP 005624766 1 547048 Proteobacteria
Yersinia pseudotuberculosis IP 31758 YP 001401700 1 349747 Proteobacteria
Desulfovibrio vulgaris RCH1 YP 005703413 1 573059 Proteobacteria
Bordetella petrii DSM 12804 YP 001633043 1 340100 Proteobacteria
Myxococcus stipitatus DSM 14675 YP 007358935 1 1278073 Proteobacteria
Syntrophus aciditrophicus SB YP 460607 1 56780 Proteobacteria
Desulfovibrio africanus str Walvis Bay YP 005051916 1 690850 Proteobacteria
Pelobacter carbinolicus DSM 2380 YP 006717550 1 338963 Proteobacteria
Shimwellia blattae DSM 4481 NBRC 105725 YP 006320560 1 630626 Proteobacteria
Xenorhabdus nematophila ATCC 19061 YP 003713557 1 406817 Proteobacteria
Xenorhabdus bovienii SS 2004 YP 003468532 1 406818 Proteobacteria
Escherichia coli E24377A YP 001462165 1 331111 Proteobacteria
Citrobacter rodentium ICC168 YP 003368136 1 637910 Proteobacteria
Shigella sonnei 53G YP 005457597 1 216599 Proteobacteria
Desulfovibrio aespoeensis Aspo 2 YP 004122140 1 643562 Proteobacteria
Polaromonas sp JS666 YP 550532 1 296591 Proteobacteria
Desulfovibrio africanus str Walvis Bay YP 005050715 1 690850 Proteobacteria
Pseudomonas syringae pv syringae B728a YP 237657 1 205918 Proteobacteria
Pseudomonas protegens CHA0 YP 007998553 1 1124983 Proteobacteria
Pseudomonas fluorescens Pf0 1 YP 346892 1 205922 Proteobacteria
Pseudomonas entomophila L48 YP 609612 1 384676 Proteobacteria
Pseudomonas brassicacearum subsp brassicacearum NFM421 YP 004352291 1 994484 Proteobacteria
Dickeya solani RNS 05 1 2A WP 057083756 1 Dsolani
Dickeya zeae DZ2Q WP 038926717 1 Dzeae
Sodalis glossinidius str morsitans YP 456032 1 343509 Proteobacteria
Sodalis glossinidius str morsitans YP 454395 1 343509 Proteobacteria
Erwinia pyrifoliae DSM 12163 YP 005802945 1 644651 Proteobacteria
Enterobacter sp 638 YP 001177305 1 399742 Proteobacteria
Shigella flexneri 5 str 8401 YP 687832 1 373384 Proteobacteria
Escherichia coli O7 K1 str CE10 YP 006145093 1 1072459 Proteobacteria
Escherichia fergusonii ATCC 35469 YP 002381805 1 585054 Proteobacteria
Escherichia coli 042 YP 006095169 1 216592 Proteobacteria
Shigella sonnei 53G YP 005457737 1 216599 Proteobacteria
Salmonella enterica subsp enterica serovar Bovismorbificans str 3114 YP 008505072 1 1320309 Proteobacteria
Salmonella enterica subsp enterica serovar Paratyphi B str SPB7 YP 001588366 1 1016998 Proteobacteria
Salmonella enterica subsp enterica serovar Schwarzengrund str CVM19633 YP 002117166 1 439843 Proteobacteria
Salmonella enterica subsp enterica serovar Schwarzengrund str CVM19633 YP 002115112 1 439843 Proteobacteria
Salmonella enterica subsp enterica serovar Enteritidis str P125109 YP 002244022 1 550537 Proteobacteria
Salmonella enterica subsp enterica serovar Typhimurium str ST4 74 YP 005242913 1 909946 Proteobacteria
Salmonella enterica subsp enterica serovar Typhimurium str 14028S YP 005237981 1 588858 Proteobacteria
Salmonella enterica subsp enterica serovar 4 5 12 i str 08 1736 YP 008266208 1 866913 Proteobacteria
Salmonella enterica subsp enterica serovar Dublin str CT 02021853 YP 002216016 1 439851 Proteobacteria

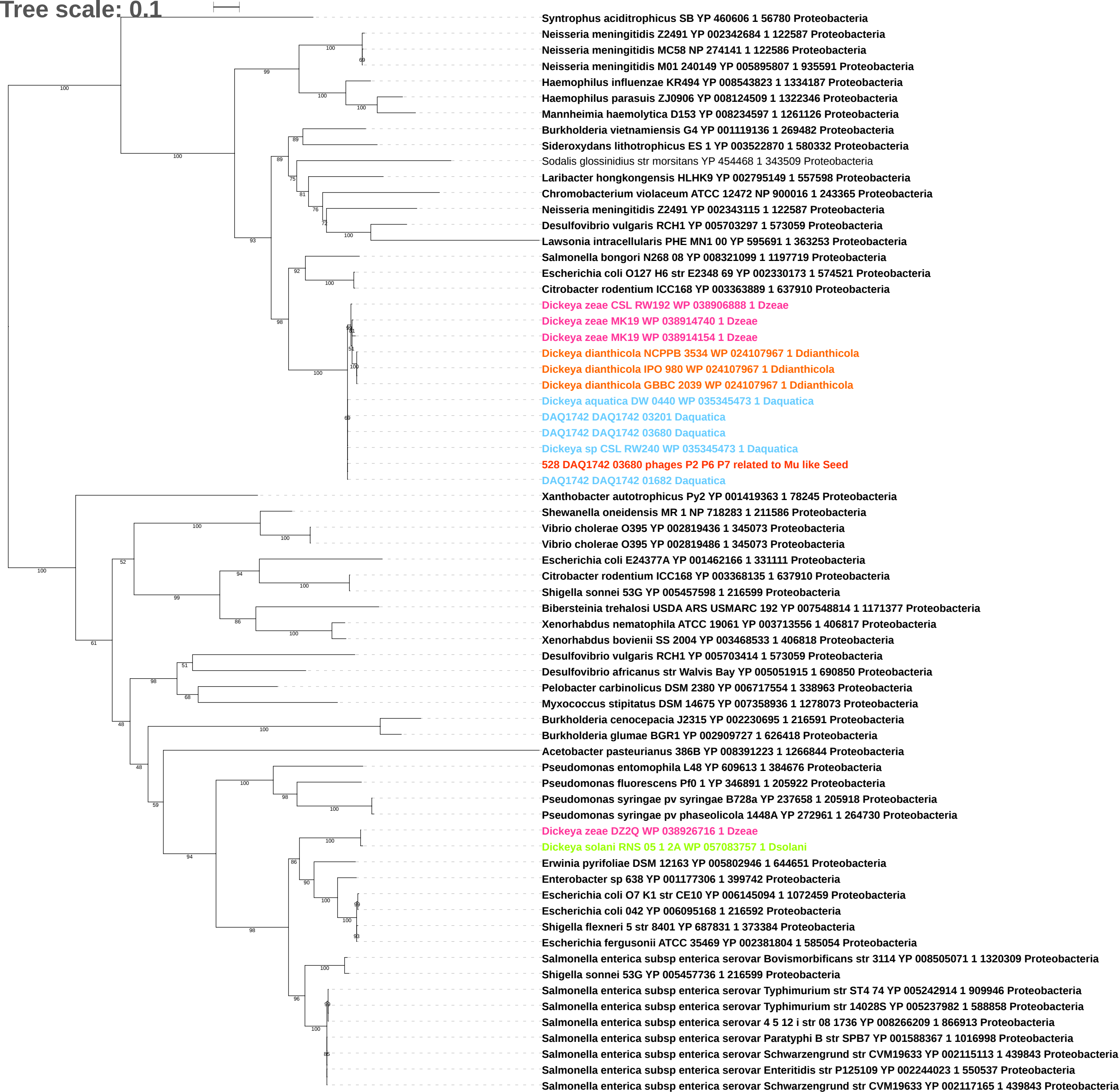
Tree scale: 0.1
Syntrophus aciditrophicus SB YP 460606 1 56780 Proteobacteria
Neisseria meningitidis Z2491 YP 002342684 1 122587 Proteobacteria
Neisseria meningitidis MC58 NP 274141 1 122586 Proteobacteria
Neisseria meningitidis M01 240149 YP 005895807 1 935591 Proteobacteria
Haemophilus influenzae KR494 YP 008543823 1 1334187 Proteobacteria
Haemophilus parasuis ZJ0906 YP 008124509 1 1322346 Proteobacteria
Mannheimia haemolytica D153 YP 008234597 1 1261126 Proteobacteria
Burkholderia vietnamiensis G4 YP 001119136 1 269482 Proteobacteria
Sideroxydans lithotrophicus ES 1 YP 003522870 1 580332 Proteobacteria
Sodalis glossinidius str morsitans YP 454468 1 343509 Proteobacteria
Laribacter hongkongensis HLHK9 YP 002795149 1 557598 Proteobacteria
Chromobacterium violaceum ATCC 12472 NP 900016 1 243365 Proteobacteria
Neisseria meningitidis Z2491 YP 002343115 1 122587 Proteobacteria
Desulfovibrio vulgaris RCH1 YP 005703297 1 573059 Proteobacteria
Lawsonia intracellularis PHE MN1 00 YP 595691 1 363253 Proteobacteria
Salmonella bongori N268 08 YP 008321099 1 1197719 Proteobacteria
Escherichia coli O127 H6 str E2348 69 YP 002330173 1 574521 Proteobacteria
Citrobacter rodentium ICC168 YP 003363889 1 637910 Proteobacteria
Dickeya zeae CSL RW192 WP 038906888 1 Dzeae
Dickeya zeae MK19 WP 038914740 1 Dzeae
Dickeya zeae MK19 WP 038914154 1 Dzeae
Dickeya dianthicola NCPPB 3534 WP 024107967 1 Ddianthicola
Dickeya dianthicola IPO 980 WP 024107967 1 Ddianthicola
Dickeya dianthicola GBBC 2039 WP 024107967 1 Ddianthicola
Dickeya aquatica DW 0440 WP 035345473 1 Daquatica
DAQ1742 DAQ1742 03201 Daquatica
DAQ1742 DAQ1742 03680 Daquatica
Dickeya sp CSL RW240 WP 035345473 1 Daquatica
528 DAQ1742 03680 phages P2 P6 P7 related to Mu like Seed
DAQ1742 DAQ1742 01682 Daquatica
Xanthobacter autotrophicus Py2 YP 001419363 1 78245 Proteobacteria
Shewanella oneidensis MR 1 NP 718283 1 211586 Proteobacteria
Vibrio cholerae O395 YP 002819436 1 345073 Proteobacteria
Vibrio cholerae O395 YP 002819486 1 345073 Proteobacteria
Escherichia coli E24377A YP 001462166 1 331111 Proteobacteria
Citrobacter rodentium ICC168 YP 003368135 1 637910 Proteobacteria
Shigella sonnei 53G YP 005457598 1 216599 Proteobacteria
Bibersteinia trehalosi USDA ARS USMARC 192 YP 007548814 1 1171377 Proteobacteria
Xenorhabdus nematophila ATCC 19061 YP 003713556 1 406817 Proteobacteria
Xenorhabdus bovienii SS 2004 YP 003468533 1 406818 Proteobacteria
Desulfovibrio vulgaris RCH1 YP 005703414 1 573059 Proteobacteria
Desulfovibrio africanus str Walvis Bay YP 005051915 1 690850 Proteobacteria
Pelobacter carbinolicus DSM 2380 YP 006717554 1 338963 Proteobacteria
Myxococcus stipitatus DSM 14675 YP 007358936 1 1278073 Proteobacteria
Burkholderia cenocepacia J2315 YP 002230695 1 216591 Proteobacteria
Burkholderia glumae BGR1 YP 002909727 1 626418 Proteobacteria
Acetobacter pasteurianus 386B YP 008391223 1 1266844 Proteobacteria
Pseudomonas entomophila L48 YP 609613 1 384676 Proteobacteria
Pseudomonas fluorescens Pf0 1 YP 346891 1 205922 Proteobacteria
Pseudomonas syringae pv syringae B728a YP 237658 1 205918 Proteobacteria
Pseudomonas syringae pv phaseolicola 1448A YP 272961 1 264730 Proteobacteria
Dickeya zeae DZ2Q WP 038926716 1 Dzeae
Dickeya solani RNS 05 1 2A WP 057083757 1 Dsolani
Erwinia pyrifoliae DSM 12163 YP 005802946 1 644651 Proteobacteria
Enterobacter sp 638 YP 001177306 1 399742 Proteobacteria
Escherichia coli O7 K1 str CE10 YP 006145094 1 1072459 Proteobacteria
Escherichia coli 042 YP 006095168 1 216592 Proteobacteria
Shigella flexneri 5 str 8401 YP 687831 1 373384 Proteobacteria
Escherichia fergusonii ATCC 35469 YP 002381804 1 585054 Proteobacteria
Salmonella enterica subsp enterica serovar Bovismorbificans str 3114 YP 008505071 1 1320309 Proteobacteria
Shigella sonnei 53G YP 005457736 1 216599 Proteobacteria
Salmonella enterica subsp enterica serovar Typhimurium str ST4 74 YP 005242914 1 909946 Proteobacteria
Salmonella enterica subsp enterica serovar Typhimurium str 14028S YP 005237982 1 588858 Proteobacteria
Salmonella enterica subsp enterica serovar 4 5 12 i str 08 1736 YP 008266209 1 866913 Proteobacteria
Salmonella enterica subsp enterica serovar Paratyphi B str SPB7 YP 001588367 1 1016998 Proteobacteria
Salmonella enterica subsp enterica serovar Schwarzengrund str CVM19633 YP 002115113 1 439843 Proteobacteria
Salmonella enterica subsp enterica serovar Enteritidis str P125109 YP 002244023 1 550537 Proteobacteria
Salmonella enterica subsp enterica serovar Schwarzengrund str CVM19633 YP 002117165 1 439843 Proteobacteria

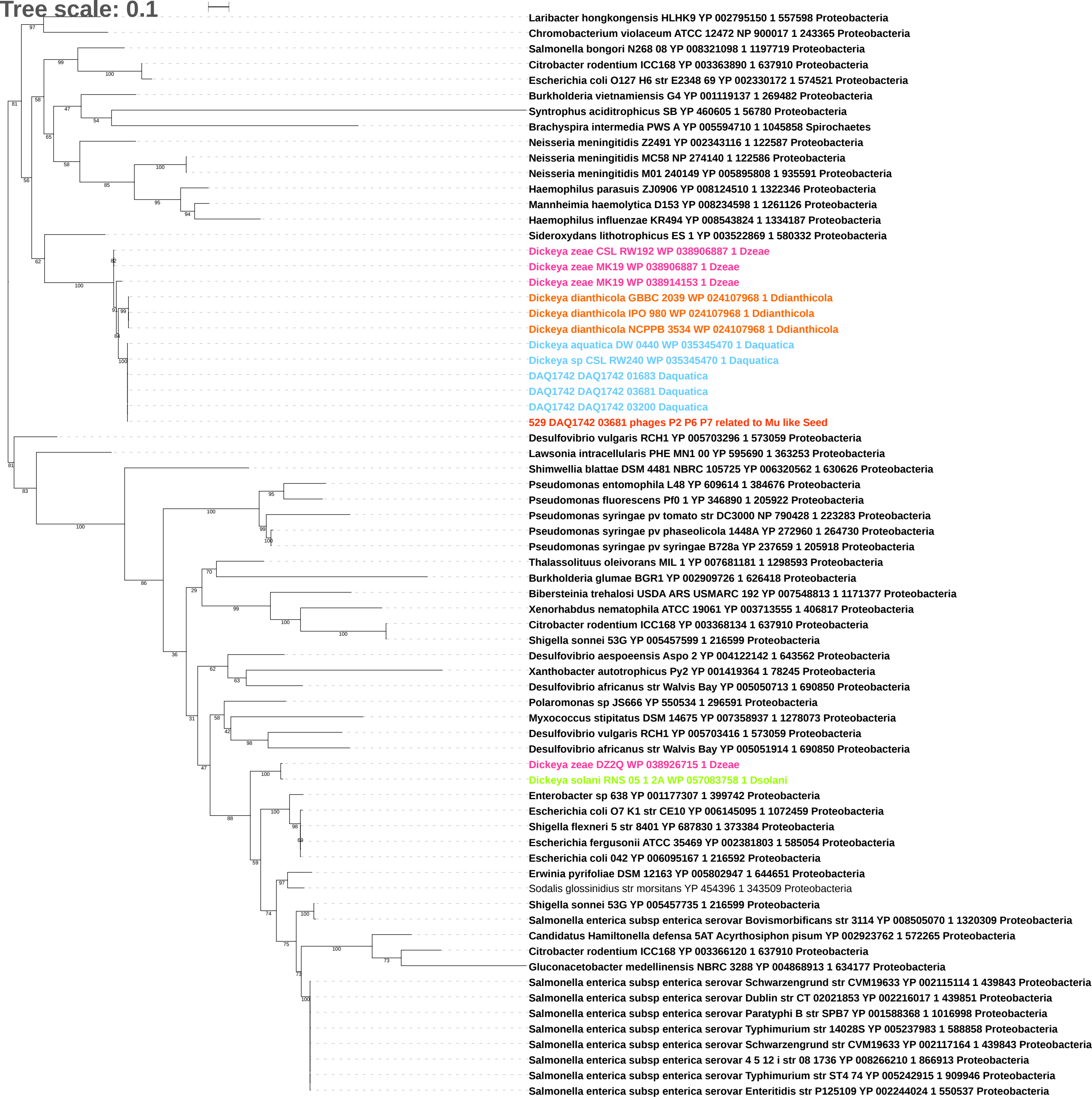
Tree scale: 0.1
Laribacter hongkongensis HLHK9 YP 002795150 1 557598 Proteobacteria
Chromobacterium violaceum ATCC 12472 NP 900017 1 243365 Proteobacteria
Salmonella bongori N268 08 YP 008321098 1 1197719 Proteobacteria
Citrobacter rodentium ICC168 YP 003363890 1 637910 Proteobacteria
Escherichia coli O127 H6 str E2348 69 YP 002330172 1 574521 Proteobacteria
Burkholderia vietnamiensis G4 YP 001119137 1 269482 Proteobacteria
Syntrophus aciditrophicus SB YP 460605 1 56780 Proteobacteria
Brachyspira intermedia PWS A YP 005594710 1 1045858 Spirochaetes
Neisseria meningitidis Z2491 YP 002343116 1 122587 Proteobacteria
Neisseria meningitidis MC58 NP 274140 1 122586 Proteobacteria
Neisseria meningitidis M01 240149 YP 005895808 1 935591 Proteobacteria
Haemophilus parasuis ZJ0906 YP 008124510 1 1322346 Proteobacteria
Mannheimia haemolytica D153 YP 008234598 1 1261126 Proteobacteria
Haemophilus influenzae KR494 YP 008543824 1 1334187 Proteobacteria
Sideroxydans lithotrophicus ES 1 YP 003522869 1 580332 Proteobacteria
Dickeya zeae CSL RW192 WP 038906887 1 Dzeae
Dickeya zeae MK19 WP 038906887 1 Dzeae
Dickeya zeae MK19 WP 038914153 1 Dzeae
Dickeya dianthicola GBBC 2039 WP 024107968 1 Ddianthicola
Dickeya dianthicola IPO 980 WP 024107968 1 Ddianthicola
Dickeya dianthicola NCPPB 3534 WP 024107968 1 Ddianthicola
Dickeya aquatica DW 0440 WP 035345470 1 Daquatica
Dickeya sp CSL RW240 WP 035345470 1 Daquatica
DAQ1742 DAQ1742 01683 Daquatica
DAQ1742 DAQ1742 03681 Daquatica
DAQ1742 DAQ1742 03200 Daquatica
529 DAQ1742 03681 phages P2 P6 P7 related to Mu like Seed
Desulfovibrio vulgaris RCH1 YP 005703296 1 573059 Proteobacteria
Lawsonia intracellularis PHE MN1 00 YP 595690 1 363253 Proteobacteria
Shimwellia blattae DSM 4481 NBRC 105725 YP 006320562 1 630626 Proteobacteria
Pseudomonas entomophila L48 YP 609614 1 384676 Proteobacteria
Pseudomonas fluorescens Pf0 1 YP 346890 1 205922 Proteobacteria
Pseudomonas syringae pv tomato str DC3000 NP 790428 1 223283 Proteobacteria
Pseudomonas syringae pv phaseolicola 1448A YP 272960 1 264730 Proteobacteria
Pseudomonas syringae pv syringae B728a YP 237659 1 205918 Proteobacteria
Thalassolituus oleivorans MIL 1 YP 007681181 1 1298593 Proteobacteria
Burkholderia glumae BGR1 YP 002909726 1 626418 Proteobacteria
Bibersteinia trehalosi USDA ARS USMARC 192 YP 007548813 1 1171377 Proteobacteria
Xenorhabdus nematophila ATCC 19061 YP 003713555 1 406817 Proteobacteria
Citrobacter rodentium ICC168 YP 003368134 1 637910 Proteobacteria
Shigella sonnei 53G YP 005457599 1 216599 Proteobacteria
Desulfovibrio aespoeensis Aspo 2 YP 004122142 1 643562 Proteobacteria
Xanthobacter autotrophicus Py2 YP 001419364 1 78245 Proteobacteria
Desulfovibrio africanus str Walvis Bay YP 005050713 1 690850 Proteobacteria
Polaromonas sp JS666 YP 550534 1 296591 Proteobacteria
Myxococcus stipitatus DSM 14675 YP 007358937 1 1278073 Proteobacteria
Desulfovibrio vulgaris RCH1 YP 005703416 1 573059 Proteobacteria
Desulfovibrio africanus str Walvis Bay YP 005051914 1 690850 Proteobacteria
Dickeya zeae DZ2Q WP 038926715 1 Dzeae
Dickeya solani RNS 05 1 2A WP 057083758 1 Dsolani
Enterobacter sp 638 YP 001177307 1 399742 Proteobacteria
Escherichia coli O7 K1 str CE10 YP 006145095 1 1072459 Proteobacteria
Shigella flexneri 5 str 8401 YP 687830 1 373384 Proteobacteria
Escherichia fergusonii ATCC 35469 YP 002381803 1 585054 Proteobacteria
Escherichia coli 042 YP 006095167 1 216592 Proteobacteria
Erwinia pyrifoliae DSM 12163 YP 005802947 1 644651 Proteobacteria
Sodalis glossinidius str morsitans YP 454396 1 343509 Proteobacteria
Shigella sonnei 53G YP 005457735 1 216599 Proteobacteria
Salmonella enterica subsp enterica serovar Bovismorbificans str 3114 YP 008505070 1 1320309 Proteobacteria
Candidatus Hamiltonella defensa 5AT Acyrthosiphon pisum YP 002923762 1 572265 Proteobacteria
Citrobacter rodentium ICC168 YP 003366120 1 637910 Proteobacteria
Gluconacetobacter medellinensis NBRC 3288 YP 004868913 1 634177 Proteobacteria
Salmonella enterica subsp enterica serovar Schwarzengrund str CVM19633 YP 002115114 1 439843 Proteobacteria
Salmonella enterica subsp enterica serovar Dublin str CT 02021853 YP 002216017 1 439851 Proteobacteria
Salmonella enterica subsp enterica serovar Paratyphi B str SPB7 YP 001588368 1 1016998 Proteobacteria
Salmonella enterica subsp enterica serovar Typhimurium str 14028S YP 005237983 1 588858 Proteobacteria
Salmonella enterica subsp enterica serovar Schwarzengrund str CVM19633 YP 002117164 1 439843 Proteobacteria
Salmonella enterica subsp enterica serovar 4 5 12 i str 08 1736 YP 008266210 1 866913 Proteobacteria
Salmonella enterica subsp enterica serovar Typhimurium str ST4 74 YP 005242915 1 909946 Proteobacteria
Salmonella enterica subsp enterica serovar Enteritidis str P125109 YP 002244024 1 550537 Proteobacteria

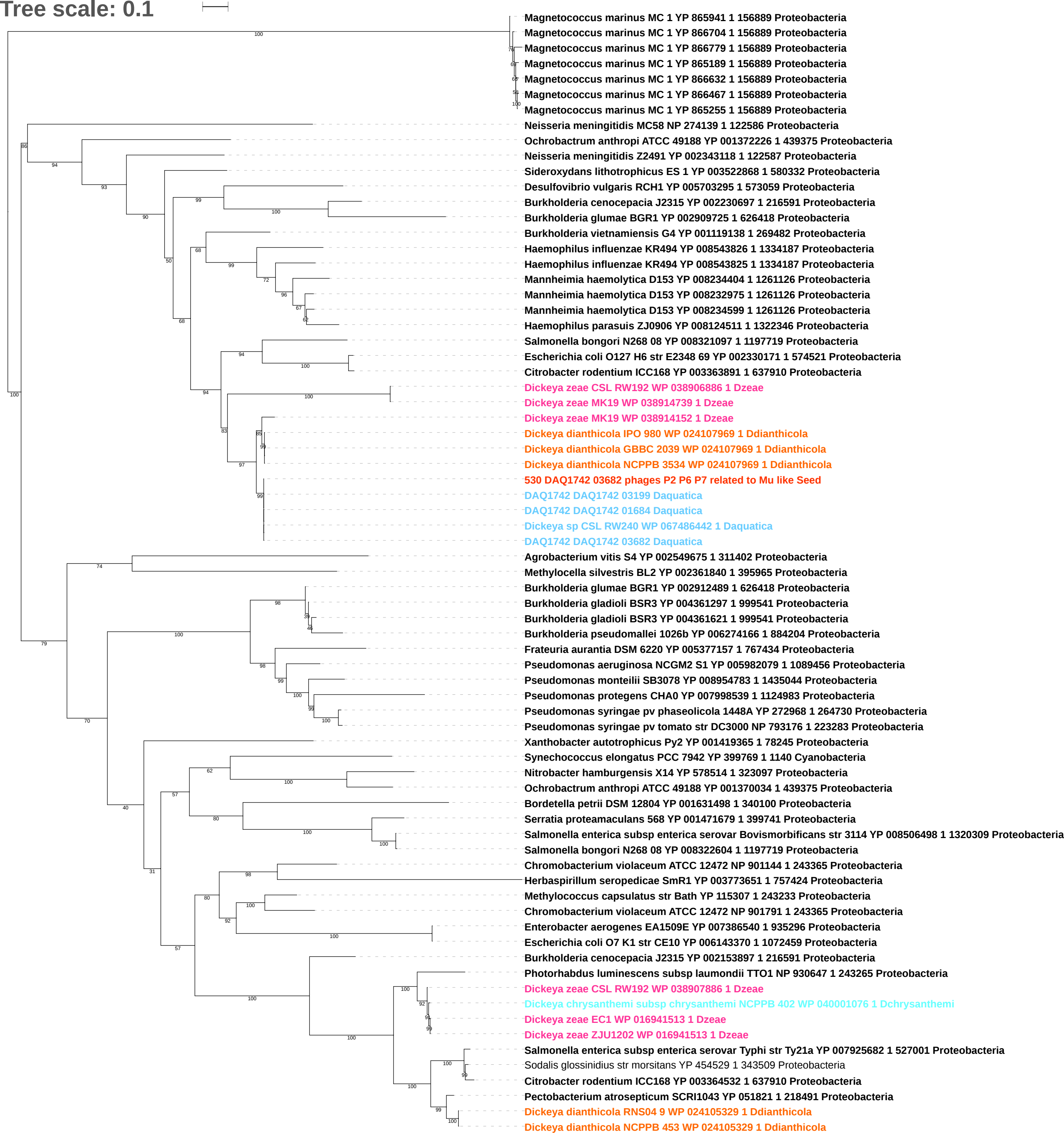

Tree scale: 0.1
Magnetococcus marinus MC 1 YP 865941 1 156889 Proteobacteria
Magnetococcus marinus MC 1 YP 866704 1 156889 Proteobacteria
Magnetococcus marinus MC 1 YP 866779 1 156889 Proteobacteria
Magnetococcus marinus MC 1 YP 865189 1 156889 Proteobacteria
Magnetococcus marinus MC 1 YP 866632 1 156889 Proteobacteria
Magnetococcus marinus MC 1 YP 866467 1 156889 Proteobacteria
Magnetococcus marinus MC 1 YP 865255 1 156889 Proteobacteria
Neisseria meningitidis MC58 NP 274139 1 122586 Proteobacteria
Ochrobactrum anthropi ATCC 49188 YP 001372226 1 439375 Proteobacteria
Neisseria meningitidis Z2491 YP 002343118 1 122587 Proteobacteria
Sideroxydans lithotrophicus ES 1 YP 003522868 1 580332 Proteobacteria
Desulfovibrio vulgaris RCH1 YP 005703295 1 573059 Proteobacteria
Burkholderia cenocepacia J2315 YP 002230697 1 216591 Proteobacteria
Burkholderia glumae BGR1 YP 002909725 1 626418 Proteobacteria
Burkholderia vietnamiensis G4 YP 001119138 1 269482 Proteobacteria
Haemophilus influenzae KR494 YP 008543826 1 1334187 Proteobacteria
Haemophilus influenzae KR494 YP 008543825 1 1334187 Proteobacteria
Mannheimia haemolytica D153 YP 008234404 1 1261126 Proteobacteria
Mannheimia haemolytica D153 YP 008232975 1 1261126 Proteobacteria
Mannheimia haemolytica D153 YP 008234599 1 1261126 Proteobacteria
Haemophilus parasuis ZJ0906 YP 008124511 1 1322346 Proteobacteria
Salmonella bongori N268 08 YP 008321097 1 1197719 Proteobacteria
Escherichia coli O127 H6 str E2348 69 YP 002330171 1 574521 Proteobacteria
Citrobacter rodentium ICC168 YP 003363891 1 637910 Proteobacteria
Dickeya zeae CSL RW192 WP 038906886 1 Dzeae
Dickeya zeae MK19 WP 038914739 1 Dzeae
Dickeya zeae MK19 WP 038914152 1 Dzeae
Dickeya dianthicola IPO 980 WP 024107969 1 Ddianthicola
Dickeya dianthicola GBBC 2039 WP 024107969 1 Ddianthicola
Dickeya dianthicola NCPPB 3534 WP 024107969 1 Ddianthicola
530 DAQ1742 03682 phages P2 P6 P7 related to Mu like Seed
DAQ1742 DAQ1742 03199 Daquatica
DAQ1742 DAQ1742 01684 Daquatica
Dickeya sp CSL RW240 WP 067486442 1 Daquatica
DAQ1742 DAQ1742 03682 Daquatica
Agrobacterium vitis S4 YP 002549675 1 311402 Proteobacteria
Methylocella silvestris BL2 YP 002361840 1 395965 Proteobacteria
Burkholderia glumae BGR1 YP 002912489 1 626418 Proteobacteria
Burkholderia gladioli BSR3 YP 004361297 1 999541 Proteobacteria
Burkholderia gladioli BSR3 YP 004361621 1 999541 Proteobacteria
Burkholderia pseudomallei 1026b YP 006274166 1 884204 Proteobacteria
Frateuria aurantia DSM 6220 YP 005377157 1 767434 Proteobacteria
Pseudomonas aeruginosa NCGM2 S1 YP 005982079 1 1089456 Proteobacteria
Pseudomonas monteilii SB3078 YP 008954783 1 1435044 Proteobacteria
Pseudomonas protegens CHA0 YP 007998539 1 1124983 Proteobacteria
Pseudomonas syringae pv phaseolicola 1448A YP 272968 1 264730 Proteobacteria
Pseudomonas syringae pv tomato str DC3000 NP 793176 1 223283 Proteobacteria
Xanthobacter autotrophicus Py2 YP 001419365 1 78245 Proteobacteria
Synechococcus elongatus PCC 7942 YP 399769 1 1140 Cyanobacteria
Nitrobacter hamburgensis X14 YP 578514 1 323097 Proteobacteria
Ochrobactrum anthropi ATCC 49188 YP 001370034 1 439375 Proteobacteria
Bordetella petrii DSM 12804 YP 001631498 1 340100 Proteobacteria
Serratia proteamaculans 568 YP 001471679 1 399741 Proteobacteria
Salmonella enterica subsp enterica serovar Bovismorbificans str 3114 YP 008506498 1 1320309 Proteobacteria
Salmonella bongori N268 08 YP 008322604 1 1197719 Proteobacteria
Chromobacterium violaceum ATCC 12472 NP 901144 1 243365 Proteobacteria
Herbaspirillum seropedicae SmR1 YP 003773651 1 757424 Proteobacteria
Methylococcus capsulatus str Bath YP 115307 1 243233 Proteobacteria
Chromobacterium violaceum ATCC 12472 NP 901791 1 243365 Proteobacteria
Enterobacter aerogenes EA1509E YP 007386540 1 935296 Proteobacteria
Escherichia coli O7 K1 str CE10 YP 006143370 1 1072459 Proteobacteria
Burkholderia cenocepacia J2315 YP 002153897 1 216591 Proteobacteria
Photorhabdus luminescens subsp laumondii TTO1 NP 930647 1 243265 Proteobacteria
Dickeya zeae CSL RW192 WP 038907886 1 Dzeae
Dickeya chrysanthemi subsp chrysanthemi NCPPB 402 WP 040001076 1 Dchrysanthemi
Dickeya zeae EC1 WP 016941513 1 Dzeae
Dickeya zeae ZJU1202 WP 016941513 1 Dzeae
Salmonella enterica subsp enterica serovar Typhi str Ty21a YP 007925682 1 527001 Proteobacteria
Sodalis glossinidius str morsitans YP 454529 1 343509 Proteobacteria
Citrobacter rodentium ICC168 YP 003364532 1 637910 Proteobacteria
Pectobacterium atrosepticum SCRI1043 YP 051821 1 218491 Proteobacteria
Dickeya dianthicola RNS04 9 WP 024105329 1 Ddianthicola
Dickeya dianthicola NCPPB 453 WP 024105329 1 Ddianthicola

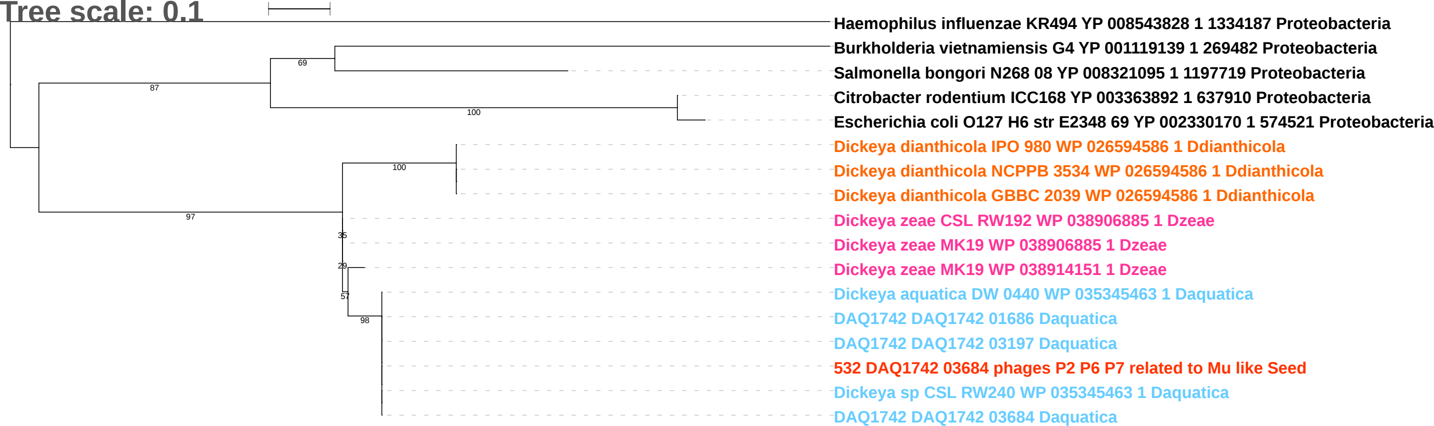

Tree scale: 0.1
Haemophilus influenzae KR494 YP 008543828 1 1334187 Proteobacteria
Burkholderia vietnamiensis G4 YP 001119139 1 269482 Proteobacteria
Salmonella bongori N268 08 YP 008321095 1 1197719 Proteobacteria
Citrobacter rodentium ICC168 YP 003363892 1 637910 Proteobacteria
Escherichia coli O127 H6 str E2348 69 YP 002330170 1 574521 Proteobacteria
Dickeya dianthicola IPO 980 WP 026594586 1 Ddianthicola
Dickeya dianthicola NCPPB 3534 WP 026594586 1 Ddianthicola
Dickeya dianthicola GBBC 2039 WP 026594586 1 Ddianthicola
Dickeya zeae CSL RW192 WP 038906885 1 Dzeae
Dickeya zeae MK19 WP 038906885 1 Dzeae
Dickeya zeae MK19 WP 038914151 1 Dzeae
Dickeya aquatica DW 0440 WP 035345463 1 Daquatica
DAQ1742 DAQ1742 01686 Daquatica
DAQ1742 DAQ1742 03197 Daquatica
532 DAQ1742 03684 phages P2 P6 P7 related to Mu like Seed
Dickeya sp CSL RW240 WP 035345463 1 Daquatica
DAQ1742 DAQ1742 03684 Daquatica
69
87
100
100
97
35
29
57
98

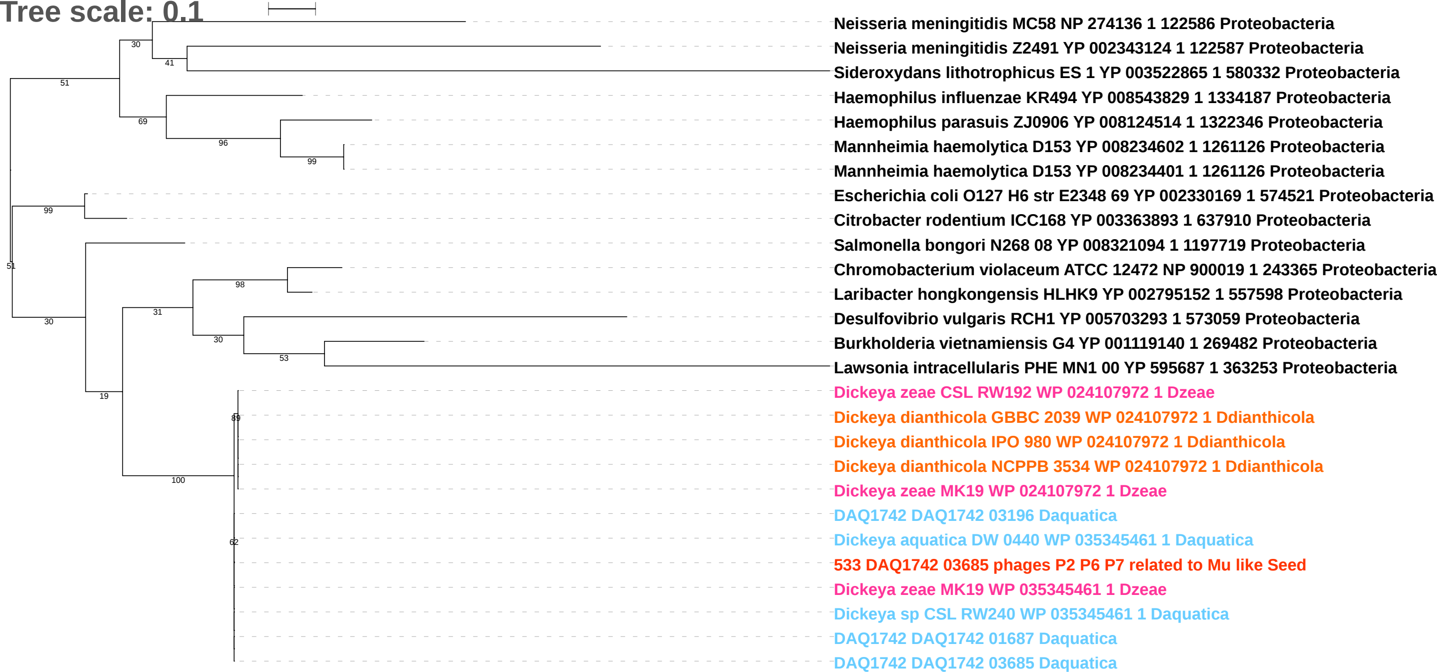
Tree scale: 0.1
Neisseria meningitidis MC58 NP 274136 1 122586 Proteobacteria
Neisseria meningitidis Z2491 YP 002343124 1 122587 Proteobacteria
Sideroxydans lithotrophicus ES 1 YP 003522865 1 580332 Proteobacteria
Haemophilus influenzae KR494 YP 008543829 1 1334187 Proteobacteria
Haemophilus parasuis ZJ0906 YP 008124514 1 1322346 Proteobacteria
Mannheimia haemolytica D153 YP 008234602 1 1261126 Proteobacteria
Mannheimia haemolytica D153 YP 008234401 1 1261126 Proteobacteria
Escherichia coli O127 H6 str E2348 69 YP 002330169 1 574521 Proteobacteria
Citrobacter rodentium ICC168 YP 003363893 1 637910 Proteobacteria
Salmonella bongori N268 08 YP 008321094 1 1197719 Proteobacteria
Chromobacterium violaceum ATCC 12472 NP 900019 1 243365 Proteobacteria
Laribacter hongkongensis HLHK9 YP 002795152 1 557598 Proteobacteria
Desulfovibrio vulgaris RCH1 YP 005703293 1 573059 Proteobacteria
Burkholderia vietnamiensis G4 YP 001119140 1 269482 Proteobacteria
Lawsonia intracellularis PHE MN1 00 YP 595687 1 363253 Proteobacteria
Dickeya zeae CSL RW192 WP 024107972 1 Dzeae
Dickeya dianthicola GBBC 2039 WP 024107972 1 Ddianthicola
Dickeya dianthicola IPO 980 WP 024107972 1 Ddianthicola
Dickeya dianthicola NCPPB 3534 WP 024107972 1 Ddianthicola
Dickeya zeae MK19 WP 024107972 1 Dzeae
DAQ1742 DAQ1742 03196 Daquatica
Dickeya aquatica DW 0440 WP 035345461 1 Daquatica
533 DAQ1742 03685 phages P2 P6 P7 related to Mu like Seed
Dickeya zeae MK19 WP 035345461 1 Dzeae
Dickeya sp CSL RW240 WP 035345461 1 Daquatica
DAQ1742 DAQ1742 01687 Daquatica
DAQ1742 DAQ1742 03685 Daquatica
30
41
51
69
96
99
99
51
30
98
31
30
53
19
89
100
62

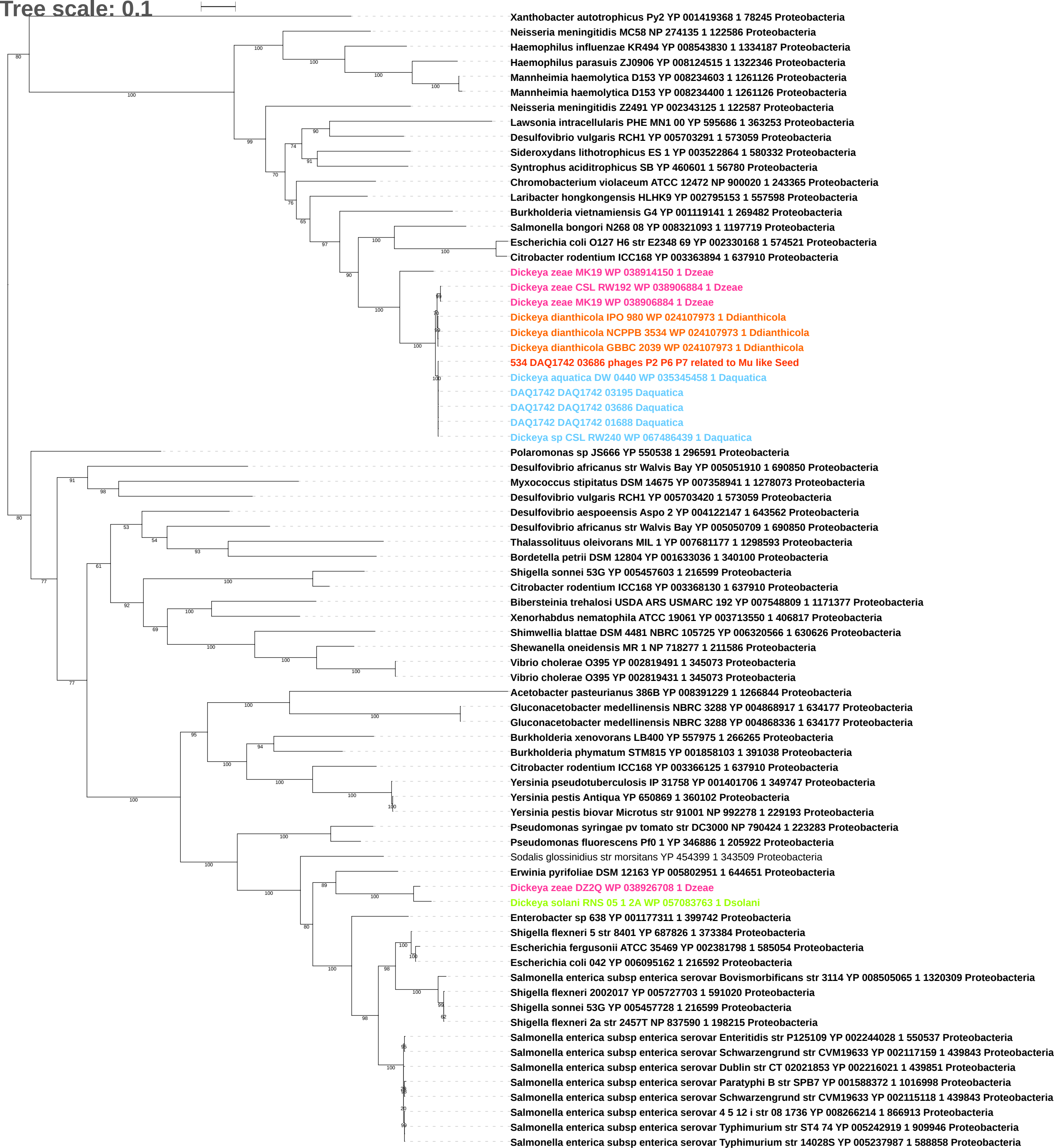

Tree scale: 0.1
Xanthobacter autotrophicus Py2 YP 001419368 1 78245 Proteobacteria
Neisseria meningitidis MC58 NP 274135 1 122586 Proteobacteria
Haemophilus influenzae KR494 YP 008543830 1 1334187 Proteobacteria
Haemophilus parasuis ZJ0906 YP 008124515 1 1322346 Proteobacteria
Mannheimia haemolytica D153 YP 008234603 1 1261126 Proteobacteria
Mannheimia haemolytica D153 YP 008234400 1 1261126 Proteobacteria
Neisseria meningitidis Z2491 YP 002343125 1 122587 Proteobacteria
Lawsonia intracellularis PHE MN1 00 YP 595686 1 363253 Proteobacteria
Desulfovibrio vulgaris RCH1 YP 005703291 1 573059 Proteobacteria
Sideroxydans lithotrophicus ES 1 YP 003522864 1 580332 Proteobacteria
Syntrophus aciditrophicus SB YP 460601 1 56780 Proteobacteria
Chromobacterium violaceum ATCC 12472 NP 900020 1 243365 Proteobacteria
Laribacter hongkongensis HLHK9 YP 002795153 1 557598 Proteobacteria
Burkholderia vietnamiensis G4 YP 001119141 1 269482 Proteobacteria
Salmonella bongori N268 08 YP 008321093 1 1197719 Proteobacteria
Escherichia coli O127 H6 str E2348 69 YP 002330168 1 574521 Proteobacteria
Citrobacter rodentium ICC168 YP 003363894 1 637910 Proteobacteria
Dickeya zeae MK19 WP 038914150 1 Dzeae
Dickeya zeae CSL RW192 WP 038906884 1 Dzeae
Dickeya zeae MK19 WP 038906884 1 Dzeae
Dickeya dianthicola IPO 980 WP 024107973 1 Ddianthicola
Dickeya dianthicola NCPPB 3534 WP 024107973 1 Ddianthicola
Dickeya dianthicola GBBC 2039 WP 024107973 1 Ddianthicola
534 DAQ1742 03686 phages P2 P6 P7 related to Mu like Seed
Dickeya aquatica DW 0440 WP 035345458 1 Daquatica
DAQ1742 DAQ1742 03195 Daquatica
DAQ1742 DAQ1742 03686 Daquatica
DAQ1742 DAQ1742 01688 Daquatica
Dickeya sp CSL RW240 WP 067486439 1 Daquatica
Polaromonas sp JS666 YP 550538 1 296591 Proteobacteria
Desulfovibrio africanus str Walvis Bay YP 005051910 1 690850 Proteobacteria
Myxococcus stipitatus DSM 14675 YP 007358941 1 1278073 Proteobacteria
Desulfovibrio vulgaris RCH1 YP 005703420 1 573059 Proteobacteria
Desulfovibrio aespoeensis Aspo 2 YP 004122147 1 643562 Proteobacteria
Desulfovibrio africanus str Walvis Bay YP 005050709 1 690850 Proteobacteria
Thalassolituus oleivorans MIL 1 YP 007681177 1 1298593 Proteobacteria
Bordetella petrii DSM 12804 YP 001633036 1 340100 Proteobacteria
Shigella sonnei 53G YP 005457603 1 216599 Proteobacteria
Citrobacter rodentium ICC168 YP 003368130 1 637910 Proteobacteria
Bibersteinia trehalosi USDA ARS USMARC 192 YP 007548809 1 1171377 Proteobacteria
Xenorhabdus nematophila ATCC 19061 YP 003713550 1 406817 Proteobacteria
Shimwellia blattae DSM 4481 NBRC 105725 YP 006320566 1 630626 Proteobacteria
Shewanella oneidensis MR 1 NP 718277 1 211586 Proteobacteria
Vibrio cholerae O395 YP 002819491 1 345073 Proteobacteria
Vibrio cholerae O395 YP 002819431 1 345073 Proteobacteria
Acetobacter pasteurianus 386B YP 008391229 1 1266844 Proteobacteria
Gluconacetobacter medellinensis NBRC 3288 YP 004868917 1 634177 Proteobacteria
Gluconacetobacter medellinensis NBRC 3288 YP 004868336 1 634177 Proteobacteria
Burkholderia xenovorans LB400 YP 557975 1 266265 Proteobacteria
Burkholderia phymatum STM815 YP 001858103 1 391038 Proteobacteria
Citrobacter rodentium ICC168 YP 003366125 1 637910 Proteobacteria
Yersinia pseudotuberculosis IP 31758 YP 001401706 1 349747 Proteobacteria
Yersinia pestis Antiqua YP 650869 1 360102 Proteobacteria
Yersinia pestis biovar Microtus str 91001 NP 992278 1 229193 Proteobacteria
Pseudomonas syringae pv tomato str DC3000 NP 790424 1 223283 Proteobacteria
Pseudomonas fluorescens Pf0 1 YP 346886 1 205922 Proteobacteria
Sodalis glossinidius str morsitans YP 454399 1 343509 Proteobacteria
Erwinia pyrifoliae DSM 12163 YP 005802951 1 644651 Proteobacteria
Dickeya zeae DZ2Q WP 038926708 1 Dzeae
Dickeya solani RNS 05 1 2A WP 057083763 1 Dsolani
Enterobacter sp 638 YP 001177311 1 399742 Proteobacteria
Shigella flexneri 5 str 8401 YP 687826 1 373384 Proteobacteria
Escherichia fergusonii ATCC 35469 YP 002381798 1 585054 Proteobacteria
Escherichia coli 042 YP 006095162 1 216592 Proteobacteria
Salmonella enterica subsp enterica serovar Bovismorbificans str 3114 YP 008505065 1 1320309 Proteobacteria
Shigella flexneri 2002017 YP 005727703 1 591020 Proteobacteria
Shigella sonnei 53G YP 005457728 1 216599 Proteobacteria
Shigella flexneri 2a str 2457T NP 837590 1 198215 Proteobacteria
Salmonella enterica subsp enterica serovar Enteritidis str P125109 YP 002244028 1 550537 Proteobacteria
Salmonella enterica subsp enterica serovar Schwarzengrund str CVM19633 YP 002117159 1 439843 Proteobacteria
Salmonella enterica subsp enterica serovar Dublin str CT 02021853 YP 002216021 1 439851 Proteobacteria
Salmonella enterica subsp enterica serovar Paratyphi B str SPB7 YP 001588372 1 1016998 Proteobacteria
Salmonella enterica subsp enterica serovar Schwarzengrund str CVM19633 YP 002115118 1 439843 Proteobacteria
Salmonella enterica subsp enterica serovar 4 5 12 i str 08 1736 YP 008266214 1 866913 Proteobacteria
Salmonella enterica subsp enterica serovar Typhimurium str ST4 74 YP 005242919 1 909946 Proteobacteria
Salmonella enterica subsp enterica serovar Typhimurium str 14028S YP 005237987 1 588858 Proteobacteria

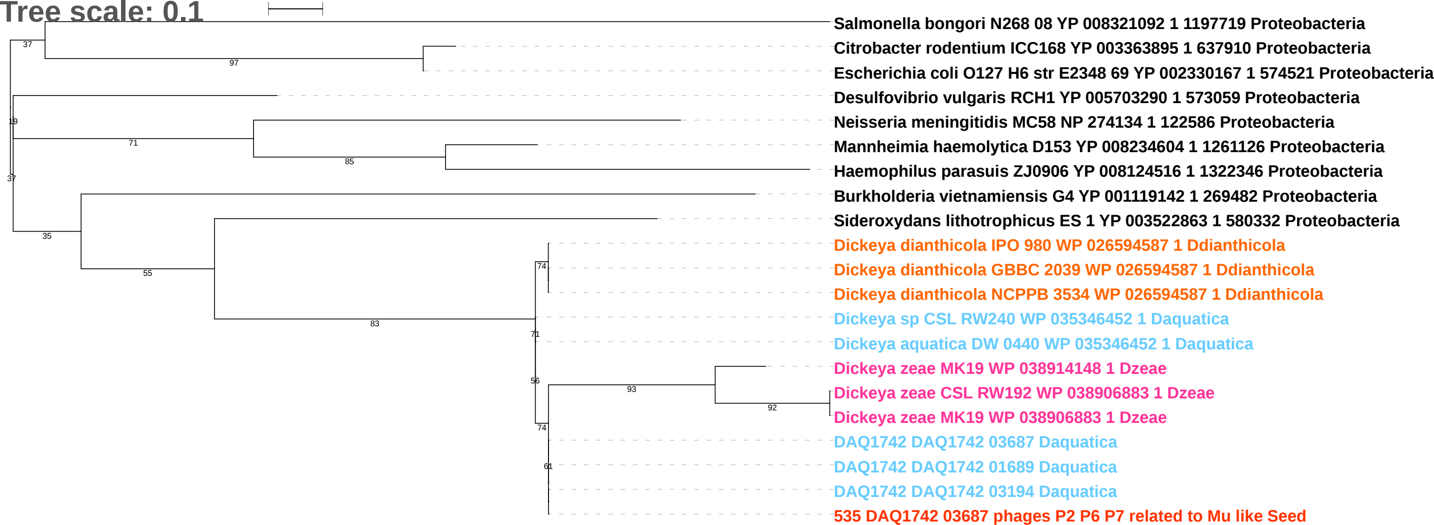
Tree scale: 0.1
Salmonella bongori N268 08 YP 008321092 1 1197719 Proteobacteria
Citrobacter rodentium ICC168 YP 003363895 1 637910 Proteobacteria
Escherichia coli O127 H6 str E2348 69 YP 002330167 1 574521 Proteobacteria
Desulfovibrio vulgaris RCH1 YP 005703290 1 573059 Proteobacteria
Neisseria meningitidis MC58 NP 274134 1 122586 Proteobacteria
Mannheimia haemolytica D153 YP 008234604 1 1261126 Proteobacteria
Haemophilus parasuis ZJ0906 YP 008124516 1 1322346 Proteobacteria
Burkholderia vietnamiensis G4 YP 001119142 1 269482 Proteobacteria
Sideroxydans lithotrophicus ES 1 YP 003522863 1 580332 Proteobacteria
Dickeya dianthicola IPO 980 WP 026594587 1 Ddianthicola
Dickeya dianthicola GBBC 2039 WP 026594587 1 Ddianthicola
Dickeya dianthicola NCPPB 3534 WP 026594587 1 Ddianthicola
Dickeya sp CSL RW240 WP 035346452 1 Daquatica
Dickeya aquatica DW 0440 WP 035346452 1 Daquatica
Dickeya zeae MK19 WP 038914148 1 Dzeae
Dickeya zeae CSL RW192 WP 038906883 1 Dzeae
Dickeya zeae MK19 WP 038906883 1 Dzeae
DAQ1742 DAQ1742 03687 Daquatica
DAQ1742 DAQ1742 01689 Daquatica
DAQ1742 DAQ1742 03194 Daquatica
535 DAQ1742 03687 phages P2 P6 P7 related to Mu like Seed

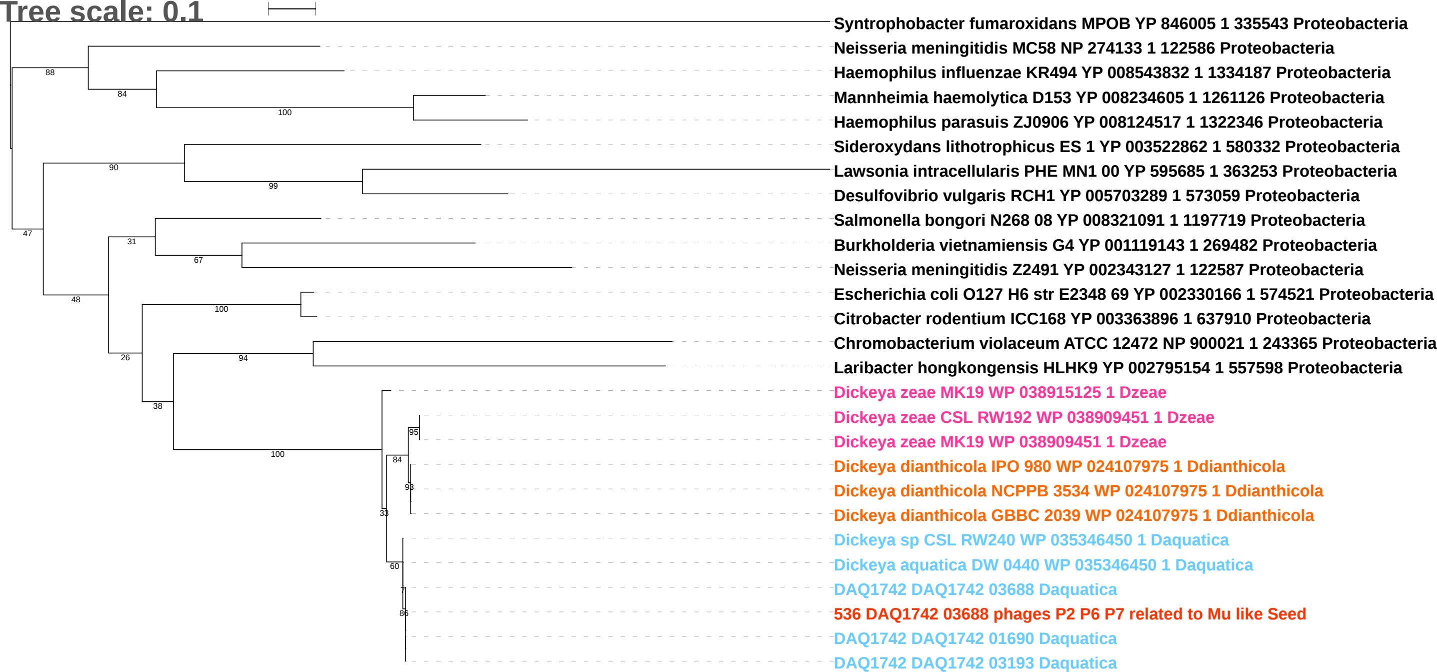
Tree scale: 0.1
Syntrophobacter fumaroxidans MPOB YP 846005 1 335543 Proteobacteria
Neisseria meningitidis MC58 NP 274133 1 122586 Proteobacteria
Haemophilus influenzae KR494 YP 008543832 1 1334187 Proteobacteria
Mannheimia haemolytica D153 YP 008234605 1 1261126 Proteobacteria
Haemophilus parasuis ZJ0906 YP 008124517 1 1322346 Proteobacteria
Sideroxydans lithotrophicus ES 1 YP 003522862 1 580332 Proteobacteria
Lawsonia intracellularis PHE MN1 00 YP 595685 1 363253 Proteobacteria
Desulfovibrio vulgaris RCH1 YP 005703289 1 573059 Proteobacteria
Salmonella bongori N268 08 YP 008321091 1 1197719 Proteobacteria
Burkholderia vietnamiensis G4 YP 001119143 1 269482 Proteobacteria
Neisseria meningitidis Z2491 YP 002343127 1 122587 Proteobacteria
Escherichia coli O127 H6 str E2348 69 YP 002330166 1 574521 Proteobacteria
Citrobacter rodentium ICC168 YP 003363896 1 637910 Proteobacteria
Chromobacterium violaceum ATCC 12472 NP 900021 1 243365 Proteobacteria
Laribacter hongkongensis HLHK9 YP 002795154 1 557598 Proteobacteria
Dickeya zeae MK19 WP 038915125 1 Dzeae
Dickeya zeae CSL RW192 WP 038909451 1 Dzeae
Dickeya zeae MK19 WP 038909451 1 Dzeae
Dickeya dianthicola IPO 980 WP 024107975 1 Ddianthicola
Dickeya dianthicola NCPPB 3534 WP 024107975 1 Ddianthicola
Dickeya dianthicola GBBC 2039 WP 024107975 1 Ddianthicola
Dickeya sp CSL RW240 WP 035346450 1 Daquatica
Dickeya aquatica DW 0440 WP 035346450 1 Daquatica
DAQ1742 DAQ1742 03688 Daquatica
536 DAQ1742 03688 phages P2 P6 P7 related to Mu like Seed
DAQ1742 DAQ1742 01690 Daquatica
DAQ1742 DAQ1742 03193 Daquatica
88
84
100
90
99
47
31
67
48
100
26
94
38
100
95
84
93
33
60
7
86

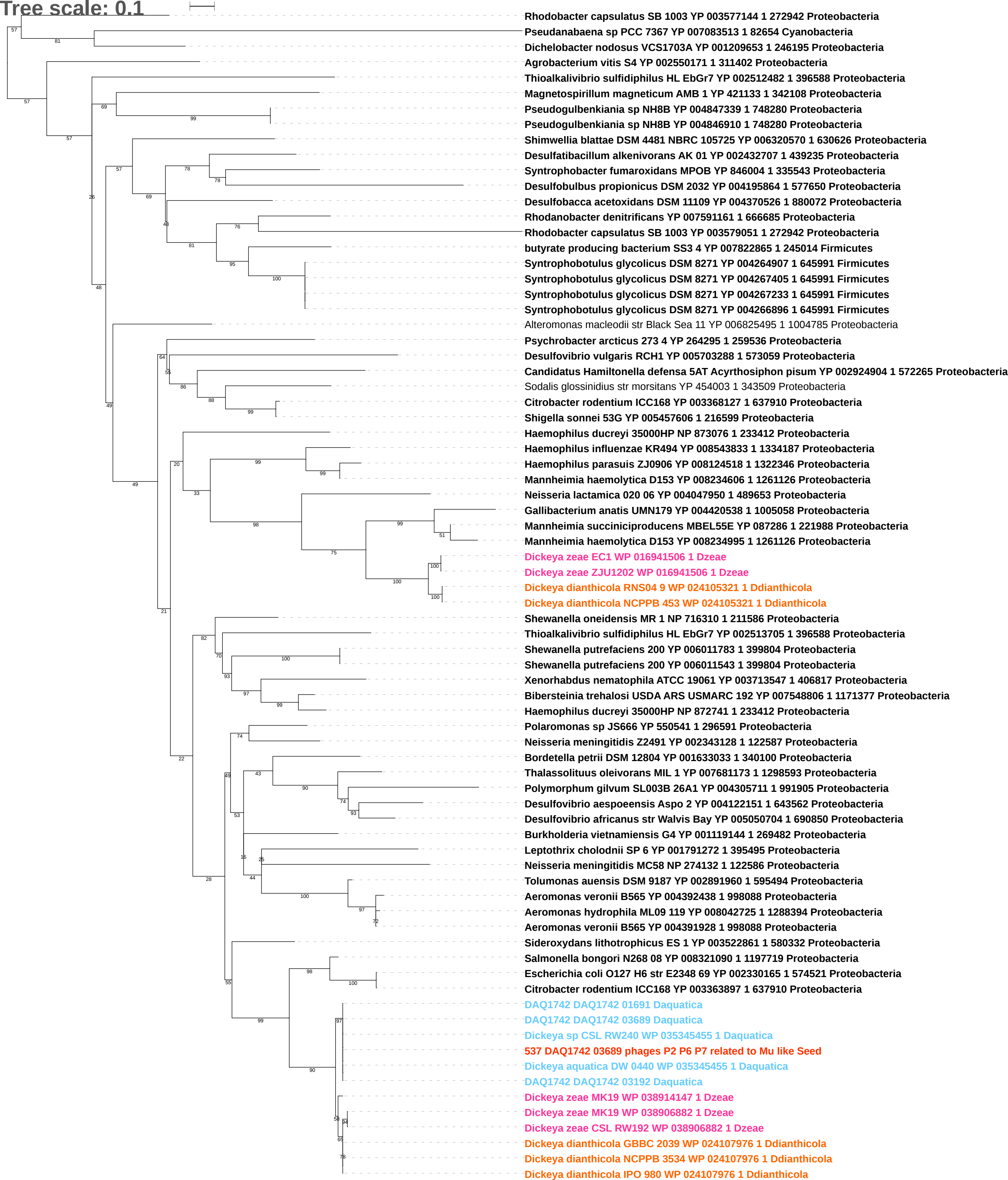
Tree scale: 0.1
Rhodobacter capsulatus SB 1003 YP 003577144 1 272942 Proteobacteria
Pseudanabaena sp PCC 7367 YP 007083513 1 82654 Cyanobacteria
Dichelobacter nodosus VCS1703A YP 001209653 1 246195 Proteobacteria
Agrobacterium vitis S4 YP 002550171 1 311402 Proteobacteria
Thioalkalivibrio sulfidiphilus HL EbGr7 YP 002512482 1 396588 Proteobacteria
Magnetospirillum magneticum AMB 1 YP 421133 1 342108 Proteobacteria
Pseudogulbenkiania sp NH8B YP 004847339 1 748280 Proteobacteria
Pseudogulbenkiania sp NH8B YP 004846910 1 748280 Proteobacteria
Shimwellia blattae DSM 4481 NBRC 105725 YP 006320570 1 630626 Proteobacteria
Desulfatibacillum alkenivorans AK 01 YP 002432707 1 439235 Proteobacteria
Syntrophobacter fumaroxidans MPOB YP 846004 1 335543 Proteobacteria
Desulfobulbus propionicus DSM 2032 YP 004195864 1 577650 Proteobacteria
Desulfobacca acetoxidans DSM 11109 YP 004370526 1 880072 Proteobacteria
Rhodanobacter denitrificans YP 007591161 1 666685 Proteobacteria
Rhodobacter capsulatus SB 1003 YP 003579051 1 272942 Proteobacteria
butyrate producing bacterium SS3 4 YP 007822865 1 245014 Firmicutes
Syntrophobotulus glycolicus DSM 8271 YP 004264907 1 645991 Firmicutes
Syntrophobotulus glycolicus DSM 8271 YP 004267405 1 645991 Firmicutes
Syntrophobotulus glycolicus DSM 8271 YP 004267233 1 645991 Firmicutes
Syntrophobotulus glycolicus DSM 8271 YP 004266896 1 645991 Firmicutes
Alteromonas macleodii str Black Sea 11 YP 006825495 1 1004785 Proteobacteria
Psychrobacter arcticus 273 4 YP 264295 1 259536 Proteobacteria
Desulfovibrio vulgaris RCH1 YP 005703288 1 573059 Proteobacteria
Candidatus Hamiltonella defensa 5AT Acyrthosiphon pisum YP 002924904 1 572265 Proteobacteria
Sodalis glossinidius str morsitans YP 454003 1 343509 Proteobacteria
Citrobacter rodentium ICC168 YP 003368127 1 637910 Proteobacteria
Shigella sonnei 53G YP 005457606 1 216599 Proteobacteria
Haemophilus ducreyi 35000HP NP 873076 1 233412 Proteobacteria
Haemophilus influenzae KR494 YP 008543833 1 1334187 Proteobacteria
Haemophilus parasuis ZJ0906 YP 008124518 1 1322346 Proteobacteria
Mannheimia haemolytica D153 YP 008234606 1 1261126 Proteobacteria
Neisseria lactamica 020 06 YP 004047950 1 489653 Proteobacteria
Gallibacterium anatis UMN179 YP 004420538 1 1005058 Proteobacteria
Mannheimia succiniciproducens MBEL55E YP 087286 1 221988 Proteobacteria
Mannheimia haemolytica D153 YP 008234995 1 1261126 Proteobacteria
Dickeya zeae EC1 WP 016941506 1 Dzeae
Dickeya zeae ZJU1202 WP 016941506 1 Dzeae
Dickeya dianthicola RNS04 9 WP 024105321 1 Ddianthicola
Dickeya dianthicola NCPPB 453 WP 024105321 1 Ddianthicola
Shewanella oneidensis MR 1 NP 716310 1 211586 Proteobacteria
Thioalkalivibrio sulfidiphilus HL EbGr7 YP 002513705 1 396588 Proteobacteria
Shewanella putrefaciens 200 YP 006011783 1 399804 Proteobacteria
Shewanella putrefaciens 200 YP 006011543 1 399804 Proteobacteria
Xenorhabdus nematophila ATCC 19061 YP 003713547 1 406817 Proteobacteria
Bibersteinia trehalosi USDA ARS USMARC 192 YP 007548806 1 1171377 Proteobacteria
Haemophilus ducreyi 35000HP NP 872741 1 233412 Proteobacteria
Polaromonas sp JS666 YP 550541 1 296591 Proteobacteria
Neisseria meningitidis Z2491 YP 002343128 1 122587 Proteobacteria
Bordetella petrii DSM 12804 YP 001633033 1 340100 Proteobacteria
Thalassolituus oleivorans MIL 1 YP 007681173 1 1298593 Proteobacteria
Polymorphum gilvum SL003B 26A1 YP 004305711 1 991905 Proteobacteria
Desulfovibrio aespoeensis Aspo 2 YP 004122151 1 643562 Proteobacteria
Desulfovibrio africanus str Walvis Bay YP 005050704 1 690850 Proteobacteria
Burkholderia vietnamiensis G4 YP 001119144 1 269482 Proteobacteria
Leptothrix cholodnii SP 6 YP 001791272 1 395495 Proteobacteria
Neisseria meningitidis MC58 NP 274132 1 122586 Proteobacteria
Tolumonas auensis DSM 9187 YP 002891960 1 595494 Proteobacteria
Aeromonas veronii B565 YP 004392438 1 998088 Proteobacteria
Aeromonas hydrophila ML09 119 YP 008042725 1 1288394 Proteobacteria
Aeromonas veronii B565 YP 004391928 1 998088 Proteobacteria
Sideroxydans lithotrophicus ES 1 YP 003522861 1 580332 Proteobacteria
Salmonella bongori N268 08 YP 008321090 1 1197719 Proteobacteria
Escherichia coli O127 H6 str E2348 69 YP 002330165 1 574521 Proteobacteria
Citrobacter rodentium ICC168 YP 003363897 1 637910 Proteobacteria
DAQ1742 DAQ1742 01691 Daquatica
DAQ1742 DAQ1742 03689 Daquatica
Dickeya sp CSL RW240 WP 035345455 1 Daquatica
537 DAQ1742 03689 phages P2 P6 P7 related to Mu like Seed
Dickeya aquatica DW 0440 WP 035345455 1 Daquatica
DAQ1742 DAQ1742 03192 Daquatica
Dickeya zeae MK19 WP 038914147 1 Dzeae
Dickeya zeae MK19 WP 038906882 1 Dzeae
Dickeya zeae CSL RW192 WP 038906882 1 Dzeae
Dickeya dianthicola GBBC 2039 WP 024107976 1 Ddianthicola
Dickeya dianthicola NCPPB 3534 WP 024107976 1 Ddianthicola
Dickeya dianthicola IPO 980 WP 024107976 1 Ddianthicola

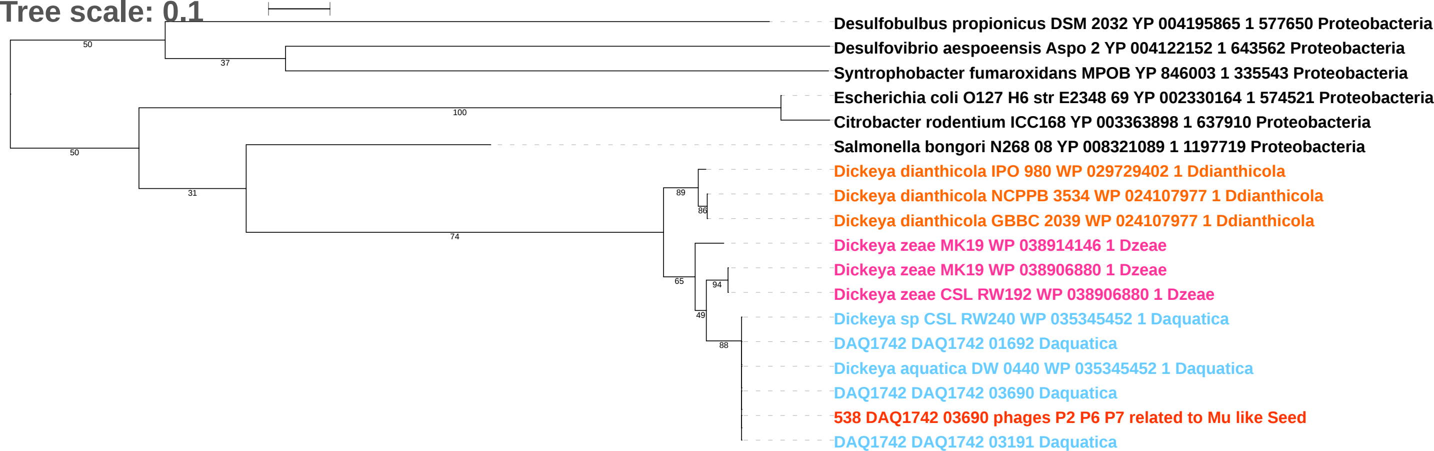

Tree scale: 0.1
50
37
100
50
31
89
86
74
65
94
49
88
Desulfobulbus propionicus DSM 2032 YP 004195865 1 577650 Proteobacteria
Desulfovibrio aespoeensis Aspo 2 YP 004122152 1 643562 Proteobacteria
Syntrophobacter fumaroxidans MPOB YP 846003 1 335543 Proteobacteria
Escherichia coli O127 H6 str E2348 69 YP 002330164 1 574521 Proteobacteria
Citrobacter rodentium ICC168 YP 003363898 1 637910 Proteobacteria
Salmonella bongori N268 08 YP 008321089 1 1197719 Proteobacteria
Dickeya dianthicola IPO 980 WP 029729402 1 Ddianthicola
Dickeya dianthicola NCPPB 3534 WP 024107977 1 Ddianthicola
Dickeya dianthicola GBBC 2039 WP 024107977 1 Ddianthicola
Dickeya zeae MK19 WP 038914146 1 Dzeae
Dickeya zeae MK19 WP 038906880 1 Dzeae
Dickeya zeae CSL RW192 WP 038906880 1 Dzeae
Dickeya sp CSL RW240 WP 035345452 1 Daquatica
DAQ1742 DAQ1742 01692 Daquatica
Dickeya aquatica DW 0440 WP 035345452 1 Daquatica
DAQ1742 DAQ1742 03690 Daquatica
538 DAQ1742 03690 phages P2 P6 P7 related to Mu like Seed
DAQ1742 DAQ1742 03191 Daquatica

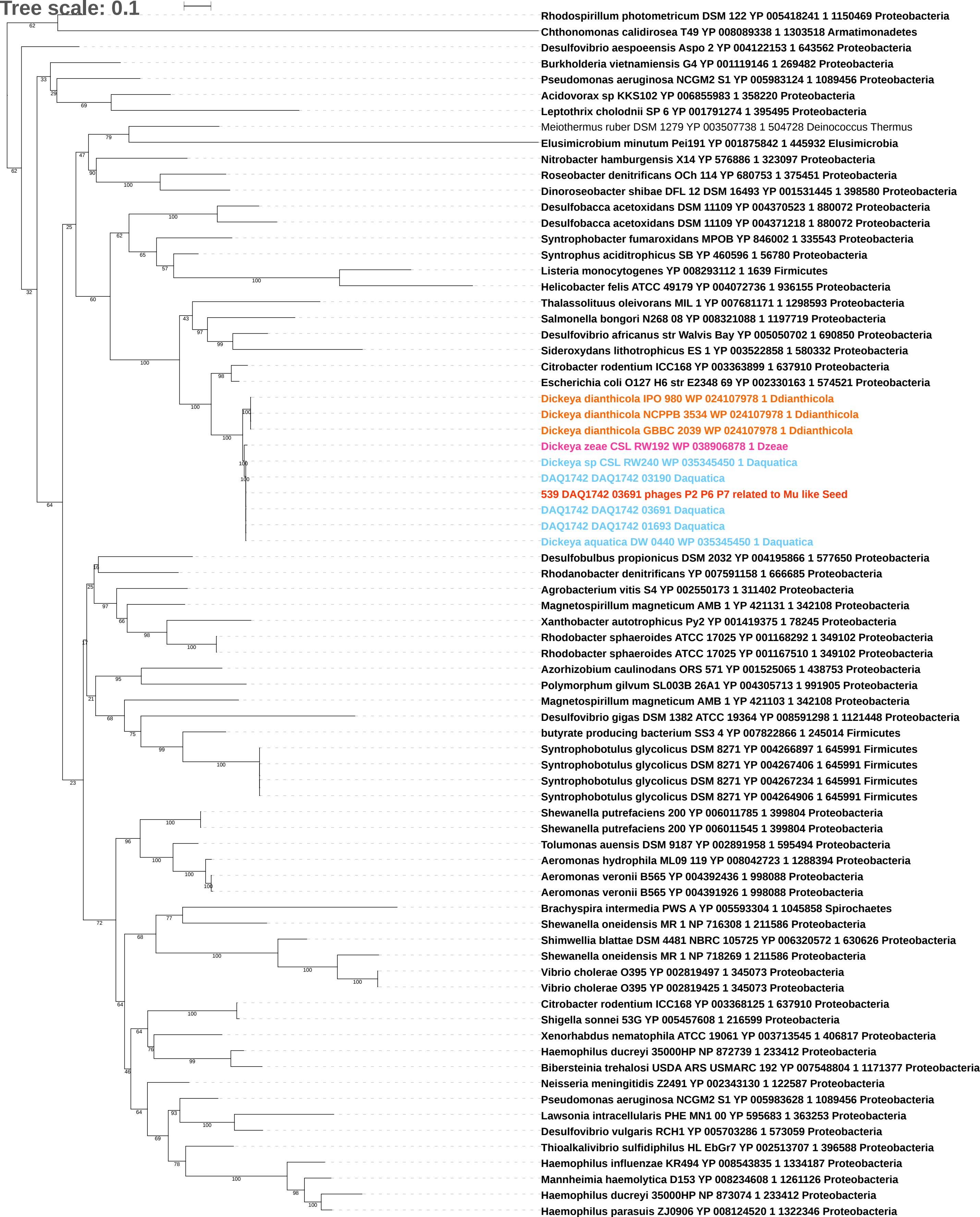
Tree scale: 0.1
Rhodospirillum photometricum DSM 122 YP 005418241 1 1150469 Proteobacteria
Chthonomonas calidirosea T49 YP 008089338 1 1303518 Armatimonadetes
Desulfovibrio aespoeensis Aspo 2 YP 004122153 1 643562 Proteobacteria
Burkholderia vietnamiensis G4 YP 001119146 1 269482 Proteobacteria
Pseudomonas aeruginosa NCGM2 S1 YP 005983124 1 1089456 Proteobacteria
Acidovorax sp KKS102 YP 006855983 1 358220 Proteobacteria
Leptothrix cholodnii SP 6 YP 001791274 1 395495 Proteobacteria
Meiothermus ruber DSM 1279 YP 003507738 1 504728 Deinococcus Thermus
Elusimicrobium minutum Pei191 YP 001875842 1 445932 Elusimicrobia
Nitrobacter hamburgensis X14 YP 576886 1 323097 Proteobacteria
Roseobacter denitrificans OCh 114 YP 680753 1 375451 Proteobacteria
Dinoroseobacter shibae DFL 12 DSM 16493 YP 001531445 1 398580 Proteobacteria
Desulfobacca acetoxidans DSM 11109 YP 004370523 1 880072 Proteobacteria
Desulfobacca acetoxidans DSM 11109 YP 004371218 1 880072 Proteobacteria
Syntrophobacter fumaroxidans MPOB YP 846002 1 335543 Proteobacteria
Syntrophus aciditrophicus SB YP 460596 1 56780 Proteobacteria
Listeria monocytogenes YP 008293112 1 1639 Firmicutes
Helicobacter felis ATCC 49179 YP 004072736 1 936155 Proteobacteria
Thalassolituus oleivorans MIL 1 YP 007681171 1 1298593 Proteobacteria
Salmonella bongori N268 08 YP 008321088 1 1197719 Proteobacteria
Desulfovibrio africanus str Walvis Bay YP 005050702 1 690850 Proteobacteria
Sideroxydans lithotrophicus ES 1 YP 003522858 1 580332 Proteobacteria
Citrobacter rodentium ICC168 YP 003363899 1 637910 Proteobacteria
Escherichia coli O127 H6 str E2348 69 YP 002330163 1 574521 Proteobacteria
Dickeya dianthicola IPO 980 WP 024107978 1 Ddianthicola
Dickeya dianthicola NCPPB 3534 WP 024107978 1 Ddianthicola
Dickeya dianthicola GBBC 2039 WP 024107978 1 Ddianthicola
Dickeya zeae CSL RW192 WP 038906878 1 Dzeae
Dickeya sp CSL RW240 WP 035345450 1 Daquatica
DAQ1742 DAQ1742 03190 Daquatica
539 DAQ1742 03691 phages P2 P6 P7 related to Mu like Seed
DAQ1742 DAQ1742 03691 Daquatica
DAQ1742 DAQ1742 01693 Daquatica
Dickeya aquatica DW 0440 WP 035345450 1 Daquatica
Desulfobulbus propionicus DSM 2032 YP 004195866 1 577650 Proteobacteria
Rhodanobacter denitrificans YP 007591158 1 666685 Proteobacteria
Agrobacterium vitis S4 YP 002550173 1 311402 Proteobacteria
Magnetospirillum magneticum AMB 1 YP 421131 1 342108 Proteobacteria
Xanthobacter autotrophicus Py2 YP 001419375 1 78245 Proteobacteria
Rhodobacter sphaeroides ATCC 17025 YP 001168292 1 349102 Proteobacteria
Rhodobacter sphaeroides ATCC 17025 YP 001167510 1 349102 Proteobacteria
Azorhizobium caulinodans ORS 571 YP 001525065 1 438753 Proteobacteria
Polymorphum gilvum SL003B 26A1 YP 004305713 1 991905 Proteobacteria
Magnetospirillum magneticum AMB 1 YP 421103 1 342108 Proteobacteria
Desulfovibrio gigas DSM 1382 ATCC 19364 YP 008591298 1 1121448 Proteobacteria
butyrate producing bacterium SS3 4 YP 007822866 1 245014 Firmicutes
Syntrophobotulus glycolicus DSM 8271 YP 004266897 1 645991 Firmicutes
Syntrophobotulus glycolicus DSM 8271 YP 004267406 1 645991 Firmicutes
Syntrophobotulus glycolicus DSM 8271 YP 004267234 1 645991 Firmicutes
Syntrophobotulus glycolicus DSM 8271 YP 004264906 1 645991 Firmicutes
Shewanella putrefaciens 200 YP 006011785 1 399804 Proteobacteria
Shewanella putrefaciens 200 YP 006011545 1 399804 Proteobacteria
Tolumonas auensis DSM 9187 YP 002891958 1 595494 Proteobacteria
Aeromonas hydrophila ML09 119 YP 008042723 1 1288394 Proteobacteria
Aeromonas veronii B565 YP 004392436 1 998088 Proteobacteria
Aeromonas veronii B565 YP 004391926 1 998088 Proteobacteria
Brachyspira intermedia PWS A YP 005593304 1 1045858 Spirochaetes
Shewanella oneidensis MR 1 NP 716308 1 211586 Proteobacteria
Shimwellia blattae DSM 4481 NBRC 105725 YP 006320572 1 630626 Proteobacteria
Shewanella oneidensis MR 1 NP 718269 1 211586 Proteobacteria
Vibrio cholerae O395 YP 002819497 1 345073 Proteobacteria
Vibrio cholerae O395 YP 002819425 1 345073 Proteobacteria
Citrobacter rodentium ICC168 YP 003368125 1 637910 Proteobacteria
Shigella sonnei 53G YP 005457608 1 216599 Proteobacteria
Xenorhabdus nematophila ATCC 19061 YP 003713545 1 406817 Proteobacteria
Haemophilus ducreyi 35000HP NP 872739 1 233412 Proteobacteria
Bibersteinia trehalosi USDA ARS USMARC 192 YP 007548804 1 1171377 Proteobacteria
Neisseria meningitidis Z2491 YP 002343130 1 122587 Proteobacteria
Pseudomonas aeruginosa NCGM2 S1 YP 005983628 1 1089456 Proteobacteria
Lawsonia intracellularis PHE MN1 00 YP 595683 1 363253 Proteobacteria
Desulfovibrio vulgaris RCH1 YP 005703286 1 573059 Proteobacteria
Thioalkalivibrio sulfidiphilus HL EbGr7 YP 002513707 1 396588 Proteobacteria
Haemophilus influenzae KR494 YP 008543835 1 1334187 Proteobacteria
Mannheimia haemolytica D153 YP 008234608 1 1261126 Proteobacteria
Haemophilus ducreyi 35000HP NP 873074 1 233412 Proteobacteria
Haemophilus parasuis ZJ0906 YP 008124520 1 1322346 Proteobacteria

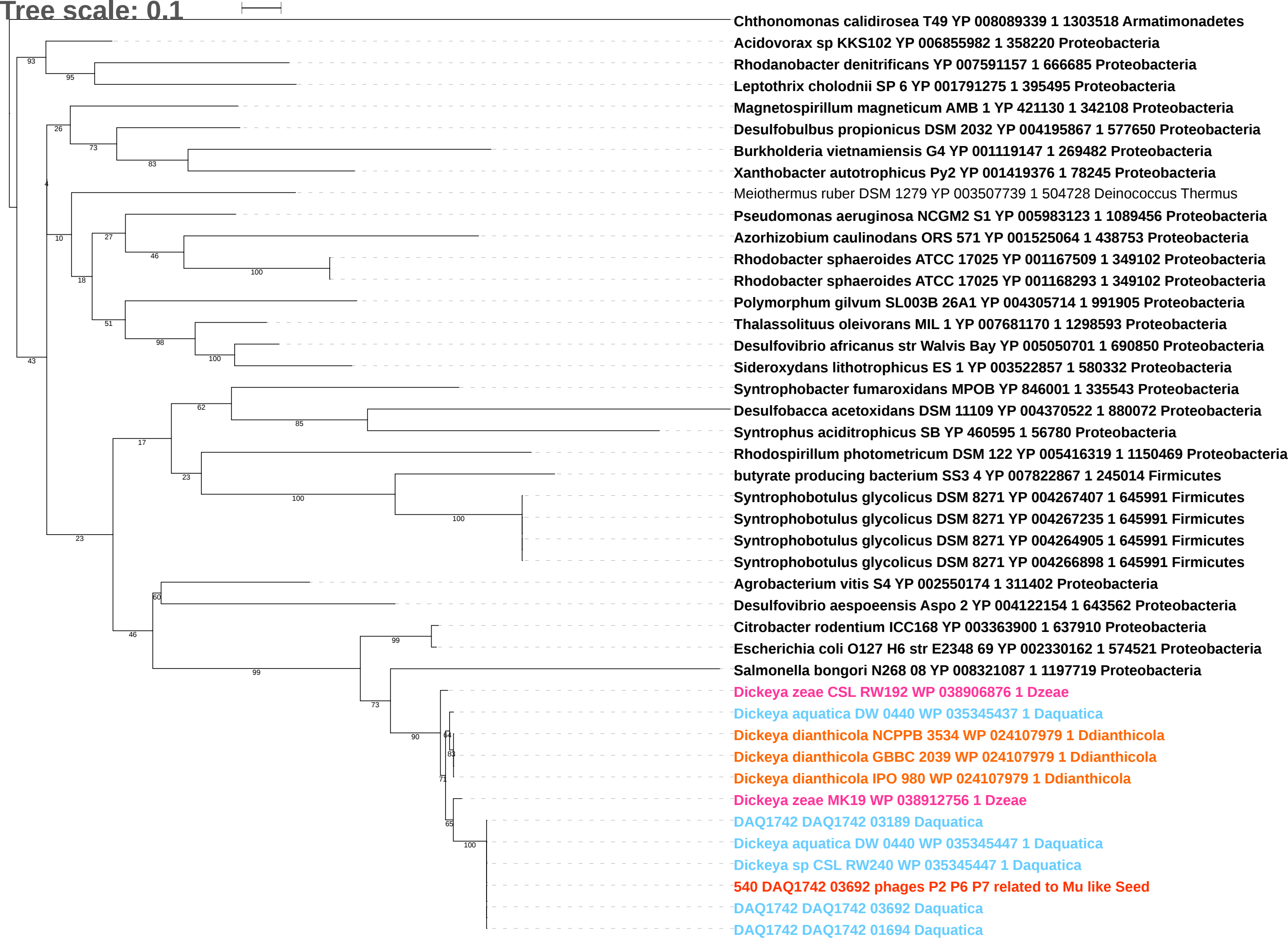
Tree scale: 0.1
Chthonomonas calidirosea T49 YP 008089339 1 1303518 Armatimonadetes
Acidovorax sp KKS102 YP 006855982 1 358220 Proteobacteria
Rhodanobacter denitrificans YP 007591157 1 666685 Proteobacteria
Leptothrix cholodnii SP 6 YP 001791275 1 395495 Proteobacteria
Magnetospirillum magneticum AMB 1 YP 421130 1 342108 Proteobacteria
Desulfobulbus propionicus DSM 2032 YP 004195867 1 577650 Proteobacteria
Burkholderia vietnamiensis G4 YP 001119147 1 269482 Proteobacteria
Xanthobacter autotrophicus Py2 YP 001419376 1 78245 Proteobacteria
Meiothermus ruber DSM 1279 YP 003507739 1 504728 Deinococcus Thermus
Pseudomonas aeruginosa NCGM2 S1 YP 005983123 1 1089456 Proteobacteria
Azorhizobium caulinodans ORS 571 YP 001525064 1 438753 Proteobacteria
Rhodobacter sphaeroides ATCC 17025 YP 001167509 1 349102 Proteobacteria
Rhodobacter sphaeroides ATCC 17025 YP 001168293 1 349102 Proteobacteria
Polymorphum gilvum SL003B 26A1 YP 004305714 1 991905 Proteobacteria
Thalassolituus oleivorans MIL 1 YP 007681170 1 1298593 Proteobacteria
Desulfovibrio africanus str Walvis Bay YP 005050701 1 690850 Proteobacteria
Sideroxydans lithotrophicus ES 1 YP 003522857 1 580332 Proteobacteria
Syntrophobacter fumaroxidans MPOB YP 846001 1 335543 Proteobacteria
Desulfobacca acetoxidans DSM 11109 YP 004370522 1 880072 Proteobacteria
Syntrophus aciditrophicus SB YP 460595 1 56780 Proteobacteria
Rhodospirillum photometricum DSM 122 YP 005416319 1 1150469 Proteobacteria
butyrate producing bacterium SS3 4 YP 007822867 1 245014 Firmicutes
Syntrophobotulus glycolicus DSM 8271 YP 004267407 1 645991 Firmicutes
Syntrophobotulus glycolicus DSM 8271 YP 004267235 1 645991 Firmicutes
Syntrophobotulus glycolicus DSM 8271 YP 004264905 1 645991 Firmicutes
Syntrophobotulus glycolicus DSM 8271 YP 004266898 1 645991 Firmicutes
Agrobacterium vitis S4 YP 002550174 1 311402 Proteobacteria
Desulfovibrio aespoeensis Aspo 2 YP 004122154 1 643562 Proteobacteria
Citrobacter rodentium ICC168 YP 003363900 1 637910 Proteobacteria
Escherichia coli O127 H6 str E2348 69 YP 002330162 1 574521 Proteobacteria
Salmonella bongori N268 08 YP 008321087 1 1197719 Proteobacteria
Dickeya zeae CSL RW192 WP 038906876 1 Dzeae
Dickeya aquatica DW 0440 WP 035345437 1 Daquatica
Dickeya dianthicola NCPPB 3534 WP 024107979 1 Ddianthicola
Dickeya dianthicola GBBC 2039 WP 024107979 1 Ddianthicola
Dickeya dianthicola IPO 980 WP 024107979 1 Ddianthicola
Dickeya zeae MK19 WP 038912756 1 Dzeae
DAQ1742 DAQ1742 03189 Daquatica
Dickeya aquatica DW 0440 WP 035345447 1 Daquatica
Dickeya sp CSL RW240 WP 035345447 1 Daquatica
540 DAQ1742 03692 phages P2 P6 P7 related to Mu like Seed
DAQ1742 DAQ1742 03692 Daquatica
DAQ1742 DAQ1742 01694 Daquatica

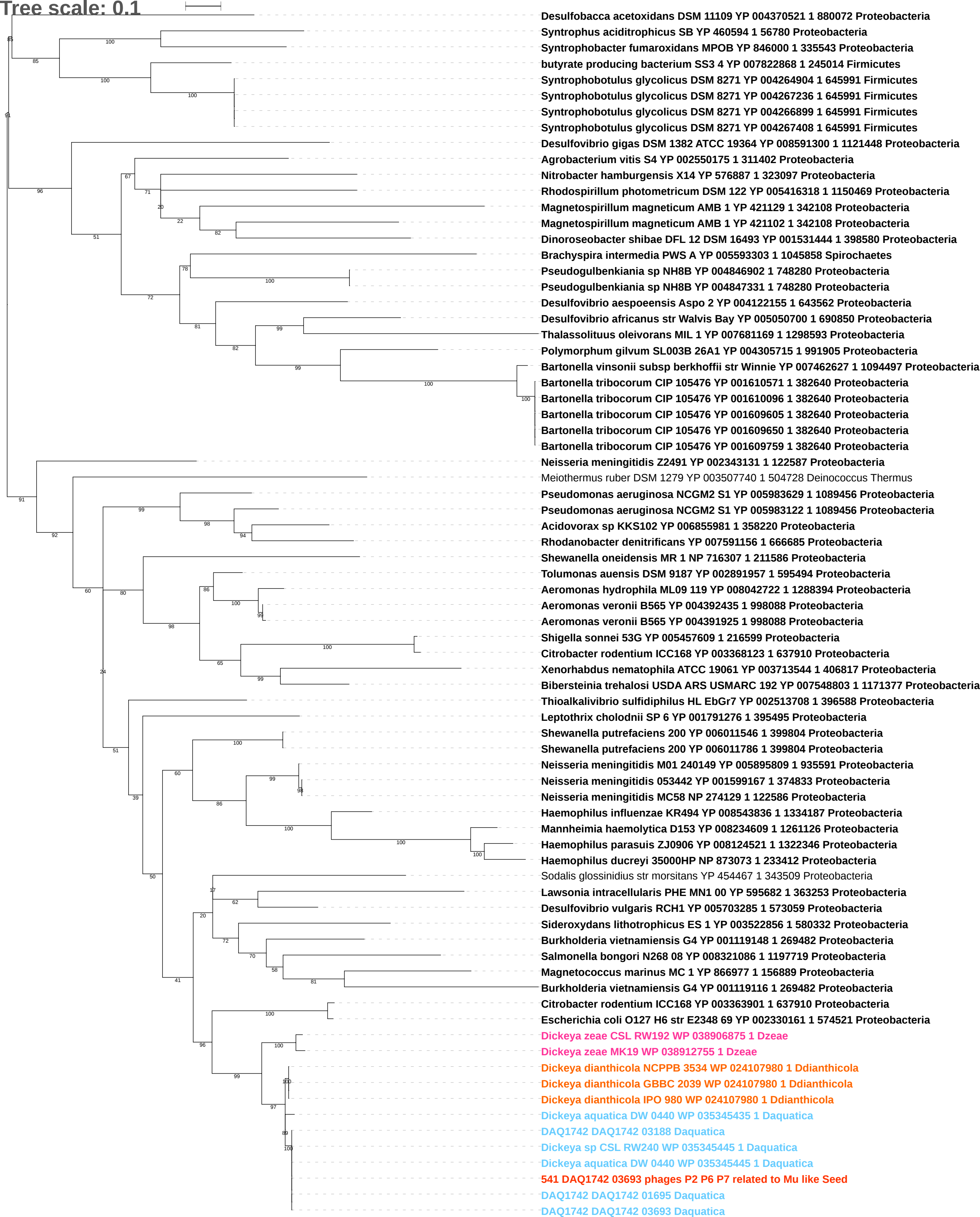
Tree scale: 0.1
Desulfobacca acetoxidans DSM 11109 YP 004370521 1 880072 Proteobacteria
Syntrophus aciditrophicus SB YP 460594 1 56780 Proteobacteria
Syntrophobacter fumaroxidans MPOB YP 846000 1 335543 Proteobacteria
butyrate producing bacterium SS3 4 YP 007822868 1 245014 Firmicutes
Syntrophobotulus glycolicus DSM 8271 YP 004264904 1 645991 Firmicutes
Syntrophobotulus glycolicus DSM 8271 YP 004267236 1 645991 Firmicutes
Syntrophobotulus glycolicus DSM 8271 YP 004266899 1 645991 Firmicutes
Syntrophobotulus glycolicus DSM 8271 YP 004267408 1 645991 Firmicutes
Desulfovibrio gigas DSM 1382 ATCC 19364 YP 008591300 1 1121448 Proteobacteria
Agrobacterium vitis S4 YP 002550175 1 311402 Proteobacteria
Nitrobacter hamburgensis X14 YP 576887 1 323097 Proteobacteria
Rhodospirillum photometricum DSM 122 YP 005416318 1 1150469 Proteobacteria
Magnetospirillum magneticum AMB 1 YP 421129 1 342108 Proteobacteria
Magnetospirillum magneticum AMB 1 YP 421102 1 342108 Proteobacteria
Dinoroseobacter shibae DFL 12 DSM 16493 YP 001531444 1 398580 Proteobacteria
Brachyspira intermedia PWS A YP 005593303 1 1045858 Spirochaetes
Pseudogulbenkiania sp NH8B YP 004846902 1 748280 Proteobacteria
Pseudogulbenkiania sp NH8B YP 004847331 1 748280 Proteobacteria
Desulfovibrio aespoeensis Aspo 2 YP 004122155 1 643562 Proteobacteria
Desulfovibrio africanus str Walvis Bay YP 005050700 1 690850 Proteobacteria
Thalassolituus oleivorans MIL 1 YP 007681169 1 1298593 Proteobacteria
Polymorphum gilvum SL003B 26A1 YP 004305715 1 991905 Proteobacteria
Bartonella vinsonii subsp berkhoffii str Winnie YP 007462627 1 1094497 Proteobacteria
Bartonella tribocorum CIP 105476 YP 001610571 1 382640 Proteobacteria
Bartonella tribocorum CIP 105476 YP 001610096 1 382640 Proteobacteria
Bartonella tribocorum CIP 105476 YP 001609605 1 382640 Proteobacteria
Bartonella tribocorum CIP 105476 YP 001609650 1 382640 Proteobacteria
Bartonella tribocorum CIP 105476 YP 001609759 1 382640 Proteobacteria
Neisseria meningitidis Z2491 YP 002343131 1 122587 Proteobacteria
Meiothermus ruber DSM 1279 YP 003507740 1 504728 Deinococcus Thermus
Pseudomonas aeruginosa NCGM2 S1 YP 005983629 1 1089456 Proteobacteria
Pseudomonas aeruginosa NCGM2 S1 YP 005983122 1 1089456 Proteobacteria
Acidovorax sp KKS102 YP 006855981 1 358220 Proteobacteria
Rhodanobacter denitrificans YP 007591156 1 666685 Proteobacteria
Shewanella oneidensis MR 1 NP 716307 1 211586 Proteobacteria
Tolumonas auensis DSM 9187 YP 002891957 1 595494 Proteobacteria
Aeromonas hydrophila ML09 119 YP 008042722 1 1288394 Proteobacteria
Aeromonas veronii B565 YP 004392435 1 998088 Proteobacteria
Aeromonas veronii B565 YP 004391925 1 998088 Proteobacteria
Shigella sonnei 53G YP 005457609 1 216599 Proteobacteria
Citrobacter rodentium ICC168 YP 003368123 1 637910 Proteobacteria
Xenorhabdus nematophila ATCC 19061 YP 003713544 1 406817 Proteobacteria
Bibersteinia trehalosi USDA ARS USMARC 192 YP 007548803 1 1171377 Proteobacteria
Thioalkalivibrio sulfidiphilus HL EbGr7 YP 002513708 1 396588 Proteobacteria
Leptothrix cholodnii SP 6 YP 001791276 1 395495 Proteobacteria
Shewanella putrefaciens 200 YP 006011546 1 399804 Proteobacteria
Shewanella putrefaciens 200 YP 006011786 1 399804 Proteobacteria
Neisseria meningitidis M01 240149 YP 005895809 1 935591 Proteobacteria
Neisseria meningitidis 053442 YP 001599167 1 374833 Proteobacteria
Neisseria meningitidis MC58 NP 274129 1 122586 Proteobacteria
Haemophilus influenzae KR494 YP 008543836 1 1334187 Proteobacteria
Mannheimia haemolytica D153 YP 008234609 1 1261126 Proteobacteria
Haemophilus parasuis ZJ0906 YP 008124521 1 1322346 Proteobacteria
Haemophilus ducreyi 35000HP NP 873073 1 233412 Proteobacteria
Sodalis glossinidius str morsitans YP 454467 1 343509 Proteobacteria
Lawsonia intracellularis PHE MN1 00 YP 595682 1 363253 Proteobacteria
Desulfovibrio vulgaris RCH1 YP 005703285 1 573059 Proteobacteria
Sideroxydans lithotrophicus ES 1 YP 003522856 1 580332 Proteobacteria
Burkholderia vietnamiensis G4 YP 001119148 1 269482 Proteobacteria
Salmonella bongori N268 08 YP 008321086 1 1197719 Proteobacteria
Magnetococcus marinus MC 1 YP 866977 1 156889 Proteobacteria
Burkholderia vietnamiensis G4 YP 001119116 1 269482 Proteobacteria
Citrobacter rodentium ICC168 YP 003363901 1 637910 Proteobacteria
Escherichia coli O127 H6 str E2348 69 YP 002330161 1 574521 Proteobacteria
Dickeya zeae CSL RW192 WP 038906875 1 Dzeae
Dickeya zeae MK19 WP 038912755 1 Dzeae
Dickeya dianthicola NCPPB 3534 WP 024107980 1 Ddianthicola
Dickeya dianthicola GBBC 2039 WP 024107980 1 Ddianthicola
Dickeya dianthicola IPO 980 WP 024107980 1 Ddianthicola
Dickeya aquatica DW 0440 WP 035345435 1 Daquatica
DAQ1742 DAQ1742 03188 Daquatica
Dickeya sp CSL RW240 WP 035345445 1 Daquatica
Dickeya aquatica DW 0440 WP 035345445 1 Daquatica
541 DAQ1742 03693 phages P2 P6 P7 related to Mu like Seed
DAQ1742 DAQ1742 01695 Daquatica
DAQ1742 DAQ1742 03693 Daquatica

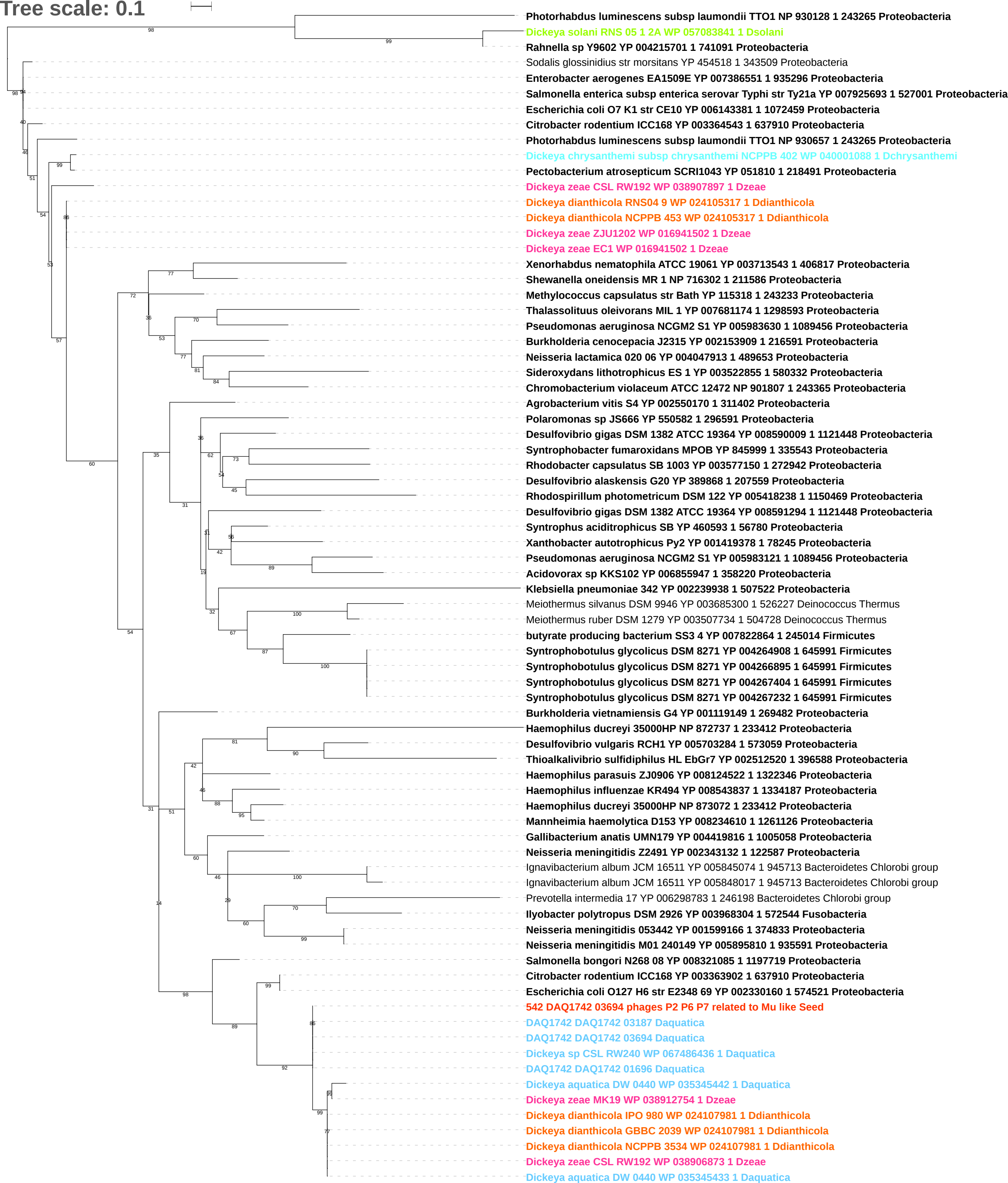
Tree scale: 0.1
Photorhabdus luminescens subsp laumondii TTO1 NP 930128 1 243265 Proteobacteria
Dickeya solani RNS 05 1 2A WP 057083841 1 Dsolani
Rahnella sp Y9602 YP 004215701 1 741091 Proteobacteria
Sodalis glossinidius str morsitans YP 454518 1 343509 Proteobacteria
Enterobacter aerogenes EA1509E YP 007386551 1 935296 Proteobacteria
Salmonella enterica subsp enterica serovar Typhi str Ty21a YP 007925693 1 527001 Proteobacteria
Escherichia coli O7 K1 str CE10 YP 006143381 1 1072459 Proteobacteria
Citrobacter rodentium ICC168 YP 003364543 1 637910 Proteobacteria
Photorhabdus luminescens subsp laumondii TTO1 NP 930657 1 243265 Proteobacteria
Dickeya chrysanthemi subsp chrysanthemi NCPPB 402 WP 040001088 1 Dchrysanthemi
Pectobacterium atrosepticum SCRI1043 YP 051810 1 218491 Proteobacteria
Dickeya zeae CSL RW192 WP 038907897 1 Dzeae
Dickeya dianthicola RNS04 9 WP 024105317 1 Ddianthicola
Dickeya dianthicola NCPPB 453 WP 024105317 1 Ddianthicola
Dickeya zeae ZJU1202 WP 016941502 1 Dzeae
Dickeya zeae EC1 WP 016941502 1 Dzeae
Xenorhabdus nematophila ATCC 19061 YP 003713543 1 406817 Proteobacteria
Shewanella oneidensis MR 1 NP 716302 1 211586 Proteobacteria
Methylococcus capsulatus str Bath YP 115318 1 243233 Proteobacteria
Thalassolituus oleivorans MIL 1 YP 007681174 1 1298593 Proteobacteria
Pseudomonas aeruginosa NCGM2 S1 YP 005983630 1 1089456 Proteobacteria
Burkholderia cenocepacia J2315 YP 002153909 1 216591 Proteobacteria
Neisseria lactamica 020 06 YP 004047913 1 489653 Proteobacteria
Sideroxydans lithotrophicus ES 1 YP 003522855 1 580332 Proteobacteria
Chromobacterium violaceum ATCC 12472 NP 901807 1 243365 Proteobacteria
Agrobacterium vitis S4 YP 002550170 1 311402 Proteobacteria
Polaromonas sp JS666 YP 550582 1 296591 Proteobacteria
Desulfovibrio gigas DSM 1382 ATCC 19364 YP 008590009 1 1121448 Proteobacteria
Syntrophobacter fumaroxidans MPOB YP 845999 1 335543 Proteobacteria
Rhodobacter capsulatus SB 1003 YP 003577150 1 272942 Proteobacteria
Desulfovibrio alaskensis G20 YP 389868 1 207559 Proteobacteria
Rhodospirillum photometricum DSM 122 YP 005418238 1 1150469 Proteobacteria
Desulfovibrio gigas DSM 1382 ATCC 19364 YP 008591294 1 1121448 Proteobacteria
Syntrophus aciditrophicus SB YP 460593 1 56780 Proteobacteria
Xanthobacter autotrophicus Py2 YP 001419378 1 78245 Proteobacteria
Pseudomonas aeruginosa NCGM2 S1 YP 005983121 1 1089456 Proteobacteria
Acidovorax sp KKS102 YP 006855947 1 358220 Proteobacteria
Klebsiella pneumoniae 342 YP 002239938 1 507522 Proteobacteria
Meiothermus silvanus DSM 9946 YP 003685300 1 526227 Deinococcus Thermus
Meiothermus ruber DSM 1279 YP 003507734 1 504728 Deinococcus Thermus
butyrate producing bacterium SS3 4 YP 007822864 1 245014 Firmicutes
Syntrophobotulus glycolicus DSM 8271 YP 004264908 1 645991 Firmicutes
Syntrophobotulus glycolicus DSM 8271 YP 004266895 1 645991 Firmicutes
Syntrophobotulus glycolicus DSM 8271 YP 004267404 1 645991 Firmicutes
Syntrophobotulus glycolicus DSM 8271 YP 004267232 1 645991 Firmicutes
Burkholderia vietnamiensis G4 YP 001119149 1 269482 Proteobacteria
Haemophilus ducreyi 35000HP NP 872737 1 233412 Proteobacteria
Desulfovibrio vulgaris RCH1 YP 005703284 1 573059 Proteobacteria
Thioalkalivibrio sulfidiphilus HL EbGr7 YP 002512520 1 396588 Proteobacteria
Haemophilus parasuis ZJ0906 YP 008124522 1 1322346 Proteobacteria
Haemophilus influenzae KR494 YP 008543837 1 1334187 Proteobacteria
Haemophilus ducreyi 35000HP NP 873072 1 233412 Proteobacteria
Mannheimia haemolytica D153 YP 008234610 1 1261126 Proteobacteria
Gallibacterium anatis UMN179 YP 004419816 1 1005058 Proteobacteria
Neisseria meningitidis Z2491 YP 002343132 1 122587 Proteobacteria
Ignavibacterium album JCM 16511 YP 005845074 1 945713 Bacteroidetes Chlorobi group
Ignavibacterium album JCM 16511 YP 005848017 1 945713 Bacteroidetes Chlorobi group
Prevotella intermedia 17 YP 006298783 1 246198 Bacteroidetes Chlorobi group
Ilyobacter polytropus DSM 2926 YP 003968304 1 572544 Fusobacteria
Neisseria meningitidis 053442 YP 001599166 1 374833 Proteobacteria
Neisseria meningitidis M01 240149 YP 005895810 1 935591 Proteobacteria
Salmonella bongori N268 08 YP 008321085 1 1197719 Proteobacteria
Citrobacter rodentium ICC168 YP 003363902 1 637910 Proteobacteria
Escherichia coli O127 H6 str E2348 69 YP 002330160 1 574521 Proteobacteria
542 DAQ1742 03694 phages P2 P6 P7 related to Mu like Seed
DAQ1742 DAQ1742 03187 Daquatica
DAQ1742 DAQ1742 03694 Daquatica
Dickeya sp CSL RW240 WP 067486436 1 Daquatica
DAQ1742 DAQ1742 01696 Daquatica
Dickeya aquatica DW 0440 WP 035345442 1 Daquatica
Dickeya zeae MK19 WP 038912754 1 Dzeae
Dickeya dianthicola IPO 980 WP 024107981 1 Ddianthicola
Dickeya dianthicola GBBC 2039 WP 024107981 1 Ddianthicola
Dickeya dianthicola NCPPB 3534 WP 024107981 1 Ddianthicola
Dickeya zeae CSL RW192 WP 038906873 1 Dzeae
Dickeya aquatica DW 0440 WP 035345433 1 Daquatica

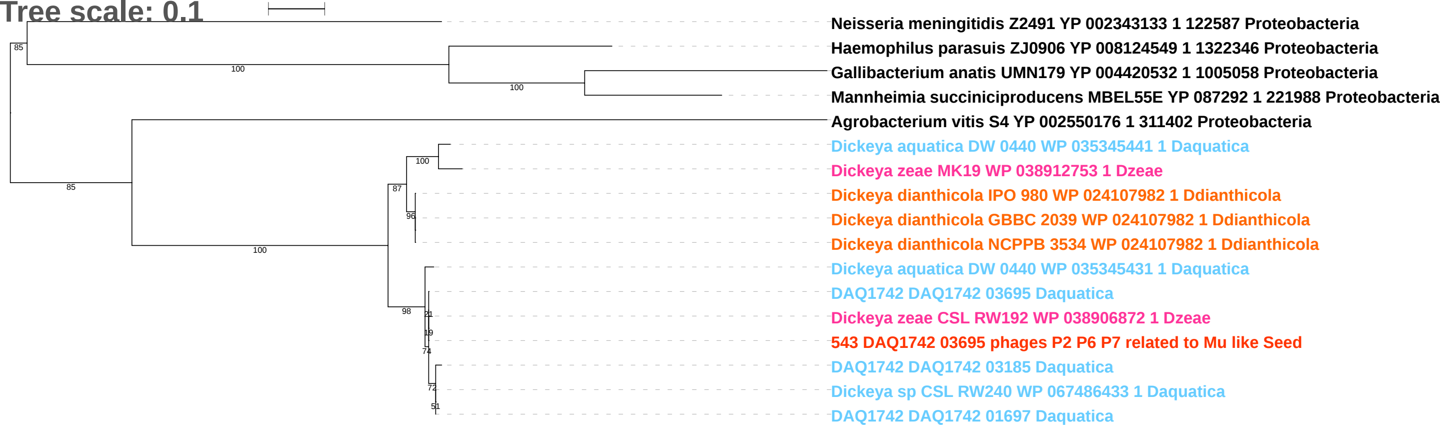

Tree scale: 0.1
Neisseria meningitidis Z2491 YP 002343133 1 122587 Proteobacteria
Haemophilus parasuis ZJ0906 YP 008124549 1 1322346 Proteobacteria
Gallibacterium anatis UMN179 YP 004420532 1 1005058 Proteobacteria
Mannheimia succiniciproducens MBEL55E YP 087292 1 221988 Proteobacteria
Agrobacterium vitis S4 YP 002550176 1 311402 Proteobacteria
Dickeya aquatica DW 0440 WP 035345441 1 Daquatica
Dickeya zeae MK19 WP 038912753 1 Dzeae
Dickeya dianthicola IPO 980 WP 024107982 1 Ddianthicola
Dickeya dianthicola GBBC 2039 WP 024107982 1 Ddianthicola
Dickeya dianthicola NCPPB 3534 WP 024107982 1 Ddianthicola
Dickeya aquatica DW 0440 WP 035345431 1 Daquatica
DAQ1742 DAQ1742 03695 Daquatica
Dickeya zeae CSL RW192 WP 038906872 1 Dzeae
543 DAQ1742 03695 phages P2 P6 P7 related to Mu like Seed
DAQ1742 DAQ1742 03185 Daquatica
Dickeya sp CSL RW240 WP 067486433 1 Daquatica
DAQ1742 DAQ1742 01697 Daquatica
85
100
100
85
100
100
87
96
98
21
19
74
72
51

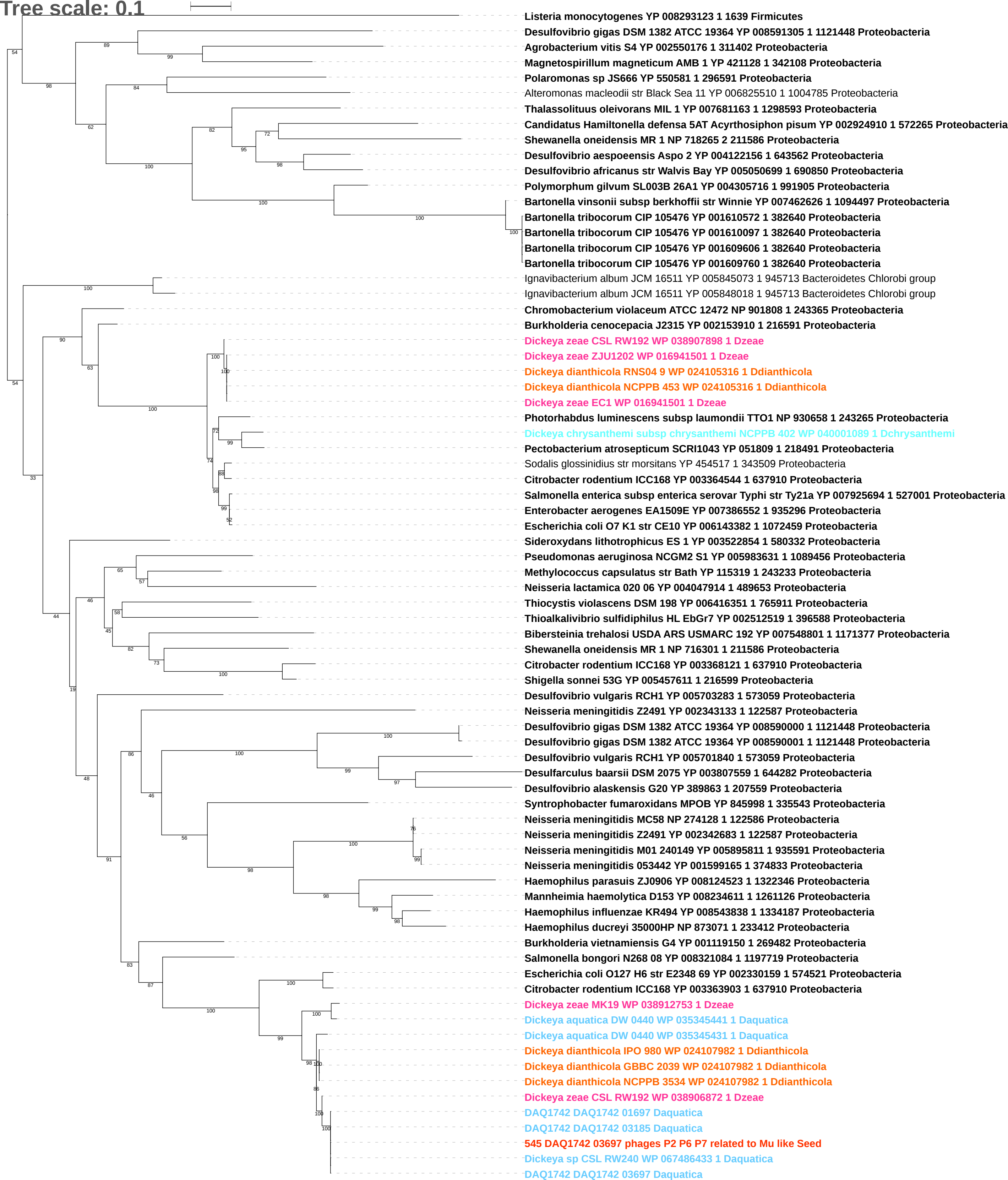
Tree scale: 0.1
Listeria monocytogenes YP 008293123 1 1639 Firmicutes
Desulfovibrio gigas DSM 1382 ATCC 19364 YP 008591305 1 1121448 Proteobacteria
Agrobacterium vitis S4 YP 002550176 1 311402 Proteobacteria
Magnetospirillum magneticum AMB 1 YP 421128 1 342108 Proteobacteria
Polaromonas sp JS666 YP 550581 1 296591 Proteobacteria
Alteromonas macleodii str Black Sea 11 YP 006825510 1 1004785 Proteobacteria
Thalassolituus oleivorans MIL 1 YP 007681163 1 1298593 Proteobacteria
Candidatus Hamiltonella defensa 5AT Acyrthosiphon pisum YP 002924910 1 572265 Proteobacteria
Shewanella oneidensis MR 1 NP 718265 2 211586 Proteobacteria
Desulfovibrio aespoeensis Aspo 2 YP 004122156 1 643562 Proteobacteria
Desulfovibrio africanus str Walvis Bay YP 005050699 1 690850 Proteobacteria
Polymorphum gilvum SL003B 26A1 YP 004305716 1 991905 Proteobacteria
Bartonella vinsonii subsp berkhoffii str Winnie YP 007462626 1 1094497 Proteobacteria
Bartonella tribocorum CIP 105476 YP 001610572 1 382640 Proteobacteria
Bartonella tribocorum CIP 105476 YP 001610097 1 382640 Proteobacteria
Bartonella tribocorum CIP 105476 YP 001609606 1 382640 Proteobacteria
Bartonella tribocorum CIP 105476 YP 001609760 1 382640 Proteobacteria
Ignavibacterium album JCM 16511 YP 005845073 1 945713 Bacteroidetes Chlorobi group
Ignavibacterium album JCM 16511 YP 005848018 1 945713 Bacteroidetes Chlorobi group
Chromobacterium violaceum ATCC 12472 NP 901808 1 243365 Proteobacteria
Burkholderia cenocepacia J2315 YP 002153910 1 216591 Proteobacteria
Dickeya zeae CSL RW192 WP 038907898 1 Dzeae
Dickeya zeae ZJU1202 WP 016941501 1 Dzeae
Dickeya dianthicola RNS04 9 WP 024105316 1 Ddianthicola
Dickeya dianthicola NCPPB 453 WP 024105316 1 Ddianthicola
Dickeya zeae EC1 WP 016941501 1 Dzeae
Photorhabdus luminescens subsp laumondii TTO1 NP 930658 1 243265 Proteobacteria
Dickeya chrysanthemi subsp chrysanthemi NCPPB 402 WP 040001089 1 Dchrysanthemi
Pectobacterium atrosepticum SCRI1043 YP 051809 1 218491 Proteobacteria
Sodalis glossinidius str morsitans YP 454517 1 343509 Proteobacteria
Citrobacter rodentium ICC168 YP 003364544 1 637910 Proteobacteria
Salmonella enterica subsp enterica serovar Typhi str Ty21a YP 007925694 1 527001 Proteobacteria
Enterobacter aerogenes EA1509E YP 007386552 1 935296 Proteobacteria
Escherichia coli O7 K1 str CE10 YP 006143382 1 1072459 Proteobacteria
Sideroxydans lithotrophicus ES 1 YP 003522854 1 580332 Proteobacteria
Pseudomonas aeruginosa NCGM2 S1 YP 005983631 1 1089456 Proteobacteria
Methylococcus capsulatus str Bath YP 115319 1 243233 Proteobacteria
Neisseria lactamica 020 06 YP 004047914 1 489653 Proteobacteria
Thiocystis violascens DSM 198 YP 006416351 1 765911 Proteobacteria
Thioalkalivibrio sulfidiphilus HL EbGr7 YP 002512519 1 396588 Proteobacteria
Bibersteinia trehalosi USDA ARS USMARC 192 YP 007548801 1 1171377 Proteobacteria
Shewanella oneidensis MR 1 NP 716301 1 211586 Proteobacteria
Citrobacter rodentium ICC168 YP 003368121 1 637910 Proteobacteria
Shigella sonnei 53G YP 005457611 1 216599 Proteobacteria
Desulfovibrio vulgaris RCH1 YP 005703283 1 573059 Proteobacteria
Neisseria meningitidis Z2491 YP 002343133 1 122587 Proteobacteria
Desulfovibrio gigas DSM 1382 ATCC 19364 YP 008590000 1 1121448 Proteobacteria
Desulfovibrio gigas DSM 1382 ATCC 19364 YP 008590001 1 1121448 Proteobacteria
Desulfovibrio vulgaris RCH1 YP 005701840 1 573059 Proteobacteria
Desulfarculus baarsii DSM 2075 YP 003807559 1 644282 Proteobacteria
Desulfovibrio alaskensis G20 YP 389863 1 207559 Proteobacteria
Syntrophobacter fumaroxidans MPOB YP 845998 1 335543 Proteobacteria
Neisseria meningitidis MC58 NP 274128 1 122586 Proteobacteria
Neisseria meningitidis Z2491 YP 002342683 1 122587 Proteobacteria
Neisseria meningitidis M01 240149 YP 005895811 1 935591 Proteobacteria
Neisseria meningitidis 053442 YP 001599165 1 374833 Proteobacteria
Haemophilus parasuis ZJ0906 YP 008124523 1 1322346 Proteobacteria
Mannheimia haemolytica D153 YP 008234611 1 1261126 Proteobacteria
Haemophilus influenzae KR494 YP 008543838 1 1334187 Proteobacteria
Haemophilus ducreyi 35000HP NP 873071 1 233412 Proteobacteria
Burkholderia vietnamiensis G4 YP 001119150 1 269482 Proteobacteria
Salmonella bongori N268 08 YP 008321084 1 1197719 Proteobacteria
Escherichia coli O127 H6 str E2348 69 YP 002330159 1 574521 Proteobacteria
Citrobacter rodentium ICC168 YP 003363903 1 637910 Proteobacteria
Dickeya zeae MK19 WP 038912753 1 Dzeae
Dickeya aquatica DW 0440 WP 035345441 1 Daquatica
Dickeya aquatica DW 0440 WP 035345431 1 Daquatica
Dickeya dianthicola IPO 980 WP 024107982 1 Ddianthicola
Dickeya dianthicola GBBC 2039 WP 024107982 1 Ddianthicola
Dickeya dianthicola NCPPB 3534 WP 024107982 1 Ddianthicola
Dickeya zeae CSL RW192 WP 038906872 1 Dzeae
DAQ1742 DAQ1742 01697 Daquatica
DAQ1742 DAQ1742 03185 Daquatica
545 DAQ1742 03697 phages P2 P6 P7 related to Mu like Seed
Dickeya sp CSL RW240 WP 067486433 1 Daquatica
DAQ1742 DAQ1742 03697 Daquatica

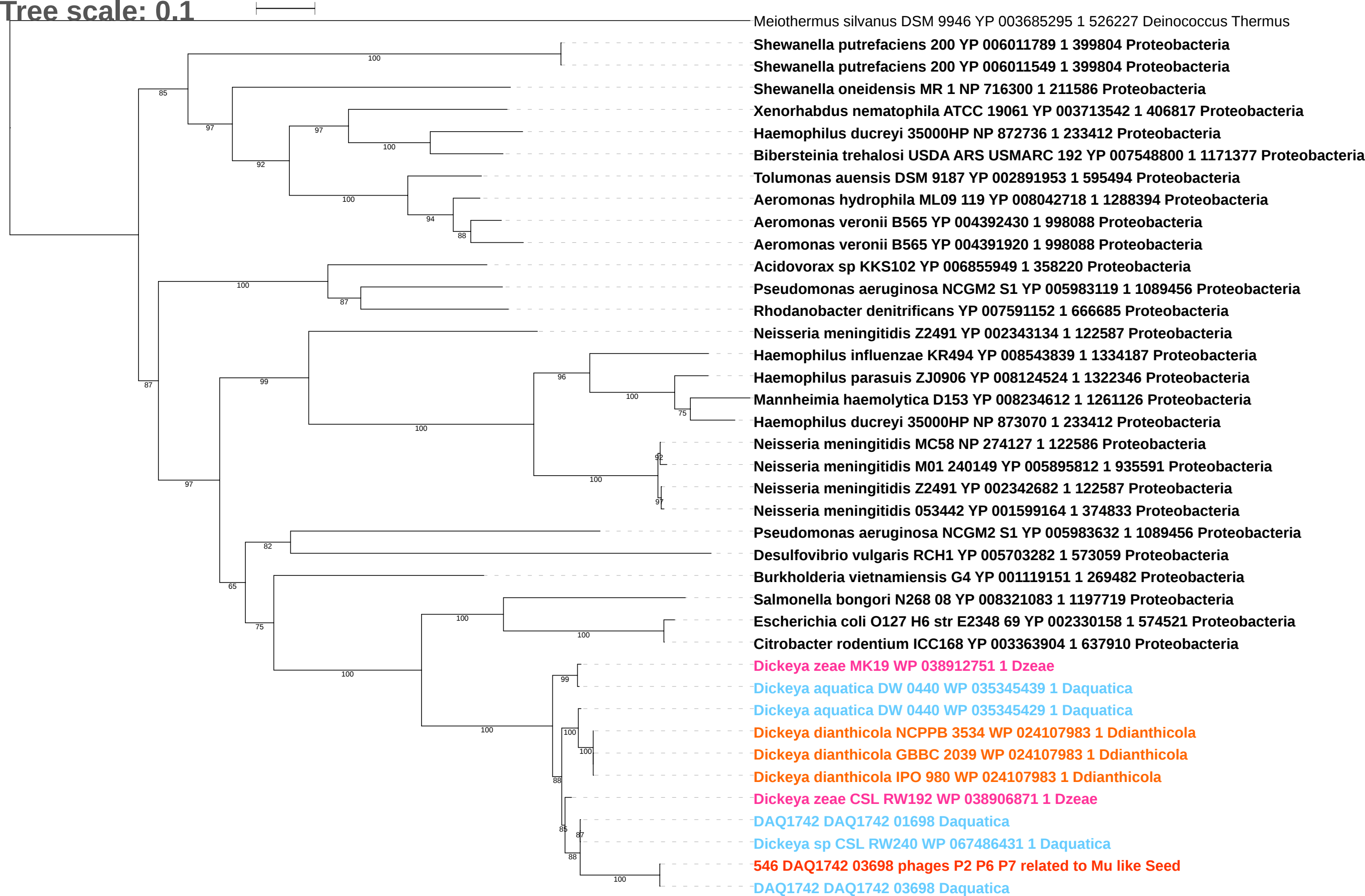

Tree scale: 0.1
Meiothermus silvanus DSM 9946 YP 003685295 1 526227 Deinococcus Thermus
Shewanella putrefaciens 200 YP 006011789 1 399804 Proteobacteria
Shewanella putrefaciens 200 YP 006011549 1 399804 Proteobacteria
Shewanella oneidensis MR 1 NP 716300 1 211586 Proteobacteria
Xenorhabdus nematophila ATCC 19061 YP 003713542 1 406817 Proteobacteria
Haemophilus ducreyi 35000HP NP 872736 1 233412 Proteobacteria
Bibersteinia trehalosi USDA ARS USMARC 192 YP 007548800 1 1171377 Proteobacteria
Tolumonas auensis DSM 9187 YP 002891953 1 595494 Proteobacteria
Aeromonas hydrophila ML09 119 YP 008042718 1 1288394 Proteobacteria
Aeromonas veronii B565 YP 004392430 1 998088 Proteobacteria
Aeromonas veronii B565 YP 004391920 1 998088 Proteobacteria
Acidovorax sp KKS102 YP 006855949 1 358220 Proteobacteria
Pseudomonas aeruginosa NCGM2 S1 YP 005983119 1 1089456 Proteobacteria
Rhodanobacter denitrificans YP 007591152 1 666685 Proteobacteria
Neisseria meningitidis Z2491 YP 002343134 1 122587 Proteobacteria
Haemophilus influenzae KR494 YP 008543839 1 1334187 Proteobacteria
Haemophilus parasuis ZJ0906 YP 008124524 1 1322346 Proteobacteria
Mannheimia haemolytica D153 YP 008234612 1 1261126 Proteobacteria
Haemophilus ducreyi 35000HP NP 873070 1 233412 Proteobacteria
Neisseria meningitidis MC58 NP 274127 1 122586 Proteobacteria
Neisseria meningitidis M01 240149 YP 005895812 1 935591 Proteobacteria
Neisseria meningitidis Z2491 YP 002342682 1 122587 Proteobacteria
Neisseria meningitidis 053442 YP 001599164 1 374833 Proteobacteria
Pseudomonas aeruginosa NCGM2 S1 YP 005983632 1 1089456 Proteobacteria
Desulfovibrio vulgaris RCH1 YP 005703282 1 573059 Proteobacteria
Burkholderia vietnamiensis G4 YP 001119151 1 269482 Proteobacteria
Salmonella bongori N268 08 YP 008321083 1 1197719 Proteobacteria
Escherichia coli O127 H6 str E2348 69 YP 002330158 1 574521 Proteobacteria
Citrobacter rodentium ICC168 YP 003363904 1 637910 Proteobacteria
Dickeya zeae MK19 WP 038912751 1 Dzeae
Dickeya aquatica DW 0440 WP 035345439 1 Daquatica
Dickeya aquatica DW 0440 WP 035345429 1 Daquatica
Dickeya dianthicola NCPPB 3534 WP 024107983 1 Ddianthicola
Dickeya dianthicola GBBC 2039 WP 024107983 1 Ddianthicola
Dickeya dianthicola IPO 980 WP 024107983 1 Ddianthicola
Dickeya zeae CSL RW192 WP 038906871 1 Dzeae
DAQ1742 DAQ1742 01698 Daquatica
Dickeya sp CSL RW240 WP 067486431 1 Daquatica
546 DAQ1742 03698 phages P2 P6 P7 related to Mu like Seed
DAQ1742 DAQ1742 03698 Daquatica

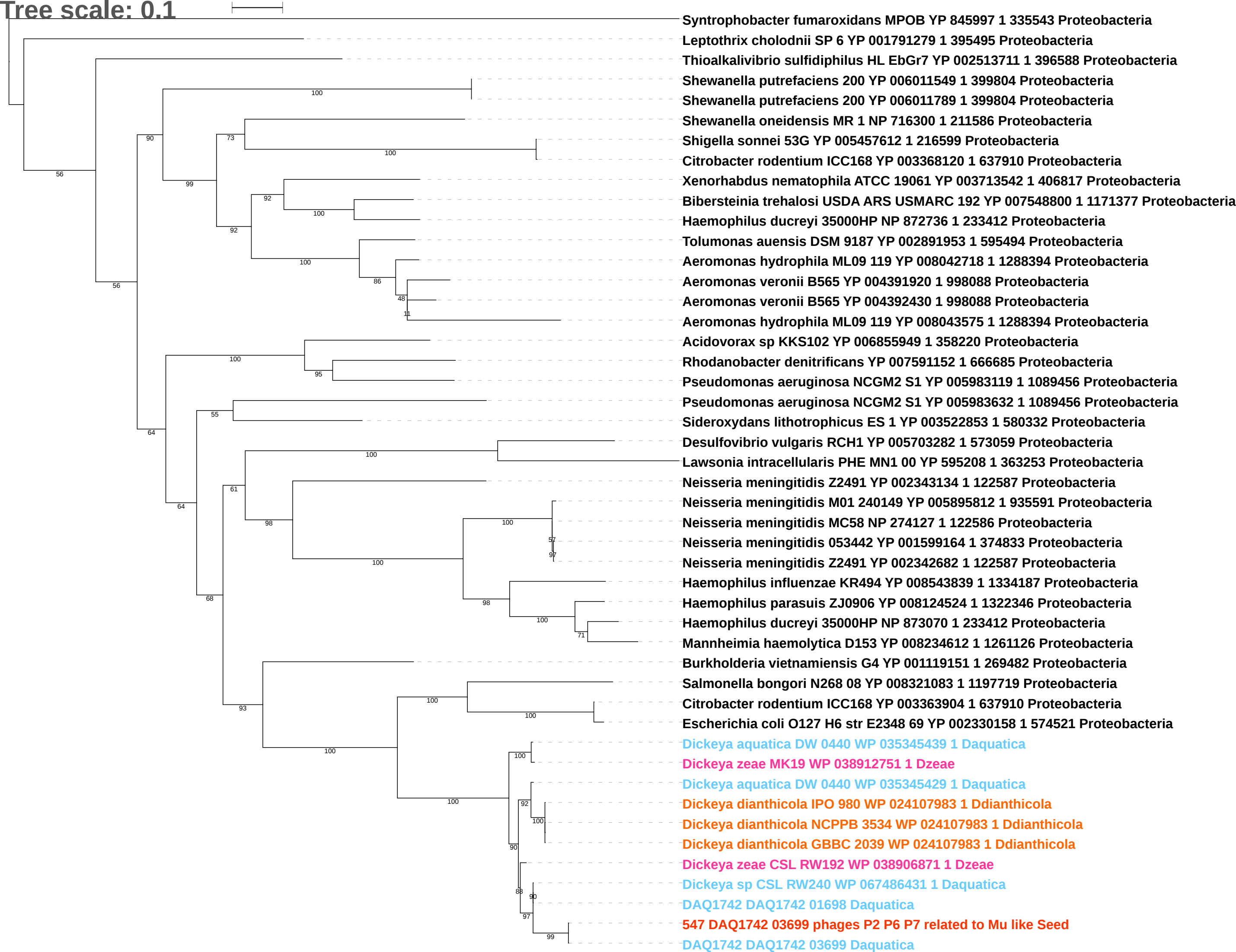

Tree scale: 0.1
Syntrophobacter fumaroxidans MPOB YP 845997 1 335543 Proteobacteria
Leptothrix cholodnii SP 6 YP 001791279 1 395495 Proteobacteria
Thioalkalivibrio sulfidiphilus HL EbGr7 YP 002513711 1 396588 Proteobacteria
Shewanella putrefaciens 200 YP 006011549 1 399804 Proteobacteria
Shewanella putrefaciens 200 YP 006011789 1 399804 Proteobacteria
Shewanella oneidensis MR 1 NP 716300 1 211586 Proteobacteria
Shigella sonnei 53G YP 005457612 1 216599 Proteobacteria
Citrobacter rodentium ICC168 YP 003368120 1 637910 Proteobacteria
Xenorhabdus nematophila ATCC 19061 YP 003713542 1 406817 Proteobacteria
Bibersteinia trehalosi USDA ARS USMARC 192 YP 007548800 1 1171377 Proteobacteria
Haemophilus ducreyi 35000HP NP 872736 1 233412 Proteobacteria
Tolumonas auensis DSM 9187 YP 002891953 1 595494 Proteobacteria
Aeromonas hydrophila ML09 119 YP 008042718 1 1288394 Proteobacteria
Aeromonas veronii B565 YP 004391920 1 998088 Proteobacteria
Aeromonas veronii B565 YP 004392430 1 998088 Proteobacteria
Aeromonas hydrophila ML09 119 YP 008043575 1 1288394 Proteobacteria
Acidovorax sp KKS102 YP 006855949 1 358220 Proteobacteria
Rhodanobacter denitrificans YP 007591152 1 666685 Proteobacteria
Pseudomonas aeruginosa NCGM2 S1 YP 005983119 1 1089456 Proteobacteria
Pseudomonas aeruginosa NCGM2 S1 YP 005983632 1 1089456 Proteobacteria
Sideroxydans lithotrophicus ES 1 YP 003522853 1 580332 Proteobacteria
Desulfovibrio vulgaris RCH1 YP 005703282 1 573059 Proteobacteria
Lawsonia intracellularis PHE MN1 00 YP 595208 1 363253 Proteobacteria
Neisseria meningitidis Z2491 YP 002343134 1 122587 Proteobacteria
Neisseria meningitidis M01 240149 YP 005895812 1 935591 Proteobacteria
Neisseria meningitidis MC58 NP 274127 1 122586 Proteobacteria
Neisseria meningitidis 053442 YP 001599164 1 374833 Proteobacteria
Neisseria meningitidis Z2491 YP 002342682 1 122587 Proteobacteria
Haemophilus influenzae KR494 YP 008543839 1 1334187 Proteobacteria
Haemophilus parasuis ZJ0906 YP 008124524 1 1322346 Proteobacteria
Haemophilus ducreyi 35000HP NP 873070 1 233412 Proteobacteria
Mannheimia haemolytica D153 YP 008234612 1 1261126 Proteobacteria
Burkholderia vietnamiensis G4 YP 001119151 1 269482 Proteobacteria
Salmonella bongori N268 08 YP 008321083 1 1197719 Proteobacteria
Citrobacter rodentium ICC168 YP 003363904 1 637910 Proteobacteria
Escherichia coli O127 H6 str E2348 69 YP 002330158 1 574521 Proteobacteria
Dickeya aquatica DW 0440 WP 035345439 1 Daquatica
Dickeya zeae MK19 WP 038912751 1 Dzeae
Dickeya aquatica DW 0440 WP 035345429 1 Daquatica
Dickeya dianthicola IPO 980 WP 024107983 1 Ddianthicola
Dickeya dianthicola NCPPB 3534 WP 024107983 1 Ddianthicola
Dickeya dianthicola GBBC 2039 WP 024107983 1 Ddianthicola
Dickeya zeae CSL RW192 WP 038906871 1 Dzeae
Dickeya sp CSL RW240 WP 067486431 1 Daquatica
DAQ1742 DAQ1742 01698 Daquatica
547 DAQ1742 03699 phages P2 P6 P7 related to Mu like Seed
DAQ1742 DAQ1742 03699 Daquatica

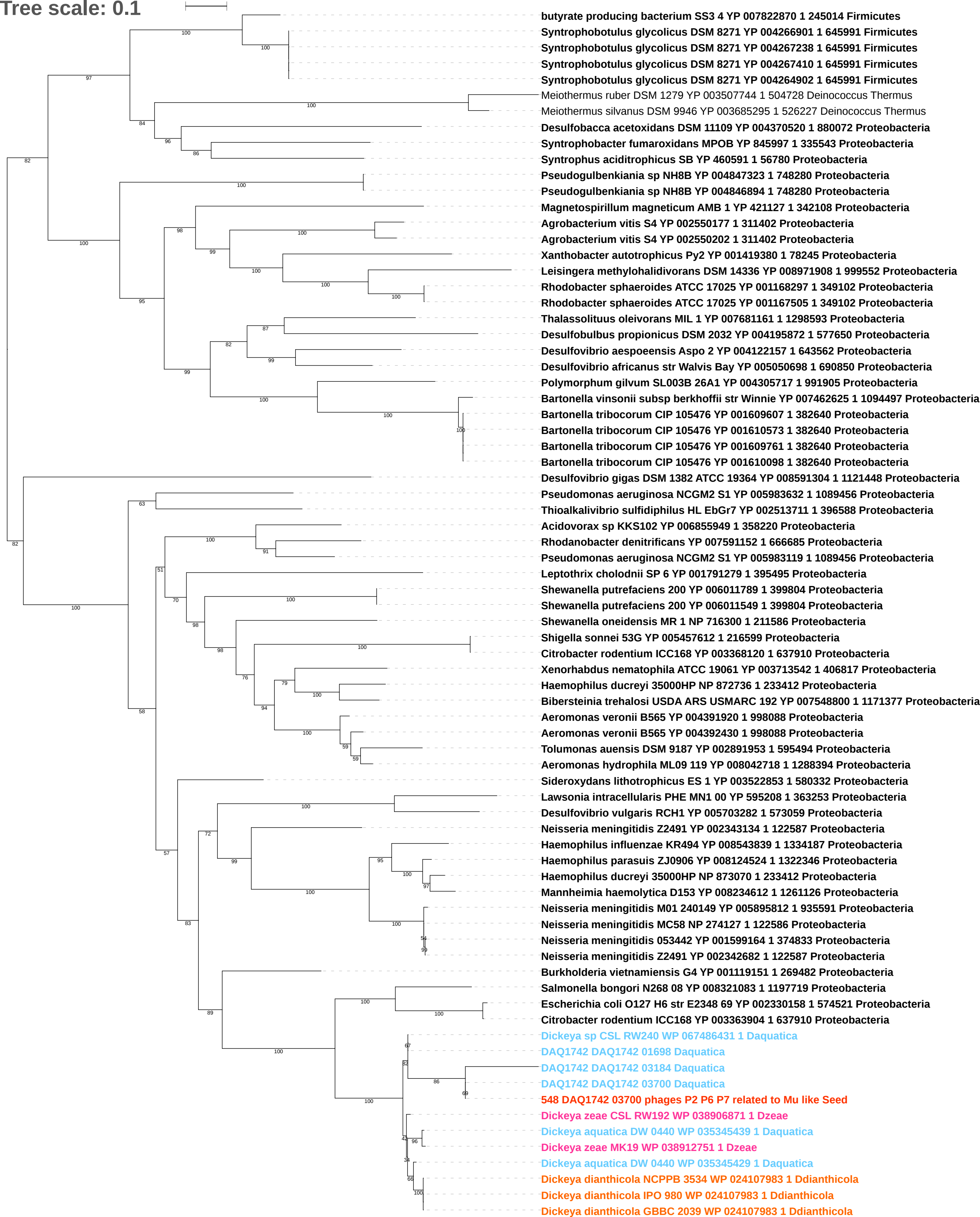
Tree scale: 0.1
butyrate producing bacterium SS3 4 YP 007822870 1 245014 Firmicutes
Syntrophobotulus glycolicus DSM 8271 YP 004266901 1 645991 Firmicutes
Syntrophobotulus glycolicus DSM 8271 YP 004267238 1 645991 Firmicutes
Syntrophobotulus glycolicus DSM 8271 YP 004267410 1 645991 Firmicutes
Syntrophobotulus glycolicus DSM 8271 YP 004264902 1 645991 Firmicutes
Meiothermus ruber DSM 1279 YP 003507744 1 504728 Deinococcus Thermus
Meiothermus silvanus DSM 9946 YP 003685295 1 526227 Deinococcus Thermus
Desulfobacca acetoxidans DSM 11109 YP 004370520 1 880072 Proteobacteria
Syntrophobacter fumaroxidans MPOB YP 845997 1 335543 Proteobacteria
Syntrophus aciditrophicus SB YP 460591 1 56780 Proteobacteria
Pseudogulbenkiania sp NH8B YP 004847323 1 748280 Proteobacteria
Pseudogulbenkiania sp NH8B YP 004846894 1 748280 Proteobacteria
Magnetospirillum magneticum AMB 1 YP 421127 1 342108 Proteobacteria
Agrobacterium vitis S4 YP 002550177 1 311402 Proteobacteria
Agrobacterium vitis S4 YP 002550202 1 311402 Proteobacteria
Xanthobacter autotrophicus Py2 YP 001419380 1 78245 Proteobacteria
Leisingera methylohalidivorans DSM 14336 YP 008971908 1 999552 Proteobacteria
Rhodobacter sphaeroides ATCC 17025 YP 001168297 1 349102 Proteobacteria
Rhodobacter sphaeroides ATCC 17025 YP 001167505 1 349102 Proteobacteria
Thalassolituus oleivorans MIL 1 YP 007681161 1 1298593 Proteobacteria
Desulfobulbus propionicus DSM 2032 YP 004195872 1 577650 Proteobacteria
Desulfovibrio aespoeensis Aspo 2 YP 004122157 1 643562 Proteobacteria
Desulfovibrio africanus str Walvis Bay YP 005050698 1 690850 Proteobacteria
Polymorphum gilvum SL003B 26A1 YP 004305717 1 991905 Proteobacteria
Bartonella vinsonii subsp berkhoffii str Winnie YP 007462625 1 1094497 Proteobacteria
Bartonella tribocorum CIP 105476 YP 001609607 1 382640 Proteobacteria
Bartonella tribocorum CIP 105476 YP 001610573 1 382640 Proteobacteria
Bartonella tribocorum CIP 105476 YP 001609761 1 382640 Proteobacteria
Bartonella tribocorum CIP 105476 YP 001610098 1 382640 Proteobacteria
Desulfovibrio gigas DSM 1382 ATCC 19364 YP 008591304 1 1121448 Proteobacteria
Pseudomonas aeruginosa NCGM2 S1 YP 005983632 1 1089456 Proteobacteria
Thioalkalivibrio sulfidiphilus HL EbGr7 YP 002513711 1 396588 Proteobacteria
Acidovorax sp KKS102 YP 006855949 1 358220 Proteobacteria
Rhodanobacter denitrificans YP 007591152 1 666685 Proteobacteria
Pseudomonas aeruginosa NCGM2 S1 YP 005983119 1 1089456 Proteobacteria
Leptothrix cholodnii SP 6 YP 001791279 1 395495 Proteobacteria
Shewanella putrefaciens 200 YP 006011789 1 399804 Proteobacteria
Shewanella putrefaciens 200 YP 006011549 1 399804 Proteobacteria
Shewanella oneidensis MR 1 NP 716300 1 211586 Proteobacteria
Shigella sonnei 53G YP 005457612 1 216599 Proteobacteria
Citrobacter rodentium ICC168 YP 003368120 1 637910 Proteobacteria
Xenorhabdus nematophila ATCC 19061 YP 003713542 1 406817 Proteobacteria
Haemophilus ducreyi 35000HP NP 872736 1 233412 Proteobacteria
Bibersteinia trehalosi USDA ARS USMARC 192 YP 007548800 1 1171377 Proteobacteria
Aeromonas veronii B565 YP 004391920 1 998088 Proteobacteria
Aeromonas veronii B565 YP 004392430 1 998088 Proteobacteria
Tolumonas auensis DSM 9187 YP 002891953 1 595494 Proteobacteria
Aeromonas hydrophila ML09 119 YP 008042718 1 1288394 Proteobacteria
Sideroxydans lithotrophicus ES 1 YP 003522853 1 580332 Proteobacteria
Lawsonia intracellularis PHE MN1 00 YP 595208 1 363253 Proteobacteria
Desulfovibrio vulgaris RCH1 YP 005703282 1 573059 Proteobacteria
Neisseria meningitidis Z2491 YP 002343134 1 122587 Proteobacteria
Haemophilus influenzae KR494 YP 008543839 1 1334187 Proteobacteria
Haemophilus parasuis ZJ0906 YP 008124524 1 1322346 Proteobacteria
Haemophilus ducreyi 35000HP NP 873070 1 233412 Proteobacteria
Mannheimia haemolytica D153 YP 008234612 1 1261126 Proteobacteria
Neisseria meningitidis M01 240149 YP 005895812 1 935591 Proteobacteria
Neisseria meningitidis MC58 NP 274127 1 122586 Proteobacteria
Neisseria meningitidis 053442 YP 001599164 1 374833 Proteobacteria
Neisseria meningitidis Z2491 YP 002342682 1 122587 Proteobacteria
Burkholderia vietnamiensis G4 YP 001119151 1 269482 Proteobacteria
Salmonella bongori N268 08 YP 008321083 1 1197719 Proteobacteria
Escherichia coli O127 H6 str E2348 69 YP 002330158 1 574521 Proteobacteria
Citrobacter rodentium ICC168 YP 003363904 1 637910 Proteobacteria
Dickeya sp CSL RW240 WP 067486431 1 Daquatica
DAQ1742 DAQ1742 01698 Daquatica
DAQ1742 DAQ1742 03184 Daquatica
DAQ1742 DAQ1742 03700 Daquatica
548 DAQ1742 03700 phages P2 P6 P7 related to Mu like Seed
Dickeya zeae CSL RW192 WP 038906871 1 Dzeae
Dickeya aquatica DW 0440 WP 035345439 1 Daquatica
Dickeya zeae MK19 WP 038912751 1 Dzeae
Dickeya aquatica DW 0440 WP 035345429 1 Daquatica
Dickeya dianthicola NCPPB 3534 WP 024107983 1 Ddianthicola
Dickeya dianthicola IPO 980 WP 024107983 1 Ddianthicola
Dickeya dianthicola GBBC 2039 WP 024107983 1 Ddianthicola

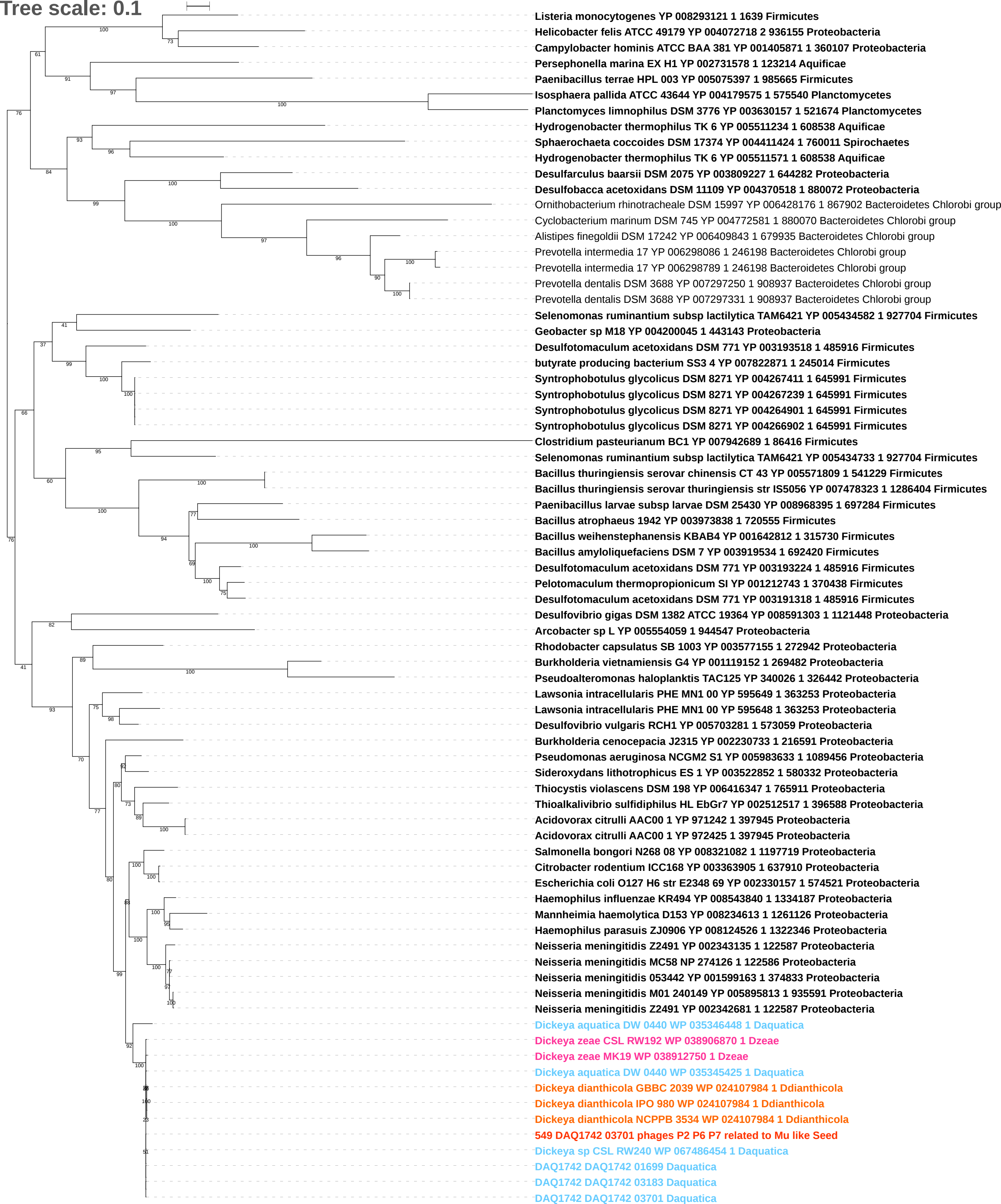
Tree scale: 0.1
Listeria monocytogenes YP 008293121 1 1639 Firmicutes
Helicobacter felis ATCC 49179 YP 004072718 2 936155 Proteobacteria
Campylobacter hominis ATCC BAA 381 YP 001405871 1 360107 Proteobacteria
Persephonella marina EX H1 YP 002731578 1 123214 Aquificae
Paenibacillus terrae HPL 003 YP 005075397 1 985665 Firmicutes
Isosphaera pallida ATCC 43644 YP 004179575 1 575540 Planctomycetes
Planctomyces limnophilus DSM 3776 YP 003630157 1 521674 Planctomycetes
Hydrogenobacter thermophilus TK 6 YP 005511234 1 608538 Aquificae
Sphaerochaeta coccoides DSM 17374 YP 004411424 1 760011 Spirochaetes
Hydrogenobacter thermophilus TK 6 YP 005511571 1 608538 Aquificae
Desulfarculus baarsii DSM 2075 YP 003809227 1 644282 Proteobacteria
Desulfobacca acetoxidans DSM 11109 YP 004370518 1 880072 Proteobacteria
Ornithobacterium rhinotracheale DSM 15997 YP 006428176 1 867902 Bacteroidetes Chlorobi group
Cyclobacterium marinum DSM 745 YP 004772581 1 880070 Bacteroidetes Chlorobi group
Alistipes finegoldii DSM 17242 YP 006409843 1 679935 Bacteroidetes Chlorobi group
Prevotella intermedia 17 YP 006298086 1 246198 Bacteroidetes Chlorobi group
Prevotella intermedia 17 YP 006298789 1 246198 Bacteroidetes Chlorobi group
Prevotella dentalis DSM 3688 YP 007297250 1 908937 Bacteroidetes Chlorobi group
Prevotella dentalis DSM 3688 YP 007297331 1 908937 Bacteroidetes Chlorobi group
Selenomonas ruminantium subsp lactilytica TAM6421 YP 005434582 1 927704 Firmicutes
Geobacter sp M18 YP 004200045 1 443143 Proteobacteria
Desulfotomaculum acetoxidans DSM 771 YP 003193518 1 485916 Firmicutes
butyrate producing bacterium SS3 4 YP 007822871 1 245014 Firmicutes
Syntrophobotulus glycolicus DSM 8271 YP 004267411 1 645991 Firmicutes
Syntrophobotulus glycolicus DSM 8271 YP 004267239 1 645991 Firmicutes
Syntrophobotulus glycolicus DSM 8271 YP 004264901 1 645991 Firmicutes
Syntrophobotulus glycolicus DSM 8271 YP 004266902 1 645991 Firmicutes
Clostridium pasteurianum BC1 YP 007942689 1 86416 Firmicutes
Selenomonas ruminantium subsp lactilytica TAM6421 YP 005434733 1 927704 Firmicutes
Bacillus thuringiensis serovar chinensis CT 43 YP 005571809 1 541229 Firmicutes
Bacillus thuringiensis serovar thuringiensis str IS5056 YP 007478323 1 1286404 Firmicutes
Paenibacillus larvae subsp larvae DSM 25430 YP 008968395 1 697284 Firmicutes
Bacillus atrophaeus 1942 YP 003973838 1 720555 Firmicutes
Bacillus weihenstephanensis KBAB4 YP 001642812 1 315730 Firmicutes
Bacillus amyloliquefaciens DSM 7 YP 003919534 1 692420 Firmicutes
Desulfotomaculum acetoxidans DSM 771 YP 003193224 1 485916 Firmicutes
Pelotomaculum thermopropionicum SI YP 001212743 1 370438 Firmicutes
Desulfotomaculum acetoxidans DSM 771 YP 003191318 1 485916 Firmicutes
Desulfovibrio gigas DSM 1382 ATCC 19364 YP 008591303 1 1121448 Proteobacteria
Arcobacter sp L YP 005554059 1 944547 Proteobacteria
Rhodobacter capsulatus SB 1003 YP 003577155 1 272942 Proteobacteria
Burkholderia vietnamiensis G4 YP 001119152 1 269482 Proteobacteria
Pseudoalteromonas haloplanktis TAC125 YP 340026 1 326442 Proteobacteria
Lawsonia intracellularis PHE MN1 00 YP 595649 1 363253 Proteobacteria
Lawsonia intracellularis PHE MN1 00 YP 595648 1 363253 Proteobacteria
Desulfovibrio vulgaris RCH1 YP 005703281 1 573059 Proteobacteria
Burkholderia cenocepacia J2315 YP 002230733 1 216591 Proteobacteria
Pseudomonas aeruginosa NCGM2 S1 YP 005983633 1 1089456 Proteobacteria
Sideroxydans lithotrophicus ES 1 YP 003522852 1 580332 Proteobacteria
Thiocystis violascens DSM 198 YP 006416347 1 765911 Proteobacteria
Thioalkalivibrio sulfidiphilus HL EbGr7 YP 002512517 1 396588 Proteobacteria
Acidovorax citrulli AAC00 1 YP 971242 1 397945 Proteobacteria
Acidovorax citrulli AAC00 1 YP 972425 1 397945 Proteobacteria
Salmonella bongori N268 08 YP 008321082 1 1197719 Proteobacteria
Citrobacter rodentium ICC168 YP 003363905 1 637910 Proteobacteria
Escherichia coli O127 H6 str E2348 69 YP 002330157 1 574521 Proteobacteria
Haemophilus influenzae KR494 YP 008543840 1 1334187 Proteobacteria
Mannheimia haemolytica D153 YP 008234613 1 1261126 Proteobacteria
Haemophilus parasuis ZJ0906 YP 008124526 1 1322346 Proteobacteria
Neisseria meningitidis Z2491 YP 002343135 1 122587 Proteobacteria
Neisseria meningitidis MC58 NP 274126 1 122586 Proteobacteria
Neisseria meningitidis 053442 YP 001599163 1 374833 Proteobacteria
Neisseria meningitidis M01 240149 YP 005895813 1 935591 Proteobacteria
Neisseria meningitidis Z2491 YP 002342681 1 122587 Proteobacteria
Dickeya aquatica DW 0440 WP 035346448 1 Daquatica
Dickeya zeae CSL RW192 WP 038906870 1 Dzeae
Dickeya zeae MK19 WP 038912750 1 Dzeae
Dickeya aquatica DW 0440 WP 035345425 1 Daquatica
Dickeya dianthicola GBBC 2039 WP 024107984 1 Ddianthicola
Dickeya dianthicola IPO 980 WP 024107984 1 Ddianthicola
Dickeya dianthicola NCPPB 3534 WP 024107984 1 Ddianthicola
549 DAQ1742 03701 phages P2 P6 P7 related to Mu like Seed
Dickeya sp CSL RW240 WP 067486454 1 Daquatica
DAQ1742 DAQ1742 01699 Daquatica
DAQ1742 DAQ1742 03183 Daquatica
DAQ1742 DAQ1742 03701 Daquatica

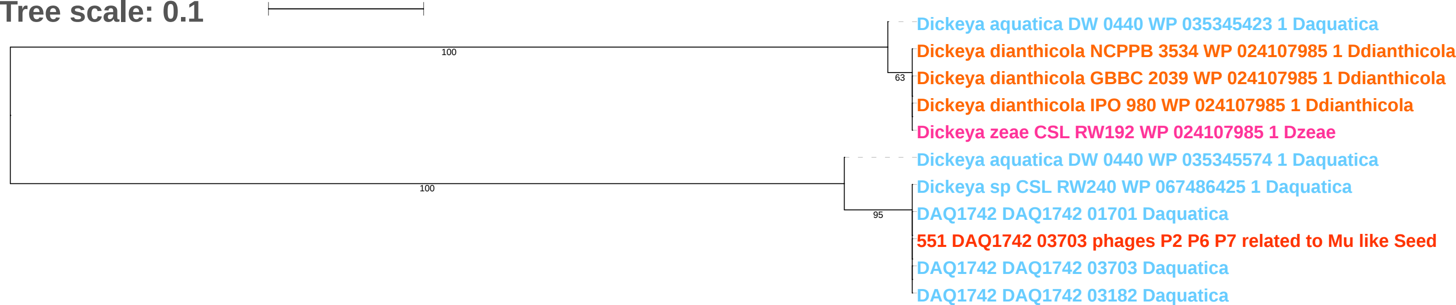

Tree scale: 0.1
100
100
63
95
Dickeya aquatica DW 0440 WP 035345423 1 Daquatica
Dickeya dianthicola NCPPB 3534 WP 024107985 1 Ddianthicola
Dickeya dianthicola GBBC 2039 WP 024107985 1 Ddianthicola
Dickeya dianthicola IPO 980 WP 024107985 1 Ddianthicola
Dickeya zeae CSL RW192 WP 024107985 1 Dzeae
Dickeya aquatica DW 0440 WP 035345574 1 Daquatica
Dickeya sp CSL RW240 WP 067486425 1 Daquatica
DAQ1742 DAQ1742 01701 Daquatica
551 DAQ1742 03703 phages P2 P6 P7 related to Mu like Seed
DAQ1742 DAQ1742 03703 Daquatica
DAQ1742 DAQ1742 03182 Daquatica

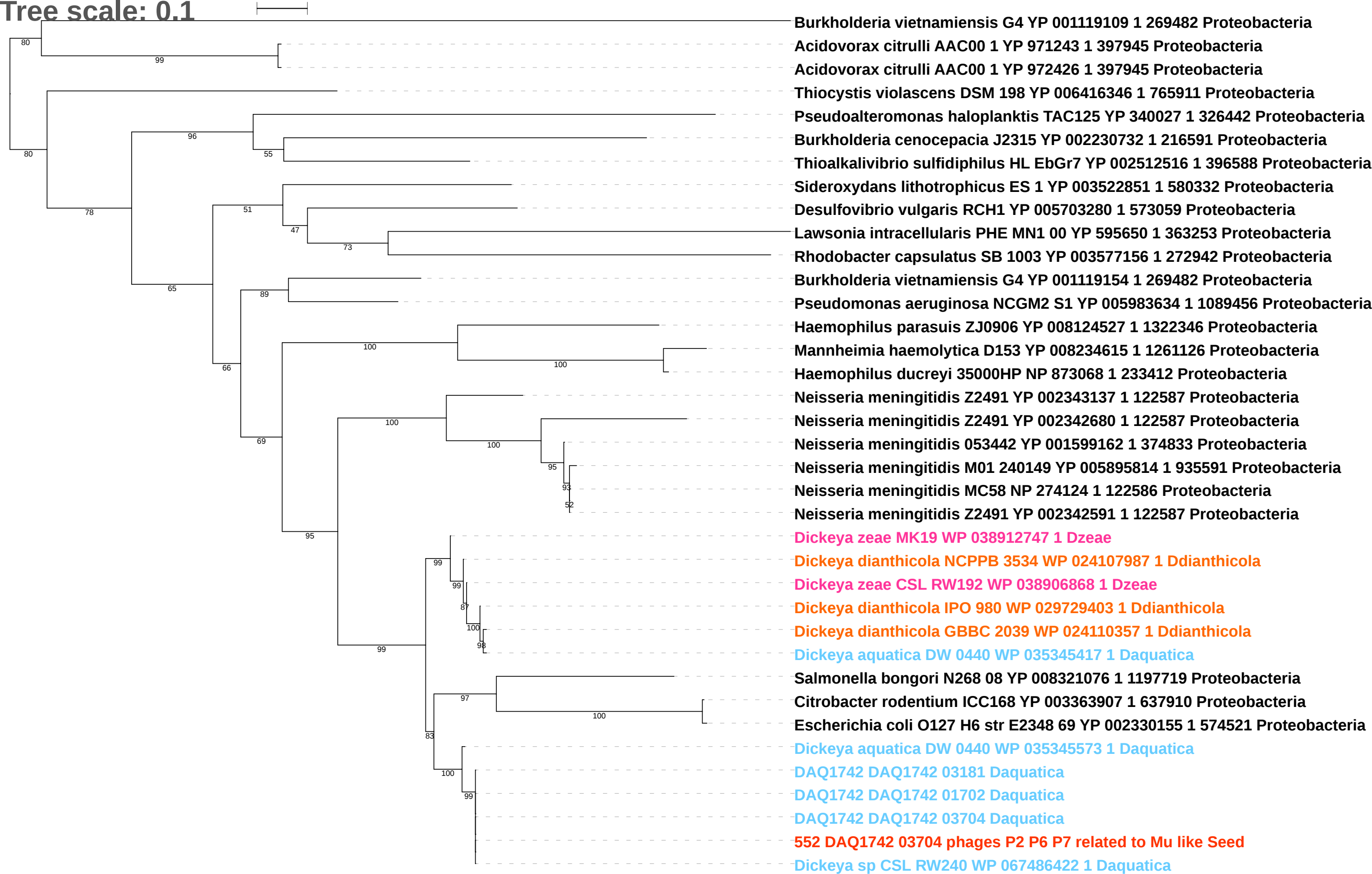

Tree scale: 0.1
80
99
80
96
55
78
51
47
73
65
89
66
100
100
69
100
100
95
93
52
95
99
99
87
100
98
99
97
100
83
100
99
Burkholderia vietnamiensis G4 YP 001119109 1 269482 Proteobacteria
Acidovorax citrulli AAC00 1 YP 971243 1 397945 Proteobacteria
Acidovorax citrulli AAC00 1 YP 972426 1 397945 Proteobacteria
Thiocystis violascens DSM 198 YP 006416346 1 765911 Proteobacteria
Pseudoalteromonas haloplanktis TAC125 YP 340027 1 326442 Proteobacteria
Burkholderia cenocepacia J2315 YP 002230732 1 216591 Proteobacteria
Thioalkalivibrio sulfidiphilus HL EbGr7 YP 002512516 1 396588 Proteobacteria
Sideroxydans lithotrophicus ES 1 YP 003522851 1 580332 Proteobacteria
Desulfovibrio vulgaris RCH1 YP 005703280 1 573059 Proteobacteria
Lawsonia intracellularis PHE MN1 00 YP 595650 1 363253 Proteobacteria
Rhodobacter capsulatus SB 1003 YP 003577156 1 272942 Proteobacteria
Burkholderia vietnamiensis G4 YP 001119154 1 269482 Proteobacteria
Pseudomonas aeruginosa NCGM2 S1 YP 005983634 1 1089456 Proteobacteria
Haemophilus parasuis ZJ0906 YP 008124527 1 1322346 Proteobacteria
Mannheimia haemolytica D153 YP 008234615 1 1261126 Proteobacteria
Haemophilus ducreyi 35000HP NP 873068 1 233412 Proteobacteria
Neisseria meningitidis Z2491 YP 002343137 1 122587 Proteobacteria
Neisseria meningitidis Z2491 YP 002342680 1 122587 Proteobacteria
Neisseria meningitidis 053442 YP 001599162 1 374833 Proteobacteria
Neisseria meningitidis M01 240149 YP 005895814 1 935591 Proteobacteria
Neisseria meningitidis MC58 NP 274124 1 122586 Proteobacteria
Neisseria meningitidis Z2491 YP 002342591 1 122587 Proteobacteria
Dickeya zeae MK19 WP 038912747 1 Dzeae
Dickeya dianthicola NCPPB 3534 WP 024107987 1 Ddianthicola
Dickeya zeae CSL RW192 WP 038906868 1 Dzeae
Dickeya dianthicola IPO 980 WP 029729403 1 Ddianthicola
Dickeya dianthicola GBBC 2039 WP 024110357 1 Ddianthicola
Dickeya aquatica DW 0440 WP 035345417 1 Daquatica
Salmonella bongori N268 08 YP 008321076 1 1197719 Proteobacteria
Citrobacter rodentium ICC168 YP 003363907 1 637910 Proteobacteria
Escherichia coli O127 H6 str E2348 69 YP 002330155 1 574521 Proteobacteria
Dickeya aquatica DW 0440 WP 035345573 1 Daquatica
DAQ1742 DAQ1742 03181 Daquatica
DAQ1742 DAQ1742 01702 Daquatica
DAQ1742 DAQ1742 03704 Daquatica
552 DAQ1742 03704 phages P2 P6 P7 related to Mu like Seed
Dickeya sp CSL RW240 WP 067486422 1 Daquatica

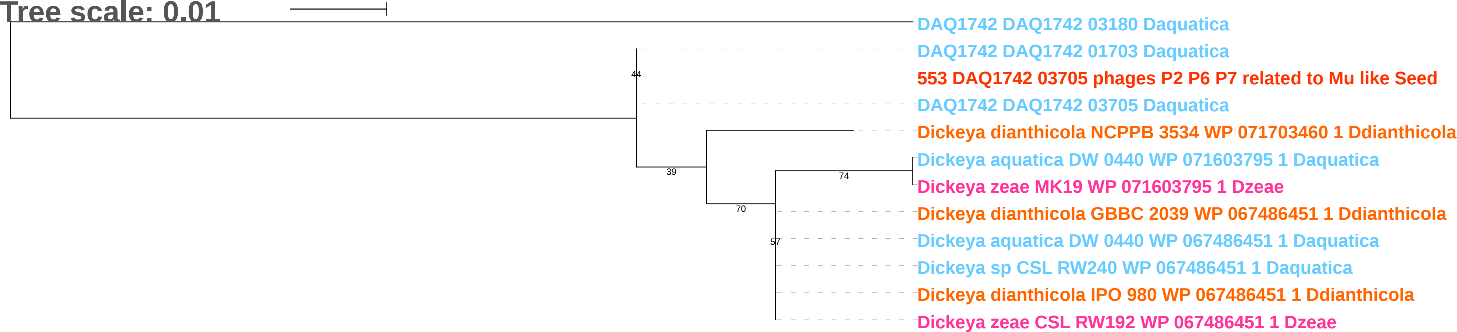

Tree scale: 0.01
DAQ1742 DAQ1742 03180 Daquatica
DAQ1742 DAQ1742 01703 Daquatica
44
553 DAQ1742 03705 phages P2 P6 P7 related to Mu like Seed
DAQ1742 DAQ1742 03705 Daquatica
Dickeya dianthicola NCPPB 3534 WP 071703460 1 Ddianthicola
39
Dickeya aquatica DW 0440 WP 071603795 1 Daquatica
74
Dickeya zeae MK19 WP 071603795 1 Dzeae
70
Dickeya dianthicola GBBC 2039 WP 067486451 1 Ddianthicola
57
Dickeya aquatica DW 0440 WP 067486451 1 Daquatica
Dickeya sp CSL RW240 WP 067486451 1 Daquatica
Dickeya dianthicola IPO 980 WP 067486451 1 Ddianthicola
Dickeya zeae CSL RW192 WP 067486451 1 Dzeae

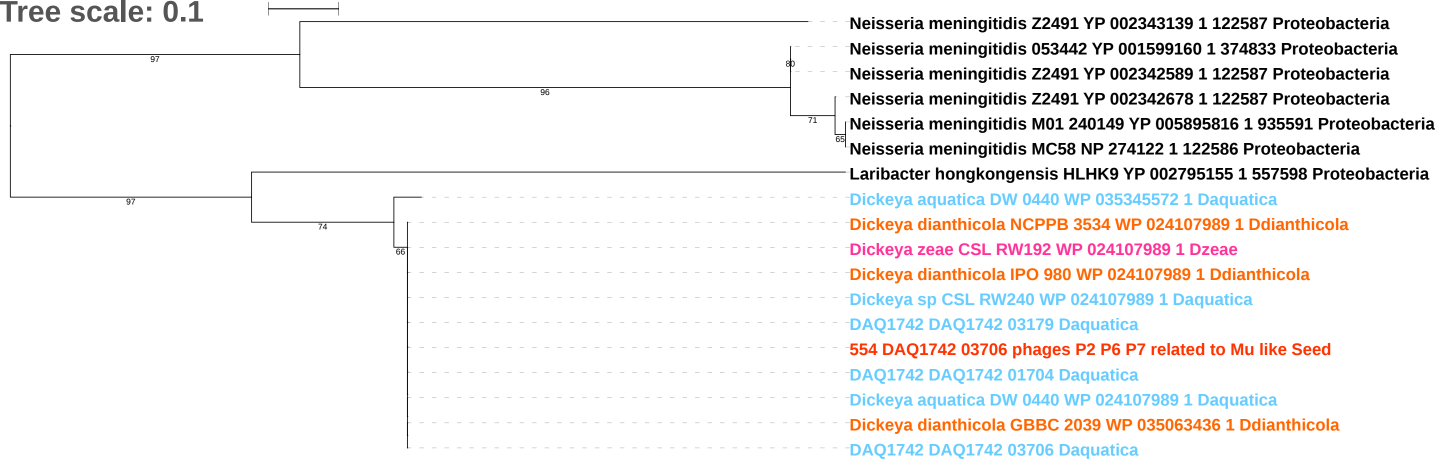
Tree scale: 0.1
Neisseria meningitidis Z2491 YP 002343139 1 122587 Proteobacteria
Neisseria meningitidis 053442 YP 001599160 1 374833 Proteobacteria
Neisseria meningitidis Z2491 YP 002342589 1 122587 Proteobacteria
Neisseria meningitidis Z2491 YP 002342678 1 122587 Proteobacteria
Neisseria meningitidis M01 240149 YP 005895816 1 935591 Proteobacteria
Neisseria meningitidis MC58 NP 274122 1 122586 Proteobacteria
Laribacter hongkongensis HLHK9 YP 002795155 1 557598 Proteobacteria
Dickeya aquatica DW 0440 WP 035345572 1 Daquatica
Dickeya dianthicola NCPPB 3534 WP 024107989 1 Ddianthicola
Dickeya zeae CSL RW192 WP 024107989 1 Dzeae
Dickeya dianthicola IPO 980 WP 024107989 1 Ddianthicola
Dickeya sp CSL RW240 WP 024107989 1 Daquatica
DAQ1742 DAQ1742 03179 Daquatica
554 DAQ1742 03706 phages P2 P6 P7 related to Mu like Seed
DAQ1742 DAQ1742 01704 Daquatica
Dickeya aquatica DW 0440 WP 024107989 1 Daquatica
Dickeya dianthicola GBBC 2039 WP 035063436 1 Ddianthicola
DAQ1742 DAQ1742 03706 Daquatica
97
96
80
71
65
97
74
66

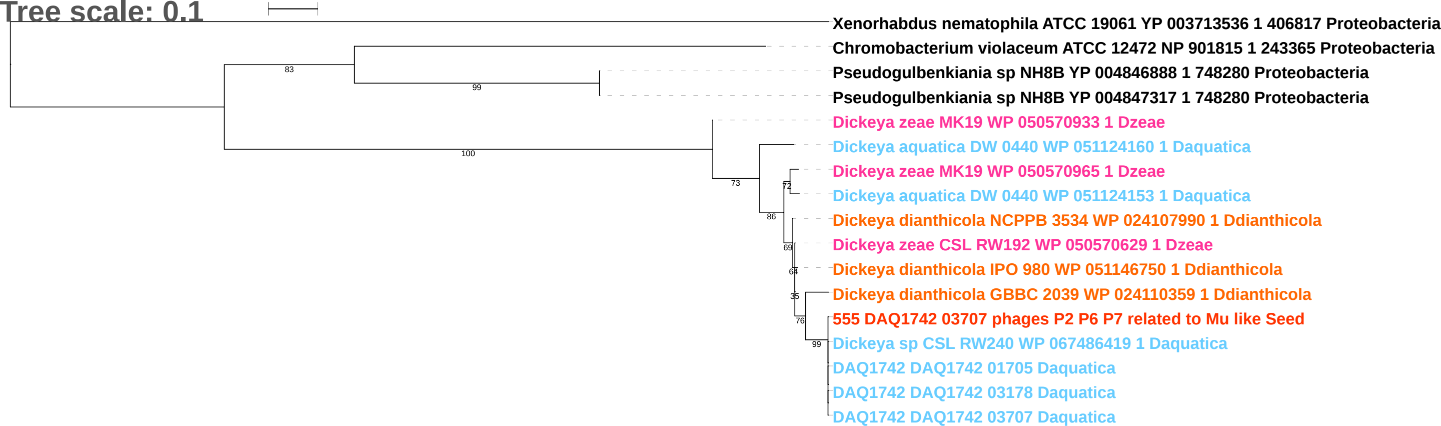
Tree scale: 0.1
Xenorhabdus nematophila ATCC 19061 YP 003713536 1 406817 Proteobacteria
Chromobacterium violaceum ATCC 12472 NP 901815 1 243365 Proteobacteria
Pseudogulbenkiania sp NH8B YP 004846888 1 748280 Proteobacteria
Pseudogulbenkiania sp NH8B YP 004847317 1 748280 Proteobacteria
Dickeya zeae MK19 WP 050570933 1 Dzeae
Dickeya aquatica DW 0440 WP 051124160 1 Daquatica
Dickeya zeae MK19 WP 050570965 1 Dzeae
Dickeya aquatica DW 0440 WP 051124153 1 Daquatica
Dickeya dianthicola NCPPB 3534 WP 024107990 1 Ddianthicola
Dickeya zeae CSL RW192 WP 050570629 1 Dzeae
Dickeya dianthicola IPO 980 WP 051146750 1 Ddianthicola
Dickeya dianthicola GBBC 2039 WP 024110359 1 Ddianthicola
555 DAQ1742 03707 phages P2 P6 P7 related to Mu like Seed
Dickeya sp CSL RW240 WP 067486419 1 Daquatica
DAQ1742 DAQ1742 01705 Daquatica
DAQ1742 DAQ1742 03178 Daquatica
DAQ1742 DAQ1742 03707 Daquatica
83
99
100
73
72
86
69
64
35
76
99

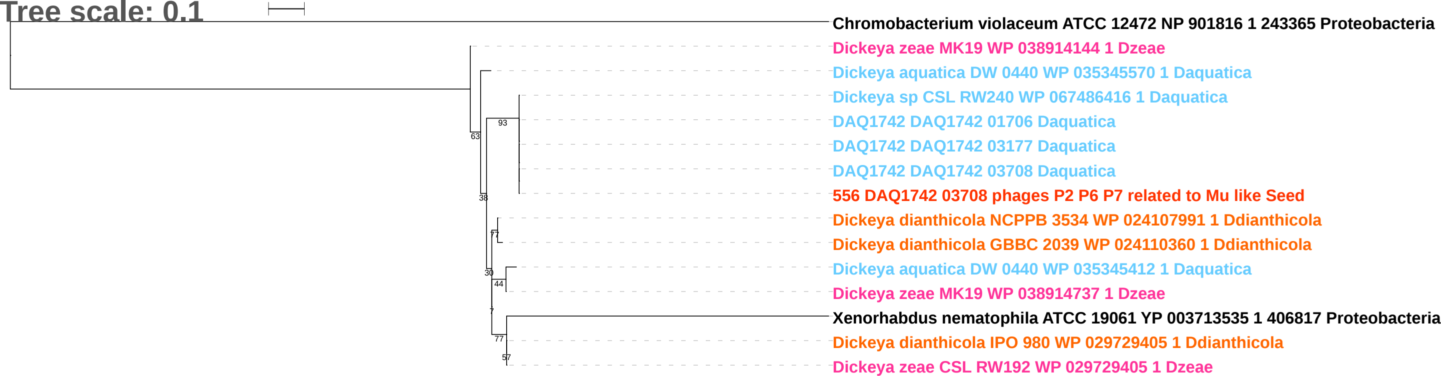
Tree scale: 0.1
Chromobacterium violaceum ATCC 12472 NP 901816 1 243365 Proteobacteria
Dickeya zeae MK19 WP 038914144 1 Dzeae
Dickeya aquatica DW 0440 WP 035345570 1 Daquatica
Dickeya sp CSL RW240 WP 067486416 1 Daquatica
DAQ1742 DAQ1742 01706 Daquatica
DAQ1742 DAQ1742 03177 Daquatica
DAQ1742 DAQ1742 03708 Daquatica
556 DAQ1742 03708 phages P2 P6 P7 related to Mu like Seed
Dickeya dianthicola NCPPB 3534 WP 024107991 1 Ddianthicola
Dickeya dianthicola GBBC 2039 WP 024110360 1 Ddianthicola
Dickeya aquatica DW 0440 WP 035345412 1 Daquatica
Dickeya zeae MK19 WP 038914737 1 Dzeae
Xenorhabdus nematophila ATCC 19061 YP 003713535 1 406817 Proteobacteria
Dickeya dianthicola IPO 980 WP 029729405 1 Ddianthicola
Dickeya zeae CSL RW192 WP 029729405 1 Dzeae
93
63
38
77
30
44
7
77
57

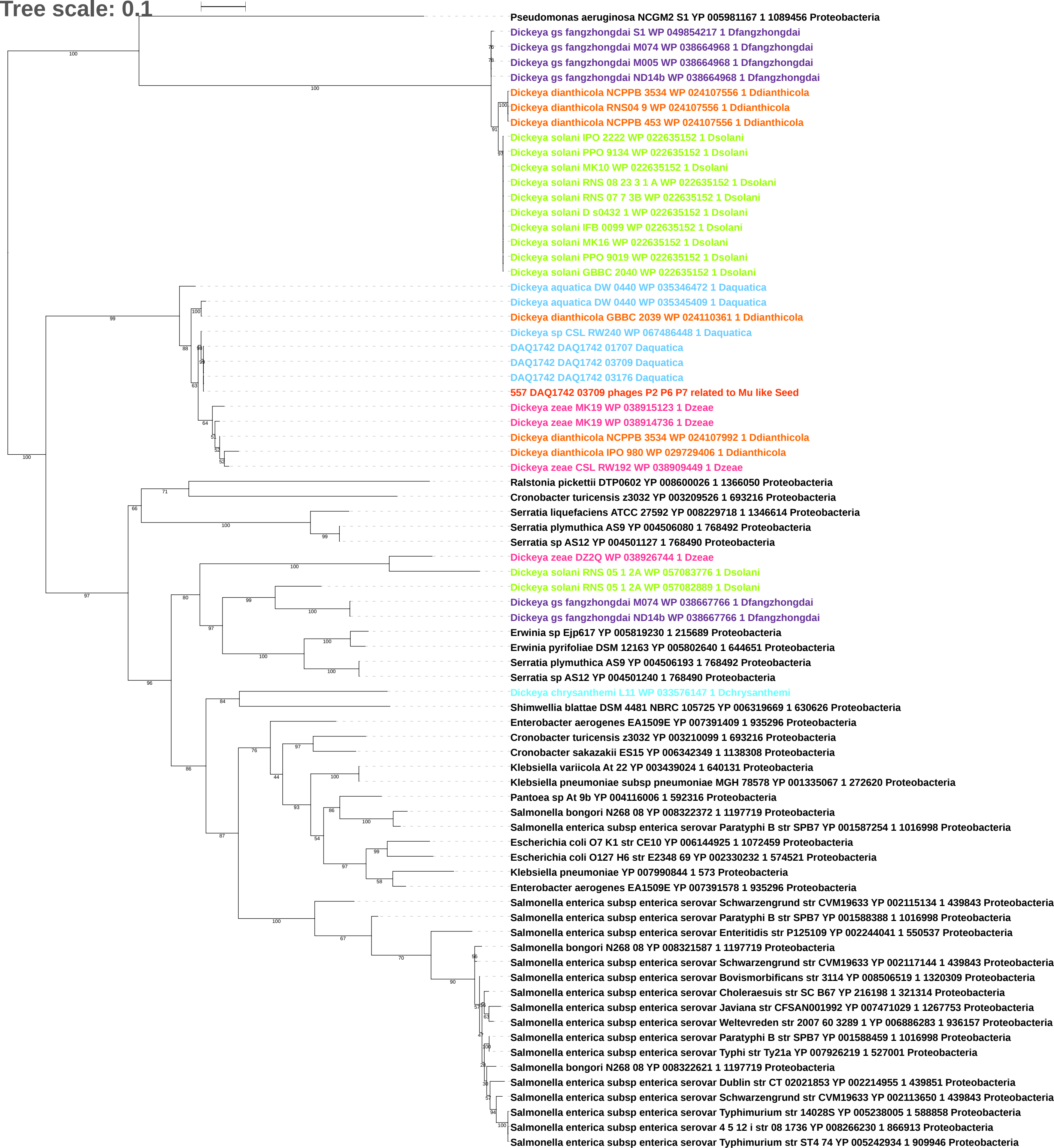
Tree scale: 0.1
Pseudomonas aeruginosa NCGM2 S1 YP 005981167 1 1089456 Proteobacteria
Dickeya gs fangzhongdai S1 WP 049854217 1 Dfangzhongdai
Dickeya gs fangzhongdai M074 WP 038664968 1 Dfangzhongdai
Dickeya gs fangzhongdai M005 WP 038664968 1 Dfangzhongdai
Dickeya gs fangzhongdai ND14b WP 038664968 1 Dfangzhongdai
Dickeya dianthicola NCPPB 3534 WP 024107556 1 Ddianthicola
Dickeya dianthicola RNS04 9 WP 024107556 1 Ddianthicola
Dickeya dianthicola NCPPB 453 WP 024107556 1 Ddianthicola
Dickeya solani IPO 2222 WP 022635152 1 Dsolani
Dickeya solani PPO 9134 WP 022635152 1 Dsolani
Dickeya solani MK10 WP 022635152 1 Dsolani
Dickeya solani RNS 08 23 3 1 A WP 022635152 1 Dsolani
Dickeya solani RNS 07 7 3B WP 022635152 1 Dsolani
Dickeya solani D s0432 1 WP 022635152 1 Dsolani
Dickeya solani IFB 0099 WP 022635152 1 Dsolani
Dickeya solani MK16 WP 022635152 1 Dsolani
Dickeya solani PPO 9019 WP 022635152 1 Dsolani
Dickeya solani GBBC 2040 WP 022635152 1 Dsolani
Dickeya aquatica DW 0440 WP 035346472 1 Daquatica
Dickeya aquatica DW 0440 WP 035345409 1 Daquatica
Dickeya dianthicola GBBC 2039 WP 024110361 1 Ddianthicola
Dickeya sp CSL RW240 WP 067486448 1 Daquatica
DAQ1742 DAQ1742 01707 Daquatica
DAQ1742 DAQ1742 03709 Daquatica
DAQ1742 DAQ1742 03176 Daquatica
557 DAQ1742 03709 phages P2 P6 P7 related to Mu like Seed
Dickeya zeae MK19 WP 038915123 1 Dzeae
Dickeya zeae MK19 WP 038914736 1 Dzeae
Dickeya dianthicola NCPPB 3534 WP 024107992 1 Ddianthicola
Dickeya dianthicola IPO 980 WP 029729406 1 Ddianthicola
Dickeya zeae CSL RW192 WP 038909449 1 Dzeae
Ralstonia pickettii DTP0602 YP 008600026 1 1366050 Proteobacteria
Cronobacter turicensis z3032 YP 003209526 1 693216 Proteobacteria
Serratia liquefaciens ATCC 27592 YP 008229718 1 1346614 Proteobacteria
Serratia plymuthica AS9 YP 004506080 1 768492 Proteobacteria
Serratia sp AS12 YP 004501127 1 768490 Proteobacteria
Dickeya zeae DZ2Q WP 038926744 1 Dzeae
Dickeya solani RNS 05 1 2A WP 057083776 1 Dsolani
Dickeya solani RNS 05 1 2A WP 057082889 1 Dsolani
Dickeya gs fangzhongdai M074 WP 038667766 1 Dfangzhongdai
Dickeya gs fangzhongdai ND14b WP 038667766 1 Dfangzhongdai
Erwinia sp Ejp617 YP 005819230 1 215689 Proteobacteria
Erwinia pyrifoliae DSM 12163 YP 005802640 1 644651 Proteobacteria
Serratia plymuthica AS9 YP 004506193 1 768492 Proteobacteria
Serratia sp AS12 YP 004501240 1 768490 Proteobacteria
Dickeya chrysanthemi L11 WP 033576147 1 Dchrysanthemi
Shimwellia blattae DSM 4481 NBRC 105725 YP 006319669 1 630626 Proteobacteria
Enterobacter aerogenes EA1509E YP 007391409 1 935296 Proteobacteria
Cronobacter turicensis z3032 YP 003210099 1 693216 Proteobacteria
Cronobacter sakazakii ES15 YP 006342349 1 1138308 Proteobacteria
Klebsiella variicola At 22 YP 003439024 1 640131 Proteobacteria
Klebsiella pneumoniae subsp pneumoniae MGH 78578 YP 001335067 1 272620 Proteobacteria
Pantoea sp At 9b YP 004116006 1 592316 Proteobacteria
Salmonella bongori N268 08 YP 008322372 1 1197719 Proteobacteria
Salmonella enterica subsp enterica serovar Paratyphi B str SPB7 YP 001587254 1 1016998 Proteobacteria
Escherichia coli O7 K1 str CE10 YP 006144925 1 1072459 Proteobacteria
Escherichia coli O127 H6 str E2348 69 YP 002330232 1 574521 Proteobacteria
Klebsiella pneumoniae YP 007990844 1 573 Proteobacteria
Enterobacter aerogenes EA1509E YP 007391578 1 935296 Proteobacteria
Salmonella enterica subsp enterica serovar Schwarzengrund str CVM19633 YP 002115134 1 439843 Proteobacteria
Salmonella enterica subsp enterica serovar Paratyphi B str SPB7 YP 001588388 1 1016998 Proteobacteria
Salmonella enterica subsp enterica serovar Enteritidis str P125109 YP 002244041 1 550537 Proteobacteria
Salmonella bongori N268 08 YP 008321587 1 1197719 Proteobacteria
Salmonella enterica subsp enterica serovar Schwarzengrund str CVM19633 YP 002117144 1 439843 Proteobacteria
Salmonella enterica subsp enterica serovar Bovismorbificans str 3114 YP 008506519 1 1320309 Proteobacteria
Salmonella enterica subsp enterica serovar Choleraesuis str SC B67 YP 216198 1 321314 Proteobacteria
Salmonella enterica subsp enterica serovar Javiana str CFSAN001992 YP 007471029 1 1267753 Proteobacteria
Salmonella enterica subsp enterica serovar Weltevreden str 2007 60 3289 1 YP 006886283 1 936157 Proteobacteria
Salmonella enterica subsp enterica serovar Paratyphi B str SPB7 YP 001588459 1 1016998 Proteobacteria
Salmonella enterica subsp enterica serovar Typhi str Ty21a YP 007926219 1 527001 Proteobacteria
Salmonella bongori N268 08 YP 008322621 1 1197719 Proteobacteria
Salmonella enterica subsp enterica serovar Dublin str CT 02021853 YP 002214955 1 439851 Proteobacteria
Salmonella enterica subsp enterica serovar Schwarzengrund str CVM19633 YP 002113650 1 439843 Proteobacteria
Salmonella enterica subsp enterica serovar Typhimurium str 14028S YP 005238005 1 588858 Proteobacteria
Salmonella enterica subsp enterica serovar 4 5 12 i str 08 1736 YP 008266230 1 866913 Proteobacteria
Salmonella enterica subsp enterica serovar Typhimurium str ST4 74 YP 005242934 1 909946 Proteobacteria

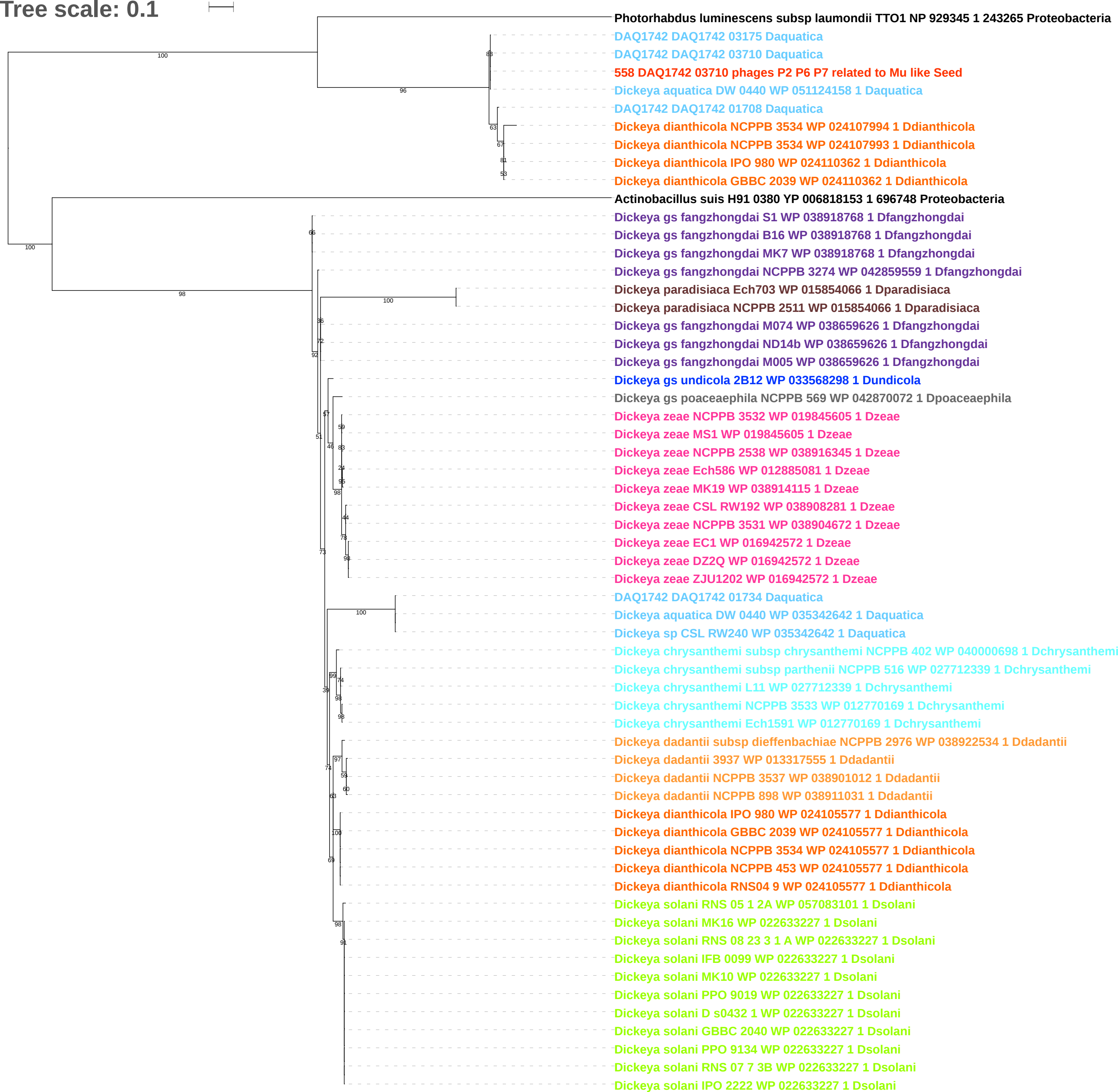
Tree scale: 0.1
Photorhabdus luminescens subsp laumondii TTO1 NP 929345 1 243265 Proteobacteria
DAQ1742 DAQ1742 03175 Daquatica
DAQ1742 DAQ1742 03710 Daquatica
558 DAQ1742 03710 phages P2 P6 P7 related to Mu like Seed
Dickeya aquatica DW 0440 WP 051124158 1 Daquatica
DAQ1742 DAQ1742 01708 Daquatica
Dickeya dianthicola NCPPB 3534 WP 024107994 1 Ddianthicola
Dickeya dianthicola NCPPB 3534 WP 024107993 1 Ddianthicola
Dickeya dianthicola IPO 980 WP 024110362 1 Ddianthicola
Dickeya dianthicola GBBC 2039 WP 024110362 1 Ddianthicola
Actinobacillus suis H91 0380 YP 006818153 1 696748 Proteobacteria
Dickeya gs fangzhongdai S1 WP 038918768 1 Dfangzhongdai
Dickeya gs fangzhongdai B16 WP 038918768 1 Dfangzhongdai
Dickeya gs fangzhongdai MK7 WP 038918768 1 Dfangzhongdai
Dickeya gs fangzhongdai NCPPB 3274 WP 042859559 1 Dfangzhongdai
Dickeya paradisiaca Ech703 WP 015854066 1 Dparadisiaca
Dickeya paradisiaca NCPPB 2511 WP 015854066 1 Dparadisiaca
Dickeya gs fangzhongdai M074 WP 038659626 1 Dfangzhongdai
Dickeya gs fangzhongdai ND14b WP 038659626 1 Dfangzhongdai
Dickeya gs fangzhongdai M005 WP 038659626 1 Dfangzhongdai
Dickeya gs undicola 2B12 WP 033568298 1 Dundicola
Dickeya gs poaceaephila NCPPB 569 WP 042870072 1 Dpoaceaephila
Dickeya zeae NCPPB 3532 WP 019845605 1 Dzeae
Dickeya zeae MS1 WP 019845605 1 Dzeae
Dickeya zeae NCPPB 2538 WP 038916345 1 Dzeae
Dickeya zeae Ech586 WP 012885081 1 Dzeae
Dickeya zeae MK19 WP 038914115 1 Dzeae
Dickeya zeae CSL RW192 WP 038908281 1 Dzeae
Dickeya zeae NCPPB 3531 WP 038904672 1 Dzeae
Dickeya zeae EC1 WP 016942572 1 Dzeae
Dickeya zeae DZ2Q WP 016942572 1 Dzeae
Dickeya zeae ZJU1202 WP 016942572 1 Dzeae
DAQ1742 DAQ1742 01734 Daquatica
Dickeya aquatica DW 0440 WP 035342642 1 Daquatica
Dickeya sp CSL RW240 WP 035342642 1 Daquatica
Dickeya chrysanthemi subsp chrysanthemi NCPPB 402 WP 040000698 1 Dchrysanthemi
Dickeya chrysanthemi subsp parthenii NCPPB 516 WP 027712339 1 Dchrysanthemi
Dickeya chrysanthemi L11 WP 027712339 1 Dchrysanthemi
Dickeya chrysanthemi NCPPB 3533 WP 012770169 1 Dchrysanthemi
Dickeya chrysanthemi Ech1591 WP 012770169 1 Dchrysanthemi
Dickeya dadantii subsp dieffenbachiae NCPPB 2976 WP 038922534 1 Ddadantii
Dickeya dadantii 3937 WP 013317555 1 Ddadantii
Dickeya dadantii NCPPB 3537 WP 038901012 1 Ddadantii
Dickeya dadantii NCPPB 898 WP 038911031 1 Ddadantii
Dickeya dianthicola IPO 980 WP 024105577 1 Ddianthicola
Dickeya dianthicola GBBC 2039 WP 024105577 1 Ddianthicola
Dickeya dianthicola NCPPB 3534 WP 024105577 1 Ddianthicola
Dickeya dianthicola NCPPB 453 WP 024105577 1 Ddianthicola
Dickeya dianthicola RNS04 9 WP 024105577 1 Ddianthicola
Dickeya solani RNS 05 1 2A WP 057083101 1 Dsolani
Dickeya solani MK16 WP 022633227 1 Dsolani
Dickeya solani RNS 08 23 3 1 A WP 022633227 1 Dsolani
Dickeya solani IFB 0099 WP 022633227 1 Dsolani
Dickeya solani MK10 WP 022633227 1 Dsolani
Dickeya solani PPO 9019 WP 022633227 1 Dsolani
Dickeya solani D s0432 1 WP 022633227 1 Dsolani
Dickeya solani GBBC 2040 WP 022633227 1 Dsolani
Dickeya solani PPO 9134 WP 022633227 1 Dsolani
Dickeya solani RNS 07 7 3B WP 022633227 1 Dsolani
Dickeya solani IPO 2222 WP 022633227 1 Dsolani

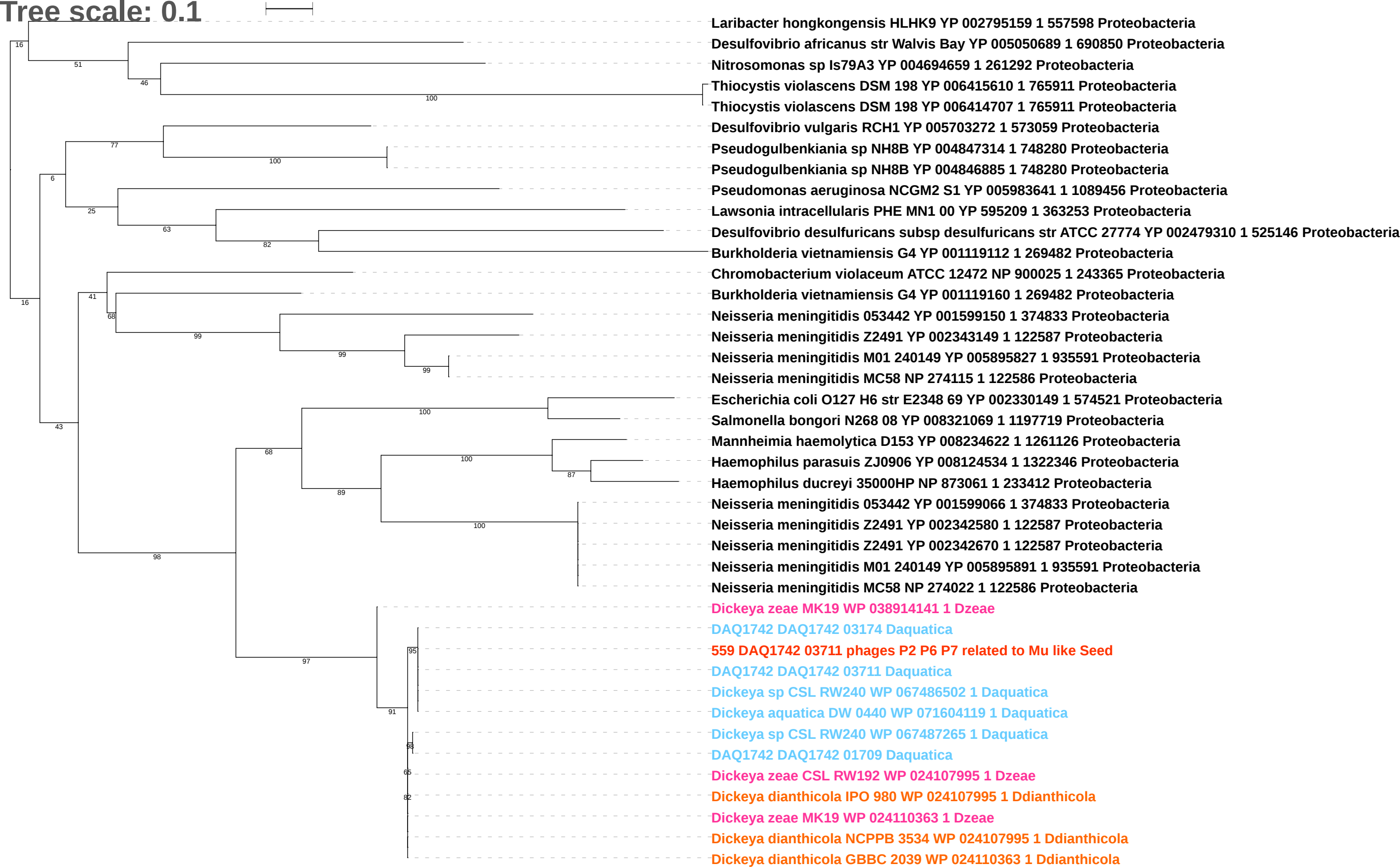
Tree scale: 0.1
Laribacter hongkongensis HLHK9 YP 002795159 1 557598 Proteobacteria
Desulfovibrio africanus str Walvis Bay YP 005050689 1 690850 Proteobacteria
Nitrosomonas sp Is79A3 YP 004694659 1 261292 Proteobacteria
Thiocystis violascens DSM 198 YP 006415610 1 765911 Proteobacteria
Thiocystis violascens DSM 198 YP 006414707 1 765911 Proteobacteria
Desulfovibrio vulgaris RCH1 YP 005703272 1 573059 Proteobacteria
Pseudogulbenkiania sp NH8B YP 004847314 1 748280 Proteobacteria
Pseudogulbenkiania sp NH8B YP 004846885 1 748280 Proteobacteria
Pseudomonas aeruginosa NCGM2 S1 YP 005983641 1 1089456 Proteobacteria
Lawsonia intracellularis PHE MN1 00 YP 595209 1 363253 Proteobacteria
Desulfovibrio desulfuricans subsp desulfuricans str ATCC 27774 YP 002479310 1 525146 Proteobacteria
Burkholderia vietnamiensis G4 YP 001119112 1 269482 Proteobacteria
Chromobacterium violaceum ATCC 12472 NP 900025 1 243365 Proteobacteria
Burkholderia vietnamiensis G4 YP 001119160 1 269482 Proteobacteria
Neisseria meningitidis 053442 YP 001599150 1 374833 Proteobacteria
Neisseria meningitidis Z2491 YP 002343149 1 122587 Proteobacteria
Neisseria meningitidis M01 240149 YP 005895827 1 935591 Proteobacteria
Neisseria meningitidis MC58 NP 274115 1 122586 Proteobacteria
Escherichia coli O127 H6 str E2348 69 YP 002330149 1 574521 Proteobacteria
Salmonella bongori N268 08 YP 008321069 1 1197719 Proteobacteria
Mannheimia haemolytica D153 YP 008234622 1 1261126 Proteobacteria
Haemophilus parasuis ZJ0906 YP 008124534 1 1322346 Proteobacteria
Haemophilus ducreyi 35000HP NP 873061 1 233412 Proteobacteria
Neisseria meningitidis 053442 YP 001599066 1 374833 Proteobacteria
Neisseria meningitidis Z2491 YP 002342580 1 122587 Proteobacteria
Neisseria meningitidis Z2491 YP 002342670 1 122587 Proteobacteria
Neisseria meningitidis M01 240149 YP 005895891 1 935591 Proteobacteria
Neisseria meningitidis MC58 NP 274022 1 122586 Proteobacteria
Dickeya zeae MK19 WP 038914141 1 Dzeae
DAQ1742 DAQ1742 03174 Daquatica
559 DAQ1742 03711 phages P2 P6 P7 related to Mu like Seed
DAQ1742 DAQ1742 03711 Daquatica
Dickeya sp CSL RW240 WP 067486502 1 Daquatica
Dickeya aquatica DW 0440 WP 071604119 1 Daquatica
Dickeya sp CSL RW240 WP 067487265 1 Daquatica
DAQ1742 DAQ1742 01709 Daquatica
Dickeya zeae CSL RW192 WP 024107995 1 Dzeae
Dickeya dianthicola IPO 980 WP 024107995 1 Ddianthicola
Dickeya zeae MK19 WP 024110363 1 Dzeae
Dickeya dianthicola NCPPB 3534 WP 024107995 1 Ddianthicola
Dickeya dianthicola GBBC 2039 WP 024110363 1 Ddianthicola

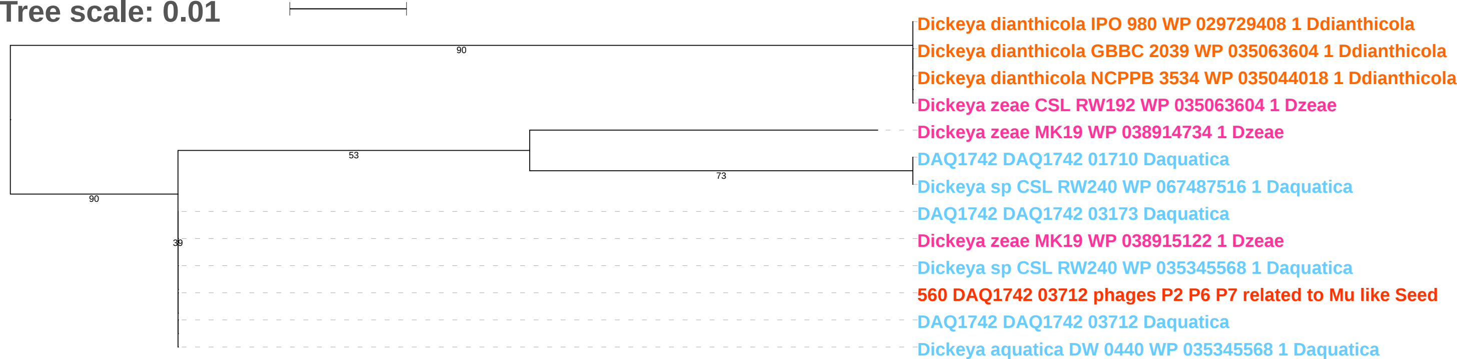
Tree scale: 0.01
90
53
73
90
39
Dickeya dianthicola IPO 980 WP 029729408 1 Ddianthicola
Dickeya dianthicola GBBC 2039 WP 035063604 1 Ddianthicola
Dickeya dianthicola NCPPB 3534 WP 035044018 1 Ddianthicola
Dickeya zeae CSL RW192 WP 035063604 1 Dzeae
Dickeya zeae MK19 WP 038914734 1 Dzeae
DAQ1742 DAQ1742 01710 Daquatica
Dickeya sp CSL RW240 WP 067487516 1 Daquatica
DAQ1742 DAQ1742 03173 Daquatica
Dickeya zeae MK19 WP 038915122 1 Dzeae
Dickeya sp CSL RW240 WP 035345568 1 Daquatica
560 DAQ1742 03712 phages P2 P6 P7 related to Mu like Seed
DAQ1742 DAQ1742 03712 Daquatica
Dickeya aquatica DW 0440 WP 035345568 1 Daquatica

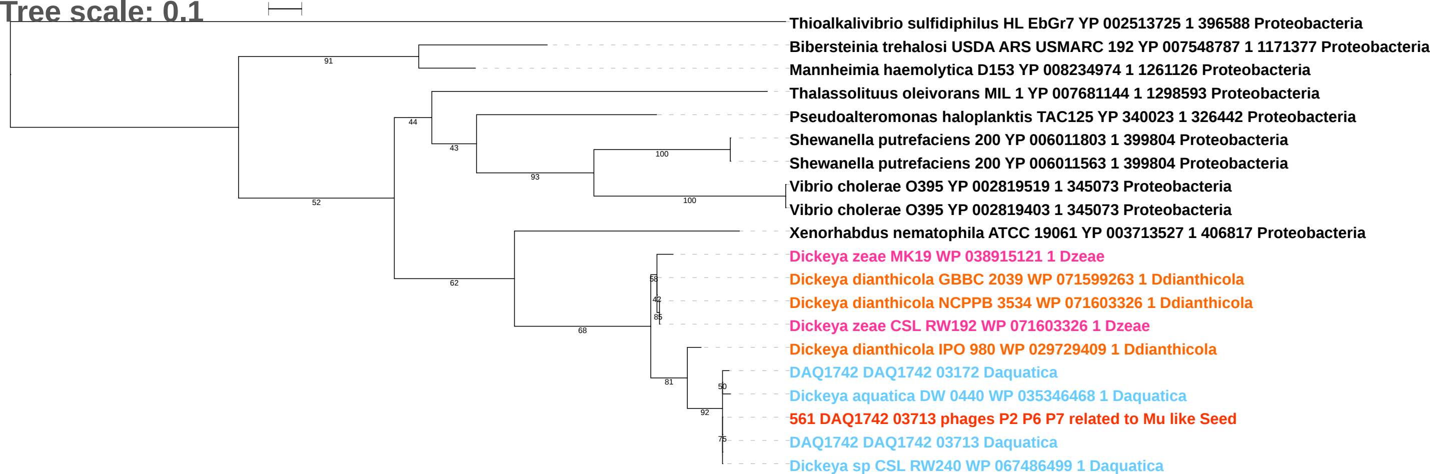
Tree scale: 0.1
Thioalkalivibrio sulfidiphilus HL EbGr7 YP 002513725 1 396588 Proteobacteria
Bibersteinia trehalosi USDA ARS USMARC 192 YP 007548787 1 1171377 Proteobacteria
Mannheimia haemolytica D153 YP 008234974 1 1261126 Proteobacteria
Thalassolituus oleivorans MIL 1 YP 007681144 1 1298593 Proteobacteria
Pseudoalteromonas haloplanktis TAC125 YP 340023 1 326442 Proteobacteria
Shewanella putrefaciens 200 YP 006011803 1 399804 Proteobacteria
Shewanella putrefaciens 200 YP 006011563 1 399804 Proteobacteria
Vibrio cholerae O395 YP 002819519 1 345073 Proteobacteria
Vibrio cholerae O395 YP 002819403 1 345073 Proteobacteria
Xenorhabdus nematophila ATCC 19061 YP 003713527 1 406817 Proteobacteria
Dickeya zeae MK19 WP 038915121 1 Dzeae
Dickeya dianthicola GBBC 2039 WP 071599263 1 Ddianthicola
Dickeya dianthicola NCPPB 3534 WP 071603326 1 Ddianthicola
Dickeya zeae CSL RW192 WP 071603326 1 Dzeae
Dickeya dianthicola IPO 980 WP 029729409 1 Ddianthicola
DAQ1742 DAQ1742 03172 Daquatica
Dickeya aquatica DW 0440 WP 035346468 1 Daquatica
561 DAQ1742 03713 phages P2 P6 P7 related to Mu like Seed
DAQ1742 DAQ1742 03713 Daquatica
Dickeya sp CSL RW240 WP 067486499 1 Daquatica
91
44
43
100
93
100
52
62
58
42
85
68
81
50
92
75

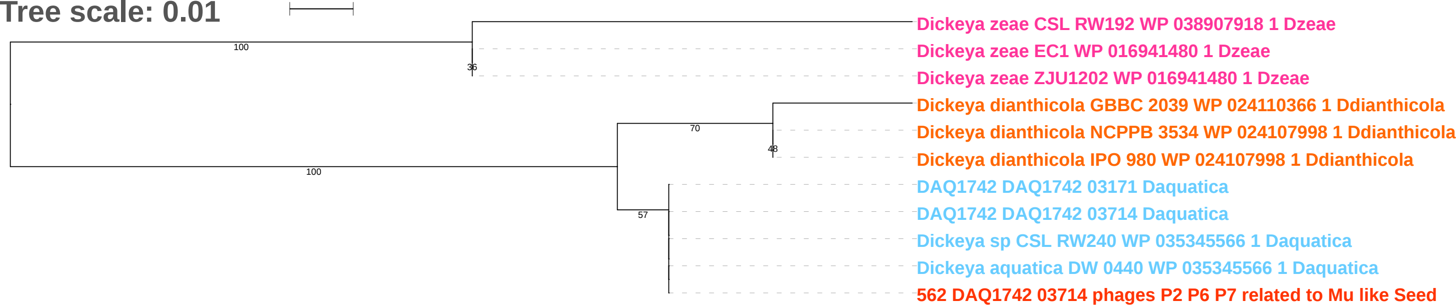
Tree scale: 0.01
100
36
100
70
48
57
Dickeya zeae CSL RW192 WP 038907918 1 Dzeae
Dickeya zeae EC1 WP 016941480 1 Dzeae
Dickeya zeae ZJU1202 WP 016941480 1 Dzeae
Dickeya dianthicola GBBC 2039 WP 024110366 1 Ddianthicola
Dickeya dianthicola NCPPB 3534 WP 024107998 1 Ddianthicola
Dickeya dianthicola IPO 980 WP 024107998 1 Ddianthicola
DAQ1742 DAQ1742 03171 Daquatica
DAQ1742 DAQ1742 03714 Daquatica
Dickeya sp CSL RW240 WP 035345566 1 Daquatica
Dickeya aquatica DW 0440 WP 035345566 1 Daquatica
562 DAQ1742 03714 phages P2 P6 P7 related to Mu like Seed

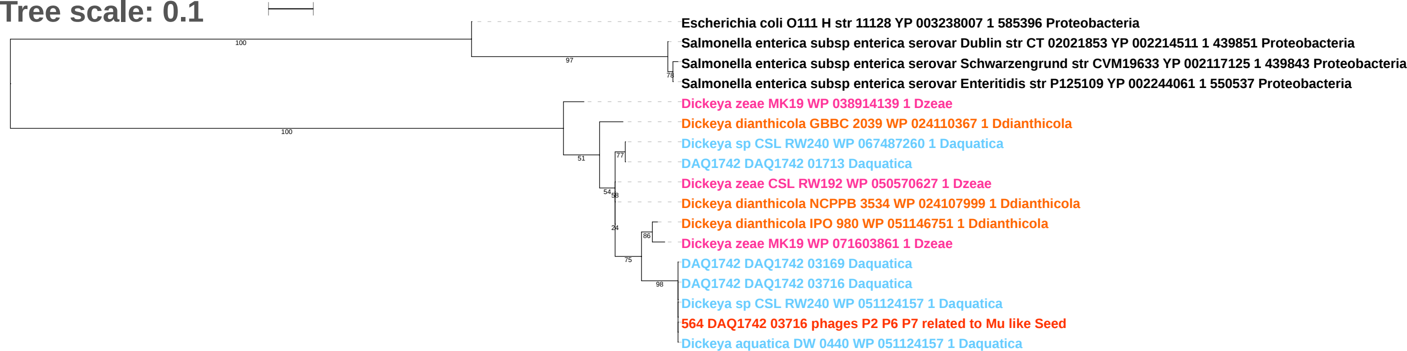
Tree scale: 0.1
Escherichia coli O111 H str 11128 YP 003238007 1 585396 Proteobacteria
Salmonella enterica subsp enterica serovar Dublin str CT 02021853 YP 002214511 1 439851 Proteobacteria
Salmonella enterica subsp enterica serovar Schwarzengrund str CVM19633 YP 002117125 1 439843 Proteobacteria
Salmonella enterica subsp enterica serovar Enteritidis str P125109 YP 002244061 1 550537 Proteobacteria
Dickeya zeae MK19 WP 038914139 1 Dzeae
Dickeya dianthicola GBBC 2039 WP 024110367 1 Ddianthicola
Dickeya sp CSL RW240 WP 067487260 1 Daquatica
DAQ1742 DAQ1742 01713 Daquatica
Dickeya zeae CSL RW192 WP 050570627 1 Dzeae
Dickeya dianthicola NCPPB 3534 WP 024107999 1 Ddianthicola
Dickeya dianthicola IPO 980 WP 051146751 1 Ddianthicola
Dickeya zeae MK19 WP 071603861 1 Dzeae
DAQ1742 DAQ1742 03169 Daquatica
DAQ1742 DAQ1742 03716 Daquatica
Dickeya sp CSL RW240 WP 051124157 1 Daquatica
564 DAQ1742 03716 phages P2 P6 P7 related to Mu like Seed
Dickeya aquatica DW 0440 WP 051124157 1 Daquatica
100
97
78
100
51
77
54
58
24
86
75
98

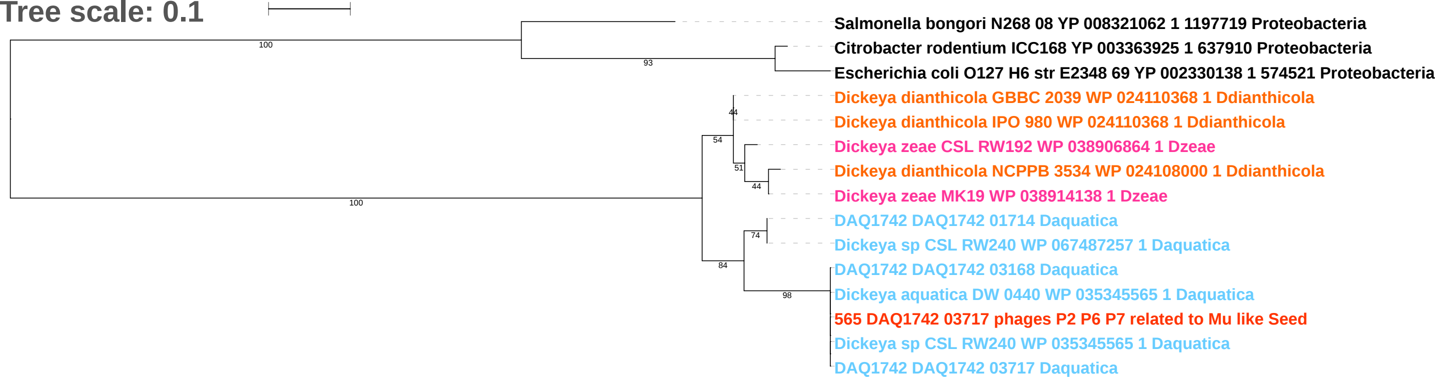

Tree scale: 0.1
Salmonella bongori N268 08 YP 008321062 1 1197719 Proteobacteria
Citrobacter rodentium ICC168 YP 003363925 1 637910 Proteobacteria
Escherichia coli O127 H6 str E2348 69 YP 002330138 1 574521 Proteobacteria
Dickeya dianthicola GBBC 2039 WP 024110368 1 Ddianthicola
Dickeya dianthicola IPO 980 WP 024110368 1 Ddianthicola
Dickeya zeae CSL RW192 WP 038906864 1 Dzeae
Dickeya dianthicola NCPPB 3534 WP 024108000 1 Ddianthicola
Dickeya zeae MK19 WP 038914138 1 Dzeae
DAQ1742 DAQ1742 01714 Daquatica
Dickeya sp CSL RW240 WP 067487257 1 Daquatica
DAQ1742 DAQ1742 03168 Daquatica
Dickeya aquatica DW 0440 WP 035345565 1 Daquatica
565 DAQ1742 03717 phages P2 P6 P7 related to Mu like Seed
Dickeya sp CSL RW240 WP 035345565 1 Daquatica
DAQ1742 DAQ1742 03717 Daquatica
100
93
44
54
51
44
100
74
84
98

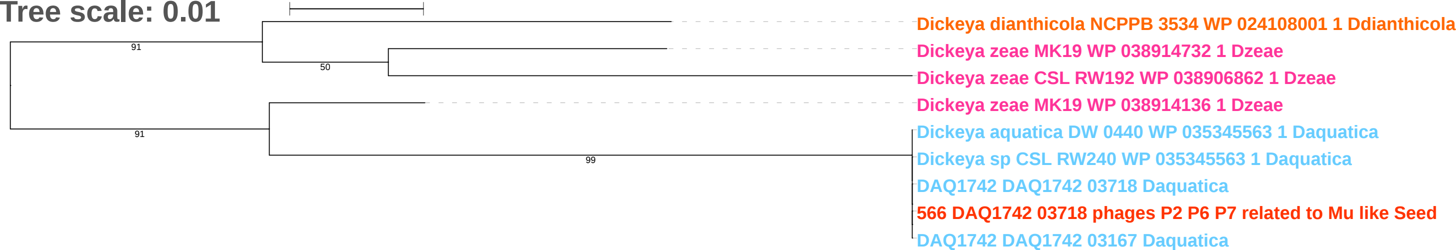
Tree scale: 0.01
91
50
91
99
Dickeya dianthicola NCPPB 3534 WP 024108001 1 Ddianthicola
Dickeya zeae MK19 WP 038914732 1 Dzeae
Dickeya zeae CSL RW192 WP 038906862 1 Dzeae
Dickeya zeae MK19 WP 038914136 1 Dzeae
Dickeya aquatica DW 0440 WP 035345563 1 Daquatica
Dickeya sp CSL RW240 WP 035345563 1 Daquatica
DAQ1742 DAQ1742 03718 Daquatica
566 DAQ1742 03718 phages P2 P6 P7 related to Mu like Seed
DAQ1742 DAQ1742 03167 Daquatica

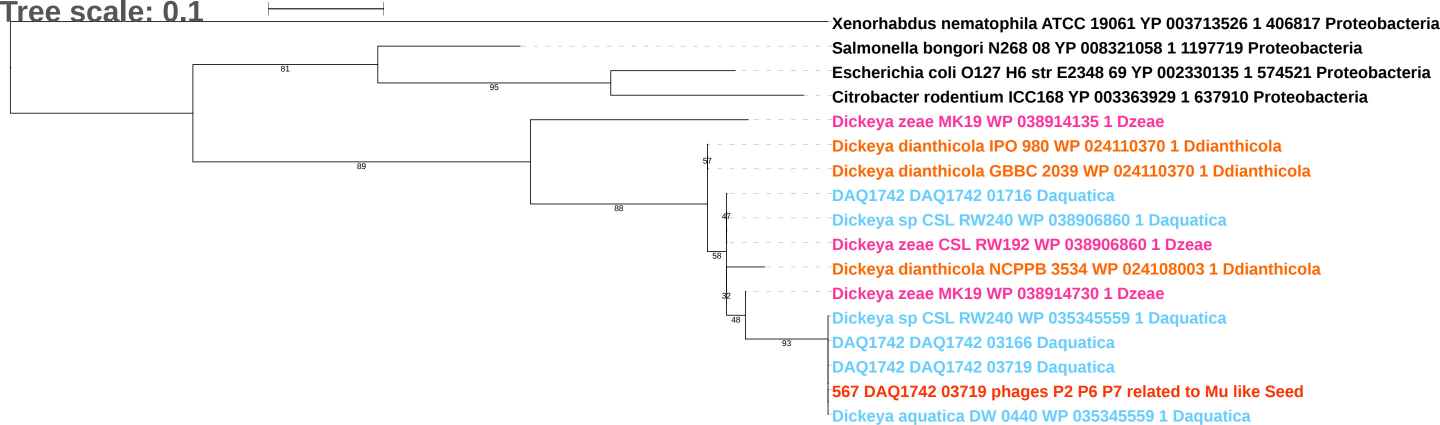
Tree scale: 0.1
Xenorhabdus nematophila ATCC 19061 YP 003713526 1 406817 Proteobacteria
Salmonella bongori N268 08 YP 008321058 1 1197719 Proteobacteria
Escherichia coli O127 H6 str E2348 69 YP 002330135 1 574521 Proteobacteria
Citrobacter rodentium ICC168 YP 003363929 1 637910 Proteobacteria
Dickeya zeae MK19 WP 038914135 1 Dzeae
Dickeya dianthicola IPO 980 WP 024110370 1 Ddianthicola
Dickeya dianthicola GBBC 2039 WP 024110370 1 Ddianthicola
DAQ1742 DAQ1742 01716 Daquatica
Dickeya sp CSL RW240 WP 038906860 1 Daquatica
Dickeya zeae CSL RW192 WP 038906860 1 Dzeae
Dickeya dianthicola NCPPB 3534 WP 024108003 1 Ddianthicola
Dickeya zeae MK19 WP 038914730 1 Dzeae
Dickeya sp CSL RW240 WP 035345559 1 Daquatica
DAQ1742 DAQ1742 03166 Daquatica
DAQ1742 DAQ1742 03719 Daquatica
567 DAQ1742 03719 phages P2 P6 P7 related to Mu like Seed
Dickeya aquatica DW 0440 WP 035345559 1 Daquatica
81
95
89
88
57
47
58
32
48
93

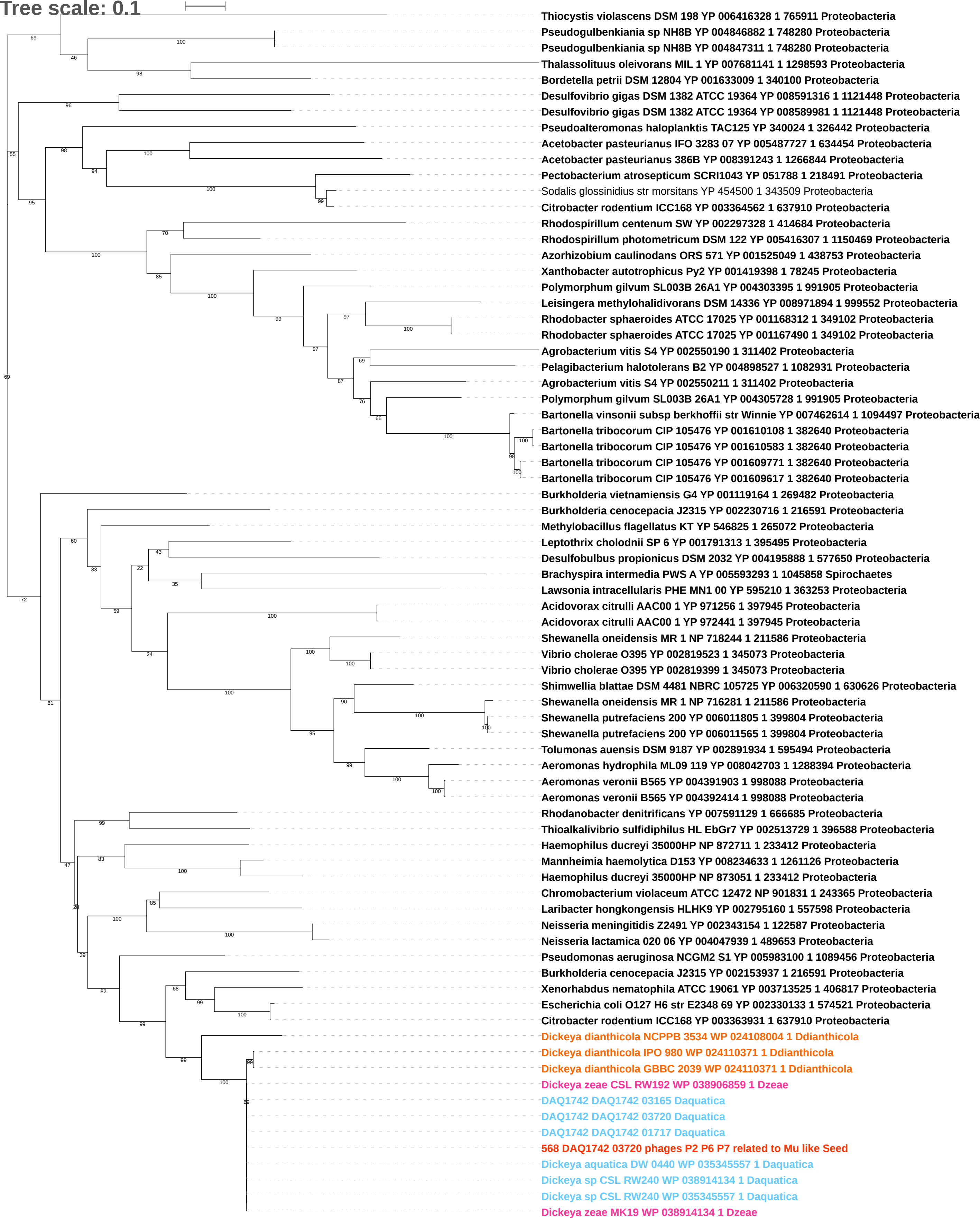
Tree scale: 0.1
Thiocystis violascens DSM 198 YP 006416328 1 765911 Proteobacteria
Pseudogulbenkiania sp NH8B YP 004846882 1 748280 Proteobacteria
Pseudogulbenkiania sp NH8B YP 004847311 1 748280 Proteobacteria
Thalassolituus oleivorans MIL 1 YP 007681141 1 1298593 Proteobacteria
Bordetella petrii DSM 12804 YP 001633009 1 340100 Proteobacteria
Desulfovibrio gigas DSM 1382 ATCC 19364 YP 008591316 1 1121448 Proteobacteria
Desulfovibrio gigas DSM 1382 ATCC 19364 YP 008589981 1 1121448 Proteobacteria
Pseudoalteromonas haloplanktis TAC125 YP 340024 1 326442 Proteobacteria
Acetobacter pasteurianus IFO 3283 07 YP 005487727 1 634454 Proteobacteria
Acetobacter pasteurianus 386B YP 008391243 1 1266844 Proteobacteria
Pectobacterium atrosepticum SCRI1043 YP 051788 1 218491 Proteobacteria
Sodalis glossinidius str morsitans YP 454500 1 343509 Proteobacteria
Citrobacter rodentium ICC168 YP 003364562 1 637910 Proteobacteria
Rhodospirillum centenum SW YP 002297328 1 414684 Proteobacteria
Rhodospirillum photometricum DSM 122 YP 005416307 1 1150469 Proteobacteria
Azorhizobium caulinodans ORS 571 YP 001525049 1 438753 Proteobacteria
Xanthobacter autotrophicus Py2 YP 001419398 1 78245 Proteobacteria
Polymorphum gilvum SL003B 26A1 YP 004303395 1 991905 Proteobacteria
Leisingera methylohalidivorans DSM 14336 YP 008971894 1 999552 Proteobacteria
Rhodobacter sphaeroides ATCC 17025 YP 001168312 1 349102 Proteobacteria
Rhodobacter sphaeroides ATCC 17025 YP 001167490 1 349102 Proteobacteria
Agrobacterium vitis S4 YP 002550190 1 311402 Proteobacteria
Pelagibacterium halotolerans B2 YP 004898527 1 1082931 Proteobacteria
Agrobacterium vitis S4 YP 002550211 1 311402 Proteobacteria
Polymorphum gilvum SL003B 26A1 YP 004305728 1 991905 Proteobacteria
Bartonella vinsonii subsp berkhoffii str Winnie YP 007462614 1 1094497 Proteobacteria
Bartonella tribocorum CIP 105476 YP 001610108 1 382640 Proteobacteria
Bartonella tribocorum CIP 105476 YP 001610583 1 382640 Proteobacteria
Bartonella tribocorum CIP 105476 YP 001609771 1 382640 Proteobacteria
Bartonella tribocorum CIP 105476 YP 001609617 1 382640 Proteobacteria
Burkholderia vietnamiensis G4 YP 001119164 1 269482 Proteobacteria
Burkholderia cenocepacia J2315 YP 002230716 1 216591 Proteobacteria
Methylobacillus flagellatus KT YP 546825 1 265072 Proteobacteria
Leptothrix cholodnii SP 6 YP 001791313 1 395495 Proteobacteria
Desulfobulbus propionicus DSM 2032 YP 004195888 1 577650 Proteobacteria
Brachyspira intermedia PWS A YP 005593293 1 1045858 Spirochaetes
Lawsonia intracellularis PHE MN1 00 YP 595210 1 363253 Proteobacteria
Acidovorax citrulli AAC00 1 YP 971256 1 397945 Proteobacteria
Acidovorax citrulli AAC00 1 YP 972441 1 397945 Proteobacteria
Shewanella oneidensis MR 1 NP 718244 1 211586 Proteobacteria
Vibrio cholerae O395 YP 002819523 1 345073 Proteobacteria
Vibrio cholerae O395 YP 002819399 1 345073 Proteobacteria
Shimwellia blattae DSM 4481 NBRC 105725 YP 006320590 1 630626 Proteobacteria
Shewanella oneidensis MR 1 NP 716281 1 211586 Proteobacteria
Shewanella putrefaciens 200 YP 006011805 1 399804 Proteobacteria
Shewanella putrefaciens 200 YP 006011565 1 399804 Proteobacteria
Tolumonas auensis DSM 9187 YP 002891934 1 595494 Proteobacteria
Aeromonas hydrophila ML09 119 YP 008042703 1 1288394 Proteobacteria
Aeromonas veronii B565 YP 004391903 1 998088 Proteobacteria
Aeromonas veronii B565 YP 004392414 1 998088 Proteobacteria
Rhodanobacter denitrificans YP 007591129 1 666685 Proteobacteria
Thioalkalivibrio sulfidiphilus HL EbGr7 YP 002513729 1 396588 Proteobacteria
Haemophilus ducreyi 35000HP NP 872711 1 233412 Proteobacteria
Mannheimia haemolytica D153 YP 008234633 1 1261126 Proteobacteria
Haemophilus ducreyi 35000HP NP 873051 1 233412 Proteobacteria
Chromobacterium violaceum ATCC 12472 NP 901831 1 243365 Proteobacteria
Laribacter hongkongensis HLHK9 YP 002795160 1 557598 Proteobacteria
Neisseria meningitidis Z2491 YP 002343154 1 122587 Proteobacteria
Neisseria lactamica 020 06 YP 004047939 1 489653 Proteobacteria
Pseudomonas aeruginosa NCGM2 S1 YP 005983100 1 1089456 Proteobacteria
Burkholderia cenocepacia J2315 YP 002153937 1 216591 Proteobacteria
Xenorhabdus nematophila ATCC 19061 YP 003713525 1 406817 Proteobacteria
Escherichia coli O127 H6 str E2348 69 YP 002330133 1 574521 Proteobacteria
Citrobacter rodentium ICC168 YP 003363931 1 637910 Proteobacteria
Dickeya dianthicola NCPPB 3534 WP 024108004 1 Ddianthicola
Dickeya dianthicola IPO 980 WP 024110371 1 Ddianthicola
Dickeya dianthicola GBBC 2039 WP 024110371 1 Ddianthicola
Dickeya zeae CSL RW192 WP 038906859 1 Dzeae
DAQ1742 DAQ1742 03165 Daquatica
DAQ1742 DAQ1742 03720 Daquatica
DAQ1742 DAQ1742 01717 Daquatica
568 DAQ1742 03720 phages P2 P6 P7 related to Mu like Seed
Dickeya aquatica DW 0440 WP 035345557 1 Daquatica
Dickeya sp CSL RW240 WP 038914134 1 Daquatica
Dickeya sp CSL RW240 WP 035345557 1 Daquatica
Dickeya zeae MK19 WP 038914134 1 Dzeae

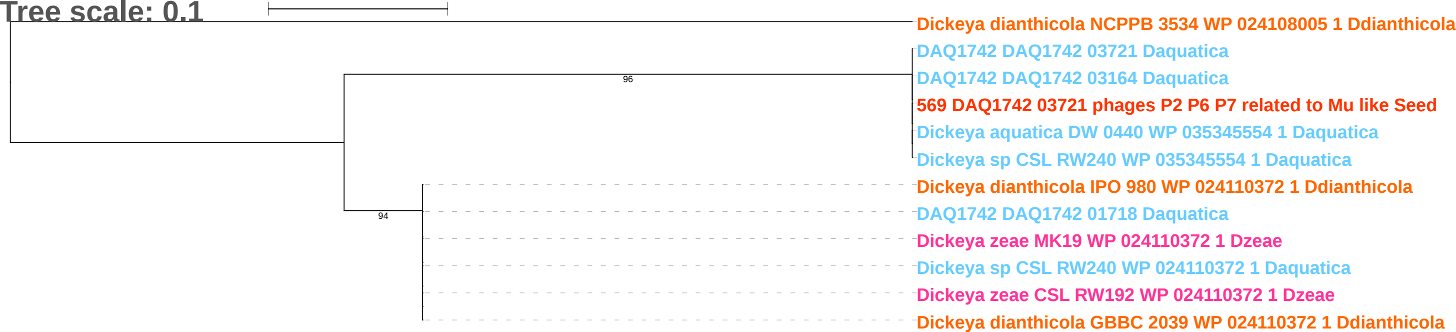
Tree scale: 0.1
Dickeya dianthicola NCPPB 3534 WP 024108005 1 Ddianthicola
DAQ1742 DAQ1742 03721 Daquatica
DAQ1742 DAQ1742 03164 Daquatica
569 DAQ1742 03721 phages P2 P6 P7 related to Mu like Seed
Dickeya aquatica DW 0440 WP 035345554 1 Daquatica
Dickeya sp CSL RW240 WP 035345554 1 Daquatica
Dickeya dianthicola IPO 980 WP 024110372 1 Ddianthicola
DAQ1742 DAQ1742 01718 Daquatica
Dickeya zeae MK19 WP 024110372 1 Dzeae
Dickeya sp CSL RW240 WP 024110372 1 Daquatica
Dickeya zeae CSL RW192 WP 024110372 1 Dzeae
Dickeya dianthicola GBBC 2039 WP 024110372 1 Ddianthicola
96
94

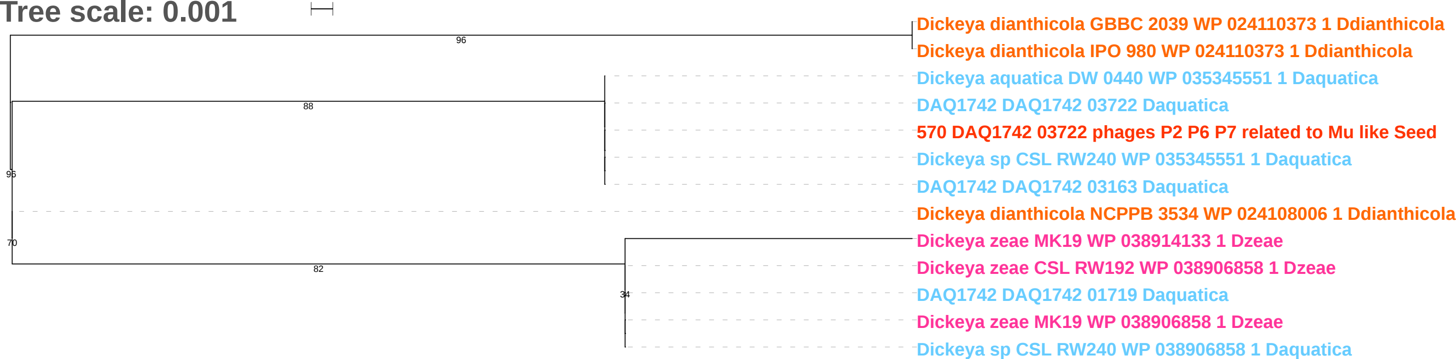
Tree scale: 0.001
96
88
96
70
82
34
Dickeya dianthicola GBBC 2039 WP 024110373 1 Ddianthicola
Dickeya dianthicola IPO 980 WP 024110373 1 Ddianthicola
Dickeya aquatica DW 0440 WP 035345551 1 Daquatica
DAQ1742 DAQ1742 03722 Daquatica
570 DAQ1742 03722 phages P2 P6 P7 related to Mu like Seed
Dickeya sp CSL RW240 WP 035345551 1 Daquatica
DAQ1742 DAQ1742 03163 Daquatica
Dickeya dianthicola NCPPB 3534 WP 024108006 1 Ddianthicola
Dickeya zeae MK19 WP 038914133 1 Dzeae
Dickeya zeae CSL RW192 WP 038906858 1 Dzeae
DAQ1742 DAQ1742 01719 Daquatica
Dickeya zeae MK19 WP 038906858 1 Dzeae
Dickeya sp CSL RW240 WP 038906858 1 Daquatica

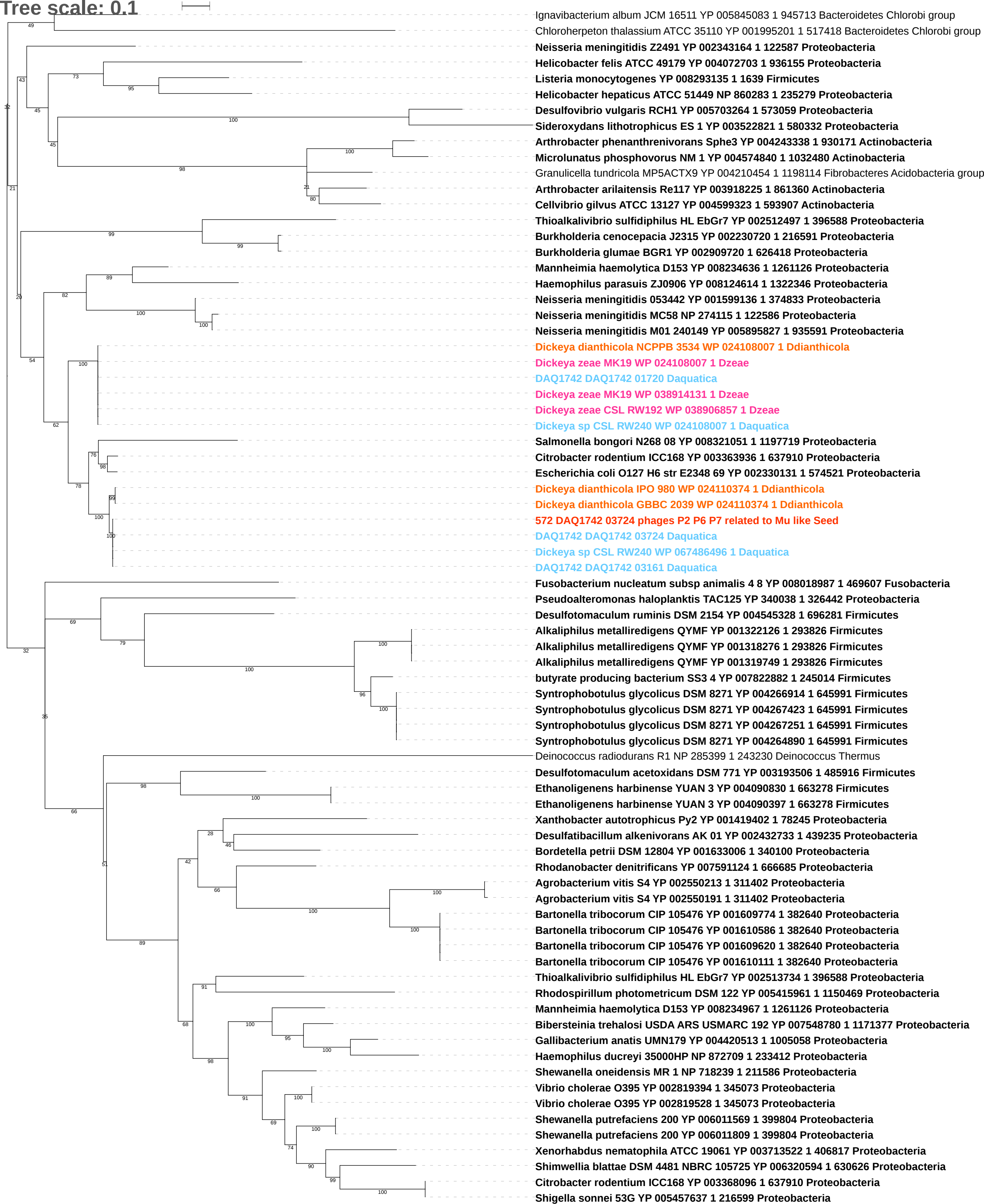
Tree scale: 0.1
Ignavibacterium album JCM 16511 YP 005845083 1 945713 Bacteroidetes Chlorobi group
Chloroherpeton thalassium ATCC 35110 YP 001995201 1 517418 Bacteroidetes Chlorobi group
Neisseria meningitidis Z2491 YP 002343164 1 122587 Proteobacteria
Helicobacter felis ATCC 49179 YP 004072703 1 936155 Proteobacteria
Listeria monocytogenes YP 008293135 1 1639 Firmicutes
Helicobacter hepaticus ATCC 51449 NP 860283 1 235279 Proteobacteria
Desulfovibrio vulgaris RCH1 YP 005703264 1 573059 Proteobacteria
Sideroxydans lithotrophicus ES 1 YP 003522821 1 580332 Proteobacteria
Arthrobacter phenanthrenivorans Sphe3 YP 004243338 1 930171 Actinobacteria
Microlunatus phosphovorus NM 1 YP 004574840 1 1032480 Actinobacteria
Granulicella tundricola MP5ACTX9 YP 004210454 1 1198114 Fibrobacteres Acidobacteria group
Arthrobacter arilaitensis Re117 YP 003918225 1 861360 Actinobacteria
Cellvibrio gilvus ATCC 13127 YP 004599323 1 593907 Actinobacteria
Thioalkalivibrio sulfidiphilus HL EbGr7 YP 002512497 1 396588 Proteobacteria
Burkholderia cenocepacia J2315 YP 002230720 1 216591 Proteobacteria
Burkholderia glumae BGR1 YP 002909720 1 626418 Proteobacteria
Mannheimia haemolytica D153 YP 008234636 1 1261126 Proteobacteria
Haemophilus parasuis ZJ0906 YP 008124614 1 1322346 Proteobacteria
Neisseria meningitidis 053442 YP 001599136 1 374833 Proteobacteria
Neisseria meningitidis MC58 NP 274115 1 122586 Proteobacteria
Neisseria meningitidis M01 240149 YP 005895827 1 935591 Proteobacteria
Dickeya dianthicola NCPPB 3534 WP 024108007 1 Ddianthicola
Dickeya zeae MK19 WP 024108007 1 Dzeae
DAQ1742 DAQ1742 01720 Daquatica
Dickeya zeae MK19 WP 038914131 1 Dzeae
Dickeya zeae CSL RW192 WP 038906857 1 Dzeae
Dickeya sp CSL RW240 WP 024108007 1 Daquatica
Salmonella bongori N268 08 YP 008321051 1 1197719 Proteobacteria
Citrobacter rodentium ICC168 YP 003363936 1 637910 Proteobacteria
Escherichia coli O127 H6 str E2348 69 YP 002330131 1 574521 Proteobacteria
Dickeya dianthicola IPO 980 WP 024110374 1 Ddianthicola
Dickeya dianthicola GBBC 2039 WP 024110374 1 Ddianthicola
572 DAQ1742 03724 phages P2 P6 P7 related to Mu like Seed
DAQ1742 DAQ1742 03724 Daquatica
Dickeya sp CSL RW240 WP 067486496 1 Daquatica
DAQ1742 DAQ1742 03161 Daquatica
Fusobacterium nucleatum subsp animalis 4 8 YP 008018987 1 469607 Fusobacteria
Pseudoalteromonas haloplanktis TAC125 YP 340038 1 326442 Proteobacteria
Desulfotomaculum ruminis DSM 2154 YP 004545328 1 696281 Firmicutes
Alkaliphilus metalliredigens QYMF YP 001322126 1 293826 Firmicutes
Alkaliphilus metalliredigens QYMF YP 001318276 1 293826 Firmicutes
Alkaliphilus metalliredigens QYMF YP 001319749 1 293826 Firmicutes
butyrate producing bacterium SS3 4 YP 007822882 1 245014 Firmicutes
Syntrophobotulus glycolicus DSM 8271 YP 004266914 1 645991 Firmicutes
Syntrophobotulus glycolicus DSM 8271 YP 004267423 1 645991 Firmicutes
Syntrophobotulus glycolicus DSM 8271 YP 004267251 1 645991 Firmicutes
Syntrophobotulus glycolicus DSM 8271 YP 004264890 1 645991 Firmicutes
Deinococcus radiodurans R1 NP 285399 1 243230 Deinococcus Thermus
Desulfotomaculum acetoxidans DSM 771 YP 003193506 1 485916 Firmicutes
Ethanoligenens harbinense YUAN 3 YP 004090830 1 663278 Firmicutes
Ethanoligenens harbinense YUAN 3 YP 004090397 1 663278 Firmicutes
Xanthobacter autotrophicus Py2 YP 001419402 1 78245 Proteobacteria
Desulfatibacillum alkenivorans AK 01 YP 002432733 1 439235 Proteobacteria
Bordetella petrii DSM 12804 YP 001633006 1 340100 Proteobacteria
Rhodanobacter denitrificans YP 007591124 1 666685 Proteobacteria
Agrobacterium vitis S4 YP 002550213 1 311402 Proteobacteria
Agrobacterium vitis S4 YP 002550191 1 311402 Proteobacteria
Bartonella tribocorum CIP 105476 YP 001609774 1 382640 Proteobacteria
Bartonella tribocorum CIP 105476 YP 001610586 1 382640 Proteobacteria
Bartonella tribocorum CIP 105476 YP 001609620 1 382640 Proteobacteria
Bartonella tribocorum CIP 105476 YP 001610111 1 382640 Proteobacteria
Thioalkalivibrio sulfidiphilus HL EbGr7 YP 002513734 1 396588 Proteobacteria
Rhodospirillum photometricum DSM 122 YP 005415961 1 1150469 Proteobacteria
Mannheimia haemolytica D153 YP 008234967 1 1261126 Proteobacteria
Bibersteinia trehalosi USDA ARS USMARC 192 YP 007548780 1 1171377 Proteobacteria
Gallibacterium anatis UMN179 YP 004420513 1 1005058 Proteobacteria
Haemophilus ducreyi 35000HP NP 872709 1 233412 Proteobacteria
Shewanella oneidensis MR 1 NP 718239 1 211586 Proteobacteria
Vibrio cholerae O395 YP 002819394 1 345073 Proteobacteria
Vibrio cholerae O395 YP 002819528 1 345073 Proteobacteria
Shewanella putrefaciens 200 YP 006011569 1 399804 Proteobacteria
Shewanella putrefaciens 200 YP 006011809 1 399804 Proteobacteria
Xenorhabdus nematophila ATCC 19061 YP 003713522 1 406817 Proteobacteria
Shimwellia blattae DSM 4481 NBRC 105725 YP 006320594 1 630626 Proteobacteria
Citrobacter rodentium ICC168 YP 003368096 1 637910 Proteobacteria
Shigella sonnei 53G YP 005457637 1 216599 Proteobacteria

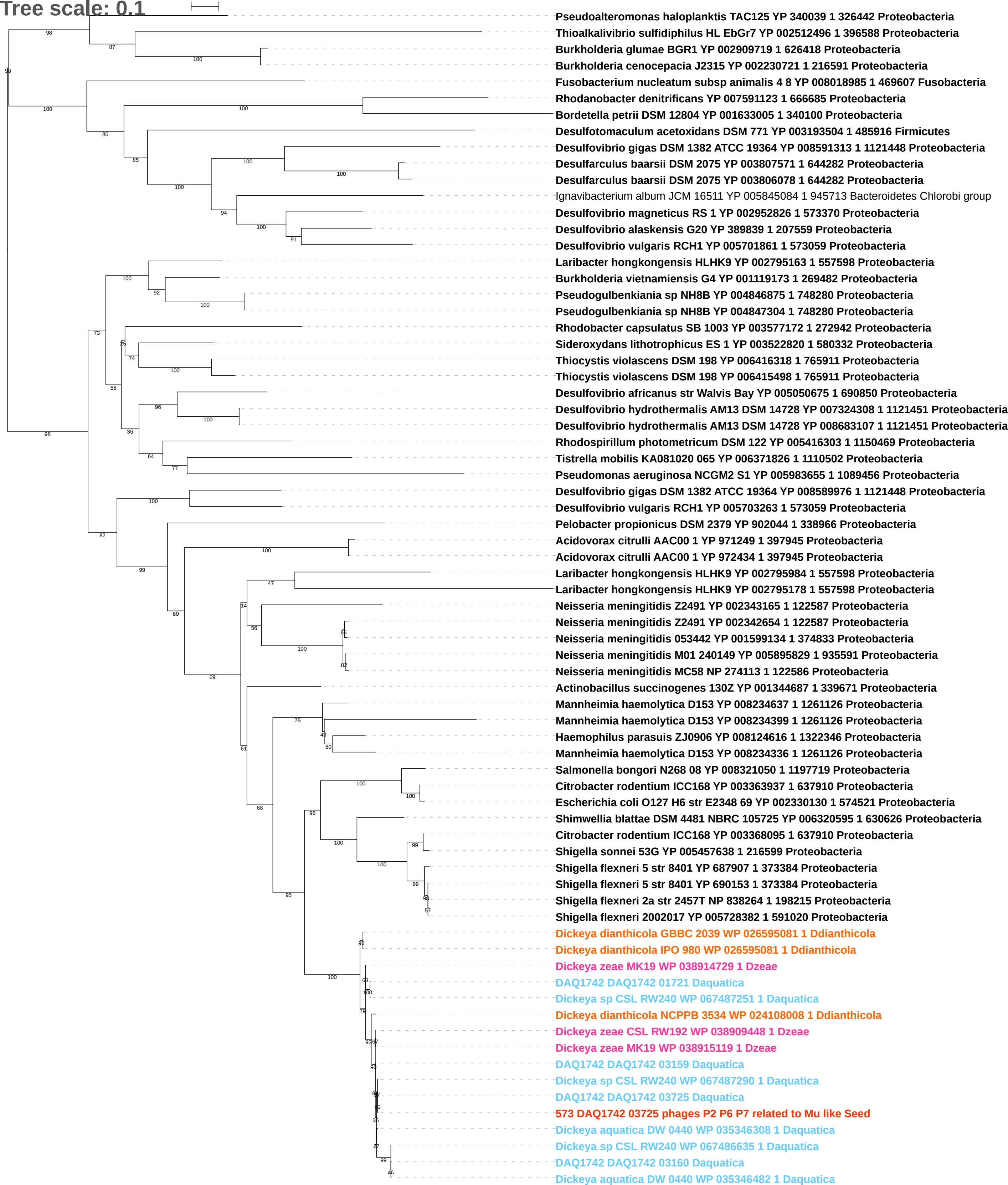
Tree scale: 0.1
Pseudoalteromonas haloplanktis TAC125 YP 340039 1 326442 Proteobacteria
Thioalkalivibrio sulfidiphilus HL EbGr7 YP 002512496 1 396588 Proteobacteria
Burkholderia glumae BGR1 YP 002909719 1 626418 Proteobacteria
Burkholderia cenocepacia J2315 YP 002230721 1 216591 Proteobacteria
Fusobacterium nucleatum subsp animalis 4 8 YP 008018985 1 469607 Fusobacteria
Rhodanobacter denitrificans YP 007591123 1 666685 Proteobacteria
Bordetella petrii DSM 12804 YP 001633005 1 340100 Proteobacteria
Desulfotomaculum acetoxidans DSM 771 YP 003193504 1 485916 Firmicutes
Desulfovibrio gigas DSM 1382 ATCC 19364 YP 008591313 1 1121448 Proteobacteria
Desulfarculus baarsii DSM 2075 YP 003807571 1 644282 Proteobacteria
Desulfarculus baarsii DSM 2075 YP 003806078 1 644282 Proteobacteria
Ignavibacterium album JCM 16511 YP 005845084 1 945713 Bacteroidetes Chlorobi group
Desulfovibrio magneticus RS 1 YP 002952826 1 573370 Proteobacteria
Desulfovibrio alaskensis G20 YP 389839 1 207559 Proteobacteria
Desulfovibrio vulgaris RCH1 YP 005701861 1 573059 Proteobacteria
Laribacter hongkongensis HLHK9 YP 002795163 1 557598 Proteobacteria
Burkholderia vietnamiensis G4 YP 001119173 1 269482 Proteobacteria
Pseudogulbenkiania sp NH8B YP 004846875 1 748280 Proteobacteria
Pseudogulbenkiania sp NH8B YP 004847304 1 748280 Proteobacteria
Rhodobacter capsulatus SB 1003 YP 003577172 1 272942 Proteobacteria
Sideroxydans lithotrophicus ES 1 YP 003522820 1 580332 Proteobacteria
Thiocystis violascens DSM 198 YP 006416318 1 765911 Proteobacteria
Thiocystis violascens DSM 198 YP 006415498 1 765911 Proteobacteria
Desulfovibrio africanus str Walvis Bay YP 005050675 1 690850 Proteobacteria
Desulfovibrio hydrothermalis AM13 DSM 14728 YP 007324308 1 1121451 Proteobacteria
Desulfovibrio hydrothermalis AM13 DSM 14728 YP 008683107 1 1121451 Proteobacteria
Rhodospirillum photometricum DSM 122 YP 005416303 1 1150469 Proteobacteria
Tistrella mobilis KA081020 065 YP 006371826 1 1110502 Proteobacteria
Pseudomonas aeruginosa NCGM2 S1 YP 005983655 1 1089456 Proteobacteria
Desulfovibrio gigas DSM 1382 ATCC 19364 YP 008589976 1 1121448 Proteobacteria
Desulfovibrio vulgaris RCH1 YP 005703263 1 573059 Proteobacteria
Pelobacter propionicus DSM 2379 YP 902044 1 338966 Proteobacteria
Acidovorax citrulli AAC00 1 YP 971249 1 397945 Proteobacteria
Acidovorax citrulli AAC00 1 YP 972434 1 397945 Proteobacteria
Laribacter hongkongensis HLHK9 YP 002795984 1 557598 Proteobacteria
Laribacter hongkongensis HLHK9 YP 002795178 1 557598 Proteobacteria
Neisseria meningitidis Z2491 YP 002343165 1 122587 Proteobacteria
Neisseria meningitidis Z2491 YP 002342654 1 122587 Proteobacteria
Neisseria meningitidis 053442 YP 001599134 1 374833 Proteobacteria
Neisseria meningitidis M01 240149 YP 005895829 1 935591 Proteobacteria
Neisseria meningitidis MC58 NP 274113 1 122586 Proteobacteria
Actinobacillus succinogenes 130Z YP 001344687 1 339671 Proteobacteria
Mannheimia haemolytica D153 YP 008234637 1 1261126 Proteobacteria
Mannheimia haemolytica D153 YP 008234399 1 1261126 Proteobacteria
Haemophilus parasuis ZJ0906 YP 008124616 1 1322346 Proteobacteria
Mannheimia haemolytica D153 YP 008234336 1 1261126 Proteobacteria
Salmonella bongori N268 08 YP 008321050 1 1197719 Proteobacteria
Citrobacter rodentium ICC168 YP 003363937 1 637910 Proteobacteria
Escherichia coli O127 H6 str E2348 69 YP 002330130 1 574521 Proteobacteria
Shimwellia blattae DSM 4481 NBRC 105725 YP 006320595 1 630626 Proteobacteria
Citrobacter rodentium ICC168 YP 003368095 1 637910 Proteobacteria
Shigella sonnei 53G YP 005457638 1 216599 Proteobacteria
Shigella flexneri 5 str 8401 YP 687907 1 373384 Proteobacteria
Shigella flexneri 5 str 8401 YP 690153 1 373384 Proteobacteria
Shigella flexneri 2a str 2457T NP 838264 1 198215 Proteobacteria
Shigella flexneri 2002017 YP 005728382 1 591020 Proteobacteria
Dickeya dianthicola GBBC 2039 WP 026595081 1 Ddianthicola
Dickeya dianthicola IPO 980 WP 026595081 1 Ddianthicola
Dickeya zeae MK19 WP 038914729 1 Dzeae
DAQ1742 DAQ1742 01721 Daquatica
Dickeya sp CSL RW240 WP 067487251 1 Daquatica
Dickeya dianthicola NCPPB 3534 WP 024108008 1 Ddianthicola
Dickeya zeae CSL RW192 WP 038909448 1 Dzeae
Dickeya zeae MK19 WP 038915119 1 Dzeae
DAQ1742 DAQ1742 03159 Daquatica
Dickeya sp CSL RW240 WP 067487290 1 Daquatica
DAQ1742 DAQ1742 03725 Daquatica
573 DAQ1742 03725 phages P2 P6 P7 related to Mu like Seed
Dickeya aquatica DW 0440 WP 035346308 1 Daquatica
Dickeya sp CSL RW240 WP 067486635 1 Daquatica
DAQ1742 DAQ1742 03160 Daquatica
Dickeya aquatica DW 0440 WP 035346482 1 Daquatica

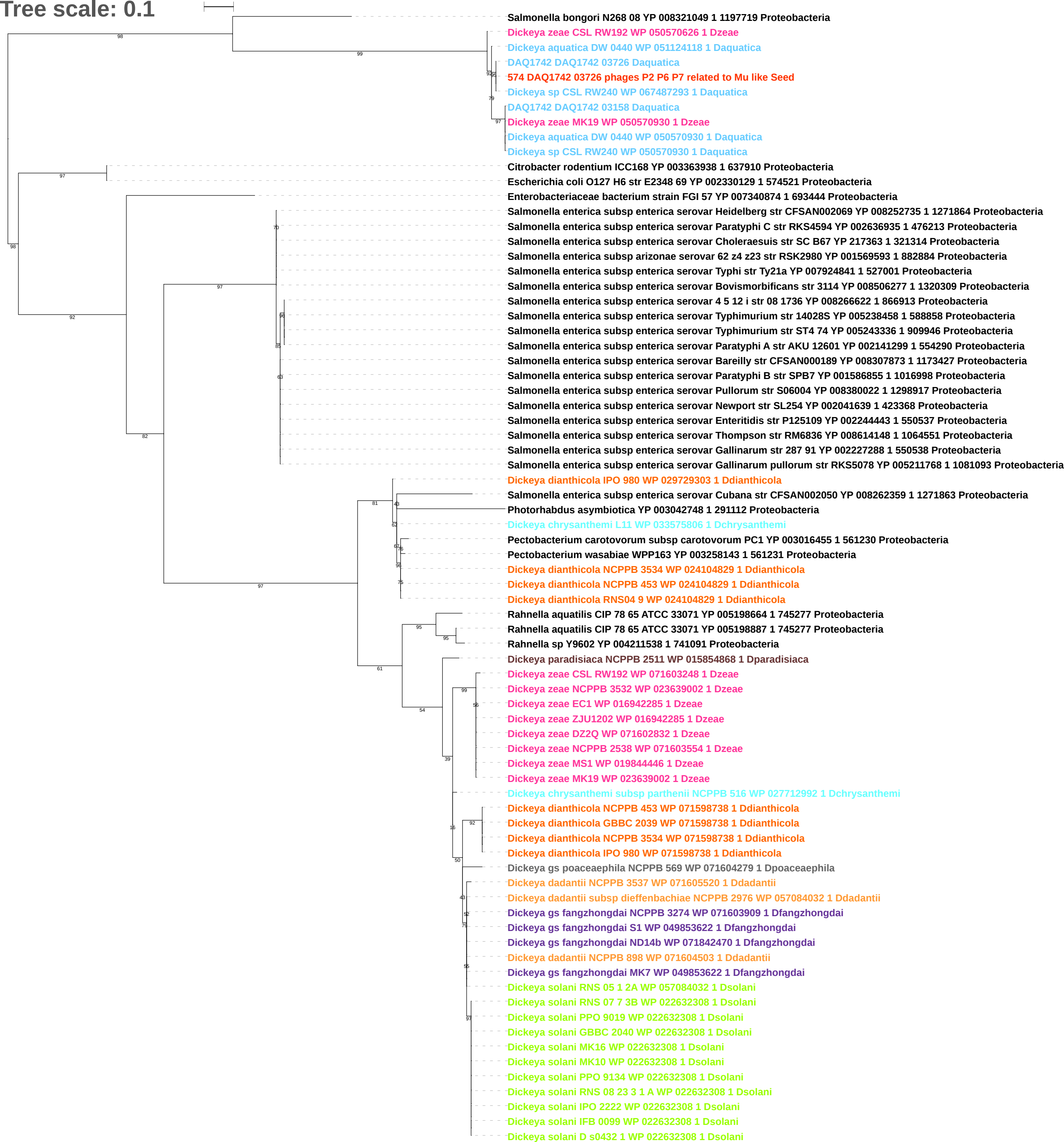
Tree scale: 0.1
Salmonella bongori N268 08 YP 008321049 1 1197719 Proteobacteria
Dickeya zeae CSL RW192 WP 050570626 1 Dzeae
Dickeya aquatica DW 0440 WP 051124118 1 Daquatica
DAQ1742 DAQ1742 03726 Daquatica
574 DAQ1742 03726 phages P2 P6 P7 related to Mu like Seed
Dickeya sp CSL RW240 WP 067487293 1 Daquatica
DAQ1742 DAQ1742 03158 Daquatica
Dickeya zeae MK19 WP 050570930 1 Dzeae
Dickeya aquatica DW 0440 WP 050570930 1 Daquatica
Dickeya sp CSL RW240 WP 050570930 1 Daquatica
Citrobacter rodentium ICC168 YP 003363938 1 637910 Proteobacteria
Escherichia coli O127 H6 str E2348 69 YP 002330129 1 574521 Proteobacteria
Enterobacteriaceae bacterium strain FGI 57 YP 007340874 1 693444 Proteobacteria
Salmonella enterica subsp enterica serovar Heidelberg str CFSAN002069 YP 008252735 1 1271864 Proteobacteria
Salmonella enterica subsp enterica serovar Paratyphi C str RKS4594 YP 002636935 1 476213 Proteobacteria
Salmonella enterica subsp enterica serovar Choleraesuis str SC B67 YP 217363 1 321314 Proteobacteria
Salmonella enterica subsp arizonae serovar 62 z4 z23 str RSK2980 YP 001569593 1 882884 Proteobacteria
Salmonella enterica subsp enterica serovar Typhi str Ty21a YP 007924841 1 527001 Proteobacteria
Salmonella enterica subsp enterica serovar Bovismorbificans str 3114 YP 008506277 1 1320309 Proteobacteria
Salmonella enterica subsp enterica serovar 4 5 12 i str 08 1736 YP 008266622 1 866913 Proteobacteria
Salmonella enterica subsp enterica serovar Typhimurium str 14028S YP 005238458 1 588858 Proteobacteria
Salmonella enterica subsp enterica serovar Typhimurium str ST4 74 YP 005243336 1 909946 Proteobacteria
Salmonella enterica subsp enterica serovar Paratyphi A str AKU 12601 YP 002141299 1 554290 Proteobacteria
Salmonella enterica subsp enterica serovar Bareilly str CFSAN000189 YP 008307873 1 1173427 Proteobacteria
Salmonella enterica subsp enterica serovar Paratyphi B str SPB7 YP 001586855 1 1016998 Proteobacteria
Salmonella enterica subsp enterica serovar Pullorum str S06004 YP 008380022 1 1298917 Proteobacteria
Salmonella enterica subsp enterica serovar Newport str SL254 YP 002041639 1 423368 Proteobacteria
Salmonella enterica subsp enterica serovar Enteritidis str P125109 YP 002244443 1 550537 Proteobacteria
Salmonella enterica subsp enterica serovar Thompson str RM6836 YP 008614148 1 1064551 Proteobacteria
Salmonella enterica subsp enterica serovar Gallinarum str 287 91 YP 002227288 1 550538 Proteobacteria
Salmonella enterica subsp enterica serovar Gallinarum pullorum str RKS5078 YP 005211768 1 1081093 Proteobacteria
Dickeya dianthicola IPO 980 WP 029729303 1 Ddianthicola
Salmonella enterica subsp enterica serovar Cubana str CFSAN002050 YP 008262359 1 1271863 Proteobacteria
Photorhabdus asymbiotica YP 003042748 1 291112 Proteobacteria
Dickeya chrysanthemi L11 WP 033575806 1 Dchrysanthemi
Pectobacterium carotovorum subsp carotovorum PC1 YP 003016455 1 561230 Proteobacteria
Pectobacterium wasabiae WPP163 YP 003258143 1 561231 Proteobacteria
Dickeya dianthicola NCPPB 3534 WP 024104829 1 Ddianthicola
Dickeya dianthicola NCPPB 453 WP 024104829 1 Ddianthicola
Dickeya dianthicola RNS04 9 WP 024104829 1 Ddianthicola
Rahnella aquatilis CIP 78 65 ATCC 33071 YP 005198664 1 745277 Proteobacteria
Rahnella aquatilis CIP 78 65 ATCC 33071 YP 005198887 1 745277 Proteobacteria
Rahnella sp Y9602 YP 004211538 1 741091 Proteobacteria
Dickeya paradisiaca NCPPB 2511 WP 015854868 1 Dparadisiaca
Dickeya zeae CSL RW192 WP 071603248 1 Dzeae
Dickeya zeae NCPPB 3532 WP 023639002 1 Dzeae
Dickeya zeae EC1 WP 016942285 1 Dzeae
Dickeya zeae ZJU1202 WP 016942285 1 Dzeae
Dickeya zeae DZ2Q WP 071602832 1 Dzeae
Dickeya zeae NCPPB 2538 WP 071603554 1 Dzeae
Dickeya zeae MS1 WP 019844446 1 Dzeae
Dickeya zeae MK19 WP 023639002 1 Dzeae
Dickeya chrysanthemi subsp parthenii NCPPB 516 WP 027712992 1 Dchrysanthemi
Dickeya dianthicola NCPPB 453 WP 071598738 1 Ddianthicola
Dickeya dianthicola GBBC 2039 WP 071598738 1 Ddianthicola
Dickeya dianthicola NCPPB 3534 WP 071598738 1 Ddianthicola
Dickeya dianthicola IPO 980 WP 071598738 1 Ddianthicola
Dickeya gs poaceaephila NCPPB 569 WP 071604279 1 Dpoaceaephila
Dickeya dadantii NCPPB 3537 WP 071605520 1 Ddadantii
Dickeya dadantii subsp dieffenbachiae NCPPB 2976 WP 057084032 1 Ddadantii
Dickeya gs fangzhongdai NCPPB 3274 WP 071603909 1 Dfangzhongdai
Dickeya gs fangzhongdai S1 WP 049853622 1 Dfangzhongdai
Dickeya gs fangzhongdai ND14b WP 071842470 1 Dfangzhongdai
Dickeya dadantii NCPPB 898 WP 071604503 1 Ddadantii
Dickeya gs fangzhongdai MK7 WP 049853622 1 Dfangzhongdai
Dickeya solani RNS 05 1 2A WP 057084032 1 Dsolani
Dickeya solani RNS 07 7 3B WP 022632308 1 Dsolani
Dickeya solani PPO 9019 WP 022632308 1 Dsolani
Dickeya solani GBBC 2040 WP 022632308 1 Dsolani
Dickeya solani MK16 WP 022632308 1 Dsolani
Dickeya solani MK10 WP 022632308 1 Dsolani
Dickeya solani PPO 9134 WP 022632308 1 Dsolani
Dickeya solani RNS 08 23 3 1 A WP 022632308 1 Dsolani
Dickeya solani IPO 2222 WP 022632308 1 Dsolani
Dickeya solani IFB 0099 WP 022632308 1 Dsolani
Dickeya solani D s0432 1 WP 022632308 1 Dsolani

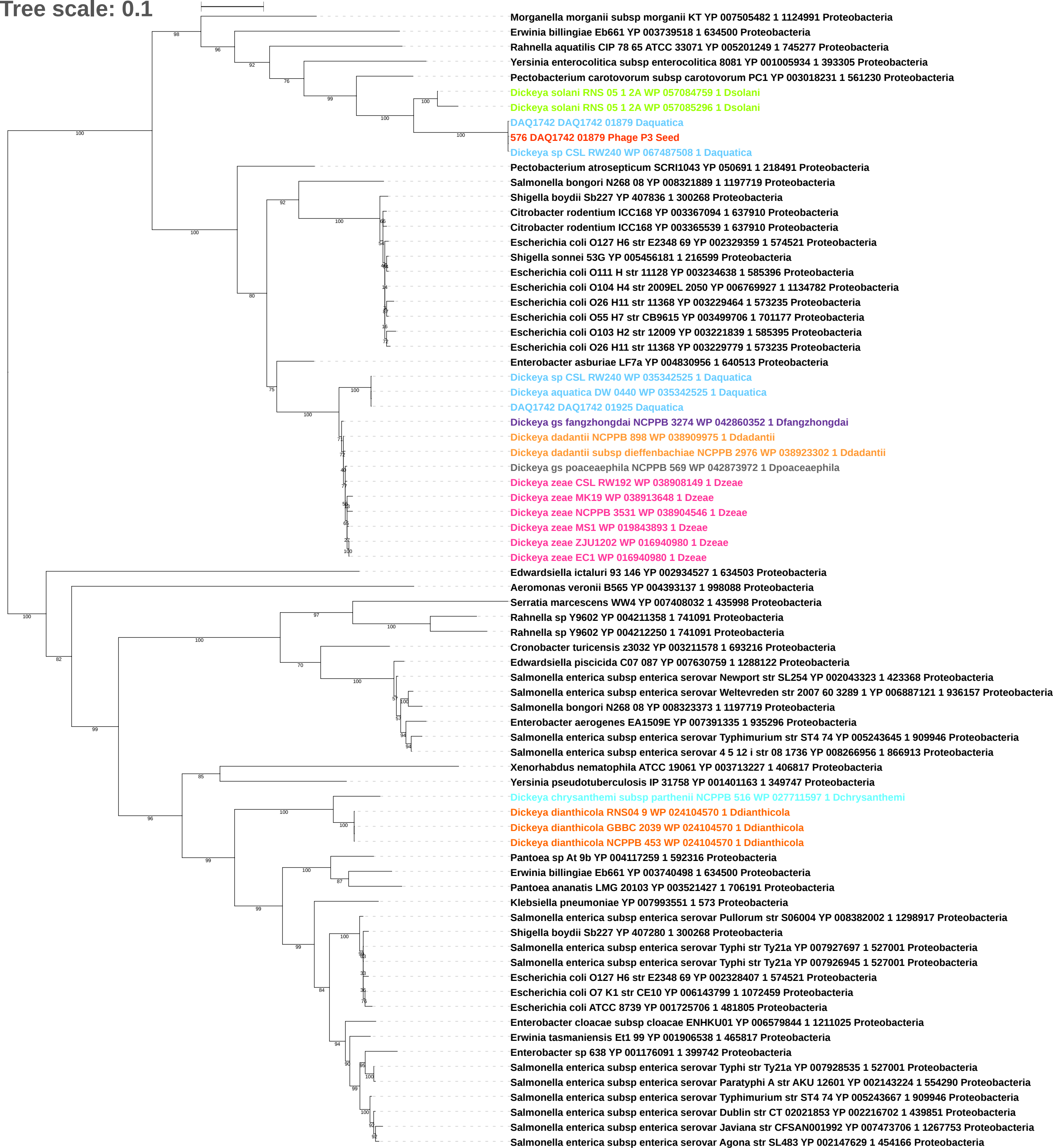
Tree scale: 0.1
Morganella morganii subsp morganii KT YP 007505482 1 1124991 Proteobacteria
Erwinia billingiae Eb661 YP 003739518 1 634500 Proteobacteria
Rahnella aquatilis CIP 78 65 ATCC 33071 YP 005201249 1 745277 Proteobacteria
Yersinia enterocolitica subsp enterocolitica 8081 YP 001005934 1 393305 Proteobacteria
Pectobacterium carotovorum subsp carotovorum PC1 YP 003018231 1 561230 Proteobacteria
Dickeya solani RNS 05 1 2A WP 057084759 1 Dsolani
Dickeya solani RNS 05 1 2A WP 057085296 1 Dsolani
DAQ1742 DAQ1742 01879 Daquatica
576 DAQ1742 01879 Phage P3 Seed
Dickeya sp CSL RW240 WP 067487508 1 Daquatica
Pectobacterium atrosepticum SCRI1043 YP 050691 1 218491 Proteobacteria
Salmonella bongori N268 08 YP 008321889 1 1197719 Proteobacteria
Shigella boydii Sb227 YP 407836 1 300268 Proteobacteria
Citrobacter rodentium ICC168 YP 003367094 1 637910 Proteobacteria
Citrobacter rodentium ICC168 YP 003365539 1 637910 Proteobacteria
Escherichia coli O127 H6 str E2348 69 YP 002329359 1 574521 Proteobacteria
Shigella sonnei 53G YP 005456181 1 216599 Proteobacteria
Escherichia coli O111 H str 11128 YP 003234638 1 585396 Proteobacteria
Escherichia coli O104 H4 str 2009EL 2050 YP 006769927 1 1134782 Proteobacteria
Escherichia coli O26 H11 str 11368 YP 003229464 1 573235 Proteobacteria
Escherichia coli O55 H7 str CB9615 YP 003499706 1 701177 Proteobacteria
Escherichia coli O103 H2 str 12009 YP 003221839 1 585395 Proteobacteria
Escherichia coli O26 H11 str 11368 YP 003229779 1 573235 Proteobacteria
Enterobacter asburiae LF7a YP 004830956 1 640513 Proteobacteria
Dickeya sp CSL RW240 WP 035342525 1 Daquatica
Dickeya aquatica DW 0440 WP 035342525 1 Daquatica
DAQ1742 DAQ1742 01925 Daquatica
Dickeya gs fangzhongdai NCPPB 3274 WP 042860352 1 Dfangzhongdai
Dickeya dadantii NCPPB 898 WP 038909975 1 Ddadantii
Dickeya dadantii subsp dieffenbachiae NCPPB 2976 WP 038923302 1 Ddadantii
Dickeya gs poaceaephila NCPPB 569 WP 042873972 1 Dpoaceaephila
Dickeya zeae CSL RW192 WP 038908149 1 Dzeae
Dickeya zeae MK19 WP 038913648 1 Dzeae
Dickeya zeae NCPPB 3531 WP 038904546 1 Dzeae
Dickeya zeae MS1 WP 019843893 1 Dzeae
Dickeya zeae ZJU1202 WP 016940980 1 Dzeae
Dickeya zeae EC1 WP 016940980 1 Dzeae
Edwardsiella ictaluri 93 146 YP 002934527 1 634503 Proteobacteria
Aeromonas veronii B565 YP 004393137 1 998088 Proteobacteria
Serratia marcescens WW4 YP 007408032 1 435998 Proteobacteria
Rahnella sp Y9602 YP 004211358 1 741091 Proteobacteria
Rahnella sp Y9602 YP 004212250 1 741091 Proteobacteria
Cronobacter turicensis z3032 YP 003211578 1 693216 Proteobacteria
Edwardsiella piscicida C07 087 YP 007630759 1 1288122 Proteobacteria
Salmonella enterica subsp enterica serovar Newport str SL254 YP 002043323 1 423368 Proteobacteria
Salmonella enterica subsp enterica serovar Weltevreden str 2007 60 3289 1 YP 006887121 1 936157 Proteobacteria
Salmonella bongori N268 08 YP 008323373 1 1197719 Proteobacteria
Enterobacter aerogenes EA1509E YP 007391335 1 935296 Proteobacteria
Salmonella enterica subsp enterica serovar Typhimurium str ST4 74 YP 005243645 1 909946 Proteobacteria
Salmonella enterica subsp enterica serovar 4 5 12 i str 08 1736 YP 008266956 1 866913 Proteobacteria
Xenorhabdus nematophila ATCC 19061 YP 003713227 1 406817 Proteobacteria
Yersinia pseudotuberculosis IP 31758 YP 001401163 1 349747 Proteobacteria
Dickeya chrysanthemi subsp parthenii NCPPB 516 WP 027711597 1 Dchrysanthemi
Dickeya dianthicola RNS04 9 WP 024104570 1 Ddianthicola
Dickeya dianthicola GBBC 2039 WP 024104570 1 Ddianthicola
Dickeya dianthicola NCPPB 453 WP 024104570 1 Ddianthicola
Pantoea sp At 9b YP 004117259 1 592316 Proteobacteria
Erwinia billingiae Eb661 YP 003740498 1 634500 Proteobacteria
Pantoea ananatis LMG 20103 YP 003521427 1 706191 Proteobacteria
Klebsiella pneumoniae YP 007993551 1 573 Proteobacteria
Salmonella enterica subsp enterica serovar Pullorum str S06004 YP 008382002 1 1298917 Proteobacteria
Shigella boydii Sb227 YP 407280 1 300268 Proteobacteria
Salmonella enterica subsp enterica serovar Typhi str Ty21a YP 007927697 1 527001 Proteobacteria
Salmonella enterica subsp enterica serovar Typhi str Ty21a YP 007926945 1 527001 Proteobacteria
Escherichia coli O127 H6 str E2348 69 YP 002328407 1 574521 Proteobacteria
Escherichia coli O7 K1 str CE10 YP 006143799 1 1072459 Proteobacteria
Escherichia coli ATCC 8739 YP 001725706 1 481805 Proteobacteria
Enterobacter cloacae subsp cloacae ENHKU01 YP 006579844 1 1211025 Proteobacteria
Erwinia tasmaniensis Et1 99 YP 001906538 1 465817 Proteobacteria
Enterobacter sp 638 YP 001176091 1 399742 Proteobacteria
Salmonella enterica subsp enterica serovar Typhi str Ty21a YP 007928535 1 527001 Proteobacteria
Salmonella enterica subsp enterica serovar Paratyphi A str AKU 12601 YP 002143224 1 554290 Proteobacteria
Salmonella enterica subsp enterica serovar Typhimurium str ST4 74 YP 005243667 1 909946 Proteobacteria
Salmonella enterica subsp enterica serovar Dublin str CT 02021853 YP 002216702 1 439851 Proteobacteria
Salmonella enterica subsp enterica serovar Javiana str CFSAN001992 YP 007473706 1 1267753 Proteobacteria
Salmonella enterica subsp enterica serovar Agona str SL483 YP 002147629 1 454166 Proteobacteria

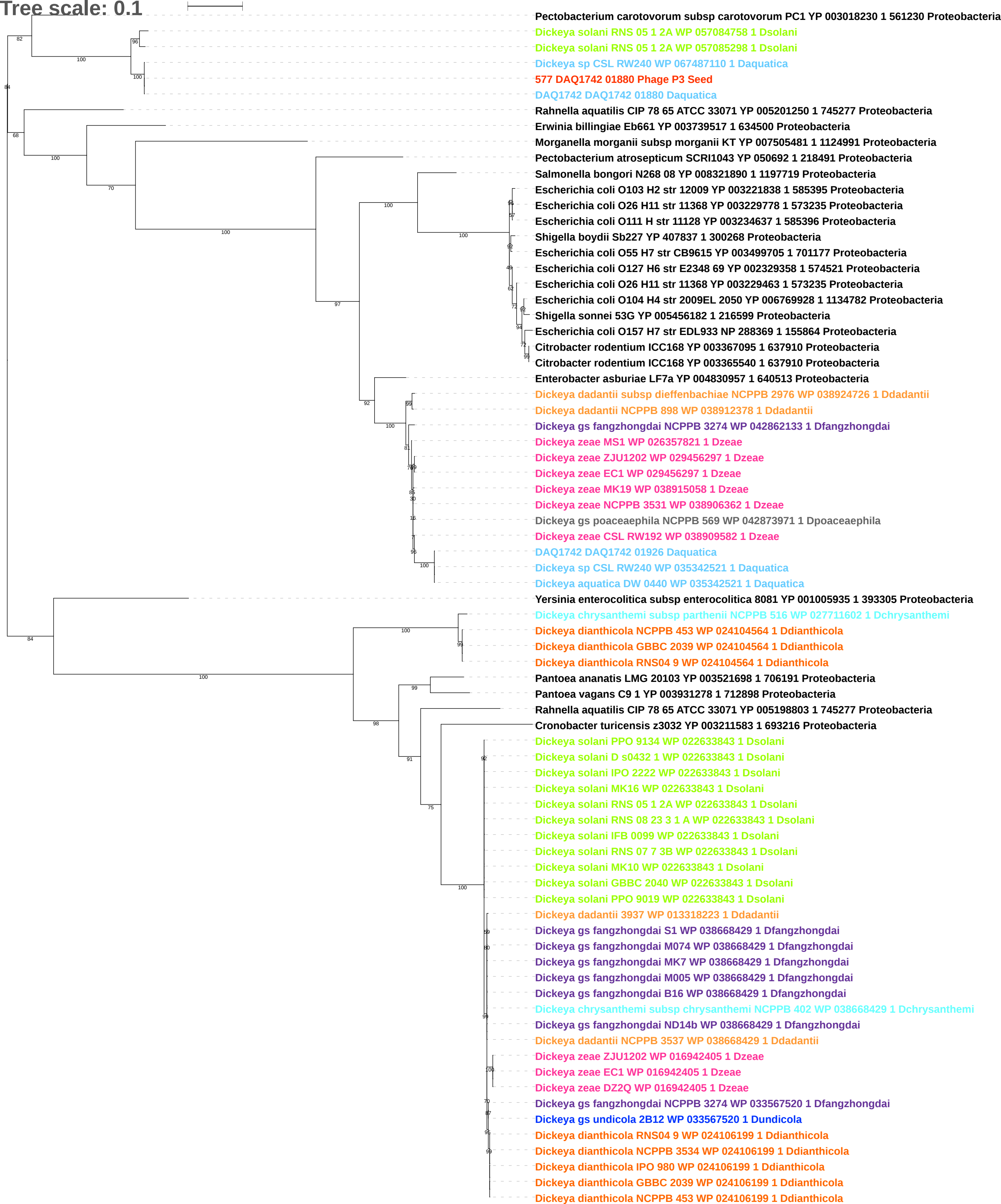
Tree scale: 0.1
Pectobacterium carotovorum subsp carotovorum PC1 YP 003018230 1 561230 Proteobacteria
Dickeya solani RNS 05 1 2A WP 057084758 1 Dsolani
Dickeya solani RNS 05 1 2A WP 057085298 1 Dsolani
Dickeya sp CSL RW240 WP 067487110 1 Daquatica
577 DAQ1742 01880 Phage P3 Seed
DAQ1742 DAQ1742 01880 Daquatica
Rahnella aquatilis CIP 78 65 ATCC 33071 YP 005201250 1 745277 Proteobacteria
Erwinia billingiae Eb661 YP 003739517 1 634500 Proteobacteria
Morganella morganii subsp morganii KT YP 007505481 1 1124991 Proteobacteria
Pectobacterium atrosepticum SCRI1043 YP 050692 1 218491 Proteobacteria
Salmonella bongori N268 08 YP 008321890 1 1197719 Proteobacteria
Escherichia coli O103 H2 str 12009 YP 003221838 1 585395 Proteobacteria
Escherichia coli O26 H11 str 11368 YP 003229778 1 573235 Proteobacteria
Escherichia coli O111 H str 11128 YP 003234637 1 585396 Proteobacteria
Shigella boydii Sb227 YP 407837 1 300268 Proteobacteria
Escherichia coli O55 H7 str CB9615 YP 003499705 1 701177 Proteobacteria
Escherichia coli O127 H6 str E2348 69 YP 002329358 1 574521 Proteobacteria
Escherichia coli O26 H11 str 11368 YP 003229463 1 573235 Proteobacteria
Escherichia coli O104 H4 str 2009EL 2050 YP 006769928 1 1134782 Proteobacteria
Shigella sonnei 53G YP 005456182 1 216599 Proteobacteria
Escherichia coli O157 H7 str EDL933 NP 288369 1 155864 Proteobacteria
Citrobacter rodentium ICC168 YP 003367095 1 637910 Proteobacteria
Citrobacter rodentium ICC168 YP 003365540 1 637910 Proteobacteria
Enterobacter asburiae LF7a YP 004830957 1 640513 Proteobacteria
Dickeya dadantii subsp dieffenbachiae NCPPB 2976 WP 038924726 1 Ddadantii
Dickeya dadantii NCPPB 898 WP 038912378 1 Ddadantii
Dickeya gs fangzhongdai NCPPB 3274 WP 042862133 1 Dfangzhongdai
Dickeya zeae MS1 WP 026357821 1 Dzeae
Dickeya zeae ZJU1202 WP 029456297 1 Dzeae
Dickeya zeae EC1 WP 029456297 1 Dzeae
Dickeya zeae MK19 WP 038915058 1 Dzeae
Dickeya zeae NCPPB 3531 WP 038906362 1 Dzeae
Dickeya gs poaceaephila NCPPB 569 WP 042873971 1 Dpoaceaephila
Dickeya zeae CSL RW192 WP 038909582 1 Dzeae
DAQ1742 DAQ1742 01926 Daquatica
Dickeya sp CSL RW240 WP 035342521 1 Daquatica
Dickeya aquatica DW 0440 WP 035342521 1 Daquatica
Yersinia enterocolitica subsp enterocolitica 8081 YP 001005935 1 393305 Proteobacteria
Dickeya chrysanthemi subsp parthenii NCPPB 516 WP 027711602 1 Dchrysanthemi
Dickeya dianthicola NCPPB 453 WP 024104564 1 Ddianthicola
Dickeya dianthicola GBBC 2039 WP 024104564 1 Ddianthicola
Dickeya dianthicola RNS04 9 WP 024104564 1 Ddianthicola
Pantoea ananatis LMG 20103 YP 003521698 1 706191 Proteobacteria
Pantoea vagans C9 1 YP 003931278 1 712898 Proteobacteria
Rahnella aquatilis CIP 78 65 ATCC 33071 YP 005198803 1 745277 Proteobacteria
Cronobacter turicensis z3032 YP 003211583 1 693216 Proteobacteria
Dickeya solani PPO 9134 WP 022633843 1 Dsolani
Dickeya solani D s0432 1 WP 022633843 1 Dsolani
Dickeya solani IPO 2222 WP 022633843 1 Dsolani
Dickeya solani MK16 WP 022633843 1 Dsolani
Dickeya solani RNS 05 1 2A WP 022633843 1 Dsolani
Dickeya solani RNS 08 23 3 1 A WP 022633843 1 Dsolani
Dickeya solani IFB 0099 WP 022633843 1 Dsolani
Dickeya solani RNS 07 7 3B WP 022633843 1 Dsolani
Dickeya solani MK10 WP 022633843 1 Dsolani
Dickeya solani GBBC 2040 WP 022633843 1 Dsolani
Dickeya solani PPO 9019 WP 022633843 1 Dsolani
Dickeya dadantii 3937 WP 013318223 1 Ddadantii
Dickeya gs fangzhongdai S1 WP 038668429 1 Dfangzhongdai
Dickeya gs fangzhongdai M074 WP 038668429 1 Dfangzhongdai
Dickeya gs fangzhongdai MK7 WP 038668429 1 Dfangzhongdai
Dickeya gs fangzhongdai M005 WP 038668429 1 Dfangzhongdai
Dickeya gs fangzhongdai B16 WP 038668429 1 Dfangzhongdai
Dickeya chrysanthemi subsp chrysanthemi NCPPB 402 WP 038668429 1 Dchrysanthemi
Dickeya gs fangzhongdai ND14b WP 038668429 1 Dfangzhongdai
Dickeya dadantii NCPPB 3537 WP 038668429 1 Ddadantii
Dickeya zeae ZJU1202 WP 016942405 1 Dzeae
Dickeya zeae EC1 WP 016942405 1 Dzeae
Dickeya zeae DZ2Q WP 016942405 1 Dzeae
Dickeya gs fangzhongdai NCPPB 3274 WP 033567520 1 Dfangzhongdai
Dickeya gs undicola 2B12 WP 033567520 1 Dundicola
Dickeya dianthicola RNS04 9 WP 024106199 1 Ddianthicola
Dickeya dianthicola NCPPB 3534 WP 024106199 1 Ddianthicola
Dickeya dianthicola IPO 980 WP 024106199 1 Ddianthicola
Dickeya dianthicola GBBC 2039 WP 024106199 1 Ddianthicola
Dickeya dianthicola NCPPB 453 WP 024106199 1 Ddianthicola

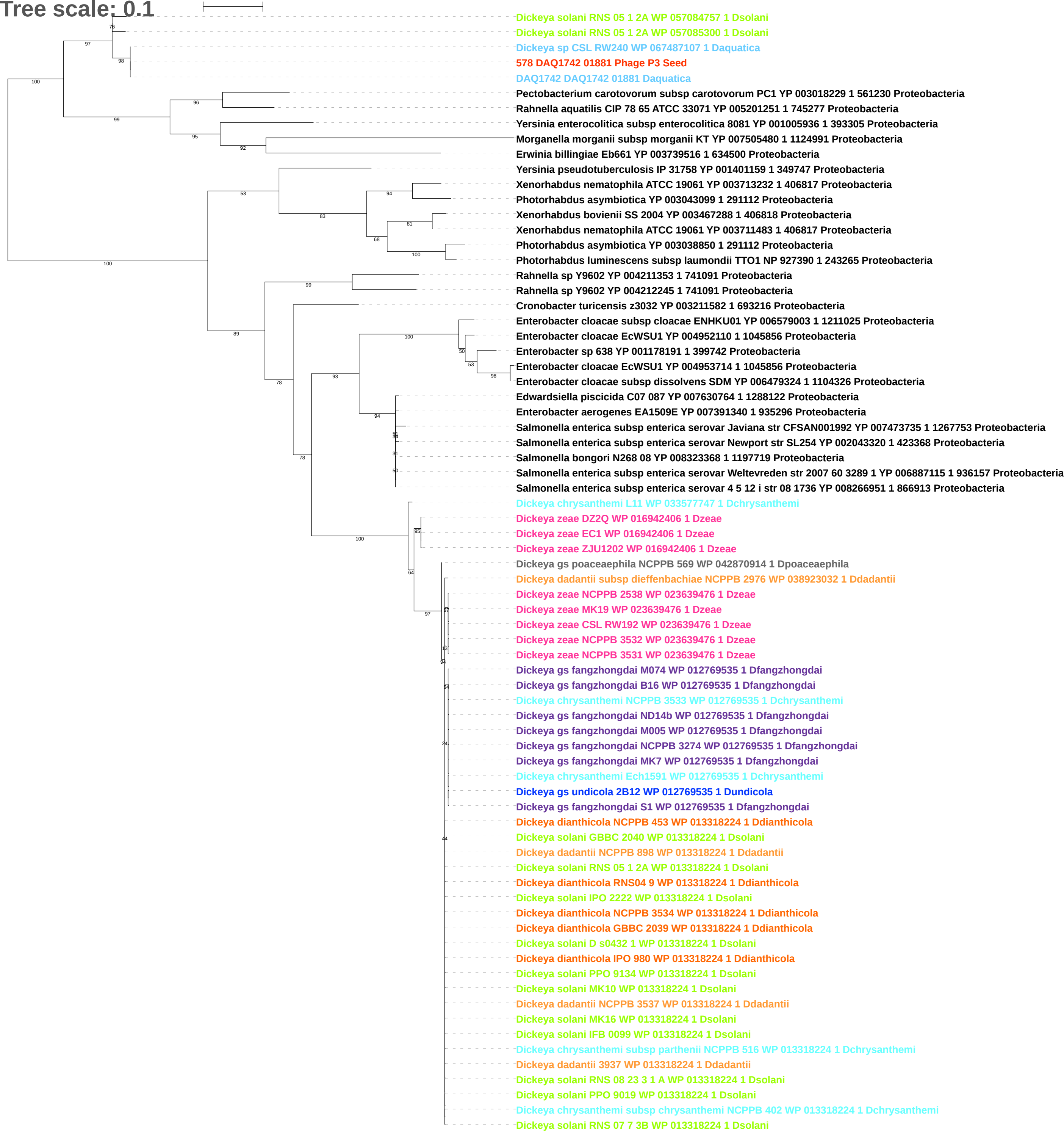
Tree scale: 0.1
Dickeya solani RNS 05 1 2A WP 057084757 1 Dsolani
Dickeya solani RNS 05 1 2A WP 057085300 1 Dsolani
Dickeya sp CSL RW240 WP 067487107 1 Daquatica
578 DAQ1742 01881 Phage P3 Seed
DAQ1742 DAQ1742 01881 Daquatica
Pectobacterium carotovorum subsp carotovorum PC1 YP 003018229 1 561230 Proteobacteria
Rahnella aquatilis CIP 78 65 ATCC 33071 YP 005201251 1 745277 Proteobacteria
Yersinia enterocolitica subsp enterocolitica 8081 YP 001005936 1 393305 Proteobacteria
Morganella morganii subsp morganii KT YP 007505480 1 1124991 Proteobacteria
Erwinia billingiae Eb661 YP 003739516 1 634500 Proteobacteria
Yersinia pseudotuberculosis IP 31758 YP 001401159 1 349747 Proteobacteria
Xenorhabdus nematophila ATCC 19061 YP 003713232 1 406817 Proteobacteria
Photorhabdus asymbiotica YP 003043099 1 291112 Proteobacteria
Xenorhabdus bovienii SS 2004 YP 003467288 1 406818 Proteobacteria
Xenorhabdus nematophila ATCC 19061 YP 003711483 1 406817 Proteobacteria
Photorhabdus asymbiotica YP 003038850 1 291112 Proteobacteria
Photorhabdus luminescens subsp laumondii TTO1 NP 927390 1 243265 Proteobacteria
Rahnella sp Y9602 YP 004211353 1 741091 Proteobacteria
Rahnella sp Y9602 YP 004212245 1 741091 Proteobacteria
Cronobacter turicensis z3032 YP 003211582 1 693216 Proteobacteria
Enterobacter cloacae subsp cloacae ENHKU01 YP 006579003 1 1211025 Proteobacteria
Enterobacter cloacae EcWSU1 YP 004952110 1 1045856 Proteobacteria
Enterobacter sp 638 YP 001178191 1 399742 Proteobacteria
Enterobacter cloacae EcWSU1 YP 004953714 1 1045856 Proteobacteria
Enterobacter cloacae subsp dissolvens SDM YP 006479324 1 1104326 Proteobacteria
Edwardsiella piscicida C07 087 YP 007630764 1 1288122 Proteobacteria
Enterobacter aerogenes EA1509E YP 007391340 1 935296 Proteobacteria
Salmonella enterica subsp enterica serovar Javiana str CFSAN001992 YP 007473735 1 1267753 Proteobacteria
Salmonella enterica subsp enterica serovar Newport str SL254 YP 002043320 1 423368 Proteobacteria
Salmonella bongori N268 08 YP 008323368 1 1197719 Proteobacteria
Salmonella enterica subsp enterica serovar Weltevreden str 2007 60 3289 1 YP 006887115 1 936157 Proteobacteria
Salmonella enterica subsp enterica serovar 4 5 12 i str 08 1736 YP 008266951 1 866913 Proteobacteria
Dickeya chrysanthemi L11 WP 033577747 1 Dchrysanthemi
Dickeya zeae DZ2Q WP 016942406 1 Dzeae
Dickeya zeae EC1 WP 016942406 1 Dzeae
Dickeya zeae ZJU1202 WP 016942406 1 Dzeae
Dickeya gs poaceaephila NCPPB 569 WP 042870914 1 Dpoaceaephila
Dickeya dadantii subsp dieffenbachiae NCPPB 2976 WP 038923032 1 Ddadantii
Dickeya zeae NCPPB 2538 WP 023639476 1 Dzeae
Dickeya zeae MK19 WP 023639476 1 Dzeae
Dickeya zeae CSL RW192 WP 023639476 1 Dzeae
Dickeya zeae NCPPB 3532 WP 023639476 1 Dzeae
Dickeya zeae NCPPB 3531 WP 023639476 1 Dzeae
Dickeya gs fangzhongdai M074 WP 012769535 1 Dfangzhongdai
Dickeya gs fangzhongdai B16 WP 012769535 1 Dfangzhongdai
Dickeya chrysanthemi NCPPB 3533 WP 012769535 1 Dchrysanthemi
Dickeya gs fangzhongdai ND14b WP 012769535 1 Dfangzhongdai
Dickeya gs fangzhongdai M005 WP 012769535 1 Dfangzhongdai
Dickeya gs fangzhongdai NCPPB 3274 WP 012769535 1 Dfangzhongdai
Dickeya gs fangzhongdai MK7 WP 012769535 1 Dfangzhongdai
Dickeya chrysanthemi Ech1591 WP 012769535 1 Dchrysanthemi
Dickeya gs undicola 2B12 WP 012769535 1 Dundicola
Dickeya gs fangzhongdai S1 WP 012769535 1 Dfangzhongdai
Dickeya dianthicola NCPPB 453 WP 013318224 1 Ddianthicola
Dickeya solani GBBC 2040 WP 013318224 1 Dsolani
Dickeya dadantii NCPPB 898 WP 013318224 1 Ddadantii
Dickeya solani RNS 05 1 2A WP 013318224 1 Dsolani
Dickeya dianthicola RNS04 9 WP 013318224 1 Ddianthicola
Dickeya solani IPO 2222 WP 013318224 1 Dsolani
Dickeya dianthicola NCPPB 3534 WP 013318224 1 Ddianthicola
Dickeya dianthicola GBBC 2039 WP 013318224 1 Ddianthicola
Dickeya solani D s0432 1 WP 013318224 1 Dsolani
Dickeya dianthicola IPO 980 WP 013318224 1 Ddianthicola
Dickeya solani PPO 9134 WP 013318224 1 Dsolani
Dickeya solani MK10 WP 013318224 1 Dsolani
Dickeya dadantii NCPPB 3537 WP 013318224 1 Ddadantii
Dickeya solani MK16 WP 013318224 1 Dsolani
Dickeya solani IFB 0099 WP 013318224 1 Dsolani
Dickeya chrysanthemi subsp parthenii NCPPB 516 WP 013318224 1 Dchrysanthemi
Dickeya dadantii 3937 WP 013318224 1 Ddadantii
Dickeya solani RNS 08 23 3 1 A WP 013318224 1 Dsolani
Dickeya solani PPO 9019 WP 013318224 1 Dsolani
Dickeya chrysanthemi subsp chrysanthemi NCPPB 402 WP 013318224 1 Dchrysanthemi
Dickeya solani RNS 07 7 3B WP 013318224 1 Dsolani

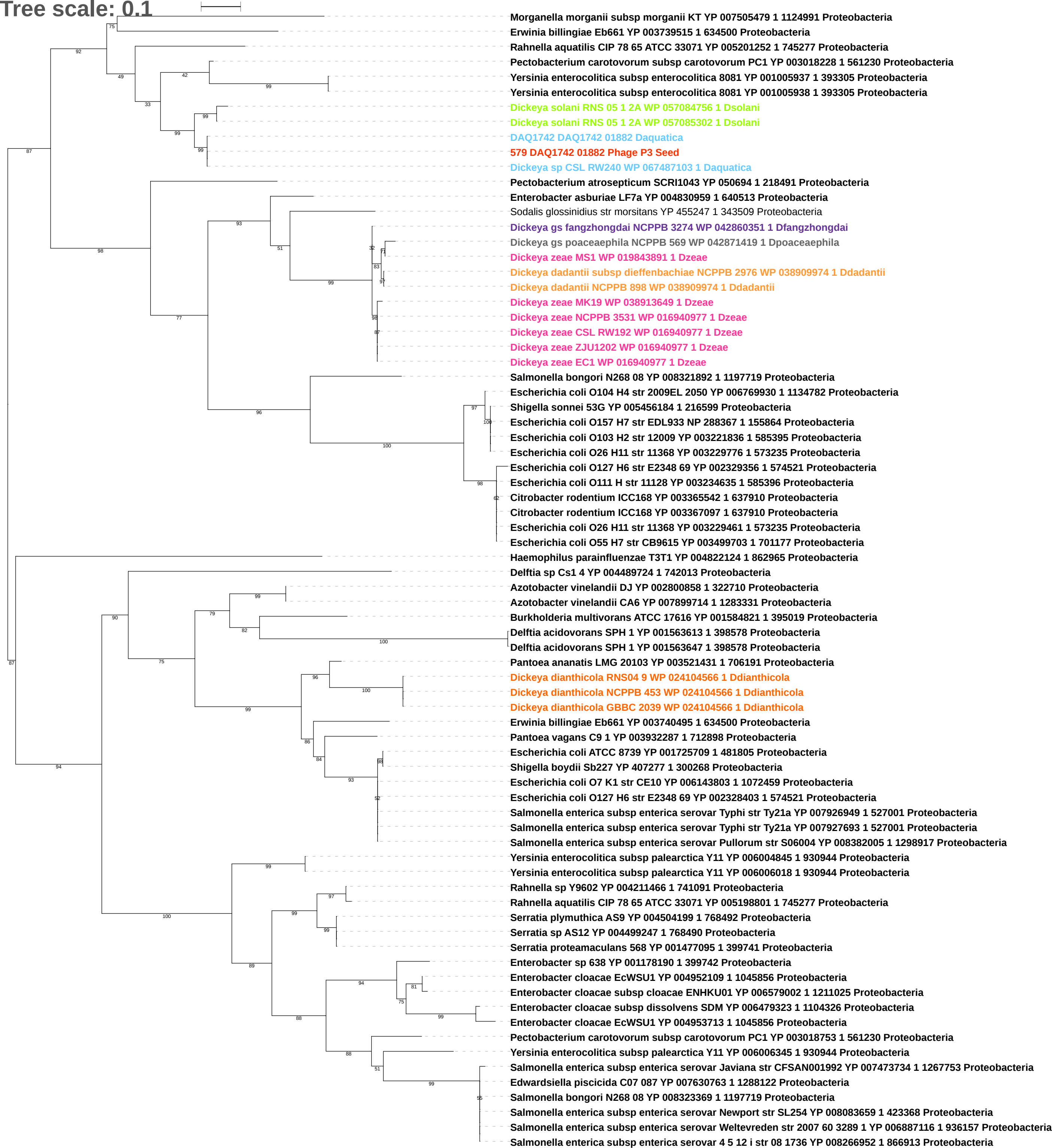
Tree scale: 0.1
Morganella morganii subsp morganii KT YP 007505479 1 1124991 Proteobacteria
Erwinia billingiae Eb661 YP 003739515 1 634500 Proteobacteria
Rahnella aquatilis CIP 78 65 ATCC 33071 YP 005201252 1 745277 Proteobacteria
Pectobacterium carotovorum subsp carotovorum PC1 YP 003018228 1 561230 Proteobacteria
Yersinia enterocolitica subsp enterocolitica 8081 YP 001005937 1 393305 Proteobacteria
Yersinia enterocolitica subsp enterocolitica 8081 YP 001005938 1 393305 Proteobacteria
Dickeya solani RNS 05 1 2A WP 057084756 1 Dsolani
Dickeya solani RNS 05 1 2A WP 057085302 1 Dsolani
DAQ1742 DAQ1742 01882 Daquatica
579 DAQ1742 01882 Phage P3 Seed
Dickeya sp CSL RW240 WP 067487103 1 Daquatica
Pectobacterium atrosepticum SCRI1043 YP 050694 1 218491 Proteobacteria
Enterobacter asburiae LF7a YP 004830959 1 640513 Proteobacteria
Sodalis glossinidius str morsitans YP 455247 1 343509 Proteobacteria
Dickeya gs fangzhongdai NCPPB 3274 WP 042860351 1 Dfangzhongdai
Dickeya gs poaceaephila NCPPB 569 WP 042871419 1 Dpoaceaephila
Dickeya zeae MS1 WP 019843891 1 Dzeae
Dickeya dadantii subsp dieffenbachiae NCPPB 2976 WP 038909974 1 Ddadantii
Dickeya dadantii NCPPB 898 WP 038909974 1 Ddadantii
Dickeya zeae MK19 WP 038913649 1 Dzeae
Dickeya zeae NCPPB 3531 WP 016940977 1 Dzeae
Dickeya zeae CSL RW192 WP 016940977 1 Dzeae
Dickeya zeae ZJU1202 WP 016940977 1 Dzeae
Dickeya zeae EC1 WP 016940977 1 Dzeae
Salmonella bongori N268 08 YP 008321892 1 1197719 Proteobacteria
Escherichia coli O104 H4 str 2009EL 2050 YP 006769930 1 1134782 Proteobacteria
Shigella sonnei 53G YP 005456184 1 216599 Proteobacteria
Escherichia coli O157 H7 str EDL933 NP 288367 1 155864 Proteobacteria
Escherichia coli O103 H2 str 12009 YP 003221836 1 585395 Proteobacteria
Escherichia coli O26 H11 str 11368 YP 003229776 1 573235 Proteobacteria
Escherichia coli O127 H6 str E2348 69 YP 002329356 1 574521 Proteobacteria
Escherichia coli O111 H str 11128 YP 003234635 1 585396 Proteobacteria
Citrobacter rodentium ICC168 YP 003365542 1 637910 Proteobacteria
Citrobacter rodentium ICC168 YP 003367097 1 637910 Proteobacteria
Escherichia coli O26 H11 str 11368 YP 003229461 1 573235 Proteobacteria
Escherichia coli O55 H7 str CB9615 YP 003499703 1 701177 Proteobacteria
Haemophilus parainfluenzae T3T1 YP 004822124 1 862965 Proteobacteria
Delftia sp Cs1 4 YP 004489724 1 742013 Proteobacteria
Azotobacter vinelandii DJ YP 002800858 1 322710 Proteobacteria
Azotobacter vinelandii CA6 YP 007899714 1 1283331 Proteobacteria
Burkholderia multivorans ATCC 17616 YP 001584821 1 395019 Proteobacteria
Delftia acidovorans SPH 1 YP 001563613 1 398578 Proteobacteria
Delftia acidovorans SPH 1 YP 001563647 1 398578 Proteobacteria
Pantoea ananatis LMG 20103 YP 003521431 1 706191 Proteobacteria
Dickeya dianthicola RNS04 9 WP 024104566 1 Ddianthicola
Dickeya dianthicola NCPPB 453 WP 024104566 1 Ddianthicola
Dickeya dianthicola GBBC 2039 WP 024104566 1 Ddianthicola
Erwinia billingiae Eb661 YP 003740495 1 634500 Proteobacteria
Pantoea vagans C9 1 YP 003932287 1 712898 Proteobacteria
Escherichia coli ATCC 8739 YP 001725709 1 481805 Proteobacteria
Shigella boydii Sb227 YP 407277 1 300268 Proteobacteria
Escherichia coli O7 K1 str CE10 YP 006143803 1 1072459 Proteobacteria
Escherichia coli O127 H6 str E2348 69 YP 002328403 1 574521 Proteobacteria
Salmonella enterica subsp enterica serovar Typhi str Ty21a YP 007926949 1 527001 Proteobacteria
Salmonella enterica subsp enterica serovar Typhi str Ty21a YP 007927693 1 527001 Proteobacteria
Salmonella enterica subsp enterica serovar Pullorum str S06004 YP 008382005 1 1298917 Proteobacteria
Yersinia enterocolitica subsp palearctica Y11 YP 006004845 1 930944 Proteobacteria
Yersinia enterocolitica subsp palearctica Y11 YP 006006018 1 930944 Proteobacteria
Rahnella sp Y9602 YP 004211466 1 741091 Proteobacteria
Rahnella aquatilis CIP 78 65 ATCC 33071 YP 005198801 1 745277 Proteobacteria
Serratia plymuthica AS9 YP 004504199 1 768492 Proteobacteria
Serratia sp AS12 YP 004499247 1 768490 Proteobacteria
Serratia proteamaculans 568 YP 001477095 1 399741 Proteobacteria
Enterobacter sp 638 YP 001178190 1 399742 Proteobacteria
Enterobacter cloacae EcWSU1 YP 004952109 1 1045856 Proteobacteria
Enterobacter cloacae subsp cloacae ENHKU01 YP 006579002 1 1211025 Proteobacteria
Enterobacter cloacae subsp dissolvens SDM YP 006479323 1 1104326 Proteobacteria
Enterobacter cloacae EcWSU1 YP 004953713 1 1045856 Proteobacteria
Pectobacterium carotovorum subsp carotovorum PC1 YP 003018753 1 561230 Proteobacteria
Yersinia enterocolitica subsp palearctica Y11 YP 006006345 1 930944 Proteobacteria
Salmonella enterica subsp enterica serovar Javiana str CFSAN001992 YP 007473734 1 1267753 Proteobacteria
Edwardsiella piscicida C07 087 YP 007630763 1 1288122 Proteobacteria
Salmonella bongori N268 08 YP 008323369 1 1197719 Proteobacteria
Salmonella enterica subsp enterica serovar Newport str SL254 YP 008083659 1 423368 Proteobacteria
Salmonella enterica subsp enterica serovar Weltevreden str 2007 60 3289 1 YP 006887116 1 936157 Proteobacteria
Salmonella enterica subsp enterica serovar 4 5 12 i str 08 1736 YP 008266952 1 866913 Proteobacteria

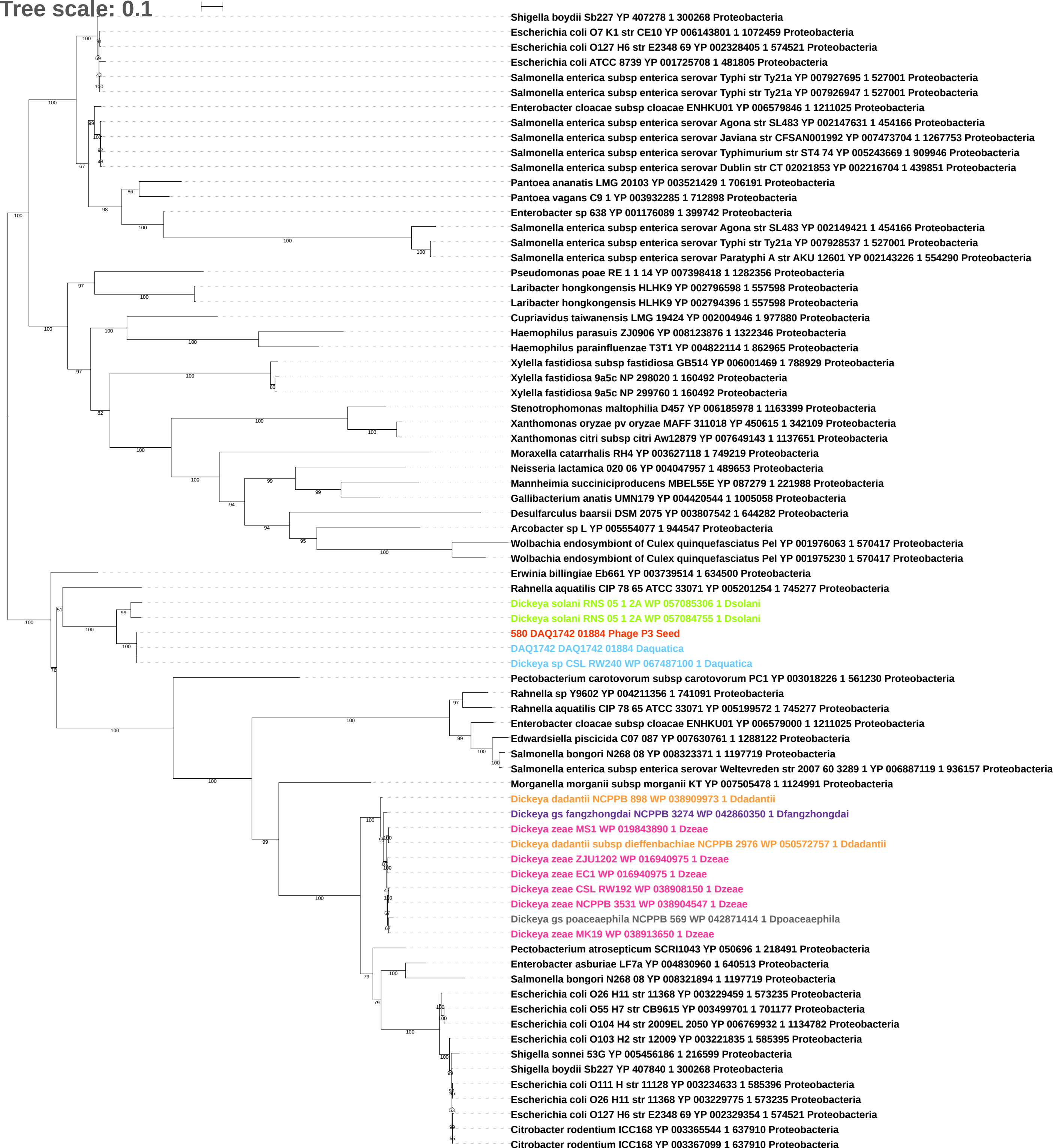
Tree scale: 0.1
Shigella boydii Sb227 YP 407278 1 300268 Proteobacteria
Escherichia coli O7 K1 str CE10 YP 006143801 1 1072459 Proteobacteria
Escherichia coli O127 H6 str E2348 69 YP 002328405 1 574521 Proteobacteria
Escherichia coli ATCC 8739 YP 001725708 1 481805 Proteobacteria
Salmonella enterica subsp enterica serovar Typhi str Ty21a YP 007927695 1 527001 Proteobacteria
Salmonella enterica subsp enterica serovar Typhi str Ty21a YP 007926947 1 527001 Proteobacteria
Enterobacter cloacae subsp cloacae ENHKU01 YP 006579846 1 1211025 Proteobacteria
Salmonella enterica subsp enterica serovar Agona str SL483 YP 002147631 1 454166 Proteobacteria
Salmonella enterica subsp enterica serovar Javiana str CFSAN001992 YP 007473704 1 1267753 Proteobacteria
Salmonella enterica subsp enterica serovar Typhimurium str ST4 74 YP 005243669 1 909946 Proteobacteria
Salmonella enterica subsp enterica serovar Dublin str CT 02021853 YP 002216704 1 439851 Proteobacteria
Pantoea ananatis LMG 20103 YP 003521429 1 706191 Proteobacteria
Pantoea vagans C9 1 YP 003932285 1 712898 Proteobacteria
Enterobacter sp 638 YP 001176089 1 399742 Proteobacteria
Salmonella enterica subsp enterica serovar Agona str SL483 YP 002149421 1 454166 Proteobacteria
Salmonella enterica subsp enterica serovar Typhi str Ty21a YP 007928537 1 527001 Proteobacteria
Salmonella enterica subsp enterica serovar Paratyphi A str AKU 12601 YP 002143226 1 554290 Proteobacteria
Pseudomonas poae RE 1 1 14 YP 007398418 1 1282356 Proteobacteria
Laribacter hongkongensis HLHK9 YP 002796598 1 557598 Proteobacteria
Laribacter hongkongensis HLHK9 YP 002794396 1 557598 Proteobacteria
Cupriavidus taiwanensis LMG 19424 YP 002004946 1 977880 Proteobacteria
Haemophilus parasuis ZJ0906 YP 008123876 1 1322346 Proteobacteria
Haemophilus parainfluenzae T3T1 YP 004822114 1 862965 Proteobacteria
Xylella fastidiosa subsp fastidiosa GB514 YP 006001469 1 788929 Proteobacteria
Xylella fastidiosa 9a5c NP 298020 1 160492 Proteobacteria
Xylella fastidiosa 9a5c NP 299760 1 160492 Proteobacteria
Stenotrophomonas maltophilia D457 YP 006185978 1 1163399 Proteobacteria
Xanthomonas oryzae pv oryzae MAFF 311018 YP 450615 1 342109 Proteobacteria
Xanthomonas citri subsp citri Aw12879 YP 007649143 1 1137651 Proteobacteria
Moraxella catarrhalis RH4 YP 003627118 1 749219 Proteobacteria
Neisseria lactamica 020 06 YP 004047957 1 489653 Proteobacteria
Mannheimia succiniciproducens MBEL55E YP 087279 1 221988 Proteobacteria
Gallibacterium anatis UMN179 YP 004420544 1 1005058 Proteobacteria
Desulfarculus baarsii DSM 2075 YP 003807542 1 644282 Proteobacteria
Arcobacter sp L YP 005554077 1 944547 Proteobacteria
Wolbachia endosymbiont of Culex quinquefasciatus Pel YP 001976063 1 570417 Proteobacteria
Wolbachia endosymbiont of Culex quinquefasciatus Pel YP 001975230 1 570417 Proteobacteria
Erwinia billingiae Eb661 YP 003739514 1 634500 Proteobacteria
Rahnella aquatilis CIP 78 65 ATCC 33071 YP 005201254 1 745277 Proteobacteria
Dickeya solani RNS 05 1 2A WP 057085306 1 Dsolani
Dickeya solani RNS 05 1 2A WP 057084755 1 Dsolani
580 DAQ1742 01884 Phage P3 Seed
DAQ1742 DAQ1742 01884 Daquatica
Dickeya sp CSL RW240 WP 067487100 1 Daquatica
Pectobacterium carotovorum subsp carotovorum PC1 YP 003018226 1 561230 Proteobacteria
Rahnella sp Y9602 YP 004211356 1 741091 Proteobacteria
Rahnella aquatilis CIP 78 65 ATCC 33071 YP 005199572 1 745277 Proteobacteria
Enterobacter cloacae subsp cloacae ENHKU01 YP 006579000 1 1211025 Proteobacteria
Edwardsiella piscicida C07 087 YP 007630761 1 1288122 Proteobacteria
Salmonella bongori N268 08 YP 008323371 1 1197719 Proteobacteria
Salmonella enterica subsp enterica serovar Weltevreden str 2007 60 3289 1 YP 006887119 1 936157 Proteobacteria
Morganella morganii subsp morganii KT YP 007505478 1 1124991 Proteobacteria
Dickeya dadantii NCPPB 898 WP 038909973 1 Ddadantii
Dickeya gs fangzhongdai NCPPB 3274 WP 042860350 1 Dfangzhongdai
Dickeya zeae MS1 WP 019843890 1 Dzeae
Dickeya dadantii subsp dieffenbachiae NCPPB 2976 WP 050572757 1 Ddadantii
Dickeya zeae ZJU1202 WP 016940975 1 Dzeae
Dickeya zeae EC1 WP 016940975 1 Dzeae
Dickeya zeae CSL RW192 WP 038908150 1 Dzeae
Dickeya zeae NCPPB 3531 WP 038904547 1 Dzeae
Dickeya gs poaceaephila NCPPB 569 WP 042871414 1 Dpoaceaephila
Dickeya zeae MK19 WP 038913650 1 Dzeae
Pectobacterium atrosepticum SCRI1043 YP 050696 1 218491 Proteobacteria
Enterobacter asburiae LF7a YP 004830960 1 640513 Proteobacteria
Salmonella bongori N268 08 YP 008321894 1 1197719 Proteobacteria
Escherichia coli O26 H11 str 11368 YP 003229459 1 573235 Proteobacteria
Escherichia coli O55 H7 str CB9615 YP 003499701 1 701177 Proteobacteria
Escherichia coli O104 H4 str 2009EL 2050 YP 006769932 1 1134782 Proteobacteria
Escherichia coli O103 H2 str 12009 YP 003221835 1 585395 Proteobacteria
Shigella sonnei 53G YP 005456186 1 216599 Proteobacteria
Shigella boydii Sb227 YP 407840 1 300268 Proteobacteria
Escherichia coli O111 H str 11128 YP 003234633 1 585396 Proteobacteria
Escherichia coli O26 H11 str 11368 YP 003229775 1 573235 Proteobacteria
Escherichia coli O127 H6 str E2348 69 YP 002329354 1 574521 Proteobacteria
Citrobacter rodentium ICC168 YP 003365544 1 637910 Proteobacteria
Citrobacter rodentium ICC168 YP 003367099 1 637910 Proteobacteria

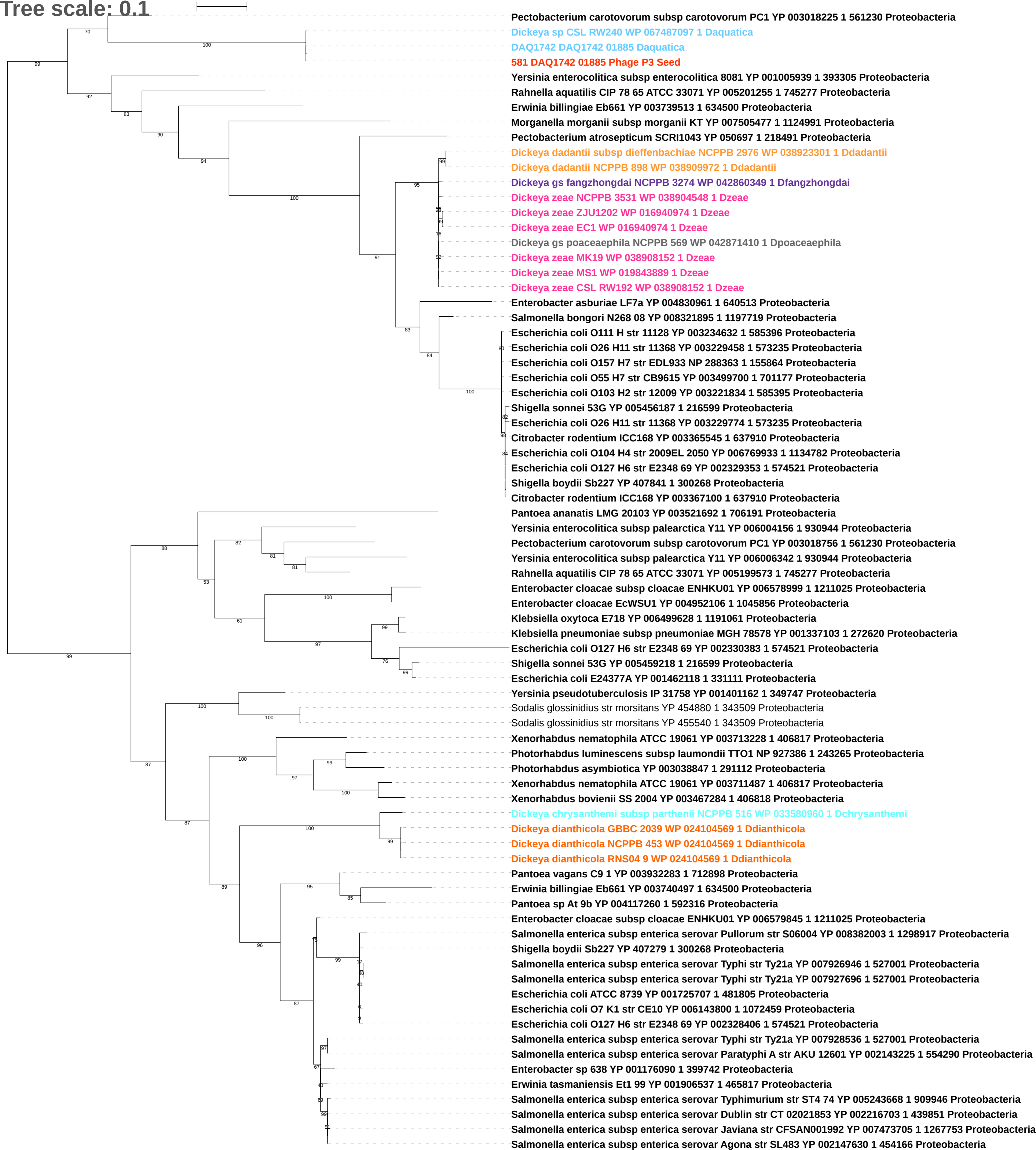
Tree scale: 0.1
Pectobacterium carotovorum subsp carotovorum PC1 YP 003018225 1 561230 Proteobacteria
Dickeya sp CSL RW240 WP 067487097 1 Daquatica
DAQ1742 DAQ1742 01885 Daquatica
581 DAQ1742 01885 Phage P3 Seed
Yersinia enterocolitica subsp enterocolitica 8081 YP 001005939 1 393305 Proteobacteria
Rahnella aquatilis CIP 78 65 ATCC 33071 YP 005201255 1 745277 Proteobacteria
Erwinia billingiae Eb661 YP 003739513 1 634500 Proteobacteria
Morganella morganii subsp morganii KT YP 007505477 1 1124991 Proteobacteria
Pectobacterium atrosepticum SCRI1043 YP 050697 1 218491 Proteobacteria
Dickeya dadantii subsp dieffenbachiae NCPPB 2976 WP 038923301 1 Ddadantii
Dickeya dadantii NCPPB 898 WP 038909972 1 Ddadantii
Dickeya gs fangzhongdai NCPPB 3274 WP 042860349 1 Dfangzhongdai
Dickeya zeae NCPPB 3531 WP 038904548 1 Dzeae
Dickeya zeae ZJU1202 WP 016940974 1 Dzeae
Dickeya zeae EC1 WP 016940974 1 Dzeae
Dickeya gs poaceaephila NCPPB 569 WP 042871410 1 Dpoaceaephila
Dickeya zeae MK19 WP 038908152 1 Dzeae
Dickeya zeae MS1 WP 019843889 1 Dzeae
Dickeya zeae CSL RW192 WP 038908152 1 Dzeae
Enterobacter asburiae LF7a YP 004830961 1 640513 Proteobacteria
Salmonella bongori N268 08 YP 008321895 1 1197719 Proteobacteria
Escherichia coli O111 H str 11128 YP 003234632 1 585396 Proteobacteria
Escherichia coli O26 H11 str 11368 YP 003229458 1 573235 Proteobacteria
Escherichia coli O157 H7 str EDL933 NP 288363 1 155864 Proteobacteria
Escherichia coli O55 H7 str CB9615 YP 003499700 1 701177 Proteobacteria
Escherichia coli O103 H2 str 12009 YP 003221834 1 585395 Proteobacteria
Shigella sonnei 53G YP 005456187 1 216599 Proteobacteria
Escherichia coli O26 H11 str 11368 YP 003229774 1 573235 Proteobacteria
Citrobacter rodentium ICC168 YP 003365545 1 637910 Proteobacteria
Escherichia coli O104 H4 str 2009EL 2050 YP 006769933 1 1134782 Proteobacteria
Escherichia coli O127 H6 str E2348 69 YP 002329353 1 574521 Proteobacteria
Shigella boydii Sb227 YP 407841 1 300268 Proteobacteria
Citrobacter rodentium ICC168 YP 003367100 1 637910 Proteobacteria
Pantoea ananatis LMG 20103 YP 003521692 1 706191 Proteobacteria
Yersinia enterocolitica subsp palearctica Y11 YP 006004156 1 930944 Proteobacteria
Pectobacterium carotovorum subsp carotovorum PC1 YP 003018756 1 561230 Proteobacteria
Yersinia enterocolitica subsp palearctica Y11 YP 006006342 1 930944 Proteobacteria
Rahnella aquatilis CIP 78 65 ATCC 33071 YP 005199573 1 745277 Proteobacteria
Enterobacter cloacae subsp cloacae ENHKU01 YP 006578999 1 1211025 Proteobacteria
Enterobacter cloacae EcWSU1 YP 004952106 1 1045856 Proteobacteria
Klebsiella oxytoca E718 YP 006499628 1 1191061 Proteobacteria
Klebsiella pneumoniae subsp pneumoniae MGH 78578 YP 001337103 1 272620 Proteobacteria
Escherichia coli O127 H6 str E2348 69 YP 002330383 1 574521 Proteobacteria
Shigella sonnei 53G YP 005459218 1 216599 Proteobacteria
Escherichia coli E24377A YP 001462118 1 331111 Proteobacteria
Yersinia pseudotuberculosis IP 31758 YP 001401162 1 349747 Proteobacteria
Sodalis glossinidius str morsitans YP 454880 1 343509 Proteobacteria
Sodalis glossinidius str morsitans YP 455540 1 343509 Proteobacteria
Xenorhabdus nematophila ATCC 19061 YP 003713228 1 406817 Proteobacteria
Photorhabdus luminescens subsp laumondii TTO1 NP 927386 1 243265 Proteobacteria
Photorhabdus asymbiotica YP 003038847 1 291112 Proteobacteria
Xenorhabdus nematophila ATCC 19061 YP 003711487 1 406817 Proteobacteria
Xenorhabdus bovienii SS 2004 YP 003467284 1 406818 Proteobacteria
Dickeya chrysanthemi subsp parthenii NCPPB 516 WP 033580960 1 Dchrysanthemi
Dickeya dianthicola GBBC 2039 WP 024104569 1 Ddianthicola
Dickeya dianthicola NCPPB 453 WP 024104569 1 Ddianthicola
Dickeya dianthicola RNS04 9 WP 024104569 1 Ddianthicola
Pantoea vagans C9 1 YP 003932283 1 712898 Proteobacteria
Erwinia billingiae Eb661 YP 003740497 1 634500 Proteobacteria
Pantoea sp At 9b YP 004117260 1 592316 Proteobacteria
Enterobacter cloacae subsp cloacae ENHKU01 YP 006579845 1 1211025 Proteobacteria
Salmonella enterica subsp enterica serovar Pullorum str S06004 YP 008382003 1 1298917 Proteobacteria
Shigella boydii Sb227 YP 407279 1 300268 Proteobacteria
Salmonella enterica subsp enterica serovar Typhi str Ty21a YP 007926946 1 527001 Proteobacteria
Salmonella enterica subsp enterica serovar Typhi str Ty21a YP 007927696 1 527001 Proteobacteria
Escherichia coli ATCC 8739 YP 001725707 1 481805 Proteobacteria
Escherichia coli O7 K1 str CE10 YP 006143800 1 1072459 Proteobacteria
Escherichia coli O127 H6 str E2348 69 YP 002328406 1 574521 Proteobacteria
Salmonella enterica subsp enterica serovar Typhi str Ty21a YP 007928536 1 527001 Proteobacteria
Salmonella enterica subsp enterica serovar Paratyphi A str AKU 12601 YP 002143225 1 554290 Proteobacteria
Enterobacter sp 638 YP 001176090 1 399742 Proteobacteria
Erwinia tasmaniensis Et1 99 YP 001906537 1 465817 Proteobacteria
Salmonella enterica subsp enterica serovar Typhimurium str ST4 74 YP 005243668 1 909946 Proteobacteria
Salmonella enterica subsp enterica serovar Dublin str CT 02021853 YP 002216703 1 439851 Proteobacteria
Salmonella enterica subsp enterica serovar Javiana str CFSAN001992 YP 007473705 1 1267753 Proteobacteria
Salmonella enterica subsp enterica serovar Agona str SL483 YP 002147630 1 454166 Proteobacteria

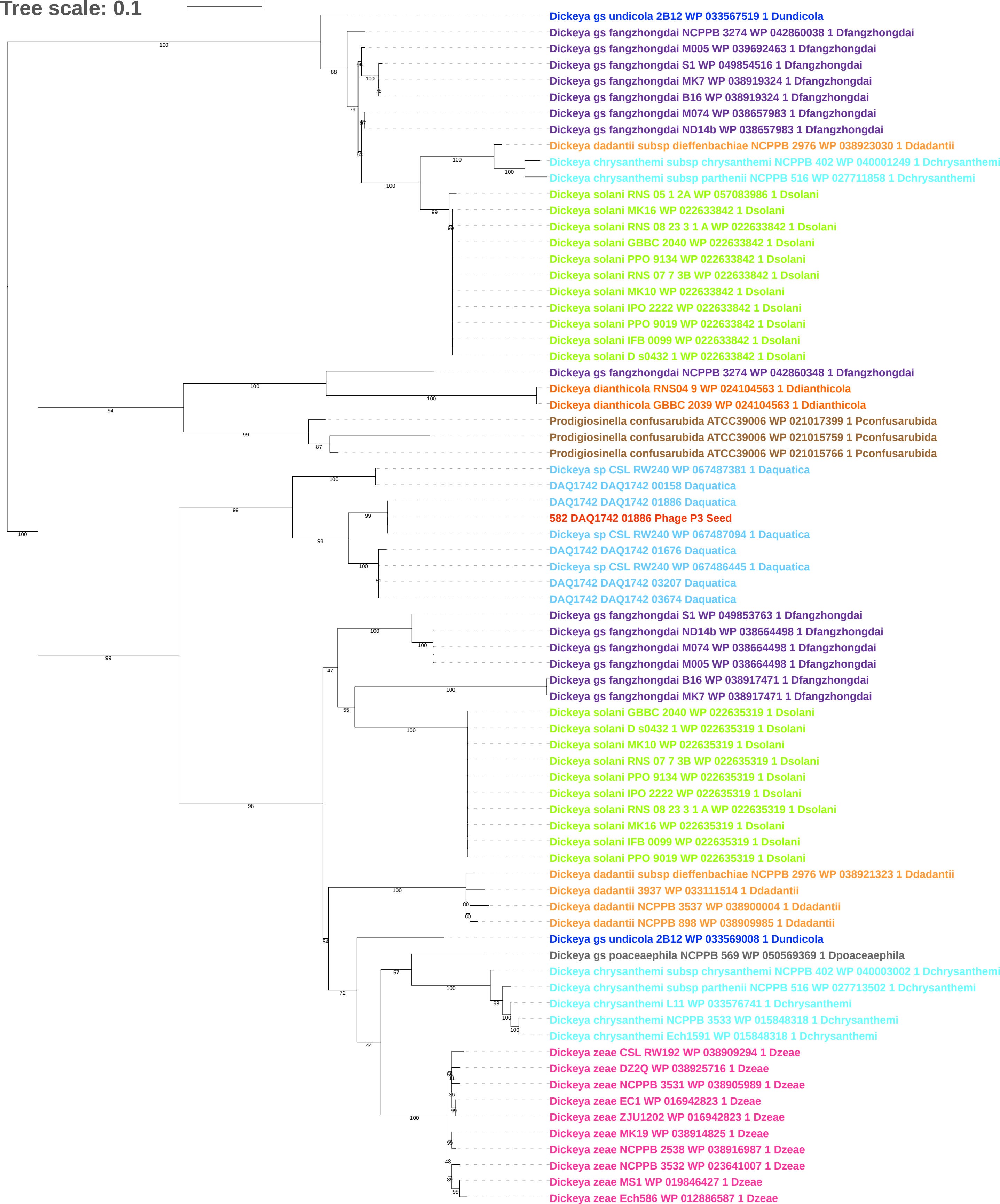
Tree scale: 0.1
Dickeya gs undicola 2B12 WP 033567519 1 Dundicola
Dickeya gs fangzhongdai NCPPB 3274 WP 042860038 1 Dfangzhongdai
Dickeya gs fangzhongdai M005 WP 039692463 1 Dfangzhongdai
Dickeya gs fangzhongdai S1 WP 049854516 1 Dfangzhongdai
Dickeya gs fangzhongdai MK7 WP 038919324 1 Dfangzhongdai
Dickeya gs fangzhongdai B16 WP 038919324 1 Dfangzhongdai
Dickeya gs fangzhongdai M074 WP 038657983 1 Dfangzhongdai
Dickeya gs fangzhongdai ND14b WP 038657983 1 Dfangzhongdai
Dickeya dadantii subsp dieffenbachiae NCPPB 2976 WP 038923030 1 Ddadantii
Dickeya chrysanthemi subsp chrysanthemi NCPPB 402 WP 040001249 1 Dchrysanthemi
Dickeya chrysanthemi subsp parthenii NCPPB 516 WP 027711858 1 Dchrysanthemi
Dickeya solani RNS 05 1 2A WP 057083986 1 Dsolani
Dickeya solani MK16 WP 022633842 1 Dsolani
Dickeya solani RNS 08 23 3 1 A WP 022633842 1 Dsolani
Dickeya solani GBBC 2040 WP 022633842 1 Dsolani
Dickeya solani PPO 9134 WP 022633842 1 Dsolani
Dickeya solani RNS 07 7 3B WP 022633842 1 Dsolani
Dickeya solani MK10 WP 022633842 1 Dsolani
Dickeya solani IPO 2222 WP 022633842 1 Dsolani
Dickeya solani PPO 9019 WP 022633842 1 Dsolani
Dickeya solani IFB 0099 WP 022633842 1 Dsolani
Dickeya solani D s0432 1 WP 022633842 1 Dsolani
Dickeya gs fangzhongdai NCPPB 3274 WP 042860348 1 Dfangzhongdai
Dickeya dianthicola RNS04 9 WP 024104563 1 Ddianthicola
Dickeya dianthicola GBBC 2039 WP 024104563 1 Ddianthicola
Prodigiosinella confusarubida ATCC39006 WP 021017399 1 Pconfusarubida
Prodigiosinella confusarubida ATCC39006 WP 021015759 1 Pconfusarubida
Prodigiosinella confusarubida ATCC39006 WP 021015766 1 Pconfusarubida
Dickeya sp CSL RW240 WP 067487381 1 Daquatica
DAQ1742 DAQ1742 00158 Daquatica
DAQ1742 DAQ1742 01886 Daquatica
582 DAQ1742 01886 Phage P3 Seed
Dickeya sp CSL RW240 WP 067487094 1 Daquatica
DAQ1742 DAQ1742 01676 Daquatica
Dickeya sp CSL RW240 WP 067486445 1 Daquatica
DAQ1742 DAQ1742 03207 Daquatica
DAQ1742 DAQ1742 03674 Daquatica
Dickeya gs fangzhongdai S1 WP 049853763 1 Dfangzhongdai
Dickeya gs fangzhongdai ND14b WP 038664498 1 Dfangzhongdai
Dickeya gs fangzhongdai M074 WP 038664498 1 Dfangzhongdai
Dickeya gs fangzhongdai M005 WP 038664498 1 Dfangzhongdai
Dickeya gs fangzhongdai B16 WP 038917471 1 Dfangzhongdai
Dickeya gs fangzhongdai MK7 WP 038917471 1 Dfangzhongdai
Dickeya solani GBBC 2040 WP 022635319 1 Dsolani
Dickeya solani D s0432 1 WP 022635319 1 Dsolani
Dickeya solani MK10 WP 022635319 1 Dsolani
Dickeya solani RNS 07 7 3B WP 022635319 1 Dsolani
Dickeya solani PPO 9134 WP 022635319 1 Dsolani
Dickeya solani IPO 2222 WP 022635319 1 Dsolani
Dickeya solani RNS 08 23 3 1 A WP 022635319 1 Dsolani
Dickeya solani MK16 WP 022635319 1 Dsolani
Dickeya solani IFB 0099 WP 022635319 1 Dsolani
Dickeya solani PPO 9019 WP 022635319 1 Dsolani
Dickeya dadantii subsp dieffenbachiae NCPPB 2976 WP 038921323 1 Ddadantii
Dickeya dadantii 3937 WP 033111514 1 Ddadantii
Dickeya dadantii NCPPB 3537 WP 038900004 1 Ddadantii
Dickeya dadantii NCPPB 898 WP 038909985 1 Ddadantii
Dickeya gs undicola 2B12 WP 033569008 1 Dundicola
Dickeya gs poaceaephila NCPPB 569 WP 050569369 1 Dpoaceaephila
Dickeya chrysanthemi subsp chrysanthemi NCPPB 402 WP 040003002 1 Dchrysanthemi
Dickeya chrysanthemi subsp parthenii NCPPB 516 WP 027713502 1 Dchrysanthemi
Dickeya chrysanthemi L11 WP 033576741 1 Dchrysanthemi
Dickeya chrysanthemi NCPPB 3533 WP 015848318 1 Dchrysanthemi
Dickeya chrysanthemi Ech1591 WP 015848318 1 Dchrysanthemi
Dickeya zeae CSL RW192 WP 038909294 1 Dzeae
Dickeya zeae DZ2Q WP 038925716 1 Dzeae
Dickeya zeae NCPPB 3531 WP 038905989 1 Dzeae
Dickeya zeae EC1 WP 016942823 1 Dzeae
Dickeya zeae ZJU1202 WP 016942823 1 Dzeae
Dickeya zeae MK19 WP 038914825 1 Dzeae
Dickeya zeae NCPPB 2538 WP 038916987 1 Dzeae
Dickeya zeae NCPPB 3532 WP 023641007 1 Dzeae
Dickeya zeae MS1 WP 019846427 1 Dzeae
Dickeya zeae Ech586 WP 012886587 1 Dzeae

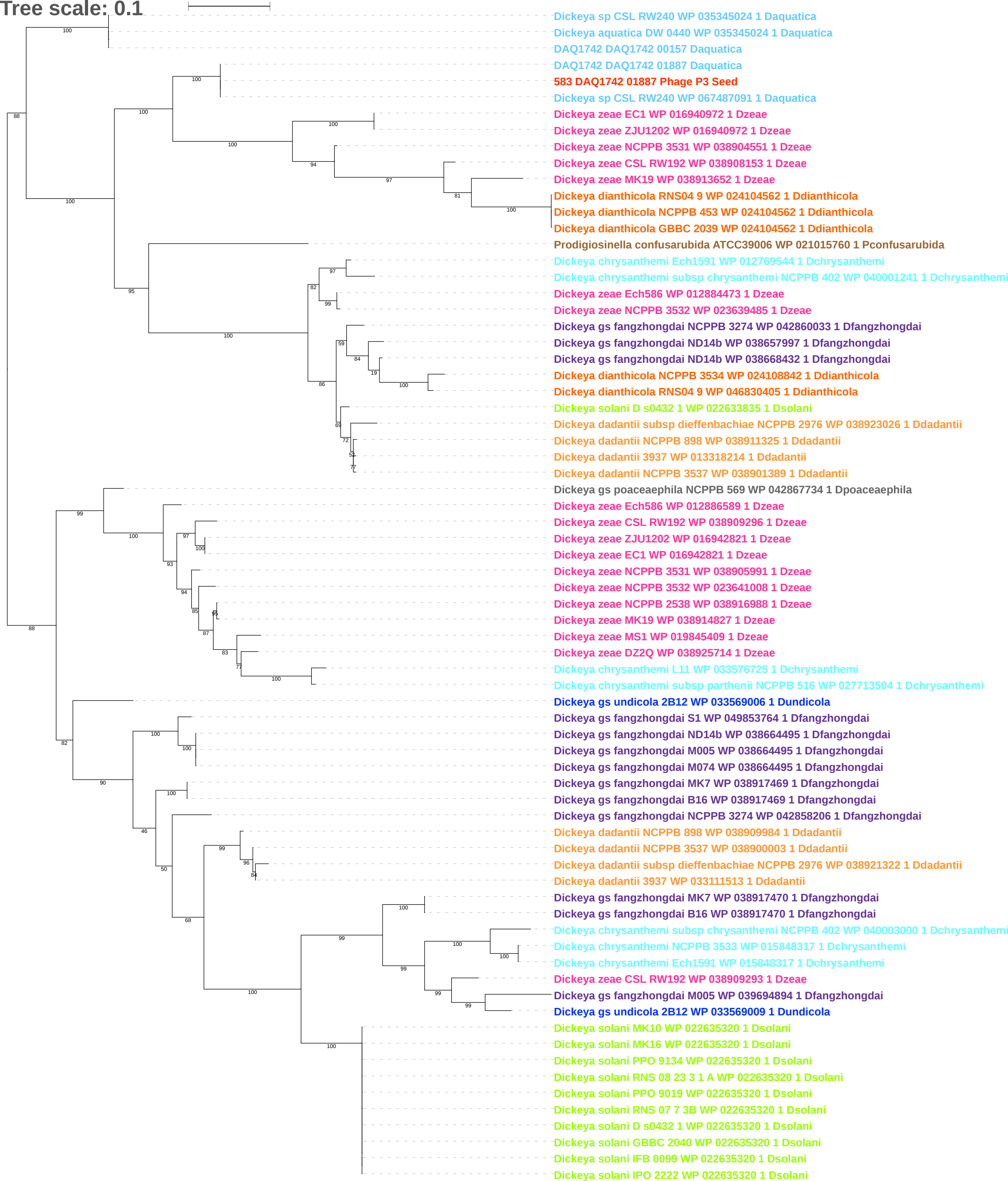
Tree scale: 0.1
Dickeya sp CSL RW240 WP 035345024 1 Daquatica
Dickeya aquatica DW 0440 WP 035345024 1 Daquatica
DAQ1742 DAQ1742 00157 Daquatica
DAQ1742 DAQ1742 01887 Daquatica
583 DAQ1742 01887 Phage P3 Seed
Dickeya sp CSL RW240 WP 067487091 1 Daquatica
Dickeya zeae EC1 WP 016940972 1 Dzeae
Dickeya zeae ZJU1202 WP 016940972 1 Dzeae
Dickeya zeae NCPPB 3531 WP 038904551 1 Dzeae
Dickeya zeae CSL RW192 WP 038908153 1 Dzeae
Dickeya zeae MK19 WP 038913652 1 Dzeae
Dickeya dianthicola RNS04 9 WP 024104562 1 Ddianthicola
Dickeya dianthicola NCPPB 453 WP 024104562 1 Ddianthicola
Dickeya dianthicola GBBC 2039 WP 024104562 1 Ddianthicola
Prodigiosinella confusarubida ATCC39006 WP 021015760 1 Pconfusarubida
Dickeya chrysanthemi Ech1591 WP 012769544 1 Dchrysanthemi
Dickeya chrysanthemi subsp chrysanthemi NCPPB 402 WP 040001241 1 Dchrysanthemi
Dickeya zeae Ech586 WP 012884473 1 Dzeae
Dickeya zeae NCPPB 3532 WP 023639485 1 Dzeae
Dickeya gs fangzhongdai NCPPB 3274 WP 042860033 1 Dfangzhongdai
Dickeya gs fangzhongdai ND14b WP 038657997 1 Dfangzhongdai
Dickeya gs fangzhongdai ND14b WP 038668432 1 Dfangzhongdai
Dickeya dianthicola NCPPB 3534 WP 024108842 1 Ddianthicola
Dickeya dianthicola RNS04 9 WP 046830405 1 Ddianthicola
Dickeya solani D s0432 1 WP 022633835 1 Dsolani
Dickeya dadantii subsp dieffenbachiae NCPPB 2976 WP 038923026 1 Ddadantii
Dickeya dadantii NCPPB 898 WP 038911325 1 Ddadantii
Dickeya dadantii 3937 WP 013318214 1 Ddadantii
Dickeya dadantii NCPPB 3537 WP 038901389 1 Ddadantii
Dickeya gs poaceaephila NCPPB 569 WP 042867734 1 Dpoaceaephila
Dickeya zeae Ech586 WP 012886589 1 Dzeae
Dickeya zeae CSL RW192 WP 038909296 1 Dzeae
Dickeya zeae ZJU1202 WP 016942821 1 Dzeae
Dickeya zeae EC1 WP 016942821 1 Dzeae
Dickeya zeae NCPPB 3531 WP 038905991 1 Dzeae
Dickeya zeae NCPPB 3532 WP 023641008 1 Dzeae
Dickeya zeae NCPPB 2538 WP 038916988 1 Dzeae
Dickeya zeae MK19 WP 038914827 1 Dzeae
Dickeya zeae MS1 WP 019845409 1 Dzeae
Dickeya zeae DZ2Q WP 038925714 1 Dzeae
Dickeya chrysanthemi L11 WP 033576725 1 Dchrysanthemi
Dickeya chrysanthemi subsp parthenii NCPPB 516 WP 027713504 1 Dchrysanthemi
Dickeya gs undicola 2B12 WP 033569006 1 Dundicola
Dickeya gs fangzhongdai S1 WP 049853764 1 Dfangzhongdai
Dickeya gs fangzhongdai ND14b WP 038664495 1 Dfangzhongdai
Dickeya gs fangzhongdai M005 WP 038664495 1 Dfangzhongdai
Dickeya gs fangzhongdai M074 WP 038664495 1 Dfangzhongdai
Dickeya gs fangzhongdai MK7 WP 038917469 1 Dfangzhongdai
Dickeya gs fangzhongdai B16 WP 038917469 1 Dfangzhongdai
Dickeya gs fangzhongdai NCPPB 3274 WP 042858206 1 Dfangzhongdai
Dickeya dadantii NCPPB 898 WP 038909984 1 Ddadantii
Dickeya dadantii NCPPB 3537 WP 038900003 1 Ddadantii
Dickeya dadantii subsp dieffenbachiae NCPPB 2976 WP 038921322 1 Ddadantii
Dickeya dadantii 3937 WP 033111513 1 Ddadantii
Dickeya gs fangzhongdai MK7 WP 038917470 1 Dfangzhongdai
Dickeya gs fangzhongdai B16 WP 038917470 1 Dfangzhongdai
Dickeya chrysanthemi subsp chrysanthemi NCPPB 402 WP 040003000 1 Dchrysanthemi
Dickeya chrysanthemi NCPPB 3533 WP 015848317 1 Dchrysanthemi
Dickeya chrysanthemi Ech1591 WP 015848317 1 Dchrysanthemi
Dickeya zeae CSL RW192 WP 038909293 1 Dzeae
Dickeya gs fangzhongdai M005 WP 039694894 1 Dfangzhongdai
Dickeya gs undicola 2B12 WP 033569009 1 Dundicola
Dickeya solani MK10 WP 022635320 1 Dsolani
Dickeya solani MK16 WP 022635320 1 Dsolani
Dickeya solani PPO 9134 WP 022635320 1 Dsolani
Dickeya solani RNS 08 23 3 1 A WP 022635320 1 Dsolani
Dickeya solani PPO 9019 WP 022635320 1 Dsolani
Dickeya solani RNS 07 7 3B WP 022635320 1 Dsolani
Dickeya solani D s0432 1 WP 022635320 1 Dsolani
Dickeya solani GBBC 2040 WP 022635320 1 Dsolani
Dickeya solani IFB 0099 WP 022635320 1 Dsolani
Dickeya solani IPO 2222 WP 022635320 1 Dsolani

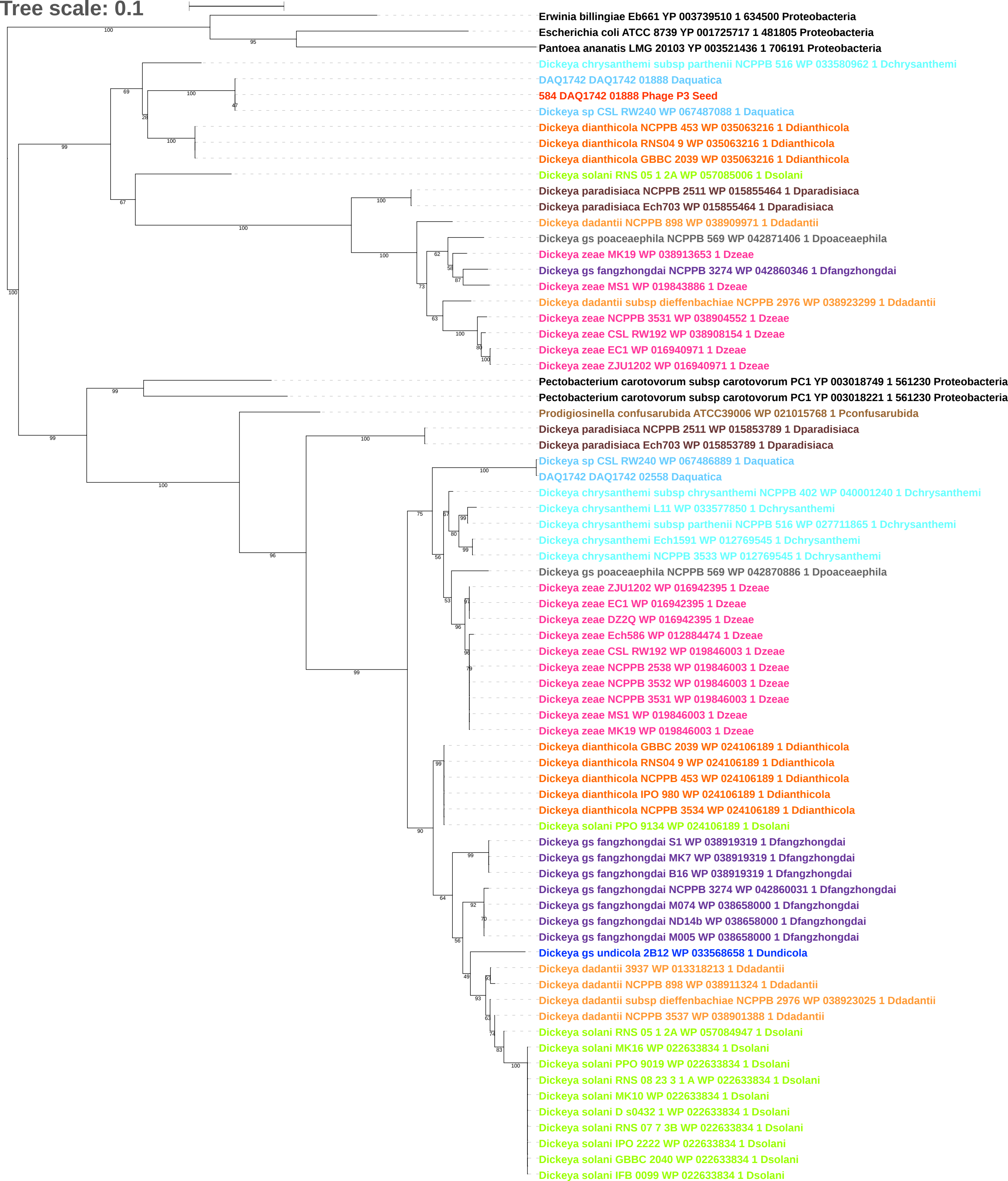
Tree scale: 0.1
Erwinia billingiae Eb661 YP 003739510 1 634500 Proteobacteria
Escherichia coli ATCC 8739 YP 001725717 1 481805 Proteobacteria
Pantoea ananatis LMG 20103 YP 003521436 1 706191 Proteobacteria
Dickeya chrysanthemi subsp parthenii NCPPB 516 WP 033580962 1 Dchrysanthemi
DAQ1742 DAQ1742 01888 Daquatica
584 DAQ1742 01888 Phage P3 Seed
Dickeya sp CSL RW240 WP 067487088 1 Daquatica
Dickeya dianthicola NCPPB 453 WP 035063216 1 Ddianthicola
Dickeya dianthicola RNS04 9 WP 035063216 1 Ddianthicola
Dickeya dianthicola GBBC 2039 WP 035063216 1 Ddianthicola
Dickeya solani RNS 05 1 2A WP 057085006 1 Dsolani
Dickeya paradisiaca NCPPB 2511 WP 015855464 1 Dparadisiaca
Dickeya paradisiaca Ech703 WP 015855464 1 Dparadisiaca
Dickeya dadantii NCPPB 898 WP 038909971 1 Ddadantii
Dickeya gs poaceaephila NCPPB 569 WP 042871406 1 Dpoaceaephila
Dickeya zeae MK19 WP 038913653 1 Dzeae
Dickeya gs fangzhongdai NCPPB 3274 WP 042860346 1 Dfangzhongdai
Dickeya zeae MS1 WP 019843886 1 Dzeae
Dickeya dadantii subsp dieffenbachiae NCPPB 2976 WP 038923299 1 Ddadantii
Dickeya zeae NCPPB 3531 WP 038904552 1 Dzeae
Dickeya zeae CSL RW192 WP 038908154 1 Dzeae
Dickeya zeae EC1 WP 016940971 1 Dzeae
Dickeya zeae ZJU1202 WP 016940971 1 Dzeae
Pectobacterium carotovorum subsp carotovorum PC1 YP 003018749 1 561230 Proteobacteria
Pectobacterium carotovorum subsp carotovorum PC1 YP 003018221 1 561230 Proteobacteria
Prodigiosinella confusarubida ATCC39006 WP 021015768 1 Pconfusarubida
Dickeya paradisiaca NCPPB 2511 WP 015853789 1 Dparadisiaca
Dickeya paradisiaca Ech703 WP 015853789 1 Dparadisiaca
Dickeya sp CSL RW240 WP 067486889 1 Daquatica
DAQ1742 DAQ1742 02558 Daquatica
Dickeya chrysanthemi subsp chrysanthemi NCPPB 402 WP 040001240 1 Dchrysanthemi
Dickeya chrysanthemi L11 WP 033577850 1 Dchrysanthemi
Dickeya chrysanthemi subsp parthenii NCPPB 516 WP 027711865 1 Dchrysanthemi
Dickeya chrysanthemi Ech1591 WP 012769545 1 Dchrysanthemi
Dickeya chrysanthemi NCPPB 3533 WP 012769545 1 Dchrysanthemi
Dickeya gs poaceaephila NCPPB 569 WP 042870886 1 Dpoaceaephila
Dickeya zeae ZJU1202 WP 016942395 1 Dzeae
Dickeya zeae EC1 WP 016942395 1 Dzeae
Dickeya zeae DZ2Q WP 016942395 1 Dzeae
Dickeya zeae Ech586 WP 012884474 1 Dzeae
Dickeya zeae CSL RW192 WP 019846003 1 Dzeae
Dickeya zeae NCPPB 2538 WP 019846003 1 Dzeae
Dickeya zeae NCPPB 3532 WP 019846003 1 Dzeae
Dickeya zeae NCPPB 3531 WP 019846003 1 Dzeae
Dickeya zeae MS1 WP 019846003 1 Dzeae
Dickeya zeae MK19 WP 019846003 1 Dzeae
Dickeya dianthicola GBBC 2039 WP 024106189 1 Ddianthicola
Dickeya dianthicola RNS04 9 WP 024106189 1 Ddianthicola
Dickeya dianthicola NCPPB 453 WP 024106189 1 Ddianthicola
Dickeya dianthicola IPO 980 WP 024106189 1 Ddianthicola
Dickeya dianthicola NCPPB 3534 WP 024106189 1 Ddianthicola
Dickeya solani PPO 9134 WP 024106189 1 Dsolani
Dickeya gs fangzhongdai S1 WP 038919319 1 Dfangzhongdai
Dickeya gs fangzhongdai MK7 WP 038919319 1 Dfangzhongdai
Dickeya gs fangzhongdai B16 WP 038919319 1 Dfangzhongdai
Dickeya gs fangzhongdai NCPPB 3274 WP 042860031 1 Dfangzhongdai
Dickeya gs fangzhongdai M074 WP 038658000 1 Dfangzhongdai
Dickeya gs fangzhongdai ND14b WP 038658000 1 Dfangzhongdai
Dickeya gs fangzhongdai M005 WP 038658000 1 Dfangzhongdai
Dickeya gs undicola 2B12 WP 033568658 1 Dundicola
Dickeya dadantii 3937 WP 013318213 1 Ddadantii
Dickeya dadantii NCPPB 898 WP 038911324 1 Ddadantii
Dickeya dadantii subsp dieffenbachiae NCPPB 2976 WP 038923025 1 Ddadantii
Dickeya dadantii NCPPB 3537 WP 038901388 1 Ddadantii
Dickeya solani RNS 05 1 2A WP 057084947 1 Dsolani
Dickeya solani MK16 WP 022633834 1 Dsolani
Dickeya solani PPO 9019 WP 022633834 1 Dsolani
Dickeya solani RNS 08 23 3 1 A WP 022633834 1 Dsolani
Dickeya solani MK10 WP 022633834 1 Dsolani
Dickeya solani D s0432 1 WP 022633834 1 Dsolani
Dickeya solani RNS 07 7 3B WP 022633834 1 Dsolani
Dickeya solani IPO 2222 WP 022633834 1 Dsolani
Dickeya solani GBBC 2040 WP 022633834 1 Dsolani
Dickeya solani IFB 0099 WP 022633834 1 Dsolani

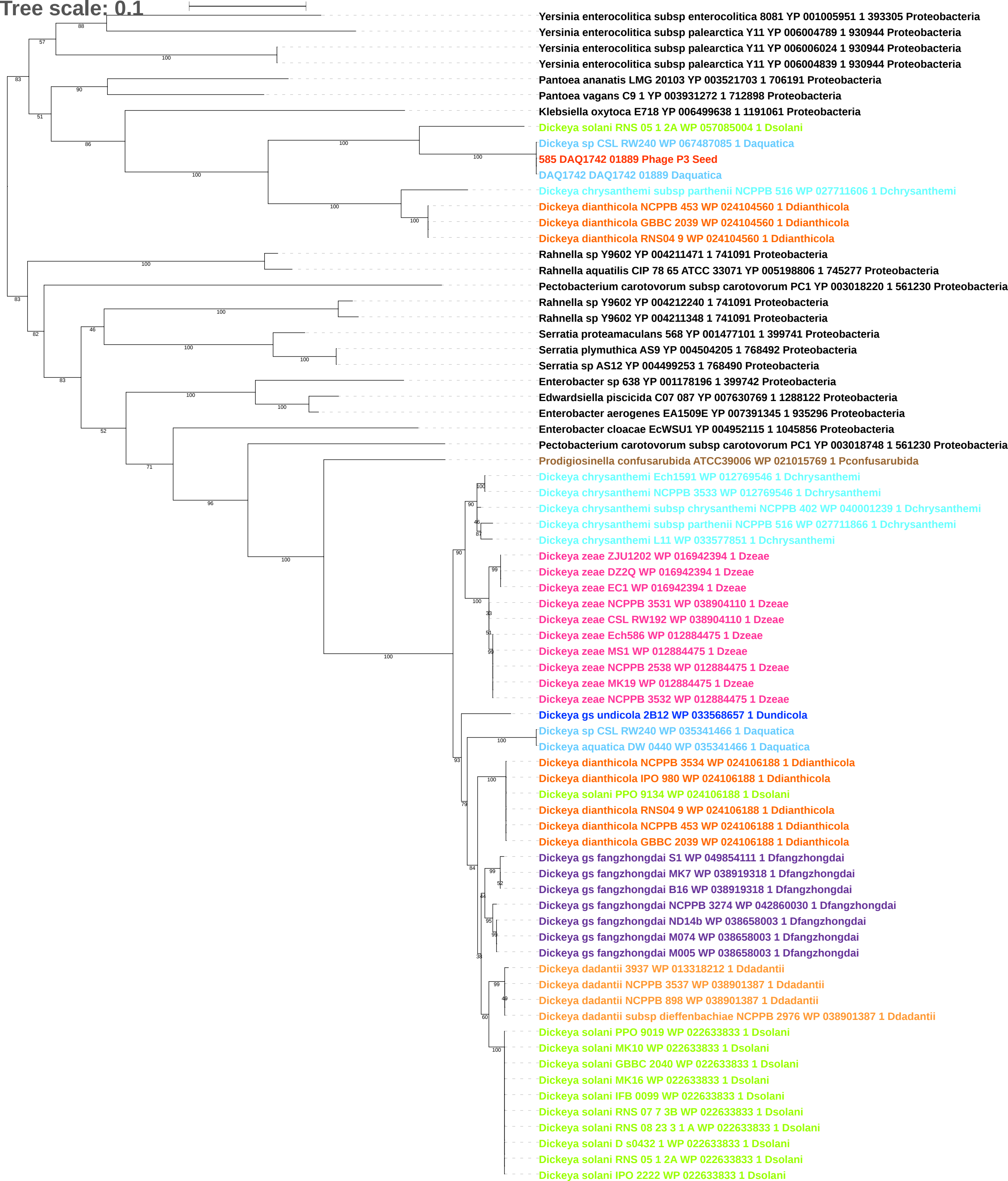
Tree scale: 0.1
Yersinia enterocolitica subsp enterocolitica 8081 YP 001005951 1 393305 Proteobacteria
Yersinia enterocolitica subsp palearctica Y11 YP 006004789 1 930944 Proteobacteria
Yersinia enterocolitica subsp palearctica Y11 YP 006006024 1 930944 Proteobacteria
Yersinia enterocolitica subsp palearctica Y11 YP 006004839 1 930944 Proteobacteria
Pantoea ananatis LMG 20103 YP 003521703 1 706191 Proteobacteria
Pantoea vagans C9 1 YP 003931272 1 712898 Proteobacteria
Klebsiella oxytoca E718 YP 006499638 1 1191061 Proteobacteria
Dickeya solani RNS 05 1 2A WP 057085004 1 Dsolani
Dickeya sp CSL RW240 WP 067487085 1 Daquatica
585 DAQ1742 01889 Phage P3 Seed
DAQ1742 DAQ1742 01889 Daquatica
Dickeya chrysanthemi subsp parthenii NCPPB 516 WP 027711606 1 Dchrysanthemi
Dickeya dianthicola NCPPB 453 WP 024104560 1 Ddianthicola
Dickeya dianthicola GBBC 2039 WP 024104560 1 Ddianthicola
Dickeya dianthicola RNS04 9 WP 024104560 1 Ddianthicola
Rahnella sp Y9602 YP 004211471 1 741091 Proteobacteria
Rahnella aquatilis CIP 78 65 ATCC 33071 YP 005198806 1 745277 Proteobacteria
Pectobacterium carotovorum subsp carotovorum PC1 YP 003018220 1 561230 Proteobacteria
Rahnella sp Y9602 YP 004212240 1 741091 Proteobacteria
Rahnella sp Y9602 YP 004211348 1 741091 Proteobacteria
Serratia proteamaculans 568 YP 001477101 1 399741 Proteobacteria
Serratia plymuthica AS9 YP 004504205 1 768492 Proteobacteria
Serratia sp AS12 YP 004499253 1 768490 Proteobacteria
Enterobacter sp 638 YP 001178196 1 399742 Proteobacteria
Edwardsiella piscicida C07 087 YP 007630769 1 1288122 Proteobacteria
Enterobacter aerogenes EA1509E YP 007391345 1 935296 Proteobacteria
Enterobacter cloacae EcWSU1 YP 004952115 1 1045856 Proteobacteria
Pectobacterium carotovorum subsp carotovorum PC1 YP 003018748 1 561230 Proteobacteria
Prodigiosinella confusarubida ATCC39006 WP 021015769 1 Pconfusarubida
Dickeya chrysanthemi Ech1591 WP 012769546 1 Dchrysanthemi
Dickeya chrysanthemi NCPPB 3533 WP 012769546 1 Dchrysanthemi
Dickeya chrysanthemi subsp chrysanthemi NCPPB 402 WP 040001239 1 Dchrysanthemi
Dickeya chrysanthemi subsp parthenii NCPPB 516 WP 027711866 1 Dchrysanthemi
Dickeya chrysanthemi L11 WP 033577851 1 Dchrysanthemi
Dickeya zeae ZJU1202 WP 016942394 1 Dzeae
Dickeya zeae DZ2Q WP 016942394 1 Dzeae
Dickeya zeae EC1 WP 016942394 1 Dzeae
Dickeya zeae NCPPB 3531 WP 038904110 1 Dzeae
Dickeya zeae CSL RW192 WP 038904110 1 Dzeae
Dickeya zeae Ech586 WP 012884475 1 Dzeae
Dickeya zeae MS1 WP 012884475 1 Dzeae
Dickeya zeae NCPPB 2538 WP 012884475 1 Dzeae
Dickeya zeae MK19 WP 012884475 1 Dzeae
Dickeya zeae NCPPB 3532 WP 012884475 1 Dzeae
Dickeya gs undicola 2B12 WP 033568657 1 Dundicola
Dickeya sp CSL RW240 WP 035341466 1 Daquatica
Dickeya aquatica DW 0440 WP 035341466 1 Daquatica
Dickeya dianthicola NCPPB 3534 WP 024106188 1 Ddianthicola
Dickeya dianthicola IPO 980 WP 024106188 1 Ddianthicola
Dickeya solani PPO 9134 WP 024106188 1 Dsolani
Dickeya dianthicola RNS04 9 WP 024106188 1 Ddianthicola
Dickeya dianthicola NCPPB 453 WP 024106188 1 Ddianthicola
Dickeya dianthicola GBBC 2039 WP 024106188 1 Ddianthicola
Dickeya gs fangzhongdai S1 WP 049854111 1 Dfangzhongdai
Dickeya gs fangzhongdai MK7 WP 038919318 1 Dfangzhongdai
Dickeya gs fangzhongdai B16 WP 038919318 1 Dfangzhongdai
Dickeya gs fangzhongdai NCPPB 3274 WP 042860030 1 Dfangzhongdai
Dickeya gs fangzhongdai ND14b WP 038658003 1 Dfangzhongdai
Dickeya gs fangzhongdai M074 WP 038658003 1 Dfangzhongdai
Dickeya gs fangzhongdai M005 WP 038658003 1 Dfangzhongdai
Dickeya dadantii 3937 WP 013318212 1 Ddadantii
Dickeya dadantii NCPPB 3537 WP 038901387 1 Ddadantii
Dickeya dadantii NCPPB 898 WP 038901387 1 Ddadantii
Dickeya dadantii subsp dieffenbachiae NCPPB 2976 WP 038901387 1 Ddadantii
Dickeya solani PPO 9019 WP 022633833 1 Dsolani
Dickeya solani MK10 WP 022633833 1 Dsolani
Dickeya solani GBBC 2040 WP 022633833 1 Dsolani
Dickeya solani MK16 WP 022633833 1 Dsolani
Dickeya solani IFB 0099 WP 022633833 1 Dsolani
Dickeya solani RNS 07 7 3B WP 022633833 1 Dsolani
Dickeya solani RNS 08 23 3 1 A WP 022633833 1 Dsolani
Dickeya solani D s0432 1 WP 022633833 1 Dsolani
Dickeya solani RNS 05 1 2A WP 022633833 1 Dsolani
Dickeya solani IPO 2222 WP 022633833 1 Dsolani

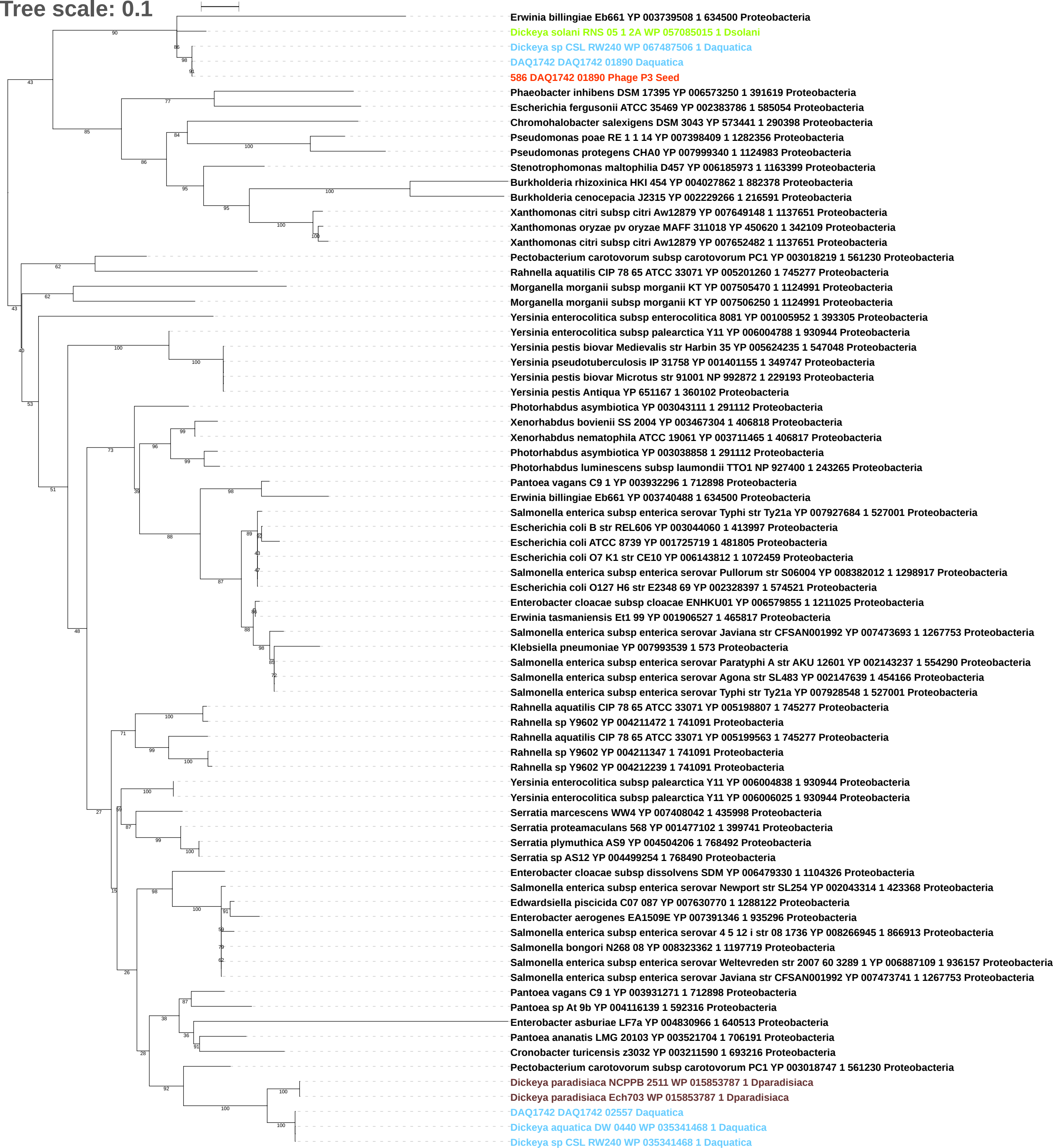

Tree scale: 0.1
Erwinia billingiae Eb661 YP 003739508 1 634500 Proteobacteria
Dickeya solani RNS 05 1 2A WP 057085015 1 Dsolani
Dickeya sp CSL RW240 WP 067487506 1 Daquatica
DAQ1742 DAQ1742 01890 Daquatica
586 DAQ1742 01890 Phage P3 Seed
Phaeobacter inhibens DSM 17395 YP 006573250 1 391619 Proteobacteria
Escherichia fergusonii ATCC 35469 YP 002383786 1 585054 Proteobacteria
Chromohalobacter salexigens DSM 3043 YP 573441 1 290398 Proteobacteria
Pseudomonas poae RE 1 1 14 YP 007398409 1 1282356 Proteobacteria
Pseudomonas protegens CHA0 YP 007999340 1 1124983 Proteobacteria
Stenotrophomonas maltophilia D457 YP 006185973 1 1163399 Proteobacteria
Burkholderia rhizoxinica HKI 454 YP 004027862 1 882378 Proteobacteria
Burkholderia cenocepacia J2315 YP 002229266 1 216591 Proteobacteria
Xanthomonas citri subsp citri Aw12879 YP 007649148 1 1137651 Proteobacteria
Xanthomonas oryzae pv oryzae MAFF 311018 YP 450620 1 342109 Proteobacteria
Xanthomonas citri subsp citri Aw12879 YP 007652482 1 1137651 Proteobacteria
Pectobacterium carotovorum subsp carotovorum PC1 YP 003018219 1 561230 Proteobacteria
Rahnella aquatilis CIP 78 65 ATCC 33071 YP 005201260 1 745277 Proteobacteria
Morganella morganii subsp morganii KT YP 007505470 1 1124991 Proteobacteria
Morganella morganii subsp morganii KT YP 007506250 1 1124991 Proteobacteria
Yersinia enterocolitica subsp enterocolitica 8081 YP 001005952 1 393305 Proteobacteria
Yersinia enterocolitica subsp palearctica Y11 YP 006004788 1 930944 Proteobacteria
Yersinia pestis biovar Medievalis str Harbin 35 YP 005624235 1 547048 Proteobacteria
Yersinia pseudotuberculosis IP 31758 YP 001401155 1 349747 Proteobacteria
Yersinia pestis biovar Microtus str 91001 NP 992872 1 229193 Proteobacteria
Yersinia pestis Antiqua YP 651167 1 360102 Proteobacteria
Photorhabdus asymbiotica YP 003043111 1 291112 Proteobacteria
Xenorhabdus bovienii SS 2004 YP 003467304 1 406818 Proteobacteria
Xenorhabdus nematophila ATCC 19061 YP 003711465 1 406817 Proteobacteria
Photorhabdus asymbiotica YP 003038858 1 291112 Proteobacteria
Photorhabdus luminescens subsp laumondii TTO1 NP 927400 1 243265 Proteobacteria
Pantoea vagans C9 1 YP 003932296 1 712898 Proteobacteria
Erwinia billingiae Eb661 YP 003740488 1 634500 Proteobacteria
Salmonella enterica subsp enterica serovar Typhi str Ty21a YP 007927684 1 527001 Proteobacteria
Escherichia coli B str REL606 YP 003044060 1 413997 Proteobacteria
Escherichia coli ATCC 8739 YP 001725719 1 481805 Proteobacteria
Escherichia coli O7 K1 str CE10 YP 006143812 1 1072459 Proteobacteria
Salmonella enterica subsp enterica serovar Pullorum str S06004 YP 008382012 1 1298917 Proteobacteria
Escherichia coli O127 H6 str E2348 69 YP 002328397 1 574521 Proteobacteria
Enterobacter cloacae subsp cloacae ENHKU01 YP 006579855 1 1211025 Proteobacteria
Erwinia tasmaniensis Et1 99 YP 001906527 1 465817 Proteobacteria
Salmonella enterica subsp enterica serovar Javiana str CFSAN001992 YP 007473693 1 1267753 Proteobacteria
Klebsiella pneumoniae YP 007993539 1 573 Proteobacteria
Salmonella enterica subsp enterica serovar Paratyphi A str AKU 12601 YP 002143237 1 554290 Proteobacteria
Salmonella enterica subsp enterica serovar Agona str SL483 YP 002147639 1 454166 Proteobacteria
Salmonella enterica subsp enterica serovar Typhi str Ty21a YP 007928548 1 527001 Proteobacteria
Rahnella aquatilis CIP 78 65 ATCC 33071 YP 005198807 1 745277 Proteobacteria
Rahnella sp Y9602 YP 004211472 1 741091 Proteobacteria
Rahnella aquatilis CIP 78 65 ATCC 33071 YP 005199563 1 745277 Proteobacteria
Rahnella sp Y9602 YP 004211347 1 741091 Proteobacteria
Rahnella sp Y9602 YP 004212239 1 741091 Proteobacteria
Yersinia enterocolitica subsp palearctica Y11 YP 006004838 1 930944 Proteobacteria
Yersinia enterocolitica subsp palearctica Y11 YP 006006025 1 930944 Proteobacteria
Serratia marcescens WW4 YP 007408042 1 435998 Proteobacteria
Serratia proteamaculans 568 YP 001477102 1 399741 Proteobacteria
Serratia plymuthica AS9 YP 004504206 1 768492 Proteobacteria
Serratia sp AS12 YP 004499254 1 768490 Proteobacteria
Enterobacter cloacae subsp dissolvens SDM YP 006479330 1 1104326 Proteobacteria
Salmonella enterica subsp enterica serovar Newport str SL254 YP 002043314 1 423368 Proteobacteria
Edwardsiella piscicida C07 087 YP 007630770 1 1288122 Proteobacteria
Enterobacter aerogenes EA1509E YP 007391346 1 935296 Proteobacteria
Salmonella enterica subsp enterica serovar 4 5 12 i str 08 1736 YP 008266945 1 866913 Proteobacteria
Salmonella bongori N268 08 YP 008323362 1 1197719 Proteobacteria
Salmonella enterica subsp enterica serovar Weltevreden str 2007 60 3289 1 YP 006887109 1 936157 Proteobacteria
Salmonella enterica subsp enterica serovar Javiana str CFSAN001992 YP 007473741 1 1267753 Proteobacteria
Pantoea vagans C9 1 YP 003931271 1 712898 Proteobacteria
Pantoea sp At 9b YP 004116139 1 592316 Proteobacteria
Enterobacter asburiae LF7a YP 004830966 1 640513 Proteobacteria
Pantoea ananatis LMG 20103 YP 003521704 1 706191 Proteobacteria
Cronobacter turicensis z3032 YP 003211590 1 693216 Proteobacteria
Pectobacterium carotovorum subsp carotovorum PC1 YP 003018747 1 561230 Proteobacteria
Dickeya paradisiaca NCPPB 2511 WP 015853787 1 Dparadisiaca
Dickeya paradisiaca Ech703 WP 015853787 1 Dparadisiaca
DAQ1742 DAQ1742 02557 Daquatica
Dickeya aquatica DW 0440 WP 035341468 1 Daquatica
Dickeya sp CSL RW240 WP 035341468 1 Daquatica

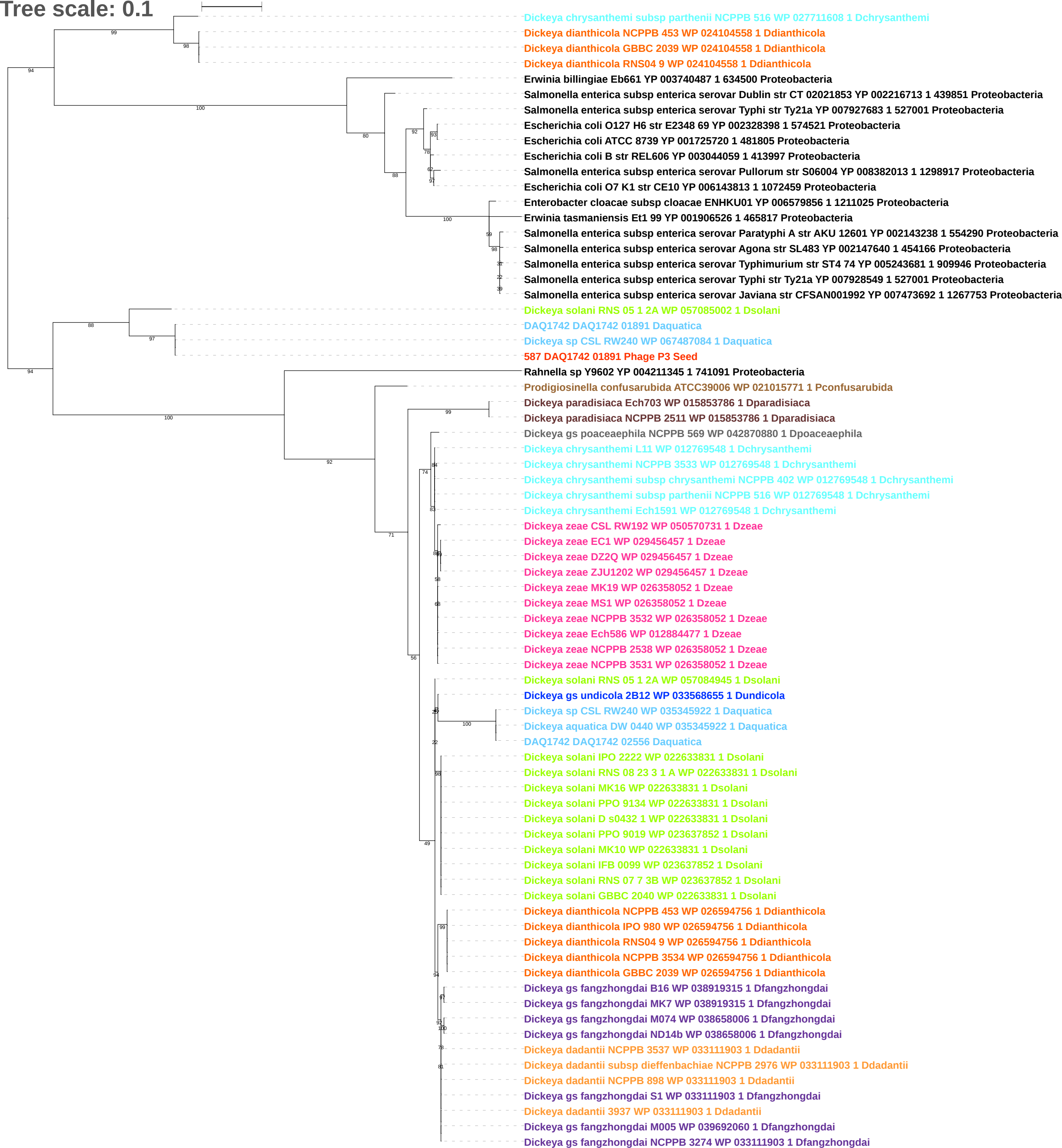
Tree scale: 0.1
Dickeya chrysanthemi subsp parthenii NCPPB 516 WP 027711608 1 Dchrysanthemi
Dickeya dianthicola NCPPB 453 WP 024104558 1 Ddianthicola
Dickeya dianthicola GBBC 2039 WP 024104558 1 Ddianthicola
Dickeya dianthicola RNS04 9 WP 024104558 1 Ddianthicola
Erwinia billingiae Eb661 YP 003740487 1 634500 Proteobacteria
Salmonella enterica subsp enterica serovar Dublin str CT 02021853 YP 002216713 1 439851 Proteobacteria
Salmonella enterica subsp enterica serovar Typhi str Ty21a YP 007927683 1 527001 Proteobacteria
Escherichia coli O127 H6 str E2348 69 YP 002328398 1 574521 Proteobacteria
Escherichia coli ATCC 8739 YP 001725720 1 481805 Proteobacteria
Escherichia coli B str REL606 YP 003044059 1 413997 Proteobacteria
Salmonella enterica subsp enterica serovar Pullorum str S06004 YP 008382013 1 1298917 Proteobacteria
Escherichia coli O7 K1 str CE10 YP 006143813 1 1072459 Proteobacteria
Enterobacter cloacae subsp cloacae ENHKU01 YP 006579856 1 1211025 Proteobacteria
Erwinia tasmaniensis Et1 99 YP 001906526 1 465817 Proteobacteria
Salmonella enterica subsp enterica serovar Paratyphi A str AKU 12601 YP 002143238 1 554290 Proteobacteria
Salmonella enterica subsp enterica serovar Agona str SL483 YP 002147640 1 454166 Proteobacteria
Salmonella enterica subsp enterica serovar Typhimurium str ST4 74 YP 005243681 1 909946 Proteobacteria
Salmonella enterica subsp enterica serovar Typhi str Ty21a YP 007928549 1 527001 Proteobacteria
Salmonella enterica subsp enterica serovar Javiana str CFSAN001992 YP 007473692 1 1267753 Proteobacteria
Dickeya solani RNS 05 1 2A WP 057085002 1 Dsolani
DAQ1742 DAQ1742 01891 Daquatica
Dickeya sp CSL RW240 WP 067487084 1 Daquatica
587 DAQ1742 01891 Phage P3 Seed
Rahnella sp Y9602 YP 004211345 1 741091 Proteobacteria
Prodigiosinella confusarubida ATCC39006 WP 021015771 1 Pconfusarubida
Dickeya paradisiaca Ech703 WP 015853786 1 Dparadisiaca
Dickeya paradisiaca NCPPB 2511 WP 015853786 1 Dparadisiaca
Dickeya gs poaceaephila NCPPB 569 WP 042870880 1 Dpoaceaephila
Dickeya chrysanthemi L11 WP 012769548 1 Dchrysanthemi
Dickeya chrysanthemi NCPPB 3533 WP 012769548 1 Dchrysanthemi
Dickeya chrysanthemi subsp chrysanthemi NCPPB 402 WP 012769548 1 Dchrysanthemi
Dickeya chrysanthemi subsp parthenii NCPPB 516 WP 012769548 1 Dchrysanthemi
Dickeya chrysanthemi Ech1591 WP 012769548 1 Dchrysanthemi
Dickeya zeae CSL RW192 WP 050570731 1 Dzeae
Dickeya zeae EC1 WP 029456457 1 Dzeae
Dickeya zeae DZ2Q WP 029456457 1 Dzeae
Dickeya zeae ZJU1202 WP 029456457 1 Dzeae
Dickeya zeae MK19 WP 026358052 1 Dzeae
Dickeya zeae MS1 WP 026358052 1 Dzeae
Dickeya zeae NCPPB 3532 WP 026358052 1 Dzeae
Dickeya zeae Ech586 WP 012884477 1 Dzeae
Dickeya zeae NCPPB 2538 WP 026358052 1 Dzeae
Dickeya zeae NCPPB 3531 WP 026358052 1 Dzeae
Dickeya solani RNS 05 1 2A WP 057084945 1 Dsolani
Dickeya gs undicola 2B12 WP 033568655 1 Dundicola
Dickeya sp CSL RW240 WP 035345922 1 Daquatica
Dickeya aquatica DW 0440 WP 035345922 1 Daquatica
DAQ1742 DAQ1742 02556 Daquatica
Dickeya solani IPO 2222 WP 022633831 1 Dsolani
Dickeya solani RNS 08 23 3 1 A WP 022633831 1 Dsolani
Dickeya solani MK16 WP 022633831 1 Dsolani
Dickeya solani PPO 9134 WP 022633831 1 Dsolani
Dickeya solani D s0432 1 WP 022633831 1 Dsolani
Dickeya solani PPO 9019 WP 023637852 1 Dsolani
Dickeya solani MK10 WP 022633831 1 Dsolani
Dickeya solani IFB 0099 WP 023637852 1 Dsolani
Dickeya solani RNS 07 7 3B WP 023637852 1 Dsolani
Dickeya solani GBBC 2040 WP 022633831 1 Dsolani
Dickeya dianthicola NCPPB 453 WP 026594756 1 Ddianthicola
Dickeya dianthicola IPO 980 WP 026594756 1 Ddianthicola
Dickeya dianthicola RNS04 9 WP 026594756 1 Ddianthicola
Dickeya dianthicola NCPPB 3534 WP 026594756 1 Ddianthicola
Dickeya dianthicola GBBC 2039 WP 026594756 1 Ddianthicola
Dickeya gs fangzhongdai B16 WP 038919315 1 Dfangzhongdai
Dickeya gs fangzhongdai MK7 WP 038919315 1 Dfangzhongdai
Dickeya gs fangzhongdai M074 WP 038658006 1 Dfangzhongdai
Dickeya gs fangzhongdai ND14b WP 038658006 1 Dfangzhongdai
Dickeya dadantii NCPPB 3537 WP 033111903 1 Ddadantii
Dickeya dadantii subsp dieffenbachiae NCPPB 2976 WP 033111903 1 Ddadantii
Dickeya dadantii NCPPB 898 WP 033111903 1 Ddadantii
Dickeya gs fangzhongdai S1 WP 033111903 1 Dfangzhongdai
Dickeya dadantii 3937 WP 033111903 1 Ddadantii
Dickeya gs fangzhongdai M005 WP 039692060 1 Dfangzhongdai
Dickeya gs fangzhongdai NCPPB 3274 WP 033111903 1 Dfangzhongdai

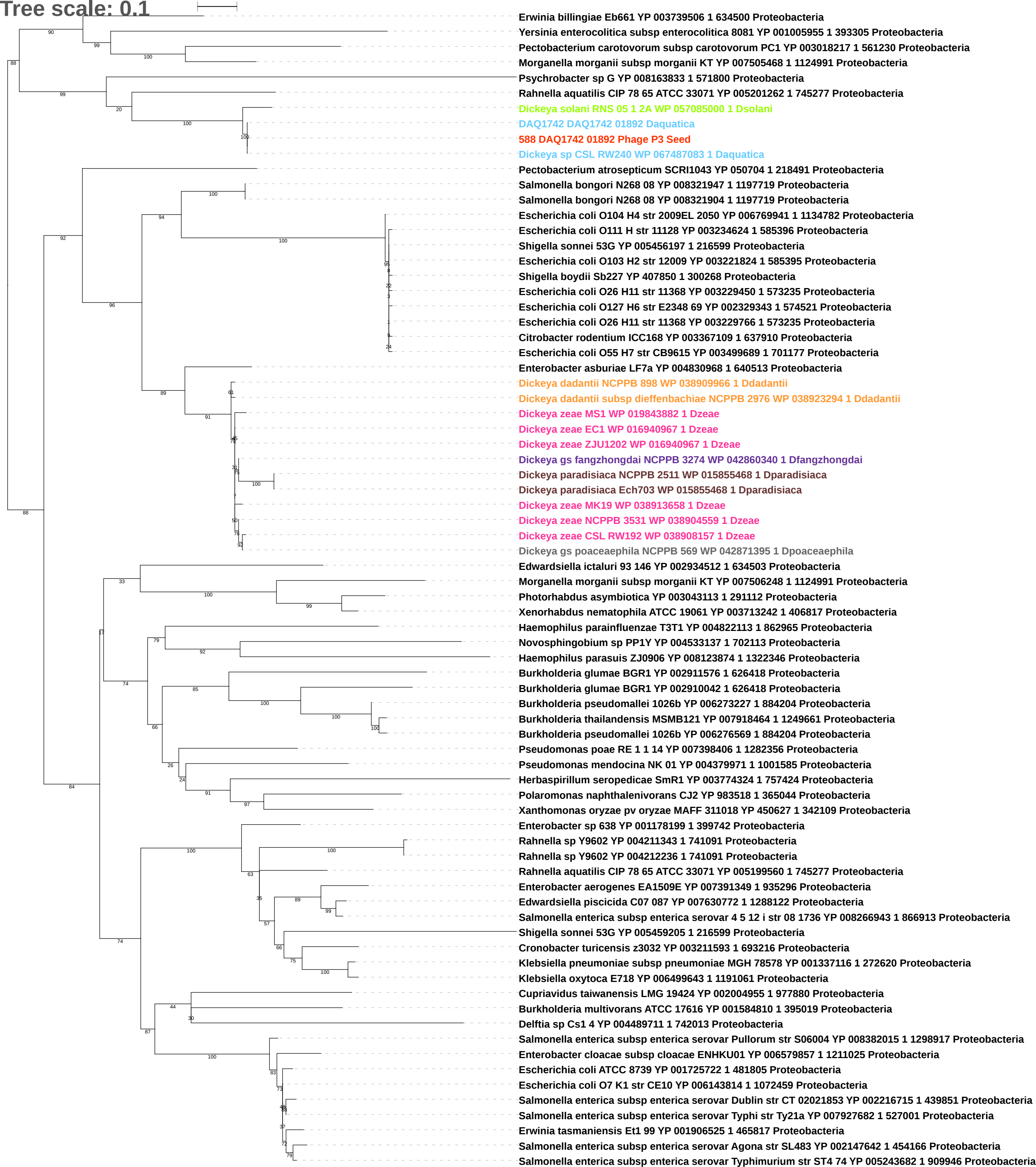
Tree scale: 0.1
Erwinia billingiae Eb661 YP 003739506 1 634500 Proteobacteria
Yersinia enterocolitica subsp enterocolitica 8081 YP 001005955 1 393305 Proteobacteria
Pectobacterium carotovorum subsp carotovorum PC1 YP 003018217 1 561230 Proteobacteria
Morganella morganii subsp morganii KT YP 007505468 1 1124991 Proteobacteria
Psychrobacter sp G YP 008163833 1 571800 Proteobacteria
Rahnella aquatilis CIP 78 65 ATCC 33071 YP 005201262 1 745277 Proteobacteria
Dickeya solani RNS 05 1 2A WP 057085000 1 Dsolani
DAQ1742 DAQ1742 01892 Daquatica
588 DAQ1742 01892 Phage P3 Seed
Dickeya sp CSL RW240 WP 067487083 1 Daquatica
Pectobacterium atrosepticum SCRI1043 YP 050704 1 218491 Proteobacteria
Salmonella bongori N268 08 YP 008321947 1 1197719 Proteobacteria
Salmonella bongori N268 08 YP 008321904 1 1197719 Proteobacteria
Escherichia coli O104 H4 str 2009EL 2050 YP 006769941 1 1134782 Proteobacteria
Escherichia coli O111 H str 11128 YP 003234624 1 585396 Proteobacteria
Shigella sonnei 53G YP 005456197 1 216599 Proteobacteria
Escherichia coli O103 H2 str 12009 YP 003221824 1 585395 Proteobacteria
Shigella boydii Sb227 YP 407850 1 300268 Proteobacteria
Escherichia coli O26 H11 str 11368 YP 003229450 1 573235 Proteobacteria
Escherichia coli O127 H6 str E2348 69 YP 002329343 1 574521 Proteobacteria
Escherichia coli O26 H11 str 11368 YP 003229766 1 573235 Proteobacteria
Citrobacter rodentium ICC168 YP 003367109 1 637910 Proteobacteria
Escherichia coli O55 H7 str CB9615 YP 003499689 1 701177 Proteobacteria
Enterobacter asburiae LF7a YP 004830968 1 640513 Proteobacteria
Dickeya dadantii NCPPB 898 WP 038909966 1 Ddadantii
Dickeya dadantii subsp dieffenbachiae NCPPB 2976 WP 038923294 1 Ddadantii
Dickeya zeae MS1 WP 019843882 1 Dzeae
Dickeya zeae EC1 WP 016940967 1 Dzeae
Dickeya zeae ZJU1202 WP 016940967 1 Dzeae
Dickeya gs fangzhongdai NCPPB 3274 WP 042860340 1 Dfangzhongdai
Dickeya paradisiaca NCPPB 2511 WP 015855468 1 Dparadisiaca
Dickeya paradisiaca Ech703 WP 015855468 1 Dparadisiaca
Dickeya zeae MK19 WP 038913658 1 Dzeae
Dickeya zeae NCPPB 3531 WP 038904559 1 Dzeae
Dickeya zeae CSL RW192 WP 038908157 1 Dzeae
Dickeya gs poaceaephila NCPPB 569 WP 042871395 1 Dpoaceaephila
Edwardsiella ictaluri 93 146 YP 002934512 1 634503 Proteobacteria
Morganella morganii subsp morganii KT YP 007506248 1 1124991 Proteobacteria
Photorhabdus asymbiotica YP 003043113 1 291112 Proteobacteria
Xenorhabdus nematophila ATCC 19061 YP 003713242 1 406817 Proteobacteria
Haemophilus parainfluenzae T3T1 YP 004822113 1 862965 Proteobacteria
Novosphingobium sp PP1Y YP 004533137 1 702113 Proteobacteria
Haemophilus parasuis ZJ0906 YP 008123874 1 1322346 Proteobacteria
Burkholderia glumae BGR1 YP 002911576 1 626418 Proteobacteria
Burkholderia glumae BGR1 YP 002910042 1 626418 Proteobacteria
Burkholderia pseudomallei 1026b YP 006273227 1 884204 Proteobacteria
Burkholderia thailandensis MSMB121 YP 007918464 1 1249661 Proteobacteria
Burkholderia pseudomallei 1026b YP 006276569 1 884204 Proteobacteria
Pseudomonas poae RE 1 1 14 YP 007398406 1 1282356 Proteobacteria
Pseudomonas mendocina NK 01 YP 004379971 1 1001585 Proteobacteria
Herbaspirillum seropedicae SmR1 YP 003774324 1 757424 Proteobacteria
Polaromonas naphthalenivorans CJ2 YP 983518 1 365044 Proteobacteria
Xanthomonas oryzae pv oryzae MAFF 311018 YP 450627 1 342109 Proteobacteria
Enterobacter sp 638 YP 001178199 1 399742 Proteobacteria
Rahnella sp Y9602 YP 004211343 1 741091 Proteobacteria
Rahnella sp Y9602 YP 004212236 1 741091 Proteobacteria
Rahnella aquatilis CIP 78 65 ATCC 33071 YP 005199560 1 745277 Proteobacteria
Enterobacter aerogenes EA1509E YP 007391349 1 935296 Proteobacteria
Edwardsiella piscicida C07 087 YP 007630772 1 1288122 Proteobacteria
Salmonella enterica subsp enterica serovar 4 5 12 i str 08 1736 YP 008266943 1 866913 Proteobacteria
Shigella sonnei 53G YP 005459205 1 216599 Proteobacteria
Cronobacter turicensis z3032 YP 003211593 1 693216 Proteobacteria
Klebsiella pneumoniae subsp pneumoniae MGH 78578 YP 001337116 1 272620 Proteobacteria
Klebsiella oxytoca E718 YP 006499643 1 1191061 Proteobacteria
Cupriavidus taiwanensis LMG 19424 YP 002004955 1 977880 Proteobacteria
Burkholderia multivorans ATCC 17616 YP 001584810 1 395019 Proteobacteria
Delftia sp Cs1 4 YP 004489711 1 742013 Proteobacteria
Salmonella enterica subsp enterica serovar Pullorum str S06004 YP 008382015 1 1298917 Proteobacteria
Enterobacter cloacae subsp cloacae ENHKU01 YP 006579857 1 1211025 Proteobacteria
Escherichia coli ATCC 8739 YP 001725722 1 481805 Proteobacteria
Escherichia coli O7 K1 str CE10 YP 006143814 1 1072459 Proteobacteria
Salmonella enterica subsp enterica serovar Dublin str CT 02021853 YP 002216715 1 439851 Proteobacteria
Salmonella enterica subsp enterica serovar Typhi str Ty21a YP 007927682 1 527001 Proteobacteria
Erwinia tasmaniensis Et1 99 YP 001906525 1 465817 Proteobacteria
Salmonella enterica subsp enterica serovar Agona str SL483 YP 002147642 1 454166 Proteobacteria
Salmonella enterica subsp enterica serovar Typhimurium str ST4 74 YP 005243682 1 909946 Proteobacteria

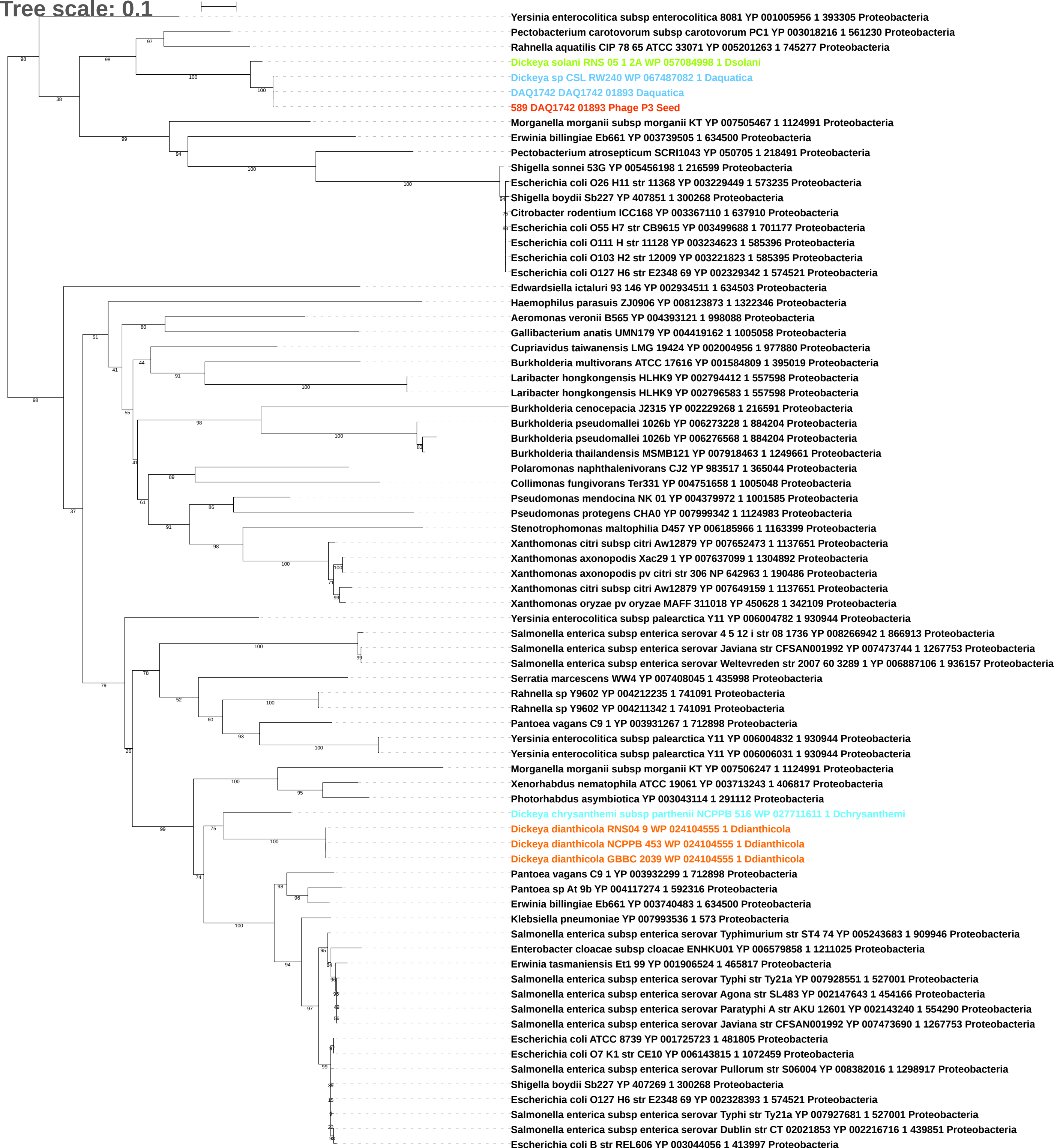
Tree scale: 0.1
Yersinia enterocolitica subsp enterocolitica 8081 YP 001005956 1 393305 Proteobacteria
Pectobacterium carotovorum subsp carotovorum PC1 YP 003018216 1 561230 Proteobacteria
Rahnella aquatilis CIP 78 65 ATCC 33071 YP 005201263 1 745277 Proteobacteria
Dickeya solani RNS 05 1 2A WP 057084998 1 Dsolani
Dickeya sp CSL RW240 WP 067487082 1 Daquatica
DAQ1742 DAQ1742 01893 Daquatica
589 DAQ1742 01893 Phage P3 Seed
Morganella morganii subsp morganii KT YP 007505467 1 1124991 Proteobacteria
Erwinia billingiae Eb661 YP 003739505 1 634500 Proteobacteria
Pectobacterium atrosepticum SCRI1043 YP 050705 1 218491 Proteobacteria
Shigella sonnei 53G YP 005456198 1 216599 Proteobacteria
Escherichia coli O26 H11 str 11368 YP 003229449 1 573235 Proteobacteria
Shigella boydii Sb227 YP 407851 1 300268 Proteobacteria
Citrobacter rodentium ICC168 YP 003367110 1 637910 Proteobacteria
Escherichia coli O55 H7 str CB9615 YP 003499688 1 701177 Proteobacteria
Escherichia coli O111 H str 11128 YP 003234623 1 585396 Proteobacteria
Escherichia coli O103 H2 str 12009 YP 003221823 1 585395 Proteobacteria
Escherichia coli O127 H6 str E2348 69 YP 002329342 1 574521 Proteobacteria
Edwardsiella ictaluri 93 146 YP 002934511 1 634503 Proteobacteria
Haemophilus parasuis ZJ0906 YP 008123873 1 1322346 Proteobacteria
Aeromonas veronii B565 YP 004393121 1 998088 Proteobacteria
Gallibacterium anatis UMN179 YP 004419162 1 1005058 Proteobacteria
Cupriavidus taiwanensis LMG 19424 YP 002004956 1 977880 Proteobacteria
Burkholderia multivorans ATCC 17616 YP 001584809 1 395019 Proteobacteria
Laribacter hongkongensis HLHK9 YP 002794412 1 557598 Proteobacteria
Laribacter hongkongensis HLHK9 YP 002796583 1 557598 Proteobacteria
Burkholderia cenocepacia J2315 YP 002229268 1 216591 Proteobacteria
Burkholderia pseudomallei 1026b YP 006273228 1 884204 Proteobacteria
Burkholderia pseudomallei 1026b YP 006276568 1 884204 Proteobacteria
Burkholderia thailandensis MSMB121 YP 007918463 1 1249661 Proteobacteria
Polaromonas naphthalenivorans CJ2 YP 983517 1 365044 Proteobacteria
Collimonas fungivorans Ter331 YP 004751658 1 1005048 Proteobacteria
Pseudomonas mendocina NK 01 YP 004379972 1 1001585 Proteobacteria
Pseudomonas protegens CHA0 YP 007999342 1 1124983 Proteobacteria
Stenotrophomonas maltophilia D457 YP 006185966 1 1163399 Proteobacteria
Xanthomonas citri subsp citri Aw12879 YP 007652473 1 1137651 Proteobacteria
Xanthomonas axonopodis Xac29 1 YP 007637099 1 1304892 Proteobacteria
Xanthomonas axonopodis pv citri str 306 NP 642963 1 190486 Proteobacteria
Xanthomonas citri subsp citri Aw12879 YP 007649159 1 1137651 Proteobacteria
Xanthomonas oryzae pv oryzae MAFF 311018 YP 450628 1 342109 Proteobacteria
Yersinia enterocolitica subsp palearctica Y11 YP 006004782 1 930944 Proteobacteria
Salmonella enterica subsp enterica serovar 4 5 12 i str 08 1736 YP 008266942 1 866913 Proteobacteria
Salmonella enterica subsp enterica serovar Javiana str CFSAN001992 YP 007473744 1 1267753 Proteobacteria
Salmonella enterica subsp enterica serovar Weltevreden str 2007 60 3289 1 YP 006887106 1 936157 Proteobacteria
Serratia marcescens WW4 YP 007408045 1 435998 Proteobacteria
Rahnella sp Y9602 YP 004212235 1 741091 Proteobacteria
Rahnella sp Y9602 YP 004211342 1 741091 Proteobacteria
Pantoea vagans C9 1 YP 003931267 1 712898 Proteobacteria
Yersinia enterocolitica subsp palearctica Y11 YP 006004832 1 930944 Proteobacteria
Yersinia enterocolitica subsp palearctica Y11 YP 006006031 1 930944 Proteobacteria
Morganella morganii subsp morganii KT YP 007506247 1 1124991 Proteobacteria
Xenorhabdus nematophila ATCC 19061 YP 003713243 1 406817 Proteobacteria
Photorhabdus asymbiotica YP 003043114 1 291112 Proteobacteria
Dickeya chrysanthemi subsp parthenii NCPPB 516 WP 027711611 1 Dchrysanthemi
Dickeya dianthicola RNS04 9 WP 024104555 1 Ddianthicola
Dickeya dianthicola NCPPB 453 WP 024104555 1 Ddianthicola
Dickeya dianthicola GBBC 2039 WP 024104555 1 Ddianthicola
Pantoea vagans C9 1 YP 003932299 1 712898 Proteobacteria
Pantoea sp At 9b YP 004117274 1 592316 Proteobacteria
Erwinia billingiae Eb661 YP 003740483 1 634500 Proteobacteria
Klebsiella pneumoniae YP 007993536 1 573 Proteobacteria
Salmonella enterica subsp enterica serovar Typhimurium str ST4 74 YP 005243683 1 909946 Proteobacteria
Enterobacter cloacae subsp cloacae ENHKU01 YP 006579858 1 1211025 Proteobacteria
Erwinia tasmaniensis Et1 99 YP 001906524 1 465817 Proteobacteria
Salmonella enterica subsp enterica serovar Typhi str Ty21a YP 007928551 1 527001 Proteobacteria
Salmonella enterica subsp enterica serovar Agona str SL483 YP 002147643 1 454166 Proteobacteria
Salmonella enterica subsp enterica serovar Paratyphi A str AKU 12601 YP 002143240 1 554290 Proteobacteria
Salmonella enterica subsp enterica serovar Javiana str CFSAN001992 YP 007473690 1 1267753 Proteobacteria
Escherichia coli ATCC 8739 YP 001725723 1 481805 Proteobacteria
Escherichia coli O7 K1 str CE10 YP 006143815 1 1072459 Proteobacteria
Salmonella enterica subsp enterica serovar Pullorum str S06004 YP 008382016 1 1298917 Proteobacteria
Shigella boydii Sb227 YP 407269 1 300268 Proteobacteria
Escherichia coli O127 H6 str E2348 69 YP 002328393 1 574521 Proteobacteria
Salmonella enterica subsp enterica serovar Typhi str Ty21a YP 007927681 1 527001 Proteobacteria
Salmonella enterica subsp enterica serovar Dublin str CT 02021853 YP 002216716 1 439851 Proteobacteria
Escherichia coli B str REL606 YP 003044056 1 413997 Proteobacteria

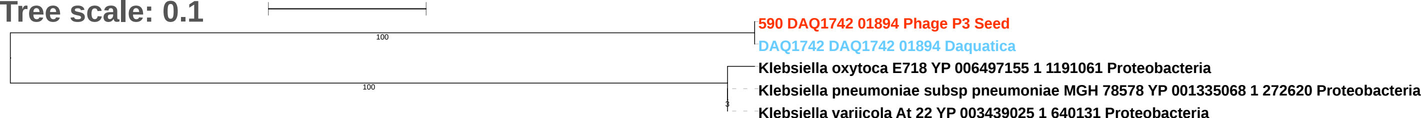
Tree scale: 0.1
590 DAQ1742 01894 Phage P3 Seed
DAQ1742 DAQ1742 01894 Daquatica
Klebsiella oxytoca E718 YP 006497155 1 1191061 Proteobacteria
Klebsiella pneumoniae subsp pneumoniae MGH 78578 YP 001335068 1 272620 Proteobacteria
Klebsiella variicola At 22 YP 003439025 1 640131 Proteobacteria
100
100
3

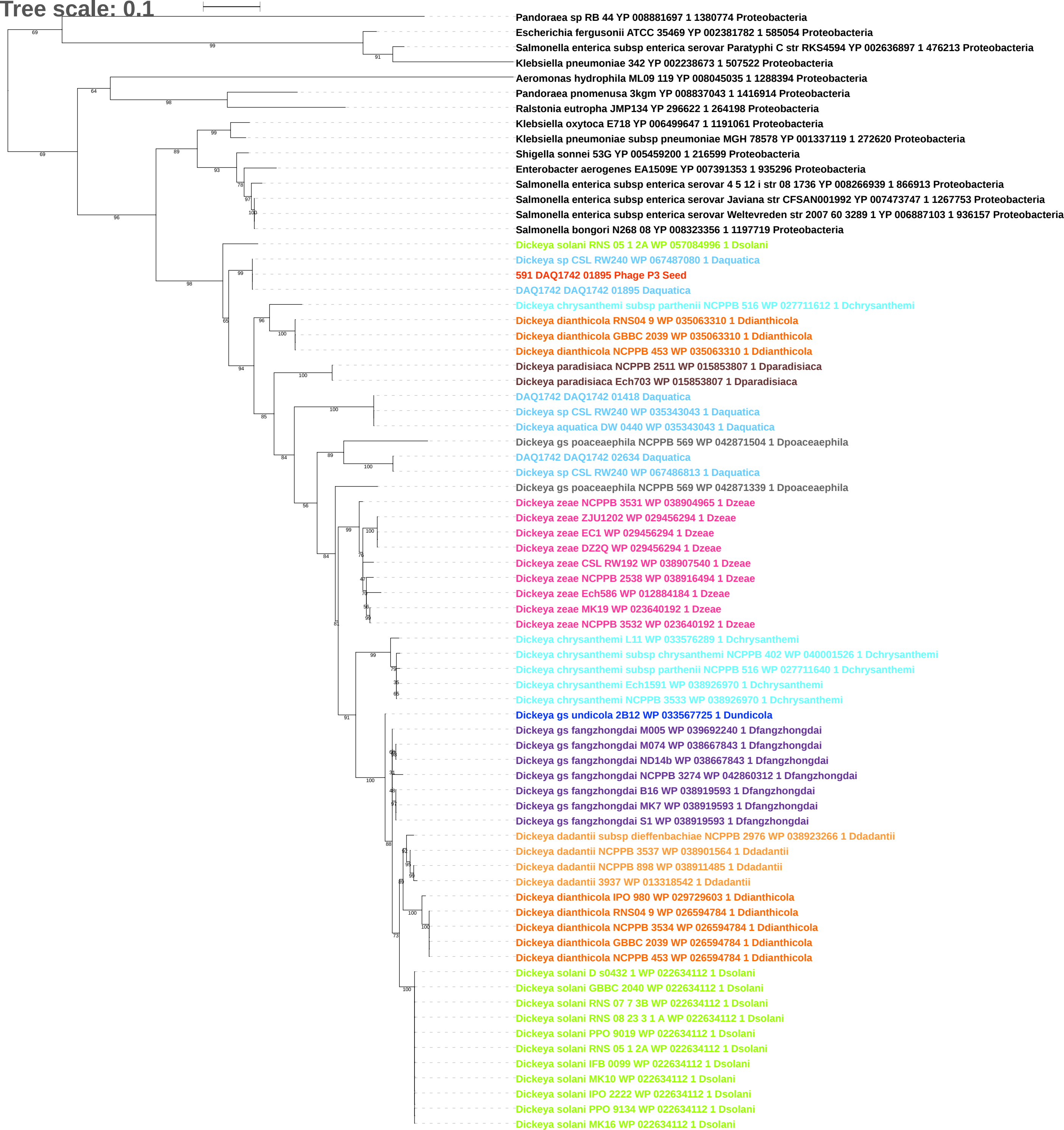
Tree scale: 0.1
Pandoraea sp RB 44 YP 008881697 1 1380774 Proteobacteria
Escherichia fergusonii ATCC 35469 YP 002381782 1 585054 Proteobacteria
Salmonella enterica subsp enterica serovar Paratyphi C str RKS4594 YP 002636897 1 476213 Proteobacteria
Klebsiella pneumoniae 342 YP 002238673 1 507522 Proteobacteria
Aeromonas hydrophila ML09 119 YP 008045035 1 1288394 Proteobacteria
Pandoraea pnomenusa 3kgm YP 008837043 1 1416914 Proteobacteria
Ralstonia eutropha JMP134 YP 296622 1 264198 Proteobacteria
Klebsiella oxytoca E718 YP 006499647 1 1191061 Proteobacteria
Klebsiella pneumoniae subsp pneumoniae MGH 78578 YP 001337119 1 272620 Proteobacteria
Shigella sonnei 53G YP 005459200 1 216599 Proteobacteria
Enterobacter aerogenes EA1509E YP 007391353 1 935296 Proteobacteria
Salmonella enterica subsp enterica serovar 4 5 12 i str 08 1736 YP 008266939 1 866913 Proteobacteria
Salmonella enterica subsp enterica serovar Javiana str CFSAN001992 YP 007473747 1 1267753 Proteobacteria
Salmonella enterica subsp enterica serovar Weltevreden str 2007 60 3289 1 YP 006887103 1 936157 Proteobacteria
Salmonella bongori N268 08 YP 008323356 1 1197719 Proteobacteria
Dickeya solani RNS 05 1 2A WP 057084996 1 Dsolani
Dickeya sp CSL RW240 WP 067487080 1 Daquatica
591 DAQ1742 01895 Phage P3 Seed
DAQ1742 DAQ1742 01895 Daquatica
Dickeya chrysanthemi subsp parthenii NCPPB 516 WP 027711612 1 Dchrysanthemi
Dickeya dianthicola RNS04 9 WP 035063310 1 Ddianthicola
Dickeya dianthicola GBBC 2039 WP 035063310 1 Ddianthicola
Dickeya dianthicola NCPPB 453 WP 035063310 1 Ddianthicola
Dickeya paradisiaca NCPPB 2511 WP 015853807 1 Dparadisiaca
Dickeya paradisiaca Ech703 WP 015853807 1 Dparadisiaca
DAQ1742 DAQ1742 01418 Daquatica
Dickeya sp CSL RW240 WP 035343043 1 Daquatica
Dickeya aquatica DW 0440 WP 035343043 1 Daquatica
Dickeya gs poaceaephila NCPPB 569 WP 042871504 1 Dpoaceaephila
DAQ1742 DAQ1742 02634 Daquatica
Dickeya sp CSL RW240 WP 067486813 1 Daquatica
Dickeya gs poaceaephila NCPPB 569 WP 042871339 1 Dpoaceaephila
Dickeya zeae NCPPB 3531 WP 038904965 1 Dzeae
Dickeya zeae ZJU1202 WP 029456294 1 Dzeae
Dickeya zeae EC1 WP 029456294 1 Dzeae
Dickeya zeae DZ2Q WP 029456294 1 Dzeae
Dickeya zeae CSL RW192 WP 038907540 1 Dzeae
Dickeya zeae NCPPB 2538 WP 038916494 1 Dzeae
Dickeya zeae Ech586 WP 012884184 1 Dzeae
Dickeya zeae MK19 WP 023640192 1 Dzeae
Dickeya zeae NCPPB 3532 WP 023640192 1 Dzeae
Dickeya chrysanthemi L11 WP 033576289 1 Dchrysanthemi
Dickeya chrysanthemi subsp chrysanthemi NCPPB 402 WP 040001526 1 Dchrysanthemi
Dickeya chrysanthemi subsp parthenii NCPPB 516 WP 027711640 1 Dchrysanthemi
Dickeya chrysanthemi Ech1591 WP 038926970 1 Dchrysanthemi
Dickeya chrysanthemi NCPPB 3533 WP 038926970 1 Dchrysanthemi
Dickeya gs undicola 2B12 WP 033567725 1 Dundicola
Dickeya gs fangzhongdai M005 WP 039692240 1 Dfangzhongdai
Dickeya gs fangzhongdai M074 WP 038667843 1 Dfangzhongdai
Dickeya gs fangzhongdai ND14b WP 038667843 1 Dfangzhongdai
Dickeya gs fangzhongdai NCPPB 3274 WP 042860312 1 Dfangzhongdai
Dickeya gs fangzhongdai B16 WP 038919593 1 Dfangzhongdai
Dickeya gs fangzhongdai MK7 WP 038919593 1 Dfangzhongdai
Dickeya gs fangzhongdai S1 WP 038919593 1 Dfangzhongdai
Dickeya dadantii subsp dieffenbachiae NCPPB 2976 WP 038923266 1 Ddadantii
Dickeya dadantii NCPPB 3537 WP 038901564 1 Ddadantii
Dickeya dadantii NCPPB 898 WP 038911485 1 Ddadantii
Dickeya dadantii 3937 WP 013318542 1 Ddadantii
Dickeya dianthicola IPO 980 WP 029729603 1 Ddianthicola
Dickeya dianthicola RNS04 9 WP 026594784 1 Ddianthicola
Dickeya dianthicola NCPPB 3534 WP 026594784 1 Ddianthicola
Dickeya dianthicola GBBC 2039 WP 026594784 1 Ddianthicola
Dickeya dianthicola NCPPB 453 WP 026594784 1 Ddianthicola
Dickeya solani D s0432 1 WP 022634112 1 Dsolani
Dickeya solani GBBC 2040 WP 022634112 1 Dsolani
Dickeya solani RNS 07 7 3B WP 022634112 1 Dsolani
Dickeya solani RNS 08 23 3 1 A WP 022634112 1 Dsolani
Dickeya solani PPO 9019 WP 022634112 1 Dsolani
Dickeya solani RNS 05 1 2A WP 022634112 1 Dsolani
Dickeya solani IFB 0099 WP 022634112 1 Dsolani
Dickeya solani MK10 WP 022634112 1 Dsolani
Dickeya solani IPO 2222 WP 022634112 1 Dsolani
Dickeya solani PPO 9134 WP 022634112 1 Dsolani
Dickeya solani MK16 WP 022634112 1 Dsolani

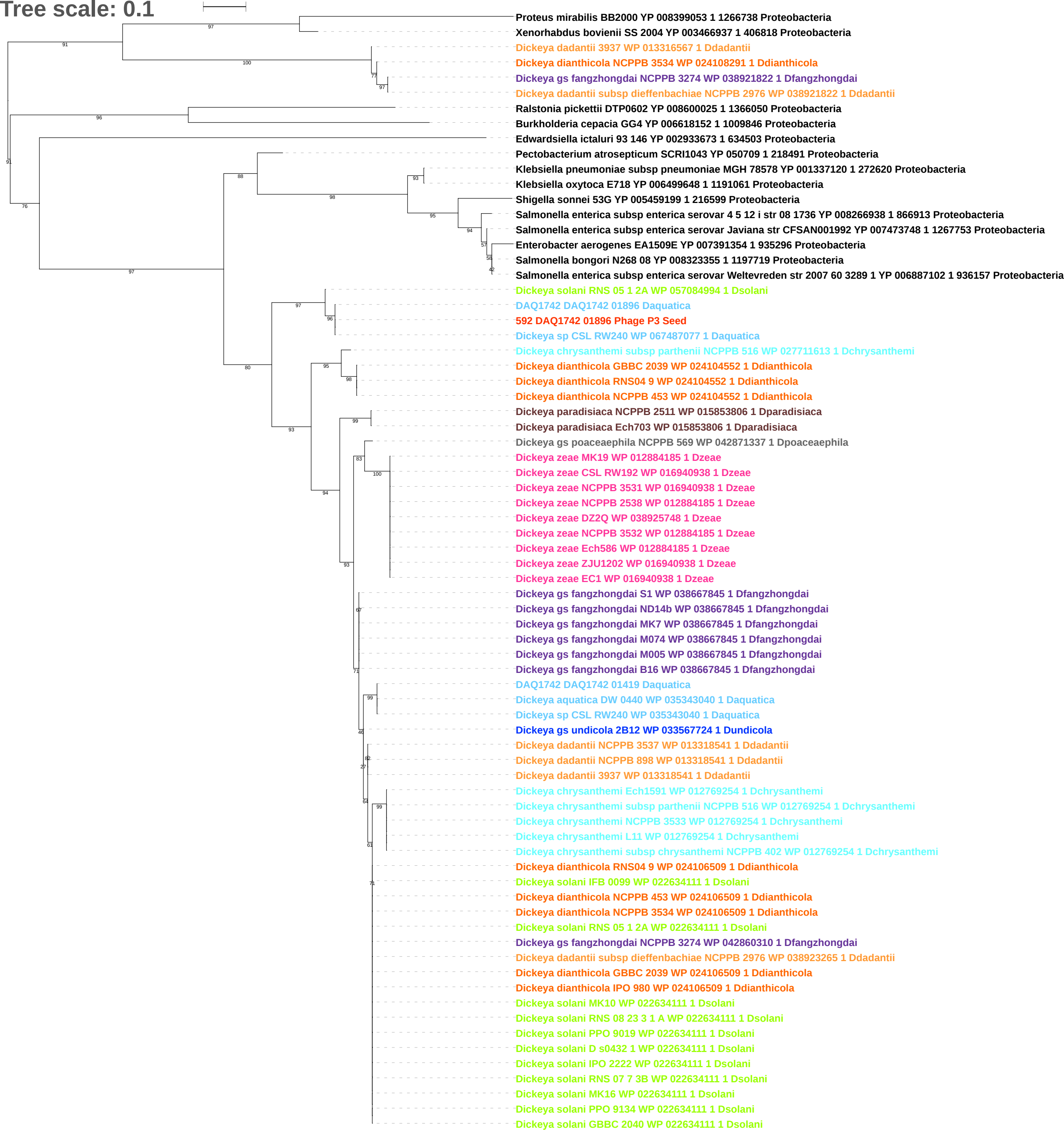
Tree scale: 0.1
Proteus mirabilis BB2000 YP 008399053 1 1266738 Proteobacteria
Xenorhabdus bovienii SS 2004 YP 003466937 1 406818 Proteobacteria
Dickeya dadantii 3937 WP 013316567 1 Ddadantii
Dickeya dianthicola NCPPB 3534 WP 024108291 1 Ddianthicola
Dickeya gs fangzhongdai NCPPB 3274 WP 038921822 1 Dfangzhongdai
Dickeya dadantii subsp dieffenbachiae NCPPB 2976 WP 038921822 1 Ddadantii
Ralstonia pickettii DTP0602 YP 008600025 1 1366050 Proteobacteria
Burkholderia cepacia GG4 YP 006618152 1 1009846 Proteobacteria
Edwardsiella ictaluri 93 146 YP 002933673 1 634503 Proteobacteria
Pectobacterium atrosepticum SCRI1043 YP 050709 1 218491 Proteobacteria
Klebsiella pneumoniae subsp pneumoniae MGH 78578 YP 001337120 1 272620 Proteobacteria
Klebsiella oxytoca E718 YP 006499648 1 1191061 Proteobacteria
Shigella sonnei 53G YP 005459199 1 216599 Proteobacteria
Salmonella enterica subsp enterica serovar 4 5 12 i str 08 1736 YP 008266938 1 866913 Proteobacteria
Salmonella enterica subsp enterica serovar Javiana str CFSAN001992 YP 007473748 1 1267753 Proteobacteria
Enterobacter aerogenes EA1509E YP 007391354 1 935296 Proteobacteria
Salmonella bongori N268 08 YP 008323355 1 1197719 Proteobacteria
Salmonella enterica subsp enterica serovar Weltevreden str 2007 60 3289 1 YP 006887102 1 936157 Proteobacteria
Dickeya solani RNS 05 1 2A WP 057084994 1 Dsolani
DAQ1742 DAQ1742 01896 Daquatica
592 DAQ1742 01896 Phage P3 Seed
Dickeya sp CSL RW240 WP 067487077 1 Daquatica
Dickeya chrysanthemi subsp parthenii NCPPB 516 WP 027711613 1 Dchrysanthemi
Dickeya dianthicola GBBC 2039 WP 024104552 1 Ddianthicola
Dickeya dianthicola RNS04 9 WP 024104552 1 Ddianthicola
Dickeya dianthicola NCPPB 453 WP 024104552 1 Ddianthicola
Dickeya paradisiaca NCPPB 2511 WP 015853806 1 Dparadisiaca
Dickeya paradisiaca Ech703 WP 015853806 1 Dparadisiaca
Dickeya gs poaceaephila NCPPB 569 WP 042871337 1 Dpoaceaephila
Dickeya zeae MK19 WP 012884185 1 Dzeae
Dickeya zeae CSL RW192 WP 016940938 1 Dzeae
Dickeya zeae NCPPB 3531 WP 016940938 1 Dzeae
Dickeya zeae NCPPB 2538 WP 012884185 1 Dzeae
Dickeya zeae DZ2Q WP 038925748 1 Dzeae
Dickeya zeae NCPPB 3532 WP 012884185 1 Dzeae
Dickeya zeae Ech586 WP 012884185 1 Dzeae
Dickeya zeae ZJU1202 WP 016940938 1 Dzeae
Dickeya zeae EC1 WP 016940938 1 Dzeae
Dickeya gs fangzhongdai S1 WP 038667845 1 Dfangzhongdai
Dickeya gs fangzhongdai ND14b WP 038667845 1 Dfangzhongdai
Dickeya gs fangzhongdai MK7 WP 038667845 1 Dfangzhongdai
Dickeya gs fangzhongdai M074 WP 038667845 1 Dfangzhongdai
Dickeya gs fangzhongdai M005 WP 038667845 1 Dfangzhongdai
Dickeya gs fangzhongdai B16 WP 038667845 1 Dfangzhongdai
DAQ1742 DAQ1742 01419 Daquatica
Dickeya aquatica DW 0440 WP 035343040 1 Daquatica
Dickeya sp CSL RW240 WP 035343040 1 Daquatica
Dickeya gs undicola 2B12 WP 033567724 1 Dundicola
Dickeya dadantii NCPPB 3537 WP 013318541 1 Ddadantii
Dickeya dadantii NCPPB 898 WP 013318541 1 Ddadantii
Dickeya dadantii 3937 WP 013318541 1 Ddadantii
Dickeya chrysanthemi Ech1591 WP 012769254 1 Dchrysanthemi
Dickeya chrysanthemi subsp parthenii NCPPB 516 WP 012769254 1 Dchrysanthemi
Dickeya chrysanthemi NCPPB 3533 WP 012769254 1 Dchrysanthemi
Dickeya chrysanthemi L11 WP 012769254 1 Dchrysanthemi
Dickeya chrysanthemi subsp chrysanthemi NCPPB 402 WP 012769254 1 Dchrysanthemi
Dickeya dianthicola RNS04 9 WP 024106509 1 Ddianthicola
Dickeya solani IFB 0099 WP 022634111 1 Dsolani
Dickeya dianthicola NCPPB 453 WP 024106509 1 Ddianthicola
Dickeya dianthicola NCPPB 3534 WP 024106509 1 Ddianthicola
Dickeya solani RNS 05 1 2A WP 022634111 1 Dsolani
Dickeya gs fangzhongdai NCPPB 3274 WP 042860310 1 Dfangzhongdai
Dickeya dadantii subsp dieffenbachiae NCPPB 2976 WP 038923265 1 Ddadantii
Dickeya dianthicola GBBC 2039 WP 024106509 1 Ddianthicola
Dickeya dianthicola IPO 980 WP 024106509 1 Ddianthicola
Dickeya solani MK10 WP 022634111 1 Dsolani
Dickeya solani RNS 08 23 3 1 A WP 022634111 1 Dsolani
Dickeya solani PPO 9019 WP 022634111 1 Dsolani
Dickeya solani D s0432 1 WP 022634111 1 Dsolani
Dickeya solani IPO 2222 WP 022634111 1 Dsolani
Dickeya solani RNS 07 7 3B WP 022634111 1 Dsolani
Dickeya solani MK16 WP 022634111 1 Dsolani
Dickeya solani PPO 9134 WP 022634111 1 Dsolani
Dickeya solani GBBC 2040 WP 022634111 1 Dsolani

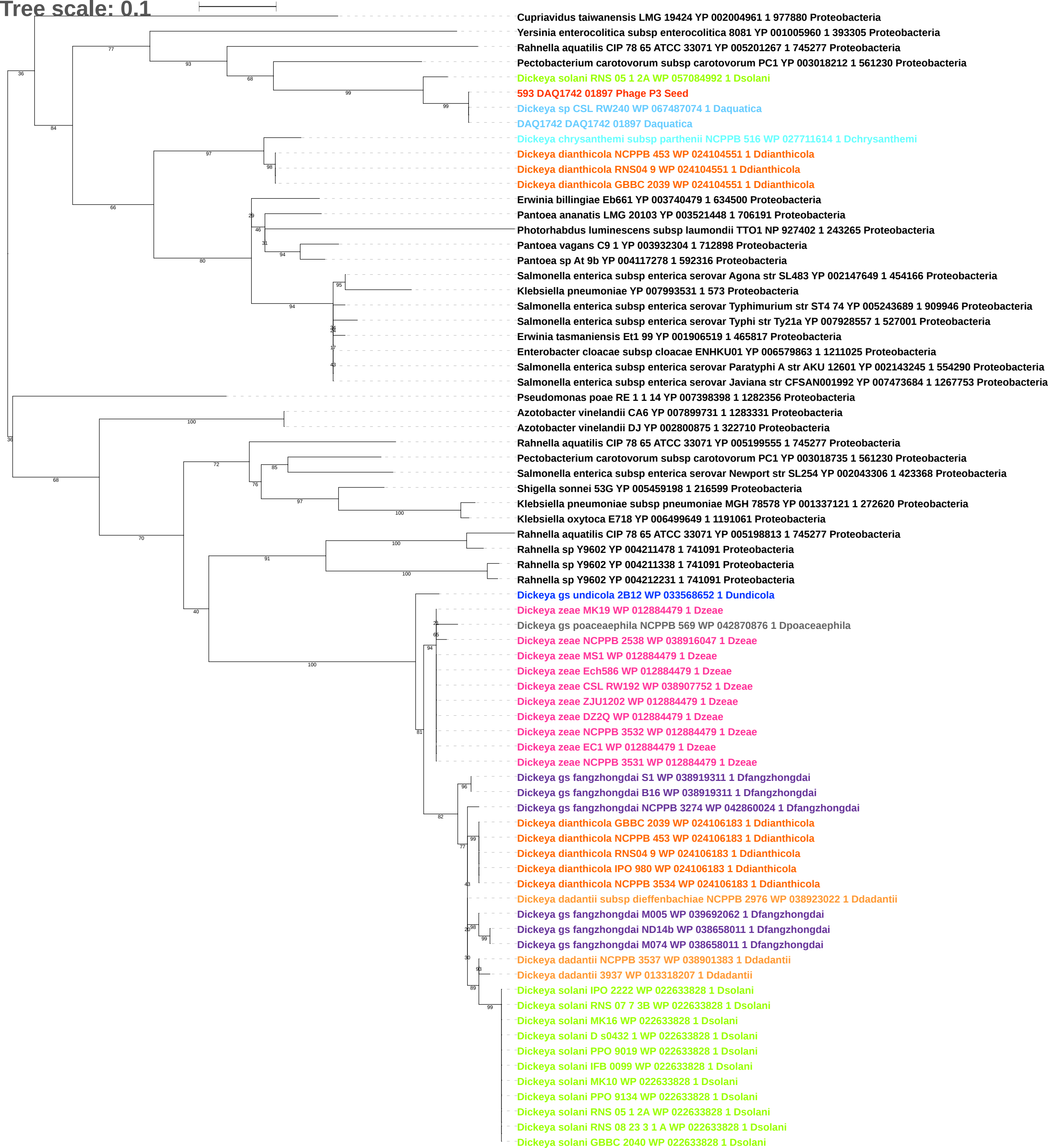
Tree scale: 0.1
Cupriavidus taiwanensis LMG 19424 YP 002004961 1 977880 Proteobacteria
Yersinia enterocolitica subsp enterocolitica 8081 YP 001005960 1 393305 Proteobacteria
Rahnella aquatilis CIP 78 65 ATCC 33071 YP 005201267 1 745277 Proteobacteria
Pectobacterium carotovorum subsp carotovorum PC1 YP 003018212 1 561230 Proteobacteria
Dickeya solani RNS 05 1 2A WP 057084992 1 Dsolani
593 DAQ1742 01897 Phage P3 Seed
Dickeya sp CSL RW240 WP 067487074 1 Daquatica
DAQ1742 DAQ1742 01897 Daquatica
Dickeya chrysanthemi subsp parthenii NCPPB 516 WP 027711614 1 Dchrysanthemi
Dickeya dianthicola NCPPB 453 WP 024104551 1 Ddianthicola
Dickeya dianthicola RNS04 9 WP 024104551 1 Ddianthicola
Dickeya dianthicola GBBC 2039 WP 024104551 1 Ddianthicola
Erwinia billingiae Eb661 YP 003740479 1 634500 Proteobacteria
Pantoea ananatis LMG 20103 YP 003521448 1 706191 Proteobacteria
Photorhabdus luminescens subsp laumondii TTO1 NP 927402 1 243265 Proteobacteria
Pantoea vagans C9 1 YP 003932304 1 712898 Proteobacteria
Pantoea sp At 9b YP 004117278 1 592316 Proteobacteria
Salmonella enterica subsp enterica serovar Agona str SL483 YP 002147649 1 454166 Proteobacteria
Klebsiella pneumoniae YP 007993531 1 573 Proteobacteria
Salmonella enterica subsp enterica serovar Typhimurium str ST4 74 YP 005243689 1 909946 Proteobacteria
Salmonella enterica subsp enterica serovar Typhi str Ty21a YP 007928557 1 527001 Proteobacteria
Erwinia tasmaniensis Et1 99 YP 001906519 1 465817 Proteobacteria
Enterobacter cloacae subsp cloacae ENHKU01 YP 006579863 1 1211025 Proteobacteria
Salmonella enterica subsp enterica serovar Paratyphi A str AKU 12601 YP 002143245 1 554290 Proteobacteria
Salmonella enterica subsp enterica serovar Javiana str CFSAN001992 YP 007473684 1 1267753 Proteobacteria
Pseudomonas poae RE 1 1 14 YP 007398398 1 1282356 Proteobacteria
Azotobacter vinelandii CA6 YP 007899731 1 1283331 Proteobacteria
Azotobacter vinelandii DJ YP 002800875 1 322710 Proteobacteria
Rahnella aquatilis CIP 78 65 ATCC 33071 YP 005199555 1 745277 Proteobacteria
Pectobacterium carotovorum subsp carotovorum PC1 YP 003018735 1 561230 Proteobacteria
Salmonella enterica subsp enterica serovar Newport str SL254 YP 002043306 1 423368 Proteobacteria
Shigella sonnei 53G YP 005459198 1 216599 Proteobacteria
Klebsiella pneumoniae subsp pneumoniae MGH 78578 YP 001337121 1 272620 Proteobacteria
Klebsiella oxytoca E718 YP 006499649 1 1191061 Proteobacteria
Rahnella aquatilis CIP 78 65 ATCC 33071 YP 005198813 1 745277 Proteobacteria
Rahnella sp Y9602 YP 004211478 1 741091 Proteobacteria
Rahnella sp Y9602 YP 004211338 1 741091 Proteobacteria
Rahnella sp Y9602 YP 004212231 1 741091 Proteobacteria
Dickeya gs undicola 2B12 WP 033568652 1 Dundicola
Dickeya zeae MK19 WP 012884479 1 Dzeae
Dickeya gs poaceaephila NCPPB 569 WP 042870876 1 Dpoaceaephila
Dickeya zeae NCPPB 2538 WP 038916047 1 Dzeae
Dickeya zeae MS1 WP 012884479 1 Dzeae
Dickeya zeae Ech586 WP 012884479 1 Dzeae
Dickeya zeae CSL RW192 WP 038907752 1 Dzeae
Dickeya zeae ZJU1202 WP 012884479 1 Dzeae
Dickeya zeae DZ2Q WP 012884479 1 Dzeae
Dickeya zeae NCPPB 3532 WP 012884479 1 Dzeae
Dickeya zeae EC1 WP 012884479 1 Dzeae
Dickeya zeae NCPPB 3531 WP 012884479 1 Dzeae
Dickeya gs fangzhongdai S1 WP 038919311 1 Dfangzhongdai
Dickeya gs fangzhongdai B16 WP 038919311 1 Dfangzhongdai
Dickeya gs fangzhongdai NCPPB 3274 WP 042860024 1 Dfangzhongdai
Dickeya dianthicola GBBC 2039 WP 024106183 1 Ddianthicola
Dickeya dianthicola NCPPB 453 WP 024106183 1 Ddianthicola
Dickeya dianthicola RNS04 9 WP 024106183 1 Ddianthicola
Dickeya dianthicola IPO 980 WP 024106183 1 Ddianthicola
Dickeya dianthicola NCPPB 3534 WP 024106183 1 Ddianthicola
Dickeya dadantii subsp dieffenbachiae NCPPB 2976 WP 038923022 1 Ddadantii
Dickeya gs fangzhongdai M005 WP 039692062 1 Dfangzhongdai
Dickeya gs fangzhongdai ND14b WP 038658011 1 Dfangzhongdai
Dickeya gs fangzhongdai M074 WP 038658011 1 Dfangzhongdai
Dickeya dadantii NCPPB 3537 WP 038901383 1 Ddadantii
Dickeya dadantii 3937 WP 013318207 1 Ddadantii
Dickeya solani IPO 2222 WP 022633828 1 Dsolani
Dickeya solani RNS 07 7 3B WP 022633828 1 Dsolani
Dickeya solani MK16 WP 022633828 1 Dsolani
Dickeya solani D s0432 1 WP 022633828 1 Dsolani
Dickeya solani PPO 9019 WP 022633828 1 Dsolani
Dickeya solani IFB 0099 WP 022633828 1 Dsolani
Dickeya solani MK10 WP 022633828 1 Dsolani
Dickeya solani PPO 9134 WP 022633828 1 Dsolani
Dickeya solani RNS 05 1 2A WP 022633828 1 Dsolani
Dickeya solani RNS 08 23 3 1 A WP 022633828 1 Dsolani
Dickeya solani GBBC 2040 WP 022633828 1 Dsolani

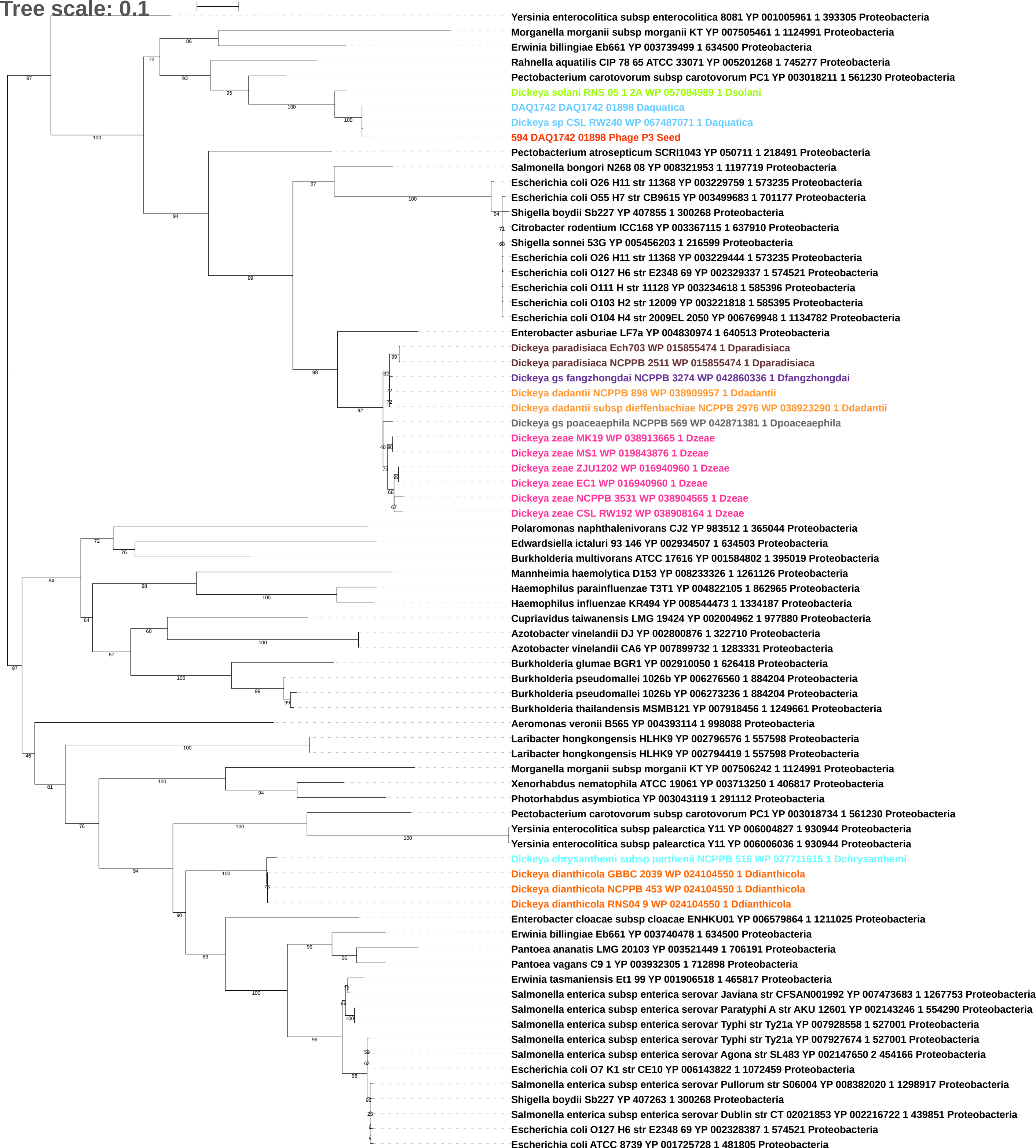
Tree scale: 0.1
Yersinia enterocolitica subsp enterocolitica 8081 YP 001005961 1 393305 Proteobacteria
Morganella morganii subsp morganii KT YP 007505461 1 1124991 Proteobacteria
Erwinia billingiae Eb661 YP 003739499 1 634500 Proteobacteria
Rahnella aquatilis CIP 78 65 ATCC 33071 YP 005201268 1 745277 Proteobacteria
Pectobacterium carotovorum subsp carotovorum PC1 YP 003018211 1 561230 Proteobacteria
Dickeya solani RNS 05 1 2A WP 057084989 1 Dsolani
DAQ1742 DAQ1742 01898 Daquatica
Dickeya sp CSL RW240 WP 067487071 1 Daquatica
594 DAQ1742 01898 Phage P3 Seed
Pectobacterium atrosepticum SCRI1043 YP 050711 1 218491 Proteobacteria
Salmonella bongori N268 08 YP 008321953 1 1197719 Proteobacteria
Escherichia coli O26 H11 str 11368 YP 003229759 1 573235 Proteobacteria
Escherichia coli O55 H7 str CB9615 YP 003499683 1 701177 Proteobacteria
Shigella boydii Sb227 YP 407855 1 300268 Proteobacteria
Citrobacter rodentium ICC168 YP 003367115 1 637910 Proteobacteria
Shigella sonnei 53G YP 005456203 1 216599 Proteobacteria
Escherichia coli O26 H11 str 11368 YP 003229444 1 573235 Proteobacteria
Escherichia coli O127 H6 str E2348 69 YP 002329337 1 574521 Proteobacteria
Escherichia coli O111 H str 11128 YP 003234618 1 585396 Proteobacteria
Escherichia coli O103 H2 str 12009 YP 003221818 1 585395 Proteobacteria
Escherichia coli O104 H4 str 2009EL 2050 YP 006769948 1 1134782 Proteobacteria
Enterobacter asburiae LF7a YP 004830974 1 640513 Proteobacteria
Dickeya paradisiaca Ech703 WP 015855474 1 Dparadisiaca
Dickeya paradisiaca NCPPB 2511 WP 015855474 1 Dparadisiaca
Dickeya gs fangzhongdai NCPPB 3274 WP 042860336 1 Dfangzhongdai
Dickeya dadantii NCPPB 898 WP 038909957 1 Ddadantii
Dickeya dadantii subsp dieffenbachiae NCPPB 2976 WP 038923290 1 Ddadantii
Dickeya gs poaceaephila NCPPB 569 WP 042871381 1 Dpoaceaephila
Dickeya zeae MK19 WP 038913665 1 Dzeae
Dickeya zeae MS1 WP 019843876 1 Dzeae
Dickeya zeae ZJU1202 WP 016940960 1 Dzeae
Dickeya zeae EC1 WP 016940960 1 Dzeae
Dickeya zeae NCPPB 3531 WP 038904565 1 Dzeae
Dickeya zeae CSL RW192 WP 038908164 1 Dzeae
Polaromonas naphthalenivorans CJ2 YP 983512 1 365044 Proteobacteria
Edwardsiella ictaluri 93 146 YP 002934507 1 634503 Proteobacteria
Burkholderia multivorans ATCC 17616 YP 001584802 1 395019 Proteobacteria
Mannheimia haemolytica D153 YP 008233326 1 1261126 Proteobacteria
Haemophilus parainfluenzae T3T1 YP 004822105 1 862965 Proteobacteria
Haemophilus influenzae KR494 YP 008544473 1 1334187 Proteobacteria
Cupriavidus taiwanensis LMG 19424 YP 002004962 1 977880 Proteobacteria
Azotobacter vinelandii DJ YP 002800876 1 322710 Proteobacteria
Azotobacter vinelandii CA6 YP 007899732 1 1283331 Proteobacteria
Burkholderia glumae BGR1 YP 002910050 1 626418 Proteobacteria
Burkholderia pseudomallei 1026b YP 006276560 1 884204 Proteobacteria
Burkholderia pseudomallei 1026b YP 006273236 1 884204 Proteobacteria
Burkholderia thailandensis MSMB121 YP 007918456 1 1249661 Proteobacteria
Aeromonas veronii B565 YP 004393114 1 998088 Proteobacteria
Laribacter hongkongensis HLHK9 YP 002796576 1 557598 Proteobacteria
Laribacter hongkongensis HLHK9 YP 002794419 1 557598 Proteobacteria
Morganella morganii subsp morganii KT YP 007506242 1 1124991 Proteobacteria
Xenorhabdus nematophila ATCC 19061 YP 003713250 1 406817 Proteobacteria
Photorhabdus asymbiotica YP 003043119 1 291112 Proteobacteria
Pectobacterium carotovorum subsp carotovorum PC1 YP 003018734 1 561230 Proteobacteria
Yersinia enterocolitica subsp palearctica Y11 YP 006004827 1 930944 Proteobacteria
Yersinia enterocolitica subsp palearctica Y11 YP 006006036 1 930944 Proteobacteria
Dickeya chrysanthemi subsp parthenii NCPPB 516 WP 027711615 1 Dchrysanthemi
Dickeya dianthicola GBBC 2039 WP 024104550 1 Ddianthicola
Dickeya dianthicola NCPPB 453 WP 024104550 1 Ddianthicola
Dickeya dianthicola RNS04 9 WP 024104550 1 Ddianthicola
Enterobacter cloacae subsp cloacae ENHKU01 YP 006579864 1 1211025 Proteobacteria
Erwinia billingiae Eb661 YP 003740478 1 634500 Proteobacteria
Pantoea ananatis LMG 20103 YP 003521449 1 706191 Proteobacteria
Pantoea vagans C9 1 YP 003932305 1 712898 Proteobacteria
Erwinia tasmaniensis Et1 99 YP 001906518 1 465817 Proteobacteria
Salmonella enterica subsp enterica serovar Javiana str CFSAN001992 YP 007473683 1 1267753 Proteobacteria
Salmonella enterica subsp enterica serovar Paratyphi A str AKU 12601 YP 002143246 1 554290 Proteobacteria
Salmonella enterica subsp enterica serovar Typhi str Ty21a YP 007928558 1 527001 Proteobacteria
Salmonella enterica subsp enterica serovar Typhi str Ty21a YP 007927674 1 527001 Proteobacteria
Salmonella enterica subsp enterica serovar Agona str SL483 YP 002147650 2 454166 Proteobacteria
Escherichia coli O7 K1 str CE10 YP 006143822 1 1072459 Proteobacteria
Salmonella enterica subsp enterica serovar Pullorum str S06004 YP 008382020 1 1298917 Proteobacteria
Shigella boydii Sb227 YP 407263 1 300268 Proteobacteria
Salmonella enterica subsp enterica serovar Dublin str CT 02021853 YP 002216722 1 439851 Proteobacteria
Escherichia coli O127 H6 str E2348 69 YP 002328387 1 574521 Proteobacteria
Escherichia coli ATCC 8739 YP 001725728 1 481805 Proteobacteria

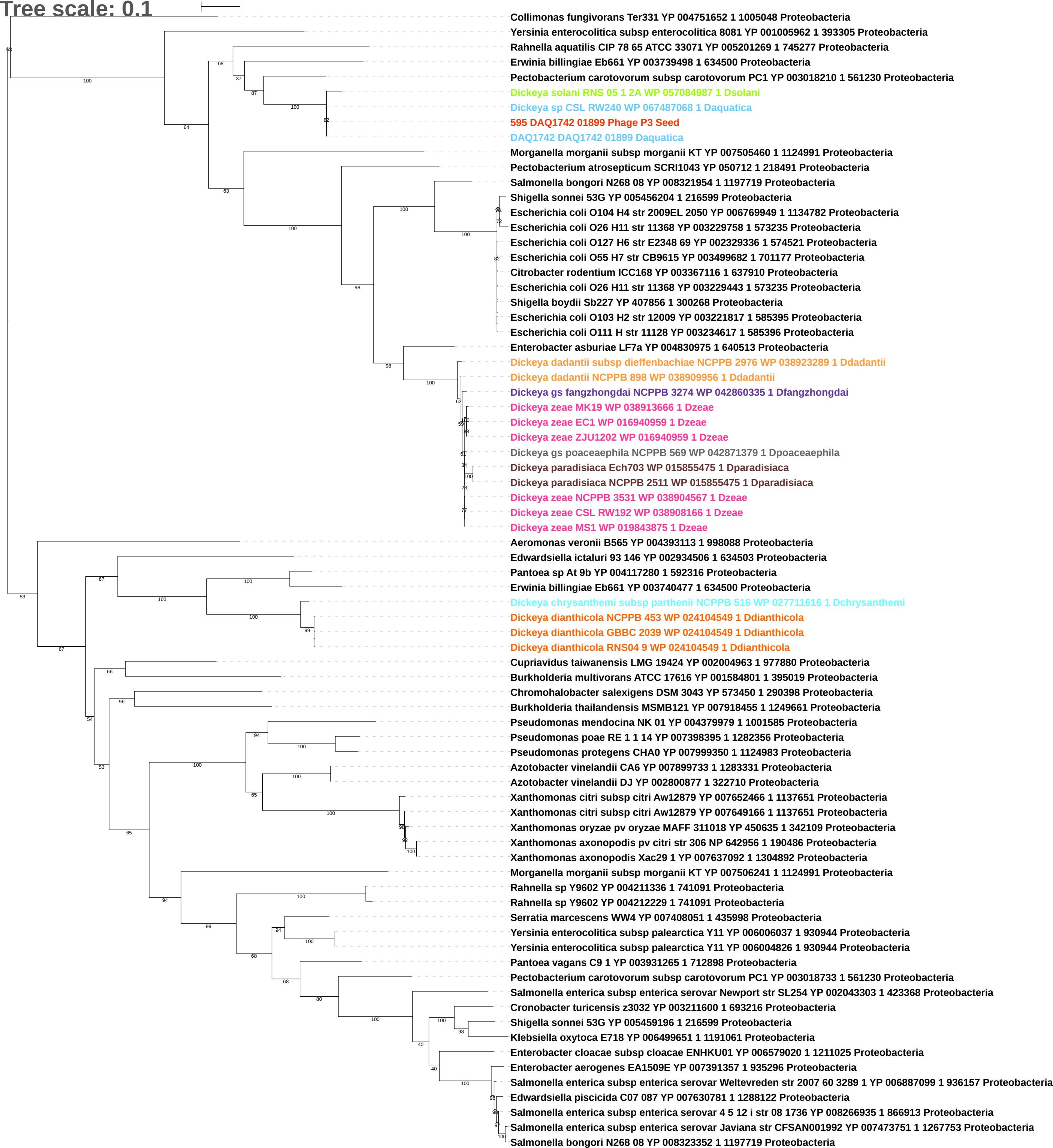
Tree scale: 0.1
Collimonas fungivorans Ter331 YP 004751652 1 1005048 Proteobacteria
Yersinia enterocolitica subsp enterocolitica 8081 YP 001005962 1 393305 Proteobacteria
Rahnella aquatilis CIP 78 65 ATCC 33071 YP 005201269 1 745277 Proteobacteria
Erwinia billingiae Eb661 YP 003739498 1 634500 Proteobacteria
Pectobacterium carotovorum subsp carotovorum PC1 YP 003018210 1 561230 Proteobacteria
Dickeya solani RNS 05 1 2A WP 057084987 1 Dsolani
Dickeya sp CSL RW240 WP 067487068 1 Daquatica
595 DAQ1742 01899 Phage P3 Seed
DAQ1742 DAQ1742 01899 Daquatica
Morganella morganii subsp morganii KT YP 007505460 1 1124991 Proteobacteria
Pectobacterium atrosepticum SCRI1043 YP 050712 1 218491 Proteobacteria
Salmonella bongori N268 08 YP 008321954 1 1197719 Proteobacteria
Shigella sonnei 53G YP 005456204 1 216599 Proteobacteria
Escherichia coli O104 H4 str 2009EL 2050 YP 006769949 1 1134782 Proteobacteria
Escherichia coli O26 H11 str 11368 YP 003229758 1 573235 Proteobacteria
Escherichia coli O127 H6 str E2348 69 YP 002329336 1 574521 Proteobacteria
Escherichia coli O55 H7 str CB9615 YP 003499682 1 701177 Proteobacteria
Citrobacter rodentium ICC168 YP 003367116 1 637910 Proteobacteria
Escherichia coli O26 H11 str 11368 YP 003229443 1 573235 Proteobacteria
Shigella boydii Sb227 YP 407856 1 300268 Proteobacteria
Escherichia coli O103 H2 str 12009 YP 003221817 1 585395 Proteobacteria
Escherichia coli O111 H str 11128 YP 003234617 1 585396 Proteobacteria
Enterobacter asburiae LF7a YP 004830975 1 640513 Proteobacteria
Dickeya dadantii subsp dieffenbachiae NCPPB 2976 WP 038923289 1 Ddadantii
Dickeya dadantii NCPPB 898 WP 038909956 1 Ddadantii
Dickeya gs fangzhongdai NCPPB 3274 WP 042860335 1 Dfangzhongdai
Dickeya zeae MK19 WP 038913666 1 Dzeae
Dickeya zeae EC1 WP 016940959 1 Dzeae
Dickeya zeae ZJU1202 WP 016940959 1 Dzeae
Dickeya gs poaceaephila NCPPB 569 WP 042871379 1 Dpoaceaephila
Dickeya paradisiaca Ech703 WP 015855475 1 Dparadisiaca
Dickeya paradisiaca NCPPB 2511 WP 015855475 1 Dparadisiaca
Dickeya zeae NCPPB 3531 WP 038904567 1 Dzeae
Dickeya zeae CSL RW192 WP 038908166 1 Dzeae
Dickeya zeae MS1 WP 019843875 1 Dzeae
Aeromonas veronii B565 YP 004393113 1 998088 Proteobacteria
Edwardsiella ictaluri 93 146 YP 002934506 1 634503 Proteobacteria
Pantoea sp At 9b YP 004117280 1 592316 Proteobacteria
Erwinia billingiae Eb661 YP 003740477 1 634500 Proteobacteria
Dickeya chrysanthemi subsp parthenii NCPPB 516 WP 027711616 1 Dchrysanthemi
Dickeya dianthicola NCPPB 453 WP 024104549 1 Ddianthicola
Dickeya dianthicola GBBC 2039 WP 024104549 1 Ddianthicola
Dickeya dianthicola RNS04 9 WP 024104549 1 Ddianthicola
Cupriavidus taiwanensis LMG 19424 YP 002004963 1 977880 Proteobacteria
Burkholderia multivorans ATCC 17616 YP 001584801 1 395019 Proteobacteria
Chromohalobacter salexigens DSM 3043 YP 573450 1 290398 Proteobacteria
Burkholderia thailandensis MSMB121 YP 007918455 1 1249661 Proteobacteria
Pseudomonas mendocina NK 01 YP 004379979 1 1001585 Proteobacteria
Pseudomonas poae RE 1 1 14 YP 007398395 1 1282356 Proteobacteria
Pseudomonas protegens CHA0 YP 007999350 1 1124983 Proteobacteria
Azotobacter vinelandii CA6 YP 007899733 1 1283331 Proteobacteria
Azotobacter vinelandii DJ YP 002800877 1 322710 Proteobacteria
Xanthomonas citri subsp citri Aw12879 YP 007652466 1 1137651 Proteobacteria
Xanthomonas citri subsp citri Aw12879 YP 007649166 1 1137651 Proteobacteria
Xanthomonas oryzae pv oryzae MAFF 311018 YP 450635 1 342109 Proteobacteria
Xanthomonas axonopodis pv citri str 306 NP 642956 1 190486 Proteobacteria
Xanthomonas axonopodis Xac29 1 YP 007637092 1 1304892 Proteobacteria
Morganella morganii subsp morganii KT YP 007506241 1 1124991 Proteobacteria
Rahnella sp Y9602 YP 004211336 1 741091 Proteobacteria
Rahnella sp Y9602 YP 004212229 1 741091 Proteobacteria
Serratia marcescens WW4 YP 007408051 1 435998 Proteobacteria
Yersinia enterocolitica subsp palearctica Y11 YP 006006037 1 930944 Proteobacteria
Yersinia enterocolitica subsp palearctica Y11 YP 006004826 1 930944 Proteobacteria
Pantoea vagans C9 1 YP 003931265 1 712898 Proteobacteria
Pectobacterium carotovorum subsp carotovorum PC1 YP 003018733 1 561230 Proteobacteria
Salmonella enterica subsp enterica serovar Newport str SL254 YP 002043303 1 423368 Proteobacteria
Cronobacter turicensis z3032 YP 003211600 1 693216 Proteobacteria
Shigella sonnei 53G YP 005459196 1 216599 Proteobacteria
Klebsiella oxytoca E718 YP 006499651 1 1191061 Proteobacteria
Enterobacter cloacae subsp cloacae ENHKU01 YP 006579020 1 1211025 Proteobacteria
Enterobacter aerogenes EA1509E YP 007391357 1 935296 Proteobacteria
Salmonella enterica subsp enterica serovar Weltevreden str 2007 60 3289 1 YP 006887099 1 936157 Proteobacteria
Edwardsiella piscicida C07 087 YP 007630781 1 1288122 Proteobacteria
Salmonella enterica subsp enterica serovar 4 5 12 i str 08 1736 YP 008266935 1 866913 Proteobacteria
Salmonella enterica subsp enterica serovar Javiana str CFSAN001992 YP 007473751 1 1267753 Proteobacteria
Salmonella bongori N268 08 YP 008323352 1 1197719 Proteobacteria

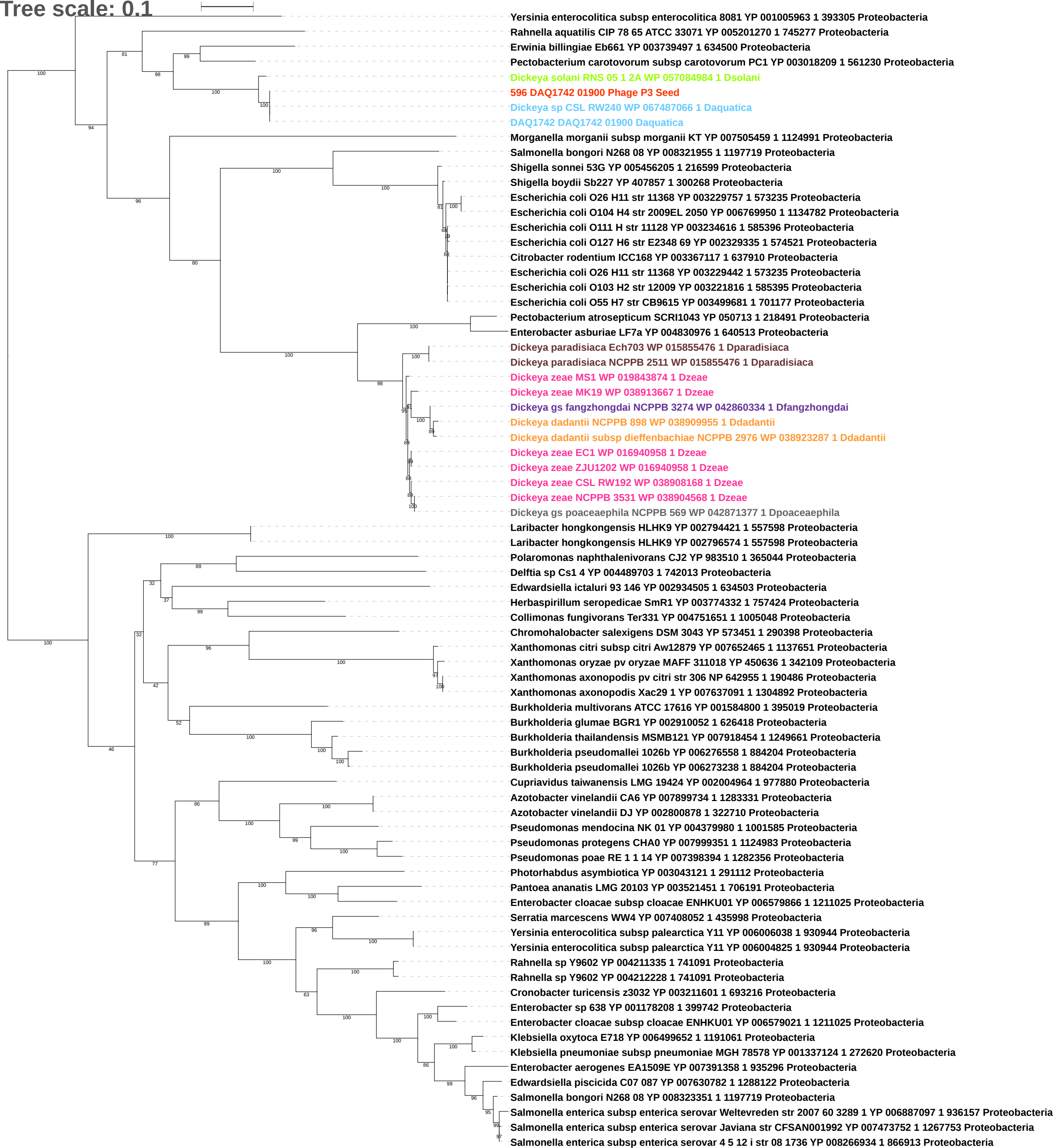
Tree scale: 0.1
Yersinia enterocolitica subsp enterocolitica 8081 YP 001005963 1 393305 Proteobacteria
Rahnella aquatilis CIP 78 65 ATCC 33071 YP 005201270 1 745277 Proteobacteria
Erwinia billingiae Eb661 YP 003739497 1 634500 Proteobacteria
Pectobacterium carotovorum subsp carotovorum PC1 YP 003018209 1 561230 Proteobacteria
Dickeya solani RNS 05 1 2A WP 057084984 1 Dsolani
596 DAQ1742 01900 Phage P3 Seed
Dickeya sp CSL RW240 WP 067487066 1 Daquatica
DAQ1742 DAQ1742 01900 Daquatica
Morganella morganii subsp morganii KT YP 007505459 1 1124991 Proteobacteria
Salmonella bongori N268 08 YP 008321955 1 1197719 Proteobacteria
Shigella sonnei 53G YP 005456205 1 216599 Proteobacteria
Shigella boydii Sb227 YP 407857 1 300268 Proteobacteria
Escherichia coli O26 H11 str 11368 YP 003229757 1 573235 Proteobacteria
Escherichia coli O104 H4 str 2009EL 2050 YP 006769950 1 1134782 Proteobacteria
Escherichia coli O111 H str 11128 YP 003234616 1 585396 Proteobacteria
Escherichia coli O127 H6 str E2348 69 YP 002329335 1 574521 Proteobacteria
Citrobacter rodentium ICC168 YP 003367117 1 637910 Proteobacteria
Escherichia coli O26 H11 str 11368 YP 003229442 1 573235 Proteobacteria
Escherichia coli O103 H2 str 12009 YP 003221816 1 585395 Proteobacteria
Escherichia coli O55 H7 str CB9615 YP 003499681 1 701177 Proteobacteria
Pectobacterium atrosepticum SCRI1043 YP 050713 1 218491 Proteobacteria
Enterobacter asburiae LF7a YP 004830976 1 640513 Proteobacteria
Dickeya paradisiaca Ech703 WP 015855476 1 Dparadisiaca
Dickeya paradisiaca NCPPB 2511 WP 015855476 1 Dparadisiaca
Dickeya zeae MS1 WP 019843874 1 Dzeae
Dickeya zeae MK19 WP 038913667 1 Dzeae
Dickeya gs fangzhongdai NCPPB 3274 WP 042860334 1 Dfangzhongdai
Dickeya dadantii NCPPB 898 WP 038909955 1 Ddadantii
Dickeya dadantii subsp dieffenbachiae NCPPB 2976 WP 038923287 1 Ddadantii
Dickeya zeae EC1 WP 016940958 1 Dzeae
Dickeya zeae ZJU1202 WP 016940958 1 Dzeae
Dickeya zeae CSL RW192 WP 038908168 1 Dzeae
Dickeya zeae NCPPB 3531 WP 038904568 1 Dzeae
Dickeya gs poaceaephila NCPPB 569 WP 042871377 1 Dpoaceaephila
Laribacter hongkongensis HLHK9 YP 002794421 1 557598 Proteobacteria
Laribacter hongkongensis HLHK9 YP 002796574 1 557598 Proteobacteria
Polaromonas naphthalenivorans CJ2 YP 983510 1 365044 Proteobacteria
Delftia sp Cs1 4 YP 004489703 1 742013 Proteobacteria
Edwardsiella ictaluri 93 146 YP 002934505 1 634503 Proteobacteria
Herbaspirillum seropedicae SmR1 YP 003774332 1 757424 Proteobacteria
Collimonas fungivorans Ter331 YP 004751651 1 1005048 Proteobacteria
Chromohalobacter salexigens DSM 3043 YP 573451 1 290398 Proteobacteria
Xanthomonas citri subsp citri Aw12879 YP 007652465 1 1137651 Proteobacteria
Xanthomonas oryzae pv oryzae MAFF 311018 YP 450636 1 342109 Proteobacteria
Xanthomonas axonopodis pv citri str 306 NP 642955 1 190486 Proteobacteria
Xanthomonas axonopodis Xac29 1 YP 007637091 1 1304892 Proteobacteria
Burkholderia multivorans ATCC 17616 YP 001584800 1 395019 Proteobacteria
Burkholderia glumae BGR1 YP 002910052 1 626418 Proteobacteria
Burkholderia thailandensis MSMB121 YP 007918454 1 1249661 Proteobacteria
Burkholderia pseudomallei 1026b YP 006276558 1 884204 Proteobacteria
Burkholderia pseudomallei 1026b YP 006273238 1 884204 Proteobacteria
Cupriavidus taiwanensis LMG 19424 YP 002004964 1 977880 Proteobacteria
Azotobacter vinelandii CA6 YP 007899734 1 1283331 Proteobacteria
Azotobacter vinelandii DJ YP 002800878 1 322710 Proteobacteria
Pseudomonas mendocina NK 01 YP 004379980 1 1001585 Proteobacteria
Pseudomonas protegens CHA0 YP 007999351 1 1124983 Proteobacteria
Pseudomonas poae RE 1 1 14 YP 007398394 1 1282356 Proteobacteria
Photorhabdus asymbiotica YP 003043121 1 291112 Proteobacteria
Pantoea ananatis LMG 20103 YP 003521451 1 706191 Proteobacteria
Enterobacter cloacae subsp cloacae ENHKU01 YP 006579866 1 1211025 Proteobacteria
Serratia marcescens WW4 YP 007408052 1 435998 Proteobacteria
Yersinia enterocolitica subsp palearctica Y11 YP 006006038 1 930944 Proteobacteria
Yersinia enterocolitica subsp palearctica Y11 YP 006004825 1 930944 Proteobacteria
Rahnella sp Y9602 YP 004211335 1 741091 Proteobacteria
Rahnella sp Y9602 YP 004212228 1 741091 Proteobacteria
Cronobacter turicensis z3032 YP 003211601 1 693216 Proteobacteria
Enterobacter sp 638 YP 001178208 1 399742 Proteobacteria
Enterobacter cloacae subsp cloacae ENHKU01 YP 006579021 1 1211025 Proteobacteria
Klebsiella oxytoca E718 YP 006499652 1 1191061 Proteobacteria
Klebsiella pneumoniae subsp pneumoniae MGH 78578 YP 001337124 1 272620 Proteobacteria
Enterobacter aerogenes EA1509E YP 007391358 1 935296 Proteobacteria
Edwardsiella piscicida C07 087 YP 007630782 1 1288122 Proteobacteria
Salmonella bongori N268 08 YP 008323351 1 1197719 Proteobacteria
Salmonella enterica subsp enterica serovar Weltevreden str 2007 60 3289 1 YP 006887097 1 936157 Proteobacteria
Salmonella enterica subsp enterica serovar Javiana str CFSAN001992 YP 007473752 1 1267753 Proteobacteria
Salmonella enterica subsp enterica serovar 4 5 12 i str 08 1736 YP 008266934 1 866913 Proteobacteria

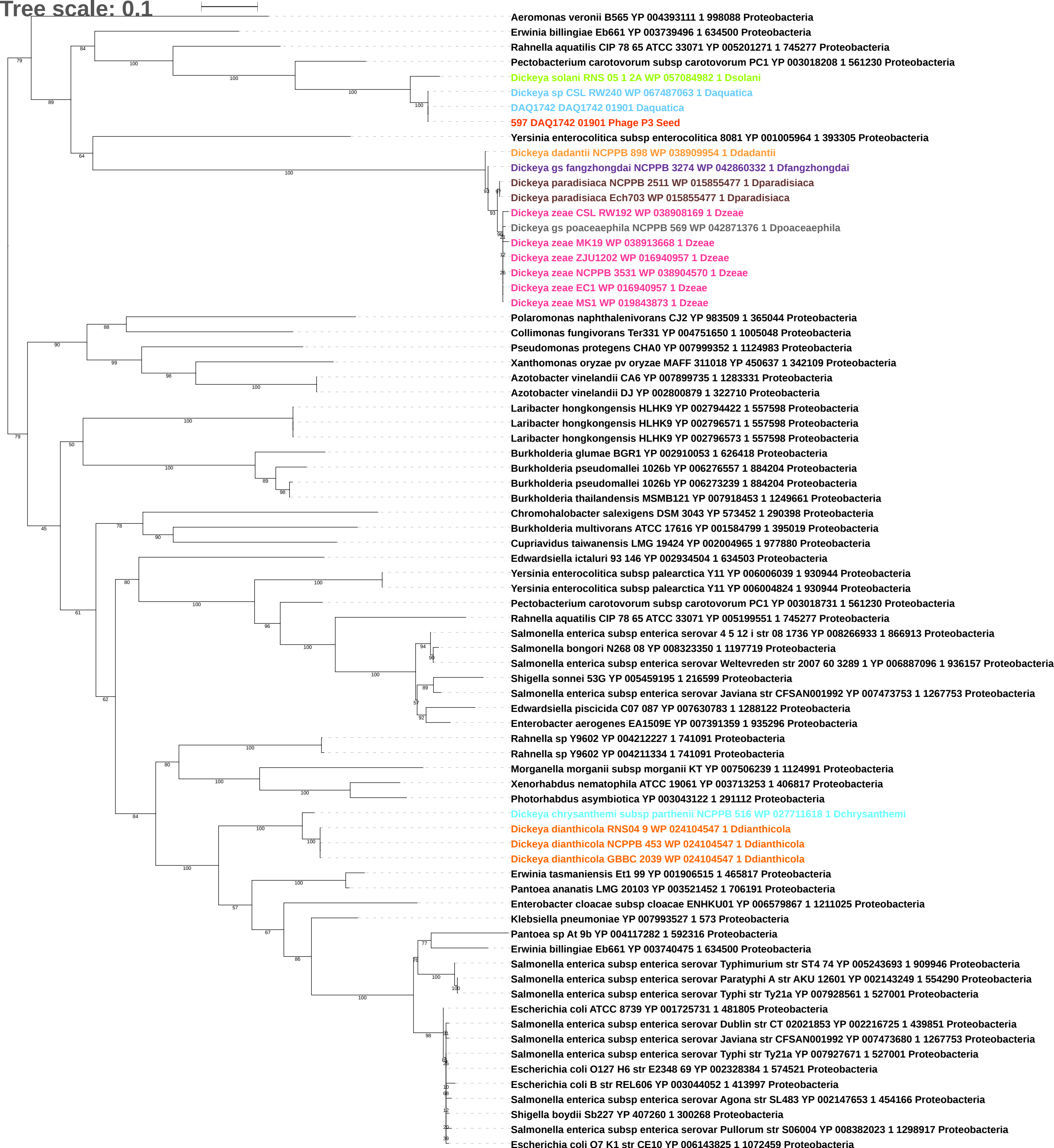
Tree scale: 0.1
Aeromonas veronii B565 YP 004393111 1 998088 Proteobacteria
Erwinia billingiae Eb661 YP 003739496 1 634500 Proteobacteria
Rahnella aquatilis CIP 78 65 ATCC 33071 YP 005201271 1 745277 Proteobacteria
Pectobacterium carotovorum subsp carotovorum PC1 YP 003018208 1 561230 Proteobacteria
Dickeya solani RNS 05 1 2A WP 057084982 1 Dsolani
Dickeya sp CSL RW240 WP 067487063 1 Daquatica
DAQ1742 DAQ1742 01901 Daquatica
597 DAQ1742 01901 Phage P3 Seed
Yersinia enterocolitica subsp enterocolitica 8081 YP 001005964 1 393305 Proteobacteria
Dickeya dadantii NCPPB 898 WP 038909954 1 Ddadantii
Dickeya gs fangzhongdai NCPPB 3274 WP 042860332 1 Dfangzhongdai
Dickeya paradisiaca NCPPB 2511 WP 015855477 1 Dparadisiaca
Dickeya paradisiaca Ech703 WP 015855477 1 Dparadisiaca
Dickeya zeae CSL RW192 WP 038908169 1 Dzeae
Dickeya gs poaceaephila NCPPB 569 WP 042871376 1 Dpoaceaephila
Dickeya zeae MK19 WP 038913668 1 Dzeae
Dickeya zeae ZJU1202 WP 016940957 1 Dzeae
Dickeya zeae NCPPB 3531 WP 038904570 1 Dzeae
Dickeya zeae EC1 WP 016940957 1 Dzeae
Dickeya zeae MS1 WP 019843873 1 Dzeae
Polaromonas naphthalenivorans CJ2 YP 983509 1 365044 Proteobacteria
Collimonas fungivorans Ter331 YP 004751650 1 1005048 Proteobacteria
Pseudomonas protegens CHA0 YP 007999352 1 1124983 Proteobacteria
Xanthomonas oryzae pv oryzae MAFF 311018 YP 450637 1 342109 Proteobacteria
Azotobacter vinelandii CA6 YP 007899735 1 1283331 Proteobacteria
Azotobacter vinelandii DJ YP 002800879 1 322710 Proteobacteria
Laribacter hongkongensis HLHK9 YP 002794422 1 557598 Proteobacteria
Laribacter hongkongensis HLHK9 YP 002796571 1 557598 Proteobacteria
Laribacter hongkongensis HLHK9 YP 002796573 1 557598 Proteobacteria
Burkholderia glumae BGR1 YP 002910053 1 626418 Proteobacteria
Burkholderia pseudomallei 1026b YP 006276557 1 884204 Proteobacteria
Burkholderia pseudomallei 1026b YP 006273239 1 884204 Proteobacteria
Burkholderia thailandensis MSMB121 YP 007918453 1 1249661 Proteobacteria
Chromohalobacter salexigens DSM 3043 YP 573452 1 290398 Proteobacteria
Burkholderia multivorans ATCC 17616 YP 001584799 1 395019 Proteobacteria
Cupriavidus taiwanensis LMG 19424 YP 002004965 1 977880 Proteobacteria
Edwardsiella ictaluri 93 146 YP 002934504 1 634503 Proteobacteria
Yersinia enterocolitica subsp palearctica Y11 YP 006006039 1 930944 Proteobacteria
Yersinia enterocolitica subsp palearctica Y11 YP 006004824 1 930944 Proteobacteria
Pectobacterium carotovorum subsp carotovorum PC1 YP 003018731 1 561230 Proteobacteria
Rahnella aquatilis CIP 78 65 ATCC 33071 YP 005199551 1 745277 Proteobacteria
Salmonella enterica subsp enterica serovar 4 5 12 i str 08 1736 YP 008266933 1 866913 Proteobacteria
Salmonella bongori N268 08 YP 008323350 1 1197719 Proteobacteria
Salmonella enterica subsp enterica serovar Weltevreden str 2007 60 3289 1 YP 006887096 1 936157 Proteobacteria
Shigella sonnei 53G YP 005459195 1 216599 Proteobacteria
Salmonella enterica subsp enterica serovar Javiana str CFSAN001992 YP 007473753 1 1267753 Proteobacteria
Edwardsiella piscicida C07 087 YP 007630783 1 1288122 Proteobacteria
Enterobacter aerogenes EA1509E YP 007391359 1 935296 Proteobacteria
Rahnella sp Y9602 YP 004212227 1 741091 Proteobacteria
Rahnella sp Y9602 YP 004211334 1 741091 Proteobacteria
Morganella morganii subsp morganii KT YP 007506239 1 1124991 Proteobacteria
Xenorhabdus nematophila ATCC 19061 YP 003713253 1 406817 Proteobacteria
Photorhabdus asymbiotica YP 003043122 1 291112 Proteobacteria
Dickeya chrysanthemi subsp parthenii NCPPB 516 WP 027711618 1 Dchrysanthemi
Dickeya dianthicola RNS04 9 WP 024104547 1 Ddianthicola
Dickeya dianthicola NCPPB 453 WP 024104547 1 Ddianthicola
Dickeya dianthicola GBBC 2039 WP 024104547 1 Ddianthicola
Erwinia tasmaniensis Et1 99 YP 001906515 1 465817 Proteobacteria
Pantoea ananatis LMG 20103 YP 003521452 1 706191 Proteobacteria
Enterobacter cloacae subsp cloacae ENHKU01 YP 006579867 1 1211025 Proteobacteria
Klebsiella pneumoniae YP 007993527 1 573 Proteobacteria
Pantoea sp At 9b YP 004117282 1 592316 Proteobacteria
Erwinia billingiae Eb661 YP 003740475 1 634500 Proteobacteria
Salmonella enterica subsp enterica serovar Typhimurium str ST4 74 YP 005243693 1 909946 Proteobacteria
Salmonella enterica subsp enterica serovar Paratyphi A str AKU 12601 YP 002143249 1 554290 Proteobacteria
Salmonella enterica subsp enterica serovar Typhi str Ty21a YP 007928561 1 527001 Proteobacteria
Escherichia coli ATCC 8739 YP 001725731 1 481805 Proteobacteria
Salmonella enterica subsp enterica serovar Dublin str CT 02021853 YP 002216725 1 439851 Proteobacteria
Salmonella enterica subsp enterica serovar Javiana str CFSAN001992 YP 007473680 1 1267753 Proteobacteria
Salmonella enterica subsp enterica serovar Typhi str Ty21a YP 007927671 1 527001 Proteobacteria
Escherichia coli O127 H6 str E2348 69 YP 002328384 1 574521 Proteobacteria
Escherichia coli B str REL606 YP 003044052 1 413997 Proteobacteria
Salmonella enterica subsp enterica serovar Agona str SL483 YP 002147653 1 454166 Proteobacteria
Shigella boydii Sb227 YP 407260 1 300268 Proteobacteria
Salmonella enterica subsp enterica serovar Pullorum str S06004 YP 008382023 1 1298917 Proteobacteria
Escherichia coli O7 K1 str CE10 YP 006143825 1 1072459 Proteobacteria

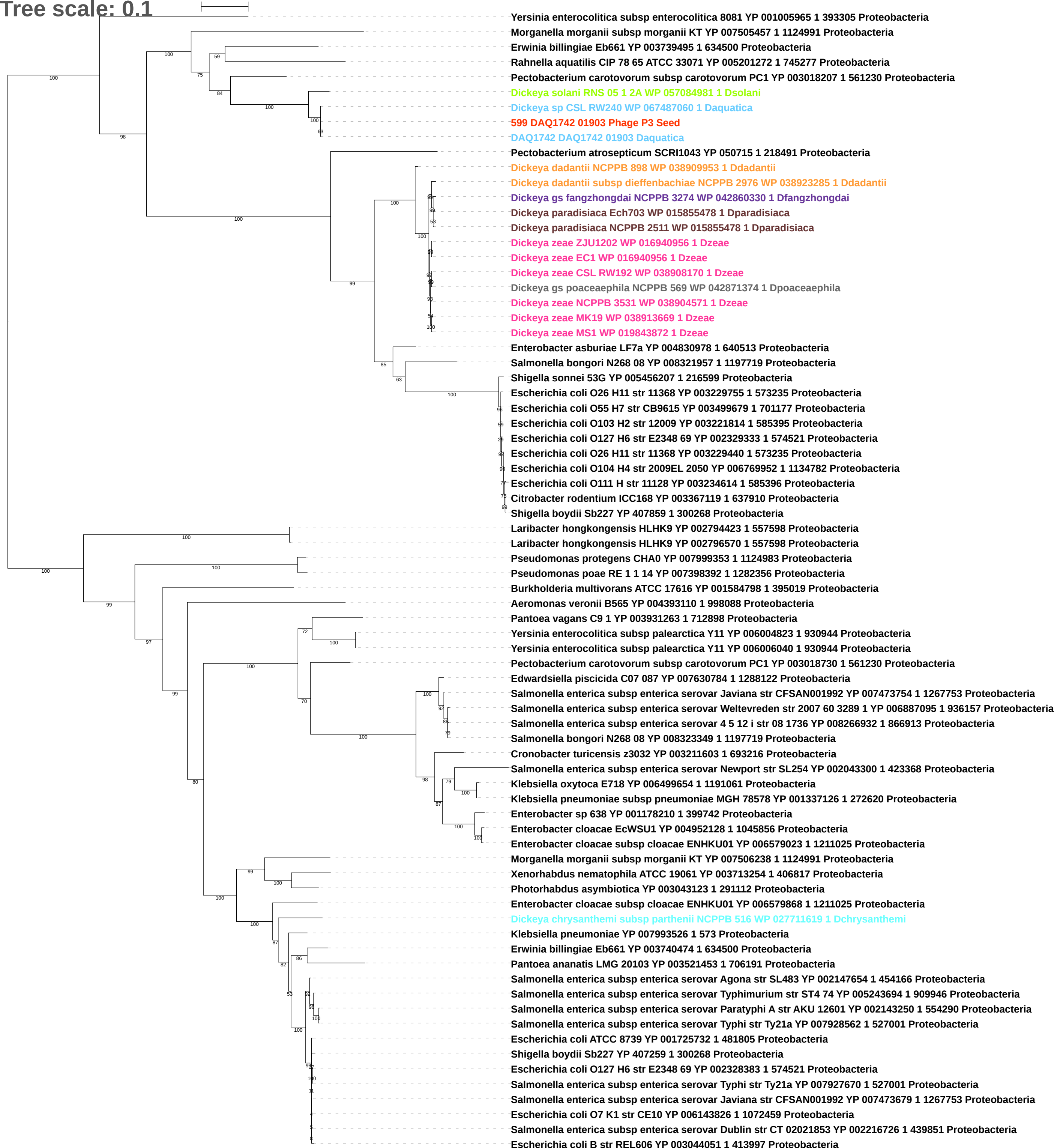
Tree scale: 0.1
Yersinia enterocolitica subsp enterocolitica 8081 YP 001005965 1 393305 Proteobacteria
Morganella morganii subsp morganii KT YP 007505457 1 1124991 Proteobacteria
Erwinia billingiae Eb661 YP 003739495 1 634500 Proteobacteria
Rahnella aquatilis CIP 78 65 ATCC 33071 YP 005201272 1 745277 Proteobacteria
Pectobacterium carotovorum subsp carotovorum PC1 YP 003018207 1 561230 Proteobacteria
Dickeya solani RNS 05 1 2A WP 057084981 1 Dsolani
Dickeya sp CSL RW240 WP 067487060 1 Daquatica
599 DAQ1742 01903 Phage P3 Seed
DAQ1742 DAQ1742 01903 Daquatica
Pectobacterium atrosepticum SCRI1043 YP 050715 1 218491 Proteobacteria
Dickeya dadantii NCPPB 898 WP 038909953 1 Ddadantii
Dickeya dadantii subsp dieffenbachiae NCPPB 2976 WP 038923285 1 Ddadantii
Dickeya gs fangzhongdai NCPPB 3274 WP 042860330 1 Dfangzhongdai
Dickeya paradisiaca Ech703 WP 015855478 1 Dparadisiaca
Dickeya paradisiaca NCPPB 2511 WP 015855478 1 Dparadisiaca
Dickeya zeae ZJU1202 WP 016940956 1 Dzeae
Dickeya zeae EC1 WP 016940956 1 Dzeae
Dickeya zeae CSL RW192 WP 038908170 1 Dzeae
Dickeya gs poaceaephila NCPPB 569 WP 042871374 1 Dpoaceaephila
Dickeya zeae NCPPB 3531 WP 038904571 1 Dzeae
Dickeya zeae MK19 WP 038913669 1 Dzeae
Dickeya zeae MS1 WP 019843872 1 Dzeae
Enterobacter asburiae LF7a YP 004830978 1 640513 Proteobacteria
Salmonella bongori N268 08 YP 008321957 1 1197719 Proteobacteria
Shigella sonnei 53G YP 005456207 1 216599 Proteobacteria
Escherichia coli O26 H11 str 11368 YP 003229755 1 573235 Proteobacteria
Escherichia coli O55 H7 str CB9615 YP 003499679 1 701177 Proteobacteria
Escherichia coli O103 H2 str 12009 YP 003221814 1 585395 Proteobacteria
Escherichia coli O127 H6 str E2348 69 YP 002329333 1 574521 Proteobacteria
Escherichia coli O26 H11 str 11368 YP 003229440 1 573235 Proteobacteria
Escherichia coli O104 H4 str 2009EL 2050 YP 006769952 1 1134782 Proteobacteria
Escherichia coli O111 H str 11128 YP 003234614 1 585396 Proteobacteria
Citrobacter rodentium ICC168 YP 003367119 1 637910 Proteobacteria
Shigella boydii Sb227 YP 407859 1 300268 Proteobacteria
Laribacter hongkongensis HLHK9 YP 002794423 1 557598 Proteobacteria
Laribacter hongkongensis HLHK9 YP 002796570 1 557598 Proteobacteria
Pseudomonas protegens CHA0 YP 007999353 1 1124983 Proteobacteria
Pseudomonas poae RE 1 1 14 YP 007398392 1 1282356 Proteobacteria
Burkholderia multivorans ATCC 17616 YP 001584798 1 395019 Proteobacteria
Aeromonas veronii B565 YP 004393110 1 998088 Proteobacteria
Pantoea vagans C9 1 YP 003931263 1 712898 Proteobacteria
Yersinia enterocolitica subsp palearctica Y11 YP 006004823 1 930944 Proteobacteria
Yersinia enterocolitica subsp palearctica Y11 YP 006006040 1 930944 Proteobacteria
Pectobacterium carotovorum subsp carotovorum PC1 YP 003018730 1 561230 Proteobacteria
Edwardsiella piscicida C07 087 YP 007630784 1 1288122 Proteobacteria
Salmonella enterica subsp enterica serovar Javiana str CFSAN001992 YP 007473754 1 1267753 Proteobacteria
Salmonella enterica subsp enterica serovar Weltevreden str 2007 60 3289 1 YP 006887095 1 936157 Proteobacteria
Salmonella enterica subsp enterica serovar 4 5 12 i str 08 1736 YP 008266932 1 866913 Proteobacteria
Salmonella bongori N268 08 YP 008323349 1 1197719 Proteobacteria
Cronobacter turicensis z3032 YP 003211603 1 693216 Proteobacteria
Salmonella enterica subsp enterica serovar Newport str SL254 YP 002043300 1 423368 Proteobacteria
Klebsiella oxytoca E718 YP 006499654 1 1191061 Proteobacteria
Klebsiella pneumoniae subsp pneumoniae MGH 78578 YP 001337126 1 272620 Proteobacteria
Enterobacter sp 638 YP 001178210 1 399742 Proteobacteria
Enterobacter cloacae EcWSU1 YP 004952128 1 1045856 Proteobacteria
Enterobacter cloacae subsp cloacae ENHKU01 YP 006579023 1 1211025 Proteobacteria
Morganella morganii subsp morganii KT YP 007506238 1 1124991 Proteobacteria
Xenorhabdus nematophila ATCC 19061 YP 003713254 1 406817 Proteobacteria
Photorhabdus asymbiotica YP 003043123 1 291112 Proteobacteria
Enterobacter cloacae subsp cloacae ENHKU01 YP 006579868 1 1211025 Proteobacteria
Dickeya chrysanthemi subsp parthenii NCPPB 516 WP 027711619 1 Dchrysanthemi
Klebsiella pneumoniae YP 007993526 1 573 Proteobacteria
Erwinia billingiae Eb661 YP 003740474 1 634500 Proteobacteria
Pantoea ananatis LMG 20103 YP 003521453 1 706191 Proteobacteria
Salmonella enterica subsp enterica serovar Agona str SL483 YP 002147654 1 454166 Proteobacteria
Salmonella enterica subsp enterica serovar Typhimurium str ST4 74 YP 005243694 1 909946 Proteobacteria
Salmonella enterica subsp enterica serovar Paratyphi A str AKU 12601 YP 002143250 1 554290 Proteobacteria
Salmonella enterica subsp enterica serovar Typhi str Ty21a YP 007928562 1 527001 Proteobacteria
Escherichia coli ATCC 8739 YP 001725732 1 481805 Proteobacteria
Shigella boydii Sb227 YP 407259 1 300268 Proteobacteria
Escherichia coli O127 H6 str E2348 69 YP 002328383 1 574521 Proteobacteria
Salmonella enterica subsp enterica serovar Typhi str Ty21a YP 007927670 1 527001 Proteobacteria
Salmonella enterica subsp enterica serovar Javiana str CFSAN001992 YP 007473679 1 1267753 Proteobacteria
Escherichia coli O7 K1 str CE10 YP 006143826 1 1072459 Proteobacteria
Salmonella enterica subsp enterica serovar Dublin str CT 02021853 YP 002216726 1 439851 Proteobacteria
Escherichia coli B str REL606 YP 003044051 1 413997 Proteobacteria

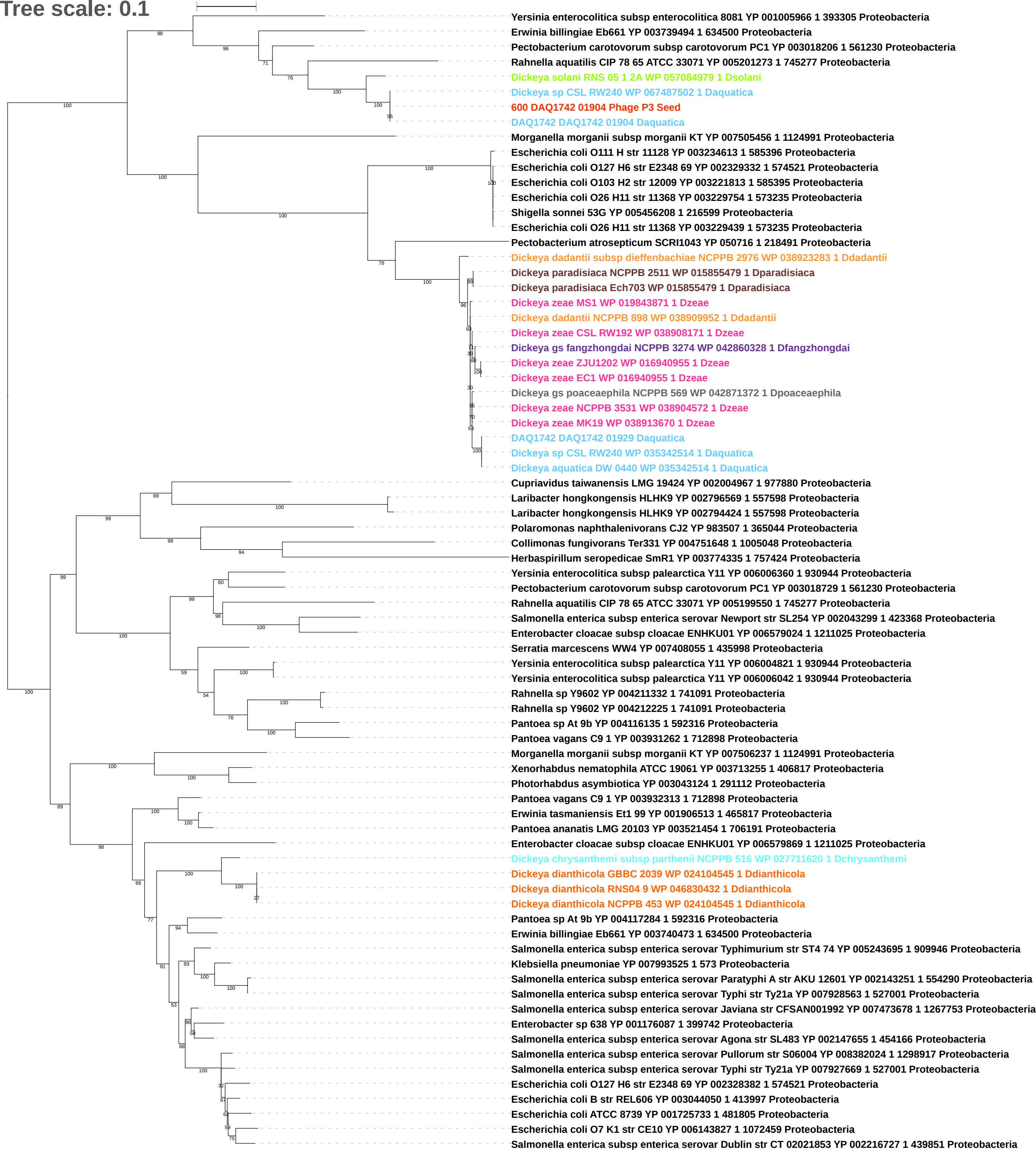
Tree scale: 0.1
Yersinia enterocolitica subsp enterocolitica 8081 YP 001005966 1 393305 Proteobacteria
Erwinia billingiae Eb661 YP 003739494 1 634500 Proteobacteria
Pectobacterium carotovorum subsp carotovorum PC1 YP 003018206 1 561230 Proteobacteria
Rahnella aquatilis CIP 78 65 ATCC 33071 YP 005201273 1 745277 Proteobacteria
Dickeya solani RNS 05 1 2A WP 057084979 1 Dsolani
Dickeya sp CSL RW240 WP 067487502 1 Daquatica
600 DAQ1742 01904 Phage P3 Seed
DAQ1742 DAQ1742 01904 Daquatica
Morganella morganii subsp morganii KT YP 007505456 1 1124991 Proteobacteria
Escherichia coli O111 H str 11128 YP 003234613 1 585396 Proteobacteria
Escherichia coli O127 H6 str E2348 69 YP 002329332 1 574521 Proteobacteria
Escherichia coli O103 H2 str 12009 YP 003221813 1 585395 Proteobacteria
Escherichia coli O26 H11 str 11368 YP 003229754 1 573235 Proteobacteria
Shigella sonnei 53G YP 005456208 1 216599 Proteobacteria
Escherichia coli O26 H11 str 11368 YP 003229439 1 573235 Proteobacteria
Pectobacterium atrosepticum SCRI1043 YP 050716 1 218491 Proteobacteria
Dickeya dadantii subsp dieffenbachiae NCPPB 2976 WP 038923283 1 Ddadantii
Dickeya paradisiaca NCPPB 2511 WP 015855479 1 Dparadisiaca
Dickeya paradisiaca Ech703 WP 015855479 1 Dparadisiaca
Dickeya zeae MS1 WP 019843871 1 Dzeae
Dickeya dadantii NCPPB 898 WP 038909952 1 Ddadantii
Dickeya zeae CSL RW192 WP 038908171 1 Dzeae
Dickeya gs fangzhongdai NCPPB 3274 WP 042860328 1 Dfangzhongdai
Dickeya zeae ZJU1202 WP 016940955 1 Dzeae
Dickeya zeae EC1 WP 016940955 1 Dzeae
Dickeya gs poaceaephila NCPPB 569 WP 042871372 1 Dpoaceaephila
Dickeya zeae NCPPB 3531 WP 038904572 1 Dzeae
Dickeya zeae MK19 WP 038913670 1 Dzeae
DAQ1742 DAQ1742 01929 Daquatica
Dickeya sp CSL RW240 WP 035342514 1 Daquatica
Dickeya aquatica DW 0440 WP 035342514 1 Daquatica
Cupriavidus taiwanensis LMG 19424 YP 002004967 1 977880 Proteobacteria
Laribacter hongkongensis HLHK9 YP 002796569 1 557598 Proteobacteria
Laribacter hongkongensis HLHK9 YP 002794424 1 557598 Proteobacteria
Polaromonas naphthalenivorans CJ2 YP 983507 1 365044 Proteobacteria
Collimonas fungivorans Ter331 YP 004751648 1 1005048 Proteobacteria
Herbaspirillum seropedicae SmR1 YP 003774335 1 757424 Proteobacteria
Yersinia enterocolitica subsp palearctica Y11 YP 006006360 1 930944 Proteobacteria
Pectobacterium carotovorum subsp carotovorum PC1 YP 003018729 1 561230 Proteobacteria
Rahnella aquatilis CIP 78 65 ATCC 33071 YP 005199550 1 745277 Proteobacteria
Salmonella enterica subsp enterica serovar Newport str SL254 YP 002043299 1 423368 Proteobacteria
Enterobacter cloacae subsp cloacae ENHKU01 YP 006579024 1 1211025 Proteobacteria
Serratia marcescens WW4 YP 007408055 1 435998 Proteobacteria
Yersinia enterocolitica subsp palearctica Y11 YP 006004821 1 930944 Proteobacteria
Yersinia enterocolitica subsp palearctica Y11 YP 006006042 1 930944 Proteobacteria
Rahnella sp Y9602 YP 004211332 1 741091 Proteobacteria
Rahnella sp Y9602 YP 004212225 1 741091 Proteobacteria
Pantoea sp At 9b YP 004116135 1 592316 Proteobacteria
Pantoea vagans C9 1 YP 003931262 1 712898 Proteobacteria
Morganella morganii subsp morganii KT YP 007506237 1 1124991 Proteobacteria
Xenorhabdus nematophila ATCC 19061 YP 003713255 1 406817 Proteobacteria
Photorhabdus asymbiotica YP 003043124 1 291112 Proteobacteria
Pantoea vagans C9 1 YP 003932313 1 712898 Proteobacteria
Erwinia tasmaniensis Et1 99 YP 001906513 1 465817 Proteobacteria
Pantoea ananatis LMG 20103 YP 003521454 1 706191 Proteobacteria
Enterobacter cloacae subsp cloacae ENHKU01 YP 006579869 1 1211025 Proteobacteria
Dickeya chrysanthemi subsp parthenii NCPPB 516 WP 027711620 1 Dchrysanthemi
Dickeya dianthicola GBBC 2039 WP 024104545 1 Ddianthicola
Dickeya dianthicola RNS04 9 WP 046830432 1 Ddianthicola
Dickeya dianthicola NCPPB 453 WP 024104545 1 Ddianthicola
Pantoea sp At 9b YP 004117284 1 592316 Proteobacteria
Erwinia billingiae Eb661 YP 003740473 1 634500 Proteobacteria
Salmonella enterica subsp enterica serovar Typhimurium str ST4 74 YP 005243695 1 909946 Proteobacteria
Klebsiella pneumoniae YP 007993525 1 573 Proteobacteria
Salmonella enterica subsp enterica serovar Paratyphi A str AKU 12601 YP 002143251 1 554290 Proteobacteria
Salmonella enterica subsp enterica serovar Typhi str Ty21a YP 007928563 1 527001 Proteobacteria
Salmonella enterica subsp enterica serovar Javiana str CFSAN001992 YP 007473678 1 1267753 Proteobacteria
Enterobacter sp 638 YP 001176087 1 399742 Proteobacteria
Salmonella enterica subsp enterica serovar Agona str SL483 YP 002147655 1 454166 Proteobacteria
Salmonella enterica subsp enterica serovar Pullorum str S06004 YP 008382024 1 1298917 Proteobacteria
Salmonella enterica subsp enterica serovar Typhi str Ty21a YP 007927669 1 527001 Proteobacteria
Escherichia coli O127 H6 str E2348 69 YP 002328382 1 574521 Proteobacteria
Escherichia coli B str REL606 YP 003044050 1 413997 Proteobacteria
Escherichia coli ATCC 8739 YP 001725733 1 481805 Proteobacteria
Escherichia coli O7 K1 str CE10 YP 006143827 1 1072459 Proteobacteria
Salmonella enterica subsp enterica serovar Dublin str CT 02021853 YP 002216727 1 439851 Proteobacteria

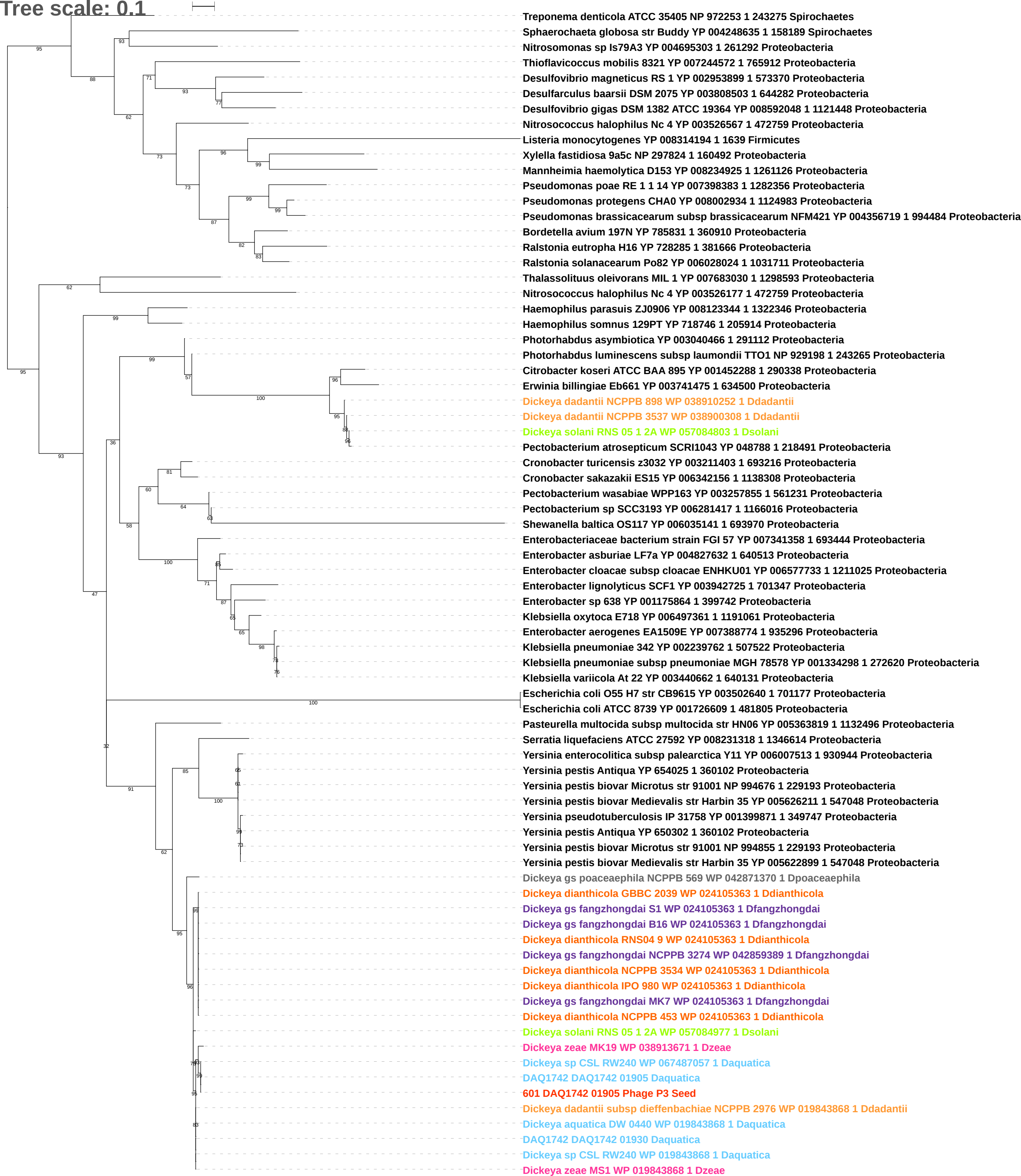
Tree scale: 0.1
Treponema denticola ATCC 35405 NP 972253 1 243275 Spirochaetes
Sphaerochaeta globosa str Buddy YP 004248635 1 158189 Spirochaetes
Nitrosomonas sp Is79A3 YP 004695303 1 261292 Proteobacteria
Thioflavicoccus mobilis 8321 YP 007244572 1 765912 Proteobacteria
Desulfovibrio magneticus RS 1 YP 002953899 1 573370 Proteobacteria
Desulfarculus baarsii DSM 2075 YP 003808503 1 644282 Proteobacteria
Desulfovibrio gigas DSM 1382 ATCC 19364 YP 008592048 1 1121448 Proteobacteria
Nitrosococcus halophilus Nc 4 YP 003526567 1 472759 Proteobacteria
Listeria monocytogenes YP 008314194 1 1639 Firmicutes
Xylella fastidiosa 9a5c NP 297824 1 160492 Proteobacteria
Mannheimia haemolytica D153 YP 008234925 1 1261126 Proteobacteria
Pseudomonas poae RE 1 1 14 YP 007398383 1 1282356 Proteobacteria
Pseudomonas protegens CHA0 YP 008002934 1 1124983 Proteobacteria
Pseudomonas brassicacearum subsp brassicacearum NFM421 YP 004356719 1 994484 Proteobacteria
Bordetella avium 197N YP 785831 1 360910 Proteobacteria
Ralstonia eutropha H16 YP 728285 1 381666 Proteobacteria
Ralstonia solanacearum Po82 YP 006028024 1 1031711 Proteobacteria
Thalassolituus oleivorans MIL 1 YP 007683030 1 1298593 Proteobacteria
Nitrosococcus halophilus Nc 4 YP 003526177 1 472759 Proteobacteria
Haemophilus parasuis ZJ0906 YP 008123344 1 1322346 Proteobacteria
Haemophilus somnus 129PT YP 718746 1 205914 Proteobacteria
Photorhabdus asymbiotica YP 003040466 1 291112 Proteobacteria
Photorhabdus luminescens subsp laumondii TTO1 NP 929198 1 243265 Proteobacteria
Citrobacter koseri ATCC BAA 895 YP 001452288 1 290338 Proteobacteria
Erwinia billingiae Eb661 YP 003741475 1 634500 Proteobacteria
Dickeya dadantii NCPPB 898 WP 038910252 1 Ddadantii
Dickeya dadantii NCPPB 3537 WP 038900308 1 Ddadantii
Dickeya solani RNS 05 1 2A WP 057084803 1 Dsolani
Pectobacterium atrosepticum SCRI1043 YP 048788 1 218491 Proteobacteria
Cronobacter turicensis z3032 YP 003211403 1 693216 Proteobacteria
Cronobacter sakazakii ES15 YP 006342156 1 1138308 Proteobacteria
Pectobacterium wasabiae WPP163 YP 003257855 1 561231 Proteobacteria
Pectobacterium sp SCC3193 YP 006281417 1 1166016 Proteobacteria
Shewanella baltica OS117 YP 006035141 1 693970 Proteobacteria
Enterobacteriaceae bacterium strain FGI 57 YP 007341358 1 693444 Proteobacteria
Enterobacter asburiae LF7a YP 004827632 1 640513 Proteobacteria
Enterobacter cloacae subsp cloacae ENHKU01 YP 006577733 1 1211025 Proteobacteria
Enterobacter lignolyticus SCF1 YP 003942725 1 701347 Proteobacteria
Enterobacter sp 638 YP 001175864 1 399742 Proteobacteria
Klebsiella oxytoca E718 YP 006497361 1 1191061 Proteobacteria
Enterobacter aerogenes EA1509E YP 007388774 1 935296 Proteobacteria
Klebsiella pneumoniae 342 YP 002239762 1 507522 Proteobacteria
Klebsiella pneumoniae subsp pneumoniae MGH 78578 YP 001334298 1 272620 Proteobacteria
Klebsiella variicola At 22 YP 003440662 1 640131 Proteobacteria
Escherichia coli O55 H7 str CB9615 YP 003502640 1 701177 Proteobacteria
Escherichia coli ATCC 8739 YP 001726609 1 481805 Proteobacteria
Pasteurella multocida subsp multocida str HN06 YP 005363819 1 1132496 Proteobacteria
Serratia liquefaciens ATCC 27592 YP 008231318 1 1346614 Proteobacteria
Yersinia enterocolitica subsp palearctica Y11 YP 006007513 1 930944 Proteobacteria
Yersinia pestis Antiqua YP 654025 1 360102 Proteobacteria
Yersinia pestis biovar Microtus str 91001 NP 994676 1 229193 Proteobacteria
Yersinia pestis biovar Medievalis str Harbin 35 YP 005626211 1 547048 Proteobacteria
Yersinia pseudotuberculosis IP 31758 YP 001399871 1 349747 Proteobacteria
Yersinia pestis Antiqua YP 650302 1 360102 Proteobacteria
Yersinia pestis biovar Microtus str 91001 NP 994855 1 229193 Proteobacteria
Yersinia pestis biovar Medievalis str Harbin 35 YP 005622899 1 547048 Proteobacteria
Dickeya gs poaceaephila NCPPB 569 WP 042871370 1 Dpoaceaephila
Dickeya dianthicola GBBC 2039 WP 024105363 1 Ddianthicola
Dickeya gs fangzhongdai S1 WP 024105363 1 Dfangzhongdai
Dickeya gs fangzhongdai B16 WP 024105363 1 Dfangzhongdai
Dickeya dianthicola RNS04 9 WP 024105363 1 Ddianthicola
Dickeya gs fangzhongdai NCPPB 3274 WP 042859389 1 Dfangzhongdai
Dickeya dianthicola NCPPB 3534 WP 024105363 1 Ddianthicola
Dickeya dianthicola IPO 980 WP 024105363 1 Ddianthicola
Dickeya gs fangzhongdai MK7 WP 024105363 1 Dfangzhongdai
Dickeya dianthicola NCPPB 453 WP 024105363 1 Ddianthicola
Dickeya solani RNS 05 1 2A WP 057084977 1 Dsolani
Dickeya zeae MK19 WP 038913671 1 Dzeae
Dickeya sp CSL RW240 WP 067487057 1 Daquatica
DAQ1742 DAQ1742 01905 Daquatica
601 DAQ1742 01905 Phage P3 Seed
Dickeya dadantii subsp dieffenbachiae NCPPB 2976 WP 019843868 1 Ddadantii
Dickeya aquatica DW 0440 WP 019843868 1 Daquatica
DAQ1742 DAQ1742 01930 Daquatica
Dickeya sp CSL RW240 WP 019843868 1 Daquatica
Dickeya zeae MS1 WP 019843868 1 Dzeae

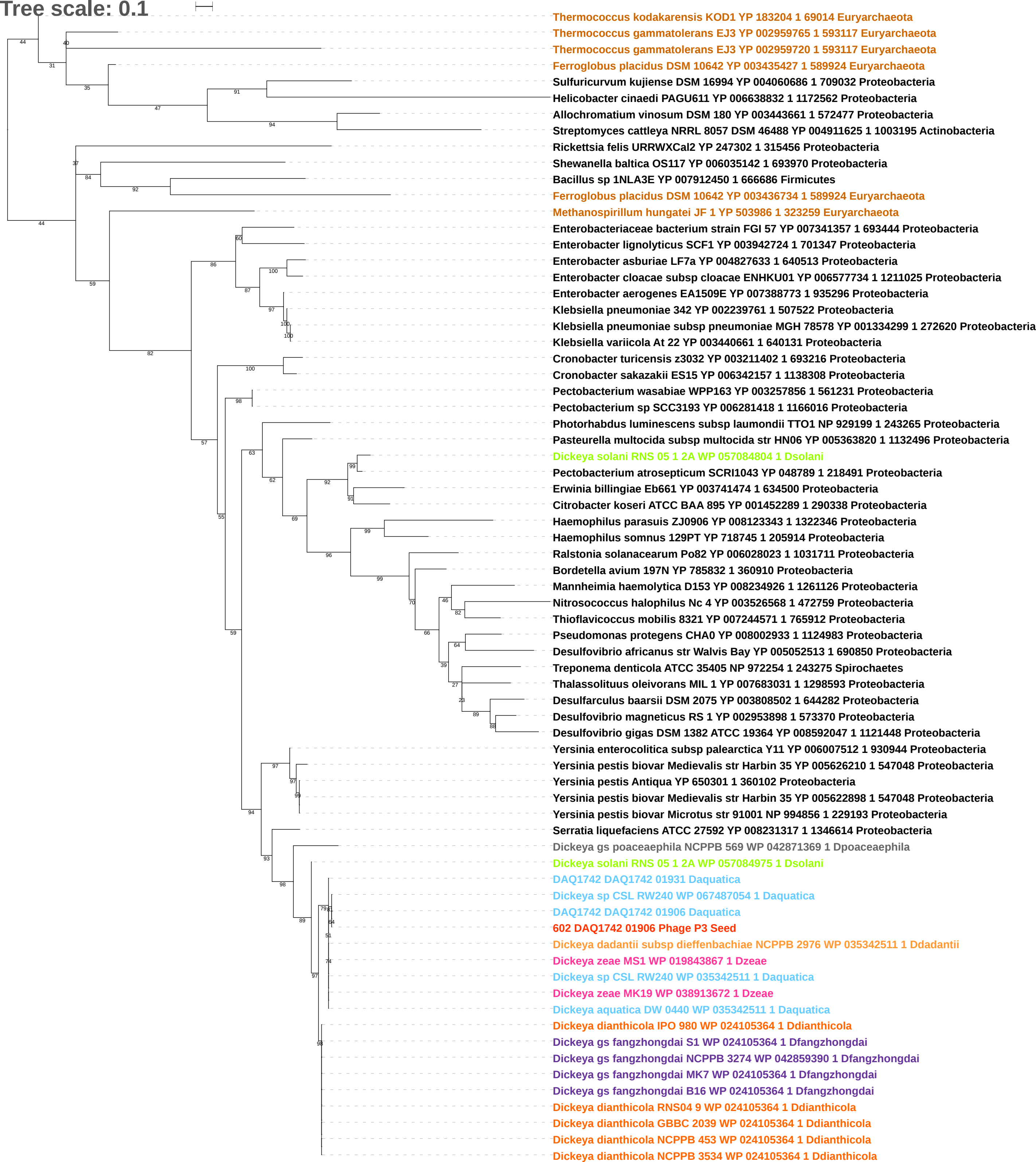

Tree scale: 0.1
Thermococcus kodakarensis KOD1 YP 183204 1 69014 Euryarchaeota
Thermococcus gammatolerans EJ3 YP 002959765 1 593117 Euryarchaeota
Thermococcus gammatolerans EJ3 YP 002959720 1 593117 Euryarchaeota
Ferroglobus placidus DSM 10642 YP 003435427 1 589924 Euryarchaeota
Sulfuricurvum kujiense DSM 16994 YP 004060686 1 709032 Proteobacteria
Helicobacter cinaedi PAGU611 YP 006638832 1 1172562 Proteobacteria
Allochromatium vinosum DSM 180 YP 003443661 1 572477 Proteobacteria
Streptomyces cattleya NRRL 8057 DSM 46488 YP 004911625 1 1003195 Actinobacteria
Rickettsia felis URRWXCal2 YP 247302 1 315456 Proteobacteria
Shewanella baltica OS117 YP 006035142 1 693970 Proteobacteria
Bacillus sp 1NLA3E YP 007912450 1 666686 Firmicutes
Ferroglobus placidus DSM 10642 YP 003436734 1 589924 Euryarchaeota
Methanospirillum hungatei JF 1 YP 503986 1 323259 Euryarchaeota
Enterobacteriaceae bacterium strain FGI 57 YP 007341357 1 693444 Proteobacteria
Enterobacter lignolyticus SCF1 YP 003942724 1 701347 Proteobacteria
Enterobacter asburiae LF7a YP 004827633 1 640513 Proteobacteria
Enterobacter cloacae subsp cloacae ENHKU01 YP 006577734 1 1211025 Proteobacteria
Enterobacter aerogenes EA1509E YP 007388773 1 935296 Proteobacteria
Klebsiella pneumoniae 342 YP 002239761 1 507522 Proteobacteria
Klebsiella pneumoniae subsp pneumoniae MGH 78578 YP 001334299 1 272620 Proteobacteria
Klebsiella variicola At 22 YP 003440661 1 640131 Proteobacteria
Cronobacter turicensis z3032 YP 003211402 1 693216 Proteobacteria
Cronobacter sakazakii ES15 YP 006342157 1 1138308 Proteobacteria
Pectobacterium wasabiae WPP163 YP 003257856 1 561231 Proteobacteria
Pectobacterium sp SCC3193 YP 006281418 1 1166016 Proteobacteria
Photorhabdus luminescens subsp laumondii TTO1 NP 929199 1 243265 Proteobacteria
Pasteurella multocida subsp multocida str HN06 YP 005363820 1 1132496 Proteobacteria
Dickeya solani RNS 05 1 2A WP 057084804 1 Dsolani
Pectobacterium atrosepticum SCRI1043 YP 048789 1 218491 Proteobacteria
Erwinia billingiae Eb661 YP 003741474 1 634500 Proteobacteria
Citrobacter koseri ATCC BAA 895 YP 001452289 1 290338 Proteobacteria
Haemophilus parasuis ZJ0906 YP 008123343 1 1322346 Proteobacteria
Haemophilus somnus 129PT YP 718745 1 205914 Proteobacteria
Ralstonia solanacearum Po82 YP 006028023 1 1031711 Proteobacteria
Bordetella avium 197N YP 785832 1 360910 Proteobacteria
Mannheimia haemolytica D153 YP 008234926 1 1261126 Proteobacteria
Nitrosococcus halophilus Nc 4 YP 003526568 1 472759 Proteobacteria
Thioflavicoccus mobilis 8321 YP 007244571 1 765912 Proteobacteria
Pseudomonas protegens CHA0 YP 008002933 1 1124983 Proteobacteria
Desulfovibrio africanus str Walvis Bay YP 005052513 1 690850 Proteobacteria
Treponema denticola ATCC 35405 NP 972254 1 243275 Spirochaetes
Thalassolituus oleivorans MIL 1 YP 007683031 1 1298593 Proteobacteria
Desulfarculus baarsii DSM 2075 YP 003808502 1 644282 Proteobacteria
Desulfovibrio magneticus RS 1 YP 002953898 1 573370 Proteobacteria
Desulfovibrio gigas DSM 1382 ATCC 19364 YP 008592047 1 1121448 Proteobacteria
Yersinia enterocolitica subsp palearctica Y11 YP 006007512 1 930944 Proteobacteria
Yersinia pestis biovar Medievalis str Harbin 35 YP 005626210 1 547048 Proteobacteria
Yersinia pestis Antiqua YP 650301 1 360102 Proteobacteria
Yersinia pestis biovar Medievalis str Harbin 35 YP 005622898 1 547048 Proteobacteria
Yersinia pestis biovar Microtus str 91001 NP 994856 1 229193 Proteobacteria
Serratia liquefaciens ATCC 27592 YP 008231317 1 1346614 Proteobacteria
Dickeya gs poaceaephila NCPPB 569 WP 042871369 1 Dpoaceaephila
Dickeya solani RNS 05 1 2A WP 057084975 1 Dsolani
DAQ1742 DAQ1742 01931 Daquatica
Dickeya sp CSL RW240 WP 067487054 1 Daquatica
DAQ1742 DAQ1742 01906 Daquatica
602 DAQ1742 01906 Phage P3 Seed
Dickeya dadantii subsp dieffenbachiae NCPPB 2976 WP 035342511 1 Ddadantii
Dickeya zeae MS1 WP 019843867 1 Dzeae
Dickeya sp CSL RW240 WP 035342511 1 Daquatica
Dickeya zeae MK19 WP 038913672 1 Dzeae
Dickeya aquatica DW 0440 WP 035342511 1 Daquatica
Dickeya dianthicola IPO 980 WP 024105364 1 Ddianthicola
Dickeya gs fangzhongdai S1 WP 024105364 1 Dfangzhongdai
Dickeya gs fangzhongdai NCPPB 3274 WP 042859390 1 Dfangzhongdai
Dickeya gs fangzhongdai MK7 WP 024105364 1 Dfangzhongdai
Dickeya gs fangzhongdai B16 WP 024105364 1 Dfangzhongdai
Dickeya dianthicola RNS04 9 WP 024105364 1 Ddianthicola
Dickeya dianthicola GBBC 2039 WP 024105364 1 Ddianthicola
Dickeya dianthicola NCPPB 453 WP 024105364 1 Ddianthicola
Dickeya dianthicola NCPPB 3534 WP 024105364 1 Ddianthicola

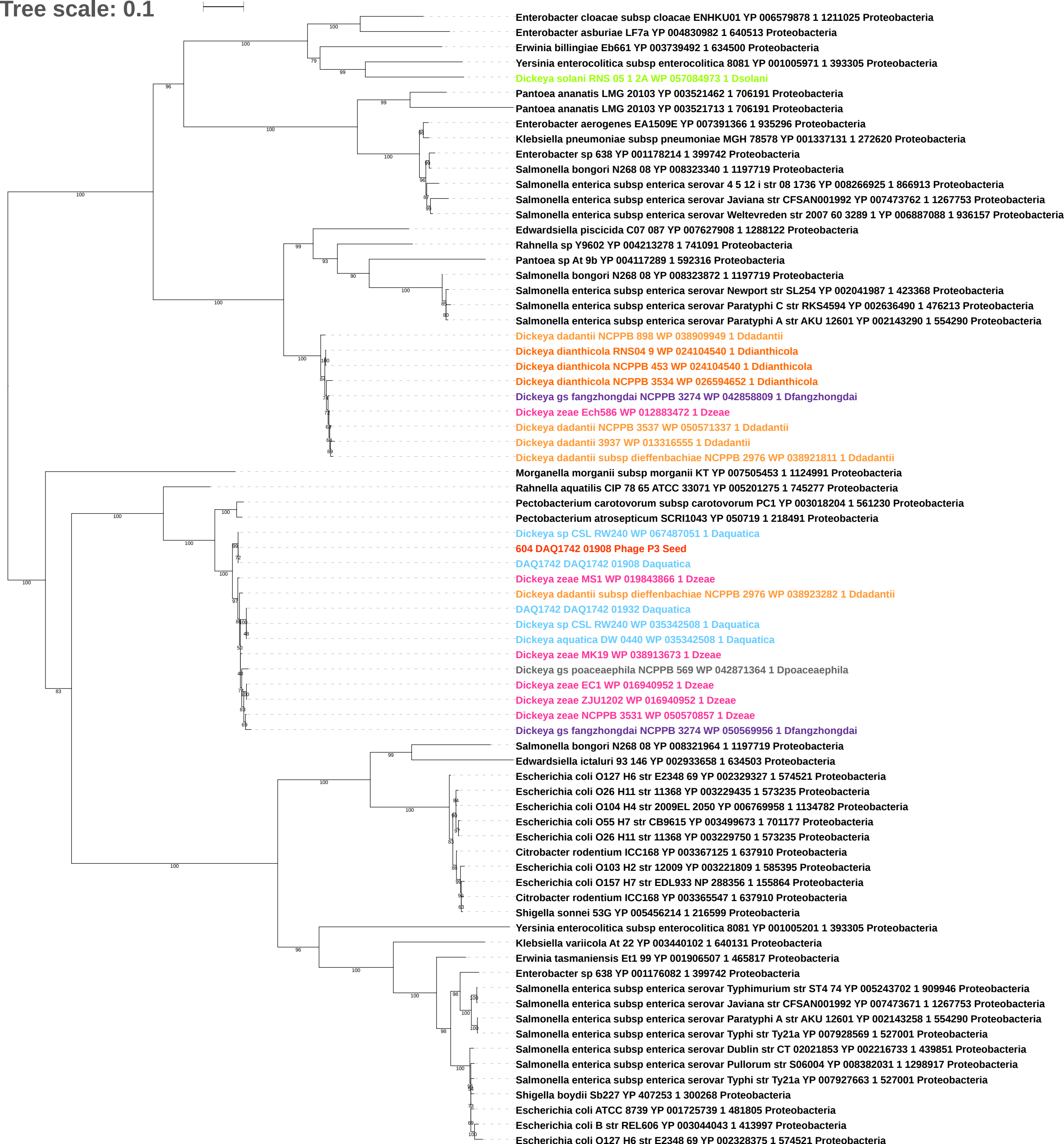
Tree scale: 0.1
Enterobacter cloacae subsp cloacae ENHKU01 YP 006579878 1 1211025 Proteobacteria
Enterobacter asburiae LF7a YP 004830982 1 640513 Proteobacteria
Erwinia billingiae Eb661 YP 003739492 1 634500 Proteobacteria
Yersinia enterocolitica subsp enterocolitica 8081 YP 001005971 1 393305 Proteobacteria
Dickeya solani RNS 05 1 2A WP 057084973 1 Dsolani
Pantoea ananatis LMG 20103 YP 003521462 1 706191 Proteobacteria
Pantoea ananatis LMG 20103 YP 003521713 1 706191 Proteobacteria
Enterobacter aerogenes EA1509E YP 007391366 1 935296 Proteobacteria
Klebsiella pneumoniae subsp pneumoniae MGH 78578 YP 001337131 1 272620 Proteobacteria
Enterobacter sp 638 YP 001178214 1 399742 Proteobacteria
Salmonella bongori N268 08 YP 008323340 1 1197719 Proteobacteria
Salmonella enterica subsp enterica serovar 4 5 12 i str 08 1736 YP 008266925 1 866913 Proteobacteria
Salmonella enterica subsp enterica serovar Javiana str CFSAN001992 YP 007473762 1 1267753 Proteobacteria
Salmonella enterica subsp enterica serovar Weltevreden str 2007 60 3289 1 YP 006887088 1 936157 Proteobacteria
Edwardsiella piscicida C07 087 YP 007627908 1 1288122 Proteobacteria
Rahnella sp Y9602 YP 004213278 1 741091 Proteobacteria
Pantoea sp At 9b YP 004117289 1 592316 Proteobacteria
Salmonella bongori N268 08 YP 008323872 1 1197719 Proteobacteria
Salmonella enterica subsp enterica serovar Newport str SL254 YP 002041987 1 423368 Proteobacteria
Salmonella enterica subsp enterica serovar Paratyphi C str RKS4594 YP 002636490 1 476213 Proteobacteria
Salmonella enterica subsp enterica serovar Paratyphi A str AKU 12601 YP 002143290 1 554290 Proteobacteria
Dickeya dadantii NCPPB 898 WP 038909949 1 Ddadantii
Dickeya dianthicola RNS04 9 WP 024104540 1 Ddianthicola
Dickeya dianthicola NCPPB 453 WP 024104540 1 Ddianthicola
Dickeya dianthicola NCPPB 3534 WP 026594652 1 Ddianthicola
Dickeya gs fangzhongdai NCPPB 3274 WP 042858809 1 Dfangzhongdai
Dickeya zeae Ech586 WP 012883472 1 Dzeae
Dickeya dadantii NCPPB 3537 WP 050571337 1 Ddadantii
Dickeya dadantii 3937 WP 013316555 1 Ddadantii
Dickeya dadantii subsp dieffenbachiae NCPPB 2976 WP 038921811 1 Ddadantii
Morganella morganii subsp morganii KT YP 007505453 1 1124991 Proteobacteria
Rahnella aquatilis CIP 78 65 ATCC 33071 YP 005201275 1 745277 Proteobacteria
Pectobacterium carotovorum subsp carotovorum PC1 YP 003018204 1 561230 Proteobacteria
Pectobacterium atrosepticum SCRI1043 YP 050719 1 218491 Proteobacteria
Dickeya sp CSL RW240 WP 067487051 1 Daquatica
604 DAQ1742 01908 Phage P3 Seed
DAQ1742 DAQ1742 01908 Daquatica
Dickeya zeae MS1 WP 019843866 1 Dzeae
Dickeya dadantii subsp dieffenbachiae NCPPB 2976 WP 038923282 1 Ddadantii
DAQ1742 DAQ1742 01932 Daquatica
Dickeya sp CSL RW240 WP 035342508 1 Daquatica
Dickeya aquatica DW 0440 WP 035342508 1 Daquatica
Dickeya zeae MK19 WP 038913673 1 Dzeae
Dickeya gs poaceaephila NCPPB 569 WP 042871364 1 Dpoaceaephila
Dickeya zeae EC1 WP 016940952 1 Dzeae
Dickeya zeae ZJU1202 WP 016940952 1 Dzeae
Dickeya zeae NCPPB 3531 WP 050570857 1 Dzeae
Dickeya gs fangzhongdai NCPPB 3274 WP 050569956 1 Dfangzhongdai
Salmonella bongori N268 08 YP 008321964 1 1197719 Proteobacteria
Edwardsiella ictaluri 93 146 YP 002933658 1 634503 Proteobacteria
Escherichia coli O127 H6 str E2348 69 YP 002329327 1 574521 Proteobacteria
Escherichia coli O26 H11 str 11368 YP 003229435 1 573235 Proteobacteria
Escherichia coli O104 H4 str 2009EL 2050 YP 006769958 1 1134782 Proteobacteria
Escherichia coli O55 H7 str CB9615 YP 003499673 1 701177 Proteobacteria
Escherichia coli O26 H11 str 11368 YP 003229750 1 573235 Proteobacteria
Citrobacter rodentium ICC168 YP 003367125 1 637910 Proteobacteria
Escherichia coli O103 H2 str 12009 YP 003221809 1 585395 Proteobacteria
Escherichia coli O157 H7 str EDL933 NP 288356 1 155864 Proteobacteria
Citrobacter rodentium ICC168 YP 003365547 1 637910 Proteobacteria
Shigella sonnei 53G YP 005456214 1 216599 Proteobacteria
Yersinia enterocolitica subsp enterocolitica 8081 YP 001005201 1 393305 Proteobacteria
Klebsiella variicola At 22 YP 003440102 1 640131 Proteobacteria
Erwinia tasmaniensis Et1 99 YP 001906507 1 465817 Proteobacteria
Enterobacter sp 638 YP 001176082 1 399742 Proteobacteria
Salmonella enterica subsp enterica serovar Typhimurium str ST4 74 YP 005243702 1 909946 Proteobacteria
Salmonella enterica subsp enterica serovar Javiana str CFSAN001992 YP 007473671 1 1267753 Proteobacteria
Salmonella enterica subsp enterica serovar Paratyphi A str AKU 12601 YP 002143258 1 554290 Proteobacteria
Salmonella enterica subsp enterica serovar Typhi str Ty21a YP 007928569 1 527001 Proteobacteria
Salmonella enterica subsp enterica serovar Dublin str CT 02021853 YP 002216733 1 439851 Proteobacteria
Salmonella enterica subsp enterica serovar Pullorum str S06004 YP 008382031 1 1298917 Proteobacteria
Salmonella enterica subsp enterica serovar Typhi str Ty21a YP 007927663 1 527001 Proteobacteria
Shigella boydii Sb227 YP 407253 1 300268 Proteobacteria
Escherichia coli ATCC 8739 YP 001725739 1 481805 Proteobacteria
Escherichia coli B str REL606 YP 003044043 1 413997 Proteobacteria
Escherichia coli O127 H6 str E2348 69 YP 002328375 1 574521 Proteobacteria

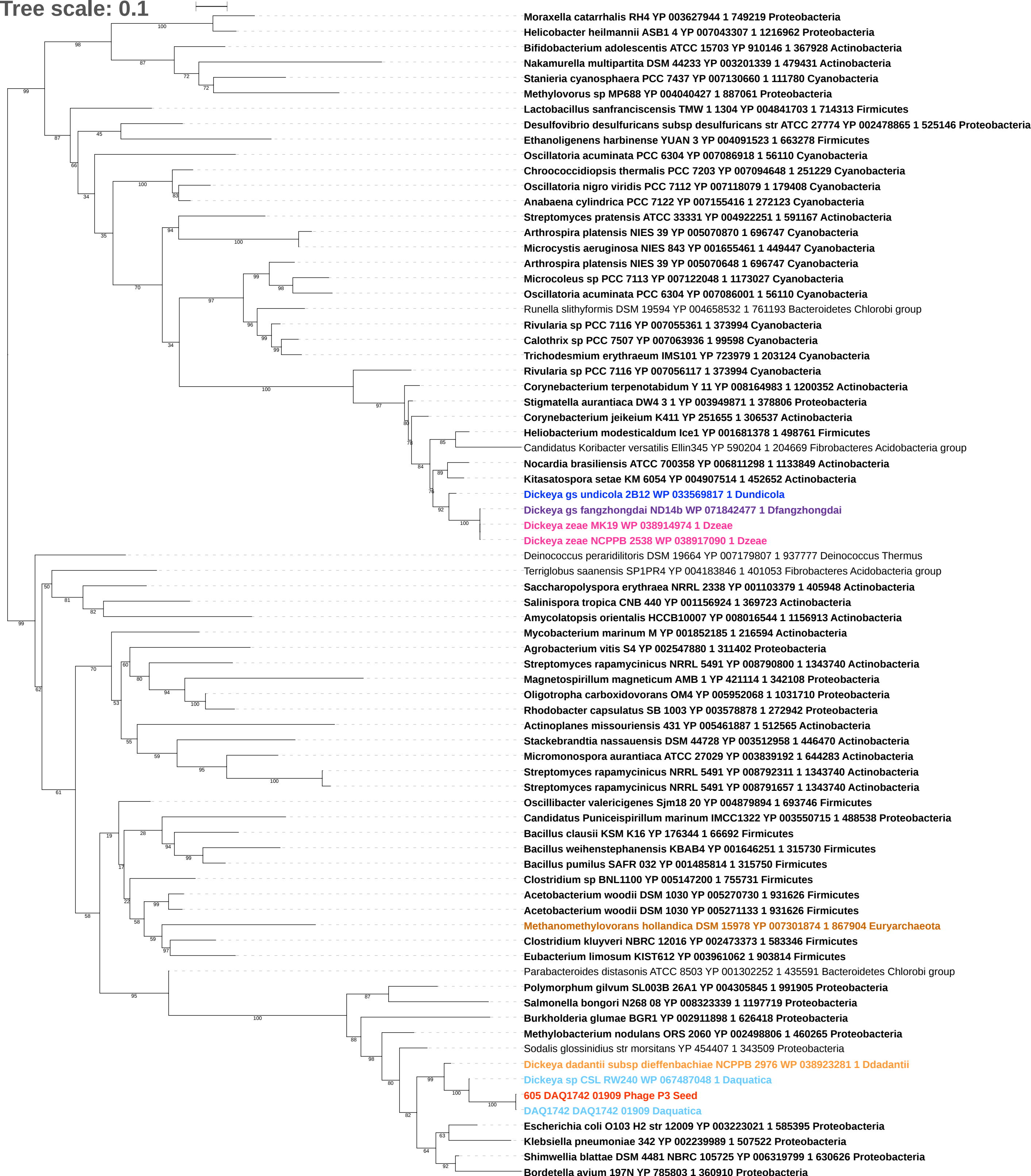
Tree scale: 0.1
Moraxella catarrhalis RH4 YP 003627944 1 749219 Proteobacteria
Helicobacter heilmannii ASB1 4 YP 007043307 1 1216962 Proteobacteria
Bifidobacterium adolescentis ATCC 15703 YP 910146 1 367928 Actinobacteria
Nakamurella multipartita DSM 44233 YP 003201339 1 479431 Actinobacteria
Stanieria cyanosphaera PCC 7437 YP 007130660 1 111780 Cyanobacteria
Methylovorus sp MP688 YP 004040427 1 887061 Proteobacteria
Lactobacillus sanfranciscensis TMW 1 1304 YP 004841703 1 714313 Firmicutes
Desulfovibrio desulfuricans subsp desulfuricans str ATCC 27774 YP 002478865 1 525146 Proteobacteria
Ethanoligenens harbinense YUAN 3 YP 004091523 1 663278 Firmicutes
Oscillatoria acuminata PCC 6304 YP 007086918 1 56110 Cyanobacteria
Chroococcidiopsis thermalis PCC 7203 YP 007094648 1 251229 Cyanobacteria
Oscillatoria nigro viridis PCC 7112 YP 007118079 1 179408 Cyanobacteria
Anabaena cylindrica PCC 7122 YP 007155416 1 272123 Cyanobacteria
Streptomyces pratensis ATCC 33331 YP 004922251 1 591167 Actinobacteria
Arthrospira platensis NIES 39 YP 005070870 1 696747 Cyanobacteria
Microcystis aeruginosa NIES 843 YP 001655461 1 449447 Cyanobacteria
Arthrospira platensis NIES 39 YP 005070648 1 696747 Cyanobacteria
Microcoleus sp PCC 7113 YP 007122048 1 1173027 Cyanobacteria
Oscillatoria acuminata PCC 6304 YP 007086001 1 56110 Cyanobacteria
Runella slithyformis DSM 19594 YP 004658532 1 761193 Bacteroidetes Chlorobi group
Rivularia sp PCC 7116 YP 007055361 1 373994 Cyanobacteria
Calothrix sp PCC 7507 YP 007063936 1 99598 Cyanobacteria
Trichodesmium erythraeum IMS101 YP 723979 1 203124 Cyanobacteria
Rivularia sp PCC 7116 YP 007056117 1 373994 Cyanobacteria
Corynebacterium terpenotabidum Y 11 YP 008164983 1 1200352 Actinobacteria
Stigmatella aurantiaca DW4 3 1 YP 003949871 1 378806 Proteobacteria
Corynebacterium jeikeium K411 YP 251655 1 306537 Actinobacteria
Heliobacterium modesticaldum Ice1 YP 001681378 1 498761 Firmicutes
Candidatus Koribacter versatilis Ellin345 YP 590204 1 204669 Fibrobacteres Acidobacteria group
Nocardia brasiliensis ATCC 700358 YP 006811298 1 1133849 Actinobacteria
Kitasatospora setae KM 6054 YP 004907514 1 452652 Actinobacteria
Dickeya gs undicola 2B12 WP 033569817 1 Dundicola
Dickeya gs fangzhongdai ND14b WP 071842477 1 Dfangzhongdai
Dickeya zeae MK19 WP 038914974 1 Dzeae
Dickeya zeae NCPPB 2538 WP 038917090 1 Dzeae
Deinococcus peraridilitoris DSM 19664 YP 007179807 1 937777 Deinococcus Thermus
Terriglobus saanensis SP1PR4 YP 004183846 1 401053 Fibrobacteres Acidobacteria group
Saccharopolyspora erythraea NRRL 2338 YP 001103379 1 405948 Actinobacteria
Salinispora tropica CNB 440 YP 001156924 1 369723 Actinobacteria
Amycolatopsis orientalis HCCB10007 YP 008016544 1 1156913 Actinobacteria
Mycobacterium marinum M YP 001852185 1 216594 Actinobacteria
Agrobacterium vitis S4 YP 002547880 1 311402 Proteobacteria
Streptomyces rapamycinicus NRRL 5491 YP 008790800 1 1343740 Actinobacteria
Magnetospirillum magneticum AMB 1 YP 421114 1 342108 Proteobacteria
Oligotropha carboxidovorans OM4 YP 005952068 1 1031710 Proteobacteria
Rhodobacter capsulatus SB 1003 YP 003578878 1 272942 Proteobacteria
Actinoplanes missouriensis 431 YP 005461887 1 512565 Actinobacteria
Stackebrandtia nassauensis DSM 44728 YP 003512958 1 446470 Actinobacteria
Micromonospora aurantiaca ATCC 27029 YP 003839192 1 644283 Actinobacteria
Streptomyces rapamycinicus NRRL 5491 YP 008792311 1 1343740 Actinobacteria
Streptomyces rapamycinicus NRRL 5491 YP 008791657 1 1343740 Actinobacteria
Oscillibacter valericigenes Sjm18 20 YP 004879894 1 693746 Firmicutes
Candidatus Puniceispirillum marinum IMCC1322 YP 003550715 1 488538 Proteobacteria
Bacillus clausii KSM K16 YP 176344 1 66692 Firmicutes
Bacillus weihenstephanensis KBAB4 YP 001646251 1 315730 Firmicutes
Bacillus pumilus SAFR 032 YP 001485814 1 315750 Firmicutes
Clostridium sp BNL1100 YP 005147200 1 755731 Firmicutes
Acetobacterium woodii DSM 1030 YP 005270730 1 931626 Firmicutes
Acetobacterium woodii DSM 1030 YP 005271133 1 931626 Firmicutes
Methanomethylovorans hollandica DSM 15978 YP 007301874 1 867904 Euryarchaeota
Clostridium kluyveri NBRC 12016 YP 002473373 1 583346 Firmicutes
Eubacterium limosum KIST612 YP 003961062 1 903814 Firmicutes
Parabacteroides distasonis ATCC 8503 YP 001302252 1 435591 Bacteroidetes Chlorobi group
Polymorphum gilvum SL003B 26A1 YP 004305845 1 991905 Proteobacteria
Salmonella bongori N268 08 YP 008323339 1 1197719 Proteobacteria
Burkholderia glumae BGR1 YP 002911898 1 626418 Proteobacteria
Methylobacterium nodulans ORS 2060 YP 002498806 1 460265 Proteobacteria
Sodalis glossinidius str morsitans YP 454407 1 343509 Proteobacteria
Dickeya dadantii subsp dieffenbachiae NCPPB 2976 WP 038923281 1 Ddadantii
Dickeya sp CSL RW240 WP 067487048 1 Daquatica
605 DAQ1742 01909 Phage P3 Seed
DAQ1742 DAQ1742 01909 Daquatica
Escherichia coli O103 H2 str 12009 YP 003223021 1 585395 Proteobacteria
Klebsiella pneumoniae 342 YP 002239989 1 507522 Proteobacteria
Shimwellia blattae DSM 4481 NBRC 105725 YP 006319799 1 630626 Proteobacteria
Bordetella avium 197N YP 785803 1 360910 Proteobacteria

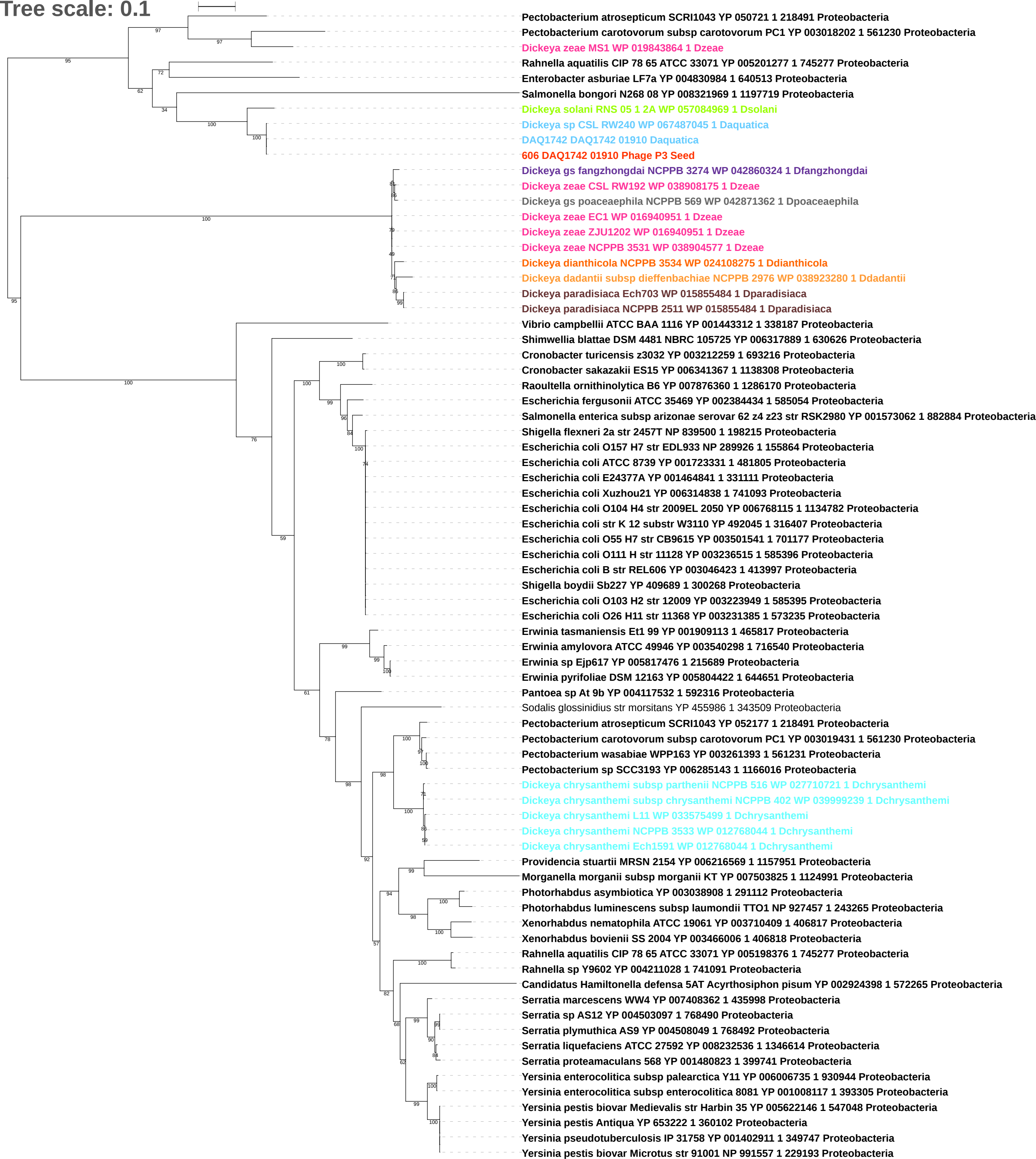
Tree scale: 0.1
Pectobacterium atrosepticum SCRI1043 YP 050721 1 218491 Proteobacteria
Pectobacterium carotovorum subsp carotovorum PC1 YP 003018202 1 561230 Proteobacteria
Dickeya zeae MS1 WP 019843864 1 Dzeae
Rahnella aquatilis CIP 78 65 ATCC 33071 YP 005201277 1 745277 Proteobacteria
Enterobacter asburiae LF7a YP 004830984 1 640513 Proteobacteria
Salmonella bongori N268 08 YP 008321969 1 1197719 Proteobacteria
Dickeya solani RNS 05 1 2A WP 057084969 1 Dsolani
Dickeya sp CSL RW240 WP 067487045 1 Daquatica
DAQ1742 DAQ1742 01910 Daquatica
606 DAQ1742 01910 Phage P3 Seed
Dickeya gs fangzhongdai NCPPB 3274 WP 042860324 1 Dfangzhongdai
Dickeya zeae CSL RW192 WP 038908175 1 Dzeae
Dickeya gs poaceaephila NCPPB 569 WP 042871362 1 Dpoaceaephila
Dickeya zeae EC1 WP 016940951 1 Dzeae
Dickeya zeae ZJU1202 WP 016940951 1 Dzeae
Dickeya zeae NCPPB 3531 WP 038904577 1 Dzeae
Dickeya dianthicola NCPPB 3534 WP 024108275 1 Ddianthicola
Dickeya dadantii subsp dieffenbachiae NCPPB 2976 WP 038923280 1 Ddadantii
Dickeya paradisiaca Ech703 WP 015855484 1 Dparadisiaca
Dickeya paradisiaca NCPPB 2511 WP 015855484 1 Dparadisiaca
Vibrio campbellii ATCC BAA 1116 YP 001443312 1 338187 Proteobacteria
Shimwellia blattae DSM 4481 NBRC 105725 YP 006317889 1 630626 Proteobacteria
Cronobacter turicensis z3032 YP 003212259 1 693216 Proteobacteria
Cronobacter sakazakii ES15 YP 006341367 1 1138308 Proteobacteria
Raoultella ornithinolytica B6 YP 007876360 1 1286170 Proteobacteria
Escherichia fergusonii ATCC 35469 YP 002384434 1 585054 Proteobacteria
Salmonella enterica subsp arizonae serovar 62 z4 z23 str RSK2980 YP 001573062 1 882884 Proteobacteria
Shigella flexneri 2a str 2457T NP 839500 1 198215 Proteobacteria
Escherichia coli O157 H7 str EDL933 NP 289926 1 155864 Proteobacteria
Escherichia coli ATCC 8739 YP 001723331 1 481805 Proteobacteria
Escherichia coli E24377A YP 001464841 1 331111 Proteobacteria
Escherichia coli Xuzhou21 YP 006314838 1 741093 Proteobacteria
Escherichia coli O104 H4 str 2009EL 2050 YP 006768115 1 1134782 Proteobacteria
Escherichia coli str K 12 substr W3110 YP 492045 1 316407 Proteobacteria
Escherichia coli O55 H7 str CB9615 YP 003501541 1 701177 Proteobacteria
Escherichia coli O111 H str 11128 YP 003236515 1 585396 Proteobacteria
Escherichia coli B str REL606 YP 003046423 1 413997 Proteobacteria
Shigella boydii Sb227 YP 409689 1 300268 Proteobacteria
Escherichia coli O103 H2 str 12009 YP 003223949 1 585395 Proteobacteria
Escherichia coli O26 H11 str 11368 YP 003231385 1 573235 Proteobacteria
Erwinia tasmaniensis Et1 99 YP 001909113 1 465817 Proteobacteria
Erwinia amylovora ATCC 49946 YP 003540298 1 716540 Proteobacteria
Erwinia sp Ejp617 YP 005817476 1 215689 Proteobacteria
Erwinia pyrifoliae DSM 12163 YP 005804422 1 644651 Proteobacteria
Pantoea sp At 9b YP 004117532 1 592316 Proteobacteria
Sodalis glossinidius str morsitans YP 455986 1 343509 Proteobacteria
Pectobacterium atrosepticum SCRI1043 YP 052177 1 218491 Proteobacteria
Pectobacterium carotovorum subsp carotovorum PC1 YP 003019431 1 561230 Proteobacteria
Pectobacterium wasabiae WPP163 YP 003261393 1 561231 Proteobacteria
Pectobacterium sp SCC3193 YP 006285143 1 1166016 Proteobacteria
Dickeya chrysanthemi subsp parthenii NCPPB 516 WP 027710721 1 Dchrysanthemi
Dickeya chrysanthemi subsp chrysanthemi NCPPB 402 WP 039999239 1 Dchrysanthemi
Dickeya chrysanthemi L11 WP 033575499 1 Dchrysanthemi
Dickeya chrysanthemi NCPPB 3533 WP 012768044 1 Dchrysanthemi
Dickeya chrysanthemi Ech1591 WP 012768044 1 Dchrysanthemi
Providencia stuartii MRSN 2154 YP 006216569 1 1157951 Proteobacteria
Morganella morganii subsp morganii KT YP 007503825 1 1124991 Proteobacteria
Photorhabdus asymbiotica YP 003038908 1 291112 Proteobacteria
Photorhabdus luminescens subsp laumondii TTO1 NP 927457 1 243265 Proteobacteria
Xenorhabdus nematophila ATCC 19061 YP 003710409 1 406817 Proteobacteria
Xenorhabdus bovienii SS 2004 YP 003466006 1 406818 Proteobacteria
Rahnella aquatilis CIP 78 65 ATCC 33071 YP 005198376 1 745277 Proteobacteria
Rahnella sp Y9602 YP 004211028 1 741091 Proteobacteria
Candidatus Hamiltonella defensa 5AT Acyrthosiphon pisum YP 002924398 1 572265 Proteobacteria
Serratia marcescens WW4 YP 007408362 1 435998 Proteobacteria
Serratia sp AS12 YP 004503097 1 768490 Proteobacteria
Serratia plymuthica AS9 YP 004508049 1 768492 Proteobacteria
Serratia liquefaciens ATCC 27592 YP 008232536 1 1346614 Proteobacteria
Serratia proteamaculans 568 YP 001480823 1 399741 Proteobacteria
Yersinia enterocolitica subsp palearctica Y11 YP 006006735 1 930944 Proteobacteria
Yersinia enterocolitica subsp enterocolitica 8081 YP 001008117 1 393305 Proteobacteria
Yersinia pestis biovar Medievalis str Harbin 35 YP 005622146 1 547048 Proteobacteria
Yersinia pestis Antiqua YP 653222 1 360102 Proteobacteria
Yersinia pseudotuberculosis IP 31758 YP 001402911 1 349747 Proteobacteria
Yersinia pestis biovar Microtus str 91001 NP 991557 1 229193 Proteobacteria

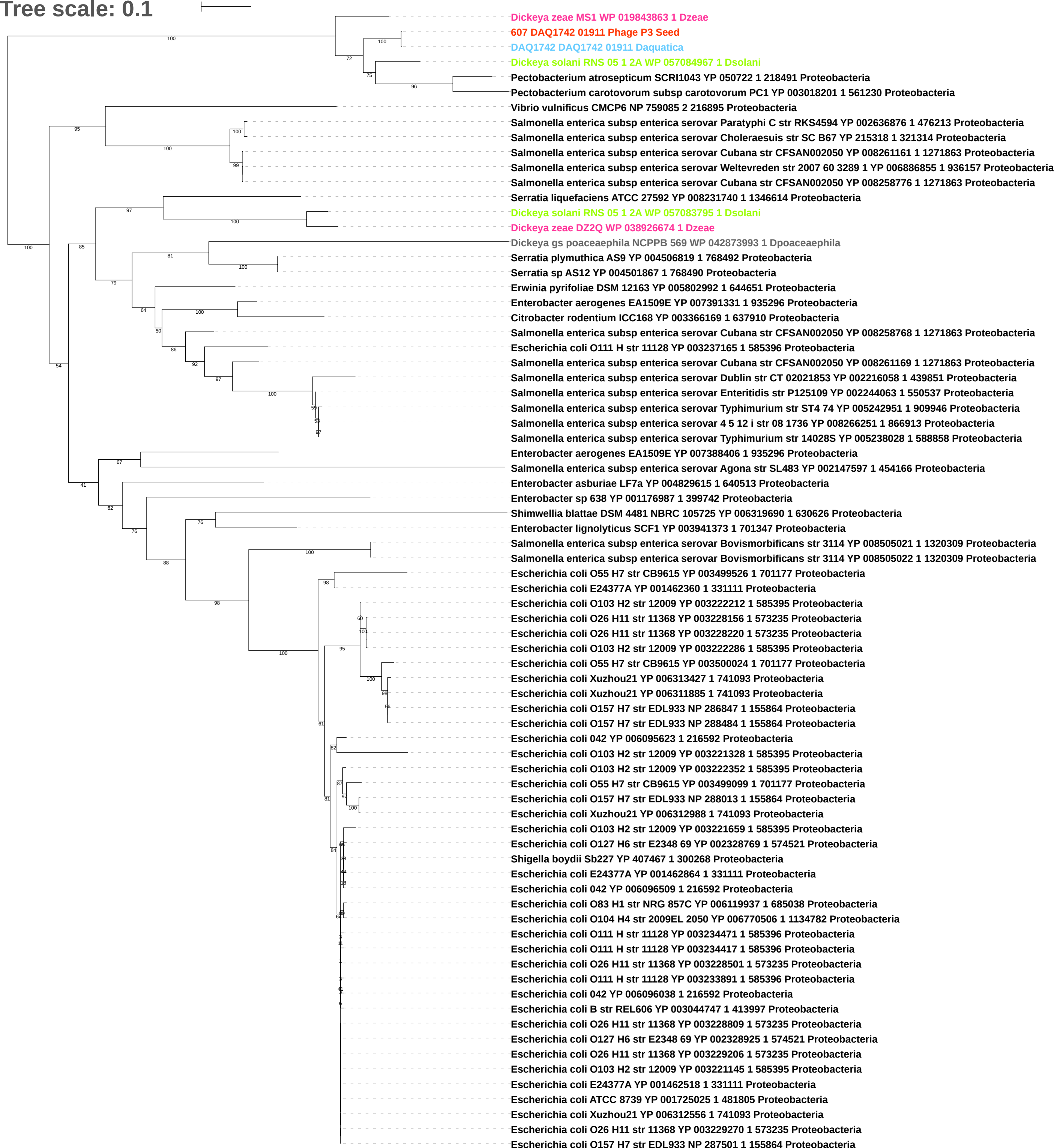
Tree scale: 0.1
Dickeya zeae MS1 WP 019843863 1 Dzeae
607 DAQ1742 01911 Phage P3 Seed
DAQ1742 DAQ1742 01911 Daquatica
Dickeya solani RNS 05 1 2A WP 057084967 1 Dsolani
Pectobacterium atrosepticum SCRI1043 YP 050722 1 218491 Proteobacteria
Pectobacterium carotovorum subsp carotovorum PC1 YP 003018201 1 561230 Proteobacteria
Vibrio vulnificus CMCP6 NP 759085 2 216895 Proteobacteria
Salmonella enterica subsp enterica serovar Paratyphi C str RKS4594 YP 002636876 1 476213 Proteobacteria
Salmonella enterica subsp enterica serovar Choleraesuis str SC B67 YP 215318 1 321314 Proteobacteria
Salmonella enterica subsp enterica serovar Cubana str CFSAN002050 YP 008261161 1 1271863 Proteobacteria
Salmonella enterica subsp enterica serovar Weltevreden str 2007 60 3289 1 YP 006886855 1 936157 Proteobacteria
Salmonella enterica subsp enterica serovar Cubana str CFSAN002050 YP 008258776 1 1271863 Proteobacteria
Serratia liquefaciens ATCC 27592 YP 008231740 1 1346614 Proteobacteria
Dickeya solani RNS 05 1 2A WP 057083795 1 Dsolani
Dickeya zeae DZ2Q WP 038926674 1 Dzeae
Dickeya gs poaceaephila NCPPB 569 WP 042873993 1 Dpoaceaephila
Serratia plymuthica AS9 YP 004506819 1 768492 Proteobacteria
Serratia sp AS12 YP 004501867 1 768490 Proteobacteria
Erwinia pyrifoliae DSM 12163 YP 005802992 1 644651 Proteobacteria
Enterobacter aerogenes EA1509E YP 007391331 1 935296 Proteobacteria
Citrobacter rodentium ICC168 YP 003366169 1 637910 Proteobacteria
Salmonella enterica subsp enterica serovar Cubana str CFSAN002050 YP 008258768 1 1271863 Proteobacteria
Escherichia coli O111 H str 11128 YP 003237165 1 585396 Proteobacteria
Salmonella enterica subsp enterica serovar Cubana str CFSAN002050 YP 008261169 1 1271863 Proteobacteria
Salmonella enterica subsp enterica serovar Dublin str CT 02021853 YP 002216058 1 439851 Proteobacteria
Salmonella enterica subsp enterica serovar Enteritidis str P125109 YP 002244063 1 550537 Proteobacteria
Salmonella enterica subsp enterica serovar Typhimurium str ST4 74 YP 005242951 1 909946 Proteobacteria
Salmonella enterica subsp enterica serovar 4 5 12 i str 08 1736 YP 008266251 1 866913 Proteobacteria
Salmonella enterica subsp enterica serovar Typhimurium str 14028S YP 005238028 1 588858 Proteobacteria
Enterobacter aerogenes EA1509E YP 007388406 1 935296 Proteobacteria
Salmonella enterica subsp enterica serovar Agona str SL483 YP 002147597 1 454166 Proteobacteria
Enterobacter asburiae LF7a YP 004829615 1 640513 Proteobacteria
Enterobacter sp 638 YP 001176987 1 399742 Proteobacteria
Shimwellia blattae DSM 4481 NBRC 105725 YP 006319690 1 630626 Proteobacteria
Enterobacter lignolyticus SCF1 YP 003941373 1 701347 Proteobacteria
Salmonella enterica subsp enterica serovar Bovismorbificans str 3114 YP 008505021 1 1320309 Proteobacteria
Salmonella enterica subsp enterica serovar Bovismorbificans str 3114 YP 008505022 1 1320309 Proteobacteria
Escherichia coli O55 H7 str CB9615 YP 003499526 1 701177 Proteobacteria
Escherichia coli E24377A YP 001462360 1 331111 Proteobacteria
Escherichia coli O103 H2 str 12009 YP 003222212 1 585395 Proteobacteria
Escherichia coli O26 H11 str 11368 YP 003228156 1 573235 Proteobacteria
Escherichia coli O26 H11 str 11368 YP 003228220 1 573235 Proteobacteria
Escherichia coli O103 H2 str 12009 YP 003222286 1 585395 Proteobacteria
Escherichia coli O55 H7 str CB9615 YP 003500024 1 701177 Proteobacteria
Escherichia coli Xuzhou21 YP 006313427 1 741093 Proteobacteria
Escherichia coli Xuzhou21 YP 006311885 1 741093 Proteobacteria
Escherichia coli O157 H7 str EDL933 NP 286847 1 155864 Proteobacteria
Escherichia coli O157 H7 str EDL933 NP 288484 1 155864 Proteobacteria
Escherichia coli 042 YP 006095623 1 216592 Proteobacteria
Escherichia coli O103 H2 str 12009 YP 003221328 1 585395 Proteobacteria
Escherichia coli O103 H2 str 12009 YP 003222352 1 585395 Proteobacteria
Escherichia coli O55 H7 str CB9615 YP 003499099 1 701177 Proteobacteria
Escherichia coli O157 H7 str EDL933 NP 288013 1 155864 Proteobacteria
Escherichia coli Xuzhou21 YP 006312988 1 741093 Proteobacteria
Escherichia coli O103 H2 str 12009 YP 003221659 1 585395 Proteobacteria
Escherichia coli O127 H6 str E2348 69 YP 002328769 1 574521 Proteobacteria
Shigella boydii Sb227 YP 407467 1 300268 Proteobacteria
Escherichia coli E24377A YP 001462864 1 331111 Proteobacteria
Escherichia coli 042 YP 006096509 1 216592 Proteobacteria
Escherichia coli O83 H1 str NRG 857C YP 006119937 1 685038 Proteobacteria
Escherichia coli O104 H4 str 2009EL 2050 YP 006770506 1 1134782 Proteobacteria
Escherichia coli O111 H str 11128 YP 003234471 1 585396 Proteobacteria
Escherichia coli O111 H str 11128 YP 003234417 1 585396 Proteobacteria
Escherichia coli O26 H11 str 11368 YP 003228501 1 573235 Proteobacteria
Escherichia coli O111 H str 11128 YP 003233891 1 585396 Proteobacteria
Escherichia coli 042 YP 006096038 1 216592 Proteobacteria
Escherichia coli B str REL606 YP 003044747 1 413997 Proteobacteria
Escherichia coli O26 H11 str 11368 YP 003228809 1 573235 Proteobacteria
Escherichia coli O127 H6 str E2348 69 YP 002328925 1 574521 Proteobacteria
Escherichia coli O26 H11 str 11368 YP 003229206 1 573235 Proteobacteria
Escherichia coli O103 H2 str 12009 YP 003221145 1 585395 Proteobacteria
Escherichia coli E24377A YP 001462518 1 331111 Proteobacteria
Escherichia coli ATCC 8739 YP 001725025 1 481805 Proteobacteria
Escherichia coli Xuzhou21 YP 006312556 1 741093 Proteobacteria
Escherichia coli O26 H11 str 11368 YP 003229270 1 573235 Proteobacteria
Escherichia coli O157 H7 str EDL933 NP 287501 1 155864 Proteobacteria

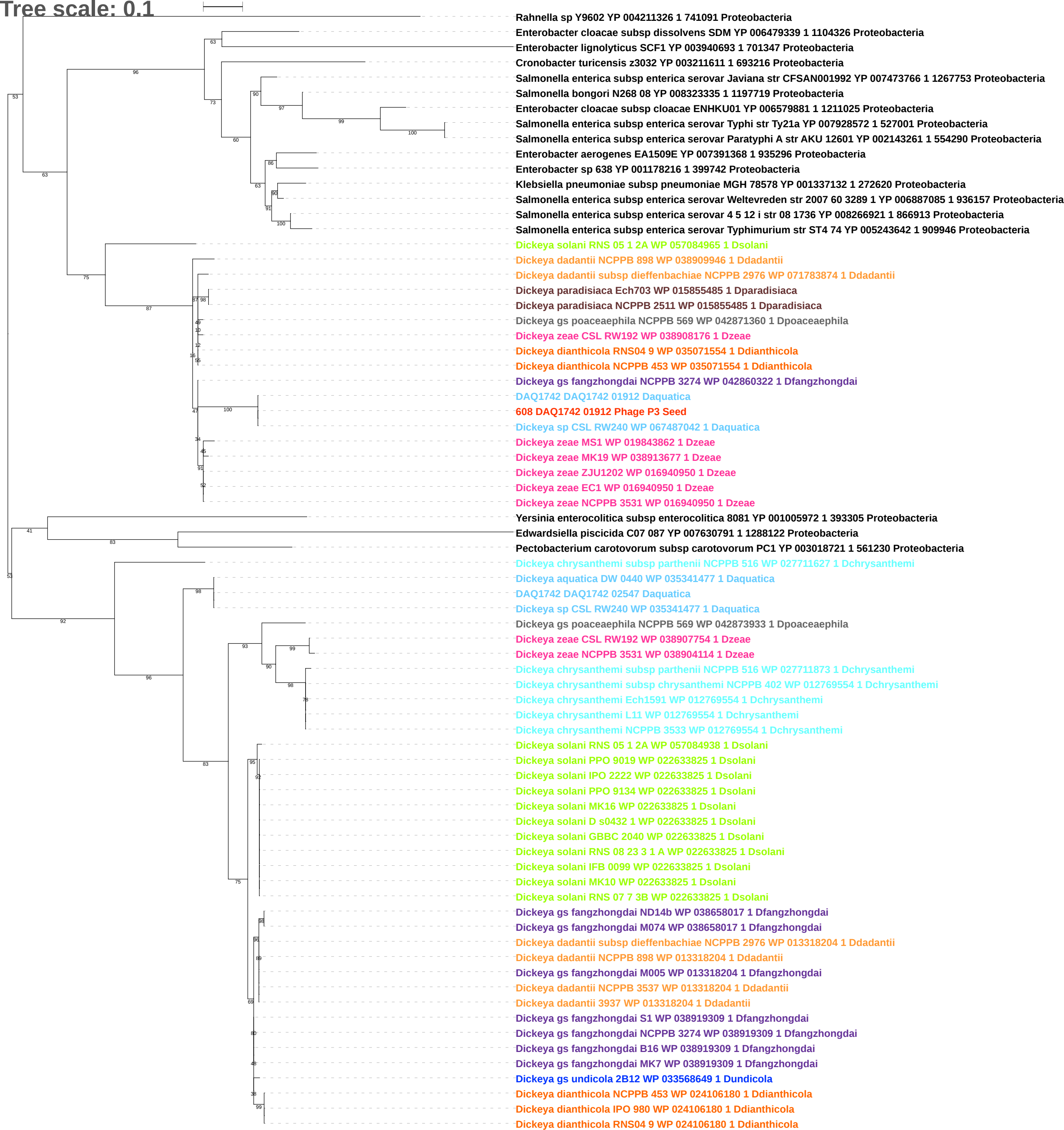
Tree scale: 0.1
Rahnella sp Y9602 YP 004211326 1 741091 Proteobacteria
Enterobacter cloacae subsp dissolvens SDM YP 006479339 1 1104326 Proteobacteria
Enterobacter lignolyticus SCF1 YP 003940693 1 701347 Proteobacteria
Cronobacter turicensis z3032 YP 003211611 1 693216 Proteobacteria
Salmonella enterica subsp enterica serovar Javiana str CFSAN001992 YP 007473766 1 1267753 Proteobacteria
Salmonella bongori N268 08 YP 008323335 1 1197719 Proteobacteria
Enterobacter cloacae subsp cloacae ENHKU01 YP 006579881 1 1211025 Proteobacteria
Salmonella enterica subsp enterica serovar Typhi str Ty21a YP 007928572 1 527001 Proteobacteria
Salmonella enterica subsp enterica serovar Paratyphi A str AKU 12601 YP 002143261 1 554290 Proteobacteria
Enterobacter aerogenes EA1509E YP 007391368 1 935296 Proteobacteria
Enterobacter sp 638 YP 001178216 1 399742 Proteobacteria
Klebsiella pneumoniae subsp pneumoniae MGH 78578 YP 001337132 1 272620 Proteobacteria
Salmonella enterica subsp enterica serovar Weltevreden str 2007 60 3289 1 YP 006887085 1 936157 Proteobacteria
Salmonella enterica subsp enterica serovar 4 5 12 i str 08 1736 YP 008266921 1 866913 Proteobacteria
Salmonella enterica subsp enterica serovar Typhimurium str ST4 74 YP 005243642 1 909946 Proteobacteria
Dickeya solani RNS 05 1 2A WP 057084965 1 Dsolani
Dickeya dadantii NCPPB 898 WP 038909946 1 Ddadantii
Dickeya dadantii subsp dieffenbachiae NCPPB 2976 WP 071783874 1 Ddadantii
Dickeya paradisiaca Ech703 WP 015855485 1 Dparadisiaca
Dickeya paradisiaca NCPPB 2511 WP 015855485 1 Dparadisiaca
Dickeya gs poaceaephila NCPPB 569 WP 042871360 1 Dpoaceaephila
Dickeya zeae CSL RW192 WP 038908176 1 Dzeae
Dickeya dianthicola RNS04 9 WP 035071554 1 Ddianthicola
Dickeya dianthicola NCPPB 453 WP 035071554 1 Ddianthicola
Dickeya gs fangzhongdai NCPPB 3274 WP 042860322 1 Dfangzhongdai
DAQ1742 DAQ1742 01912 Daquatica
608 DAQ1742 01912 Phage P3 Seed
Dickeya sp CSL RW240 WP 067487042 1 Daquatica
Dickeya zeae MS1 WP 019843862 1 Dzeae
Dickeya zeae MK19 WP 038913677 1 Dzeae
Dickeya zeae ZJU1202 WP 016940950 1 Dzeae
Dickeya zeae EC1 WP 016940950 1 Dzeae
Dickeya zeae NCPPB 3531 WP 016940950 1 Dzeae
Yersinia enterocolitica subsp enterocolitica 8081 YP 001005972 1 393305 Proteobacteria
Edwardsiella piscicida C07 087 YP 007630791 1 1288122 Proteobacteria
Pectobacterium carotovorum subsp carotovorum PC1 YP 003018721 1 561230 Proteobacteria
Dickeya chrysanthemi subsp parthenii NCPPB 516 WP 027711627 1 Dchrysanthemi
Dickeya aquatica DW 0440 WP 035341477 1 Daquatica
DAQ1742 DAQ1742 02547 Daquatica
Dickeya sp CSL RW240 WP 035341477 1 Daquatica
Dickeya gs poaceaephila NCPPB 569 WP 042873933 1 Dpoaceaephila
Dickeya zeae CSL RW192 WP 038907754 1 Dzeae
Dickeya zeae NCPPB 3531 WP 038904114 1 Dzeae
Dickeya chrysanthemi subsp parthenii NCPPB 516 WP 027711873 1 Dchrysanthemi
Dickeya chrysanthemi subsp chrysanthemi NCPPB 402 WP 012769554 1 Dchrysanthemi
Dickeya chrysanthemi Ech1591 WP 012769554 1 Dchrysanthemi
Dickeya chrysanthemi L11 WP 012769554 1 Dchrysanthemi
Dickeya chrysanthemi NCPPB 3533 WP 012769554 1 Dchrysanthemi
Dickeya solani RNS 05 1 2A WP 057084938 1 Dsolani
Dickeya solani PPO 9019 WP 022633825 1 Dsolani
Dickeya solani IPO 2222 WP 022633825 1 Dsolani
Dickeya solani PPO 9134 WP 022633825 1 Dsolani
Dickeya solani MK16 WP 022633825 1 Dsolani
Dickeya solani D s0432 1 WP 022633825 1 Dsolani
Dickeya solani GBBC 2040 WP 022633825 1 Dsolani
Dickeya solani RNS 08 23 3 1 A WP 022633825 1 Dsolani
Dickeya solani IFB 0099 WP 022633825 1 Dsolani
Dickeya solani MK10 WP 022633825 1 Dsolani
Dickeya solani RNS 07 7 3B WP 022633825 1 Dsolani
Dickeya gs fangzhongdai ND14b WP 038658017 1 Dfangzhongdai
Dickeya gs fangzhongdai M074 WP 038658017 1 Dfangzhongdai
Dickeya dadantii subsp dieffenbachiae NCPPB 2976 WP 013318204 1 Ddadantii
Dickeya dadantii NCPPB 898 WP 013318204 1 Ddadantii
Dickeya gs fangzhongdai M005 WP 013318204 1 Dfangzhongdai
Dickeya dadantii NCPPB 3537 WP 013318204 1 Ddadantii
Dickeya dadantii 3937 WP 013318204 1 Ddadantii
Dickeya gs fangzhongdai S1 WP 038919309 1 Dfangzhongdai
Dickeya gs fangzhongdai NCPPB 3274 WP 038919309 1 Dfangzhongdai
Dickeya gs fangzhongdai B16 WP 038919309 1 Dfangzhongdai
Dickeya gs fangzhongdai MK7 WP 038919309 1 Dfangzhongdai
Dickeya gs undicola 2B12 WP 033568649 1 Dundicola
Dickeya dianthicola NCPPB 453 WP 024106180 1 Ddianthicola
Dickeya dianthicola IPO 980 WP 024106180 1 Ddianthicola
Dickeya dianthicola RNS04 9 WP 024106180 1 Ddianthicola

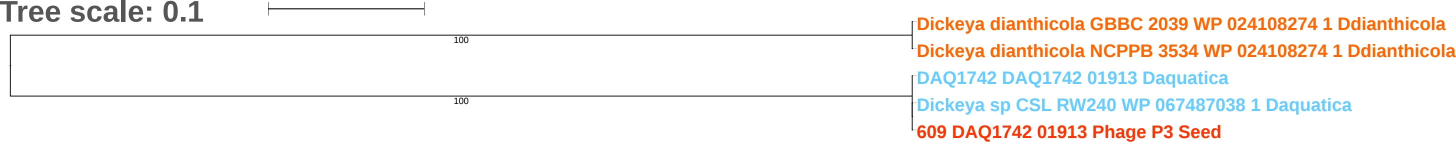
Tree scale: 0.1
100
100
Dickeya dianthicola GBBC 2039 WP 024108274 1 Ddianthicola
Dickeya dianthicola NCPPB 3534 WP 024108274 1 Ddianthicola
DAQ1742 DAQ1742 01913 Daquatica
Dickeya sp CSL RW240 WP 067487038 1 Daquatica
609 DAQ1742 01913 Phage P3 Seed

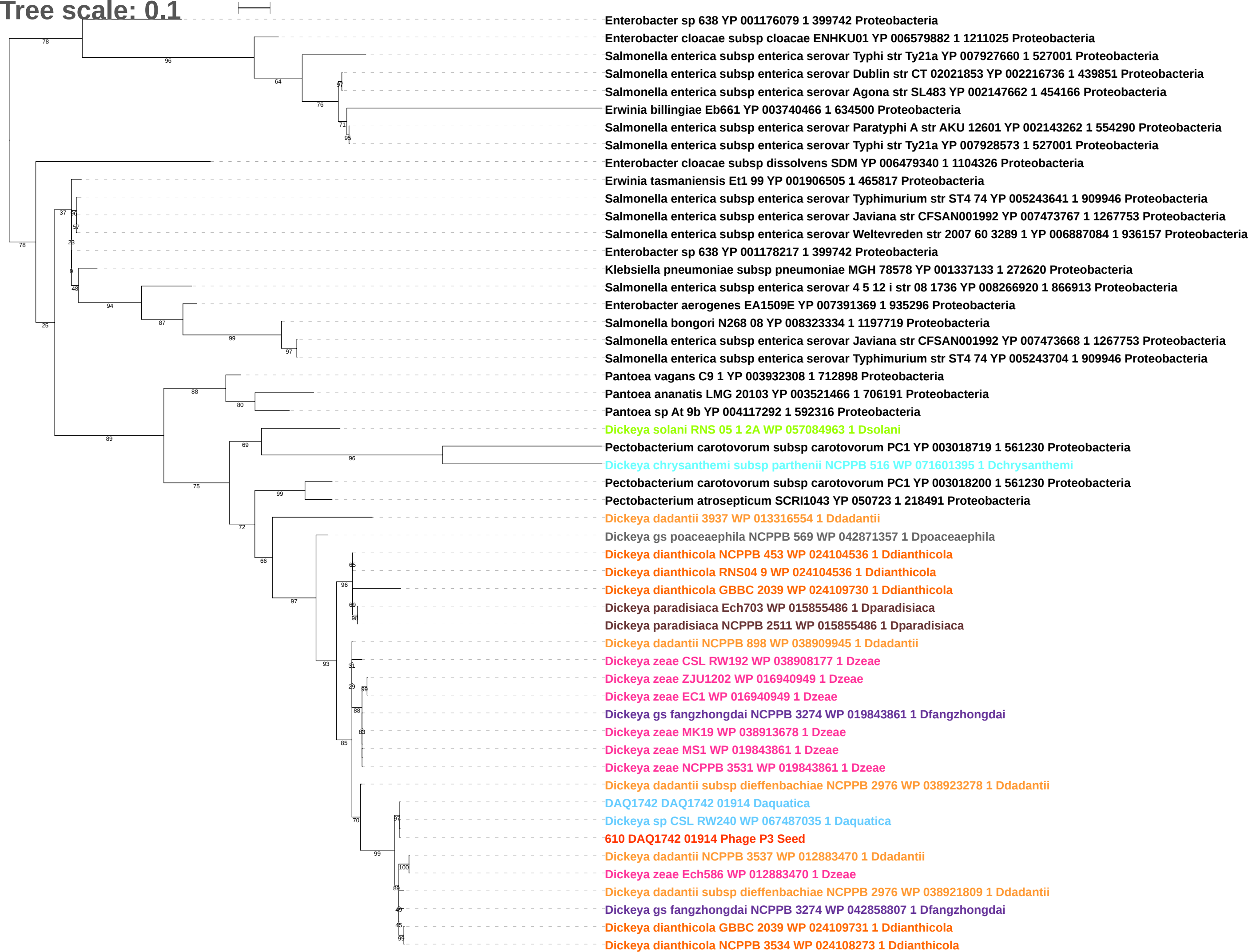

Tree scale: 0.1
Enterobacter sp 638 YP 001176079 1 399742 Proteobacteria
Enterobacter cloacae subsp cloacae ENHKU01 YP 006579882 1 1211025 Proteobacteria
Salmonella enterica subsp enterica serovar Typhi str Ty21a YP 007927660 1 527001 Proteobacteria
Salmonella enterica subsp enterica serovar Dublin str CT 02021853 YP 002216736 1 439851 Proteobacteria
Salmonella enterica subsp enterica serovar Agona str SL483 YP 002147662 1 454166 Proteobacteria
Erwinia billingiae Eb661 YP 003740466 1 634500 Proteobacteria
Salmonella enterica subsp enterica serovar Paratyphi A str AKU 12601 YP 002143262 1 554290 Proteobacteria
Salmonella enterica subsp enterica serovar Typhi str Ty21a YP 007928573 1 527001 Proteobacteria
Enterobacter cloacae subsp dissolvens SDM YP 006479340 1 1104326 Proteobacteria
Erwinia tasmaniensis Et1 99 YP 001906505 1 465817 Proteobacteria
Salmonella enterica subsp enterica serovar Typhimurium str ST4 74 YP 005243641 1 909946 Proteobacteria
Salmonella enterica subsp enterica serovar Javiana str CFSAN001992 YP 007473767 1 1267753 Proteobacteria
Salmonella enterica subsp enterica serovar Weltevreden str 2007 60 3289 1 YP 006887084 1 936157 Proteobacteria
Enterobacter sp 638 YP 001178217 1 399742 Proteobacteria
Klebsiella pneumoniae subsp pneumoniae MGH 78578 YP 001337133 1 272620 Proteobacteria
Salmonella enterica subsp enterica serovar 4 5 12 i str 08 1736 YP 008266920 1 866913 Proteobacteria
Enterobacter aerogenes EA1509E YP 007391369 1 935296 Proteobacteria
Salmonella bongori N268 08 YP 008323334 1 1197719 Proteobacteria
Salmonella enterica subsp enterica serovar Javiana str CFSAN001992 YP 007473668 1 1267753 Proteobacteria
Salmonella enterica subsp enterica serovar Typhimurium str ST4 74 YP 005243704 1 909946 Proteobacteria
Pantoea vagans C9 1 YP 003932308 1 712898 Proteobacteria
Pantoea ananatis LMG 20103 YP 003521466 1 706191 Proteobacteria
Pantoea sp At 9b YP 004117292 1 592316 Proteobacteria
Dickeya solani RNS 05 1 2A WP 057084963 1 Dsolani
Pectobacterium carotovorum subsp carotovorum PC1 YP 003018719 1 561230 Proteobacteria
Dickeya chrysanthemi subsp parthenii NCPPB 516 WP 071601395 1 Dchrysanthemi
Pectobacterium carotovorum subsp carotovorum PC1 YP 003018200 1 561230 Proteobacteria
Pectobacterium atrosepticum SCRI1043 YP 050723 1 218491 Proteobacteria
Dickeya dadantii 3937 WP 013316554 1 Ddadantii
Dickeya gs poaceaephila NCPPB 569 WP 042871357 1 Dpoaceaephila
Dickeya dianthicola NCPPB 453 WP 024104536 1 Ddianthicola
Dickeya dianthicola RNS04 9 WP 024104536 1 Ddianthicola
Dickeya dianthicola GBBC 2039 WP 024109730 1 Ddianthicola
Dickeya paradisiaca Ech703 WP 015855486 1 Dparadisiaca
Dickeya paradisiaca NCPPB 2511 WP 015855486 1 Dparadisiaca
Dickeya dadantii NCPPB 898 WP 038909945 1 Ddadantii
Dickeya zeae CSL RW192 WP 038908177 1 Dzeae
Dickeya zeae ZJU1202 WP 016940949 1 Dzeae
Dickeya zeae EC1 WP 016940949 1 Dzeae
Dickeya gs fangzhongdai NCPPB 3274 WP 019843861 1 Dfangzhongdai
Dickeya zeae MK19 WP 038913678 1 Dzeae
Dickeya zeae MS1 WP 019843861 1 Dzeae
Dickeya zeae NCPPB 3531 WP 019843861 1 Dzeae
Dickeya dadantii subsp dieffenbachiae NCPPB 2976 WP 038923278 1 Ddadantii
DAQ1742 DAQ1742 01914 Daquatica
Dickeya sp CSL RW240 WP 067487035 1 Daquatica
610 DAQ1742 01914 Phage P3 Seed
Dickeya dadantii NCPPB 3537 WP 012883470 1 Ddadantii
Dickeya zeae Ech586 WP 012883470 1 Dzeae
Dickeya dadantii subsp dieffenbachiae NCPPB 2976 WP 038921809 1 Ddadantii
Dickeya gs fangzhongdai NCPPB 3274 WP 042858807 1 Dfangzhongdai
Dickeya dianthicola GBBC 2039 WP 024109731 1 Ddianthicola
Dickeya dianthicola NCPPB 3534 WP 024108273 1 Ddianthicola

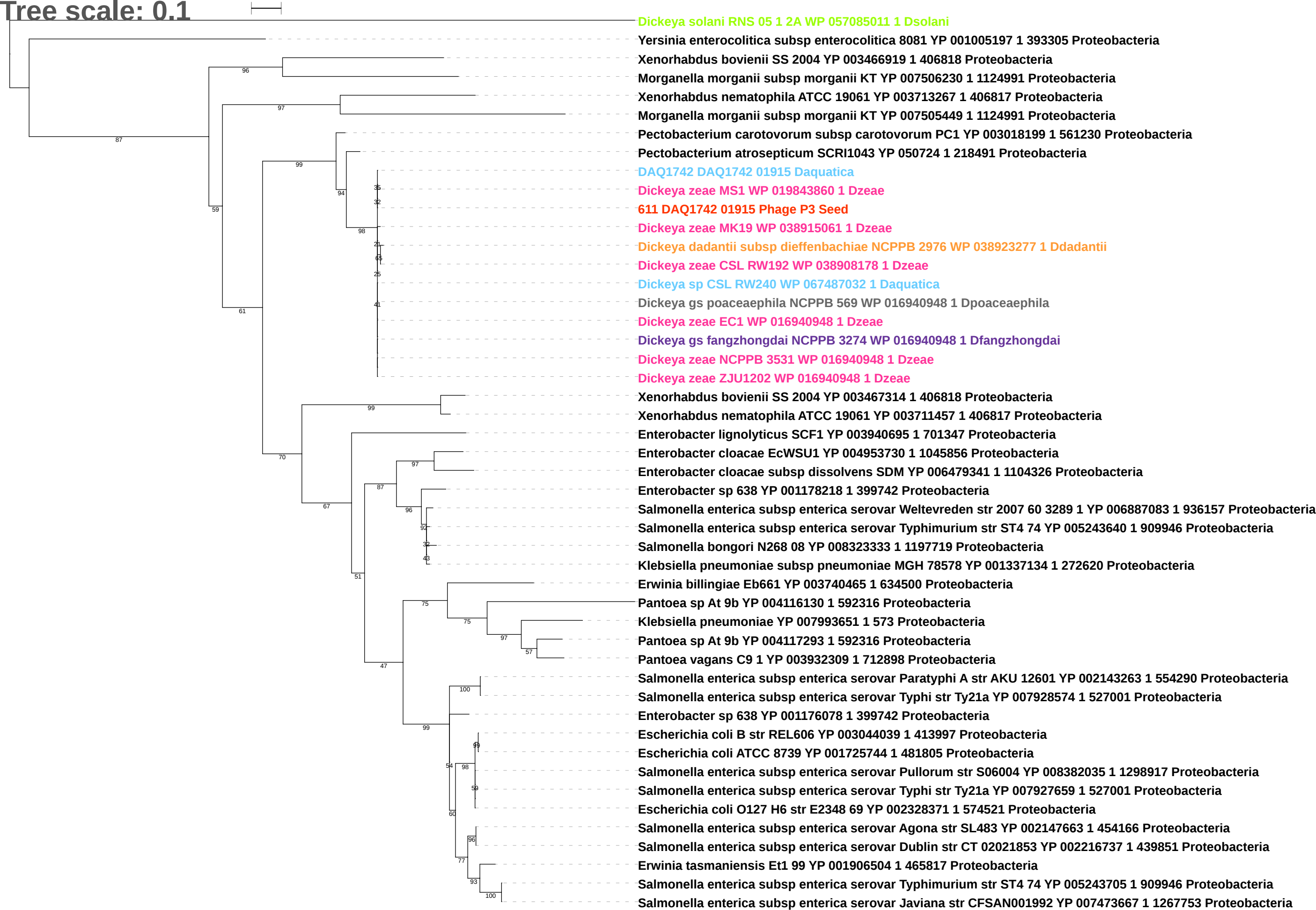
Tree scale: 0.1
Dickeya solani RNS 05 1 2A WP 057085011 1 Dsolani
Yersinia enterocolitica subsp enterocolitica 8081 YP 001005197 1 393305 Proteobacteria
Xenorhabdus bovienii SS 2004 YP 003466919 1 406818 Proteobacteria
Morganella morganii subsp morganii KT YP 007506230 1 1124991 Proteobacteria
Xenorhabdus nematophila ATCC 19061 YP 003713267 1 406817 Proteobacteria
Morganella morganii subsp morganii KT YP 007505449 1 1124991 Proteobacteria
Pectobacterium carotovorum subsp carotovorum PC1 YP 003018199 1 561230 Proteobacteria
Pectobacterium atrosepticum SCRI1043 YP 050724 1 218491 Proteobacteria
DAQ1742 DAQ1742 01915 Daquatica
Dickeya zeae MS1 WP 019843860 1 Dzeae
611 DAQ1742 01915 Phage P3 Seed
Dickeya zeae MK19 WP 038915061 1 Dzeae
Dickeya dadantii subsp dieffenbachiae NCPPB 2976 WP 038923277 1 Ddadantii
Dickeya zeae CSL RW192 WP 038908178 1 Dzeae
Dickeya sp CSL RW240 WP 067487032 1 Daquatica
Dickeya gs poaceaephila NCPPB 569 WP 016940948 1 Dpoaceaephila
Dickeya zeae EC1 WP 016940948 1 Dzeae
Dickeya gs fangzhongdai NCPPB 3274 WP 016940948 1 Dfangzhongdai
Dickeya zeae NCPPB 3531 WP 016940948 1 Dzeae
Dickeya zeae ZJU1202 WP 016940948 1 Dzeae
Xenorhabdus bovienii SS 2004 YP 003467314 1 406818 Proteobacteria
Xenorhabdus nematophila ATCC 19061 YP 003711457 1 406817 Proteobacteria
Enterobacter lignolyticus SCF1 YP 003940695 1 701347 Proteobacteria
Enterobacter cloacae EcWSU1 YP 004953730 1 1045856 Proteobacteria
Enterobacter cloacae subsp dissolvens SDM YP 006479341 1 1104326 Proteobacteria
Enterobacter sp 638 YP 001178218 1 399742 Proteobacteria
Salmonella enterica subsp enterica serovar Weltevreden str 2007 60 3289 1 YP 006887083 1 936157 Proteobacteria
Salmonella enterica subsp enterica serovar Typhimurium str ST4 74 YP 005243640 1 909946 Proteobacteria
Salmonella bongori N268 08 YP 008323333 1 1197719 Proteobacteria
Klebsiella pneumoniae subsp pneumoniae MGH 78578 YP 001337134 1 272620 Proteobacteria
Erwinia billingiae Eb661 YP 003740465 1 634500 Proteobacteria
Pantoea sp At 9b YP 004116130 1 592316 Proteobacteria
Klebsiella pneumoniae YP 007993651 1 573 Proteobacteria
Pantoea sp At 9b YP 004117293 1 592316 Proteobacteria
Pantoea vagans C9 1 YP 003932309 1 712898 Proteobacteria
Salmonella enterica subsp enterica serovar Paratyphi A str AKU 12601 YP 002143263 1 554290 Proteobacteria
Salmonella enterica subsp enterica serovar Typhi str Ty21a YP 007928574 1 527001 Proteobacteria
Enterobacter sp 638 YP 001176078 1 399742 Proteobacteria
Escherichia coli B str REL606 YP 003044039 1 413997 Proteobacteria
Escherichia coli ATCC 8739 YP 001725744 1 481805 Proteobacteria
Salmonella enterica subsp enterica serovar Pullorum str S06004 YP 008382035 1 1298917 Proteobacteria
Salmonella enterica subsp enterica serovar Typhi str Ty21a YP 007927659 1 527001 Proteobacteria
Escherichia coli O127 H6 str E2348 69 YP 002328371 1 574521 Proteobacteria
Salmonella enterica subsp enterica serovar Agona str SL483 YP 002147663 1 454166 Proteobacteria
Salmonella enterica subsp enterica serovar Dublin str CT 02021853 YP 002216737 1 439851 Proteobacteria
Erwinia tasmaniensis Et1 99 YP 001906504 1 465817 Proteobacteria
Salmonella enterica subsp enterica serovar Typhimurium str ST4 74 YP 005243705 1 909946 Proteobacteria
Salmonella enterica subsp enterica serovar Javiana str CFSAN001992 YP 007473667 1 1267753 Proteobacteria

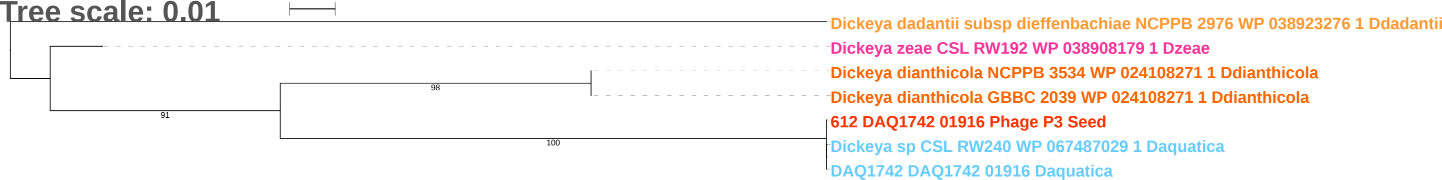

Tree scale: 0.01
Dickeya dadantii subsp dieffenbachiae NCPPB 2976 WP 038923276 1 Ddadantii
Dickeya zeae CSL RW192 WP 038908179 1 Dzeae
Dickeya dianthicola NCPPB 3534 WP 024108271 1 Ddianthicola
Dickeya dianthicola GBBC 2039 WP 024108271 1 Ddianthicola
612 DAQ1742 01916 Phage P3 Seed
Dickeya sp CSL RW240 WP 067487029 1 Daquatica
DAQ1742 DAQ1742 01916 Daquatica
98
91
100

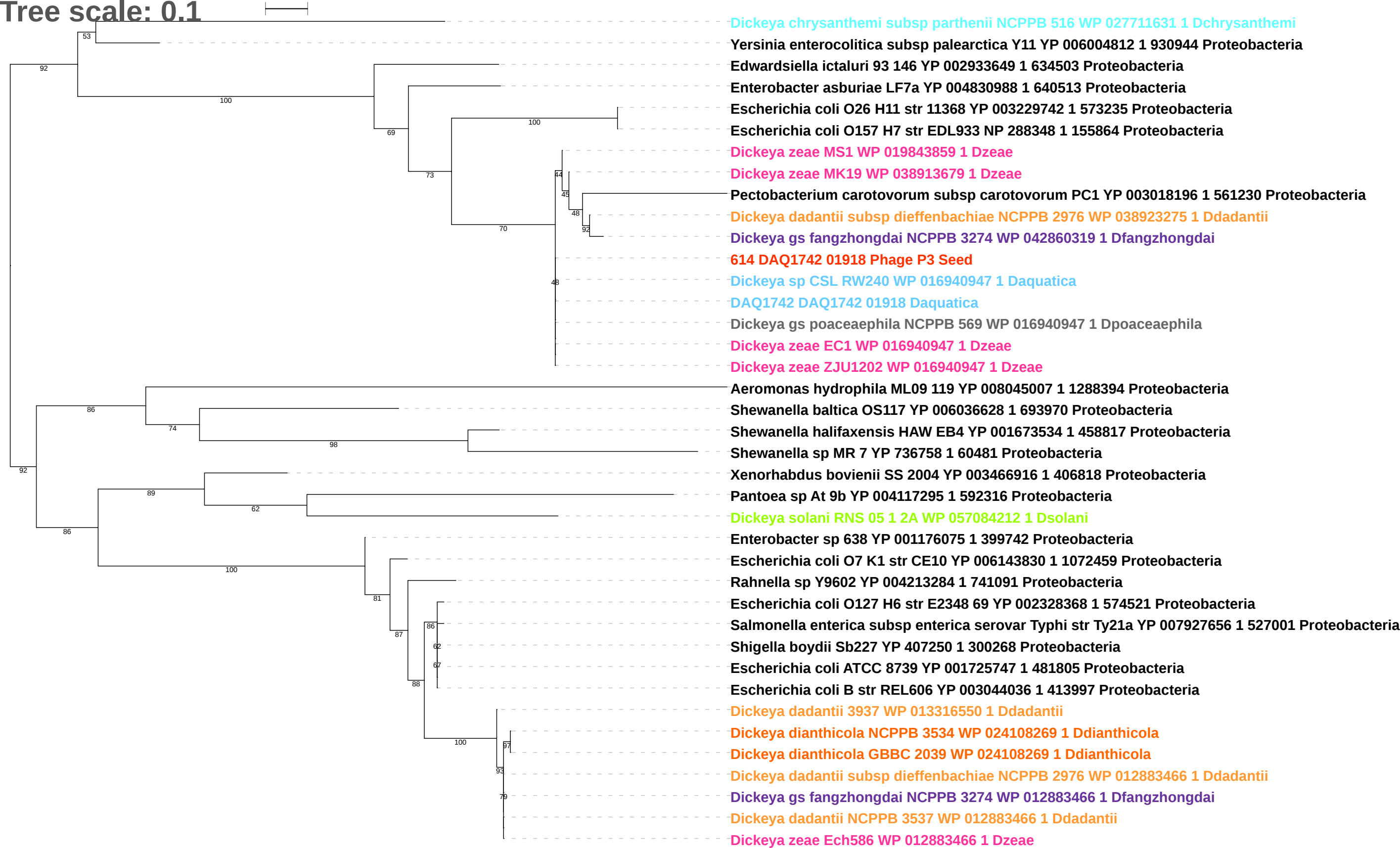
Tree scale: 0.1
Dickeya chrysanthemi subsp parthenii NCPPB 516 WP 027711631 1 Dchrysanthemi
Yersinia enterocolitica subsp palearctica Y11 YP 006004812 1 930944 Proteobacteria
Edwardsiella ictaluri 93 146 YP 002933649 1 634503 Proteobacteria
Enterobacter asburiae LF7a YP 004830988 1 640513 Proteobacteria
Escherichia coli O26 H11 str 11368 YP 003229742 1 573235 Proteobacteria
Escherichia coli O157 H7 str EDL933 NP 288348 1 155864 Proteobacteria
Dickeya zeae MS1 WP 019843859 1 Dzeae
Dickeya zeae MK19 WP 038913679 1 Dzeae
Pectobacterium carotovorum subsp carotovorum PC1 YP 003018196 1 561230 Proteobacteria
Dickeya dadantii subsp dieffenbachiae NCPPB 2976 WP 038923275 1 Ddadantii
Dickeya gs fangzhongdai NCPPB 3274 WP 042860319 1 Dfangzhongdai
614 DAQ1742 01918 Phage P3 Seed
Dickeya sp CSL RW240 WP 016940947 1 Daquatica
DAQ1742 DAQ1742 01918 Daquatica
Dickeya gs poaceaephila NCPPB 569 WP 016940947 1 Dpoaceaephila
Dickeya zeae EC1 WP 016940947 1 Dzeae
Dickeya zeae ZJU1202 WP 016940947 1 Dzeae
Aeromonas hydrophila ML09 119 YP 008045007 1 1288394 Proteobacteria
Shewanella baltica OS117 YP 006036628 1 693970 Proteobacteria
Shewanella halifaxensis HAW EB4 YP 001673534 1 458817 Proteobacteria
Shewanella sp MR 7 YP 736758 1 60481 Proteobacteria
Xenorhabdus bovienii SS 2004 YP 003466916 1 406818 Proteobacteria
Pantoea sp At 9b YP 004117295 1 592316 Proteobacteria
Dickeya solani RNS 05 1 2A WP 057084212 1 Dsolani
Enterobacter sp 638 YP 001176075 1 399742 Proteobacteria
Escherichia coli O7 K1 str CE10 YP 006143830 1 1072459 Proteobacteria
Rahnella sp Y9602 YP 004213284 1 741091 Proteobacteria
Escherichia coli O127 H6 str E2348 69 YP 002328368 1 574521 Proteobacteria
Salmonella enterica subsp enterica serovar Typhi str Ty21a YP 007927656 1 527001 Proteobacteria
Shigella boydii Sb227 YP 407250 1 300268 Proteobacteria
Escherichia coli ATCC 8739 YP 001725747 1 481805 Proteobacteria
Escherichia coli B str REL606 YP 003044036 1 413997 Proteobacteria
Dickeya dadantii 3937 WP 013316550 1 Ddadantii
Dickeya dianthicola NCPPB 3534 WP 024108269 1 Ddianthicola
Dickeya dianthicola GBBC 2039 WP 024108269 1 Ddianthicola
Dickeya dadantii subsp dieffenbachiae NCPPB 2976 WP 012883466 1 Ddadantii
Dickeya gs fangzhongdai NCPPB 3274 WP 012883466 1 Dfangzhongdai
Dickeya dadantii NCPPB 3537 WP 012883466 1 Ddadantii
Dickeya zeae Ech586 WP 012883466 1 Dzeae

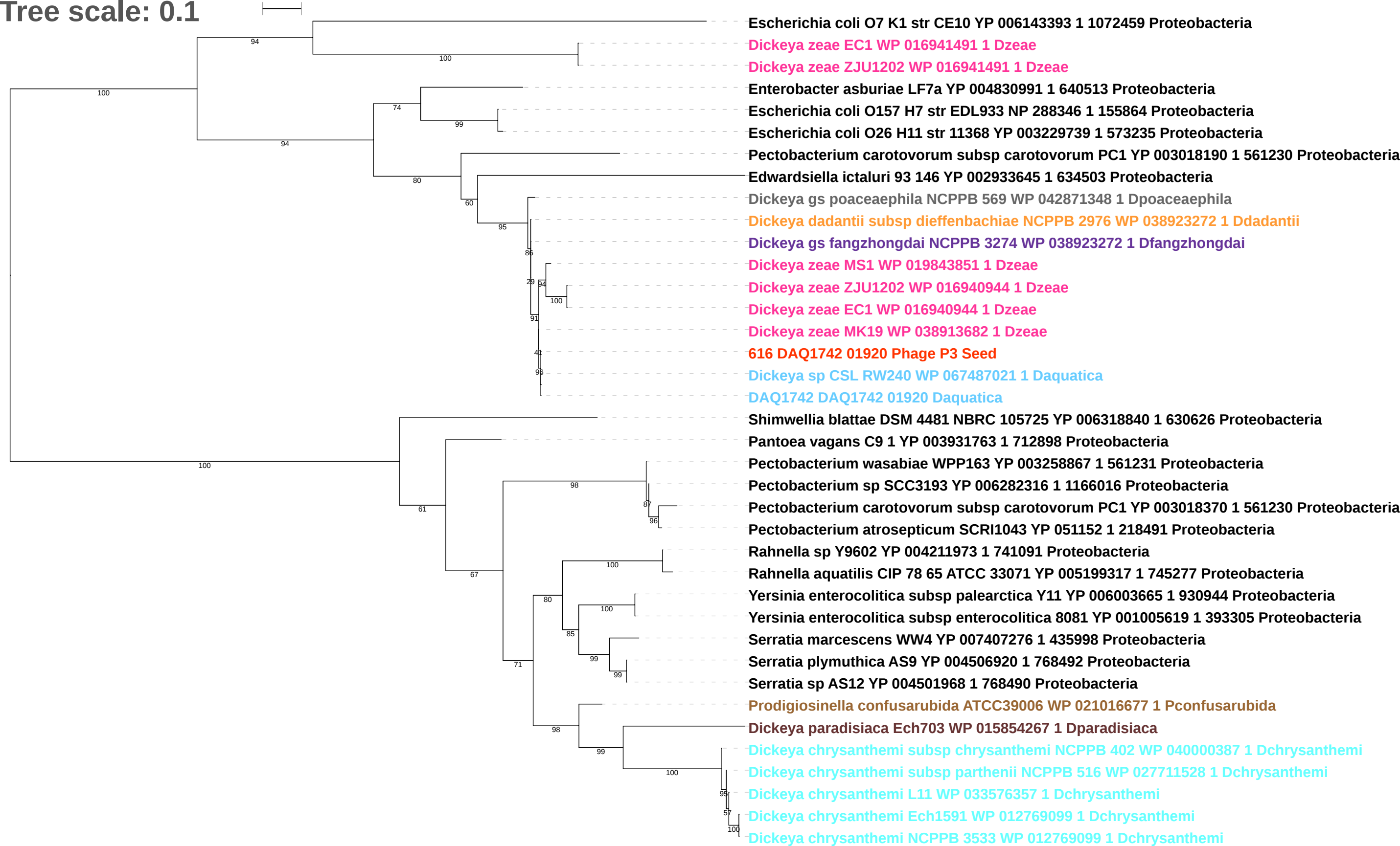
Tree scale: 0.1
Escherichia coli O7 K1 str CE10 YP 006143393 1 1072459 Proteobacteria
Dickeya zeae EC1 WP 016941491 1 Dzeae
Dickeya zeae ZJU1202 WP 016941491 1 Dzeae
Enterobacter asburiae LF7a YP 004830991 1 640513 Proteobacteria
Escherichia coli O157 H7 str EDL933 NP 288346 1 155864 Proteobacteria
Escherichia coli O26 H11 str 11368 YP 003229739 1 573235 Proteobacteria
Pectobacterium carotovorum subsp carotovorum PC1 YP 003018190 1 561230 Proteobacteria
Edwardsiella ictaluri 93 146 YP 002933645 1 634503 Proteobacteria
Dickeya gs poaceaephila NCPPB 569 WP 042871348 1 Dpoaceaephila
Dickeya dadantii subsp dieffenbachiae NCPPB 2976 WP 038923272 1 Ddadantii
Dickeya gs fangzhongdai NCPPB 3274 WP 038923272 1 Dfangzhongdai
Dickeya zeae MS1 WP 019843851 1 Dzeae
Dickeya zeae ZJU1202 WP 016940944 1 Dzeae
Dickeya zeae EC1 WP 016940944 1 Dzeae
Dickeya zeae MK19 WP 038913682 1 Dzeae
616 DAQ1742 01920 Phage P3 Seed
Dickeya sp CSL RW240 WP 067487021 1 Daquatica
DAQ1742 DAQ1742 01920 Daquatica
Shimwellia blattae DSM 4481 NBRC 105725 YP 006318840 1 630626 Proteobacteria
Pantoea vagans C9 1 YP 003931763 1 712898 Proteobacteria
Pectobacterium wasabiae WPP163 YP 003258867 1 561231 Proteobacteria
Pectobacterium sp SCC3193 YP 006282316 1 1166016 Proteobacteria
Pectobacterium carotovorum subsp carotovorum PC1 YP 003018370 1 561230 Proteobacteria
Pectobacterium atrosepticum SCRI1043 YP 051152 1 218491 Proteobacteria
Rahnella sp Y9602 YP 004211973 1 741091 Proteobacteria
Rahnella aquatilis CIP 78 65 ATCC 33071 YP 005199317 1 745277 Proteobacteria
Yersinia enterocolitica subsp palearctica Y11 YP 006003665 1 930944 Proteobacteria
Yersinia enterocolitica subsp enterocolitica 8081 YP 001005619 1 393305 Proteobacteria
Serratia marcescens WW4 YP 007407276 1 435998 Proteobacteria
Serratia plymuthica AS9 YP 004506920 1 768492 Proteobacteria
Serratia sp AS12 YP 004501968 1 768490 Proteobacteria
Prodigiosinella confusarubida ATCC39006 WP 021016677 1 Pconfusarubida
Dickeya paradisiaca Ech703 WP 015854267 1 Dparadisiaca
Dickeya chrysanthemi subsp chrysanthemi NCPPB 402 WP 040000387 1 Dchrysanthemi
Dickeya chrysanthemi subsp parthenii NCPPB 516 WP 027711528 1 Dchrysanthemi
Dickeya chrysanthemi L11 WP 033576357 1 Dchrysanthemi
Dickeya chrysanthemi Ech1591 WP 012769099 1 Dchrysanthemi
Dickeya chrysanthemi NCPPB 3533 WP 012769099 1 Dchrysanthemi

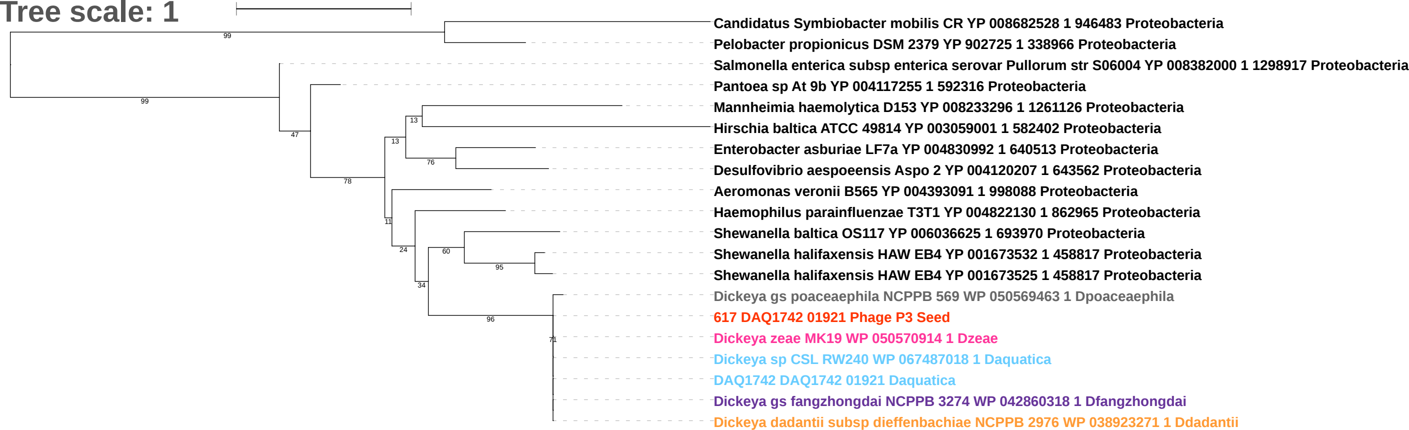

Tree scale: 1
Candidatus Symbiobacter mobilis CR YP 008682528 1 946483 Proteobacteria
Pelobacter propionicus DSM 2379 YP 902725 1 338966 Proteobacteria
Salmonella enterica subsp enterica serovar Pullorum str S06004 YP 008382000 1 1298917 Proteobacteria
Pantoea sp At 9b YP 004117255 1 592316 Proteobacteria
Mannheimia haemolytica D153 YP 008233296 1 1261126 Proteobacteria
Hirschia baltica ATCC 49814 YP 003059001 1 582402 Proteobacteria
Enterobacter asburiae LF7a YP 004830992 1 640513 Proteobacteria
Desulfovibrio aespoeensis Aspo 2 YP 004120207 1 643562 Proteobacteria
Aeromonas veronii B565 YP 004393091 1 998088 Proteobacteria
Haemophilus parainfluenzae T3T1 YP 004822130 1 862965 Proteobacteria
Shewanella baltica OS117 YP 006036625 1 693970 Proteobacteria
Shewanella halifaxensis HAW EB4 YP 001673532 1 458817 Proteobacteria
Shewanella halifaxensis HAW EB4 YP 001673525 1 458817 Proteobacteria
Dickeya gs poaceaephila NCPPB 569 WP 050569463 1 Dpoaceaephila
617 DAQ1742 01921 Phage P3 Seed
Dickeya zeae MK19 WP 050570914 1 Dzeae
Dickeya sp CSL RW240 WP 067487018 1 Daquatica
DAQ1742 DAQ1742 01921 Daquatica
Dickeya gs fangzhongdai NCPPB 3274 WP 042860318 1 Dfangzhongdai
Dickeya dadantii subsp dieffenbachiae NCPPB 2976 WP 038923271 1 Ddadantii
99
99
47
13
13
76
78
11
24
60
95
34
96
71

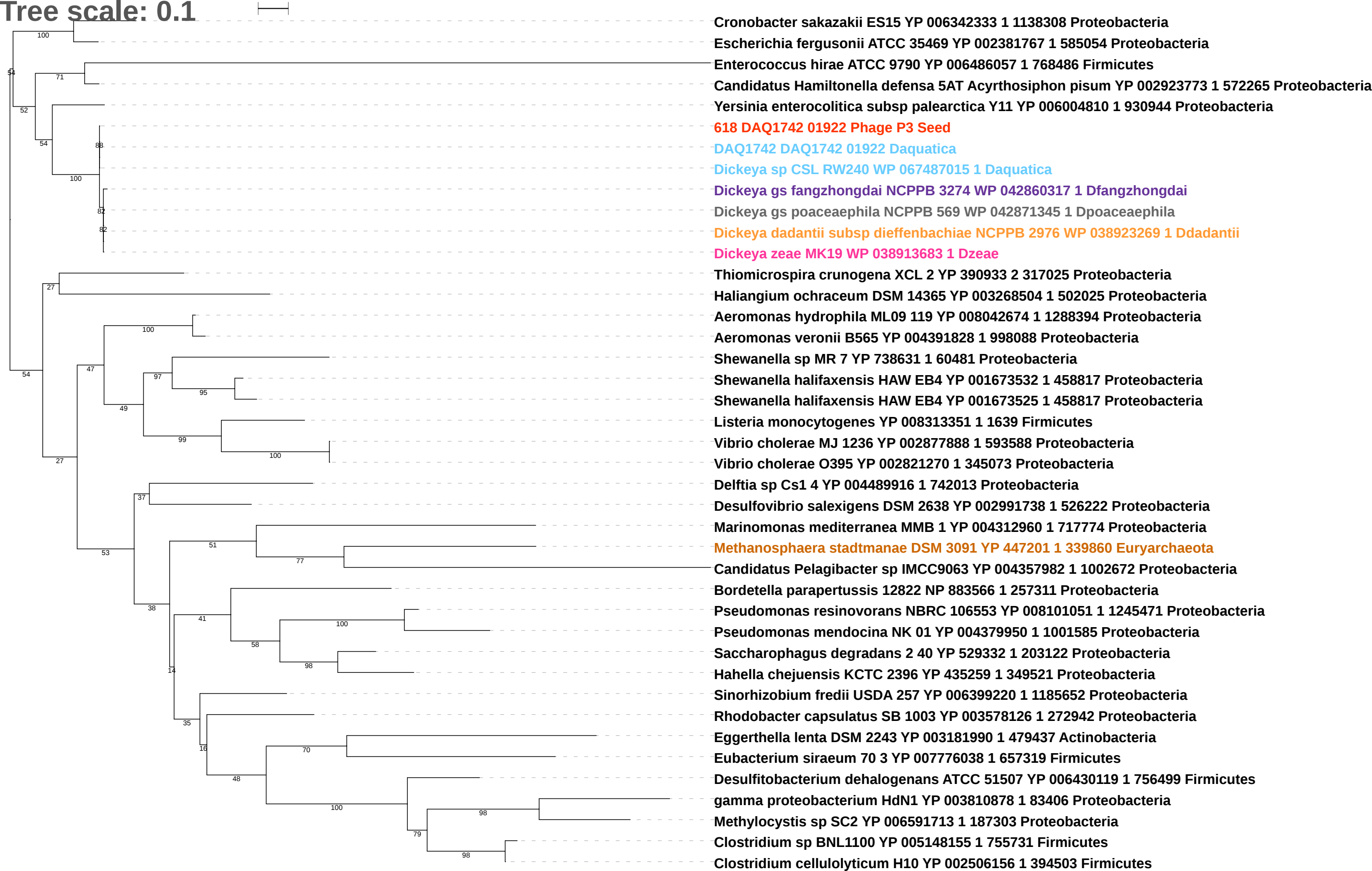
Tree scale: 0.1
Cronobacter sakazakii ES15 YP 006342333 1 1138308 Proteobacteria
Escherichia fergusonii ATCC 35469 YP 002381767 1 585054 Proteobacteria
Enterococcus hirae ATCC 9790 YP 006486057 1 768486 Firmicutes
Candidatus Hamiltonella defensa 5AT Acyrthosiphon pisum YP 002923773 1 572265 Proteobacteria
Yersinia enterocolitica subsp palearctica Y11 YP 006004810 1 930944 Proteobacteria
618 DAQ1742 01922 Phage P3 Seed
DAQ1742 DAQ1742 01922 Daquatica
Dickeya sp CSL RW240 WP 067487015 1 Daquatica
Dickeya gs fangzhongdai NCPPB 3274 WP 042860317 1 Dfangzhongdai
Dickeya gs poaceaephila NCPPB 569 WP 042871345 1 Dpoaceaephila
Dickeya dadantii subsp dieffenbachiae NCPPB 2976 WP 038923269 1 Ddadantii
Dickeya zeae MK19 WP 038913683 1 Dzeae
Thiomicrospira crunogena XCL 2 YP 390933 2 317025 Proteobacteria
Haliangium ochraceum DSM 14365 YP 003268504 1 502025 Proteobacteria
Aeromonas hydrophila ML09 119 YP 008042674 1 1288394 Proteobacteria
Aeromonas veronii B565 YP 004391828 1 998088 Proteobacteria
Shewanella sp MR 7 YP 738631 1 60481 Proteobacteria
Shewanella halifaxensis HAW EB4 YP 001673532 1 458817 Proteobacteria
Shewanella halifaxensis HAW EB4 YP 001673525 1 458817 Proteobacteria
Listeria monocytogenes YP 008313351 1 1639 Firmicutes
Vibrio cholerae MJ 1236 YP 002877888 1 593588 Proteobacteria
Vibrio cholerae O395 YP 002821270 1 345073 Proteobacteria
Delftia sp Cs1 4 YP 004489916 1 742013 Proteobacteria
Desulfovibrio salexigens DSM 2638 YP 002991738 1 526222 Proteobacteria
Marinomonas mediterranea MMB 1 YP 004312960 1 717774 Proteobacteria
Methanosphaera stadtmanae DSM 3091 YP 447201 1 339860 Euryarchaeota
Candidatus Pelagibacter sp IMCC9063 YP 004357982 1 1002672 Proteobacteria
Bordetella parapertussis 12822 NP 883566 1 257311 Proteobacteria
Pseudomonas resinovorans NBRC 106553 YP 008101051 1 1245471 Proteobacteria
Pseudomonas mendocina NK 01 YP 004379950 1 1001585 Proteobacteria
Saccharophagus degradans 2 40 YP 529332 1 203122 Proteobacteria
Hahella chejuensis KCTC 2396 YP 435259 1 349521 Proteobacteria
Sinorhizobium fredii USDA 257 YP 006399220 1 1185652 Proteobacteria
Rhodobacter capsulatus SB 1003 YP 003578126 1 272942 Proteobacteria
Eggerthella lenta DSM 2243 YP 003181990 1 479437 Actinobacteria
Eubacterium siraeum 70 3 YP 007776038 1 657319 Firmicutes
Desulfitobacterium dehalogenans ATCC 51507 YP 006430119 1 756499 Firmicutes
gamma proteobacterium HdN1 YP 003810878 1 83406 Proteobacteria
Methylocystis sp SC2 YP 006591713 1 187303 Proteobacteria
Clostridium sp BNL1100 YP 005148155 1 755731 Firmicutes
Clostridium cellulolyticum H10 YP 002506156 1 394503 Firmicutes

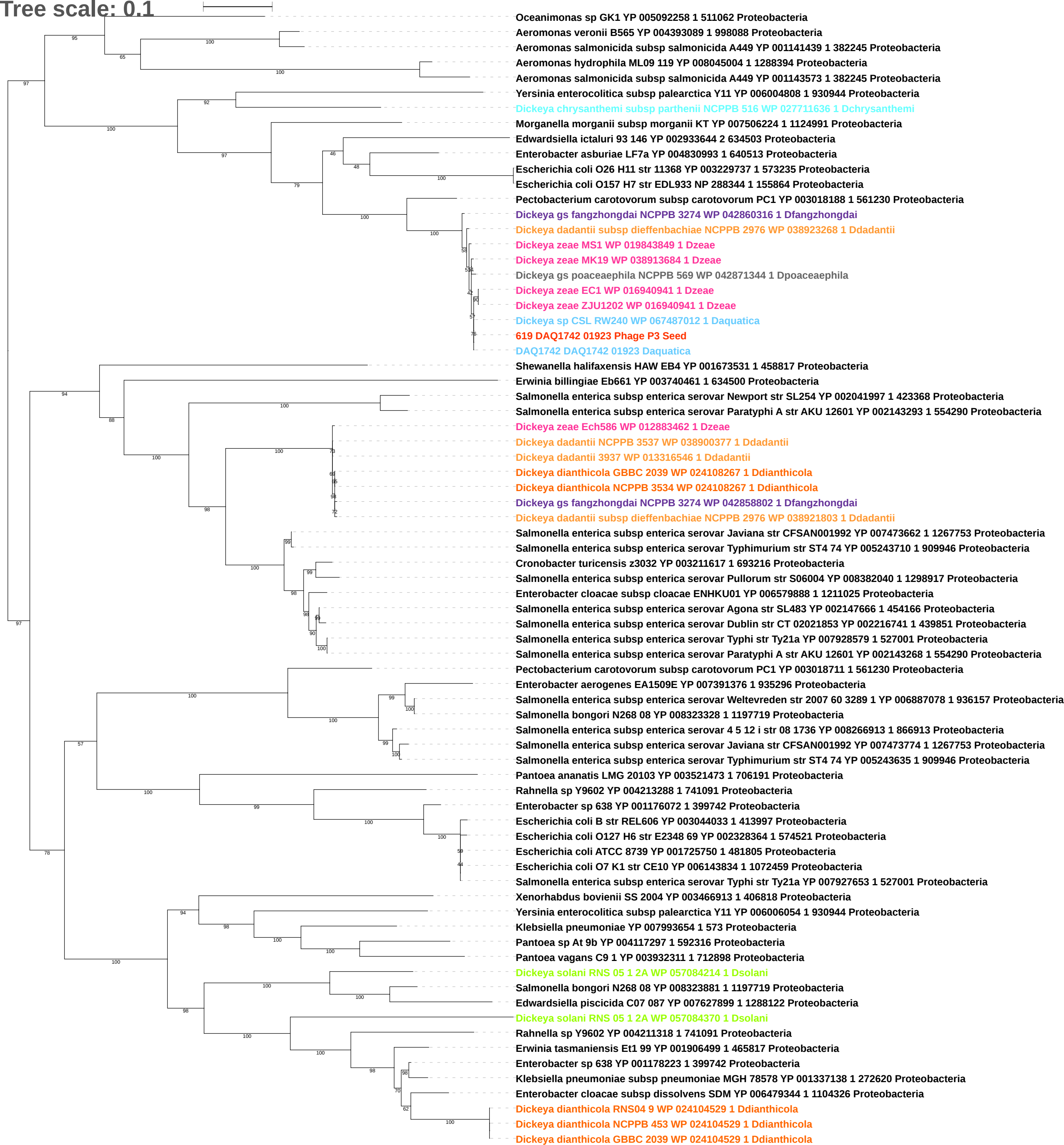
Tree scale: 0.1
Oceanimonas sp GK1 YP 005092258 1 511062 Proteobacteria
Aeromonas veronii B565 YP 004393089 1 998088 Proteobacteria
Aeromonas salmonicida subsp salmonicida A449 YP 001141439 1 382245 Proteobacteria
Aeromonas hydrophila ML09 119 YP 008045004 1 1288394 Proteobacteria
Aeromonas salmonicida subsp salmonicida A449 YP 001143573 1 382245 Proteobacteria
Yersinia enterocolitica subsp palearctica Y11 YP 006004808 1 930944 Proteobacteria
Dickeya chrysanthemi subsp parthenii NCPPB 516 WP 027711636 1 Dchrysanthemi
Morganella morganii subsp morganii KT YP 007506224 1 1124991 Proteobacteria
Edwardsiella ictaluri 93 146 YP 002933644 2 634503 Proteobacteria
Enterobacter asburiae LF7a YP 004830993 1 640513 Proteobacteria
Escherichia coli O26 H11 str 11368 YP 003229737 1 573235 Proteobacteria
Escherichia coli O157 H7 str EDL933 NP 288344 1 155864 Proteobacteria
Pectobacterium carotovorum subsp carotovorum PC1 YP 003018188 1 561230 Proteobacteria
Dickeya gs fangzhongdai NCPPB 3274 WP 042860316 1 Dfangzhongdai
Dickeya dadantii subsp dieffenbachiae NCPPB 2976 WP 038923268 1 Ddadantii
Dickeya zeae MS1 WP 019843849 1 Dzeae
Dickeya zeae MK19 WP 038913684 1 Dzeae
Dickeya gs poaceaephila NCPPB 569 WP 042871344 1 Dpoaceaephila
Dickeya zeae EC1 WP 016940941 1 Dzeae
Dickeya zeae ZJU1202 WP 016940941 1 Dzeae
Dickeya sp CSL RW240 WP 067487012 1 Daquatica
619 DAQ1742 01923 Phage P3 Seed
DAQ1742 DAQ1742 01923 Daquatica
Shewanella halifaxensis HAW EB4 YP 001673531 1 458817 Proteobacteria
Erwinia billingiae Eb661 YP 003740461 1 634500 Proteobacteria
Salmonella enterica subsp enterica serovar Newport str SL254 YP 002041997 1 423368 Proteobacteria
Salmonella enterica subsp enterica serovar Paratyphi A str AKU 12601 YP 002143293 1 554290 Proteobacteria
Dickeya zeae Ech586 WP 012883462 1 Dzeae
Dickeya dadantii NCPPB 3537 WP 038900377 1 Ddadantii
Dickeya dadantii 3937 WP 013316546 1 Ddadantii
Dickeya dianthicola GBBC 2039 WP 024108267 1 Ddianthicola
Dickeya dianthicola NCPPB 3534 WP 024108267 1 Ddianthicola
Dickeya gs fangzhongdai NCPPB 3274 WP 042858802 1 Dfangzhongdai
Dickeya dadantii subsp dieffenbachiae NCPPB 2976 WP 038921803 1 Ddadantii
Salmonella enterica subsp enterica serovar Javiana str CFSAN001992 YP 007473662 1 1267753 Proteobacteria
Salmonella enterica subsp enterica serovar Typhimurium str ST4 74 YP 005243710 1 909946 Proteobacteria
Cronobacter turicensis z3032 YP 003211617 1 693216 Proteobacteria
Salmonella enterica subsp enterica serovar Pullorum str S06004 YP 008382040 1 1298917 Proteobacteria
Enterobacter cloacae subsp cloacae ENHKU01 YP 006579888 1 1211025 Proteobacteria
Salmonella enterica subsp enterica serovar Agona str SL483 YP 002147666 1 454166 Proteobacteria
Salmonella enterica subsp enterica serovar Dublin str CT 02021853 YP 002216741 1 439851 Proteobacteria
Salmonella enterica subsp enterica serovar Typhi str Ty21a YP 007928579 1 527001 Proteobacteria
Salmonella enterica subsp enterica serovar Paratyphi A str AKU 12601 YP 002143268 1 554290 Proteobacteria
Pectobacterium carotovorum subsp carotovorum PC1 YP 003018711 1 561230 Proteobacteria
Enterobacter aerogenes EA1509E YP 007391376 1 935296 Proteobacteria
Salmonella enterica subsp enterica serovar Weltevreden str 2007 60 3289 1 YP 006887078 1 936157 Proteobacteria
Salmonella bongori N268 08 YP 008323328 1 1197719 Proteobacteria
Salmonella enterica subsp enterica serovar 4 5 12 i str 08 1736 YP 008266913 1 866913 Proteobacteria
Salmonella enterica subsp enterica serovar Javiana str CFSAN001992 YP 007473774 1 1267753 Proteobacteria
Salmonella enterica subsp enterica serovar Typhimurium str ST4 74 YP 005243635 1 909946 Proteobacteria
Pantoea ananatis LMG 20103 YP 003521473 1 706191 Proteobacteria
Rahnella sp Y9602 YP 004213288 1 741091 Proteobacteria
Enterobacter sp 638 YP 001176072 1 399742 Proteobacteria
Escherichia coli B str REL606 YP 003044033 1 413997 Proteobacteria
Escherichia coli O127 H6 str E2348 69 YP 002328364 1 574521 Proteobacteria
Escherichia coli ATCC 8739 YP 001725750 1 481805 Proteobacteria
Escherichia coli O7 K1 str CE10 YP 006143834 1 1072459 Proteobacteria
Salmonella enterica subsp enterica serovar Typhi str Ty21a YP 007927653 1 527001 Proteobacteria
Xenorhabdus bovienii SS 2004 YP 003466913 1 406818 Proteobacteria
Yersinia enterocolitica subsp palearctica Y11 YP 006006054 1 930944 Proteobacteria
Klebsiella pneumoniae YP 007993654 1 573 Proteobacteria
Pantoea sp At 9b YP 004117297 1 592316 Proteobacteria
Pantoea vagans C9 1 YP 003932311 1 712898 Proteobacteria
Dickeya solani RNS 05 1 2A WP 057084214 1 Dsolani
Salmonella bongori N268 08 YP 008323881 1 1197719 Proteobacteria
Edwardsiella piscicida C07 087 YP 007627899 1 1288122 Proteobacteria
Dickeya solani RNS 05 1 2A WP 057084370 1 Dsolani
Rahnella sp Y9602 YP 004211318 1 741091 Proteobacteria
Erwinia tasmaniensis Et1 99 YP 001906499 1 465817 Proteobacteria
Enterobacter sp 638 YP 001178223 1 399742 Proteobacteria
Klebsiella pneumoniae subsp pneumoniae MGH 78578 YP 001337138 1 272620 Proteobacteria
Enterobacter cloacae subsp dissolvens SDM YP 006479344 1 1104326 Proteobacteria
Dickeya dianthicola RNS04 9 WP 024104529 1 Ddianthicola
Dickeya dianthicola NCPPB 453 WP 024104529 1 Ddianthicola
Dickeya dianthicola GBBC 2039 WP 024104529 1 Ddianthicola

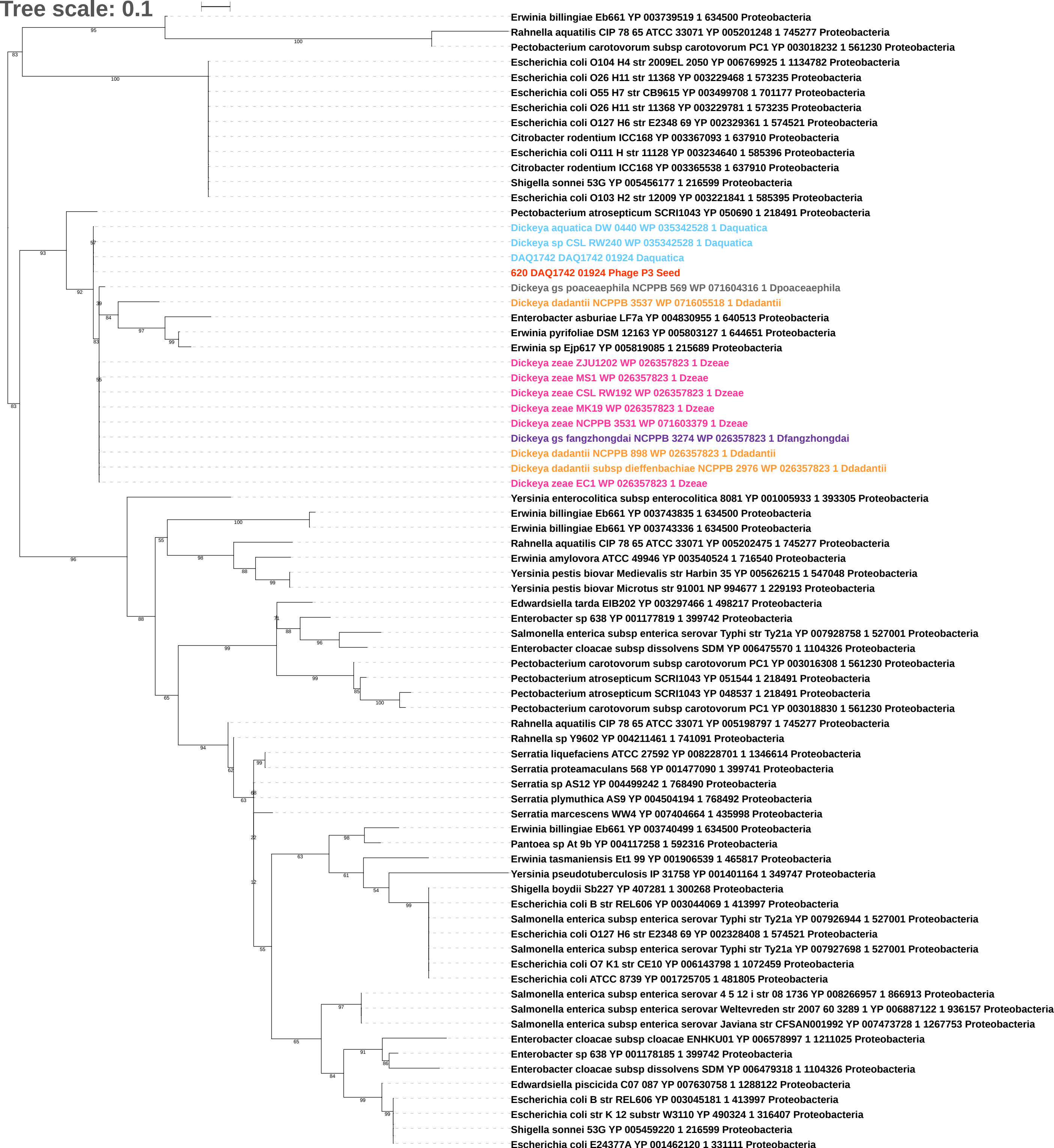
Tree scale: 0.1
Erwinia billingiae Eb661 YP 003739519 1 634500 Proteobacteria
Rahnella aquatilis CIP 78 65 ATCC 33071 YP 005201248 1 745277 Proteobacteria
Pectobacterium carotovorum subsp carotovorum PC1 YP 003018232 1 561230 Proteobacteria
Escherichia coli O104 H4 str 2009EL 2050 YP 006769925 1 1134782 Proteobacteria
Escherichia coli O26 H11 str 11368 YP 003229468 1 573235 Proteobacteria
Escherichia coli O55 H7 str CB9615 YP 003499708 1 701177 Proteobacteria
Escherichia coli O26 H11 str 11368 YP 003229781 1 573235 Proteobacteria
Escherichia coli O127 H6 str E2348 69 YP 002329361 1 574521 Proteobacteria
Citrobacter rodentium ICC168 YP 003367093 1 637910 Proteobacteria
Escherichia coli O111 H str 11128 YP 003234640 1 585396 Proteobacteria
Citrobacter rodentium ICC168 YP 003365538 1 637910 Proteobacteria
Shigella sonnei 53G YP 005456177 1 216599 Proteobacteria
Escherichia coli O103 H2 str 12009 YP 003221841 1 585395 Proteobacteria
Pectobacterium atrosepticum SCRI1043 YP 050690 1 218491 Proteobacteria
Dickeya aquatica DW 0440 WP 035342528 1 Daquatica
Dickeya sp CSL RW240 WP 035342528 1 Daquatica
DAQ1742 DAQ1742 01924 Daquatica
620 DAQ1742 01924 Phage P3 Seed
Dickeya gs poaceaephila NCPPB 569 WP 071604316 1 Dpoaceaephila
Dickeya dadantii NCPPB 3537 WP 071605518 1 Ddadantii
Enterobacter asburiae LF7a YP 004830955 1 640513 Proteobacteria
Erwinia pyrifoliae DSM 12163 YP 005803127 1 644651 Proteobacteria
Erwinia sp Ejp617 YP 005819085 1 215689 Proteobacteria
Dickeya zeae ZJU1202 WP 026357823 1 Dzeae
Dickeya zeae MS1 WP 026357823 1 Dzeae
Dickeya zeae CSL RW192 WP 026357823 1 Dzeae
Dickeya zeae MK19 WP 026357823 1 Dzeae
Dickeya zeae NCPPB 3531 WP 071603379 1 Dzeae
Dickeya gs fangzhongdai NCPPB 3274 WP 026357823 1 Dfangzhongdai
Dickeya dadantii NCPPB 898 WP 026357823 1 Ddadantii
Dickeya dadantii subsp dieffenbachiae NCPPB 2976 WP 026357823 1 Ddadantii
Dickeya zeae EC1 WP 026357823 1 Dzeae
Yersinia enterocolitica subsp enterocolitica 8081 YP 001005933 1 393305 Proteobacteria
Erwinia billingiae Eb661 YP 003743835 1 634500 Proteobacteria
Erwinia billingiae Eb661 YP 003743336 1 634500 Proteobacteria
Rahnella aquatilis CIP 78 65 ATCC 33071 YP 005202475 1 745277 Proteobacteria
Erwinia amylovora ATCC 49946 YP 003540524 1 716540 Proteobacteria
Yersinia pestis biovar Medievalis str Harbin 35 YP 005626215 1 547048 Proteobacteria
Yersinia pestis biovar Microtus str 91001 NP 994677 1 229193 Proteobacteria
Edwardsiella tarda EIB202 YP 003297466 1 498217 Proteobacteria
Enterobacter sp 638 YP 001177819 1 399742 Proteobacteria
Salmonella enterica subsp enterica serovar Typhi str Ty21a YP 007928758 1 527001 Proteobacteria
Enterobacter cloacae subsp dissolvens SDM YP 006475570 1 1104326 Proteobacteria
Pectobacterium carotovorum subsp carotovorum PC1 YP 003016308 1 561230 Proteobacteria
Pectobacterium atrosepticum SCRI1043 YP 051544 1 218491 Proteobacteria
Pectobacterium atrosepticum SCRI1043 YP 048537 1 218491 Proteobacteria
Pectobacterium carotovorum subsp carotovorum PC1 YP 003018830 1 561230 Proteobacteria
Rahnella aquatilis CIP 78 65 ATCC 33071 YP 005198797 1 745277 Proteobacteria
Rahnella sp Y9602 YP 004211461 1 741091 Proteobacteria
Serratia liquefaciens ATCC 27592 YP 008228701 1 1346614 Proteobacteria
Serratia proteamaculans 568 YP 001477090 1 399741 Proteobacteria
Serratia sp AS12 YP 004499242 1 768490 Proteobacteria
Serratia plymuthica AS9 YP 004504194 1 768492 Proteobacteria
Serratia marcescens WW4 YP 007404664 1 435998 Proteobacteria
Erwinia billingiae Eb661 YP 003740499 1 634500 Proteobacteria
Pantoea sp At 9b YP 004117258 1 592316 Proteobacteria
Erwinia tasmaniensis Et1 99 YP 001906539 1 465817 Proteobacteria
Yersinia pseudotuberculosis IP 31758 YP 001401164 1 349747 Proteobacteria
Shigella boydii Sb227 YP 407281 1 300268 Proteobacteria
Escherichia coli B str REL606 YP 003044069 1 413997 Proteobacteria
Salmonella enterica subsp enterica serovar Typhi str Ty21a YP 007926944 1 527001 Proteobacteria
Escherichia coli O127 H6 str E2348 69 YP 002328408 1 574521 Proteobacteria
Salmonella enterica subsp enterica serovar Typhi str Ty21a YP 007927698 1 527001 Proteobacteria
Escherichia coli O7 K1 str CE10 YP 006143798 1 1072459 Proteobacteria
Escherichia coli ATCC 8739 YP 001725705 1 481805 Proteobacteria
Salmonella enterica subsp enterica serovar 4 5 12 i str 08 1736 YP 008266957 1 866913 Proteobacteria
Salmonella enterica subsp enterica serovar Weltevreden str 2007 60 3289 1 YP 006887122 1 936157 Proteobacteria
Salmonella enterica subsp enterica serovar Javiana str CFSAN001992 YP 007473728 1 1267753 Proteobacteria
Enterobacter cloacae subsp cloacae ENHKU01 YP 006578997 1 1211025 Proteobacteria
Enterobacter sp 638 YP 001178185 1 399742 Proteobacteria
Enterobacter cloacae subsp dissolvens SDM YP 006479318 1 1104326 Proteobacteria
Edwardsiella piscicida C07 087 YP 007630758 1 1288122 Proteobacteria
Escherichia coli B str REL606 YP 003045181 1 413997 Proteobacteria
Escherichia coli str K 12 substr W3110 YP 490324 1 316407 Proteobacteria
Shigella sonnei 53G YP 005459220 1 216599 Proteobacteria
Escherichia coli E24377A YP 001462120 1 331111 Proteobacteria

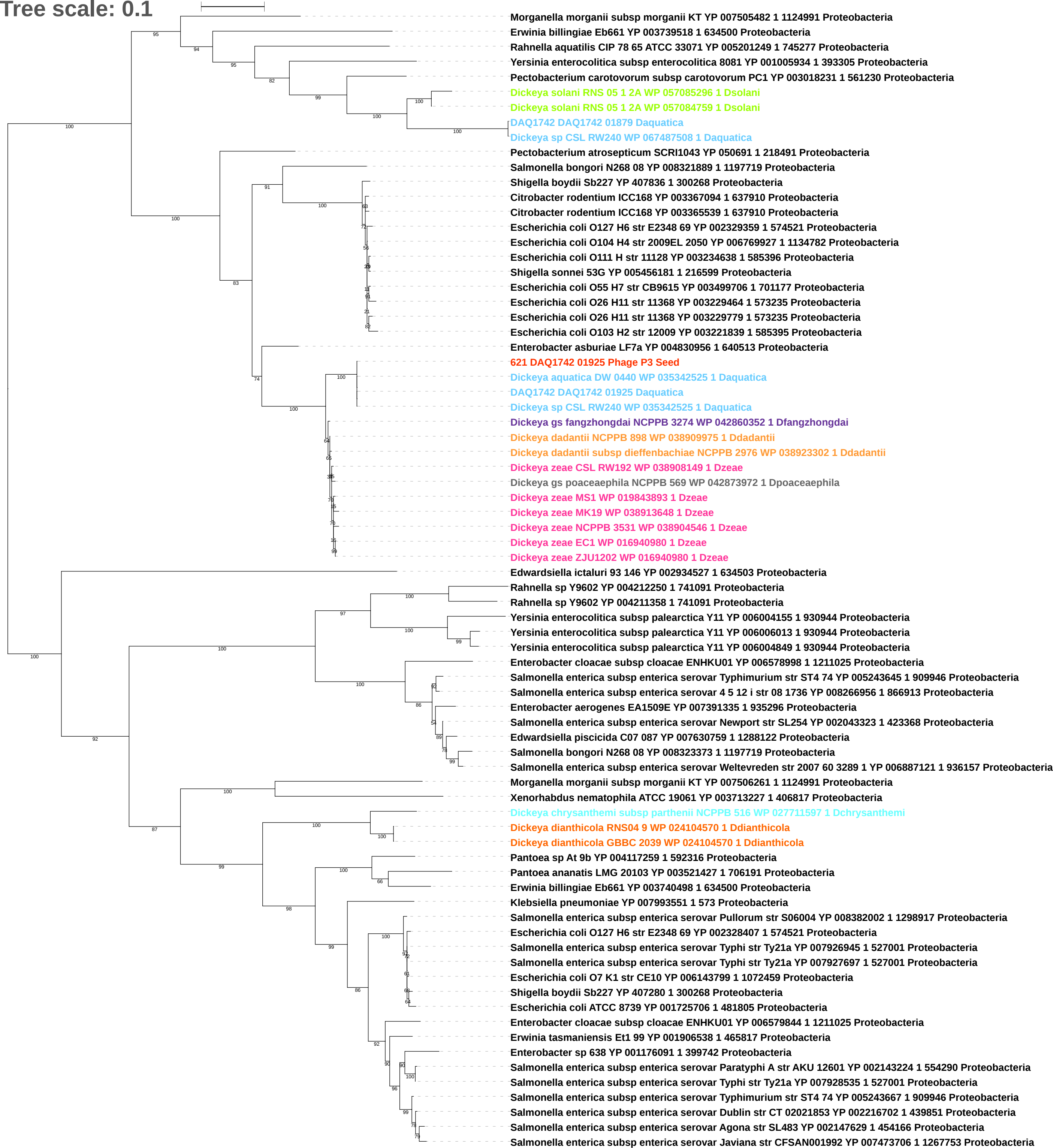

Tree scale: 0.1
Morganella morganii subsp morganii KT YP 007505482 1 1124991 Proteobacteria
Erwinia billingiae Eb661 YP 003739518 1 634500 Proteobacteria
Rahnella aquatilis CIP 78 65 ATCC 33071 YP 005201249 1 745277 Proteobacteria
Yersinia enterocolitica subsp enterocolitica 8081 YP 001005934 1 393305 Proteobacteria
Pectobacterium carotovorum subsp carotovorum PC1 YP 003018231 1 561230 Proteobacteria
Dickeya solani RNS 05 1 2A WP 057085296 1 Dsolani
Dickeya solani RNS 05 1 2A WP 057084759 1 Dsolani
DAQ1742 DAQ1742 01879 Daquatica
Dickeya sp CSL RW240 WP 067487508 1 Daquatica
Pectobacterium atrosepticum SCRI1043 YP 050691 1 218491 Proteobacteria
Salmonella bongori N268 08 YP 008321889 1 1197719 Proteobacteria
Shigella boydii Sb227 YP 407836 1 300268 Proteobacteria
Citrobacter rodentium ICC168 YP 003367094 1 637910 Proteobacteria
Citrobacter rodentium ICC168 YP 003365539 1 637910 Proteobacteria
Escherichia coli O127 H6 str E2348 69 YP 002329359 1 574521 Proteobacteria
Escherichia coli O104 H4 str 2009EL 2050 YP 006769927 1 1134782 Proteobacteria
Escherichia coli O111 H str 11128 YP 003234638 1 585396 Proteobacteria
Shigella sonnei 53G YP 005456181 1 216599 Proteobacteria
Escherichia coli O55 H7 str CB9615 YP 003499706 1 701177 Proteobacteria
Escherichia coli O26 H11 str 11368 YP 003229464 1 573235 Proteobacteria
Escherichia coli O26 H11 str 11368 YP 003229779 1 573235 Proteobacteria
Escherichia coli O103 H2 str 12009 YP 003221839 1 585395 Proteobacteria
Enterobacter asburiae LF7a YP 004830956 1 640513 Proteobacteria
621 DAQ1742 01925 Phage P3 Seed
Dickeya aquatica DW 0440 WP 035342525 1 Daquatica
DAQ1742 DAQ1742 01925 Daquatica
Dickeya sp CSL RW240 WP 035342525 1 Daquatica
Dickeya gs fangzhongdai NCPPB 3274 WP 042860352 1 Dfangzhongdai
Dickeya dadantii NCPPB 898 WP 038909975 1 Ddadantii
Dickeya dadantii subsp dieffenbachiae NCPPB 2976 WP 038923302 1 Ddadantii
Dickeya zeae CSL RW192 WP 038908149 1 Dzeae
Dickeya gs poaceaephila NCPPB 569 WP 042873972 1 Dpoaceaephila
Dickeya zeae MS1 WP 019843893 1 Dzeae
Dickeya zeae MK19 WP 038913648 1 Dzeae
Dickeya zeae NCPPB 3531 WP 038904546 1 Dzeae
Dickeya zeae EC1 WP 016940980 1 Dzeae
Dickeya zeae ZJU1202 WP 016940980 1 Dzeae
Edwardsiella ictaluri 93 146 YP 002934527 1 634503 Proteobacteria
Rahnella sp Y9602 YP 004212250 1 741091 Proteobacteria
Rahnella sp Y9602 YP 004211358 1 741091 Proteobacteria
Yersinia enterocolitica subsp palearctica Y11 YP 006004155 1 930944 Proteobacteria
Yersinia enterocolitica subsp palearctica Y11 YP 006006013 1 930944 Proteobacteria
Yersinia enterocolitica subsp palearctica Y11 YP 006004849 1 930944 Proteobacteria
Enterobacter cloacae subsp cloacae ENHKU01 YP 006578998 1 1211025 Proteobacteria
Salmonella enterica subsp enterica serovar Typhimurium str ST4 74 YP 005243645 1 909946 Proteobacteria
Salmonella enterica subsp enterica serovar 4 5 12 i str 08 1736 YP 008266956 1 866913 Proteobacteria
Enterobacter aerogenes EA1509E YP 007391335 1 935296 Proteobacteria
Salmonella enterica subsp enterica serovar Newport str SL254 YP 002043323 1 423368 Proteobacteria
Edwardsiella piscicida C07 087 YP 007630759 1 1288122 Proteobacteria
Salmonella bongori N268 08 YP 008323373 1 1197719 Proteobacteria
Salmonella enterica subsp enterica serovar Weltevreden str 2007 60 3289 1 YP 006887121 1 936157 Proteobacteria
Morganella morganii subsp morganii KT YP 007506261 1 1124991 Proteobacteria
Xenorhabdus nematophila ATCC 19061 YP 003713227 1 406817 Proteobacteria
Dickeya chrysanthemi subsp parthenii NCPPB 516 WP 027711597 1 Dchrysanthemi
Dickeya dianthicola RNS04 9 WP 024104570 1 Ddianthicola
Dickeya dianthicola GBBC 2039 WP 024104570 1 Ddianthicola
Pantoea sp At 9b YP 004117259 1 592316 Proteobacteria
Pantoea ananatis LMG 20103 YP 003521427 1 706191 Proteobacteria
Erwinia billingiae Eb661 YP 003740498 1 634500 Proteobacteria
Klebsiella pneumoniae YP 007993551 1 573 Proteobacteria
Salmonella enterica subsp enterica serovar Pullorum str S06004 YP 008382002 1 1298917 Proteobacteria
Escherichia coli O127 H6 str E2348 69 YP 002328407 1 574521 Proteobacteria
Salmonella enterica subsp enterica serovar Typhi str Ty21a YP 007926945 1 527001 Proteobacteria
Salmonella enterica subsp enterica serovar Typhi str Ty21a YP 007927697 1 527001 Proteobacteria
Escherichia coli O7 K1 str CE10 YP 006143799 1 1072459 Proteobacteria
Shigella boydii Sb227 YP 407280 1 300268 Proteobacteria
Escherichia coli ATCC 8739 YP 001725706 1 481805 Proteobacteria
Enterobacter cloacae subsp cloacae ENHKU01 YP 006579844 1 1211025 Proteobacteria
Erwinia tasmaniensis Et1 99 YP 001906538 1 465817 Proteobacteria
Enterobacter sp 638 YP 001176091 1 399742 Proteobacteria
Salmonella enterica subsp enterica serovar Paratyphi A str AKU 12601 YP 002143224 1 554290 Proteobacteria
Salmonella enterica subsp enterica serovar Typhi str Ty21a YP 007928535 1 527001 Proteobacteria
Salmonella enterica subsp enterica serovar Typhimurium str ST4 74 YP 005243667 1 909946 Proteobacteria
Salmonella enterica subsp enterica serovar Dublin str CT 02021853 YP 002216702 1 439851 Proteobacteria
Salmonella enterica subsp enterica serovar Agona str SL483 YP 002147629 1 454166 Proteobacteria
Salmonella enterica subsp enterica serovar Javiana str CFSAN001992 YP 007473706 1 1267753 Proteobacteria

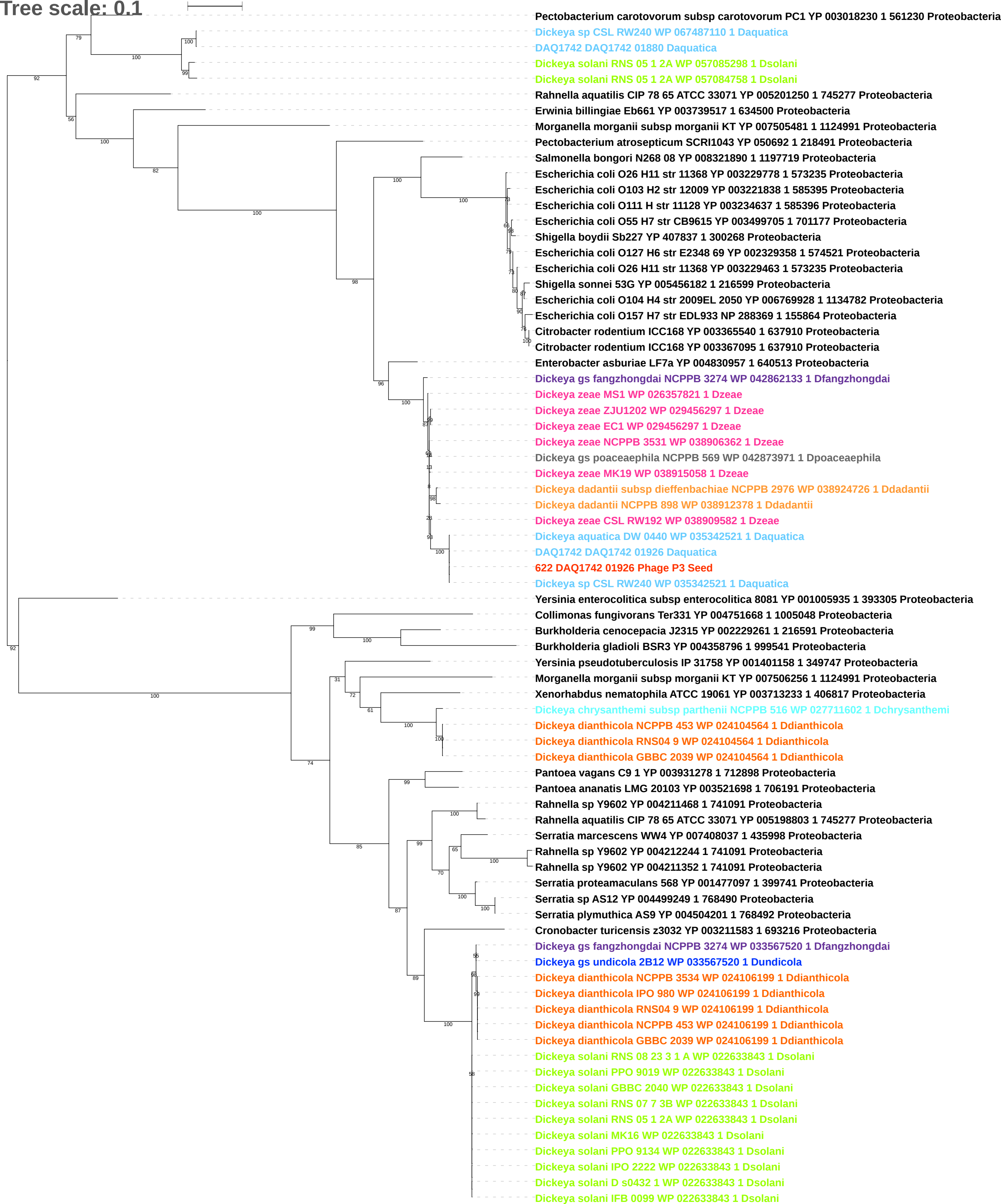
Tree scale: 0.1
Pectobacterium carotovorum subsp carotovorum PC1 YP 003018230 1 561230 Proteobacteria
Dickeya sp CSL RW240 WP 067487110 1 Daquatica
DAQ1742 DAQ1742 01880 Daquatica
Dickeya solani RNS 05 1 2A WP 057085298 1 Dsolani
Dickeya solani RNS 05 1 2A WP 057084758 1 Dsolani
Rahnella aquatilis CIP 78 65 ATCC 33071 YP 005201250 1 745277 Proteobacteria
Erwinia billingiae Eb661 YP 003739517 1 634500 Proteobacteria
Morganella morganii subsp morganii KT YP 007505481 1 1124991 Proteobacteria
Pectobacterium atrosepticum SCRI1043 YP 050692 1 218491 Proteobacteria
Salmonella bongori N268 08 YP 008321890 1 1197719 Proteobacteria
Escherichia coli O26 H11 str 11368 YP 003229778 1 573235 Proteobacteria
Escherichia coli O103 H2 str 12009 YP 003221838 1 585395 Proteobacteria
Escherichia coli O111 H str 11128 YP 003234637 1 585396 Proteobacteria
Escherichia coli O55 H7 str CB9615 YP 003499705 1 701177 Proteobacteria
Shigella boydii Sb227 YP 407837 1 300268 Proteobacteria
Escherichia coli O127 H6 str E2348 69 YP 002329358 1 574521 Proteobacteria
Escherichia coli O26 H11 str 11368 YP 003229463 1 573235 Proteobacteria
Shigella sonnei 53G YP 005456182 1 216599 Proteobacteria
Escherichia coli O104 H4 str 2009EL 2050 YP 006769928 1 1134782 Proteobacteria
Escherichia coli O157 H7 str EDL933 NP 288369 1 155864 Proteobacteria
Citrobacter rodentium ICC168 YP 003365540 1 637910 Proteobacteria
Citrobacter rodentium ICC168 YP 003367095 1 637910 Proteobacteria
Enterobacter asburiae LF7a YP 004830957 1 640513 Proteobacteria
Dickeya gs fangzhongdai NCPPB 3274 WP 042862133 1 Dfangzhongdai
Dickeya zeae MS1 WP 026357821 1 Dzeae
Dickeya zeae ZJU1202 WP 029456297 1 Dzeae
Dickeya zeae EC1 WP 029456297 1 Dzeae
Dickeya zeae NCPPB 3531 WP 038906362 1 Dzeae
Dickeya gs poaceaephila NCPPB 569 WP 042873971 1 Dpoaceaephila
Dickeya zeae MK19 WP 038915058 1 Dzeae
Dickeya dadantii subsp dieffenbachiae NCPPB 2976 WP 038924726 1 Ddadantii
Dickeya dadantii NCPPB 898 WP 038912378 1 Ddadantii
Dickeya zeae CSL RW192 WP 038909582 1 Dzeae
Dickeya aquatica DW 0440 WP 035342521 1 Daquatica
DAQ1742 DAQ1742 01926 Daquatica
622 DAQ1742 01926 Phage P3 Seed
Dickeya sp CSL RW240 WP 035342521 1 Daquatica
Yersinia enterocolitica subsp enterocolitica 8081 YP 001005935 1 393305 Proteobacteria
Collimonas fungivorans Ter331 YP 004751668 1 1005048 Proteobacteria
Burkholderia cenocepacia J2315 YP 002229261 1 216591 Proteobacteria
Burkholderia gladioli BSR3 YP 004358796 1 999541 Proteobacteria
Yersinia pseudotuberculosis IP 31758 YP 001401158 1 349747 Proteobacteria
Morganella morganii subsp morganii KT YP 007506256 1 1124991 Proteobacteria
Xenorhabdus nematophila ATCC 19061 YP 003713233 1 406817 Proteobacteria
Dickeya chrysanthemi subsp parthenii NCPPB 516 WP 027711602 1 Dchrysanthemi
Dickeya dianthicola NCPPB 453 WP 024104564 1 Ddianthicola
Dickeya dianthicola RNS04 9 WP 024104564 1 Ddianthicola
Dickeya dianthicola GBBC 2039 WP 024104564 1 Ddianthicola
Pantoea vagans C9 1 YP 003931278 1 712898 Proteobacteria
Pantoea ananatis LMG 20103 YP 003521698 1 706191 Proteobacteria
Rahnella sp Y9602 YP 004211468 1 741091 Proteobacteria
Rahnella aquatilis CIP 78 65 ATCC 33071 YP 005198803 1 745277 Proteobacteria
Serratia marcescens WW4 YP 007408037 1 435998 Proteobacteria
Rahnella sp Y9602 YP 004212244 1 741091 Proteobacteria
Rahnella sp Y9602 YP 004211352 1 741091 Proteobacteria
Serratia proteamaculans 568 YP 001477097 1 399741 Proteobacteria
Serratia sp AS12 YP 004499249 1 768490 Proteobacteria
Serratia plymuthica AS9 YP 004504201 1 768492 Proteobacteria
Cronobacter turicensis z3032 YP 003211583 1 693216 Proteobacteria
Dickeya gs fangzhongdai NCPPB 3274 WP 033567520 1 Dfangzhongdai
Dickeya gs undicola 2B12 WP 033567520 1 Dundicola
Dickeya dianthicola NCPPB 3534 WP 024106199 1 Ddianthicola
Dickeya dianthicola IPO 980 WP 024106199 1 Ddianthicola
Dickeya dianthicola RNS04 9 WP 024106199 1 Ddianthicola
Dickeya dianthicola NCPPB 453 WP 024106199 1 Ddianthicola
Dickeya dianthicola GBBC 2039 WP 024106199 1 Ddianthicola
Dickeya solani RNS 08 23 3 1 A WP 022633843 1 Dsolani
Dickeya solani PPO 9019 WP 022633843 1 Dsolani
Dickeya solani GBBC 2040 WP 022633843 1 Dsolani
Dickeya solani RNS 07 7 3B WP 022633843 1 Dsolani
Dickeya solani RNS 05 1 2A WP 022633843 1 Dsolani
Dickeya solani MK16 WP 022633843 1 Dsolani
Dickeya solani PPO 9134 WP 022633843 1 Dsolani
Dickeya solani IPO 2222 WP 022633843 1 Dsolani
Dickeya solani D s0432 1 WP 022633843 1 Dsolani
Dickeya solani IFB 0099 WP 022633843 1 Dsolani

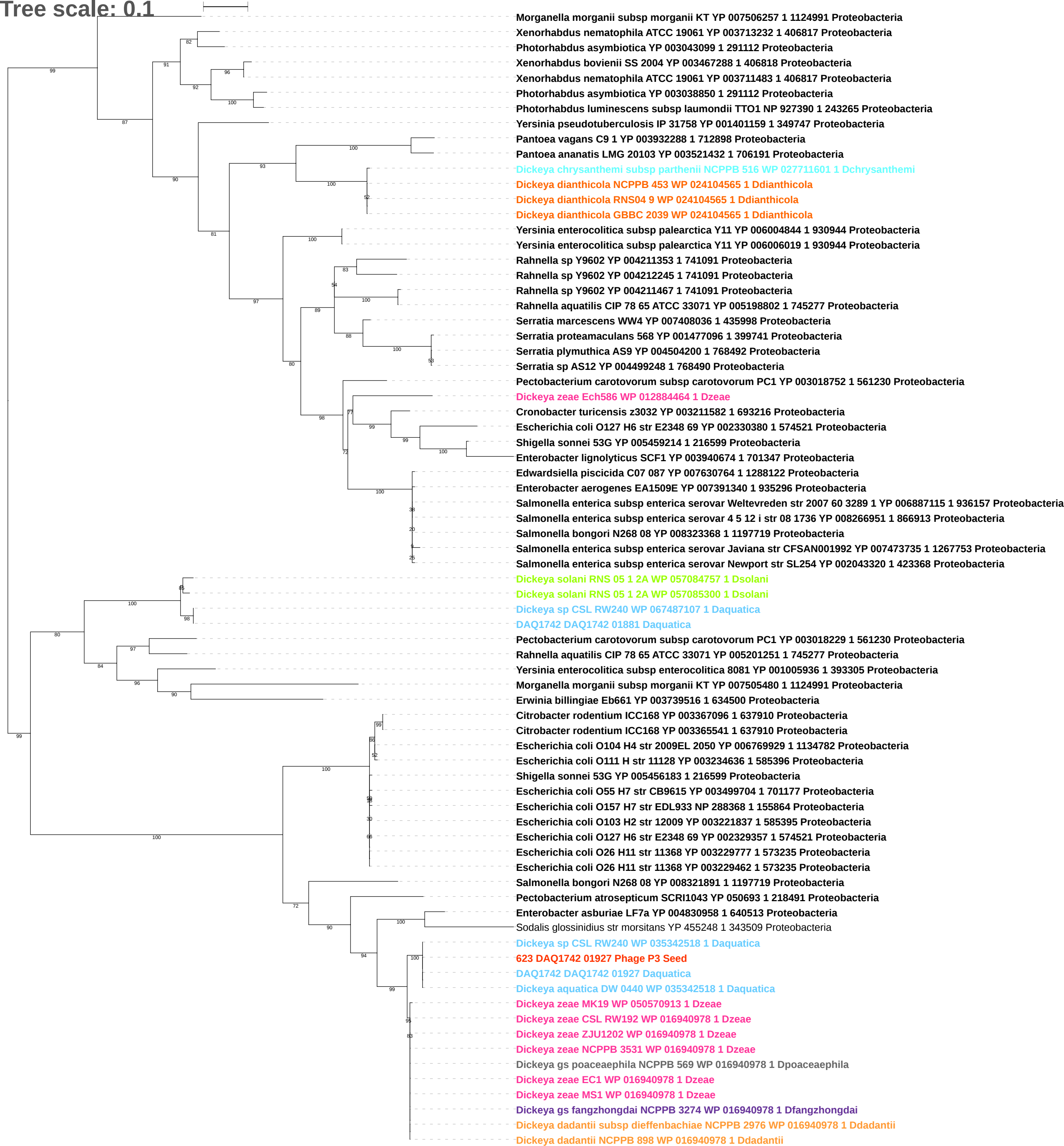
Tree scale: 0.1
Morganella morganii subsp morganii KT YP 007506257 1 1124991 Proteobacteria
Xenorhabdus nematophila ATCC 19061 YP 003713232 1 406817 Proteobacteria
Photorhabdus asymbiotica YP 003043099 1 291112 Proteobacteria
Xenorhabdus bovienii SS 2004 YP 003467288 1 406818 Proteobacteria
Xenorhabdus nematophila ATCC 19061 YP 003711483 1 406817 Proteobacteria
Photorhabdus asymbiotica YP 003038850 1 291112 Proteobacteria
Photorhabdus luminescens subsp laumondii TTO1 NP 927390 1 243265 Proteobacteria
Yersinia pseudotuberculosis IP 31758 YP 001401159 1 349747 Proteobacteria
Pantoea vagans C9 1 YP 003932288 1 712898 Proteobacteria
Pantoea ananatis LMG 20103 YP 003521432 1 706191 Proteobacteria
Dickeya chrysanthemi subsp parthenii NCPPB 516 WP 027711601 1 Dchrysanthemi
Dickeya dianthicola NCPPB 453 WP 024104565 1 Ddianthicola
Dickeya dianthicola RNS04 9 WP 024104565 1 Ddianthicola
Dickeya dianthicola GBBC 2039 WP 024104565 1 Ddianthicola
Yersinia enterocolitica subsp palearctica Y11 YP 006004844 1 930944 Proteobacteria
Yersinia enterocolitica subsp palearctica Y11 YP 006006019 1 930944 Proteobacteria
Rahnella sp Y9602 YP 004211353 1 741091 Proteobacteria
Rahnella sp Y9602 YP 004212245 1 741091 Proteobacteria
Rahnella sp Y9602 YP 004211467 1 741091 Proteobacteria
Rahnella aquatilis CIP 78 65 ATCC 33071 YP 005198802 1 745277 Proteobacteria
Serratia marcescens WW4 YP 007408036 1 435998 Proteobacteria
Serratia proteamaculans 568 YP 001477096 1 399741 Proteobacteria
Serratia plymuthica AS9 YP 004504200 1 768492 Proteobacteria
Serratia sp AS12 YP 004499248 1 768490 Proteobacteria
Pectobacterium carotovorum subsp carotovorum PC1 YP 003018752 1 561230 Proteobacteria
Dickeya zeae Ech586 WP 012884464 1 Dzeae
Cronobacter turicensis z3032 YP 003211582 1 693216 Proteobacteria
Escherichia coli O127 H6 str E2348 69 YP 002330380 1 574521 Proteobacteria
Shigella sonnei 53G YP 005459214 1 216599 Proteobacteria
Enterobacter lignolyticus SCF1 YP 003940674 1 701347 Proteobacteria
Edwardsiella piscicida C07 087 YP 007630764 1 1288122 Proteobacteria
Enterobacter aerogenes EA1509E YP 007391340 1 935296 Proteobacteria
Salmonella enterica subsp enterica serovar Weltevreden str 2007 60 3289 1 YP 006887115 1 936157 Proteobacteria
Salmonella enterica subsp enterica serovar 4 5 12 i str 08 1736 YP 008266951 1 866913 Proteobacteria
Salmonella bongori N268 08 YP 008323368 1 1197719 Proteobacteria
Salmonella enterica subsp enterica serovar Javiana str CFSAN001992 YP 007473735 1 1267753 Proteobacteria
Salmonella enterica subsp enterica serovar Newport str SL254 YP 002043320 1 423368 Proteobacteria
Dickeya solani RNS 05 1 2A WP 057084757 1 Dsolani
Dickeya solani RNS 05 1 2A WP 057085300 1 Dsolani
Dickeya sp CSL RW240 WP 067487107 1 Daquatica
DAQ1742 DAQ1742 01881 Daquatica
Pectobacterium carotovorum subsp carotovorum PC1 YP 003018229 1 561230 Proteobacteria
Rahnella aquatilis CIP 78 65 ATCC 33071 YP 005201251 1 745277 Proteobacteria
Yersinia enterocolitica subsp enterocolitica 8081 YP 001005936 1 393305 Proteobacteria
Morganella morganii subsp morganii KT YP 007505480 1 1124991 Proteobacteria
Erwinia billingiae Eb661 YP 003739516 1 634500 Proteobacteria
Citrobacter rodentium ICC168 YP 003367096 1 637910 Proteobacteria
Citrobacter rodentium ICC168 YP 003365541 1 637910 Proteobacteria
Escherichia coli O104 H4 str 2009EL 2050 YP 006769929 1 1134782 Proteobacteria
Escherichia coli O111 H str 11128 YP 003234636 1 585396 Proteobacteria
Shigella sonnei 53G YP 005456183 1 216599 Proteobacteria
Escherichia coli O55 H7 str CB9615 YP 003499704 1 701177 Proteobacteria
Escherichia coli O157 H7 str EDL933 NP 288368 1 155864 Proteobacteria
Escherichia coli O103 H2 str 12009 YP 003221837 1 585395 Proteobacteria
Escherichia coli O127 H6 str E2348 69 YP 002329357 1 574521 Proteobacteria
Escherichia coli O26 H11 str 11368 YP 003229777 1 573235 Proteobacteria
Escherichia coli O26 H11 str 11368 YP 003229462 1 573235 Proteobacteria
Salmonella bongori N268 08 YP 008321891 1 1197719 Proteobacteria
Pectobacterium atrosepticum SCRI1043 YP 050693 1 218491 Proteobacteria
Enterobacter asburiae LF7a YP 004830958 1 640513 Proteobacteria
Sodalis glossinidius str morsitans YP 455248 1 343509 Proteobacteria
Dickeya sp CSL RW240 WP 035342518 1 Daquatica
623 DAQ1742 01927 Phage P3 Seed
DAQ1742 DAQ1742 01927 Daquatica
Dickeya aquatica DW 0440 WP 035342518 1 Daquatica
Dickeya zeae MK19 WP 050570913 1 Dzeae
Dickeya zeae CSL RW192 WP 016940978 1 Dzeae
Dickeya zeae ZJU1202 WP 016940978 1 Dzeae
Dickeya zeae NCPPB 3531 WP 016940978 1 Dzeae
Dickeya gs poaceaephila NCPPB 569 WP 016940978 1 Dpoaceaephila
Dickeya zeae EC1 WP 016940978 1 Dzeae
Dickeya zeae MS1 WP 016940978 1 Dzeae
Dickeya gs fangzhongdai NCPPB 3274 WP 016940978 1 Dfangzhongdai
Dickeya dadantii subsp dieffenbachiae NCPPB 2976 WP 016940978 1 Ddadantii
Dickeya dadantii NCPPB 898 WP 016940978 1 Ddadantii

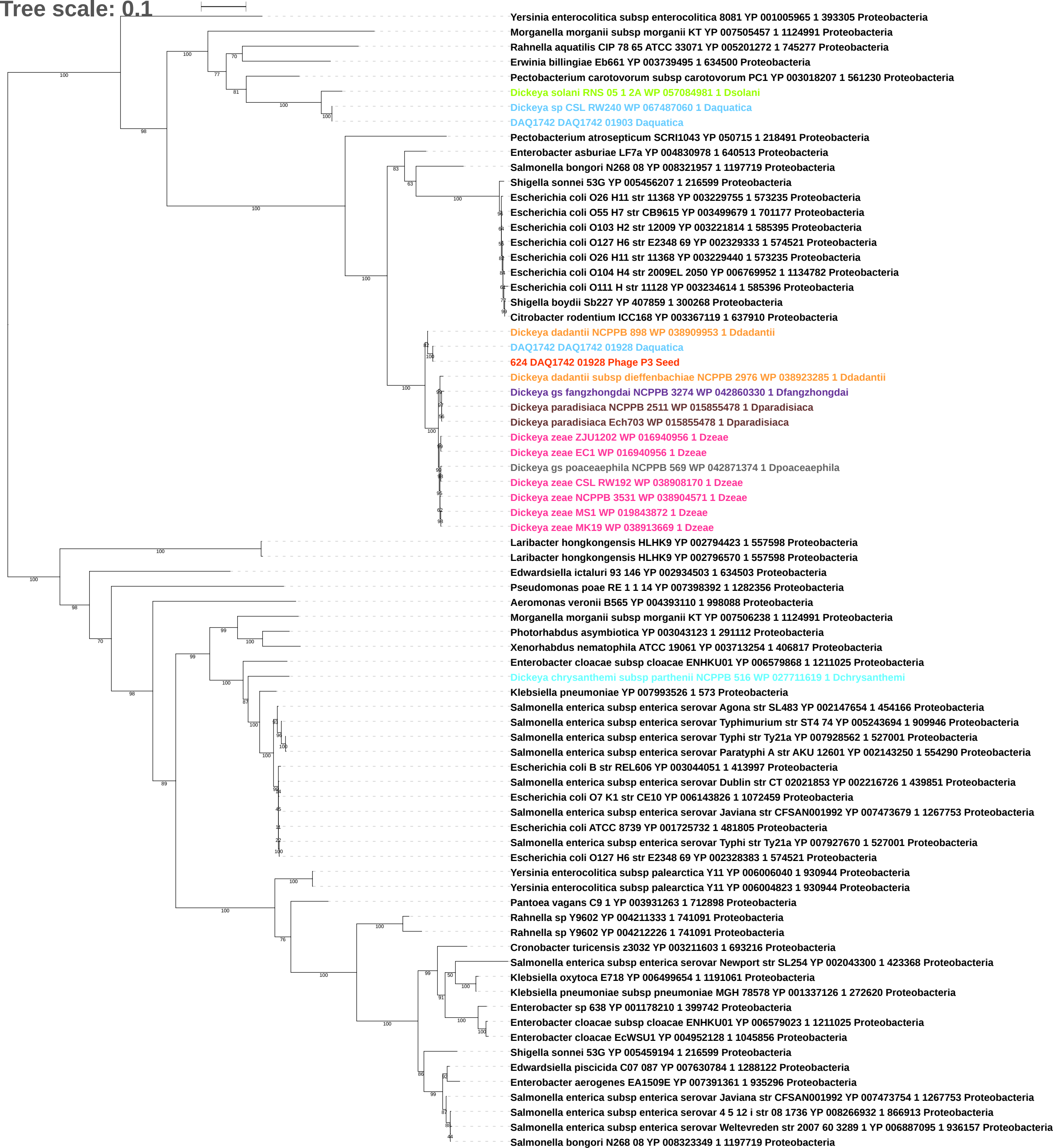
Tree scale: 0.1
Yersinia enterocolitica subsp enterocolitica 8081 YP 001005965 1 393305 Proteobacteria
Morganella morganii subsp morganii KT YP 007505457 1 1124991 Proteobacteria
Rahnella aquatilis CIP 78 65 ATCC 33071 YP 005201272 1 745277 Proteobacteria
Erwinia billingiae Eb661 YP 003739495 1 634500 Proteobacteria
Pectobacterium carotovorum subsp carotovorum PC1 YP 003018207 1 561230 Proteobacteria
Dickeya solani RNS 05 1 2A WP 057084981 1 Dsolani
Dickeya sp CSL RW240 WP 067487060 1 Daquatica
DAQ1742 DAQ1742 01903 Daquatica
Pectobacterium atrosepticum SCRI1043 YP 050715 1 218491 Proteobacteria
Enterobacter asburiae LF7a YP 004830978 1 640513 Proteobacteria
Salmonella bongori N268 08 YP 008321957 1 1197719 Proteobacteria
Shigella sonnei 53G YP 005456207 1 216599 Proteobacteria
Escherichia coli O26 H11 str 11368 YP 003229755 1 573235 Proteobacteria
Escherichia coli O55 H7 str CB9615 YP 003499679 1 701177 Proteobacteria
Escherichia coli O103 H2 str 12009 YP 003221814 1 585395 Proteobacteria
Escherichia coli O127 H6 str E2348 69 YP 002329333 1 574521 Proteobacteria
Escherichia coli O26 H11 str 11368 YP 003229440 1 573235 Proteobacteria
Escherichia coli O104 H4 str 2009EL 2050 YP 006769952 1 1134782 Proteobacteria
Escherichia coli O111 H str 11128 YP 003234614 1 585396 Proteobacteria
Shigella boydii Sb227 YP 407859 1 300268 Proteobacteria
Citrobacter rodentium ICC168 YP 003367119 1 637910 Proteobacteria
Dickeya dadantii NCPPB 898 WP 038909953 1 Ddadantii
DAQ1742 DAQ1742 01928 Daquatica
624 DAQ1742 01928 Phage P3 Seed
Dickeya dadantii subsp dieffenbachiae NCPPB 2976 WP 038923285 1 Ddadantii
Dickeya gs fangzhongdai NCPPB 3274 WP 042860330 1 Dfangzhongdai
Dickeya paradisiaca NCPPB 2511 WP 015855478 1 Dparadisiaca
Dickeya paradisiaca Ech703 WP 015855478 1 Dparadisiaca
Dickeya zeae ZJU1202 WP 016940956 1 Dzeae
Dickeya zeae EC1 WP 016940956 1 Dzeae
Dickeya gs poaceaephila NCPPB 569 WP 042871374 1 Dpoaceaephila
Dickeya zeae CSL RW192 WP 038908170 1 Dzeae
Dickeya zeae NCPPB 3531 WP 038904571 1 Dzeae
Dickeya zeae MS1 WP 019843872 1 Dzeae
Dickeya zeae MK19 WP 038913669 1 Dzeae
Laribacter hongkongensis HLHK9 YP 002794423 1 557598 Proteobacteria
Laribacter hongkongensis HLHK9 YP 002796570 1 557598 Proteobacteria
Edwardsiella ictaluri 93 146 YP 002934503 1 634503 Proteobacteria
Pseudomonas poae RE 1 1 14 YP 007398392 1 1282356 Proteobacteria
Aeromonas veronii B565 YP 004393110 1 998088 Proteobacteria
Morganella morganii subsp morganii KT YP 007506238 1 1124991 Proteobacteria
Photorhabdus asymbiotica YP 003043123 1 291112 Proteobacteria
Xenorhabdus nematophila ATCC 19061 YP 003713254 1 406817 Proteobacteria
Enterobacter cloacae subsp cloacae ENHKU01 YP 006579868 1 1211025 Proteobacteria
Dickeya chrysanthemi subsp parthenii NCPPB 516 WP 027711619 1 Dchrysanthemi
Klebsiella pneumoniae YP 007993526 1 573 Proteobacteria
Salmonella enterica subsp enterica serovar Agona str SL483 YP 002147654 1 454166 Proteobacteria
Salmonella enterica subsp enterica serovar Typhimurium str ST4 74 YP 005243694 1 909946 Proteobacteria
Salmonella enterica subsp enterica serovar Typhi str Ty21a YP 007928562 1 527001 Proteobacteria
Salmonella enterica subsp enterica serovar Paratyphi A str AKU 12601 YP 002143250 1 554290 Proteobacteria
Escherichia coli B str REL606 YP 003044051 1 413997 Proteobacteria
Salmonella enterica subsp enterica serovar Dublin str CT 02021853 YP 002216726 1 439851 Proteobacteria
Escherichia coli O7 K1 str CE10 YP 006143826 1 1072459 Proteobacteria
Salmonella enterica subsp enterica serovar Javiana str CFSAN001992 YP 007473679 1 1267753 Proteobacteria
Escherichia coli ATCC 8739 YP 001725732 1 481805 Proteobacteria
Salmonella enterica subsp enterica serovar Typhi str Ty21a YP 007927670 1 527001 Proteobacteria
Escherichia coli O127 H6 str E2348 69 YP 002328383 1 574521 Proteobacteria
Yersinia enterocolitica subsp palearctica Y11 YP 006006040 1 930944 Proteobacteria
Yersinia enterocolitica subsp palearctica Y11 YP 006004823 1 930944 Proteobacteria
Pantoea vagans C9 1 YP 003931263 1 712898 Proteobacteria
Rahnella sp Y9602 YP 004211333 1 741091 Proteobacteria
Rahnella sp Y9602 YP 004212226 1 741091 Proteobacteria
Cronobacter turicensis z3032 YP 003211603 1 693216 Proteobacteria
Salmonella enterica subsp enterica serovar Newport str SL254 YP 002043300 1 423368 Proteobacteria
Klebsiella oxytoca E718 YP 006499654 1 1191061 Proteobacteria
Klebsiella pneumoniae subsp pneumoniae MGH 78578 YP 001337126 1 272620 Proteobacteria
Enterobacter sp 638 YP 001178210 1 399742 Proteobacteria
Enterobacter cloacae subsp cloacae ENHKU01 YP 006579023 1 1211025 Proteobacteria
Enterobacter cloacae EcWSU1 YP 004952128 1 1045856 Proteobacteria
Shigella sonnei 53G YP 005459194 1 216599 Proteobacteria
Edwardsiella piscicida C07 087 YP 007630784 1 1288122 Proteobacteria
Enterobacter aerogenes EA1509E YP 007391361 1 935296 Proteobacteria
Salmonella enterica subsp enterica serovar Javiana str CFSAN001992 YP 007473754 1 1267753 Proteobacteria
Salmonella enterica subsp enterica serovar 4 5 12 i str 08 1736 YP 008266932 1 866913 Proteobacteria
Salmonella enterica subsp enterica serovar Weltevreden str 2007 60 3289 1 YP 006887095 1 936157 Proteobacteria
Salmonella bongori N268 08 YP 008323349 1 1197719 Proteobacteria

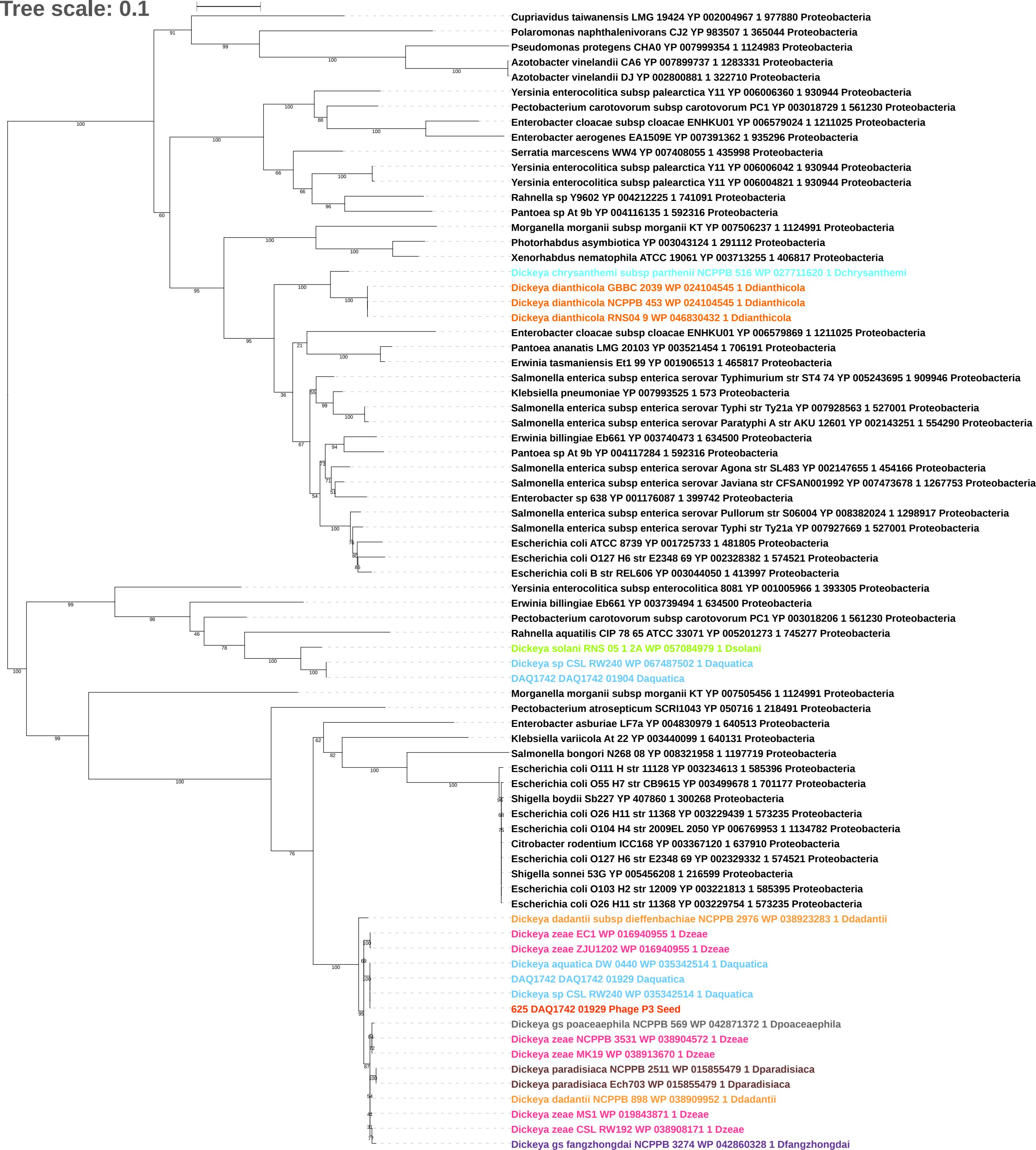
Tree scale: 0.1
Cupriavidus taiwanensis LMG 19424 YP 002004967 1 977880 Proteobacteria
Polaromonas naphthalenivorans CJ2 YP 983507 1 365044 Proteobacteria
Pseudomonas protegens CHA0 YP 007999354 1 1124983 Proteobacteria
Azotobacter vinelandii CA6 YP 007899737 1 1283331 Proteobacteria
Azotobacter vinelandii DJ YP 002800881 1 322710 Proteobacteria
Yersinia enterocolitica subsp palearctica Y11 YP 006006360 1 930944 Proteobacteria
Pectobacterium carotovorum subsp carotovorum PC1 YP 003018729 1 561230 Proteobacteria
Enterobacter cloacae subsp cloacae ENHKU01 YP 006579024 1 1211025 Proteobacteria
Enterobacter aerogenes EA1509E YP 007391362 1 935296 Proteobacteria
Serratia marcescens WW4 YP 007408055 1 435998 Proteobacteria
Yersinia enterocolitica subsp palearctica Y11 YP 006006042 1 930944 Proteobacteria
Yersinia enterocolitica subsp palearctica Y11 YP 006004821 1 930944 Proteobacteria
Rahnella sp Y9602 YP 004212225 1 741091 Proteobacteria
Pantoea sp At 9b YP 004116135 1 592316 Proteobacteria
Morganella morganii subsp morganii KT YP 007506237 1 1124991 Proteobacteria
Photorhabdus asymbiotica YP 003043124 1 291112 Proteobacteria
Xenorhabdus nematophila ATCC 19061 YP 003713255 1 406817 Proteobacteria
Dickeya chrysanthemi subsp parthenii NCPPB 516 WP 027711620 1 Dchrysanthemi
Dickeya dianthicola GBBC 2039 WP 024104545 1 Ddianthicola
Dickeya dianthicola NCPPB 453 WP 024104545 1 Ddianthicola
Dickeya dianthicola RNS04 9 WP 046830432 1 Ddianthicola
Enterobacter cloacae subsp cloacae ENHKU01 YP 006579869 1 1211025 Proteobacteria
Pantoea ananatis LMG 20103 YP 003521454 1 706191 Proteobacteria
Erwinia tasmaniensis Et1 99 YP 001906513 1 465817 Proteobacteria
Salmonella enterica subsp enterica serovar Typhimurium str ST4 74 YP 005243695 1 909946 Proteobacteria
Klebsiella pneumoniae YP 007993525 1 573 Proteobacteria
Salmonella enterica subsp enterica serovar Typhi str Ty21a YP 007928563 1 527001 Proteobacteria
Salmonella enterica subsp enterica serovar Paratyphi A str AKU 12601 YP 002143251 1 554290 Proteobacteria
Erwinia billingiae Eb661 YP 003740473 1 634500 Proteobacteria
Pantoea sp At 9b YP 004117284 1 592316 Proteobacteria
Salmonella enterica subsp enterica serovar Agona str SL483 YP 002147655 1 454166 Proteobacteria
Salmonella enterica subsp enterica serovar Javiana str CFSAN001992 YP 007473678 1 1267753 Proteobacteria
Enterobacter sp 638 YP 001176087 1 399742 Proteobacteria
Salmonella enterica subsp enterica serovar Pullorum str S06004 YP 008382024 1 1298917 Proteobacteria
Salmonella enterica subsp enterica serovar Typhi str Ty21a YP 007927669 1 527001 Proteobacteria
Escherichia coli ATCC 8739 YP 001725733 1 481805 Proteobacteria
Escherichia coli O127 H6 str E2348 69 YP 002328382 1 574521 Proteobacteria
Escherichia coli B str REL606 YP 003044050 1 413997 Proteobacteria
Yersinia enterocolitica subsp enterocolitica 8081 YP 001005966 1 393305 Proteobacteria
Erwinia billingiae Eb661 YP 003739494 1 634500 Proteobacteria
Pectobacterium carotovorum subsp carotovorum PC1 YP 003018206 1 561230 Proteobacteria
Rahnella aquatilis CIP 78 65 ATCC 33071 YP 005201273 1 745277 Proteobacteria
Dickeya solani RNS 05 1 2A WP 057084979 1 Dsolani
Dickeya sp CSL RW240 WP 067487502 1 Daquatica
DAQ1742 DAQ1742 01904 Daquatica
Morganella morganii subsp morganii KT YP 007505456 1 1124991 Proteobacteria
Pectobacterium atrosepticum SCRI1043 YP 050716 1 218491 Proteobacteria
Enterobacter asburiae LF7a YP 004830979 1 640513 Proteobacteria
Klebsiella variicola At 22 YP 003440099 1 640131 Proteobacteria
Salmonella bongori N268 08 YP 008321958 1 1197719 Proteobacteria
Escherichia coli O111 H str 11128 YP 003234613 1 585396 Proteobacteria
Escherichia coli O55 H7 str CB9615 YP 003499678 1 701177 Proteobacteria
Shigella boydii Sb227 YP 407860 1 300268 Proteobacteria
Escherichia coli O26 H11 str 11368 YP 003229439 1 573235 Proteobacteria
Escherichia coli O104 H4 str 2009EL 2050 YP 006769953 1 1134782 Proteobacteria
Citrobacter rodentium ICC168 YP 003367120 1 637910 Proteobacteria
Escherichia coli O127 H6 str E2348 69 YP 002329332 1 574521 Proteobacteria
Shigella sonnei 53G YP 005456208 1 216599 Proteobacteria
Escherichia coli O103 H2 str 12009 YP 003221813 1 585395 Proteobacteria
Escherichia coli O26 H11 str 11368 YP 003229754 1 573235 Proteobacteria
Dickeya dadantii subsp dieffenbachiae NCPPB 2976 WP 038923283 1 Ddadantii
Dickeya zeae EC1 WP 016940955 1 Dzeae
Dickeya zeae ZJU1202 WP 016940955 1 Dzeae
Dickeya aquatica DW 0440 WP 035342514 1 Daquatica
DAQ1742 DAQ1742 01929 Daquatica
Dickeya sp CSL RW240 WP 035342514 1 Daquatica
625 DAQ1742 01929 Phage P3 Seed
Dickeya gs poaceaephila NCPPB 569 WP 042871372 1 Dpoaceaephila
Dickeya zeae NCPPB 3531 WP 038904572 1 Dzeae
Dickeya zeae MK19 WP 038913670 1 Dzeae
Dickeya paradisiaca NCPPB 2511 WP 015855479 1 Dparadisiaca
Dickeya paradisiaca Ech703 WP 015855479 1 Dparadisiaca
Dickeya dadantii NCPPB 898 WP 038909952 1 Ddadantii
Dickeya zeae MS1 WP 019843871 1 Dzeae
Dickeya zeae CSL RW192 WP 038908171 1 Dzeae
Dickeya gs fangzhongdai NCPPB 3274 WP 042860328 1 Dfangzhongdai

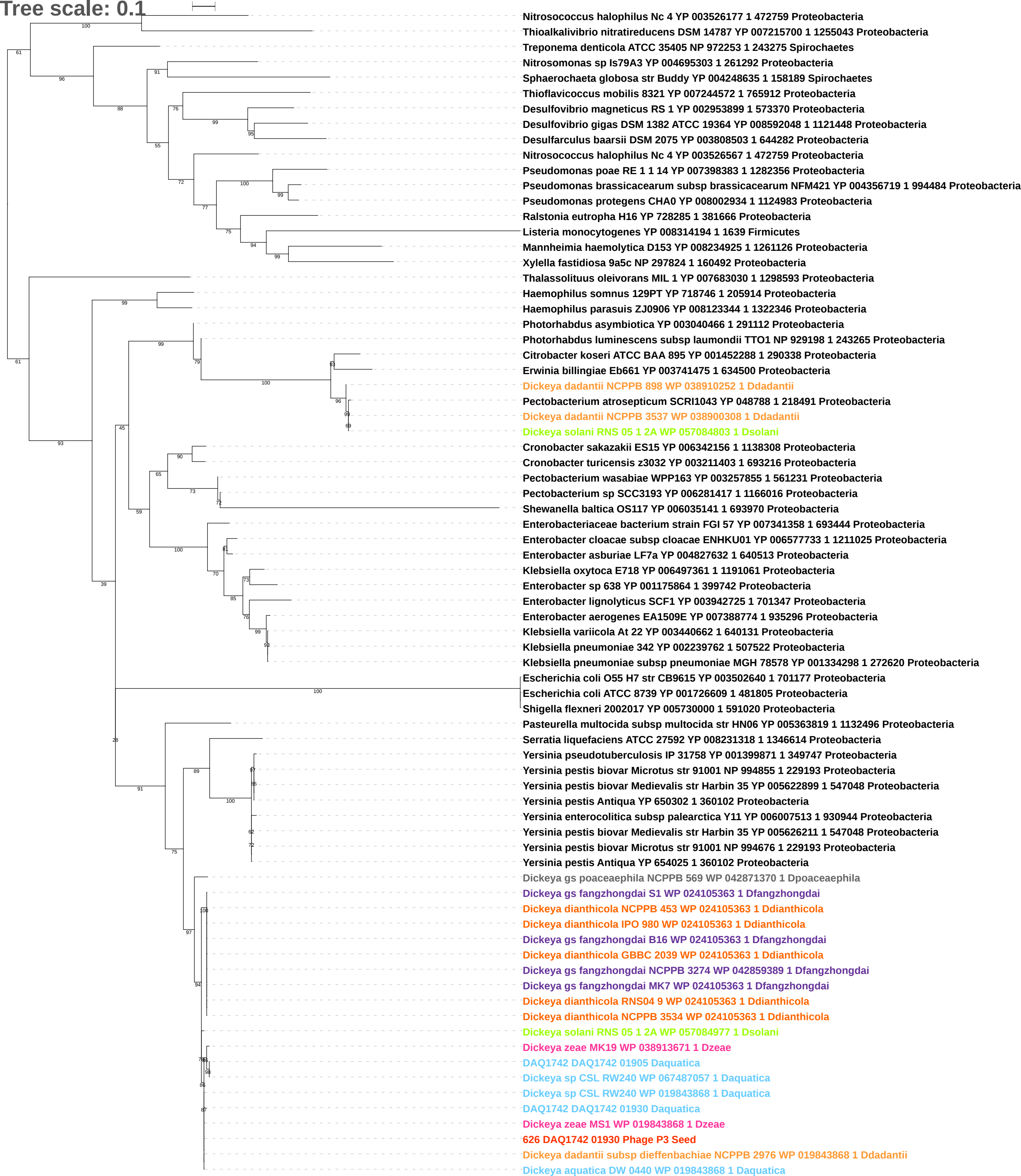
Tree scale: 0.1
Nitrosococcus halophilus Nc 4 YP 003526177 1 472759 Proteobacteria
Thioalkalivibrio nitratireducens DSM 14787 YP 007215700 1 1255043 Proteobacteria
Treponema denticola ATCC 35405 NP 972253 1 243275 Spirochaetes
Nitrosomonas sp Is79A3 YP 004695303 1 261292 Proteobacteria
Sphaerochaeta globosa str Buddy YP 004248635 1 158189 Spirochaetes
Thioflavicoccus mobilis 8321 YP 007244572 1 765912 Proteobacteria
Desulfovibrio magneticus RS 1 YP 002953899 1 573370 Proteobacteria
Desulfovibrio gigas DSM 1382 ATCC 19364 YP 008592048 1 1121448 Proteobacteria
Desulfarculus baarsii DSM 2075 YP 003808503 1 644282 Proteobacteria
Nitrosococcus halophilus Nc 4 YP 003526567 1 472759 Proteobacteria
Pseudomonas poae RE 1 1 14 YP 007398383 1 1282356 Proteobacteria
Pseudomonas brassicacearum subsp brassicacearum NFM421 YP 004356719 1 994484 Proteobacteria
Pseudomonas protegens CHA0 YP 008002934 1 1124983 Proteobacteria
Ralstonia eutropha H16 YP 728285 1 381666 Proteobacteria
Listeria monocytogenes YP 008314194 1 1639 Firmicutes
Mannheimia haemolytica D153 YP 008234925 1 1261126 Proteobacteria
Xylella fastidiosa 9a5c NP 297824 1 160492 Proteobacteria
Thalassolituus oleivorans MIL 1 YP 007683030 1 1298593 Proteobacteria
Haemophilus somnus 129PT YP 718746 1 205914 Proteobacteria
Haemophilus parasuis ZJ0906 YP 008123344 1 1322346 Proteobacteria
Photorhabdus asymbiotica YP 003040466 1 291112 Proteobacteria
Photorhabdus luminescens subsp laumondii TTO1 NP 929198 1 243265 Proteobacteria
Citrobacter koseri ATCC BAA 895 YP 001452288 1 290338 Proteobacteria
Erwinia billingiae Eb661 YP 003741475 1 634500 Proteobacteria
Dickeya dadantii NCPPB 898 WP 038910252 1 Ddadantii
Pectobacterium atrosepticum SCRI1043 YP 048788 1 218491 Proteobacteria
Dickeya dadantii NCPPB 3537 WP 038900308 1 Ddadantii
Dickeya solani RNS 05 1 2A WP 057084803 1 Dsolani
Cronobacter sakazakii ES15 YP 006342156 1 1138308 Proteobacteria
Cronobacter turicensis z3032 YP 003211403 1 693216 Proteobacteria
Pectobacterium wasabiae WPP163 YP 003257855 1 561231 Proteobacteria
Pectobacterium sp SCC3193 YP 006281417 1 1166016 Proteobacteria
Shewanella baltica OS117 YP 006035141 1 693970 Proteobacteria
Enterobacteriaceae bacterium strain FGI 57 YP 007341358 1 693444 Proteobacteria
Enterobacter cloacae subsp cloacae ENHKU01 YP 006577733 1 1211025 Proteobacteria
Enterobacter asburiae LF7a YP 004827632 1 640513 Proteobacteria
Klebsiella oxytoca E718 YP 006497361 1 1191061 Proteobacteria
Enterobacter sp 638 YP 001175864 1 399742 Proteobacteria
Enterobacter lignolyticus SCF1 YP 003942725 1 701347 Proteobacteria
Enterobacter aerogenes EA1509E YP 007388774 1 935296 Proteobacteria
Klebsiella variicola At 22 YP 003440662 1 640131 Proteobacteria
Klebsiella pneumoniae 342 YP 002239762 1 507522 Proteobacteria
Klebsiella pneumoniae subsp pneumoniae MGH 78578 YP 001334298 1 272620 Proteobacteria
Escherichia coli O55 H7 str CB9615 YP 003502640 1 701177 Proteobacteria
Escherichia coli ATCC 8739 YP 001726609 1 481805 Proteobacteria
Shigella flexneri 2002017 YP 005730000 1 591020 Proteobacteria
Pasteurella multocida subsp multocida str HN06 YP 005363819 1 1132496 Proteobacteria
Serratia liquefaciens ATCC 27592 YP 008231318 1 1346614 Proteobacteria
Yersinia pseudotuberculosis IP 31758 YP 001399871 1 349747 Proteobacteria
Yersinia pestis biovar Microtus str 91001 NP 994855 1 229193 Proteobacteria
Yersinia pestis biovar Medievalis str Harbin 35 YP 005622899 1 547048 Proteobacteria
Yersinia pestis Antiqua YP 650302 1 360102 Proteobacteria
Yersinia enterocolitica subsp palearctica Y11 YP 006007513 1 930944 Proteobacteria
Yersinia pestis biovar Medievalis str Harbin 35 YP 005626211 1 547048 Proteobacteria
Yersinia pestis biovar Microtus str 91001 NP 994676 1 229193 Proteobacteria
Yersinia pestis Antiqua YP 654025 1 360102 Proteobacteria
Dickeya gs poaceaephila NCPPB 569 WP 042871370 1 Dpoaceaephila
Dickeya gs fangzhongdai S1 WP 024105363 1 Dfangzhongdai
Dickeya dianthicola NCPPB 453 WP 024105363 1 Ddianthicola
Dickeya dianthicola IPO 980 WP 024105363 1 Ddianthicola
Dickeya gs fangzhongdai B16 WP 024105363 1 Dfangzhongdai
Dickeya dianthicola GBBC 2039 WP 024105363 1 Ddianthicola
Dickeya gs fangzhongdai NCPPB 3274 WP 042859389 1 Dfangzhongdai
Dickeya gs fangzhongdai MK7 WP 024105363 1 Dfangzhongdai
Dickeya dianthicola RNS04 9 WP 024105363 1 Ddianthicola
Dickeya dianthicola NCPPB 3534 WP 024105363 1 Ddianthicola
Dickeya solani RNS 05 1 2A WP 057084977 1 Dsolani
Dickeya zeae MK19 WP 038913671 1 Dzeae
DAQ1742 DAQ1742 01905 Daquatica
Dickeya sp CSL RW240 WP 067487057 1 Daquatica
Dickeya sp CSL RW240 WP 019843868 1 Daquatica
DAQ1742 DAQ1742 01930 Daquatica
Dickeya zeae MS1 WP 019843868 1 Dzeae
626 DAQ1742 01930 Phage P3 Seed
Dickeya dadantii subsp dieffenbachiae NCPPB 2976 WP 019843868 1 Ddadantii
Dickeya aquatica DW 0440 WP 019843868 1 Daquatica

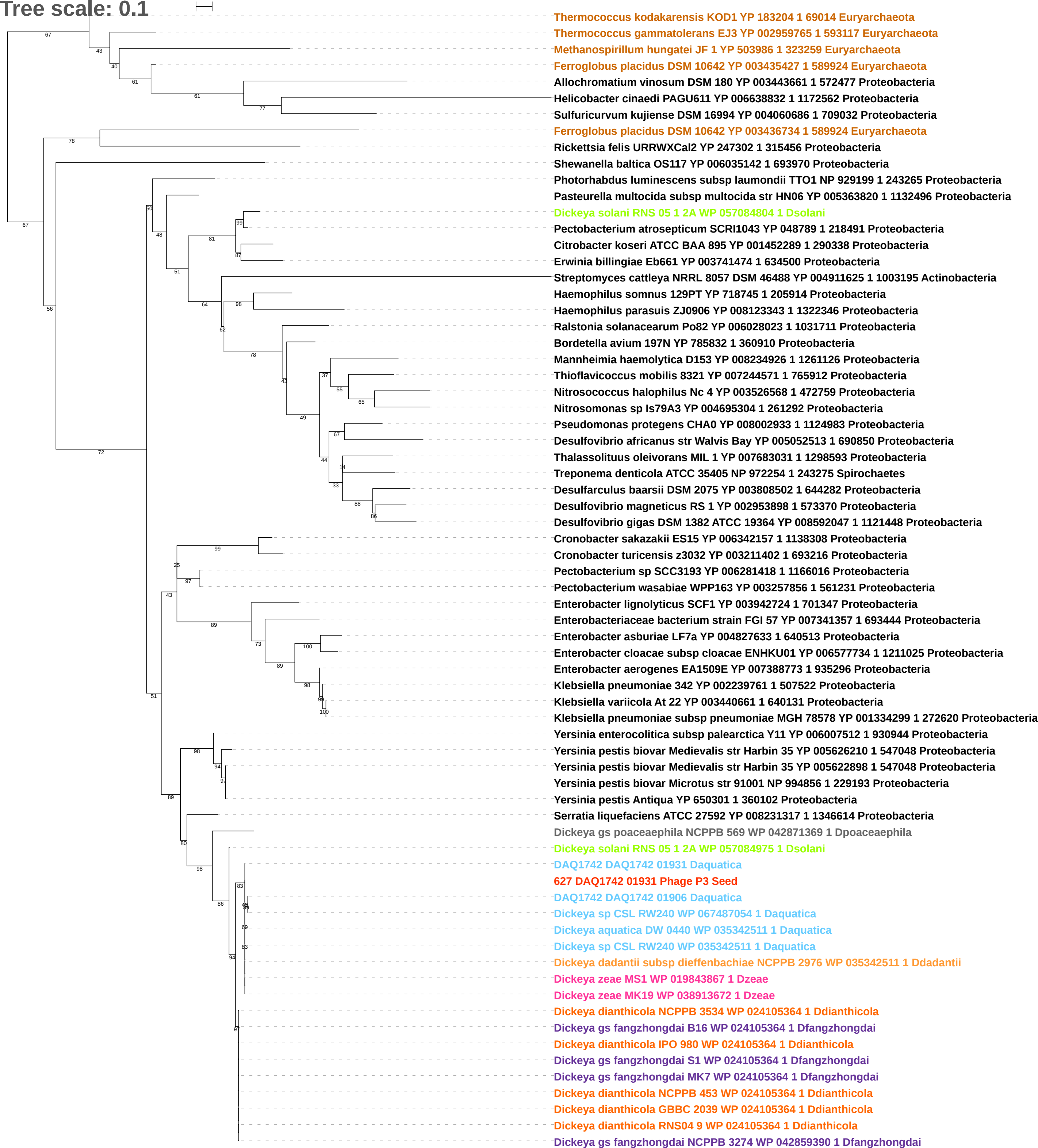
Tree scale: 0.1
Thermococcus kodakarensis KOD1 YP 183204 1 69014 Euryarchaeota
Thermococcus gammatolerans EJ3 YP 002959765 1 593117 Euryarchaeota
Methanospirillum hungatei JF 1 YP 503986 1 323259 Euryarchaeota
Ferroglobus placidus DSM 10642 YP 003435427 1 589924 Euryarchaeota
Allochromatium vinosum DSM 180 YP 003443661 1 572477 Proteobacteria
Helicobacter cinaedi PAGU611 YP 006638832 1 1172562 Proteobacteria
Sulfuricurvum kujiense DSM 16994 YP 004060686 1 709032 Proteobacteria
Ferroglobus placidus DSM 10642 YP 003436734 1 589924 Euryarchaeota
Rickettsia felis URRWXCal2 YP 247302 1 315456 Proteobacteria
Shewanella baltica OS117 YP 006035142 1 693970 Proteobacteria
Photorhabdus luminescens subsp laumondii TTO1 NP 929199 1 243265 Proteobacteria
Pasteurella multocida subsp multocida str HN06 YP 005363820 1 1132496 Proteobacteria
Dickeya solani RNS 05 1 2A WP 057084804 1 Dsolani
Pectobacterium atrosepticum SCRI1043 YP 048789 1 218491 Proteobacteria
Citrobacter koseri ATCC BAA 895 YP 001452289 1 290338 Proteobacteria
Erwinia billingiae Eb661 YP 003741474 1 634500 Proteobacteria
Streptomyces cattleya NRRL 8057 DSM 46488 YP 004911625 1 1003195 Actinobacteria
Haemophilus somnus 129PT YP 718745 1 205914 Proteobacteria
Haemophilus parasuis ZJ0906 YP 008123343 1 1322346 Proteobacteria
Ralstonia solanacearum Po82 YP 006028023 1 1031711 Proteobacteria
Bordetella avium 197N YP 785832 1 360910 Proteobacteria
Mannheimia haemolytica D153 YP 008234926 1 1261126 Proteobacteria
Thioflavicoccus mobilis 8321 YP 007244571 1 765912 Proteobacteria
Nitrosococcus halophilus Nc 4 YP 003526568 1 472759 Proteobacteria
Nitrosomonas sp Is79A3 YP 004695304 1 261292 Proteobacteria
Pseudomonas protegens CHA0 YP 008002933 1 1124983 Proteobacteria
Desulfovibrio africanus str Walvis Bay YP 005052513 1 690850 Proteobacteria
Thalassolituus oleivorans MIL 1 YP 007683031 1 1298593 Proteobacteria
Treponema denticola ATCC 35405 NP 972254 1 243275 Spirochaetes
Desulfarculus baarsii DSM 2075 YP 003808502 1 644282 Proteobacteria
Desulfovibrio magneticus RS 1 YP 002953898 1 573370 Proteobacteria
Desulfovibrio gigas DSM 1382 ATCC 19364 YP 008592047 1 1121448 Proteobacteria
Cronobacter sakazakii ES15 YP 006342157 1 1138308 Proteobacteria
Cronobacter turicensis z3032 YP 003211402 1 693216 Proteobacteria
Pectobacterium sp SCC3193 YP 006281418 1 1166016 Proteobacteria
Pectobacterium wasabiae WPP163 YP 003257856 1 561231 Proteobacteria
Enterobacter lignolyticus SCF1 YP 003942724 1 701347 Proteobacteria
Enterobacteriaceae bacterium strain FGI 57 YP 007341357 1 693444 Proteobacteria
Enterobacter asburiae LF7a YP 004827633 1 640513 Proteobacteria
Enterobacter cloacae subsp cloacae ENHKU01 YP 006577734 1 1211025 Proteobacteria
Enterobacter aerogenes EA1509E YP 007388773 1 935296 Proteobacteria
Klebsiella pneumoniae 342 YP 002239761 1 507522 Proteobacteria
Klebsiella variicola At 22 YP 003440661 1 640131 Proteobacteria
Klebsiella pneumoniae subsp pneumoniae MGH 78578 YP 001334299 1 272620 Proteobacteria
Yersinia enterocolitica subsp palearctica Y11 YP 006007512 1 930944 Proteobacteria
Yersinia pestis biovar Medievalis str Harbin 35 YP 005626210 1 547048 Proteobacteria
Yersinia pestis biovar Medievalis str Harbin 35 YP 005622898 1 547048 Proteobacteria
Yersinia pestis biovar Microtus str 91001 NP 994856 1 229193 Proteobacteria
Yersinia pestis Antiqua YP 650301 1 360102 Proteobacteria
Serratia liquefaciens ATCC 27592 YP 008231317 1 1346614 Proteobacteria
Dickeya gs poaceaephila NCPPB 569 WP 042871369 1 Dpoaceaephila
Dickeya solani RNS 05 1 2A WP 057084975 1 Dsolani
DAQ1742 DAQ1742 01931 Daquatica
627 DAQ1742 01931 Phage P3 Seed
DAQ1742 DAQ1742 01906 Daquatica
Dickeya sp CSL RW240 WP 067487054 1 Daquatica
Dickeya aquatica DW 0440 WP 035342511 1 Daquatica
Dickeya sp CSL RW240 WP 035342511 1 Daquatica
Dickeya dadantii subsp dieffenbachiae NCPPB 2976 WP 035342511 1 Ddadantii
Dickeya zeae MS1 WP 019843867 1 Dzeae
Dickeya zeae MK19 WP 038913672 1 Dzeae
Dickeya dianthicola NCPPB 3534 WP 024105364 1 Ddianthicola
Dickeya gs fangzhongdai B16 WP 024105364 1 Dfangzhongdai
Dickeya dianthicola IPO 980 WP 024105364 1 Ddianthicola
Dickeya gs fangzhongdai S1 WP 024105364 1 Dfangzhongdai
Dickeya gs fangzhongdai MK7 WP 024105364 1 Dfangzhongdai
Dickeya dianthicola NCPPB 453 WP 024105364 1 Ddianthicola
Dickeya dianthicola GBBC 2039 WP 024105364 1 Ddianthicola
Dickeya dianthicola RNS04 9 WP 024105364 1 Ddianthicola
Dickeya gs fangzhongdai NCPPB 3274 WP 042859390 1 Dfangzhongdai

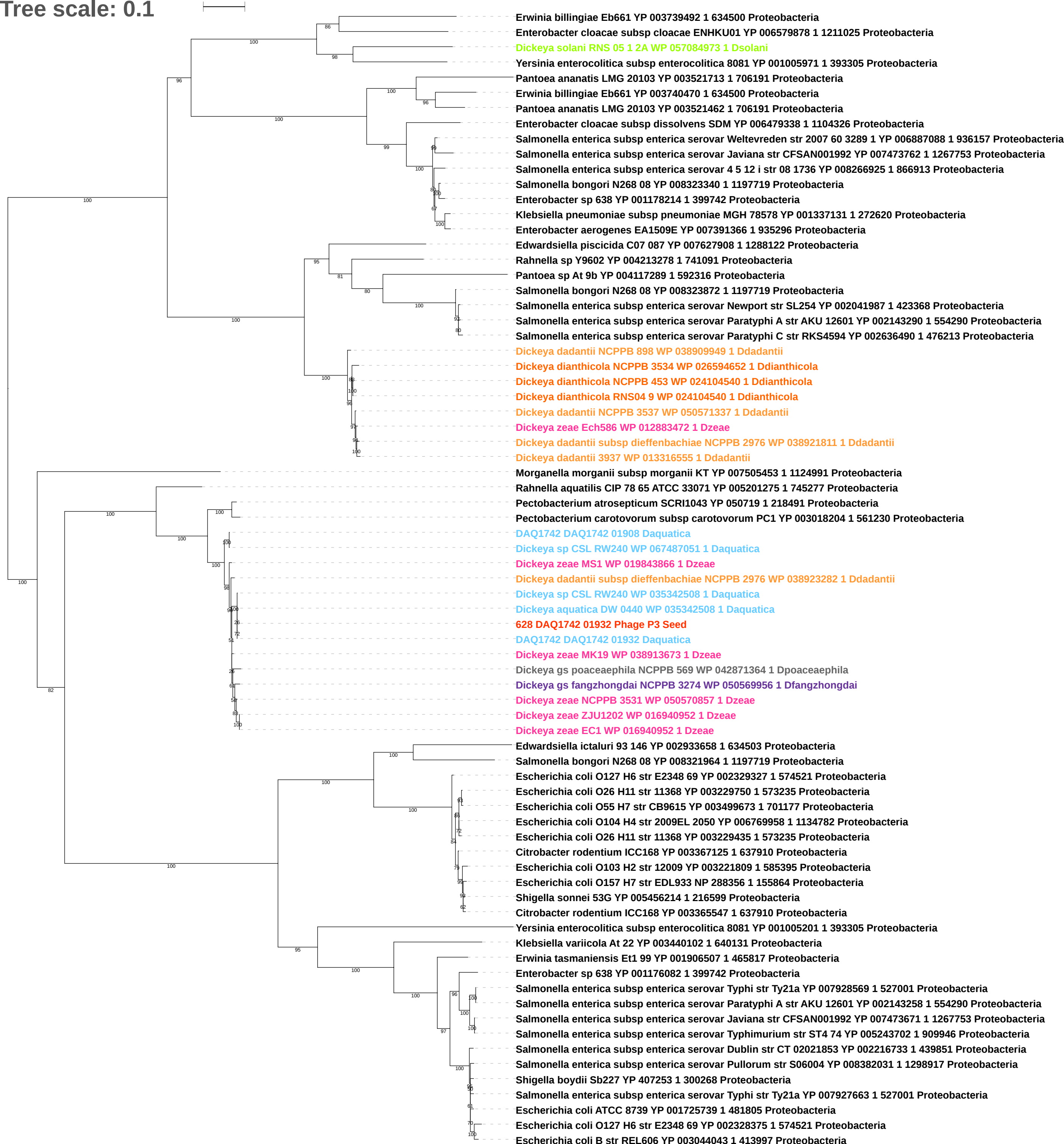
Tree scale: 0.1
Erwinia billingiae Eb661 YP 003739492 1 634500 Proteobacteria
Enterobacter cloacae subsp cloacae ENHKU01 YP 006579878 1 1211025 Proteobacteria
Dickeya solani RNS 05 1 2A WP 057084973 1 Dsolani
Yersinia enterocolitica subsp enterocolitica 8081 YP 001005971 1 393305 Proteobacteria
Pantoea ananatis LMG 20103 YP 003521713 1 706191 Proteobacteria
Erwinia billingiae Eb661 YP 003740470 1 634500 Proteobacteria
Pantoea ananatis LMG 20103 YP 003521462 1 706191 Proteobacteria
Enterobacter cloacae subsp dissolvens SDM YP 006479338 1 1104326 Proteobacteria
Salmonella enterica subsp enterica serovar Weltevreden str 2007 60 3289 1 YP 006887088 1 936157 Proteobacteria
Salmonella enterica subsp enterica serovar Javiana str CFSAN001992 YP 007473762 1 1267753 Proteobacteria
Salmonella enterica subsp enterica serovar 4 5 12 i str 08 1736 YP 008266925 1 866913 Proteobacteria
Salmonella bongori N268 08 YP 008323340 1 1197719 Proteobacteria
Enterobacter sp 638 YP 001178214 1 399742 Proteobacteria
Klebsiella pneumoniae subsp pneumoniae MGH 78578 YP 001337131 1 272620 Proteobacteria
Enterobacter aerogenes EA1509E YP 007391366 1 935296 Proteobacteria
Edwardsiella piscicida C07 087 YP 007627908 1 1288122 Proteobacteria
Rahnella sp Y9602 YP 004213278 1 741091 Proteobacteria
Pantoea sp At 9b YP 004117289 1 592316 Proteobacteria
Salmonella bongori N268 08 YP 008323872 1 1197719 Proteobacteria
Salmonella enterica subsp enterica serovar Newport str SL254 YP 002041987 1 423368 Proteobacteria
Salmonella enterica subsp enterica serovar Paratyphi A str AKU 12601 YP 002143290 1 554290 Proteobacteria
Salmonella enterica subsp enterica serovar Paratyphi C str RKS4594 YP 002636490 1 476213 Proteobacteria
Dickeya dadantii NCPPB 898 WP 038909949 1 Ddadantii
Dickeya dianthicola NCPPB 3534 WP 026594652 1 Ddianthicola
Dickeya dianthicola NCPPB 453 WP 024104540 1 Ddianthicola
Dickeya dianthicola RNS04 9 WP 024104540 1 Ddianthicola
Dickeya dadantii NCPPB 3537 WP 050571337 1 Ddadantii
Dickeya zeae Ech586 WP 012883472 1 Dzeae
Dickeya dadantii subsp dieffenbachiae NCPPB 2976 WP 038921811 1 Ddadantii
Dickeya dadantii 3937 WP 013316555 1 Ddadantii
Morganella morganii subsp morganii KT YP 007505453 1 1124991 Proteobacteria
Rahnella aquatilis CIP 78 65 ATCC 33071 YP 005201275 1 745277 Proteobacteria
Pectobacterium atrosepticum SCRI1043 YP 050719 1 218491 Proteobacteria
Pectobacterium carotovorum subsp carotovorum PC1 YP 003018204 1 561230 Proteobacteria
DAQ1742 DAQ1742 01908 Daquatica
Dickeya sp CSL RW240 WP 067487051 1 Daquatica
Dickeya zeae MS1 WP 019843866 1 Dzeae
Dickeya dadantii subsp dieffenbachiae NCPPB 2976 WP 038923282 1 Ddadantii
Dickeya sp CSL RW240 WP 035342508 1 Daquatica
Dickeya aquatica DW 0440 WP 035342508 1 Daquatica
628 DAQ1742 01932 Phage P3 Seed
DAQ1742 DAQ1742 01932 Daquatica
Dickeya zeae MK19 WP 038913673 1 Dzeae
Dickeya gs poaceaephila NCPPB 569 WP 042871364 1 Dpoaceaephila
Dickeya gs fangzhongdai NCPPB 3274 WP 050569956 1 Dfangzhongdai
Dickeya zeae NCPPB 3531 WP 050570857 1 Dzeae
Dickeya zeae ZJU1202 WP 016940952 1 Dzeae
Dickeya zeae EC1 WP 016940952 1 Dzeae
Edwardsiella ictaluri 93 146 YP 002933658 1 634503 Proteobacteria
Salmonella bongori N268 08 YP 008321964 1 1197719 Proteobacteria
Escherichia coli O127 H6 str E2348 69 YP 002329327 1 574521 Proteobacteria
Escherichia coli O26 H11 str 11368 YP 003229750 1 573235 Proteobacteria
Escherichia coli O55 H7 str CB9615 YP 003499673 1 701177 Proteobacteria
Escherichia coli O104 H4 str 2009EL 2050 YP 006769958 1 1134782 Proteobacteria
Escherichia coli O26 H11 str 11368 YP 003229435 1 573235 Proteobacteria
Citrobacter rodentium ICC168 YP 003367125 1 637910 Proteobacteria
Escherichia coli O103 H2 str 12009 YP 003221809 1 585395 Proteobacteria
Escherichia coli O157 H7 str EDL933 NP 288356 1 155864 Proteobacteria
Shigella sonnei 53G YP 005456214 1 216599 Proteobacteria
Citrobacter rodentium ICC168 YP 003365547 1 637910 Proteobacteria
Yersinia enterocolitica subsp enterocolitica 8081 YP 001005201 1 393305 Proteobacteria
Klebsiella variicola At 22 YP 003440102 1 640131 Proteobacteria
Erwinia tasmaniensis Et1 99 YP 001906507 1 465817 Proteobacteria
Enterobacter sp 638 YP 001176082 1 399742 Proteobacteria
Salmonella enterica subsp enterica serovar Typhi str Ty21a YP 007928569 1 527001 Proteobacteria
Salmonella enterica subsp enterica serovar Paratyphi A str AKU 12601 YP 002143258 1 554290 Proteobacteria
Salmonella enterica subsp enterica serovar Javiana str CFSAN001992 YP 007473671 1 1267753 Proteobacteria
Salmonella enterica subsp enterica serovar Typhimurium str ST4 74 YP 005243702 1 909946 Proteobacteria
Salmonella enterica subsp enterica serovar Dublin str CT 02021853 YP 002216733 1 439851 Proteobacteria
Salmonella enterica subsp enterica serovar Pullorum str S06004 YP 008382031 1 1298917 Proteobacteria
Shigella boydii Sb227 YP 407253 1 300268 Proteobacteria
Salmonella enterica subsp enterica serovar Typhi str Ty21a YP 007927663 1 527001 Proteobacteria
Escherichia coli ATCC 8739 YP 001725739 1 481805 Proteobacteria
Escherichia coli O127 H6 str E2348 69 YP 002328375 1 574521 Proteobacteria
Escherichia coli B str REL606 YP 003044043 1 413997 Proteobacteria

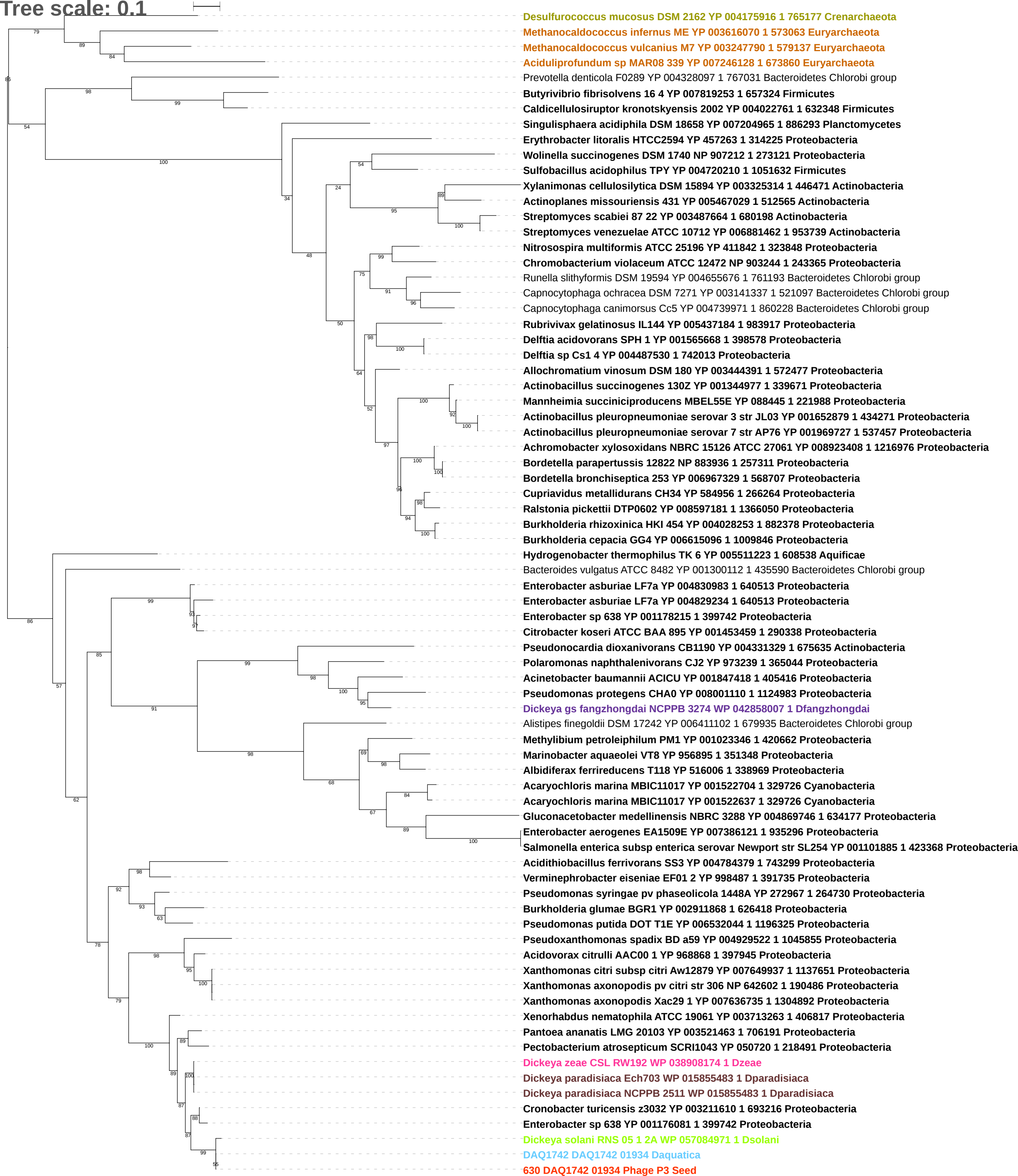

Tree scale: 0.1
Desulfurococcus mucosus DSM 2162 YP 004175916 1 765177 Crenarchaeota
Methanocaldococcus infernus ME YP 003616070 1 573063 Euryarchaeota
Methanocaldococcus vulcanius M7 YP 003247790 1 579137 Euryarchaeota
Aciduliprofundum sp MAR08 339 YP 007246128 1 673860 Euryarchaeota
Prevotella denticola F0289 YP 004328097 1 767031 Bacteroidetes Chlorobi group
Butyrivibrio fibrisolvens 16 4 YP 007819253 1 657324 Firmicutes
Caldicellulosiruptor kronotskyensis 2002 YP 004022761 1 632348 Firmicutes
Singulisphaera acidiphila DSM 18658 YP 007204965 1 886293 Planctomycetes
Erythrobacter litoralis HTCC2594 YP 457263 1 314225 Proteobacteria
Wolinella succinogenes DSM 1740 NP 907212 1 273121 Proteobacteria
Sulfobacillus acidophilus TPY YP 004720210 1 1051632 Firmicutes
Xylanimonas cellulosilytica DSM 15894 YP 003325314 1 446471 Actinobacteria
Actinoplanes missouriensis 431 YP 005467029 1 512565 Actinobacteria
Streptomyces scabiei 87 22 YP 003487664 1 680198 Actinobacteria
Streptomyces venezuelae ATCC 10712 YP 006881462 1 953739 Actinobacteria
Nitrosospira multiformis ATCC 25196 YP 411842 1 323848 Proteobacteria
Chromobacterium violaceum ATCC 12472 NP 903244 1 243365 Proteobacteria
Runella slithyformis DSM 19594 YP 004655676 1 761193 Bacteroidetes Chlorobi group
Capnocytophaga ochracea DSM 7271 YP 003141337 1 521097 Bacteroidetes Chlorobi group
Capnocytophaga canimorsus Cc5 YP 004739971 1 860228 Bacteroidetes Chlorobi group
Rubrivivax gelatinosus IL144 YP 005437184 1 983917 Proteobacteria
Delftia acidovorans SPH 1 YP 001565668 1 398578 Proteobacteria
Delftia sp Cs1 4 YP 004487530 1 742013 Proteobacteria
Allochromatium vinosum DSM 180 YP 003444391 1 572477 Proteobacteria
Actinobacillus succinogenes 130Z YP 001344977 1 339671 Proteobacteria
Mannheimia succiniciproducens MBEL55E YP 088445 1 221988 Proteobacteria
Actinobacillus pleuropneumoniae serovar 3 str JL03 YP 001652879 1 434271 Proteobacteria
Actinobacillus pleuropneumoniae serovar 7 str AP76 YP 001969727 1 537457 Proteobacteria
Achromobacter xylosoxidans NBRC 15126 ATCC 27061 YP 008923408 1 1216976 Proteobacteria
Bordetella parapertussis 12822 NP 883936 1 257311 Proteobacteria
Bordetella bronchiseptica 253 YP 006967329 1 568707 Proteobacteria
Cupriavidus metallidurans CH34 YP 584956 1 266264 Proteobacteria
Ralstonia pickettii DTP0602 YP 008597181 1 1366050 Proteobacteria
Burkholderia rhizoxinica HKI 454 YP 004028253 1 882378 Proteobacteria
Burkholderia cepacia GG4 YP 006615096 1 1009846 Proteobacteria
Hydrogenobacter thermophilus TK 6 YP 005511223 1 608538 Aquificae
Bacteroides vulgatus ATCC 8482 YP 001300112 1 435590 Bacteroidetes Chlorobi group
Enterobacter asburiae LF7a YP 004830983 1 640513 Proteobacteria
Enterobacter asburiae LF7a YP 004829234 1 640513 Proteobacteria
Enterobacter sp 638 YP 001178215 1 399742 Proteobacteria
Citrobacter koseri ATCC BAA 895 YP 001453459 1 290338 Proteobacteria
Pseudonocardia dioxanivorans CB1190 YP 004331329 1 675635 Actinobacteria
Polaromonas naphthalenivorans CJ2 YP 973239 1 365044 Proteobacteria
Acinetobacter baumannii ACICU YP 001847418 1 405416 Proteobacteria
Pseudomonas protegens CHA0 YP 008001110 1 1124983 Proteobacteria
Dickeya gs fangzhongdai NCPPB 3274 WP 042858007 1 Dfangzhongdai
Alistipes finegoldii DSM 17242 YP 006411102 1 679935 Bacteroidetes Chlorobi group
Methylibium petroleiphilum PM1 YP 001023346 1 420662 Proteobacteria
Marinobacter aquaeolei VT8 YP 956895 1 351348 Proteobacteria
Albidiferax ferrireducens T118 YP 516006 1 338969 Proteobacteria
Acaryochloris marina MBIC11017 YP 001522704 1 329726 Cyanobacteria
Acaryochloris marina MBIC11017 YP 001522637 1 329726 Cyanobacteria
Gluconacetobacter medellinensis NBRC 3288 YP 004869746 1 634177 Proteobacteria
Enterobacter aerogenes EA1509E YP 007386121 1 935296 Proteobacteria
Salmonella enterica subsp enterica serovar Newport str SL254 YP 001101885 1 423368 Proteobacteria
Acidithiobacillus ferrivorans SS3 YP 004784379 1 743299 Proteobacteria
Verminephrobacter eiseniae EF01 2 YP 998487 1 391735 Proteobacteria
Pseudomonas syringae pv phaseolicola 1448A YP 272967 1 264730 Proteobacteria
Burkholderia glumae BGR1 YP 002911868 1 626418 Proteobacteria
Pseudomonas putida DOT T1E YP 006532044 1 1196325 Proteobacteria
Pseudoxanthomonas spadix BD a59 YP 004929522 1 1045855 Proteobacteria
Acidovorax citrulli AAC00 1 YP 968868 1 397945 Proteobacteria
Xanthomonas citri subsp citri Aw12879 YP 007649937 1 1137651 Proteobacteria
Xanthomonas axonopodis pv citri str 306 NP 642602 1 190486 Proteobacteria
Xanthomonas axonopodis Xac29 1 YP 007636735 1 1304892 Proteobacteria
Xenorhabdus nematophila ATCC 19061 YP 003713263 1 406817 Proteobacteria
Pantoea ananatis LMG 20103 YP 003521463 1 706191 Proteobacteria
Pectobacterium atrosepticum SCRI1043 YP 050720 1 218491 Proteobacteria
Dickeya zeae CSL RW192 WP 038908174 1 Dzeae
Dickeya paradisiaca Ech703 WP 015855483 1 Dparadisiaca
Dickeya paradisiaca NCPPB 2511 WP 015855483 1 Dparadisiaca
Cronobacter turicensis z3032 YP 003211610 1 693216 Proteobacteria
Enterobacter sp 638 YP 001176081 1 399742 Proteobacteria
Dickeya solani RNS 05 1 2A WP 057084971 1 Dsolani
DAQ1742 DAQ1742 01934 Daquatica
630 DAQ1742 01934 Phage P3 Seed

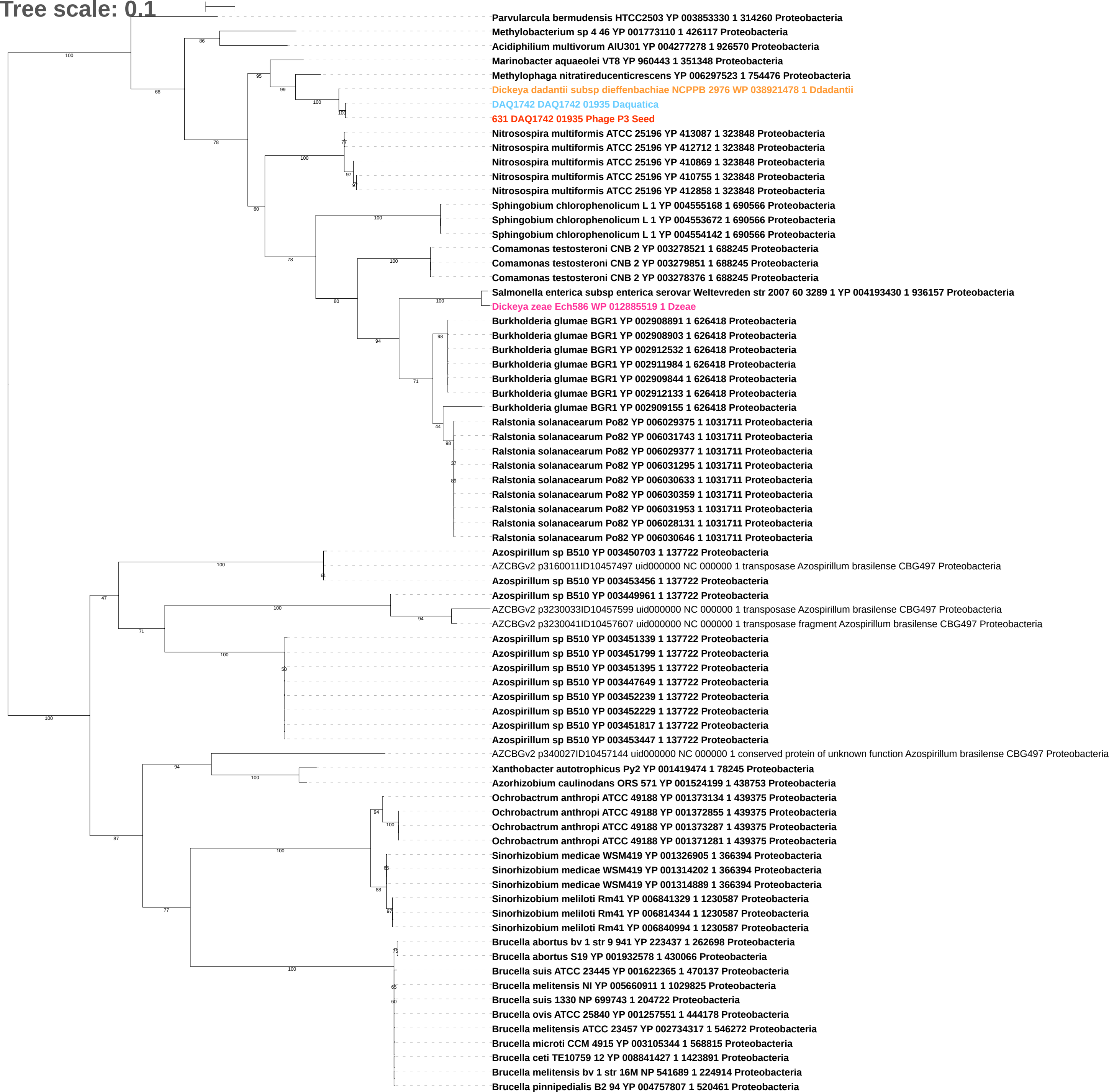

Tree scale: 0.1
Parvularcula bermudensis HTCC2503 YP 003853330 1 314260 Proteobacteria
Methylobacterium sp 4 46 YP 001773110 1 426117 Proteobacteria
Acidiphilium multivorum AIU301 YP 004277278 1 926570 Proteobacteria
Marinobacter aquaeolei VT8 YP 960443 1 351348 Proteobacteria
Methylophaga nitratireducenticrescens YP 006297523 1 754476 Proteobacteria
Dickeya dadantii subsp dieffenbachiae NCPPB 2976 WP 038921478 1 Ddadantii
DAQ1742 DAQ1742 01935 Daquatica
631 DAQ1742 01935 Phage P3 Seed
Nitrosospira multiformis ATCC 25196 YP 413087 1 323848 Proteobacteria
Nitrosospira multiformis ATCC 25196 YP 412712 1 323848 Proteobacteria
Nitrosospira multiformis ATCC 25196 YP 410869 1 323848 Proteobacteria
Nitrosospira multiformis ATCC 25196 YP 410755 1 323848 Proteobacteria
Nitrosospira multiformis ATCC 25196 YP 412858 1 323848 Proteobacteria
Sphingobium chlorophenolicum L 1 YP 004555168 1 690566 Proteobacteria
Sphingobium chlorophenolicum L 1 YP 004553672 1 690566 Proteobacteria
Sphingobium chlorophenolicum L 1 YP 004554142 1 690566 Proteobacteria
Comamonas testosteroni CNB 2 YP 003278521 1 688245 Proteobacteria
Comamonas testosteroni CNB 2 YP 003279851 1 688245 Proteobacteria
Comamonas testosteroni CNB 2 YP 003278376 1 688245 Proteobacteria
Salmonella enterica subsp enterica serovar Weltevreden str 2007 60 3289 1 YP 004193430 1 936157 Proteobacteria
Dickeya zeae Ech586 WP 012885519 1 Dzeae
Burkholderia glumae BGR1 YP 002908891 1 626418 Proteobacteria
Burkholderia glumae BGR1 YP 002908903 1 626418 Proteobacteria
Burkholderia glumae BGR1 YP 002912532 1 626418 Proteobacteria
Burkholderia glumae BGR1 YP 002911984 1 626418 Proteobacteria
Burkholderia glumae BGR1 YP 002909844 1 626418 Proteobacteria
Burkholderia glumae BGR1 YP 002912133 1 626418 Proteobacteria
Burkholderia glumae BGR1 YP 002909155 1 626418 Proteobacteria
Ralstonia solanacearum Po82 YP 006029375 1 1031711 Proteobacteria
Ralstonia solanacearum Po82 YP 006031743 1 1031711 Proteobacteria
Ralstonia solanacearum Po82 YP 006029377 1 1031711 Proteobacteria
Ralstonia solanacearum Po82 YP 006031295 1 1031711 Proteobacteria
Ralstonia solanacearum Po82 YP 006030633 1 1031711 Proteobacteria
Ralstonia solanacearum Po82 YP 006030359 1 1031711 Proteobacteria
Ralstonia solanacearum Po82 YP 006031953 1 1031711 Proteobacteria
Ralstonia solanacearum Po82 YP 006028131 1 1031711 Proteobacteria
Ralstonia solanacearum Po82 YP 006030646 1 1031711 Proteobacteria
Azospirillum sp B510 YP 003450703 1 137722 Proteobacteria
AZCBGv2 p3160011ID10457497 uid000000 NC 000000 1 transposase Azospirillum brasilense CBG497 Proteobacteria
Azospirillum sp B510 YP 003453456 1 137722 Proteobacteria
Azospirillum sp B510 YP 003449961 1 137722 Proteobacteria
AZCBGv2 p3230033ID10457599 uid000000 NC 000000 1 transposase Azospirillum brasilense CBG497 Proteobacteria
AZCBGv2 p3230041ID10457607 uid000000 NC 000000 1 transposase fragment Azospirillum brasilense CBG497 Proteobacteria
Azospirillum sp B510 YP 003451339 1 137722 Proteobacteria
Azospirillum sp B510 YP 003451799 1 137722 Proteobacteria
Azospirillum sp B510 YP 003451395 1 137722 Proteobacteria
Azospirillum sp B510 YP 003447649 1 137722 Proteobacteria
Azospirillum sp B510 YP 003452239 1 137722 Proteobacteria
Azospirillum sp B510 YP 003452229 1 137722 Proteobacteria
Azospirillum sp B510 YP 003451817 1 137722 Proteobacteria
Azospirillum sp B510 YP 003453447 1 137722 Proteobacteria
AZCBGv2 p340027ID10457144 uid000000 NC 000000 1 conserved protein of unknown function Azospirillum brasilense CBG497 Proteobacteria
Xanthobacter autotrophicus Py2 YP 001419474 1 78245 Proteobacteria
Azorhizobium caulinodans ORS 571 YP 001524199 1 438753 Proteobacteria
Ochrobactrum anthropi ATCC 49188 YP 001373134 1 439375 Proteobacteria
Ochrobactrum anthropi ATCC 49188 YP 001372855 1 439375 Proteobacteria
Ochrobactrum anthropi ATCC 49188 YP 001373287 1 439375 Proteobacteria
Ochrobactrum anthropi ATCC 49188 YP 001371281 1 439375 Proteobacteria
Sinorhizobium medicae WSM419 YP 001326905 1 366394 Proteobacteria
Sinorhizobium medicae WSM419 YP 001314202 1 366394 Proteobacteria
Sinorhizobium medicae WSM419 YP 001314889 1 366394 Proteobacteria
Sinorhizobium meliloti Rm41 YP 006841329 1 1230587 Proteobacteria
Sinorhizobium meliloti Rm41 YP 006814344 1 1230587 Proteobacteria
Sinorhizobium meliloti Rm41 YP 006840994 1 1230587 Proteobacteria
Brucella abortus bv 1 str 9 941 YP 223437 1 262698 Proteobacteria
Brucella abortus S19 YP 001932578 1 430066 Proteobacteria
Brucella suis ATCC 23445 YP 001622365 1 470137 Proteobacteria
Brucella melitensis NI YP 005660911 1 1029825 Proteobacteria
Brucella suis 1330 NP 699743 1 204722 Proteobacteria
Brucella ovis ATCC 25840 YP 001257551 1 444178 Proteobacteria
Brucella melitensis ATCC 23457 YP 002734317 1 546272 Proteobacteria
Brucella microti CCM 4915 YP 003105344 1 568815 Proteobacteria
Brucella ceti TE10759 12 YP 008841427 1 1423891 Proteobacteria
Brucella melitensis bv 1 str 16M NP 541689 1 224914 Proteobacteria
Brucella pinnipedialis B2 94 YP 004757807 1 520461 Proteobacteria

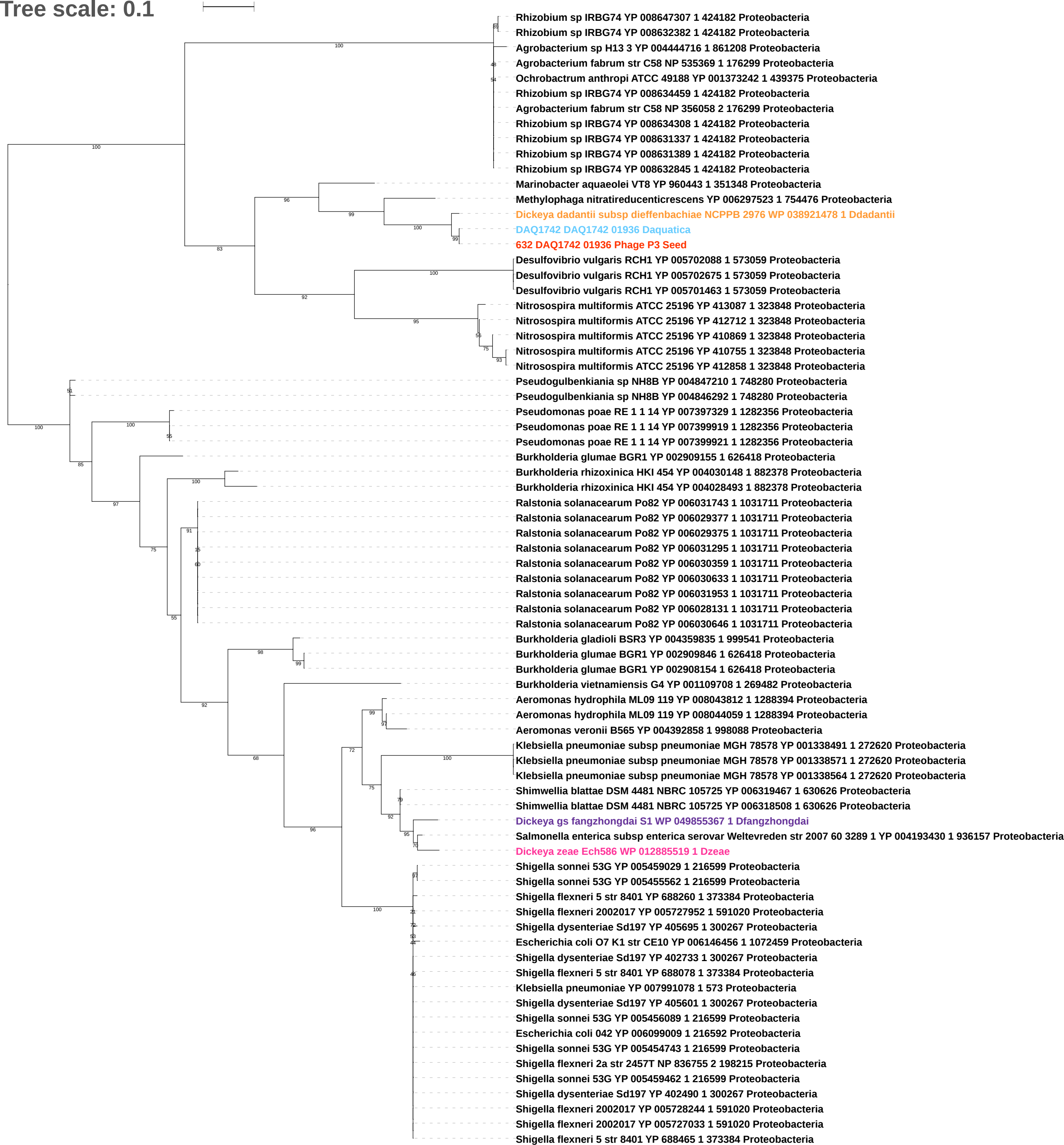
Tree scale: 0.1
Rhizobium sp IRBG74 YP 008647307 1 424182 Proteobacteria
Rhizobium sp IRBG74 YP 008632382 1 424182 Proteobacteria
Agrobacterium sp H13 3 YP 004444716 1 861208 Proteobacteria
Agrobacterium fabrum str C58 NP 535369 1 176299 Proteobacteria
Ochrobactrum anthropi ATCC 49188 YP 001373242 1 439375 Proteobacteria
Rhizobium sp IRBG74 YP 008634459 1 424182 Proteobacteria
Agrobacterium fabrum str C58 NP 356058 2 176299 Proteobacteria
Rhizobium sp IRBG74 YP 008634308 1 424182 Proteobacteria
Rhizobium sp IRBG74 YP 008631337 1 424182 Proteobacteria
Rhizobium sp IRBG74 YP 008631389 1 424182 Proteobacteria
Rhizobium sp IRBG74 YP 008632845 1 424182 Proteobacteria
Marinobacter aquaeolei VT8 YP 960443 1 351348 Proteobacteria
Methylophaga nitratireducenticrescens YP 006297523 1 754476 Proteobacteria
Dickeya dadantii subsp dieffenbachiae NCPPB 2976 WP 038921478 1 Ddadantii
DAQ1742 DAQ1742 01936 Daquatica
632 DAQ1742 01936 Phage P3 Seed
Desulfovibrio vulgaris RCH1 YP 005702088 1 573059 Proteobacteria
Desulfovibrio vulgaris RCH1 YP 005702675 1 573059 Proteobacteria
Desulfovibrio vulgaris RCH1 YP 005701463 1 573059 Proteobacteria
Nitrosospira multiformis ATCC 25196 YP 413087 1 323848 Proteobacteria
Nitrosospira multiformis ATCC 25196 YP 412712 1 323848 Proteobacteria
Nitrosospira multiformis ATCC 25196 YP 410869 1 323848 Proteobacteria
Nitrosospira multiformis ATCC 25196 YP 410755 1 323848 Proteobacteria
Nitrosospira multiformis ATCC 25196 YP 412858 1 323848 Proteobacteria
Pseudogulbenkiania sp NH8B YP 004847210 1 748280 Proteobacteria
Pseudogulbenkiania sp NH8B YP 004846292 1 748280 Proteobacteria
Pseudomonas poae RE 1 1 14 YP 007397329 1 1282356 Proteobacteria
Pseudomonas poae RE 1 1 14 YP 007399919 1 1282356 Proteobacteria
Pseudomonas poae RE 1 1 14 YP 007399921 1 1282356 Proteobacteria
Burkholderia glumae BGR1 YP 002909155 1 626418 Proteobacteria
Burkholderia rhizoxinica HKI 454 YP 004030148 1 882378 Proteobacteria
Burkholderia rhizoxinica HKI 454 YP 004028493 1 882378 Proteobacteria
Ralstonia solanacearum Po82 YP 006031743 1 1031711 Proteobacteria
Ralstonia solanacearum Po82 YP 006029377 1 1031711 Proteobacteria
Ralstonia solanacearum Po82 YP 006029375 1 1031711 Proteobacteria
Ralstonia solanacearum Po82 YP 006031295 1 1031711 Proteobacteria
Ralstonia solanacearum Po82 YP 006030359 1 1031711 Proteobacteria
Ralstonia solanacearum Po82 YP 006030633 1 1031711 Proteobacteria
Ralstonia solanacearum Po82 YP 006031953 1 1031711 Proteobacteria
Ralstonia solanacearum Po82 YP 006028131 1 1031711 Proteobacteria
Ralstonia solanacearum Po82 YP 006030646 1 1031711 Proteobacteria
Burkholderia gladioli BSR3 YP 004359835 1 999541 Proteobacteria
Burkholderia glumae BGR1 YP 002909846 1 626418 Proteobacteria
Burkholderia glumae BGR1 YP 002908154 1 626418 Proteobacteria
Burkholderia vietnamiensis G4 YP 001109708 1 269482 Proteobacteria
Aeromonas hydrophila ML09 119 YP 008043812 1 1288394 Proteobacteria
Aeromonas hydrophila ML09 119 YP 008044059 1 1288394 Proteobacteria
Aeromonas veronii B565 YP 004392858 1 998088 Proteobacteria
Klebsiella pneumoniae subsp pneumoniae MGH 78578 YP 001338491 1 272620 Proteobacteria
Klebsiella pneumoniae subsp pneumoniae MGH 78578 YP 001338571 1 272620 Proteobacteria
Klebsiella pneumoniae subsp pneumoniae MGH 78578 YP 001338564 1 272620 Proteobacteria
Shimwellia blattae DSM 4481 NBRC 105725 YP 006319467 1 630626 Proteobacteria
Shimwellia blattae DSM 4481 NBRC 105725 YP 006318508 1 630626 Proteobacteria
Dickeya gs fangzhongdai S1 WP 049855367 1 Dfangzhongdai
Salmonella enterica subsp enterica serovar Weltevreden str 2007 60 3289 1 YP 004193430 1 936157 Proteobacteria
Dickeya zeae Ech586 WP 012885519 1 Dzeae
Shigella sonnei 53G YP 005459029 1 216599 Proteobacteria
Shigella sonnei 53G YP 005455562 1 216599 Proteobacteria
Shigella flexneri 5 str 8401 YP 688260 1 373384 Proteobacteria
Shigella flexneri 2002017 YP 005727952 1 591020 Proteobacteria
Shigella dysenteriae Sd197 YP 405695 1 300267 Proteobacteria
Escherichia coli O7 K1 str CE10 YP 006146456 1 1072459 Proteobacteria
Shigella dysenteriae Sd197 YP 402733 1 300267 Proteobacteria
Shigella flexneri 5 str 8401 YP 688078 1 373384 Proteobacteria
Klebsiella pneumoniae YP 007991078 1 573 Proteobacteria
Shigella dysenteriae Sd197 YP 405601 1 300267 Proteobacteria
Shigella sonnei 53G YP 005456089 1 216599 Proteobacteria
Escherichia coli 042 YP 006099009 1 216592 Proteobacteria
Shigella sonnei 53G YP 005454743 1 216599 Proteobacteria
Shigella flexneri 2a str 2457T NP 836755 2 198215 Proteobacteria
Shigella sonnei 53G YP 005459462 1 216599 Proteobacteria
Shigella dysenteriae Sd197 YP 402490 1 300267 Proteobacteria
Shigella flexneri 2002017 YP 005728244 1 591020 Proteobacteria
Shigella flexneri 2002017 YP 005727033 1 591020 Proteobacteria
Shigella flexneri 5 str 8401 YP 688465 1 373384 Proteobacteria

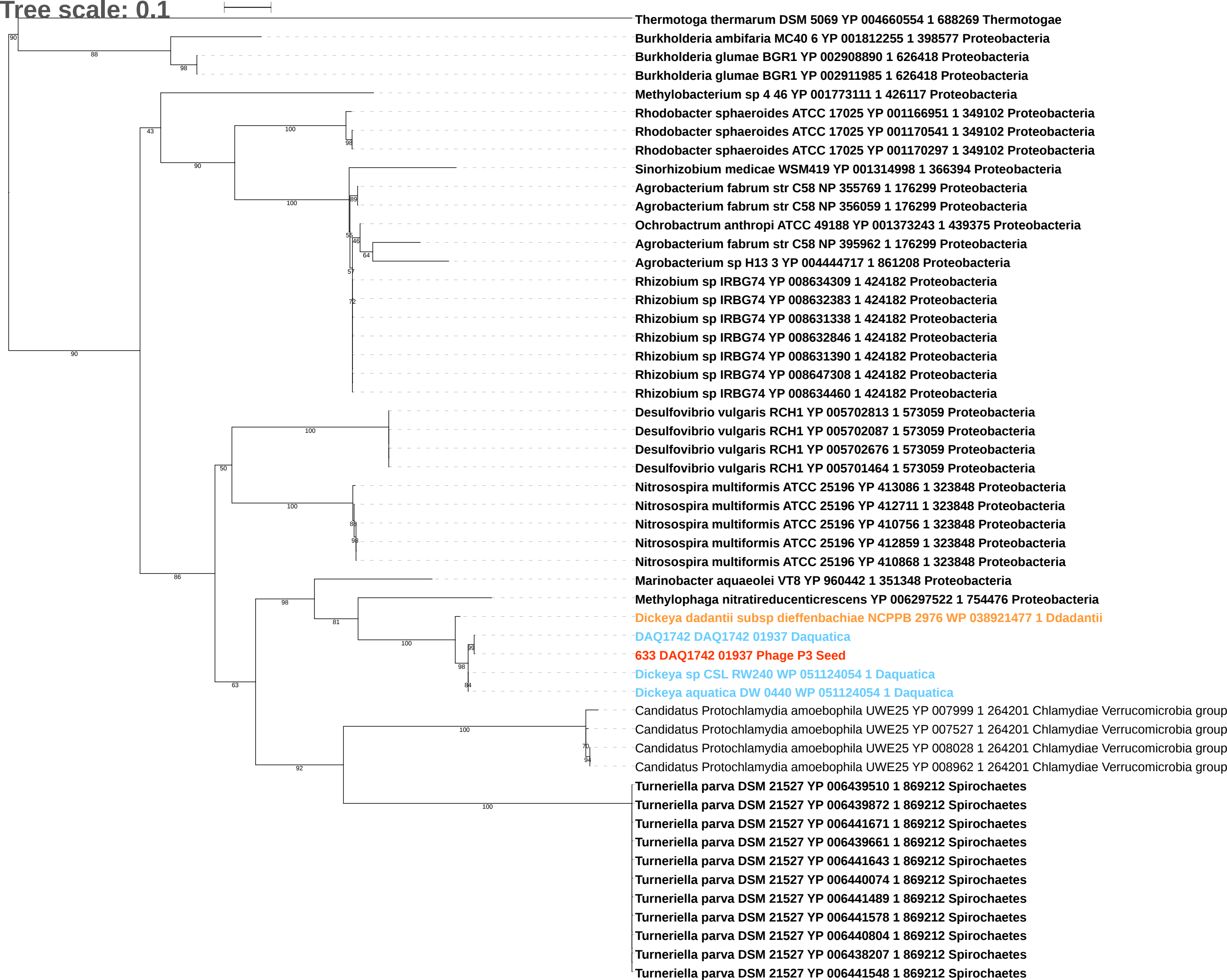
Tree scale: 0.1
Thermotoga thermarum DSM 5069 YP 004660554 1 688269 Thermotogae
Burkholderia ambifaria MC40 6 YP 001812255 1 398577 Proteobacteria
Burkholderia glumae BGR1 YP 002908890 1 626418 Proteobacteria
Burkholderia glumae BGR1 YP 002911985 1 626418 Proteobacteria
Methylobacterium sp 4 46 YP 001773111 1 426117 Proteobacteria
Rhodobacter sphaeroides ATCC 17025 YP 001166951 1 349102 Proteobacteria
Rhodobacter sphaeroides ATCC 17025 YP 001170541 1 349102 Proteobacteria
Rhodobacter sphaeroides ATCC 17025 YP 001170297 1 349102 Proteobacteria
Sinorhizobium medicae WSM419 YP 001314998 1 366394 Proteobacteria
Agrobacterium fabrum str C58 NP 355769 1 176299 Proteobacteria
Agrobacterium fabrum str C58 NP 356059 1 176299 Proteobacteria
Ochrobactrum anthropi ATCC 49188 YP 001373243 1 439375 Proteobacteria
Agrobacterium fabrum str C58 NP 395962 1 176299 Proteobacteria
Agrobacterium sp H13 3 YP 004444717 1 861208 Proteobacteria
Rhizobium sp IRBG74 YP 008634309 1 424182 Proteobacteria
Rhizobium sp IRBG74 YP 008632383 1 424182 Proteobacteria
Rhizobium sp IRBG74 YP 008631338 1 424182 Proteobacteria
Rhizobium sp IRBG74 YP 008632846 1 424182 Proteobacteria
Rhizobium sp IRBG74 YP 008631390 1 424182 Proteobacteria
Rhizobium sp IRBG74 YP 008647308 1 424182 Proteobacteria
Rhizobium sp IRBG74 YP 008634460 1 424182 Proteobacteria
Desulfovibrio vulgaris RCH1 YP 005702813 1 573059 Proteobacteria
Desulfovibrio vulgaris RCH1 YP 005702087 1 573059 Proteobacteria
Desulfovibrio vulgaris RCH1 YP 005702676 1 573059 Proteobacteria
Desulfovibrio vulgaris RCH1 YP 005701464 1 573059 Proteobacteria
Nitrosospira multiformis ATCC 25196 YP 413086 1 323848 Proteobacteria
Nitrosospira multiformis ATCC 25196 YP 412711 1 323848 Proteobacteria
Nitrosospira multiformis ATCC 25196 YP 410756 1 323848 Proteobacteria
Nitrosospira multiformis ATCC 25196 YP 412859 1 323848 Proteobacteria
Nitrosospira multiformis ATCC 25196 YP 410868 1 323848 Proteobacteria
Marinobacter aquaeolei VT8 YP 960442 1 351348 Proteobacteria
Methylophaga nitratireducenticrescens YP 006297522 1 754476 Proteobacteria
Dickeya dadantii subsp dieffenbachiae NCPPB 2976 WP 038921477 1 Ddadantii
DAQ1742 DAQ1742 01937 Daquatica
633 DAQ1742 01937 Phage P3 Seed
Dickeya sp CSL RW240 WP 051124054 1 Daquatica
Dickeya aquatica DW 0440 WP 051124054 1 Daquatica
Candidatus Protochlamydia amoebophila UWE25 YP 007999 1 264201 Chlamydiae Verrucomicrobia group
Candidatus Protochlamydia amoebophila UWE25 YP 007527 1 264201 Chlamydiae Verrucomicrobia group
Candidatus Protochlamydia amoebophila UWE25 YP 008028 1 264201 Chlamydiae Verrucomicrobia group
Candidatus Protochlamydia amoebophila UWE25 YP 008962 1 264201 Chlamydiae Verrucomicrobia group
Turneriella parva DSM 21527 YP 006439510 1 869212 Spirochaetes
Turneriella parva DSM 21527 YP 006439872 1 869212 Spirochaetes
Turneriella parva DSM 21527 YP 006441671 1 869212 Spirochaetes
Turneriella parva DSM 21527 YP 006439661 1 869212 Spirochaetes
Turneriella parva DSM 21527 YP 006441643 1 869212 Spirochaetes
Turneriella parva DSM 21527 YP 006440074 1 869212 Spirochaetes
Turneriella parva DSM 21527 YP 006441489 1 869212 Spirochaetes
Turneriella parva DSM 21527 YP 006441578 1 869212 Spirochaetes
Turneriella parva DSM 21527 YP 006440804 1 869212 Spirochaetes
Turneriella parva DSM 21527 YP 006438207 1 869212 Spirochaetes
Turneriella parva DSM 21527 YP 006441548 1 869212 Spirochaetes

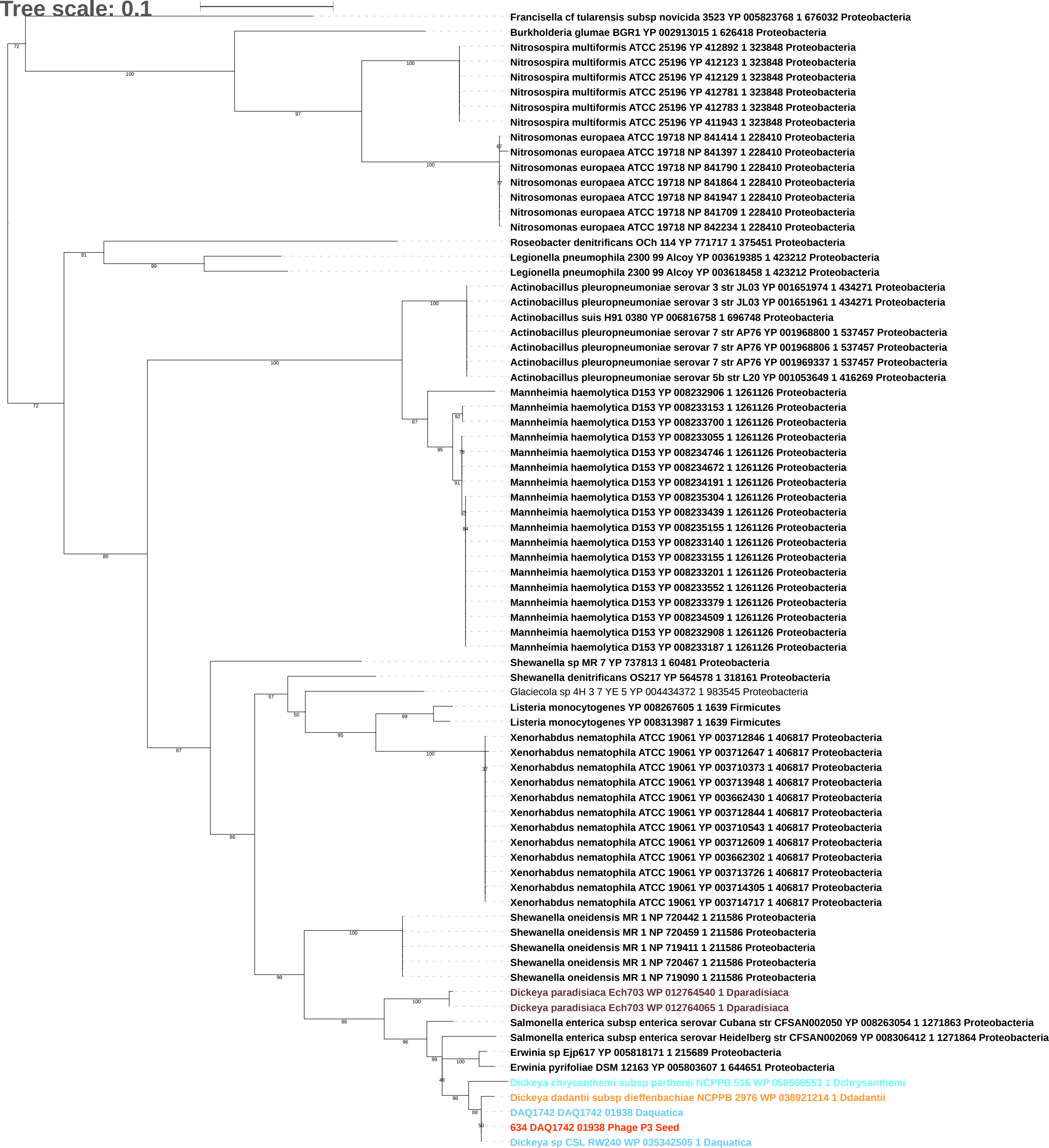
Tree scale: 0.1
Francisella cf tularensis subsp novicida 3523 YP 005823768 1 676032 Proteobacteria
Burkholderia glumae BGR1 YP 002913015 1 626418 Proteobacteria
Nitrosospira multiformis ATCC 25196 YP 412892 1 323848 Proteobacteria
Nitrosospira multiformis ATCC 25196 YP 412123 1 323848 Proteobacteria
Nitrosospira multiformis ATCC 25196 YP 412129 1 323848 Proteobacteria
Nitrosospira multiformis ATCC 25196 YP 412781 1 323848 Proteobacteria
Nitrosospira multiformis ATCC 25196 YP 412783 1 323848 Proteobacteria
Nitrosospira multiformis ATCC 25196 YP 411943 1 323848 Proteobacteria
Nitrosomonas europaea ATCC 19718 NP 841414 1 228410 Proteobacteria
Nitrosomonas europaea ATCC 19718 NP 841397 1 228410 Proteobacteria
Nitrosomonas europaea ATCC 19718 NP 841790 1 228410 Proteobacteria
Nitrosomonas europaea ATCC 19718 NP 841864 1 228410 Proteobacteria
Nitrosomonas europaea ATCC 19718 NP 841947 1 228410 Proteobacteria
Nitrosomonas europaea ATCC 19718 NP 841709 1 228410 Proteobacteria
Nitrosomonas europaea ATCC 19718 NP 842234 1 228410 Proteobacteria
Roseobacter denitrificans OCh 114 YP 771717 1 375451 Proteobacteria
Legionella pneumophila 2300 99 Alcoy YP 003619385 1 423212 Proteobacteria
Legionella pneumophila 2300 99 Alcoy YP 003618458 1 423212 Proteobacteria
Actinobacillus pleuropneumoniae serovar 3 str JL03 YP 001651974 1 434271 Proteobacteria
Actinobacillus pleuropneumoniae serovar 3 str JL03 YP 001651961 1 434271 Proteobacteria
Actinobacillus suis H91 0380 YP 006816758 1 696748 Proteobacteria
Actinobacillus pleuropneumoniae serovar 7 str AP76 YP 001968800 1 537457 Proteobacteria
Actinobacillus pleuropneumoniae serovar 7 str AP76 YP 001968806 1 537457 Proteobacteria
Actinobacillus pleuropneumoniae serovar 7 str AP76 YP 001969337 1 537457 Proteobacteria
Actinobacillus pleuropneumoniae serovar 5b str L20 YP 001053649 1 416269 Proteobacteria
Mannheimia haemolytica D153 YP 008232906 1 1261126 Proteobacteria
Mannheimia haemolytica D153 YP 008233153 1 1261126 Proteobacteria
Mannheimia haemolytica D153 YP 008233700 1 1261126 Proteobacteria
Mannheimia haemolytica D153 YP 008233055 1 1261126 Proteobacteria
Mannheimia haemolytica D153 YP 008234746 1 1261126 Proteobacteria
Mannheimia haemolytica D153 YP 008234672 1 1261126 Proteobacteria
Mannheimia haemolytica D153 YP 008234191 1 1261126 Proteobacteria
Mannheimia haemolytica D153 YP 008235304 1 1261126 Proteobacteria
Mannheimia haemolytica D153 YP 008233439 1 1261126 Proteobacteria
Mannheimia haemolytica D153 YP 008235155 1 1261126 Proteobacteria
Mannheimia haemolytica D153 YP 008233140 1 1261126 Proteobacteria
Mannheimia haemolytica D153 YP 008233155 1 1261126 Proteobacteria
Mannheimia haemolytica D153 YP 008233201 1 1261126 Proteobacteria
Mannheimia haemolytica D153 YP 008233552 1 1261126 Proteobacteria
Mannheimia haemolytica D153 YP 008233379 1 1261126 Proteobacteria
Mannheimia haemolytica D153 YP 008234509 1 1261126 Proteobacteria
Mannheimia haemolytica D153 YP 008232908 1 1261126 Proteobacteria
Mannheimia haemolytica D153 YP 008233187 1 1261126 Proteobacteria
Shewanella sp MR 7 YP 737813 1 60481 Proteobacteria
Shewanella denitrificans OS217 YP 564578 1 318161 Proteobacteria
Glaciecola sp 4H 3 7 YE 5 YP 004434372 1 983545 Proteobacteria
Listeria monocytogenes YP 008267605 1 1639 Firmicutes
Listeria monocytogenes YP 008313987 1 1639 Firmicutes
Xenorhabdus nematophila ATCC 19061 YP 003712846 1 406817 Proteobacteria
Xenorhabdus nematophila ATCC 19061 YP 003712647 1 406817 Proteobacteria
Xenorhabdus nematophila ATCC 19061 YP 003710373 1 406817 Proteobacteria
Xenorhabdus nematophila ATCC 19061 YP 003713948 1 406817 Proteobacteria
Xenorhabdus nematophila ATCC 19061 YP 003662430 1 406817 Proteobacteria
Xenorhabdus nematophila ATCC 19061 YP 003712844 1 406817 Proteobacteria
Xenorhabdus nematophila ATCC 19061 YP 003710543 1 406817 Proteobacteria
Xenorhabdus nematophila ATCC 19061 YP 003712609 1 406817 Proteobacteria
Xenorhabdus nematophila ATCC 19061 YP 003662302 1 406817 Proteobacteria
Xenorhabdus nematophila ATCC 19061 YP 003713726 1 406817 Proteobacteria
Xenorhabdus nematophila ATCC 19061 YP 003714305 1 406817 Proteobacteria
Xenorhabdus nematophila ATCC 19061 YP 003714717 1 406817 Proteobacteria
Shewanella oneidensis MR 1 NP 720442 1 211586 Proteobacteria
Shewanella oneidensis MR 1 NP 720459 1 211586 Proteobacteria
Shewanella oneidensis MR 1 NP 719411 1 211586 Proteobacteria
Shewanella oneidensis MR 1 NP 720467 1 211586 Proteobacteria
Shewanella oneidensis MR 1 NP 719090 1 211586 Proteobacteria
Dickeya paradisiaca Ech703 WP 012764540 1 Dparadisiaca
Dickeya paradisiaca Ech703 WP 012764065 1 Dparadisiaca
Salmonella enterica subsp enterica serovar Cubana str CFSAN002050 YP 008263054 1 1271863 Proteobacteria
Salmonella enterica subsp enterica serovar Heidelberg str CFSAN002069 YP 008306412 1 1271864 Proteobacteria
Erwinia sp Ejp617 YP 005818171 1 215689 Proteobacteria
Erwinia pyrifoliae DSM 12163 YP 005803607 1 644651 Proteobacteria
Dickeya chrysanthemi subsp parthenii NCPPB 516 WP 050568553 1 Dchrysanthemi
Dickeya dadantii subsp dieffenbachiae NCPPB 2976 WP 038921214 1 Ddadantii
DAQ1742 DAQ1742 01938 Daquatica
634 DAQ1742 01938 Phage P3 Seed
Dickeya sp CSL RW240 WP 035342505 1 Daquatica

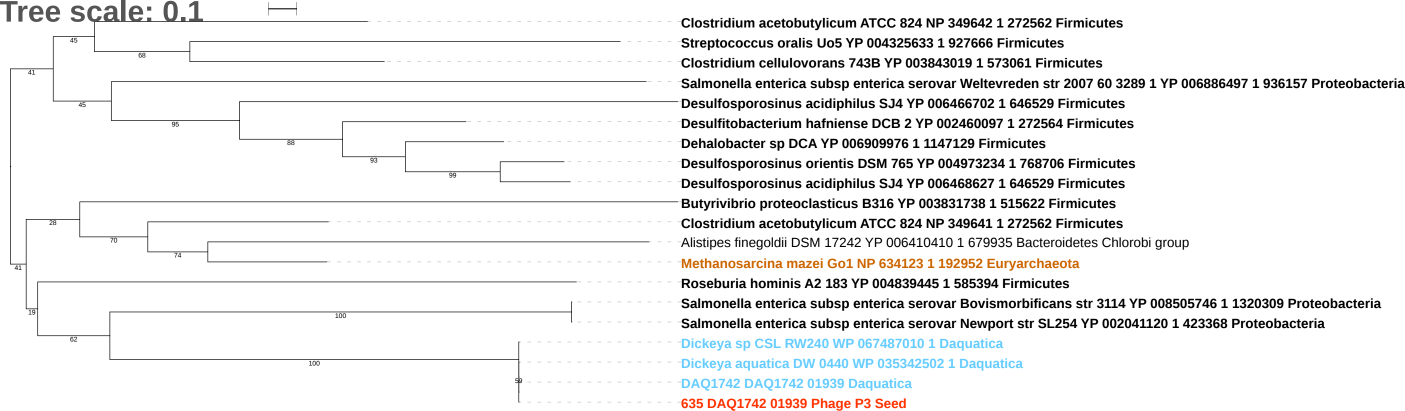
Tree scale: 0.1
Clostridium acetobutylicum ATCC 824 NP 349642 1 272562 Firmicutes
Streptococcus oralis Uo5 YP 004325633 1 927666 Firmicutes
Clostridium cellulovorans 743B YP 003843019 1 573061 Firmicutes
Salmonella enterica subsp enterica serovar Weltevreden str 2007 60 3289 1 YP 006886497 1 936157 Proteobacteria
Desulfosporosinus acidiphilus SJ4 YP 006466702 1 646529 Firmicutes
Desulfitobacterium hafniense DCB 2 YP 002460097 1 272564 Firmicutes
Dehalobacter sp DCA YP 006909976 1 1147129 Firmicutes
Desulfosporosinus orientis DSM 765 YP 004973234 1 768706 Firmicutes
Desulfosporosinus acidiphilus SJ4 YP 006468627 1 646529 Firmicutes
Butyrivibrio proteoclasticus B316 YP 003831738 1 515622 Firmicutes
Clostridium acetobutylicum ATCC 824 NP 349641 1 272562 Firmicutes
Alistipes finegoldii DSM 17242 YP 006410410 1 679935 Bacteroidetes Chlorobi group
Methanosarcina mazei Go1 NP 634123 1 192952 Euryarchaeota
Roseburia hominis A2 183 YP 004839445 1 585394 Firmicutes
Salmonella enterica subsp enterica serovar Bovismorbificans str 3114 YP 008505746 1 1320309 Proteobacteria
Salmonella enterica subsp enterica serovar Newport str SL254 YP 002041120 1 423368 Proteobacteria
Dickeya sp CSL RW240 WP 067487010 1 Daquatica
Dickeya aquatica DW 0440 WP 035342502 1 Daquatica
DAQ1742 DAQ1742 01939 Daquatica
635 DAQ1742 01939 Phage P3 Seed
45
68
41
45
95
88
93
99
28
70
74
41
19
100
62
100
59

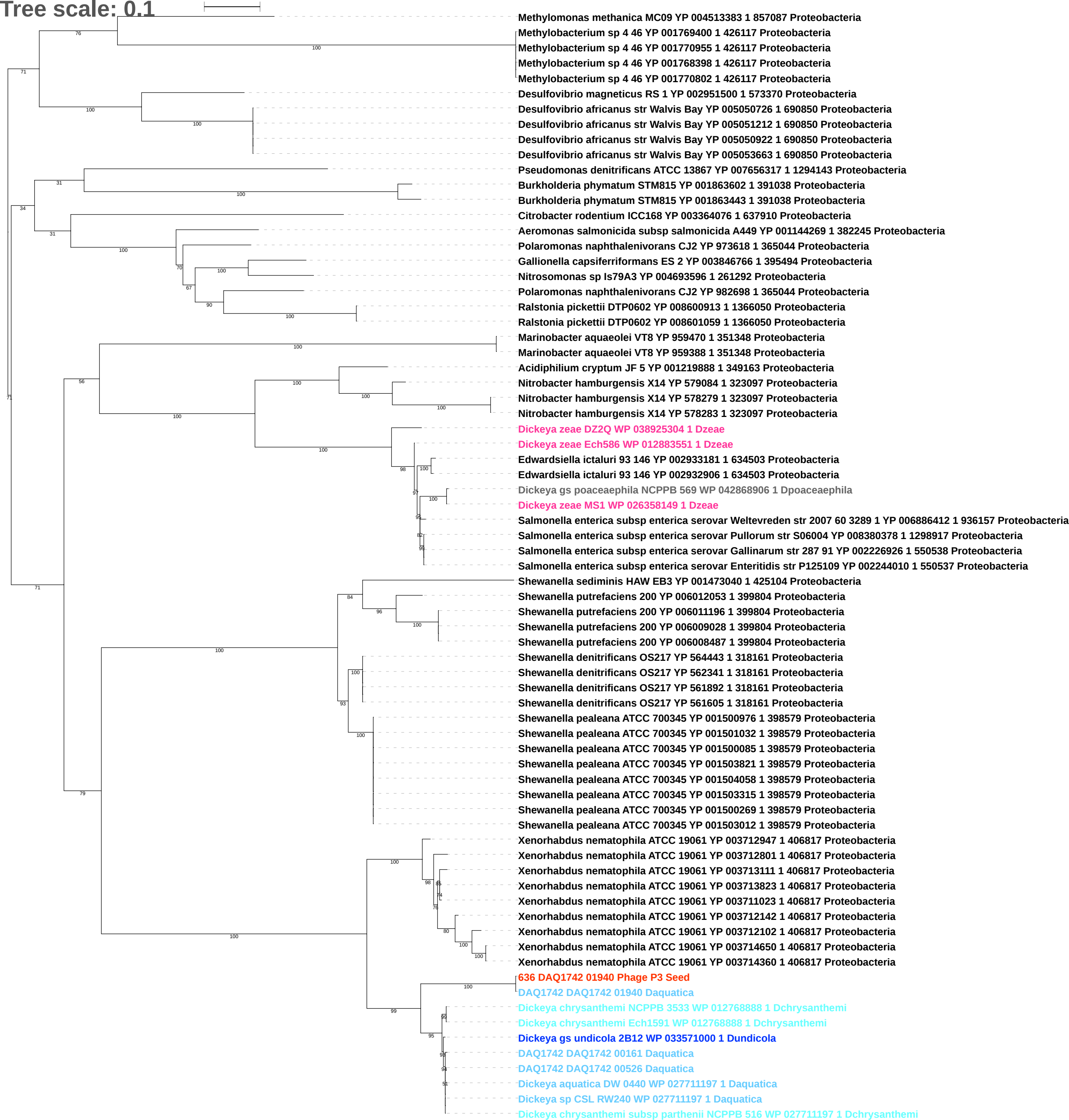
Tree scale: 0.1
Methylomonas methanica MC09 YP 004513383 1 857087 Proteobacteria
Methylobacterium sp 4 46 YP 001769400 1 426117 Proteobacteria
Methylobacterium sp 4 46 YP 001770955 1 426117 Proteobacteria
Methylobacterium sp 4 46 YP 001768398 1 426117 Proteobacteria
Methylobacterium sp 4 46 YP 001770802 1 426117 Proteobacteria
Desulfovibrio magneticus RS 1 YP 002951500 1 573370 Proteobacteria
Desulfovibrio africanus str Walvis Bay YP 005050726 1 690850 Proteobacteria
Desulfovibrio africanus str Walvis Bay YP 005051212 1 690850 Proteobacteria
Desulfovibrio africanus str Walvis Bay YP 005050922 1 690850 Proteobacteria
Desulfovibrio africanus str Walvis Bay YP 005053663 1 690850 Proteobacteria
Pseudomonas denitrificans ATCC 13867 YP 007656317 1 1294143 Proteobacteria
Burkholderia phymatum STM815 YP 001863602 1 391038 Proteobacteria
Burkholderia phymatum STM815 YP 001863443 1 391038 Proteobacteria
Citrobacter rodentium ICC168 YP 003364076 1 637910 Proteobacteria
Aeromonas salmonicida subsp salmonicida A449 YP 001144269 1 382245 Proteobacteria
Polaromonas naphthalenivorans CJ2 YP 973618 1 365044 Proteobacteria
Gallionella capsiferriformans ES 2 YP 003846766 1 395494 Proteobacteria
Nitrosomonas sp Is79A3 YP 004693596 1 261292 Proteobacteria
Polaromonas naphthalenivorans CJ2 YP 982698 1 365044 Proteobacteria
Ralstonia pickettii DTP0602 YP 008600913 1 1366050 Proteobacteria
Ralstonia pickettii DTP0602 YP 008601059 1 1366050 Proteobacteria
Marinobacter aquaeolei VT8 YP 959470 1 351348 Proteobacteria
Marinobacter aquaeolei VT8 YP 959388 1 351348 Proteobacteria
Acidiphilium cryptum JF 5 YP 001219888 1 349163 Proteobacteria
Nitrobacter hamburgensis X14 YP 579084 1 323097 Proteobacteria
Nitrobacter hamburgensis X14 YP 578279 1 323097 Proteobacteria
Nitrobacter hamburgensis X14 YP 578283 1 323097 Proteobacteria
Dickeya zeae DZ2Q WP 038925304 1 Dzeae
Dickeya zeae Ech586 WP 012883551 1 Dzeae
Edwardsiella ictaluri 93 146 YP 002933181 1 634503 Proteobacteria
Edwardsiella ictaluri 93 146 YP 002932906 1 634503 Proteobacteria
Dickeya gs poaceaephila NCPPB 569 WP 042868906 1 Dpoaceaephila
Dickeya zeae MS1 WP 026358149 1 Dzeae
Salmonella enterica subsp enterica serovar Weltevreden str 2007 60 3289 1 YP 006886412 1 936157 Proteobacteria
Salmonella enterica subsp enterica serovar Pullorum str S06004 YP 008380378 1 1298917 Proteobacteria
Salmonella enterica subsp enterica serovar Gallinarum str 287 91 YP 002226926 1 550538 Proteobacteria
Salmonella enterica subsp enterica serovar Enteritidis str P125109 YP 002244010 1 550537 Proteobacteria
Shewanella sediminis HAW EB3 YP 001473040 1 425104 Proteobacteria
Shewanella putrefaciens 200 YP 006012053 1 399804 Proteobacteria
Shewanella putrefaciens 200 YP 006011196 1 399804 Proteobacteria
Shewanella putrefaciens 200 YP 006009028 1 399804 Proteobacteria
Shewanella putrefaciens 200 YP 006008487 1 399804 Proteobacteria
Shewanella denitrificans OS217 YP 564443 1 318161 Proteobacteria
Shewanella denitrificans OS217 YP 562341 1 318161 Proteobacteria
Shewanella denitrificans OS217 YP 561892 1 318161 Proteobacteria
Shewanella denitrificans OS217 YP 561605 1 318161 Proteobacteria
Shewanella pealeana ATCC 700345 YP 001500976 1 398579 Proteobacteria
Shewanella pealeana ATCC 700345 YP 001501032 1 398579 Proteobacteria
Shewanella pealeana ATCC 700345 YP 001500085 1 398579 Proteobacteria
Shewanella pealeana ATCC 700345 YP 001503821 1 398579 Proteobacteria
Shewanella pealeana ATCC 700345 YP 001504058 1 398579 Proteobacteria
Shewanella pealeana ATCC 700345 YP 001503315 1 398579 Proteobacteria
Shewanella pealeana ATCC 700345 YP 001500269 1 398579 Proteobacteria
Shewanella pealeana ATCC 700345 YP 001503012 1 398579 Proteobacteria
Xenorhabdus nematophila ATCC 19061 YP 003712947 1 406817 Proteobacteria
Xenorhabdus nematophila ATCC 19061 YP 003712801 1 406817 Proteobacteria
Xenorhabdus nematophila ATCC 19061 YP 003713111 1 406817 Proteobacteria
Xenorhabdus nematophila ATCC 19061 YP 003713823 1 406817 Proteobacteria
Xenorhabdus nematophila ATCC 19061 YP 003711023 1 406817 Proteobacteria
Xenorhabdus nematophila ATCC 19061 YP 003712142 1 406817 Proteobacteria
Xenorhabdus nematophila ATCC 19061 YP 003712102 1 406817 Proteobacteria
Xenorhabdus nematophila ATCC 19061 YP 003714650 1 406817 Proteobacteria
Xenorhabdus nematophila ATCC 19061 YP 003714360 1 406817 Proteobacteria
636 DAQ1742 01940 Phage P3 Seed
DAQ1742 DAQ1742 01940 Daquatica
Dickeya chrysanthemi NCPPB 3533 WP 012768888 1 Dchrysanthemi
Dickeya chrysanthemi Ech1591 WP 012768888 1 Dchrysanthemi
Dickeya gs undicola 2B12 WP 033571000 1 Dundicola
DAQ1742 DAQ1742 00161 Daquatica
DAQ1742 DAQ1742 00526 Daquatica
Dickeya aquatica DW 0440 WP 027711197 1 Daquatica
Dickeya sp CSL RW240 WP 027711197 1 Daquatica
Dickeya chrysanthemi subsp parthenii NCPPB 516 WP 027711197 1 Dchrysanthemi

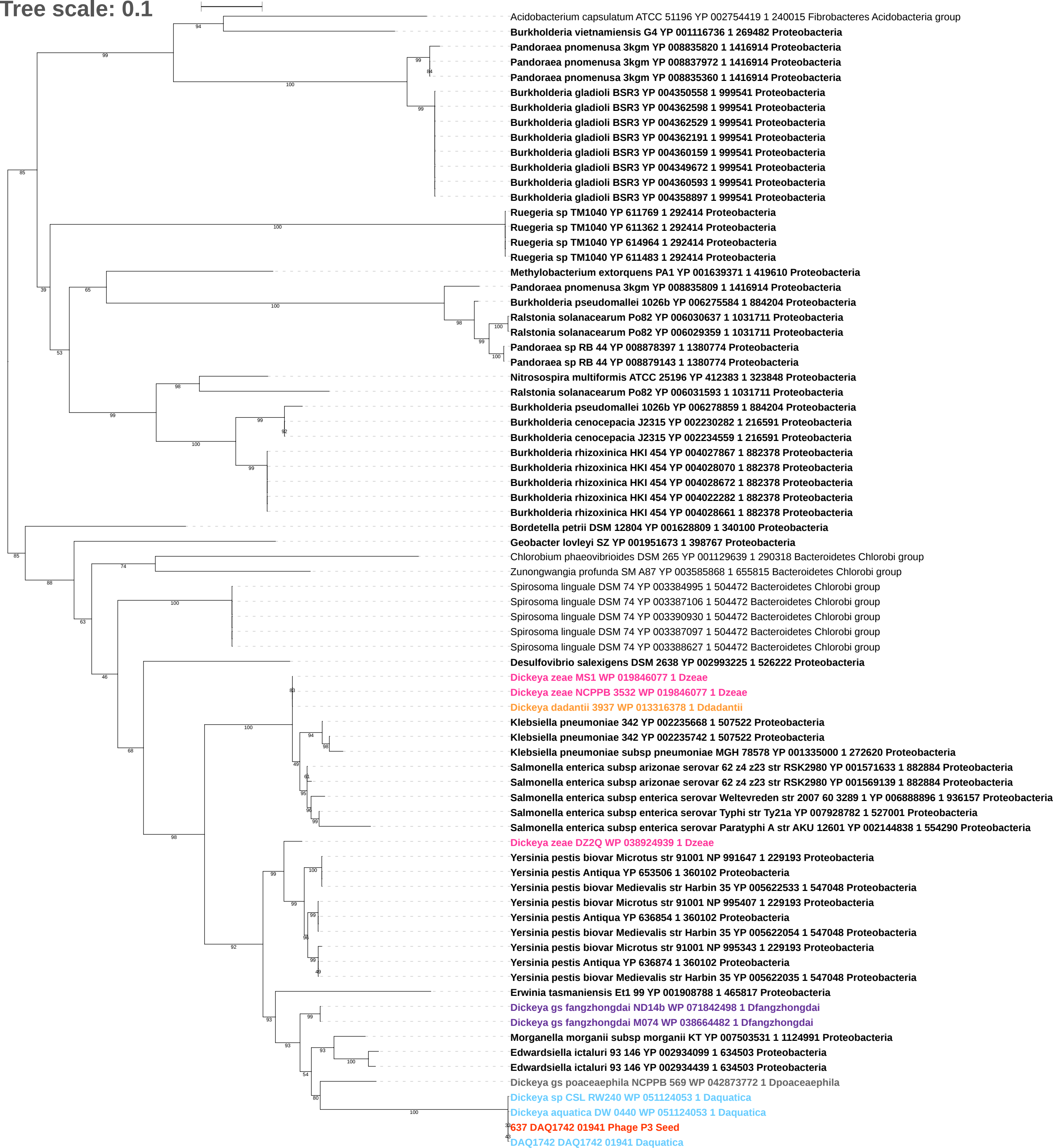
Tree scale: 0.1
Acidobacterium capsulatum ATCC 51196 YP 002754419 1 240015 Fibrobacteres Acidobacteria group
Burkholderia vietnamiensis G4 YP 001116736 1 269482 Proteobacteria
Pandoraea pnomenusa 3kgm YP 008835820 1 1416914 Proteobacteria
Pandoraea pnomenusa 3kgm YP 008837972 1 1416914 Proteobacteria
Pandoraea pnomenusa 3kgm YP 008835360 1 1416914 Proteobacteria
Burkholderia gladioli BSR3 YP 004350558 1 999541 Proteobacteria
Burkholderia gladioli BSR3 YP 004362598 1 999541 Proteobacteria
Burkholderia gladioli BSR3 YP 004362529 1 999541 Proteobacteria
Burkholderia gladioli BSR3 YP 004362191 1 999541 Proteobacteria
Burkholderia gladioli BSR3 YP 004360159 1 999541 Proteobacteria
Burkholderia gladioli BSR3 YP 004349672 1 999541 Proteobacteria
Burkholderia gladioli BSR3 YP 004360593 1 999541 Proteobacteria
Burkholderia gladioli BSR3 YP 004358897 1 999541 Proteobacteria
Ruegeria sp TM1040 YP 611769 1 292414 Proteobacteria
Ruegeria sp TM1040 YP 611362 1 292414 Proteobacteria
Ruegeria sp TM1040 YP 614964 1 292414 Proteobacteria
Ruegeria sp TM1040 YP 611483 1 292414 Proteobacteria
Methylobacterium extorquens PA1 YP 001639371 1 419610 Proteobacteria
Pandoraea pnomenusa 3kgm YP 008835809 1 1416914 Proteobacteria
Burkholderia pseudomallei 1026b YP 006275584 1 884204 Proteobacteria
Ralstonia solanacearum Po82 YP 006030637 1 1031711 Proteobacteria
Ralstonia solanacearum Po82 YP 006029359 1 1031711 Proteobacteria
Pandoraea sp RB 44 YP 008878397 1 1380774 Proteobacteria
Pandoraea sp RB 44 YP 008879143 1 1380774 Proteobacteria
Nitrosospira multiformis ATCC 25196 YP 412383 1 323848 Proteobacteria
Ralstonia solanacearum Po82 YP 006031593 1 1031711 Proteobacteria
Burkholderia pseudomallei 1026b YP 006278859 1 884204 Proteobacteria
Burkholderia cenocepacia J2315 YP 002230282 1 216591 Proteobacteria
Burkholderia cenocepacia J2315 YP 002234559 1 216591 Proteobacteria
Burkholderia rhizoxinica HKI 454 YP 004027867 1 882378 Proteobacteria
Burkholderia rhizoxinica HKI 454 YP 004028070 1 882378 Proteobacteria
Burkholderia rhizoxinica HKI 454 YP 004028672 1 882378 Proteobacteria
Burkholderia rhizoxinica HKI 454 YP 004022282 1 882378 Proteobacteria
Burkholderia rhizoxinica HKI 454 YP 004028661 1 882378 Proteobacteria
Bordetella petrii DSM 12804 YP 001628809 1 340100 Proteobacteria
Geobacter lovleyi SZ YP 001951673 1 398767 Proteobacteria
Chlorobium phaeovibrioides DSM 265 YP 001129639 1 290318 Bacteroidetes Chlorobi group
Zunongwangia profunda SM A87 YP 003585868 1 655815 Bacteroidetes Chlorobi group
Spirosoma linguale DSM 74 YP 003384995 1 504472 Bacteroidetes Chlorobi group
Spirosoma linguale DSM 74 YP 003387106 1 504472 Bacteroidetes Chlorobi group
Spirosoma linguale DSM 74 YP 003390930 1 504472 Bacteroidetes Chlorobi group
Spirosoma linguale DSM 74 YP 003387097 1 504472 Bacteroidetes Chlorobi group
Spirosoma linguale DSM 74 YP 003388627 1 504472 Bacteroidetes Chlorobi group
Desulfovibrio salexigens DSM 2638 YP 002993225 1 526222 Proteobacteria
Dickeya zeae MS1 WP 019846077 1 Dzeae
Dickeya zeae NCPPB 3532 WP 019846077 1 Dzeae
Dickeya dadantii 3937 WP 013316378 1 Ddadantii
Klebsiella pneumoniae 342 YP 002235668 1 507522 Proteobacteria
Klebsiella pneumoniae 342 YP 002235742 1 507522 Proteobacteria
Klebsiella pneumoniae subsp pneumoniae MGH 78578 YP 001335000 1 272620 Proteobacteria
Salmonella enterica subsp arizonae serovar 62 z4 z23 str RSK2980 YP 001571633 1 882884 Proteobacteria
Salmonella enterica subsp arizonae serovar 62 z4 z23 str RSK2980 YP 001569139 1 882884 Proteobacteria
Salmonella enterica subsp enterica serovar Weltevreden str 2007 60 3289 1 YP 006888896 1 936157 Proteobacteria
Salmonella enterica subsp enterica serovar Typhi str Ty21a YP 007928782 1 527001 Proteobacteria
Salmonella enterica subsp enterica serovar Paratyphi A str AKU 12601 YP 002144838 1 554290 Proteobacteria
Dickeya zeae DZ2Q WP 038924939 1 Dzeae
Yersinia pestis biovar Microtus str 91001 NP 991647 1 229193 Proteobacteria
Yersinia pestis Antiqua YP 653506 1 360102 Proteobacteria
Yersinia pestis biovar Medievalis str Harbin 35 YP 005622533 1 547048 Proteobacteria
Yersinia pestis biovar Microtus str 91001 NP 995407 1 229193 Proteobacteria
Yersinia pestis Antiqua YP 636854 1 360102 Proteobacteria
Yersinia pestis biovar Medievalis str Harbin 35 YP 005622054 1 547048 Proteobacteria
Yersinia pestis biovar Microtus str 91001 NP 995343 1 229193 Proteobacteria
Yersinia pestis Antiqua YP 636874 1 360102 Proteobacteria
Yersinia pestis biovar Medievalis str Harbin 35 YP 005622035 1 547048 Proteobacteria
Erwinia tasmaniensis Et1 99 YP 001908788 1 465817 Proteobacteria
Dickeya gs fangzhongdai ND14b WP 071842498 1 Dfangzhongdai
Dickeya gs fangzhongdai M074 WP 038664482 1 Dfangzhongdai
Morganella morganii subsp morganii KT YP 007503531 1 1124991 Proteobacteria
Edwardsiella ictaluri 93 146 YP 002934099 1 634503 Proteobacteria
Edwardsiella ictaluri 93 146 YP 002934439 1 634503 Proteobacteria
Dickeya gs poaceaephila NCPPB 569 WP 042873772 1 Dpoaceaephila
Dickeya sp CSL RW240 WP 051124053 1 Daquatica
Dickeya aquatica DW 0440 WP 051124053 1 Daquatica
637 DAQ1742 01941 Phage P3 Seed
DAQ1742 DAQ1742 01941 Daquatica

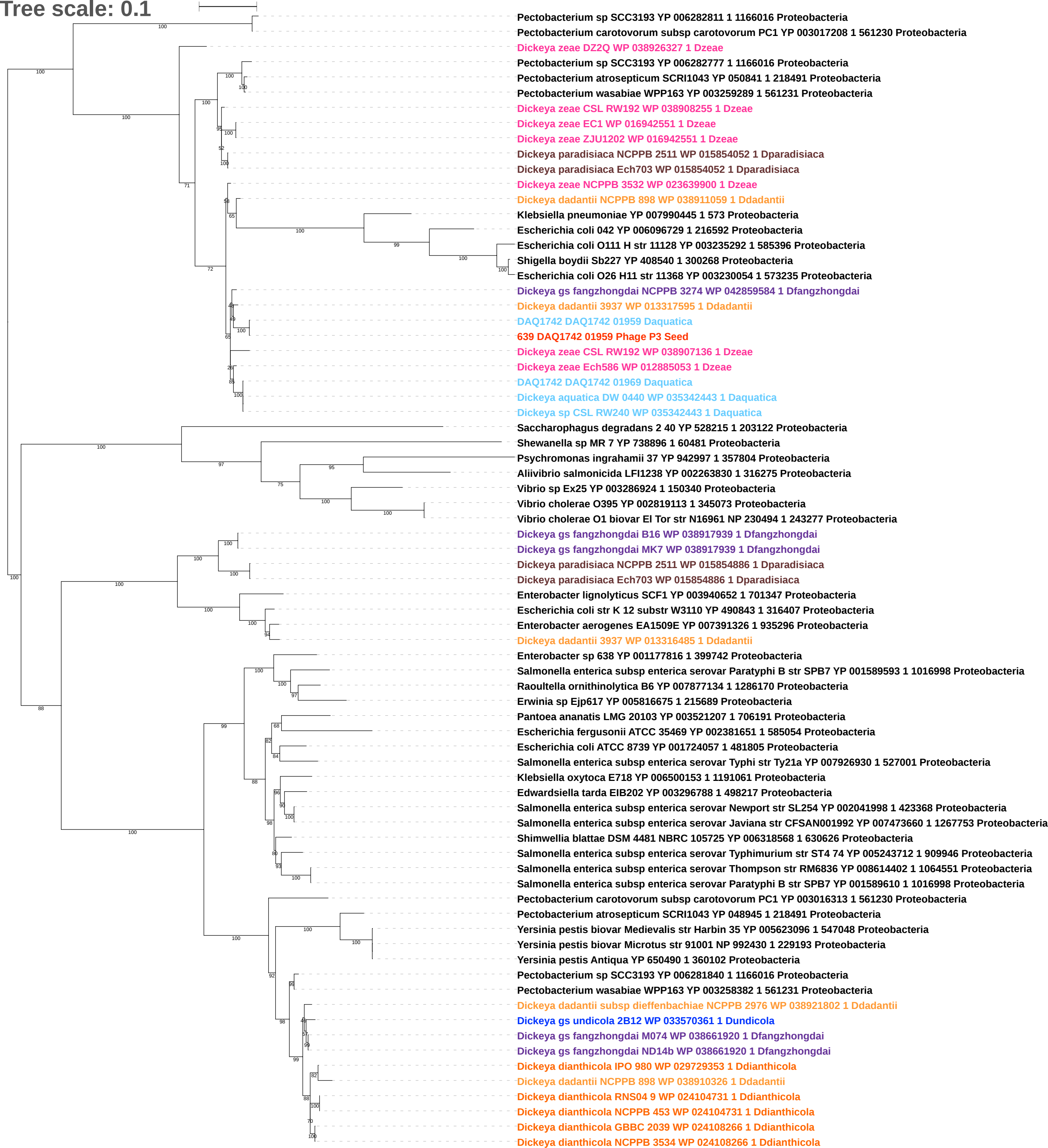
Tree scale: 0.1
Pectobacterium sp SCC3193 YP 006282811 1 1166016 Proteobacteria
Pectobacterium carotovorum subsp carotovorum PC1 YP 003017208 1 561230 Proteobacteria
Dickeya zeae DZ2Q WP 038926327 1 Dzeae
Pectobacterium sp SCC3193 YP 006282777 1 1166016 Proteobacteria
Pectobacterium atrosepticum SCRI1043 YP 050841 1 218491 Proteobacteria
Pectobacterium wasabiae WPP163 YP 003259289 1 561231 Proteobacteria
Dickeya zeae CSL RW192 WP 038908255 1 Dzeae
Dickeya zeae EC1 WP 016942551 1 Dzeae
Dickeya zeae ZJU1202 WP 016942551 1 Dzeae
Dickeya paradisiaca NCPPB 2511 WP 015854052 1 Dparadisiaca
Dickeya paradisiaca Ech703 WP 015854052 1 Dparadisiaca
Dickeya zeae NCPPB 3532 WP 023639900 1 Dzeae
Dickeya dadantii NCPPB 898 WP 038911059 1 Ddadantii
Klebsiella pneumoniae YP 007990445 1 573 Proteobacteria
Escherichia coli 042 YP 006096729 1 216592 Proteobacteria
Escherichia coli O111 H str 11128 YP 003235292 1 585396 Proteobacteria
Shigella boydii Sb227 YP 408540 1 300268 Proteobacteria
Escherichia coli O26 H11 str 11368 YP 003230054 1 573235 Proteobacteria
Dickeya gs fangzhongdai NCPPB 3274 WP 042859584 1 Dfangzhongdai
Dickeya dadantii 3937 WP 013317595 1 Ddadantii
DAQ1742 DAQ1742 01959 Daquatica
639 DAQ1742 01959 Phage P3 Seed
Dickeya zeae CSL RW192 WP 038907136 1 Dzeae
Dickeya zeae Ech586 WP 012885053 1 Dzeae
DAQ1742 DAQ1742 01969 Daquatica
Dickeya aquatica DW 0440 WP 035342443 1 Daquatica
Dickeya sp CSL RW240 WP 035342443 1 Daquatica
Saccharophagus degradans 2 40 YP 528215 1 203122 Proteobacteria
Shewanella sp MR 7 YP 738896 1 60481 Proteobacteria
Psychromonas ingrahamii 37 YP 942997 1 357804 Proteobacteria
Aliivibrio salmonicida LFI1238 YP 002263830 1 316275 Proteobacteria
Vibrio sp Ex25 YP 003286924 1 150340 Proteobacteria
Vibrio cholerae O395 YP 002819113 1 345073 Proteobacteria
Vibrio cholerae O1 biovar El Tor str N16961 NP 230494 1 243277 Proteobacteria
Dickeya gs fangzhongdai B16 WP 038917939 1 Dfangzhongdai
Dickeya gs fangzhongdai MK7 WP 038917939 1 Dfangzhongdai
Dickeya paradisiaca NCPPB 2511 WP 015854886 1 Dparadisiaca
Dickeya paradisiaca Ech703 WP 015854886 1 Dparadisiaca
Enterobacter lignolyticus SCF1 YP 003940652 1 701347 Proteobacteria
Escherichia coli str K 12 substr W3110 YP 490843 1 316407 Proteobacteria
Enterobacter aerogenes EA1509E YP 007391326 1 935296 Proteobacteria
Dickeya dadantii 3937 WP 013316485 1 Ddadantii
Enterobacter sp 638 YP 001177816 1 399742 Proteobacteria
Salmonella enterica subsp enterica serovar Paratyphi B str SPB7 YP 001589593 1 1016998 Proteobacteria
Raoultella ornithinolytica B6 YP 007877134 1 1286170 Proteobacteria
Erwinia sp Ejp617 YP 005816675 1 215689 Proteobacteria
Pantoea ananatis LMG 20103 YP 003521207 1 706191 Proteobacteria
Escherichia fergusonii ATCC 35469 YP 002381651 1 585054 Proteobacteria
Escherichia coli ATCC 8739 YP 001724057 1 481805 Proteobacteria
Salmonella enterica subsp enterica serovar Typhi str Ty21a YP 007926930 1 527001 Proteobacteria
Klebsiella oxytoca E718 YP 006500153 1 1191061 Proteobacteria
Edwardsiella tarda EIB202 YP 003296788 1 498217 Proteobacteria
Salmonella enterica subsp enterica serovar Newport str SL254 YP 002041998 1 423368 Proteobacteria
Salmonella enterica subsp enterica serovar Javiana str CFSAN001992 YP 007473660 1 1267753 Proteobacteria
Shimwellia blattae DSM 4481 NBRC 105725 YP 006318568 1 630626 Proteobacteria
Salmonella enterica subsp enterica serovar Typhimurium str ST4 74 YP 005243712 1 909946 Proteobacteria
Salmonella enterica subsp enterica serovar Thompson str RM6836 YP 008614402 1 1064551 Proteobacteria
Salmonella enterica subsp enterica serovar Paratyphi B str SPB7 YP 001589610 1 1016998 Proteobacteria
Pectobacterium carotovorum subsp carotovorum PC1 YP 003016313 1 561230 Proteobacteria
Pectobacterium atrosepticum SCRI1043 YP 048945 1 218491 Proteobacteria
Yersinia pestis biovar Medievalis str Harbin 35 YP 005623096 1 547048 Proteobacteria
Yersinia pestis biovar Microtus str 91001 NP 992430 1 229193 Proteobacteria
Yersinia pestis Antiqua YP 650490 1 360102 Proteobacteria
Pectobacterium sp SCC3193 YP 006281840 1 1166016 Proteobacteria
Pectobacterium wasabiae WPP163 YP 003258382 1 561231 Proteobacteria
Dickeya dadantii subsp dieffenbachiae NCPPB 2976 WP 038921802 1 Ddadantii
Dickeya gs undicola 2B12 WP 033570361 1 Dundicola
Dickeya gs fangzhongdai M074 WP 038661920 1 Dfangzhongdai
Dickeya gs fangzhongdai ND14b WP 038661920 1 Dfangzhongdai
Dickeya dianthicola IPO 980 WP 029729353 1 Ddianthicola
Dickeya dadantii NCPPB 898 WP 038910326 1 Ddadantii
Dickeya dianthicola RNS04 9 WP 024104731 1 Ddianthicola
Dickeya dianthicola NCPPB 453 WP 024104731 1 Ddianthicola
Dickeya dianthicola GBBC 2039 WP 024108266 1 Ddianthicola
Dickeya dianthicola NCPPB 3534 WP 024108266 1 Ddianthicola

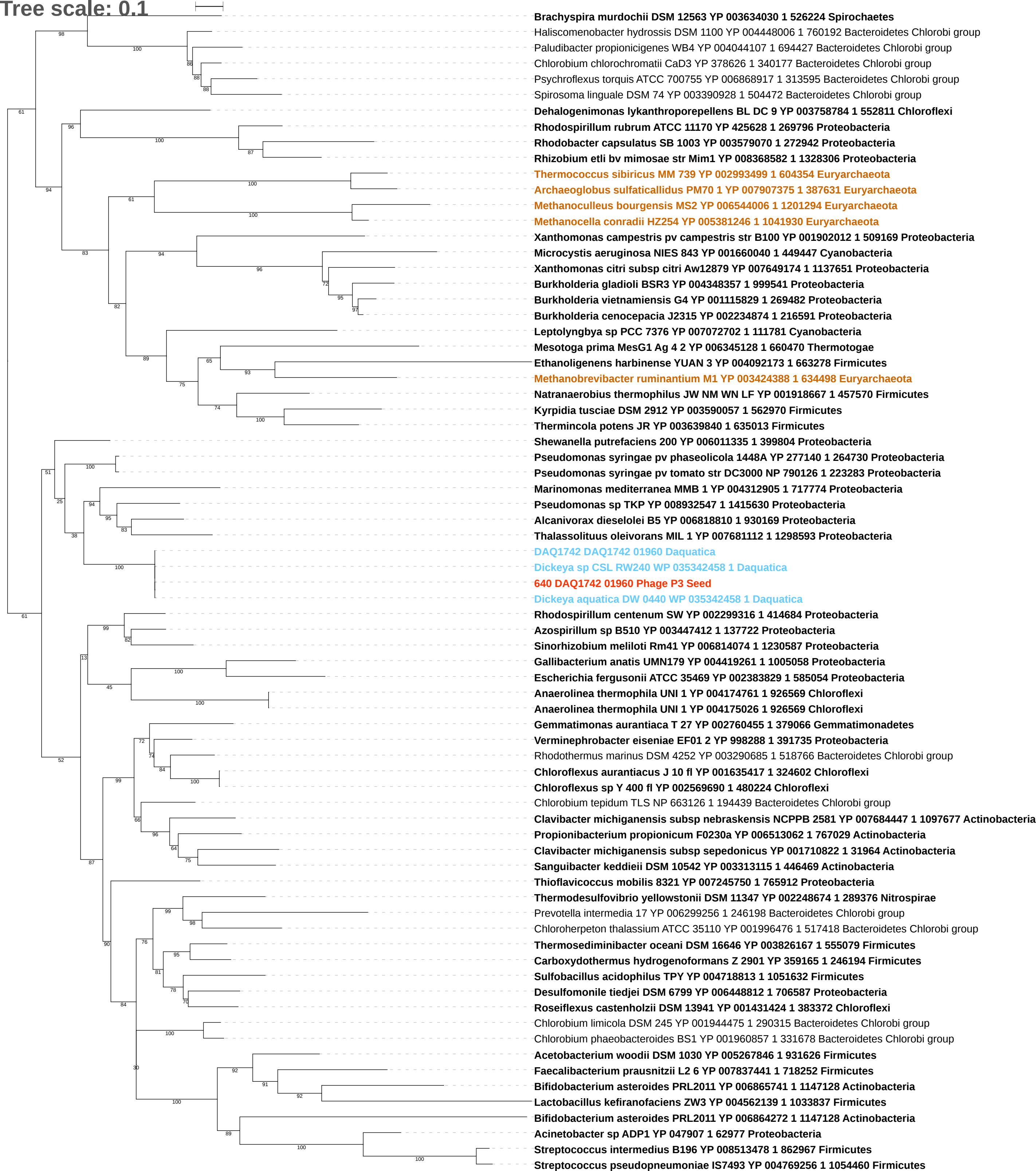
Tree scale: 0.1
Brachyspira murdochii DSM 12563 YP 003634030 1 526224 Spirochaetes
Haliscomenobacter hydrossis DSM 1100 YP 004448006 1 760192 Bacteroidetes Chlorobi group
Paludibacter propionicigenes WB4 YP 004044107 1 694427 Bacteroidetes Chlorobi group
Chlorobium chlorochromatii CaD3 YP 378626 1 340177 Bacteroidetes Chlorobi group
Psychroflexus torquis ATCC 700755 YP 006868917 1 313595 Bacteroidetes Chlorobi group
Spirosoma linguale DSM 74 YP 003390928 1 504472 Bacteroidetes Chlorobi group
Dehalogenimonas lykanthroporepellens BL DC 9 YP 003758784 1 552811 Chloroflexi
Rhodospirillum rubrum ATCC 11170 YP 425628 1 269796 Proteobacteria
Rhodobacter capsulatus SB 1003 YP 003579070 1 272942 Proteobacteria
Rhizobium etli bv mimosae str Mim1 YP 008368582 1 1328306 Proteobacteria
Thermococcus sibiricus MM 739 YP 002993499 1 604354 Euryarchaeota
Archaeoglobus sulfaticallidus PM70 1 YP 007907375 1 387631 Euryarchaeota
Methanoculleus bourgensis MS2 YP 006544006 1 1201294 Euryarchaeota
Methanocella conradii HZ254 YP 005381246 1 1041930 Euryarchaeota
Xanthomonas campestris pv campestris str B100 YP 001902012 1 509169 Proteobacteria
Microcystis aeruginosa NIES 843 YP 001660040 1 449447 Cyanobacteria
Xanthomonas citri subsp citri Aw12879 YP 007649174 1 1137651 Proteobacteria
Burkholderia gladioli BSR3 YP 004348357 1 999541 Proteobacteria
Burkholderia vietnamiensis G4 YP 001115829 1 269482 Proteobacteria
Burkholderia cenocepacia J2315 YP 002234874 1 216591 Proteobacteria
Leptolyngbya sp PCC 7376 YP 007072702 1 111781 Cyanobacteria
Mesotoga prima MesG1 Ag 4 2 YP 006345128 1 660470 Thermotogae
Ethanoligenens harbinense YUAN 3 YP 004092173 1 663278 Firmicutes
Methanobrevibacter ruminantium M1 YP 003424388 1 634498 Euryarchaeota
Natranaerobius thermophilus JW NM WN LF YP 001918667 1 457570 Firmicutes
Kyrpidia tusciae DSM 2912 YP 003590057 1 562970 Firmicutes
Thermincola potens JR YP 003639840 1 635013 Firmicutes
Shewanella putrefaciens 200 YP 006011335 1 399804 Proteobacteria
Pseudomonas syringae pv phaseolicola 1448A YP 277140 1 264730 Proteobacteria
Pseudomonas syringae pv tomato str DC3000 NP 790126 1 223283 Proteobacteria
Marinomonas mediterranea MMB 1 YP 004312905 1 717774 Proteobacteria
Pseudomonas sp TKP YP 008932547 1 1415630 Proteobacteria
Alcanivorax dieselolei B5 YP 006818810 1 930169 Proteobacteria
Thalassolituus oleivorans MIL 1 YP 007681112 1 1298593 Proteobacteria
DAQ1742 DAQ1742 01960 Daquatica
Dickeya sp CSL RW240 WP 035342458 1 Daquatica
640 DAQ1742 01960 Phage P3 Seed
Dickeya aquatica DW 0440 WP 035342458 1 Daquatica
Rhodospirillum centenum SW YP 002299316 1 414684 Proteobacteria
Azospirillum sp B510 YP 003447412 1 137722 Proteobacteria
Sinorhizobium meliloti Rm41 YP 006814074 1 1230587 Proteobacteria
Gallibacterium anatis UMN179 YP 004419261 1 1005058 Proteobacteria
Escherichia fergusonii ATCC 35469 YP 002383829 1 585054 Proteobacteria
Anaerolinea thermophila UNI 1 YP 004174761 1 926569 Chloroflexi
Anaerolinea thermophila UNI 1 YP 004175026 1 926569 Chloroflexi
Gemmatimonas aurantiaca T 27 YP 002760455 1 379066 Gemmatimonadetes
Verminephrobacter eiseniae EF01 2 YP 998288 1 391735 Proteobacteria
Rhodothermus marinus DSM 4252 YP 003290685 1 518766 Bacteroidetes Chlorobi group
Chloroflexus aurantiacus J 10 fl YP 001635417 1 324602 Chloroflexi
Chloroflexus sp Y 400 fl YP 002569690 1 480224 Chloroflexi
Chlorobium tepidum TLS NP 663126 1 194439 Bacteroidetes Chlorobi group
Clavibacter michiganensis subsp nebraskensis NCPPB 2581 YP 007684447 1 1097677 Actinobacteria
Propionibacterium propionicum F0230a YP 006513062 1 767029 Actinobacteria
Clavibacter michiganensis subsp sepedonicus YP 001710822 1 31964 Actinobacteria
Sanguibacter keddieii DSM 10542 YP 003313115 1 446469 Actinobacteria
Thioflavicoccus mobilis 8321 YP 007245750 1 765912 Proteobacteria
Thermodesulfovibrio yellowstonii DSM 11347 YP 002248674 1 289376 Nitrospirae
Prevotella intermedia 17 YP 006299256 1 246198 Bacteroidetes Chlorobi group
Chloroherpeton thalassium ATCC 35110 YP 001996476 1 517418 Bacteroidetes Chlorobi group
Thermosediminibacter oceani DSM 16646 YP 003826167 1 555079 Firmicutes
Carboxydothermus hydrogenoformans Z 2901 YP 359165 1 246194 Firmicutes
Sulfobacillus acidophilus TPY YP 004718813 1 1051632 Firmicutes
Desulfomonile tiedjei DSM 6799 YP 006448812 1 706587 Proteobacteria
Roseiflexus castenholzii DSM 13941 YP 001431424 1 383372 Chloroflexi
Chlorobium limicola DSM 245 YP 001944475 1 290315 Bacteroidetes Chlorobi group
Chlorobium phaeobacteroides BS1 YP 001960857 1 331678 Bacteroidetes Chlorobi group
Acetobacterium woodii DSM 1030 YP 005267846 1 931626 Firmicutes
Faecalibacterium prausnitzii L2 6 YP 007837441 1 718252 Firmicutes
Bifidobacterium asteroides PRL2011 YP 006865741 1 1147128 Actinobacteria
Lactobacillus kefiranofaciens ZW3 YP 004562139 1 1033837 Firmicutes
Bifidobacterium asteroides PRL2011 YP 006864272 1 1147128 Actinobacteria
Acinetobacter sp ADP1 YP 047907 1 62977 Proteobacteria
Streptococcus intermedius B196 YP 008513478 1 862967 Firmicutes
Streptococcus pseudopneumoniae IS7493 YP 004769256 1 1054460 Firmicutes

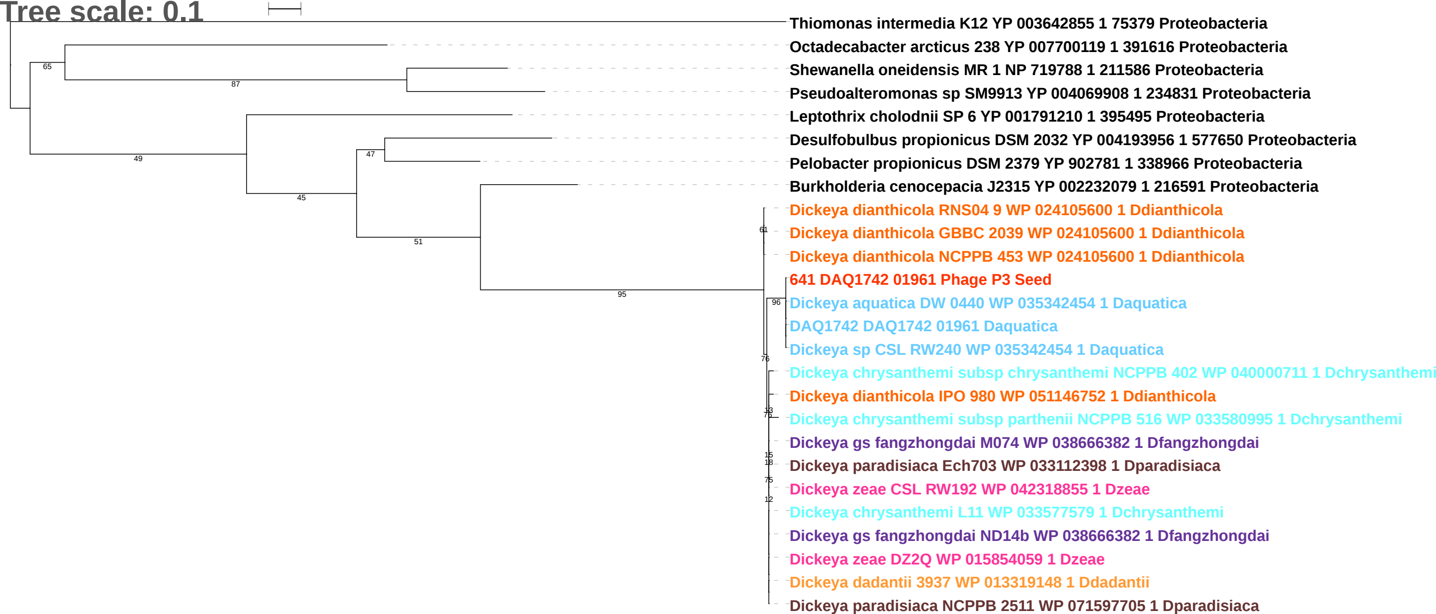
Tree scale: 0.1
Thiomonas intermedia K12 YP 003642855 1 75379 Proteobacteria
Octadecabacter arcticus 238 YP 007700119 1 391616 Proteobacteria
Shewanella oneidensis MR 1 NP 719788 1 211586 Proteobacteria
Pseudoalteromonas sp SM9913 YP 004069908 1 234831 Proteobacteria
Leptothrix cholodnii SP 6 YP 001791210 1 395495 Proteobacteria
Desulfobulbus propionicus DSM 2032 YP 004193956 1 577650 Proteobacteria
Pelobacter propionicus DSM 2379 YP 902781 1 338966 Proteobacteria
Burkholderia cenocepacia J2315 YP 002232079 1 216591 Proteobacteria
Dickeya dianthicola RNS04 9 WP 024105600 1 Ddianthicola
Dickeya dianthicola GBBC 2039 WP 024105600 1 Ddianthicola
Dickeya dianthicola NCPPB 453 WP 024105600 1 Ddianthicola
641 DAQ1742 01961 Phage P3 Seed
Dickeya aquatica DW 0440 WP 035342454 1 Daquatica
DAQ1742 DAQ1742 01961 Daquatica
Dickeya sp CSL RW240 WP 035342454 1 Daquatica
Dickeya chrysanthemi subsp chrysanthemi NCPPB 402 WP 040000711 1 Dchrysanthemi
Dickeya dianthicola IPO 980 WP 051146752 1 Ddianthicola
Dickeya chrysanthemi subsp parthenii NCPPB 516 WP 033580995 1 Dchrysanthemi
Dickeya gs fangzhongdai M074 WP 038666382 1 Dfangzhongdai
Dickeya paradisiaca Ech703 WP 033112398 1 Dparadisiaca
Dickeya zeae CSL RW192 WP 042318855 1 Dzeae
Dickeya chrysanthemi L11 WP 033577579 1 Dchrysanthemi
Dickeya gs fangzhongdai ND14b WP 038666382 1 Dfangzhongdai
Dickeya zeae DZ2Q WP 015854059 1 Dzeae
Dickeya dadantii 3937 WP 013319148 1 Ddadantii
Dickeya paradisiaca NCPPB 2511 WP 071597705 1 Dparadisiaca
65
87
49
45
47
51
95
61
96
76

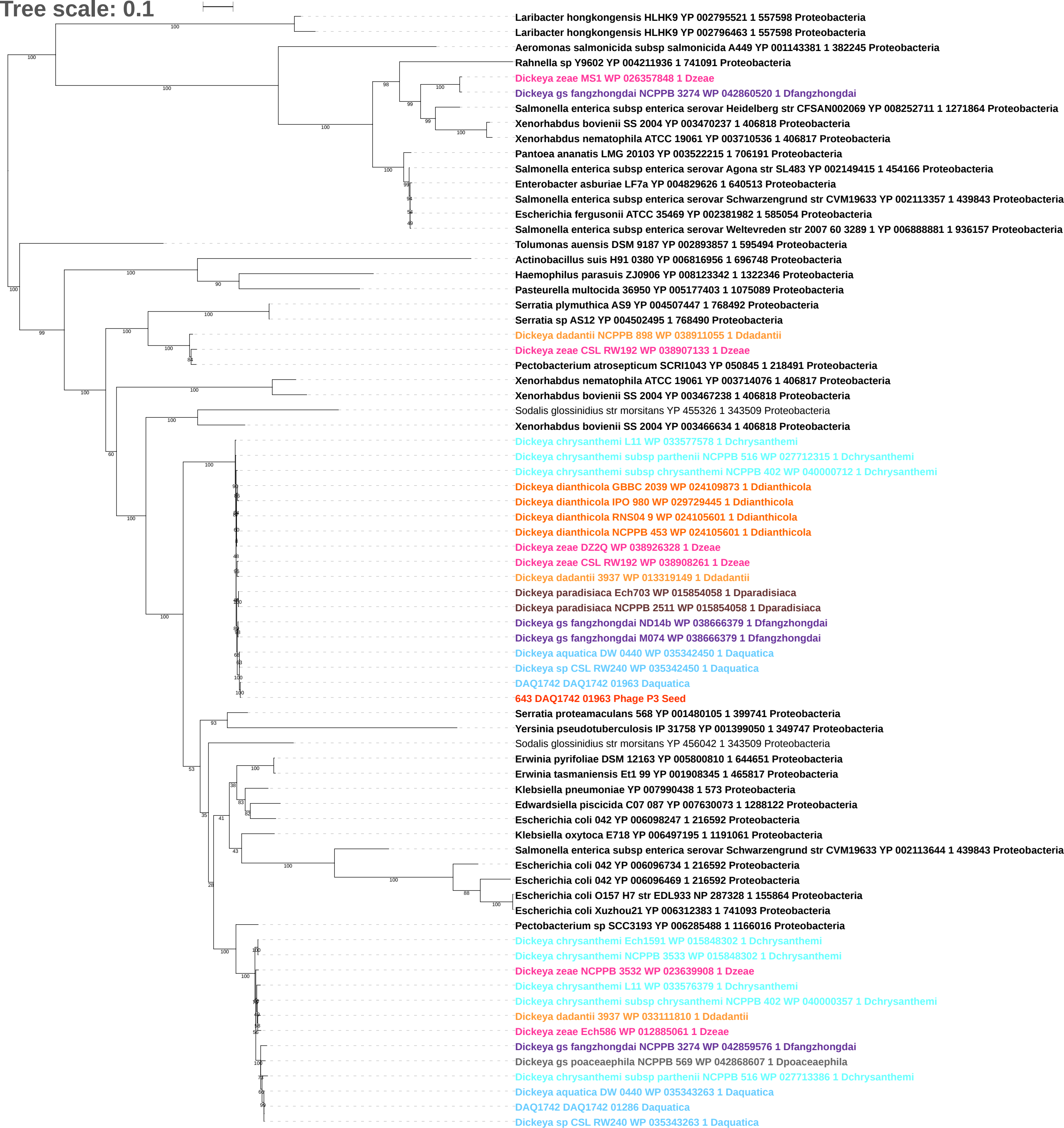
Tree scale: 0.1
Laribacter hongkongensis HLHK9 YP 002795521 1 557598 Proteobacteria
Laribacter hongkongensis HLHK9 YP 002796463 1 557598 Proteobacteria
Aeromonas salmonicida subsp salmonicida A449 YP 001143381 1 382245 Proteobacteria
Rahnella sp Y9602 YP 004211936 1 741091 Proteobacteria
Dickeya zeae MS1 WP 026357848 1 Dzeae
Dickeya gs fangzhongdai NCPPB 3274 WP 042860520 1 Dfangzhongdai
Salmonella enterica subsp enterica serovar Heidelberg str CFSAN002069 YP 008252711 1 1271864 Proteobacteria
Xenorhabdus bovienii SS 2004 YP 003470237 1 406818 Proteobacteria
Xenorhabdus nematophila ATCC 19061 YP 003710536 1 406817 Proteobacteria
Pantoea ananatis LMG 20103 YP 003522215 1 706191 Proteobacteria
Salmonella enterica subsp enterica serovar Agona str SL483 YP 002149415 1 454166 Proteobacteria
Enterobacter asburiae LF7a YP 004829626 1 640513 Proteobacteria
Salmonella enterica subsp enterica serovar Schwarzengrund str CVM19633 YP 002113357 1 439843 Proteobacteria
Escherichia fergusonii ATCC 35469 YP 002381982 1 585054 Proteobacteria
Salmonella enterica subsp enterica serovar Weltevreden str 2007 60 3289 1 YP 006888881 1 936157 Proteobacteria
Tolumonas auensis DSM 9187 YP 002893857 1 595494 Proteobacteria
Actinobacillus suis H91 0380 YP 006816956 1 696748 Proteobacteria
Haemophilus parasuis ZJ0906 YP 008123342 1 1322346 Proteobacteria
Pasteurella multocida 36950 YP 005177403 1 1075089 Proteobacteria
Serratia plymuthica AS9 YP 004507447 1 768492 Proteobacteria
Serratia sp AS12 YP 004502495 1 768490 Proteobacteria
Dickeya dadantii NCPPB 898 WP 038911055 1 Ddadantii
Dickeya zeae CSL RW192 WP 038907133 1 Dzeae
Pectobacterium atrosepticum SCRI1043 YP 050845 1 218491 Proteobacteria
Xenorhabdus nematophila ATCC 19061 YP 003714076 1 406817 Proteobacteria
Xenorhabdus bovienii SS 2004 YP 003467238 1 406818 Proteobacteria
Sodalis glossinidius str morsitans YP 455326 1 343509 Proteobacteria
Xenorhabdus bovienii SS 2004 YP 003466634 1 406818 Proteobacteria
Dickeya chrysanthemi L11 WP 033577578 1 Dchrysanthemi
Dickeya chrysanthemi subsp parthenii NCPPB 516 WP 027712315 1 Dchrysanthemi
Dickeya chrysanthemi subsp chrysanthemi NCPPB 402 WP 040000712 1 Dchrysanthemi
Dickeya dianthicola GBBC 2039 WP 024109873 1 Ddianthicola
Dickeya dianthicola IPO 980 WP 029729445 1 Ddianthicola
Dickeya dianthicola RNS04 9 WP 024105601 1 Ddianthicola
Dickeya dianthicola NCPPB 453 WP 024105601 1 Ddianthicola
Dickeya zeae DZ2Q WP 038926328 1 Dzeae
Dickeya zeae CSL RW192 WP 038908261 1 Dzeae
Dickeya dadantii 3937 WP 013319149 1 Ddadantii
Dickeya paradisiaca Ech703 WP 015854058 1 Dparadisiaca
Dickeya paradisiaca NCPPB 2511 WP 015854058 1 Dparadisiaca
Dickeya gs fangzhongdai ND14b WP 038666379 1 Dfangzhongdai
Dickeya gs fangzhongdai M074 WP 038666379 1 Dfangzhongdai
Dickeya aquatica DW 0440 WP 035342450 1 Daquatica
Dickeya sp CSL RW240 WP 035342450 1 Daquatica
DAQ1742 DAQ1742 01963 Daquatica
643 DAQ1742 01963 Phage P3 Seed
Serratia proteamaculans 568 YP 001480105 1 399741 Proteobacteria
Yersinia pseudotuberculosis IP 31758 YP 001399050 1 349747 Proteobacteria
Sodalis glossinidius str morsitans YP 456042 1 343509 Proteobacteria
Erwinia pyrifoliae DSM 12163 YP 005800810 1 644651 Proteobacteria
Erwinia tasmaniensis Et1 99 YP 001908345 1 465817 Proteobacteria
Klebsiella pneumoniae YP 007990438 1 573 Proteobacteria
Edwardsiella piscicida C07 087 YP 007630073 1 1288122 Proteobacteria
Escherichia coli 042 YP 006098247 1 216592 Proteobacteria
Klebsiella oxytoca E718 YP 006497195 1 1191061 Proteobacteria
Salmonella enterica subsp enterica serovar Schwarzengrund str CVM19633 YP 002113644 1 439843 Proteobacteria
Escherichia coli 042 YP 006096734 1 216592 Proteobacteria
Escherichia coli 042 YP 006096469 1 216592 Proteobacteria
Escherichia coli O157 H7 str EDL933 NP 287328 1 155864 Proteobacteria
Escherichia coli Xuzhou21 YP 006312383 1 741093 Proteobacteria
Pectobacterium sp SCC3193 YP 006285488 1 1166016 Proteobacteria
Dickeya chrysanthemi Ech1591 WP 015848302 1 Dchrysanthemi
Dickeya chrysanthemi NCPPB 3533 WP 015848302 1 Dchrysanthemi
Dickeya zeae NCPPB 3532 WP 023639908 1 Dzeae
Dickeya chrysanthemi L11 WP 033576379 1 Dchrysanthemi
Dickeya chrysanthemi subsp chrysanthemi NCPPB 402 WP 040000357 1 Dchrysanthemi
Dickeya dadantii 3937 WP 033111810 1 Ddadantii
Dickeya zeae Ech586 WP 012885061 1 Dzeae
Dickeya gs fangzhongdai NCPPB 3274 WP 042859576 1 Dfangzhongdai
Dickeya gs poaceaephila NCPPB 569 WP 042868607 1 Dpoaceaephila
Dickeya chrysanthemi subsp parthenii NCPPB 516 WP 027713386 1 Dchrysanthemi
Dickeya aquatica DW 0440 WP 035343263 1 Daquatica
DAQ1742 DAQ1742 01286 Daquatica
Dickeya sp CSL RW240 WP 035343263 1 Daquatica

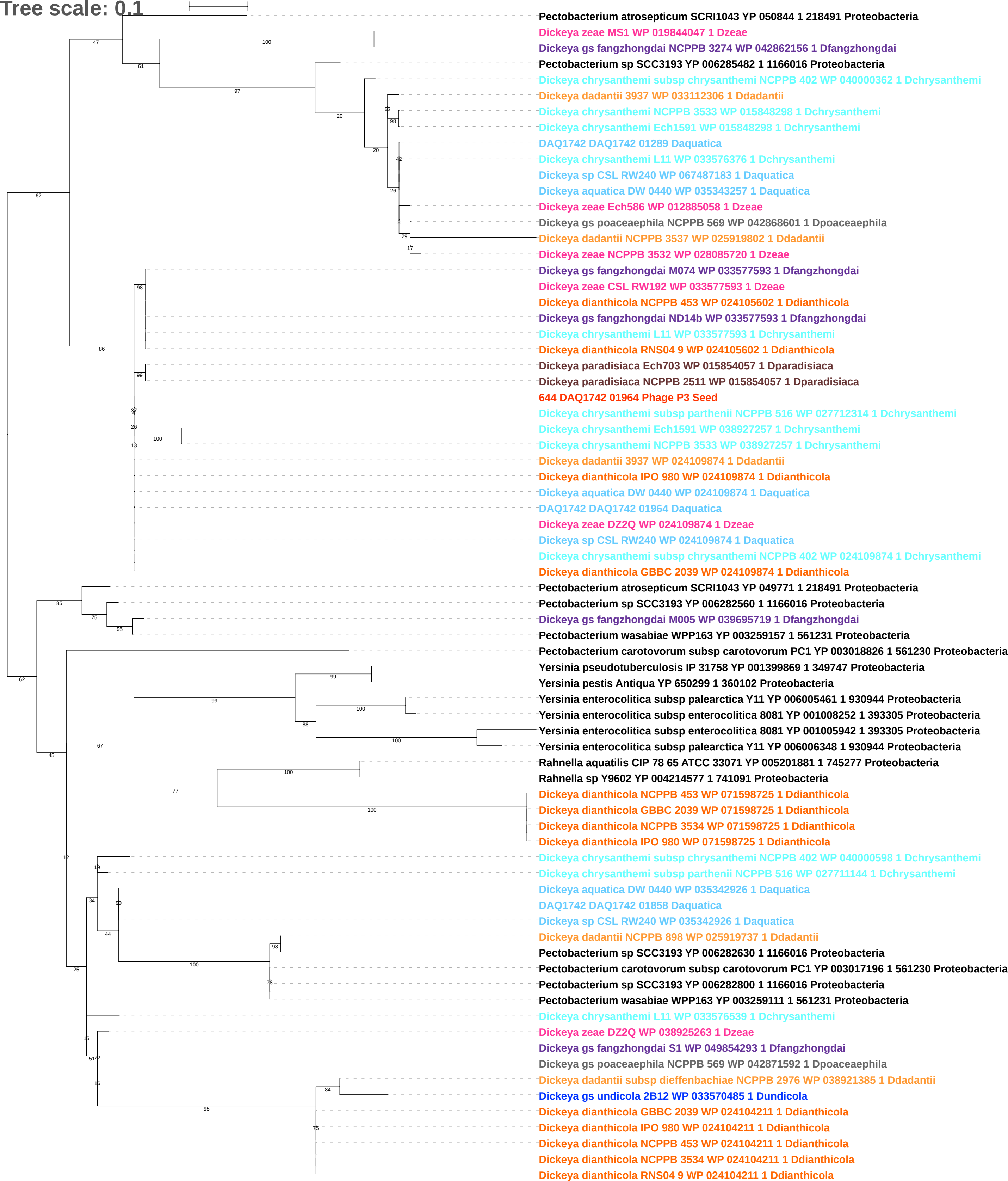
Tree scale: 0.1
Pectobacterium atrosepticum SCRI1043 YP 050844 1 218491 Proteobacteria
Dickeya zeae MS1 WP 019844047 1 Dzeae
Dickeya gs fangzhongdai NCPPB 3274 WP 042862156 1 Dfangzhongdai
Pectobacterium sp SCC3193 YP 006285482 1 1166016 Proteobacteria
Dickeya chrysanthemi subsp chrysanthemi NCPPB 402 WP 040000362 1 Dchrysanthemi
Dickeya dadantii 3937 WP 033112306 1 Ddadantii
Dickeya chrysanthemi NCPPB 3533 WP 015848298 1 Dchrysanthemi
Dickeya chrysanthemi Ech1591 WP 015848298 1 Dchrysanthemi
DAQ1742 DAQ1742 01289 Daquatica
Dickeya chrysanthemi L11 WP 033576376 1 Dchrysanthemi
Dickeya sp CSL RW240 WP 067487183 1 Daquatica
Dickeya aquatica DW 0440 WP 035343257 1 Daquatica
Dickeya zeae Ech586 WP 012885058 1 Dzeae
Dickeya gs poaceaephila NCPPB 569 WP 042868601 1 Dpoaceaephila
Dickeya dadantii NCPPB 3537 WP 025919802 1 Ddadantii
Dickeya zeae NCPPB 3532 WP 028085720 1 Dzeae
Dickeya gs fangzhongdai M074 WP 033577593 1 Dfangzhongdai
Dickeya zeae CSL RW192 WP 033577593 1 Dzeae
Dickeya dianthicola NCPPB 453 WP 024105602 1 Ddianthicola
Dickeya gs fangzhongdai ND14b WP 033577593 1 Dfangzhongdai
Dickeya chrysanthemi L11 WP 033577593 1 Dchrysanthemi
Dickeya dianthicola RNS04 9 WP 024105602 1 Ddianthicola
Dickeya paradisiaca Ech703 WP 015854057 1 Dparadisiaca
Dickeya paradisiaca NCPPB 2511 WP 015854057 1 Dparadisiaca
644 DAQ1742 01964 Phage P3 Seed
Dickeya chrysanthemi subsp parthenii NCPPB 516 WP 027712314 1 Dchrysanthemi
Dickeya chrysanthemi Ech1591 WP 038927257 1 Dchrysanthemi
Dickeya chrysanthemi NCPPB 3533 WP 038927257 1 Dchrysanthemi
Dickeya dadantii 3937 WP 024109874 1 Ddadantii
Dickeya dianthicola IPO 980 WP 024109874 1 Ddianthicola
Dickeya aquatica DW 0440 WP 024109874 1 Daquatica
DAQ1742 DAQ1742 01964 Daquatica
Dickeya zeae DZ2Q WP 024109874 1 Dzeae
Dickeya sp CSL RW240 WP 024109874 1 Daquatica
Dickeya chrysanthemi subsp chrysanthemi NCPPB 402 WP 024109874 1 Dchrysanthemi
Dickeya dianthicola GBBC 2039 WP 024109874 1 Ddianthicola
Pectobacterium atrosepticum SCRI1043 YP 049771 1 218491 Proteobacteria
Pectobacterium sp SCC3193 YP 006282560 1 1166016 Proteobacteria
Dickeya gs fangzhongdai M005 WP 039695719 1 Dfangzhongdai
Pectobacterium wasabiae WPP163 YP 003259157 1 561231 Proteobacteria
Pectobacterium carotovorum subsp carotovorum PC1 YP 003018826 1 561230 Proteobacteria
Yersinia pseudotuberculosis IP 31758 YP 001399869 1 349747 Proteobacteria
Yersinia pestis Antiqua YP 650299 1 360102 Proteobacteria
Yersinia enterocolitica subsp palearctica Y11 YP 006005461 1 930944 Proteobacteria
Yersinia enterocolitica subsp enterocolitica 8081 YP 001008252 1 393305 Proteobacteria
Yersinia enterocolitica subsp enterocolitica 8081 YP 001005942 1 393305 Proteobacteria
Yersinia enterocolitica subsp palearctica Y11 YP 006006348 1 930944 Proteobacteria
Rahnella aquatilis CIP 78 65 ATCC 33071 YP 005201881 1 745277 Proteobacteria
Rahnella sp Y9602 YP 004214577 1 741091 Proteobacteria
Dickeya dianthicola NCPPB 453 WP 071598725 1 Ddianthicola
Dickeya dianthicola GBBC 2039 WP 071598725 1 Ddianthicola
Dickeya dianthicola NCPPB 3534 WP 071598725 1 Ddianthicola
Dickeya dianthicola IPO 980 WP 071598725 1 Ddianthicola
Dickeya chrysanthemi subsp chrysanthemi NCPPB 402 WP 040000598 1 Dchrysanthemi
Dickeya chrysanthemi subsp parthenii NCPPB 516 WP 027711144 1 Dchrysanthemi
Dickeya aquatica DW 0440 WP 035342926 1 Daquatica
DAQ1742 DAQ1742 01858 Daquatica
Dickeya sp CSL RW240 WP 035342926 1 Daquatica
Dickeya dadantii NCPPB 898 WP 025919737 1 Ddadantii
Pectobacterium sp SCC3193 YP 006282630 1 1166016 Proteobacteria
Pectobacterium carotovorum subsp carotovorum PC1 YP 003017196 1 561230 Proteobacteria
Pectobacterium sp SCC3193 YP 006282800 1 1166016 Proteobacteria
Pectobacterium wasabiae WPP163 YP 003259111 1 561231 Proteobacteria
Dickeya chrysanthemi L11 WP 033576539 1 Dchrysanthemi
Dickeya zeae DZ2Q WP 038925263 1 Dzeae
Dickeya gs fangzhongdai S1 WP 049854293 1 Dfangzhongdai
Dickeya gs poaceaephila NCPPB 569 WP 042871592 1 Dpoaceaephila
Dickeya dadantii subsp dieffenbachiae NCPPB 2976 WP 038921385 1 Ddadantii
Dickeya gs undicola 2B12 WP 033570485 1 Dundicola
Dickeya dianthicola GBBC 2039 WP 024104211 1 Ddianthicola
Dickeya dianthicola IPO 980 WP 024104211 1 Ddianthicola
Dickeya dianthicola NCPPB 453 WP 024104211 1 Ddianthicola
Dickeya dianthicola NCPPB 3534 WP 024104211 1 Ddianthicola
Dickeya dianthicola RNS04 9 WP 024104211 1 Ddianthicola

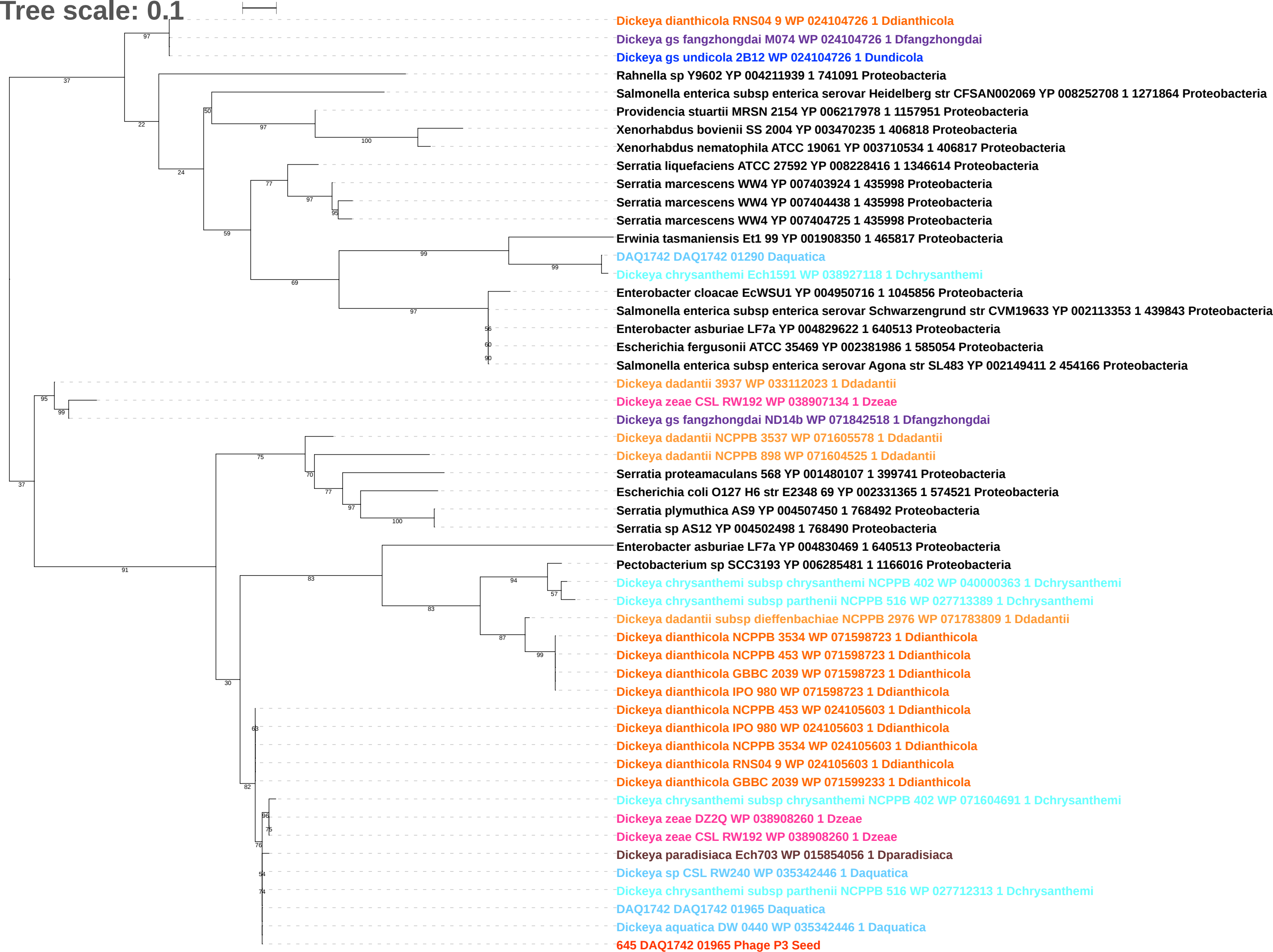
Tree scale: 0.1
Dickeya dianthicola RNS04 9 WP 024104726 1 Ddianthicola
Dickeya gs fangzhongdai M074 WP 024104726 1 Dfangzhongdai
Dickeya gs undicola 2B12 WP 024104726 1 Dundicola
Rahnella sp Y9602 YP 004211939 1 741091 Proteobacteria
Salmonella enterica subsp enterica serovar Heidelberg str CFSAN002069 YP 008252708 1 1271864 Proteobacteria
Providencia stuartii MRSN 2154 YP 006217978 1 1157951 Proteobacteria
Xenorhabdus bovienii SS 2004 YP 003470235 1 406818 Proteobacteria
Xenorhabdus nematophila ATCC 19061 YP 003710534 1 406817 Proteobacteria
Serratia liquefaciens ATCC 27592 YP 008228416 1 1346614 Proteobacteria
Serratia marcescens WW4 YP 007403924 1 435998 Proteobacteria
Serratia marcescens WW4 YP 007404438 1 435998 Proteobacteria
Serratia marcescens WW4 YP 007404725 1 435998 Proteobacteria
Erwinia tasmaniensis Et1 99 YP 001908350 1 465817 Proteobacteria
DAQ1742 DAQ1742 01290 Daquatica
Dickeya chrysanthemi Ech1591 WP 038927118 1 Dchrysanthemi
Enterobacter cloacae EcWSU1 YP 004950716 1 1045856 Proteobacteria
Salmonella enterica subsp enterica serovar Schwarzengrund str CVM19633 YP 002113353 1 439843 Proteobacteria
Enterobacter asburiae LF7a YP 004829622 1 640513 Proteobacteria
Escherichia fergusonii ATCC 35469 YP 002381986 1 585054 Proteobacteria
Salmonella enterica subsp enterica serovar Agona str SL483 YP 002149411 2 454166 Proteobacteria
Dickeya dadantii 3937 WP 033112023 1 Ddadantii
Dickeya zeae CSL RW192 WP 038907134 1 Dzeae
Dickeya gs fangzhongdai ND14b WP 071842518 1 Dfangzhongdai
Dickeya dadantii NCPPB 3537 WP 071605578 1 Ddadantii
Dickeya dadantii NCPPB 898 WP 071604525 1 Ddadantii
Serratia proteamaculans 568 YP 001480107 1 399741 Proteobacteria
Escherichia coli O127 H6 str E2348 69 YP 002331365 1 574521 Proteobacteria
Serratia plymuthica AS9 YP 004507450 1 768492 Proteobacteria
Serratia sp AS12 YP 004502498 1 768490 Proteobacteria
Enterobacter asburiae LF7a YP 004830469 1 640513 Proteobacteria
Pectobacterium sp SCC3193 YP 006285481 1 1166016 Proteobacteria
Dickeya chrysanthemi subsp chrysanthemi NCPPB 402 WP 040000363 1 Dchrysanthemi
Dickeya chrysanthemi subsp parthenii NCPPB 516 WP 027713389 1 Dchrysanthemi
Dickeya dadantii subsp dieffenbachiae NCPPB 2976 WP 071783809 1 Ddadantii
Dickeya dianthicola NCPPB 3534 WP 071598723 1 Ddianthicola
Dickeya dianthicola NCPPB 453 WP 071598723 1 Ddianthicola
Dickeya dianthicola GBBC 2039 WP 071598723 1 Ddianthicola
Dickeya dianthicola IPO 980 WP 071598723 1 Ddianthicola
Dickeya dianthicola NCPPB 453 WP 024105603 1 Ddianthicola
Dickeya dianthicola IPO 980 WP 024105603 1 Ddianthicola
Dickeya dianthicola NCPPB 3534 WP 024105603 1 Ddianthicola
Dickeya dianthicola RNS04 9 WP 024105603 1 Ddianthicola
Dickeya dianthicola GBBC 2039 WP 071599233 1 Ddianthicola
Dickeya chrysanthemi subsp chrysanthemi NCPPB 402 WP 071604691 1 Dchrysanthemi
Dickeya zeae DZ2Q WP 038908260 1 Dzeae
Dickeya zeae CSL RW192 WP 038908260 1 Dzeae
Dickeya paradisiaca Ech703 WP 015854056 1 Dparadisiaca
Dickeya sp CSL RW240 WP 035342446 1 Daquatica
Dickeya chrysanthemi subsp parthenii NCPPB 516 WP 027712313 1 Dchrysanthemi
DAQ1742 DAQ1742 01965 Daquatica
Dickeya aquatica DW 0440 WP 035342446 1 Daquatica
645 DAQ1742 01965 Phage P3 Seed

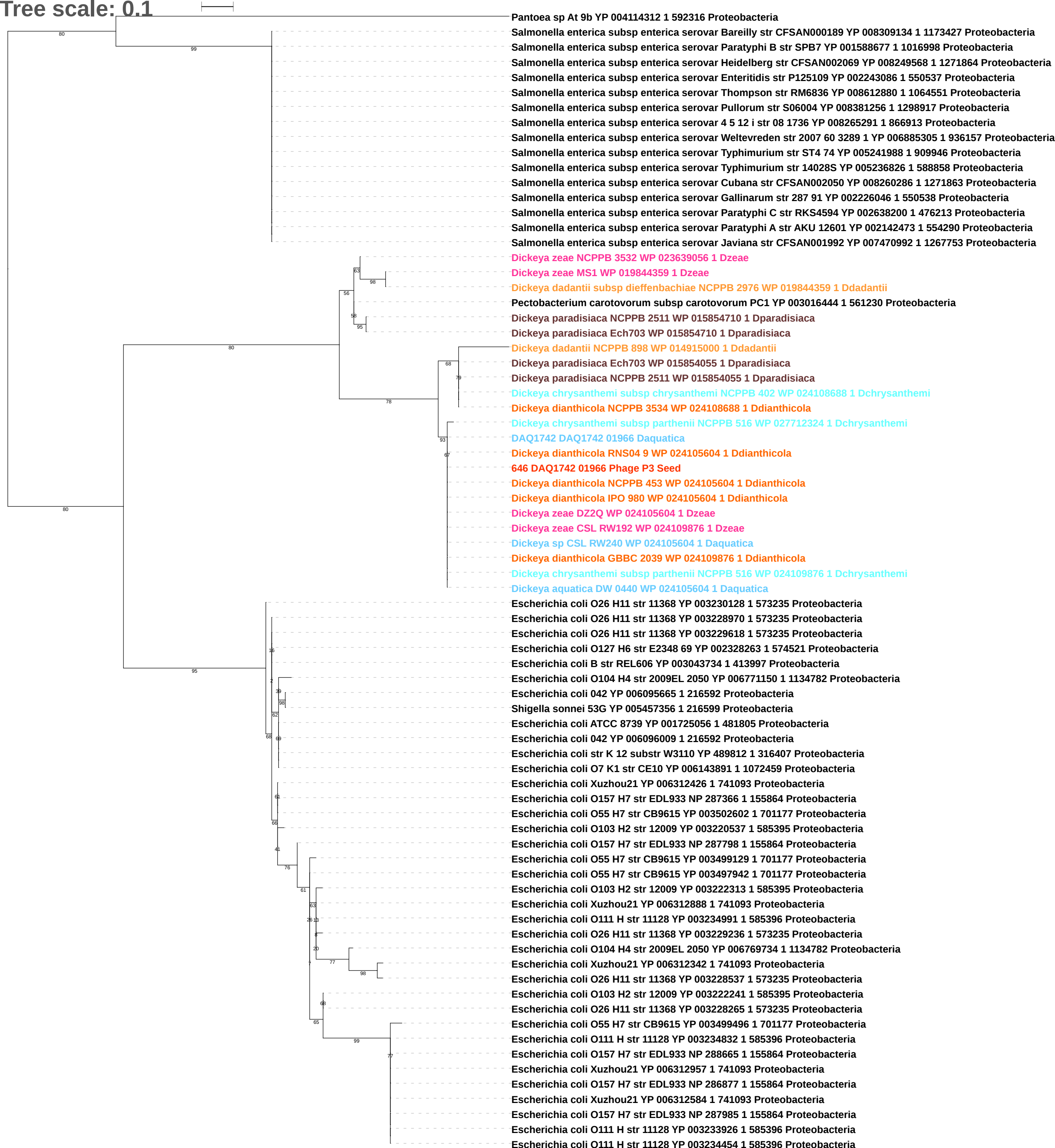
Tree scale: 0.1
Pantoea sp At 9b YP 004114312 1 592316 Proteobacteria
Salmonella enterica subsp enterica serovar Bareilly str CFSAN000189 YP 008309134 1 1173427 Proteobacteria
Salmonella enterica subsp enterica serovar Paratyphi B str SPB7 YP 001588677 1 1016998 Proteobacteria
Salmonella enterica subsp enterica serovar Heidelberg str CFSAN002069 YP 008249568 1 1271864 Proteobacteria
Salmonella enterica subsp enterica serovar Enteritidis str P125109 YP 002243086 1 550537 Proteobacteria
Salmonella enterica subsp enterica serovar Thompson str RM6836 YP 008612880 1 1064551 Proteobacteria
Salmonella enterica subsp enterica serovar Pullorum str S06004 YP 008381256 1 1298917 Proteobacteria
Salmonella enterica subsp enterica serovar 4 5 12 i str 08 1736 YP 008265291 1 866913 Proteobacteria
Salmonella enterica subsp enterica serovar Weltevreden str 2007 60 3289 1 YP 006885305 1 936157 Proteobacteria
Salmonella enterica subsp enterica serovar Typhimurium str ST4 74 YP 005241988 1 909946 Proteobacteria
Salmonella enterica subsp enterica serovar Typhimurium str 14028S YP 005236826 1 588858 Proteobacteria
Salmonella enterica subsp enterica serovar Cubana str CFSAN002050 YP 008260286 1 1271863 Proteobacteria
Salmonella enterica subsp enterica serovar Gallinarum str 287 91 YP 002226046 1 550538 Proteobacteria
Salmonella enterica subsp enterica serovar Paratyphi C str RKS4594 YP 002638200 1 476213 Proteobacteria
Salmonella enterica subsp enterica serovar Paratyphi A str AKU 12601 YP 002142473 1 554290 Proteobacteria
Salmonella enterica subsp enterica serovar Javiana str CFSAN001992 YP 007470992 1 1267753 Proteobacteria
Dickeya zeae NCPPB 3532 WP 023639056 1 Dzeae
Dickeya zeae MS1 WP 019844359 1 Dzeae
Dickeya dadantii subsp dieffenbachiae NCPPB 2976 WP 019844359 1 Ddadantii
Pectobacterium carotovorum subsp carotovorum PC1 YP 003016444 1 561230 Proteobacteria
Dickeya paradisiaca NCPPB 2511 WP 015854710 1 Dparadisiaca
Dickeya paradisiaca Ech703 WP 015854710 1 Dparadisiaca
Dickeya dadantii NCPPB 898 WP 014915000 1 Ddadantii
Dickeya paradisiaca Ech703 WP 015854055 1 Dparadisiaca
Dickeya paradisiaca NCPPB 2511 WP 015854055 1 Dparadisiaca
Dickeya chrysanthemi subsp chrysanthemi NCPPB 402 WP 024108688 1 Dchrysanthemi
Dickeya dianthicola NCPPB 3534 WP 024108688 1 Ddianthicola
Dickeya chrysanthemi subsp parthenii NCPPB 516 WP 027712324 1 Dchrysanthemi
DAQ1742 DAQ1742 01966 Daquatica
Dickeya dianthicola RNS04 9 WP 024105604 1 Ddianthicola
646 DAQ1742 01966 Phage P3 Seed
Dickeya dianthicola NCPPB 453 WP 024105604 1 Ddianthicola
Dickeya dianthicola IPO 980 WP 024105604 1 Ddianthicola
Dickeya zeae DZ2Q WP 024105604 1 Dzeae
Dickeya zeae CSL RW192 WP 024109876 1 Dzeae
Dickeya sp CSL RW240 WP 024105604 1 Daquatica
Dickeya dianthicola GBBC 2039 WP 024109876 1 Ddianthicola
Dickeya chrysanthemi subsp parthenii NCPPB 516 WP 024109876 1 Dchrysanthemi
Dickeya aquatica DW 0440 WP 024105604 1 Daquatica
Escherichia coli O26 H11 str 11368 YP 003230128 1 573235 Proteobacteria
Escherichia coli O26 H11 str 11368 YP 003228970 1 573235 Proteobacteria
Escherichia coli O26 H11 str 11368 YP 003229618 1 573235 Proteobacteria
Escherichia coli O127 H6 str E2348 69 YP 002328263 1 574521 Proteobacteria
Escherichia coli B str REL606 YP 003043734 1 413997 Proteobacteria
Escherichia coli O104 H4 str 2009EL 2050 YP 006771150 1 1134782 Proteobacteria
Escherichia coli 042 YP 006095665 1 216592 Proteobacteria
Shigella sonnei 53G YP 005457356 1 216599 Proteobacteria
Escherichia coli ATCC 8739 YP 001725056 1 481805 Proteobacteria
Escherichia coli 042 YP 006096009 1 216592 Proteobacteria
Escherichia coli str K 12 substr W3110 YP 489812 1 316407 Proteobacteria
Escherichia coli O7 K1 str CE10 YP 006143891 1 1072459 Proteobacteria
Escherichia coli Xuzhou21 YP 006312426 1 741093 Proteobacteria
Escherichia coli O157 H7 str EDL933 NP 287366 1 155864 Proteobacteria
Escherichia coli O55 H7 str CB9615 YP 003502602 1 701177 Proteobacteria
Escherichia coli O103 H2 str 12009 YP 003220537 1 585395 Proteobacteria
Escherichia coli O157 H7 str EDL933 NP 287798 1 155864 Proteobacteria
Escherichia coli O55 H7 str CB9615 YP 003499129 1 701177 Proteobacteria
Escherichia coli O55 H7 str CB9615 YP 003497942 1 701177 Proteobacteria
Escherichia coli O103 H2 str 12009 YP 003222313 1 585395 Proteobacteria
Escherichia coli Xuzhou21 YP 006312888 1 741093 Proteobacteria
Escherichia coli O111 H str 11128 YP 003234991 1 585396 Proteobacteria
Escherichia coli O26 H11 str 11368 YP 003229236 1 573235 Proteobacteria
Escherichia coli O104 H4 str 2009EL 2050 YP 006769734 1 1134782 Proteobacteria
Escherichia coli Xuzhou21 YP 006312342 1 741093 Proteobacteria
Escherichia coli O26 H11 str 11368 YP 003228537 1 573235 Proteobacteria
Escherichia coli O103 H2 str 12009 YP 003222241 1 585395 Proteobacteria
Escherichia coli O26 H11 str 11368 YP 003228265 1 573235 Proteobacteria
Escherichia coli O55 H7 str CB9615 YP 003499496 1 701177 Proteobacteria
Escherichia coli O111 H str 11128 YP 003234832 1 585396 Proteobacteria
Escherichia coli O157 H7 str EDL933 NP 288665 1 155864 Proteobacteria
Escherichia coli Xuzhou21 YP 006312957 1 741093 Proteobacteria
Escherichia coli O157 H7 str EDL933 NP 286877 1 155864 Proteobacteria
Escherichia coli Xuzhou21 YP 006312584 1 741093 Proteobacteria
Escherichia coli O157 H7 str EDL933 NP 287985 1 155864 Proteobacteria
Escherichia coli O111 H str 11128 YP 003233926 1 585396 Proteobacteria
Escherichia coli O111 H str 11128 YP 003234454 1 585396 Proteobacteria

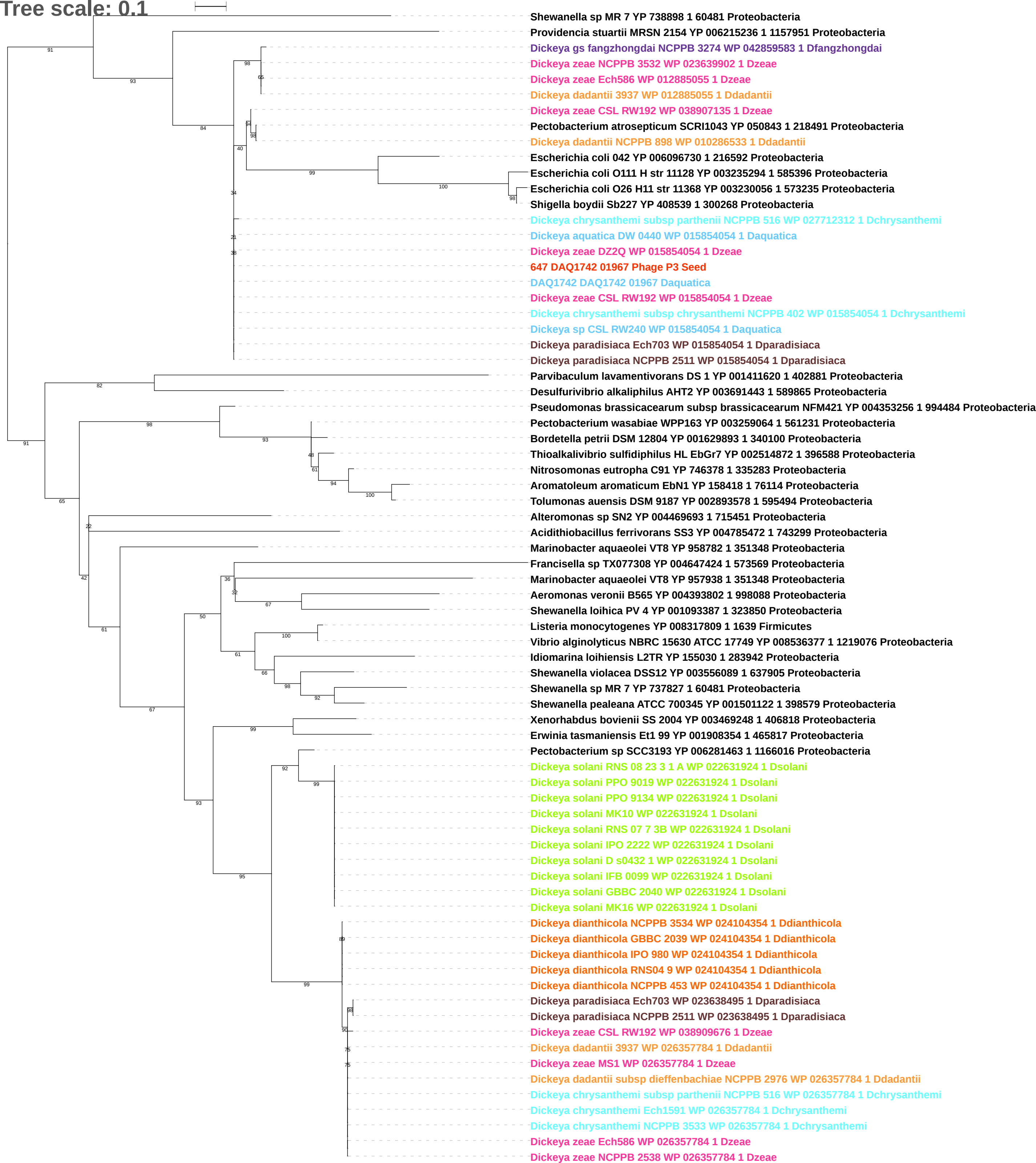

Tree scale: 0.1
Shewanella sp MR 7 YP 738898 1 60481 Proteobacteria
Providencia stuartii MRSN 2154 YP 006215236 1 1157951 Proteobacteria
Dickeya gs fangzhongdai NCPPB 3274 WP 042859583 1 Dfangzhongdai
Dickeya zeae NCPPB 3532 WP 023639902 1 Dzeae
Dickeya zeae Ech586 WP 012885055 1 Dzeae
Dickeya dadantii 3937 WP 012885055 1 Ddadantii
Dickeya zeae CSL RW192 WP 038907135 1 Dzeae
Pectobacterium atrosepticum SCRI1043 YP 050843 1 218491 Proteobacteria
Dickeya dadantii NCPPB 898 WP 010286533 1 Ddadantii
Escherichia coli 042 YP 006096730 1 216592 Proteobacteria
Escherichia coli O111 H str 11128 YP 003235294 1 585396 Proteobacteria
Escherichia coli O26 H11 str 11368 YP 003230056 1 573235 Proteobacteria
Shigella boydii Sb227 YP 408539 1 300268 Proteobacteria
Dickeya chrysanthemi subsp parthenii NCPPB 516 WP 027712312 1 Dchrysanthemi
Dickeya aquatica DW 0440 WP 015854054 1 Daquatica
Dickeya zeae DZ2Q WP 015854054 1 Dzeae
647 DAQ1742 01967 Phage P3 Seed
DAQ1742 DAQ1742 01967 Daquatica
Dickeya zeae CSL RW192 WP 015854054 1 Dzeae
Dickeya chrysanthemi subsp chrysanthemi NCPPB 402 WP 015854054 1 Dchrysanthemi
Dickeya sp CSL RW240 WP 015854054 1 Daquatica
Dickeya paradisiaca Ech703 WP 015854054 1 Dparadisiaca
Dickeya paradisiaca NCPPB 2511 WP 015854054 1 Dparadisiaca
Parvibaculum lavamentivorans DS 1 YP 001411620 1 402881 Proteobacteria
Desulfurivibrio alkaliphilus AHT2 YP 003691443 1 589865 Proteobacteria
Pseudomonas brassicacearum subsp brassicacearum NFM421 YP 004353256 1 994484 Proteobacteria
Pectobacterium wasabiae WPP163 YP 003259064 1 561231 Proteobacteria
Bordetella petrii DSM 12804 YP 001629893 1 340100 Proteobacteria
Thioalkalivibrio sulfidiphilus HL EbGr7 YP 002514872 1 396588 Proteobacteria
Nitrosomonas eutropha C91 YP 746378 1 335283 Proteobacteria
Aromatoleum aromaticum EbN1 YP 158418 1 76114 Proteobacteria
Tolumonas auensis DSM 9187 YP 002893578 1 595494 Proteobacteria
Alteromonas sp SN2 YP 004469693 1 715451 Proteobacteria
Acidithiobacillus ferrivorans SS3 YP 004785472 1 743299 Proteobacteria
Marinobacter aquaeolei VT8 YP 958782 1 351348 Proteobacteria
Francisella sp TX077308 YP 004647424 1 573569 Proteobacteria
Marinobacter aquaeolei VT8 YP 957938 1 351348 Proteobacteria
Aeromonas veronii B565 YP 004393802 1 998088 Proteobacteria
Shewanella loihica PV 4 YP 001093387 1 323850 Proteobacteria
Listeria monocytogenes YP 008317809 1 1639 Firmicutes
Vibrio alginolyticus NBRC 15630 ATCC 17749 YP 008536377 1 1219076 Proteobacteria
Idiomarina loihiensis L2TR YP 155030 1 283942 Proteobacteria
Shewanella violacea DSS12 YP 003556089 1 637905 Proteobacteria
Shewanella sp MR 7 YP 737827 1 60481 Proteobacteria
Shewanella pealeana ATCC 700345 YP 001501122 1 398579 Proteobacteria
Xenorhabdus bovienii SS 2004 YP 003469248 1 406818 Proteobacteria
Erwinia tasmaniensis Et1 99 YP 001908354 1 465817 Proteobacteria
Pectobacterium sp SCC3193 YP 006281463 1 1166016 Proteobacteria
Dickeya solani RNS 08 23 3 1 A WP 022631924 1 Dsolani
Dickeya solani PPO 9019 WP 022631924 1 Dsolani
Dickeya solani PPO 9134 WP 022631924 1 Dsolani
Dickeya solani MK10 WP 022631924 1 Dsolani
Dickeya solani RNS 07 7 3B WP 022631924 1 Dsolani
Dickeya solani IPO 2222 WP 022631924 1 Dsolani
Dickeya solani D s0432 1 WP 022631924 1 Dsolani
Dickeya solani IFB 0099 WP 022631924 1 Dsolani
Dickeya solani GBBC 2040 WP 022631924 1 Dsolani
Dickeya solani MK16 WP 022631924 1 Dsolani
Dickeya dianthicola NCPPB 3534 WP 024104354 1 Ddianthicola
Dickeya dianthicola GBBC 2039 WP 024104354 1 Ddianthicola
Dickeya dianthicola IPO 980 WP 024104354 1 Ddianthicola
Dickeya dianthicola RNS04 9 WP 024104354 1 Ddianthicola
Dickeya dianthicola NCPPB 453 WP 024104354 1 Ddianthicola
Dickeya paradisiaca Ech703 WP 023638495 1 Dparadisiaca
Dickeya paradisiaca NCPPB 2511 WP 023638495 1 Dparadisiaca
Dickeya zeae CSL RW192 WP 038909676 1 Dzeae
Dickeya dadantii 3937 WP 026357784 1 Ddadantii
Dickeya zeae MS1 WP 026357784 1 Dzeae
Dickeya dadantii subsp dieffenbachiae NCPPB 2976 WP 026357784 1 Ddadantii
Dickeya chrysanthemi subsp parthenii NCPPB 516 WP 026357784 1 Dchrysanthemi
Dickeya chrysanthemi Ech1591 WP 026357784 1 Dchrysanthemi
Dickeya chrysanthemi NCPPB 3533 WP 026357784 1 Dchrysanthemi
Dickeya zeae Ech586 WP 026357784 1 Dzeae
Dickeya zeae NCPPB 2538 WP 026357784 1 Dzeae

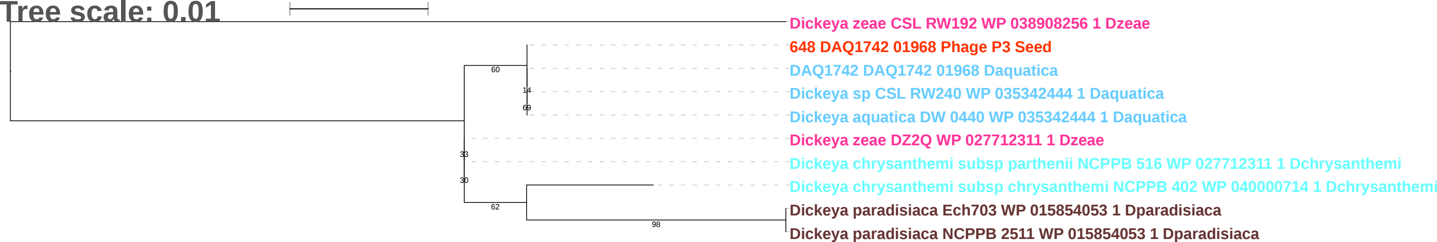
Tree scale: 0.01
Dickeya zeae CSL RW192 WP 038908256 1 Dzeae
648 DAQ1742 01968 Phage P3 Seed
DAQ1742 DAQ1742 01968 Daquatica
Dickeya sp CSL RW240 WP 035342444 1 Daquatica
Dickeya aquatica DW 0440 WP 035342444 1 Daquatica
Dickeya zeae DZ2Q WP 027712311 1 Dzeae
Dickeya chrysanthemi subsp parthenii NCPPB 516 WP 027712311 1 Dchrysanthemi
Dickeya chrysanthemi subsp chrysanthemi NCPPB 402 WP 040000714 1 Dchrysanthemi
Dickeya paradisiaca Ech703 WP 015854053 1 Dparadisiaca
Dickeya paradisiaca NCPPB 2511 WP 015854053 1 Dparadisiaca
60
14
69
33
30
62
98

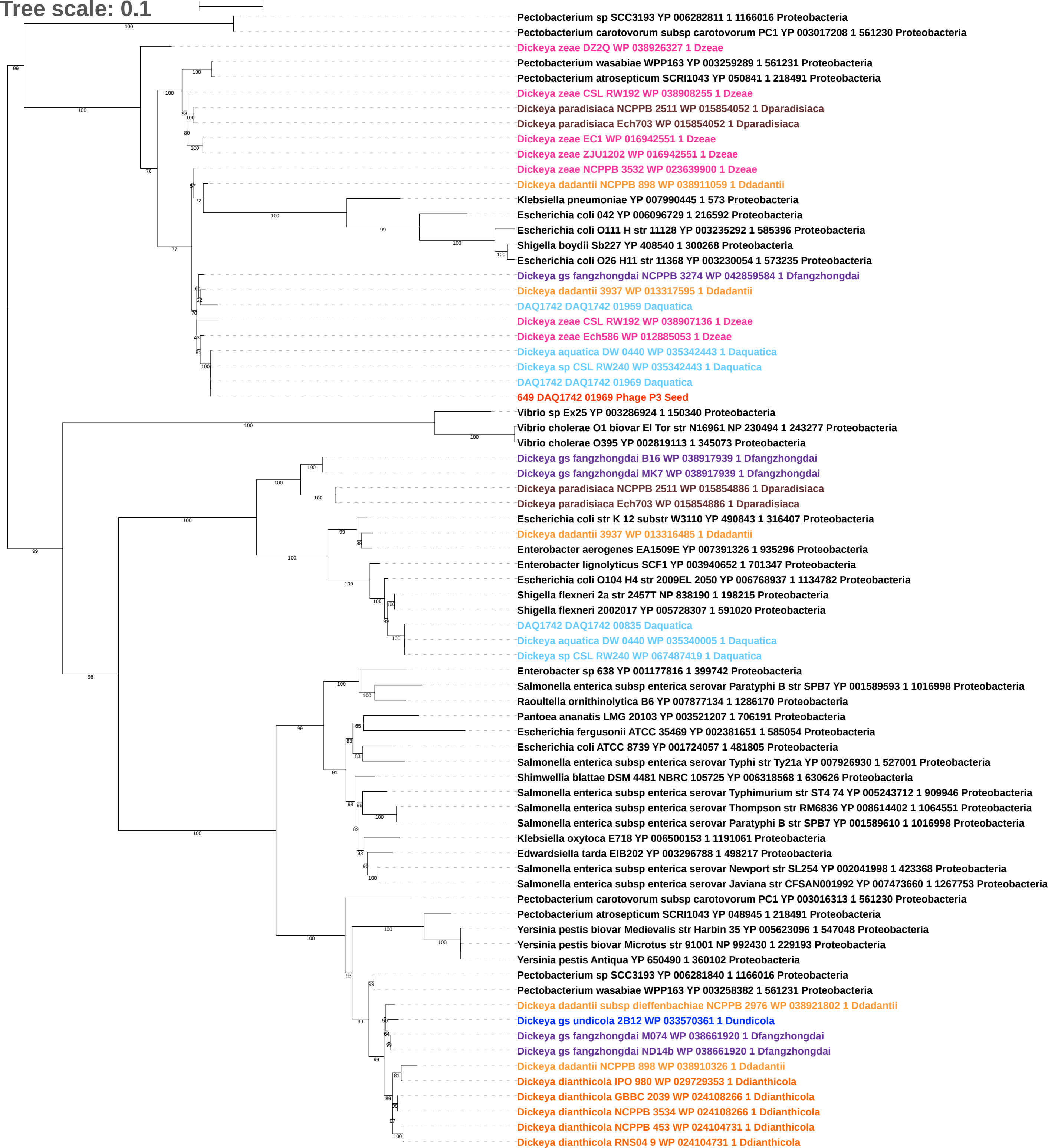
Tree scale: 0.1
Pectobacterium sp SCC3193 YP 006282811 1 1166016 Proteobacteria
Pectobacterium carotovorum subsp carotovorum PC1 YP 003017208 1 561230 Proteobacteria
Dickeya zeae DZ2Q WP 038926327 1 Dzeae
Pectobacterium wasabiae WPP163 YP 003259289 1 561231 Proteobacteria
Pectobacterium atrosepticum SCRI1043 YP 050841 1 218491 Proteobacteria
Dickeya zeae CSL RW192 WP 038908255 1 Dzeae
Dickeya paradisiaca NCPPB 2511 WP 015854052 1 Dparadisiaca
Dickeya paradisiaca Ech703 WP 015854052 1 Dparadisiaca
Dickeya zeae EC1 WP 016942551 1 Dzeae
Dickeya zeae ZJU1202 WP 016942551 1 Dzeae
Dickeya zeae NCPPB 3532 WP 023639900 1 Dzeae
Dickeya dadantii NCPPB 898 WP 038911059 1 Ddadantii
Klebsiella pneumoniae YP 007990445 1 573 Proteobacteria
Escherichia coli 042 YP 006096729 1 216592 Proteobacteria
Escherichia coli O111 H str 11128 YP 003235292 1 585396 Proteobacteria
Shigella boydii Sb227 YP 408540 1 300268 Proteobacteria
Escherichia coli O26 H11 str 11368 YP 003230054 1 573235 Proteobacteria
Dickeya gs fangzhongdai NCPPB 3274 WP 042859584 1 Dfangzhongdai
Dickeya dadantii 3937 WP 013317595 1 Ddadantii
DAQ1742 DAQ1742 01959 Daquatica
Dickeya zeae CSL RW192 WP 038907136 1 Dzeae
Dickeya zeae Ech586 WP 012885053 1 Dzeae
Dickeya aquatica DW 0440 WP 035342443 1 Daquatica
Dickeya sp CSL RW240 WP 035342443 1 Daquatica
DAQ1742 DAQ1742 01969 Daquatica
649 DAQ1742 01969 Phage P3 Seed
Vibrio sp Ex25 YP 003286924 1 150340 Proteobacteria
Vibrio cholerae O1 biovar El Tor str N16961 NP 230494 1 243277 Proteobacteria
Vibrio cholerae O395 YP 002819113 1 345073 Proteobacteria
Dickeya gs fangzhongdai B16 WP 038917939 1 Dfangzhongdai
Dickeya gs fangzhongdai MK7 WP 038917939 1 Dfangzhongdai
Dickeya paradisiaca NCPPB 2511 WP 015854886 1 Dparadisiaca
Dickeya paradisiaca Ech703 WP 015854886 1 Dparadisiaca
Escherichia coli str K 12 substr W3110 YP 490843 1 316407 Proteobacteria
Dickeya dadantii 3937 WP 013316485 1 Ddadantii
Enterobacter aerogenes EA1509E YP 007391326 1 935296 Proteobacteria
Enterobacter lignolyticus SCF1 YP 003940652 1 701347 Proteobacteria
Escherichia coli O104 H4 str 2009EL 2050 YP 006768937 1 1134782 Proteobacteria
Shigella flexneri 2a str 2457T NP 838190 1 198215 Proteobacteria
Shigella flexneri 2002017 YP 005728307 1 591020 Proteobacteria
DAQ1742 DAQ1742 00835 Daquatica
Dickeya aquatica DW 0440 WP 035340005 1 Daquatica
Dickeya sp CSL RW240 WP 067487419 1 Daquatica
Enterobacter sp 638 YP 001177816 1 399742 Proteobacteria
Salmonella enterica subsp enterica serovar Paratyphi B str SPB7 YP 001589593 1 1016998 Proteobacteria
Raoultella ornithinolytica B6 YP 007877134 1 1286170 Proteobacteria
Pantoea ananatis LMG 20103 YP 003521207 1 706191 Proteobacteria
Escherichia fergusonii ATCC 35469 YP 002381651 1 585054 Proteobacteria
Escherichia coli ATCC 8739 YP 001724057 1 481805 Proteobacteria
Salmonella enterica subsp enterica serovar Typhi str Ty21a YP 007926930 1 527001 Proteobacteria
Shimwellia blattae DSM 4481 NBRC 105725 YP 006318568 1 630626 Proteobacteria
Salmonella enterica subsp enterica serovar Typhimurium str ST4 74 YP 005243712 1 909946 Proteobacteria
Salmonella enterica subsp enterica serovar Thompson str RM6836 YP 008614402 1 1064551 Proteobacteria
Salmonella enterica subsp enterica serovar Paratyphi B str SPB7 YP 001589610 1 1016998 Proteobacteria
Klebsiella oxytoca E718 YP 006500153 1 1191061 Proteobacteria
Edwardsiella tarda EIB202 YP 003296788 1 498217 Proteobacteria
Salmonella enterica subsp enterica serovar Newport str SL254 YP 002041998 1 423368 Proteobacteria
Salmonella enterica subsp enterica serovar Javiana str CFSAN001992 YP 007473660 1 1267753 Proteobacteria
Pectobacterium carotovorum subsp carotovorum PC1 YP 003016313 1 561230 Proteobacteria
Pectobacterium atrosepticum SCRI1043 YP 048945 1 218491 Proteobacteria
Yersinia pestis biovar Medievalis str Harbin 35 YP 005623096 1 547048 Proteobacteria
Yersinia pestis biovar Microtus str 91001 NP 992430 1 229193 Proteobacteria
Yersinia pestis Antiqua YP 650490 1 360102 Proteobacteria
Pectobacterium sp SCC3193 YP 006281840 1 1166016 Proteobacteria
Pectobacterium wasabiae WPP163 YP 003258382 1 561231 Proteobacteria
Dickeya dadantii subsp dieffenbachiae NCPPB 2976 WP 038921802 1 Ddadantii
Dickeya gs undicola 2B12 WP 033570361 1 Dundicola
Dickeya gs fangzhongdai M074 WP 038661920 1 Dfangzhongdai
Dickeya gs fangzhongdai ND14b WP 038661920 1 Dfangzhongdai
Dickeya dadantii NCPPB 898 WP 038910326 1 Ddadantii
Dickeya dianthicola IPO 980 WP 029729353 1 Ddianthicola
Dickeya dianthicola GBBC 2039 WP 024108266 1 Ddianthicola
Dickeya dianthicola NCPPB 3534 WP 024108266 1 Ddianthicola
Dickeya dianthicola NCPPB 453 WP 024104731 1 Ddianthicola
Dickeya dianthicola RNS04 9 WP 024104731 1 Ddianthicola

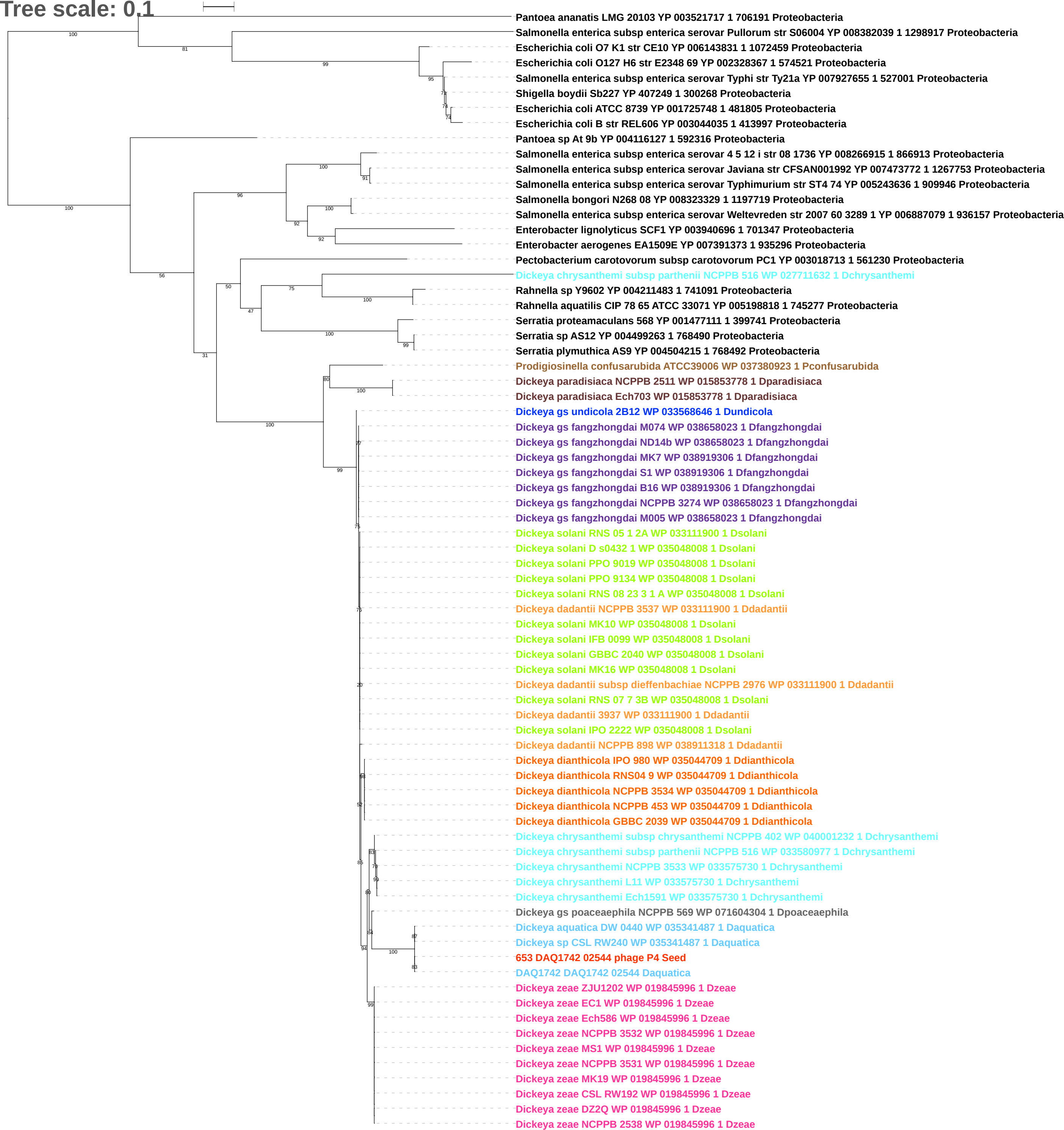
Tree scale: 0.1
Pantoea ananatis LMG 20103 YP 003521717 1 706191 Proteobacteria
Salmonella enterica subsp enterica serovar Pullorum str S06004 YP 008382039 1 1298917 Proteobacteria
Escherichia coli O7 K1 str CE10 YP 006143831 1 1072459 Proteobacteria
Escherichia coli O127 H6 str E2348 69 YP 002328367 1 574521 Proteobacteria
Salmonella enterica subsp enterica serovar Typhi str Ty21a YP 007927655 1 527001 Proteobacteria
Shigella boydii Sb227 YP 407249 1 300268 Proteobacteria
Escherichia coli ATCC 8739 YP 001725748 1 481805 Proteobacteria
Escherichia coli B str REL606 YP 003044035 1 413997 Proteobacteria
Pantoea sp At 9b YP 004116127 1 592316 Proteobacteria
Salmonella enterica subsp enterica serovar 4 5 12 i str 08 1736 YP 008266915 1 866913 Proteobacteria
Salmonella enterica subsp enterica serovar Javiana str CFSAN001992 YP 007473772 1 1267753 Proteobacteria
Salmonella enterica subsp enterica serovar Typhimurium str ST4 74 YP 005243636 1 909946 Proteobacteria
Salmonella bongori N268 08 YP 008323329 1 1197719 Proteobacteria
Salmonella enterica subsp enterica serovar Weltevreden str 2007 60 3289 1 YP 006887079 1 936157 Proteobacteria
Enterobacter lignolyticus SCF1 YP 003940696 1 701347 Proteobacteria
Enterobacter aerogenes EA1509E YP 007391373 1 935296 Proteobacteria
Pectobacterium carotovorum subsp carotovorum PC1 YP 003018713 1 561230 Proteobacteria
Dickeya chrysanthemi subsp parthenii NCPPB 516 WP 027711632 1 Dchrysanthemi
Rahnella sp Y9602 YP 004211483 1 741091 Proteobacteria
Rahnella aquatilis CIP 78 65 ATCC 33071 YP 005198818 1 745277 Proteobacteria
Serratia proteamaculans 568 YP 001477111 1 399741 Proteobacteria
Serratia sp AS12 YP 004499263 1 768490 Proteobacteria
Serratia plymuthica AS9 YP 004504215 1 768492 Proteobacteria
Prodigiosinella confusarubida ATCC39006 WP 037380923 1 Pconfusarubida
Dickeya paradisiaca NCPPB 2511 WP 015853778 1 Dparadisiaca
Dickeya paradisiaca Ech703 WP 015853778 1 Dparadisiaca
Dickeya gs undicola 2B12 WP 033568646 1 Dundicola
Dickeya gs fangzhongdai M074 WP 038658023 1 Dfangzhongdai
Dickeya gs fangzhongdai ND14b WP 038658023 1 Dfangzhongdai
Dickeya gs fangzhongdai MK7 WP 038919306 1 Dfangzhongdai
Dickeya gs fangzhongdai S1 WP 038919306 1 Dfangzhongdai
Dickeya gs fangzhongdai B16 WP 038919306 1 Dfangzhongdai
Dickeya gs fangzhongdai NCPPB 3274 WP 038658023 1 Dfangzhongdai
Dickeya gs fangzhongdai M005 WP 038658023 1 Dfangzhongdai
Dickeya solani RNS 05 1 2A WP 033111900 1 Dsolani
Dickeya solani D s0432 1 WP 035048008 1 Dsolani
Dickeya solani PPO 9019 WP 035048008 1 Dsolani
Dickeya solani PPO 9134 WP 035048008 1 Dsolani
Dickeya solani RNS 08 23 3 1 A WP 035048008 1 Dsolani
Dickeya dadantii NCPPB 3537 WP 033111900 1 Ddadantii
Dickeya solani MK10 WP 035048008 1 Dsolani
Dickeya solani IFB 0099 WP 035048008 1 Dsolani
Dickeya solani GBBC 2040 WP 035048008 1 Dsolani
Dickeya solani MK16 WP 035048008 1 Dsolani
Dickeya dadantii subsp dieffenbachiae NCPPB 2976 WP 033111900 1 Ddadantii
Dickeya solani RNS 07 7 3B WP 035048008 1 Dsolani
Dickeya dadantii 3937 WP 033111900 1 Ddadantii
Dickeya solani IPO 2222 WP 035048008 1 Dsolani
Dickeya dadantii NCPPB 898 WP 038911318 1 Ddadantii
Dickeya dianthicola IPO 980 WP 035044709 1 Ddianthicola
Dickeya dianthicola RNS04 9 WP 035044709 1 Ddianthicola
Dickeya dianthicola NCPPB 3534 WP 035044709 1 Ddianthicola
Dickeya dianthicola NCPPB 453 WP 035044709 1 Ddianthicola
Dickeya dianthicola GBBC 2039 WP 035044709 1 Ddianthicola
Dickeya chrysanthemi subsp chrysanthemi NCPPB 402 WP 040001232 1 Dchrysanthemi
Dickeya chrysanthemi subsp parthenii NCPPB 516 WP 033580977 1 Dchrysanthemi
Dickeya chrysanthemi NCPPB 3533 WP 033575730 1 Dchrysanthemi
Dickeya chrysanthemi L11 WP 033575730 1 Dchrysanthemi
Dickeya chrysanthemi Ech1591 WP 033575730 1 Dchrysanthemi
Dickeya gs poaceaephila NCPPB 569 WP 071604304 1 Dpoaceaephila
Dickeya aquatica DW 0440 WP 035341487 1 Daquatica
Dickeya sp CSL RW240 WP 035341487 1 Daquatica
653 DAQ1742 02544 phage P4 Seed
DAQ1742 DAQ1742 02544 Daquatica
Dickeya zeae ZJU1202 WP 019845996 1 Dzeae
Dickeya zeae EC1 WP 019845996 1 Dzeae
Dickeya zeae Ech586 WP 019845996 1 Dzeae
Dickeya zeae NCPPB 3532 WP 019845996 1 Dzeae
Dickeya zeae MS1 WP 019845996 1 Dzeae
Dickeya zeae NCPPB 3531 WP 019845996 1 Dzeae
Dickeya zeae MK19 WP 019845996 1 Dzeae
Dickeya zeae CSL RW192 WP 019845996 1 Dzeae
Dickeya zeae DZ2Q WP 019845996 1 Dzeae
Dickeya zeae NCPPB 2538 WP 019845996 1 Dzeae

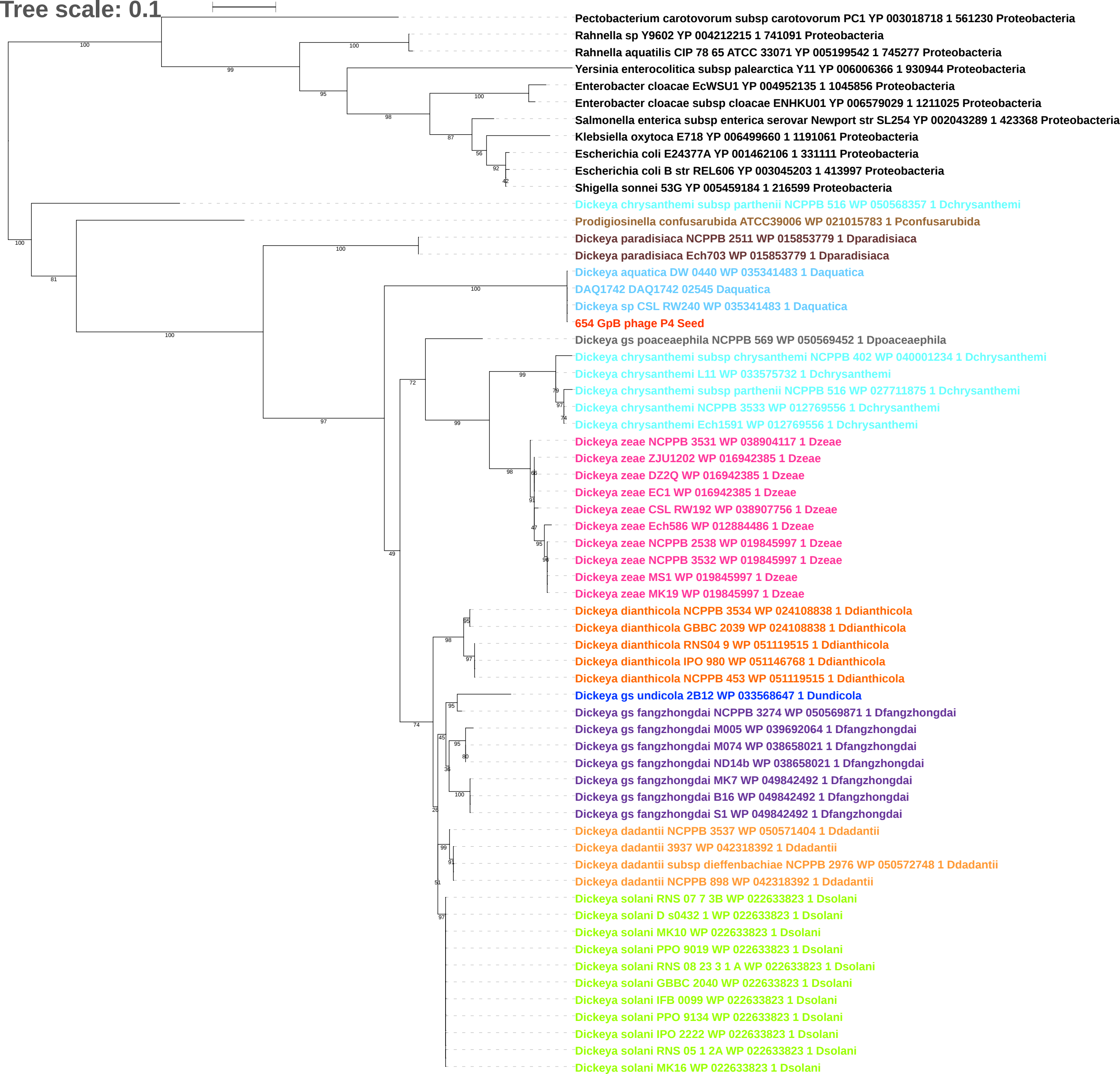

Tree scale: 0.1
Pectobacterium carotovorum subsp carotovorum PC1 YP 003018718 1 561230 Proteobacteria
Rahnella sp Y9602 YP 004212215 1 741091 Proteobacteria
Rahnella aquatilis CIP 78 65 ATCC 33071 YP 005199542 1 745277 Proteobacteria
Yersinia enterocolitica subsp palearctica Y11 YP 006006366 1 930944 Proteobacteria
Enterobacter cloacae EcWSU1 YP 004952135 1 1045856 Proteobacteria
Enterobacter cloacae subsp cloacae ENHKU01 YP 006579029 1 1211025 Proteobacteria
Salmonella enterica subsp enterica serovar Newport str SL254 YP 002043289 1 423368 Proteobacteria
Klebsiella oxytoca E718 YP 006499660 1 1191061 Proteobacteria
Escherichia coli E24377A YP 001462106 1 331111 Proteobacteria
Escherichia coli B str REL606 YP 003045203 1 413997 Proteobacteria
Shigella sonnei 53G YP 005459184 1 216599 Proteobacteria
Dickeya chrysanthemi subsp parthenii NCPPB 516 WP 050568357 1 Dchrysanthemi
Prodigiosinella confusarubida ATCC39006 WP 021015783 1 Pconfusarubida
Dickeya paradisiaca NCPPB 2511 WP 015853779 1 Dparadisiaca
Dickeya paradisiaca Ech703 WP 015853779 1 Dparadisiaca
Dickeya aquatica DW 0440 WP 035341483 1 Daquatica
DAQ1742 DAQ1742 02545 Daquatica
Dickeya sp CSL RW240 WP 035341483 1 Daquatica
654 GpB phage P4 Seed
Dickeya gs poaceaephila NCPPB 569 WP 050569452 1 Dpoaceaephila
Dickeya chrysanthemi subsp chrysanthemi NCPPB 402 WP 040001234 1 Dchrysanthemi
Dickeya chrysanthemi L11 WP 033575732 1 Dchrysanthemi
Dickeya chrysanthemi subsp parthenii NCPPB 516 WP 027711875 1 Dchrysanthemi
Dickeya chrysanthemi NCPPB 3533 WP 012769556 1 Dchrysanthemi
Dickeya chrysanthemi Ech1591 WP 012769556 1 Dchrysanthemi
Dickeya zeae NCPPB 3531 WP 038904117 1 Dzeae
Dickeya zeae ZJU1202 WP 016942385 1 Dzeae
Dickeya zeae DZ2Q WP 016942385 1 Dzeae
Dickeya zeae EC1 WP 016942385 1 Dzeae
Dickeya zeae CSL RW192 WP 038907756 1 Dzeae
Dickeya zeae Ech586 WP 012884486 1 Dzeae
Dickeya zeae NCPPB 2538 WP 019845997 1 Dzeae
Dickeya zeae NCPPB 3532 WP 019845997 1 Dzeae
Dickeya zeae MS1 WP 019845997 1 Dzeae
Dickeya zeae MK19 WP 019845997 1 Dzeae
Dickeya dianthicola NCPPB 3534 WP 024108838 1 Ddianthicola
Dickeya dianthicola GBBC 2039 WP 024108838 1 Ddianthicola
Dickeya dianthicola RNS04 9 WP 051119515 1 Ddianthicola
Dickeya dianthicola IPO 980 WP 051146768 1 Ddianthicola
Dickeya dianthicola NCPPB 453 WP 051119515 1 Ddianthicola
Dickeya gs undicola 2B12 WP 033568647 1 Dundicola
Dickeya gs fangzhongdai NCPPB 3274 WP 050569871 1 Dfangzhongdai
Dickeya gs fangzhongdai M005 WP 039692064 1 Dfangzhongdai
Dickeya gs fangzhongdai M074 WP 038658021 1 Dfangzhongdai
Dickeya gs fangzhongdai ND14b WP 038658021 1 Dfangzhongdai
Dickeya gs fangzhongdai MK7 WP 049842492 1 Dfangzhongdai
Dickeya gs fangzhongdai B16 WP 049842492 1 Dfangzhongdai
Dickeya gs fangzhongdai S1 WP 049842492 1 Dfangzhongdai
Dickeya dadantii NCPPB 3537 WP 050571404 1 Ddadantii
Dickeya dadantii 3937 WP 042318392 1 Ddadantii
Dickeya dadantii subsp dieffenbachiae NCPPB 2976 WP 050572748 1 Ddadantii
Dickeya dadantii NCPPB 898 WP 042318392 1 Ddadantii
Dickeya solani RNS 07 7 3B WP 022633823 1 Dsolani
Dickeya solani D s0432 1 WP 022633823 1 Dsolani
Dickeya solani MK10 WP 022633823 1 Dsolani
Dickeya solani PPO 9019 WP 022633823 1 Dsolani
Dickeya solani RNS 08 23 3 1 A WP 022633823 1 Dsolani
Dickeya solani GBBC 2040 WP 022633823 1 Dsolani
Dickeya solani IFB 0099 WP 022633823 1 Dsolani
Dickeya solani PPO 9134 WP 022633823 1 Dsolani
Dickeya solani IPO 2222 WP 022633823 1 Dsolani
Dickeya solani RNS 05 1 2A WP 022633823 1 Dsolani
Dickeya solani MK16 WP 022633823 1 Dsolani

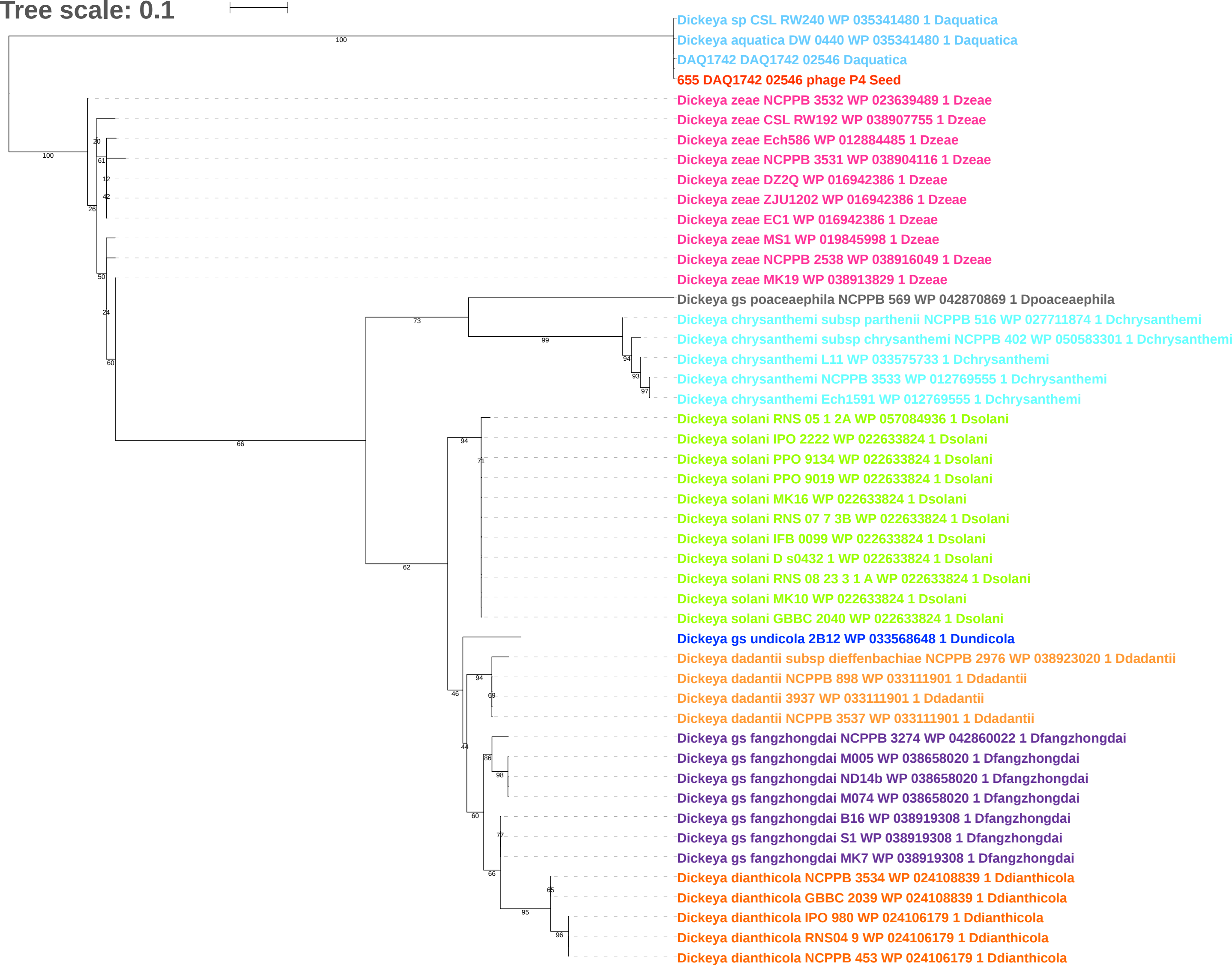
Tree scale: 0.1
Dickeya sp CSL RW240 WP 035341480 1 Daquatica
Dickeya aquatica DW 0440 WP 035341480 1 Daquatica
DAQ1742 DAQ1742 02546 Daquatica
655 DAQ1742 02546 phage P4 Seed
Dickeya zeae NCPPB 3532 WP 023639489 1 Dzeae
Dickeya zeae CSL RW192 WP 038907755 1 Dzeae
Dickeya zeae Ech586 WP 012884485 1 Dzeae
Dickeya zeae NCPPB 3531 WP 038904116 1 Dzeae
Dickeya zeae DZ2Q WP 016942386 1 Dzeae
Dickeya zeae ZJU1202 WP 016942386 1 Dzeae
Dickeya zeae EC1 WP 016942386 1 Dzeae
Dickeya zeae MS1 WP 019845998 1 Dzeae
Dickeya zeae NCPPB 2538 WP 038916049 1 Dzeae
Dickeya zeae MK19 WP 038913829 1 Dzeae
Dickeya gs poaceaephila NCPPB 569 WP 042870869 1 Dpoaceaephila
Dickeya chrysanthemi subsp parthenii NCPPB 516 WP 027711874 1 Dchrysanthemi
Dickeya chrysanthemi subsp chrysanthemi NCPPB 402 WP 050583301 1 Dchrysanthemi
Dickeya chrysanthemi L11 WP 033575733 1 Dchrysanthemi
Dickeya chrysanthemi NCPPB 3533 WP 012769555 1 Dchrysanthemi
Dickeya chrysanthemi Ech1591 WP 012769555 1 Dchrysanthemi
Dickeya solani RNS 05 1 2A WP 057084936 1 Dsolani
Dickeya solani IPO 2222 WP 022633824 1 Dsolani
Dickeya solani PPO 9134 WP 022633824 1 Dsolani
Dickeya solani PPO 9019 WP 022633824 1 Dsolani
Dickeya solani MK16 WP 022633824 1 Dsolani
Dickeya solani RNS 07 7 3B WP 022633824 1 Dsolani
Dickeya solani IFB 0099 WP 022633824 1 Dsolani
Dickeya solani D s0432 1 WP 022633824 1 Dsolani
Dickeya solani RNS 08 23 3 1 A WP 022633824 1 Dsolani
Dickeya solani MK10 WP 022633824 1 Dsolani
Dickeya solani GBBC 2040 WP 022633824 1 Dsolani
Dickeya gs undicola 2B12 WP 033568648 1 Dundicola
Dickeya dadantii subsp dieffenbachiae NCPPB 2976 WP 038923020 1 Ddadantii
Dickeya dadantii NCPPB 898 WP 033111901 1 Ddadantii
Dickeya dadantii 3937 WP 033111901 1 Ddadantii
Dickeya dadantii NCPPB 3537 WP 033111901 1 Ddadantii
Dickeya gs fangzhongdai NCPPB 3274 WP 042860022 1 Dfangzhongdai
Dickeya gs fangzhongdai M005 WP 038658020 1 Dfangzhongdai
Dickeya gs fangzhongdai ND14b WP 038658020 1 Dfangzhongdai
Dickeya gs fangzhongdai M074 WP 038658020 1 Dfangzhongdai
Dickeya gs fangzhongdai B16 WP 038919308 1 Dfangzhongdai
Dickeya gs fangzhongdai S1 WP 038919308 1 Dfangzhongdai
Dickeya gs fangzhongdai MK7 WP 038919308 1 Dfangzhongdai
Dickeya dianthicola NCPPB 3534 WP 024108839 1 Ddianthicola
Dickeya dianthicola GBBC 2039 WP 024108839 1 Ddianthicola
Dickeya dianthicola IPO 980 WP 024106179 1 Ddianthicola
Dickeya dianthicola RNS04 9 WP 024106179 1 Ddianthicola
Dickeya dianthicola NCPPB 453 WP 024106179 1 Ddianthicola

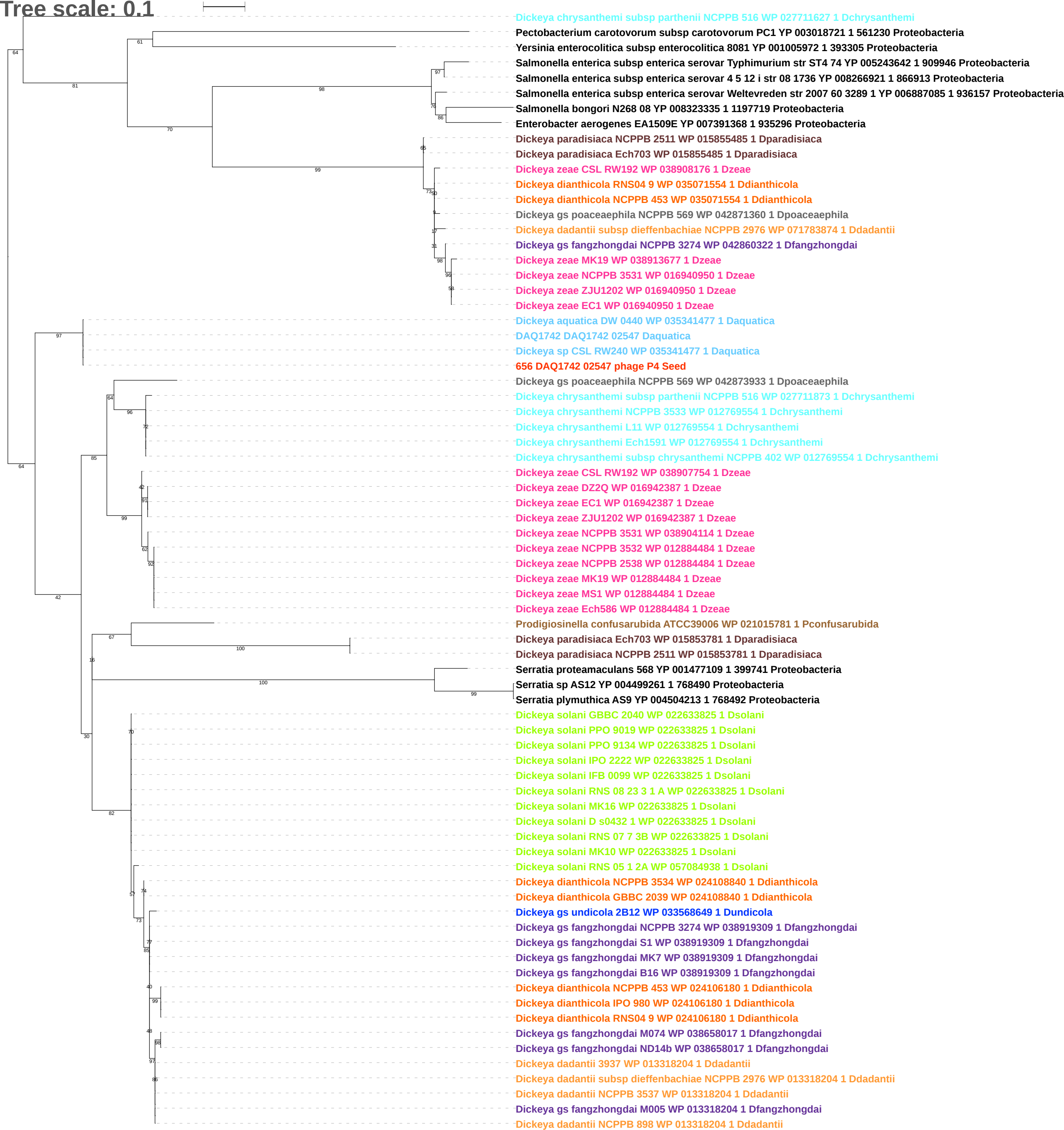
Tree scale: 0.1
Dickeya chrysanthemi subsp parthenii NCPPB 516 WP 027711627 1 Dchrysanthemi
Pectobacterium carotovorum subsp carotovorum PC1 YP 003018721 1 561230 Proteobacteria
Yersinia enterocolitica subsp enterocolitica 8081 YP 001005972 1 393305 Proteobacteria
Salmonella enterica subsp enterica serovar Typhimurium str ST4 74 YP 005243642 1 909946 Proteobacteria
Salmonella enterica subsp enterica serovar 4 5 12 i str 08 1736 YP 008266921 1 866913 Proteobacteria
Salmonella enterica subsp enterica serovar Weltevreden str 2007 60 3289 1 YP 006887085 1 936157 Proteobacteria
Salmonella bongori N268 08 YP 008323335 1 1197719 Proteobacteria
Enterobacter aerogenes EA1509E YP 007391368 1 935296 Proteobacteria
Dickeya paradisiaca NCPPB 2511 WP 015855485 1 Dparadisiaca
Dickeya paradisiaca Ech703 WP 015855485 1 Dparadisiaca
Dickeya zeae CSL RW192 WP 038908176 1 Dzeae
Dickeya dianthicola RNS04 9 WP 035071554 1 Ddianthicola
Dickeya dianthicola NCPPB 453 WP 035071554 1 Ddianthicola
Dickeya gs poaceaephila NCPPB 569 WP 042871360 1 Dpoaceaephila
Dickeya dadantii subsp dieffenbachiae NCPPB 2976 WP 071783874 1 Ddadantii
Dickeya gs fangzhongdai NCPPB 3274 WP 042860322 1 Dfangzhongdai
Dickeya zeae MK19 WP 038913677 1 Dzeae
Dickeya zeae NCPPB 3531 WP 016940950 1 Dzeae
Dickeya zeae ZJU1202 WP 016940950 1 Dzeae
Dickeya zeae EC1 WP 016940950 1 Dzeae
Dickeya aquatica DW 0440 WP 035341477 1 Daquatica
DAQ1742 DAQ1742 02547 Daquatica
Dickeya sp CSL RW240 WP 035341477 1 Daquatica
656 DAQ1742 02547 phage P4 Seed
Dickeya gs poaceaephila NCPPB 569 WP 042873933 1 Dpoaceaephila
Dickeya chrysanthemi subsp parthenii NCPPB 516 WP 027711873 1 Dchrysanthemi
Dickeya chrysanthemi NCPPB 3533 WP 012769554 1 Dchrysanthemi
Dickeya chrysanthemi L11 WP 012769554 1 Dchrysanthemi
Dickeya chrysanthemi Ech1591 WP 012769554 1 Dchrysanthemi
Dickeya chrysanthemi subsp chrysanthemi NCPPB 402 WP 012769554 1 Dchrysanthemi
Dickeya zeae CSL RW192 WP 038907754 1 Dzeae
Dickeya zeae DZ2Q WP 016942387 1 Dzeae
Dickeya zeae EC1 WP 016942387 1 Dzeae
Dickeya zeae ZJU1202 WP 016942387 1 Dzeae
Dickeya zeae NCPPB 3531 WP 038904114 1 Dzeae
Dickeya zeae NCPPB 3532 WP 012884484 1 Dzeae
Dickeya zeae NCPPB 2538 WP 012884484 1 Dzeae
Dickeya zeae MK19 WP 012884484 1 Dzeae
Dickeya zeae MS1 WP 012884484 1 Dzeae
Dickeya zeae Ech586 WP 012884484 1 Dzeae
Prodigiosinella confusarubida ATCC39006 WP 021015781 1 Pconfusarubida
Dickeya paradisiaca Ech703 WP 015853781 1 Dparadisiaca
Dickeya paradisiaca NCPPB 2511 WP 015853781 1 Dparadisiaca
Serratia proteamaculans 568 YP 001477109 1 399741 Proteobacteria
Serratia sp AS12 YP 004499261 1 768490 Proteobacteria
Serratia plymuthica AS9 YP 004504213 1 768492 Proteobacteria
Dickeya solani GBBC 2040 WP 022633825 1 Dsolani
Dickeya solani PPO 9019 WP 022633825 1 Dsolani
Dickeya solani PPO 9134 WP 022633825 1 Dsolani
Dickeya solani IPO 2222 WP 022633825 1 Dsolani
Dickeya solani IFB 0099 WP 022633825 1 Dsolani
Dickeya solani RNS 08 23 3 1 A WP 022633825 1 Dsolani
Dickeya solani MK16 WP 022633825 1 Dsolani
Dickeya solani D s0432 1 WP 022633825 1 Dsolani
Dickeya solani RNS 07 7 3B WP 022633825 1 Dsolani
Dickeya solani MK10 WP 022633825 1 Dsolani
Dickeya solani RNS 05 1 2A WP 057084938 1 Dsolani
Dickeya dianthicola NCPPB 3534 WP 024108840 1 Ddianthicola
Dickeya dianthicola GBBC 2039 WP 024108840 1 Ddianthicola
Dickeya gs undicola 2B12 WP 033568649 1 Dundicola
Dickeya gs fangzhongdai NCPPB 3274 WP 038919309 1 Dfangzhongdai
Dickeya gs fangzhongdai S1 WP 038919309 1 Dfangzhongdai
Dickeya gs fangzhongdai MK7 WP 038919309 1 Dfangzhongdai
Dickeya gs fangzhongdai B16 WP 038919309 1 Dfangzhongdai
Dickeya dianthicola NCPPB 453 WP 024106180 1 Ddianthicola
Dickeya dianthicola IPO 980 WP 024106180 1 Ddianthicola
Dickeya dianthicola RNS04 9 WP 024106180 1 Ddianthicola
Dickeya gs fangzhongdai M074 WP 038658017 1 Dfangzhongdai
Dickeya gs fangzhongdai ND14b WP 038658017 1 Dfangzhongdai
Dickeya dadantii 3937 WP 013318204 1 Ddadantii
Dickeya dadantii subsp dieffenbachiae NCPPB 2976 WP 013318204 1 Ddadantii
Dickeya dadantii NCPPB 3537 WP 013318204 1 Ddadantii
Dickeya gs fangzhongdai M005 WP 013318204 1 Dfangzhongdai
Dickeya dadantii NCPPB 898 WP 013318204 1 Ddadantii

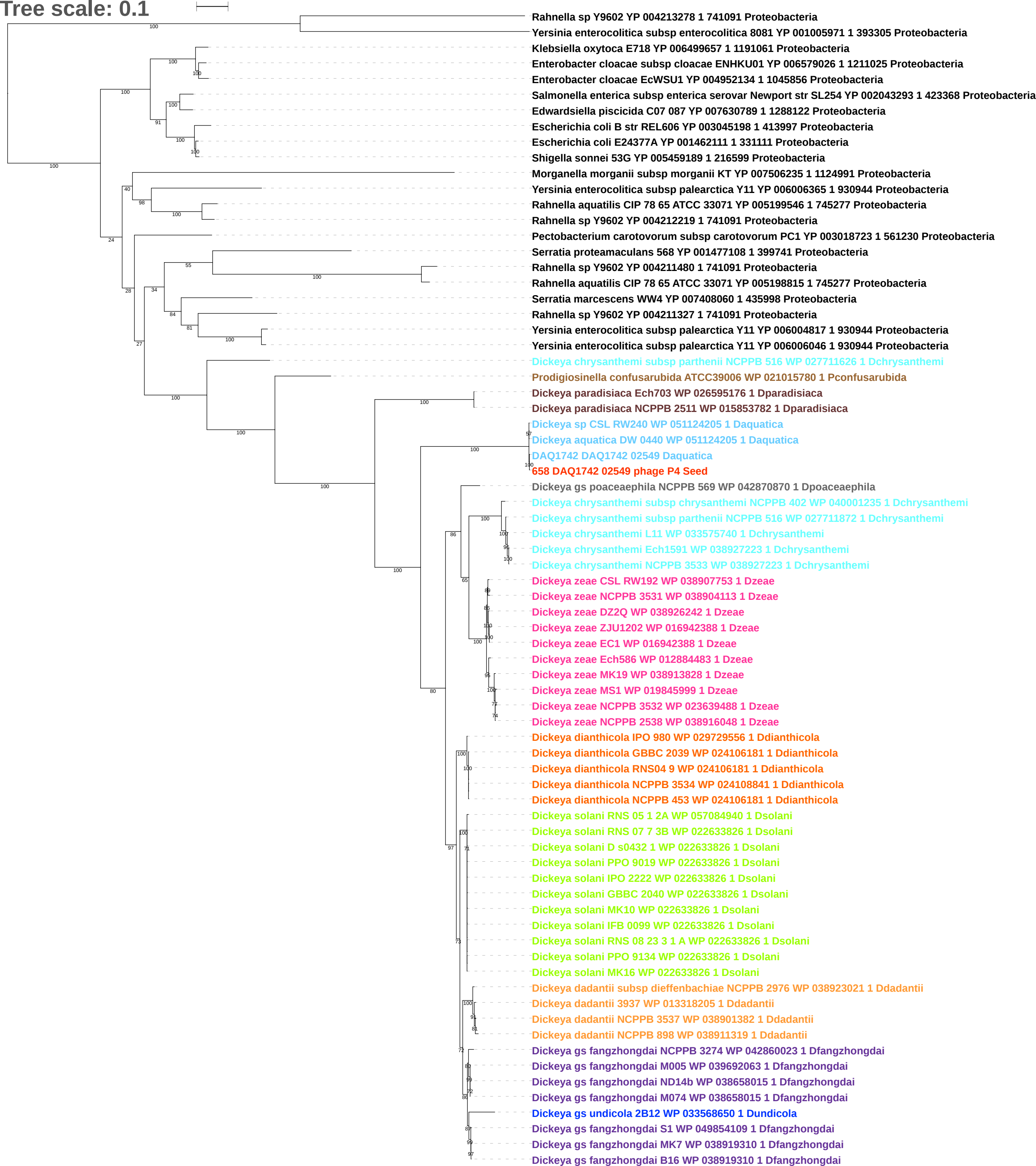
Tree scale: 0.1
Rahnella sp Y9602 YP 004213278 1 741091 Proteobacteria
Yersinia enterocolitica subsp enterocolitica 8081 YP 001005971 1 393305 Proteobacteria
Klebsiella oxytoca E718 YP 006499657 1 1191061 Proteobacteria
Enterobacter cloacae subsp cloacae ENHKU01 YP 006579026 1 1211025 Proteobacteria
Enterobacter cloacae EcWSU1 YP 004952134 1 1045856 Proteobacteria
Salmonella enterica subsp enterica serovar Newport str SL254 YP 002043293 1 423368 Proteobacteria
Edwardsiella piscicida C07 087 YP 007630789 1 1288122 Proteobacteria
Escherichia coli B str REL606 YP 003045198 1 413997 Proteobacteria
Escherichia coli E24377A YP 001462111 1 331111 Proteobacteria
Shigella sonnei 53G YP 005459189 1 216599 Proteobacteria
Morganella morganii subsp morganii KT YP 007506235 1 1124991 Proteobacteria
Yersinia enterocolitica subsp palearctica Y11 YP 006006365 1 930944 Proteobacteria
Rahnella aquatilis CIP 78 65 ATCC 33071 YP 005199546 1 745277 Proteobacteria
Rahnella sp Y9602 YP 004212219 1 741091 Proteobacteria
Pectobacterium carotovorum subsp carotovorum PC1 YP 003018723 1 561230 Proteobacteria
Serratia proteamaculans 568 YP 001477108 1 399741 Proteobacteria
Rahnella sp Y9602 YP 004211480 1 741091 Proteobacteria
Rahnella aquatilis CIP 78 65 ATCC 33071 YP 005198815 1 745277 Proteobacteria
Serratia marcescens WW4 YP 007408060 1 435998 Proteobacteria
Rahnella sp Y9602 YP 004211327 1 741091 Proteobacteria
Yersinia enterocolitica subsp palearctica Y11 YP 006004817 1 930944 Proteobacteria
Yersinia enterocolitica subsp palearctica Y11 YP 006006046 1 930944 Proteobacteria
Dickeya chrysanthemi subsp parthenii NCPPB 516 WP 027711626 1 Dchrysanthemi
Prodigiosinella confusarubida ATCC39006 WP 021015780 1 Pconfusarubida
Dickeya paradisiaca Ech703 WP 026595176 1 Dparadisiaca
Dickeya paradisiaca NCPPB 2511 WP 015853782 1 Dparadisiaca
Dickeya sp CSL RW240 WP 051124205 1 Daquatica
Dickeya aquatica DW 0440 WP 051124205 1 Daquatica
DAQ1742 DAQ1742 02549 Daquatica
658 DAQ1742 02549 phage P4 Seed
Dickeya gs poaceaephila NCPPB 569 WP 042870870 1 Dpoaceaephila
Dickeya chrysanthemi subsp chrysanthemi NCPPB 402 WP 040001235 1 Dchrysanthemi
Dickeya chrysanthemi subsp parthenii NCPPB 516 WP 027711872 1 Dchrysanthemi
Dickeya chrysanthemi L11 WP 033575740 1 Dchrysanthemi
Dickeya chrysanthemi Ech1591 WP 038927223 1 Dchrysanthemi
Dickeya chrysanthemi NCPPB 3533 WP 038927223 1 Dchrysanthemi
Dickeya zeae CSL RW192 WP 038907753 1 Dzeae
Dickeya zeae NCPPB 3531 WP 038904113 1 Dzeae
Dickeya zeae DZ2Q WP 038926242 1 Dzeae
Dickeya zeae ZJU1202 WP 016942388 1 Dzeae
Dickeya zeae EC1 WP 016942388 1 Dzeae
Dickeya zeae Ech586 WP 012884483 1 Dzeae
Dickeya zeae MK19 WP 038913828 1 Dzeae
Dickeya zeae MS1 WP 019845999 1 Dzeae
Dickeya zeae NCPPB 3532 WP 023639488 1 Dzeae
Dickeya zeae NCPPB 2538 WP 038916048 1 Dzeae
Dickeya dianthicola IPO 980 WP 029729556 1 Ddianthicola
Dickeya dianthicola GBBC 2039 WP 024106181 1 Ddianthicola
Dickeya dianthicola RNS04 9 WP 024106181 1 Ddianthicola
Dickeya dianthicola NCPPB 3534 WP 024108841 1 Ddianthicola
Dickeya dianthicola NCPPB 453 WP 024106181 1 Ddianthicola
Dickeya solani RNS 05 1 2A WP 057084940 1 Dsolani
Dickeya solani RNS 07 7 3B WP 022633826 1 Dsolani
Dickeya solani D s0432 1 WP 022633826 1 Dsolani
Dickeya solani PPO 9019 WP 022633826 1 Dsolani
Dickeya solani IPO 2222 WP 022633826 1 Dsolani
Dickeya solani GBBC 2040 WP 022633826 1 Dsolani
Dickeya solani MK10 WP 022633826 1 Dsolani
Dickeya solani IFB 0099 WP 022633826 1 Dsolani
Dickeya solani RNS 08 23 3 1 A WP 022633826 1 Dsolani
Dickeya solani PPO 9134 WP 022633826 1 Dsolani
Dickeya solani MK16 WP 022633826 1 Dsolani
Dickeya dadantii subsp dieffenbachiae NCPPB 2976 WP 038923021 1 Ddadantii
Dickeya dadantii 3937 WP 013318205 1 Ddadantii
Dickeya dadantii NCPPB 3537 WP 038901382 1 Ddadantii
Dickeya dadantii NCPPB 898 WP 038911319 1 Ddadantii
Dickeya gs fangzhongdai NCPPB 3274 WP 042860023 1 Dfangzhongdai
Dickeya gs fangzhongdai M005 WP 039692063 1 Dfangzhongdai
Dickeya gs fangzhongdai ND14b WP 038658015 1 Dfangzhongdai
Dickeya gs fangzhongdai M074 WP 038658015 1 Dfangzhongdai
Dickeya gs undicola 2B12 WP 033568650 1 Dundicola
Dickeya gs fangzhongdai S1 WP 049854109 1 Dfangzhongdai
Dickeya gs fangzhongdai MK7 WP 038919310 1 Dfangzhongdai
Dickeya gs fangzhongdai B16 WP 038919310 1 Dfangzhongdai

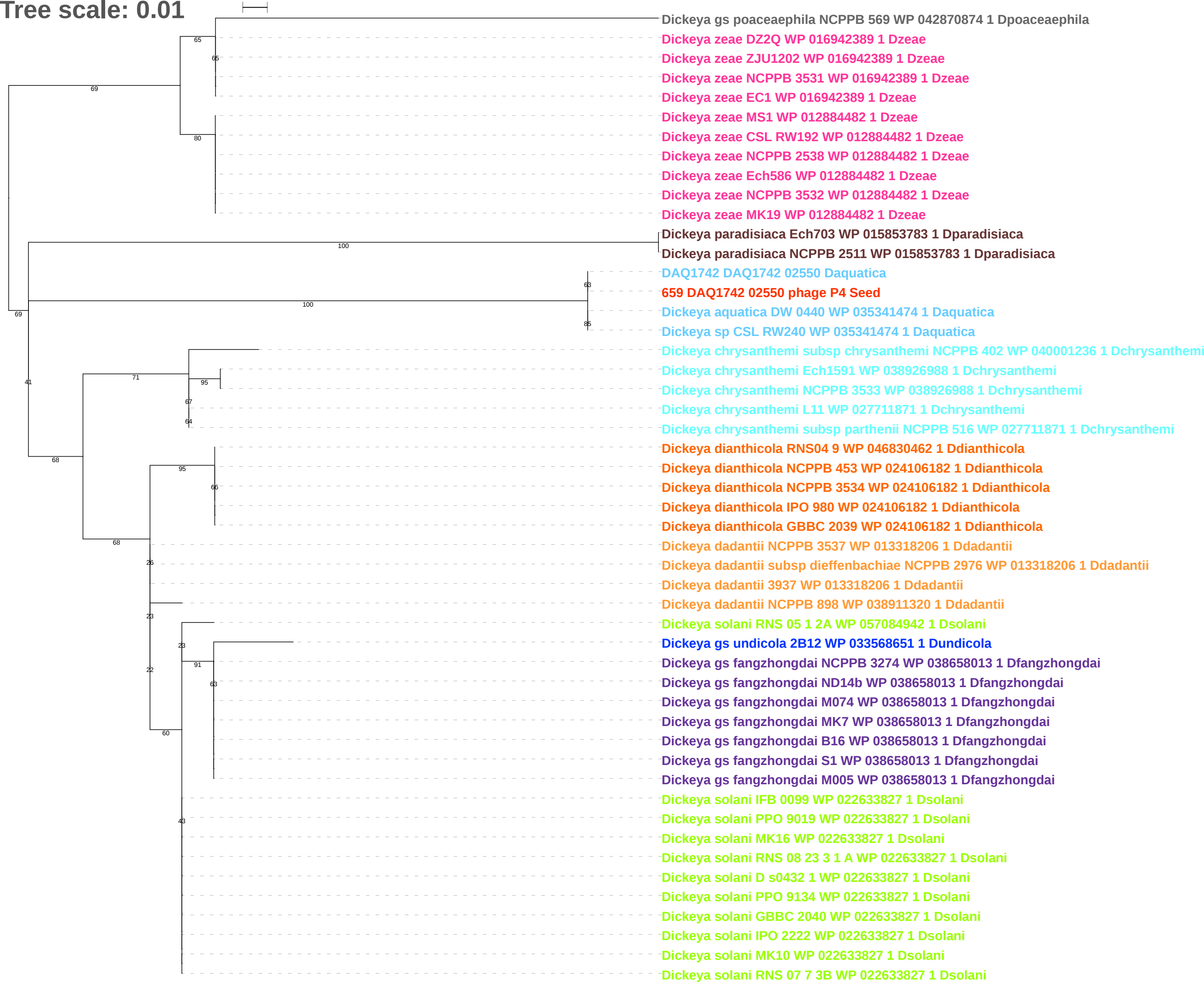

Tree scale: 0.01
Dickeya gs poaceaephila NCPPB 569 WP 042870874 1 Dpoaceaephila
Dickeya zeae DZ2Q WP 016942389 1 Dzeae
Dickeya zeae ZJU1202 WP 016942389 1 Dzeae
Dickeya zeae NCPPB 3531 WP 016942389 1 Dzeae
Dickeya zeae EC1 WP 016942389 1 Dzeae
Dickeya zeae MS1 WP 012884482 1 Dzeae
Dickeya zeae CSL RW192 WP 012884482 1 Dzeae
Dickeya zeae NCPPB 2538 WP 012884482 1 Dzeae
Dickeya zeae Ech586 WP 012884482 1 Dzeae
Dickeya zeae NCPPB 3532 WP 012884482 1 Dzeae
Dickeya zeae MK19 WP 012884482 1 Dzeae
Dickeya paradisiaca Ech703 WP 015853783 1 Dparadisiaca
Dickeya paradisiaca NCPPB 2511 WP 015853783 1 Dparadisiaca
DAQ1742 DAQ1742 02550 Daquatica
659 DAQ1742 02550 phage P4 Seed
Dickeya aquatica DW 0440 WP 035341474 1 Daquatica
Dickeya sp CSL RW240 WP 035341474 1 Daquatica
Dickeya chrysanthemi subsp chrysanthemi NCPPB 402 WP 040001236 1 Dchrysanthemi
Dickeya chrysanthemi Ech1591 WP 038926988 1 Dchrysanthemi
Dickeya chrysanthemi NCPPB 3533 WP 038926988 1 Dchrysanthemi
Dickeya chrysanthemi L11 WP 027711871 1 Dchrysanthemi
Dickeya chrysanthemi subsp parthenii NCPPB 516 WP 027711871 1 Dchrysanthemi
Dickeya dianthicola RNS04 9 WP 046830462 1 Ddianthicola
Dickeya dianthicola NCPPB 453 WP 024106182 1 Ddianthicola
Dickeya dianthicola NCPPB 3534 WP 024106182 1 Ddianthicola
Dickeya dianthicola IPO 980 WP 024106182 1 Ddianthicola
Dickeya dianthicola GBBC 2039 WP 024106182 1 Ddianthicola
Dickeya dadantii NCPPB 3537 WP 013318206 1 Ddadantii
Dickeya dadantii subsp dieffenbachiae NCPPB 2976 WP 013318206 1 Ddadantii
Dickeya dadantii 3937 WP 013318206 1 Ddadantii
Dickeya dadantii NCPPB 898 WP 038911320 1 Ddadantii
Dickeya solani RNS 05 1 2A WP 057084942 1 Dsolani
Dickeya gs undicola 2B12 WP 033568651 1 Dundicola
Dickeya gs fangzhongdai NCPPB 3274 WP 038658013 1 Dfangzhongdai
Dickeya gs fangzhongdai ND14b WP 038658013 1 Dfangzhongdai
Dickeya gs fangzhongdai M074 WP 038658013 1 Dfangzhongdai
Dickeya gs fangzhongdai MK7 WP 038658013 1 Dfangzhongdai
Dickeya gs fangzhongdai B16 WP 038658013 1 Dfangzhongdai
Dickeya gs fangzhongdai S1 WP 038658013 1 Dfangzhongdai
Dickeya gs fangzhongdai M005 WP 038658013 1 Dfangzhongdai
Dickeya solani IFB 0099 WP 022633827 1 Dsolani
Dickeya solani PPO 9019 WP 022633827 1 Dsolani
Dickeya solani MK16 WP 022633827 1 Dsolani
Dickeya solani RNS 08 23 3 1 A WP 022633827 1 Dsolani
Dickeya solani D s0432 1 WP 022633827 1 Dsolani
Dickeya solani PPO 9134 WP 022633827 1 Dsolani
Dickeya solani GBBC 2040 WP 022633827 1 Dsolani
Dickeya solani IPO 2222 WP 022633827 1 Dsolani
Dickeya solani MK10 WP 022633827 1 Dsolani
Dickeya solani RNS 07 7 3B WP 022633827 1 Dsolani

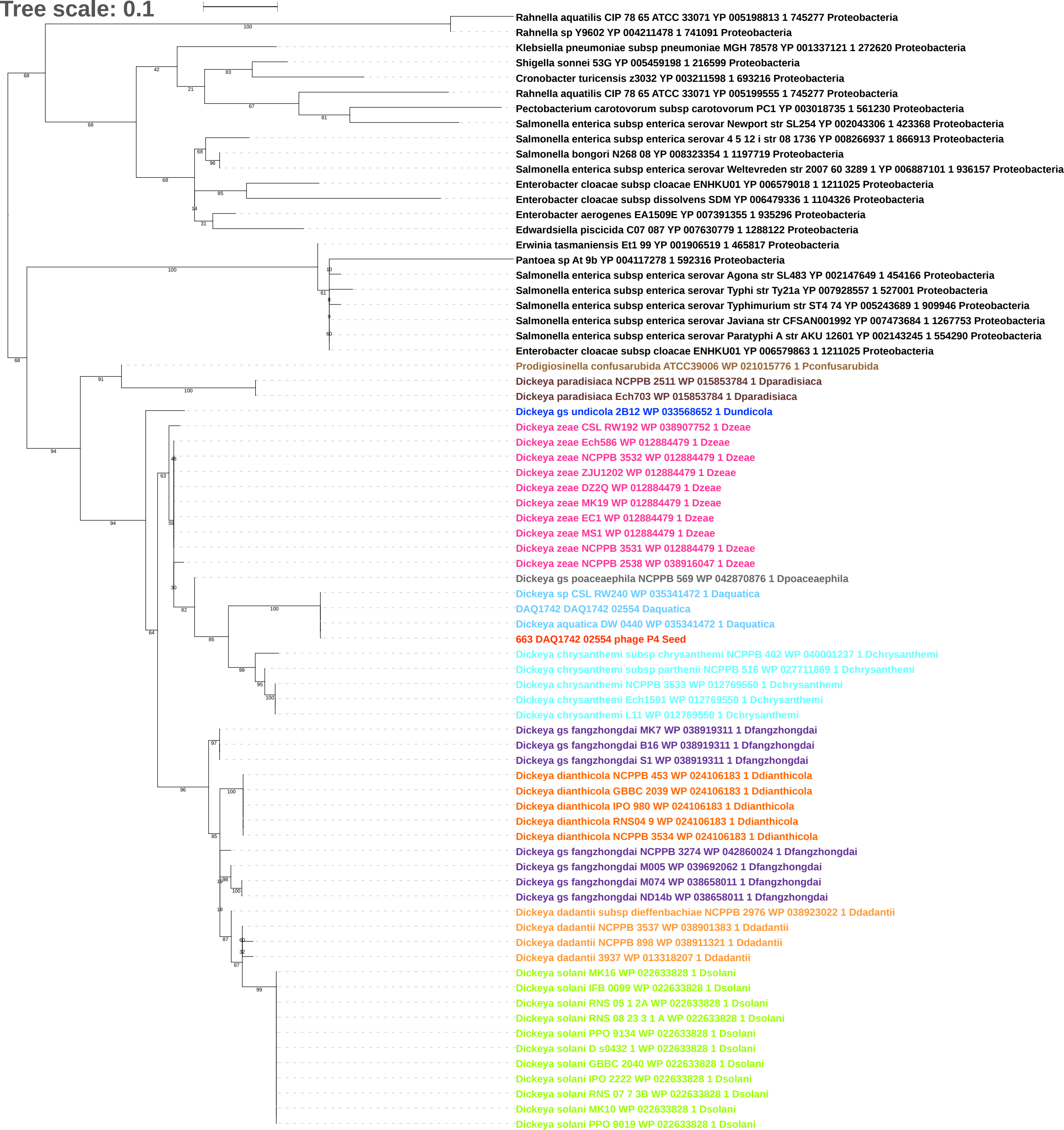

Tree scale: 0.1
Rahnella aquatilis CIP 78 65 ATCC 33071 YP 005198813 1 745277 Proteobacteria
Rahnella sp Y9602 YP 004211478 1 741091 Proteobacteria
Klebsiella pneumoniae subsp pneumoniae MGH 78578 YP 001337121 1 272620 Proteobacteria
Shigella sonnei 53G YP 005459198 1 216599 Proteobacteria
Cronobacter turicensis z3032 YP 003211598 1 693216 Proteobacteria
Rahnella aquatilis CIP 78 65 ATCC 33071 YP 005199555 1 745277 Proteobacteria
Pectobacterium carotovorum subsp carotovorum PC1 YP 003018735 1 561230 Proteobacteria
Salmonella enterica subsp enterica serovar Newport str SL254 YP 002043306 1 423368 Proteobacteria
Salmonella enterica subsp enterica serovar 4 5 12 i str 08 1736 YP 008266937 1 866913 Proteobacteria
Salmonella bongori N268 08 YP 008323354 1 1197719 Proteobacteria
Salmonella enterica subsp enterica serovar Weltevreden str 2007 60 3289 1 YP 006887101 1 936157 Proteobacteria
Enterobacter cloacae subsp cloacae ENHKU01 YP 006579018 1 1211025 Proteobacteria
Enterobacter cloacae subsp dissolvens SDM YP 006479336 1 1104326 Proteobacteria
Enterobacter aerogenes EA1509E YP 007391355 1 935296 Proteobacteria
Edwardsiella piscicida C07 087 YP 007630779 1 1288122 Proteobacteria
Erwinia tasmaniensis Et1 99 YP 001906519 1 465817 Proteobacteria
Pantoea sp At 9b YP 004117278 1 592316 Proteobacteria
Salmonella enterica subsp enterica serovar Agona str SL483 YP 002147649 1 454166 Proteobacteria
Salmonella enterica subsp enterica serovar Typhi str Ty21a YP 007928557 1 527001 Proteobacteria
Salmonella enterica subsp enterica serovar Typhimurium str ST4 74 YP 005243689 1 909946 Proteobacteria
Salmonella enterica subsp enterica serovar Javiana str CFSAN001992 YP 007473684 1 1267753 Proteobacteria
Salmonella enterica subsp enterica serovar Paratyphi A str AKU 12601 YP 002143245 1 554290 Proteobacteria
Enterobacter cloacae subsp cloacae ENHKU01 YP 006579863 1 1211025 Proteobacteria
Prodigiosinella confusarubida ATCC39006 WP 021015776 1 Pconfusarubida
Dickeya paradisiaca NCPPB 2511 WP 015853784 1 Dparadisiaca
Dickeya paradisiaca Ech703 WP 015853784 1 Dparadisiaca
Dickeya gs undicola 2B12 WP 033568652 1 Dundicola
Dickeya zeae CSL RW192 WP 038907752 1 Dzeae
Dickeya zeae Ech586 WP 012884479 1 Dzeae
Dickeya zeae NCPPB 3532 WP 012884479 1 Dzeae
Dickeya zeae ZJU1202 WP 012884479 1 Dzeae
Dickeya zeae DZ2Q WP 012884479 1 Dzeae
Dickeya zeae MK19 WP 012884479 1 Dzeae
Dickeya zeae EC1 WP 012884479 1 Dzeae
Dickeya zeae MS1 WP 012884479 1 Dzeae
Dickeya zeae NCPPB 3531 WP 012884479 1 Dzeae
Dickeya zeae NCPPB 2538 WP 038916047 1 Dzeae
Dickeya gs poaceaephila NCPPB 569 WP 042870876 1 Dpoaceaephila
Dickeya sp CSL RW240 WP 035341472 1 Daquatica
DAQ1742 DAQ1742 02554 Daquatica
Dickeya aquatica DW 0440 WP 035341472 1 Daquatica
663 DAQ1742 02554 phage P4 Seed
Dickeya chrysanthemi subsp chrysanthemi NCPPB 402 WP 040001237 1 Dchrysanthemi
Dickeya chrysanthemi subsp parthenii NCPPB 516 WP 027711869 1 Dchrysanthemi
Dickeya chrysanthemi NCPPB 3533 WP 012769550 1 Dchrysanthemi
Dickeya chrysanthemi Ech1591 WP 012769550 1 Dchrysanthemi
Dickeya chrysanthemi L11 WP 012769550 1 Dchrysanthemi
Dickeya gs fangzhongdai MK7 WP 038919311 1 Dfangzhongdai
Dickeya gs fangzhongdai B16 WP 038919311 1 Dfangzhongdai
Dickeya gs fangzhongdai S1 WP 038919311 1 Dfangzhongdai
Dickeya dianthicola NCPPB 453 WP 024106183 1 Ddianthicola
Dickeya dianthicola GBBC 2039 WP 024106183 1 Ddianthicola
Dickeya dianthicola IPO 980 WP 024106183 1 Ddianthicola
Dickeya dianthicola RNS04 9 WP 024106183 1 Ddianthicola
Dickeya dianthicola NCPPB 3534 WP 024106183 1 Ddianthicola
Dickeya gs fangzhongdai NCPPB 3274 WP 042860024 1 Dfangzhongdai
Dickeya gs fangzhongdai M005 WP 039692062 1 Dfangzhongdai
Dickeya gs fangzhongdai M074 WP 038658011 1 Dfangzhongdai
Dickeya gs fangzhongdai ND14b WP 038658011 1 Dfangzhongdai
Dickeya dadantii subsp dieffenbachiae NCPPB 2976 WP 038923022 1 Ddadantii
Dickeya dadantii NCPPB 3537 WP 038901383 1 Ddadantii
Dickeya dadantii NCPPB 898 WP 038911321 1 Ddadantii
Dickeya dadantii 3937 WP 013318207 1 Ddadantii
Dickeya solani MK16 WP 022633828 1 Dsolani
Dickeya solani IFB 0099 WP 022633828 1 Dsolani
Dickeya solani RNS 05 1 2A WP 022633828 1 Dsolani
Dickeya solani RNS 08 23 3 1 A WP 022633828 1 Dsolani
Dickeya solani PPO 9134 WP 022633828 1 Dsolani
Dickeya solani D s0432 1 WP 022633828 1 Dsolani
Dickeya solani GBBC 2040 WP 022633828 1 Dsolani
Dickeya solani IPO 2222 WP 022633828 1 Dsolani
Dickeya solani RNS 07 7 3B WP 022633828 1 Dsolani
Dickeya solani MK10 WP 022633828 1 Dsolani
Dickeya solani PPO 9019 WP 022633828 1 Dsolani

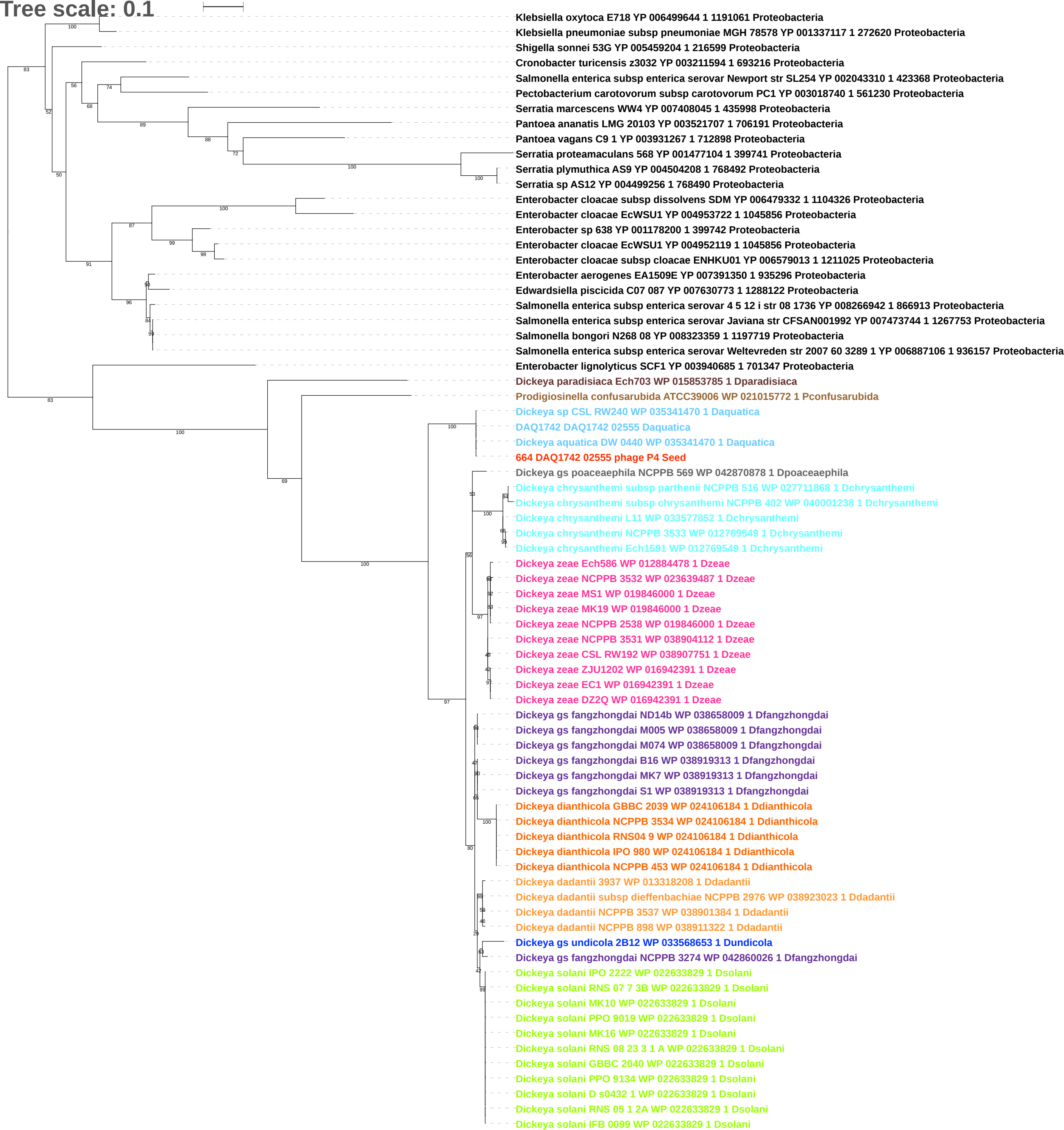
Tree scale: 0.1
Klebsiella oxytoca E718 YP 006499644 1 1191061 Proteobacteria
Klebsiella pneumoniae subsp pneumoniae MGH 78578 YP 001337117 1 272620 Proteobacteria
Shigella sonnei 53G YP 005459204 1 216599 Proteobacteria
Cronobacter turicensis z3032 YP 003211594 1 693216 Proteobacteria
Salmonella enterica subsp enterica serovar Newport str SL254 YP 002043310 1 423368 Proteobacteria
Pectobacterium carotovorum subsp carotovorum PC1 YP 003018740 1 561230 Proteobacteria
Serratia marcescens WW4 YP 007408045 1 435998 Proteobacteria
Pantoea ananatis LMG 20103 YP 003521707 1 706191 Proteobacteria
Pantoea vagans C9 1 YP 003931267 1 712898 Proteobacteria
Serratia proteamaculans 568 YP 001477104 1 399741 Proteobacteria
Serratia plymuthica AS9 YP 004504208 1 768492 Proteobacteria
Serratia sp AS12 YP 004499256 1 768490 Proteobacteria
Enterobacter cloacae subsp dissolvens SDM YP 006479332 1 1104326 Proteobacteria
Enterobacter cloacae EcWSU1 YP 004953722 1 1045856 Proteobacteria
Enterobacter sp 638 YP 001178200 1 399742 Proteobacteria
Enterobacter cloacae EcWSU1 YP 004952119 1 1045856 Proteobacteria
Enterobacter cloacae subsp cloacae ENHKU01 YP 006579013 1 1211025 Proteobacteria
Enterobacter aerogenes EA1509E YP 007391350 1 935296 Proteobacteria
Edwardsiella piscicida C07 087 YP 007630773 1 1288122 Proteobacteria
Salmonella enterica subsp enterica serovar 4 5 12 i str 08 1736 YP 008266942 1 866913 Proteobacteria
Salmonella enterica subsp enterica serovar Javiana str CFSAN001992 YP 007473744 1 1267753 Proteobacteria
Salmonella bongori N268 08 YP 008323359 1 1197719 Proteobacteria
Salmonella enterica subsp enterica serovar Weltevreden str 2007 60 3289 1 YP 006887106 1 936157 Proteobacteria
Enterobacter lignolyticus SCF1 YP 003940685 1 701347 Proteobacteria
Dickeya paradisiaca Ech703 WP 015853785 1 Dparadisiaca
Prodigiosinella confusarubida ATCC39006 WP 021015772 1 Pconfusarubida
Dickeya sp CSL RW240 WP 035341470 1 Daquatica
DAQ1742 DAQ1742 02555 Daquatica
Dickeya aquatica DW 0440 WP 035341470 1 Daquatica
664 DAQ1742 02555 phage P4 Seed
Dickeya gs poaceaephila NCPPB 569 WP 042870878 1 Dpoaceaephila
Dickeya chrysanthemi subsp parthenii NCPPB 516 WP 027711868 1 Dchrysanthemi
Dickeya chrysanthemi subsp chrysanthemi NCPPB 402 WP 040001238 1 Dchrysanthemi
Dickeya chrysanthemi L11 WP 033577852 1 Dchrysanthemi
Dickeya chrysanthemi NCPPB 3533 WP 012769549 1 Dchrysanthemi
Dickeya chrysanthemi Ech1591 WP 012769549 1 Dchrysanthemi
Dickeya zeae Ech586 WP 012884478 1 Dzeae
Dickeya zeae NCPPB 3532 WP 023639487 1 Dzeae
Dickeya zeae MS1 WP 019846000 1 Dzeae
Dickeya zeae MK19 WP 019846000 1 Dzeae
Dickeya zeae NCPPB 2538 WP 019846000 1 Dzeae
Dickeya zeae NCPPB 3531 WP 038904112 1 Dzeae
Dickeya zeae CSL RW192 WP 038907751 1 Dzeae
Dickeya zeae ZJU1202 WP 016942391 1 Dzeae
Dickeya zeae EC1 WP 016942391 1 Dzeae
Dickeya zeae DZ2Q WP 016942391 1 Dzeae
Dickeya gs fangzhongdai ND14b WP 038658009 1 Dfangzhongdai
Dickeya gs fangzhongdai M005 WP 038658009 1 Dfangzhongdai
Dickeya gs fangzhongdai M074 WP 038658009 1 Dfangzhongdai
Dickeya gs fangzhongdai B16 WP 038919313 1 Dfangzhongdai
Dickeya gs fangzhongdai MK7 WP 038919313 1 Dfangzhongdai
Dickeya gs fangzhongdai S1 WP 038919313 1 Dfangzhongdai
Dickeya dianthicola GBBC 2039 WP 024106184 1 Ddianthicola
Dickeya dianthicola NCPPB 3534 WP 024106184 1 Ddianthicola
Dickeya dianthicola RNS04 9 WP 024106184 1 Ddianthicola
Dickeya dianthicola IPO 980 WP 024106184 1 Ddianthicola
Dickeya dianthicola NCPPB 453 WP 024106184 1 Ddianthicola
Dickeya dadantii 3937 WP 013318208 1 Ddadantii
Dickeya dadantii subsp dieffenbachiae NCPPB 2976 WP 038923023 1 Ddadantii
Dickeya dadantii NCPPB 3537 WP 038901384 1 Ddadantii
Dickeya dadantii NCPPB 898 WP 038911322 1 Ddadantii
Dickeya gs undicola 2B12 WP 033568653 1 Dundicola
Dickeya gs fangzhongdai NCPPB 3274 WP 042860026 1 Dfangzhongdai
Dickeya solani IPO 2222 WP 022633829 1 Dsolani
Dickeya solani RNS 07 7 3B WP 022633829 1 Dsolani
Dickeya solani MK10 WP 022633829 1 Dsolani
Dickeya solani PPO 9019 WP 022633829 1 Dsolani
Dickeya solani MK16 WP 022633829 1 Dsolani
Dickeya solani RNS 08 23 3 1 A WP 022633829 1 Dsolani
Dickeya solani GBBC 2040 WP 022633829 1 Dsolani
Dickeya solani PPO 9134 WP 022633829 1 Dsolani
Dickeya solani D s0432 1 WP 022633829 1 Dsolani
Dickeya solani RNS 05 1 2A WP 022633829 1 Dsolani
Dickeya solani IFB 0099 WP 022633829 1 Dsolani

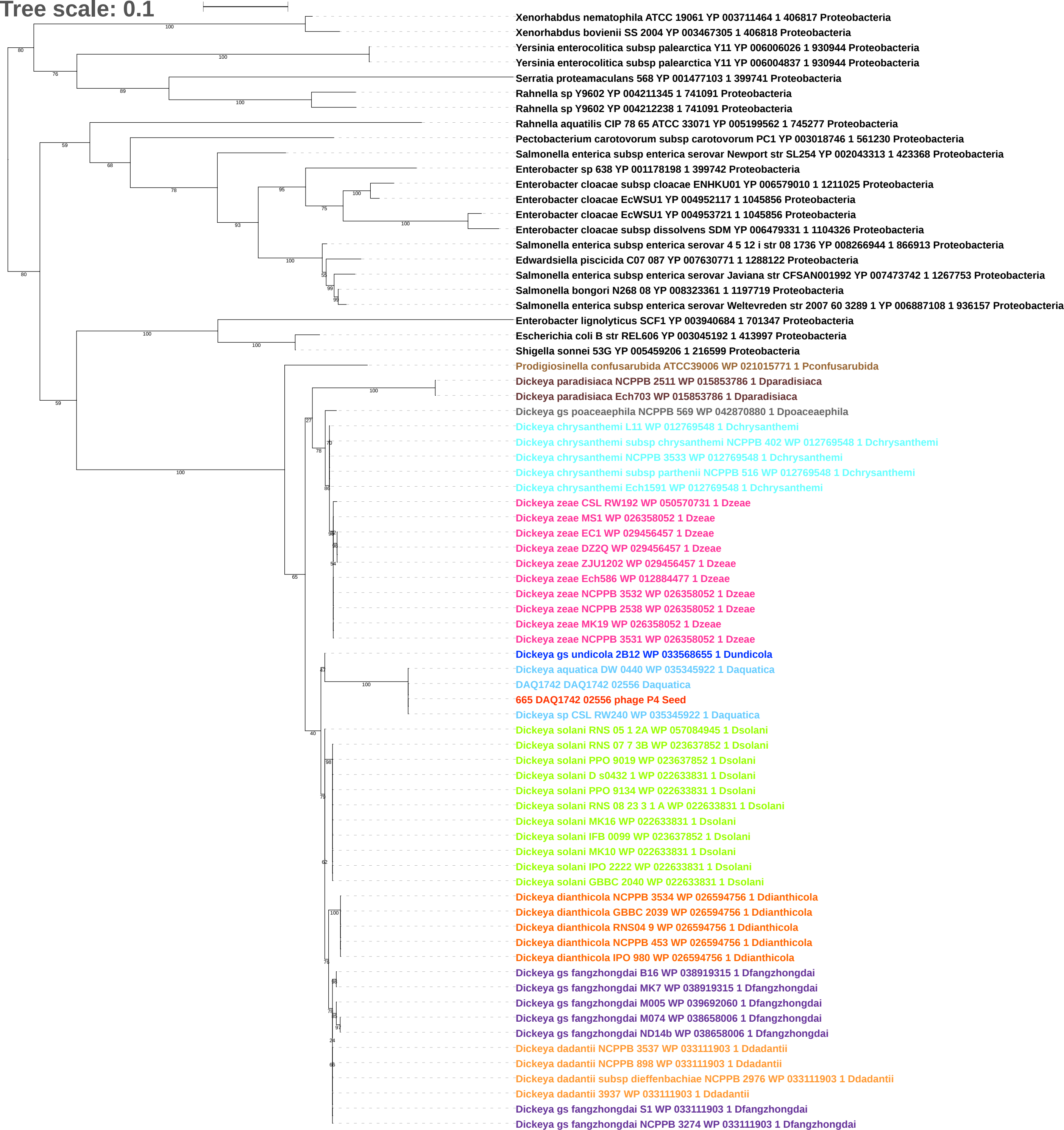
Tree scale: 0.1
Xenorhabdus nematophila ATCC 19061 YP 003711464 1 406817 Proteobacteria
Xenorhabdus bovienii SS 2004 YP 003467305 1 406818 Proteobacteria
Yersinia enterocolitica subsp palearctica Y11 YP 006006026 1 930944 Proteobacteria
Yersinia enterocolitica subsp palearctica Y11 YP 006004837 1 930944 Proteobacteria
Serratia proteamaculans 568 YP 001477103 1 399741 Proteobacteria
Rahnella sp Y9602 YP 004211345 1 741091 Proteobacteria
Rahnella sp Y9602 YP 004212238 1 741091 Proteobacteria
Rahnella aquatilis CIP 78 65 ATCC 33071 YP 005199562 1 745277 Proteobacteria
Pectobacterium carotovorum subsp carotovorum PC1 YP 003018746 1 561230 Proteobacteria
Salmonella enterica subsp enterica serovar Newport str SL254 YP 002043313 1 423368 Proteobacteria
Enterobacter sp 638 YP 001178198 1 399742 Proteobacteria
Enterobacter cloacae subsp cloacae ENHKU01 YP 006579010 1 1211025 Proteobacteria
Enterobacter cloacae EcWSU1 YP 004952117 1 1045856 Proteobacteria
Enterobacter cloacae EcWSU1 YP 004953721 1 1045856 Proteobacteria
Enterobacter cloacae subsp dissolvens SDM YP 006479331 1 1104326 Proteobacteria
Salmonella enterica subsp enterica serovar 4 5 12 i str 08 1736 YP 008266944 1 866913 Proteobacteria
Edwardsiella piscicida C07 087 YP 007630771 1 1288122 Proteobacteria
Salmonella enterica subsp enterica serovar Javiana str CFSAN001992 YP 007473742 1 1267753 Proteobacteria
Salmonella bongori N268 08 YP 008323361 1 1197719 Proteobacteria
Salmonella enterica subsp enterica serovar Weltevreden str 2007 60 3289 1 YP 006887108 1 936157 Proteobacteria
Enterobacter lignolyticus SCF1 YP 003940684 1 701347 Proteobacteria
Escherichia coli B str REL606 YP 003045192 1 413997 Proteobacteria
Shigella sonnei 53G YP 005459206 1 216599 Proteobacteria
Prodigiosinella confusarubida ATCC39006 WP 021015771 1 Pconfusarubida
Dickeya paradisiaca NCPPB 2511 WP 015853786 1 Dparadisiaca
Dickeya paradisiaca Ech703 WP 015853786 1 Dparadisiaca
Dickeya gs poaceaephila NCPPB 569 WP 042870880 1 Dpoaceaephila
Dickeya chrysanthemi L11 WP 012769548 1 Dchrysanthemi
Dickeya chrysanthemi subsp chrysanthemi NCPPB 402 WP 012769548 1 Dchrysanthemi
Dickeya chrysanthemi NCPPB 3533 WP 012769548 1 Dchrysanthemi
Dickeya chrysanthemi subsp parthenii NCPPB 516 WP 012769548 1 Dchrysanthemi
Dickeya chrysanthemi Ech1591 WP 012769548 1 Dchrysanthemi
Dickeya zeae CSL RW192 WP 050570731 1 Dzeae
Dickeya zeae MS1 WP 026358052 1 Dzeae
Dickeya zeae EC1 WP 029456457 1 Dzeae
Dickeya zeae DZ2Q WP 029456457 1 Dzeae
Dickeya zeae ZJU1202 WP 029456457 1 Dzeae
Dickeya zeae Ech586 WP 012884477 1 Dzeae
Dickeya zeae NCPPB 3532 WP 026358052 1 Dzeae
Dickeya zeae NCPPB 2538 WP 026358052 1 Dzeae
Dickeya zeae MK19 WP 026358052 1 Dzeae
Dickeya zeae NCPPB 3531 WP 026358052 1 Dzeae
Dickeya gs undicola 2B12 WP 033568655 1 Dundicola
Dickeya aquatica DW 0440 WP 035345922 1 Daquatica
DAQ1742 DAQ1742 02556 Daquatica
665 DAQ1742 02556 phage P4 Seed
Dickeya sp CSL RW240 WP 035345922 1 Daquatica
Dickeya solani RNS 05 1 2A WP 057084945 1 Dsolani
Dickeya solani RNS 07 7 3B WP 023637852 1 Dsolani
Dickeya solani PPO 9019 WP 023637852 1 Dsolani
Dickeya solani D s0432 1 WP 022633831 1 Dsolani
Dickeya solani PPO 9134 WP 022633831 1 Dsolani
Dickeya solani RNS 08 23 3 1 A WP 022633831 1 Dsolani
Dickeya solani MK16 WP 022633831 1 Dsolani
Dickeya solani IFB 0099 WP 023637852 1 Dsolani
Dickeya solani MK10 WP 022633831 1 Dsolani
Dickeya solani IPO 2222 WP 022633831 1 Dsolani
Dickeya solani GBBC 2040 WP 022633831 1 Dsolani
Dickeya dianthicola NCPPB 3534 WP 026594756 1 Ddianthicola
Dickeya dianthicola GBBC 2039 WP 026594756 1 Ddianthicola
Dickeya dianthicola RNS04 9 WP 026594756 1 Ddianthicola
Dickeya dianthicola NCPPB 453 WP 026594756 1 Ddianthicola
Dickeya dianthicola IPO 980 WP 026594756 1 Ddianthicola
Dickeya gs fangzhongdai B16 WP 038919315 1 Dfangzhongdai
Dickeya gs fangzhongdai MK7 WP 038919315 1 Dfangzhongdai
Dickeya gs fangzhongdai M005 WP 039692060 1 Dfangzhongdai
Dickeya gs fangzhongdai M074 WP 038658006 1 Dfangzhongdai
Dickeya gs fangzhongdai ND14b WP 038658006 1 Dfangzhongdai
Dickeya dadantii NCPPB 3537 WP 033111903 1 Ddadantii
Dickeya dadantii NCPPB 898 WP 033111903 1 Ddadantii
Dickeya dadantii subsp dieffenbachiae NCPPB 2976 WP 033111903 1 Ddadantii
Dickeya dadantii 3937 WP 033111903 1 Ddadantii
Dickeya gs fangzhongdai S1 WP 033111903 1 Dfangzhongdai
Dickeya gs fangzhongdai NCPPB 3274 WP 033111903 1 Dfangzhongdai

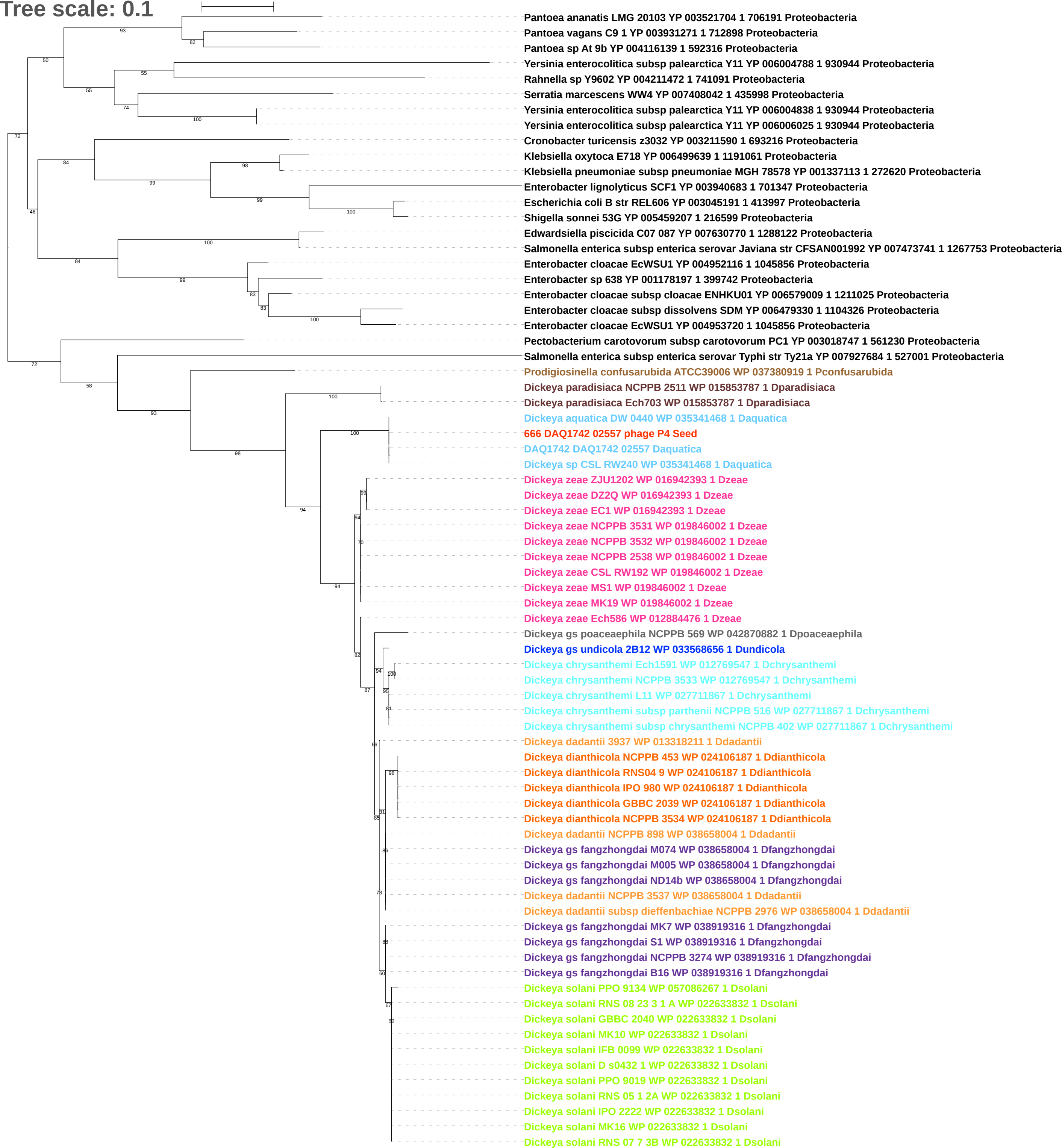
Tree scale: 0.1
Pantoea ananatis LMG 20103 YP 003521704 1 706191 Proteobacteria
Pantoea vagans C9 1 YP 003931271 1 712898 Proteobacteria
Pantoea sp At 9b YP 004116139 1 592316 Proteobacteria
Yersinia enterocolitica subsp palearctica Y11 YP 006004788 1 930944 Proteobacteria
Rahnella sp Y9602 YP 004211472 1 741091 Proteobacteria
Serratia marcescens WW4 YP 007408042 1 435998 Proteobacteria
Yersinia enterocolitica subsp palearctica Y11 YP 006004838 1 930944 Proteobacteria
Yersinia enterocolitica subsp palearctica Y11 YP 006006025 1 930944 Proteobacteria
Cronobacter turicensis z3032 YP 003211590 1 693216 Proteobacteria
Klebsiella oxytoca E718 YP 006499639 1 1191061 Proteobacteria
Klebsiella pneumoniae subsp pneumoniae MGH 78578 YP 001337113 1 272620 Proteobacteria
Enterobacter lignolyticus SCF1 YP 003940683 1 701347 Proteobacteria
Escherichia coli B str REL606 YP 003045191 1 413997 Proteobacteria
Shigella sonnei 53G YP 005459207 1 216599 Proteobacteria
Edwardsiella piscicida C07 087 YP 007630770 1 1288122 Proteobacteria
Salmonella enterica subsp enterica serovar Javiana str CFSAN001992 YP 007473741 1 1267753 Proteobacteria
Enterobacter cloacae EcWSU1 YP 004952116 1 1045856 Proteobacteria
Enterobacter sp 638 YP 001178197 1 399742 Proteobacteria
Enterobacter cloacae subsp cloacae ENHKU01 YP 006579009 1 1211025 Proteobacteria
Enterobacter cloacae subsp dissolvens SDM YP 006479330 1 1104326 Proteobacteria
Enterobacter cloacae EcWSU1 YP 004953720 1 1045856 Proteobacteria
Pectobacterium carotovorum subsp carotovorum PC1 YP 003018747 1 561230 Proteobacteria
Salmonella enterica subsp enterica serovar Typhi str Ty21a YP 007927684 1 527001 Proteobacteria
Prodigiosinella confusarubida ATCC39006 WP 037380919 1 Pconfusarubida
Dickeya paradisiaca NCPPB 2511 WP 015853787 1 Dparadisiaca
Dickeya paradisiaca Ech703 WP 015853787 1 Dparadisiaca
Dickeya aquatica DW 0440 WP 035341468 1 Daquatica
666 DAQ1742 02557 phage P4 Seed
DAQ1742 DAQ1742 02557 Daquatica
Dickeya sp CSL RW240 WP 035341468 1 Daquatica
Dickeya zeae ZJU1202 WP 016942393 1 Dzeae
Dickeya zeae DZ2Q WP 016942393 1 Dzeae
Dickeya zeae EC1 WP 016942393 1 Dzeae
Dickeya zeae NCPPB 3531 WP 019846002 1 Dzeae
Dickeya zeae NCPPB 3532 WP 019846002 1 Dzeae
Dickeya zeae NCPPB 2538 WP 019846002 1 Dzeae
Dickeya zeae CSL RW192 WP 019846002 1 Dzeae
Dickeya zeae MS1 WP 019846002 1 Dzeae
Dickeya zeae MK19 WP 019846002 1 Dzeae
Dickeya zeae Ech586 WP 012884476 1 Dzeae
Dickeya gs poaceaephila NCPPB 569 WP 042870882 1 Dpoaceaephila
Dickeya gs undicola 2B12 WP 033568656 1 Dundicola
Dickeya chrysanthemi Ech1591 WP 012769547 1 Dchrysanthemi
Dickeya chrysanthemi NCPPB 3533 WP 012769547 1 Dchrysanthemi
Dickeya chrysanthemi L11 WP 027711867 1 Dchrysanthemi
Dickeya chrysanthemi subsp parthenii NCPPB 516 WP 027711867 1 Dchrysanthemi
Dickeya chrysanthemi subsp chrysanthemi NCPPB 402 WP 027711867 1 Dchrysanthemi
Dickeya dadantii 3937 WP 013318211 1 Ddadantii
Dickeya dianthicola NCPPB 453 WP 024106187 1 Ddianthicola
Dickeya dianthicola RNS04 9 WP 024106187 1 Ddianthicola
Dickeya dianthicola IPO 980 WP 024106187 1 Ddianthicola
Dickeya dianthicola GBBC 2039 WP 024106187 1 Ddianthicola
Dickeya dianthicola NCPPB 3534 WP 024106187 1 Ddianthicola
Dickeya dadantii NCPPB 898 WP 038658004 1 Ddadantii
Dickeya gs fangzhongdai M074 WP 038658004 1 Dfangzhongdai
Dickeya gs fangzhongdai M005 WP 038658004 1 Dfangzhongdai
Dickeya gs fangzhongdai ND14b WP 038658004 1 Dfangzhongdai
Dickeya dadantii NCPPB 3537 WP 038658004 1 Ddadantii
Dickeya dadantii subsp dieffenbachiae NCPPB 2976 WP 038658004 1 Ddadantii
Dickeya gs fangzhongdai MK7 WP 038919316 1 Dfangzhongdai
Dickeya gs fangzhongdai S1 WP 038919316 1 Dfangzhongdai
Dickeya gs fangzhongdai NCPPB 3274 WP 038919316 1 Dfangzhongdai
Dickeya gs fangzhongdai B16 WP 038919316 1 Dfangzhongdai
Dickeya solani PPO 9134 WP 057086267 1 Dsolani
Dickeya solani RNS 08 23 3 1 A WP 022633832 1 Dsolani
Dickeya solani GBBC 2040 WP 022633832 1 Dsolani
Dickeya solani MK10 WP 022633832 1 Dsolani
Dickeya solani IFB 0099 WP 022633832 1 Dsolani
Dickeya solani D s0432 1 WP 022633832 1 Dsolani
Dickeya solani PPO 9019 WP 022633832 1 Dsolani
Dickeya solani RNS 05 1 2A WP 022633832 1 Dsolani
Dickeya solani IPO 2222 WP 022633832 1 Dsolani
Dickeya solani MK16 WP 022633832 1 Dsolani
Dickeya solani RNS 07 7 3B WP 022633832 1 Dsolani

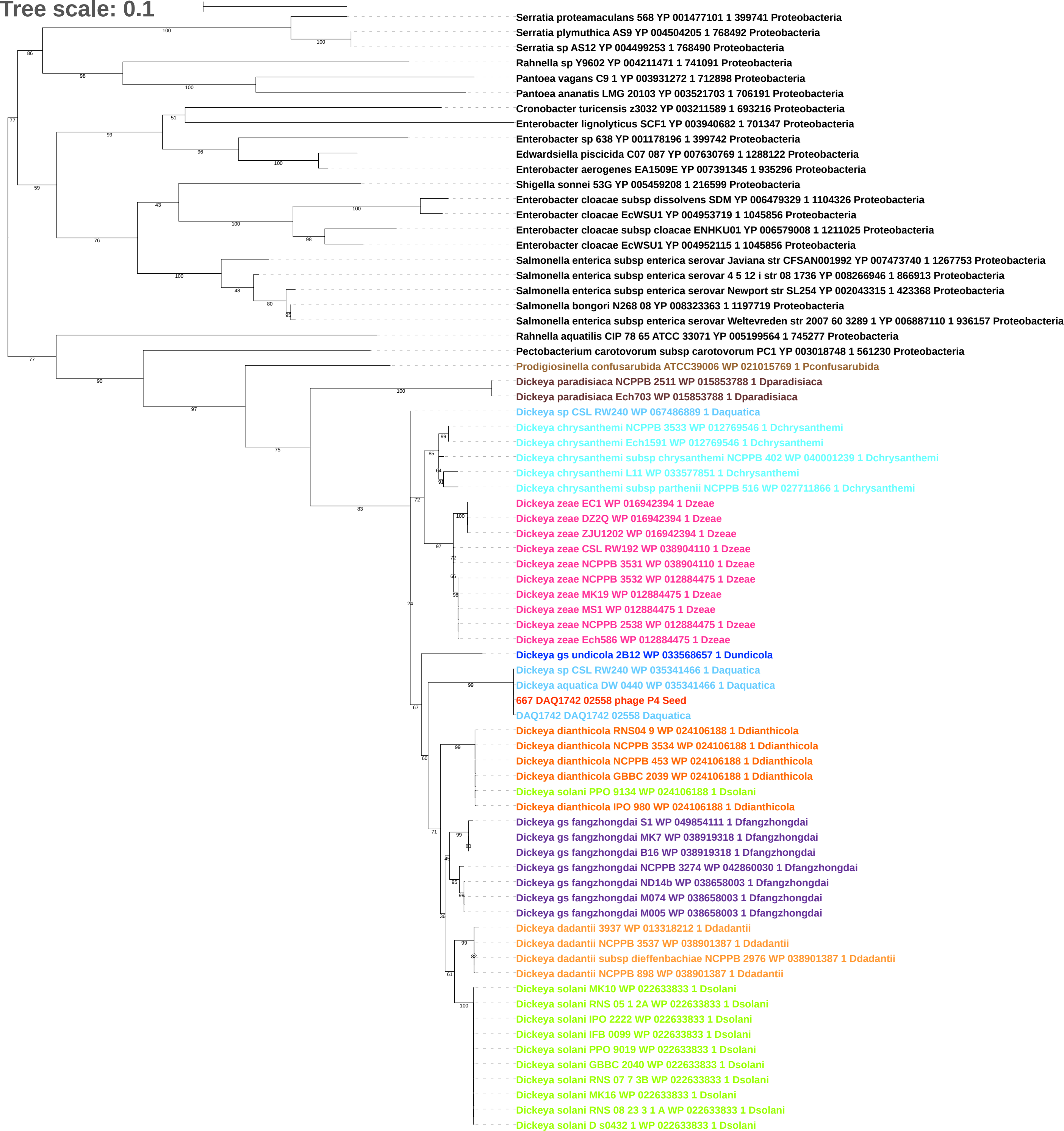

Tree scale: 0.1
Serratia proteamaculans 568 YP 001477101 1 399741 Proteobacteria
Serratia plymuthica AS9 YP 004504205 1 768492 Proteobacteria
Serratia sp AS12 YP 004499253 1 768490 Proteobacteria
Rahnella sp Y9602 YP 004211471 1 741091 Proteobacteria
Pantoea vagans C9 1 YP 003931272 1 712898 Proteobacteria
Pantoea ananatis LMG 20103 YP 003521703 1 706191 Proteobacteria
Cronobacter turicensis z3032 YP 003211589 1 693216 Proteobacteria
Enterobacter lignolyticus SCF1 YP 003940682 1 701347 Proteobacteria
Enterobacter sp 638 YP 001178196 1 399742 Proteobacteria
Edwardsiella piscicida C07 087 YP 007630769 1 1288122 Proteobacteria
Enterobacter aerogenes EA1509E YP 007391345 1 935296 Proteobacteria
Shigella sonnei 53G YP 005459208 1 216599 Proteobacteria
Enterobacter cloacae subsp dissolvens SDM YP 006479329 1 1104326 Proteobacteria
Enterobacter cloacae EcWSU1 YP 004953719 1 1045856 Proteobacteria
Enterobacter cloacae subsp cloacae ENHKU01 YP 006579008 1 1211025 Proteobacteria
Enterobacter cloacae EcWSU1 YP 004952115 1 1045856 Proteobacteria
Salmonella enterica subsp enterica serovar Javiana str CFSAN001992 YP 007473740 1 1267753 Proteobacteria
Salmonella enterica subsp enterica serovar 4 5 12 i str 08 1736 YP 008266946 1 866913 Proteobacteria
Salmonella enterica subsp enterica serovar Newport str SL254 YP 002043315 1 423368 Proteobacteria
Salmonella bongori N268 08 YP 008323363 1 1197719 Proteobacteria
Salmonella enterica subsp enterica serovar Weltevreden str 2007 60 3289 1 YP 006887110 1 936157 Proteobacteria
Rahnella aquatilis CIP 78 65 ATCC 33071 YP 005199564 1 745277 Proteobacteria
Pectobacterium carotovorum subsp carotovorum PC1 YP 003018748 1 561230 Proteobacteria
Prodigiosinella confusarubida ATCC39006 WP 021015769 1 Pconfusarubida
Dickeya paradisiaca NCPPB 2511 WP 015853788 1 Dparadisiaca
Dickeya paradisiaca Ech703 WP 015853788 1 Dparadisiaca
Dickeya sp CSL RW240 WP 067486889 1 Daquatica
Dickeya chrysanthemi NCPPB 3533 WP 012769546 1 Dchrysanthemi
Dickeya chrysanthemi Ech1591 WP 012769546 1 Dchrysanthemi
Dickeya chrysanthemi subsp chrysanthemi NCPPB 402 WP 040001239 1 Dchrysanthemi
Dickeya chrysanthemi L11 WP 033577851 1 Dchrysanthemi
Dickeya chrysanthemi subsp parthenii NCPPB 516 WP 027711866 1 Dchrysanthemi
Dickeya zeae EC1 WP 016942394 1 Dzeae
Dickeya zeae DZ2Q WP 016942394 1 Dzeae
Dickeya zeae ZJU1202 WP 016942394 1 Dzeae
Dickeya zeae CSL RW192 WP 038904110 1 Dzeae
Dickeya zeae NCPPB 3531 WP 038904110 1 Dzeae
Dickeya zeae NCPPB 3532 WP 012884475 1 Dzeae
Dickeya zeae MK19 WP 012884475 1 Dzeae
Dickeya zeae MS1 WP 012884475 1 Dzeae
Dickeya zeae NCPPB 2538 WP 012884475 1 Dzeae
Dickeya zeae Ech586 WP 012884475 1 Dzeae
Dickeya gs undicola 2B12 WP 033568657 1 Dundicola
Dickeya sp CSL RW240 WP 035341466 1 Daquatica
Dickeya aquatica DW 0440 WP 035341466 1 Daquatica
667 DAQ1742 02558 phage P4 Seed
DAQ1742 DAQ1742 02558 Daquatica
Dickeya dianthicola RNS04 9 WP 024106188 1 Ddianthicola
Dickeya dianthicola NCPPB 3534 WP 024106188 1 Ddianthicola
Dickeya dianthicola NCPPB 453 WP 024106188 1 Ddianthicola
Dickeya dianthicola GBBC 2039 WP 024106188 1 Ddianthicola
Dickeya solani PPO 9134 WP 024106188 1 Dsolani
Dickeya dianthicola IPO 980 WP 024106188 1 Ddianthicola
Dickeya gs fangzhongdai S1 WP 049854111 1 Dfangzhongdai
Dickeya gs fangzhongdai MK7 WP 038919318 1 Dfangzhongdai
Dickeya gs fangzhongdai B16 WP 038919318 1 Dfangzhongdai
Dickeya gs fangzhongdai NCPPB 3274 WP 042860030 1 Dfangzhongdai
Dickeya gs fangzhongdai ND14b WP 038658003 1 Dfangzhongdai
Dickeya gs fangzhongdai M074 WP 038658003 1 Dfangzhongdai
Dickeya gs fangzhongdai M005 WP 038658003 1 Dfangzhongdai
Dickeya dadantii 3937 WP 013318212 1 Ddadantii
Dickeya dadantii NCPPB 3537 WP 038901387 1 Ddadantii
Dickeya dadantii subsp dieffenbachiae NCPPB 2976 WP 038901387 1 Ddadantii
Dickeya dadantii NCPPB 898 WP 038901387 1 Ddadantii
Dickeya solani MK10 WP 022633833 1 Dsolani
Dickeya solani RNS 05 1 2A WP 022633833 1 Dsolani
Dickeya solani IPO 2222 WP 022633833 1 Dsolani
Dickeya solani IFB 0099 WP 022633833 1 Dsolani
Dickeya solani PPO 9019 WP 022633833 1 Dsolani
Dickeya solani GBBC 2040 WP 022633833 1 Dsolani
Dickeya solani RNS 07 7 3B WP 022633833 1 Dsolani
Dickeya solani MK16 WP 022633833 1 Dsolani
Dickeya solani RNS 08 23 3 1 A WP 022633833 1 Dsolani
Dickeya solani D s0432 1 WP 022633833 1 Dsolani

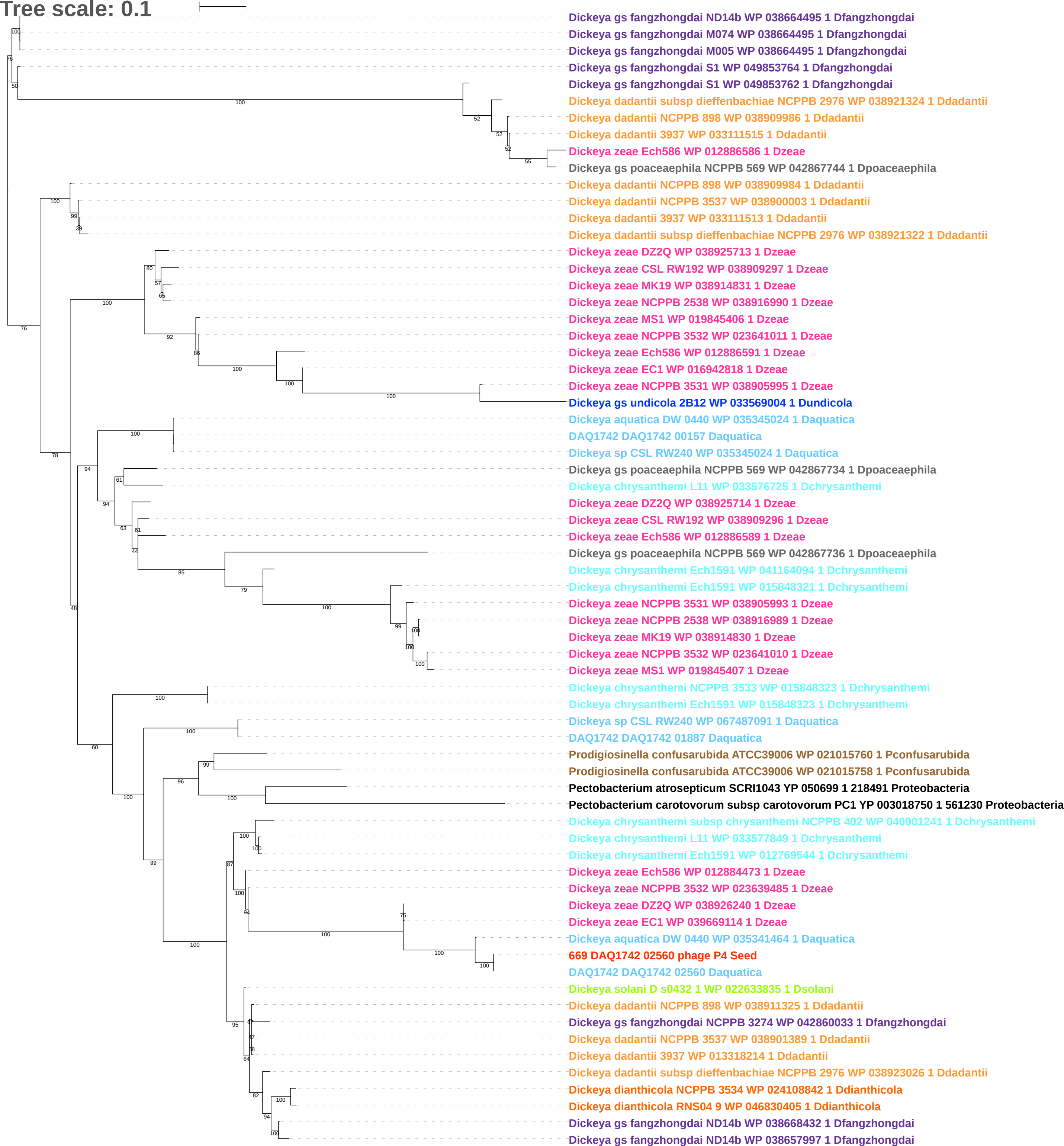

Tree scale: 0.1
Dickeya gs fangzhongdai ND14b WP 038664495 1 Dfangzhongdai
Dickeya gs fangzhongdai M074 WP 038664495 1 Dfangzhongdai
Dickeya gs fangzhongdai M005 WP 038664495 1 Dfangzhongdai
Dickeya gs fangzhongdai S1 WP 049853764 1 Dfangzhongdai
Dickeya gs fangzhongdai S1 WP 049853762 1 Dfangzhongdai
Dickeya dadantii subsp dieffenbachiae NCPPB 2976 WP 038921324 1 Ddadantii
Dickeya dadantii NCPPB 898 WP 038909986 1 Ddadantii
Dickeya dadantii 3937 WP 033111515 1 Ddadantii
Dickeya zeae Ech586 WP 012886586 1 Dzeae
Dickeya gs poaceaephila NCPPB 569 WP 042867744 1 Dpoaceaephila
Dickeya dadantii NCPPB 898 WP 038909984 1 Ddadantii
Dickeya dadantii NCPPB 3537 WP 038900003 1 Ddadantii
Dickeya dadantii 3937 WP 033111513 1 Ddadantii
Dickeya dadantii subsp dieffenbachiae NCPPB 2976 WP 038921322 1 Ddadantii
Dickeya zeae DZ2Q WP 038925713 1 Dzeae
Dickeya zeae CSL RW192 WP 038909297 1 Dzeae
Dickeya zeae MK19 WP 038914831 1 Dzeae
Dickeya zeae NCPPB 2538 WP 038916990 1 Dzeae
Dickeya zeae MS1 WP 019845406 1 Dzeae
Dickeya zeae NCPPB 3532 WP 023641011 1 Dzeae
Dickeya zeae Ech586 WP 012886591 1 Dzeae
Dickeya zeae EC1 WP 016942818 1 Dzeae
Dickeya zeae NCPPB 3531 WP 038905995 1 Dzeae
Dickeya gs undicola 2B12 WP 033569004 1 Dundicola
Dickeya aquatica DW 0440 WP 035345024 1 Daquatica
DAQ1742 DAQ1742 00157 Daquatica
Dickeya sp CSL RW240 WP 035345024 1 Daquatica
Dickeya gs poaceaephila NCPPB 569 WP 042867734 1 Dpoaceaephila
Dickeya chrysanthemi L11 WP 033576725 1 Dchrysanthemi
Dickeya zeae DZ2Q WP 038925714 1 Dzeae
Dickeya zeae CSL RW192 WP 038909296 1 Dzeae
Dickeya zeae Ech586 WP 012886589 1 Dzeae
Dickeya gs poaceaephila NCPPB 569 WP 042867736 1 Dpoaceaephila
Dickeya chrysanthemi Ech1591 WP 041164094 1 Dchrysanthemi
Dickeya chrysanthemi Ech1591 WP 015848321 1 Dchrysanthemi
Dickeya zeae NCPPB 3531 WP 038905993 1 Dzeae
Dickeya zeae NCPPB 2538 WP 038916989 1 Dzeae
Dickeya zeae MK19 WP 038914830 1 Dzeae
Dickeya zeae NCPPB 3532 WP 023641010 1 Dzeae
Dickeya zeae MS1 WP 019845407 1 Dzeae
Dickeya chrysanthemi NCPPB 3533 WP 015848323 1 Dchrysanthemi
Dickeya chrysanthemi Ech1591 WP 015848323 1 Dchrysanthemi
Dickeya sp CSL RW240 WP 067487091 1 Daquatica
DAQ1742 DAQ1742 01887 Daquatica
Prodigiosinella confusarubida ATCC39006 WP 021015760 1 Pconfusarubida
Prodigiosinella confusarubida ATCC39006 WP 021015758 1 Pconfusarubida
Pectobacterium atrosepticum SCRI1043 YP 050699 1 218491 Proteobacteria
Pectobacterium carotovorum subsp carotovorum PC1 YP 003018750 1 561230 Proteobacteria
Dickeya chrysanthemi subsp chrysanthemi NCPPB 402 WP 040001241 1 Dchrysanthemi
Dickeya chrysanthemi L11 WP 033577849 1 Dchrysanthemi
Dickeya chrysanthemi Ech1591 WP 012769544 1 Dchrysanthemi
Dickeya zeae Ech586 WP 012884473 1 Dzeae
Dickeya zeae NCPPB 3532 WP 023639485 1 Dzeae
Dickeya zeae DZ2Q WP 038926240 1 Dzeae
Dickeya zeae EC1 WP 039669114 1 Dzeae
Dickeya aquatica DW 0440 WP 035341464 1 Daquatica
669 DAQ1742 02560 phage P4 Seed
DAQ1742 DAQ1742 02560 Daquatica
Dickeya solani D s0432 1 WP 022633835 1 Dsolani
Dickeya dadantii NCPPB 898 WP 038911325 1 Ddadantii
Dickeya gs fangzhongdai NCPPB 3274 WP 042860033 1 Dfangzhongdai
Dickeya dadantii NCPPB 3537 WP 038901389 1 Ddadantii
Dickeya dadantii 3937 WP 013318214 1 Ddadantii
Dickeya dadantii subsp dieffenbachiae NCPPB 2976 WP 038923026 1 Ddadantii
Dickeya dianthicola NCPPB 3534 WP 024108842 1 Ddianthicola
Dickeya dianthicola RNS04 9 WP 046830405 1 Ddianthicola
Dickeya gs fangzhongdai ND14b WP 038668432 1 Dfangzhongdai
Dickeya gs fangzhongdai ND14b WP 038657997 1 Dfangzhongdai

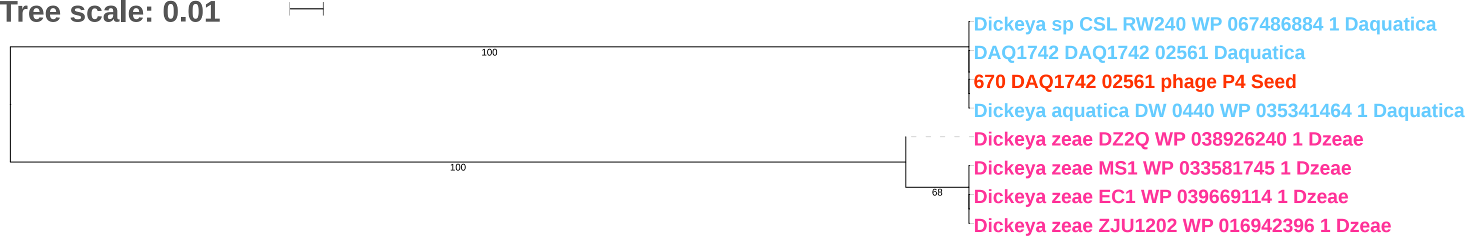
Tree scale: 0.01
100
100
68
Dickeya sp CSL RW240 WP 067486884 1 Daquatica
DAQ1742 DAQ1742 02561 Daquatica
670 DAQ1742 02561 phage P4 Seed
Dickeya aquatica DW 0440 WP 035341464 1 Daquatica
Dickeya zeae DZ2Q WP 038926240 1 Dzeae
Dickeya zeae MS1 WP 033581745 1 Dzeae
Dickeya zeae EC1 WP 039669114 1 Dzeae
Dickeya zeae ZJU1202 WP 016942396 1 Dzeae

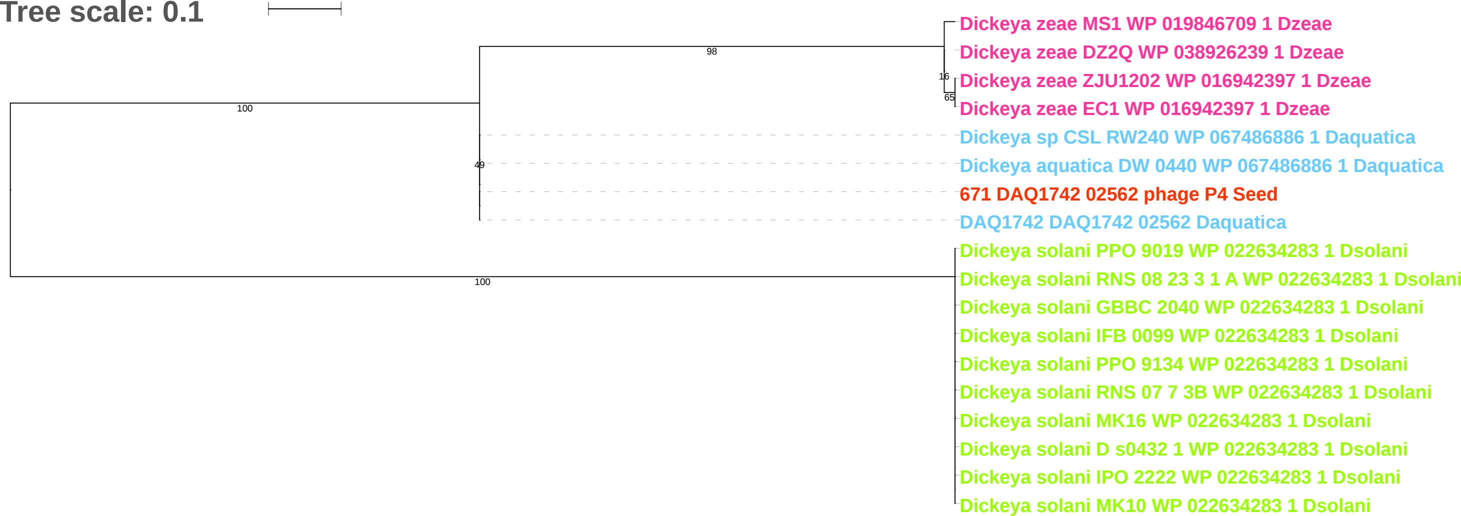
Tree scale: 0.1
Dickeya zeae MS1 WP 019846709 1 Dzeae
Dickeya zeae DZ2Q WP 038926239 1 Dzeae
Dickeya zeae ZJU1202 WP 016942397 1 Dzeae
Dickeya zeae EC1 WP 016942397 1 Dzeae
Dickeya sp CSL RW240 WP 067486886 1 Daquatica
Dickeya aquatica DW 0440 WP 067486886 1 Daquatica
671 DAQ1742 02562 phage P4 Seed
DAQ1742 DAQ1742 02562 Daquatica
Dickeya solani PPO 9019 WP 022634283 1 Dsolani
Dickeya solani RNS 08 23 3 1 A WP 022634283 1 Dsolani
Dickeya solani GBBC 2040 WP 022634283 1 Dsolani
Dickeya solani IFB 0099 WP 022634283 1 Dsolani
Dickeya solani PPO 9134 WP 022634283 1 Dsolani
Dickeya solani RNS 07 7 3B WP 022634283 1 Dsolani
Dickeya solani MK16 WP 022634283 1 Dsolani
Dickeya solani D s0432 1 WP 022634283 1 Dsolani
Dickeya solani IPO 2222 WP 022634283 1 Dsolani
Dickeya solani MK10 WP 022634283 1 Dsolani
98
100
16
65
49
100

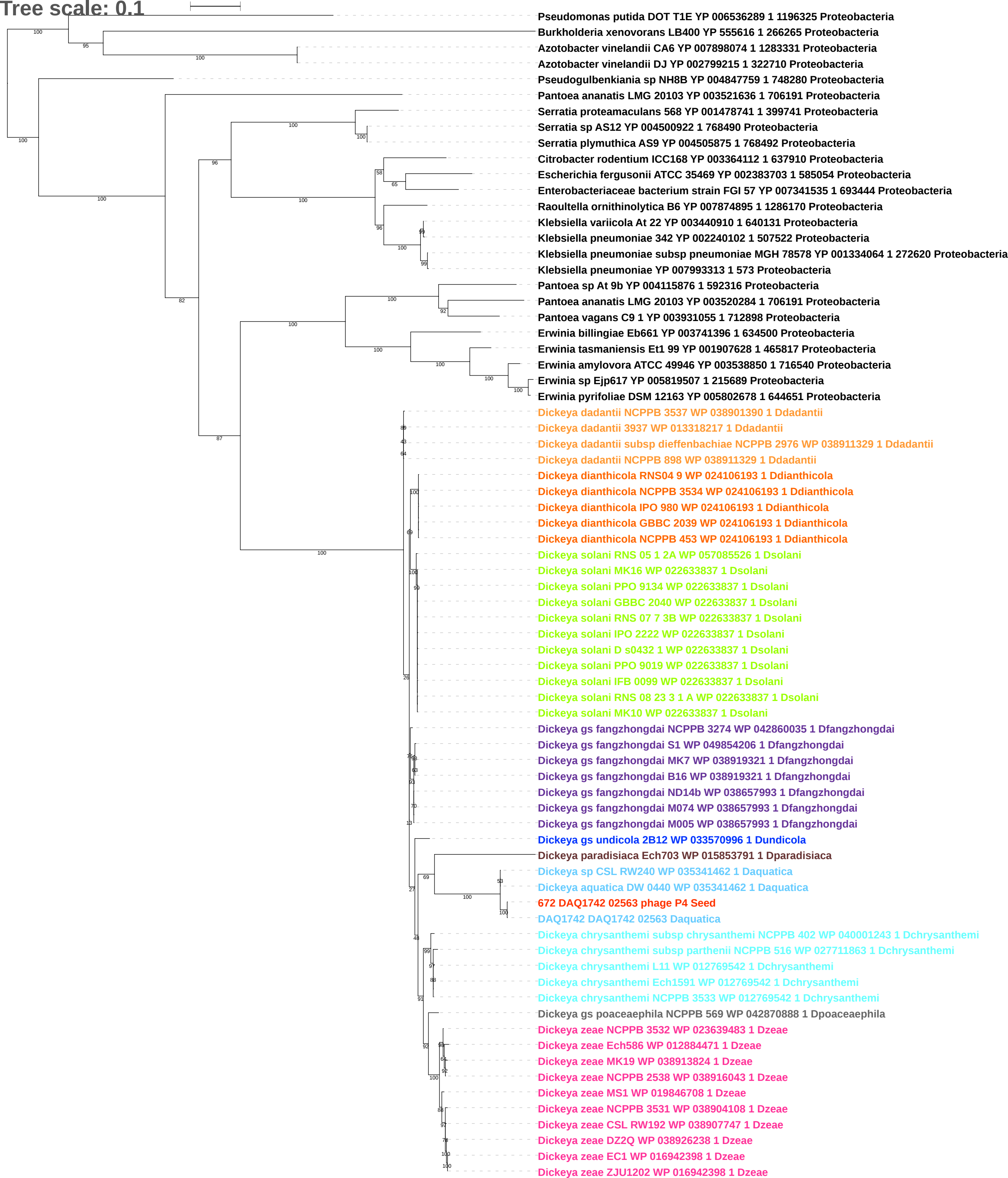
Tree scale: 0.1
Pseudomonas putida DOT T1E YP 006536289 1 1196325 Proteobacteria
Burkholderia xenovorans LB400 YP 555616 1 266265 Proteobacteria
Azotobacter vinelandii CA6 YP 007898074 1 1283331 Proteobacteria
Azotobacter vinelandii DJ YP 002799215 1 322710 Proteobacteria
Pseudogulbenkiania sp NH8B YP 004847759 1 748280 Proteobacteria
Pantoea ananatis LMG 20103 YP 003521636 1 706191 Proteobacteria
Serratia proteamaculans 568 YP 001478741 1 399741 Proteobacteria
Serratia sp AS12 YP 004500922 1 768490 Proteobacteria
Serratia plymuthica AS9 YP 004505875 1 768492 Proteobacteria
Citrobacter rodentium ICC168 YP 003364112 1 637910 Proteobacteria
Escherichia fergusonii ATCC 35469 YP 002383703 1 585054 Proteobacteria
Enterobacteriaceae bacterium strain FGI 57 YP 007341535 1 693444 Proteobacteria
Raoultella ornithinolytica B6 YP 007874895 1 1286170 Proteobacteria
Klebsiella variicola At 22 YP 003440910 1 640131 Proteobacteria
Klebsiella pneumoniae 342 YP 002240102 1 507522 Proteobacteria
Klebsiella pneumoniae subsp pneumoniae MGH 78578 YP 001334064 1 272620 Proteobacteria
Klebsiella pneumoniae YP 007993313 1 573 Proteobacteria
Pantoea sp At 9b YP 004115876 1 592316 Proteobacteria
Pantoea ananatis LMG 20103 YP 003520284 1 706191 Proteobacteria
Pantoea vagans C9 1 YP 003931055 1 712898 Proteobacteria
Erwinia billingiae Eb661 YP 003741396 1 634500 Proteobacteria
Erwinia tasmaniensis Et1 99 YP 001907628 1 465817 Proteobacteria
Erwinia amylovora ATCC 49946 YP 003538850 1 716540 Proteobacteria
Erwinia sp Ejp617 YP 005819507 1 215689 Proteobacteria
Erwinia pyrifoliae DSM 12163 YP 005802678 1 644651 Proteobacteria
Dickeya dadantii NCPPB 3537 WP 038901390 1 Ddadantii
Dickeya dadantii 3937 WP 013318217 1 Ddadantii
Dickeya dadantii subsp dieffenbachiae NCPPB 2976 WP 038911329 1 Ddadantii
Dickeya dadantii NCPPB 898 WP 038911329 1 Ddadantii
Dickeya dianthicola RNS04 9 WP 024106193 1 Ddianthicola
Dickeya dianthicola NCPPB 3534 WP 024106193 1 Ddianthicola
Dickeya dianthicola IPO 980 WP 024106193 1 Ddianthicola
Dickeya dianthicola GBBC 2039 WP 024106193 1 Ddianthicola
Dickeya dianthicola NCPPB 453 WP 024106193 1 Ddianthicola
Dickeya solani RNS 05 1 2A WP 057085526 1 Dsolani
Dickeya solani MK16 WP 022633837 1 Dsolani
Dickeya solani PPO 9134 WP 022633837 1 Dsolani
Dickeya solani GBBC 2040 WP 022633837 1 Dsolani
Dickeya solani RNS 07 7 3B WP 022633837 1 Dsolani
Dickeya solani IPO 2222 WP 022633837 1 Dsolani
Dickeya solani D s0432 1 WP 022633837 1 Dsolani
Dickeya solani PPO 9019 WP 022633837 1 Dsolani
Dickeya solani IFB 0099 WP 022633837 1 Dsolani
Dickeya solani RNS 08 23 3 1 A WP 022633837 1 Dsolani
Dickeya solani MK10 WP 022633837 1 Dsolani
Dickeya gs fangzhongdai NCPPB 3274 WP 042860035 1 Dfangzhongdai
Dickeya gs fangzhongdai S1 WP 049854206 1 Dfangzhongdai
Dickeya gs fangzhongdai MK7 WP 038919321 1 Dfangzhongdai
Dickeya gs fangzhongdai B16 WP 038919321 1 Dfangzhongdai
Dickeya gs fangzhongdai ND14b WP 038657993 1 Dfangzhongdai
Dickeya gs fangzhongdai M074 WP 038657993 1 Dfangzhongdai
Dickeya gs fangzhongdai M005 WP 038657993 1 Dfangzhongdai
Dickeya gs undicola 2B12 WP 033570996 1 Dundicola
Dickeya paradisiaca Ech703 WP 015853791 1 Dparadisiaca
Dickeya sp CSL RW240 WP 035341462 1 Daquatica
Dickeya aquatica DW 0440 WP 035341462 1 Daquatica
672 DAQ1742 02563 phage P4 Seed
DAQ1742 DAQ1742 02563 Daquatica
Dickeya chrysanthemi subsp chrysanthemi NCPPB 402 WP 040001243 1 Dchrysanthemi
Dickeya chrysanthemi subsp parthenii NCPPB 516 WP 027711863 1 Dchrysanthemi
Dickeya chrysanthemi L11 WP 012769542 1 Dchrysanthemi
Dickeya chrysanthemi Ech1591 WP 012769542 1 Dchrysanthemi
Dickeya chrysanthemi NCPPB 3533 WP 012769542 1 Dchrysanthemi
Dickeya gs poaceaephila NCPPB 569 WP 042870888 1 Dpoaceaephila
Dickeya zeae NCPPB 3532 WP 023639483 1 Dzeae
Dickeya zeae Ech586 WP 012884471 1 Dzeae
Dickeya zeae MK19 WP 038913824 1 Dzeae
Dickeya zeae NCPPB 2538 WP 038916043 1 Dzeae
Dickeya zeae MS1 WP 019846708 1 Dzeae
Dickeya zeae NCPPB 3531 WP 038904108 1 Dzeae
Dickeya zeae CSL RW192 WP 038907747 1 Dzeae
Dickeya zeae DZ2Q WP 038926238 1 Dzeae
Dickeya zeae EC1 WP 016942398 1 Dzeae
Dickeya zeae ZJU1202 WP 016942398 1 Dzeae

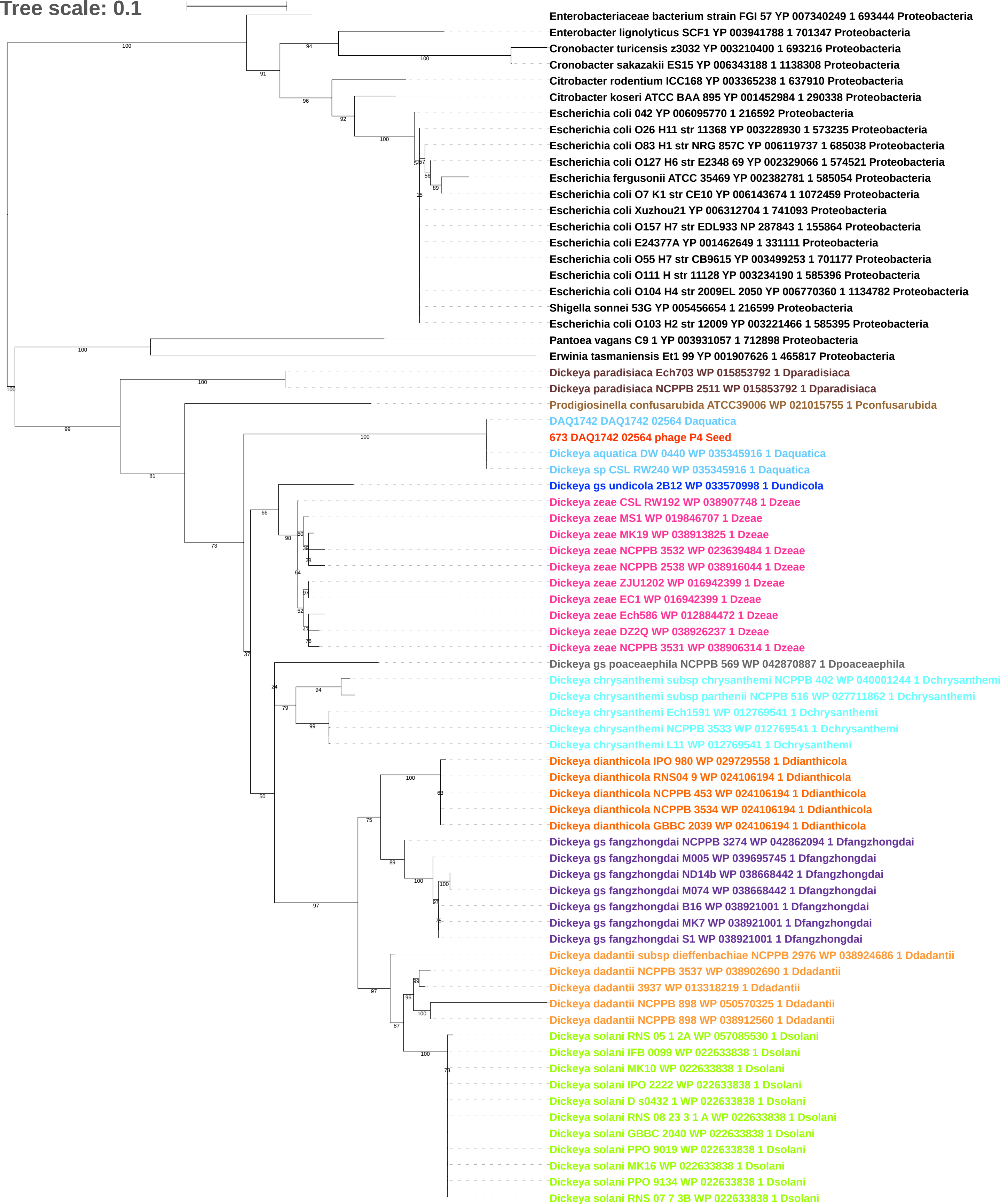
Tree scale: 0.1
Enterobacteriaceae bacterium strain FGI 57 YP 007340249 1 693444 Proteobacteria
Enterobacter lignolyticus SCF1 YP 003941788 1 701347 Proteobacteria
Cronobacter turicensis z3032 YP 003210400 1 693216 Proteobacteria
Cronobacter sakazakii ES15 YP 006343188 1 1138308 Proteobacteria
Citrobacter rodentium ICC168 YP 003365238 1 637910 Proteobacteria
Citrobacter koseri ATCC BAA 895 YP 001452984 1 290338 Proteobacteria
Escherichia coli 042 YP 006095770 1 216592 Proteobacteria
Escherichia coli O26 H11 str 11368 YP 003228930 1 573235 Proteobacteria
Escherichia coli O83 H1 str NRG 857C YP 006119737 1 685038 Proteobacteria
Escherichia coli O127 H6 str E2348 69 YP 002329066 1 574521 Proteobacteria
Escherichia fergusonii ATCC 35469 YP 002382781 1 585054 Proteobacteria
Escherichia coli O7 K1 str CE10 YP 006143674 1 1072459 Proteobacteria
Escherichia coli Xuzhou21 YP 006312704 1 741093 Proteobacteria
Escherichia coli O157 H7 str EDL933 NP 287843 1 155864 Proteobacteria
Escherichia coli E24377A YP 001462649 1 331111 Proteobacteria
Escherichia coli O55 H7 str CB9615 YP 003499253 1 701177 Proteobacteria
Escherichia coli O111 H str 11128 YP 003234190 1 585396 Proteobacteria
Escherichia coli O104 H4 str 2009EL 2050 YP 006770360 1 1134782 Proteobacteria
Shigella sonnei 53G YP 005456654 1 216599 Proteobacteria
Escherichia coli O103 H2 str 12009 YP 003221466 1 585395 Proteobacteria
Pantoea vagans C9 1 YP 003931057 1 712898 Proteobacteria
Erwinia tasmaniensis Et1 99 YP 001907626 1 465817 Proteobacteria
Dickeya paradisiaca Ech703 WP 015853792 1 Dparadisiaca
Dickeya paradisiaca NCPPB 2511 WP 015853792 1 Dparadisiaca
Prodigiosinella confusarubida ATCC39006 WP 021015755 1 Pconfusarubida
DAQ1742 DAQ1742 02564 Daquatica
673 DAQ1742 02564 phage P4 Seed
Dickeya aquatica DW 0440 WP 035345916 1 Daquatica
Dickeya sp CSL RW240 WP 035345916 1 Daquatica
Dickeya gs undicola 2B12 WP 033570998 1 Dundicola
Dickeya zeae CSL RW192 WP 038907748 1 Dzeae
Dickeya zeae MS1 WP 019846707 1 Dzeae
Dickeya zeae MK19 WP 038913825 1 Dzeae
Dickeya zeae NCPPB 3532 WP 023639484 1 Dzeae
Dickeya zeae NCPPB 2538 WP 038916044 1 Dzeae
Dickeya zeae ZJU1202 WP 016942399 1 Dzeae
Dickeya zeae EC1 WP 016942399 1 Dzeae
Dickeya zeae Ech586 WP 012884472 1 Dzeae
Dickeya zeae DZ2Q WP 038926237 1 Dzeae
Dickeya zeae NCPPB 3531 WP 038906314 1 Dzeae
Dickeya gs poaceaephila NCPPB 569 WP 042870887 1 Dpoaceaephila
Dickeya chrysanthemi subsp chrysanthemi NCPPB 402 WP 040001244 1 Dchrysanthemi
Dickeya chrysanthemi subsp parthenii NCPPB 516 WP 027711862 1 Dchrysanthemi
Dickeya chrysanthemi Ech1591 WP 012769541 1 Dchrysanthemi
Dickeya chrysanthemi NCPPB 3533 WP 012769541 1 Dchrysanthemi
Dickeya chrysanthemi L11 WP 012769541 1 Dchrysanthemi
Dickeya dianthicola IPO 980 WP 029729558 1 Ddianthicola
Dickeya dianthicola RNS04 9 WP 024106194 1 Ddianthicola
Dickeya dianthicola NCPPB 453 WP 024106194 1 Ddianthicola
Dickeya dianthicola NCPPB 3534 WP 024106194 1 Ddianthicola
Dickeya dianthicola GBBC 2039 WP 024106194 1 Ddianthicola
Dickeya gs fangzhongdai NCPPB 3274 WP 042862094 1 Dfangzhongdai
Dickeya gs fangzhongdai M005 WP 039695745 1 Dfangzhongdai
Dickeya gs fangzhongdai ND14b WP 038668442 1 Dfangzhongdai
Dickeya gs fangzhongdai M074 WP 038668442 1 Dfangzhongdai
Dickeya gs fangzhongdai B16 WP 038921001 1 Dfangzhongdai
Dickeya gs fangzhongdai MK7 WP 038921001 1 Dfangzhongdai
Dickeya gs fangzhongdai S1 WP 038921001 1 Dfangzhongdai
Dickeya dadantii subsp dieffenbachiae NCPPB 2976 WP 038924686 1 Ddadantii
Dickeya dadantii NCPPB 3537 WP 038902690 1 Ddadantii
Dickeya dadantii 3937 WP 013318219 1 Ddadantii
Dickeya dadantii NCPPB 898 WP 050570325 1 Ddadantii
Dickeya dadantii NCPPB 898 WP 038912560 1 Ddadantii
Dickeya solani RNS 05 1 2A WP 057085530 1 Dsolani
Dickeya solani IFB 0099 WP 022633838 1 Dsolani
Dickeya solani MK10 WP 022633838 1 Dsolani
Dickeya solani IPO 2222 WP 022633838 1 Dsolani
Dickeya solani D s0432 1 WP 022633838 1 Dsolani
Dickeya solani RNS 08 23 3 1 A WP 022633838 1 Dsolani
Dickeya solani GBBC 2040 WP 022633838 1 Dsolani
Dickeya solani PPO 9019 WP 022633838 1 Dsolani
Dickeya solani MK16 WP 022633838 1 Dsolani
Dickeya solani PPO 9134 WP 022633838 1 Dsolani
Dickeya solani RNS 07 7 3B WP 022633838 1 Dsolani

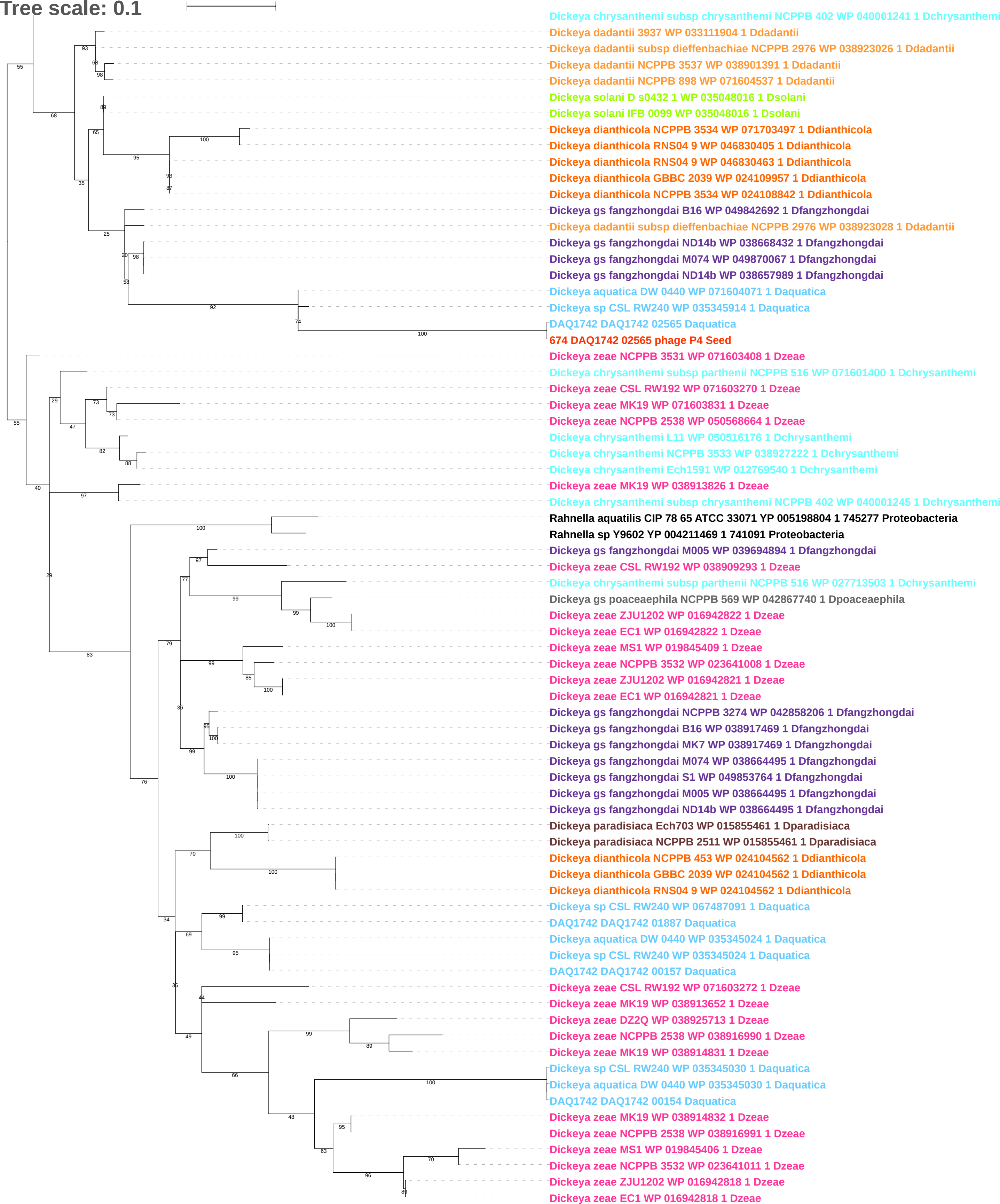

Tree scale: 0.1
Dickeya chrysanthemi subsp chrysanthemi NCPPB 402 WP 040001241 1 Dchrysanthemi
Dickeya dadantii 3937 WP 033111904 1 Ddadantii
Dickeya dadantii subsp dieffenbachiae NCPPB 2976 WP 038923026 1 Ddadantii
Dickeya dadantii NCPPB 3537 WP 038901391 1 Ddadantii
Dickeya dadantii NCPPB 898 WP 071604537 1 Ddadantii
Dickeya solani D s0432 1 WP 035048016 1 Dsolani
Dickeya solani IFB 0099 WP 035048016 1 Dsolani
Dickeya dianthicola NCPPB 3534 WP 071703497 1 Ddianthicola
Dickeya dianthicola RNS04 9 WP 046830405 1 Ddianthicola
Dickeya dianthicola RNS04 9 WP 046830463 1 Ddianthicola
Dickeya dianthicola GBBC 2039 WP 024109957 1 Ddianthicola
Dickeya dianthicola NCPPB 3534 WP 024108842 1 Ddianthicola
Dickeya gs fangzhongdai B16 WP 049842692 1 Dfangzhongdai
Dickeya dadantii subsp dieffenbachiae NCPPB 2976 WP 038923028 1 Ddadantii
Dickeya gs fangzhongdai ND14b WP 038668432 1 Dfangzhongdai
Dickeya gs fangzhongdai M074 WP 049870067 1 Dfangzhongdai
Dickeya gs fangzhongdai ND14b WP 038657989 1 Dfangzhongdai
Dickeya aquatica DW 0440 WP 071604071 1 Daquatica
Dickeya sp CSL RW240 WP 035345914 1 Daquatica
DAQ1742 DAQ1742 02565 Daquatica
674 DAQ1742 02565 phage P4 Seed
Dickeya zeae NCPPB 3531 WP 071603408 1 Dzeae
Dickeya chrysanthemi subsp parthenii NCPPB 516 WP 071601400 1 Dchrysanthemi
Dickeya zeae CSL RW192 WP 071603270 1 Dzeae
Dickeya zeae MK19 WP 071603831 1 Dzeae
Dickeya zeae NCPPB 2538 WP 050568664 1 Dzeae
Dickeya chrysanthemi L11 WP 050516176 1 Dchrysanthemi
Dickeya chrysanthemi NCPPB 3533 WP 038927222 1 Dchrysanthemi
Dickeya chrysanthemi Ech1591 WP 012769540 1 Dchrysanthemi
Dickeya zeae MK19 WP 038913826 1 Dzeae
Dickeya chrysanthemi subsp chrysanthemi NCPPB 402 WP 040001245 1 Dchrysanthemi
Rahnella aquatilis CIP 78 65 ATCC 33071 YP 005198804 1 745277 Proteobacteria
Rahnella sp Y9602 YP 004211469 1 741091 Proteobacteria
Dickeya gs fangzhongdai M005 WP 039694894 1 Dfangzhongdai
Dickeya zeae CSL RW192 WP 038909293 1 Dzeae
Dickeya chrysanthemi subsp parthenii NCPPB 516 WP 027713503 1 Dchrysanthemi
Dickeya gs poaceaephila NCPPB 569 WP 042867740 1 Dpoaceaephila
Dickeya zeae ZJU1202 WP 016942822 1 Dzeae
Dickeya zeae EC1 WP 016942822 1 Dzeae
Dickeya zeae MS1 WP 019845409 1 Dzeae
Dickeya zeae NCPPB 3532 WP 023641008 1 Dzeae
Dickeya zeae ZJU1202 WP 016942821 1 Dzeae
Dickeya zeae EC1 WP 016942821 1 Dzeae
Dickeya gs fangzhongdai NCPPB 3274 WP 042858206 1 Dfangzhongdai
Dickeya gs fangzhongdai B16 WP 038917469 1 Dfangzhongdai
Dickeya gs fangzhongdai MK7 WP 038917469 1 Dfangzhongdai
Dickeya gs fangzhongdai M074 WP 038664495 1 Dfangzhongdai
Dickeya gs fangzhongdai S1 WP 049853764 1 Dfangzhongdai
Dickeya gs fangzhongdai M005 WP 038664495 1 Dfangzhongdai
Dickeya gs fangzhongdai ND14b WP 038664495 1 Dfangzhongdai
Dickeya paradisiaca Ech703 WP 015855461 1 Dparadisiaca
Dickeya paradisiaca NCPPB 2511 WP 015855461 1 Dparadisiaca
Dickeya dianthicola NCPPB 453 WP 024104562 1 Ddianthicola
Dickeya dianthicola GBBC 2039 WP 024104562 1 Ddianthicola
Dickeya dianthicola RNS04 9 WP 024104562 1 Ddianthicola
Dickeya sp CSL RW240 WP 067487091 1 Daquatica
DAQ1742 DAQ1742 01887 Daquatica
Dickeya aquatica DW 0440 WP 035345024 1 Daquatica
Dickeya sp CSL RW240 WP 035345024 1 Daquatica
DAQ1742 DAQ1742 00157 Daquatica
Dickeya zeae CSL RW192 WP 071603272 1 Dzeae
Dickeya zeae MK19 WP 038913652 1 Dzeae
Dickeya zeae DZ2Q WP 038925713 1 Dzeae
Dickeya zeae NCPPB 2538 WP 038916990 1 Dzeae
Dickeya zeae MK19 WP 038914831 1 Dzeae
Dickeya sp CSL RW240 WP 035345030 1 Daquatica
Dickeya aquatica DW 0440 WP 035345030 1 Daquatica
DAQ1742 DAQ1742 00154 Daquatica
Dickeya zeae MK19 WP 038914832 1 Dzeae
Dickeya zeae NCPPB 2538 WP 038916991 1 Dzeae
Dickeya zeae MS1 WP 019845406 1 Dzeae
Dickeya zeae NCPPB 3532 WP 023641011 1 Dzeae
Dickeya zeae ZJU1202 WP 016942818 1 Dzeae
Dickeya zeae EC1 WP 016942818 1 Dzeae

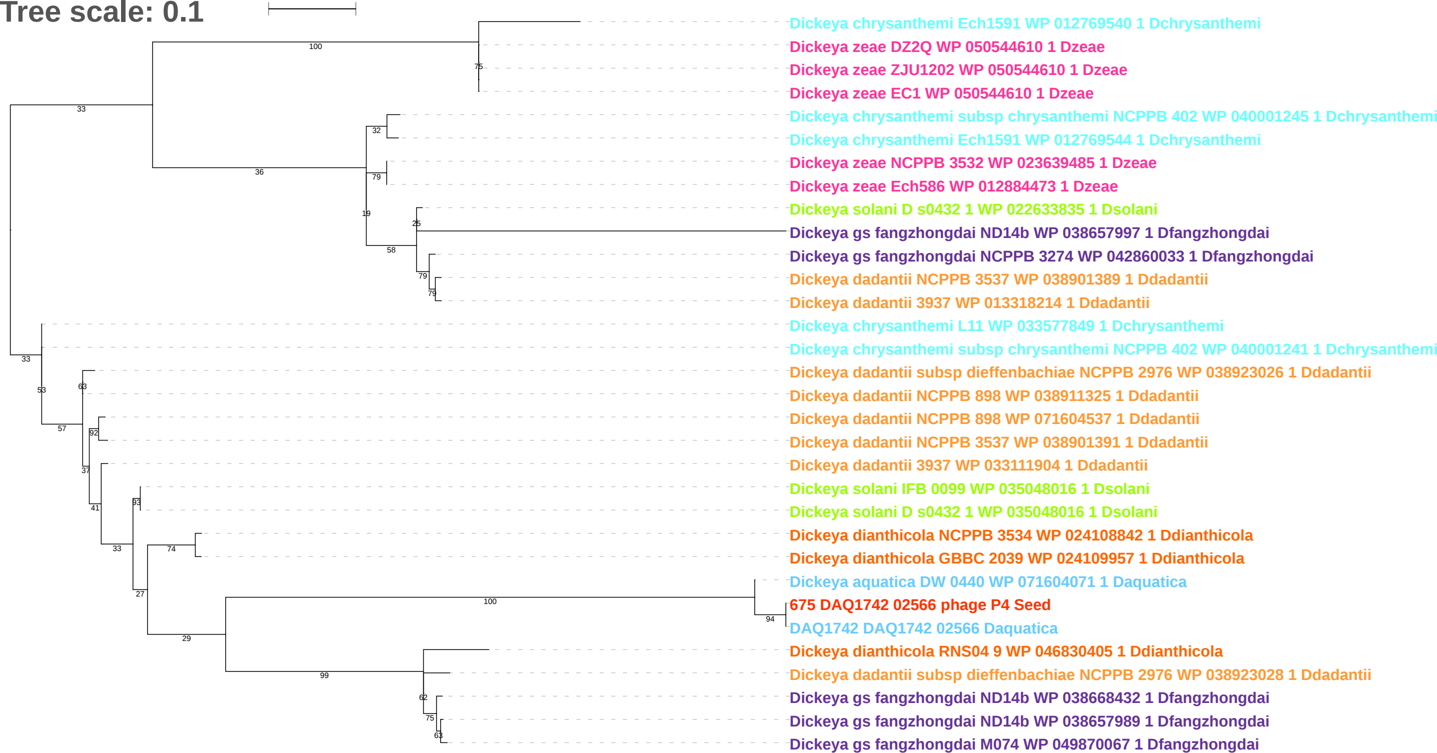

Tree scale: 0.1
Dickeya chrysanthemi Ech1591 WP 012769540 1 Dchrysanthemi
Dickeya zeae DZ2Q WP 050544610 1 Dzeae
Dickeya zeae ZJU1202 WP 050544610 1 Dzeae
Dickeya zeae EC1 WP 050544610 1 Dzeae
Dickeya chrysanthemi subsp chrysanthemi NCPPB 402 WP 040001245 1 Dchrysanthemi
Dickeya chrysanthemi Ech1591 WP 012769544 1 Dchrysanthemi
Dickeya zeae NCPPB 3532 WP 023639485 1 Dzeae
Dickeya zeae Ech586 WP 012884473 1 Dzeae
Dickeya solani D s0432 1 WP 022633835 1 Dsolani
Dickeya gs fangzhongdai ND14b WP 038657997 1 Dfangzhongdai
Dickeya gs fangzhongdai NCPPB 3274 WP 042860033 1 Dfangzhongdai
Dickeya dadantii NCPPB 3537 WP 038901389 1 Ddadantii
Dickeya dadantii 3937 WP 013318214 1 Ddadantii
Dickeya chrysanthemi L11 WP 033577849 1 Dchrysanthemi
Dickeya chrysanthemi subsp chrysanthemi NCPPB 402 WP 040001241 1 Dchrysanthemi
Dickeya dadantii subsp dieffenbachiae NCPPB 2976 WP 038923026 1 Ddadantii
Dickeya dadantii NCPPB 898 WP 038911325 1 Ddadantii
Dickeya dadantii NCPPB 898 WP 071604537 1 Ddadantii
Dickeya dadantii NCPPB 3537 WP 038901391 1 Ddadantii
Dickeya dadantii 3937 WP 033111904 1 Ddadantii
Dickeya solani IFB 0099 WP 035048016 1 Dsolani
Dickeya solani D s0432 1 WP 035048016 1 Dsolani
Dickeya dianthicola NCPPB 3534 WP 024108842 1 Ddianthicola
Dickeya dianthicola GBBC 2039 WP 024109957 1 Ddianthicola
Dickeya aquatica DW 0440 WP 071604071 1 Daquatica
675 DAQ1742 02566 phage P4 Seed
DAQ1742 DAQ1742 02566 Daquatica
Dickeya dianthicola RNS04 9 WP 046830405 1 Ddianthicola
Dickeya dadantii subsp dieffenbachiae NCPPB 2976 WP 038923028 1 Ddadantii
Dickeya gs fangzhongdai ND14b WP 038668432 1 Dfangzhongdai
Dickeya gs fangzhongdai ND14b WP 038657989 1 Dfangzhongdai
Dickeya gs fangzhongdai M074 WP 049870067 1 Dfangzhongdai

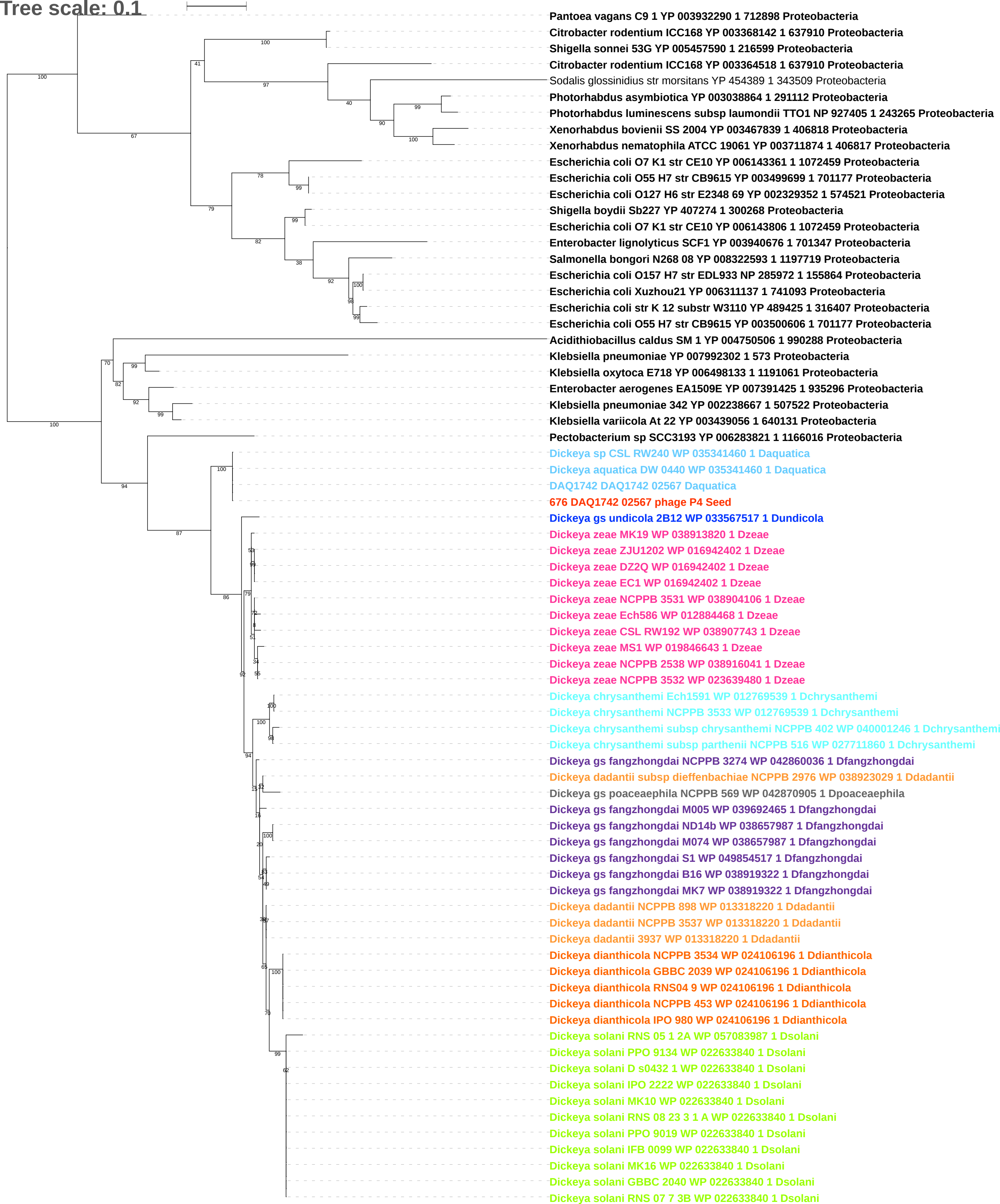
Tree scale: 0.1
Pantoea vagans C9 1 YP 003932290 1 712898 Proteobacteria
Citrobacter rodentium ICC168 YP 003368142 1 637910 Proteobacteria
Shigella sonnei 53G YP 005457590 1 216599 Proteobacteria
Citrobacter rodentium ICC168 YP 003364518 1 637910 Proteobacteria
Sodalis glossinidius str morsitans YP 454389 1 343509 Proteobacteria
Photorhabdus asymbiotica YP 003038864 1 291112 Proteobacteria
Photorhabdus luminescens subsp laumondii TTO1 NP 927405 1 243265 Proteobacteria
Xenorhabdus bovienii SS 2004 YP 003467839 1 406818 Proteobacteria
Xenorhabdus nematophila ATCC 19061 YP 003711874 1 406817 Proteobacteria
Escherichia coli O7 K1 str CE10 YP 006143361 1 1072459 Proteobacteria
Escherichia coli O55 H7 str CB9615 YP 003499699 1 701177 Proteobacteria
Escherichia coli O127 H6 str E2348 69 YP 002329352 1 574521 Proteobacteria
Shigella boydii Sb227 YP 407274 1 300268 Proteobacteria
Escherichia coli O7 K1 str CE10 YP 006143806 1 1072459 Proteobacteria
Enterobacter lignolyticus SCF1 YP 003940676 1 701347 Proteobacteria
Salmonella bongori N268 08 YP 008322593 1 1197719 Proteobacteria
Escherichia coli O157 H7 str EDL933 NP 285972 1 155864 Proteobacteria
Escherichia coli Xuzhou21 YP 006311137 1 741093 Proteobacteria
Escherichia coli str K 12 substr W3110 YP 489425 1 316407 Proteobacteria
Escherichia coli O55 H7 str CB9615 YP 003500606 1 701177 Proteobacteria
Acidithiobacillus caldus SM 1 YP 004750506 1 990288 Proteobacteria
Klebsiella pneumoniae YP 007992302 1 573 Proteobacteria
Klebsiella oxytoca E718 YP 006498133 1 1191061 Proteobacteria
Enterobacter aerogenes EA1509E YP 007391425 1 935296 Proteobacteria
Klebsiella pneumoniae 342 YP 002238667 1 507522 Proteobacteria
Klebsiella variicola At 22 YP 003439056 1 640131 Proteobacteria
Pectobacterium sp SCC3193 YP 006283821 1 1166016 Proteobacteria
Dickeya sp CSL RW240 WP 035341460 1 Daquatica
Dickeya aquatica DW 0440 WP 035341460 1 Daquatica
DAQ1742 DAQ1742 02567 Daquatica
676 DAQ1742 02567 phage P4 Seed
Dickeya gs undicola 2B12 WP 033567517 1 Dundicola
Dickeya zeae MK19 WP 038913820 1 Dzeae
Dickeya zeae ZJU1202 WP 016942402 1 Dzeae
Dickeya zeae DZ2Q WP 016942402 1 Dzeae
Dickeya zeae EC1 WP 016942402 1 Dzeae
Dickeya zeae NCPPB 3531 WP 038904106 1 Dzeae
Dickeya zeae Ech586 WP 012884468 1 Dzeae
Dickeya zeae CSL RW192 WP 038907743 1 Dzeae
Dickeya zeae MS1 WP 019846643 1 Dzeae
Dickeya zeae NCPPB 2538 WP 038916041 1 Dzeae
Dickeya zeae NCPPB 3532 WP 023639480 1 Dzeae
Dickeya chrysanthemi Ech1591 WP 012769539 1 Dchrysanthemi
Dickeya chrysanthemi NCPPB 3533 WP 012769539 1 Dchrysanthemi
Dickeya chrysanthemi subsp chrysanthemi NCPPB 402 WP 040001246 1 Dchrysanthemi
Dickeya chrysanthemi subsp parthenii NCPPB 516 WP 027711860 1 Dchrysanthemi
Dickeya gs fangzhongdai NCPPB 3274 WP 042860036 1 Dfangzhongdai
Dickeya dadantii subsp dieffenbachiae NCPPB 2976 WP 038923029 1 Ddadantii
Dickeya gs poaceaephila NCPPB 569 WP 042870905 1 Dpoaceaephila
Dickeya gs fangzhongdai M005 WP 039692465 1 Dfangzhongdai
Dickeya gs fangzhongdai ND14b WP 038657987 1 Dfangzhongdai
Dickeya gs fangzhongdai M074 WP 038657987 1 Dfangzhongdai
Dickeya gs fangzhongdai S1 WP 049854517 1 Dfangzhongdai
Dickeya gs fangzhongdai B16 WP 038919322 1 Dfangzhongdai
Dickeya gs fangzhongdai MK7 WP 038919322 1 Dfangzhongdai
Dickeya dadantii NCPPB 898 WP 013318220 1 Ddadantii
Dickeya dadantii NCPPB 3537 WP 013318220 1 Ddadantii
Dickeya dadantii 3937 WP 013318220 1 Ddadantii
Dickeya dianthicola NCPPB 3534 WP 024106196 1 Ddianthicola
Dickeya dianthicola GBBC 2039 WP 024106196 1 Ddianthicola
Dickeya dianthicola RNS04 9 WP 024106196 1 Ddianthicola
Dickeya dianthicola NCPPB 453 WP 024106196 1 Ddianthicola
Dickeya dianthicola IPO 980 WP 024106196 1 Ddianthicola
Dickeya solani RNS 05 1 2A WP 057083987 1 Dsolani
Dickeya solani PPO 9134 WP 022633840 1 Dsolani
Dickeya solani D s0432 1 WP 022633840 1 Dsolani
Dickeya solani IPO 2222 WP 022633840 1 Dsolani
Dickeya solani MK10 WP 022633840 1 Dsolani
Dickeya solani RNS 08 23 3 1 A WP 022633840 1 Dsolani
Dickeya solani PPO 9019 WP 022633840 1 Dsolani
Dickeya solani IFB 0099 WP 022633840 1 Dsolani
Dickeya solani MK16 WP 022633840 1 Dsolani
Dickeya solani GBBC 2040 WP 022633840 1 Dsolani
Dickeya solani RNS 07 7 3B WP 022633840 1 Dsolani

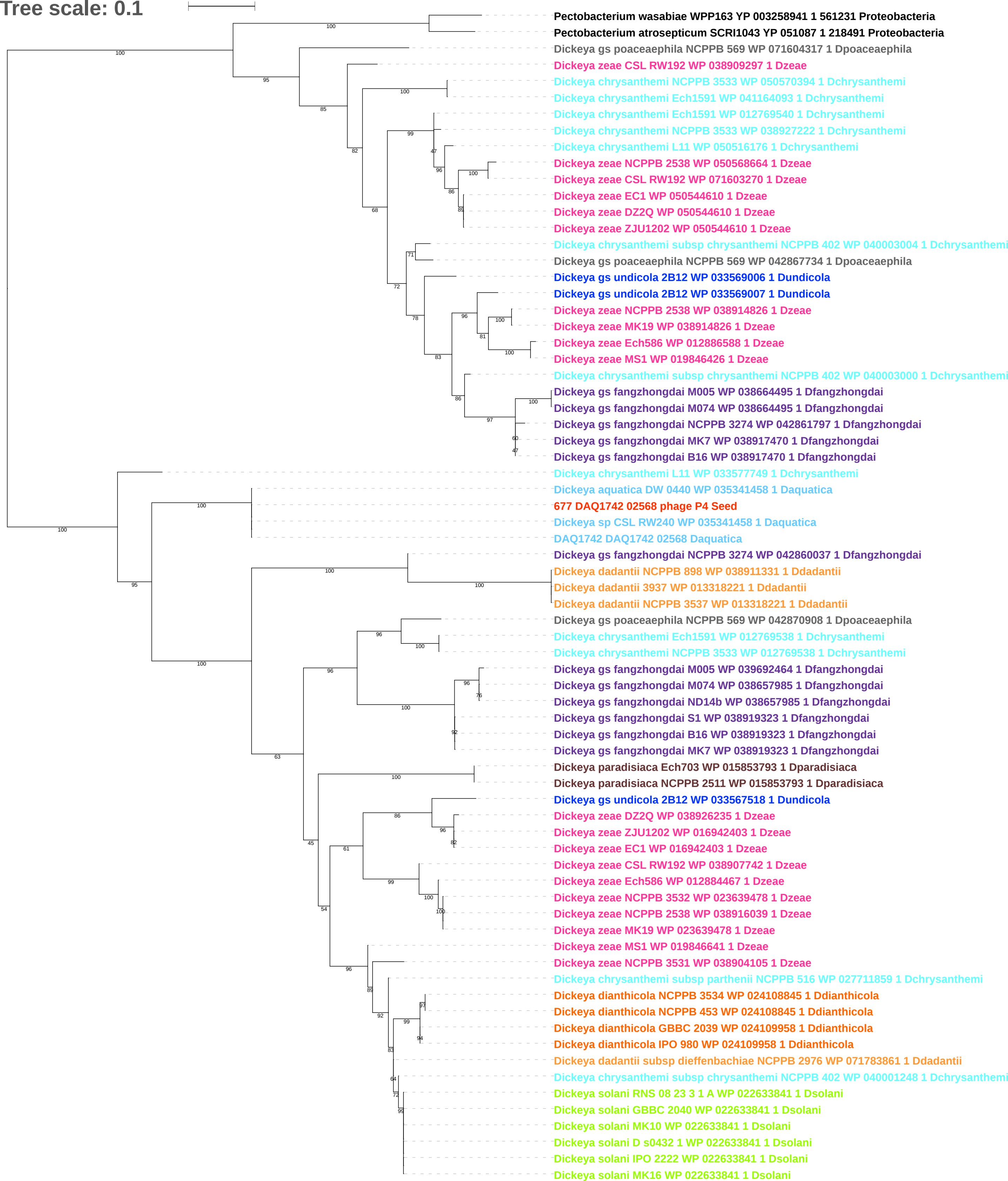
Tree scale: 0.1
Pectobacterium wasabiae WPP163 YP 003258941 1 561231 Proteobacteria
Pectobacterium atrosepticum SCRI1043 YP 051087 1 218491 Proteobacteria
Dickeya gs poaceaephila NCPPB 569 WP 071604317 1 Dpoaceaephila
Dickeya zeae CSL RW192 WP 038909297 1 Dzeae
Dickeya chrysanthemi NCPPB 3533 WP 050570394 1 Dchrysanthemi
Dickeya chrysanthemi Ech1591 WP 041164093 1 Dchrysanthemi
Dickeya chrysanthemi Ech1591 WP 012769540 1 Dchrysanthemi
Dickeya chrysanthemi NCPPB 3533 WP 038927222 1 Dchrysanthemi
Dickeya chrysanthemi L11 WP 050516176 1 Dchrysanthemi
Dickeya zeae NCPPB 2538 WP 050568664 1 Dzeae
Dickeya zeae CSL RW192 WP 071603270 1 Dzeae
Dickeya zeae EC1 WP 050544610 1 Dzeae
Dickeya zeae DZ2Q WP 050544610 1 Dzeae
Dickeya zeae ZJU1202 WP 050544610 1 Dzeae
Dickeya chrysanthemi subsp chrysanthemi NCPPB 402 WP 040003004 1 Dchrysanthemi
Dickeya gs poaceaephila NCPPB 569 WP 042867734 1 Dpoaceaephila
Dickeya gs undicola 2B12 WP 033569006 1 Dundicola
Dickeya gs undicola 2B12 WP 033569007 1 Dundicola
Dickeya zeae NCPPB 2538 WP 038914826 1 Dzeae
Dickeya zeae MK19 WP 038914826 1 Dzeae
Dickeya zeae Ech586 WP 012886588 1 Dzeae
Dickeya zeae MS1 WP 019846426 1 Dzeae
Dickeya chrysanthemi subsp chrysanthemi NCPPB 402 WP 040003000 1 Dchrysanthemi
Dickeya gs fangzhongdai M005 WP 038664495 1 Dfangzhongdai
Dickeya gs fangzhongdai M074 WP 038664495 1 Dfangzhongdai
Dickeya gs fangzhongdai NCPPB 3274 WP 042861797 1 Dfangzhongdai
Dickeya gs fangzhongdai MK7 WP 038917470 1 Dfangzhongdai
Dickeya gs fangzhongdai B16 WP 038917470 1 Dfangzhongdai
Dickeya chrysanthemi L11 WP 033577749 1 Dchrysanthemi
Dickeya aquatica DW 0440 WP 035341458 1 Daquatica
677 DAQ1742 02568 phage P4 Seed
Dickeya sp CSL RW240 WP 035341458 1 Daquatica
DAQ1742 DAQ1742 02568 Daquatica
Dickeya gs fangzhongdai NCPPB 3274 WP 042860037 1 Dfangzhongdai
Dickeya dadantii NCPPB 898 WP 038911331 1 Ddadantii
Dickeya dadantii 3937 WP 013318221 1 Ddadantii
Dickeya dadantii NCPPB 3537 WP 013318221 1 Ddadantii
Dickeya gs poaceaephila NCPPB 569 WP 042870908 1 Dpoaceaephila
Dickeya chrysanthemi Ech1591 WP 012769538 1 Dchrysanthemi
Dickeya chrysanthemi NCPPB 3533 WP 012769538 1 Dchrysanthemi
Dickeya gs fangzhongdai M005 WP 039692464 1 Dfangzhongdai
Dickeya gs fangzhongdai M074 WP 038657985 1 Dfangzhongdai
Dickeya gs fangzhongdai ND14b WP 038657985 1 Dfangzhongdai
Dickeya gs fangzhongdai S1 WP 038919323 1 Dfangzhongdai
Dickeya gs fangzhongdai B16 WP 038919323 1 Dfangzhongdai
Dickeya gs fangzhongdai MK7 WP 038919323 1 Dfangzhongdai
Dickeya paradisiaca Ech703 WP 015853793 1 Dparadisiaca
Dickeya paradisiaca NCPPB 2511 WP 015853793 1 Dparadisiaca
Dickeya gs undicola 2B12 WP 033567518 1 Dundicola
Dickeya zeae DZ2Q WP 038926235 1 Dzeae
Dickeya zeae ZJU1202 WP 016942403 1 Dzeae
Dickeya zeae EC1 WP 016942403 1 Dzeae
Dickeya zeae CSL RW192 WP 038907742 1 Dzeae
Dickeya zeae Ech586 WP 012884467 1 Dzeae
Dickeya zeae NCPPB 3532 WP 023639478 1 Dzeae
Dickeya zeae NCPPB 2538 WP 038916039 1 Dzeae
Dickeya zeae MK19 WP 023639478 1 Dzeae
Dickeya zeae MS1 WP 019846641 1 Dzeae
Dickeya zeae NCPPB 3531 WP 038904105 1 Dzeae
Dickeya chrysanthemi subsp parthenii NCPPB 516 WP 027711859 1 Dchrysanthemi
Dickeya dianthicola NCPPB 3534 WP 024108845 1 Ddianthicola
Dickeya dianthicola NCPPB 453 WP 024108845 1 Ddianthicola
Dickeya dianthicola GBBC 2039 WP 024109958 1 Ddianthicola
Dickeya dianthicola IPO 980 WP 024109958 1 Ddianthicola
Dickeya dadantii subsp dieffenbachiae NCPPB 2976 WP 071783861 1 Ddadantii
Dickeya chrysanthemi subsp chrysanthemi NCPPB 402 WP 040001248 1 Dchrysanthemi
Dickeya solani RNS 08 23 3 1 A WP 022633841 1 Dsolani
Dickeya solani GBBC 2040 WP 022633841 1 Dsolani
Dickeya solani MK10 WP 022633841 1 Dsolani
Dickeya solani D s0432 1 WP 022633841 1 Dsolani
Dickeya solani IPO 2222 WP 022633841 1 Dsolani
Dickeya solani MK16 WP 022633841 1 Dsolani

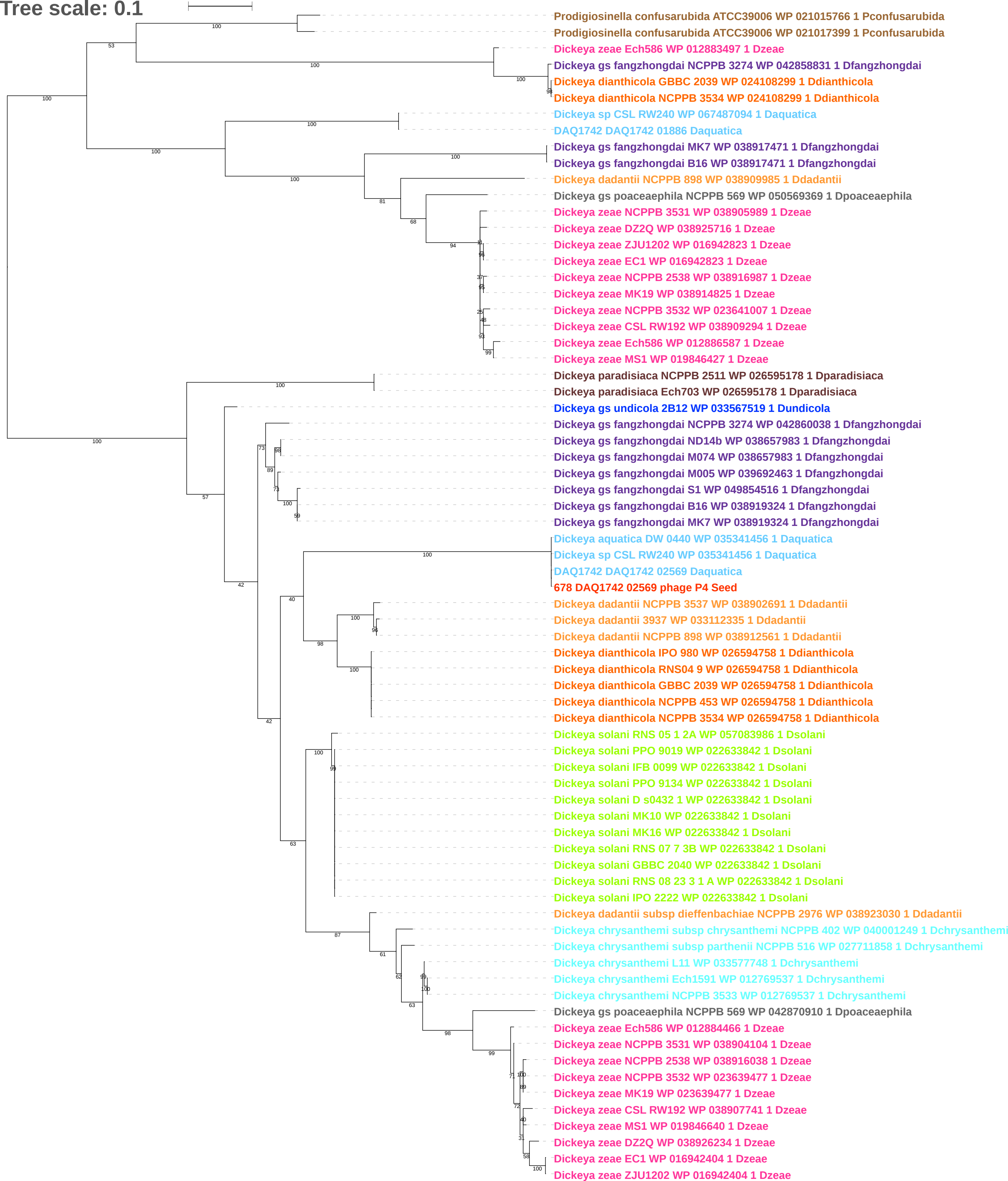
Tree scale: 0.1
Prodigiosinella confusarubida ATCC39006 WP 021015766 1 Pconfusarubida
Prodigiosinella confusarubida ATCC39006 WP 021017399 1 Pconfusarubida
Dickeya zeae Ech586 WP 012883497 1 Dzeae
Dickeya gs fangzhongdai NCPPB 3274 WP 042858831 1 Dfangzhongdai
Dickeya dianthicola GBBC 2039 WP 024108299 1 Ddianthicola
Dickeya dianthicola NCPPB 3534 WP 024108299 1 Ddianthicola
Dickeya sp CSL RW240 WP 067487094 1 Daquatica
DAQ1742 DAQ1742 01886 Daquatica
Dickeya gs fangzhongdai MK7 WP 038917471 1 Dfangzhongdai
Dickeya gs fangzhongdai B16 WP 038917471 1 Dfangzhongdai
Dickeya dadantii NCPPB 898 WP 038909985 1 Ddadantii
Dickeya gs poaceaephila NCPPB 569 WP 050569369 1 Dpoaceaephila
Dickeya zeae NCPPB 3531 WP 038905989 1 Dzeae
Dickeya zeae DZ2Q WP 038925716 1 Dzeae
Dickeya zeae ZJU1202 WP 016942823 1 Dzeae
Dickeya zeae EC1 WP 016942823 1 Dzeae
Dickeya zeae NCPPB 2538 WP 038916987 1 Dzeae
Dickeya zeae MK19 WP 038914825 1 Dzeae
Dickeya zeae NCPPB 3532 WP 023641007 1 Dzeae
Dickeya zeae CSL RW192 WP 038909294 1 Dzeae
Dickeya zeae Ech586 WP 012886587 1 Dzeae
Dickeya zeae MS1 WP 019846427 1 Dzeae
Dickeya paradisiaca NCPPB 2511 WP 026595178 1 Dparadisiaca
Dickeya paradisiaca Ech703 WP 026595178 1 Dparadisiaca
Dickeya gs undicola 2B12 WP 033567519 1 Dundicola
Dickeya gs fangzhongdai NCPPB 3274 WP 042860038 1 Dfangzhongdai
Dickeya gs fangzhongdai ND14b WP 038657983 1 Dfangzhongdai
Dickeya gs fangzhongdai M074 WP 038657983 1 Dfangzhongdai
Dickeya gs fangzhongdai M005 WP 039692463 1 Dfangzhongdai
Dickeya gs fangzhongdai S1 WP 049854516 1 Dfangzhongdai
Dickeya gs fangzhongdai B16 WP 038919324 1 Dfangzhongdai
Dickeya gs fangzhongdai MK7 WP 038919324 1 Dfangzhongdai
Dickeya aquatica DW 0440 WP 035341456 1 Daquatica
Dickeya sp CSL RW240 WP 035341456 1 Daquatica
DAQ1742 DAQ1742 02569 Daquatica
678 DAQ1742 02569 phage P4 Seed
Dickeya dadantii NCPPB 3537 WP 038902691 1 Ddadantii
Dickeya dadantii 3937 WP 033112335 1 Ddadantii
Dickeya dadantii NCPPB 898 WP 038912561 1 Ddadantii
Dickeya dianthicola IPO 980 WP 026594758 1 Ddianthicola
Dickeya dianthicola RNS04 9 WP 026594758 1 Ddianthicola
Dickeya dianthicola GBBC 2039 WP 026594758 1 Ddianthicola
Dickeya dianthicola NCPPB 453 WP 026594758 1 Ddianthicola
Dickeya dianthicola NCPPB 3534 WP 026594758 1 Ddianthicola
Dickeya solani RNS 05 1 2A WP 057083986 1 Dsolani
Dickeya solani PPO 9019 WP 022633842 1 Dsolani
Dickeya solani IFB 0099 WP 022633842 1 Dsolani
Dickeya solani PPO 9134 WP 022633842 1 Dsolani
Dickeya solani D s0432 1 WP 022633842 1 Dsolani
Dickeya solani MK10 WP 022633842 1 Dsolani
Dickeya solani MK16 WP 022633842 1 Dsolani
Dickeya solani RNS 07 7 3B WP 022633842 1 Dsolani
Dickeya solani GBBC 2040 WP 022633842 1 Dsolani
Dickeya solani RNS 08 23 3 1 A WP 022633842 1 Dsolani
Dickeya solani IPO 2222 WP 022633842 1 Dsolani
Dickeya dadantii subsp dieffenbachiae NCPPB 2976 WP 038923030 1 Ddadantii
Dickeya chrysanthemi subsp chrysanthemi NCPPB 402 WP 040001249 1 Dchrysanthemi
Dickeya chrysanthemi subsp parthenii NCPPB 516 WP 027711858 1 Dchrysanthemi
Dickeya chrysanthemi L11 WP 033577748 1 Dchrysanthemi
Dickeya chrysanthemi Ech1591 WP 012769537 1 Dchrysanthemi
Dickeya chrysanthemi NCPPB 3533 WP 012769537 1 Dchrysanthemi
Dickeya gs poaceaephila NCPPB 569 WP 042870910 1 Dpoaceaephila
Dickeya zeae Ech586 WP 012884466 1 Dzeae
Dickeya zeae NCPPB 3531 WP 038904104 1 Dzeae
Dickeya zeae NCPPB 2538 WP 038916038 1 Dzeae
Dickeya zeae NCPPB 3532 WP 023639477 1 Dzeae
Dickeya zeae MK19 WP 023639477 1 Dzeae
Dickeya zeae CSL RW192 WP 038907741 1 Dzeae
Dickeya zeae MS1 WP 019846640 1 Dzeae
Dickeya zeae DZ2Q WP 038926234 1 Dzeae
Dickeya zeae EC1 WP 016942404 1 Dzeae
Dickeya zeae ZJU1202 WP 016942404 1 Dzeae

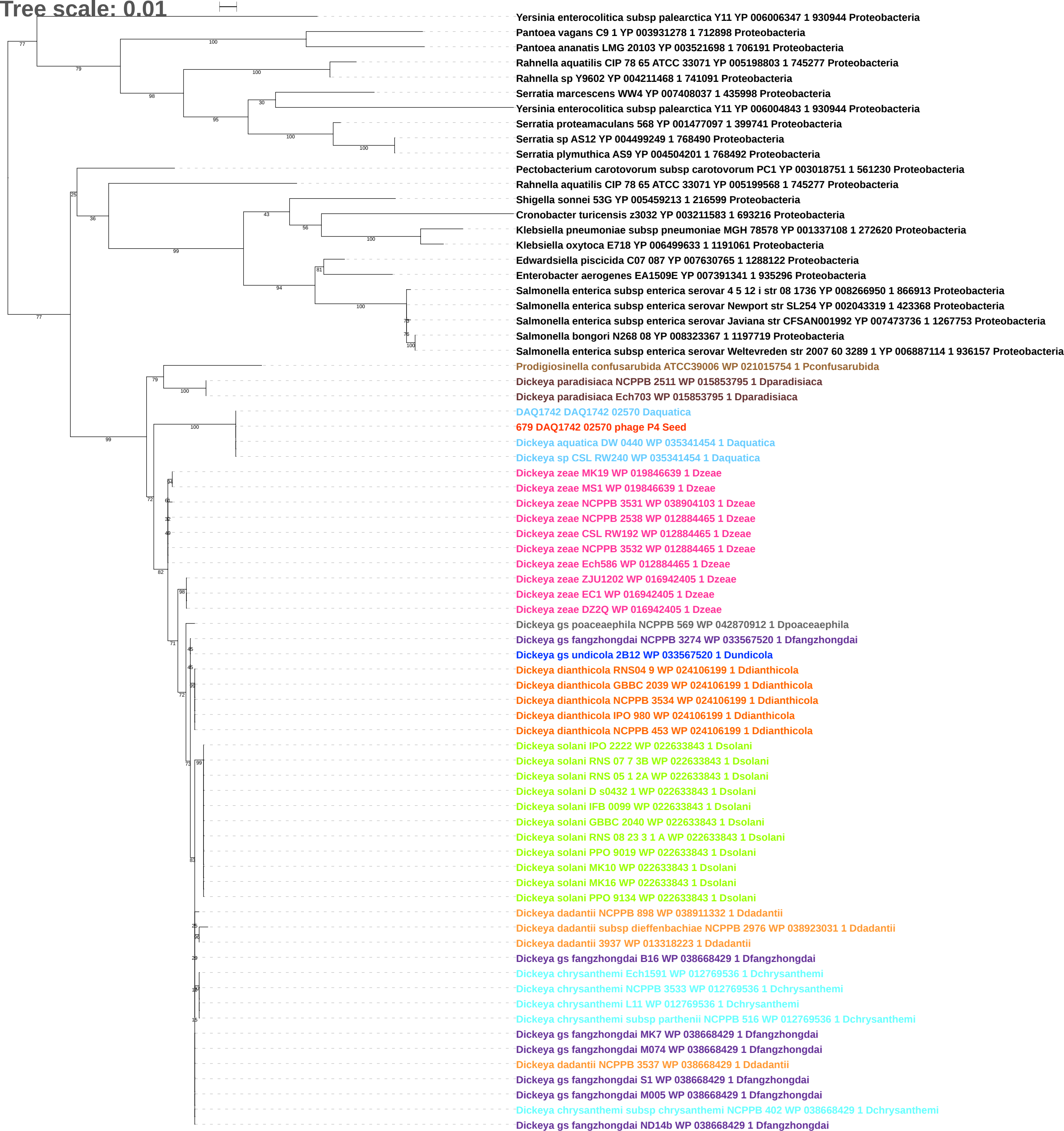
Tree scale: 0.01
Yersinia enterocolitica subsp palearctica Y11 YP 006006347 1 930944 Proteobacteria
Pantoea vagans C9 1 YP 003931278 1 712898 Proteobacteria
Pantoea ananatis LMG 20103 YP 003521698 1 706191 Proteobacteria
Rahnella aquatilis CIP 78 65 ATCC 33071 YP 005198803 1 745277 Proteobacteria
Rahnella sp Y9602 YP 004211468 1 741091 Proteobacteria
Serratia marcescens WW4 YP 007408037 1 435998 Proteobacteria
Yersinia enterocolitica subsp palearctica Y11 YP 006004843 1 930944 Proteobacteria
Serratia proteamaculans 568 YP 001477097 1 399741 Proteobacteria
Serratia sp AS12 YP 004499249 1 768490 Proteobacteria
Serratia plymuthica AS9 YP 004504201 1 768492 Proteobacteria
Pectobacterium carotovorum subsp carotovorum PC1 YP 003018751 1 561230 Proteobacteria
Rahnella aquatilis CIP 78 65 ATCC 33071 YP 005199568 1 745277 Proteobacteria
Shigella sonnei 53G YP 005459213 1 216599 Proteobacteria
Cronobacter turicensis z3032 YP 003211583 1 693216 Proteobacteria
Klebsiella pneumoniae subsp pneumoniae MGH 78578 YP 001337108 1 272620 Proteobacteria
Klebsiella oxytoca E718 YP 006499633 1 1191061 Proteobacteria
Edwardsiella piscicida C07 087 YP 007630765 1 1288122 Proteobacteria
Enterobacter aerogenes EA1509E YP 007391341 1 935296 Proteobacteria
Salmonella enterica subsp enterica serovar 4 5 12 i str 08 1736 YP 008266950 1 866913 Proteobacteria
Salmonella enterica subsp enterica serovar Newport str SL254 YP 002043319 1 423368 Proteobacteria
Salmonella enterica subsp enterica serovar Javiana str CFSAN001992 YP 007473736 1 1267753 Proteobacteria
Salmonella bongori N268 08 YP 008323367 1 1197719 Proteobacteria
Salmonella enterica subsp enterica serovar Weltevreden str 2007 60 3289 1 YP 006887114 1 936157 Proteobacteria
Prodigiosinella confusarubida ATCC39006 WP 021015754 1 Pconfusarubida
Dickeya paradisiaca NCPPB 2511 WP 015853795 1 Dparadisiaca
Dickeya paradisiaca Ech703 WP 015853795 1 Dparadisiaca
DAQ1742 DAQ1742 02570 Daquatica
679 DAQ1742 02570 phage P4 Seed
Dickeya aquatica DW 0440 WP 035341454 1 Daquatica
Dickeya sp CSL RW240 WP 035341454 1 Daquatica
Dickeya zeae MK19 WP 019846639 1 Dzeae
Dickeya zeae MS1 WP 019846639 1 Dzeae
Dickeya zeae NCPPB 3531 WP 038904103 1 Dzeae
Dickeya zeae NCPPB 2538 WP 012884465 1 Dzeae
Dickeya zeae CSL RW192 WP 012884465 1 Dzeae
Dickeya zeae NCPPB 3532 WP 012884465 1 Dzeae
Dickeya zeae Ech586 WP 012884465 1 Dzeae
Dickeya zeae ZJU1202 WP 016942405 1 Dzeae
Dickeya zeae EC1 WP 016942405 1 Dzeae
Dickeya zeae DZ2Q WP 016942405 1 Dzeae
Dickeya gs poaceaephila NCPPB 569 WP 042870912 1 Dpoaceaephila
Dickeya gs fangzhongdai NCPPB 3274 WP 033567520 1 Dfangzhongdai
Dickeya gs undicola 2B12 WP 033567520 1 Dundicola
Dickeya dianthicola RNS04 9 WP 024106199 1 Ddianthicola
Dickeya dianthicola GBBC 2039 WP 024106199 1 Ddianthicola
Dickeya dianthicola NCPPB 3534 WP 024106199 1 Ddianthicola
Dickeya dianthicola IPO 980 WP 024106199 1 Ddianthicola
Dickeya dianthicola NCPPB 453 WP 024106199 1 Ddianthicola
Dickeya solani IPO 2222 WP 022633843 1 Dsolani
Dickeya solani RNS 07 7 3B WP 022633843 1 Dsolani
Dickeya solani RNS 05 1 2A WP 022633843 1 Dsolani
Dickeya solani D s0432 1 WP 022633843 1 Dsolani
Dickeya solani IFB 0099 WP 022633843 1 Dsolani
Dickeya solani GBBC 2040 WP 022633843 1 Dsolani
Dickeya solani RNS 08 23 3 1 A WP 022633843 1 Dsolani
Dickeya solani PPO 9019 WP 022633843 1 Dsolani
Dickeya solani MK10 WP 022633843 1 Dsolani
Dickeya solani MK16 WP 022633843 1 Dsolani
Dickeya solani PPO 9134 WP 022633843 1 Dsolani
Dickeya dadantii NCPPB 898 WP 038911332 1 Ddadantii
Dickeya dadantii subsp dieffenbachiae NCPPB 2976 WP 038923031 1 Ddadantii
Dickeya dadantii 3937 WP 013318223 1 Ddadantii
Dickeya gs fangzhongdai B16 WP 038668429 1 Dfangzhongdai
Dickeya chrysanthemi Ech1591 WP 012769536 1 Dchrysanthemi
Dickeya chrysanthemi NCPPB 3533 WP 012769536 1 Dchrysanthemi
Dickeya chrysanthemi L11 WP 012769536 1 Dchrysanthemi
Dickeya chrysanthemi subsp parthenii NCPPB 516 WP 012769536 1 Dchrysanthemi
Dickeya gs fangzhongdai MK7 WP 038668429 1 Dfangzhongdai
Dickeya gs fangzhongdai M074 WP 038668429 1 Dfangzhongdai
Dickeya dadantii NCPPB 3537 WP 038668429 1 Ddadantii
Dickeya gs fangzhongdai S1 WP 038668429 1 Dfangzhongdai
Dickeya gs fangzhongdai M005 WP 038668429 1 Dfangzhongdai
Dickeya chrysanthemi subsp chrysanthemi NCPPB 402 WP 038668429 1 Dchrysanthemi
Dickeya gs fangzhongdai ND14b WP 038668429 1 Dfangzhongdai

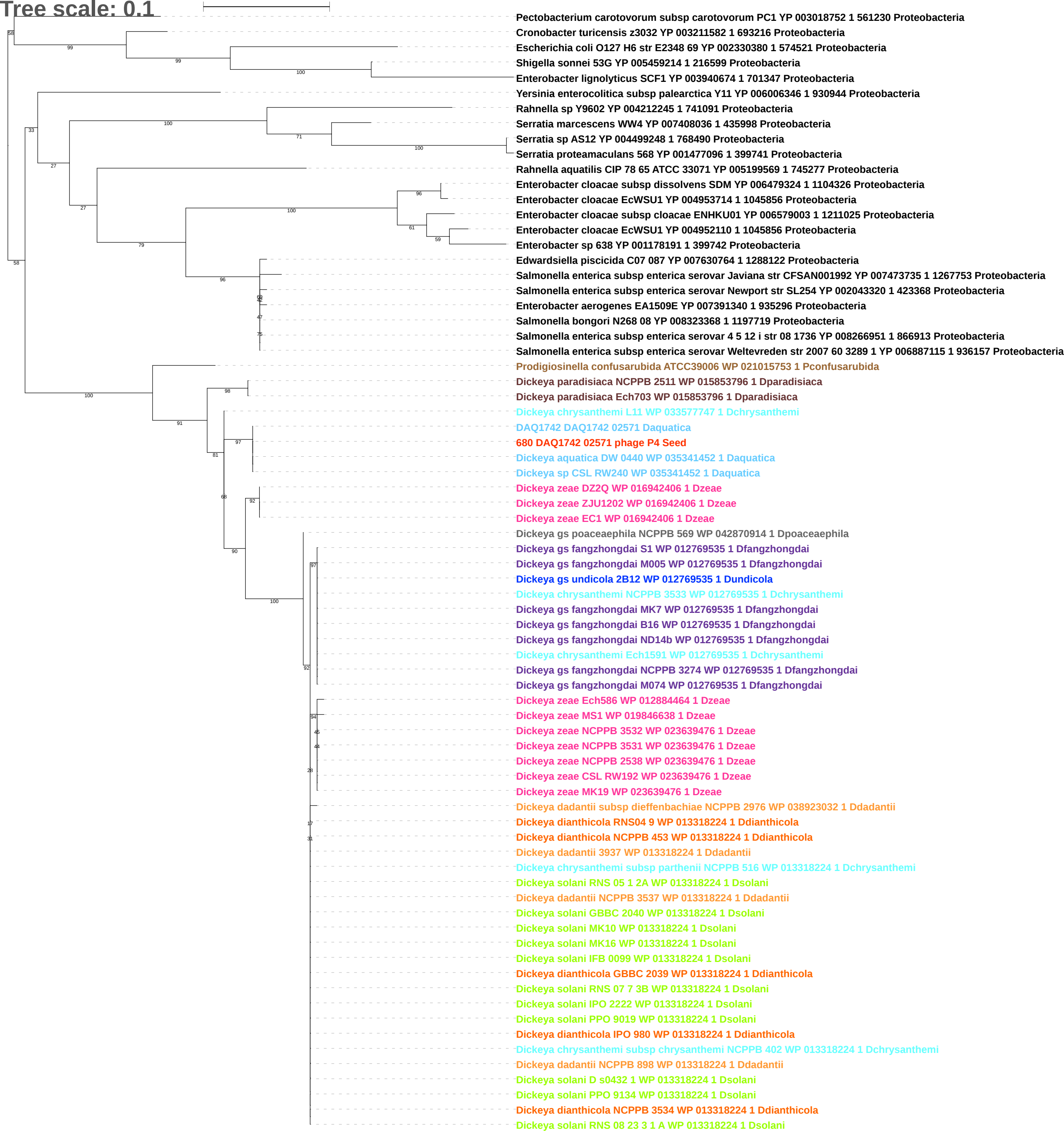
Tree scale: 0.1
Pectobacterium carotovorum subsp carotovorum PC1 YP 003018752 1 561230 Proteobacteria
Cronobacter turicensis z3032 YP 003211582 1 693216 Proteobacteria
Escherichia coli O127 H6 str E2348 69 YP 002330380 1 574521 Proteobacteria
Shigella sonnei 53G YP 005459214 1 216599 Proteobacteria
Enterobacter lignolyticus SCF1 YP 003940674 1 701347 Proteobacteria
Yersinia enterocolitica subsp palearctica Y11 YP 006006346 1 930944 Proteobacteria
Rahnella sp Y9602 YP 004212245 1 741091 Proteobacteria
Serratia marcescens WW4 YP 007408036 1 435998 Proteobacteria
Serratia sp AS12 YP 004499248 1 768490 Proteobacteria
Serratia proteamaculans 568 YP 001477096 1 399741 Proteobacteria
Rahnella aquatilis CIP 78 65 ATCC 33071 YP 005199569 1 745277 Proteobacteria
Enterobacter cloacae subsp dissolvens SDM YP 006479324 1 1104326 Proteobacteria
Enterobacter cloacae EcWSU1 YP 004953714 1 1045856 Proteobacteria
Enterobacter cloacae subsp cloacae ENHKU01 YP 006579003 1 1211025 Proteobacteria
Enterobacter cloacae EcWSU1 YP 004952110 1 1045856 Proteobacteria
Enterobacter sp 638 YP 001178191 1 399742 Proteobacteria
Edwardsiella piscicida C07 087 YP 007630764 1 1288122 Proteobacteria
Salmonella enterica subsp enterica serovar Javiana str CFSAN001992 YP 007473735 1 1267753 Proteobacteria
Salmonella enterica subsp enterica serovar Newport str SL254 YP 002043320 1 423368 Proteobacteria
Enterobacter aerogenes EA1509E YP 007391340 1 935296 Proteobacteria
Salmonella bongori N268 08 YP 008323368 1 1197719 Proteobacteria
Salmonella enterica subsp enterica serovar 4 5 12 i str 08 1736 YP 008266951 1 866913 Proteobacteria
Salmonella enterica subsp enterica serovar Weltevreden str 2007 60 3289 1 YP 006887115 1 936157 Proteobacteria
Prodigiosinella confusarubida ATCC39006 WP 021015753 1 Pconfusarubida
Dickeya paradisiaca NCPPB 2511 WP 015853796 1 Dparadisiaca
Dickeya paradisiaca Ech703 WP 015853796 1 Dparadisiaca
Dickeya chrysanthemi L11 WP 033577747 1 Dchrysanthemi
DAQ1742 DAQ1742 02571 Daquatica
680 DAQ1742 02571 phage P4 Seed
Dickeya aquatica DW 0440 WP 035341452 1 Daquatica
Dickeya sp CSL RW240 WP 035341452 1 Daquatica
Dickeya zeae DZ2Q WP 016942406 1 Dzeae
Dickeya zeae ZJU1202 WP 016942406 1 Dzeae
Dickeya zeae EC1 WP 016942406 1 Dzeae
Dickeya gs poaceaephila NCPPB 569 WP 042870914 1 Dpoaceaephila
Dickeya gs fangzhongdai S1 WP 012769535 1 Dfangzhongdai
Dickeya gs fangzhongdai M005 WP 012769535 1 Dfangzhongdai
Dickeya gs undicola 2B12 WP 012769535 1 Dundicola
Dickeya chrysanthemi NCPPB 3533 WP 012769535 1 Dchrysanthemi
Dickeya gs fangzhongdai MK7 WP 012769535 1 Dfangzhongdai
Dickeya gs fangzhongdai B16 WP 012769535 1 Dfangzhongdai
Dickeya gs fangzhongdai ND14b WP 012769535 1 Dfangzhongdai
Dickeya chrysanthemi Ech1591 WP 012769535 1 Dchrysanthemi
Dickeya gs fangzhongdai NCPPB 3274 WP 012769535 1 Dfangzhongdai
Dickeya gs fangzhongdai M074 WP 012769535 1 Dfangzhongdai
Dickeya zeae Ech586 WP 012884464 1 Dzeae
Dickeya zeae MS1 WP 019846638 1 Dzeae
Dickeya zeae NCPPB 3532 WP 023639476 1 Dzeae
Dickeya zeae NCPPB 3531 WP 023639476 1 Dzeae
Dickeya zeae NCPPB 2538 WP 023639476 1 Dzeae
Dickeya zeae CSL RW192 WP 023639476 1 Dzeae
Dickeya zeae MK19 WP 023639476 1 Dzeae
Dickeya dadantii subsp dieffenbachiae NCPPB 2976 WP 038923032 1 Ddadantii
Dickeya dianthicola RNS04 9 WP 013318224 1 Ddianthicola
Dickeya dianthicola NCPPB 453 WP 013318224 1 Ddianthicola
Dickeya dadantii 3937 WP 013318224 1 Ddadantii
Dickeya chrysanthemi subsp parthenii NCPPB 516 WP 013318224 1 Dchrysanthemi
Dickeya solani RNS 05 1 2A WP 013318224 1 Dsolani
Dickeya dadantii NCPPB 3537 WP 013318224 1 Ddadantii
Dickeya solani GBBC 2040 WP 013318224 1 Dsolani
Dickeya solani MK10 WP 013318224 1 Dsolani
Dickeya solani MK16 WP 013318224 1 Dsolani
Dickeya solani IFB 0099 WP 013318224 1 Dsolani
Dickeya dianthicola GBBC 2039 WP 013318224 1 Ddianthicola
Dickeya solani RNS 07 7 3B WP 013318224 1 Dsolani
Dickeya solani IPO 2222 WP 013318224 1 Dsolani
Dickeya solani PPO 9019 WP 013318224 1 Dsolani
Dickeya dianthicola IPO 980 WP 013318224 1 Ddianthicola
Dickeya chrysanthemi subsp chrysanthemi NCPPB 402 WP 013318224 1 Dchrysanthemi
Dickeya dadantii NCPPB 898 WP 013318224 1 Ddadantii
Dickeya solani D s0432 1 WP 013318224 1 Dsolani
Dickeya solani PPO 9134 WP 013318224 1 Dsolani
Dickeya dianthicola NCPPB 3534 WP 013318224 1 Ddianthicola
Dickeya solani RNS 08 23 3 1 A WP 013318224 1 Dsolani

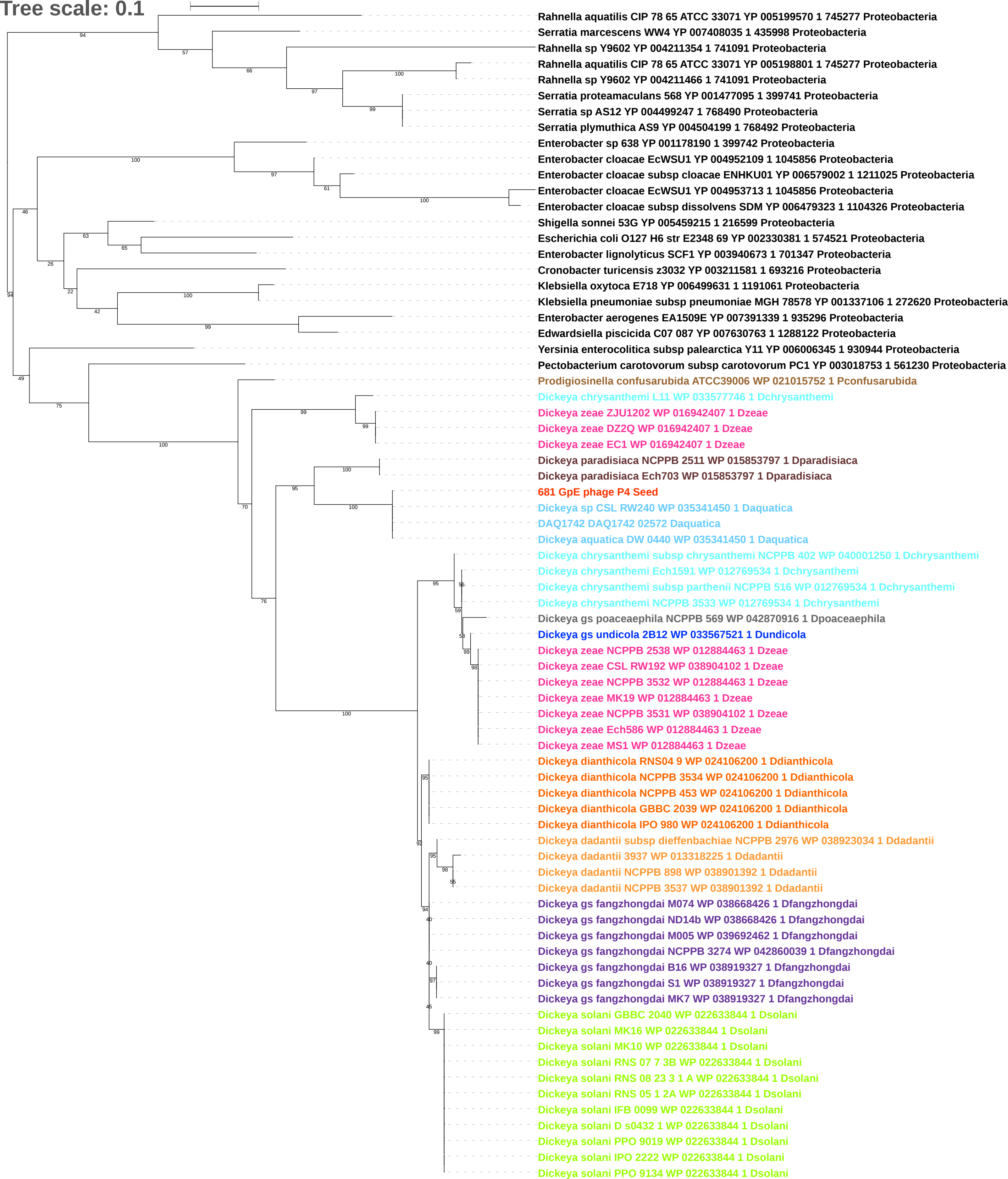
Tree scale: 0.1
Rahnella aquatilis CIP 78 65 ATCC 33071 YP 005199570 1 745277 Proteobacteria
Serratia marcescens WW4 YP 007408035 1 435998 Proteobacteria
Rahnella sp Y9602 YP 004211354 1 741091 Proteobacteria
Rahnella aquatilis CIP 78 65 ATCC 33071 YP 005198801 1 745277 Proteobacteria
Rahnella sp Y9602 YP 004211466 1 741091 Proteobacteria
Serratia proteamaculans 568 YP 001477095 1 399741 Proteobacteria
Serratia sp AS12 YP 004499247 1 768490 Proteobacteria
Serratia plymuthica AS9 YP 004504199 1 768492 Proteobacteria
Enterobacter sp 638 YP 001178190 1 399742 Proteobacteria
Enterobacter cloacae EcWSU1 YP 004952109 1 1045856 Proteobacteria
Enterobacter cloacae subsp cloacae ENHKU01 YP 006579002 1 1211025 Proteobacteria
Enterobacter cloacae EcWSU1 YP 004953713 1 1045856 Proteobacteria
Enterobacter cloacae subsp dissolvens SDM YP 006479323 1 1104326 Proteobacteria
Shigella sonnei 53G YP 005459215 1 216599 Proteobacteria
Escherichia coli O127 H6 str E2348 69 YP 002330381 1 574521 Proteobacteria
Enterobacter lignolyticus SCF1 YP 003940673 1 701347 Proteobacteria
Cronobacter turicensis z3032 YP 003211581 1 693216 Proteobacteria
Klebsiella oxytoca E718 YP 006499631 1 1191061 Proteobacteria
Klebsiella pneumoniae subsp pneumoniae MGH 78578 YP 001337106 1 272620 Proteobacteria
Enterobacter aerogenes EA1509E YP 007391339 1 935296 Proteobacteria
Edwardsiella piscicida C07 087 YP 007630763 1 1288122 Proteobacteria
Yersinia enterocolitica subsp palearctica Y11 YP 006006345 1 930944 Proteobacteria
Pectobacterium carotovorum subsp carotovorum PC1 YP 003018753 1 561230 Proteobacteria
Prodigiosinella confusarubida ATCC39006 WP 021015752 1 Pconfusarubida
Dickeya chrysanthemi L11 WP 033577746 1 Dchrysanthemi
Dickeya zeae ZJU1202 WP 016942407 1 Dzeae
Dickeya zeae DZ2Q WP 016942407 1 Dzeae
Dickeya zeae EC1 WP 016942407 1 Dzeae
Dickeya paradisiaca NCPPB 2511 WP 015853797 1 Dparadisiaca
Dickeya paradisiaca Ech703 WP 015853797 1 Dparadisiaca
681 GpE phage P4 Seed
Dickeya sp CSL RW240 WP 035341450 1 Daquatica
DAQ1742 DAQ1742 02572 Daquatica
Dickeya aquatica DW 0440 WP 035341450 1 Daquatica
Dickeya chrysanthemi subsp chrysanthemi NCPPB 402 WP 040001250 1 Dchrysanthemi
Dickeya chrysanthemi Ech1591 WP 012769534 1 Dchrysanthemi
Dickeya chrysanthemi subsp parthenii NCPPB 516 WP 012769534 1 Dchrysanthemi
Dickeya chrysanthemi NCPPB 3533 WP 012769534 1 Dchrysanthemi
Dickeya gs poaceaephila NCPPB 569 WP 042870916 1 Dpoaceaephila
Dickeya gs undicola 2B12 WP 033567521 1 Dundicola
Dickeya zeae NCPPB 2538 WP 012884463 1 Dzeae
Dickeya zeae CSL RW192 WP 038904102 1 Dzeae
Dickeya zeae NCPPB 3532 WP 012884463 1 Dzeae
Dickeya zeae MK19 WP 012884463 1 Dzeae
Dickeya zeae NCPPB 3531 WP 038904102 1 Dzeae
Dickeya zeae Ech586 WP 012884463 1 Dzeae
Dickeya zeae MS1 WP 012884463 1 Dzeae
Dickeya dianthicola RNS04 9 WP 024106200 1 Ddianthicola
Dickeya dianthicola NCPPB 3534 WP 024106200 1 Ddianthicola
Dickeya dianthicola NCPPB 453 WP 024106200 1 Ddianthicola
Dickeya dianthicola GBBC 2039 WP 024106200 1 Ddianthicola
Dickeya dianthicola IPO 980 WP 024106200 1 Ddianthicola
Dickeya dadantii subsp dieffenbachiae NCPPB 2976 WP 038923034 1 Ddadantii
Dickeya dadantii 3937 WP 013318225 1 Ddadantii
Dickeya dadantii NCPPB 898 WP 038901392 1 Ddadantii
Dickeya dadantii NCPPB 3537 WP 038901392 1 Ddadantii
Dickeya gs fangzhongdai M074 WP 038668426 1 Dfangzhongdai
Dickeya gs fangzhongdai ND14b WP 038668426 1 Dfangzhongdai
Dickeya gs fangzhongdai M005 WP 039692462 1 Dfangzhongdai
Dickeya gs fangzhongdai NCPPB 3274 WP 042860039 1 Dfangzhongdai
Dickeya gs fangzhongdai B16 WP 038919327 1 Dfangzhongdai
Dickeya gs fangzhongdai S1 WP 038919327 1 Dfangzhongdai
Dickeya gs fangzhongdai MK7 WP 038919327 1 Dfangzhongdai
Dickeya solani GBBC 2040 WP 022633844 1 Dsolani
Dickeya solani MK16 WP 022633844 1 Dsolani
Dickeya solani MK10 WP 022633844 1 Dsolani
Dickeya solani RNS 07 7 3B WP 022633844 1 Dsolani
Dickeya solani RNS 08 23 3 1 A WP 022633844 1 Dsolani
Dickeya solani RNS 05 1 2A WP 022633844 1 Dsolani
Dickeya solani IFB 0099 WP 022633844 1 Dsolani
Dickeya solani D s0432 1 WP 022633844 1 Dsolani
Dickeya solani PPO 9019 WP 022633844 1 Dsolani
Dickeya solani IPO 2222 WP 022633844 1 Dsolani
Dickeya solani PPO 9134 WP 022633844 1 Dsolani

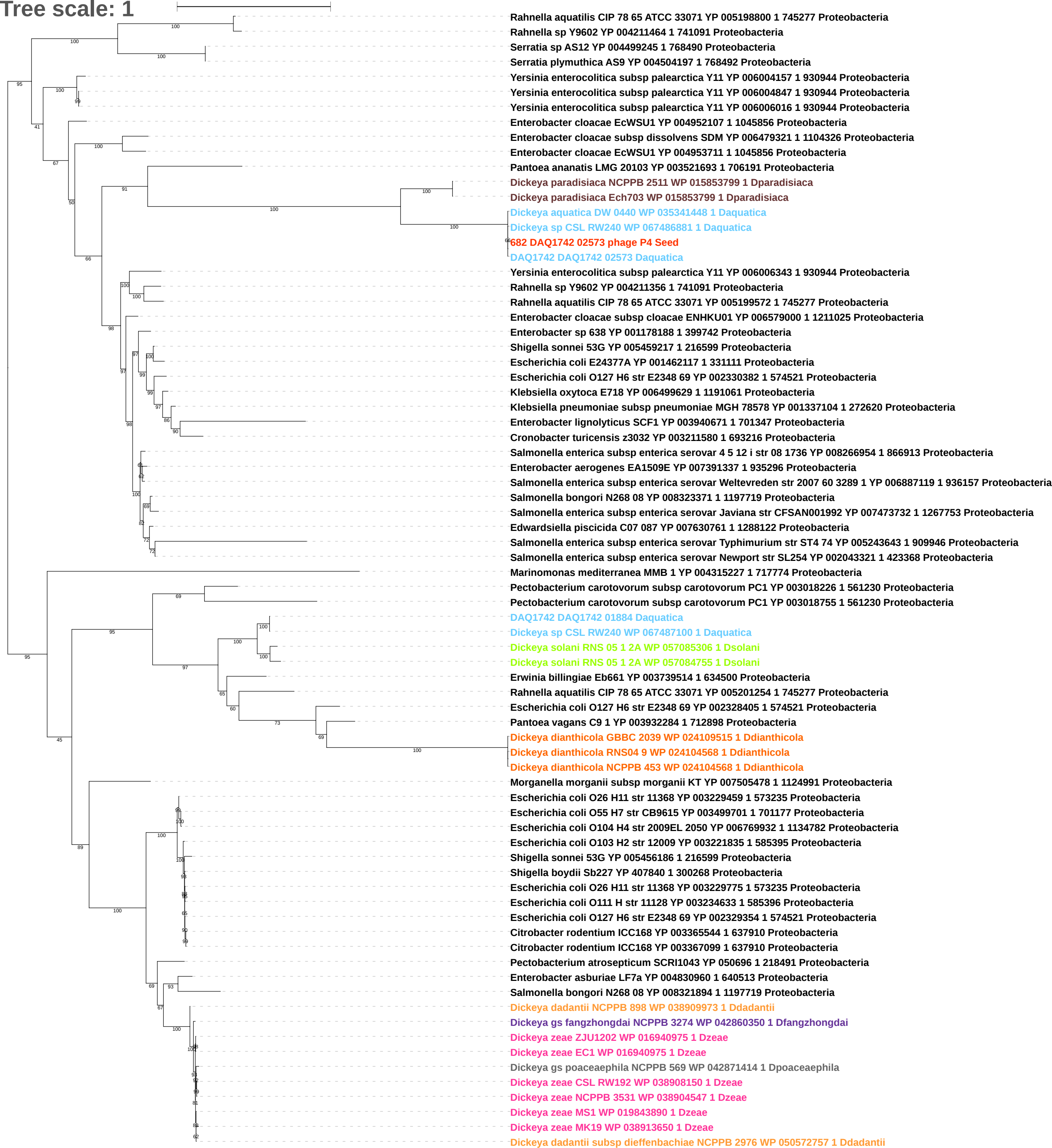
Tree scale: 1
Rahnella aquatilis CIP 78 65 ATCC 33071 YP 005198800 1 745277 Proteobacteria
Rahnella sp Y9602 YP 004211464 1 741091 Proteobacteria
Serratia sp AS12 YP 004499245 1 768490 Proteobacteria
Serratia plymuthica AS9 YP 004504197 1 768492 Proteobacteria
Yersinia enterocolitica subsp palearctica Y11 YP 006004157 1 930944 Proteobacteria
Yersinia enterocolitica subsp palearctica Y11 YP 006004847 1 930944 Proteobacteria
Yersinia enterocolitica subsp palearctica Y11 YP 006006016 1 930944 Proteobacteria
Enterobacter cloacae EcWSU1 YP 004952107 1 1045856 Proteobacteria
Enterobacter cloacae subsp dissolvens SDM YP 006479321 1 1104326 Proteobacteria
Enterobacter cloacae EcWSU1 YP 004953711 1 1045856 Proteobacteria
Pantoea ananatis LMG 20103 YP 003521693 1 706191 Proteobacteria
Dickeya paradisiaca NCPPB 2511 WP 015853799 1 Dparadisiaca
Dickeya paradisiaca Ech703 WP 015853799 1 Dparadisiaca
Dickeya aquatica DW 0440 WP 035341448 1 Daquatica
Dickeya sp CSL RW240 WP 067486881 1 Daquatica
682 DAQ1742 02573 phage P4 Seed
DAQ1742 DAQ1742 02573 Daquatica
Yersinia enterocolitica subsp palearctica Y11 YP 006006343 1 930944 Proteobacteria
Rahnella sp Y9602 YP 004211356 1 741091 Proteobacteria
Rahnella aquatilis CIP 78 65 ATCC 33071 YP 005199572 1 745277 Proteobacteria
Enterobacter cloacae subsp cloacae ENHKU01 YP 006579000 1 1211025 Proteobacteria
Enterobacter sp 638 YP 001178188 1 399742 Proteobacteria
Shigella sonnei 53G YP 005459217 1 216599 Proteobacteria
Escherichia coli E24377A YP 001462117 1 331111 Proteobacteria
Escherichia coli O127 H6 str E2348 69 YP 002330382 1 574521 Proteobacteria
Klebsiella oxytoca E718 YP 006499629 1 1191061 Proteobacteria
Klebsiella pneumoniae subsp pneumoniae MGH 78578 YP 001337104 1 272620 Proteobacteria
Enterobacter lignolyticus SCF1 YP 003940671 1 701347 Proteobacteria
Cronobacter turicensis z3032 YP 003211580 1 693216 Proteobacteria
Salmonella enterica subsp enterica serovar 4 5 12 i str 08 1736 YP 008266954 1 866913 Proteobacteria
Enterobacter aerogenes EA1509E YP 007391337 1 935296 Proteobacteria
Salmonella enterica subsp enterica serovar Weltevreden str 2007 60 3289 1 YP 006887119 1 936157 Proteobacteria
Salmonella bongori N268 08 YP 008323371 1 1197719 Proteobacteria
Salmonella enterica subsp enterica serovar Javiana str CFSAN001992 YP 007473732 1 1267753 Proteobacteria
Edwardsiella piscicida C07 087 YP 007630761 1 1288122 Proteobacteria
Salmonella enterica subsp enterica serovar Typhimurium str ST4 74 YP 005243643 1 909946 Proteobacteria
Salmonella enterica subsp enterica serovar Newport str SL254 YP 002043321 1 423368 Proteobacteria
Marinomonas mediterranea MMB 1 YP 004315227 1 717774 Proteobacteria
Pectobacterium carotovorum subsp carotovorum PC1 YP 003018226 1 561230 Proteobacteria
Pectobacterium carotovorum subsp carotovorum PC1 YP 003018755 1 561230 Proteobacteria
DAQ1742 DAQ1742 01884 Daquatica
Dickeya sp CSL RW240 WP 067487100 1 Daquatica
Dickeya solani RNS 05 1 2A WP 057085306 1 Dsolani
Dickeya solani RNS 05 1 2A WP 057084755 1 Dsolani
Erwinia billingiae Eb661 YP 003739514 1 634500 Proteobacteria
Rahnella aquatilis CIP 78 65 ATCC 33071 YP 005201254 1 745277 Proteobacteria
Escherichia coli O127 H6 str E2348 69 YP 002328405 1 574521 Proteobacteria
Pantoea vagans C9 1 YP 003932284 1 712898 Proteobacteria
Dickeya dianthicola GBBC 2039 WP 024109515 1 Ddianthicola
Dickeya dianthicola RNS04 9 WP 024104568 1 Ddianthicola
Dickeya dianthicola NCPPB 453 WP 024104568 1 Ddianthicola
Morganella morganii subsp morganii KT YP 007505478 1 1124991 Proteobacteria
Escherichia coli O26 H11 str 11368 YP 003229459 1 573235 Proteobacteria
Escherichia coli O55 H7 str CB9615 YP 003499701 1 701177 Proteobacteria
Escherichia coli O104 H4 str 2009EL 2050 YP 006769932 1 1134782 Proteobacteria
Escherichia coli O103 H2 str 12009 YP 003221835 1 585395 Proteobacteria
Shigella sonnei 53G YP 005456186 1 216599 Proteobacteria
Shigella boydii Sb227 YP 407840 1 300268 Proteobacteria
Escherichia coli O26 H11 str 11368 YP 003229775 1 573235 Proteobacteria
Escherichia coli O111 H str 11128 YP 003234633 1 585396 Proteobacteria
Escherichia coli O127 H6 str E2348 69 YP 002329354 1 574521 Proteobacteria
Citrobacter rodentium ICC168 YP 003365544 1 637910 Proteobacteria
Citrobacter rodentium ICC168 YP 003367099 1 637910 Proteobacteria
Pectobacterium atrosepticum SCRI1043 YP 050696 1 218491 Proteobacteria
Enterobacter asburiae LF7a YP 004830960 1 640513 Proteobacteria
Salmonella bongori N268 08 YP 008321894 1 1197719 Proteobacteria
Dickeya dadantii NCPPB 898 WP 038909973 1 Ddadantii
Dickeya gs fangzhongdai NCPPB 3274 WP 042860350 1 Dfangzhongdai
Dickeya zeae ZJU1202 WP 016940975 1 Dzeae
Dickeya zeae EC1 WP 016940975 1 Dzeae
Dickeya gs poaceaephila NCPPB 569 WP 042871414 1 Dpoaceaephila
Dickeya zeae CSL RW192 WP 038908150 1 Dzeae
Dickeya zeae NCPPB 3531 WP 038904547 1 Dzeae
Dickeya zeae MS1 WP 019843890 1 Dzeae
Dickeya zeae MK19 WP 038913650 1 Dzeae
Dickeya dadantii subsp dieffenbachiae NCPPB 2976 WP 050572757 1 Ddadantii

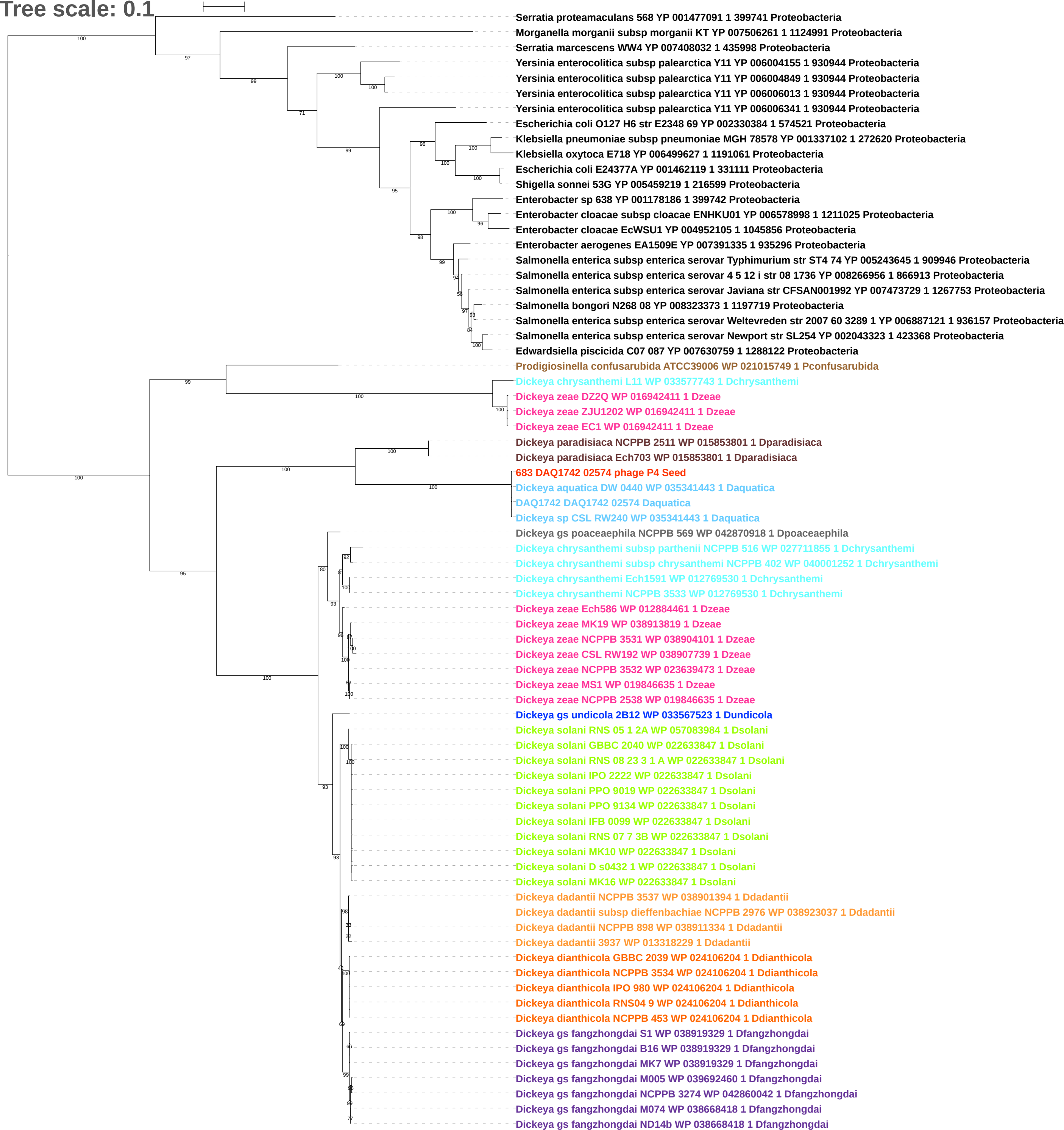
Tree scale: 0.1
Serratia proteamaculans 568 YP 001477091 1 399741 Proteobacteria
Morganella morganii subsp morganii KT YP 007506261 1 1124991 Proteobacteria
Serratia marcescens WW4 YP 007408032 1 435998 Proteobacteria
Yersinia enterocolitica subsp palearctica Y11 YP 006004155 1 930944 Proteobacteria
Yersinia enterocolitica subsp palearctica Y11 YP 006004849 1 930944 Proteobacteria
Yersinia enterocolitica subsp palearctica Y11 YP 006006013 1 930944 Proteobacteria
Yersinia enterocolitica subsp palearctica Y11 YP 006006341 1 930944 Proteobacteria
Escherichia coli O127 H6 str E2348 69 YP 002330384 1 574521 Proteobacteria
Klebsiella pneumoniae subsp pneumoniae MGH 78578 YP 001337102 1 272620 Proteobacteria
Klebsiella oxytoca E718 YP 006499627 1 1191061 Proteobacteria
Escherichia coli E24377A YP 001462119 1 331111 Proteobacteria
Shigella sonnei 53G YP 005459219 1 216599 Proteobacteria
Enterobacter sp 638 YP 001178186 1 399742 Proteobacteria
Enterobacter cloacae subsp cloacae ENHKU01 YP 006578998 1 1211025 Proteobacteria
Enterobacter cloacae EcWSU1 YP 004952105 1 1045856 Proteobacteria
Enterobacter aerogenes EA1509E YP 007391335 1 935296 Proteobacteria
Salmonella enterica subsp enterica serovar Typhimurium str ST4 74 YP 005243645 1 909946 Proteobacteria
Salmonella enterica subsp enterica serovar 4 5 12 i str 08 1736 YP 008266956 1 866913 Proteobacteria
Salmonella enterica subsp enterica serovar Javiana str CFSAN001992 YP 007473729 1 1267753 Proteobacteria
Salmonella bongori N268 08 YP 008323373 1 1197719 Proteobacteria
Salmonella enterica subsp enterica serovar Weltevreden str 2007 60 3289 1 YP 006887121 1 936157 Proteobacteria
Salmonella enterica subsp enterica serovar Newport str SL254 YP 002043323 1 423368 Proteobacteria
Edwardsiella piscicida C07 087 YP 007630759 1 1288122 Proteobacteria
Prodigiosinella confusarubida ATCC39006 WP 021015749 1 Pconfusarubida
Dickeya chrysanthemi L11 WP 033577743 1 Dchrysanthemi
Dickeya zeae DZ2Q WP 016942411 1 Dzeae
Dickeya zeae ZJU1202 WP 016942411 1 Dzeae
Dickeya zeae EC1 WP 016942411 1 Dzeae
Dickeya paradisiaca NCPPB 2511 WP 015853801 1 Dparadisiaca
Dickeya paradisiaca Ech703 WP 015853801 1 Dparadisiaca
683 DAQ1742 02574 phage P4 Seed
Dickeya aquatica DW 0440 WP 035341443 1 Daquatica
DAQ1742 DAQ1742 02574 Daquatica
Dickeya sp CSL RW240 WP 035341443 1 Daquatica
Dickeya gs poaceaephila NCPPB 569 WP 042870918 1 Dpoaceaephila
Dickeya chrysanthemi subsp parthenii NCPPB 516 WP 027711855 1 Dchrysanthemi
Dickeya chrysanthemi subsp chrysanthemi NCPPB 402 WP 040001252 1 Dchrysanthemi
Dickeya chrysanthemi Ech1591 WP 012769530 1 Dchrysanthemi
Dickeya chrysanthemi NCPPB 3533 WP 012769530 1 Dchrysanthemi
Dickeya zeae Ech586 WP 012884461 1 Dzeae
Dickeya zeae MK19 WP 038913819 1 Dzeae
Dickeya zeae NCPPB 3531 WP 038904101 1 Dzeae
Dickeya zeae CSL RW192 WP 038907739 1 Dzeae
Dickeya zeae NCPPB 3532 WP 023639473 1 Dzeae
Dickeya zeae MS1 WP 019846635 1 Dzeae
Dickeya zeae NCPPB 2538 WP 019846635 1 Dzeae
Dickeya gs undicola 2B12 WP 033567523 1 Dundicola
Dickeya solani RNS 05 1 2A WP 057083984 1 Dsolani
Dickeya solani GBBC 2040 WP 022633847 1 Dsolani
Dickeya solani RNS 08 23 3 1 A WP 022633847 1 Dsolani
Dickeya solani IPO 2222 WP 022633847 1 Dsolani
Dickeya solani PPO 9019 WP 022633847 1 Dsolani
Dickeya solani PPO 9134 WP 022633847 1 Dsolani
Dickeya solani IFB 0099 WP 022633847 1 Dsolani
Dickeya solani RNS 07 7 3B WP 022633847 1 Dsolani
Dickeya solani MK10 WP 022633847 1 Dsolani
Dickeya solani D s0432 1 WP 022633847 1 Dsolani
Dickeya solani MK16 WP 022633847 1 Dsolani
Dickeya dadantii NCPPB 3537 WP 038901394 1 Ddadantii
Dickeya dadantii subsp dieffenbachiae NCPPB 2976 WP 038923037 1 Ddadantii
Dickeya dadantii NCPPB 898 WP 038911334 1 Ddadantii
Dickeya dadantii 3937 WP 013318229 1 Ddadantii
Dickeya dianthicola GBBC 2039 WP 024106204 1 Ddianthicola
Dickeya dianthicola NCPPB 3534 WP 024106204 1 Ddianthicola
Dickeya dianthicola IPO 980 WP 024106204 1 Ddianthicola
Dickeya dianthicola RNS04 9 WP 024106204 1 Ddianthicola
Dickeya dianthicola NCPPB 453 WP 024106204 1 Ddianthicola
Dickeya gs fangzhongdai S1 WP 038919329 1 Dfangzhongdai
Dickeya gs fangzhongdai B16 WP 038919329 1 Dfangzhongdai
Dickeya gs fangzhongdai MK7 WP 038919329 1 Dfangzhongdai
Dickeya gs fangzhongdai M005 WP 039692460 1 Dfangzhongdai
Dickeya gs fangzhongdai NCPPB 3274 WP 042860042 1 Dfangzhongdai
Dickeya gs fangzhongdai M074 WP 038668418 1 Dfangzhongdai
Dickeya gs fangzhongdai ND14b WP 038668418 1 Dfangzhongdai

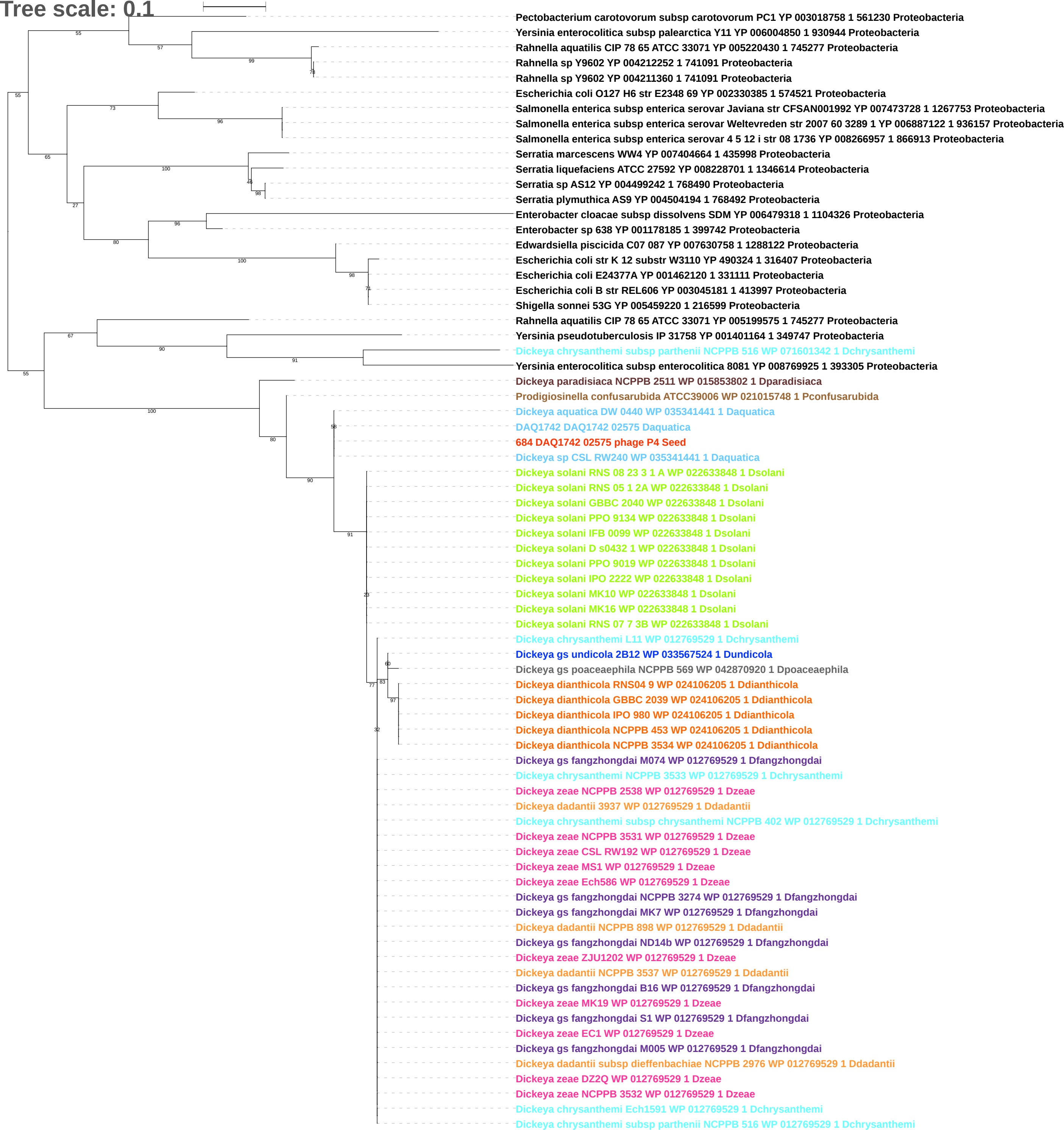
Tree scale: 0.1
Pectobacterium carotovorum subsp carotovorum PC1 YP 003018758 1 561230 Proteobacteria
Yersinia enterocolitica subsp palearctica Y11 YP 006004850 1 930944 Proteobacteria
Rahnella aquatilis CIP 78 65 ATCC 33071 YP 005220430 1 745277 Proteobacteria
Rahnella sp Y9602 YP 004212252 1 741091 Proteobacteria
Rahnella sp Y9602 YP 004211360 1 741091 Proteobacteria
Escherichia coli O127 H6 str E2348 69 YP 002330385 1 574521 Proteobacteria
Salmonella enterica subsp enterica serovar Javiana str CFSAN001992 YP 007473728 1 1267753 Proteobacteria
Salmonella enterica subsp enterica serovar Weltevreden str 2007 60 3289 1 YP 006887122 1 936157 Proteobacteria
Salmonella enterica subsp enterica serovar 4 5 12 i str 08 1736 YP 008266957 1 866913 Proteobacteria
Serratia marcescens WW4 YP 007404664 1 435998 Proteobacteria
Serratia liquefaciens ATCC 27592 YP 008228701 1 1346614 Proteobacteria
Serratia sp AS12 YP 004499242 1 768490 Proteobacteria
Serratia plymuthica AS9 YP 004504194 1 768492 Proteobacteria
Enterobacter cloacae subsp dissolvens SDM YP 006479318 1 1104326 Proteobacteria
Enterobacter sp 638 YP 001178185 1 399742 Proteobacteria
Edwardsiella piscicida C07 087 YP 007630758 1 1288122 Proteobacteria
Escherichia coli str K 12 substr W3110 YP 490324 1 316407 Proteobacteria
Escherichia coli E24377A YP 001462120 1 331111 Proteobacteria
Escherichia coli B str REL606 YP 003045181 1 413997 Proteobacteria
Shigella sonnei 53G YP 005459220 1 216599 Proteobacteria
Rahnella aquatilis CIP 78 65 ATCC 33071 YP 005199575 1 745277 Proteobacteria
Yersinia pseudotuberculosis IP 31758 YP 001401164 1 349747 Proteobacteria
Dickeya chrysanthemi subsp parthenii NCPPB 516 WP 071601342 1 Dchrysanthemi
Yersinia enterocolitica subsp enterocolitica 8081 YP 008769925 1 393305 Proteobacteria
Dickeya paradisiaca NCPPB 2511 WP 015853802 1 Dparadisiaca
Prodigiosinella confusarubida ATCC39006 WP 021015748 1 Pconfusarubida
Dickeya aquatica DW 0440 WP 035341441 1 Daquatica
DAQ1742 DAQ1742 02575 Daquatica
684 DAQ1742 02575 phage P4 Seed
Dickeya sp CSL RW240 WP 035341441 1 Daquatica
Dickeya solani RNS 08 23 3 1 A WP 022633848 1 Dsolani
Dickeya solani RNS 05 1 2A WP 022633848 1 Dsolani
Dickeya solani GBBC 2040 WP 022633848 1 Dsolani
Dickeya solani PPO 9134 WP 022633848 1 Dsolani
Dickeya solani IFB 0099 WP 022633848 1 Dsolani
Dickeya solani D s0432 1 WP 022633848 1 Dsolani
Dickeya solani PPO 9019 WP 022633848 1 Dsolani
Dickeya solani IPO 2222 WP 022633848 1 Dsolani
Dickeya solani MK10 WP 022633848 1 Dsolani
Dickeya solani MK16 WP 022633848 1 Dsolani
Dickeya solani RNS 07 7 3B WP 022633848 1 Dsolani
Dickeya chrysanthemi L11 WP 012769529 1 Dchrysanthemi
Dickeya gs undicola 2B12 WP 033567524 1 Dundicola
Dickeya gs poaceaephila NCPPB 569 WP 042870920 1 Dpoaceaephila
Dickeya dianthicola RNS04 9 WP 024106205 1 Ddianthicola
Dickeya dianthicola GBBC 2039 WP 024106205 1 Ddianthicola
Dickeya dianthicola IPO 980 WP 024106205 1 Ddianthicola
Dickeya dianthicola NCPPB 453 WP 024106205 1 Ddianthicola
Dickeya dianthicola NCPPB 3534 WP 024106205 1 Ddianthicola
Dickeya gs fangzhongdai M074 WP 012769529 1 Dfangzhongdai
Dickeya chrysanthemi NCPPB 3533 WP 012769529 1 Dchrysanthemi
Dickeya zeae NCPPB 2538 WP 012769529 1 Dzeae
Dickeya dadantii 3937 WP 012769529 1 Ddadantii
Dickeya chrysanthemi subsp chrysanthemi NCPPB 402 WP 012769529 1 Dchrysanthemi
Dickeya zeae NCPPB 3531 WP 012769529 1 Dzeae
Dickeya zeae CSL RW192 WP 012769529 1 Dzeae
Dickeya zeae MS1 WP 012769529 1 Dzeae
Dickeya zeae Ech586 WP 012769529 1 Dzeae
Dickeya gs fangzhongdai NCPPB 3274 WP 012769529 1 Dfangzhongdai
Dickeya gs fangzhongdai MK7 WP 012769529 1 Dfangzhongdai
Dickeya dadantii NCPPB 898 WP 012769529 1 Ddadantii
Dickeya gs fangzhongdai ND14b WP 012769529 1 Dfangzhongdai
Dickeya zeae ZJU1202 WP 012769529 1 Dzeae
Dickeya dadantii NCPPB 3537 WP 012769529 1 Ddadantii
Dickeya gs fangzhongdai B16 WP 012769529 1 Dfangzhongdai
Dickeya zeae MK19 WP 012769529 1 Dzeae
Dickeya gs fangzhongdai S1 WP 012769529 1 Dfangzhongdai
Dickeya zeae EC1 WP 012769529 1 Dzeae
Dickeya gs fangzhongdai M005 WP 012769529 1 Dfangzhongdai
Dickeya dadantii subsp dieffenbachiae NCPPB 2976 WP 012769529 1 Ddadantii
Dickeya zeae DZ2Q WP 012769529 1 Dzeae
Dickeya zeae NCPPB 3532 WP 012769529 1 Dzeae
Dickeya chrysanthemi Ech1591 WP 012769529 1 Dchrysanthemi
Dickeya chrysanthemi subsp parthenii NCPPB 516 WP 012769529 1 Dchrysanthemi

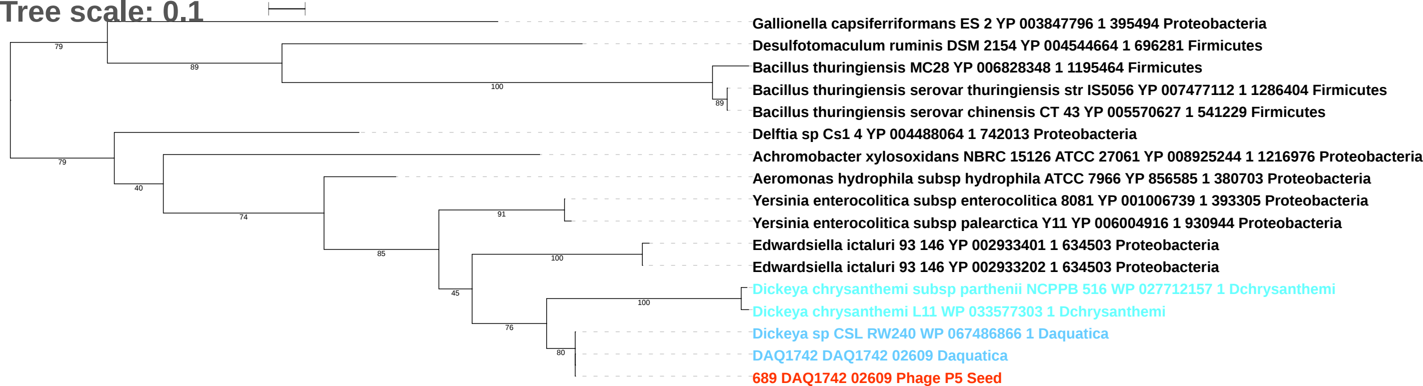
Tree scale: 0.1
Gallionella capsiferriformans ES 2 YP 003847796 1 395494 Proteobacteria
Desulfotomaculum ruminis DSM 2154 YP 004544664 1 696281 Firmicutes
Bacillus thuringiensis MC28 YP 006828348 1 1195464 Firmicutes
Bacillus thuringiensis serovar thuringiensis str IS5056 YP 007477112 1 1286404 Firmicutes
Bacillus thuringiensis serovar chinensis CT 43 YP 005570627 1 541229 Firmicutes
Delftia sp Cs1 4 YP 004488064 1 742013 Proteobacteria
Achromobacter xylosoxidans NBRC 15126 ATCC 27061 YP 008925244 1 1216976 Proteobacteria
Aeromonas hydrophila subsp hydrophila ATCC 7966 YP 856585 1 380703 Proteobacteria
Yersinia enterocolitica subsp enterocolitica 8081 YP 001006739 1 393305 Proteobacteria
Yersinia enterocolitica subsp palearctica Y11 YP 006004916 1 930944 Proteobacteria
Edwardsiella ictaluri 93 146 YP 002933401 1 634503 Proteobacteria
Edwardsiella ictaluri 93 146 YP 002933202 1 634503 Proteobacteria
Dickeya chrysanthemi subsp parthenii NCPPB 516 WP 027712157 1 Dchrysanthemi
Dickeya chrysanthemi L11 WP 033577303 1 Dchrysanthemi
Dickeya sp CSL RW240 WP 067486866 1 Daquatica
DAQ1742 DAQ1742 02609 Daquatica
689 DAQ1742 02609 Phage P5 Seed
79
89
100
89
79
40
74
91
85
100
45
100
76
80

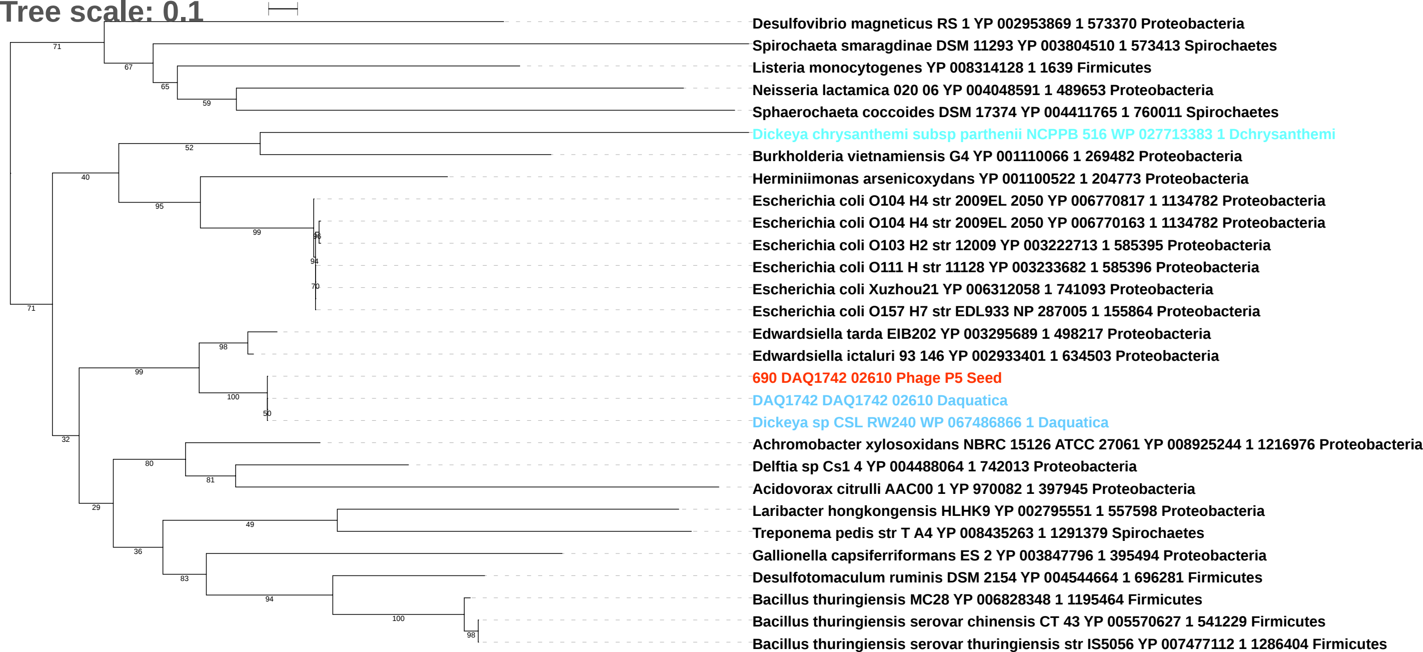

Tree scale: 0.1
Desulfovibrio magneticus RS 1 YP 002953869 1 573370 Proteobacteria
Spirochaeta smaragdinae DSM 11293 YP 003804510 1 573413 Spirochaetes
Listeria monocytogenes YP 008314128 1 1639 Firmicutes
Neisseria lactamica 020 06 YP 004048591 1 489653 Proteobacteria
Sphaerochaeta coccoides DSM 17374 YP 004411765 1 760011 Spirochaetes
Dickeya chrysanthemi subsp parthenii NCPPB 516 WP 027713383 1 Dchrysanthemi
Burkholderia vietnamiensis G4 YP 001110066 1 269482 Proteobacteria
Herminiimonas arsenicoxydans YP 001100522 1 204773 Proteobacteria
Escherichia coli O104 H4 str 2009EL 2050 YP 006770817 1 1134782 Proteobacteria
Escherichia coli O104 H4 str 2009EL 2050 YP 006770163 1 1134782 Proteobacteria
Escherichia coli O103 H2 str 12009 YP 003222713 1 585395 Proteobacteria
Escherichia coli O111 H str 11128 YP 003233682 1 585396 Proteobacteria
Escherichia coli Xuzhou21 YP 006312058 1 741093 Proteobacteria
Escherichia coli O157 H7 str EDL933 NP 287005 1 155864 Proteobacteria
Edwardsiella tarda EIB202 YP 003295689 1 498217 Proteobacteria
Edwardsiella ictaluri 93 146 YP 002933401 1 634503 Proteobacteria
690 DAQ1742 02610 Phage P5 Seed
DAQ1742 DAQ1742 02610 Daquatica
Dickeya sp CSL RW240 WP 067486866 1 Daquatica
Achromobacter xylosoxidans NBRC 15126 ATCC 27061 YP 008925244 1 1216976 Proteobacteria
Delftia sp Cs1 4 YP 004488064 1 742013 Proteobacteria
Acidovorax citrulli AAC00 1 YP 970082 1 397945 Proteobacteria
Laribacter hongkongensis HLHK9 YP 002795551 1 557598 Proteobacteria
Treponema pedis str T A4 YP 008435263 1 1291379 Spirochaetes
Gallionella capsiferriformans ES 2 YP 003847796 1 395494 Proteobacteria
Desulfotomaculum ruminis DSM 2154 YP 004544664 1 696281 Firmicutes
Bacillus thuringiensis MC28 YP 006828348 1 1195464 Firmicutes
Bacillus thuringiensis serovar chinensis CT 43 YP 005570627 1 541229 Firmicutes
Bacillus thuringiensis serovar thuringiensis str IS5056 YP 007477112 1 1286404 Firmicutes
71
67
65
59
52
40
95
99
96
94
70
71
98
99
100
50
32
80
81
29
49
36
83
94
100
98

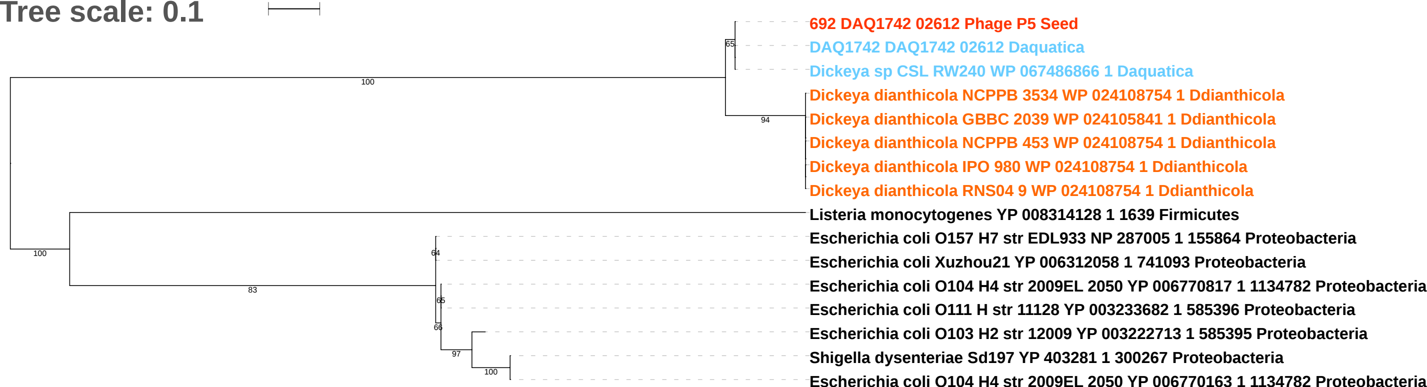

Tree scale: 0.1
692 DAQ1742 02612 Phage P5 Seed
DAQ1742 DAQ1742 02612 Daquatica
Dickeya sp CSL RW240 WP 067486866 1 Daquatica
Dickeya dianthicola NCPPB 3534 WP 024108754 1 Ddianthicola
Dickeya dianthicola GBBC 2039 WP 024105841 1 Ddianthicola
Dickeya dianthicola NCPPB 453 WP 024108754 1 Ddianthicola
Dickeya dianthicola IPO 980 WP 024108754 1 Ddianthicola
Dickeya dianthicola RNS04 9 WP 024108754 1 Ddianthicola
Listeria monocytogenes YP 008314128 1 1639 Firmicutes
Escherichia coli O157 H7 str EDL933 NP 287005 1 155864 Proteobacteria
Escherichia coli Xuzhou21 YP 006312058 1 741093 Proteobacteria
Escherichia coli O104 H4 str 2009EL 2050 YP 006770817 1 1134782 Proteobacteria
Escherichia coli O111 H str 11128 YP 003233682 1 585396 Proteobacteria
Escherichia coli O103 H2 str 12009 YP 003222713 1 585395 Proteobacteria
Shigella dysenteriae Sd197 YP 403281 1 300267 Proteobacteria
Escherichia coli O104 H4 str 2009EL 2050 YP 006770163 1 1134782 Proteobacteria
100
65
94
100
83
64
65
66
97
100

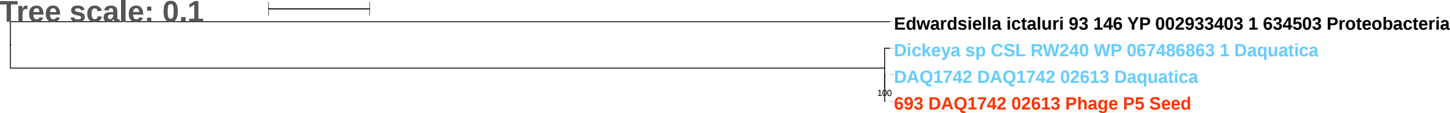
Tree scale: 0.1
Edwardsiella ictaluri 93 146 YP 002933403 1 634503 Proteobacteria
Dickeya sp CSL RW240 WP 067486863 1 Daquatica
DAQ1742 DAQ1742 02613 Daquatica
100
693 DAQ1742 02613 Phage P5 Seed

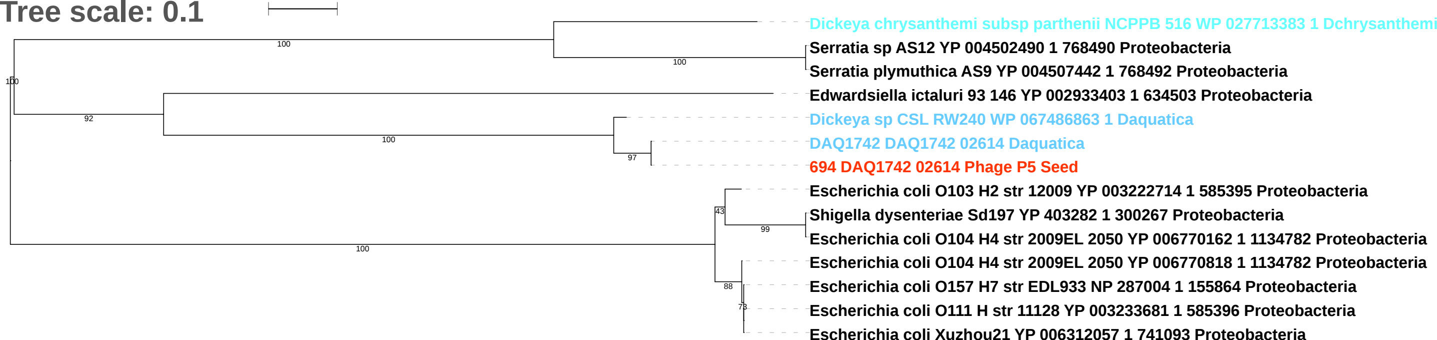
Tree scale: 0.1
Dickeya chrysanthemi subsp parthenii NCPPB 516 WP 027713383 1 Dchrysanthemi
Serratia sp AS12 YP 004502490 1 768490 Proteobacteria
Serratia plymuthica AS9 YP 004507442 1 768492 Proteobacteria
Edwardsiella ictaluri 93 146 YP 002933403 1 634503 Proteobacteria
Dickeya sp CSL RW240 WP 067486863 1 Daquatica
DAQ1742 DAQ1742 02614 Daquatica
694 DAQ1742 02614 Phage P5 Seed
Escherichia coli O103 H2 str 12009 YP 003222714 1 585395 Proteobacteria
Shigella dysenteriae Sd197 YP 403282 1 300267 Proteobacteria
Escherichia coli O104 H4 str 2009EL 2050 YP 006770162 1 1134782 Proteobacteria
Escherichia coli O104 H4 str 2009EL 2050 YP 006770818 1 1134782 Proteobacteria
Escherichia coli O157 H7 str EDL933 NP 287004 1 155864 Proteobacteria
Escherichia coli O111 H str 11128 YP 003233681 1 585396 Proteobacteria
Escherichia coli Xuzhou21 YP 006312057 1 741093 Proteobacteria
100
100
100
92
100
97
43
99
100
88
73

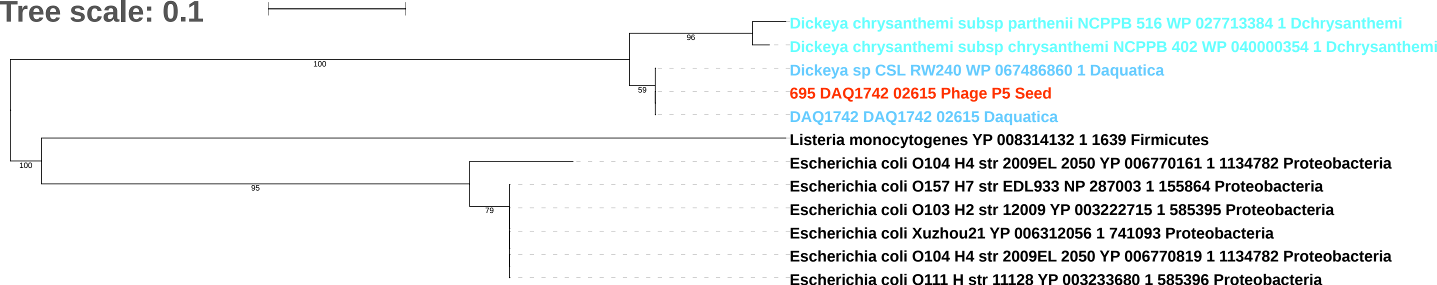

Tree scale: 0.1
Dickeya chrysanthemi subsp parthenii NCPPB 516 WP 027713384 1 Dchrysanthemi
Dickeya chrysanthemi subsp chrysanthemi NCPPB 402 WP 040000354 1 Dchrysanthemi
Dickeya sp CSL RW240 WP 067486860 1 Daquatica
695 DAQ1742 02615 Phage P5 Seed
DAQ1742 DAQ1742 02615 Daquatica
Listeria monocytogenes YP 008314132 1 1639 Firmicutes
Escherichia coli O104 H4 str 2009EL 2050 YP 006770161 1 1134782 Proteobacteria
Escherichia coli O157 H7 str EDL933 NP 287003 1 155864 Proteobacteria
Escherichia coli O103 H2 str 12009 YP 003222715 1 585395 Proteobacteria
Escherichia coli Xuzhou21 YP 006312056 1 741093 Proteobacteria
Escherichia coli O104 H4 str 2009EL 2050 YP 006770819 1 1134782 Proteobacteria
Escherichia coli O111 H str 11128 YP 003233680 1 585396 Proteobacteria
96
100
59
100
95
79

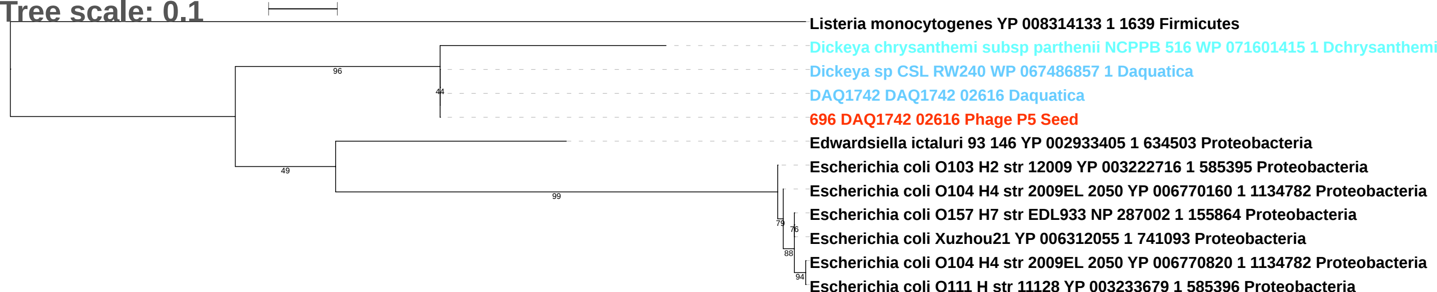
Tree scale: 0.1
Listeria monocytogenes YP 008314133 1 1639 Firmicutes
Dickeya chrysanthemi subsp parthenii NCPPB 516 WP 071601415 1 Dchrysanthemi
Dickeya sp CSL RW240 WP 067486857 1 Daquatica
DAQ1742 DAQ1742 02616 Daquatica
696 DAQ1742 02616 Phage P5 Seed
Edwardsiella ictaluri 93 146 YP 002933405 1 634503 Proteobacteria
Escherichia coli O103 H2 str 12009 YP 003222716 1 585395 Proteobacteria
Escherichia coli O104 H4 str 2009EL 2050 YP 006770160 1 1134782 Proteobacteria
Escherichia coli O157 H7 str EDL933 NP 287002 1 155864 Proteobacteria
Escherichia coli Xuzhou21 YP 006312055 1 741093 Proteobacteria
Escherichia coli O104 H4 str 2009EL 2050 YP 006770820 1 1134782 Proteobacteria
Escherichia coli O111 H str 11128 YP 003233679 1 585396 Proteobacteria
96
44
49
99
79
76
88
94

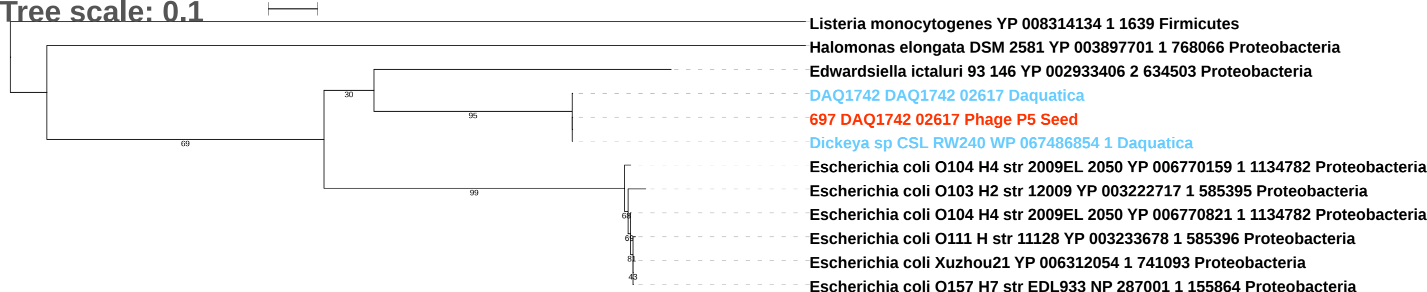
Tree scale: 0.1
Listeria monocytogenes YP 008314134 1 1639 Firmicutes
Halomonas elongata DSM 2581 YP 003897701 1 768066 Proteobacteria
Edwardsiella ictaluri 93 146 YP 002933406 2 634503 Proteobacteria
DAQ1742 DAQ1742 02617 Daquatica
697 DAQ1742 02617 Phage P5 Seed
Dickeya sp CSL RW240 WP 067486854 1 Daquatica
Escherichia coli O104 H4 str 2009EL 2050 YP 006770159 1 1134782 Proteobacteria
Escherichia coli O103 H2 str 12009 YP 003222717 1 585395 Proteobacteria
Escherichia coli O104 H4 str 2009EL 2050 YP 006770821 1 1134782 Proteobacteria
Escherichia coli O111 H str 11128 YP 003233678 1 585396 Proteobacteria
Escherichia coli Xuzhou21 YP 006312054 1 741093 Proteobacteria
Escherichia coli O157 H7 str EDL933 NP 287001 1 155864 Proteobacteria
30
95
69
99
68
69
81
43

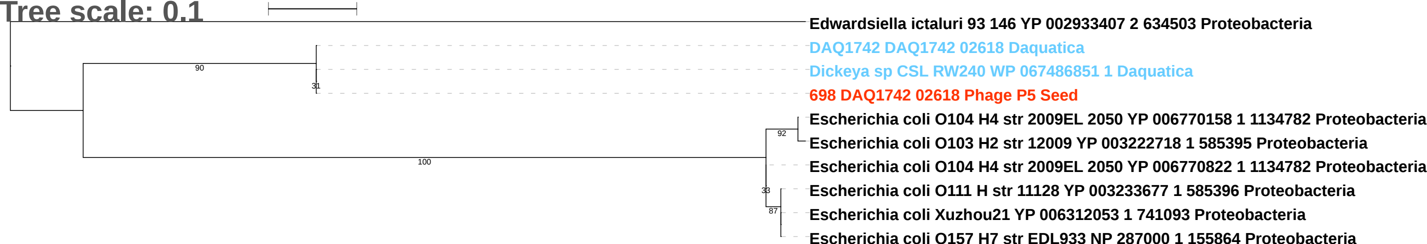
Tree scale: 0.1
Edwardsiella ictaluri 93 146 YP 002933407 2 634503 Proteobacteria
DAQ1742 DAQ1742 02618 Daquatica
Dickeya sp CSL RW240 WP 067486851 1 Daquatica
698 DAQ1742 02618 Phage P5 Seed
Escherichia coli O104 H4 str 2009EL 2050 YP 006770158 1 1134782 Proteobacteria
Escherichia coli O103 H2 str 12009 YP 003222718 1 585395 Proteobacteria
Escherichia coli O104 H4 str 2009EL 2050 YP 006770822 1 1134782 Proteobacteria
Escherichia coli O111 H str 11128 YP 003233677 1 585396 Proteobacteria
Escherichia coli Xuzhou21 YP 006312053 1 741093 Proteobacteria
Escherichia coli O157 H7 str EDL933 NP 287000 1 155864 Proteobacteria
90
31
92
100
33
87

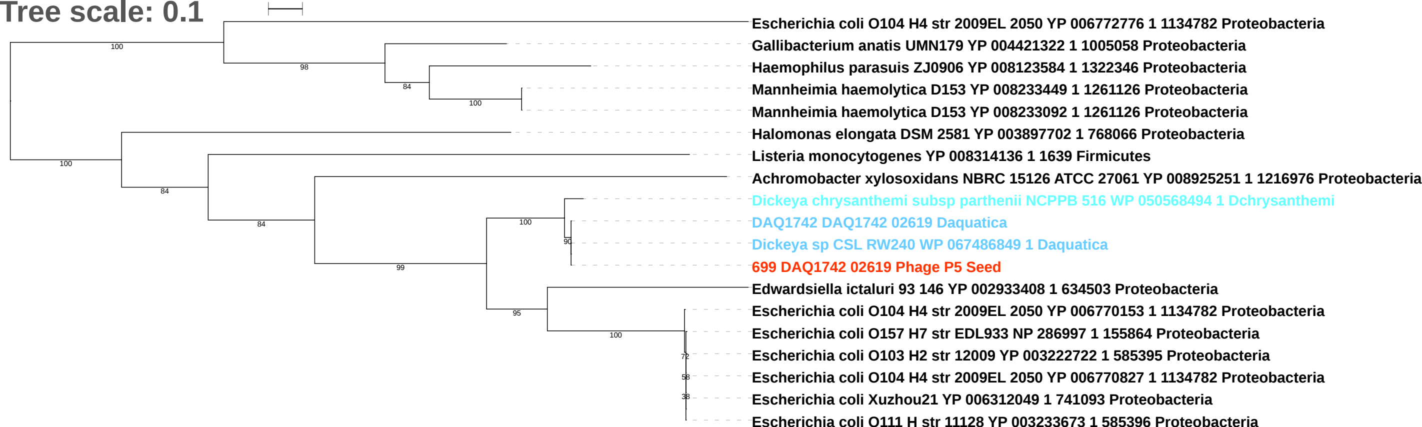
Tree scale: 0.1
Escherichia coli O104 H4 str 2009EL 2050 YP 006772776 1 1134782 Proteobacteria
Gallibacterium anatis UMN179 YP 004421322 1 1005058 Proteobacteria
Haemophilus parasuis ZJ0906 YP 008123584 1 1322346 Proteobacteria
Mannheimia haemolytica D153 YP 008233449 1 1261126 Proteobacteria
Mannheimia haemolytica D153 YP 008233092 1 1261126 Proteobacteria
Halomonas elongata DSM 2581 YP 003897702 1 768066 Proteobacteria
Listeria monocytogenes YP 008314136 1 1639 Firmicutes
Achromobacter xylosoxidans NBRC 15126 ATCC 27061 YP 008925251 1 1216976 Proteobacteria
Dickeya chrysanthemi subsp parthenii NCPPB 516 WP 050568494 1 Dchrysanthemi
DAQ1742 DAQ1742 02619 Daquatica
Dickeya sp CSL RW240 WP 067486849 1 Daquatica
699 DAQ1742 02619 Phage P5 Seed
Edwardsiella ictaluri 93 146 YP 002933408 1 634503 Proteobacteria
Escherichia coli O104 H4 str 2009EL 2050 YP 006770153 1 1134782 Proteobacteria
Escherichia coli O157 H7 str EDL933 NP 286997 1 155864 Proteobacteria
Escherichia coli O103 H2 str 12009 YP 003222722 1 585395 Proteobacteria
Escherichia coli O104 H4 str 2009EL 2050 YP 006770827 1 1134782 Proteobacteria
Escherichia coli Xuzhou21 YP 006312049 1 741093 Proteobacteria
Escherichia coli O111 H str 11128 YP 003233673 1 585396 Proteobacteria
100
98
84
100
100
84
84
99
100
90
95
100
72
58
38

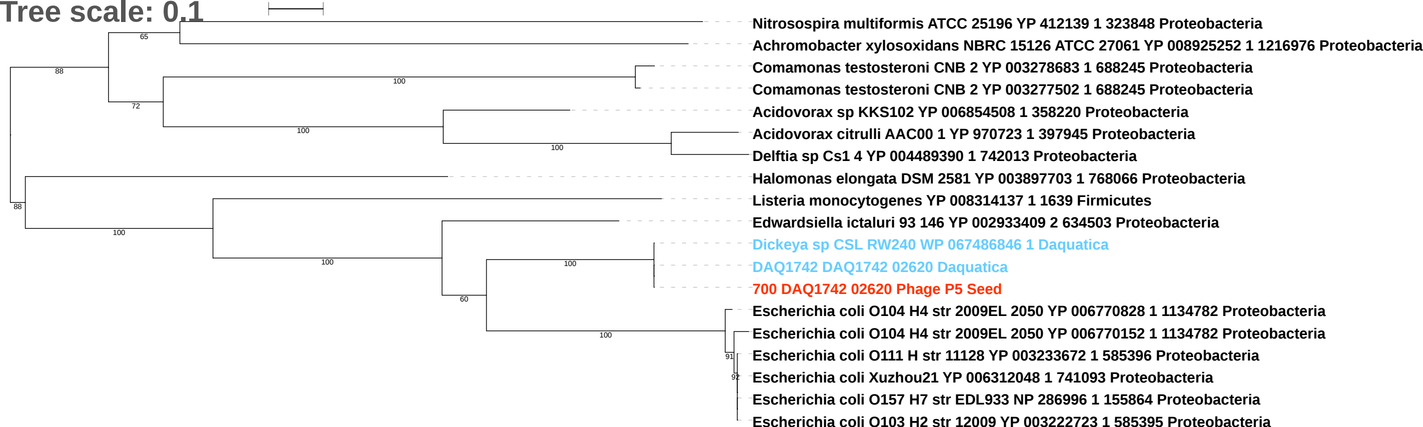
Tree scale: 0.1
Nitrosospira multiformis ATCC 25196 YP 412139 1 323848 Proteobacteria
Achromobacter xylosoxidans NBRC 15126 ATCC 27061 YP 008925252 1 1216976 Proteobacteria
Comamonas testosteroni CNB 2 YP 003278683 1 688245 Proteobacteria
Comamonas testosteroni CNB 2 YP 003277502 1 688245 Proteobacteria
Acidovorax sp KKS102 YP 006854508 1 358220 Proteobacteria
Acidovorax citrulli AAC00 1 YP 970723 1 397945 Proteobacteria
Delftia sp Cs1 4 YP 004489390 1 742013 Proteobacteria
Halomonas elongata DSM 2581 YP 003897703 1 768066 Proteobacteria
Listeria monocytogenes YP 008314137 1 1639 Firmicutes
Edwardsiella ictaluri 93 146 YP 002933409 2 634503 Proteobacteria
Dickeya sp CSL RW240 WP 067486846 1 Daquatica
DAQ1742 DAQ1742 02620 Daquatica
700 DAQ1742 02620 Phage P5 Seed
Escherichia coli O104 H4 str 2009EL 2050 YP 006770828 1 1134782 Proteobacteria
Escherichia coli O104 H4 str 2009EL 2050 YP 006770152 1 1134782 Proteobacteria
Escherichia coli O111 H str 11128 YP 003233672 1 585396 Proteobacteria
Escherichia coli Xuzhou21 YP 006312048 1 741093 Proteobacteria
Escherichia coli O157 H7 str EDL933 NP 286996 1 155864 Proteobacteria
Escherichia coli O103 H2 str 12009 YP 003222723 1 585395 Proteobacteria
65
88
100
72
100
100
88
100
100
100
60
100
91
92

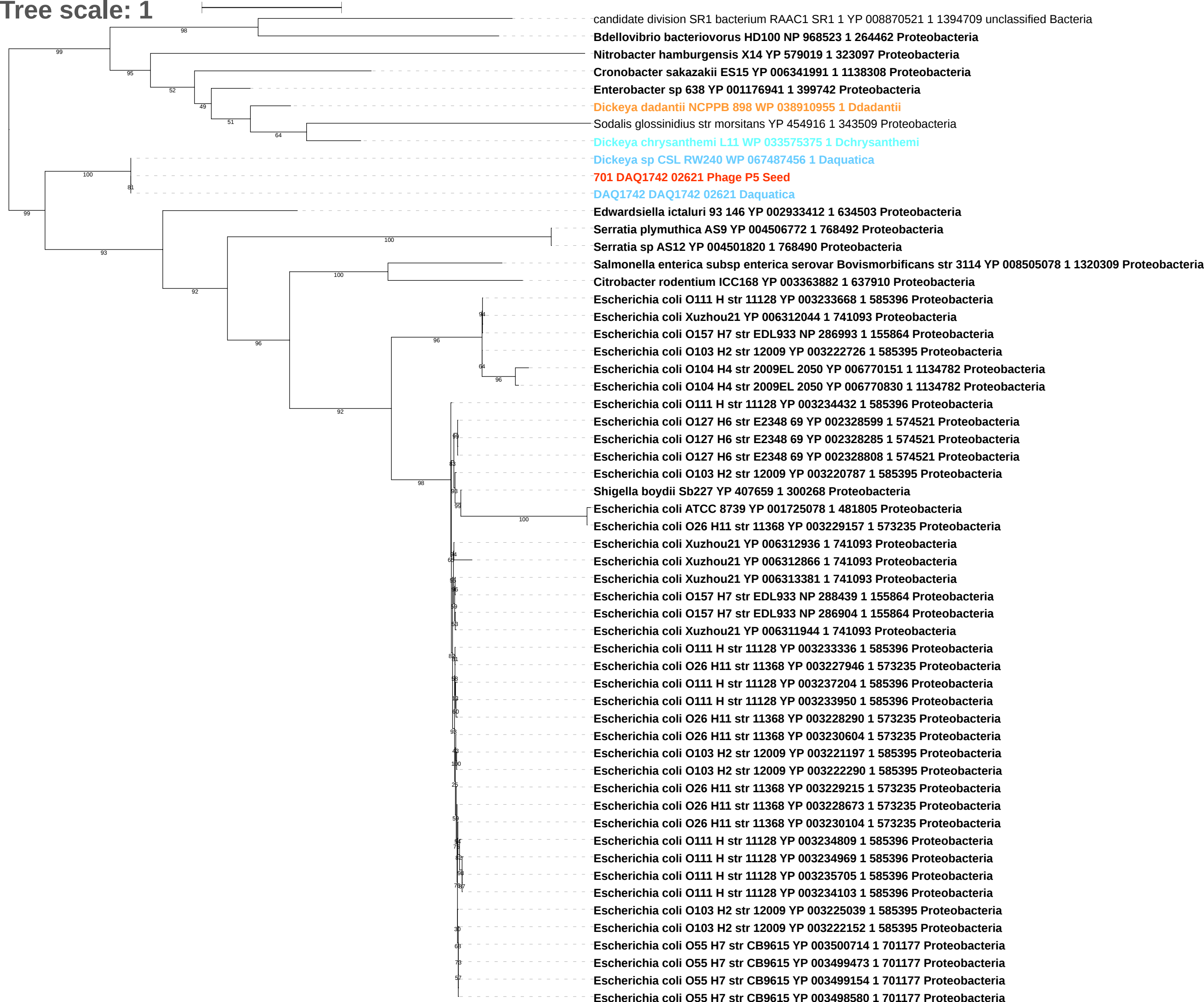
Tree scale: 1
candidate division SR1 bacterium RAAC1 SR1 1 YP 008870521 1 1394709 unclassified Bacteria
Bdellovibrio bacteriovorus HD100 NP 968523 1 264462 Proteobacteria
Nitrobacter hamburgensis X14 YP 579019 1 323097 Proteobacteria
Cronobacter sakazakii ES15 YP 006341991 1 1138308 Proteobacteria
Enterobacter sp 638 YP 001176941 1 399742 Proteobacteria
Dickeya dadantii NCPPB 898 WP 038910955 1 Ddadantii
Sodalis glossinidius str morsitans YP 454916 1 343509 Proteobacteria
Dickeya chrysanthemi L11 WP 033575375 1 Dchrysanthemi
Dickeya sp CSL RW240 WP 067487456 1 Daquatica
701 DAQ1742 02621 Phage P5 Seed
DAQ1742 DAQ1742 02621 Daquatica
Edwardsiella ictaluri 93 146 YP 002933412 1 634503 Proteobacteria
Serratia plymuthica AS9 YP 004506772 1 768492 Proteobacteria
Serratia sp AS12 YP 004501820 1 768490 Proteobacteria
Salmonella enterica subsp enterica serovar Bovismorbificans str 3114 YP 008505078 1 1320309 Proteobacteria
Citrobacter rodentium ICC168 YP 003363882 1 637910 Proteobacteria
Escherichia coli O111 H str 11128 YP 003233668 1 585396 Proteobacteria
Escherichia coli Xuzhou21 YP 006312044 1 741093 Proteobacteria
Escherichia coli O157 H7 str EDL933 NP 286993 1 155864 Proteobacteria
Escherichia coli O103 H2 str 12009 YP 003222726 1 585395 Proteobacteria
Escherichia coli O104 H4 str 2009EL 2050 YP 006770151 1 1134782 Proteobacteria
Escherichia coli O104 H4 str 2009EL 2050 YP 006770830 1 1134782 Proteobacteria
Escherichia coli O111 H str 11128 YP 003234432 1 585396 Proteobacteria
Escherichia coli O127 H6 str E2348 69 YP 002328599 1 574521 Proteobacteria
Escherichia coli O127 H6 str E2348 69 YP 002328285 1 574521 Proteobacteria
Escherichia coli O127 H6 str E2348 69 YP 002328808 1 574521 Proteobacteria
Escherichia coli O103 H2 str 12009 YP 003220787 1 585395 Proteobacteria
Shigella boydii Sb227 YP 407659 1 300268 Proteobacteria
Escherichia coli ATCC 8739 YP 001725078 1 481805 Proteobacteria
Escherichia coli O26 H11 str 11368 YP 003229157 1 573235 Proteobacteria
Escherichia coli Xuzhou21 YP 006312936 1 741093 Proteobacteria
Escherichia coli Xuzhou21 YP 006312866 1 741093 Proteobacteria
Escherichia coli Xuzhou21 YP 006313381 1 741093 Proteobacteria
Escherichia coli O157 H7 str EDL933 NP 288439 1 155864 Proteobacteria
Escherichia coli O157 H7 str EDL933 NP 286904 1 155864 Proteobacteria
Escherichia coli Xuzhou21 YP 006311944 1 741093 Proteobacteria
Escherichia coli O111 H str 11128 YP 003233336 1 585396 Proteobacteria
Escherichia coli O26 H11 str 11368 YP 003227946 1 573235 Proteobacteria
Escherichia coli O111 H str 11128 YP 003237204 1 585396 Proteobacteria
Escherichia coli O111 H str 11128 YP 003233950 1 585396 Proteobacteria
Escherichia coli O26 H11 str 11368 YP 003228290 1 573235 Proteobacteria
Escherichia coli O26 H11 str 11368 YP 003230604 1 573235 Proteobacteria
Escherichia coli O103 H2 str 12009 YP 003221197 1 585395 Proteobacteria
Escherichia coli O103 H2 str 12009 YP 003222290 1 585395 Proteobacteria
Escherichia coli O26 H11 str 11368 YP 003229215 1 573235 Proteobacteria
Escherichia coli O26 H11 str 11368 YP 003228673 1 573235 Proteobacteria
Escherichia coli O26 H11 str 11368 YP 003230104 1 573235 Proteobacteria
Escherichia coli O111 H str 11128 YP 003234809 1 585396 Proteobacteria
Escherichia coli O111 H str 11128 YP 003234969 1 585396 Proteobacteria
Escherichia coli O111 H str 11128 YP 003235705 1 585396 Proteobacteria
Escherichia coli O111 H str 11128 YP 003234103 1 585396 Proteobacteria
Escherichia coli O103 H2 str 12009 YP 003225039 1 585395 Proteobacteria
Escherichia coli O103 H2 str 12009 YP 003222152 1 585395 Proteobacteria
Escherichia coli O55 H7 str CB9615 YP 003500714 1 701177 Proteobacteria
Escherichia coli O55 H7 str CB9615 YP 003499473 1 701177 Proteobacteria
Escherichia coli O55 H7 str CB9615 YP 003499154 1 701177 Proteobacteria
Escherichia coli O55 H7 str CB9615 YP 003498580 1 701177 Proteobacteria

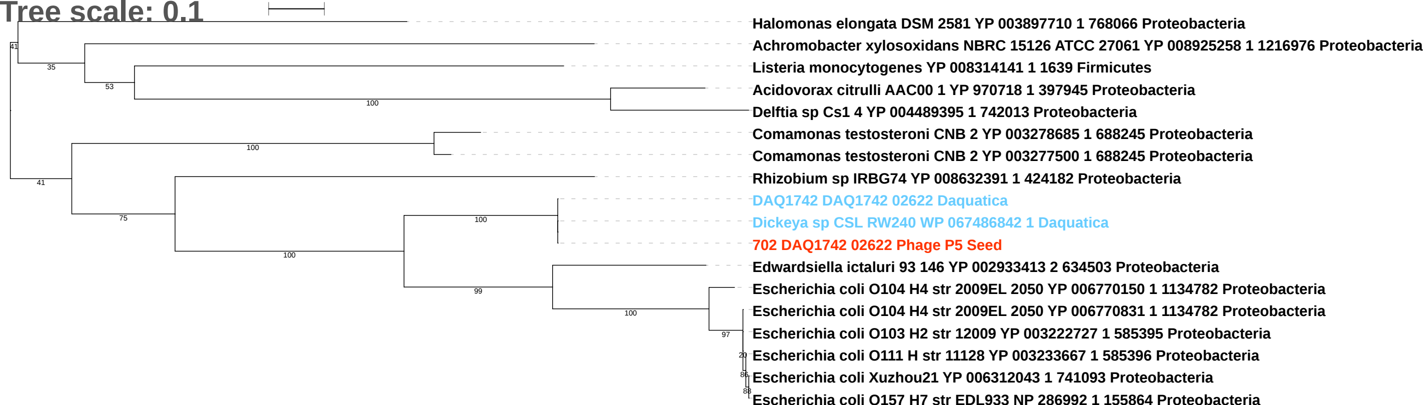
Tree scale: 0.1
Halomonas elongata DSM 2581 YP 003897710 1 768066 Proteobacteria
Achromobacter xylosoxidans NBRC 15126 ATCC 27061 YP 008925258 1 1216976 Proteobacteria
Listeria monocytogenes YP 008314141 1 1639 Firmicutes
Acidovorax citrulli AAC00 1 YP 970718 1 397945 Proteobacteria
Delftia sp Cs1 4 YP 004489395 1 742013 Proteobacteria
Comamonas testosteroni CNB 2 YP 003278685 1 688245 Proteobacteria
Comamonas testosteroni CNB 2 YP 003277500 1 688245 Proteobacteria
Rhizobium sp IRBG74 YP 008632391 1 424182 Proteobacteria
DAQ1742 DAQ1742 02622 Daquatica
Dickeya sp CSL RW240 WP 067486842 1 Daquatica
702 DAQ1742 02622 Phage P5 Seed
Edwardsiella ictaluri 93 146 YP 002933413 2 634503 Proteobacteria
Escherichia coli O104 H4 str 2009EL 2050 YP 006770150 1 1134782 Proteobacteria
Escherichia coli O104 H4 str 2009EL 2050 YP 006770831 1 1134782 Proteobacteria
Escherichia coli O103 H2 str 12009 YP 003222727 1 585395 Proteobacteria
Escherichia coli O111 H str 11128 YP 003233667 1 585396 Proteobacteria
Escherichia coli Xuzhou21 YP 006312043 1 741093 Proteobacteria
Escherichia coli O157 H7 str EDL933 NP 286992 1 155864 Proteobacteria
41
35
53
100
100
41
75
100
100
99
100
97
20
86
88

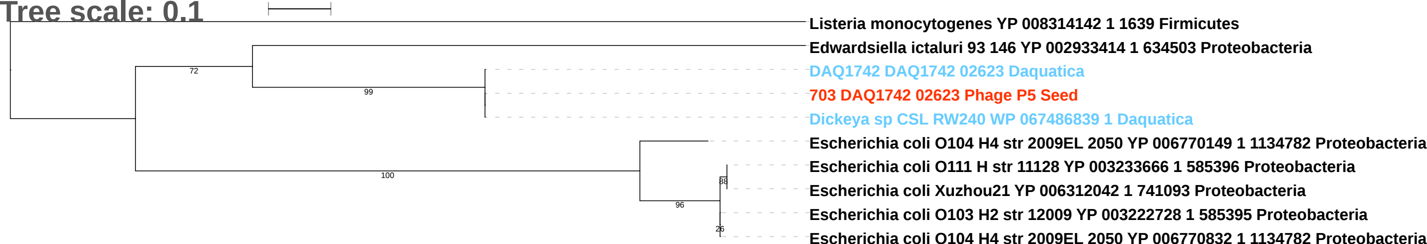
Tree scale: 0.1
Listeria monocytogenes YP 008314142 1 1639 Firmicutes
Edwardsiella ictaluri 93 146 YP 002933414 1 634503 Proteobacteria
DAQ1742 DAQ1742 02623 Daquatica
703 DAQ1742 02623 Phage P5 Seed
Dickeya sp CSL RW240 WP 067486839 1 Daquatica
Escherichia coli O104 H4 str 2009EL 2050 YP 006770149 1 1134782 Proteobacteria
Escherichia coli O111 H str 11128 YP 003233666 1 585396 Proteobacteria
Escherichia coli Xuzhou21 YP 006312042 1 741093 Proteobacteria
Escherichia coli O103 H2 str 12009 YP 003222728 1 585395 Proteobacteria
Escherichia coli O104 H4 str 2009EL 2050 YP 006770832 1 1134782 Proteobacteria
72
99
100
96
88
26

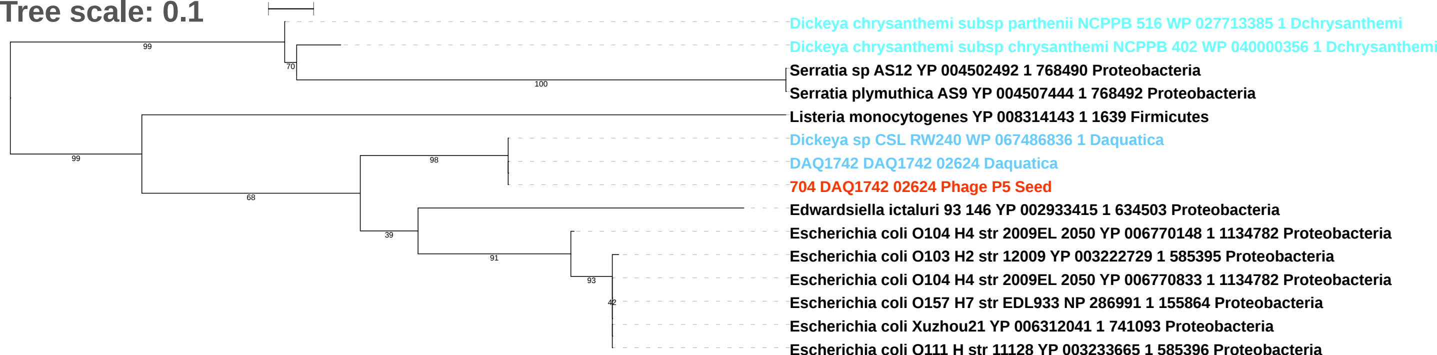

Tree scale: 0.1
Dickeya chrysanthemi subsp parthenii NCPPB 516 WP 027713385 1 Dchrysanthemi
Dickeya chrysanthemi subsp chrysanthemi NCPPB 402 WP 040000356 1 Dchrysanthemi
Serratia sp AS12 YP 004502492 1 768490 Proteobacteria
Serratia plymuthica AS9 YP 004507444 1 768492 Proteobacteria
Listeria monocytogenes YP 008314143 1 1639 Firmicutes
Dickeya sp CSL RW240 WP 067486836 1 Daquatica
DAQ1742 DAQ1742 02624 Daquatica
704 DAQ1742 02624 Phage P5 Seed
Edwardsiella ictaluri 93 146 YP 002933415 1 634503 Proteobacteria
Escherichia coli O104 H4 str 2009EL 2050 YP 006770148 1 1134782 Proteobacteria
Escherichia coli O103 H2 str 12009 YP 003222729 1 585395 Proteobacteria
Escherichia coli O104 H4 str 2009EL 2050 YP 006770833 1 1134782 Proteobacteria
Escherichia coli O157 H7 str EDL933 NP 286991 1 155864 Proteobacteria
Escherichia coli Xuzhou21 YP 006312041 1 741093 Proteobacteria
Escherichia coli O111 H str 11128 YP 003233665 1 585396 Proteobacteria
99
70
100
99
98
68
39
91
93
42

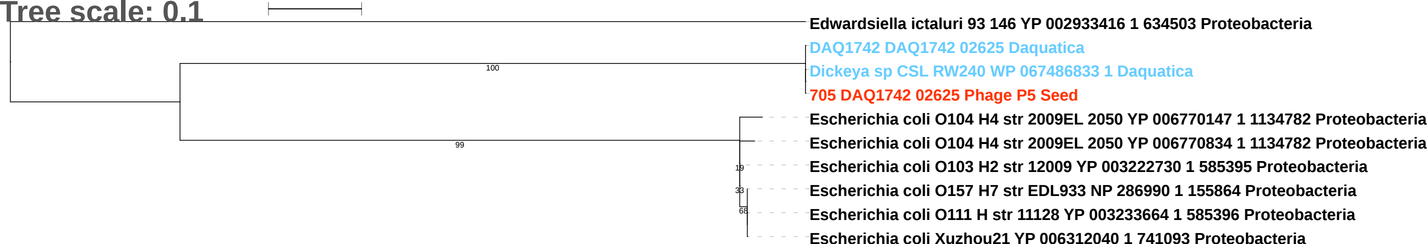
Tree scale: 0.1
Edwardsiella ictaluri 93 146 YP 002933416 1 634503 Proteobacteria
DAQ1742 DAQ1742 02625 Daquatica
Dickeya sp CSL RW240 WP 067486833 1 Daquatica
705 DAQ1742 02625 Phage P5 Seed
Escherichia coli O104 H4 str 2009EL 2050 YP 006770147 1 1134782 Proteobacteria
Escherichia coli O104 H4 str 2009EL 2050 YP 006770834 1 1134782 Proteobacteria
Escherichia coli O103 H2 str 12009 YP 003222730 1 585395 Proteobacteria
Escherichia coli O157 H7 str EDL933 NP 286990 1 155864 Proteobacteria
Escherichia coli O111 H str 11128 YP 003233664 1 585396 Proteobacteria
Escherichia coli Xuzhou21 YP 006312040 1 741093 Proteobacteria
100
99
19
33
68

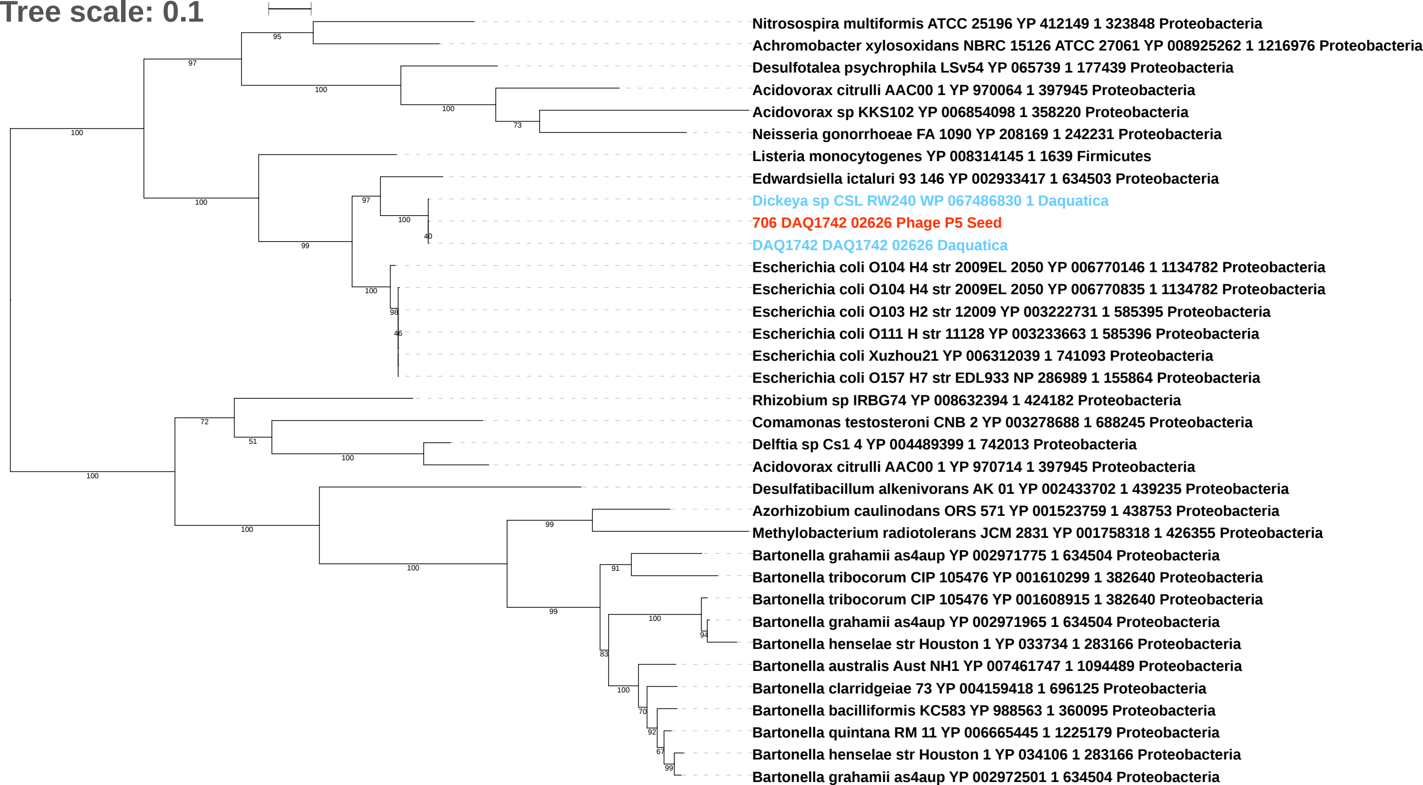
Tree scale: 0.1
Nitrosospira multiformis ATCC 25196 YP 412149 1 323848 Proteobacteria
Achromobacter xylosoxidans NBRC 15126 ATCC 27061 YP 008925262 1 1216976 Proteobacteria
Desulfotalea psychrophila LSv54 YP 065739 1 177439 Proteobacteria
Acidovorax citrulli AAC00 1 YP 970064 1 397945 Proteobacteria
Acidovorax sp KKS102 YP 006854098 1 358220 Proteobacteria
Neisseria gonorrhoeae FA 1090 YP 208169 1 242231 Proteobacteria
Listeria monocytogenes YP 008314145 1 1639 Firmicutes
Edwardsiella ictaluri 93 146 YP 002933417 1 634503 Proteobacteria
Dickeya sp CSL RW240 WP 067486830 1 Daquatica
706 DAQ1742 02626 Phage P5 Seed
DAQ1742 DAQ1742 02626 Daquatica
Escherichia coli O104 H4 str 2009EL 2050 YP 006770146 1 1134782 Proteobacteria
Escherichia coli O104 H4 str 2009EL 2050 YP 006770835 1 1134782 Proteobacteria
Escherichia coli O103 H2 str 12009 YP 003222731 1 585395 Proteobacteria
Escherichia coli O111 H str 11128 YP 003233663 1 585396 Proteobacteria
Escherichia coli Xuzhou21 YP 006312039 1 741093 Proteobacteria
Escherichia coli O157 H7 str EDL933 NP 286989 1 155864 Proteobacteria
Rhizobium sp IRBG74 YP 008632394 1 424182 Proteobacteria
Comamonas testosteroni CNB 2 YP 003278688 1 688245 Proteobacteria
Delftia sp Cs1 4 YP 004489399 1 742013 Proteobacteria
Acidovorax citrulli AAC00 1 YP 970714 1 397945 Proteobacteria
Desulfatibacillum alkenivorans AK 01 YP 002433702 1 439235 Proteobacteria
Azorhizobium caulinodans ORS 571 YP 001523759 1 438753 Proteobacteria
Methylobacterium radiotolerans JCM 2831 YP 001758318 1 426355 Proteobacteria
Bartonella grahamii as4aup YP 002971775 1 634504 Proteobacteria
Bartonella tribocorum CIP 105476 YP 001610299 1 382640 Proteobacteria
Bartonella tribocorum CIP 105476 YP 001608915 1 382640 Proteobacteria
Bartonella grahamii as4aup YP 002971965 1 634504 Proteobacteria
Bartonella henselae str Houston 1 YP 033734 1 283166 Proteobacteria
Bartonella australis Aust NH1 YP 007461747 1 1094489 Proteobacteria
Bartonella clarridgeiae 73 YP 004159418 1 696125 Proteobacteria
Bartonella bacilliformis KC583 YP 988563 1 360095 Proteobacteria
Bartonella quintana RM 11 YP 006665445 1 1225179 Proteobacteria
Bartonella henselae str Houston 1 YP 034106 1 283166 Proteobacteria
Bartonella grahamii as4aup YP 002972501 1 634504 Proteobacteria

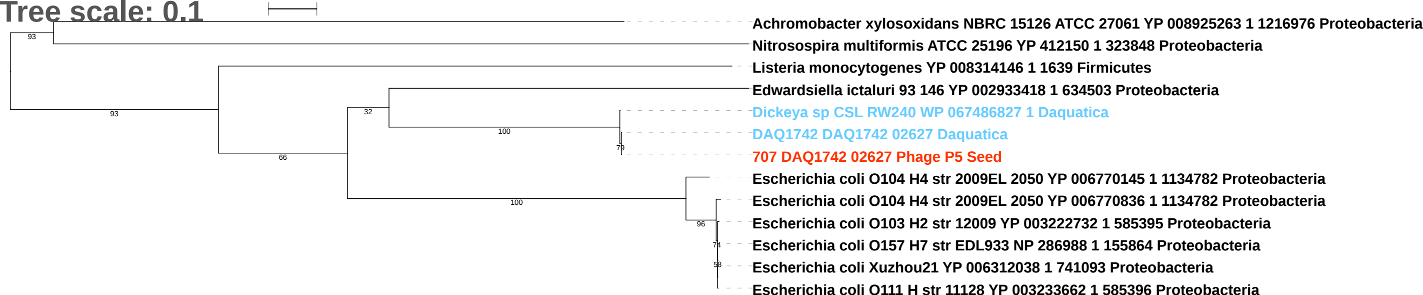
Tree scale: 0.1
Achromobacter xylosoxidans NBRC 15126 ATCC 27061 YP 008925263 1 1216976 Proteobacteria
Nitrosospira multiformis ATCC 25196 YP 412150 1 323848 Proteobacteria
Listeria monocytogenes YP 008314146 1 1639 Firmicutes
Edwardsiella ictaluri 93 146 YP 002933418 1 634503 Proteobacteria
Dickeya sp CSL RW240 WP 067486827 1 Daquatica
DAQ1742 DAQ1742 02627 Daquatica
707 DAQ1742 02627 Phage P5 Seed
Escherichia coli O104 H4 str 2009EL 2050 YP 006770145 1 1134782 Proteobacteria
Escherichia coli O104 H4 str 2009EL 2050 YP 006770836 1 1134782 Proteobacteria
Escherichia coli O103 H2 str 12009 YP 003222732 1 585395 Proteobacteria
Escherichia coli O157 H7 str EDL933 NP 286988 1 155864 Proteobacteria
Escherichia coli Xuzhou21 YP 006312038 1 741093 Proteobacteria
Escherichia coli O111 H str 11128 YP 003233662 1 585396 Proteobacteria
93
93
32
100
79
66
100
96
74
58

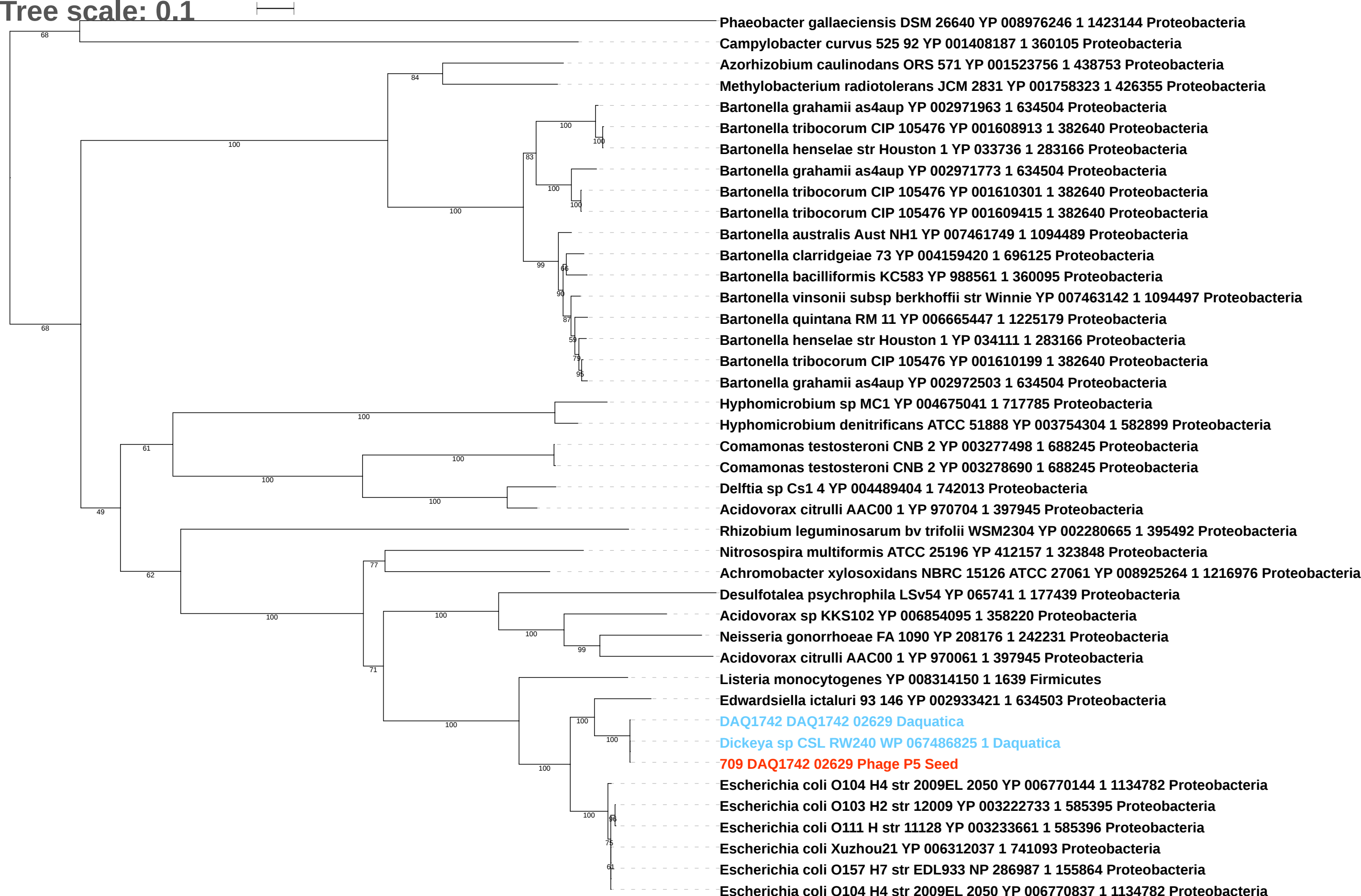
Tree scale: 0.1
Phaeobacter gallaeciensis DSM 26640 YP 008976246 1 1423144 Proteobacteria
Campylobacter curvus 525 92 YP 001408187 1 360105 Proteobacteria
Azorhizobium caulinodans ORS 571 YP 001523756 1 438753 Proteobacteria
Methylobacterium radiotolerans JCM 2831 YP 001758323 1 426355 Proteobacteria
Bartonella grahamii as4aup YP 002971963 1 634504 Proteobacteria
Bartonella tribocorum CIP 105476 YP 001608913 1 382640 Proteobacteria
Bartonella henselae str Houston 1 YP 033736 1 283166 Proteobacteria
Bartonella grahamii as4aup YP 002971773 1 634504 Proteobacteria
Bartonella tribocorum CIP 105476 YP 001610301 1 382640 Proteobacteria
Bartonella tribocorum CIP 105476 YP 001609415 1 382640 Proteobacteria
Bartonella australis Aust NH1 YP 007461749 1 1094489 Proteobacteria
Bartonella clarridgeiae 73 YP 004159420 1 696125 Proteobacteria
Bartonella bacilliformis KC583 YP 988561 1 360095 Proteobacteria
Bartonella vinsonii subsp berkhoffii str Winnie YP 007463142 1 1094497 Proteobacteria
Bartonella quintana RM 11 YP 006665447 1 1225179 Proteobacteria
Bartonella henselae str Houston 1 YP 034111 1 283166 Proteobacteria
Bartonella tribocorum CIP 105476 YP 001610199 1 382640 Proteobacteria
Bartonella grahamii as4aup YP 002972503 1 634504 Proteobacteria
Hyphomicrobium sp MC1 YP 004675041 1 717785 Proteobacteria
Hyphomicrobium denitrificans ATCC 51888 YP 003754304 1 582899 Proteobacteria
Comamonas testosteroni CNB 2 YP 003277498 1 688245 Proteobacteria
Comamonas testosteroni CNB 2 YP 003278690 1 688245 Proteobacteria
Delftia sp Cs1 4 YP 004489404 1 742013 Proteobacteria
Acidovorax citrulli AAC00 1 YP 970704 1 397945 Proteobacteria
Rhizobium leguminosarum bv trifolii WSM2304 YP 002280665 1 395492 Proteobacteria
Nitrosospira multiformis ATCC 25196 YP 412157 1 323848 Proteobacteria
Achromobacter xylosoxidans NBRC 15126 ATCC 27061 YP 008925264 1 1216976 Proteobacteria
Desulfotalea psychrophila LSv54 YP 065741 1 177439 Proteobacteria
Acidovorax sp KKS102 YP 006854095 1 358220 Proteobacteria
Neisseria gonorrhoeae FA 1090 YP 208176 1 242231 Proteobacteria
Acidovorax citrulli AAC00 1 YP 970061 1 397945 Proteobacteria
Listeria monocytogenes YP 008314150 1 1639 Firmicutes
Edwardsiella ictaluri 93 146 YP 002933421 1 634503 Proteobacteria
DAQ1742 DAQ1742 02629 Daquatica
Dickeya sp CSL RW240 WP 067486825 1 Daquatica
709 DAQ1742 02629 Phage P5 Seed
Escherichia coli O104 H4 str 2009EL 2050 YP 006770144 1 1134782 Proteobacteria
Escherichia coli O103 H2 str 12009 YP 003222733 1 585395 Proteobacteria
Escherichia coli O111 H str 11128 YP 003233661 1 585396 Proteobacteria
Escherichia coli Xuzhou21 YP 006312037 1 741093 Proteobacteria
Escherichia coli O157 H7 str EDL933 NP 286987 1 155864 Proteobacteria
Escherichia coli O104 H4 str 2009EL 2050 YP 006770837 1 1134782 Proteobacteria

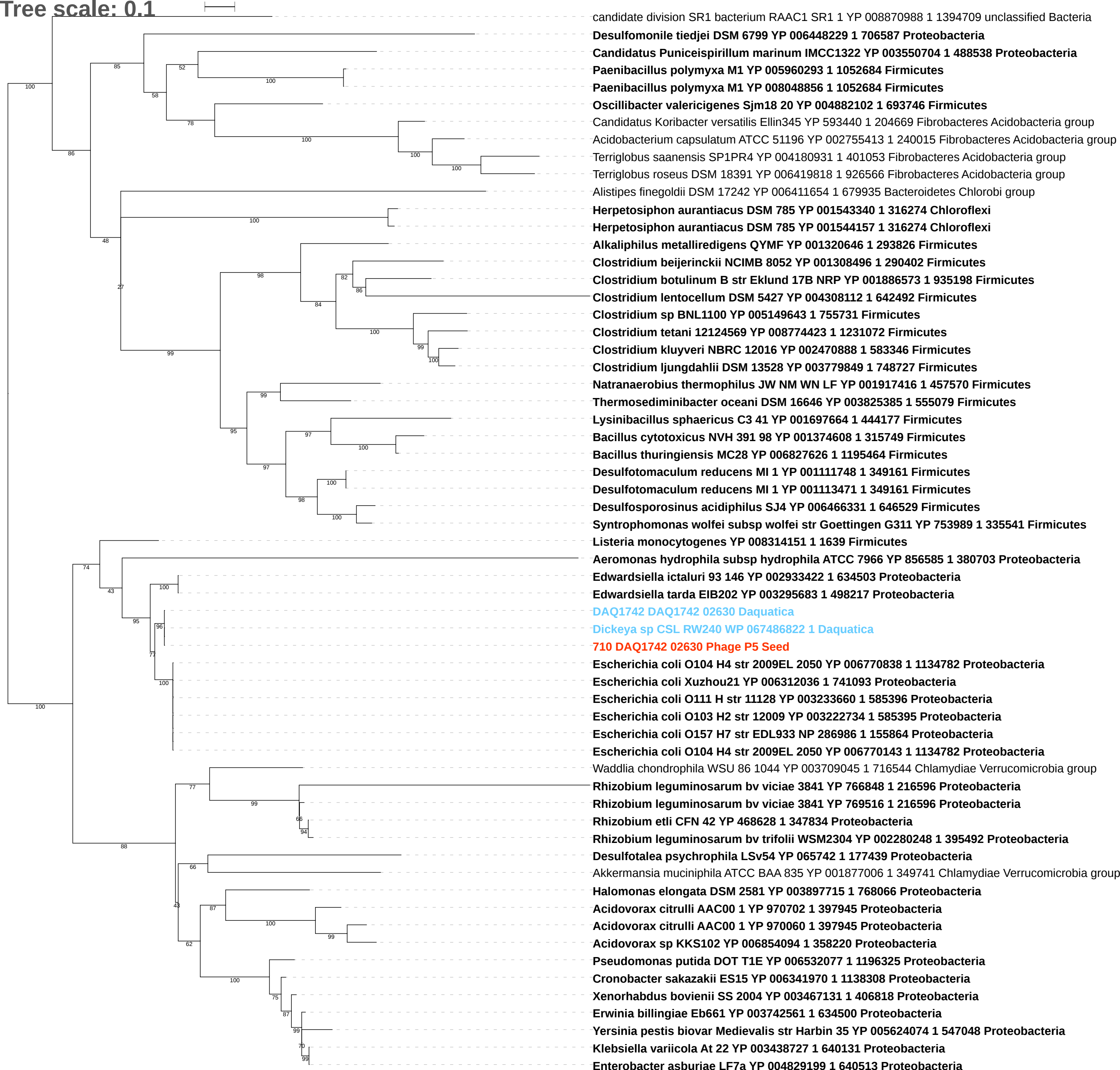
Tree scale: 0.1
candidate division SR1 bacterium RAAC1 SR1 1 YP 008870988 1 1394709 unclassified Bacteria
Desulfomonile tiedjei DSM 6799 YP 006448229 1 706587 Proteobacteria
Candidatus Puniceispirillum marinum IMCC1322 YP 003550704 1 488538 Proteobacteria
Paenibacillus polymyxa M1 YP 005960293 1 1052684 Firmicutes
Paenibacillus polymyxa M1 YP 008048856 1 1052684 Firmicutes
Oscillibacter valericigenes Sjm18 20 YP 004882102 1 693746 Firmicutes
Candidatus Koribacter versatilis Ellin345 YP 593440 1 204669 Fibrobacteres Acidobacteria group
Acidobacterium capsulatum ATCC 51196 YP 002755413 1 240015 Fibrobacteres Acidobacteria group
Terriglobus saanensis SP1PR4 YP 004180931 1 401053 Fibrobacteres Acidobacteria group
Terriglobus roseus DSM 18391 YP 006419818 1 926566 Fibrobacteres Acidobacteria group
Alistipes finegoldii DSM 17242 YP 006411654 1 679935 Bacteroidetes Chlorobi group
Herpetosiphon aurantiacus DSM 785 YP 001543340 1 316274 Chloroflexi
Herpetosiphon aurantiacus DSM 785 YP 001544157 1 316274 Chloroflexi
Alkaliphilus metalliredigens QYMF YP 001320646 1 293826 Firmicutes
Clostridium beijerinckii NCIMB 8052 YP 001308496 1 290402 Firmicutes
Clostridium botulinum B str Eklund 17B NRP YP 001886573 1 935198 Firmicutes
Clostridium lentocellum DSM 5427 YP 004308112 1 642492 Firmicutes
Clostridium sp BNL1100 YP 005149643 1 755731 Firmicutes
Clostridium tetani 12124569 YP 008774423 1 1231072 Firmicutes
Clostridium kluyveri NBRC 12016 YP 002470888 1 583346 Firmicutes
Clostridium ljungdahlii DSM 13528 YP 003779849 1 748727 Firmicutes
Natranaerobius thermophilus JW NM WN LF YP 001917416 1 457570 Firmicutes
Thermosediminibacter oceani DSM 16646 YP 003825385 1 555079 Firmicutes
Lysinibacillus sphaericus C3 41 YP 001697664 1 444177 Firmicutes
Bacillus cytotoxicus NVH 391 98 YP 001374608 1 315749 Firmicutes
Bacillus thuringiensis MC28 YP 006827626 1 1195464 Firmicutes
Desulfotomaculum reducens MI 1 YP 001111748 1 349161 Firmicutes
Desulfotomaculum reducens MI 1 YP 001113471 1 349161 Firmicutes
Desulfosporosinus acidiphilus SJ4 YP 006466331 1 646529 Firmicutes
Syntrophomonas wolfei subsp wolfei str Goettingen G311 YP 753989 1 335541 Firmicutes
Listeria monocytogenes YP 008314151 1 1639 Firmicutes
Aeromonas hydrophila subsp hydrophila ATCC 7966 YP 856585 1 380703 Proteobacteria
Edwardsiella ictaluri 93 146 YP 002933422 1 634503 Proteobacteria
Edwardsiella tarda EIB202 YP 003295683 1 498217 Proteobacteria
DAQ1742 DAQ1742 02630 Daquatica
Dickeya sp CSL RW240 WP 067486822 1 Daquatica
710 DAQ1742 02630 Phage P5 Seed
Escherichia coli O104 H4 str 2009EL 2050 YP 006770838 1 1134782 Proteobacteria
Escherichia coli Xuzhou21 YP 006312036 1 741093 Proteobacteria
Escherichia coli O111 H str 11128 YP 003233660 1 585396 Proteobacteria
Escherichia coli O103 H2 str 12009 YP 003222734 1 585395 Proteobacteria
Escherichia coli O157 H7 str EDL933 NP 286986 1 155864 Proteobacteria
Escherichia coli O104 H4 str 2009EL 2050 YP 006770143 1 1134782 Proteobacteria
Waddlia chondrophila WSU 86 1044 YP 003709045 1 716544 Chlamydiae Verrucomicrobia group
Rhizobium leguminosarum bv viciae 3841 YP 766848 1 216596 Proteobacteria
Rhizobium leguminosarum bv viciae 3841 YP 769516 1 216596 Proteobacteria
Rhizobium etli CFN 42 YP 468628 1 347834 Proteobacteria
Rhizobium leguminosarum bv trifolii WSM2304 YP 002280248 1 395492 Proteobacteria
Desulfotalea psychrophila LSv54 YP 065742 1 177439 Proteobacteria
Akkermansia muciniphila ATCC BAA 835 YP 001877006 1 349741 Chlamydiae Verrucomicrobia group
Halomonas elongata DSM 2581 YP 003897715 1 768066 Proteobacteria
Acidovorax citrulli AAC00 1 YP 970702 1 397945 Proteobacteria
Acidovorax citrulli AAC00 1 YP 970060 1 397945 Proteobacteria
Acidovorax sp KKS102 YP 006854094 1 358220 Proteobacteria
Pseudomonas putida DOT T1E YP 006532077 1 1196325 Proteobacteria
Cronobacter sakazakii ES15 YP 006341970 1 1138308 Proteobacteria
Xenorhabdus bovienii SS 2004 YP 003467131 1 406818 Proteobacteria
Erwinia billingiae Eb661 YP 003742561 1 634500 Proteobacteria
Yersinia pestis biovar Medievalis str Harbin 35 YP 005624074 1 547048 Proteobacteria
Klebsiella variicola At 22 YP 003438727 1 640131 Proteobacteria
Enterobacter asburiae LF7a YP 004829199 1 640513 Proteobacteria

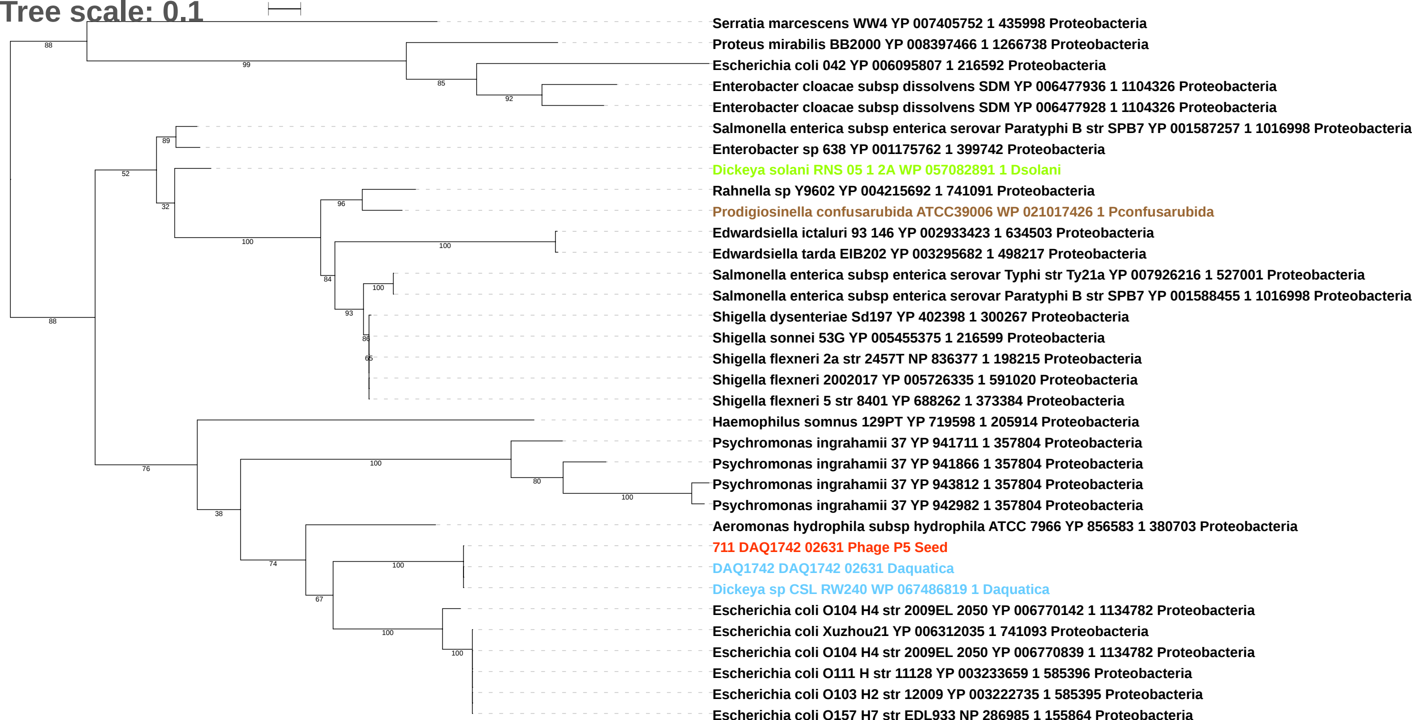
Tree scale: 0.1
Serratia marcescens WW4 YP 007405752 1 435998 Proteobacteria
Proteus mirabilis BB2000 YP 008397466 1 1266738 Proteobacteria
Escherichia coli 042 YP 006095807 1 216592 Proteobacteria
Enterobacter cloacae subsp dissolvens SDM YP 006477936 1 1104326 Proteobacteria
Enterobacter cloacae subsp dissolvens SDM YP 006477928 1 1104326 Proteobacteria
Salmonella enterica subsp enterica serovar Paratyphi B str SPB7 YP 001587257 1 1016998 Proteobacteria
Enterobacter sp 638 YP 001175762 1 399742 Proteobacteria
Dickeya solani RNS 05 1 2A WP 057082891 1 Dsolani
Rahnella sp Y9602 YP 004215692 1 741091 Proteobacteria
Prodigiosinella confusarubida ATCC39006 WP 021017426 1 Pconfusarubida
Edwardsiella ictaluri 93 146 YP 002933423 1 634503 Proteobacteria
Edwardsiella tarda EIB202 YP 003295682 1 498217 Proteobacteria
Salmonella enterica subsp enterica serovar Typhi str Ty21a YP 007926216 1 527001 Proteobacteria
Salmonella enterica subsp enterica serovar Paratyphi B str SPB7 YP 001588455 1 1016998 Proteobacteria
Shigella dysenteriae Sd197 YP 402398 1 300267 Proteobacteria
Shigella sonnei 53G YP 005455375 1 216599 Proteobacteria
Shigella flexneri 2a str 2457T NP 836377 1 198215 Proteobacteria
Shigella flexneri 2002017 YP 005726335 1 591020 Proteobacteria
Shigella flexneri 5 str 8401 YP 688262 1 373384 Proteobacteria
Haemophilus somnus 129PT YP 719598 1 205914 Proteobacteria
Psychromonas ingrahamii 37 YP 941711 1 357804 Proteobacteria
Psychromonas ingrahamii 37 YP 941866 1 357804 Proteobacteria
Psychromonas ingrahamii 37 YP 943812 1 357804 Proteobacteria
Psychromonas ingrahamii 37 YP 942982 1 357804 Proteobacteria
Aeromonas hydrophila subsp hydrophila ATCC 7966 YP 856583 1 380703 Proteobacteria
711 DAQ1742 02631 Phage P5 Seed
DAQ1742 DAQ1742 02631 Daquatica
Dickeya sp CSL RW240 WP 067486819 1 Daquatica
Escherichia coli O104 H4 str 2009EL 2050 YP 006770142 1 1134782 Proteobacteria
Escherichia coli Xuzhou21 YP 006312035 1 741093 Proteobacteria
Escherichia coli O104 H4 str 2009EL 2050 YP 006770839 1 1134782 Proteobacteria
Escherichia coli O111 H str 11128 YP 003233659 1 585396 Proteobacteria
Escherichia coli O103 H2 str 12009 YP 003222735 1 585395 Proteobacteria
Escherichia coli O157 H7 str EDL933 NP 286985 1 155864 Proteobacteria

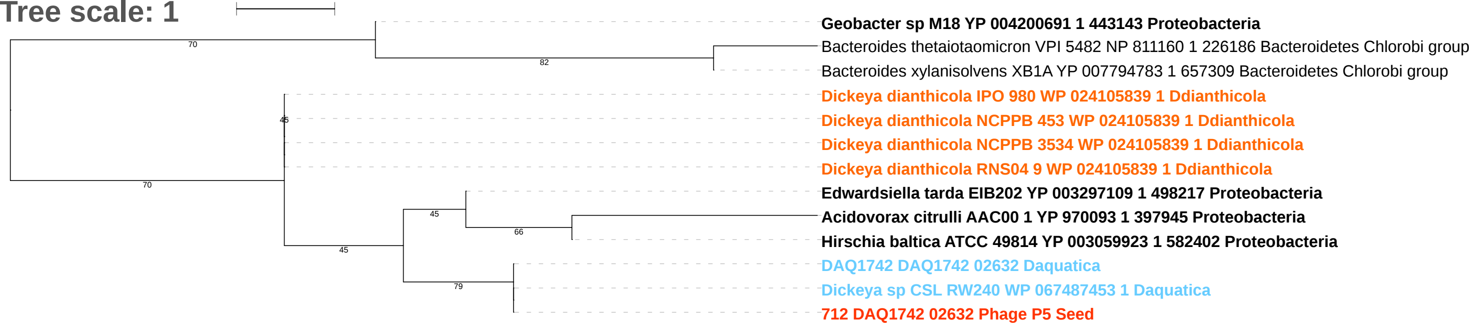

Tree scale: 1
70
82
45
70
45
66
45
79
Geobacter sp M18 YP 004200691 1 443143 Proteobacteria
Bacteroides thetaiotaomicron VPI 5482 NP 811160 1 226186 Bacteroidetes Chlorobi group
Bacteroides xylanisolvens XB1A YP 007794783 1 657309 Bacteroidetes Chlorobi group
Dickeya dianthicola IPO 980 WP 024105839 1 Ddianthicola
Dickeya dianthicola NCPPB 453 WP 024105839 1 Ddianthicola
Dickeya dianthicola NCPPB 3534 WP 024105839 1 Ddianthicola
Dickeya dianthicola RNS04 9 WP 024105839 1 Ddianthicola
Edwardsiella tarda EIB202 YP 003297109 1 498217 Proteobacteria
Acidovorax citrulli AAC00 1 YP 970093 1 397945 Proteobacteria
Hirschia baltica ATCC 49814 YP 003059923 1 582402 Proteobacteria
DAQ1742 DAQ1742 02632 Daquatica
Dickeya sp CSL RW240 WP 067487453 1 Daquatica
712 DAQ1742 02632 Phage P5 Seed

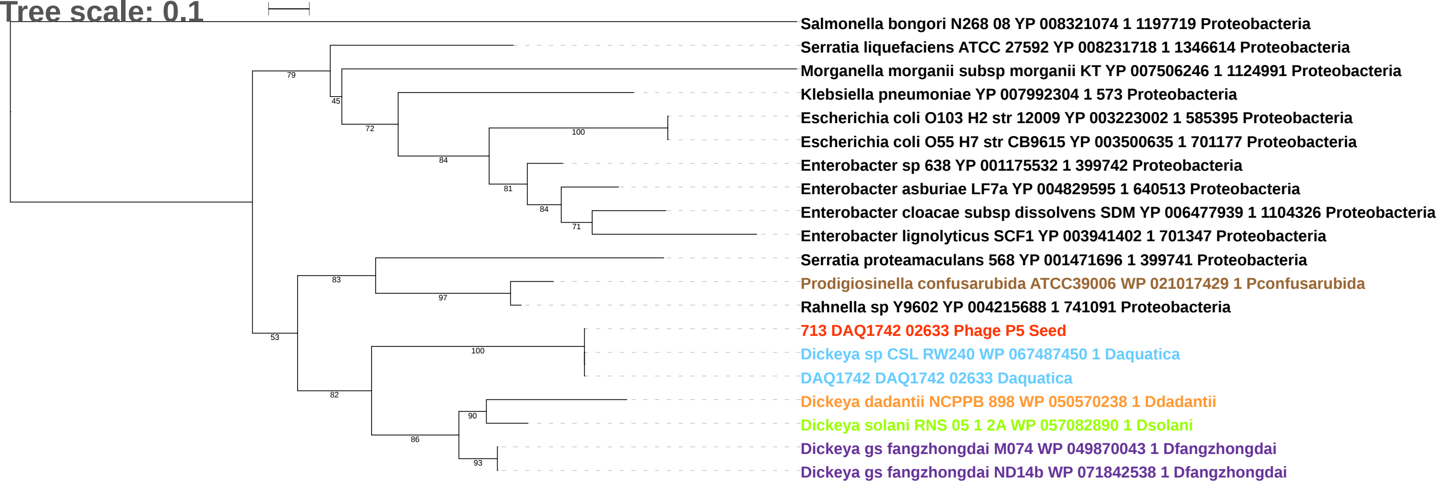
Tree scale: 0.1
Salmonella bongori N268 08 YP 008321074 1 1197719 Proteobacteria
Serratia liquefaciens ATCC 27592 YP 008231718 1 1346614 Proteobacteria
Morganella morganii subsp morganii KT YP 007506246 1 1124991 Proteobacteria
Klebsiella pneumoniae YP 007992304 1 573 Proteobacteria
Escherichia coli O103 H2 str 12009 YP 003223002 1 585395 Proteobacteria
Escherichia coli O55 H7 str CB9615 YP 003500635 1 701177 Proteobacteria
Enterobacter sp 638 YP 001175532 1 399742 Proteobacteria
Enterobacter asburiae LF7a YP 004829595 1 640513 Proteobacteria
Enterobacter cloacae subsp dissolvens SDM YP 006477939 1 1104326 Proteobacteria
Enterobacter lignolyticus SCF1 YP 003941402 1 701347 Proteobacteria
Serratia proteamaculans 568 YP 001471696 1 399741 Proteobacteria
Prodigiosinella confusarubida ATCC39006 WP 021017429 1 Pconfusarubida
Rahnella sp Y9602 YP 004215688 1 741091 Proteobacteria
713 DAQ1742 02633 Phage P5 Seed
Dickeya sp CSL RW240 WP 067487450 1 Daquatica
DAQ1742 DAQ1742 02633 Daquatica
Dickeya dadantii NCPPB 898 WP 050570238 1 Ddadantii
Dickeya solani RNS 05 1 2A WP 057082890 1 Dsolani
Dickeya gs fangzhongdai M074 WP 049870043 1 Dfangzhongdai
Dickeya gs fangzhongdai ND14b WP 071842538 1 Dfangzhongdai
79
45
72
100
84
81
84
71
83
97
53
100
82
90
86
93

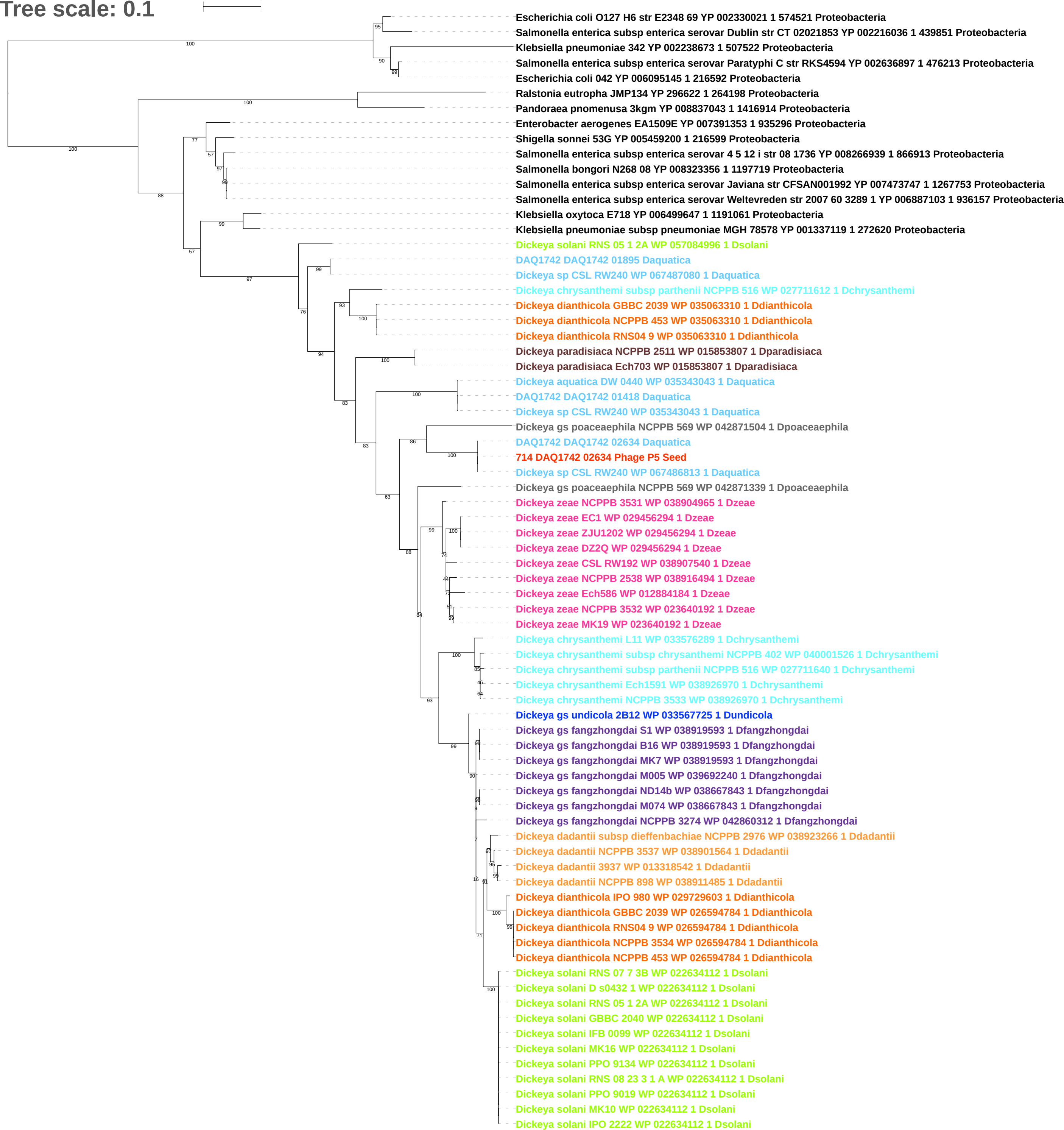
Tree scale: 0.1
Escherichia coli O127 H6 str E2348 69 YP 002330021 1 574521 Proteobacteria
Salmonella enterica subsp enterica serovar Dublin str CT 02021853 YP 002216036 1 439851 Proteobacteria
Klebsiella pneumoniae 342 YP 002238673 1 507522 Proteobacteria
Salmonella enterica subsp enterica serovar Paratyphi C str RKS4594 YP 002636897 1 476213 Proteobacteria
Escherichia coli 042 YP 006095145 1 216592 Proteobacteria
Ralstonia eutropha JMP134 YP 296622 1 264198 Proteobacteria
Pandoraea pnomenusa 3kgm YP 008837043 1 1416914 Proteobacteria
Enterobacter aerogenes EA1509E YP 007391353 1 935296 Proteobacteria
Shigella sonnei 53G YP 005459200 1 216599 Proteobacteria
Salmonella enterica subsp enterica serovar 4 5 12 i str 08 1736 YP 008266939 1 866913 Proteobacteria
Salmonella bongori N268 08 YP 008323356 1 1197719 Proteobacteria
Salmonella enterica subsp enterica serovar Javiana str CFSAN001992 YP 007473747 1 1267753 Proteobacteria
Salmonella enterica subsp enterica serovar Weltevreden str 2007 60 3289 1 YP 006887103 1 936157 Proteobacteria
Klebsiella oxytoca E718 YP 006499647 1 1191061 Proteobacteria
Klebsiella pneumoniae subsp pneumoniae MGH 78578 YP 001337119 1 272620 Proteobacteria
Dickeya solani RNS 05 1 2A WP 057084996 1 Dsolani
DAQ1742 DAQ1742 01895 Daquatica
Dickeya sp CSL RW240 WP 067487080 1 Daquatica
Dickeya chrysanthemi subsp parthenii NCPPB 516 WP 027711612 1 Dchrysanthemi
Dickeya dianthicola GBBC 2039 WP 035063310 1 Ddianthicola
Dickeya dianthicola NCPPB 453 WP 035063310 1 Ddianthicola
Dickeya dianthicola RNS04 9 WP 035063310 1 Ddianthicola
Dickeya paradisiaca NCPPB 2511 WP 015853807 1 Dparadisiaca
Dickeya paradisiaca Ech703 WP 015853807 1 Dparadisiaca
Dickeya aquatica DW 0440 WP 035343043 1 Daquatica
DAQ1742 DAQ1742 01418 Daquatica
Dickeya sp CSL RW240 WP 035343043 1 Daquatica
Dickeya gs poaceaephila NCPPB 569 WP 042871504 1 Dpoaceaephila
DAQ1742 DAQ1742 02634 Daquatica
714 DAQ1742 02634 Phage P5 Seed
Dickeya sp CSL RW240 WP 067486813 1 Daquatica
Dickeya gs poaceaephila NCPPB 569 WP 042871339 1 Dpoaceaephila
Dickeya zeae NCPPB 3531 WP 038904965 1 Dzeae
Dickeya zeae EC1 WP 029456294 1 Dzeae
Dickeya zeae ZJU1202 WP 029456294 1 Dzeae
Dickeya zeae DZ2Q WP 029456294 1 Dzeae
Dickeya zeae CSL RW192 WP 038907540 1 Dzeae
Dickeya zeae NCPPB 2538 WP 038916494 1 Dzeae
Dickeya zeae Ech586 WP 012884184 1 Dzeae
Dickeya zeae NCPPB 3532 WP 023640192 1 Dzeae
Dickeya zeae MK19 WP 023640192 1 Dzeae
Dickeya chrysanthemi L11 WP 033576289 1 Dchrysanthemi
Dickeya chrysanthemi subsp chrysanthemi NCPPB 402 WP 040001526 1 Dchrysanthemi
Dickeya chrysanthemi subsp parthenii NCPPB 516 WP 027711640 1 Dchrysanthemi
Dickeya chrysanthemi Ech1591 WP 038926970 1 Dchrysanthemi
Dickeya chrysanthemi NCPPB 3533 WP 038926970 1 Dchrysanthemi
Dickeya gs undicola 2B12 WP 033567725 1 Dundicola
Dickeya gs fangzhongdai S1 WP 038919593 1 Dfangzhongdai
Dickeya gs fangzhongdai B16 WP 038919593 1 Dfangzhongdai
Dickeya gs fangzhongdai MK7 WP 038919593 1 Dfangzhongdai
Dickeya gs fangzhongdai M005 WP 039692240 1 Dfangzhongdai
Dickeya gs fangzhongdai ND14b WP 038667843 1 Dfangzhongdai
Dickeya gs fangzhongdai M074 WP 038667843 1 Dfangzhongdai
Dickeya gs fangzhongdai NCPPB 3274 WP 042860312 1 Dfangzhongdai
Dickeya dadantii subsp dieffenbachiae NCPPB 2976 WP 038923266 1 Ddadantii
Dickeya dadantii NCPPB 3537 WP 038901564 1 Ddadantii
Dickeya dadantii 3937 WP 013318542 1 Ddadantii
Dickeya dadantii NCPPB 898 WP 038911485 1 Ddadantii
Dickeya dianthicola IPO 980 WP 029729603 1 Ddianthicola
Dickeya dianthicola GBBC 2039 WP 026594784 1 Ddianthicola
Dickeya dianthicola RNS04 9 WP 026594784 1 Ddianthicola
Dickeya dianthicola NCPPB 3534 WP 026594784 1 Ddianthicola
Dickeya dianthicola NCPPB 453 WP 026594784 1 Ddianthicola
Dickeya solani RNS 07 7 3B WP 022634112 1 Dsolani
Dickeya solani D s0432 1 WP 022634112 1 Dsolani
Dickeya solani RNS 05 1 2A WP 022634112 1 Dsolani
Dickeya solani GBBC 2040 WP 022634112 1 Dsolani
Dickeya solani IFB 0099 WP 022634112 1 Dsolani
Dickeya solani MK16 WP 022634112 1 Dsolani
Dickeya solani PPO 9134 WP 022634112 1 Dsolani
Dickeya solani RNS 08 23 3 1 A WP 022634112 1 Dsolani
Dickeya solani PPO 9019 WP 022634112 1 Dsolani
Dickeya solani MK10 WP 022634112 1 Dsolani
Dickeya solani IPO 2222 WP 022634112 1 Dsolani

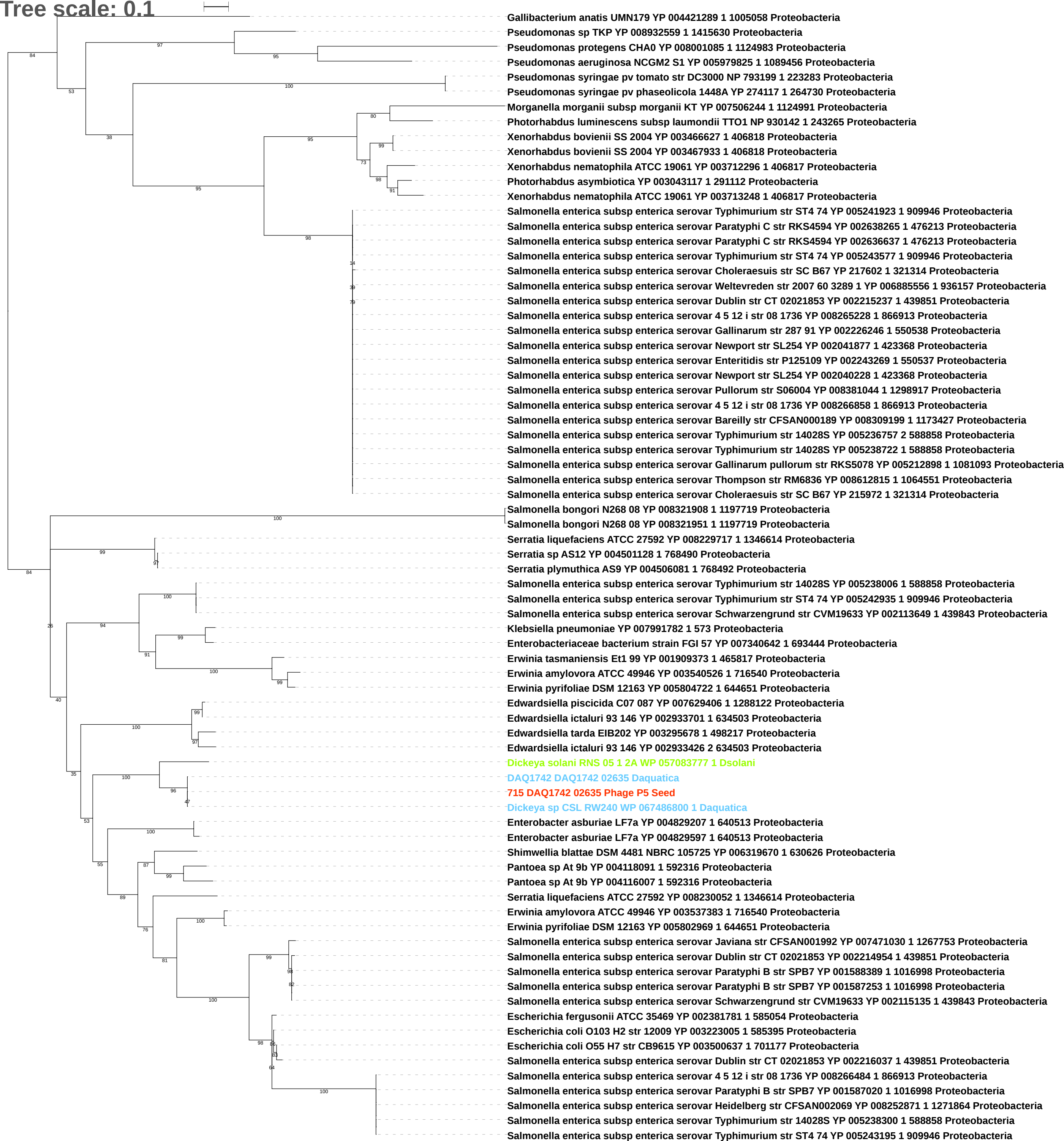
Tree scale: 0.1
Gallibacterium anatis UMN179 YP 004421289 1 1005058 Proteobacteria
Pseudomonas sp TKP YP 008932559 1 1415630 Proteobacteria
Pseudomonas protegens CHA0 YP 008001085 1 1124983 Proteobacteria
Pseudomonas aeruginosa NCGM2 S1 YP 005979825 1 1089456 Proteobacteria
Pseudomonas syringae pv tomato str DC3000 NP 793199 1 223283 Proteobacteria
Pseudomonas syringae pv phaseolicola 1448A YP 274117 1 264730 Proteobacteria
Morganella morganii subsp morganii KT YP 007506244 1 1124991 Proteobacteria
Photorhabdus luminescens subsp laumondii TTO1 NP 930142 1 243265 Proteobacteria
Xenorhabdus bovienii SS 2004 YP 003466627 1 406818 Proteobacteria
Xenorhabdus bovienii SS 2004 YP 003467933 1 406818 Proteobacteria
Xenorhabdus nematophila ATCC 19061 YP 003712296 1 406817 Proteobacteria
Photorhabdus asymbiotica YP 003043117 1 291112 Proteobacteria
Xenorhabdus nematophila ATCC 19061 YP 003713248 1 406817 Proteobacteria
Salmonella enterica subsp enterica serovar Typhimurium str ST4 74 YP 005241923 1 909946 Proteobacteria
Salmonella enterica subsp enterica serovar Paratyphi C str RKS4594 YP 002638265 1 476213 Proteobacteria
Salmonella enterica subsp enterica serovar Paratyphi C str RKS4594 YP 002636637 1 476213 Proteobacteria
Salmonella enterica subsp enterica serovar Typhimurium str ST4 74 YP 005243577 1 909946 Proteobacteria
Salmonella enterica subsp enterica serovar Choleraesuis str SC B67 YP 217602 1 321314 Proteobacteria
Salmonella enterica subsp enterica serovar Weltevreden str 2007 60 3289 1 YP 006885556 1 936157 Proteobacteria
Salmonella enterica subsp enterica serovar Dublin str CT 02021853 YP 002215237 1 439851 Proteobacteria
Salmonella enterica subsp enterica serovar 4 5 12 i str 08 1736 YP 008265228 1 866913 Proteobacteria
Salmonella enterica subsp enterica serovar Gallinarum str 287 91 YP 002226246 1 550538 Proteobacteria
Salmonella enterica subsp enterica serovar Newport str SL254 YP 002041877 1 423368 Proteobacteria
Salmonella enterica subsp enterica serovar Enteritidis str P125109 YP 002243269 1 550537 Proteobacteria
Salmonella enterica subsp enterica serovar Newport str SL254 YP 002040228 1 423368 Proteobacteria
Salmonella enterica subsp enterica serovar Pullorum str S06004 YP 008381044 1 1298917 Proteobacteria
Salmonella enterica subsp enterica serovar 4 5 12 i str 08 1736 YP 008266858 1 866913 Proteobacteria
Salmonella enterica subsp enterica serovar Bareilly str CFSAN000189 YP 008309199 1 1173427 Proteobacteria
Salmonella enterica subsp enterica serovar Typhimurium str 14028S YP 005236757 2 588858 Proteobacteria
Salmonella enterica subsp enterica serovar Typhimurium str 14028S YP 005238722 1 588858 Proteobacteria
Salmonella enterica subsp enterica serovar Gallinarum pullorum str RKS5078 YP 005212898 1 1081093 Proteobacteria
Salmonella enterica subsp enterica serovar Thompson str RM6836 YP 008612815 1 1064551 Proteobacteria
Salmonella enterica subsp enterica serovar Choleraesuis str SC B67 YP 215972 1 321314 Proteobacteria
Salmonella bongori N268 08 YP 008321908 1 1197719 Proteobacteria
Salmonella bongori N268 08 YP 008321951 1 1197719 Proteobacteria
Serratia liquefaciens ATCC 27592 YP 008229717 1 1346614 Proteobacteria
Serratia sp AS12 YP 004501128 1 768490 Proteobacteria
Serratia plymuthica AS9 YP 004506081 1 768492 Proteobacteria
Salmonella enterica subsp enterica serovar Typhimurium str 14028S YP 005238006 1 588858 Proteobacteria
Salmonella enterica subsp enterica serovar Typhimurium str ST4 74 YP 005242935 1 909946 Proteobacteria
Salmonella enterica subsp enterica serovar Schwarzengrund str CVM19633 YP 002113649 1 439843 Proteobacteria
Klebsiella pneumoniae YP 007991782 1 573 Proteobacteria
Enterobacteriaceae bacterium strain FGI 57 YP 007340642 1 693444 Proteobacteria
Erwinia tasmaniensis Et1 99 YP 001909373 1 465817 Proteobacteria
Erwinia amylovora ATCC 49946 YP 003540526 1 716540 Proteobacteria
Erwinia pyrifoliae DSM 12163 YP 005804722 1 644651 Proteobacteria
Edwardsiella piscicida C07 087 YP 007629406 1 1288122 Proteobacteria
Edwardsiella ictaluri 93 146 YP 002933701 1 634503 Proteobacteria
Edwardsiella tarda EIB202 YP 003295678 1 498217 Proteobacteria
Edwardsiella ictaluri 93 146 YP 002933426 2 634503 Proteobacteria
Dickeya solani RNS 05 1 2A WP 057083777 1 Dsolani
DAQ1742 DAQ1742 02635 Daquatica
715 DAQ1742 02635 Phage P5 Seed
Dickeya sp CSL RW240 WP 067486800 1 Daquatica
Enterobacter asburiae LF7a YP 004829207 1 640513 Proteobacteria
Enterobacter asburiae LF7a YP 004829597 1 640513 Proteobacteria
Shimwellia blattae DSM 4481 NBRC 105725 YP 006319670 1 630626 Proteobacteria
Pantoea sp At 9b YP 004118091 1 592316 Proteobacteria
Pantoea sp At 9b YP 004116007 1 592316 Proteobacteria
Serratia liquefaciens ATCC 27592 YP 008230052 1 1346614 Proteobacteria
Erwinia amylovora ATCC 49946 YP 003537383 1 716540 Proteobacteria
Erwinia pyrifoliae DSM 12163 YP 005802969 1 644651 Proteobacteria
Salmonella enterica subsp enterica serovar Javiana str CFSAN001992 YP 007471030 1 1267753 Proteobacteria
Salmonella enterica subsp enterica serovar Dublin str CT 02021853 YP 002214954 1 439851 Proteobacteria
Salmonella enterica subsp enterica serovar Paratyphi B str SPB7 YP 001588389 1 1016998 Proteobacteria
Salmonella enterica subsp enterica serovar Paratyphi B str SPB7 YP 001587253 1 1016998 Proteobacteria
Salmonella enterica subsp enterica serovar Schwarzengrund str CVM19633 YP 002115135 1 439843 Proteobacteria
Escherichia fergusonii ATCC 35469 YP 002381781 1 585054 Proteobacteria
Escherichia coli O103 H2 str 12009 YP 003223005 1 585395 Proteobacteria
Escherichia coli O55 H7 str CB9615 YP 003500637 1 701177 Proteobacteria
Salmonella enterica subsp enterica serovar Dublin str CT 02021853 YP 002216037 1 439851 Proteobacteria
Salmonella enterica subsp enterica serovar 4 5 12 i str 08 1736 YP 008266484 1 866913 Proteobacteria
Salmonella enterica subsp enterica serovar Paratyphi B str SPB7 YP 001587020 1 1016998 Proteobacteria
Salmonella enterica subsp enterica serovar Heidelberg str CFSAN002069 YP 008252871 1 1271864 Proteobacteria
Salmonella enterica subsp enterica serovar Typhimurium str 14028S YP 005238300 1 588858 Proteobacteria
Salmonella enterica subsp enterica serovar Typhimurium str ST4 74 YP 005243195 1 909946 Proteobacteria

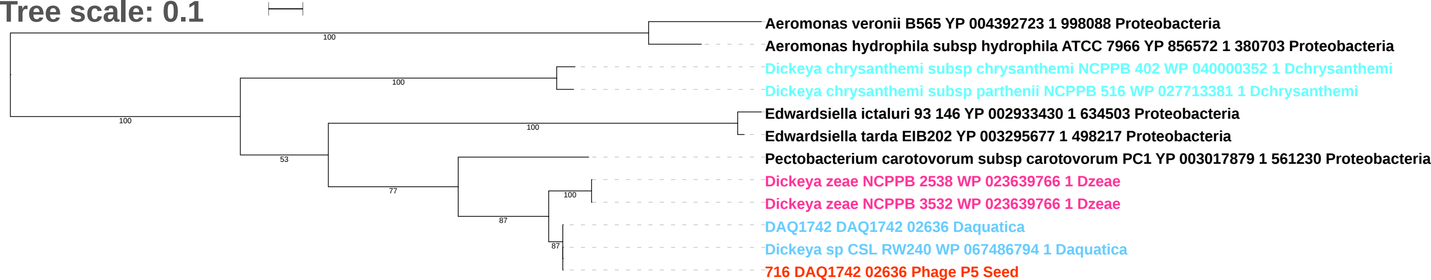

Tree scale: 0.1
100
100
100
100
53
77
100
87
87
Aeromonas veronii B565 YP 004392723 1 998088 Proteobacteria
Aeromonas hydrophila subsp hydrophila ATCC 7966 YP 856572 1 380703 Proteobacteria
Dickeya chrysanthemi subsp chrysanthemi NCPPB 402 WP 040000352 1 Dchrysanthemi
Dickeya chrysanthemi subsp parthenii NCPPB 516 WP 027713381 1 Dchrysanthemi
Edwardsiella ictaluri 93 146 YP 002933430 1 634503 Proteobacteria
Edwardsiella tarda EIB202 YP 003295677 1 498217 Proteobacteria
Pectobacterium carotovorum subsp carotovorum PC1 YP 003017879 1 561230 Proteobacteria
Dickeya zeae NCPPB 2538 WP 023639766 1 Dzeae
Dickeya zeae NCPPB 3532 WP 023639766 1 Dzeae
DAQ1742 DAQ1742 02636 Daquatica
Dickeya sp CSL RW240 WP 067486794 1 Daquatica
716 DAQ1742 02636 Phage P5 Seed

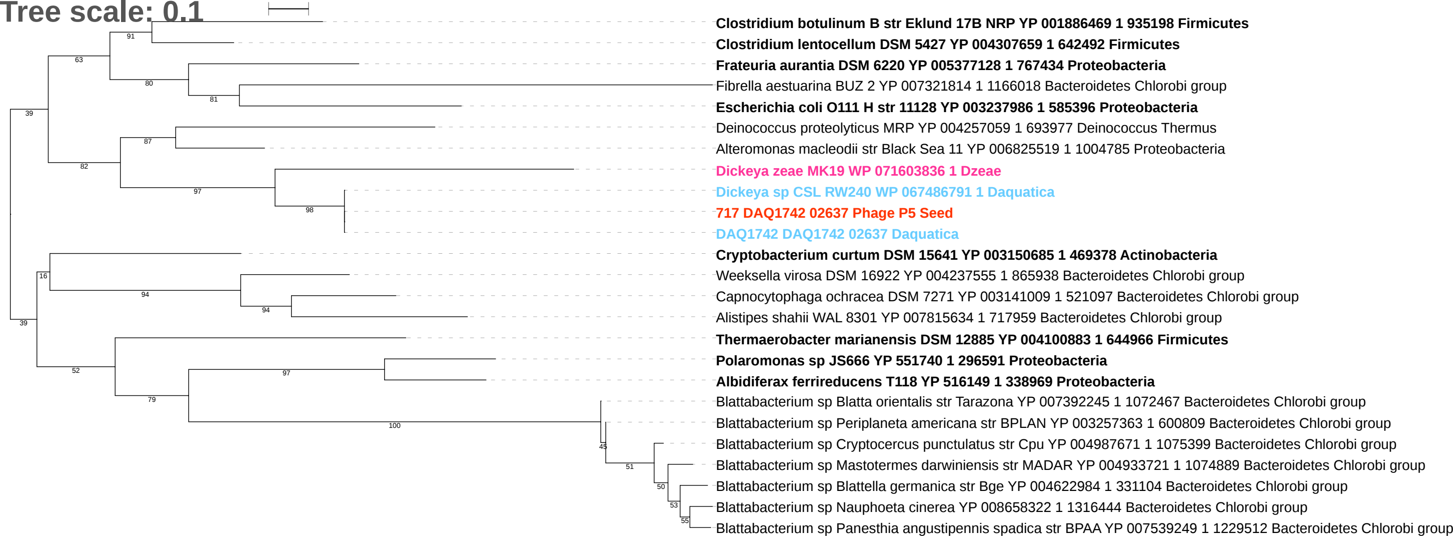
Tree scale: 0.1
Clostridium botulinum B str Eklund 17B NRP YP 001886469 1 935198 Firmicutes
Clostridium lentocellum DSM 5427 YP 004307659 1 642492 Firmicutes
Frateuria aurantia DSM 6220 YP 005377128 1 767434 Proteobacteria
Fibrella aestuarina BUZ 2 YP 007321814 1 1166018 Bacteroidetes Chlorobi group
Escherichia coli O111 H str 11128 YP 003237986 1 585396 Proteobacteria
Deinococcus proteolyticus MRP YP 004257059 1 693977 Deinococcus Thermus
Alteromonas macleodii str Black Sea 11 YP 006825519 1 1004785 Proteobacteria
Dickeya zeae MK19 WP 071603836 1 Dzeae
Dickeya sp CSL RW240 WP 067486791 1 Daquatica
717 DAQ1742 02637 Phage P5 Seed
DAQ1742 DAQ1742 02637 Daquatica
Cryptobacterium curtum DSM 15641 YP 003150685 1 469378 Actinobacteria
Weeksella virosa DSM 16922 YP 004237555 1 865938 Bacteroidetes Chlorobi group
Capnocytophaga ochracea DSM 7271 YP 003141009 1 521097 Bacteroidetes Chlorobi group
Alistipes shahii WAL 8301 YP 007815634 1 717959 Bacteroidetes Chlorobi group
Thermaerobacter marianensis DSM 12885 YP 004100883 1 644966 Firmicutes
Polaromonas sp JS666 YP 551740 1 296591 Proteobacteria
Albidiferax ferrireducens T118 YP 516149 1 338969 Proteobacteria
Blattabacterium sp Blatta orientalis str Tarazona YP 007392245 1 1072467 Bacteroidetes Chlorobi group
Blattabacterium sp Periplaneta americana str BPLAN YP 003257363 1 600809 Bacteroidetes Chlorobi group
Blattabacterium sp Cryptocercus punctulatus str Cpu YP 004987671 1 1075399 Bacteroidetes Chlorobi group
Blattabacterium sp Mastotermes darwiniensis str MADAR YP 004933721 1 1074889 Bacteroidetes Chlorobi group
Blattabacterium sp Blattella germanica str Bge YP 004622984 1 331104 Bacteroidetes Chlorobi group
Blattabacterium sp Nauphoeta cinerea YP 008658322 1 1316444 Bacteroidetes Chlorobi group
Blattabacterium sp Panesthia angustipennis spadica str BPAA YP 007539249 1 1229512 Bacteroidetes Chlorobi group

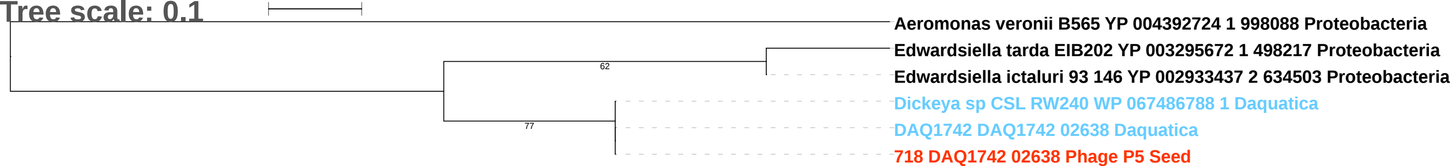
Tree scale: 0.1
Aeromonas veronii B565 YP 004392724 1 998088 Proteobacteria
Edwardsiella tarda EIB202 YP 003295672 1 498217 Proteobacteria
Edwardsiella ictaluri 93 146 YP 002933437 2 634503 Proteobacteria
Dickeya sp CSL RW240 WP 067486788 1 Daquatica
DAQ1742 DAQ1742 02638 Daquatica
718 DAQ1742 02638 Phage P5 Seed
62
77

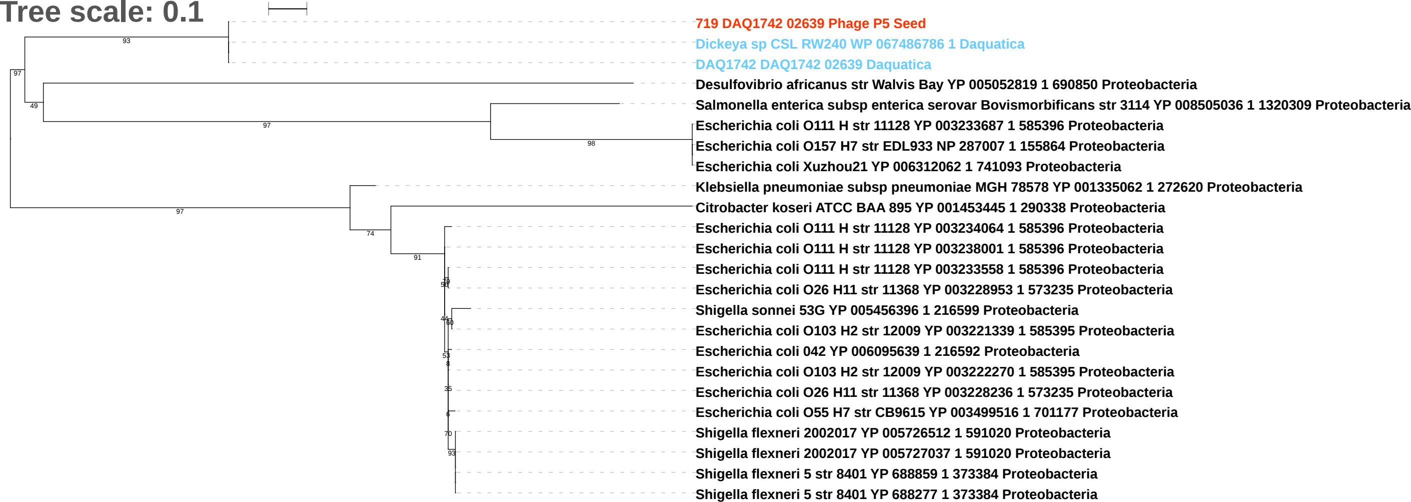
Tree scale: 0.1
719 DAQ1742 02639 Phage P5 Seed
Dickeya sp CSL RW240 WP 067486786 1 Daquatica
DAQ1742 DAQ1742 02639 Daquatica
Desulfovibrio africanus str Walvis Bay YP 005052819 1 690850 Proteobacteria
Salmonella enterica subsp enterica serovar Bovismorbificans str 3114 YP 008505036 1 1320309 Proteobacteria
Escherichia coli O111 H str 11128 YP 003233687 1 585396 Proteobacteria
Escherichia coli O157 H7 str EDL933 NP 287007 1 155864 Proteobacteria
Escherichia coli Xuzhou21 YP 006312062 1 741093 Proteobacteria
Klebsiella pneumoniae subsp pneumoniae MGH 78578 YP 001335062 1 272620 Proteobacteria
Citrobacter koseri ATCC BAA 895 YP 001453445 1 290338 Proteobacteria
Escherichia coli O111 H str 11128 YP 003234064 1 585396 Proteobacteria
Escherichia coli O111 H str 11128 YP 003238001 1 585396 Proteobacteria
Escherichia coli O111 H str 11128 YP 003233558 1 585396 Proteobacteria
Escherichia coli O26 H11 str 11368 YP 003228953 1 573235 Proteobacteria
Shigella sonnei 53G YP 005456396 1 216599 Proteobacteria
Escherichia coli O103 H2 str 12009 YP 003221339 1 585395 Proteobacteria
Escherichia coli 042 YP 006095639 1 216592 Proteobacteria
Escherichia coli O103 H2 str 12009 YP 003222270 1 585395 Proteobacteria
Escherichia coli O26 H11 str 11368 YP 003228236 1 573235 Proteobacteria
Escherichia coli O55 H7 str CB9615 YP 003499516 1 701177 Proteobacteria
Shigella flexneri 2002017 YP 005726512 1 591020 Proteobacteria
Shigella flexneri 2002017 YP 005727037 1 591020 Proteobacteria
Shigella flexneri 5 str 8401 YP 688859 1 373384 Proteobacteria
Shigella flexneri 5 str 8401 YP 688277 1 373384 Proteobacteria

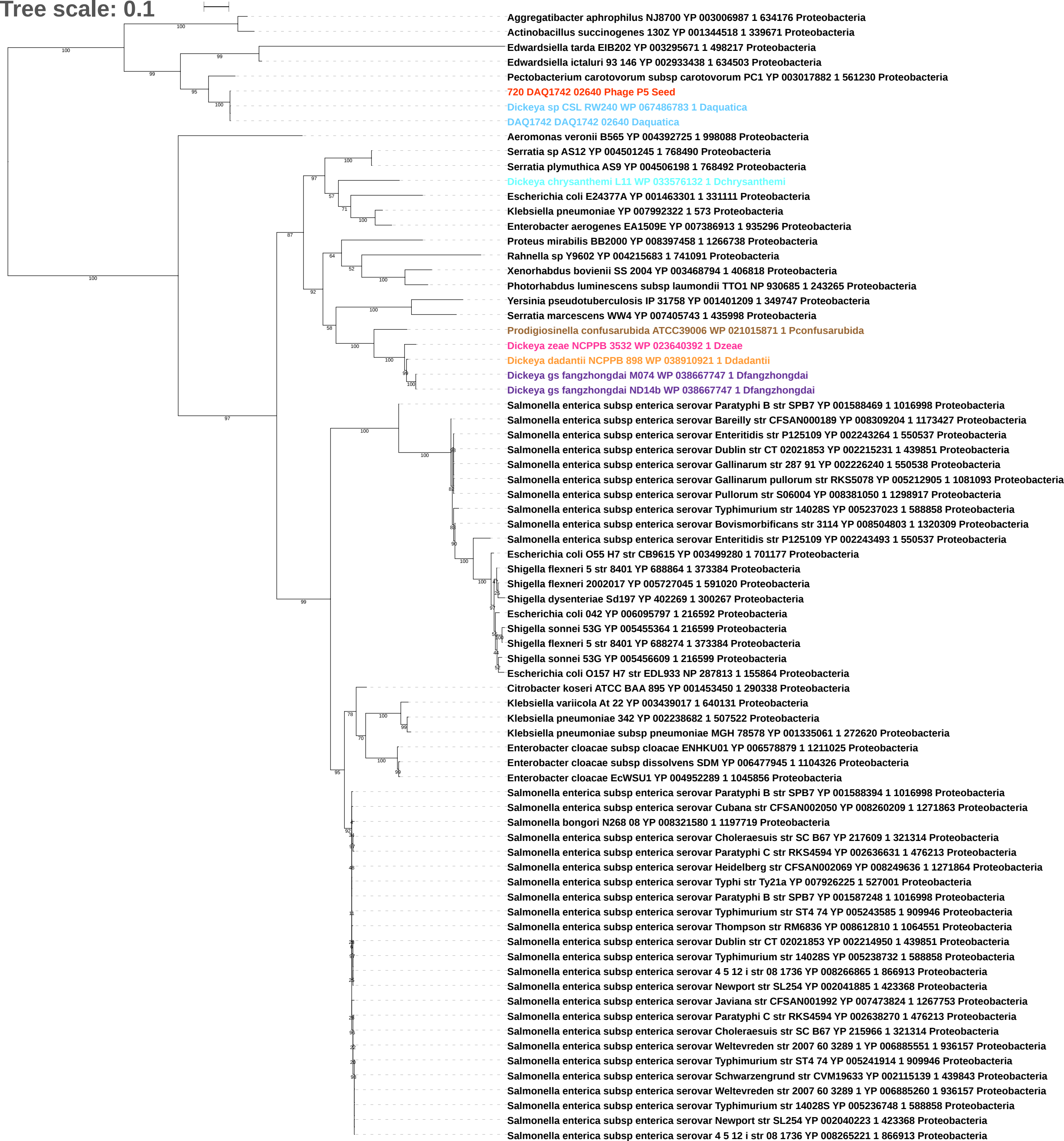

Tree scale: 0.1
Aggregatibacter aphrophilus NJ8700 YP 003006987 1 634176 Proteobacteria
Actinobacillus succinogenes 130Z YP 001344518 1 339671 Proteobacteria
Edwardsiella tarda EIB202 YP 003295671 1 498217 Proteobacteria
Edwardsiella ictaluri 93 146 YP 002933438 1 634503 Proteobacteria
Pectobacterium carotovorum subsp carotovorum PC1 YP 003017882 1 561230 Proteobacteria
720 DAQ1742 02640 Phage P5 Seed
Dickeya sp CSL RW240 WP 067486783 1 Daquatica
DAQ1742 DAQ1742 02640 Daquatica
Aeromonas veronii B565 YP 004392725 1 998088 Proteobacteria
Serratia sp AS12 YP 004501245 1 768490 Proteobacteria
Serratia plymuthica AS9 YP 004506198 1 768492 Proteobacteria
Dickeya chrysanthemi L11 WP 033576132 1 Dchrysanthemi
Escherichia coli E24377A YP 001463301 1 331111 Proteobacteria
Klebsiella pneumoniae YP 007992322 1 573 Proteobacteria
Enterobacter aerogenes EA1509E YP 007386913 1 935296 Proteobacteria
Proteus mirabilis BB2000 YP 008397458 1 1266738 Proteobacteria
Rahnella sp Y9602 YP 004215683 1 741091 Proteobacteria
Xenorhabdus bovienii SS 2004 YP 003468794 1 406818 Proteobacteria
Photorhabdus luminescens subsp laumondii TTO1 NP 930685 1 243265 Proteobacteria
Yersinia pseudotuberculosis IP 31758 YP 001401209 1 349747 Proteobacteria
Serratia marcescens WW4 YP 007405743 1 435998 Proteobacteria
Prodigiosinella confusarubida ATCC39006 WP 021015871 1 Pconfusarubida
Dickeya zeae NCPPB 3532 WP 023640392 1 Dzeae
Dickeya dadantii NCPPB 898 WP 038910921 1 Ddadantii
Dickeya gs fangzhongdai M074 WP 038667747 1 Dfangzhongdai
Dickeya gs fangzhongdai ND14b WP 038667747 1 Dfangzhongdai
Salmonella enterica subsp enterica serovar Paratyphi B str SPB7 YP 001588469 1 1016998 Proteobacteria
Salmonella enterica subsp enterica serovar Bareilly str CFSAN000189 YP 008309204 1 1173427 Proteobacteria
Salmonella enterica subsp enterica serovar Enteritidis str P125109 YP 002243264 1 550537 Proteobacteria
Salmonella enterica subsp enterica serovar Dublin str CT 02021853 YP 002215231 1 439851 Proteobacteria
Salmonella enterica subsp enterica serovar Gallinarum str 287 91 YP 002226240 1 550538 Proteobacteria
Salmonella enterica subsp enterica serovar Gallinarum pullorum str RKS5078 YP 005212905 1 1081093 Proteobacteria
Salmonella enterica subsp enterica serovar Pullorum str S06004 YP 008381050 1 1298917 Proteobacteria
Salmonella enterica subsp enterica serovar Typhimurium str 14028S YP 005237023 1 588858 Proteobacteria
Salmonella enterica subsp enterica serovar Bovismorbificans str 3114 YP 008504803 1 1320309 Proteobacteria
Salmonella enterica subsp enterica serovar Enteritidis str P125109 YP 002243493 1 550537 Proteobacteria
Escherichia coli O55 H7 str CB9615 YP 003499280 1 701177 Proteobacteria
Shigella flexneri 5 str 8401 YP 688864 1 373384 Proteobacteria
Shigella flexneri 2002017 YP 005727045 1 591020 Proteobacteria
Shigella dysenteriae Sd197 YP 402269 1 300267 Proteobacteria
Escherichia coli 042 YP 006095797 1 216592 Proteobacteria
Shigella sonnei 53G YP 005455364 1 216599 Proteobacteria
Shigella flexneri 5 str 8401 YP 688274 1 373384 Proteobacteria
Shigella sonnei 53G YP 005456609 1 216599 Proteobacteria
Escherichia coli O157 H7 str EDL933 NP 287813 1 155864 Proteobacteria
Citrobacter koseri ATCC BAA 895 YP 001453450 1 290338 Proteobacteria
Klebsiella variicola At 22 YP 003439017 1 640131 Proteobacteria
Klebsiella pneumoniae 342 YP 002238682 1 507522 Proteobacteria
Klebsiella pneumoniae subsp pneumoniae MGH 78578 YP 001335061 1 272620 Proteobacteria
Enterobacter cloacae subsp cloacae ENHKU01 YP 006578879 1 1211025 Proteobacteria
Enterobacter cloacae subsp dissolvens SDM YP 006477945 1 1104326 Proteobacteria
Enterobacter cloacae EcWSU1 YP 004952289 1 1045856 Proteobacteria
Salmonella enterica subsp enterica serovar Paratyphi B str SPB7 YP 001588394 1 1016998 Proteobacteria
Salmonella enterica subsp enterica serovar Cubana str CFSAN002050 YP 008260209 1 1271863 Proteobacteria
Salmonella bongori N268 08 YP 008321580 1 1197719 Proteobacteria
Salmonella enterica subsp enterica serovar Choleraesuis str SC B67 YP 217609 1 321314 Proteobacteria
Salmonella enterica subsp enterica serovar Paratyphi C str RKS4594 YP 002636631 1 476213 Proteobacteria
Salmonella enterica subsp enterica serovar Heidelberg str CFSAN002069 YP 008249636 1 1271864 Proteobacteria
Salmonella enterica subsp enterica serovar Typhi str Ty21a YP 007926225 1 527001 Proteobacteria
Salmonella enterica subsp enterica serovar Paratyphi B str SPB7 YP 001587248 1 1016998 Proteobacteria
Salmonella enterica subsp enterica serovar Typhimurium str ST4 74 YP 005243585 1 909946 Proteobacteria
Salmonella enterica subsp enterica serovar Thompson str RM6836 YP 008612810 1 1064551 Proteobacteria
Salmonella enterica subsp enterica serovar Dublin str CT 02021853 YP 002214950 1 439851 Proteobacteria
Salmonella enterica subsp enterica serovar Typhimurium str 14028S YP 005238732 1 588858 Proteobacteria
Salmonella enterica subsp enterica serovar 4 5 12 i str 08 1736 YP 008266865 1 866913 Proteobacteria
Salmonella enterica subsp enterica serovar Newport str SL254 YP 002041885 1 423368 Proteobacteria
Salmonella enterica subsp enterica serovar Javiana str CFSAN001992 YP 007473824 1 1267753 Proteobacteria
Salmonella enterica subsp enterica serovar Paratyphi C str RKS4594 YP 002638270 1 476213 Proteobacteria
Salmonella enterica subsp enterica serovar Choleraesuis str SC B67 YP 215966 1 321314 Proteobacteria
Salmonella enterica subsp enterica serovar Weltevreden str 2007 60 3289 1 YP 006885551 1 936157 Proteobacteria
Salmonella enterica subsp enterica serovar Typhimurium str ST4 74 YP 005241914 1 909946 Proteobacteria
Salmonella enterica subsp enterica serovar Schwarzengrund str CVM19633 YP 002115139 1 439843 Proteobacteria
Salmonella enterica subsp enterica serovar Weltevreden str 2007 60 3289 1 YP 006885260 1 936157 Proteobacteria
Salmonella enterica subsp enterica serovar Typhimurium str 14028S YP 005236748 1 588858 Proteobacteria
Salmonella enterica subsp enterica serovar Newport str SL254 YP 002040223 1 423368 Proteobacteria
Salmonella enterica subsp enterica serovar 4 5 12 i str 08 1736 YP 008265221 1 866913 Proteobacteria

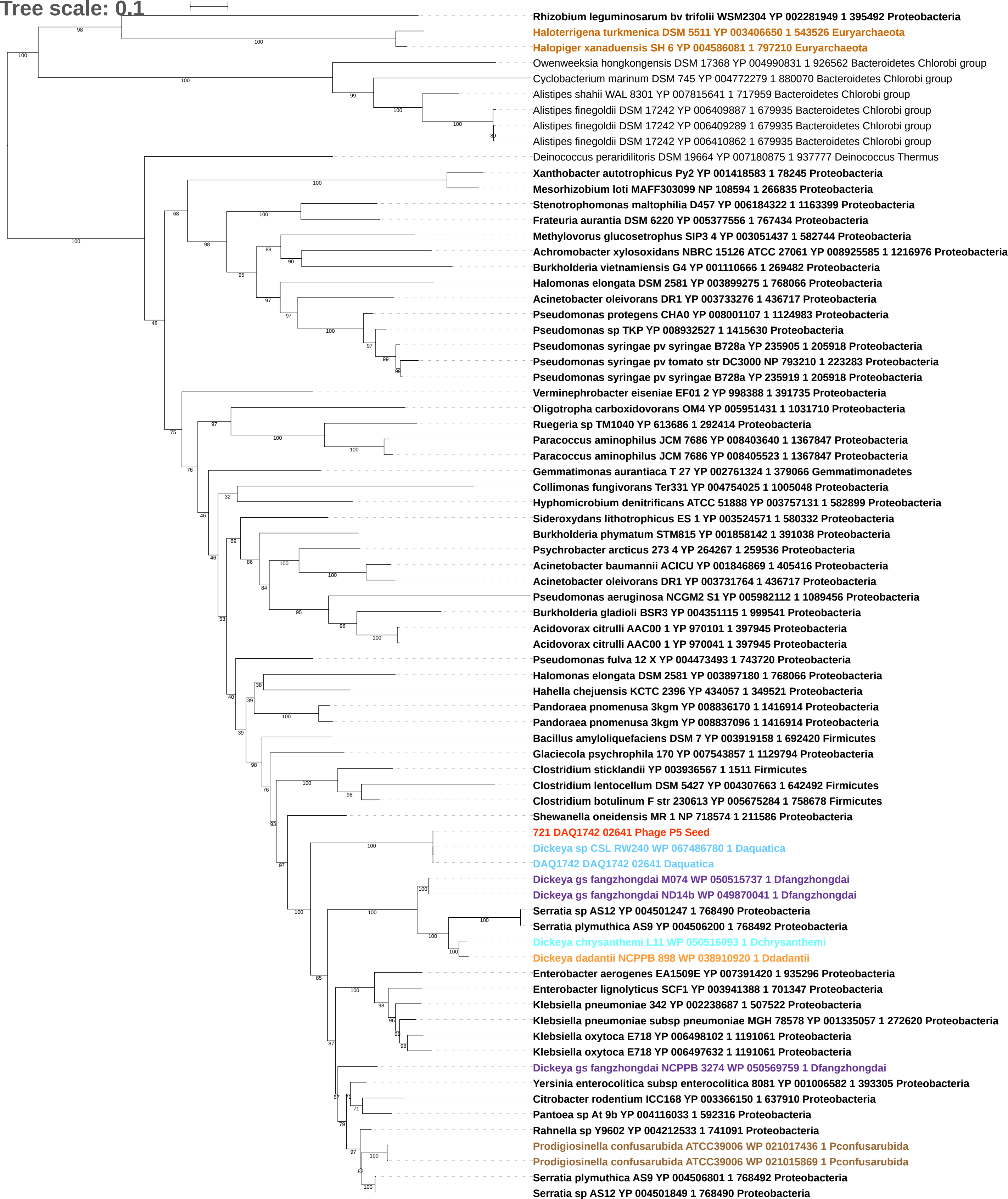

Tree scale: 0.1
Rhizobium leguminosarum bv trifolii WSM2304 YP 002281949 1 395492 Proteobacteria
Haloterrigena turkmenica DSM 5511 YP 003406650 1 543526 Euryarchaeota
Halopiger xanaduensis SH 6 YP 004586081 1 797210 Euryarchaeota
Owenweeksia hongkongensis DSM 17368 YP 004990831 1 926562 Bacteroidetes Chlorobi group
Cyclobacterium marinum DSM 745 YP 004772279 1 880070 Bacteroidetes Chlorobi group
Alistipes shahii WAL 8301 YP 007815641 1 717959 Bacteroidetes Chlorobi group
Alistipes finegoldii DSM 17242 YP 006409887 1 679935 Bacteroidetes Chlorobi group
Alistipes finegoldii DSM 17242 YP 006409289 1 679935 Bacteroidetes Chlorobi group
Alistipes finegoldii DSM 17242 YP 006410862 1 679935 Bacteroidetes Chlorobi group
Deinococcus perariditoris DSM 19664 YP 007180875 1 937777 Deinococcus Thermus
Xanthobacter autotrophicus Py2 YP 001418583 1 78245 Proteobacteria
Mesorhizobium loti MAFF303099 NP 108594 1 266835 Proteobacteria
Stenotrophomonas maltophilia D457 YP 006184322 1 1163399 Proteobacteria
Frateuria aurantia DSM 6220 YP 005377556 1 767434 Proteobacteria
Methylovorus glucosetrophus SIP3 4 YP 003051437 1 582744 Proteobacteria
Achromobacter xylosoxidans NBRC 15126 ATCC 27061 YP 008925585 1 1216976 Proteobacteria
Burkholderia vietnamiensis G4 YP 001110666 1 269482 Proteobacteria
Halomonas elongata DSM 2581 YP 003899275 1 768066 Proteobacteria
Acinetobacter oleivorans DR1 YP 003733276 1 436717 Proteobacteria
Pseudomonas protegens CHA0 YP 008001107 1 1124983 Proteobacteria
Pseudomonas sp TKP YP 008932527 1 1415630 Proteobacteria
Pseudomonas syringae pv syringae B728a YP 235905 1 205918 Proteobacteria
Pseudomonas syringae pv tomato str DC3000 NP 793210 1 223283 Proteobacteria
Pseudomonas syringae pv syringae B728a YP 235919 1 205918 Proteobacteria
Verminephrobacter eiseniae EF01 2 YP 998388 1 391735 Proteobacteria
Oligotropha carboxidovorans OM4 YP 005951431 1 1031710 Proteobacteria
Ruegeria sp TM1040 YP 613686 1 292414 Proteobacteria
Paracoccus aminophilus JCM 7686 YP 008403640 1 1367847 Proteobacteria
Paracoccus aminophilus JCM 7686 YP 008405523 1 1367847 Proteobacteria
Gemmatimonas aurantiaca T 27 YP 002761324 1 379066 Gemmatimonadetes
Collimonas fungivorans Ter331 YP 004754025 1 1005048 Proteobacteria
Hyphomicrobium denitrificans ATCC 51888 YP 003757131 1 582899 Proteobacteria
Sideroxydans lithotrophicus ES 1 YP 003524571 1 580332 Proteobacteria
Burkholderia phymatum STM815 YP 001858142 1 391038 Proteobacteria
Psychrobacter arcticus 273 4 YP 264267 1 259536 Proteobacteria
Acinetobacter baumannii ACICU YP 001846869 1 405416 Proteobacteria
Acinetobacter oleivorans DR1 YP 003731764 1 436717 Proteobacteria
Pseudomonas aeruginosa NCGM2 S1 YP 005982112 1 1089456 Proteobacteria
Burkholderia gladioli BSR3 YP 004351115 1 999541 Proteobacteria
Acidovorax citrulli AAC00 1 YP 970101 1 397945 Proteobacteria
Acidovorax citrulli AAC00 1 YP 970041 1 397945 Proteobacteria
Pseudomonas fulva 12 X YP 004473493 1 743720 Proteobacteria
Halomonas elongata DSM 2581 YP 003897180 1 768066 Proteobacteria
Hahella chejuensis KCTC 2396 YP 434057 1 349521 Proteobacteria
Pandoraea pnomenusa 3kgm YP 008836170 1 1416914 Proteobacteria
Pandoraea pnomenusa 3kgm YP 008837096 1 1416914 Proteobacteria
Bacillus amyloliquefaciens DSM 7 YP 003919158 1 692420 Firmicutes
Glaciecola psychrophila 170 YP 007543857 1 1129794 Proteobacteria
Clostridium sticklandii YP 003936567 1 1511 Firmicutes
Clostridium lentocellum DSM 5427 YP 004307663 1 642492 Firmicutes
Clostridium botulinum F str 230613 YP 005675284 1 758678 Firmicutes
Shewanella oneidensis MR 1 NP 718574 1 211586 Proteobacteria
721 DAQ1742 02641 Phage P5 Seed
Dickeya sp CSL RW240 WP 067486780 1 Daquatica
DAQ1742 DAQ1742 02641 Daquatica
Dickeya gs fangzhongdai M074 WP 050515737 1 Dfangzhongdai
Dickeya gs fangzhongdai ND14b WP 049870041 1 Dfangzhongdai
Serratia sp AS12 YP 004501247 1 768490 Proteobacteria
Serratia plymuthica AS9 YP 004506200 1 768492 Proteobacteria
Dickeya chrysanthemi L11 WP 050516093 1 Dchrysanthemi
Dickeya dadantii NCPPB 898 WP 038910920 1 Ddadantii
Enterobacter aerogenes EA1509E YP 007391420 1 935296 Proteobacteria
Enterobacter lignolyticus SCF1 YP 003941388 1 701347 Proteobacteria
Klebsiella pneumoniae 342 YP 002238687 1 507522 Proteobacteria
Klebsiella pneumoniae subsp pneumoniae MGH 78578 YP 001335057 1 272620 Proteobacteria
Klebsiella oxytoca E718 YP 006498102 1 1191061 Proteobacteria
Klebsiella oxytoca E718 YP 006497632 1 1191061 Proteobacteria
Dickeya gs fangzhongdai NCPPB 3274 WP 050569759 1 Dfangzhongdai
Yersinia enterocolitica subsp enterocolitica 8081 YP 001006582 1 393305 Proteobacteria
Citrobacter rodentium ICC168 YP 003366150 1 637910 Proteobacteria
Pantoea sp At 9b YP 004116033 1 592316 Proteobacteria
Rahnella sp Y9602 YP 004212533 1 741091 Proteobacteria
Prodigiosinella confusarubida ATCC39006 WP 021017436 1 Pconfusarubida
Prodigiosinella confusarubida ATCC39006 WP 021015869 1 Pconfusarubida
Serratia plymuthica AS9 YP 004506801 1 768492 Proteobacteria
Serratia sp AS12 YP 004501849 1 768490 Proteobacteria

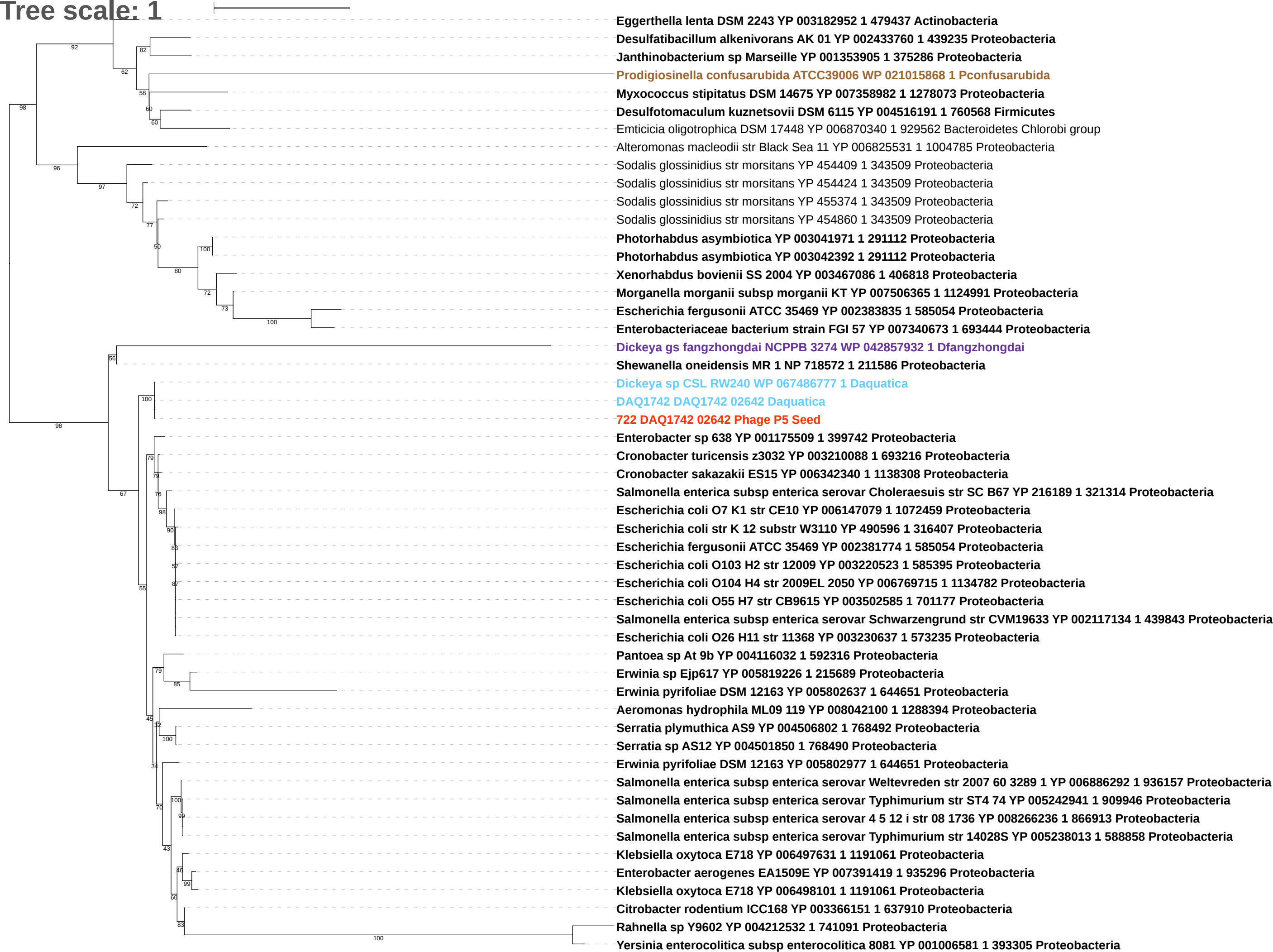
Tree scale: 1
Eggerthella lenta DSM 2243 YP 003182952 1 479437 Actinobacteria
Desulfatibacillum alkenivorans AK 01 YP 002433760 1 439235 Proteobacteria
Janthinobacterium sp Marseille YP 001353905 1 375286 Proteobacteria
Prodigiosinella confusarubida ATCC39006 WP 021015868 1 Pconfusarubida
Myxococcus stipitatus DSM 14675 YP 007358982 1 1278073 Proteobacteria
Desulfotomaculum kuznetsovii DSM 6115 YP 004516191 1 760568 Firmicutes
Emticicia oligotrophica DSM 17448 YP 006870340 1 929562 Bacteroidetes Chlorobi group
Alteromonas macleodii str Black Sea 11 YP 006825531 1 1004785 Proteobacteria
Sodalis glossinidius str morsitans YP 454409 1 343509 Proteobacteria
Sodalis glossinidius str morsitans YP 454424 1 343509 Proteobacteria
Sodalis glossinidius str morsitans YP 455374 1 343509 Proteobacteria
Sodalis glossinidius str morsitans YP 454860 1 343509 Proteobacteria
Photorhabdus asymbiotica YP 003041971 1 291112 Proteobacteria
Photorhabdus asymbiotica YP 003042392 1 291112 Proteobacteria
Xenorhabdus bovienii SS 2004 YP 003467086 1 406818 Proteobacteria
Morganella morganii subsp morganii KT YP 007506365 1 1124991 Proteobacteria
Escherichia fergusonii ATCC 35469 YP 002383835 1 585054 Proteobacteria
Enterobacteriaceae bacterium strain FGI 57 YP 007340673 1 693444 Proteobacteria
Dickeya gs fangzhongdai NCPPB 3274 WP 042857932 1 Dfangzhongdai
Shewanella oneidensis MR 1 NP 718572 1 211586 Proteobacteria
Dickeya sp CSL RW240 WP 067486777 1 Daquatica
DAQ1742 DAQ1742 02642 Daquatica
722 DAQ1742 02642 Phage P5 Seed
Enterobacter sp 638 YP 001175509 1 399742 Proteobacteria
Cronobacter turicensis z3032 YP 003210088 1 693216 Proteobacteria
Cronobacter sakazakii ES15 YP 006342340 1 1138308 Proteobacteria
Salmonella enterica subsp enterica serovar Choleraesuis str SC B67 YP 216189 1 321314 Proteobacteria
Escherichia coli O7 K1 str CE10 YP 006147079 1 1072459 Proteobacteria
Escherichia coli str K 12 substr W3110 YP 490596 1 316407 Proteobacteria
Escherichia fergusonii ATCC 35469 YP 002381774 1 585054 Proteobacteria
Escherichia coli O103 H2 str 12009 YP 003220523 1 585395 Proteobacteria
Escherichia coli O104 H4 str 2009EL 2050 YP 006769715 1 1134782 Proteobacteria
Escherichia coli O55 H7 str CB9615 YP 003502585 1 701177 Proteobacteria
Salmonella enterica subsp enterica serovar Schwarzengrund str CVM19633 YP 002117134 1 439843 Proteobacteria
Escherichia coli O26 H11 str 11368 YP 003230637 1 573235 Proteobacteria
Pantoea sp At 9b YP 004116032 1 592316 Proteobacteria
Erwinia sp Ejp617 YP 005819226 1 215689 Proteobacteria
Erwinia pyrifoliae DSM 12163 YP 005802637 1 644651 Proteobacteria
Aeromonas hydrophila ML09 119 YP 008042100 1 1288394 Proteobacteria
Serratia plymuthica AS9 YP 004506802 1 768492 Proteobacteria
Serratia sp AS12 YP 004501850 1 768490 Proteobacteria
Erwinia pyrifoliae DSM 12163 YP 005802977 1 644651 Proteobacteria
Salmonella enterica subsp enterica serovar Weltevreden str 2007 60 3289 1 YP 006886292 1 936157 Proteobacteria
Salmonella enterica subsp enterica serovar Typhimurium str ST4 74 YP 005242941 1 909946 Proteobacteria
Salmonella enterica subsp enterica serovar 4 5 12 i str 08 1736 YP 008266236 1 866913 Proteobacteria
Salmonella enterica subsp enterica serovar Typhimurium str 14028S YP 005238013 1 588858 Proteobacteria
Klebsiella oxytoca E718 YP 006497631 1 1191061 Proteobacteria
Enterobacter aerogenes EA1509E YP 007391419 1 935296 Proteobacteria
Klebsiella oxytoca E718 YP 006498101 1 1191061 Proteobacteria
Citrobacter rodentium ICC168 YP 003366151 1 637910 Proteobacteria
Rahnella sp Y9602 YP 004212532 1 741091 Proteobacteria
Yersinia enterocolitica subsp enterocolitica 8081 YP 001006581 1 393305 Proteobacteria

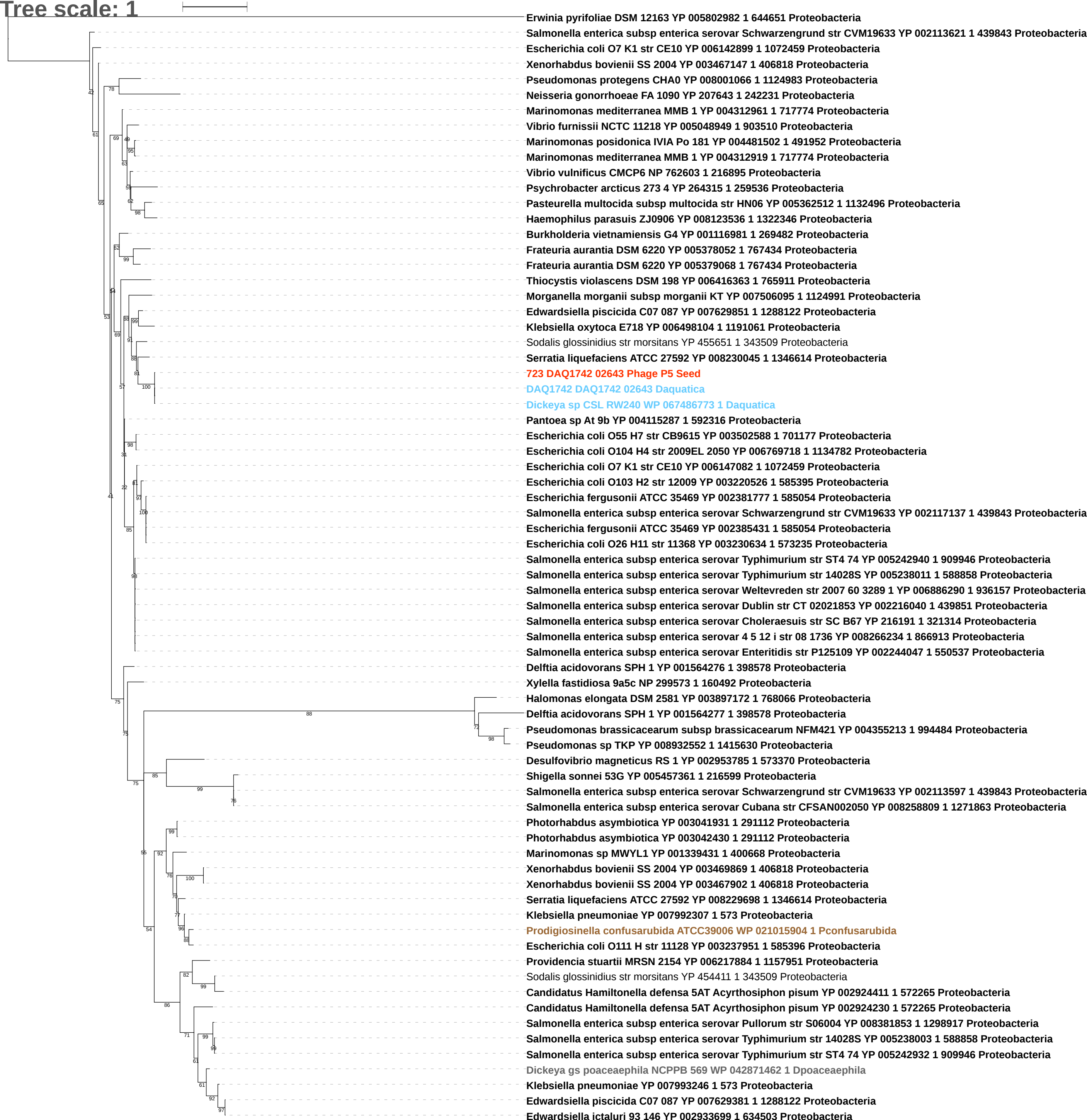
Tree scale: 1
Erwinia pyrifoliae DSM 12163 YP 005802982 1 644651 Proteobacteria
Salmonella enterica subsp enterica serovar Schwarzengrund str CVM19633 YP 002113621 1 439843 Proteobacteria
Escherichia coli O7 K1 str CE10 YP 006142899 1 1072459 Proteobacteria
Xenorhabdus bovienii SS 2004 YP 003467147 1 406818 Proteobacteria
Pseudomonas protegens CHA0 YP 008001066 1 1124983 Proteobacteria
Neisseria gonorrhoeae FA 1090 YP 207643 1 242231 Proteobacteria
Marinomonas mediterranea MMB 1 YP 004312961 1 717774 Proteobacteria
Vibrio furnissii NCTC 11218 YP 005048949 1 903510 Proteobacteria
Marinomonas posidonica IVIA Po 181 YP 004481502 1 491952 Proteobacteria
Marinomonas mediterranea MMB 1 YP 004312919 1 717774 Proteobacteria
Vibrio vulnificus CMCP6 NP 762603 1 216895 Proteobacteria
Psychrobacter arcticus 273 4 YP 264315 1 259536 Proteobacteria
Pasteurella multocida subsp multocida str HN06 YP 005362512 1 1132496 Proteobacteria
Haemophilus parasuis ZJ0906 YP 008123536 1 1322346 Proteobacteria
Burkholderia vietnamiensis G4 YP 001116981 1 269482 Proteobacteria
Frateuria aurantia DSM 6220 YP 005378052 1 767434 Proteobacteria
Frateuria aurantia DSM 6220 YP 005379068 1 767434 Proteobacteria
Thiocystis violascens DSM 198 YP 006416363 1 765911 Proteobacteria
Morganella morganii subsp morganii KT YP 007506095 1 1124991 Proteobacteria
Edwardsiella piscicida C07 087 YP 007629851 1 1288122 Proteobacteria
Klebsiella oxytoca E718 YP 006498104 1 1191061 Proteobacteria
Sodalis glossinidius str morsitans YP 455651 1 343509 Proteobacteria
Serratia liquefaciens ATCC 27592 YP 008230045 1 1346614 Proteobacteria
723 DAQ1742 02643 Phage P5 Seed
DAQ1742 DAQ1742 02643 Daquatica
Dickeya sp CSL RW240 WP 067486773 1 Daquatica
Pantoea sp At 9b YP 004115287 1 592316 Proteobacteria
Escherichia coli O55 H7 str CB9615 YP 003502588 1 701177 Proteobacteria
Escherichia coli O104 H4 str 2009EL 2050 YP 006769718 1 1134782 Proteobacteria
Escherichia coli O7 K1 str CE10 YP 006147082 1 1072459 Proteobacteria
Escherichia coli O103 H2 str 12009 YP 003220526 1 585395 Proteobacteria
Escherichia fergusonii ATCC 35469 YP 002381777 1 585054 Proteobacteria
Salmonella enterica subsp enterica serovar Schwarzengrund str CVM19633 YP 002117137 1 439843 Proteobacteria
Escherichia fergusonii ATCC 35469 YP 002385431 1 585054 Proteobacteria
Escherichia coli O26 H11 str 11368 YP 003230634 1 573235 Proteobacteria
Salmonella enterica subsp enterica serovar Typhimurium str ST4 74 YP 005242940 1 909946 Proteobacteria
Salmonella enterica subsp enterica serovar Typhimurium str 14028S YP 005238011 1 588858 Proteobacteria
Salmonella enterica subsp enterica serovar Weltevreden str 2007 60 3289 1 YP 006886290 1 936157 Proteobacteria
Salmonella enterica subsp enterica serovar Dublin str CT 02021853 YP 002216040 1 439851 Proteobacteria
Salmonella enterica subsp enterica serovar Choleraesuis str SC B67 YP 216191 1 321314 Proteobacteria
Salmonella enterica subsp enterica serovar 4 5 12 i str 08 1736 YP 008266234 1 866913 Proteobacteria
Salmonella enterica subsp enterica serovar Enteritidis str P125109 YP 002244047 1 550537 Proteobacteria
Delftia acidovorans SPH 1 YP 001564276 1 398578 Proteobacteria
Xylella fastidiosa 9a5c NP 299573 1 160492 Proteobacteria
Halomonas elongata DSM 2581 YP 003897172 1 768066 Proteobacteria
Delftia acidovorans SPH 1 YP 001564277 1 398578 Proteobacteria
Pseudomonas brassicacearum subsp brassicacearum NFM421 YP 004355213 1 994484 Proteobacteria
Pseudomonas sp TKP YP 008932552 1 1415630 Proteobacteria
Desulfovibrio magneticus RS 1 YP 002953785 1 573370 Proteobacteria
Shigella sonnei 53G YP 005457361 1 216599 Proteobacteria
Salmonella enterica subsp enterica serovar Schwarzengrund str CVM19633 YP 002113597 1 439843 Proteobacteria
Salmonella enterica subsp enterica serovar Cubana str CFSAN002050 YP 008258809 1 1271863 Proteobacteria
Photorhabdus asymbiotica YP 003041931 1 291112 Proteobacteria
Photorhabdus asymbiotica YP 003042430 1 291112 Proteobacteria
Marinomonas sp MWYL1 YP 001339431 1 400668 Proteobacteria
Xenorhabdus bovienii SS 2004 YP 003469869 1 406818 Proteobacteria
Xenorhabdus bovienii SS 2004 YP 003467902 1 406818 Proteobacteria
Serratia liquefaciens ATCC 27592 YP 008229698 1 1346614 Proteobacteria
Klebsiella pneumoniae YP 007992307 1 573 Proteobacteria
Prodigiosinella confusarubida ATCC39006 WP 021015904 1 Pconfusarubida
Escherichia coli O111 H str 11128 YP 003237951 1 585396 Proteobacteria
Providencia stuartii MRSN 2154 YP 006217884 1 1157951 Proteobacteria
Sodalis glossinidius str morsitans YP 454411 1 343509 Proteobacteria
Candidatus Hamiltonella defensa 5AT Acyrthosiphon pisum YP 002924411 1 572265 Proteobacteria
Candidatus Hamiltonella defensa 5AT Acyrthosiphon pisum YP 002924230 1 572265 Proteobacteria
Salmonella enterica subsp enterica serovar Pullorum str S06004 YP 008381853 1 1298917 Proteobacteria
Salmonella enterica subsp enterica serovar Typhimurium str 14028S YP 005238003 1 588858 Proteobacteria
Salmonella enterica subsp enterica serovar Typhimurium str ST4 74 YP 005242932 1 909946 Proteobacteria
Dickeya gs poaceaephila NCPPB 569 WP 042871462 1 Dpoaceaephila
Klebsiella pneumoniae YP 007993246 1 573 Proteobacteria
Edwardsiella piscicida C07 087 YP 007629381 1 1288122 Proteobacteria
Edwardsiella ictaluri 93 146 YP 002933699 1 634503 Proteobacteria

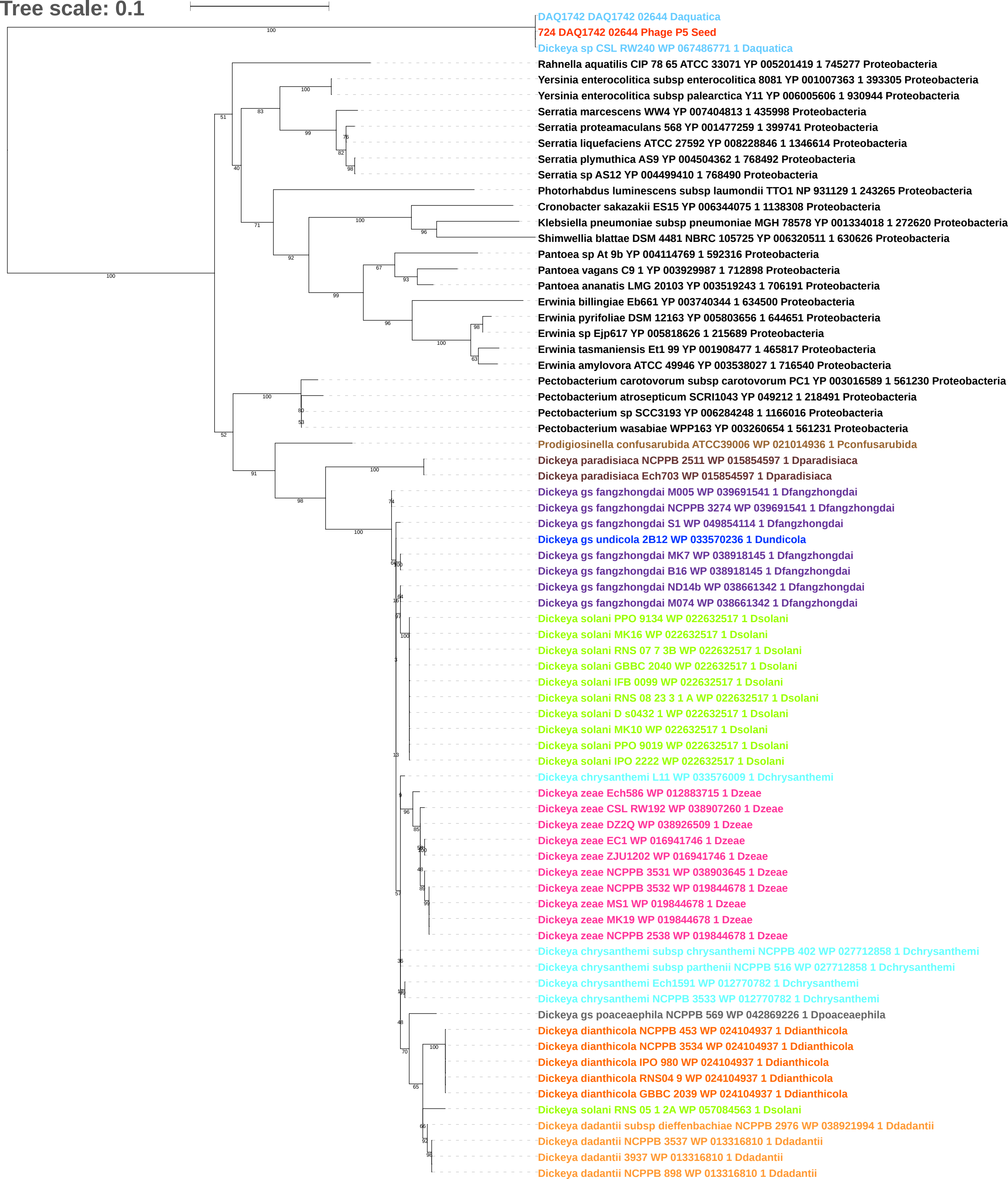
Tree scale: 0.1
DAQ1742 DAQ1742 02644 Daquatica
724 DAQ1742 02644 Phage P5 Seed
Dickeya sp CSL RW240 WP 067486771 1 Daquatica
Rahnella aquatilis CIP 78 65 ATCC 33071 YP 005201419 1 745277 Proteobacteria
Yersinia enterocolitica subsp enterocolitica 8081 YP 001007363 1 393305 Proteobacteria
Yersinia enterocolitica subsp palearctica Y11 YP 006005606 1 930944 Proteobacteria
Serratia marcescens WW4 YP 007404813 1 435998 Proteobacteria
Serratia proteamaculans 568 YP 001477259 1 399741 Proteobacteria
Serratia liquefaciens ATCC 27592 YP 008228846 1 1346614 Proteobacteria
Serratia plymuthica AS9 YP 004504362 1 768492 Proteobacteria
Serratia sp AS12 YP 004499410 1 768490 Proteobacteria
Photorhabdus luminescens subsp laumondii TTO1 NP 931129 1 243265 Proteobacteria
Cronobacter sakazakii ES15 YP 006344075 1 1138308 Proteobacteria
Klebsiella pneumoniae subsp pneumoniae MGH 78578 YP 001334018 1 272620 Proteobacteria
Shimwellia blattae DSM 4481 NBRC 105725 YP 006320511 1 630626 Proteobacteria
Pantoea sp At 9b YP 004114769 1 592316 Proteobacteria
Pantoea vagans C9 1 YP 003929987 1 712898 Proteobacteria
Pantoea ananatis LMG 20103 YP 003519243 1 706191 Proteobacteria
Erwinia billingiae Eb661 YP 003740344 1 634500 Proteobacteria
Erwinia pyrifoliae DSM 12163 YP 005803656 1 644651 Proteobacteria
Erwinia sp Ejp617 YP 005818626 1 215689 Proteobacteria
Erwinia tasmaniensis Et1 99 YP 001908477 1 465817 Proteobacteria
Erwinia amylovora ATCC 49946 YP 003538027 1 716540 Proteobacteria
Pectobacterium carotovorum subsp carotovorum PC1 YP 003016589 1 561230 Proteobacteria
Pectobacterium atrosepticum SCRI1043 YP 049212 1 218491 Proteobacteria
Pectobacterium sp SCC3193 YP 006284248 1 1166016 Proteobacteria
Pectobacterium wasabiae WPP163 YP 003260654 1 561231 Proteobacteria
Prodigiosinella confusarubida ATCC39006 WP 021014936 1 Pconfusarubida
Dickeya paradisiaca NCPPB 2511 WP 015854597 1 Dparadisiaca
Dickeya paradisiaca Ech703 WP 015854597 1 Dparadisiaca
Dickeya gs fangzhongdai M005 WP 039691541 1 Dfangzhongdai
Dickeya gs fangzhongdai NCPPB 3274 WP 039691541 1 Dfangzhongdai
Dickeya gs fangzhongdai S1 WP 049854114 1 Dfangzhongdai
Dickeya gs undicola 2B12 WP 033570236 1 Dundicola
Dickeya gs fangzhongdai MK7 WP 038918145 1 Dfangzhongdai
Dickeya gs fangzhongdai B16 WP 038918145 1 Dfangzhongdai
Dickeya gs fangzhongdai ND14b WP 038661342 1 Dfangzhongdai
Dickeya gs fangzhongdai M074 WP 038661342 1 Dfangzhongdai
Dickeya solani PPO 9134 WP 022632517 1 Dsolani
Dickeya solani MK16 WP 022632517 1 Dsolani
Dickeya solani RNS 07 7 3B WP 022632517 1 Dsolani
Dickeya solani GBBC 2040 WP 022632517 1 Dsolani
Dickeya solani IFB 0099 WP 022632517 1 Dsolani
Dickeya solani RNS 08 23 3 1 A WP 022632517 1 Dsolani
Dickeya solani D s0432 1 WP 022632517 1 Dsolani
Dickeya solani MK10 WP 022632517 1 Dsolani
Dickeya solani PPO 9019 WP 022632517 1 Dsolani
Dickeya solani IPO 2222 WP 022632517 1 Dsolani
Dickeya chrysanthemi L11 WP 033576009 1 Dchrysanthemi
Dickeya zeae Ech586 WP 012883715 1 Dzeae
Dickeya zeae CSL RW192 WP 038907260 1 Dzeae
Dickeya zeae DZ2Q WP 038926509 1 Dzeae
Dickeya zeae EC1 WP 016941746 1 Dzeae
Dickeya zeae ZJU1202 WP 016941746 1 Dzeae
Dickeya zeae NCPPB 3531 WP 038903645 1 Dzeae
Dickeya zeae NCPPB 3532 WP 019844678 1 Dzeae
Dickeya zeae MS1 WP 019844678 1 Dzeae
Dickeya zeae MK19 WP 019844678 1 Dzeae
Dickeya zeae NCPPB 2538 WP 019844678 1 Dzeae
Dickeya chrysanthemi subsp chrysanthemi NCPPB 402 WP 027712858 1 Dchrysanthemi
Dickeya chrysanthemi subsp parthenii NCPPB 516 WP 027712858 1 Dchrysanthemi
Dickeya chrysanthemi Ech1591 WP 012770782 1 Dchrysanthemi
Dickeya chrysanthemi NCPPB 3533 WP 012770782 1 Dchrysanthemi
Dickeya gs poaceaephila NCPPB 569 WP 042869226 1 Dpoaceaephila
Dickeya dianthicola NCPPB 453 WP 024104937 1 Ddianthicola
Dickeya dianthicola NCPPB 3534 WP 024104937 1 Ddianthicola
Dickeya dianthicola IPO 980 WP 024104937 1 Ddianthicola
Dickeya dianthicola RNS04 9 WP 024104937 1 Ddianthicola
Dickeya dianthicola GBBC 2039 WP 024104937 1 Ddianthicola
Dickeya solani RNS 05 1 2A WP 057084563 1 Dsolani
Dickeya dadantii subsp dieffenbachiae NCPPB 2976 WP 038921994 1 Ddadantii
Dickeya dadantii NCPPB 3537 WP 013316810 1 Ddadantii
Dickeya dadantii 3937 WP 013316810 1 Ddadantii
Dickeya dadantii NCPPB 898 WP 013316810 1 Ddadantii

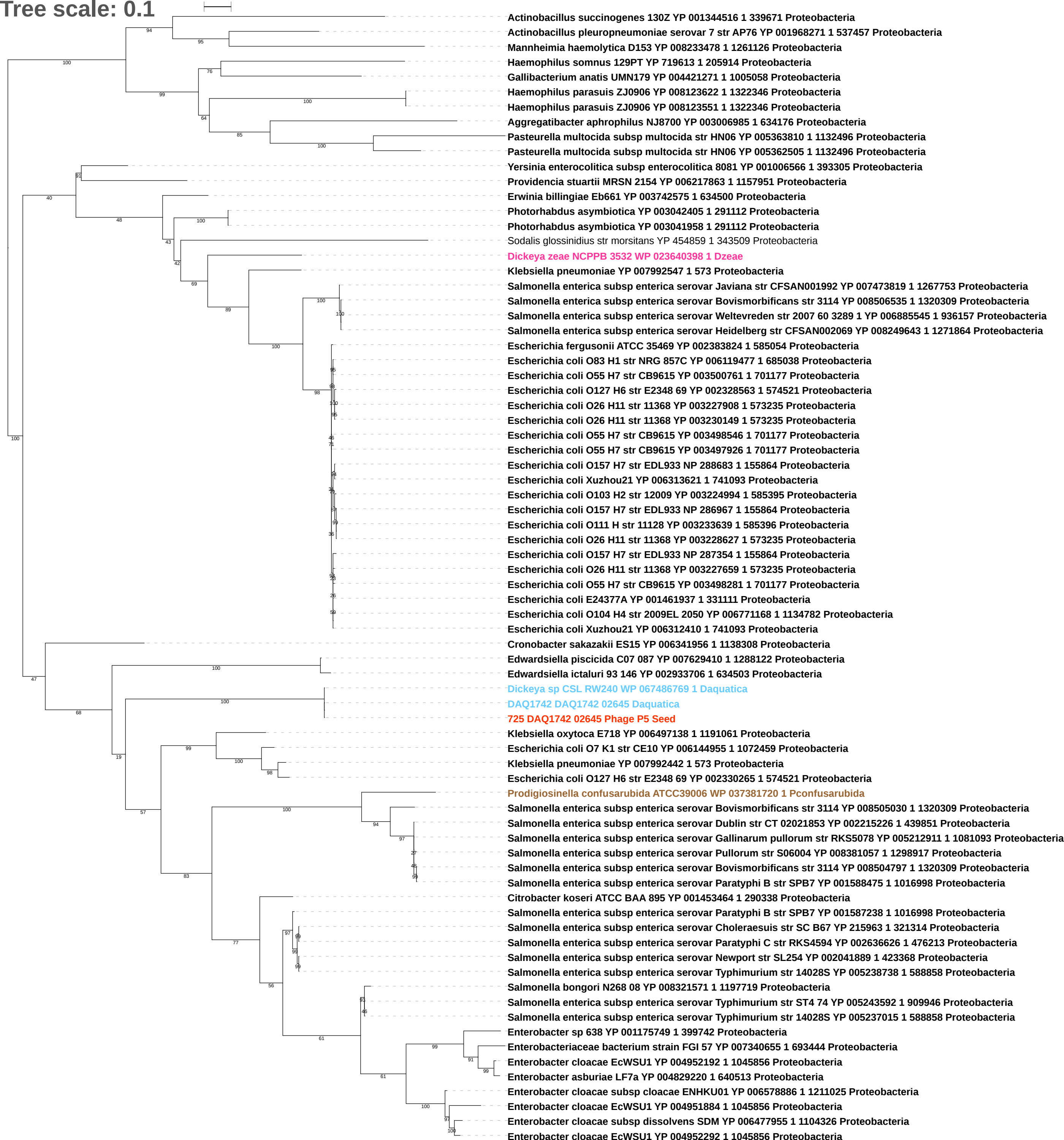
Tree scale: 0.1
Actinobacillus succinogenes 130Z YP 001344516 1 339671 Proteobacteria
Actinobacillus pleuropneumoniae serovar 7 str AP76 YP 001968271 1 537457 Proteobacteria
Mannheimia haemolytica D153 YP 008233478 1 1261126 Proteobacteria
Haemophilus somnus 129PT YP 719613 1 205914 Proteobacteria
Gallibacterium anatis UMN179 YP 004421271 1 1005058 Proteobacteria
Haemophilus parasuis ZJ0906 YP 008123622 1 1322346 Proteobacteria
Haemophilus parasuis ZJ0906 YP 008123551 1 1322346 Proteobacteria
Aggregatibacter aphrophilus NJ8700 YP 003006985 1 634176 Proteobacteria
Pasteurella multocida subsp multocida str HN06 YP 005363810 1 1132496 Proteobacteria
Pasteurella multocida subsp multocida str HN06 YP 005362505 1 1132496 Proteobacteria
Yersinia enterocolitica subsp enterocolitica 8081 YP 001006566 1 393305 Proteobacteria
Providencia stuartii MRSN 2154 YP 006217863 1 1157951 Proteobacteria
Erwinia billingiae Eb661 YP 003742575 1 634500 Proteobacteria
Photorhabdus asymbiotica YP 003042405 1 291112 Proteobacteria
Photorhabdus asymbiotica YP 003041958 1 291112 Proteobacteria
Sodalis glossinidius str morsitans YP 454859 1 343509 Proteobacteria
Dickeya zeae NCPPB 3532 WP 023640398 1 Dzeae
Klebsiella pneumoniae YP 007992547 1 573 Proteobacteria
Salmonella enterica subsp enterica serovar Javiana str CFSAN001992 YP 007473819 1 1267753 Proteobacteria
Salmonella enterica subsp enterica serovar Bovismorbificans str 3114 YP 008506535 1 1320309 Proteobacteria
Salmonella enterica subsp enterica serovar Weltevreden str 2007 60 3289 1 YP 006885545 1 936157 Proteobacteria
Salmonella enterica subsp enterica serovar Heidelberg str CFSAN002069 YP 008249643 1 1271864 Proteobacteria
Escherichia fergusonii ATCC 35469 YP 002383824 1 585054 Proteobacteria
Escherichia coli O83 H1 str NRG 857C YP 006119477 1 685038 Proteobacteria
Escherichia coli O55 H7 str CB9615 YP 003500761 1 701177 Proteobacteria
Escherichia coli O127 H6 str E2348 69 YP 002328563 1 574521 Proteobacteria
Escherichia coli O26 H11 str 11368 YP 003227908 1 573235 Proteobacteria
Escherichia coli O26 H11 str 11368 YP 003230149 1 573235 Proteobacteria
Escherichia coli O55 H7 str CB9615 YP 003498546 1 701177 Proteobacteria
Escherichia coli O55 H7 str CB9615 YP 003497926 1 701177 Proteobacteria
Escherichia coli O157 H7 str EDL933 NP 288683 1 155864 Proteobacteria
Escherichia coli Xuzhou21 YP 006313621 1 741093 Proteobacteria
Escherichia coli O103 H2 str 12009 YP 003224994 1 585395 Proteobacteria
Escherichia coli O157 H7 str EDL933 NP 286967 1 155864 Proteobacteria
Escherichia coli O111 H str 11128 YP 003233639 1 585396 Proteobacteria
Escherichia coli O26 H11 str 11368 YP 003228627 1 573235 Proteobacteria
Escherichia coli O157 H7 str EDL933 NP 287354 1 155864 Proteobacteria
Escherichia coli O26 H11 str 11368 YP 003227659 1 573235 Proteobacteria
Escherichia coli O55 H7 str CB9615 YP 003498281 1 701177 Proteobacteria
Escherichia coli E24377A YP 001461937 1 331111 Proteobacteria
Escherichia coli O104 H4 str 2009EL 2050 YP 006771168 1 1134782 Proteobacteria
Escherichia coli Xuzhou21 YP 006312410 1 741093 Proteobacteria
Cronobacter sakazakii ES15 YP 006341956 1 1138308 Proteobacteria
Edwardsiella piscicida C07 087 YP 007629410 1 1288122 Proteobacteria
Edwardsiella ictaluri 93 146 YP 002933706 1 634503 Proteobacteria
Dickeya sp CSL RW240 WP 067486769 1 Daquatica
DAQ1742 DAQ1742 02645 Daquatica
725 DAQ1742 02645 Phage P5 Seed
Klebsiella oxytoca E718 YP 006497138 1 1191061 Proteobacteria
Escherichia coli O7 K1 str CE10 YP 006144955 1 1072459 Proteobacteria
Klebsiella pneumoniae YP 007992442 1 573 Proteobacteria
Escherichia coli O127 H6 str E2348 69 YP 002330265 1 574521 Proteobacteria
Prodigiosinella confusarubida ATCC39006 WP 037381720 1 Pconfusarubida
Salmonella enterica subsp enterica serovar Bovismorbificans str 3114 YP 008505030 1 1320309 Proteobacteria
Salmonella enterica subsp enterica serovar Dublin str CT 02021853 YP 002215226 1 439851 Proteobacteria
Salmonella enterica subsp enterica serovar Gallinarum pullorum str RKS5078 YP 005212911 1 1081093 Proteobacteria
Salmonella enterica subsp enterica serovar Pullorum str S06004 YP 008381057 1 1298917 Proteobacteria
Salmonella enterica subsp enterica serovar Bovismorbificans str 3114 YP 008504797 1 1320309 Proteobacteria
Salmonella enterica subsp enterica serovar Paratyphi B str SPB7 YP 001588475 1 1016998 Proteobacteria
Citrobacter koseri ATCC BAA 895 YP 001453464 1 290338 Proteobacteria
Salmonella enterica subsp enterica serovar Paratyphi B str SPB7 YP 001587238 1 1016998 Proteobacteria
Salmonella enterica subsp enterica serovar Choleraesuis str SC B67 YP 215963 1 321314 Proteobacteria
Salmonella enterica subsp enterica serovar Paratyphi C str RKS4594 YP 002636626 1 476213 Proteobacteria
Salmonella enterica subsp enterica serovar Newport str SL254 YP 002041889 1 423368 Proteobacteria
Salmonella enterica subsp enterica serovar Typhimurium str 14028S YP 005238738 1 588858 Proteobacteria
Salmonella bongori N268 08 YP 008321571 1 1197719 Proteobacteria
Salmonella enterica subsp enterica serovar Typhimurium str ST4 74 YP 005243592 1 909946 Proteobacteria
Salmonella enterica subsp enterica serovar Typhimurium str 14028S YP 005237015 1 588858 Proteobacteria
Enterobacter sp 638 YP 001175749 1 399742 Proteobacteria
Enterobacteriaceae bacterium strain FGI 57 YP 007340655 1 693444 Proteobacteria
Enterobacter cloacae EcWSU1 YP 004952192 1 1045856 Proteobacteria
Enterobacter asburiae LF7a YP 004829220 1 640513 Proteobacteria
Enterobacter cloacae subsp cloacae ENHKU01 YP 006578886 1 1211025 Proteobacteria
Enterobacter cloacae EcWSU1 YP 004951884 1 1045856 Proteobacteria
Enterobacter cloacae subsp dissolvens SDM YP 006477955 1 1104326 Proteobacteria
Enterobacter cloacae EcWSU1 YP 004952292 1 1045856 Proteobacteria

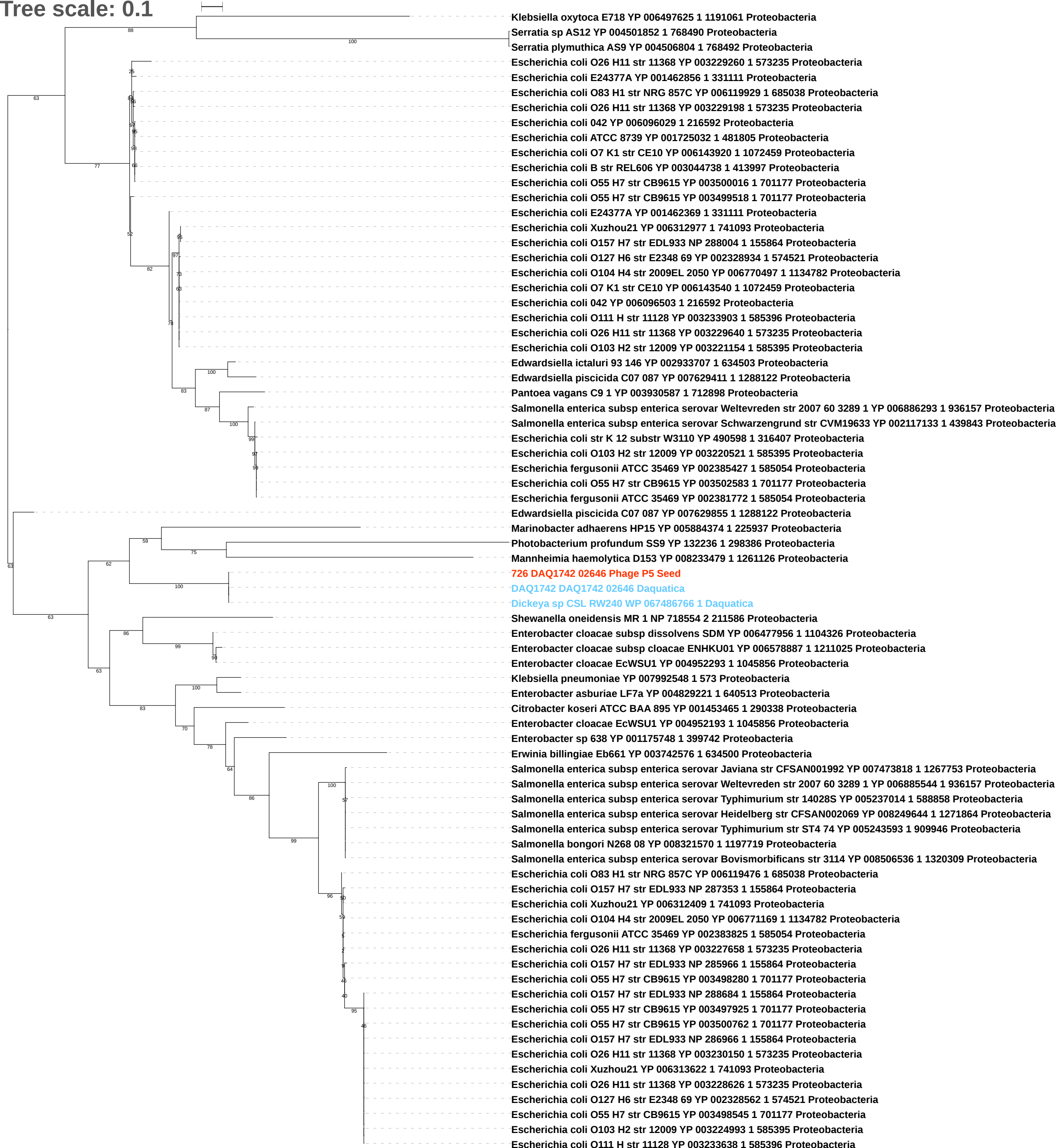
Tree scale: 0.1
Klebsiella oxytoca E718 YP 006497625 1 1191061 Proteobacteria
Serratia sp AS12 YP 004501852 1 768490 Proteobacteria
Serratia plymuthica AS9 YP 004506804 1 768492 Proteobacteria
Escherichia coli O26 H11 str 11368 YP 003229260 1 573235 Proteobacteria
Escherichia coli E24377A YP 001462856 1 331111 Proteobacteria
Escherichia coli O83 H1 str NRG 857C YP 006119929 1 685038 Proteobacteria
Escherichia coli O26 H11 str 11368 YP 003229198 1 573235 Proteobacteria
Escherichia coli 042 YP 006096029 1 216592 Proteobacteria
Escherichia coli ATCC 8739 YP 001725032 1 481805 Proteobacteria
Escherichia coli O7 K1 str CE10 YP 006143920 1 1072459 Proteobacteria
Escherichia coli B str REL606 YP 003044738 1 413997 Proteobacteria
Escherichia coli O55 H7 str CB9615 YP 003500016 1 701177 Proteobacteria
Escherichia coli O55 H7 str CB9615 YP 003499518 1 701177 Proteobacteria
Escherichia coli E24377A YP 001462369 1 331111 Proteobacteria
Escherichia coli Xuzhou21 YP 006312977 1 741093 Proteobacteria
Escherichia coli O157 H7 str EDL933 NP 288004 1 155864 Proteobacteria
Escherichia coli O127 H6 str E2348 69 YP 002328934 1 574521 Proteobacteria
Escherichia coli O104 H4 str 2009EL 2050 YP 006770497 1 1134782 Proteobacteria
Escherichia coli O7 K1 str CE10 YP 006143540 1 1072459 Proteobacteria
Escherichia coli 042 YP 006096503 1 216592 Proteobacteria
Escherichia coli O111 H str 11128 YP 003233903 1 585396 Proteobacteria
Escherichia coli O26 H11 str 11368 YP 003229640 1 573235 Proteobacteria
Escherichia coli O103 H2 str 12009 YP 003221154 1 585395 Proteobacteria
Edwardsiella ictaluri 93 146 YP 002933707 1 634503 Proteobacteria
Edwardsiella piscicida C07 087 YP 007629411 1 1288122 Proteobacteria
Pantoea vagans C9 1 YP 003930587 1 712898 Proteobacteria
Salmonella enterica subsp enterica serovar Weltevreden str 2007 60 3289 1 YP 006886293 1 936157 Proteobacteria
Salmonella enterica subsp enterica serovar Schwarzengrund str CVM19633 YP 002117133 1 439843 Proteobacteria
Escherichia coli str K 12 substr W3110 YP 490598 1 316407 Proteobacteria
Escherichia coli O103 H2 str 12009 YP 003220521 1 585395 Proteobacteria
Escherichia fergusonii ATCC 35469 YP 002385427 1 585054 Proteobacteria
Escherichia coli O55 H7 str CB9615 YP 003502583 1 701177 Proteobacteria
Escherichia fergusonii ATCC 35469 YP 002381772 1 585054 Proteobacteria
Edwardsiella piscicida C07 087 YP 007629855 1 1288122 Proteobacteria
Marinobacter adhaerens HP15 YP 005884374 1 225937 Proteobacteria
Photobacterium profundum SS9 YP 132236 1 298386 Proteobacteria
Mannheimia haemolytica D153 YP 008233479 1 1261126 Proteobacteria
726 DAQ1742 02646 Phage P5 Seed
DAQ1742 DAQ1742 02646 Daquatica
Dickeya sp CSL RW240 WP 067486766 1 Daquatica
Shewanella oneidensis MR 1 NP 718554 2 211586 Proteobacteria
Enterobacter cloacae subsp dissolvens SDM YP 006477956 1 1104326 Proteobacteria
Enterobacter cloacae subsp cloacae ENHKU01 YP 006578887 1 1211025 Proteobacteria
Enterobacter cloacae EcWSU1 YP 004952293 1 1045856 Proteobacteria
Klebsiella pneumoniae YP 007992548 1 573 Proteobacteria
Enterobacter asburiae LF7a YP 004829221 1 640513 Proteobacteria
Citrobacter koseri ATCC BAA 895 YP 001453465 1 290338 Proteobacteria
Enterobacter cloacae EcWSU1 YP 004952193 1 1045856 Proteobacteria
Enterobacter sp 638 YP 001175748 1 399742 Proteobacteria
Erwinia billingiae Eb661 YP 003742576 1 634500 Proteobacteria
Salmonella enterica subsp enterica serovar Javiana str CFSAN001992 YP 007473818 1 1267753 Proteobacteria
Salmonella enterica subsp enterica serovar Weltevreden str 2007 60 3289 1 YP 006885544 1 936157 Proteobacteria
Salmonella enterica subsp enterica serovar Typhimurium str 14028S YP 005237014 1 588858 Proteobacteria
Salmonella enterica subsp enterica serovar Heidelberg str CFSAN002069 YP 008249644 1 1271864 Proteobacteria
Salmonella enterica subsp enterica serovar Typhimurium str ST4 74 YP 005243593 1 909946 Proteobacteria
Salmonella bongori N268 08 YP 008321570 1 1197719 Proteobacteria
Salmonella enterica subsp enterica serovar Bovismorbificans str 3114 YP 008506536 1 1320309 Proteobacteria
Escherichia coli O83 H1 str NRG 857C YP 006119476 1 685038 Proteobacteria
Escherichia coli O157 H7 str EDL933 NP 287353 1 155864 Proteobacteria
Escherichia coli Xuzhou21 YP 006312409 1 741093 Proteobacteria
Escherichia coli O104 H4 str 2009EL 2050 YP 006771169 1 1134782 Proteobacteria
Escherichia fergusonii ATCC 35469 YP 002383825 1 585054 Proteobacteria
Escherichia coli O26 H11 str 11368 YP 003227658 1 573235 Proteobacteria
Escherichia coli O157 H7 str EDL933 NP 285966 1 155864 Proteobacteria
Escherichia coli O55 H7 str CB9615 YP 003498280 1 701177 Proteobacteria
Escherichia coli O157 H7 str EDL933 NP 288684 1 155864 Proteobacteria
Escherichia coli O55 H7 str CB9615 YP 003497925 1 701177 Proteobacteria
Escherichia coli O55 H7 str CB9615 YP 003500762 1 701177 Proteobacteria
Escherichia coli O157 H7 str EDL933 NP 286966 1 155864 Proteobacteria
Escherichia coli O26 H11 str 11368 YP 003230150 1 573235 Proteobacteria
Escherichia coli Xuzhou21 YP 006313622 1 741093 Proteobacteria
Escherichia coli O26 H11 str 11368 YP 003228626 1 573235 Proteobacteria
Escherichia coli O127 H6 str E2348 69 YP 002328562 1 574521 Proteobacteria
Escherichia coli O55 H7 str CB9615 YP 003498545 1 701177 Proteobacteria
Escherichia coli O103 H2 str 12009 YP 003224993 1 585395 Proteobacteria
Escherichia coli O111 H str 11128 YP 003233638 1 585396 Proteobacteria

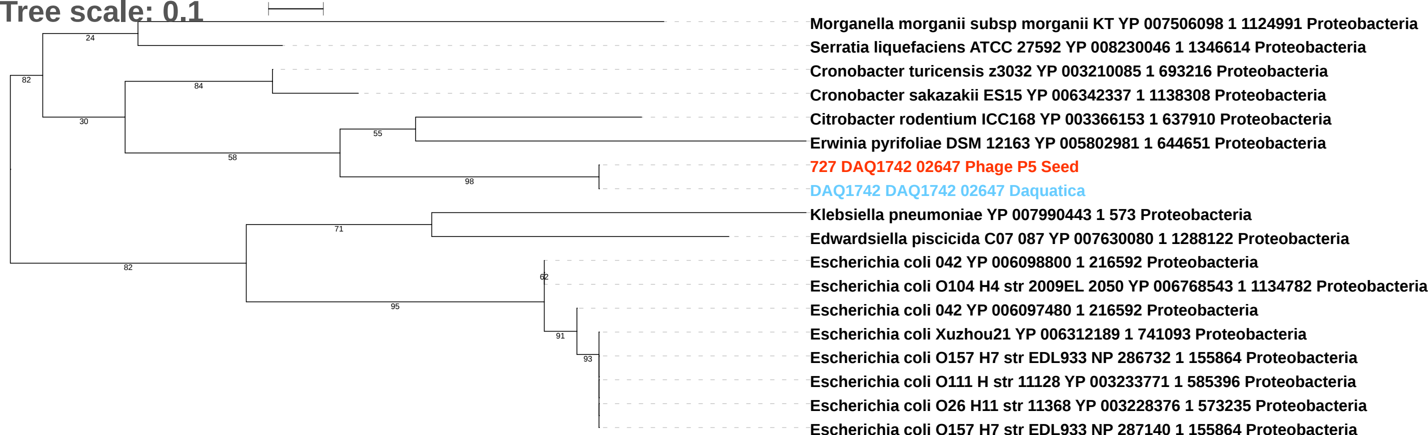

Tree scale: 0.1
Morganella morganii subsp morganii KT YP 007506098 1 1124991 Proteobacteria
Serratia liquefaciens ATCC 27592 YP 008230046 1 1346614 Proteobacteria
Cronobacter turicensis z3032 YP 003210085 1 693216 Proteobacteria
Cronobacter sakazakii ES15 YP 006342337 1 1138308 Proteobacteria
Citrobacter rodentium ICC168 YP 003366153 1 637910 Proteobacteria
Erwinia pyrifoliae DSM 12163 YP 005802981 1 644651 Proteobacteria
727 DAQ1742 02647 Phage P5 Seed
DAQ1742 DAQ1742 02647 Daquatica
Klebsiella pneumoniae YP 007990443 1 573 Proteobacteria
Edwardsiella piscicida C07 087 YP 007630080 1 1288122 Proteobacteria
Escherichia coli 042 YP 006098800 1 216592 Proteobacteria
Escherichia coli O104 H4 str 2009EL 2050 YP 006768543 1 1134782 Proteobacteria
Escherichia coli 042 YP 006097480 1 216592 Proteobacteria
Escherichia coli Xuzhou21 YP 006312189 1 741093 Proteobacteria
Escherichia coli O157 H7 str EDL933 NP 286732 1 155864 Proteobacteria
Escherichia coli O111 H str 11128 YP 003233771 1 585396 Proteobacteria
Escherichia coli O26 H11 str 11368 YP 003228376 1 573235 Proteobacteria
Escherichia coli O157 H7 str EDL933 NP 287140 1 155864 Proteobacteria
24
82
84
30
55
58
98
71
82
62
95
91
93

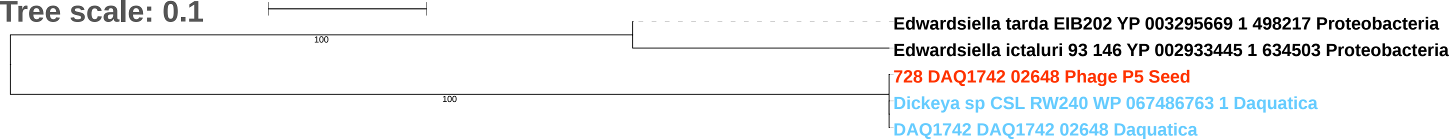
Tree scale: 0.1
100
100
Edwardsiella tarda EIB202 YP 003295669 1 498217 Proteobacteria
Edwardsiella ictaluri 93 146 YP 002933445 1 634503 Proteobacteria
728 DAQ1742 02648 Phage P5 Seed
Dickeya sp CSL RW240 WP 067486763 1 Daquatica
DAQ1742 DAQ1742 02648 Daquatica

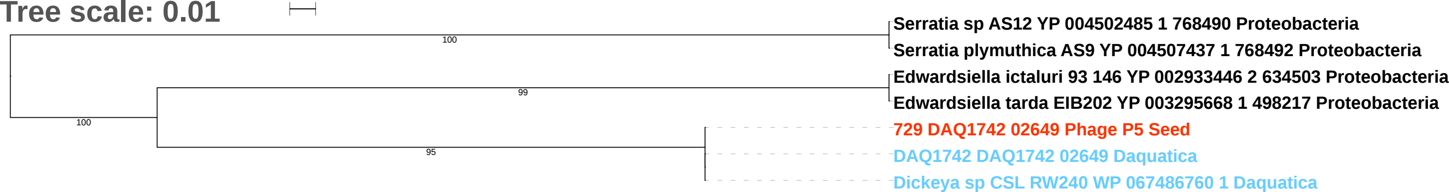
Tree scale: 0.01
Serratia sp AS12 YP 004502485 1 768490 Proteobacteria
Serratia plymuthica AS9 YP 004507437 1 768492 Proteobacteria
Edwardsiella ictaluri 93 146 YP 002933446 2 634503 Proteobacteria
Edwardsiella tarda EIB202 YP 003295668 1 498217 Proteobacteria
729 DAQ1742 02649 Phage P5 Seed
DAQ1742 DAQ1742 02649 Daquatica
Dickeya sp CSL RW240 WP 067486760 1 Daquatica
100
99
100
95

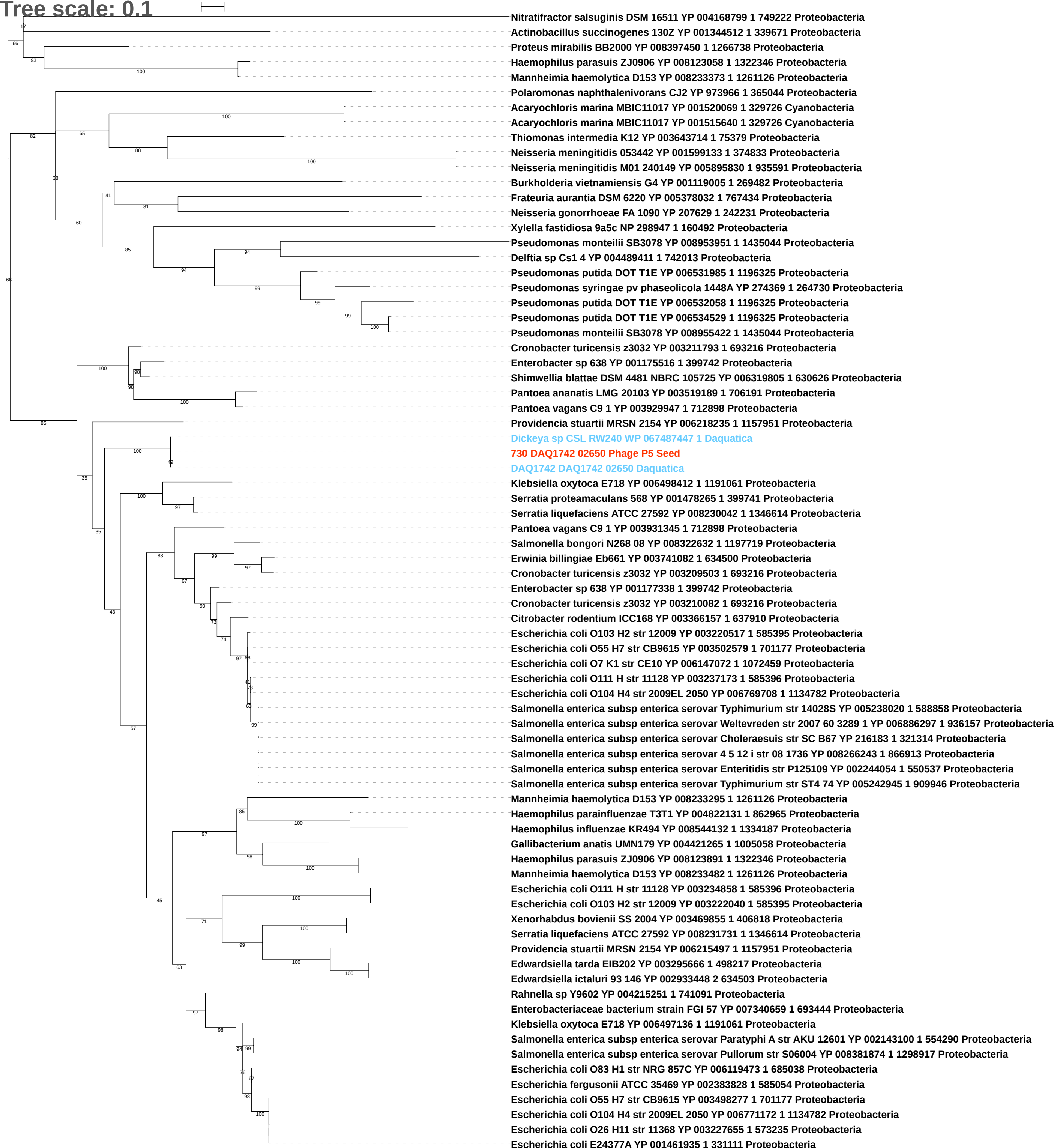
Tree scale: 0.1
Nitratifractor salsuginis DSM 16511 YP 004168799 1 749222 Proteobacteria
Actinobacillus succinogenes 130Z YP 001344512 1 339671 Proteobacteria
Proteus mirabilis BB2000 YP 008397450 1 1266738 Proteobacteria
Haemophilus parasuis ZJ0906 YP 008123058 1 1322346 Proteobacteria
Mannheimia haemolytica D153 YP 008233373 1 1261126 Proteobacteria
Polaromonas naphthalenivorans CJ2 YP 973966 1 365044 Proteobacteria
Acaryochloris marina MBIC11017 YP 001520069 1 329726 Cyanobacteria
Acaryochloris marina MBIC11017 YP 001515640 1 329726 Cyanobacteria
Thiomonas intermedia K12 YP 003643714 1 75379 Proteobacteria
Neisseria meningitidis 053442 YP 001599133 1 374833 Proteobacteria
Neisseria meningitidis M01 240149 YP 005895830 1 935591 Proteobacteria
Burkholderia vietnamiensis G4 YP 001119005 1 269482 Proteobacteria
Frateuria aurantia DSM 6220 YP 005378032 1 767434 Proteobacteria
Neisseria gonorrhoeae FA 1090 YP 207629 1 242231 Proteobacteria
Xylella fastidiosa 9a5c NP 298947 1 160492 Proteobacteria
Pseudomonas monteilii SB3078 YP 008953951 1 1435044 Proteobacteria
Delftia sp Cs1 4 YP 004489411 1 742013 Proteobacteria
Pseudomonas putida DOT T1E YP 006531985 1 1196325 Proteobacteria
Pseudomonas syringae pv phaseolicola 1448A YP 274369 1 264730 Proteobacteria
Pseudomonas putida DOT T1E YP 006532058 1 1196325 Proteobacteria
Pseudomonas putida DOT T1E YP 006534529 1 1196325 Proteobacteria
Pseudomonas monteilii SB3078 YP 008955422 1 1435044 Proteobacteria
Cronobacter turicensis z3032 YP 003211793 1 693216 Proteobacteria
Enterobacter sp 638 YP 001175516 1 399742 Proteobacteria
Shimwellia blattae DSM 4481 NBRC 105725 YP 006319805 1 630626 Proteobacteria
Pantoea ananatis LMG 20103 YP 003519189 1 706191 Proteobacteria
Pantoea vagans C9 1 YP 003929947 1 712898 Proteobacteria
Providencia stuartii MRSN 2154 YP 006218235 1 1157951 Proteobacteria
Dickeya sp CSL RW240 WP 067487447 1 Daquatica
730 DAQ1742 02650 Phage P5 Seed
DAQ1742 DAQ1742 02650 Daquatica
Klebsiella oxytoca E718 YP 006498412 1 1191061 Proteobacteria
Serratia proteamaculans 568 YP 001478265 1 399741 Proteobacteria
Serratia liquefaciens ATCC 27592 YP 008230042 1 1346614 Proteobacteria
Pantoea vagans C9 1 YP 003931345 1 712898 Proteobacteria
Salmonella bongori N268 08 YP 008322632 1 1197719 Proteobacteria
Erwinia billingiae Eb661 YP 003741082 1 634500 Proteobacteria
Cronobacter turicensis z3032 YP 003209503 1 693216 Proteobacteria
Enterobacter sp 638 YP 001177338 1 399742 Proteobacteria
Cronobacter turicensis z3032 YP 003210082 1 693216 Proteobacteria
Citrobacter rodentium ICC168 YP 003366157 1 637910 Proteobacteria
Escherichia coli O103 H2 str 12009 YP 003220517 1 585395 Proteobacteria
Escherichia coli O55 H7 str CB9615 YP 003502579 1 701177 Proteobacteria
Escherichia coli O7 K1 str CE10 YP 006147072 1 1072459 Proteobacteria
Escherichia coli O111 H str 11128 YP 003237173 1 585396 Proteobacteria
Escherichia coli O104 H4 str 2009EL 2050 YP 006769708 1 1134782 Proteobacteria
Salmonella enterica subsp enterica serovar Typhimurium str 14028S YP 005238020 1 588858 Proteobacteria
Salmonella enterica subsp enterica serovar Weltevreden str 2007 60 3289 1 YP 006886297 1 936157 Proteobacteria
Salmonella enterica subsp enterica serovar Choleraesuis str SC B67 YP 216183 1 321314 Proteobacteria
Salmonella enterica subsp enterica serovar 4 5 12 i str 08 1736 YP 008266243 1 866913 Proteobacteria
Salmonella enterica subsp enterica serovar Enteritidis str P125109 YP 002244054 1 550537 Proteobacteria
Salmonella enterica subsp enterica serovar Typhimurium str ST4 74 YP 005242945 1 909946 Proteobacteria
Mannheimia haemolytica D153 YP 008233295 1 1261126 Proteobacteria
Haemophilus parainfluenzae T3T1 YP 004822131 1 862965 Proteobacteria
Haemophilus influenzae KR494 YP 008544132 1 1334187 Proteobacteria
Gallibacterium anatis UMN179 YP 004421265 1 1005058 Proteobacteria
Haemophilus parasuis ZJ0906 YP 008123891 1 1322346 Proteobacteria
Mannheimia haemolytica D153 YP 008233482 1 1261126 Proteobacteria
Escherichia coli O111 H str 11128 YP 003234858 1 585396 Proteobacteria
Escherichia coli O103 H2 str 12009 YP 003222040 1 585395 Proteobacteria
Xenorhabdus bovienii SS 2004 YP 003469855 1 406818 Proteobacteria
Serratia liquefaciens ATCC 27592 YP 008231731 1 1346614 Proteobacteria
Providencia stuartii MRSN 2154 YP 006215497 1 1157951 Proteobacteria
Edwardsiella tarda EIB202 YP 003295666 1 498217 Proteobacteria
Edwardsiella ictaluri 93 146 YP 002933448 2 634503 Proteobacteria
Rahnella sp Y9602 YP 004215251 1 741091 Proteobacteria
Enterobacteriaceae bacterium strain FGI 57 YP 007340659 1 693444 Proteobacteria
Klebsiella oxytoca E718 YP 006497136 1 1191061 Proteobacteria
Salmonella enterica subsp enterica serovar Paratyphi A str AKU 12601 YP 002143100 1 554290 Proteobacteria
Salmonella enterica subsp enterica serovar Pullorum str S06004 YP 008381874 1 1298917 Proteobacteria
Escherichia coli O83 H1 str NRG 857C YP 006119473 1 685038 Proteobacteria
Escherichia fergusonii ATCC 35469 YP 002383828 1 585054 Proteobacteria
Escherichia coli O55 H7 str CB9615 YP 003498277 1 701177 Proteobacteria
Escherichia coli O104 H4 str 2009EL 2050 YP 006771172 1 1134782 Proteobacteria
Escherichia coli O26 H11 str 11368 YP 003227655 1 573235 Proteobacteria
Escherichia coli E24377A YP 001461935 1 331111 Proteobacteria

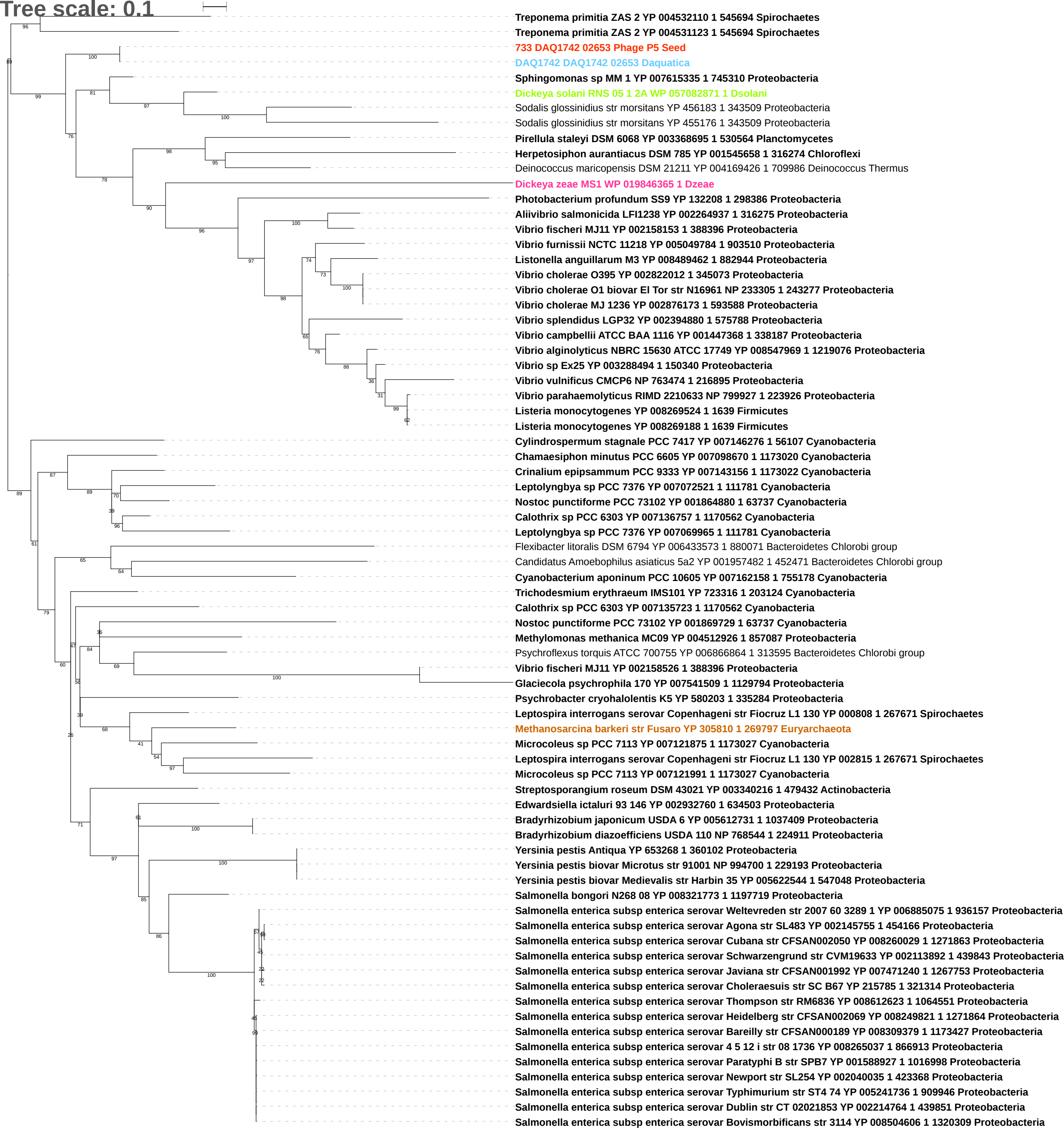
Tree scale: 0.1
Treponema primitia ZAS 2 YP 004532110 1 545694 Spirochaetes
Treponema primitia ZAS 2 YP 004531123 1 545694 Spirochaetes
733 DAQ1742 02653 Phage P5 Seed
DAQ1742 DAQ1742 02653 Daquatica
Sphingomonas sp MM 1 YP 007615335 1 745310 Proteobacteria
Dickeya solani RNS 05 1 2A WP 057082871 1 Dsolani
Sodalis glossinidius str morsitans YP 456183 1 343509 Proteobacteria
Sodalis glossinidius str morsitans YP 455176 1 343509 Proteobacteria
Pirellula staleyi DSM 6068 YP 003368695 1 530564 Planctomycetes
Herpetosiphon aurantiacus DSM 785 YP 001545658 1 316274 Chloroflexi
Deinococcus maricopensis DSM 21211 YP 004169426 1 709986 Deinococcus Thermus
Dickeya zeae MS1 WP 019846365 1 Dzeae
Photobacterium profundum SS9 YP 132208 1 298386 Proteobacteria
Aliivibrio salmonicida LFI1238 YP 002264937 1 316275 Proteobacteria
Vibrio fischeri MJ11 YP 002158153 1 388396 Proteobacteria
Vibrio furnissii NCTC 11218 YP 005049784 1 903510 Proteobacteria
Listonella anguillarum M3 YP 008489462 1 882944 Proteobacteria
Vibrio cholerae O395 YP 002822012 1 345073 Proteobacteria
Vibrio cholerae O1 biovar El Tor str N16961 NP 233305 1 243277 Proteobacteria
Vibrio cholerae MJ 1236 YP 002876173 1 593588 Proteobacteria
Vibrio splendidus LGP32 YP 002394880 1 575788 Proteobacteria
Vibrio campbellii ATCC BAA 1116 YP 001447368 1 338187 Proteobacteria
Vibrio alginolyticus NBRC 15630 ATCC 17749 YP 008547969 1 1219076 Proteobacteria
Vibrio sp Ex25 YP 003288494 1 150340 Proteobacteria
Vibrio vulnificus CMCP6 NP 763474 1 216895 Proteobacteria
Vibrio parahaemolyticus RIMD 2210633 NP 799927 1 223926 Proteobacteria
Listeria monocytogenes YP 008269524 1 1639 Firmicutes
Listeria monocytogenes YP 008269188 1 1639 Firmicutes
Cylindrospermum stagnale PCC 7417 YP 007146276 1 56107 Cyanobacteria
Chamaesiphon minutus PCC 6605 YP 007098670 1 1173020 Cyanobacteria
Crinalium epipsammum PCC 9333 YP 007143156 1 1173022 Cyanobacteria
Leptolyngbya sp PCC 7376 YP 007072521 1 111781 Cyanobacteria
Nostoc punctiforme PCC 73102 YP 001864880 1 63737 Cyanobacteria
Calothrix sp PCC 6303 YP 007136757 1 1170562 Cyanobacteria
Leptolyngbya sp PCC 7376 YP 007069965 1 111781 Cyanobacteria
Flexibacter litoralis DSM 6794 YP 006433573 1 880071 Bacteroidetes Chlorobi group
Candidatus Amoebophilus asiaticus 5a2 YP 001957482 1 452471 Bacteroidetes Chlorobi group
Cyanobacterium aponinum PCC 10605 YP 007162158 1 755178 Cyanobacteria
Trichodesmium erythraeum IMS101 YP 723316 1 203124 Cyanobacteria
Calothrix sp PCC 6303 YP 007135723 1 1170562 Cyanobacteria
Nostoc punctiforme PCC 73102 YP 001869729 1 63737 Cyanobacteria
Methylomonas methanica MC09 YP 004512926 1 857087 Proteobacteria
Psychroflexus torquis ATCC 700755 YP 006866864 1 313595 Bacteroidetes Chlorobi group
Vibrio fischeri MJ11 YP 002158526 1 388396 Proteobacteria
Glaciecola psychrophila 170 YP 007541509 1 1129794 Proteobacteria
Psychrobacter cryohalolentis K5 YP 580203 1 335284 Proteobacteria
Leptospira interrogans serovar Copenhageni str Fiocruz L1 130 YP 000808 1 267671 Spirochaetes
Methanosarcina barkeri str Fusaro YP 305810 1 269797 Euryarchaeota
Microcoleus sp PCC 7113 YP 007121875 1 1173027 Cyanobacteria
Leptospira interrogans serovar Copenhageni str Fiocruz L1 130 YP 002815 1 267671 Spirochaetes
Microcoleus sp PCC 7113 YP 007121991 1 1173027 Cyanobacteria
Streptosporangium roseum DSM 43021 YP 003340216 1 479432 Actinobacteria
Edwardsiella ictaluri 93 146 YP 002932760 1 634503 Proteobacteria
Bradyrhizobium japonicum USDA 6 YP 005612731 1 1037409 Proteobacteria
Bradyrhizobium diazoefficiens USDA 110 NP 768544 1 224911 Proteobacteria
Yersinia pestis Antiqua YP 653268 1 360102 Proteobacteria
Yersinia pestis biovar Microtus str 91001 NP 994700 1 229193 Proteobacteria
Yersinia pestis biovar Medievalis str Harbin 35 YP 005622544 1 547048 Proteobacteria
Salmonella bongori N268 08 YP 008321773 1 1197719 Proteobacteria
Salmonella enterica subsp enterica serovar Weltevreden str 2007 60 3289 1 YP 006885075 1 936157 Proteobacteria
Salmonella enterica subsp enterica serovar Agona str SL483 YP 002145755 1 454166 Proteobacteria
Salmonella enterica subsp enterica serovar Cubana str CFSAN002050 YP 008260029 1 1271863 Proteobacteria
Salmonella enterica subsp enterica serovar Schwarzengrund str CVM19633 YP 002113892 1 439843 Proteobacteria
Salmonella enterica subsp enterica serovar Javiana str CFSAN001992 YP 007471240 1 1267753 Proteobacteria
Salmonella enterica subsp enterica serovar Choleraesuis str SC B67 YP 215785 1 321314 Proteobacteria
Salmonella enterica subsp enterica serovar Thompson str RM6836 YP 008612623 1 1064551 Proteobacteria
Salmonella enterica subsp enterica serovar Heidelberg str CFSAN002069 YP 008249821 1 1271864 Proteobacteria
Salmonella enterica subsp enterica serovar Bareilly str CFSAN000189 YP 008309379 1 1173427 Proteobacteria
Salmonella enterica subsp enterica serovar 4 5 12 i str 08 1736 YP 008265037 1 866913 Proteobacteria
Salmonella enterica subsp enterica serovar Paratyphi B str SPB7 YP 001588927 1 1016998 Proteobacteria
Salmonella enterica subsp enterica serovar Newport str SL254 YP 002040035 1 423368 Proteobacteria
Salmonella enterica subsp enterica serovar Typhimurium str ST4 74 YP 005241736 1 909946 Proteobacteria
Salmonella enterica subsp enterica serovar Dublin str CT 02021853 YP 002214764 1 439851 Proteobacteria
Salmonella enterica subsp enterica serovar Bovismorbificans str 3114 YP 008504606 1 1320309 Proteobacteria

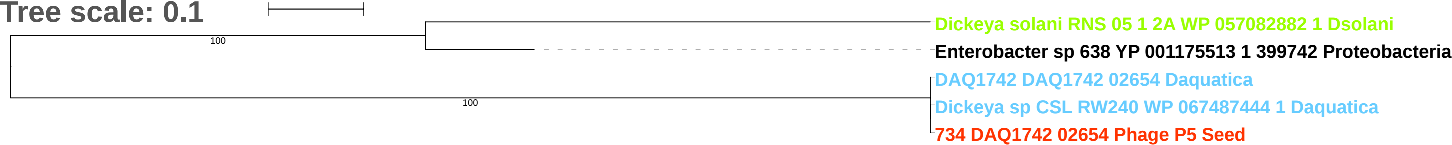
Tree scale: 0.1
100
100
Dickeya solani RNS 05 1 2A WP 057082882 1 Dsolani
Enterobacter sp 638 YP 001175513 1 399742 Proteobacteria
DAQ1742 DAQ1742 02654 Daquatica
Dickeya sp CSL RW240 WP 067487444 1 Daquatica
734 DAQ1742 02654 Phage P5 Seed

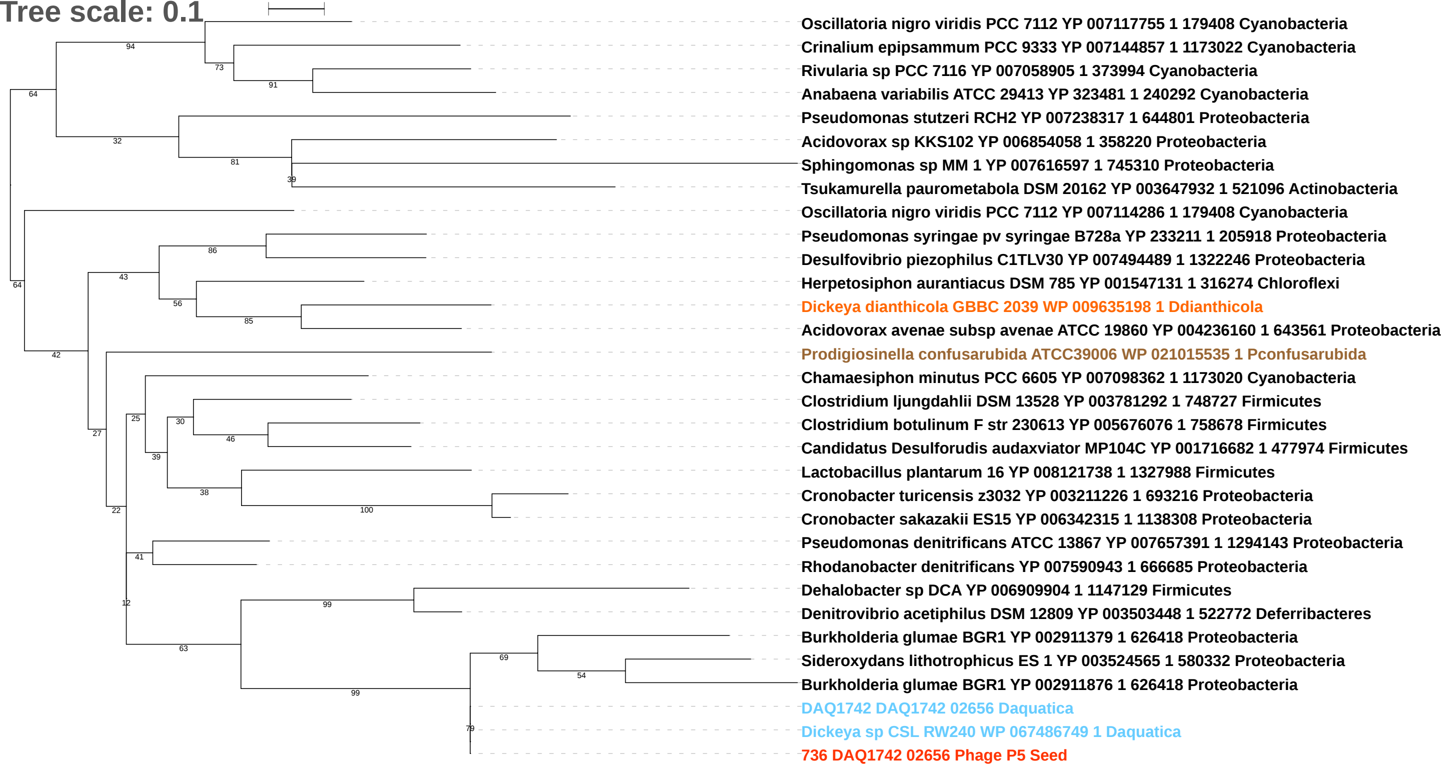
Tree scale: 0.1
Oscillatoria nigro viridis PCC 7112 YP 007117755 1 179408 Cyanobacteria
Crinalium epipsammum PCC 9333 YP 007144857 1 1173022 Cyanobacteria
Rivularia sp PCC 7116 YP 007058905 1 373994 Cyanobacteria
Anabaena variabilis ATCC 29413 YP 323481 1 240292 Cyanobacteria
Pseudomonas stutzeri RCH2 YP 007238317 1 644801 Proteobacteria
Acidovorax sp KKS102 YP 006854058 1 358220 Proteobacteria
Sphingomonas sp MM 1 YP 007616597 1 745310 Proteobacteria
Tsukamurella paurometabola DSM 20162 YP 003647932 1 521096 Actinobacteria
Oscillatoria nigro viridis PCC 7112 YP 007114286 1 179408 Cyanobacteria
Pseudomonas syringae pv syringae B728a YP 233211 1 205918 Proteobacteria
Desulfovibrio piezophilus C1TLV30 YP 007494489 1 1322246 Proteobacteria
Herpetosiphon aurantiacus DSM 785 YP 001547131 1 316274 Chloroflexi
Dickeya dianthicola GBBC 2039 WP 009635198 1 Ddianthicola
Acidovorax avenae subsp avenae ATCC 19860 YP 004236160 1 643561 Proteobacteria
Prodigiosinella confusarubida ATCC39006 WP 021015535 1 Pconfusarubida
Chamaesiphon minutus PCC 6605 YP 007098362 1 1173020 Cyanobacteria
Clostridium ljungdahlii DSM 13528 YP 003781292 1 748727 Firmicutes
Clostridium botulinum F str 230613 YP 005676076 1 758678 Firmicutes
Candidatus Desulforudis audaxviator MP104C YP 001716682 1 477974 Firmicutes
Lactobacillus plantarum 16 YP 008121738 1 1327988 Firmicutes
Cronobacter turicensis z3032 YP 003211226 1 693216 Proteobacteria
Cronobacter sakazakii ES15 YP 006342315 1 1138308 Proteobacteria
Pseudomonas denitrificans ATCC 13867 YP 007657391 1 1294143 Proteobacteria
Rhodanobacter denitrificans YP 007590943 1 666685 Proteobacteria
Dehalobacter sp DCA YP 006909904 1 1147129 Firmicutes
Denitrovibrio acetiphilus DSM 12809 YP 003503448 1 522772 Deferribacteres
Burkholderia glumae BGR1 YP 002911379 1 626418 Proteobacteria
Sideroxydans lithotrophicus ES 1 YP 003524565 1 580332 Proteobacteria
Burkholderia glumae BGR1 YP 002911876 1 626418 Proteobacteria
DAQ1742 DAQ1742 02656 Daquatica
Dickeya sp CSL RW240 WP 067486749 1 Daquatica
736 DAQ1742 02656 Phage P5 Seed

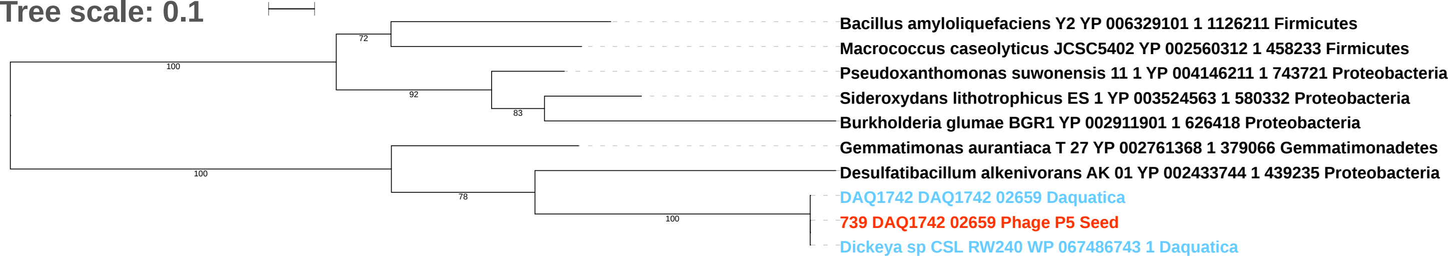
Tree scale: 0.1
Bacillus amyloliquefaciens Y2 YP 006329101 1 1126211 Firmicutes
Macrococcus caseolyticus JCSC5402 YP 002560312 1 458233 Firmicutes
Pseudoxanthomonas suwonensis 11 1 YP 004146211 1 743721 Proteobacteria
Sideroxydans lithotrophicus ES 1 YP 003524563 1 580332 Proteobacteria
Burkholderia glumae BGR1 YP 002911901 1 626418 Proteobacteria
Gemmatimonas aurantiaca T 27 YP 002761368 1 379066 Gemmatimonadetes
Desulfatibacillum alkenivorans AK 01 YP 002433744 1 439235 Proteobacteria
DAQ1742 DAQ1742 02659 Daquatica
739 DAQ1742 02659 Phage P5 Seed
Dickeya sp CSL RW240 WP 067486743 1 Daquatica
72
100
92
83
100
78
100

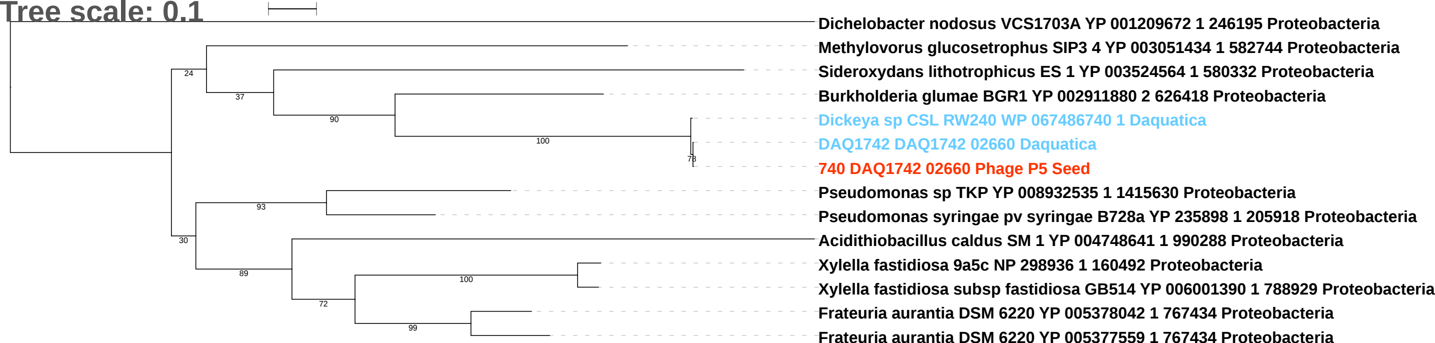
Tree scale: 0.1
Dichelobacter nodosus VCS1703A YP 001209672 1 246195 Proteobacteria
Methylovorus glucosetrophus SIP3 4 YP 003051434 1 582744 Proteobacteria
Sideroxydans lithotrophicus ES 1 YP 003524564 1 580332 Proteobacteria
Burkholderia glumae BGR1 YP 002911880 2 626418 Proteobacteria
Dickeya sp CSL RW240 WP 067486740 1 Daquatica
DAQ1742 DAQ1742 02660 Daquatica
740 DAQ1742 02660 Phage P5 Seed
Pseudomonas sp TKP YP 008932535 1 1415630 Proteobacteria
Pseudomonas syringae pv syringae B728a YP 235898 1 205918 Proteobacteria
Acidithiobacillus caldus SM 1 YP 004748641 1 990288 Proteobacteria
Xylella fastidiosa 9a5c NP 298936 1 160492 Proteobacteria
Xylella fastidiosa subsp fastidiosa GB514 YP 006001390 1 788929 Proteobacteria
Frateuria aurantia DSM 6220 YP 005378042 1 767434 Proteobacteria
Frateuria aurantia DSM 6220 YP 005377559 1 767434 Proteobacteria
24
37
90
100
78
93
30
89
100
72
99

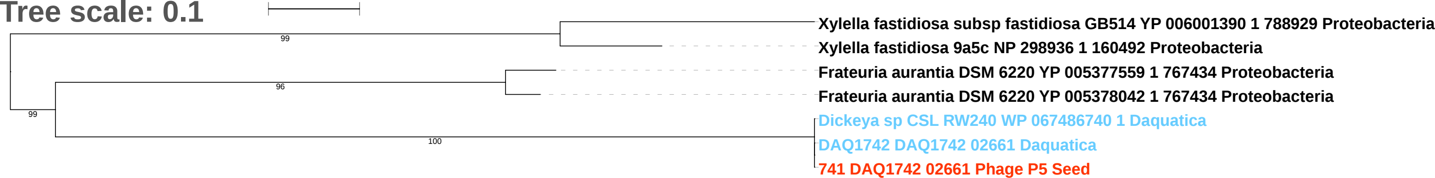

Tree scale: 0.1
Xylella fastidiosa subsp fastidiosa GB514 YP 006001390 1 788929 Proteobacteria
Xylella fastidiosa 9a5c NP 298936 1 160492 Proteobacteria
Frateuria aurantia DSM 6220 YP 005377559 1 767434 Proteobacteria
Frateuria aurantia DSM 6220 YP 005378042 1 767434 Proteobacteria
Dickeya sp CSL RW240 WP 067486740 1 Daquatica
DAQ1742 DAQ1742 02661 Daquatica
741 DAQ1742 02661 Phage P5 Seed
99
96
99
100

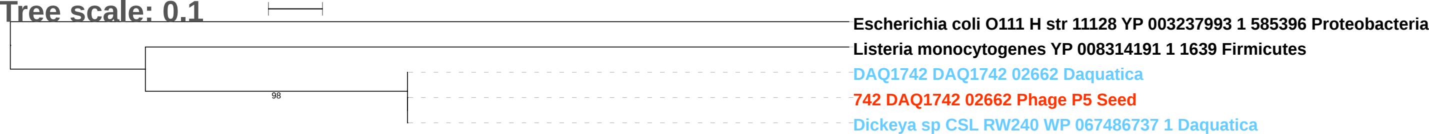
Tree scale: 0.1
Escherichia coli O111 H str 11128 YP 003237993 1 585396 Proteobacteria
Listeria monocytogenes YP 008314191 1 1639 Firmicutes
DAQ1742 DAQ1742 02662 Daquatica
742 DAQ1742 02662 Phage P5 Seed
Dickeya sp CSL RW240 WP 067486737 1 Daquatica
98

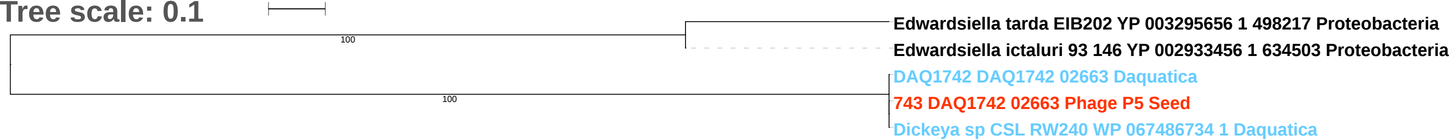
Tree scale: 0.1
100
100
Edwardsiella tarda EIB202 YP 003295656 1 498217 Proteobacteria
Edwardsiella ictaluri 93 146 YP 002933456 1 634503 Proteobacteria
DAQ1742 DAQ1742 02663 Daquatica
743 DAQ1742 02663 Phage P5 Seed
Dickeya sp CSL RW240 WP 067486734 1 Daquatica

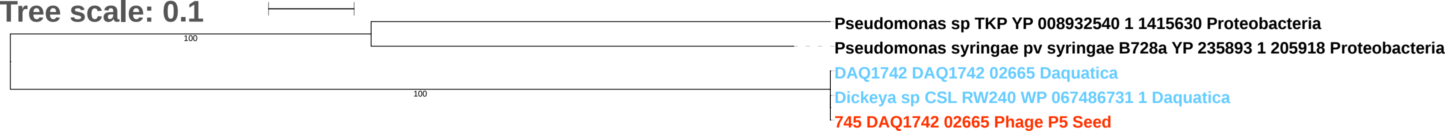
Tree scale: 0.1
100
100
Pseudomonas sp TKP YP 008932540 1 1415630 Proteobacteria
Pseudomonas syringae pv syringae B728a YP 235893 1 205918 Proteobacteria
DAQ1742 DAQ1742 02665 Daquatica
Dickeya sp CSL RW240 WP 067486731 1 Daquatica
745 DAQ1742 02665 Phage P5 Seed

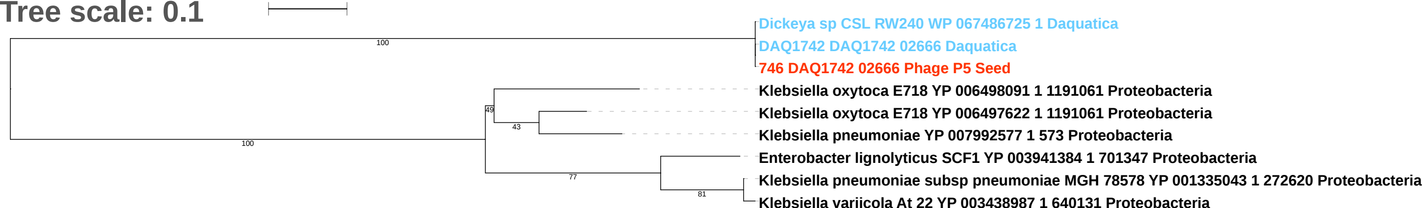

Tree scale: 0.1
Dickeya sp CSL RW240 WP 067486725 1 Daquatica
DAQ1742 DAQ1742 02666 Daquatica
746 DAQ1742 02666 Phage P5 Seed
Klebsiella oxytoca E718 YP 006498091 1 1191061 Proteobacteria
Klebsiella oxytoca E718 YP 006497622 1 1191061 Proteobacteria
Klebsiella pneumoniae YP 007992577 1 573 Proteobacteria
Enterobacter lignolyticus SCF1 YP 003941384 1 701347 Proteobacteria
Klebsiella pneumoniae subsp pneumoniae MGH 78578 YP 001335043 1 272620 Proteobacteria
Klebsiella variicola At 22 YP 003438987 1 640131 Proteobacteria
100
100
49
43
77
81

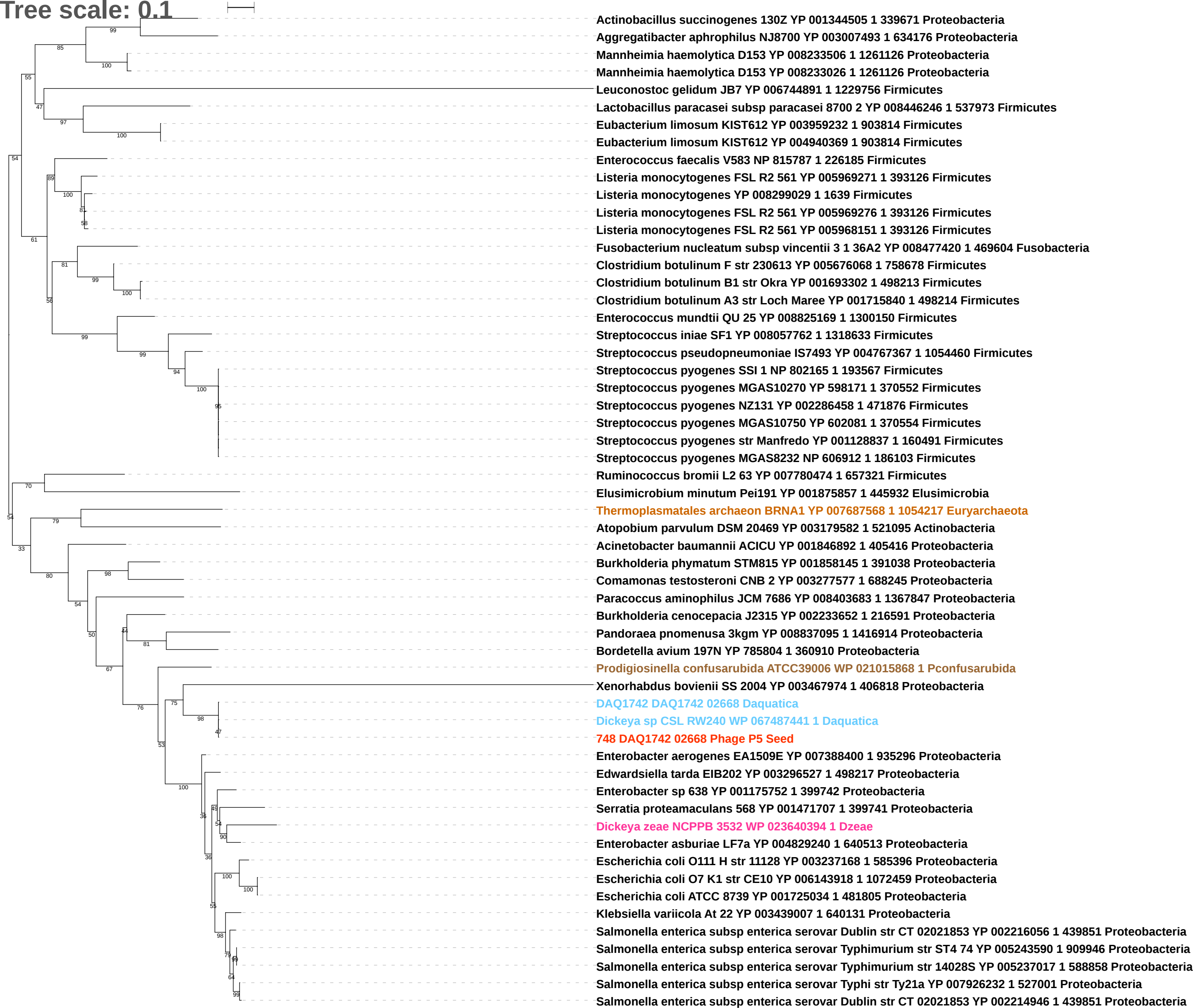
Tree scale: 0.1
Actinobacillus succinogenes 130Z YP 001344505 1 339671 Proteobacteria
Aggregatibacter aphrophilus NJ8700 YP 003007493 1 634176 Proteobacteria
Mannheimia haemolytica D153 YP 008233506 1 1261126 Proteobacteria
Mannheimia haemolytica D153 YP 008233026 1 1261126 Proteobacteria
Leuconostoc gelidum JB7 YP 006744891 1 1229756 Firmicutes
Lactobacillus paracasei subsp paracasei 8700 2 YP 008446246 1 537973 Firmicutes
Eubacterium limosum KIST612 YP 003959232 1 903814 Firmicutes
Eubacterium limosum KIST612 YP 004940369 1 903814 Firmicutes
Enterococcus faecalis V583 NP 815787 1 226185 Firmicutes
Listeria monocytogenes FSL R2 561 YP 005969271 1 393126 Firmicutes
Listeria monocytogenes YP 008299029 1 1639 Firmicutes
Listeria monocytogenes FSL R2 561 YP 005969276 1 393126 Firmicutes
Listeria monocytogenes FSL R2 561 YP 005968151 1 393126 Firmicutes
Fusobacterium nucleatum subsp vincentii 3 1 36A2 YP 008477420 1 469604 Fusobacteria
Clostridium botulinum F str 230613 YP 005676068 1 758678 Firmicutes
Clostridium botulinum B1 str Okra YP 001693302 1 498213 Firmicutes
Clostridium botulinum A3 str Loch Maree YP 001715840 1 498214 Firmicutes
Enterococcus mundtii QU 25 YP 008825169 1 1300150 Firmicutes
Streptococcus iniae SF1 YP 008057762 1 1318633 Firmicutes
Streptococcus pseudopneumoniae IS7493 YP 004767367 1 1054460 Firmicutes
Streptococcus pyogenes SSI 1 NP 802165 1 193567 Firmicutes
Streptococcus pyogenes MGAS10270 YP 598171 1 370552 Firmicutes
Streptococcus pyogenes NZ131 YP 002286458 1 471876 Firmicutes
Streptococcus pyogenes MGAS10750 YP 602081 1 370554 Firmicutes
Streptococcus pyogenes str Manfredo YP 001128837 1 160491 Firmicutes
Streptococcus pyogenes MGAS8232 NP 606912 1 186103 Firmicutes
Ruminococcus bromii L2 63 YP 007780474 1 657321 Firmicutes
Elusimicrobium minutum Pei191 YP 001875857 1 445932 Elusimicrobia
Thermoplasmatales archaeon BRNA1 YP 007687568 1 1054217 Euryarchaeota
Atopobium parvulum DSM 20469 YP 003179582 1 521095 Actinobacteria
Acinetobacter baumannii ACICU YP 001846892 1 405416 Proteobacteria
Burkholderia phymatum STM815 YP 001858145 1 391038 Proteobacteria
Comamonas testosteroni CNB 2 YP 003277577 1 688245 Proteobacteria
Paracoccus aminophilus JCM 7686 YP 008403683 1 1367847 Proteobacteria
Burkholderia cenocepacia J2315 YP 002233652 1 216591 Proteobacteria
Pandoraea pnomenusa 3kgm YP 008837095 1 1416914 Proteobacteria
Bordetella avium 197N YP 785804 1 360910 Proteobacteria
Prodigiosinella confusarubida ATCC39006 WP 021015868 1 Pconfusarubida
Xenorhabdus bovienii SS 2004 YP 003467974 1 406818 Proteobacteria
DAQ1742 DAQ1742 02668 Daquatica
Dickeya sp CSL RW240 WP 067487441 1 Daquatica
748 DAQ1742 02668 Phage P5 Seed
Enterobacter aerogenes EA1509E YP 007388400 1 935296 Proteobacteria
Edwardsiella tarda EIB202 YP 003296527 1 498217 Proteobacteria
Enterobacter sp 638 YP 001175752 1 399742 Proteobacteria
Serratia proteamaculans 568 YP 001471707 1 399741 Proteobacteria
Dickeya zeae NCPPB 3532 WP 023640394 1 Dzeae
Enterobacter asburiae LF7a YP 004829240 1 640513 Proteobacteria
Escherichia coli O111 H str 11128 YP 003237168 1 585396 Proteobacteria
Escherichia coli O7 K1 str CE10 YP 006143918 1 1072459 Proteobacteria
Escherichia coli ATCC 8739 YP 001725034 1 481805 Proteobacteria
Klebsiella variicola At 22 YP 003439007 1 640131 Proteobacteria
Salmonella enterica subsp enterica serovar Dublin str CT 02021853 YP 002216056 1 439851 Proteobacteria
Salmonella enterica subsp enterica serovar Typhimurium str ST4 74 YP 005243590 1 909946 Proteobacteria
Salmonella enterica subsp enterica serovar Typhimurium str 14028S YP 005237017 1 588858 Proteobacteria
Salmonella enterica subsp enterica serovar Typhi str Ty21a YP 007926232 1 527001 Proteobacteria
Salmonella enterica subsp enterica serovar Dublin str CT 02021853 YP 002214946 1 439851 Proteobacteria

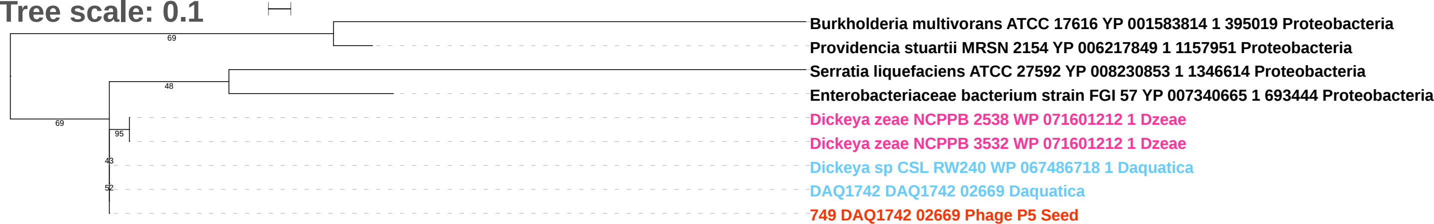
Tree scale: 0.1
Burkholderia multivorans ATCC 17616 YP 001583814 1 395019 Proteobacteria
Providencia stuartii MRSN 2154 YP 006217849 1 1157951 Proteobacteria
Serratia liquefaciens ATCC 27592 YP 008230853 1 1346614 Proteobacteria
Enterobacteriaceae bacterium strain FGI 57 YP 007340665 1 693444 Proteobacteria
Dickeya zeae NCPPB 2538 WP 071601212 1 Dzeae
Dickeya zeae NCPPB 3532 WP 071601212 1 Dzeae
Dickeya sp CSL RW240 WP 067486718 1 Daquatica
DAQ1742 DAQ1742 02669 Daquatica
749 DAQ1742 02669 Phage P5 Seed
69
69
48
95
43
52

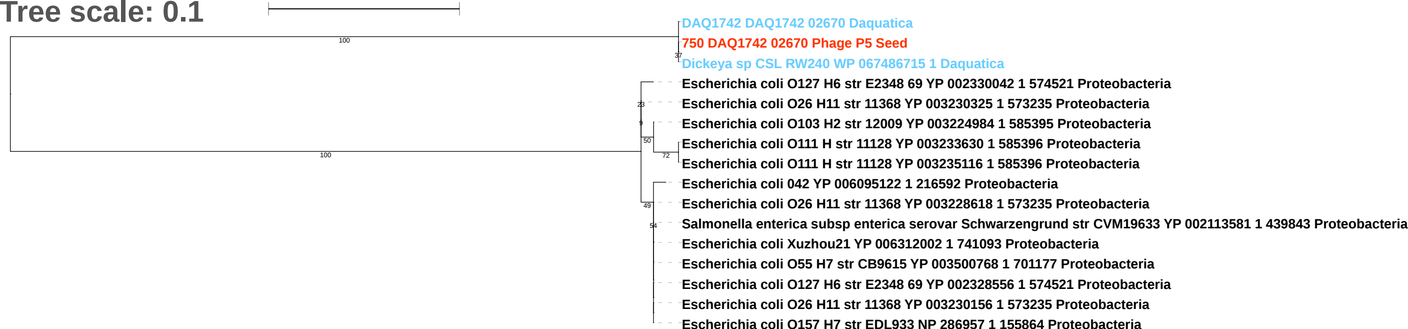

Tree scale: 0.1
DAQ1742 DAQ1742 02670 Daquatica
750 DAQ1742 02670 Phage P5 Seed
Dickeya sp CSL RW240 WP 067486715 1 Daquatica
Escherichia coli O127 H6 str E2348 69 YP 002330042 1 574521 Proteobacteria
Escherichia coli O26 H11 str 11368 YP 003230325 1 573235 Proteobacteria
Escherichia coli O103 H2 str 12009 YP 003224984 1 585395 Proteobacteria
Escherichia coli O111 H str 11128 YP 003233630 1 585396 Proteobacteria
Escherichia coli O111 H str 11128 YP 003235116 1 585396 Proteobacteria
Escherichia coli 042 YP 006095122 1 216592 Proteobacteria
Escherichia coli O26 H11 str 11368 YP 003228618 1 573235 Proteobacteria
Salmonella enterica subsp enterica serovar Schwarzengrund str CVM19633 YP 002113581 1 439843 Proteobacteria
Escherichia coli Xuzhou21 YP 006312002 1 741093 Proteobacteria
Escherichia coli O55 H7 str CB9615 YP 003500768 1 701177 Proteobacteria
Escherichia coli O127 H6 str E2348 69 YP 002328556 1 574521 Proteobacteria
Escherichia coli O26 H11 str 11368 YP 003230156 1 573235 Proteobacteria
Escherichia coli O157 H7 str EDL933 NP 286957 1 155864 Proteobacteria
100
100
37
23
9
50
72
49
54

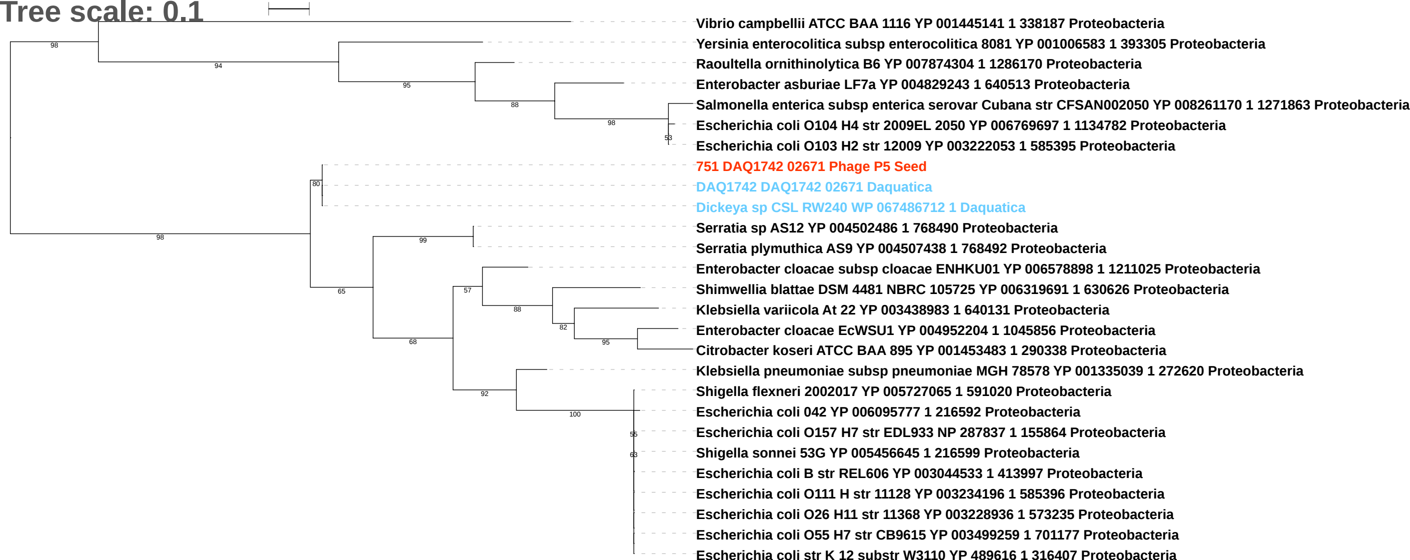
Tree scale: 0.1
Vibrio campbellii ATCC BAA 1116 YP 001445141 1 338187 Proteobacteria
Yersinia enterocolitica subsp enterocolitica 8081 YP 001006583 1 393305 Proteobacteria
Raoultella ornithinolytica B6 YP 007874304 1 1286170 Proteobacteria
Enterobacter asburiae LF7a YP 004829243 1 640513 Proteobacteria
Salmonella enterica subsp enterica serovar Cubana str CFSAN002050 YP 008261170 1 1271863 Proteobacteria
Escherichia coli O104 H4 str 2009EL 2050 YP 006769697 1 1134782 Proteobacteria
Escherichia coli O103 H2 str 12009 YP 003222053 1 585395 Proteobacteria
751 DAQ1742 02671 Phage P5 Seed
DAQ1742 DAQ1742 02671 Daquatica
Dickeya sp CSL RW240 WP 067486712 1 Daquatica
Serratia sp AS12 YP 004502486 1 768490 Proteobacteria
Serratia plymuthica AS9 YP 004507438 1 768492 Proteobacteria
Enterobacter cloacae subsp cloacae ENHKU01 YP 006578898 1 1211025 Proteobacteria
Shimwellia blattae DSM 4481 NBRC 105725 YP 006319691 1 630626 Proteobacteria
Klebsiella variicola At 22 YP 003438983 1 640131 Proteobacteria
Enterobacter cloacae EcWSU1 YP 004952204 1 1045856 Proteobacteria
Citrobacter koseri ATCC BAA 895 YP 001453483 1 290338 Proteobacteria
Klebsiella pneumoniae subsp pneumoniae MGH 78578 YP 001335039 1 272620 Proteobacteria
Shigella flexneri 2002017 YP 005727065 1 591020 Proteobacteria
Escherichia coli 042 YP 006095777 1 216592 Proteobacteria
Escherichia coli O157 H7 str EDL933 NP 287837 1 155864 Proteobacteria
Shigella sonnei 53G YP 005456645 1 216599 Proteobacteria
Escherichia coli B str REL606 YP 003044533 1 413997 Proteobacteria
Escherichia coli O111 H str 11128 YP 003234196 1 585396 Proteobacteria
Escherichia coli O26 H11 str 11368 YP 003228936 1 573235 Proteobacteria
Escherichia coli O55 H7 str CB9615 YP 003499259 1 701177 Proteobacteria
Escherichia coli str K 12 substr W3110 YP 489616 1 316407 Proteobacteria
98
94
95
88
98
53
80
98
99
65
57
88
82
95
68
92
100
55
63

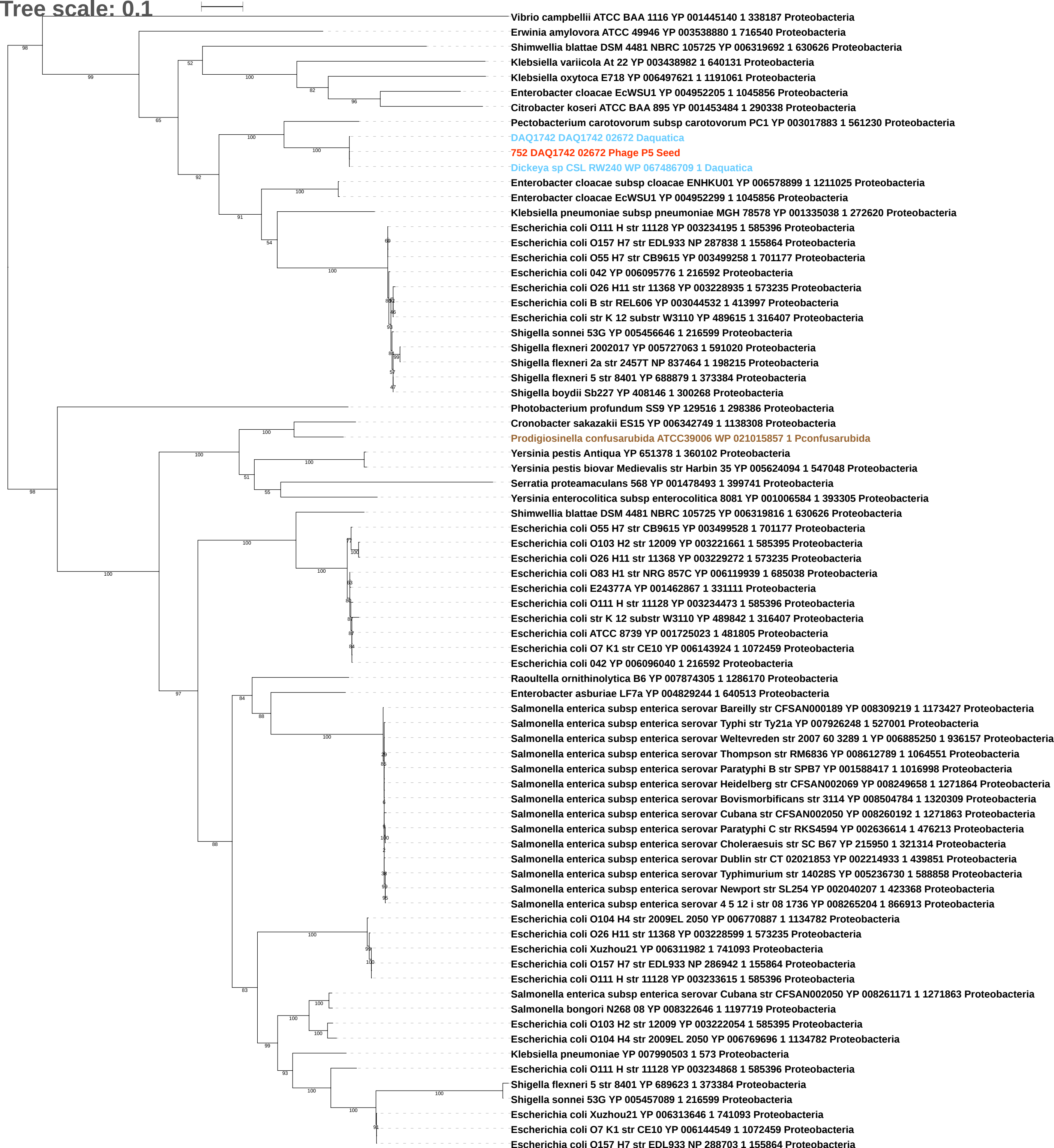

Tree scale: 0.1
Vibrio campbellii ATCC BAA 1116 YP 001445140 1 338187 Proteobacteria
Erwinia amylovora ATCC 49946 YP 003538880 1 716540 Proteobacteria
Shimwellia blattae DSM 4481 NBRC 105725 YP 006319692 1 630626 Proteobacteria
Klebsiella variicola At 22 YP 003438982 1 640131 Proteobacteria
Klebsiella oxytoca E718 YP 006497621 1 1191061 Proteobacteria
Enterobacter cloacae EcWSU1 YP 004952205 1 1045856 Proteobacteria
Citrobacter koseri ATCC BAA 895 YP 001453484 1 290338 Proteobacteria
Pectobacterium carotovorum subsp carotovorum PC1 YP 003017883 1 561230 Proteobacteria
DAQ1742 DAQ1742 02672 Daquatica
752 DAQ1742 02672 Phage P5 Seed
Dickeya sp CSL RW240 WP 067486709 1 Daquatica
Enterobacter cloacae subsp cloacae ENHKU01 YP 006578899 1 1211025 Proteobacteria
Enterobacter cloacae EcWSU1 YP 004952299 1 1045856 Proteobacteria
Klebsiella pneumoniae subsp pneumoniae MGH 78578 YP 001335038 1 272620 Proteobacteria
Escherichia coli O111 H str 11128 YP 003234195 1 585396 Proteobacteria
Escherichia coli O157 H7 str EDL933 NP 287838 1 155864 Proteobacteria
Escherichia coli O55 H7 str CB9615 YP 003499258 1 701177 Proteobacteria
Escherichia coli 042 YP 006095776 1 216592 Proteobacteria
Escherichia coli O26 H11 str 11368 YP 003228935 1 573235 Proteobacteria
Escherichia coli B str REL606 YP 003044532 1 413997 Proteobacteria
Escherichia coli str K 12 substr W3110 YP 489615 1 316407 Proteobacteria
Shigella sonnei 53G YP 005456646 1 216599 Proteobacteria
Shigella flexneri 2002017 YP 005727063 1 591020 Proteobacteria
Shigella flexneri 2a str 2457T NP 837464 1 198215 Proteobacteria
Shigella flexneri 5 str 8401 YP 688879 1 373384 Proteobacteria
Shigella boydii Sb227 YP 408146 1 300268 Proteobacteria
Photobacterium profundum SS9 YP 129516 1 298386 Proteobacteria
Cronobacter sakazakii ES15 YP 006342749 1 1138308 Proteobacteria
Prodigiosinella confusarubida ATCC39006 WP 021015857 1 Pconfusarubida
Yersinia pestis Antiqua YP 651378 1 360102 Proteobacteria
Yersinia pestis biovar Medievalis str Harbin 35 YP 005624094 1 547048 Proteobacteria
Serratia proteamaculans 568 YP 001478493 1 399741 Proteobacteria
Yersinia enterocolitica subsp enterocolitica 8081 YP 001006584 1 393305 Proteobacteria
Shimwellia blattae DSM 4481 NBRC 105725 YP 006319816 1 630626 Proteobacteria
Escherichia coli O55 H7 str CB9615 YP 003499528 1 701177 Proteobacteria
Escherichia coli O103 H2 str 12009 YP 003221661 1 585395 Proteobacteria
Escherichia coli O26 H11 str 11368 YP 003229272 1 573235 Proteobacteria
Escherichia coli O83 H1 str NRG 857C YP 006119939 1 685038 Proteobacteria
Escherichia coli E24377A YP 001462867 1 331111 Proteobacteria
Escherichia coli O111 H str 11128 YP 003234473 1 585396 Proteobacteria
Escherichia coli str K 12 substr W3110 YP 489842 1 316407 Proteobacteria
Escherichia coli ATCC 8739 YP 001725023 1 481805 Proteobacteria
Escherichia coli O7 K1 str CE10 YP 006143924 1 1072459 Proteobacteria
Escherichia coli 042 YP 006096040 1 216592 Proteobacteria
Raoultella ornithinolytica B6 YP 007874305 1 1286170 Proteobacteria
Enterobacter asburiae LF7a YP 004829244 1 640513 Proteobacteria
Salmonella enterica subsp enterica serovar Bareilly str CFSAN000189 YP 008309219 1 1173427 Proteobacteria
Salmonella enterica subsp enterica serovar Typhi str Ty21a YP 007926248 1 527001 Proteobacteria
Salmonella enterica subsp enterica serovar Weltevreden str 2007 60 3289 1 YP 006885250 1 936157 Proteobacteria
Salmonella enterica subsp enterica serovar Thompson str RM6836 YP 008612789 1 1064551 Proteobacteria
Salmonella enterica subsp enterica serovar Paratyphi B str SPB7 YP 001588417 1 1016998 Proteobacteria
Salmonella enterica subsp enterica serovar Heidelberg str CFSAN002069 YP 008249658 1 1271864 Proteobacteria
Salmonella enterica subsp enterica serovar Bovismorbificans str 3114 YP 008504784 1 1320309 Proteobacteria
Salmonella enterica subsp enterica serovar Cubana str CFSAN002050 YP 008260192 1 1271863 Proteobacteria
Salmonella enterica subsp enterica serovar Paratyphi C str RKS4594 YP 002636614 1 476213 Proteobacteria
Salmonella enterica subsp enterica serovar Choleraesuis str SC B67 YP 215950 1 321314 Proteobacteria
Salmonella enterica subsp enterica serovar Dublin str CT 02021853 YP 002214933 1 439851 Proteobacteria
Salmonella enterica subsp enterica serovar Typhimurium str 14028S YP 005236730 1 588858 Proteobacteria
Salmonella enterica subsp enterica serovar Newport str SL254 YP 002040207 1 423368 Proteobacteria
Salmonella enterica subsp enterica serovar 4 5 12 i str 08 1736 YP 008265204 1 866913 Proteobacteria
Escherichia coli O104 H4 str 2009EL 2050 YP 006770887 1 1134782 Proteobacteria
Escherichia coli O26 H11 str 11368 YP 003228599 1 573235 Proteobacteria
Escherichia coli Xuzhou21 YP 006311982 1 741093 Proteobacteria
Escherichia coli O157 H7 str EDL933 NP 286942 1 155864 Proteobacteria
Escherichia coli O111 H str 11128 YP 003233615 1 585396 Proteobacteria
Salmonella enterica subsp enterica serovar Cubana str CFSAN002050 YP 008261171 1 1271863 Proteobacteria
Salmonella bongori N268 08 YP 008322646 1 1197719 Proteobacteria
Escherichia coli O103 H2 str 12009 YP 003222054 1 585395 Proteobacteria
Escherichia coli O104 H4 str 2009EL 2050 YP 006769696 1 1134782 Proteobacteria
Klebsiella pneumoniae YP 007990503 1 573 Proteobacteria
Escherichia coli O111 H str 11128 YP 003234868 1 585396 Proteobacteria
Shigella flexneri 5 str 8401 YP 689623 1 373384 Proteobacteria
Shigella sonnei 53G YP 005457089 1 216599 Proteobacteria
Escherichia coli Xuzhou21 YP 006313646 1 741093 Proteobacteria
Escherichia coli O7 K1 str CE10 YP 006144549 1 1072459 Proteobacteria
Escherichia coli O157 H7 str EDL933 NP 288703 1 155864 Proteobacteria

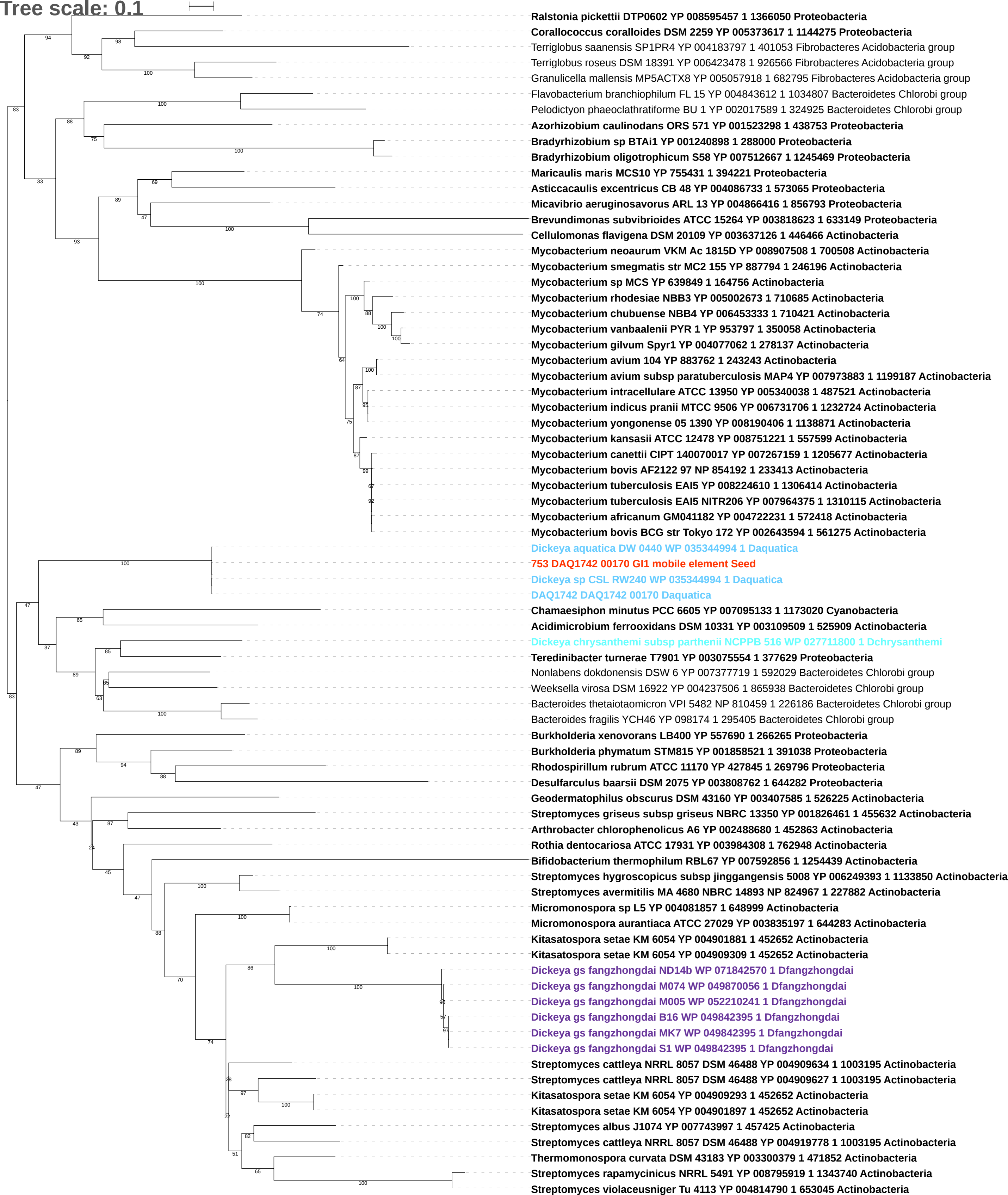
Tree scale: 0.1
Ralstonia pickettii DTP0602 YP 008595457 1 1366050 Proteobacteria
Corallococcus coralloides DSM 2259 YP 005373617 1 1144275 Proteobacteria
Terriglobus saanensis SP1PR4 YP 004183797 1 401053 Fibrobacteres Acidobacteria group
Terriglobus roseus DSM 18391 YP 006423478 1 926566 Fibrobacteres Acidobacteria group
Granulicella mallensis MP5ACTX8 YP 005057918 1 682795 Fibrobacteres Acidobacteria group
Flavobacterium branchiophilum FL 15 YP 004843612 1 1034807 Bacteroidetes Chlorobi group
Pelodictyon phaeoclathratiforme BU 1 YP 002017589 1 324925 Bacteroidetes Chlorobi group
Azorhizobium caulinodans ORS 571 YP 001523298 1 438753 Proteobacteria
Bradyrhizobium sp BTAi1 YP 001240898 1 288000 Proteobacteria
Bradyrhizobium oligotrophicum S58 YP 007512667 1 1245469 Proteobacteria
Maricaulis maris MCS10 YP 755431 1 394221 Proteobacteria
Asticcacaulis excentricus CB 48 YP 004086733 1 573065 Proteobacteria
Micavibrio aeruginosavorus ARL 13 YP 004866416 1 856793 Proteobacteria
Brevundimonas subvibrioides ATCC 15264 YP 003818623 1 633149 Proteobacteria
Cellulomonas flavigena DSM 20109 YP 003637126 1 446466 Actinobacteria
Mycobacterium neoaurum VKM Ac 1815D YP 008907508 1 700508 Actinobacteria
Mycobacterium smegmatis str MC2 155 YP 887794 1 246196 Actinobacteria
Mycobacterium sp MCS YP 639849 1 164756 Actinobacteria
Mycobacterium rhodesiae NBB3 YP 005002673 1 710685 Actinobacteria
Mycobacterium chubuense NBB4 YP 006453333 1 710421 Actinobacteria
Mycobacterium vanbaalenii PYR 1 YP 953797 1 350058 Actinobacteria
Mycobacterium gilvum Spyr1 YP 004077062 1 278137 Actinobacteria
Mycobacterium avium 104 YP 883762 1 243243 Actinobacteria
Mycobacterium avium subsp paratuberculosis MAP4 YP 007973883 1 1199187 Actinobacteria
Mycobacterium intracellulare ATCC 13950 YP 005340038 1 487521 Actinobacteria
Mycobacterium indicus pranii MTCC 9506 YP 006731706 1 1232724 Actinobacteria
Mycobacterium yongonense 05 1390 YP 008190406 1 1138871 Actinobacteria
Mycobacterium kansasii ATCC 12478 YP 008751221 1 557599 Actinobacteria
Mycobacterium canettii CIPT 140070017 YP 007267159 1 1205677 Actinobacteria
Mycobacterium bovis AF2122 97 NP 854192 1 233413 Actinobacteria
Mycobacterium tuberculosis EAI5 YP 008224610 1 1306414 Actinobacteria
Mycobacterium tuberculosis EAI5 NITR206 YP 007964375 1 1310115 Actinobacteria
Mycobacterium africanum GM041182 YP 004722231 1 572418 Actinobacteria
Mycobacterium bovis BCG str Tokyo 172 YP 002643594 1 561275 Actinobacteria
Dickeya aquatica DW 0440 WP 035344994 1 Daquatica
753 DAQ1742 00170 GI1 mobile element Seed
Dickeya sp CSL RW240 WP 035344994 1 Daquatica
DAQ1742 DAQ1742 00170 Daquatica
Chamaesiphon minutus PCC 6605 YP 007095133 1 1173020 Cyanobacteria
Acidimicrobium ferrooxidans DSM 10331 YP 003109509 1 525909 Actinobacteria
Dickeya chrysanthemi subsp parthenii NCPPB 516 WP 027711800 1 Dchrysanthemi
Teredinibacter turnerae T7901 YP 003075554 1 377629 Proteobacteria
Nonlabens dokdonensis DSW 6 YP 007377719 1 592029 Bacteroidetes Chlorobi group
Weeksella virosa DSM 16922 YP 004237506 1 865938 Bacteroidetes Chlorobi group
Bacteroides thetaiotaomicron VPI 5482 NP 810459 1 226186 Bacteroidetes Chlorobi group
Bacteroides fragilis YCH46 YP 098174 1 295405 Bacteroidetes Chlorobi group
Burkholderia xenovorans LB400 YP 557690 1 266265 Proteobacteria
Burkholderia phymatum STM815 YP 001858521 1 391038 Proteobacteria
Rhodospirillum rubrum ATCC 11170 YP 427845 1 269796 Proteobacteria
Desulfarculus baarsii DSM 2075 YP 003808762 1 644282 Proteobacteria
Geodermatophilus obscurus DSM 43160 YP 003407585 1 526225 Actinobacteria
Streptomyces griseus subsp griseus NBRC 13350 YP 001826461 1 455632 Actinobacteria
Arthrobacter chlorophenolicus A6 YP 002488680 1 452863 Actinobacteria
Rothia dentocariosa ATCC 17931 YP 003984308 1 762948 Actinobacteria
Bifidobacterium thermophilum RBL67 YP 007592856 1 1254439 Actinobacteria
Streptomyces hygroscopicus subsp jinggangensis 5008 YP 006249393 1 1133850 Actinobacteria
Streptomyces avermitilis MA 4680 NBRC 14893 NP 824967 1 227882 Actinobacteria
Micromonospora sp L5 YP 004081857 1 648999 Actinobacteria
Micromonospora aurantiaca ATCC 27029 YP 003835197 1 644283 Actinobacteria
Kitasatospora setae KM 6054 YP 004901881 1 452652 Actinobacteria
Kitasatospora setae KM 6054 YP 004909309 1 452652 Actinobacteria
Dickeya gs fangzhongdai ND14b WP 071842570 1 Dfangzhongdai
Dickeya gs fangzhongdai M074 WP 049870056 1 Dfangzhongdai
Dickeya gs fangzhongdai M005 WP 052210241 1 Dfangzhongdai
Dickeya gs fangzhongdai B16 WP 049842395 1 Dfangzhongdai
Dickeya gs fangzhongdai MK7 WP 049842395 1 Dfangzhongdai
Dickeya gs fangzhongdai S1 WP 049842395 1 Dfangzhongdai
Streptomyces cattleya NRRL 8057 DSM 46488 YP 004909634 1 1003195 Actinobacteria
Streptomyces cattleya NRRL 8057 DSM 46488 YP 004909627 1 1003195 Actinobacteria
Kitasatospora setae KM 6054 YP 004909293 1 452652 Actinobacteria
Kitasatospora setae KM 6054 YP 004901897 1 452652 Actinobacteria
Streptomyces albus J1074 YP 007743997 1 457425 Actinobacteria
Streptomyces cattleya NRRL 8057 DSM 46488 YP 004919778 1 1003195 Actinobacteria
Thermomonospora curvata DSM 43183 YP 003300379 1 471852 Actinobacteria
Streptomyces rapamycinicus NRRL 5491 YP 008795919 1 1343740 Actinobacteria
Streptomyces violaceusniger Tu 4113 YP 004814790 1 653045 Actinobacteria

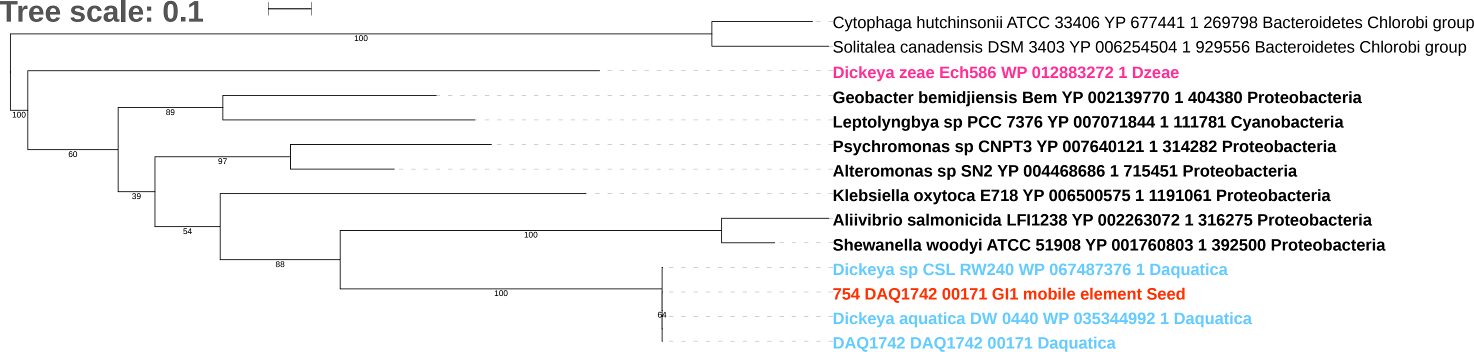
Tree scale: 0.1
Cytophaga hutchinsonii ATCC 33406 YP 677441 1 269798 Bacteroidetes Chlorobi group
Solitalea canadensis DSM 3403 YP 006254504 1 929556 Bacteroidetes Chlorobi group
Dickeya zeae Ech586 WP 012883272 1 Dzeae
Geobacter bemidjiensis Bem YP 002139770 1 404380 Proteobacteria
Leptolyngbya sp PCC 7376 YP 007071844 1 111781 Cyanobacteria
Psychromonas sp CNPT3 YP 007640121 1 314282 Proteobacteria
Alteromonas sp SN2 YP 004468686 1 715451 Proteobacteria
Klebsiella oxytoca E718 YP 006500575 1 1191061 Proteobacteria
Aliivibrio salmonicida LFI1238 YP 002263072 1 316275 Proteobacteria
Shewanella woodyi ATCC 51908 YP 001760803 1 392500 Proteobacteria
Dickeya sp CSL RW240 WP 067487376 1 Daquatica
754 DAQ1742 00171 GI1 mobile element Seed
Dickeya aquatica DW 0440 WP 035344992 1 Daquatica
DAQ1742 DAQ1742 00171 Daquatica
100
100
89
60
97
39
54
100
88
100
64

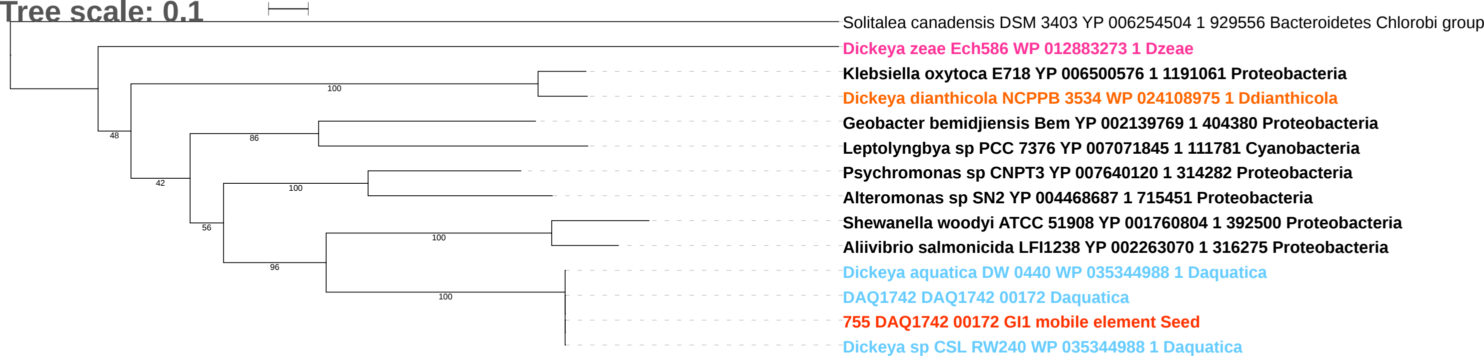
Tree scale: 0.1
Solitalea canadensis DSM 3403 YP 006254504 1 929556 Bacteroidetes Chlorobi group
Dickeya zeae Ech586 WP 012883273 1 Dzeae
Klebsiella oxytoca E718 YP 006500576 1 1191061 Proteobacteria
Dickeya dianthicola NCPPB 3534 WP 024108975 1 Ddianthicola
Geobacter bemidjiensis Bem YP 002139769 1 404380 Proteobacteria
Leptolyngbya sp PCC 7376 YP 007071845 1 111781 Cyanobacteria
Psychromonas sp CNPT3 YP 007640120 1 314282 Proteobacteria
Alteromonas sp SN2 YP 004468687 1 715451 Proteobacteria
Shewanella woodyi ATCC 51908 YP 001760804 1 392500 Proteobacteria
Aliivibrio salmonicida LFI1238 YP 002263070 1 316275 Proteobacteria
Dickeya aquatica DW 0440 WP 035344988 1 Daquatica
DAQ1742 DAQ1742 00172 Daquatica
755 DAQ1742 00172 GI1 mobile element Seed
Dickeya sp CSL RW240 WP 035344988 1 Daquatica
100
48
86
42
100
56
100
96
100

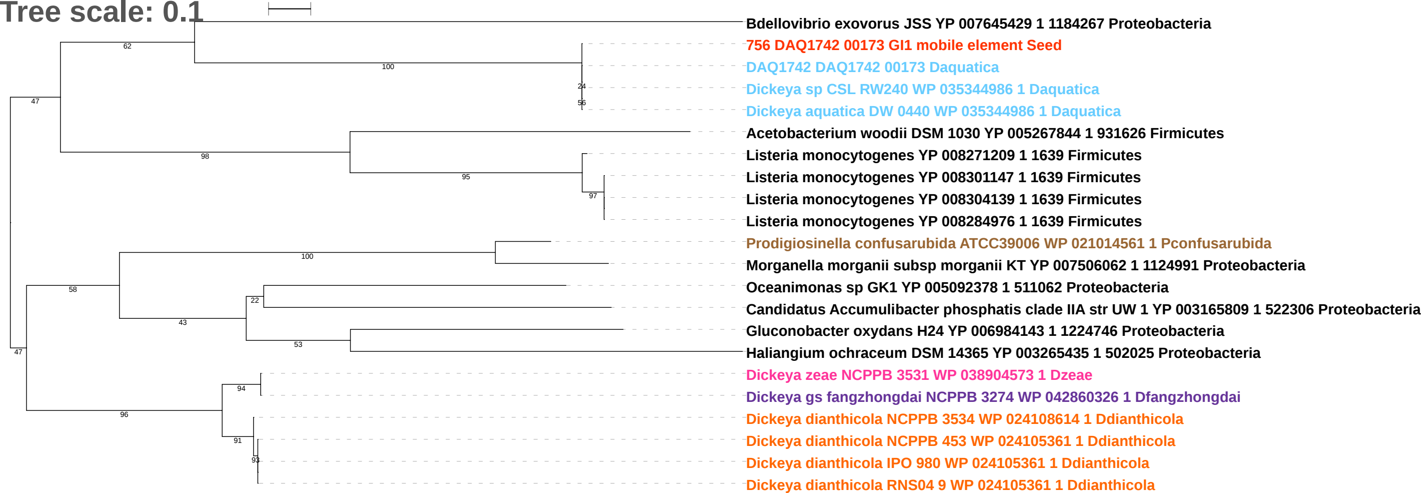

Tree scale: 0.1
Bdellovibrio exovorus JSS YP 007645429 1 1184267 Proteobacteria
756 DAQ1742 00173 GI1 mobile element Seed
DAQ1742 DAQ1742 00173 Daquatica
Dickeya sp CSL RW240 WP 035344986 1 Daquatica
Dickeya aquatica DW 0440 WP 035344986 1 Daquatica
Acetobacterium woodii DSM 1030 YP 005267844 1 931626 Firmicutes
Listeria monocytogenes YP 008271209 1 1639 Firmicutes
Listeria monocytogenes YP 008301147 1 1639 Firmicutes
Listeria monocytogenes YP 008304139 1 1639 Firmicutes
Listeria monocytogenes YP 008284976 1 1639 Firmicutes
Prodigiosinella confusarubida ATCC39006 WP 021014561 1 Pconfusarubida
Morganella morganii subsp morganii KT YP 007506062 1 1124991 Proteobacteria
Oceanimonas sp GK1 YP 005092378 1 511062 Proteobacteria
Candidatus Accumulibacter phosphatis clade IIA str UW 1 YP 003165809 1 522306 Proteobacteria
Gluconobacter oxydans H24 YP 006984143 1 1224746 Proteobacteria
Haliangium ochraceum DSM 14365 YP 003265435 1 502025 Proteobacteria
Dickeya zeae NCPPB 3531 WP 038904573 1 Dzeae
Dickeya gs fangzhongdai NCPPB 3274 WP 042860326 1 Dfangzhongdai
Dickeya dianthicola NCPPB 3534 WP 024108614 1 Ddianthicola
Dickeya dianthicola NCPPB 453 WP 024105361 1 Ddianthicola
Dickeya dianthicola IPO 980 WP 024105361 1 Ddianthicola
Dickeya dianthicola RNS04 9 WP 024105361 1 Ddianthicola
62
100
24
56
47
98
95
97
100
58
22
43
53
47
94
96
91
93

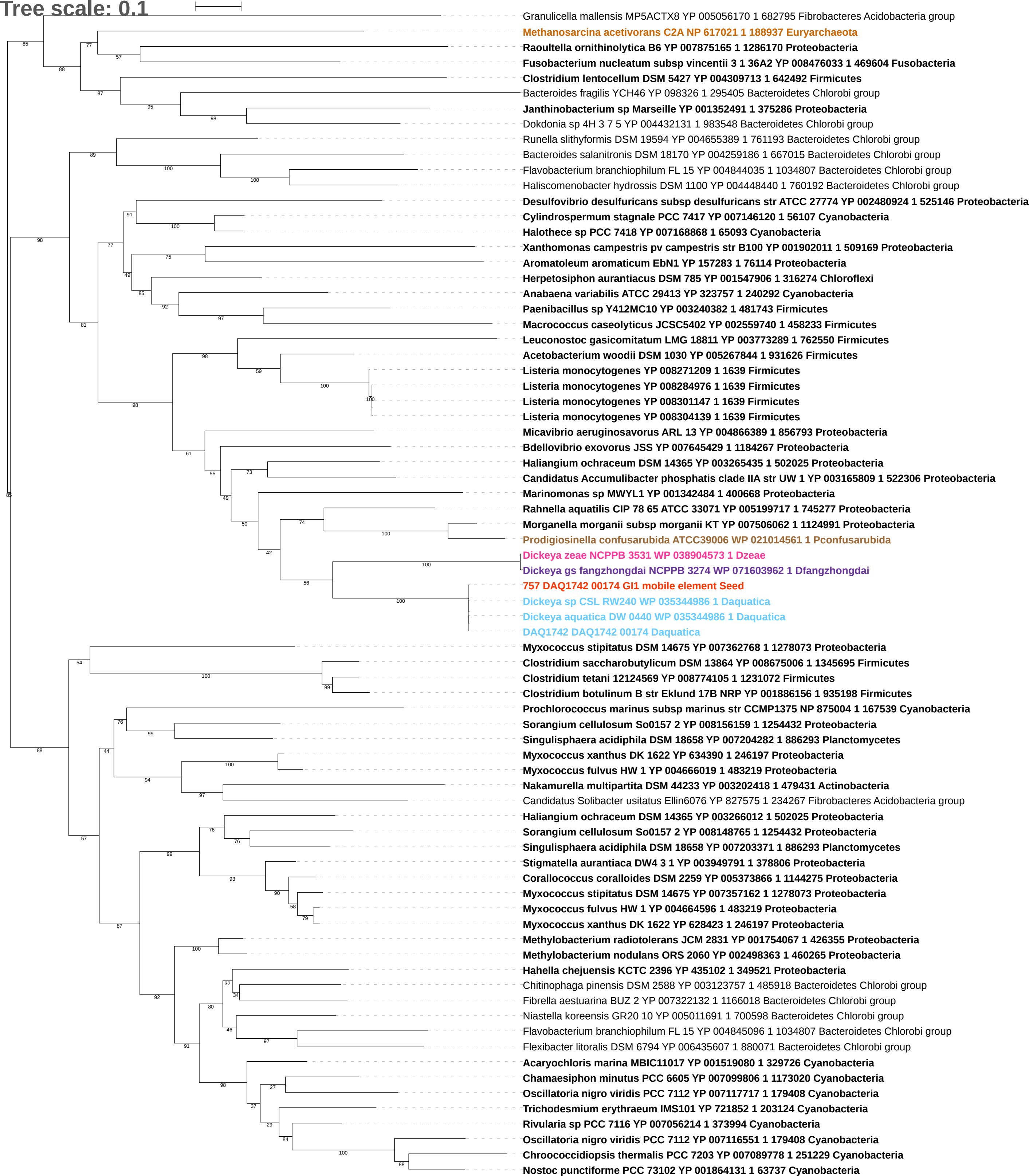
Tree scale: 0.1
Granulicella mallensis MP5ACTX8 YP 005056170 1 682795 Fibrobacteres Acidobacteria group
Methanosarcina acetivorans C2A NP 617021 1 188937 Euryarchaeota
Raoultella ornithinolytica B6 YP 007875165 1 1286170 Proteobacteria
Fusobacterium nucleatum subsp vincentii 3 1 36A2 YP 008476033 1 469604 Fusobacteria
Clostridium lentocellum DSM 5427 YP 004309713 1 642492 Firmicutes
Bacteroides fragilis YCH46 YP 098326 1 295405 Bacteroidetes Chlorobi group
Janthinobacterium sp Marseille YP 001352491 1 375286 Proteobacteria
Dokdonia sp 4H 3 7 5 YP 004432131 1 983548 Bacteroidetes Chlorobi group
Runella slithyformis DSM 19594 YP 004655389 1 761193 Bacteroidetes Chlorobi group
Bacteroides salanitronis DSM 18170 YP 004259186 1 667015 Bacteroidetes Chlorobi group
Flavobacterium branchiophilum FL 15 YP 004844035 1 1034807 Bacteroidetes Chlorobi group
Haliscomenobacter hydrossis DSM 1100 YP 004448440 1 760192 Bacteroidetes Chlorobi group
Desulfovibrio desulfuricans subsp desulfuricans str ATCC 27774 YP 002480924 1 525146 Proteobacteria
Cylindrospermum stagnale PCC 7417 YP 007146120 1 56107 Cyanobacteria
Halothece sp PCC 7418 YP 007168868 1 65093 Cyanobacteria
Xanthomonas campestris pv campestris str B100 YP 001902011 1 509169 Proteobacteria
Aromatoleum aromaticum EbN1 YP 157283 1 76114 Proteobacteria
Herpetosiphon aurantiacus DSM 785 YP 001547906 1 316274 Chloroflexi
Anabaena variabilis ATCC 29413 YP 323757 1 240292 Cyanobacteria
Paenibacillus sp Y412MC10 YP 003240382 1 481743 Firmicutes
Macrococcus caseolyticus JCSC5402 YP 002559740 1 458233 Firmicutes
Leuconostoc gasicomitatum LMG 18811 YP 003773289 1 762550 Firmicutes
Acetobacterium woodii DSM 1030 YP 005267844 1 931626 Firmicutes
Listeria monocytogenes YP 008271209 1 1639 Firmicutes
Listeria monocytogenes YP 008284976 1 1639 Firmicutes
Listeria monocytogenes YP 008301147 1 1639 Firmicutes
Listeria monocytogenes YP 008304139 1 1639 Firmicutes
Micavibrio aeruginosavorus ARL 13 YP 004866389 1 856793 Proteobacteria
Bdellovibrio exovorus JSS YP 007645429 1 1184267 Proteobacteria
Haliangium ochraceum DSM 14365 YP 003265435 1 502025 Proteobacteria
Candidatus Accumulibacter phosphatis clade IIA str UW 1 YP 003165809 1 522306 Proteobacteria
Marinomonas sp MWYL1 YP 001342484 1 400668 Proteobacteria
Rahnella aquatilis CIP 78 65 ATCC 33071 YP 005199717 1 745277 Proteobacteria
Morganella morganii subsp morganii KT YP 007506062 1 1124991 Proteobacteria
Prodigiosinella confusarubida ATCC39006 WP 021014561 1 Pconfusarubida
Dickeya zeae NCPPB 3531 WP 038904573 1 Dzeae
Dickeya gs fangzhongdai NCPPB 3274 WP 071603962 1 Dfangzhongdai
757 DAQ1742 00174 GI1 mobile element Seed
Dickeya sp CSL RW240 WP 035344986 1 Daquatica
Dickeya aquatica DW 0440 WP 035344986 1 Daquatica
DAQ1742 DAQ1742 00174 Daquatica
Myxococcus stipitatus DSM 14675 YP 007362768 1 1278073 Proteobacteria
Clostridium saccharobutylicum DSM 13864 YP 008675006 1 1345695 Firmicutes
Clostridium tetani 12124569 YP 008774105 1 1231072 Firmicutes
Clostridium botulinum B str Eklund 17B NRP YP 001886156 1 935198 Firmicutes
Prochlorococcus marinus subsp marinus str CCMP1375 NP 875004 1 167539 Cyanobacteria
Sorangium cellulosum So0157 2 YP 008156159 1 1254432 Proteobacteria
Singulisphaera acidiphila DSM 18658 YP 007204282 1 886293 Planctomycetes
Myxococcus xanthus DK 1622 YP 634390 1 246197 Proteobacteria
Myxococcus fulvus HW 1 YP 004666019 1 483219 Proteobacteria
Nakamurella multipartita DSM 44233 YP 003202418 1 479431 Actinobacteria
Candidatus Solibacter usitatus Ellin6076 YP 827575 1 234267 Fibrobacteres Acidobacteria group
Haliangium ochraceum DSM 14365 YP 003266012 1 502025 Proteobacteria
Sorangium cellulosum So0157 2 YP 008148765 1 1254432 Proteobacteria
Singulisphaera acidiphila DSM 18658 YP 007203371 1 886293 Planctomycetes
Stigmatella aurantiaca DW4 3 1 YP 003949791 1 378806 Proteobacteria
Corallococcus coralloides DSM 2259 YP 005373866 1 1144275 Proteobacteria
Myxococcus stipitatus DSM 14675 YP 007357162 1 1278073 Proteobacteria
Myxococcus fulvus HW 1 YP 004664596 1 483219 Proteobacteria
Myxococcus xanthus DK 1622 YP 628423 1 246197 Proteobacteria
Methylobacterium radiotolerans JCM 2831 YP 001754067 1 426355 Proteobacteria
Methylobacterium nodulans ORS 2060 YP 002498363 1 460265 Proteobacteria
Hahella chejuensis KCTC 2396 YP 435102 1 349521 Proteobacteria
Chitinophaga pinensis DSM 2588 YP 003123757 1 485918 Bacteroidetes Chlorobi group
Fibrella aestuarina BUZ 2 YP 007322132 1 1166018 Bacteroidetes Chlorobi group
Niastella koreensis GR20 10 YP 005011691 1 700598 Bacteroidetes Chlorobi group
Flavobacterium branchiophilum FL 15 YP 004845096 1 1034807 Bacteroidetes Chlorobi group
Flexibacter litoralis DSM 6794 YP 006435607 1 880071 Bacteroidetes Chlorobi group
Acaryochloris marina MBIC11017 YP 001519080 1 329726 Cyanobacteria
Chamaesiphon minutus PCC 6605 YP 007099806 1 1173020 Cyanobacteria
Oscillatoria nigro viridis PCC 7112 YP 007117717 1 179408 Cyanobacteria
Trichodesmium erythraeum IMS101 YP 721852 1 203124 Cyanobacteria
Rivularia sp PCC 7116 YP 007056214 1 373994 Cyanobacteria
Oscillatoria nigro viridis PCC 7112 YP 007116551 1 179408 Cyanobacteria
Chroococcidiopsis thermalis PCC 7203 YP 007089778 1 251229 Cyanobacteria
Nostoc punctiforme PCC 73102 YP 001864131 1 63737 Cyanobacteria

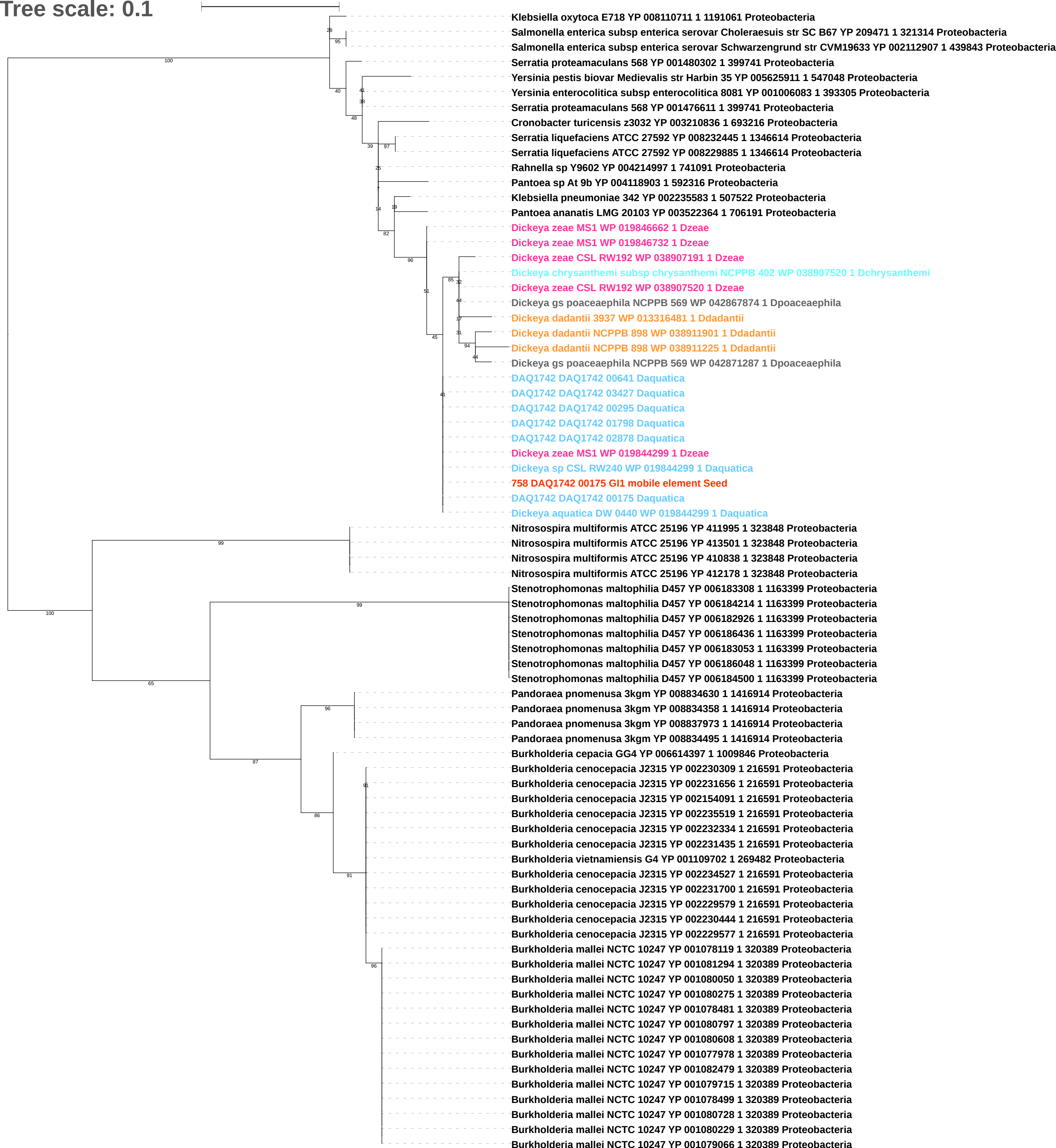
Tree scale: 0.1
Klebsiella oxytoca E718 YP 008110711 1 1191061 Proteobacteria
Salmonella enterica subsp enterica serovar Choleraesuis str SC B67 YP 209471 1 321314 Proteobacteria
Salmonella enterica subsp enterica serovar Schwarzengrund str CVM19633 YP 002112907 1 439843 Proteobacteria
Serratia proteamaculans 568 YP 001480302 1 399741 Proteobacteria
Yersinia pestis biovar Medievalis str Harbin 35 YP 005625911 1 547048 Proteobacteria
Yersinia enterocolitica subsp enterocolitica 8081 YP 001006083 1 393305 Proteobacteria
Serratia proteamaculans 568 YP 001476611 1 399741 Proteobacteria
Cronobacter turicensis z3032 YP 003210836 1 693216 Proteobacteria
Serratia liquefaciens ATCC 27592 YP 008232445 1 1346614 Proteobacteria
Serratia liquefaciens ATCC 27592 YP 008229885 1 1346614 Proteobacteria
Rahnella sp Y9602 YP 004214997 1 741091 Proteobacteria
Pantoea sp At 9b YP 004118903 1 592316 Proteobacteria
Klebsiella pneumoniae 342 YP 002235583 1 507522 Proteobacteria
Pantoea ananatis LMG 20103 YP 003522364 1 706191 Proteobacteria
Dickeya zeae MS1 WP 019846662 1 Dzeae
Dickeya zeae MS1 WP 019846732 1 Dzeae
Dickeya zeae CSL RW192 WP 038907191 1 Dzeae
Dickeya chrysanthemi subsp chrysanthemi NCPPB 402 WP 038907520 1 Dchrysanthemi
Dickeya zeae CSL RW192 WP 038907520 1 Dzeae
Dickeya gs poaceaephila NCPPB 569 WP 042867874 1 Dpoaceaephila
Dickeya dadantii 3937 WP 013316481 1 Ddadantii
Dickeya dadantii NCPPB 898 WP 038911901 1 Ddadantii
Dickeya dadantii NCPPB 898 WP 038911225 1 Ddadantii
Dickeya gs poaceaephila NCPPB 569 WP 042871287 1 Dpoaceaephila
DAQ1742 DAQ1742 00641 Daquatica
DAQ1742 DAQ1742 03427 Daquatica
DAQ1742 DAQ1742 00295 Daquatica
DAQ1742 DAQ1742 01798 Daquatica
DAQ1742 DAQ1742 02878 Daquatica
Dickeya zeae MS1 WP 019844299 1 Dzeae
Dickeya sp CSL RW240 WP 019844299 1 Daquatica
758 DAQ1742 00175 GI1 mobile element Seed
DAQ1742 DAQ1742 00175 Daquatica
Dickeya aquatica DW 0440 WP 019844299 1 Daquatica
Nitrosospira multiformis ATCC 25196 YP 411995 1 323848 Proteobacteria
Nitrosospira multiformis ATCC 25196 YP 413501 1 323848 Proteobacteria
Nitrosospira multiformis ATCC 25196 YP 410838 1 323848 Proteobacteria
Nitrosospira multiformis ATCC 25196 YP 412178 1 323848 Proteobacteria
Stenotrophomonas maltophilia D457 YP 006183308 1 1163399 Proteobacteria
Stenotrophomonas maltophilia D457 YP 006184214 1 1163399 Proteobacteria
Stenotrophomonas maltophilia D457 YP 006182926 1 1163399 Proteobacteria
Stenotrophomonas maltophilia D457 YP 006186436 1 1163399 Proteobacteria
Stenotrophomonas maltophilia D457 YP 006183053 1 1163399 Proteobacteria
Stenotrophomonas maltophilia D457 YP 006186048 1 1163399 Proteobacteria
Stenotrophomonas maltophilia D457 YP 006184500 1 1163399 Proteobacteria
Pandoraea pnomenusa 3kgm YP 008834630 1 1416914 Proteobacteria
Pandoraea pnomenusa 3kgm YP 008834358 1 1416914 Proteobacteria
Pandoraea pnomenusa 3kgm YP 008837973 1 1416914 Proteobacteria
Pandoraea pnomenusa 3kgm YP 008834495 1 1416914 Proteobacteria
Burkholderia cepacia GG4 YP 006614397 1 1009846 Proteobacteria
Burkholderia cenocepacia J2315 YP 002230309 1 216591 Proteobacteria
Burkholderia cenocepacia J2315 YP 002231656 1 216591 Proteobacteria
Burkholderia cenocepacia J2315 YP 002154091 1 216591 Proteobacteria
Burkholderia cenocepacia J2315 YP 002235519 1 216591 Proteobacteria
Burkholderia cenocepacia J2315 YP 002232334 1 216591 Proteobacteria
Burkholderia cenocepacia J2315 YP 002231435 1 216591 Proteobacteria
Burkholderia vietnamiensis G4 YP 001109702 1 269482 Proteobacteria
Burkholderia cenocepacia J2315 YP 002234527 1 216591 Proteobacteria
Burkholderia cenocepacia J2315 YP 002231700 1 216591 Proteobacteria
Burkholderia cenocepacia J2315 YP 002229579 1 216591 Proteobacteria
Burkholderia cenocepacia J2315 YP 002230444 1 216591 Proteobacteria
Burkholderia cenocepacia J2315 YP 002229577 1 216591 Proteobacteria
Burkholderia mallei NCTC 10247 YP 001078119 1 320389 Proteobacteria
Burkholderia mallei NCTC 10247 YP 001081294 1 320389 Proteobacteria
Burkholderia mallei NCTC 10247 YP 001080050 1 320389 Proteobacteria
Burkholderia mallei NCTC 10247 YP 001080275 1 320389 Proteobacteria
Burkholderia mallei NCTC 10247 YP 001078481 1 320389 Proteobacteria
Burkholderia mallei NCTC 10247 YP 001080797 1 320389 Proteobacteria
Burkholderia mallei NCTC 10247 YP 001080608 1 320389 Proteobacteria
Burkholderia mallei NCTC 10247 YP 001077978 1 320389 Proteobacteria
Burkholderia mallei NCTC 10247 YP 001082479 1 320389 Proteobacteria
Burkholderia mallei NCTC 10247 YP 001079715 1 320389 Proteobacteria
Burkholderia mallei NCTC 10247 YP 001078499 1 320389 Proteobacteria
Burkholderia mallei NCTC 10247 YP 001080728 1 320389 Proteobacteria
Burkholderia mallei NCTC 10247 YP 001080229 1 320389 Proteobacteria
Burkholderia mallei NCTC 10247 YP 001079066 1 320389 Proteobacteria

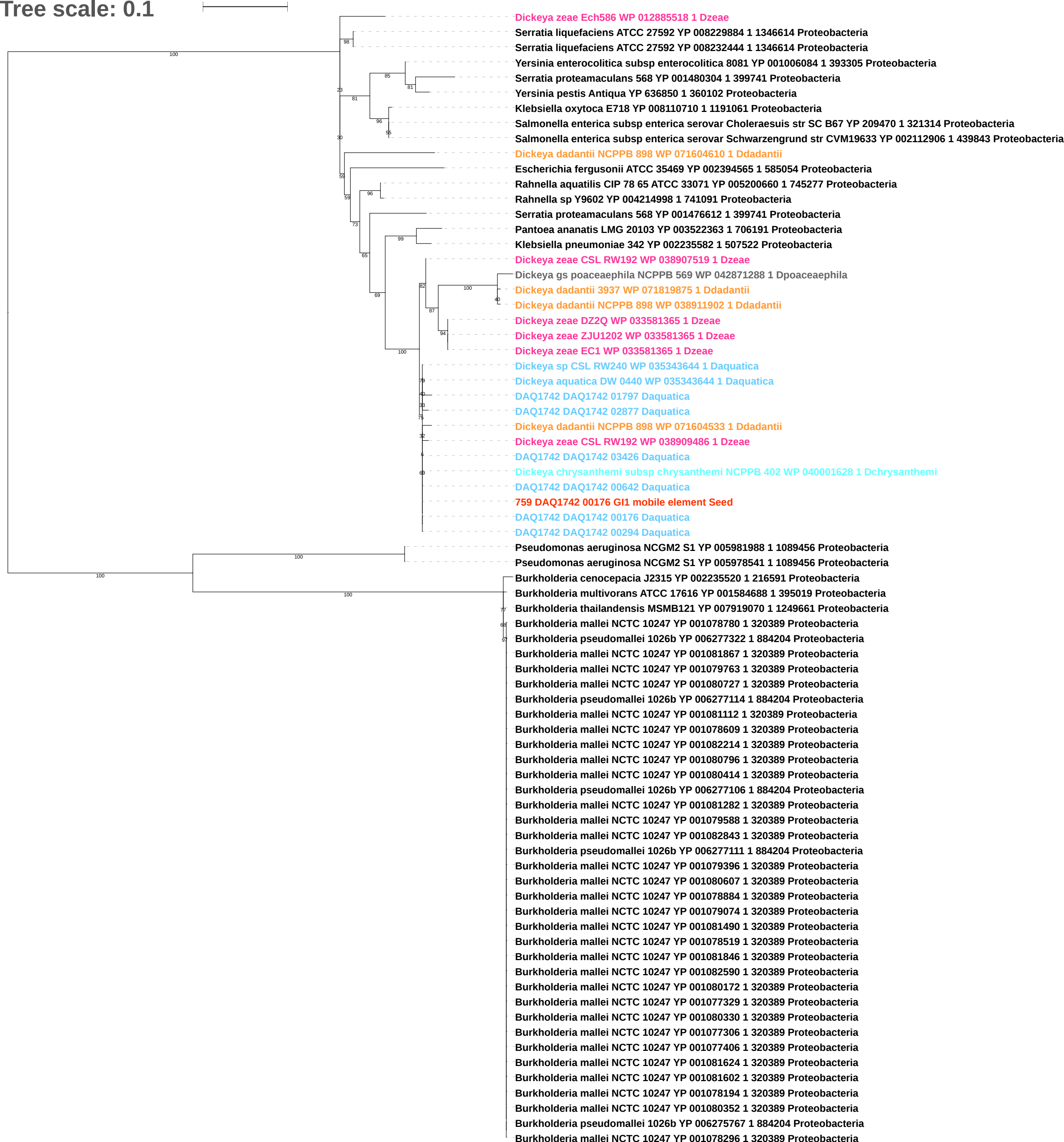
Tree scale: 0.1
Dickeya zeae Ech586 WP 012885518 1 Dzeae
Serratia liquefaciens ATCC 27592 YP 008229884 1 1346614 Proteobacteria
Serratia liquefaciens ATCC 27592 YP 008232444 1 1346614 Proteobacteria
Yersinia enterocolitica subsp enterocolitica 8081 YP 001006084 1 393305 Proteobacteria
Serratia proteamaculans 568 YP 001480304 1 399741 Proteobacteria
Yersinia pestis Antiqua YP 636850 1 360102 Proteobacteria
Klebsiella oxytoca E718 YP 008110710 1 1191061 Proteobacteria
Salmonella enterica subsp enterica serovar Choleraesuis str SC B67 YP 209470 1 321314 Proteobacteria
Salmonella enterica subsp enterica serovar Schwarzengrund str CVM19633 YP 002112906 1 439843 Proteobacteria
Dickeya dadantii NCPPB 898 WP 071604610 1 Ddadantii
Escherichia fergusonii ATCC 35469 YP 002394565 1 585054 Proteobacteria
Rahnella aquatilis CIP 78 65 ATCC 33071 YP 005200660 1 745277 Proteobacteria
Rahnella sp Y9602 YP 004214998 1 741091 Proteobacteria
Serratia proteamaculans 568 YP 001476612 1 399741 Proteobacteria
Pantoea ananatis LMG 20103 YP 003522363 1 706191 Proteobacteria
Klebsiella pneumoniae 342 YP 002235582 1 507522 Proteobacteria
Dickeya zeae CSL RW192 WP 038907519 1 Dzeae
Dickeya gs poaceaephila NCPPB 569 WP 042871288 1 Dpoaceaephila
Dickeya dadantii 3937 WP 071819875 1 Ddadantii
Dickeya dadantii NCPPB 898 WP 038911902 1 Ddadantii
Dickeya zeae DZ2Q WP 033581365 1 Dzeae
Dickeya zeae ZJU1202 WP 033581365 1 Dzeae
Dickeya zeae EC1 WP 033581365 1 Dzeae
Dickeya sp CSL RW240 WP 035343644 1 Daquatica
Dickeya aquatica DW 0440 WP 035343644 1 Daquatica
DAQ1742 DAQ1742 01797 Daquatica
DAQ1742 DAQ1742 02877 Daquatica
Dickeya dadantii NCPPB 898 WP 071604533 1 Ddadantii
Dickeya zeae CSL RW192 WP 038909486 1 Dzeae
DAQ1742 DAQ1742 03426 Daquatica
Dickeya chrysanthemi subsp chrysanthemi NCPPB 402 WP 040001628 1 Dchrysanthemi
DAQ1742 DAQ1742 00642 Daquatica
759 DAQ1742 00176 GI1 mobile element Seed
DAQ1742 DAQ1742 00176 Daquatica
DAQ1742 DAQ1742 00294 Daquatica
Pseudomonas aeruginosa NCGM2 S1 YP 005981988 1 1089456 Proteobacteria
Pseudomonas aeruginosa NCGM2 S1 YP 005978541 1 1089456 Proteobacteria
Burkholderia cenocepacia J2315 YP 002235520 1 216591 Proteobacteria
Burkholderia multivorans ATCC 17616 YP 001584688 1 395019 Proteobacteria
Burkholderia thailandensis MSMB121 YP 007919070 1 1249661 Proteobacteria
Burkholderia mallei NCTC 10247 YP 001078780 1 320389 Proteobacteria
Burkholderia pseudomallei 1026b YP 006277322 1 884204 Proteobacteria
Burkholderia mallei NCTC 10247 YP 001081867 1 320389 Proteobacteria
Burkholderia mallei NCTC 10247 YP 001079763 1 320389 Proteobacteria
Burkholderia mallei NCTC 10247 YP 001080727 1 320389 Proteobacteria
Burkholderia pseudomallei 1026b YP 006277114 1 884204 Proteobacteria
Burkholderia mallei NCTC 10247 YP 001081112 1 320389 Proteobacteria
Burkholderia mallei NCTC 10247 YP 001078609 1 320389 Proteobacteria
Burkholderia mallei NCTC 10247 YP 001082214 1 320389 Proteobacteria
Burkholderia mallei NCTC 10247 YP 001080796 1 320389 Proteobacteria
Burkholderia mallei NCTC 10247 YP 001080414 1 320389 Proteobacteria
Burkholderia pseudomallei 1026b YP 006277106 1 884204 Proteobacteria
Burkholderia mallei NCTC 10247 YP 001081282 1 320389 Proteobacteria
Burkholderia mallei NCTC 10247 YP 001079588 1 320389 Proteobacteria
Burkholderia mallei NCTC 10247 YP 001082843 1 320389 Proteobacteria
Burkholderia pseudomallei 1026b YP 006277111 1 884204 Proteobacteria
Burkholderia mallei NCTC 10247 YP 001079396 1 320389 Proteobacteria
Burkholderia mallei NCTC 10247 YP 001080607 1 320389 Proteobacteria
Burkholderia mallei NCTC 10247 YP 001078884 1 320389 Proteobacteria
Burkholderia mallei NCTC 10247 YP 001079074 1 320389 Proteobacteria
Burkholderia mallei NCTC 10247 YP 001081490 1 320389 Proteobacteria
Burkholderia mallei NCTC 10247 YP 001078519 1 320389 Proteobacteria
Burkholderia mallei NCTC 10247 YP 001081846 1 320389 Proteobacteria
Burkholderia mallei NCTC 10247 YP 001082590 1 320389 Proteobacteria
Burkholderia mallei NCTC 10247 YP 001080172 1 320389 Proteobacteria
Burkholderia mallei NCTC 10247 YP 001077329 1 320389 Proteobacteria
Burkholderia mallei NCTC 10247 YP 001080330 1 320389 Proteobacteria
Burkholderia mallei NCTC 10247 YP 001077306 1 320389 Proteobacteria
Burkholderia mallei NCTC 10247 YP 001077406 1 320389 Proteobacteria
Burkholderia mallei NCTC 10247 YP 001081624 1 320389 Proteobacteria
Burkholderia mallei NCTC 10247 YP 001081602 1 320389 Proteobacteria
Burkholderia mallei NCTC 10247 YP 001078194 1 320389 Proteobacteria
Burkholderia mallei NCTC 10247 YP 001080352 1 320389 Proteobacteria
Burkholderia pseudomallei 1026b YP 006275767 1 884204 Proteobacteria
Burkholderia mallei NCTC 10247 YP 001078296 1 320389 Proteobacteria

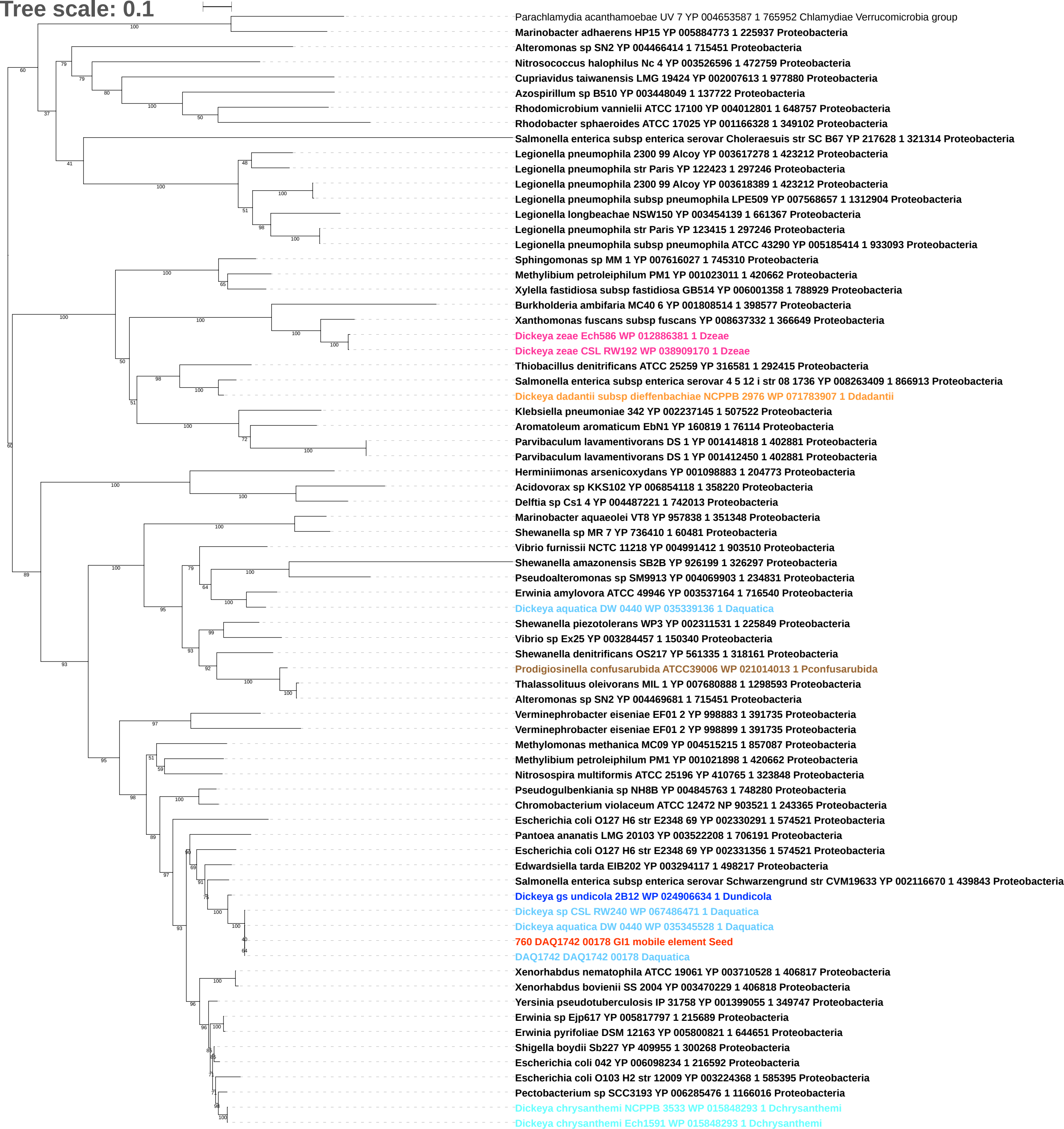
Tree scale: 0.1
Parachlamydia acanthamoebae UV 7 YP 004653587 1 765952 Chlamydiae Verrucomicrobia group
Marinobacter adhaerens HP15 YP 005884773 1 225937 Proteobacteria
Alteromonas sp SN2 YP 004466414 1 715451 Proteobacteria
Nitrosococcus halophilus Nc 4 YP 003526596 1 472759 Proteobacteria
Cupriavidus taiwanensis LMG 19424 YP 002007613 1 977880 Proteobacteria
Azospirillum sp B510 YP 003448049 1 137722 Proteobacteria
Rhodomicrobium vannielii ATCC 17100 YP 004012801 1 648757 Proteobacteria
Rhodobacter sphaeroides ATCC 17025 YP 001166328 1 349102 Proteobacteria
Salmonella enterica subsp enterica serovar Choleraesuis str SC B67 YP 217628 1 321314 Proteobacteria
Legionella pneumophila 2300 99 Alcoy YP 003617278 1 423212 Proteobacteria
Legionella pneumophila str Paris YP 122423 1 297246 Proteobacteria
Legionella pneumophila 2300 99 Alcoy YP 003618389 1 423212 Proteobacteria
Legionella pneumophila subsp pneumophila LPE509 YP 007568657 1 1312904 Proteobacteria
Legionella longbeachae NSW150 YP 003454139 1 661367 Proteobacteria
Legionella pneumophila str Paris YP 123415 1 297246 Proteobacteria
Legionella pneumophila subsp pneumophila ATCC 43290 YP 005185414 1 933093 Proteobacteria
Sphingomonas sp MM 1 YP 007616027 1 745310 Proteobacteria
Methylibium petroleiphilum PM1 YP 001023011 1 420662 Proteobacteria
Xylella fastidiosa subsp fastidiosa GB514 YP 006001358 1 788929 Proteobacteria
Burkholderia ambifaria MC40 6 YP 001808514 1 398577 Proteobacteria
Xanthomonas fuscans subsp fuscans YP 008637332 1 366649 Proteobacteria
Dickeya zeae Ech586 WP 012886381 1 Dzeae
Dickeya zeae CSL RW192 WP 038909170 1 Dzeae
Thiobacillus denitrificans ATCC 25259 YP 316581 1 292415 Proteobacteria
Salmonella enterica subsp enterica serovar 4 5 12 i str 08 1736 YP 008263409 1 866913 Proteobacteria
Dickeya dadantii subsp dieffenbachiae NCPPB 2976 WP 071783907 1 Ddadantii
Klebsiella pneumoniae 342 YP 002237145 1 507522 Proteobacteria
Aromatoleum aromaticum EbN1 YP 160819 1 76114 Proteobacteria
Parvibaculum lavamentivorans DS 1 YP 001414818 1 402881 Proteobacteria
Parvibaculum lavamentivorans DS 1 YP 001412450 1 402881 Proteobacteria
Herminiimonas arsenicoxydans YP 001098883 1 204773 Proteobacteria
Acidovorax sp KKS102 YP 006854118 1 358220 Proteobacteria
Delftia sp Cs1 4 YP 004487221 1 742013 Proteobacteria
Marinobacter aquaeolei VT8 YP 957838 1 351348 Proteobacteria
Shewanella sp MR 7 YP 736410 1 60481 Proteobacteria
Vibrio furnissii NCTC 11218 YP 004991412 1 903510 Proteobacteria
Shewanella amazonensis SB2B YP 926199 1 326297 Proteobacteria
Pseudoalteromonas sp SM9913 YP 004069903 1 234831 Proteobacteria
Erwinia amylovora ATCC 49946 YP 003537164 1 716540 Proteobacteria
Dickeya aquatica DW 0440 WP 035339136 1 Daquatica
Shewanella piezotolerans WP3 YP 002311531 1 225849 Proteobacteria
Vibrio sp Ex25 YP 003284457 1 150340 Proteobacteria
Shewanella denitrificans OS217 YP 561335 1 318161 Proteobacteria
Prodigiosinella confusarubida ATCC39006 WP 021014013 1 Pconfusarubida
Thalassolituus oleivorans MIL 1 YP 007680888 1 1298593 Proteobacteria
Alteromonas sp SN2 YP 004469681 1 715451 Proteobacteria
Verminephrobacter eiseniae EF01 2 YP 998883 1 391735 Proteobacteria
Verminephrobacter eiseniae EF01 2 YP 998899 1 391735 Proteobacteria
Methylomonas methanica MC09 YP 004515215 1 857087 Proteobacteria
Methylibium petroleiphilum PM1 YP 001021898 1 420662 Proteobacteria
Nitrosospira multiformis ATCC 25196 YP 410765 1 323848 Proteobacteria
Pseudogulbenkiania sp NH8B YP 004845763 1 748280 Proteobacteria
Chromobacterium violaceum ATCC 12472 NP 903521 1 243365 Proteobacteria
Escherichia coli O127 H6 str E2348 69 YP 002330291 1 574521 Proteobacteria
Pantoea ananatis LMG 20103 YP 003522208 1 706191 Proteobacteria
Escherichia coli O127 H6 str E2348 69 YP 002331356 1 574521 Proteobacteria
Edwardsiella tarda EIB202 YP 003294117 1 498217 Proteobacteria
Salmonella enterica subsp enterica serovar Schwarzengrund str CVM19633 YP 002116670 1 439843 Proteobacteria
Dickeya gs undicola 2B12 WP 024906634 1 Dundicola
Dickeya sp CSL RW240 WP 067486471 1 Daquatica
Dickeya aquatica DW 0440 WP 035345528 1 Daquatica
760 DAQ1742 00178 GI1 mobile element Seed
DAQ1742 DAQ1742 00178 Daquatica
Xenorhabdus nematophila ATCC 19061 YP 003710528 1 406817 Proteobacteria
Xenorhabdus bovienii SS 2004 YP 003470229 1 406818 Proteobacteria
Yersinia pseudotuberculosis IP 31758 YP 001399055 1 349747 Proteobacteria
Erwinia sp Ejp617 YP 005817797 1 215689 Proteobacteria
Erwinia pyrifoliae DSM 12163 YP 005800821 1 644651 Proteobacteria
Shigella boydii Sb227 YP 409955 1 300268 Proteobacteria
Escherichia coli 042 YP 006098234 1 216592 Proteobacteria
Escherichia coli O103 H2 str 12009 YP 003224368 1 585395 Proteobacteria
Pectobacterium sp SCC3193 YP 006285476 1 1166016 Proteobacteria
Dickeya chrysanthemi NCPPB 3533 WP 015848293 1 Dchrysanthemi
Dickeya chrysanthemi Ech1591 WP 015848293 1 Dchrysanthemi

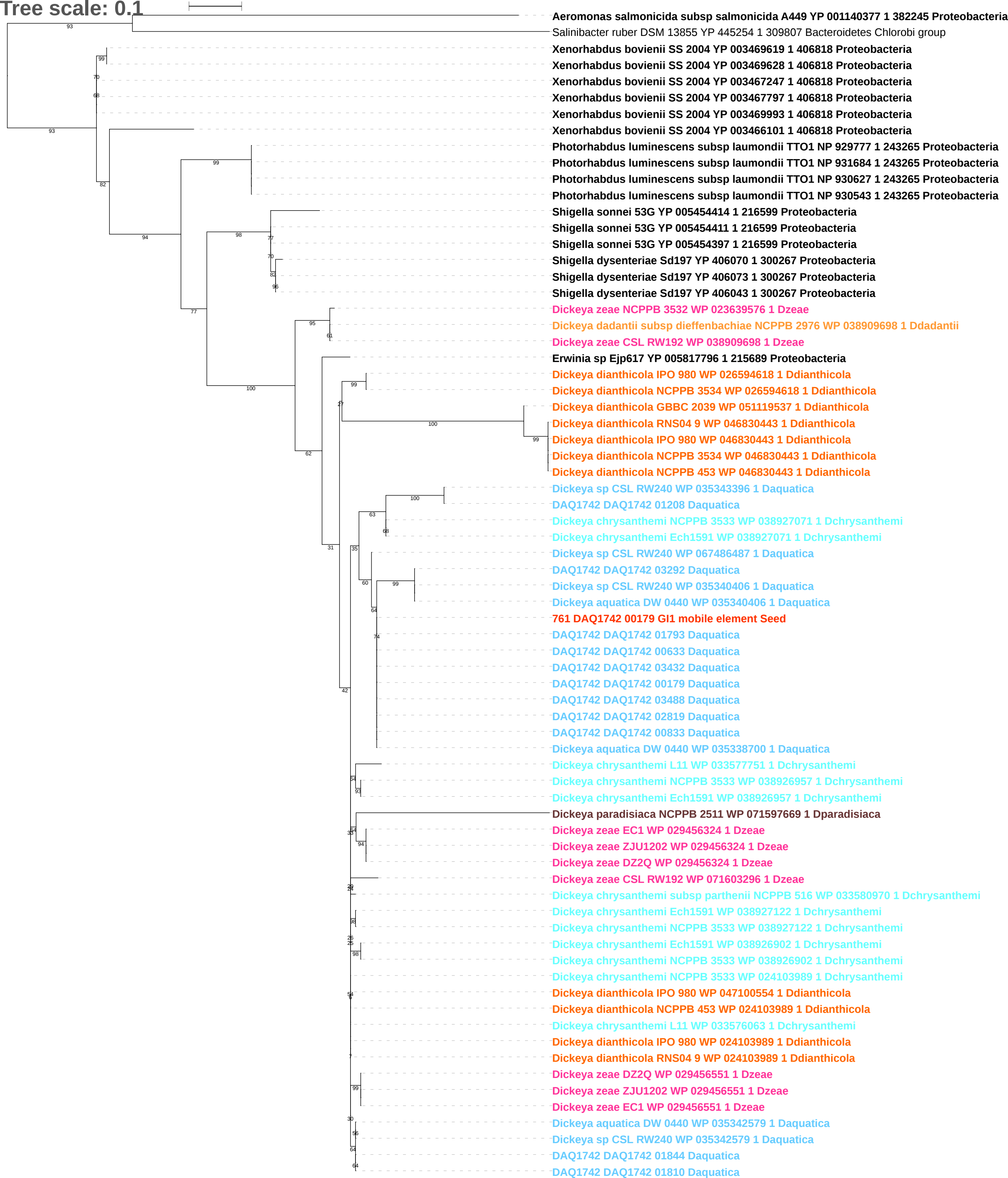
Tree scale: 0.1
Aeromonas salmonicida subsp salmonicida A449 YP 001140377 1 382245 Proteobacteria
Salinibacter ruber DSM 13855 YP 445254 1 309807 Bacteroidetes Chlorobi group
Xenorhabdus bovienii SS 2004 YP 003469619 1 406818 Proteobacteria
Xenorhabdus bovienii SS 2004 YP 003469628 1 406818 Proteobacteria
Xenorhabdus bovienii SS 2004 YP 003467247 1 406818 Proteobacteria
Xenorhabdus bovienii SS 2004 YP 003467797 1 406818 Proteobacteria
Xenorhabdus bovienii SS 2004 YP 003469993 1 406818 Proteobacteria
Xenorhabdus bovienii SS 2004 YP 003466101 1 406818 Proteobacteria
Photorhabdus luminescens subsp laumondii TTO1 NP 929777 1 243265 Proteobacteria
Photorhabdus luminescens subsp laumondii TTO1 NP 931684 1 243265 Proteobacteria
Photorhabdus luminescens subsp laumondii TTO1 NP 930627 1 243265 Proteobacteria
Photorhabdus luminescens subsp laumondii TTO1 NP 930543 1 243265 Proteobacteria
Shigella sonnei 53G YP 005454414 1 216599 Proteobacteria
Shigella sonnei 53G YP 005454411 1 216599 Proteobacteria
Shigella sonnei 53G YP 005454397 1 216599 Proteobacteria
Shigella dysenteriae Sd197 YP 406070 1 300267 Proteobacteria
Shigella dysenteriae Sd197 YP 406073 1 300267 Proteobacteria
Shigella dysenteriae Sd197 YP 406043 1 300267 Proteobacteria
Dickeya zeae NCPPB 3532 WP 023639576 1 Dzeae
Dickeya dadantii subsp dieffenbachiae NCPPB 2976 WP 038909698 1 Ddadantii
Dickeya zeae CSL RW192 WP 038909698 1 Dzeae
Erwinia sp Ejp617 YP 005817796 1 215689 Proteobacteria
Dickeya dianthicola IPO 980 WP 026594618 1 Ddianthicola
Dickeya dianthicola NCPPB 3534 WP 026594618 1 Ddianthicola
Dickeya dianthicola GBBC 2039 WP 051119537 1 Ddianthicola
Dickeya dianthicola RNS04 9 WP 046830443 1 Ddianthicola
Dickeya dianthicola IPO 980 WP 046830443 1 Ddianthicola
Dickeya dianthicola NCPPB 3534 WP 046830443 1 Ddianthicola
Dickeya dianthicola NCPPB 453 WP 046830443 1 Ddianthicola
Dickeya sp CSL RW240 WP 035343396 1 Daquatica
DAQ1742 DAQ1742 01208 Daquatica
Dickeya chrysanthemi NCPPB 3533 WP 038927071 1 Dchrysanthemi
Dickeya chrysanthemi Ech1591 WP 038927071 1 Dchrysanthemi
Dickeya sp CSL RW240 WP 067486487 1 Daquatica
DAQ1742 DAQ1742 03292 Daquatica
Dickeya sp CSL RW240 WP 035340406 1 Daquatica
Dickeya aquatica DW 0440 WP 035340406 1 Daquatica
761 DAQ1742 00179 GI1 mobile element Seed
DAQ1742 DAQ1742 01793 Daquatica
DAQ1742 DAQ1742 00633 Daquatica
DAQ1742 DAQ1742 03432 Daquatica
DAQ1742 DAQ1742 00179 Daquatica
DAQ1742 DAQ1742 03488 Daquatica
DAQ1742 DAQ1742 02819 Daquatica
DAQ1742 DAQ1742 00833 Daquatica
Dickeya aquatica DW 0440 WP 035338700 1 Daquatica
Dickeya chrysanthemi L11 WP 033577751 1 Dchrysanthemi
Dickeya chrysanthemi NCPPB 3533 WP 038926957 1 Dchrysanthemi
Dickeya chrysanthemi Ech1591 WP 038926957 1 Dchrysanthemi
Dickeya paradisiaca NCPPB 2511 WP 071597669 1 Dparadisiaca
Dickeya zeae EC1 WP 029456324 1 Dzeae
Dickeya zeae ZJU1202 WP 029456324 1 Dzeae
Dickeya zeae DZ2Q WP 029456324 1 Dzeae
Dickeya zeae CSL RW192 WP 071603296 1 Dzeae
Dickeya chrysanthemi subsp parthenii NCPPB 516 WP 033580970 1 Dchrysanthemi
Dickeya chrysanthemi Ech1591 WP 038927122 1 Dchrysanthemi
Dickeya chrysanthemi NCPPB 3533 WP 038927122 1 Dchrysanthemi
Dickeya chrysanthemi Ech1591 WP 038926902 1 Dchrysanthemi
Dickeya chrysanthemi NCPPB 3533 WP 038926902 1 Dchrysanthemi
Dickeya chrysanthemi NCPPB 3533 WP 024103989 1 Dchrysanthemi
Dickeya dianthicola IPO 980 WP 047100554 1 Ddianthicola
Dickeya dianthicola NCPPB 453 WP 024103989 1 Ddianthicola
Dickeya chrysanthemi L11 WP 033576063 1 Dchrysanthemi
Dickeya dianthicola IPO 980 WP 024103989 1 Ddianthicola
Dickeya dianthicola RNS04 9 WP 024103989 1 Ddianthicola
Dickeya zeae DZ2Q WP 029456551 1 Dzeae
Dickeya zeae ZJU1202 WP 029456551 1 Dzeae
Dickeya zeae EC1 WP 029456551 1 Dzeae
Dickeya aquatica DW 0440 WP 035342579 1 Daquatica
Dickeya sp CSL RW240 WP 035342579 1 Daquatica
DAQ1742 DAQ1742 01844 Daquatica
DAQ1742 DAQ1742 01810 Daquatica

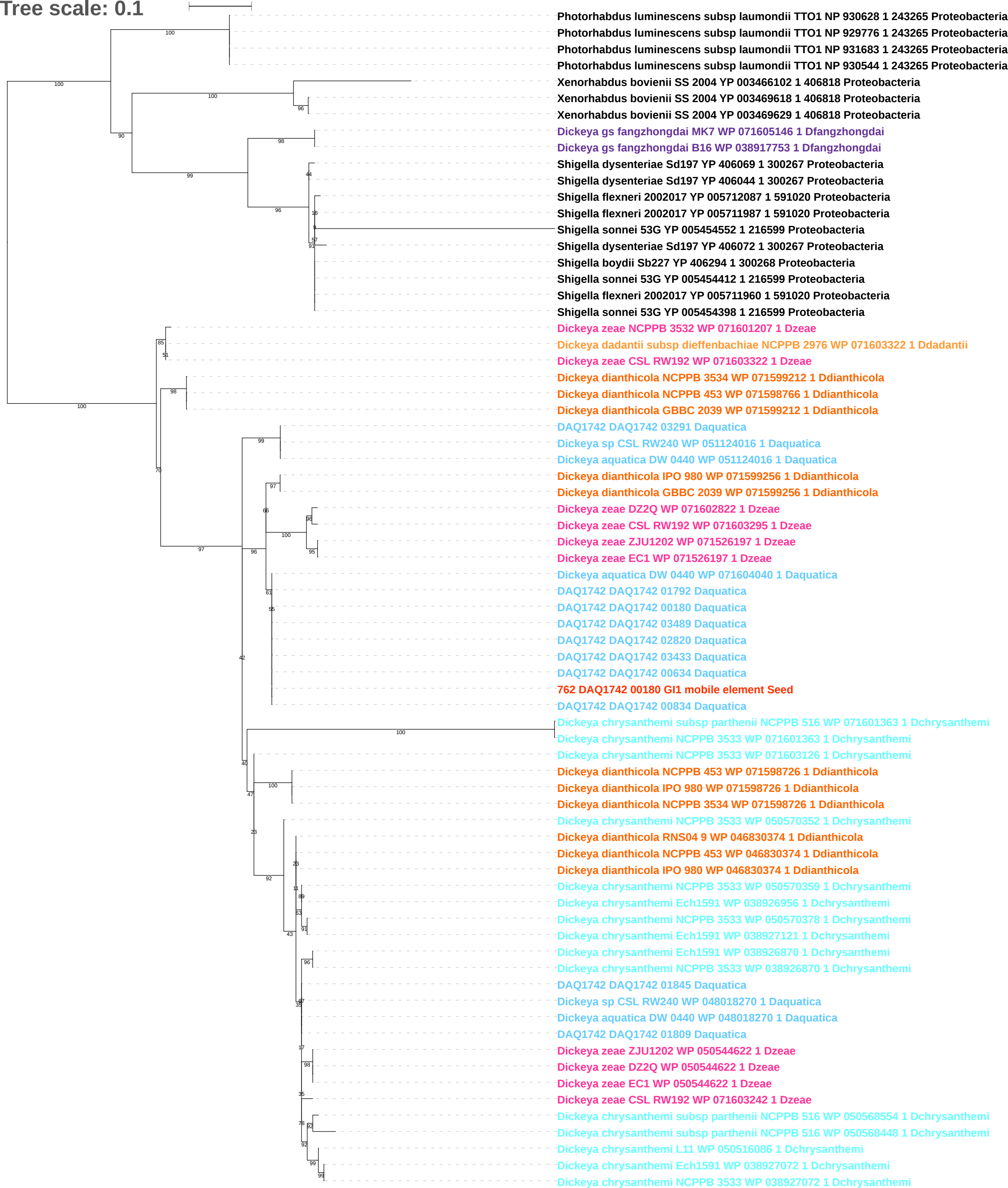

Tree scale: 0.1
Photorhabdus luminescens subsp laumondii TTO1 NP 930628 1 243265 Proteobacteria
Photorhabdus luminescens subsp laumondii TTO1 NP 929776 1 243265 Proteobacteria
Photorhabdus luminescens subsp laumondii TTO1 NP 931683 1 243265 Proteobacteria
Photorhabdus luminescens subsp laumondii TTO1 NP 930544 1 243265 Proteobacteria
Xenorhabdus bovienii SS 2004 YP 003466102 1 406818 Proteobacteria
Xenorhabdus bovienii SS 2004 YP 003469618 1 406818 Proteobacteria
Xenorhabdus bovienii SS 2004 YP 003469629 1 406818 Proteobacteria
Dickeya gs fangzhongdai MK7 WP 071605146 1 Dfangzhongdai
Dickeya gs fangzhongdai B16 WP 038917753 1 Dfangzhongdai
Shigella dysenteriae Sd197 YP 406069 1 300267 Proteobacteria
Shigella dysenteriae Sd197 YP 406044 1 300267 Proteobacteria
Shigella flexneri 2002017 YP 005712087 1 591020 Proteobacteria
Shigella flexneri 2002017 YP 005711987 1 591020 Proteobacteria
Shigella sonnei 53G YP 005454552 1 216599 Proteobacteria
Shigella dysenteriae Sd197 YP 406072 1 300267 Proteobacteria
Shigella boydii Sb227 YP 406294 1 300268 Proteobacteria
Shigella sonnei 53G YP 005454412 1 216599 Proteobacteria
Shigella flexneri 2002017 YP 005711960 1 591020 Proteobacteria
Shigella sonnei 53G YP 005454398 1 216599 Proteobacteria
Dickeya zeae NCPPB 3532 WP 071601207 1 Dzeae
Dickeya dadantii subsp dieffenbachiae NCPPB 2976 WP 071603322 1 Ddadantii
Dickeya zeae CSL RW192 WP 071603322 1 Dzeae
Dickeya dianthicola NCPPB 3534 WP 071599212 1 Ddianthicola
Dickeya dianthicola NCPPB 453 WP 071598766 1 Ddianthicola
Dickeya dianthicola GBBC 2039 WP 071599212 1 Ddianthicola
DAQ1742 DAQ1742 03291 Daquatica
Dickeya sp CSL RW240 WP 051124016 1 Daquatica
Dickeya aquatica DW 0440 WP 051124016 1 Daquatica
Dickeya dianthicola IPO 980 WP 071599256 1 Ddianthicola
Dickeya dianthicola GBBC 2039 WP 071599256 1 Ddianthicola
Dickeya zeae DZ2Q WP 071602822 1 Dzeae
Dickeya zeae CSL RW192 WP 071603295 1 Dzeae
Dickeya zeae ZJU1202 WP 071526197 1 Dzeae
Dickeya zeae EC1 WP 071526197 1 Dzeae
Dickeya aquatica DW 0440 WP 071604040 1 Daquatica
DAQ1742 DAQ1742 01792 Daquatica
DAQ1742 DAQ1742 00180 Daquatica
DAQ1742 DAQ1742 03489 Daquatica
DAQ1742 DAQ1742 02820 Daquatica
DAQ1742 DAQ1742 03433 Daquatica
DAQ1742 DAQ1742 00634 Daquatica
762 DAQ1742 00180 GI1 mobile element Seed
DAQ1742 DAQ1742 00834 Daquatica
Dickeya chrysanthemi subsp parthenii NCPPB 516 WP 071601363 1 Dchrysanthemi
Dickeya chrysanthemi NCPPB 3533 WP 071601363 1 Dchrysanthemi
Dickeya chrysanthemi NCPPB 3533 WP 071603126 1 Dchrysanthemi
Dickeya dianthicola NCPPB 453 WP 071598726 1 Ddianthicola
Dickeya dianthicola IPO 980 WP 071598726 1 Ddianthicola
Dickeya dianthicola NCPPB 3534 WP 071598726 1 Ddianthicola
Dickeya chrysanthemi NCPPB 3533 WP 050570352 1 Dchrysanthemi
Dickeya dianthicola RNS04 9 WP 046830374 1 Ddianthicola
Dickeya dianthicola NCPPB 453 WP 046830374 1 Ddianthicola
Dickeya dianthicola IPO 980 WP 046830374 1 Ddianthicola
Dickeya chrysanthemi NCPPB 3533 WP 050570359 1 Dchrysanthemi
Dickeya chrysanthemi Ech1591 WP 038926956 1 Dchrysanthemi
Dickeya chrysanthemi NCPPB 3533 WP 050570378 1 Dchrysanthemi
Dickeya chrysanthemi Ech1591 WP 038927121 1 Dchrysanthemi
Dickeya chrysanthemi Ech1591 WP 038926870 1 Dchrysanthemi
Dickeya chrysanthemi NCPPB 3533 WP 038926870 1 Dchrysanthemi
DAQ1742 DAQ1742 01845 Daquatica
Dickeya sp CSL RW240 WP 048018270 1 Daquatica
Dickeya aquatica DW 0440 WP 048018270 1 Daquatica
DAQ1742 DAQ1742 01809 Daquatica
Dickeya zeae ZJU1202 WP 050544622 1 Dzeae
Dickeya zeae DZ2Q WP 050544622 1 Dzeae
Dickeya zeae EC1 WP 050544622 1 Dzeae
Dickeya zeae CSL RW192 WP 071603242 1 Dzeae
Dickeya chrysanthemi subsp parthenii NCPPB 516 WP 050568554 1 Dchrysanthemi
Dickeya chrysanthemi subsp parthenii NCPPB 516 WP 050568448 1 Dchrysanthemi
Dickeya chrysanthemi L11 WP 050516086 1 Dchrysanthemi
Dickeya chrysanthemi Ech1591 WP 038927072 1 Dchrysanthemi
Dickeya chrysanthemi NCPPB 3533 WP 038927072 1 Dchrysanthemi

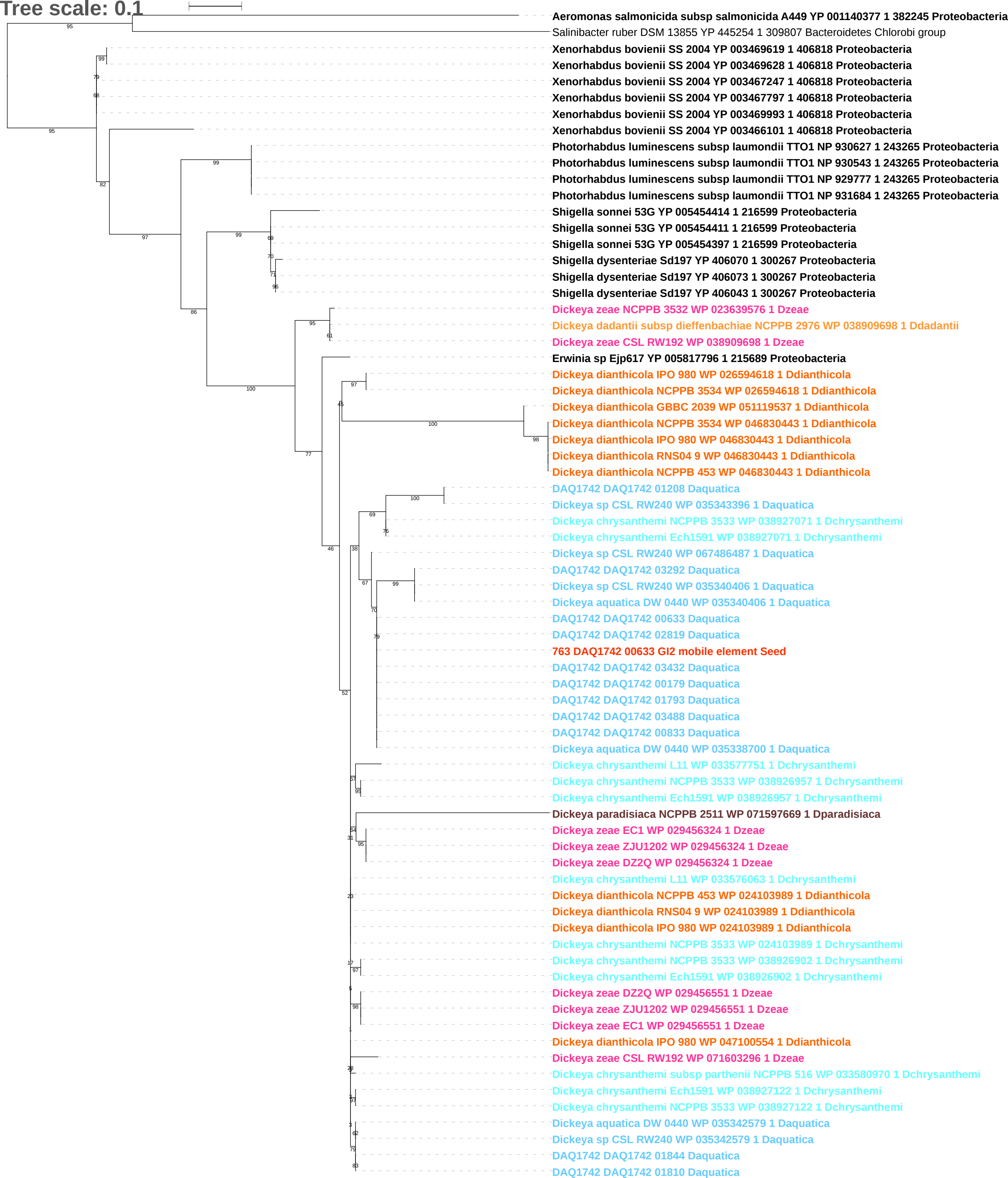
Tree scale: 0.1
Aeromonas salmonicida subsp salmonicida A449 YP 001140377 1 382245 Proteobacteria
Salinibacter ruber DSM 13855 YP 445254 1 309807 Bacteroidetes Chlorobi group
Xenorhabdus bovienii SS 2004 YP 003469619 1 406818 Proteobacteria
Xenorhabdus bovienii SS 2004 YP 003469628 1 406818 Proteobacteria
Xenorhabdus bovienii SS 2004 YP 003467247 1 406818 Proteobacteria
Xenorhabdus bovienii SS 2004 YP 003467797 1 406818 Proteobacteria
Xenorhabdus bovienii SS 2004 YP 003469993 1 406818 Proteobacteria
Xenorhabdus bovienii SS 2004 YP 003466101 1 406818 Proteobacteria
Photorhabdus luminescens subsp laumondii TTO1 NP 930627 1 243265 Proteobacteria
Photorhabdus luminescens subsp laumondii TTO1 NP 930543 1 243265 Proteobacteria
Photorhabdus luminescens subsp laumondii TTO1 NP 929777 1 243265 Proteobacteria
Photorhabdus luminescens subsp laumondii TTO1 NP 931684 1 243265 Proteobacteria
Shigella sonnei 53G YP 005454414 1 216599 Proteobacteria
Shigella sonnei 53G YP 005454411 1 216599 Proteobacteria
Shigella sonnei 53G YP 005454397 1 216599 Proteobacteria
Shigella dysenteriae Sd197 YP 406070 1 300267 Proteobacteria
Shigella dysenteriae Sd197 YP 406073 1 300267 Proteobacteria
Shigella dysenteriae Sd197 YP 406043 1 300267 Proteobacteria
Dickeya zeae NCPPB 3532 WP 023639576 1 Dzeae
Dickeya dadantii subsp dieffenbachiae NCPPB 2976 WP 038909698 1 Ddadantii
Dickeya zeae CSL RW192 WP 038909698 1 Dzeae
Erwinia sp Ejp617 YP 005817796 1 215689 Proteobacteria
Dickeya dianthicola IPO 980 WP 026594618 1 Ddianthicola
Dickeya dianthicola NCPPB 3534 WP 026594618 1 Ddianthicola
Dickeya dianthicola GBBC 2039 WP 051119537 1 Ddianthicola
Dickeya dianthicola NCPPB 3534 WP 046830443 1 Ddianthicola
Dickeya dianthicola IPO 980 WP 046830443 1 Ddianthicola
Dickeya dianthicola RNS04 9 WP 046830443 1 Ddianthicola
Dickeya dianthicola NCPPB 453 WP 046830443 1 Ddianthicola
DAQ1742 DAQ1742 01208 Daquatica
Dickeya sp CSL RW240 WP 035343396 1 Daquatica
Dickeya chrysanthemi NCPPB 3533 WP 038927071 1 Dchrysanthemi
Dickeya chrysanthemi Ech1591 WP 038927071 1 Dchrysanthemi
Dickeya sp CSL RW240 WP 067486487 1 Daquatica
DAQ1742 DAQ1742 03292 Daquatica
Dickeya sp CSL RW240 WP 035340406 1 Daquatica
Dickeya aquatica DW 0440 WP 035340406 1 Daquatica
DAQ1742 DAQ1742 00633 Daquatica
DAQ1742 DAQ1742 02819 Daquatica
763 DAQ1742 00633 GI2 mobile element Seed
DAQ1742 DAQ1742 03432 Daquatica
DAQ1742 DAQ1742 00179 Daquatica
DAQ1742 DAQ1742 01793 Daquatica
DAQ1742 DAQ1742 03488 Daquatica
DAQ1742 DAQ1742 00833 Daquatica
Dickeya aquatica DW 0440 WP 035338700 1 Daquatica
Dickeya chrysanthemi L11 WP 033577751 1 Dchrysanthemi
Dickeya chrysanthemi NCPPB 3533 WP 038926957 1 Dchrysanthemi
Dickeya chrysanthemi Ech1591 WP 038926957 1 Dchrysanthemi
Dickeya paradisiaca NCPPB 2511 WP 071597669 1 Dparadisiaca
Dickeya zeae EC1 WP 029456324 1 Dzeae
Dickeya zeae ZJU1202 WP 029456324 1 Dzeae
Dickeya zeae DZ2Q WP 029456324 1 Dzeae
Dickeya chrysanthemi L11 WP 033576063 1 Dchrysanthemi
Dickeya dianthicola NCPPB 453 WP 024103989 1 Ddianthicola
Dickeya dianthicola RNS04 9 WP 024103989 1 Ddianthicola
Dickeya dianthicola IPO 980 WP 024103989 1 Ddianthicola
Dickeya chrysanthemi NCPPB 3533 WP 024103989 1 Dchrysanthemi
Dickeya chrysanthemi NCPPB 3533 WP 038926902 1 Dchrysanthemi
Dickeya chrysanthemi Ech1591 WP 038926902 1 Dchrysanthemi
Dickeya zeae DZ2Q WP 029456551 1 Dzeae
Dickeya zeae ZJU1202 WP 029456551 1 Dzeae
Dickeya zeae EC1 WP 029456551 1 Dzeae
Dickeya dianthicola IPO 980 WP 047100554 1 Ddianthicola
Dickeya zeae CSL RW192 WP 071603296 1 Dzeae
Dickeya chrysanthemi subsp parthenii NCPPB 516 WP 033580970 1 Dchrysanthemi
Dickeya chrysanthemi Ech1591 WP 038927122 1 Dchrysanthemi
Dickeya chrysanthemi NCPPB 3533 WP 038927122 1 Dchrysanthemi
Dickeya aquatica DW 0440 WP 035342579 1 Daquatica
Dickeya sp CSL RW240 WP 035342579 1 Daquatica
DAQ1742 DAQ1742 01844 Daquatica
DAQ1742 DAQ1742 01810 Daquatica

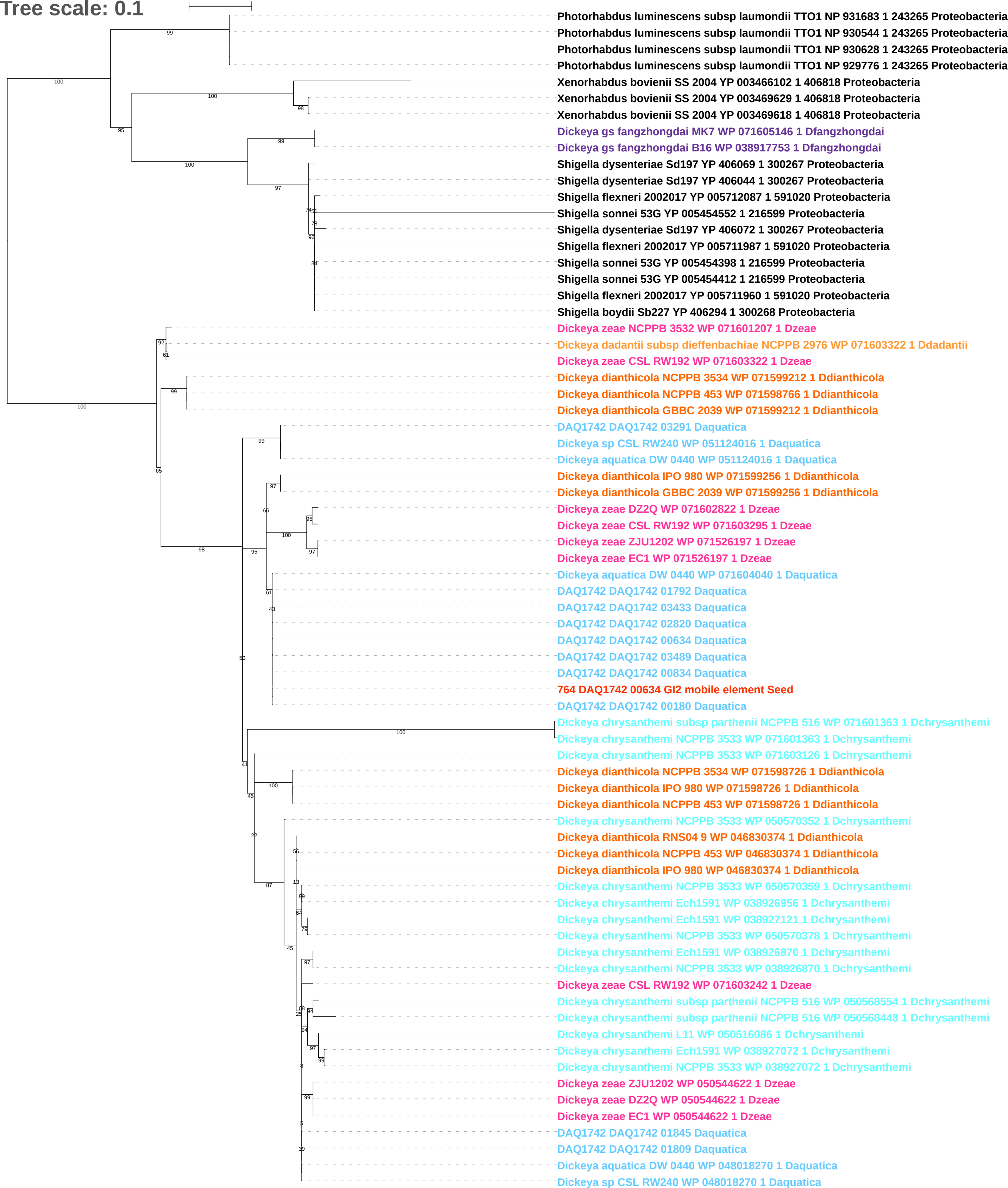
Tree scale: 0.1
Photorhabdus luminescens subsp laumondii TTO1 NP 931683 1 243265 Proteobacteria
Photorhabdus luminescens subsp laumondii TTO1 NP 930544 1 243265 Proteobacteria
Photorhabdus luminescens subsp laumondii TTO1 NP 930628 1 243265 Proteobacteria
Photorhabdus luminescens subsp laumondii TTO1 NP 929776 1 243265 Proteobacteria
Xenorhabdus bovienii SS 2004 YP 003466102 1 406818 Proteobacteria
Xenorhabdus bovienii SS 2004 YP 003469629 1 406818 Proteobacteria
Xenorhabdus bovienii SS 2004 YP 003469618 1 406818 Proteobacteria
Dickeya gs fangzhongdai MK7 WP 071605146 1 Dfangzhongdai
Dickeya gs fangzhongdai B16 WP 038917753 1 Dfangzhongdai
Shigella dysenteriae Sd197 YP 406069 1 300267 Proteobacteria
Shigella dysenteriae Sd197 YP 406044 1 300267 Proteobacteria
Shigella flexneri 2002017 YP 005712087 1 591020 Proteobacteria
Shigella sonnei 53G YP 005454552 1 216599 Proteobacteria
Shigella dysenteriae Sd197 YP 406072 1 300267 Proteobacteria
Shigella flexneri 2002017 YP 005711987 1 591020 Proteobacteria
Shigella sonnei 53G YP 005454398 1 216599 Proteobacteria
Shigella sonnei 53G YP 005454412 1 216599 Proteobacteria
Shigella flexneri 2002017 YP 005711960 1 591020 Proteobacteria
Shigella boydii Sb227 YP 406294 1 300268 Proteobacteria
Dickeya zeae NCPPB 3532 WP 071601207 1 Dzeae
Dickeya dadantii subsp dieffenbachiae NCPPB 2976 WP 071603322 1 Ddadantii
Dickeya zeae CSL RW192 WP 071603322 1 Dzeae
Dickeya dianthicola NCPPB 3534 WP 071599212 1 Ddianthicola
Dickeya dianthicola NCPPB 453 WP 071598766 1 Ddianthicola
Dickeya dianthicola GBBC 2039 WP 071599212 1 Ddianthicola
DAQ1742 DAQ1742 03291 Daquatica
Dickeya sp CSL RW240 WP 051124016 1 Daquatica
Dickeya aquatica DW 0440 WP 051124016 1 Daquatica
Dickeya dianthicola IPO 980 WP 071599256 1 Ddianthicola
Dickeya dianthicola GBBC 2039 WP 071599256 1 Ddianthicola
Dickeya zeae DZ2Q WP 071602822 1 Dzeae
Dickeya zeae CSL RW192 WP 071603295 1 Dzeae
Dickeya zeae ZJU1202 WP 071526197 1 Dzeae
Dickeya zeae EC1 WP 071526197 1 Dzeae
Dickeya aquatica DW 0440 WP 071604040 1 Daquatica
DAQ1742 DAQ1742 01792 Daquatica
DAQ1742 DAQ1742 03433 Daquatica
DAQ1742 DAQ1742 02820 Daquatica
DAQ1742 DAQ1742 00634 Daquatica
DAQ1742 DAQ1742 03489 Daquatica
DAQ1742 DAQ1742 00834 Daquatica
764 DAQ1742 00634 GI2 mobile element Seed
DAQ1742 DAQ1742 00180 Daquatica
Dickeya chrysanthemi subsp parthenii NCPPB 516 WP 071601363 1 Dchrysanthemi
Dickeya chrysanthemi NCPPB 3533 WP 071601363 1 Dchrysanthemi
Dickeya chrysanthemi NCPPB 3533 WP 071603126 1 Dchrysanthemi
Dickeya dianthicola NCPPB 3534 WP 071598726 1 Ddianthicola
Dickeya dianthicola IPO 980 WP 071598726 1 Ddianthicola
Dickeya dianthicola NCPPB 453 WP 071598726 1 Ddianthicola
Dickeya chrysanthemi NCPPB 3533 WP 050570352 1 Dchrysanthemi
Dickeya dianthicola RNS04 9 WP 046830374 1 Ddianthicola
Dickeya dianthicola NCPPB 453 WP 046830374 1 Ddianthicola
Dickeya dianthicola IPO 980 WP 046830374 1 Ddianthicola
Dickeya chrysanthemi NCPPB 3533 WP 050570359 1 Dchrysanthemi
Dickeya chrysanthemi Ech1591 WP 038926956 1 Dchrysanthemi
Dickeya chrysanthemi Ech1591 WP 038927121 1 Dchrysanthemi
Dickeya chrysanthemi NCPPB 3533 WP 050570378 1 Dchrysanthemi
Dickeya chrysanthemi Ech1591 WP 038926870 1 Dchrysanthemi
Dickeya chrysanthemi NCPPB 3533 WP 038926870 1 Dchrysanthemi
Dickeya zeae CSL RW192 WP 071603242 1 Dzeae
Dickeya chrysanthemi subsp parthenii NCPPB 516 WP 050568554 1 Dchrysanthemi
Dickeya chrysanthemi subsp parthenii NCPPB 516 WP 050568448 1 Dchrysanthemi
Dickeya chrysanthemi L11 WP 050516086 1 Dchrysanthemi
Dickeya chrysanthemi Ech1591 WP 038927072 1 Dchrysanthemi
Dickeya chrysanthemi NCPPB 3533 WP 038927072 1 Dchrysanthemi
Dickeya zeae ZJU1202 WP 050544622 1 Dzeae
Dickeya zeae DZ2Q WP 050544622 1 Dzeae
Dickeya zeae EC1 WP 050544622 1 Dzeae
DAQ1742 DAQ1742 01845 Daquatica
DAQ1742 DAQ1742 01809 Daquatica
Dickeya aquatica DW 0440 WP 048018270 1 Daquatica
Dickeya sp CSL RW240 WP 048018270 1 Daquatica

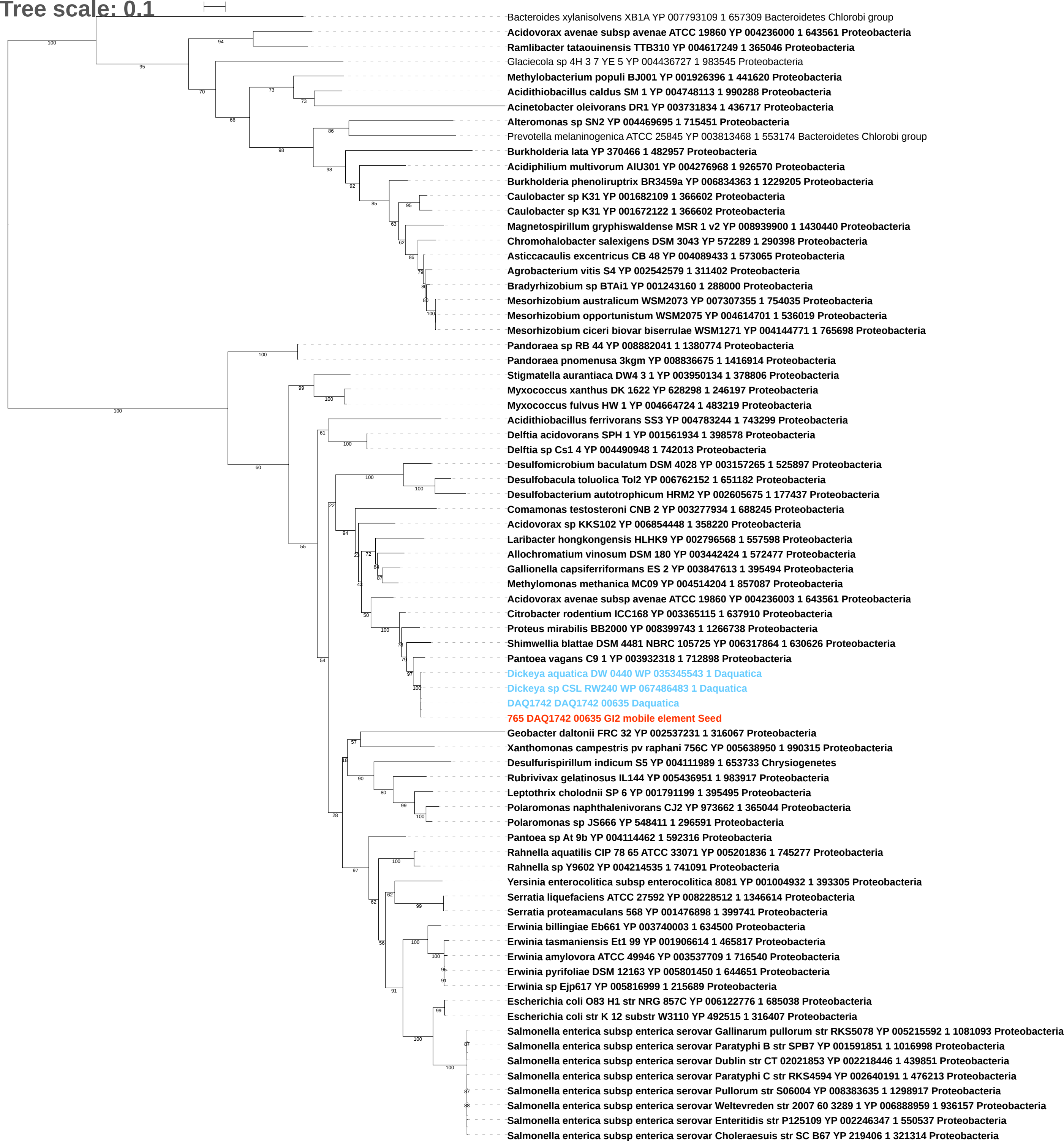
Tree scale: 0.1
Bacteroides xylanisolvens XB1A YP 007793109 1 657309 Bacteroidetes Chlorobi group
Acidovorax avenae subsp avenae ATCC 19860 YP 004236000 1 643561 Proteobacteria
Ramlibacter tataouinensis TTB310 YP 004617249 1 365046 Proteobacteria
Glaciecola sp 4H 3 7 YE 5 YP 004436727 1 983545 Proteobacteria
Methylobacterium populi BJ001 YP 001926396 1 441620 Proteobacteria
Acidithiobacillus caldus SM 1 YP 004748113 1 990288 Proteobacteria
Acinetobacter oleivorans DR1 YP 003731834 1 436717 Proteobacteria
Alteromonas sp SN2 YP 004469695 1 715451 Proteobacteria
Prevotella melaninogenica ATCC 25845 YP 003813468 1 553174 Bacteroidetes Chlorobi group
Burkholderia lata YP 370466 1 482957 Proteobacteria
Acidiphilium multivorum AIU301 YP 004276968 1 926570 Proteobacteria
Burkholderia phenoliruptrix BR3459a YP 006834363 1 1229205 Proteobacteria
Caulobacter sp K31 YP 001682109 1 366602 Proteobacteria
Caulobacter sp K31 YP 001672122 1 366602 Proteobacteria
Magnetospirillum gryphiswaldense MSR 1 v2 YP 008939900 1 1430440 Proteobacteria
Chromohalobacter salexigens DSM 3043 YP 572289 1 290398 Proteobacteria
Asticcacaulis excentricus CB 48 YP 004089433 1 573065 Proteobacteria
Agrobacterium vitis S4 YP 002542579 1 311402 Proteobacteria
Bradyrhizobium sp BTAi1 YP 001243160 1 288000 Proteobacteria
Mesorhizobium australicum WSM2073 YP 007307355 1 754035 Proteobacteria
Mesorhizobium opportunistum WSM2075 YP 004614701 1 536019 Proteobacteria
Mesorhizobium ciceri biovar biserrulae WSM1271 YP 004144771 1 765698 Proteobacteria
Pandoraea sp RB 44 YP 008882041 1 1380774 Proteobacteria
Pandoraea pnomenusa 3kgm YP 008836675 1 1416914 Proteobacteria
Stigmatella aurantiaca DW4 3 1 YP 003950134 1 378806 Proteobacteria
Myxococcus xanthus DK 1622 YP 628298 1 246197 Proteobacteria
Myxococcus fulvus HW 1 YP 004664724 1 483219 Proteobacteria
Acidithiobacillus ferrivorans SS3 YP 004783244 1 743299 Proteobacteria
Delftia acidovorans SPH 1 YP 001561934 1 398578 Proteobacteria
Delftia sp Cs1 4 YP 004490948 1 742013 Proteobacteria
Desulfomicrobium baculatum DSM 4028 YP 003157265 1 525897 Proteobacteria
Desulfobacula toluolica Tol2 YP 006762152 1 651182 Proteobacteria
Desulfobacterium autotrophicum HRM2 YP 002605675 1 177437 Proteobacteria
Comamonas testosteroni CNB 2 YP 003277934 1 688245 Proteobacteria
Acidovorax sp KKS102 YP 006854448 1 358220 Proteobacteria
Laribacter hongkongensis HLHK9 YP 002796568 1 557598 Proteobacteria
Allochromatium vinosum DSM 180 YP 003442424 1 572477 Proteobacteria
Gallionella capsiferriformans ES 2 YP 003847613 1 395494 Proteobacteria
Methylomonas methanica MC09 YP 004514204 1 857087 Proteobacteria
Acidovorax avenae subsp avenae ATCC 19860 YP 004236003 1 643561 Proteobacteria
Citrobacter rodentium ICC168 YP 003365115 1 637910 Proteobacteria
Proteus mirabilis BB2000 YP 008399743 1 1266738 Proteobacteria
Shimwellia blattae DSM 4481 NBRC 105725 YP 006317864 1 630626 Proteobacteria
Pantoea vagans C9 1 YP 003932318 1 712898 Proteobacteria
Dickeya aquatica DW 0440 WP 035345543 1 Daquatica
Dickeya sp CSL RW240 WP 067486483 1 Daquatica
DAQ1742 DAQ1742 00635 Daquatica
765 DAQ1742 00635 GI2 mobile element Seed
Geobacter daltonii FRC 32 YP 002537231 1 316067 Proteobacteria
Xanthomonas campestris pv raphani 756C YP 005638950 1 990315 Proteobacteria
Desulfurispirillum indicum S5 YP 004111989 1 653733 Chrysiogenetes
Rubrivivax gelatinosus IL144 YP 005436951 1 983917 Proteobacteria
Leptothrix cholodnii SP 6 YP 001791199 1 395495 Proteobacteria
Polaromonas naphthalenivorans CJ2 YP 973662 1 365044 Proteobacteria
Polaromonas sp JS666 YP 548411 1 296591 Proteobacteria
Pantoea sp At 9b YP 004114462 1 592316 Proteobacteria
Rahnella aquatilis CIP 78 65 ATCC 33071 YP 005201836 1 745277 Proteobacteria
Rahnella sp Y9602 YP 004214535 1 741091 Proteobacteria
Yersinia enterocolitica subsp enterocolitica 8081 YP 001004932 1 393305 Proteobacteria
Serratia liquefaciens ATCC 27592 YP 008228512 1 1346614 Proteobacteria
Serratia proteamaculans 568 YP 001476898 1 399741 Proteobacteria
Erwinia billingiae Eb661 YP 003740003 1 634500 Proteobacteria
Erwinia tasmaniensis Et1 99 YP 001906614 1 465817 Proteobacteria
Erwinia amylovora ATCC 49946 YP 003537709 1 716540 Proteobacteria
Erwinia pyrifoliae DSM 12163 YP 005801450 1 644651 Proteobacteria
Erwinia sp Ejp617 YP 005816999 1 215689 Proteobacteria
Escherichia coli O83 H1 str NRG 857C YP 006122776 1 685038 Proteobacteria
Escherichia coli str K 12 substr W3110 YP 492515 1 316407 Proteobacteria
Salmonella enterica subsp enterica serovar Gallinarum pullorum str RKS5078 YP 005215592 1 1081093 Proteobacteria
Salmonella enterica subsp enterica serovar Paratyphi B str SPB7 YP 001591851 1 1016998 Proteobacteria
Salmonella enterica subsp enterica serovar Dublin str CT 02021853 YP 002218446 1 439851 Proteobacteria
Salmonella enterica subsp enterica serovar Paratyphi C str RKS4594 YP 002640191 1 476213 Proteobacteria
Salmonella enterica subsp enterica serovar Pullorum str S06004 YP 008383635 1 1298917 Proteobacteria
Salmonella enterica subsp enterica serovar Weltevreden str 2007 60 3289 1 YP 006888959 1 936157 Proteobacteria
Salmonella enterica subsp enterica serovar Enteritidis str P125109 YP 002246347 1 550537 Proteobacteria
Salmonella enterica subsp enterica serovar Choleraesuis str SC B67 YP 219406 1 321314 Proteobacteria

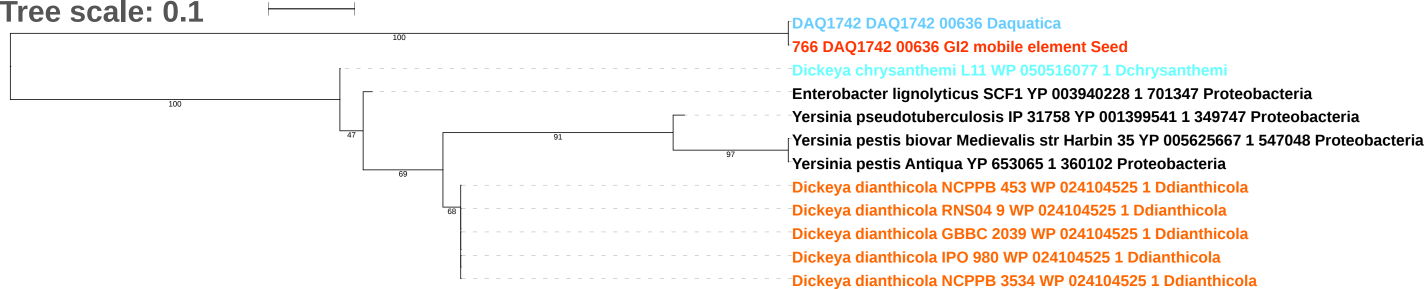
Tree scale: 0.1
DAQ1742 DAQ1742 00636 Daquatica
766 DAQ1742 00636 GI2 mobile element Seed
Dickeya chrysanthemi L11 WP 050516077 1 Dchrysanthemi
Enterobacter lignolyticus SCF1 YP 003940228 1 701347 Proteobacteria
Yersinia pseudotuberculosis IP 31758 YP 001399541 1 349747 Proteobacteria
Yersinia pestis biovar Medievalis str Harbin 35 YP 005625667 1 547048 Proteobacteria
Yersinia pestis Antiqua YP 653065 1 360102 Proteobacteria
Dickeya dianthicola NCPPB 453 WP 024104525 1 Ddianthicola
Dickeya dianthicola RNS04 9 WP 024104525 1 Ddianthicola
Dickeya dianthicola GBBC 2039 WP 024104525 1 Ddianthicola
Dickeya dianthicola IPO 980 WP 024104525 1 Ddianthicola
Dickeya dianthicola NCPPB 3534 WP 024104525 1 Ddianthicola
100
100
47
91
97
69
68

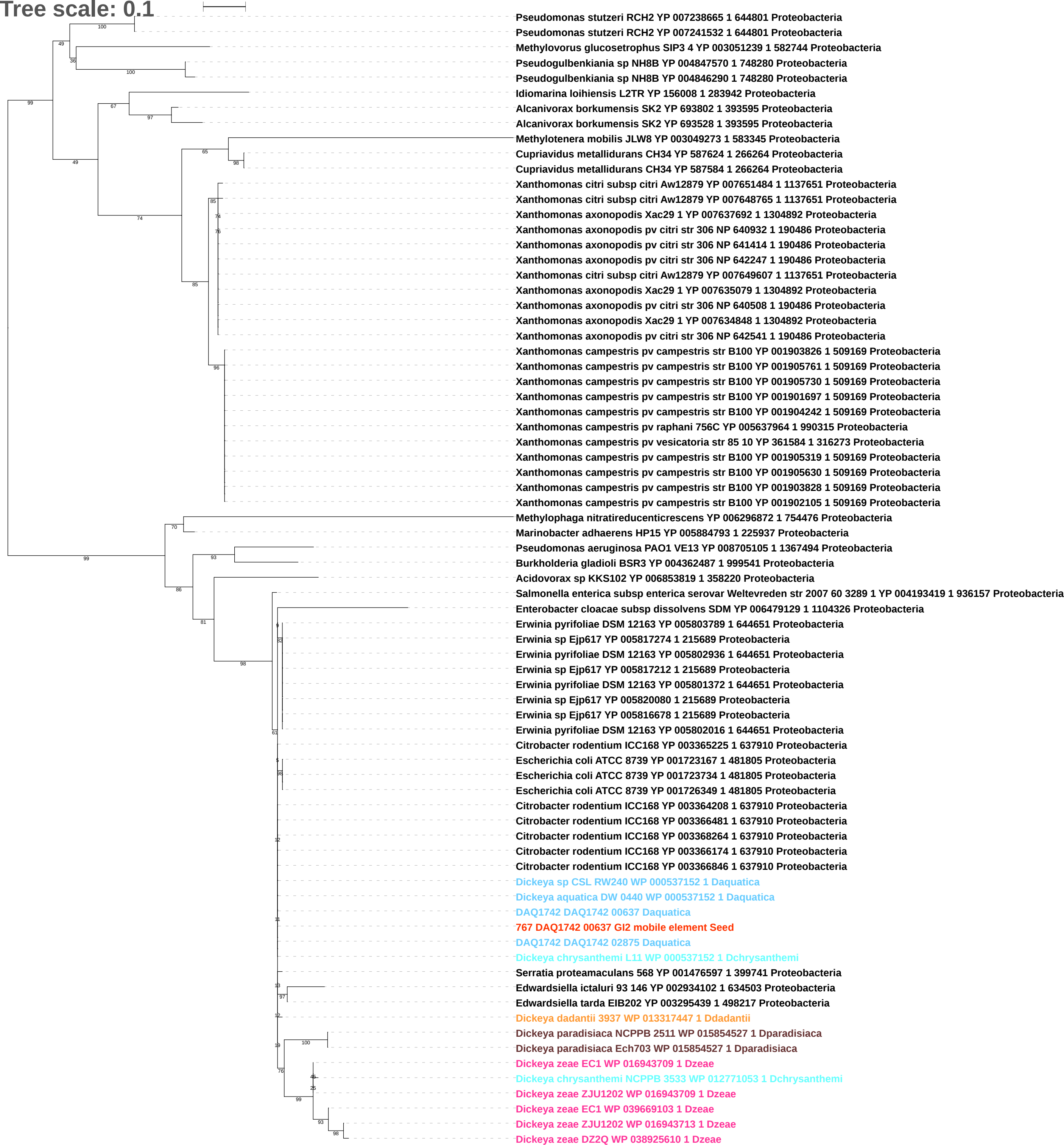

Tree scale: 0.1
Pseudomonas stutzeri RCH2 YP 007238665 1 644801 Proteobacteria
Pseudomonas stutzeri RCH2 YP 007241532 1 644801 Proteobacteria
Methylovorus glucosetrophus SIP3 4 YP 003051239 1 582744 Proteobacteria
Pseudogulbenkiania sp NH8B YP 004847570 1 748280 Proteobacteria
Pseudogulbenkiania sp NH8B YP 004846290 1 748280 Proteobacteria
Idiomarina loihiensis L2TR YP 156008 1 283942 Proteobacteria
Alcanivorax borkumensis SK2 YP 693802 1 393595 Proteobacteria
Alcanivorax borkumensis SK2 YP 693528 1 393595 Proteobacteria
Methylotenera mobilis JLW8 YP 003049273 1 583345 Proteobacteria
Cupriavidus metallidurans CH34 YP 587624 1 266264 Proteobacteria
Cupriavidus metallidurans CH34 YP 587584 1 266264 Proteobacteria
Xanthomonas citri subsp citri Aw12879 YP 007651484 1 1137651 Proteobacteria
Xanthomonas citri subsp citri Aw12879 YP 007648765 1 1137651 Proteobacteria
Xanthomonas axonopodis Xac29 1 YP 007637692 1 1304892 Proteobacteria
Xanthomonas axonopodis pv citri str 306 NP 640932 1 190486 Proteobacteria
Xanthomonas axonopodis pv citri str 306 NP 641414 1 190486 Proteobacteria
Xanthomonas axonopodis pv citri str 306 NP 642247 1 190486 Proteobacteria
Xanthomonas citri subsp citri Aw12879 YP 007649607 1 1137651 Proteobacteria
Xanthomonas axonopodis Xac29 1 YP 007635079 1 1304892 Proteobacteria
Xanthomonas axonopodis pv citri str 306 NP 640508 1 190486 Proteobacteria
Xanthomonas axonopodis Xac29 1 YP 007634848 1 1304892 Proteobacteria
Xanthomonas axonopodis pv citri str 306 NP 642541 1 190486 Proteobacteria
Xanthomonas campestris pv campestris str B100 YP 001903826 1 509169 Proteobacteria
Xanthomonas campestris pv campestris str B100 YP 001905761 1 509169 Proteobacteria
Xanthomonas campestris pv campestris str B100 YP 001905730 1 509169 Proteobacteria
Xanthomonas campestris pv campestris str B100 YP 001901697 1 509169 Proteobacteria
Xanthomonas campestris pv campestris str B100 YP 001904242 1 509169 Proteobacteria
Xanthomonas campestris pv raphani 756C YP 005637964 1 990315 Proteobacteria
Xanthomonas campestris pv vesicatoria str 85 10 YP 361584 1 316273 Proteobacteria
Xanthomonas campestris pv campestris str B100 YP 001905319 1 509169 Proteobacteria
Xanthomonas campestris pv campestris str B100 YP 001905630 1 509169 Proteobacteria
Xanthomonas campestris pv campestris str B100 YP 001903828 1 509169 Proteobacteria
Xanthomonas campestris pv campestris str B100 YP 001902105 1 509169 Proteobacteria
Methylophaga nitratireducenticrescens YP 006296872 1 754476 Proteobacteria
Marinobacter adhaerens HP15 YP 005884793 1 225937 Proteobacteria
Pseudomonas aeruginosa PAO1 VE13 YP 008705105 1 1367494 Proteobacteria
Burkholderia gladioli BSR3 YP 004362487 1 999541 Proteobacteria
Acidovorax sp KKS102 YP 006853819 1 358220 Proteobacteria
Salmonella enterica subsp enterica serovar Weltevreden str 2007 60 3289 1 YP 004193419 1 936157 Proteobacteria
Enterobacter cloacae subsp dissolvens SDM YP 006479129 1 1104326 Proteobacteria
Erwinia pyrifoliae DSM 12163 YP 005803789 1 644651 Proteobacteria
Erwinia sp Ejp617 YP 005817274 1 215689 Proteobacteria
Erwinia pyrifoliae DSM 12163 YP 005802936 1 644651 Proteobacteria
Erwinia sp Ejp617 YP 005817212 1 215689 Proteobacteria
Erwinia pyrifoliae DSM 12163 YP 005801372 1 644651 Proteobacteria
Erwinia sp Ejp617 YP 005820080 1 215689 Proteobacteria
Erwinia sp Ejp617 YP 005816678 1 215689 Proteobacteria
Erwinia pyrifoliae DSM 12163 YP 005802016 1 644651 Proteobacteria
Citrobacter rodentium ICC168 YP 003365225 1 637910 Proteobacteria
Escherichia coli ATCC 8739 YP 001723167 1 481805 Proteobacteria
Escherichia coli ATCC 8739 YP 001723734 1 481805 Proteobacteria
Escherichia coli ATCC 8739 YP 001726349 1 481805 Proteobacteria
Citrobacter rodentium ICC168 YP 003364208 1 637910 Proteobacteria
Citrobacter rodentium ICC168 YP 003366481 1 637910 Proteobacteria
Citrobacter rodentium ICC168 YP 003368264 1 637910 Proteobacteria
Citrobacter rodentium ICC168 YP 003366174 1 637910 Proteobacteria
Citrobacter rodentium ICC168 YP 003366846 1 637910 Proteobacteria
Dickeya sp CSL RW240 WP 000537152 1 Daquatica
Dickeya aquatica DW 0440 WP 000537152 1 Daquatica
DAQ1742 DAQ1742 00637 Daquatica
767 DAQ1742 00637 GI2 mobile element Seed
DAQ1742 DAQ1742 02875 Daquatica
Dickeya chrysanthemi L11 WP 000537152 1 Dchrysanthemi
Serratia proteamaculans 568 YP 001476597 1 399741 Proteobacteria
Edwardsiella ictaluri 93 146 YP 002934102 1 634503 Proteobacteria
Edwardsiella tarda EIB202 YP 003295439 1 498217 Proteobacteria
Dickeya dadantii 3937 WP 013317447 1 Ddadantii
Dickeya paradisiaca NCPPB 2511 WP 015854527 1 Dparadisiaca
Dickeya paradisiaca Ech703 WP 015854527 1 Dparadisiaca
Dickeya zeae EC1 WP 016943709 1 Dzeae
Dickeya chrysanthemi NCPPB 3533 WP 012771053 1 Dchrysanthemi
Dickeya zeae ZJU1202 WP 016943709 1 Dzeae
Dickeya zeae EC1 WP 039669103 1 Dzeae
Dickeya zeae ZJU1202 WP 016943713 1 Dzeae
Dickeya zeae DZ2Q WP 038925610 1 Dzeae

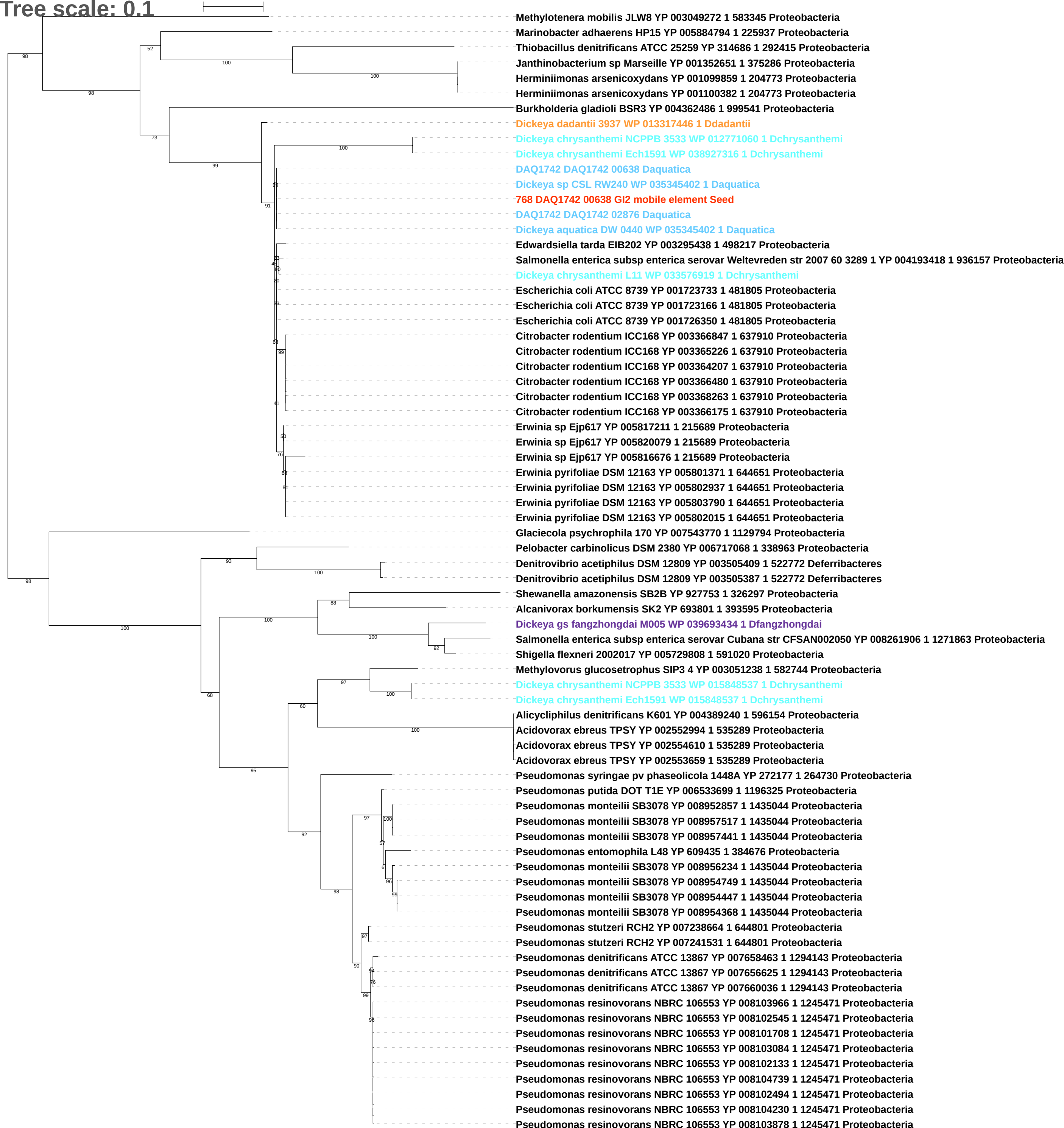
Tree scale: 0.1
Methylotenera mobilis JLW8 YP 003049272 1 583345 Proteobacteria
Marinobacter adhaerens HP15 YP 005884794 1 225937 Proteobacteria
Thiobacillus denitrificans ATCC 25259 YP 314686 1 292415 Proteobacteria
Janthinobacterium sp Marseille YP 001352651 1 375286 Proteobacteria
Herminiimonas arsenicoxydans YP 001099859 1 204773 Proteobacteria
Herminiimonas arsenicoxydans YP 001100382 1 204773 Proteobacteria
Burkholderia gladioli BSR3 YP 004362486 1 999541 Proteobacteria
Dickeya dadantii 3937 WP 013317446 1 Ddadantii
Dickeya chrysanthemi NCPPB 3533 WP 012771060 1 Dchrysanthemi
Dickeya chrysanthemi Ech1591 WP 038927316 1 Dchrysanthemi
DAQ1742 DAQ1742 00638 Daquatica
Dickeya sp CSL RW240 WP 035345402 1 Daquatica
768 DAQ1742 00638 GI2 mobile element Seed
DAQ1742 DAQ1742 02876 Daquatica
Dickeya aquatica DW 0440 WP 035345402 1 Daquatica
Edwardsiella tarda EIB202 YP 003295438 1 498217 Proteobacteria
Salmonella enterica subsp enterica serovar Weltevreden str 2007 60 3289 1 YP 004193418 1 936157 Proteobacteria
Dickeya chrysanthemi L11 WP 033576919 1 Dchrysanthemi
Escherichia coli ATCC 8739 YP 001723733 1 481805 Proteobacteria
Escherichia coli ATCC 8739 YP 001723166 1 481805 Proteobacteria
Escherichia coli ATCC 8739 YP 001726350 1 481805 Proteobacteria
Citrobacter rodentium ICC168 YP 003366847 1 637910 Proteobacteria
Citrobacter rodentium ICC168 YP 003365226 1 637910 Proteobacteria
Citrobacter rodentium ICC168 YP 003364207 1 637910 Proteobacteria
Citrobacter rodentium ICC168 YP 003366480 1 637910 Proteobacteria
Citrobacter rodentium ICC168 YP 003368263 1 637910 Proteobacteria
Citrobacter rodentium ICC168 YP 003366175 1 637910 Proteobacteria
Erwinia sp Ejp617 YP 005817211 1 215689 Proteobacteria
Erwinia sp Ejp617 YP 005820079 1 215689 Proteobacteria
Erwinia sp Ejp617 YP 005816676 1 215689 Proteobacteria
Erwinia pyrifoliae DSM 12163 YP 005801371 1 644651 Proteobacteria
Erwinia pyrifoliae DSM 12163 YP 005802937 1 644651 Proteobacteria
Erwinia pyrifoliae DSM 12163 YP 005803790 1 644651 Proteobacteria
Erwinia pyrifoliae DSM 12163 YP 005802015 1 644651 Proteobacteria
Glaciecola psychrophila 170 YP 007543770 1 1129794 Proteobacteria
Pelobacter carbinolicus DSM 2380 YP 006717068 1 338963 Proteobacteria
Denitrovibrio acetiphilus DSM 12809 YP 003505409 1 522772 Deferribacteres
Denitrovibrio acetiphilus DSM 12809 YP 003505387 1 522772 Deferribacteres
Shewanella amazonensis SB2B YP 927753 1 326297 Proteobacteria
Alcanivorax borkumensis SK2 YP 693801 1 393595 Proteobacteria
Dickeya gs fangzhongdai M005 WP 039693434 1 Dfangzhongdai
Salmonella enterica subsp enterica serovar Cubana str CFSAN002050 YP 008261906 1 1271863 Proteobacteria
Shigella flexneri 2002017 YP 005729808 1 591020 Proteobacteria
Methylovorus glucosetrophus SIP3 4 YP 003051238 1 582744 Proteobacteria
Dickeya chrysanthemi NCPPB 3533 WP 015848537 1 Dchrysanthemi
Dickeya chrysanthemi Ech1591 WP 015848537 1 Dchrysanthemi
Alicycliphilus denitrificans K601 YP 004389240 1 596154 Proteobacteria
Acidovorax ebreus TPSY YP 002552994 1 535289 Proteobacteria
Acidovorax ebreus TPSY YP 002554610 1 535289 Proteobacteria
Acidovorax ebreus TPSY YP 002553659 1 535289 Proteobacteria
Pseudomonas syringae pv phaseolicola 1448A YP 272177 1 264730 Proteobacteria
Pseudomonas putida DOT T1E YP 006533699 1 1196325 Proteobacteria
Pseudomonas monteilii SB3078 YP 008952857 1 1435044 Proteobacteria
Pseudomonas monteilii SB3078 YP 008957517 1 1435044 Proteobacteria
Pseudomonas monteilii SB3078 YP 008957441 1 1435044 Proteobacteria
Pseudomonas entomophila L48 YP 609435 1 384676 Proteobacteria
Pseudomonas monteilii SB3078 YP 008956234 1 1435044 Proteobacteria
Pseudomonas monteilii SB3078 YP 008954749 1 1435044 Proteobacteria
Pseudomonas monteilii SB3078 YP 008954447 1 1435044 Proteobacteria
Pseudomonas monteilii SB3078 YP 008954368 1 1435044 Proteobacteria
Pseudomonas stutzeri RCH2 YP 007238664 1 644801 Proteobacteria
Pseudomonas stutzeri RCH2 YP 007241531 1 644801 Proteobacteria
Pseudomonas denitrificans ATCC 13867 YP 007658463 1 1294143 Proteobacteria
Pseudomonas denitrificans ATCC 13867 YP 007656625 1 1294143 Proteobacteria
Pseudomonas denitrificans ATCC 13867 YP 007660036 1 1294143 Proteobacteria
Pseudomonas resinovorans NBRC 106553 YP 008103966 1 1245471 Proteobacteria
Pseudomonas resinovorans NBRC 106553 YP 008102545 1 1245471 Proteobacteria
Pseudomonas resinovorans NBRC 106553 YP 008101708 1 1245471 Proteobacteria
Pseudomonas resinovorans NBRC 106553 YP 008103084 1 1245471 Proteobacteria
Pseudomonas resinovorans NBRC 106553 YP 008102133 1 1245471 Proteobacteria
Pseudomonas resinovorans NBRC 106553 YP 008104739 1 1245471 Proteobacteria
Pseudomonas resinovorans NBRC 106553 YP 008102494 1 1245471 Proteobacteria
Pseudomonas resinovorans NBRC 106553 YP 008104230 1 1245471 Proteobacteria
Pseudomonas resinovorans NBRC 106553 YP 008103878 1 1245471 Proteobacteria

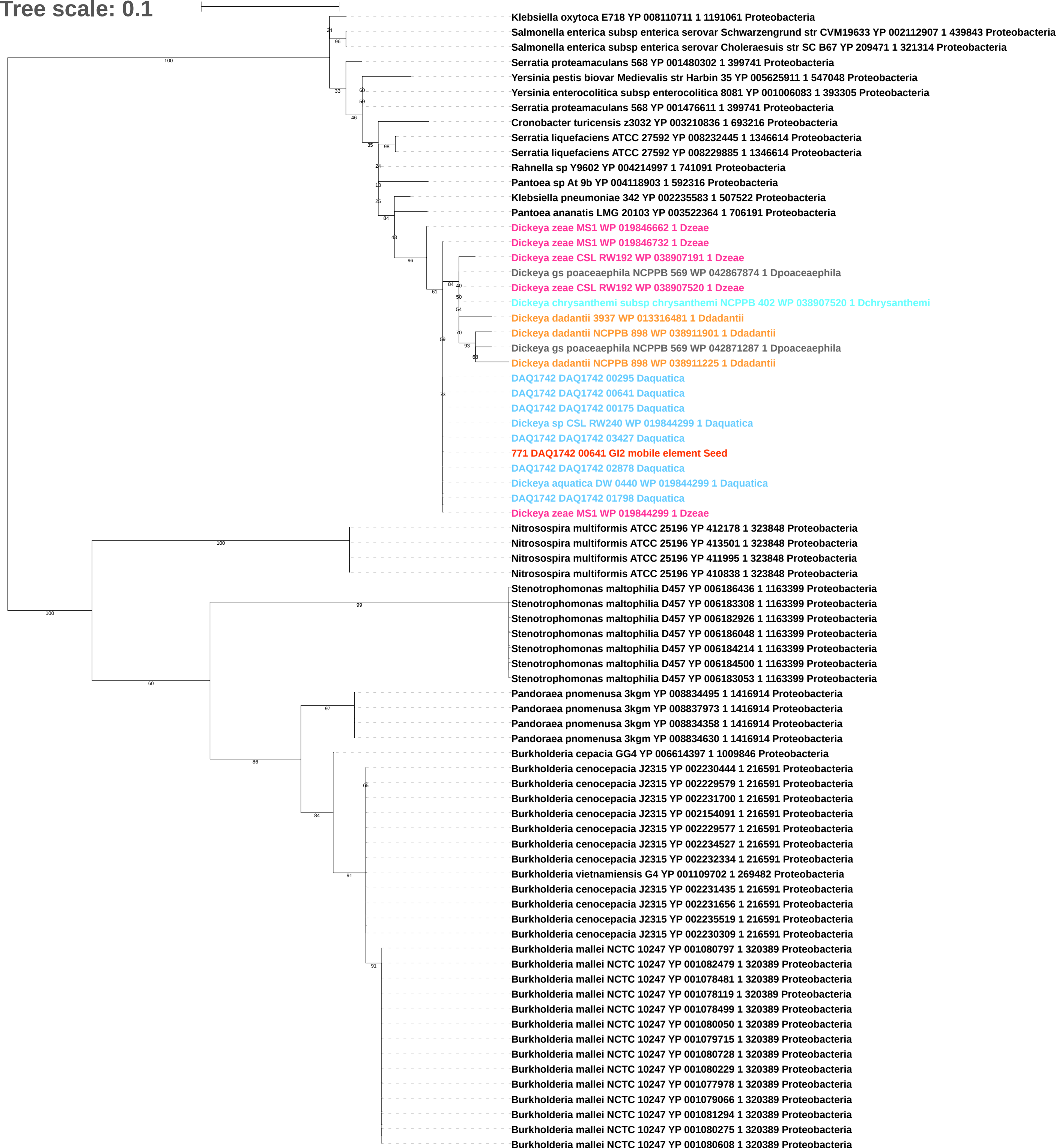
Tree scale: 0.1
Klebsiella oxytoca E718 YP 008110711 1 1191061 Proteobacteria
Salmonella enterica subsp enterica serovar Schwarzengrund str CVM19633 YP 002112907 1 439843 Proteobacteria
Salmonella enterica subsp enterica serovar Choleraesuis str SC B67 YP 209471 1 321314 Proteobacteria
Serratia proteamaculans 568 YP 001480302 1 399741 Proteobacteria
Yersinia pestis biovar Medievalis str Harbin 35 YP 005625911 1 547048 Proteobacteria
Yersinia enterocolitica subsp enterocolitica 8081 YP 001006083 1 393305 Proteobacteria
Serratia proteamaculans 568 YP 001476611 1 399741 Proteobacteria
Cronobacter turicensis z3032 YP 003210836 1 693216 Proteobacteria
Serratia liquefaciens ATCC 27592 YP 008232445 1 1346614 Proteobacteria
Serratia liquefaciens ATCC 27592 YP 008229885 1 1346614 Proteobacteria
Rahnella sp Y9602 YP 004214997 1 741091 Proteobacteria
Pantoea sp At 9b YP 004118903 1 592316 Proteobacteria
Klebsiella pneumoniae 342 YP 002235583 1 507522 Proteobacteria
Pantoea ananatis LMG 20103 YP 003522364 1 706191 Proteobacteria
Dickeya zeae MS1 WP 019846662 1 Dzeae
Dickeya zeae MS1 WP 019846732 1 Dzeae
Dickeya zeae CSL RW192 WP 038907191 1 Dzeae
Dickeya gs poaceaephila NCPPB 569 WP 042867874 1 Dpoaceaephila
Dickeya zeae CSL RW192 WP 038907520 1 Dzeae
Dickeya chrysanthemi subsp chrysanthemi NCPPB 402 WP 038907520 1 Dchrysanthemi
Dickeya dadantii 3937 WP 013316481 1 Ddadantii
Dickeya dadantii NCPPB 898 WP 038911901 1 Ddadantii
Dickeya gs poaceaephila NCPPB 569 WP 042871287 1 Dpoaceaephila
Dickeya dadantii NCPPB 898 WP 038911225 1 Ddadantii
DAQ1742 DAQ1742 00295 Daquatica
DAQ1742 DAQ1742 00641 Daquatica
DAQ1742 DAQ1742 00175 Daquatica
Dickeya sp CSL RW240 WP 019844299 1 Daquatica
DAQ1742 DAQ1742 03427 Daquatica
771 DAQ1742 00641 GI2 mobile element Seed
DAQ1742 DAQ1742 02878 Daquatica
Dickeya aquatica DW 0440 WP 019844299 1 Daquatica
DAQ1742 DAQ1742 01798 Daquatica
Dickeya zeae MS1 WP 019844299 1 Dzeae
Nitrosospira multiformis ATCC 25196 YP 412178 1 323848 Proteobacteria
Nitrosospira multiformis ATCC 25196 YP 413501 1 323848 Proteobacteria
Nitrosospira multiformis ATCC 25196 YP 411995 1 323848 Proteobacteria
Nitrosospira multiformis ATCC 25196 YP 410838 1 323848 Proteobacteria
Stenotrophomonas maltophilia D457 YP 006186436 1 1163399 Proteobacteria
Stenotrophomonas maltophilia D457 YP 006183308 1 1163399 Proteobacteria
Stenotrophomonas maltophilia D457 YP 006182926 1 1163399 Proteobacteria
Stenotrophomonas maltophilia D457 YP 006186048 1 1163399 Proteobacteria
Stenotrophomonas maltophilia D457 YP 006184214 1 1163399 Proteobacteria
Stenotrophomonas maltophilia D457 YP 006184500 1 1163399 Proteobacteria
Stenotrophomonas maltophilia D457 YP 006183053 1 1163399 Proteobacteria
Pandoraea pnomenusa 3kgm YP 008834495 1 1416914 Proteobacteria
Pandoraea pnomenusa 3kgm YP 008837973 1 1416914 Proteobacteria
Pandoraea pnomenusa 3kgm YP 008834358 1 1416914 Proteobacteria
Pandoraea pnomenusa 3kgm YP 008834630 1 1416914 Proteobacteria
Burkholderia cepacia GG4 YP 006614397 1 1009846 Proteobacteria
Burkholderia cenocepacia J2315 YP 002230444 1 216591 Proteobacteria
Burkholderia cenocepacia J2315 YP 002229579 1 216591 Proteobacteria
Burkholderia cenocepacia J2315 YP 002231700 1 216591 Proteobacteria
Burkholderia cenocepacia J2315 YP 002154091 1 216591 Proteobacteria
Burkholderia cenocepacia J2315 YP 002229577 1 216591 Proteobacteria
Burkholderia cenocepacia J2315 YP 002234527 1 216591 Proteobacteria
Burkholderia cenocepacia J2315 YP 002232334 1 216591 Proteobacteria
Burkholderia vietnamiensis G4 YP 001109702 1 269482 Proteobacteria
Burkholderia cenocepacia J2315 YP 002231435 1 216591 Proteobacteria
Burkholderia cenocepacia J2315 YP 002231656 1 216591 Proteobacteria
Burkholderia cenocepacia J2315 YP 002235519 1 216591 Proteobacteria
Burkholderia cenocepacia J2315 YP 002230309 1 216591 Proteobacteria
Burkholderia mallei NCTC 10247 YP 001080797 1 320389 Proteobacteria
Burkholderia mallei NCTC 10247 YP 001082479 1 320389 Proteobacteria
Burkholderia mallei NCTC 10247 YP 001078481 1 320389 Proteobacteria
Burkholderia mallei NCTC 10247 YP 001078119 1 320389 Proteobacteria
Burkholderia mallei NCTC 10247 YP 001078499 1 320389 Proteobacteria
Burkholderia mallei NCTC 10247 YP 001080050 1 320389 Proteobacteria
Burkholderia mallei NCTC 10247 YP 001079715 1 320389 Proteobacteria
Burkholderia mallei NCTC 10247 YP 001080728 1 320389 Proteobacteria
Burkholderia mallei NCTC 10247 YP 001080229 1 320389 Proteobacteria
Burkholderia mallei NCTC 10247 YP 001077978 1 320389 Proteobacteria
Burkholderia mallei NCTC 10247 YP 001079066 1 320389 Proteobacteria
Burkholderia mallei NCTC 10247 YP 001081294 1 320389 Proteobacteria
Burkholderia mallei NCTC 10247 YP 001080275 1 320389 Proteobacteria
Burkholderia mallei NCTC 10247 YP 001080608 1 320389 Proteobacteria

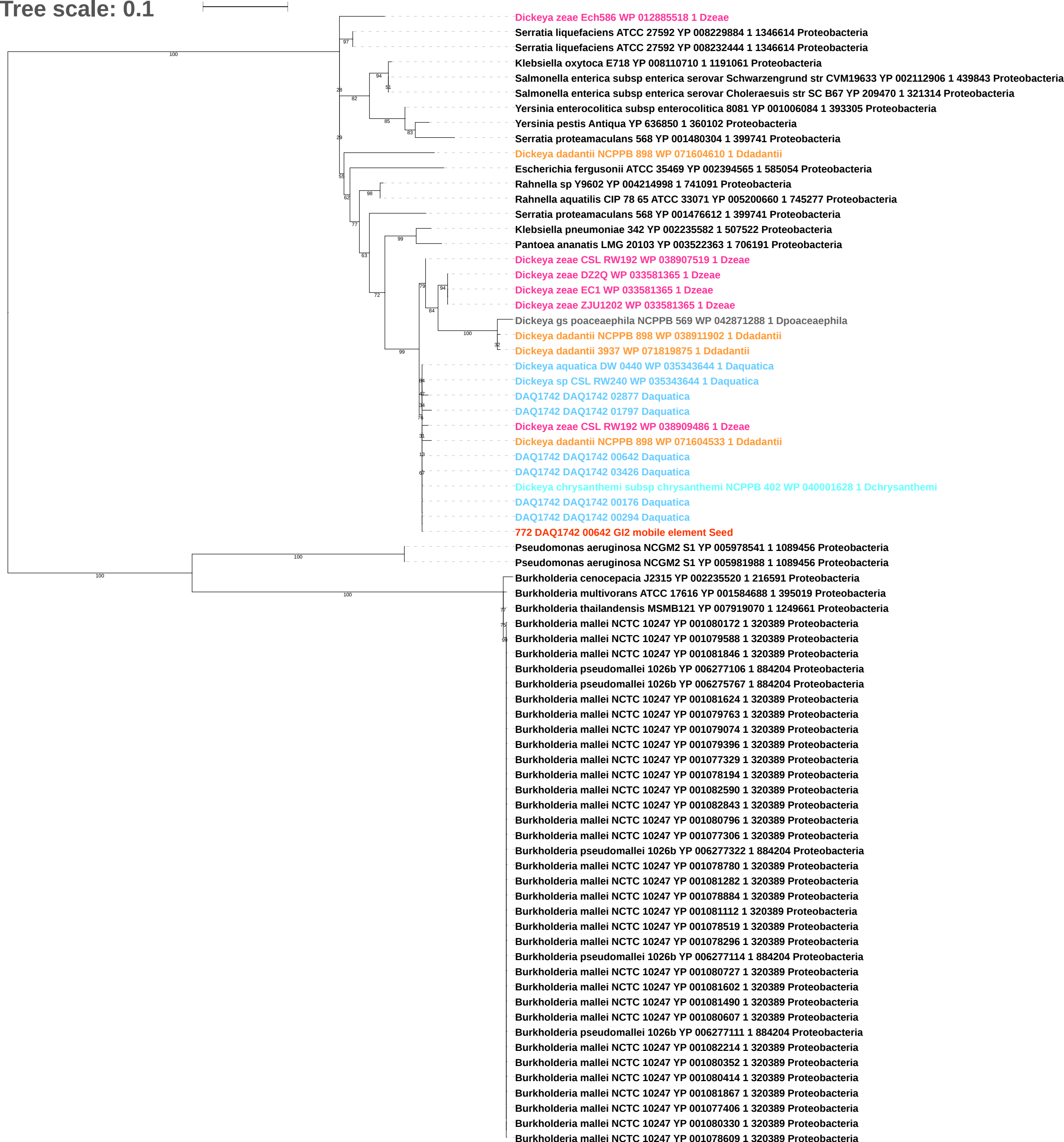
Tree scale: 0.1
Dickeya zeae Ech586 WP 012885518 1 Dzeae
Serratia liquefaciens ATCC 27592 YP 008229884 1 1346614 Proteobacteria
Serratia liquefaciens ATCC 27592 YP 008232444 1 1346614 Proteobacteria
Klebsiella oxytoca E718 YP 008110710 1 1191061 Proteobacteria
Salmonella enterica subsp enterica serovar Schwarzengrund str CVM19633 YP 002112906 1 439843 Proteobacteria
Salmonella enterica subsp enterica serovar Choleraesuis str SC B67 YP 209470 1 321314 Proteobacteria
Yersinia enterocolitica subsp enterocolitica 8081 YP 001006084 1 393305 Proteobacteria
Yersinia pestis Antiqua YP 636850 1 360102 Proteobacteria
Serratia proteamaculans 568 YP 001480304 1 399741 Proteobacteria
Dickeya dadantii NCPPB 898 WP 071604610 1 Ddadantii
Escherichia fergusonii ATCC 35469 YP 002394565 1 585054 Proteobacteria
Rahnella sp Y9602 YP 004214998 1 741091 Proteobacteria
Rahnella aquatilis CIP 78 65 ATCC 33071 YP 005200660 1 745277 Proteobacteria
Serratia proteamaculans 568 YP 001476612 1 399741 Proteobacteria
Klebsiella pneumoniae 342 YP 002235582 1 507522 Proteobacteria
Pantoea ananatis LMG 20103 YP 003522363 1 706191 Proteobacteria
Dickeya zeae CSL RW192 WP 038907519 1 Dzeae
Dickeya zeae DZ2Q WP 033581365 1 Dzeae
Dickeya zeae EC1 WP 033581365 1 Dzeae
Dickeya zeae ZJU1202 WP 033581365 1 Dzeae
Dickeya gs poaceaephila NCPPB 569 WP 042871288 1 Dpoaceaephila
Dickeya dadantii NCPPB 898 WP 038911902 1 Ddadantii
Dickeya dadantii 3937 WP 071819875 1 Ddadantii
Dickeya aquatica DW 0440 WP 035343644 1 Daquatica
Dickeya sp CSL RW240 WP 035343644 1 Daquatica
DAQ1742 DAQ1742 02877 Daquatica
DAQ1742 DAQ1742 01797 Daquatica
Dickeya zeae CSL RW192 WP 038909486 1 Dzeae
Dickeya dadantii NCPPB 898 WP 071604533 1 Ddadantii
DAQ1742 DAQ1742 00642 Daquatica
DAQ1742 DAQ1742 03426 Daquatica
Dickeya chrysanthemi subsp chrysanthemi NCPPB 402 WP 040001628 1 Dchrysanthemi
DAQ1742 DAQ1742 00176 Daquatica
DAQ1742 DAQ1742 00294 Daquatica
772 DAQ1742 00642 GI2 mobile element Seed
Pseudomonas aeruginosa NCGM2 S1 YP 005978541 1 1089456 Proteobacteria
Pseudomonas aeruginosa NCGM2 S1 YP 005981988 1 1089456 Proteobacteria
Burkholderia cenocepacia J2315 YP 002235520 1 216591 Proteobacteria
Burkholderia multivorans ATCC 17616 YP 001584688 1 395019 Proteobacteria
Burkholderia thailandensis MSMB121 YP 007919070 1 1249661 Proteobacteria
Burkholderia mallei NCTC 10247 YP 001080172 1 320389 Proteobacteria
Burkholderia mallei NCTC 10247 YP 001079588 1 320389 Proteobacteria
Burkholderia mallei NCTC 10247 YP 001081846 1 320389 Proteobacteria
Burkholderia pseudomallei 1026b YP 006277106 1 884204 Proteobacteria
Burkholderia pseudomallei 1026b YP 006275767 1 884204 Proteobacteria
Burkholderia mallei NCTC 10247 YP 001081624 1 320389 Proteobacteria
Burkholderia mallei NCTC 10247 YP 001079763 1 320389 Proteobacteria
Burkholderia mallei NCTC 10247 YP 001079074 1 320389 Proteobacteria
Burkholderia mallei NCTC 10247 YP 001079396 1 320389 Proteobacteria
Burkholderia mallei NCTC 10247 YP 001077329 1 320389 Proteobacteria
Burkholderia mallei NCTC 10247 YP 001078194 1 320389 Proteobacteria
Burkholderia mallei NCTC 10247 YP 001082590 1 320389 Proteobacteria
Burkholderia mallei NCTC 10247 YP 001082843 1 320389 Proteobacteria
Burkholderia mallei NCTC 10247 YP 001080796 1 320389 Proteobacteria
Burkholderia mallei NCTC 10247 YP 001077306 1 320389 Proteobacteria
Burkholderia pseudomallei 1026b YP 006277322 1 884204 Proteobacteria
Burkholderia mallei NCTC 10247 YP 001078780 1 320389 Proteobacteria
Burkholderia mallei NCTC 10247 YP 001081282 1 320389 Proteobacteria
Burkholderia mallei NCTC 10247 YP 001078884 1 320389 Proteobacteria
Burkholderia mallei NCTC 10247 YP 001081112 1 320389 Proteobacteria
Burkholderia mallei NCTC 10247 YP 001078519 1 320389 Proteobacteria
Burkholderia mallei NCTC 10247 YP 001078296 1 320389 Proteobacteria
Burkholderia pseudomallei 1026b YP 006277114 1 884204 Proteobacteria
Burkholderia mallei NCTC 10247 YP 001080727 1 320389 Proteobacteria
Burkholderia mallei NCTC 10247 YP 001081602 1 320389 Proteobacteria
Burkholderia mallei NCTC 10247 YP 001081490 1 320389 Proteobacteria
Burkholderia mallei NCTC 10247 YP 001080607 1 320389 Proteobacteria
Burkholderia pseudomallei 1026b YP 006277111 1 884204 Proteobacteria
Burkholderia mallei NCTC 10247 YP 001082214 1 320389 Proteobacteria
Burkholderia mallei NCTC 10247 YP 001080352 1 320389 Proteobacteria
Burkholderia mallei NCTC 10247 YP 001080414 1 320389 Proteobacteria
Burkholderia mallei NCTC 10247 YP 001081867 1 320389 Proteobacteria
Burkholderia mallei NCTC 10247 YP 001077406 1 320389 Proteobacteria
Burkholderia mallei NCTC 10247 YP 001080330 1 320389 Proteobacteria
Burkholderia mallei NCTC 10247 YP 001078609 1 320389 Proteobacteria

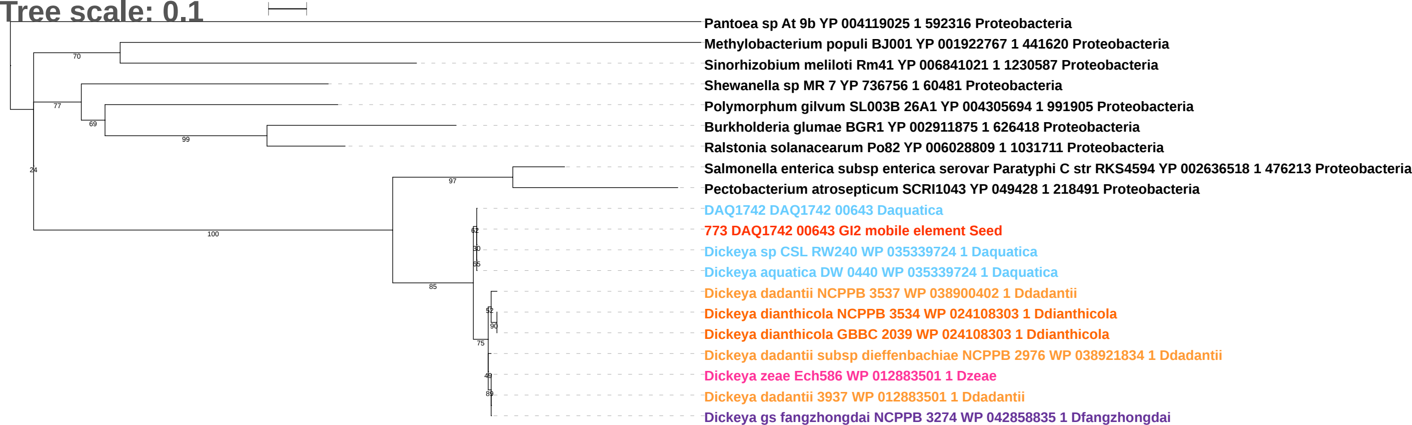

Tree scale: 0.1
Pantoea sp At 9b YP 004119025 1 592316 Proteobacteria
Methylobacterium populi BJ001 YP 001922767 1 441620 Proteobacteria
Sinorhizobium meliloti Rm41 YP 006841021 1 1230587 Proteobacteria
Shewanella sp MR 7 YP 736756 1 60481 Proteobacteria
Polymorphum gilvum SL003B 26A1 YP 004305694 1 991905 Proteobacteria
Burkholderia glumae BGR1 YP 002911875 1 626418 Proteobacteria
Ralstonia solanacearum Po82 YP 006028809 1 1031711 Proteobacteria
Salmonella enterica subsp enterica serovar Paratyphi C str RKS4594 YP 002636518 1 476213 Proteobacteria
Pectobacterium atrosepticum SCRI1043 YP 049428 1 218491 Proteobacteria
DAQ1742 DAQ1742 00643 Daquatica
773 DAQ1742 00643 GI2 mobile element Seed
Dickeya sp CSL RW240 WP 035339724 1 Daquatica
Dickeya aquatica DW 0440 WP 035339724 1 Daquatica
Dickeya dadantii NCPPB 3537 WP 038900402 1 Ddadantii
Dickeya dianthicola NCPPB 3534 WP 024108303 1 Ddianthicola
Dickeya dianthicola GBBC 2039 WP 024108303 1 Ddianthicola
Dickeya dadantii subsp dieffenbachiae NCPPB 2976 WP 038921834 1 Ddadantii
Dickeya zeae Ech586 WP 012883501 1 Dzeae
Dickeya dadantii 3937 WP 012883501 1 Ddadantii
Dickeya gs fangzhongdai NCPPB 3274 WP 042858835 1 Dfangzhongdai
70
77
69
99
24
97
100
62
30
65
85
52
90
75
49
89

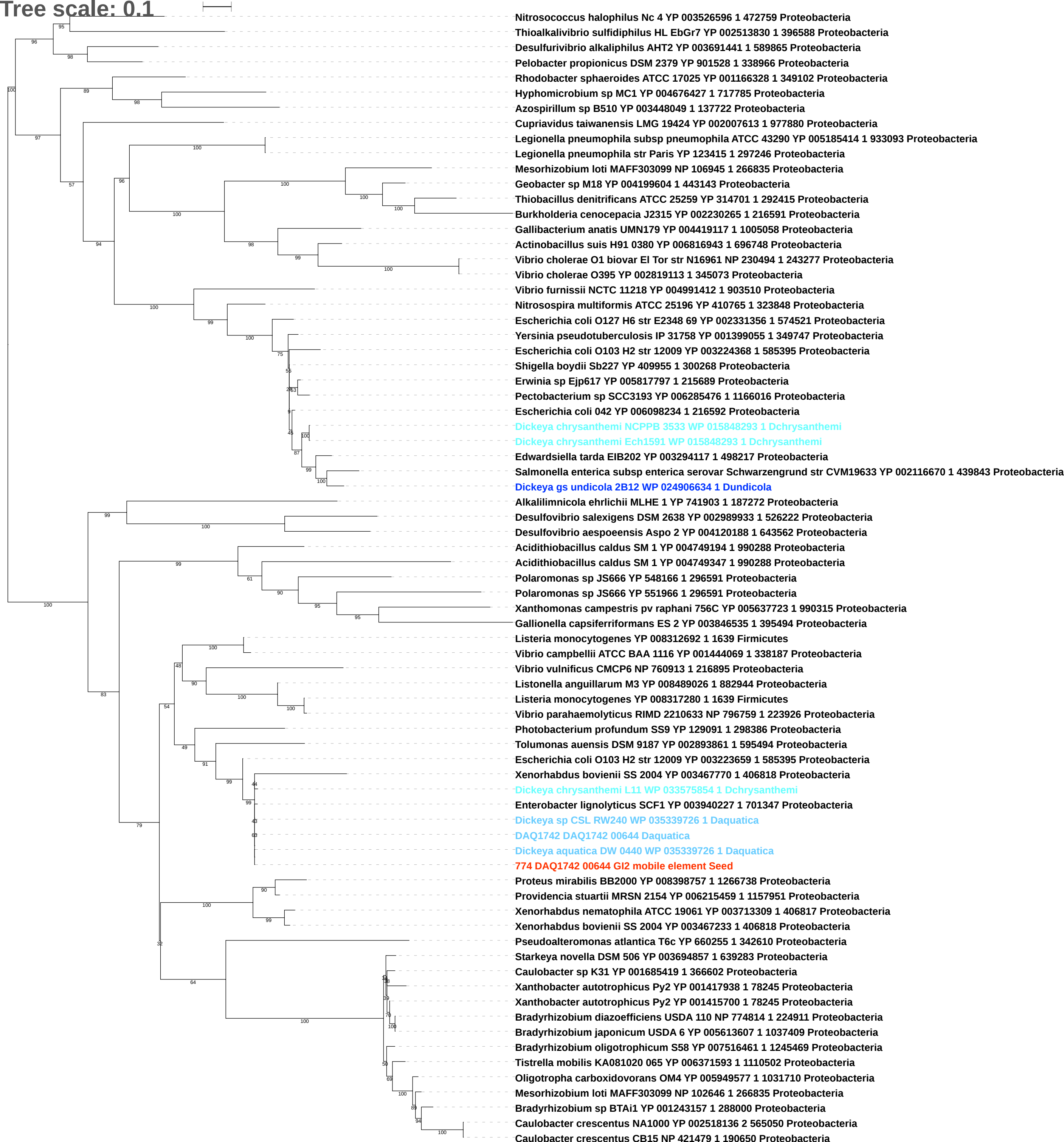
Tree scale: 0.1
Nitrosococcus halophilus Nc 4 YP 003526596 1 472759 Proteobacteria
Thioalkalivibrio sulfidiphilus HL EbGr7 YP 002513830 1 396588 Proteobacteria
Desulfurivibrio alkaliphilus AHT2 YP 003691441 1 589865 Proteobacteria
Pelobacter propionicus DSM 2379 YP 901528 1 338966 Proteobacteria
Rhodobacter sphaeroides ATCC 17025 YP 001166328 1 349102 Proteobacteria
Hyphomicrobium sp MC1 YP 004676427 1 717785 Proteobacteria
Azospirillum sp B510 YP 003448049 1 137722 Proteobacteria
Cupriavidus taiwanensis LMG 19424 YP 002007613 1 977880 Proteobacteria
Legionella pneumophila subsp pneumophila ATCC 43290 YP 005185414 1 933093 Proteobacteria
Legionella pneumophila str Paris YP 123415 1 297246 Proteobacteria
Mesorhizobium loti MAFF303099 NP 106945 1 266835 Proteobacteria
Geobacter sp M18 YP 004199604 1 443143 Proteobacteria
Thiobacillus denitrificans ATCC 25259 YP 314701 1 292415 Proteobacteria
Burkholderia cenocepacia J2315 YP 002230265 1 216591 Proteobacteria
Gallibacterium anatis UMN179 YP 004419117 1 1005058 Proteobacteria
Actinobacillus suis H91 0380 YP 006816943 1 696748 Proteobacteria
Vibrio cholerae O1 biovar El Tor str N16961 NP 230494 1 243277 Proteobacteria
Vibrio cholerae O395 YP 002819113 1 345073 Proteobacteria
Vibrio furnissii NCTC 11218 YP 004991412 1 903510 Proteobacteria
Nitrosospira multiformis ATCC 25196 YP 410765 1 323848 Proteobacteria
Escherichia coli O127 H6 str E2348 69 YP 002331356 1 574521 Proteobacteria
Yersinia pseudotuberculosis IP 31758 YP 001399055 1 349747 Proteobacteria
Escherichia coli O103 H2 str 12009 YP 003224368 1 585395 Proteobacteria
Shigella boydii Sb227 YP 409955 1 300268 Proteobacteria
Erwinia sp Ejp617 YP 005817797 1 215689 Proteobacteria
Pectobacterium sp SCC3193 YP 006285476 1 1166016 Proteobacteria
Escherichia coli 042 YP 006098234 1 216592 Proteobacteria
Dickeya chrysanthemi NCPPB 3533 WP 015848293 1 Dchrysanthemi
Dickeya chrysanthemi Ech1591 WP 015848293 1 Dchrysanthemi
Edwardsiella tarda EIB202 YP 003294117 1 498217 Proteobacteria
Salmonella enterica subsp enterica serovar Schwarzengrund str CVM19633 YP 002116670 1 439843 Proteobacteria
Dickeya gs undicola 2B12 WP 024906634 1 Dundicola
Alkalilimnicola ehrlichii MLHE 1 YP 741903 1 187272 Proteobacteria
Desulfovibrio salexigens DSM 2638 YP 002989933 1 526222 Proteobacteria
Desulfovibrio aespoeensis Aspo 2 YP 004120188 1 643562 Proteobacteria
Acidithiobacillus caldus SM 1 YP 004749194 1 990288 Proteobacteria
Acidithiobacillus caldus SM 1 YP 004749347 1 990288 Proteobacteria
Polaromonas sp JS666 YP 548166 1 296591 Proteobacteria
Polaromonas sp JS666 YP 551966 1 296591 Proteobacteria
Xanthomonas campestris pv raphani 756C YP 005637723 1 990315 Proteobacteria
Gallionella capsiferriformans ES 2 YP 003846535 1 395494 Proteobacteria
Listeria monocytogenes YP 008312692 1 1639 Firmicutes
Vibrio campbellii ATCC BAA 1116 YP 001444069 1 338187 Proteobacteria
Vibrio vulnificus CMCP6 NP 760913 1 216895 Proteobacteria
Listonella anguillarum M3 YP 008489026 1 882944 Proteobacteria
Listeria monocytogenes YP 008317280 1 1639 Firmicutes
Vibrio parahaemolyticus RIMD 2210633 NP 796759 1 223926 Proteobacteria
Photobacterium profundum SS9 YP 129091 1 298386 Proteobacteria
Tolumonas auensis DSM 9187 YP 002893861 1 595494 Proteobacteria
Escherichia coli O103 H2 str 12009 YP 003223659 1 585395 Proteobacteria
Xenorhabdus bovienii SS 2004 YP 003467770 1 406818 Proteobacteria
Dickeya chrysanthemi L11 WP 033575854 1 Dchrysanthemi
Enterobacter lignolyticus SCF1 YP 003940227 1 701347 Proteobacteria
Dickeya sp CSL RW240 WP 035339726 1 Daquatica
DAQ1742 DAQ1742 00644 Daquatica
Dickeya aquatica DW 0440 WP 035339726 1 Daquatica
774 DAQ1742 00644 GI2 mobile element Seed
Proteus mirabilis BB2000 YP 008398757 1 1266738 Proteobacteria
Providencia stuartii MRSN 2154 YP 006215459 1 1157951 Proteobacteria
Xenorhabdus nematophila ATCC 19061 YP 003713309 1 406817 Proteobacteria
Xenorhabdus bovienii SS 2004 YP 003467233 1 406818 Proteobacteria
Pseudoalteromonas atlantica T6c YP 660255 1 342610 Proteobacteria
Starkeya novella DSM 506 YP 003694857 1 639283 Proteobacteria
Caulobacter sp K31 YP 001685419 1 366602 Proteobacteria
Xanthobacter autotrophicus Py2 YP 001417938 1 78245 Proteobacteria
Xanthobacter autotrophicus Py2 YP 001415700 1 78245 Proteobacteria
Bradyrhizobium diazoefficiens USDA 110 NP 774814 1 224911 Proteobacteria
Bradyrhizobium japonicum USDA 6 YP 005613607 1 1037409 Proteobacteria
Bradyrhizobium oligotrophicum S58 YP 007516461 1 1245469 Proteobacteria
Tistrella mobilis KA081020 065 YP 006371593 1 1110502 Proteobacteria
Oligotropha carboxidovorans OM4 YP 005949577 1 1031710 Proteobacteria
Mesorhizobium loti MAFF303099 NP 102646 1 266835 Proteobacteria
Bradyrhizobium sp BTAi1 YP 001243157 1 288000 Proteobacteria
Caulobacter crescentus NA1000 YP 002518136 2 565050 Proteobacteria
Caulobacter crescentus CB15 NP 421479 1 190650 Proteobacteria

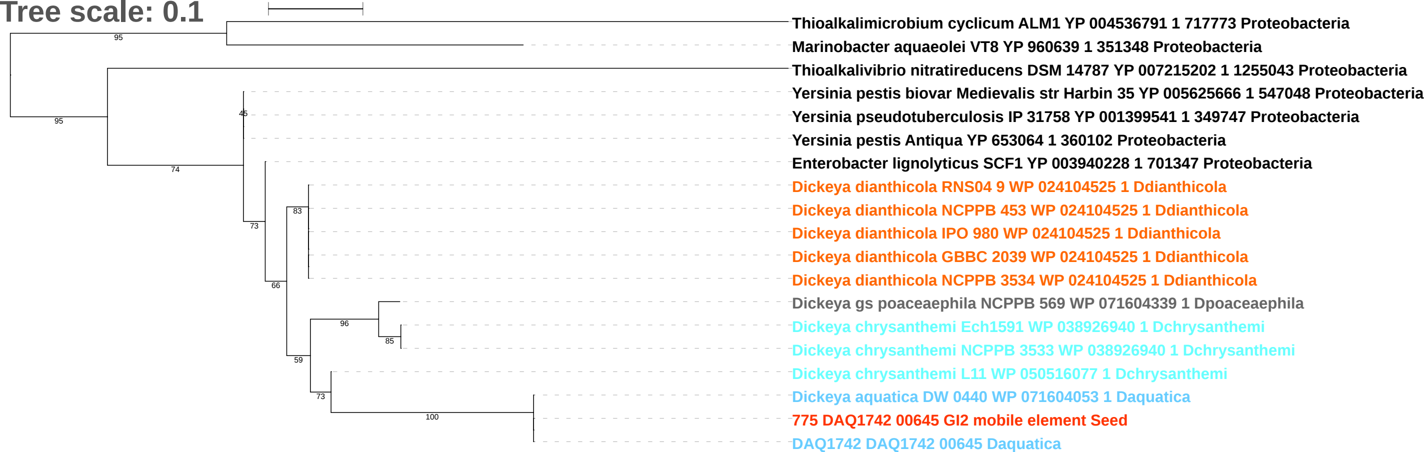
Tree scale: 0.1
Thioalkalimicrobium cyclicum ALM1 YP 004536791 1 717773 Proteobacteria
Marinobacter aquaeolei VT8 YP 960639 1 351348 Proteobacteria
Thioalkalivibrio nitratireducens DSM 14787 YP 007215202 1 1255043 Proteobacteria
Yersinia pestis biovar Medievalis str Harbin 35 YP 005625666 1 547048 Proteobacteria
Yersinia pseudotuberculosis IP 31758 YP 001399541 1 349747 Proteobacteria
Yersinia pestis Antiqua YP 653064 1 360102 Proteobacteria
Enterobacter lignolyticus SCF1 YP 003940228 1 701347 Proteobacteria
Dickeya dianthicola RNS04 9 WP 024104525 1 Ddianthicola
Dickeya dianthicola NCPPB 453 WP 024104525 1 Ddianthicola
Dickeya dianthicola IPO 980 WP 024104525 1 Ddianthicola
Dickeya dianthicola GBBC 2039 WP 024104525 1 Ddianthicola
Dickeya dianthicola NCPPB 3534 WP 024104525 1 Ddianthicola
Dickeya gs poaceaephila NCPPB 569 WP 071604339 1 Dpoaceaephila
Dickeya chrysanthemi Ech1591 WP 038926940 1 Dchrysanthemi
Dickeya chrysanthemi NCPPB 3533 WP 038926940 1 Dchrysanthemi
Dickeya chrysanthemi L11 WP 050516077 1 Dchrysanthemi
Dickeya aquatica DW 0440 WP 071604053 1 Daquatica
775 DAQ1742 00645 GI2 mobile element Seed
DAQ1742 DAQ1742 00645 Daquatica
95
95
45
74
73
83
66
96
85
59
73
100

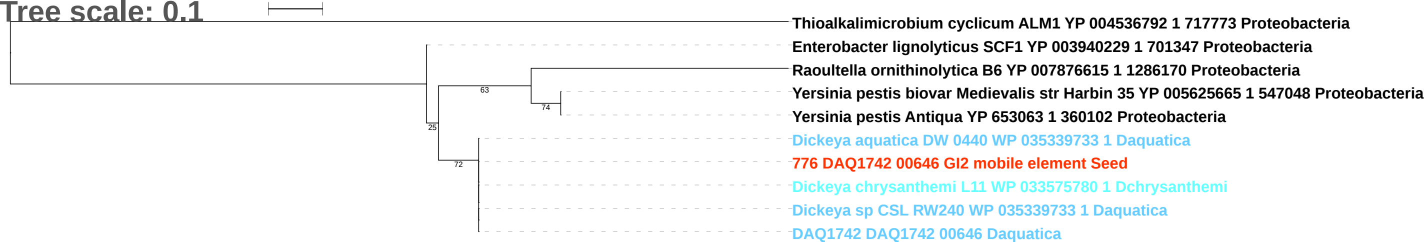
Tree scale: 0.1
Thioalkalimicrobium cyclicum ALM1 YP 004536792 1 717773 Proteobacteria
Enterobacter lignolyticus SCF1 YP 003940229 1 701347 Proteobacteria
Raoultella ornithinolytica B6 YP 007876615 1 1286170 Proteobacteria
Yersinia pestis biovar Medievalis str Harbin 35 YP 005625665 1 547048 Proteobacteria
Yersinia pestis Antiqua YP 653063 1 360102 Proteobacteria
Dickeya aquatica DW 0440 WP 035339733 1 Daquatica
776 DAQ1742 00646 GI2 mobile element Seed
Dickeya chrysanthemi L11 WP 033575780 1 Dchrysanthemi
Dickeya sp CSL RW240 WP 035339733 1 Daquatica
DAQ1742 DAQ1742 00646 Daquatica
63
74
25
72

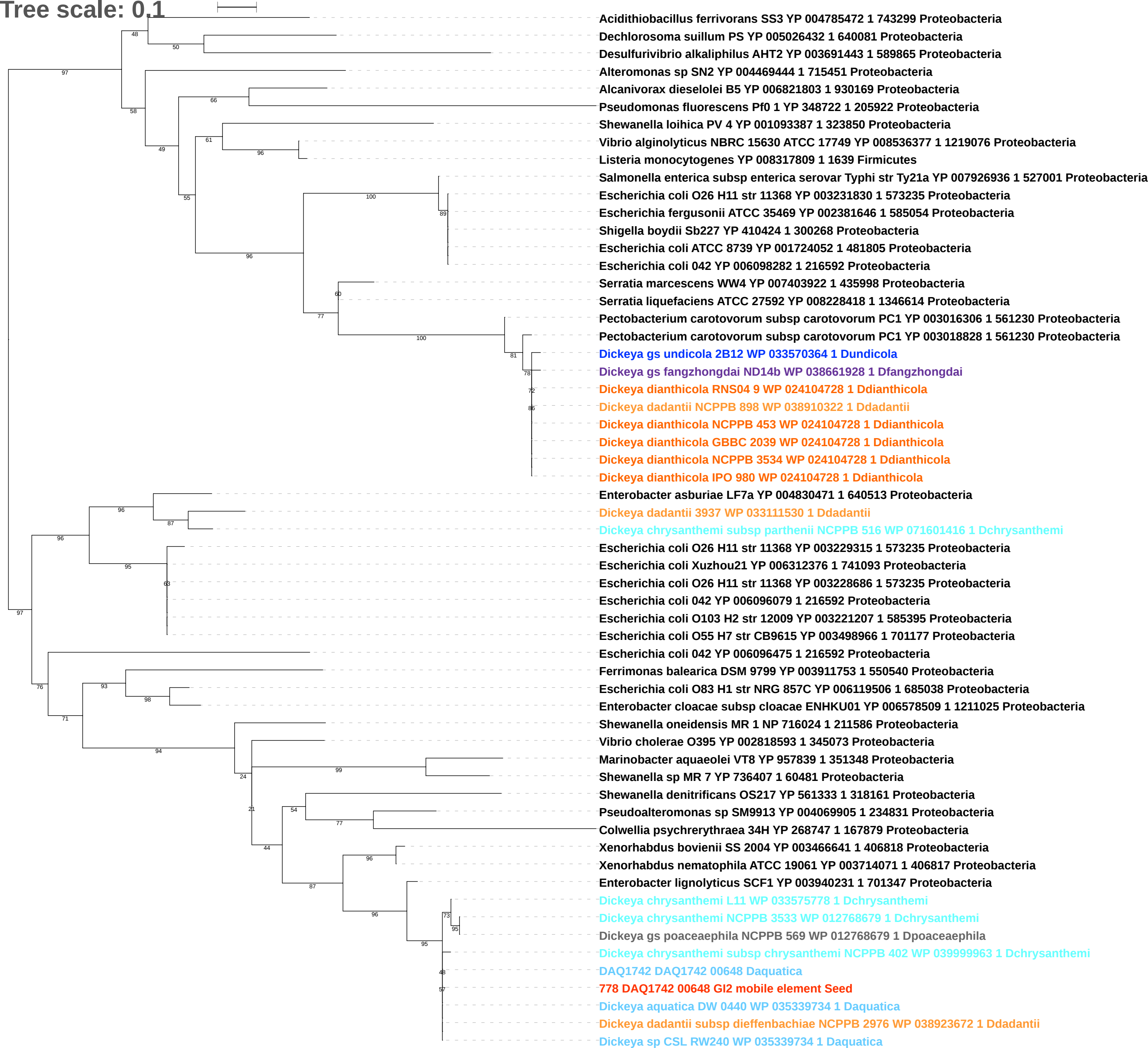
Tree scale: 0.1
Acidithiobacillus ferrivorans SS3 YP 004785472 1 743299 Proteobacteria
Dechlorosoma suillum PS YP 005026432 1 640081 Proteobacteria
Desulfurivibrio alkaliphilus AHT2 YP 003691443 1 589865 Proteobacteria
Alteromonas sp SN2 YP 004469444 1 715451 Proteobacteria
Alcanivorax dieselolei B5 YP 006821803 1 930169 Proteobacteria
Pseudomonas fluorescens Pf0 1 YP 348722 1 205922 Proteobacteria
Shewanella loihica PV 4 YP 001093387 1 323850 Proteobacteria
Vibrio alginolyticus NBRC 15630 ATCC 17749 YP 008536377 1 1219076 Proteobacteria
Listeria monocytogenes YP 008317809 1 1639 Firmicutes
Salmonella enterica subsp enterica serovar Typhi str Ty21a YP 007926936 1 527001 Proteobacteria
Escherichia coli O26 H11 str 11368 YP 003231830 1 573235 Proteobacteria
Escherichia fergusonii ATCC 35469 YP 002381646 1 585054 Proteobacteria
Shigella boydii Sb227 YP 410424 1 300268 Proteobacteria
Escherichia coli ATCC 8739 YP 001724052 1 481805 Proteobacteria
Escherichia coli 042 YP 006098282 1 216592 Proteobacteria
Serratia marcescens WW4 YP 007403922 1 435998 Proteobacteria
Serratia liquefaciens ATCC 27592 YP 008228418 1 1346614 Proteobacteria
Pectobacterium carotovorum subsp carotovorum PC1 YP 003016306 1 561230 Proteobacteria
Pectobacterium carotovorum subsp carotovorum PC1 YP 003018828 1 561230 Proteobacteria
Dickeya gs undicola 2B12 WP 033570364 1 Dundicola
Dickeya gs fangzhongdai ND14b WP 038661928 1 Dfangzhongdai
Dickeya dianthicola RNS04 9 WP 024104728 1 Ddianthicola
Dickeya dadantii NCPPB 898 WP 038910322 1 Ddadantii
Dickeya dianthicola NCPPB 453 WP 024104728 1 Ddianthicola
Dickeya dianthicola GBBC 2039 WP 024104728 1 Ddianthicola
Dickeya dianthicola NCPPB 3534 WP 024104728 1 Ddianthicola
Dickeya dianthicola IPO 980 WP 024104728 1 Ddianthicola
Enterobacter asburiae LF7a YP 004830471 1 640513 Proteobacteria
Dickeya dadantii 3937 WP 033111530 1 Ddadantii
Dickeya chrysanthemi subsp parthenii NCPPB 516 WP 071601416 1 Dchrysanthemi
Escherichia coli O26 H11 str 11368 YP 003229315 1 573235 Proteobacteria
Escherichia coli Xuzhou21 YP 006312376 1 741093 Proteobacteria
Escherichia coli O26 H11 str 11368 YP 003228686 1 573235 Proteobacteria
Escherichia coli 042 YP 006096079 1 216592 Proteobacteria
Escherichia coli O103 H2 str 12009 YP 003221207 1 585395 Proteobacteria
Escherichia coli O55 H7 str CB9615 YP 003498966 1 701177 Proteobacteria
Escherichia coli 042 YP 006096475 1 216592 Proteobacteria
Ferrimonas balearica DSM 9799 YP 003911753 1 550540 Proteobacteria
Escherichia coli O83 H1 str NRG 857C YP 006119506 1 685038 Proteobacteria
Enterobacter cloacae subsp cloacae ENHKU01 YP 006578509 1 1211025 Proteobacteria
Shewanella oneidensis MR 1 NP 716024 1 211586 Proteobacteria
Vibrio cholerae O395 YP 002818593 1 345073 Proteobacteria
Marinobacter aquaeolei VT8 YP 957839 1 351348 Proteobacteria
Shewanella sp MR 7 YP 736407 1 60481 Proteobacteria
Shewanella denitrificans OS217 YP 561333 1 318161 Proteobacteria
Pseudoalteromonas sp SM9913 YP 004069905 1 234831 Proteobacteria
Colwellia psychrerythraea 34H YP 268747 1 167879 Proteobacteria
Xenorhabdus bovienii SS 2004 YP 003466641 1 406818 Proteobacteria
Xenorhabdus nematophila ATCC 19061 YP 003714071 1 406817 Proteobacteria
Enterobacter lignolyticus SCF1 YP 003940231 1 701347 Proteobacteria
Dickeya chrysanthemi L11 WP 033575778 1 Dchrysanthemi
Dickeya chrysanthemi NCPPB 3533 WP 012768679 1 Dchrysanthemi
Dickeya gs poaceaephila NCPPB 569 WP 012768679 1 Dpoaceaephila
Dickeya chrysanthemi subsp chrysanthemi NCPPB 402 WP 039999963 1 Dchrysanthemi
DAQ1742 DAQ1742 00648 Daquatica
778 DAQ1742 00648 GI2 mobile element Seed
Dickeya aquatica DW 0440 WP 035339734 1 Daquatica
Dickeya dadantii subsp dieffenbachiae NCPPB 2976 WP 038923672 1 Ddadantii
Dickeya sp CSL RW240 WP 035339734 1 Daquatica

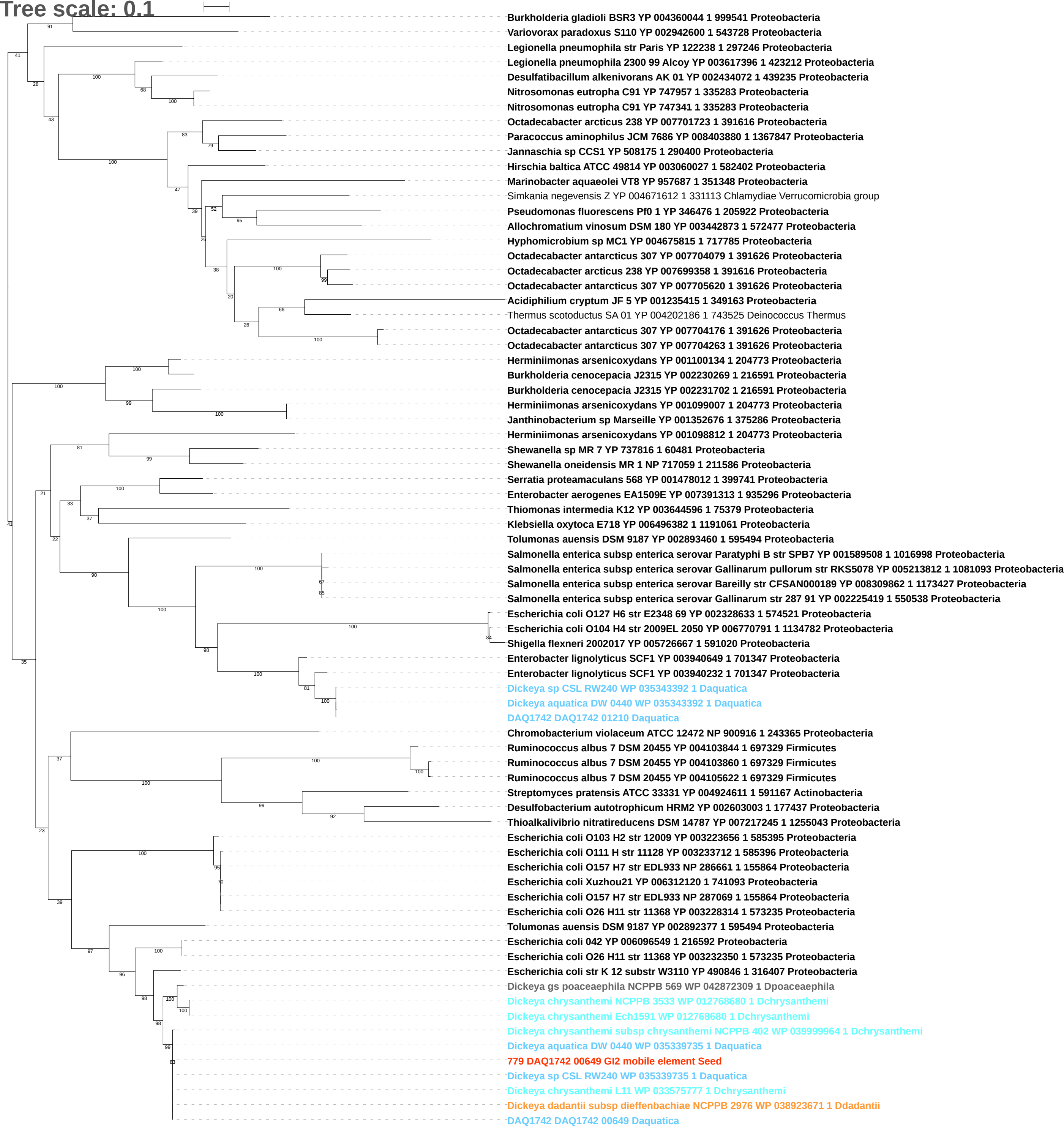
Tree scale: 0.1
Burkholderia gladioli BSR3 YP 004360044 1 999541 Proteobacteria
Variovorax paradoxus S110 YP 002942600 1 543728 Proteobacteria
Legionella pneumophila str Paris YP 122238 1 297246 Proteobacteria
Legionella pneumophila 2300 99 Alcoy YP 003617396 1 423212 Proteobacteria
Desulfatibacillum alkenivorans AK 01 YP 002434072 1 439235 Proteobacteria
Nitrosomonas eutropha C91 YP 747957 1 335283 Proteobacteria
Nitrosomonas eutropha C91 YP 747341 1 335283 Proteobacteria
Octadecabacter arcticus 238 YP 007701723 1 391616 Proteobacteria
Paracoccus aminophilus JCM 7686 YP 008403880 1 1367847 Proteobacteria
Jannaschia sp CCS1 YP 508175 1 290400 Proteobacteria
Hirschia baltica ATCC 49814 YP 003060027 1 582402 Proteobacteria
Marinobacter aquaeolei VT8 YP 957687 1 351348 Proteobacteria
Simkania negevensis Z YP 004671612 1 331113 Chlamydiae Verrucomicrobia group
Pseudomonas fluorescens Pf0 1 YP 346476 1 205922 Proteobacteria
Allochromatium vinosum DSM 180 YP 003442873 1 572477 Proteobacteria
Hyphomicrobium sp MC1 YP 004675815 1 717785 Proteobacteria
Octadecabacter antarcticus 307 YP 007704079 1 391626 Proteobacteria
Octadecabacter arcticus 238 YP 007699358 1 391616 Proteobacteria
Octadecabacter antarcticus 307 YP 007705620 1 391626 Proteobacteria
Acidiphilium cryptum JF 5 YP 001235415 1 349163 Proteobacteria
Thermus scotoductus SA 01 YP 004202186 1 743525 Deinococcus Thermus
Octadecabacter antarcticus 307 YP 007704176 1 391626 Proteobacteria
Octadecabacter antarcticus 307 YP 007704263 1 391626 Proteobacteria
Herminiimonas arsenicoxydans YP 001100134 1 204773 Proteobacteria
Burkholderia cenocepacia J2315 YP 002230269 1 216591 Proteobacteria
Burkholderia cenocepacia J2315 YP 002231702 1 216591 Proteobacteria
Herminiimonas arsenicoxydans YP 001099007 1 204773 Proteobacteria
Janthinobacterium sp Marseille YP 001352676 1 375286 Proteobacteria
Herminiimonas arsenicoxydans YP 001098812 1 204773 Proteobacteria
Shewanella sp MR 7 YP 737816 1 60481 Proteobacteria
Shewanella oneidensis MR 1 NP 717059 1 211586 Proteobacteria
Serratia proteamaculans 568 YP 001478012 1 399741 Proteobacteria
Enterobacter aerogenes EA1509E YP 007391313 1 935296 Proteobacteria
Thiomonas intermedia K12 YP 003644596 1 75379 Proteobacteria
Klebsiella oxytoca E718 YP 006496382 1 1191061 Proteobacteria
Tolumonas auensis DSM 9187 YP 002893460 1 595494 Proteobacteria
Salmonella enterica subsp enterica serovar Paratyphi B str SPB7 YP 001589508 1 1016998 Proteobacteria
Salmonella enterica subsp enterica serovar Gallinarum pullorum str RKS5078 YP 005213812 1 1081093 Proteobacteria
Salmonella enterica subsp enterica serovar Bareilly str CFSAN000189 YP 008309862 1 1173427 Proteobacteria
Salmonella enterica subsp enterica serovar Gallinarum str 287 91 YP 002225419 1 550538 Proteobacteria
Escherichia coli O127 H6 str E2348 69 YP 002328633 1 574521 Proteobacteria
Escherichia coli O104 H4 str 2009EL 2050 YP 006770791 1 1134782 Proteobacteria
Shigella flexneri 2002017 YP 005726667 1 591020 Proteobacteria
Enterobacter lignolyticus SCF1 YP 003940649 1 701347 Proteobacteria
Enterobacter lignolyticus SCF1 YP 003940232 1 701347 Proteobacteria
Dickeya sp CSL RW240 WP 035343392 1 Daquatica
Dickeya aquatica DW 0440 WP 035343392 1 Daquatica
DAQ1742 DAQ1742 01210 Daquatica
Chromobacterium violaceum ATCC 12472 NP 900916 1 243365 Proteobacteria
Ruminococcus albus 7 DSM 20455 YP 004103844 1 697329 Firmicutes
Ruminococcus albus 7 DSM 20455 YP 004103860 1 697329 Firmicutes
Ruminococcus albus 7 DSM 20455 YP 004105622 1 697329 Firmicutes
Streptomyces pratensis ATCC 33331 YP 004924611 1 591167 Actinobacteria
Desulfobacterium autotrophicum HRM2 YP 002603003 1 177437 Proteobacteria
Thioalkalivibrio nitratireducens DSM 14787 YP 007217245 1 1255043 Proteobacteria
Escherichia coli O103 H2 str 12009 YP 003223656 1 585395 Proteobacteria
Escherichia coli O111 H str 11128 YP 003233712 1 585396 Proteobacteria
Escherichia coli O157 H7 str EDL933 NP 286661 1 155864 Proteobacteria
Escherichia coli Xuzhou21 YP 006312120 1 741093 Proteobacteria
Escherichia coli O157 H7 str EDL933 NP 287069 1 155864 Proteobacteria
Escherichia coli O26 H11 str 11368 YP 003228314 1 573235 Proteobacteria
Tolumonas auensis DSM 9187 YP 002892377 1 595494 Proteobacteria
Escherichia coli 042 YP 006096549 1 216592 Proteobacteria
Escherichia coli O26 H11 str 11368 YP 003232350 1 573235 Proteobacteria
Escherichia coli str K 12 substr W3110 YP 490846 1 316407 Proteobacteria
Dickeya gs poaceaephila NCPPB 569 WP 042872309 1 Dpoaceaephila
Dickeya chrysanthemi NCPPB 3533 WP 012768680 1 Dchrysanthemi
Dickeya chrysanthemi Ech1591 WP 012768680 1 Dchrysanthemi
Dickeya chrysanthemi subsp chrysanthemi NCPPB 402 WP 039999964 1 Dchrysanthemi
Dickeya aquatica DW 0440 WP 035339735 1 Daquatica
779 DAQ1742 00649 GI2 mobile element Seed
Dickeya sp CSL RW240 WP 035339735 1 Daquatica
Dickeya chrysanthemi L11 WP 033575777 1 Dchrysanthemi
Dickeya dadantii subsp dieffenbachiae NCPPB 2976 WP 038923671 1 Ddadantii
DAQ1742 DAQ1742 00649 Daquatica

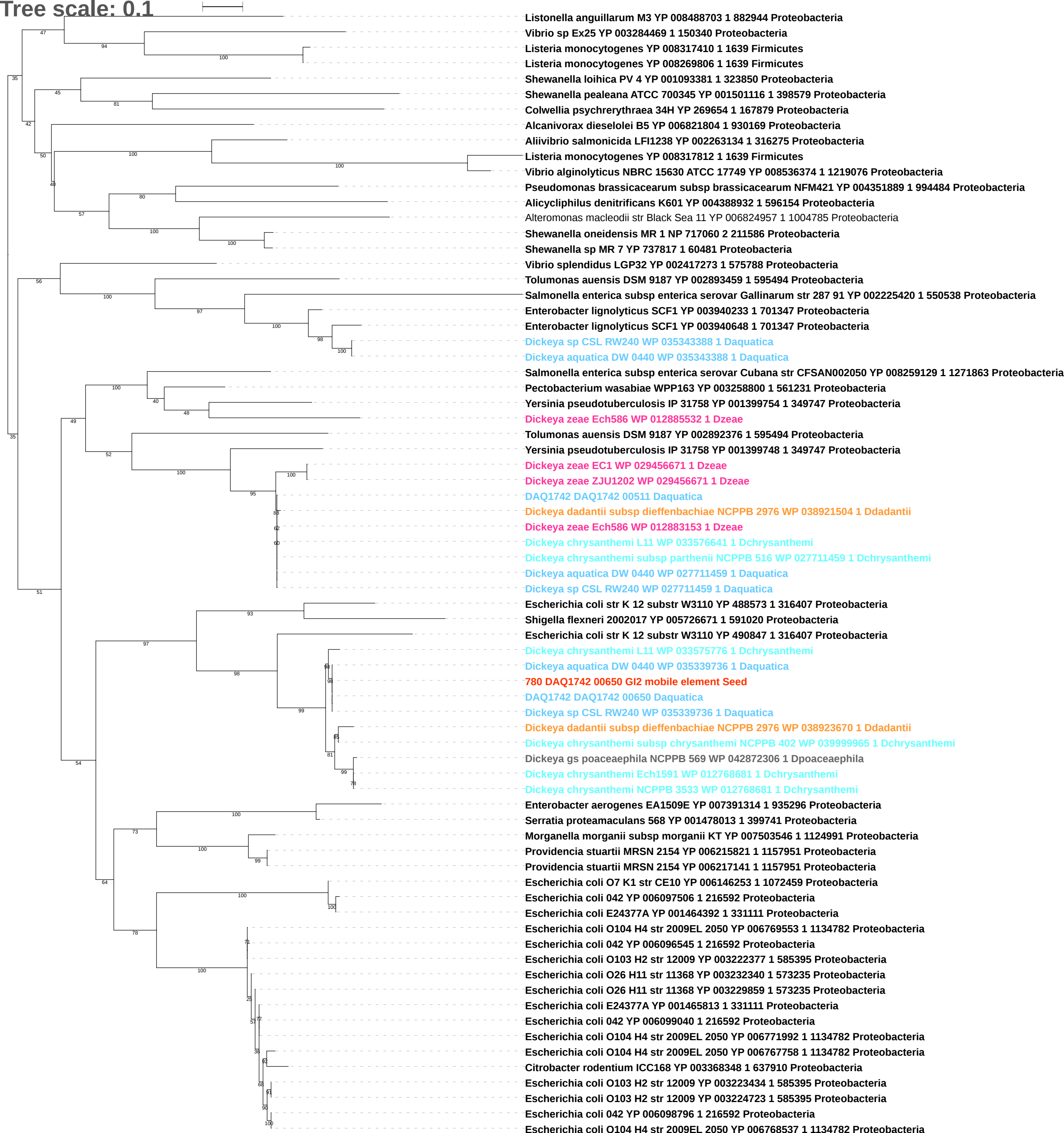

Tree scale: 0.1
Listonella anguillarum M3 YP 008488703 1 882944 Proteobacteria
Vibrio sp Ex25 YP 003284469 1 150340 Proteobacteria
Listeria monocytogenes YP 008317410 1 1639 Firmicutes
Listeria monocytogenes YP 008269806 1 1639 Firmicutes
Shewanella loihica PV 4 YP 001093381 1 323850 Proteobacteria
Shewanella pealeana ATCC 700345 YP 001501116 1 398579 Proteobacteria
Colwellia psychrerythraea 34H YP 269654 1 167879 Proteobacteria
Alcanivorax dieselolei B5 YP 006821804 1 930169 Proteobacteria
Aliivibrio salmonicida LFI1238 YP 002263134 1 316275 Proteobacteria
Listeria monocytogenes YP 008317812 1 1639 Firmicutes
Vibrio alginolyticus NBRC 15630 ATCC 17749 YP 008536374 1 1219076 Proteobacteria
Pseudomonas brassicacearum subsp brassicacearum NFM421 YP 004351889 1 994484 Proteobacteria
Alicycliphilus denitrificans K601 YP 004388932 1 596154 Proteobacteria
Alteromonas macleodii str Black Sea 11 YP 006824957 1 1004785 Proteobacteria
Shewanella oneidensis MR 1 NP 717060 2 211586 Proteobacteria
Shewanella sp MR 7 YP 737817 1 60481 Proteobacteria
Vibrio splendidus LGP32 YP 002417273 1 575788 Proteobacteria
Tolumonas auensis DSM 9187 YP 002893459 1 595494 Proteobacteria
Salmonella enterica subsp enterica serovar Gallinarum str 287 91 YP 002225420 1 550538 Proteobacteria
Enterobacter lignolyticus SCF1 YP 003940233 1 701347 Proteobacteria
Enterobacter lignolyticus SCF1 YP 003940648 1 701347 Proteobacteria
Dickeya sp CSL RW240 WP 035343388 1 Daquatica
Dickeya aquatica DW 0440 WP 035343388 1 Daquatica
Salmonella enterica subsp enterica serovar Cubana str CFSAN002050 YP 008259129 1 1271863 Proteobacteria
Pectobacterium wasabiae WPP163 YP 003258800 1 561231 Proteobacteria
Yersinia pseudotuberculosis IP 31758 YP 001399754 1 349747 Proteobacteria
Dickeya zeae Ech586 WP 012885532 1 Dzeae
Tolumonas auensis DSM 9187 YP 002892376 1 595494 Proteobacteria
Yersinia pseudotuberculosis IP 31758 YP 001399748 1 349747 Proteobacteria
Dickeya zeae EC1 WP 029456671 1 Dzeae
Dickeya zeae ZJU1202 WP 029456671 1 Dzeae
DAQ1742 DAQ1742 00511 Daquatica
Dickeya dadantii subsp dieffenbachiae NCPPB 2976 WP 038921504 1 Ddadantii
Dickeya zeae Ech586 WP 012883153 1 Dzeae
Dickeya chrysanthemi L11 WP 033576641 1 Dchrysanthemi
Dickeya chrysanthemi subsp parthenii NCPPB 516 WP 027711459 1 Dchrysanthemi
Dickeya aquatica DW 0440 WP 027711459 1 Daquatica
Dickeya sp CSL RW240 WP 027711459 1 Daquatica
Escherichia coli str K 12 substr W3110 YP 488573 1 316407 Proteobacteria
Shigella flexneri 2002017 YP 005726671 1 591020 Proteobacteria
Escherichia coli str K 12 substr W3110 YP 490847 1 316407 Proteobacteria
Dickeya chrysanthemi L11 WP 033575776 1 Dchrysanthemi
Dickeya aquatica DW 0440 WP 035339736 1 Daquatica
780 DAQ1742 00650 GI2 mobile element Seed
DAQ1742 DAQ1742 00650 Daquatica
Dickeya sp CSL RW240 WP 035339736 1 Daquatica
Dickeya dadantii subsp dieffenbachiae NCPPB 2976 WP 038923670 1 Ddadantii
Dickeya chrysanthemi subsp chrysanthemi NCPPB 402 WP 039999965 1 Dchrysanthemi
Dickeya gs poaceaephila NCPPB 569 WP 042872306 1 Dpoaceaephila
Dickeya chrysanthemi Ech1591 WP 012768681 1 Dchrysanthemi
Dickeya chrysanthemi NCPPB 3533 WP 012768681 1 Dchrysanthemi
Enterobacter aerogenes EA1509E YP 007391314 1 935296 Proteobacteria
Serratia proteamaculans 568 YP 001478013 1 399741 Proteobacteria
Morganella morganii subsp morganii KT YP 007503546 1 1124991 Proteobacteria
Providencia stuartii MRSN 2154 YP 006215821 1 1157951 Proteobacteria
Providencia stuartii MRSN 2154 YP 006217141 1 1157951 Proteobacteria
Escherichia coli O7 K1 str CE10 YP 006146253 1 1072459 Proteobacteria
Escherichia coli 042 YP 006097506 1 216592 Proteobacteria
Escherichia coli E24377A YP 001464392 1 331111 Proteobacteria
Escherichia coli O104 H4 str 2009EL 2050 YP 006769553 1 1134782 Proteobacteria
Escherichia coli 042 YP 006096545 1 216592 Proteobacteria
Escherichia coli O103 H2 str 12009 YP 003222377 1 585395 Proteobacteria
Escherichia coli O26 H11 str 11368 YP 003232340 1 573235 Proteobacteria
Escherichia coli O26 H11 str 11368 YP 003229859 1 573235 Proteobacteria
Escherichia coli E24377A YP 001465813 1 331111 Proteobacteria
Escherichia coli 042 YP 006099040 1 216592 Proteobacteria
Escherichia coli O104 H4 str 2009EL 2050 YP 006771992 1 1134782 Proteobacteria
Escherichia coli O104 H4 str 2009EL 2050 YP 006767758 1 1134782 Proteobacteria
Citrobacter rodentium ICC168 YP 003368348 1 637910 Proteobacteria
Escherichia coli O103 H2 str 12009 YP 003223434 1 585395 Proteobacteria
Escherichia coli O103 H2 str 12009 YP 003224723 1 585395 Proteobacteria
Escherichia coli 042 YP 006098796 1 216592 Proteobacteria
Escherichia coli O104 H4 str 2009EL 2050 YP 006768537 1 1134782 Proteobacteria

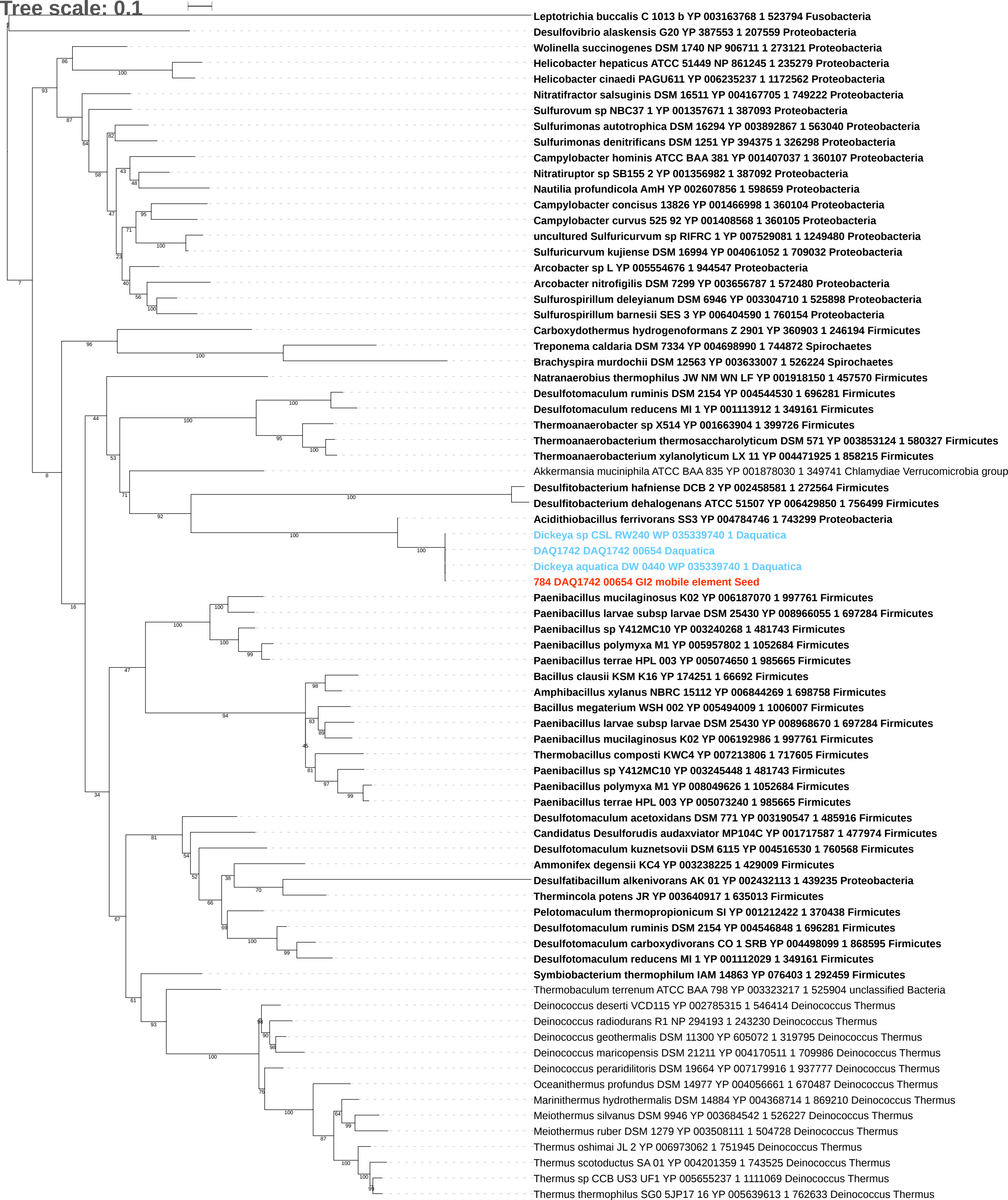
Tree scale: 0.1
Leptotrichia buccalis C 1013 b YP 003163768 1 523794 Fusobacteria
Desulfovibrio alaskensis G20 YP 387553 1 207559 Proteobacteria
Wolinella succinogenes DSM 1740 NP 906711 1 273121 Proteobacteria
Helicobacter hepaticus ATCC 51449 NP 861245 1 235279 Proteobacteria
Helicobacter cinaedi PAGU611 YP 006235237 1 1172562 Proteobacteria
Nitratifractor salsuginis DSM 16511 YP 004167705 1 749222 Proteobacteria
Sulfurovum sp NBC37 1 YP 001357671 1 387093 Proteobacteria
Sulfurimonas autotrophica DSM 16294 YP 003892867 1 563040 Proteobacteria
Sulfurimonas denitrificans DSM 1251 YP 394375 1 326298 Proteobacteria
Campylobacter hominis ATCC BAA 381 YP 001407037 1 360107 Proteobacteria
Nitratiruptor sp SB155 2 YP 001356982 1 387092 Proteobacteria
Nautilia profundicola AmH YP 002607856 1 598659 Proteobacteria
Campylobacter concisus 13826 YP 001466998 1 360104 Proteobacteria
Campylobacter curvus 525 92 YP 001408568 1 360105 Proteobacteria
uncultured Sulfuricurvum sp RIFRC 1 YP 007529081 1 1249480 Proteobacteria
Sulfuricurvum kujiense DSM 16994 YP 004061052 1 709032 Proteobacteria
Arcobacter sp L YP 005554676 1 944547 Proteobacteria
Arcobacter nitrofigilis DSM 7299 YP 003656787 1 572480 Proteobacteria
Sulfurospirillum deleyianum DSM 6946 YP 003304710 1 525898 Proteobacteria
Sulfurospirillum barnesii SES 3 YP 006404590 1 760154 Proteobacteria
Carboxydothermus hydrogenoformans Z 2901 YP 360903 1 246194 Firmicutes
Treponema caldaria DSM 7334 YP 004698990 1 744872 Spirochaetes
Brachyspira murdochii DSM 12563 YP 003633007 1 526224 Spirochaetes
Natranaerobius thermophilus JW NM WN LF YP 001918150 1 457570 Firmicutes
Desulfotomaculum ruminis DSM 2154 YP 004544530 1 696281 Firmicutes
Desulfotomaculum reducens MI 1 YP 001113912 1 349161 Firmicutes
Thermoanaerobacter sp X514 YP 001663904 1 399726 Firmicutes
Thermoanaerobacterium thermosaccharolyticum DSM 571 YP 003853124 1 580327 Firmicutes
Thermoanaerobacterium xylanolyticum LX 11 YP 004471925 1 858215 Firmicutes
Akkermansia muciniphila ATCC BAA 835 YP 001878030 1 349741 Chlamydiae Verrucomicrobia group
Desulfitobacterium hafniense DCB 2 YP 002458581 1 272564 Firmicutes
Desulfitobacterium dehalogenans ATCC 51507 YP 006429850 1 756499 Firmicutes
Acidithiobacillus ferrivorans SS3 YP 004784746 1 743299 Proteobacteria
Dickeya sp CSL RW240 WP 035339740 1 Daquatica
DAQ1742 DAQ1742 00654 Daquatica
Dickeya aquatica DW 0440 WP 035339740 1 Daquatica
784 DAQ1742 00654 GI2 mobile element Seed
Paenibacillus mucilaginosus K02 YP 006187070 1 997761 Firmicutes
Paenibacillus larvae subsp larvae DSM 25430 YP 008966055 1 697284 Firmicutes
Paenibacillus sp Y412MC10 YP 003240268 1 481743 Firmicutes
Paenibacillus polymyxa M1 YP 005957802 1 1052684 Firmicutes
Paenibacillus terrae HPL 003 YP 005074650 1 985665 Firmicutes
Bacillus clausii KSM K16 YP 174251 1 66692 Firmicutes
Amphibacillus xylanus NBRC 15112 YP 006844269 1 698758 Firmicutes
Bacillus megaterium WSH 002 YP 005494009 1 1006007 Firmicutes
Paenibacillus larvae subsp larvae DSM 25430 YP 008968670 1 697284 Firmicutes
Paenibacillus mucilaginosus K02 YP 006192986 1 997761 Firmicutes
Thermobacillus composti KWC4 YP 007213806 1 717605 Firmicutes
Paenibacillus sp Y412MC10 YP 003245448 1 481743 Firmicutes
Paenibacillus polymyxa M1 YP 008049626 1 1052684 Firmicutes
Paenibacillus terrae HPL 003 YP 005073240 1 985665 Firmicutes
Desulfotomaculum acetoxidans DSM 771 YP 003190547 1 485916 Firmicutes
Candidatus Desulforudis audaxviator MP104C YP 001717587 1 477974 Firmicutes
Desulfotomaculum kuznetsovii DSM 6115 YP 004516530 1 760568 Firmicutes
Ammonifex degensii KC4 YP 003238225 1 429009 Firmicutes
Desulfatibacillum alkenivorans AK 01 YP 002432113 1 439235 Proteobacteria
Thermincola potens JR YP 003640917 1 635013 Firmicutes
Pelotomaculum thermopropionicum SI YP 001212422 1 370438 Firmicutes
Desulfotomaculum ruminis DSM 2154 YP 004546848 1 696281 Firmicutes
Desulfotomaculum carboxydivorans CO 1 SRB YP 004498099 1 868595 Firmicutes
Desulfotomaculum reducens MI 1 YP 001112029 1 349161 Firmicutes
Symbiobacterium thermophilum IAM 14863 YP 076403 1 292459 Firmicutes
Thermobaculum terrenum ATCC BAA 798 YP 003323217 1 525904 unclassified Bacteria
Deinococcus deserti VCD115 YP 002785315 1 546414 Deinococcus Thermus
Deinococcus radiodurans R1 NP 294193 1 243230 Deinococcus Thermus
Deinococcus geothermalis DSM 11300 YP 605072 1 319795 Deinococcus Thermus
Deinococcus maricopensis DSM 21211 YP 004170511 1 709986 Deinococcus Thermus
Deinococcus peraridilitoris DSM 19664 YP 007179916 1 937777 Deinococcus Thermus
Oceanithermus profundus DSM 14977 YP 004056661 1 670487 Deinococcus Thermus
Marinithermus hydrothermalis DSM 14884 YP 004368714 1 869210 Deinococcus Thermus
Meiothermus silvanus DSM 9946 YP 003684542 1 526227 Deinococcus Thermus
Meiothermus ruber DSM 1279 YP 003508111 1 504728 Deinococcus Thermus
Thermus oshimai JL 2 YP 006973062 1 751945 Deinococcus Thermus
Thermus scotoductus SA 01 YP 004201359 1 743525 Deinococcus Thermus
Thermus sp CCB US3 UF1 YP 005655237 1 1111069 Deinococcus Thermus
Thermus thermophilus SG0 5JP17 16 YP 005639613 1 762633 Deinococcus Thermus

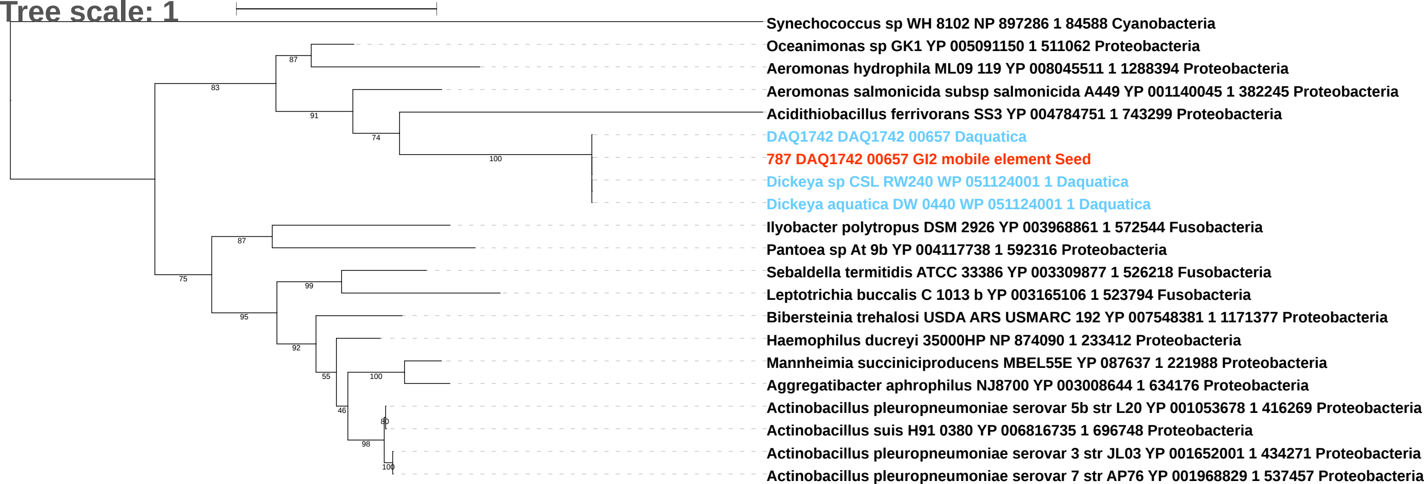
Tree scale: 1
Synechococcus sp WH 8102 NP 897286 1 84588 Cyanobacteria
Oceanimonas sp GK1 YP 005091150 1 511062 Proteobacteria
Aeromonas hydrophila ML09 119 YP 008045511 1 1288394 Proteobacteria
Aeromonas salmonicida subsp salmonicida A449 YP 001140045 1 382245 Proteobacteria
Acidithiobacillus ferrivorans SS3 YP 004784751 1 743299 Proteobacteria
DAQ1742 DAQ1742 00657 Daquatica
787 DAQ1742 00657 GI2 mobile element Seed
Dickeya sp CSL RW240 WP 051124001 1 Daquatica
Dickeya aquatica DW 0440 WP 051124001 1 Daquatica
Ilyobacter polytropus DSM 2926 YP 003968861 1 572544 Fusobacteria
Pantoea sp At 9b YP 004117738 1 592316 Proteobacteria
Sebaldella termitidis ATCC 33386 YP 003309877 1 526218 Fusobacteria
Leptotrichia buccalis C 1013 b YP 003165106 1 523794 Fusobacteria
Bibersteinia trehalosi USDA ARS USMARC 192 YP 007548381 1 1171377 Proteobacteria
Haemophilus ducreyi 35000HP NP 874090 1 233412 Proteobacteria
Mannheimia succiniciproducens MBEL55E YP 087637 1 221988 Proteobacteria
Aggregatibacter aphrophilus NJ8700 YP 003008644 1 634176 Proteobacteria
Actinobacillus pleuropneumoniae serovar 5b str L20 YP 001053678 1 416269 Proteobacteria
Actinobacillus suis H91 0380 YP 006816735 1 696748 Proteobacteria
Actinobacillus pleuropneumoniae serovar 3 str JL03 YP 001652001 1 434271 Proteobacteria
Actinobacillus pleuropneumoniae serovar 7 str AP76 YP 001968829 1 537457 Proteobacteria
87
83
91
74
100
87
75
99
95
92
55
100
46
80
98
100

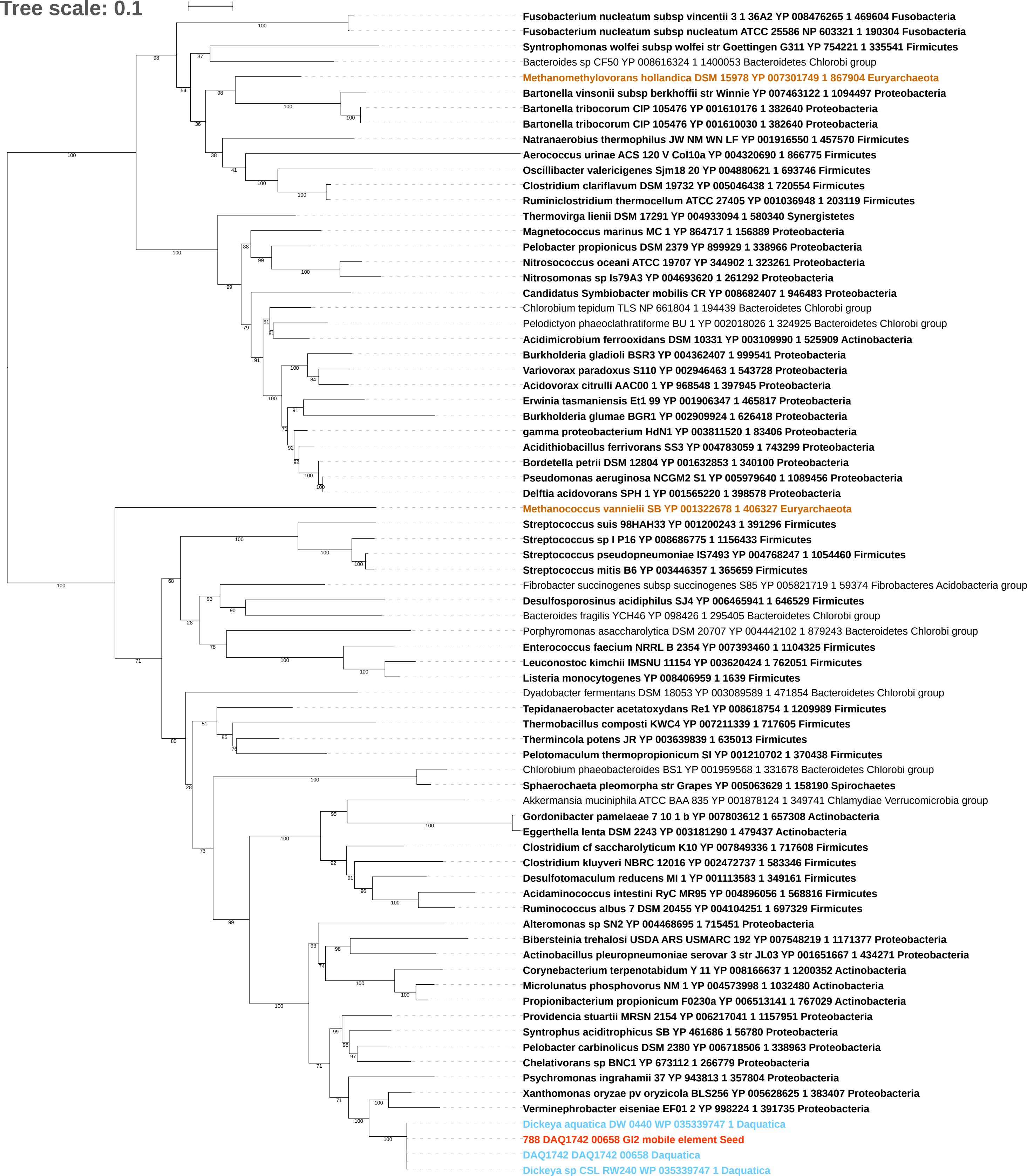
Tree scale: 0.1
Fusobacterium nucleatum subsp vincentii 3 1 36A2 YP 008476265 1 469604 Fusobacteria
Fusobacterium nucleatum subsp nucleatum ATCC 25586 NP 603321 1 190304 Fusobacteria
Syntrophomonas wolfei subsp wolfei str Goettingen G311 YP 754221 1 335541 Firmicutes
Bacteroides sp CF50 YP 008616324 1 1400053 Bacteroidetes Chlorobi group
Methanomethylovorans hollandica DSM 15978 YP 007301749 1 867904 Euryarchaeota
Bartonella vinsonii subsp berkhoffii str Winnie YP 007463122 1 1094497 Proteobacteria
Bartonella tribocorum CIP 105476 YP 001610176 1 382640 Proteobacteria
Bartonella tribocorum CIP 105476 YP 001610030 1 382640 Proteobacteria
Natranaerobius thermophilus JW NM WN LF YP 001916550 1 457570 Firmicutes
Aerococcus urinae ACS 120 V Col10a YP 004320690 1 866775 Firmicutes
Oscillibacter valericigenes Sjm18 20 YP 004880621 1 693746 Firmicutes
Clostridium clariflavum DSM 19732 YP 005046438 1 720554 Firmicutes
Ruminiclostridium thermocellum ATCC 27405 YP 001036948 1 203119 Firmicutes
Thermovirga lienii DSM 17291 YP 004933094 1 580340 Synergistetes
Magnetococcus marinus MC 1 YP 864717 1 156889 Proteobacteria
Pelobacter propionicus DSM 2379 YP 899929 1 338966 Proteobacteria
Nitrosococcus oceani ATCC 19707 YP 344902 1 323261 Proteobacteria
Nitrosomonas sp Is79A3 YP 004693620 1 261292 Proteobacteria
Candidatus Symbiobacter mobilis CR YP 008682407 1 946483 Proteobacteria
Chlorobium tepidum TLS NP 661804 1 194439 Bacteroidetes Chlorobi group
Pelodictyon phaeoclathratiforme BU 1 YP 002018026 1 324925 Bacteroidetes Chlorobi group
Acidimicrobium ferrooxidans DSM 10331 YP 003109990 1 525909 Actinobacteria
Burkholderia gladioli BSR3 YP 004362407 1 999541 Proteobacteria
Variovorax paradoxus S110 YP 002946463 1 543728 Proteobacteria
Acidovorax citrulli AAC00 1 YP 968548 1 397945 Proteobacteria
Erwinia tasmaniensis Et1 99 YP 001906347 1 465817 Proteobacteria
Burkholderia glumae BGR1 YP 002909924 1 626418 Proteobacteria
gamma proteobacterium HdN1 YP 003811520 1 83406 Proteobacteria
Acidithiobacillus ferrivorans SS3 YP 004783059 1 743299 Proteobacteria
Bordetella petrii DSM 12804 YP 001632853 1 340100 Proteobacteria
Pseudomonas aeruginosa NCGM2 S1 YP 005979640 1 1089456 Proteobacteria
Delftia acidovorans SPH 1 YP 001565220 1 398578 Proteobacteria
Methanococcus vannielii SB YP 001322678 1 406327 Euryarchaeota
Streptococcus suis 98HAH33 YP 001200243 1 391296 Firmicutes
Streptococcus sp I P16 YP 008686775 1 1156433 Firmicutes
Streptococcus pseudopneumoniae IS7493 YP 004768247 1 1054460 Firmicutes
Streptococcus mitis B6 YP 003446357 1 365659 Firmicutes
Fibrobacter succinogenes subsp succinogenes S85 YP 005821719 1 59374 Fibrobacteres Acidobacteria group
Desulfosporosinus acidiphilus SJ4 YP 006465941 1 646529 Firmicutes
Bacteroides fragilis YCH46 YP 098426 1 295405 Bacteroidetes Chlorobi group
Porphyromonas asaccharolytica DSM 20707 YP 004442102 1 879243 Bacteroidetes Chlorobi group
Enterococcus faecium NRRL B 2354 YP 007393460 1 1104325 Firmicutes
Leuconostoc kimchii IMSNU 11154 YP 003620424 1 762051 Firmicutes
Listeria monocytogenes YP 008406959 1 1639 Firmicutes
Dyadobacter fermentans DSM 18053 YP 003089589 1 471854 Bacteroidetes Chlorobi group
Tepidanaerobacter acetatoxydans Re1 YP 008618754 1 1209989 Firmicutes
Thermobacillus composti KWC4 YP 007211339 1 717605 Firmicutes
Thermincola potens JR YP 003639839 1 635013 Firmicutes
Pelotomaculum thermopropionicum SI YP 001210702 1 370438 Firmicutes
Chlorobium phaeobacteroides BS1 YP 001959568 1 331678 Bacteroidetes Chlorobi group
Sphaerochaeta pleomorpha str Grapes YP 005063629 1 158190 Spirochaetes
Akkermansia muciniphila ATCC BAA 835 YP 001878124 1 349741 Chlamydiae Verrucomicrobia group
Gordonibacter pamelaeae 7 10 1 b YP 007803612 1 657308 Actinobacteria
Eggerthella lenta DSM 2243 YP 003181290 1 479437 Actinobacteria
Clostridium cf saccharolyticum K10 YP 007849336 1 717608 Firmicutes
Clostridium kluyveri NBRC 12016 YP 002472737 1 583346 Firmicutes
Desulfotomaculum reducens MI 1 YP 001113583 1 349161 Firmicutes
Acidaminococcus intestini RyC MR95 YP 004896056 1 568816 Firmicutes
Ruminococcus albus 7 DSM 20455 YP 004104251 1 697329 Firmicutes
Alteromonas sp SN2 YP 004468695 1 715451 Proteobacteria
Bibersteinia trehalosi USDA ARS USMARC 192 YP 007548219 1 1171377 Proteobacteria
Actinobacillus pleuropneumoniae serovar 3 str JL03 YP 001651667 1 434271 Proteobacteria
Corynebacterium terpenotabidum Y 11 YP 008166637 1 1200352 Actinobacteria
Microlunatus phosphovorus NM 1 YP 004573998 1 1032480 Actinobacteria
Propionibacterium propionicum F0230a YP 006513141 1 767029 Actinobacteria
Providencia stuartii MRSN 2154 YP 006217041 1 1157951 Proteobacteria
Syntrophus aciditrophicus SB YP 461686 1 56780 Proteobacteria
Pelobacter carbinolicus DSM 2380 YP 006718506 1 338963 Proteobacteria
Chelativorans sp BNC1 YP 673112 1 266779 Proteobacteria
Psychromonas ingrahamii 37 YP 943813 1 357804 Proteobacteria
Xanthomonas oryzae pv oryzicola BLS256 YP 005628625 1 383407 Proteobacteria
Verminephrobacter eiseniae EF01 2 YP 998224 1 391735 Proteobacteria
Dickeya aquatica DW 0440 WP 035339747 1 Daquatica
788 DAQ1742 00658 GI2 mobile element Seed
DAQ1742 DAQ1742 00658 Daquatica
Dickeya sp CSL RW240 WP 035339747 1 Daquatica

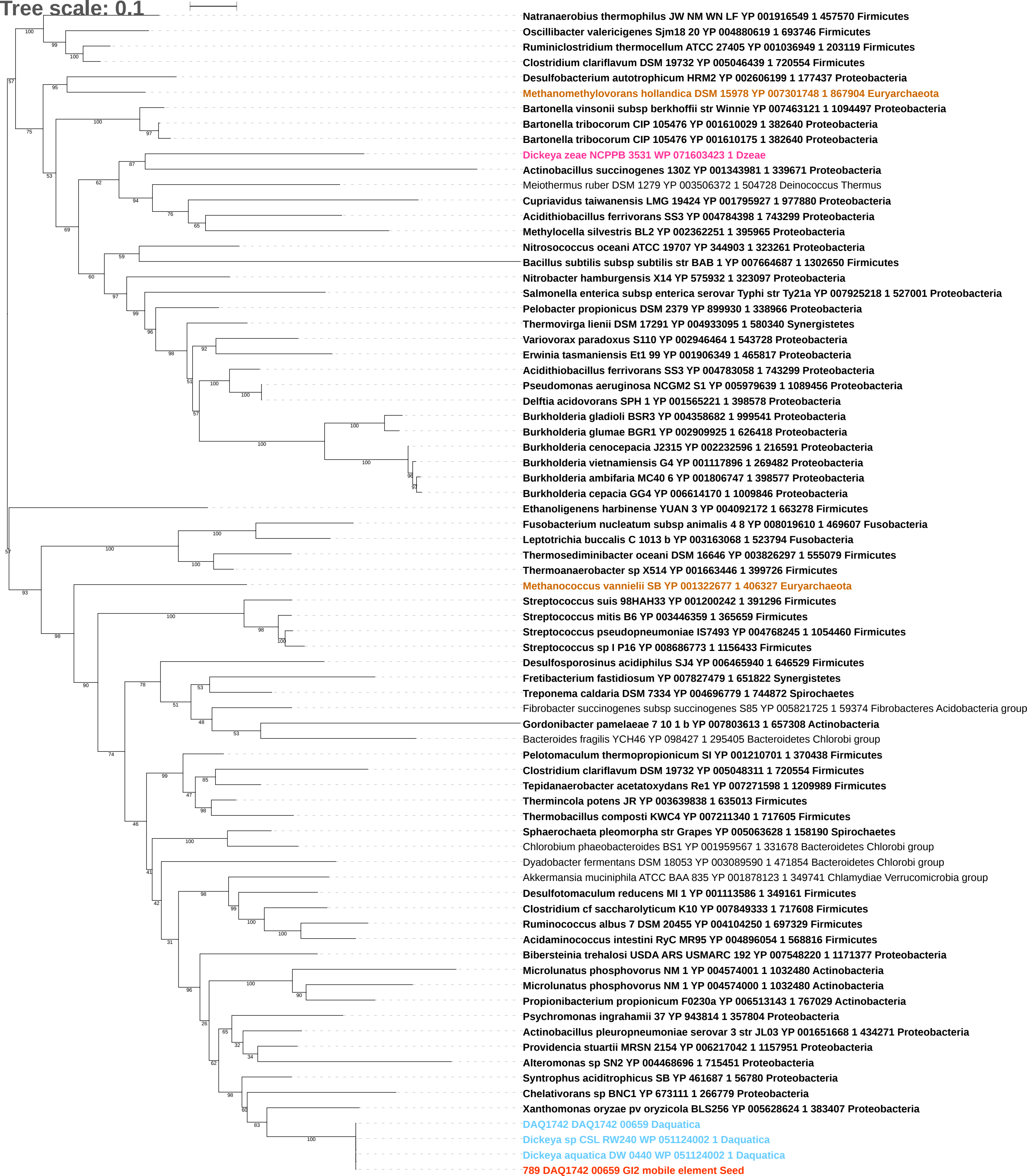
Tree scale: 0.1
Natranaerobius thermophilus JW NM WN LF YP 001916549 1 457570 Firmicutes
Oscillibacter valericigenes Sjm18 20 YP 004880619 1 693746 Firmicutes
Ruminiclostridium thermocellum ATCC 27405 YP 001036949 1 203119 Firmicutes
Clostridium clariflavum DSM 19732 YP 005046439 1 720554 Firmicutes
Desulfobacterium autotrophicum HRM2 YP 002606199 1 177437 Proteobacteria
Methanomethylovorans hollandica DSM 15978 YP 007301748 1 867904 Euryarchaeota
Bartonella vinsonii subsp berkhoffii str Winnie YP 007463121 1 1094497 Proteobacteria
Bartonella tribocorum CIP 105476 YP 001610029 1 382640 Proteobacteria
Bartonella tribocorum CIP 105476 YP 001610175 1 382640 Proteobacteria
Dickeya zeae NCPPB 3531 WP 071603423 1 Dzeae
Actinobacillus succinogenes 130Z YP 001343981 1 339671 Proteobacteria
Meiothermus ruber DSM 1279 YP 003506372 1 504728 Deinococcus Thermus
Cupriavidus taiwanensis LMG 19424 YP 001795927 1 977880 Proteobacteria
Acidithiobacillus ferrivorans SS3 YP 004784398 1 743299 Proteobacteria
Methylocella silvestris BL2 YP 002362251 1 395965 Proteobacteria
Nitrosococcus oceani ATCC 19707 YP 344903 1 323261 Proteobacteria
Bacillus subtilis subsp subtilis str BAB 1 YP 007664687 1 1302650 Firmicutes
Nitrobacter hamburgensis X14 YP 575932 1 323097 Proteobacteria
Salmonella enterica subsp enterica serovar Typhi str Ty21a YP 007925218 1 527001 Proteobacteria
Pelobacter propionicus DSM 2379 YP 899930 1 338966 Proteobacteria
Thermovirga lienii DSM 17291 YP 004933095 1 580340 Synergistetes
Variovorax paradoxus S110 YP 002946464 1 543728 Proteobacteria
Erwinia tasmaniensis Et1 99 YP 001906349 1 465817 Proteobacteria
Acidithiobacillus ferrivorans SS3 YP 004783058 1 743299 Proteobacteria
Pseudomonas aeruginosa NCGM2 S1 YP 005979639 1 1089456 Proteobacteria
Delftia acidovorans SPH 1 YP 001565221 1 398578 Proteobacteria
Burkholderia gladioli BSR3 YP 004358682 1 999541 Proteobacteria
Burkholderia glumae BGR1 YP 002909925 1 626418 Proteobacteria
Burkholderia cenocepacia J2315 YP 002232596 1 216591 Proteobacteria
Burkholderia vietnamiensis G4 YP 001117896 1 269482 Proteobacteria
Burkholderia ambifaria MC40 6 YP 001806747 1 398577 Proteobacteria
Burkholderia cepacia GG4 YP 006614170 1 1009846 Proteobacteria
Ethanoligenens harbinense YUAN 3 YP 004092172 1 663278 Firmicutes
Fusobacterium nucleatum subsp animalis 4 8 YP 008019610 1 469607 Fusobacteria
Leptotrichia buccalis C 1013 b YP 003163068 1 523794 Fusobacteria
Thermosediminibacter oceani DSM 16646 YP 003826297 1 555079 Firmicutes
Thermoanaerobacter sp X514 YP 001663446 1 399726 Firmicutes
Methanococcus vannielii SB YP 001322677 1 406327 Euryarchaeota
Streptococcus suis 98HAH33 YP 001200242 1 391296 Firmicutes
Streptococcus mitis B6 YP 003446359 1 365659 Firmicutes
Streptococcus pseudopneumoniae IS7493 YP 004768245 1 1054460 Firmicutes
Streptococcus sp I P16 YP 008686773 1 1156433 Firmicutes
Desulfosporosinus acidiphilus SJ4 YP 006465940 1 646529 Firmicutes
Fretibacterium fastidiosum YP 007827479 1 651822 Synergistetes
Treponema caldaria DSM 7334 YP 004696779 1 744872 Spirochaetes
Fibrobacter succinogenes subsp succinogenes S85 YP 005821725 1 59374 Fibrobacteres Acidobacteria group
Gordonibacter pamelaeae 7 10 1 b YP 007803613 1 657308 Actinobacteria
Bacteroides fragilis YCH46 YP 098427 1 295405 Bacteroidetes Chlorobi group
Pelotomaculum thermopropionicum SI YP 001210701 1 370438 Firmicutes
Clostridium clariflavum DSM 19732 YP 005048311 1 720554 Firmicutes
Tepidanaerobacter acetatoxydans Re1 YP 007271598 1 1209989 Firmicutes
Thermincola potens JR YP 003639838 1 635013 Firmicutes
Thermobacillus composti KWC4 YP 007211340 1 717605 Firmicutes
Sphaerochaeta pleomorpha str Grapes YP 005063628 1 158190 Spirochaetes
Chlorobium phaeobacteroides BS1 YP 001959567 1 331678 Bacteroidetes Chlorobi group
Dyadobacter fermentans DSM 18053 YP 003089590 1 471854 Bacteroidetes Chlorobi group
Akkermansia muciniphila ATCC BAA 835 YP 001878123 1 349741 Chlamydiae Verrucomicrobia group
Desulfotomaculum reducens MI 1 YP 001113586 1 349161 Firmicutes
Clostridium cf saccharolyticum K10 YP 007849333 1 717608 Firmicutes
Ruminococcus albus 7 DSM 20455 YP 004104250 1 697329 Firmicutes
Acidaminococcus intestini RyC MR95 YP 004896054 1 568816 Firmicutes
Bibersteinia trehalosi USDA ARS USMARC 192 YP 007548220 1 1171377 Proteobacteria
Microlunatus phosphovorus NM 1 YP 004574001 1 1032480 Actinobacteria
Microlunatus phosphovorus NM 1 YP 004574000 1 1032480 Actinobacteria
Propionibacterium propionicum F0230a YP 006513143 1 767029 Actinobacteria
Psychromonas ingrahamii 37 YP 943814 1 357804 Proteobacteria
Actinobacillus pleuropneumoniae serovar 3 str JL03 YP 001651668 1 434271 Proteobacteria
Providencia stuartii MRSN 2154 YP 006217042 1 1157951 Proteobacteria
Alteromonas sp SN2 YP 004468696 1 715451 Proteobacteria
Syntrophus aciditrophicus SB YP 461687 1 56780 Proteobacteria
Chelativorans sp BNC1 YP 673111 1 266779 Proteobacteria
Xanthomonas oryzae pv oryzicola BLS256 YP 005628624 1 383407 Proteobacteria
DAQ1742 DAQ1742 00659 Daquatica
Dickeya sp CSL RW240 WP 051124002 1 Daquatica
Dickeya aquatica DW 0440 WP 051124002 1 Daquatica
789 DAQ1742 00659 GI2 mobile element Seed

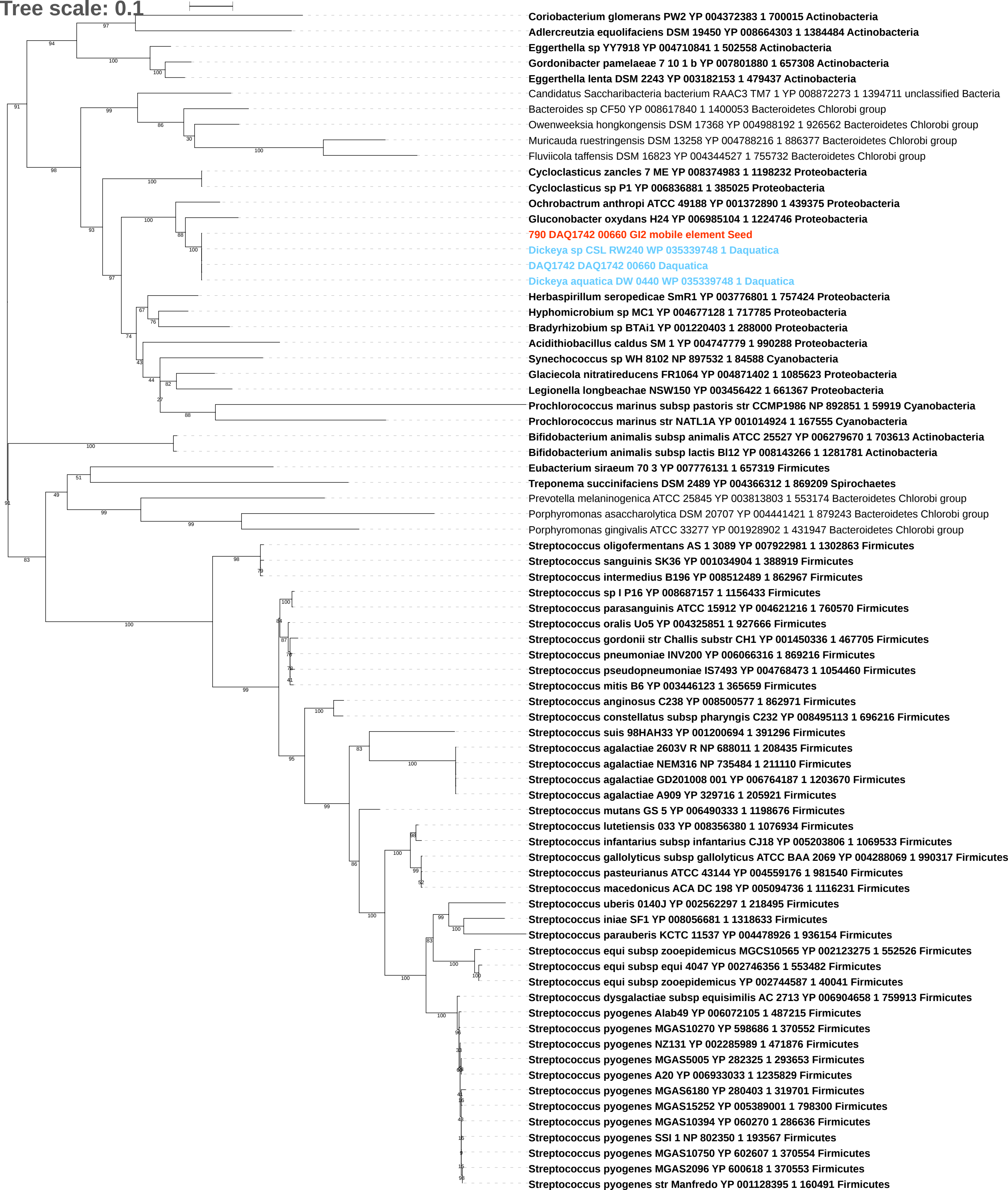
Tree scale: 0.1
Coriobacterium glomerans PW2 YP 004372383 1 700015 Actinobacteria
Adlercreutzia equolifaciens DSM 19450 YP 008664303 1 1384484 Actinobacteria
Eggerthella sp YY7918 YP 004710841 1 502558 Actinobacteria
Gordonibacter pamelaeae 7 10 1 b YP 007801880 1 657308 Actinobacteria
Eggerthella lenta DSM 2243 YP 003182153 1 479437 Actinobacteria
Candidatus Saccharibacteria bacterium RAAC3 TM7 1 YP 008872273 1 1394711 unclassified Bacteria
Bacteroides sp CF50 YP 008617840 1 1400053 Bacteroidetes Chlorobi group
Owenweeksia hongkongensis DSM 17368 YP 004988192 1 926562 Bacteroidetes Chlorobi group
Muricauda ruestringensis DSM 13258 YP 004788216 1 886377 Bacteroidetes Chlorobi group
Fluviicola taffensis DSM 16823 YP 004344527 1 755732 Bacteroidetes Chlorobi group
Cycloclasticus zancles 7 ME YP 008374983 1 1198232 Proteobacteria
Cycloclasticus sp P1 YP 006836881 1 385025 Proteobacteria
Ochrobactrum anthropi ATCC 49188 YP 001372890 1 439375 Proteobacteria
Gluconobacter oxydans H24 YP 006985104 1 1224746 Proteobacteria
790 DAQ1742 00660 GI2 mobile element Seed
Dickeya sp CSL RW240 WP 035339748 1 Daquatica
DAQ1742 DAQ1742 00660 Daquatica
Dickeya aquatica DW 0440 WP 035339748 1 Daquatica
Herbaspirillum seropedicae SmR1 YP 003776801 1 757424 Proteobacteria
Hyphomicrobium sp MC1 YP 004677128 1 717785 Proteobacteria
Bradyrhizobium sp BTAi1 YP 001220403 1 288000 Proteobacteria
Acidithiobacillus caldus SM 1 YP 004747779 1 990288 Proteobacteria
Synechococcus sp WH 8102 NP 897532 1 84588 Cyanobacteria
Glaciecola nitratireducens FR1064 YP 004871402 1 1085623 Proteobacteria
Legionella longbeachae NSW150 YP 003456422 1 661367 Proteobacteria
Prochlorococcus marinus subsp pastoris str CCMP1986 NP 892851 1 59919 Cyanobacteria
Prochlorococcus marinus str NATL1A YP 001014924 1 167555 Cyanobacteria
Bifidobacterium animalis subsp animalis ATCC 25527 YP 006279670 1 703613 Actinobacteria
Bifidobacterium animalis subsp lactis Bl12 YP 008143266 1 1281781 Actinobacteria
Eubacterium siraeum 70 3 YP 007776131 1 657319 Firmicutes
Treponema succinifaciens DSM 2489 YP 004366312 1 869209 Spirochaetes
Prevotella melaninogenica ATCC 25845 YP 003813803 1 553174 Bacteroidetes Chlorobi group
Porphyromonas asaccharolytica DSM 20707 YP 004441421 1 879243 Bacteroidetes Chlorobi group
Porphyromonas gingivalis ATCC 33277 YP 001928902 1 431947 Bacteroidetes Chlorobi group
Streptococcus oligofermentans AS 1 3089 YP 007922981 1 1302863 Firmicutes
Streptococcus sanguinis SK36 YP 001034904 1 388919 Firmicutes
Streptococcus intermedius B196 YP 008512489 1 862967 Firmicutes
Streptococcus sp I P16 YP 008687157 1 1156433 Firmicutes
Streptococcus parasanguinis ATCC 15912 YP 004621216 1 760570 Firmicutes
Streptococcus oralis Uo5 YP 004325851 1 927666 Firmicutes
Streptococcus gordonii str Challis substr CH1 YP 001450336 1 467705 Firmicutes
Streptococcus pneumoniae INV200 YP 006066316 1 869216 Firmicutes
Streptococcus pseudopneumoniae IS7493 YP 004768473 1 1054460 Firmicutes
Streptococcus mitis B6 YP 003446123 1 365659 Firmicutes
Streptococcus anginosus C238 YP 008500577 1 862971 Firmicutes
Streptococcus constellatus subsp pharyngis C232 YP 008495113 1 696216 Firmicutes
Streptococcus suis 98HAH33 YP 001200694 1 391296 Firmicutes
Streptococcus agalactiae 2603V R NP 688011 1 208435 Firmicutes
Streptococcus agalactiae NEM316 NP 735484 1 211110 Firmicutes
Streptococcus agalactiae GD201008 001 YP 006764187 1 1203670 Firmicutes
Streptococcus agalactiae A909 YP 329716 1 205921 Firmicutes
Streptococcus mutans GS 5 YP 006490333 1 1198676 Firmicutes
Streptococcus lutetiensis 033 YP 008356380 1 1076934 Firmicutes
Streptococcus infantarius subsp infantarius CJ18 YP 005203806 1 1069533 Firmicutes
Streptococcus gallolyticus subsp gallolyticus ATCC BAA 2069 YP 004288069 1 990317 Firmicutes
Streptococcus pasteurianus ATCC 43144 YP 004559176 1 981540 Firmicutes
Streptococcus macedonicus ACA DC 198 YP 005094736 1 1116231 Firmicutes
Streptococcus uberis 0140J YP 002562297 1 218495 Firmicutes
Streptococcus iniae SF1 YP 008056681 1 1318633 Firmicutes
Streptococcus parauberis KCTC 11537 YP 004478926 1 936154 Firmicutes
Streptococcus equi subsp zooepidemicus MGCS10565 YP 002123275 1 552526 Firmicutes
Streptococcus equi subsp equi 4047 YP 002746356 1 553482 Firmicutes
Streptococcus equi subsp zooepidemicus YP 002744587 1 40041 Firmicutes
Streptococcus dysgalactiae subsp equisimilis AC 2713 YP 006904658 1 759913 Firmicutes
Streptococcus pyogenes Alab49 YP 006072105 1 487215 Firmicutes
Streptococcus pyogenes MGAS10270 YP 598686 1 370552 Firmicutes
Streptococcus pyogenes NZ131 YP 002285989 1 471876 Firmicutes
Streptococcus pyogenes MGAS5005 YP 282325 1 293653 Firmicutes
Streptococcus pyogenes A20 YP 006933033 1 1235829 Firmicutes
Streptococcus pyogenes MGAS6180 YP 280403 1 319701 Firmicutes
Streptococcus pyogenes MGAS15252 YP 005389001 1 798300 Firmicutes
Streptococcus pyogenes MGAS10394 YP 060270 1 286636 Firmicutes
Streptococcus pyogenes SSI 1 NP 802350 1 193567 Firmicutes
Streptococcus pyogenes MGAS10750 YP 602607 1 370554 Firmicutes
Streptococcus pyogenes MGAS2096 YP 600618 1 370553 Firmicutes
Streptococcus pyogenes str Manfredo YP 001128395 1 160491 Firmicutes

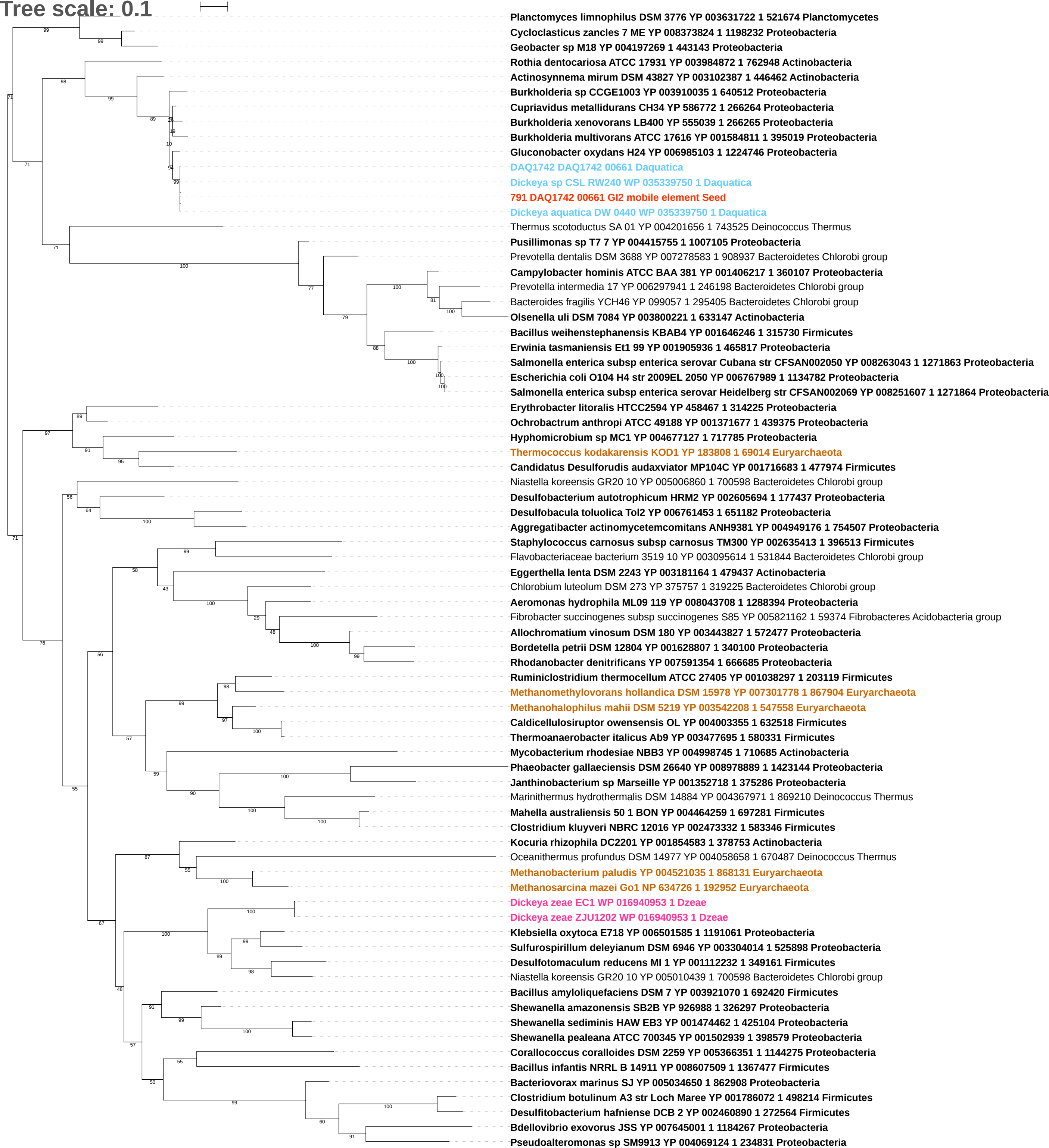
Tree scale: 0.1
Planctomyces limnophilus DSM 3776 YP 003631722 1 521674 Planctomycetes
Cycloclasticus zancles 7 ME YP 008373824 1 1198232 Proteobacteria
Geobacter sp M18 YP 004197269 1 443143 Proteobacteria
Rothia dentocariosa ATCC 17931 YP 003984872 1 762948 Actinobacteria
Actinosynnema mirum DSM 43827 YP 003102387 1 446462 Actinobacteria
Burkholderia sp CCGE1003 YP 003910035 1 640512 Proteobacteria
Cupriavidus metallidurans CH34 YP 586772 1 266264 Proteobacteria
Burkholderia xenovorans LB400 YP 555039 1 266265 Proteobacteria
Burkholderia multivorans ATCC 17616 YP 001584811 1 395019 Proteobacteria
Gluconobacter oxydans H24 YP 006985103 1 1224746 Proteobacteria
DAQ1742 DAQ1742 00661 Daquatica
Dickeya sp CSL RW240 WP 035339750 1 Daquatica
791 DAQ1742 00661 GI2 mobile element Seed
Dickeya aquatica DW 0440 WP 035339750 1 Daquatica
Thermus scotoductus SA 01 YP 004201656 1 743525 Deinococcus Thermus
Pusillimonas sp T7 7 YP 004415755 1 1007105 Proteobacteria
Prevotella dentalis DSM 3688 YP 007278583 1 908937 Bacteroidetes Chlorobi group
Campylobacter hominis ATCC BAA 381 YP 001406217 1 360107 Proteobacteria
Prevotella intermedia 17 YP 006297941 1 246198 Bacteroidetes Chlorobi group
Bacteroides fragilis YCH46 YP 099057 1 295405 Bacteroidetes Chlorobi group
Olsenella uli DSM 7084 YP 003800221 1 633147 Actinobacteria
Bacillus weihenstephanensis KBAB4 YP 001646246 1 315730 Firmicutes
Erwinia tasmaniensis Et1 99 YP 001905936 1 465817 Proteobacteria
Salmonella enterica subsp enterica serovar Cubana str CFSAN002050 YP 008263043 1 1271863 Proteobacteria
Escherichia coli O104 H4 str 2009EL 2050 YP 006767989 1 1134782 Proteobacteria
Salmonella enterica subsp enterica serovar Heidelberg str CFSAN002069 YP 008251607 1 1271864 Proteobacteria
Erythrobacter litoralis HTCC2594 YP 458467 1 314225 Proteobacteria
Ochrobactrum anthropi ATCC 49188 YP 001371677 1 439375 Proteobacteria
Hyphomicrobium sp MC1 YP 004677127 1 717785 Proteobacteria
Thermococcus kodakarensis KOD1 YP 183808 1 69014 Euryarchaeota
Candidatus Desulforudis audaxviator MP104C YP 001716683 1 477974 Firmicutes
Niastella koreensis GR20 10 YP 005006860 1 700598 Bacteroidetes Chlorobi group
Desulfobacterium autotrophicum HRM2 YP 002605694 1 177437 Proteobacteria
Desulfobacula toluolica Tol2 YP 006761453 1 651182 Proteobacteria
Aggregatibacter actinomycetemcomitans ANH9381 YP 004949176 1 754507 Proteobacteria
Staphylococcus carnosus subsp carnosus TM300 YP 002635413 1 396513 Firmicutes
Flavobacteriaceae bacterium 3519 10 YP 003095614 1 531844 Bacteroidetes Chlorobi group
Eggerthella lenta DSM 2243 YP 003181164 1 479437 Actinobacteria
Chlorobium luteolum DSM 273 YP 375757 1 319225 Bacteroidetes Chlorobi group
Aeromonas hydrophila ML09 119 YP 008043708 1 1288394 Proteobacteria
Fibrobacter succinogenes subsp succinogenes S85 YP 005821162 1 59374 Fibrobacteres Acidobacteria group
Allochromatium vinosum DSM 180 YP 003443827 1 572477 Proteobacteria
Bordetella petrii DSM 12804 YP 001628807 1 340100 Proteobacteria
Rhodanobacter denitrificans YP 007591354 1 666685 Proteobacteria
Ruminiclostridium thermocellum ATCC 27405 YP 001038297 1 203119 Firmicutes
Methanomethylovorans hollandica DSM 15978 YP 007301778 1 867904 Euryarchaeota
Methanohalophilus mahii DSM 5219 YP 003542208 1 547558 Euryarchaeota
Caldicellulosiruptor owensensis OL YP 004003355 1 632518 Firmicutes
Thermoanaerobacter italicus Ab9 YP 003477695 1 580331 Firmicutes
Mycobacterium rhodesiae NBB3 YP 004998745 1 710685 Actinobacteria
Phaeobacter gallaeciensis DSM 26640 YP 008978889 1 1423144 Proteobacteria
Janthinobacterium sp Marseille YP 001352718 1 375286 Proteobacteria
Marinithermus hydrothermalis DSM 14884 YP 004367971 1 869210 Deinococcus Thermus
Mahella australiensis 50 1 BON YP 004464259 1 697281 Firmicutes
Clostridium kluyveri NBRC 12016 YP 002473332 1 583346 Firmicutes
Kocuria rhizophila DC2201 YP 001854583 1 378753 Actinobacteria
Oceanithermus profundus DSM 14977 YP 004058658 1 670487 Deinococcus Thermus
Methanobacterium paludis YP 004521035 1 868131 Euryarchaeota
Methanosarcina mazei Go1 NP 634726 1 192952 Euryarchaeota
Dickeya zeae EC1 WP 016940953 1 Dzeae
Dickeya zeae ZJU1202 WP 016940953 1 Dzeae
Klebsiella oxytoca E718 YP 006501585 1 1191061 Proteobacteria
Sulfurospirillum deleyianum DSM 6946 YP 003304014 1 525898 Proteobacteria
Desulfotomaculum reducens MI 1 YP 001112232 1 349161 Firmicutes
Niastella koreensis GR20 10 YP 005010439 1 700598 Bacteroidetes Chlorobi group
Bacillus amyloliquefaciens DSM 7 YP 003921070 1 692420 Firmicutes
Shewanella amazonensis SB2B YP 926988 1 326297 Proteobacteria
Shewanella sediminis HAW EB3 YP 001474462 1 425104 Proteobacteria
Shewanella pealeana ATCC 700345 YP 001502939 1 398579 Proteobacteria
Corallococcus coralloides DSM 2259 YP 005366351 1 1144275 Proteobacteria
Bacillus infantis NRRL B 14911 YP 008607509 1 1367477 Firmicutes
Bacteriovorax marinus SJ YP 005034650 1 862908 Proteobacteria
Clostridium botulinum A3 str Loch Maree YP 001786072 1 498214 Firmicutes
Desulfitobacterium hafniense DCB 2 YP 002460890 1 272564 Firmicutes
Bdellovibrio exovorus JSS YP 007645001 1 1184267 Proteobacteria
Pseudoalteromonas sp SM9913 YP 004069124 1 234831 Proteobacteria

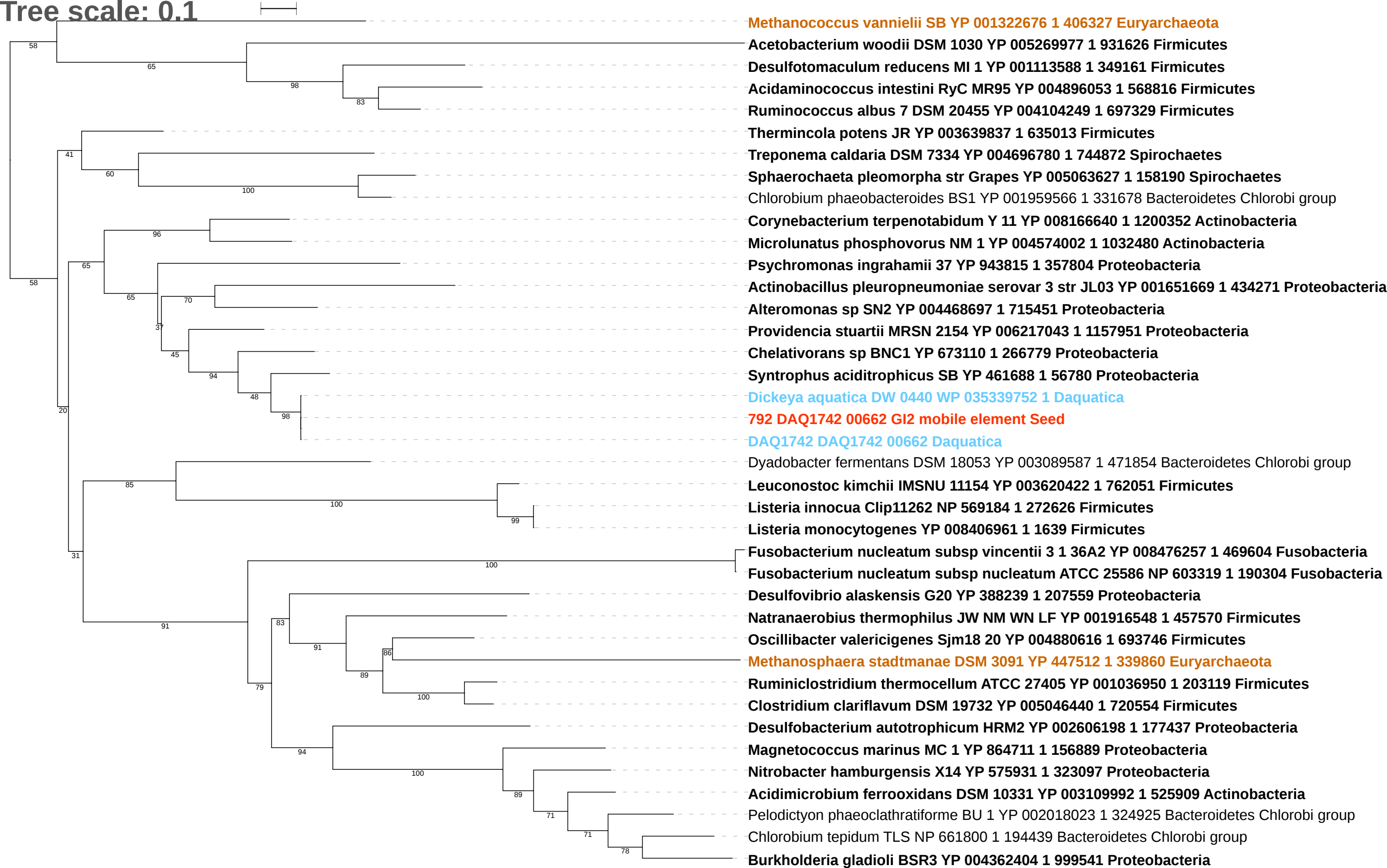
Tree scale: 0.1
Methanococcus vannielii SB YP 001322676 1 406327 Euryarchaeota
Acetobacterium woodii DSM 1030 YP 005269977 1 931626 Firmicutes
Desulfotomaculum reducens MI 1 YP 001113588 1 349161 Firmicutes
Acidaminococcus intestini RyC MR95 YP 004896053 1 568816 Firmicutes
Ruminococcus albus 7 DSM 20455 YP 004104249 1 697329 Firmicutes
Thermincola potens JR YP 003639837 1 635013 Firmicutes
Treponema caldaria DSM 7334 YP 004696780 1 744872 Spirochaetes
Sphaerochaeta pleomorpha str Grapes YP 005063627 1 158190 Spirochaetes
Chlorobium phaeobacteroides BS1 YP 001959566 1 331678 Bacteroidetes Chlorobi group
Corynebacterium terpenotabidum Y 11 YP 008166640 1 1200352 Actinobacteria
Microlunatus phosphovorus NM 1 YP 004574002 1 1032480 Actinobacteria
Psychromonas ingrahamii 37 YP 943815 1 357804 Proteobacteria
Actinobacillus pleuropneumoniae serovar 3 str JL03 YP 001651669 1 434271 Proteobacteria
Alteromonas sp SN2 YP 004468697 1 715451 Proteobacteria
Providencia stuartii MRSN 2154 YP 006217043 1 1157951 Proteobacteria
Chelativorans sp BNC1 YP 673110 1 266779 Proteobacteria
Syntrophus aciditrophicus SB YP 461688 1 56780 Proteobacteria
Dickeya aquatica DW 0440 WP 035339752 1 Daquatica
792 DAQ1742 00662 GI2 mobile element Seed
DAQ1742 DAQ1742 00662 Daquatica
Dyadobacter fermentans DSM 18053 YP 003089587 1 471854 Bacteroidetes Chlorobi group
Leuconostoc kimchii IMSNU 11154 YP 003620422 1 762051 Firmicutes
Listeria innocua Clip11262 NP 569184 1 272626 Firmicutes
Listeria monocytogenes YP 008406961 1 1639 Firmicutes
Fusobacterium nucleatum subsp vincentii 3 1 36A2 YP 008476257 1 469604 Fusobacteria
Fusobacterium nucleatum subsp nucleatum ATCC 25586 NP 603319 1 190304 Fusobacteria
Desulfovibrio alaskensis G20 YP 388239 1 207559 Proteobacteria
Natranaerobius thermophilus JW NM WN LF YP 001916548 1 457570 Firmicutes
Oscillibacter valericigenes Sjm18 20 YP 004880616 1 693746 Firmicutes
Methanosphaera stadtmanae DSM 3091 YP 447512 1 339860 Euryarchaeota
Ruminiclostridium thermocellum ATCC 27405 YP 001036950 1 203119 Firmicutes
Clostridium clariflavum DSM 19732 YP 005046440 1 720554 Firmicutes
Desulfobacterium autotrophicum HRM2 YP 002606198 1 177437 Proteobacteria
Magnetococcus marinus MC 1 YP 864711 1 156889 Proteobacteria
Nitrobacter hamburgensis X14 YP 575931 1 323097 Proteobacteria
Acidimicrobium ferrooxidans DSM 10331 YP 003109992 1 525909 Actinobacteria
Pelodictyon phaeoclathratiforme BU 1 YP 002018023 1 324925 Bacteroidetes Chlorobi group
Chlorobium tepidum TLS NP 661800 1 194439 Bacteroidetes Chlorobi group
Burkholderia gladioli BSR3 YP 004362404 1 999541 Proteobacteria

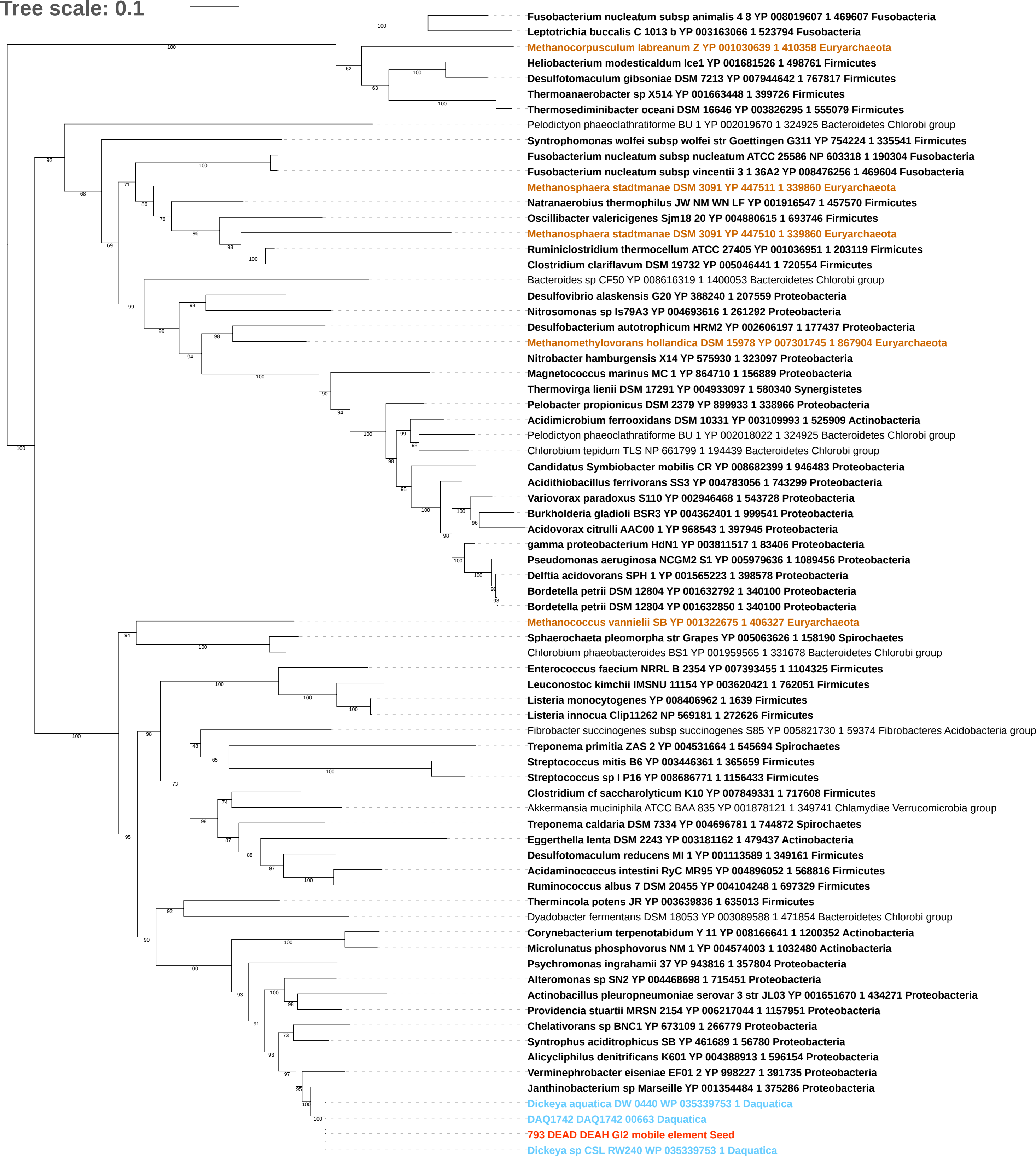
Tree scale: 0.1
Fusobacterium nucleatum subsp animalis 4 8 YP 008019607 1 469607 Fusobacteria
Leptotrichia buccalis C 1013 b YP 003163066 1 523794 Fusobacteria
Methanocorpusculum labreanum Z YP 001030639 1 410358 Euryarchaeota
Heliobacterium modesticaldum Ice1 YP 001681526 1 498761 Firmicutes
Desulfotomaculum gibsoniae DSM 7213 YP 007944642 1 767817 Firmicutes
Thermoanaerobacter sp X514 YP 001663448 1 399726 Firmicutes
Thermosediminibacter oceani DSM 16646 YP 003826295 1 555079 Firmicutes
Pelodictyon phaeoclathratiforme BU 1 YP 002019670 1 324925 Bacteroidetes Chlorobi group
Syntrophomonas wolfei subsp wolfei str Goettingen G311 YP 754224 1 335541 Firmicutes
Fusobacterium nucleatum subsp nucleatum ATCC 25586 NP 603318 1 190304 Fusobacteria
Fusobacterium nucleatum subsp vincentii 3 1 36A2 YP 008476256 1 469604 Fusobacteria
Methanosphaera stadtmanae DSM 3091 YP 447511 1 339860 Euryarchaeota
Natranaerobius thermophilus JW NM WN LF YP 001916547 1 457570 Firmicutes
Oscillibacter valericigenes Sjm18 20 YP 004880615 1 693746 Firmicutes
Methanosphaera stadtmanae DSM 3091 YP 447510 1 339860 Euryarchaeota
Ruminiclostridium thermocellum ATCC 27405 YP 001036951 1 203119 Firmicutes
Clostridium clariflavum DSM 19732 YP 005046441 1 720554 Firmicutes
Bacteroides sp CF50 YP 008616319 1 1400053 Bacteroidetes Chlorobi group
Desulfovibrio alaskensis G20 YP 388240 1 207559 Proteobacteria
Nitrosomonas sp Is79A3 YP 004693616 1 261292 Proteobacteria
Desulfobacterium autotrophicum HRM2 YP 002606197 1 177437 Proteobacteria
Methanomethylovorans hollandica DSM 15978 YP 007301745 1 867904 Euryarchaeota
Nitrobacter hamburgensis X14 YP 575930 1 323097 Proteobacteria
Magnetococcus marinus MC 1 YP 864710 1 156889 Proteobacteria
Thermovirga lienii DSM 17291 YP 004933097 1 580340 Synergistetes
Pelobacter propionicus DSM 2379 YP 899933 1 338966 Proteobacteria
Acidimicrobium ferrooxidans DSM 10331 YP 003109993 1 525909 Actinobacteria
Pelodictyon phaeoclathratiforme BU 1 YP 002018022 1 324925 Bacteroidetes Chlorobi group
Chlorobium tepidum TLS NP 661799 1 194439 Bacteroidetes Chlorobi group
Candidatus Symbiobacter mobilis CR YP 008682399 1 946483 Proteobacteria
Acidithiobacillus ferrivorans SS3 YP 004783056 1 743299 Proteobacteria
Variovorax paradoxus S110 YP 002946468 1 543728 Proteobacteria
Burkholderia gladioli BSR3 YP 004362401 1 999541 Proteobacteria
Acidovorax citrulli AAC00 1 YP 968543 1 397945 Proteobacteria
gamma proteobacterium HdN1 YP 003811517 1 83406 Proteobacteria
Pseudomonas aeruginosa NCGM2 S1 YP 005979636 1 1089456 Proteobacteria
Delftia acidovorans SPH 1 YP 001565223 1 398578 Proteobacteria
Bordetella petrii DSM 12804 YP 001632792 1 340100 Proteobacteria
Bordetella petrii DSM 12804 YP 001632850 1 340100 Proteobacteria
Methanococcus vannielii SB YP 001322675 1 406327 Euryarchaeota
Sphaerochaeta pleomorpha str Grapes YP 005063626 1 158190 Spirochaetes
Chlorobium phaeobacteroides BS1 YP 001959565 1 331678 Bacteroidetes Chlorobi group
Enterococcus faecium NRRL B 2354 YP 007393455 1 1104325 Firmicutes
Leuconostoc kimchii IMSNU 11154 YP 003620421 1 762051 Firmicutes
Listeria monocytogenes YP 008406962 1 1639 Firmicutes
Listeria innocua Clip11262 NP 569181 1 272626 Firmicutes
Fibrobacter succinogenes subsp succinogenes S85 YP 005821730 1 59374 Fibrobacteres Acidobacteria group
Treponema primitia ZAS 2 YP 004531664 1 545694 Spirochaetes
Streptococcus mitis B6 YP 003446361 1 365659 Firmicutes
Streptococcus sp I P16 YP 008686771 1 1156433 Firmicutes
Clostridium cf saccharolyticum K10 YP 007849331 1 717608 Firmicutes
Akkermansia muciniphila ATCC BAA 835 YP 001878121 1 349741 Chlamydiae Verrucomicrobia group
Treponema caldaria DSM 7334 YP 004696781 1 744872 Spirochaetes
Eggerthella lenta DSM 2243 YP 003181162 1 479437 Actinobacteria
Desulfotomaculum reducens MI 1 YP 001113589 1 349161 Firmicutes
Acidaminococcus intestini RyC MR95 YP 004896052 1 568816 Firmicutes
Ruminococcus albus 7 DSM 20455 YP 004104248 1 697329 Firmicutes
Thermincola potens JR YP 003639836 1 635013 Firmicutes
Dyadobacter fermentans DSM 18053 YP 003089588 1 471854 Bacteroidetes Chlorobi group
Corynebacterium terpenotabidum Y 11 YP 008166641 1 1200352 Actinobacteria
Microlunatus phosphovorus NM 1 YP 004574003 1 1032480 Actinobacteria
Psychromonas ingrahamii 37 YP 943816 1 357804 Proteobacteria
Alteromonas sp SN2 YP 004468698 1 715451 Proteobacteria
Actinobacillus pleuropneumoniae serovar 3 str JL03 YP 001651670 1 434271 Proteobacteria
Providencia stuartii MRSN 2154 YP 006217044 1 1157951 Proteobacteria
Chelativorans sp BNC1 YP 673109 1 266779 Proteobacteria
Syntrophus aciditrophicus SB YP 461689 1 56780 Proteobacteria
Alicycliphilus denitrificans K601 YP 004388913 1 596154 Proteobacteria
Verminephrobacter eiseniae EF01 2 YP 998227 1 391735 Proteobacteria
Janthinobacterium sp Marseille YP 001354484 1 375286 Proteobacteria
Dickeya aquatica DW 0440 WP 035339753 1 Daquatica
DAQ1742 DAQ1742 00663 Daquatica
793 DEAD DEAH GI2 mobile element Seed
Dickeya sp CSL RW240 WP 035339753 1 Daquatica

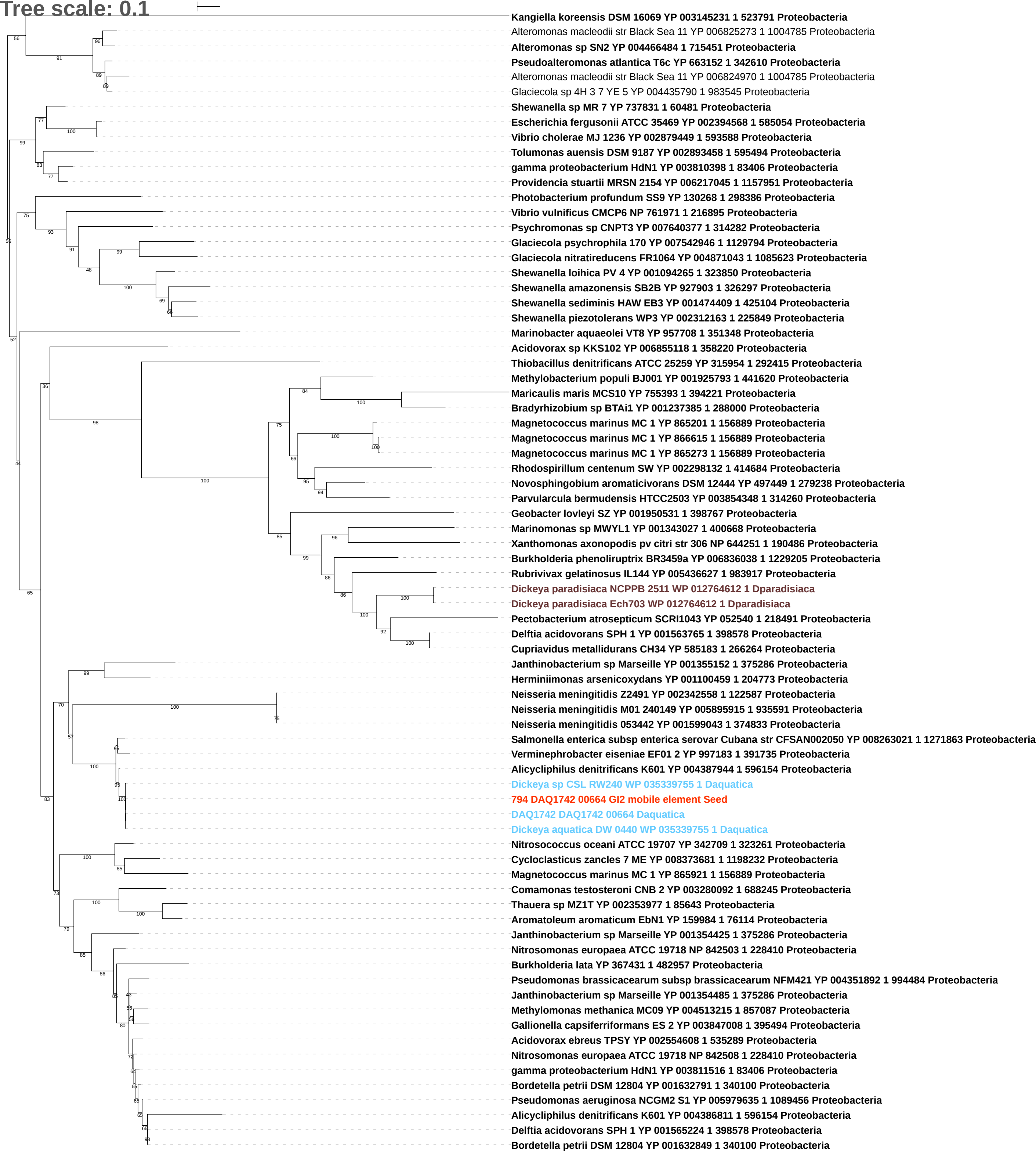
Tree scale: 0.1
Kangiella koreensis DSM 16069 YP 003145231 1 523791 Proteobacteria
Alteromonas macleodii str Black Sea 11 YP 006825273 1 1004785 Proteobacteria
Alteromonas sp SN2 YP 004466484 1 715451 Proteobacteria
Pseudoalteromonas atlantica T6c YP 663152 1 342610 Proteobacteria
Alteromonas macleodii str Black Sea 11 YP 006824970 1 1004785 Proteobacteria
Glaciecola sp 4H 3 7 YE 5 YP 004435790 1 983545 Proteobacteria
Shewanella sp MR 7 YP 737831 1 60481 Proteobacteria
Escherichia fergusonii ATCC 35469 YP 002394568 1 585054 Proteobacteria
Vibrio cholerae MJ 1236 YP 002879449 1 593588 Proteobacteria
Tolumonas auensis DSM 9187 YP 002893458 1 595494 Proteobacteria
gamma proteobacterium HdN1 YP 003810398 1 83406 Proteobacteria
Providencia stuartii MRSN 2154 YP 006217045 1 1157951 Proteobacteria
Photobacterium profundum SS9 YP 130268 1 298386 Proteobacteria
Vibrio vulnificus CMCP6 NP 761971 1 216895 Proteobacteria
Psychromonas sp CNPT3 YP 007640377 1 314282 Proteobacteria
Glaciecola psychrophila 170 YP 007542946 1 1129794 Proteobacteria
Glaciecola nitratireducens FR1064 YP 004871043 1 1085623 Proteobacteria
Shewanella loihica PV 4 YP 001094265 1 323850 Proteobacteria
Shewanella amazonensis SB2B YP 927903 1 326297 Proteobacteria
Shewanella sediminis HAW EB3 YP 001474409 1 425104 Proteobacteria
Shewanella piezotolerans WP3 YP 002312163 1 225849 Proteobacteria
Marinobacter aquaeolei VT8 YP 957708 1 351348 Proteobacteria
Acidovorax sp KKS102 YP 006855118 1 358220 Proteobacteria
Thiobacillus denitrificans ATCC 25259 YP 315954 1 292415 Proteobacteria
Methylobacterium populi BJ001 YP 001925793 1 441620 Proteobacteria
Maricaulis maris MCS10 YP 755393 1 394221 Proteobacteria
Bradyrhizobium sp BTAi1 YP 001237385 1 288000 Proteobacteria
Magnetococcus marinus MC 1 YP 865201 1 156889 Proteobacteria
Magnetococcus marinus MC 1 YP 866615 1 156889 Proteobacteria
Magnetococcus marinus MC 1 YP 865273 1 156889 Proteobacteria
Rhodospirillum centenum SW YP 002298132 1 414684 Proteobacteria
Novosphingobium aromaticivorans DSM 12444 YP 497449 1 279238 Proteobacteria
Parvularcula bermudensis HTCC2503 YP 003854348 1 314260 Proteobacteria
Geobacter lovleyi SZ YP 001950531 1 398767 Proteobacteria
Marinomonas sp MWYL1 YP 001343027 1 400668 Proteobacteria
Xanthomonas axonopodis pv citri str 306 NP 644251 1 190486 Proteobacteria
Burkholderia phenoliruptrix BR3459a YP 006836038 1 1229205 Proteobacteria
Rubrivivax gelatinosus IL144 YP 005436627 1 983917 Proteobacteria
Dickeya paradisiaca NCPPB 2511 WP 012764612 1 Dparadisiaca
Dickeya paradisiaca Ech703 WP 012764612 1 Dparadisiaca
Pectobacterium atrosepticum SCRI1043 YP 052540 1 218491 Proteobacteria
Delftia acidovorans SPH 1 YP 001563765 1 398578 Proteobacteria
Cupriavidus metallidurans CH34 YP 585183 1 266264 Proteobacteria
Janthinobacterium sp Marseille YP 001355152 1 375286 Proteobacteria
Herminiimonas arsenicoxydans YP 001100459 1 204773 Proteobacteria
Neisseria meningitidis Z2491 YP 002342558 1 122587 Proteobacteria
Neisseria meningitidis M01 240149 YP 005895915 1 935591 Proteobacteria
Neisseria meningitidis 053442 YP 001599043 1 374833 Proteobacteria
Salmonella enterica subsp enterica serovar Cubana str CFSAN002050 YP 008263021 1 1271863 Proteobacteria
Verminephrobacter eiseniae EF01 2 YP 997183 1 391735 Proteobacteria
Alicycliphilus denitrificans K601 YP 004387944 1 596154 Proteobacteria
Dickeya sp CSL RW240 WP 035339755 1 Daquatica
794 DAQ1742 00664 GI2 mobile element Seed
DAQ1742 DAQ1742 00664 Daquatica
Dickeya aquatica DW 0440 WP 035339755 1 Daquatica
Nitrosococcus oceani ATCC 19707 YP 342709 1 323261 Proteobacteria
Cycloclasticus zancles 7 ME YP 008373681 1 1198232 Proteobacteria
Magnetococcus marinus MC 1 YP 865921 1 156889 Proteobacteria
Comamonas testosteroni CNB 2 YP 003280092 1 688245 Proteobacteria
Thauera sp MZ1T YP 002353977 1 85643 Proteobacteria
Aromatoleum aromaticum EbN1 YP 159984 1 76114 Proteobacteria
Janthinobacterium sp Marseille YP 001354425 1 375286 Proteobacteria
Nitrosomonas europaea ATCC 19718 NP 842503 1 228410 Proteobacteria
Burkholderia lata YP 367431 1 482957 Proteobacteria
Pseudomonas brassicacearum subsp brassicacearum NFM421 YP 004351892 1 994484 Proteobacteria
Janthinobacterium sp Marseille YP 001354485 1 375286 Proteobacteria
Methylomonas methanica MC09 YP 004513215 1 857087 Proteobacteria
Gallionella capsiferriformans ES 2 YP 003847008 1 395494 Proteobacteria
Acidovorax ebreus TPSY YP 002554608 1 535289 Proteobacteria
Nitrosomonas europaea ATCC 19718 NP 842508 1 228410 Proteobacteria
gamma proteobacterium HdN1 YP 003811516 1 83406 Proteobacteria
Bordetella petrii DSM 12804 YP 001632791 1 340100 Proteobacteria
Pseudomonas aeruginosa NCGM2 S1 YP 005979635 1 1089456 Proteobacteria
Alicycliphilus denitrificans K601 YP 004386811 1 596154 Proteobacteria
Delftia acidovorans SPH 1 YP 001565224 1 398578 Proteobacteria
Bordetella petrii DSM 12804 YP 001632849 1 340100 Proteobacteria

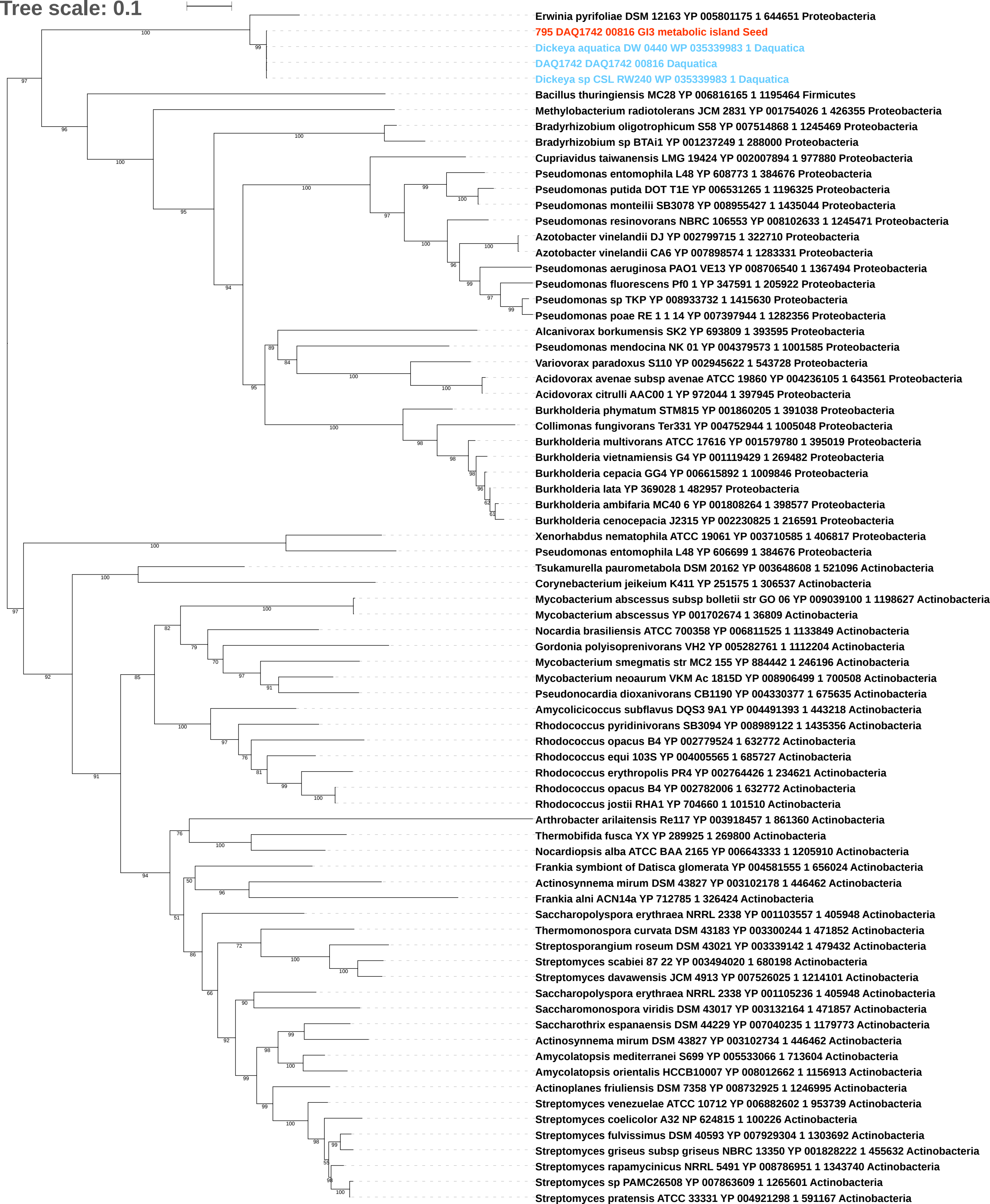
Tree scale: 0.1
Erwinia pyrifoliae DSM 12163 YP 005801175 1 644651 Proteobacteria
795 DAQ1742 00816 GI3 metabolic island Seed
Dickeya aquatica DW 0440 WP 035339983 1 Daquatica
DAQ1742 DAQ1742 00816 Daquatica
Dickeya sp CSL RW240 WP 035339983 1 Daquatica
Bacillus thuringiensis MC28 YP 006816165 1 1195464 Firmicutes
Methylobacterium radiotolerans JCM 2831 YP 001754026 1 426355 Proteobacteria
Bradyrhizobium oligotrophicum S58 YP 007514868 1 1245469 Proteobacteria
Bradyrhizobium sp BTAi1 YP 001237249 1 288000 Proteobacteria
Cupriavidus taiwanensis LMG 19424 YP 002007894 1 977880 Proteobacteria
Pseudomonas entomophila L48 YP 608773 1 384676 Proteobacteria
Pseudomonas putida DOT T1E YP 006531265 1 1196325 Proteobacteria
Pseudomonas monteilii SB3078 YP 008955427 1 1435044 Proteobacteria
Pseudomonas resinovorans NBRC 106553 YP 008102633 1 1245471 Proteobacteria
Azotobacter vinelandii DJ YP 002799715 1 322710 Proteobacteria
Azotobacter vinelandii CA6 YP 007898574 1 1283331 Proteobacteria
Pseudomonas aeruginosa PAO1 VE13 YP 008706540 1 1367494 Proteobacteria
Pseudomonas fluorescens Pf0 1 YP 347591 1 205922 Proteobacteria
Pseudomonas sp TKP YP 008933732 1 1415630 Proteobacteria
Pseudomonas poae RE 1 1 14 YP 007397944 1 1282356 Proteobacteria
Alcanivorax borkumensis SK2 YP 693809 1 393595 Proteobacteria
Pseudomonas mendocina NK 01 YP 004379573 1 1001585 Proteobacteria
Variovorax paradoxus S110 YP 002945622 1 543728 Proteobacteria
Acidovorax avenae subsp avenae ATCC 19860 YP 004236105 1 643561 Proteobacteria
Acidovorax citrulli AAC00 1 YP 972044 1 397945 Proteobacteria
Burkholderia phymatum STM815 YP 001860205 1 391038 Proteobacteria
Collimonas fungivorans Ter331 YP 004752944 1 1005048 Proteobacteria
Burkholderia multivorans ATCC 17616 YP 001579780 1 395019 Proteobacteria
Burkholderia vietnamiensis G4 YP 001119429 1 269482 Proteobacteria
Burkholderia cepacia GG4 YP 006615892 1 1009846 Proteobacteria
Burkholderia lata YP 369028 1 482957 Proteobacteria
Burkholderia ambifaria MC40 6 YP 001808264 1 398577 Proteobacteria
Burkholderia cenocepacia J2315 YP 002230825 1 216591 Proteobacteria
Xenorhabdus nematophila ATCC 19061 YP 003710585 1 406817 Proteobacteria
Pseudomonas entomophila L48 YP 606699 1 384676 Proteobacteria
Tsukamurella paurometabola DSM 20162 YP 003648608 1 521096 Actinobacteria
Corynebacterium jeikeium K411 YP 251575 1 306537 Actinobacteria
Mycobacterium abscessus subsp bolletii str GO 06 YP 009039100 1 1198627 Actinobacteria
Mycobacterium abscessus YP 001702674 1 36809 Actinobacteria
Nocardia brasiliensis ATCC 700358 YP 006811525 1 1133849 Actinobacteria
Gordonia polyisoprenivorans VH2 YP 005282761 1 1112204 Actinobacteria
Mycobacterium smegmatis str MC2 155 YP 884442 1 246196 Actinobacteria
Mycobacterium neoaurum VKM Ac 1815D YP 008906499 1 700508 Actinobacteria
Pseudonocardia dioxanivorans CB1190 YP 004330377 1 675635 Actinobacteria
Amycolicicoccus subflavus DQS3 9A1 YP 004491393 1 443218 Actinobacteria
Rhodococcus pyridinivorans SB3094 YP 008989122 1 1435356 Actinobacteria
Rhodococcus opacus B4 YP 002779524 1 632772 Actinobacteria
Rhodococcus equi 103S YP 004005565 1 685727 Actinobacteria
Rhodococcus erythropolis PR4 YP 002764426 1 234621 Actinobacteria
Rhodococcus opacus B4 YP 002782006 1 632772 Actinobacteria
Rhodococcus jostii RHA1 YP 704660 1 101510 Actinobacteria
Arthrobacter arilaitensis Re117 YP 003918457 1 861360 Actinobacteria
Thermobifida fusca YX YP 289925 1 269800 Actinobacteria
Nocardiopsis alba ATCC BAA 2165 YP 006643333 1 1205910 Actinobacteria
Frankia symbiont of Datisca glomerata YP 004581555 1 656024 Actinobacteria
Actinosynnema mirum DSM 43827 YP 003102178 1 446462 Actinobacteria
Frankia alni ACN14a YP 712785 1 326424 Actinobacteria
Saccharopolyspora erythraea NRRL 2338 YP 001103557 1 405948 Actinobacteria
Thermomonospora curvata DSM 43183 YP 003300244 1 471852 Actinobacteria
Streptosporangium roseum DSM 43021 YP 003339142 1 479432 Actinobacteria
Streptomyces scabiei 87 22 YP 003494020 1 680198 Actinobacteria
Streptomyces davawensis JCM 4913 YP 007526025 1 1214101 Actinobacteria
Saccharopolyspora erythraea NRRL 2338 YP 001105236 1 405948 Actinobacteria
Saccharomonospora viridis DSM 43017 YP 003132164 1 471857 Actinobacteria
Saccharothrix espanaensis DSM 44229 YP 007040235 1 1179773 Actinobacteria
Actinosynnema mirum DSM 43827 YP 003102734 1 446462 Actinobacteria
Amycolatopsis mediterranei S699 YP 005533066 1 713604 Actinobacteria
Amycolatopsis orientalis HCCB10007 YP 008012662 1 1156913 Actinobacteria
Actinoplanes friuliensis DSM 7358 YP 008732925 1 1246995 Actinobacteria
Streptomyces venezuelae ATCC 10712 YP 006882602 1 953739 Actinobacteria
Streptomyces coelicolor A32 NP 624815 1 100226 Actinobacteria
Streptomyces fulvissimus DSM 40593 YP 007929304 1 1303692 Actinobacteria
Streptomyces griseus subsp griseus NBRC 13350 YP 001828222 1 455632 Actinobacteria
Streptomyces rapamycinicus NRRL 5491 YP 008786951 1 1343740 Actinobacteria
Streptomyces sp PAMC26508 YP 007863609 1 1265601 Actinobacteria
Streptomyces pratensis ATCC 33331 YP 004921298 1 591167 Actinobacteria

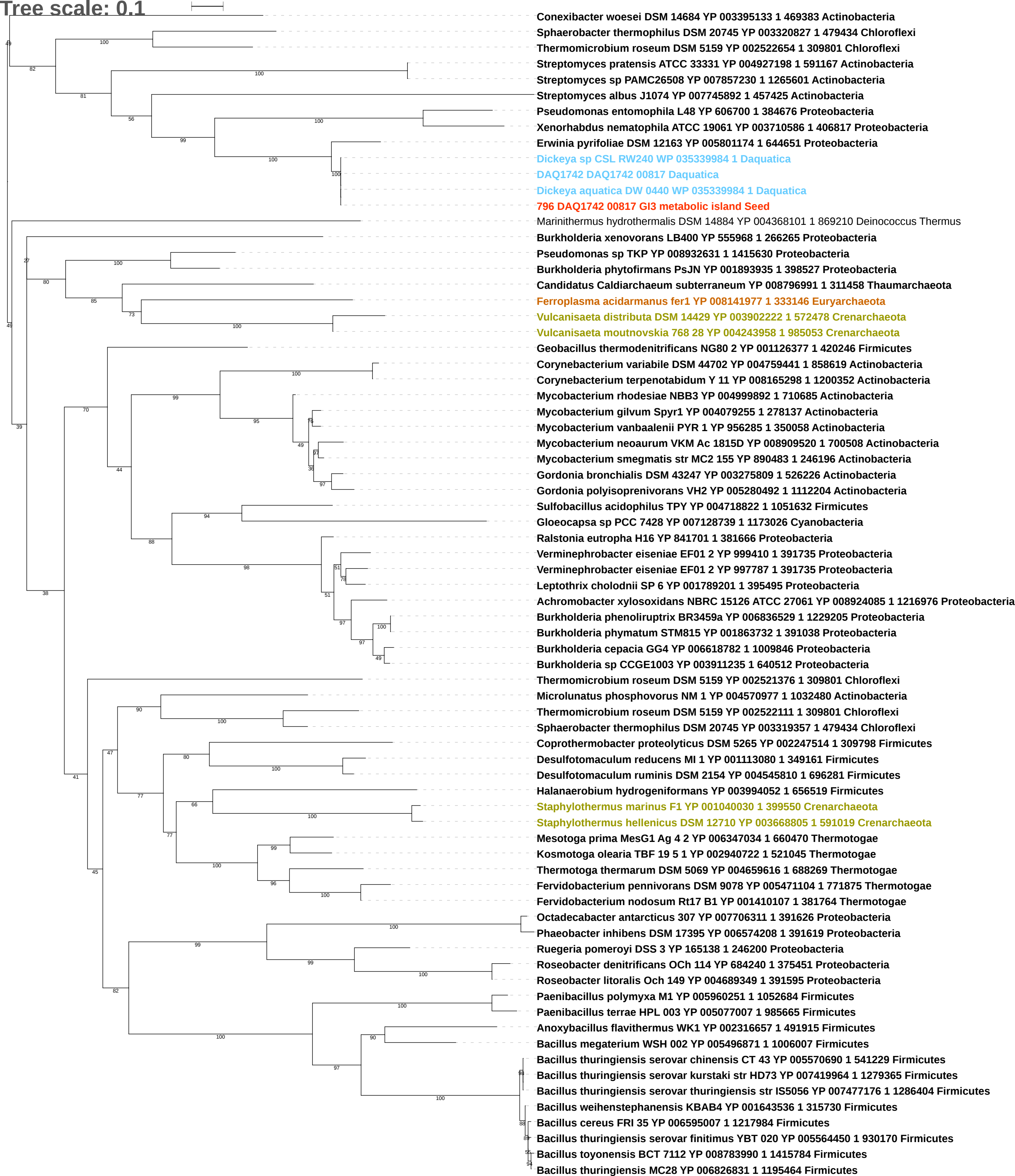
Tree scale: 0.1
Conexibacter woesei DSM 14684 YP 003395133 1 469383 Actinobacteria
Sphaerobacter thermophilus DSM 20745 YP 003320827 1 479434 Chloroflexi
Thermomicrobium roseum DSM 5159 YP 002522654 1 309801 Chloroflexi
Streptomyces pratensis ATCC 33331 YP 004927198 1 591167 Actinobacteria
Streptomyces sp PAMC26508 YP 007857230 1 1265601 Actinobacteria
Streptomyces albus J1074 YP 007745892 1 457425 Actinobacteria
Pseudomonas entomophila L48 YP 606700 1 384676 Proteobacteria
Xenorhabdus nematophila ATCC 19061 YP 003710586 1 406817 Proteobacteria
Erwinia pyrifoliae DSM 12163 YP 005801174 1 644651 Proteobacteria
Dickeya sp CSL RW240 WP 035339984 1 Daquatica
DAQ1742 DAQ1742 00817 Daquatica
Dickeya aquatica DW 0440 WP 035339984 1 Daquatica
796 DAQ1742 00817 GI3 metabolic island Seed
Marinithermus hydrothermalis DSM 14884 YP 004368101 1 869210 Deinococcus Thermus
Burkholderia xenovorans LB400 YP 555968 1 266265 Proteobacteria
Pseudomonas sp TKP YP 008932631 1 1415630 Proteobacteria
Burkholderia phytofirmans PsJN YP 001893935 1 398527 Proteobacteria
Candidatus Caldiarchaeum subterraneum YP 008796991 1 311458 Thaumarchaeota
Ferroplasma acidarmanus fer1 YP 008141977 1 333146 Euryarchaeota
Vulcanisaeta distributa DSM 14429 YP 003902222 1 572478 Crenarchaeota
Vulcanisaeta moutnovskia 768 28 YP 004243958 1 985053 Crenarchaeota
Geobacillus thermodenitrificans NG80 2 YP 001126377 1 420246 Firmicutes
Corynebacterium variabile DSM 44702 YP 004759441 1 858619 Actinobacteria
Corynebacterium terpenotabidum Y 11 YP 008165298 1 1200352 Actinobacteria
Mycobacterium rhodesiae NBB3 YP 004999892 1 710685 Actinobacteria
Mycobacterium gilvum Spyr1 YP 004079255 1 278137 Actinobacteria
Mycobacterium vanbaalenii PYR 1 YP 956285 1 350058 Actinobacteria
Mycobacterium neoaurum VKM Ac 1815D YP 008909520 1 700508 Actinobacteria
Mycobacterium smegmatis str MC2 155 YP 890483 1 246196 Actinobacteria
Gordonia bronchialis DSM 43247 YP 003275809 1 526226 Actinobacteria
Gordonia polyisoprenivorans VH2 YP 005280492 1 1112204 Actinobacteria
Sulfobacillus acidophilus TPY YP 004718822 1 1051632 Firmicutes
Gloeocapsa sp PCC 7428 YP 007128739 1 1173026 Cyanobacteria
Ralstonia eutropha H16 YP 841701 1 381666 Proteobacteria
Verminephrobacter eiseniae EF01 2 YP 999410 1 391735 Proteobacteria
Verminephrobacter eiseniae EF01 2 YP 997787 1 391735 Proteobacteria
Leptothrix cholodnii SP 6 YP 001789201 1 395495 Proteobacteria
Achromobacter xylosoxidans NBRC 15126 ATCC 27061 YP 008924085 1 1216976 Proteobacteria
Burkholderia phenoliruptrix BR3459a YP 006836529 1 1229205 Proteobacteria
Burkholderia phymatum STM815 YP 001863732 1 391038 Proteobacteria
Burkholderia cepacia GG4 YP 006618782 1 1009846 Proteobacteria
Burkholderia sp CCGE1003 YP 003911235 1 640512 Proteobacteria
Thermomicrobium roseum DSM 5159 YP 002521376 1 309801 Chloroflexi
Microlunatus phosphovorus NM 1 YP 004570977 1 1032480 Actinobacteria
Thermomicrobium roseum DSM 5159 YP 002522111 1 309801 Chloroflexi
Sphaerobacter thermophilus DSM 20745 YP 003319357 1 479434 Chloroflexi
Coprothermobacter proteolyticus DSM 5265 YP 002247514 1 309798 Firmicutes
Desulfotomaculum reducens MI 1 YP 001113080 1 349161 Firmicutes
Desulfotomaculum ruminis DSM 2154 YP 004545810 1 696281 Firmicutes
Halanaerobium hydrogeniformans YP 003994052 1 656519 Firmicutes
Staphylothermus marinus F1 YP 001040030 1 399550 Crenarchaeota
Staphylothermus hellenicus DSM 12710 YP 003668805 1 591019 Crenarchaeota
Mesotoga prima MesG1 Ag 4 2 YP 006347034 1 660470 Thermotogae
Kosmotoga olearia TBF 19 5 1 YP 002940722 1 521045 Thermotogae
Thermotoga thermarum DSM 5069 YP 004659616 1 688269 Thermotogae
Fervidobacterium pennivorans DSM 9078 YP 005471104 1 771875 Thermotogae
Fervidobacterium nodosum Rt17 B1 YP 001410107 1 381764 Thermotogae
Octadecabacter antarcticus 307 YP 007706311 1 391626 Proteobacteria
Phaeobacter inhibens DSM 17395 YP 006574208 1 391619 Proteobacteria
Ruegeria pomeroyi DSS 3 YP 165138 1 246200 Proteobacteria
Roseobacter denitrificans OCh 114 YP 684240 1 375451 Proteobacteria
Roseobacter litoralis Och 149 YP 004689349 1 391595 Proteobacteria
Paenibacillus polymyxa M1 YP 005960251 1 1052684 Firmicutes
Paenibacillus terrae HPL 003 YP 005077007 1 985665 Firmicutes
Anoxybacillus flavithermus WK1 YP 002316657 1 491915 Firmicutes
Bacillus megaterium WSH 002 YP 005496871 1 1006007 Firmicutes
Bacillus thuringiensis serovar chinensis CT 43 YP 005570690 1 541229 Firmicutes
Bacillus thuringiensis serovar kurstaki str HD73 YP 007419964 1 1279365 Firmicutes
Bacillus thuringiensis serovar thuringiensis str IS5056 YP 007477176 1 1286404 Firmicutes
Bacillus weihenstephanensis KBAB4 YP 001643536 1 315730 Firmicutes
Bacillus cereus FRI 35 YP 006595007 1 1217984 Firmicutes
Bacillus thuringiensis serovar finitimus YBT 020 YP 005564450 1 930170 Firmicutes
Bacillus toyonensis BCT 7112 YP 008783990 1 1415784 Firmicutes
Bacillus thuringiensis MC28 YP 006826831 1 1195464 Firmicutes

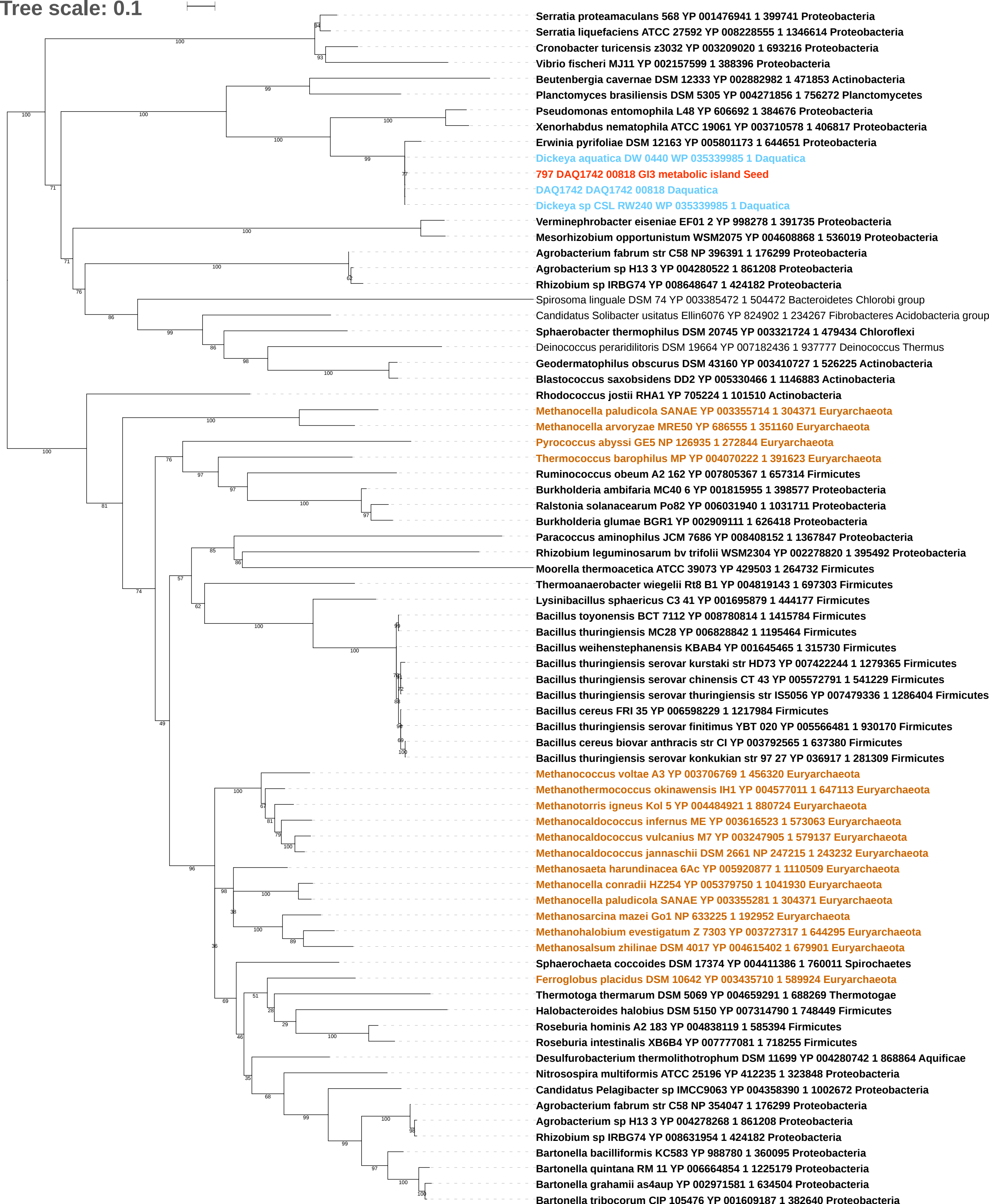
Tree scale: 0.1
Serratia proteamaculans 568 YP 001476941 1 399741 Proteobacteria
Serratia liquefaciens ATCC 27592 YP 008228555 1 1346614 Proteobacteria
Cronobacter turicensis z3032 YP 003209020 1 693216 Proteobacteria
Vibrio fischeri MJ11 YP 002157599 1 388396 Proteobacteria
Beutenbergia cavernae DSM 12333 YP 002882982 1 471853 Actinobacteria
Planctomyces brasiliensis DSM 5305 YP 004271856 1 756272 Planctomycetes
Pseudomonas entomophila L48 YP 606692 1 384676 Proteobacteria
Xenorhabdus nematophila ATCC 19061 YP 003710578 1 406817 Proteobacteria
Erwinia pyrifoliae DSM 12163 YP 005801173 1 644651 Proteobacteria
Dickeya aquatica DW 0440 WP 035339985 1 Daquatica
797 DAQ1742 00818 GI3 metabolic island Seed
DAQ1742 DAQ1742 00818 Daquatica
Dickeya sp CSL RW240 WP 035339985 1 Daquatica
Verminephrobacter eiseniae EF01 2 YP 998278 1 391735 Proteobacteria
Mesorhizobium opportunistum WSM2075 YP 004608868 1 536019 Proteobacteria
Agrobacterium fabrum str C58 NP 396391 1 176299 Proteobacteria
Agrobacterium sp H13 3 YP 004280522 1 861208 Proteobacteria
Rhizobium sp IRBG74 YP 008648647 1 424182 Proteobacteria
Spirosoma linguale DSM 74 YP 003385472 1 504472 Bacteroidetes Chlorobi group
Candidatus Solibacter usitatus Ellin6076 YP 824902 1 234267 Fibrobacteres Acidobacteria group
Sphaerobacter thermophilus DSM 20745 YP 003321724 1 479434 Chloroflexi
Deinococcus peraridilitoris DSM 19664 YP 007182436 1 937777 Deinococcus Thermus
Geodermatophilus obscurus DSM 43160 YP 003410727 1 526225 Actinobacteria
Blastococcus saxobsidens DD2 YP 005330466 1 1146883 Actinobacteria
Rhodococcus jostii RHA1 YP 705224 1 101510 Actinobacteria
Methanocella paludicola SANAE YP 003355714 1 304371 Euryarchaeota
Methanocella arvoryzae MRE50 YP 686555 1 351160 Euryarchaeota
Pyrococcus abyssi GE5 NP 126935 1 272844 Euryarchaeota
Thermococcus barophilus MP YP 004070222 1 391623 Euryarchaeota
Ruminococcus obeum A2 162 YP 007805367 1 657314 Firmicutes
Burkholderia ambifaria MC40 6 YP 001815955 1 398577 Proteobacteria
Ralstonia solanacearum Po82 YP 006031940 1 1031711 Proteobacteria
Burkholderia glumae BGR1 YP 002909111 1 626418 Proteobacteria
Paracoccus aminophilus JCM 7686 YP 008408152 1 1367847 Proteobacteria
Rhizobium leguminosarum bv trifolii WSM2304 YP 002278820 1 395492 Proteobacteria
Moorella thermoacetica ATCC 39073 YP 429503 1 264732 Firmicutes
Thermoanaerobacter wiegelii Rt8 B1 YP 004819143 1 697303 Firmicutes
Lysinibacillus sphaericus C3 41 YP 001695879 1 444177 Firmicutes
Bacillus toyonensis BCT 7112 YP 008780814 1 1415784 Firmicutes
Bacillus thuringiensis MC28 YP 006828842 1 1195464 Firmicutes
Bacillus weihenstephanensis KBAB4 YP 001645465 1 315730 Firmicutes
Bacillus thuringiensis serovar kurstaki str HD73 YP 007422244 1 1279365 Firmicutes
Bacillus thuringiensis serovar chinensis CT 43 YP 005572791 1 541229 Firmicutes
Bacillus thuringiensis serovar thuringiensis str IS5056 YP 007479336 1 1286404 Firmicutes
Bacillus cereus FRI 35 YP 006598229 1 1217984 Firmicutes
Bacillus thuringiensis serovar finitimus YBT 020 YP 005566481 1 930170 Firmicutes
Bacillus cereus biovar anthracis str CI YP 003792565 1 637380 Firmicutes
Bacillus thuringiensis serovar konkukian str 97 27 YP 036917 1 281309 Firmicutes
Methanococcus voltae A3 YP 003706769 1 456320 Euryarchaeota
Methanothermococcus okinawensis IH1 YP 004577011 1 647113 Euryarchaeota
Methanotorris igneus Kol 5 YP 004484921 1 880724 Euryarchaeota
Methanocaldococcus infernus ME YP 003616523 1 573063 Euryarchaeota
Methanocaldococcus vulcanius M7 YP 003247905 1 579137 Euryarchaeota
Methanocaldococcus jannaschii DSM 2661 NP 247215 1 243232 Euryarchaeota
Methanosaeta harundinacea 6Ac YP 005920877 1 1110509 Euryarchaeota
Methanocella conradii HZ254 YP 005379750 1 1041930 Euryarchaeota
Methanocella paludicola SANAE YP 003355281 1 304371 Euryarchaeota
Methanosarcina mazei Go1 NP 633225 1 192952 Euryarchaeota
Methanohalobium evestigatum Z 7303 YP 003727317 1 644295 Euryarchaeota
Methanosalsum zhilinae DSM 4017 YP 004615402 1 679901 Euryarchaeota
Sphaerochaeta coccoides DSM 17374 YP 004411386 1 760011 Spirochaetes
Ferroglobus placidus DSM 10642 YP 003435710 1 589924 Euryarchaeota
Thermotoga thermarum DSM 5069 YP 004659291 1 688269 Thermotogae
Halobacteroides halobius DSM 5150 YP 007314790 1 748449 Firmicutes
Roseburia hominis A2 183 YP 004838119 1 585394 Firmicutes
Roseburia intestinalis XB6B4 YP 007777081 1 718255 Firmicutes
Desulfurobacterium thermolithotrophum DSM 11699 YP 004280742 1 868864 Aquificae
Nitrosospira multiformis ATCC 25196 YP 412235 1 323848 Proteobacteria
Candidatus Pelagibacter sp IMCC9063 YP 004358390 1 1002672 Proteobacteria
Agrobacterium fabrum str C58 NP 354047 1 176299 Proteobacteria
Agrobacterium sp H13 3 YP 004278268 1 861208 Proteobacteria
Rhizobium sp IRBG74 YP 008631954 1 424182 Proteobacteria
Bartonella bacilliformis KC583 YP 988780 1 360095 Proteobacteria
Bartonella quintana RM 11 YP 006664854 1 1225179 Proteobacteria
Bartonella grahamii as4aup YP 002971581 1 634504 Proteobacteria
Bartonella tribocorum CIP 105476 YP 001609187 1 382640 Proteobacteria

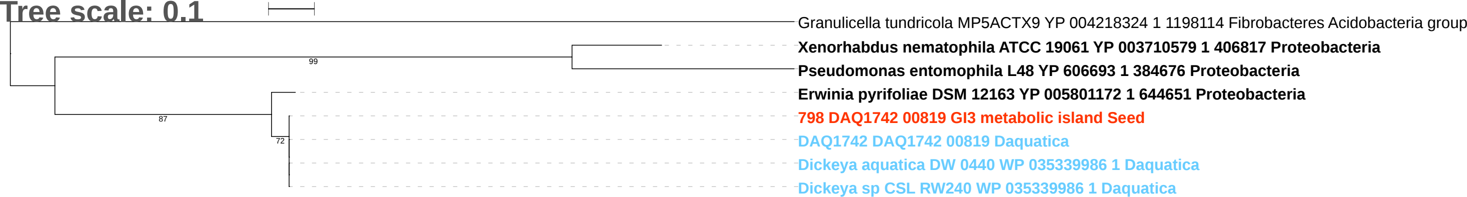

Tree scale: 0.1
Granulicella tundricola MP5ACTX9 YP 004218324 1 1198114 Fibrobacteres Acidobacteria group
Xenorhabdus nematophila ATCC 19061 YP 003710579 1 406817 Proteobacteria
Pseudomonas entomophila L48 YP 606693 1 384676 Proteobacteria
Erwinia pyrifoliae DSM 12163 YP 005801172 1 644651 Proteobacteria
798 DAQ1742 00819 GI3 metabolic island Seed
DAQ1742 DAQ1742 00819 Daquatica
Dickeya aquatica DW 0440 WP 035339986 1 Daquatica
Dickeya sp CSL RW240 WP 035339986 1 Daquatica
99
87
72

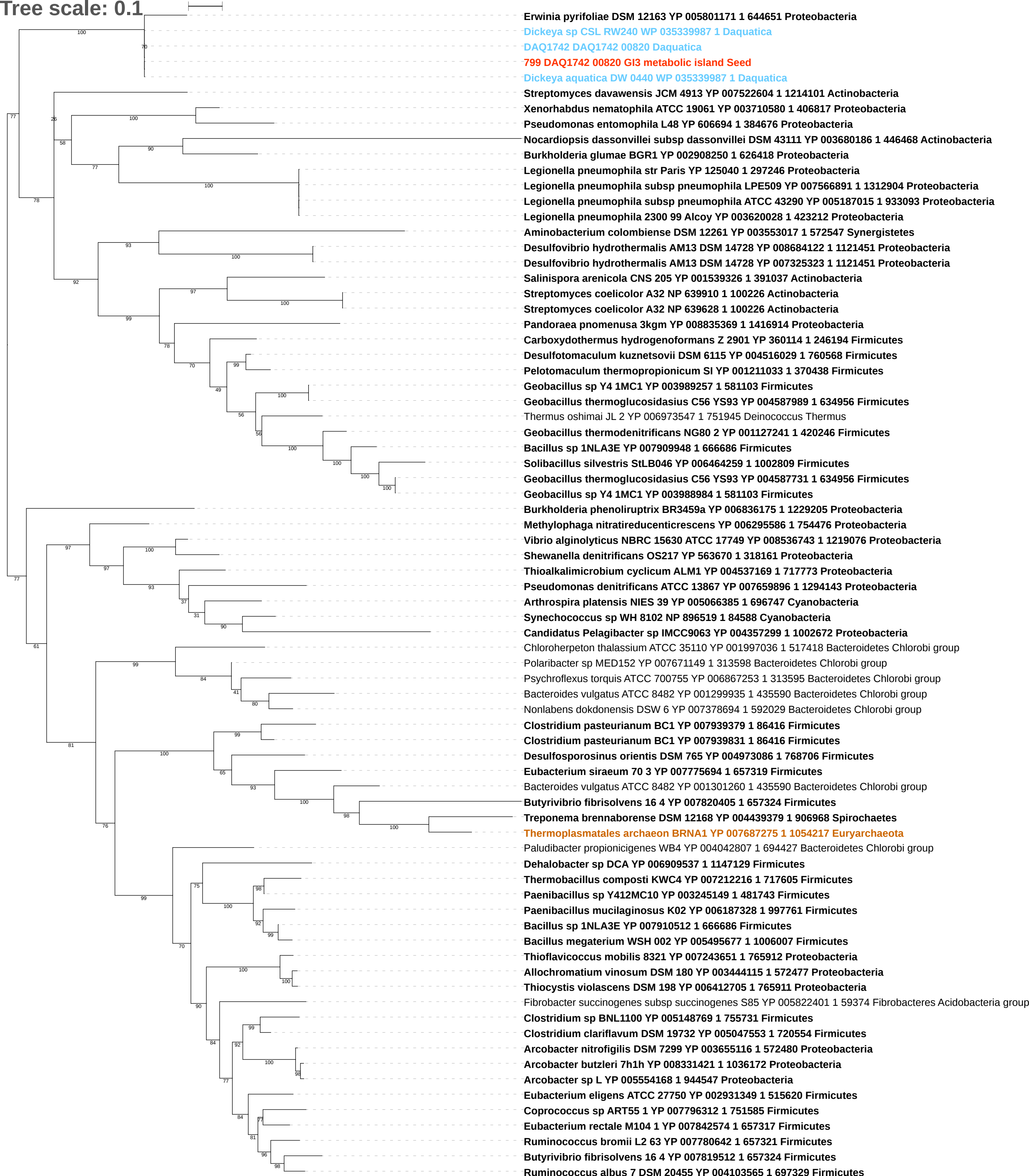
Tree scale: 0.1
Erwinia pyrifoliae DSM 12163 YP 005801171 1 644651 Proteobacteria
Dickeya sp CSL RW240 WP 035339987 1 Daquatica
DAQ1742 DAQ1742 00820 Daquatica
799 DAQ1742 00820 GI3 metabolic island Seed
Dickeya aquatica DW 0440 WP 035339987 1 Daquatica
Streptomyces davawensis JCM 4913 YP 007522604 1 1214101 Actinobacteria
Xenorhabdus nematophila ATCC 19061 YP 003710580 1 406817 Proteobacteria
Pseudomonas entomophila L48 YP 606694 1 384676 Proteobacteria
Nocardiopsis dassonvillei subsp dassonvillei DSM 43111 YP 003680186 1 446468 Actinobacteria
Burkholderia glumae BGR1 YP 002908250 1 626418 Proteobacteria
Legionella pneumophila str Paris YP 125040 1 297246 Proteobacteria
Legionella pneumophila subsp pneumophila LPE509 YP 007566891 1 1312904 Proteobacteria
Legionella pneumophila subsp pneumophila ATCC 43290 YP 005187015 1 933093 Proteobacteria
Legionella pneumophila 2300 99 Alcoy YP 003620028 1 423212 Proteobacteria
Aminobacterium colombiense DSM 12261 YP 003553017 1 572547 Synergistetes
Desulfovibrio hydrothermalis AM13 DSM 14728 YP 008684122 1 1121451 Proteobacteria
Desulfovibrio hydrothermalis AM13 DSM 14728 YP 007325323 1 1121451 Proteobacteria
Salinispora arenicola CNS 205 YP 001539326 1 391037 Actinobacteria
Streptomyces coelicolor A32 NP 639910 1 100226 Actinobacteria
Streptomyces coelicolor A32 NP 639628 1 100226 Actinobacteria
Pandoraea pnomenusa 3kgm YP 008835369 1 1416914 Proteobacteria
Carboxydothermus hydrogenoformans Z 2901 YP 360114 1 246194 Firmicutes
Desulfotomaculum kuznetsovii DSM 6115 YP 004516029 1 760568 Firmicutes
Pelotomaculum thermopropionicum SI YP 001211033 1 370438 Firmicutes
Geobacillus sp Y4 1MC1 YP 003989257 1 581103 Firmicutes
Geobacillus thermoglucosidasius C56 YS93 YP 004587989 1 634956 Firmicutes
Thermus oshimai JL 2 YP 006973547 1 751945 Deinococcus Thermus
Geobacillus thermodenitrificans NG80 2 YP 001127241 1 420246 Firmicutes
Bacillus sp 1NLA3E YP 007909948 1 666686 Firmicutes
Solibacillus silvestris StLB046 YP 006464259 1 1002809 Firmicutes
Geobacillus thermoglucosidasius C56 YS93 YP 004587731 1 634956 Firmicutes
Geobacillus sp Y4 1MC1 YP 003988984 1 581103 Firmicutes
Burkholderia phenoliruptrix BR3459a YP 006836175 1 1229205 Proteobacteria
Methylophaga nitratireducenticrescens YP 006295586 1 754476 Proteobacteria
Vibrio alginolyticus NBRC 15630 ATCC 17749 YP 008536743 1 1219076 Proteobacteria
Shewanella denitrificans OS217 YP 563670 1 318161 Proteobacteria
Thioalkalimicrobium cyclicum ALM1 YP 004537169 1 717773 Proteobacteria
Pseudomonas denitrificans ATCC 13867 YP 007659896 1 1294143 Proteobacteria
Arthrospira platensis NIES 39 YP 005066385 1 696747 Cyanobacteria
Synechococcus sp WH 8102 NP 896519 1 84588 Cyanobacteria
Candidatus Pelagibacter sp IMCC9063 YP 004357299 1 1002672 Proteobacteria
Chloroherpeton thalassium ATCC 35110 YP 001997036 1 517418 Bacteroidetes Chlorobi group
Polaribacter sp MED152 YP 007671149 1 313598 Bacteroidetes Chlorobi group
Psychroflexus torquis ATCC 700755 YP 006867253 1 313595 Bacteroidetes Chlorobi group
Bacteroides vulgatus ATCC 8482 YP 001299935 1 435590 Bacteroidetes Chlorobi group
Nonlabens dokdonensis DSW 6 YP 007378694 1 592029 Bacteroidetes Chlorobi group
Clostridium pasteurianum BC1 YP 007939379 1 86416 Firmicutes
Clostridium pasteurianum BC1 YP 007939831 1 86416 Firmicutes
Desulfosporosinus orientis DSM 765 YP 004973086 1 768706 Firmicutes
Eubacterium siraeum 70 3 YP 007775694 1 657319 Firmicutes
Bacteroides vulgatus ATCC 8482 YP 001301260 1 435590 Bacteroidetes Chlorobi group
Butyrivibrio fibrisolvens 16 4 YP 007820405 1 657324 Firmicutes
Treponema brennaborense DSM 12168 YP 004439379 1 906968 Spirochaetes
Thermoplasmatales archaeon BRNA1 YP 007687275 1 1054217 Euryarchaeota
Paludibacter propionicigenes WB4 YP 004042807 1 694427 Bacteroidetes Chlorobi group
Dehalobacter sp DCA YP 006909537 1 1147129 Firmicutes
Thermobacillus composti KWC4 YP 007212216 1 717605 Firmicutes
Paenibacillus sp Y412MC10 YP 003245149 1 481743 Firmicutes
Paenibacillus mucilaginosus K02 YP 006187328 1 997761 Firmicutes
Bacillus sp 1NLA3E YP 007910512 1 666686 Firmicutes
Bacillus megaterium WSH 002 YP 005495677 1 1006007 Firmicutes
Thioflavicoccus mobilis 8321 YP 007243651 1 765912 Proteobacteria
Allochromatium vinosum DSM 180 YP 003444115 1 572477 Proteobacteria
Thiocystis violascens DSM 198 YP 006412705 1 765911 Proteobacteria
Fibrobacter succinogenes subsp succinogenes S85 YP 005822401 1 59374 Fibrobacteres Acidobacteria group
Clostridium sp BNL1100 YP 005148769 1 755731 Firmicutes
Clostridium clariflavum DSM 19732 YP 005047553 1 720554 Firmicutes
Arcobacter nitrofigilis DSM 7299 YP 003655116 1 572480 Proteobacteria
Arcobacter butzleri 7h1h YP 008331421 1 1036172 Proteobacteria
Arcobacter sp L YP 005554168 1 944547 Proteobacteria
Eubacterium eligens ATCC 27750 YP 002931349 1 515620 Firmicutes
Coprococcus sp ART55 1 YP 007796312 1 751585 Firmicutes
Eubacterium rectale M104 1 YP 007842574 1 657317 Firmicutes
Ruminococcus bromii L2 63 YP 007780642 1 657321 Firmicutes
Butyrivibrio fibrisolvens 16 4 YP 007819512 1 657324 Firmicutes
Ruminococcus albus 7 DSM 20455 YP 004103565 1 697329 Firmicutes

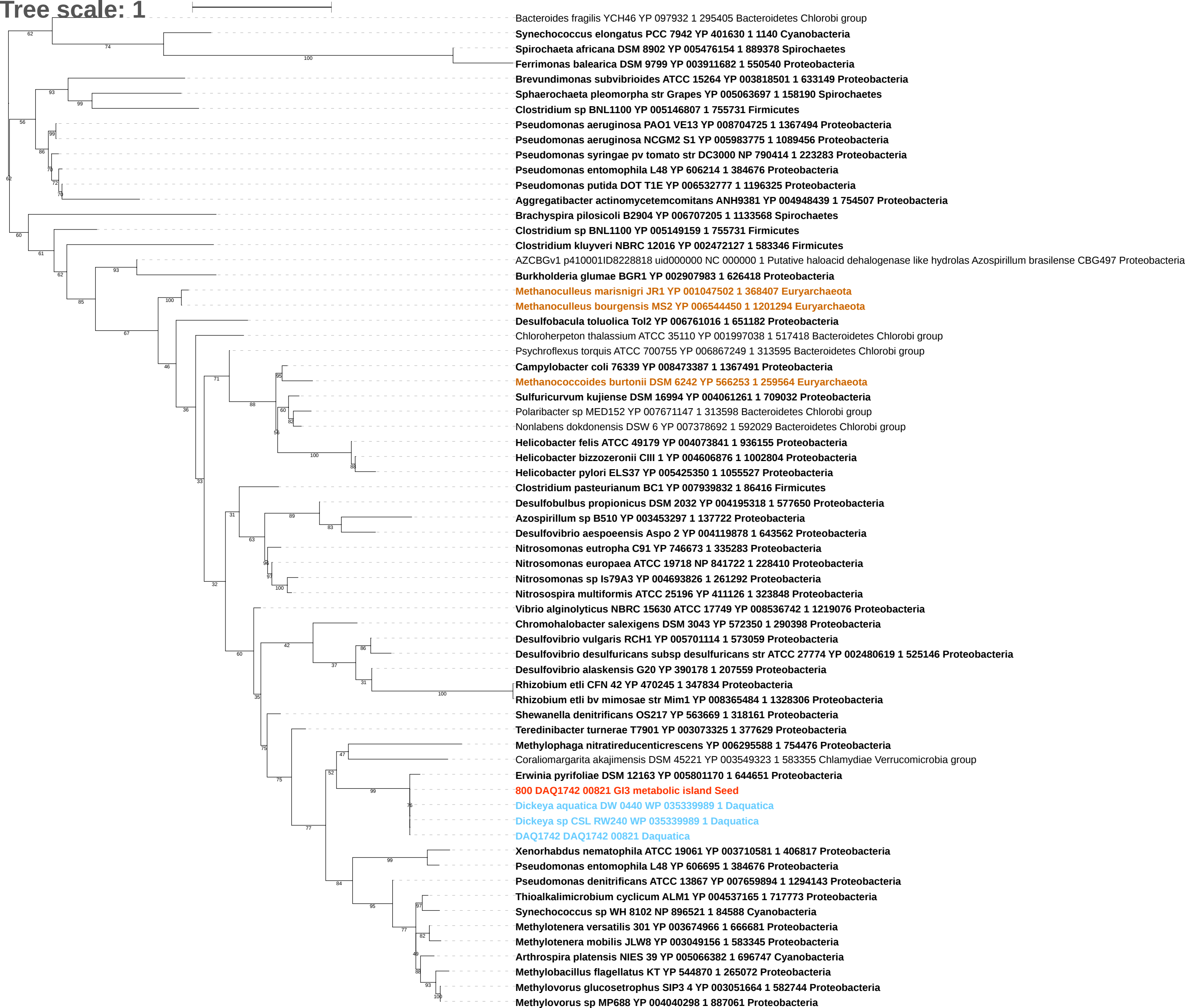
Tree scale: 1
Bacteroides fragilis YCH46 YP 097932 1 295405 Bacteroidetes Chlorobi group
Synechococcus elongatus PCC 7942 YP 401630 1 1140 Cyanobacteria
Spirochaeta africana DSM 8902 YP 005476154 1 889378 Spirochaetes
Ferrimonas balearica DSM 9799 YP 003911682 1 550540 Proteobacteria
Brevundimonas subvibrioides ATCC 15264 YP 003818501 1 633149 Proteobacteria
Sphaerochaeta pleomorpha str Grapes YP 005063697 1 158190 Spirochaetes
Clostridium sp BNL1100 YP 005146807 1 755731 Firmicutes
Pseudomonas aeruginosa PAO1 VE13 YP 008704725 1 1367494 Proteobacteria
Pseudomonas aeruginosa NCGM2 S1 YP 005983775 1 1089456 Proteobacteria
Pseudomonas syringae pv tomato str DC3000 NP 790414 1 223283 Proteobacteria
Pseudomonas entomophila L48 YP 606214 1 384676 Proteobacteria
Pseudomonas putida DOT T1E YP 006532777 1 1196325 Proteobacteria
Aggregatibacter actinomycetemcomitans ANH9381 YP 004948439 1 754507 Proteobacteria
Brachyspira pilosicoli B2904 YP 006707205 1 1133568 Spirochaetes
Clostridium sp BNL1100 YP 005149159 1 755731 Firmicutes
Clostridium kluyveri NBRC 12016 YP 002472127 1 583346 Firmicutes
AZCBGv1 p410001ID8228818 uid000000 NC 000000 1 Putative haloacid dehalogenase like hydrolas Azospirillum brasilense CBG497 Proteobacteria
Burkholderia glumae BGR1 YP 002907983 1 626418 Proteobacteria
Methanoculleus marisnigri JR1 YP 001047502 1 368407 Euryarchaeota
Methanoculleus bourgensis MS2 YP 006544450 1 1201294 Euryarchaeota
Desulfobacula toluolica Tol2 YP 006761016 1 651182 Proteobacteria
Chloroherpeton thalassium ATCC 35110 YP 001997038 1 517418 Bacteroidetes Chlorobi group
Psychroflexus torquis ATCC 700755 YP 006867249 1 313595 Bacteroidetes Chlorobi group
Campylobacter coli 76339 YP 008473387 1 1367491 Proteobacteria
Methanococcoides burtonii DSM 6242 YP 566253 1 259564 Euryarchaeota
Sulfuricurvum kujiense DSM 16994 YP 004061261 1 709032 Proteobacteria
Polaribacter sp MED152 YP 007671147 1 313598 Bacteroidetes Chlorobi group
Nonlabens dokdonensis DSW 6 YP 007378692 1 592029 Bacteroidetes Chlorobi group
Helicobacter felis ATCC 49179 YP 004073841 1 936155 Proteobacteria
Helicobacter bizzozeronii CIII 1 YP 004606876 1 1002804 Proteobacteria
Helicobacter pylori ELS37 YP 005425350 1 1055527 Proteobacteria
Clostridium pasteurianum BC1 YP 007939832 1 86416 Firmicutes
Desulfobulbus propionicus DSM 2032 YP 004195318 1 577650 Proteobacteria
Azospirillum sp B510 YP 003453297 1 137722 Proteobacteria
Desulfovibrio aespoeensis Aspo 2 YP 004119878 1 643562 Proteobacteria
Nitrosomonas eutropha C91 YP 746673 1 335283 Proteobacteria
Nitrosomonas europaea ATCC 19718 NP 841722 1 228410 Proteobacteria
Nitrosomonas sp Is79A3 YP 004693826 1 261292 Proteobacteria
Nitrosospira multiformis ATCC 25196 YP 411126 1 323848 Proteobacteria
Vibrio alginolyticus NBRC 15630 ATCC 17749 YP 008536742 1 1219076 Proteobacteria
Chromohalobacter salexigens DSM 3043 YP 572350 1 290398 Proteobacteria
Desulfovibrio vulgaris RCH1 YP 005701114 1 573059 Proteobacteria
Desulfovibrio desulfuricans subsp desulfuricans str ATCC 27774 YP 002480619 1 525146 Proteobacteria
Desulfovibrio alaskensis G20 YP 390178 1 207559 Proteobacteria
Rhizobium etli CFN 42 YP 470245 1 347834 Proteobacteria
Rhizobium etli bv mimosae str Mim1 YP 008365484 1 1328306 Proteobacteria
Shewanella denitrificans OS217 YP 563669 1 318161 Proteobacteria
Teredinibacter turnerae T7901 YP 003073325 1 377629 Proteobacteria
Methylophaga nitratireducenticrescens YP 006295588 1 754476 Proteobacteria
Coraliomargarita akajimensis DSM 45221 YP 003549323 1 583355 Chlamydiae Verrucomicrobia group
Erwinia pyrifoliae DSM 12163 YP 005801170 1 644651 Proteobacteria
800 DAQ1742 00821 GI3 metabolic island Seed
Dickeya aquatica DW 0440 WP 035339989 1 Daquatica
Dickeya sp CSL RW240 WP 035339989 1 Daquatica
DAQ1742 DAQ1742 00821 Daquatica
Xenorhabdus nematophila ATCC 19061 YP 003710581 1 406817 Proteobacteria
Pseudomonas entomophila L48 YP 606695 1 384676 Proteobacteria
Pseudomonas denitrificans ATCC 13867 YP 007659894 1 1294143 Proteobacteria
Thioalkalimicrobium cyclicum ALM1 YP 004537165 1 717773 Proteobacteria
Synechococcus sp WH 8102 NP 896521 1 84588 Cyanobacteria
Methylotenera versatilis 301 YP 003674966 1 666681 Proteobacteria
Methylotenera mobilis JLW8 YP 003049156 1 583345 Proteobacteria
Arthrospira platensis NIES 39 YP 005066382 1 696747 Cyanobacteria
Methylobacillus flagellatus KT YP 544870 1 265072 Proteobacteria
Methylovorus glucosetrophus SIP3 4 YP 003051664 1 582744 Proteobacteria
Methylovorus sp MP688 YP 004040298 1 887061 Proteobacteria

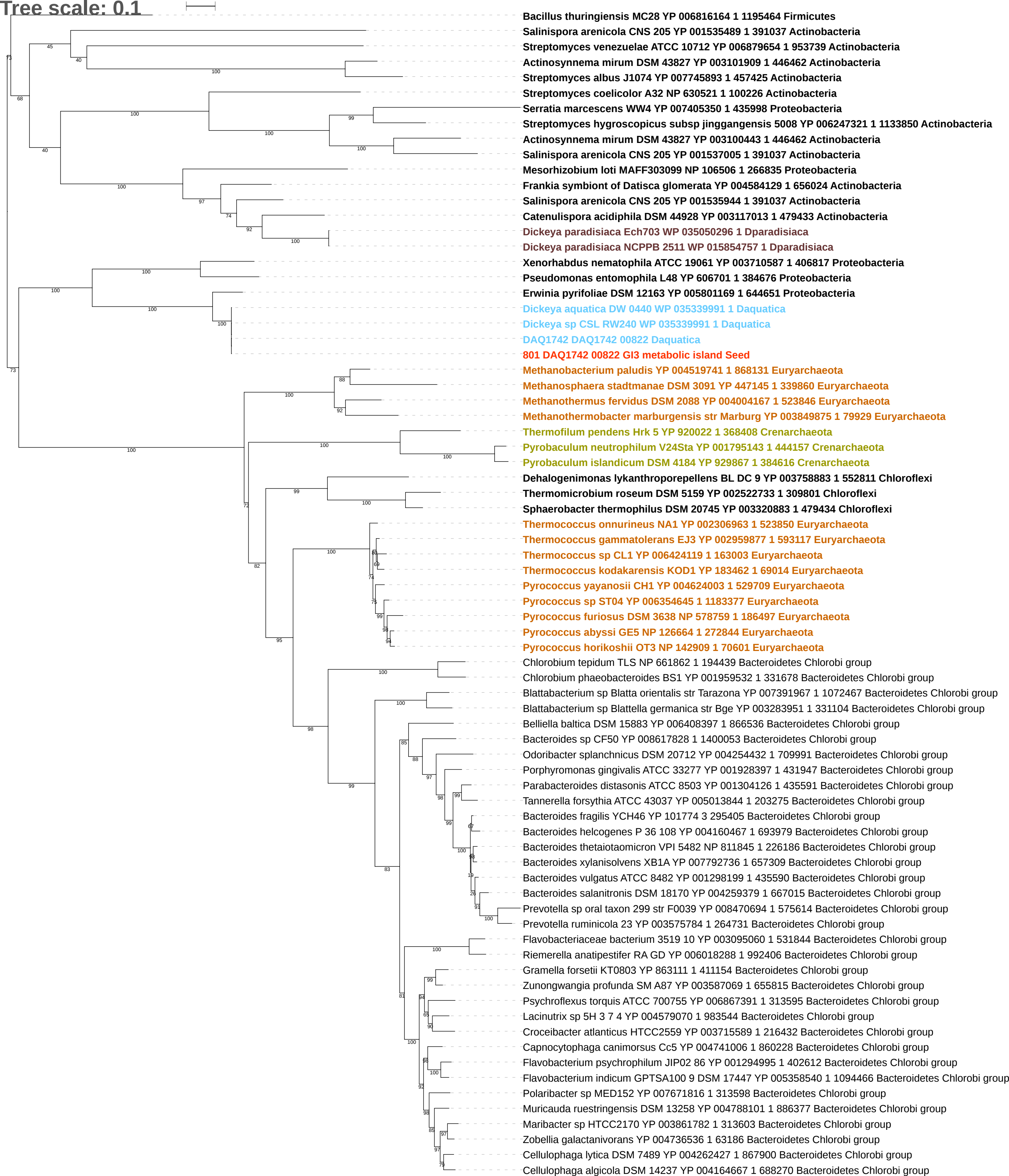
Tree scale: 0.1
Bacillus thuringiensis MC28 YP 006816164 1 1195464 Firmicutes
Salinispora arenicola CNS 205 YP 001535489 1 391037 Actinobacteria
Streptomyces venezuelae ATCC 10712 YP 006879654 1 953739 Actinobacteria
Actinosynnema mirum DSM 43827 YP 003101909 1 446462 Actinobacteria
Streptomyces albus J1074 YP 007745893 1 457425 Actinobacteria
Streptomyces coelicolor A32 NP 630521 1 100226 Actinobacteria
Serratia marcescens WW4 YP 007405350 1 435998 Proteobacteria
Streptomyces hygroscopicus subsp jinggangensis 5008 YP 006247321 1 1133850 Actinobacteria
Actinosynnema mirum DSM 43827 YP 003100443 1 446462 Actinobacteria
Salinispora arenicola CNS 205 YP 001537005 1 391037 Actinobacteria
Mesorhizobium loti MAFF303099 NP 106506 1 266835 Proteobacteria
Frankia symbiont of Datisca glomerata YP 004584129 1 656024 Actinobacteria
Salinispora arenicola CNS 205 YP 001535944 1 391037 Actinobacteria
Catenulispora acidiphila DSM 44928 YP 003117013 1 479433 Actinobacteria
Dickeya paradisiaca Ech703 WP 035050296 1 Dparadisiaca
Dickeya paradisiaca NCPPB 2511 WP 015854757 1 Dparadisiaca
Xenorhabdus nematophila ATCC 19061 YP 003710587 1 406817 Proteobacteria
Pseudomonas entomophila L48 YP 606701 1 384676 Proteobacteria
Erwinia pyrifoliae DSM 12163 YP 005801169 1 644651 Proteobacteria
Dickeya aquatica DW 0440 WP 035339991 1 Daquatica
Dickeya sp CSL RW240 WP 035339991 1 Daquatica
DAQ1742 DAQ1742 00822 Daquatica
801 DAQ1742 00822 GI3 metabolic island Seed
Methanobacterium paludis YP 004519741 1 868131 Euryarchaeota
Methanosphaera stadtmanae DSM 3091 YP 447145 1 339860 Euryarchaeota
Methanothermus fervidus DSM 2088 YP 004004167 1 523846 Euryarchaeota
Methanothermobacter marburgensis str Marburg YP 003849875 1 79929 Euryarchaeota
Thermofilum pendens Hrk 5 YP 920022 1 368408 Crenarchaeota
Pyrobaculum neutrophilum V24Sta YP 001795143 1 444157 Crenarchaeota
Pyrobaculum islandicum DSM 4184 YP 929867 1 384616 Crenarchaeota
Dehalogenimonas lykanthroporepellens BL DC 9 YP 003758883 1 552811 Chloroflexi
Thermomicrobium roseum DSM 5159 YP 002522733 1 309801 Chloroflexi
Sphaerobacter thermophilus DSM 20745 YP 003320883 1 479434 Chloroflexi
Thermococcus onnurineus NA1 YP 002306963 1 523850 Euryarchaeota
Thermococcus gammatolerans EJ3 YP 002959877 1 593117 Euryarchaeota
Thermococcus sp CL1 YP 006424119 1 163003 Euryarchaeota
Thermococcus kodakarensis KOD1 YP 183462 1 69014 Euryarchaeota
Pyrococcus yayanosii CH1 YP 004624003 1 529709 Euryarchaeota
Pyrococcus sp ST04 YP 006354645 1 1183377 Euryarchaeota
Pyrococcus furiosus DSM 3638 NP 578759 1 186497 Euryarchaeota
Pyrococcus abyssi GE5 NP 126664 1 272844 Euryarchaeota
Pyrococcus horikoshii OT3 NP 142909 1 70601 Euryarchaeota
Chlorobium tepidum TLS NP 661862 1 194439 Bacteroidetes Chlorobi group
Chlorobium phaeobacteroides BS1 YP 001959532 1 331678 Bacteroidetes Chlorobi group
Blattabacterium sp Blatta orientalis str Tarazona YP 007391967 1 1072467 Bacteroidetes Chlorobi group
Blattabacterium sp Blattella germanica str Bge YP 003283951 1 331104 Bacteroidetes Chlorobi group
Belliella baltica DSM 15883 YP 006408397 1 866536 Bacteroidetes Chlorobi group
Bacteroides sp CF50 YP 008617828 1 1400053 Bacteroidetes Chlorobi group
Odoribacter splanchnicus DSM 20712 YP 004254432 1 709991 Bacteroidetes Chlorobi group
Porphyromonas gingivalis ATCC 33277 YP 001928397 1 431947 Bacteroidetes Chlorobi group
Parabacteroides distasonis ATCC 8503 YP 001304126 1 435591 Bacteroidetes Chlorobi group
Tannerella forsythia ATCC 43037 YP 005013844 1 203275 Bacteroidetes Chlorobi group
Bacteroides fragilis YCH46 YP 101774 3 295405 Bacteroidetes Chlorobi group
Bacteroides helcogenes P 36 108 YP 004160467 1 693979 Bacteroidetes Chlorobi group
Bacteroides thetaiotaomicron VPI 5482 NP 811845 1 226186 Bacteroidetes Chlorobi group
Bacteroides xylanisolvens XB1A YP 007792736 1 657309 Bacteroidetes Chlorobi group
Bacteroides vulgatus ATCC 8482 YP 001298199 1 435590 Bacteroidetes Chlorobi group
Bacteroides salanitronis DSM 18170 YP 004259379 1 667015 Bacteroidetes Chlorobi group
Prevotella sp oral taxon 299 str F0039 YP 008470694 1 575614 Bacteroidetes Chlorobi group
Prevotella ruminicola 23 YP 003575784 1 264731 Bacteroidetes Chlorobi group
Flavobacteriaceae bacterium 3519 10 YP 003095060 1 531844 Bacteroidetes Chlorobi group
Riemerella anatipestifer RA GD YP 006018288 1 992406 Bacteroidetes Chlorobi group
Gramella forsetii KT0803 YP 863111 1 411154 Bacteroidetes Chlorobi group
Zunongwangia profunda SM A87 YP 003587069 1 655815 Bacteroidetes Chlorobi group
Psychroflexus torquis ATCC 700755 YP 006867391 1 313595 Bacteroidetes Chlorobi group
Lacinutrix sp 5H 3 7 4 YP 004579070 1 983544 Bacteroidetes Chlorobi group
Croceibacter atlanticus HTCC2559 YP 003715589 1 216432 Bacteroidetes Chlorobi group
Capnocytophaga canimorsus Cc5 YP 004741006 1 860228 Bacteroidetes Chlorobi group
Flavobacterium psychrophilum JIP02 86 YP 001294995 1 402612 Bacteroidetes Chlorobi group
Flavobacterium indicum GPTSA100 9 DSM 17447 YP 005358540 1 1094466 Bacteroidetes Chlorobi group
Polaribacter sp MED152 YP 007671816 1 313598 Bacteroidetes Chlorobi group
Muricauda ruestringensis DSM 13258 YP 004788101 1 886377 Bacteroidetes Chlorobi group
Maribacter sp HTCC2170 YP 003861782 1 313603 Bacteroidetes Chlorobi group
Zobellia galactanivorans YP 004736536 1 63186 Bacteroidetes Chlorobi group
Cellulophaga lytica DSM 7489 YP 004262427 1 867900 Bacteroidetes Chlorobi group
Cellulophaga algicola DSM 14237 YP 004164667 1 688270 Bacteroidetes Chlorobi group

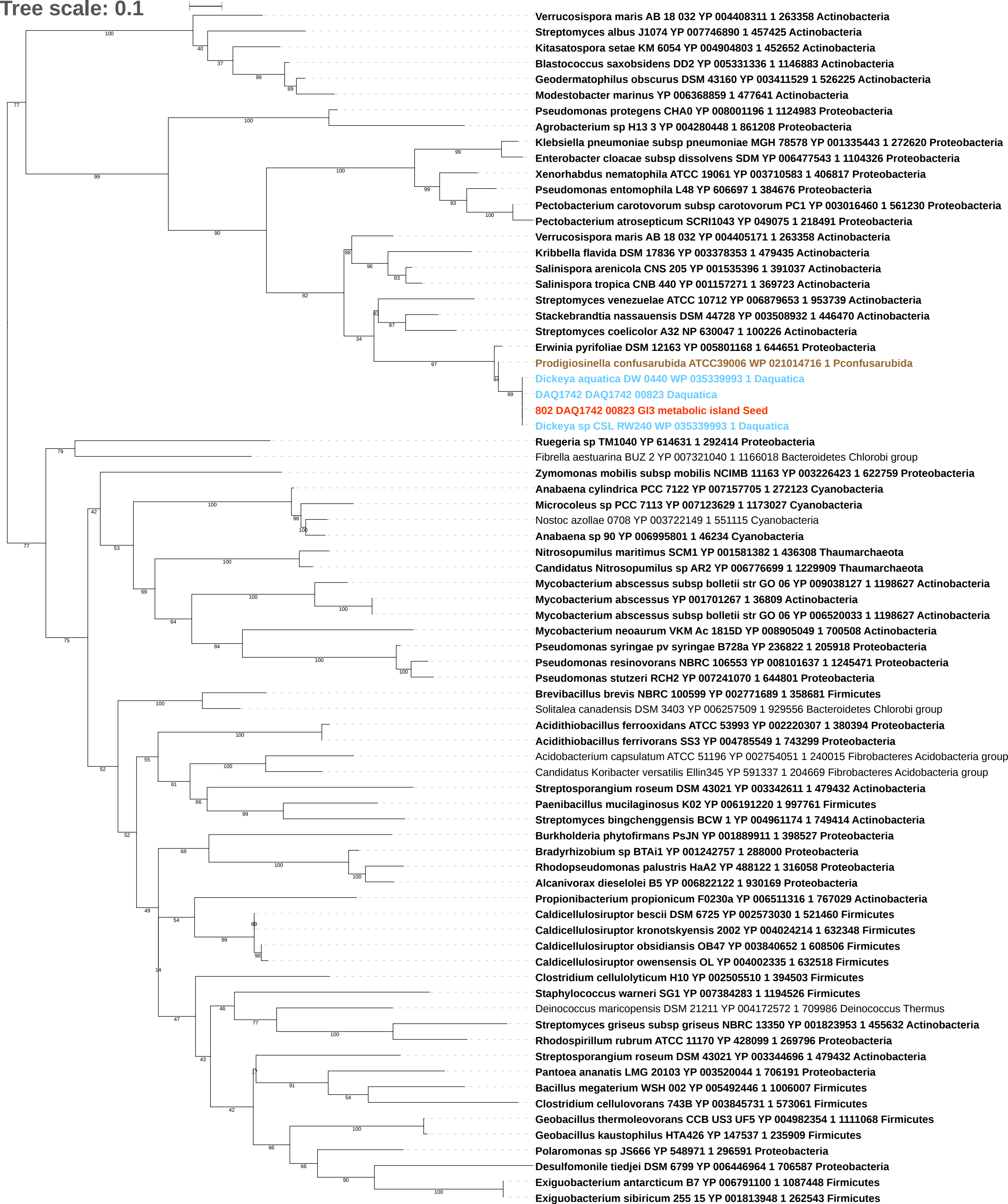
Tree scale: 0.1
Verrucosispora maris AB 18 032 YP 004408311 1 263358 Actinobacteria
Streptomyces albus J1074 YP 007746890 1 457425 Actinobacteria
Kitasatospora setae KM 6054 YP 004904803 1 452652 Actinobacteria
Blastococcus saxobsidens DD2 YP 005331336 1 1146883 Actinobacteria
Geodermatophilus obscurus DSM 43160 YP 003411529 1 526225 Actinobacteria
Modestobacter marinus YP 006368859 1 477641 Actinobacteria
Pseudomonas protegens CHA0 YP 008001196 1 1124983 Proteobacteria
Agrobacterium sp H13 3 YP 004280448 1 861208 Proteobacteria
Klebsiella pneumoniae subsp pneumoniae MGH 78578 YP 001335443 1 272620 Proteobacteria
Enterobacter cloacae subsp dissolvens SDM YP 006477543 1 1104326 Proteobacteria
Xenorhabdus nematophila ATCC 19061 YP 003710583 1 406817 Proteobacteria
Pseudomonas entomophila L48 YP 606697 1 384676 Proteobacteria
Pectobacterium carotovorum subsp carotovorum PC1 YP 003016460 1 561230 Proteobacteria
Pectobacterium atrosepticum SCRI1043 YP 049075 1 218491 Proteobacteria
Verrucosispora maris AB 18 032 YP 004405171 1 263358 Actinobacteria
Kribbella flavida DSM 17836 YP 003378353 1 479435 Actinobacteria
Salinispora arenicola CNS 205 YP 001535396 1 391037 Actinobacteria
Salinispora tropica CNB 440 YP 001157271 1 369723 Actinobacteria
Streptomyces venezuelae ATCC 10712 YP 006879653 1 953739 Actinobacteria
Stackebrandtia nassauensis DSM 44728 YP 003508932 1 446470 Actinobacteria
Streptomyces coelicolor A32 NP 630047 1 100226 Actinobacteria
Erwinia pyrifoliae DSM 12163 YP 005801168 1 644651 Proteobacteria
Prodigiosinella confusarubida ATCC39006 WP 021014716 1 Pconfusarubida
Dickeya aquatica DW 0440 WP 035339993 1 Daquatica
DAQ1742 DAQ1742 00823 Daquatica
802 DAQ1742 00823 GI3 metabolic island Seed
Dickeya sp CSL RW240 WP 035339993 1 Daquatica
Ruegeria sp TM1040 YP 614631 1 292414 Proteobacteria
Fibrella aestuarina BUZ 2 YP 007321040 1 1166018 Bacteroidetes Chlorobi group
Zymomonas mobilis subsp mobilis NCIMB 11163 YP 003226423 1 622759 Proteobacteria
Anabaena cylindrica PCC 7122 YP 007157705 1 272123 Cyanobacteria
Microcoleus sp PCC 7113 YP 007123629 1 1173027 Cyanobacteria
Nostoc azollae 0708 YP 003722149 1 551115 Cyanobacteria
Anabaena sp 90 YP 006995801 1 46234 Cyanobacteria
Nitrosopumilus maritimus SCM1 YP 001581382 1 436308 Thaumarchaeota
Candidatus Nitrosopumilus sp AR2 YP 006776699 1 1229909 Thaumarchaeota
Mycobacterium abscessus subsp bolletii str GO 06 YP 009038127 1 1198627 Actinobacteria
Mycobacterium abscessus YP 001701267 1 36809 Actinobacteria
Mycobacterium abscessus subsp bolletii str GO 06 YP 006520033 1 1198627 Actinobacteria
Mycobacterium neoaurum VKM Ac 1815D YP 008905049 1 700508 Actinobacteria
Pseudomonas syringae pv syringae B728a YP 236822 1 205918 Proteobacteria
Pseudomonas resinovorans NBRC 106553 YP 008101637 1 1245471 Proteobacteria
Pseudomonas stutzeri RCH2 YP 007241070 1 644801 Proteobacteria
Brevibacillus brevis NBRC 100599 YP 002771689 1 358681 Firmicutes
Solitalea canadensis DSM 3403 YP 006257509 1 929556 Bacteroidetes Chlorobi group
Acidithiobacillus ferrooxidans ATCC 53993 YP 002220307 1 380394 Proteobacteria
Acidithiobacillus ferrivorans SS3 YP 004785549 1 743299 Proteobacteria
Acidobacterium capsulatum ATCC 51196 YP 002754051 1 240015 Fibrobacteres Acidobacteria group
Candidatus Koribacter versatilis Ellin345 YP 591337 1 204669 Fibrobacteres Acidobacteria group
Streptosporangium roseum DSM 43021 YP 003342611 1 479432 Actinobacteria
Paenibacillus mucilaginosus K02 YP 006191220 1 997761 Firmicutes
Streptomyces bingchenggensis BCW 1 YP 004961174 1 749414 Actinobacteria
Burkholderia phytofirmans PsJN YP 001889911 1 398527 Proteobacteria
Bradyrhizobium sp BTAi1 YP 001242757 1 288000 Proteobacteria
Rhodopseudomonas palustris HaA2 YP 488122 1 316058 Proteobacteria
Alcanivorax dieselolei B5 YP 006822122 1 930169 Proteobacteria
Propionibacterium propionicum F0230a YP 006511316 1 767029 Actinobacteria
Caldicellulosiruptor bescii DSM 6725 YP 002573030 1 521460 Firmicutes
Caldicellulosiruptor kronotskyensis 2002 YP 004024214 1 632348 Firmicutes
Caldicellulosiruptor obsidiansis OB47 YP 003840652 1 608506 Firmicutes
Caldicellulosiruptor owensensis OL YP 004002335 1 632518 Firmicutes
Clostridium cellulolyticum H10 YP 002505510 1 394503 Firmicutes
Staphylococcus warneri SG1 YP 007384283 1 1194526 Firmicutes
Deinococcus maricopensis DSM 21211 YP 004172572 1 709986 Deinococcus Thermus
Streptomyces griseus subsp griseus NBRC 13350 YP 001823953 1 455632 Actinobacteria
Rhodospirillum rubrum ATCC 11170 YP 428099 1 269796 Proteobacteria
Streptosporangium roseum DSM 43021 YP 003344696 1 479432 Actinobacteria
Pantoea ananatis LMG 20103 YP 003520044 1 706191 Proteobacteria
Bacillus megaterium WSH 002 YP 005492446 1 1006007 Firmicutes
Clostridium cellulovorans 743B YP 003845731 1 573061 Firmicutes
Geobacillus thermoleovorans CCB US3 UF5 YP 004982354 1 1111068 Firmicutes
Geobacillus kaustophilus HTA426 YP 147537 1 235909 Firmicutes
Polaromonas sp JS666 YP 548971 1 296591 Proteobacteria
Desulfomonile tiedjei DSM 6799 YP 006446964 1 706587 Proteobacteria
Exiguobacterium antarcticum B7 YP 006791100 1 1087448 Firmicutes
Exiguobacterium sibiricum 255 15 YP 001813948 1 262543 Firmicutes

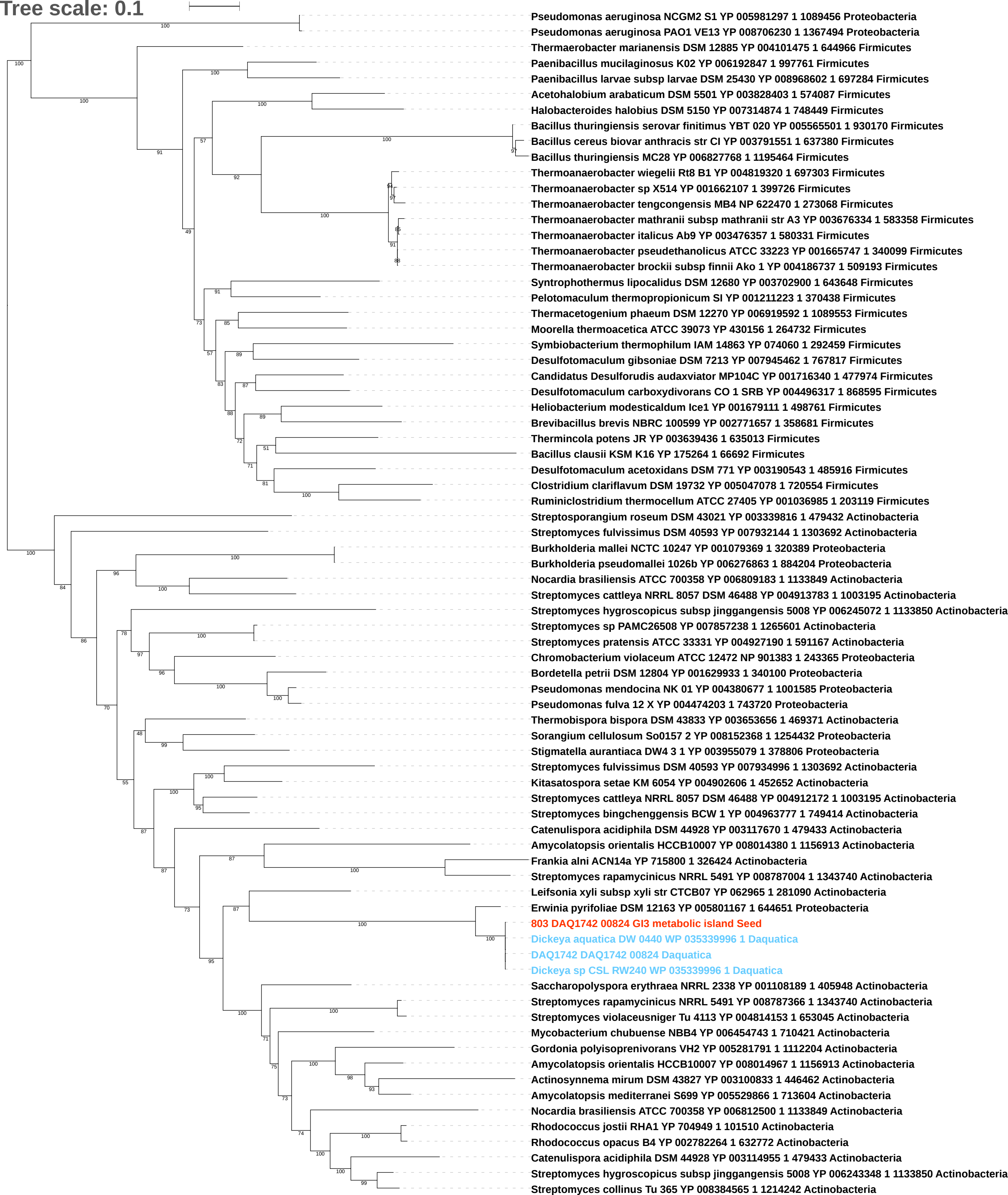
Tree scale: 0.1
Pseudomonas aeruginosa NCGM2 S1 YP 005981297 1 1089456 Proteobacteria
Pseudomonas aeruginosa PAO1 VE13 YP 008706230 1 1367494 Proteobacteria
Thermaerobacter marianensis DSM 12885 YP 004101475 1 644966 Firmicutes
Paenibacillus mucilaginosus K02 YP 006192847 1 997761 Firmicutes
Paenibacillus larvae subsp larvae DSM 25430 YP 008968602 1 697284 Firmicutes
Acetohalobium arabaticum DSM 5501 YP 003828403 1 574087 Firmicutes
Halobacteroides halobius DSM 5150 YP 007314874 1 748449 Firmicutes
Bacillus thuringiensis serovar finitimus YBT 020 YP 005565501 1 930170 Firmicutes
Bacillus cereus biovar anthracis str CI YP 003791551 1 637380 Firmicutes
Bacillus thuringiensis MC28 YP 006827768 1 1195464 Firmicutes
Thermoanaerobacter wiegelii Rt8 B1 YP 004819320 1 697303 Firmicutes
Thermoanaerobacter sp X514 YP 001662107 1 399726 Firmicutes
Thermoanaerobacter tengcongensis MB4 NP 622470 1 273068 Firmicutes
Thermoanaerobacter mathranii subsp mathranii str A3 YP 003676334 1 583358 Firmicutes
Thermoanaerobacter italicus Ab9 YP 003476357 1 580331 Firmicutes
Thermoanaerobacter pseudethanolicus ATCC 33223 YP 001665747 1 340099 Firmicutes
Thermoanaerobacter brockii subsp finnii Ako 1 YP 004186737 1 509193 Firmicutes
Syntrophothermus lipocalidus DSM 12680 YP 003702900 1 643648 Firmicutes
Pelotomaculum thermopropionicum SI YP 001211223 1 370438 Firmicutes
Thermacetogenium phaeum DSM 12270 YP 006919592 1 1089553 Firmicutes
Moorella thermoacetica ATCC 39073 YP 430156 1 264732 Firmicutes
Symbiobacterium thermophilum IAM 14863 YP 074060 1 292459 Firmicutes
Desulfotomaculum gibsoniae DSM 7213 YP 007945462 1 767817 Firmicutes
Candidatus Desulforudis audaxviator MP104C YP 001716340 1 477974 Firmicutes
Desulfotomaculum carboxydivorans CO 1 SRB YP 004496317 1 868595 Firmicutes
Heliobacterium modesticaldum Ice1 YP 001679111 1 498761 Firmicutes
Brevibacillus brevis NBRC 100599 YP 002771657 1 358681 Firmicutes
Thermincola potens JR YP 003639436 1 635013 Firmicutes
Bacillus clausii KSM K16 YP 175264 1 66692 Firmicutes
Desulfotomaculum acetoxidans DSM 771 YP 003190543 1 485916 Firmicutes
Clostridium clariflavum DSM 19732 YP 005047078 1 720554 Firmicutes
Ruminiclostridium thermocellum ATCC 27405 YP 001036985 1 203119 Firmicutes
Streptosporangium roseum DSM 43021 YP 003339816 1 479432 Actinobacteria
Streptomyces fulvissimus DSM 40593 YP 007932144 1 1303692 Actinobacteria
Burkholderia mallei NCTC 10247 YP 001079369 1 320389 Proteobacteria
Burkholderia pseudomallei 1026b YP 006276863 1 884204 Proteobacteria
Nocardia brasiliensis ATCC 700358 YP 006809183 1 1133849 Actinobacteria
Streptomyces cattleya NRRL 8057 DSM 46488 YP 004913783 1 1003195 Actinobacteria
Streptomyces hygroscopicus subsp jinggangensis 5008 YP 006245072 1 1133850 Actinobacteria
Streptomyces sp PAMC26508 YP 007857238 1 1265601 Actinobacteria
Streptomyces pratensis ATCC 33331 YP 004927190 1 591167 Actinobacteria
Chromobacterium violaceum ATCC 12472 NP 901383 1 243365 Proteobacteria
Bordetella petrii DSM 12804 YP 001629933 1 340100 Proteobacteria
Pseudomonas mendocina NK 01 YP 004380677 1 1001585 Proteobacteria
Pseudomonas fulva 12 X YP 004474203 1 743720 Proteobacteria
Thermobispora bispora DSM 43833 YP 003653656 1 469371 Actinobacteria
Sorangium cellulosum So0157 2 YP 008152368 1 1254432 Proteobacteria
Stigmatella aurantiaca DW4 3 1 YP 003955079 1 378806 Proteobacteria
Streptomyces fulvissimus DSM 40593 YP 007934996 1 1303692 Actinobacteria
Kitasatospora setae KM 6054 YP 004902606 1 452652 Actinobacteria
Streptomyces cattleya NRRL 8057 DSM 46488 YP 004912172 1 1003195 Actinobacteria
Streptomyces bingchenggensis BCW 1 YP 004963777 1 749414 Actinobacteria
Catenulispora acidiphila DSM 44928 YP 003117670 1 479433 Actinobacteria
Amycolatopsis orientalis HCCB10007 YP 008014380 1 1156913 Actinobacteria
Frankia alni ACN14a YP 715800 1 326424 Actinobacteria
Streptomyces rapamycinicus NRRL 5491 YP 008787004 1 1343740 Actinobacteria
Leifsonia xyli subsp xyli str CTCB07 YP 062965 1 281090 Actinobacteria
Erwinia pyrifoliae DSM 12163 YP 005801167 1 644651 Proteobacteria
803 DAQ1742 00824 GI3 metabolic island Seed
Dickeya aquatica DW 0440 WP 035339996 1 Daquatica
DAQ1742 DAQ1742 00824 Daquatica
Dickeya sp CSL RW240 WP 035339996 1 Daquatica
Saccharopolyspora erythraea NRRL 2338 YP 001108189 1 405948 Actinobacteria
Streptomyces rapamycinicus NRRL 5491 YP 008787366 1 1343740 Actinobacteria
Streptomyces violaceusniger Tu 4113 YP 004814153 1 653045 Actinobacteria
Mycobacterium chubuense NBB4 YP 006454743 1 710421 Actinobacteria
Gordonia polyisoprenivorans VH2 YP 005281791 1 1112204 Actinobacteria
Amycolatopsis orientalis HCCB10007 YP 008014967 1 1156913 Actinobacteria
Actinosynnema mirum DSM 43827 YP 003100833 1 446462 Actinobacteria
Amycolatopsis mediterranei S699 YP 005529866 1 713604 Actinobacteria
Nocardia brasiliensis ATCC 700358 YP 006812500 1 1133849 Actinobacteria
Rhodococcus jostii RHA1 YP 704949 1 101510 Actinobacteria
Rhodococcus opacus B4 YP 002782264 1 632772 Actinobacteria
Catenulispora acidiphila DSM 44928 YP 003114955 1 479433 Actinobacteria
Streptomyces hygroscopicus subsp jinggangensis 5008 YP 006243348 1 1133850 Actinobacteria
Streptomyces collinus Tu 365 YP 008384565 1 1214242 Actinobacteria

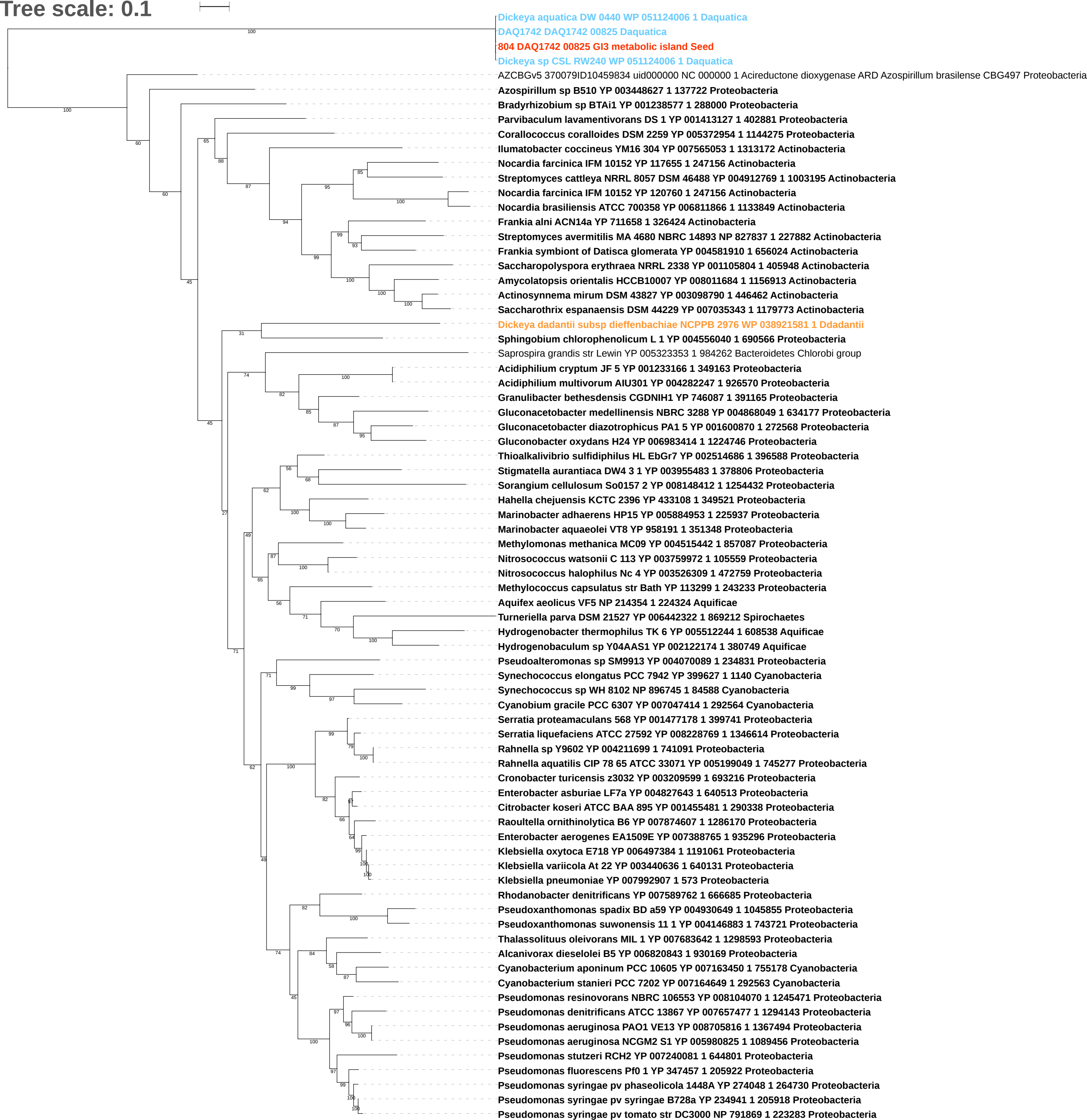
Tree scale: 0.1
Dickeya aquatica DW 0440 WP 051124006 1 Daquatica
DAQ1742 DAQ1742 00825 Daquatica
804 DAQ1742 00825 GI3 metabolic island Seed
Dickeya sp CSL RW240 WP 051124006 1 Daquatica
AZCBGv5 370079ID10459834 uid000000 NC 000000 1 Acireductone dioxygenase ARD Azospirillum brasilense CBG497 Proteobacteria
Azospirillum sp B510 YP 003448627 1 137722 Proteobacteria
Bradyrhizobium sp BTAi1 YP 001238577 1 288000 Proteobacteria
Parvibaculum lavamentivorans DS 1 YP 001413127 1 402881 Proteobacteria
Corallococcus coralloides DSM 2259 YP 005372954 1 1144275 Proteobacteria
Ilumatobacter coccineus YM16 304 YP 007565053 1 1313172 Actinobacteria
Nocardia farcinica IFM 10152 YP 117655 1 247156 Actinobacteria
Streptomyces cattleya NRRL 8057 DSM 46488 YP 004912769 1 1003195 Actinobacteria
Nocardia farcinica IFM 10152 YP 120760 1 247156 Actinobacteria
Nocardia brasiliensis ATCC 700358 YP 006811866 1 1133849 Actinobacteria
Frankia alni ACN14a YP 711658 1 326424 Actinobacteria
Streptomyces avermitilis MA 4680 NBRC 14893 NP 827837 1 227882 Actinobacteria
Frankia symbiont of Datisca glomerata YP 004581910 1 656024 Actinobacteria
Saccharopolyspora erythraea NRRL 2338 YP 001105804 1 405948 Actinobacteria
Amycolatopsis orientalis HCCB10007 YP 008011684 1 1156913 Actinobacteria
Actinosynnema mirum DSM 43827 YP 003098790 1 446462 Actinobacteria
Saccharothrix espanaensis DSM 44229 YP 007035343 1 1179773 Actinobacteria
Dickeya dadantii subsp dieffenbachiae NCPPB 2976 WP 038921581 1 Ddadantii
Sphingobium chlorophenolicum L 1 YP 004556040 1 690566 Proteobacteria
Saprospira grandis str Lewin YP 005323353 1 984262 Bacteroidetes Chlorobi group
Acidiphilium cryptum JF 5 YP 001233166 1 349163 Proteobacteria
Acidiphilium multivorum AIU301 YP 004282247 1 926570 Proteobacteria
Granulibacter bethesdensis CGDNIH1 YP 746087 1 391165 Proteobacteria
Gluconacetobacter medellinensis NBRC 3288 YP 004868049 1 634177 Proteobacteria
Gluconacetobacter diazotrophicus PA1 5 YP 001600870 1 272568 Proteobacteria
Gluconobacter oxydans H24 YP 006983414 1 1224746 Proteobacteria
Thioalkalivibrio sulfidiphilus HL EbGr7 YP 002514686 1 396588 Proteobacteria
Stigmatella aurantiaca DW4 3 1 YP 003955483 1 378806 Proteobacteria
Sorangium cellulosum So0157 2 YP 008148412 1 1254432 Proteobacteria
Hahella chejuensis KCTC 2396 YP 433108 1 349521 Proteobacteria
Marinobacter adhaerens HP15 YP 005884953 1 225937 Proteobacteria
Marinobacter aquaeolei VT8 YP 958191 1 351348 Proteobacteria
Methylomonas methanica MC09 YP 004515442 1 857087 Proteobacteria
Nitrosococcus watsonii C 113 YP 003759972 1 105559 Proteobacteria
Nitrosococcus halophilus Nc 4 YP 003526309 1 472759 Proteobacteria
Methylococcus capsulatus str Bath YP 113299 1 243233 Proteobacteria
Aquifex aeolicus VF5 NP 214354 1 224324 Aquificae
Turneriella parva DSM 21527 YP 006442322 1 869212 Spirochaetes
Hydrogenobacter thermophilus TK 6 YP 005512244 1 608538 Aquificae
Hydrogenobaculum sp Y04AAS1 YP 002122174 1 380749 Aquificae
Pseudoalteromonas sp SM9913 YP 004070089 1 234831 Proteobacteria
Synechococcus elongatus PCC 7942 YP 399627 1 1140 Cyanobacteria
Synechococcus sp WH 8102 NP 896745 1 84588 Cyanobacteria
Cyanobium gracile PCC 6307 YP 007047414 1 292564 Cyanobacteria
Serratia proteamaculans 568 YP 001477178 1 399741 Proteobacteria
Serratia liquefaciens ATCC 27592 YP 008228769 1 1346614 Proteobacteria
Rahnella sp Y9602 YP 004211699 1 741091 Proteobacteria
Rahnella aquatilis CIP 78 65 ATCC 33071 YP 005199049 1 745277 Proteobacteria
Cronobacter turicensis z3032 YP 003209599 1 693216 Proteobacteria
Enterobacter asburiae LF7a YP 004827643 1 640513 Proteobacteria
Citrobacter koseri ATCC BAA 895 YP 001455481 1 290338 Proteobacteria
Raoultella ornithinolytica B6 YP 007874607 1 1286170 Proteobacteria
Enterobacter aerogenes EA1509E YP 007388765 1 935296 Proteobacteria
Klebsiella oxytoca E718 YP 006497384 1 1191061 Proteobacteria
Klebsiella variicola At 22 YP 003440636 1 640131 Proteobacteria
Klebsiella pneumoniae YP 007992907 1 573 Proteobacteria
Rhodanobacter denitrificans YP 007589762 1 666685 Proteobacteria
Pseudoxanthomonas spadix BD a59 YP 004930649 1 1045855 Proteobacteria
Pseudoxanthomonas suwonensis 11 1 YP 004146883 1 743721 Proteobacteria
Thalassolituus oleivorans MIL 1 YP 007683642 1 1298593 Proteobacteria
Alcanivorax dieselolei B5 YP 006820843 1 930169 Proteobacteria
Cyanobacterium aponinum PCC 10605 YP 007163450 1 755178 Cyanobacteria
Cyanobacterium stanieri PCC 7202 YP 007164649 1 292563 Cyanobacteria
Pseudomonas resinovorans NBRC 106553 YP 008104070 1 1245471 Proteobacteria
Pseudomonas denitrificans ATCC 13867 YP 007657477 1 1294143 Proteobacteria
Pseudomonas aeruginosa PAO1 VE13 YP 008705816 1 1367494 Proteobacteria
Pseudomonas aeruginosa NCGM2 S1 YP 005980825 1 1089456 Proteobacteria
Pseudomonas stutzeri RCH2 YP 007240081 1 644801 Proteobacteria
Pseudomonas fluorescens Pf0 1 YP 347457 1 205922 Proteobacteria
Pseudomonas syringae pv phaseolicola 1448A YP 274048 1 264730 Proteobacteria
Pseudomonas syringae pv syringae B728a YP 234941 1 205918 Proteobacteria
Pseudomonas syringae pv tomato str DC3000 NP 791869 1 223283 Proteobacteria

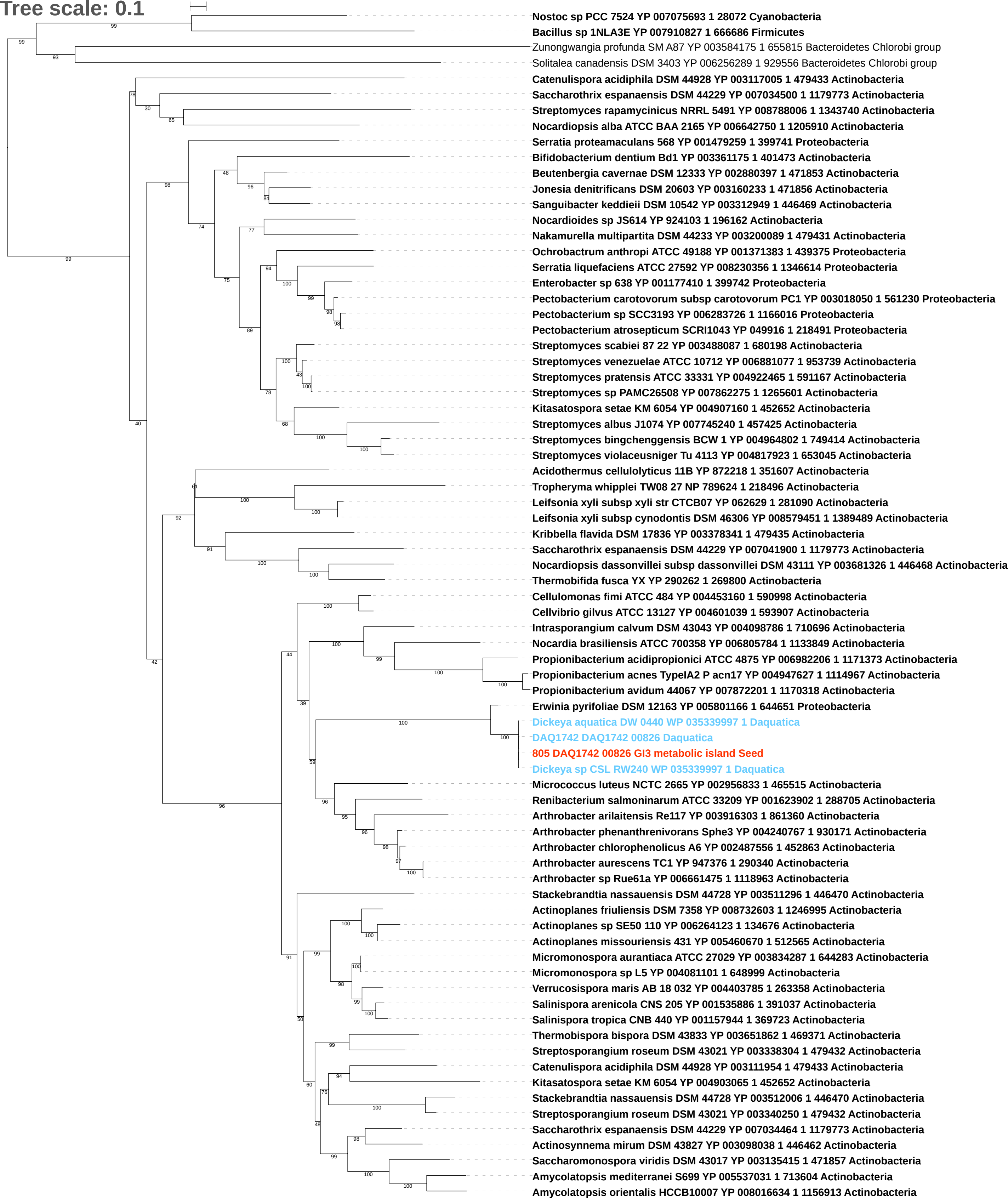
Tree scale: 0.1
Nostoc sp PCC 7524 YP 007075693 1 28072 Cyanobacteria
Bacillus sp 1NLA3E YP 007910827 1 666686 Firmicutes
Zunongwangia profunda SM A87 YP 003584175 1 655815 Bacteroidetes Chlorobi group
Solitalea canadensis DSM 3403 YP 006256289 1 929556 Bacteroidetes Chlorobi group
Catenulispora acidiphila DSM 44928 YP 003117005 1 479433 Actinobacteria
Saccharothrix espanaensis DSM 44229 YP 007034500 1 1179773 Actinobacteria
Streptomyces rapamycinicus NRRL 5491 YP 008788006 1 1343740 Actinobacteria
Nocardiopsis alba ATCC BAA 2165 YP 006642750 1 1205910 Actinobacteria
Serratia proteamaculans 568 YP 001479259 1 399741 Proteobacteria
Bifidobacterium dentium Bd1 YP 003361175 1 401473 Actinobacteria
Beutenbergia cavernae DSM 12333 YP 002880397 1 471853 Actinobacteria
Jonesia denitrificans DSM 20603 YP 003160233 1 471856 Actinobacteria
Sanguibacter keddieii DSM 10542 YP 003312949 1 446469 Actinobacteria
Nocardioides sp JS614 YP 924103 1 196162 Actinobacteria
Nakamurella multipartita DSM 44233 YP 003200089 1 479431 Actinobacteria
Ochrobactrum anthropi ATCC 49188 YP 001371383 1 439375 Proteobacteria
Serratia liquefaciens ATCC 27592 YP 008230356 1 1346614 Proteobacteria
Enterobacter sp 638 YP 001177410 1 399742 Proteobacteria
Pectobacterium carotovorum subsp carotovorum PC1 YP 003018050 1 561230 Proteobacteria
Pectobacterium sp SCC3193 YP 006283726 1 1166016 Proteobacteria
Pectobacterium atrosepticum SCRI1043 YP 049916 1 218491 Proteobacteria
Streptomyces scabiei 87 22 YP 003488087 1 680198 Actinobacteria
Streptomyces venezuelae ATCC 10712 YP 006881077 1 953739 Actinobacteria
Streptomyces pratensis ATCC 33331 YP 004922465 1 591167 Actinobacteria
Streptomyces sp PAMC26508 YP 007862275 1 1265601 Actinobacteria
Kitasatospora setae KM 6054 YP 004907160 1 452652 Actinobacteria
Streptomyces albus J1074 YP 007745240 1 457425 Actinobacteria
Streptomyces bingchenggensis BCW 1 YP 004964802 1 749414 Actinobacteria
Streptomyces violaceusniger Tu 4113 YP 004817923 1 653045 Actinobacteria
Acidothermus cellulolyticus 11B YP 872218 1 351607 Actinobacteria
Tropheryma whipplei TW08 27 NP 789624 1 218496 Actinobacteria
Leifsonia xyli subsp xyli str CTCB07 YP 062629 1 281090 Actinobacteria
Leifsonia xyli subsp cynodontis DSM 46306 YP 008579451 1 1389489 Actinobacteria
Kribbella flavida DSM 17836 YP 003378341 1 479435 Actinobacteria
Saccharothrix espanaensis DSM 44229 YP 007041900 1 1179773 Actinobacteria
Nocardiopsis dassonvillei subsp dassonvillei DSM 43111 YP 003681326 1 446468 Actinobacteria
Thermobifida fusca YX YP 290262 1 269800 Actinobacteria
Cellulomonas fimi ATCC 484 YP 004453160 1 590998 Actinobacteria
Cellvibrio gilvus ATCC 13127 YP 004601039 1 593907 Actinobacteria
Intrasporangium calvum DSM 43043 YP 004098786 1 710696 Actinobacteria
Nocardia brasiliensis ATCC 700358 YP 006805784 1 1133849 Actinobacteria
Propionibacterium acidipropionici ATCC 4875 YP 006982206 1 1171373 Actinobacteria
Propionibacterium acnes TypeIA2 P acn17 YP 004947627 1 1114967 Actinobacteria
Propionibacterium avidum 44067 YP 007872201 1 1170318 Actinobacteria
Erwinia pyrifoliae DSM 12163 YP 005801166 1 644651 Proteobacteria
Dickeya aquatica DW 0440 WP 035339997 1 Daquatica
DAQ1742 DAQ1742 00826 Daquatica
805 DAQ1742 00826 GI3 metabolic island Seed
Dickeya sp CSL RW240 WP 035339997 1 Daquatica
Micrococcus luteus NCTC 2665 YP 002956833 1 465515 Actinobacteria
Renibacterium salmoninarum ATCC 33209 YP 001623902 1 288705 Actinobacteria
Arthrobacter arilaitensis Re117 YP 003916303 1 861360 Actinobacteria
Arthrobacter phenanthrenivorans Sphe3 YP 004240767 1 930171 Actinobacteria
Arthrobacter chlorophenolicus A6 YP 002487556 1 452863 Actinobacteria
Arthrobacter aurescens TC1 YP 947376 1 290340 Actinobacteria
Arthrobacter sp Rue61a YP 006661475 1 1118963 Actinobacteria
Stackebrandtia nassauensis DSM 44728 YP 003511296 1 446470 Actinobacteria
Actinoplanes friuliensis DSM 7358 YP 008732603 1 1246995 Actinobacteria
Actinoplanes sp SE50 110 YP 006264123 1 134676 Actinobacteria
Actinoplanes missouriensis 431 YP 005460670 1 512565 Actinobacteria
Micromonospora aurantiaca ATCC 27029 YP 003834287 1 644283 Actinobacteria
Micromonospora sp L5 YP 004081101 1 648999 Actinobacteria
Verrucosispora maris AB 18 032 YP 004403785 1 263358 Actinobacteria
Salinispora arenicola CNS 205 YP 001535886 1 391037 Actinobacteria
Salinispora tropica CNB 440 YP 001157944 1 369723 Actinobacteria
Thermobispora bispora DSM 43833 YP 003651862 1 469371 Actinobacteria
Streptosporangium roseum DSM 43021 YP 003338304 1 479432 Actinobacteria
Catenulispora acidiphila DSM 44928 YP 003111954 1 479433 Actinobacteria
Kitasatospora setae KM 6054 YP 004903065 1 452652 Actinobacteria
Stackebrandtia nassauensis DSM 44728 YP 003512006 1 446470 Actinobacteria
Streptosporangium roseum DSM 43021 YP 003340250 1 479432 Actinobacteria
Saccharothrix espanaensis DSM 44229 YP 007034464 1 1179773 Actinobacteria
Actinosynnema mirum DSM 43827 YP 003098038 1 446462 Actinobacteria
Saccharomonospora viridis DSM 43017 YP 003135415 1 471857 Actinobacteria
Amycolatopsis mediterranei S699 YP 005537031 1 713604 Actinobacteria
Amycolatopsis orientalis HCCB10007 YP 008016634 1 1156913 Actinobacteria

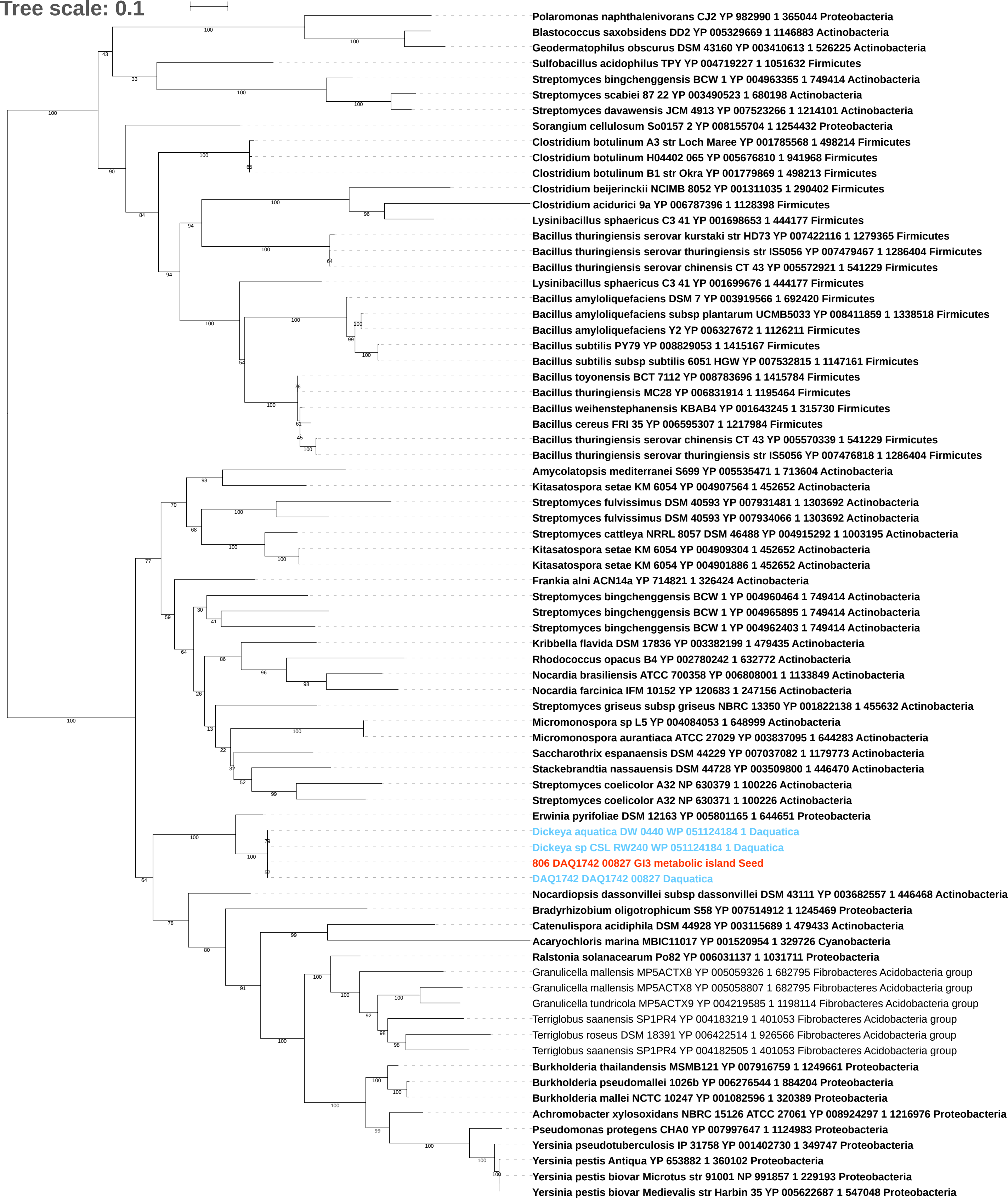
Tree scale: 0.1
Polaromonas naphthalenivorans CJ2 YP 982990 1 365044 Proteobacteria
Blastococcus saxobsidens DD2 YP 005329669 1 1146883 Actinobacteria
Geodermatophilus obscurus DSM 43160 YP 003410613 1 526225 Actinobacteria
Sulfobacillus acidophilus TPY YP 004719227 1 1051632 Firmicutes
Streptomyces bingchenggensis BCW 1 YP 004963355 1 749414 Actinobacteria
Streptomyces scabiei 87 22 YP 003490523 1 680198 Actinobacteria
Streptomyces davawensis JCM 4913 YP 007523266 1 1214101 Actinobacteria
Sorangium cellulosum So0157 2 YP 008155704 1 1254432 Proteobacteria
Clostridium botulinum A3 str Loch Maree YP 001785568 1 498214 Firmicutes
Clostridium botulinum H04402 065 YP 005676810 1 941968 Firmicutes
Clostridium botulinum B1 str Okra YP 001779869 1 498213 Firmicutes
Clostridium beijerinckii NCIMB 8052 YP 001311035 1 290402 Firmicutes
Clostridium acidurici 9a YP 006787396 1 1128398 Firmicutes
Lysinibacillus sphaericus C3 41 YP 001698653 1 444177 Firmicutes
Bacillus thuringiensis serovar kurstaki str HD73 YP 007422116 1 1279365 Firmicutes
Bacillus thuringiensis serovar thuringiensis str IS5056 YP 007479467 1 1286404 Firmicutes
Bacillus thuringiensis serovar chinensis CT 43 YP 005572921 1 541229 Firmicutes
Lysinibacillus sphaericus C3 41 YP 001699676 1 444177 Firmicutes
Bacillus amyloliquefaciens DSM 7 YP 003919566 1 692420 Firmicutes
Bacillus amyloliquefaciens subsp plantarum UCMB5033 YP 008411859 1 1338518 Firmicutes
Bacillus amyloliquefaciens Y2 YP 006327672 1 1126211 Firmicutes
Bacillus subtilis PY79 YP 008829053 1 1415167 Firmicutes
Bacillus subtilis subsp subtilis 6051 HGW YP 007532815 1 1147161 Firmicutes
Bacillus toyonensis BCT 7112 YP 008783696 1 1415784 Firmicutes
Bacillus thuringiensis MC28 YP 006831914 1 1195464 Firmicutes
Bacillus weihenstephanensis KBAB4 YP 001643245 1 315730 Firmicutes
Bacillus cereus FRI 35 YP 006595307 1 1217984 Firmicutes
Bacillus thuringiensis serovar chinensis CT 43 YP 005570339 1 541229 Firmicutes
Bacillus thuringiensis serovar thuringiensis str IS5056 YP 007476818 1 1286404 Firmicutes
Amycolatopsis mediterranei S699 YP 005535471 1 713604 Actinobacteria
Kitasatospora setae KM 6054 YP 004907564 1 452652 Actinobacteria
Streptomyces fulvissimus DSM 40593 YP 007931481 1 1303692 Actinobacteria
Streptomyces fulvissimus DSM 40593 YP 007934066 1 1303692 Actinobacteria
Streptomyces cattleya NRRL 8057 DSM 46488 YP 004915292 1 1003195 Actinobacteria
Kitasatospora setae KM 6054 YP 004909304 1 452652 Actinobacteria
Kitasatospora setae KM 6054 YP 004901886 1 452652 Actinobacteria
Frankia alni ACN14a YP 714821 1 326424 Actinobacteria
Streptomyces bingchenggensis BCW 1 YP 004960464 1 749414 Actinobacteria
Streptomyces bingchenggensis BCW 1 YP 004965895 1 749414 Actinobacteria
Streptomyces bingchenggensis BCW 1 YP 004962403 1 749414 Actinobacteria
Kribbella flavida DSM 17836 YP 003382199 1 479435 Actinobacteria
Rhodococcus opacus B4 YP 002780242 1 632772 Actinobacteria
Nocardia brasiliensis ATCC 700358 YP 006808001 1 1133849 Actinobacteria
Nocardia farcinica IFM 10152 YP 120683 1 247156 Actinobacteria
Streptomyces griseus subsp griseus NBRC 13350 YP 001822138 1 455632 Actinobacteria
Micromonospora sp L5 YP 004084053 1 648999 Actinobacteria
Micromonospora aurantiaca ATCC 27029 YP 003837095 1 644283 Actinobacteria
Saccharothrix espanaensis DSM 44229 YP 007037082 1 1179773 Actinobacteria
Stackebrandtia nassauensis DSM 44728 YP 003509800 1 446470 Actinobacteria
Streptomyces coelicolor A32 NP 630379 1 100226 Actinobacteria
Streptomyces coelicolor A32 NP 630371 1 100226 Actinobacteria
Erwinia pyrifoliae DSM 12163 YP 005801165 1 644651 Proteobacteria
Dickeya aquatica DW 0440 WP 051124184 1 Daquatica
Dickeya sp CSL RW240 WP 051124184 1 Daquatica
806 DAQ1742 00827 GI3 metabolic island Seed
DAQ1742 DAQ1742 00827 Daquatica
Nocardiopsis dassonvillei subsp dassonvillei DSM 43111 YP 003682557 1 446468 Actinobacteria
Bradyrhizobium oligotrophicum S58 YP 007514912 1 1245469 Proteobacteria
Catenulispora acidiphila DSM 44928 YP 003115689 1 479433 Actinobacteria
Acaryochloris marina MBIC11017 YP 001520954 1 329726 Cyanobacteria
Ralstonia solanacearum Po82 YP 006031137 1 1031711 Proteobacteria
Granulicella mallensis MP5ACTX8 YP 005059326 1 682795 Fibrobacteres Acidobacteria group
Granulicella mallensis MP5ACTX8 YP 005058807 1 682795 Fibrobacteres Acidobacteria group
Granulicella tundricola MP5ACTX9 YP 004219585 1 1198114 Fibrobacteres Acidobacteria group
Terriglobus saanensis SP1PR4 YP 004183219 1 401053 Fibrobacteres Acidobacteria group
Terriglobus roseus DSM 18391 YP 006422514 1 926566 Fibrobacteres Acidobacteria group
Terriglobus saanensis SP1PR4 YP 004182505 1 401053 Fibrobacteres Acidobacteria group
Burkholderia thailandensis MSMB121 YP 007916759 1 1249661 Proteobacteria
Burkholderia pseudomallei 1026b YP 006276544 1 884204 Proteobacteria
Burkholderia mallei NCTC 10247 YP 001082596 1 320389 Proteobacteria
Achromobacter xylosoxidans NBRC 15126 ATCC 27061 YP 008924297 1 1216976 Proteobacteria
Pseudomonas protegens CHA0 YP 007997647 1 1124983 Proteobacteria
Yersinia pseudotuberculosis IP 31758 YP 001402730 1 349747 Proteobacteria
Yersinia pestis Antiqua YP 653882 1 360102 Proteobacteria
Yersinia pestis biovar Microtus str 91001 NP 991857 1 229193 Proteobacteria
Yersinia pestis biovar Medievalis str Harbin 35 YP 005622687 1 547048 Proteobacteria

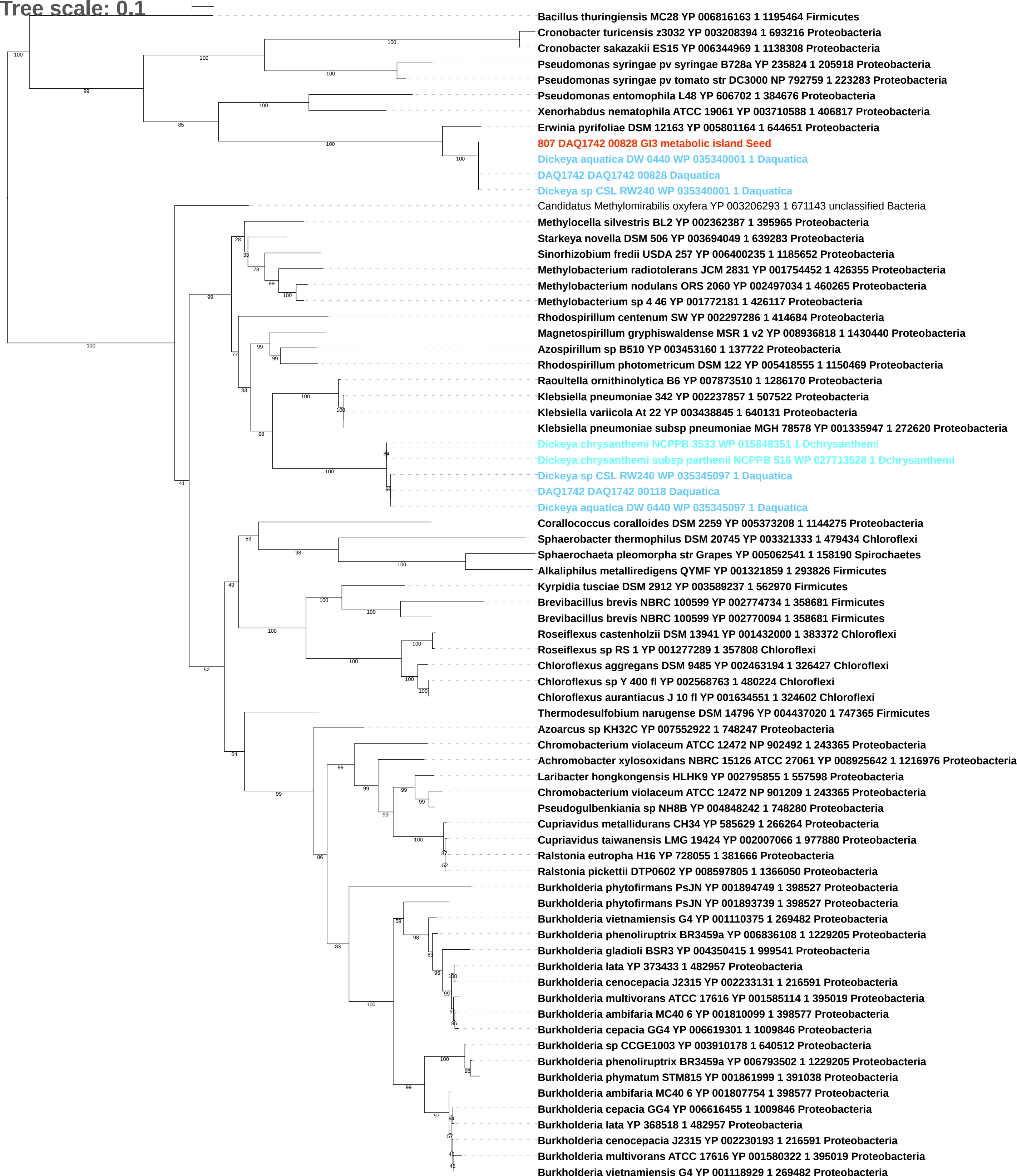

Tree scale: 0.1
Bacillus thuringiensis MC28 YP 006816163 1 1195464 Firmicutes
Cronobacter turicensis z3032 YP 003208394 1 693216 Proteobacteria
Cronobacter sakazakii ES15 YP 006344969 1 1138308 Proteobacteria
Pseudomonas syringae pv syringae B728a YP 235824 1 205918 Proteobacteria
Pseudomonas syringae pv tomato str DC3000 NP 792759 1 223283 Proteobacteria
Pseudomonas entomophila L48 YP 606702 1 384676 Proteobacteria
Xenorhabdus nematophila ATCC 19061 YP 003710588 1 406817 Proteobacteria
Erwinia pyrifoliae DSM 12163 YP 005801164 1 644651 Proteobacteria
807 DAQ1742 00828 GI3 metabolic island Seed
Dickeya aquatica DW 0440 WP 035340001 1 Daquatica
DAQ1742 DAQ1742 00828 Daquatica
Dickeya sp CSL RW240 WP 035340001 1 Daquatica
Candidatus Methylomirabilis oxyfera YP 003206293 1 671143 unclassified Bacteria
Methylocella silvestris BL2 YP 002362387 1 395965 Proteobacteria
Starkeya novella DSM 506 YP 003694049 1 639283 Proteobacteria
Sinorhizobium fredii USDA 257 YP 006400235 1 1185652 Proteobacteria
Methylobacterium radiotolerans JCM 2831 YP 001754452 1 426355 Proteobacteria
Methylobacterium nodulans ORS 2060 YP 002497034 1 460265 Proteobacteria
Methylobacterium sp 4 46 YP 001772181 1 426117 Proteobacteria
Rhodospirillum centenum SW YP 002297286 1 414684 Proteobacteria
Magnetospirillum gryphiswaldense MSR 1 v2 YP 008936818 1 1430440 Proteobacteria
Azospirillum sp B510 YP 003453160 1 137722 Proteobacteria
Rhodospirillum photometricum DSM 122 YP 005418555 1 1150469 Proteobacteria
Raoultella ornithinolytica B6 YP 007873510 1 1286170 Proteobacteria
Klebsiella pneumoniae 342 YP 002237857 1 507522 Proteobacteria
Klebsiella variicola At 22 YP 003438845 1 640131 Proteobacteria
Klebsiella pneumoniae subsp pneumoniae MGH 78578 YP 001335947 1 272620 Proteobacteria
Dickeya chrysanthemi NCPPB 3533 WP 015848351 1 Dchrysanthemi
Dickeya chrysanthemi subsp parthenii NCPPB 516 WP 027713528 1 Dchrysanthemi
Dickeya sp CSL RW240 WP 035345097 1 Daquatica
DAQ1742 DAQ1742 00118 Daquatica
Dickeya aquatica DW 0440 WP 035345097 1 Daquatica
Corallococcus coralloides DSM 2259 YP 005373208 1 1144275 Proteobacteria
Sphaerobacter thermophilus DSM 20745 YP 003321333 1 479434 Chloroflexi
Sphaerochaeta pleomorpha str Grapes YP 005062541 1 158190 Spirochaetes
Alkaliphilus metalliredigens QYMF YP 001321859 1 293826 Firmicutes
Kyrpidia tusciae DSM 2912 YP 003589237 1 562970 Firmicutes
Brevibacillus brevis NBRC 100599 YP 002774734 1 358681 Firmicutes
Brevibacillus brevis NBRC 100599 YP 002770094 1 358681 Firmicutes
Roseiflexus castenholzii DSM 13941 YP 001432000 1 383372 Chloroflexi
Roseiflexus sp RS 1 YP 001277289 1 357808 Chloroflexi
Chloroflexus aggregans DSM 9485 YP 002463194 1 326427 Chloroflexi
Chloroflexus sp Y 400 fl YP 002568763 1 480224 Chloroflexi
Chloroflexus aurantiacus J 10 fl YP 001634551 1 324602 Chloroflexi
Thermodesulfobium narugense DSM 14796 YP 004437020 1 747365 Firmicutes
Azoarcus sp KH32C YP 007552922 1 748247 Proteobacteria
Chromobacterium violaceum ATCC 12472 NP 902492 1 243365 Proteobacteria
Achromobacter xylosoxidans NBRC 15126 ATCC 27061 YP 008925642 1 1216976 Proteobacteria
Laribacter hongkongensis HLHK9 YP 002795855 1 557598 Proteobacteria
Chromobacterium violaceum ATCC 12472 NP 901209 1 243365 Proteobacteria
Pseudogulbenkiania sp NH8B YP 004848242 1 748280 Proteobacteria
Cupriavidus metallidurans CH34 YP 585629 1 266264 Proteobacteria
Cupriavidus taiwanensis LMG 19424 YP 002007066 1 977880 Proteobacteria
Ralstonia eutropha H16 YP 728055 1 381666 Proteobacteria
Ralstonia pickettii DTP0602 YP 008597805 1 1366050 Proteobacteria
Burkholderia phytofirmans PsJN YP 001894749 1 398527 Proteobacteria
Burkholderia phytofirmans PsJN YP 001893739 1 398527 Proteobacteria
Burkholderia vietnamiensis G4 YP 001110375 1 269482 Proteobacteria
Burkholderia phenoliruptrix BR3459a YP 006836108 1 1229205 Proteobacteria
Burkholderia gladioli BSR3 YP 004350415 1 999541 Proteobacteria
Burkholderia lata YP 373433 1 482957 Proteobacteria
Burkholderia cenocepacia J2315 YP 002233131 1 216591 Proteobacteria
Burkholderia multivorans ATCC 17616 YP 001585114 1 395019 Proteobacteria
Burkholderia ambifaria MC40 6 YP 001810099 1 398577 Proteobacteria
Burkholderia cepacia GG4 YP 006619301 1 1009846 Proteobacteria
Burkholderia sp CCGE1003 YP 003910178 1 640512 Proteobacteria
Burkholderia phenoliruptrix BR3459a YP 006793502 1 1229205 Proteobacteria
Burkholderia phymatum STM815 YP 001861999 1 391038 Proteobacteria
Burkholderia ambifaria MC40 6 YP 001807754 1 398577 Proteobacteria
Burkholderia cepacia GG4 YP 006616455 1 1009846 Proteobacteria
Burkholderia lata YP 368518 1 482957 Proteobacteria
Burkholderia cenocepacia J2315 YP 002230193 1 216591 Proteobacteria
Burkholderia multivorans ATCC 17616 YP 001580322 1 395019 Proteobacteria
Burkholderia vietnamiensis G4 YP 001118929 1 269482 Proteobacteria

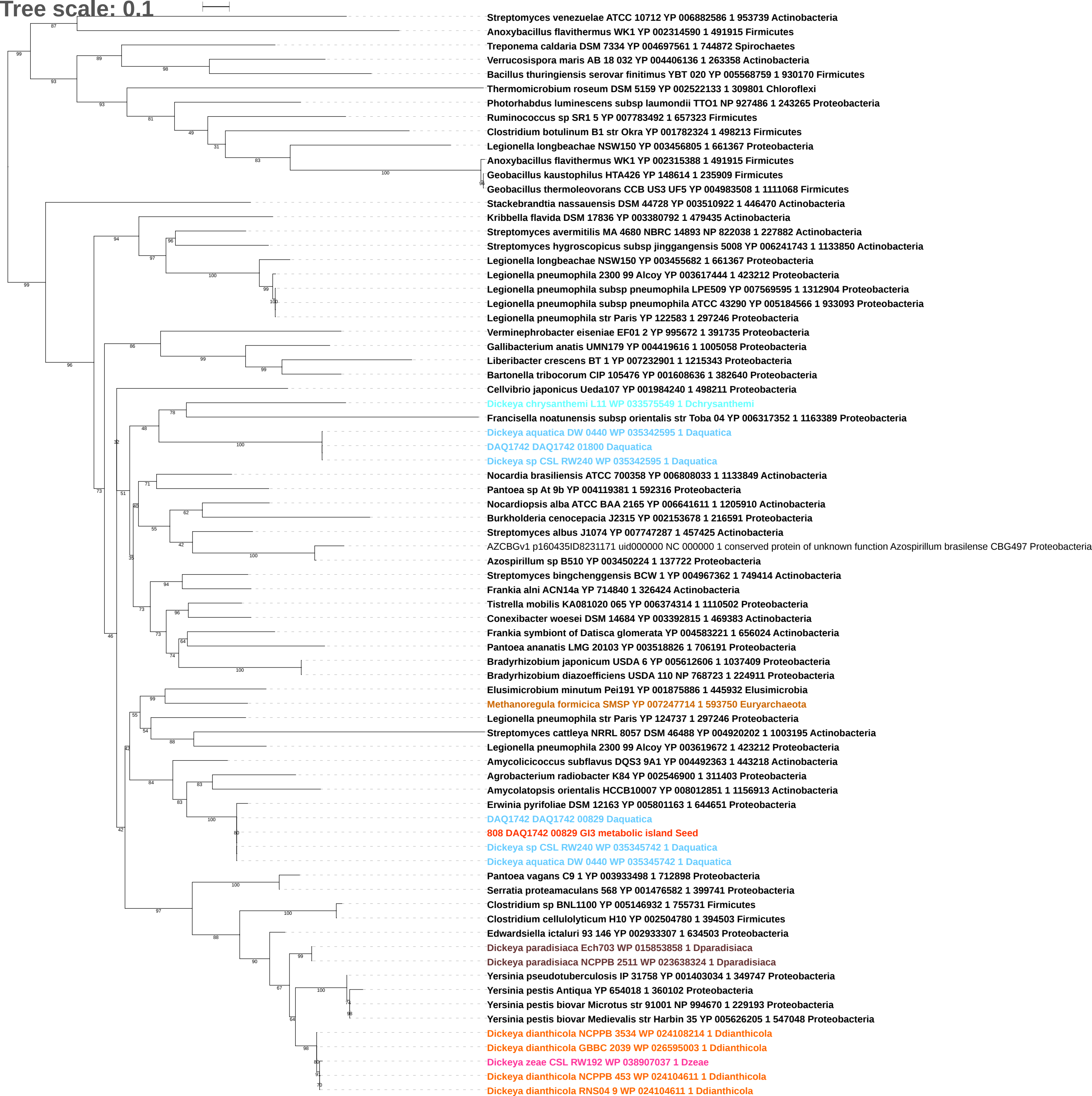
Tree scale: 0.1
Streptomyces venezuelae ATCC 10712 YP 006882586 1 953739 Actinobacteria
Anoxybacillus flavithermus WK1 YP 002314590 1 491915 Firmicutes
Treponema caldaria DSM 7334 YP 004697561 1 744872 Spirochaetes
Verrucosispora maris AB 18 032 YP 004406136 1 263358 Actinobacteria
Bacillus thuringiensis serovar finitimus YBT 020 YP 005568759 1 930170 Firmicutes
Thermomicrobium roseum DSM 5159 YP 002522133 1 309801 Chloroflexi
Photorhabdus luminescens subsp laumondii TTO1 NP 927486 1 243265 Proteobacteria
Ruminococcus sp SR1 5 YP 007783492 1 657323 Firmicutes
Clostridium botulinum B1 str Okra YP 001782324 1 498213 Firmicutes
Legionella longbeachae NSW150 YP 003456805 1 661367 Proteobacteria
Anoxybacillus flavithermus WK1 YP 002315388 1 491915 Firmicutes
Geobacillus kaustophilus HTA426 YP 148614 1 235909 Firmicutes
Geobacillus thermoleovorans CCB US3 UF5 YP 004983508 1 1111068 Firmicutes
Stackebrandtia nassauensis DSM 44728 YP 003510922 1 446470 Actinobacteria
Kribbella flavida DSM 17836 YP 003380792 1 479435 Actinobacteria
Streptomyces avermitilis MA 4680 NBRC 14893 NP 822038 1 227882 Actinobacteria
Streptomyces hygroscopicus subsp jinggangensis 5008 YP 006241743 1 1133850 Actinobacteria
Legionella longbeachae NSW150 YP 003455682 1 661367 Proteobacteria
Legionella pneumophila 2300 99 Alcoy YP 003617444 1 423212 Proteobacteria
Legionella pneumophila subsp pneumophila LPE509 YP 007569595 1 1312904 Proteobacteria
Legionella pneumophila subsp pneumophila ATCC 43290 YP 005184566 1 933093 Proteobacteria
Legionella pneumophila str Paris YP 122583 1 297246 Proteobacteria
Verminephrobacter eiseniae EF01 2 YP 995672 1 391735 Proteobacteria
Gallibacterium anatis UMN179 YP 004419616 1 1005058 Proteobacteria
Liberibacter crescens BT 1 YP 007232901 1 1215343 Proteobacteria
Bartonella tribocorum CIP 105476 YP 001608636 1 382640 Proteobacteria
Cellvibrio japonicus Ueda107 YP 001984240 1 498211 Proteobacteria
Dickeya chrysanthemi L11 WP 033575549 1 Dchrysanthemi
Francisella noatunensis subsp orientalis str Toba 04 YP 006317352 1 1163389 Proteobacteria
Dickeya aquatica DW 0440 WP 035342595 1 Daquatica
DAQ1742 DAQ1742 01800 Daquatica
Dickeya sp CSL RW240 WP 035342595 1 Daquatica
Nocardia brasiliensis ATCC 700358 YP 006808033 1 1133849 Actinobacteria
Pantoea sp At 9b YP 004119381 1 592316 Proteobacteria
Nocardiopsis alba ATCC BAA 2165 YP 006641611 1 1205910 Actinobacteria
Burkholderia cenocepacia J2315 YP 002153678 1 216591 Proteobacteria
Streptomyces albus J1074 YP 007747287 1 457425 Actinobacteria
AZCBGv1 p160435ID8231171 uid000000 NC 000000 1 conserved protein of unknown function Azospirillum brasilense CBG497 Proteobacteria
Azospirillum sp B510 YP 003450224 1 137722 Proteobacteria
Streptomyces bingchenggensis BCW 1 YP 004967362 1 749414 Actinobacteria
Frankia alni ACN14a YP 714840 1 326424 Actinobacteria
Tistrella mobilis KA081020 065 YP 006374314 1 1110502 Proteobacteria
Conexibacter woesei DSM 14684 YP 003392815 1 469383 Actinobacteria
Frankia symbiont of Datisca glomerata YP 004583221 1 656024 Actinobacteria
Pantoea ananatis LMG 20103 YP 003518826 1 706191 Proteobacteria
Bradyrhizobium japonicum USDA 6 YP 005612606 1 1037409 Proteobacteria
Bradyrhizobium diazoefficiens USDA 110 NP 768723 1 224911 Proteobacteria
Elusimicrobium minutum Pei191 YP 001875886 1 445932 Elusimicrobia
Methanoregula formicica SMSP YP 007247714 1 593750 Euryarchaeota
Legionella pneumophila str Paris YP 124737 1 297246 Proteobacteria
Streptomyces cattleya NRRL 8057 DSM 46488 YP 004920202 1 1003195 Actinobacteria
Legionella pneumophila 2300 99 Alcoy YP 003619672 1 423212 Proteobacteria
Amycolicicoccus subflavus DQS3 9A1 YP 004492363 1 443218 Actinobacteria
Agrobacterium radiobacter K84 YP 002546900 1 311403 Proteobacteria
Amycolatopsis orientalis HCCB10007 YP 008012851 1 1156913 Actinobacteria
Erwinia pyrifoliae DSM 12163 YP 005801163 1 644651 Proteobacteria
DAQ1742 DAQ1742 00829 Daquatica
808 DAQ1742 00829 GI3 metabolic island Seed
Dickeya sp CSL RW240 WP 035345742 1 Daquatica
Dickeya aquatica DW 0440 WP 035345742 1 Daquatica
Pantoea vagans C9 1 YP 003933498 1 712898 Proteobacteria
Serratia proteamaculans 568 YP 001476582 1 399741 Proteobacteria
Clostridium sp BNL1100 YP 005146932 1 755731 Firmicutes
Clostridium cellulolyticum H10 YP 002504780 1 394503 Firmicutes
Edwardsiella ictaluri 93 146 YP 002933307 1 634503 Proteobacteria
Dickeya paradisiaca Ech703 WP 015853858 1 Dparadisiaca
Dickeya paradisiaca NCPPB 2511 WP 023638324 1 Dparadisiaca
Yersinia pseudotuberculosis IP 31758 YP 001403034 1 349747 Proteobacteria
Yersinia pestis Antiqua YP 654018 1 360102 Proteobacteria
Yersinia pestis biovar Microtus str 91001 NP 994670 1 229193 Proteobacteria
Yersinia pestis biovar Medievalis str Harbin 35 YP 005626205 1 547048 Proteobacteria
Dickeya dianthicola NCPPB 3534 WP 024108214 1 Ddianthicola
Dickeya dianthicola GBBC 2039 WP 026595003 1 Ddianthicola
Dickeya zeae CSL RW192 WP 038907037 1 Dzeae
Dickeya dianthicola NCPPB 453 WP 024104611 1 Ddianthicola
Dickeya dianthicola RNS04 9 WP 024104611 1 Ddianthicola

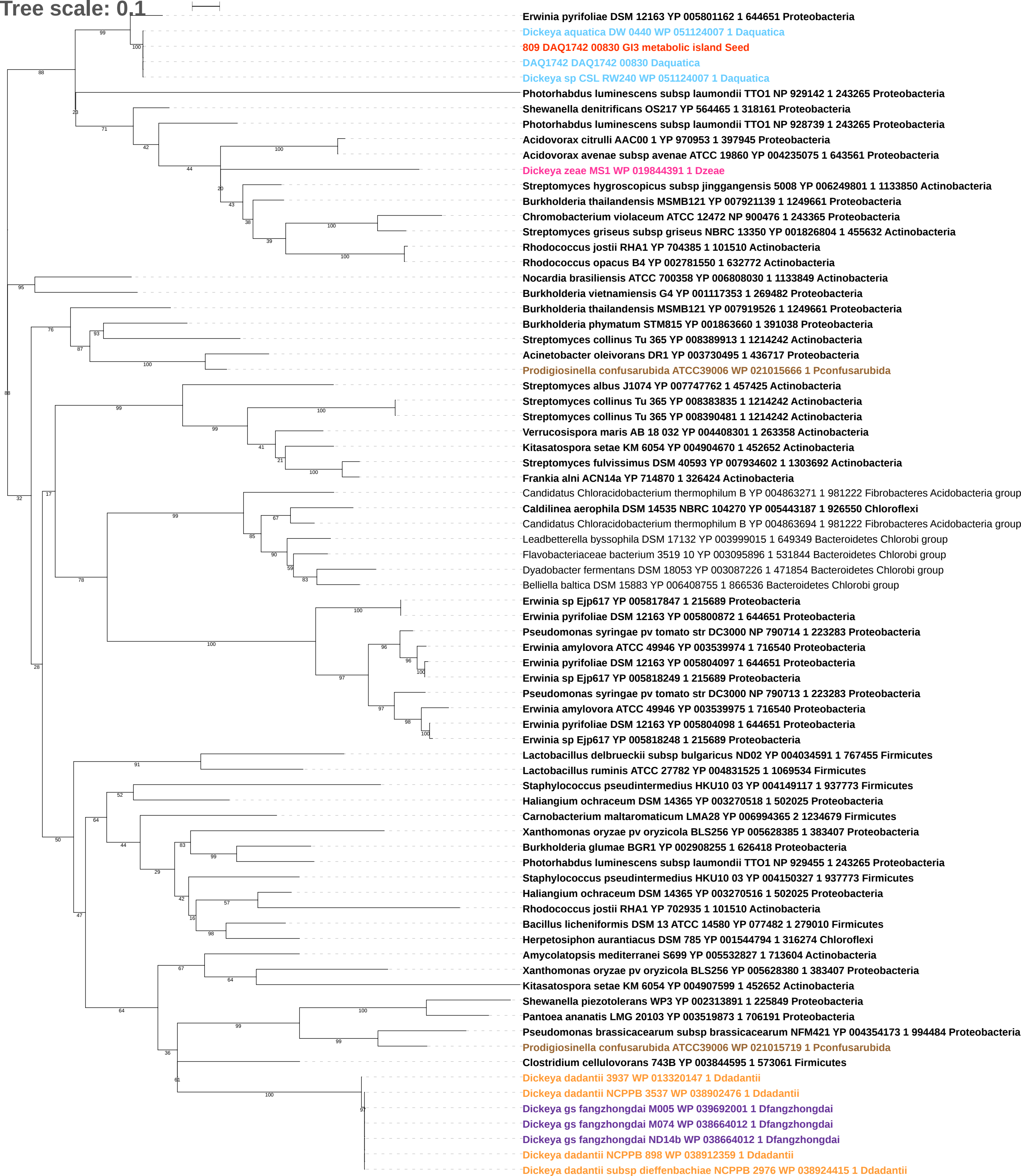
Tree scale: 0.1
Erwinia pyrifoliae DSM 12163 YP 005801162 1 644651 Proteobacteria
Dickeya aquatica DW 0440 WP 051124007 1 Daquatica
809 DAQ1742 00830 GI3 metabolic island Seed
DAQ1742 DAQ1742 00830 Daquatica
Dickeya sp CSL RW240 WP 051124007 1 Daquatica
Photorhabdus luminescens subsp laumondii TTO1 NP 929142 1 243265 Proteobacteria
Shewanella denitrificans OS217 YP 564465 1 318161 Proteobacteria
Photorhabdus luminescens subsp laumondii TTO1 NP 928739 1 243265 Proteobacteria
Acidovorax citrulli AAC00 1 YP 970953 1 397945 Proteobacteria
Acidovorax avenae subsp avenae ATCC 19860 YP 004235075 1 643561 Proteobacteria
Dickeya zeae MS1 WP 019844391 1 Dzeae
Streptomyces hygroscopicus subsp jinggangensis 5008 YP 006249801 1 1133850 Actinobacteria
Burkholderia thailandensis MSMB121 YP 007921139 1 1249661 Proteobacteria
Chromobacterium violaceum ATCC 12472 NP 900476 1 243365 Proteobacteria
Streptomyces griseus subsp griseus NBRC 13350 YP 001826804 1 455632 Actinobacteria
Rhodococcus jostii RHA1 YP 704385 1 101510 Actinobacteria
Rhodococcus opacus B4 YP 002781550 1 632772 Actinobacteria
Nocardia brasiliensis ATCC 700358 YP 006808030 1 1133849 Actinobacteria
Burkholderia vietnamiensis G4 YP 001117353 1 269482 Proteobacteria
Burkholderia thailandensis MSMB121 YP 007919526 1 1249661 Proteobacteria
Burkholderia phymatum STM815 YP 001863660 1 391038 Proteobacteria
Streptomyces collinus Tu 365 YP 008389913 1 1214242 Actinobacteria
Acinetobacter oleivorans DR1 YP 003730495 1 436717 Proteobacteria
Prodigiosinella confusarubida ATCC39006 WP 021015666 1 Pconfusarubida
Streptomyces albus J1074 YP 007747762 1 457425 Actinobacteria
Streptomyces collinus Tu 365 YP 008383835 1 1214242 Actinobacteria
Streptomyces collinus Tu 365 YP 008390481 1 1214242 Actinobacteria
Verrucosispora maris AB 18 032 YP 004408301 1 263358 Actinobacteria
Kitasatospora setae KM 6054 YP 004904670 1 452652 Actinobacteria
Streptomyces fulvissimus DSM 40593 YP 007934602 1 1303692 Actinobacteria
Frankia alni ACN14a YP 714870 1 326424 Actinobacteria
Candidatus Chloracidobacterium thermophilum B YP 004863271 1 981222 Fibrobacteres Acidobacteria group
Caldilinea aerophila DSM 14535 NBRC 104270 YP 005443187 1 926550 Chloroflexi
Candidatus Chloracidobacterium thermophilum B YP 004863694 1 981222 Fibrobacteres Acidobacteria group
Leadbetterella byssophila DSM 17132 YP 003999015 1 649349 Bacteroidetes Chlorobi group
Flavobacteriaceae bacterium 3519 10 YP 003095896 1 531844 Bacteroidetes Chlorobi group
Dyadobacter fermentans DSM 18053 YP 003087226 1 471854 Bacteroidetes Chlorobi group
Belliella baltica DSM 15883 YP 006408755 1 866536 Bacteroidetes Chlorobi group
Erwinia sp Ejp617 YP 005817847 1 215689 Proteobacteria
Erwinia pyrifoliae DSM 12163 YP 005800872 1 644651 Proteobacteria
Pseudomonas syringae pv tomato str DC3000 NP 790714 1 223283 Proteobacteria
Erwinia amylovora ATCC 49946 YP 003539974 1 716540 Proteobacteria
Erwinia pyrifoliae DSM 12163 YP 005804097 1 644651 Proteobacteria
Erwinia sp Ejp617 YP 005818249 1 215689 Proteobacteria
Pseudomonas syringae pv tomato str DC3000 NP 790713 1 223283 Proteobacteria
Erwinia amylovora ATCC 49946 YP 003539975 1 716540 Proteobacteria
Erwinia pyrifoliae DSM 12163 YP 005804098 1 644651 Proteobacteria
Erwinia sp Ejp617 YP 005818248 1 215689 Proteobacteria
Lactobacillus delbrueckii subsp bulgaricus ND02 YP 004034591 1 767455 Firmicutes
Lactobacillus ruminis ATCC 27782 YP 004831525 1 1069534 Firmicutes
Staphylococcus pseudintermedius HKU10 03 YP 004149117 1 937773 Firmicutes
Haliangium ochraceum DSM 14365 YP 003270518 1 502025 Proteobacteria
Carnobacterium maltaromaticum LMA28 YP 006994365 2 1234679 Firmicutes
Xanthomonas oryzae pv oryzicola BLS256 YP 005628385 1 383407 Proteobacteria
Burkholderia glumae BGR1 YP 002908255 1 626418 Proteobacteria
Photorhabdus luminescens subsp laumondii TTO1 NP 929455 1 243265 Proteobacteria
Staphylococcus pseudintermedius HKU10 03 YP 004150327 1 937773 Firmicutes
Haliangium ochraceum DSM 14365 YP 003270516 1 502025 Proteobacteria
Rhodococcus jostii RHA1 YP 702935 1 101510 Actinobacteria
Bacillus licheniformis DSM 13 ATCC 14580 YP 077482 1 279010 Firmicutes
Herpetosiphon aurantiacus DSM 785 YP 001544794 1 316274 Chloroflexi
Amycolatopsis mediterranei S699 YP 005532827 1 713604 Actinobacteria
Xanthomonas oryzae pv oryzicola BLS256 YP 005628380 1 383407 Proteobacteria
Kitasatospora setae KM 6054 YP 004907599 1 452652 Actinobacteria
Shewanella piezotolerans WP3 YP 002313891 1 225849 Proteobacteria
Pantoea ananatis LMG 20103 YP 003519873 1 706191 Proteobacteria
Pseudomonas brassicacearum subsp brassicacearum NFM421 YP 004354173 1 994484 Proteobacteria
Prodigiosinella confusarubida ATCC39006 WP 021015719 1 Pconfusarubida
Clostridium cellulovorans 743B YP 003844595 1 573061 Firmicutes
Dickeya dadantii 3937 WP 013320147 1 Ddadantii
Dickeya dadantii NCPPB 3537 WP 038902476 1 Ddadantii
Dickeya gs fangzhongdai M005 WP 039692001 1 Dfangzhongdai
Dickeya gs fangzhongdai M074 WP 038664012 1 Dfangzhongdai
Dickeya gs fangzhongdai ND14b WP 038664012 1 Dfangzhongdai
Dickeya dadantii NCPPB 898 WP 038912359 1 Ddadantii
Dickeya dadantii subsp dieffenbachiae NCPPB 2976 WP 038924415 1 Ddadantii

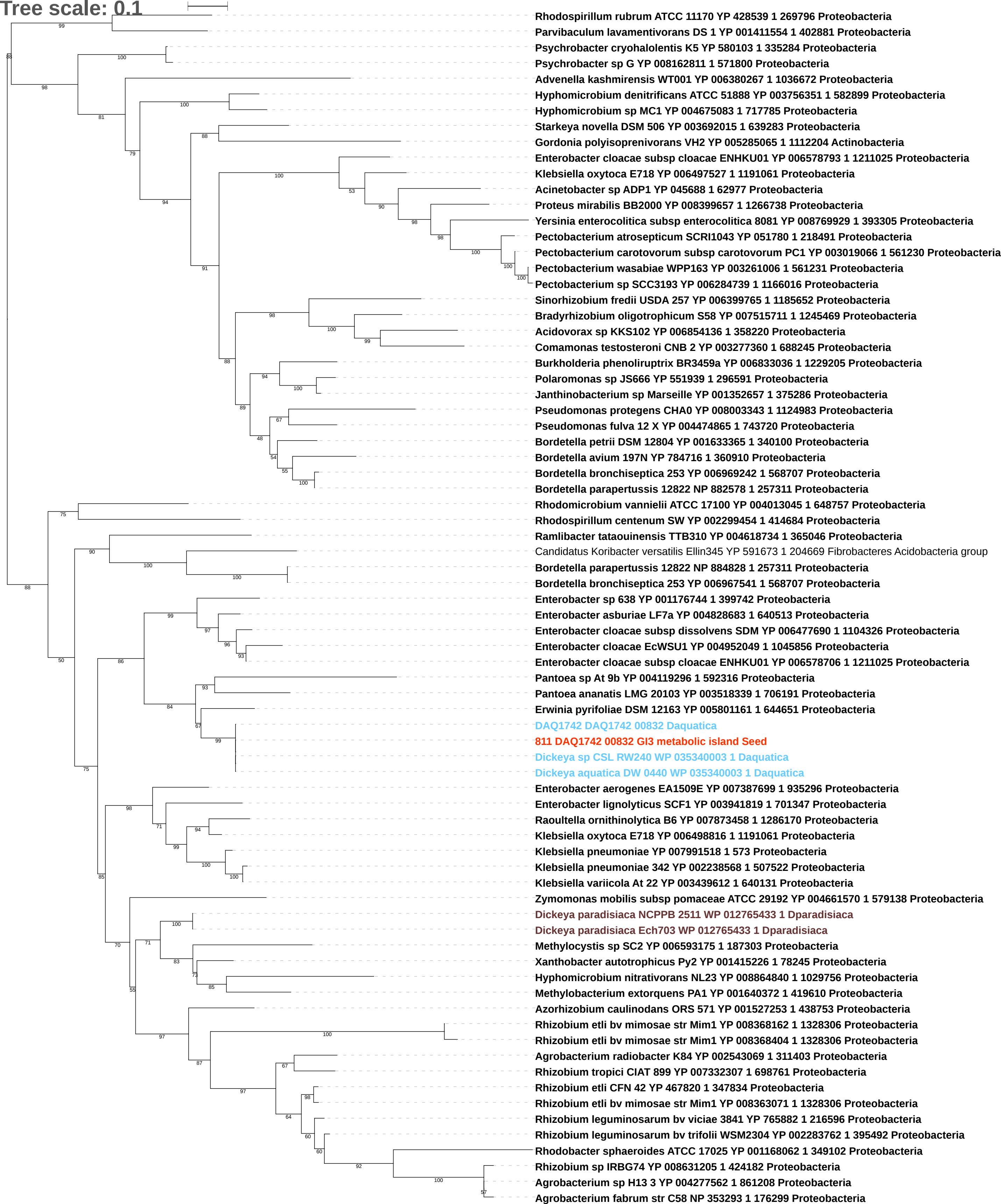
Tree scale: 0.1
Rhodospirillum rubrum ATCC 11170 YP 428539 1 269796 Proteobacteria
Parvibaculum lavamentivorans DS 1 YP 001411554 1 402881 Proteobacteria
Psychrobacter cryohalolentis K5 YP 580103 1 335284 Proteobacteria
Psychrobacter sp G YP 008162811 1 571800 Proteobacteria
Advenella kashmirensis WT001 YP 006380267 1 1036672 Proteobacteria
Hyphomicrobium denitrificans ATCC 51888 YP 003756351 1 582899 Proteobacteria
Hyphomicrobium sp MC1 YP 004675083 1 717785 Proteobacteria
Starkeya novella DSM 506 YP 003692015 1 639283 Proteobacteria
Gordonia polyisoprenivorans VH2 YP 005285065 1 1112204 Actinobacteria
Enterobacter cloacae subsp cloacae ENHKU01 YP 006578793 1 1211025 Proteobacteria
Klebsiella oxytoca E718 YP 006497527 1 1191061 Proteobacteria
Acinetobacter sp ADP1 YP 045688 1 62977 Proteobacteria
Proteus mirabilis BB2000 YP 008399657 1 1266738 Proteobacteria
Yersinia enterocolitica subsp enterocolitica 8081 YP 008769929 1 393305 Proteobacteria
Pectobacterium atrosepticum SCRI1043 YP 051780 1 218491 Proteobacteria
Pectobacterium carotovorum subsp carotovorum PC1 YP 003019066 1 561230 Proteobacteria
Pectobacterium wasabiae WPP163 YP 003261006 1 561231 Proteobacteria
Pectobacterium sp SCC3193 YP 006284739 1 1166016 Proteobacteria
Sinorhizobium fredii USDA 257 YP 006399765 1 1185652 Proteobacteria
Bradyrhizobium oligotrophicum S58 YP 007515711 1 1245469 Proteobacteria
Acidovorax sp KKS102 YP 006854136 1 358220 Proteobacteria
Comamonas testosteroni CNB 2 YP 003277360 1 688245 Proteobacteria
Burkholderia phenoliruptrix BR3459a YP 006833036 1 1229205 Proteobacteria
Polaromonas sp JS666 YP 551939 1 296591 Proteobacteria
Janthinobacterium sp Marseille YP 001352657 1 375286 Proteobacteria
Pseudomonas protegens CHA0 YP 008003343 1 1124983 Proteobacteria
Pseudomonas fulva 12 X YP 004474865 1 743720 Proteobacteria
Bordetella petrii DSM 12804 YP 001633365 1 340100 Proteobacteria
Bordetella avium 197N YP 784716 1 360910 Proteobacteria
Bordetella bronchiseptica 253 YP 006969242 1 568707 Proteobacteria
Bordetella parapertussis 12822 NP 882578 1 257311 Proteobacteria
Rhodomicrobium vannielii ATCC 17100 YP 004013045 1 648757 Proteobacteria
Rhodospirillum centenum SW YP 002299454 1 414684 Proteobacteria
Ramlibacter tataouinensis TTB310 YP 004618734 1 365046 Proteobacteria
Candidatus Koribacter versatilis Ellin345 YP 591673 1 204669 Fibrobacteres Acidobacteria group
Bordetella parapertussis 12822 NP 884828 1 257311 Proteobacteria
Bordetella bronchiseptica 253 YP 006967541 1 568707 Proteobacteria
Enterobacter sp 638 YP 001176744 1 399742 Proteobacteria
Enterobacter asburiae LF7a YP 004828683 1 640513 Proteobacteria
Enterobacter cloacae subsp dissolvens SDM YP 006477690 1 1104326 Proteobacteria
Enterobacter cloacae EcWSU1 YP 004952049 1 1045856 Proteobacteria
Enterobacter cloacae subsp cloacae ENHKU01 YP 006578706 1 1211025 Proteobacteria
Pantoea sp At 9b YP 004119296 1 592316 Proteobacteria
Pantoea ananatis LMG 20103 YP 003518339 1 706191 Proteobacteria
Erwinia pyrifoliae DSM 12163 YP 005801161 1 644651 Proteobacteria
DAQ1742 DAQ1742 00832 Daquatica
811 DAQ1742 00832 GI3 metabolic island Seed
Dickeya sp CSL RW240 WP 035340003 1 Daquatica
Dickeya aquatica DW 0440 WP 035340003 1 Daquatica
Enterobacter aerogenes EA1509E YP 007387699 1 935296 Proteobacteria
Enterobacter lignolyticus SCF1 YP 003941819 1 701347 Proteobacteria
Raoultella ornithinolytica B6 YP 007873458 1 1286170 Proteobacteria
Klebsiella oxytoca E718 YP 006498816 1 1191061 Proteobacteria
Klebsiella pneumoniae YP 007991518 1 573 Proteobacteria
Klebsiella pneumoniae 342 YP 002238568 1 507522 Proteobacteria
Klebsiella variicola At 22 YP 003439612 1 640131 Proteobacteria
Zymomonas mobilis subsp pomaceae ATCC 29192 YP 004661570 1 579138 Proteobacteria
Dickeya paradisiaca NCPPB 2511 WP 012765433 1 Dparadisiaca
Dickeya paradisiaca Ech703 WP 012765433 1 Dparadisiaca
Methylocystis sp SC2 YP 006593175 1 187303 Proteobacteria
Xanthobacter autotrophicus Py2 YP 001415226 1 78245 Proteobacteria
Hyphomicrobium nitrativorans NL23 YP 008864840 1 1029756 Proteobacteria
Methylobacterium extorquens PA1 YP 001640372 1 419610 Proteobacteria
Azorhizobium caulinodans ORS 571 YP 001527253 1 438753 Proteobacteria
Rhizobium etli bv mimosae str Mim1 YP 008368162 1 1328306 Proteobacteria
Rhizobium etli bv mimosae str Mim1 YP 008368404 1 1328306 Proteobacteria
Agrobacterium radiobacter K84 YP 002543069 1 311403 Proteobacteria
Rhizobium tropici CIAT 899 YP 007332307 1 698761 Proteobacteria
Rhizobium etli CFN 42 YP 467820 1 347834 Proteobacteria
Rhizobium etli bv mimosae str Mim1 YP 008363071 1 1328306 Proteobacteria
Rhizobium leguminosarum bv viciae 3841 YP 765882 1 216596 Proteobacteria
Rhizobium leguminosarum bv trifolii WSM2304 YP 002283762 1 395492 Proteobacteria
Rhodobacter sphaeroides ATCC 17025 YP 001168062 1 349102 Proteobacteria
Rhizobium sp IRBG74 YP 008631205 1 424182 Proteobacteria
Agrobacterium sp H13 3 YP 004277562 1 861208 Proteobacteria
Agrobacterium fabrum str C58 NP 353293 1 176299 Proteobacteria

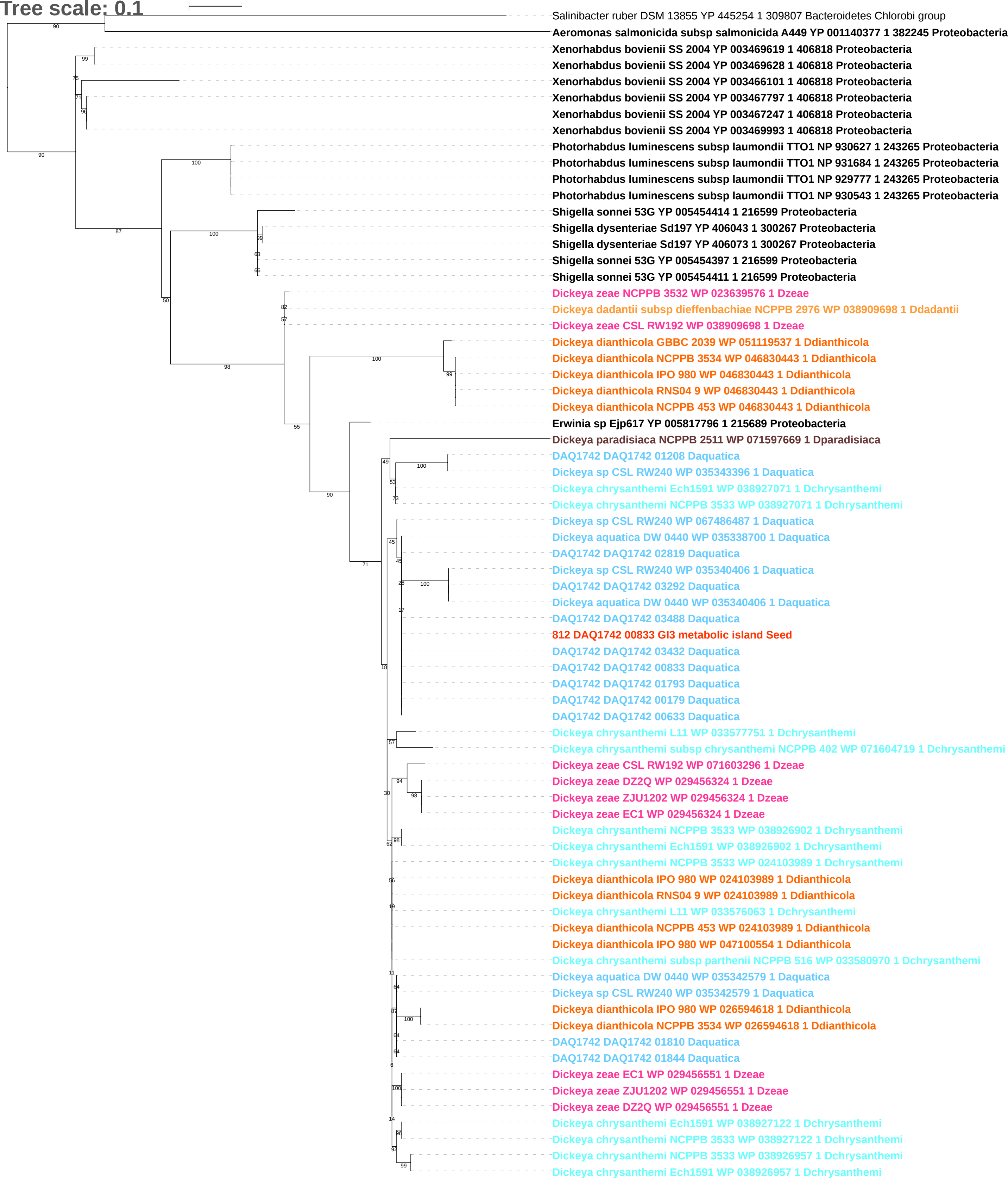
Tree scale: 0.1
Salinibacter ruber DSM 13855 YP 445254 1 309807 Bacteroidetes Chlorobi group
Aeromonas salmonicida subsp salmonicida A449 YP 001140377 1 382245 Proteobacteria
Xenorhabdus bovienii SS 2004 YP 003469619 1 406818 Proteobacteria
Xenorhabdus bovienii SS 2004 YP 003469628 1 406818 Proteobacteria
Xenorhabdus bovienii SS 2004 YP 003466101 1 406818 Proteobacteria
Xenorhabdus bovienii SS 2004 YP 003467797 1 406818 Proteobacteria
Xenorhabdus bovienii SS 2004 YP 003467247 1 406818 Proteobacteria
Xenorhabdus bovienii SS 2004 YP 003469993 1 406818 Proteobacteria
Photorhabdus luminescens subsp laumondii TTO1 NP 930627 1 243265 Proteobacteria
Photorhabdus luminescens subsp laumondii TTO1 NP 931684 1 243265 Proteobacteria
Photorhabdus luminescens subsp laumondii TTO1 NP 929777 1 243265 Proteobacteria
Photorhabdus luminescens subsp laumondii TTO1 NP 930543 1 243265 Proteobacteria
Shigella sonnei 53G YP 005454414 1 216599 Proteobacteria
Shigella dysenteriae Sd197 YP 406043 1 300267 Proteobacteria
Shigella dysenteriae Sd197 YP 406073 1 300267 Proteobacteria
Shigella sonnei 53G YP 005454397 1 216599 Proteobacteria
Shigella sonnei 53G YP 005454411 1 216599 Proteobacteria
Dickeya zeae NCPPB 3532 WP 023639576 1 Dzeae
Dickeya dadantii subsp dieffenbachiae NCPPB 2976 WP 038909698 1 Ddadantii
Dickeya zeae CSL RW192 WP 038909698 1 Dzeae
Dickeya dianthicola GBBC 2039 WP 051119537 1 Ddianthicola
Dickeya dianthicola NCPPB 3534 WP 046830443 1 Ddianthicola
Dickeya dianthicola IPO 980 WP 046830443 1 Ddianthicola
Dickeya dianthicola RNS04 9 WP 046830443 1 Ddianthicola
Dickeya dianthicola NCPPB 453 WP 046830443 1 Ddianthicola
Erwinia sp Ejp617 YP 005817796 1 215689 Proteobacteria
Dickeya paradisiaca NCPPB 2511 WP 071597669 1 Dparadisiaca
DAQ1742 DAQ1742 01208 Daquatica
Dickeya sp CSL RW240 WP 035343396 1 Daquatica
Dickeya chrysanthemi Ech1591 WP 038927071 1 Dchrysanthemi
Dickeya chrysanthemi NCPPB 3533 WP 038927071 1 Dchrysanthemi
Dickeya sp CSL RW240 WP 067486487 1 Daquatica
Dickeya aquatica DW 0440 WP 035338700 1 Daquatica
DAQ1742 DAQ1742 02819 Daquatica
Dickeya sp CSL RW240 WP 035340406 1 Daquatica
DAQ1742 DAQ1742 03292 Daquatica
Dickeya aquatica DW 0440 WP 035340406 1 Daquatica
DAQ1742 DAQ1742 03488 Daquatica
812 DAQ1742 00833 GI3 metabolic island Seed
DAQ1742 DAQ1742 03432 Daquatica
DAQ1742 DAQ1742 00833 Daquatica
DAQ1742 DAQ1742 01793 Daquatica
DAQ1742 DAQ1742 00179 Daquatica
DAQ1742 DAQ1742 00633 Daquatica
Dickeya chrysanthemi L11 WP 033577751 1 Dchrysanthemi
Dickeya chrysanthemi subsp chrysanthemi NCPPB 402 WP 071604719 1 Dchrysanthemi
Dickeya zeae CSL RW192 WP 071603296 1 Dzeae
Dickeya zeae DZ2Q WP 029456324 1 Dzeae
Dickeya zeae ZJU1202 WP 029456324 1 Dzeae
Dickeya zeae EC1 WP 029456324 1 Dzeae
Dickeya chrysanthemi NCPPB 3533 WP 038926902 1 Dchrysanthemi
Dickeya chrysanthemi Ech1591 WP 038926902 1 Dchrysanthemi
Dickeya chrysanthemi NCPPB 3533 WP 024103989 1 Dchrysanthemi
Dickeya dianthicola IPO 980 WP 024103989 1 Ddianthicola
Dickeya dianthicola RNS04 9 WP 024103989 1 Ddianthicola
Dickeya chrysanthemi L11 WP 033576063 1 Dchrysanthemi
Dickeya dianthicola NCPPB 453 WP 024103989 1 Ddianthicola
Dickeya dianthicola IPO 980 WP 047100554 1 Ddianthicola
Dickeya chrysanthemi subsp parthenii NCPPB 516 WP 033580970 1 Dchrysanthemi
Dickeya aquatica DW 0440 WP 035342579 1 Daquatica
Dickeya sp CSL RW240 WP 035342579 1 Daquatica
Dickeya dianthicola IPO 980 WP 026594618 1 Ddianthicola
Dickeya dianthicola NCPPB 3534 WP 026594618 1 Ddianthicola
DAQ1742 DAQ1742 01810 Daquatica
DAQ1742 DAQ1742 01844 Daquatica
Dickeya zeae EC1 WP 029456551 1 Dzeae
Dickeya zeae ZJU1202 WP 029456551 1 Dzeae
Dickeya zeae DZ2Q WP 029456551 1 Dzeae
Dickeya chrysanthemi Ech1591 WP 038927122 1 Dchrysanthemi
Dickeya chrysanthemi NCPPB 3533 WP 038927122 1 Dchrysanthemi
Dickeya chrysanthemi NCPPB 3533 WP 038926957 1 Dchrysanthemi
Dickeya chrysanthemi Ech1591 WP 038926957 1 Dchrysanthemi

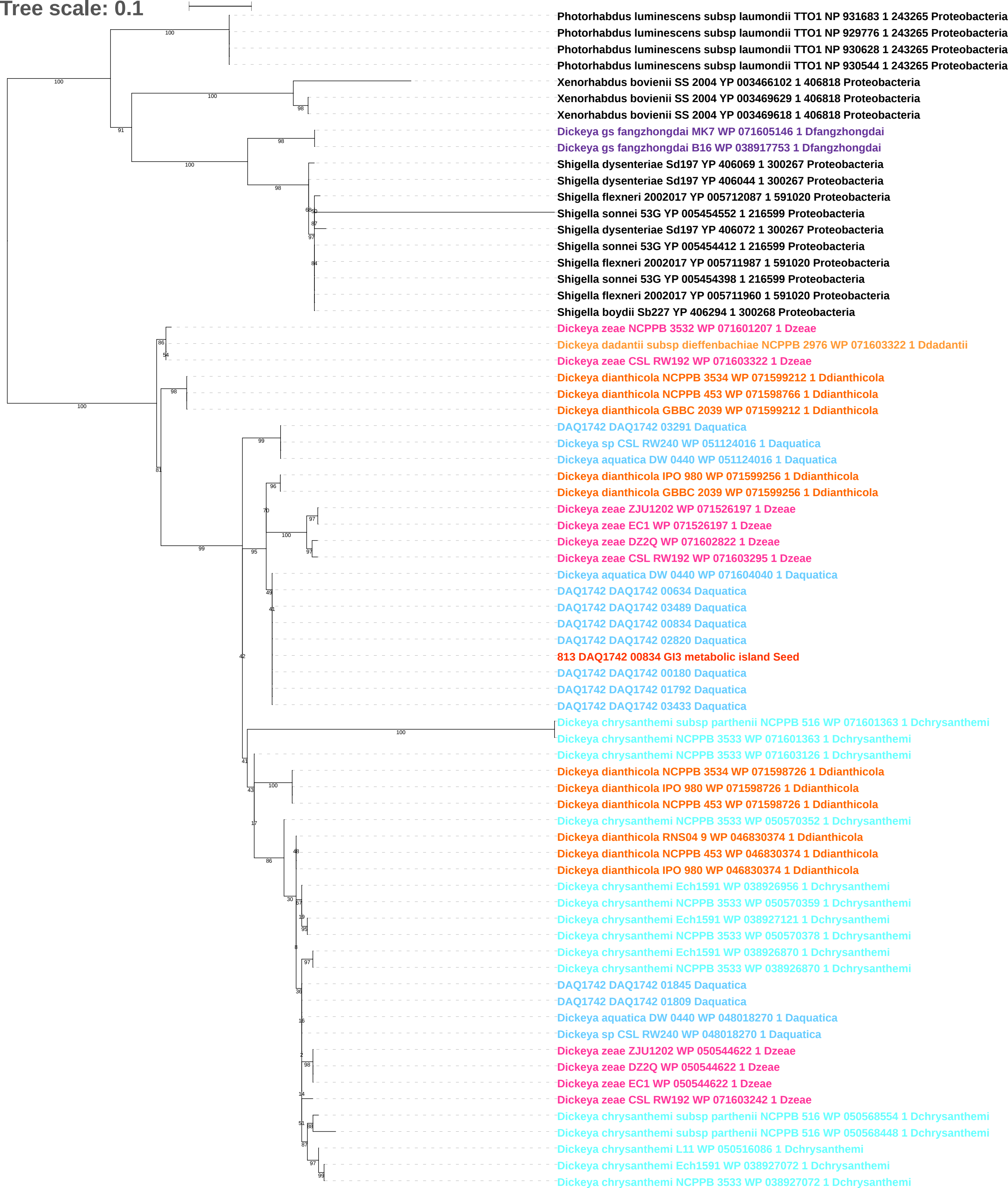
Tree scale: 0.1
Photorhabdus luminescens subsp laumondii TTO1 NP 931683 1 243265 Proteobacteria
Photorhabdus luminescens subsp laumondii TTO1 NP 929776 1 243265 Proteobacteria
Photorhabdus luminescens subsp laumondii TTO1 NP 930628 1 243265 Proteobacteria
Photorhabdus luminescens subsp laumondii TTO1 NP 930544 1 243265 Proteobacteria
Xenorhabdus bovienii SS 2004 YP 003466102 1 406818 Proteobacteria
Xenorhabdus bovienii SS 2004 YP 003469629 1 406818 Proteobacteria
Xenorhabdus bovienii SS 2004 YP 003469618 1 406818 Proteobacteria
Dickeya gs fangzhongdai MK7 WP 071605146 1 Dfangzhongdai
Dickeya gs fangzhongdai B16 WP 038917753 1 Dfangzhongdai
Shigella dysenteriae Sd197 YP 406069 1 300267 Proteobacteria
Shigella dysenteriae Sd197 YP 406044 1 300267 Proteobacteria
Shigella flexneri 2002017 YP 005712087 1 591020 Proteobacteria
Shigella sonnei 53G YP 005454552 1 216599 Proteobacteria
Shigella dysenteriae Sd197 YP 406072 1 300267 Proteobacteria
Shigella sonnei 53G YP 005454412 1 216599 Proteobacteria
Shigella flexneri 2002017 YP 005711987 1 591020 Proteobacteria
Shigella sonnei 53G YP 005454398 1 216599 Proteobacteria
Shigella flexneri 2002017 YP 005711960 1 591020 Proteobacteria
Shigella boydii Sb227 YP 406294 1 300268 Proteobacteria
Dickeya zeae NCPPB 3532 WP 071601207 1 Dzeae
Dickeya dadantii subsp dieffenbachiae NCPPB 2976 WP 071603322 1 Ddadantii
Dickeya zeae CSL RW192 WP 071603322 1 Dzeae
Dickeya dianthicola NCPPB 3534 WP 071599212 1 Ddianthicola
Dickeya dianthicola NCPPB 453 WP 071598766 1 Ddianthicola
Dickeya dianthicola GBBC 2039 WP 071599212 1 Ddianthicola
DAQ1742 DAQ1742 03291 Daquatica
Dickeya sp CSL RW240 WP 051124016 1 Daquatica
Dickeya aquatica DW 0440 WP 051124016 1 Daquatica
Dickeya dianthicola IPO 980 WP 071599256 1 Ddianthicola
Dickeya dianthicola GBBC 2039 WP 071599256 1 Ddianthicola
Dickeya zeae ZJU1202 WP 071526197 1 Dzeae
Dickeya zeae EC1 WP 071526197 1 Dzeae
Dickeya zeae DZ2Q WP 071602822 1 Dzeae
Dickeya zeae CSL RW192 WP 071603295 1 Dzeae
Dickeya aquatica DW 0440 WP 071604040 1 Daquatica
DAQ1742 DAQ1742 00634 Daquatica
DAQ1742 DAQ1742 03489 Daquatica
DAQ1742 DAQ1742 00834 Daquatica
DAQ1742 DAQ1742 02820 Daquatica
813 DAQ1742 00834 GI3 metabolic island Seed
DAQ1742 DAQ1742 00180 Daquatica
DAQ1742 DAQ1742 01792 Daquatica
DAQ1742 DAQ1742 03433 Daquatica
Dickeya chrysanthemi subsp parthenii NCPPB 516 WP 071601363 1 Dchrysanthemi
Dickeya chrysanthemi NCPPB 3533 WP 071601363 1 Dchrysanthemi
Dickeya chrysanthemi NCPPB 3533 WP 071603126 1 Dchrysanthemi
Dickeya dianthicola NCPPB 3534 WP 071598726 1 Ddianthicola
Dickeya dianthicola IPO 980 WP 071598726 1 Ddianthicola
Dickeya dianthicola NCPPB 453 WP 071598726 1 Ddianthicola
Dickeya chrysanthemi NCPPB 3533 WP 050570352 1 Dchrysanthemi
Dickeya dianthicola RNS04 9 WP 046830374 1 Ddianthicola
Dickeya dianthicola NCPPB 453 WP 046830374 1 Ddianthicola
Dickeya dianthicola IPO 980 WP 046830374 1 Ddianthicola
Dickeya chrysanthemi Ech1591 WP 038926956 1 Dchrysanthemi
Dickeya chrysanthemi NCPPB 3533 WP 050570359 1 Dchrysanthemi
Dickeya chrysanthemi Ech1591 WP 038927121 1 Dchrysanthemi
Dickeya chrysanthemi NCPPB 3533 WP 050570378 1 Dchrysanthemi
Dickeya chrysanthemi Ech1591 WP 038926870 1 Dchrysanthemi
Dickeya chrysanthemi NCPPB 3533 WP 038926870 1 Dchrysanthemi
DAQ1742 DAQ1742 01845 Daquatica
DAQ1742 DAQ1742 01809 Daquatica
Dickeya aquatica DW 0440 WP 048018270 1 Daquatica
Dickeya sp CSL RW240 WP 048018270 1 Daquatica
Dickeya zeae ZJU1202 WP 050544622 1 Dzeae
Dickeya zeae DZ2Q WP 050544622 1 Dzeae
Dickeya zeae EC1 WP 050544622 1 Dzeae
Dickeya zeae CSL RW192 WP 071603242 1 Dzeae
Dickeya chrysanthemi subsp parthenii NCPPB 516 WP 050568554 1 Dchrysanthemi
Dickeya chrysanthemi subsp parthenii NCPPB 516 WP 050568448 1 Dchrysanthemi
Dickeya chrysanthemi L11 WP 050516086 1 Dchrysanthemi
Dickeya chrysanthemi Ech1591 WP 038927072 1 Dchrysanthemi
Dickeya chrysanthemi NCPPB 3533 WP 038927072 1 Dchrysanthemi

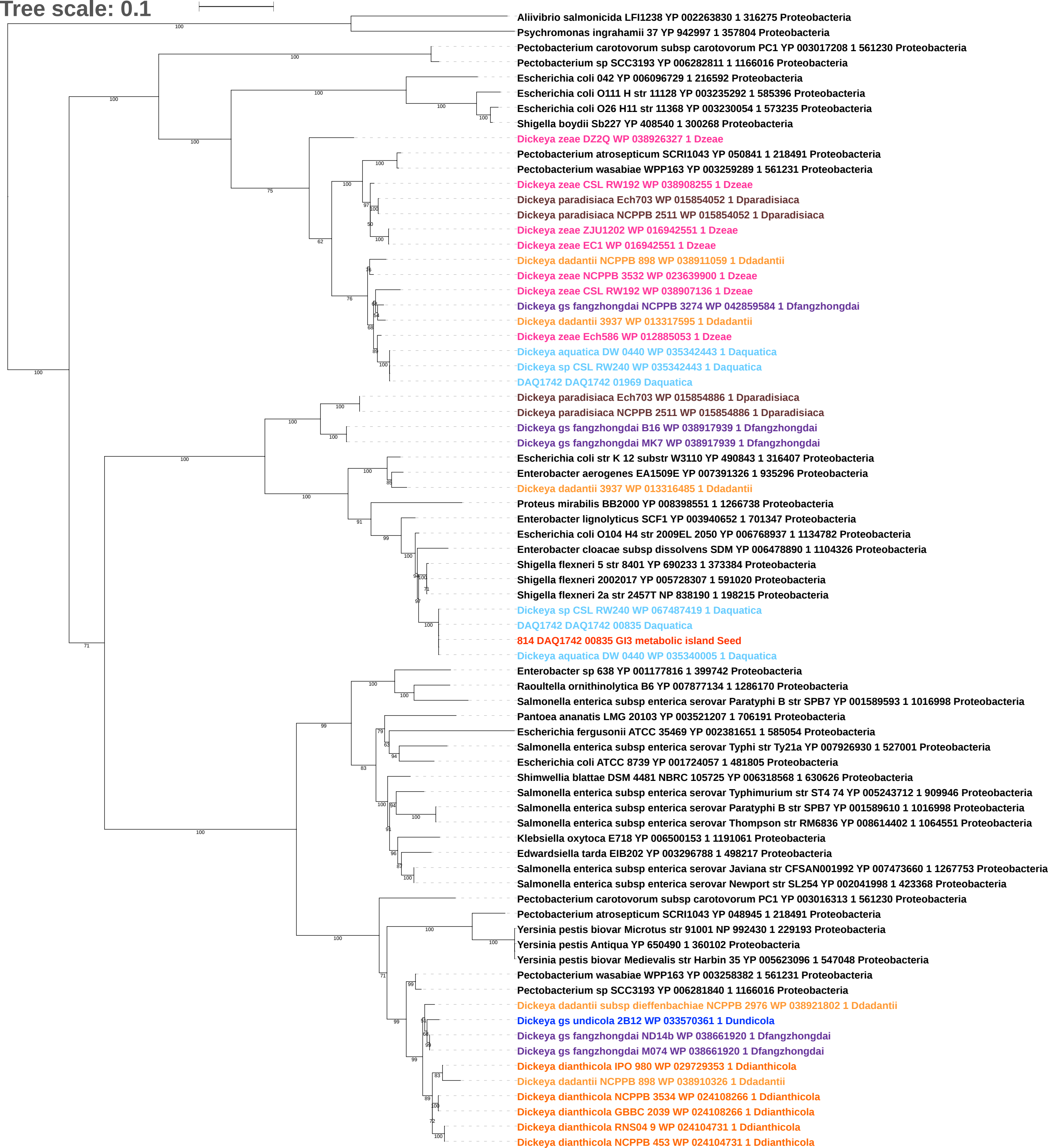

Tree scale: 0.1
Aliivibrio salmonicida LFI1238 YP 002263830 1 316275 Proteobacteria
Psychromonas ingrahamii 37 YP 942997 1 357804 Proteobacteria
Pectobacterium carotovorum subsp carotovorum PC1 YP 003017208 1 561230 Proteobacteria
Pectobacterium sp SCC3193 YP 006282811 1 1166016 Proteobacteria
Escherichia coli 042 YP 006096729 1 216592 Proteobacteria
Escherichia coli O111 H str 11128 YP 003235292 1 585396 Proteobacteria
Escherichia coli O26 H11 str 11368 YP 003230054 1 573235 Proteobacteria
Shigella boydii Sb227 YP 408540 1 300268 Proteobacteria
Dickeya zeae DZ2Q WP 038926327 1 Dzeae
Pectobacterium atrosepticum SCRI1043 YP 050841 1 218491 Proteobacteria
Pectobacterium wasabiae WPP163 YP 003259289 1 561231 Proteobacteria
Dickeya zeae CSL RW192 WP 038908255 1 Dzeae
Dickeya paradisiaca Ech703 WP 015854052 1 Dparadisiaca
Dickeya paradisiaca NCPPB 2511 WP 015854052 1 Dparadisiaca
Dickeya zeae ZJU1202 WP 016942551 1 Dzeae
Dickeya zeae EC1 WP 016942551 1 Dzeae
Dickeya dadantii NCPPB 898 WP 038911059 1 Ddadantii
Dickeya zeae NCPPB 3532 WP 023639900 1 Dzeae
Dickeya zeae CSL RW192 WP 038907136 1 Dzeae
Dickeya gs fangzhongdai NCPPB 3274 WP 042859584 1 Dfangzhongdai
Dickeya dadantii 3937 WP 013317595 1 Ddadantii
Dickeya zeae Ech586 WP 012885053 1 Dzeae
Dickeya aquatica DW 0440 WP 035342443 1 Daquatica
Dickeya sp CSL RW240 WP 035342443 1 Daquatica
DAQ1742 DAQ1742 01969 Daquatica
Dickeya paradisiaca Ech703 WP 015854886 1 Dparadisiaca
Dickeya paradisiaca NCPPB 2511 WP 015854886 1 Dparadisiaca
Dickeya gs fangzhongdai B16 WP 038917939 1 Dfangzhongdai
Dickeya gs fangzhongdai MK7 WP 038917939 1 Dfangzhongdai
Escherichia coli str K 12 substr W3110 YP 490843 1 316407 Proteobacteria
Enterobacter aerogenes EA1509E YP 007391326 1 935296 Proteobacteria
Dickeya dadantii 3937 WP 013316485 1 Ddadantii
Proteus mirabilis BB2000 YP 008398551 1 1266738 Proteobacteria
Enterobacter lignolyticus SCF1 YP 003940652 1 701347 Proteobacteria
Escherichia coli O104 H4 str 2009EL 2050 YP 006768937 1 1134782 Proteobacteria
Enterobacter cloacae subsp dissolvens SDM YP 006478890 1 1104326 Proteobacteria
Shigella flexneri 5 str 8401 YP 690233 1 373384 Proteobacteria
Shigella flexneri 2002017 YP 005728307 1 591020 Proteobacteria
Shigella flexneri 2a str 2457T NP 838190 1 198215 Proteobacteria
Dickeya sp CSL RW240 WP 067487419 1 Daquatica
DAQ1742 DAQ1742 00835 Daquatica
814 DAQ1742 00835 GI3 metabolic island Seed
Dickeya aquatica DW 0440 WP 035340005 1 Daquatica
Enterobacter sp 638 YP 001177816 1 399742 Proteobacteria
Raoultella ornithinolytica B6 YP 007877134 1 1286170 Proteobacteria
Salmonella enterica subsp enterica serovar Paratyphi B str SPB7 YP 001589593 1 1016998 Proteobacteria
Pantoea ananatis LMG 20103 YP 003521207 1 706191 Proteobacteria
Escherichia fergusonii ATCC 35469 YP 002381651 1 585054 Proteobacteria
Salmonella enterica subsp enterica serovar Typhi str Ty21a YP 007926930 1 527001 Proteobacteria
Escherichia coli ATCC 8739 YP 001724057 1 481805 Proteobacteria
Shimwellia blattae DSM 4481 NBRC 105725 YP 006318568 1 630626 Proteobacteria
Salmonella enterica subsp enterica serovar Typhimurium str ST4 74 YP 005243712 1 909946 Proteobacteria
Salmonella enterica subsp enterica serovar Paratyphi B str SPB7 YP 001589610 1 1016998 Proteobacteria
Salmonella enterica subsp enterica serovar Thompson str RM6836 YP 008614402 1 1064551 Proteobacteria
Klebsiella oxytoca E718 YP 006500153 1 1191061 Proteobacteria
Edwardsiella tarda EIB202 YP 003296788 1 498217 Proteobacteria
Salmonella enterica subsp enterica serovar Javiana str CFSAN001992 YP 007473660 1 1267753 Proteobacteria
Salmonella enterica subsp enterica serovar Newport str SL254 YP 002041998 1 423368 Proteobacteria
Pectobacterium carotovorum subsp carotovorum PC1 YP 003016313 1 561230 Proteobacteria
Pectobacterium atrosepticum SCRI1043 YP 048945 1 218491 Proteobacteria
Yersinia pestis biovar Microtus str 91001 NP 992430 1 229193 Proteobacteria
Yersinia pestis Antiqua YP 650490 1 360102 Proteobacteria
Yersinia pestis biovar Medievalis str Harbin 35 YP 005623096 1 547048 Proteobacteria
Pectobacterium wasabiae WPP163 YP 003258382 1 561231 Proteobacteria
Pectobacterium sp SCC3193 YP 006281840 1 1166016 Proteobacteria
Dickeya dadantii subsp dieffenbachiae NCPPB 2976 WP 038921802 1 Ddadantii
Dickeya gs undicola 2B12 WP 033570361 1 Dundicola
Dickeya gs fangzhongdai ND14b WP 038661920 1 Dfangzhongdai
Dickeya gs fangzhongdai M074 WP 038661920 1 Dfangzhongdai
Dickeya dianthicola IPO 980 WP 029729353 1 Ddianthicola
Dickeya dadantii NCPPB 898 WP 038910326 1 Ddadantii
Dickeya dianthicola NCPPB 3534 WP 024108266 1 Ddianthicola
Dickeya dianthicola GBBC 2039 WP 024108266 1 Ddianthicola
Dickeya dianthicola RNS04 9 WP 024104731 1 Ddianthicola
Dickeya dianthicola NCPPB 453 WP 024104731 1 Ddianthicola

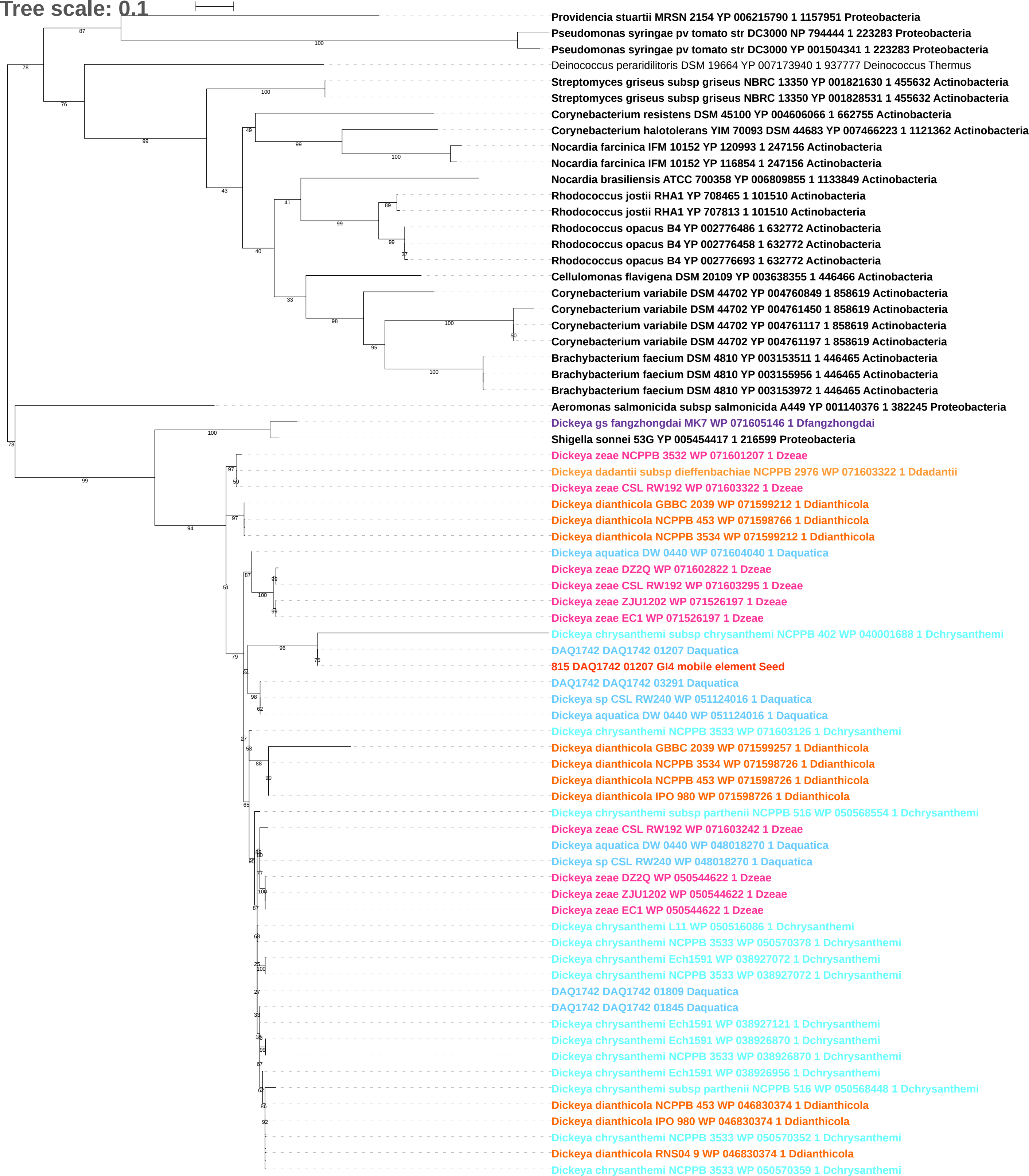
Tree scale: 0.1
Providencia stuartii MRSN 2154 YP 006215790 1 1157951 Proteobacteria
Pseudomonas syringae pv tomato str DC3000 NP 794444 1 223283 Proteobacteria
Pseudomonas syringae pv tomato str DC3000 YP 001504341 1 223283 Proteobacteria
Deinococcus peraridilitoris DSM 19664 YP 007173940 1 937777 Deinococcus Thermus
Streptomyces griseus subsp griseus NBRC 13350 YP 001821630 1 455632 Actinobacteria
Streptomyces griseus subsp griseus NBRC 13350 YP 001828531 1 455632 Actinobacteria
Corynebacterium resistens DSM 45100 YP 004606066 1 662755 Actinobacteria
Corynebacterium halotolerans YIM 70093 DSM 44683 YP 007466223 1 1121362 Actinobacteria
Nocardia farcinica IFM 10152 YP 120993 1 247156 Actinobacteria
Nocardia farcinica IFM 10152 YP 116854 1 247156 Actinobacteria
Nocardia brasiliensis ATCC 700358 YP 006809855 1 1133849 Actinobacteria
Rhodococcus jostii RHA1 YP 708465 1 101510 Actinobacteria
Rhodococcus jostii RHA1 YP 707813 1 101510 Actinobacteria
Rhodococcus opacus B4 YP 002776486 1 632772 Actinobacteria
Rhodococcus opacus B4 YP 002776458 1 632772 Actinobacteria
Rhodococcus opacus B4 YP 002776693 1 632772 Actinobacteria
Cellulomonas flavigena DSM 20109 YP 003638355 1 446466 Actinobacteria
Corynebacterium variabile DSM 44702 YP 004760849 1 858619 Actinobacteria
Corynebacterium variabile DSM 44702 YP 004761450 1 858619 Actinobacteria
Corynebacterium variabile DSM 44702 YP 004761117 1 858619 Actinobacteria
Corynebacterium variabile DSM 44702 YP 004761197 1 858619 Actinobacteria
Brachybacterium faecium DSM 4810 YP 003153511 1 446465 Actinobacteria
Brachybacterium faecium DSM 4810 YP 003155956 1 446465 Actinobacteria
Brachybacterium faecium DSM 4810 YP 003153972 1 446465 Actinobacteria
Aeromonas salmonicida subsp salmonicida A449 YP 001140376 1 382245 Proteobacteria
Dickeya gs fangzhongdai MK7 WP 071605146 1 Dfangzhongdai
Shigella sonnei 53G YP 005454417 1 216599 Proteobacteria
Dickeya zeae NCPPB 3532 WP 071601207 1 Dzeae
Dickeya dadantii subsp dieffenbachiae NCPPB 2976 WP 071603322 1 Ddadantii
Dickeya zeae CSL RW192 WP 071603322 1 Dzeae
Dickeya dianthicola GBBC 2039 WP 071599212 1 Ddianthicola
Dickeya dianthicola NCPPB 453 WP 071598766 1 Ddianthicola
Dickeya dianthicola NCPPB 3534 WP 071599212 1 Ddianthicola
Dickeya aquatica DW 0440 WP 071604040 1 Daquatica
Dickeya zeae DZ2Q WP 071602822 1 Dzeae
Dickeya zeae CSL RW192 WP 071603295 1 Dzeae
Dickeya zeae ZJU1202 WP 071526197 1 Dzeae
Dickeya zeae EC1 WP 071526197 1 Dzeae
Dickeya chrysanthemi subsp chrysanthemi NCPPB 402 WP 040001688 1 Dchrysanthemi
DAQ1742 DAQ1742 01207 Daquatica
815 DAQ1742 01207 GI4 mobile element Seed
DAQ1742 DAQ1742 03291 Daquatica
Dickeya sp CSL RW240 WP 051124016 1 Daquatica
Dickeya aquatica DW 0440 WP 051124016 1 Daquatica
Dickeya chrysanthemi NCPPB 3533 WP 071603126 1 Dchrysanthemi
Dickeya dianthicola GBBC 2039 WP 071599257 1 Ddianthicola
Dickeya dianthicola NCPPB 3534 WP 071598726 1 Ddianthicola
Dickeya dianthicola NCPPB 453 WP 071598726 1 Ddianthicola
Dickeya dianthicola IPO 980 WP 071598726 1 Ddianthicola
Dickeya chrysanthemi subsp parthenii NCPPB 516 WP 050568554 1 Dchrysanthemi
Dickeya zeae CSL RW192 WP 071603242 1 Dzeae
Dickeya aquatica DW 0440 WP 048018270 1 Daquatica
Dickeya sp CSL RW240 WP 048018270 1 Daquatica
Dickeya zeae DZ2Q WP 050544622 1 Dzeae
Dickeya zeae ZJU1202 WP 050544622 1 Dzeae
Dickeya zeae EC1 WP 050544622 1 Dzeae
Dickeya chrysanthemi L11 WP 050516086 1 Dchrysanthemi
Dickeya chrysanthemi NCPPB 3533 WP 050570378 1 Dchrysanthemi
Dickeya chrysanthemi Ech1591 WP 038927072 1 Dchrysanthemi
Dickeya chrysanthemi NCPPB 3533 WP 038927072 1 Dchrysanthemi
DAQ1742 DAQ1742 01809 Daquatica
DAQ1742 DAQ1742 01845 Daquatica
Dickeya chrysanthemi Ech1591 WP 038927121 1 Dchrysanthemi
Dickeya chrysanthemi Ech1591 WP 038926870 1 Dchrysanthemi
Dickeya chrysanthemi NCPPB 3533 WP 038926870 1 Dchrysanthemi
Dickeya chrysanthemi Ech1591 WP 038926956 1 Dchrysanthemi
Dickeya chrysanthemi subsp parthenii NCPPB 516 WP 050568448 1 Dchrysanthemi
Dickeya dianthicola NCPPB 453 WP 046830374 1 Ddianthicola
Dickeya dianthicola IPO 980 WP 046830374 1 Ddianthicola
Dickeya chrysanthemi NCPPB 3533 WP 050570352 1 Dchrysanthemi
Dickeya dianthicola RNS04 9 WP 046830374 1 Ddianthicola
Dickeya chrysanthemi NCPPB 3533 WP 050570359 1 Dchrysanthemi

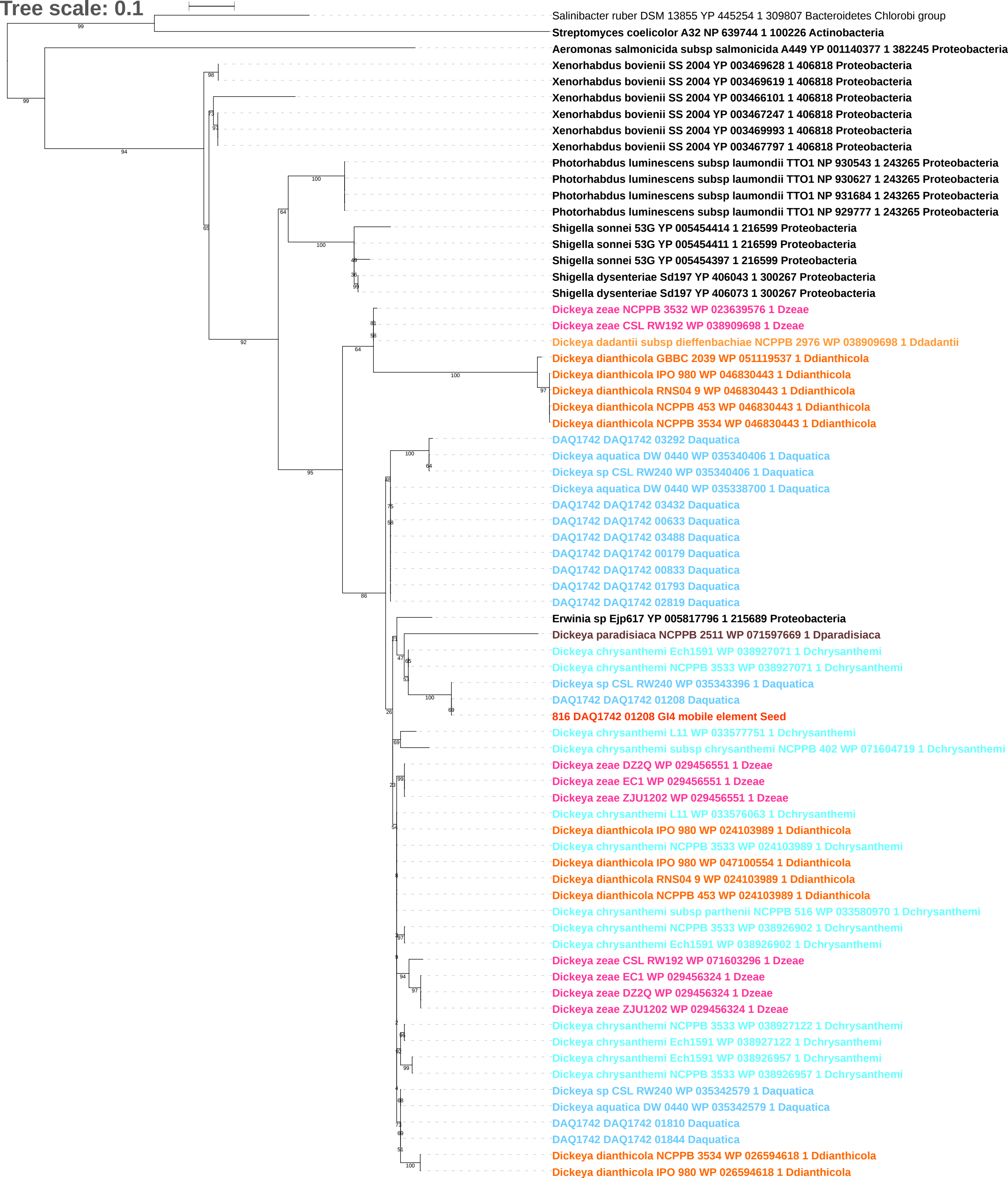
Tree scale: 0.1
Salinibacter ruber DSM 13855 YP 445254 1 309807 Bacteroidetes Chlorobi group
Streptomyces coelicolor A32 NP 639744 1 100226 Actinobacteria
Aeromonas salmonicida subsp salmonicida A449 YP 001140377 1 382245 Proteobacteria
Xenorhabdus bovienii SS 2004 YP 003469628 1 406818 Proteobacteria
Xenorhabdus bovienii SS 2004 YP 003469619 1 406818 Proteobacteria
Xenorhabdus bovienii SS 2004 YP 003466101 1 406818 Proteobacteria
Xenorhabdus bovienii SS 2004 YP 003467247 1 406818 Proteobacteria
Xenorhabdus bovienii SS 2004 YP 003469993 1 406818 Proteobacteria
Xenorhabdus bovienii SS 2004 YP 003467797 1 406818 Proteobacteria
Photorhabdus luminescens subsp laumondii TTO1 NP 930543 1 243265 Proteobacteria
Photorhabdus luminescens subsp laumondii TTO1 NP 930627 1 243265 Proteobacteria
Photorhabdus luminescens subsp laumondii TTO1 NP 931684 1 243265 Proteobacteria
Photorhabdus luminescens subsp laumondii TTO1 NP 929777 1 243265 Proteobacteria
Shigella sonnei 53G YP 005454414 1 216599 Proteobacteria
Shigella sonnei 53G YP 005454411 1 216599 Proteobacteria
Shigella sonnei 53G YP 005454397 1 216599 Proteobacteria
Shigella dysenteriae Sd197 YP 406043 1 300267 Proteobacteria
Shigella dysenteriae Sd197 YP 406073 1 300267 Proteobacteria
Dickeya zeae NCPPB 3532 WP 023639576 1 Dzeae
Dickeya zeae CSL RW192 WP 038909698 1 Dzeae
Dickeya dadantii subsp dieffenbachiae NCPPB 2976 WP 038909698 1 Ddadantii
Dickeya dianthicola GBBC 2039 WP 051119537 1 Ddianthicola
Dickeya dianthicola IPO 980 WP 046830443 1 Ddianthicola
Dickeya dianthicola RNS04 9 WP 046830443 1 Ddianthicola
Dickeya dianthicola NCPPB 453 WP 046830443 1 Ddianthicola
Dickeya dianthicola NCPPB 3534 WP 046830443 1 Ddianthicola
DAQ1742 DAQ1742 03292 Daquatica
Dickeya aquatica DW 0440 WP 035340406 1 Daquatica
Dickeya sp CSL RW240 WP 035340406 1 Daquatica
Dickeya aquatica DW 0440 WP 035338700 1 Daquatica
DAQ1742 DAQ1742 03432 Daquatica
DAQ1742 DAQ1742 00633 Daquatica
DAQ1742 DAQ1742 03488 Daquatica
DAQ1742 DAQ1742 00179 Daquatica
DAQ1742 DAQ1742 00833 Daquatica
DAQ1742 DAQ1742 01793 Daquatica
DAQ1742 DAQ1742 02819 Daquatica
Erwinia sp Ejp617 YP 005817796 1 215689 Proteobacteria
Dickeya paradisiaca NCPPB 2511 WP 071597669 1 Dparadisiaca
Dickeya chrysanthemi Ech1591 WP 038927071 1 Dchrysanthemi
Dickeya chrysanthemi NCPPB 3533 WP 038927071 1 Dchrysanthemi
Dickeya sp CSL RW240 WP 035343396 1 Daquatica
DAQ1742 DAQ1742 01208 Daquatica
816 DAQ1742 01208 GI4 mobile element Seed
Dickeya chrysanthemi L11 WP 033577751 1 Dchrysanthemi
Dickeya chrysanthemi subsp chrysanthemi NCPPB 402 WP 071604719 1 Dchrysanthemi
Dickeya zeae DZ2Q WP 029456551 1 Dzeae
Dickeya zeae EC1 WP 029456551 1 Dzeae
Dickeya zeae ZJU1202 WP 029456551 1 Dzeae
Dickeya chrysanthemi L11 WP 033576063 1 Dchrysanthemi
Dickeya dianthicola IPO 980 WP 024103989 1 Ddianthicola
Dickeya chrysanthemi NCPPB 3533 WP 024103989 1 Dchrysanthemi
Dickeya dianthicola IPO 980 WP 047100554 1 Ddianthicola
Dickeya dianthicola RNS04 9 WP 024103989 1 Ddianthicola
Dickeya dianthicola NCPPB 453 WP 024103989 1 Ddianthicola
Dickeya chrysanthemi subsp parthenii NCPPB 516 WP 033580970 1 Dchrysanthemi
Dickeya chrysanthemi NCPPB 3533 WP 038926902 1 Dchrysanthemi
Dickeya chrysanthemi Ech1591 WP 038926902 1 Dchrysanthemi
Dickeya zeae CSL RW192 WP 071603296 1 Dzeae
Dickeya zeae EC1 WP 029456324 1 Dzeae
Dickeya zeae DZ2Q WP 029456324 1 Dzeae
Dickeya zeae ZJU1202 WP 029456324 1 Dzeae
Dickeya chrysanthemi NCPPB 3533 WP 038927122 1 Dchrysanthemi
Dickeya chrysanthemi Ech1591 WP 038927122 1 Dchrysanthemi
Dickeya chrysanthemi Ech1591 WP 038926957 1 Dchrysanthemi
Dickeya chrysanthemi NCPPB 3533 WP 038926957 1 Dchrysanthemi
Dickeya sp CSL RW240 WP 035342579 1 Daquatica
Dickeya aquatica DW 0440 WP 035342579 1 Daquatica
DAQ1742 DAQ1742 01810 Daquatica
DAQ1742 DAQ1742 01844 Daquatica
Dickeya dianthicola NCPPB 3534 WP 026594618 1 Ddianthicola
Dickeya dianthicola IPO 980 WP 026594618 1 Ddianthicola

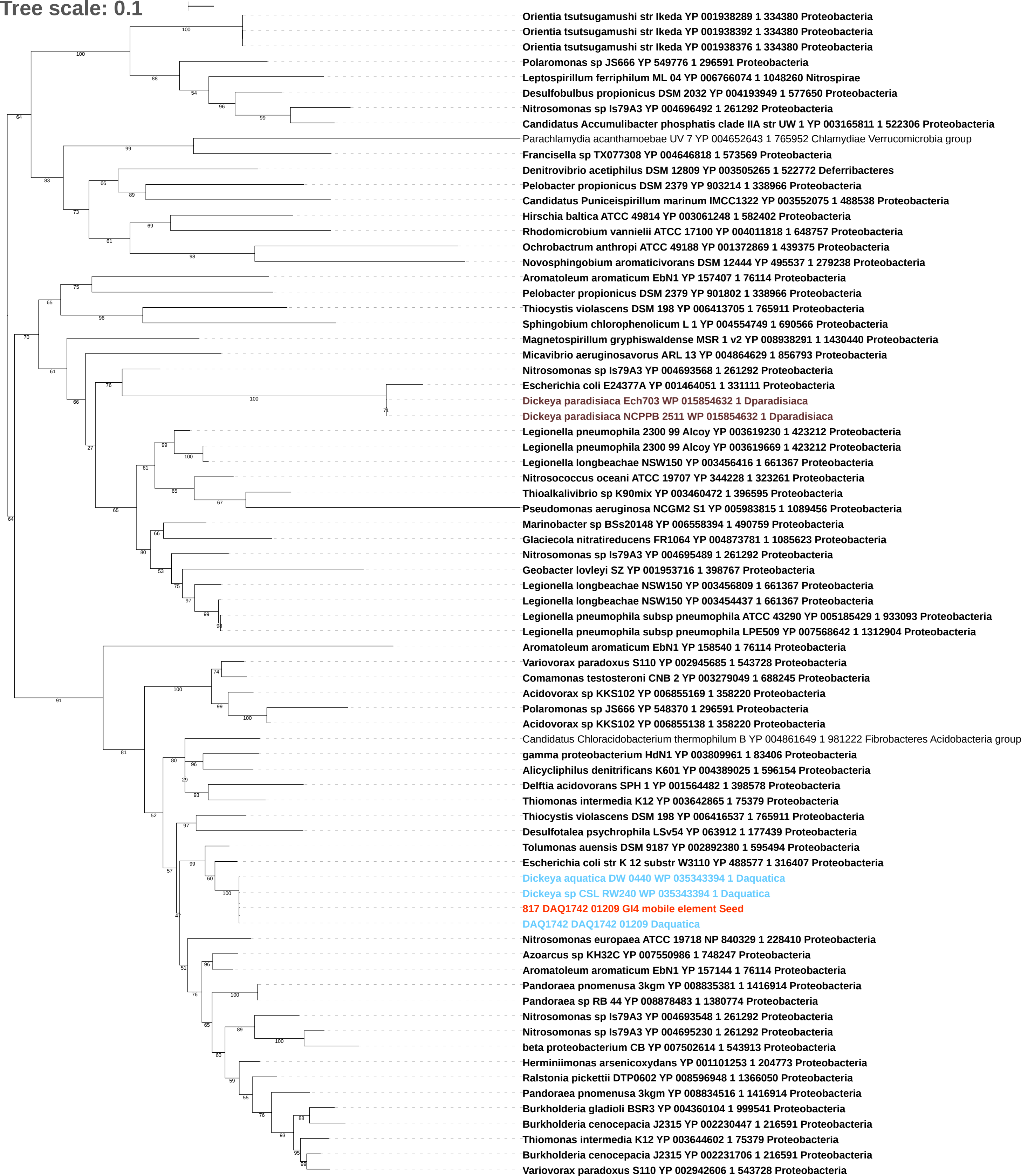

Tree scale: 0.1
Orientia tsutsugamushi str Ikeda YP 001938289 1 334380 Proteobacteria
Orientia tsutsugamushi str Ikeda YP 001938392 1 334380 Proteobacteria
Orientia tsutsugamushi str Ikeda YP 001938376 1 334380 Proteobacteria
Polaromonas sp JS666 YP 549776 1 296591 Proteobacteria
Leptospirillum ferriphilum ML 04 YP 006766074 1 1048260 Nitrospirae
Desulfobulbus propionicus DSM 2032 YP 004193949 1 577650 Proteobacteria
Nitrosomonas sp Is79A3 YP 004696492 1 261292 Proteobacteria
Candidatus Accumulibacter phosphatis clade IIA str UW 1 YP 003165811 1 522306 Proteobacteria
Parachlamydia acanthamoebae UV 7 YP 004652643 1 765952 Chlamydiae Verrucomicrobia group
Francisella sp TX077308 YP 004646818 1 573569 Proteobacteria
Denitrovibrio acetiphilus DSM 12809 YP 003505265 1 522772 Deferribacteres
Pelobacter propionicus DSM 2379 YP 903214 1 338966 Proteobacteria
Candidatus Puniceispirillum marinum IMCC1322 YP 003552075 1 488538 Proteobacteria
Hirschia baltica ATCC 49814 YP 003061248 1 582402 Proteobacteria
Rhodomicrobium vannielii ATCC 17100 YP 004011818 1 648757 Proteobacteria
Ochrobactrum anthropi ATCC 49188 YP 001372869 1 439375 Proteobacteria
Novosphingobium aromaticivorans DSM 12444 YP 495537 1 279238 Proteobacteria
Aromatoleum aromaticum EbN1 YP 157407 1 76114 Proteobacteria
Pelobacter propionicus DSM 2379 YP 901802 1 338966 Proteobacteria
Thiocystis violascens DSM 198 YP 006413705 1 765911 Proteobacteria
Sphingobium chlorophenolicum L 1 YP 004554749 1 690566 Proteobacteria
Magnetospirillum gryphiswaldense MSR 1 v2 YP 008938291 1 1430440 Proteobacteria
Micavibrio aeruginosavorus ARL 13 YP 004864629 1 856793 Proteobacteria
Nitrosomonas sp Is79A3 YP 004693568 1 261292 Proteobacteria
Escherichia coli E24377A YP 001464051 1 331111 Proteobacteria
Dickeya paradisiaca Ech703 WP 015854632 1 Dparadisiaca
Dickeya paradisiaca NCPPB 2511 WP 015854632 1 Dparadisiaca
Legionella pneumophila 2300 99 Alcoy YP 003619230 1 423212 Proteobacteria
Legionella pneumophila 2300 99 Alcoy YP 003619669 1 423212 Proteobacteria
Legionella longbeachae NSW150 YP 003456416 1 661367 Proteobacteria
Nitrosococcus oceani ATCC 19707 YP 344228 1 323261 Proteobacteria
Thioalkalivibrio sp K90mix YP 003460472 1 396595 Proteobacteria
Pseudomonas aeruginosa NCGM2 S1 YP 005983815 1 1089456 Proteobacteria
Marinobacter sp BSs20148 YP 006558394 1 490759 Proteobacteria
Glaciecola nitratireducens FR1064 YP 004873781 1 1085623 Proteobacteria
Nitrosomonas sp Is79A3 YP 004695489 1 261292 Proteobacteria
Geobacter lovleyi SZ YP 001953716 1 398767 Proteobacteria
Legionella longbeachae NSW150 YP 003456809 1 661367 Proteobacteria
Legionella longbeachae NSW150 YP 003454437 1 661367 Proteobacteria
Legionella pneumophila subsp pneumophila ATCC 43290 YP 005185429 1 933093 Proteobacteria
Legionella pneumophila subsp pneumophila LPE509 YP 007568642 1 1312904 Proteobacteria
Aromatoleum aromaticum EbN1 YP 158540 1 76114 Proteobacteria
Variovorax paradoxus S110 YP 002945685 1 543728 Proteobacteria
Comamonas testosteroni CNB 2 YP 003279049 1 688245 Proteobacteria
Acidovorax sp KKS102 YP 006855169 1 358220 Proteobacteria
Polaromonas sp JS666 YP 548370 1 296591 Proteobacteria
Acidovorax sp KKS102 YP 006855138 1 358220 Proteobacteria
Candidatus Chloracidobacterium thermophilum B YP 004861649 1 981222 Fibrobacteres Acidobacteria group
gamma proteobacterium HdN1 YP 003809961 1 83406 Proteobacteria
Alicycliphilus denitrificans K601 YP 004389025 1 596154 Proteobacteria
Delftia acidovorans SPH 1 YP 001564482 1 398578 Proteobacteria
Thiomonas intermedia K12 YP 003642865 1 75379 Proteobacteria
Thiocystis violascens DSM 198 YP 006416537 1 765911 Proteobacteria
Desulfotalea psychrophila LSv54 YP 063912 1 177439 Proteobacteria
Tolumonas auensis DSM 9187 YP 002892380 1 595494 Proteobacteria
Escherichia coli str K 12 substr W3110 YP 488577 1 316407 Proteobacteria
Dickeya aquatica DW 0440 WP 035343394 1 Daquatica
Dickeya sp CSL RW240 WP 035343394 1 Daquatica
817 DAQ1742 01209 GI4 mobile element Seed
DAQ1742 DAQ1742 01209 Daquatica
Nitrosomonas europaea ATCC 19718 NP 840329 1 228410 Proteobacteria
Azoarcus sp KH32C YP 007550986 1 748247 Proteobacteria
Aromatoleum aromaticum EbN1 YP 157144 1 76114 Proteobacteria
Pandoraea pnomenusa 3kgm YP 008835381 1 1416914 Proteobacteria
Pandoraea sp RB 44 YP 008878483 1 1380774 Proteobacteria
Nitrosomonas sp Is79A3 YP 004693548 1 261292 Proteobacteria
Nitrosomonas sp Is79A3 YP 004695230 1 261292 Proteobacteria
beta proteobacterium CB YP 007502614 1 543913 Proteobacteria
Herminiimonas arsenicoxydans YP 001101253 1 204773 Proteobacteria
Ralstonia pickettii DTP0602 YP 008596948 1 1366050 Proteobacteria
Pandoraea pnomenusa 3kgm YP 008834516 1 1416914 Proteobacteria
Burkholderia gladioli BSR3 YP 004360104 1 999541 Proteobacteria
Burkholderia cenocepacia J2315 YP 002230447 1 216591 Proteobacteria
Thiomonas intermedia K12 YP 003644602 1 75379 Proteobacteria
Burkholderia cenocepacia J2315 YP 002231706 1 216591 Proteobacteria
Variovorax paradoxus S110 YP 002942606 1 543728 Proteobacteria

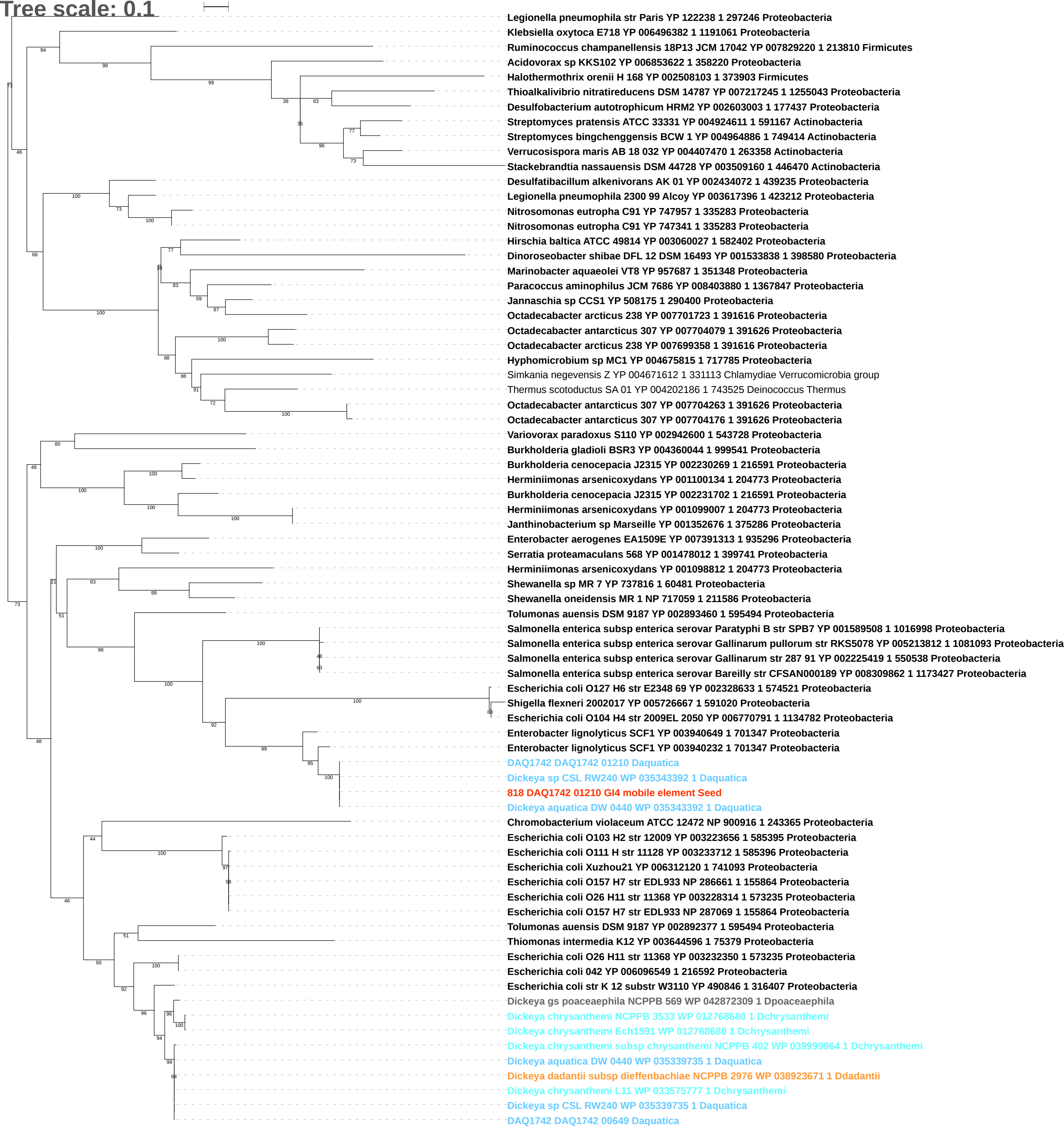
Tree scale: 0.1
Legionella pneumophila str Paris YP 122238 1 297246 Proteobacteria
Klebsiella oxytoca E718 YP 006496382 1 1191061 Proteobacteria
Ruminococcus champanellensis 18P13 JCM 17042 YP 007829220 1 213810 Firmicutes
Acidovorax sp KKS102 YP 006853622 1 358220 Proteobacteria
Halothermothrix orenii H 168 YP 002508103 1 373903 Firmicutes
Thioalkalivibrio nitratireducens DSM 14787 YP 007217245 1 1255043 Proteobacteria
Desulfobacterium autotrophicum HRM2 YP 002603003 1 177437 Proteobacteria
Streptomyces pratensis ATCC 33331 YP 004924611 1 591167 Actinobacteria
Streptomyces bingchenggensis BCW 1 YP 004964886 1 749414 Actinobacteria
Verrucosispora maris AB 18 032 YP 004407470 1 263358 Actinobacteria
Stackebrandtia nassauensis DSM 44728 YP 003509160 1 446470 Actinobacteria
Desulfatibacillum alkenivorans AK 01 YP 002434072 1 439235 Proteobacteria
Legionella pneumophila 2300 99 Alcoy YP 003617396 1 423212 Proteobacteria
Nitrosomonas eutropha C91 YP 747957 1 335283 Proteobacteria
Nitrosomonas eutropha C91 YP 747341 1 335283 Proteobacteria
Hirschia baltica ATCC 49814 YP 003060027 1 582402 Proteobacteria
Dinoroseobacter shibae DFL 12 DSM 16493 YP 001533838 1 398580 Proteobacteria
Marinobacter aquaeolei VT8 YP 957687 1 351348 Proteobacteria
Paracoccus aminophilus JCM 7686 YP 008403880 1 1367847 Proteobacteria
Jannaschia sp CCS1 YP 508175 1 290400 Proteobacteria
Octadecabacter arcticus 238 YP 007701723 1 391616 Proteobacteria
Octadecabacter antarcticus 307 YP 007704079 1 391626 Proteobacteria
Octadecabacter arcticus 238 YP 007699358 1 391616 Proteobacteria
Hyphomicrobium sp MC1 YP 004675815 1 717785 Proteobacteria
Simkania negevensis Z YP 004671612 1 331113 Chlamydiae Verrucomicrobia group
Thermus scotoductus SA 01 YP 004202186 1 743525 Deinococcus Thermus
Octadecabacter antarcticus 307 YP 007704263 1 391626 Proteobacteria
Octadecabacter antarcticus 307 YP 007704176 1 391626 Proteobacteria
Variovorax paradoxus S110 YP 002942600 1 543728 Proteobacteria
Burkholderia gladioli BSR3 YP 004360044 1 999541 Proteobacteria
Burkholderia cenocepacia J2315 YP 002230269 1 216591 Proteobacteria
Herminiimonas arsenicoxydans YP 001100134 1 204773 Proteobacteria
Burkholderia cenocepacia J2315 YP 002231702 1 216591 Proteobacteria
Herminiimonas arsenicoxydans YP 001099007 1 204773 Proteobacteria
Janthinobacterium sp Marseille YP 001352676 1 375286 Proteobacteria
Enterobacter aerogenes EA1509E YP 007391313 1 935296 Proteobacteria
Serratia proteamaculans 568 YP 001478012 1 399741 Proteobacteria
Herminiimonas arsenicoxydans YP 001098812 1 204773 Proteobacteria
Shewanella sp MR 7 YP 737816 1 60481 Proteobacteria
Shewanella oneidensis MR 1 NP 717059 1 211586 Proteobacteria
Tolumonas auensis DSM 9187 YP 002893460 1 595494 Proteobacteria
Salmonella enterica subsp enterica serovar Paratyphi B str SPB7 YP 001589508 1 1016998 Proteobacteria
Salmonella enterica subsp enterica serovar Gallinarum pullorum str RKS5078 YP 005213812 1 1081093 Proteobacteria
Salmonella enterica subsp enterica serovar Gallinarum str 287 91 YP 002225419 1 550538 Proteobacteria
Salmonella enterica subsp enterica serovar Bareilly str CFSAN000189 YP 008309862 1 1173427 Proteobacteria
Escherichia coli O127 H6 str E2348 69 YP 002328633 1 574521 Proteobacteria
Shigella flexneri 2002017 YP 005726667 1 591020 Proteobacteria
Escherichia coli O104 H4 str 2009EL 2050 YP 006770791 1 1134782 Proteobacteria
Enterobacter lignolyticus SCF1 YP 003940649 1 701347 Proteobacteria
Enterobacter lignolyticus SCF1 YP 003940232 1 701347 Proteobacteria
DAQ1742 DAQ1742 01210 Daquatica
Dickeya sp CSL RW240 WP 035343392 1 Daquatica
818 DAQ1742 01210 GI4 mobile element Seed
Dickeya aquatica DW 0440 WP 035343392 1 Daquatica
Chromobacterium violaceum ATCC 12472 NP 900916 1 243365 Proteobacteria
Escherichia coli O103 H2 str 12009 YP 003223656 1 585395 Proteobacteria
Escherichia coli O111 H str 11128 YP 003233712 1 585396 Proteobacteria
Escherichia coli Xuzhou21 YP 006312120 1 741093 Proteobacteria
Escherichia coli O157 H7 str EDL933 NP 286661 1 155864 Proteobacteria
Escherichia coli O26 H11 str 11368 YP 003228314 1 573235 Proteobacteria
Escherichia coli O157 H7 str EDL933 NP 287069 1 155864 Proteobacteria
Tolumonas auensis DSM 9187 YP 002892377 1 595494 Proteobacteria
Thiomonas intermedia K12 YP 003644596 1 75379 Proteobacteria
Escherichia coli O26 H11 str 11368 YP 003232350 1 573235 Proteobacteria
Escherichia coli 042 YP 006096549 1 216592 Proteobacteria
Escherichia coli str K 12 substr W3110 YP 490846 1 316407 Proteobacteria
Dickeya gs poaceaephila NCPPB 569 WP 042872309 1 Dpoaceaephila
Dickeya chrysanthemi NCPPB 3533 WP 012768680 1 Dchrysanthemi
Dickeya chrysanthemi Ech1591 WP 012768680 1 Dchrysanthemi
Dickeya chrysanthemi subsp chrysanthemi NCPPB 402 WP 039999964 1 Dchrysanthemi
Dickeya aquatica DW 0440 WP 035339735 1 Daquatica
Dickeya dadantii subsp dieffenbachiae NCPPB 2976 WP 038923671 1 Ddadantii
Dickeya chrysanthemi L11 WP 033575777 1 Dchrysanthemi
Dickeya sp CSL RW240 WP 035339735 1 Daquatica
DAQ1742 DAQ1742 00649 Daquatica

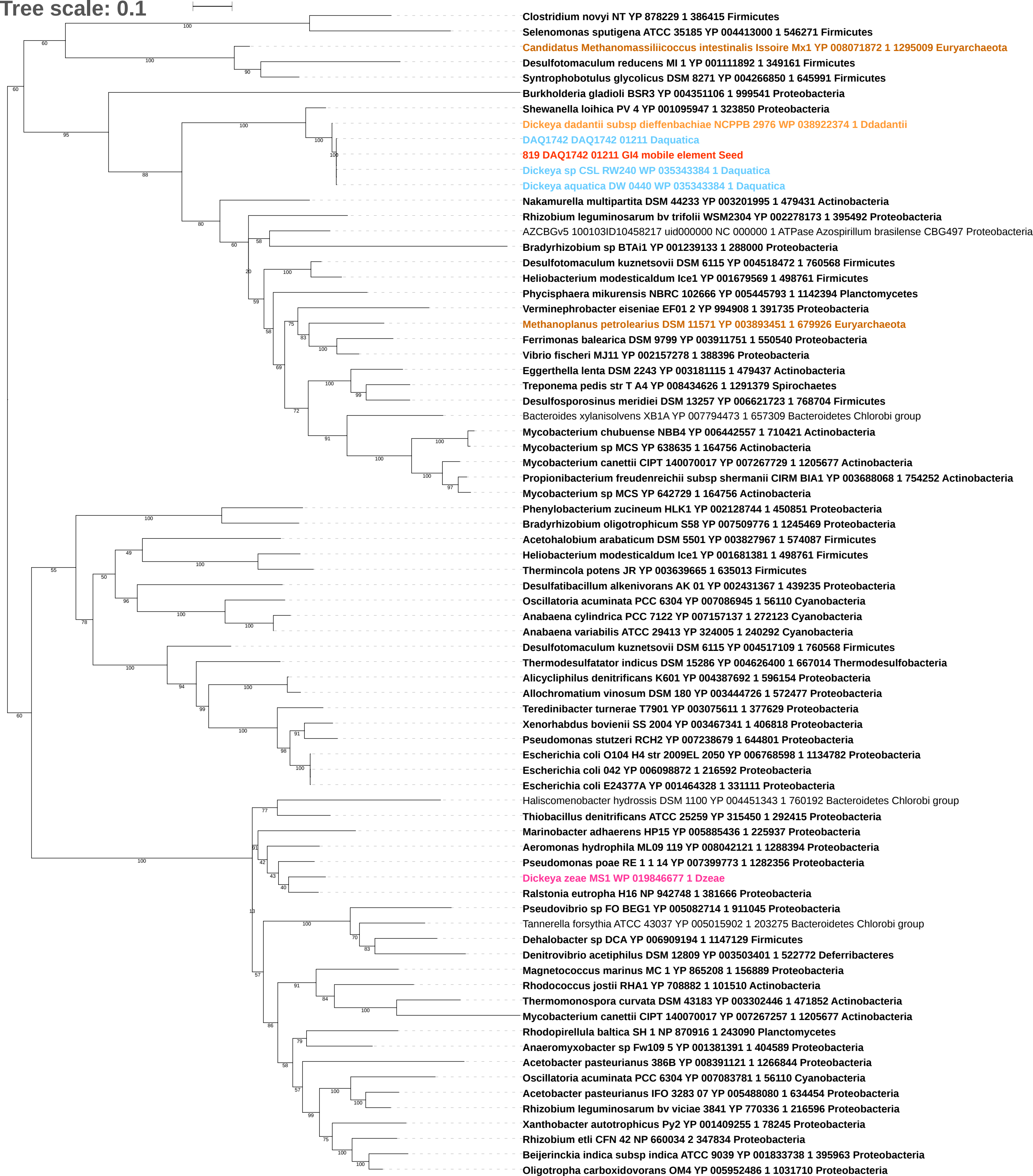
Tree scale: 0.1
Clostridium novyi NT YP 878229 1 386415 Firmicutes
Selenomonas sputigena ATCC 35185 YP 004413000 1 546271 Firmicutes
Candidatus Methanomassiliicoccus intestinalis Issoire Mx1 YP 008071872 1 1295009 Euryarchaeota
Desulfotomaculum reducens MI 1 YP 001111892 1 349161 Firmicutes
Syntrophobotulus glycolicus DSM 8271 YP 004266850 1 645991 Firmicutes
Burkholderia gladioli BSR3 YP 004351106 1 999541 Proteobacteria
Shewanella loihica PV 4 YP 001095947 1 323850 Proteobacteria
Dickeya dadantii subsp dieffenbachiae NCPPB 2976 WP 038922374 1 Ddadantii
DAQ1742 DAQ1742 01211 Daquatica
819 DAQ1742 01211 GI4 mobile element Seed
Dickeya sp CSL RW240 WP 035343384 1 Daquatica
Dickeya aquatica DW 0440 WP 035343384 1 Daquatica
Nakamurella multipartita DSM 44233 YP 003201995 1 479431 Actinobacteria
Rhizobium leguminosarum bv trifolii WSM2304 YP 002278173 1 395492 Proteobacteria
AZCBGv5 100103ID10458217 uid000000 NC 000000 1 ATPase Azospirillum brasilense CBG497 Proteobacteria
Bradyrhizobium sp BTAi1 YP 001239133 1 288000 Proteobacteria
Desulfotomaculum kuznetsovii DSM 6115 YP 004518472 1 760568 Firmicutes
Heliobacterium modesticaldum Ice1 YP 001679569 1 498761 Firmicutes
Phycisphaera mikurensis NBRC 102666 YP 005445793 1 1142394 Planctomycetes
Verminephrobacter eiseniae EF01 2 YP 994908 1 391735 Proteobacteria
Methanoplanus petrolearius DSM 11571 YP 003893451 1 679926 Euryarchaeota
Ferrimonas balearica DSM 9799 YP 003911751 1 550540 Proteobacteria
Vibrio fischeri MJ11 YP 002157278 1 388396 Proteobacteria
Eggerthella lenta DSM 2243 YP 003181115 1 479437 Actinobacteria
Treponema pedis str T A4 YP 008434626 1 1291379 Spirochaetes
Desulfosporosinus meridiei DSM 13257 YP 006621723 1 768704 Firmicutes
Bacteroides xylanisolvens XB1A YP 007794473 1 657309 Bacteroidetes Chlorobi group
Mycobacterium chubuense NBB4 YP 006442557 1 710421 Actinobacteria
Mycobacterium sp MCS YP 638635 1 164756 Actinobacteria
Mycobacterium canettii CIPT 140070017 YP 007267729 1 1205677 Actinobacteria
Propionibacterium freudenreichii subsp shermanii CIRM BIA1 YP 003688068 1 754252 Actinobacteria
Mycobacterium sp MCS YP 642729 1 164756 Actinobacteria
Phenylobacterium zucineum HLK1 YP 002128744 1 450851 Proteobacteria
Bradyrhizobium oligotrophicum S58 YP 007509776 1 1245469 Proteobacteria
Acetohalobium arabaticum DSM 5501 YP 003827967 1 574087 Firmicutes
Heliobacterium modesticaldum Ice1 YP 001681381 1 498761 Firmicutes
Thermincola potens JR YP 003639665 1 635013 Firmicutes
Desulfatibacillum alkenivorans AK 01 YP 002431367 1 439235 Proteobacteria
Oscillatoria acuminata PCC 6304 YP 007086945 1 56110 Cyanobacteria
Anabaena cylindrica PCC 7122 YP 007157137 1 272123 Cyanobacteria
Anabaena variabilis ATCC 29413 YP 324005 1 240292 Cyanobacteria
Desulfotomaculum kuznetsovii DSM 6115 YP 004517109 1 760568 Firmicutes
Thermodesulfatator indicus DSM 15286 YP 004626400 1 667014 Thermodesulfobacteria
Alicycliphilus denitrificans K601 YP 004387692 1 596154 Proteobacteria
Allochromatium vinosum DSM 180 YP 003444726 1 572477 Proteobacteria
Teredinibacter turnerae T7901 YP 003075611 1 377629 Proteobacteria
Xenorhabdus bovienii SS 2004 YP 003467341 1 406818 Proteobacteria
Pseudomonas stutzeri RCH2 YP 007238679 1 644801 Proteobacteria
Escherichia coli O104 H4 str 2009EL 2050 YP 006768598 1 1134782 Proteobacteria
Escherichia coli 042 YP 006098872 1 216592 Proteobacteria
Escherichia coli E24377A YP 001464328 1 331111 Proteobacteria
Haliscomenobacter hydrossis DSM 1100 YP 004451343 1 760192 Bacteroidetes Chlorobi group
Thiobacillus denitrificans ATCC 25259 YP 315450 1 292415 Proteobacteria
Marinobacter adhaerens HP15 YP 005885436 1 225937 Proteobacteria
Aeromonas hydrophila ML09 119 YP 008042121 1 1288394 Proteobacteria
Pseudomonas poae RE 1 1 14 YP 007399773 1 1282356 Proteobacteria
Dickeya zeae MS1 WP 019846677 1 Dzeae
Ralstonia eutropha H16 NP 942748 1 381666 Proteobacteria
Pseudovibrio sp FO BEG1 YP 005082714 1 911045 Proteobacteria
Tannerella forsythia ATCC 43037 YP 005015902 1 203275 Bacteroidetes Chlorobi group
Dehalobacter sp DCA YP 006909194 1 1147129 Firmicutes
Denitrovibrio acetiphilus DSM 12809 YP 003503401 1 522772 Deferribacteres
Magnetococcus marinus MC 1 YP 865208 1 156889 Proteobacteria
Rhodococcus jostii RHA1 YP 708882 1 101510 Actinobacteria
Thermomonospora curvata DSM 43183 YP 003302446 1 471852 Actinobacteria
Mycobacterium canettii CIPT 140070017 YP 007267257 1 1205677 Actinobacteria
Rhodopirellula baltica SH 1 NP 870916 1 243090 Planctomycetes
Anaeromyxobacter sp Fw109 5 YP 001381391 1 404589 Proteobacteria
Acetobacter pasteurianus 386B YP 008391121 1 1266844 Proteobacteria
Oscillatoria acuminata PCC 6304 YP 007083781 1 56110 Cyanobacteria
Acetobacter pasteurianus IFO 3283 07 YP 005488080 1 634454 Proteobacteria
Rhizobium leguminosarum bv viciae 3841 YP 770336 1 216596 Proteobacteria
Xanthobacter autotrophicus Py2 YP 001409255 1 78245 Proteobacteria
Rhizobium etli CFN 42 NP 660034 2 347834 Proteobacteria
Beijerinckia indica subsp indica ATCC 9039 YP 001833738 1 395963 Proteobacteria
Oligotropha carboxidovorans OM4 YP 005952486 1 1031710 Proteobacteria

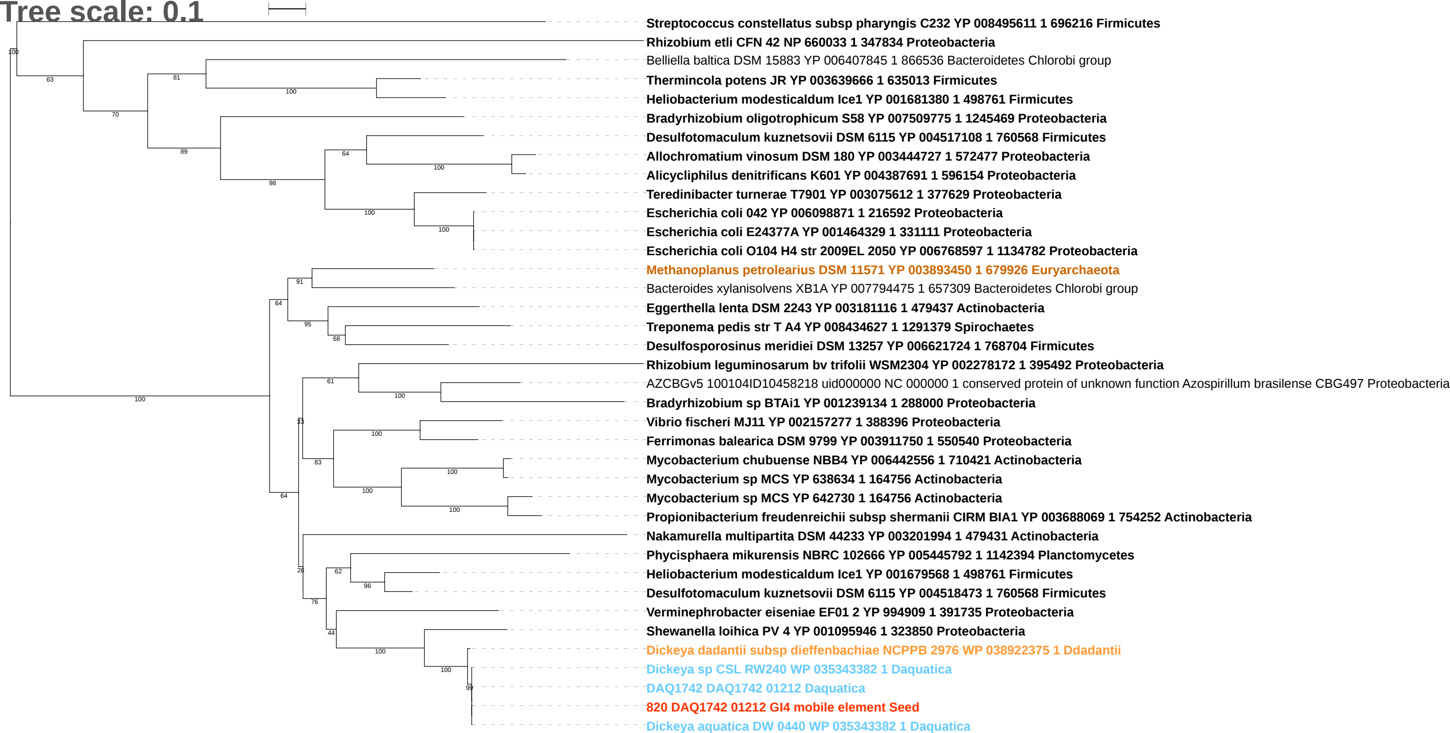
Tree scale: 0.1
Streptococcus constellatus subsp pharyngis C232 YP 008495611 1 696216 Firmicutes
Rhizobium etli CFN 42 NP 660033 1 347834 Proteobacteria
Belliella baltica DSM 15883 YP 006407845 1 866536 Bacteroidetes Chlorobi group
Thermincola potens JR YP 003639666 1 635013 Firmicutes
Heliobacterium modesticaldum Ice1 YP 001681380 1 498761 Firmicutes
Bradyrhizobium oligotrophicum S58 YP 007509775 1 1245469 Proteobacteria
Desulfotomaculum kuznetsovii DSM 6115 YP 004517108 1 760568 Firmicutes
Allochromatium vinosum DSM 180 YP 003444727 1 572477 Proteobacteria
Alicycliphilus denitrificans K601 YP 004387691 1 596154 Proteobacteria
Teredinibacter turnerae T7901 YP 003075612 1 377629 Proteobacteria
Escherichia coli 042 YP 006098871 1 216592 Proteobacteria
Escherichia coli E24377A YP 001464329 1 331111 Proteobacteria
Escherichia coli O104 H4 str 2009EL 2050 YP 006768597 1 1134782 Proteobacteria
Methanoplanus petrolearius DSM 11571 YP 003893450 1 679926 Euryarchaeota
Bacteroides xylanisolvens XB1A YP 007794475 1 657309 Bacteroidetes Chlorobi group
Eggerthella lenta DSM 2243 YP 003181116 1 479437 Actinobacteria
Treponema pedis str T A4 YP 008434627 1 1291379 Spirochaetes
Desulfosporosinus meridiei DSM 13257 YP 006621724 1 768704 Firmicutes
Rhizobium leguminosarum bv trifolii WSM2304 YP 002278172 1 395492 Proteobacteria
AZCBGv5 100104ID10458218 uid000000 NC 000000 1 conserved protein of unknown function Azospirillum brasilense CBG497 Proteobacteria
Bradyrhizobium sp BTAi1 YP 001239134 1 288000 Proteobacteria
Vibrio fischeri MJ11 YP 002157277 1 388396 Proteobacteria
Ferrimonas balearica DSM 9799 YP 003911750 1 550540 Proteobacteria
Mycobacterium chubuense NBB4 YP 006442556 1 710421 Actinobacteria
Mycobacterium sp MCS YP 638634 1 164756 Actinobacteria
Mycobacterium sp MCS YP 642730 1 164756 Actinobacteria
Propionibacterium freudenreichii subsp shermanii CIRM BIA1 YP 003688069 1 754252 Actinobacteria
Nakamurella multipartita DSM 44233 YP 003201994 1 479431 Actinobacteria
Phycisphaera mikurensis NBRC 102666 YP 005445792 1 1142394 Planctomycetes
Heliobacterium modesticaldum Ice1 YP 001679568 1 498761 Firmicutes
Desulfotomaculum kuznetsovii DSM 6115 YP 004518473 1 760568 Firmicutes
Verminephrobacter eiseniae EF01 2 YP 994909 1 391735 Proteobacteria
Shewanella loihica PV 4 YP 001095946 1 323850 Proteobacteria
Dickeya dadantii subsp dieffenbachiae NCPPB 2976 WP 038922375 1 Ddadantii
Dickeya sp CSL RW240 WP 035343382 1 Daquatica
DAQ1742 DAQ1742 01212 Daquatica
820 DAQ1742 01212 GI4 mobile element Seed
Dickeya aquatica DW 0440 WP 035343382 1 Daquatica

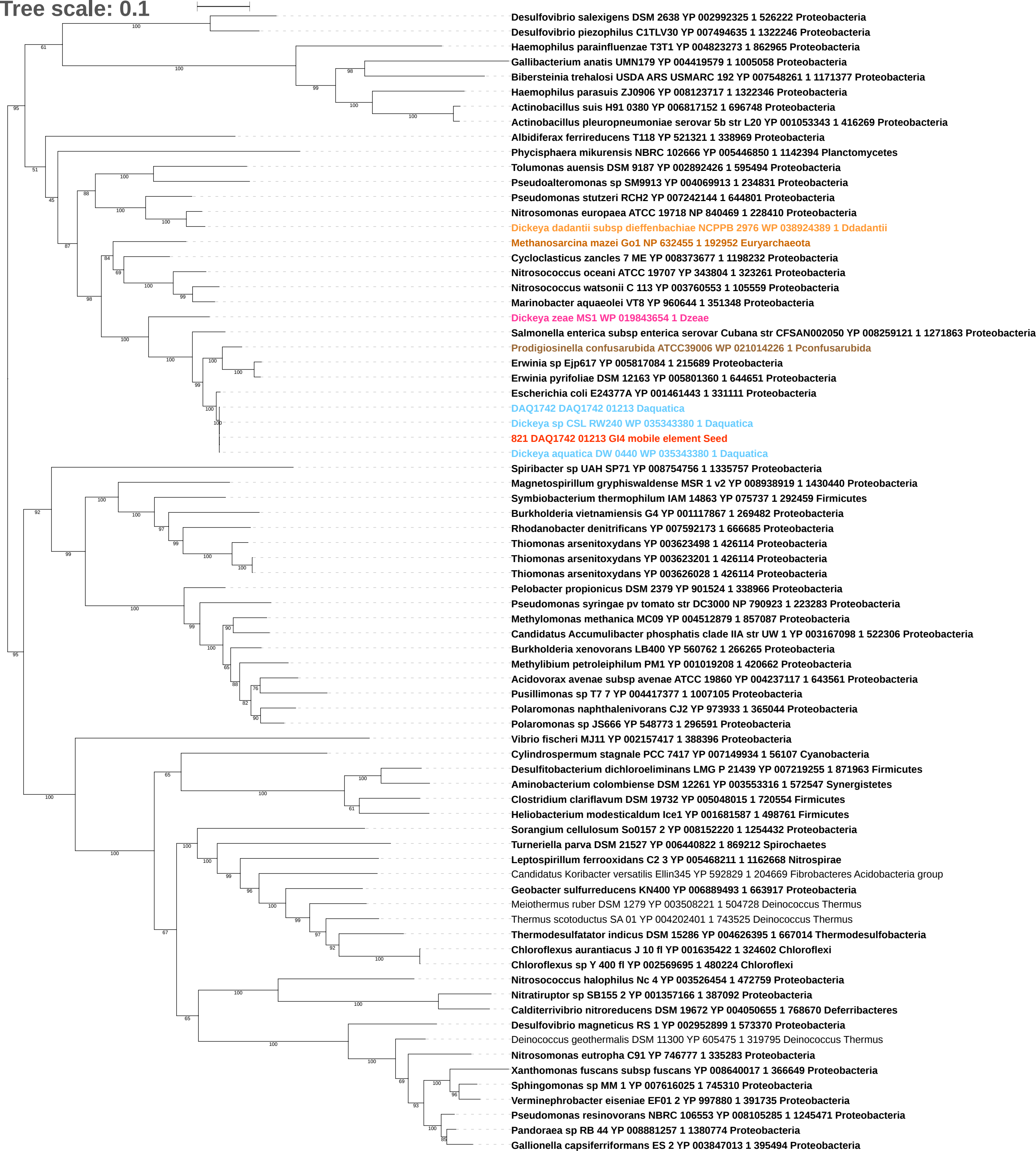
Tree scale: 0.1
Desulfovibrio salexigens DSM 2638 YP 002992325 1 526222 Proteobacteria
Desulfovibrio piezophilus C1TLV30 YP 007494635 1 1322246 Proteobacteria
Haemophilus parainfluenzae T3T1 YP 004823273 1 862965 Proteobacteria
Gallibacterium anatis UMN179 YP 004419579 1 1005058 Proteobacteria
Bibersteinia trehalosi USDA ARS USMARC 192 YP 007548261 1 1171377 Proteobacteria
Haemophilus parasuis ZJ0906 YP 008123717 1 1322346 Proteobacteria
Actinobacillus suis H91 0380 YP 006817152 1 696748 Proteobacteria
Actinobacillus pleuropneumoniae serovar 5b str L20 YP 001053343 1 416269 Proteobacteria
Albidiferax ferrireducens T118 YP 521321 1 338969 Proteobacteria
Phycisphaera mikurensis NBRC 102666 YP 005446850 1 1142394 Planctomycetes
Tolumonas auensis DSM 9187 YP 002892426 1 595494 Proteobacteria
Pseudoalteromonas sp SM9913 YP 004069913 1 234831 Proteobacteria
Pseudomonas stutzeri RCH2 YP 007242144 1 644801 Proteobacteria
Nitrosomonas europaea ATCC 19718 NP 840469 1 228410 Proteobacteria
Dickeya dadantii subsp dieffenbachiae NCPPB 2976 WP 038924389 1 Ddadantii
Methanosarcina mazei Go1 NP 632455 1 192952 Euryarchaeota
Cycloclasticus zancles 7 ME YP 008373677 1 1198232 Proteobacteria
Nitrosococcus oceani ATCC 19707 YP 343804 1 323261 Proteobacteria
Nitrosococcus watsonii C 113 YP 003760553 1 105559 Proteobacteria
Marinobacter aquaeolei VT8 YP 960644 1 351348 Proteobacteria
Dickeya zeae MS1 WP 019843654 1 Dzeae
Salmonella enterica subsp enterica serovar Cubana str CFSAN002050 YP 008259121 1 1271863 Proteobacteria
Prodigiosinella confusarubida ATCC39006 WP 021014226 1 Pconfusarubida
Erwinia sp Ejp617 YP 005817084 1 215689 Proteobacteria
Erwinia pyrifoliae DSM 12163 YP 005801360 1 644651 Proteobacteria
Escherichia coli E24377A YP 001461443 1 331111 Proteobacteria
DAQ1742 DAQ1742 01213 Daquatica
Dickeya sp CSL RW240 WP 035343380 1 Daquatica
821 DAQ1742 01213 GI4 mobile element Seed
Dickeya aquatica DW 0440 WP 035343380 1 Daquatica
Spiribacter sp UAH SP71 YP 008754756 1 1335757 Proteobacteria
Magnetospirillum gryphiswaldense MSR 1 v2 YP 008938919 1 1430440 Proteobacteria
Symbiobacterium thermophilum IAM 14863 YP 075737 1 292459 Firmicutes
Burkholderia vietnamiensis G4 YP 001117867 1 269482 Proteobacteria
Rhodanobacter denitrificans YP 007592173 1 666685 Proteobacteria
Thiomonas arsenitoxydans YP 003623498 1 426114 Proteobacteria
Thiomonas arsenitoxydans YP 003623201 1 426114 Proteobacteria
Thiomonas arsenitoxydans YP 003626028 1 426114 Proteobacteria
Pelobacter propionicus DSM 2379 YP 901524 1 338966 Proteobacteria
Pseudomonas syringae pv tomato str DC3000 NP 790923 1 223283 Proteobacteria
Methylomonas methanica MC09 YP 004512879 1 857087 Proteobacteria
Candidatus Accumulibacter phosphatis clade IIA str UW 1 YP 003167098 1 522306 Proteobacteria
Burkholderia xenovorans LB400 YP 560762 1 266265 Proteobacteria
Methylibium petroleiphilum PM1 YP 001019208 1 420662 Proteobacteria
Acidovorax avenae subsp avenae ATCC 19860 YP 004237117 1 643561 Proteobacteria
Pusillimonas sp T7 7 YP 004417377 1 1007105 Proteobacteria
Polaromonas naphthalenivorans CJ2 YP 973933 1 365044 Proteobacteria
Polaromonas sp JS666 YP 548773 1 296591 Proteobacteria
Vibrio fischeri MJ11 YP 002157417 1 388396 Proteobacteria
Cylindrospermum stagnale PCC 7417 YP 007149934 1 56107 Cyanobacteria
Desulfitobacterium dichloroeliminans LMG P 21439 YP 007219255 1 871963 Firmicutes
Aminobacterium colombiense DSM 12261 YP 003553316 1 572547 Synergistetes
Clostridium clariflavum DSM 19732 YP 005048015 1 720554 Firmicutes
Heliobacterium modesticaldum Ice1 YP 001681587 1 498761 Firmicutes
Sorangium cellulosum So0157 2 YP 008152220 1 1254432 Proteobacteria
Turneriella parva DSM 21527 YP 006440822 1 869212 Spirochaetes
Leptospirillum ferrooxidans C2 3 YP 005468211 1 1162668 Nitrospirae
Candidatus Koribacter versatilis Ellin345 YP 592829 1 204669 Fibrobacteres Acidobacteria group
Geobacter sulfurreducens KN400 YP 006889493 1 663917 Proteobacteria
Meiothermus ruber DSM 1279 YP 003508221 1 504728 Deinococcus Thermus
Thermus scotoductus SA 01 YP 004202401 1 743525 Deinococcus Thermus
Thermodesulfatator indicus DSM 15286 YP 004626395 1 667014 Thermodesulfobacteria
Chloroflexus aurantiacus J 10 fl YP 001635422 1 324602 Chloroflexi
Chloroflexus sp Y 400 fl YP 002569695 1 480224 Chloroflexi
Nitrosococcus halophilus Nc 4 YP 003526454 1 472759 Proteobacteria
Nitratiruptor sp SB155 2 YP 001357166 1 387092 Proteobacteria
Calditerrivibrio nitroreducens DSM 19672 YP 004050655 1 768670 Deferribacteres
Desulfovibrio magneticus RS 1 YP 002952899 1 573370 Proteobacteria
Deinococcus geothermalis DSM 11300 YP 605475 1 319795 Deinococcus Thermus
Nitrosomonas eutropha C91 YP 746777 1 335283 Proteobacteria
Xanthomonas fuscans subsp fuscans YP 008640017 1 366649 Proteobacteria
Sphingomonas sp MM 1 YP 007616025 1 745310 Proteobacteria
Verminephrobacter eiseniae EF01 2 YP 997880 1 391735 Proteobacteria
Pseudomonas resinovorans NBRC 106553 YP 008105285 1 1245471 Proteobacteria
Pandoraea sp RB 44 YP 008881257 1 1380774 Proteobacteria
Gallionella capsiferriformans ES 2 YP 003847013 1 395494 Proteobacteria

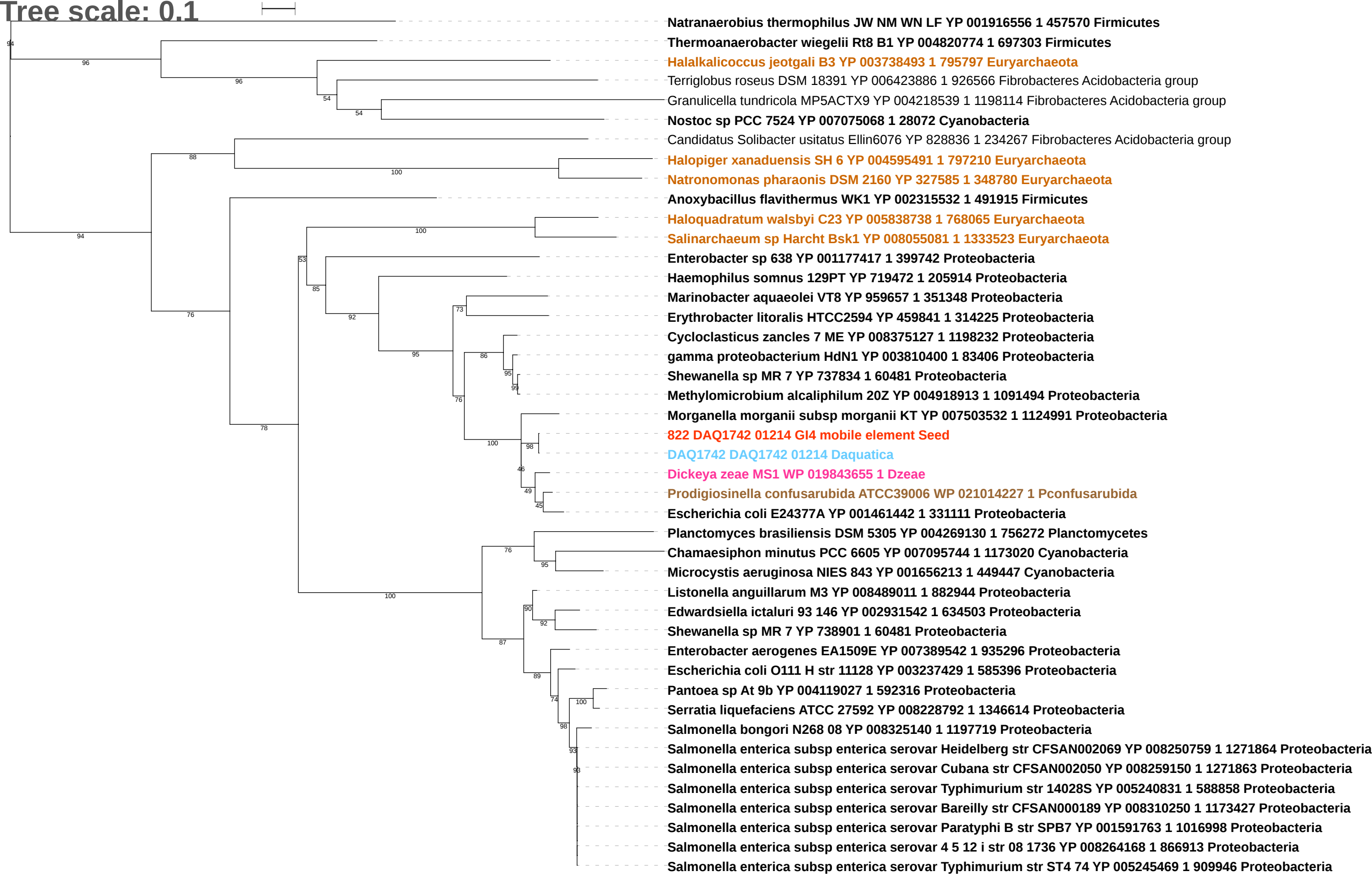
Tree scale: 0.1
Natranaerobius thermophilus JW NM WN LF YP 001916556 1 457570 Firmicutes
Thermoanaerobacter wiegelii Rt8 B1 YP 004820774 1 697303 Firmicutes
Halalkalicoccus jeotgali B3 YP 003738493 1 795797 Euryarchaeota
Terriglobus roseus DSM 18391 YP 006423886 1 926566 Fibrobacteres Acidobacteria group
Granulicella tundricola MP5ACTX9 YP 004218539 1 1198114 Fibrobacteres Acidobacteria group
Nostoc sp PCC 7524 YP 007075068 1 28072 Cyanobacteria
Candidatus Solibacter usitatus Ellin6076 YP 828836 1 234267 Fibrobacteres Acidobacteria group
Halopiger xanaduensis SH 6 YP 004595491 1 797210 Euryarchaeota
Natronomonas pharaonis DSM 2160 YP 327585 1 348780 Euryarchaeota
Anoxybacillus flavithermus WK1 YP 002315532 1 491915 Firmicutes
Haloquadratum walsbyi C23 YP 005838738 1 768065 Euryarchaeota
Salinarchaeum sp Harcht Bsk1 YP 008055081 1 1333523 Euryarchaeota
Enterobacter sp 638 YP 001177417 1 399742 Proteobacteria
Haemophilus somnus 129PT YP 719472 1 205914 Proteobacteria
Marinobacter aquaeolei VT8 YP 959657 1 351348 Proteobacteria
Erythrobacter litoralis HTCC2594 YP 459841 1 314225 Proteobacteria
Cycloclasticus zancles 7 ME YP 008375127 1 1198232 Proteobacteria
gamma proteobacterium HdN1 YP 003810400 1 83406 Proteobacteria
Shewanella sp MR 7 YP 737834 1 60481 Proteobacteria
Methylomicrobium alcaliphilum 20Z YP 004918913 1 1091494 Proteobacteria
Morganella morganii subsp morganii KT YP 007503532 1 1124991 Proteobacteria
822 DAQ1742 01214 GI4 mobile element Seed
DAQ1742 DAQ1742 01214 Daquatica
Dickeya zeae MS1 WP 019843655 1 Dzeae
Prodigiosinella confusarubida ATCC39006 WP 021014227 1 Pconfusarubida
Escherichia coli E24377A YP 001461442 1 331111 Proteobacteria
Planctomyces brasiliensis DSM 5305 YP 004269130 1 756272 Planctomycetes
Chamaesiphon minutus PCC 6605 YP 007095744 1 1173020 Cyanobacteria
Microcystis aeruginosa NIES 843 YP 001656213 1 449447 Cyanobacteria
Listonella anguillarum M3 YP 008489011 1 882944 Proteobacteria
Edwardsiella ictaluri 93 146 YP 002931542 1 634503 Proteobacteria
Shewanella sp MR 7 YP 738901 1 60481 Proteobacteria
Enterobacter aerogenes EA1509E YP 007389542 1 935296 Proteobacteria
Escherichia coli O111 H str 11128 YP 003237429 1 585396 Proteobacteria
Pantoea sp At 9b YP 004119027 1 592316 Proteobacteria
Serratia liquefaciens ATCC 27592 YP 008228792 1 1346614 Proteobacteria
Salmonella bongori N268 08 YP 008325140 1 1197719 Proteobacteria
Salmonella enterica subsp enterica serovar Heidelberg str CFSAN002069 YP 008250759 1 1271864 Proteobacteria
Salmonella enterica subsp enterica serovar Cubana str CFSAN002050 YP 008259150 1 1271863 Proteobacteria
Salmonella enterica subsp enterica serovar Typhimurium str 14028S YP 005240831 1 588858 Proteobacteria
Salmonella enterica subsp enterica serovar Bareilly str CFSAN000189 YP 008310250 1 1173427 Proteobacteria
Salmonella enterica subsp enterica serovar Paratyphi B str SPB7 YP 001591763 1 1016998 Proteobacteria
Salmonella enterica subsp enterica serovar 4 5 12 i str 08 1736 YP 008264168 1 866913 Proteobacteria
Salmonella enterica subsp enterica serovar Typhimurium str ST4 74 YP 005245469 1 909946 Proteobacteria

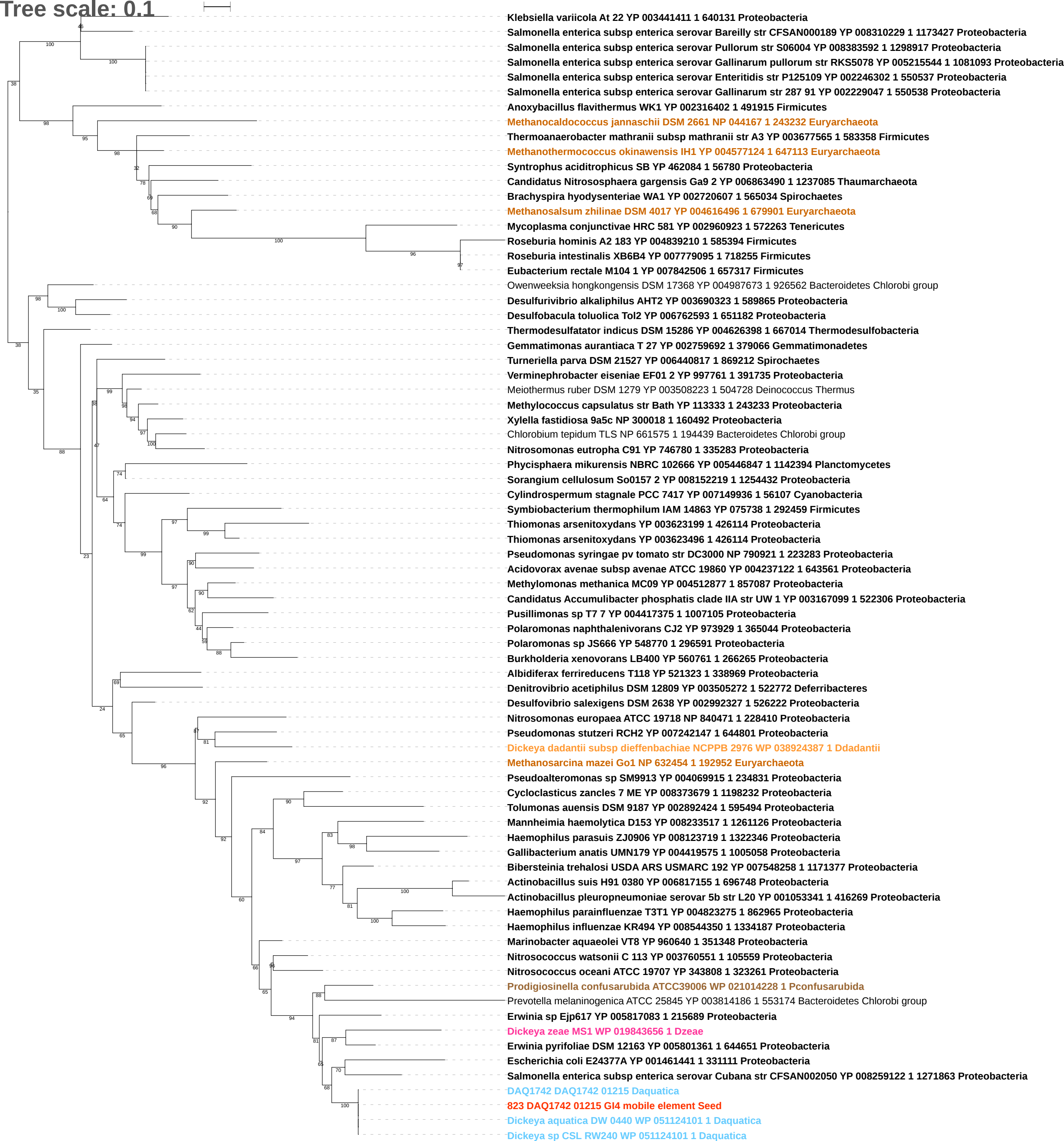
Tree scale: 0.1
Klebsiella variicola At 22 YP 003441411 1 640131 Proteobacteria
Salmonella enterica subsp enterica serovar Bareilly str CFSAN000189 YP 008310229 1 1173427 Proteobacteria
Salmonella enterica subsp enterica serovar Pullorum str S06004 YP 008383592 1 1298917 Proteobacteria
Salmonella enterica subsp enterica serovar Gallinarum pullorum str RKS5078 YP 005215544 1 1081093 Proteobacteria
Salmonella enterica subsp enterica serovar Enteritidis str P125109 YP 002246302 1 550537 Proteobacteria
Salmonella enterica subsp enterica serovar Gallinarum str 287 91 YP 002229047 1 550538 Proteobacteria
Anoxybacillus flavithermus WK1 YP 002316402 1 491915 Firmicutes
Methanocaldococcus jannaschii DSM 2661 NP 044167 1 243232 Euryarchaeota
Thermoanaerobacter mathranii subsp mathranii str A3 YP 003677565 1 583358 Firmicutes
Methanothermococcus okinawensis IH1 YP 004577124 1 647113 Euryarchaeota
Syntrophus aciditrophicus SB YP 462084 1 56780 Proteobacteria
Candidatus Nitrososphaera gargensis Ga9 2 YP 006863490 1 1237085 Thaumarchaeota
Brachyspira hyodysenteriae WA1 YP 002720607 1 565034 Spirochaetes
Methanosalsum zhilinae DSM 4017 YP 004616496 1 679901 Euryarchaeota
Mycoplasma conjunctivae HRC 581 YP 002960923 1 572263 Tenericutes
Roseburia hominis A2 183 YP 004839210 1 585394 Firmicutes
Roseburia intestinalis XB6B4 YP 007779095 1 718255 Firmicutes
Eubacterium rectale M104 1 YP 007842506 1 657317 Firmicutes
Owenweeksia hongkongensis DSM 17368 YP 004987673 1 926562 Bacteroidetes Chlorobi group
Desulfurivibrio alkaliphilus AHT2 YP 003690323 1 589865 Proteobacteria
Desulfobacula toluolica Tol2 YP 006762593 1 651182 Proteobacteria
Thermodesulfatator indicus DSM 15286 YP 004626398 1 667014 Thermodesulfobacteria
Gemmatimonas aurantiaca T 27 YP 002759692 1 379066 Gemmatimonadetes
Turneriella parva DSM 21527 YP 006440817 1 869212 Spirochaetes
Verminephrobacter eiseniae EF01 2 YP 997761 1 391735 Proteobacteria
Meiothermus ruber DSM 1279 YP 003508223 1 504728 Deinococcus Thermus
Methylococcus capsulatus str Bath YP 113333 1 243233 Proteobacteria
Xylella fastidiosa 9a5c NP 300018 1 160492 Proteobacteria
Chlorobium tepidum TLS NP 661575 1 194439 Bacteroidetes Chlorobi group
Nitrosomonas eutropha C91 YP 746780 1 335283 Proteobacteria
Phycisphaera mikurensis NBRC 102666 YP 005446847 1 1142394 Planctomycetes
Sorangium cellulosum So0157 2 YP 008152219 1 1254432 Proteobacteria
Cylindrospermum stagnale PCC 7417 YP 007149936 1 56107 Cyanobacteria
Symbiobacterium thermophilum IAM 14863 YP 075738 1 292459 Firmicutes
Thiomonas arsenitoxydans YP 003623199 1 426114 Proteobacteria
Thiomonas arsenitoxydans YP 003623496 1 426114 Proteobacteria
Pseudomonas syringae pv tomato str DC3000 NP 790921 1 223283 Proteobacteria
Acidovorax avenae subsp avenae ATCC 19860 YP 004237122 1 643561 Proteobacteria
Methylomonas methanica MC09 YP 004512877 1 857087 Proteobacteria
Candidatus Accumulibacter phosphatis clade IIA str UW 1 YP 003167099 1 522306 Proteobacteria
Pusillimonas sp T7 7 YP 004417375 1 1007105 Proteobacteria
Polaromonas naphthalenivorans CJ2 YP 973929 1 365044 Proteobacteria
Polaromonas sp JS666 YP 548770 1 296591 Proteobacteria
Burkholderia xenovorans LB400 YP 560761 1 266265 Proteobacteria
Albidiferax ferrireducens T118 YP 521323 1 338969 Proteobacteria
Denitrovibrio acetiphilus DSM 12809 YP 003505272 1 522772 Deferribacteres
Desulfovibrio salexigens DSM 2638 YP 002992327 1 526222 Proteobacteria
Nitrosomonas europaea ATCC 19718 NP 840471 1 228410 Proteobacteria
Pseudomonas stutzeri RCH2 YP 007242147 1 644801 Proteobacteria
Dickeya dadantii subsp dieffenbachiae NCPPB 2976 WP 038924387 1 Ddadantii
Methanosarcina mazei Go1 NP 632454 1 192952 Euryarchaeota
Pseudoalteromonas sp SM9913 YP 004069915 1 234831 Proteobacteria
Cycloclasticus zancles 7 ME YP 008373679 1 1198232 Proteobacteria
Tolumonas auensis DSM 9187 YP 002892424 1 595494 Proteobacteria
Mannheimia haemolytica D153 YP 008233517 1 1261126 Proteobacteria
Haemophilus parasuis ZJ0906 YP 008123719 1 1322346 Proteobacteria
Gallibacterium anatis UMN179 YP 004419575 1 1005058 Proteobacteria
Bibersteinia trehalosi USDA ARS USMARC 192 YP 007548258 1 1171377 Proteobacteria
Actinobacillus suis H91 0380 YP 006817155 1 696748 Proteobacteria
Actinobacillus pleuropneumoniae serovar 5b str L20 YP 001053341 1 416269 Proteobacteria
Haemophilus parainfluenzae T3T1 YP 004823275 1 862965 Proteobacteria
Haemophilus influenzae KR494 YP 008544350 1 1334187 Proteobacteria
Marinobacter aquaeolei VT8 YP 960640 1 351348 Proteobacteria
Nitrosococcus watsonii C 113 YP 003760551 1 105559 Proteobacteria
Nitrosococcus oceani ATCC 19707 YP 343808 1 323261 Proteobacteria
Prodigiosinella confusarubida ATCC39006 WP 021014228 1 Pconfusarubida
Prevotella melaninogenica ATCC 25845 YP 003814186 1 553174 Bacteroidetes Chlorobi group
Erwinia sp Ejp617 YP 005817083 1 215689 Proteobacteria
Dickeya zeae MS1 WP 019843656 1 Dzeae
Erwinia pyrifoliae DSM 12163 YP 005801361 1 644651 Proteobacteria
Escherichia coli E24377A YP 001461441 1 331111 Proteobacteria
Salmonella enterica subsp enterica serovar Cubana str CFSAN002050 YP 008259122 1 1271863 Proteobacteria
DAQ1742 DAQ1742 01215 Daquatica
823 DAQ1742 01215 GI4 mobile element Seed
Dickeya aquatica DW 0440 WP 051124101 1 Daquatica
Dickeya sp CSL RW240 WP 051124101 1 Daquatica

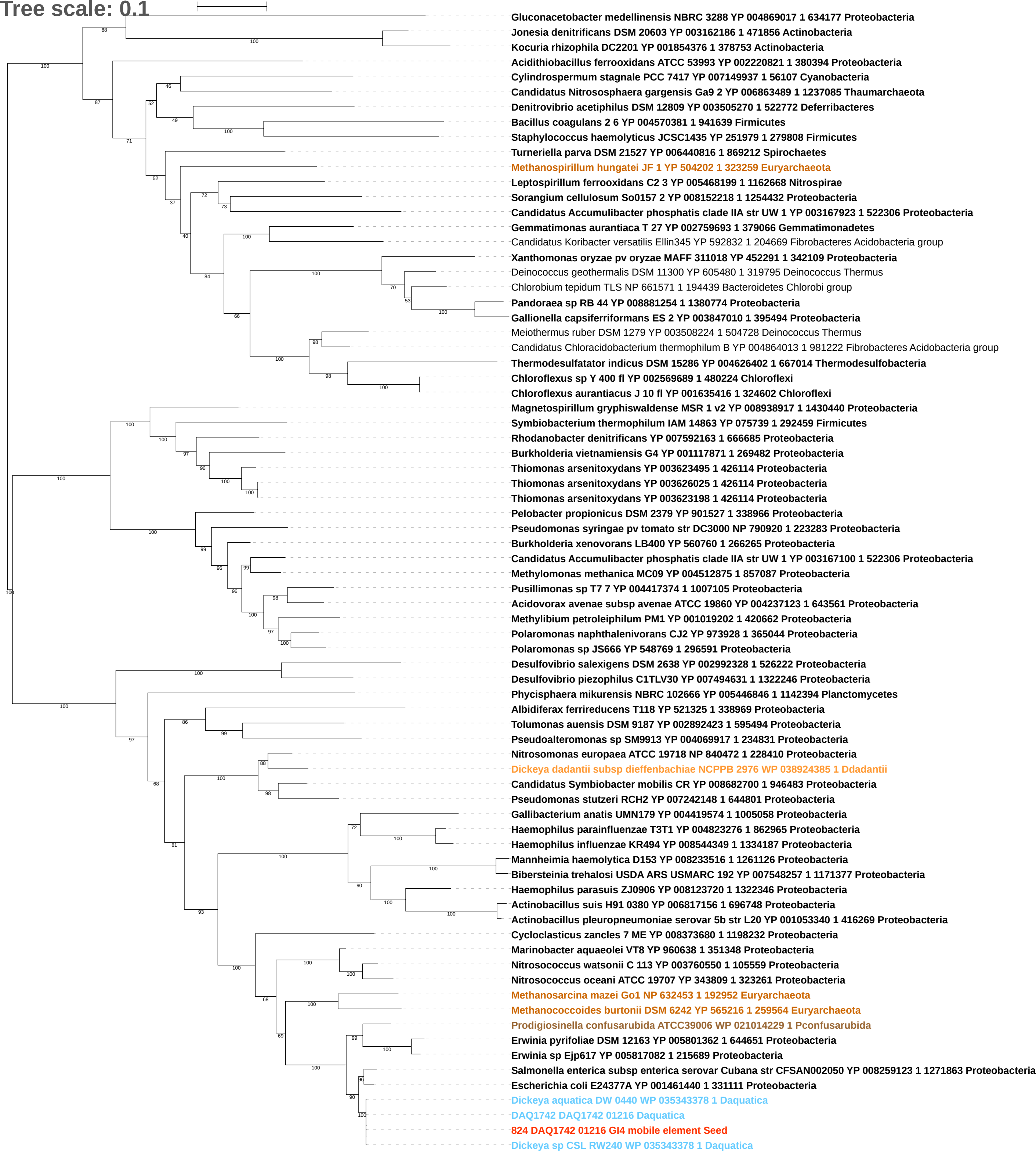
Tree scale: 0.1
Gluconacetobacter medellinensis NBRC 3288 YP 004869017 1 634177 Proteobacteria
Jonesia denitrificans DSM 20603 YP 003162186 1 471856 Actinobacteria
Kocuria rhizophila DC2201 YP 001854376 1 378753 Actinobacteria
Acidithiobacillus ferrooxidans ATCC 53993 YP 002220821 1 380394 Proteobacteria
Cylindrospermum stagnale PCC 7417 YP 007149937 1 56107 Cyanobacteria
Candidatus Nitrososphaera gargensis Ga9 2 YP 006863489 1 1237085 Thaumarchaeota
Denitrovibrio acetiphilus DSM 12809 YP 003505270 1 522772 Deferribacteres
Bacillus coagulans 2 6 YP 004570381 1 941639 Firmicutes
Staphylococcus haemolyticus JCSC1435 YP 251979 1 279808 Firmicutes
Turneriella parva DSM 21527 YP 006440816 1 869212 Spirochaetes
Methanospirillum hungatei JF 1 YP 504202 1 323259 Euryarchaeota
Leptospirillum ferrooxidans C2 3 YP 005468199 1 1162668 Nitrospirae
Sorangium cellulosum So0157 2 YP 008152218 1 1254432 Proteobacteria
Candidatus Accumulibacter phosphatis clade IIA str UW 1 YP 003167923 1 522306 Proteobacteria
Gemmatimonas aurantiaca T 27 YP 002759693 1 379066 Gemmatimonadetes
Candidatus Koribacter versatilis Ellin345 YP 592832 1 204669 Fibrobacteres Acidobacteria group
Xanthomonas oryzae pv oryzae MAFF 311018 YP 452291 1 342109 Proteobacteria
Deinococcus geothermalis DSM 11300 YP 605480 1 319795 Deinococcus Thermus
Chlorobium tepidum TLS NP 661571 1 194439 Bacteroidetes Chlorobi group
Pandoraea sp RB 44 YP 008881254 1 1380774 Proteobacteria
Gallionella capsiferriformans ES 2 YP 003847010 1 395494 Proteobacteria
Meiothermus ruber DSM 1279 YP 003508224 1 504728 Deinococcus Thermus
Candidatus Chloracidobacterium thermophilum B YP 004864013 1 981222 Fibrobacteres Acidobacteria group
Thermodesulfatator indicus DSM 15286 YP 004626402 1 667014 Thermodesulfobacteria
Chloroflexus sp Y 400 fl YP 002569689 1 480224 Chloroflexi
Chloroflexus aurantiacus J 10 fl YP 001635416 1 324602 Chloroflexi
Magnetospirillum gryphiswaldense MSR 1 v2 YP 008938917 1 1430440 Proteobacteria
Symbiobacterium thermophilum IAM 14863 YP 075739 1 292459 Firmicutes
Rhodanobacter denitrificans YP 007592163 1 666685 Proteobacteria
Burkholderia vietnamiensis G4 YP 001117871 1 269482 Proteobacteria
Thiomonas arsenitoxydans YP 003623495 1 426114 Proteobacteria
Thiomonas arsenitoxydans YP 003626025 1 426114 Proteobacteria
Thiomonas arsenitoxydans YP 003623198 1 426114 Proteobacteria
Pelobacter propionicus DSM 2379 YP 901527 1 338966 Proteobacteria
Pseudomonas syringae pv tomato str DC3000 NP 790920 1 223283 Proteobacteria
Burkholderia xenovorans LB400 YP 560760 1 266265 Proteobacteria
Candidatus Accumulibacter phosphatis clade IIA str UW 1 YP 003167100 1 522306 Proteobacteria
Methylomonas methanica MC09 YP 004512875 1 857087 Proteobacteria
Pusillimonas sp T7 7 YP 004417374 1 1007105 Proteobacteria
Acidovorax avenae subsp avenae ATCC 19860 YP 004237123 1 643561 Proteobacteria
Methylibium petroleiphilum PM1 YP 001019202 1 420662 Proteobacteria
Polaromonas naphthalenivorans CJ2 YP 973928 1 365044 Proteobacteria
Polaromonas sp JS666 YP 548769 1 296591 Proteobacteria
Desulfovibrio salexigens DSM 2638 YP 002992328 1 526222 Proteobacteria
Desulfovibrio piezophilus C1TLV30 YP 007494631 1 1322246 Proteobacteria
Phycisphaera mikurensis NBRC 102666 YP 005446846 1 1142394 Planctomycetes
Albidiferax ferrireducens T118 YP 521325 1 338969 Proteobacteria
Tolumonas auensis DSM 9187 YP 002892423 1 595494 Proteobacteria
Pseudoalteromonas sp SM9913 YP 004069917 1 234831 Proteobacteria
Nitrosomonas europaea ATCC 19718 NP 840472 1 228410 Proteobacteria
Dickeya dadantii subsp dieffenbachiae NCPPB 2976 WP 038924385 1 Ddadantii
Candidatus Symbiobacter mobilis CR YP 008682700 1 946483 Proteobacteria
Pseudomonas stutzeri RCH2 YP 007242148 1 644801 Proteobacteria
Gallibacterium anatis UMN179 YP 004419574 1 1005058 Proteobacteria
Haemophilus parainfluenzae T3T1 YP 004823276 1 862965 Proteobacteria
Haemophilus influenzae KR494 YP 008544349 1 1334187 Proteobacteria
Mannheimia haemolytica D153 YP 008233516 1 1261126 Proteobacteria
Bibersteinia trehalosi USDA ARS USMARC 192 YP 007548257 1 1171377 Proteobacteria
Haemophilus parasuis ZJ0906 YP 008123720 1 1322346 Proteobacteria
Actinobacillus suis H91 0380 YP 006817156 1 696748 Proteobacteria
Actinobacillus pleuropneumoniae serovar 5b str L20 YP 001053340 1 416269 Proteobacteria
Cycloclasticus zancles 7 ME YP 008373680 1 1198232 Proteobacteria
Marinobacter aquaeolei VT8 YP 960638 1 351348 Proteobacteria
Nitrosococcus watsonii C 113 YP 003760550 1 105559 Proteobacteria
Nitrosococcus oceani ATCC 19707 YP 343809 1 323261 Proteobacteria
Methanosarcina mazei Go1 NP 632453 1 192952 Euryarchaeota
Methanococcoides burtonii DSM 6242 YP 565216 1 259564 Euryarchaeota
Prodigiosinella confusarubida ATCC39006 WP 021014229 1 Pconfusarubida
Erwinia pyrifoliae DSM 12163 YP 005801362 1 644651 Proteobacteria
Erwinia sp Ejp617 YP 005817082 1 215689 Proteobacteria
Salmonella enterica subsp enterica serovar Cubana str CFSAN002050 YP 008259123 1 1271863 Proteobacteria
Escherichia coli E24377A YP 001461440 1 331111 Proteobacteria
Dickeya aquatica DW 0440 WP 035343378 1 Daquatica
DAQ1742 DAQ1742 01216 Daquatica
824 DAQ1742 01216 GI4 mobile element Seed
Dickeya sp CSL RW240 WP 035343378 1 Daquatica

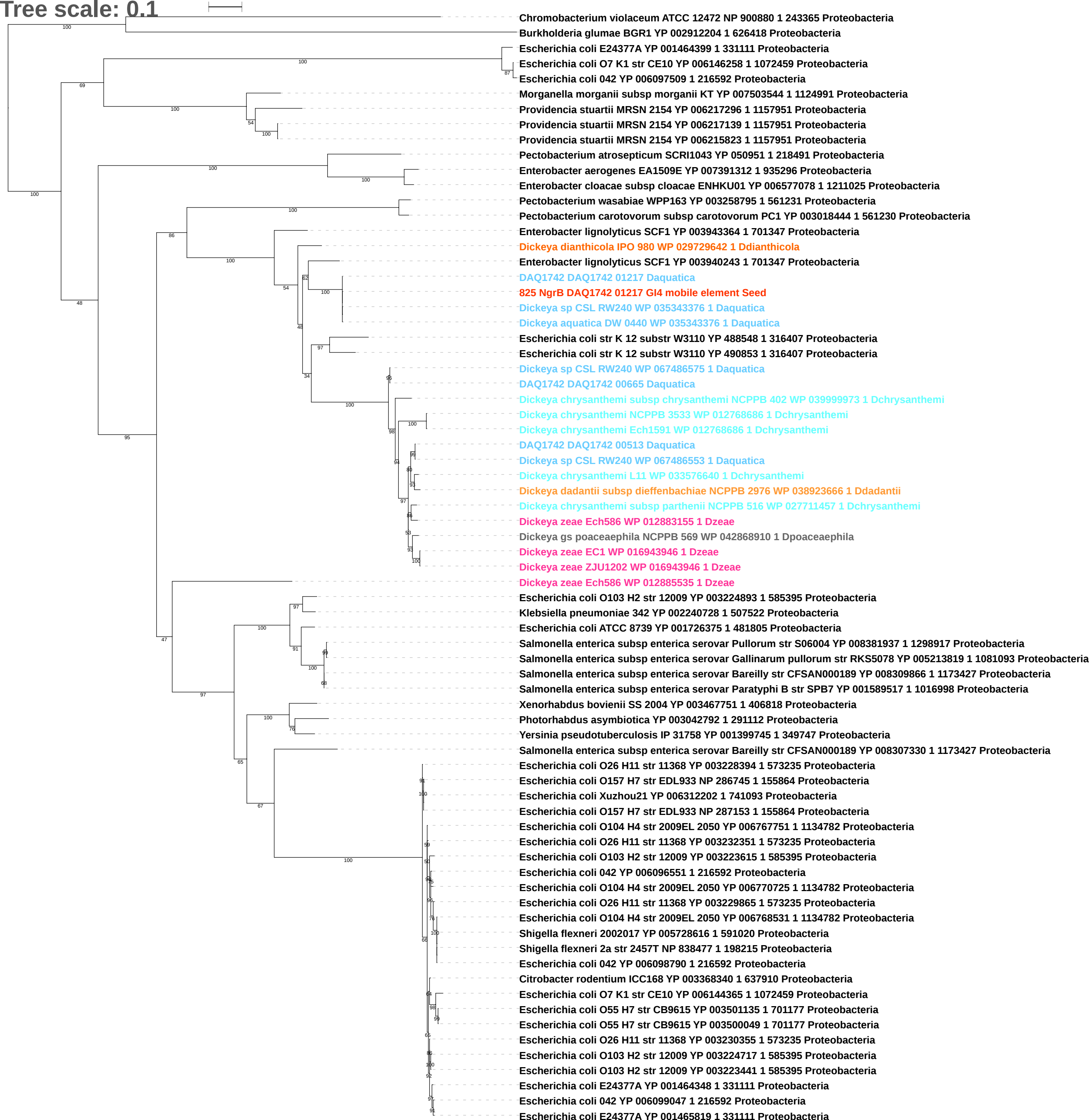
Tree scale: 0.1
Chromobacterium violaceum ATCC 12472 NP 900880 1 243365 Proteobacteria
Burkholderia glumae BGR1 YP 002912204 1 626418 Proteobacteria
Escherichia coli E24377A YP 001464399 1 331111 Proteobacteria
Escherichia coli O7 K1 str CE10 YP 006146258 1 1072459 Proteobacteria
Escherichia coli 042 YP 006097509 1 216592 Proteobacteria
Morganella morganii subsp morganii KT YP 007503544 1 1124991 Proteobacteria
Providencia stuartii MRSN 2154 YP 006217296 1 1157951 Proteobacteria
Providencia stuartii MRSN 2154 YP 006217139 1 1157951 Proteobacteria
Providencia stuartii MRSN 2154 YP 006215823 1 1157951 Proteobacteria
Pectobacterium atrosepticum SCRI1043 YP 050951 1 218491 Proteobacteria
Enterobacter aerogenes EA1509E YP 007391312 1 935296 Proteobacteria
Enterobacter cloacae subsp cloacae ENHKU01 YP 006577078 1 1211025 Proteobacteria
Pectobacterium wasabiae WPP163 YP 003258795 1 561231 Proteobacteria
Pectobacterium carotovorum subsp carotovorum PC1 YP 003018444 1 561230 Proteobacteria
Enterobacter lignolyticus SCF1 YP 003943364 1 701347 Proteobacteria
Dickeya dianthicola IPO 980 WP 029729642 1 Ddianthicola
Enterobacter lignolyticus SCF1 YP 003940243 1 701347 Proteobacteria
DAQ1742 DAQ1742 01217 Daquatica
825 NgrB DAQ1742 01217 GI4 mobile element Seed
Dickeya sp CSL RW240 WP 035343376 1 Daquatica
Dickeya aquatica DW 0440 WP 035343376 1 Daquatica
Escherichia coli str K 12 substr W3110 YP 488548 1 316407 Proteobacteria
Escherichia coli str K 12 substr W3110 YP 490853 1 316407 Proteobacteria
Dickeya sp CSL RW240 WP 067486575 1 Daquatica
DAQ1742 DAQ1742 00665 Daquatica
Dickeya chrysanthemi subsp chrysanthemi NCPPB 402 WP 039999973 1 Dchrysanthemi
Dickeya chrysanthemi NCPPB 3533 WP 012768686 1 Dchrysanthemi
Dickeya chrysanthemi Ech1591 WP 012768686 1 Dchrysanthemi
DAQ1742 DAQ1742 00513 Daquatica
Dickeya sp CSL RW240 WP 067486553 1 Daquatica
Dickeya chrysanthemi L11 WP 033576640 1 Dchrysanthemi
Dickeya dadantii subsp dieffenbachiae NCPPB 2976 WP 038923666 1 Ddadantii
Dickeya chrysanthemi subsp parthenii NCPPB 516 WP 027711457 1 Dchrysanthemi
Dickeya zeae Ech586 WP 012883155 1 Dzeae
Dickeya gs poaceaephila NCPPB 569 WP 042868910 1 Dpoaceaephila
Dickeya zeae EC1 WP 016943946 1 Dzeae
Dickeya zeae ZJU1202 WP 016943946 1 Dzeae
Dickeya zeae Ech586 WP 012885535 1 Dzeae
Escherichia coli O103 H2 str 12009 YP 003224893 1 585395 Proteobacteria
Klebsiella pneumoniae 342 YP 002240728 1 507522 Proteobacteria
Escherichia coli ATCC 8739 YP 001726375 1 481805 Proteobacteria
Salmonella enterica subsp enterica serovar Pullorum str S06004 YP 008381937 1 1298917 Proteobacteria
Salmonella enterica subsp enterica serovar Gallinarum pullorum str RKS5078 YP 005213819 1 1081093 Proteobacteria
Salmonella enterica subsp enterica serovar Bareilly str CFSAN000189 YP 008309866 1 1173427 Proteobacteria
Salmonella enterica subsp enterica serovar Paratyphi B str SPB7 YP 001589517 1 1016998 Proteobacteria
Xenorhabdus bovienii SS 2004 YP 003467751 1 406818 Proteobacteria
Photorhabdus asymbiotica YP 003042792 1 291112 Proteobacteria
Yersinia pseudotuberculosis IP 31758 YP 001399745 1 349747 Proteobacteria
Salmonella enterica subsp enterica serovar Bareilly str CFSAN000189 YP 008307330 1 1173427 Proteobacteria
Escherichia coli O26 H11 str 11368 YP 003228394 1 573235 Proteobacteria
Escherichia coli O157 H7 str EDL933 NP 286745 1 155864 Proteobacteria
Escherichia coli Xuzhou21 YP 006312202 1 741093 Proteobacteria
Escherichia coli O157 H7 str EDL933 NP 287153 1 155864 Proteobacteria
Escherichia coli O104 H4 str 2009EL 2050 YP 006767751 1 1134782 Proteobacteria
Escherichia coli O26 H11 str 11368 YP 003232351 1 573235 Proteobacteria
Escherichia coli O103 H2 str 12009 YP 003223615 1 585395 Proteobacteria
Escherichia coli 042 YP 006096551 1 216592 Proteobacteria
Escherichia coli O104 H4 str 2009EL 2050 YP 006770725 1 1134782 Proteobacteria
Escherichia coli O26 H11 str 11368 YP 003229865 1 573235 Proteobacteria
Escherichia coli O104 H4 str 2009EL 2050 YP 006768531 1 1134782 Proteobacteria
Shigella flexneri 2002017 YP 005728616 1 591020 Proteobacteria
Shigella flexneri 2a str 2457T NP 838477 1 198215 Proteobacteria
Escherichia coli 042 YP 006098790 1 216592 Proteobacteria
Citrobacter rodentium ICC168 YP 003368340 1 637910 Proteobacteria
Escherichia coli O7 K1 str CE10 YP 006144365 1 1072459 Proteobacteria
Escherichia coli O55 H7 str CB9615 YP 003501135 1 701177 Proteobacteria
Escherichia coli O55 H7 str CB9615 YP 003500049 1 701177 Proteobacteria
Escherichia coli O26 H11 str 11368 YP 003230355 1 573235 Proteobacteria
Escherichia coli O103 H2 str 12009 YP 003224717 1 585395 Proteobacteria
Escherichia coli O103 H2 str 12009 YP 003223441 1 585395 Proteobacteria
Escherichia coli E24377A YP 001464348 1 331111 Proteobacteria
Escherichia coli 042 YP 006099047 1 216592 Proteobacteria
Escherichia coli E24377A YP 001465819 1 331111 Proteobacteria

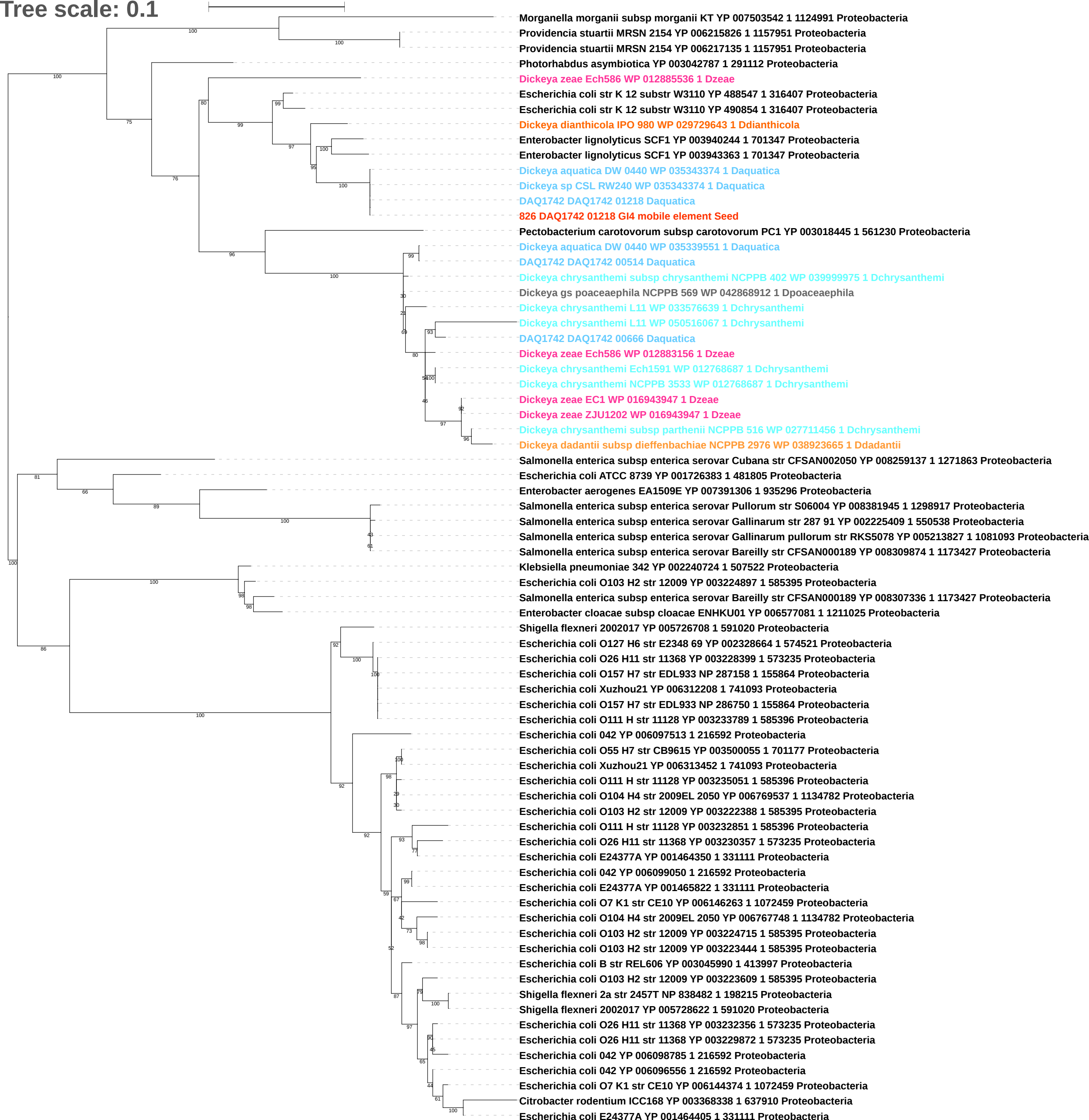
Tree scale: 0.1
Morganella morganii subsp morganii KT YP 007503542 1 1124991 Proteobacteria
Providencia stuartii MRSN 2154 YP 006215826 1 1157951 Proteobacteria
Providencia stuartii MRSN 2154 YP 006217135 1 1157951 Proteobacteria
Photorhabdus asymbiotica YP 003042787 1 291112 Proteobacteria
Dickeya zeae Ech586 WP 012885536 1 Dzeae
Escherichia coli str K 12 substr W3110 YP 488547 1 316407 Proteobacteria
Escherichia coli str K 12 substr W3110 YP 490854 1 316407 Proteobacteria
Dickeya dianthicola IPO 980 WP 029729643 1 Ddianthicola
Enterobacter lignolyticus SCF1 YP 003940244 1 701347 Proteobacteria
Enterobacter lignolyticus SCF1 YP 003943363 1 701347 Proteobacteria
Dickeya aquatica DW 0440 WP 035343374 1 Daquatica
Dickeya sp CSL RW240 WP 035343374 1 Daquatica
DAQ1742 DAQ1742 01218 Daquatica
826 DAQ1742 01218 GI4 mobile element Seed
Pectobacterium carotovorum subsp carotovorum PC1 YP 003018445 1 561230 Proteobacteria
Dickeya aquatica DW 0440 WP 035339551 1 Daquatica
DAQ1742 DAQ1742 00514 Daquatica
Dickeya chrysanthemi subsp chrysanthemi NCPPB 402 WP 039999975 1 Dchrysanthemi
Dickeya gs poaceaephila NCPPB 569 WP 042868912 1 Dpoaceaephila
Dickeya chrysanthemi L11 WP 033576639 1 Dchrysanthemi
Dickeya chrysanthemi L11 WP 050516067 1 Dchrysanthemi
DAQ1742 DAQ1742 00666 Daquatica
Dickeya zeae Ech586 WP 012883156 1 Dzeae
Dickeya chrysanthemi Ech1591 WP 012768687 1 Dchrysanthemi
Dickeya chrysanthemi NCPPB 3533 WP 012768687 1 Dchrysanthemi
Dickeya zeae EC1 WP 016943947 1 Dzeae
Dickeya zeae ZJU1202 WP 016943947 1 Dzeae
Dickeya chrysanthemi subsp parthenii NCPPB 516 WP 027711456 1 Dchrysanthemi
Dickeya dadantii subsp dieffenbachiae NCPPB 2976 WP 038923665 1 Ddadantii
Salmonella enterica subsp enterica serovar Cubana str CFSAN002050 YP 008259137 1 1271863 Proteobacteria
Escherichia coli ATCC 8739 YP 001726383 1 481805 Proteobacteria
Enterobacter aerogenes EA1509E YP 007391306 1 935296 Proteobacteria
Salmonella enterica subsp enterica serovar Pullorum str S06004 YP 008381945 1 1298917 Proteobacteria
Salmonella enterica subsp enterica serovar Gallinarum str 287 91 YP 002225409 1 550538 Proteobacteria
Salmonella enterica subsp enterica serovar Gallinarum pullorum str RKS5078 YP 005213827 1 1081093 Proteobacteria
Salmonella enterica subsp enterica serovar Bareilly str CFSAN000189 YP 008309874 1 1173427 Proteobacteria
Klebsiella pneumoniae 342 YP 002240724 1 507522 Proteobacteria
Escherichia coli O103 H2 str 12009 YP 003224897 1 585395 Proteobacteria
Salmonella enterica subsp enterica serovar Bareilly str CFSAN000189 YP 008307336 1 1173427 Proteobacteria
Enterobacter cloacae subsp cloacae ENHKU01 YP 006577081 1 1211025 Proteobacteria
Shigella flexneri 2002017 YP 005726708 1 591020 Proteobacteria
Escherichia coli O127 H6 str E2348 69 YP 002328664 1 574521 Proteobacteria
Escherichia coli O26 H11 str 11368 YP 003228399 1 573235 Proteobacteria
Escherichia coli O157 H7 str EDL933 NP 287158 1 155864 Proteobacteria
Escherichia coli Xuzhou21 YP 006312208 1 741093 Proteobacteria
Escherichia coli O157 H7 str EDL933 NP 286750 1 155864 Proteobacteria
Escherichia coli O111 H str 11128 YP 003233789 1 585396 Proteobacteria
Escherichia coli 042 YP 006097513 1 216592 Proteobacteria
Escherichia coli O55 H7 str CB9615 YP 003500055 1 701177 Proteobacteria
Escherichia coli Xuzhou21 YP 006313452 1 741093 Proteobacteria
Escherichia coli O111 H str 11128 YP 003235051 1 585396 Proteobacteria
Escherichia coli O104 H4 str 2009EL 2050 YP 006769537 1 1134782 Proteobacteria
Escherichia coli O103 H2 str 12009 YP 003222388 1 585395 Proteobacteria
Escherichia coli O111 H str 11128 YP 003232851 1 585396 Proteobacteria
Escherichia coli O26 H11 str 11368 YP 003230357 1 573235 Proteobacteria
Escherichia coli E24377A YP 001464350 1 331111 Proteobacteria
Escherichia coli 042 YP 006099050 1 216592 Proteobacteria
Escherichia coli E24377A YP 001465822 1 331111 Proteobacteria
Escherichia coli O7 K1 str CE10 YP 006146263 1 1072459 Proteobacteria
Escherichia coli O104 H4 str 2009EL 2050 YP 006767748 1 1134782 Proteobacteria
Escherichia coli O103 H2 str 12009 YP 003224715 1 585395 Proteobacteria
Escherichia coli O103 H2 str 12009 YP 003223444 1 585395 Proteobacteria
Escherichia coli B str REL606 YP 003045990 1 413997 Proteobacteria
Escherichia coli O103 H2 str 12009 YP 003223609 1 585395 Proteobacteria
Shigella flexneri 2a str 2457T NP 838482 1 198215 Proteobacteria
Shigella flexneri 2002017 YP 005728622 1 591020 Proteobacteria
Escherichia coli O26 H11 str 11368 YP 003232356 1 573235 Proteobacteria
Escherichia coli O26 H11 str 11368 YP 003229872 1 573235 Proteobacteria
Escherichia coli 042 YP 006098785 1 216592 Proteobacteria
Escherichia coli 042 YP 006096556 1 216592 Proteobacteria
Escherichia coli O7 K1 str CE10 YP 006144374 1 1072459 Proteobacteria
Citrobacter rodentium ICC168 YP 003368338 1 637910 Proteobacteria
Escherichia coli E24377A YP 001464405 1 331111 Proteobacteria

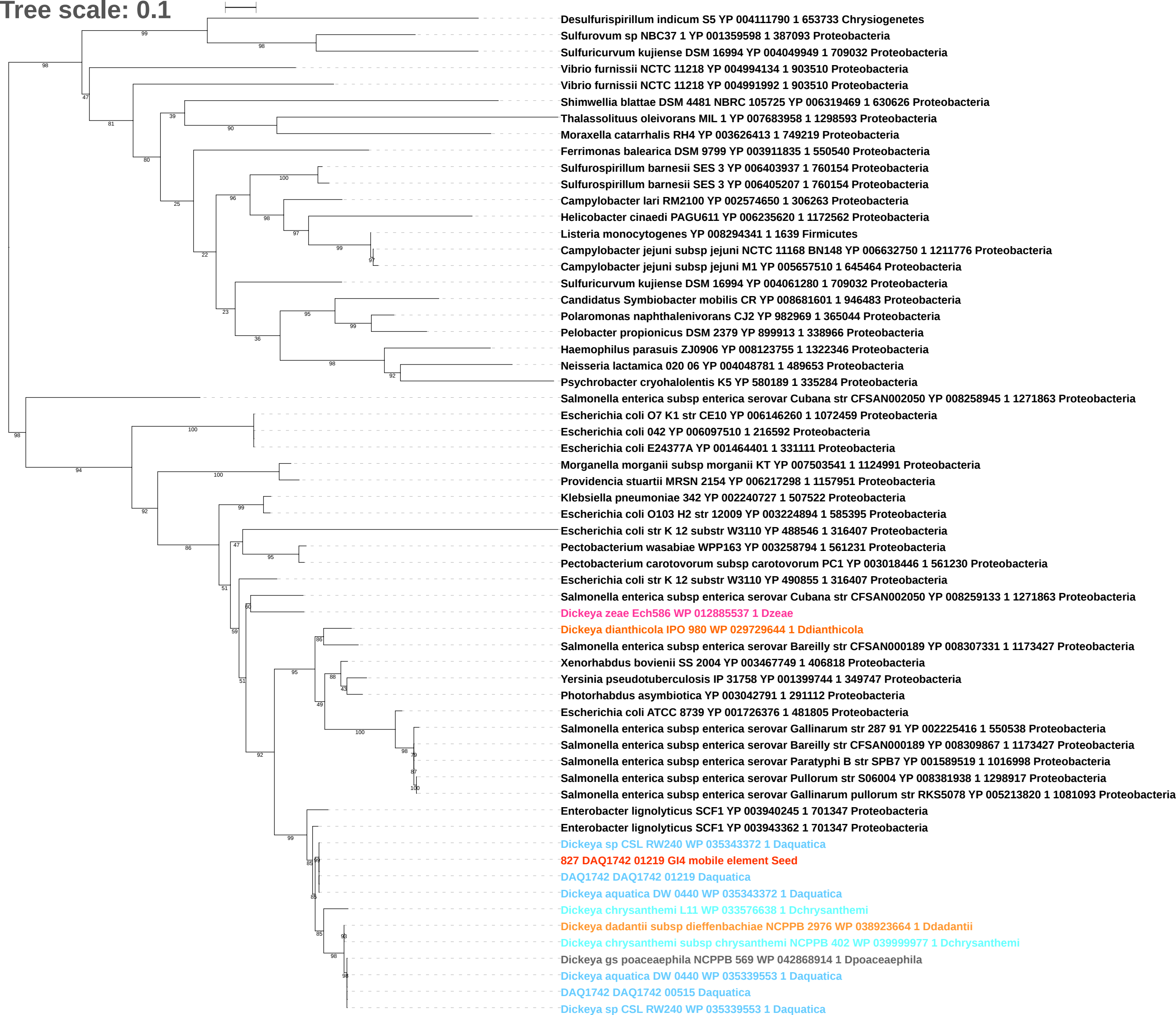
Tree scale: 0.1
Desulfurispirillum indicum S5 YP 004111790 1 653733 Chrysiogenetes
Sulfurovum sp NBC37 1 YP 001359598 1 387093 Proteobacteria
Sulfuricurvum kujiense DSM 16994 YP 004049949 1 709032 Proteobacteria
Vibrio furnissii NCTC 11218 YP 004994134 1 903510 Proteobacteria
Vibrio furnissii NCTC 11218 YP 004991992 1 903510 Proteobacteria
Shimwellia blattae DSM 4481 NBRC 105725 YP 006319469 1 630626 Proteobacteria
Thalassolituus oleivorans MIL 1 YP 007683958 1 1298593 Proteobacteria
Moraxella catarrhalis RH4 YP 003626413 1 749219 Proteobacteria
Ferrimonas balearica DSM 9799 YP 003911835 1 550540 Proteobacteria
Sulfurospirillum barnesii SES 3 YP 006403937 1 760154 Proteobacteria
Sulfurospirillum barnesii SES 3 YP 006405207 1 760154 Proteobacteria
Campylobacter lari RM2100 YP 002574650 1 306263 Proteobacteria
Helicobacter cinaedi PAGU611 YP 006235620 1 1172562 Proteobacteria
Listeria monocytogenes YP 008294341 1 1639 Firmicutes
Campylobacter jejuni subsp jejuni NCTC 11168 BN148 YP 006632750 1 1211776 Proteobacteria
Campylobacter jejuni subsp jejuni M1 YP 005657510 1 645464 Proteobacteria
Sulfuricurvum kujiense DSM 16994 YP 004061280 1 709032 Proteobacteria
Candidatus Symbiobacter mobilis CR YP 008681601 1 946483 Proteobacteria
Polaromonas naphthalenivorans CJ2 YP 982969 1 365044 Proteobacteria
Pelobacter propionicus DSM 2379 YP 899913 1 338966 Proteobacteria
Haemophilus parasuis ZJ0906 YP 008123755 1 1322346 Proteobacteria
Neisseria lactamica 020 06 YP 004048781 1 489653 Proteobacteria
Psychrobacter cryohalolentis K5 YP 580189 1 335284 Proteobacteria
Salmonella enterica subsp enterica serovar Cubana str CFSAN002050 YP 008258945 1 1271863 Proteobacteria
Escherichia coli O7 K1 str CE10 YP 006146260 1 1072459 Proteobacteria
Escherichia coli 042 YP 006097510 1 216592 Proteobacteria
Escherichia coli E24377A YP 001464401 1 331111 Proteobacteria
Morganella morganii subsp morganii KT YP 007503541 1 1124991 Proteobacteria
Providencia stuartii MRSN 2154 YP 006217298 1 1157951 Proteobacteria
Klebsiella pneumoniae 342 YP 002240727 1 507522 Proteobacteria
Escherichia coli O103 H2 str 12009 YP 003224894 1 585395 Proteobacteria
Escherichia coli str K 12 substr W3110 YP 488546 1 316407 Proteobacteria
Pectobacterium wasabiae WPP163 YP 003258794 1 561231 Proteobacteria
Pectobacterium carotovorum subsp carotovorum PC1 YP 003018446 1 561230 Proteobacteria
Escherichia coli str K 12 substr W3110 YP 490855 1 316407 Proteobacteria
Salmonella enterica subsp enterica serovar Cubana str CFSAN002050 YP 008259133 1 1271863 Proteobacteria
Dickeya zeae Ech586 WP 012885537 1 Dzeae
Dickeya dianthicola IPO 980 WP 029729644 1 Ddianthicola
Salmonella enterica subsp enterica serovar Bareilly str CFSAN000189 YP 008307331 1 1173427 Proteobacteria
Xenorhabdus bovienii SS 2004 YP 003467749 1 406818 Proteobacteria
Yersinia pseudotuberculosis IP 31758 YP 001399744 1 349747 Proteobacteria
Photorhabdus asymbiotica YP 003042791 1 291112 Proteobacteria
Escherichia coli ATCC 8739 YP 001726376 1 481805 Proteobacteria
Salmonella enterica subsp enterica serovar Gallinarum str 287 91 YP 002225416 1 550538 Proteobacteria
Salmonella enterica subsp enterica serovar Bareilly str CFSAN000189 YP 008309867 1 1173427 Proteobacteria
Salmonella enterica subsp enterica serovar Paratyphi B str SPB7 YP 001589519 1 1016998 Proteobacteria
Salmonella enterica subsp enterica serovar Pullorum str S06004 YP 008381938 1 1298917 Proteobacteria
Salmonella enterica subsp enterica serovar Gallinarum pullorum str RKS5078 YP 005213820 1 1081093 Proteobacteria
Enterobacter lignolyticus SCF1 YP 003940245 1 701347 Proteobacteria
Enterobacter lignolyticus SCF1 YP 003943362 1 701347 Proteobacteria
Dickeya sp CSL RW240 WP 035343372 1 Daquatica
827 DAQ1742 01219 GI4 mobile element Seed
DAQ1742 DAQ1742 01219 Daquatica
Dickeya aquatica DW 0440 WP 035343372 1 Daquatica
Dickeya chrysanthemi L11 WP 033576638 1 Dchrysanthemi
Dickeya dadantii subsp dieffenbachiae NCPPB 2976 WP 038923664 1 Ddadantii
Dickeya chrysanthemi subsp chrysanthemi NCPPB 402 WP 039999977 1 Dchrysanthemi
Dickeya gs poaceaephila NCPPB 569 WP 042868914 1 Dpoaceaephila
Dickeya aquatica DW 0440 WP 035339553 1 Daquatica
DAQ1742 DAQ1742 00515 Daquatica
Dickeya sp CSL RW240 WP 035339553 1 Daquatica

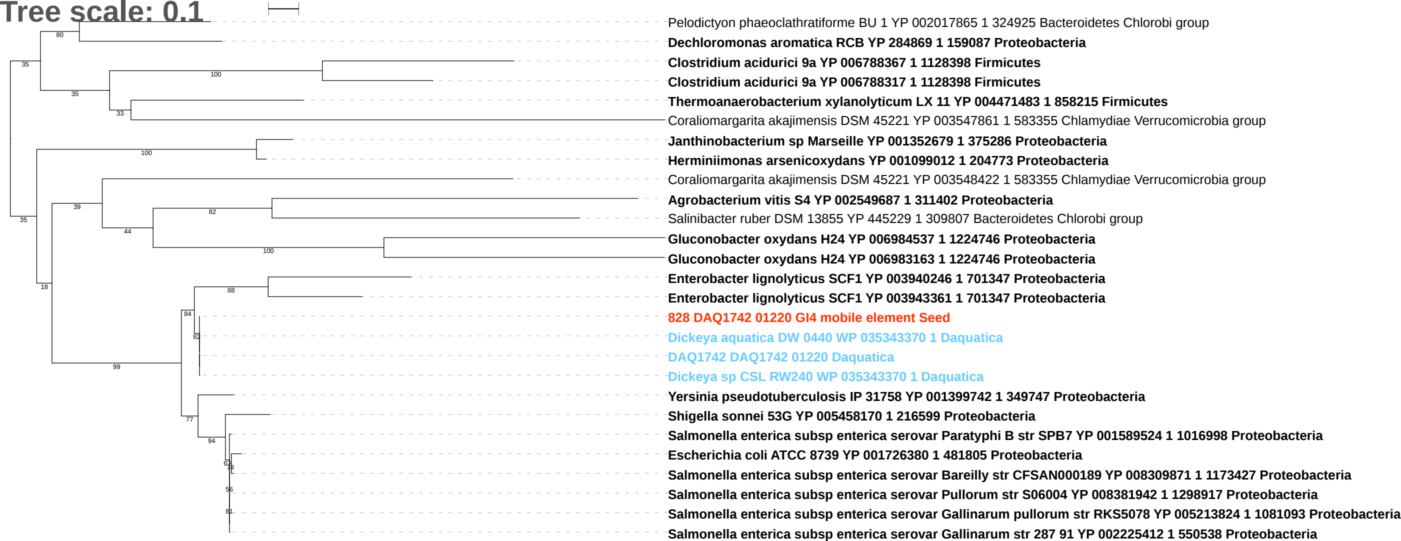

Tree scale: 0.1
Pelodictyon phaeoclathratiforme BU 1 YP 002017865 1 324925 Bacteroidetes Chlorobi group
Dechloromonas aromatica RCB YP 284869 1 159087 Proteobacteria
Clostridium acidurici 9a YP 006788367 1 1128398 Firmicutes
Clostridium acidurici 9a YP 006788317 1 1128398 Firmicutes
Thermoanaerobacterium xylanolyticum LX 11 YP 004471483 1 858215 Firmicutes
Coraliomargarita akajimensis DSM 45221 YP 003547861 1 583355 Chlamydiae Verrucomicrobia group
Janthinobacterium sp Marseille YP 001352679 1 375286 Proteobacteria
Herminiimonas arsenicoxydans YP 001099012 1 204773 Proteobacteria
Coraliomargarita akajimensis DSM 45221 YP 003548422 1 583355 Chlamydiae Verrucomicrobia group
Agrobacterium vitis S4 YP 002549687 1 311402 Proteobacteria
Salinibacter ruber DSM 13855 YP 445229 1 309807 Bacteroidetes Chlorobi group
Gluconobacter oxydans H24 YP 006984537 1 1224746 Proteobacteria
Gluconobacter oxydans H24 YP 006983163 1 1224746 Proteobacteria
Enterobacter lignolyticus SCF1 YP 003940246 1 701347 Proteobacteria
Enterobacter lignolyticus SCF1 YP 003943361 1 701347 Proteobacteria
828 DAQ1742 01220 GI4 mobile element Seed
Dickeya aquatica DW 0440 WP 035343370 1 Daquatica
DAQ1742 DAQ1742 01220 Daquatica
Dickeya sp CSL RW240 WP 035343370 1 Daquatica
Yersinia pseudotuberculosis IP 31758 YP 001399742 1 349747 Proteobacteria
Shigella sonnei 53G YP 005458170 1 216599 Proteobacteria
Salmonella enterica subsp enterica serovar Paratyphi B str SPB7 YP 001589524 1 1016998 Proteobacteria
Escherichia coli ATCC 8739 YP 001726380 1 481805 Proteobacteria
Salmonella enterica subsp enterica serovar Bareilly str CFSAN000189 YP 008309871 1 1173427 Proteobacteria
Salmonella enterica subsp enterica serovar Pullorum str S06004 YP 008381942 1 1298917 Proteobacteria
Salmonella enterica subsp enterica serovar Gallinarum pullorum str RKS5078 YP 005213824 1 1081093 Proteobacteria
Salmonella enterica subsp enterica serovar Gallinarum str 287 91 YP 002225412 1 550538 Proteobacteria

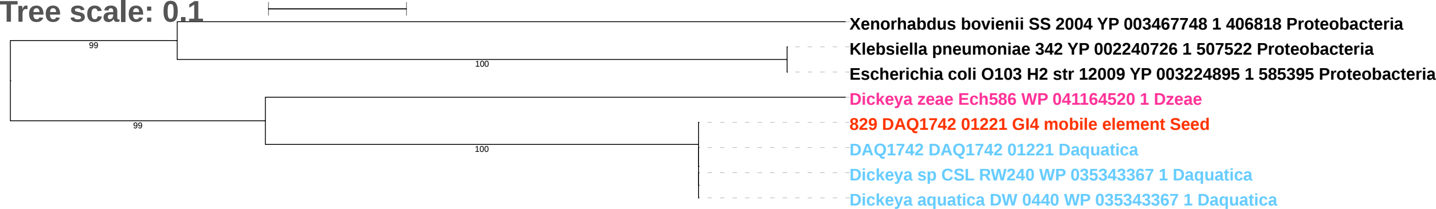
Tree scale: 0.1
Xenorhabdus bovienii SS 2004 YP 003467748 1 406818 Proteobacteria
Klebsiella pneumoniae 342 YP 002240726 1 507522 Proteobacteria
Escherichia coli O103 H2 str 12009 YP 003224895 1 585395 Proteobacteria
Dickeya zeae Ech586 WP 041164520 1 Dzeae
829 DAQ1742 01221 GI4 mobile element Seed
DAQ1742 DAQ1742 01221 Daquatica
Dickeya sp CSL RW240 WP 035343367 1 Daquatica
Dickeya aquatica DW 0440 WP 035343367 1 Daquatica
99
100
99
100

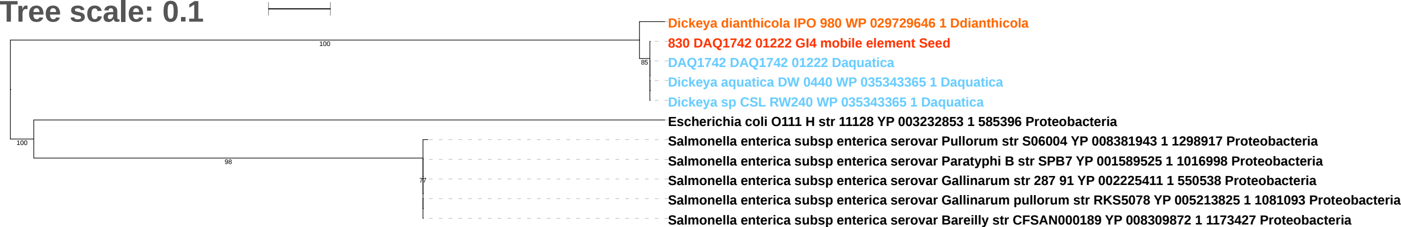
Tree scale: 0.1
Dickeya dianthicola IPO 980 WP 029729646 1 Ddianthicola
830 DAQ1742 01222 GI4 mobile element Seed
DAQ1742 DAQ1742 01222 Daquatica
Dickeya aquatica DW 0440 WP 035343365 1 Daquatica
Dickeya sp CSL RW240 WP 035343365 1 Daquatica
Escherichia coli O111 H str 11128 YP 003232853 1 585396 Proteobacteria
Salmonella enterica subsp enterica serovar Pullorum str S06004 YP 008381943 1 1298917 Proteobacteria
Salmonella enterica subsp enterica serovar Paratyphi B str SPB7 YP 001589525 1 1016998 Proteobacteria
Salmonella enterica subsp enterica serovar Gallinarum str 287 91 YP 002225411 1 550538 Proteobacteria
Salmonella enterica subsp enterica serovar Gallinarum pullorum str RKS5078 YP 005213825 1 1081093 Proteobacteria
Salmonella enterica subsp enterica serovar Bareilly str CFSAN000189 YP 008309872 1 1173427 Proteobacteria
100
85
100
98
77

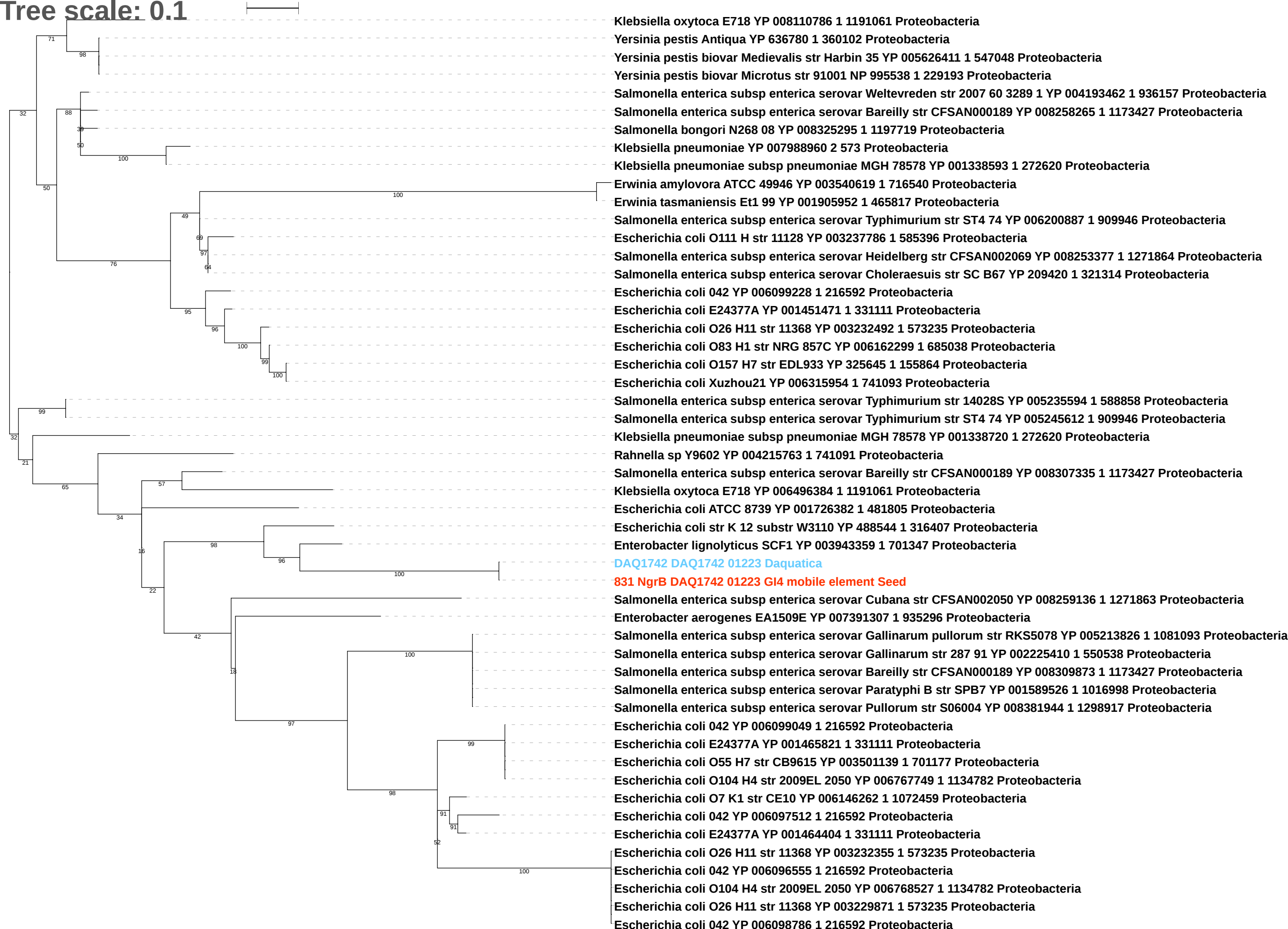
Tree scale: 0.1
Klebsiella oxytoca E718 YP 008110786 1 1191061 Proteobacteria
Yersinia pestis Antiqua YP 636780 1 360102 Proteobacteria
Yersinia pestis biovar Medievalis str Harbin 35 YP 005626411 1 547048 Proteobacteria
Yersinia pestis biovar Microtus str 91001 NP 995538 1 229193 Proteobacteria
Salmonella enterica subsp enterica serovar Weltevreden str 2007 60 3289 1 YP 004193462 1 936157 Proteobacteria
Salmonella enterica subsp enterica serovar Bareilly str CFSAN000189 YP 008258265 1 1173427 Proteobacteria
Salmonella bongori N268 08 YP 008325295 1 1197719 Proteobacteria
Klebsiella pneumoniae YP 007988960 2 573 Proteobacteria
Klebsiella pneumoniae subsp pneumoniae MGH 78578 YP 001338593 1 272620 Proteobacteria
Erwinia amylovora ATCC 49946 YP 003540619 1 716540 Proteobacteria
Erwinia tasmaniensis Et1 99 YP 001905952 1 465817 Proteobacteria
Salmonella enterica subsp enterica serovar Typhimurium str ST4 74 YP 006200887 1 909946 Proteobacteria
Escherichia coli O111 H str 11128 YP 003237786 1 585396 Proteobacteria
Salmonella enterica subsp enterica serovar Heidelberg str CFSAN002069 YP 008253377 1 1271864 Proteobacteria
Salmonella enterica subsp enterica serovar Choleraesuis str SC B67 YP 209420 1 321314 Proteobacteria
Escherichia coli 042 YP 006099228 1 216592 Proteobacteria
Escherichia coli E24377A YP 001451471 1 331111 Proteobacteria
Escherichia coli O26 H11 str 11368 YP 003232492 1 573235 Proteobacteria
Escherichia coli O83 H1 str NRG 857C YP 006162299 1 685038 Proteobacteria
Escherichia coli O157 H7 str EDL933 YP 325645 1 155864 Proteobacteria
Escherichia coli Xuzhou21 YP 006315954 1 741093 Proteobacteria
Salmonella enterica subsp enterica serovar Typhimurium str 14028S YP 005235594 1 588858 Proteobacteria
Salmonella enterica subsp enterica serovar Typhimurium str ST4 74 YP 005245612 1 909946 Proteobacteria
Klebsiella pneumoniae subsp pneumoniae MGH 78578 YP 001338720 1 272620 Proteobacteria
Rahnella sp Y9602 YP 004215763 1 741091 Proteobacteria
Salmonella enterica subsp enterica serovar Bareilly str CFSAN000189 YP 008307335 1 1173427 Proteobacteria
Klebsiella oxytoca E718 YP 006496384 1 1191061 Proteobacteria
Escherichia coli ATCC 8739 YP 001726382 1 481805 Proteobacteria
Escherichia coli str K 12 substr W3110 YP 488544 1 316407 Proteobacteria
Enterobacter lignolyticus SCF1 YP 003943359 1 701347 Proteobacteria
DAQ1742 DAQ1742 01223 Daquatica
831 NgrB DAQ1742 01223 GI4 mobile element Seed
Salmonella enterica subsp enterica serovar Cubana str CFSAN002050 YP 008259136 1 1271863 Proteobacteria
Enterobacter aerogenes EA1509E YP 007391307 1 935296 Proteobacteria
Salmonella enterica subsp enterica serovar Gallinarum pullorum str RKS5078 YP 005213826 1 1081093 Proteobacteria
Salmonella enterica subsp enterica serovar Gallinarum str 287 91 YP 002225410 1 550538 Proteobacteria
Salmonella enterica subsp enterica serovar Bareilly str CFSAN000189 YP 008309873 1 1173427 Proteobacteria
Salmonella enterica subsp enterica serovar Paratyphi B str SPB7 YP 001589526 1 1016998 Proteobacteria
Salmonella enterica subsp enterica serovar Pullorum str S06004 YP 008381944 1 1298917 Proteobacteria
Escherichia coli 042 YP 006099049 1 216592 Proteobacteria
Escherichia coli E24377A YP 001465821 1 331111 Proteobacteria
Escherichia coli O55 H7 str CB9615 YP 003501139 1 701177 Proteobacteria
Escherichia coli O104 H4 str 2009EL 2050 YP 006767749 1 1134782 Proteobacteria
Escherichia coli O7 K1 str CE10 YP 006146262 1 1072459 Proteobacteria
Escherichia coli 042 YP 006097512 1 216592 Proteobacteria
Escherichia coli E24377A YP 001464404 1 331111 Proteobacteria
Escherichia coli O26 H11 str 11368 YP 003232355 1 573235 Proteobacteria
Escherichia coli 042 YP 006096555 1 216592 Proteobacteria
Escherichia coli O104 H4 str 2009EL 2050 YP 006768527 1 1134782 Proteobacteria
Escherichia coli O26 H11 str 11368 YP 003229871 1 573235 Proteobacteria
Escherichia coli 042 YP 006098786 1 216592 Proteobacteria

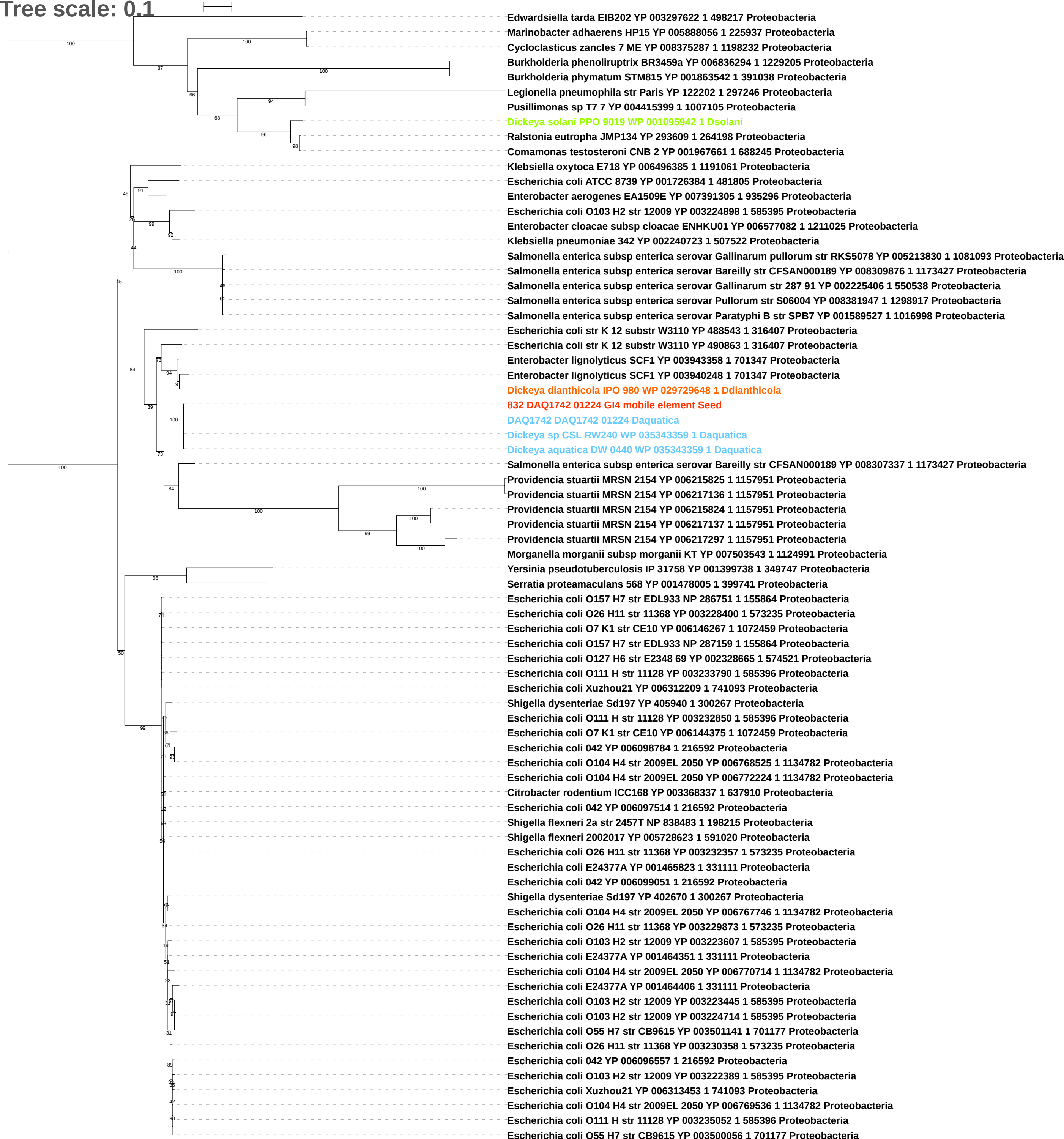
Tree scale: 0.1
Edwardsiella tarda EIB202 YP 003297622 1 498217 Proteobacteria
Marinobacter adhaerens HP15 YP 005888056 1 225937 Proteobacteria
Cycloclasticus zancles 7 ME YP 008375287 1 1198232 Proteobacteria
Burkholderia phenoliruptrix BR3459a YP 006836294 1 1229205 Proteobacteria
Burkholderia phymatum STM815 YP 001863542 1 391038 Proteobacteria
Legionella pneumophila str Paris YP 122202 1 297246 Proteobacteria
Pusillimonas sp T7 7 YP 004415399 1 1007105 Proteobacteria
Dickeya solani PPO 9019 WP 001095942 1 Dsolani
Ralstonia eutropha JMP134 YP 293609 1 264198 Proteobacteria
Comamonas testosteroni CNB 2 YP 001967661 1 688245 Proteobacteria
Klebsiella oxytoca E718 YP 006496385 1 1191061 Proteobacteria
Escherichia coli ATCC 8739 YP 001726384 1 481805 Proteobacteria
Enterobacter aerogenes EA1509E YP 007391305 1 935296 Proteobacteria
Escherichia coli O103 H2 str 12009 YP 003224898 1 585395 Proteobacteria
Enterobacter cloacae subsp cloacae ENHKU01 YP 006577082 1 1211025 Proteobacteria
Klebsiella pneumoniae 342 YP 002240723 1 507522 Proteobacteria
Salmonella enterica subsp enterica serovar Gallinarum pullorum str RKS5078 YP 005213830 1 1081093 Proteobacteria
Salmonella enterica subsp enterica serovar Bareilly str CFSAN000189 YP 008309876 1 1173427 Proteobacteria
Salmonella enterica subsp enterica serovar Gallinarum str 287 91 YP 002225406 1 550538 Proteobacteria
Salmonella enterica subsp enterica serovar Pullorum str S06004 YP 008381947 1 1298917 Proteobacteria
Salmonella enterica subsp enterica serovar Paratyphi B str SPB7 YP 001589527 1 1016998 Proteobacteria
Escherichia coli str K 12 substr W3110 YP 488543 1 316407 Proteobacteria
Escherichia coli str K 12 substr W3110 YP 490863 1 316407 Proteobacteria
Enterobacter lignolyticus SCF1 YP 003943358 1 701347 Proteobacteria
Enterobacter lignolyticus SCF1 YP 003940248 1 701347 Proteobacteria
Dickeya dianthicola IPO 980 WP 029729648 1 Ddianthicola
832 DAQ1742 01224 GI4 mobile element Seed
DAQ1742 DAQ1742 01224 Daquatica
Dickeya sp CSL RW240 WP 035343359 1 Daquatica
Dickeya aquatica DW 0440 WP 035343359 1 Daquatica
Salmonella enterica subsp enterica serovar Bareilly str CFSAN000189 YP 008307337 1 1173427 Proteobacteria
Providencia stuartii MRSN 2154 YP 006215825 1 1157951 Proteobacteria
Providencia stuartii MRSN 2154 YP 006217136 1 1157951 Proteobacteria
Providencia stuartii MRSN 2154 YP 006215824 1 1157951 Proteobacteria
Providencia stuartii MRSN 2154 YP 006217137 1 1157951 Proteobacteria
Providencia stuartii MRSN 2154 YP 006217297 1 1157951 Proteobacteria
Morganella morganii subsp morganii KT YP 007503543 1 1124991 Proteobacteria
Yersinia pseudotuberculosis IP 31758 YP 001399738 1 349747 Proteobacteria
Serratia proteamaculans 568 YP 001478005 1 399741 Proteobacteria
Escherichia coli O157 H7 str EDL933 NP 286751 1 155864 Proteobacteria
Escherichia coli O26 H11 str 11368 YP 003228400 1 573235 Proteobacteria
Escherichia coli O7 K1 str CE10 YP 006146267 1 1072459 Proteobacteria
Escherichia coli O157 H7 str EDL933 NP 287159 1 155864 Proteobacteria
Escherichia coli O127 H6 str E2348 69 YP 002328665 1 574521 Proteobacteria
Escherichia coli O111 H str 11128 YP 003233790 1 585396 Proteobacteria
Escherichia coli Xuzhou21 YP 006312209 1 741093 Proteobacteria
Shigella dysenteriae Sd197 YP 405940 1 300267 Proteobacteria
Escherichia coli O111 H str 11128 YP 003232850 1 585396 Proteobacteria
Escherichia coli O7 K1 str CE10 YP 006144375 1 1072459 Proteobacteria
Escherichia coli 042 YP 006098784 1 216592 Proteobacteria
Escherichia coli O104 H4 str 2009EL 2050 YP 006768525 1 1134782 Proteobacteria
Escherichia coli O104 H4 str 2009EL 2050 YP 006772224 1 1134782 Proteobacteria
Citrobacter rodentium ICC168 YP 003368337 1 637910 Proteobacteria
Escherichia coli 042 YP 006097514 1 216592 Proteobacteria
Shigella flexneri 2a str 2457T NP 838483 1 198215 Proteobacteria
Shigella flexneri 2002017 YP 005728623 1 591020 Proteobacteria
Escherichia coli O26 H11 str 11368 YP 003232357 1 573235 Proteobacteria
Escherichia coli E24377A YP 001465823 1 331111 Proteobacteria
Escherichia coli 042 YP 006099051 1 216592 Proteobacteria
Shigella dysenteriae Sd197 YP 402670 1 300267 Proteobacteria
Escherichia coli O104 H4 str 2009EL 2050 YP 006767746 1 1134782 Proteobacteria
Escherichia coli O26 H11 str 11368 YP 003229873 1 573235 Proteobacteria
Escherichia coli O103 H2 str 12009 YP 003223607 1 585395 Proteobacteria
Escherichia coli E24377A YP 001464351 1 331111 Proteobacteria
Escherichia coli O104 H4 str 2009EL 2050 YP 006770714 1 1134782 Proteobacteria
Escherichia coli E24377A YP 001464406 1 331111 Proteobacteria
Escherichia coli O103 H2 str 12009 YP 003223445 1 585395 Proteobacteria
Escherichia coli O103 H2 str 12009 YP 003224714 1 585395 Proteobacteria
Escherichia coli O55 H7 str CB9615 YP 003501141 1 701177 Proteobacteria
Escherichia coli O26 H11 str 11368 YP 003230358 1 573235 Proteobacteria
Escherichia coli 042 YP 006096557 1 216592 Proteobacteria
Escherichia coli O103 H2 str 12009 YP 003222389 1 585395 Proteobacteria
Escherichia coli Xuzhou21 YP 006313453 1 741093 Proteobacteria
Escherichia coli O104 H4 str 2009EL 2050 YP 006769536 1 1134782 Proteobacteria
Escherichia coli O111 H str 11128 YP 003235052 1 585396 Proteobacteria
Escherichia coli O55 H7 str CB9615 YP 003500056 1 701177 Proteobacteria

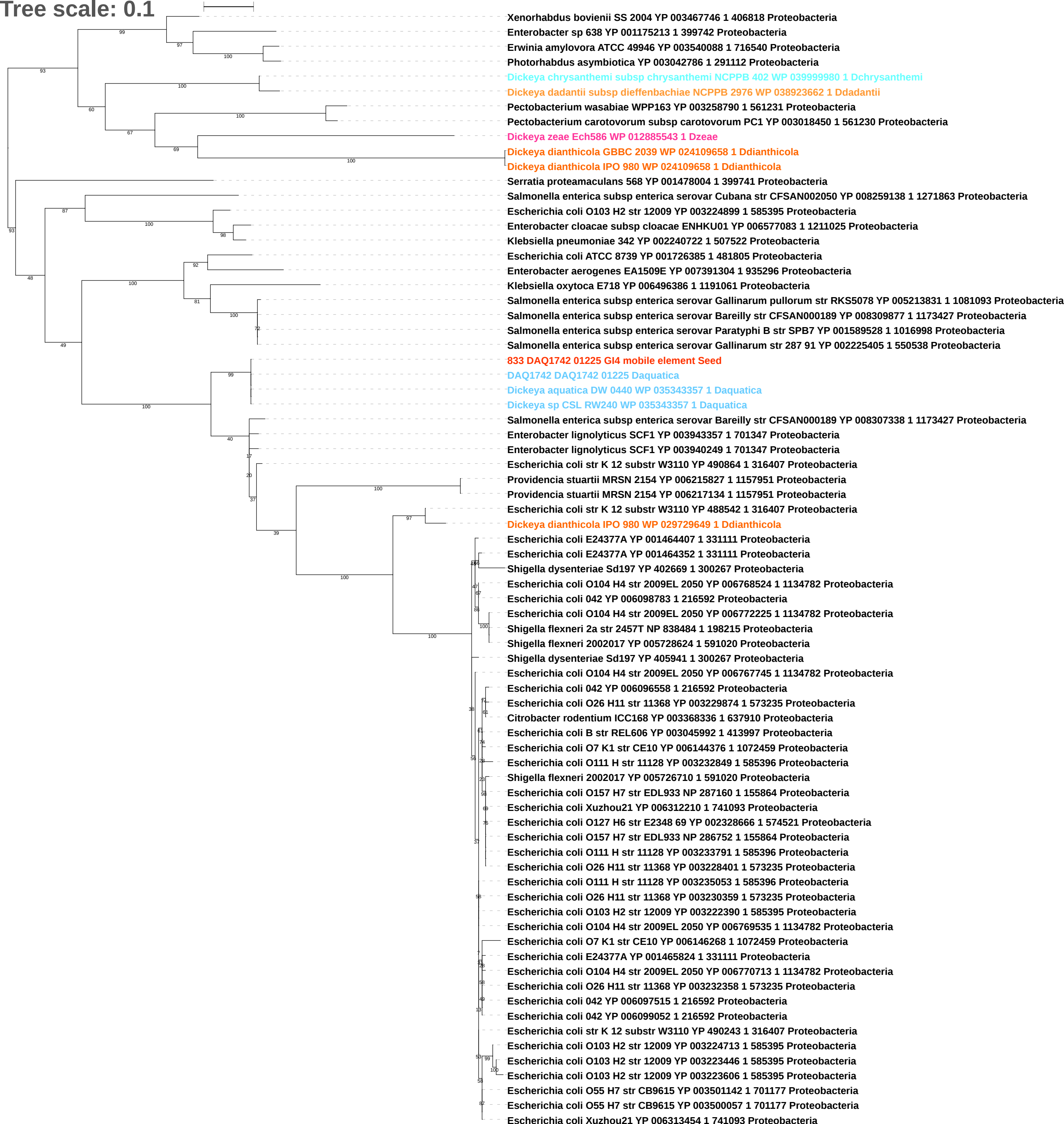
Tree scale: 0.1
Xenorhabdus bovienii SS 2004 YP 003467746 1 406818 Proteobacteria
Enterobacter sp 638 YP 001175213 1 399742 Proteobacteria
Erwinia amylovora ATCC 49946 YP 003540088 1 716540 Proteobacteria
Photorhabdus asymbiotica YP 003042786 1 291112 Proteobacteria
Dickeya chrysanthemi subsp chrysanthemi NCPPB 402 WP 039999980 1 Dchrysanthemi
Dickeya dadantii subsp dieffenbachiae NCPPB 2976 WP 038923662 1 Ddadantii
Pectobacterium wasabiae WPP163 YP 003258790 1 561231 Proteobacteria
Pectobacterium carotovorum subsp carotovorum PC1 YP 003018450 1 561230 Proteobacteria
Dickeya zeae Ech586 WP 012885543 1 Dzeae
Dickeya dianthicola GBBC 2039 WP 024109658 1 Ddianthicola
Dickeya dianthicola IPO 980 WP 024109658 1 Ddianthicola
Serratia proteamaculans 568 YP 001478004 1 399741 Proteobacteria
Salmonella enterica subsp enterica serovar Cubana str CFSAN002050 YP 008259138 1 1271863 Proteobacteria
Escherichia coli O103 H2 str 12009 YP 003224899 1 585395 Proteobacteria
Enterobacter cloacae subsp cloacae ENHKU01 YP 006577083 1 1211025 Proteobacteria
Klebsiella pneumoniae 342 YP 002240722 1 507522 Proteobacteria
Escherichia coli ATCC 8739 YP 001726385 1 481805 Proteobacteria
Enterobacter aerogenes EA1509E YP 007391304 1 935296 Proteobacteria
Klebsiella oxytoca E718 YP 006496386 1 1191061 Proteobacteria
Salmonella enterica subsp enterica serovar Gallinarum pullorum str RKS5078 YP 005213831 1 1081093 Proteobacteria
Salmonella enterica subsp enterica serovar Bareilly str CFSAN000189 YP 008309877 1 1173427 Proteobacteria
Salmonella enterica subsp enterica serovar Paratyphi B str SPB7 YP 001589528 1 1016998 Proteobacteria
Salmonella enterica subsp enterica serovar Gallinarum str 287 91 YP 002225405 1 550538 Proteobacteria
833 DAQ1742 01225 GI4 mobile element Seed
DAQ1742 DAQ1742 01225 Daquatica
Dickeya aquatica DW 0440 WP 035343357 1 Daquatica
Dickeya sp CSL RW240 WP 035343357 1 Daquatica
Salmonella enterica subsp enterica serovar Bareilly str CFSAN000189 YP 008307338 1 1173427 Proteobacteria
Enterobacter lignolyticus SCF1 YP 003943357 1 701347 Proteobacteria
Enterobacter lignolyticus SCF1 YP 003940249 1 701347 Proteobacteria
Escherichia coli str K 12 substr W3110 YP 490864 1 316407 Proteobacteria
Providencia stuartii MRSN 2154 YP 006215827 1 1157951 Proteobacteria
Providencia stuartii MRSN 2154 YP 006217134 1 1157951 Proteobacteria
Escherichia coli str K 12 substr W3110 YP 488542 1 316407 Proteobacteria
Dickeya dianthicola IPO 980 WP 029729649 1 Ddianthicola
Escherichia coli E24377A YP 001464407 1 331111 Proteobacteria
Escherichia coli E24377A YP 001464352 1 331111 Proteobacteria
Shigella dysenteriae Sd197 YP 402669 1 300267 Proteobacteria
Escherichia coli O104 H4 str 2009EL 2050 YP 006768524 1 1134782 Proteobacteria
Escherichia coli 042 YP 006098783 1 216592 Proteobacteria
Escherichia coli O104 H4 str 2009EL 2050 YP 006772225 1 1134782 Proteobacteria
Shigella flexneri 2a str 2457T NP 838484 1 198215 Proteobacteria
Shigella flexneri 2002017 YP 005728624 1 591020 Proteobacteria
Shigella dysenteriae Sd197 YP 405941 1 300267 Proteobacteria
Escherichia coli O104 H4 str 2009EL 2050 YP 006767745 1 1134782 Proteobacteria
Escherichia coli 042 YP 006096558 1 216592 Proteobacteria
Escherichia coli O26 H11 str 11368 YP 003229874 1 573235 Proteobacteria
Citrobacter rodentium ICC168 YP 003368336 1 637910 Proteobacteria
Escherichia coli B str REL606 YP 003045992 1 413997 Proteobacteria
Escherichia coli O7 K1 str CE10 YP 006144376 1 1072459 Proteobacteria
Escherichia coli O111 H str 11128 YP 003232849 1 585396 Proteobacteria
Shigella flexneri 2002017 YP 005726710 1 591020 Proteobacteria
Escherichia coli O157 H7 str EDL933 NP 287160 1 155864 Proteobacteria
Escherichia coli Xuzhou21 YP 006312210 1 741093 Proteobacteria
Escherichia coli O127 H6 str E2348 69 YP 002328666 1 574521 Proteobacteria
Escherichia coli O157 H7 str EDL933 NP 286752 1 155864 Proteobacteria
Escherichia coli O111 H str 11128 YP 003233791 1 585396 Proteobacteria
Escherichia coli O26 H11 str 11368 YP 003228401 1 573235 Proteobacteria
Escherichia coli O111 H str 11128 YP 003235053 1 585396 Proteobacteria
Escherichia coli O26 H11 str 11368 YP 003230359 1 573235 Proteobacteria
Escherichia coli O103 H2 str 12009 YP 003222390 1 585395 Proteobacteria
Escherichia coli O104 H4 str 2009EL 2050 YP 006769535 1 1134782 Proteobacteria
Escherichia coli O7 K1 str CE10 YP 006146268 1 1072459 Proteobacteria
Escherichia coli E24377A YP 001465824 1 331111 Proteobacteria
Escherichia coli O104 H4 str 2009EL 2050 YP 006770713 1 1134782 Proteobacteria
Escherichia coli O26 H11 str 11368 YP 003232358 1 573235 Proteobacteria
Escherichia coli 042 YP 006097515 1 216592 Proteobacteria
Escherichia coli 042 YP 006099052 1 216592 Proteobacteria
Escherichia coli str K 12 substr W3110 YP 490243 1 316407 Proteobacteria
Escherichia coli O103 H2 str 12009 YP 003224713 1 585395 Proteobacteria
Escherichia coli O103 H2 str 12009 YP 003223446 1 585395 Proteobacteria
Escherichia coli O103 H2 str 12009 YP 003223606 1 585395 Proteobacteria
Escherichia coli O55 H7 str CB9615 YP 003501142 1 701177 Proteobacteria
Escherichia coli O55 H7 str CB9615 YP 003500057 1 701177 Proteobacteria
Escherichia coli Xuzhou21 YP 006313454 1 741093 Proteobacteria

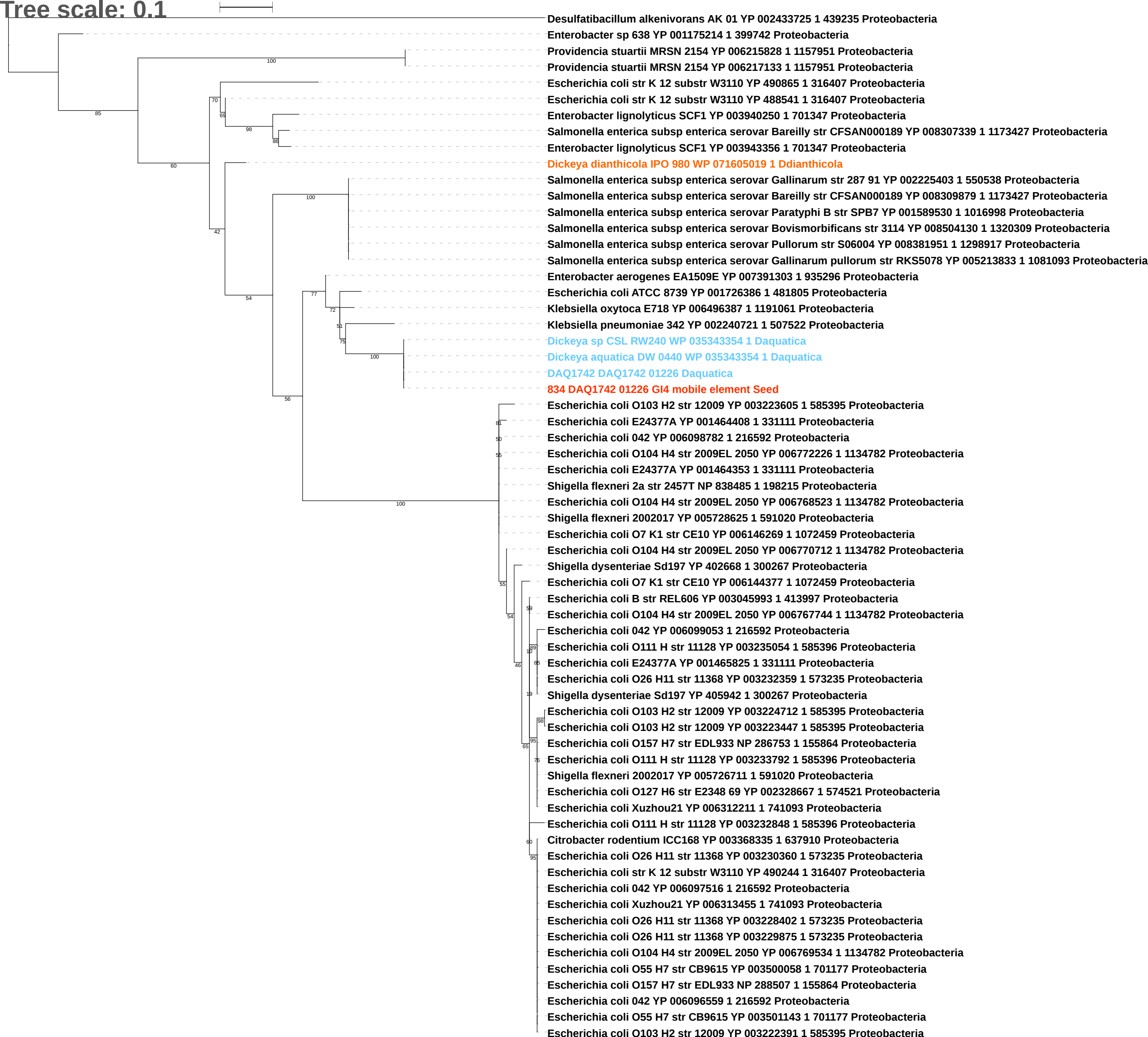
Tree scale: 0.1
Desulfatibacillum alkenivorans AK 01 YP 002433725 1 439235 Proteobacteria
Enterobacter sp 638 YP 001175214 1 399742 Proteobacteria
Providencia stuartii MRSN 2154 YP 006215828 1 1157951 Proteobacteria
Providencia stuartii MRSN 2154 YP 006217133 1 1157951 Proteobacteria
Escherichia coli str K 12 substr W3110 YP 490865 1 316407 Proteobacteria
Escherichia coli str K 12 substr W3110 YP 488541 1 316407 Proteobacteria
Enterobacter lignolyticus SCF1 YP 003940250 1 701347 Proteobacteria
Salmonella enterica subsp enterica serovar Bareilly str CFSAN000189 YP 008307339 1 1173427 Proteobacteria
Enterobacter lignolyticus SCF1 YP 003943356 1 701347 Proteobacteria
Dickeya dianthicola IPO 980 WP 071605019 1 Ddianthicola
Salmonella enterica subsp enterica serovar Gallinarum str 287 91 YP 002225403 1 550538 Proteobacteria
Salmonella enterica subsp enterica serovar Bareilly str CFSAN000189 YP 008309879 1 1173427 Proteobacteria
Salmonella enterica subsp enterica serovar Paratyphi B str SPB7 YP 001589530 1 1016998 Proteobacteria
Salmonella enterica subsp enterica serovar Bovismorbificans str 3114 YP 008504130 1 1320309 Proteobacteria
Salmonella enterica subsp enterica serovar Pullorum str S06004 YP 008381951 1 1298917 Proteobacteria
Salmonella enterica subsp enterica serovar Gallinarum pullorum str RKS5078 YP 005213833 1 1081093 Proteobacteria
Enterobacter aerogenes EA1509E YP 007391303 1 935296 Proteobacteria
Escherichia coli ATCC 8739 YP 001726386 1 481805 Proteobacteria
Klebsiella oxytoca E718 YP 006496387 1 1191061 Proteobacteria
Klebsiella pneumoniae 342 YP 002240721 1 507522 Proteobacteria
Dickeya sp CSL RW240 WP 035343354 1 Daquatica
Dickeya aquatica DW 0440 WP 035343354 1 Daquatica
DAQ1742 DAQ1742 01226 Daquatica
834 DAQ1742 01226 GI4 mobile element Seed
Escherichia coli O103 H2 str 12009 YP 003223605 1 585395 Proteobacteria
Escherichia coli E24377A YP 001464408 1 331111 Proteobacteria
Escherichia coli 042 YP 006098782 1 216592 Proteobacteria
Escherichia coli O104 H4 str 2009EL 2050 YP 006772226 1 1134782 Proteobacteria
Escherichia coli E24377A YP 001464353 1 331111 Proteobacteria
Shigella flexneri 2a str 2457T NP 838485 1 198215 Proteobacteria
Escherichia coli O104 H4 str 2009EL 2050 YP 006768523 1 1134782 Proteobacteria
Shigella flexneri 2002017 YP 005728625 1 591020 Proteobacteria
Escherichia coli O7 K1 str CE10 YP 006146269 1 1072459 Proteobacteria
Escherichia coli O104 H4 str 2009EL 2050 YP 006770712 1 1134782 Proteobacteria
Shigella dysenteriae Sd197 YP 402668 1 300267 Proteobacteria
Escherichia coli O7 K1 str CE10 YP 006144377 1 1072459 Proteobacteria
Escherichia coli B str REL606 YP 003045993 1 413997 Proteobacteria
Escherichia coli O104 H4 str 2009EL 2050 YP 006767744 1 1134782 Proteobacteria
Escherichia coli 042 YP 006099053 1 216592 Proteobacteria
Escherichia coli O111 H str 11128 YP 003235054 1 585396 Proteobacteria
Escherichia coli E24377A YP 001465825 1 331111 Proteobacteria
Escherichia coli O26 H11 str 11368 YP 003232359 1 573235 Proteobacteria
Shigella dysenteriae Sd197 YP 405942 1 300267 Proteobacteria
Escherichia coli O103 H2 str 12009 YP 003224712 1 585395 Proteobacteria
Escherichia coli O103 H2 str 12009 YP 003223447 1 585395 Proteobacteria
Escherichia coli O157 H7 str EDL933 NP 286753 1 155864 Proteobacteria
Escherichia coli O111 H str 11128 YP 003233792 1 585396 Proteobacteria
Shigella flexneri 2002017 YP 005726711 1 591020 Proteobacteria
Escherichia coli O127 H6 str E2348 69 YP 002328667 1 574521 Proteobacteria
Escherichia coli Xuzhou21 YP 006312211 1 741093 Proteobacteria
Escherichia coli O111 H str 11128 YP 003232848 1 585396 Proteobacteria
Citrobacter rodentium ICC168 YP 003368335 1 637910 Proteobacteria
Escherichia coli O26 H11 str 11368 YP 003230360 1 573235 Proteobacteria
Escherichia coli str K 12 substr W3110 YP 490244 1 316407 Proteobacteria
Escherichia coli 042 YP 006097516 1 216592 Proteobacteria
Escherichia coli Xuzhou21 YP 006313455 1 741093 Proteobacteria
Escherichia coli O26 H11 str 11368 YP 003228402 1 573235 Proteobacteria
Escherichia coli O26 H11 str 11368 YP 003229875 1 573235 Proteobacteria
Escherichia coli O104 H4 str 2009EL 2050 YP 006769534 1 1134782 Proteobacteria
Escherichia coli O55 H7 str CB9615 YP 003500058 1 701177 Proteobacteria
Escherichia coli O157 H7 str EDL933 NP 288507 1 155864 Proteobacteria
Escherichia coli 042 YP 006096559 1 216592 Proteobacteria
Escherichia coli O55 H7 str CB9615 YP 003501143 1 701177 Proteobacteria
Escherichia coli O103 H2 str 12009 YP 003222391 1 585395 Proteobacteria

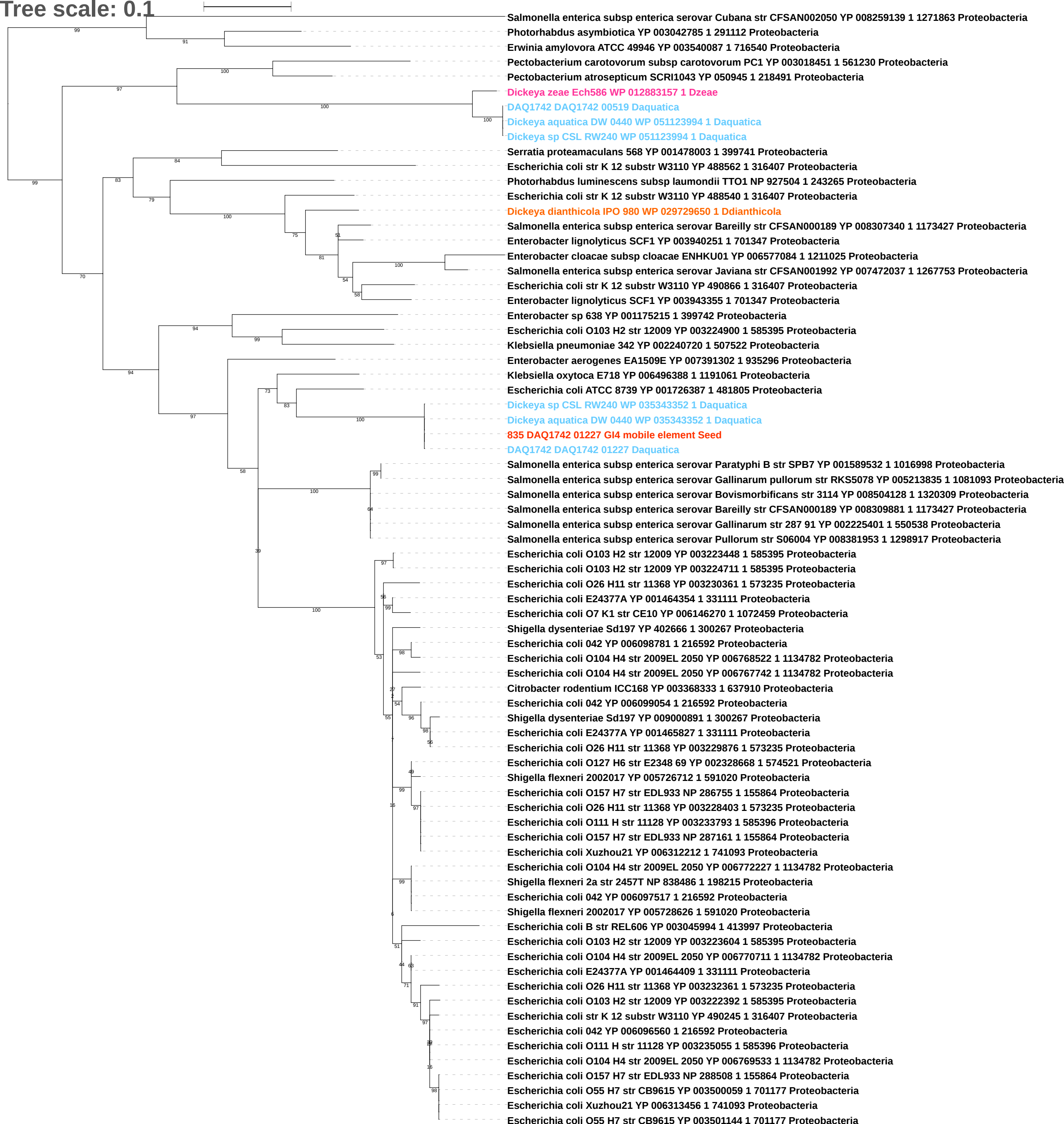

Tree scale: 0.1
Salmonella enterica subsp enterica serovar Cubana str CFSAN002050 YP 008259139 1 1271863 Proteobacteria
Photorhabdus asymbiotica YP 003042785 1 291112 Proteobacteria
Erwinia amylovora ATCC 49946 YP 003540087 1 716540 Proteobacteria
Pectobacterium carotovorum subsp carotovorum PC1 YP 003018451 1 561230 Proteobacteria
Pectobacterium atrosepticum SCRI1043 YP 050945 1 218491 Proteobacteria
Dickeya zeae Ech586 WP 012883157 1 Dzeae
DAQ1742 DAQ1742 00519 Daquatica
Dickeya aquatica DW 0440 WP 051123994 1 Daquatica
Dickeya sp CSL RW240 WP 051123994 1 Daquatica
Serratia proteamaculans 568 YP 001478003 1 399741 Proteobacteria
Escherichia coli str K 12 substr W3110 YP 488562 1 316407 Proteobacteria
Photorhabdus luminescens subsp laumondii TTO1 NP 927504 1 243265 Proteobacteria
Escherichia coli str K 12 substr W3110 YP 488540 1 316407 Proteobacteria
Dickeya dianthicola IPO 980 WP 029729650 1 Ddianthicola
Salmonella enterica subsp enterica serovar Bareilly str CFSAN000189 YP 008307340 1 1173427 Proteobacteria
Enterobacter lignolyticus SCF1 YP 003940251 1 701347 Proteobacteria
Enterobacter cloacae subsp cloacae ENHKU01 YP 006577084 1 1211025 Proteobacteria
Salmonella enterica subsp enterica serovar Javiana str CFSAN001992 YP 007472037 1 1267753 Proteobacteria
Escherichia coli str K 12 substr W3110 YP 490866 1 316407 Proteobacteria
Enterobacter lignolyticus SCF1 YP 003943355 1 701347 Proteobacteria
Enterobacter sp 638 YP 001175215 1 399742 Proteobacteria
Escherichia coli O103 H2 str 12009 YP 003224900 1 585395 Proteobacteria
Klebsiella pneumoniae 342 YP 002240720 1 507522 Proteobacteria
Enterobacter aerogenes EA1509E YP 007391302 1 935296 Proteobacteria
Klebsiella oxytoca E718 YP 006496388 1 1191061 Proteobacteria
Escherichia coli ATCC 8739 YP 001726387 1 481805 Proteobacteria
Dickeya sp CSL RW240 WP 035343352 1 Daquatica
Dickeya aquatica DW 0440 WP 035343352 1 Daquatica
835 DAQ1742 01227 GI4 mobile element Seed
DAQ1742 DAQ1742 01227 Daquatica
Salmonella enterica subsp enterica serovar Paratyphi B str SPB7 YP 001589532 1 1016998 Proteobacteria
Salmonella enterica subsp enterica serovar Gallinarum pullorum str RKS5078 YP 005213835 1 1081093 Proteobacteria
Salmonella enterica subsp enterica serovar Bovismorbificans str 3114 YP 008504128 1 1320309 Proteobacteria
Salmonella enterica subsp enterica serovar Bareilly str CFSAN000189 YP 008309881 1 1173427 Proteobacteria
Salmonella enterica subsp enterica serovar Gallinarum str 287 91 YP 002225401 1 550538 Proteobacteria
Salmonella enterica subsp enterica serovar Pullorum str S06004 YP 008381953 1 1298917 Proteobacteria
Escherichia coli O103 H2 str 12009 YP 003223448 1 585395 Proteobacteria
Escherichia coli O103 H2 str 12009 YP 003224711 1 585395 Proteobacteria
Escherichia coli O26 H11 str 11368 YP 003230361 1 573235 Proteobacteria
Escherichia coli E24377A YP 001464354 1 331111 Proteobacteria
Escherichia coli O7 K1 str CE10 YP 006146270 1 1072459 Proteobacteria
Shigella dysenteriae Sd197 YP 402666 1 300267 Proteobacteria
Escherichia coli 042 YP 006098781 1 216592 Proteobacteria
Escherichia coli O104 H4 str 2009EL 2050 YP 006768522 1 1134782 Proteobacteria
Escherichia coli O104 H4 str 2009EL 2050 YP 006767742 1 1134782 Proteobacteria
Citrobacter rodentium ICC168 YP 003368333 1 637910 Proteobacteria
Escherichia coli 042 YP 006099054 1 216592 Proteobacteria
Shigella dysenteriae Sd197 YP 009000891 1 300267 Proteobacteria
Escherichia coli E24377A YP 001465827 1 331111 Proteobacteria
Escherichia coli O26 H11 str 11368 YP 003229876 1 573235 Proteobacteria
Escherichia coli O127 H6 str E2348 69 YP 002328668 1 574521 Proteobacteria
Shigella flexneri 2002017 YP 005726712 1 591020 Proteobacteria
Escherichia coli O157 H7 str EDL933 NP 286755 1 155864 Proteobacteria
Escherichia coli O26 H11 str 11368 YP 003228403 1 573235 Proteobacteria
Escherichia coli O111 H str 11128 YP 003233793 1 585396 Proteobacteria
Escherichia coli O157 H7 str EDL933 NP 287161 1 155864 Proteobacteria
Escherichia coli Xuzhou21 YP 006312212 1 741093 Proteobacteria
Escherichia coli O104 H4 str 2009EL 2050 YP 006772227 1 1134782 Proteobacteria
Shigella flexneri 2a str 2457T NP 838486 1 198215 Proteobacteria
Escherichia coli 042 YP 006097517 1 216592 Proteobacteria
Shigella flexneri 2002017 YP 005728626 1 591020 Proteobacteria
Escherichia coli B str REL606 YP 003045994 1 413997 Proteobacteria
Escherichia coli O103 H2 str 12009 YP 003223604 1 585395 Proteobacteria
Escherichia coli O104 H4 str 2009EL 2050 YP 006770711 1 1134782 Proteobacteria
Escherichia coli E24377A YP 001464409 1 331111 Proteobacteria
Escherichia coli O26 H11 str 11368 YP 003232361 1 573235 Proteobacteria
Escherichia coli O103 H2 str 12009 YP 003222392 1 585395 Proteobacteria
Escherichia coli str K 12 substr W3110 YP 490245 1 316407 Proteobacteria
Escherichia coli 042 YP 006096560 1 216592 Proteobacteria
Escherichia coli O111 H str 11128 YP 003235055 1 585396 Proteobacteria
Escherichia coli O104 H4 str 2009EL 2050 YP 006769533 1 1134782 Proteobacteria
Escherichia coli O157 H7 str EDL933 NP 288508 1 155864 Proteobacteria
Escherichia coli O55 H7 str CB9615 YP 003500059 1 701177 Proteobacteria
Escherichia coli Xuzhou21 YP 006313456 1 741093 Proteobacteria
Escherichia coli O55 H7 str CB9615 YP 003501144 1 701177 Proteobacteria

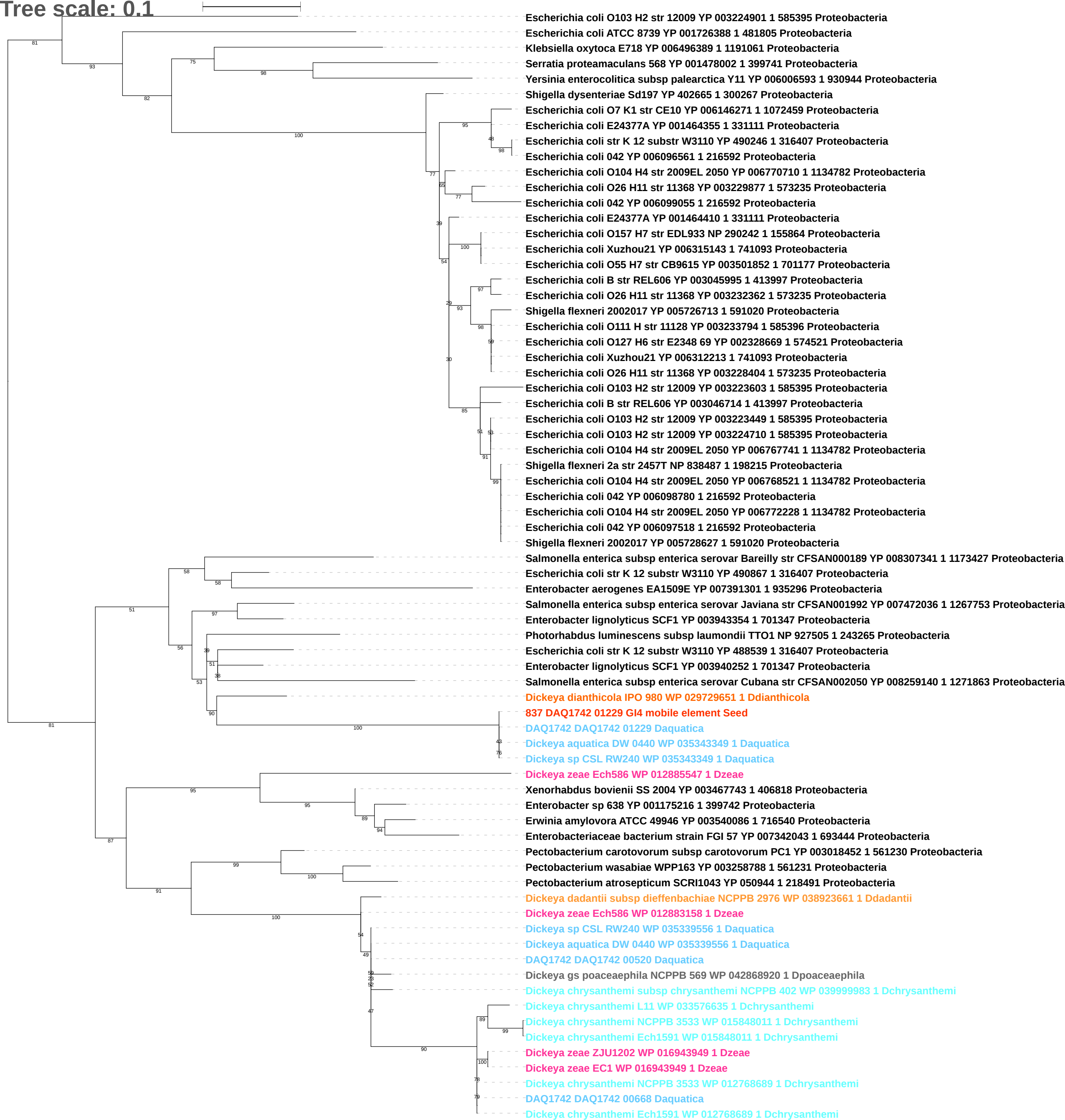
Tree scale: 0.1
Escherichia coli O103 H2 str 12009 YP 003224901 1 585395 Proteobacteria
Escherichia coli ATCC 8739 YP 001726388 1 481805 Proteobacteria
Klebsiella oxytoca E718 YP 006496389 1 1191061 Proteobacteria
Serratia proteamaculans 568 YP 001478002 1 399741 Proteobacteria
Yersinia enterocolitica subsp palearctica Y11 YP 006006593 1 930944 Proteobacteria
Shigella dysenteriae Sd197 YP 402665 1 300267 Proteobacteria
Escherichia coli O7 K1 str CE10 YP 006146271 1 1072459 Proteobacteria
Escherichia coli E24377A YP 001464355 1 331111 Proteobacteria
Escherichia coli str K 12 substr W3110 YP 490246 1 316407 Proteobacteria
Escherichia coli 042 YP 006096561 1 216592 Proteobacteria
Escherichia coli O104 H4 str 2009EL 2050 YP 006770710 1 1134782 Proteobacteria
Escherichia coli O26 H11 str 11368 YP 003229877 1 573235 Proteobacteria
Escherichia coli 042 YP 006099055 1 216592 Proteobacteria
Escherichia coli E24377A YP 001464410 1 331111 Proteobacteria
Escherichia coli O157 H7 str EDL933 NP 290242 1 155864 Proteobacteria
Escherichia coli Xuzhou21 YP 006315143 1 741093 Proteobacteria
Escherichia coli O55 H7 str CB9615 YP 003501852 1 701177 Proteobacteria
Escherichia coli B str REL606 YP 003045995 1 413997 Proteobacteria
Escherichia coli O26 H11 str 11368 YP 003232362 1 573235 Proteobacteria
Shigella flexneri 2002017 YP 005726713 1 591020 Proteobacteria
Escherichia coli O111 H str 11128 YP 003233794 1 585396 Proteobacteria
Escherichia coli O127 H6 str E2348 69 YP 002328669 1 574521 Proteobacteria
Escherichia coli Xuzhou21 YP 006312213 1 741093 Proteobacteria
Escherichia coli O26 H11 str 11368 YP 003228404 1 573235 Proteobacteria
Escherichia coli O103 H2 str 12009 YP 003223603 1 585395 Proteobacteria
Escherichia coli B str REL606 YP 003046714 1 413997 Proteobacteria
Escherichia coli O103 H2 str 12009 YP 003223449 1 585395 Proteobacteria
Escherichia coli O103 H2 str 12009 YP 003224710 1 585395 Proteobacteria
Escherichia coli O104 H4 str 2009EL 2050 YP 006767741 1 1134782 Proteobacteria
Shigella flexneri 2a str 2457T NP 838487 1 198215 Proteobacteria
Escherichia coli O104 H4 str 2009EL 2050 YP 006768521 1 1134782 Proteobacteria
Escherichia coli 042 YP 006098780 1 216592 Proteobacteria
Escherichia coli O104 H4 str 2009EL 2050 YP 006772228 1 1134782 Proteobacteria
Escherichia coli 042 YP 006097518 1 216592 Proteobacteria
Shigella flexneri 2002017 YP 005728627 1 591020 Proteobacteria
Salmonella enterica subsp enterica serovar Bareilly str CFSAN000189 YP 008307341 1 1173427 Proteobacteria
Escherichia coli str K 12 substr W3110 YP 490867 1 316407 Proteobacteria
Enterobacter aerogenes EA1509E YP 007391301 1 935296 Proteobacteria
Salmonella enterica subsp enterica serovar Javiana str CFSAN001992 YP 007472036 1 1267753 Proteobacteria
Enterobacter lignolyticus SCF1 YP 003943354 1 701347 Proteobacteria
Photorhabdus luminescens subsp laumondii TTO1 NP 927505 1 243265 Proteobacteria
Escherichia coli str K 12 substr W3110 YP 488539 1 316407 Proteobacteria
Enterobacter lignolyticus SCF1 YP 003940252 1 701347 Proteobacteria
Salmonella enterica subsp enterica serovar Cubana str CFSAN002050 YP 008259140 1 1271863 Proteobacteria
Dickeya dianthicola IPO 980 WP 029729651 1 Ddianthicola
837 DAQ1742 01229 GI4 mobile element Seed
DAQ1742 DAQ1742 01229 Daquatica
Dickeya aquatica DW 0440 WP 035343349 1 Daquatica
Dickeya sp CSL RW240 WP 035343349 1 Daquatica
Dickeya zeae Ech586 WP 012885547 1 Dzeae
Xenorhabdus bovienii SS 2004 YP 003467743 1 406818 Proteobacteria
Enterobacter sp 638 YP 001175216 1 399742 Proteobacteria
Erwinia amylovora ATCC 49946 YP 003540086 1 716540 Proteobacteria
Enterobacteriaceae bacterium strain FGI 57 YP 007342043 1 693444 Proteobacteria
Pectobacterium carotovorum subsp carotovorum PC1 YP 003018452 1 561230 Proteobacteria
Pectobacterium wasabiae WPP163 YP 003258788 1 561231 Proteobacteria
Pectobacterium atrosepticum SCRI1043 YP 050944 1 218491 Proteobacteria
Dickeya dadantii subsp dieffenbachiae NCPPB 2976 WP 038923661 1 Ddadantii
Dickeya zeae Ech586 WP 012883158 1 Dzeae
Dickeya sp CSL RW240 WP 035339556 1 Daquatica
Dickeya aquatica DW 0440 WP 035339556 1 Daquatica
DAQ1742 DAQ1742 00520 Daquatica
Dickeya gs poaceaephila NCPPB 569 WP 042868920 1 Dpoaceaephila
Dickeya chrysanthemi subsp chrysanthemi NCPPB 402 WP 039999983 1 Dchrysanthemi
Dickeya chrysanthemi L11 WP 033576635 1 Dchrysanthemi
Dickeya chrysanthemi NCPPB 3533 WP 015848011 1 Dchrysanthemi
Dickeya chrysanthemi Ech1591 WP 015848011 1 Dchrysanthemi
Dickeya zeae ZJU1202 WP 016943949 1 Dzeae
Dickeya zeae EC1 WP 016943949 1 Dzeae
Dickeya chrysanthemi NCPPB 3533 WP 012768689 1 Dchrysanthemi
DAQ1742 DAQ1742 00668 Daquatica
Dickeya chrysanthemi Ech1591 WP 012768689 1 Dchrysanthemi

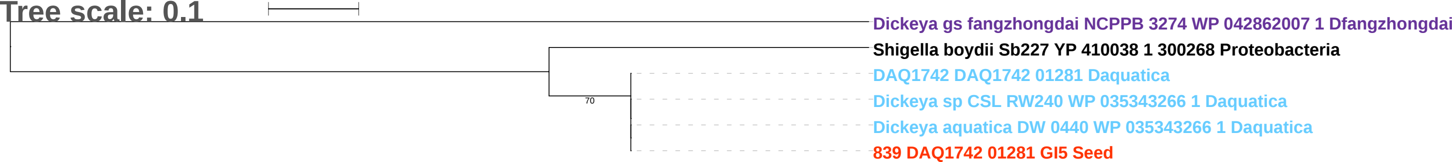

Tree scale: 0.1
Dickeya gs fangzhongdai NCPPB 3274 WP 042862007 1 Dfangzhongdai
Shigella boydii Sb227 YP 410038 1 300268 Proteobacteria
70
DAQ1742 DAQ1742 01281 Daquatica
Dickeya sp CSL RW240 WP 035343266 1 Daquatica
Dickeya aquatica DW 0440 WP 035343266 1 Daquatica
839 DAQ1742 01281 GI5 Seed

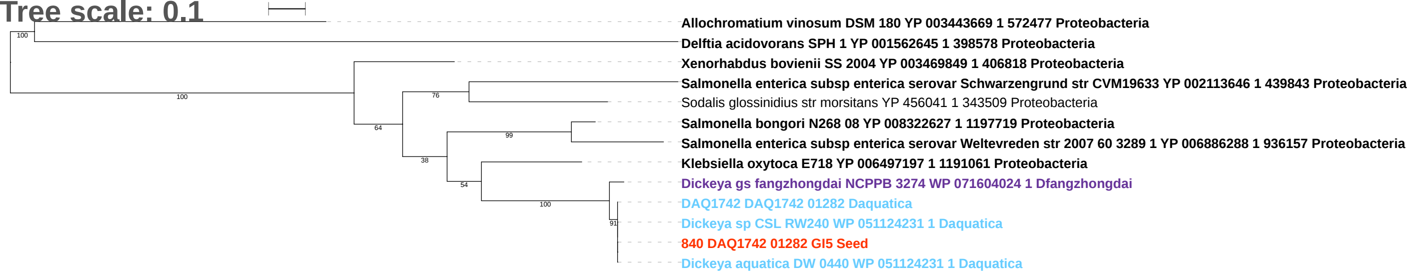

Tree scale: 0.1
Allochromatium vinosum DSM 180 YP 003443669 1 572477 Proteobacteria
Delftia acidovorans SPH 1 YP 001562645 1 398578 Proteobacteria
Xenorhabdus bovienii SS 2004 YP 003469849 1 406818 Proteobacteria
Salmonella enterica subsp enterica serovar Schwarzengrund str CVM19633 YP 002113646 1 439843 Proteobacteria
Sodalis glossinidius str morsitans YP 456041 1 343509 Proteobacteria
Salmonella bongori N268 08 YP 008322627 1 1197719 Proteobacteria
Salmonella enterica subsp enterica serovar Weltevreden str 2007 60 3289 1 YP 006886288 1 936157 Proteobacteria
Klebsiella oxytoca E718 YP 006497197 1 1191061 Proteobacteria
Dickeya gs fangzhongdai NCPPB 3274 WP 071604024 1 Dfangzhongdai
DAQ1742 DAQ1742 01282 Daquatica
Dickeya sp CSL RW240 WP 051124231 1 Daquatica
840 DAQ1742 01282 GI5 Seed
Dickeya aquatica DW 0440 WP 051124231 1 Daquatica
100
100
76
64
99
38
54
100
91

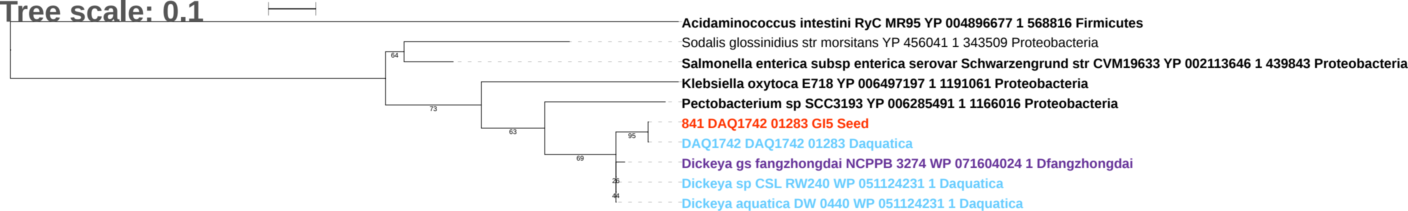

Tree scale: 0.1
Acidaminococcus intestini RyC MR95 YP 004896677 1 568816 Firmicutes
Sodalis glossinidius str morsitans YP 456041 1 343509 Proteobacteria
Salmonella enterica subsp enterica serovar Schwarzengrund str CVM19633 YP 002113646 1 439843 Proteobacteria
Klebsiella oxytoca E718 YP 006497197 1 1191061 Proteobacteria
Pectobacterium sp SCC3193 YP 006285491 1 1166016 Proteobacteria
841 DAQ1742 01283 GI5 Seed
DAQ1742 DAQ1742 01283 Daquatica
Dickeya gs fangzhongdai NCPPB 3274 WP 071604024 1 Dfangzhongdai
Dickeya sp CSL RW240 WP 051124231 1 Daquatica
Dickeya aquatica DW 0440 WP 051124231 1 Daquatica
64
73
63
95
69
26
44

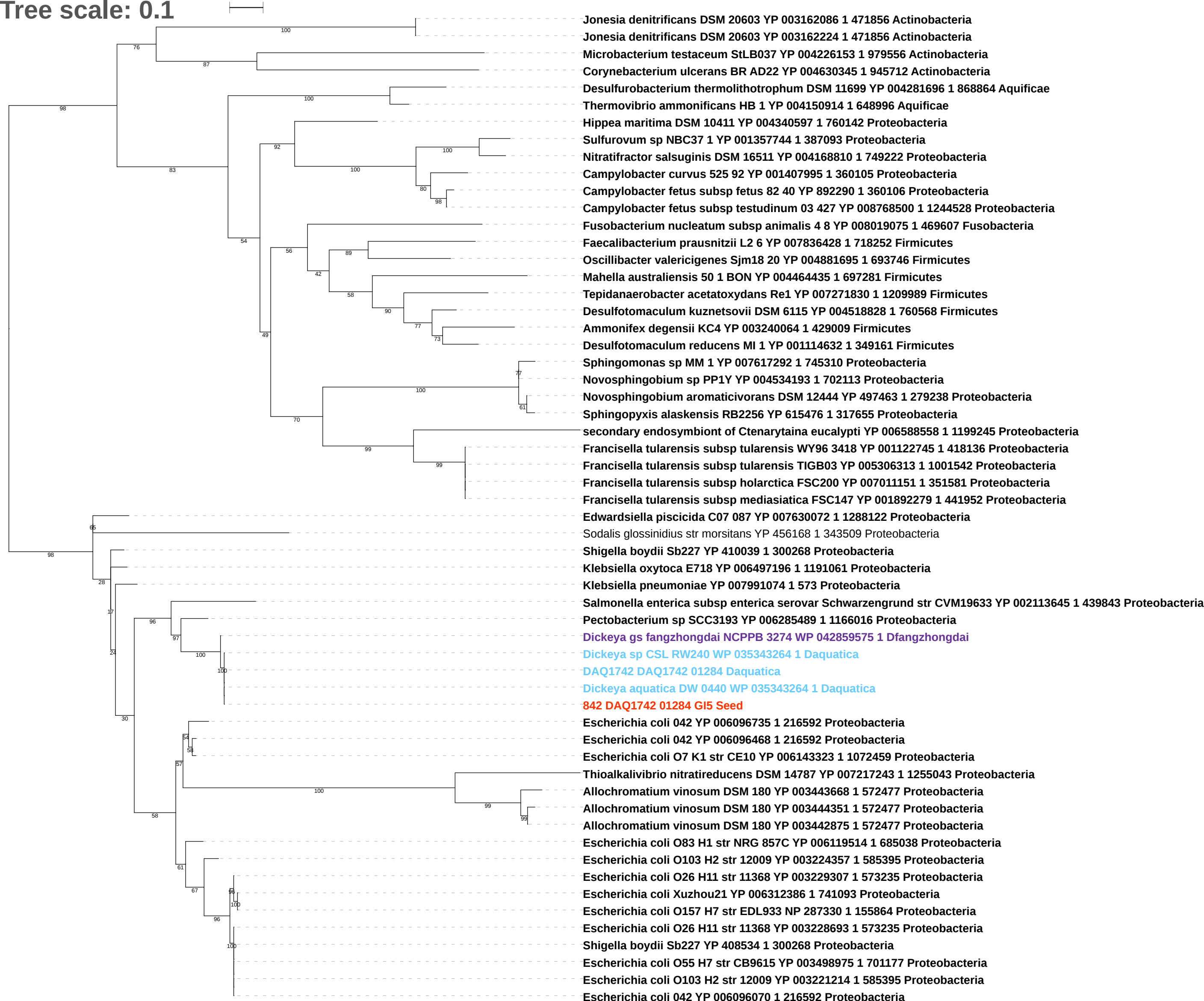
Tree scale: 0.1
Jonesia denitrificans DSM 20603 YP 003162086 1 471856 Actinobacteria
Jonesia denitrificans DSM 20603 YP 003162224 1 471856 Actinobacteria
Microbacterium testaceum StLB037 YP 004226153 1 979556 Actinobacteria
Corynebacterium ulcerans BR AD22 YP 004630345 1 945712 Actinobacteria
Desulfurobacterium thermolithotrophum DSM 11699 YP 004281696 1 868864 Aquificae
Thermovibrio ammonificans HB 1 YP 004150914 1 648996 Aquificae
Hippea maritima DSM 10411 YP 004340597 1 760142 Proteobacteria
Sulfurovum sp NBC37 1 YP 001357744 1 387093 Proteobacteria
Nitratifractor salsuginis DSM 16511 YP 004168810 1 749222 Proteobacteria
Campylobacter curvus 525 92 YP 001407995 1 360105 Proteobacteria
Campylobacter fetus subsp fetus 82 40 YP 892290 1 360106 Proteobacteria
Campylobacter fetus subsp testudinum 03 427 YP 008768500 1 1244528 Proteobacteria
Fusobacterium nucleatum subsp animalis 4 8 YP 008019075 1 469607 Fusobacteria
Faecalibacterium prausnitzii L2 6 YP 007836428 1 718252 Firmicutes
Oscillibacter valericigenes Sjm18 20 YP 004881695 1 693746 Firmicutes
Mahella australiensis 50 1 BON YP 004464435 1 697281 Firmicutes
Tepidanaerobacter acetatoxydans Re1 YP 007271830 1 1209989 Firmicutes
Desulfotomaculum kuznetsovii DSM 6115 YP 004518828 1 760568 Firmicutes
Ammonifex degensii KC4 YP 003240064 1 429009 Firmicutes
Desulfotomaculum reducens MI 1 YP 001114632 1 349161 Firmicutes
Sphingomonas sp MM 1 YP 007617292 1 745310 Proteobacteria
Novosphingobium sp PP1Y YP 004534193 1 702113 Proteobacteria
Novosphingobium aromaticivorans DSM 12444 YP 497463 1 279238 Proteobacteria
Sphingopyxis alaskensis RB2256 YP 615476 1 317655 Proteobacteria
secondary endosymbiont of Ctenarytaina eucalypti YP 006588558 1 1199245 Proteobacteria
Francisella tularensis subsp tularensis WY96 3418 YP 001122745 1 418136 Proteobacteria
Francisella tularensis subsp tularensis TIGB03 YP 005306313 1 1001542 Proteobacteria
Francisella tularensis subsp holarctica FSC200 YP 007011151 1 351581 Proteobacteria
Francisella tularensis subsp mediasiatica FSC147 YP 001892279 1 441952 Proteobacteria
Edwardsiella piscicida C07 087 YP 007630072 1 1288122 Proteobacteria
Sodalis glossinidius str morsitans YP 456168 1 343509 Proteobacteria
Shigella boydii Sb227 YP 410039 1 300268 Proteobacteria
Klebsiella oxytoca E718 YP 006497196 1 1191061 Proteobacteria
Klebsiella pneumoniae YP 007991074 1 573 Proteobacteria
Salmonella enterica subsp enterica serovar Schwarzengrund str CVM19633 YP 002113645 1 439843 Proteobacteria
Pectobacterium sp SCC3193 YP 006285489 1 1166016 Proteobacteria
Dickeya gs fangzhongdai NCPPB 3274 WP 042859575 1 Dfangzhongdai
Dickeya sp CSL RW240 WP 035343264 1 Daquatica
DAQ1742 DAQ1742 01284 Daquatica
Dickeya aquatica DW 0440 WP 035343264 1 Daquatica
842 DAQ1742 01284 GI5 Seed
Escherichia coli 042 YP 006096735 1 216592 Proteobacteria
Escherichia coli 042 YP 006096468 1 216592 Proteobacteria
Escherichia coli O7 K1 str CE10 YP 006143323 1 1072459 Proteobacteria
Thioalkalivibrio nitratireducens DSM 14787 YP 007217243 1 1255043 Proteobacteria
Allochromatium vinosum DSM 180 YP 003443668 1 572477 Proteobacteria
Allochromatium vinosum DSM 180 YP 003444351 1 572477 Proteobacteria
Allochromatium vinosum DSM 180 YP 003442875 1 572477 Proteobacteria
Escherichia coli O83 H1 str NRG 857C YP 006119514 1 685038 Proteobacteria
Escherichia coli O103 H2 str 12009 YP 003224357 1 585395 Proteobacteria
Escherichia coli O26 H11 str 11368 YP 003229307 1 573235 Proteobacteria
Escherichia coli Xuzhou21 YP 006312386 1 741093 Proteobacteria
Escherichia coli O157 H7 str EDL933 NP 287330 1 155864 Proteobacteria
Escherichia coli O26 H11 str 11368 YP 003228693 1 573235 Proteobacteria
Shigella boydii Sb227 YP 408534 1 300268 Proteobacteria
Escherichia coli O55 H7 str CB9615 YP 003498975 1 701177 Proteobacteria
Escherichia coli O103 H2 str 12009 YP 003221214 1 585395 Proteobacteria
Escherichia coli 042 YP 006096070 1 216592 Proteobacteria

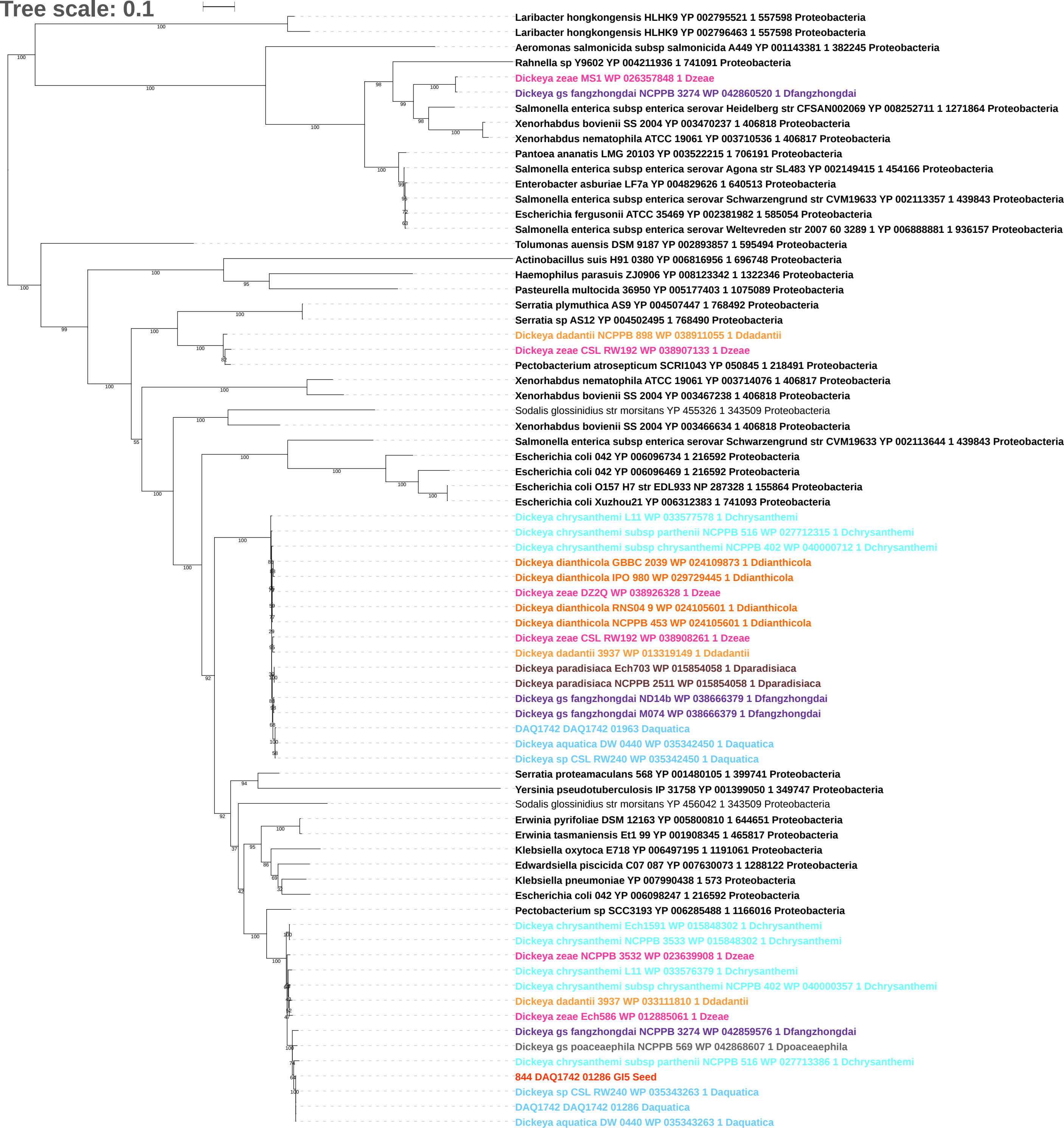
Tree scale: 0.1
Laribacter hongkongensis HLHK9 YP 002795521 1 557598 Proteobacteria
Laribacter hongkongensis HLHK9 YP 002796463 1 557598 Proteobacteria
Aeromonas salmonicida subsp salmonicida A449 YP 001143381 1 382245 Proteobacteria
Rahnella sp Y9602 YP 004211936 1 741091 Proteobacteria
Dickeya zeae MS1 WP 026357848 1 Dzeae
Dickeya gs fangzhongdai NCPPB 3274 WP 042860520 1 Dfangzhongdai
Salmonella enterica subsp enterica serovar Heidelberg str CFSAN002069 YP 008252711 1 1271864 Proteobacteria
Xenorhabdus bovienii SS 2004 YP 003470237 1 406818 Proteobacteria
Xenorhabdus nematophila ATCC 19061 YP 003710536 1 406817 Proteobacteria
Pantoea ananatis LMG 20103 YP 003522215 1 706191 Proteobacteria
Salmonella enterica subsp enterica serovar Agona str SL483 YP 002149415 1 454166 Proteobacteria
Enterobacter asburiae LF7a YP 004829626 1 640513 Proteobacteria
Salmonella enterica subsp enterica serovar Schwarzengrund str CVM19633 YP 002113357 1 439843 Proteobacteria
Escherichia fergusonii ATCC 35469 YP 002381982 1 585054 Proteobacteria
Salmonella enterica subsp enterica serovar Weltevreden str 2007 60 3289 1 YP 006888881 1 936157 Proteobacteria
Tolumonas auensis DSM 9187 YP 002893857 1 595494 Proteobacteria
Actinobacillus suis H91 0380 YP 006816956 1 696748 Proteobacteria
Haemophilus parasuis ZJ0906 YP 008123342 1 1322346 Proteobacteria
Pasteurella multocida 36950 YP 005177403 1 1075089 Proteobacteria
Serratia plymuthica AS9 YP 004507447 1 768492 Proteobacteria
Serratia sp AS12 YP 004502495 1 768490 Proteobacteria
Dickeya dadantii NCPPB 898 WP 038911055 1 Ddadantii
Dickeya zeae CSL RW192 WP 038907133 1 Dzeae
Pectobacterium atrosepticum SCRI1043 YP 050845 1 218491 Proteobacteria
Xenorhabdus nematophila ATCC 19061 YP 003714076 1 406817 Proteobacteria
Xenorhabdus bovienii SS 2004 YP 003467238 1 406818 Proteobacteria
Sodalis glossinidius str morsitans YP 455326 1 343509 Proteobacteria
Xenorhabdus bovienii SS 2004 YP 003466634 1 406818 Proteobacteria
Salmonella enterica subsp enterica serovar Schwarzengrund str CVM19633 YP 002113644 1 439843 Proteobacteria
Escherichia coli 042 YP 006096734 1 216592 Proteobacteria
Escherichia coli 042 YP 006096469 1 216592 Proteobacteria
Escherichia coli O157 H7 str EDL933 NP 287328 1 155864 Proteobacteria
Escherichia coli Xuzhou21 YP 006312383 1 741093 Proteobacteria
Dickeya chrysanthemi L11 WP 033577578 1 Dchrysanthemi
Dickeya chrysanthemi subsp parthenii NCPPB 516 WP 027712315 1 Dchrysanthemi
Dickeya chrysanthemi subsp chrysanthemi NCPPB 402 WP 040000712 1 Dchrysanthemi
Dickeya dianthicola GBBC 2039 WP 024109873 1 Ddianthicola
Dickeya dianthicola IPO 980 WP 029729445 1 Ddianthicola
Dickeya zeae DZ2Q WP 038926328 1 Dzeae
Dickeya dianthicola RNS04 9 WP 024105601 1 Ddianthicola
Dickeya dianthicola NCPPB 453 WP 024105601 1 Ddianthicola
Dickeya zeae CSL RW192 WP 038908261 1 Dzeae
Dickeya dadantii 3937 WP 013319149 1 Ddadantii
Dickeya paradisiaca Ech703 WP 015854058 1 Dparadisiaca
Dickeya paradisiaca NCPPB 2511 WP 015854058 1 Dparadisiaca
Dickeya gs fangzhongdai ND14b WP 038666379 1 Dfangzhongdai
Dickeya gs fangzhongdai M074 WP 038666379 1 Dfangzhongdai
DAQ1742 DAQ1742 01963 Daquatica
Dickeya aquatica DW 0440 WP 035342450 1 Daquatica
Dickeya sp CSL RW240 WP 035342450 1 Daquatica
Serratia proteamaculans 568 YP 001480105 1 399741 Proteobacteria
Yersinia pseudotuberculosis IP 31758 YP 001399050 1 349747 Proteobacteria
Sodalis glossinidius str morsitans YP 456042 1 343509 Proteobacteria
Erwinia pyrifoliae DSM 12163 YP 005800810 1 644651 Proteobacteria
Erwinia tasmaniensis Et1 99 YP 001908345 1 465817 Proteobacteria
Klebsiella oxytoca E718 YP 006497195 1 1191061 Proteobacteria
Edwardsiella piscicida C07 087 YP 007630073 1 1288122 Proteobacteria
Klebsiella pneumoniae YP 007990438 1 573 Proteobacteria
Escherichia coli 042 YP 006098247 1 216592 Proteobacteria
Pectobacterium sp SCC3193 YP 006285488 1 1166016 Proteobacteria
Dickeya chrysanthemi Ech1591 WP 015848302 1 Dchrysanthemi
Dickeya chrysanthemi NCPPB 3533 WP 015848302 1 Dchrysanthemi
Dickeya zeae NCPPB 3532 WP 023639908 1 Dzeae
Dickeya chrysanthemi L11 WP 033576379 1 Dchrysanthemi
Dickeya chrysanthemi subsp chrysanthemi NCPPB 402 WP 040000357 1 Dchrysanthemi
Dickeya dadantii 3937 WP 033111810 1 Ddadantii
Dickeya zeae Ech586 WP 012885061 1 Dzeae
Dickeya gs fangzhongdai NCPPB 3274 WP 042859576 1 Dfangzhongdai
Dickeya gs poaceaephila NCPPB 569 WP 042868607 1 Dpoaceaephila
Dickeya chrysanthemi subsp parthenii NCPPB 516 WP 027713386 1 Dchrysanthemi
844 DAQ1742 01286 GI5 Seed
Dickeya sp CSL RW240 WP 035343263 1 Daquatica
DAQ1742 DAQ1742 01286 Daquatica
Dickeya aquatica DW 0440 WP 035343263 1 Daquatica

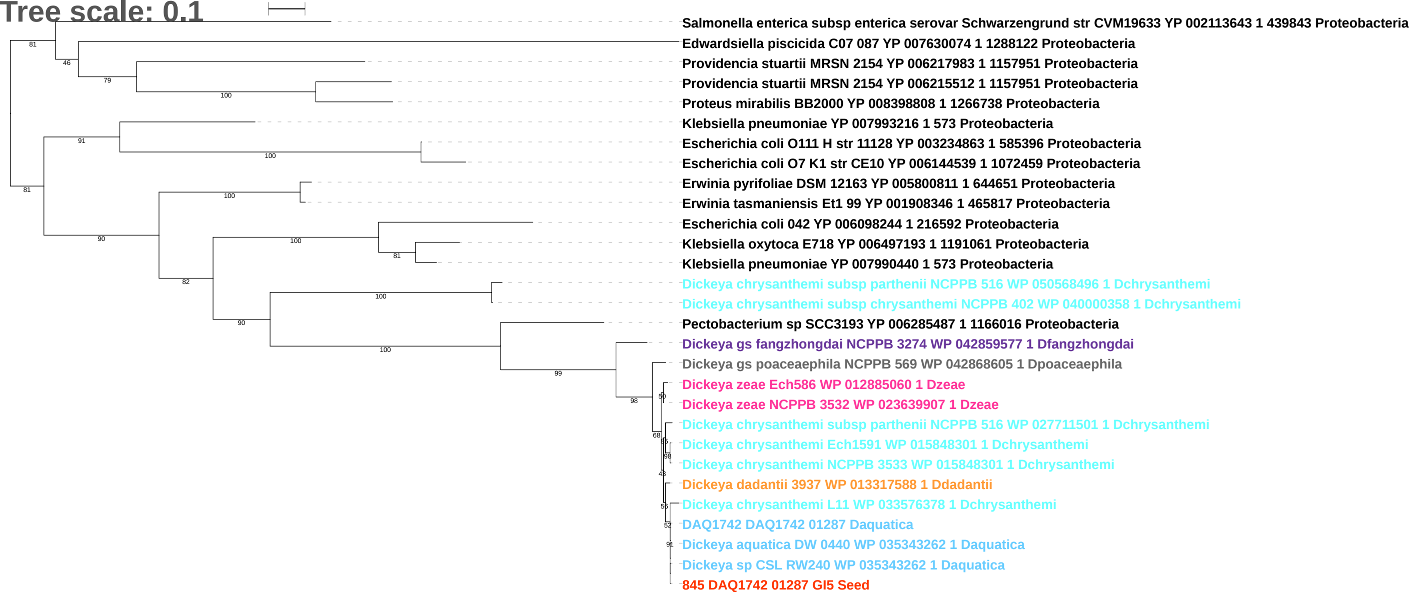

Tree scale: 0.1
Salmonella enterica subsp enterica serovar Schwarzengrund str CVM19633 YP 002113643 1 439843 Proteobacteria
Edwardsiella piscicida C07 087 YP 007630074 1 1288122 Proteobacteria
Providencia stuartii MRSN 2154 YP 006217983 1 1157951 Proteobacteria
Providencia stuartii MRSN 2154 YP 006215512 1 1157951 Proteobacteria
Proteus mirabilis BB2000 YP 008398808 1 1266738 Proteobacteria
Klebsiella pneumoniae YP 007993216 1 573 Proteobacteria
Escherichia coli O111 H str 11128 YP 003234863 1 585396 Proteobacteria
Escherichia coli O7 K1 str CE10 YP 006144539 1 1072459 Proteobacteria
Erwinia pyrifoliae DSM 12163 YP 005800811 1 644651 Proteobacteria
Erwinia tasmaniensis Et1 99 YP 001908346 1 465817 Proteobacteria
Escherichia coli 042 YP 006098244 1 216592 Proteobacteria
Klebsiella oxytoca E718 YP 006497193 1 1191061 Proteobacteria
Klebsiella pneumoniae YP 007990440 1 573 Proteobacteria
Dickeya chrysanthemi subsp parthenii NCPPB 516 WP 050568496 1 Dchrysanthemi
Dickeya chrysanthemi subsp chrysanthemi NCPPB 402 WP 040000358 1 Dchrysanthemi
Pectobacterium sp SCC3193 YP 006285487 1 1166016 Proteobacteria
Dickeya gs fangzhongdai NCPPB 3274 WP 042859577 1 Dfangzhongdai
Dickeya gs poaceaephila NCPPB 569 WP 042868605 1 Dpoaceaephila
Dickeya zeae Ech586 WP 012885060 1 Dzeae
Dickeya zeae NCPPB 3532 WP 023639907 1 Dzeae
Dickeya chrysanthemi subsp parthenii NCPPB 516 WP 027711501 1 Dchrysanthemi
Dickeya chrysanthemi Ech1591 WP 015848301 1 Dchrysanthemi
Dickeya chrysanthemi NCPPB 3533 WP 015848301 1 Dchrysanthemi
Dickeya dadantii 3937 WP 013317588 1 Ddadantii
Dickeya chrysanthemi L11 WP 033576378 1 Dchrysanthemi
DAQ1742 DAQ1742 01287 Daquatica
Dickeya aquatica DW 0440 WP 035343262 1 Daquatica
Dickeya sp CSL RW240 WP 035343262 1 Daquatica
845 DAQ1742 01287 GI5 Seed

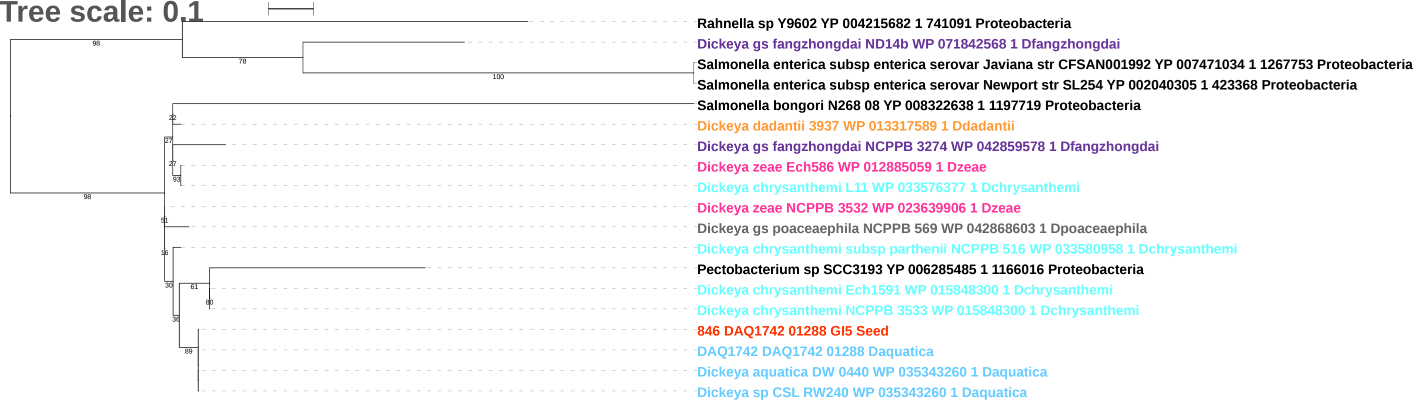
Tree scale: 0.1
Rahnella sp Y9602 YP 004215682 1 741091 Proteobacteria
Dickeya gs fangzhongdai ND14b WP 071842568 1 Dfangzhongdai
Salmonella enterica subsp enterica serovar Javiana str CFSAN001992 YP 007471034 1 1267753 Proteobacteria
Salmonella enterica subsp enterica serovar Newport str SL254 YP 002040305 1 423368 Proteobacteria
Salmonella bongori N268 08 YP 008322638 1 1197719 Proteobacteria
Dickeya dadantii 3937 WP 013317589 1 Ddadantii
Dickeya gs fangzhongdai NCPPB 3274 WP 042859578 1 Dfangzhongdai
Dickeya zeae Ech586 WP 012885059 1 Dzeae
Dickeya chrysanthemi L11 WP 033576377 1 Dchrysanthemi
Dickeya zeae NCPPB 3532 WP 023639906 1 Dzeae
Dickeya gs poaceaephila NCPPB 569 WP 042868603 1 Dpoaceaephila
Dickeya chrysanthemi subsp parthenii NCPPB 516 WP 033580958 1 Dchrysanthemi
Pectobacterium sp SCC3193 YP 006285485 1 1166016 Proteobacteria
Dickeya chrysanthemi Ech1591 WP 015848300 1 Dchrysanthemi
Dickeya chrysanthemi NCPPB 3533 WP 015848300 1 Dchrysanthemi
846 DAQ1742 01288 GI5 Seed
DAQ1742 DAQ1742 01288 Daquatica
Dickeya aquatica DW 0440 WP 035343260 1 Daquatica
Dickeya sp CSL RW240 WP 035343260 1 Daquatica
98
78
100
22
27
27
93
98
51
16
30
61
80
36
89

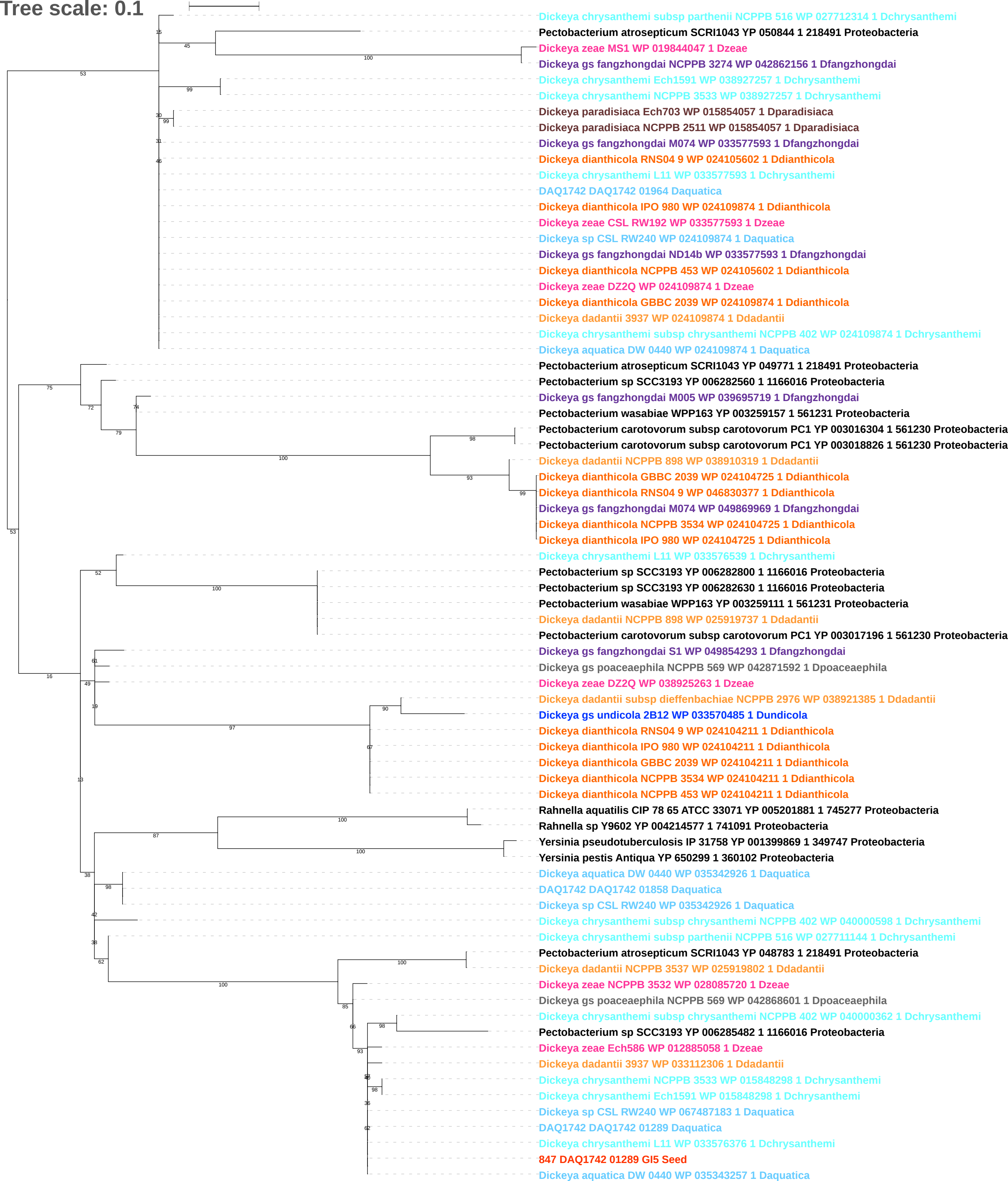
Tree scale: 0.1
Dickeya chrysanthemi subsp parthenii NCPPB 516 WP 027712314 1 Dchrysanthemi
Pectobacterium atrosepticum SCRI1043 YP 050844 1 218491 Proteobacteria
Dickeya zeae MS1 WP 019844047 1 Dzeae
Dickeya gs fangzhongdai NCPPB 3274 WP 042862156 1 Dfangzhongdai
Dickeya chrysanthemi Ech1591 WP 038927257 1 Dchrysanthemi
Dickeya chrysanthemi NCPPB 3533 WP 038927257 1 Dchrysanthemi
Dickeya paradisiaca Ech703 WP 015854057 1 Dparadisiaca
Dickeya paradisiaca NCPPB 2511 WP 015854057 1 Dparadisiaca
Dickeya gs fangzhongdai M074 WP 033577593 1 Dfangzhongdai
Dickeya dianthicola RNS04 9 WP 024105602 1 Ddianthicola
Dickeya chrysanthemi L11 WP 033577593 1 Dchrysanthemi
DAQ1742 DAQ1742 01964 Daquatica
Dickeya dianthicola IPO 980 WP 024109874 1 Ddianthicola
Dickeya zeae CSL RW192 WP 033577593 1 Dzeae
Dickeya sp CSL RW240 WP 024109874 1 Daquatica
Dickeya gs fangzhongdai ND14b WP 033577593 1 Dfangzhongdai
Dickeya dianthicola NCPPB 453 WP 024105602 1 Ddianthicola
Dickeya zeae DZ2Q WP 024109874 1 Dzeae
Dickeya dianthicola GBBC 2039 WP 024109874 1 Ddianthicola
Dickeya dadantii 3937 WP 024109874 1 Ddadantii
Dickeya chrysanthemi subsp chrysanthemi NCPPB 402 WP 024109874 1 Dchrysanthemi
Dickeya aquatica DW 0440 WP 024109874 1 Daquatica
Pectobacterium atrosepticum SCRI1043 YP 049771 1 218491 Proteobacteria
Pectobacterium sp SCC3193 YP 006282560 1 1166016 Proteobacteria
Dickeya gs fangzhongdai M005 WP 039695719 1 Dfangzhongdai
Pectobacterium wasabiae WPP163 YP 003259157 1 561231 Proteobacteria
Pectobacterium carotovorum subsp carotovorum PC1 YP 003016304 1 561230 Proteobacteria
Pectobacterium carotovorum subsp carotovorum PC1 YP 003018826 1 561230 Proteobacteria
Dickeya dadantii NCPPB 898 WP 038910319 1 Ddadantii
Dickeya dianthicola GBBC 2039 WP 024104725 1 Ddianthicola
Dickeya dianthicola RNS04 9 WP 046830377 1 Ddianthicola
Dickeya gs fangzhongdai M074 WP 049869969 1 Dfangzhongdai
Dickeya dianthicola NCPPB 3534 WP 024104725 1 Ddianthicola
Dickeya dianthicola IPO 980 WP 024104725 1 Ddianthicola
Dickeya chrysanthemi L11 WP 033576539 1 Dchrysanthemi
Pectobacterium sp SCC3193 YP 006282800 1 1166016 Proteobacteria
Pectobacterium sp SCC3193 YP 006282630 1 1166016 Proteobacteria
Pectobacterium wasabiae WPP163 YP 003259111 1 561231 Proteobacteria
Dickeya dadantii NCPPB 898 WP 025919737 1 Ddadantii
Pectobacterium carotovorum subsp carotovorum PC1 YP 003017196 1 561230 Proteobacteria
Dickeya gs fangzhongdai S1 WP 049854293 1 Dfangzhongdai
Dickeya gs poaceaephila NCPPB 569 WP 042871592 1 Dpoaceaephila
Dickeya zeae DZ2Q WP 038925263 1 Dzeae
Dickeya dadantii subsp dieffenbachiae NCPPB 2976 WP 038921385 1 Ddadantii
Dickeya gs undicola 2B12 WP 033570485 1 Dundicola
Dickeya dianthicola RNS04 9 WP 024104211 1 Ddianthicola
Dickeya dianthicola IPO 980 WP 024104211 1 Ddianthicola
Dickeya dianthicola GBBC 2039 WP 024104211 1 Ddianthicola
Dickeya dianthicola NCPPB 3534 WP 024104211 1 Ddianthicola
Dickeya dianthicola NCPPB 453 WP 024104211 1 Ddianthicola
Rahnella aquatilis CIP 78 65 ATCC 33071 YP 005201881 1 745277 Proteobacteria
Rahnella sp Y9602 YP 004214577 1 741091 Proteobacteria
Yersinia pseudotuberculosis IP 31758 YP 001399869 1 349747 Proteobacteria
Yersinia pestis Antiqua YP 650299 1 360102 Proteobacteria
Dickeya aquatica DW 0440 WP 035342926 1 Daquatica
DAQ1742 DAQ1742 01858 Daquatica
Dickeya sp CSL RW240 WP 035342926 1 Daquatica
Dickeya chrysanthemi subsp chrysanthemi NCPPB 402 WP 040000598 1 Dchrysanthemi
Dickeya chrysanthemi subsp parthenii NCPPB 516 WP 027711144 1 Dchrysanthemi
Pectobacterium atrosepticum SCRI1043 YP 048783 1 218491 Proteobacteria
Dickeya dadantii NCPPB 3537 WP 025919802 1 Ddadantii
Dickeya zeae NCPPB 3532 WP 028085720 1 Dzeae
Dickeya gs poaceaephila NCPPB 569 WP 042868601 1 Dpoaceaephila
Dickeya chrysanthemi subsp chrysanthemi NCPPB 402 WP 040000362 1 Dchrysanthemi
Pectobacterium sp SCC3193 YP 006285482 1 1166016 Proteobacteria
Dickeya zeae Ech586 WP 012885058 1 Dzeae
Dickeya dadantii 3937 WP 033112306 1 Ddadantii
Dickeya chrysanthemi NCPPB 3533 WP 015848298 1 Dchrysanthemi
Dickeya chrysanthemi Ech1591 WP 015848298 1 Dchrysanthemi
Dickeya sp CSL RW240 WP 067487183 1 Daquatica
DAQ1742 DAQ1742 01289 Daquatica
Dickeya chrysanthemi L11 WP 033576376 1 Dchrysanthemi
847 DAQ1742 01289 GI5 Seed
Dickeya aquatica DW 0440 WP 035343257 1 Daquatica

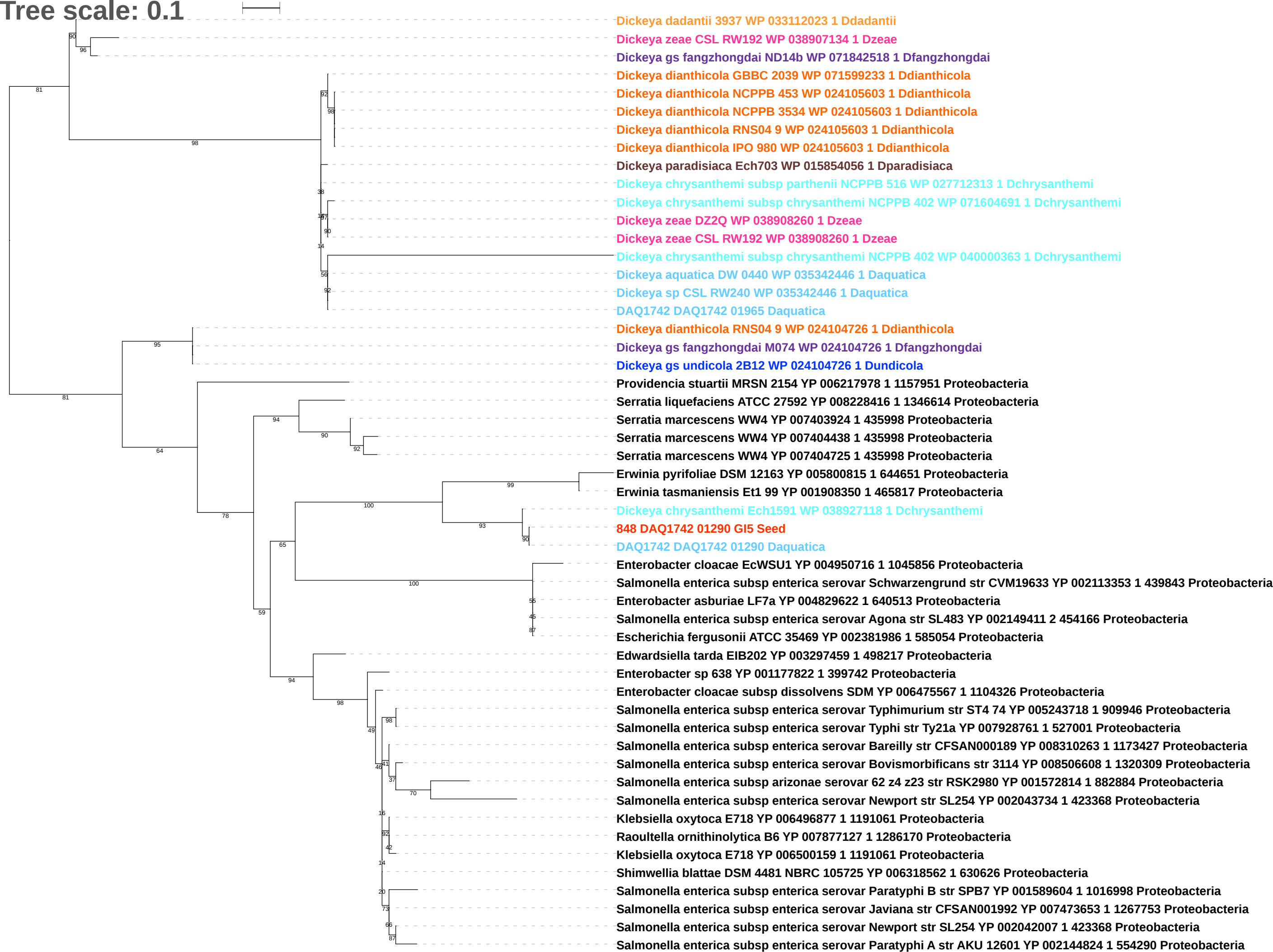
Tree scale: 0.1
Dickeya dadantii 3937 WP 033112023 1 Ddadantii
Dickeya zeae CSL RW192 WP 038907134 1 Dzeae
Dickeya gs fangzhongdai ND14b WP 071842518 1 Dfangzhongdai
Dickeya dianthicola GBBC 2039 WP 071599233 1 Ddianthicola
Dickeya dianthicola NCPPB 453 WP 024105603 1 Ddianthicola
Dickeya dianthicola NCPPB 3534 WP 024105603 1 Ddianthicola
Dickeya dianthicola RNS04 9 WP 024105603 1 Ddianthicola
Dickeya dianthicola IPO 980 WP 024105603 1 Ddianthicola
Dickeya paradisiaca Ech703 WP 015854056 1 Dparadisiaca
Dickeya chrysanthemi subsp parthenii NCPPB 516 WP 027712313 1 Dchrysanthemi
Dickeya chrysanthemi subsp chrysanthemi NCPPB 402 WP 071604691 1 Dchrysanthemi
Dickeya zeae DZ2Q WP 038908260 1 Dzeae
Dickeya zeae CSL RW192 WP 038908260 1 Dzeae
Dickeya chrysanthemi subsp chrysanthemi NCPPB 402 WP 040000363 1 Dchrysanthemi
Dickeya aquatica DW 0440 WP 035342446 1 Daquatica
Dickeya sp CSL RW240 WP 035342446 1 Daquatica
DAQ1742 DAQ1742 01965 Daquatica
Dickeya dianthicola RNS04 9 WP 024104726 1 Ddianthicola
Dickeya gs fangzhongdai M074 WP 024104726 1 Dfangzhongdai
Dickeya gs undicola 2B12 WP 024104726 1 Dundicola
Providencia stuartii MRSN 2154 YP 006217978 1 1157951 Proteobacteria
Serratia liquefaciens ATCC 27592 YP 008228416 1 1346614 Proteobacteria
Serratia marcescens WW4 YP 007403924 1 435998 Proteobacteria
Serratia marcescens WW4 YP 007404438 1 435998 Proteobacteria
Serratia marcescens WW4 YP 007404725 1 435998 Proteobacteria
Erwinia pyrifoliae DSM 12163 YP 005800815 1 644651 Proteobacteria
Erwinia tasmaniensis Et1 99 YP 001908350 1 465817 Proteobacteria
Dickeya chrysanthemi Ech1591 WP 038927118 1 Dchrysanthemi
848 DAQ1742 01290 GI5 Seed
DAQ1742 DAQ1742 01290 Daquatica
Enterobacter cloacae EcWSU1 YP 004950716 1 1045856 Proteobacteria
Salmonella enterica subsp enterica serovar Schwarzengrund str CVM19633 YP 002113353 1 439843 Proteobacteria
Enterobacter asburiae LF7a YP 004829622 1 640513 Proteobacteria
Salmonella enterica subsp enterica serovar Agona str SL483 YP 002149411 2 454166 Proteobacteria
Escherichia fergusonii ATCC 35469 YP 002381986 1 585054 Proteobacteria
Edwardsiella tarda EIB202 YP 003297459 1 498217 Proteobacteria
Enterobacter sp 638 YP 001177822 1 399742 Proteobacteria
Enterobacter cloacae subsp dissolvens SDM YP 006475567 1 1104326 Proteobacteria
Salmonella enterica subsp enterica serovar Typhimurium str ST4 74 YP 005243718 1 909946 Proteobacteria
Salmonella enterica subsp enterica serovar Typhi str Ty21a YP 007928761 1 527001 Proteobacteria
Salmonella enterica subsp enterica serovar Bareilly str CFSAN000189 YP 008310263 1 1173427 Proteobacteria
Salmonella enterica subsp enterica serovar Bovismorbificans str 3114 YP 008506608 1 1320309 Proteobacteria
Salmonella enterica subsp arizonae serovar 62 z4 z23 str RSK2980 YP 001572814 1 882884 Proteobacteria
Salmonella enterica subsp enterica serovar Newport str SL254 YP 002043734 1 423368 Proteobacteria
Klebsiella oxytoca E718 YP 006496877 1 1191061 Proteobacteria
Raoultella ornithinolytica B6 YP 007877127 1 1286170 Proteobacteria
Klebsiella oxytoca E718 YP 006500159 1 1191061 Proteobacteria
Shimwellia blattae DSM 4481 NBRC 105725 YP 006318562 1 630626 Proteobacteria
Salmonella enterica subsp enterica serovar Paratyphi B str SPB7 YP 001589604 1 1016998 Proteobacteria
Salmonella enterica subsp enterica serovar Javiana str CFSAN001992 YP 007473653 1 1267753 Proteobacteria
Salmonella enterica subsp enterica serovar Newport str SL254 YP 002042007 1 423368 Proteobacteria
Salmonella enterica subsp enterica serovar Paratyphi A str AKU 12601 YP 002144824 1 554290 Proteobacteria

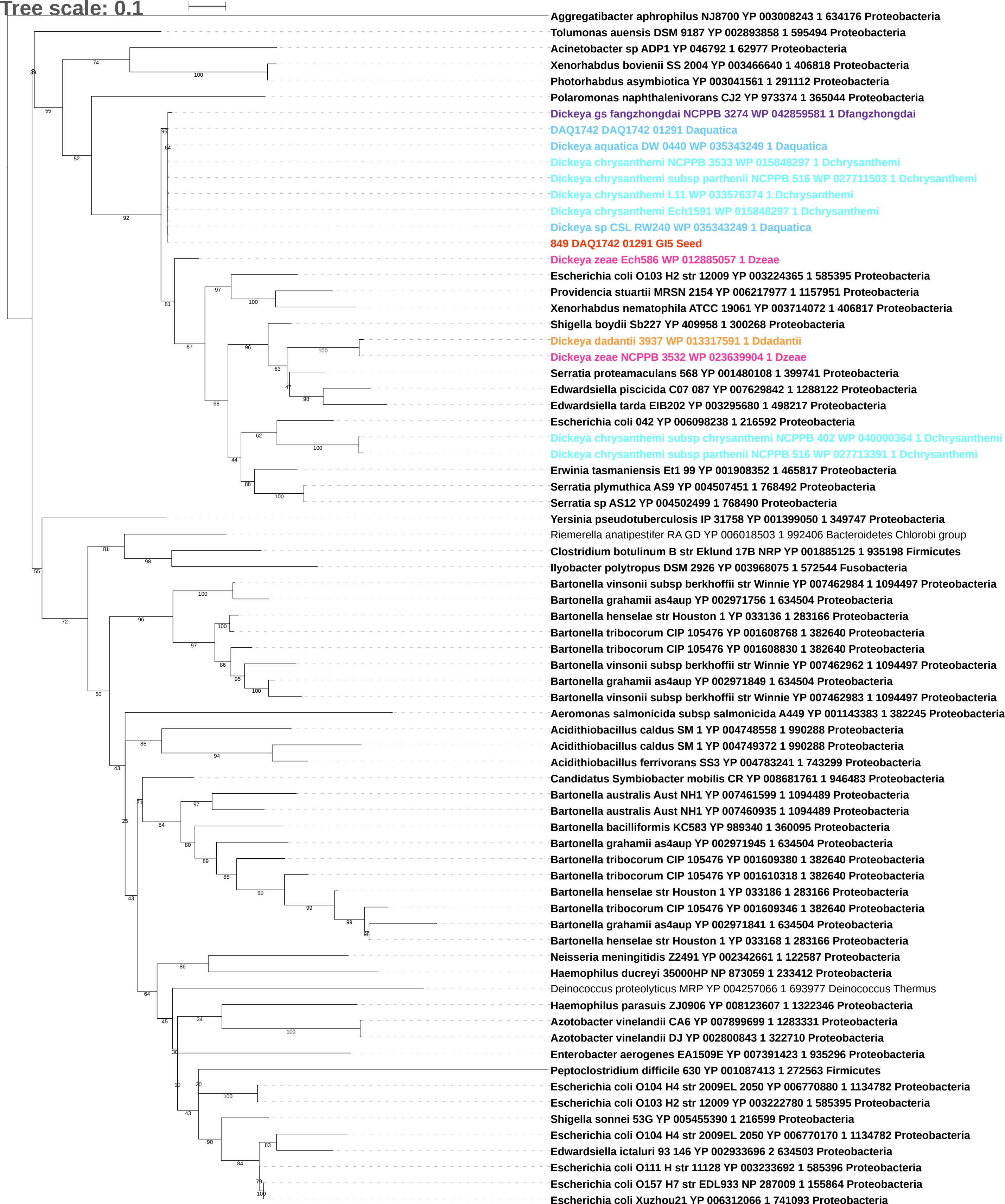
Tree scale: 0.1
Aggregatibacter aphrophilus NJ8700 YP 003008243 1 634176 Proteobacteria
Tolumonas auensis DSM 9187 YP 002893858 1 595494 Proteobacteria
Acinetobacter sp ADP1 YP 046792 1 62977 Proteobacteria
Xenorhabdus bovienii SS 2004 YP 003466640 1 406818 Proteobacteria
Photorhabdus asymbiotica YP 003041561 1 291112 Proteobacteria
Polaromonas naphthalenivorans CJ2 YP 973374 1 365044 Proteobacteria
Dickeya gs fangzhongdai NCPPB 3274 WP 042859581 1 Dfangzhongdai
DAQ1742 DAQ1742 01291 Daquatica
Dickeya aquatica DW 0440 WP 035343249 1 Daquatica
Dickeya chrysanthemi NCPPB 3533 WP 015848297 1 Dchrysanthemi
Dickeya chrysanthemi subsp parthenii NCPPB 516 WP 027711503 1 Dchrysanthemi
Dickeya chrysanthemi L11 WP 033576374 1 Dchrysanthemi
Dickeya chrysanthemi Ech1591 WP 015848297 1 Dchrysanthemi
Dickeya sp CSL RW240 WP 035343249 1 Daquatica
849 DAQ1742 01291 GI5 Seed
Dickeya zeae Ech586 WP 012885057 1 Dzeae
Escherichia coli O103 H2 str 12009 YP 003224365 1 585395 Proteobacteria
Providencia stuartii MRSN 2154 YP 006217977 1 1157951 Proteobacteria
Xenorhabdus nematophila ATCC 19061 YP 003714072 1 406817 Proteobacteria
Shigella boydii Sb227 YP 409958 1 300268 Proteobacteria
Dickeya dadantii 3937 WP 013317591 1 Ddadantii
Dickeya zeae NCPPB 3532 WP 023639904 1 Dzeae
Serratia proteamaculans 568 YP 001480108 1 399741 Proteobacteria
Edwardsiella piscicida C07 087 YP 007629842 1 1288122 Proteobacteria
Edwardsiella tarda EIB202 YP 003295680 1 498217 Proteobacteria
Escherichia coli 042 YP 006098238 1 216592 Proteobacteria
Dickeya chrysanthemi subsp chrysanthemi NCPPB 402 WP 040000364 1 Dchrysanthemi
Dickeya chrysanthemi subsp parthenii NCPPB 516 WP 027713391 1 Dchrysanthemi
Erwinia tasmaniensis Et1 99 YP 001908352 1 465817 Proteobacteria
Serratia plymuthica AS9 YP 004507451 1 768492 Proteobacteria
Serratia sp AS12 YP 004502499 1 768490 Proteobacteria
Yersinia pseudotuberculosis IP 31758 YP 001399050 1 349747 Proteobacteria
Riemerella anatipestifer RA GD YP 006018503 1 992406 Bacteroidetes Chlorobi group
Clostridium botulinum B str Eklund 17B NRP YP 001885125 1 935198 Firmicutes
Ilyobacter polytropus DSM 2926 YP 003968075 1 572544 Fusobacteria
Bartonella vinsonii subsp berkhoffii str Winnie YP 007462984 1 1094497 Proteobacteria
Bartonella grahamii as4aup YP 002971756 1 634504 Proteobacteria
Bartonella henselae str Houston 1 YP 033136 1 283166 Proteobacteria
Bartonella tribocorum CIP 105476 YP 001608768 1 382640 Proteobacteria
Bartonella tribocorum CIP 105476 YP 001608830 1 382640 Proteobacteria
Bartonella vinsonii subsp berkhoffii str Winnie YP 007462962 1 1094497 Proteobacteria
Bartonella grahamii as4aup YP 002971849 1 634504 Proteobacteria
Bartonella vinsonii subsp berkhoffii str Winnie YP 007462983 1 1094497 Proteobacteria
Aeromonas salmonicida subsp salmonicida A449 YP 001143383 1 382245 Proteobacteria
Acidithiobacillus caldus SM 1 YP 004748558 1 990288 Proteobacteria
Acidithiobacillus caldus SM 1 YP 004749372 1 990288 Proteobacteria
Acidithiobacillus ferrivorans SS3 YP 004783241 1 743299 Proteobacteria
Candidatus Symbiobacter mobilis CR YP 008681761 1 946483 Proteobacteria
Bartonella australis Aust NH1 YP 007461599 1 1094489 Proteobacteria
Bartonella australis Aust NH1 YP 007460935 1 1094489 Proteobacteria
Bartonella bacilliformis KC583 YP 989340 1 360095 Proteobacteria
Bartonella grahamii as4aup YP 002971945 1 634504 Proteobacteria
Bartonella tribocorum CIP 105476 YP 001609380 1 382640 Proteobacteria
Bartonella tribocorum CIP 105476 YP 001610318 1 382640 Proteobacteria
Bartonella henselae str Houston 1 YP 033186 1 283166 Proteobacteria
Bartonella tribocorum CIP 105476 YP 001609346 1 382640 Proteobacteria
Bartonella grahamii as4aup YP 002971841 1 634504 Proteobacteria
Bartonella henselae str Houston 1 YP 033168 1 283166 Proteobacteria
Neisseria meningitidis Z2491 YP 002342661 1 122587 Proteobacteria
Haemophilus ducreyi 35000HP NP 873059 1 233412 Proteobacteria
Deinococcus proteolyticus MRP YP 004257066 1 693977 Deinococcus Thermus
Haemophilus parasuis ZJ0906 YP 008123607 1 1322346 Proteobacteria
Azotobacter vinelandii CA6 YP 007899699 1 1283331 Proteobacteria
Azotobacter vinelandii DJ YP 002800843 1 322710 Proteobacteria
Enterobacter aerogenes EA1509E YP 007391423 1 935296 Proteobacteria
Peptoclostridium difficile 630 YP 001087413 1 272563 Firmicutes
Escherichia coli O104 H4 str 2009EL 2050 YP 006770880 1 1134782 Proteobacteria
Escherichia coli O103 H2 str 12009 YP 003222780 1 585395 Proteobacteria
Shigella sonnei 53G YP 005455390 1 216599 Proteobacteria
Escherichia coli O104 H4 str 2009EL 2050 YP 006770170 1 1134782 Proteobacteria
Edwardsiella ictaluri 93 146 YP 002933696 2 634503 Proteobacteria
Escherichia coli O111 H str 11128 YP 003233692 1 585396 Proteobacteria
Escherichia coli O157 H7 str EDL933 NP 287009 1 155864 Proteobacteria
Escherichia coli Xuzhou21 YP 006312066 1 741093 Proteobacteria

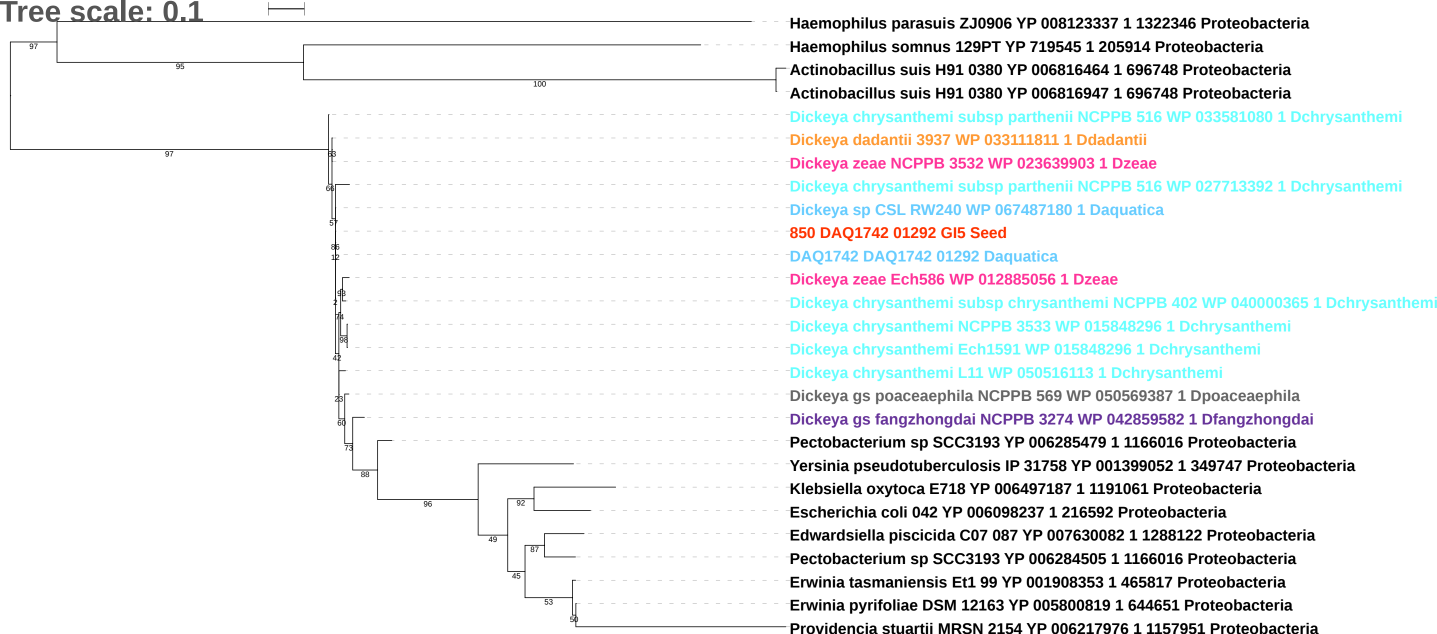
Tree scale: 0.1
Haemophilus parasuis ZJ0906 YP 008123337 1 1322346 Proteobacteria
Haemophilus somnus 129PT YP 719545 1 205914 Proteobacteria
Actinobacillus suis H91 0380 YP 006816464 1 696748 Proteobacteria
Actinobacillus suis H91 0380 YP 006816947 1 696748 Proteobacteria
Dickeya chrysanthemi subsp parthenii NCPPB 516 WP 033581080 1 Dchrysanthemi
Dickeya dadantii 3937 WP 033111811 1 Ddadantii
Dickeya zeae NCPPB 3532 WP 023639903 1 Dzeae
Dickeya chrysanthemi subsp parthenii NCPPB 516 WP 027713392 1 Dchrysanthemi
Dickeya sp CSL RW240 WP 067487180 1 Daquatica
850 DAQ1742 01292 GI5 Seed
DAQ1742 DAQ1742 01292 Daquatica
Dickeya zeae Ech586 WP 012885056 1 Dzeae
Dickeya chrysanthemi subsp chrysanthemi NCPPB 402 WP 040000365 1 Dchrysanthemi
Dickeya chrysanthemi NCPPB 3533 WP 015848296 1 Dchrysanthemi
Dickeya chrysanthemi Ech1591 WP 015848296 1 Dchrysanthemi
Dickeya chrysanthemi L11 WP 050516113 1 Dchrysanthemi
Dickeya gs poaceaephila NCPPB 569 WP 050569387 1 Dpoaceaephila
Dickeya gs fangzhongdai NCPPB 3274 WP 042859582 1 Dfangzhongdai
Pectobacterium sp SCC3193 YP 006285479 1 1166016 Proteobacteria
Yersinia pseudotuberculosis IP 31758 YP 001399052 1 349747 Proteobacteria
Klebsiella oxytoca E718 YP 006497187 1 1191061 Proteobacteria
Escherichia coli 042 YP 006098237 1 216592 Proteobacteria
Edwardsiella piscicida C07 087 YP 007630082 1 1288122 Proteobacteria
Pectobacterium sp SCC3193 YP 006284505 1 1166016 Proteobacteria
Erwinia tasmaniensis Et1 99 YP 001908353 1 465817 Proteobacteria
Erwinia pyrifoliae DSM 12163 YP 005800819 1 644651 Proteobacteria
Providencia stuartii MRSN 2154 YP 006217976 1 1157951 Proteobacteria

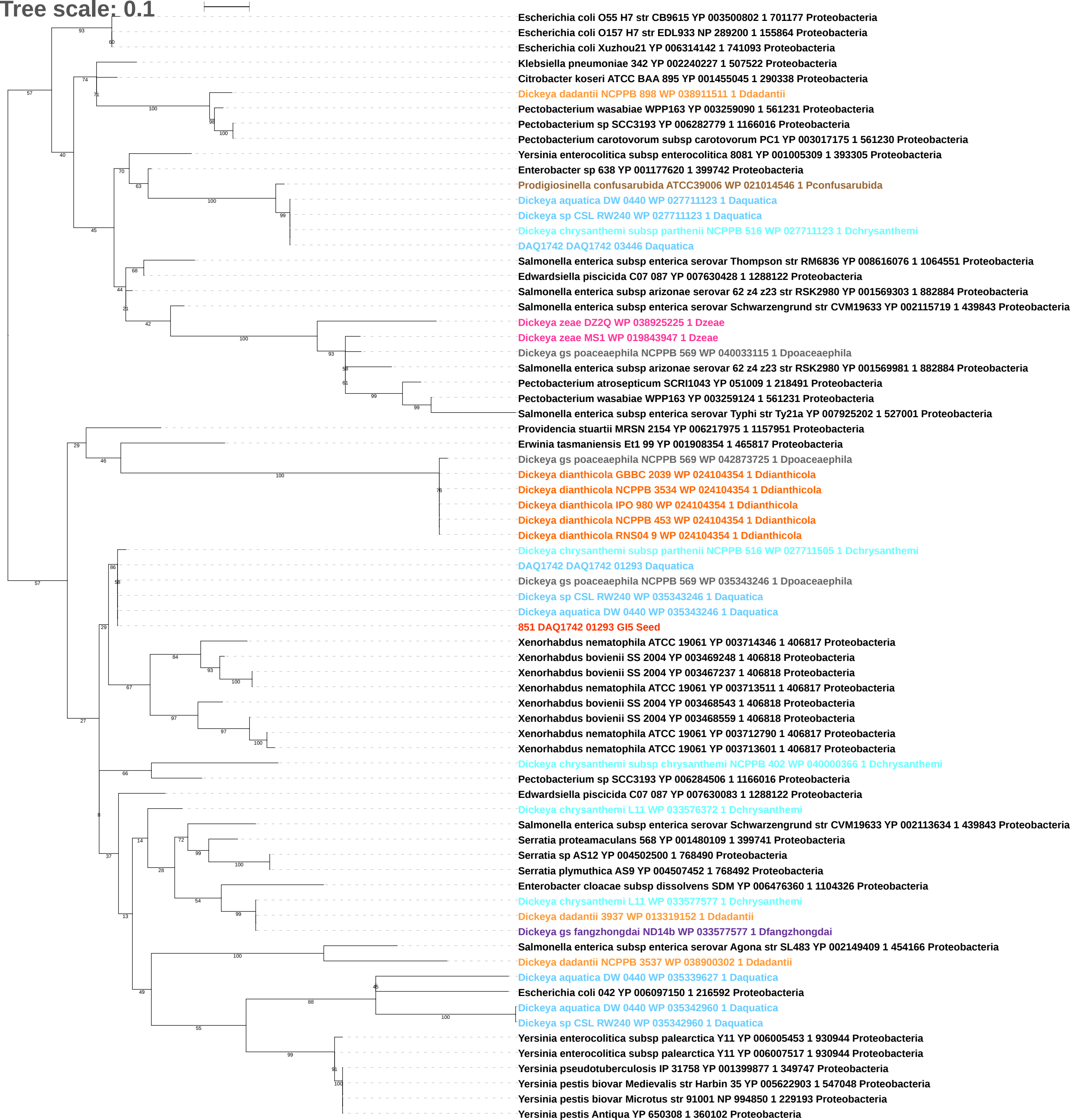
Tree scale: 0.1
Escherichia coli O55 H7 str CB9615 YP 003500802 1 701177 Proteobacteria
Escherichia coli O157 H7 str EDL933 NP 289200 1 155864 Proteobacteria
Escherichia coli Xuzhou21 YP 006314142 1 741093 Proteobacteria
Klebsiella pneumoniae 342 YP 002240227 1 507522 Proteobacteria
Citrobacter koseri ATCC BAA 895 YP 001455045 1 290338 Proteobacteria
Dickeya dadantii NCPPB 898 WP 038911511 1 Ddadantii
Pectobacterium wasabiae WPP163 YP 003259090 1 561231 Proteobacteria
Pectobacterium sp SCC3193 YP 006282779 1 1166016 Proteobacteria
Pectobacterium carotovorum subsp carotovorum PC1 YP 003017175 1 561230 Proteobacteria
Yersinia enterocolitica subsp enterocolitica 8081 YP 001005309 1 393305 Proteobacteria
Enterobacter sp 638 YP 001177620 1 399742 Proteobacteria
Prodigiosinella confusarubida ATCC39006 WP 021014546 1 Pconfusarubida
Dickeya aquatica DW 0440 WP 027711123 1 Daquatica
Dickeya sp CSL RW240 WP 027711123 1 Daquatica
Dickeya chrysanthemi subsp parthenii NCPPB 516 WP 027711123 1 Dchrysanthemi
DAQ1742 DAQ1742 03446 Daquatica
Salmonella enterica subsp enterica serovar Thompson str RM6836 YP 008616076 1 1064551 Proteobacteria
Edwardsiella piscicida C07 087 YP 007630428 1 1288122 Proteobacteria
Salmonella enterica subsp arizonae serovar 62 z4 z23 str RSK2980 YP 001569303 1 882884 Proteobacteria
Salmonella enterica subsp enterica serovar Schwarzengrund str CVM19633 YP 002115719 1 439843 Proteobacteria
Dickeya zeae DZ2Q WP 038925225 1 Dzeae
Dickeya zeae MS1 WP 019843947 1 Dzeae
Dickeya gs poaceaephila NCPPB 569 WP 040033115 1 Dpoaceaephila
Salmonella enterica subsp arizonae serovar 62 z4 z23 str RSK2980 YP 001569981 1 882884 Proteobacteria
Pectobacterium atrosepticum SCRI1043 YP 051009 1 218491 Proteobacteria
Pectobacterium wasabiae WPP163 YP 003259124 1 561231 Proteobacteria
Salmonella enterica subsp enterica serovar Typhi str Ty21a YP 007925202 1 527001 Proteobacteria
Providencia stuartii MRSN 2154 YP 006217975 1 1157951 Proteobacteria
Erwinia tasmaniensis Et1 99 YP 001908354 1 465817 Proteobacteria
Dickeya gs poaceaephila NCPPB 569 WP 042873725 1 Dpoaceaephila
Dickeya dianthicola GBBC 2039 WP 024104354 1 Ddianthicola
Dickeya dianthicola NCPPB 3534 WP 024104354 1 Ddianthicola
Dickeya dianthicola IPO 980 WP 024104354 1 Ddianthicola
Dickeya dianthicola NCPPB 453 WP 024104354 1 Ddianthicola
Dickeya dianthicola RNS04 9 WP 024104354 1 Ddianthicola
Dickeya chrysanthemi subsp parthenii NCPPB 516 WP 027711505 1 Dchrysanthemi
DAQ1742 DAQ1742 01293 Daquatica
Dickeya gs poaceaephila NCPPB 569 WP 035343246 1 Dpoaceaephila
Dickeya sp CSL RW240 WP 035343246 1 Daquatica
Dickeya aquatica DW 0440 WP 035343246 1 Daquatica
851 DAQ1742 01293 GI5 Seed
Xenorhabdus nematophila ATCC 19061 YP 003714346 1 406817 Proteobacteria
Xenorhabdus bovienii SS 2004 YP 003469248 1 406818 Proteobacteria
Xenorhabdus bovienii SS 2004 YP 003467237 1 406818 Proteobacteria
Xenorhabdus nematophila ATCC 19061 YP 003713511 1 406817 Proteobacteria
Xenorhabdus bovienii SS 2004 YP 003468543 1 406818 Proteobacteria
Xenorhabdus bovienii SS 2004 YP 003468559 1 406818 Proteobacteria
Xenorhabdus nematophila ATCC 19061 YP 003712790 1 406817 Proteobacteria
Xenorhabdus nematophila ATCC 19061 YP 003713601 1 406817 Proteobacteria
Dickeya chrysanthemi subsp chrysanthemi NCPPB 402 WP 040000366 1 Dchrysanthemi
Pectobacterium sp SCC3193 YP 006284506 1 1166016 Proteobacteria
Edwardsiella piscicida C07 087 YP 007630083 1 1288122 Proteobacteria
Dickeya chrysanthemi L11 WP 033576372 1 Dchrysanthemi
Salmonella enterica subsp enterica serovar Schwarzengrund str CVM19633 YP 002113634 1 439843 Proteobacteria
Serratia proteamaculans 568 YP 001480109 1 399741 Proteobacteria
Serratia sp AS12 YP 004502500 1 768490 Proteobacteria
Serratia plymuthica AS9 YP 004507452 1 768492 Proteobacteria
Enterobacter cloacae subsp dissolvens SDM YP 006476360 1 1104326 Proteobacteria
Dickeya chrysanthemi L11 WP 033577577 1 Dchrysanthemi
Dickeya dadantii 3937 WP 013319152 1 Ddadantii
Dickeya gs fangzhongdai ND14b WP 033577577 1 Dfangzhongdai
Salmonella enterica subsp enterica serovar Agona str SL483 YP 002149409 1 454166 Proteobacteria
Dickeya dadantii NCPPB 3537 WP 038900302 1 Ddadantii
Dickeya aquatica DW 0440 WP 035339627 1 Daquatica
Escherichia coli 042 YP 006097150 1 216592 Proteobacteria
Dickeya aquatica DW 0440 WP 035342960 1 Daquatica
Dickeya sp CSL RW240 WP 035342960 1 Daquatica
Yersinia enterocolitica subsp palearctica Y11 YP 006005453 1 930944 Proteobacteria
Yersinia enterocolitica subsp palearctica Y11 YP 006007517 1 930944 Proteobacteria
Yersinia pseudotuberculosis IP 31758 YP 001399877 1 349747 Proteobacteria
Yersinia pestis biovar Medievalis str Harbin 35 YP 005622903 1 547048 Proteobacteria
Yersinia pestis biovar Microtus str 91001 NP 994850 1 229193 Proteobacteria
Yersinia pestis Antiqua YP 650308 1 360102 Proteobacteria

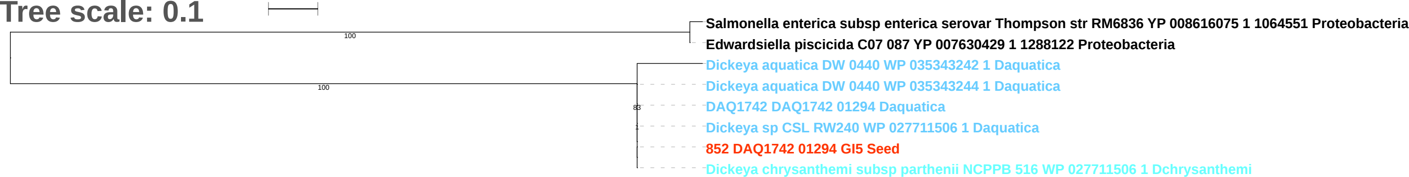
Tree scale: 0.1
100
100
83
Salmonella enterica subsp enterica serovar Thompson str RM6836 YP 008616075 1 1064551 Proteobacteria
Edwardsiella piscicida C07 087 YP 007630429 1 1288122 Proteobacteria
Dickeya aquatica DW 0440 WP 035343242 1 Daquatica
Dickeya aquatica DW 0440 WP 035343244 1 Daquatica
DAQ1742 DAQ1742 01294 Daquatica
Dickeya sp CSL RW240 WP 027711506 1 Daquatica
852 DAQ1742 01294 GI5 Seed
Dickeya chrysanthemi subsp parthenii NCPPB 516 WP 027711506 1 Dchrysanthemi

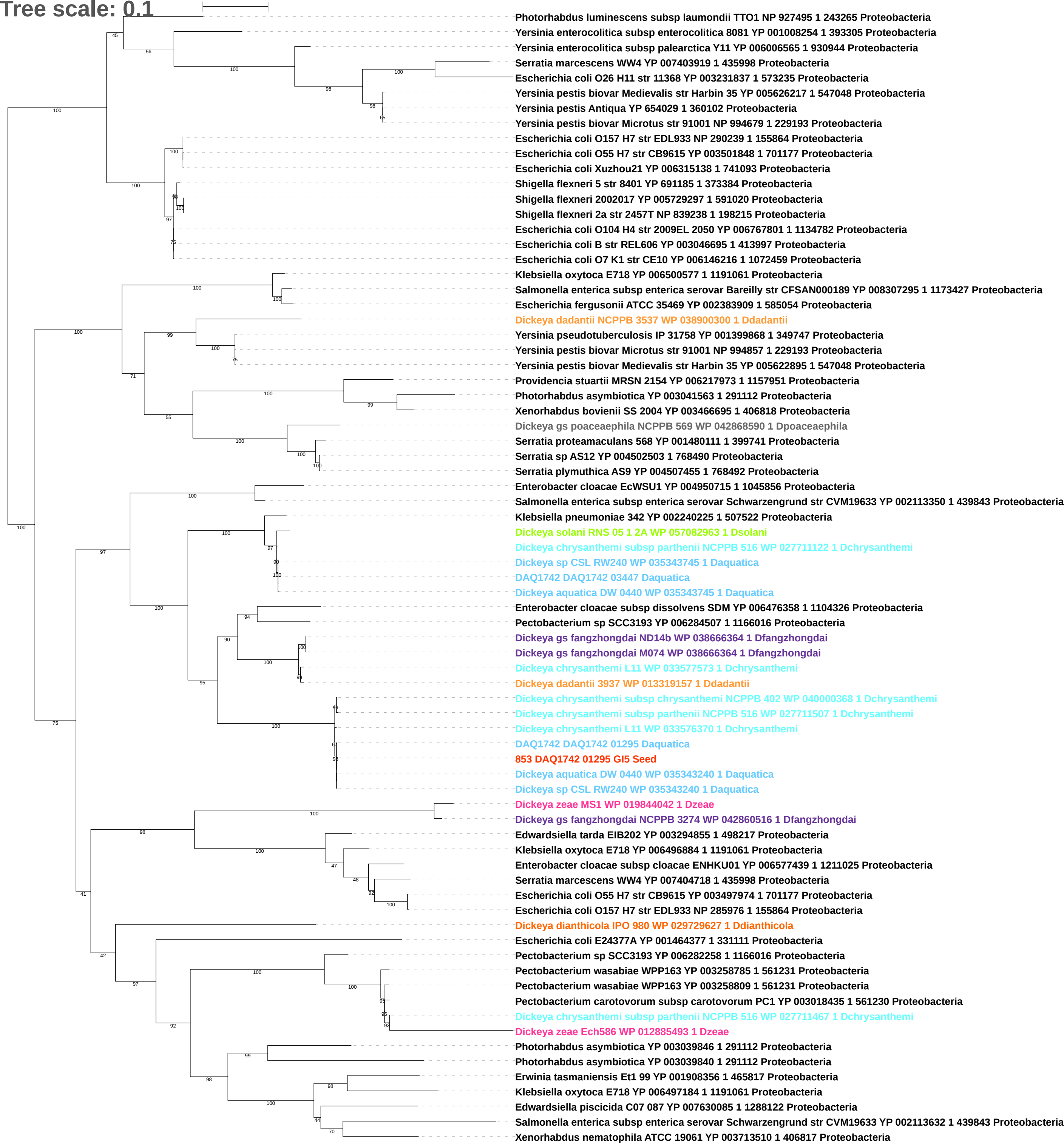
Tree scale: 0.1
Photorhabdus luminescens subsp laumondii TTO1 NP 927495 1 243265 Proteobacteria
Yersinia enterocolitica subsp enterocolitica 8081 YP 001008254 1 393305 Proteobacteria
Yersinia enterocolitica subsp palearctica Y11 YP 006006565 1 930944 Proteobacteria
Serratia marcescens WW4 YP 007403919 1 435998 Proteobacteria
Escherichia coli O26 H11 str 11368 YP 003231837 1 573235 Proteobacteria
Yersinia pestis biovar Medievalis str Harbin 35 YP 005626217 1 547048 Proteobacteria
Yersinia pestis Antiqua YP 654029 1 360102 Proteobacteria
Yersinia pestis biovar Microtus str 91001 NP 994679 1 229193 Proteobacteria
Escherichia coli O157 H7 str EDL933 NP 290239 1 155864 Proteobacteria
Escherichia coli O55 H7 str CB9615 YP 003501848 1 701177 Proteobacteria
Escherichia coli Xuzhou21 YP 006315138 1 741093 Proteobacteria
Shigella flexneri 5 str 8401 YP 691185 1 373384 Proteobacteria
Shigella flexneri 2002017 YP 005729297 1 591020 Proteobacteria
Shigella flexneri 2a str 2457T NP 839238 1 198215 Proteobacteria
Escherichia coli O104 H4 str 2009EL 2050 YP 006767801 1 1134782 Proteobacteria
Escherichia coli B str REL606 YP 003046695 1 413997 Proteobacteria
Escherichia coli O7 K1 str CE10 YP 006146216 1 1072459 Proteobacteria
Klebsiella oxytoca E718 YP 006500577 1 1191061 Proteobacteria
Salmonella enterica subsp enterica serovar Bareilly str CFSAN000189 YP 008307295 1 1173427 Proteobacteria
Escherichia fergusonii ATCC 35469 YP 002383909 1 585054 Proteobacteria
Dickeya dadantii NCPPB 3537 WP 038900300 1 Ddadantii
Yersinia pseudotuberculosis IP 31758 YP 001399868 1 349747 Proteobacteria
Yersinia pestis biovar Microtus str 91001 NP 994857 1 229193 Proteobacteria
Yersinia pestis biovar Medievalis str Harbin 35 YP 005622895 1 547048 Proteobacteria
Providencia stuartii MRSN 2154 YP 006217973 1 1157951 Proteobacteria
Photorhabdus asymbiotica YP 003041563 1 291112 Proteobacteria
Xenorhabdus bovienii SS 2004 YP 003466695 1 406818 Proteobacteria
Dickeya gs poaceaephila NCPPB 569 WP 042868590 1 Dpoaceaephila
Serratia proteamaculans 568 YP 001480111 1 399741 Proteobacteria
Serratia sp AS12 YP 004502503 1 768490 Proteobacteria
Serratia plymuthica AS9 YP 004507455 1 768492 Proteobacteria
Enterobacter cloacae EcWSU1 YP 004950715 1 1045856 Proteobacteria
Salmonella enterica subsp enterica serovar Schwarzengrund str CVM19633 YP 002113350 1 439843 Proteobacteria
Klebsiella pneumoniae 342 YP 002240225 1 507522 Proteobacteria
Dickeya solani RNS 05 1 2A WP 057082963 1 Dsolani
Dickeya chrysanthemi subsp parthenii NCPPB 516 WP 027711122 1 Dchrysanthemi
Dickeya sp CSL RW240 WP 035343745 1 Daquatica
DAQ1742 DAQ1742 03447 Daquatica
Dickeya aquatica DW 0440 WP 035343745 1 Daquatica
Enterobacter cloacae subsp dissolvens SDM YP 006476358 1 1104326 Proteobacteria
Pectobacterium sp SCC3193 YP 006284507 1 1166016 Proteobacteria
Dickeya gs fangzhongdai ND14b WP 038666364 1 Dfangzhongdai
Dickeya gs fangzhongdai M074 WP 038666364 1 Dfangzhongdai
Dickeya chrysanthemi L11 WP 033577573 1 Dchrysanthemi
Dickeya dadantii 3937 WP 013319157 1 Ddadantii
Dickeya chrysanthemi subsp chrysanthemi NCPPB 402 WP 040000368 1 Dchrysanthemi
Dickeya chrysanthemi subsp parthenii NCPPB 516 WP 027711507 1 Dchrysanthemi
Dickeya chrysanthemi L11 WP 033576370 1 Dchrysanthemi
DAQ1742 DAQ1742 01295 Daquatica
853 DAQ1742 01295 GI5 Seed
Dickeya aquatica DW 0440 WP 035343240 1 Daquatica
Dickeya sp CSL RW240 WP 035343240 1 Daquatica
Dickeya zeae MS1 WP 019844042 1 Dzeae
Dickeya gs fangzhongdai NCPPB 3274 WP 042860516 1 Dfangzhongdai
Edwardsiella tarda EIB202 YP 003294855 1 498217 Proteobacteria
Klebsiella oxytoca E718 YP 006496884 1 1191061 Proteobacteria
Enterobacter cloacae subsp cloacae ENHKU01 YP 006577439 1 1211025 Proteobacteria
Serratia marcescens WW4 YP 007404718 1 435998 Proteobacteria
Escherichia coli O55 H7 str CB9615 YP 003497974 1 701177 Proteobacteria
Escherichia coli O157 H7 str EDL933 NP 285976 1 155864 Proteobacteria
Dickeya dianthicola IPO 980 WP 029729627 1 Ddianthicola
Escherichia coli E24377A YP 001464377 1 331111 Proteobacteria
Pectobacterium sp SCC3193 YP 006282258 1 1166016 Proteobacteria
Pectobacterium wasabiae WPP163 YP 003258785 1 561231 Proteobacteria
Pectobacterium wasabiae WPP163 YP 003258809 1 561231 Proteobacteria
Pectobacterium carotovorum subsp carotovorum PC1 YP 003018435 1 561230 Proteobacteria
Dickeya chrysanthemi subsp parthenii NCPPB 516 WP 027711467 1 Dchrysanthemi
Dickeya zeae Ech586 WP 012885493 1 Dzeae
Photorhabdus asymbiotica YP 003039846 1 291112 Proteobacteria
Photorhabdus asymbiotica YP 003039840 1 291112 Proteobacteria
Erwinia tasmaniensis Et1 99 YP 001908356 1 465817 Proteobacteria
Klebsiella oxytoca E718 YP 006497184 1 1191061 Proteobacteria
Edwardsiella piscicida C07 087 YP 007630085 1 1288122 Proteobacteria
Salmonella enterica subsp enterica serovar Schwarzengrund str CVM19633 YP 002113632 1 439843 Proteobacteria
Xenorhabdus nematophila ATCC 19061 YP 003713510 1 406817 Proteobacteria

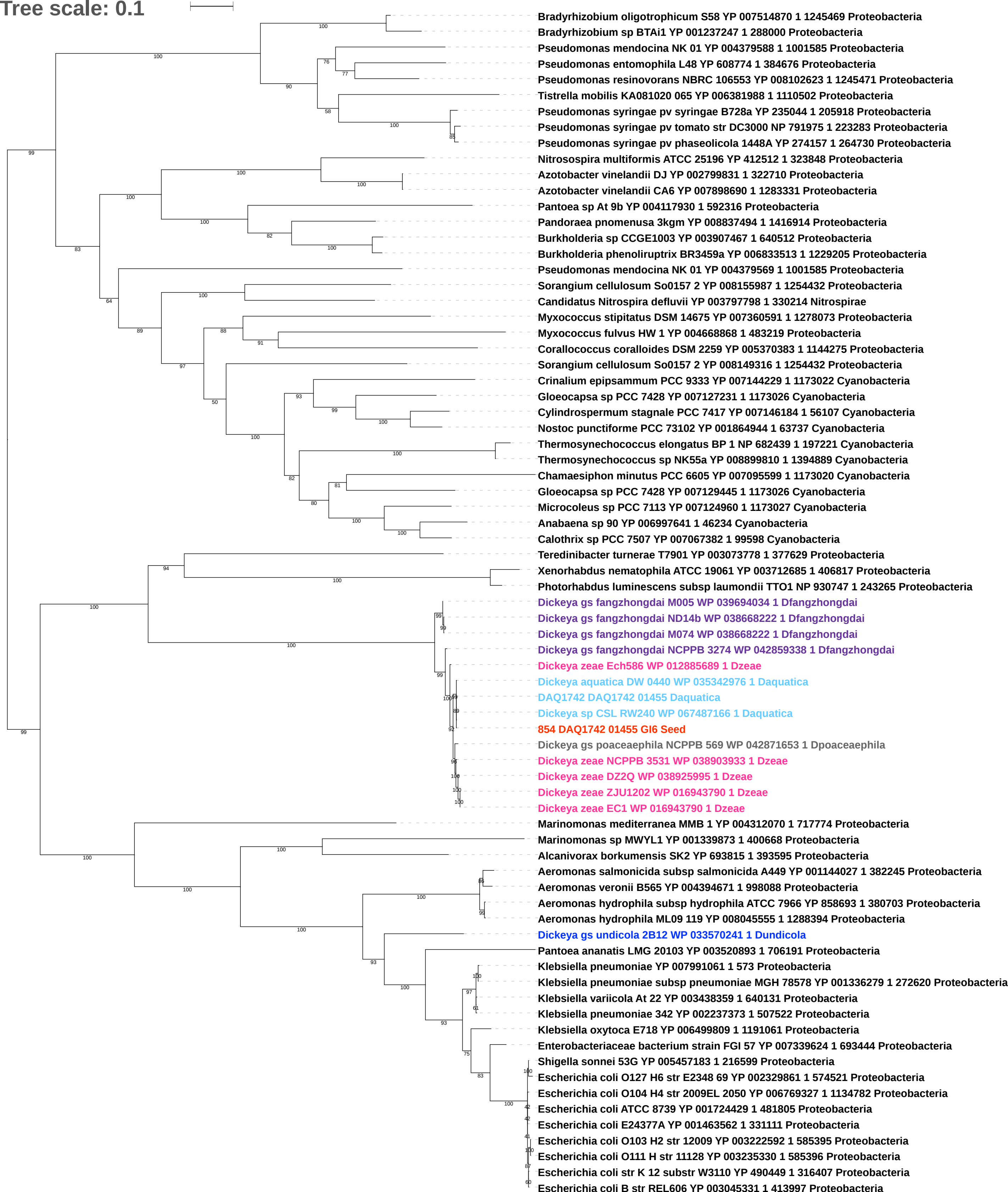
Tree scale: 0.1
Bradyrhizobium oligotrophicum S58 YP 007514870 1 1245469 Proteobacteria
Bradyrhizobium sp BTAi1 YP 001237247 1 288000 Proteobacteria
Pseudomonas mendocina NK 01 YP 004379588 1 1001585 Proteobacteria
Pseudomonas entomophila L48 YP 608774 1 384676 Proteobacteria
Pseudomonas resinovorans NBRC 106553 YP 008102623 1 1245471 Proteobacteria
Tistrella mobilis KA081020 065 YP 006381988 1 1110502 Proteobacteria
Pseudomonas syringae pv syringae B728a YP 235044 1 205918 Proteobacteria
Pseudomonas syringae pv tomato str DC3000 NP 791975 1 223283 Proteobacteria
Pseudomonas syringae pv phaseolicola 1448A YP 274157 1 264730 Proteobacteria
Nitrosospira multiformis ATCC 25196 YP 412512 1 323848 Proteobacteria
Azotobacter vinelandii DJ YP 002799831 1 322710 Proteobacteria
Azotobacter vinelandii CA6 YP 007898690 1 1283331 Proteobacteria
Pantoea sp At 9b YP 004117930 1 592316 Proteobacteria
Pandoraea pnomenusa 3kgm YP 008837494 1 1416914 Proteobacteria
Burkholderia sp CCGE1003 YP 003907467 1 640512 Proteobacteria
Burkholderia phenoliruptrix BR3459a YP 006833513 1 1229205 Proteobacteria
Pseudomonas mendocina NK 01 YP 004379569 1 1001585 Proteobacteria
Sorangium cellulosum So0157 2 YP 008155987 1 1254432 Proteobacteria
Candidatus Nitrospira defluvii YP 003797798 1 330214 Nitrospirae
Myxococcus stipitatus DSM 14675 YP 007360591 1 1278073 Proteobacteria
Myxococcus fulvus HW 1 YP 004668868 1 483219 Proteobacteria
Corallococcus coralloides DSM 2259 YP 005370383 1 1144275 Proteobacteria
Sorangium cellulosum So0157 2 YP 008149316 1 1254432 Proteobacteria
Crinalium epipsammum PCC 9333 YP 007144229 1 1173022 Cyanobacteria
Gloeocapsa sp PCC 7428 YP 007127231 1 1173026 Cyanobacteria
Cylindrospermum stagnale PCC 7417 YP 007146184 1 56107 Cyanobacteria
Nostoc punctiforme PCC 73102 YP 001864944 1 63737 Cyanobacteria
Thermosynechococcus elongatus BP 1 NP 682439 1 197221 Cyanobacteria
Thermosynechococcus sp NK55a YP 008899810 1 1394889 Cyanobacteria
Chamaesiphon minutus PCC 6605 YP 007095599 1 1173020 Cyanobacteria
Gloeocapsa sp PCC 7428 YP 007129445 1 1173026 Cyanobacteria
Microcoleus sp PCC 7113 YP 007124960 1 1173027 Cyanobacteria
Anabaena sp 90 YP 006997641 1 46234 Cyanobacteria
Calothrix sp PCC 7507 YP 007067382 1 99598 Cyanobacteria
Teredinibacter turnerae T7901 YP 003073778 1 377629 Proteobacteria
Xenorhabdus nematophila ATCC 19061 YP 003712685 1 406817 Proteobacteria
Photorhabdus luminescens subsp laumondii TTO1 NP 930747 1 243265 Proteobacteria
Dickeya gs fangzhongdai M005 WP 039694034 1 Dfangzhongdai
Dickeya gs fangzhongdai ND14b WP 038668222 1 Dfangzhongdai
Dickeya gs fangzhongdai M074 WP 038668222 1 Dfangzhongdai
Dickeya gs fangzhongdai NCPPB 3274 WP 042859338 1 Dfangzhongdai
Dickeya zeae Ech586 WP 012885689 1 Dzeae
Dickeya aquatica DW 0440 WP 035342976 1 Daquatica
DAQ1742 DAQ1742 01455 Daquatica
Dickeya sp CSL RW240 WP 067487166 1 Daquatica
854 DAQ1742 01455 GI6 Seed
Dickeya gs poaceaephila NCPPB 569 WP 042871653 1 Dpoaceaephila
Dickeya zeae NCPPB 3531 WP 038903933 1 Dzeae
Dickeya zeae DZ2Q WP 038925995 1 Dzeae
Dickeya zeae ZJU1202 WP 016943790 1 Dzeae
Dickeya zeae EC1 WP 016943790 1 Dzeae
Marinomonas mediterranea MMB 1 YP 004312070 1 717774 Proteobacteria
Marinomonas sp MWYL1 YP 001339873 1 400668 Proteobacteria
Alcanivorax borkumensis SK2 YP 693815 1 393595 Proteobacteria
Aeromonas salmonicida subsp salmonicida A449 YP 001144027 1 382245 Proteobacteria
Aeromonas veronii B565 YP 004394671 1 998088 Proteobacteria
Aeromonas hydrophila subsp hydrophila ATCC 7966 YP 858693 1 380703 Proteobacteria
Aeromonas hydrophila ML09 119 YP 008045555 1 1288394 Proteobacteria
Dickeya gs undicola 2B12 WP 033570241 1 Dundicola
Pantoea ananatis LMG 20103 YP 003520893 1 706191 Proteobacteria
Klebsiella pneumoniae YP 007991061 1 573 Proteobacteria
Klebsiella pneumoniae subsp pneumoniae MGH 78578 YP 001336279 1 272620 Proteobacteria
Klebsiella variicola At 22 YP 003438359 1 640131 Proteobacteria
Klebsiella pneumoniae 342 YP 002237373 1 507522 Proteobacteria
Klebsiella oxytoca E718 YP 006499809 1 1191061 Proteobacteria
Enterobacteriaceae bacterium strain FGI 57 YP 007339624 1 693444 Proteobacteria
Shigella sonnei 53G YP 005457183 1 216599 Proteobacteria
Escherichia coli O127 H6 str E2348 69 YP 002329861 1 574521 Proteobacteria
Escherichia coli O104 H4 str 2009EL 2050 YP 006769327 1 1134782 Proteobacteria
Escherichia coli ATCC 8739 YP 001724429 1 481805 Proteobacteria
Escherichia coli E24377A YP 001463562 1 331111 Proteobacteria
Escherichia coli O103 H2 str 12009 YP 003222592 1 585395 Proteobacteria
Escherichia coli O111 H str 11128 YP 003235330 1 585396 Proteobacteria
Escherichia coli str K 12 substr W3110 YP 490449 1 316407 Proteobacteria
Escherichia coli B str REL606 YP 003045331 1 413997 Proteobacteria

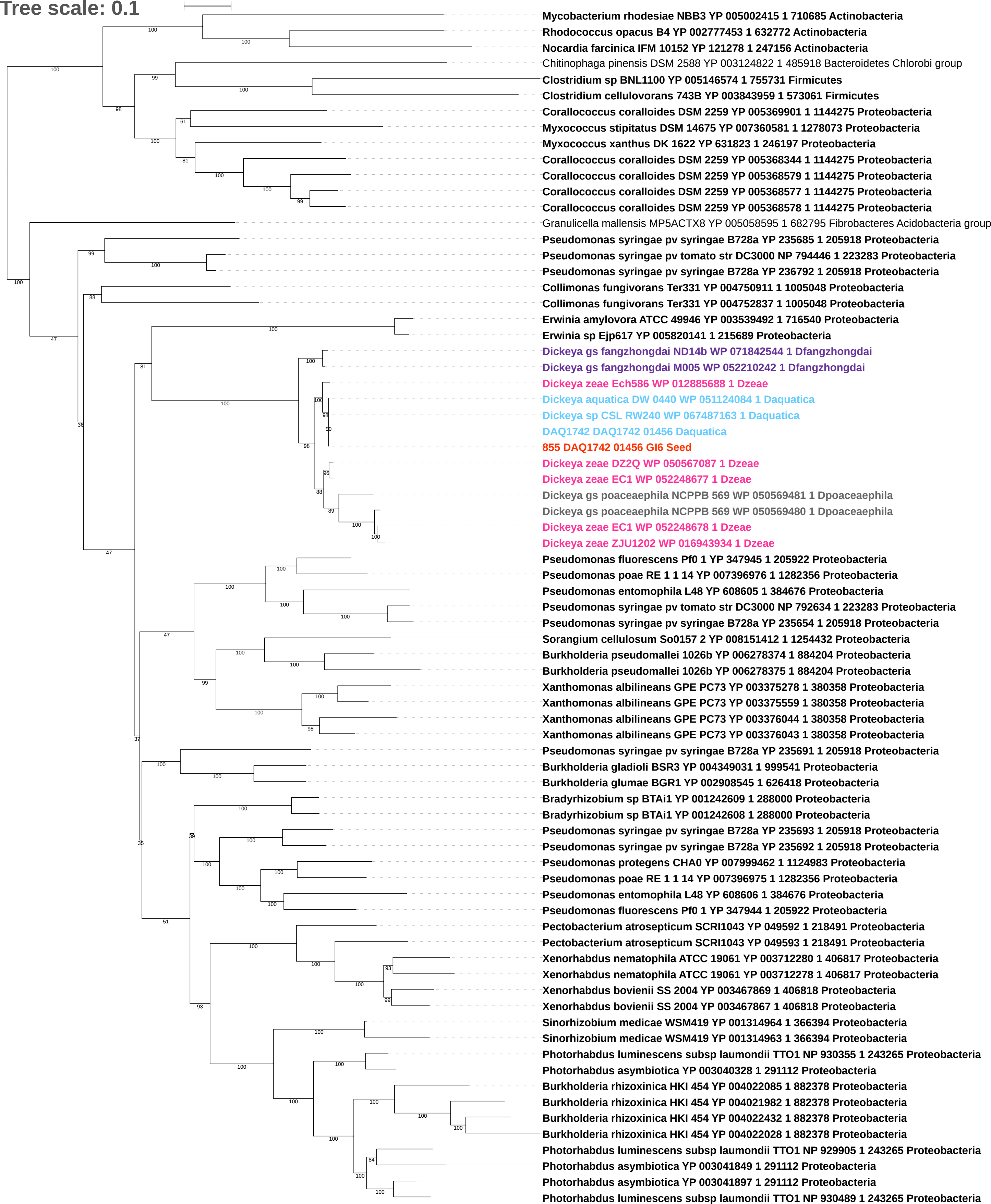
Tree scale: 0.1
Mycobacterium rhodesiae NBB3 YP 005002415 1 710685 Actinobacteria
Rhodococcus opacus B4 YP 002777453 1 632772 Actinobacteria
Nocardia farcinica IFM 10152 YP 121278 1 247156 Actinobacteria
Chitinophaga pinensis DSM 2588 YP 003124822 1 485918 Bacteroidetes Chlorobi group
Clostridium sp BNL1100 YP 005146574 1 755731 Firmicutes
Clostridium cellulovorans 743B YP 003843959 1 573061 Firmicutes
Corallococcus coralloides DSM 2259 YP 005369901 1 1144275 Proteobacteria
Myxococcus stipitatus DSM 14675 YP 007360581 1 1278073 Proteobacteria
Myxococcus xanthus DK 1622 YP 631823 1 246197 Proteobacteria
Corallococcus coralloides DSM 2259 YP 005368344 1 1144275 Proteobacteria
Corallococcus coralloides DSM 2259 YP 005368579 1 1144275 Proteobacteria
Corallococcus coralloides DSM 2259 YP 005368577 1 1144275 Proteobacteria
Corallococcus coralloides DSM 2259 YP 005368578 1 1144275 Proteobacteria
Granulicella mallensis MP5ACTX8 YP 005058595 1 682795 Fibrobacteres Acidobacteria group
Pseudomonas syringae pv syringae B728a YP 235685 1 205918 Proteobacteria
Pseudomonas syringae pv tomato str DC3000 NP 794446 1 223283 Proteobacteria
Pseudomonas syringae pv syringae B728a YP 236792 1 205918 Proteobacteria
Collimonas fungivorans Ter331 YP 004750911 1 1005048 Proteobacteria
Collimonas fungivorans Ter331 YP 004752837 1 1005048 Proteobacteria
Erwinia amylovora ATCC 49946 YP 003539492 1 716540 Proteobacteria
Erwinia sp Ejp617 YP 005820141 1 215689 Proteobacteria
Dickeya gs fangzhongdai ND14b WP 071842544 1 Dfangzhongdai
Dickeya gs fangzhongdai M005 WP 052210242 1 Dfangzhongdai
Dickeya zeae Ech586 WP 012885688 1 Dzeae
Dickeya aquatica DW 0440 WP 051124084 1 Daquatica
Dickeya sp CSL RW240 WP 067487163 1 Daquatica
DAQ1742 DAQ1742 01456 Daquatica
855 DAQ1742 01456 GI6 Seed
Dickeya zeae DZ2Q WP 050567087 1 Dzeae
Dickeya zeae EC1 WP 052248677 1 Dzeae
Dickeya gs poaceaephila NCPPB 569 WP 050569481 1 Dpoaceaephila
Dickeya gs poaceaephila NCPPB 569 WP 050569480 1 Dpoaceaephila
Dickeya zeae EC1 WP 052248678 1 Dzeae
Dickeya zeae ZJU1202 WP 016943934 1 Dzeae
Pseudomonas fluorescens Pf0 1 YP 347945 1 205922 Proteobacteria
Pseudomonas poae RE 1 1 14 YP 007396976 1 1282356 Proteobacteria
Pseudomonas entomophila L48 YP 608605 1 384676 Proteobacteria
Pseudomonas syringae pv tomato str DC3000 NP 792634 1 223283 Proteobacteria
Pseudomonas syringae pv syringae B728a YP 235654 1 205918 Proteobacteria
Sorangium cellulosum So0157 2 YP 008151412 1 1254432 Proteobacteria
Burkholderia pseudomallei 1026b YP 006278374 1 884204 Proteobacteria
Burkholderia pseudomallei 1026b YP 006278375 1 884204 Proteobacteria
Xanthomonas albilineans GPE PC73 YP 003375278 1 380358 Proteobacteria
Xanthomonas albilineans GPE PC73 YP 003375559 1 380358 Proteobacteria
Xanthomonas albilineans GPE PC73 YP 003376044 1 380358 Proteobacteria
Xanthomonas albilineans GPE PC73 YP 003376043 1 380358 Proteobacteria
Pseudomonas syringae pv syringae B728a YP 235691 1 205918 Proteobacteria
Burkholderia gladioli BSR3 YP 004349031 1 999541 Proteobacteria
Burkholderia glumae BGR1 YP 002908545 1 626418 Proteobacteria
Bradyrhizobium sp BTAi1 YP 001242609 1 288000 Proteobacteria
Bradyrhizobium sp BTAi1 YP 001242608 1 288000 Proteobacteria
Pseudomonas syringae pv syringae B728a YP 235693 1 205918 Proteobacteria
Pseudomonas syringae pv syringae B728a YP 235692 1 205918 Proteobacteria
Pseudomonas protegens CHA0 YP 007999462 1 1124983 Proteobacteria
Pseudomonas poae RE 1 1 14 YP 007396975 1 1282356 Proteobacteria
Pseudomonas entomophila L48 YP 608606 1 384676 Proteobacteria
Pseudomonas fluorescens Pf0 1 YP 347944 1 205922 Proteobacteria
Pectobacterium atrosepticum SCRI1043 YP 049592 1 218491 Proteobacteria
Pectobacterium atrosepticum SCRI1043 YP 049593 1 218491 Proteobacteria
Xenorhabdus nematophila ATCC 19061 YP 003712280 1 406817 Proteobacteria
Xenorhabdus nematophila ATCC 19061 YP 003712278 1 406817 Proteobacteria
Xenorhabdus bovienii SS 2004 YP 003467869 1 406818 Proteobacteria
Xenorhabdus bovienii SS 2004 YP 003467867 1 406818 Proteobacteria
Sinorhizobium medicae WSM419 YP 001314964 1 366394 Proteobacteria
Sinorhizobium medicae WSM419 YP 001314963 1 366394 Proteobacteria
Photorhabdus luminescens subsp laumondii TTO1 NP 930355 1 243265 Proteobacteria
Photorhabdus asymbiotica YP 003040328 1 291112 Proteobacteria
Burkholderia rhizoxinica HKI 454 YP 004022085 1 882378 Proteobacteria
Burkholderia rhizoxinica HKI 454 YP 004021982 1 882378 Proteobacteria
Burkholderia rhizoxinica HKI 454 YP 004022432 1 882378 Proteobacteria
Burkholderia rhizoxinica HKI 454 YP 004022028 1 882378 Proteobacteria
Photorhabdus luminescens subsp laumondii TTO1 NP 929905 1 243265 Proteobacteria
Photorhabdus asymbiotica YP 003041849 1 291112 Proteobacteria
Photorhabdus asymbiotica YP 003041897 1 291112 Proteobacteria
Photorhabdus luminescens subsp laumondii TTO1 NP 930489 1 243265 Proteobacteria

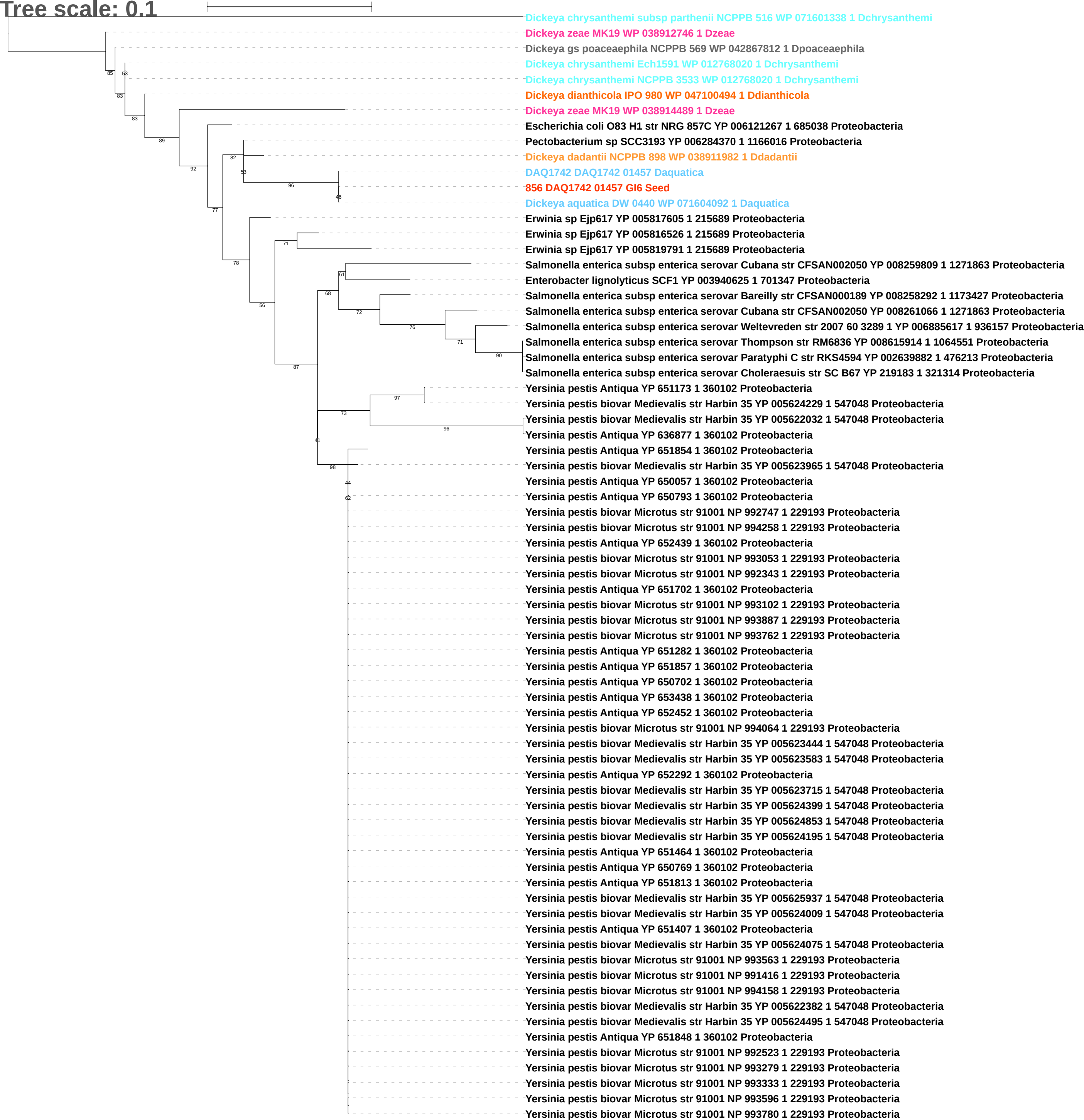

Tree scale: 0.1
Dickeya chrysanthemi subsp parthenii NCPPB 516 WP 071601338 1 Dchrysanthemi
Dickeya zeae MK19 WP 038912746 1 Dzeae
Dickeya gs poaceaephila NCPPB 569 WP 042867812 1 Dpoaceaephila
Dickeya chrysanthemi Ech1591 WP 012768020 1 Dchrysanthemi
Dickeya chrysanthemi NCPPB 3533 WP 012768020 1 Dchrysanthemi
Dickeya dianthicola IPO 980 WP 047100494 1 Ddianthicola
Dickeya zeae MK19 WP 038914489 1 Dzeae
Escherichia coli O83 H1 str NRG 857C YP 006121267 1 685038 Proteobacteria
Pectobacterium sp SCC3193 YP 006284370 1 1166016 Proteobacteria
Dickeya dadantii NCPPB 898 WP 038911982 1 Ddadantii
DAQ1742 DAQ1742 01457 Daquatica
856 DAQ1742 01457 GI6 Seed
Dickeya aquatica DW 0440 WP 071604092 1 Daquatica
Erwinia sp Ejp617 YP 005817605 1 215689 Proteobacteria
Erwinia sp Ejp617 YP 005816526 1 215689 Proteobacteria
Erwinia sp Ejp617 YP 005819791 1 215689 Proteobacteria
Salmonella enterica subsp enterica serovar Cubana str CFSAN002050 YP 008259809 1 1271863 Proteobacteria
Enterobacter lignolyticus SCF1 YP 003940625 1 701347 Proteobacteria
Salmonella enterica subsp enterica serovar Bareilly str CFSAN000189 YP 008258292 1 1173427 Proteobacteria
Salmonella enterica subsp enterica serovar Cubana str CFSAN002050 YP 008261066 1 1271863 Proteobacteria
Salmonella enterica subsp enterica serovar Weltevreden str 2007 60 3289 1 YP 006885617 1 936157 Proteobacteria
Salmonella enterica subsp enterica serovar Thompson str RM6836 YP 008615914 1 1064551 Proteobacteria
Salmonella enterica subsp enterica serovar Paratyphi C str RKS4594 YP 002639882 1 476213 Proteobacteria
Salmonella enterica subsp enterica serovar Choleraesuis str SC B67 YP 219183 1 321314 Proteobacteria
Yersinia pestis Antiqua YP 651173 1 360102 Proteobacteria
Yersinia pestis biovar Medievalis str Harbin 35 YP 005624229 1 547048 Proteobacteria
Yersinia pestis biovar Medievalis str Harbin 35 YP 005622032 1 547048 Proteobacteria
Yersinia pestis Antiqua YP 636877 1 360102 Proteobacteria
Yersinia pestis Antiqua YP 651854 1 360102 Proteobacteria
Yersinia pestis biovar Medievalis str Harbin 35 YP 005623965 1 547048 Proteobacteria
Yersinia pestis Antiqua YP 650057 1 360102 Proteobacteria
Yersinia pestis Antiqua YP 650793 1 360102 Proteobacteria
Yersinia pestis biovar Microtus str 91001 NP 992747 1 229193 Proteobacteria
Yersinia pestis biovar Microtus str 91001 NP 994258 1 229193 Proteobacteria
Yersinia pestis Antiqua YP 652439 1 360102 Proteobacteria
Yersinia pestis biovar Microtus str 91001 NP 993053 1 229193 Proteobacteria
Yersinia pestis biovar Microtus str 91001 NP 992343 1 229193 Proteobacteria
Yersinia pestis Antiqua YP 651702 1 360102 Proteobacteria
Yersinia pestis biovar Microtus str 91001 NP 993102 1 229193 Proteobacteria
Yersinia pestis biovar Microtus str 91001 NP 993887 1 229193 Proteobacteria
Yersinia pestis biovar Microtus str 91001 NP 993762 1 229193 Proteobacteria
Yersinia pestis Antiqua YP 651282 1 360102 Proteobacteria
Yersinia pestis Antiqua YP 651857 1 360102 Proteobacteria
Yersinia pestis Antiqua YP 650702 1 360102 Proteobacteria
Yersinia pestis Antiqua YP 653438 1 360102 Proteobacteria
Yersinia pestis Antiqua YP 652452 1 360102 Proteobacteria
Yersinia pestis biovar Microtus str 91001 NP 994064 1 229193 Proteobacteria
Yersinia pestis biovar Medievalis str Harbin 35 YP 005623444 1 547048 Proteobacteria
Yersinia pestis biovar Medievalis str Harbin 35 YP 005623583 1 547048 Proteobacteria
Yersinia pestis Antiqua YP 652292 1 360102 Proteobacteria
Yersinia pestis biovar Medievalis str Harbin 35 YP 005623715 1 547048 Proteobacteria
Yersinia pestis biovar Medievalis str Harbin 35 YP 005624399 1 547048 Proteobacteria
Yersinia pestis biovar Medievalis str Harbin 35 YP 005624853 1 547048 Proteobacteria
Yersinia pestis biovar Medievalis str Harbin 35 YP 005624195 1 547048 Proteobacteria
Yersinia pestis Antiqua YP 651464 1 360102 Proteobacteria
Yersinia pestis Antiqua YP 650769 1 360102 Proteobacteria
Yersinia pestis Antiqua YP 651813 1 360102 Proteobacteria
Yersinia pestis biovar Medievalis str Harbin 35 YP 005625937 1 547048 Proteobacteria
Yersinia pestis biovar Medievalis str Harbin 35 YP 005624009 1 547048 Proteobacteria
Yersinia pestis Antiqua YP 651407 1 360102 Proteobacteria
Yersinia pestis biovar Medievalis str Harbin 35 YP 005624075 1 547048 Proteobacteria
Yersinia pestis biovar Microtus str 91001 NP 993563 1 229193 Proteobacteria
Yersinia pestis biovar Microtus str 91001 NP 991416 1 229193 Proteobacteria
Yersinia pestis biovar Microtus str 91001 NP 994158 1 229193 Proteobacteria
Yersinia pestis biovar Medievalis str Harbin 35 YP 005622382 1 547048 Proteobacteria
Yersinia pestis biovar Medievalis str Harbin 35 YP 005624495 1 547048 Proteobacteria
Yersinia pestis Antiqua YP 651848 1 360102 Proteobacteria
Yersinia pestis biovar Microtus str 91001 NP 992523 1 229193 Proteobacteria
Yersinia pestis biovar Microtus str 91001 NP 993279 1 229193 Proteobacteria
Yersinia pestis biovar Microtus str 91001 NP 993333 1 229193 Proteobacteria
Yersinia pestis biovar Microtus str 91001 NP 993596 1 229193 Proteobacteria
Yersinia pestis biovar Microtus str 91001 NP 993780 1 229193 Proteobacteria

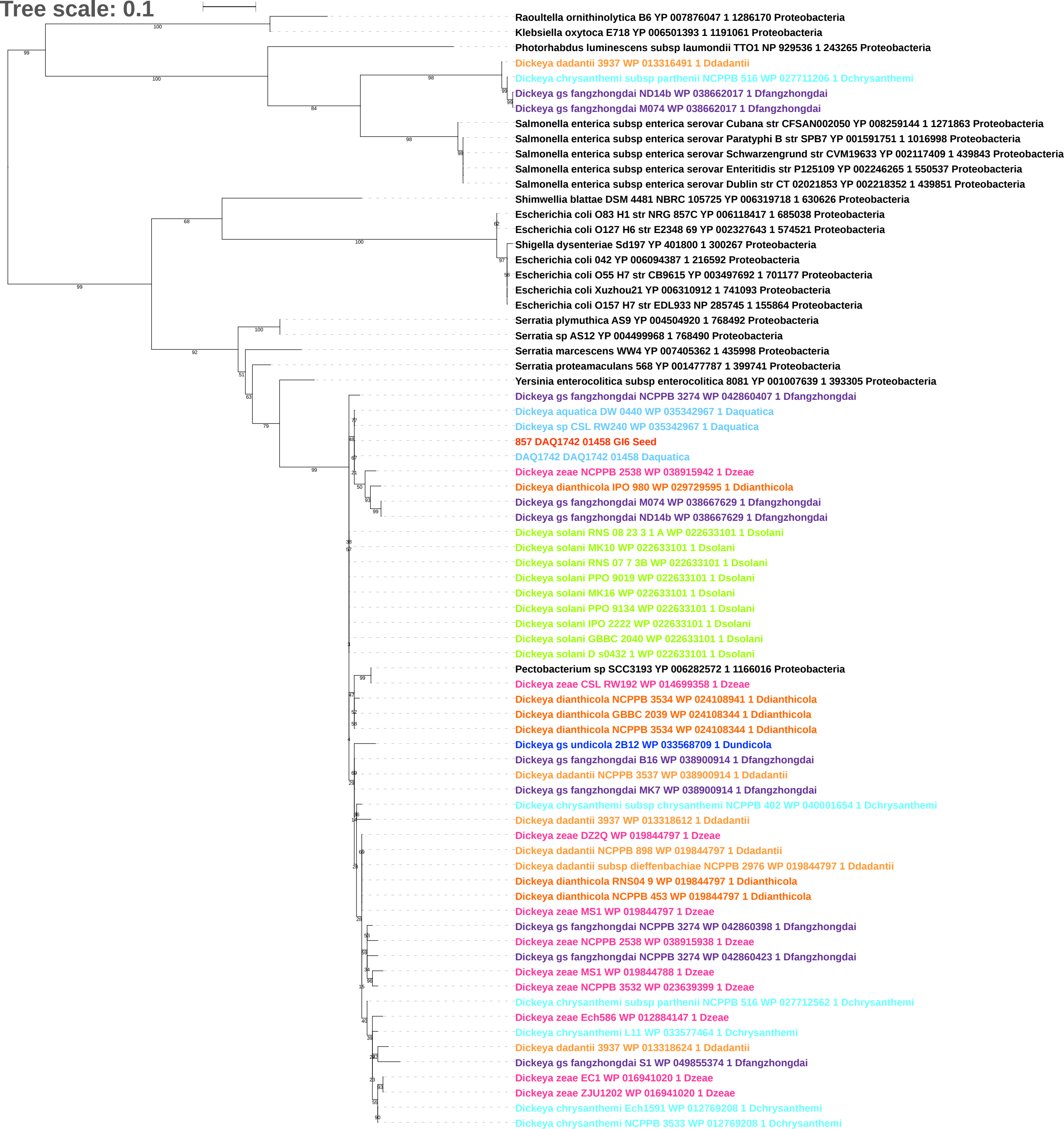
Tree scale: 0.1
Raoultella ornithinolytica B6 YP 007876047 1 1286170 Proteobacteria
Klebsiella oxytoca E718 YP 006501393 1 1191061 Proteobacteria
Photorhabdus luminescens subsp laumondii TTO1 NP 929536 1 243265 Proteobacteria
Dickeya dadantii 3937 WP 013316491 1 Ddadantii
Dickeya chrysanthemi subsp parthenii NCPPB 516 WP 027711206 1 Dchrysanthemi
Dickeya gs fangzhongdai ND14b WP 038662017 1 Dfangzhongdai
Dickeya gs fangzhongdai M074 WP 038662017 1 Dfangzhongdai
Salmonella enterica subsp enterica serovar Cubana str CFSAN002050 YP 008259144 1 1271863 Proteobacteria
Salmonella enterica subsp enterica serovar Paratyphi B str SPB7 YP 001591751 1 1016998 Proteobacteria
Salmonella enterica subsp enterica serovar Schwarzengrund str CVM19633 YP 002117409 1 439843 Proteobacteria
Salmonella enterica subsp enterica serovar Enteritidis str P125109 YP 002246265 1 550537 Proteobacteria
Salmonella enterica subsp enterica serovar Dublin str CT 02021853 YP 002218352 1 439851 Proteobacteria
Shimwellia blattae DSM 4481 NBRC 105725 YP 006319718 1 630626 Proteobacteria
Escherichia coli O83 H1 str NRG 857C YP 006118417 1 685038 Proteobacteria
Escherichia coli O127 H6 str E2348 69 YP 002327643 1 574521 Proteobacteria
Shigella dysenteriae Sd197 YP 401800 1 300267 Proteobacteria
Escherichia coli 042 YP 006094387 1 216592 Proteobacteria
Escherichia coli O55 H7 str CB9615 YP 003497692 1 701177 Proteobacteria
Escherichia coli Xuzhou21 YP 006310912 1 741093 Proteobacteria
Escherichia coli O157 H7 str EDL933 NP 285745 1 155864 Proteobacteria
Serratia plymuthica AS9 YP 004504920 1 768492 Proteobacteria
Serratia sp AS12 YP 004499968 1 768490 Proteobacteria
Serratia marcescens WW4 YP 007405362 1 435998 Proteobacteria
Serratia proteamaculans 568 YP 001477787 1 399741 Proteobacteria
Yersinia enterocolitica subsp enterocolitica 8081 YP 001007639 1 393305 Proteobacteria
Dickeya gs fangzhongdai NCPPB 3274 WP 042860407 1 Dfangzhongdai
Dickeya aquatica DW 0440 WP 035342967 1 Daquatica
Dickeya sp CSL RW240 WP 035342967 1 Daquatica
857 DAQ1742 01458 GI6 Seed
DAQ1742 DAQ1742 01458 Daquatica
Dickeya zeae NCPPB 2538 WP 038915942 1 Dzeae
Dickeya dianthicola IPO 980 WP 029729595 1 Ddianthicola
Dickeya gs fangzhongdai M074 WP 038667629 1 Dfangzhongdai
Dickeya gs fangzhongdai ND14b WP 038667629 1 Dfangzhongdai
Dickeya solani RNS 08 23 3 1 A WP 022633101 1 Dsolani
Dickeya solani MK10 WP 022633101 1 Dsolani
Dickeya solani RNS 07 7 3B WP 022633101 1 Dsolani
Dickeya solani PPO 9019 WP 022633101 1 Dsolani
Dickeya solani MK16 WP 022633101 1 Dsolani
Dickeya solani PPO 9134 WP 022633101 1 Dsolani
Dickeya solani IPO 2222 WP 022633101 1 Dsolani
Dickeya solani GBBC 2040 WP 022633101 1 Dsolani
Dickeya solani D s0432 1 WP 022633101 1 Dsolani
Pectobacterium sp SCC3193 YP 006282572 1 1166016 Proteobacteria
Dickeya zeae CSL RW192 WP 014699358 1 Dzeae
Dickeya dianthicola NCPPB 3534 WP 024108941 1 Ddianthicola
Dickeya dianthicola GBBC 2039 WP 024108344 1 Ddianthicola
Dickeya dianthicola NCPPB 3534 WP 024108344 1 Ddianthicola
Dickeya gs undicola 2B12 WP 033568709 1 Dundicola
Dickeya gs fangzhongdai B16 WP 038900914 1 Dfangzhongdai
Dickeya dadantii NCPPB 3537 WP 038900914 1 Ddadantii
Dickeya gs fangzhongdai MK7 WP 038900914 1 Dfangzhongdai
Dickeya chrysanthemi subsp chrysanthemi NCPPB 402 WP 040001654 1 Dchrysanthemi
Dickeya dadantii 3937 WP 013318612 1 Ddadantii
Dickeya zeae DZ2Q WP 019844797 1 Dzeae
Dickeya dadantii NCPPB 898 WP 019844797 1 Ddadantii
Dickeya dadantii subsp dieffenbachiae NCPPB 2976 WP 019844797 1 Ddadantii
Dickeya dianthicola RNS04 9 WP 019844797 1 Ddianthicola
Dickeya dianthicola NCPPB 453 WP 019844797 1 Ddianthicola
Dickeya zeae MS1 WP 019844797 1 Dzeae
Dickeya gs fangzhongdai NCPPB 3274 WP 042860398 1 Dfangzhongdai
Dickeya zeae NCPPB 2538 WP 038915938 1 Dzeae
Dickeya gs fangzhongdai NCPPB 3274 WP 042860423 1 Dfangzhongdai
Dickeya zeae MS1 WP 019844788 1 Dzeae
Dickeya zeae NCPPB 3532 WP 023639399 1 Dzeae
Dickeya chrysanthemi subsp parthenii NCPPB 516 WP 027712562 1 Dchrysanthemi
Dickeya zeae Ech586 WP 012884147 1 Dzeae
Dickeya chrysanthemi L11 WP 033577464 1 Dchrysanthemi
Dickeya dadantii 3937 WP 013318624 1 Ddadantii
Dickeya gs fangzhongdai S1 WP 049855374 1 Dfangzhongdai
Dickeya zeae EC1 WP 016941020 1 Dzeae
Dickeya zeae ZJU1202 WP 016941020 1 Dzeae
Dickeya chrysanthemi Ech1591 WP 012769208 1 Dchrysanthemi
Dickeya chrysanthemi NCPPB 3533 WP 012769208 1 Dchrysanthemi

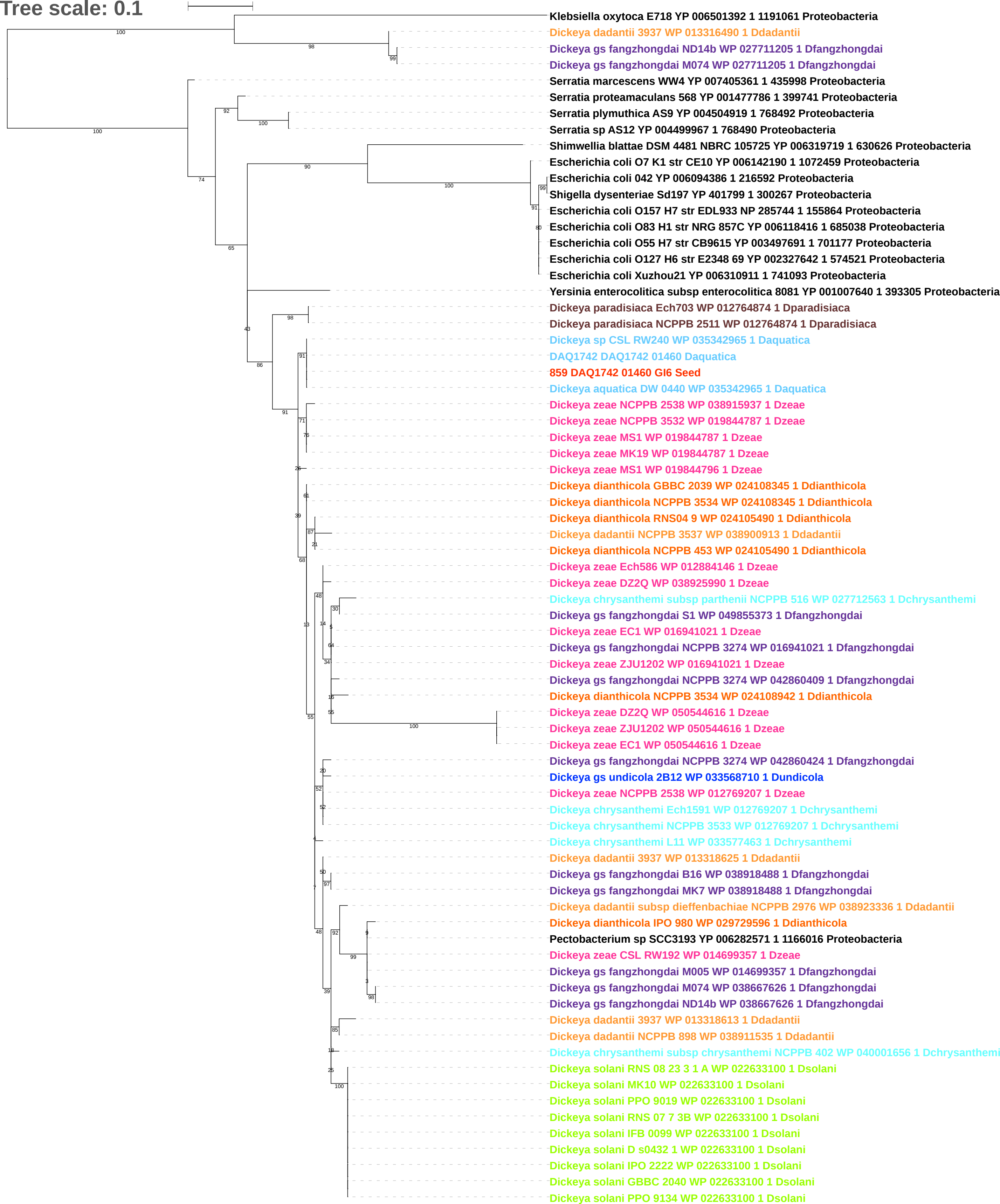
Tree scale: 0.1
Klebsiella oxytoca E718 YP 006501392 1 1191061 Proteobacteria
Dickeya dadantii 3937 WP 013316490 1 Ddadantii
Dickeya gs fangzhongdai ND14b WP 027711205 1 Dfangzhongdai
Dickeya gs fangzhongdai M074 WP 027711205 1 Dfangzhongdai
Serratia marcescens WW4 YP 007405361 1 435998 Proteobacteria
Serratia proteamaculans 568 YP 001477786 1 399741 Proteobacteria
Serratia plymuthica AS9 YP 004504919 1 768492 Proteobacteria
Serratia sp AS12 YP 004499967 1 768490 Proteobacteria
Shimwellia blattae DSM 4481 NBRC 105725 YP 006319719 1 630626 Proteobacteria
Escherichia coli O7 K1 str CE10 YP 006142190 1 1072459 Proteobacteria
Escherichia coli 042 YP 003094386 1 216592 Proteobacteria
Shigella dysenteriae Sd197 YP 401799 1 300267 Proteobacteria
Escherichia coli O157 H7 str EDL933 NP 285744 1 155864 Proteobacteria
Escherichia coli O83 H1 str NRG 857C YP 006118416 1 685038 Proteobacteria
Escherichia coli O55 H7 str CB9615 YP 003497691 1 701177 Proteobacteria
Escherichia coli O127 H6 str E2348 69 YP 002327642 1 574521 Proteobacteria
Escherichia coli Xuzhou21 YP 006310911 1 741093 Proteobacteria
Yersinia enterocolitica subsp enterocolitica 8081 YP 001007640 1 393305 Proteobacteria
Dickeya paradisiaca Ech703 WP 012764874 1 Dparadisiaca
Dickeya paradisiaca NCPPB 2511 WP 012764874 1 Dparadisiaca
Dickeya sp CSL RW240 WP 035342965 1 Daquatica
DAQ1742 DAQ1742 01460 Daquatica
859 DAQ1742 01460 GI6 Seed
Dickeya aquatica DW 0440 WP 035342965 1 Daquatica
Dickeya zeae NCPPB 2538 WP 038915937 1 Dzeae
Dickeya zeae NCPPB 3532 WP 019844787 1 Dzeae
Dickeya zeae MS1 WP 019844787 1 Dzeae
Dickeya zeae MK19 WP 019844787 1 Dzeae
Dickeya zeae MS1 WP 019844796 1 Dzeae
Dickeya dianthicola GBBC 2039 WP 024108345 1 Ddianthicola
Dickeya dianthicola NCPPB 3534 WP 024108345 1 Ddianthicola
Dickeya dianthicola RNS04 9 WP 024105490 1 Ddianthicola
Dickeya dadantii NCPPB 3537 WP 038900913 1 Ddadantii
Dickeya dianthicola NCPPB 453 WP 024105490 1 Ddianthicola
Dickeya zeae Ech586 WP 012884146 1 Dzeae
Dickeya zeae DZ2Q WP 038925990 1 Dzeae
Dickeya chrysanthemi subsp parthenii NCPPB 516 WP 027712563 1 Dchrysanthemi
Dickeya gs fangzhongdai S1 WP 049855373 1 Dfangzhongdai
Dickeya zeae EC1 WP 016941021 1 Dzeae
Dickeya gs fangzhongdai NCPPB 3274 WP 016941021 1 Dfangzhongdai
Dickeya zeae ZJU1202 WP 016941021 1 Dzeae
Dickeya gs fangzhongdai NCPPB 3274 WP 042860409 1 Dfangzhongdai
Dickeya dianthicola NCPPB 3534 WP 024108942 1 Ddianthicola
Dickeya zeae DZ2Q WP 050544616 1 Dzeae
Dickeya zeae ZJU1202 WP 050544616 1 Dzeae
Dickeya zeae EC1 WP 050544616 1 Dzeae
Dickeya gs fangzhongdai NCPPB 3274 WP 042860424 1 Dfangzhongdai
Dickeya gs undicola 2B12 WP 033568710 1 Dundicola
Dickeya zeae NCPPB 2538 WP 012769207 1 Dzeae
Dickeya chrysanthemi Ech1591 WP 012769207 1 Dchrysanthemi
Dickeya chrysanthemi NCPPB 3533 WP 012769207 1 Dchrysanthemi
Dickeya chrysanthemi L11 WP 033577463 1 Dchrysanthemi
Dickeya dadantii 3937 WP 013318625 1 Ddadantii
Dickeya gs fangzhongdai B16 WP 038918488 1 Dfangzhongdai
Dickeya gs fangzhongdai MK7 WP 038918488 1 Dfangzhongdai
Dickeya dadantii subsp dieffenbachiae NCPPB 2976 WP 038923336 1 Ddadantii
Dickeya dianthicola IPO 980 WP 029729596 1 Ddianthicola
Pectobacterium sp SCC3193 YP 006282571 1 1166016 Proteobacteria
Dickeya zeae CSL RW192 WP 014699357 1 Dzeae
Dickeya gs fangzhongdai M005 WP 014699357 1 Dfangzhongdai
Dickeya gs fangzhongdai M074 WP 038667626 1 Dfangzhongdai
Dickeya gs fangzhongdai ND14b WP 038667626 1 Dfangzhongdai
Dickeya dadantii 3937 WP 013318613 1 Ddadantii
Dickeya dadantii NCPPB 898 WP 038911535 1 Ddadantii
Dickeya chrysanthemi subsp chrysanthemi NCPPB 402 WP 040001656 1 Dchrysanthemi
Dickeya solani RNS 08 23 3 1 A WP 022633100 1 Dsolani
Dickeya solani MK10 WP 022633100 1 Dsolani
Dickeya solani PPO 9019 WP 022633100 1 Dsolani
Dickeya solani RNS 07 7 3B WP 022633100 1 Dsolani
Dickeya solani IFB 0099 WP 022633100 1 Dsolani
Dickeya solani D s0432 1 WP 022633100 1 Dsolani
Dickeya solani IPO 2222 WP 022633100 1 Dsolani
Dickeya solani GBBC 2040 WP 022633100 1 Dsolani
Dickeya solani PPO 9134 WP 022633100 1 Dsolani

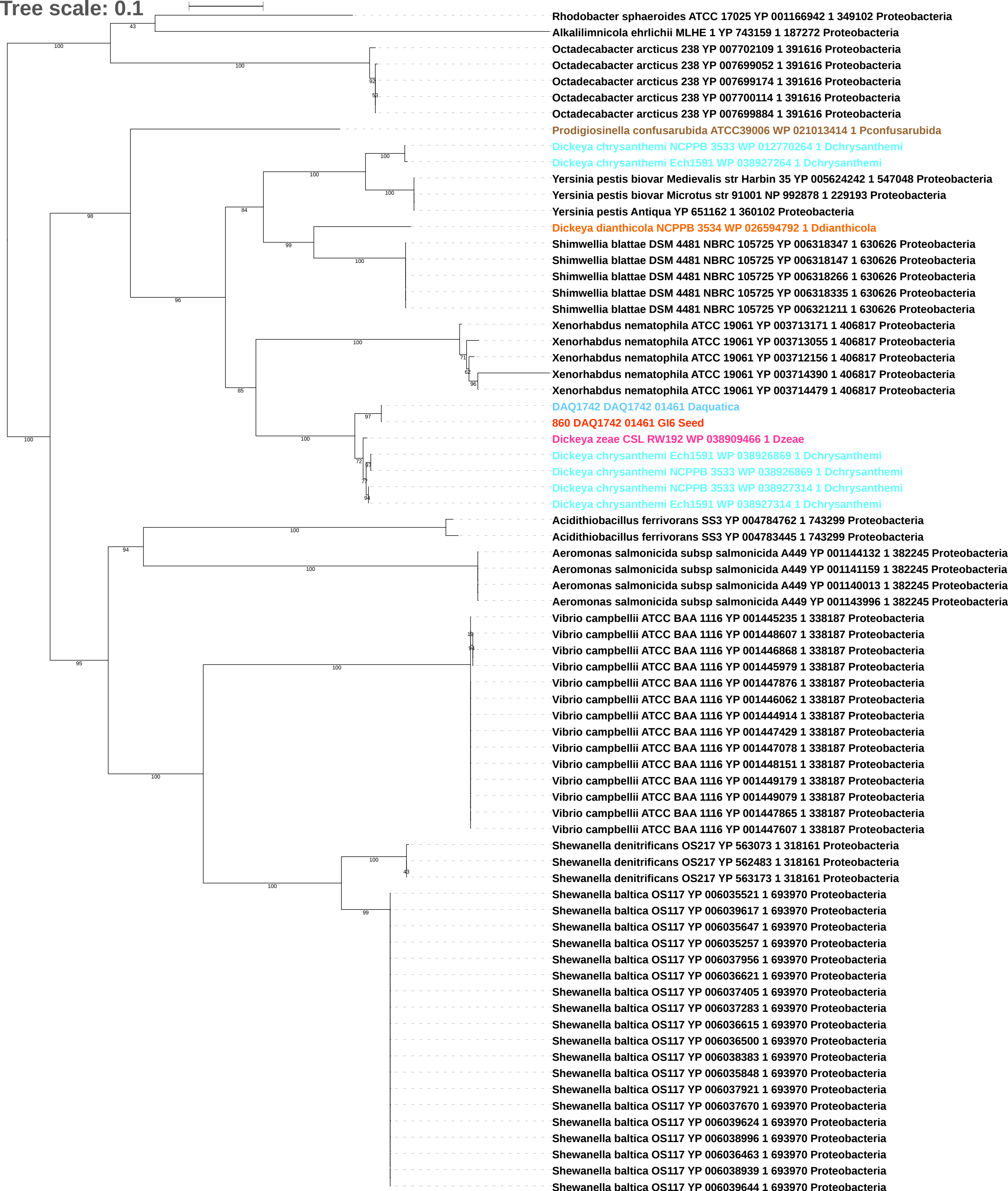
Tree scale: 0.1
Rhodobacter sphaeroides ATCC 17025 YP 001166942 1 349102 Proteobacteria
Alkalilimnicola ehrlichii MLHE 1 YP 743159 1 187272 Proteobacteria
Octadecabacter arcticus 238 YP 007702109 1 391616 Proteobacteria
Octadecabacter arcticus 238 YP 007699052 1 391616 Proteobacteria
Octadecabacter arcticus 238 YP 007699174 1 391616 Proteobacteria
Octadecabacter arcticus 238 YP 007700114 1 391616 Proteobacteria
Octadecabacter arcticus 238 YP 007699884 1 391616 Proteobacteria
Prodigiosinella confusarubida ATCC39006 WP 021013414 1 Pconfusarubida
Dickeya chrysanthemi NCPPB 3533 WP 012770264 1 Dchrysanthemi
Dickeya chrysanthemi Ech1591 WP 038927264 1 Dchrysanthemi
Yersinia pestis biovar Medievalis str Harbin 35 YP 005624242 1 547048 Proteobacteria
Yersinia pestis biovar Microtus str 91001 NP 992878 1 229193 Proteobacteria
Yersinia pestis Antiqua YP 651162 1 360102 Proteobacteria
Dickeya dianthicola NCPPB 3534 WP 026594792 1 Ddianthicola
Shimwellia blattae DSM 4481 NBRC 105725 YP 006318347 1 630626 Proteobacteria
Shimwellia blattae DSM 4481 NBRC 105725 YP 006318147 1 630626 Proteobacteria
Shimwellia blattae DSM 4481 NBRC 105725 YP 006318266 1 630626 Proteobacteria
Shimwellia blattae DSM 4481 NBRC 105725 YP 006318335 1 630626 Proteobacteria
Shimwellia blattae DSM 4481 NBRC 105725 YP 006321211 1 630626 Proteobacteria
Xenorhabdus nematophila ATCC 19061 YP 003713171 1 406817 Proteobacteria
Xenorhabdus nematophila ATCC 19061 YP 003713055 1 406817 Proteobacteria
Xenorhabdus nematophila ATCC 19061 YP 003712156 1 406817 Proteobacteria
Xenorhabdus nematophila ATCC 19061 YP 003714390 1 406817 Proteobacteria
Xenorhabdus nematophila ATCC 19061 YP 003714479 1 406817 Proteobacteria
DAQ1742 DAQ1742 01461 Daquatica
860 DAQ1742 01461 GI6 Seed
Dickeya zeae CSL RW192 WP 038909466 1 Dzeae
Dickeya chrysanthemi Ech1591 WP 038926869 1 Dchrysanthemi
Dickeya chrysanthemi NCPPB 3533 WP 038926869 1 Dchrysanthemi
Dickeya chrysanthemi NCPPB 3533 WP 038927314 1 Dchrysanthemi
Dickeya chrysanthemi Ech1591 WP 038927314 1 Dchrysanthemi
Acidithiobacillus ferrivorans SS3 YP 004784762 1 743299 Proteobacteria
Acidithiobacillus ferrivorans SS3 YP 004783445 1 743299 Proteobacteria
Aeromonas salmonicida subsp salmonicida A449 YP 001144132 1 382245 Proteobacteria
Aeromonas salmonicida subsp salmonicida A449 YP 001141159 1 382245 Proteobacteria
Aeromonas salmonicida subsp salmonicida A449 YP 001140013 1 382245 Proteobacteria
Aeromonas salmonicida subsp salmonicida A449 YP 001143996 1 382245 Proteobacteria
Vibrio campbellii ATCC BAA 1116 YP 001445235 1 338187 Proteobacteria
Vibrio campbellii ATCC BAA 1116 YP 001448607 1 338187 Proteobacteria
Vibrio campbellii ATCC BAA 1116 YP 001446868 1 338187 Proteobacteria
Vibrio campbellii ATCC BAA 1116 YP 001445979 1 338187 Proteobacteria
Vibrio campbellii ATCC BAA 1116 YP 001447876 1 338187 Proteobacteria
Vibrio campbellii ATCC BAA 1116 YP 001446062 1 338187 Proteobacteria
Vibrio campbellii ATCC BAA 1116 YP 001444914 1 338187 Proteobacteria
Vibrio campbellii ATCC BAA 1116 YP 001447429 1 338187 Proteobacteria
Vibrio campbellii ATCC BAA 1116 YP 001447078 1 338187 Proteobacteria
Vibrio campbellii ATCC BAA 1116 YP 001448151 1 338187 Proteobacteria
Vibrio campbellii ATCC BAA 1116 YP 001449179 1 338187 Proteobacteria
Vibrio campbellii ATCC BAA 1116 YP 001449079 1 338187 Proteobacteria
Vibrio campbellii ATCC BAA 1116 YP 001447865 1 338187 Proteobacteria
Vibrio campbellii ATCC BAA 1116 YP 001447607 1 338187 Proteobacteria
Shewanella denitrificans OS217 YP 563073 1 318161 Proteobacteria
Shewanella denitrificans OS217 YP 562483 1 318161 Proteobacteria
Shewanella denitrificans OS217 YP 563173 1 318161 Proteobacteria
Shewanella baltica OS117 YP 006035521 1 693970 Proteobacteria
Shewanella baltica OS117 YP 006039617 1 693970 Proteobacteria
Shewanella baltica OS117 YP 006035647 1 693970 Proteobacteria
Shewanella baltica OS117 YP 006035257 1 693970 Proteobacteria
Shewanella baltica OS117 YP 006037956 1 693970 Proteobacteria
Shewanella baltica OS117 YP 006036621 1 693970 Proteobacteria
Shewanella baltica OS117 YP 006037405 1 693970 Proteobacteria
Shewanella baltica OS117 YP 006037283 1 693970 Proteobacteria
Shewanella baltica OS117 YP 006036615 1 693970 Proteobacteria
Shewanella baltica OS117 YP 006036500 1 693970 Proteobacteria
Shewanella baltica OS117 YP 006038383 1 693970 Proteobacteria
Shewanella baltica OS117 YP 006035848 1 693970 Proteobacteria
Shewanella baltica OS117 YP 006037921 1 693970 Proteobacteria
Shewanella baltica OS117 YP 006037670 1 693970 Proteobacteria
Shewanella baltica OS117 YP 006039624 1 693970 Proteobacteria
Shewanella baltica OS117 YP 006038996 1 693970 Proteobacteria
Shewanella baltica OS117 YP 006036463 1 693970 Proteobacteria
Shewanella baltica OS117 YP 006038939 1 693970 Proteobacteria
Shewanella baltica OS117 YP 006039644 1 693970 Proteobacteria

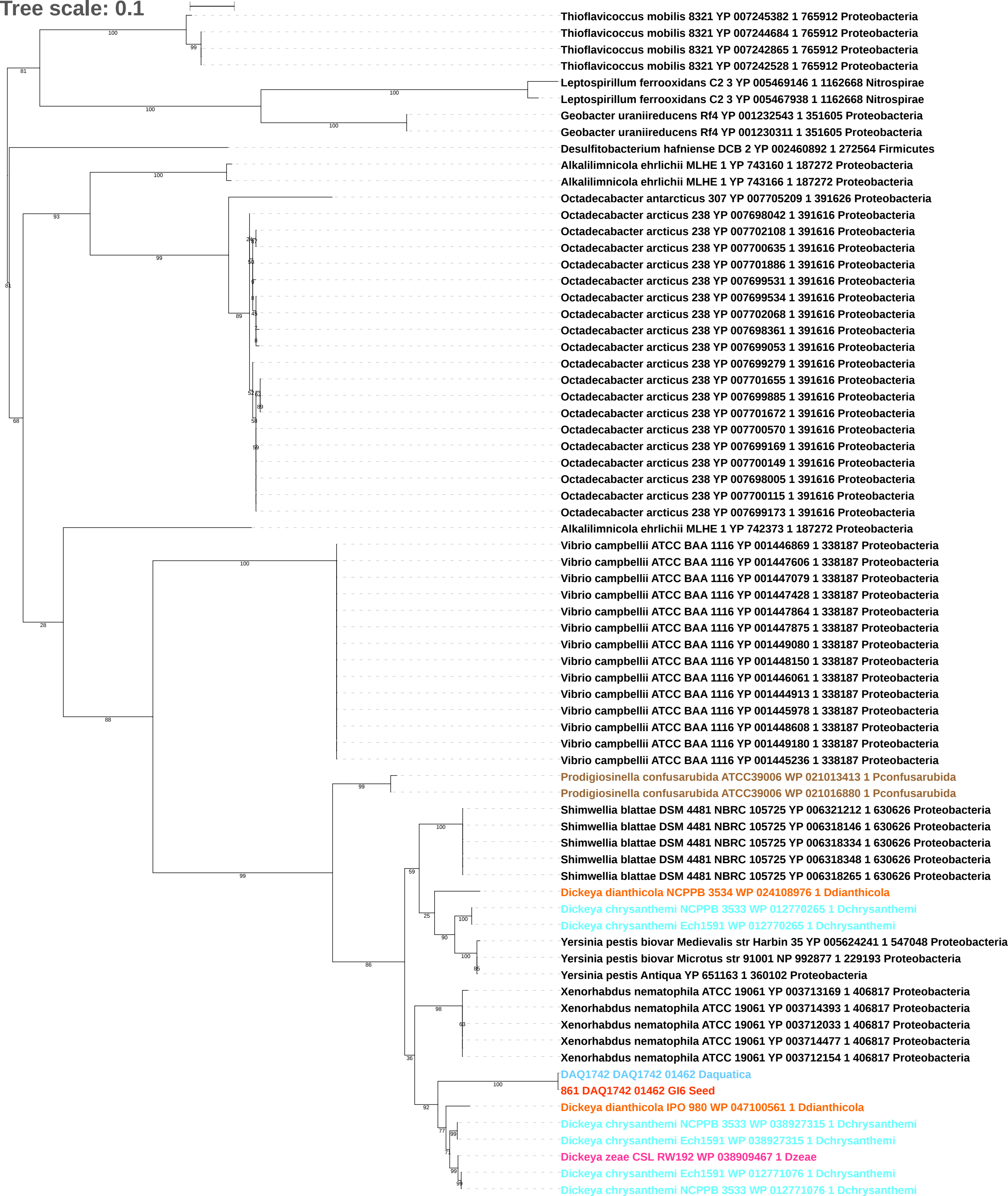
Tree scale: 0.1
Thioflavicoccus mobilis 8321 YP 007245382 1 765912 Proteobacteria
Thioflavicoccus mobilis 8321 YP 007244684 1 765912 Proteobacteria
Thioflavicoccus mobilis 8321 YP 007242865 1 765912 Proteobacteria
Thioflavicoccus mobilis 8321 YP 007242528 1 765912 Proteobacteria
Leptospirillum ferrooxidans C2 3 YP 005469146 1 1162668 Nitrospirae
Leptospirillum ferrooxidans C2 3 YP 005467938 1 1162668 Nitrospirae
Geobacter uraniireducens Rf4 YP 001232543 1 351605 Proteobacteria
Geobacter uraniireducens Rf4 YP 001230311 1 351605 Proteobacteria
Desulfitobacterium hafniense DCB 2 YP 002460892 1 272564 Firmicutes
Alkalilimnicola ehrlichii MLHE 1 YP 743160 1 187272 Proteobacteria
Alkalilimnicola ehrlichii MLHE 1 YP 743166 1 187272 Proteobacteria
Octadecabacter antarcticus 307 YP 007705209 1 391626 Proteobacteria
Octadecabacter arcticus 238 YP 007698042 1 391616 Proteobacteria
Octadecabacter arcticus 238 YP 007702108 1 391616 Proteobacteria
Octadecabacter arcticus 238 YP 007700635 1 391616 Proteobacteria
Octadecabacter arcticus 238 YP 007701886 1 391616 Proteobacteria
Octadecabacter arcticus 238 YP 007699531 1 391616 Proteobacteria
Octadecabacter arcticus 238 YP 007699534 1 391616 Proteobacteria
Octadecabacter arcticus 238 YP 007702068 1 391616 Proteobacteria
Octadecabacter arcticus 238 YP 007698361 1 391616 Proteobacteria
Octadecabacter arcticus 238 YP 007699053 1 391616 Proteobacteria
Octadecabacter arcticus 238 YP 007699279 1 391616 Proteobacteria
Octadecabacter arcticus 238 YP 007701655 1 391616 Proteobacteria
Octadecabacter arcticus 238 YP 007699885 1 391616 Proteobacteria
Octadecabacter arcticus 238 YP 007701672 1 391616 Proteobacteria
Octadecabacter arcticus 238 YP 007700570 1 391616 Proteobacteria
Octadecabacter arcticus 238 YP 007699169 1 391616 Proteobacteria
Octadecabacter arcticus 238 YP 007700149 1 391616 Proteobacteria
Octadecabacter arcticus 238 YP 007698005 1 391616 Proteobacteria
Octadecabacter arcticus 238 YP 007700115 1 391616 Proteobacteria
Octadecabacter arcticus 238 YP 007699173 1 391616 Proteobacteria
Alkalilimnicola ehrlichii MLHE 1 YP 742373 1 187272 Proteobacteria
Vibrio campbellii ATCC BAA 1116 YP 001446869 1 338187 Proteobacteria
Vibrio campbellii ATCC BAA 1116 YP 001447606 1 338187 Proteobacteria
Vibrio campbellii ATCC BAA 1116 YP 001447079 1 338187 Proteobacteria
Vibrio campbellii ATCC BAA 1116 YP 001447428 1 338187 Proteobacteria
Vibrio campbellii ATCC BAA 1116 YP 001447864 1 338187 Proteobacteria
Vibrio campbellii ATCC BAA 1116 YP 001447875 1 338187 Proteobacteria
Vibrio campbellii ATCC BAA 1116 YP 001449080 1 338187 Proteobacteria
Vibrio campbellii ATCC BAA 1116 YP 001448150 1 338187 Proteobacteria
Vibrio campbellii ATCC BAA 1116 YP 001446061 1 338187 Proteobacteria
Vibrio campbellii ATCC BAA 1116 YP 001444913 1 338187 Proteobacteria
Vibrio campbellii ATCC BAA 1116 YP 001445978 1 338187 Proteobacteria
Vibrio campbellii ATCC BAA 1116 YP 001448608 1 338187 Proteobacteria
Vibrio campbellii ATCC BAA 1116 YP 001449180 1 338187 Proteobacteria
Vibrio campbellii ATCC BAA 1116 YP 001445236 1 338187 Proteobacteria
Prodigiosinella confusarubida ATCC39006 WP 021013413 1 Pconfusarubida
Prodigiosinella confusarubida ATCC39006 WP 021016880 1 Pconfusarubida
Shimwellia blattae DSM 4481 NBRC 105725 YP 006321212 1 630626 Proteobacteria
Shimwellia blattae DSM 4481 NBRC 105725 YP 006318146 1 630626 Proteobacteria
Shimwellia blattae DSM 4481 NBRC 105725 YP 006318334 1 630626 Proteobacteria
Shimwellia blattae DSM 4481 NBRC 105725 YP 006318348 1 630626 Proteobacteria
Shimwellia blattae DSM 4481 NBRC 105725 YP 006318265 1 630626 Proteobacteria
Dickeya dianthicola NCPPB 3534 WP 024108976 1 Ddianthicola
Dickeya chrysanthemi NCPPB 3533 WP 012770265 1 Dchrysanthemi
Dickeya chrysanthemi Ech1591 WP 012770265 1 Dchrysanthemi
Yersinia pestis biovar Medievalis str Harbin 35 YP 005624241 1 547048 Proteobacteria
Yersinia pestis biovar Microtus str 91001 NP 992877 1 229193 Proteobacteria
Yersinia pestis Antiqua YP 651163 1 360102 Proteobacteria
Xenorhabdus nematophila ATCC 19061 YP 003713169 1 406817 Proteobacteria
Xenorhabdus nematophila ATCC 19061 YP 003714393 1 406817 Proteobacteria
Xenorhabdus nematophila ATCC 19061 YP 003712033 1 406817 Proteobacteria
Xenorhabdus nematophila ATCC 19061 YP 003714477 1 406817 Proteobacteria
Xenorhabdus nematophila ATCC 19061 YP 003712154 1 406817 Proteobacteria
DAQ1742 DAQ1742 01462 Daquatica
861 DAQ1742 01462 GI6 Seed
Dickeya dianthicola IPO 980 WP 047100561 1 Ddianthicola
Dickeya chrysanthemi NCPPB 3533 WP 038927315 1 Dchrysanthemi
Dickeya chrysanthemi Ech1591 WP 038927315 1 Dchrysanthemi
Dickeya zeae CSL RW192 WP 038909467 1 Dzeae
Dickeya chrysanthemi Ech1591 WP 012771076 1 Dchrysanthemi
Dickeya chrysanthemi NCPPB 3533 WP 012771076 1 Dchrysanthemi

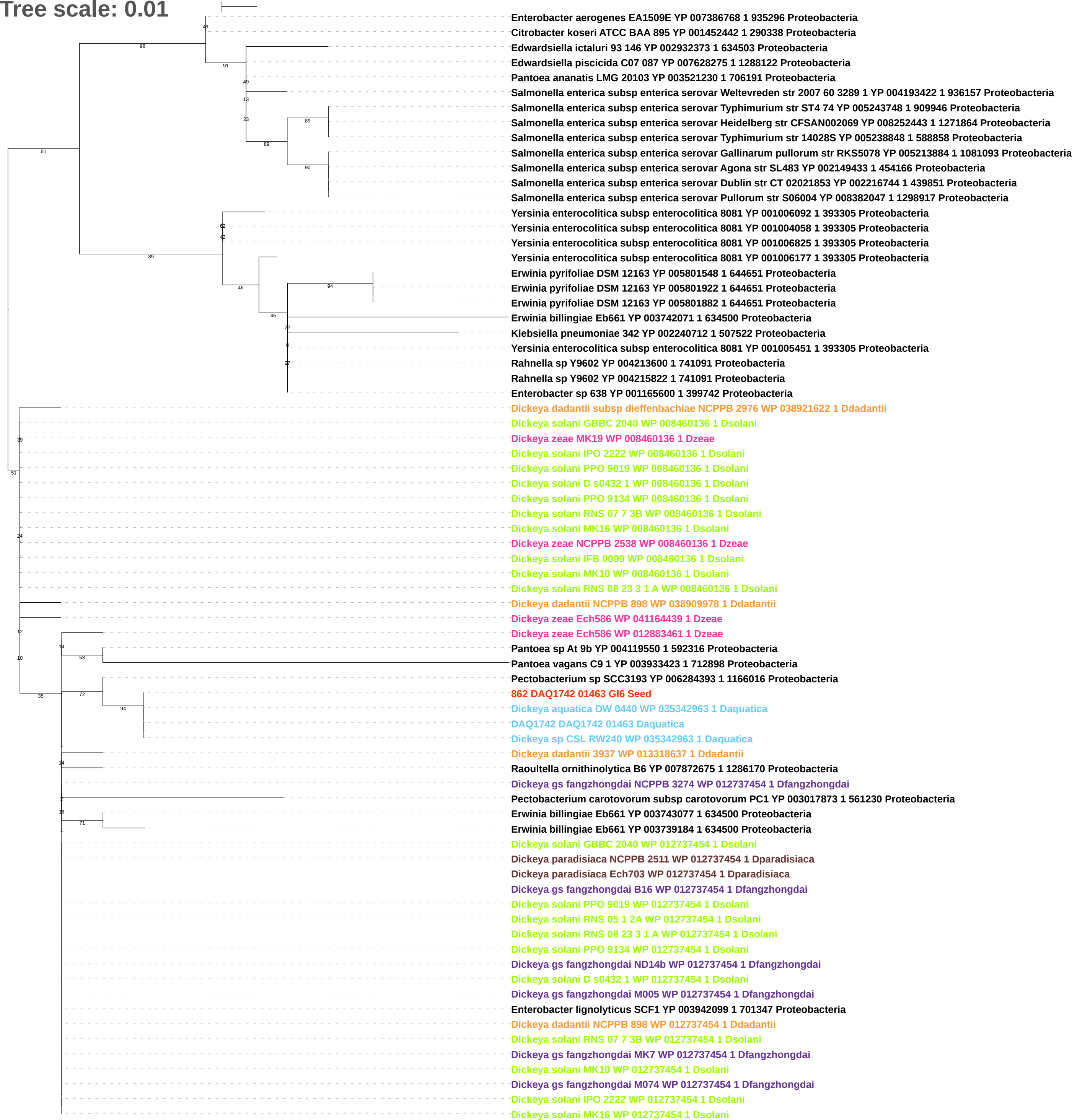
Tree scale: 0.01
Enterobacter aerogenes EA1509E YP 007386768 1 935296 Proteobacteria
Citrobacter koseri ATCC BAA 895 YP 001452442 1 290338 Proteobacteria
Edwardsiella ictaluri 93 146 YP 002932373 1 634503 Proteobacteria
Edwardsiella piscicida C07 087 YP 007628275 1 1288122 Proteobacteria
Pantoea ananatis LMG 20103 YP 003521230 1 706191 Proteobacteria
Salmonella enterica subsp enterica serovar Weltevreden str 2007 60 3289 1 YP 004193422 1 936157 Proteobacteria
Salmonella enterica subsp enterica serovar Typhimurium str ST4 74 YP 005243748 1 909946 Proteobacteria
Salmonella enterica subsp enterica serovar Heidelberg str CFSAN002069 YP 008252443 1 1271864 Proteobacteria
Salmonella enterica subsp enterica serovar Typhimurium str 14028S YP 005238848 1 588858 Proteobacteria
Salmonella enterica subsp enterica serovar Gallinarum pullorum str RKS5078 YP 005213884 1 1081093 Proteobacteria
Salmonella enterica subsp enterica serovar Agona str SL483 YP 002149433 1 454166 Proteobacteria
Salmonella enterica subsp enterica serovar Dublin str CT 02021853 YP 002216744 1 439851 Proteobacteria
Salmonella enterica subsp enterica serovar Pullorum str S06004 YP 008382047 1 1298917 Proteobacteria
Yersinia enterocolitica subsp enterocolitica 8081 YP 001006092 1 393305 Proteobacteria
Yersinia enterocolitica subsp enterocolitica 8081 YP 001004058 1 393305 Proteobacteria
Yersinia enterocolitica subsp enterocolitica 8081 YP 001006825 1 393305 Proteobacteria
Yersinia enterocolitica subsp enterocolitica 8081 YP 001006177 1 393305 Proteobacteria
Erwinia pyrifoliae DSM 12163 YP 005801548 1 644651 Proteobacteria
Erwinia pyrifoliae DSM 12163 YP 005801922 1 644651 Proteobacteria
Erwinia pyrifoliae DSM 12163 YP 005801882 1 644651 Proteobacteria
Erwinia billingiae Eb661 YP 003742071 1 634500 Proteobacteria
Klebsiella pneumoniae 342 YP 002240712 1 507522 Proteobacteria
Yersinia enterocolitica subsp enterocolitica 8081 YP 001005451 1 393305 Proteobacteria
Rahnella sp Y9602 YP 004213600 1 741091 Proteobacteria
Rahnella sp Y9602 YP 004215822 1 741091 Proteobacteria
Enterobacter sp 638 YP 001165600 1 399742 Proteobacteria
Dickeya dadantii subsp dieffenbachiae NCPPB 2976 WP 038921622 1 Ddadantii
Dickeya solani GBBC 2040 WP 008460136 1 Dsolani
Dickeya zeae MK19 WP 008460136 1 Dzeae
Dickeya solani IPO 2222 WP 008460136 1 Dsolani
Dickeya solani PPO 9019 WP 008460136 1 Dsolani
Dickeya solani D s0432 1 WP 008460136 1 Dsolani
Dickeya solani PPO 9134 WP 008460136 1 Dsolani
Dickeya solani RNS 07 7 3B WP 008460136 1 Dsolani
Dickeya solani MK16 WP 008460136 1 Dsolani
Dickeya zeae NCPPB 2538 WP 008460136 1 Dzeae
Dickeya solani IFB 0099 WP 008460136 1 Dsolani
Dickeya solani MK10 WP 008460136 1 Dsolani
Dickeya solani RNS 08 23 3 1 A WP 008460136 1 Dsolani
Dickeya dadantii NCPPB 898 WP 038909978 1 Ddadantii
Dickeya zeae Ech586 WP 041164439 1 Dzeae
Dickeya zeae Ech586 WP 012883461 1 Dzeae
Pantoea sp At 9b YP 004119550 1 592316 Proteobacteria
Pantoea vagans C9 1 YP 003933423 1 712898 Proteobacteria
Pectobacterium sp SCC3193 YP 006284393 1 1166016 Proteobacteria
862 DAQ1742 01463 GI6 Seed
Dickeya aquatica DW 0440 WP 035342963 1 Daquatica
DAQ1742 DAQ1742 01463 Daquatica
Dickeya sp CSL RW240 WP 035342963 1 Daquatica
Dickeya dadantii 3937 WP 013318637 1 Ddadantii
Raoultella ornithinolytica B6 YP 007872675 1 1286170 Proteobacteria
Dickeya gs fangzhongdai NCPPB 3274 WP 012737454 1 Dfangzhongdai
Pectobacterium carotovorum subsp carotovorum PC1 YP 003017873 1 561230 Proteobacteria
Erwinia billingiae Eb661 YP 003743077 1 634500 Proteobacteria
Erwinia billingiae Eb661 YP 003739184 1 634500 Proteobacteria
Dickeya solani GBBC 2040 WP 012737454 1 Dsolani
Dickeya paradisiaca NCPPB 2511 WP 012737454 1 Dparadisiaca
Dickeya paradisiaca Ech703 WP 012737454 1 Dparadisiaca
Dickeya gs fangzhongdai B16 WP 012737454 1 Dfangzhongdai
Dickeya solani PPO 9019 WP 012737454 1 Dsolani
Dickeya solani RNS 05 1 2A WP 012737454 1 Dsolani
Dickeya solani RNS 08 23 3 1 A WP 012737454 1 Dsolani
Dickeya solani PPO 9134 WP 012737454 1 Dsolani
Dickeya gs fangzhongdai ND14b WP 012737454 1 Dfangzhongdai
Dickeya solani D s0432 1 WP 012737454 1 Dsolani
Dickeya gs fangzhongdai M005 WP 012737454 1 Dfangzhongdai
Enterobacter lignolyticus SCF1 YP 003942099 1 701347 Proteobacteria
Dickeya dadantii NCPPB 898 WP 012737454 1 Ddadantii
Dickeya solani RNS 07 7 3B WP 012737454 1 Dsolani
Dickeya gs fangzhongdai MK7 WP 012737454 1 Dfangzhongdai
Dickeya solani MK10 WP 012737454 1 Dsolani
Dickeya gs fangzhongdai M074 WP 012737454 1 Dfangzhongdai
Dickeya solani IPO 2222 WP 012737454 1 Dsolani
Dickeya solani MK16 WP 012737454 1 Dsolani

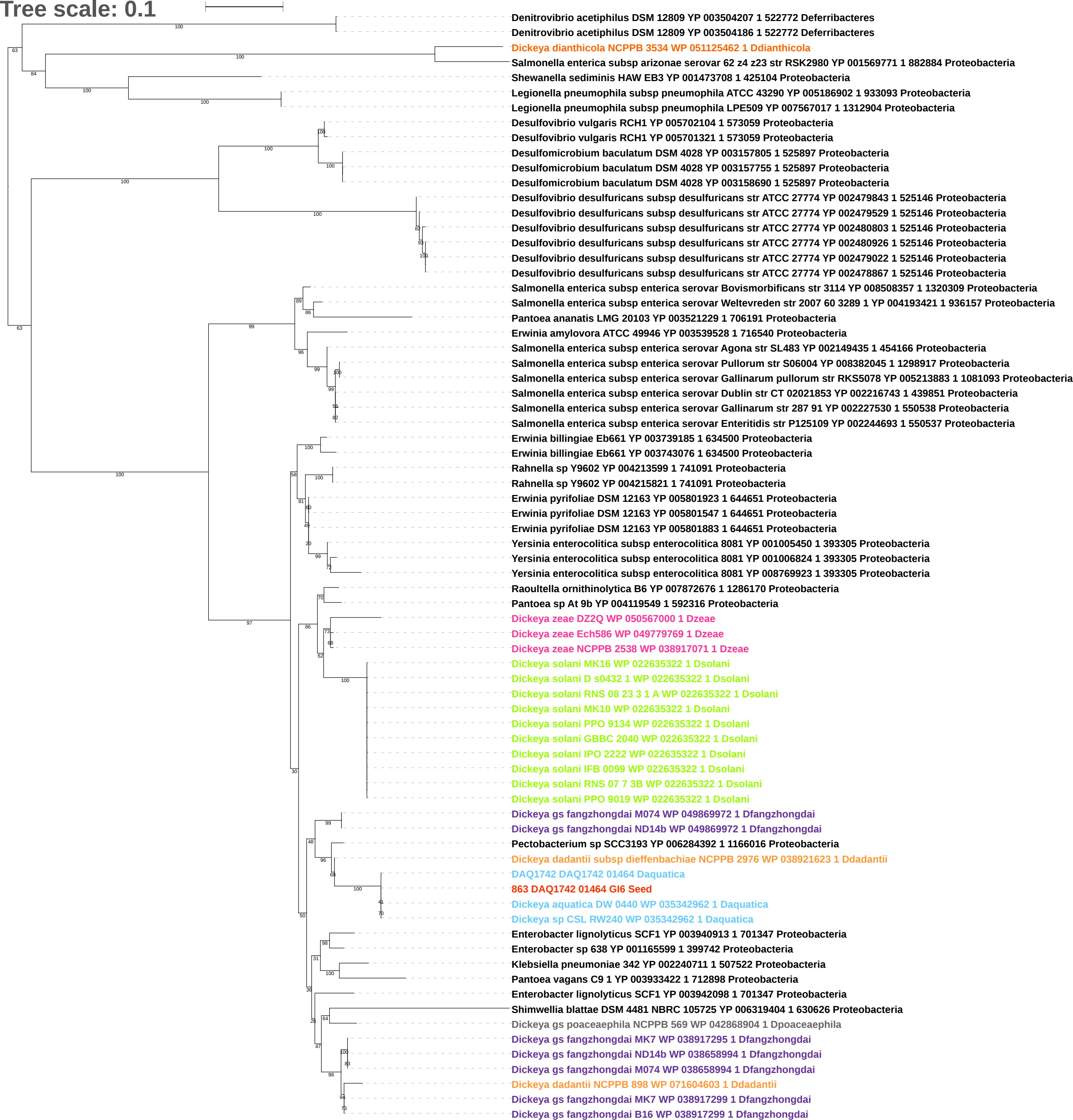
Tree scale: 0.1
Denitrovibrio acetiphilus DSM 12809 YP 003504207 1 522772 Deferribacteres
Denitrovibrio acetiphilus DSM 12809 YP 003504186 1 522772 Deferribacteres
Dickeya dianthicola NCPPB 3534 WP 051125462 1 Ddianthicola
Salmonella enterica subsp arizonae serovar 62 z4 z23 str RSK2980 YP 001569771 1 882884 Proteobacteria
Shewanella sediminis HAW EB3 YP 001473708 1 425104 Proteobacteria
Legionella pneumophila subsp pneumophila ATCC 43290 YP 005186902 1 933093 Proteobacteria
Legionella pneumophila subsp pneumophila LPE509 YP 007567017 1 1312904 Proteobacteria
Desulfovibrio vulgaris RCH1 YP 005702104 1 573059 Proteobacteria
Desulfovibrio vulgaris RCH1 YP 005701321 1 573059 Proteobacteria
Desulfomicrobium baculatum DSM 4028 YP 003157805 1 525897 Proteobacteria
Desulfomicrobium baculatum DSM 4028 YP 003157755 1 525897 Proteobacteria
Desulfomicrobium baculatum DSM 4028 YP 003158690 1 525897 Proteobacteria
Desulfovibrio desulfuricans subsp desulfuricans str ATCC 27774 YP 002479843 1 525146 Proteobacteria
Desulfovibrio desulfuricans subsp desulfuricans str ATCC 27774 YP 002479529 1 525146 Proteobacteria
Desulfovibrio desulfuricans subsp desulfuricans str ATCC 27774 YP 002480803 1 525146 Proteobacteria
Desulfovibrio desulfuricans subsp desulfuricans str ATCC 27774 YP 002480926 1 525146 Proteobacteria
Desulfovibrio desulfuricans subsp desulfuricans str ATCC 27774 YP 002479022 1 525146 Proteobacteria
Desulfovibrio desulfuricans subsp desulfuricans str ATCC 27774 YP 002478867 1 525146 Proteobacteria
Salmonella enterica subsp enterica serovar Bovismorbificans str 3114 YP 008508357 1 1320309 Proteobacteria
Salmonella enterica subsp enterica serovar Weltevreden str 2007 60 3289 1 YP 004193421 1 936157 Proteobacteria
Pantoea ananatis LMG 20103 YP 003521229 1 706191 Proteobacteria
Erwinia amylovora ATCC 49946 YP 003539528 1 716540 Proteobacteria
Salmonella enterica subsp enterica serovar Agona str SL483 YP 002149435 1 454166 Proteobacteria
Salmonella enterica subsp enterica serovar Pullorum str S06004 YP 008382045 1 1298917 Proteobacteria
Salmonella enterica subsp enterica serovar Gallinarum pullorum str RKS5078 YP 005213883 1 1081093 Proteobacteria
Salmonella enterica subsp enterica serovar Dublin str CT 02021853 YP 002216743 1 439851 Proteobacteria
Salmonella enterica subsp enterica serovar Gallinarum str 287 91 YP 002227530 1 550538 Proteobacteria
Salmonella enterica subsp enterica serovar Enteritidis str P125109 YP 002244693 1 550537 Proteobacteria
Erwinia billingiae Eb661 YP 003739185 1 634500 Proteobacteria
Erwinia billingiae Eb661 YP 003743076 1 634500 Proteobacteria
Rahnella sp Y9602 YP 004213599 1 741091 Proteobacteria
Rahnella sp Y9602 YP 004215821 1 741091 Proteobacteria
Erwinia pyrifoliae DSM 12163 YP 005801923 1 644651 Proteobacteria
Erwinia pyrifoliae DSM 12163 YP 005801547 1 644651 Proteobacteria
Erwinia pyrifoliae DSM 12163 YP 005801883 1 644651 Proteobacteria
Yersinia enterocolitica subsp enterocolitica 8081 YP 001005450 1 393305 Proteobacteria
Yersinia enterocolitica subsp enterocolitica 8081 YP 001006824 1 393305 Proteobacteria
Yersinia enterocolitica subsp enterocolitica 8081 YP 008769923 1 393305 Proteobacteria
Raoultella ornithinolytica B6 YP 007872676 1 1286170 Proteobacteria
Pantoea sp At 9b YP 004119549 1 592316 Proteobacteria
Dickeya zeae DZ2Q WP 050567000 1 Dzeae
Dickeya zeae Ech586 WP 049779769 1 Dzeae
Dickeya zeae NCPPB 2538 WP 038917071 1 Dzeae
Dickeya solani MK16 WP 022635322 1 Dsolani
Dickeya solani D s0432 1 WP 022635322 1 Dsolani
Dickeya solani RNS 08 23 3 1 A WP 022635322 1 Dsolani
Dickeya solani MK10 WP 022635322 1 Dsolani
Dickeya solani PPO 9134 WP 022635322 1 Dsolani
Dickeya solani GBBC 2040 WP 022635322 1 Dsolani
Dickeya solani IPO 2222 WP 022635322 1 Dsolani
Dickeya solani IFB 0099 WP 022635322 1 Dsolani
Dickeya solani RNS 07 7 3B WP 022635322 1 Dsolani
Dickeya solani PPO 9019 WP 022635322 1 Dsolani
Dickeya gs fangzhongdai M074 WP 049869972 1 Dfangzhongdai
Dickeya gs fangzhongdai ND14b WP 049869972 1 Dfangzhongdai
Pectobacterium sp SCC3193 YP 006284392 1 1166016 Proteobacteria
Dickeya dadantii subsp dieffenbachiae NCPPB 2976 WP 038921623 1 Ddadantii
DAQ1742 DAQ1742 01464 Daquatica
863 DAQ1742 01464 GI6 Seed
Dickeya aquatica DW 0440 WP 035342962 1 Daquatica
Dickeya sp CSL RW240 WP 035342962 1 Daquatica
Enterobacter lignolyticus SCF1 YP 003940913 1 701347 Proteobacteria
Enterobacter sp 638 YP 001165599 1 399742 Proteobacteria
Klebsiella pneumoniae 342 YP 002240711 1 507522 Proteobacteria
Pantoea vagans C9 1 YP 003933422 1 712898 Proteobacteria
Enterobacter lignolyticus SCF1 YP 003942098 1 701347 Proteobacteria
Shimwellia blattae DSM 4481 NBRC 105725 YP 006319404 1 630626 Proteobacteria
Dickeya gs poaceaephila NCPPB 569 WP 042868904 1 Dpoaceaephila
Dickeya gs fangzhongdai MK7 WP 038917295 1 Dfangzhongdai
Dickeya gs fangzhongdai ND14b WP 038658994 1 Dfangzhongdai
Dickeya gs fangzhongdai M074 WP 038658994 1 Dfangzhongdai
Dickeya dadantii NCPPB 898 WP 071604603 1 Ddadantii
Dickeya gs fangzhongdai MK7 WP 038917299 1 Dfangzhongdai
Dickeya gs fangzhongdai B16 WP 038917299 1 Dfangzhongdai

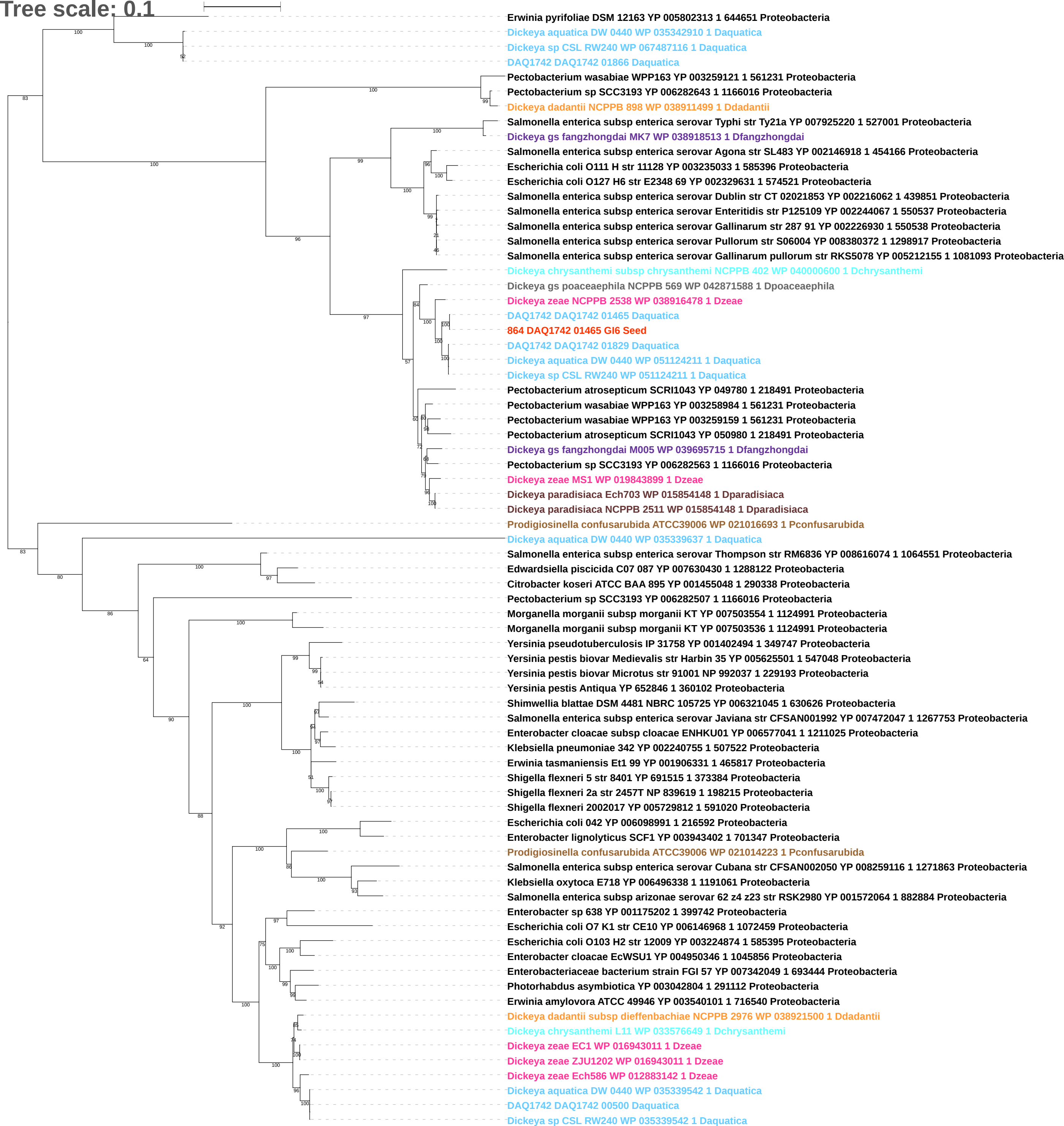
Tree scale: 0.1
Erwinia pyrifoliae DSM 12163 YP 005802313 1 644651 Proteobacteria
Dickeya aquatica DW 0440 WP 035342910 1 Daquatica
Dickeya sp CSL RW240 WP 067487116 1 Daquatica
DAQ1742 DAQ1742 01866 Daquatica
Pectobacterium wasabiae WPP163 YP 003259121 1 561231 Proteobacteria
Pectobacterium sp SCC3193 YP 006282643 1 1166016 Proteobacteria
Dickeya dadantii NCPPB 898 WP 038911499 1 Ddadantii
Salmonella enterica subsp enterica serovar Typhi str Ty21a YP 007925220 1 527001 Proteobacteria
Dickeya gs fangzhongdai MK7 WP 038918513 1 Dfangzhongdai
Salmonella enterica subsp enterica serovar Agona str SL483 YP 002146918 1 454166 Proteobacteria
Escherichia coli O111 H str 11128 YP 003235033 1 585396 Proteobacteria
Escherichia coli O127 H6 str E2348 69 YP 002329631 1 574521 Proteobacteria
Salmonella enterica subsp enterica serovar Dublin str CT 02021853 YP 002216062 1 439851 Proteobacteria
Salmonella enterica subsp enterica serovar Enteritidis str P125109 YP 002244067 1 550537 Proteobacteria
Salmonella enterica subsp enterica serovar Gallinarum str 287 91 YP 002226930 1 550538 Proteobacteria
Salmonella enterica subsp enterica serovar Pullorum str S06004 YP 008380372 1 1298917 Proteobacteria
Salmonella enterica subsp enterica serovar Gallinarum pullorum str RKS5078 YP 005212155 1 1081093 Proteobacteria
Dickeya chrysanthemi subsp chrysanthemi NCPPB 402 WP 040000600 1 Dchrysanthemi
Dickeya gs poaceaephila NCPPB 569 WP 042871588 1 Dpoaceaephila
Dickeya zeae NCPPB 2538 WP 038916478 1 Dzeae
DAQ1742 DAQ1742 01465 Daquatica
864 DAQ1742 01465 GI6 Seed
DAQ1742 DAQ1742 01829 Daquatica
Dickeya aquatica DW 0440 WP 051124211 1 Daquatica
Dickeya sp CSL RW240 WP 051124211 1 Daquatica
Pectobacterium atrosepticum SCRI1043 YP 049780 1 218491 Proteobacteria
Pectobacterium wasabiae WPP163 YP 003258984 1 561231 Proteobacteria
Pectobacterium wasabiae WPP163 YP 003259159 1 561231 Proteobacteria
Pectobacterium atrosepticum SCRI1043 YP 050980 1 218491 Proteobacteria
Dickeya gs fangzhongdai M005 WP 039695715 1 Dfangzhongdai
Pectobacterium sp SCC3193 YP 006282563 1 1166016 Proteobacteria
Dickeya zeae MS1 WP 019843899 1 Dzeae
Dickeya paradisiaca Ech703 WP 015854148 1 Dparadisiaca
Dickeya paradisiaca NCPPB 2511 WP 015854148 1 Dparadisiaca
Prodigiosinella confusarubida ATCC39006 WP 021016693 1 Pconfusarubida
Dickeya aquatica DW 0440 WP 035339637 1 Daquatica
Salmonella enterica subsp enterica serovar Thompson str RM6836 YP 008616074 1 1064551 Proteobacteria
Edwardsiella piscicida C07 087 YP 007630430 1 1288122 Proteobacteria
Citrobacter koseri ATCC BAA 895 YP 001455048 1 290338 Proteobacteria
Pectobacterium sp SCC3193 YP 006282507 1 1166016 Proteobacteria
Morganella morganii subsp morganii KT YP 007503554 1 1124991 Proteobacteria
Morganella morganii subsp morganii KT YP 007503536 1 1124991 Proteobacteria
Yersinia pseudotuberculosis IP 31758 YP 001402494 1 349747 Proteobacteria
Yersinia pestis biovar Medievalis str Harbin 35 YP 005625501 1 547048 Proteobacteria
Yersinia pestis biovar Microtus str 91001 NP 992037 1 229193 Proteobacteria
Yersinia pestis Antiqua YP 652846 1 360102 Proteobacteria
Shimwellia blattae DSM 4481 NBRC 105725 YP 006321045 1 630626 Proteobacteria
Salmonella enterica subsp enterica serovar Javiana str CFSAN001992 YP 007472047 1 1267753 Proteobacteria
Enterobacter cloacae subsp cloacae ENHKU01 YP 006577041 1 1211025 Proteobacteria
Klebsiella pneumoniae 342 YP 002240755 1 507522 Proteobacteria
Erwinia tasmaniensis Et1 99 YP 001906331 1 465817 Proteobacteria
Shigella flexneri 5 str 8401 YP 691515 1 373384 Proteobacteria
Shigella flexneri 2a str 2457T NP 839619 1 198215 Proteobacteria
Shigella flexneri 2002017 YP 005729812 1 591020 Proteobacteria
Escherichia coli 042 YP 006098991 1 216592 Proteobacteria
Enterobacter lignolyticus SCF1 YP 003943402 1 701347 Proteobacteria
Prodigiosinella confusarubida ATCC39006 WP 021014223 1 Pconfusarubida
Salmonella enterica subsp enterica serovar Cubana str CFSAN002050 YP 008259116 1 1271863 Proteobacteria
Klebsiella oxytoca E718 YP 006496338 1 1191061 Proteobacteria
Salmonella enterica subsp arizonae serovar 62 z4 z23 str RSK2980 YP 001572064 1 882884 Proteobacteria
Enterobacter sp 638 YP 001175202 1 399742 Proteobacteria
Escherichia coli O7 K1 str CE10 YP 006146968 1 1072459 Proteobacteria
Escherichia coli O103 H2 str 12009 YP 003224874 1 585395 Proteobacteria
Enterobacter cloacae EcWSU1 YP 004950346 1 1045856 Proteobacteria
Enterobacteriaceae bacterium strain FGI 57 YP 007342049 1 693444 Proteobacteria
Photorhabdus asymbiotica YP 003042804 1 291112 Proteobacteria
Erwinia amylovora ATCC 49946 YP 003540101 1 716540 Proteobacteria
Dickeya dadantii subsp dieffenbachiae NCPPB 2976 WP 038921500 1 Ddadantii
Dickeya chrysanthemi L11 WP 033576649 1 Dchrysanthemi
Dickeya zeae EC1 WP 016943011 1 Dzeae
Dickeya zeae ZJU1202 WP 016943011 1 Dzeae
Dickeya zeae Ech586 WP 012883142 1 Dzeae
Dickeya aquatica DW 0440 WP 035339542 1 Daquatica
DAQ1742 DAQ1742 00500 Daquatica
Dickeya sp CSL RW240 WP 035339542 1 Daquatica

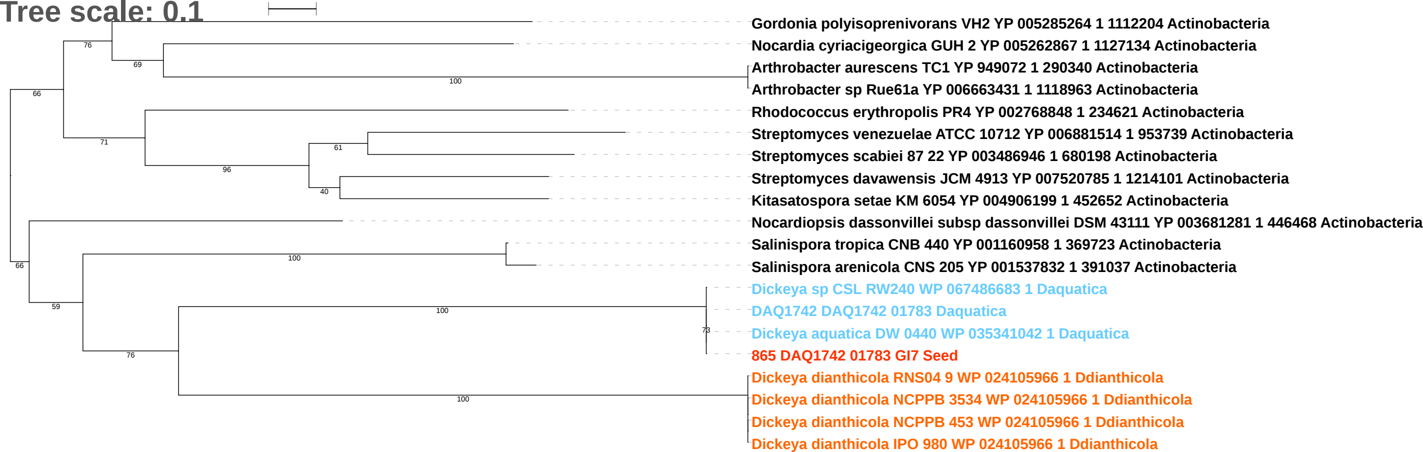
Tree scale: 0.1
Gordonia polyisoprenivorans VH2 YP 005285264 1 1112204 Actinobacteria
Nocardia cyriacigeorgica GUH 2 YP 005262867 1 1127134 Actinobacteria
Arthrobacter aurescens TC1 YP 949072 1 290340 Actinobacteria
Arthrobacter sp Rue61a YP 006663431 1 1118963 Actinobacteria
Rhodococcus erythropolis PR4 YP 002768848 1 234621 Actinobacteria
Streptomyces venezuelae ATCC 10712 YP 006881514 1 953739 Actinobacteria
Streptomyces scabiei 87 22 YP 003486946 1 680198 Actinobacteria
Streptomyces davawensis JCM 4913 YP 007520785 1 1214101 Actinobacteria
Kitasatospora setae KM 6054 YP 004906199 1 452652 Actinobacteria
Nocardiopsis dassonvillei subsp dassonvillei DSM 43111 YP 003681281 1 446468 Actinobacteria
Salinispora tropica CNB 440 YP 001160958 1 369723 Actinobacteria
Salinispora arenicola CNS 205 YP 001537832 1 391037 Actinobacteria
Dickeya sp CSL RW240 WP 067486683 1 Daquatica
DAQ1742 DAQ1742 01783 Daquatica
Dickeya aquatica DW 0440 WP 035341042 1 Daquatica
865 DAQ1742 01783 GI7 Seed
Dickeya dianthicola RNS04 9 WP 024105966 1 Ddianthicola
Dickeya dianthicola NCPPB 3534 WP 024105966 1 Ddianthicola
Dickeya dianthicola NCPPB 453 WP 024105966 1 Ddianthicola
Dickeya dianthicola IPO 980 WP 024105966 1 Ddianthicola
76
69
100
66
71
61
96
40
100
66
59
100
73
76
100

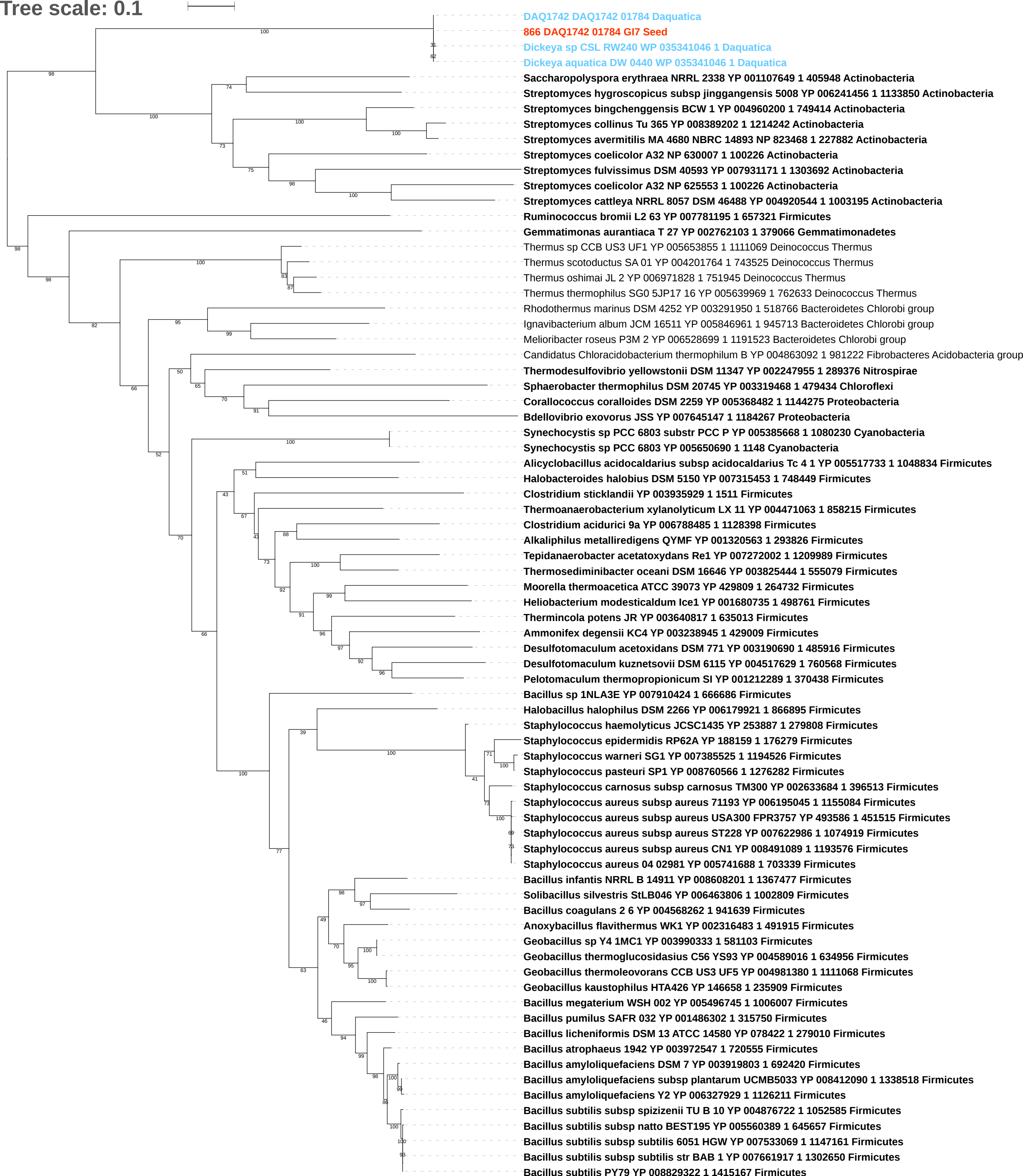
Tree scale: 0.1
DAQ1742 DAQ1742 01784 Daquatica
866 DAQ1742 01784 GI7 Seed
Dickeya sp CSL RW240 WP 035341046 1 Daquatica
Dickeya aquatica DW 0440 WP 035341046 1 Daquatica
Saccharopolyspora erythraea NRRL 2338 YP 001107649 1 405948 Actinobacteria
Streptomyces hygroscopicus subsp jinggangensis 5008 YP 006241456 1 1133850 Actinobacteria
Streptomyces bingchenggensis BCW 1 YP 004960200 1 749414 Actinobacteria
Streptomyces collinus Tu 365 YP 008389202 1 1214242 Actinobacteria
Streptomyces avermitilis MA 4680 NBRC 14893 NP 823468 1 227882 Actinobacteria
Streptomyces coelicolor A32 NP 630007 1 100226 Actinobacteria
Streptomyces fulvissimus DSM 40593 YP 007931171 1 1303692 Actinobacteria
Streptomyces coelicolor A32 NP 625553 1 100226 Actinobacteria
Streptomyces cattleya NRRL 8057 DSM 46488 YP 004920544 1 1003195 Actinobacteria
Ruminococcus bromii L2 63 YP 007781195 1 657321 Firmicutes
Gemmatimonas aurantiaca T 27 YP 002762103 1 379066 Gemmatimonadetes
Thermus sp CCB US3 UF1 YP 005653855 1 1111069 Deinococcus Thermus
Thermus scotoductus SA 01 YP 004201764 1 743525 Deinococcus Thermus
Thermus oshimai JL 2 YP 006971828 1 751945 Deinococcus Thermus
Thermus thermophilus SG0 5JP17 16 YP 005639969 1 762633 Deinococcus Thermus
Rhodothermus marinus DSM 4252 YP 003291950 1 518766 Bacteroidetes Chlorobi group
Ignavibacterium album JCM 16511 YP 005846961 1 945713 Bacteroidetes Chlorobi group
Melioribacter roseus P3M 2 YP 006528699 1 1191523 Bacteroidetes Chlorobi group
Candidatus Chloracidobacterium thermophilum B YP 004863092 1 981222 Fibrobacteres Acidobacteria group
Thermodesulfovibrio yellowstonii DSM 11347 YP 002247955 1 289376 Nitrospirae
Sphaerobacter thermophilus DSM 20745 YP 003319468 1 479434 Chloroflexi
Corallococcus coralloides DSM 2259 YP 005368482 1 1144275 Proteobacteria
Bdellovibrio exovorus JSS YP 007645147 1 1184267 Proteobacteria
Synechocystis sp PCC 6803 substr PCC P YP 005385668 1 1080230 Cyanobacteria
Synechocystis sp PCC 6803 YP 005650690 1 1148 Cyanobacteria
Alicyclobacillus acidocaldarius subsp acidocaldarius Tc 4 1 YP 005517733 1 1048834 Firmicutes
Halobacteroides halobius DSM 5150 YP 007315453 1 748449 Firmicutes
Clostridium sticklandii YP 003935929 1 1511 Firmicutes
Thermoanaerobacterium xylanolyticum LX 11 YP 004471063 1 858215 Firmicutes
Clostridium acidurici 9a YP 006788485 1 1128398 Firmicutes
Alkaliphilus metalliredigens QYMF YP 001320563 1 293826 Firmicutes
Tepidanaerobacter acetatoxydans Re1 YP 007272002 1 1209989 Firmicutes
Thermosediminibacter oceani DSM 16646 YP 003825444 1 555079 Firmicutes
Moorella thermoacetica ATCC 39073 YP 429809 1 264732 Firmicutes
Heliobacterium modesticaldum Ice1 YP 001680735 1 498761 Firmicutes
Thermincola potens JR YP 003640817 1 635013 Firmicutes
Ammonifex degensii KC4 YP 003238945 1 429009 Firmicutes
Desulfotomaculum acetoxidans DSM 771 YP 003190690 1 485916 Firmicutes
Desulfotomaculum kuznetsovii DSM 6115 YP 004517629 1 760568 Firmicutes
Pelotomaculum thermopropionicum SI YP 001212289 1 370438 Firmicutes
Bacillus sp 1NLA3E YP 007910424 1 666686 Firmicutes
Halobacillus halophilus DSM 2266 YP 006179921 1 866895 Firmicutes
Staphylococcus haemolyticus JCSC1435 YP 253887 1 279808 Firmicutes
Staphylococcus epidermidis RP62A YP 188159 1 176279 Firmicutes
Staphylococcus warneri SG1 YP 007385525 1 1194526 Firmicutes
Staphylococcus pasteuri SP1 YP 008760566 1 1276282 Firmicutes
Staphylococcus carnosus subsp carnosus TM300 YP 002633684 1 396513 Firmicutes
Staphylococcus aureus subsp aureus 71193 YP 006195045 1 1155084 Firmicutes
Staphylococcus aureus subsp aureus USA300 FPR3757 YP 493586 1 451515 Firmicutes
Staphylococcus aureus subsp aureus ST228 YP 007622986 1 1074919 Firmicutes
Staphylococcus aureus subsp aureus CN1 YP 008491089 1 1193576 Firmicutes
Staphylococcus aureus 04 02981 YP 005741688 1 703339 Firmicutes
Bacillus infantis NRRL B 14911 YP 008608201 1 1367477 Firmicutes
Solibacillus silvestris StLB046 YP 006463806 1 1002809 Firmicutes
Bacillus coagulans 2 6 YP 004568262 1 941639 Firmicutes
Anoxybacillus flavithermus WK1 YP 002316483 1 491915 Firmicutes
Geobacillus sp Y4 1MC1 YP 003990333 1 581103 Firmicutes
Geobacillus thermoglucosidasius C56 YS93 YP 004589016 1 634956 Firmicutes
Geobacillus thermoleovorans CCB US3 UF5 YP 004981380 1 1111068 Firmicutes
Geobacillus kaustophilus HTA426 YP 146658 1 235909 Firmicutes
Bacillus megaterium WSH 002 YP 005496745 1 1006007 Firmicutes
Bacillus pumilus SAFR 032 YP 001486302 1 315750 Firmicutes
Bacillus licheniformis DSM 13 ATCC 14580 YP 078422 1 279010 Firmicutes
Bacillus atrophaeus 1942 YP 003972547 1 720555 Firmicutes
Bacillus amyloliquefaciens DSM 7 YP 003919803 1 692420 Firmicutes
Bacillus amyloliquefaciens subsp plantarum UCMB5033 YP 008412090 1 1338518 Firmicutes
Bacillus amyloliquefaciens Y2 YP 006327929 1 1126211 Firmicutes
Bacillus subtilis subsp spizizenii TU B 10 YP 004876722 1 1052585 Firmicutes
Bacillus subtilis subsp natto BEST195 YP 005560389 1 645657 Firmicutes
Bacillus subtilis subsp subtilis 6051 HGW YP 007533069 1 1147161 Firmicutes
Bacillus subtilis subsp subtilis str BAB 1 YP 007661917 1 1302650 Firmicutes
Bacillus subtilis PY79 YP 008829322 1 1415167 Firmicutes

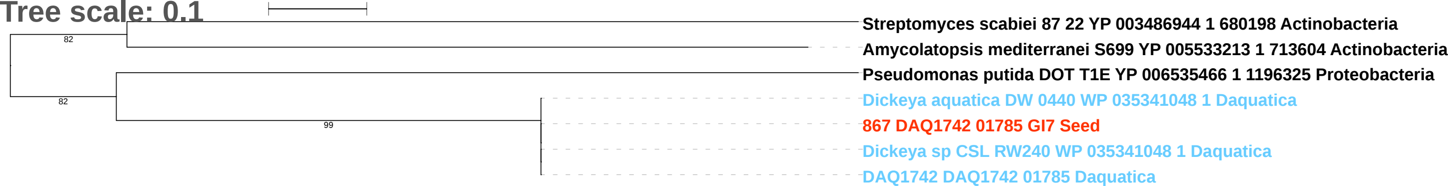
Tree scale: 0.1
Streptomyces scabiei 87 22 YP 003486944 1 680198 Actinobacteria
Amycolatopsis mediterranei S699 YP 005533213 1 713604 Actinobacteria
Pseudomonas putida DOT T1E YP 006535466 1 1196325 Proteobacteria
Dickeya aquatica DW 0440 WP 035341048 1 Daquatica
867 DAQ1742 01785 GI7 Seed
Dickeya sp CSL RW240 WP 035341048 1 Daquatica
DAQ1742 DAQ1742 01785 Daquatica
82
82
99

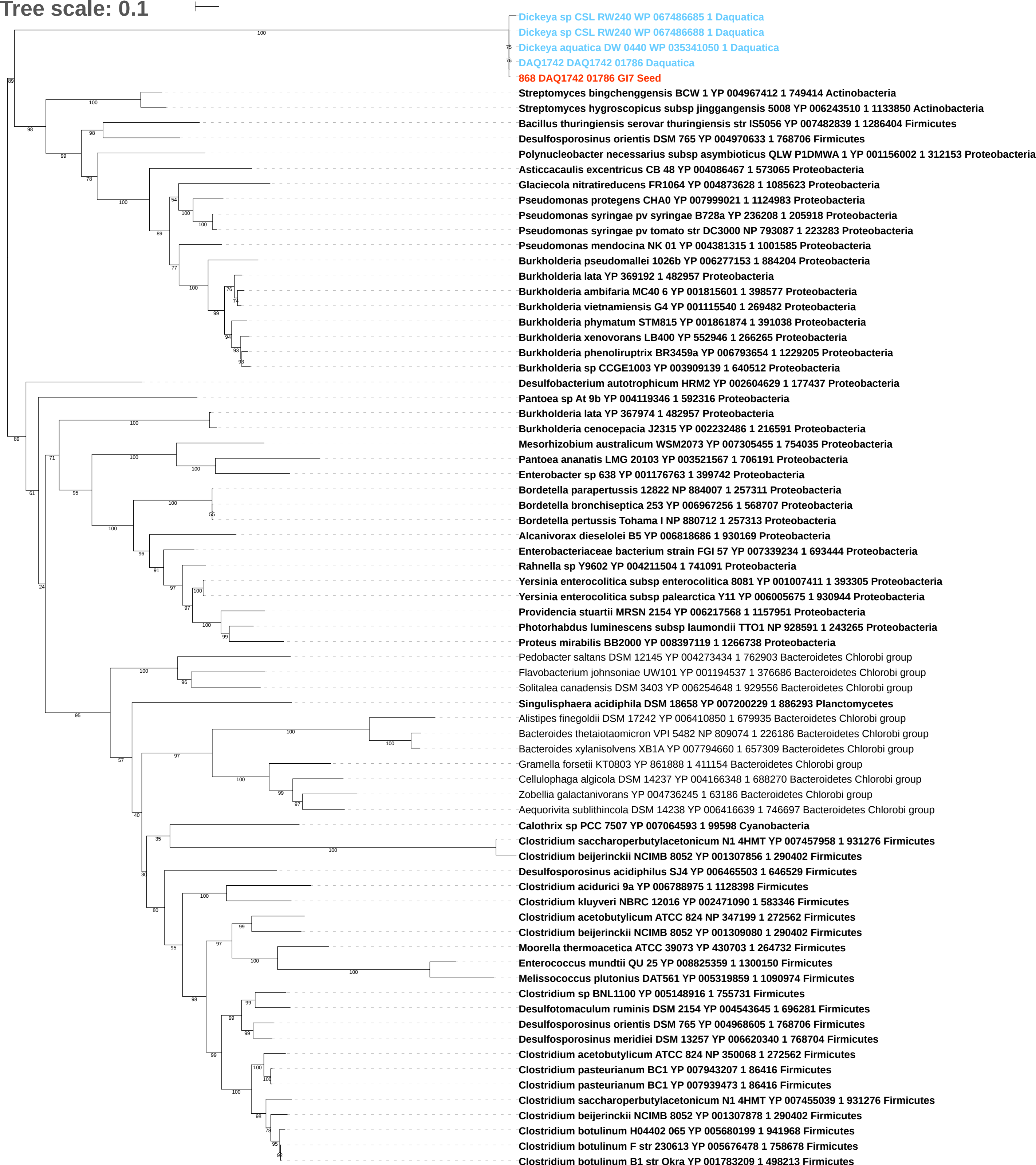

Tree scale: 0.1
Dickeya sp CSL RW240 WP 067486685 1 Daquatica
Dickeya sp CSL RW240 WP 067486688 1 Daquatica
Dickeya aquatica DW 0440 WP 035341050 1 Daquatica
DAQ1742 DAQ1742 01786 Daquatica
868 DAQ1742 01786 GI7 Seed
Streptomyces bingchenggensis BCW 1 YP 004967412 1 749414 Actinobacteria
Streptomyces hygroscopicus subsp jinggangensis 5008 YP 006243510 1 1133850 Actinobacteria
Bacillus thuringiensis serovar thuringiensis str IS5056 YP 007482839 1 1286404 Firmicutes
Desulfosporosinus orientis DSM 765 YP 004970633 1 768706 Firmicutes
Polynucleobacter necessarius subsp asymbioticus QLW P1DMWA 1 YP 001156002 1 312153 Proteobacteria
Asticcacaulis excentricus CB 48 YP 004086467 1 573065 Proteobacteria
Glaciecola nitratireducens FR1064 YP 004873628 1 1085623 Proteobacteria
Pseudomonas protegens CHA0 YP 007999021 1 1124983 Proteobacteria
Pseudomonas syringae pv syringae B728a YP 236208 1 205918 Proteobacteria
Pseudomonas syringae pv tomato str DC3000 NP 793087 1 223283 Proteobacteria
Pseudomonas mendocina NK 01 YP 004381315 1 1001585 Proteobacteria
Burkholderia pseudomallei 1026b YP 006277153 1 884204 Proteobacteria
Burkholderia lata YP 369192 1 482957 Proteobacteria
Burkholderia ambifaria MC40 6 YP 001815601 1 398577 Proteobacteria
Burkholderia vietnamiensis G4 YP 001115540 1 269482 Proteobacteria
Burkholderia phymatum STM815 YP 001861874 1 391038 Proteobacteria
Burkholderia xenovorans LB400 YP 552946 1 266265 Proteobacteria
Burkholderia phenoliruptrix BR3459a YP 006793654 1 1229205 Proteobacteria
Burkholderia sp CCGE1003 YP 003909139 1 640512 Proteobacteria
Desulfobacterium autotrophicum HRM2 YP 002604629 1 177437 Proteobacteria
Pantoea sp At 9b YP 004119346 1 592316 Proteobacteria
Burkholderia lata YP 367974 1 482957 Proteobacteria
Burkholderia cenocepacia J2315 YP 002232486 1 216591 Proteobacteria
Mesorhizobium australicum WSM2073 YP 007305455 1 754035 Proteobacteria
Pantoea ananatis LMG 20103 YP 003521567 1 706191 Proteobacteria
Enterobacter sp 638 YP 001176763 1 399742 Proteobacteria
Bordetella parapertussis 12822 NP 884007 1 257311 Proteobacteria
Bordetella bronchiseptica 253 YP 006967256 1 568707 Proteobacteria
Bordetella pertussis Tohama I NP 880712 1 257313 Proteobacteria
Alcanivorax dieselolei B5 YP 006818686 1 930169 Proteobacteria
Enterobacteriaceae bacterium strain FGI 57 YP 007339234 1 693444 Proteobacteria
Rahnella sp Y9602 YP 004211504 1 741091 Proteobacteria
Yersinia enterocolitica subsp enterocolitica 8081 YP 001007411 1 393305 Proteobacteria
Yersinia enterocolitica subsp palearctica Y11 YP 006005675 1 930944 Proteobacteria
Providencia stuartii MRSN 2154 YP 006217568 1 1157951 Proteobacteria
Photorhabdus luminescens subsp laumondii TTO1 NP 928591 1 243265 Proteobacteria
Proteus mirabilis BB2000 YP 008397119 1 1266738 Proteobacteria
Pedobacter saltans DSM 12145 YP 004273434 1 762903 Bacteroidetes Chlorobi group
Flavobacterium johnsoniae UW101 YP 001194537 1 376686 Bacteroidetes Chlorobi group
Solitalea canadensis DSM 3403 YP 006254648 1 929556 Bacteroidetes Chlorobi group
Singulisphaera acidiphila DSM 18658 YP 007200229 1 886293 Planctomycetes
Alistipes finegoldii DSM 17242 YP 006410850 1 679935 Bacteroidetes Chlorobi group
Bacteroides thetaiotaomicron VPI 5482 NP 809074 1 226186 Bacteroidetes Chlorobi group
Bacteroides xylanisolvens XB1A YP 007794660 1 657309 Bacteroidetes Chlorobi group
Gramella forsetii KT0803 YP 861888 1 411154 Bacteroidetes Chlorobi group
Cellulophaga algicola DSM 14237 YP 004166348 1 688270 Bacteroidetes Chlorobi group
Zobellia galactanivorans YP 004736245 1 63186 Bacteroidetes Chlorobi group
Aequorivita sublithincola DSM 14238 YP 006416639 1 746697 Bacteroidetes Chlorobi group
Calothrix sp PCC 7507 YP 007064593 1 99598 Cyanobacteria
Clostridium saccharoperbutylacetonicum N1 4HMT YP 007457958 1 931276 Firmicutes
Clostridium beijerinckii NCIMB 8052 YP 001307856 1 290402 Firmicutes
Desulfosporosinus acidiphilus SJ4 YP 006465503 1 646529 Firmicutes
Clostridium acidurici 9a YP 006788975 1 1128398 Firmicutes
Clostridium kluyveri NBRC 12016 YP 002471090 1 583346 Firmicutes
Clostridium acetobutylicum ATCC 824 NP 347199 1 272562 Firmicutes
Clostridium beijerinckii NCIMB 8052 YP 001309080 1 290402 Firmicutes
Moorella thermoacetica ATCC 39073 YP 430703 1 264732 Firmicutes
Enterococcus mundtii QU 25 YP 008825359 1 1300150 Firmicutes
Melissococcus plutonius DAT561 YP 005319859 1 1090974 Firmicutes
Clostridium sp BNL1100 YP 005148916 1 755731 Firmicutes
Desulfotomaculum ruminis DSM 2154 YP 004543645 1 696281 Firmicutes
Desulfosporosinus orientis DSM 765 YP 004968605 1 768706 Firmicutes
Desulfosporosinus meridiei DSM 13257 YP 006620340 1 768704 Firmicutes
Clostridium acetobutylicum ATCC 824 NP 350068 1 272562 Firmicutes
Clostridium pasteurianum BC1 YP 007943207 1 86416 Firmicutes
Clostridium pasteurianum BC1 YP 007939473 1 86416 Firmicutes
Clostridium saccharoperbutylacetonicum N1 4HMT YP 007455039 1 931276 Firmicutes
Clostridium beijerinckii NCIMB 8052 YP 001307878 1 290402 Firmicutes
Clostridium botulinum H04402 065 YP 005680199 1 941968 Firmicutes
Clostridium botulinum F str 230613 YP 005676478 1 758678 Firmicutes
Clostridium botulinum B1 str Okra YP 001783209 1 498213 Firmicutes

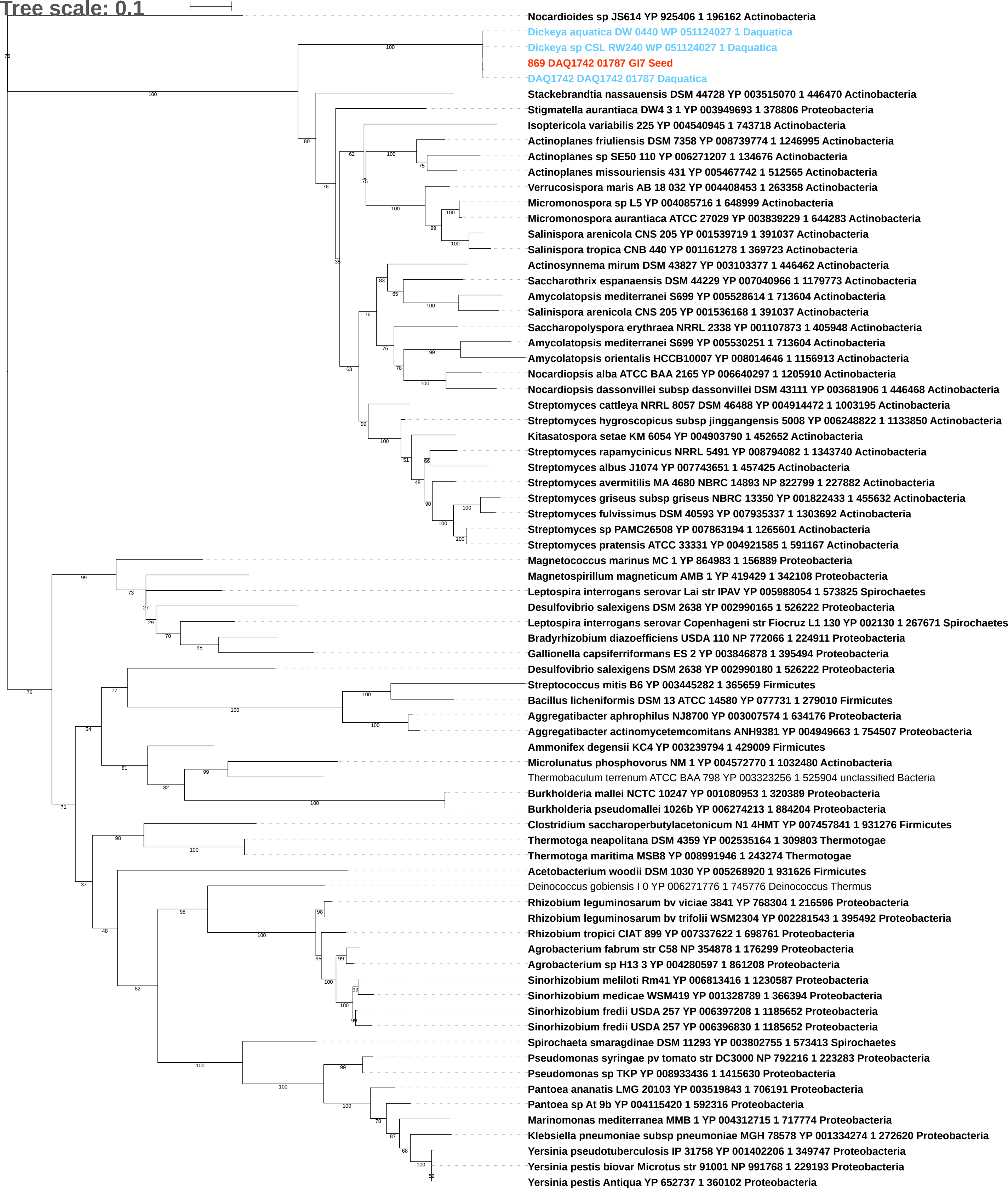
Tree scale: 0.1
Nocardioides sp JS614 YP 925406 1 196162 Actinobacteria
Dickeya aquatica DW 0440 WP 051124027 1 Daquatica
Dickeya sp CSL RW240 WP 051124027 1 Daquatica
869 DAQ1742 01787 GI7 Seed
DAQ1742 DAQ1742 01787 Daquatica
Stackebrandtia nassauensis DSM 44728 YP 003515070 1 446470 Actinobacteria
Stigmatella aurantiaca DW4 3 1 YP 003949693 1 378806 Proteobacteria
Isoptericola variabilis 225 YP 004540945 1 743718 Actinobacteria
Actinoplanes friuliensis DSM 7358 YP 008739774 1 1246995 Actinobacteria
Actinoplanes sp SE50 110 YP 006271207 1 134676 Actinobacteria
Actinoplanes missouriensis 431 YP 005467742 1 512565 Actinobacteria
Verrucosispora maris AB 18 032 YP 004408453 1 263358 Actinobacteria
Micromonospora sp L5 YP 004085716 1 648999 Actinobacteria
Micromonospora aurantiaca ATCC 27029 YP 003839229 1 644283 Actinobacteria
Salinispora arenicola CNS 205 YP 001539719 1 391037 Actinobacteria
Salinispora tropica CNB 440 YP 001161278 1 369723 Actinobacteria
Actinosynnema mirum DSM 43827 YP 003103377 1 446462 Actinobacteria
Saccharothrix espanaensis DSM 44229 YP 007040966 1 1179773 Actinobacteria
Amycolatopsis mediterranei S699 YP 005528614 1 713604 Actinobacteria
Salinispora arenicola CNS 205 YP 001536168 1 391037 Actinobacteria
Saccharopolyspora erythraea NRRL 2338 YP 001107873 1 405948 Actinobacteria
Amycolatopsis mediterranei S699 YP 005530251 1 713604 Actinobacteria
Amycolatopsis orientalis HCCB10007 YP 008014646 1 1156913 Actinobacteria
Nocardiopsis alba ATCC BAA 2165 YP 006640297 1 1205910 Actinobacteria
Nocardiopsis dassonvillei subsp dassonvillei DSM 43111 YP 003681906 1 446468 Actinobacteria
Streptomyces cattleya NRRL 8057 DSM 46488 YP 004914472 1 1003195 Actinobacteria
Streptomyces hygroscopicus subsp jinggangensis 5008 YP 006248822 1 1133850 Actinobacteria
Kitasatospora setae KM 6054 YP 004903790 1 452652 Actinobacteria
Streptomyces rapamycinicus NRRL 5491 YP 008794082 1 1343740 Actinobacteria
Streptomyces albus J1074 YP 007743651 1 457425 Actinobacteria
Streptomyces avermitilis MA 4680 NBRC 14893 NP 822799 1 227882 Actinobacteria
Streptomyces griseus subsp griseus NBRC 13350 YP 001822433 1 455632 Actinobacteria
Streptomyces fulvissimus DSM 40593 YP 007935337 1 1303692 Actinobacteria
Streptomyces sp PAMC26508 YP 007863194 1 1265601 Actinobacteria
Streptomyces pratensis ATCC 33331 YP 004921585 1 591167 Actinobacteria
Magnetococcus marinus MC 1 YP 864983 1 156889 Proteobacteria
Magnetospirillum magneticum AMB 1 YP 419429 1 342108 Proteobacteria
Leptospira interrogans serovar Lai str IPAV YP 005988054 1 573825 Spirochaetes
Desulfovibrio salexigens DSM 2638 YP 002990165 1 526222 Proteobacteria
Leptospira interrogans serovar Copenhageni str Fiocruz L1 130 YP 002130 1 267671 Spirochaetes
Bradyrhizobium diazoefficiens USDA 110 NP 772066 1 224911 Proteobacteria
Gallionella capsiferriformans ES 2 YP 003846878 1 395494 Proteobacteria
Desulfovibrio salexigens DSM 2638 YP 002990180 1 526222 Proteobacteria
Streptococcus mitis B6 YP 003445282 1 365659 Firmicutes
Bacillus licheniformis DSM 13 ATCC 14580 YP 077731 1 279010 Firmicutes
Aggregatibacter aphrophilus NJ8700 YP 003007574 1 634176 Proteobacteria
Aggregatibacter actinomycetemcomitans ANH9381 YP 004949663 1 754507 Proteobacteria
Ammonifex degensii KC4 YP 003239794 1 429009 Firmicutes
Microlunatus phosphovorus NM 1 YP 004572770 1 1032480 Actinobacteria
Thermobaculum terrenum ATCC BAA 798 YP 003323256 1 525904 unclassified Bacteria
Burkholderia mallei NCTC 10247 YP 001080953 1 320389 Proteobacteria
Burkholderia pseudomallei 1026b YP 006274213 1 884204 Proteobacteria
Clostridium saccharoperbutylacetonicum N1 4HMT YP 007457841 1 931276 Firmicutes
Thermotoga neapolitana DSM 4359 YP 002535164 1 309803 Thermotogae
Thermotoga maritima MSB8 YP 008991946 1 243274 Thermotogae
Acetobacterium woodii DSM 1030 YP 005268920 1 931626 Firmicutes
Deinococcus gobiensis I 0 YP 006271776 1 745776 Deinococcus Thermus
Rhizobium leguminosarum bv viciae 3841 YP 768304 1 216596 Proteobacteria
Rhizobium leguminosarum bv trifolii WSM2304 YP 002281543 1 395492 Proteobacteria
Rhizobium tropici CIAT 899 YP 007337622 1 698761 Proteobacteria
Agrobacterium fabrum str C58 NP 354878 1 176299 Proteobacteria
Agrobacterium sp H13 3 YP 004280597 1 861208 Proteobacteria
Sinorhizobium meliloti Rm41 YP 006813416 1 1230587 Proteobacteria
Sinorhizobium medicae WSM419 YP 001328789 1 366394 Proteobacteria
Sinorhizobium fredii USDA 257 YP 006397208 1 1185652 Proteobacteria
Sinorhizobium fredii USDA 257 YP 006396830 1 1185652 Proteobacteria
Spirochaeta smaragdinae DSM 11293 YP 003802755 1 573413 Spirochaetes
Pseudomonas syringae pv tomato str DC3000 NP 792216 1 223283 Proteobacteria
Pseudomonas sp TKP YP 008933436 1 1415630 Proteobacteria
Pantoea ananatis LMG 20103 YP 003519843 1 706191 Proteobacteria
Pantoea sp At 9b YP 004115420 1 592316 Proteobacteria
Marinomonas mediterranea MMB 1 YP 004312715 1 717774 Proteobacteria
Klebsiella pneumoniae subsp pneumoniae MGH 78578 YP 001334274 1 272620 Proteobacteria
Yersinia pseudotuberculosis IP 31758 YP 001402206 1 349747 Proteobacteria
Yersinia pestis biovar Microtus str 91001 NP 991768 1 229193 Proteobacteria
Yersinia pestis Antiqua YP 652737 1 360102 Proteobacteria

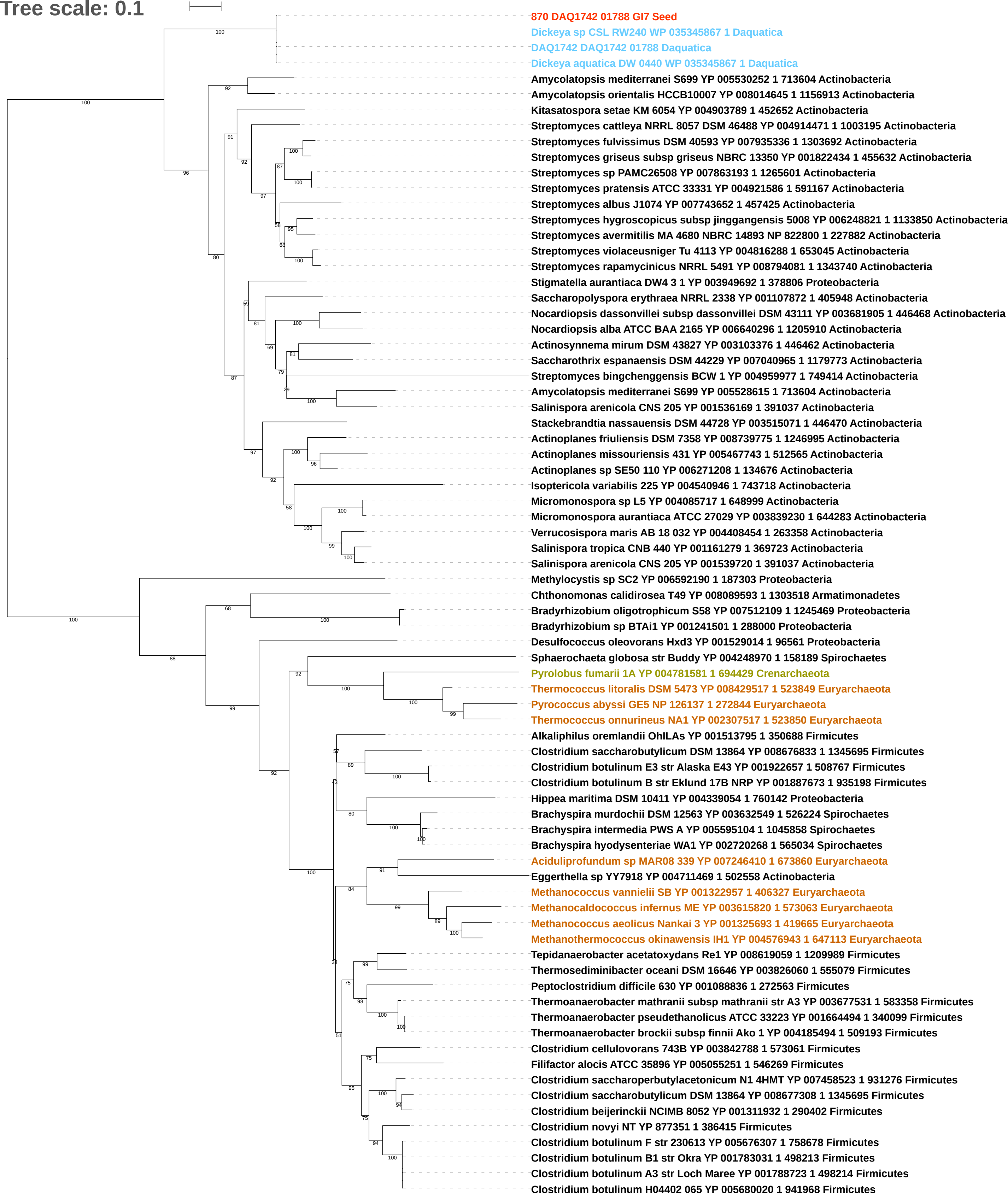

Tree scale: 0.1
870 DAQ1742 01788 GI7 Seed
Dickeya sp CSL RW240 WP 035345867 1 Daquatica
DAQ1742 DAQ1742 01788 Daquatica
Dickeya aquatica DW 0440 WP 035345867 1 Daquatica
Amycolatopsis mediterranei S699 YP 005530252 1 713604 Actinobacteria
Amycolatopsis orientalis HCCB10007 YP 008014645 1 1156913 Actinobacteria
Kitasatospora setae KM 6054 YP 004903789 1 452652 Actinobacteria
Streptomyces cattleya NRRL 8057 DSM 46488 YP 004914471 1 1003195 Actinobacteria
Streptomyces fulvissimus DSM 40593 YP 007935336 1 1303692 Actinobacteria
Streptomyces griseus subsp griseus NBRC 13350 YP 001822434 1 455632 Actinobacteria
Streptomyces sp PAMC26508 YP 007863193 1 1265601 Actinobacteria
Streptomyces pratensis ATCC 33331 YP 004921586 1 591167 Actinobacteria
Streptomyces albus J1074 YP 007743652 1 457425 Actinobacteria
Streptomyces hygroscopicus subsp jinggangensis 5008 YP 006248821 1 1133850 Actinobacteria
Streptomyces avermitilis MA 4680 NBRC 14893 NP 822800 1 227882 Actinobacteria
Streptomyces violaceusniger Tu 4113 YP 004816288 1 653045 Actinobacteria
Streptomyces rapamycinicus NRRL 5491 YP 008794081 1 1343740 Actinobacteria
Stigmatella aurantiaca DW4 3 1 YP 003949692 1 378806 Proteobacteria
Saccharopolyspora erythraea NRRL 2338 YP 001107872 1 405948 Actinobacteria
Nocardiopsis dassonvillei subsp dassonvillei DSM 43111 YP 003681905 1 446468 Actinobacteria
Nocardiopsis alba ATCC BAA 2165 YP 006640296 1 1205910 Actinobacteria
Actinosynnema mirum DSM 43827 YP 003103376 1 446462 Actinobacteria
Saccharothrix espanaensis DSM 44229 YP 007040965 1 1179773 Actinobacteria
Streptomyces bingchenggensis BCW 1 YP 004959977 1 749414 Actinobacteria
Amycolatopsis mediterranei S699 YP 005528615 1 713604 Actinobacteria
Salinispora arenicola CNS 205 YP 001536169 1 391037 Actinobacteria
Stackebrandtia nassauensis DSM 44728 YP 003515071 1 446470 Actinobacteria
Actinoplanes friuliensis DSM 7358 YP 008739775 1 1246995 Actinobacteria
Actinoplanes missouriensis 431 YP 005467743 1 512565 Actinobacteria
Actinoplanes sp SE50 110 YP 006271208 1 134676 Actinobacteria
Isoptericola variabilis 225 YP 004540946 1 743718 Actinobacteria
Micromonospora sp L5 YP 004085717 1 648999 Actinobacteria
Micromonospora aurantiaca ATCC 27029 YP 003839230 1 644283 Actinobacteria
Verrucosispora maris AB 18 032 YP 004408454 1 263358 Actinobacteria
Salinispora tropica CNB 440 YP 001161279 1 369723 Actinobacteria
Salinispora arenicola CNS 205 YP 001539720 1 391037 Actinobacteria
Methylocystis sp SC2 YP 006592190 1 187303 Proteobacteria
Chthonomonas calidirosea T49 YP 008089593 1 1303518 Armatimonadetes
Bradyrhizobium oligotrophicum S58 YP 007512109 1 1245469 Proteobacteria
Bradyrhizobium sp BTAi1 YP 001241501 1 288000 Proteobacteria
Desulfococcus oleovorans Hxd3 YP 001529014 1 96561 Proteobacteria
Sphaerochaeta globosa str Buddy YP 004248970 1 158189 Spirochaetes
Pyrolobus fumarii 1A YP 004781581 1 694429 Crenarchaeota
Thermococcus litoralis DSM 5473 YP 008429517 1 523849 Euryarchaeota
Pyrococcus abyssi GE5 NP 126137 1 272844 Euryarchaeota
Thermococcus onnurineus NA1 YP 002307517 1 523850 Euryarchaeota
Alkaliphilus oremlandii OhILAs YP 001513795 1 350688 Firmicutes
Clostridium saccharobutylicum DSM 13864 YP 008676833 1 1345695 Firmicutes
Clostridium botulinum E3 str Alaska E43 YP 001922657 1 508767 Firmicutes
Clostridium botulinum B str Eklund 17B NRP YP 001887673 1 935198 Firmicutes
Hippea maritima DSM 10411 YP 004339054 1 760142 Proteobacteria
Brachyspira murdochii DSM 12563 YP 003632549 1 526224 Spirochaetes
Brachyspira intermedia PWS A YP 005595104 1 1045858 Spirochaetes
Brachyspira hyodysenteriae WA1 YP 002720268 1 565034 Spirochaetes
Aciduliprofundum sp MAR08 339 YP 007246410 1 673860 Euryarchaeota
Eggerthella sp YY7918 YP 004711469 1 502558 Actinobacteria
Methanococcus vannielii SB YP 001322957 1 406327 Euryarchaeota
Methanocaldococcus infernus ME YP 003615820 1 573063 Euryarchaeota
Methanococcus aeolicus Nankai 3 YP 001325693 1 419665 Euryarchaeota
Methanothermococcus okinawensis IH1 YP 004576943 1 647113 Euryarchaeota
Tepidanaerobacter acetatoxydans Re1 YP 008619059 1 1209989 Firmicutes
Thermosediminibacter oceani DSM 16646 YP 003826060 1 555079 Firmicutes
Peptoclostridium difficile 630 YP 001088836 1 272563 Firmicutes
Thermoanaerobacter mathranii subsp mathranii str A3 YP 003677531 1 583358 Firmicutes
Thermoanaerobacter pseudethanolicus ATCC 33223 YP 001664494 1 340099 Firmicutes
Thermoanaerobacter brockii subsp finnii Ako 1 YP 004185494 1 509193 Firmicutes
Clostridium cellulovorans 743B YP 003842788 1 573061 Firmicutes
Filifactor alocis ATCC 35896 YP 005055251 1 546269 Firmicutes
Clostridium saccharoperbutylacetonicum N1 4HMT YP 007458523 1 931276 Firmicutes
Clostridium saccharobutylicum DSM 13864 YP 008677308 1 1345695 Firmicutes
Clostridium beijerinckii NCIMB 8052 YP 001311932 1 290402 Firmicutes
Clostridium novyi NT YP 877351 1 386415 Firmicutes
Clostridium botulinum F str 230613 YP 005676307 1 758678 Firmicutes
Clostridium botulinum B1 str Okra YP 001783031 1 498213 Firmicutes
Clostridium botulinum A3 str Loch Maree YP 001788723 1 498214 Firmicutes
Clostridium botulinum H04402 065 YP 005680020 1 941968 Firmicutes

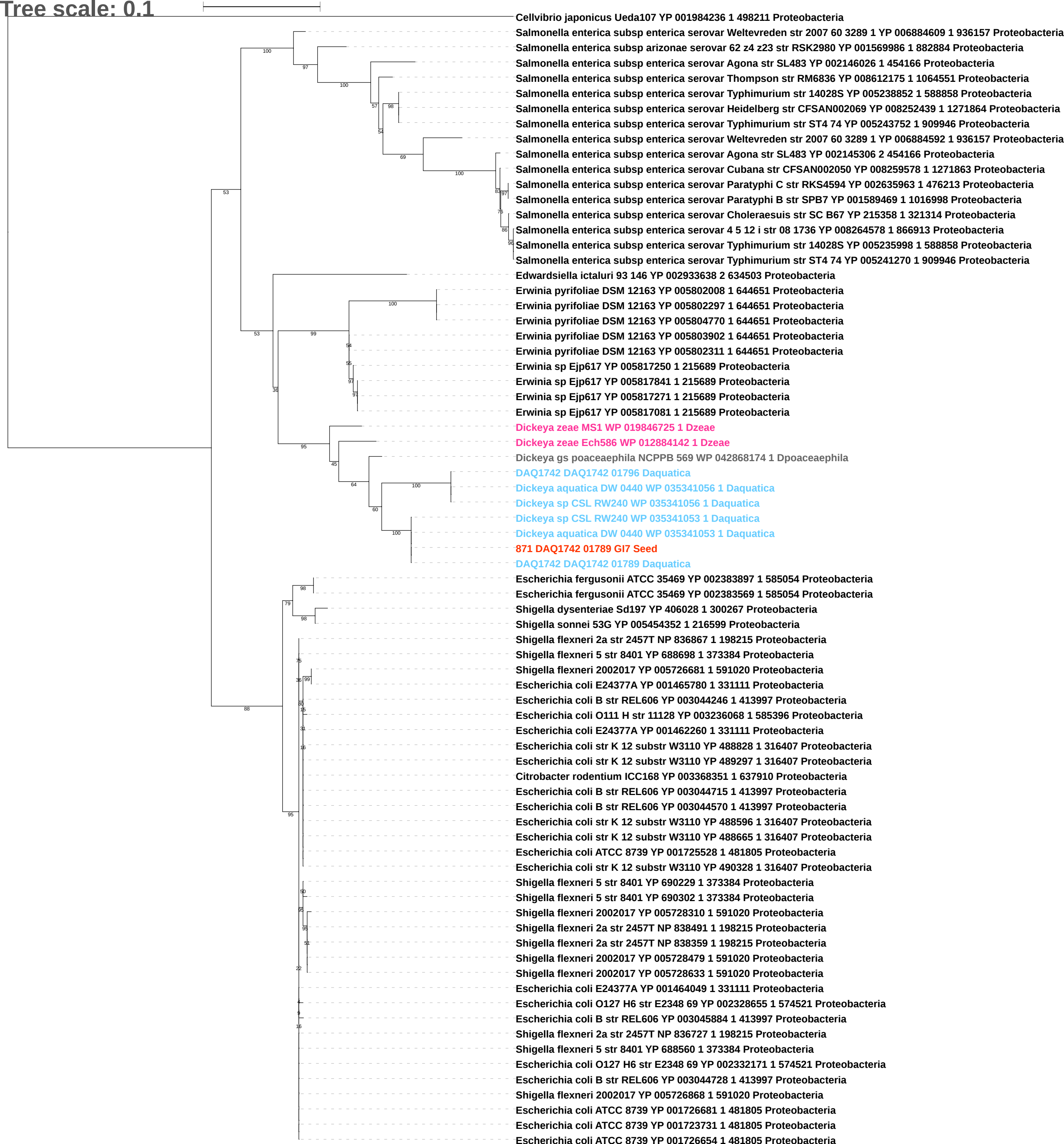
Tree scale: 0.1
Cellvibrio japonicus Ueda107 YP 001984236 1 498211 Proteobacteria
Salmonella enterica subsp enterica serovar Weltevreden str 2007 60 3289 1 YP 006884609 1 936157 Proteobacteria
Salmonella enterica subsp arizonae serovar 62 z4 z23 str RSK2980 YP 001569986 1 882884 Proteobacteria
Salmonella enterica subsp enterica serovar Agona str SL483 YP 002146026 1 454166 Proteobacteria
Salmonella enterica subsp enterica serovar Thompson str RM6836 YP 008612175 1 1064551 Proteobacteria
Salmonella enterica subsp enterica serovar Typhimurium str 14028S YP 005238852 1 588858 Proteobacteria
Salmonella enterica subsp enterica serovar Heidelberg str CFSAN002069 YP 008252439 1 1271864 Proteobacteria
Salmonella enterica subsp enterica serovar Typhimurium str ST4 74 YP 005243752 1 909946 Proteobacteria
Salmonella enterica subsp enterica serovar Weltevreden str 2007 60 3289 1 YP 006884592 1 936157 Proteobacteria
Salmonella enterica subsp enterica serovar Agona str SL483 YP 002145306 2 454166 Proteobacteria
Salmonella enterica subsp enterica serovar Cubana str CFSAN002050 YP 008259578 1 1271863 Proteobacteria
Salmonella enterica subsp enterica serovar Paratyphi C str RKS4594 YP 002635963 1 476213 Proteobacteria
Salmonella enterica subsp enterica serovar Paratyphi B str SPB7 YP 001589469 1 1016998 Proteobacteria
Salmonella enterica subsp enterica serovar Choleraesuis str SC B67 YP 215358 1 321314 Proteobacteria
Salmonella enterica subsp enterica serovar 4 5 12 i str 08 1736 YP 008264578 1 866913 Proteobacteria
Salmonella enterica subsp enterica serovar Typhimurium str 14028S YP 005235998 1 588858 Proteobacteria
Salmonella enterica subsp enterica serovar Typhimurium str ST4 74 YP 005241270 1 909946 Proteobacteria
Edwardsiella ictaluri 93 146 YP 002933638 2 634503 Proteobacteria
Erwinia pyrifoliae DSM 12163 YP 005802008 1 644651 Proteobacteria
Erwinia pyrifoliae DSM 12163 YP 005802297 1 644651 Proteobacteria
Erwinia pyrifoliae DSM 12163 YP 005804770 1 644651 Proteobacteria
Erwinia pyrifoliae DSM 12163 YP 005803902 1 644651 Proteobacteria
Erwinia pyrifoliae DSM 12163 YP 005802311 1 644651 Proteobacteria
Erwinia sp Ejp617 YP 005817250 1 215689 Proteobacteria
Erwinia sp Ejp617 YP 005817841 1 215689 Proteobacteria
Erwinia sp Ejp617 YP 005817271 1 215689 Proteobacteria
Erwinia sp Ejp617 YP 005817081 1 215689 Proteobacteria
Dickeya zeae MS1 WP 019846725 1 Dzeae
Dickeya zeae Ech586 WP 012884142 1 Dzeae
Dickeya gs poaceaephila NCPPB 569 WP 042868174 1 Dpoaceaephila
DAQ1742 DAQ1742 01796 Daquatica
Dickeya aquatica DW 0440 WP 035341056 1 Daquatica
Dickeya sp CSL RW240 WP 035341056 1 Daquatica
Dickeya sp CSL RW240 WP 035341053 1 Daquatica
Dickeya aquatica DW 0440 WP 035341053 1 Daquatica
871 DAQ1742 01789 GI7 Seed
DAQ1742 DAQ1742 01789 Daquatica
Escherichia fergusonii ATCC 35469 YP 002383897 1 585054 Proteobacteria
Escherichia fergusonii ATCC 35469 YP 002383569 1 585054 Proteobacteria
Shigella dysenteriae Sd197 YP 406028 1 300267 Proteobacteria
Shigella sonnei 53G YP 005454352 1 216599 Proteobacteria
Shigella flexneri 2a str 2457T NP 836867 1 198215 Proteobacteria
Shigella flexneri 5 str 8401 YP 688698 1 373384 Proteobacteria
Shigella flexneri 2002017 YP 005726681 1 591020 Proteobacteria
Escherichia coli E24377A YP 001465780 1 331111 Proteobacteria
Escherichia coli B str REL606 YP 003044246 1 413997 Proteobacteria
Escherichia coli O111 H str 11128 YP 003236068 1 585396 Proteobacteria
Escherichia coli E24377A YP 001462260 1 331111 Proteobacteria
Escherichia coli str K 12 substr W3110 YP 488828 1 316407 Proteobacteria
Escherichia coli str K 12 substr W3110 YP 489297 1 316407 Proteobacteria
Citrobacter rodentium ICC168 YP 003368351 1 637910 Proteobacteria
Escherichia coli B str REL606 YP 003044715 1 413997 Proteobacteria
Escherichia coli B str REL606 YP 003044570 1 413997 Proteobacteria
Escherichia coli str K 12 substr W3110 YP 488596 1 316407 Proteobacteria
Escherichia coli str K 12 substr W3110 YP 488665 1 316407 Proteobacteria
Escherichia coli ATCC 8739 YP 001725528 1 481805 Proteobacteria
Escherichia coli str K 12 substr W3110 YP 490328 1 316407 Proteobacteria
Shigella flexneri 5 str 8401 YP 690229 1 373384 Proteobacteria
Shigella flexneri 5 str 8401 YP 690302 1 373384 Proteobacteria
Shigella flexneri 2002017 YP 005728310 1 591020 Proteobacteria
Shigella flexneri 2a str 2457T NP 838491 1 198215 Proteobacteria
Shigella flexneri 2a str 2457T NP 838359 1 198215 Proteobacteria
Shigella flexneri 2002017 YP 005728479 1 591020 Proteobacteria
Shigella flexneri 2002017 YP 005728633 1 591020 Proteobacteria
Escherichia coli E24377A YP 001464049 1 331111 Proteobacteria
Escherichia coli O127 H6 str E2348 69 YP 002328655 1 574521 Proteobacteria
Escherichia coli B str REL606 YP 003045884 1 413997 Proteobacteria
Shigella flexneri 2a str 2457T NP 836727 1 198215 Proteobacteria
Shigella flexneri 5 str 8401 YP 688560 1 373384 Proteobacteria
Escherichia coli O127 H6 str E2348 69 YP 002332171 1 574521 Proteobacteria
Escherichia coli B str REL606 YP 003044728 1 413997 Proteobacteria
Shigella flexneri 2002017 YP 005726868 1 591020 Proteobacteria
Escherichia coli ATCC 8739 YP 001726681 1 481805 Proteobacteria
Escherichia coli ATCC 8739 YP 001723731 1 481805 Proteobacteria
Escherichia coli ATCC 8739 YP 001726654 1 481805 Proteobacteria

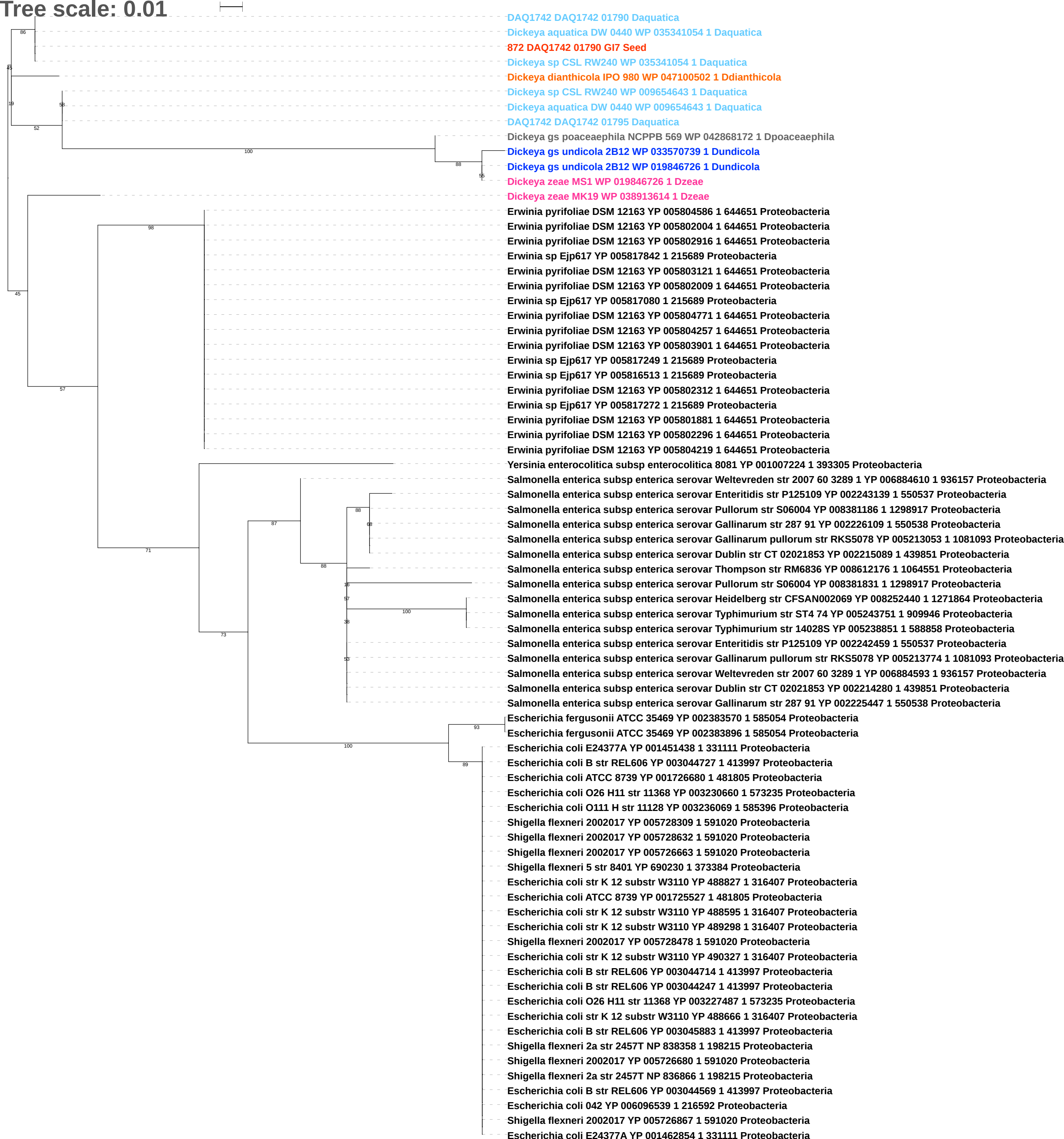

Tree scale: 0.01
DAQ1742 DAQ1742 01790 Daquatica
Dickeya aquatica DW 0440 WP 035341054 1 Daquatica
872 DAQ1742 01790 GI7 Seed
Dickeya sp CSL RW240 WP 035341054 1 Daquatica
Dickeya dianthicola IPO 980 WP 047100502 1 Ddianthicola
Dickeya sp CSL RW240 WP 009654643 1 Daquatica
Dickeya aquatica DW 0440 WP 009654643 1 Daquatica
DAQ1742 DAQ1742 01795 Daquatica
Dickeya gs poaceaephila NCPPB 569 WP 042868172 1 Dpoaceaephila
Dickeya gs undicola 2B12 WP 033570739 1 Dundicola
Dickeya gs undicola 2B12 WP 019846726 1 Dundicola
Dickeya zeae MS1 WP 019846726 1 Dzeae
Dickeya zeae MK19 WP 038913614 1 Dzeae
Erwinia pyrifoliae DSM 12163 YP 005804586 1 644651 Proteobacteria
Erwinia pyrifoliae DSM 12163 YP 005802004 1 644651 Proteobacteria
Erwinia pyrifoliae DSM 12163 YP 005802916 1 644651 Proteobacteria
Erwinia sp Ejp617 YP 005817842 1 215689 Proteobacteria
Erwinia pyrifoliae DSM 12163 YP 005803121 1 644651 Proteobacteria
Erwinia pyrifoliae DSM 12163 YP 005802009 1 644651 Proteobacteria
Erwinia sp Ejp617 YP 005817080 1 215689 Proteobacteria
Erwinia pyrifoliae DSM 12163 YP 005804771 1 644651 Proteobacteria
Erwinia pyrifoliae DSM 12163 YP 005804257 1 644651 Proteobacteria
Erwinia pyrifoliae DSM 12163 YP 005803901 1 644651 Proteobacteria
Erwinia sp Ejp617 YP 005817249 1 215689 Proteobacteria
Erwinia sp Ejp617 YP 005816513 1 215689 Proteobacteria
Erwinia pyrifoliae DSM 12163 YP 005802312 1 644651 Proteobacteria
Erwinia sp Ejp617 YP 005817272 1 215689 Proteobacteria
Erwinia pyrifoliae DSM 12163 YP 005801881 1 644651 Proteobacteria
Erwinia pyrifoliae DSM 12163 YP 005802296 1 644651 Proteobacteria
Erwinia pyrifoliae DSM 12163 YP 005804219 1 644651 Proteobacteria
Yersinia enterocolitica subsp enterocolitica 8081 YP 001007224 1 393305 Proteobacteria
Salmonella enterica subsp enterica serovar Weltevreden str 2007 60 3289 1 YP 006884610 1 936157 Proteobacteria
Salmonella enterica subsp enterica serovar Enteritidis str P125109 YP 002243139 1 550537 Proteobacteria
Salmonella enterica subsp enterica serovar Pullorum str S06004 YP 008381186 1 1298917 Proteobacteria
Salmonella enterica subsp enterica serovar Gallinarum str 287 91 YP 002226109 1 550538 Proteobacteria
Salmonella enterica subsp enterica serovar Gallinarum pullorum str RKS5078 YP 005213053 1 1081093 Proteobacteria
Salmonella enterica subsp enterica serovar Dublin str CT 02021853 YP 002215089 1 439851 Proteobacteria
Salmonella enterica subsp enterica serovar Thompson str RM6836 YP 008612176 1 1064551 Proteobacteria
Salmonella enterica subsp enterica serovar Pullorum str S06004 YP 008381831 1 1298917 Proteobacteria
Salmonella enterica subsp enterica serovar Heidelberg str CFSAN002069 YP 008252440 1 1271864 Proteobacteria
Salmonella enterica subsp enterica serovar Typhimurium str ST4 74 YP 005243751 1 909946 Proteobacteria
Salmonella enterica subsp enterica serovar Typhimurium str 14028S YP 005238851 1 588858 Proteobacteria
Salmonella enterica subsp enterica serovar Enteritidis str P125109 YP 002242459 1 550537 Proteobacteria
Salmonella enterica subsp enterica serovar Gallinarum pullorum str RKS5078 YP 005213774 1 1081093 Proteobacteria
Salmonella enterica subsp enterica serovar Weltevreden str 2007 60 3289 1 YP 006884593 1 936157 Proteobacteria
Salmonella enterica subsp enterica serovar Dublin str CT 02021853 YP 002214280 1 439851 Proteobacteria
Salmonella enterica subsp enterica serovar Gallinarum str 287 91 YP 002225447 1 550538 Proteobacteria
Escherichia fergusonii ATCC 35469 YP 002383570 1 585054 Proteobacteria
Escherichia fergusonii ATCC 35469 YP 002383896 1 585054 Proteobacteria
Escherichia coli E24377A YP 001451438 1 331111 Proteobacteria
Escherichia coli B str REL606 YP 003044727 1 413997 Proteobacteria
Escherichia coli ATCC 8739 YP 001726680 1 481805 Proteobacteria
Escherichia coli O26 H11 str 11368 YP 003230660 1 573235 Proteobacteria
Escherichia coli O111 H str 11128 YP 003236069 1 585396 Proteobacteria
Shigella flexneri 2002017 YP 005728309 1 591020 Proteobacteria
Shigella flexneri 2002017 YP 005728632 1 591020 Proteobacteria
Shigella flexneri 2002017 YP 005726663 1 591020 Proteobacteria
Shigella flexneri 5 str 8401 YP 690230 1 373384 Proteobacteria
Escherichia coli str K 12 substr W3110 YP 488827 1 316407 Proteobacteria
Escherichia coli ATCC 8739 YP 001725527 1 481805 Proteobacteria
Escherichia coli str K 12 substr W3110 YP 488595 1 316407 Proteobacteria
Escherichia coli str K 12 substr W3110 YP 489298 1 316407 Proteobacteria
Shigella flexneri 2002017 YP 005728478 1 591020 Proteobacteria
Escherichia coli str K 12 substr W3110 YP 490327 1 316407 Proteobacteria
Escherichia coli B str REL606 YP 003044714 1 413997 Proteobacteria
Escherichia coli B str REL606 YP 003044247 1 413997 Proteobacteria
Escherichia coli O26 H11 str 11368 YP 003227487 1 573235 Proteobacteria
Escherichia coli str K 12 substr W3110 YP 488666 1 316407 Proteobacteria
Escherichia coli B str REL606 YP 003045883 1 413997 Proteobacteria
Shigella flexneri 2a str 2457T NP 838358 1 198215 Proteobacteria
Shigella flexneri 2002017 YP 005726680 1 591020 Proteobacteria
Shigella flexneri 2a str 2457T NP 836866 1 198215 Proteobacteria
Escherichia coli B str REL606 YP 003044569 1 413997 Proteobacteria
Escherichia coli 042 YP 006096539 1 216592 Proteobacteria
Shigella flexneri 2002017 YP 005726867 1 591020 Proteobacteria
Escherichia coli E24377A YP 001462854 1 331111 Proteobacteria

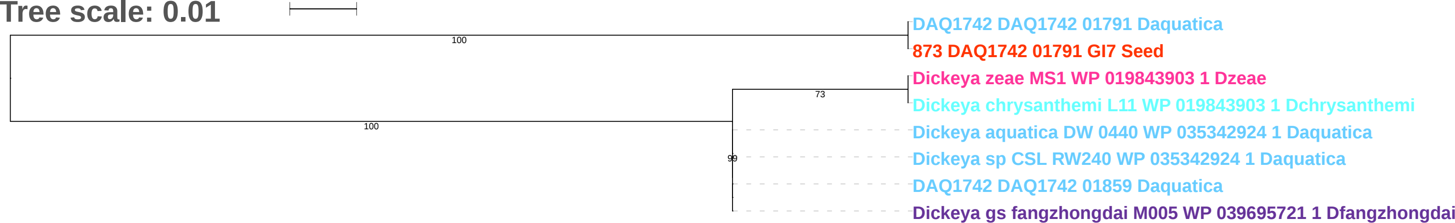

Tree scale: 0.01
100
100
73
99
DAQ1742 DAQ1742 01791 Daquatica
873 DAQ1742 01791 GI7 Seed
Dickeya zeae MS1 WP 019843903 1 Dzeae
Dickeya chrysanthemi L11 WP 019843903 1 Dchrysanthemi
Dickeya aquatica DW 0440 WP 035342924 1 Daquatica
Dickeya sp CSL RW240 WP 035342924 1 Daquatica
DAQ1742 DAQ1742 01859 Daquatica
Dickeya gs fangzhongdai M005 WP 039695721 1 Dfangzhongdai

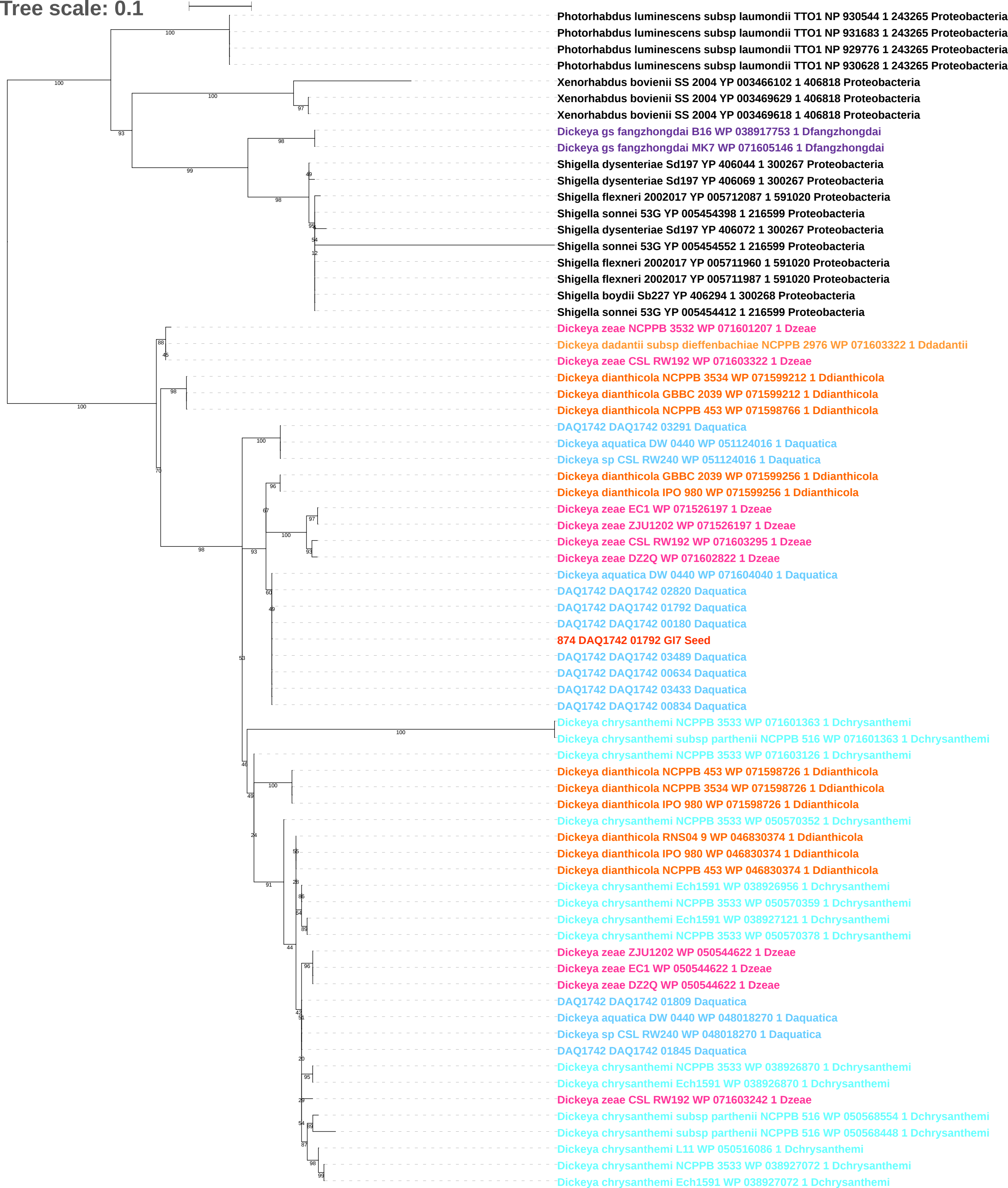
Tree scale: 0.1
Photorhabdus luminescens subsp laumondii TTO1 NP 930544 1 243265 Proteobacteria
Photorhabdus luminescens subsp laumondii TTO1 NP 931683 1 243265 Proteobacteria
Photorhabdus luminescens subsp laumondii TTO1 NP 929776 1 243265 Proteobacteria
Photorhabdus luminescens subsp laumondii TTO1 NP 930628 1 243265 Proteobacteria
Xenorhabdus bovienii SS 2004 YP 003466102 1 406818 Proteobacteria
Xenorhabdus bovienii SS 2004 YP 003469629 1 406818 Proteobacteria
Xenorhabdus bovienii SS 2004 YP 003469618 1 406818 Proteobacteria
Dickeya gs fangzhongdai B16 WP 038917753 1 Dfangzhongdai
Dickeya gs fangzhongdai MK7 WP 071605146 1 Dfangzhongdai
Shigella dysenteriae Sd197 YP 406044 1 300267 Proteobacteria
Shigella dysenteriae Sd197 YP 406069 1 300267 Proteobacteria
Shigella flexneri 2002017 YP 005712087 1 591020 Proteobacteria
Shigella sonnei 53G YP 005454398 1 216599 Proteobacteria
Shigella dysenteriae Sd197 YP 406072 1 300267 Proteobacteria
Shigella sonnei 53G YP 005454552 1 216599 Proteobacteria
Shigella flexneri 2002017 YP 005711960 1 591020 Proteobacteria
Shigella flexneri 2002017 YP 005711987 1 591020 Proteobacteria
Shigella boydii Sb227 YP 406294 1 300268 Proteobacteria
Shigella sonnei 53G YP 005454412 1 216599 Proteobacteria
Dickeya zeae NCPPB 3532 WP 071601207 1 Dzeae
Dickeya dadantii subsp dieffenbachiae NCPPB 2976 WP 071603322 1 Ddadantii
Dickeya zeae CSL RW192 WP 071603322 1 Dzeae
Dickeya dianthicola NCPPB 3534 WP 071599212 1 Ddianthicola
Dickeya dianthicola GBBC 2039 WP 071599212 1 Ddianthicola
Dickeya dianthicola NCPPB 453 WP 071598766 1 Ddianthicola
DAQ1742 DAQ1742 03291 Daquatica
Dickeya aquatica DW 0440 WP 051124016 1 Daquatica
Dickeya sp CSL RW240 WP 051124016 1 Daquatica
Dickeya dianthicola GBBC 2039 WP 071599256 1 Ddianthicola
Dickeya dianthicola IPO 980 WP 071599256 1 Ddianthicola
Dickeya zeae EC1 WP 071526197 1 Dzeae
Dickeya zeae ZJU1202 WP 071526197 1 Dzeae
Dickeya zeae CSL RW192 WP 071603295 1 Dzeae
Dickeya zeae DZ2Q WP 071602822 1 Dzeae
Dickeya aquatica DW 0440 WP 071604040 1 Daquatica
DAQ1742 DAQ1742 02820 Daquatica
DAQ1742 DAQ1742 01792 Daquatica
DAQ1742 DAQ1742 00180 Daquatica
874 DAQ1742 01792 GI7 Seed
DAQ1742 DAQ1742 03489 Daquatica
DAQ1742 DAQ1742 00634 Daquatica
DAQ1742 DAQ1742 03433 Daquatica
DAQ1742 DAQ1742 00834 Daquatica
Dickeya chrysanthemi NCPPB 3533 WP 071601363 1 Dchrysanthemi
Dickeya chrysanthemi subsp parthenii NCPPB 516 WP 071601363 1 Dchrysanthemi
Dickeya chrysanthemi NCPPB 3533 WP 071603126 1 Dchrysanthemi
Dickeya dianthicola NCPPB 453 WP 071598726 1 Ddianthicola
Dickeya dianthicola NCPPB 3534 WP 071598726 1 Ddianthicola
Dickeya dianthicola IPO 980 WP 071598726 1 Ddianthicola
Dickeya chrysanthemi NCPPB 3533 WP 050570352 1 Dchrysanthemi
Dickeya dianthicola RNS04 9 WP 046830374 1 Ddianthicola
Dickeya dianthicola IPO 980 WP 046830374 1 Ddianthicola
Dickeya dianthicola NCPPB 453 WP 046830374 1 Ddianthicola
Dickeya chrysanthemi Ech1591 WP 038926956 1 Dchrysanthemi
Dickeya chrysanthemi NCPPB 3533 WP 050570359 1 Dchrysanthemi
Dickeya chrysanthemi Ech1591 WP 038927121 1 Dchrysanthemi
Dickeya chrysanthemi NCPPB 3533 WP 050570378 1 Dchrysanthemi
Dickeya zeae ZJU1202 WP 050544622 1 Dzeae
Dickeya zeae EC1 WP 050544622 1 Dzeae
Dickeya zeae DZ2Q WP 050544622 1 Dzeae
DAQ1742 DAQ1742 01809 Daquatica
Dickeya aquatica DW 0440 WP 048018270 1 Daquatica
Dickeya sp CSL RW240 WP 048018270 1 Daquatica
DAQ1742 DAQ1742 01845 Daquatica
Dickeya chrysanthemi NCPPB 3533 WP 038926870 1 Dchrysanthemi
Dickeya chrysanthemi Ech1591 WP 038926870 1 Dchrysanthemi
Dickeya zeae CSL RW192 WP 071603242 1 Dzeae
Dickeya chrysanthemi subsp parthenii NCPPB 516 WP 050568554 1 Dchrysanthemi
Dickeya chrysanthemi subsp parthenii NCPPB 516 WP 050568448 1 Dchrysanthemi
Dickeya chrysanthemi L11 WP 050516086 1 Dchrysanthemi
Dickeya chrysanthemi NCPPB 3533 WP 038927072 1 Dchrysanthemi
Dickeya chrysanthemi Ech1591 WP 038927072 1 Dchrysanthemi

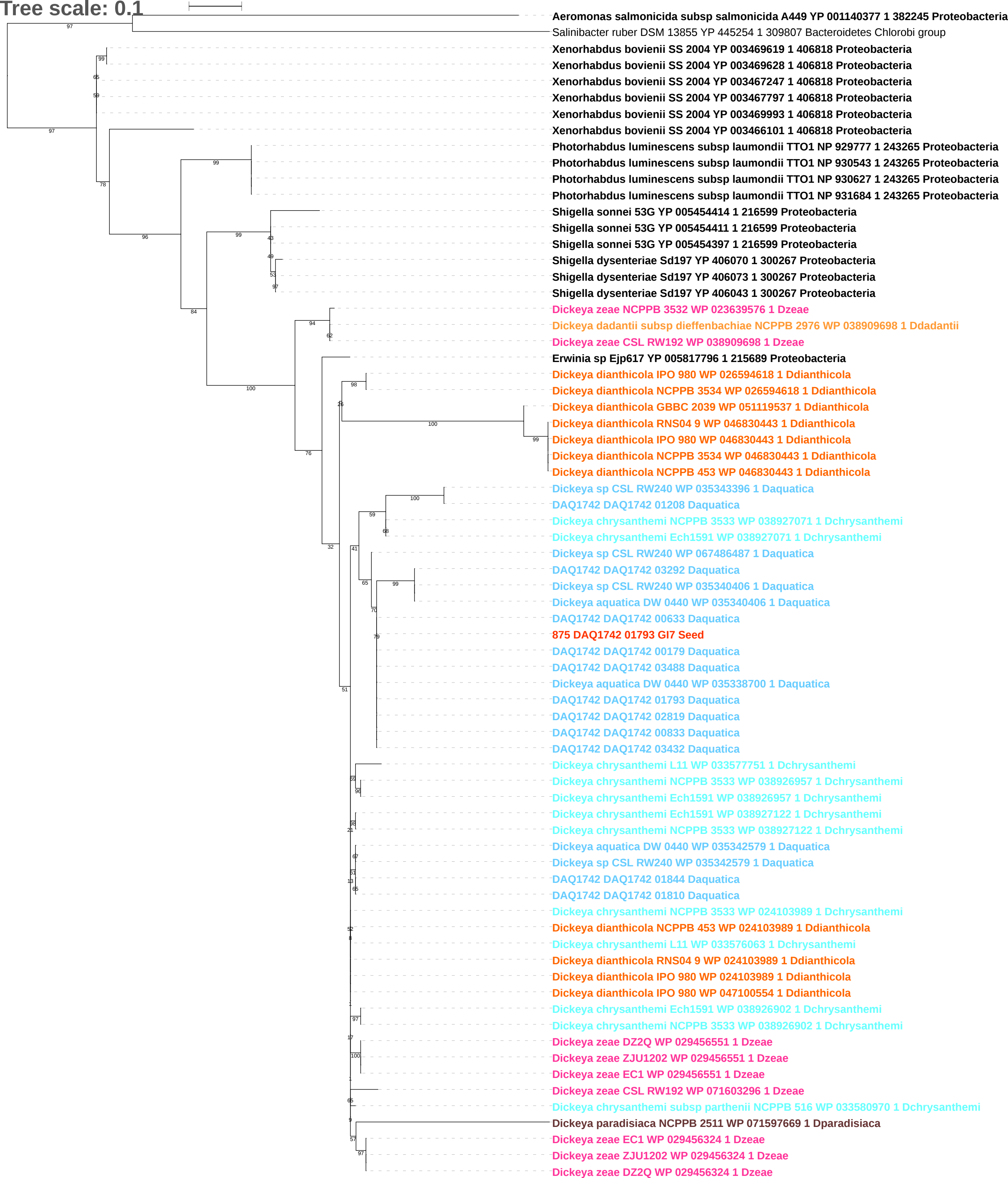

Tree scale: 0.1
Aeromonas salmonicida subsp salmonicida A449 YP 001140377 1 382245 Proteobacteria
Salinibacter ruber DSM 13855 YP 445254 1 309807 Bacteroidetes Chlorobi group
Xenorhabdus bovienii SS 2004 YP 003469619 1 406818 Proteobacteria
Xenorhabdus bovienii SS 2004 YP 003469628 1 406818 Proteobacteria
Xenorhabdus bovienii SS 2004 YP 003467247 1 406818 Proteobacteria
Xenorhabdus bovienii SS 2004 YP 003467797 1 406818 Proteobacteria
Xenorhabdus bovienii SS 2004 YP 003469993 1 406818 Proteobacteria
Xenorhabdus bovienii SS 2004 YP 003466101 1 406818 Proteobacteria
Photorhabdus luminescens subsp laumondii TTO1 NP 929777 1 243265 Proteobacteria
Photorhabdus luminescens subsp laumondii TTO1 NP 930543 1 243265 Proteobacteria
Photorhabdus luminescens subsp laumondii TTO1 NP 930627 1 243265 Proteobacteria
Photorhabdus luminescens subsp laumondii TTO1 NP 931684 1 243265 Proteobacteria
Shigella sonnei 53G YP 005454414 1 216599 Proteobacteria
Shigella sonnei 53G YP 005454411 1 216599 Proteobacteria
Shigella sonnei 53G YP 005454397 1 216599 Proteobacteria
Shigella dysenteriae Sd197 YP 406070 1 300267 Proteobacteria
Shigella dysenteriae Sd197 YP 406073 1 300267 Proteobacteria
Shigella dysenteriae Sd197 YP 406043 1 300267 Proteobacteria
Dickeya zeae NCPPB 3532 WP 023639576 1 Dzeae
Dickeya dadantii subsp dieffenbachiae NCPPB 2976 WP 038909698 1 Ddadantii
Dickeya zeae CSL RW192 WP 038909698 1 Dzeae
Erwinia sp Ejp617 YP 005817796 1 215689 Proteobacteria
Dickeya dianthicola IPO 980 WP 026594618 1 Ddianthicola
Dickeya dianthicola NCPPB 3534 WP 026594618 1 Ddianthicola
Dickeya dianthicola GBBC 2039 WP 051119537 1 Ddianthicola
Dickeya dianthicola RNS04 9 WP 046830443 1 Ddianthicola
Dickeya dianthicola IPO 980 WP 046830443 1 Ddianthicola
Dickeya dianthicola NCPPB 3534 WP 046830443 1 Ddianthicola
Dickeya dianthicola NCPPB 453 WP 046830443 1 Ddianthicola
Dickeya sp CSL RW240 WP 035343396 1 Daquatica
DAQ1742 DAQ1742 01208 Daquatica
Dickeya chrysanthemi NCPPB 3533 WP 038927071 1 Dchrysanthemi
Dickeya chrysanthemi Ech1591 WP 038927071 1 Dchrysanthemi
Dickeya sp CSL RW240 WP 067486487 1 Daquatica
DAQ1742 DAQ1742 03292 Daquatica
Dickeya sp CSL RW240 WP 035340406 1 Daquatica
Dickeya aquatica DW 0440 WP 035340406 1 Daquatica
DAQ1742 DAQ1742 00633 Daquatica
875 DAQ1742 01793 GI7 Seed
DAQ1742 DAQ1742 00179 Daquatica
DAQ1742 DAQ1742 03488 Daquatica
Dickeya aquatica DW 0440 WP 035338700 1 Daquatica
DAQ1742 DAQ1742 01793 Daquatica
DAQ1742 DAQ1742 02819 Daquatica
DAQ1742 DAQ1742 00833 Daquatica
DAQ1742 DAQ1742 03432 Daquatica
Dickeya chrysanthemi L11 WP 033577751 1 Dchrysanthemi
Dickeya chrysanthemi NCPPB 3533 WP 038926957 1 Dchrysanthemi
Dickeya chrysanthemi Ech1591 WP 038926957 1 Dchrysanthemi
Dickeya chrysanthemi Ech1591 WP 038927122 1 Dchrysanthemi
Dickeya chrysanthemi NCPPB 3533 WP 038927122 1 Dchrysanthemi
Dickeya aquatica DW 0440 WP 035342579 1 Daquatica
Dickeya sp CSL RW240 WP 035342579 1 Daquatica
DAQ1742 DAQ1742 01844 Daquatica
DAQ1742 DAQ1742 01810 Daquatica
Dickeya chrysanthemi NCPPB 3533 WP 024103989 1 Dchrysanthemi
Dickeya dianthicola NCPPB 453 WP 024103989 1 Ddianthicola
Dickeya chrysanthemi L11 WP 033576063 1 Dchrysanthemi
Dickeya dianthicola RNS04 9 WP 024103989 1 Ddianthicola
Dickeya dianthicola IPO 980 WP 024103989 1 Ddianthicola
Dickeya dianthicola IPO 980 WP 047100554 1 Ddianthicola
Dickeya chrysanthemi Ech1591 WP 038926902 1 Dchrysanthemi
Dickeya chrysanthemi NCPPB 3533 WP 038926902 1 Dchrysanthemi
Dickeya zeae DZ2Q WP 029456551 1 Dzeae
Dickeya zeae ZJU1202 WP 029456551 1 Dzeae
Dickeya zeae EC1 WP 029456551 1 Dzeae
Dickeya zeae CSL RW192 WP 071603296 1 Dzeae
Dickeya chrysanthemi subsp parthenii NCPPB 516 WP 033580970 1 Dchrysanthemi
Dickeya paradisiaca NCPPB 2511 WP 071597669 1 Dparadisiaca
Dickeya zeae EC1 WP 029456324 1 Dzeae
Dickeya zeae ZJU1202 WP 029456324 1 Dzeae
Dickeya zeae DZ2Q WP 029456324 1 Dzeae

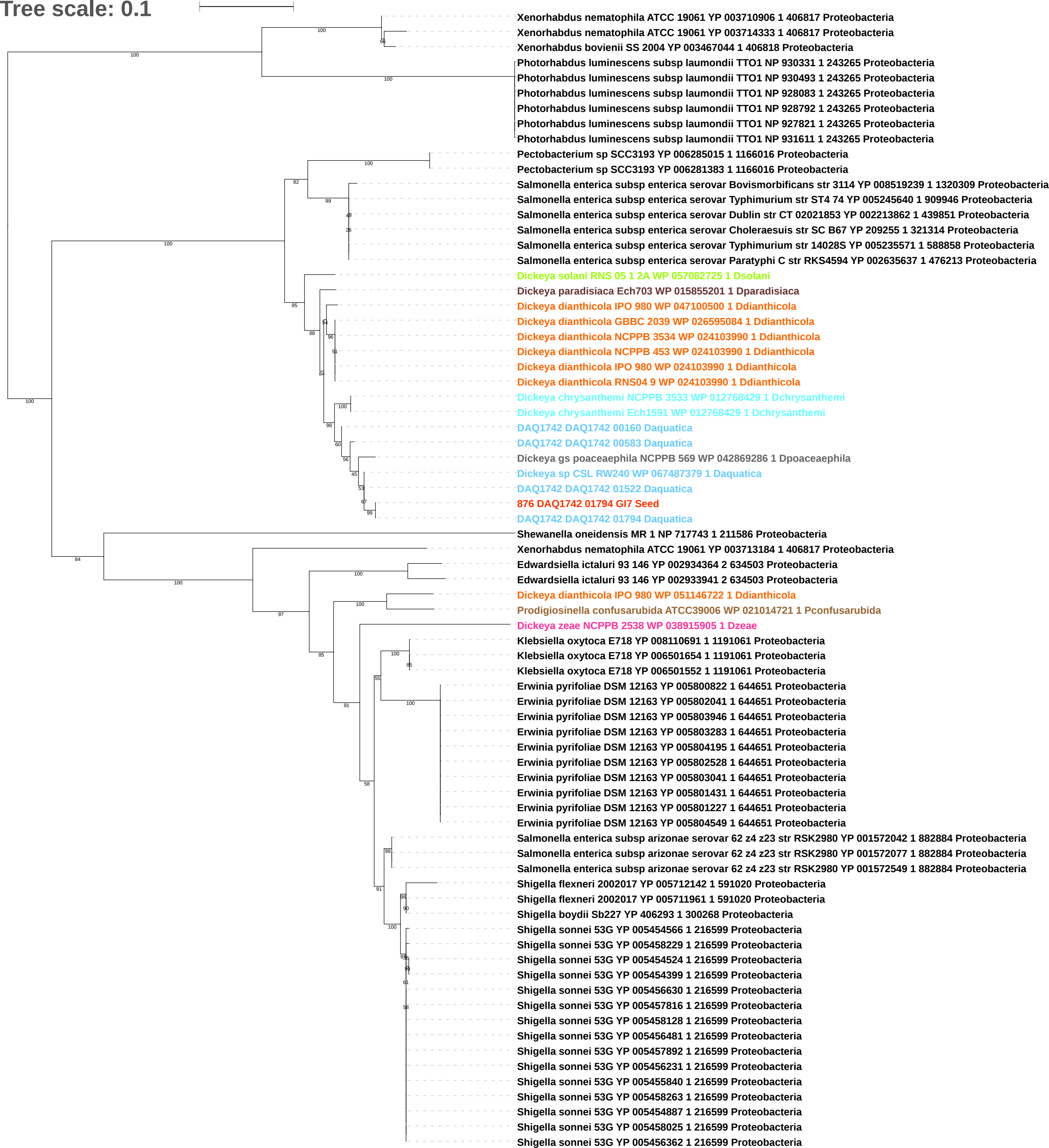
Tree scale: 0.1
Xenorhabdus nematophila ATCC 19061 YP 003710906 1 406817 Proteobacteria
Xenorhabdus nematophila ATCC 19061 YP 003714333 1 406817 Proteobacteria
Xenorhabdus bovienii SS 2004 YP 003467044 1 406818 Proteobacteria
Photorhabdus luminescens subsp laumondii TTO1 NP 930331 1 243265 Proteobacteria
Photorhabdus luminescens subsp laumondii TTO1 NP 930493 1 243265 Proteobacteria
Photorhabdus luminescens subsp laumondii TTO1 NP 928083 1 243265 Proteobacteria
Photorhabdus luminescens subsp laumondii TTO1 NP 928792 1 243265 Proteobacteria
Photorhabdus luminescens subsp laumondii TTO1 NP 927821 1 243265 Proteobacteria
Photorhabdus luminescens subsp laumondii TTO1 NP 931611 1 243265 Proteobacteria
Pectobacterium sp SCC3193 YP 006285015 1 1166016 Proteobacteria
Pectobacterium sp SCC3193 YP 006281383 1 1166016 Proteobacteria
Salmonella enterica subsp enterica serovar Bovismorbificans str 3114 YP 008519239 1 1320309 Proteobacteria
Salmonella enterica subsp enterica serovar Typhimurium str ST4 74 YP 005245640 1 909946 Proteobacteria
Salmonella enterica subsp enterica serovar Dublin str CT 02021853 YP 002213862 1 439851 Proteobacteria
Salmonella enterica subsp enterica serovar Choleraesuis str SC B67 YP 209255 1 321314 Proteobacteria
Salmonella enterica subsp enterica serovar Typhimurium str 14028S YP 005235571 1 588858 Proteobacteria
Salmonella enterica subsp enterica serovar Paratyphi C str RKS4594 YP 002635637 1 476213 Proteobacteria
Dickeya solani RNS 05 1 2A WP 057082725 1 Dsolani
Dickeya paradisiaca Ech703 WP 015855201 1 Dparadisiaca
Dickeya dianthicola IPO 980 WP 047100500 1 Ddianthicola
Dickeya dianthicola GBBC 2039 WP 026595084 1 Ddianthicola
Dickeya dianthicola NCPPB 3534 WP 024103990 1 Ddianthicola
Dickeya dianthicola NCPPB 453 WP 024103990 1 Ddianthicola
Dickeya dianthicola IPO 980 WP 024103990 1 Ddianthicola
Dickeya dianthicola RNS04 9 WP 024103990 1 Ddianthicola
Dickeya chrysanthemi NCPPB 3533 WP 012768429 1 Dchrysanthemi
Dickeya chrysanthemi Ech1591 WP 012768429 1 Dchrysanthemi
DAQ1742 DAQ1742 00160 Daquatica
DAQ1742 DAQ1742 00583 Daquatica
Dickeya gs poaceaephila NCPPB 569 WP 042869286 1 Dpoaceaephila
Dickeya sp CSL RW240 WP 067487379 1 Daquatica
DAQ1742 DAQ1742 01522 Daquatica
876 DAQ1742 01794 GI7 Seed
DAQ1742 DAQ1742 01794 Daquatica
Shewanella oneidensis MR 1 NP 717743 1 211586 Proteobacteria
Xenorhabdus nematophila ATCC 19061 YP 003713184 1 406817 Proteobacteria
Edwardsiella ictaluri 93 146 YP 002934364 2 634503 Proteobacteria
Edwardsiella ictaluri 93 146 YP 002933941 2 634503 Proteobacteria
Dickeya dianthicola IPO 980 WP 051146722 1 Ddianthicola
Prodigiosinella confusarubida ATCC39006 WP 021014721 1 Pconfusarubida
Dickeya zeae NCPPB 2538 WP 038915905 1 Dzeae
Klebsiella oxytoca E718 YP 008110691 1 1191061 Proteobacteria
Klebsiella oxytoca E718 YP 006501654 1 1191061 Proteobacteria
Klebsiella oxytoca E718 YP 006501552 1 1191061 Proteobacteria
Erwinia pyrifoliae DSM 12163 YP 005800822 1 644651 Proteobacteria
Erwinia pyrifoliae DSM 12163 YP 005802041 1 644651 Proteobacteria
Erwinia pyrifoliae DSM 12163 YP 005803946 1 644651 Proteobacteria
Erwinia pyrifoliae DSM 12163 YP 005803283 1 644651 Proteobacteria
Erwinia pyrifoliae DSM 12163 YP 005804195 1 644651 Proteobacteria
Erwinia pyrifoliae DSM 12163 YP 005802528 1 644651 Proteobacteria
Erwinia pyrifoliae DSM 12163 YP 005803041 1 644651 Proteobacteria
Erwinia pyrifoliae DSM 12163 YP 005801431 1 644651 Proteobacteria
Erwinia pyrifoliae DSM 12163 YP 005801227 1 644651 Proteobacteria
Erwinia pyrifoliae DSM 12163 YP 005804549 1 644651 Proteobacteria
Salmonella enterica subsp arizonae serovar 62 z4 z23 str RSK2980 YP 001572042 1 882884 Proteobacteria
Salmonella enterica subsp arizonae serovar 62 z4 z23 str RSK2980 YP 001572077 1 882884 Proteobacteria
Salmonella enterica subsp arizonae serovar 62 z4 z23 str RSK2980 YP 001572549 1 882884 Proteobacteria
Shigella flexneri 2002017 YP 005712142 1 591020 Proteobacteria
Shigella flexneri 2002017 YP 005711961 1 591020 Proteobacteria
Shigella boydii Sb227 YP 406293 1 300268 Proteobacteria
Shigella sonnei 53G YP 005454566 1 216599 Proteobacteria
Shigella sonnei 53G YP 005458229 1 216599 Proteobacteria
Shigella sonnei 53G YP 005454524 1 216599 Proteobacteria
Shigella sonnei 53G YP 005454399 1 216599 Proteobacteria
Shigella sonnei 53G YP 005456630 1 216599 Proteobacteria
Shigella sonnei 53G YP 005457816 1 216599 Proteobacteria
Shigella sonnei 53G YP 005458128 1 216599 Proteobacteria
Shigella sonnei 53G YP 005456481 1 216599 Proteobacteria
Shigella sonnei 53G YP 005457892 1 216599 Proteobacteria
Shigella sonnei 53G YP 005456231 1 216599 Proteobacteria
Shigella sonnei 53G YP 005455840 1 216599 Proteobacteria
Shigella sonnei 53G YP 005458263 1 216599 Proteobacteria
Shigella sonnei 53G YP 005454887 1 216599 Proteobacteria
Shigella sonnei 53G YP 005458025 1 216599 Proteobacteria
Shigella sonnei 53G YP 005456362 1 216599 Proteobacteria

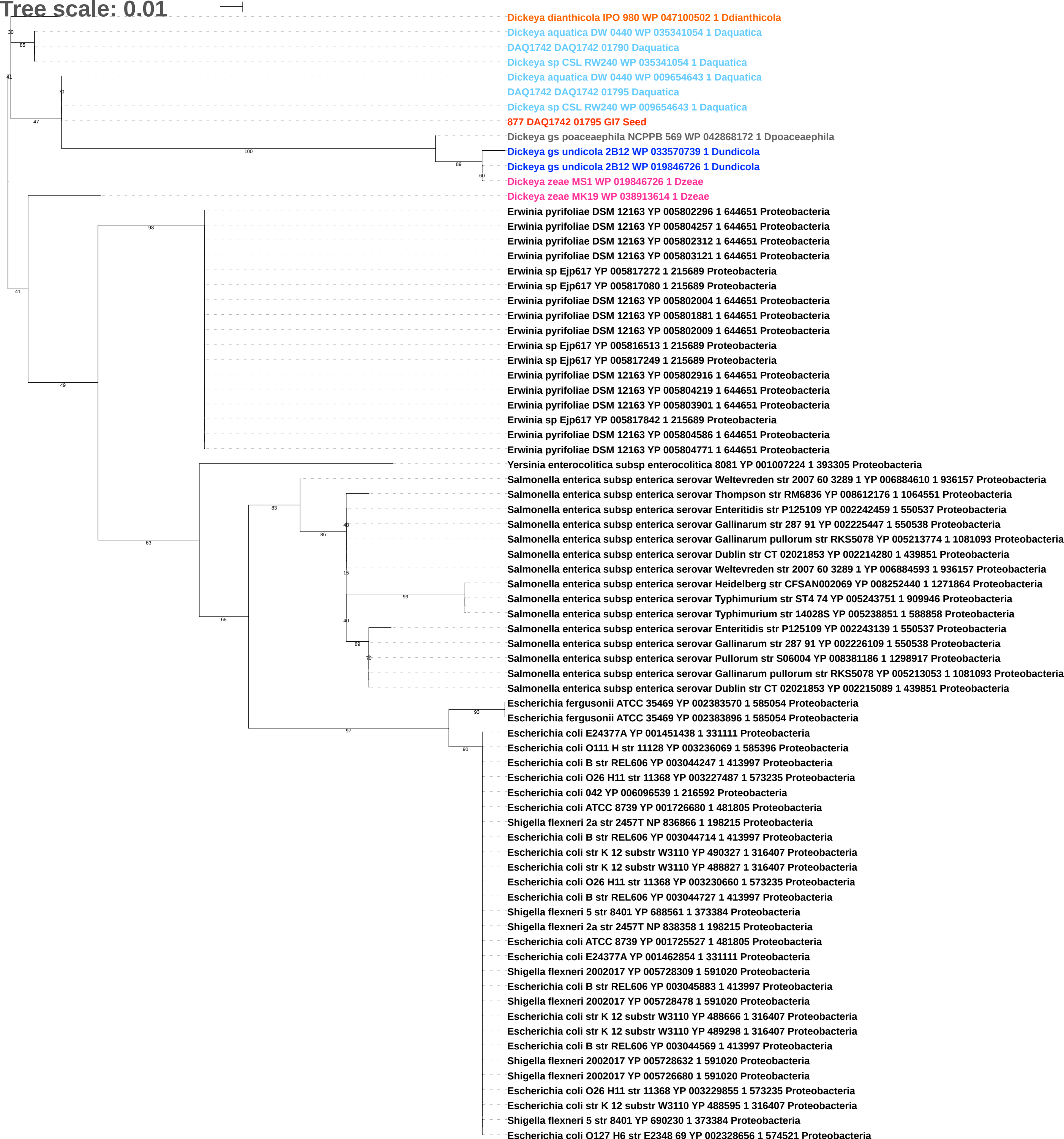
Tree scale: 0.01
Dickeya dianthicola IPO 980 WP 047100502 1 Ddianthicola
Dickeya aquatica DW 0440 WP 035341054 1 Daquatica
DAQ1742 DAQ1742 01790 Daquatica
Dickeya sp CSL RW240 WP 035341054 1 Daquatica
Dickeya aquatica DW 0440 WP 009654643 1 Daquatica
DAQ1742 DAQ1742 01795 Daquatica
Dickeya sp CSL RW240 WP 009654643 1 Daquatica
877 DAQ1742 01795 GI7 Seed
Dickeya gs poaceaephila NCPPB 569 WP 042868172 1 Dpoaceaephila
Dickeya gs undicola 2B12 WP 033570739 1 Dundicola
Dickeya gs undicola 2B12 WP 019846726 1 Dundicola
Dickeya zeae MS1 WP 019846726 1 Dzeae
Dickeya zeae MK19 WP 038913614 1 Dzeae
Erwinia pyrifoliae DSM 12163 YP 005802296 1 644651 Proteobacteria
Erwinia pyrifoliae DSM 12163 YP 005804257 1 644651 Proteobacteria
Erwinia pyrifoliae DSM 12163 YP 005802312 1 644651 Proteobacteria
Erwinia pyrifoliae DSM 12163 YP 005803121 1 644651 Proteobacteria
Erwinia sp Ejp617 YP 005817272 1 215689 Proteobacteria
Erwinia sp Ejp617 YP 005817080 1 215689 Proteobacteria
Erwinia pyrifoliae DSM 12163 YP 005802004 1 644651 Proteobacteria
Erwinia pyrifoliae DSM 12163 YP 005801881 1 644651 Proteobacteria
Erwinia pyrifoliae DSM 12163 YP 005802009 1 644651 Proteobacteria
Erwinia sp Ejp617 YP 005816513 1 215689 Proteobacteria
Erwinia sp Ejp617 YP 005817249 1 215689 Proteobacteria
Erwinia pyrifoliae DSM 12163 YP 005802916 1 644651 Proteobacteria
Erwinia pyrifoliae DSM 12163 YP 005804219 1 644651 Proteobacteria
Erwinia pyrifoliae DSM 12163 YP 005803901 1 644651 Proteobacteria
Erwinia sp Ejp617 YP 005817842 1 215689 Proteobacteria
Erwinia pyrifoliae DSM 12163 YP 005804586 1 644651 Proteobacteria
Erwinia pyrifoliae DSM 12163 YP 005804771 1 644651 Proteobacteria
Yersinia enterocolitica subsp enterocolitica 8081 YP 001007224 1 393305 Proteobacteria
Salmonella enterica subsp enterica serovar Weltevreden str 2007 60 3289 1 YP 006884610 1 936157 Proteobacteria
Salmonella enterica subsp enterica serovar Thompson str RM6836 YP 008612176 1 1064551 Proteobacteria
Salmonella enterica subsp enterica serovar Enteritidis str P125109 YP 002242459 1 550537 Proteobacteria
Salmonella enterica subsp enterica serovar Gallinarum str 287 91 YP 002225447 1 550538 Proteobacteria
Salmonella enterica subsp enterica serovar Gallinarum pullorum str RKS5078 YP 005213774 1 1081093 Proteobacteria
Salmonella enterica subsp enterica serovar Dublin str CT 02021853 YP 002214280 1 439851 Proteobacteria
Salmonella enterica subsp enterica serovar Weltevreden str 2007 60 3289 1 YP 006884593 1 936157 Proteobacteria
Salmonella enterica subsp enterica serovar Heidelberg str CFSAN002069 YP 008252440 1 1271864 Proteobacteria
Salmonella enterica subsp enterica serovar Typhimurium str ST4 74 YP 005243751 1 909946 Proteobacteria
Salmonella enterica subsp enterica serovar Typhimurium str 14028S YP 005238851 1 588858 Proteobacteria
Salmonella enterica subsp enterica serovar Enteritidis str P125109 YP 002243139 1 550537 Proteobacteria
Salmonella enterica subsp enterica serovar Gallinarum str 287 91 YP 002226109 1 550538 Proteobacteria
Salmonella enterica subsp enterica serovar Pullorum str S06004 YP 008381186 1 1298917 Proteobacteria
Salmonella enterica subsp enterica serovar Gallinarum pullorum str RKS5078 YP 005213053 1 1081093 Proteobacteria
Salmonella enterica subsp enterica serovar Dublin str CT 02021853 YP 002215089 1 439851 Proteobacteria
Escherichia fergusonii ATCC 35469 YP 002383570 1 585054 Proteobacteria
Escherichia fergusonii ATCC 35469 YP 002383896 1 585054 Proteobacteria
Escherichia coli E24377A YP 001451438 1 331111 Proteobacteria
Escherichia coli O111 H str 11128 YP 003236069 1 585396 Proteobacteria
Escherichia coli B str REL606 YP 003044247 1 413997 Proteobacteria
Escherichia coli O26 H11 str 11368 YP 003227487 1 573235 Proteobacteria
Escherichia coli 042 YP 006096539 1 216592 Proteobacteria
Escherichia coli ATCC 8739 YP 001726680 1 481805 Proteobacteria
Shigella flexneri 2a str 2457T NP 836866 1 198215 Proteobacteria
Escherichia coli B str REL606 YP 003044714 1 413997 Proteobacteria
Escherichia coli str K 12 substr W3110 YP 490327 1 316407 Proteobacteria
Escherichia coli str K 12 substr W3110 YP 488827 1 316407 Proteobacteria
Escherichia coli O26 H11 str 11368 YP 003230660 1 573235 Proteobacteria
Escherichia coli B str REL606 YP 003044727 1 413997 Proteobacteria
Shigella flexneri 5 str 8401 YP 688561 1 373384 Proteobacteria
Shigella flexneri 2a str 2457T NP 838358 1 198215 Proteobacteria
Escherichia coli ATCC 8739 YP 001725527 1 481805 Proteobacteria
Escherichia coli E24377A YP 001462854 1 331111 Proteobacteria
Shigella flexneri 2002017 YP 005728309 1 591020 Proteobacteria
Escherichia coli B str REL606 YP 003045883 1 413997 Proteobacteria
Shigella flexneri 2002017 YP 005728478 1 591020 Proteobacteria
Escherichia coli str K 12 substr W3110 YP 488666 1 316407 Proteobacteria
Escherichia coli str K 12 substr W3110 YP 489298 1 316407 Proteobacteria
Escherichia coli B str REL606 YP 003044569 1 413997 Proteobacteria
Shigella flexneri 2002017 YP 005728632 1 591020 Proteobacteria
Shigella flexneri 2002017 YP 005726680 1 591020 Proteobacteria
Escherichia coli O26 H11 str 11368 YP 003229855 1 573235 Proteobacteria
Escherichia coli str K 12 substr W3110 YP 488595 1 316407 Proteobacteria
Shigella flexneri 5 str 8401 YP 690230 1 373384 Proteobacteria
Escherichia coli O127 H6 str E2348 69 YP 002328656 1 574521 Proteobacteria

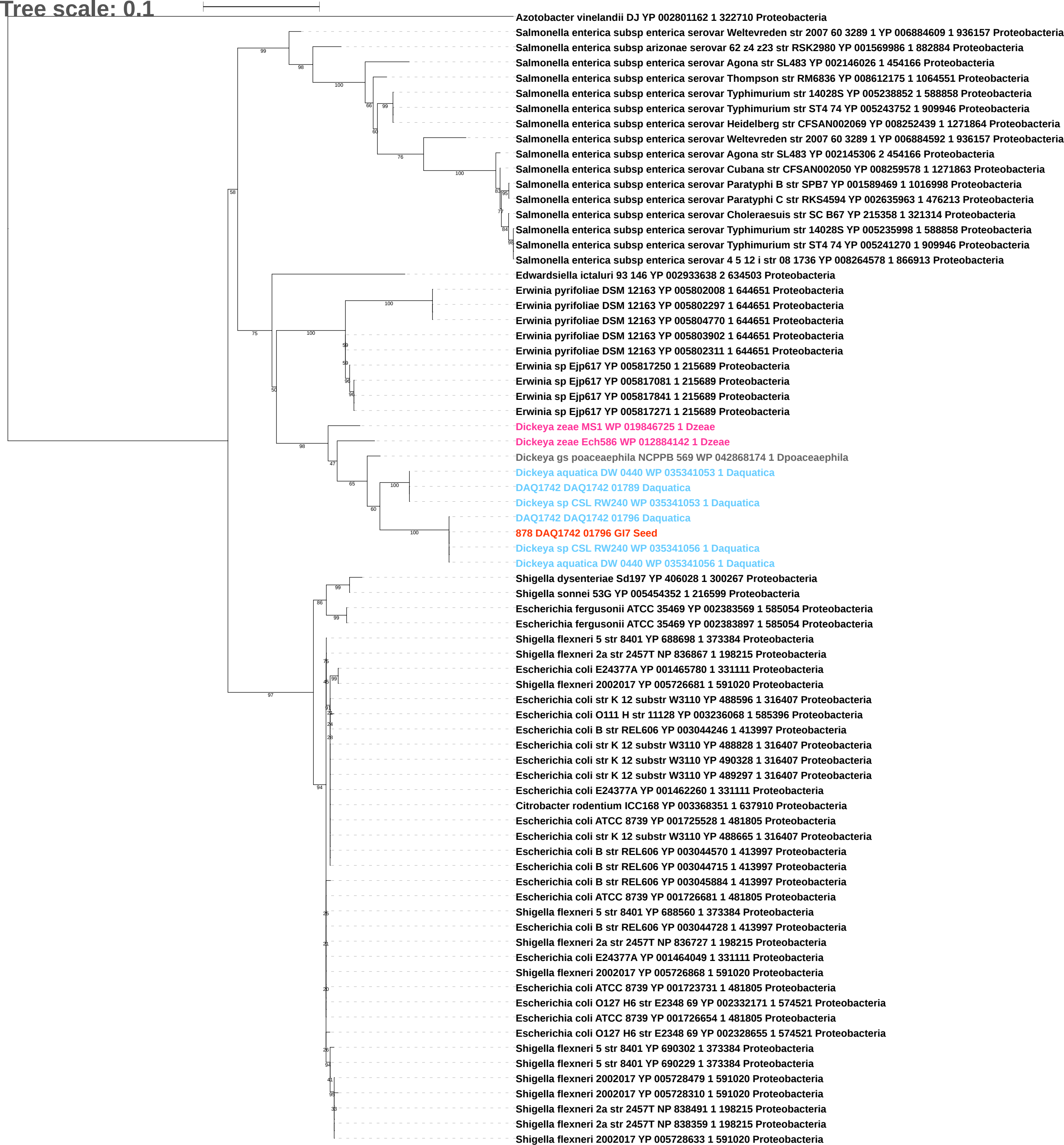

Tree scale: 0.1
Azotobacter vinelandii DJ YP 002801162 1 322710 Proteobacteria
Salmonella enterica subsp enterica serovar Weltevreden str 2007 60 3289 1 YP 006884609 1 936157 Proteobacteria
Salmonella enterica subsp arizonae serovar 62 z4 z23 str RSK2980 YP 001569986 1 882884 Proteobacteria
Salmonella enterica subsp enterica serovar Agona str SL483 YP 002146026 1 454166 Proteobacteria
Salmonella enterica subsp enterica serovar Thompson str RM6836 YP 008612175 1 1064551 Proteobacteria
Salmonella enterica subsp enterica serovar Typhimurium str 14028S YP 005238852 1 588858 Proteobacteria
Salmonella enterica subsp enterica serovar Typhimurium str ST4 74 YP 005243752 1 909946 Proteobacteria
Salmonella enterica subsp enterica serovar Heidelberg str CFSAN002069 YP 008252439 1 1271864 Proteobacteria
Salmonella enterica subsp enterica serovar Weltevreden str 2007 60 3289 1 YP 006884592 1 936157 Proteobacteria
Salmonella enterica subsp enterica serovar Agona str SL483 YP 002145306 2 454166 Proteobacteria
Salmonella enterica subsp enterica serovar Cubana str CFSAN002050 YP 008259578 1 1271863 Proteobacteria
Salmonella enterica subsp enterica serovar Paratyphi B str SPB7 YP 001589469 1 1016998 Proteobacteria
Salmonella enterica subsp enterica serovar Paratyphi C str RKS4594 YP 002635963 1 476213 Proteobacteria
Salmonella enterica subsp enterica serovar Choleraesuis str SC B67 YP 215358 1 321314 Proteobacteria
Salmonella enterica subsp enterica serovar Typhimurium str 14028S YP 005235998 1 588858 Proteobacteria
Salmonella enterica subsp enterica serovar Typhimurium str ST4 74 YP 005241270 1 909946 Proteobacteria
Salmonella enterica subsp enterica serovar 4 5 12 i str 08 1736 YP 008264578 1 866913 Proteobacteria
Edwardsiella ictaluri 93 146 YP 002933638 2 634503 Proteobacteria
Erwinia pyrifoliae DSM 12163 YP 005802008 1 644651 Proteobacteria
Erwinia pyrifoliae DSM 12163 YP 005802297 1 644651 Proteobacteria
Erwinia pyrifoliae DSM 12163 YP 005804770 1 644651 Proteobacteria
Erwinia pyrifoliae DSM 12163 YP 005803902 1 644651 Proteobacteria
Erwinia pyrifoliae DSM 12163 YP 005802311 1 644651 Proteobacteria
Erwinia sp Ejp617 YP 005817250 1 215689 Proteobacteria
Erwinia sp Ejp617 YP 005817081 1 215689 Proteobacteria
Erwinia sp Ejp617 YP 005817841 1 215689 Proteobacteria
Erwinia sp Ejp617 YP 005817271 1 215689 Proteobacteria
Dickeya zeae MS1 WP 019846725 1 Dzeae
Dickeya zeae Ech586 WP 012884142 1 Dzeae
Dickeya gs poaceaephila NCPPB 569 WP 042868174 1 Dpoaceaephila
Dickeya aquatica DW 0440 WP 035341053 1 Daquatica
DAQ1742 DAQ1742 01789 Daquatica
Dickeya sp CSL RW240 WP 035341053 1 Daquatica
DAQ1742 DAQ1742 01796 Daquatica
878 DAQ1742 01796 GI7 Seed
Dickeya sp CSL RW240 WP 035341056 1 Daquatica
Dickeya aquatica DW 0440 WP 035341056 1 Daquatica
Shigella dysenteriae Sd197 YP 406028 1 300267 Proteobacteria
Shigella sonnei 53G YP 005454352 1 216599 Proteobacteria
Escherichia fergusonii ATCC 35469 YP 002383569 1 585054 Proteobacteria
Escherichia fergusonii ATCC 35469 YP 002383897 1 585054 Proteobacteria
Shigella flexneri 5 str 8401 YP 688698 1 373384 Proteobacteria
Shigella flexneri 2a str 2457T NP 836867 1 198215 Proteobacteria
Escherichia coli E24377A YP 001465780 1 331111 Proteobacteria
Shigella flexneri 2002017 YP 005726681 1 591020 Proteobacteria
Escherichia coli str K 12 substr W3110 YP 488596 1 316407 Proteobacteria
Escherichia coli O111 H str 11128 YP 003236068 1 585396 Proteobacteria
Escherichia coli B str REL606 YP 003044246 1 413997 Proteobacteria
Escherichia coli str K 12 substr W3110 YP 488828 1 316407 Proteobacteria
Escherichia coli str K 12 substr W3110 YP 490328 1 316407 Proteobacteria
Escherichia coli str K 12 substr W3110 YP 489297 1 316407 Proteobacteria
Escherichia coli E24377A YP 001462260 1 331111 Proteobacteria
Citrobacter rodentium ICC168 YP 003368351 1 637910 Proteobacteria
Escherichia coli ATCC 8739 YP 001725528 1 481805 Proteobacteria
Escherichia coli str K 12 substr W3110 YP 488665 1 316407 Proteobacteria
Escherichia coli B str REL606 YP 003044570 1 413997 Proteobacteria
Escherichia coli B str REL606 YP 003044715 1 413997 Proteobacteria
Escherichia coli B str REL606 YP 003045884 1 413997 Proteobacteria
Escherichia coli ATCC 8739 YP 001726681 1 481805 Proteobacteria
Shigella flexneri 5 str 8401 YP 688560 1 373384 Proteobacteria
Escherichia coli B str REL606 YP 003044728 1 413997 Proteobacteria
Shigella flexneri 2a str 2457T NP 836727 1 198215 Proteobacteria
Escherichia coli E24377A YP 001464049 1 331111 Proteobacteria
Shigella flexneri 2002017 YP 005726868 1 591020 Proteobacteria
Escherichia coli ATCC 8739 YP 001723731 1 481805 Proteobacteria
Escherichia coli O127 H6 str E2348 69 YP 002332171 1 574521 Proteobacteria
Escherichia coli ATCC 8739 YP 001726654 1 481805 Proteobacteria
Escherichia coli O127 H6 str E2348 69 YP 002328655 1 574521 Proteobacteria
Shigella flexneri 5 str 8401 YP 690302 1 373384 Proteobacteria
Shigella flexneri 5 str 8401 YP 690229 1 373384 Proteobacteria
Shigella flexneri 2002017 YP 005728479 1 591020 Proteobacteria
Shigella flexneri 2002017 YP 005728310 1 591020 Proteobacteria
Shigella flexneri 2a str 2457T NP 838491 1 198215 Proteobacteria
Shigella flexneri 2a str 2457T NP 838359 1 198215 Proteobacteria
Shigella flexneri 2002017 YP 005728633 1 591020 Proteobacteria

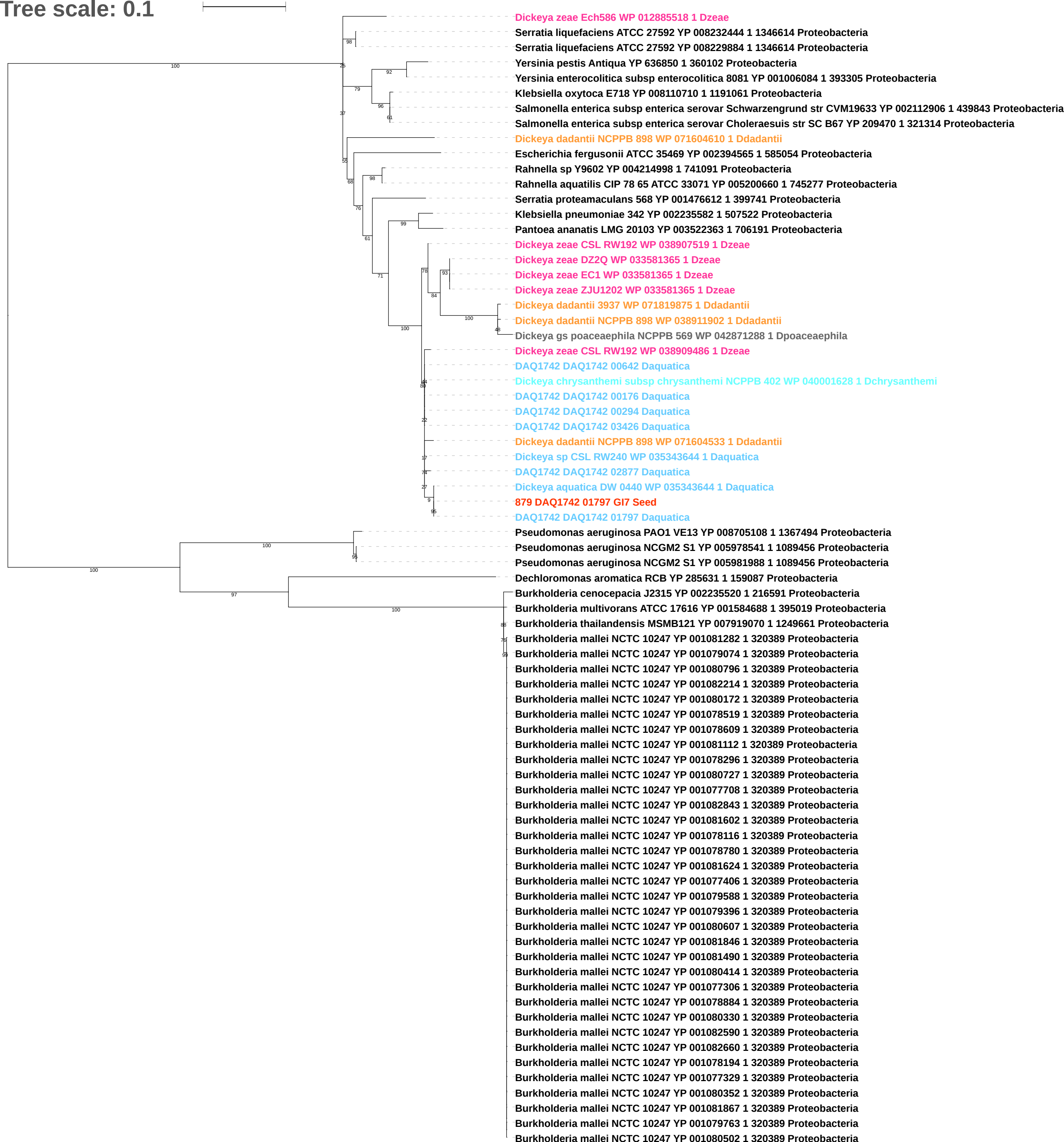
Tree scale: 0.1
Dickeya zeae Ech586 WP 012885518 1 Dzeae
Serratia liquefaciens ATCC 27592 YP 008232444 1 1346614 Proteobacteria
Serratia liquefaciens ATCC 27592 YP 008229884 1 1346614 Proteobacteria
Yersinia pestis Antiqua YP 636850 1 360102 Proteobacteria
Yersinia enterocolitica subsp enterocolitica 8081 YP 001006084 1 393305 Proteobacteria
Klebsiella oxytoca E718 YP 008110710 1 1191061 Proteobacteria
Salmonella enterica subsp enterica serovar Schwarzengrund str CVM19633 YP 002112906 1 439843 Proteobacteria
Salmonella enterica subsp enterica serovar Choleraesuis str SC B67 YP 209470 1 321314 Proteobacteria
Dickeya dadantii NCPPB 898 WP 071604610 1 Ddadantii
Escherichia fergusonii ATCC 35469 YP 002394565 1 585054 Proteobacteria
Rahnella sp Y9602 YP 004214998 1 741091 Proteobacteria
Rahnella aquatilis CIP 78 65 ATCC 33071 YP 005200660 1 745277 Proteobacteria
Serratia proteamaculans 568 YP 001476612 1 399741 Proteobacteria
Klebsiella pneumoniae 342 YP 002235582 1 507522 Proteobacteria
Pantoea ananatis LMG 20103 YP 003522363 1 706191 Proteobacteria
Dickeya zeae CSL RW192 WP 038907519 1 Dzeae
Dickeya zeae DZ2Q WP 033581365 1 Dzeae
Dickeya zeae EC1 WP 033581365 1 Dzeae
Dickeya zeae ZJU1202 WP 033581365 1 Dzeae
Dickeya dadantii 3937 WP 071819875 1 Ddadantii
Dickeya dadantii NCPPB 898 WP 038911902 1 Ddadantii
Dickeya gs poaceaephila NCPPB 569 WP 042871288 1 Dpoaceaephila
Dickeya zeae CSL RW192 WP 038909486 1 Dzeae
DAQ1742 DAQ1742 00642 Daquatica
Dickeya chrysanthemi subsp chrysanthemi NCPPB 402 WP 040001628 1 Dchrysanthemi
DAQ1742 DAQ1742 00176 Daquatica
DAQ1742 DAQ1742 00294 Daquatica
DAQ1742 DAQ1742 03426 Daquatica
Dickeya dadantii NCPPB 898 WP 071604533 1 Ddadantii
Dickeya sp CSL RW240 WP 035343644 1 Daquatica
DAQ1742 DAQ1742 02877 Daquatica
Dickeya aquatica DW 0440 WP 035343644 1 Daquatica
879 DAQ1742 01797 GI7 Seed
DAQ1742 DAQ1742 01797 Daquatica
Pseudomonas aeruginosa PAO1 VE13 YP 008705108 1 1367494 Proteobacteria
Pseudomonas aeruginosa NCGM2 S1 YP 005978541 1 1089456 Proteobacteria
Pseudomonas aeruginosa NCGM2 S1 YP 005981988 1 1089456 Proteobacteria
Dechloromonas aromatica RCB YP 285631 1 159087 Proteobacteria
Burkholderia cenocepacia J2315 YP 002235520 1 216591 Proteobacteria
Burkholderia multivorans ATCC 17616 YP 001584688 1 395019 Proteobacteria
Burkholderia thailandensis MSMB121 YP 007919070 1 1249661 Proteobacteria
Burkholderia mallei NCTC 10247 YP 001081282 1 320389 Proteobacteria
Burkholderia mallei NCTC 10247 YP 001079074 1 320389 Proteobacteria
Burkholderia mallei NCTC 10247 YP 001080796 1 320389 Proteobacteria
Burkholderia mallei NCTC 10247 YP 001082214 1 320389 Proteobacteria
Burkholderia mallei NCTC 10247 YP 001080172 1 320389 Proteobacteria
Burkholderia mallei NCTC 10247 YP 001078519 1 320389 Proteobacteria
Burkholderia mallei NCTC 10247 YP 001078609 1 320389 Proteobacteria
Burkholderia mallei NCTC 10247 YP 001081112 1 320389 Proteobacteria
Burkholderia mallei NCTC 10247 YP 001078296 1 320389 Proteobacteria
Burkholderia mallei NCTC 10247 YP 001080727 1 320389 Proteobacteria
Burkholderia mallei NCTC 10247 YP 001077708 1 320389 Proteobacteria
Burkholderia mallei NCTC 10247 YP 001082843 1 320389 Proteobacteria
Burkholderia mallei NCTC 10247 YP 001081602 1 320389 Proteobacteria
Burkholderia mallei NCTC 10247 YP 001078116 1 320389 Proteobacteria
Burkholderia mallei NCTC 10247 YP 001078780 1 320389 Proteobacteria
Burkholderia mallei NCTC 10247 YP 001081624 1 320389 Proteobacteria
Burkholderia mallei NCTC 10247 YP 001077406 1 320389 Proteobacteria
Burkholderia mallei NCTC 10247 YP 001079588 1 320389 Proteobacteria
Burkholderia mallei NCTC 10247 YP 001079396 1 320389 Proteobacteria
Burkholderia mallei NCTC 10247 YP 001080607 1 320389 Proteobacteria
Burkholderia mallei NCTC 10247 YP 001081846 1 320389 Proteobacteria
Burkholderia mallei NCTC 10247 YP 001081490 1 320389 Proteobacteria
Burkholderia mallei NCTC 10247 YP 001080414 1 320389 Proteobacteria
Burkholderia mallei NCTC 10247 YP 001077306 1 320389 Proteobacteria
Burkholderia mallei NCTC 10247 YP 001078884 1 320389 Proteobacteria
Burkholderia mallei NCTC 10247 YP 001080330 1 320389 Proteobacteria
Burkholderia mallei NCTC 10247 YP 001082590 1 320389 Proteobacteria
Burkholderia mallei NCTC 10247 YP 001082660 1 320389 Proteobacteria
Burkholderia mallei NCTC 10247 YP 001078194 1 320389 Proteobacteria
Burkholderia mallei NCTC 10247 YP 001077329 1 320389 Proteobacteria
Burkholderia mallei NCTC 10247 YP 001080352 1 320389 Proteobacteria
Burkholderia mallei NCTC 10247 YP 001081867 1 320389 Proteobacteria
Burkholderia mallei NCTC 10247 YP 001079763 1 320389 Proteobacteria
Burkholderia mallei NCTC 10247 YP 001080502 1 320389 Proteobacteria

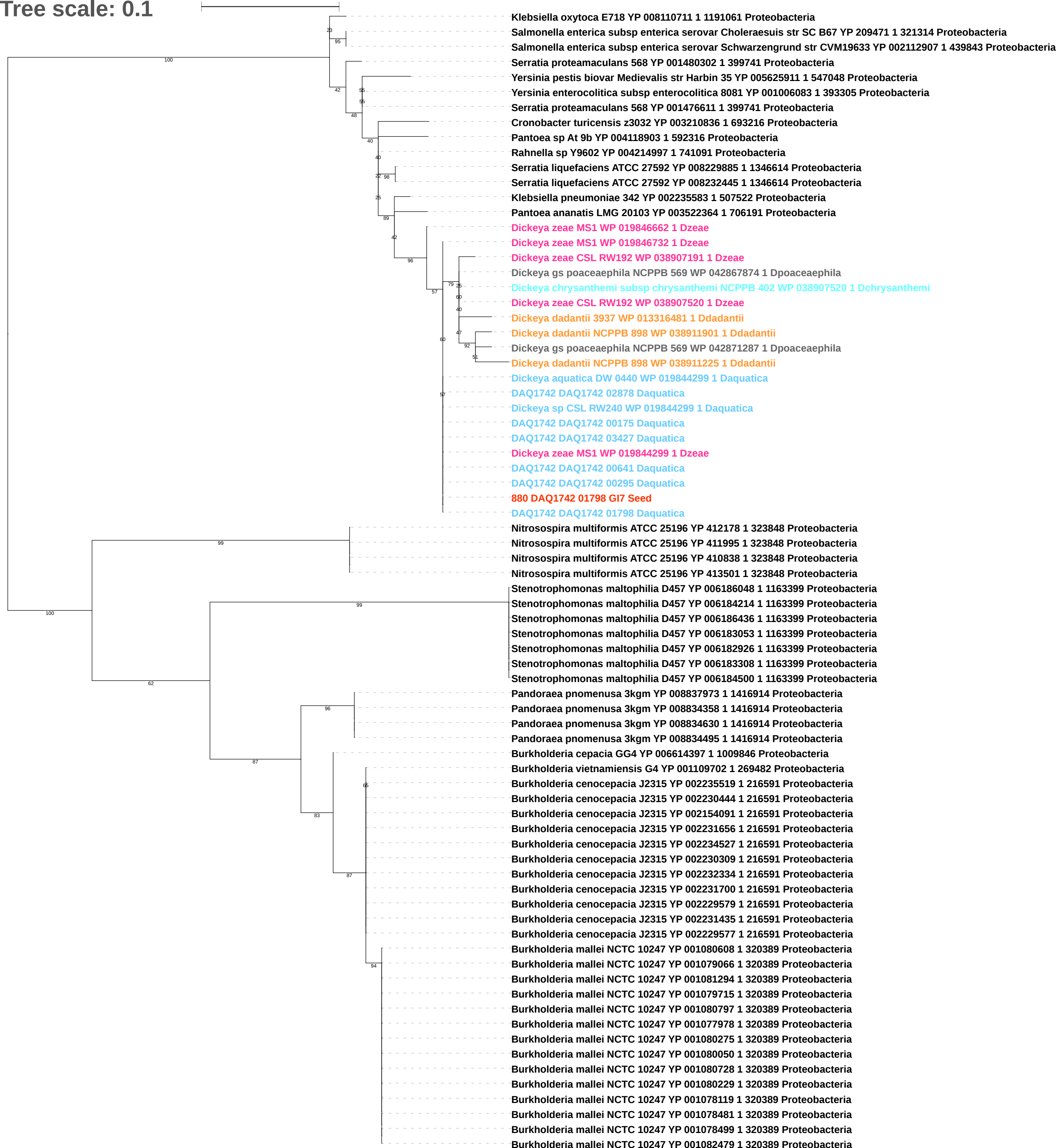
Tree scale: 0.1
Klebsiella oxytoca E718 YP 008110711 1 1191061 Proteobacteria
Salmonella enterica subsp enterica serovar Choleraesuis str SC B67 YP 209471 1 321314 Proteobacteria
Salmonella enterica subsp enterica serovar Schwarzengrund str CVM19633 YP 002112907 1 439843 Proteobacteria
Serratia proteamaculans 568 YP 001480302 1 399741 Proteobacteria
Yersinia pestis biovar Medievalis str Harbin 35 YP 005625911 1 547048 Proteobacteria
Yersinia enterocolitica subsp enterocolitica 8081 YP 001006083 1 393305 Proteobacteria
Serratia proteamaculans 568 YP 001476611 1 399741 Proteobacteria
Cronobacter turicensis z3032 YP 003210836 1 693216 Proteobacteria
Pantoea sp At 9b YP 004118903 1 592316 Proteobacteria
Rahnella sp Y9602 YP 004214997 1 741091 Proteobacteria
Serratia liquefaciens ATCC 27592 YP 008229885 1 1346614 Proteobacteria
Serratia liquefaciens ATCC 27592 YP 008232445 1 1346614 Proteobacteria
Klebsiella pneumoniae 342 YP 002235583 1 507522 Proteobacteria
Pantoea ananatis LMG 20103 YP 003522364 1 706191 Proteobacteria
Dickeya zeae MS1 WP 019846662 1 Dzeae
Dickeya zeae MS1 WP 019846732 1 Dzeae
Dickeya zeae CSL RW192 WP 038907191 1 Dzeae
Dickeya gs poaceaephila NCPPB 569 WP 042867874 1 Dpoaceaephila
Dickeya chrysanthemi subsp chrysanthemi NCPPB 402 WP 038907520 1 Dchrysanthemi
Dickeya zeae CSL RW192 WP 038907520 1 Dzeae
Dickeya dadantii 3937 WP 013316481 1 Ddadantii
Dickeya dadantii NCPPB 898 WP 038911901 1 Ddadantii
Dickeya gs poaceaephila NCPPB 569 WP 042871287 1 Dpoaceaephila
Dickeya dadantii NCPPB 898 WP 038911225 1 Ddadantii
Dickeya aquatica DW 0440 WP 019844299 1 Daquatica
DAQ1742 DAQ1742 02878 Daquatica
Dickeya sp CSL RW240 WP 019844299 1 Daquatica
DAQ1742 DAQ1742 00175 Daquatica
DAQ1742 DAQ1742 03427 Daquatica
Dickeya zeae MS1 WP 019844299 1 Dzeae
DAQ1742 DAQ1742 00641 Daquatica
DAQ1742 DAQ1742 00295 Daquatica
880 DAQ1742 01798 GI7 Seed
DAQ1742 DAQ1742 01798 Daquatica
Nitrosospira multiformis ATCC 25196 YP 412178 1 323848 Proteobacteria
Nitrosospira multiformis ATCC 25196 YP 411995 1 323848 Proteobacteria
Nitrosospira multiformis ATCC 25196 YP 410838 1 323848 Proteobacteria
Nitrosospira multiformis ATCC 25196 YP 413501 1 323848 Proteobacteria
Stenotrophomonas maltophilia D457 YP 006186048 1 1163399 Proteobacteria
Stenotrophomonas maltophilia D457 YP 006184214 1 1163399 Proteobacteria
Stenotrophomonas maltophilia D457 YP 006186436 1 1163399 Proteobacteria
Stenotrophomonas maltophilia D457 YP 006183053 1 1163399 Proteobacteria
Stenotrophomonas maltophilia D457 YP 006182926 1 1163399 Proteobacteria
Stenotrophomonas maltophilia D457 YP 006183308 1 1163399 Proteobacteria
Stenotrophomonas maltophilia D457 YP 006184500 1 1163399 Proteobacteria
Pandoraea pnomenusa 3kgm YP 008837973 1 1416914 Proteobacteria
Pandoraea pnomenusa 3kgm YP 008834358 1 1416914 Proteobacteria
Pandoraea pnomenusa 3kgm YP 008834630 1 1416914 Proteobacteria
Pandoraea pnomenusa 3kgm YP 008834495 1 1416914 Proteobacteria
Burkholderia cepacia GG4 YP 006614397 1 1009846 Proteobacteria
Burkholderia vietnamiensis G4 YP 001109702 1 269482 Proteobacteria
Burkholderia cenocepacia J2315 YP 002235519 1 216591 Proteobacteria
Burkholderia cenocepacia J2315 YP 002230444 1 216591 Proteobacteria
Burkholderia cenocepacia J2315 YP 002154091 1 216591 Proteobacteria
Burkholderia cenocepacia J2315 YP 002231656 1 216591 Proteobacteria
Burkholderia cenocepacia J2315 YP 002234527 1 216591 Proteobacteria
Burkholderia cenocepacia J2315 YP 002230309 1 216591 Proteobacteria
Burkholderia cenocepacia J2315 YP 002232334 1 216591 Proteobacteria
Burkholderia cenocepacia J2315 YP 002231700 1 216591 Proteobacteria
Burkholderia cenocepacia J2315 YP 002229579 1 216591 Proteobacteria
Burkholderia cenocepacia J2315 YP 002231435 1 216591 Proteobacteria
Burkholderia cenocepacia J2315 YP 002229577 1 216591 Proteobacteria
Burkholderia mallei NCTC 10247 YP 001080608 1 320389 Proteobacteria
Burkholderia mallei NCTC 10247 YP 001079066 1 320389 Proteobacteria
Burkholderia mallei NCTC 10247 YP 001081294 1 320389 Proteobacteria
Burkholderia mallei NCTC 10247 YP 001079715 1 320389 Proteobacteria
Burkholderia mallei NCTC 10247 YP 001080797 1 320389 Proteobacteria
Burkholderia mallei NCTC 10247 YP 001077978 1 320389 Proteobacteria
Burkholderia mallei NCTC 10247 YP 001080275 1 320389 Proteobacteria
Burkholderia mallei NCTC 10247 YP 001080050 1 320389 Proteobacteria
Burkholderia mallei NCTC 10247 YP 001080728 1 320389 Proteobacteria
Burkholderia mallei NCTC 10247 YP 001080229 1 320389 Proteobacteria
Burkholderia mallei NCTC 10247 YP 001078119 1 320389 Proteobacteria
Burkholderia mallei NCTC 10247 YP 001078481 1 320389 Proteobacteria
Burkholderia mallei NCTC 10247 YP 001078499 1 320389 Proteobacteria
Burkholderia mallei NCTC 10247 YP 001082479 1 320389 Proteobacteria

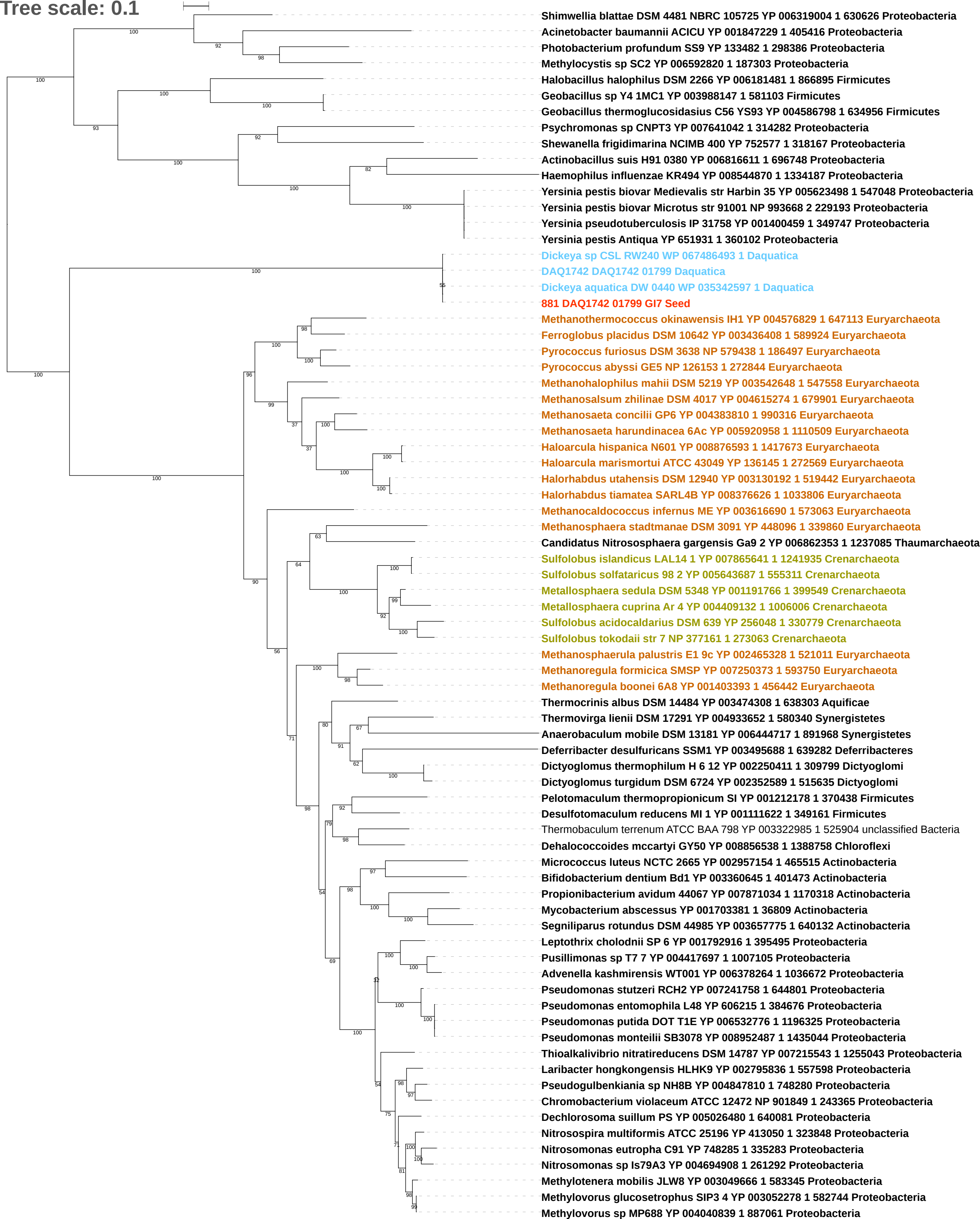
Tree scale: 0.1
Shimwellia blattae DSM 4481 NBRC 105725 YP 006319004 1 630626 Proteobacteria
Acinetobacter baumannii ACICU YP 001847229 1 405416 Proteobacteria
Photobacterium profundum SS9 YP 133482 1 298386 Proteobacteria
Methylocystis sp SC2 YP 006592820 1 187303 Proteobacteria
Halobacillus halophilus DSM 2266 YP 006181481 1 866895 Firmicutes
Geobacillus sp Y4 1MC1 YP 003988147 1 581103 Firmicutes
Geobacillus thermoglucosidasius C56 YS93 YP 004586798 1 634956 Firmicutes
Psychromonas sp CNPT3 YP 007641042 1 314282 Proteobacteria
Shewanella frigidimarina NCIMB 400 YP 752577 1 318167 Proteobacteria
Actinobacillus suis H91 0380 YP 006816611 1 696748 Proteobacteria
Haemophilus influenzae KR494 YP 008544870 1 1334187 Proteobacteria
Yersinia pestis biovar Medievalis str Harbin 35 YP 005623498 1 547048 Proteobacteria
Yersinia pestis biovar Microtus str 91001 NP 993668 2 229193 Proteobacteria
Yersinia pseudotuberculosis IP 31758 YP 001400459 1 349747 Proteobacteria
Yersinia pestis Antiqua YP 651931 1 360102 Proteobacteria
Dickeya sp CSL RW240 WP 067486493 1 Daquatica
DAQ1742 DAQ1742 01799 Daquatica
Dickeya aquatica DW 0440 WP 035342597 1 Daquatica
881 DAQ1742 01799 GI7 Seed
Methanothermococcus okinawensis IH1 YP 004576829 1 647113 Euryarchaeota
Ferroglobus placidus DSM 10642 YP 003436408 1 589924 Euryarchaeota
Pyrococcus furiosus DSM 3638 NP 579438 1 186497 Euryarchaeota
Pyrococcus abyssi GE5 NP 126153 1 272844 Euryarchaeota
Methanohalophilus mahii DSM 5219 YP 003542648 1 547558 Euryarchaeota
Methanosalsum zhilinae DSM 4017 YP 004615274 1 679901 Euryarchaeota
Methanosaeta concilii GP6 YP 004383810 1 990316 Euryarchaeota
Methanosaeta harundinacea 6Ac YP 005920958 1 1110509 Euryarchaeota
Haloarcula hispanica N601 YP 008876593 1 1417673 Euryarchaeota
Haloarcula marismortui ATCC 43049 YP 136145 1 272569 Euryarchaeota
Halorhabdus utahensis DSM 12940 YP 003130192 1 519442 Euryarchaeota
Halorhabdus tiamatea SARL4B YP 008376626 1 1033806 Euryarchaeota
Methanocaldococcus infernus ME YP 003616690 1 573063 Euryarchaeota
Methanosphaera stadtmanae DSM 3091 YP 448096 1 339860 Euryarchaeota
Candidatus Nitrososphaera gargensis Ga9 2 YP 006862353 1 1237085 Thaumarchaeota
Sulfolobus islandicus LAL14 1 YP 007865641 1 1241935 Crenarchaeota
Sulfolobus solfataricus 98 2 YP 005643687 1 555311 Crenarchaeota
Metallosphaera sedula DSM 5348 YP 001191766 1 399549 Crenarchaeota
Metallosphaera cuprina Ar 4 YP 004409132 1 1006006 Crenarchaeota
Sulfolobus acidocaldarius DSM 639 YP 256048 1 330779 Crenarchaeota
Sulfolobus tokodaii str 7 NP 377161 1 273063 Crenarchaeota
Methanosphaerula palustris E1 9c YP 002465328 1 521011 Euryarchaeota
Methanoregula formicica SMSP YP 007250373 1 593750 Euryarchaeota
Methanoregula boonei 6A8 YP 001403393 1 456442 Euryarchaeota
Thermocrinis albus DSM 14484 YP 003474308 1 638303 Aquificae
Thermovirga lienii DSM 17291 YP 004933652 1 580340 Synergistetes
Anaerobaculum mobile DSM 13181 YP 006444717 1 891968 Synergistetes
Deferribacter desulfuricans SSM1 YP 003495688 1 639282 Deferribacteres
Dictyoglomus thermophilum H 6 12 YP 002250411 1 309799 Dictyoglomi
Dictyoglomus turgidum DSM 6724 YP 002352589 1 515635 Dictyoglomi
Pelotomaculum thermopropionicum SI YP 001212178 1 370438 Firmicutes
Desulfotomaculum reducens MI 1 YP 001111622 1 349161 Firmicutes
Thermobaculum terrenum ATCC BAA 798 YP 003322985 1 525904 unclassified Bacteria
Dehalococcoides mccartyi GY50 YP 008856538 1 1388758 Chloroflexi
Micrococcus luteus NCTC 2665 YP 002957154 1 465515 Actinobacteria
Bifidobacterium dentium Bd1 YP 003360645 1 401473 Actinobacteria
Propionibacterium avidum 44067 YP 007871034 1 1170318 Actinobacteria
Mycobacterium abscessus YP 001703381 1 36809 Actinobacteria
Segniliparus rotundus DSM 44985 YP 003657775 1 640132 Actinobacteria
Leptothrix cholodnii SP 6 YP 001792916 1 395495 Proteobacteria
Pusillimonas sp T7 7 YP 004417697 1 1007105 Proteobacteria
Advenella kashmirensis WT001 YP 006378264 1 1036672 Proteobacteria
Pseudomonas stutzeri RCH2 YP 007241758 1 644801 Proteobacteria
Pseudomonas entomophila L48 YP 606215 1 384676 Proteobacteria
Pseudomonas putida DOT T1E YP 006532776 1 1196325 Proteobacteria
Pseudomonas monteilii SB3078 YP 008952487 1 1435044 Proteobacteria
Thioalkalivibrio nitratireducens DSM 14787 YP 007215543 1 1255043 Proteobacteria
Laribacter hongkongensis HLHK9 YP 002795836 1 557598 Proteobacteria
Pseudogulbenkiania sp NH8B YP 004847810 1 748280 Proteobacteria
Chromobacterium violaceum ATCC 12472 NP 901849 1 243365 Proteobacteria
Dechlorosoma suillum PS YP 005026480 1 640081 Proteobacteria
Nitrosospira multiformis ATCC 25196 YP 413050 1 323848 Proteobacteria
Nitrosomonas eutropha C91 YP 748285 1 335283 Proteobacteria
Nitrosomonas sp Is79A3 YP 004694908 1 261292 Proteobacteria
Methylotenera mobilis JLW8 YP 003049666 1 583345 Proteobacteria
Methylovorus glucosetrophus SIP3 4 YP 003052278 1 582744 Proteobacteria
Methylovorus sp MP688 YP 004040839 1 887061 Proteobacteria

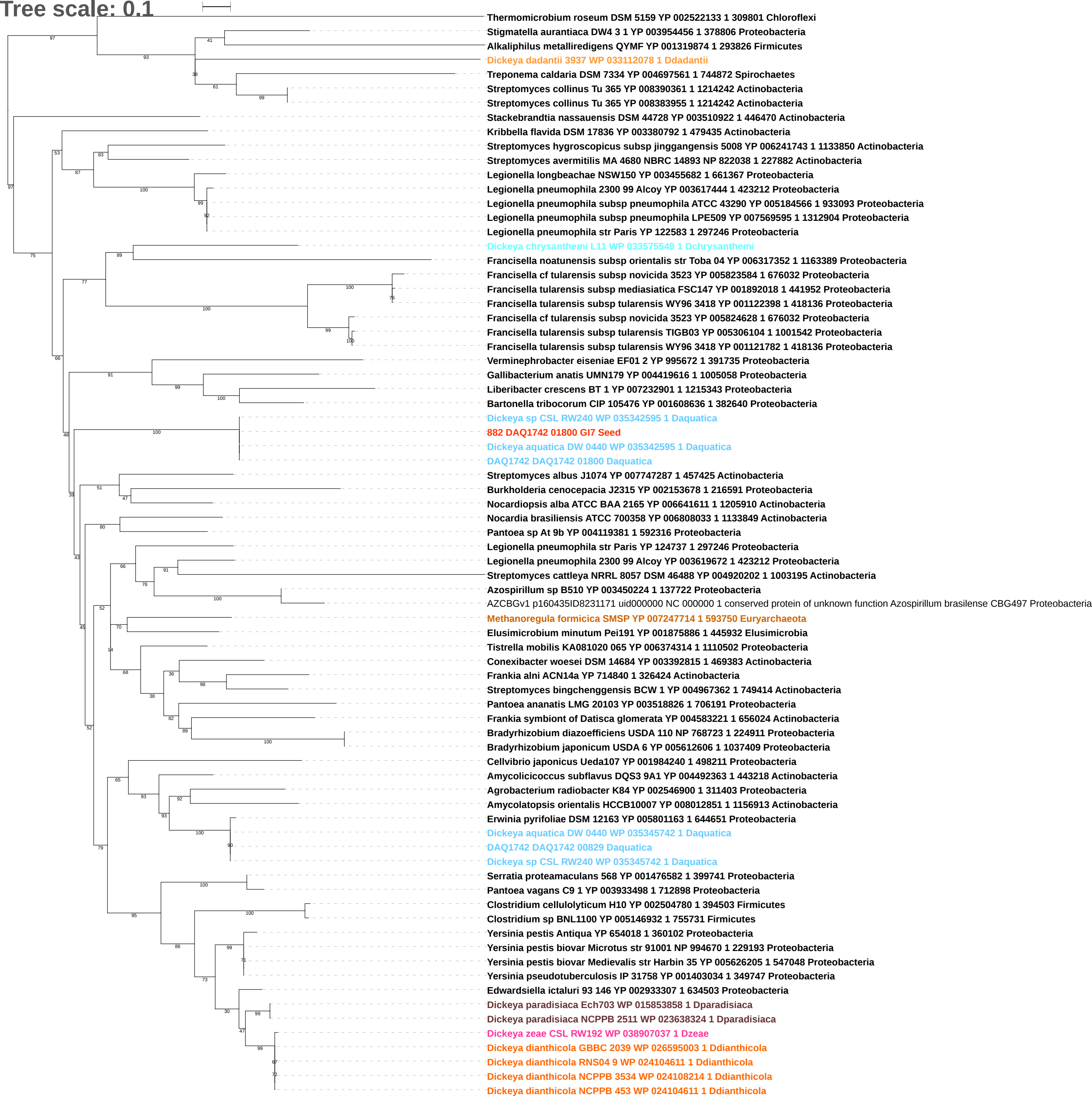
Tree scale: 0.1
Thermomicrobium roseum DSM 5159 YP 002522133 1 309801 Chloroflexi
Stigmatella aurantiaca DW4 3 1 YP 003954456 1 378806 Proteobacteria
Alkaliphilus metalliredigens QYMF YP 001319874 1 293826 Firmicutes
Dickeya dadantii 3937 WP 033112078 1 Ddadantii
Treponema caldaria DSM 7334 YP 004697561 1 744872 Spirochaetes
Streptomyces collinus Tu 365 YP 008390361 1 1214242 Actinobacteria
Streptomyces collinus Tu 365 YP 008383955 1 1214242 Actinobacteria
Stackebrandtia nassauensis DSM 44728 YP 003510922 1 446470 Actinobacteria
Kribbella flavida DSM 17836 YP 003380792 1 479435 Actinobacteria
Streptomyces hygroscopicus subsp jinggangensis 5008 YP 006241743 1 1133850 Actinobacteria
Streptomyces avermitilis MA 4680 NBRC 14893 NP 822038 1 227882 Actinobacteria
Legionella longbeachae NSW150 YP 003455682 1 661367 Proteobacteria
Legionella pneumophila 2300 99 Alcoy YP 003617444 1 423212 Proteobacteria
Legionella pneumophila subsp pneumophila ATCC 43290 YP 005184566 1 933093 Proteobacteria
Legionella pneumophila subsp pneumophila LPE509 YP 007569595 1 1312904 Proteobacteria
Legionella pneumophila str Paris YP 122583 1 297246 Proteobacteria
Dickeya chrysanthemi L11 WP 033575549 1 Dchrysanthemi
Francisella noatunensis subsp orientalis str Toba 04 YP 006317352 1 1163389 Proteobacteria
Francisella cf tularensis subsp novicida 3523 YP 005823584 1 676032 Proteobacteria
Francisella tularensis subsp mediasiatica FSC147 YP 001892018 1 441952 Proteobacteria
Francisella tularensis subsp tularensis WY96 3418 YP 001122398 1 418136 Proteobacteria
Francisella cf tularensis subsp novicida 3523 YP 005824628 1 676032 Proteobacteria
Francisella tularensis subsp tularensis TIGB03 YP 005306104 1 1001542 Proteobacteria
Francisella tularensis subsp tularensis WY96 3418 YP 001121782 1 418136 Proteobacteria
Verminephrobacter eiseniae EF01 2 YP 995672 1 391735 Proteobacteria
Gallibacterium anatis UMN179 YP 004419616 1 1005058 Proteobacteria
Liberibacter crescens BT 1 YP 007232901 1 1215343 Proteobacteria
Bartonella tribocorum CIP 105476 YP 001608636 1 382640 Proteobacteria
Dickeya sp CSL RW240 WP 035342595 1 Daquatica
882 DAQ1742 01800 GI7 Seed
Dickeya aquatica DW 0440 WP 035342595 1 Daquatica
DAQ1742 DAQ1742 01800 Daquatica
Streptomyces albus J1074 YP 007747287 1 457425 Actinobacteria
Burkholderia cenocepacia J2315 YP 002153678 1 216591 Proteobacteria
Nocardiopsis alba ATCC BAA 2165 YP 006641611 1 1205910 Actinobacteria
Nocardia brasiliensis ATCC 700358 YP 006808033 1 1133849 Actinobacteria
Pantoea sp At 9b YP 004119381 1 592316 Proteobacteria
Legionella pneumophila str Paris YP 124737 1 297246 Proteobacteria
Legionella pneumophila 2300 99 Alcoy YP 003619672 1 423212 Proteobacteria
Streptomyces cattleya NRRL 8057 DSM 46488 YP 004920202 1 1003195 Actinobacteria
Azospirillum sp B510 YP 003450224 1 137722 Proteobacteria
AZCBGv1 p160435ID8231171 uid000000 NC 000000 1 conserved protein of unknown function Azospirillum brasilense CBG497 Proteobacteria
Methanoregula formicica SMSP YP 007247714 1 593750 Euryarchaeota
Elusimicrobium minutum Pei191 YP 001875886 1 445932 Elusimicrobia
Tistrella mobilis KA081020 065 YP 006374314 1 1110502 Proteobacteria
Conexibacter woesei DSM 14684 YP 003392815 1 469383 Actinobacteria
Frankia alni ACN14a YP 714840 1 326424 Actinobacteria
Streptomyces bingchenggensis BCW 1 YP 004967362 1 749414 Actinobacteria
Pantoea ananatis LMG 20103 YP 003518826 1 706191 Proteobacteria
Frankia symbiont of Datisca glomerata YP 004583221 1 656024 Actinobacteria
Bradyrhizobium diazoefficiens USDA 110 NP 768723 1 224911 Proteobacteria
Bradyrhizobium japonicum USDA 6 YP 005612606 1 1037409 Proteobacteria
Cellvibrio japonicus Ueda107 YP 001984240 1 498211 Proteobacteria
Amycolicicoccus subflavus DQS3 9A1 YP 004492363 1 443218 Actinobacteria
Agrobacterium radiobacter K84 YP 002546900 1 311403 Proteobacteria
Amycolatopsis orientalis HCCB10007 YP 008012851 1 1156913 Actinobacteria
Erwinia pyrifoliae DSM 12163 YP 005801163 1 644651 Proteobacteria
Dickeya aquatica DW 0440 WP 035345742 1 Daquatica
DAQ1742 DAQ1742 00829 Daquatica
Dickeya sp CSL RW240 WP 035345742 1 Daquatica
Serratia proteamaculans 568 YP 001476582 1 399741 Proteobacteria
Pantoea vagans C9 1 YP 003933498 1 712898 Proteobacteria
Clostridium cellulolyticum H10 YP 002504780 1 394503 Firmicutes
Clostridium sp BNL1100 YP 005146932 1 755731 Firmicutes
Yersinia pestis Antiqua YP 654018 1 360102 Proteobacteria
Yersinia pestis biovar Microtus str 91001 NP 994670 1 229193 Proteobacteria
Yersinia pestis biovar Medievalis str Harbin 35 YP 005626205 1 547048 Proteobacteria
Yersinia pseudotuberculosis IP 31758 YP 001403034 1 349747 Proteobacteria
Edwardsiella ictaluri 93 146 YP 002933307 1 634503 Proteobacteria
Dickeya paradisiaca Ech703 WP 015853858 1 Dparadisiaca
Dickeya paradisiaca NCPPB 2511 WP 023638324 1 Dparadisiaca
Dickeya zeae CSL RW192 WP 038907037 1 Dzeae
Dickeya dianthicola GBBC 2039 WP 026595003 1 Ddianthicola
Dickeya dianthicola RNS04 9 WP 024104611 1 Ddianthicola
Dickeya dianthicola NCPPB 3534 WP 024108214 1 Ddianthicola
Dickeya dianthicola NCPPB 453 WP 024104611 1 Ddianthicola

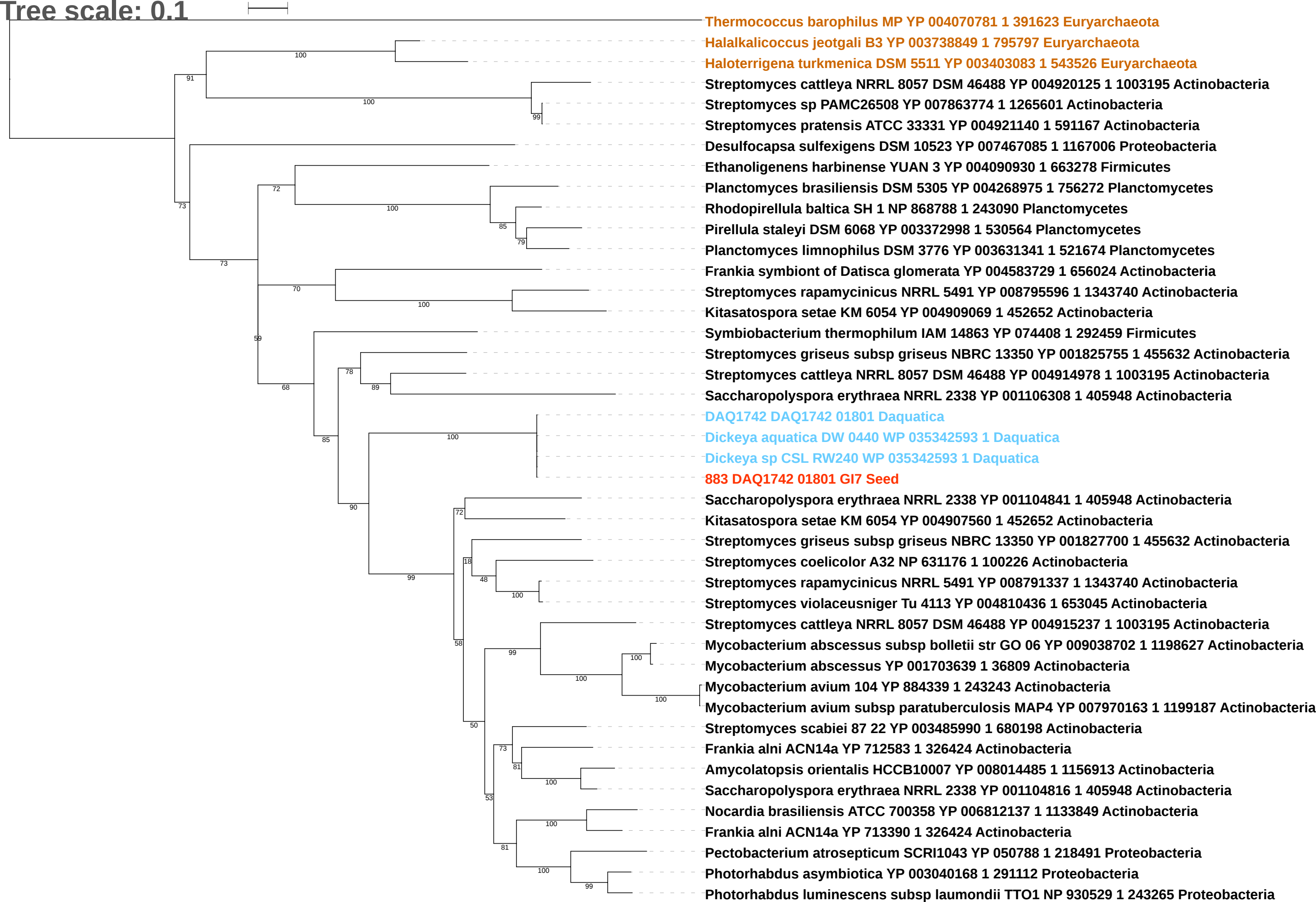
Tree scale: 0.1
Thermococcus barophilus MP YP 004070781 1 391623 Euryarchaeota
Halalkalicoccus jeotgali B3 YP 003738849 1 795797 Euryarchaeota
Haloterrigena turkmenica DSM 5511 YP 003403083 1 543526 Euryarchaeota
Streptomyces cattleya NRRL 8057 DSM 46488 YP 004920125 1 1003195 Actinobacteria
Streptomyces sp PAMC26508 YP 007863774 1 1265601 Actinobacteria
Streptomyces pratensis ATCC 33331 YP 004921140 1 591167 Actinobacteria
Desulfocapsa sulfexigens DSM 10523 YP 007467085 1 1167006 Proteobacteria
Ethanoligenens harbinense YUAN 3 YP 004090930 1 663278 Firmicutes
Planctomyces brasiliensis DSM 5305 YP 004268975 1 756272 Planctomycetes
Rhodopirellula baltica SH 1 NP 868788 1 243090 Planctomycetes
Pirellula staleyi DSM 6068 YP 003372998 1 530564 Planctomycetes
Planctomyces limnophilus DSM 3776 YP 003631341 1 521674 Planctomycetes
Frankia symbiont of Datisca glomerata YP 004583729 1 656024 Actinobacteria
Streptomyces rapamycinicus NRRL 5491 YP 008795596 1 1343740 Actinobacteria
Kitasatospora setae KM 6054 YP 004909069 1 452652 Actinobacteria
Symbiobacterium thermophilum IAM 14863 YP 074408 1 292459 Firmicutes
Streptomyces griseus subsp griseus NBRC 13350 YP 001825755 1 455632 Actinobacteria
Streptomyces cattleya NRRL 8057 DSM 46488 YP 004914978 1 1003195 Actinobacteria
Saccharopolyspora erythraea NRRL 2338 YP 001106308 1 405948 Actinobacteria
DAQ1742 DAQ1742 01801 Daquatica
Dickeya aquatica DW 0440 WP 035342593 1 Daquatica
Dickeya sp CSL RW240 WP 035342593 1 Daquatica
883 DAQ1742 01801 GI7 Seed
Saccharopolyspora erythraea NRRL 2338 YP 001104841 1 405948 Actinobacteria
Kitasatospora setae KM 6054 YP 004907560 1 452652 Actinobacteria
Streptomyces griseus subsp griseus NBRC 13350 YP 001827700 1 455632 Actinobacteria
Streptomyces coelicolor A32 NP 631176 1 100226 Actinobacteria
Streptomyces rapamycinicus NRRL 5491 YP 008791337 1 1343740 Actinobacteria
Streptomyces violaceusniger Tu 4113 YP 004810436 1 653045 Actinobacteria
Streptomyces cattleya NRRL 8057 DSM 46488 YP 004915237 1 1003195 Actinobacteria
Mycobacterium abscessus subsp bolletii str GO 06 YP 009038702 1 1198627 Actinobacteria
Mycobacterium abscessus YP 001703639 1 36809 Actinobacteria
Mycobacterium avium 104 YP 884339 1 243243 Actinobacteria
Mycobacterium avium subsp paratuberculosis MAP4 YP 007970163 1 1199187 Actinobacteria
Streptomyces scabiei 87 22 YP 003485990 1 680198 Actinobacteria
Frankia alni ACN14a YP 712583 1 326424 Actinobacteria
Amycolatopsis orientalis HCCB10007 YP 008014485 1 1156913 Actinobacteria
Saccharopolyspora erythraea NRRL 2338 YP 001104816 1 405948 Actinobacteria
Nocardia brasiliensis ATCC 700358 YP 006812137 1 1133849 Actinobacteria
Frankia alni ACN14a YP 713390 1 326424 Actinobacteria
Pectobacterium atrosepticum SCRI1043 YP 050788 1 218491 Proteobacteria
Photorhabdus asymbiotica YP 003040168 1 291112 Proteobacteria
Photorhabdus luminescens subsp laumondii TTO1 NP 930529 1 243265 Proteobacteria

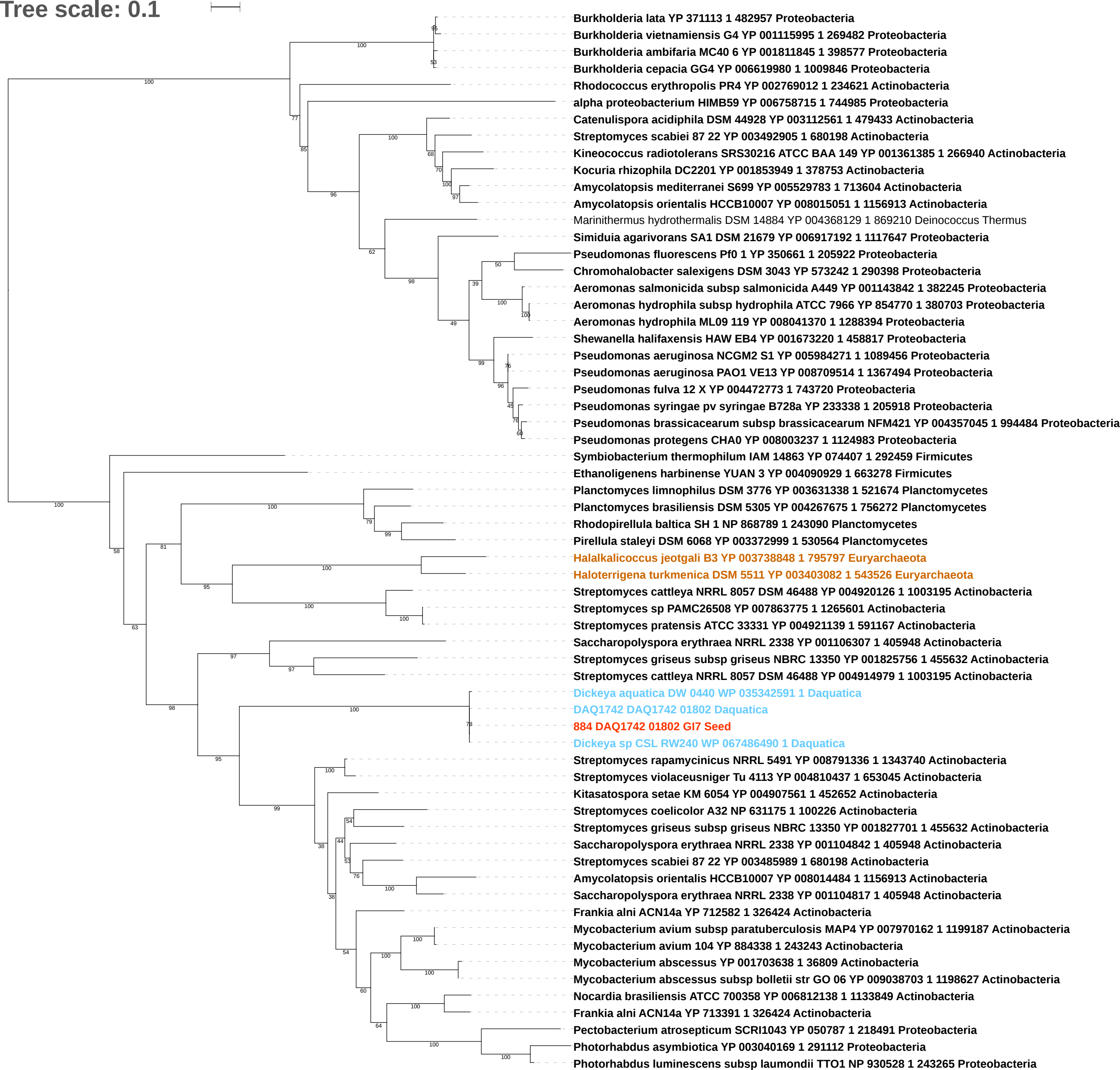
Tree scale: 0.1
Burkholderia lata YP 371113 1 482957 Proteobacteria
Burkholderia vietnamiensis G4 YP 001115995 1 269482 Proteobacteria
Burkholderia ambifaria MC40 6 YP 001811845 1 398577 Proteobacteria
Burkholderia cepacia GG4 YP 006619980 1 1009846 Proteobacteria
Rhodococcus erythropolis PR4 YP 002769012 1 234621 Actinobacteria
alpha proteobacterium HIMB59 YP 006758715 1 744985 Proteobacteria
Catenulispora acidiphila DSM 44928 YP 003112561 1 479433 Actinobacteria
Streptomyces scabiei 87 22 YP 003492905 1 680198 Actinobacteria
Kineococcus radiotolerans SRS30216 ATCC BAA 149 YP 001361385 1 266940 Actinobacteria
Kocuria rhizophila DC2201 YP 001853949 1 378753 Actinobacteria
Amycolatopsis mediterranei S699 YP 005529783 1 713604 Actinobacteria
Amycolatopsis orientalis HCCB10007 YP 008015051 1 1156913 Actinobacteria
Marinithermus hydrothermalis DSM 14884 YP 004368129 1 869210 Deinococcus Thermus
Simiduia agarivorans SA1 DSM 21679 YP 006917192 1 1117647 Proteobacteria
Pseudomonas fluorescens Pf0 1 YP 350661 1 205922 Proteobacteria
Chromohalobacter salexigens DSM 3043 YP 573242 1 290398 Proteobacteria
Aeromonas salmonicida subsp salmonicida A449 YP 001143842 1 382245 Proteobacteria
Aeromonas hydrophila subsp hydrophila ATCC 7966 YP 854770 1 380703 Proteobacteria
Aeromonas hydrophila ML09 119 YP 008041370 1 1288394 Proteobacteria
Shewanella halifaxensis HAW EB4 YP 001673220 1 458817 Proteobacteria
Pseudomonas aeruginosa NCGM2 S1 YP 005984271 1 1089456 Proteobacteria
Pseudomonas aeruginosa PAO1 VE13 YP 008709514 1 1367494 Proteobacteria
Pseudomonas fulva 12 X YP 004472773 1 743720 Proteobacteria
Pseudomonas syringae pv syringae B728a YP 233338 1 205918 Proteobacteria
Pseudomonas brassicacearum subsp brassicacearum NFM421 YP 004357045 1 994484 Proteobacteria
Pseudomonas protegens CHA0 YP 008003237 1 1124983 Proteobacteria
Symbiobacterium thermophilum IAM 14863 YP 074407 1 292459 Firmicutes
Ethanoligenens harbinense YUAN 3 YP 004090929 1 663278 Firmicutes
Planctomyces limnophilus DSM 3776 YP 003631338 1 521674 Planctomycetes
Planctomyces brasiliensis DSM 5305 YP 004267675 1 756272 Planctomycetes
Rhodopirellula baltica SH 1 NP 868789 1 243090 Planctomycetes
Pirellula staleyi DSM 6068 YP 003372999 1 530564 Planctomycetes
Halalkalicoccus jeotgali B3 YP 003738848 1 795797 Euryarchaeota
Haloterrigena turkmenica DSM 5511 YP 003403082 1 543526 Euryarchaeota
Streptomyces cattleya NRRL 8057 DSM 46488 YP 004920126 1 1003195 Actinobacteria
Streptomyces sp PAMC26508 YP 007863775 1 1265601 Actinobacteria
Streptomyces pratensis ATCC 33331 YP 004921139 1 591167 Actinobacteria
Saccharopolyspora erythraea NRRL 2338 YP 001106307 1 405948 Actinobacteria
Streptomyces griseus subsp griseus NBRC 13350 YP 001825756 1 455632 Actinobacteria
Streptomyces cattleya NRRL 8057 DSM 46488 YP 004914979 1 1003195 Actinobacteria
Dickeya aquatica DW 0440 WP 035342591 1 Daquatica
DAQ1742 DAQ1742 01802 Daquatica
884 DAQ1742 01802 GI7 Seed
Dickeya sp CSL RW240 WP 067486490 1 Daquatica
Streptomyces rapamycinicus NRRL 5491 YP 008791336 1 1343740 Actinobacteria
Streptomyces violaceusniger Tu 4113 YP 004810437 1 653045 Actinobacteria
Kitasatospora setae KM 6054 YP 004907561 1 452652 Actinobacteria
Streptomyces coelicolor A32 NP 631175 1 100226 Actinobacteria
Streptomyces griseus subsp griseus NBRC 13350 YP 001827701 1 455632 Actinobacteria
Saccharopolyspora erythraea NRRL 2338 YP 001104842 1 405948 Actinobacteria
Streptomyces scabiei 87 22 YP 003485989 1 680198 Actinobacteria
Amycolatopsis orientalis HCCB10007 YP 008014484 1 1156913 Actinobacteria
Saccharopolyspora erythraea NRRL 2338 YP 001104817 1 405948 Actinobacteria
Frankia alni ACN14a YP 712582 1 326424 Actinobacteria
Mycobacterium avium subsp paratuberculosis MAP4 YP 007970162 1 1199187 Actinobacteria
Mycobacterium avium 104 YP 884338 1 243243 Actinobacteria
Mycobacterium abscessus YP 001703638 1 36809 Actinobacteria
Mycobacterium abscessus subsp bolletii str GO 06 YP 009038703 1 1198627 Actinobacteria
Nocardia brasiliensis ATCC 700358 YP 006812138 1 1133849 Actinobacteria
Frankia alni ACN14a YP 713391 1 326424 Actinobacteria
Pectobacterium atrosepticum SCRI1043 YP 050787 1 218491 Proteobacteria
Photorhabdus asymbiotica YP 003040169 1 291112 Proteobacteria
Photorhabdus luminescens subsp laumondii TTO1 NP 930528 1 243265 Proteobacteria

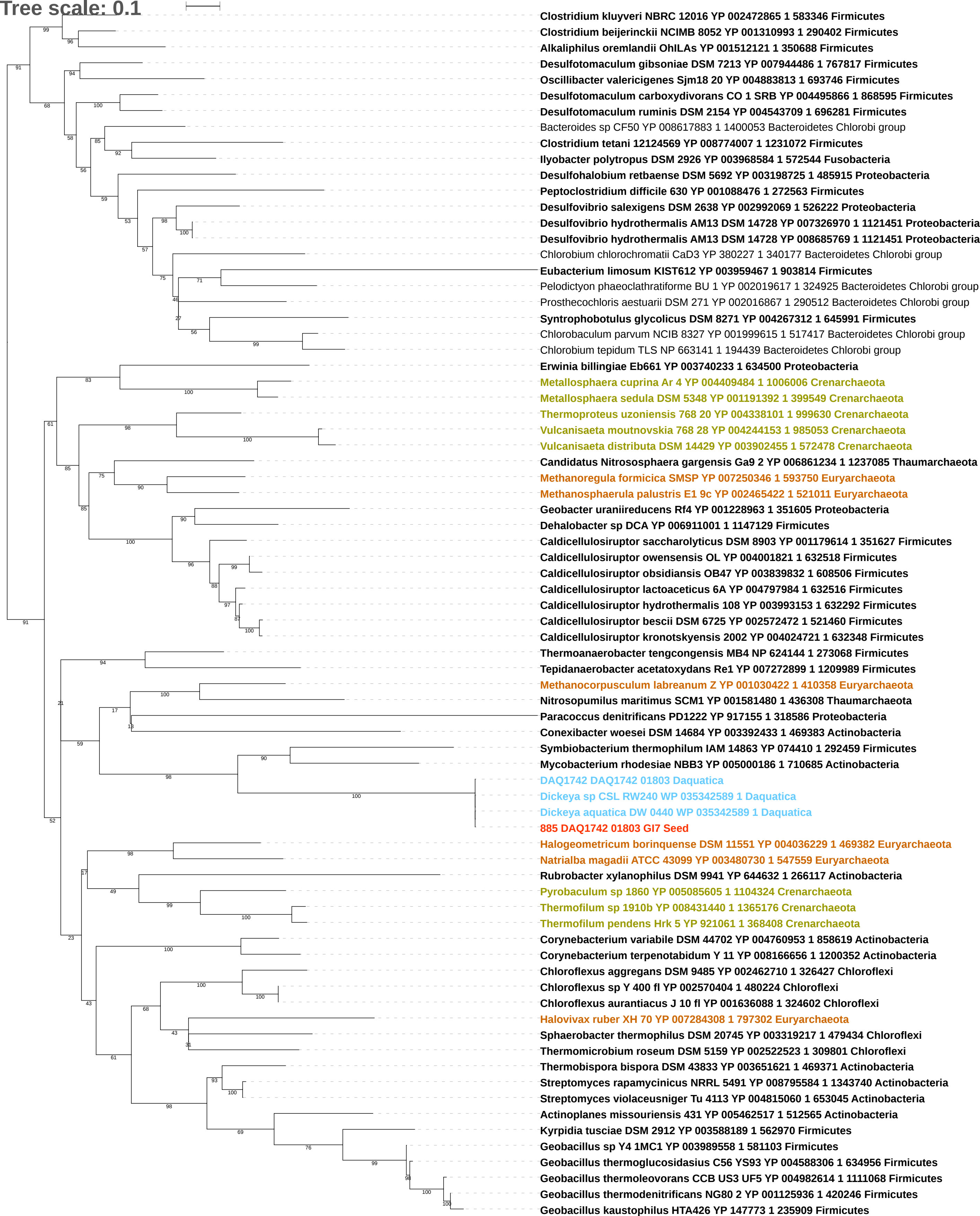
Tree scale: 0.1
Clostridium kluyveri NBRC 12016 YP 002472865 1 583346 Firmicutes
Clostridium beijerinckii NCIMB 8052 YP 001310993 1 290402 Firmicutes
Alkaliphilus oremlandii OhILAs YP 001512121 1 350688 Firmicutes
Desulfotomaculum gibsoniae DSM 7213 YP 007944486 1 767817 Firmicutes
Oscillibacter valericigenes Sjm18 20 YP 004883813 1 693746 Firmicutes
Desulfotomaculum carboxydivorans CO 1 SRB YP 004495866 1 868595 Firmicutes
Desulfotomaculum ruminis DSM 2154 YP 004543709 1 696281 Firmicutes
Bacteroides sp CF50 YP 008617883 1 1400053 Bacteroidetes Chlorobi group
Clostridium tetani 12124569 YP 008774007 1 1231072 Firmicutes
Ilyobacter polytropus DSM 2926 YP 003968584 1 572544 Fusobacteria
Desulfohalobium retbaense DSM 5692 YP 003198725 1 485915 Proteobacteria
Peptoclostridium difficile 630 YP 001088476 1 272563 Firmicutes
Desulfovibrio salexigens DSM 2638 YP 002992069 1 526222 Proteobacteria
Desulfovibrio hydrothermalis AM13 DSM 14728 YP 007326970 1 1121451 Proteobacteria
Desulfovibrio hydrothermalis AM13 DSM 14728 YP 008685769 1 1121451 Proteobacteria
Chlorobium chlorochromatii CaD3 YP 380227 1 340177 Bacteroidetes Chlorobi group
Eubacterium limosum KIST612 YP 003959467 1 903814 Firmicutes
Pelodictyon phaeoclathratiforme BU 1 YP 002019617 1 324925 Bacteroidetes Chlorobi group
Prosthecochloris aestuarii DSM 271 YP 002016867 1 290512 Bacteroidetes Chlorobi group
Syntrophobotulus glycolicus DSM 8271 YP 004267312 1 645991 Firmicutes
Chlorobaculum parvum NCIB 8327 YP 001999615 1 517417 Bacteroidetes Chlorobi group
Chlorobium tepidum TLS NP 663141 1 194439 Bacteroidetes Chlorobi group
Erwinia billingiae Eb661 YP 003740233 1 634500 Proteobacteria
Metallosphaera cuprina Ar 4 YP 004409484 1 1006006 Crenarchaeota
Metallosphaera sedula DSM 5348 YP 001191392 1 399549 Crenarchaeota
Thermoproteus uzoniensis 768 20 YP 004338101 1 999630 Crenarchaeota
Vulcanisaeta moutnovskia 768 28 YP 004244153 1 985053 Crenarchaeota
Vulcanisaeta distributa DSM 14429 YP 003902455 1 572478 Crenarchaeota
Candidatus Nitrososphaera gargensis Ga9 2 YP 006861234 1 1237085 Thaumarchaeota
Methanoregula formicica SMSP YP 007250346 1 593750 Euryarchaeota
Methanosphaerula palustris E1 9c YP 002465422 1 521011 Euryarchaeota
Geobacter uraniireducens Rf4 YP 001228963 1 351605 Proteobacteria
Dehalobacter sp DCA YP 006911001 1 1147129 Firmicutes
Caldicellulosiruptor saccharolyticus DSM 8903 YP 001179614 1 351627 Firmicutes
Caldicellulosiruptor owensensis OL YP 004001821 1 632518 Firmicutes
Caldicellulosiruptor obsidiansis OB47 YP 003839832 1 608506 Firmicutes
Caldicellulosiruptor lactoaceticus 6A YP 004797984 1 632516 Firmicutes
Caldicellulosiruptor hydrothermalis 108 YP 003993153 1 632292 Firmicutes
Caldicellulosiruptor bescii DSM 6725 YP 002572472 1 521460 Firmicutes
Caldicellulosiruptor kronotskyensis 2002 YP 004024721 1 632348 Firmicutes
Thermoanaerobacter tengcongensis MB4 NP 624144 1 273068 Firmicutes
Tepidanaerobacter acetatoxydans Re1 YP 007272899 1 1209989 Firmicutes
Methanocorpusculum labreanum Z YP 001030422 1 410358 Euryarchaeota
Nitrosopumilus maritimus SCM1 YP 001581480 1 436308 Thaumarchaeota
Paracoccus denitrificans PD1222 YP 917155 1 318586 Proteobacteria
Conexibacter woesei DSM 14684 YP 003392433 1 469383 Actinobacteria
Symbiobacterium thermophilum IAM 14863 YP 074410 1 292459 Firmicutes
Mycobacterium rhodesiae NBB3 YP 005000186 1 710685 Actinobacteria
DAQ1742 DAQ1742 01803 Daquatica
Dickeya sp CSL RW240 WP 035342589 1 Daquatica
Dickeya aquatica DW 0440 WP 035342589 1 Daquatica
885 DAQ1742 01803 GI7 Seed
Halogeometricum borinquense DSM 11551 YP 004036229 1 469382 Euryarchaeota
Natrialba magadii ATCC 43099 YP 003480730 1 547559 Euryarchaeota
Rubrobacter xylanophilus DSM 9941 YP 644632 1 266117 Actinobacteria
Pyrobaculum sp 1860 YP 005085605 1 1104324 Crenarchaeota
Thermofilum sp 1910b YP 008431440 1 1365176 Crenarchaeota
Thermofilum pendens Hrk 5 YP 921061 1 368408 Crenarchaeota
Corynebacterium variabile DSM 44702 YP 004760953 1 858619 Actinobacteria
Corynebacterium terpenotabidum Y 11 YP 008166656 1 1200352 Actinobacteria
Chloroflexus aggregans DSM 9485 YP 002462710 1 326427 Chloroflexi
Chloroflexus sp Y 400 fl YP 002570404 1 480224 Chloroflexi
Chloroflexus aurantiacus J 10 fl YP 001636088 1 324602 Chloroflexi
Halovivax ruber XH 70 YP 007284308 1 797302 Euryarchaeota
Sphaerobacter thermophilus DSM 20745 YP 003319217 1 479434 Chloroflexi
Thermomicrobium roseum DSM 5159 YP 002522523 1 309801 Chloroflexi
Thermobispora bispora DSM 43833 YP 003651621 1 469371 Actinobacteria
Streptomyces rapamycinicus NRRL 5491 YP 008795584 1 1343740 Actinobacteria
Streptomyces violaceusniger Tu 4113 YP 004815060 1 653045 Actinobacteria
Actinoplanes missouriensis 431 YP 005462517 1 512565 Actinobacteria
Kyrpidia tusciae DSM 2912 YP 003588189 1 562970 Firmicutes
Geobacillus sp Y4 1MC1 YP 003989558 1 581103 Firmicutes
Geobacillus thermoglucosidasius C56 YS93 YP 004588306 1 634956 Firmicutes
Geobacillus thermoleovorans CCB US3 UF5 YP 004982614 1 1111068 Firmicutes
Geobacillus thermodenitrificans NG80 2 YP 001125936 1 420246 Firmicutes
Geobacillus kaustophilus HTA426 YP 147773 1 235909 Firmicutes

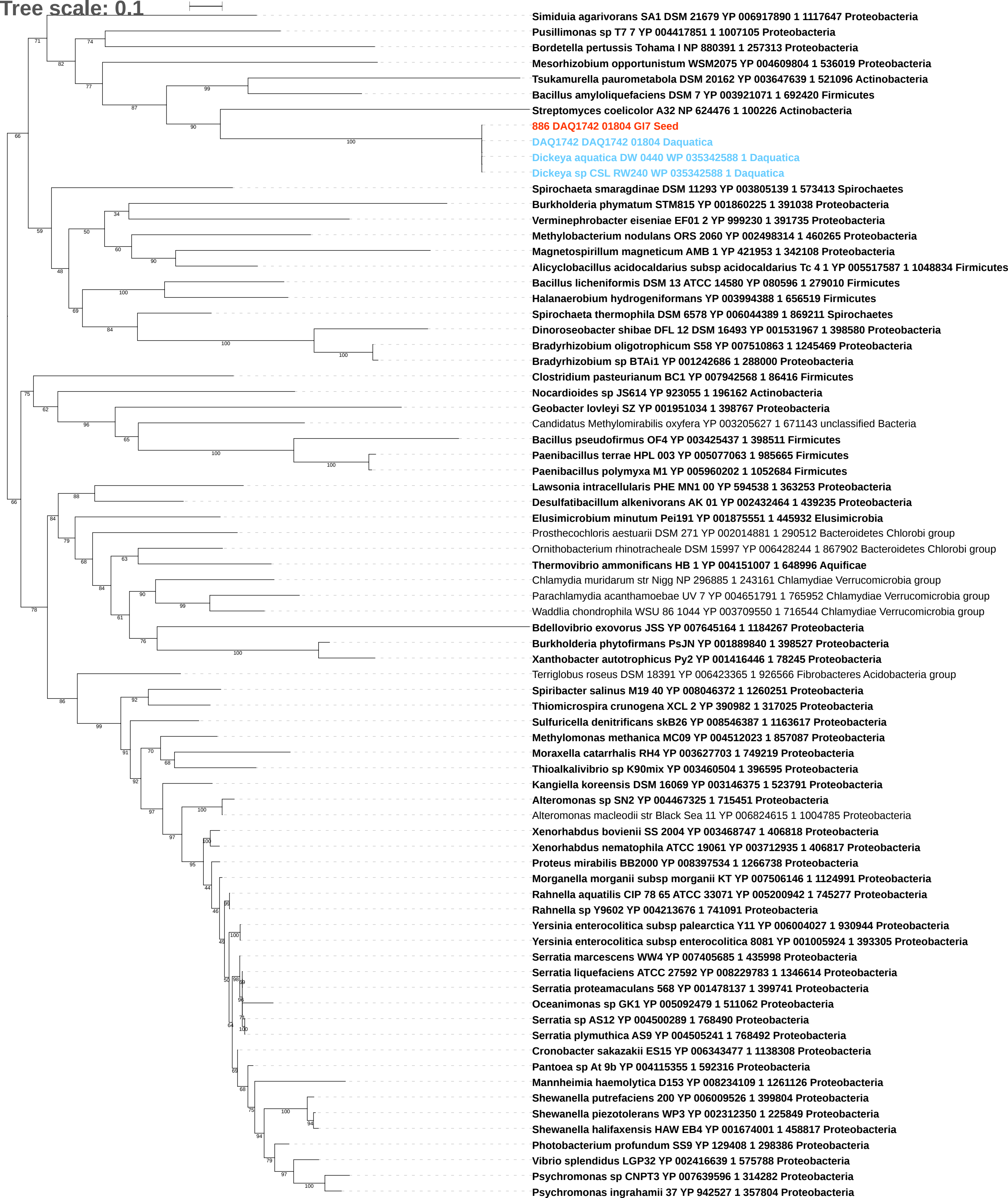
Tree scale: 0.1
Simiduia agarivorans SA1 DSM 21679 YP 006917890 1 1117647 Proteobacteria
Pusillimonas sp T7 7 YP 004417851 1 1007105 Proteobacteria
Bordetella pertussis Tohama I NP 880391 1 257313 Proteobacteria
Mesorhizobium opportunistum WSM2075 YP 004609804 1 536019 Proteobacteria
Tsukamurella paurometabola DSM 20162 YP 003647639 1 521096 Actinobacteria
Bacillus amyloliquefaciens DSM 7 YP 003921071 1 692420 Firmicutes
Streptomyces coelicolor A32 NP 624476 1 100226 Actinobacteria
886 DAQ1742 01804 GI7 Seed
DAQ1742 DAQ1742 01804 Daquatica
Dickeya aquatica DW 0440 WP 035342588 1 Daquatica
Dickeya sp CSL RW240 WP 035342588 1 Daquatica
Spirochaeta smaragdinae DSM 11293 YP 003805139 1 573413 Spirochaetes
Burkholderia phymatum STM815 YP 001860225 1 391038 Proteobacteria
Verminephrobacter eiseniae EF01 2 YP 999230 1 391735 Proteobacteria
Methylobacterium nodulans ORS 2060 YP 002498314 1 460265 Proteobacteria
Magnetospirillum magneticum AMB 1 YP 421953 1 342108 Proteobacteria
Alicyclobacillus acidocaldarius subsp acidocaldarius Tc 4 1 YP 005517587 1 1048834 Firmicutes
Bacillus licheniformis DSM 13 ATCC 14580 YP 080596 1 279010 Firmicutes
Halanaerobium hydrogeniformans YP 003994388 1 656519 Firmicutes
Spirochaeta thermophila DSM 6578 YP 006044389 1 869211 Spirochaetes
Dinoroseobacter shibae DFL 12 DSM 16493 YP 001531967 1 398580 Proteobacteria
Bradyrhizobium oligotrophicum S58 YP 007510863 1 1245469 Proteobacteria
Bradyrhizobium sp BTAi1 YP 001242686 1 288000 Proteobacteria
Clostridium pasteurianum BC1 YP 007942568 1 86416 Firmicutes
Nocardioides sp JS614 YP 923055 1 196162 Actinobacteria
Geobacter lovleyi SZ YP 001951034 1 398767 Proteobacteria
Candidatus Methylomirabilis oxyfera YP 003205627 1 671143 unclassified Bacteria
Bacillus pseudofirmus OF4 YP 003425437 1 398511 Firmicutes
Paenibacillus terrae HPL 003 YP 005077063 1 985665 Firmicutes
Paenibacillus polymyxa M1 YP 005960202 1 1052684 Firmicutes
Lawsonia intracellularis PHE MN1 00 YP 594538 1 363253 Proteobacteria
Desulfatibacillum alkenivorans AK 01 YP 002432464 1 439235 Proteobacteria
Elusimicrobium minutum Pei191 YP 001875551 1 445932 Elusimicrobia
Prosthecochloris aestuarii DSM 271 YP 002014881 1 290512 Bacteroidetes Chlorobi group
Ornithobacterium rhinotracheale DSM 15997 YP 006428244 1 867902 Bacteroidetes Chlorobi group
Thermovibrio ammonificans HB 1 YP 004151007 1 648996 Aquificae
Chlamydia muridarum str Nigg NP 296885 1 243161 Chlamydiae Verrucomicrobia group
Parachlamydia acanthamoebae UV 7 YP 004651791 1 765952 Chlamydiae Verrucomicrobia group
Waddlia chondrophila WSU 86 1044 YP 003709550 1 716544 Chlamydiae Verrucomicrobia group
Bdellovibrio exovorus JSS YP 007645164 1 1184267 Proteobacteria
Burkholderia phytofirmans PsJN YP 001889840 1 398527 Proteobacteria
Xanthobacter autotrophicus Py2 YP 001416446 1 78245 Proteobacteria
Terriglobus roseus DSM 18391 YP 006423365 1 926566 Fibrobacteres Acidobacteria group
Spiribacter salinus M19 40 YP 008046372 1 1260251 Proteobacteria
Thiomicrospira crunogena XCL 2 YP 390982 1 317025 Proteobacteria
Sulfuricella denitrificans skB26 YP 008546387 1 1163617 Proteobacteria
Methylomonas methanica MC09 YP 004512023 1 857087 Proteobacteria
Moraxella catarrhalis RH4 YP 003627703 1 749219 Proteobacteria
Thioalkalivibrio sp K90mix YP 003460504 1 396595 Proteobacteria
Kangiella koreensis DSM 16069 YP 003146375 1 523791 Proteobacteria
Alteromonas sp SN2 YP 004467325 1 715451 Proteobacteria
Alteromonas macleodii str Black Sea 11 YP 006824615 1 1004785 Proteobacteria
Xenorhabdus bovienii SS 2004 YP 003468747 1 406818 Proteobacteria
Xenorhabdus nematophila ATCC 19061 YP 003712935 1 406817 Proteobacteria
Proteus mirabilis BB2000 YP 008397534 1 1266738 Proteobacteria
Morganella morganii subsp morganii KT YP 007506146 1 1124991 Proteobacteria
Rahnella aquatilis CIP 78 65 ATCC 33071 YP 005200942 1 745277 Proteobacteria
Rahnella sp Y9602 YP 004213676 1 741091 Proteobacteria
Yersinia enterocolitica subsp palearctica Y11 YP 006004027 1 930944 Proteobacteria
Yersinia enterocolitica subsp enterocolitica 8081 YP 001005924 1 393305 Proteobacteria
Serratia marcescens WW4 YP 007405685 1 435998 Proteobacteria
Serratia liquefaciens ATCC 27592 YP 008229783 1 1346614 Proteobacteria
Serratia proteamaculans 568 YP 001478137 1 399741 Proteobacteria
Oceanimonas sp GK1 YP 005092479 1 511062 Proteobacteria
Serratia sp AS12 YP 004500289 1 768490 Proteobacteria
Serratia plymuthica AS9 YP 004505241 1 768492 Proteobacteria
Cronobacter sakazakii ES15 YP 006343477 1 1138308 Proteobacteria
Pantoea sp At 9b YP 004115355 1 592316 Proteobacteria
Mannheimia haemolytica D153 YP 008234109 1 1261126 Proteobacteria
Shewanella putrefaciens 200 YP 006009526 1 399804 Proteobacteria
Shewanella piezotolerans WP3 YP 002312350 1 225849 Proteobacteria
Shewanella halifaxensis HAW EB4 YP 001674001 1 458817 Proteobacteria
Photobacterium profundum SS9 YP 129408 1 298386 Proteobacteria
Vibrio splendidus LGP32 YP 002416639 1 575788 Proteobacteria
Psychromonas sp CNPT3 YP 007639596 1 314282 Proteobacteria
Psychromonas ingrahamii 37 YP 942527 1 357804 Proteobacteria

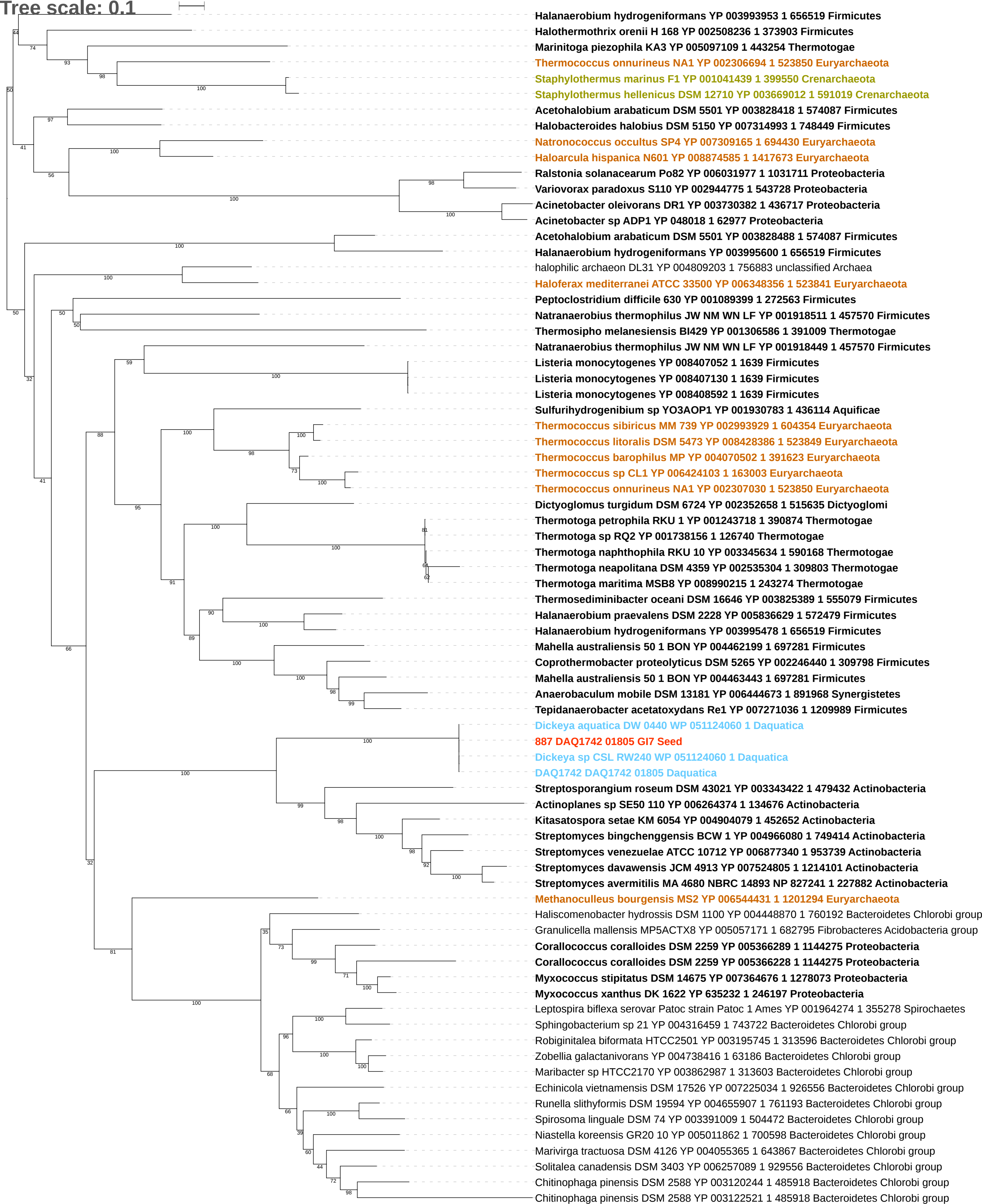
Tree scale: 0.1
Halanaerobium hydrogeniformans YP 003993953 1 656519 Firmicutes
Halothermothrix orenii H 168 YP 002508236 1 373903 Firmicutes
Marinitoga piezophila KA3 YP 005097109 1 443254 Thermotogae
Thermococcus onnurineus NA1 YP 002306694 1 523850 Euryarchaeota
Staphylothermus marinus F1 YP 001041439 1 399550 Crenarchaeota
Staphylothermus hellenicus DSM 12710 YP 003669012 1 591019 Crenarchaeota
Acetohalobium arabaticum DSM 5501 YP 003828418 1 574087 Firmicutes
Halobacteroides halobius DSM 5150 YP 007314993 1 748449 Firmicutes
Natronococcus occultus SP4 YP 007309165 1 694430 Euryarchaeota
Haloarcula hispanica N601 YP 008874585 1 1417673 Euryarchaeota
Ralstonia solanacearum Po82 YP 006031977 1 1031711 Proteobacteria
Variovorax paradoxus S110 YP 002944775 1 543728 Proteobacteria
Acinetobacter oleivorans DR1 YP 003730382 1 436717 Proteobacteria
Acinetobacter sp ADP1 YP 048018 1 62977 Proteobacteria
Acetohalobium arabaticum DSM 5501 YP 003828488 1 574087 Firmicutes
Halanaerobium hydrogeniformans YP 003995600 1 656519 Firmicutes
halophilic archaeon DL31 YP 004809203 1 756883 unclassified Archaea
Haloferax mediterranei ATCC 33500 YP 006348356 1 523841 Euryarchaeota
Peptoclostridium difficile 630 YP 001089399 1 272563 Firmicutes
Natranaerobius thermophilus JW NM WN LF YP 001918511 1 457570 Firmicutes
Thermosipho melanesiensis BI429 YP 001306586 1 391009 Thermotogae
Natranaerobius thermophilus JW NM WN LF YP 001918449 1 457570 Firmicutes
Listeria monocytogenes YP 008407052 1 1639 Firmicutes
Listeria monocytogenes YP 008407130 1 1639 Firmicutes
Listeria monocytogenes YP 008408592 1 1639 Firmicutes
Sulfurihydrogenibium sp YO3AOP1 YP 001930783 1 436114 Aquificae
Thermococcus sibiricus MM 739 YP 002993929 1 604354 Euryarchaeota
Thermococcus litoralis DSM 5473 YP 008428386 1 523849 Euryarchaeota
Thermococcus barophilus MP YP 004070502 1 391623 Euryarchaeota
Thermococcus sp CL1 YP 006424103 1 163003 Euryarchaeota
Thermococcus onnurineus NA1 YP 002307030 1 523850 Euryarchaeota
Dictyoglomus turgidum DSM 6724 YP 002352658 1 515635 Dictyoglomi
Thermotoga petrophila RKU 1 YP 001243718 1 390874 Thermotogae
Thermotoga sp RQ2 YP 001738156 1 126740 Thermotogae
Thermotoga naphthophila RKU 10 YP 003345634 1 590168 Thermotogae
Thermotoga neapolitana DSM 4359 YP 002535304 1 309803 Thermotogae
Thermotoga maritima MSB8 YP 008990215 1 243274 Thermotogae
Thermosediminibacter oceani DSM 16646 YP 003825389 1 555079 Firmicutes
Halanaerobium praevalens DSM 2228 YP 005836629 1 572479 Firmicutes
Halanaerobium hydrogeniformans YP 003995478 1 656519 Firmicutes
Mahella australiensis 50 1 BON YP 004462199 1 697281 Firmicutes
Coprothermobacter proteolyticus DSM 5265 YP 002246440 1 309798 Firmicutes
Mahella australiensis 50 1 BON YP 004463443 1 697281 Firmicutes
Anaerobaculum mobile DSM 13181 YP 006444673 1 891968 Synergistetes
Tepidanaerobacter acetatoxydans Re1 YP 007271036 1 1209989 Firmicutes
Dickeya aquatica DW 0440 WP 051124060 1 Daquatica
887 DAQ1742 01805 GI7 Seed
Dickeya sp CSL RW240 WP 051124060 1 Daquatica
DAQ1742 DAQ1742 01805 Daquatica
Streptosporangium roseum DSM 43021 YP 003343422 1 479432 Actinobacteria
Actinoplanes sp SE50 110 YP 006264374 1 134676 Actinobacteria
Kitasatospora setae KM 6054 YP 004904079 1 452652 Actinobacteria
Streptomyces bingchenggensis BCW 1 YP 004966080 1 749414 Actinobacteria
Streptomyces venezuelae ATCC 10712 YP 006877340 1 953739 Actinobacteria
Streptomyces davawensis JCM 4913 YP 007524805 1 1214101 Actinobacteria
Streptomyces avermitilis MA 4680 NBRC 14893 NP 827241 1 227882 Actinobacteria
Methanoculleus bourgensis MS2 YP 006544431 1 1201294 Euryarchaeota
Haliscomenobacter hydrossis DSM 1100 YP 004448870 1 760192 Bacteroidetes Chlorobi group
Granulicella mallensis MP5ACTX8 YP 005057171 1 682795 Fibrobacteres Acidobacteria group
Corallococcus coralloides DSM 2259 YP 005366289 1 1144275 Proteobacteria
Corallococcus coralloides DSM 2259 YP 005366228 1 1144275 Proteobacteria
Myxococcus stipitatus DSM 14675 YP 007364676 1 1278073 Proteobacteria
Myxococcus xanthus DK 1622 YP 635232 1 246197 Proteobacteria
Leptospira biflexa serovar Patoc strain Patoc 1 Ames YP 001964274 1 355278 Spirochaetes
Sphingobacterium sp 21 YP 004316459 1 743722 Bacteroidetes Chlorobi group
Robiginitalea biformata HTCC2501 YP 003195745 1 313596 Bacteroidetes Chlorobi group
Zobellia galactanivorans YP 004738416 1 63186 Bacteroidetes Chlorobi group
Maribacter sp HTCC2170 YP 003862987 1 313603 Bacteroidetes Chlorobi group
Echinicola vietnamensis DSM 17526 YP 007225034 1 926556 Bacteroidetes Chlorobi group
Runella slithyformis DSM 19594 YP 004655907 1 761193 Bacteroidetes Chlorobi group
Spirosoma linguale DSM 74 YP 003391009 1 504472 Bacteroidetes Chlorobi group
Niastella koreensis GR20 10 YP 005011862 1 700598 Bacteroidetes Chlorobi group
Marivirga tractuosa DSM 4126 YP 004055365 1 643867 Bacteroidetes Chlorobi group
Solitalea canadensis DSM 3403 YP 006257089 1 929556 Bacteroidetes Chlorobi group
Chitinophaga pinensis DSM 2588 YP 003120244 1 485918 Bacteroidetes Chlorobi group
Chitinophaga pinensis DSM 2588 YP 003122521 1 485918 Bacteroidetes Chlorobi group

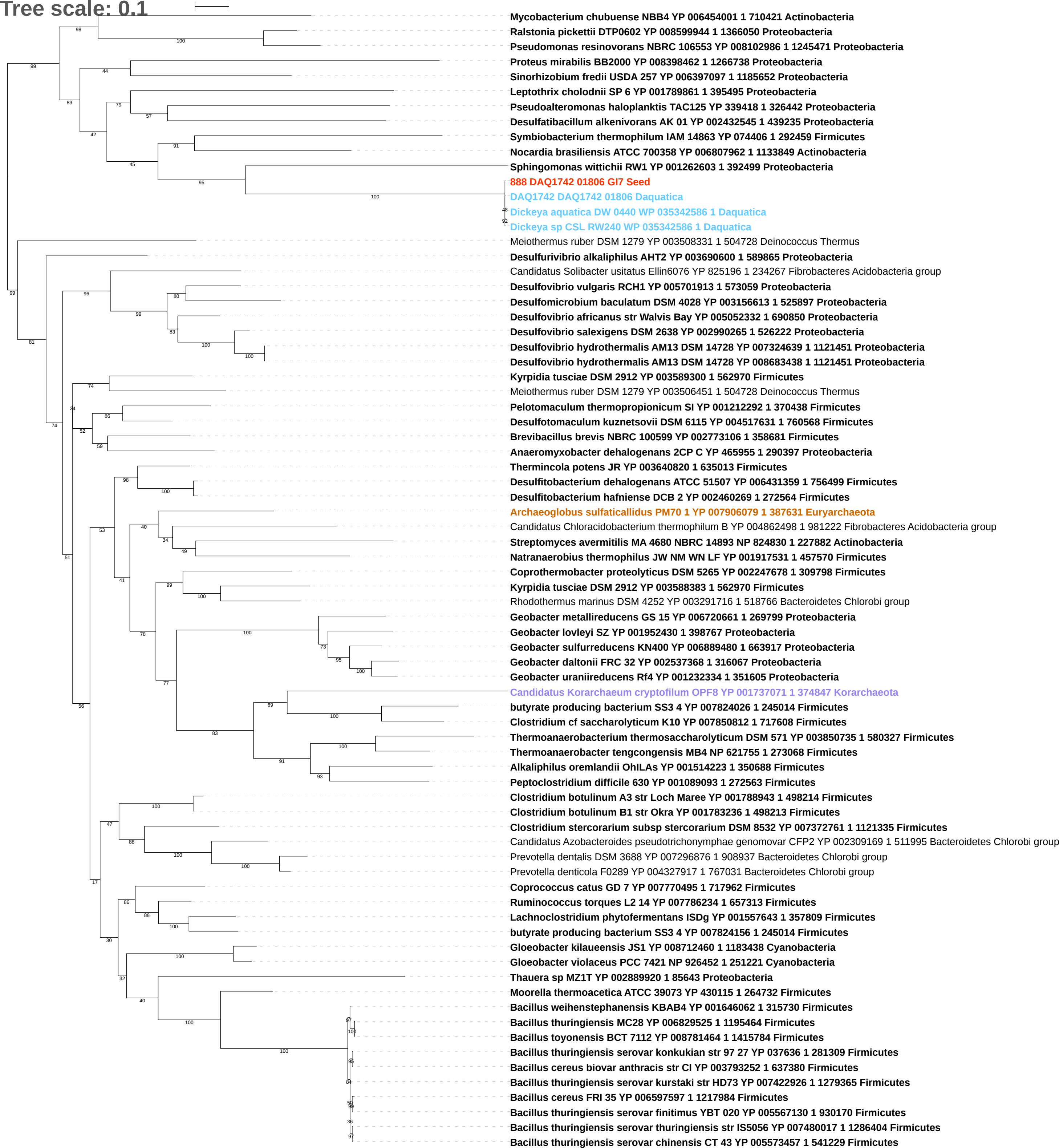

Tree scale: 0.1
Mycobacterium chubuense NBB4 YP 006454001 1 710421 Actinobacteria
Ralstonia pickettii DTP0602 YP 008599944 1 1366050 Proteobacteria
Pseudomonas resinovorans NBRC 106553 YP 008102986 1 1245471 Proteobacteria
Proteus mirabilis BB2000 YP 008398462 1 1266738 Proteobacteria
Sinorhizobium fredii USDA 257 YP 006397097 1 1185652 Proteobacteria
Leptothrix cholodnii SP 6 YP 001789861 1 395495 Proteobacteria
Pseudoalteromonas haloplanktis TAC125 YP 339418 1 326442 Proteobacteria
Desulfatibacillum alkenivorans AK 01 YP 002432545 1 439235 Proteobacteria
Symbiobacterium thermophilum IAM 14863 YP 074406 1 292459 Firmicutes
Nocardia brasiliensis ATCC 700358 YP 006807962 1 1133849 Actinobacteria
Sphingomonas wittichii RW1 YP 001262603 1 392499 Proteobacteria
888 DAQ1742 01806 GI7 Seed
DAQ1742 DAQ1742 01806 Daquatica
Dickeya aquatica DW 0440 WP 035342586 1 Daquatica
Dickeya sp CSL RW240 WP 035342586 1 Daquatica
Meiothermus ruber DSM 1279 YP 003508331 1 504728 Deinococcus Thermus
Desulfurivibrio alkaliphilus AHT2 YP 003690600 1 589865 Proteobacteria
Candidatus Solibacter usitatus Ellin6076 YP 825196 1 234267 Fibrobacteres Acidobacteria group
Desulfovibrio vulgaris RCH1 YP 005701913 1 573059 Proteobacteria
Desulfomicrobium baculatum DSM 4028 YP 003156613 1 525897 Proteobacteria
Desulfovibrio africanus str Walvis Bay YP 005052332 1 690850 Proteobacteria
Desulfovibrio salexigens DSM 2638 YP 002990265 1 526222 Proteobacteria
Desulfovibrio hydrothermalis AM13 DSM 14728 YP 007324639 1 1121451 Proteobacteria
Desulfovibrio hydrothermalis AM13 DSM 14728 YP 008683438 1 1121451 Proteobacteria
Kyrpidia tusciae DSM 2912 YP 003589300 1 562970 Firmicutes
Meiothermus ruber DSM 1279 YP 003506451 1 504728 Deinococcus Thermus
Pelotomaculum thermopropionicum SI YP 001212292 1 370438 Firmicutes
Desulfotomaculum kuznetsovii DSM 6115 YP 004517631 1 760568 Firmicutes
Brevibacillus brevis NBRC 100599 YP 002773106 1 358681 Firmicutes
Anaeromyxobacter dehalogenans 2CP C YP 465955 1 290397 Proteobacteria
Thermincola potens JR YP 003640820 1 635013 Firmicutes
Desulfitobacterium dehalogenans ATCC 51507 YP 006431359 1 756499 Firmicutes
Desulfitobacterium hafniense DCB 2 YP 002460269 1 272564 Firmicutes
Archaeoglobus sulfaticallidus PM70 1 YP 007906079 1 387631 Euryarchaeota
Candidatus Chloracidobacterium thermophilum B YP 004862498 1 981222 Fibrobacteres Acidobacteria group
Streptomyces avermitilis MA 4680 NBRC 14893 NP 824830 1 227882 Actinobacteria
Natranaerobius thermophilus JW NM WN LF YP 001917531 1 457570 Firmicutes
Coprothermobacter proteolyticus DSM 5265 YP 002247678 1 309798 Firmicutes
Kyrpidia tusciae DSM 2912 YP 003588383 1 562970 Firmicutes
Rhodothermus marinus DSM 4252 YP 003291716 1 518766 Bacteroidetes Chlorobi group
Geobacter metallireducens GS 15 YP 006720661 1 269799 Proteobacteria
Geobacter lovleyi SZ YP 001952430 1 398767 Proteobacteria
Geobacter sulfurreducens KN400 YP 006889480 1 663917 Proteobacteria
Geobacter daltonii FRC 32 YP 002537368 1 316067 Proteobacteria
Geobacter uraniireducens Rf4 YP 001232334 1 351605 Proteobacteria
Candidatus Korarchaeum cryptofilum OPF8 YP 001737071 1 374847 Korarchaeota
butyrate producing bacterium SS3 4 YP 007824026 1 245014 Firmicutes
Clostridium cf saccharolyticum K10 YP 007850812 1 717608 Firmicutes
Thermoanaerobacterium thermosaccharolyticum DSM 571 YP 003850735 1 580327 Firmicutes
Thermoanaerobacter tengcongensis MB4 NP 621755 1 273068 Firmicutes
Alkaliphilus oremlandii OhILAs YP 001514223 1 350688 Firmicutes
Peptoclostridium difficile 630 YP 001089093 1 272563 Firmicutes
Clostridium botulinum A3 str Loch Maree YP 001788943 1 498214 Firmicutes
Clostridium botulinum B1 str Okra YP 001783236 1 498213 Firmicutes
Clostridium stercorarium subsp stercorarium DSM 8532 YP 007372761 1 1121335 Firmicutes
Candidatus Azobacteroides pseudotrichonymphae genomovar CFP2 YP 002309169 1 511995 Bacteroidetes Chlorobi group
Prevotella dentalis DSM 3688 YP 007296876 1 908937 Bacteroidetes Chlorobi group
Prevotella denticola F0289 YP 004327917 1 767031 Bacteroidetes Chlorobi group
Coprococcus catus GD 7 YP 007770495 1 717962 Firmicutes
Ruminococcus torques L2 14 YP 007786234 1 657313 Firmicutes
Lachnoclostridium phytofermentans ISDg YP 001557643 1 357809 Firmicutes
butyrate producing bacterium SS3 4 YP 007824156 1 245014 Firmicutes
Gloeobacter kilaueensis JS1 YP 008712460 1 1183438 Cyanobacteria
Gloeobacter violaceus PCC 7421 NP 926452 1 251221 Cyanobacteria
Thauera sp MZ1T YP 002889920 1 85643 Proteobacteria
Moorella thermoacetica ATCC 39073 YP 430115 1 264732 Firmicutes
Bacillus weihenstephanensis KBAB4 YP 001646062 1 315730 Firmicutes
Bacillus thuringiensis MC28 YP 006829525 1 1195464 Firmicutes
Bacillus toyonensis BCT 7112 YP 008781464 1 1415784 Firmicutes
Bacillus thuringiensis serovar konkukian str 97 27 YP 037636 1 281309 Firmicutes
Bacillus cereus biovar anthracis str CI YP 003793252 1 637380 Firmicutes
Bacillus thuringiensis serovar kurstaki str HD73 YP 007422926 1 1279365 Firmicutes
Bacillus cereus FRI 35 YP 006597597 1 1217984 Firmicutes
Bacillus thuringiensis serovar finitimus YBT 020 YP 005567130 1 930170 Firmicutes
Bacillus thuringiensis serovar thuringiensis str IS5056 YP 007480017 1 1286404 Firmicutes
Bacillus thuringiensis serovar chinensis CT 43 YP 005573457 1 541229 Firmicutes

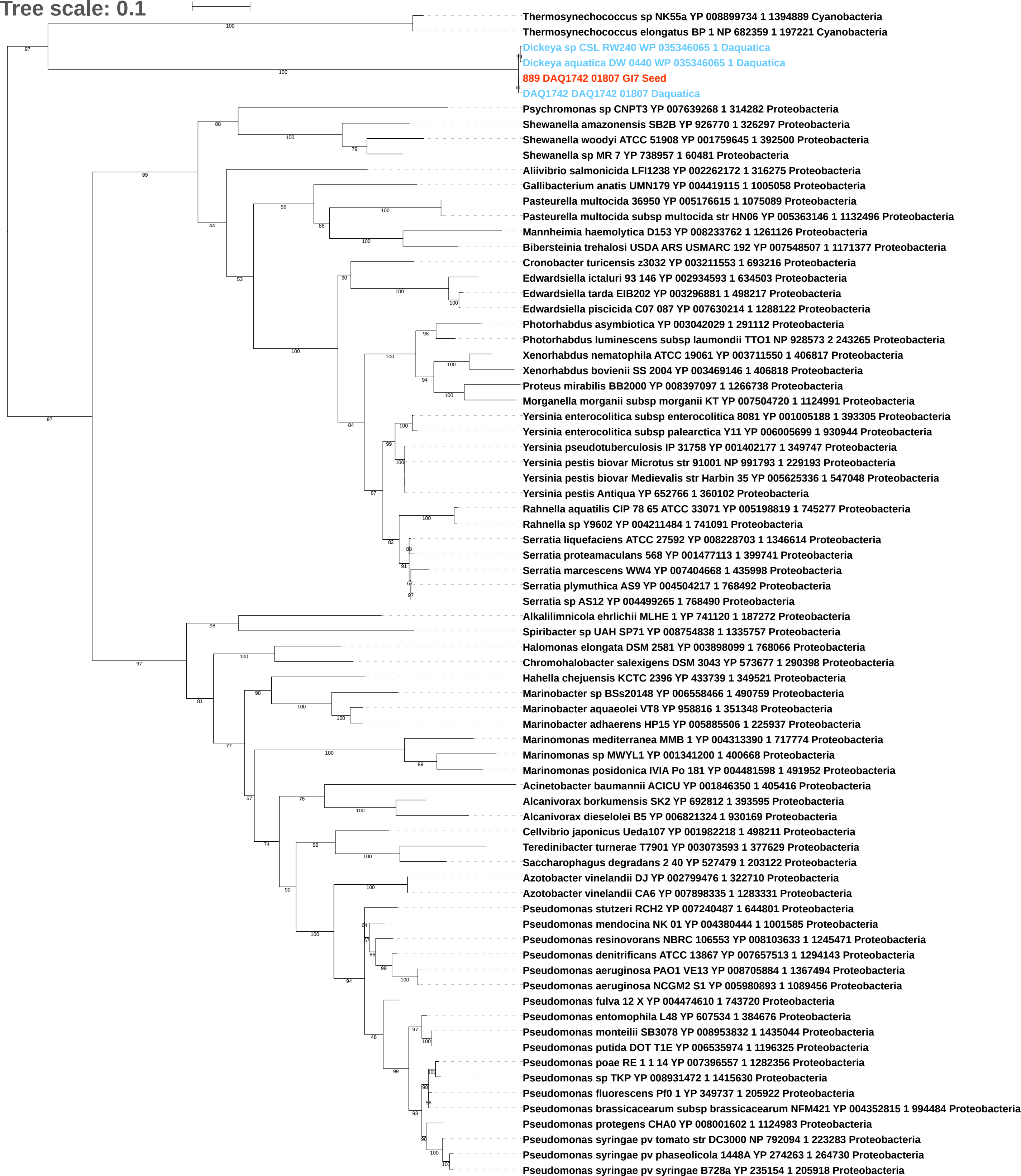
Tree scale: 0.1
Thermosynechococcus sp NK55a YP 008899734 1 1394889 Cyanobacteria
Thermosynechococcus elongatus BP 1 NP 682359 1 197221 Cyanobacteria
Dickeya sp CSL RW240 WP 035346065 1 Daquatica
Dickeya aquatica DW 0440 WP 035346065 1 Daquatica
889 DAQ1742 01807 GI7 Seed
DAQ1742 DAQ1742 01807 Daquatica
Psychromonas sp CNPT3 YP 007639268 1 314282 Proteobacteria
Shewanella amazonensis SB2B YP 926770 1 326297 Proteobacteria
Shewanella woodyi ATCC 51908 YP 001759645 1 392500 Proteobacteria
Shewanella sp MR 7 YP 738957 1 60481 Proteobacteria
Aliivibrio salmonicida LFI1238 YP 002262172 1 316275 Proteobacteria
Gallibacterium anatis UMN179 YP 004419115 1 1005058 Proteobacteria
Pasteurella multocida 36950 YP 005176615 1 1075089 Proteobacteria
Pasteurella multocida subsp multocida str HN06 YP 005363146 1 1132496 Proteobacteria
Mannheimia haemolytica D153 YP 008233762 1 1261126 Proteobacteria
Bibersteinia trehalosi USDA ARS USMARC 192 YP 007548507 1 1171377 Proteobacteria
Cronobacter turicensis z3032 YP 003211553 1 693216 Proteobacteria
Edwardsiella ictaluri 93 146 YP 002934593 1 634503 Proteobacteria
Edwardsiella tarda EIB202 YP 003296881 1 498217 Proteobacteria
Edwardsiella piscicida C07 087 YP 007630214 1 1288122 Proteobacteria
Photorhabdus asymbiotica YP 003042029 1 291112 Proteobacteria
Photorhabdus luminescens subsp laumondii TTO1 NP 928573 2 243265 Proteobacteria
Xenorhabdus nematophila ATCC 19061 YP 003711550 1 406817 Proteobacteria
Xenorhabdus bovienii SS 2004 YP 003469146 1 406818 Proteobacteria
Proteus mirabilis BB2000 YP 008397097 1 1266738 Proteobacteria
Morganella morganii subsp morganii KT YP 007504720 1 1124991 Proteobacteria
Yersinia enterocolitica subsp enterocolitica 8081 YP 001005188 1 393305 Proteobacteria
Yersinia enterocolitica subsp palearctica Y11 YP 006005699 1 930944 Proteobacteria
Yersinia pseudotuberculosis IP 31758 YP 001402177 1 349747 Proteobacteria
Yersinia pestis biovar Microtus str 91001 NP 991793 1 229193 Proteobacteria
Yersinia pestis biovar Medievalis str Harbin 35 YP 005625336 1 547048 Proteobacteria
Yersinia pestis Antiqua YP 652766 1 360102 Proteobacteria
Rahnella aquatilis CIP 78 65 ATCC 33071 YP 005198819 1 745277 Proteobacteria
Rahnella sp Y9602 YP 004211484 1 741091 Proteobacteria
Serratia liquefaciens ATCC 27592 YP 008228703 1 1346614 Proteobacteria
Serratia proteamaculans 568 YP 001477113 1 399741 Proteobacteria
Serratia marcescens WW4 YP 007404668 1 435998 Proteobacteria
Serratia plymuthica AS9 YP 004504217 1 768492 Proteobacteria
Serratia sp AS12 YP 004499265 1 768490 Proteobacteria
Alkalilimnicola ehrlichii MLHE 1 YP 741120 1 187272 Proteobacteria
Spiribacter sp UAH SP71 YP 008754838 1 1335757 Proteobacteria
Halomonas elongata DSM 2581 YP 003898099 1 768066 Proteobacteria
Chromohalobacter salexigens DSM 3043 YP 573677 1 290398 Proteobacteria
Hahella chejuensis KCTC 2396 YP 433739 1 349521 Proteobacteria
Marinobacter sp BSs20148 YP 006558466 1 490759 Proteobacteria
Marinobacter aquaeolei VT8 YP 958816 1 351348 Proteobacteria
Marinobacter adhaerens HP15 YP 005885506 1 225937 Proteobacteria
Marinomonas mediterranea MMB 1 YP 004313390 1 717774 Proteobacteria
Marinomonas sp MWYL1 YP 001341200 1 400668 Proteobacteria
Marinomonas posidonica IVIA Po 181 YP 004481598 1 491952 Proteobacteria
Acinetobacter baumannii ACICU YP 001846350 1 405416 Proteobacteria
Alcanivorax borkumensis SK2 YP 692812 1 393595 Proteobacteria
Alcanivorax dieselolei B5 YP 006821324 1 930169 Proteobacteria
Cellvibrio japonicus Ueda107 YP 001982218 1 498211 Proteobacteria
Teredinibacter turnerae T7901 YP 003073593 1 377629 Proteobacteria
Saccharophagus degradans 2 40 YP 527479 1 203122 Proteobacteria
Azotobacter vinelandii DJ YP 002799476 1 322710 Proteobacteria
Azotobacter vinelandii CA6 YP 007898335 1 1283331 Proteobacteria
Pseudomonas stutzeri RCH2 YP 007240487 1 644801 Proteobacteria
Pseudomonas mendocina NK 01 YP 004380444 1 1001585 Proteobacteria
Pseudomonas resinovorans NBRC 106553 YP 008103633 1 1245471 Proteobacteria
Pseudomonas denitrificans ATCC 13867 YP 007657513 1 1294143 Proteobacteria
Pseudomonas aeruginosa PAO1 VE13 YP 008705884 1 1367494 Proteobacteria
Pseudomonas aeruginosa NCGM2 S1 YP 005980893 1 1089456 Proteobacteria
Pseudomonas fulva 12 X YP 004474610 1 743720 Proteobacteria
Pseudomonas entomophila L48 YP 607534 1 384676 Proteobacteria
Pseudomonas monteilii SB3078 YP 008953832 1 1435044 Proteobacteria
Pseudomonas putida DOT T1E YP 006535974 1 1196325 Proteobacteria
Pseudomonas poae RE 1 1 14 YP 007396557 1 1282356 Proteobacteria
Pseudomonas sp TKP YP 008931472 1 1415630 Proteobacteria
Pseudomonas fluorescens Pf0 1 YP 349737 1 205922 Proteobacteria
Pseudomonas brassicacearum subsp brassicacearum NFM421 YP 004352815 1 994484 Proteobacteria
Pseudomonas protegens CHA0 YP 008001602 1 1124983 Proteobacteria
Pseudomonas syringae pv tomato str DC3000 NP 792094 1 223283 Proteobacteria
Pseudomonas syringae pv phaseolicola 1448A YP 274263 1 264730 Proteobacteria
Pseudomonas syringae pv syringae B728a YP 235154 1 205918 Proteobacteria

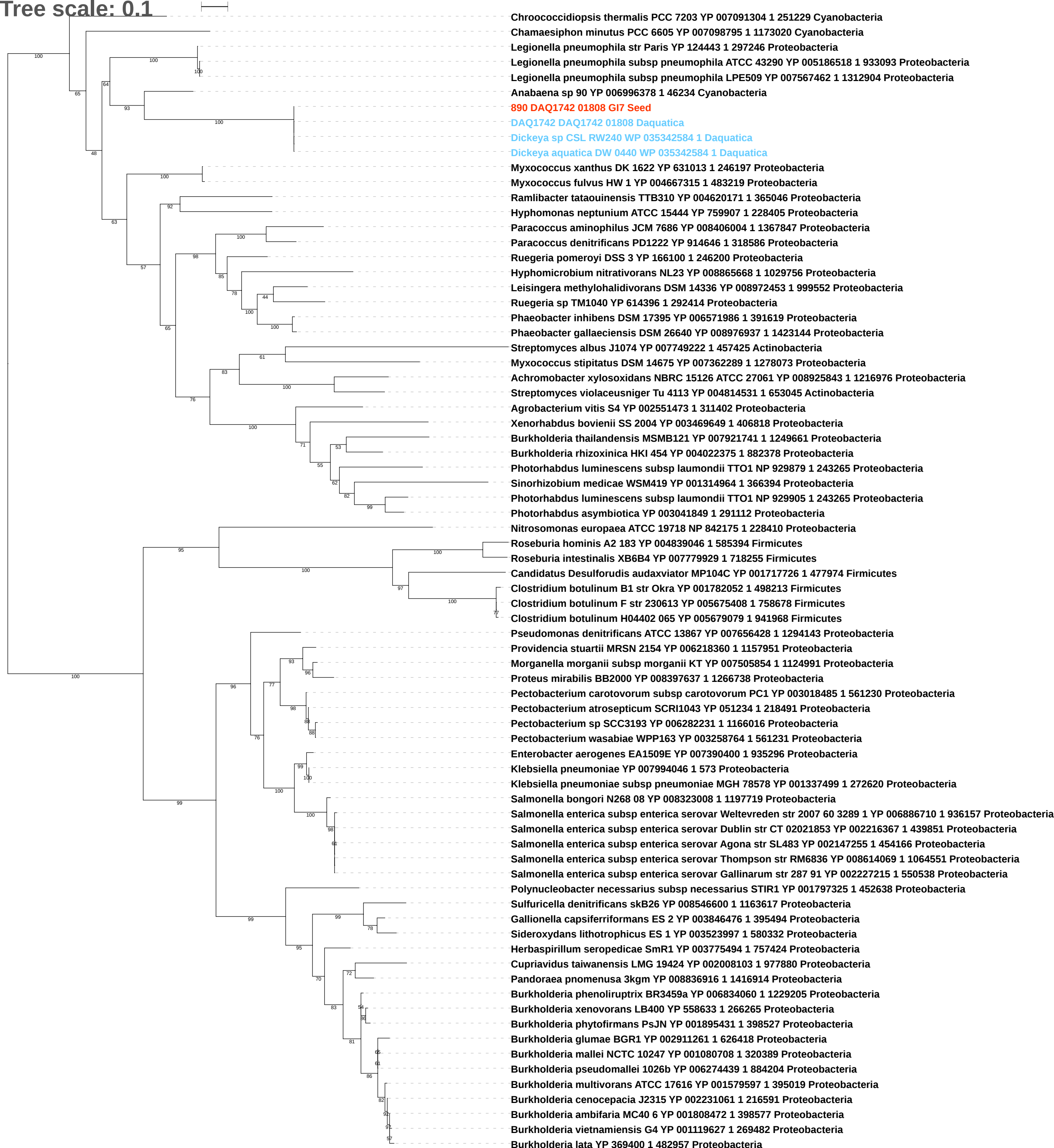
Tree scale: 0.1
Chroococcidiopsis thermalis PCC 7203 YP 007091304 1 251229 Cyanobacteria
Chamaesiphon minutus PCC 6605 YP 007098795 1 1173020 Cyanobacteria
Legionella pneumophila str Paris YP 124443 1 297246 Proteobacteria
Legionella pneumophila subsp pneumophila ATCC 43290 YP 005186518 1 933093 Proteobacteria
Legionella pneumophila subsp pneumophila LPE509 YP 007567462 1 1312904 Proteobacteria
Anabaena sp 90 YP 006996378 1 46234 Cyanobacteria
890 DAQ1742 01808 GI7 Seed
DAQ1742 DAQ1742 01808 Daquatica
Dickeya sp CSL RW240 WP 035342584 1 Daquatica
Dickeya aquatica DW 0440 WP 035342584 1 Daquatica
Myxococcus xanthus DK 1622 YP 631013 1 246197 Proteobacteria
Myxococcus fulvus HW 1 YP 004667315 1 483219 Proteobacteria
Ramlibacter tataouinensis TTB310 YP 004620171 1 365046 Proteobacteria
Hyphomonas neptunium ATCC 15444 YP 759907 1 228405 Proteobacteria
Paracoccus aminophilus JCM 7686 YP 008406004 1 1367847 Proteobacteria
Paracoccus denitrificans PD1222 YP 914646 1 318586 Proteobacteria
Ruegeria pomeroyi DSS 3 YP 166100 1 246200 Proteobacteria
Hyphomicrobium nitrativorans NL23 YP 008865668 1 1029756 Proteobacteria
Leisingera methylohalidivorans DSM 14336 YP 008972453 1 999552 Proteobacteria
Ruegeria sp TM1040 YP 614396 1 292414 Proteobacteria
Phaeobacter inhibens DSM 17395 YP 006571986 1 391619 Proteobacteria
Phaeobacter gallaeciensis DSM 26640 YP 008976937 1 1423144 Proteobacteria
Streptomyces albus J1074 YP 007749222 1 457425 Actinobacteria
Myxococcus stipitatus DSM 14675 YP 007362289 1 1278073 Proteobacteria
Achromobacter xylosoxidans NBRC 15126 ATCC 27061 YP 008925843 1 1216976 Proteobacteria
Streptomyces violaceusniger Tu 4113 YP 004814531 1 653045 Actinobacteria
Agrobacterium vitis S4 YP 002551473 1 311402 Proteobacteria
Xenorhabdus bovienii SS 2004 YP 003469649 1 406818 Proteobacteria
Burkholderia thailandensis MSMB121 YP 007921741 1 1249661 Proteobacteria
Burkholderia rhizoxinica HKI 454 YP 004022375 1 882378 Proteobacteria
Photorhabdus luminescens subsp laumondii TTO1 NP 929879 1 243265 Proteobacteria
Sinorhizobium medicae WSM419 YP 001314964 1 366394 Proteobacteria
Photorhabdus luminescens subsp laumondii TTO1 NP 929905 1 243265 Proteobacteria
Photorhabdus asymbiotica YP 003041849 1 291112 Proteobacteria
Nitrosomonas europaea ATCC 19718 NP 842175 1 228410 Proteobacteria
Roseburia hominis A2 183 YP 004839046 1 585394 Firmicutes
Roseburia intestinalis XB6B4 YP 007779929 1 718255 Firmicutes
Candidatus Desulforudis audaxviator MP104C YP 001717726 1 477974 Firmicutes
Clostridium botulinum B1 str Okra YP 001782052 1 498213 Firmicutes
Clostridium botulinum F str 230613 YP 005675408 1 758678 Firmicutes
Clostridium botulinum H04402 065 YP 005679079 1 941968 Firmicutes
Pseudomonas denitrificans ATCC 13867 YP 007656428 1 1294143 Proteobacteria
Providencia stuartii MRSN 2154 YP 006218360 1 1157951 Proteobacteria
Morganella morganii subsp morganii KT YP 007505854 1 1124991 Proteobacteria
Proteus mirabilis BB2000 YP 008397637 1 1266738 Proteobacteria
Pectobacterium carotovorum subsp carotovorum PC1 YP 003018485 1 561230 Proteobacteria
Pectobacterium atrosepticum SCRI1043 YP 051234 1 218491 Proteobacteria
Pectobacterium sp SCC3193 YP 006282231 1 1166016 Proteobacteria
Pectobacterium wasabiae WPP163 YP 003258764 1 561231 Proteobacteria
Enterobacter aerogenes EA1509E YP 007390400 1 935296 Proteobacteria
Klebsiella pneumoniae YP 007994046 1 573 Proteobacteria
Klebsiella pneumoniae subsp pneumoniae MGH 78578 YP 001337499 1 272620 Proteobacteria
Salmonella bongori N268 08 YP 008323008 1 1197719 Proteobacteria
Salmonella enterica subsp enterica serovar Weltevreden str 2007 60 3289 1 YP 006886710 1 936157 Proteobacteria
Salmonella enterica subsp enterica serovar Dublin str CT 02021853 YP 002216367 1 439851 Proteobacteria
Salmonella enterica subsp enterica serovar Agona str SL483 YP 002147255 1 454166 Proteobacteria
Salmonella enterica subsp enterica serovar Thompson str RM6836 YP 008614069 1 1064551 Proteobacteria
Salmonella enterica subsp enterica serovar Gallinarum str 287 91 YP 002227215 1 550538 Proteobacteria
Polynucleobacter necessarius subsp necessarius STIR1 YP 001797325 1 452638 Proteobacteria
Sulfuricella denitrificans skB26 YP 008546600 1 1163617 Proteobacteria
Gallionella capsiferriformans ES 2 YP 003846476 1 395494 Proteobacteria
Sideroxydans lithotrophicus ES 1 YP 003523997 1 580332 Proteobacteria
Herbaspirillum seropedicae SmR1 YP 003775494 1 757424 Proteobacteria
Cupriavidus taiwanensis LMG 19424 YP 002008103 1 977880 Proteobacteria
Pandoraea pnomenusa 3kgm YP 008836916 1 1416914 Proteobacteria
Burkholderia phenoliruptrix BR3459a YP 006834060 1 1229205 Proteobacteria
Burkholderia xenovorans LB400 YP 558633 1 266265 Proteobacteria
Burkholderia phytofirmans PsJN YP 001895431 1 398527 Proteobacteria
Burkholderia glumae BGR1 YP 002911261 1 626418 Proteobacteria
Burkholderia mallei NCTC 10247 YP 001080708 1 320389 Proteobacteria
Burkholderia pseudomallei 1026b YP 006274439 1 884204 Proteobacteria
Burkholderia multivorans ATCC 17616 YP 001579597 1 395019 Proteobacteria
Burkholderia cenocepacia J2315 YP 002231061 1 216591 Proteobacteria
Burkholderia ambifaria MC40 6 YP 001808472 1 398577 Proteobacteria
Burkholderia vietnamiensis G4 YP 001119627 1 269482 Proteobacteria
Burkholderia lata YP 369400 1 482957 Proteobacteria

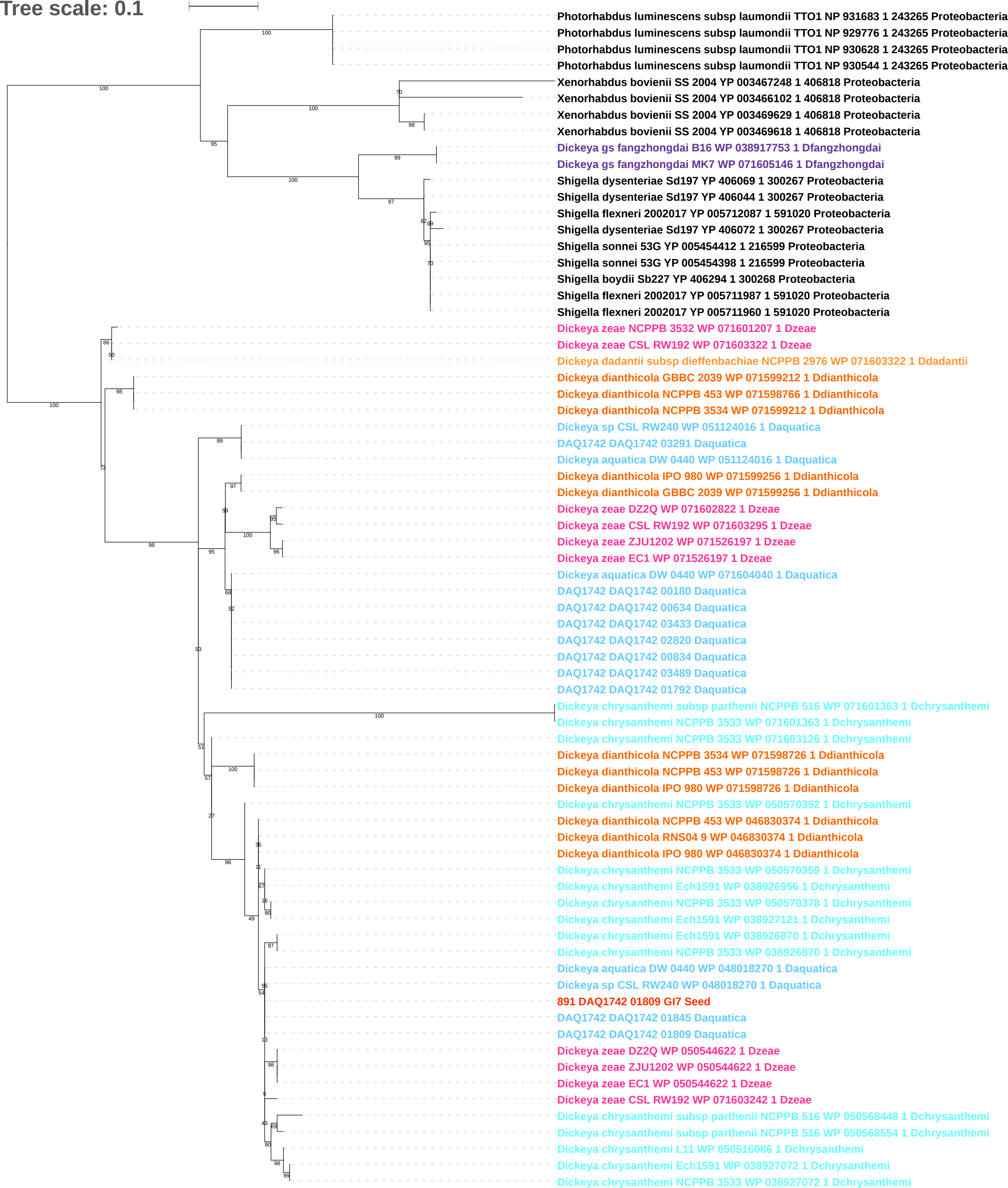
Tree scale: 0.1
Photorhabdus luminescens subsp laumondii TTO1 NP 931683 1 243265 Proteobacteria
Photorhabdus luminescens subsp laumondii TTO1 NP 929776 1 243265 Proteobacteria
Photorhabdus luminescens subsp laumondii TTO1 NP 930628 1 243265 Proteobacteria
Photorhabdus luminescens subsp laumondii TTO1 NP 930544 1 243265 Proteobacteria
Xenorhabdus bovienii SS 2004 YP 003467248 1 406818 Proteobacteria
Xenorhabdus bovienii SS 2004 YP 003466102 1 406818 Proteobacteria
Xenorhabdus bovienii SS 2004 YP 003469629 1 406818 Proteobacteria
Xenorhabdus bovienii SS 2004 YP 003469618 1 406818 Proteobacteria
Dickeya gs fangzhongdai B16 WP 038917753 1 Dfangzhongdai
Dickeya gs fangzhongdai MK7 WP 071605146 1 Dfangzhongdai
Shigella dysenteriae Sd197 YP 406069 1 300267 Proteobacteria
Shigella dysenteriae Sd197 YP 406044 1 300267 Proteobacteria
Shigella flexneri 2002017 YP 005712087 1 591020 Proteobacteria
Shigella dysenteriae Sd197 YP 406072 1 300267 Proteobacteria
Shigella sonnei 53G YP 005454412 1 216599 Proteobacteria
Shigella sonnei 53G YP 005454398 1 216599 Proteobacteria
Shigella boydii Sb227 YP 406294 1 300268 Proteobacteria
Shigella flexneri 2002017 YP 005711987 1 591020 Proteobacteria
Shigella flexneri 2002017 YP 005711960 1 591020 Proteobacteria
Dickeya zeae NCPPB 3532 WP 071601207 1 Dzeae
Dickeya zeae CSL RW192 WP 071603322 1 Dzeae
Dickeya dadantii subsp dieffenbachiae NCPPB 2976 WP 071603322 1 Ddadantii
Dickeya dianthicola GBBC 2039 WP 071599212 1 Ddianthicola
Dickeya dianthicola NCPPB 453 WP 071598766 1 Ddianthicola
Dickeya dianthicola NCPPB 3534 WP 071599212 1 Ddianthicola
Dickeya sp CSL RW240 WP 051124016 1 Daquatica
DAQ1742 DAQ1742 03291 Daquatica
Dickeya aquatica DW 0440 WP 051124016 1 Daquatica
Dickeya dianthicola IPO 980 WP 071599256 1 Ddianthicola
Dickeya dianthicola GBBC 2039 WP 071599256 1 Ddianthicola
Dickeya zeae DZ2Q WP 071602822 1 Dzeae
Dickeya zeae CSL RW192 WP 071603295 1 Dzeae
Dickeya zeae ZJU1202 WP 071526197 1 Dzeae
Dickeya zeae EC1 WP 071526197 1 Dzeae
Dickeya aquatica DW 0440 WP 071604040 1 Daquatica
DAQ1742 DAQ1742 00180 Daquatica
DAQ1742 DAQ1742 00634 Daquatica
DAQ1742 DAQ1742 03433 Daquatica
DAQ1742 DAQ1742 02820 Daquatica
DAQ1742 DAQ1742 00834 Daquatica
DAQ1742 DAQ1742 03489 Daquatica
DAQ1742 DAQ1742 01792 Daquatica
Dickeya chrysanthemi subsp parthenii NCPPB 516 WP 071601363 1 Dchrysanthemi
Dickeya chrysanthemi NCPPB 3533 WP 071601363 1 Dchrysanthemi
Dickeya chrysanthemi NCPPB 3533 WP 071603126 1 Dchrysanthemi
Dickeya dianthicola NCPPB 3534 WP 071598726 1 Ddianthicola
Dickeya dianthicola NCPPB 453 WP 071598726 1 Ddianthicola
Dickeya dianthicola IPO 980 WP 071598726 1 Ddianthicola
Dickeya chrysanthemi NCPPB 3533 WP 050570352 1 Dchrysanthemi
Dickeya dianthicola NCPPB 453 WP 046830374 1 Ddianthicola
Dickeya dianthicola RNS04 9 WP 046830374 1 Ddianthicola
Dickeya dianthicola IPO 980 WP 046830374 1 Ddianthicola
Dickeya chrysanthemi NCPPB 3533 WP 050570359 1 Dchrysanthemi
Dickeya chrysanthemi Ech1591 WP 038926956 1 Dchrysanthemi
Dickeya chrysanthemi NCPPB 3533 WP 050570378 1 Dchrysanthemi
Dickeya chrysanthemi Ech1591 WP 038927121 1 Dchrysanthemi
Dickeya chrysanthemi Ech1591 WP 038926870 1 Dchrysanthemi
Dickeya chrysanthemi NCPPB 3533 WP 038926870 1 Dchrysanthemi
Dickeya aquatica DW 0440 WP 048018270 1 Daquatica
Dickeya sp CSL RW240 WP 048018270 1 Daquatica
891 DAQ1742 01809 GI7 Seed
DAQ1742 DAQ1742 01845 Daquatica
DAQ1742 DAQ1742 01809 Daquatica
Dickeya zeae DZ2Q WP 050544622 1 Dzeae
Dickeya zeae ZJU1202 WP 050544622 1 Dzeae
Dickeya zeae EC1 WP 050544622 1 Dzeae
Dickeya zeae CSL RW192 WP 071603242 1 Dzeae
Dickeya chrysanthemi subsp parthenii NCPPB 516 WP 050568448 1 Dchrysanthemi
Dickeya chrysanthemi subsp parthenii NCPPB 516 WP 050568554 1 Dchrysanthemi
Dickeya chrysanthemi L11 WP 050516086 1 Dchrysanthemi
Dickeya chrysanthemi Ech1591 WP 038927072 1 Dchrysanthemi
Dickeya chrysanthemi NCPPB 3533 WP 038927072 1 Dchrysanthemi

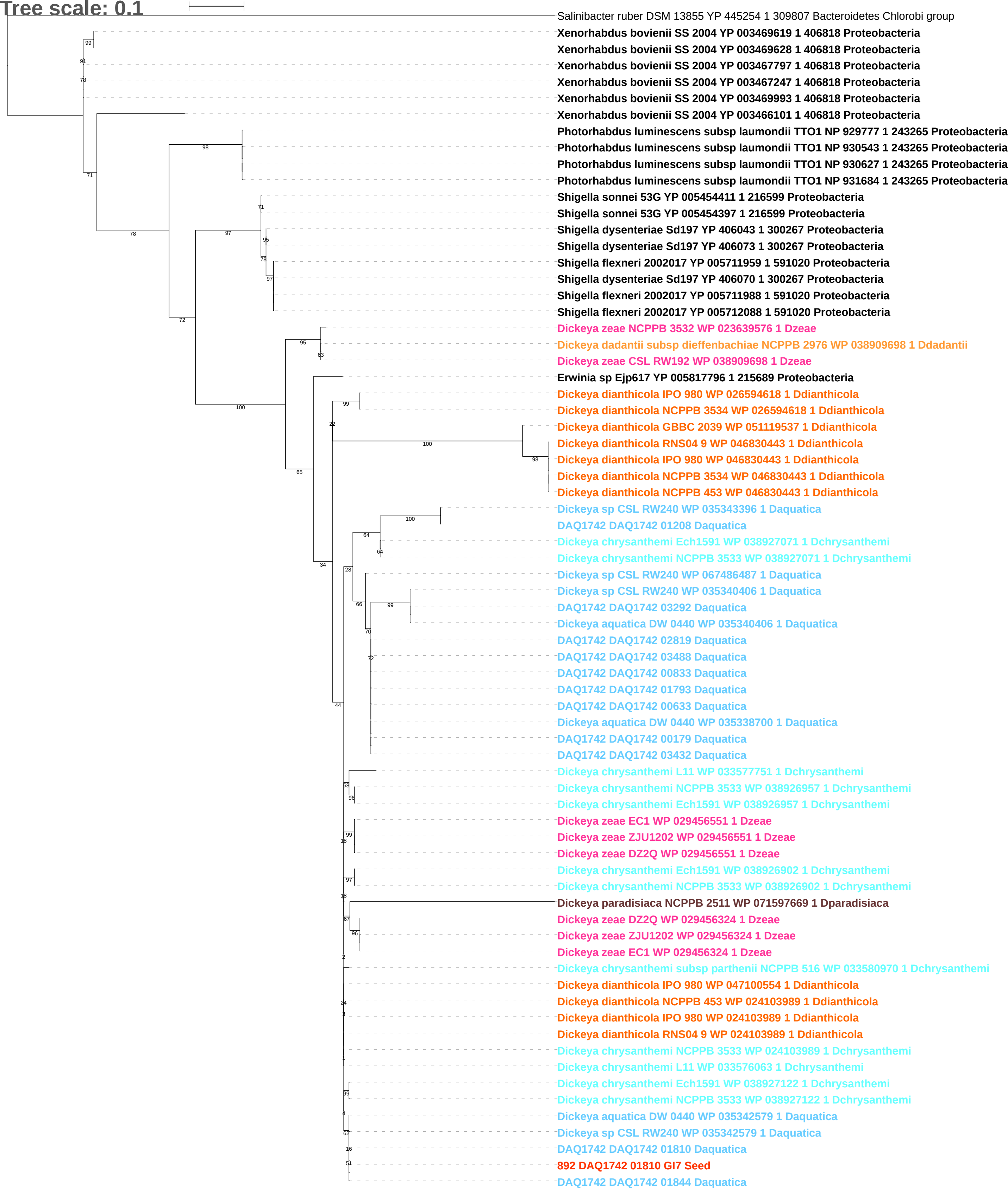

Tree scale: 0.1
Salinibacter ruber DSM 13855 YP 445254 1 309807 Bacteroidetes Chlorobi group
Xenorhabdus bovienii SS 2004 YP 003469619 1 406818 Proteobacteria
Xenorhabdus bovienii SS 2004 YP 003469628 1 406818 Proteobacteria
Xenorhabdus bovienii SS 2004 YP 003467797 1 406818 Proteobacteria
Xenorhabdus bovienii SS 2004 YP 003467247 1 406818 Proteobacteria
Xenorhabdus bovienii SS 2004 YP 003469993 1 406818 Proteobacteria
Xenorhabdus bovienii SS 2004 YP 003466101 1 406818 Proteobacteria
Photorhabdus luminescens subsp laumondii TTO1 NP 929777 1 243265 Proteobacteria
Photorhabdus luminescens subsp laumondii TTO1 NP 930543 1 243265 Proteobacteria
Photorhabdus luminescens subsp laumondii TTO1 NP 930627 1 243265 Proteobacteria
Photorhabdus luminescens subsp laumondii TTO1 NP 931684 1 243265 Proteobacteria
Shigella sonnei 53G YP 005454411 1 216599 Proteobacteria
Shigella sonnei 53G YP 005454397 1 216599 Proteobacteria
Shigella dysenteriae Sd197 YP 406043 1 300267 Proteobacteria
Shigella dysenteriae Sd197 YP 406073 1 300267 Proteobacteria
Shigella flexneri 2002017 YP 005711959 1 591020 Proteobacteria
Shigella dysenteriae Sd197 YP 406070 1 300267 Proteobacteria
Shigella flexneri 2002017 YP 005711988 1 591020 Proteobacteria
Shigella flexneri 2002017 YP 005712088 1 591020 Proteobacteria
Dickeya zeae NCPPB 3532 WP 023639576 1 Dzeae
Dickeya dadantii subsp dieffenbachiae NCPPB 2976 WP 038909698 1 Ddadantii
Dickeya zeae CSL RW192 WP 038909698 1 Dzeae
Erwinia sp Ejp617 YP 005817796 1 215689 Proteobacteria
Dickeya dianthicola IPO 980 WP 026594618 1 Ddianthicola
Dickeya dianthicola NCPPB 3534 WP 026594618 1 Ddianthicola
Dickeya dianthicola GBBC 2039 WP 051119537 1 Ddianthicola
Dickeya dianthicola RNS04 9 WP 046830443 1 Ddianthicola
Dickeya dianthicola IPO 980 WP 046830443 1 Ddianthicola
Dickeya dianthicola NCPPB 3534 WP 046830443 1 Ddianthicola
Dickeya dianthicola NCPPB 453 WP 046830443 1 Ddianthicola
Dickeya sp CSL RW240 WP 035343396 1 Daquatica
DAQ1742 DAQ1742 01208 Daquatica
Dickeya chrysanthemi Ech1591 WP 038927071 1 Dchrysanthemi
Dickeya chrysanthemi NCPPB 3533 WP 038927071 1 Dchrysanthemi
Dickeya sp CSL RW240 WP 067486487 1 Daquatica
Dickeya sp CSL RW240 WP 035340406 1 Daquatica
DAQ1742 DAQ1742 03292 Daquatica
Dickeya aquatica DW 0440 WP 035340406 1 Daquatica
DAQ1742 DAQ1742 02819 Daquatica
DAQ1742 DAQ1742 03488 Daquatica
DAQ1742 DAQ1742 00833 Daquatica
DAQ1742 DAQ1742 01793 Daquatica
DAQ1742 DAQ1742 00633 Daquatica
Dickeya aquatica DW 0440 WP 035338700 1 Daquatica
DAQ1742 DAQ1742 00179 Daquatica
DAQ1742 DAQ1742 03432 Daquatica
Dickeya chrysanthemi L11 WP 033577751 1 Dchrysanthemi
Dickeya chrysanthemi NCPPB 3533 WP 038926957 1 Dchrysanthemi
Dickeya chrysanthemi Ech1591 WP 038926957 1 Dchrysanthemi
Dickeya zeae EC1 WP 029456551 1 Dzeae
Dickeya zeae ZJU1202 WP 029456551 1 Dzeae
Dickeya zeae DZ2Q WP 029456551 1 Dzeae
Dickeya chrysanthemi Ech1591 WP 038926902 1 Dchrysanthemi
Dickeya chrysanthemi NCPPB 3533 WP 038926902 1 Dchrysanthemi
Dickeya paradisiaca NCPPB 2511 WP 071597669 1 Dparadisiaca
Dickeya zeae DZ2Q WP 029456324 1 Dzeae
Dickeya zeae ZJU1202 WP 029456324 1 Dzeae
Dickeya zeae EC1 WP 029456324 1 Dzeae
Dickeya chrysanthemi subsp parthenii NCPPB 516 WP 033580970 1 Dchrysanthemi
Dickeya dianthicola IPO 980 WP 047100554 1 Ddianthicola
Dickeya dianthicola NCPPB 453 WP 024103989 1 Ddianthicola
Dickeya dianthicola IPO 980 WP 024103989 1 Ddianthicola
Dickeya dianthicola RNS04 9 WP 024103989 1 Ddianthicola
Dickeya chrysanthemi NCPPB 3533 WP 024103989 1 Dchrysanthemi
Dickeya chrysanthemi L11 WP 033576063 1 Dchrysanthemi
Dickeya chrysanthemi Ech1591 WP 038927122 1 Dchrysanthemi
Dickeya chrysanthemi NCPPB 3533 WP 038927122 1 Dchrysanthemi
Dickeya aquatica DW 0440 WP 035342579 1 Daquatica
Dickeya sp CSL RW240 WP 035342579 1 Daquatica
DAQ1742 DAQ1742 01810 Daquatica
892 DAQ1742 01810 GI7 Seed
DAQ1742 DAQ1742 01844 Daquatica

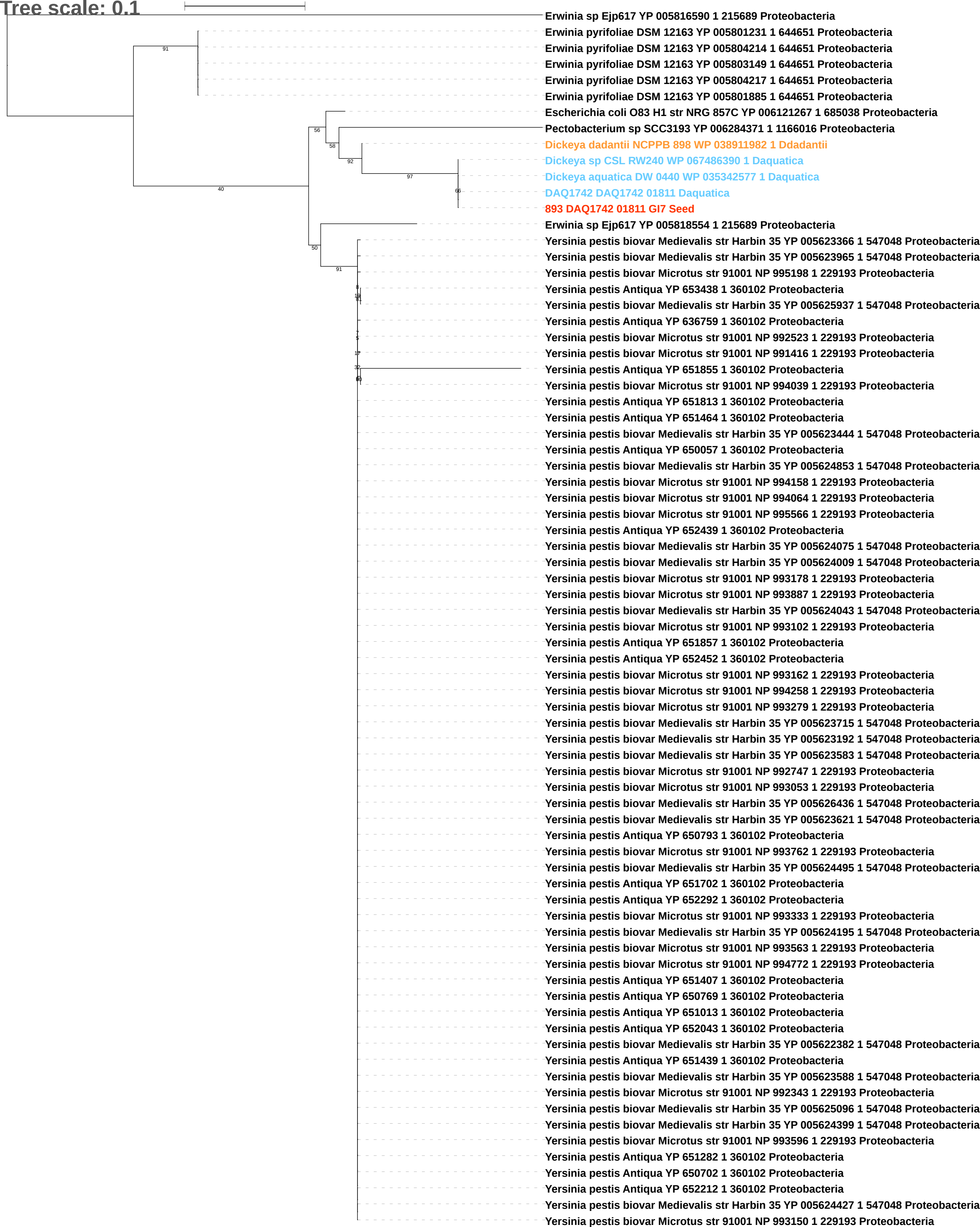
Tree scale: 0.1
Erwinia sp Ejp617 YP 005816590 1 215689 Proteobacteria
Erwinia pyrifoliae DSM 12163 YP 005801231 1 644651 Proteobacteria
Erwinia pyrifoliae DSM 12163 YP 005804214 1 644651 Proteobacteria
Erwinia pyrifoliae DSM 12163 YP 005803149 1 644651 Proteobacteria
Erwinia pyrifoliae DSM 12163 YP 005804217 1 644651 Proteobacteria
Erwinia pyrifoliae DSM 12163 YP 005801885 1 644651 Proteobacteria
Escherichia coli O83 H1 str NRG 857C YP 006121267 1 685038 Proteobacteria
Pectobacterium sp SCC3193 YP 006284371 1 1166016 Proteobacteria
Dickeya dadantii NCPPB 898 WP 038911982 1 Ddadantii
Dickeya sp CSL RW240 WP 067486390 1 Daquatica
Dickeya aquatica DW 0440 WP 035342577 1 Daquatica
DAQ1742 DAQ1742 01811 Daquatica
893 DAQ1742 01811 GI7 Seed
Erwinia sp Ejp617 YP 005818554 1 215689 Proteobacteria
Yersinia pestis biovar Medievalis str Harbin 35 YP 005623366 1 547048 Proteobacteria
Yersinia pestis biovar Medievalis str Harbin 35 YP 005623965 1 547048 Proteobacteria
Yersinia pestis biovar Microtus str 91001 NP 995198 1 229193 Proteobacteria
Yersinia pestis Antiqua YP 653438 1 360102 Proteobacteria
Yersinia pestis biovar Medievalis str Harbin 35 YP 005625937 1 547048 Proteobacteria
Yersinia pestis Antiqua YP 636759 1 360102 Proteobacteria
Yersinia pestis biovar Microtus str 91001 NP 992523 1 229193 Proteobacteria
Yersinia pestis biovar Microtus str 91001 NP 991416 1 229193 Proteobacteria
Yersinia pestis Antiqua YP 651855 1 360102 Proteobacteria
Yersinia pestis biovar Microtus str 91001 NP 994039 1 229193 Proteobacteria
Yersinia pestis Antiqua YP 651813 1 360102 Proteobacteria
Yersinia pestis Antiqua YP 651464 1 360102 Proteobacteria
Yersinia pestis biovar Medievalis str Harbin 35 YP 005623444 1 547048 Proteobacteria
Yersinia pestis Antiqua YP 650057 1 360102 Proteobacteria
Yersinia pestis biovar Medievalis str Harbin 35 YP 005624853 1 547048 Proteobacteria
Yersinia pestis biovar Microtus str 91001 NP 994158 1 229193 Proteobacteria
Yersinia pestis biovar Microtus str 91001 NP 994064 1 229193 Proteobacteria
Yersinia pestis biovar Microtus str 91001 NP 995566 1 229193 Proteobacteria
Yersinia pestis Antiqua YP 652439 1 360102 Proteobacteria
Yersinia pestis biovar Medievalis str Harbin 35 YP 005624075 1 547048 Proteobacteria
Yersinia pestis biovar Medievalis str Harbin 35 YP 005624009 1 547048 Proteobacteria
Yersinia pestis biovar Microtus str 91001 NP 993178 1 229193 Proteobacteria
Yersinia pestis biovar Microtus str 91001 NP 993887 1 229193 Proteobacteria
Yersinia pestis biovar Medievalis str Harbin 35 YP 005624043 1 547048 Proteobacteria
Yersinia pestis biovar Microtus str 91001 NP 993102 1 229193 Proteobacteria
Yersinia pestis Antiqua YP 651857 1 360102 Proteobacteria
Yersinia pestis Antiqua YP 652452 1 360102 Proteobacteria
Yersinia pestis biovar Microtus str 91001 NP 993162 1 229193 Proteobacteria
Yersinia pestis biovar Microtus str 91001 NP 994258 1 229193 Proteobacteria
Yersinia pestis biovar Microtus str 91001 NP 993279 1 229193 Proteobacteria
Yersinia pestis biovar Medievalis str Harbin 35 YP 005623715 1 547048 Proteobacteria
Yersinia pestis biovar Medievalis str Harbin 35 YP 005623192 1 547048 Proteobacteria
Yersinia pestis biovar Medievalis str Harbin 35 YP 005623583 1 547048 Proteobacteria
Yersinia pestis biovar Microtus str 91001 NP 992747 1 229193 Proteobacteria
Yersinia pestis biovar Microtus str 91001 NP 993053 1 229193 Proteobacteria
Yersinia pestis biovar Medievalis str Harbin 35 YP 005626436 1 547048 Proteobacteria
Yersinia pestis biovar Medievalis str Harbin 35 YP 005623621 1 547048 Proteobacteria
Yersinia pestis Antiqua YP 650793 1 360102 Proteobacteria
Yersinia pestis biovar Microtus str 91001 NP 993762 1 229193 Proteobacteria
Yersinia pestis biovar Medievalis str Harbin 35 YP 005624495 1 547048 Proteobacteria
Yersinia pestis Antiqua YP 651702 1 360102 Proteobacteria
Yersinia pestis Antiqua YP 652292 1 360102 Proteobacteria
Yersinia pestis biovar Microtus str 91001 NP 993333 1 229193 Proteobacteria
Yersinia pestis biovar Medievalis str Harbin 35 YP 005624195 1 547048 Proteobacteria
Yersinia pestis biovar Microtus str 91001 NP 993563 1 229193 Proteobacteria
Yersinia pestis biovar Microtus str 91001 NP 994772 1 229193 Proteobacteria
Yersinia pestis Antiqua YP 651407 1 360102 Proteobacteria
Yersinia pestis Antiqua YP 650769 1 360102 Proteobacteria
Yersinia pestis Antiqua YP 651013 1 360102 Proteobacteria
Yersinia pestis Antiqua YP 652043 1 360102 Proteobacteria
Yersinia pestis biovar Medievalis str Harbin 35 YP 005622382 1 547048 Proteobacteria
Yersinia pestis Antiqua YP 651439 1 360102 Proteobacteria
Yersinia pestis biovar Medievalis str Harbin 35 YP 005623588 1 547048 Proteobacteria
Yersinia pestis biovar Microtus str 91001 NP 992343 1 229193 Proteobacteria
Yersinia pestis biovar Medievalis str Harbin 35 YP 005625096 1 547048 Proteobacteria
Yersinia pestis biovar Medievalis str Harbin 35 YP 005624399 1 547048 Proteobacteria
Yersinia pestis biovar Microtus str 91001 NP 993596 1 229193 Proteobacteria
Yersinia pestis Antiqua YP 651282 1 360102 Proteobacteria
Yersinia pestis Antiqua YP 650702 1 360102 Proteobacteria
Yersinia pestis Antiqua YP 652212 1 360102 Proteobacteria
Yersinia pestis biovar Medievalis str Harbin 35 YP 005624427 1 547048 Proteobacteria
Yersinia pestis biovar Microtus str 91001 NP 993150 1 229193 Proteobacteria

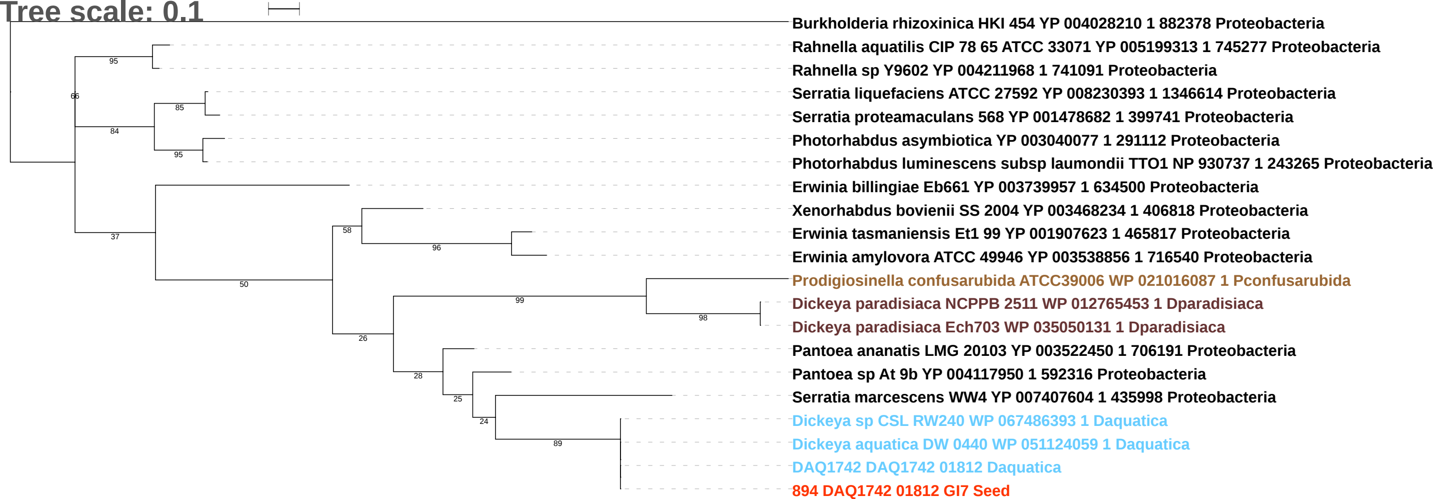
Tree scale: 0.1
Burkholderia rhizoxinica HKI 454 YP 004028210 1 882378 Proteobacteria
Rahnella aquatilis CIP 78 65 ATCC 33071 YP 005199313 1 745277 Proteobacteria
Rahnella sp Y9602 YP 004211968 1 741091 Proteobacteria
Serratia liquefaciens ATCC 27592 YP 008230393 1 1346614 Proteobacteria
Serratia proteamaculans 568 YP 001478682 1 399741 Proteobacteria
Photorhabdus asymbiotica YP 003040077 1 291112 Proteobacteria
Photorhabdus luminescens subsp laumondii TTO1 NP 930737 1 243265 Proteobacteria
Erwinia billingiae Eb661 YP 003739957 1 634500 Proteobacteria
Xenorhabdus bovienii SS 2004 YP 003468234 1 406818 Proteobacteria
Erwinia tasmaniensis Et1 99 YP 001907623 1 465817 Proteobacteria
Erwinia amylovora ATCC 49946 YP 003538856 1 716540 Proteobacteria
Prodigiosinella confusarubida ATCC39006 WP 021016087 1 Pconfusarubida
Dickeya paradisiaca NCPPB 2511 WP 012765453 1 Dparadisiaca
Dickeya paradisiaca Ech703 WP 035050131 1 Dparadisiaca
Pantoea ananatis LMG 20103 YP 003522450 1 706191 Proteobacteria
Pantoea sp At 9b YP 004117950 1 592316 Proteobacteria
Serratia marcescens WW4 YP 007407604 1 435998 Proteobacteria
Dickeya sp CSL RW240 WP 067486393 1 Daquatica
Dickeya aquatica DW 0440 WP 051124059 1 Daquatica
DAQ1742 DAQ1742 01812 Daquatica
894 DAQ1742 01812 GI7 Seed
95
66
85
84
95
37
58
96
50
99
98
26
28
25
24
89

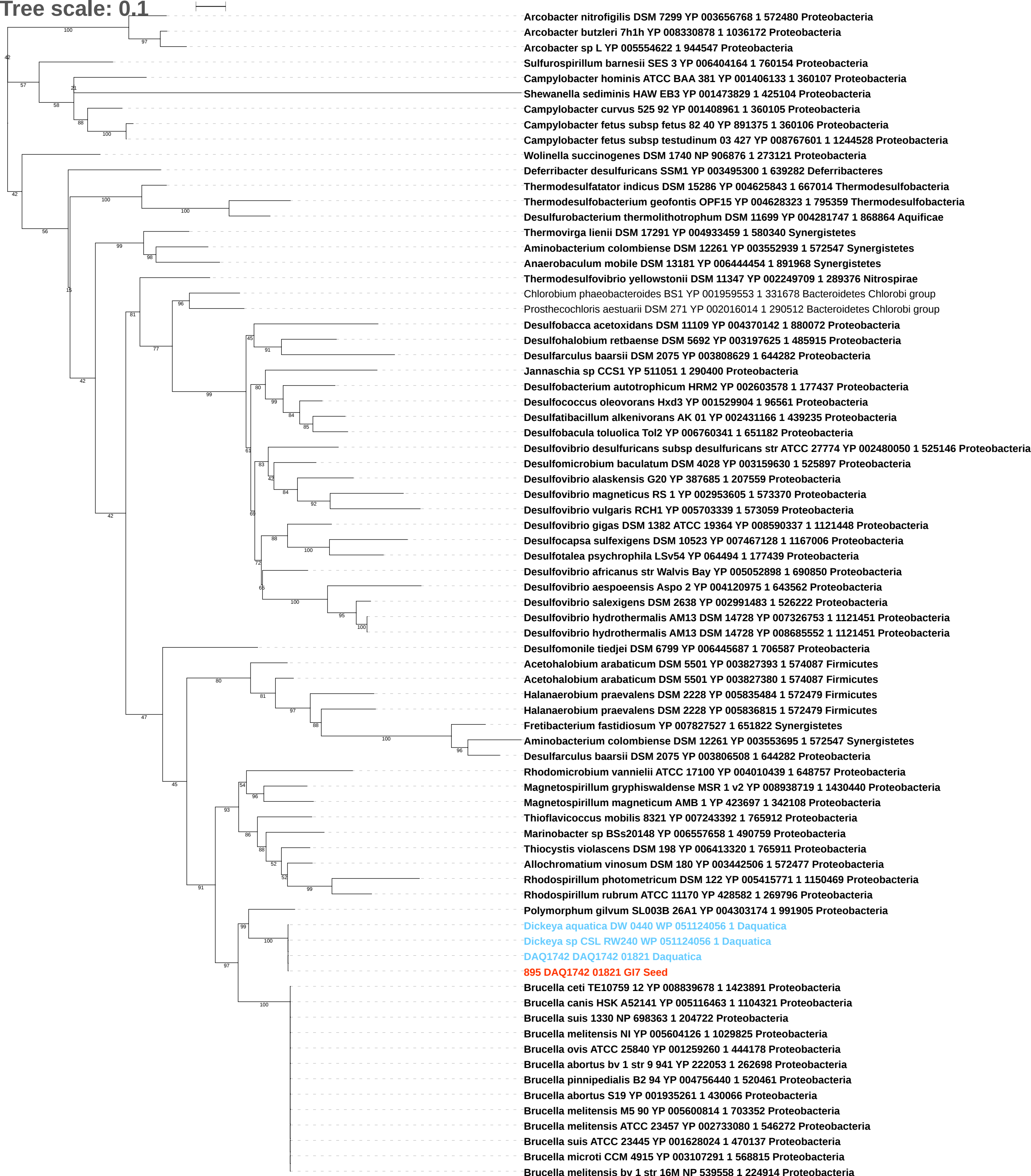
Tree scale: 0.1
Arcobacter nitrofigilis DSM 7299 YP 003656768 1 572480 Proteobacteria
Arcobacter butzleri 7h1h YP 008330878 1 1036172 Proteobacteria
Arcobacter sp L YP 005554622 1 944547 Proteobacteria
Sulfurospirillum barnesii SES 3 YP 006404164 1 760154 Proteobacteria
Campylobacter hominis ATCC BAA 381 YP 001406133 1 360107 Proteobacteria
Shewanella sediminis HAW EB3 YP 001473829 1 425104 Proteobacteria
Campylobacter curvus 525 92 YP 001408961 1 360105 Proteobacteria
Campylobacter fetus subsp fetus 82 40 YP 891375 1 360106 Proteobacteria
Campylobacter fetus subsp testudinum 03 427 YP 008767601 1 1244528 Proteobacteria
Wolinella succinogenes DSM 1740 NP 906876 1 273121 Proteobacteria
Deferribacter desulfuricans SSM1 YP 003495300 1 639282 Deferribacteres
Thermodesulfatator indicus DSM 15286 YP 004625843 1 667014 Thermodesulfobacteria
Thermodesulfobacterium geofontis OPF15 YP 004628323 1 795359 Thermodesulfobacteria
Desulfurobacterium thermolithotrophum DSM 11699 YP 004281747 1 868864 Aquificae
Thermovirga lienii DSM 17291 YP 004933459 1 580340 Synergistetes
Aminobacterium colombiense DSM 12261 YP 003552939 1 572547 Synergistetes
Anaerobaculum mobile DSM 13181 YP 006444454 1 891968 Synergistetes
Thermodesulfovibrio yellowstonii DSM 11347 YP 002249709 1 289376 Nitrospirae
Chlorobium phaeobacteroides BS1 YP 001959553 1 331678 Bacteroidetes Chlorobi group
Prosthecochloris aestuarii DSM 271 YP 002016014 1 290512 Bacteroidetes Chlorobi group
Desulfobacca acetoxidans DSM 11109 YP 004370142 1 880072 Proteobacteria
Desulfohalobium retbaense DSM 5692 YP 003197625 1 485915 Proteobacteria
Desulfarculus baarsii DSM 2075 YP 003808629 1 644282 Proteobacteria
Jannaschia sp CCS1 YP 511051 1 290400 Proteobacteria
Desulfobacterium autotrophicum HRM2 YP 002603578 1 177437 Proteobacteria
Desulfococcus oleovorans Hxd3 YP 001529904 1 96561 Proteobacteria
Desulfatibacillum alkenivorans AK 01 YP 002431166 1 439235 Proteobacteria
Desulfobacula toluolica Tol2 YP 006760341 1 651182 Proteobacteria
Desulfovibrio desulfuricans subsp desulfuricans str ATCC 27774 YP 002480050 1 525146 Proteobacteria
Desulfomicrobium baculatum DSM 4028 YP 003159630 1 525897 Proteobacteria
Desulfovibrio alaskensis G20 YP 387685 1 207559 Proteobacteria
Desulfovibrio magneticus RS 1 YP 002953605 1 573370 Proteobacteria
Desulfovibrio vulgaris RCH1 YP 005703339 1 573059 Proteobacteria
Desulfovibrio gigas DSM 1382 ATCC 19364 YP 008590337 1 1121448 Proteobacteria
Desulfocapsa sulfexigens DSM 10523 YP 007467128 1 1167006 Proteobacteria
Desulfotalea psychrophila LSv54 YP 064494 1 177439 Proteobacteria
Desulfovibrio africanus str Walvis Bay YP 005052898 1 690850 Proteobacteria
Desulfovibrio aespoeensis Aspo 2 YP 004120975 1 643562 Proteobacteria
Desulfovibrio salexigens DSM 2638 YP 002991483 1 526222 Proteobacteria
Desulfovibrio hydrothermalis AM13 DSM 14728 YP 007326753 1 1121451 Proteobacteria
Desulfovibrio hydrothermalis AM13 DSM 14728 YP 008685552 1 1121451 Proteobacteria
Desulfomonile tiedjei DSM 6799 YP 006445687 1 706587 Proteobacteria
Acetohalobium arabaticum DSM 5501 YP 003827393 1 574087 Firmicutes
Acetohalobium arabaticum DSM 5501 YP 003827380 1 574087 Firmicutes
Halanaerobium praevalens DSM 2228 YP 005835484 1 572479 Firmicutes
Halanaerobium praevalens DSM 2228 YP 005836815 1 572479 Firmicutes
Fretibacterium fastidiosum YP 007827527 1 651822 Synergistetes
Aminobacterium colombiense DSM 12261 YP 003553695 1 572547 Synergistetes
Desulfarculus baarsii DSM 2075 YP 003806508 1 644282 Proteobacteria
Rhodomicrobium vannielii ATCC 17100 YP 004010439 1 648757 Proteobacteria
Magnetospirillum gryphiswaldense MSR 1 v2 YP 008938719 1 1430440 Proteobacteria
Magnetospirillum magneticum AMB 1 YP 423697 1 342108 Proteobacteria
Thioflavicoccus mobilis 8321 YP 007243392 1 765912 Proteobacteria
Marinobacter sp BSs20148 YP 006557658 1 490759 Proteobacteria
Thiocystis violascens DSM 198 YP 006413320 1 765911 Proteobacteria
Allochromatium vinosum DSM 180 YP 003442506 1 572477 Proteobacteria
Rhodospirillum photometricum DSM 122 YP 005415771 1 1150469 Proteobacteria
Rhodospirillum rubrum ATCC 11170 YP 428582 1 269796 Proteobacteria
Polymorphum gilvum SL003B 26A1 YP 004303174 1 991905 Proteobacteria
Dickeya aquatica DW 0440 WP 051124056 1 Daquatica
Dickeya sp CSL RW240 WP 051124056 1 Daquatica
DAQ1742 DAQ1742 01821 Daquatica
895 DAQ1742 01821 GI7 Seed
Brucella ceti TE10759 12 YP 008839678 1 1423891 Proteobacteria
Brucella canis HSK A52141 YP 005116463 1 1104321 Proteobacteria
Brucella suis 1330 NP 698363 1 204722 Proteobacteria
Brucella melitensis NI YP 005604126 1 1029825 Proteobacteria
Brucella ovis ATCC 25840 YP 001259260 1 444178 Proteobacteria
Brucella abortus bv 1 str 9 941 YP 222053 1 262698 Proteobacteria
Brucella pinnipedialis B2 94 YP 004756440 1 520461 Proteobacteria
Brucella abortus S19 YP 001935261 1 430066 Proteobacteria
Brucella melitensis M5 90 YP 005600814 1 703352 Proteobacteria
Brucella melitensis ATCC 23457 YP 002733080 1 546272 Proteobacteria
Brucella suis ATCC 23445 YP 001628024 1 470137 Proteobacteria
Brucella microti CCM 4915 YP 003107291 1 568815 Proteobacteria
Brucella melitensis bv 1 str 16M NP 539558 1 224914 Proteobacteria

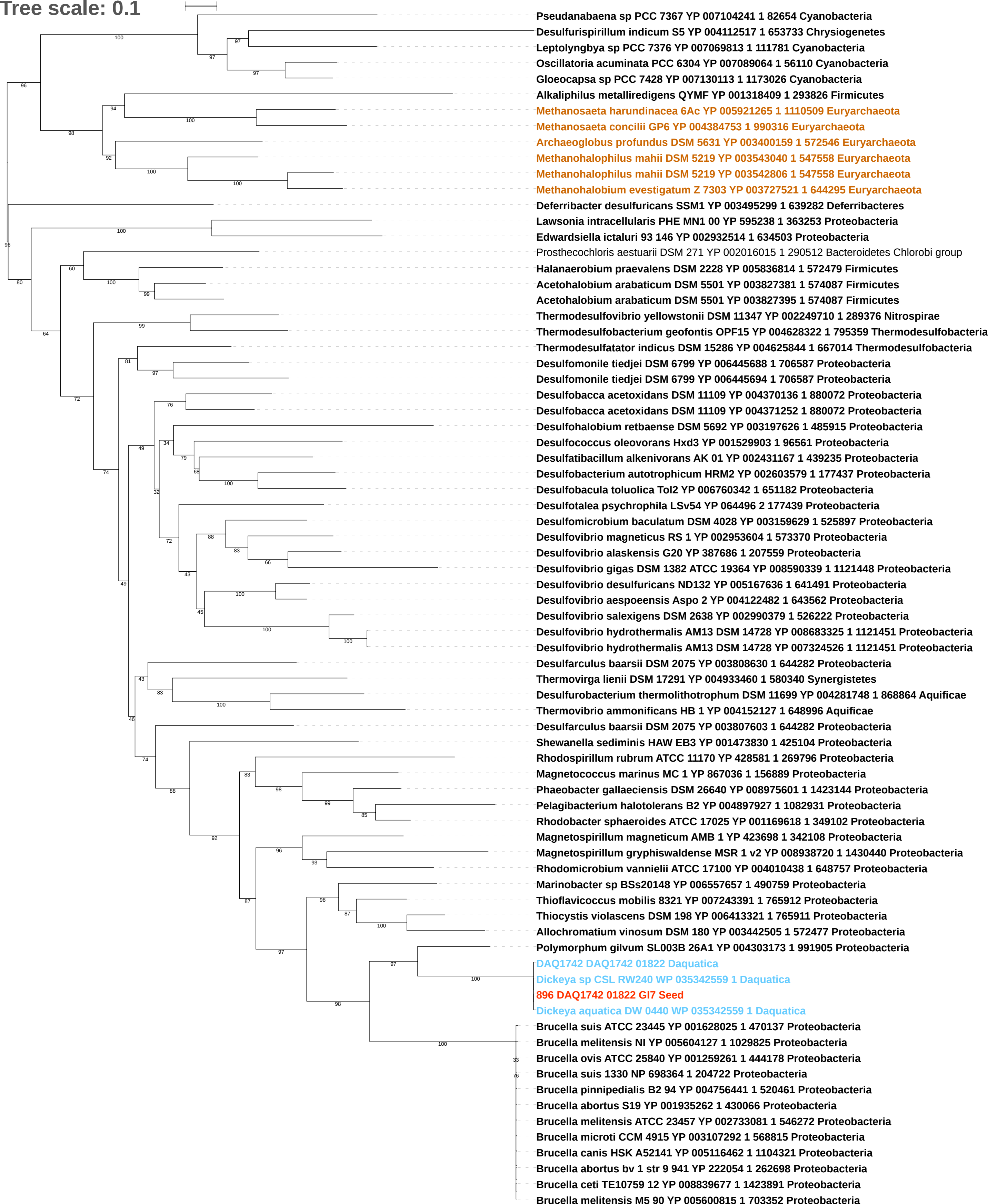
Tree scale: 0.1
Pseudanabaena sp PCC 7367 YP 007104241 1 82654 Cyanobacteria
Desulfurispirillum indicum S5 YP 004112517 1 653733 Chrysiogenetes
Leptolyngbya sp PCC 7376 YP 007069813 1 111781 Cyanobacteria
Oscillatoria acuminata PCC 6304 YP 007089064 1 56110 Cyanobacteria
Gloeocapsa sp PCC 7428 YP 007130113 1 1173026 Cyanobacteria
Alkaliphilus metalliredigens QYMF YP 001318409 1 293826 Firmicutes
Methanosaeta harundinacea 6Ac YP 005921265 1 1110509 Euryarchaeota
Methanosaeta concilii GP6 YP 004384753 1 990316 Euryarchaeota
Archaeoglobus profundus DSM 5631 YP 003400159 1 572546 Euryarchaeota
Methanohalophilus mahii DSM 5219 YP 003543040 1 547558 Euryarchaeota
Methanohalophilus mahii DSM 5219 YP 003542806 1 547558 Euryarchaeota
Methanohalobium evestigatum Z 7303 YP 003727521 1 644295 Euryarchaeota
Deferribacter desulfuricans SSM1 YP 003495299 1 639282 Deferribacteres
Lawsonia intracellularis PHE MN1 00 YP 595238 1 363253 Proteobacteria
Edwardsiella ictaluri 93 146 YP 002932514 1 634503 Proteobacteria
Prosthecochloris aestuarii DSM 271 YP 002016015 1 290512 Bacteroidetes Chlorobi group
Halanaerobium praevalens DSM 2228 YP 005836814 1 572479 Firmicutes
Acetohalobium arabaticum DSM 5501 YP 003827381 1 574087 Firmicutes
Acetohalobium arabaticum DSM 5501 YP 003827395 1 574087 Firmicutes
Thermodesulfovibrio yellowstonii DSM 11347 YP 002249710 1 289376 Nitrospirae
Thermodesulfobacterium geofontis OPF15 YP 004628322 1 795359 Thermodesulfobacteria
Thermodesulfatator indicus DSM 15286 YP 004625844 1 667014 Thermodesulfobacteria
Desulfomonile tiedjei DSM 6799 YP 006445688 1 706587 Proteobacteria
Desulfomonile tiedjei DSM 6799 YP 006445694 1 706587 Proteobacteria
Desulfobacca acetoxidans DSM 11109 YP 004370136 1 880072 Proteobacteria
Desulfobacca acetoxidans DSM 11109 YP 004371252 1 880072 Proteobacteria
Desulfohalobium retbaense DSM 5692 YP 003197626 1 485915 Proteobacteria
Desulfococcus oleovorans Hxd3 YP 001529903 1 96561 Proteobacteria
Desulfatibacillum alkenivorans AK 01 YP 002431167 1 439235 Proteobacteria
Desulfobacterium autotrophicum HRM2 YP 002603579 1 177437 Proteobacteria
Desulfobacula toluolica Tol2 YP 006760342 1 651182 Proteobacteria
Desulfotalea psychrophila LSv54 YP 064496 2 177439 Proteobacteria
Desulfomicrobium baculatum DSM 4028 YP 003159629 1 525897 Proteobacteria
Desulfovibrio magneticus RS 1 YP 002953604 1 573370 Proteobacteria
Desulfovibrio alaskensis G20 YP 387686 1 207559 Proteobacteria
Desulfovibrio gigas DSM 1382 ATCC 19364 YP 008590339 1 1121448 Proteobacteria
Desulfovibrio desulfuricans ND132 YP 005167636 1 641491 Proteobacteria
Desulfovibrio aespoeensis Aspo 2 YP 004122482 1 643562 Proteobacteria
Desulfovibrio salexigens DSM 2638 YP 002990379 1 526222 Proteobacteria
Desulfovibrio hydrothermalis AM13 DSM 14728 YP 008683325 1 1121451 Proteobacteria
Desulfovibrio hydrothermalis AM13 DSM 14728 YP 007324526 1 1121451 Proteobacteria
Desulfarculus baarsii DSM 2075 YP 003808630 1 644282 Proteobacteria
Thermovirga lienii DSM 17291 YP 004933460 1 580340 Synergistetes
Desulfurobacterium thermolithotrophum DSM 11699 YP 004281748 1 868864 Aquificae
Thermovibrio ammonificans HB 1 YP 004152127 1 648996 Aquificae
Desulfarculus baarsii DSM 2075 YP 003807603 1 644282 Proteobacteria
Shewanella sediminis HAW EB3 YP 001473830 1 425104 Proteobacteria
Rhodospirillum rubrum ATCC 11170 YP 428581 1 269796 Proteobacteria
Magnetococcus marinus MC 1 YP 867036 1 156889 Proteobacteria
Phaeobacter gallaeciensis DSM 26640 YP 008975601 1 1423144 Proteobacteria
Pelagibacterium halotolerans B2 YP 004897927 1 1082931 Proteobacteria
Rhodobacter sphaeroides ATCC 17025 YP 001169618 1 349102 Proteobacteria
Magnetospirillum magneticum AMB 1 YP 423698 1 342108 Proteobacteria
Magnetospirillum gryphiswaldense MSR 1 v2 YP 008938720 1 1430440 Proteobacteria
Rhodomicrobium vannielii ATCC 17100 YP 004010438 1 648757 Proteobacteria
Marinobacter sp BSs20148 YP 006557657 1 490759 Proteobacteria
Thioflavicoccus mobilis 8321 YP 007243391 1 765912 Proteobacteria
Thiocystis violascens DSM 198 YP 006413321 1 765911 Proteobacteria
Allochromatium vinosum DSM 180 YP 003442505 1 572477 Proteobacteria
Polymorphum gilvum SL003B 26A1 YP 004303173 1 991905 Proteobacteria
DAQ1742 DAQ1742 01822 Daquatica
Dickeya sp CSL RW240 WP 035342559 1 Daquatica
896 DAQ1742 01822 GI7 Seed
Dickeya aquatica DW 0440 WP 035342559 1 Daquatica
Brucella suis ATCC 23445 YP 001628025 1 470137 Proteobacteria
Brucella melitensis NI YP 005604127 1 1029825 Proteobacteria
Brucella ovis ATCC 25840 YP 001259261 1 444178 Proteobacteria
Brucella suis 1330 NP 698364 1 204722 Proteobacteria
Brucella pinnipedialis B2 94 YP 004756441 1 520461 Proteobacteria
Brucella abortus S19 YP 001935262 1 430066 Proteobacteria
Brucella melitensis ATCC 23457 YP 002733081 1 546272 Proteobacteria
Brucella microti CCM 4915 YP 003107292 1 568815 Proteobacteria
Brucella canis HSK A52141 YP 005116462 1 1104321 Proteobacteria
Brucella abortus bv 1 str 9 941 YP 222054 1 262698 Proteobacteria
Brucella ceti TE10759 12 YP 008839677 1 1423891 Proteobacteria
Brucella melitensis M5 90 YP 005600815 1 703352 Proteobacteria

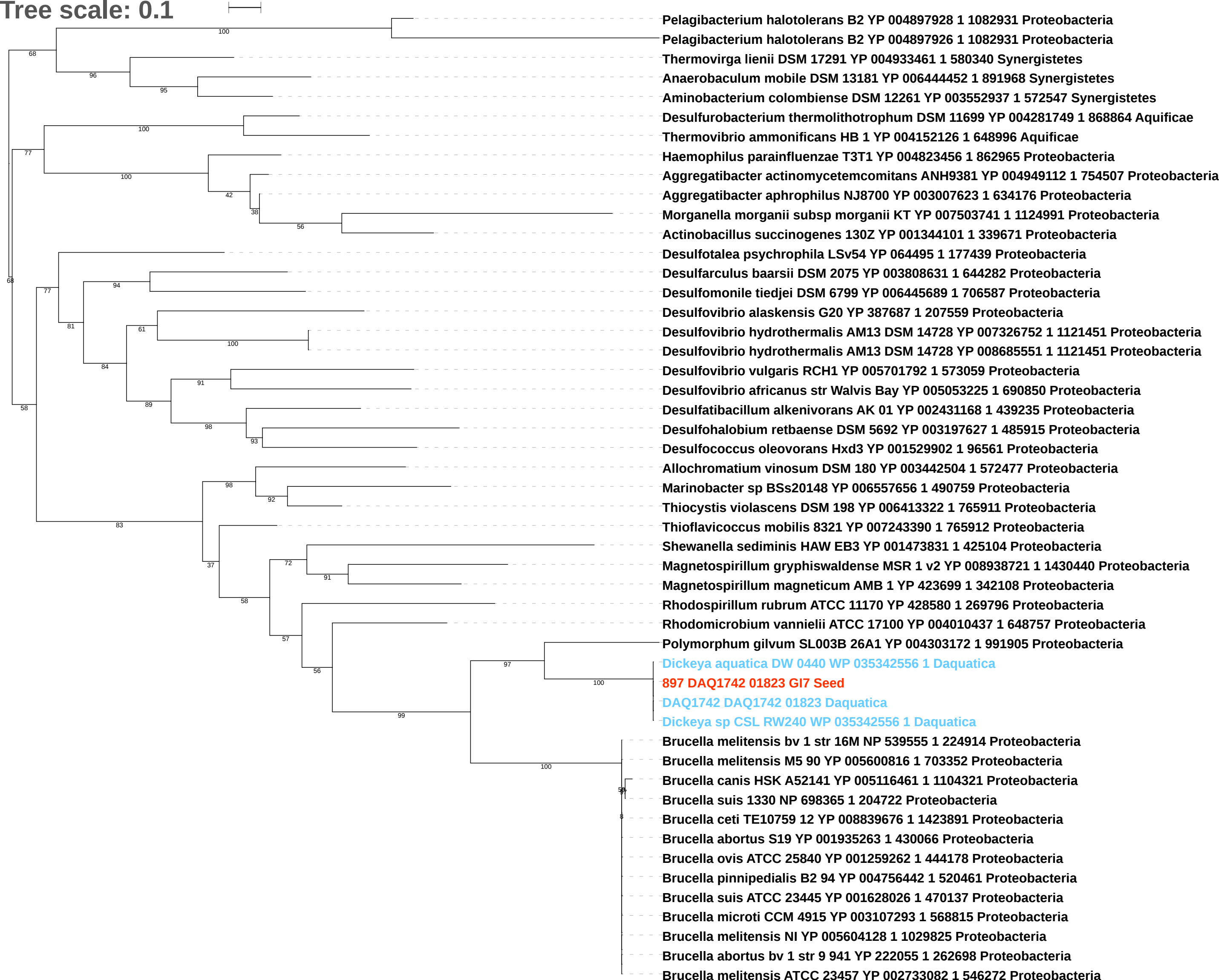
Tree scale: 0.1
Pelagibacterium halotolerans B2 YP 004897928 1 1082931 Proteobacteria
Pelagibacterium halotolerans B2 YP 004897926 1 1082931 Proteobacteria
Thermovirga lienii DSM 17291 YP 004933461 1 580340 Synergistetes
Anaerobaculum mobile DSM 13181 YP 006444452 1 891968 Synergistetes
Aminobacterium colombiense DSM 12261 YP 003552937 1 572547 Synergistetes
Desulfurobacterium thermolithotrophum DSM 11699 YP 004281749 1 868864 Aquificae
Thermovibrio ammonificans HB 1 YP 004152126 1 648996 Aquificae
Haemophilus parainfluenzae T3T1 YP 004823456 1 862965 Proteobacteria
Aggregatibacter actinomycetemcomitans ANH9381 YP 004949112 1 754507 Proteobacteria
Aggregatibacter aphrophilus NJ8700 YP 003007623 1 634176 Proteobacteria
Morganella morganii subsp morganii KT YP 007503741 1 1124991 Proteobacteria
Actinobacillus succinogenes 130Z YP 001344101 1 339671 Proteobacteria
Desulfotalea psychrophila LSv54 YP 064495 1 177439 Proteobacteria
Desulfarculus baarsii DSM 2075 YP 003808631 1 644282 Proteobacteria
Desulfomonile tiedjei DSM 6799 YP 006445689 1 706587 Proteobacteria
Desulfovibrio alaskensis G20 YP 387687 1 207559 Proteobacteria
Desulfovibrio hydrothermalis AM13 DSM 14728 YP 007326752 1 1121451 Proteobacteria
Desulfovibrio hydrothermalis AM13 DSM 14728 YP 008685551 1 1121451 Proteobacteria
Desulfovibrio vulgaris RCH1 YP 005701792 1 573059 Proteobacteria
Desulfovibrio africanus str Walvis Bay YP 005053225 1 690850 Proteobacteria
Desulfatibacillum alkenivorans AK 01 YP 002431168 1 439235 Proteobacteria
Desulfohalobium retbaense DSM 5692 YP 003197627 1 485915 Proteobacteria
Desulfococcus oleovorans Hxd3 YP 001529902 1 96561 Proteobacteria
Allochromatium vinosum DSM 180 YP 003442504 1 572477 Proteobacteria
Marinobacter sp BSs20148 YP 006557656 1 490759 Proteobacteria
Thiocystis violascens DSM 198 YP 006413322 1 765911 Proteobacteria
Thioflavicoccus mobilis 8321 YP 007243390 1 765912 Proteobacteria
Shewanella sediminis HAW EB3 YP 001473831 1 425104 Proteobacteria
Magnetospirillum gryphiswaldense MSR 1 v2 YP 008938721 1 1430440 Proteobacteria
Magnetospirillum magneticum AMB 1 YP 423699 1 342108 Proteobacteria
Rhodospirillum rubrum ATCC 11170 YP 428580 1 269796 Proteobacteria
Rhodomicrobium vannielii ATCC 17100 YP 004010437 1 648757 Proteobacteria
Polymorphum gilvum SL003B 26A1 YP 004303172 1 991905 Proteobacteria
Dickeya aquatica DW 0440 WP 035342556 1 Daquatica
897 DAQ1742 01823 GI7 Seed
DAQ1742 DAQ1742 01823 Daquatica
Dickeya sp CSL RW240 WP 035342556 1 Daquatica
Brucella melitensis bv 1 str 16M NP 539555 1 224914 Proteobacteria
Brucella melitensis M5 90 YP 005600816 1 703352 Proteobacteria
Brucella canis HSK A52141 YP 005116461 1 1104321 Proteobacteria
Brucella suis 1330 NP 698365 1 204722 Proteobacteria
Brucella ceti TE10759 12 YP 008839676 1 1423891 Proteobacteria
Brucella abortus S19 YP 001935263 1 430066 Proteobacteria
Brucella ovis ATCC 25840 YP 001259262 1 444178 Proteobacteria
Brucella pinnipedialis B2 94 YP 004756442 1 520461 Proteobacteria
Brucella suis ATCC 23445 YP 001628026 1 470137 Proteobacteria
Brucella microti CCM 4915 YP 003107293 1 568815 Proteobacteria
Brucella melitensis NI YP 005604128 1 1029825 Proteobacteria
Brucella abortus bv 1 str 9 941 YP 222055 1 262698 Proteobacteria
Brucella melitensis ATCC 23457 YP 002733082 1 546272 Proteobacteria

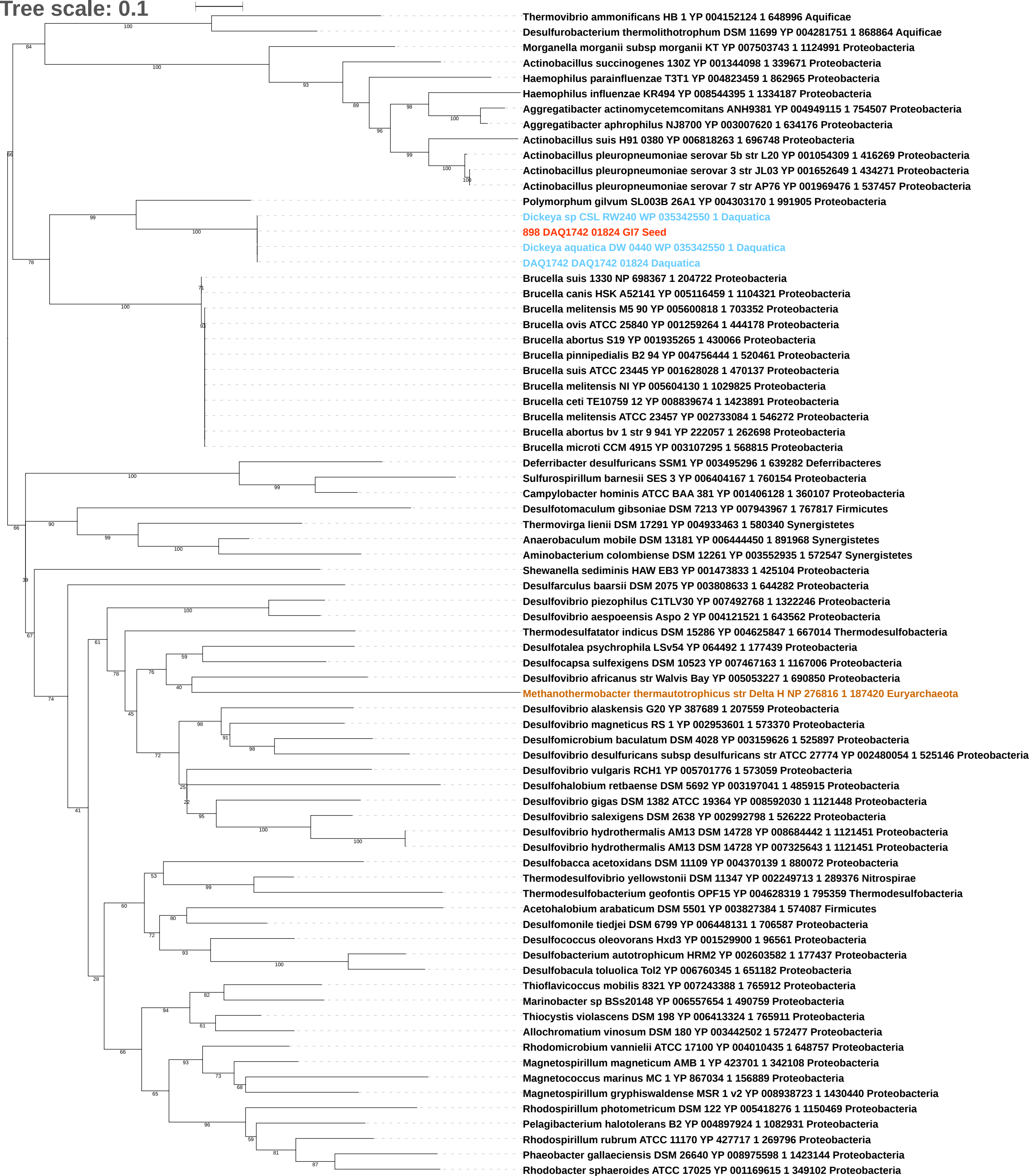
Tree scale: 0.1
Thermovibrio ammonificans HB 1 YP 004152124 1 648996 Aquificae
Desulfurobacterium thermolithotrophum DSM 11699 YP 004281751 1 868864 Aquificae
Morganella morganii subsp morganii KT YP 007503743 1 1124991 Proteobacteria
Actinobacillus succinogenes 130Z YP 001344098 1 339671 Proteobacteria
Haemophilus parainfluenzae T3T1 YP 004823459 1 862965 Proteobacteria
Haemophilus influenzae KR494 YP 008544395 1 1334187 Proteobacteria
Aggregatibacter actinomycetemcomitans ANH9381 YP 004949115 1 754507 Proteobacteria
Aggregatibacter aphrophilus NJ8700 YP 003007620 1 634176 Proteobacteria
Actinobacillus suis H91 0380 YP 006818263 1 696748 Proteobacteria
Actinobacillus pleuropneumoniae serovar 5b str L20 YP 001054309 1 416269 Proteobacteria
Actinobacillus pleuropneumoniae serovar 3 str JL03 YP 001652649 1 434271 Proteobacteria
Actinobacillus pleuropneumoniae serovar 7 str AP76 YP 001969476 1 537457 Proteobacteria
Polymorphum gilvum SL003B 26A1 YP 004303170 1 991905 Proteobacteria
Dickeya sp CSL RW240 WP 035342550 1 Daquatica
898 DAQ1742 01824 GI7 Seed
Dickeya aquatica DW 0440 WP 035342550 1 Daquatica
DAQ1742 DAQ1742 01824 Daquatica
Brucella suis 1330 NP 698367 1 204722 Proteobacteria
Brucella canis HSK A52141 YP 005116459 1 1104321 Proteobacteria
Brucella melitensis M5 90 YP 005600818 1 703352 Proteobacteria
Brucella ovis ATCC 25840 YP 001259264 1 444178 Proteobacteria
Brucella abortus S19 YP 001935265 1 430066 Proteobacteria
Brucella pinnipedialis B2 94 YP 004756444 1 520461 Proteobacteria
Brucella suis ATCC 23445 YP 001628028 1 470137 Proteobacteria
Brucella melitensis NI YP 005604130 1 1029825 Proteobacteria
Brucella ceti TE10759 12 YP 008839674 1 1423891 Proteobacteria
Brucella melitensis ATCC 23457 YP 002733084 1 546272 Proteobacteria
Brucella abortus bv 1 str 9 941 YP 222057 1 262698 Proteobacteria
Brucella microti CCM 4915 YP 003107295 1 568815 Proteobacteria
Deferribacter desulfuricans SSM1 YP 003495296 1 639282 Deferribacteres
Sulfurospirillum barnesii SES 3 YP 006404167 1 760154 Proteobacteria
Campylobacter hominis ATCC BAA 381 YP 001406128 1 360107 Proteobacteria
Desulfotomaculum gibsoniae DSM 7213 YP 007943967 1 767817 Firmicutes
Thermovirga lienii DSM 17291 YP 004933463 1 580340 Synergistetes
Anaerobaculum mobile DSM 13181 YP 006444450 1 891968 Synergistetes
Aminobacterium colombiense DSM 12261 YP 003552935 1 572547 Synergistetes
Shewanella sediminis HAW EB3 YP 001473833 1 425104 Proteobacteria
Desulfarculus baarsii DSM 2075 YP 003808633 1 644282 Proteobacteria
Desulfovibrio piezophilus C1TLV30 YP 007492768 1 1322246 Proteobacteria
Desulfovibrio aespoeensis Aspo 2 YP 004121521 1 643562 Proteobacteria
Thermodesulfatator indicus DSM 15286 YP 004625847 1 667014 Thermodesulfobacteria
Desulfotalea psychrophila LSv54 YP 064492 1 177439 Proteobacteria
Desulfocapsa sulfexigens DSM 10523 YP 007467163 1 1167006 Proteobacteria
Desulfovibrio africanus str Walvis Bay YP 005053227 1 690850 Proteobacteria
Methanothermobacter thermautotrophicus str Delta H NP 276816 1 187420 Euryarchaeota
Desulfovibrio alaskensis G20 YP 387689 1 207559 Proteobacteria
Desulfovibrio magneticus RS 1 YP 002953601 1 573370 Proteobacteria
Desulfomicrobium baculatum DSM 4028 YP 003159626 1 525897 Proteobacteria
Desulfovibrio desulfuricans subsp desulfuricans str ATCC 27774 YP 002480054 1 525146 Proteobacteria
Desulfovibrio vulgaris RCH1 YP 005701776 1 573059 Proteobacteria
Desulfohalobium retbaense DSM 5692 YP 003197041 1 485915 Proteobacteria
Desulfovibrio gigas DSM 1382 ATCC 19364 YP 008592030 1 1121448 Proteobacteria
Desulfovibrio salexigens DSM 2638 YP 002992798 1 526222 Proteobacteria
Desulfovibrio hydrothermalis AM13 DSM 14728 YP 008684442 1 1121451 Proteobacteria
Desulfovibrio hydrothermalis AM13 DSM 14728 YP 007325643 1 1121451 Proteobacteria
Desulfobacca acetoxidans DSM 11109 YP 004370139 1 880072 Proteobacteria
Thermodesulfovibrio yellowstonii DSM 11347 YP 002249713 1 289376 Nitrospirae
Thermodesulfobacterium geofontis OPF15 YP 004628319 1 795359 Thermodesulfobacteria
Acetohalobium arabaticum DSM 5501 YP 003827384 1 574087 Firmicutes
Desulfomonile tiedjei DSM 6799 YP 006448131 1 706587 Proteobacteria
Desulfococcus oleovorans Hxd3 YP 001529900 1 96561 Proteobacteria
Desulfobacterium autotrophicum HRM2 YP 002603582 1 177437 Proteobacteria
Desulfobacula toluolica Tol2 YP 006760345 1 651182 Proteobacteria
Thioflavicoccus mobilis 8321 YP 007243388 1 765912 Proteobacteria
Marinobacter sp BSs20148 YP 006557654 1 490759 Proteobacteria
Thiocystis violascens DSM 198 YP 006413324 1 765911 Proteobacteria
Allochromatium vinosum DSM 180 YP 003442502 1 572477 Proteobacteria
Rhodomicrobium vannielii ATCC 17100 YP 004010435 1 648757 Proteobacteria
Magnetospirillum magneticum AMB 1 YP 423701 1 342108 Proteobacteria
Magnetococcus marinus MC 1 YP 867034 1 156889 Proteobacteria
Magnetospirillum gryphiswaldense MSR 1 v2 YP 008938723 1 1430440 Proteobacteria
Rhodospirillum photometricum DSM 122 YP 005418276 1 1150469 Proteobacteria
Pelagibacterium halotolerans B2 YP 004897924 1 1082931 Proteobacteria
Rhodospirillum rubrum ATCC 11170 YP 427717 1 269796 Proteobacteria
Phaeobacter gallaeciensis DSM 26640 YP 008975598 1 1423144 Proteobacteria
Rhodobacter sphaeroides ATCC 17025 YP 001169615 1 349102 Proteobacteria

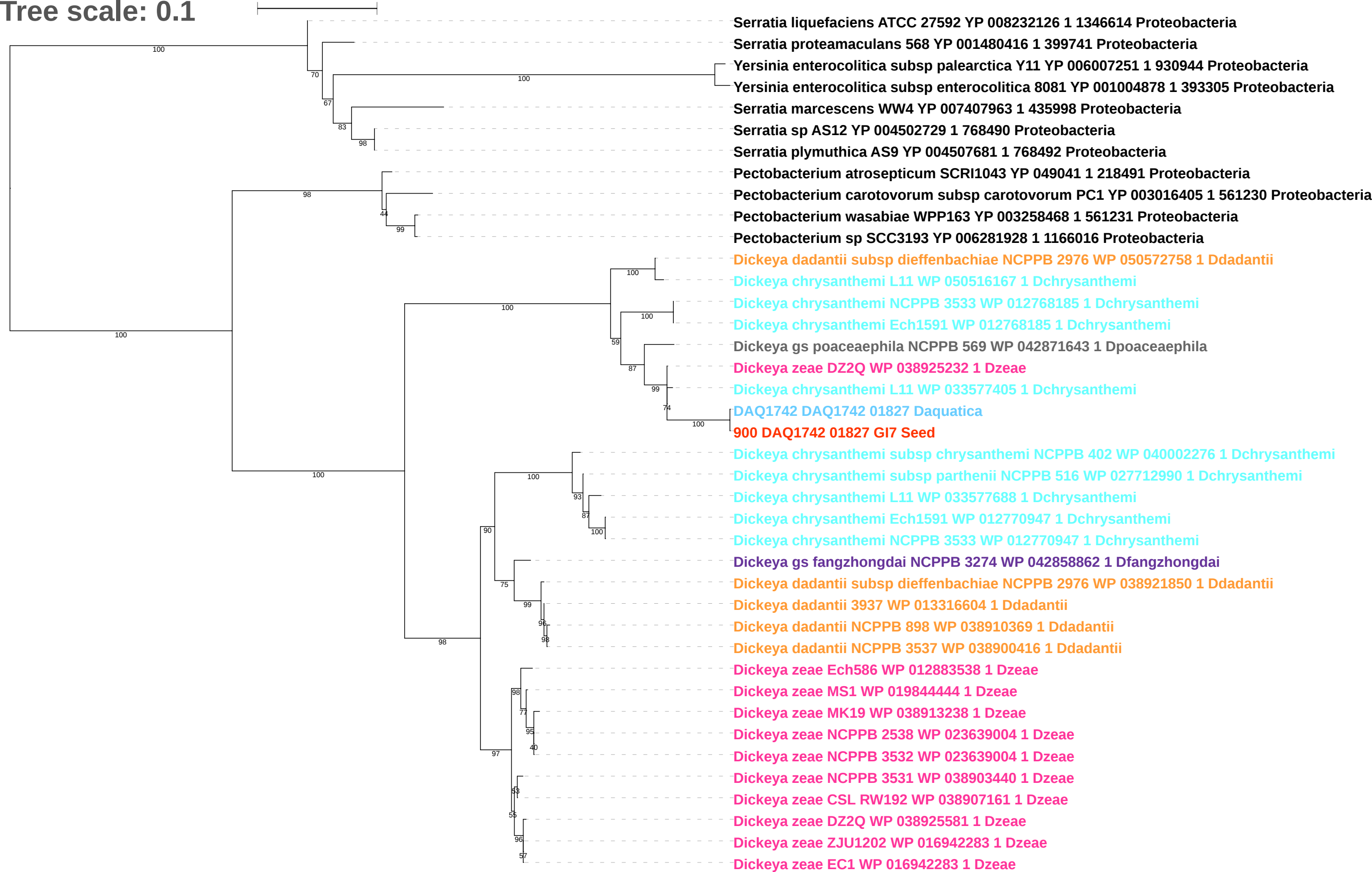
Tree scale: 0.1
Serratia liquefaciens ATCC 27592 YP 008232126 1 1346614 Proteobacteria
Serratia proteamaculans 568 YP 001480416 1 399741 Proteobacteria
Yersinia enterocolitica subsp palearctica Y11 YP 006007251 1 930944 Proteobacteria
Yersinia enterocolitica subsp enterocolitica 8081 YP 001004878 1 393305 Proteobacteria
Serratia marcescens WW4 YP 007407963 1 435998 Proteobacteria
Serratia sp AS12 YP 004502729 1 768490 Proteobacteria
Serratia plymuthica AS9 YP 004507681 1 768492 Proteobacteria
Pectobacterium atrosepticum SCRI1043 YP 049041 1 218491 Proteobacteria
Pectobacterium carotovorum subsp carotovorum PC1 YP 003016405 1 561230 Proteobacteria
Pectobacterium wasabiae WPP163 YP 003258468 1 561231 Proteobacteria
Pectobacterium sp SCC3193 YP 006281928 1 1166016 Proteobacteria
Dickeya dadantii subsp dieffenbachiae NCPPB 2976 WP 050572758 1 Ddadantii
Dickeya chrysanthemi L11 WP 050516167 1 Dchrysanthemi
Dickeya chrysanthemi NCPPB 3533 WP 012768185 1 Dchrysanthemi
Dickeya chrysanthemi Ech1591 WP 012768185 1 Dchrysanthemi
Dickeya gs poaceaephila NCPPB 569 WP 042871643 1 Dpoaceaephila
Dickeya zeae DZ2Q WP 038925232 1 Dzeae
Dickeya chrysanthemi L11 WP 033577405 1 Dchrysanthemi
DAQ1742 DAQ1742 01827 Daquatica
900 DAQ1742 01827 GI7 Seed
Dickeya chrysanthemi subsp chrysanthemi NCPPB 402 WP 040002276 1 Dchrysanthemi
Dickeya chrysanthemi subsp parthenii NCPPB 516 WP 027712990 1 Dchrysanthemi
Dickeya chrysanthemi L11 WP 033577688 1 Dchrysanthemi
Dickeya chrysanthemi Ech1591 WP 012770947 1 Dchrysanthemi
Dickeya chrysanthemi NCPPB 3533 WP 012770947 1 Dchrysanthemi
Dickeya gs fangzhongdai NCPPB 3274 WP 042858862 1 Dfangzhongdai
Dickeya dadantii subsp dieffenbachiae NCPPB 2976 WP 038921850 1 Ddadantii
Dickeya dadantii 3937 WP 013316604 1 Ddadantii
Dickeya dadantii NCPPB 898 WP 038910369 1 Ddadantii
Dickeya dadantii NCPPB 3537 WP 038900416 1 Ddadantii
Dickeya zeae Ech586 WP 012883538 1 Dzeae
Dickeya zeae MS1 WP 019844444 1 Dzeae
Dickeya zeae MK19 WP 038913238 1 Dzeae
Dickeya zeae NCPPB 2538 WP 023639004 1 Dzeae
Dickeya zeae NCPPB 3532 WP 023639004 1 Dzeae
Dickeya zeae NCPPB 3531 WP 038903440 1 Dzeae
Dickeya zeae CSL RW192 WP 038907161 1 Dzeae
Dickeya zeae DZ2Q WP 038925581 1 Dzeae
Dickeya zeae ZJU1202 WP 016942283 1 Dzeae
Dickeya zeae EC1 WP 016942283 1 Dzeae

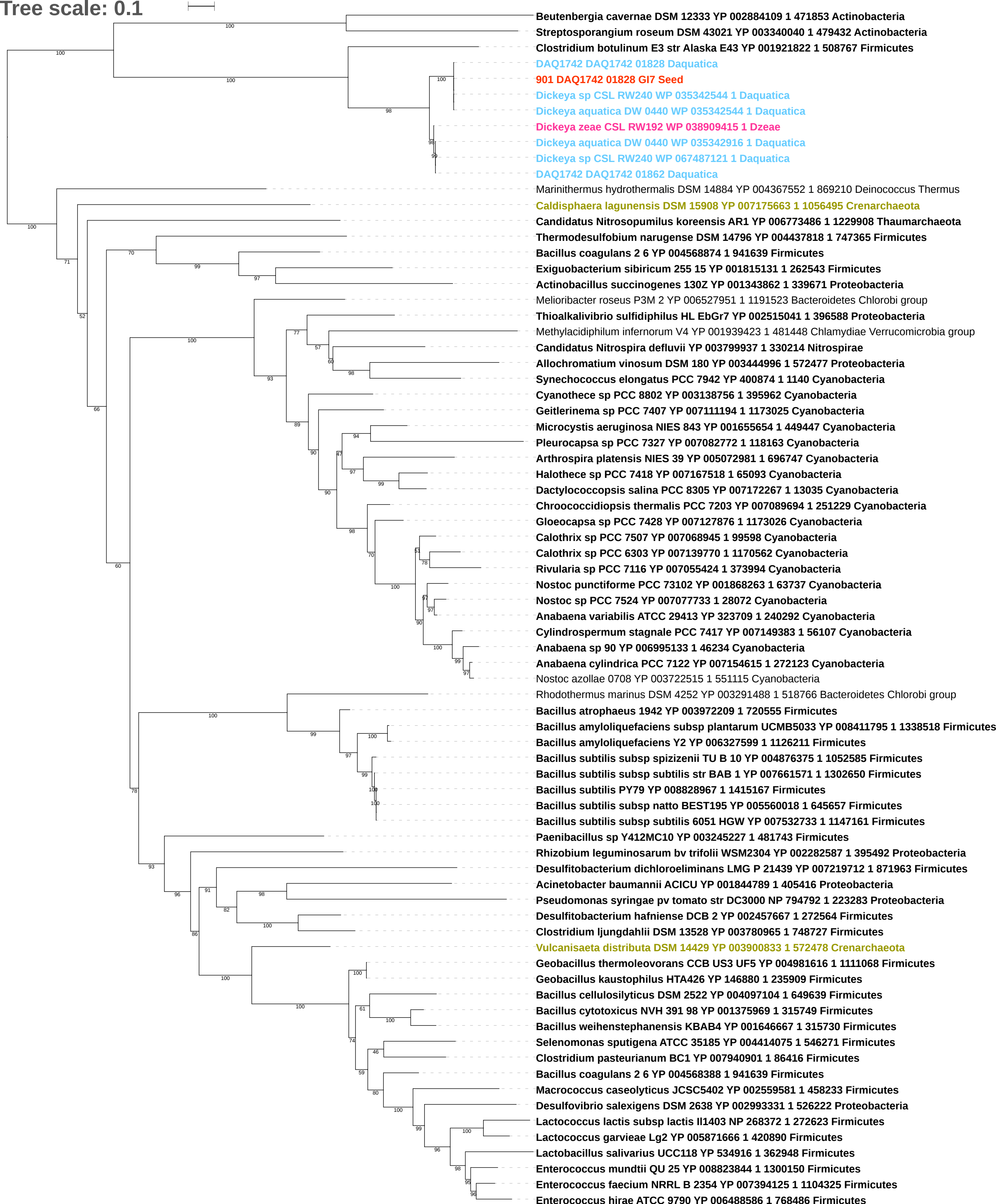

Tree scale: 0.1
Beutenbergia cavernae DSM 12333 YP 002884109 1 471853 Actinobacteria
Streptosporangium roseum DSM 43021 YP 003340040 1 479432 Actinobacteria
Clostridium botulinum E3 str Alaska E43 YP 001921822 1 508767 Firmicutes
DAQ1742 DAQ1742 01828 Daquatica
901 DAQ1742 01828 GI7 Seed
Dickeya sp CSL RW240 WP 035342544 1 Daquatica
Dickeya aquatica DW 0440 WP 035342544 1 Daquatica
Dickeya zeae CSL RW192 WP 038909415 1 Dzeae
Dickeya aquatica DW 0440 WP 035342916 1 Daquatica
Dickeya sp CSL RW240 WP 067487121 1 Daquatica
DAQ1742 DAQ1742 01862 Daquatica
Marinithermus hydrothermalis DSM 14884 YP 004367552 1 869210 Deinococcus Thermus
Caldisphaera lagunensis DSM 15908 YP 007175663 1 1056495 Crenarchaeota
Candidatus Nitrosopumilus koreensis AR1 YP 006773486 1 1229908 Thaumarchaeota
Thermodesulfobium narugense DSM 14796 YP 004437818 1 747365 Firmicutes
Bacillus coagulans 2 6 YP 004568874 1 941639 Firmicutes
Exiguobacterium sibiricum 255 15 YP 001815131 1 262543 Firmicutes
Actinobacillus succinogenes 130Z YP 001343862 1 339671 Proteobacteria
Melioribacter roseus P3M 2 YP 006527951 1 1191523 Bacteroidetes Chlorobi group
Thioalkalivibrio sulfidiphilus HL EbGr7 YP 002515041 1 396588 Proteobacteria
Methylacidiphilum infernorum V4 YP 001939423 1 481448 Chlamydiae Verrucomicrobia group
Candidatus Nitrospira defluvii YP 003799937 1 330214 Nitrospirae
Allochromatium vinosum DSM 180 YP 003444996 1 572477 Proteobacteria
Synechococcus elongatus PCC 7942 YP 400874 1 1140 Cyanobacteria
Cyanothece sp PCC 8802 YP 003138756 1 395962 Cyanobacteria
Geitlerinema sp PCC 7407 YP 007111194 1 1173025 Cyanobacteria
Microcystis aeruginosa NIES 843 YP 001655654 1 449447 Cyanobacteria
Pleurocapsa sp PCC 7327 YP 007082772 1 118163 Cyanobacteria
Arthrospira platensis NIES 39 YP 005072981 1 696747 Cyanobacteria
Halothece sp PCC 7418 YP 007167518 1 65093 Cyanobacteria
Dactylococcopsis salina PCC 8305 YP 007172267 1 13035 Cyanobacteria
Chroococcidiopsis thermalis PCC 7203 YP 007089694 1 251229 Cyanobacteria
Gloeocapsa sp PCC 7428 YP 007127876 1 1173026 Cyanobacteria
Calothrix sp PCC 7507 YP 007068945 1 99598 Cyanobacteria
Calothrix sp PCC 6303 YP 007139770 1 1170562 Cyanobacteria
Rivularia sp PCC 7116 YP 007055424 1 373994 Cyanobacteria
Nostoc punctiforme PCC 73102 YP 001868263 1 63737 Cyanobacteria
Nostoc sp PCC 7524 YP 007077733 1 28072 Cyanobacteria
Anabaena variabilis ATCC 29413 YP 323709 1 240292 Cyanobacteria
Cylindrospermum stagnale PCC 7417 YP 007149383 1 56107 Cyanobacteria
Anabaena sp 90 YP 006995133 1 46234 Cyanobacteria
Anabaena cylindrica PCC 7122 YP 007154615 1 272123 Cyanobacteria
Nostoc azollae 0708 YP 003722515 1 551115 Cyanobacteria
Rhodothermus marinus DSM 4252 YP 003291488 1 518766 Bacteroidetes Chlorobi group
Bacillus atrophaeus 1942 YP 003972209 1 720555 Firmicutes
Bacillus amyloliquefaciens subsp plantarum UCMB5033 YP 008411795 1 1338518 Firmicutes
Bacillus amyloliquefaciens Y2 YP 006327599 1 1126211 Firmicutes
Bacillus subtilis subsp spizizenii TU B 10 YP 004876375 1 1052585 Firmicutes
Bacillus subtilis subsp subtilis str BAB 1 YP 007661571 1 1302650 Firmicutes
Bacillus subtilis PY79 YP 008828967 1 1415167 Firmicutes
Bacillus subtilis subsp natto BEST195 YP 005560018 1 645657 Firmicutes
Bacillus subtilis subsp subtilis 6051 HGW YP 007532733 1 1147161 Firmicutes
Paenibacillus sp Y412MC10 YP 003245227 1 481743 Firmicutes
Rhizobium leguminosarum bv trifolii WSM2304 YP 002282587 1 395492 Proteobacteria
Desulfitobacterium dichloroeliminans LMG P 21439 YP 007219712 1 871963 Firmicutes
Acinetobacter baumannii ACICU YP 001844789 1 405416 Proteobacteria
Pseudomonas syringae pv tomato str DC3000 NP 794792 1 223283 Proteobacteria
Desulfitobacterium hafniense DCB 2 YP 002457667 1 272564 Firmicutes
Clostridium ljungdahlii DSM 13528 YP 003780965 1 748727 Firmicutes
Vulcanisaeta distributa DSM 14429 YP 003900833 1 572478 Crenarchaeota
Geobacillus thermoleovorans CCB US3 UF5 YP 004981616 1 1111068 Firmicutes
Geobacillus kaustophilus HTA426 YP 146880 1 235909 Firmicutes
Bacillus cellulosilyticus DSM 2522 YP 004097104 1 649639 Firmicutes
Bacillus cytotoxicus NVH 391 98 YP 001375969 1 315749 Firmicutes
Bacillus weihenstephanensis KBAB4 YP 001646667 1 315730 Firmicutes
Selenomonas sputigena ATCC 35185 YP 004414075 1 546271 Firmicutes
Clostridium pasteurianum BC1 YP 007940901 1 86416 Firmicutes
Bacillus coagulans 2 6 YP 004568388 1 941639 Firmicutes
Macrococcus caseolyticus JCSC5402 YP 002559581 1 458233 Firmicutes
Desulfovibrio salexigens DSM 2638 YP 002993331 1 526222 Proteobacteria
Lactococcus lactis subsp lactis Il1403 NP 268372 1 272623 Firmicutes
Lactococcus garvieae Lg2 YP 005871666 1 420890 Firmicutes
Lactobacillus salivarius UCC118 YP 534916 1 362948 Firmicutes
Enterococcus mundtii QU 25 YP 008823844 1 1300150 Firmicutes
Enterococcus faecium NRRL B 2354 YP 007394125 1 1104325 Firmicutes
Enterococcus hirae ATCC 9790 YP 006488586 1 768486 Firmicutes

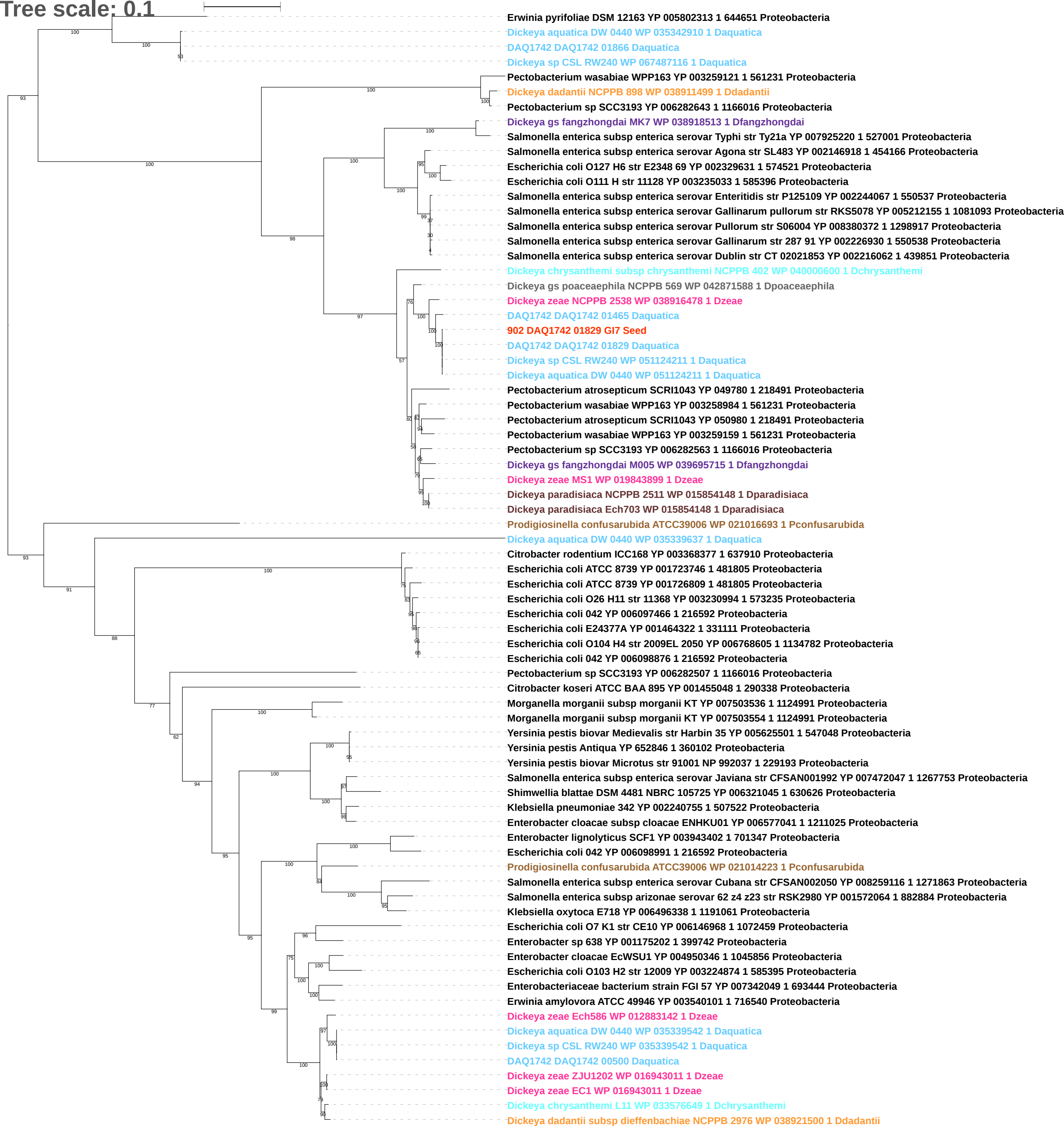
Tree scale: 0.1
Erwinia pyrifoliae DSM 12163 YP 005802313 1 644651 Proteobacteria
Dickeya aquatica DW 0440 WP 035342910 1 Daquatica
DAQ1742 DAQ1742 01866 Daquatica
Dickeya sp CSL RW240 WP 067487116 1 Daquatica
Pectobacterium wasabiae WPP163 YP 003259121 1 561231 Proteobacteria
Dickeya dadantii NCPPB 898 WP 038911499 1 Ddadantii
Pectobacterium sp SCC3193 YP 006282643 1 1166016 Proteobacteria
Dickeya gs fangzhongdai MK7 WP 038918513 1 Dfangzhongdai
Salmonella enterica subsp enterica serovar Typhi str Ty21a YP 007925220 1 527001 Proteobacteria
Salmonella enterica subsp enterica serovar Agona str SL483 YP 002146918 1 454166 Proteobacteria
Escherichia coli O127 H6 str E2348 69 YP 002329631 1 574521 Proteobacteria
Escherichia coli O111 H str 11128 YP 003235033 1 585396 Proteobacteria
Salmonella enterica subsp enterica serovar Enteritidis str P125109 YP 002244067 1 550537 Proteobacteria
Salmonella enterica subsp enterica serovar Gallinarum pullorum str RKS5078 YP 005212155 1 1081093 Proteobacteria
Salmonella enterica subsp enterica serovar Pullorum str S06004 YP 008380372 1 1298917 Proteobacteria
Salmonella enterica subsp enterica serovar Gallinarum str 287 91 YP 002226930 1 550538 Proteobacteria
Salmonella enterica subsp enterica serovar Dublin str CT 02021853 YP 002216062 1 439851 Proteobacteria
Dickeya chrysanthemi subsp chrysanthemi NCPPB 402 WP 040000600 1 Dchrysanthemi
Dickeya gs poaceaephila NCPPB 569 WP 042871588 1 Dpoaceaephila
Dickeya zeae NCPPB 2538 WP 038916478 1 Dzeae
DAQ1742 DAQ1742 01465 Daquatica
902 DAQ1742 01829 GI7 Seed
DAQ1742 DAQ1742 01829 Daquatica
Dickeya sp CSL RW240 WP 051124211 1 Daquatica
Dickeya aquatica DW 0440 WP 051124211 1 Daquatica
Pectobacterium atrosepticum SCRI1043 YP 049780 1 218491 Proteobacteria
Pectobacterium wasabiae WPP163 YP 003258984 1 561231 Proteobacteria
Pectobacterium atrosepticum SCRI1043 YP 050980 1 218491 Proteobacteria
Pectobacterium wasabiae WPP163 YP 003259159 1 561231 Proteobacteria
Pectobacterium sp SCC3193 YP 006282563 1 1166016 Proteobacteria
Dickeya gs fangzhongdai M005 WP 039695715 1 Dfangzhongdai
Dickeya zeae MS1 WP 019843899 1 Dzeae
Dickeya paradisiaca NCPPB 2511 WP 015854148 1 Dparadisiaca
Dickeya paradisiaca Ech703 WP 015854148 1 Dparadisiaca
Prodigiosinella confusarubida ATCC39006 WP 021016693 1 Pconfusarubida
Dickeya aquatica DW 0440 WP 035339637 1 Daquatica
Citrobacter rodentium ICC168 YP 003368377 1 637910 Proteobacteria
Escherichia coli ATCC 8739 YP 001723746 1 481805 Proteobacteria
Escherichia coli ATCC 8739 YP 001726809 1 481805 Proteobacteria
Escherichia coli O26 H11 str 11368 YP 003230994 1 573235 Proteobacteria
Escherichia coli 042 YP 006097466 1 216592 Proteobacteria
Escherichia coli E24377A YP 001464322 1 331111 Proteobacteria
Escherichia coli O104 H4 str 2009EL 2050 YP 006768605 1 1134782 Proteobacteria
Escherichia coli 042 YP 006098876 1 216592 Proteobacteria
Pectobacterium sp SCC3193 YP 006282507 1 1166016 Proteobacteria
Citrobacter koseri ATCC BAA 895 YP 001455048 1 290338 Proteobacteria
Morganella morganii subsp morganii KT YP 007503536 1 1124991 Proteobacteria
Morganella morganii subsp morganii KT YP 007503554 1 1124991 Proteobacteria
Yersinia pestis biovar Medievalis str Harbin 35 YP 005625501 1 547048 Proteobacteria
Yersinia pestis Antiqua YP 652846 1 360102 Proteobacteria
Yersinia pestis biovar Microtus str 91001 NP 992037 1 229193 Proteobacteria
Salmonella enterica subsp enterica serovar Javiana str CFSAN001992 YP 007472047 1 1267753 Proteobacteria
Shimwellia blattae DSM 4481 NBRC 105725 YP 006321045 1 630626 Proteobacteria
Klebsiella pneumoniae 342 YP 002240755 1 507522 Proteobacteria
Enterobacter cloacae subsp cloacae ENHKU01 YP 006577041 1 1211025 Proteobacteria
Enterobacter lignolyticus SCF1 YP 003943402 1 701347 Proteobacteria
Escherichia coli 042 YP 006098991 1 216592 Proteobacteria
Prodigiosinella confusarubida ATCC39006 WP 021014223 1 Pconfusarubida
Salmonella enterica subsp enterica serovar Cubana str CFSAN002050 YP 008259116 1 1271863 Proteobacteria
Salmonella enterica subsp arizonae serovar 62 z4 z23 str RSK2980 YP 001572064 1 882884 Proteobacteria
Klebsiella oxytoca E718 YP 006496338 1 1191061 Proteobacteria
Escherichia coli O7 K1 str CE10 YP 006146968 1 1072459 Proteobacteria
Enterobacter sp 638 YP 001175202 1 399742 Proteobacteria
Enterobacter cloacae EcWSU1 YP 004950346 1 1045856 Proteobacteria
Escherichia coli O103 H2 str 12009 YP 003224874 1 585395 Proteobacteria
Enterobacteriaceae bacterium strain FGI 57 YP 007342049 1 693444 Proteobacteria
Erwinia amylovora ATCC 49946 YP 003540101 1 716540 Proteobacteria
Dickeya zeae Ech586 WP 012883142 1 Dzeae
Dickeya aquatica DW 0440 WP 035339542 1 Daquatica
Dickeya sp CSL RW240 WP 035339542 1 Daquatica
DAQ1742 DAQ1742 00500 Daquatica
Dickeya zeae ZJU1202 WP 016943011 1 Dzeae
Dickeya zeae EC1 WP 016943011 1 Dzeae
Dickeya chrysanthemi L11 WP 033576649 1 Dchrysanthemi
Dickeya dadantii subsp dieffenbachiae NCPPB 2976 WP 038921500 1 Ddadantii

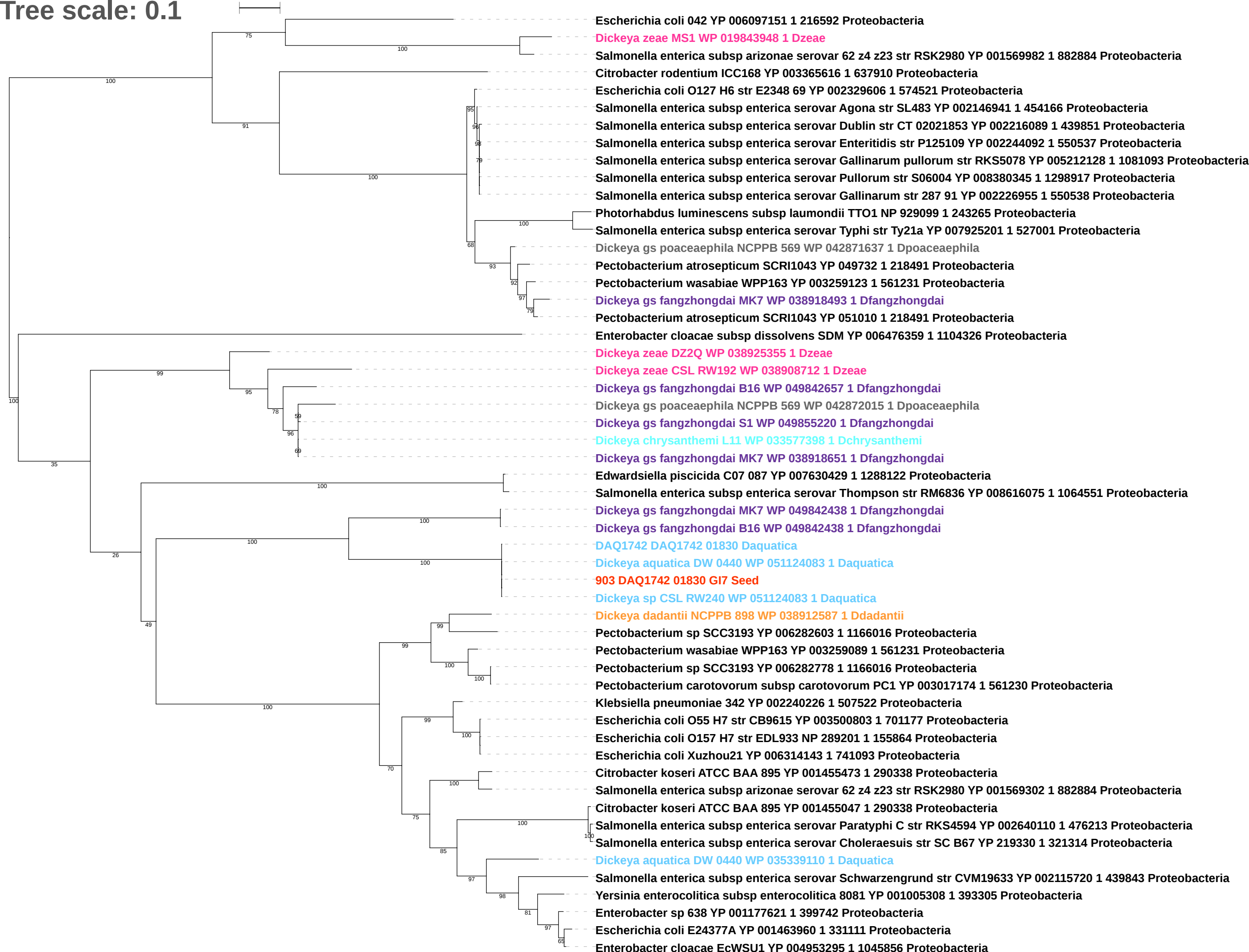
Tree scale: 0.1
Escherichia coli 042 YP 006097151 1 216592 Proteobacteria
Dickeya zeae MS1 WP 019843948 1 Dzeae
Salmonella enterica subsp arizonae serovar 62 z4 z23 str RSK2980 YP 001569982 1 882884 Proteobacteria
Citrobacter rodentium ICC168 YP 003365616 1 637910 Proteobacteria
Escherichia coli O127 H6 str E2348 69 YP 002329606 1 574521 Proteobacteria
Salmonella enterica subsp enterica serovar Agona str SL483 YP 002146941 1 454166 Proteobacteria
Salmonella enterica subsp enterica serovar Dublin str CT 02021853 YP 002216089 1 439851 Proteobacteria
Salmonella enterica subsp enterica serovar Enteritidis str P125109 YP 002244092 1 550537 Proteobacteria
Salmonella enterica subsp enterica serovar Gallinarum pullorum str RKS5078 YP 005212128 1 1081093 Proteobacteria
Salmonella enterica subsp enterica serovar Pullorum str S06004 YP 008380345 1 1298917 Proteobacteria
Salmonella enterica subsp enterica serovar Gallinarum str 287 91 YP 002226955 1 550538 Proteobacteria
Photorhabdus luminescens subsp laumondii TTO1 NP 929099 1 243265 Proteobacteria
Salmonella enterica subsp enterica serovar Typhi str Ty21a YP 007925201 1 527001 Proteobacteria
Dickeya gs poaceaephila NCPPB 569 WP 042871637 1 Dpoaceaephila
Pectobacterium atrosepticum SCRI1043 YP 049732 1 218491 Proteobacteria
Pectobacterium wasabiae WPP163 YP 003259123 1 561231 Proteobacteria
Dickeya gs fangzhongdai MK7 WP 038918493 1 Dfangzhongdai
Pectobacterium atrosepticum SCRI1043 YP 051010 1 218491 Proteobacteria
Enterobacter cloacae subsp dissolvens SDM YP 006476359 1 1104326 Proteobacteria
Dickeya zeae DZ2Q WP 038925355 1 Dzeae
Dickeya zeae CSL RW192 WP 038908712 1 Dzeae
Dickeya gs fangzhongdai B16 WP 049842657 1 Dfangzhongdai
Dickeya gs poaceaephila NCPPB 569 WP 042872015 1 Dpoaceaephila
Dickeya gs fangzhongdai S1 WP 049855220 1 Dfangzhongdai
Dickeya chrysanthemi L11 WP 033577398 1 Dchrysanthemi
Dickeya gs fangzhongdai MK7 WP 038918651 1 Dfangzhongdai
Edwardsiella piscicida C07 087 YP 007630429 1 1288122 Proteobacteria
Salmonella enterica subsp enterica serovar Thompson str RM6836 YP 008616075 1 1064551 Proteobacteria
Dickeya gs fangzhongdai MK7 WP 049842438 1 Dfangzhongdai
Dickeya gs fangzhongdai B16 WP 049842438 1 Dfangzhongdai
DAQ1742 DAQ1742 01830 Daquatica
Dickeya aquatica DW 0440 WP 051124083 1 Daquatica
903 DAQ1742 01830 GI7 Seed
Dickeya sp CSL RW240 WP 051124083 1 Daquatica
Dickeya dadantii NCPPB 898 WP 038912587 1 Ddadantii
Pectobacterium sp SCC3193 YP 006282603 1 1166016 Proteobacteria
Pectobacterium wasabiae WPP163 YP 003259089 1 561231 Proteobacteria
Pectobacterium sp SCC3193 YP 006282778 1 1166016 Proteobacteria
Pectobacterium carotovorum subsp carotovorum PC1 YP 003017174 1 561230 Proteobacteria
Klebsiella pneumoniae 342 YP 002240226 1 507522 Proteobacteria
Escherichia coli O55 H7 str CB9615 YP 003500803 1 701177 Proteobacteria
Escherichia coli O157 H7 str EDL933 NP 289201 1 155864 Proteobacteria
Escherichia coli Xuzhou21 YP 006314143 1 741093 Proteobacteria
Citrobacter koseri ATCC BAA 895 YP 001455473 1 290338 Proteobacteria
Salmonella enterica subsp arizonae serovar 62 z4 z23 str RSK2980 YP 001569302 1 882884 Proteobacteria
Citrobacter koseri ATCC BAA 895 YP 001455047 1 290338 Proteobacteria
Salmonella enterica subsp enterica serovar Paratyphi C str RKS4594 YP 002640110 1 476213 Proteobacteria
Salmonella enterica subsp enterica serovar Choleraesuis str SC B67 YP 219330 1 321314 Proteobacteria
Dickeya aquatica DW 0440 WP 035339110 1 Daquatica
Salmonella enterica subsp enterica serovar Schwarzengrund str CVM19633 YP 002115720 1 439843 Proteobacteria
Yersinia enterocolitica subsp enterocolitica 8081 YP 001005308 1 393305 Proteobacteria
Enterobacter sp 638 YP 001177621 1 399742 Proteobacteria
Escherichia coli E24377A YP 001463960 1 331111 Proteobacteria
Enterobacter cloacae EcWSU1 YP 004953295 1 1045856 Proteobacteria

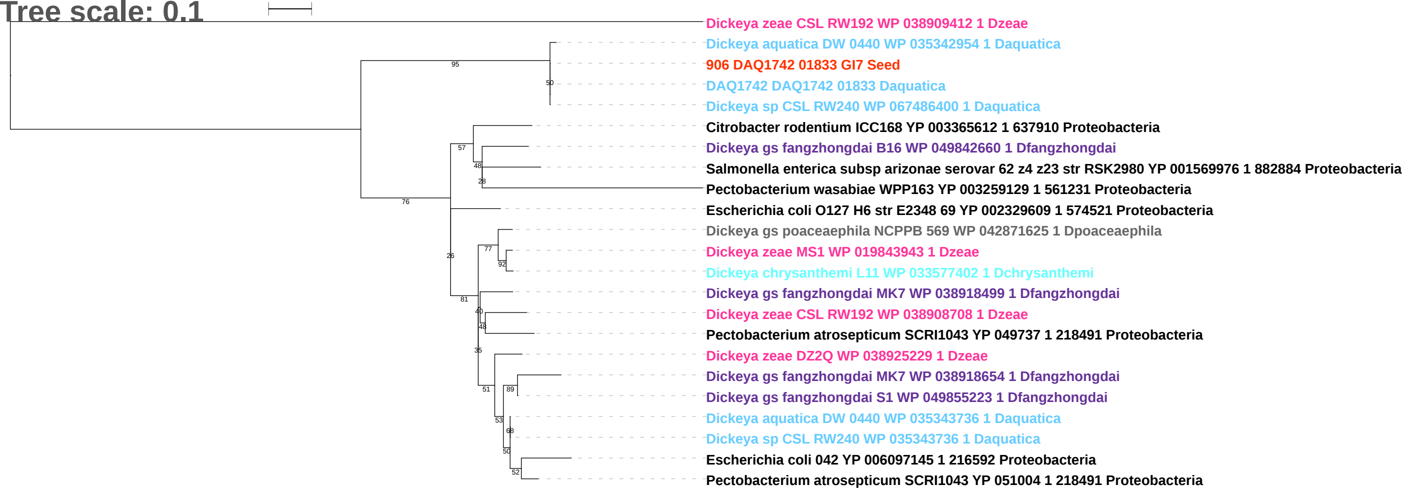
Tree scale: 0.1
Dickeya zeae CSL RW192 WP 038909412 1 Dzeae
Dickeya aquatica DW 0440 WP 035342954 1 Daquatica
906 DAQ1742 01833 GI7 Seed
DAQ1742 DAQ1742 01833 Daquatica
Dickeya sp CSL RW240 WP 067486400 1 Daquatica
Citrobacter rodentium ICC168 YP 003365612 1 637910 Proteobacteria
Dickeya gs fangzhongdai B16 WP 049842660 1 Dfangzhongdai
Salmonella enterica subsp arizonae serovar 62 z4 z23 str RSK2980 YP 001569976 1 882884 Proteobacteria
Pectobacterium wasabiae WPP163 YP 003259129 1 561231 Proteobacteria
Escherichia coli O127 H6 str E2348 69 YP 002329609 1 574521 Proteobacteria
Dickeya gs poaceaephila NCPPB 569 WP 042871625 1 Dpoaceaephila
Dickeya zeae MS1 WP 019843943 1 Dzeae
Dickeya chrysanthemi L11 WP 033577402 1 Dchrysanthemi
Dickeya gs fangzhongdai MK7 WP 038918499 1 Dfangzhongdai
Dickeya zeae CSL RW192 WP 038908708 1 Dzeae
Pectobacterium atrosepticum SCRI1043 YP 049737 1 218491 Proteobacteria
Dickeya zeae DZ2Q WP 038925229 1 Dzeae
Dickeya gs fangzhongdai MK7 WP 038918654 1 Dfangzhongdai
Dickeya gs fangzhongdai S1 WP 049855223 1 Dfangzhongdai
Dickeya aquatica DW 0440 WP 035343736 1 Daquatica
Dickeya sp CSL RW240 WP 035343736 1 Daquatica
Escherichia coli 042 YP 006097145 1 216592 Proteobacteria
Pectobacterium atrosepticum SCRI1043 YP 051004 1 218491 Proteobacteria

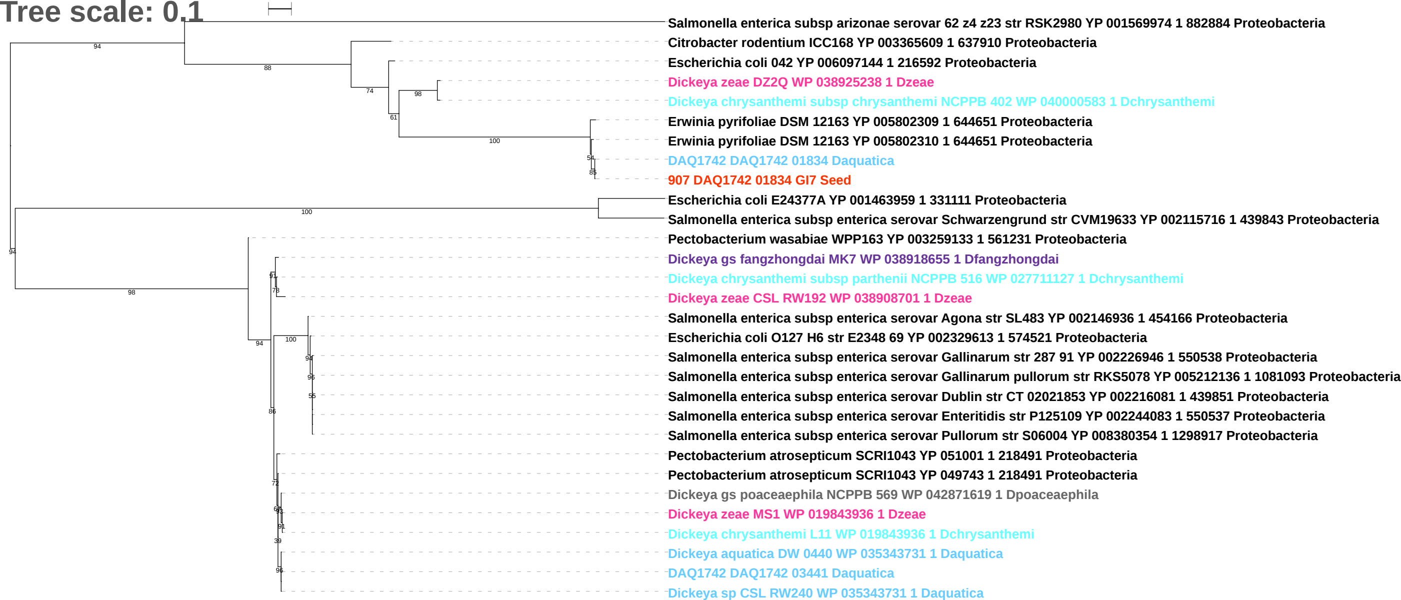
Tree scale: 0.1
Salmonella enterica subsp arizonae serovar 62 z4 z23 str RSK2980 YP 001569974 1 882884 Proteobacteria
Citrobacter rodentium ICC168 YP 003365609 1 637910 Proteobacteria
Escherichia coli 042 YP 006097144 1 216592 Proteobacteria
Dickeya zeae DZ2Q WP 038925238 1 Dzeae
Dickeya chrysanthemi subsp chrysanthemi NCPPB 402 WP 040000583 1 Dchrysanthemi
Erwinia pyrifoliae DSM 12163 YP 005802309 1 644651 Proteobacteria
Erwinia pyrifoliae DSM 12163 YP 005802310 1 644651 Proteobacteria
DAQ1742 DAQ1742 01834 Daquatica
907 DAQ1742 01834 GI7 Seed
Escherichia coli E24377A YP 001463959 1 331111 Proteobacteria
Salmonella enterica subsp enterica serovar Schwarzengrund str CVM19633 YP 002115716 1 439843 Proteobacteria
Pectobacterium wasabiae WPP163 YP 003259133 1 561231 Proteobacteria
Dickeya gs fangzhongdai MK7 WP 038918655 1 Dfangzhongdai
Dickeya chrysanthemi subsp parthenii NCPPB 516 WP 027711127 1 Dchrysanthemi
Dickeya zeae CSL RW192 WP 038908701 1 Dzeae
Salmonella enterica subsp enterica serovar Agona str SL483 YP 002146936 1 454166 Proteobacteria
Escherichia coli O127 H6 str E2348 69 YP 002329613 1 574521 Proteobacteria
Salmonella enterica subsp enterica serovar Gallinarum str 287 91 YP 002226946 1 550538 Proteobacteria
Salmonella enterica subsp enterica serovar Gallinarum pullorum str RKS5078 YP 005212136 1 1081093 Proteobacteria
Salmonella enterica subsp enterica serovar Dublin str CT 02021853 YP 002216081 1 439851 Proteobacteria
Salmonella enterica subsp enterica serovar Enteritidis str P125109 YP 002244083 1 550537 Proteobacteria
Salmonella enterica subsp enterica serovar Pullorum str S06004 YP 008380354 1 1298917 Proteobacteria
Pectobacterium atrosepticum SCRI1043 YP 051001 1 218491 Proteobacteria
Pectobacterium atrosepticum SCRI1043 YP 049743 1 218491 Proteobacteria
Dickeya gs poaceaephila NCPPB 569 WP 042871619 1 Dpoaceaephila
Dickeya zeae MS1 WP 019843936 1 Dzeae
Dickeya chrysanthemi L11 WP 019843936 1 Dchrysanthemi
Dickeya aquatica DW 0440 WP 035343731 1 Daquatica
DAQ1742 DAQ1742 03441 Daquatica
Dickeya sp CSL RW240 WP 035343731 1 Daquatica

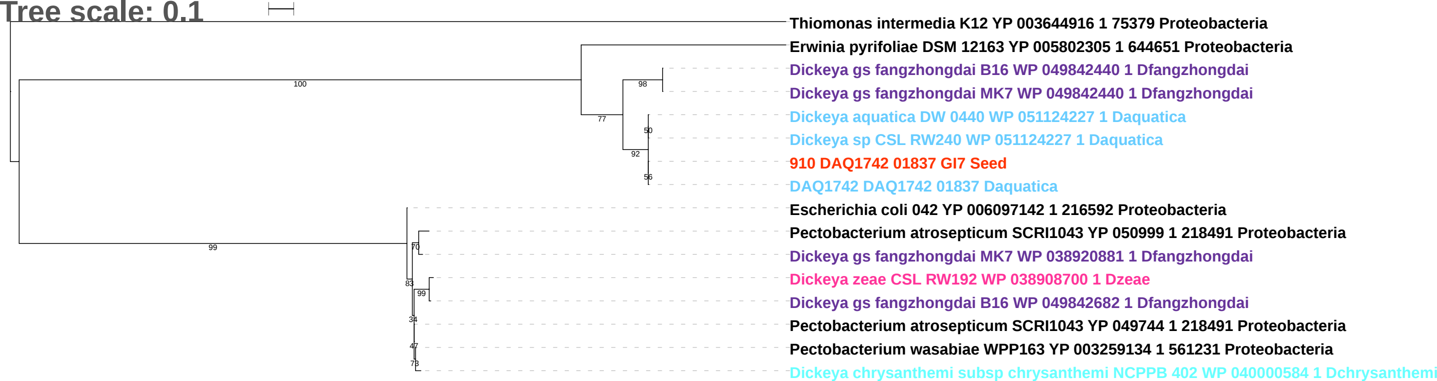

Tree scale: 0.1
Thiomonas intermedia K12 YP 003644916 1 75379 Proteobacteria
Erwinia pyrifoliae DSM 12163 YP 005802305 1 644651 Proteobacteria
Dickeya gs fangzhongdai B16 WP 049842440 1 Dfangzhongdai
Dickeya gs fangzhongdai MK7 WP 049842440 1 Dfangzhongdai
Dickeya aquatica DW 0440 WP 051124227 1 Daquatica
Dickeya sp CSL RW240 WP 051124227 1 Daquatica
910 DAQ1742 01837 GI7 Seed
DAQ1742 DAQ1742 01837 Daquatica
Escherichia coli 042 YP 006097142 1 216592 Proteobacteria
Pectobacterium atrosepticum SCRI1043 YP 050999 1 218491 Proteobacteria
Dickeya gs fangzhongdai MK7 WP 038920881 1 Dfangzhongdai
Dickeya zeae CSL RW192 WP 038908700 1 Dzeae
Dickeya gs fangzhongdai B16 WP 049842682 1 Dfangzhongdai
Pectobacterium atrosepticum SCRI1043 YP 049744 1 218491 Proteobacteria
Pectobacterium wasabiae WPP163 YP 003259134 1 561231 Proteobacteria
Dickeya chrysanthemi subsp chrysanthemi NCPPB 402 WP 040000584 1 Dchrysanthemi
100
98
77
50
92
56
99
70
83
99
34
47
73

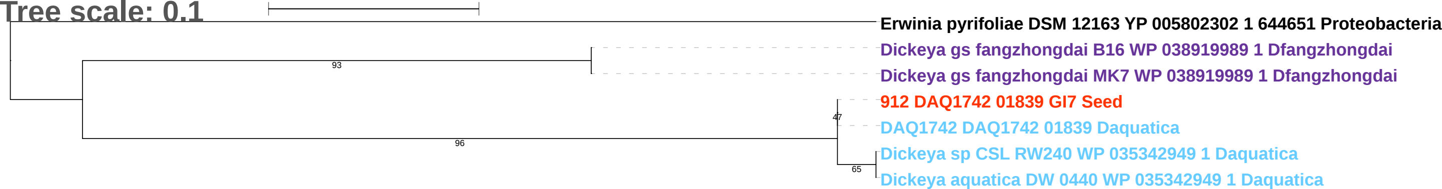

Tree scale: 0.1
Erwinia pyrifoliae DSM 12163 YP 005802302 1 644651 Proteobacteria
Dickeya gs fangzhongdai B16 WP 038919989 1 Dfangzhongdai
Dickeya gs fangzhongdai MK7 WP 038919989 1 Dfangzhongdai
912 DAQ1742 01839 GI7 Seed
DAQ1742 DAQ1742 01839 Daquatica
Dickeya sp CSL RW240 WP 035342949 1 Daquatica
Dickeya aquatica DW 0440 WP 035342949 1 Daquatica
93
47
96
65

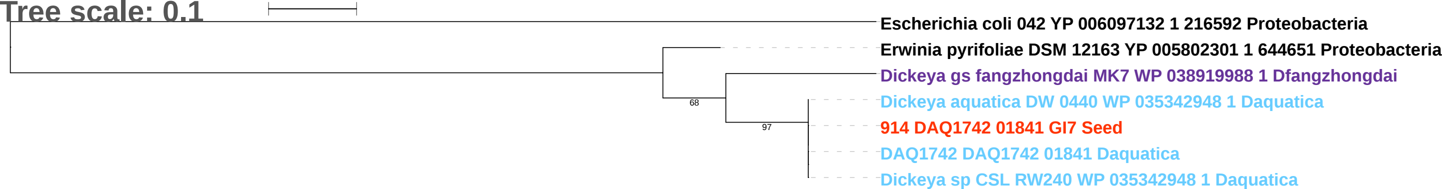
Tree scale: 0.1
Escherichia coli 042 YP 006097132 1 216592 Proteobacteria
Erwinia pyrifoliae DSM 12163 YP 005802301 1 644651 Proteobacteria
Dickeya gs fangzhongdai MK7 WP 038919988 1 Dfangzhongdai
Dickeya aquatica DW 0440 WP 035342948 1 Daquatica
914 DAQ1742 01841 GI7 Seed
DAQ1742 DAQ1742 01841 Daquatica
Dickeya sp CSL RW240 WP 035342948 1 Daquatica
68
97

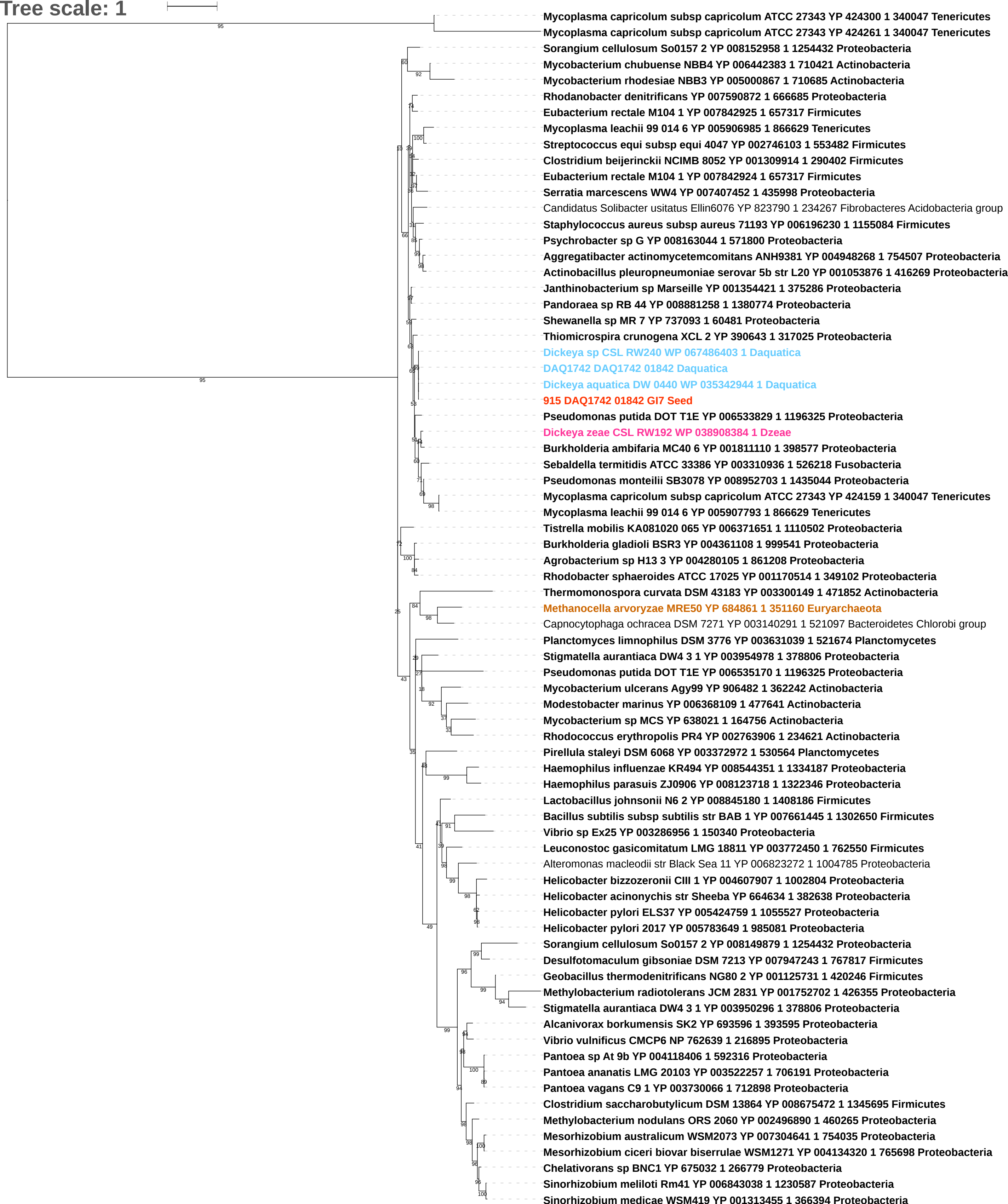
Tree scale: 1
Mycoplasma capricolum subsp capricolum ATCC 27343 YP 424300 1 340047 Tenericutes
Mycoplasma capricolum subsp capricolum ATCC 27343 YP 424261 1 340047 Tenericutes
Sorangium cellulosum So0157 2 YP 008152958 1 1254432 Proteobacteria
Mycobacterium chubuense NBB4 YP 006442383 1 710421 Actinobacteria
Mycobacterium rhodesiae NBB3 YP 005000867 1 710685 Actinobacteria
Rhodanobacter denitrificans YP 007590872 1 666685 Proteobacteria
Eubacterium rectale M104 1 YP 007842925 1 657317 Firmicutes
Mycoplasma leachii 99 014 6 YP 005906985 1 866629 Tenericutes
Streptococcus equi subsp equi 4047 YP 002746103 1 553482 Firmicutes
Clostridium beijerinckii NCIMB 8052 YP 001309914 1 290402 Firmicutes
Eubacterium rectale M104 1 YP 007842924 1 657317 Firmicutes
Serratia marcescens WW4 YP 007407452 1 435998 Proteobacteria
Candidatus Solibacter usitatus Ellin6076 YP 823790 1 234267 Fibrobacteres Acidobacteria group
Staphylococcus aureus subsp aureus 71193 YP 006196230 1 1155084 Firmicutes
Psychrobacter sp G YP 008163044 1 571800 Proteobacteria
Aggregatibacter actinomycetemcomitans ANH9381 YP 004948268 1 754507 Proteobacteria
Actinobacillus pleuropneumoniae serovar 5b str L20 YP 001053876 1 416269 Proteobacteria
Janthinobacterium sp Marseille YP 001354421 1 375286 Proteobacteria
Pandoraea sp RB 44 YP 008881258 1 1380774 Proteobacteria
Shewanella sp MR 7 YP 737093 1 60481 Proteobacteria
Thiomicrospira crunogena XCL 2 YP 390643 1 317025 Proteobacteria
Dickeya sp CSL RW240 WP 067486403 1 Daquatica
DAQ1742 DAQ1742 01842 Daquatica
Dickeya aquatica DW 0440 WP 035342944 1 Daquatica
915 DAQ1742 01842 GI7 Seed
Pseudomonas putida DOT T1E YP 006533829 1 1196325 Proteobacteria
Dickeya zeae CSL RW192 WP 038908384 1 Dzeae
Burkholderia ambifaria MC40 6 YP 001811110 1 398577 Proteobacteria
Sebaldella termitidis ATCC 33386 YP 003310936 1 526218 Fusobacteria
Pseudomonas monteilii SB3078 YP 008952703 1 1435044 Proteobacteria
Mycoplasma capricolum subsp capricolum ATCC 27343 YP 424159 1 340047 Tenericutes
Mycoplasma leachii 99 014 6 YP 005907793 1 866629 Tenericutes
Tistrella mobilis KA081020 065 YP 006371651 1 1110502 Proteobacteria
Burkholderia gladioli BSR3 YP 004361108 1 999541 Proteobacteria
Agrobacterium sp H13 3 YP 004280105 1 861208 Proteobacteria
Rhodobacter sphaeroides ATCC 17025 YP 001170514 1 349102 Proteobacteria
Thermomonospora curvata DSM 43183 YP 003300149 1 471852 Actinobacteria
Methanocella arvoryzae MRE50 YP 684861 1 351160 Euryarchaeota
Capnocytophaga ochracea DSM 7271 YP 003140291 1 521097 Bacteroidetes Chlorobi group
Planctomyces limnophilus DSM 3776 YP 003631039 1 521674 Planctomycetes
Stigmatella aurantiaca DW4 3 1 YP 003954978 1 378806 Proteobacteria
Pseudomonas putida DOT T1E YP 006535170 1 1196325 Proteobacteria
Mycobacterium ulcerans Agy99 YP 906482 1 362242 Actinobacteria
Modestobacter marinus YP 006368109 1 477641 Actinobacteria
Mycobacterium sp MCS YP 638021 1 164756 Actinobacteria
Rhodococcus erythropolis PR4 YP 002763906 1 234621 Actinobacteria
Pirellula staleyi DSM 6068 YP 003372972 1 530564 Planctomycetes
Haemophilus influenzae KR494 YP 008544351 1 1334187 Proteobacteria
Haemophilus parasuis ZJ0906 YP 008123718 1 1322346 Proteobacteria
Lactobacillus johnsonii N6 2 YP 008845180 1 1408186 Firmicutes
Bacillus subtilis subsp subtilis str BAB 1 YP 007661445 1 1302650 Firmicutes
Vibrio sp Ex25 YP 003286956 1 150340 Proteobacteria
Leuconostoc gasicomitatum LMG 18811 YP 003772450 1 762550 Firmicutes
Alteromonas macleodii str Black Sea 11 YP 006823272 1 1004785 Proteobacteria
Helicobacter bizzozeronii CIII 1 YP 004607907 1 1002804 Proteobacteria
Helicobacter acinonychis str Sheeba YP 664634 1 382638 Proteobacteria
Helicobacter pylori ELS37 YP 005424759 1 1055527 Proteobacteria
Helicobacter pylori 2017 YP 005783649 1 985081 Proteobacteria
Sorangium cellulosum So0157 2 YP 008149879 1 1254432 Proteobacteria
Desulfotomaculum gibsoniae DSM 7213 YP 007947243 1 767817 Firmicutes
Geobacillus thermodenitrificans NG80 2 YP 001125731 1 420246 Firmicutes
Methylobacterium radiotolerans JCM 2831 YP 001752702 1 426355 Proteobacteria
Stigmatella aurantiaca DW4 3 1 YP 003950296 1 378806 Proteobacteria
Alcanivorax borkumensis SK2 YP 693596 1 393595 Proteobacteria
Vibrio vulnificus CMCP6 NP 762639 1 216895 Proteobacteria
Pantoea sp At 9b YP 004118406 1 592316 Proteobacteria
Pantoea ananatis LMG 20103 YP 003522257 1 706191 Proteobacteria
Pantoea vagans C9 1 YP 003730066 1 712898 Proteobacteria
Clostridium saccharobutylicum DSM 13864 YP 008675472 1 1345695 Firmicutes
Methylobacterium nodulans ORS 2060 YP 002496890 1 460265 Proteobacteria
Mesorhizobium australicum WSM2073 YP 007304641 1 754035 Proteobacteria
Mesorhizobium ciceri biovar biserrulae WSM1271 YP 004134320 1 765698 Proteobacteria
Chelativorans sp BNC1 YP 675032 1 266779 Proteobacteria
Sinorhizobium meliloti Rm41 YP 006843038 1 1230587 Proteobacteria
Sinorhizobium medicae WSM419 YP 001313455 1 366394 Proteobacteria

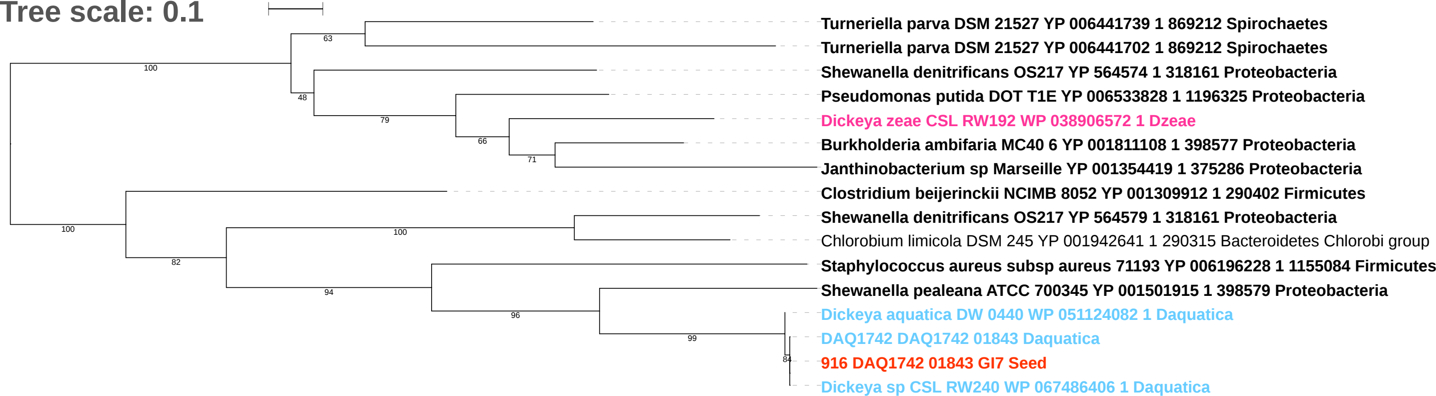
Tree scale: 0.1
Turneriella parva DSM 21527 YP 006441739 1 869212 Spirochaetes
Turneriella parva DSM 21527 YP 006441702 1 869212 Spirochaetes
Shewanella denitrificans OS217 YP 564574 1 318161 Proteobacteria
Pseudomonas putida DOT T1E YP 006533828 1 1196325 Proteobacteria
Dickeya zeae CSL RW192 WP 038906572 1 Dzeae
Burkholderia ambifaria MC40 6 YP 001811108 1 398577 Proteobacteria
Janthinobacterium sp Marseille YP 001354419 1 375286 Proteobacteria
Clostridium beijerinckii NCIMB 8052 YP 001309912 1 290402 Firmicutes
Shewanella denitrificans OS217 YP 564579 1 318161 Proteobacteria
Chlorobium limicola DSM 245 YP 001942641 1 290315 Bacteroidetes Chlorobi group
Staphylococcus aureus subsp aureus 71193 YP 006196228 1 1155084 Firmicutes
Shewanella pealeana ATCC 700345 YP 001501915 1 398579 Proteobacteria
Dickeya aquatica DW 0440 WP 051124082 1 Daquatica
DAQ1742 DAQ1742 01843 Daquatica
916 DAQ1742 01843 GI7 Seed
Dickeya sp CSL RW240 WP 067486406 1 Daquatica
63
100
48
79
66
71
100
100
82
94
96
99
84

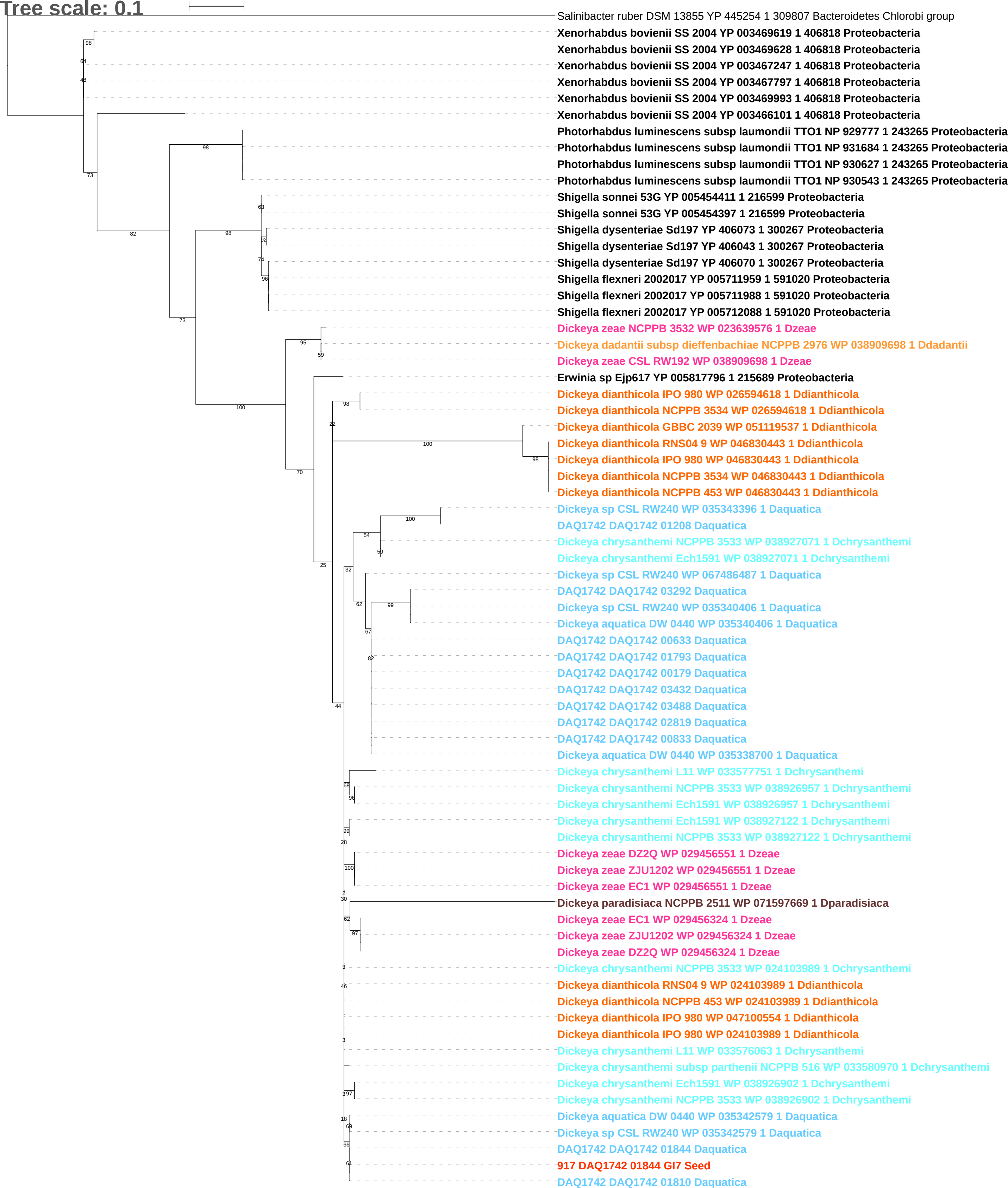
Tree scale: 0.1
Salinibacter ruber DSM 13855 YP 445254 1 309807 Bacteroidetes Chlorobi group
Xenorhabdus bovienii SS 2004 YP 003469619 1 406818 Proteobacteria
Xenorhabdus bovienii SS 2004 YP 003469628 1 406818 Proteobacteria
Xenorhabdus bovienii SS 2004 YP 003467247 1 406818 Proteobacteria
Xenorhabdus bovienii SS 2004 YP 003467797 1 406818 Proteobacteria
Xenorhabdus bovienii SS 2004 YP 003469993 1 406818 Proteobacteria
Xenorhabdus bovienii SS 2004 YP 003466101 1 406818 Proteobacteria
Photorhabdus luminescens subsp laumondii TTO1 NP 929777 1 243265 Proteobacteria
Photorhabdus luminescens subsp laumondii TTO1 NP 931684 1 243265 Proteobacteria
Photorhabdus luminescens subsp laumondii TTO1 NP 930627 1 243265 Proteobacteria
Photorhabdus luminescens subsp laumondii TTO1 NP 930543 1 243265 Proteobacteria
Shigella sonnei 53G YP 005454411 1 216599 Proteobacteria
Shigella sonnei 53G YP 005454397 1 216599 Proteobacteria
Shigella dysenteriae Sd197 YP 406073 1 300267 Proteobacteria
Shigella dysenteriae Sd197 YP 406043 1 300267 Proteobacteria
Shigella dysenteriae Sd197 YP 406070 1 300267 Proteobacteria
Shigella flexneri 2002017 YP 005711959 1 591020 Proteobacteria
Shigella flexneri 2002017 YP 005711988 1 591020 Proteobacteria
Shigella flexneri 2002017 YP 005712088 1 591020 Proteobacteria
Dickeya zeae NCPPB 3532 WP 023639576 1 Dzeae
Dickeya dadantii subsp dieffenbachiae NCPPB 2976 WP 038909698 1 Ddadantii
Dickeya zeae CSL RW192 WP 038909698 1 Dzeae
Erwinia sp Ejp617 YP 005817796 1 215689 Proteobacteria
Dickeya dianthicola IPO 980 WP 026594618 1 Ddianthicola
Dickeya dianthicola NCPPB 3534 WP 026594618 1 Ddianthicola
Dickeya dianthicola GBBC 2039 WP 051119537 1 Ddianthicola
Dickeya dianthicola RNS04 9 WP 046830443 1 Ddianthicola
Dickeya dianthicola IPO 980 WP 046830443 1 Ddianthicola
Dickeya dianthicola NCPPB 3534 WP 046830443 1 Ddianthicola
Dickeya dianthicola NCPPB 453 WP 046830443 1 Ddianthicola
Dickeya sp CSL RW240 WP 035343396 1 Daquatica
DAQ1742 DAQ1742 01208 Daquatica
Dickeya chrysanthemi NCPPB 3533 WP 038927071 1 Dchrysanthemi
Dickeya chrysanthemi Ech1591 WP 038927071 1 Dchrysanthemi
Dickeya sp CSL RW240 WP 067486487 1 Daquatica
DAQ1742 DAQ1742 03292 Daquatica
Dickeya sp CSL RW240 WP 035340406 1 Daquatica
Dickeya aquatica DW 0440 WP 035340406 1 Daquatica
DAQ1742 DAQ1742 00633 Daquatica
DAQ1742 DAQ1742 01793 Daquatica
DAQ1742 DAQ1742 00179 Daquatica
DAQ1742 DAQ1742 03432 Daquatica
DAQ1742 DAQ1742 03488 Daquatica
DAQ1742 DAQ1742 02819 Daquatica
DAQ1742 DAQ1742 00833 Daquatica
Dickeya aquatica DW 0440 WP 035338700 1 Daquatica
Dickeya chrysanthemi L11 WP 033577751 1 Dchrysanthemi
Dickeya chrysanthemi NCPPB 3533 WP 038926957 1 Dchrysanthemi
Dickeya chrysanthemi Ech1591 WP 038926957 1 Dchrysanthemi
Dickeya chrysanthemi Ech1591 WP 038927122 1 Dchrysanthemi
Dickeya chrysanthemi NCPPB 3533 WP 038927122 1 Dchrysanthemi
Dickeya zeae DZ2Q WP 029456551 1 Dzeae
Dickeya zeae ZJU1202 WP 029456551 1 Dzeae
Dickeya zeae EC1 WP 029456551 1 Dzeae
Dickeya paradisiaca NCPPB 2511 WP 071597669 1 Dparadisiaca
Dickeya zeae EC1 WP 029456324 1 Dzeae
Dickeya zeae ZJU1202 WP 029456324 1 Dzeae
Dickeya zeae DZ2Q WP 029456324 1 Dzeae
Dickeya chrysanthemi NCPPB 3533 WP 024103989 1 Dchrysanthemi
Dickeya dianthicola RNS04 9 WP 024103989 1 Ddianthicola
Dickeya dianthicola NCPPB 453 WP 024103989 1 Ddianthicola
Dickeya dianthicola IPO 980 WP 047100554 1 Ddianthicola
Dickeya dianthicola IPO 980 WP 024103989 1 Ddianthicola
Dickeya chrysanthemi L11 WP 033576063 1 Dchrysanthemi
Dickeya chrysanthemi subsp parthenii NCPPB 516 WP 033580970 1 Dchrysanthemi
Dickeya chrysanthemi Ech1591 WP 038926902 1 Dchrysanthemi
Dickeya chrysanthemi NCPPB 3533 WP 038926902 1 Dchrysanthemi
Dickeya aquatica DW 0440 WP 035342579 1 Daquatica
Dickeya sp CSL RW240 WP 035342579 1 Daquatica
DAQ1742 DAQ1742 01844 Daquatica
917 DAQ1742 01844 GI7 Seed
DAQ1742 DAQ1742 01810 Daquatica

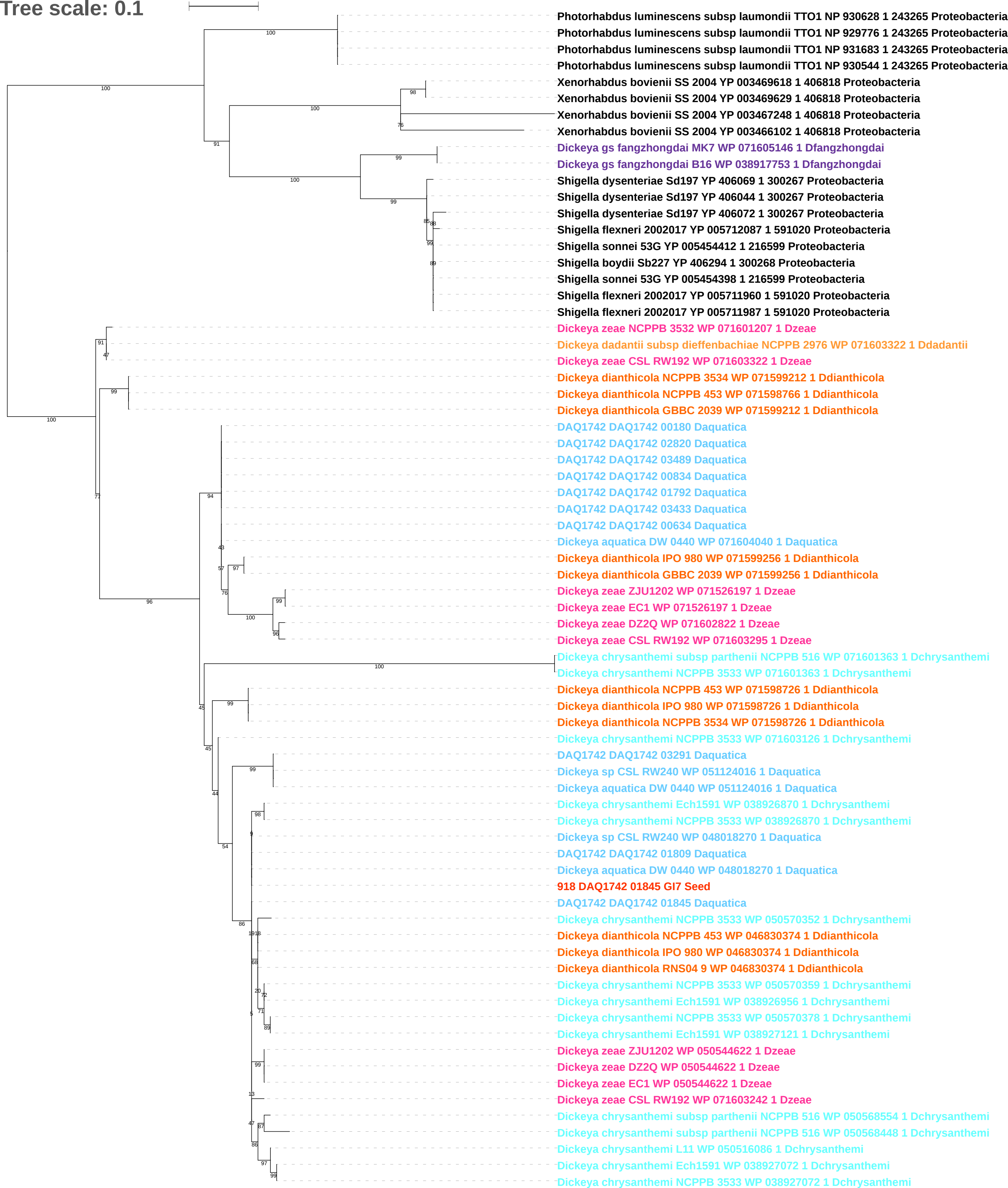
Tree scale: 0.1
Photorhabdus luminescens subsp laumondii TTO1 NP 930628 1 243265 Proteobacteria
Photorhabdus luminescens subsp laumondii TTO1 NP 929776 1 243265 Proteobacteria
Photorhabdus luminescens subsp laumondii TTO1 NP 931683 1 243265 Proteobacteria
Photorhabdus luminescens subsp laumondii TTO1 NP 930544 1 243265 Proteobacteria
Xenorhabdus bovienii SS 2004 YP 003469618 1 406818 Proteobacteria
Xenorhabdus bovienii SS 2004 YP 003469629 1 406818 Proteobacteria
Xenorhabdus bovienii SS 2004 YP 003467248 1 406818 Proteobacteria
Xenorhabdus bovienii SS 2004 YP 003466102 1 406818 Proteobacteria
Dickeya gs fangzhongdai MK7 WP 071605146 1 Dfangzhongdai
Dickeya gs fangzhongdai B16 WP 038917753 1 Dfangzhongdai
Shigella dysenteriae Sd197 YP 406069 1 300267 Proteobacteria
Shigella dysenteriae Sd197 YP 406044 1 300267 Proteobacteria
Shigella dysenteriae Sd197 YP 406072 1 300267 Proteobacteria
Shigella flexneri 2002017 YP 005712087 1 591020 Proteobacteria
Shigella sonnei 53G YP 005454412 1 216599 Proteobacteria
Shigella boydii Sb227 YP 406294 1 300268 Proteobacteria
Shigella sonnei 53G YP 005454398 1 216599 Proteobacteria
Shigella flexneri 2002017 YP 005711960 1 591020 Proteobacteria
Shigella flexneri 2002017 YP 005711987 1 591020 Proteobacteria
Dickeya zeae NCPPB 3532 WP 071601207 1 Dzeae
Dickeya dadantii subsp dieffenbachiae NCPPB 2976 WP 071603322 1 Ddadantii
Dickeya zeae CSL RW192 WP 071603322 1 Dzeae
Dickeya dianthicola NCPPB 3534 WP 071599212 1 Ddianthicola
Dickeya dianthicola NCPPB 453 WP 071598766 1 Ddianthicola
Dickeya dianthicola GBBC 2039 WP 071599212 1 Ddianthicola
DAQ1742 DAQ1742 00180 Daquatica
DAQ1742 DAQ1742 02820 Daquatica
DAQ1742 DAQ1742 03489 Daquatica
DAQ1742 DAQ1742 00834 Daquatica
DAQ1742 DAQ1742 01792 Daquatica
DAQ1742 DAQ1742 03433 Daquatica
DAQ1742 DAQ1742 00634 Daquatica
Dickeya aquatica DW 0440 WP 071604040 1 Daquatica
Dickeya dianthicola IPO 980 WP 071599256 1 Ddianthicola
Dickeya dianthicola GBBC 2039 WP 071599256 1 Ddianthicola
Dickeya zeae ZJU1202 WP 071526197 1 Dzeae
Dickeya zeae EC1 WP 071526197 1 Dzeae
Dickeya zeae DZ2Q WP 071602822 1 Dzeae
Dickeya zeae CSL RW192 WP 071603295 1 Dzeae
Dickeya chrysanthemi subsp parthenii NCPPB 516 WP 071601363 1 Dchrysanthemi
Dickeya chrysanthemi NCPPB 3533 WP 071601363 1 Dchrysanthemi
Dickeya dianthicola NCPPB 453 WP 071598726 1 Ddianthicola
Dickeya dianthicola IPO 980 WP 071598726 1 Ddianthicola
Dickeya dianthicola NCPPB 3534 WP 071598726 1 Ddianthicola
Dickeya chrysanthemi NCPPB 3533 WP 071603126 1 Dchrysanthemi
DAQ1742 DAQ1742 03291 Daquatica
Dickeya sp CSL RW240 WP 051124016 1 Daquatica
Dickeya aquatica DW 0440 WP 051124016 1 Daquatica
Dickeya chrysanthemi Ech1591 WP 038926870 1 Dchrysanthemi
Dickeya chrysanthemi NCPPB 3533 WP 038926870 1 Dchrysanthemi
Dickeya sp CSL RW240 WP 048018270 1 Daquatica
DAQ1742 DAQ1742 01809 Daquatica
Dickeya aquatica DW 0440 WP 048018270 1 Daquatica
918 DAQ1742 01845 GI7 Seed
DAQ1742 DAQ1742 01845 Daquatica
Dickeya chrysanthemi NCPPB 3533 WP 050570352 1 Dchrysanthemi
Dickeya dianthicola NCPPB 453 WP 046830374 1 Ddianthicola
Dickeya dianthicola IPO 980 WP 046830374 1 Ddianthicola
Dickeya dianthicola RNS04 9 WP 046830374 1 Ddianthicola
Dickeya chrysanthemi NCPPB 3533 WP 050570359 1 Dchrysanthemi
Dickeya chrysanthemi Ech1591 WP 038926956 1 Dchrysanthemi
Dickeya chrysanthemi NCPPB 3533 WP 050570378 1 Dchrysanthemi
Dickeya chrysanthemi Ech1591 WP 038927121 1 Dchrysanthemi
Dickeya zeae ZJU1202 WP 050544622 1 Dzeae
Dickeya zeae DZ2Q WP 050544622 1 Dzeae
Dickeya zeae EC1 WP 050544622 1 Dzeae
Dickeya zeae CSL RW192 WP 071603242 1 Dzeae
Dickeya chrysanthemi subsp parthenii NCPPB 516 WP 050568554 1 Dchrysanthemi
Dickeya chrysanthemi subsp parthenii NCPPB 516 WP 050568448 1 Dchrysanthemi
Dickeya chrysanthemi L11 WP 050516086 1 Dchrysanthemi
Dickeya chrysanthemi Ech1591 WP 038927072 1 Dchrysanthemi
Dickeya chrysanthemi NCPPB 3533 WP 038927072 1 Dchrysanthemi

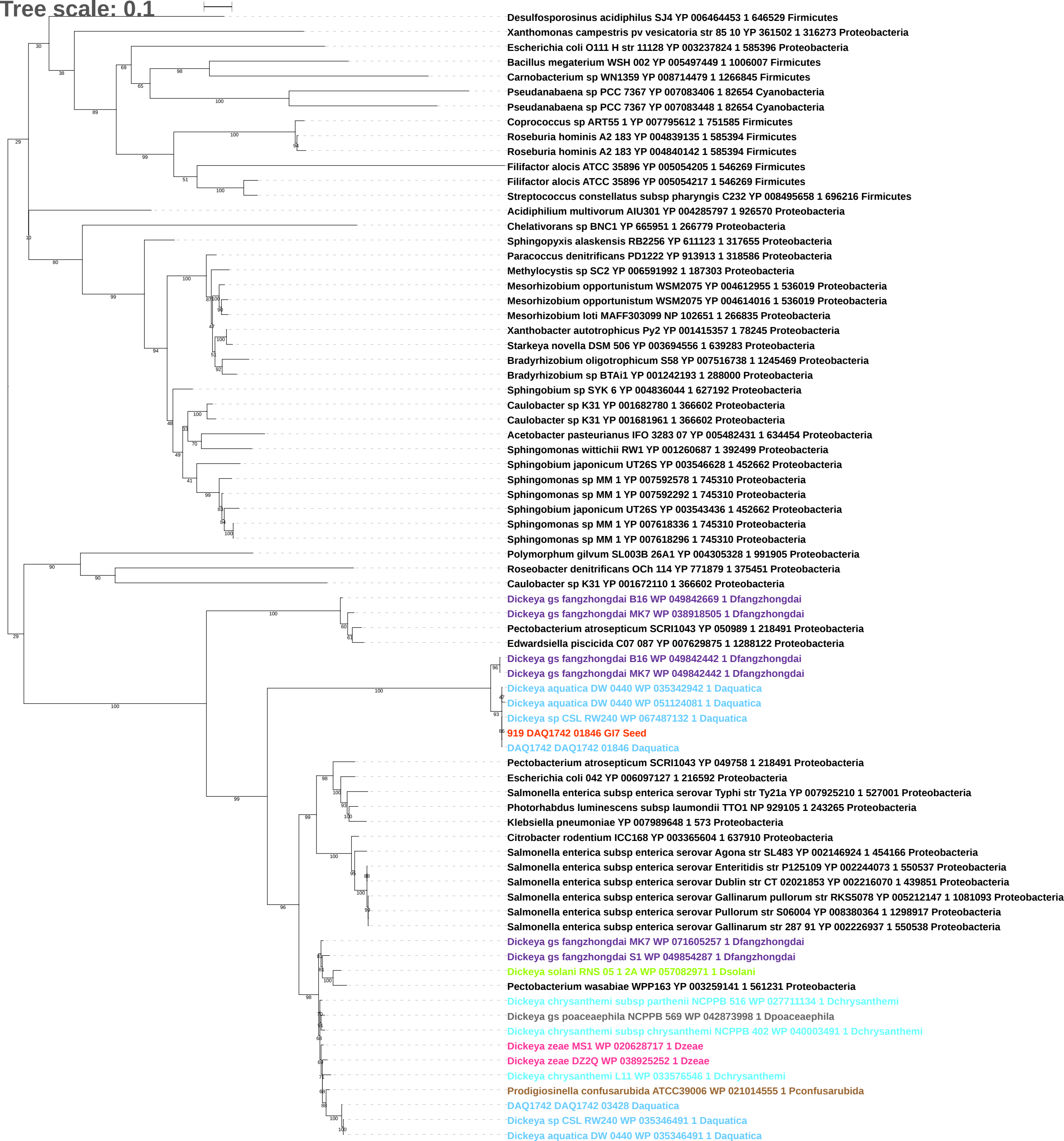
Tree scale: 0.1
Desulfosporosinus acidiphilus SJ4 YP 006464453 1 646529 Firmicutes
Xanthomonas campestris pv vesicatoria str 85 10 YP 361502 1 316273 Proteobacteria
Escherichia coli O111 H str 11128 YP 003237824 1 585396 Proteobacteria
Bacillus megaterium WSH 002 YP 005497449 1 1006007 Firmicutes
Carnobacterium sp WN1359 YP 008714479 1 1266845 Firmicutes
Pseudanabaena sp PCC 7367 YP 007083406 1 82654 Cyanobacteria
Pseudanabaena sp PCC 7367 YP 007083448 1 82654 Cyanobacteria
Coprococcus sp ART55 1 YP 007795612 1 751585 Firmicutes
Roseburia hominis A2 183 YP 004839135 1 585394 Firmicutes
Roseburia hominis A2 183 YP 004840142 1 585394 Firmicutes
Filifactor alocis ATCC 35896 YP 005054205 1 546269 Firmicutes
Filifactor alocis ATCC 35896 YP 005054217 1 546269 Firmicutes
Streptococcus constellatus subsp pharyngis C232 YP 008495658 1 696216 Firmicutes
Acidiphilium multivorum AIU301 YP 004285797 1 926570 Proteobacteria
Chelativorans sp BNC1 YP 665951 1 266779 Proteobacteria
Sphingopyxis alaskensis RB2256 YP 611123 1 317655 Proteobacteria
Paracoccus denitrificans PD1222 YP 913913 1 318586 Proteobacteria
Methylocystis sp SC2 YP 006591992 1 187303 Proteobacteria
Mesorhizobium opportunistum WSM2075 YP 004612955 1 536019 Proteobacteria
Mesorhizobium opportunistum WSM2075 YP 004614016 1 536019 Proteobacteria
Mesorhizobium loti MAFF303099 NP 102651 1 266835 Proteobacteria
Xanthobacter autotrophicus Py2 YP 001415357 1 78245 Proteobacteria
Starkeya novella DSM 506 YP 003694556 1 639283 Proteobacteria
Bradyrhizobium oligotrophicum S58 YP 007516738 1 1245469 Proteobacteria
Bradyrhizobium sp BTAi1 YP 001242193 1 288000 Proteobacteria
Sphingobium sp SYK 6 YP 004836044 1 627192 Proteobacteria
Caulobacter sp K31 YP 001682780 1 366602 Proteobacteria
Caulobacter sp K31 YP 001681961 1 366602 Proteobacteria
Acetobacter pasteurianus IFO 3283 07 YP 005482431 1 634454 Proteobacteria
Sphingomonas wittichii RW1 YP 001260687 1 392499 Proteobacteria
Sphingobium japonicum UT26S YP 003546628 1 452662 Proteobacteria
Sphingomonas sp MM 1 YP 007592578 1 745310 Proteobacteria
Sphingomonas sp MM 1 YP 007592292 1 745310 Proteobacteria
Sphingobium japonicum UT26S YP 003543436 1 452662 Proteobacteria
Sphingomonas sp MM 1 YP 007618336 1 745310 Proteobacteria
Sphingomonas sp MM 1 YP 007618296 1 745310 Proteobacteria
Polymorphum gilvum SL003B 26A1 YP 004305328 1 991905 Proteobacteria
Roseobacter denitrificans OCh 114 YP 771879 1 375451 Proteobacteria
Caulobacter sp K31 YP 001672110 1 366602 Proteobacteria
Dickeya gs fangzhongdai B16 WP 049842669 1 Dfangzhongdai
Dickeya gs fangzhongdai MK7 WP 038918505 1 Dfangzhongdai
Pectobacterium atrosepticum SCRI1043 YP 050989 1 218491 Proteobacteria
Edwardsiella piscicida C07 087 YP 007629875 1 1288122 Proteobacteria
Dickeya gs fangzhongdai B16 WP 049842442 1 Dfangzhongdai
Dickeya gs fangzhongdai MK7 WP 049842442 1 Dfangzhongdai
Dickeya aquatica DW 0440 WP 035342942 1 Daquatica
Dickeya aquatica DW 0440 WP 051124081 1 Daquatica
Dickeya sp CSL RW240 WP 067487132 1 Daquatica
919 DAQ1742 01846 GI7 Seed
DAQ1742 DAQ1742 01846 Daquatica
Pectobacterium atrosepticum SCRI1043 YP 049758 1 218491 Proteobacteria
Escherichia coli 042 YP 006097127 1 216592 Proteobacteria
Salmonella enterica subsp enterica serovar Typhi str Ty21a YP 007925210 1 527001 Proteobacteria
Photorhabdus luminescens subsp laumondii TTO1 NP 929105 1 243265 Proteobacteria
Klebsiella pneumoniae YP 007989648 1 573 Proteobacteria
Citrobacter rodentium ICC168 YP 003365604 1 637910 Proteobacteria
Salmonella enterica subsp enterica serovar Agona str SL483 YP 002146924 1 454166 Proteobacteria
Salmonella enterica subsp enterica serovar Enteritidis str P125109 YP 002244073 1 550537 Proteobacteria
Salmonella enterica subsp enterica serovar Dublin str CT 02021853 YP 002216070 1 439851 Proteobacteria
Salmonella enterica subsp enterica serovar Gallinarum pullorum str RKS5078 YP 005212147 1 1081093 Proteobacteria
Salmonella enterica subsp enterica serovar Pullorum str S06004 YP 008380364 1 1298917 Proteobacteria
Salmonella enterica subsp enterica serovar Gallinarum str 287 91 YP 002226937 1 550538 Proteobacteria
Dickeya gs fangzhongdai MK7 WP 071605257 1 Dfangzhongdai
Dickeya gs fangzhongdai S1 WP 049854287 1 Dfangzhongdai
Dickeya solani RNS 05 1 2A WP 057082971 1 Dsolani
Pectobacterium wasabiae WPP163 YP 003259141 1 561231 Proteobacteria
Dickeya chrysanthemi subsp parthenii NCPPB 516 WP 027711134 1 Dchrysanthemi
Dickeya gs poaceaephila NCPPB 569 WP 042873998 1 Dpoaceaephila
Dickeya chrysanthemi subsp chrysanthemi NCPPB 402 WP 040003491 1 Dchrysanthemi
Dickeya zeae MS1 WP 020628717 1 Dzeae
Dickeya zeae DZ2Q WP 038925252 1 Dzeae
Dickeya chrysanthemi L11 WP 033576546 1 Dchrysanthemi
Prodigiosinella confusarubida ATCC39006 WP 021014555 1 Pconfusarubida
DAQ1742 DAQ1742 03428 Daquatica
Dickeya sp CSL RW240 WP 035346491 1 Daquatica
Dickeya aquatica DW 0440 WP 035346491 1 Daquatica

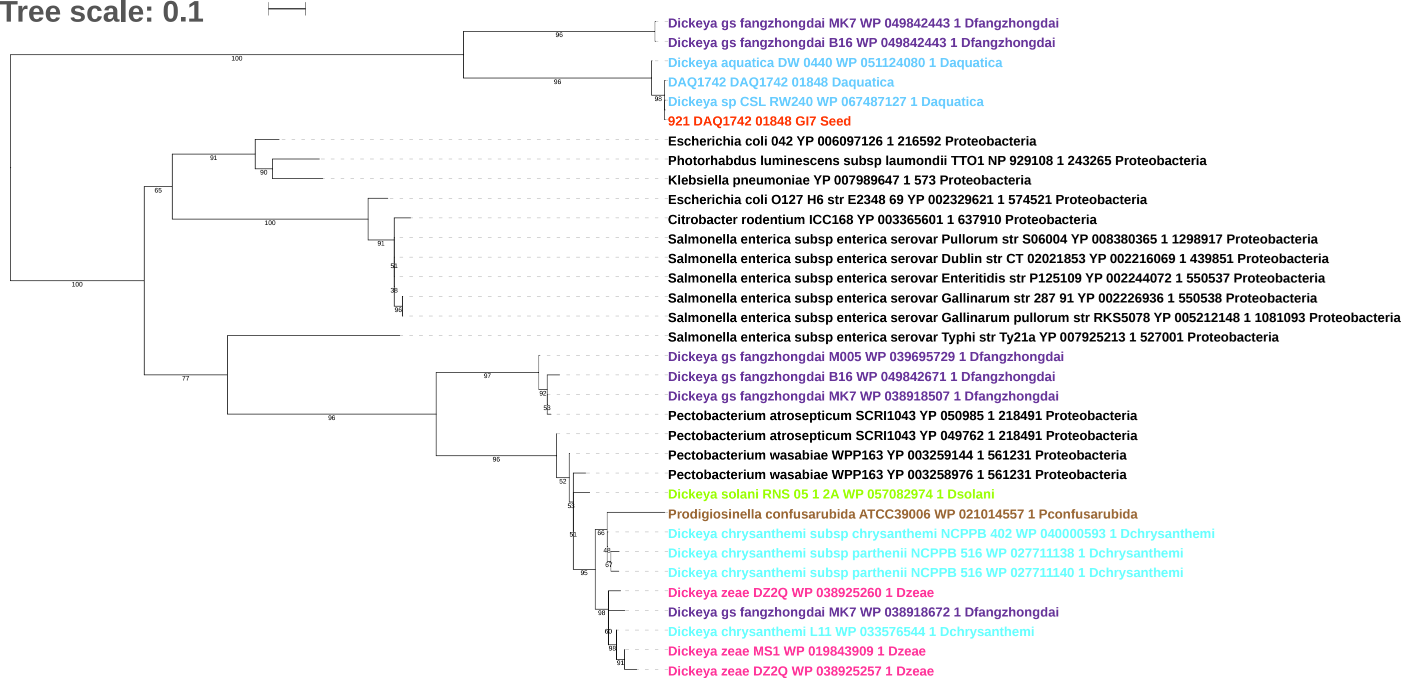

Tree scale: 0.1
Dickeya gs fangzhongdai MK7 WP 049842443 1 Dfangzhongdai
Dickeya gs fangzhongdai B16 WP 049842443 1 Dfangzhongdai
Dickeya aquatica DW 0440 WP 051124080 1 Daquatica
DAQ1742 DAQ1742 01848 Daquatica
Dickeya sp CSL RW240 WP 067487127 1 Daquatica
921 DAQ1742 01848 GI7 Seed
Escherichia coli 042 YP 006097126 1 216592 Proteobacteria
Photorhabdus luminescens subsp laumondii TTO1 NP 929108 1 243265 Proteobacteria
Klebsiella pneumoniae YP 007989647 1 573 Proteobacteria
Escherichia coli O127 H6 str E2348 69 YP 002329621 1 574521 Proteobacteria
Citrobacter rodentium ICC168 YP 003365601 1 637910 Proteobacteria
Salmonella enterica subsp enterica serovar Pullorum str S06004 YP 008380365 1 1298917 Proteobacteria
Salmonella enterica subsp enterica serovar Dublin str CT 02021853 YP 002216069 1 439851 Proteobacteria
Salmonella enterica subsp enterica serovar Enteritidis str P125109 YP 002244072 1 550537 Proteobacteria
Salmonella enterica subsp enterica serovar Gallinarum str 287 91 YP 002226936 1 550538 Proteobacteria
Salmonella enterica subsp enterica serovar Gallinarum pullorum str RKS5078 YP 005212148 1 1081093 Proteobacteria
Salmonella enterica subsp enterica serovar Typhi str Ty21a YP 007925213 1 527001 Proteobacteria
Dickeya gs fangzhongdai M005 WP 039695729 1 Dfangzhongdai
Dickeya gs fangzhongdai B16 WP 049842671 1 Dfangzhongdai
Dickeya gs fangzhongdai MK7 WP 038918507 1 Dfangzhongdai
Pectobacterium atrosepticum SCRI1043 YP 050985 1 218491 Proteobacteria
Pectobacterium atrosepticum SCRI1043 YP 049762 1 218491 Proteobacteria
Pectobacterium wasabiae WPP163 YP 003259144 1 561231 Proteobacteria
Pectobacterium wasabiae WPP163 YP 003258976 1 561231 Proteobacteria
Dickeya solani RNS 05 1 2A WP 057082974 1 Dsolani
Prodigiosinella confusarubida ATCC39006 WP 021014557 1 Pconfusarubida
Dickeya chrysanthemi subsp chrysanthemi NCPPB 402 WP 040000593 1 Dchrysanthemi
Dickeya chrysanthemi subsp parthenii NCPPB 516 WP 027711138 1 Dchrysanthemi
Dickeya chrysanthemi subsp parthenii NCPPB 516 WP 027711140 1 Dchrysanthemi
Dickeya zeae DZ2Q WP 038925260 1 Dzeae
Dickeya gs fangzhongdai MK7 WP 038918672 1 Dfangzhongdai
Dickeya chrysanthemi L11 WP 033576544 1 Dchrysanthemi
Dickeya zeae MS1 WP 019843909 1 Dzeae
Dickeya zeae DZ2Q WP 038925257 1 Dzeae

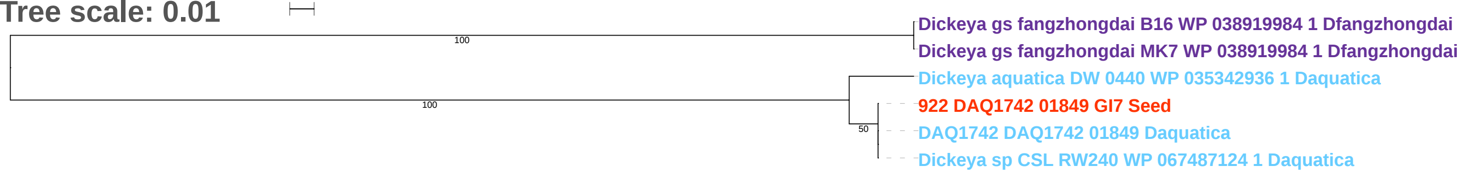

Tree scale: 0.01
100
100
50
Dickeya gs fangzhongdai B16 WP 038919984 1 Dfangzhongdai
Dickeya gs fangzhongdai MK7 WP 038919984 1 Dfangzhongdai
Dickeya aquatica DW 0440 WP 035342936 1 Daquatica
922 DAQ1742 01849 GI7 Seed
DAQ1742 DAQ1742 01849 Daquatica
Dickeya sp CSL RW240 WP 067487124 1 Daquatica

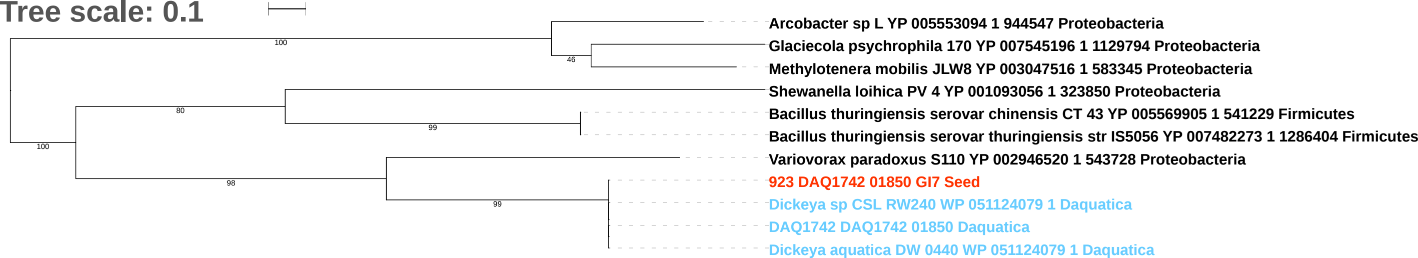
Tree scale: 0.1
Arcobacter sp L YP 005553094 1 944547 Proteobacteria
Glaciecola psychrophila 170 YP 007545196 1 1129794 Proteobacteria
Methylotenera mobilis JLW8 YP 003047516 1 583345 Proteobacteria
Shewanella loihica PV 4 YP 001093056 1 323850 Proteobacteria
Bacillus thuringiensis serovar chinensis CT 43 YP 005569905 1 541229 Firmicutes
Bacillus thuringiensis serovar thuringiensis str IS5056 YP 007482273 1 1286404 Firmicutes
Variovorax paradoxus S110 YP 002946520 1 543728 Proteobacteria
923 DAQ1742 01850 GI7 Seed
Dickeya sp CSL RW240 WP 051124079 1 Daquatica
DAQ1742 DAQ1742 01850 Daquatica
Dickeya aquatica DW 0440 WP 051124079 1 Daquatica
100
46
80
99
100
98
99

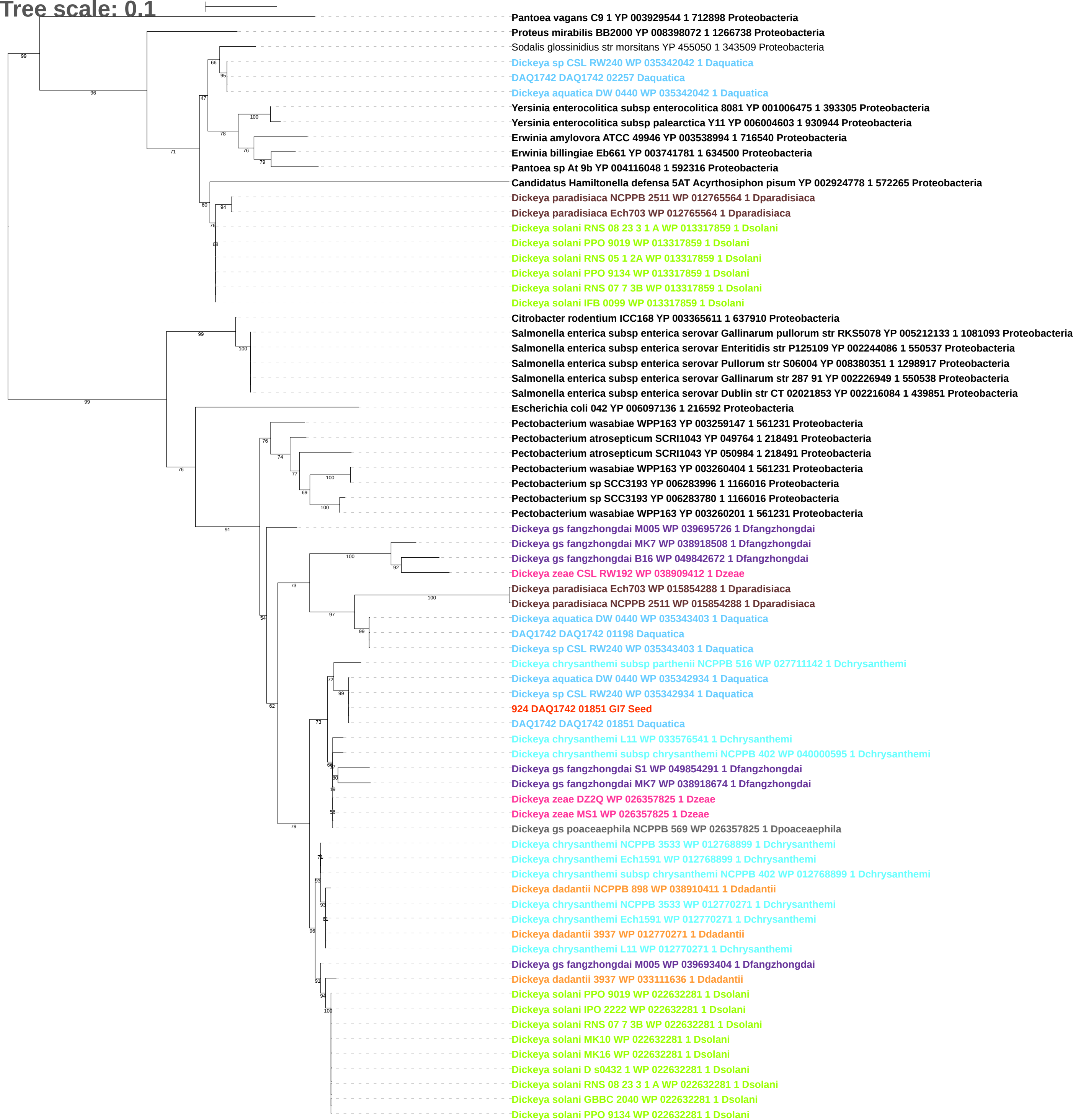

Tree scale: 0.1
Pantoea vagans C9 1 YP 003929544 1 712898 Proteobacteria
Proteus mirabilis BB2000 YP 008398072 1 1266738 Proteobacteria
Sodalis glossinidius str morsitans YP 455050 1 343509 Proteobacteria
Dickeya sp CSL RW240 WP 035342042 1 Daquatica
DAQ1742 DAQ1742 02257 Daquatica
Dickeya aquatica DW 0440 WP 035342042 1 Daquatica
Yersinia enterocolitica subsp enterocolitica 8081 YP 001006475 1 393305 Proteobacteria
Yersinia enterocolitica subsp palearctica Y11 YP 006004603 1 930944 Proteobacteria
Erwinia amylovora ATCC 49946 YP 003538994 1 716540 Proteobacteria
Erwinia billingiae Eb661 YP 003741781 1 634500 Proteobacteria
Pantoea sp At 9b YP 004116048 1 592316 Proteobacteria
Candidatus Hamiltonella defensa 5AT Acyrthosiphon pisum YP 002924778 1 572265 Proteobacteria
Dickeya paradisiaca NCPPB 2511 WP 012765564 1 Dparadisiaca
Dickeya paradisiaca Ech703 WP 012765564 1 Dparadisiaca
Dickeya solani RNS 08 23 3 1 A WP 013317859 1 Dsolani
Dickeya solani PPO 9019 WP 013317859 1 Dsolani
Dickeya solani RNS 05 1 2A WP 013317859 1 Dsolani
Dickeya solani PPO 9134 WP 013317859 1 Dsolani
Dickeya solani RNS 07 7 3B WP 013317859 1 Dsolani
Dickeya solani IFB 0099 WP 013317859 1 Dsolani
Citrobacter rodentium ICC168 YP 003365611 1 637910 Proteobacteria
Salmonella enterica subsp enterica serovar Gallinarum pullorum str RKS5078 YP 005212133 1 1081093 Proteobacteria
Salmonella enterica subsp enterica serovar Enteritidis str P125109 YP 002244086 1 550537 Proteobacteria
Salmonella enterica subsp enterica serovar Pullorum str S06004 YP 008380351 1 1298917 Proteobacteria
Salmonella enterica subsp enterica serovar Gallinarum str 287 91 YP 002226949 1 550538 Proteobacteria
Salmonella enterica subsp enterica serovar Dublin str CT 02021853 YP 002216084 1 439851 Proteobacteria
Escherichia coli 042 YP 006097136 1 216592 Proteobacteria
Pectobacterium wasabiae WPP163 YP 003259147 1 561231 Proteobacteria
Pectobacterium atrosepticum SCRI1043 YP 049764 1 218491 Proteobacteria
Pectobacterium atrosepticum SCRI1043 YP 050984 1 218491 Proteobacteria
Pectobacterium wasabiae WPP163 YP 003260404 1 561231 Proteobacteria
Pectobacterium sp SCC3193 YP 006283996 1 1166016 Proteobacteria
Pectobacterium sp SCC3193 YP 006283780 1 1166016 Proteobacteria
Pectobacterium wasabiae WPP163 YP 003260201 1 561231 Proteobacteria
Dickeya gs fangzhongdai M005 WP 039695726 1 Dfangzhongdai
Dickeya gs fangzhongdai MK7 WP 038918508 1 Dfangzhongdai
Dickeya gs fangzhongdai B16 WP 049842672 1 Dfangzhongdai
Dickeya zeae CSL RW192 WP 038909412 1 Dzeae
Dickeya paradisiaca Ech703 WP 015854288 1 Dparadisiaca
Dickeya paradisiaca NCPPB 2511 WP 015854288 1 Dparadisiaca
Dickeya aquatica DW 0440 WP 035343403 1 Daquatica
DAQ1742 DAQ1742 01198 Daquatica
Dickeya sp CSL RW240 WP 035343403 1 Daquatica
Dickeya chrysanthemi subsp parthenii NCPPB 516 WP 027711142 1 Dchrysanthemi
Dickeya aquatica DW 0440 WP 035342934 1 Daquatica
Dickeya sp CSL RW240 WP 035342934 1 Daquatica
924 DAQ1742 01851 GI7 Seed
DAQ1742 DAQ1742 01851 Daquatica
Dickeya chrysanthemi L11 WP 033576541 1 Dchrysanthemi
Dickeya chrysanthemi subsp chrysanthemi NCPPB 402 WP 040000595 1 Dchrysanthemi
Dickeya gs fangzhongdai S1 WP 049854291 1 Dfangzhongdai
Dickeya gs fangzhongdai MK7 WP 038918674 1 Dfangzhongdai
Dickeya zeae DZ2Q WP 026357825 1 Dzeae
Dickeya zeae MS1 WP 026357825 1 Dzeae
Dickeya gs poaceaephila NCPPB 569 WP 026357825 1 Dpoaceaephila
Dickeya chrysanthemi NCPPB 3533 WP 012768899 1 Dchrysanthemi
Dickeya chrysanthemi Ech1591 WP 012768899 1 Dchrysanthemi
Dickeya chrysanthemi subsp chrysanthemi NCPPB 402 WP 012768899 1 Dchrysanthemi
Dickeya dadantii NCPPB 898 WP 038910411 1 Ddadantii
Dickeya chrysanthemi NCPPB 3533 WP 012770271 1 Dchrysanthemi
Dickeya chrysanthemi Ech1591 WP 012770271 1 Dchrysanthemi
Dickeya dadantii 3937 WP 012770271 1 Ddadantii
Dickeya chrysanthemi L11 WP 012770271 1 Dchrysanthemi
Dickeya gs fangzhongdai M005 WP 039693404 1 Dfangzhongdai
Dickeya dadantii 3937 WP 033111636 1 Ddadantii
Dickeya solani PPO 9019 WP 022632281 1 Dsolani
Dickeya solani IPO 2222 WP 022632281 1 Dsolani
Dickeya solani RNS 07 7 3B WP 022632281 1 Dsolani
Dickeya solani MK10 WP 022632281 1 Dsolani
Dickeya solani MK16 WP 022632281 1 Dsolani
Dickeya solani D s0432 1 WP 022632281 1 Dsolani
Dickeya solani RNS 08 23 3 1 A WP 022632281 1 Dsolani
Dickeya solani GBBC 2040 WP 022632281 1 Dsolani
Dickeya solani PPO 9134 WP 022632281 1 Dsolani

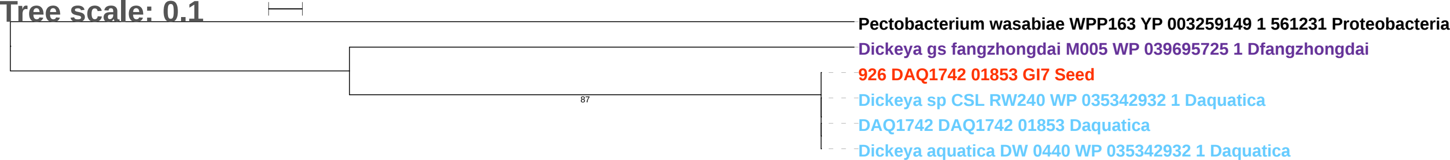

Tree scale: 0.1
Pectobacterium wasabiae WPP163 YP 003259149 1 561231 Proteobacteria
Dickeya gs fangzhongdai M005 WP 039695725 1 Dfangzhongdai
926 DAQ1742 01853 GI7 Seed
Dickeya sp CSL RW240 WP 035342932 1 Daquatica
DAQ1742 DAQ1742 01853 Daquatica
Dickeya aquatica DW 0440 WP 035342932 1 Daquatica
87

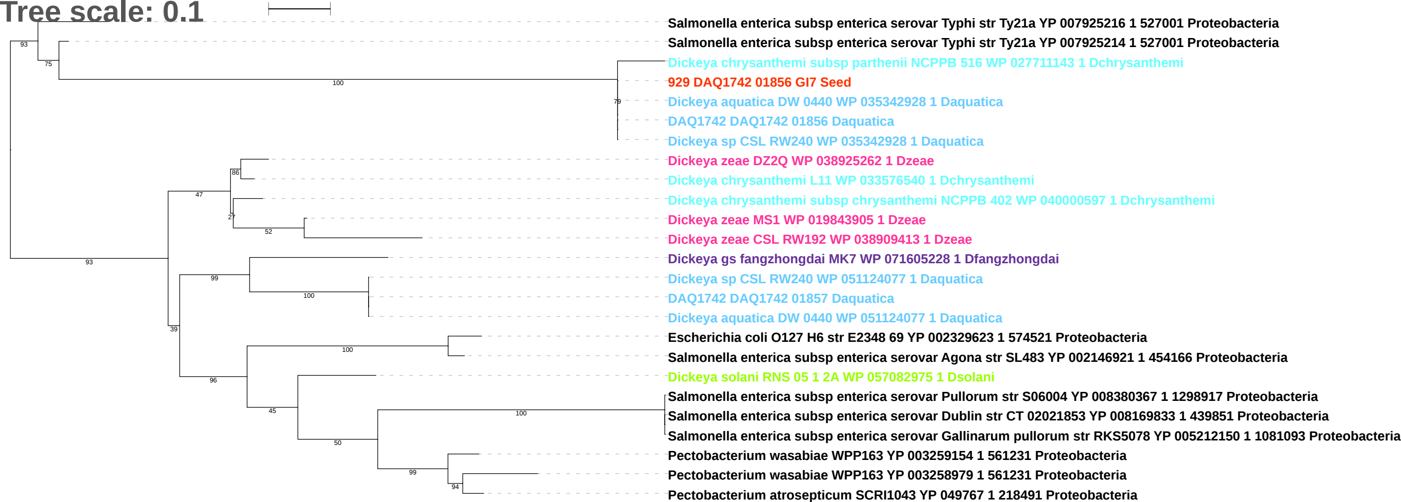
Tree scale: 0.1
Salmonella enterica subsp enterica serovar Typhi str Ty21a YP 007925216 1 527001 Proteobacteria
Salmonella enterica subsp enterica serovar Typhi str Ty21a YP 007925214 1 527001 Proteobacteria
Dickeya chrysanthemi subsp parthenii NCPPB 516 WP 027711143 1 Dchrysanthemi
929 DAQ1742 01856 GI7 Seed
Dickeya aquatica DW 0440 WP 035342928 1 Daquatica
DAQ1742 DAQ1742 01856 Daquatica
Dickeya sp CSL RW240 WP 035342928 1 Daquatica
Dickeya zeae DZ2Q WP 038925262 1 Dzeae
Dickeya chrysanthemi L11 WP 033576540 1 Dchrysanthemi
Dickeya chrysanthemi subsp chrysanthemi NCPPB 402 WP 040000597 1 Dchrysanthemi
Dickeya zeae MS1 WP 019843905 1 Dzeae
Dickeya zeae CSL RW192 WP 038909413 1 Dzeae
Dickeya gs fangzhongdai MK7 WP 071605228 1 Dfangzhongdai
Dickeya sp CSL RW240 WP 051124077 1 Daquatica
DAQ1742 DAQ1742 01857 Daquatica
Dickeya aquatica DW 0440 WP 051124077 1 Daquatica
Escherichia coli O127 H6 str E2348 69 YP 002329623 1 574521 Proteobacteria
Salmonella enterica subsp enterica serovar Agona str SL483 YP 002146921 1 454166 Proteobacteria
Dickeya solani RNS 05 1 2A WP 057082975 1 Dsolani
Salmonella enterica subsp enterica serovar Pullorum str S06004 YP 008380367 1 1298917 Proteobacteria
Salmonella enterica subsp enterica serovar Dublin str CT 02021853 YP 008169833 1 439851 Proteobacteria
Salmonella enterica subsp enterica serovar Gallinarum pullorum str RKS5078 YP 005212150 1 1081093 Proteobacteria
Pectobacterium wasabiae WPP163 YP 003259154 1 561231 Proteobacteria
Pectobacterium wasabiae WPP163 YP 003258979 1 561231 Proteobacteria
Pectobacterium atrosepticum SCRI1043 YP 049767 1 218491 Proteobacteria

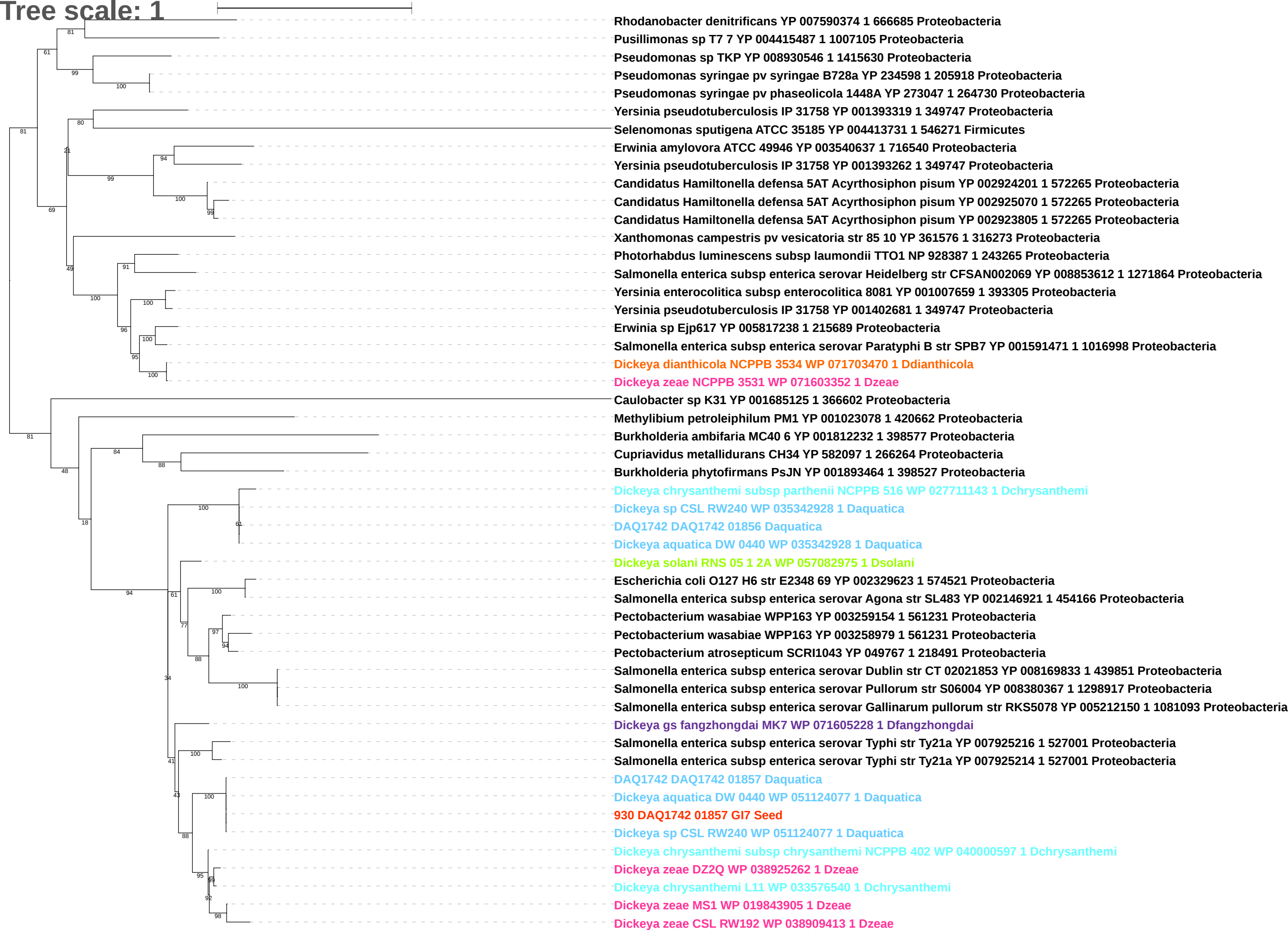
Tree scale: 1
Rhodanobacter denitrificans YP 007590374 1 666685 Proteobacteria
Pusillimonas sp T7 7 YP 004415487 1 1007105 Proteobacteria
Pseudomonas sp TKP YP 008930546 1 1415630 Proteobacteria
Pseudomonas syringae pv syringae B728a YP 234598 1 205918 Proteobacteria
Pseudomonas syringae pv phaseolicola 1448A YP 273047 1 264730 Proteobacteria
Yersinia pseudotuberculosis IP 31758 YP 001393319 1 349747 Proteobacteria
Selenomonas sputigena ATCC 35185 YP 004413731 1 546271 Firmicutes
Erwinia amylovora ATCC 49946 YP 003540637 1 716540 Proteobacteria
Yersinia pseudotuberculosis IP 31758 YP 001393262 1 349747 Proteobacteria
Candidatus Hamiltonella defensa 5AT Acyrthosiphon pisum YP 002924201 1 572265 Proteobacteria
Candidatus Hamiltonella defensa 5AT Acyrthosiphon pisum YP 002925070 1 572265 Proteobacteria
Candidatus Hamiltonella defensa 5AT Acyrthosiphon pisum YP 002923805 1 572265 Proteobacteria
Xanthomonas campestris pv vesicatoria str 85 10 YP 361576 1 316273 Proteobacteria
Photorhabdus luminescens subsp laumondii TTO1 NP 928387 1 243265 Proteobacteria
Salmonella enterica subsp enterica serovar Heidelberg str CFSAN002069 YP 008853612 1 1271864 Proteobacteria
Yersinia enterocolitica subsp enterocolitica 8081 YP 001007659 1 393305 Proteobacteria
Yersinia pseudotuberculosis IP 31758 YP 001402681 1 349747 Proteobacteria
Erwinia sp Ejp617 YP 005817238 1 215689 Proteobacteria
Salmonella enterica subsp enterica serovar Paratyphi B str SPB7 YP 001591471 1 1016998 Proteobacteria
Dickeya dianthicola NCPPB 3534 WP 071703470 1 Ddianthicola
Dickeya zeae NCPPB 3531 WP 071603352 1 Dzeae
Caulobacter sp K31 YP 001685125 1 366602 Proteobacteria
Methylibium petroleiphilum PM1 YP 001023078 1 420662 Proteobacteria
Burkholderia ambifaria MC40 6 YP 001812232 1 398577 Proteobacteria
Cupriavidus metallidurans CH34 YP 582097 1 266264 Proteobacteria
Burkholderia phytofirmans PsJN YP 001893464 1 398527 Proteobacteria
Dickeya chrysanthemi subsp parthenii NCPPB 516 WP 027711143 1 Dchrysanthemi
Dickeya sp CSL RW240 WP 035342928 1 Daquatica
DAQ1742 DAQ1742 01856 Daquatica
Dickeya aquatica DW 0440 WP 035342928 1 Daquatica
Dickeya solani RNS 05 1 2A WP 057082975 1 Dsolani
Escherichia coli O127 H6 str E2348 69 YP 002329623 1 574521 Proteobacteria
Salmonella enterica subsp enterica serovar Agona str SL483 YP 002146921 1 454166 Proteobacteria
Pectobacterium wasabiae WPP163 YP 003259154 1 561231 Proteobacteria
Pectobacterium wasabiae WPP163 YP 003258979 1 561231 Proteobacteria
Pectobacterium atrosepticum SCRI1043 YP 049767 1 218491 Proteobacteria
Salmonella enterica subsp enterica serovar Dublin str CT 02021853 YP 008169833 1 439851 Proteobacteria
Salmonella enterica subsp enterica serovar Pullorum str S06004 YP 008380367 1 1298917 Proteobacteria
Salmonella enterica subsp enterica serovar Gallinarum pullorum str RKS5078 YP 005212150 1 1081093 Proteobacteria
Dickeya gs fangzhongdai MK7 WP 071605228 1 Dfangzhongdai
Salmonella enterica subsp enterica serovar Typhi str Ty21a YP 007925216 1 527001 Proteobacteria
Salmonella enterica subsp enterica serovar Typhi str Ty21a YP 007925214 1 527001 Proteobacteria
DAQ1742 DAQ1742 01857 Daquatica
Dickeya aquatica DW 0440 WP 051124077 1 Daquatica
930 DAQ1742 01857 GI7 Seed
Dickeya sp CSL RW240 WP 051124077 1 Daquatica
Dickeya chrysanthemi subsp chrysanthemi NCPPB 402 WP 040000597 1 Dchrysanthemi
Dickeya zeae DZ2Q WP 038925262 1 Dzeae
Dickeya chrysanthemi L11 WP 033576540 1 Dchrysanthemi
Dickeya zeae MS1 WP 019843905 1 Dzeae
Dickeya zeae CSL RW192 WP 038909413 1 Dzeae

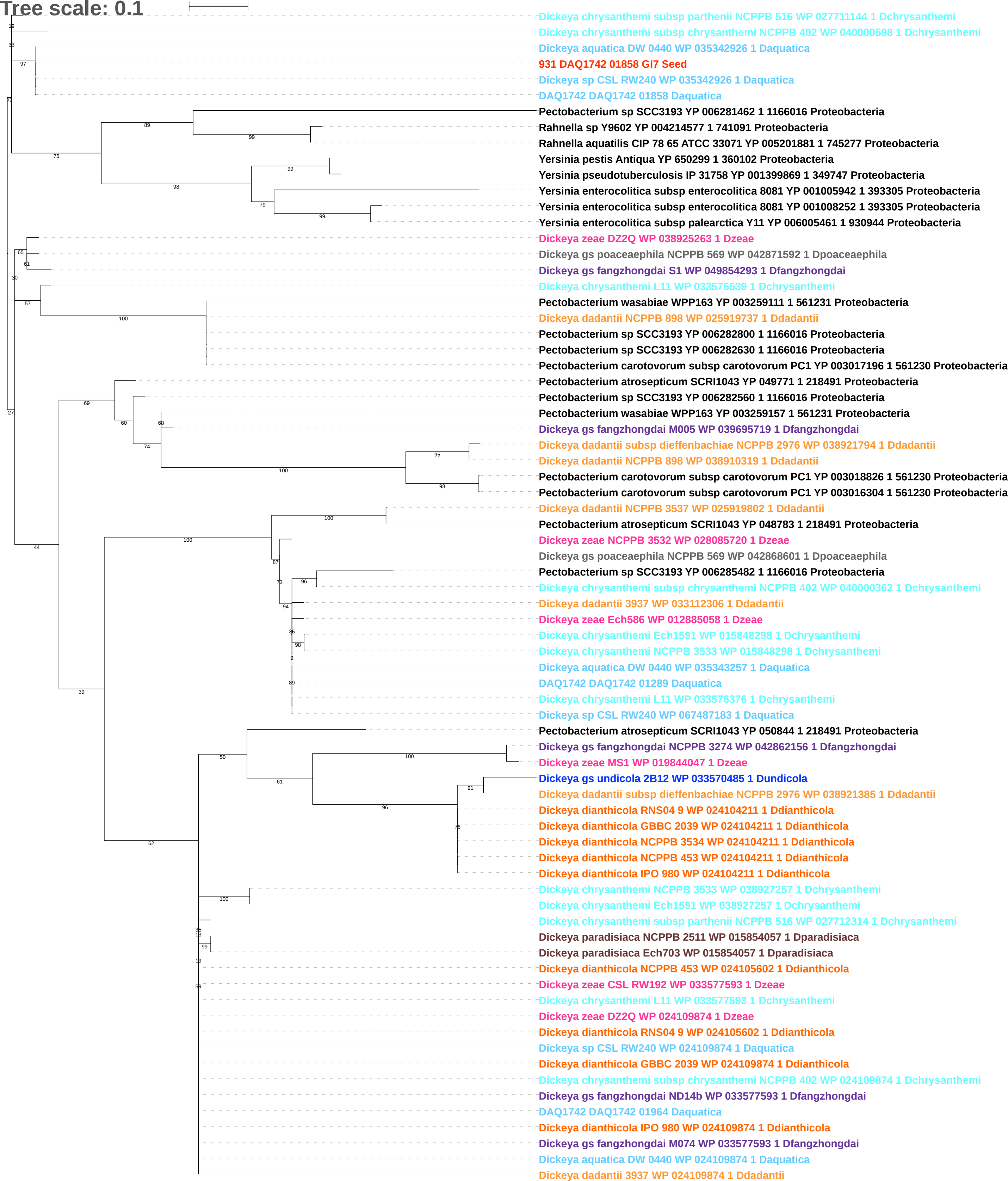
Tree scale: 0.1
Dickeya chrysanthemi subsp parthenii NCPPB 516 WP 027711144 1 Dchrysanthemi
Dickeya chrysanthemi subsp chrysanthemi NCPPB 402 WP 040000598 1 Dchrysanthemi
Dickeya aquatica DW 0440 WP 035342926 1 Daquatica
931 DAQ1742 01858 GI7 Seed
Dickeya sp CSL RW240 WP 035342926 1 Daquatica
DAQ1742 DAQ1742 01858 Daquatica
Pectobacterium sp SCC3193 YP 006281462 1 1166016 Proteobacteria
Rahnella sp Y9602 YP 004214577 1 741091 Proteobacteria
Rahnella aquatilis CIP 78 65 ATCC 33071 YP 005201881 1 745277 Proteobacteria
Yersinia pestis Antiqua YP 650299 1 360102 Proteobacteria
Yersinia pseudotuberculosis IP 31758 YP 001399869 1 349747 Proteobacteria
Yersinia enterocolitica subsp enterocolitica 8081 YP 001005942 1 393305 Proteobacteria
Yersinia enterocolitica subsp enterocolitica 8081 YP 001008252 1 393305 Proteobacteria
Yersinia enterocolitica subsp palearctica Y11 YP 006005461 1 930944 Proteobacteria
Dickeya zeae DZ2Q WP 038925263 1 Dzeae
Dickeya gs poaceaephila NCPPB 569 WP 042871592 1 Dpoaceaephila
Dickeya gs fangzhongdai S1 WP 049854293 1 Dfangzhongdai
Dickeya chrysanthemi L11 WP 033576539 1 Dchrysanthemi
Pectobacterium wasabiae WPP163 YP 003259111 1 561231 Proteobacteria
Dickeya dadantii NCPPB 898 WP 025919737 1 Ddadantii
Pectobacterium sp SCC3193 YP 006282800 1 1166016 Proteobacteria
Pectobacterium sp SCC3193 YP 006282630 1 1166016 Proteobacteria
Pectobacterium carotovorum subsp carotovorum PC1 YP 003017196 1 561230 Proteobacteria
Pectobacterium atrosepticum SCRI1043 YP 049771 1 218491 Proteobacteria
Pectobacterium sp SCC3193 YP 006282560 1 1166016 Proteobacteria
Pectobacterium wasabiae WPP163 YP 003259157 1 561231 Proteobacteria
Dickeya gs fangzhongdai M005 WP 039695719 1 Dfangzhongdai
Dickeya dadantii subsp dieffenbachiae NCPPB 2976 WP 038921794 1 Ddadantii
Dickeya dadantii NCPPB 898 WP 038910319 1 Ddadantii
Pectobacterium carotovorum subsp carotovorum PC1 YP 003018826 1 561230 Proteobacteria
Pectobacterium carotovorum subsp carotovorum PC1 YP 003016304 1 561230 Proteobacteria
Dickeya dadantii NCPPB 3537 WP 025919802 1 Ddadantii
Pectobacterium atrosepticum SCRI1043 YP 048783 1 218491 Proteobacteria
Dickeya zeae NCPPB 3532 WP 028085720 1 Dzeae
Dickeya gs poaceaephila NCPPB 569 WP 042868601 1 Dpoaceaephila
Pectobacterium sp SCC3193 YP 006285482 1 1166016 Proteobacteria
Dickeya chrysanthemi subsp chrysanthemi NCPPB 402 WP 040000362 1 Dchrysanthemi
Dickeya dadantii 3937 WP 033112306 1 Ddadantii
Dickeya zeae Ech586 WP 012885058 1 Dzeae
Dickeya chrysanthemi Ech1591 WP 015848298 1 Dchrysanthemi
Dickeya chrysanthemi NCPPB 3533 WP 015848298 1 Dchrysanthemi
Dickeya aquatica DW 0440 WP 035343257 1 Daquatica
DAQ1742 DAQ1742 01289 Daquatica
Dickeya chrysanthemi L11 WP 033576376 1 Dchrysanthemi
Dickeya sp CSL RW240 WP 067487183 1 Daquatica
Pectobacterium atrosepticum SCRI1043 YP 050844 1 218491 Proteobacteria
Dickeya gs fangzhongdai NCPPB 3274 WP 042862156 1 Dfangzhongdai
Dickeya zeae MS1 WP 019844047 1 Dzeae
Dickeya gs undicola 2B12 WP 033570485 1 Dundicola
Dickeya dadantii subsp dieffenbachiae NCPPB 2976 WP 038921385 1 Ddadantii
Dickeya dianthicola RNS04 9 WP 024104211 1 Ddianthicola
Dickeya dianthicola GBBC 2039 WP 024104211 1 Ddianthicola
Dickeya dianthicola NCPPB 3534 WP 024104211 1 Ddianthicola
Dickeya dianthicola NCPPB 453 WP 024104211 1 Ddianthicola
Dickeya dianthicola IPO 980 WP 024104211 1 Ddianthicola
Dickeya chrysanthemi NCPPB 3533 WP 038927257 1 Dchrysanthemi
Dickeya chrysanthemi Ech1591 WP 038927257 1 Dchrysanthemi
Dickeya chrysanthemi subsp parthenii NCPPB 516 WP 027712314 1 Dchrysanthemi
Dickeya paradisiaca NCPPB 2511 WP 015854057 1 Dparadisiaca
Dickeya paradisiaca Ech703 WP 015854057 1 Dparadisiaca
Dickeya dianthicola NCPPB 453 WP 024105602 1 Ddianthicola
Dickeya zeae CSL RW192 WP 033577593 1 Dzeae
Dickeya chrysanthemi L11 WP 033577593 1 Dchrysanthemi
Dickeya zeae DZ2Q WP 024109874 1 Dzeae
Dickeya dianthicola RNS04 9 WP 024105602 1 Ddianthicola
Dickeya sp CSL RW240 WP 024109874 1 Daquatica
Dickeya dianthicola GBBC 2039 WP 024109874 1 Ddianthicola
Dickeya chrysanthemi subsp chrysanthemi NCPPB 402 WP 024109874 1 Dchrysanthemi
Dickeya gs fangzhongdai ND14b WP 033577593 1 Dfangzhongdai
DAQ1742 DAQ1742 01964 Daquatica
Dickeya dianthicola IPO 980 WP 024109874 1 Ddianthicola
Dickeya gs fangzhongdai M074 WP 033577593 1 Dfangzhongdai
Dickeya aquatica DW 0440 WP 024109874 1 Daquatica
Dickeya dadantii 3937 WP 024109874 1 Ddadantii

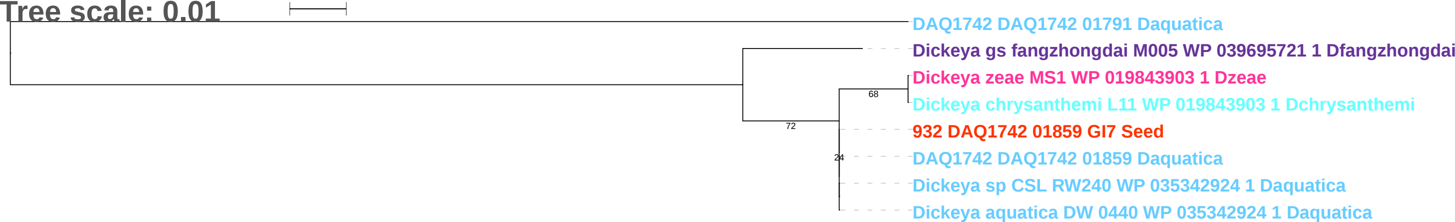
Tree scale: 0.01
DAQ1742 DAQ1742 01791 Daquatica
Dickeya gs fangzhongdai M005 WP 039695721 1 Dfangzhongdai
Dickeya zeae MS1 WP 019843903 1 Dzeae
Dickeya chrysanthemi L11 WP 019843903 1 Dchrysanthemi
932 DAQ1742 01859 GI7 Seed
DAQ1742 DAQ1742 01859 Daquatica
Dickeya sp CSL RW240 WP 035342924 1 Daquatica
Dickeya aquatica DW 0440 WP 035342924 1 Daquatica
68
72
24

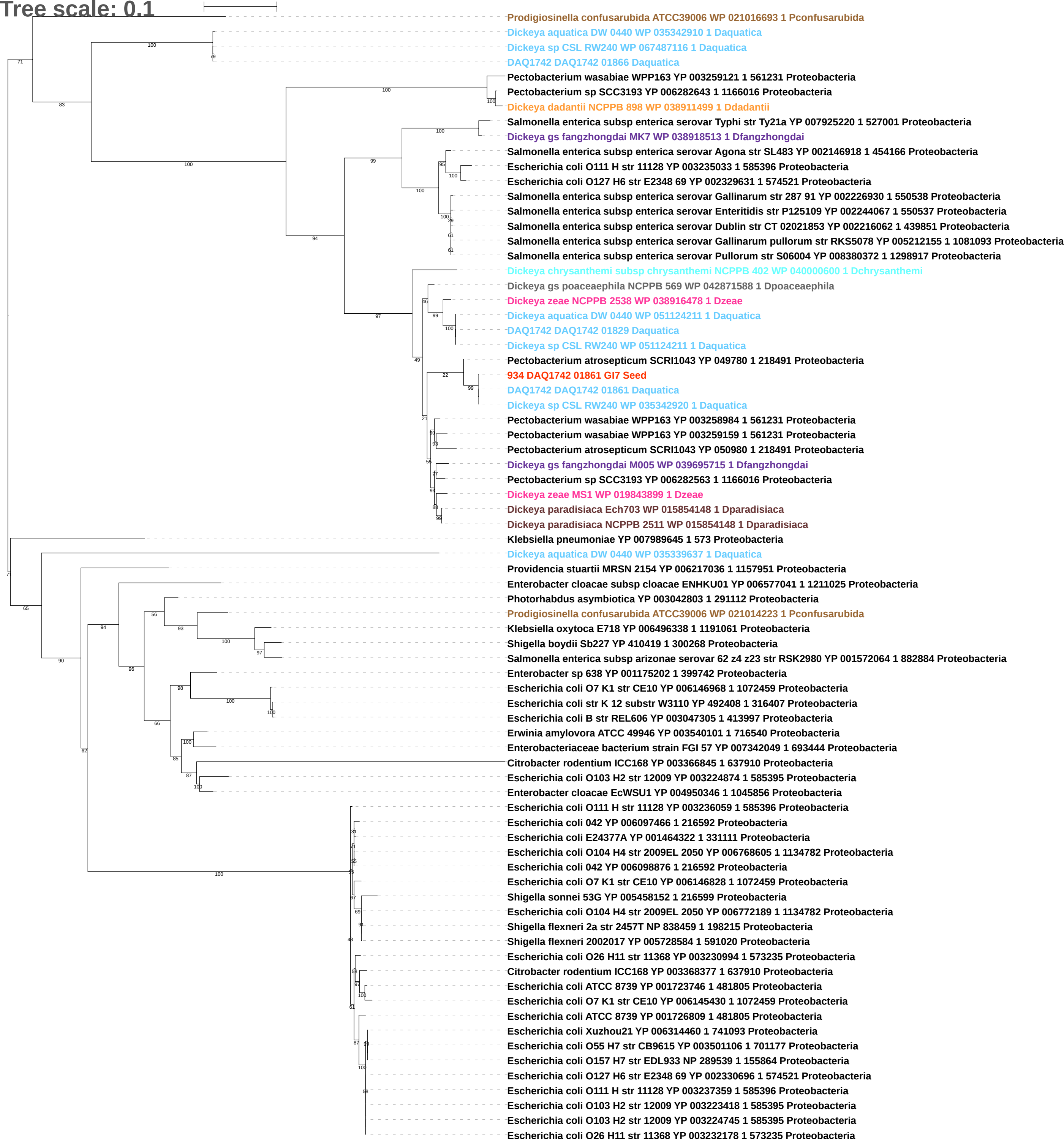
Tree scale: 0.1
Prodigiosinella confusarubida ATCC39006 WP 021016693 1 Pconfusarubida
Dickeya aquatica DW 0440 WP 035342910 1 Daquatica
Dickeya sp CSL RW240 WP 067487116 1 Daquatica
DAQ1742 DAQ1742 01866 Daquatica
Pectobacterium wasabiae WPP163 YP 003259121 1 561231 Proteobacteria
Pectobacterium sp SCC3193 YP 006282643 1 1166016 Proteobacteria
Dickeya dadantii NCPPB 898 WP 038911499 1 Ddadantii
Salmonella enterica subsp enterica serovar Typhi str Ty21a YP 007925220 1 527001 Proteobacteria
Dickeya gs fangzhongdai MK7 WP 038918513 1 Dfangzhongdai
Salmonella enterica subsp enterica serovar Agona str SL483 YP 002146918 1 454166 Proteobacteria
Escherichia coli O111 H str 11128 YP 003235033 1 585396 Proteobacteria
Escherichia coli O127 H6 str E2348 69 YP 002329631 1 574521 Proteobacteria
Salmonella enterica subsp enterica serovar Gallinarum str 287 91 YP 002226930 1 550538 Proteobacteria
Salmonella enterica subsp enterica serovar Enteritidis str P125109 YP 002244067 1 550537 Proteobacteria
Salmonella enterica subsp enterica serovar Dublin str CT 02021853 YP 002216062 1 439851 Proteobacteria
Salmonella enterica subsp enterica serovar Gallinarum pullorum str RKS5078 YP 005212155 1 1081093 Proteobacteria
Salmonella enterica subsp enterica serovar Pullorum str S06004 YP 008380372 1 1298917 Proteobacteria
Dickeya chrysanthemi subsp chrysanthemi NCPPB 402 WP 040000600 1 Dchrysanthemi
Dickeya gs poaceaephila NCPPB 569 WP 042871588 1 Dpoaceaephila
Dickeya zeae NCPPB 2538 WP 038916478 1 Dzeae
Dickeya aquatica DW 0440 WP 051124211 1 Daquatica
DAQ1742 DAQ1742 01829 Daquatica
Dickeya sp CSL RW240 WP 051124211 1 Daquatica
Pectobacterium atrosepticum SCRI1043 YP 049780 1 218491 Proteobacteria
934 DAQ1742 01861 GI7 Seed
DAQ1742 DAQ1742 01861 Daquatica
Dickeya sp CSL RW240 WP 035342920 1 Daquatica
Pectobacterium wasabiae WPP163 YP 003258984 1 561231 Proteobacteria
Pectobacterium wasabiae WPP163 YP 003259159 1 561231 Proteobacteria
Pectobacterium atrosepticum SCRI1043 YP 050980 1 218491 Proteobacteria
Dickeya gs fangzhongdai M005 WP 039695715 1 Dfangzhongdai
Pectobacterium sp SCC3193 YP 006282563 1 1166016 Proteobacteria
Dickeya zeae MS1 WP 019843899 1 Dzeae
Dickeya paradisiaca Ech703 WP 015854148 1 Dparadisiaca
Dickeya paradisiaca NCPPB 2511 WP 015854148 1 Dparadisiaca
Klebsiella pneumoniae YP 007989645 1 573 Proteobacteria
Dickeya aquatica DW 0440 WP 035339637 1 Daquatica
Providencia stuartii MRSN 2154 YP 006217036 1 1157951 Proteobacteria
Enterobacter cloacae subsp cloacae ENHKU01 YP 006577041 1 1211025 Proteobacteria
Photorhabdus asymbiotica YP 003042803 1 291112 Proteobacteria
Prodigiosinella confusarubida ATCC39006 WP 021014223 1 Pconfusarubida
Klebsiella oxytoca E718 YP 006496338 1 1191061 Proteobacteria
Shigella boydii Sb227 YP 410419 1 300268 Proteobacteria
Salmonella enterica subsp arizonae serovar 62 z4 z23 str RSK2980 YP 001572064 1 882884 Proteobacteria
Enterobacter sp 638 YP 001175202 1 399742 Proteobacteria
Escherichia coli O7 K1 str CE10 YP 006146968 1 1072459 Proteobacteria
Escherichia coli str K 12 substr W3110 YP 492408 1 316407 Proteobacteria
Escherichia coli B str REL606 YP 003047305 1 413997 Proteobacteria
Erwinia amylovora ATCC 49946 YP 003540101 1 716540 Proteobacteria
Enterobacteriaceae bacterium strain FGI 57 YP 007342049 1 693444 Proteobacteria
Citrobacter rodentium ICC168 YP 003366845 1 637910 Proteobacteria
Escherichia coli O103 H2 str 12009 YP 003224874 1 585395 Proteobacteria
Enterobacter cloacae EcWSU1 YP 004950346 1 1045856 Proteobacteria
Escherichia coli O111 H str 11128 YP 003236059 1 585396 Proteobacteria
Escherichia coli 042 YP 006097466 1 216592 Proteobacteria
Escherichia coli E24377A YP 001464322 1 331111 Proteobacteria
Escherichia coli O104 H4 str 2009EL 2050 YP 006768605 1 1134782 Proteobacteria
Escherichia coli 042 YP 006098876 1 216592 Proteobacteria
Escherichia coli O7 K1 str CE10 YP 006146828 1 1072459 Proteobacteria
Shigella sonnei 53G YP 005458152 1 216599 Proteobacteria
Escherichia coli O104 H4 str 2009EL 2050 YP 006772189 1 1134782 Proteobacteria
Shigella flexneri 2a str 2457T NP 838459 1 198215 Proteobacteria
Shigella flexneri 2002017 YP 005728584 1 591020 Proteobacteria
Escherichia coli O26 H11 str 11368 YP 003230994 1 573235 Proteobacteria
Citrobacter rodentium ICC168 YP 003368377 1 637910 Proteobacteria
Escherichia coli ATCC 8739 YP 001723746 1 481805 Proteobacteria
Escherichia coli O7 K1 str CE10 YP 006145430 1 1072459 Proteobacteria
Escherichia coli ATCC 8739 YP 001726809 1 481805 Proteobacteria
Escherichia coli Xuzhou21 YP 006314460 1 741093 Proteobacteria
Escherichia coli O55 H7 str CB9615 YP 003501106 1 701177 Proteobacteria
Escherichia coli O157 H7 str EDL933 NP 289539 1 155864 Proteobacteria
Escherichia coli O127 H6 str E2348 69 YP 002330696 1 574521 Proteobacteria
Escherichia coli O111 H str 11128 YP 003237359 1 585396 Proteobacteria
Escherichia coli O103 H2 str 12009 YP 003223418 1 585395 Proteobacteria
Escherichia coli O103 H2 str 12009 YP 003224745 1 585395 Proteobacteria
Escherichia coli O26 H11 str 11368 YP 003232178 1 573235 Proteobacteria

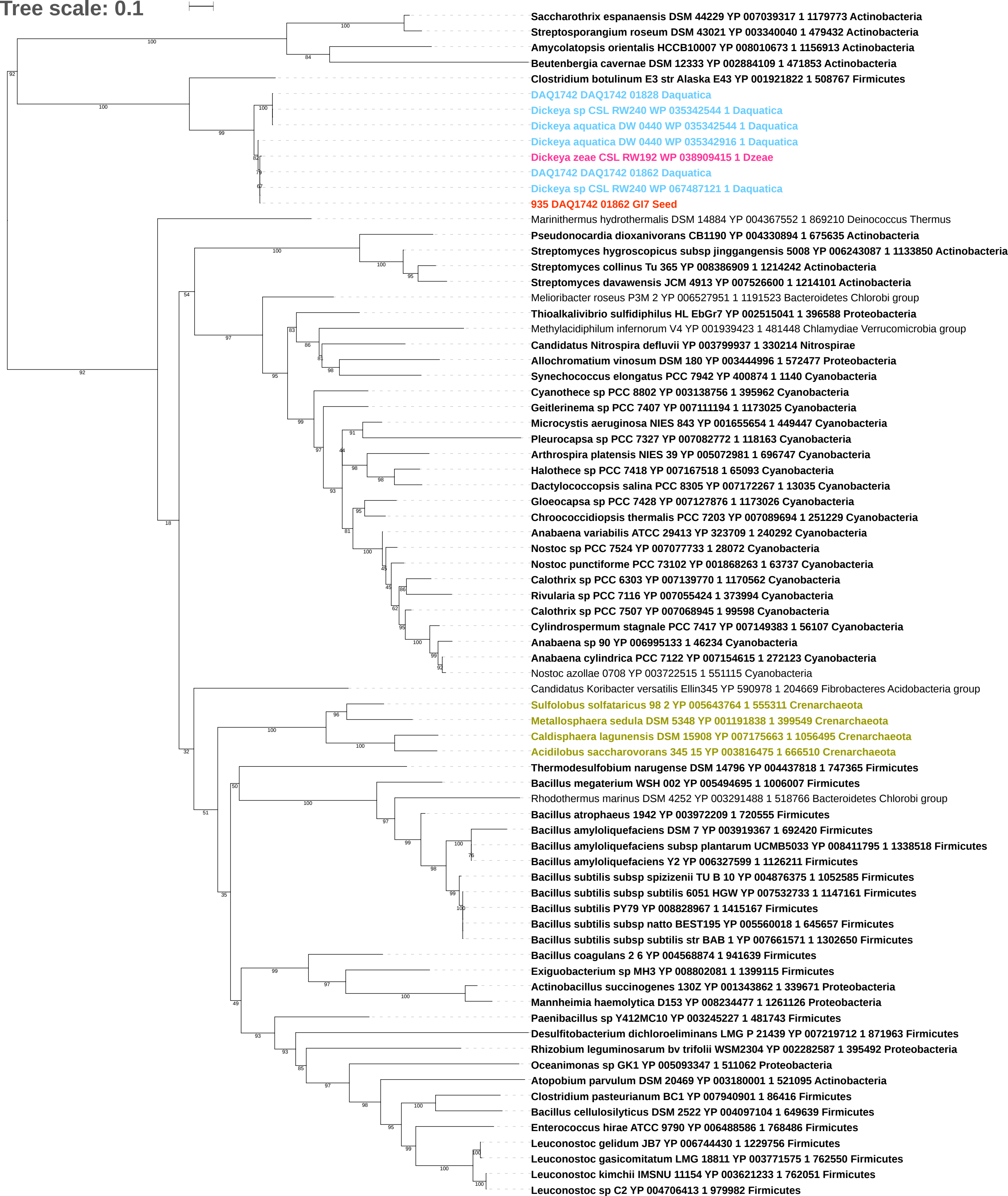

Tree scale: 0.1
Saccharothrix espanaensis DSM 44229 YP 007039317 1 1179773 Actinobacteria
Streptosporangium roseum DSM 43021 YP 003340040 1 479432 Actinobacteria
Amycolatopsis orientalis HCCB10007 YP 008010673 1 1156913 Actinobacteria
Beutenbergia cavernae DSM 12333 YP 002884109 1 471853 Actinobacteria
Clostridium botulinum E3 str Alaska E43 YP 001921822 1 508767 Firmicutes
DAQ1742 DAQ1742 01828 Daquatica
Dickeya sp CSL RW240 WP 035342544 1 Daquatica
Dickeya aquatica DW 0440 WP 035342544 1 Daquatica
Dickeya aquatica DW 0440 WP 035342916 1 Daquatica
Dickeya zeae CSL RW192 WP 038909415 1 Dzeae
DAQ1742 DAQ1742 01862 Daquatica
Dickeya sp CSL RW240 WP 067487121 1 Daquatica
935 DAQ1742 01862 GI7 Seed
Marinithermus hydrothermalis DSM 14884 YP 004367552 1 869210 Deinococcus Thermus
Pseudonocardia dioxanivorans CB1190 YP 004330894 1 675635 Actinobacteria
Streptomyces hygroscopicus subsp jinggangensis 5008 YP 006243087 1 1133850 Actinobacteria
Streptomyces collinus Tu 365 YP 008386909 1 1214242 Actinobacteria
Streptomyces davawensis JCM 4913 YP 007526600 1 1214101 Actinobacteria
Melioribacter roseus P3M 2 YP 006527951 1 1191523 Bacteroidetes Chlorobi group
Thioalkalivibrio sulfidiphilus HL EbGr7 YP 002515041 1 396588 Proteobacteria
Methylacidiphilum infernorum V4 YP 001939423 1 481448 Chlamydiae Verrucomicrobia group
Candidatus Nitrospira defluvii YP 003799937 1 330214 Nitrospirae
Allochromatium vinosum DSM 180 YP 003444996 1 572477 Proteobacteria
Synechococcus elongatus PCC 7942 YP 400874 1 1140 Cyanobacteria
Cyanothece sp PCC 8802 YP 003138756 1 395962 Cyanobacteria
Geitlerinema sp PCC 7407 YP 007111194 1 1173025 Cyanobacteria
Microcystis aeruginosa NIES 843 YP 001655654 1 449447 Cyanobacteria
Pleurocapsa sp PCC 7327 YP 007082772 1 118163 Cyanobacteria
Arthrospira platensis NIES 39 YP 005072981 1 696747 Cyanobacteria
Halothece sp PCC 7418 YP 007167518 1 65093 Cyanobacteria
Dactylococcopsis salina PCC 8305 YP 007172267 1 13035 Cyanobacteria
Gloeocapsa sp PCC 7428 YP 007127876 1 1173026 Cyanobacteria
Chroococcidiopsis thermalis PCC 7203 YP 007089694 1 251229 Cyanobacteria
Anabaena variabilis ATCC 29413 YP 323709 1 240292 Cyanobacteria
Nostoc sp PCC 7524 YP 007077733 1 28072 Cyanobacteria
Nostoc punctiforme PCC 73102 YP 001868263 1 63737 Cyanobacteria
Calothrix sp PCC 6303 YP 007139770 1 1170562 Cyanobacteria
Rivularia sp PCC 7116 YP 007055424 1 373994 Cyanobacteria
Calothrix sp PCC 7507 YP 007068945 1 99598 Cyanobacteria
Cylindrospermum stagnale PCC 7417 YP 007149383 1 56107 Cyanobacteria
Anabaena sp 90 YP 006995133 1 46234 Cyanobacteria
Anabaena cylindrica PCC 7122 YP 007154615 1 272123 Cyanobacteria
Nostoc azollae 0708 YP 003722515 1 551115 Cyanobacteria
Candidatus Koribacter versatilis Ellin345 YP 590978 1 204669 Fibrobacteres Acidobacteria group
Sulfolobus solfataricus 98 2 YP 005643764 1 555311 Crenarchaeota
Metallosphaera sedula DSM 5348 YP 001191838 1 399549 Crenarchaeota
Caldisphaera lagunensis DSM 15908 YP 007175663 1 1056495 Crenarchaeota
Acidilobus saccharovorans 345 15 YP 003816475 1 666510 Crenarchaeota
Thermodesulfobium narugense DSM 14796 YP 004437818 1 747365 Firmicutes
Bacillus megaterium WSH 002 YP 005494695 1 1006007 Firmicutes
Rhodothermus marinus DSM 4252 YP 003291488 1 518766 Bacteroidetes Chlorobi group
Bacillus atrophaeus 1942 YP 003972209 1 720555 Firmicutes
Bacillus amyloliquefaciens DSM 7 YP 003919367 1 692420 Firmicutes
Bacillus amyloliquefaciens subsp plantarum UCMB5033 YP 008411795 1 1338518 Firmicutes
Bacillus amyloliquefaciens Y2 YP 006327599 1 1126211 Firmicutes
Bacillus subtilis subsp spizizenii TU B 10 YP 004876375 1 1052585 Firmicutes
Bacillus subtilis subsp subtilis 6051 HGW YP 007532733 1 1147161 Firmicutes
Bacillus subtilis PY79 YP 008828967 1 1415167 Firmicutes
Bacillus subtilis subsp natto BEST195 YP 005560018 1 645657 Firmicutes
Bacillus subtilis subsp subtilis str BAB 1 YP 007661571 1 1302650 Firmicutes
Bacillus coagulans 2 6 YP 004568874 1 941639 Firmicutes
Exiguobacterium sp MH3 YP 008802081 1 1399115 Firmicutes
Actinobacillus succinogenes 130Z YP 001343862 1 339671 Proteobacteria
Mannheimia haemolytica D153 YP 008234477 1 1261126 Proteobacteria
Paenibacillus sp Y412MC10 YP 003245227 1 481743 Firmicutes
Desulfitobacterium dichloroeliminans LMG P 21439 YP 007219712 1 871963 Firmicutes
Rhizobium leguminosarum bv trifolii WSM2304 YP 002282587 1 395492 Proteobacteria
Oceanimonas sp GK1 YP 005093347 1 511062 Proteobacteria
Atopobium parvulum DSM 20469 YP 003180001 1 521095 Actinobacteria
Clostridium pasteurianum BC1 YP 007940901 1 86416 Firmicutes
Bacillus cellulosilyticus DSM 2522 YP 004097104 1 649639 Firmicutes
Enterococcus hirae ATCC 9790 YP 006488586 1 768486 Firmicutes
Leuconostoc gelidum JB7 YP 006744430 1 1229756 Firmicutes
Leuconostoc gasicomitatum LMG 18811 YP 003771575 1 762550 Firmicutes
Leuconostoc kimchii IMSNU 11154 YP 003621233 1 762051 Firmicutes
Leuconostoc sp C2 YP 004706413 1 979982 Firmicutes

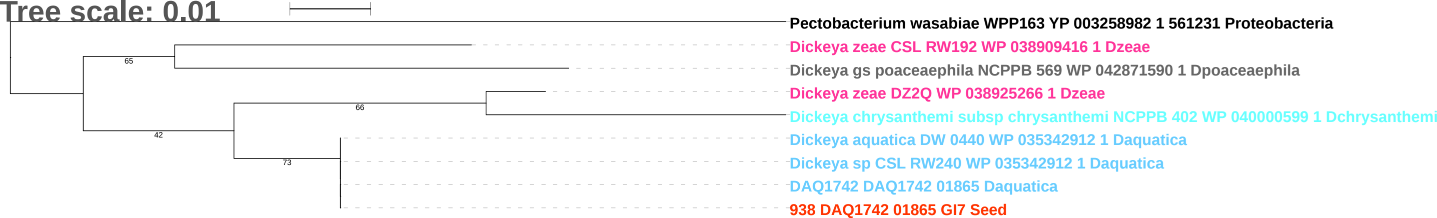

Tree scale: 0.01
Pectobacterium wasabiae WPP163 YP 003258982 1 561231 Proteobacteria
Dickeya zeae CSL RW192 WP 038909416 1 Dzeae
Dickeya gs poaceaephila NCPPB 569 WP 042871590 1 Dpoaceaephila
Dickeya zeae DZ2Q WP 038925266 1 Dzeae
Dickeya chrysanthemi subsp chrysanthemi NCPPB 402 WP 040000599 1 Dchrysanthemi
Dickeya aquatica DW 0440 WP 035342912 1 Daquatica
Dickeya sp CSL RW240 WP 035342912 1 Daquatica
DAQ1742 DAQ1742 01865 Daquatica
938 DAQ1742 01865 GI7 Seed
65
66
42
73

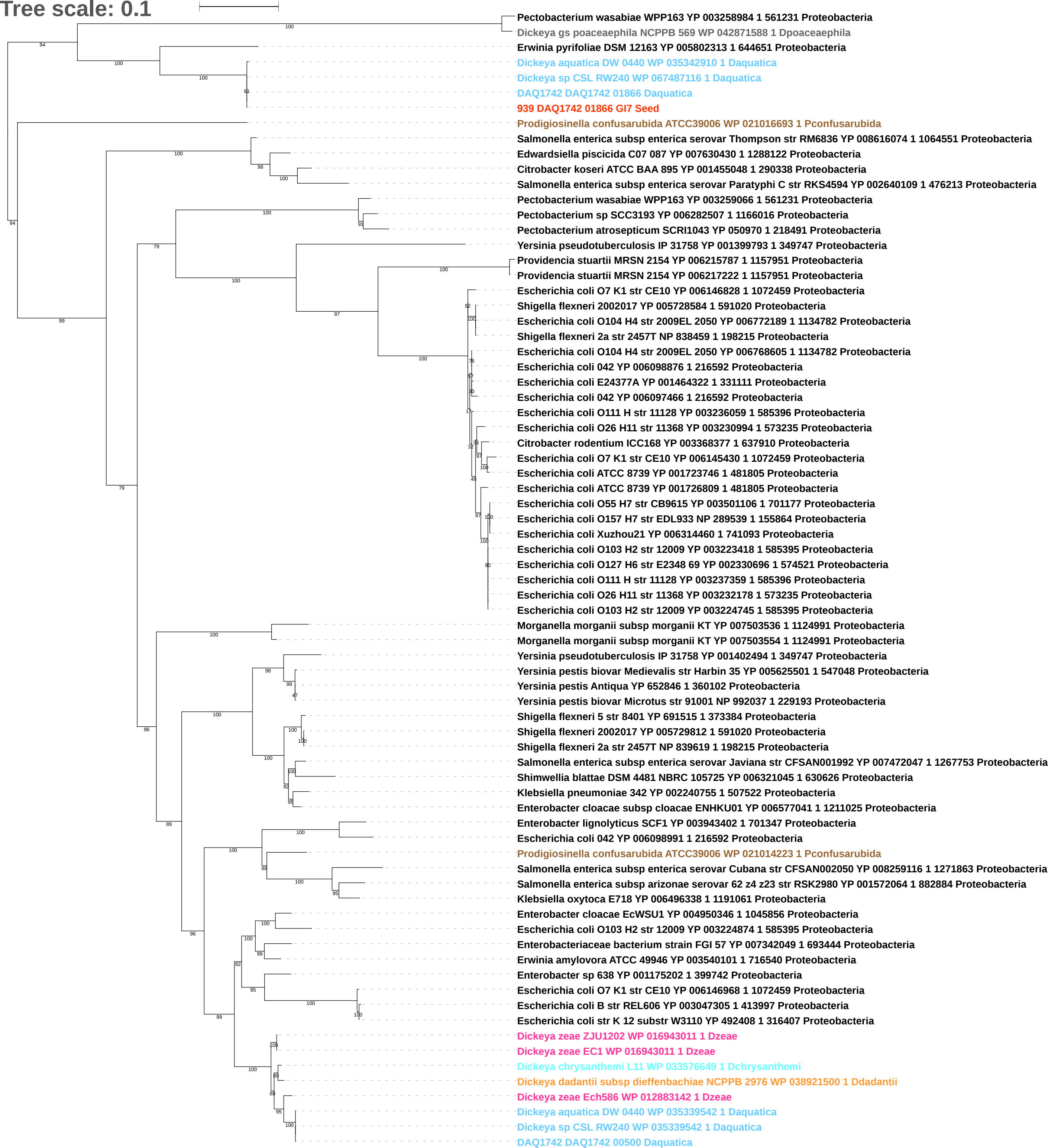
Tree scale: 0.1
Pectobacterium wasabiae WPP163 YP 003258984 1 561231 Proteobacteria
Dickeya gs poaceaephila NCPPB 569 WP 042871588 1 Dpoaceaephila
Erwinia pyrifoliae DSM 12163 YP 005802313 1 644651 Proteobacteria
Dickeya aquatica DW 0440 WP 035342910 1 Daquatica
Dickeya sp CSL RW240 WP 067487116 1 Daquatica
DAQ1742 DAQ1742 01866 Daquatica
939 DAQ1742 01866 GI7 Seed
Prodigiosinella confusarubida ATCC39006 WP 021016693 1 Pconfusarubida
Salmonella enterica subsp enterica serovar Thompson str RM6836 YP 008616074 1 1064551 Proteobacteria
Edwardsiella piscicida C07 087 YP 007630430 1 1288122 Proteobacteria
Citrobacter koseri ATCC BAA 895 YP 001455048 1 290338 Proteobacteria
Salmonella enterica subsp enterica serovar Paratyphi C str RKS4594 YP 002640109 1 476213 Proteobacteria
Pectobacterium wasabiae WPP163 YP 003259066 1 561231 Proteobacteria
Pectobacterium sp SCC3193 YP 006282507 1 1166016 Proteobacteria
Pectobacterium atrosepticum SCRI1043 YP 050970 1 218491 Proteobacteria
Yersinia pseudotuberculosis IP 31758 YP 001399793 1 349747 Proteobacteria
Providencia stuartii MRSN 2154 YP 006215787 1 1157951 Proteobacteria
Providencia stuartii MRSN 2154 YP 006217222 1 1157951 Proteobacteria
Escherichia coli O7 K1 str CE10 YP 006146828 1 1072459 Proteobacteria
Shigella flexneri 2002017 YP 005728584 1 591020 Proteobacteria
Escherichia coli O104 H4 str 2009EL 2050 YP 006772189 1 1134782 Proteobacteria
Shigella flexneri 2a str 2457T NP 838459 1 198215 Proteobacteria
Escherichia coli O104 H4 str 2009EL 2050 YP 006768605 1 1134782 Proteobacteria
Escherichia coli 042 YP 006098876 1 216592 Proteobacteria
Escherichia coli E24377A YP 001464322 1 331111 Proteobacteria
Escherichia coli 042 YP 006097466 1 216592 Proteobacteria
Escherichia coli O111 H str 11128 YP 003236059 1 585396 Proteobacteria
Escherichia coli O26 H11 str 11368 YP 003230994 1 573235 Proteobacteria
Citrobacter rodentium ICC168 YP 003368377 1 637910 Proteobacteria
Escherichia coli O7 K1 str CE10 YP 006145430 1 1072459 Proteobacteria
Escherichia coli ATCC 8739 YP 001723746 1 481805 Proteobacteria
Escherichia coli ATCC 8739 YP 001726809 1 481805 Proteobacteria
Escherichia coli O55 H7 str CB9615 YP 003501106 1 701177 Proteobacteria
Escherichia coli O157 H7 str EDL933 NP 289539 1 155864 Proteobacteria
Escherichia coli Xuzhou21 YP 006314460 1 741093 Proteobacteria
Escherichia coli O103 H2 str 12009 YP 003223418 1 585395 Proteobacteria
Escherichia coli O127 H6 str E2348 69 YP 002330696 1 574521 Proteobacteria
Escherichia coli O111 H str 11128 YP 003237359 1 585396 Proteobacteria
Escherichia coli O26 H11 str 11368 YP 003232178 1 573235 Proteobacteria
Escherichia coli O103 H2 str 12009 YP 003224745 1 585395 Proteobacteria
Morganella morganii subsp morganii KT YP 007503536 1 1124991 Proteobacteria
Morganella morganii subsp morganii KT YP 007503554 1 1124991 Proteobacteria
Yersinia pseudotuberculosis IP 31758 YP 001402494 1 349747 Proteobacteria
Yersinia pestis biovar Medievalis str Harbin 35 YP 005625501 1 547048 Proteobacteria
Yersinia pestis Antiqua YP 652846 1 360102 Proteobacteria
Yersinia pestis biovar Microtus str 91001 NP 992037 1 229193 Proteobacteria
Shigella flexneri 5 str 8401 YP 691515 1 373384 Proteobacteria
Shigella flexneri 2002017 YP 005729812 1 591020 Proteobacteria
Shigella flexneri 2a str 2457T NP 839619 1 198215 Proteobacteria
Salmonella enterica subsp enterica serovar Javiana str CFSAN001992 YP 007472047 1 1267753 Proteobacteria
Shimwellia blattae DSM 4481 NBRC 105725 YP 006321045 1 630626 Proteobacteria
Klebsiella pneumoniae 342 YP 002240755 1 507522 Proteobacteria
Enterobacter cloacae subsp cloacae ENHKU01 YP 006577041 1 1211025 Proteobacteria
Enterobacter lignolyticus SCF1 YP 003943402 1 701347 Proteobacteria
Escherichia coli 042 YP 006098991 1 216592 Proteobacteria
Prodigiosinella confusarubida ATCC39006 WP 021014223 1 Pconfusarubida
Salmonella enterica subsp enterica serovar Cubana str CFSAN002050 YP 008259116 1 1271863 Proteobacteria
Salmonella enterica subsp arizonae serovar 62 z4 z23 str RSK2980 YP 001572064 1 882884 Proteobacteria
Klebsiella oxytoca E718 YP 006496338 1 1191061 Proteobacteria
Enterobacter cloacae EcWSU1 YP 004950346 1 1045856 Proteobacteria
Escherichia coli O103 H2 str 12009 YP 003224874 1 585395 Proteobacteria
Enterobacteriaceae bacterium strain FGI 57 YP 007342049 1 693444 Proteobacteria
Erwinia amylovora ATCC 49946 YP 003540101 1 716540 Proteobacteria
Enterobacter sp 638 YP 001175202 1 399742 Proteobacteria
Escherichia coli O7 K1 str CE10 YP 006146968 1 1072459 Proteobacteria
Escherichia coli B str REL606 YP 003047305 1 413997 Proteobacteria
Escherichia coli str K 12 substr W3110 YP 492408 1 316407 Proteobacteria
Dickeya zeae ZJU1202 WP 016943011 1 Dzeae
Dickeya zeae EC1 WP 016943011 1 Dzeae
Dickeya chrysanthemi L11 WP 033576649 1 Dchrysanthemi
Dickeya dadantii subsp dieffenbachiae NCPPB 2976 WP 038921500 1 Ddadantii
Dickeya zeae Ech586 WP 012883142 1 Dzeae
Dickeya aquatica DW 0440 WP 035339542 1 Daquatica
Dickeya sp CSL RW240 WP 035339542 1 Daquatica
DAQ1742 DAQ1742 00500 Daquatica

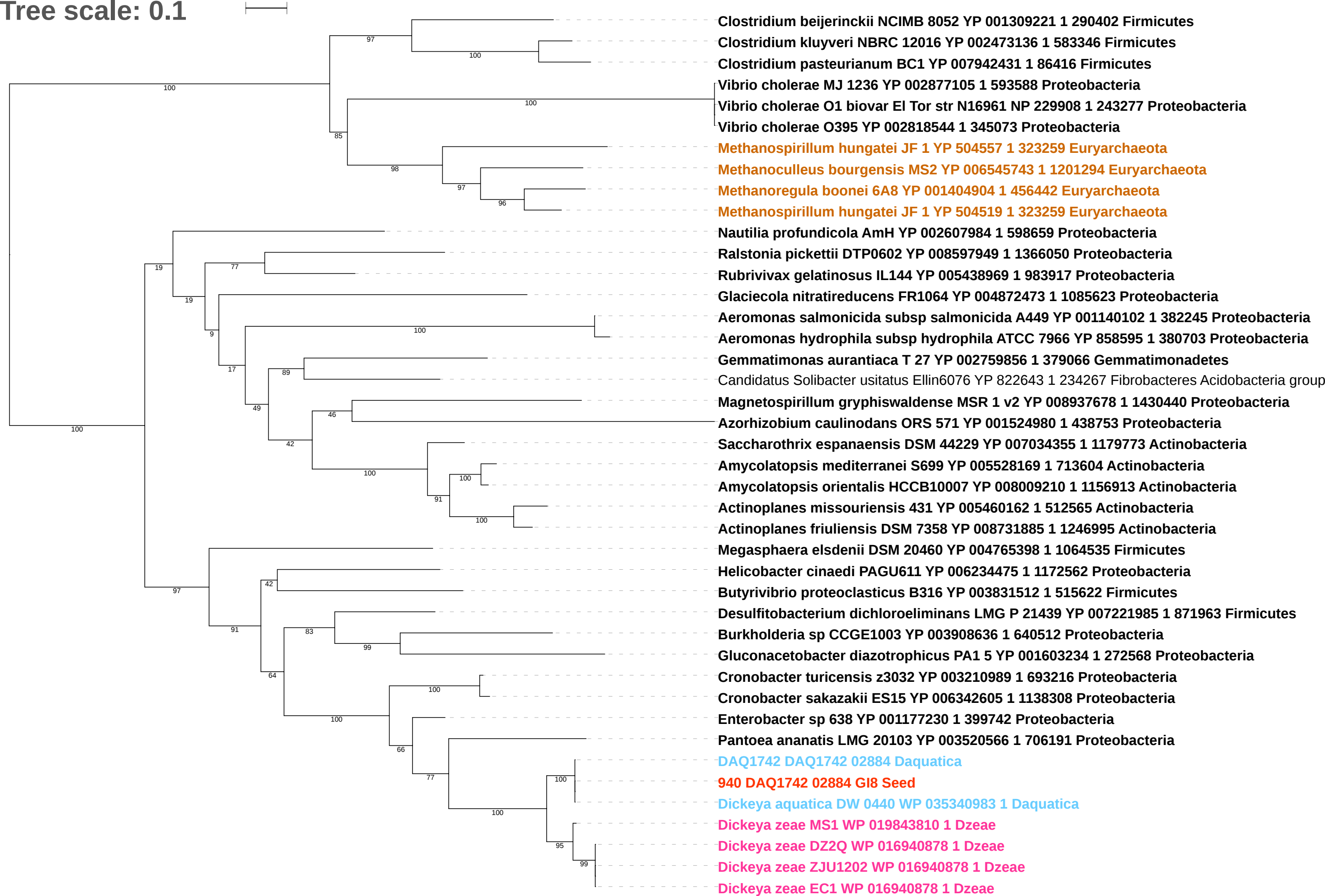
Tree scale: 0.1
Clostridium beijerinckii NCIMB 8052 YP 001309221 1 290402 Firmicutes
Clostridium kluyveri NBRC 12016 YP 002473136 1 583346 Firmicutes
Clostridium pasteurianum BC1 YP 007942431 1 86416 Firmicutes
Vibrio cholerae MJ 1236 YP 002877105 1 593588 Proteobacteria
Vibrio cholerae O1 biovar El Tor str N16961 NP 229908 1 243277 Proteobacteria
Vibrio cholerae O395 YP 002818544 1 345073 Proteobacteria
Methanospirillum hungatei JF 1 YP 504557 1 323259 Euryarchaeota
Methanoculleus bourgensis MS2 YP 006545743 1 1201294 Euryarchaeota
Methanoregula boonei 6A8 YP 001404904 1 456442 Euryarchaeota
Methanospirillum hungatei JF 1 YP 504519 1 323259 Euryarchaeota
Nautilia profundicola AmH YP 002607984 1 598659 Proteobacteria
Ralstonia pickettii DTP0602 YP 008597949 1 1366050 Proteobacteria
Rubrivivax gelatinosus IL144 YP 005438969 1 983917 Proteobacteria
Glaciecola nitratireducens FR1064 YP 004872473 1 1085623 Proteobacteria
Aeromonas salmonicida subsp salmonicida A449 YP 001140102 1 382245 Proteobacteria
Aeromonas hydrophila subsp hydrophila ATCC 7966 YP 858595 1 380703 Proteobacteria
Gemmatimonas aurantiaca T 27 YP 002759856 1 379066 Gemmatimonadetes
Candidatus Solibacter usitatus Ellin6076 YP 822643 1 234267 Fibrobacteres Acidobacteria group
Magnetospirillum gryphiswaldense MSR 1 v2 YP 008937678 1 1430440 Proteobacteria
Azorhizobium caulinodans ORS 571 YP 001524980 1 438753 Proteobacteria
Saccharothrix espanaensis DSM 44229 YP 007034355 1 1179773 Actinobacteria
Amycolatopsis mediterranei S699 YP 005528169 1 713604 Actinobacteria
Amycolatopsis orientalis HCCB10007 YP 008009210 1 1156913 Actinobacteria
Actinoplanes missouriensis 431 YP 005460162 1 512565 Actinobacteria
Actinoplanes friuliensis DSM 7358 YP 008731885 1 1246995 Actinobacteria
Megasphaera elsdenii DSM 20460 YP 004765398 1 1064535 Firmicutes
Helicobacter cinaedi PAGU611 YP 006234475 1 1172562 Proteobacteria
Butyrivibrio proteoclasticus B316 YP 003831512 1 515622 Firmicutes
Desulfitobacterium dichloroeliminans LMG P 21439 YP 007221985 1 871963 Firmicutes
Burkholderia sp CCGE1003 YP 003908636 1 640512 Proteobacteria
Gluconacetobacter diazotrophicus PA1 5 YP 001603234 1 272568 Proteobacteria
Cronobacter turicensis z3032 YP 003210989 1 693216 Proteobacteria
Cronobacter sakazakii ES15 YP 006342605 1 1138308 Proteobacteria
Enterobacter sp 638 YP 001177230 1 399742 Proteobacteria
Pantoea ananatis LMG 20103 YP 003520566 1 706191 Proteobacteria
DAQ1742 DAQ1742 02884 Daquatica
940 DAQ1742 02884 GI8 Seed
Dickeya aquatica DW 0440 WP 035340983 1 Daquatica
Dickeya zeae MS1 WP 019843810 1 Dzeae
Dickeya zeae DZ2Q WP 016940878 1 Dzeae
Dickeya zeae ZJU1202 WP 016940878 1 Dzeae
Dickeya zeae EC1 WP 016940878 1 Dzeae

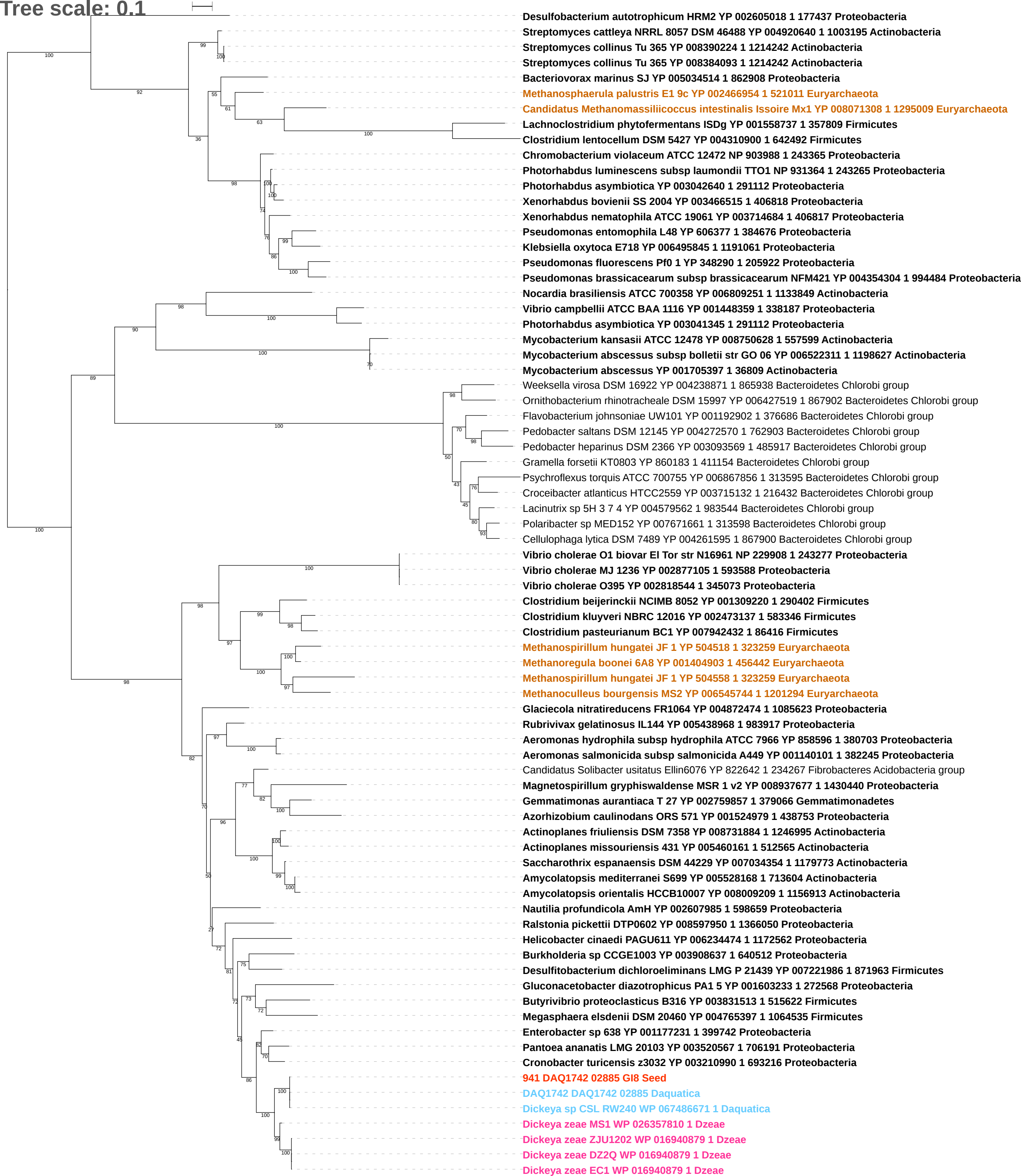
Tree scale: 0.1
Desulfobacterium autotrophicum HRM2 YP 002605018 1 177437 Proteobacteria
Streptomyces cattleya NRRL 8057 DSM 46488 YP 004920640 1 1003195 Actinobacteria
Streptomyces collinus Tu 365 YP 008390224 1 1214242 Actinobacteria
Streptomyces collinus Tu 365 YP 008384093 1 1214242 Actinobacteria
Bacteriovorax marinus SJ YP 005034514 1 862908 Proteobacteria
Methanosphaerula palustris E1 9c YP 002466954 1 521011 Euryarchaeota
Candidatus Methanomassiliicoccus intestinalis Issoire Mx1 YP 008071308 1 1295009 Euryarchaeota
Lachnoclostridium phytofermentans ISDg YP 001558737 1 357809 Firmicutes
Clostridium lentocellum DSM 5427 YP 004310900 1 642492 Firmicutes
Chromobacterium violaceum ATCC 12472 NP 903988 1 243365 Proteobacteria
Photorhabdus luminescens subsp laumondii TTO1 NP 931364 1 243265 Proteobacteria
Photorhabdus asymbiotica YP 003042640 1 291112 Proteobacteria
Xenorhabdus bovienii SS 2004 YP 003466515 1 406818 Proteobacteria
Xenorhabdus nematophila ATCC 19061 YP 003714684 1 406817 Proteobacteria
Pseudomonas entomophila L48 YP 606377 1 384676 Proteobacteria
Klebsiella oxytoca E718 YP 006495845 1 1191061 Proteobacteria
Pseudomonas fluorescens Pf0 1 YP 348290 1 205922 Proteobacteria
Pseudomonas brassicacearum subsp brassicacearum NFM421 YP 004354304 1 994484 Proteobacteria
Nocardia brasiliensis ATCC 700358 YP 006809251 1 1133849 Actinobacteria
Vibrio campbellii ATCC BAA 1116 YP 001448359 1 338187 Proteobacteria
Photorhabdus asymbiotica YP 003041345 1 291112 Proteobacteria
Mycobacterium kansasii ATCC 12478 YP 008750628 1 557599 Actinobacteria
Mycobacterium abscessus subsp bolletii str GO 06 YP 006522311 1 1198627 Actinobacteria
Mycobacterium abscessus YP 001705397 1 36809 Actinobacteria
Weeksella virosa DSM 16922 YP 004238871 1 865938 Bacteroidetes Chlorobi group
Ornithobacterium rhinotracheale DSM 15997 YP 006427519 1 867902 Bacteroidetes Chlorobi group
Flavobacterium johnsoniae UW101 YP 001192902 1 376686 Bacteroidetes Chlorobi group
Pedobacter saltans DSM 12145 YP 004272570 1 762903 Bacteroidetes Chlorobi group
Pedobacter heparinus DSM 2366 YP 003093569 1 485917 Bacteroidetes Chlorobi group
Gramella forsetii KT0803 YP 860183 1 411154 Bacteroidetes Chlorobi group
Psychroflexus torquis ATCC 700755 YP 006867856 1 313595 Bacteroidetes Chlorobi group
Croceibacter atlanticus HTCC2559 YP 003715132 1 216432 Bacteroidetes Chlorobi group
Lacinutrix sp 5H 3 7 4 YP 004579562 1 983544 Bacteroidetes Chlorobi group
Polaribacter sp MED152 YP 007671661 1 313598 Bacteroidetes Chlorobi group
Cellulophaga lytica DSM 7489 YP 004261595 1 867900 Bacteroidetes Chlorobi group
Vibrio cholerae O1 biovar El Tor str N16961 NP 229908 1 243277 Proteobacteria
Vibrio cholerae MJ 1236 YP 002877105 1 593588 Proteobacteria
Vibrio cholerae O395 YP 002818544 1 345073 Proteobacteria
Clostridium beijerinckii NCIMB 8052 YP 001309220 1 290402 Firmicutes
Clostridium kluyveri NBRC 12016 YP 002473137 1 583346 Firmicutes
Clostridium pasteurianum BC1 YP 007942432 1 86416 Firmicutes
Methanospirillum hungatei JF 1 YP 504518 1 323259 Euryarchaeota
Methanoregula boonei 6A8 YP 001404903 1 456442 Euryarchaeota
Methanospirillum hungatei JF 1 YP 504558 1 323259 Euryarchaeota
Methanoculleus bourgensis MS2 YP 006545744 1 1201294 Euryarchaeota
Glaciecola nitratireducens FR1064 YP 004872474 1 1085623 Proteobacteria
Rubrivivax gelatinosus IL144 YP 005438968 1 983917 Proteobacteria
Aeromonas hydrophila subsp hydrophila ATCC 7966 YP 858596 1 380703 Proteobacteria
Aeromonas salmonicida subsp salmonicida A449 YP 001140101 1 382245 Proteobacteria
Candidatus Solibacter usitatus Ellin6076 YP 822642 1 234267 Fibrobacteres Acidobacteria group
Magnetospirillum gryphiswaldense MSR 1 v2 YP 008937677 1 1430440 Proteobacteria
Gemmatimonas aurantiaca T 27 YP 002759857 1 379066 Gemmatimonadetes
Azorhizobium caulinodans ORS 571 YP 001524979 1 438753 Proteobacteria
Actinoplanes friuliensis DSM 7358 YP 008731884 1 1246995 Actinobacteria
Actinoplanes missouriensis 431 YP 005460161 1 512565 Actinobacteria
Saccharothrix espanaensis DSM 44229 YP 007034354 1 1179773 Actinobacteria
Amycolatopsis mediterranei S699 YP 005528168 1 713604 Actinobacteria
Amycolatopsis orientalis HCCB10007 YP 008009209 1 1156913 Actinobacteria
Nautilia profundicola AmH YP 002607985 1 598659 Proteobacteria
Ralstonia pickettii DTP0602 YP 008597950 1 1366050 Proteobacteria
Helicobacter cinaedi PAGU611 YP 006234474 1 1172562 Proteobacteria
Burkholderia sp CCGE1003 YP 003908637 1 640512 Proteobacteria
Desulfitobacterium dichloroeliminans LMG P 21439 YP 007221986 1 871963 Firmicutes
Gluconacetobacter diazotrophicus PA1 5 YP 001603233 1 272568 Proteobacteria
Butyrivibrio proteoclasticus B316 YP 003831513 1 515622 Firmicutes
Megasphaera elsdenii DSM 20460 YP 004765397 1 1064535 Firmicutes
Enterobacter sp 638 YP 001177231 1 399742 Proteobacteria
Pantoea ananatis LMG 20103 YP 003520567 1 706191 Proteobacteria
Cronobacter turicensis z3032 YP 003210990 1 693216 Proteobacteria
941 DAQ1742 02885 GI8 Seed
DAQ1742 DAQ1742 02885 Daquatica
Dickeya sp CSL RW240 WP 067486671 1 Daquatica
Dickeya zeae MS1 WP 026357810 1 Dzeae
Dickeya zeae ZJU1202 WP 016940879 1 Dzeae
Dickeya zeae DZ2Q WP 016940879 1 Dzeae
Dickeya zeae EC1 WP 016940879 1 Dzeae

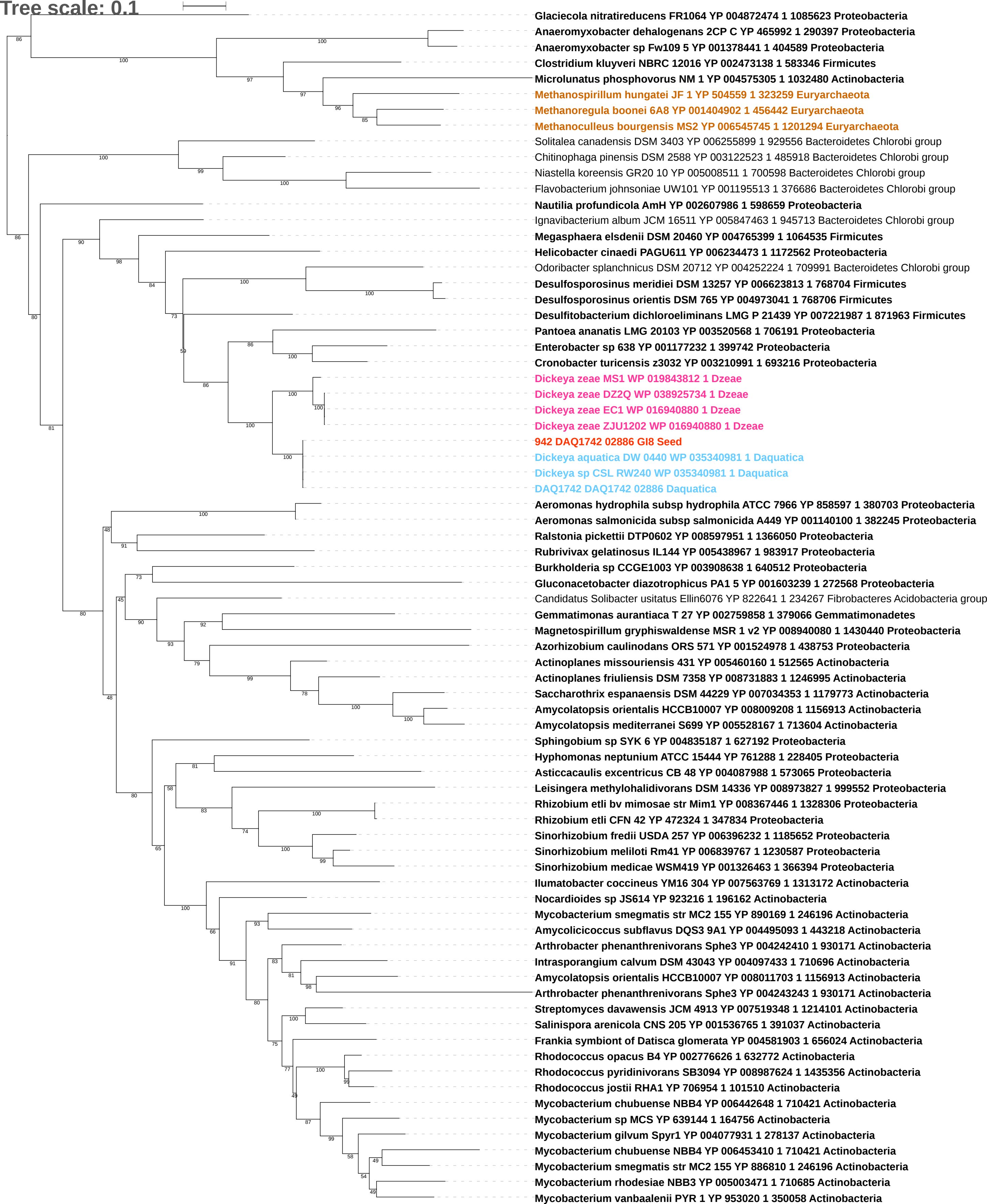
Tree scale: 0.1
Glaciecola nitratireducens FR1064 YP 004872474 1 1085623 Proteobacteria
Anaeromyxobacter dehalogenans 2CP C YP 465992 1 290397 Proteobacteria
Anaeromyxobacter sp Fw109 5 YP 001378441 1 404589 Proteobacteria
Clostridium kluyveri NBRC 12016 YP 002473138 1 583346 Firmicutes
Microlunatus phosphovorus NM 1 YP 004575305 1 1032480 Actinobacteria
Methanospirillum hungatei JF 1 YP 504559 1 323259 Euryarchaeota
Methanoregula boonei 6A8 YP 001404902 1 456442 Euryarchaeota
Methanoculleus bourgensis MS2 YP 006545745 1 1201294 Euryarchaeota
Solitalea canadensis DSM 3403 YP 006255899 1 929556 Bacteroidetes Chlorobi group
Chitinophaga pinensis DSM 2588 YP 003122523 1 485918 Bacteroidetes Chlorobi group
Niastella koreensis GR20 10 YP 005008511 1 700598 Bacteroidetes Chlorobi group
Flavobacterium johnsoniae UW101 YP 001195513 1 376686 Bacteroidetes Chlorobi group
Nautilia profundicola AmH YP 002607986 1 598659 Proteobacteria
Ignavibacterium album JCM 16511 YP 005847463 1 945713 Bacteroidetes Chlorobi group
Megasphaera elsdenii DSM 20460 YP 004765399 1 1064535 Firmicutes
Helicobacter cinaedi PAGU611 YP 006234473 1 1172562 Proteobacteria
Odoribacter splanchnicus DSM 20712 YP 004252224 1 709991 Bacteroidetes Chlorobi group
Desulfosporosinus meridiei DSM 13257 YP 006623813 1 768704 Firmicutes
Desulfosporosinus orientis DSM 765 YP 004973041 1 768706 Firmicutes
Desulfitobacterium dichloroeliminans LMG P 21439 YP 007221987 1 871963 Firmicutes
Pantoea ananatis LMG 20103 YP 003520568 1 706191 Proteobacteria
Enterobacter sp 638 YP 001177232 1 399742 Proteobacteria
Cronobacter turicensis z3032 YP 003210991 1 693216 Proteobacteria
Dickeya zeae MS1 WP 019843812 1 Dzeae
Dickeya zeae DZ2Q WP 038925734 1 Dzeae
Dickeya zeae EC1 WP 016940880 1 Dzeae
Dickeya zeae ZJU1202 WP 016940880 1 Dzeae
942 DAQ1742 02886 GI8 Seed
Dickeya aquatica DW 0440 WP 035340981 1 Daquatica
Dickeya sp CSL RW240 WP 035340981 1 Daquatica
DAQ1742 DAQ1742 02886 Daquatica
Aeromonas hydrophila subsp hydrophila ATCC 7966 YP 858597 1 380703 Proteobacteria
Aeromonas salmonicida subsp salmonicida A449 YP 001140100 1 382245 Proteobacteria
Ralstonia pickettii DTP0602 YP 008597951 1 1366050 Proteobacteria
Rubrivivax gelatinosus IL144 YP 005438967 1 983917 Proteobacteria
Burkholderia sp CCGE1003 YP 003908638 1 640512 Proteobacteria
Gluconacetobacter diazotrophicus PA1 5 YP 001603239 1 272568 Proteobacteria
Candidatus Solibacter usitatus Ellin6076 YP 822641 1 234267 Fibrobacteres Acidobacteria group
Gemmatimonas aurantiaca T 27 YP 002759858 1 379066 Gemmatimonadetes
Magnetospirillum gryphiswaldense MSR 1 v2 YP 008940080 1 1430440 Proteobacteria
Azorhizobium caulinodans ORS 571 YP 001524978 1 438753 Proteobacteria
Actinoplanes missouriensis 431 YP 005460160 1 512565 Actinobacteria
Actinoplanes friuliensis DSM 7358 YP 008731883 1 1246995 Actinobacteria
Saccharothrix espanaensis DSM 44229 YP 007034353 1 1179773 Actinobacteria
Amycolatopsis orientalis HCCB10007 YP 008009208 1 1156913 Actinobacteria
Amycolatopsis mediterranei S699 YP 005528167 1 713604 Actinobacteria
Sphingobium sp SYK 6 YP 004835187 1 627192 Proteobacteria
Hyphomonas neptunium ATCC 15444 YP 761288 1 228405 Proteobacteria
Asticcacaulis excentricus CB 48 YP 004087988 1 573065 Proteobacteria
Leisingera methylohalidivorans DSM 14336 YP 008973827 1 999552 Proteobacteria
Rhizobium etli bv mimosae str Mim1 YP 008367446 1 1328306 Proteobacteria
Rhizobium etli CFN 42 YP 472324 1 347834 Proteobacteria
Sinorhizobium fredii USDA 257 YP 006396232 1 1185652 Proteobacteria
Sinorhizobium meliloti Rm41 YP 006839767 1 1230587 Proteobacteria
Sinorhizobium medicae WSM419 YP 001326463 1 366394 Proteobacteria
Ilumatobacter coccineus YM16 304 YP 007563769 1 1313172 Actinobacteria
Nocardioides sp JS614 YP 923216 1 196162 Actinobacteria
Mycobacterium smegmatis str MC2 155 YP 890169 1 246196 Actinobacteria
Amycolicicoccus subflavus DQS3 9A1 YP 004495093 1 443218 Actinobacteria
Arthrobacter phenanthrenivorans Sphe3 YP 004242410 1 930171 Actinobacteria
Intrasporangium calvum DSM 43043 YP 004097433 1 710696 Actinobacteria
Amycolatopsis orientalis HCCB10007 YP 008011703 1 1156913 Actinobacteria
Arthrobacter phenanthrenivorans Sphe3 YP 004243243 1 930171 Actinobacteria
Streptomyces davawensis JCM 4913 YP 007519348 1 1214101 Actinobacteria
Salinispora arenicola CNS 205 YP 001536765 1 391037 Actinobacteria
Frankia symbiont of Datisca glomerata YP 004581903 1 656024 Actinobacteria
Rhodococcus opacus B4 YP 002776626 1 632772 Actinobacteria
Rhodococcus pyridinivorans SB3094 YP 008987624 1 1435356 Actinobacteria
Rhodococcus jostii RHA1 YP 706954 1 101510 Actinobacteria
Mycobacterium chubuense NBB4 YP 006442648 1 710421 Actinobacteria
Mycobacterium sp MCS YP 639144 1 164756 Actinobacteria
Mycobacterium gilvum Spyr1 YP 004077931 1 278137 Actinobacteria
Mycobacterium chubuense NBB4 YP 006453410 1 710421 Actinobacteria
Mycobacterium smegmatis str MC2 155 YP 886810 1 246196 Actinobacteria
Mycobacterium rhodesiae NBB3 YP 005003471 1 710685 Actinobacteria
Mycobacterium vanbaalenii PYR 1 YP 953020 1 350058 Actinobacteria

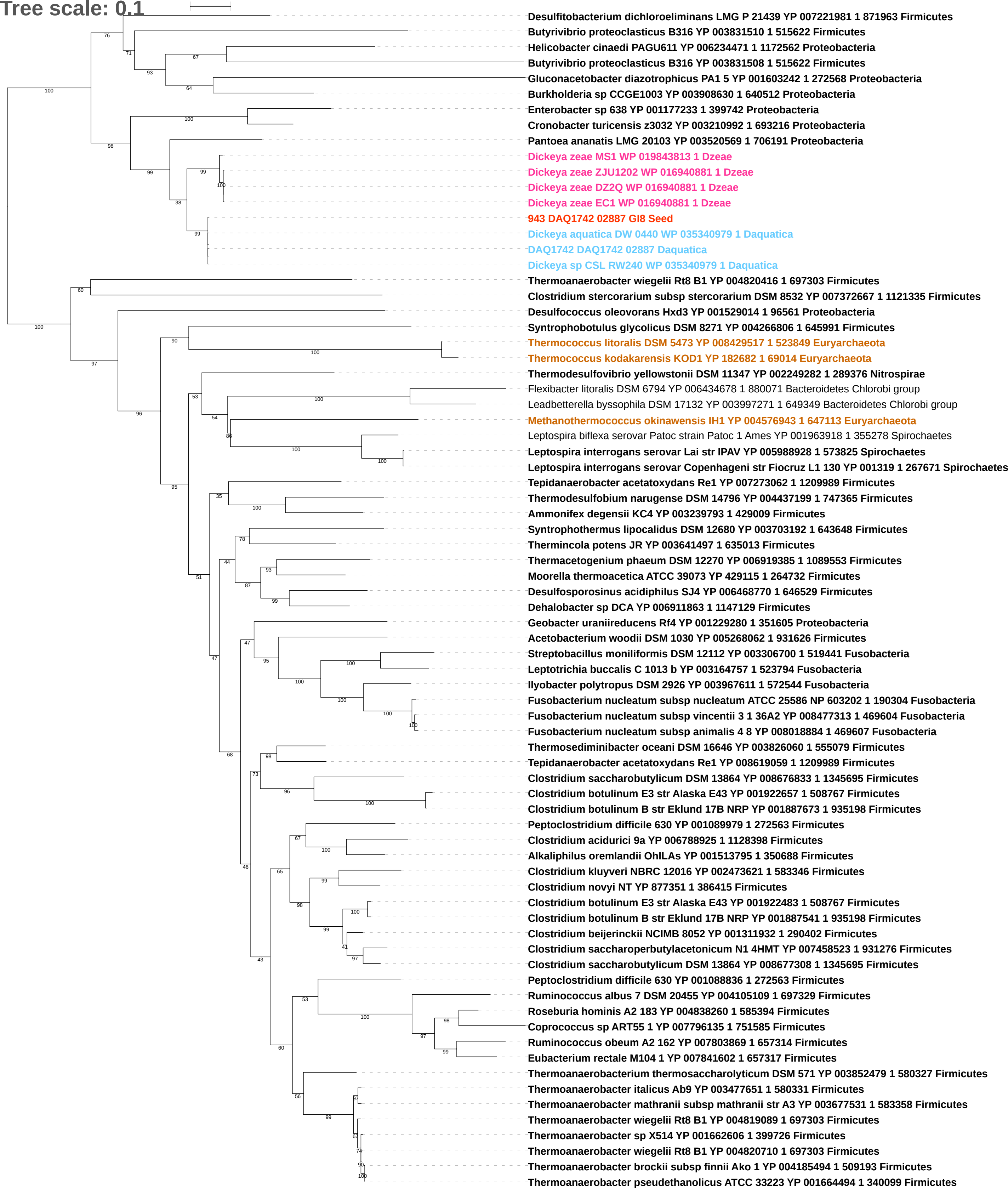
Tree scale: 0.1
Desulfitobacterium dichloroeliminans LMG P 21439 YP 007221981 1 871963 Firmicutes
Butyrivibrio proteoclasticus B316 YP 003831510 1 515622 Firmicutes
Helicobacter cinaedi PAGU611 YP 006234471 1 1172562 Proteobacteria
Butyrivibrio proteoclasticus B316 YP 003831508 1 515622 Firmicutes
Gluconacetobacter diazotrophicus PA1 5 YP 001603242 1 272568 Proteobacteria
Burkholderia sp CCGE1003 YP 003908630 1 640512 Proteobacteria
Enterobacter sp 638 YP 001177233 1 399742 Proteobacteria
Cronobacter turicensis z3032 YP 003210992 1 693216 Proteobacteria
Pantoea ananatis LMG 20103 YP 003520569 1 706191 Proteobacteria
Dickeya zeae MS1 WP 019843813 1 Dzeae
Dickeya zeae ZJU1202 WP 016940881 1 Dzeae
Dickeya zeae DZ2Q WP 016940881 1 Dzeae
Dickeya zeae EC1 WP 016940881 1 Dzeae
943 DAQ1742 02887 GI8 Seed
Dickeya aquatica DW 0440 WP 035340979 1 Daquatica
DAQ1742 DAQ1742 02887 Daquatica
Dickeya sp CSL RW240 WP 035340979 1 Daquatica
Thermoanaerobacter wiegelii Rt8 B1 YP 004820416 1 697303 Firmicutes
Clostridium stercorarium subsp stercorarium DSM 8532 YP 007372667 1 1121335 Firmicutes
Desulfococcus oleovorans Hxd3 YP 001529014 1 96561 Proteobacteria
Syntrophobotulus glycolicus DSM 8271 YP 004266806 1 645991 Firmicutes
Thermococcus litoralis DSM 5473 YP 008429517 1 523849 Euryarchaeota
Thermococcus kodakarensis KOD1 YP 182682 1 69014 Euryarchaeota
Thermodesulfovibrio yellowstonii DSM 11347 YP 002249282 1 289376 Nitrospirae
Flexibacter litoralis DSM 6794 YP 006434678 1 880071 Bacteroidetes Chlorobi group
Leadbetterella byssophila DSM 17132 YP 003997271 1 649349 Bacteroidetes Chlorobi group
Methanothermococcus okinawensis IH1 YP 004576943 1 647113 Euryarchaeota
Leptospira biflexa serovar Patoc strain Patoc 1 Ames YP 001963918 1 355278 Spirochaetes
Leptospira interrogans serovar Lai str IPAV YP 005988928 1 573825 Spirochaetes
Leptospira interrogans serovar Copenhageni str Fiocruz L1 130 YP 001319 1 267671 Spirochaetes
Tepidanaerobacter acetatoxydans Re1 YP 007273062 1 1209989 Firmicutes
Thermodesulfobium narugense DSM 14796 YP 004437199 1 747365 Firmicutes
Ammonifex degensii KC4 YP 003239793 1 429009 Firmicutes
Syntrophothermus lipocalidus DSM 12680 YP 003703192 1 643648 Firmicutes
Thermincola potens JR YP 003641497 1 635013 Firmicutes
Thermacetogenium phaeum DSM 12270 YP 006919385 1 1089553 Firmicutes
Moorella thermoacetica ATCC 39073 YP 429115 1 264732 Firmicutes
Desulfosporosinus acidiphilus SJ4 YP 006468770 1 646529 Firmicutes
Dehalobacter sp DCA YP 006911863 1 1147129 Firmicutes
Geobacter uraniireducens Rf4 YP 001229280 1 351605 Proteobacteria
Acetobacterium woodii DSM 1030 YP 005268062 1 931626 Firmicutes
Streptobacillus moniliformis DSM 12112 YP 003306700 1 519441 Fusobacteria
Leptotrichia buccalis C 1013 b YP 003164757 1 523794 Fusobacteria
Ilyobacter polytropus DSM 2926 YP 003967611 1 572544 Fusobacteria
Fusobacterium nucleatum subsp nucleatum ATCC 25586 NP 603202 1 190304 Fusobacteria
Fusobacterium nucleatum subsp vincentii 3 1 36A2 YP 008477313 1 469604 Fusobacteria
Fusobacterium nucleatum subsp animalis 4 8 YP 008018884 1 469607 Fusobacteria
Thermosediminibacter oceani DSM 16646 YP 003826060 1 555079 Firmicutes
Tepidanaerobacter acetatoxydans Re1 YP 008619059 1 1209989 Firmicutes
Clostridium saccharobutylicum DSM 13864 YP 008676833 1 1345695 Firmicutes
Clostridium botulinum E3 str Alaska E43 YP 001922657 1 508767 Firmicutes
Clostridium botulinum B str Eklund 17B NRP YP 001887673 1 935198 Firmicutes
Peptoclostridium difficile 630 YP 001089979 1 272563 Firmicutes
Clostridium acidurici 9a YP 006788925 1 1128398 Firmicutes
Alkaliphilus oremlandii OhILAs YP 001513795 1 350688 Firmicutes
Clostridium kluyveri NBRC 12016 YP 002473621 1 583346 Firmicutes
Clostridium novyi NT YP 877351 1 386415 Firmicutes
Clostridium botulinum E3 str Alaska E43 YP 001922483 1 508767 Firmicutes
Clostridium botulinum B str Eklund 17B NRP YP 001887541 1 935198 Firmicutes
Clostridium beijerinckii NCIMB 8052 YP 001311932 1 290402 Firmicutes
Clostridium saccharoperbutylacetonicum N1 4HMT YP 007458523 1 931276 Firmicutes
Clostridium saccharobutylicum DSM 13864 YP 008677308 1 1345695 Firmicutes
Peptoclostridium difficile 630 YP 001088836 1 272563 Firmicutes
Ruminococcus albus 7 DSM 20455 YP 004105109 1 697329 Firmicutes
Roseburia hominis A2 183 YP 004838260 1 585394 Firmicutes
Coprococcus sp ART55 1 YP 007796135 1 751585 Firmicutes
Ruminococcus obeum A2 162 YP 007803869 1 657314 Firmicutes
Eubacterium rectale M104 1 YP 007841602 1 657317 Firmicutes
Thermoanaerobacterium thermosaccharolyticum DSM 571 YP 003852479 1 580327 Firmicutes
Thermoanaerobacter italicus Ab9 YP 003477651 1 580331 Firmicutes
Thermoanaerobacter mathranii subsp mathranii str A3 YP 003677531 1 583358 Firmicutes
Thermoanaerobacter wiegelii Rt8 B1 YP 004819089 1 697303 Firmicutes
Thermoanaerobacter sp X514 YP 001662606 1 399726 Firmicutes
Thermoanaerobacter wiegelii Rt8 B1 YP 004820710 1 697303 Firmicutes
Thermoanaerobacter brockii subsp finnii Ako 1 YP 004185494 1 509193 Firmicutes
Thermoanaerobacter pseudethanolicus ATCC 33223 YP 001664494 1 340099 Firmicutes

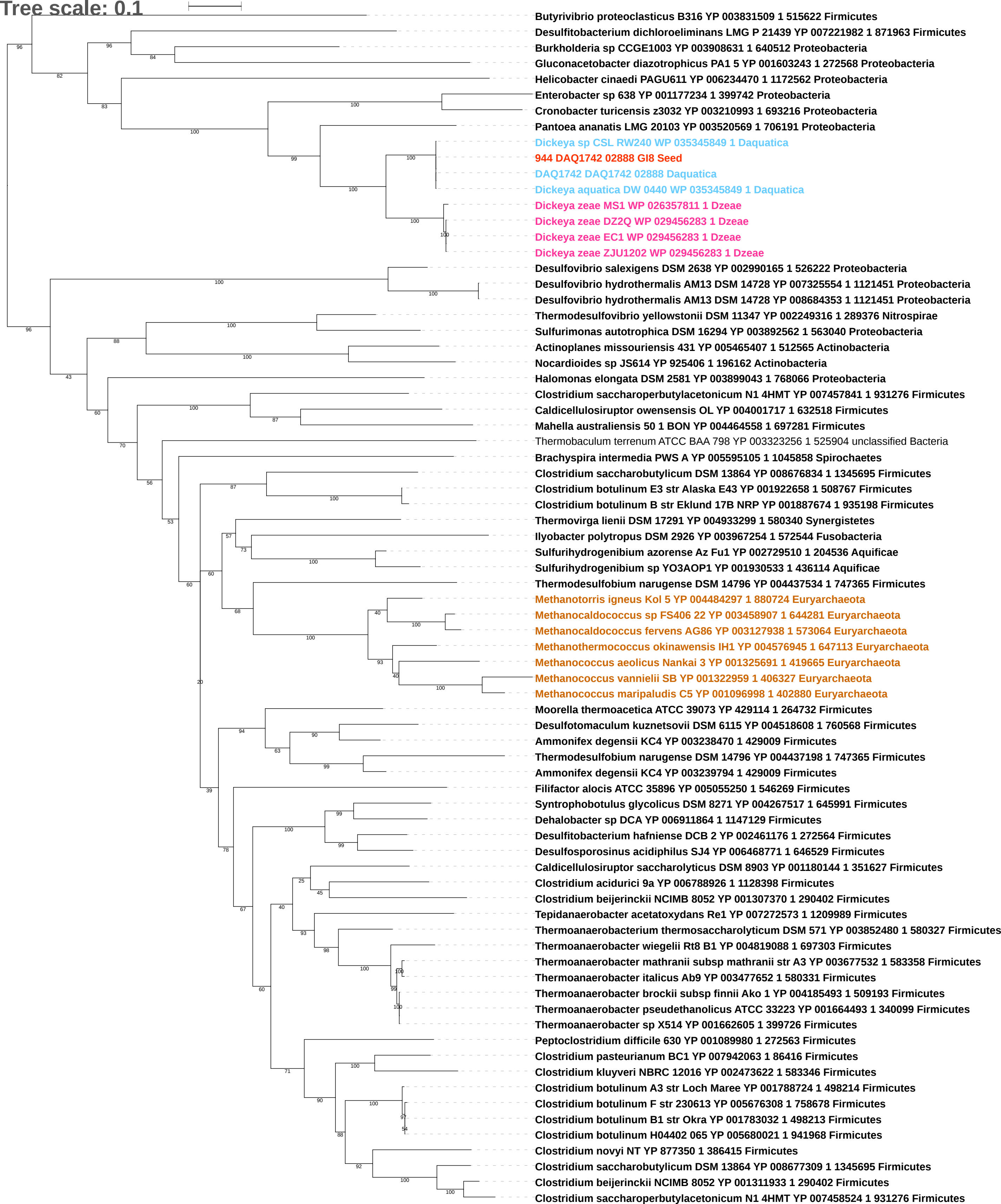
Tree scale: 0.1
Butyrivibrio proteoclasticus B316 YP 003831509 1 515622 Firmicutes
Desulfitobacterium dichloroeliminans LMG P 21439 YP 007221982 1 871963 Firmicutes
Burkholderia sp CCGE1003 YP 003908631 1 640512 Proteobacteria
Gluconacetobacter diazotrophicus PA1 5 YP 001603243 1 272568 Proteobacteria
Helicobacter cinaedi PAGU611 YP 006234470 1 1172562 Proteobacteria
Enterobacter sp 638 YP 001177234 1 399742 Proteobacteria
Cronobacter turicensis z3032 YP 003210993 1 693216 Proteobacteria
Pantoea ananatis LMG 20103 YP 003520569 1 706191 Proteobacteria
Dickeya sp CSL RW240 WP 035345849 1 Daquatica
944 DAQ1742 02888 GI8 Seed
DAQ1742 DAQ1742 02888 Daquatica
Dickeya aquatica DW 0440 WP 035345849 1 Daquatica
Dickeya zeae MS1 WP 026357811 1 Dzeae
Dickeya zeae DZ2Q WP 029456283 1 Dzeae
Dickeya zeae EC1 WP 029456283 1 Dzeae
Dickeya zeae ZJU1202 WP 029456283 1 Dzeae
Desulfovibrio salexigens DSM 2638 YP 002990165 1 526222 Proteobacteria
Desulfovibrio hydrothermalis AM13 DSM 14728 YP 007325554 1 1121451 Proteobacteria
Desulfovibrio hydrothermalis AM13 DSM 14728 YP 008684353 1 1121451 Proteobacteria
Thermodesulfovibrio yellowstonii DSM 11347 YP 002249316 1 289376 Nitrospirae
Sulfurimonas autotrophica DSM 16294 YP 003892562 1 563040 Proteobacteria
Actinoplanes missouriensis 431 YP 005465407 1 512565 Actinobacteria
Nocardioides sp JS614 YP 925406 1 196162 Actinobacteria
Halomonas elongata DSM 2581 YP 003899043 1 768066 Proteobacteria
Clostridium saccharoperbutylacetonicum N1 4HMT YP 007457841 1 931276 Firmicutes
Caldicellulosiruptor owensensis OL YP 004001717 1 632518 Firmicutes
Mahella australiensis 50 1 BON YP 004464558 1 697281 Firmicutes
Thermobaculum terrenum ATCC BAA 798 YP 003323256 1 525904 unclassified Bacteria
Brachyspira intermedia PWS A YP 005595105 1 1045858 Spirochaetes
Clostridium saccharobutylicum DSM 13864 YP 008676834 1 1345695 Firmicutes
Clostridium botulinum E3 str Alaska E43 YP 001922658 1 508767 Firmicutes
Clostridium botulinum B str Eklund 17B NRP YP 001887674 1 935198 Firmicutes
Thermovirga lienii DSM 17291 YP 004933299 1 580340 Synergistetes
Ilyobacter polytropus DSM 2926 YP 003967254 1 572544 Fusobacteria
Sulfurihydrogenibium azorense Az Fu1 YP 002729510 1 204536 Aquificae
Sulfurihydrogenibium sp YO3AOP1 YP 001930533 1 436114 Aquificae
Thermodesulfobium narugense DSM 14796 YP 004437534 1 747365 Firmicutes
Methanotorris igneus Kol 5 YP 004484297 1 880724 Euryarchaeota
Methanocaldococcus sp FS406 22 YP 003458907 1 644281 Euryarchaeota
Methanocaldococcus fervens AG86 YP 003127938 1 573064 Euryarchaeota
Methanothermococcus okinawensis IH1 YP 004576945 1 647113 Euryarchaeota
Methanococcus aeolicus Nankai 3 YP 001325691 1 419665 Euryarchaeota
Methanococcus vannielii SB YP 001322959 1 406327 Euryarchaeota
Methanococcus maripaludis C5 YP 001096998 1 402880 Euryarchaeota
Moorella thermoacetica ATCC 39073 YP 429114 1 264732 Firmicutes
Desulfotomaculum kuznetsovii DSM 6115 YP 004518608 1 760568 Firmicutes
Ammonifex degensii KC4 YP 003238470 1 429009 Firmicutes
Thermodesulfobium narugense DSM 14796 YP 004437198 1 747365 Firmicutes
Ammonifex degensii KC4 YP 003239794 1 429009 Firmicutes
Filifactor alocis ATCC 35896 YP 005055250 1 546269 Firmicutes
Syntrophobotulus glycolicus DSM 8271 YP 004267517 1 645991 Firmicutes
Dehalobacter sp DCA YP 006911864 1 1147129 Firmicutes
Desulfitobacterium hafniense DCB 2 YP 002461176 1 272564 Firmicutes
Desulfosporosinus acidiphilus SJ4 YP 006468771 1 646529 Firmicutes
Caldicellulosiruptor saccharolyticus DSM 8903 YP 001180144 1 351627 Firmicutes
Clostridium acidurici 9a YP 006788926 1 1128398 Firmicutes
Clostridium beijerinckii NCIMB 8052 YP 001307370 1 290402 Firmicutes
Tepidanaerobacter acetatoxydans Re1 YP 007272573 1 1209989 Firmicutes
Thermoanaerobacterium thermosaccharolyticum DSM 571 YP 003852480 1 580327 Firmicutes
Thermoanaerobacter wiegelii Rt8 B1 YP 004819088 1 697303 Firmicutes
Thermoanaerobacter mathranii subsp mathranii str A3 YP 003677532 1 583358 Firmicutes
Thermoanaerobacter italicus Ab9 YP 003477652 1 580331 Firmicutes
Thermoanaerobacter brockii subsp finnii Ako 1 YP 004185493 1 509193 Firmicutes
Thermoanaerobacter pseudethanolicus ATCC 33223 YP 001664493 1 340099 Firmicutes
Thermoanaerobacter sp X514 YP 001662605 1 399726 Firmicutes
Peptoclostridium difficile 630 YP 001089980 1 272563 Firmicutes
Clostridium pasteurianum BC1 YP 007942063 1 86416 Firmicutes
Clostridium kluyveri NBRC 12016 YP 002473622 1 583346 Firmicutes
Clostridium botulinum A3 str Loch Maree YP 001788724 1 498214 Firmicutes
Clostridium botulinum F str 230613 YP 005676308 1 758678 Firmicutes
Clostridium botulinum B1 str Okra YP 001783032 1 498213 Firmicutes
Clostridium botulinum H04402 065 YP 005680021 1 941968 Firmicutes
Clostridium novyi NT YP 877350 1 386415 Firmicutes
Clostridium saccharobutylicum DSM 13864 YP 008677309 1 1345695 Firmicutes
Clostridium beijerinckii NCIMB 8052 YP 001311933 1 290402 Firmicutes
Clostridium saccharoperbutylacetonicum N1 4HMT YP 007458524 1 931276 Firmicutes

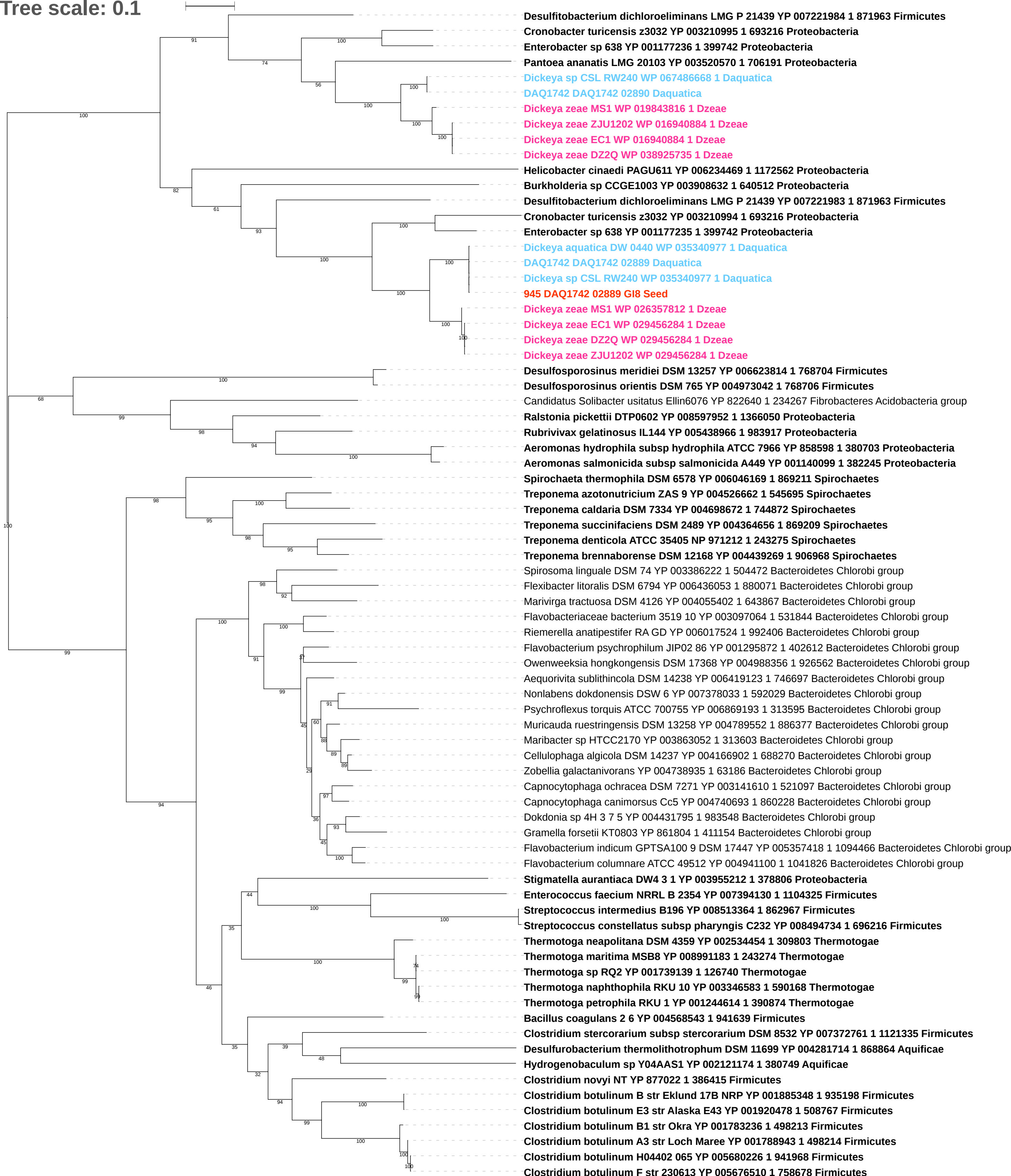
Tree scale: 0.1
Desulfitobacterium dichloroeliminans LMG P 21439 YP 007221984 1 871963 Firmicutes
Cronobacter turicensis z3032 YP 003210995 1 693216 Proteobacteria
Enterobacter sp 638 YP 001177236 1 399742 Proteobacteria
Pantoea ananatis LMG 20103 YP 003520570 1 706191 Proteobacteria
Dickeya sp CSL RW240 WP 067486668 1 Daquatica
DAQ1742 DAQ1742 02890 Daquatica
Dickeya zeae MS1 WP 019843816 1 Dzeae
Dickeya zeae ZJU1202 WP 016940884 1 Dzeae
Dickeya zeae EC1 WP 016940884 1 Dzeae
Dickeya zeae DZ2Q WP 038925735 1 Dzeae
Helicobacter cinaedi PAGU611 YP 006234469 1 1172562 Proteobacteria
Burkholderia sp CCGE1003 YP 003908632 1 640512 Proteobacteria
Desulfitobacterium dichloroeliminans LMG P 21439 YP 007221983 1 871963 Firmicutes
Cronobacter turicensis z3032 YP 003210994 1 693216 Proteobacteria
Enterobacter sp 638 YP 001177235 1 399742 Proteobacteria
Dickeya aquatica DW 0440 WP 035340977 1 Daquatica
DAQ1742 DAQ1742 02889 Daquatica
Dickeya sp CSL RW240 WP 035340977 1 Daquatica
945 DAQ1742 02889 GI8 Seed
Dickeya zeae MS1 WP 026357812 1 Dzeae
Dickeya zeae EC1 WP 029456284 1 Dzeae
Dickeya zeae DZ2Q WP 029456284 1 Dzeae
Dickeya zeae ZJU1202 WP 029456284 1 Dzeae
Desulfosporosinus meridiei DSM 13257 YP 006623814 1 768704 Firmicutes
Desulfosporosinus orientis DSM 765 YP 004973042 1 768706 Firmicutes
Candidatus Solibacter usitatus Ellin6076 YP 822640 1 234267 Fibrobacteres Acidobacteria group
Ralstonia pickettii DTP0602 YP 008597952 1 1366050 Proteobacteria
Rubrivivax gelatinosus IL144 YP 005438966 1 983917 Proteobacteria
Aeromonas hydrophila subsp hydrophila ATCC 7966 YP 858598 1 380703 Proteobacteria
Aeromonas salmonicida subsp salmonicida A449 YP 001140099 1 382245 Proteobacteria
Spirochaeta thermophila DSM 6578 YP 006046169 1 869211 Spirochaetes
Treponema azotonutricium ZAS 9 YP 004526662 1 545695 Spirochaetes
Treponema caldaria DSM 7334 YP 004698672 1 744872 Spirochaetes
Treponema succinifaciens DSM 2489 YP 004364656 1 869209 Spirochaetes
Treponema denticola ATCC 35405 NP 971212 1 243275 Spirochaetes
Treponema brennaborense DSM 12168 YP 004439269 1 906968 Spirochaetes
Spirosoma linguale DSM 74 YP 003386222 1 504472 Bacteroidetes Chlorobi group
Flexibacter litoralis DSM 6794 YP 006436053 1 880071 Bacteroidetes Chlorobi group
Marivirga tractuosa DSM 4126 YP 004055402 1 643867 Bacteroidetes Chlorobi group
Flavobacteriaceae bacterium 3519 10 YP 003097064 1 531844 Bacteroidetes Chlorobi group
Riemerella anatipestifer RA GD YP 006017524 1 992406 Bacteroidetes Chlorobi group
Flavobacterium psychrophilum JIP02 86 YP 001295872 1 402612 Bacteroidetes Chlorobi group
Owenweeksia hongkongensis DSM 17368 YP 004988356 1 926562 Bacteroidetes Chlorobi group
Aequorivita sublithincola DSM 14238 YP 006419123 1 746697 Bacteroidetes Chlorobi group
Nonlabens dokdonensis DSW 6 YP 007378033 1 592029 Bacteroidetes Chlorobi group
Psychroflexus torquis ATCC 700755 YP 006869193 1 313595 Bacteroidetes Chlorobi group
Muricauda ruestringensis DSM 13258 YP 004789552 1 886377 Bacteroidetes Chlorobi group
Maribacter sp HTCC2170 YP 003863052 1 313603 Bacteroidetes Chlorobi group
Cellulophaga algicola DSM 14237 YP 004166902 1 688270 Bacteroidetes Chlorobi group
Zobellia galactanivorans YP 004738935 1 63186 Bacteroidetes Chlorobi group
Capnocytophaga ochracea DSM 7271 YP 003141610 1 521097 Bacteroidetes Chlorobi group
Capnocytophaga canimorsus Cc5 YP 004740693 1 860228 Bacteroidetes Chlorobi group
Dokdonia sp 4H 3 7 5 YP 004431795 1 983548 Bacteroidetes Chlorobi group
Gramella forsetii KT0803 YP 861804 1 411154 Bacteroidetes Chlorobi group
Flavobacterium indicum GPTSA100 9 DSM 17447 YP 005357418 1 1094466 Bacteroidetes Chlorobi group
Flavobacterium columnare ATCC 49512 YP 004941100 1 1041826 Bacteroidetes Chlorobi group
Stigmatella aurantiaca DW4 3 1 YP 003955212 1 378806 Proteobacteria
Enterococcus faecium NRRL B 2354 YP 007394130 1 1104325 Firmicutes
Streptococcus intermedius B196 YP 008513364 1 862967 Firmicutes
Streptococcus constellatus subsp pharyngis C232 YP 008494734 1 696216 Firmicutes
Thermotoga neapolitana DSM 4359 YP 002534454 1 309803 Thermotogae
Thermotoga maritima MSB8 YP 008991183 1 243274 Thermotogae
Thermotoga sp RQ2 YP 001739139 1 126740 Thermotogae
Thermotoga naphthophila RKU 10 YP 003346583 1 590168 Thermotogae
Thermotoga petrophila RKU 1 YP 001244614 1 390874 Thermotogae
Bacillus coagulans 2 6 YP 004568543 1 941639 Firmicutes
Clostridium stercorarium subsp stercorarium DSM 8532 YP 007372761 1 1121335 Firmicutes
Desulfurobacterium thermolithotrophum DSM 11699 YP 004281714 1 868864 Aquificae
Hydrogenobaculum sp Y04AAS1 YP 002121174 1 380749 Aquificae
Clostridium novyi NT YP 877022 1 386415 Firmicutes
Clostridium botulinum B str Eklund 17B NRP YP 001885348 1 935198 Firmicutes
Clostridium botulinum E3 str Alaska E43 YP 001920478 1 508767 Firmicutes
Clostridium botulinum B1 str Okra YP 001783236 1 498213 Firmicutes
Clostridium botulinum A3 str Loch Maree YP 001788943 1 498214 Firmicutes
Clostridium botulinum H04402 065 YP 005680226 1 941968 Firmicutes
Clostridium botulinum F str 230613 YP 005676510 1 758678 Firmicutes

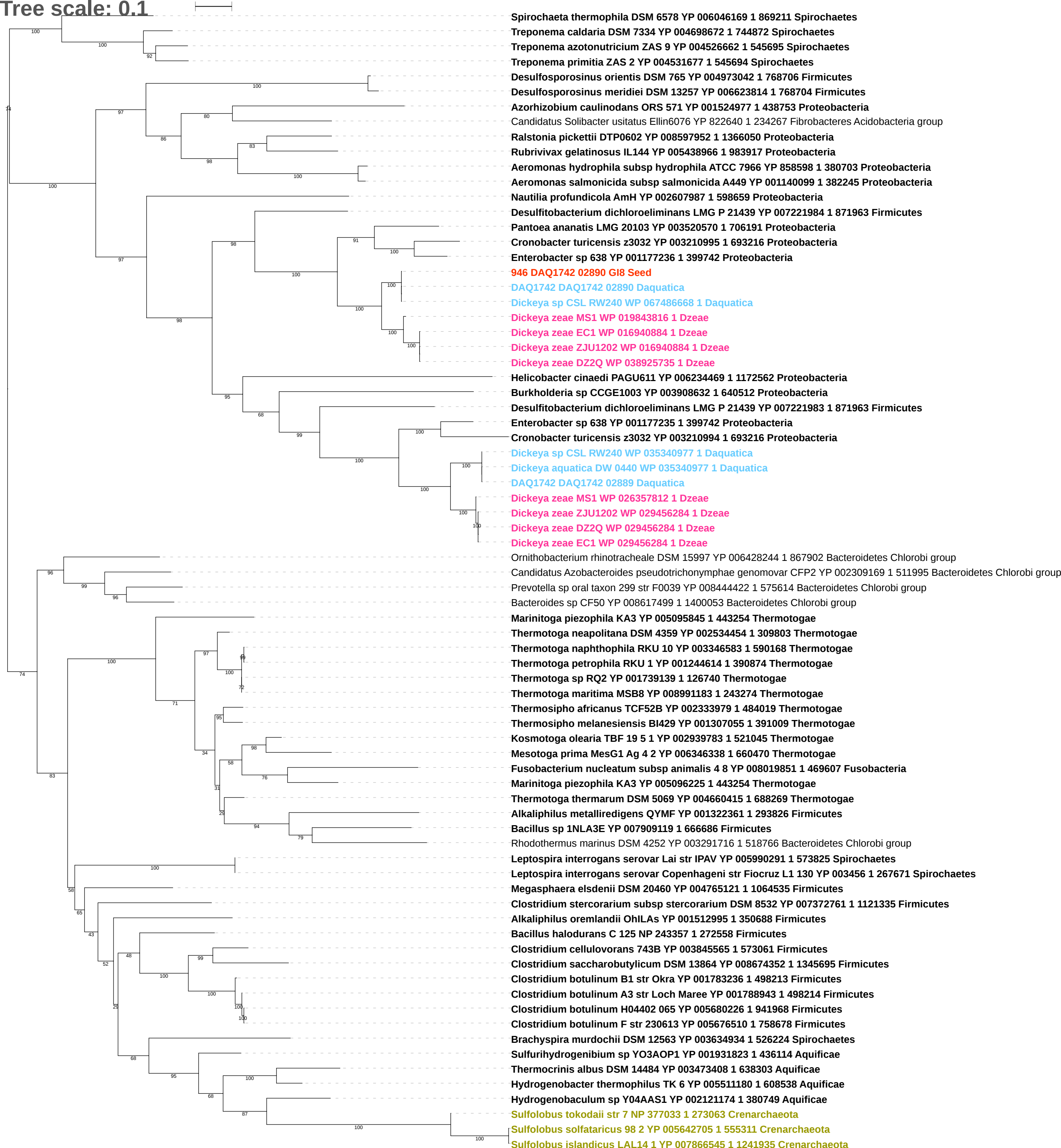

Tree scale: 0.1
Spirochaeta thermophila DSM 6578 YP 006046169 1 869211 Spirochaetes
Treponema caldaria DSM 7334 YP 004698672 1 744872 Spirochaetes
Treponema azotonutricium ZAS 9 YP 004526662 1 545695 Spirochaetes
Treponema primitia ZAS 2 YP 004531677 1 545694 Spirochaetes
Desulfosporosinus orientis DSM 765 YP 004973042 1 768706 Firmicutes
Desulfosporosinus meridiei DSM 13257 YP 006623814 1 768704 Firmicutes
Azorhizobium caulinodans ORS 571 YP 001524977 1 438753 Proteobacteria
Candidatus Solibacter usitatus Ellin6076 YP 822640 1 234267 Fibrobacteres Acidobacteria group
Ralstonia pickettii DTP0602 YP 008597952 1 1366050 Proteobacteria
Rubrivivax gelatinosus IL144 YP 005438966 1 983917 Proteobacteria
Aeromonas hydrophila subsp hydrophila ATCC 7966 YP 858598 1 380703 Proteobacteria
Aeromonas salmonicida subsp salmonicida A449 YP 001140099 1 382245 Proteobacteria
Nautilia profundicola AmH YP 002607987 1 598659 Proteobacteria
Desulfitobacterium dichloroeliminans LMG P 21439 YP 007221984 1 871963 Firmicutes
Pantoea ananatis LMG 20103 YP 003520570 1 706191 Proteobacteria
Cronobacter turicensis z3032 YP 003210995 1 693216 Proteobacteria
Enterobacter sp 638 YP 001177236 1 399742 Proteobacteria
946 DAQ1742 02890 GI8 Seed
DAQ1742 DAQ1742 02890 Daquatica
Dickeya sp CSL RW240 WP 067486668 1 Daquatica
Dickeya zeae MS1 WP 019843816 1 Dzeae
Dickeya zeae EC1 WP 016940884 1 Dzeae
Dickeya zeae ZJU1202 WP 016940884 1 Dzeae
Dickeya zeae DZ2Q WP 038925735 1 Dzeae
Helicobacter cinaedi PAGU611 YP 006234469 1 1172562 Proteobacteria
Burkholderia sp CCGE1003 YP 003908632 1 640512 Proteobacteria
Desulfitobacterium dichloroeliminans LMG P 21439 YP 007221983 1 871963 Firmicutes
Enterobacter sp 638 YP 001177235 1 399742 Proteobacteria
Cronobacter turicensis z3032 YP 003210994 1 693216 Proteobacteria
Dickeya sp CSL RW240 WP 035340977 1 Daquatica
Dickeya aquatica DW 0440 WP 035340977 1 Daquatica
DAQ1742 DAQ1742 02889 Daquatica
Dickeya zeae MS1 WP 026357812 1 Dzeae
Dickeya zeae ZJU1202 WP 029456284 1 Dzeae
Dickeya zeae DZ2Q WP 029456284 1 Dzeae
Dickeya zeae EC1 WP 029456284 1 Dzeae
Ornithobacterium rhinotracheale DSM 15997 YP 006428244 1 867902 Bacteroidetes Chlorobi group
Candidatus Azobacteroides pseudotrichonymphae genomovar CFP2 YP 002309169 1 511995 Bacteroidetes Chlorobi group
Prevotella sp oral taxon 299 str F0039 YP 008444422 1 575614 Bacteroidetes Chlorobi group
Bacteroides sp CF50 YP 008617499 1 1400053 Bacteroidetes Chlorobi group
Marinitoga piezophila KA3 YP 005095845 1 443254 Thermotogae
Thermotoga neapolitana DSM 4359 YP 002534454 1 309803 Thermotogae
Thermotoga naphthophila RKU 10 YP 003346583 1 590168 Thermotogae
Thermotoga petrophila RKU 1 YP 001244614 1 390874 Thermotogae
Thermotoga sp RQ2 YP 001739139 1 126740 Thermotogae
Thermotoga maritima MSB8 YP 008991183 1 243274 Thermotogae
Thermosipho africanus TCF52B YP 002333979 1 484019 Thermotogae
Thermosipho melanesiensis BI429 YP 001307055 1 391009 Thermotogae
Kosmotoga olearia TBF 19 5 1 YP 002939783 1 521045 Thermotogae
Mesotoga prima MesG1 Ag 4 2 YP 006346338 1 660470 Thermotogae
Fusobacterium nucleatum subsp animalis 4 8 YP 008019851 1 469607 Fusobacteria
Marinitoga piezophila KA3 YP 005096225 1 443254 Thermotogae
Thermotoga thermarum DSM 5069 YP 004660415 1 688269 Thermotogae
Alkaliphilus metalliredigens QYMF YP 001322361 1 293826 Firmicutes
Bacillus sp 1NLA3E YP 007909119 1 666686 Firmicutes
Rhodothermus marinus DSM 4252 YP 003291716 1 518766 Bacteroidetes Chlorobi group
Leptospira interrogans serovar Lai str IPAV YP 005990291 1 573825 Spirochaetes
Leptospira interrogans serovar Copenhageni str Fiocruz L1 130 YP 003456 1 267671 Spirochaetes
Megasphaera elsdenii DSM 20460 YP 004765121 1 1064535 Firmicutes
Clostridium stercorarium subsp stercorarium DSM 8532 YP 007372761 1 1121335 Firmicutes
Alkaliphilus oremlandii OhILAs YP 001512995 1 350688 Firmicutes
Bacillus halodurans C 125 NP 243357 1 272558 Firmicutes
Clostridium cellulovorans 743B YP 003845565 1 573061 Firmicutes
Clostridium saccharobutylicum DSM 13864 YP 008674352 1 1345695 Firmicutes
Clostridium botulinum B1 str Okra YP 001783236 1 498213 Firmicutes
Clostridium botulinum A3 str Loch Maree YP 001788943 1 498214 Firmicutes
Clostridium botulinum H04402 065 YP 005680226 1 941968 Firmicutes
Clostridium botulinum F str 230613 YP 005676510 1 758678 Firmicutes
Brachyspira murdochii DSM 12563 YP 003634934 1 526224 Spirochaetes
Sulfurihydrogenibium sp YO3AOP1 YP 001931823 1 436114 Aquificae
Thermocrinis albus DSM 14484 YP 003473408 1 638303 Aquificae
Hydrogenobacter thermophilus TK 6 YP 005511180 1 608538 Aquificae
Hydrogenobaculum sp Y04AAS1 YP 002121174 1 380749 Aquificae
Sulfolobus tokodaii str 7 NP 377033 1 273063 Crenarchaeota
Sulfolobus solfataricus 98 2 YP 005642705 1 555311 Crenarchaeota
Sulfolobus islandicus LAL14 1 YP 007866545 1 1241935 Crenarchaeota

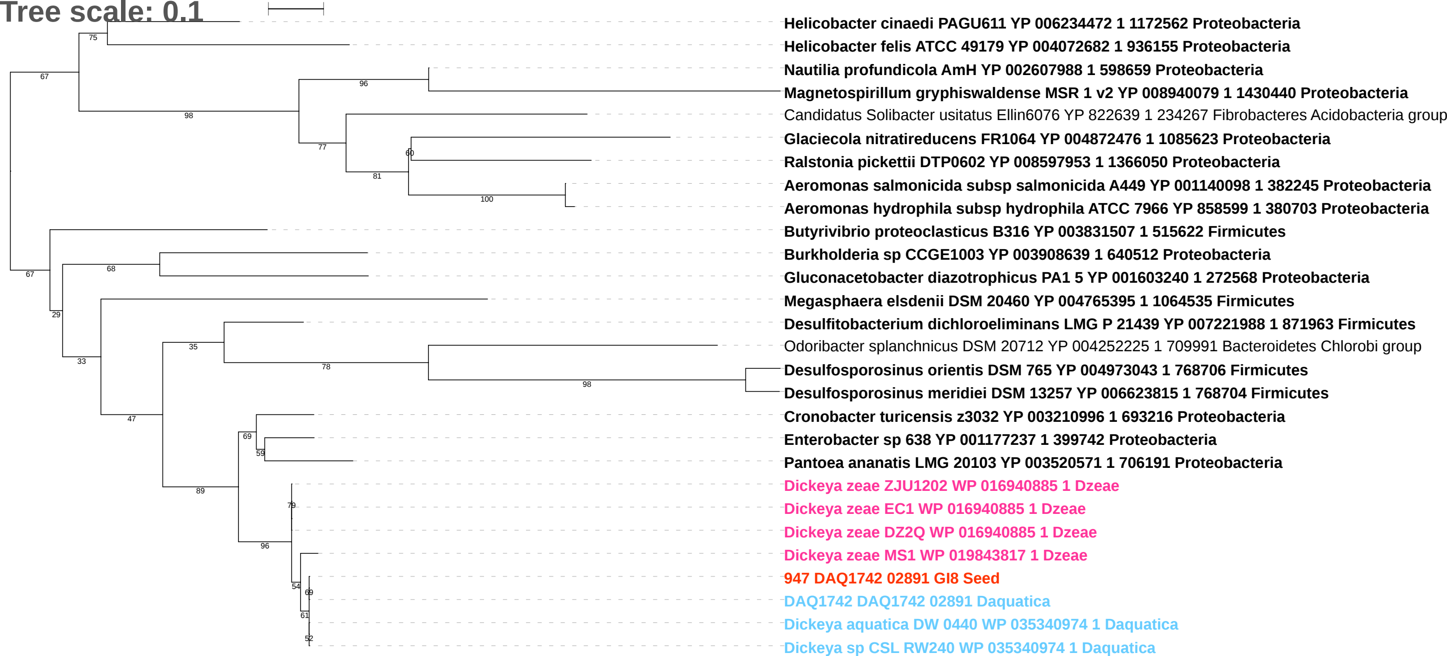
Tree scale: 0.1
Helicobacter cinaedi PAGU611 YP 006234472 1 1172562 Proteobacteria
Helicobacter felis ATCC 49179 YP 004072682 1 936155 Proteobacteria
Nautilia profundicola AmH YP 002607988 1 598659 Proteobacteria
Magnetospirillum gryphiswaldense MSR 1 v2 YP 008940079 1 1430440 Proteobacteria
Candidatus Solibacter usitatus Ellin6076 YP 822639 1 234267 Fibrobacteres Acidobacteria group
Glaciecola nitratireducens FR1064 YP 004872476 1 1085623 Proteobacteria
Ralstonia pickettii DTP0602 YP 008597953 1 1366050 Proteobacteria
Aeromonas salmonicida subsp salmonicida A449 YP 001140098 1 382245 Proteobacteria
Aeromonas hydrophila subsp hydrophila ATCC 7966 YP 858599 1 380703 Proteobacteria
Butyrivibrio proteoclasticus B316 YP 003831507 1 515622 Firmicutes
Burkholderia sp CCGE1003 YP 003908639 1 640512 Proteobacteria
Gluconacetobacter diazotrophicus PA1 5 YP 001603240 1 272568 Proteobacteria
Megasphaera elsdenii DSM 20460 YP 004765395 1 1064535 Firmicutes
Desulfitobacterium dichloroeliminans LMG P 21439 YP 007221988 1 871963 Firmicutes
Odoribacter splanchnicus DSM 20712 YP 004252225 1 709991 Bacteroidetes Chlorobi group
Desulfosporosinus orientis DSM 765 YP 004973043 1 768706 Firmicutes
Desulfosporosinus meridiei DSM 13257 YP 006623815 1 768704 Firmicutes
Cronobacter turicensis z3032 YP 003210996 1 693216 Proteobacteria
Enterobacter sp 638 YP 001177237 1 399742 Proteobacteria
Pantoea ananatis LMG 20103 YP 003520571 1 706191 Proteobacteria
Dickeya zeae ZJU1202 WP 016940885 1 Dzeae
Dickeya zeae EC1 WP 016940885 1 Dzeae
Dickeya zeae DZ2Q WP 016940885 1 Dzeae
Dickeya zeae MS1 WP 019843817 1 Dzeae
947 DAQ1742 02891 GI8 Seed
DAQ1742 DAQ1742 02891 Daquatica
Dickeya aquatica DW 0440 WP 035340974 1 Daquatica
Dickeya sp CSL RW240 WP 035340974 1 Daquatica

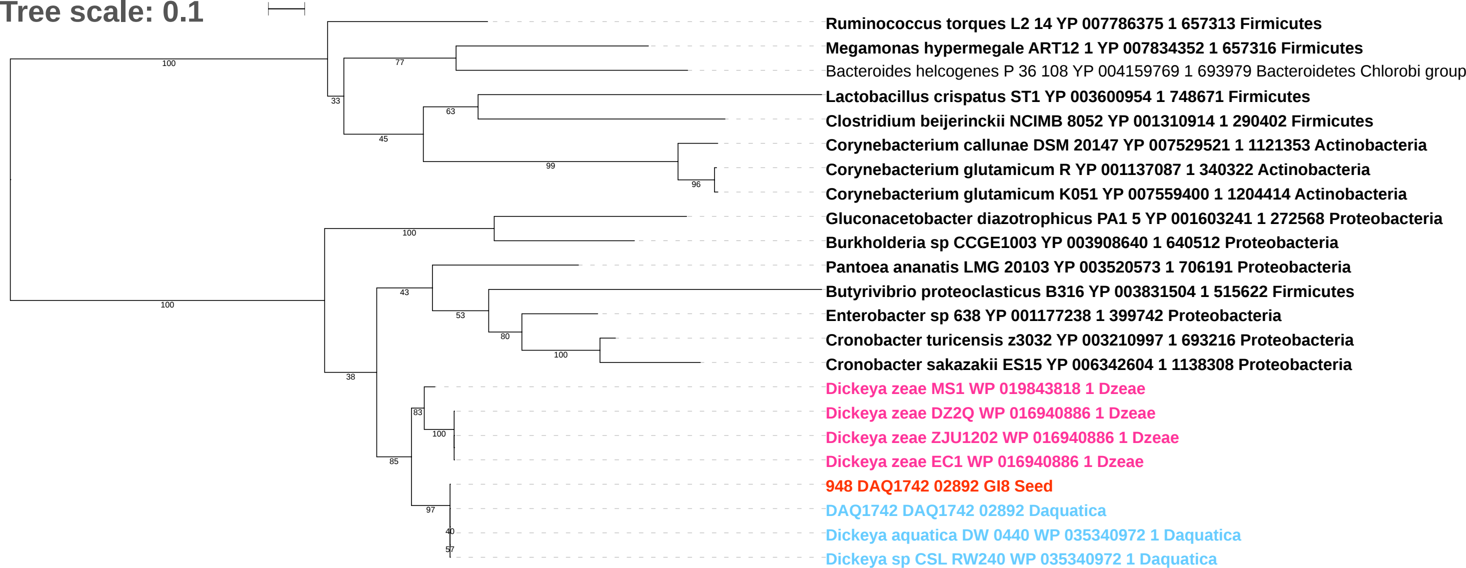
Tree scale: 0.1
Ruminococcus torques L2 14 YP 007786375 1 657313 Firmicutes
Megamonas hypermegale ART12 1 YP 007834352 1 657316 Firmicutes
Bacteroides helcogenes P 36 108 YP 004159769 1 693979 Bacteroidetes Chlorobi group
Lactobacillus crispatus ST1 YP 003600954 1 748671 Firmicutes
Clostridium beijerinckii NCIMB 8052 YP 001310914 1 290402 Firmicutes
Corynebacterium callunae DSM 20147 YP 007529521 1 1121353 Actinobacteria
Corynebacterium glutamicum R YP 001137087 1 340322 Actinobacteria
Corynebacterium glutamicum K051 YP 007559400 1 1204414 Actinobacteria
Gluconacetobacter diazotrophicus PA1 5 YP 001603241 1 272568 Proteobacteria
Burkholderia sp CCGE1003 YP 003908640 1 640512 Proteobacteria
Pantoea ananatis LMG 20103 YP 003520573 1 706191 Proteobacteria
Butyrivibrio proteoclasticus B316 YP 003831504 1 515622 Firmicutes
Enterobacter sp 638 YP 001177238 1 399742 Proteobacteria
Cronobacter turicensis z3032 YP 003210997 1 693216 Proteobacteria
Cronobacter sakazakii ES15 YP 006342604 1 1138308 Proteobacteria
Dickeya zeae MS1 WP 019843818 1 Dzeae
Dickeya zeae DZ2Q WP 016940886 1 Dzeae
Dickeya zeae ZJU1202 WP 016940886 1 Dzeae
Dickeya zeae EC1 WP 016940886 1 Dzeae
948 DAQ1742 02892 GI8 Seed
DAQ1742 DAQ1742 02892 Daquatica
Dickeya aquatica DW 0440 WP 035340972 1 Daquatica
Dickeya sp CSL RW240 WP 035340972 1 Daquatica
100
77
33
63
45
99
96
100
100
43
53
80
100
38
83
100
85
97
40
57

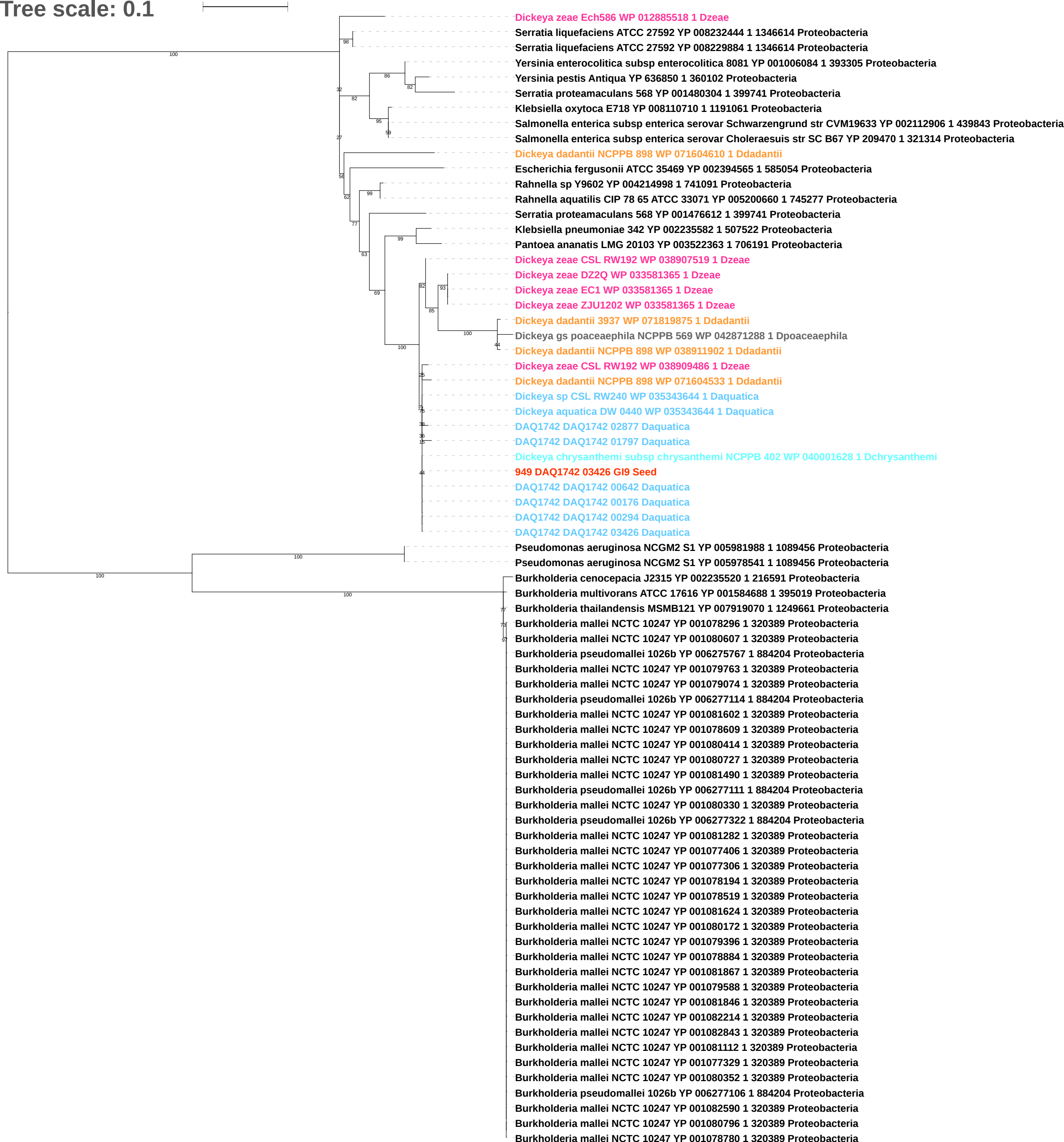
Tree scale: 0.1
Dickeya zeae Ech586 WP 012885518 1 Dzeae
Serratia liquefaciens ATCC 27592 YP 008232444 1 1346614 Proteobacteria
Serratia liquefaciens ATCC 27592 YP 008229884 1 1346614 Proteobacteria
Yersinia enterocolitica subsp enterocolitica 8081 YP 001006084 1 393305 Proteobacteria
Yersinia pestis Antiqua YP 636850 1 360102 Proteobacteria
Serratia proteamaculans 568 YP 001480304 1 399741 Proteobacteria
Klebsiella oxytoca E718 YP 008110710 1 1191061 Proteobacteria
Salmonella enterica subsp enterica serovar Schwarzengrund str CVM19633 YP 002112906 1 439843 Proteobacteria
Salmonella enterica subsp enterica serovar Choleraesuis str SC B67 YP 209470 1 321314 Proteobacteria
Dickeya dadantii NCPPB 898 WP 071604610 1 Ddadantii
Escherichia fergusonii ATCC 35469 YP 002394565 1 585054 Proteobacteria
Rahnella sp Y9602 YP 004214998 1 741091 Proteobacteria
Rahnella aquatilis CIP 78 65 ATCC 33071 YP 005200660 1 745277 Proteobacteria
Serratia proteamaculans 568 YP 001476612 1 399741 Proteobacteria
Klebsiella pneumoniae 342 YP 002235582 1 507522 Proteobacteria
Pantoea ananatis LMG 20103 YP 003522363 1 706191 Proteobacteria
Dickeya zeae CSL RW192 WP 038907519 1 Dzeae
Dickeya zeae DZ2Q WP 033581365 1 Dzeae
Dickeya zeae EC1 WP 033581365 1 Dzeae
Dickeya zeae ZJU1202 WP 033581365 1 Dzeae
Dickeya dadantii 3937 WP 071819875 1 Ddadantii
Dickeya gs poaceaephila NCPPB 569 WP 042871288 1 Dpoaceaephila
Dickeya dadantii NCPPB 898 WP 038911902 1 Ddadantii
Dickeya zeae CSL RW192 WP 038909486 1 Dzeae
Dickeya dadantii NCPPB 898 WP 071604533 1 Ddadantii
Dickeya sp CSL RW240 WP 035343644 1 Daquatica
Dickeya aquatica DW 0440 WP 035343644 1 Daquatica
DAQ1742 DAQ1742 02877 Daquatica
DAQ1742 DAQ1742 01797 Daquatica
Dickeya chrysanthemi subsp chrysanthemi NCPPB 402 WP 040001628 1 Dchrysanthemi
949 DAQ1742 03426 GI9 Seed
DAQ1742 DAQ1742 00642 Daquatica
DAQ1742 DAQ1742 00176 Daquatica
DAQ1742 DAQ1742 00294 Daquatica
DAQ1742 DAQ1742 03426 Daquatica
Pseudomonas aeruginosa NCGM2 S1 YP 005981988 1 1089456 Proteobacteria
Pseudomonas aeruginosa NCGM2 S1 YP 005978541 1 1089456 Proteobacteria
Burkholderia cenocepacia J2315 YP 002235520 1 216591 Proteobacteria
Burkholderia multivorans ATCC 17616 YP 001584688 1 395019 Proteobacteria
Burkholderia thailandensis MSMB121 YP 007919070 1 1249661 Proteobacteria
Burkholderia mallei NCTC 10247 YP 001078296 1 320389 Proteobacteria
Burkholderia mallei NCTC 10247 YP 001080607 1 320389 Proteobacteria
Burkholderia pseudomallei 1026b YP 006275767 1 884204 Proteobacteria
Burkholderia mallei NCTC 10247 YP 001079763 1 320389 Proteobacteria
Burkholderia mallei NCTC 10247 YP 001079074 1 320389 Proteobacteria
Burkholderia pseudomallei 1026b YP 006277114 1 884204 Proteobacteria
Burkholderia mallei NCTC 10247 YP 001081602 1 320389 Proteobacteria
Burkholderia mallei NCTC 10247 YP 001078609 1 320389 Proteobacteria
Burkholderia mallei NCTC 10247 YP 001080414 1 320389 Proteobacteria
Burkholderia mallei NCTC 10247 YP 001080727 1 320389 Proteobacteria
Burkholderia mallei NCTC 10247 YP 001081490 1 320389 Proteobacteria
Burkholderia pseudomallei 1026b YP 006277111 1 884204 Proteobacteria
Burkholderia mallei NCTC 10247 YP 001080330 1 320389 Proteobacteria
Burkholderia pseudomallei 1026b YP 006277322 1 884204 Proteobacteria
Burkholderia mallei NCTC 10247 YP 001081282 1 320389 Proteobacteria
Burkholderia mallei NCTC 10247 YP 001077406 1 320389 Proteobacteria
Burkholderia mallei NCTC 10247 YP 001077306 1 320389 Proteobacteria
Burkholderia mallei NCTC 10247 YP 001078194 1 320389 Proteobacteria
Burkholderia mallei NCTC 10247 YP 001078519 1 320389 Proteobacteria
Burkholderia mallei NCTC 10247 YP 001081624 1 320389 Proteobacteria
Burkholderia mallei NCTC 10247 YP 001080172 1 320389 Proteobacteria
Burkholderia mallei NCTC 10247 YP 001079396 1 320389 Proteobacteria
Burkholderia mallei NCTC 10247 YP 001078884 1 320389 Proteobacteria
Burkholderia mallei NCTC 10247 YP 001081867 1 320389 Proteobacteria
Burkholderia mallei NCTC 10247 YP 001079588 1 320389 Proteobacteria
Burkholderia mallei NCTC 10247 YP 001081846 1 320389 Proteobacteria
Burkholderia mallei NCTC 10247 YP 001082214 1 320389 Proteobacteria
Burkholderia mallei NCTC 10247 YP 001082843 1 320389 Proteobacteria
Burkholderia mallei NCTC 10247 YP 001081112 1 320389 Proteobacteria
Burkholderia mallei NCTC 10247 YP 001077329 1 320389 Proteobacteria
Burkholderia mallei NCTC 10247 YP 001080352 1 320389 Proteobacteria
Burkholderia pseudomallei 1026b YP 006277106 1 884204 Proteobacteria
Burkholderia mallei NCTC 10247 YP 001082590 1 320389 Proteobacteria
Burkholderia mallei NCTC 10247 YP 001080796 1 320389 Proteobacteria
Burkholderia mallei NCTC 10247 YP 001078780 1 320389 Proteobacteria

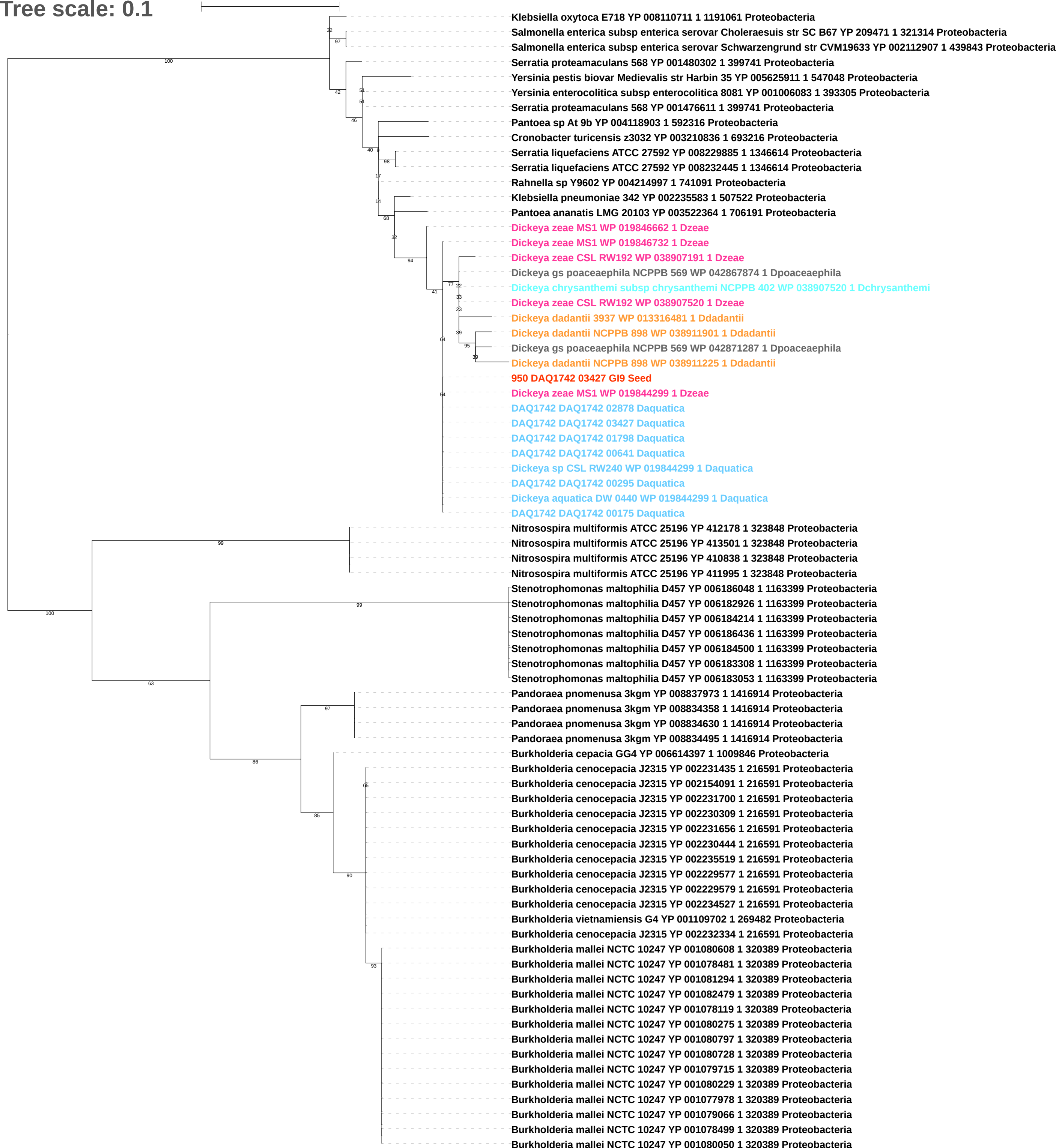

Tree scale: 0.1
Klebsiella oxytoca E718 YP 008110711 1 1191061 Proteobacteria
Salmonella enterica subsp enterica serovar Choleraesuis str SC B67 YP 209471 1 321314 Proteobacteria
Salmonella enterica subsp enterica serovar Schwarzengrund str CVM19633 YP 002112907 1 439843 Proteobacteria
Serratia proteamaculans 568 YP 001480302 1 399741 Proteobacteria
Yersinia pestis biovar Medievalis str Harbin 35 YP 005625911 1 547048 Proteobacteria
Yersinia enterocolitica subsp enterocolitica 8081 YP 001006083 1 393305 Proteobacteria
Serratia proteamaculans 568 YP 001476611 1 399741 Proteobacteria
Pantoea sp At 9b YP 004118903 1 592316 Proteobacteria
Cronobacter turicensis z3032 YP 003210836 1 693216 Proteobacteria
Serratia liquefaciens ATCC 27592 YP 008229885 1 1346614 Proteobacteria
Serratia liquefaciens ATCC 27592 YP 008232445 1 1346614 Proteobacteria
Rahnella sp Y9602 YP 004214997 1 741091 Proteobacteria
Klebsiella pneumoniae 342 YP 002235583 1 507522 Proteobacteria
Pantoea ananatis LMG 20103 YP 003522364 1 706191 Proteobacteria
Dickeya zeae MS1 WP 019846662 1 Dzeae
Dickeya zeae MS1 WP 019846732 1 Dzeae
Dickeya zeae CSL RW192 WP 038907191 1 Dzeae
Dickeya gs poaceaephila NCPPB 569 WP 042867874 1 Dpoaceaephila
Dickeya chrysanthemi subsp chrysanthemi NCPPB 402 WP 038907520 1 Dchrysanthemi
Dickeya zeae CSL RW192 WP 038907520 1 Dzeae
Dickeya dadantii 3937 WP 013316481 1 Ddadantii
Dickeya dadantii NCPPB 898 WP 038911901 1 Ddadantii
Dickeya gs poaceaephila NCPPB 569 WP 042871287 1 Dpoaceaephila
Dickeya dadantii NCPPB 898 WP 038911225 1 Ddadantii
950 DAQ1742 03427 GI9 Seed
Dickeya zeae MS1 WP 019844299 1 Dzeae
DAQ1742 DAQ1742 02878 Daquatica
DAQ1742 DAQ1742 03427 Daquatica
DAQ1742 DAQ1742 01798 Daquatica
DAQ1742 DAQ1742 00641 Daquatica
Dickeya sp CSL RW240 WP 019844299 1 Daquatica
DAQ1742 DAQ1742 00295 Daquatica
Dickeya aquatica DW 0440 WP 019844299 1 Daquatica
DAQ1742 DAQ1742 00175 Daquatica
Nitrosospira multiformis ATCC 25196 YP 412178 1 323848 Proteobacteria
Nitrosospira multiformis ATCC 25196 YP 413501 1 323848 Proteobacteria
Nitrosospira multiformis ATCC 25196 YP 410838 1 323848 Proteobacteria
Nitrosospira multiformis ATCC 25196 YP 411995 1 323848 Proteobacteria
Stenotrophomonas maltophilia D457 YP 006186048 1 1163399 Proteobacteria
Stenotrophomonas maltophilia D457 YP 006182926 1 1163399 Proteobacteria
Stenotrophomonas maltophilia D457 YP 006184214 1 1163399 Proteobacteria
Stenotrophomonas maltophilia D457 YP 006186436 1 1163399 Proteobacteria
Stenotrophomonas maltophilia D457 YP 006184500 1 1163399 Proteobacteria
Stenotrophomonas maltophilia D457 YP 006183308 1 1163399 Proteobacteria
Stenotrophomonas maltophilia D457 YP 006183053 1 1163399 Proteobacteria
Pandoraea pnomenusa 3kgm YP 008837973 1 1416914 Proteobacteria
Pandoraea pnomenusa 3kgm YP 008834358 1 1416914 Proteobacteria
Pandoraea pnomenusa 3kgm YP 008834630 1 1416914 Proteobacteria
Pandoraea pnomenusa 3kgm YP 008834495 1 1416914 Proteobacteria
Burkholderia cepacia GG4 YP 006614397 1 1009846 Proteobacteria
Burkholderia cenocepacia J2315 YP 002231435 1 216591 Proteobacteria
Burkholderia cenocepacia J2315 YP 002154091 1 216591 Proteobacteria
Burkholderia cenocepacia J2315 YP 002231700 1 216591 Proteobacteria
Burkholderia cenocepacia J2315 YP 002230309 1 216591 Proteobacteria
Burkholderia cenocepacia J2315 YP 002231656 1 216591 Proteobacteria
Burkholderia cenocepacia J2315 YP 002230444 1 216591 Proteobacteria
Burkholderia cenocepacia J2315 YP 002235519 1 216591 Proteobacteria
Burkholderia cenocepacia J2315 YP 002229577 1 216591 Proteobacteria
Burkholderia cenocepacia J2315 YP 002229579 1 216591 Proteobacteria
Burkholderia cenocepacia J2315 YP 002234527 1 216591 Proteobacteria
Burkholderia vietnamiensis G4 YP 001109702 1 269482 Proteobacteria
Burkholderia cenocepacia J2315 YP 002232334 1 216591 Proteobacteria
Burkholderia mallei NCTC 10247 YP 001080608 1 320389 Proteobacteria
Burkholderia mallei NCTC 10247 YP 001078481 1 320389 Proteobacteria
Burkholderia mallei NCTC 10247 YP 001081294 1 320389 Proteobacteria
Burkholderia mallei NCTC 10247 YP 001082479 1 320389 Proteobacteria
Burkholderia mallei NCTC 10247 YP 001078119 1 320389 Proteobacteria
Burkholderia mallei NCTC 10247 YP 001080275 1 320389 Proteobacteria
Burkholderia mallei NCTC 10247 YP 001080797 1 320389 Proteobacteria
Burkholderia mallei NCTC 10247 YP 001080728 1 320389 Proteobacteria
Burkholderia mallei NCTC 10247 YP 001079715 1 320389 Proteobacteria
Burkholderia mallei NCTC 10247 YP 001080229 1 320389 Proteobacteria
Burkholderia mallei NCTC 10247 YP 001077978 1 320389 Proteobacteria
Burkholderia mallei NCTC 10247 YP 001079066 1 320389 Proteobacteria
Burkholderia mallei NCTC 10247 YP 001078499 1 320389 Proteobacteria
Burkholderia mallei NCTC 10247 YP 001080050 1 320389 Proteobacteria

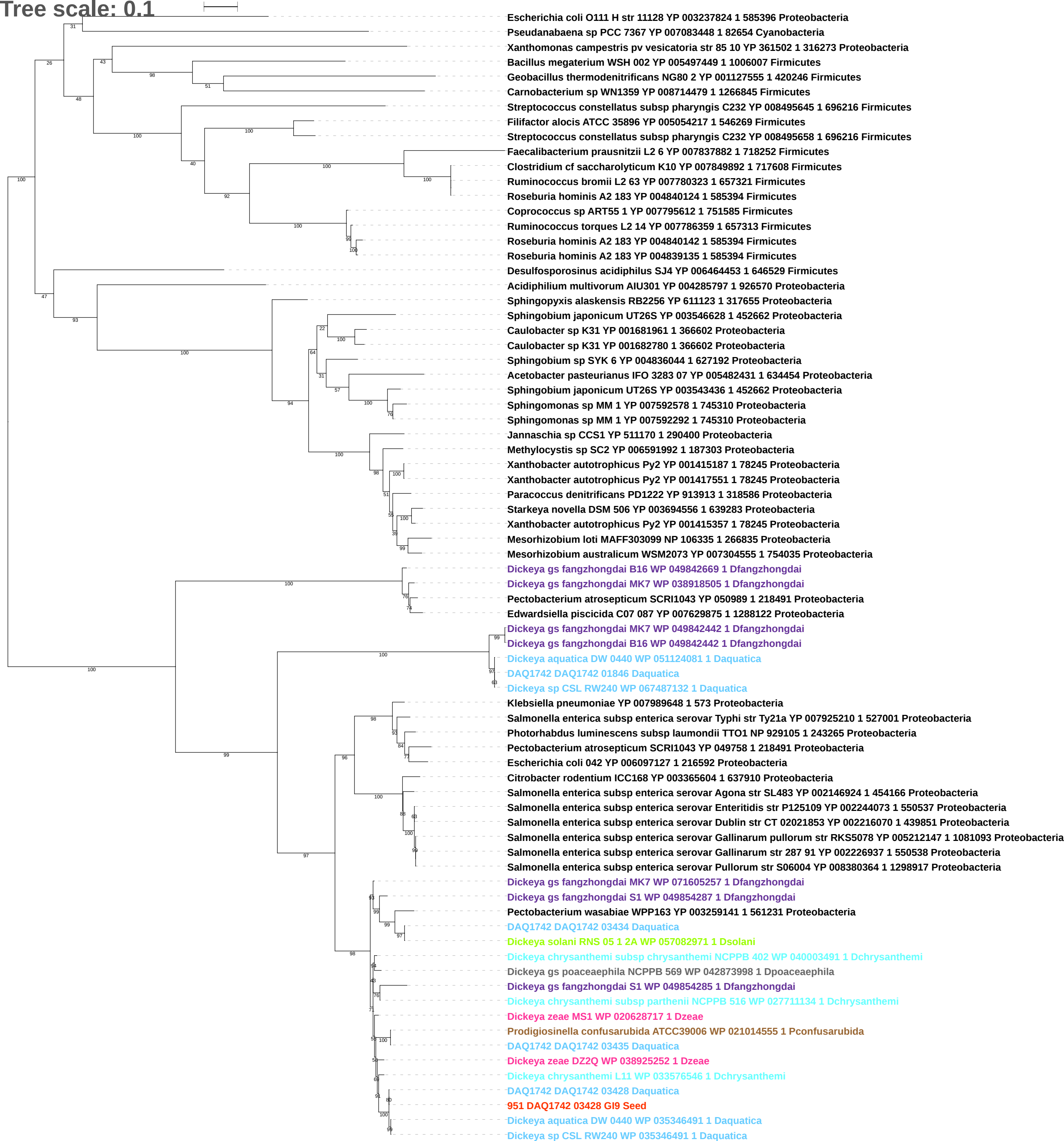
Tree scale: 0.1
Escherichia coli O111 H str 11128 YP 003237824 1 585396 Proteobacteria
Pseudanabaena sp PCC 7367 YP 007083448 1 82654 Cyanobacteria
Xanthomonas campestris pv vesicatoria str 85 10 YP 361502 1 316273 Proteobacteria
Bacillus megaterium WSH 002 YP 005497449 1 1006007 Firmicutes
Geobacillus thermodenitrificans NG80 2 YP 001127555 1 420246 Firmicutes
Carnobacterium sp WN1359 YP 008714479 1 1266845 Firmicutes
Streptococcus constellatus subsp pharyngis C232 YP 008495645 1 696216 Firmicutes
Filifactor alocis ATCC 35896 YP 005054217 1 546269 Firmicutes
Streptococcus constellatus subsp pharyngis C232 YP 008495658 1 696216 Firmicutes
Faecalibacterium prausnitzii L2 6 YP 007837882 1 718252 Firmicutes
Clostridium cf saccharolyticum K10 YP 007849892 1 717608 Firmicutes
Ruminococcus bromii L2 63 YP 007780323 1 657321 Firmicutes
Roseburia hominis A2 183 YP 004840124 1 585394 Firmicutes
Coprococcus sp ART55 1 YP 007795612 1 751585 Firmicutes
Ruminococcus torques L2 14 YP 007786359 1 657313 Firmicutes
Roseburia hominis A2 183 YP 004840142 1 585394 Firmicutes
Roseburia hominis A2 183 YP 004839135 1 585394 Firmicutes
Desulfosporosinus acidiphilus SJ4 YP 006464453 1 646529 Firmicutes
Acidiphilium multivorum AIU301 YP 004285797 1 926570 Proteobacteria
Sphingopyxis alaskensis RB2256 YP 611123 1 317655 Proteobacteria
Sphingobium japonicum UT26S YP 003546628 1 452662 Proteobacteria
Caulobacter sp K31 YP 001681961 1 366602 Proteobacteria
Caulobacter sp K31 YP 001682780 1 366602 Proteobacteria
Sphingobium sp SYK 6 YP 004836044 1 627192 Proteobacteria
Acetobacter pasteurianus IFO 3283 07 YP 005482431 1 634454 Proteobacteria
Sphingobium japonicum UT26S YP 003543436 1 452662 Proteobacteria
Sphingomonas sp MM 1 YP 007592578 1 745310 Proteobacteria
Sphingomonas sp MM 1 YP 007592292 1 745310 Proteobacteria
Jannaschia sp CCS1 YP 511170 1 290400 Proteobacteria
Methylocystis sp SC2 YP 006591992 1 187303 Proteobacteria
Xanthobacter autotrophicus Py2 YP 001415187 1 78245 Proteobacteria
Xanthobacter autotrophicus Py2 YP 001417551 1 78245 Proteobacteria
Paracoccus denitrificans PD1222 YP 913913 1 318586 Proteobacteria
Starkeya novella DSM 506 YP 003694556 1 639283 Proteobacteria
Xanthobacter autotrophicus Py2 YP 001415357 1 78245 Proteobacteria
Mesorhizobium loti MAFF303099 NP 106335 1 266835 Proteobacteria
Mesorhizobium australicum WSM2073 YP 007304555 1 754035 Proteobacteria
Dickeya gs fangzhongdai B16 WP 049842669 1 Dfangzhongdai
Dickeya gs fangzhongdai MK7 WP 038918505 1 Dfangzhongdai
Pectobacterium atrosepticum SCRI1043 YP 050989 1 218491 Proteobacteria
Edwardsiella piscicida C07 087 YP 007629875 1 1288122 Proteobacteria
Dickeya gs fangzhongdai MK7 WP 049842442 1 Dfangzhongdai
Dickeya gs fangzhongdai B16 WP 049842442 1 Dfangzhongdai
Dickeya aquatica DW 0440 WP 051124081 1 Daquatica
DAQ1742 DAQ1742 01846 Daquatica
Dickeya sp CSL RW240 WP 067487132 1 Daquatica
Klebsiella pneumoniae YP 007989648 1 573 Proteobacteria
Salmonella enterica subsp enterica serovar Typhi str Ty21a YP 007925210 1 527001 Proteobacteria
Photorhabdus luminescens subsp laumondii TTO1 NP 929105 1 243265 Proteobacteria
Pectobacterium atrosepticum SCRI1043 YP 049758 1 218491 Proteobacteria
Escherichia coli 042 YP 006097127 1 216592 Proteobacteria
Citrobacter rodentium ICC168 YP 003365604 1 637910 Proteobacteria
Salmonella enterica subsp enterica serovar Agona str SL483 YP 002146924 1 454166 Proteobacteria
Salmonella enterica subsp enterica serovar Enteritidis str P125109 YP 002244073 1 550537 Proteobacteria
Salmonella enterica subsp enterica serovar Dublin str CT 02021853 YP 002216070 1 439851 Proteobacteria
Salmonella enterica subsp enterica serovar Gallinarum pullorum str RKS5078 YP 005212147 1 1081093 Proteobacteria
Salmonella enterica subsp enterica serovar Gallinarum str 287 91 YP 002226937 1 550538 Proteobacteria
Salmonella enterica subsp enterica serovar Pullorum str S06004 YP 008380364 1 1298917 Proteobacteria
Dickeya gs fangzhongdai MK7 WP 071605257 1 Dfangzhongdai
Dickeya gs fangzhongdai S1 WP 049854287 1 Dfangzhongdai
Pectobacterium wasabiae WPP163 YP 003259141 1 561231 Proteobacteria
DAQ1742 DAQ1742 03434 Daquatica
Dickeya solani RNS 05 1 2A WP 057082971 1 Dsolani
Dickeya chrysanthemi subsp chrysanthemi NCPPB 402 WP 040003491 1 Dchrysanthemi
Dickeya gs poaceaephila NCPPB 569 WP 042873998 1 Dpoaceaephila
Dickeya gs fangzhongdai S1 WP 049854285 1 Dfangzhongdai
Dickeya chrysanthemi subsp parthenii NCPPB 516 WP 027711134 1 Dchrysanthemi
Dickeya zeae MS1 WP 020628717 1 Dzeae
Prodigiosinella confusarubida ATCC39006 WP 021014555 1 Pconfusarubida
DAQ1742 DAQ1742 03435 Daquatica
Dickeya zeae DZ2Q WP 038925252 1 Dzeae
Dickeya chrysanthemi L11 WP 033576546 1 Dchrysanthemi
DAQ1742 DAQ1742 03428 Daquatica
951 DAQ1742 03428 GI9 Seed
Dickeya aquatica DW 0440 WP 035346491 1 Daquatica
Dickeya sp CSL RW240 WP 035346491 1 Daquatica

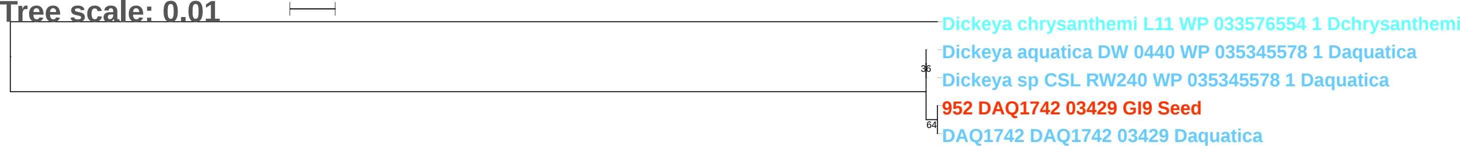
Tree scale: 0.01
Dickeya chrysanthemi L11 WP 033576554 1 Dchrysanthemi
Dickeya aquatica DW 0440 WP 035345578 1 Daquatica
36
Dickeya sp CSL RW240 WP 035345578 1 Daquatica
952 DAQ1742 03429 GI9 Seed
64
DAQ1742 DAQ1742 03429 Daquatica

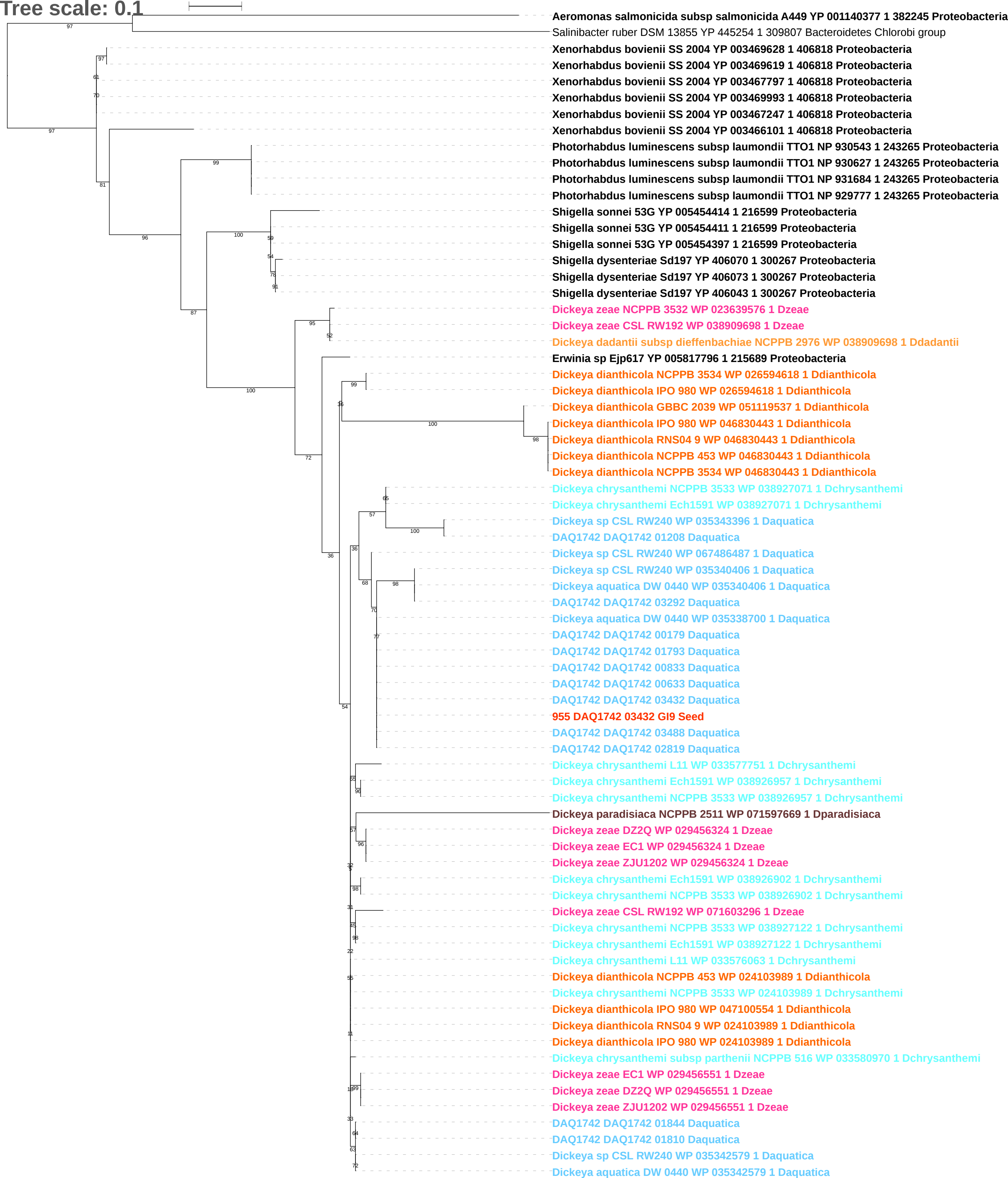
Tree scale: 0.1
Aeromonas salmonicida subsp salmonicida A449 YP 001140377 1 382245 Proteobacteria
Salinibacter ruber DSM 13855 YP 445254 1 309807 Bacteroidetes Chlorobi group
Xenorhabdus bovienii SS 2004 YP 003469628 1 406818 Proteobacteria
Xenorhabdus bovienii SS 2004 YP 003469619 1 406818 Proteobacteria
Xenorhabdus bovienii SS 2004 YP 003467797 1 406818 Proteobacteria
Xenorhabdus bovienii SS 2004 YP 003469993 1 406818 Proteobacteria
Xenorhabdus bovienii SS 2004 YP 003467247 1 406818 Proteobacteria
Xenorhabdus bovienii SS 2004 YP 003466101 1 406818 Proteobacteria
Photorhabdus luminescens subsp laumondii TTO1 NP 930543 1 243265 Proteobacteria
Photorhabdus luminescens subsp laumondii TTO1 NP 930627 1 243265 Proteobacteria
Photorhabdus luminescens subsp laumondii TTO1 NP 931684 1 243265 Proteobacteria
Photorhabdus luminescens subsp laumondii TTO1 NP 929777 1 243265 Proteobacteria
Shigella sonnei 53G YP 005454414 1 216599 Proteobacteria
Shigella sonnei 53G YP 005454411 1 216599 Proteobacteria
Shigella sonnei 53G YP 005454397 1 216599 Proteobacteria
Shigella dysenteriae Sd197 YP 406070 1 300267 Proteobacteria
Shigella dysenteriae Sd197 YP 406073 1 300267 Proteobacteria
Shigella dysenteriae Sd197 YP 406043 1 300267 Proteobacteria
Dickeya zeae NCPPB 3532 WP 023639576 1 Dzeae
Dickeya zeae CSL RW192 WP 038909698 1 Dzeae
Dickeya dadantii subsp dieffenbachiae NCPPB 2976 WP 038909698 1 Ddadantii
Erwinia sp Ejp617 YP 005817796 1 215689 Proteobacteria
Dickeya dianthicola NCPPB 3534 WP 026594618 1 Ddianthicola
Dickeya dianthicola IPO 980 WP 026594618 1 Ddianthicola
Dickeya dianthicola GBBC 2039 WP 051119537 1 Ddianthicola
Dickeya dianthicola IPO 980 WP 046830443 1 Ddianthicola
Dickeya dianthicola RNS04 9 WP 046830443 1 Ddianthicola
Dickeya dianthicola NCPPB 453 WP 046830443 1 Ddianthicola
Dickeya dianthicola NCPPB 3534 WP 046830443 1 Ddianthicola
Dickeya chrysanthemi NCPPB 3533 WP 038927071 1 Dchrysanthemi
Dickeya chrysanthemi Ech1591 WP 038927071 1 Dchrysanthemi
Dickeya sp CSL RW240 WP 035343396 1 Daquatica
DAQ1742 DAQ1742 01208 Daquatica
Dickeya sp CSL RW240 WP 067486487 1 Daquatica
Dickeya sp CSL RW240 WP 035340406 1 Daquatica
Dickeya aquatica DW 0440 WP 035340406 1 Daquatica
DAQ1742 DAQ1742 03292 Daquatica
Dickeya aquatica DW 0440 WP 035338700 1 Daquatica
DAQ1742 DAQ1742 00179 Daquatica
DAQ1742 DAQ1742 01793 Daquatica
DAQ1742 DAQ1742 00833 Daquatica
DAQ1742 DAQ1742 00633 Daquatica
DAQ1742 DAQ1742 03432 Daquatica
955 DAQ1742 03432 GI9 Seed
DAQ1742 DAQ1742 03488 Daquatica
DAQ1742 DAQ1742 02819 Daquatica
Dickeya chrysanthemi L11 WP 033577751 1 Dchrysanthemi
Dickeya chrysanthemi Ech1591 WP 038926957 1 Dchrysanthemi
Dickeya chrysanthemi NCPPB 3533 WP 038926957 1 Dchrysanthemi
Dickeya paradisiaca NCPPB 2511 WP 071597669 1 Dparadisiaca
Dickeya zeae DZ2Q WP 029456324 1 Dzeae
Dickeya zeae EC1 WP 029456324 1 Dzeae
Dickeya zeae ZJU1202 WP 029456324 1 Dzeae
Dickeya chrysanthemi Ech1591 WP 038926902 1 Dchrysanthemi
Dickeya chrysanthemi NCPPB 3533 WP 038926902 1 Dchrysanthemi
Dickeya zeae CSL RW192 WP 071603296 1 Dzeae
Dickeya chrysanthemi NCPPB 3533 WP 038927122 1 Dchrysanthemi
Dickeya chrysanthemi Ech1591 WP 038927122 1 Dchrysanthemi
Dickeya chrysanthemi L11 WP 033576063 1 Dchrysanthemi
Dickeya dianthicola NCPPB 453 WP 024103989 1 Ddianthicola
Dickeya chrysanthemi NCPPB 3533 WP 024103989 1 Dchrysanthemi
Dickeya dianthicola IPO 980 WP 047100554 1 Ddianthicola
Dickeya dianthicola RNS04 9 WP 024103989 1 Ddianthicola
Dickeya dianthicola IPO 980 WP 024103989 1 Ddianthicola
Dickeya chrysanthemi subsp parthenii NCPPB 516 WP 033580970 1 Dchrysanthemi
Dickeya zeae EC1 WP 029456551 1 Dzeae
Dickeya zeae DZ2Q WP 029456551 1 Dzeae
Dickeya zeae ZJU1202 WP 029456551 1 Dzeae
DAQ1742 DAQ1742 01844 Daquatica
DAQ1742 DAQ1742 01810 Daquatica
Dickeya sp CSL RW240 WP 035342579 1 Daquatica
Dickeya aquatica DW 0440 WP 035342579 1 Daquatica

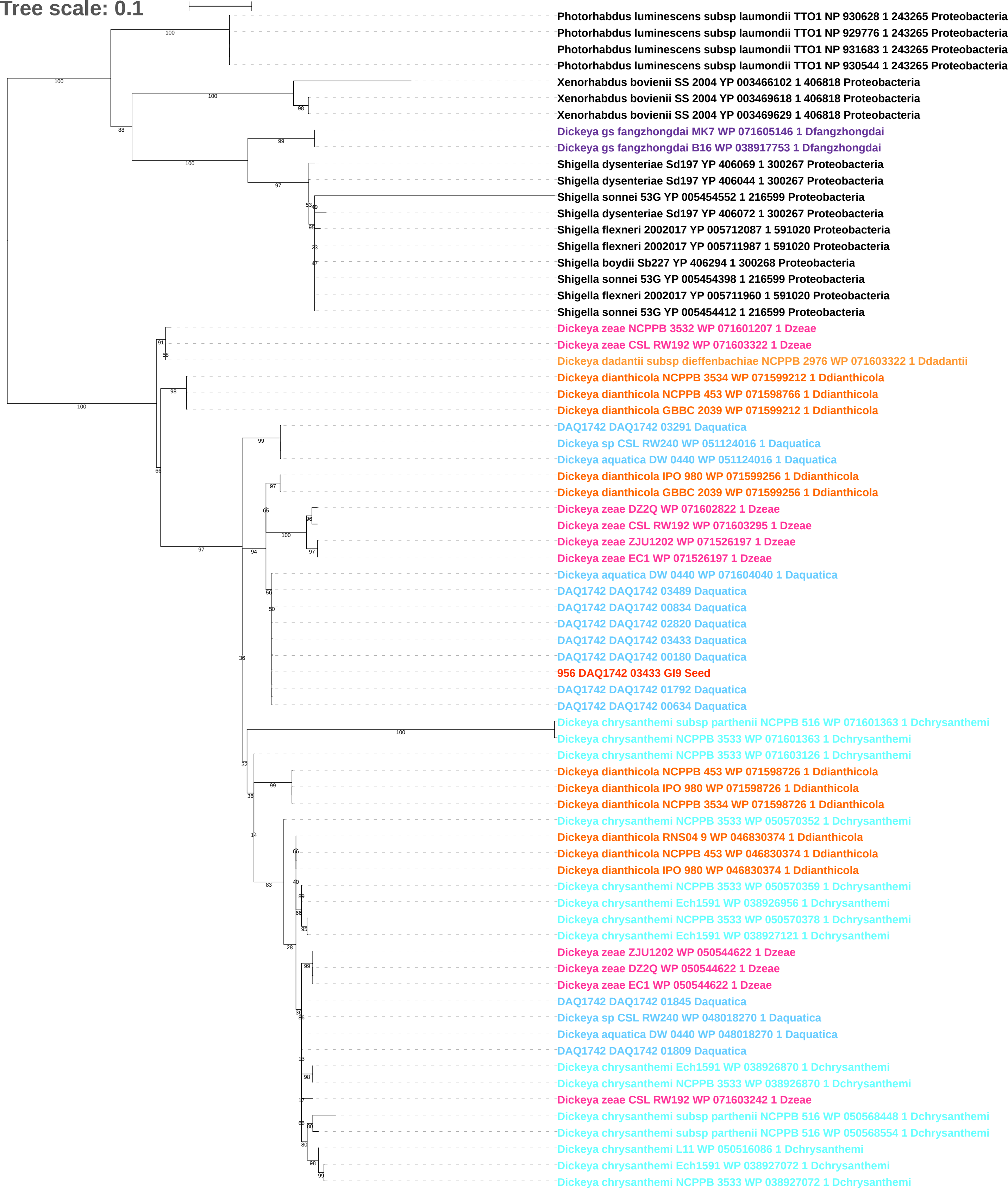

Tree scale: 0.1
Photorhabdus luminescens subsp laumondii TTO1 NP 930628 1 243265 Proteobacteria
Photorhabdus luminescens subsp laumondii TTO1 NP 929776 1 243265 Proteobacteria
Photorhabdus luminescens subsp laumondii TTO1 NP 931683 1 243265 Proteobacteria
Photorhabdus luminescens subsp laumondii TTO1 NP 930544 1 243265 Proteobacteria
Xenorhabdus bovienii SS 2004 YP 003466102 1 406818 Proteobacteria
Xenorhabdus bovienii SS 2004 YP 003469618 1 406818 Proteobacteria
Xenorhabdus bovienii SS 2004 YP 003469629 1 406818 Proteobacteria
Dickeya gs fangzhongdai MK7 WP 071605146 1 Dfangzhongdai
Dickeya gs fangzhongdai B16 WP 038917753 1 Dfangzhongdai
Shigella dysenteriae Sd197 YP 406069 1 300267 Proteobacteria
Shigella dysenteriae Sd197 YP 406044 1 300267 Proteobacteria
Shigella sonnei 53G YP 005454552 1 216599 Proteobacteria
Shigella dysenteriae Sd197 YP 406072 1 300267 Proteobacteria
Shigella flexneri 2002017 YP 005712087 1 591020 Proteobacteria
Shigella flexneri 2002017 YP 005711987 1 591020 Proteobacteria
Shigella boydii Sb227 YP 406294 1 300268 Proteobacteria
Shigella sonnei 53G YP 005454398 1 216599 Proteobacteria
Shigella flexneri 2002017 YP 005711960 1 591020 Proteobacteria
Shigella sonnei 53G YP 005454412 1 216599 Proteobacteria
Dickeya zeae NCPPB 3532 WP 071601207 1 Dzeae
Dickeya zeae CSL RW192 WP 071603322 1 Dzeae
Dickeya dadantii subsp dieffenbachiae NCPPB 2976 WP 071603322 1 Ddadantii
Dickeya dianthicola NCPPB 3534 WP 071599212 1 Ddianthicola
Dickeya dianthicola NCPPB 453 WP 071598766 1 Ddianthicola
Dickeya dianthicola GBBC 2039 WP 071599212 1 Ddianthicola
DAQ1742 DAQ1742 03291 Daquatica
Dickeya sp CSL RW240 WP 051124016 1 Daquatica
Dickeya aquatica DW 0440 WP 051124016 1 Daquatica
Dickeya dianthicola IPO 980 WP 071599256 1 Ddianthicola
Dickeya dianthicola GBBC 2039 WP 071599256 1 Ddianthicola
Dickeya zeae DZ2Q WP 071602822 1 Dzeae
Dickeya zeae CSL RW192 WP 071603295 1 Dzeae
Dickeya zeae ZJU1202 WP 071526197 1 Dzeae
Dickeya zeae EC1 WP 071526197 1 Dzeae
Dickeya aquatica DW 0440 WP 071604040 1 Daquatica
DAQ1742 DAQ1742 03489 Daquatica
DAQ1742 DAQ1742 00834 Daquatica
DAQ1742 DAQ1742 02820 Daquatica
DAQ1742 DAQ1742 03433 Daquatica
DAQ1742 DAQ1742 00180 Daquatica
956 DAQ1742 03433 GI9 Seed
DAQ1742 DAQ1742 01792 Daquatica
DAQ1742 DAQ1742 00634 Daquatica
Dickeya chrysanthemi subsp parthenii NCPPB 516 WP 071601363 1 Dchrysanthemi
Dickeya chrysanthemi NCPPB 3533 WP 071601363 1 Dchrysanthemi
Dickeya chrysanthemi NCPPB 3533 WP 071603126 1 Dchrysanthemi
Dickeya dianthicola NCPPB 453 WP 071598726 1 Ddianthicola
Dickeya dianthicola IPO 980 WP 071598726 1 Ddianthicola
Dickeya dianthicola NCPPB 3534 WP 071598726 1 Ddianthicola
Dickeya chrysanthemi NCPPB 3533 WP 050570352 1 Dchrysanthemi
Dickeya dianthicola RNS04 9 WP 046830374 1 Ddianthicola
Dickeya dianthicola NCPPB 453 WP 046830374 1 Ddianthicola
Dickeya dianthicola IPO 980 WP 046830374 1 Ddianthicola
Dickeya chrysanthemi NCPPB 3533 WP 050570359 1 Dchrysanthemi
Dickeya chrysanthemi Ech1591 WP 038926956 1 Dchrysanthemi
Dickeya chrysanthemi NCPPB 3533 WP 050570378 1 Dchrysanthemi
Dickeya chrysanthemi Ech1591 WP 038927121 1 Dchrysanthemi
Dickeya zeae ZJU1202 WP 050544622 1 Dzeae
Dickeya zeae DZ2Q WP 050544622 1 Dzeae
Dickeya zeae EC1 WP 050544622 1 Dzeae
DAQ1742 DAQ1742 01845 Daquatica
Dickeya sp CSL RW240 WP 048018270 1 Daquatica
Dickeya aquatica DW 0440 WP 048018270 1 Daquatica
DAQ1742 DAQ1742 01809 Daquatica
Dickeya chrysanthemi Ech1591 WP 038926870 1 Dchrysanthemi
Dickeya chrysanthemi NCPPB 3533 WP 038926870 1 Dchrysanthemi
Dickeya zeae CSL RW192 WP 071603242 1 Dzeae
Dickeya chrysanthemi subsp parthenii NCPPB 516 WP 050568448 1 Dchrysanthemi
Dickeya chrysanthemi subsp parthenii NCPPB 516 WP 050568554 1 Dchrysanthemi
Dickeya chrysanthemi L11 WP 050516086 1 Dchrysanthemi
Dickeya chrysanthemi Ech1591 WP 038927072 1 Dchrysanthemi
Dickeya chrysanthemi NCPPB 3533 WP 038927072 1 Dchrysanthemi

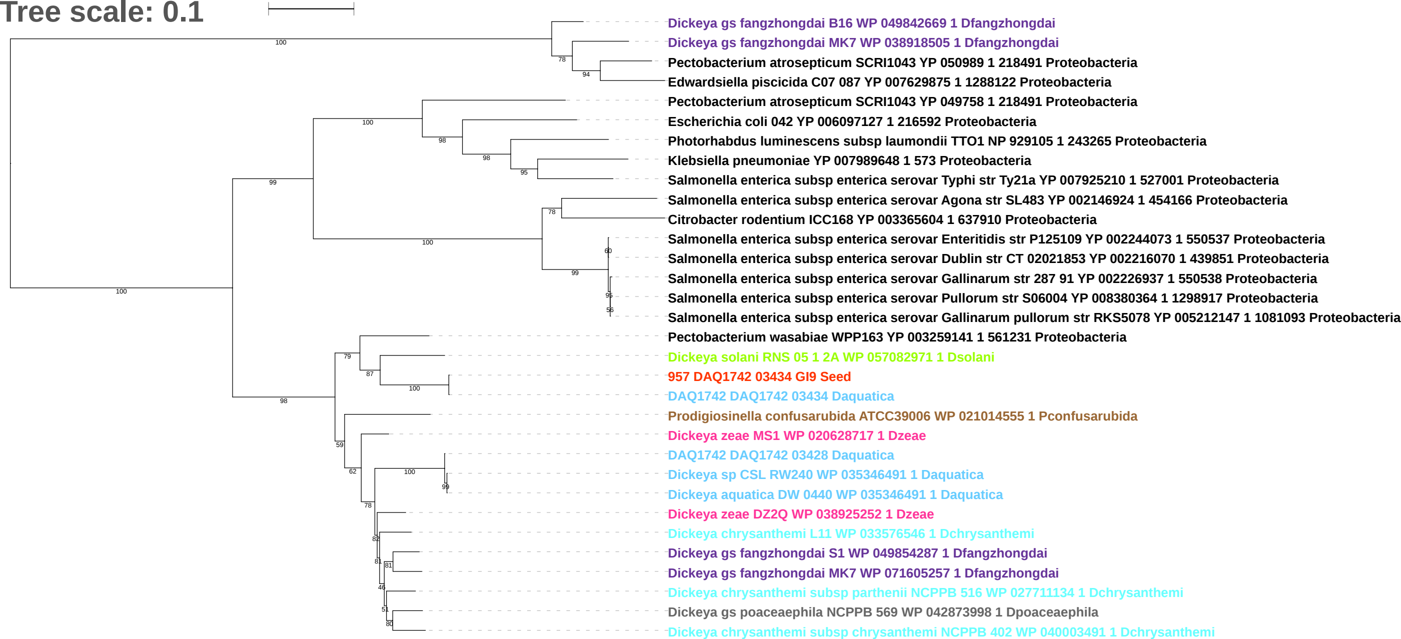
Tree scale: 0.1
Dickeya gs fangzhongdai B16 WP 049842669 1 Dfangzhongdai
Dickeya gs fangzhongdai MK7 WP 038918505 1 Dfangzhongdai
Pectobacterium atrosepticum SCRI1043 YP 050989 1 218491 Proteobacteria
Edwardsiella piscicida C07 087 YP 007629875 1 1288122 Proteobacteria
Pectobacterium atrosepticum SCRI1043 YP 049758 1 218491 Proteobacteria
Escherichia coli 042 YP 006097127 1 216592 Proteobacteria
Photorhabdus luminescens subsp laumondii TTO1 NP 929105 1 243265 Proteobacteria
Klebsiella pneumoniae YP 007989648 1 573 Proteobacteria
Salmonella enterica subsp enterica serovar Typhi str Ty21a YP 007925210 1 527001 Proteobacteria
Salmonella enterica subsp enterica serovar Agona str SL483 YP 002146924 1 454166 Proteobacteria
Citrobacter rodentium ICC168 YP 003365604 1 637910 Proteobacteria
Salmonella enterica subsp enterica serovar Enteritidis str P125109 YP 002244073 1 550537 Proteobacteria
Salmonella enterica subsp enterica serovar Dublin str CT 02021853 YP 002216070 1 439851 Proteobacteria
Salmonella enterica subsp enterica serovar Gallinarum str 287 91 YP 002226937 1 550538 Proteobacteria
Salmonella enterica subsp enterica serovar Pullorum str S06004 YP 008380364 1 1298917 Proteobacteria
Salmonella enterica subsp enterica serovar Gallinarum pullorum str RKS5078 YP 005212147 1 1081093 Proteobacteria
Pectobacterium wasabiae WPP163 YP 003259141 1 561231 Proteobacteria
Dickeya solani RNS 05 1 2A WP 057082971 1 Dsolani
957 DAQ1742 03434 GI9 Seed
DAQ1742 DAQ1742 03434 Daquatica
Prodigiosinella confusarubida ATCC39006 WP 021014555 1 Pconfusarubida
Dickeya zeae MS1 WP 020628717 1 Dzeae
DAQ1742 DAQ1742 03428 Daquatica
Dickeya sp CSL RW240 WP 035346491 1 Daquatica
Dickeya aquatica DW 0440 WP 035346491 1 Daquatica
Dickeya zeae DZ2Q WP 038925252 1 Dzeae
Dickeya chrysanthemi L11 WP 033576546 1 Dchrysanthemi
Dickeya gs fangzhongdai S1 WP 049854287 1 Dfangzhongdai
Dickeya gs fangzhongdai MK7 WP 071605257 1 Dfangzhongdai
Dickeya chrysanthemi subsp parthenii NCPPB 516 WP 027711134 1 Dchrysanthemi
Dickeya gs poaceaephila NCPPB 569 WP 042873998 1 Dpoaceaephila
Dickeya chrysanthemi subsp chrysanthemi NCPPB 402 WP 040003491 1 Dchrysanthemi

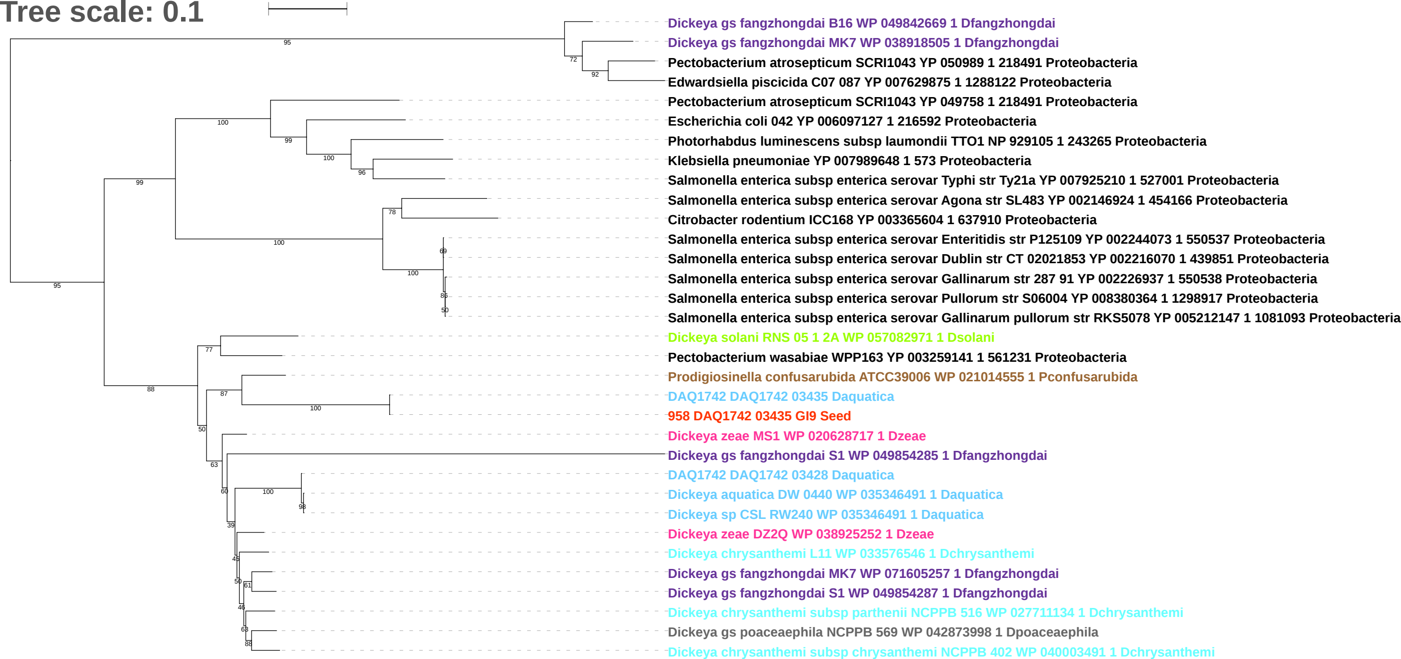
Tree scale: 0.1
Dickeya gs fangzhongdai B16 WP 049842669 1 Dfangzhongdai
Dickeya gs fangzhongdai MK7 WP 038918505 1 Dfangzhongdai
Pectobacterium atrosepticum SCRI1043 YP 050989 1 218491 Proteobacteria
Edwardsiella piscicida C07 087 YP 007629875 1 1288122 Proteobacteria
Pectobacterium atrosepticum SCRI1043 YP 049758 1 218491 Proteobacteria
Escherichia coli 042 YP 006097127 1 216592 Proteobacteria
Photorhabdus luminescens subsp laumondii TTO1 NP 929105 1 243265 Proteobacteria
Klebsiella pneumoniae YP 007989648 1 573 Proteobacteria
Salmonella enterica subsp enterica serovar Typhi str Ty21a YP 007925210 1 527001 Proteobacteria
Salmonella enterica subsp enterica serovar Agona str SL483 YP 002146924 1 454166 Proteobacteria
Citrobacter rodentium ICC168 YP 003365604 1 637910 Proteobacteria
Salmonella enterica subsp enterica serovar Enteritidis str P125109 YP 002244073 1 550537 Proteobacteria
Salmonella enterica subsp enterica serovar Dublin str CT 02021853 YP 002216070 1 439851 Proteobacteria
Salmonella enterica subsp enterica serovar Gallinarum str 287 91 YP 002226937 1 550538 Proteobacteria
Salmonella enterica subsp enterica serovar Pullorum str S06004 YP 008380364 1 1298917 Proteobacteria
Salmonella enterica subsp enterica serovar Gallinarum pullorum str RKS5078 YP 005212147 1 1081093 Proteobacteria
Dickeya solani RNS 05 1 2A WP 057082971 1 Dsolani
Pectobacterium wasabiae WPP163 YP 003259141 1 561231 Proteobacteria
Prodigiosinella confusarubida ATCC39006 WP 021014555 1 Pconfusarubida
DAQ1742 DAQ1742 03435 Daquatica
958 DAQ1742 03435 GI9 Seed
Dickeya zeae MS1 WP 020628717 1 Dzeae
Dickeya gs fangzhongdai S1 WP 049854285 1 Dfangzhongdai
DAQ1742 DAQ1742 03428 Daquatica
Dickeya aquatica DW 0440 WP 035346491 1 Daquatica
Dickeya sp CSL RW240 WP 035346491 1 Daquatica
Dickeya zeae DZ2Q WP 038925252 1 Dzeae
Dickeya chrysanthemi L11 WP 033576546 1 Dchrysanthemi
Dickeya gs fangzhongdai MK7 WP 071605257 1 Dfangzhongdai
Dickeya gs fangzhongdai S1 WP 049854287 1 Dfangzhongdai
Dickeya chrysanthemi subsp parthenii NCPPB 516 WP 027711134 1 Dchrysanthemi
Dickeya gs poaceaephila NCPPB 569 WP 042873998 1 Dpoaceaephila
Dickeya chrysanthemi subsp chrysanthemi NCPPB 402 WP 040003491 1 Dchrysanthemi

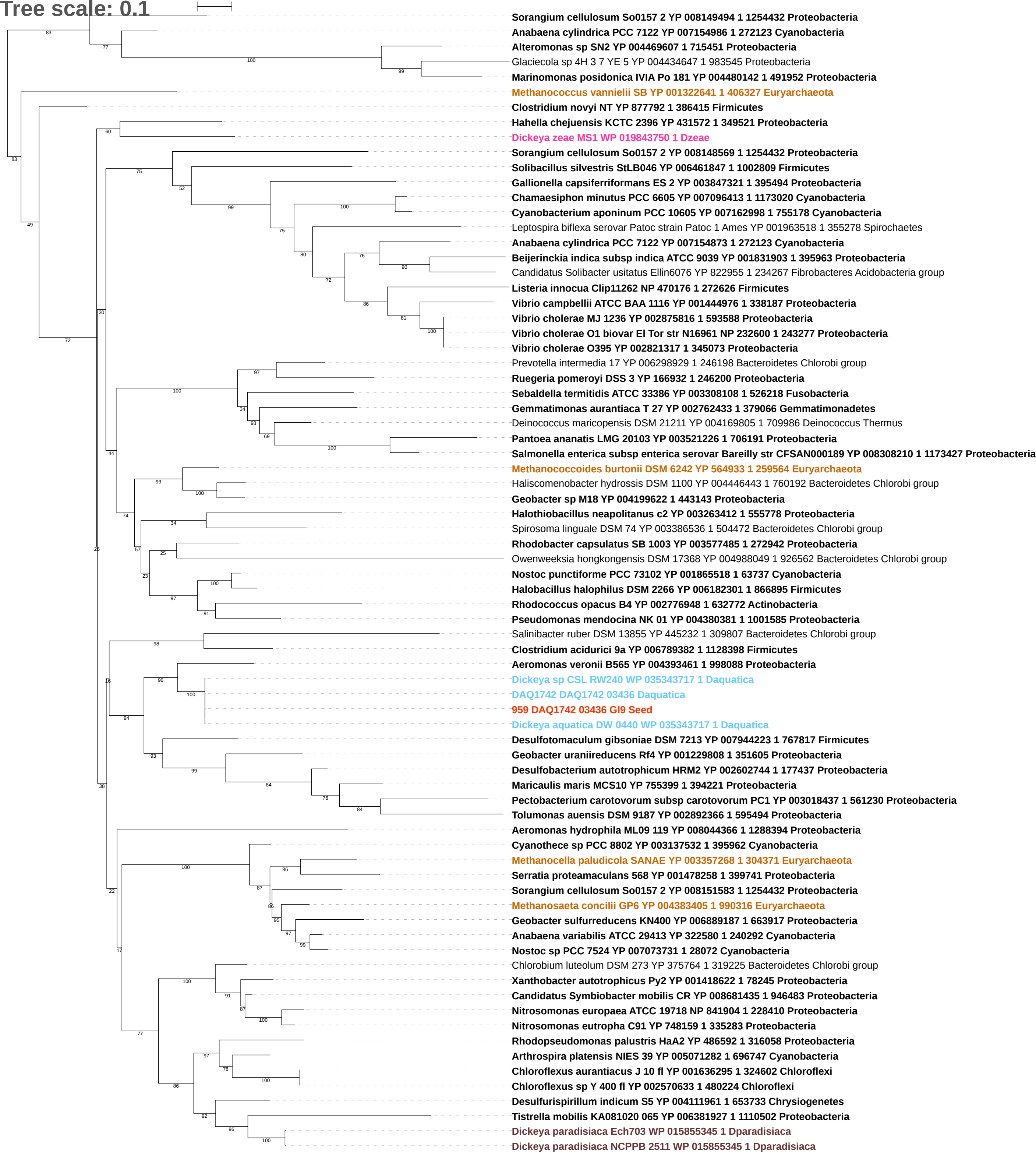

Tree scale: 0.1
Sorangium cellulosum So0157 2 YP 008149494 1 1254432 Proteobacteria
Anabaena cylindrica PCC 7122 YP 007154986 1 272123 Cyanobacteria
Alteromonas sp SN2 YP 004469607 1 715451 Proteobacteria
Glaciecola sp 4H 3 7 YE 5 YP 004434647 1 983545 Proteobacteria
Marinomonas posidonica IVIA Po 181 YP 004480142 1 491952 Proteobacteria
Methanococcus vannielii SB YP 001322641 1 406327 Euryarchaeota
Clostridium novyi NT YP 877792 1 386415 Firmicutes
Hahella chejuensis KCTC 2396 YP 431572 1 349521 Proteobacteria
Dickeya zeae MS1 WP 019843750 1 Dzeae
Sorangium cellulosum So0157 2 YP 008148569 1 1254432 Proteobacteria
Solibacillus silvestris StLB046 YP 006461847 1 1002809 Firmicutes
Gallionella capsiferriformans ES 2 YP 003847321 1 395494 Proteobacteria
Chamaesiphon minutus PCC 6605 YP 007096413 1 1173020 Cyanobacteria
Cyanobacterium aponinum PCC 10605 YP 007162998 1 755178 Cyanobacteria
Leptospira biflexa serovar Patoc strain Patoc 1 Ames YP 001963518 1 355278 Spirochaetes
Anabaena cylindrica PCC 7122 YP 007154873 1 272123 Cyanobacteria
Beijerinckia indica subsp indica ATCC 9039 YP 001831903 1 395963 Proteobacteria
Candidatus Solibacter usitatus Ellin6076 YP 822955 1 234267 Fibrobacteres Acidobacteria group
Listeria innocua Clip11262 NP 470176 1 272626 Firmicutes
Vibrio campbellii ATCC BAA 1116 YP 001444976 1 338187 Proteobacteria
Vibrio cholerae MJ 1236 YP 002875816 1 593588 Proteobacteria
Vibrio cholerae O1 biovar El Tor str N16961 NP 232600 1 243277 Proteobacteria
Vibrio cholerae O395 YP 002821317 1 345073 Proteobacteria
Prevotella intermedia 17 YP 006298929 1 246198 Bacteroidetes Chlorobi group
Ruegeria pomeroyi DSS 3 YP 166932 1 246200 Proteobacteria
Sebaldella termitidis ATCC 33386 YP 003308108 1 526218 Fusobacteria
Gemmatimonas aurantiaca T 27 YP 002762433 1 379066 Gemmatimonadetes
Deinococcus maricopensis DSM 21211 YP 004169805 1 709986 Deinococcus Thermus
Pantoea ananatis LMG 20103 YP 003521226 1 706191 Proteobacteria
Salmonella enterica subsp enterica serovar Bareilly str CFSAN000189 YP 008308210 1 1173427 Proteobacteria
Methanococcoides burtonii DSM 6242 YP 564933 1 259564 Euryarchaeota
Haliscomenobacter hydrossis DSM 1100 YP 004446443 1 760192 Bacteroidetes Chlorobi group
Geobacter sp M18 YP 004199622 1 443143 Proteobacteria
Halothiobacillus neapolitanus c2 YP 003263412 1 555778 Proteobacteria
Spirosoma linguale DSM 74 YP 003386536 1 504472 Bacteroidetes Chlorobi group
Rhodobacter capsulatus SB 1003 YP 003577485 1 272942 Proteobacteria
Owenweeksia hongkongensis DSM 17368 YP 004988049 1 926562 Bacteroidetes Chlorobi group
Nostoc punctiforme PCC 73102 YP 001865518 1 63737 Cyanobacteria
Halobacillus halophilus DSM 2266 YP 006182301 1 866895 Firmicutes
Rhodococcus opacus B4 YP 002776948 1 632772 Actinobacteria
Pseudomonas mendocina NK 01 YP 004380381 1 1001585 Proteobacteria
Salinibacter ruber DSM 13855 YP 445232 1 309807 Bacteroidetes Chlorobi group
Clostridium acidurici 9a YP 006789382 1 1128398 Firmicutes
Aeromonas veronii B565 YP 004393461 1 998088 Proteobacteria
Dickeya sp CSL RW240 WP 035343717 1 Daquatica
DAQ1742 DAQ1742 03436 Daquatica
959 DAQ1742 03436 GI9 Seed
Dickeya aquatica DW 0440 WP 035343717 1 Daquatica
Desulfotomaculum gibsoniae DSM 7213 YP 007944223 1 767817 Firmicutes
Geobacter uraniireducens Rf4 YP 001229808 1 351605 Proteobacteria
Desulfobacterium autotrophicum HRM2 YP 002602744 1 177437 Proteobacteria
Maricaulis maris MCS10 YP 755399 1 394221 Proteobacteria
Pectobacterium carotovorum subsp carotovorum PC1 YP 003018437 1 561230 Proteobacteria
Tolumonas auensis DSM 9187 YP 002892366 1 595494 Proteobacteria
Aeromonas hydrophila ML09 119 YP 008044366 1 1288394 Proteobacteria
Cyanothece sp PCC 8802 YP 003137532 1 395962 Cyanobacteria
Methanocella paludicola SANAE YP 003357268 1 304371 Euryarchaeota
Serratia proteamaculans 568 YP 001478258 1 399741 Proteobacteria
Sorangium cellulosum So0157 2 YP 008151583 1 1254432 Proteobacteria
Methanosaeta concilii GP6 YP 004383405 1 990316 Euryarchaeota
Geobacter sulfurreducens KN400 YP 006889187 1 663917 Proteobacteria
Anabaena variabilis ATCC 29413 YP 322580 1 240292 Cyanobacteria
Nostoc sp PCC 7524 YP 007073731 1 28072 Cyanobacteria
Chlorobium luteolum DSM 273 YP 375764 1 319225 Bacteroidetes Chlorobi group
Xanthobacter autotrophicus Py2 YP 001418622 1 78245 Proteobacteria
Candidatus Symbiobacter mobilis CR YP 008681435 1 946483 Proteobacteria
Nitrosomonas europaea ATCC 19718 NP 841904 1 228410 Proteobacteria
Nitrosomonas eutropha C91 YP 748159 1 335283 Proteobacteria
Rhodopseudomonas palustris HaA2 YP 486592 1 316058 Proteobacteria
Arthrospira platensis NIES 39 YP 005071282 1 696747 Cyanobacteria
Chloroflexus aurantiacus J 10 fl YP 001636295 1 324602 Chloroflexi
Chloroflexus sp Y 400 fl YP 002570633 1 480224 Chloroflexi
Desulfurispirillum indicum S5 YP 004111961 1 653733 Chrysiogenetes
Tistrella mobilis KA081020 065 YP 006381927 1 1110502 Proteobacteria
Dickeya paradisiaca Ech703 WP 015855345 1 Dparadisiaca
Dickeya paradisiaca NCPPB 2511 WP 015855345 1 Dparadisiaca

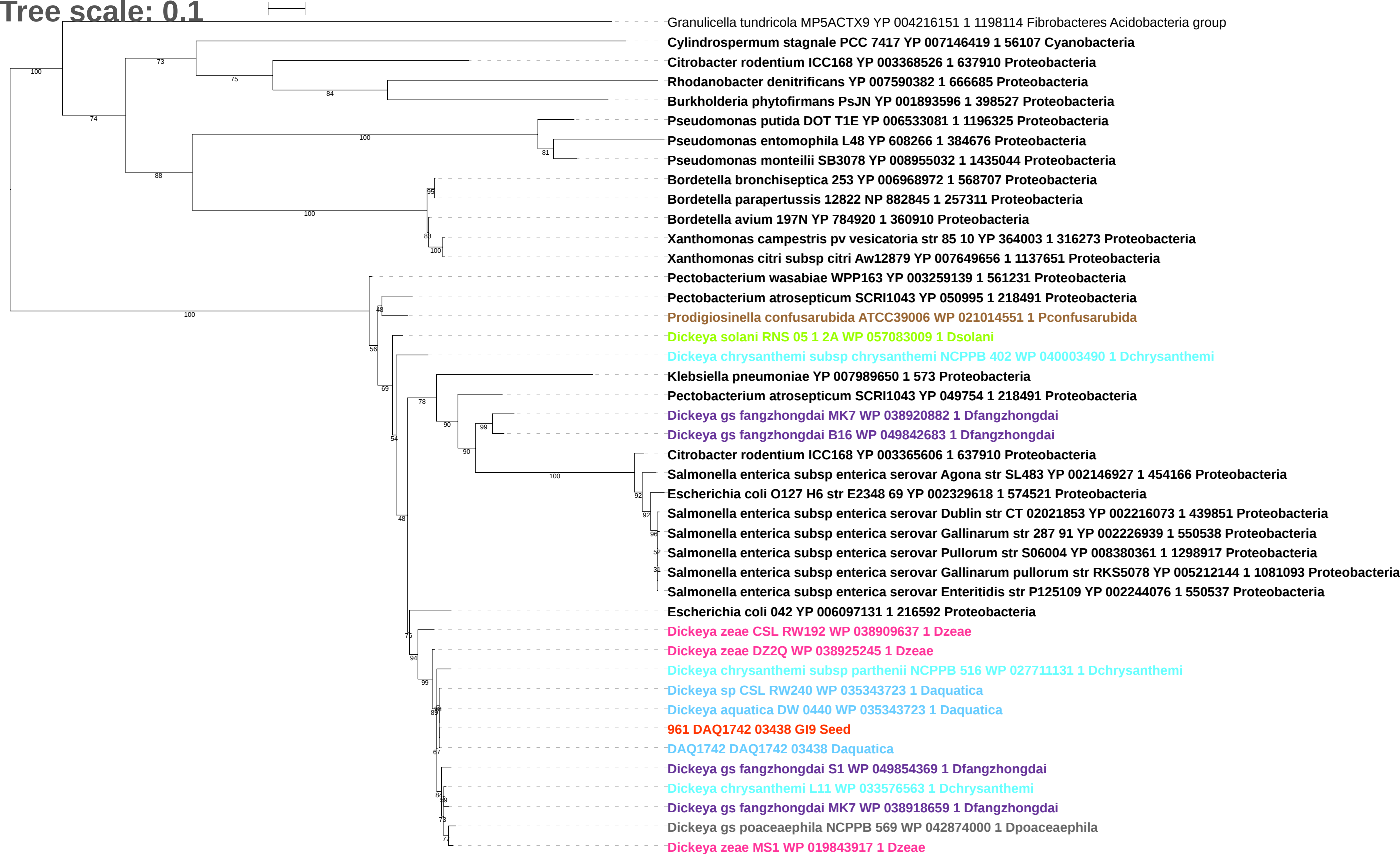
Tree scale: 0.1
Granulicella tundricola MP5ACTX9 YP 004216151 1 1198114 Fibrobacteres Acidobacteria group
Cylindrospermum stagnale PCC 7417 YP 007146419 1 56107 Cyanobacteria
Citrobacter rodentium ICC168 YP 003368526 1 637910 Proteobacteria
Rhodanobacter denitrificans YP 007590382 1 666685 Proteobacteria
Burkholderia phytofirmans PsJN YP 001893596 1 398527 Proteobacteria
Pseudomonas putida DOT T1E YP 006533081 1 1196325 Proteobacteria
Pseudomonas entomophila L48 YP 608266 1 384676 Proteobacteria
Pseudomonas monteilii SB3078 YP 008955032 1 1435044 Proteobacteria
Bordetella bronchiseptica 253 YP 006968972 1 568707 Proteobacteria
Bordetella parapertussis 12822 NP 882845 1 257311 Proteobacteria
Bordetella avium 197N YP 784920 1 360910 Proteobacteria
Xanthomonas campestris pv vesicatoria str 85 10 YP 364003 1 316273 Proteobacteria
Xanthomonas citri subsp citri Aw12879 YP 007649656 1 1137651 Proteobacteria
Pectobacterium wasabiae WPP163 YP 003259139 1 561231 Proteobacteria
Pectobacterium atrosepticum SCRI1043 YP 050995 1 218491 Proteobacteria
Prodigiosinella confusarubida ATCC39006 WP 021014551 1 Pconfusarubida
Dickeya solani RNS 05 1 2A WP 057083009 1 Dsolani
Dickeya chrysanthemi subsp chrysanthemi NCPPB 402 WP 040003490 1 Dchrysanthemi
Klebsiella pneumoniae YP 007989650 1 573 Proteobacteria
Pectobacterium atrosepticum SCRI1043 YP 049754 1 218491 Proteobacteria
Dickeya gs fangzhongdai MK7 WP 038920882 1 Dfangzhongdai
Dickeya gs fangzhongdai B16 WP 049842683 1 Dfangzhongdai
Citrobacter rodentium ICC168 YP 003365606 1 637910 Proteobacteria
Salmonella enterica subsp enterica serovar Agona str SL483 YP 002146927 1 454166 Proteobacteria
Escherichia coli O127 H6 str E2348 69 YP 002329618 1 574521 Proteobacteria
Salmonella enterica subsp enterica serovar Dublin str CT 02021853 YP 002216073 1 439851 Proteobacteria
Salmonella enterica subsp enterica serovar Gallinarum str 287 91 YP 002226939 1 550538 Proteobacteria
Salmonella enterica subsp enterica serovar Pullorum str S06004 YP 008380361 1 1298917 Proteobacteria
Salmonella enterica subsp enterica serovar Gallinarum pullorum str RKS5078 YP 005212144 1 1081093 Proteobacteria
Salmonella enterica subsp enterica serovar Enteritidis str P125109 YP 002244076 1 550537 Proteobacteria
Escherichia coli 042 YP 006097131 1 216592 Proteobacteria
Dickeya zeae CSL RW192 WP 038909637 1 Dzeae
Dickeya zeae DZ2Q WP 038925245 1 Dzeae
Dickeya chrysanthemi subsp parthenii NCPPB 516 WP 027711131 1 Dchrysanthemi
Dickeya sp CSL RW240 WP 035343723 1 Daquatica
Dickeya aquatica DW 0440 WP 035343723 1 Daquatica
961 DAQ1742 03438 GI9 Seed
DAQ1742 DAQ1742 03438 Daquatica
Dickeya gs fangzhongdai S1 WP 049854369 1 Dfangzhongdai
Dickeya chrysanthemi L11 WP 033576563 1 Dchrysanthemi
Dickeya gs fangzhongdai MK7 WP 038918659 1 Dfangzhongdai
Dickeya gs poaceaephila NCPPB 569 WP 042874000 1 Dpoaceaephila
Dickeya zeae MS1 WP 019843917 1 Dzeae

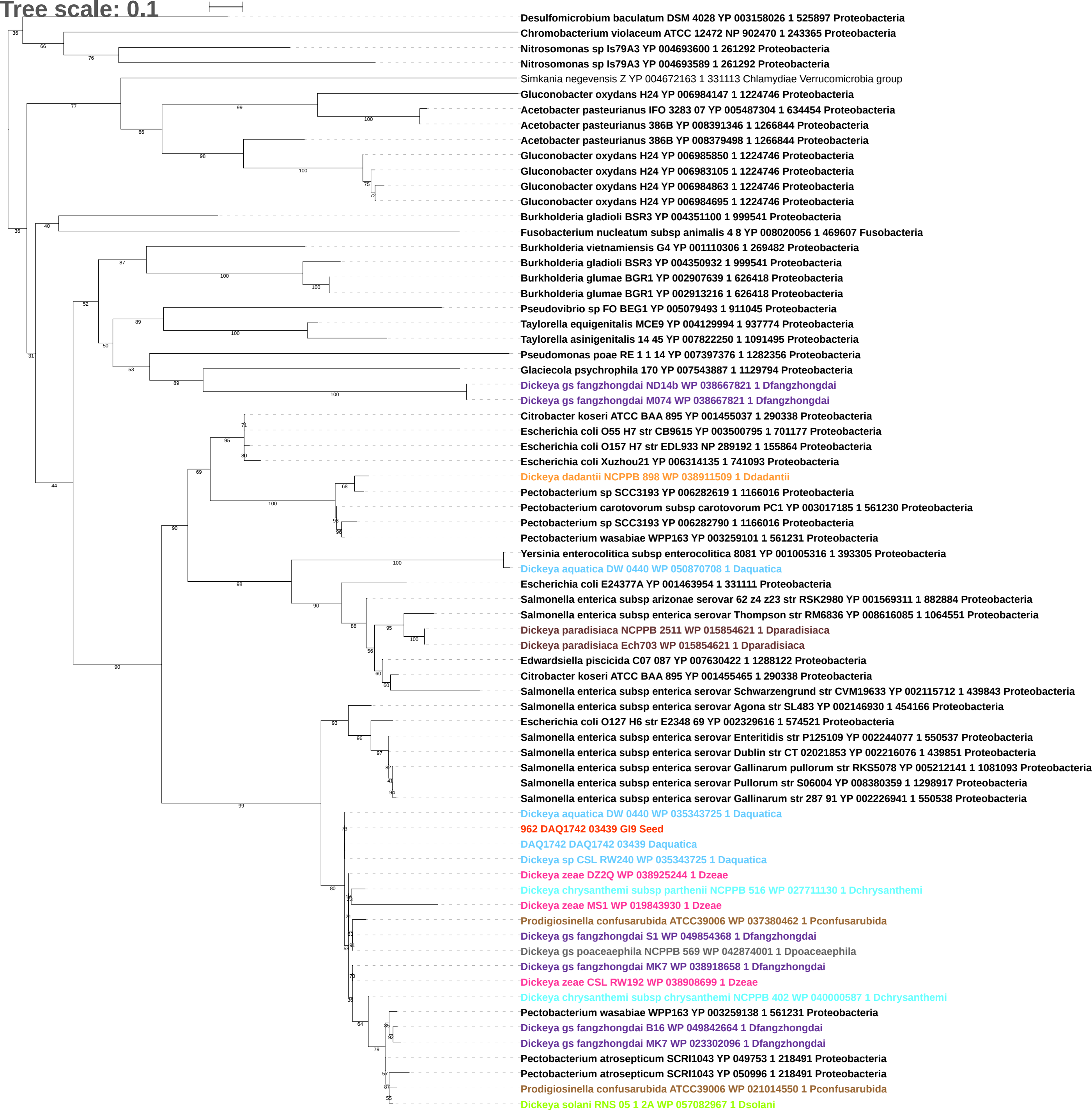
Tree scale: 0.1
Desulfomicrobium baculatum DSM 4028 YP 003158026 1 525897 Proteobacteria
Chromobacterium violaceum ATCC 12472 NP 902470 1 243365 Proteobacteria
Nitrosomonas sp Is79A3 YP 004693600 1 261292 Proteobacteria
Nitrosomonas sp Is79A3 YP 004693589 1 261292 Proteobacteria
Simkania negevensis Z YP 004672163 1 331113 Chlamydiae Verrucomicrobia group
Gluconobacter oxydans H24 YP 006984147 1 1224746 Proteobacteria
Acetobacter pasteurianus IFO 3283 07 YP 005487304 1 634454 Proteobacteria
Acetobacter pasteurianus 386B YP 008391346 1 1266844 Proteobacteria
Acetobacter pasteurianus 386B YP 008379498 1 1266844 Proteobacteria
Gluconobacter oxydans H24 YP 006985850 1 1224746 Proteobacteria
Gluconobacter oxydans H24 YP 006983105 1 1224746 Proteobacteria
Gluconobacter oxydans H24 YP 006984863 1 1224746 Proteobacteria
Gluconobacter oxydans H24 YP 006984695 1 1224746 Proteobacteria
Burkholderia gladioli BSR3 YP 004351100 1 999541 Proteobacteria
Fusobacterium nucleatum subsp animalis 4 8 YP 008020056 1 469607 Fusobacteria
Burkholderia vietnamiensis G4 YP 001110306 1 269482 Proteobacteria
Burkholderia gladioli BSR3 YP 004350932 1 999541 Proteobacteria
Burkholderia glumae BGR1 YP 002907639 1 626418 Proteobacteria
Burkholderia glumae BGR1 YP 002913216 1 626418 Proteobacteria
Pseudovibrio sp FO BEG1 YP 005079493 1 911045 Proteobacteria
Taylorella equigenitalis MCE9 YP 004129994 1 937774 Proteobacteria
Taylorella asinigenitalis 14 45 YP 007822250 1 1091495 Proteobacteria
Pseudomonas poae RE 1 1 14 YP 007397376 1 1282356 Proteobacteria
Glaciecola psychrophila 170 YP 007543887 1 1129794 Proteobacteria
Dickeya gs fangzhongdai ND14b WP 038667821 1 Dfangzhongdai
Dickeya gs fangzhongdai M074 WP 038667821 1 Dfangzhongdai
Citrobacter koseri ATCC BAA 895 YP 001455037 1 290338 Proteobacteria
Escherichia coli O55 H7 str CB9615 YP 003500795 1 701177 Proteobacteria
Escherichia coli O157 H7 str EDL933 NP 289192 1 155864 Proteobacteria
Escherichia coli Xuzhou21 YP 006314135 1 741093 Proteobacteria
Dickeya dadantii NCPPB 898 WP 038911509 1 Ddadantii
Pectobacterium sp SCC3193 YP 006282619 1 1166016 Proteobacteria
Pectobacterium carotovorum subsp carotovorum PC1 YP 003017185 1 561230 Proteobacteria
Pectobacterium sp SCC3193 YP 006282790 1 1166016 Proteobacteria
Pectobacterium wasabiae WPP163 YP 003259101 1 561231 Proteobacteria
Yersinia enterocolitica subsp enterocolitica 8081 YP 001005316 1 393305 Proteobacteria
Dickeya aquatica DW 0440 WP 050870708 1 Daquatica
Escherichia coli E24377A YP 001463954 1 331111 Proteobacteria
Salmonella enterica subsp arizonae serovar 62 z4 z23 str RSK2980 YP 001569311 1 882884 Proteobacteria
Salmonella enterica subsp enterica serovar Thompson str RM6836 YP 008616085 1 1064551 Proteobacteria
Dickeya paradisiaca NCPPB 2511 WP 015854621 1 Dparadisiaca
Dickeya paradisiaca Ech703 WP 015854621 1 Dparadisiaca
Edwardsiella piscicida C07 087 YP 007630422 1 1288122 Proteobacteria
Citrobacter koseri ATCC BAA 895 YP 001455465 1 290338 Proteobacteria
Salmonella enterica subsp enterica serovar Schwarzengrund str CVM19633 YP 002115712 1 439843 Proteobacteria
Salmonella enterica subsp enterica serovar Agona str SL483 YP 002146930 1 454166 Proteobacteria
Escherichia coli O127 H6 str E2348 69 YP 002329616 1 574521 Proteobacteria
Salmonella enterica subsp enterica serovar Enteritidis str P125109 YP 002244077 1 550537 Proteobacteria
Salmonella enterica subsp enterica serovar Dublin str CT 02021853 YP 002216076 1 439851 Proteobacteria
Salmonella enterica subsp enterica serovar Gallinarum pullorum str RKS5078 YP 005212141 1 1081093 Proteobacteria
Salmonella enterica subsp enterica serovar Pullorum str S06004 YP 008380359 1 1298917 Proteobacteria
Salmonella enterica subsp enterica serovar Gallinarum str 287 91 YP 002226941 1 550538 Proteobacteria
Dickeya aquatica DW 0440 WP 035343725 1 Daquatica
962 DAQ1742 03439 GI9 Seed
DAQ1742 DAQ1742 03439 Daquatica
Dickeya sp CSL RW240 WP 035343725 1 Daquatica
Dickeya zeae DZ2Q WP 038925244 1 Dzeae
Dickeya chrysanthemi subsp parthenii NCPPB 516 WP 027711130 1 Dchrysanthemi
Dickeya zeae MS1 WP 019843930 1 Dzeae
Prodigiosinella confusarubida ATCC39006 WP 037380462 1 Pconfusarubida
Dickeya gs fangzhongdai S1 WP 049854368 1 Dfangzhongdai
Dickeya gs poaceaephila NCPPB 569 WP 042874001 1 Dpoaceaephila
Dickeya gs fangzhongdai MK7 WP 038918658 1 Dfangzhongdai
Dickeya zeae CSL RW192 WP 038908699 1 Dzeae
Dickeya chrysanthemi subsp chrysanthemi NCPPB 402 WP 040000587 1 Dchrysanthemi
Pectobacterium wasabiae WPP163 YP 003259138 1 561231 Proteobacteria
Dickeya gs fangzhongdai B16 WP 049842664 1 Dfangzhongdai
Dickeya gs fangzhongdai MK7 WP 023302096 1 Dfangzhongdai
Pectobacterium atrosepticum SCRI1043 YP 049753 1 218491 Proteobacteria
Pectobacterium atrosepticum SCRI1043 YP 050996 1 218491 Proteobacteria
Prodigiosinella confusarubida ATCC39006 WP 021014550 1 Pconfusarubida
Dickeya solani RNS 05 1 2A WP 057082967 1 Dsolani

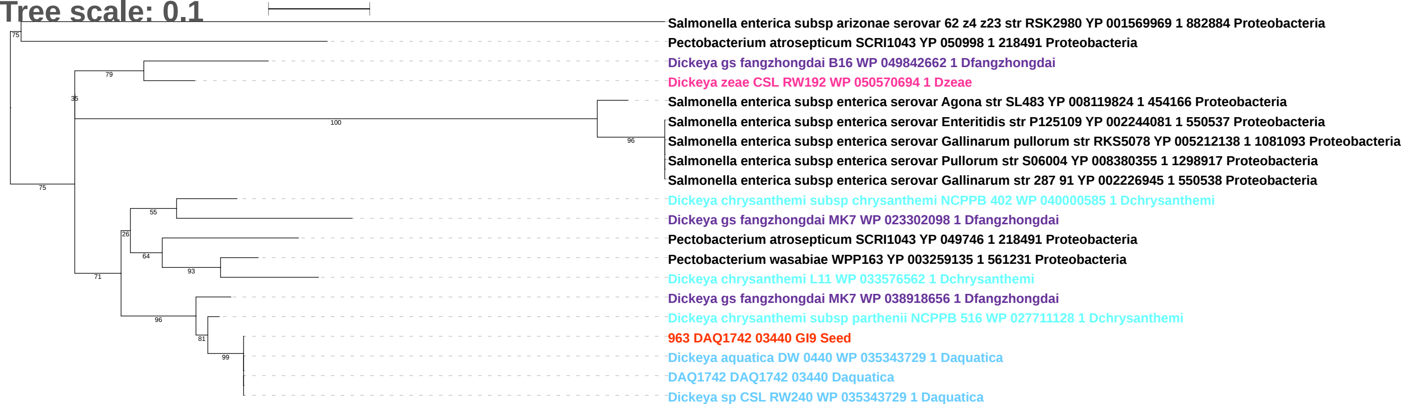
Tree scale: 0.1
Salmonella enterica subsp arizonae serovar 62 z4 z23 str RSK2980 YP 001569969 1 882884 Proteobacteria
Pectobacterium atrosepticum SCRI1043 YP 050998 1 218491 Proteobacteria
Dickeya gs fangzhongdai B16 WP 049842662 1 Dfangzhongdai
Dickeya zeae CSL RW192 WP 050570694 1 Dzeae
Salmonella enterica subsp enterica serovar Agona str SL483 YP 008119824 1 454166 Proteobacteria
Salmonella enterica subsp enterica serovar Enteritidis str P125109 YP 002244081 1 550537 Proteobacteria
Salmonella enterica subsp enterica serovar Gallinarum pullorum str RKS5078 YP 005212138 1 1081093 Proteobacteria
Salmonella enterica subsp enterica serovar Pullorum str S06004 YP 008380355 1 1298917 Proteobacteria
Salmonella enterica subsp enterica serovar Gallinarum str 287 91 YP 002226945 1 550538 Proteobacteria
Dickeya chrysanthemi subsp chrysanthemi NCPPB 402 WP 040000585 1 Dchrysanthemi
Dickeya gs fangzhongdai MK7 WP 023302098 1 Dfangzhongdai
Pectobacterium atrosepticum SCRI1043 YP 049746 1 218491 Proteobacteria
Pectobacterium wasabiae WPP163 YP 003259135 1 561231 Proteobacteria
Dickeya chrysanthemi L11 WP 033576562 1 Dchrysanthemi
Dickeya gs fangzhongdai MK7 WP 038918656 1 Dfangzhongdai
Dickeya chrysanthemi subsp parthenii NCPPB 516 WP 027711128 1 Dchrysanthemi
963 DAQ1742 03440 GI9 Seed
Dickeya aquatica DW 0440 WP 035343729 1 Daquatica
DAQ1742 DAQ1742 03440 Daquatica
Dickeya sp CSL RW240 WP 035343729 1 Daquatica
75
79
35
100
96
75
55
26
64
93
71
96
81
99

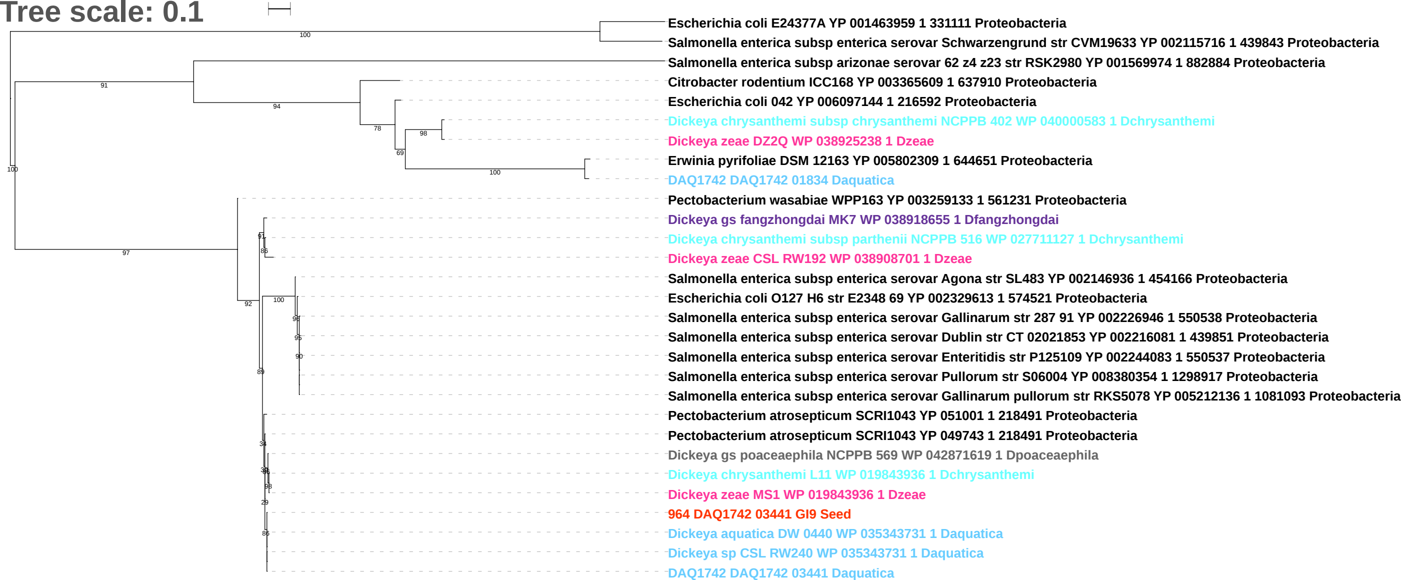
Tree scale: 0.1
Escherichia coli E24377A YP 001463959 1 331111 Proteobacteria
Salmonella enterica subsp enterica serovar Schwarzengrund str CVM19633 YP 002115716 1 439843 Proteobacteria
Salmonella enterica subsp arizonae serovar 62 z4 z23 str RSK2980 YP 001569974 1 882884 Proteobacteria
Citrobacter rodentium ICC168 YP 003365609 1 637910 Proteobacteria
Escherichia coli 042 YP 006097144 1 216592 Proteobacteria
Dickeya chrysanthemi subsp chrysanthemi NCPPB 402 WP 040000583 1 Dchrysanthemi
Dickeya zeae DZ2Q WP 038925238 1 Dzeae
Erwinia pyrifoliae DSM 12163 YP 005802309 1 644651 Proteobacteria
DAQ1742 DAQ1742 01834 Daquatica
Pectobacterium wasabiae WPP163 YP 003259133 1 561231 Proteobacteria
Dickeya gs fangzhongdai MK7 WP 038918655 1 Dfangzhongdai
Dickeya chrysanthemi subsp parthenii NCPPB 516 WP 027711127 1 Dchrysanthemi
Dickeya zeae CSL RW192 WP 038908701 1 Dzeae
Salmonella enterica subsp enterica serovar Agona str SL483 YP 002146936 1 454166 Proteobacteria
Escherichia coli O127 H6 str E2348 69 YP 002329613 1 574521 Proteobacteria
Salmonella enterica subsp enterica serovar Gallinarum str 287 91 YP 002226946 1 550538 Proteobacteria
Salmonella enterica subsp enterica serovar Dublin str CT 02021853 YP 002216081 1 439851 Proteobacteria
Salmonella enterica subsp enterica serovar Enteritidis str P125109 YP 002244083 1 550537 Proteobacteria
Salmonella enterica subsp enterica serovar Pullorum str S06004 YP 008380354 1 1298917 Proteobacteria
Salmonella enterica subsp enterica serovar Gallinarum pullorum str RKS5078 YP 005212136 1 1081093 Proteobacteria
Pectobacterium atrosepticum SCRI1043 YP 051001 1 218491 Proteobacteria
Pectobacterium atrosepticum SCRI1043 YP 049743 1 218491 Proteobacteria
Dickeya gs poaceaephila NCPPB 569 WP 042871619 1 Dpoaceaephila
Dickeya chrysanthemi L11 WP 019843936 1 Dchrysanthemi
Dickeya zeae MS1 WP 019843936 1 Dzeae
964 DAQ1742 03441 GI9 Seed
Dickeya aquatica DW 0440 WP 035343731 1 Daquatica
Dickeya sp CSL RW240 WP 035343731 1 Daquatica
DAQ1742 DAQ1742 03441 Daquatica

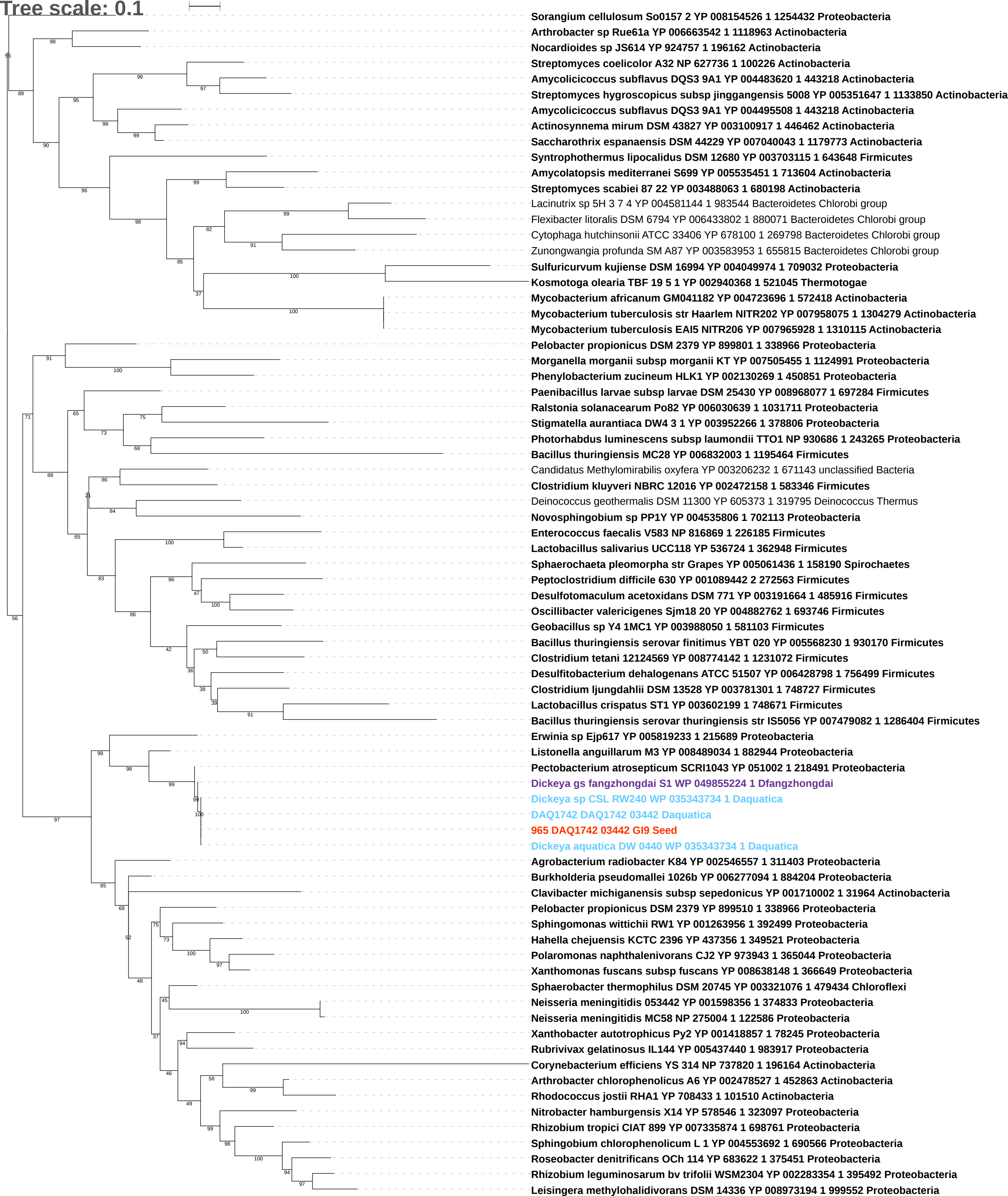
Tree scale: 0.1
Sorangium cellulosum So0157 2 YP 008154526 1 1254432 Proteobacteria
Arthrobacter sp Rue61a YP 006663542 1 1118963 Actinobacteria
Nocardioides sp JS614 YP 924757 1 196162 Actinobacteria
Streptomyces coelicolor A32 NP 627736 1 100226 Actinobacteria
Amycolicicoccus subflavus DQS3 9A1 YP 004483620 1 443218 Actinobacteria
Streptomyces hygroscopicus subsp jinggangensis 5008 YP 005351647 1 1133850 Actinobacteria
Amycolicicoccus subflavus DQS3 9A1 YP 004495508 1 443218 Actinobacteria
Actinosynnema mirum DSM 43827 YP 003100917 1 446462 Actinobacteria
Saccharothrix espanaensis DSM 44229 YP 007040043 1 1179773 Actinobacteria
Syntrophothermus lipocalidus DSM 12680 YP 003703115 1 643648 Firmicutes
Amycolatopsis mediterranei S699 YP 005535451 1 713604 Actinobacteria
Streptomyces scabiei 87 22 YP 003488063 1 680198 Actinobacteria
Lacinutrix sp 5H 3 7 4 YP 004581144 1 983544 Bacteroidetes Chlorobi group
Flexibacter litoralis DSM 6794 YP 006433802 1 880071 Bacteroidetes Chlorobi group
Cytophaga hutchinsonii ATCC 33406 YP 678100 1 269798 Bacteroidetes Chlorobi group
Zunongwangia profunda SM A87 YP 003583953 1 655815 Bacteroidetes Chlorobi group
Sulfuricurvum kujiense DSM 16994 YP 004049974 1 709032 Proteobacteria
Kosmotoga olearia TBF 19 5 1 YP 002940368 1 521045 Thermotogae
Mycobacterium africanum GM041182 YP 004723696 1 572418 Actinobacteria
Mycobacterium tuberculosis str Haarlem NITR202 YP 007958075 1 1304279 Actinobacteria
Mycobacterium tuberculosis EAI5 NITR206 YP 007965928 1 1310115 Actinobacteria
Pelobacter propionicus DSM 2379 YP 899801 1 338966 Proteobacteria
Morganella morganii subsp morganii KT YP 007505455 1 1124991 Proteobacteria
Phenylobacterium zucineum HLK1 YP 002130269 1 450851 Proteobacteria
Paenibacillus larvae subsp larvae DSM 25430 YP 008968077 1 697284 Firmicutes
Ralstonia solanacearum Po82 YP 006030639 1 1031711 Proteobacteria
Stigmatella aurantiaca DW4 3 1 YP 003952266 1 378806 Proteobacteria
Photorhabdus luminescens subsp laumondii TTO1 NP 930686 1 243265 Proteobacteria
Bacillus thuringiensis MC28 YP 006832003 1 1195464 Firmicutes
Candidatus Methylomirabilis oxyfera YP 003206232 1 671143 unclassified Bacteria
Clostridium kluyveri NBRC 12016 YP 002472158 1 583346 Firmicutes
Deinococcus geothermalis DSM 11300 YP 605373 1 319795 Deinococcus Thermus
Novosphingobium sp PP1Y YP 004535806 1 702113 Proteobacteria
Enterococcus faecalis V583 NP 816869 1 226185 Firmicutes
Lactobacillus salivarius UCC118 YP 536724 1 362948 Firmicutes
Sphaerochaeta pleomorpha str Grapes YP 005061436 1 158190 Spirochaetes
Peptoclostridium difficile 630 YP 001089442 2 272563 Firmicutes
Desulfotomaculum acetoxidans DSM 771 YP 003191664 1 485916 Firmicutes
Oscillibacter valericigenes Sjm18 20 YP 004882762 1 693746 Firmicutes
Geobacillus sp Y4 1MC1 YP 003988050 1 581103 Firmicutes
Bacillus thuringiensis serovar finitimus YBT 020 YP 005568230 1 930170 Firmicutes
Clostridium tetani 12124569 YP 008774142 1 1231072 Firmicutes
Desulfitobacterium dehalogenans ATCC 51507 YP 006428798 1 756499 Firmicutes
Clostridium ljungdahlii DSM 13528 YP 003781301 1 748727 Firmicutes
Lactobacillus crispatus ST1 YP 003602199 1 748671 Firmicutes
Bacillus thuringiensis serovar thuringiensis str IS5056 YP 007479082 1 1286404 Firmicutes
Erwinia sp Ejp617 YP 005819233 1 215689 Proteobacteria
Listonella anguillarum M3 YP 008489034 1 882944 Proteobacteria
Pectobacterium atrosepticum SCRI1043 YP 051002 1 218491 Proteobacteria
Dickeya gs fangzhongdai S1 WP 049855224 1 Dfangzhongdai
Dickeya sp CSL RW240 WP 035343734 1 Daquatica
DAQ1742 DAQ1742 03442 Daquatica
965 DAQ1742 03442 GI9 Seed
Dickeya aquatica DW 0440 WP 035343734 1 Daquatica
Agrobacterium radiobacter K84 YP 002546557 1 311403 Proteobacteria
Burkholderia pseudomallei 1026b YP 006277094 1 884204 Proteobacteria
Clavibacter michiganensis subsp sepedonicus YP 001710002 1 31964 Actinobacteria
Pelobacter propionicus DSM 2379 YP 899510 1 338966 Proteobacteria
Sphingomonas wittichii RW1 YP 001263956 1 392499 Proteobacteria
Hahella chejuensis KCTC 2396 YP 437356 1 349521 Proteobacteria
Polaromonas naphthalenivorans CJ2 YP 973943 1 365044 Proteobacteria
Xanthomonas fuscans subsp fuscans YP 008638148 1 366649 Proteobacteria
Sphaerobacter thermophilus DSM 20745 YP 003321076 1 479434 Chloroflexi
Neisseria meningitidis 053442 YP 001598356 1 374833 Proteobacteria
Neisseria meningitidis MC58 NP 275004 1 122586 Proteobacteria
Xanthobacter autotrophicus Py2 YP 001418857 1 78245 Proteobacteria
Rubrivivax gelatinosus IL144 YP 005437440 1 983917 Proteobacteria
Corynebacterium efficiens YS 314 NP 737820 1 196164 Actinobacteria
Arthrobacter chlorophenolicus A6 YP 002478527 1 452863 Actinobacteria
Rhodococcus jostii RHA1 YP 708433 1 101510 Actinobacteria
Nitrobacter hamburgensis X14 YP 578546 1 323097 Proteobacteria
Rhizobium tropici CIAT 899 YP 007335874 1 698761 Proteobacteria
Sphingobium chlorophenolicum L 1 YP 004553692 1 690566 Proteobacteria
Roseobacter denitrificans OCh 114 YP 683622 1 375451 Proteobacteria
Rhizobium leguminosarum bv trifolii WSM2304 YP 002283354 1 395492 Proteobacteria
Leisingera methylohalidivorans DSM 14336 YP 008973194 1 999552 Proteobacteria

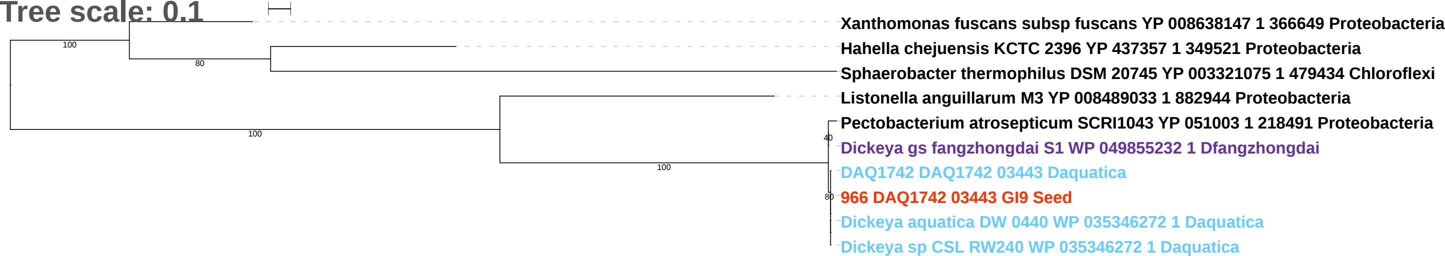

Tree scale: 0.1
Xanthomonas fuscans subsp fuscans YP 008638147 1 366649 Proteobacteria
Hahella chejuensis KCTC 2396 YP 437357 1 349521 Proteobacteria
Sphaerobacter thermophilus DSM 20745 YP 003321075 1 479434 Chloroflexi
Listonella anguillarum M3 YP 008489033 1 882944 Proteobacteria
Pectobacterium atrosepticum SCRI1043 YP 051003 1 218491 Proteobacteria
Dickeya gs fangzhongdai S1 WP 049855232 1 Dfangzhongdai
DAQ1742 DAQ1742 03443 Daquatica
966 DAQ1742 03443 GI9 Seed
Dickeya aquatica DW 0440 WP 035346272 1 Daquatica
Dickeya sp CSL RW240 WP 035346272 1 Daquatica
100
80
100
100
40
80

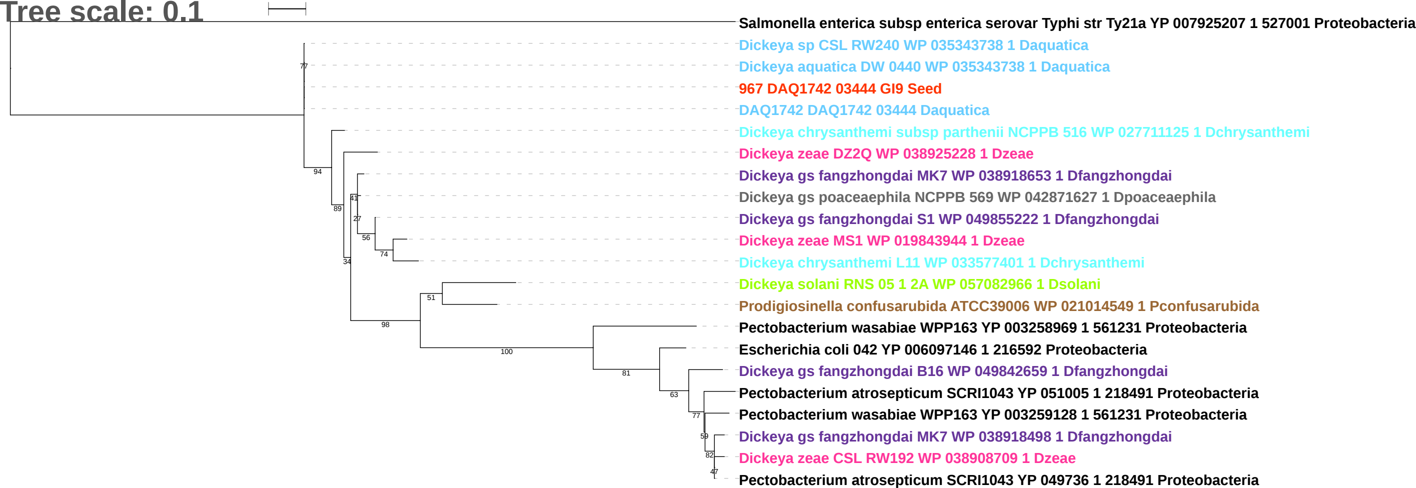
Tree scale: 0.1
Salmonella enterica subsp enterica serovar Typhi str Ty21a YP 007925207 1 527001 Proteobacteria
Dickeya sp CSL RW240 WP 035343738 1 Daquatica
Dickeya aquatica DW 0440 WP 035343738 1 Daquatica
967 DAQ1742 03444 GI9 Seed
DAQ1742 DAQ1742 03444 Daquatica
Dickeya chrysanthemi subsp parthenii NCPPB 516 WP 027711125 1 Dchrysanthemi
Dickeya zeae DZ2Q WP 038925228 1 Dzeae
Dickeya gs fangzhongdai MK7 WP 038918653 1 Dfangzhongdai
Dickeya gs poaceaephila NCPPB 569 WP 042871627 1 Dpoaceaephila
Dickeya gs fangzhongdai S1 WP 049855222 1 Dfangzhongdai
Dickeya zeae MS1 WP 019843944 1 Dzeae
Dickeya chrysanthemi L11 WP 033577401 1 Dchrysanthemi
Dickeya solani RNS 05 1 2A WP 057082966 1 Dsolani
Prodigiosinella confusarubida ATCC39006 WP 021014549 1 Pconfusarubida
Pectobacterium wasabiae WPP163 YP 003258969 1 561231 Proteobacteria
Escherichia coli 042 YP 006097146 1 216592 Proteobacteria
Dickeya gs fangzhongdai B16 WP 049842659 1 Dfangzhongdai
Pectobacterium atrosepticum SCRI1043 YP 051005 1 218491 Proteobacteria
Pectobacterium wasabiae WPP163 YP 003259128 1 561231 Proteobacteria
Dickeya gs fangzhongdai MK7 WP 038918498 1 Dfangzhongdai
Dickeya zeae CSL RW192 WP 038908709 1 Dzeae
Pectobacterium atrosepticum SCRI1043 YP 049736 1 218491 Proteobacteria
77
94
89
41
27
56
74
34
51
98
100
81
63
77
59
82
47

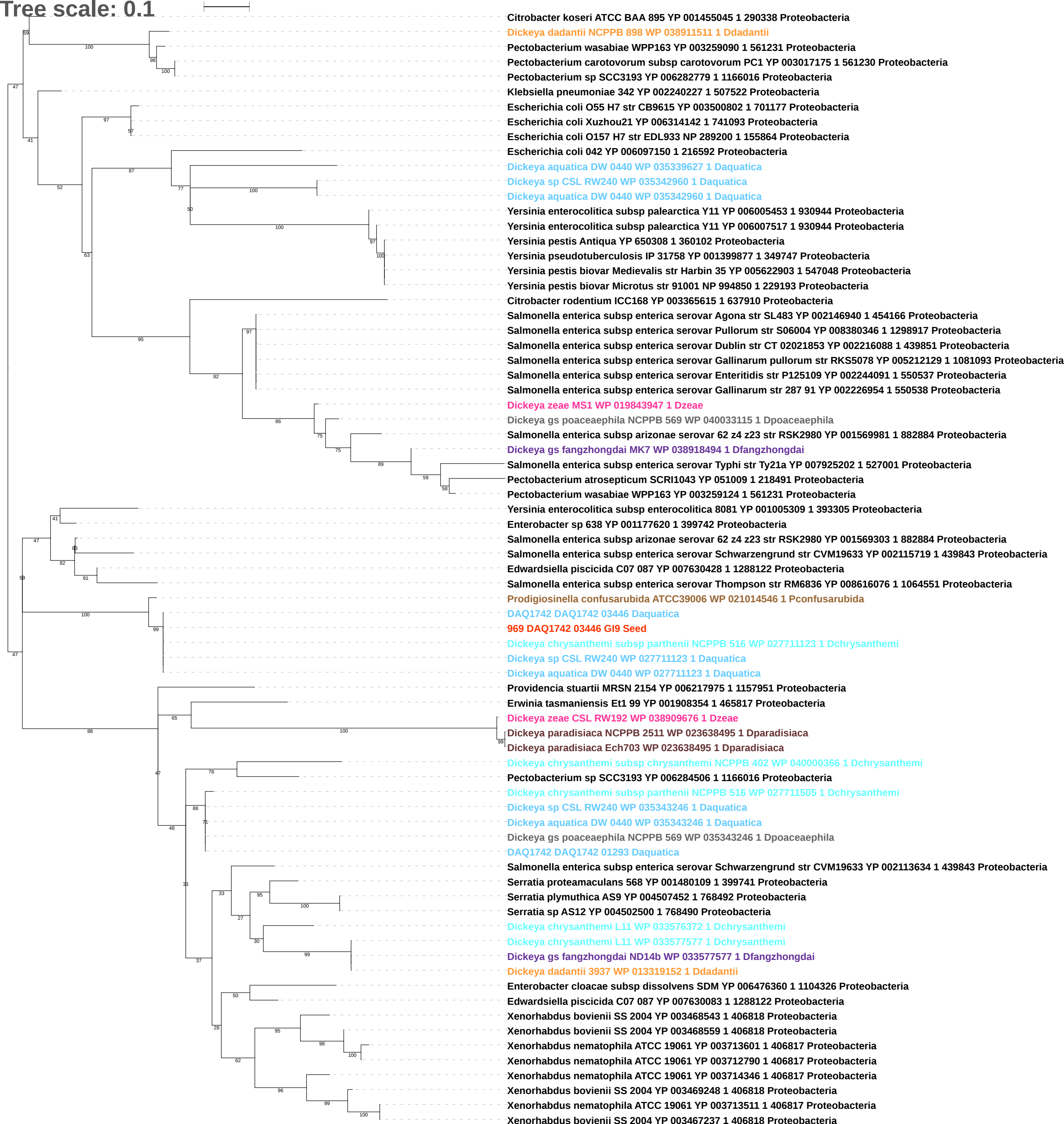
Tree scale: 0.1
Citrobacter koseri ATCC BAA 895 YP 001455045 1 290338 Proteobacteria
Dickeya dadantii NCPPB 898 WP 038911511 1 Ddadantii
Pectobacterium wasabiae WPP163 YP 003259090 1 561231 Proteobacteria
Pectobacterium carotovorum subsp carotovorum PC1 YP 003017175 1 561230 Proteobacteria
Pectobacterium sp SCC3193 YP 006282779 1 1166016 Proteobacteria
Klebsiella pneumoniae 342 YP 002240227 1 507522 Proteobacteria
Escherichia coli O55 H7 str CB9615 YP 003500802 1 701177 Proteobacteria
Escherichia coli Xuzhou21 YP 006314142 1 741093 Proteobacteria
Escherichia coli O157 H7 str EDL933 NP 289200 1 155864 Proteobacteria
Escherichia coli 042 YP 006097150 1 216592 Proteobacteria
Dickeya aquatica DW 0440 WP 035339627 1 Daquatica
Dickeya sp CSL RW240 WP 035342960 1 Daquatica
Dickeya aquatica DW 0440 WP 035342960 1 Daquatica
Yersinia enterocolitica subsp palearctica Y11 YP 006005453 1 930944 Proteobacteria
Yersinia enterocolitica subsp palearctica Y11 YP 006007517 1 930944 Proteobacteria
Yersinia pestis Antiqua YP 650308 1 360102 Proteobacteria
Yersinia pseudotuberculosis IP 31758 YP 001399877 1 349747 Proteobacteria
Yersinia pestis biovar Medievalis str Harbin 35 YP 005622903 1 547048 Proteobacteria
Yersinia pestis biovar Microtus str 91001 NP 994850 1 229193 Proteobacteria
Citrobacter rodentium ICC168 YP 003365615 1 637910 Proteobacteria
Salmonella enterica subsp enterica serovar Agona str SL483 YP 002146940 1 454166 Proteobacteria
Salmonella enterica subsp enterica serovar Pullorum str S06004 YP 008380346 1 1298917 Proteobacteria
Salmonella enterica subsp enterica serovar Dublin str CT 02021853 YP 002216088 1 439851 Proteobacteria
Salmonella enterica subsp enterica serovar Gallinarum pullorum str RKS5078 YP 005212129 1 1081093 Proteobacteria
Salmonella enterica subsp enterica serovar Enteritidis str P125109 YP 002244091 1 550537 Proteobacteria
Salmonella enterica subsp enterica serovar Gallinarum str 287 91 YP 002226954 1 550538 Proteobacteria
Dickeya zeae MS1 WP 019843947 1 Dzeae
Dickeya gs poaceaephila NCPPB 569 WP 040033115 1 Dpoaceaephila
Salmonella enterica subsp arizonae serovar 62 z4 z23 str RSK2980 YP 001569981 1 882884 Proteobacteria
Dickeya gs fangzhongdai MK7 WP 038918494 1 Dfangzhongdai
Salmonella enterica subsp enterica serovar Typhi str Ty21a YP 007925202 1 527001 Proteobacteria
Pectobacterium atrosepticum SCRI1043 YP 051009 1 218491 Proteobacteria
Pectobacterium wasabiae WPP163 YP 003259124 1 561231 Proteobacteria
Yersinia enterocolitica subsp enterocolitica 8081 YP 001005309 1 393305 Proteobacteria
Enterobacter sp 638 YP 001177620 1 399742 Proteobacteria
Salmonella enterica subsp arizonae serovar 62 z4 z23 str RSK2980 YP 001569303 1 882884 Proteobacteria
Salmonella enterica subsp enterica serovar Schwarzengrund str CVM19633 YP 002115719 1 439843 Proteobacteria
Edwardsiella piscicida C07 087 YP 007630428 1 1288122 Proteobacteria
Salmonella enterica subsp enterica serovar Thompson str RM6836 YP 008616076 1 1064551 Proteobacteria
Prodigiosinella confusarubida ATCC39006 WP 021014546 1 Pconfusarubida
DAQ1742 DAQ1742 03446 Daquatica
969 DAQ1742 03446 GI9 Seed
Dickeya chrysanthemi subsp parthenii NCPPB 516 WP 027711123 1 Dchrysanthemi
Dickeya sp CSL RW240 WP 027711123 1 Daquatica
Dickeya aquatica DW 0440 WP 027711123 1 Daquatica
Providencia stuartii MRSN 2154 YP 006217975 1 1157951 Proteobacteria
Erwinia tasmaniensis Et1 99 YP 001908354 1 465817 Proteobacteria
Dickeya zeae CSL RW192 WP 038909676 1 Dzeae
Dickeya paradisiaca NCPPB 2511 WP 023638495 1 Dparadisiaca
Dickeya paradisiaca Ech703 WP 023638495 1 Dparadisiaca
Dickeya chrysanthemi subsp chrysanthemi NCPPB 402 WP 040000366 1 Dchrysanthemi
Pectobacterium sp SCC3193 YP 006284506 1 1166016 Proteobacteria
Dickeya chrysanthemi subsp parthenii NCPPB 516 WP 027711505 1 Dchrysanthemi
Dickeya sp CSL RW240 WP 035343246 1 Daquatica
Dickeya aquatica DW 0440 WP 035343246 1 Daquatica
Dickeya gs poaceaephila NCPPB 569 WP 035343246 1 Dpoaceaephila
DAQ1742 DAQ1742 01293 Daquatica
Salmonella enterica subsp enterica serovar Schwarzengrund str CVM19633 YP 002113634 1 439843 Proteobacteria
Serratia proteamaculans 568 YP 001480109 1 399741 Proteobacteria
Serratia plymuthica AS9 YP 004507452 1 768492 Proteobacteria
Serratia sp AS12 YP 004502500 1 768490 Proteobacteria
Dickeya chrysanthemi L11 WP 033576372 1 Dchrysanthemi
Dickeya chrysanthemi L11 WP 033577577 1 Dchrysanthemi
Dickeya gs fangzhongdai ND14b WP 033577577 1 Dfangzhongdai
Dickeya dadantii 3937 WP 013319152 1 Ddadantii
Enterobacter cloacae subsp dissolvens SDM YP 006476360 1 1104326 Proteobacteria
Edwardsiella piscicida C07 087 YP 007630083 1 1288122 Proteobacteria
Xenorhabdus bovienii SS 2004 YP 003468543 1 406818 Proteobacteria
Xenorhabdus bovienii SS 2004 YP 003468559 1 406818 Proteobacteria
Xenorhabdus nematophila ATCC 19061 YP 003713601 1 406817 Proteobacteria
Xenorhabdus nematophila ATCC 19061 YP 003712790 1 406817 Proteobacteria
Xenorhabdus nematophila ATCC 19061 YP 003714346 1 406817 Proteobacteria
Xenorhabdus bovienii SS 2004 YP 003469248 1 406818 Proteobacteria
Xenorhabdus nematophila ATCC 19061 YP 003713511 1 406817 Proteobacteria
Xenorhabdus bovienii SS 2004 YP 003467237 1 406818 Proteobacteria

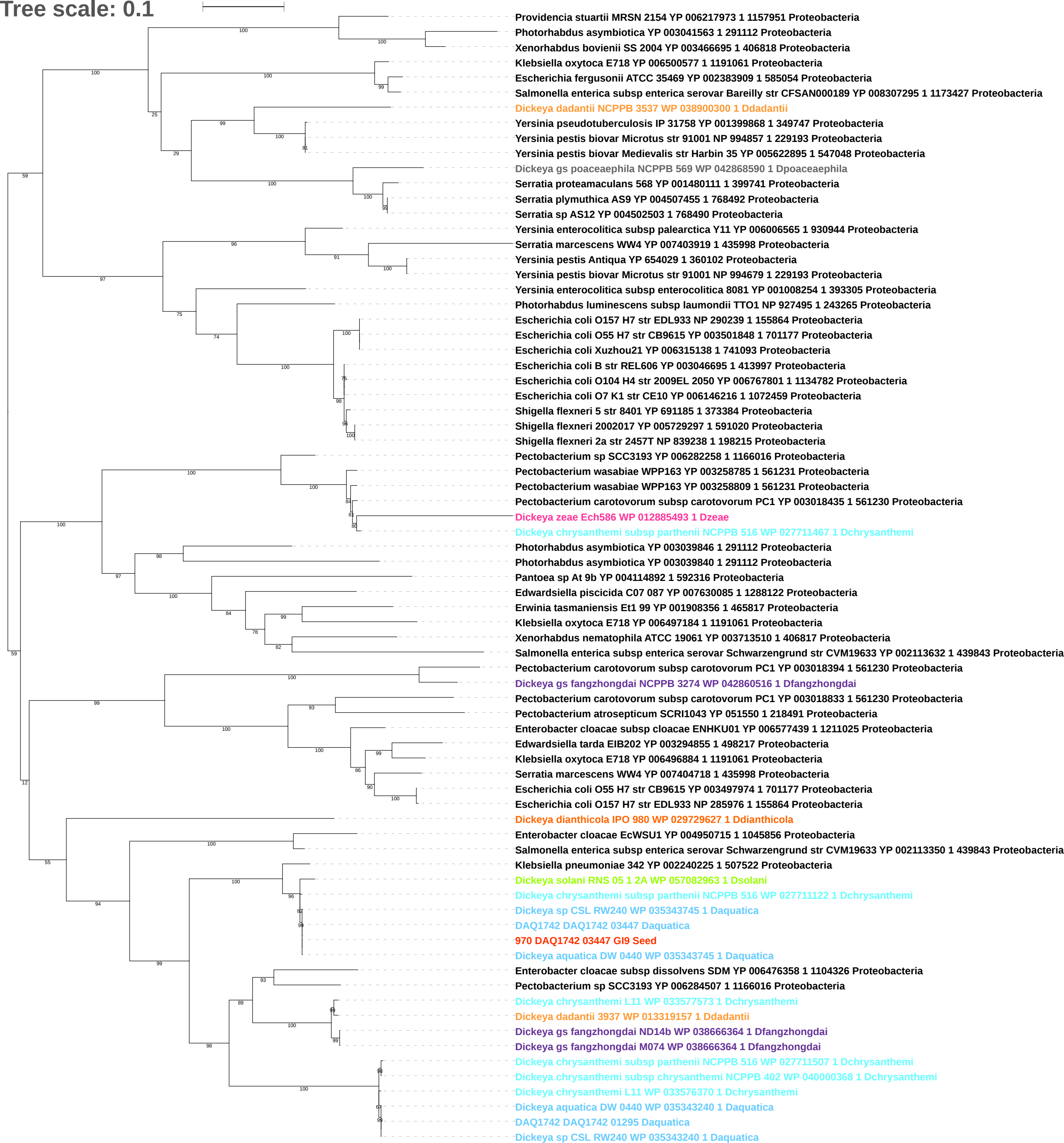
Tree scale: 0.1
Providencia stuartii MRSN 2154 YP 006217973 1 1157951 Proteobacteria
Photorhabdus asymbiotica YP 003041563 1 291112 Proteobacteria
Xenorhabdus bovienii SS 2004 YP 003466695 1 406818 Proteobacteria
Klebsiella oxytoca E718 YP 006500577 1 1191061 Proteobacteria
Escherichia fergusonii ATCC 35469 YP 002383909 1 585054 Proteobacteria
Salmonella enterica subsp enterica serovar Bareilly str CFSAN000189 YP 008307295 1 1173427 Proteobacteria
Dickeya dadantii NCPPB 3537 WP 038900300 1 Ddadantii
Yersinia pseudotuberculosis IP 31758 YP 001399868 1 349747 Proteobacteria
Yersinia pestis biovar Microtus str 91001 NP 994857 1 229193 Proteobacteria
Yersinia pestis biovar Medievalis str Harbin 35 YP 005622895 1 547048 Proteobacteria
Dickeya gs poaceaephila NCPPB 569 WP 042868590 1 Dpoaceaephila
Serratia proteamaculans 568 YP 001480111 1 399741 Proteobacteria
Serratia plymuthica AS9 YP 004507455 1 768492 Proteobacteria
Serratia sp AS12 YP 004502503 1 768490 Proteobacteria
Yersinia enterocolitica subsp palearctica Y11 YP 006006565 1 930944 Proteobacteria
Serratia marcescens WW4 YP 007403919 1 435998 Proteobacteria
Yersinia pestis Antiqua YP 654029 1 360102 Proteobacteria
Yersinia pestis biovar Microtus str 91001 NP 994679 1 229193 Proteobacteria
Yersinia enterocolitica subsp enterocolitica 8081 YP 001008254 1 393305 Proteobacteria
Photorhabdus luminescens subsp laumondii TTO1 NP 927495 1 243265 Proteobacteria
Escherichia coli O157 H7 str EDL933 NP 290239 1 155864 Proteobacteria
Escherichia coli O55 H7 str CB9615 YP 003501848 1 701177 Proteobacteria
Escherichia coli Xuzhou21 YP 006315138 1 741093 Proteobacteria
Escherichia coli B str REL606 YP 003046695 1 413997 Proteobacteria
Escherichia coli O104 H4 str 2009EL 2050 YP 006767801 1 1134782 Proteobacteria
Escherichia coli O7 K1 str CE10 YP 006146216 1 1072459 Proteobacteria
Shigella flexneri 5 str 8401 YP 691185 1 373384 Proteobacteria
Shigella flexneri 2002017 YP 005729297 1 591020 Proteobacteria
Shigella flexneri 2a str 2457T NP 839238 1 198215 Proteobacteria
Pectobacterium sp SCC3193 YP 006282258 1 1166016 Proteobacteria
Pectobacterium wasabiae WPP163 YP 003258785 1 561231 Proteobacteria
Pectobacterium wasabiae WPP163 YP 003258809 1 561231 Proteobacteria
Pectobacterium carotovorum subsp carotovorum PC1 YP 003018435 1 561230 Proteobacteria
Dickeya zeae Ech586 WP 012885493 1 Dzeae
Dickeya chrysanthemi subsp parthenii NCPPB 516 WP 027711467 1 Dchrysanthemi
Photorhabdus asymbiotica YP 003039846 1 291112 Proteobacteria
Photorhabdus asymbiotica YP 003039840 1 291112 Proteobacteria
Pantoea sp At 9b YP 004114892 1 592316 Proteobacteria
Edwardsiella piscicida C07 087 YP 007630085 1 1288122 Proteobacteria
Erwinia tasmaniensis Et1 99 YP 001908356 1 465817 Proteobacteria
Klebsiella oxytoca E718 YP 006497184 1 1191061 Proteobacteria
Xenorhabdus nematophila ATCC 19061 YP 003713510 1 406817 Proteobacteria
Salmonella enterica subsp enterica serovar Schwarzengrund str CVM19633 YP 002113632 1 439843 Proteobacteria
Pectobacterium carotovorum subsp carotovorum PC1 YP 003018394 1 561230 Proteobacteria
Dickeya gs fangzhongdai NCPPB 3274 WP 042860516 1 Dfangzhongdai
Pectobacterium carotovorum subsp carotovorum PC1 YP 003018833 1 561230 Proteobacteria
Pectobacterium atrosepticum SCRI1043 YP 051550 1 218491 Proteobacteria
Enterobacter cloacae subsp cloacae ENHKU01 YP 006577439 1 1211025 Proteobacteria
Edwardsiella tarda EIB202 YP 003294855 1 498217 Proteobacteria
Klebsiella oxytoca E718 YP 006496884 1 1191061 Proteobacteria
Serratia marcescens WW4 YP 007404718 1 435998 Proteobacteria
Escherichia coli O55 H7 str CB9615 YP 003497974 1 701177 Proteobacteria
Escherichia coli O157 H7 str EDL933 NP 285976 1 155864 Proteobacteria
Dickeya dianthicola IPO 980 WP 029729627 1 Ddianthicola
Enterobacter cloacae EcWSU1 YP 004950715 1 1045856 Proteobacteria
Salmonella enterica subsp enterica serovar Schwarzengrund str CVM19633 YP 002113350 1 439843 Proteobacteria
Klebsiella pneumoniae 342 YP 002240225 1 507522 Proteobacteria
Dickeya solani RNS 05 1 2A WP 057082963 1 Dsolani
Dickeya chrysanthemi subsp parthenii NCPPB 516 WP 027711122 1 Dchrysanthemi
Dickeya sp CSL RW240 WP 035343745 1 Daquatica
DAQ1742 DAQ1742 03447 Daquatica
970 DAQ1742 03447 GI9 Seed
Dickeya aquatica DW 0440 WP 035343745 1 Daquatica
Enterobacter cloacae subsp dissolvens SDM YP 006476358 1 1104326 Proteobacteria
Pectobacterium sp SCC3193 YP 006284507 1 1166016 Proteobacteria
Dickeya chrysanthemi L11 WP 033577573 1 Dchrysanthemi
Dickeya dadantii 3937 WP 013319157 1 Ddadantii
Dickeya gs fangzhongdai ND14b WP 038666364 1 Dfangzhongdai
Dickeya gs fangzhongdai M074 WP 038666364 1 Dfangzhongdai
Dickeya chrysanthemi subsp parthenii NCPPB 516 WP 027711507 1 Dchrysanthemi
Dickeya chrysanthemi subsp chrysanthemi NCPPB 402 WP 040000368 1 Dchrysanthemi
Dickeya chrysanthemi L11 WP 033576370 1 Dchrysanthemi
Dickeya aquatica DW 0440 WP 035343240 1 Daquatica
DAQ1742 DAQ1742 01295 Daquatica
Dickeya sp CSL RW240 WP 035343240 1 Daquatica

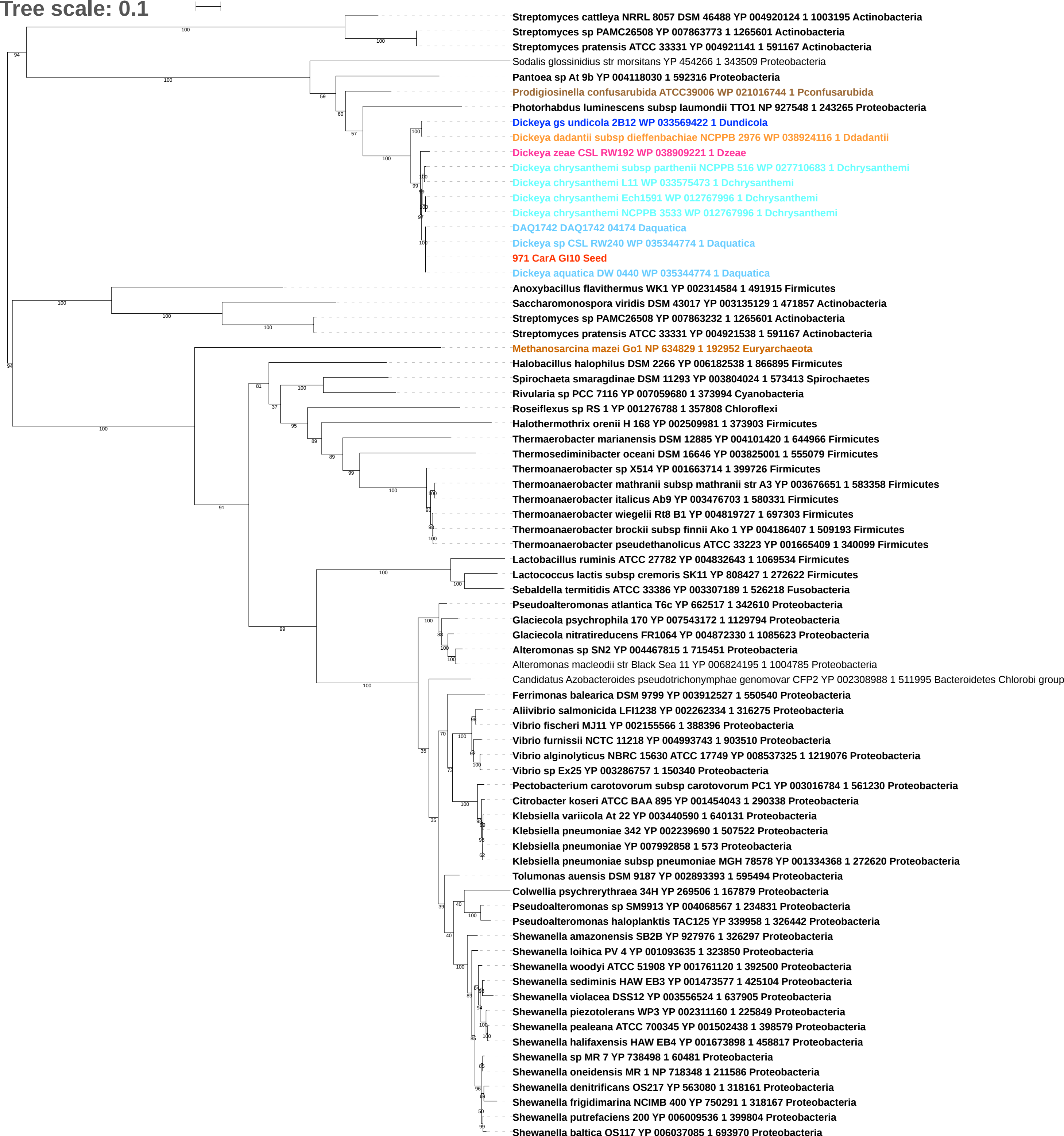
Tree scale: 0.1
Streptomyces cattleya NRRL 8057 DSM 46488 YP 004920124 1 1003195 Actinobacteria
Streptomyces sp PAMC26508 YP 007863773 1 1265601 Actinobacteria
Streptomyces pratensis ATCC 33331 YP 004921141 1 591167 Actinobacteria
Sodalis glossinidius str morsitans YP 454266 1 343509 Proteobacteria
Pantoea sp At 9b YP 004118030 1 592316 Proteobacteria
Prodigiosinella confusarubida ATCC39006 WP 021016744 1 Pconfusarubida
Photorhabdus luminescens subsp laumondii TTO1 NP 927548 1 243265 Proteobacteria
Dickeya gs undicola 2B12 WP 033569422 1 Dundicola
Dickeya dadantii subsp dieffenbachiae NCPPB 2976 WP 038924116 1 Ddadantii
Dickeya zeae CSL RW192 WP 038909221 1 Dzeae
Dickeya chrysanthemi subsp parthenii NCPPB 516 WP 027710683 1 Dchrysanthemi
Dickeya chrysanthemi L11 WP 033575473 1 Dchrysanthemi
Dickeya chrysanthemi Ech1591 WP 012767996 1 Dchrysanthemi
Dickeya chrysanthemi NCPPB 3533 WP 012767996 1 Dchrysanthemi
DAQ1742 DAQ1742 04174 Daquatica
Dickeya sp CSL RW240 WP 035344774 1 Daquatica
971 CarA GI10 Seed
Dickeya aquatica DW 0440 WP 035344774 1 Daquatica
Anoxybacillus flavithermus WK1 YP 002314584 1 491915 Firmicutes
Saccharomonospora viridis DSM 43017 YP 003135129 1 471857 Actinobacteria
Streptomyces sp PAMC26508 YP 007863232 1 1265601 Actinobacteria
Streptomyces pratensis ATCC 33331 YP 004921538 1 591167 Actinobacteria
Methanosarcina mazei Go1 NP 634829 1 192952 Euryarchaeota
Halobacillus halophilus DSM 2266 YP 006182538 1 866895 Firmicutes
Spirochaeta smaragdinae DSM 11293 YP 003804024 1 573413 Spirochaetes
Rivularia sp PCC 7116 YP 007059680 1 373994 Cyanobacteria
Roseiflexus sp RS 1 YP 001276788 1 357808 Chloroflexi
Halothermothrix orenii H 168 YP 002509981 1 373903 Firmicutes
Thermaerobacter marianensis DSM 12885 YP 004101420 1 644966 Firmicutes
Thermosediminibacter oceani DSM 16646 YP 003825001 1 555079 Firmicutes
Thermoanaerobacter sp X514 YP 001663714 1 399726 Firmicutes
Thermoanaerobacter mathranii subsp mathranii str A3 YP 003676651 1 583358 Firmicutes
Thermoanaerobacter italicus Ab9 YP 003476703 1 580331 Firmicutes
Thermoanaerobacter wiegelii Rt8 B1 YP 004819727 1 697303 Firmicutes
Thermoanaerobacter brockii subsp finnii Ako 1 YP 004186407 1 509193 Firmicutes
Thermoanaerobacter pseudethanolicus ATCC 33223 YP 001665409 1 340099 Firmicutes
Lactobacillus ruminis ATCC 27782 YP 004832643 1 1069534 Firmicutes
Lactococcus lactis subsp cremoris SK11 YP 808427 1 272622 Firmicutes
Sebaldella termitidis ATCC 33386 YP 003307189 1 526218 Fusobacteria
Pseudoalteromonas atlantica T6c YP 662517 1 342610 Proteobacteria
Glaciecola psychrophila 170 YP 007543172 1 1129794 Proteobacteria
Glaciecola nitratireducens FR1064 YP 004872330 1 1085623 Proteobacteria
Alteromonas sp SN2 YP 004467815 1 715451 Proteobacteria
Alteromonas macleodii str Black Sea 11 YP 006824195 1 1004785 Proteobacteria
Candidatus Azobacteroides pseudotrichonymphae genomovar CFP2 YP 002308988 1 511995 Bacteroidetes Chlorobi group
Ferrimonas balearica DSM 9799 YP 003912527 1 550540 Proteobacteria
Aliivibrio salmonicida LFI1238 YP 002262334 1 316275 Proteobacteria
Vibrio fischeri MJ11 YP 002155566 1 388396 Proteobacteria
Vibrio furnissii NCTC 11218 YP 004993743 1 903510 Proteobacteria
Vibrio alginolyticus NBRC 15630 ATCC 17749 YP 008537325 1 1219076 Proteobacteria
Vibrio sp Ex25 YP 003286757 1 150340 Proteobacteria
Pectobacterium carotovorum subsp carotovorum PC1 YP 003016784 1 561230 Proteobacteria
Citrobacter koseri ATCC BAA 895 YP 001454043 1 290338 Proteobacteria
Klebsiella variicola At 22 YP 003440590 1 640131 Proteobacteria
Klebsiella pneumoniae 342 YP 002239690 1 507522 Proteobacteria
Klebsiella pneumoniae YP 007992858 1 573 Proteobacteria
Klebsiella pneumoniae subsp pneumoniae MGH 78578 YP 001334368 1 272620 Proteobacteria
Tolumonas auensis DSM 9187 YP 002893393 1 595494 Proteobacteria
Colwellia psychrerythraea 34H YP 269506 1 167879 Proteobacteria
Pseudoalteromonas sp SM9913 YP 004068567 1 234831 Proteobacteria
Pseudoalteromonas haloplanktis TAC125 YP 339958 1 326442 Proteobacteria
Shewanella amazonensis SB2B YP 927976 1 326297 Proteobacteria
Shewanella loihica PV 4 YP 001093635 1 323850 Proteobacteria
Shewanella woodyi ATCC 51908 YP 001761120 1 392500 Proteobacteria
Shewanella sediminis HAW EB3 YP 001473577 1 425104 Proteobacteria
Shewanella violacea DSS12 YP 003556524 1 637905 Proteobacteria
Shewanella piezotolerans WP3 YP 002311160 1 225849 Proteobacteria
Shewanella pealeana ATCC 700345 YP 001502438 1 398579 Proteobacteria
Shewanella halifaxensis HAW EB4 YP 001673898 1 458817 Proteobacteria
Shewanella sp MR 7 YP 738498 1 60481 Proteobacteria
Shewanella oneidensis MR 1 NP 718348 1 211586 Proteobacteria
Shewanella denitrificans OS217 YP 563080 1 318161 Proteobacteria
Shewanella frigidimarina NCIMB 400 YP 750291 1 318167 Proteobacteria
Shewanella putrefaciens 200 YP 006009536 1 399804 Proteobacteria
Shewanella baltica OS117 YP 006037085 1 693970 Proteobacteria

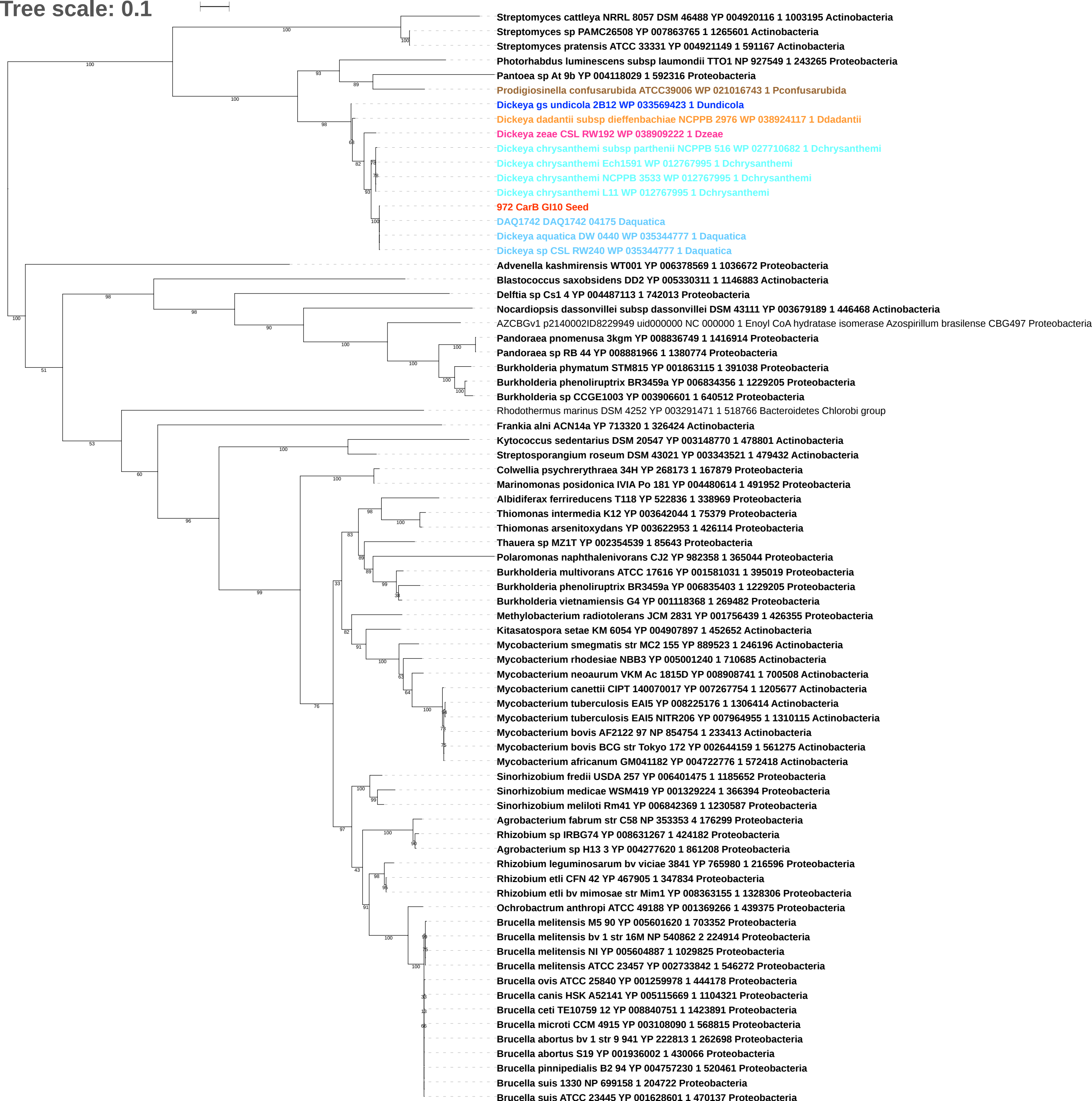
Tree scale: 0.1
Streptomyces cattleya NRRL 8057 DSM 46488 YP 004920116 1 1003195 Actinobacteria
Streptomyces sp PAMC26508 YP 007863765 1 1265601 Actinobacteria
Streptomyces pratensis ATCC 33331 YP 004921149 1 591167 Actinobacteria
Photorhabdus luminescens subsp laumondii TTO1 NP 927549 1 243265 Proteobacteria
Pantoea sp At 9b YP 004118029 1 592316 Proteobacteria
Prodigiosinella confusarubida ATCC39006 WP 021016743 1 Pconfusarubida
Dickeya gs undicola 2B12 WP 033569423 1 Dundicola
Dickeya dadantii subsp dieffenbachiae NCPPB 2976 WP 038924117 1 Ddadantii
Dickeya zeae CSL RW192 WP 038909222 1 Dzeae
Dickeya chrysanthemi subsp parthenii NCPPB 516 WP 027710682 1 Dchrysanthemi
Dickeya chrysanthemi Ech1591 WP 012767995 1 Dchrysanthemi
Dickeya chrysanthemi NCPPB 3533 WP 012767995 1 Dchrysanthemi
Dickeya chrysanthemi L11 WP 012767995 1 Dchrysanthemi
972 CarB GI10 Seed
DAQ1742 DAQ1742 04175 Daquatica
Dickeya aquatica DW 0440 WP 035344777 1 Daquatica
Dickeya sp CSL RW240 WP 035344777 1 Daquatica
Advenella kashmirensis WT001 YP 006378569 1 1036672 Proteobacteria
Blastococcus saxobsidens DD2 YP 005330311 1 1146883 Actinobacteria
Delftia sp Cs1 4 YP 004487113 1 742013 Proteobacteria
Nocardiopsis dassonvillei subsp dassonvillei DSM 43111 YP 003679189 1 446468 Actinobacteria
AZCBGv1 p2140002ID8229949 uid000000 NC 000000 1 Enoyl CoA hydratase isomerase Azospirillum brasilense CBG497 Proteobacteria
Pandoraea pnomenusa 3kgm YP 008836749 1 1416914 Proteobacteria
Pandoraea sp RB 44 YP 008881966 1 1380774 Proteobacteria
Burkholderia phymatum STM815 YP 001863115 1 391038 Proteobacteria
Burkholderia phenoliruptrix BR3459a YP 006834356 1 1229205 Proteobacteria
Burkholderia sp CCGE1003 YP 003906601 1 640512 Proteobacteria
Rhodothermus marinus DSM 4252 YP 003291471 1 518766 Bacteroidetes Chlorobi group
Frankia alni ACN14a YP 713320 1 326424 Actinobacteria
Kytococcus sedentarius DSM 20547 YP 003148770 1 478801 Actinobacteria
Streptosporangium roseum DSM 43021 YP 003343521 1 479432 Actinobacteria
Colwellia psychrerythraea 34H YP 268173 1 167879 Proteobacteria
Marinomonas posidonica IVIA Po 181 YP 004480614 1 491952 Proteobacteria
Albidiferax ferrireducens T118 YP 522836 1 338969 Proteobacteria
Thiomonas intermedia K12 YP 003642044 1 75379 Proteobacteria
Thiomonas arsenitoxydans YP 003622953 1 426114 Proteobacteria
Thauera sp MZ1T YP 002354539 1 85643 Proteobacteria
Polaromonas naphthalenivorans CJ2 YP 982358 1 365044 Proteobacteria
Burkholderia multivorans ATCC 17616 YP 001581031 1 395019 Proteobacteria
Burkholderia phenoliruptrix BR3459a YP 006835403 1 1229205 Proteobacteria
Burkholderia vietnamiensis G4 YP 001118368 1 269482 Proteobacteria
Methylobacterium radiotolerans JCM 2831 YP 001756439 1 426355 Proteobacteria
Kitasatospora setae KM 6054 YP 004907897 1 452652 Actinobacteria
Mycobacterium smegmatis str MC2 155 YP 889523 1 246196 Actinobacteria
Mycobacterium rhodesiae NBB3 YP 005001240 1 710685 Actinobacteria
Mycobacterium neoaurum VKM Ac 1815D YP 008908741 1 700508 Actinobacteria
Mycobacterium canettii CIPT 140070017 YP 007267754 1 1205677 Actinobacteria
Mycobacterium tuberculosis EAI5 YP 008225176 1 1306414 Actinobacteria
Mycobacterium tuberculosis EAI5 NITR206 YP 007964955 1 1310115 Actinobacteria
Mycobacterium bovis AF2122 97 NP 854754 1 233413 Actinobacteria
Mycobacterium bovis BCG str Tokyo 172 YP 002644159 1 561275 Actinobacteria
Mycobacterium africanum GM041182 YP 004722776 1 572418 Actinobacteria
Sinorhizobium fredii USDA 257 YP 006401475 1 1185652 Proteobacteria
Sinorhizobium medicae WSM419 YP 001329224 1 366394 Proteobacteria
Sinorhizobium meliloti Rm41 YP 006842369 1 1230587 Proteobacteria
Agrobacterium fabrum str C58 NP 353353 4 176299 Proteobacteria
Rhizobium sp IRBG74 YP 008631267 1 424182 Proteobacteria
Agrobacterium sp H13 3 YP 004277620 1 861208 Proteobacteria
Rhizobium leguminosarum bv viciae 3841 YP 765980 1 216596 Proteobacteria
Rhizobium etli CFN 42 YP 467905 1 347834 Proteobacteria
Rhizobium etli bv mimosae str Mim1 YP 008363155 1 1328306 Proteobacteria
Ochrobactrum anthropi ATCC 49188 YP 001369266 1 439375 Proteobacteria
Brucella melitensis M5 90 YP 005601620 1 703352 Proteobacteria
Brucella melitensis bv 1 str 16M NP 540862 2 224914 Proteobacteria
Brucella melitensis NI YP 005604887 1 1029825 Proteobacteria
Brucella melitensis ATCC 23457 YP 002733842 1 546272 Proteobacteria
Brucella ovis ATCC 25840 YP 001259978 1 444178 Proteobacteria
Brucella canis HSK A52141 YP 005115669 1 1104321 Proteobacteria
Brucella ceti TE10759 12 YP 008840751 1 1423891 Proteobacteria
Brucella microti CCM 4915 YP 003108090 1 568815 Proteobacteria
Brucella abortus bv 1 str 9 941 YP 222813 1 262698 Proteobacteria
Brucella abortus S19 YP 001936002 1 430066 Proteobacteria
Brucella pinnipedialis B2 94 YP 004757230 1 520461 Proteobacteria
Brucella suis 1330 NP 699158 1 204722 Proteobacteria
Brucella suis ATCC 23445 YP 001628601 1 470137 Proteobacteria

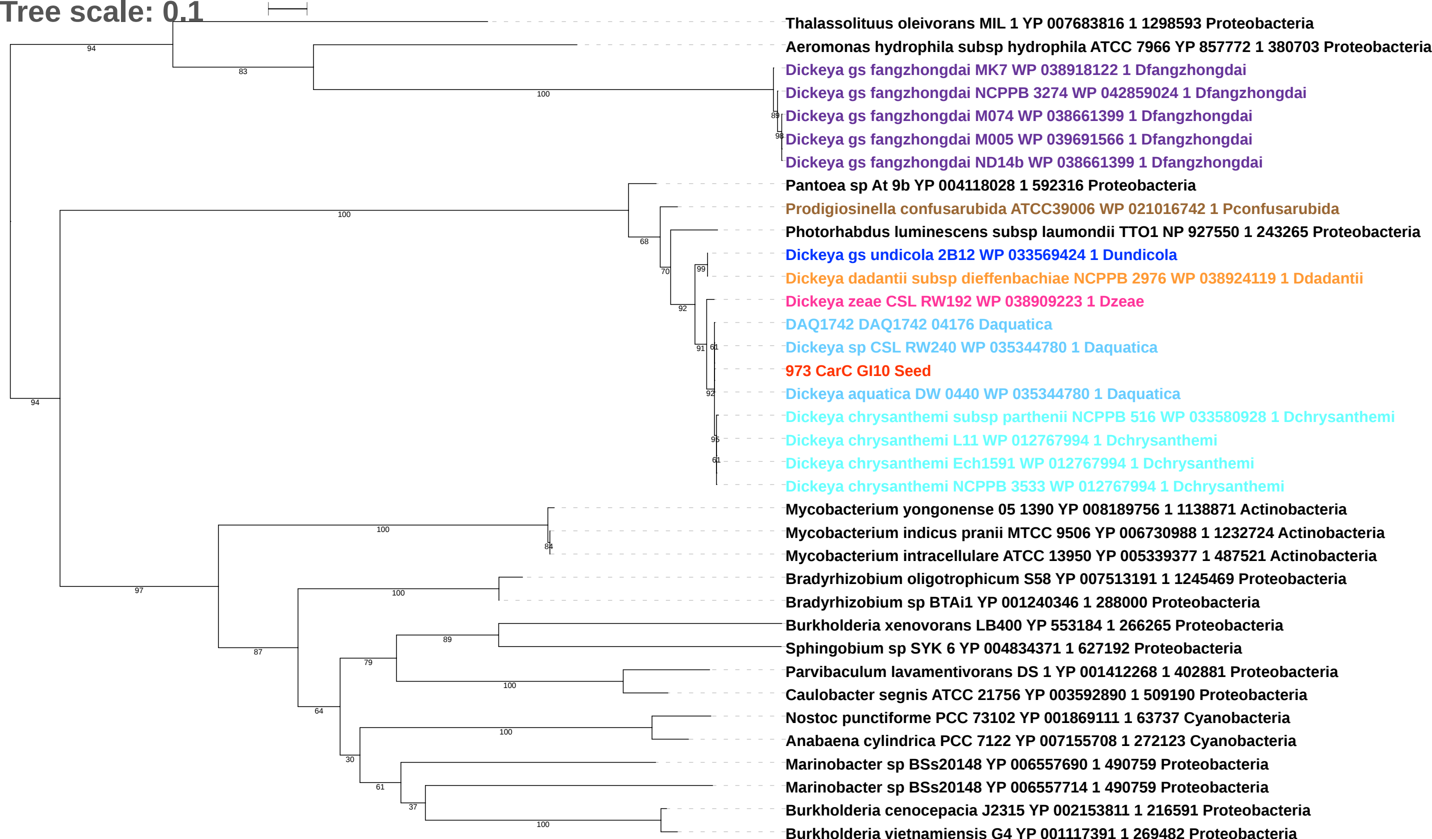
Tree scale: 0.1
Thalassolituus oleivorans MIL 1 YP 007683816 1 1298593 Proteobacteria
Aeromonas hydrophila subsp hydrophila ATCC 7966 YP 857772 1 380703 Proteobacteria
Dickeya gs fangzhongdai MK7 WP 038918122 1 Dfangzhongdai
Dickeya gs fangzhongdai NCPPB 3274 WP 042859024 1 Dfangzhongdai
Dickeya gs fangzhongdai M074 WP 038661399 1 Dfangzhongdai
Dickeya gs fangzhongdai M005 WP 039691566 1 Dfangzhongdai
Dickeya gs fangzhongdai ND14b WP 038661399 1 Dfangzhongdai
Pantoea sp At 9b YP 004118028 1 592316 Proteobacteria
Prodigiosinella confusarubida ATCC39006 WP 021016742 1 Pconfusarubida
Photorhabdus luminescens subsp laumondii TTO1 NP 927550 1 243265 Proteobacteria
Dickeya gs undicola 2B12 WP 033569424 1 Dundicola
Dickeya dadantii subsp dieffenbachiae NCPPB 2976 WP 038924119 1 Ddadantii
Dickeya zeae CSL RW192 WP 038909223 1 Dzeae
DAQ1742 DAQ1742 04176 Daquatica
Dickeya sp CSL RW240 WP 035344780 1 Daquatica
973 CarC GI10 Seed
Dickeya aquatica DW 0440 WP 035344780 1 Daquatica
Dickeya chrysanthemi subsp parthenii NCPPB 516 WP 033580928 1 Dchrysanthemi
Dickeya chrysanthemi L11 WP 012767994 1 Dchrysanthemi
Dickeya chrysanthemi Ech1591 WP 012767994 1 Dchrysanthemi
Dickeya chrysanthemi NCPPB 3533 WP 012767994 1 Dchrysanthemi
Mycobacterium yongonense 05 1390 YP 008189756 1 1138871 Actinobacteria
Mycobacterium indicus pranii MTCC 9506 YP 006730988 1 1232724 Actinobacteria
Mycobacterium intracellulare ATCC 13950 YP 005339377 1 487521 Actinobacteria
Bradyrhizobium oligotrophicum S58 YP 007513191 1 1245469 Proteobacteria
Bradyrhizobium sp BTAi1 YP 001240346 1 288000 Proteobacteria
Burkholderia xenovorans LB400 YP 553184 1 266265 Proteobacteria
Sphingobium sp SYK 6 YP 004834371 1 627192 Proteobacteria
Parvibaculum lavamentivorans DS 1 YP 001412268 1 402881 Proteobacteria
Caulobacter segnis ATCC 21756 YP 003592890 1 509190 Proteobacteria
Nostoc punctiforme PCC 73102 YP 001869111 1 63737 Cyanobacteria
Anabaena cylindrica PCC 7122 YP 007155708 1 272123 Cyanobacteria
Marinobacter sp BSs20148 YP 006557690 1 490759 Proteobacteria
Marinobacter sp BSs20148 YP 006557714 1 490759 Proteobacteria
Burkholderia cenocepacia J2315 YP 002153811 1 216591 Proteobacteria
Burkholderia vietnamiensis G4 YP 001117391 1 269482 Proteobacteria

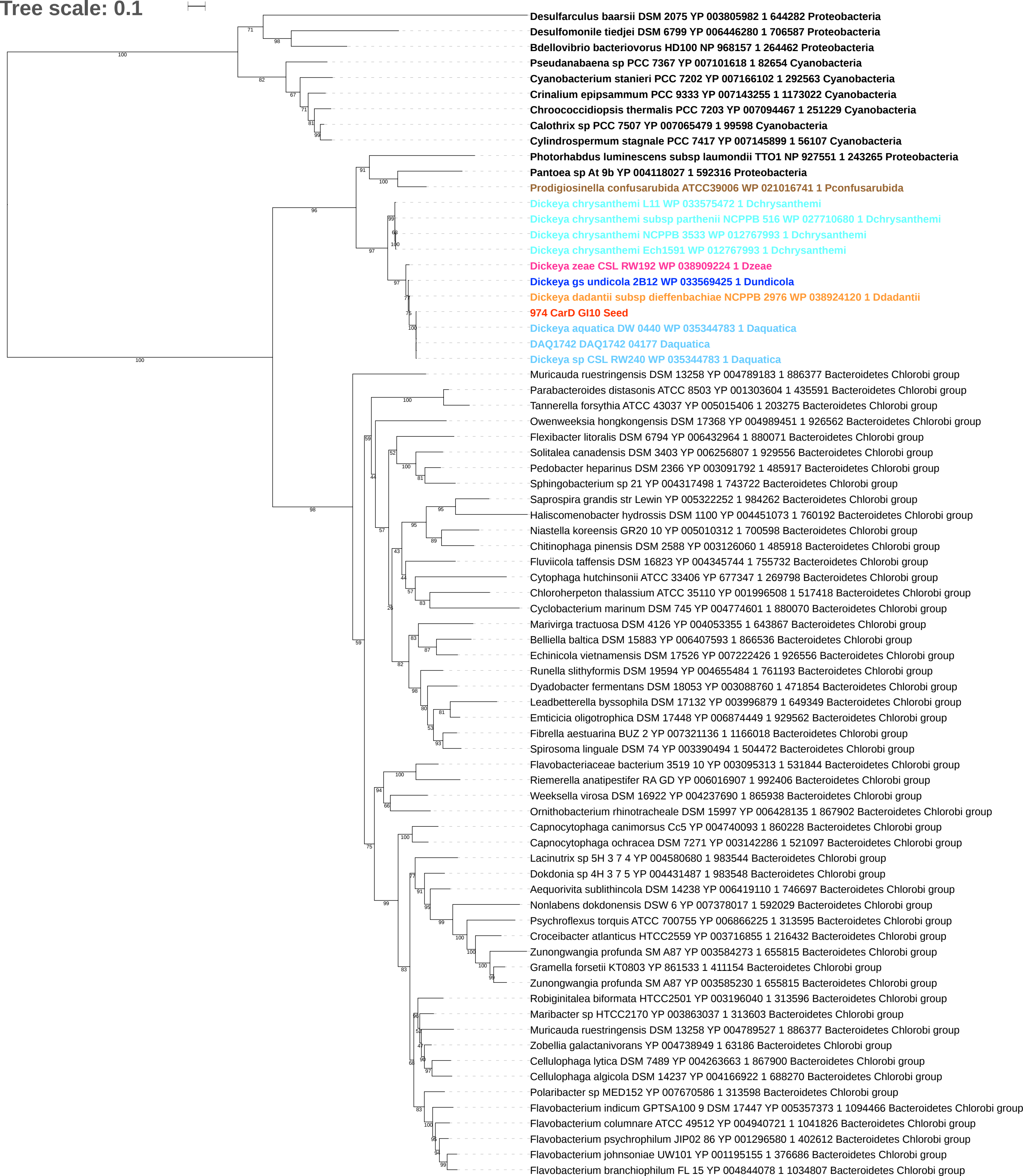
Tree scale: 0.1
Desulfarculus baarsii DSM 2075 YP 003805982 1 644282 Proteobacteria
Desulfomonile tiedjei DSM 6799 YP 006446280 1 706587 Proteobacteria
Bdellovibrio bacteriovorus HD100 NP 968157 1 264462 Proteobacteria
Pseudanabaena sp PCC 7367 YP 007101618 1 82654 Cyanobacteria
Cyanobacterium stanieri PCC 7202 YP 007166102 1 292563 Cyanobacteria
Crinalium epipsammum PCC 9333 YP 007143255 1 1173022 Cyanobacteria
Chroococcidiopsis thermalis PCC 7203 YP 007094467 1 251229 Cyanobacteria
Calothrix sp PCC 7507 YP 007065479 1 99598 Cyanobacteria
Cylindrospermum stagnale PCC 7417 YP 007145899 1 56107 Cyanobacteria
Photorhabdus luminescens subsp laumondii TTO1 NP 927551 1 243265 Proteobacteria
Pantoea sp At 9b YP 004118027 1 592316 Proteobacteria
Prodigiosinella confusarubida ATCC39006 WP 021016741 1 Pconfusarubida
Dickeya chrysanthemi L11 WP 033575472 1 Dchrysanthemi
Dickeya chrysanthemi subsp parthenii NCPPB 516 WP 027710680 1 Dchrysanthemi
Dickeya chrysanthemi NCPPB 3533 WP 012767993 1 Dchrysanthemi
Dickeya chrysanthemi Ech1591 WP 012767993 1 Dchrysanthemi
Dickeya zeae CSL RW192 WP 038909224 1 Dzeae
Dickeya gs undicola 2B12 WP 033569425 1 Dundicola
Dickeya dadantii subsp dieffenbachiae NCPPB 2976 WP 038924120 1 Ddadantii
974 CarD GI10 Seed
Dickeya aquatica DW 0440 WP 035344783 1 Daquatica
DAQ1742 DAQ1742 04177 Daquatica
Dickeya sp CSL RW240 WP 035344783 1 Daquatica
Muricauda ruestringensis DSM 13258 YP 004789183 1 886377 Bacteroidetes Chlorobi group
Parabacteroides distasonis ATCC 8503 YP 001303604 1 435591 Bacteroidetes Chlorobi group
Tannerella forsythia ATCC 43037 YP 005015406 1 203275 Bacteroidetes Chlorobi group
Owenweeksia hongkongensis DSM 17368 YP 004989451 1 926562 Bacteroidetes Chlorobi group
Flexibacter litoralis DSM 6794 YP 006432964 1 880071 Bacteroidetes Chlorobi group
Solitalea canadensis DSM 3403 YP 006256807 1 929556 Bacteroidetes Chlorobi group
Pedobacter heparinus DSM 2366 YP 003091792 1 485917 Bacteroidetes Chlorobi group
Sphingobacterium sp 21 YP 004317498 1 743722 Bacteroidetes Chlorobi group
Saprospira grandis str Lewin YP 005322252 1 984262 Bacteroidetes Chlorobi group
Haliscomenobacter hydrossis DSM 1100 YP 004451073 1 760192 Bacteroidetes Chlorobi group
Niastella koreensis GR20 10 YP 005010312 1 700598 Bacteroidetes Chlorobi group
Chitinophaga pinensis DSM 2588 YP 003126060 1 485918 Bacteroidetes Chlorobi group
Fluviicola taffensis DSM 16823 YP 004345744 1 755732 Bacteroidetes Chlorobi group
Cytophaga hutchinsonii ATCC 33406 YP 677347 1 269798 Bacteroidetes Chlorobi group
Chloroherpeton thalassium ATCC 35110 YP 001996508 1 517418 Bacteroidetes Chlorobi group
Cyclobacterium marinum DSM 745 YP 004774601 1 880070 Bacteroidetes Chlorobi group
Marivirga tractuosa DSM 4126 YP 004053355 1 643867 Bacteroidetes Chlorobi group
Belliella baltica DSM 15883 YP 006407593 1 866536 Bacteroidetes Chlorobi group
Echinicola vietnamensis DSM 17526 YP 007222426 1 926556 Bacteroidetes Chlorobi group
Runella slithyformis DSM 19594 YP 004655484 1 761193 Bacteroidetes Chlorobi group
Dyadobacter fermentans DSM 18053 YP 003088760 1 471854 Bacteroidetes Chlorobi group
Leadbetterella byssophila DSM 17132 YP 003996879 1 649349 Bacteroidetes Chlorobi group
Emticicia oligotrophica DSM 17448 YP 006874449 1 929562 Bacteroidetes Chlorobi group
Fibrella aestuarina BUZ 2 YP 007321136 1 1166018 Bacteroidetes Chlorobi group
Spirosoma linguale DSM 74 YP 003390494 1 504472 Bacteroidetes Chlorobi group
Flavobacteriaceae bacterium 3519 10 YP 003095313 1 531844 Bacteroidetes Chlorobi group
Riemerella anatipestifer RA GD YP 006016907 1 992406 Bacteroidetes Chlorobi group
Weeksella virosa DSM 16922 YP 004237690 1 865938 Bacteroidetes Chlorobi group
Ornithobacterium rhinotracheale DSM 15997 YP 006428135 1 867902 Bacteroidetes Chlorobi group
Capnocytophaga canimorsus Cc5 YP 004740093 1 860228 Bacteroidetes Chlorobi group
Capnocytophaga ochracea DSM 7271 YP 003142286 1 521097 Bacteroidetes Chlorobi group
Lacinutrix sp 5H 3 7 4 YP 004580680 1 983544 Bacteroidetes Chlorobi group
Dokdonia sp 4H 3 7 5 YP 004431487 1 983548 Bacteroidetes Chlorobi group
Aequorivita sublithincola DSM 14238 YP 006419110 1 746697 Bacteroidetes Chlorobi group
Nonlabens dokdonensis DSW 6 YP 007378017 1 592029 Bacteroidetes Chlorobi group
Psychroflexus torquis ATCC 700755 YP 006866225 1 313595 Bacteroidetes Chlorobi group
Croceibacter atlanticus HTCC2559 YP 003716855 1 216432 Bacteroidetes Chlorobi group
Zunongwangia profunda SM A87 YP 003584273 1 655815 Bacteroidetes Chlorobi group
Gramella forsetii KT0803 YP 861533 1 411154 Bacteroidetes Chlorobi group
Zunongwangia profunda SM A87 YP 003585230 1 655815 Bacteroidetes Chlorobi group
Robiginitalea biformata HTCC2501 YP 003196040 1 313596 Bacteroidetes Chlorobi group
Maribacter sp HTCC2170 YP 003863037 1 313603 Bacteroidetes Chlorobi group
Muricauda ruestringensis DSM 13258 YP 004789527 1 886377 Bacteroidetes Chlorobi group
Zobellia galactanivorans YP 004738949 1 63186 Bacteroidetes Chlorobi group
Cellulophaga lytica DSM 7489 YP 004263663 1 867900 Bacteroidetes Chlorobi group
Cellulophaga algicola DSM 14237 YP 004166922 1 688270 Bacteroidetes Chlorobi group
Polaribacter sp MED152 YP 007670586 1 313598 Bacteroidetes Chlorobi group
Flavobacterium indicum GPTSA100 9 DSM 17447 YP 005357373 1 1094466 Bacteroidetes Chlorobi group
Flavobacterium columnare ATCC 49512 YP 004940721 1 1041826 Bacteroidetes Chlorobi group
Flavobacterium psychrophilum JIP02 86 YP 001296580 1 402612 Bacteroidetes Chlorobi group
Flavobacterium johnsoniae UW101 YP 001195155 1 376686 Bacteroidetes Chlorobi group
Flavobacterium branchiophilum FL 15 YP 004844078 1 1034807 Bacteroidetes Chlorobi group

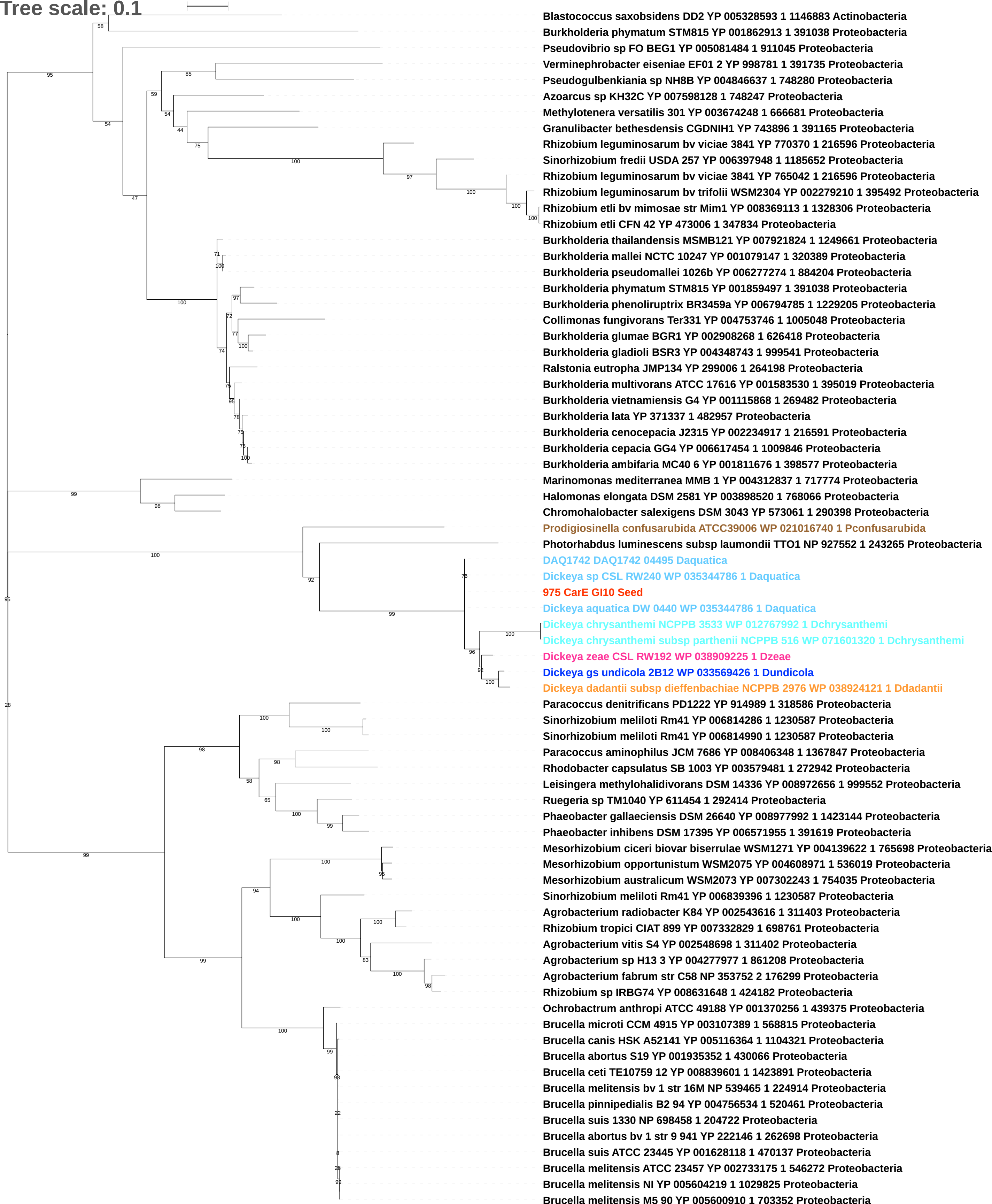
Tree scale: 0.1
Blastococcus saxobsidens DD2 YP 005328593 1 1146883 Actinobacteria
Burkholderia phymatum STM815 YP 001862913 1 391038 Proteobacteria
Pseudovibrio sp FO BEG1 YP 005081484 1 911045 Proteobacteria
Verminephrobacter eiseniae EF01 2 YP 998781 1 391735 Proteobacteria
Pseudogulbenkiania sp NH8B YP 004846637 1 748280 Proteobacteria
Azoarcus sp KH32C YP 007598128 1 748247 Proteobacteria
Methylotenera versatilis 301 YP 003674248 1 666681 Proteobacteria
Granulibacter bethesdensis CGDNIH1 YP 743896 1 391165 Proteobacteria
Rhizobium leguminosarum bv viciae 3841 YP 770370 1 216596 Proteobacteria
Sinorhizobium fredii USDA 257 YP 006397948 1 1185652 Proteobacteria
Rhizobium leguminosarum bv viciae 3841 YP 765042 1 216596 Proteobacteria
Rhizobium leguminosarum bv trifolii WSM2304 YP 002279210 1 395492 Proteobacteria
Rhizobium etli bv mimosae str Mim1 YP 008369113 1 1328306 Proteobacteria
Rhizobium etli CFN 42 YP 473006 1 347834 Proteobacteria
Burkholderia thailandensis MSMB121 YP 007921824 1 1249661 Proteobacteria
Burkholderia mallei NCTC 10247 YP 001079147 1 320389 Proteobacteria
Burkholderia pseudomallei 1026b YP 006277274 1 884204 Proteobacteria
Burkholderia phymatum STM815 YP 001859497 1 391038 Proteobacteria
Burkholderia phenoliruptrix BR3459a YP 006794785 1 1229205 Proteobacteria
Collimonas fungivorans Ter331 YP 004753746 1 1005048 Proteobacteria
Burkholderia glumae BGR1 YP 002908268 1 626418 Proteobacteria
Burkholderia gladioli BSR3 YP 004348743 1 999541 Proteobacteria
Ralstonia eutropha JMP134 YP 299006 1 264198 Proteobacteria
Burkholderia multivorans ATCC 17616 YP 001583530 1 395019 Proteobacteria
Burkholderia vietnamiensis G4 YP 001115868 1 269482 Proteobacteria
Burkholderia lata YP 371337 1 482957 Proteobacteria
Burkholderia cenocepacia J2315 YP 002234917 1 216591 Proteobacteria
Burkholderia cepacia GG4 YP 006617454 1 1009846 Proteobacteria
Burkholderia ambifaria MC40 6 YP 001811676 1 398577 Proteobacteria
Marinomonas mediterranea MMB 1 YP 004312837 1 717774 Proteobacteria
Halomonas elongata DSM 2581 YP 003898520 1 768066 Proteobacteria
Chromohalobacter salexigens DSM 3043 YP 573061 1 290398 Proteobacteria
Prodigiosinella confusarubida ATCC39006 WP 021016740 1 Pconfusarubida
Photorhabdus luminescens subsp laumondii TTO1 NP 927552 1 243265 Proteobacteria
DAQ1742 DAQ1742 04495 Daquatica
Dickeya sp CSL RW240 WP 035344786 1 Daquatica
975 CarE GI10 Seed
Dickeya aquatica DW 0440 WP 035344786 1 Daquatica
Dickeya chrysanthemi NCPPB 3533 WP 012767992 1 Dchrysanthemi
Dickeya chrysanthemi subsp parthenii NCPPB 516 WP 071601320 1 Dchrysanthemi
Dickeya zeae CSL RW192 WP 038909225 1 Dzeae
Dickeya gs undicola 2B12 WP 033569426 1 Dundicola
Dickeya dadantii subsp dieffenbachiae NCPPB 2976 WP 038924121 1 Ddadantii
Paracoccus denitrificans PD1222 YP 914989 1 318586 Proteobacteria
Sinorhizobium meliloti Rm41 YP 006814286 1 1230587 Proteobacteria
Sinorhizobium meliloti Rm41 YP 006814990 1 1230587 Proteobacteria
Paracoccus aminophilus JCM 7686 YP 008406348 1 1367847 Proteobacteria
Rhodobacter capsulatus SB 1003 YP 003579481 1 272942 Proteobacteria
Leisingera methylohalidivorans DSM 14336 YP 008972656 1 999552 Proteobacteria
Ruegeria sp TM1040 YP 611454 1 292414 Proteobacteria
Phaeobacter gallaeciensis DSM 26640 YP 008977992 1 1423144 Proteobacteria
Phaeobacter inhibens DSM 17395 YP 006571955 1 391619 Proteobacteria
Mesorhizobium ciceri biovar biserrulae WSM1271 YP 004139622 1 765698 Proteobacteria
Mesorhizobium opportunistum WSM2075 YP 004608971 1 536019 Proteobacteria
Mesorhizobium australicum WSM2073 YP 007302243 1 754035 Proteobacteria
Sinorhizobium meliloti Rm41 YP 006839396 1 1230587 Proteobacteria
Agrobacterium radiobacter K84 YP 002543616 1 311403 Proteobacteria
Rhizobium tropici CIAT 899 YP 007332829 1 698761 Proteobacteria
Agrobacterium vitis S4 YP 002548698 1 311402 Proteobacteria
Agrobacterium sp H13 3 YP 004277977 1 861208 Proteobacteria
Agrobacterium fabrum str C58 NP 353752 2 176299 Proteobacteria
Rhizobium sp IRBG74 YP 008631648 1 424182 Proteobacteria
Ochrobactrum anthropi ATCC 49188 YP 001370256 1 439375 Proteobacteria
Brucella microti CCM 4915 YP 003107389 1 568815 Proteobacteria
Brucella canis HSK A52141 YP 005116364 1 1104321 Proteobacteria
Brucella abortus S19 YP 001935352 1 430066 Proteobacteria
Brucella ceti TE10759 12 YP 008839601 1 1423891 Proteobacteria
Brucella melitensis bv 1 str 16M NP 539465 1 224914 Proteobacteria
Brucella pinnipedialis B2 94 YP 004756534 1 520461 Proteobacteria
Brucella suis 1330 NP 698458 1 204722 Proteobacteria
Brucella abortus bv 1 str 9 941 YP 222146 1 262698 Proteobacteria
Brucella suis ATCC 23445 YP 001628118 1 470137 Proteobacteria
Brucella melitensis ATCC 23457 YP 002733175 1 546272 Proteobacteria
Brucella melitensis NI YP 005604219 1 1029825 Proteobacteria
Brucella melitensis M5 90 YP 005600910 1 703352 Proteobacteria

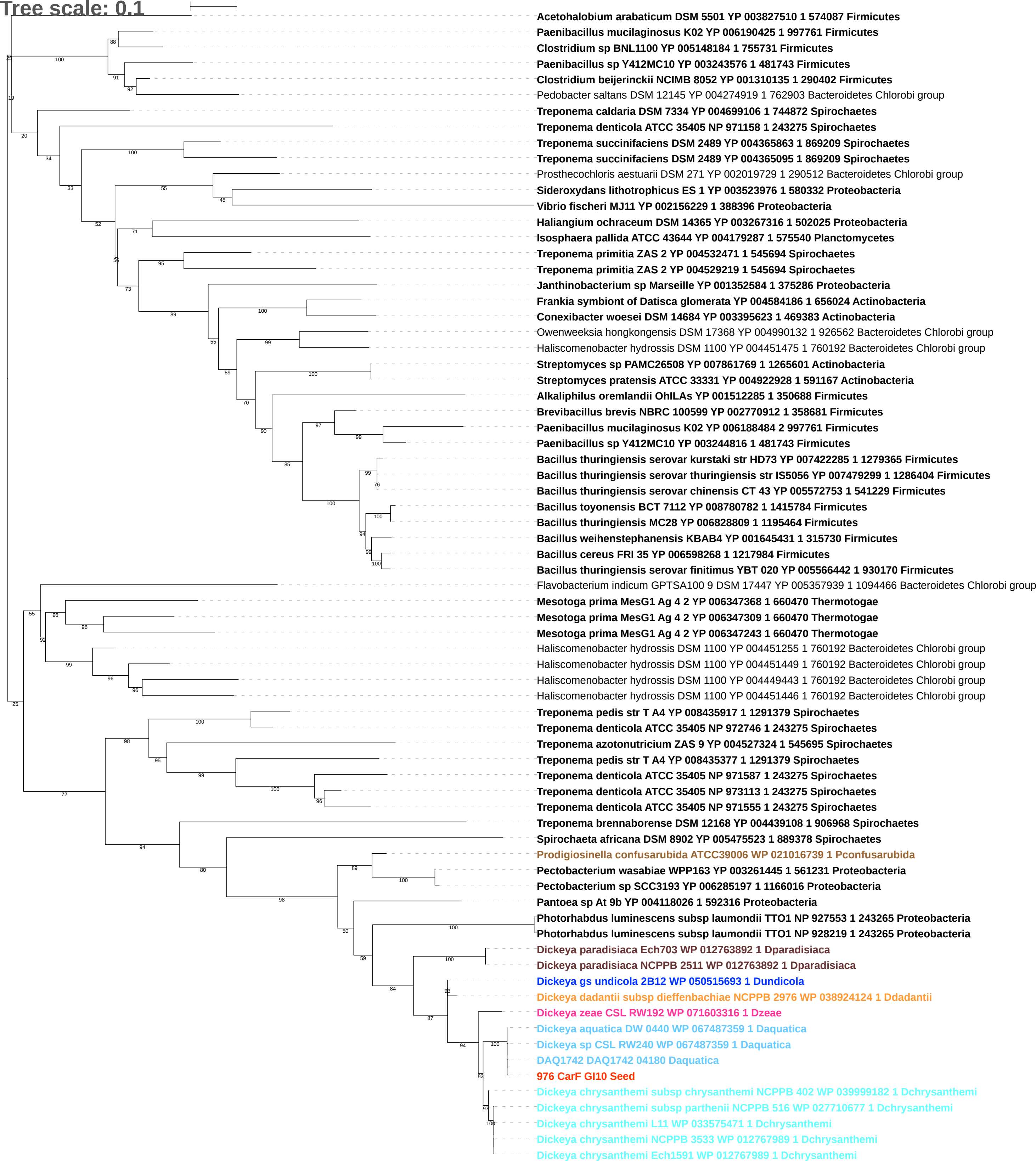
Tree scale: 0.1
Acetohalobium arabaticum DSM 5501 YP 003827510 1 574087 Firmicutes
Paenibacillus mucilaginosus K02 YP 006190425 1 997761 Firmicutes
Clostridium sp BNL1100 YP 005148184 1 755731 Firmicutes
Paenibacillus sp Y412MC10 YP 003243576 1 481743 Firmicutes
Clostridium beijerinckii NCIMB 8052 YP 001310135 1 290402 Firmicutes
Pedobacter saltans DSM 12145 YP 004274919 1 762903 Bacteroidetes Chlorobi group
Treponema caldaria DSM 7334 YP 004699106 1 744872 Spirochaetes
Treponema denticola ATCC 35405 NP 971158 1 243275 Spirochaetes
Treponema succinifaciens DSM 2489 YP 004365863 1 869209 Spirochaetes
Treponema succinifaciens DSM 2489 YP 004365095 1 869209 Spirochaetes
Prosthecochloris aestuarii DSM 271 YP 002019729 1 290512 Bacteroidetes Chlorobi group
Sideroxydans lithotrophicus ES 1 YP 003523976 1 580332 Proteobacteria
Vibrio fischeri MJ11 YP 002156229 1 388396 Proteobacteria
Haliangium ochraceum DSM 14365 YP 003267316 1 502025 Proteobacteria
Isosphaera pallida ATCC 43644 YP 004179287 1 575540 Planctomycetes
Treponema primitia ZAS 2 YP 004532471 1 545694 Spirochaetes
Treponema primitia ZAS 2 YP 004529219 1 545694 Spirochaetes
Janthinobacterium sp Marseille YP 001352584 1 375286 Proteobacteria
Frankia symbiont of Datisca glomerata YP 004584186 1 656024 Actinobacteria
Conexibacter woesei DSM 14684 YP 003395623 1 469383 Actinobacteria
Owenweeksia hongkongensis DSM 17368 YP 004990132 1 926562 Bacteroidetes Chlorobi group
Haliscomenobacter hydrossis DSM 1100 YP 004451475 1 760192 Bacteroidetes Chlorobi group
Streptomyces sp PAMC26508 YP 007861769 1 1265601 Actinobacteria
Streptomyces pratensis ATCC 33331 YP 004922928 1 591167 Actinobacteria
Alkaliphilus oremlandii OhILAs YP 001512285 1 350688 Firmicutes
Brevibacillus brevis NBRC 100599 YP 002770912 1 358681 Firmicutes
Paenibacillus mucilaginosus K02 YP 006188484 2 997761 Firmicutes
Paenibacillus sp Y412MC10 YP 003244816 1 481743 Firmicutes
Bacillus thuringiensis serovar kurstaki str HD73 YP 007422285 1 1279365 Firmicutes
Bacillus thuringiensis serovar thuringiensis str IS5056 YP 007479299 1 1286404 Firmicutes
Bacillus thuringiensis serovar chinensis CT 43 YP 005572753 1 541229 Firmicutes
Bacillus toyonensis BCT 7112 YP 008780782 1 1415784 Firmicutes
Bacillus thuringiensis MC28 YP 006828809 1 1195464 Firmicutes
Bacillus weihenstephanensis KBAB4 YP 001645431 1 315730 Firmicutes
Bacillus cereus FRI 35 YP 006598268 1 1217984 Firmicutes
Bacillus thuringiensis serovar finitimus YBT 020 YP 005566442 1 930170 Firmicutes
Flavobacterium indicum GPTSA100 9 DSM 17447 YP 005357939 1 1094466 Bacteroidetes Chlorobi group
Mesotoga prima MesG1 Ag 4 2 YP 006347368 1 660470 Thermotogae
Mesotoga prima MesG1 Ag 4 2 YP 006347309 1 660470 Thermotogae
Mesotoga prima MesG1 Ag 4 2 YP 006347243 1 660470 Thermotogae
Haliscomenobacter hydrossis DSM 1100 YP 004451255 1 760192 Bacteroidetes Chlorobi group
Haliscomenobacter hydrossis DSM 1100 YP 004451449 1 760192 Bacteroidetes Chlorobi group
Haliscomenobacter hydrossis DSM 1100 YP 004449443 1 760192 Bacteroidetes Chlorobi group
Haliscomenobacter hydrossis DSM 1100 YP 004451446 1 760192 Bacteroidetes Chlorobi group
Treponema pedis str T A4 YP 008435917 1 1291379 Spirochaetes
Treponema denticola ATCC 35405 NP 972746 1 243275 Spirochaetes
Treponema azotonutricium ZAS 9 YP 004527324 1 545695 Spirochaetes
Treponema pedis str T A4 YP 008435377 1 1291379 Spirochaetes
Treponema denticola ATCC 35405 NP 971587 1 243275 Spirochaetes
Treponema denticola ATCC 35405 NP 973113 1 243275 Spirochaetes
Treponema denticola ATCC 35405 NP 971555 1 243275 Spirochaetes
Treponema brennaborense DSM 12168 YP 004439108 1 906968 Spirochaetes
Spirochaeta africana DSM 8902 YP 005475523 1 889378 Spirochaetes
Prodigiosinella confusarubida ATCC39006 WP 021016739 1 Pconfusarubida
Pectobacterium wasabiae WPP163 YP 003261445 1 561231 Proteobacteria
Pectobacterium sp SCC3193 YP 006285197 1 1166016 Proteobacteria
Pantoea sp At 9b YP 004118026 1 592316 Proteobacteria
Photorhabdus luminescens subsp laumondii TTO1 NP 927553 1 243265 Proteobacteria
Photorhabdus luminescens subsp laumondii TTO1 NP 928219 1 243265 Proteobacteria
Dickeya paradisiaca Ech703 WP 012763892 1 Dparadisiaca
Dickeya paradisiaca NCPPB 2511 WP 012763892 1 Dparadisiaca
Dickeya gs undicola 2B12 WP 050515693 1 Dundicola
Dickeya dadantii subsp dieffenbachiae NCPPB 2976 WP 038924124 1 Ddadantii
Dickeya zeae CSL RW192 WP 071603316 1 Dzeae
Dickeya aquatica DW 0440 WP 067487359 1 Daquatica
Dickeya sp CSL RW240 WP 067487359 1 Daquatica
DAQ1742 DAQ1742 04180 Daquatica
976 CarF GI10 Seed
Dickeya chrysanthemi subsp chrysanthemi NCPPB 402 WP 039999182 1 Dchrysanthemi
Dickeya chrysanthemi subsp parthenii NCPPB 516 WP 027710677 1 Dchrysanthemi
Dickeya chrysanthemi L11 WP 033575471 1 Dchrysanthemi
Dickeya chrysanthemi NCPPB 3533 WP 012767989 1 Dchrysanthemi
Dickeya chrysanthemi Ech1591 WP 012767989 1 Dchrysanthemi

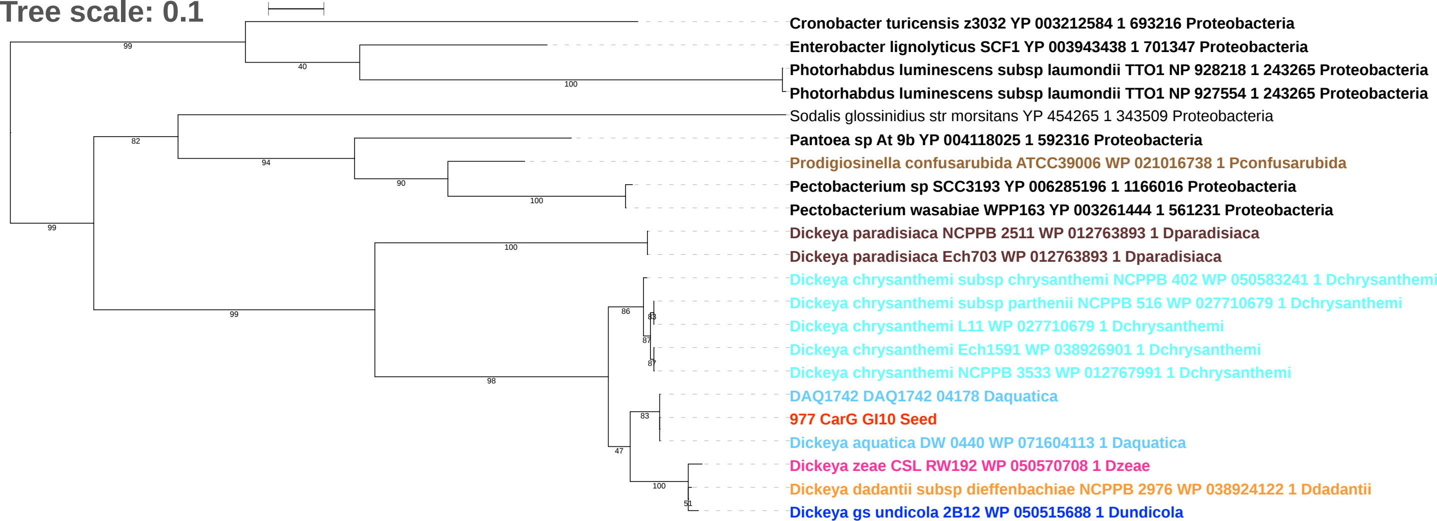

Tree scale: 0.1
Cronobacter turicensis z3032 YP 003212584 1 693216 Proteobacteria
Enterobacter lignolyticus SCF1 YP 003943438 1 701347 Proteobacteria
Photorhabdus luminescens subsp laumondii TTO1 NP 928218 1 243265 Proteobacteria
Photorhabdus luminescens subsp laumondii TTO1 NP 927554 1 243265 Proteobacteria
Sodalis glossinidius str morsitans YP 454265 1 343509 Proteobacteria
Pantoea sp At 9b YP 004118025 1 592316 Proteobacteria
Prodigiosinella confusarubida ATCC39006 WP 021016738 1 Pconfusarubida
Pectobacterium sp SCC3193 YP 006285196 1 1166016 Proteobacteria
Pectobacterium wasabiae WPP163 YP 003261444 1 561231 Proteobacteria
Dickeya paradisiaca NCPPB 2511 WP 012763893 1 Dparadisiaca
Dickeya paradisiaca Ech703 WP 012763893 1 Dparadisiaca
Dickeya chrysanthemi subsp chrysanthemi NCPPB 402 WP 050583241 1 Dchrysanthemi
Dickeya chrysanthemi subsp parthenii NCPPB 516 WP 027710679 1 Dchrysanthemi
Dickeya chrysanthemi L11 WP 027710679 1 Dchrysanthemi
Dickeya chrysanthemi Ech1591 WP 038926901 1 Dchrysanthemi
Dickeya chrysanthemi NCPPB 3533 WP 012767991 1 Dchrysanthemi
DAQ1742 DAQ1742 04178 Daquatica
977 CarG GI10 Seed
Dickeya aquatica DW 0440 WP 071604113 1 Daquatica
Dickeya zeae CSL RW192 WP 050570708 1 Dzeae
Dickeya dadantii subsp dieffenbachiae NCPPB 2976 WP 038924122 1 Ddadantii
Dickeya gs undicola 2B12 WP 050515688 1 Dundicola
99
40
100
82
94
90
100
99
100
86
83
87
87
99
98
83
47
100
51

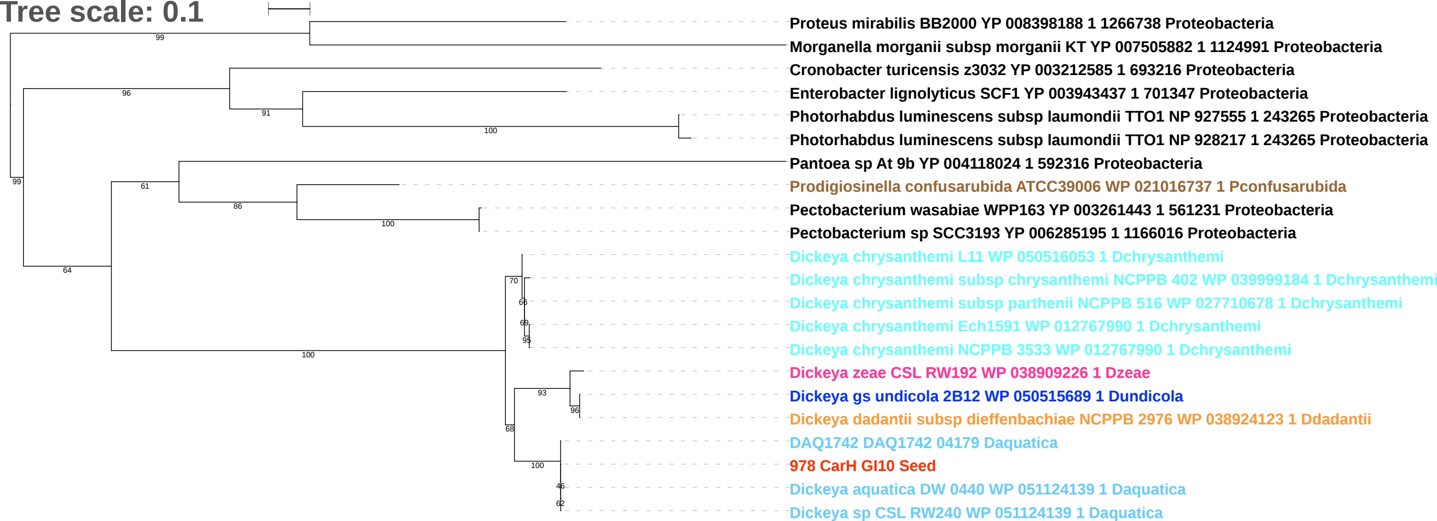
Tree scale: 0.1
Proteus mirabilis BB2000 YP 008398188 1 1266738 Proteobacteria
Morganella morganii subsp morganii KT YP 007505882 1 1124991 Proteobacteria
Cronobacter turicensis z3032 YP 003212585 1 693216 Proteobacteria
Enterobacter lignolyticus SCF1 YP 003943437 1 701347 Proteobacteria
Photorhabdus luminescens subsp laumondii TTO1 NP 927555 1 243265 Proteobacteria
Photorhabdus luminescens subsp laumondii TTO1 NP 928217 1 243265 Proteobacteria
Pantoea sp At 9b YP 004118024 1 592316 Proteobacteria
Prodigiosinella confusarubida ATCC39006 WP 021016737 1 Pconfusarubida
Pectobacterium wasabiae WPP163 YP 003261443 1 561231 Proteobacteria
Pectobacterium sp SCC3193 YP 006285195 1 1166016 Proteobacteria
Dickeya chrysanthemi L11 WP 050516053 1 Dchrysanthemi
Dickeya chrysanthemi subsp chrysanthemi NCPPB 402 WP 039999184 1 Dchrysanthemi
Dickeya chrysanthemi subsp parthenii NCPPB 516 WP 027710678 1 Dchrysanthemi
Dickeya chrysanthemi Ech1591 WP 012767990 1 Dchrysanthemi
Dickeya chrysanthemi NCPPB 3533 WP 012767990 1 Dchrysanthemi
Dickeya zeae CSL RW192 WP 038909226 1 Dzeae
Dickeya gs undicola 2B12 WP 050515689 1 Dundicola
Dickeya dadantii subsp dieffenbachiae NCPPB 2976 WP 038924123 1 Ddadantii
DAQ1742 DAQ1742 04179 Daquatica
978 CarH GI10 Seed
Dickeya aquatica DW 0440 WP 051124139 1 Daquatica
Dickeya sp CSL RW240 WP 051124139 1 Daquatica
